# Distinct roles for MNK1 and MNK2 in social and cognitive behavior through kinase-specific regulation of the synaptic proteome and phosphoproteome

**DOI:** 10.1101/2024.12.18.628637

**Authors:** Rosalba Olga Proce, Erika Uddström, Maria Steinecker, Souhaila Wüsthoff, Luiz Gustavo Teixeira Alves, Oliver Popp, Katie Maxwell, Lucie Hortmann, Philipp Mertins, Markus Landthaler, Daria Bunina, Hanna Hörnberg

**Affiliations:** Molecular and cellular basis of behavior, Max Delbrück Center for Molecular Medicine in the Helmholtz Association (MDC), Berlin, Germany; Systems biology of cardiovascular and neuronal pathologies, Max Delbrück Center for Molecular Medicine in the Helmholtz Association (MDC), Berlin, Germany; RNA Biology and Posttranscriptional Regulation, Max Delbrück Center for Molecular Medicine in the Helmholtz Association (MDC), Berlin, Germany; Proteomics, Max Delbrück Center for Molecular Medicine in the Helmholtz Association (MDC), Berlin, Germany; Institute for Biology, Humboldt University, Berlin, Germany

**Author notes:** Corresponding author: Hanna Hörnberg, PhD, Cellular and molecular mechanisms of behavior, Max Delbrück Center for Molecular Medicine in the Helmholtz association (MDC), Robert-Rössle-Straße, 10 13125 Berlin.

## Abstract

Local mRNA translation is required for adaptive changes in the synaptic proteome. The mitogen-activated protein kinase (MAPK) interacting protein kinases 1 and 2 (MNK1 and MNK2) have emerged as key regulators of neuronal translation, primarily through phosphorylation of the eukaryotic initiation factor 4E (eIF4E). The therapeutic benefits of targeting the MNKs are being investigated for nervous system conditions and disorders that affect translation, including autism, pain, and cancer. However, it is still unclear if MNK1 and MNK2 regulate distinct aspects of neuronal translation and how the activity of each kinase is associated with the synaptic and behavioral features linked to MNK signaling. To examine the individual neurobiological functions of each kinase, we used knockout mice lacking either MNK1 or MNK2. We found that the knockout of MNK1 and MNK2 leads to different social and cognitive behavioral profiles and distinct alterations of the cortical synaptic proteome, transcriptome, and phosphoproteome. Loss of MNK1 was associated with an increase in ribosomal protein expression, whereas deletion of MNK2 decreased the expression and phosphorylation of synaptic proteins. Together, our findings provide evidence for a high degree of functional specialization of the MNKs in synaptic compartments and suggest that pharmacological inhibition of individual MNK may provide more specific targets for neurological disorders.

## Introduction

Precise regulation of mRNA translation allows cells to modulate protein expression levels globally and to restrict protein localization to a specific time or cellular compartment[1]. In neurons, spatiotemporal regulation of mRNA translation is necessary for learning and memory. Part of this translation occurs at the synapse, where local translation contributes to adaptive changes in the synaptic proteome[2–6]. As synapses are space-restricted and reside far away from the soma, recent evidence suggests that translation may be regulated differently at the synapse compared to the soma[1, 7].

Translation of most mRNAs is regulated at the step of initiation by the formation of a pre-initiation complex that involves recognition and binding of the m^7^GTP cap structure by the cap-binding eukaryotic initiation factor 4E (eIF4E)[8]. eIF4E activity is modulated by phosphorylation of a single residue, Ser209, by the serine/threonine kinases mitogen-activated protein kinase (MAPK) interacting protein kinases 1 and 2 (MNK 1 and 2)[9, 10]. These non-essential kinases are mainly activated by p38 and the extracellular signal-regulated kinase (ERK)/MAPK pathways, and eIF4E is one of the few MNK targets validated *in vivo*[9, 11]. Both MNK1 and MNK2 are expressed in the brain[12], and the MNKs have been implicated in a range of adaptive behaviors, including memory formation, social behavior, and depressive-like behavior[13–16]. Most of these functions have been attributed to MNKs phosphorylation of eIF4E. For example, MNKs regulate long-term potentiation (LTP) in the dental gyrus via eIF4E phosphorylation-mediated release of cytoplasmic FMR1-interacting protein 1 (CYFIP1) from the cap[17–19]. Recently, MNKs were found to also regulate neuronal translation via phosphorylation of the brain-specific synaptic Ras GTPase-activating protein 1 (Syngap1), which acts upstream of mTOR-dependent translation[15]. MNK-Syngap1 signaling modulates hippocampal learning and memory independent of eIF4E phosphorylation, indicating that the MNKs can regulate synaptic plasticity via distinct mechanisms that lead to the translation of largely independent pools of mRNAs[15, 16].

Because of their status as the sole kinases phosphorylating eIF4E, there has been considerable interest in targeting the MNKs for the treatment of nervous system disorders that affect translation, including neurodevelopmental conditions and neuropathic pain[10, 13, 14, 20, 21]. To further develop the MNKs as drug targets, an unanswered question is to what extent there is any functional specialization of the different MNK proteins in the nervous system. Although structurally similar, MNK1 and MNK2 are known to differ in their activity and have distinct roles in some biological processes[22–25]. While both kinases phosphorylate eIF4E under basal conditions, studies suggest that MNK1 is the main kinase that regulates eIF4E phosphorylation in response to signaling, whereas MNK2 is constitutively active[11, 17].

In addition, MNK2, but not MNK1, can modulate translation via inhibition of eIF4G activation and crosstalk with the mTOR pathway[26–30]. Studies using single-knockout mice suggest that MNK1 may be the main isoform in the brain[17, 18]. However, transcriptomic data shows that MNK1 and MNK2 are broadly expressed in the mouse and human brain[12, 31], and data from ribosome sequencing suggests that translation of MNK1 and MNK2 is enriched in the neuropil compared to the somata of neurons[32]. Whether there is any functional specialization of the MNK proteins in neurons and synapses has yet to be established.

Here, we set out to characterize the individual neurobiological functions of MNK1 and MNK2. We find that mice lacking MNK1 or MNK2 have distinct behavioral profiles with different social and cognitive phenotypes. Deletion of MNK1 and MNK2 have strikingly different effects on the synaptic proteome, phosphoproteome, and transcriptome, suggesting that each kinase has separate functions at the synapse. Overall, our finding suggests that individual targeting of the MNKs should be considered when developing therapeutic approaches for disorders affecting the nervous system.

## Results

### Deletion of MNK1 or MNK2 causes specific behavioral phenotypes

To examine the individual functions of the MNKs, we took advantage of knockout mice lacking either MNK1 or MNK2 and examined their behavioral phenotype. We performed several behavioral tests covering social, cognitive, and exploratory behaviors (Figure 1A). Social interaction, habituation, and novelty preference were examined using the five-trial social habituation/recognition test and the social olfaction test. In the social habituation/recognition test, an unfamiliar sex- and age-matched conspecific is presented to the test mouse for four trials to examine social interest and habituation. On the fifth trial, a novel mouse is introduced to examine social recognition. We found that mice lacking MNK1 exhibited a reduced interest in the novel mouse on the fifth trial compared to wild-type mice (Figure 1B-C). MNK1^KO^ mice also showed a tendency to reduced interest in social odors, but not nonsocial odors, in the social odor recognition task (Figure 1D). MNK2^KO^ mice did not differ in their total social interaction, habituation, or social recognition compared to wild-type mice (Figure 1B-D). We next assessed object interaction and memory in two separate tests: the five-trial object habituation/recognition task and the novel object task. MNK1^KO^ mice had significantly reduced interaction with the first object in the habituation/recognition task compared to wild-type mice, and significantly reduced interest in the novel object presented on the 5^th^ trial (Figure 1E-F). The reduced object recognition in MNK1^KO^ mice was also seen in the novel object task (Figure 1G-H). Interestingly, mice lacking MNK2 showed increased interest in the novel object compared to wild-type mice in the object habituation/recognition task (Figure 1E-F). In the open field test, both MNK1 and MNK2 knockout mice showed a slight decrease in distance traveled compared to wild-type mice, which was most pronounced during the first minutes of the test, but no change in time spent in the center (Figure 1I-J). Mice from both sexes were used, and we observed no main effect of sex in any test (Supplementary Figure 1).

**Figure 1.**
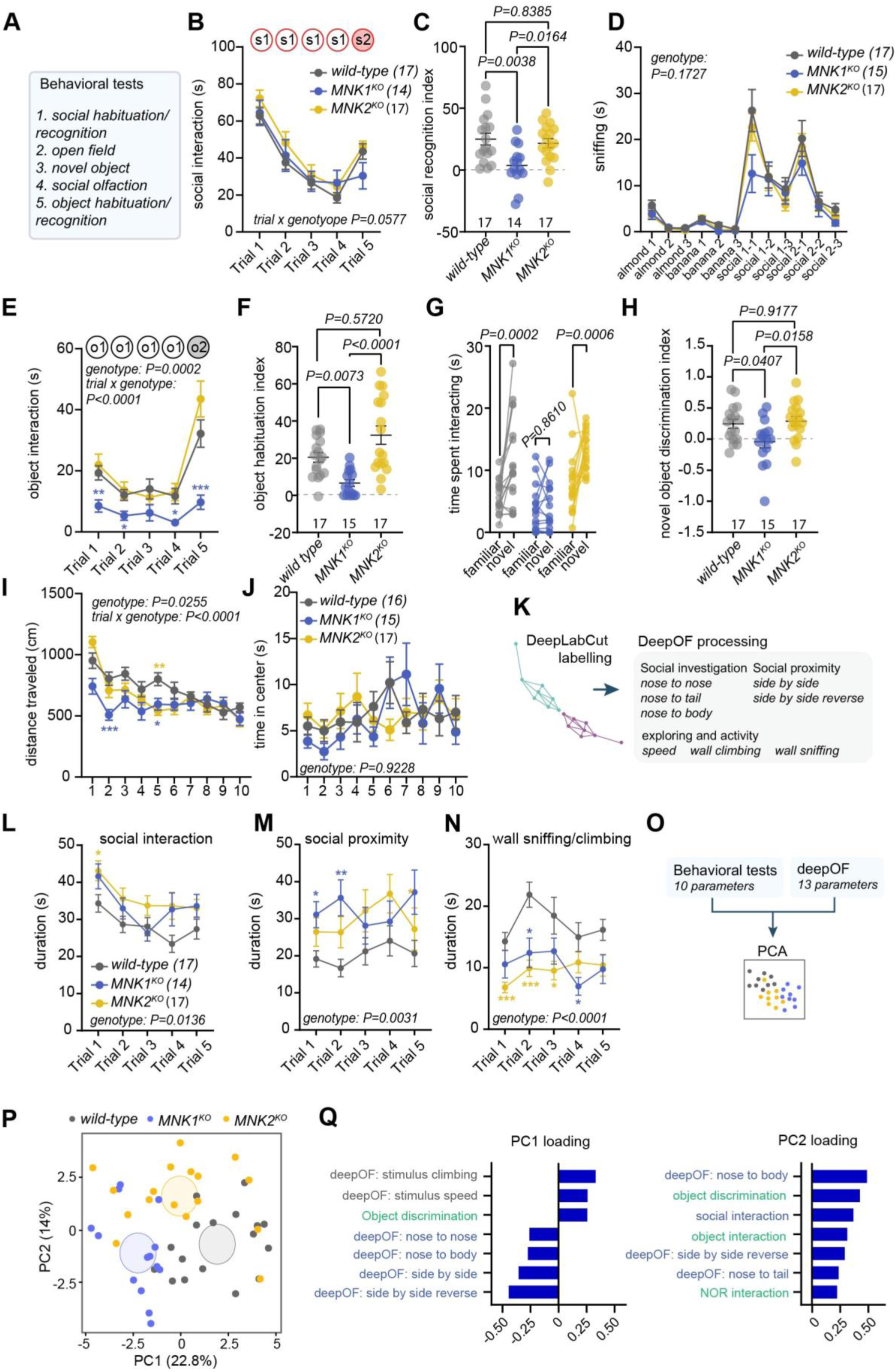
Mice lacking MNK1 or MNK2 have distinct behavioral profiles. (A) Order of behavioral tests. (B-C) Mean social interaction time (B) and recognition index (C) in the social habituation/recognition test for wild-type, MNK1^KO,^ and MNK2^KO^ mice. s1=first social stimulus, s2=second social stimulus. (D) Time spent sniffing each odor in the social olfaction habituation test. (E-F) Mean object interaction time (E) and recognition index (F) in the object habituation/recognition test. o1=first object stimulus, o2=second object stimulus. (G-H) Time spent interacting with the familiar and novel object (G) and novel object discrimination index (H) in the novel object recognition test. (I-J) Distance traveled (I) and time spent in center (J) per minute in the open field test. (K) Schematic of deep social profiling using DeepLabCut and deepOF. (L-N). Time spent in social interaction (sum of nose to nose, nose to tail, nose to body, L), social proximity (sum of side by side by side and side by side reverse, M), and wall climbing and exploring (sum of wall climbing and wall sniffing, N) in the five-trial social habituation/recognition test as quantified by deepOF. (O) Schematic of the outcome measures included in the PCA analysis. (P) PCA of all 23 behavioral parameters and (Q) loading for PC1 and PC2. Error bars show s.e.m. P values: *<0.5, **<0.01, ***<0.001 relative to wild-type. Significance was determined by two-way ANOVA followed by Tukey’s post-hoc test for B, D, E, and G, one-way ANOVA followed by Tukey’s post-hoc test for C, H, Kruskal-Wallis test followed by Dunn’s multiple comparison test for F, mixed-effect model followed by Tukey’s post-hoc test for I, J, L, M, N.

While examining the videos from the social habituation/recognition test, we noticed differences in the types of social interactions observed in the MNK1 and MNK2 knockout mice. To quantify this further, we used deepOF, an open-source system for deep social phenotyping of two freely interacting mice[34]. We first trained DeepLabCut[45] to recognize both animals during the two-minute-long videos and then analyzed all trials using the deepOF supervised pipeline (Figure 1K). We found a slight but significantly higher overall social interaction in the MNK2^KO^ mice primarily driven by increased nose-to-body contact (Figure 1L, Supplementary Figure 2A-C). Both MNK1^KO^ and MNK2^KO^ mice spent significantly more time in close proximity to the stimulus mice without directly interacting (Figure 1M, Supplementary Figure 2D-E), and significantly less time exploring the edges of the cage (Figure 1N, Supplementary Figure 2F-G). There was no difference in the social behavior of the stimulus animal or the overall speed of the MNK1^KO^ and MNK2^KO^ mice during the test (Supplementary Figure 2H-M). Taken together, this detailed social behavioral classification captures robust differences in specific types of social behaviors in both MNK1 and MNK2 knockout mice, suggesting that loss of either kinase affects the animal’s social profile.

To capture the full behavioral profile of each animal, we combined information from all behavioral tests and analyzed the parameters using a PCA. We collected 10 output measures from the classical tests and 13 from the deepOF analysis (Figure 1O-P, all parameters listed in Supplementary Table 1). The first two components explained 36.7% of the variance. The first principal component (PC1) separated all three genotypes and was mainly driven by social behaviors, whereas the second component primarily correlated with object interaction and recognition (Supplementary Figure 2N). Specifically, across all tests, MNK2^KO^ mice engaged in more social behaviors, whereas MNK1^KO^ mice had reduced object interest and recognition and spent more time in close social proximity (Figure 1Q). These results suggest that loss of MNK1 is primarily responsible for the cognitive phenotypes seen in the MNK1/2^DKO^ mice[15], whereas both MNK1 and MNK2 affect social behaviors and exploratory behavior in different ways.

### MNK1 and MNK2 are expressed in overlapping neuronal cell types in cortex and hippocampus

A possible explanation for the different behavioral phenotypes in the MNK1^KO^ and MNK2^KO^ mice is that the MNKs are expressed in distinct neuronal populations. Transcriptomic studies suggest that MNK1 and MNK2 are expressed throughout the brain in mice, humans, and pigs[12, 31]. To further characterize their expression, we took advantage of published RNA-sequencing data from ribosome-bound mRNAs (RiboTRAP) from multiple neuronal cell types in the cortex and hippocampus[46]. *Mknk1* and *Mknk2* mRNAs were broadly expressed throughout all examined cell types, with slightly higher expression of *Mknk2* (Figure 2A-B)[46], consistent with previous gene expression data[12, 31]. To further assess MNK brain expression, we performed fluorescent in situ hybridization (FiSH) using probes against *Mknk1* and *Mknk2* in cortex and hippocampus. In the cortex, we found that *Mknk1* and *Mknk2* were expressed in almost all neurons, with most excitatory (*vGlut1* positive) and inhibitory (*Gad1* positive) neurons expressing both *Mknk1* and *Mknk2* (Figure 2C-D). A similar expression pattern was found in hippocampus (Figure 2E-F). Taken together, these results suggest that MNK1 and MNK2 are expressed in largely overlapping neuronal populations throughout cortex and hippocampus in mice.

**Figure 2.**
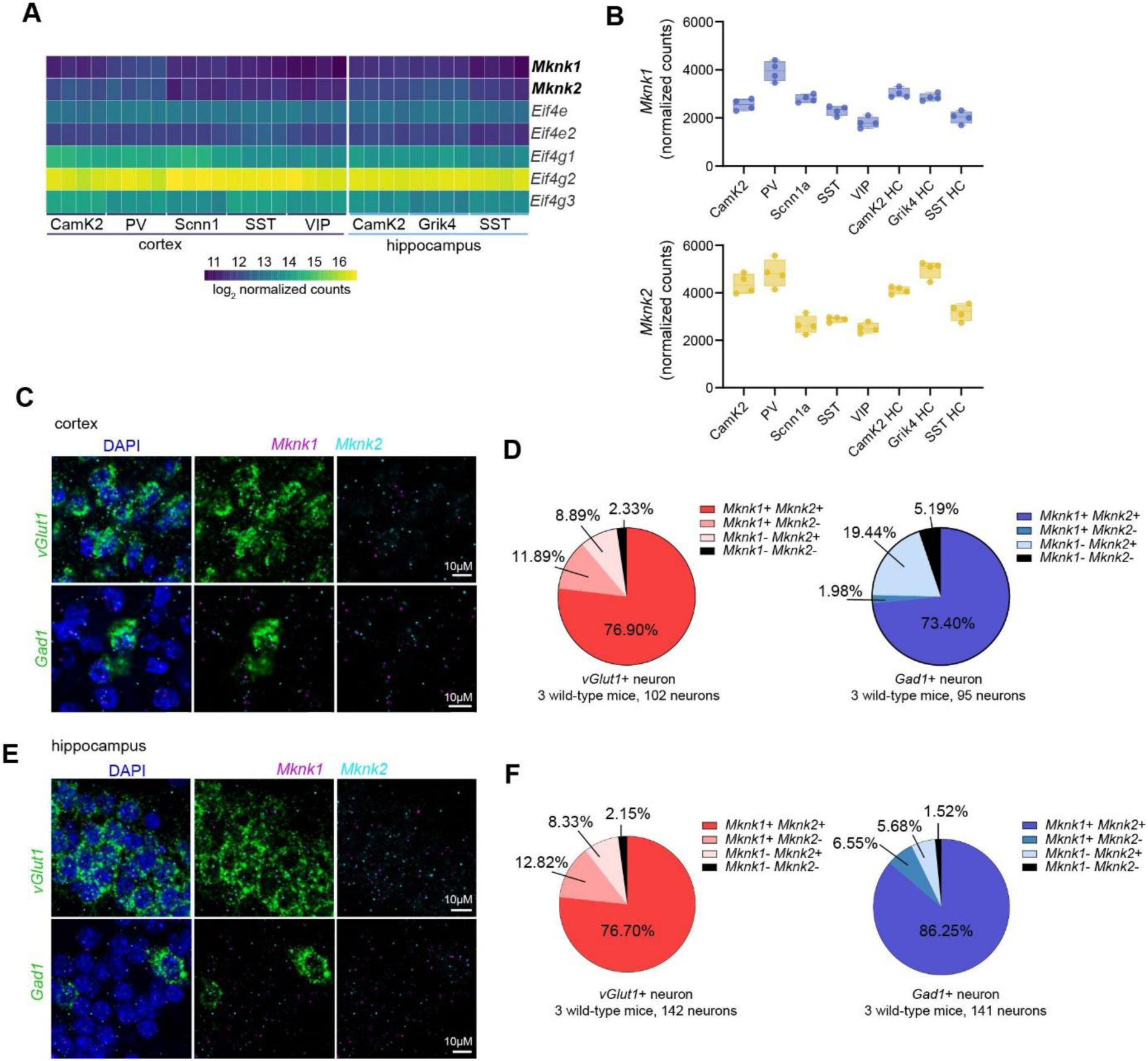
MNK1 and MNK2 have overlapping expressing patterns in cortex and hippocampus. (A-B) Heatmap (A) and boxplot (B) of *Mknk1* and *Mknk2* across different neuronal populations in cortex and hippocampus. *Eif4e* and *Eif4g* isoforms are shown for comparison in the heatmap. The mRNA expression was measured using Ribo-TRAP sequencing from Furlanis et al. (2019). (C, E) Representative fluorescent in situ hybridization (FiSH) images in cortex (C) and hippocampus (E) using probes for *vGlut1* (green), *Gad1* (green), *Mknk1* (magenta), and *Mknk2* (cyan). DAPI is in blue. (D, F) Pie-chart showing the distribution of *Mknk1* and *Mknk2* in *vGlut1* (left) and *Gad1* (right) positive neurons in cortex (D) and (F) hippocampus. The total number of neurons is listed under each pie chart. The percentage of each neuronal population is an average of three mice. Camk2= calcium/calmodulin-dependent protein kinase II positive neurons, PV= Parvalbumin-positive interneurons, Scnn1a= sodium channel, nonvoltage-gated 1α positive spiny stellate and star pyramid layer 4 (L4) neurons, SST= somatostatin-positive interneurons, VIP= vasointestinal peptide-positive interneurons, Grik4=glutamate receptor, ionotropic, kainate 4-positive interneurons, HC=hippocampus.

### Proteomic analysis of MNK1 and MNK2 knockout mice show different effects on the synaptic proteome

The specific behavioral phenotypes of MNK1 and MNK2 knockout mice suggest that the MNKs may regulate distinct aspects of neuronal translation. To start investigating how MNK1 and MNK2 affect protein expression, we performed tandem mass tag (TMT)-based mass spectrometry on the whole homogenate and isolated synaptoneurosomes from cortex of MNK1^KO^, MNK2^KO^, and MNK1/2^DKO^ mice compared to wild-type mice (Figure 3A). In cortex, we found that loss of either MNK1, MNK2, or both kinases had a similar effect on the proteome, with a significant correlation in protein log fold change (logFC) relative to wild-type between all genotypes (Figure 3B). Consistent with the proteomic profile of mice treated with an MNK inhibitor[14], few significantly differentially expressed proteins were identified in the cortical proteome for any genotype (adjusted p-value<0.05, Supplementary Table 2). We used gene set enrichment analysis (GSEA) to identify molecular pathways with altered expression between genotypes (FDR<0.25, Supplementary Table 2). Overall, loss of MNK1 or MNK2 affected similar pathways in cortex (Figure 3C-D, Supplementary Table 2). Hierarchical clustering identified two main clusters, with cluster 1 enriched in pathways related to translation and cluster 2 enriched in pathways associated with the extracellular matrix (Figure 3D, Supplementary Table 2), consistent with previous studies using MNK1/2^DKO^ mice[15].

**Figure 3.**
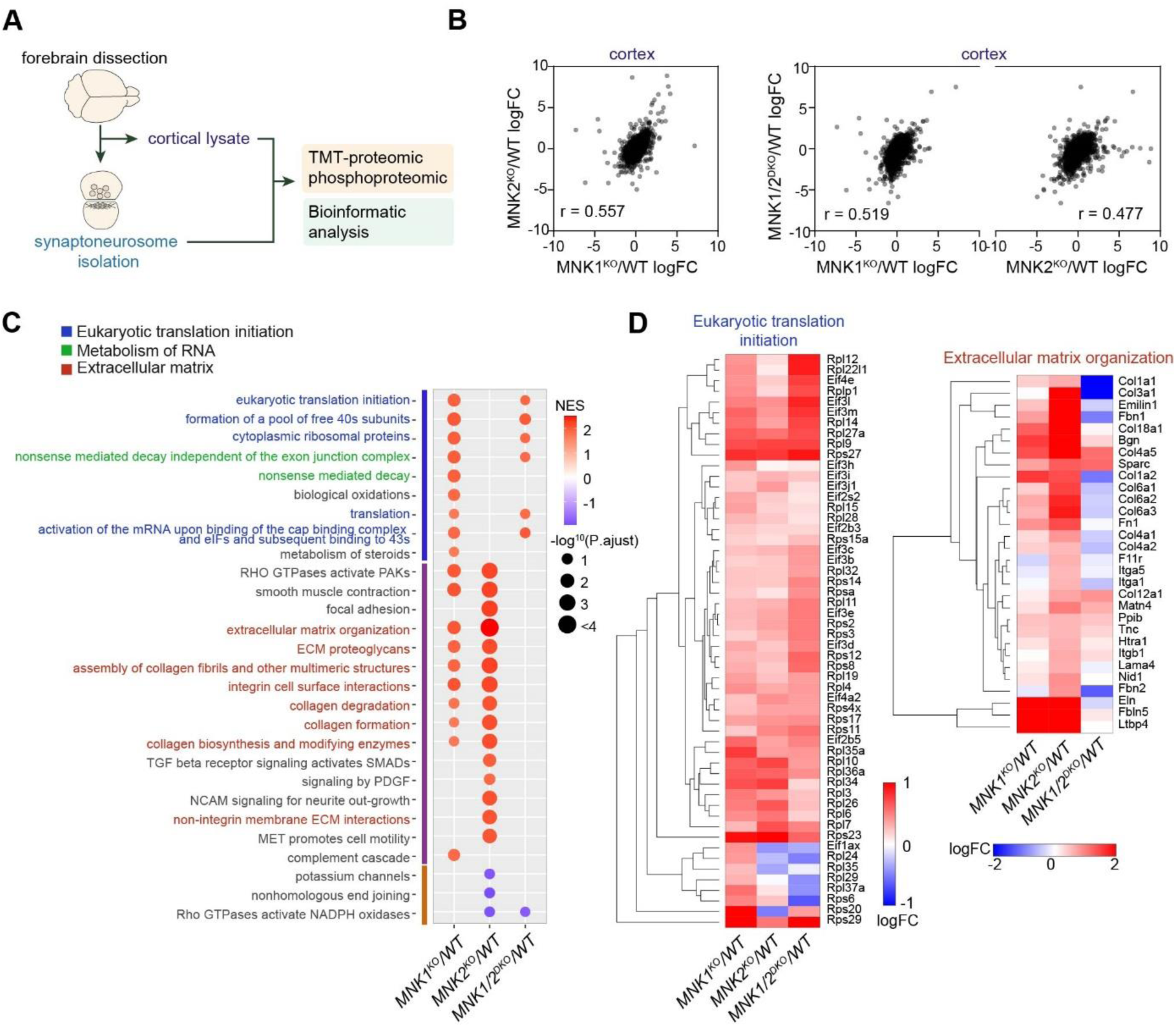
Proteomic analysis of MNK1 and MNK2 knockout mice. (A) Schematic of the experimental procedure. (B) Pearson correlation of the cortical proteome Log fold change (LogFC, relative to wild-type) in MNK1^KO^, MNK2^KO^, and MNK1/2^DKO^. (C) Dot plot of selected significant enriched (FDR<0.25) canonical pathways of Gene set enrichment analysis (GSEA) comparing MNK1^KO^, MNK2^KO,^ and MNK1/2^DKO^ cortical proteome relative to wild-type. Only significant pathways are shown per genotype. (D) Heatmap showing logFC of the core enrichment proteins in the eukaryotic translation initiation (left) and extracellular matrix organization (right) signaling pathways. WT= wild-type, NES= normalized enrichment score.

Synaptic dysfunction is a hallmark of many neurodevelopmental conditions. Synaptic activity can induce eIF4E phosphorylation, and MNK1 has previously been implicated in synaptic translation[15, 17, 18]. Therefore, we next examined how knockout of the MNKs affected the synaptic proteome. We isolated synaptoneurosomes using a protocol that enriches both the pre-and postsynaptic compartments[5, 47], and confirmed enrichment by comparing the proteome of cortical homogenate to the synaptoneurosome fraction in wild-type animals (Supplementary Figure 3). In stark contrast to the cortical proteome, comparing the protein LFC relative to wild-type in the synaptoneurosome fractions between the different genotypes showed a very low correlation between MNK1^KO^ and MNK2^KO^ mice (Figure 4A, Supplementary Figure 4A). The protein expression of MNK1/2^DKO^ mice was highly correlated with mice lacking MNK2 but not MNK1, suggesting that the majority of altered protein expression in synaptoneurosomes from MNK1/2^DKO^ mice is driven by the loss of MNK2. Of note, the low correlation between MNK1^KO^ and MNK2^KO^ mice synaptoneurosomes appears to be driven by MNK2 deletion differentially affecting the cortical and synaptoneurosome proteome, whereas deletion of MNK1 causes similar proteomic changes in both fractions (Supplementary Figure 4A).

**Figure 4.**
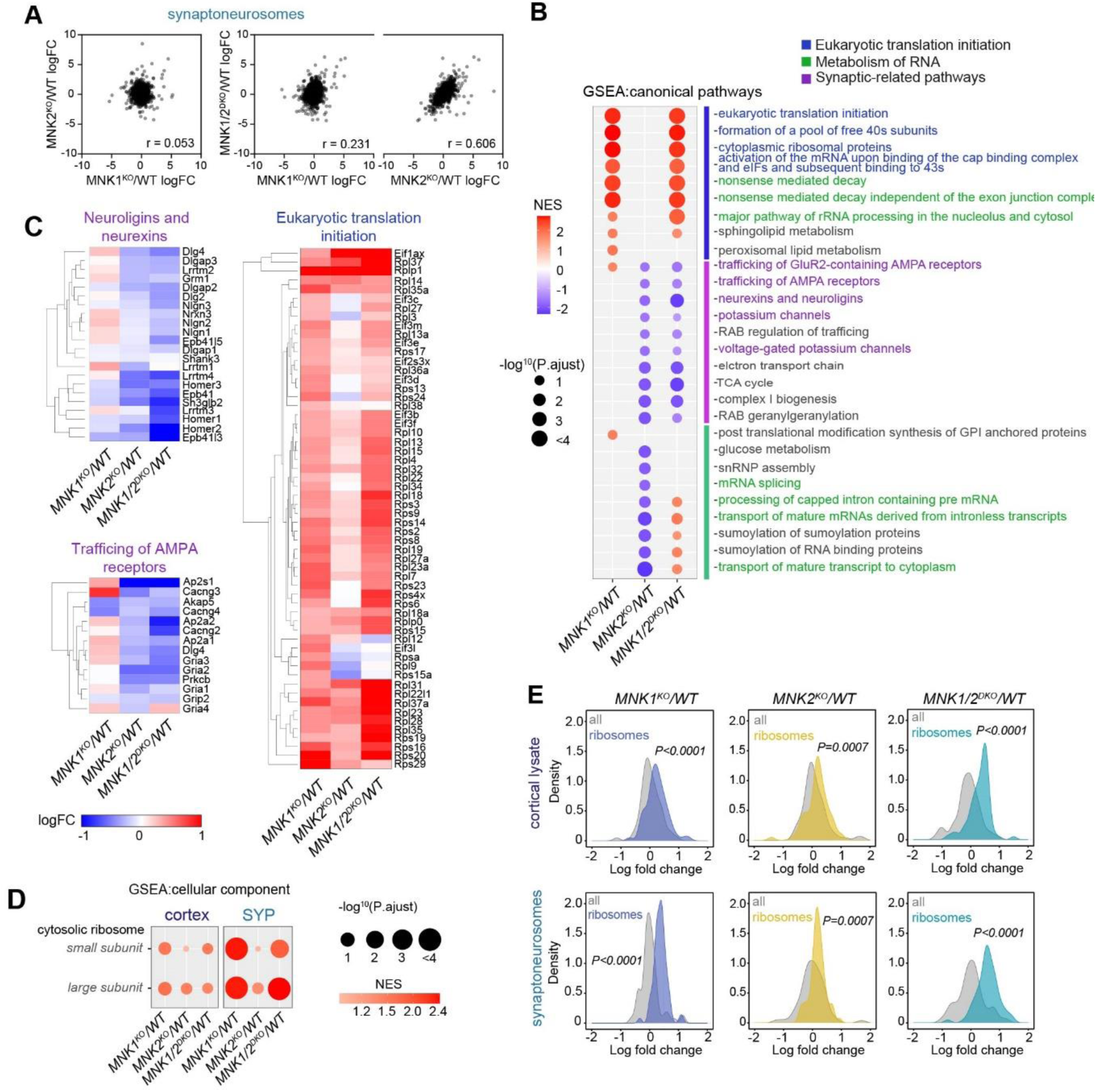
Distinct contribution of MNK1 and MNK2 to the synaptic proteome. (A) Pearson correlation of the synaptoneurosome proteome logFC relative to wild-type in MNK1^KO^ versus MNK2^KO^ (left), and of MNK1^KO^ and MNK2^KO^ versus MNK1/2^DKO^ (right). (B) Dot plot of selected significantly enriched (FDR<0.25) canonical pathways of GSEA comparing MNK1^KO^, MNK2^KO^, and MNK1/2^DKO^ synaptoneurosome proteome to wild-type. Only significant pathways are shown for each genotype. (C) Heatmap showing logFC relative to wild-type of the core enrichment proteins in the neurexins and neuroligins, trafficking of AMPA receptors, and eukaryotic translation initiation signaling pathways. (D) GSEA of cytosolic ribosome cellular component GO term enrichment for MNK1^KO^, MNK2^KO,^ and MNK1/2^DKO^ relative to wild-type. (E) Density plots of logFC ribosomal protein abundance compared to all proteins in the cortical (top) and synaptoneurosome (bottom) proteome for all genotypes relative to wild-type. P-values in E were calculated using two-sided Kolmogorov-Smirnov test.

GSEA identified three major clusters of pathways with altered abundance (Figure 4B). The first cluster consisted of pathways related to translation and RNA metabolism. These were highly enriched in MNK1^KO^ mice synaptoneurosomes, whereas synapse and RNA metabolism pathways in the second and third clusters were de-enriched in MNK2^KO^ mice (Figure 4B-C, Supplementary Table 3). Examination of the core proteins accounting for the enrichment in the translation and RNA metabolism pathways in cluster one identified ribosomal proteins as the main group of protein overexpressed in MNK1^KO^ mice (Figure 3C), and a comparison between cortex and synaptoneurosomes showed that this enrichment was more pronounced at the synapse (Figure 4D, Supplementary Table 3). To examine if this enrichment affected ribosomal proteins as a group or was specific to a subset of ribosomal proteins, we analyzed the expression of all ribosomal subunits identified in our dataset. We found a significant upregulation of both small and large ribosomal subunits that was more pronounced in synaptoneurosomes from mice lacking MNK1 (Figure 4E). Increased expression of ribosomal proteins was also found in wild-type mice treated with an MNK inhibitor (Supplementary Figure 4B)[14]. Together, these results strongly suggest that MNK1 inhibition or deletion elevates ribosomal protein expression.

### Comparison of mRNA and protein reveals translational and transcriptional upregulation of ribosomal genes in MNK1^KO^ mice

The differences in protein expression at the synapse of MNK1^KO^ and MNK2^KO^ mice could be caused by an altered abundance of local mRNAs, post-transcriptional modification, or changes in transport. To examine if mRNA abundance contributed to the proteomic differences, we performed RNA sequencing (RNA-Seq) on isolated cortical synaptoneurosomes from MNK1^KO^, MNK2^KO^, and wild-type mice. We identified 621 mRNAs significantly altered in synaptoneurosomes from MNK1^KO^ mice and 118 from MNK2^KO^ mice (Supplementary Figure 5A-B, Supplementary Table 4), suggesting that loss of MNK1 has a larger effect on the synapse-enriched transcriptome compared to loss of MNK2. In agreement with the proteomic dataset, the mRNA log fold change relative to wild-type showed a low correlation between MNK1^KO^ and MNK2^KO^ mice (Supplementary Figure 5C), suggesting that loss of MNK1 or MNK2 also has a largely distinct effect on the synaptic transcriptome.

To examine the differences between protein and mRNA expression, we performed a multi-omics integration using the log fold change of genes identified in all datasets as input and K-means clustering to identify co-regulated groups of mRNAs and proteins. Unsupervised hierarchical clustering showed that mRNA expression changes were more similar in both knockouts compared to their proteome changes. The MNK2^KO^ proteome clustering separately from other samples, pointing towards a stronger effect of post-transcriptional regulation in MNK2^KO^ mice (Figure 5A). In general, the 20 identified clusters showed low co-regulation between mRNA and protein for both genotypes, with most clusters showing anti-correlated expression for mRNAs and proteins (Figure 5A, Supplementary Table 4). We focused on a subset of clusters that were strongly correlated or anti-correlated for each genotype and performed gene-ontology (GO) analysis on each cluster (Figure 5B). Cluster 2 and 11 showed opposing mRNA expression profiles between MNK1^KO^ and MNK2^KO^ mice, with decreased mRNA expression in MNK1^KO^ mice but increased in MNK2^KO^ mice. Pathways related to behavior, cognition and learning, axon guidance, and ion transport were significantly enriched in these clusters. The change in mRNA expression was anti-correlated with protein expression, suggesting that the effect on protein expression is caused by altered translational regulation rather than mRNA abundance (Figure 5A-B, Supplementary Figure 5D).

**Figure 5.**
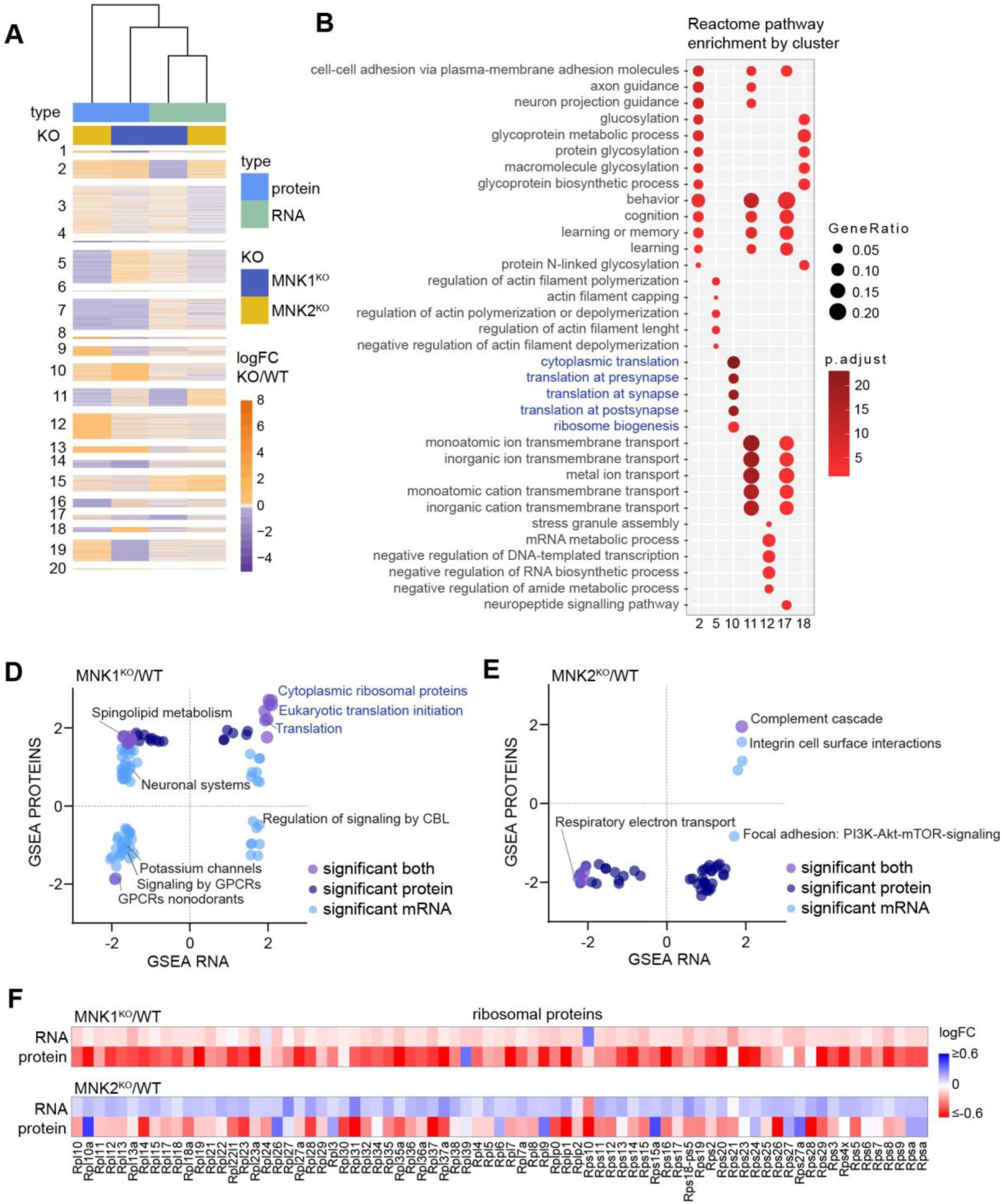
MNK1 and MNK2 knockout have different effects on the synaptic transcriptome. (A) Heatmap showing K-means clustering of protein and mRNA logFC values in MNK1^KO^ and MNK2^KO^ mice relative to wild-type. A total of 8294 genes are shown that are detected on both mRNA and protein levels. (B) Gene ontology pathway enrichment analysis (from the Reactome database) of the genes in each cluster in (A) using all detected genes as a background. Top five significantly overrepresented Reactome pathways per cluster is shown, sorted by the adjusted p-value. (C-D) Overlap of GSEA canonical pathways terms significant in either the RNA-Seq or proteomic datasets or both for synaptoneurosome from (C) MNK1 and (D) MNK2 knockout mice. (E) Heatmap showing logFC of ribosomal subunit mRNA and protein from synaptoneurosomes from MNK1^KO^ mice (top) and MNK2^KO^ mice (bottom), relative to wild-type mice.

The multi-omic integration identified one cluster (cluster 10) with strongly increased protein expression in MNK1^KO^ mice (Figure 5A). GO analysis showed an enrichment of categories involved with synaptic translation and ribosome biogenesis in this cluster (Figure 5B), consistent with our proteomic GSEA results. To further compare the transcriptomic and proteomic datasets, we performed GSEA on the transcriptomic dataset and compared gene sets significantly altered in either the transcriptome, proteome, or both (Figure 5C-D, Supplementary Table 4). Interestingly, pathways related to translation were highly enriched for both protein and mRNA in synaptoneurosomes from MNK1^KO^ mice. In agreement with the proteomic dataset, the core enriched genes in these pathways were primarily ribosomal proteins, and analysis of individual ribosomal subunits confirmed an increase in mRNA and protein expression for almost all ribosomal subunits in synaptoneurosomes from MNK1^KO^ mice (Figure 5E). These results suggest that although the majority of changes in protein expression in both MNK1 and MNK2 knockout mice is driven by post-transcriptional regulation, the increased expression of ribosomal proteins in MNK1^KO^ mice can be at least partially explained by an increase in ribosomal subunit mRNAs at the synapse.

### MNK1 and MNK2 have specific effects on the neuronal phosphoproteome

The behavioral, proteomic, and transcriptomic datasets suggest that MNK1 and MNK2 have specific functions in the nervous system and control distinct cellular processes at the synapse. To examine potential mechanisms of how the MNKs exert their function, we first focused on the activity of their substrates. The MNKs can regulate neuronal translation via phosphorylation of eIF4E or Syngap1[11, 15](Figure 6A). Mice with a mutation that prevents eIF4E phosphorylation show increased translation of ribosomal proteins[48], suggesting that MNK1-dependent eIF4E phosphorylation may regulate ribosomal protein expression. However, we observed no significant difference in eIF4E phosphorylation in cortical lysate or synaptoneurosomes in MNK1 and MNK2 knockout mice (Figure 6B-C, Supplementary Figure 6A-B). As expected, no eIF4E phosphorylation was seen in the MNK1/2^DKO^ mice (Figure 6B-C, Supplementary Figure 6A-B), and no difference was found in eIF4E protein levels (Supplementary Figure 6C-D). MNK1/2 can phosphorylate Syngap1 on S788, which promotes protein synthesis and increases phosphorylation of ribosomal protein S6 (rpS6)[15]. We found a decrease in rpS6 phosphorylation in cortical lysate in MNK1^KO^ mice and in synaptoneurosomes from both MNK1^KO^ and MNK2^KO^ mice (Figure 6D-E)[15]. Phosphorylation of the mTOR substrate 4E-binding protein1 (4EBP1) was unaffected (Supplementary Figure 6E), and there was no difference in the upstream signaling pathway ERK1/2 (Supplementary Figure 6F-G).

**Figure 6.**
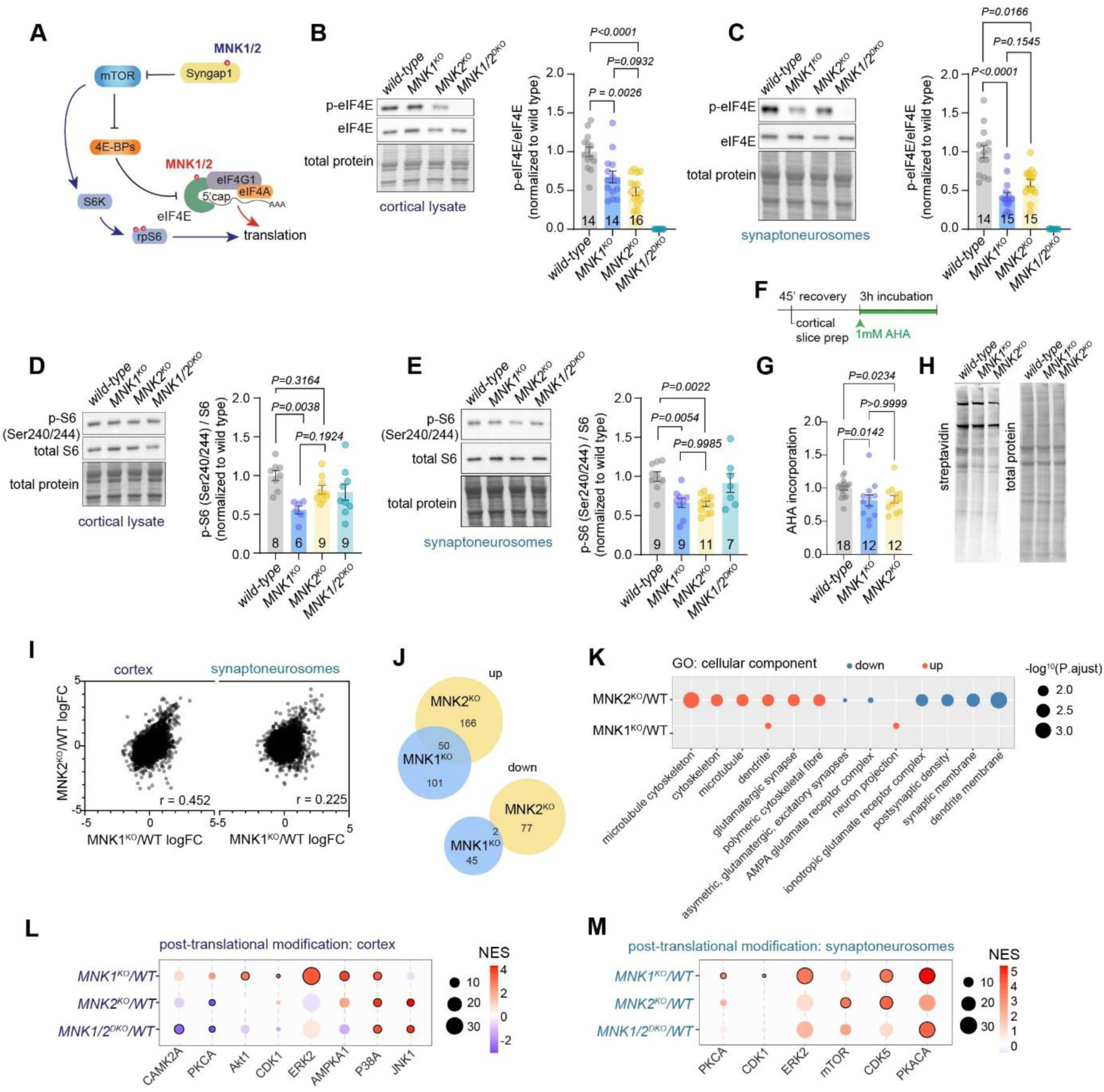
Phosphoproteomic characterization of MNK1 and MNK2 knockout mice. (A) Schematic of MNK1/2 regulation of mRNA translation via phosphorylation of Syngap1 (blue) and eIF4E (red). (B-C) Representative western blot (left) and quantification (right) of p-eIF4E compared to total eIF4E from (B) cortical lysates and (C) synaptoneurosomes from wild-type, MNK1^KO^, and MNK2^KO^ mice. MNK1/2^DKO^ mice are included for validation. (D-E) Representative western blot (left) and quantification (right) of p-rpS6 compared to total rpS6 from (D) cortical lysates and (E) synaptoneurosomes from wild-type, MNK1^KO^, MNK2^KO^, and MNK1/2^DKO^ mice. (F) Timeline of AHA incubation in brain slices. (G) Quantification and (H) representative image of AHA incorporation in cortical brain slices from wild-type, MNK1^KO,^ and MNK2^KO^ mice. (I) Alterations in MNK1^KO^ and MNK2^KO^ cortical and synaptoneurosomes phosphoproteome relative to wild-type. r was determined by Pearson correlation. (J) Venn diagram showing the overlap of phosphosites increased or decreased (>1.2logFC compared to wild-type) in synaptoneurosome phosphoproteome from MNK1^KO^ and MNK2^KO^ mice. (K) Bubble plot of cellular component GO terms enriched in MNK1^KO^ and MNK2^KO^ synaptoneurosome phosphoproteome. Red: increased, blue: decreased. (L-M) Bubble plot of selected phosphosite-specific signatures in cortex (L) and synaptoneurosomes (M) as determined by PTM-SEA. Significantly enriched kinase signatures (FDR<0.05) are marked with black circle, and size corresponds to the number of observed phosphosites. All error bars are s.e.m. Significance was determined by one-way ANOVA followed by Tukey’s multiple comparison test for B, D, E, and Kruskal-Wallis test followed by Dunn’s multiple comparison test for C, G.

We next examined if the rate of protein synthesis was altered in mice lacking MNK1 or MNK2. MNK1/2^DKO^ mice have no change in protein synthesis rate[11], but acute inhibition of the MNKs reduces the rate of protein synthesis in neurons[14]. We used incorporation of the noncanonical amino acid azidohomoalanine (AHA) to examine how the loss of MNK1 or MNK2 affected protein synthesis in cortical brain slices and found a small but significant reduction of translation rate in both MNK1 and MNK2 knockout mice (Figure 6F-H). It is possible that a small but significant decrease in MNK1 protein expression in cortex from MNK2^KO^ mice affects these results (Supplementary Figure 6H-I). None of the antibodies we tested were specific for MNK2, and neither MNK1 nor MNK2 was detected in the proteome from either cortex or synaptoneurosomes. Therefore, we could not clarify whether MNK2 expression was similarly altered in MNK1^KO^ mice.

Next, we performed phosphoproteomics to identify possible differences in other signaling pathways (Supplementary Table 5). Similar to the proteomic dataset, there was a high correlation between the phosphosite abundance changes relative to wild-type between MNK1^KO^ and MNK2^KO^ mice in the cortical homogenate but not in synaptoneurosomes (Figure 6I, Supplementary Figure 7A). To determine which cellular functions were affected by MNK1 or MNK2 deletion, we performed a gene ontology (GO) analysis of increased and decreased phosphosites. Using a cutoff value of LFC<1.2, we found multiple altered phosphosites, most of which were unique for each genotype (Figure 6J, Supplementary Table 5). Interestingly, proteins related to synaptic function, particularly synaptic membranes, were overrepresented in proteins with decreased phosphorylation in MNK2^KO^ mice, whereas phosphorylation of proteins associated with the microtubule and cytoskeleton was increased (Figure 6K). These results are consistent with the proteomic dataset and suggest that MNK2 deletion causes a decrease in both synaptic protein expression and phosphorylation, a change not seen in MNK1^KO^ mice.

To further explore how the MNKs affect signaling pathways, we performed a post-translational modifications signature enrichment analysis (PTM-SEA). We found several pathways significantly altered in both cortex and synaptoneurosomes (Figure 6L-M, Supplementary Figure 7B, Supplementary Table 5), including a decrease in Camk2a signaling, previously identified in synaptosomes from MNK1/2^DKO^ mice[15]. Hierarchical clustering showed that pathway changes were more similar between the cortical and synaptic fractions in MNK1^KO^ mice compared to MNK2^KO^ mice. The cortical fraction from MNK2^KO^ and MNK1/2^KO^ clustered separately from other samples, pointing to a different role for MNK2 depending on location (Supplementary Figure 7B). We focused on the synaptoneurosomes, where 4 PTM pathways were significantly altered in MNK1^KO^ mice and 2 in MNK2^KO^ mice. These pathways included an increase in CDK1 and cAMP-dependent protein kinase catalytic subunit alpha (PKACA) signaling in MNK1^KO^ mice, and an increase in mTOR signaling in MNK2^KO^ mice (Figure 6M, Supplementary Figure 7C). Together, this data suggests that MNK1 and MNK2 regulate distinct signaling pathways at the synapse.

## Discussion

To further develop the MNKs as drug targets, it is essential to better understand each kinase specific role in the nervous system. Using a multi-omic approach combined with detailed behavioral analysis, our study provides novel mechanistic insight into the isoform-specific function of the MNKs in the brain. We demonstrate that loss of MNK1 and MNK2 differentially affects the synaptic proteome and causes distinct social and cognitive behavioral phenotypes. Our results add to the numerous studies that suggest a degree of functional specification for the MNK proteins [17, 24, 27, 28], and indicate that it may be preferential to target each kinase individually.

Comparing the proteome between the whole cortex and the synaptic compartment allowed us to examine the location-specific effects of MNK deletion. We found that MNK1 and MNK2 have partially overlapping functions in cortex but distinct roles at the synapse, and that this difference is driven by a location-specific effect of MNK2. This finding is somewhat surprising, given that the phosphorylation levels of eIF4E in synaptic fractions suggest that both kinases do not significantly differ in their baseline activity in the synaptic compartment. MNK2 is alternatively spliced into two isoforms with somewhat specialized functions [49], and one possible explanation for the location-specific effect of MNK2 deletion is that each splice isoform localizes to distinct neuronal compartments. Another possibility is that MNK2 has different activators or targets different substrates depending on its location. However, further studies are needed to explore this.

MNK inhibition has previously been shown to alter ribosomal protein expression [14, 15, 48, 50], and we here identify MNK1 as the kinase responsible for this change. Interestingly, the shift in ribosomal protein expression was more pronounced at the synapse, suggesting the intriguing possibility that MNK1 may be of particular importance for synaptic translation. The increased abundance of ribosomal proteins at the synapse is supported by the overexpression of ribosomal subunit mRNAs in the synaptic fraction of MNK1^KO^ mice, which suggests that the increase in ribosomal proteins is caused by on-site translation rather than a shift in ribosomal protein stability. The change in ribosomal protein expression is coupled with altered social and object memory in MNK1^KO^ mice and altered spatial memory in MNK1/2^DKO^ mice[15]. These results are consistent with previous work showing that altered ribosome expression is linked to memory dysfunction and altered long-term depression (LTD)[14, 51], and add to a growing body of research suggesting that precise regulation of local ribosomal protein expression may be necessary to support synaptic plasticity and memory [4, 52–54]. Indeed, the lack of cognitive behavioral phenotype in MNK2^KO^ mice despite the reduction of synaptic proteins could be related to the slight increase in MNK1 expression and unchanged levels of synaptic ribosomal proteins in these mice.

It is likely that the increase in ribosomal protein expression is at least partially driven by reduced eIF4E phosphorylation, as increased translation of ribosomal proteins has previously been found in eIF4E phosphorylation mutant mice and cells treated with MNK inhibitors[48, 50]. However, the MNK’s impact on translation is complex and is not only dependent on their kinase activity. For example, MNK1 was recently found to interact directly with ribosomal proteins and members of the eIF complex[29], whereas MNK2 can negatively regulate translation via direct interaction with eIF4G and inhibition of mTOR[30]. Although we did not identify any changes in eIF4G phosphorylation in our dataset, our pathway analysis identified an increase in mTOR signaling in mice lacking MNK2, including increased phosphorylation of mTORC1 component regulatory-associated protein of mTOR (Raptor) and Larp1. As mTOR and Larp1 are key regulators of ribosome production[55], we cannot rule out that MNK2 can act as an enhancer of ribosome protein expression via interaction with the mTOR pathway, perhaps even via direct interaction with Raptor[29]. This could explain the slight reduction of ribosomal subunit mRNAs in MNK2^KO^ mice, although more work is needed to test this.

Taken together, our results suggest a model where MNK1 regulates ribosomal protein expression via eIF4E phosphorylation at the synapse, whereas MNK2 regulates the translation of a pool of mRNAs that include synaptic proteins via other, possible mTOR-dependent mechanisms, although the exact mechanisms remain to be determined. This model is supported by the fact that both ribosomal and synaptic proteins are altered in the MNK1/2^DKO^ mice. Overall, our work may help clarify each kinase’s individual contribution to the therapeutic effects of MNK inhibitors and suggests that targeting MNK1 or MNK2 could differentially affect synaptic function.

## Materials and methods

### Mice

Wild-type, MNK1^KO^, MNK2^KO^, and MNK1/2^DKO^ mice of both sexes were used for this study. The MNK mice were obtained from RIKEN (RBRC01512, RBRC01513, RBRC01514), and were kept on a C57BL/6j background. Animals were weaned at P21-P23 and group-housed (2-5 mice per cage) under a 12h light–dark cycle with food and water ad libitum. All experiments were performed during the light cycle. All experiments were carried out in accordance with European animal welfare law and were approved by the Berlin Landesamt für Gesundheit und Soziales (LAGeSo).

### Behavior

All animals were juveniles (postnatal day 26 – 33) at the start of the first behavior, and both sexes were used for all behavioral tests. The order of the tests was: social habituation/recognition test, open field, novel object test, social olfaction test, and object habituation/recognition test.

#### Social or object habituation/ recognition task

This test was performed as previously described[14]. A fresh home cage without grid, food, and water was used for the experimental cage. The animals were acclimated to the cage for 30 min before the start of the test. For the first trial, a novel same-sex mouse (stimulus mouse: C57BL/6 juvenile mice, P21-P28) or an object (dice or toy car) was placed into the cage for 2 min, and the mice were left to freely interact. This was repeated for 4 consecutive trials with 5 min between trial intervals. On the 5th trial, a novel mouse (littermate to the stimulus mouse) or a novel object (dice or toy car) was introduced for 2 min. For the social stimulus, the interaction was scored when the experimental mouse initiated the action and when the nose of the animal was oriented toward the social stimulus mouse only. For the object stimulus, interaction was scored when the nose of the mouse was oriented 1 cm or less toward the object. The interaction time was used to calculate the recognition index as: (Interaction trial 5) - (Interaction trial 4). One animal (MNK1^KO^) was excluded from the social habituation/recognition task because of aggressive behavior.

#### Open field

The animal was placed in a 50cmx50cm open field arena (ActiMot system, TSE), and its movements were monitored for 10 minutes using the ActiMot automated tracking system. The arena was cleaned with 70% ethanol between trials. One animal (wild-type mouse) was excluded for technical reasons as the system failed to record the movement of the animal, and for two MNK1^KO^ animals, the system failed to track the last three minutes.

#### Novel object recognition

The day after the open field test, the animals were placed back into the same arena (ActiMot system, TSE) containg two identical objects (glass flasks) for 5 minutes. After one hour, short-term memory was tested by exposing the animals to a familiar object (glass flask) and a novel object (lego blocks of similar size to the glass flask) for 5 minutes. Investigation of the object was considered when the mouse’s nose was sniffing less than a centimeter from or touching the object. The discrimination ratio was calculated as follows: (Time spent investigating novel object+ familiar object)/(total time investigating).

#### Social olfaction test

The test animals were placed in a fresh home cage with a grid without food and water. A cotton swab was attached to the grid, and the animal was left to acclimate to the environment for 30 min. Odor habituation and recognition were tested using a cotton swab soaked with an odor. Each odor was presented three times for two minutes, with one minute in between trials. The odors were: water, banana, and almond, and social odors collected by dragging the cotton swab through a dirty cage from sex-matched wild-type mice. Water was used as a habituation and is not shown in the figure. Time spent sniffing the swab was manually scored, with the observers blinded to the genotype. Sniffing was scored when the nose was within 2 cm of the cotton swab.

#### deepOF

Recordings of the social habituation/recognition task were acquired through GoPro Hero9 cameras. The videos were processed through DeepLabCut (v2.2.2). A multi-animal project was created in which 8 body parts per animal were labeled (nose, left ear, right ear, body center, left side, right side, tail base, tail end) according to the protocol in Nath et al. [33]. The tracked data of the mice was processed with DeepOF. We used the supervised annotation analysis as shown in Borders et al. [34], but we modified the parameter for close contact to 1cm. For the PCA analysis, the mean of trials 1-5 was used for all behavioral parameters. Tracking information was missing from one MNK2^KO^ mice for trial five, so the mean of trials 1-4 was used for the PCA for that animal.

#### Z-score calculation

Z-scoring was used to normalize each behavioral test against the mean of the control. The Z-score for each behavior was calculated as shown below, where X: every observation, µ: mean of the control group, and σ: standard deviation of the control group.

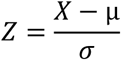

### Fluorescent in situ hybridization

Fluorescent *in situ* hybridization was performed for *Mknk1, Mknk2, Slc17a7,* and *Gad1* by using RNAscope® Multiplex Fluorescent Reagent Kit v2 assay (Advanced Cell Diagnostics, 323136) with probes Mm-Mknk1-C3 mRNA (Advanced Cell Diagnostics, 1270101-C3), Mm-Mknk2-C2 mRNA (Advanced Cell Diagnostics, 1270111-C2), Mm-Gad1-Mus mRNA (Advanced Cell Diagnostics, 400951), and Mm-Slc17a7-O2 17 (Advanced Cell Diagnostics, 501101). Briefly, snap-frozen brains were cut into 16μm thick slices with a cryostat. The brain slices were put on the glass coverslips (Thermo Fisher Scientific, 10149870, 25×75×1mm) and kept at −70°C until further proceeding. The samples were rinsed with PBS (2x) and, after that, fixed with 4% PFA in 1M PBS for 30 minutes at RT, followed by washing with PBS (2x). Dehydration was performed using an ethanol gradient of 50%, 70%, and 2×100%. Hybridization of probes and subsequent signal amplification was carried out using hydrogen peroxide and protease III, followed by the reagents in the Multiplex Fluorescent Detection Reagents v2 kits (Advanced Cell Diagnostics (ACD); Hayward, CA) as described in the manufacturer’s protocol. The fluorophores used to detect the proves were: TSA Vivid Fluorophore kit 520 (Tocris Bioscience, 7523/1, 1:1500), TSA Vivid Fluorophore kit 570 (Tocris Bioscience, 7526/1, 1:1500) and TSA Vivid Fluorophore kit 650 (Tocris Bioscience, 7527/1,1:1500). The slices were stained with DAPI (Advanced Cell Diagnostics, 323136) and mounted using ProLong Glass Antifade Mountant (Thermo Fisher Scientific, P36982). Images were obtained on the IXplore Spin Confocal Imaging Microscope (Olympus) using an x60 oil immersion objective. The images were analyzed using ImageJ by acquiring the mean intensity of *Mknk1* and *Mknk2* signals in *Gad1* or *Slc17a7* positive cells in the cortical and hippocampal area. Three wild-type mice were used. Percentages were assessed for all three mice independently and then averaged for the pie chart.

### BONCAT

400 μm thick coronal slices were cut on a vibratome in ice-cold cutting solution (NaCl 87 mM, NaHCO3 25 mM KCl 2.5 mM, NaH2PO4 1.25 mM, sucrose 75 mM, CaCl2 0.5, MgCl2 7mM, glucose 10 mM, equilibrated with 95% O2/ 5% CO2). Slices were immediately transferred to a storage chamber containing artificial cerebral spinal fluid (ACSF, NaCl 125 mM, NaHCO3 25 mM, KCl 2.5 m), NaH2PO4 1.25 mM, MgCl2 2 mM, CaCl2 2.5 mM, glucose 1 mM, pH 7.4, constantly bubbled with 95% O2 and 5% CO2). Slices were maintained at 32°C in ACSF for 45 min and then moved to an incubation chamber and incubated for an additional 3h with 1mM AHA (Jena Bioscience, CLK-AA005). At the end of incubation, slices were snap-frozen in liquid nitrogen and stored at −70°C. Slices were lysed in lysis buffer (1xPBS, 0.5% SDS + protease inhibitor) 12x per mg with a dunce homogenizer and centrifuged for 15min 14 000g at 4°C. BCA-Assay (Pierce Protein BCA-Assay Kit) was used to determine the protein concentration. 500µg of protein was diluted in the lysis buffer to a final volume of 150µl. The samples were sonicated for 30sec with a Hielscher ultrasonicator, then treated with 20mM Iodoacetamide for 1h at RT in the dark. To perform the click-reaction, the following reagents were added to the samples and briefly vortexed after each addition: 127µM TBTA, 3.75mM copper sulfate (Jena Bioscience, CLK-MI004-50), 100µm PEG4-biotine alkyne (Jena Bioscience, CLK-TA0105-25), 1mM TCEP and adjust with 1xPBS + protease inhibitor to final volume of 400µl. The samples were incubated for 2h at RT in the dark during constant rotation. After incubation, proteins were extracted using the methanol-chloroform method. The final pellet was resolubilized in 150µl RIPA (NaCl 150mM, Tx100 1%, Sodium deoxycholate 0.5%, Tris pH8 50mM, SDS 0.1%) and sonicated for 30sec. AHA incorporation was measured by western blotting using an anti-Streptavidin IRDye 800CW antibody (LI-COR, 926-32230, 1:2000). The results were normalized to total protein concentration using MEMCODE Reversible Protein Stain Kit (Pierce, # PIER24580).

### Synaponeurosome isolation

Cortex was rapidly dissected and homogenized in oxygenated Krebs buffer (NaCl 118.5mM, CaCl_2_ 2.5mM, KH_2_PO_4_ 1.18mM, MgSO_4_ 1.18mM, MgCl_2_ 3.8mM, NaHCO_3_ 24.9mM, Glucose 212.7mM in DEPC treated H_2_O) with protease and phosphatase inhibitors in a dunce grinder. An aliquot of the homogenate was taken from the whole lysate and snap-frozen in liquid nitrogen. The rest of the homogenate was passed through 2x 100µm pre-wet nylon filter (Merck Millipore, NY1H02500) using an 18G needle, followed by a second filtration with a 5µm pre-wet filter (Merck Millipore, NY0502500). The lysate was centrifuged for 10min 1000g 4°C. The synaptoneurosome pellet was washed 1x in 500µl Krebs buffer. The resulting pellet was snap-frozen in liquid nitrogen and stored at −70°C until further use.

### Western Blot and AlphaLISA immunoassay

Brain tissue was homogenized in RIPA buffer (NaCl 150mM, Tx100 1%, Sodium deoxycholate 0.5%, Tris pH8 50mM, SDS 0.1%), and complete protease and phosphatase inhibitors. The synaptoneurosome pellet was either lysed in AlphaLISA lysis buffer (PBS 1x, EDTA 5mM, Tx100 1%, protease and phosphatase inhibitors), or RIPA buffer with protease and phosphatase inhibitors. The S1 fraction from the synaptoneurosome isolation was further diluted in RIPA buffer with protease and phosphatase inhibitors. For all western blot samples, protein concentration was measured with BCA-Assay (Pierce Protein BCA-Assay Kit), and diluted to an equal concentration in RIPA-buffer and 4xLDS sample buffer (mpage, MPSB-10ml). Immunoblotting was done with HRP-conjugated secondary antibodies and WesternBright Chemilumineszenz Substrat (Biozym #541020, 541004). The following primary antibodies were used: p-4EBP1 (Cell Signaling, #2855S, 1:1000), 4EBP1 (Cell Signaling, #9644S, 1:1000). p-eIF4E (Abcam, ab76256 1:500), eIF4E (Cell Signaling, #9742S, 1:2000), p-ERK1/2 (Cell Signaling, #4370, 1:500), ERK1/2 (Cell Signaling, #4695, 1:500), MNK1 (Cell Signaling, #2195, 1:500), p-S6 (Cell Signaling, #5364 1:1000), S6 (Cell Signaling, #2317 1:1000). Secondary antibodies were anti-Rabbit IgG (Cell Signaling, #7074 1:2000) and anti-mouse IgG (Cell Signaling, #7076 1:2000). Membranes were stained with MEMCODE Reversible Protein Stain Kit (Pierce, # PIER24580) to visualize total protein concentration. Signals were acquired using an image analyzer (Bio-Rad, ChemiDoc MP Imaging System), and images were analyzed using ImageJ and Biorad Image lab software. The total protein concentration was used as a loading control for all experiments.

eIF4E phosphorylation was measured using the AlphaLISA SureFire Ultra p-eIF4E (Ser209) Assay Kits (PerkinElmer) according to the manufacturer’s protocol. AlphaLISA signals were measured using a Tecan SPARK plate reader in the recommended settings.

### TMT mass spectrometry

Global proteomes and phosphoproteomes of cortical homogenate and synaptoneurosomes were analyzed using TMT (Thermo Fisher Scientific) isobaric labels combined with deep fractionation, as described in Mertins et al., 2018[35]. Briefly, synaptoneurosome pellets and whole homogenate were lysed in SDS buffer (25mM HEPES, 2% SDS, protease, and phosphatase inhibitors) and boiled at 95°C for 3 minutes. Peptides were cleaned up and digested with trypsin using the SP3 protocol as previously described[36]. An amount of 100 µg of each peptide sample was subjected to TMTpro 18-plex (Thermo Fisher Scientific) labeling with randomized channel assignments. Quantitation across two TMT plexes was achieved by including an internal reference derived from a mixture of all samples. Samples were fractionated using an UltiMate 3000 Systems (Thermo Fisher Scientific) into 24 fractions for proteome analysis and 12 fractions for phosphoproteome analysis. For phosphoproteome analysis, the peptides were subjected to phosphopeptide enrichment using an AssayMAP Bravo Protein Sample Prep Platform (Agilent Technologies). All samples were measured on an Exploris 480 orbitrap mass spectrometer (Thermo Fisher Scientific) connected to an EASY-nLC system 1200 system (Thermo Fisher Scientific). 4-week old male and female mice (two from each sex per genotype) were used for all TMT experiments.

For analysis, MaxQuant version 2.1.4.0[37] was used, employing MS2-based reporter ion quantitation. Carbamidomethylation was set as a fixed modification and acetylated N-termini as well as oxidized methionine as variable modifications. For phosphoproteomics analysis, phosphorylation on serine, threonine and tyrosine was enabled as a variable modification. A PIF filter was applied with a threshold value of 0.5. For database search, a Uniprot mouse database (2022-03) plus common contaminants were used. Proteins with less than 2 peptides were excluded from the analysis. Corrected log2-transformed reporter ion intensities were normalized to the internal reference samples and further normalized using median-MAD normalization before applying two-sample moderated t-tests (limma)[38]. P-values were adjusted using the Benjamini-Hochberg procedure. The data was further analysed using Protigy 1.1.5 (https://github.com/broadinstitute/protigy/), using two-sample mod t-test without group-wise normalization. An adjusted p-value <0.05 was used for significance. Visualization of proteomic data was done using Protigy v1.1.5 (PCA, heatmap of protein expression), GraphPad Prism 10.2.1 and in R studio (2023.12.1, R version 4.4.4) using the package corrplot (logFC correlation). For visualization of specific phosphosite intensities with multiplicities, a mean was used.

### Protein gene ontology and gene set enrichment analysis

Gene set enrichment analysis (GSEA) was performed using GSEA v4.3.2[39] with 10 000 permutations. GO terms were collected from the mouse MSigDB database for canonical pathways or cellular components. LogFC of all proteins were used as gene ranks. Significance was set as FDR<0.25, and only GO terms significant in MNK1^KO^ or MNK2^KO^ were used for further analysis. Hierarchical clustering was done by calculating the euclidean distance between the NES in R studio using the packages pheatmap and dichromt, with the method set as ward.d2. The number of clusters was determined using Nbclust. ggplot2 was used for GSEA visualization, and the heatmaps showing logFC of the individual GO terms were done using Morpheus[40]. To compare the distribution of ribosomal proteins, a random set of the same number of proteins as ribosomal proteins was generated from the total protein expression. PTM-SEA was performed using ssGSRA2.0 in R studio, with the mouse PTMsigDB signature set[41], with logFC as the rank input list. The number of permutations was set to 10 000. Gene ontology was performed on phosphosites with a logFC value above 1.2 or below −1.2 using Enrichr[42], with all proteins with detected phosphosites set as background. Significance was set to p.ajust<0.05. P-value was adjusted using the Benjamini-Hochberg method. SynGO analyses were performed using the SynGO web page (version 1.2) with default settings[43]. ggplot2 was used for visualization. Venn diagrams were created using BioVenn[44].

### RNA-Sequencing and enrichment analysis

Samples for mRNA-seq experiment were collected from 3 four-week-old male mice for each knockout and wild-type genotypes, used as biological replicates in the subsequent analyses. Frozen synaptoneurosomes were resuspended in 600 mL TRIzol using a 22G syringe and were centrifuged for 10min 13 000g 4°C. Total RNA was isolated using the Direct-zol RNA microprep kit (Zymo Research, #R2062) and treated with DNAse, following the manufacturers’ instructions. Total RNA samples were quantified using a Qubit Fluorometer, and RNA integrity was checked on a TapeStation (Agilent). Double-indexed stranded mRNA-Seq libraries were prepared using the ILMN Stranded mRNA Library Prep Kit (Illumina, #20040534), starting from 250 ng of input material according to the manufacturer’s instructions. Libraries were equimolarly pooled based on Qubit concentration measurements and TapeStation size distributions. The loading concentration of the pool was determined using a qPCR assay (Roche, #7960573001). Libraries were then sequenced on the Illumina NovaSeq X Plus platform using PE100 sequencing mode, with a target of 50 million reads per library. RNA-seq reads were adapter trimmed using Trimmomatic v.0.39 and aligned to the mouse genome (mm10) with STAR aligner version 2.7.8a (https://github.com/alexdobin/STAR) using default parameters. Gene counts were produced with a featureCounts function from the subread v.2.0.3 and Ensembl GRCm39.109 annotation of all mouse genes. Differential expression analysis was performed using an R package DESeq2 v.1.44.0 for each mutant independently. Significantly differential genes were identified with an adjusted p-value threshold of 0.05. Gene ontology and gene set enrichment analyses were performed using R package clusterProfiler v.4.10.1 and canonical pathway database m2.cp.v2023.1.Mm.symbols for mouse genes. For GSEA comparison with the protein dataset, significance was set to the same adjusted p-value of <0.25, and only pathways present in both datasets were compared. Volcano plots were generated using EnhancedVolcano v.1.20.0 R package. K-means clustering was performed using stats base package in R v.4.3.2 with 20 clusters since the predicted number of clusters (3) did not achieve satisfactory group splitting by visual assessment.

### Statistical analysis

Statistical analyses were performed in R studio or with the GraphPad Prism software. All animals were used for further analysis unless otherwise stated. Normality was tested using the Shapiro-Wilk test. When normally distributed, the data were analyzed with one-way ANOVA or repeated measures (RM) ANOVA, followed by Tukey’s post-hoc test. Grouped data with missing values were analyzed using a Mixed effect model followed by Tukey’s multiple comparison test. 3-way ANOVA was calculated using the R package ezANOVA(ez). Correlation was assessed using Pearson correlation. For non-parametric tests, Mann-Whitney test or Kruskal-Wallis test followed by Dunn’s multiple comparison test were used. Outliers were identified using Rout Q=0.1%, this lead to the removal of one MNK1/2^DKO^ sample from Figure 6D. Differences in frequency distribution were assessed using the Kolmogorov-Smirnov test. Principal component analysis was done using the packages factominer and factoextra in the R environment. Missing values (open field from 1 wild-type and last three minutes for 2 MNK1^KO^ mice) were imputed using the median from the same genotype for the PCA. Data are represented as the mean ± s.e.m. and significance was set at P<0.05. Detailed statistical information for all figures is shown in Supplementary Table 6.

## Data availability

The mass spectrometry proteomics data have been deposited to the ProteomeXchange Consortium via the PRIDE[56] partner repository with identifier PXD058409. The RNA sequencing data will be deposited to the EBI repository and is available on request.

## Acknowledgments

We thank the Genomics technology platform and the proteomic technology platform of the Max Delbrück Center for RNA sequencing and mass-spectrometry, and the animal caretakers for their support with the mice. We thank the Scientific Computing Team of the MDC for the support with the data analysis on the HPC cluster. We are grateful to members of the Hörnberg lab for fruitful discussions. This work was funded by the Deutsche Forschungsgemeinschaft (DFG, German Research Foundation) under Germanýs Excellence Strategy – EXC-2049 – 390688087 (H.H).

## Supplementary Material

**Supplementary Figure 1.**
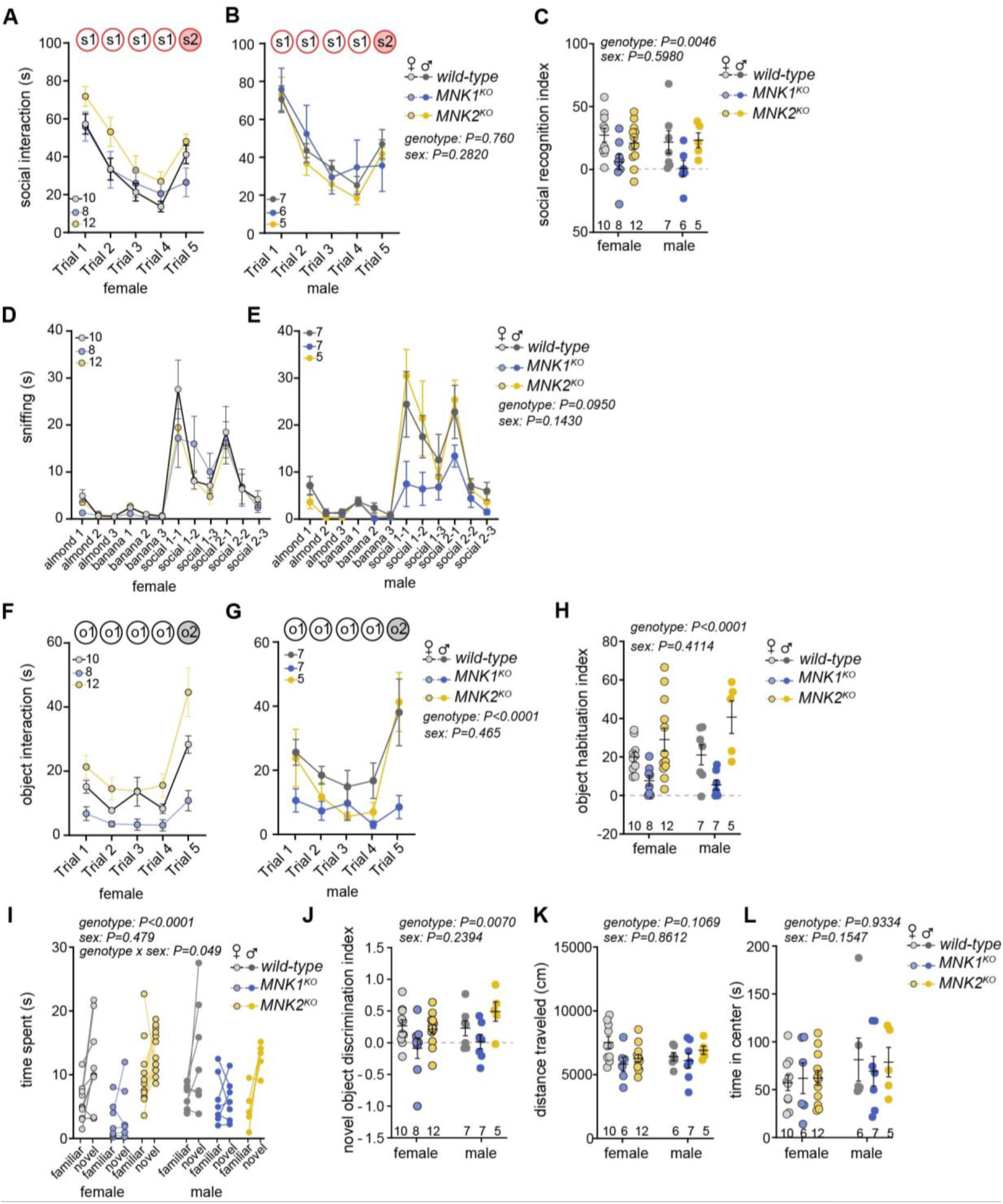
Sex-specific analysis of the behavioral phenotypes in MNK1^KO^ and MNK2^KO^ mice. (A-C) Mean social interaction time for females (A) and males (B), and recognition index (C) in the social habituation/recognition test in wild-type, MNK1^KO^ and MNK2^KO^ mice. s1=first social stimulus, s2=second social stimulus. (D-E) Time sniffing for females (D) and males (E) in the social olfaction habituation test. (F-H) Mean object interaction time for females (E) and males (G), and recognition index (H) in the object habituation/recognition test. o1=first object stimulus, o2=second object stimulus. (I-J) Time spent interacting with the familiar and novel object (I) and novel object discrimination index (J) in the novel object recognition test in male and female mice. (K-L) Total distance traveled (K) and time spent in center (L) in the open field test. Error bars show s.e.m. P values: *<0.5, **<0.01, ***<0.001 relative to wild-type. Significance was determined by three-way ANOVA for A, B, D, E, F, G, I, and two-way ANOVA for C, H, J, K, L.

**Supplementary Figure 2.**
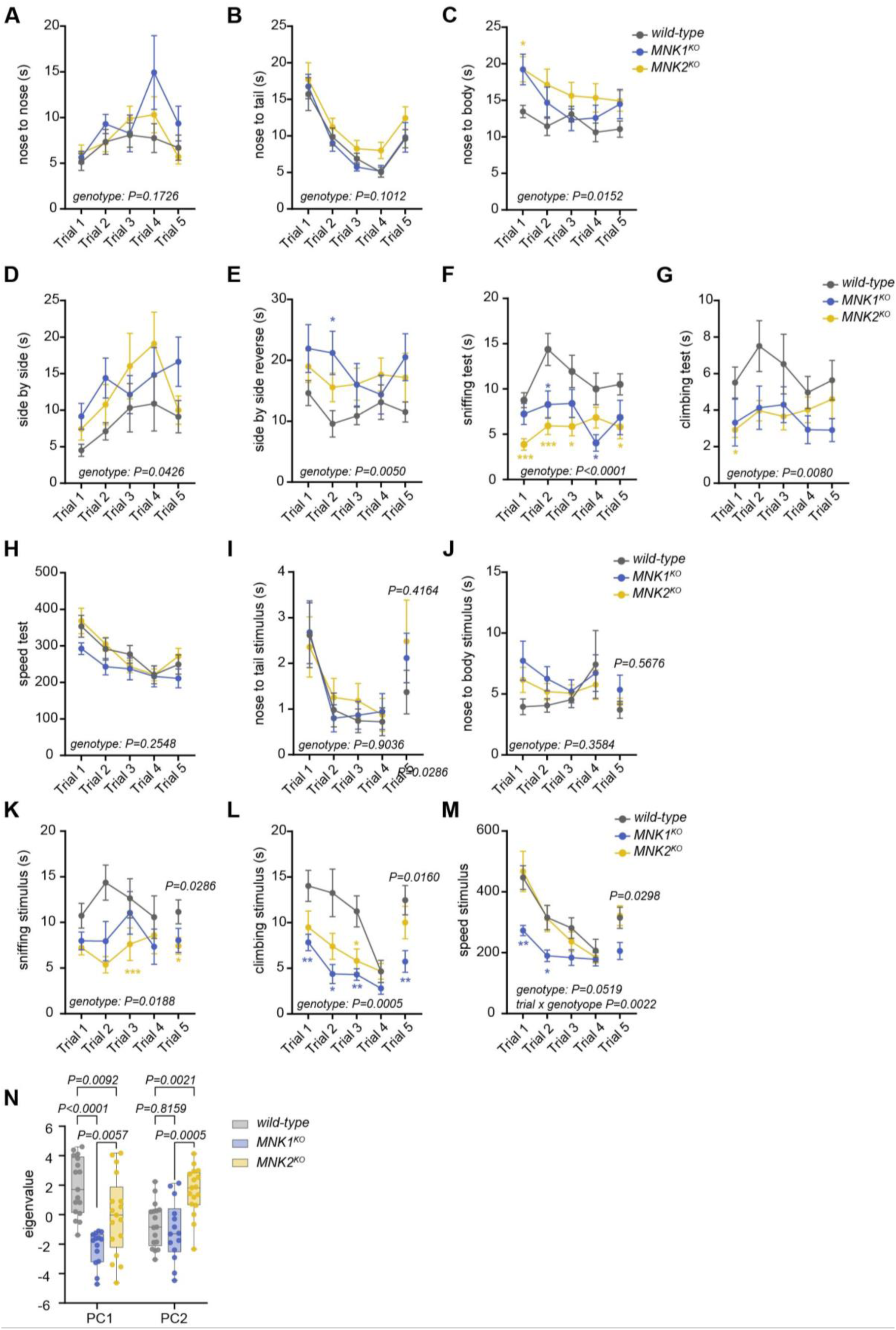
Detailed social phenotyping using deepOF shows social phenotypes in MNK1^KO^ and MNK2^KO^ mice. (A-M) DeepOF analysis of social and individual behaviors in the five-trial social habituation/dishabituation test in wild-type, MNK1^KO^, and MNK2^KO^ mice. (A-C) Social interaction from the test mouse towards the stimulus mouse covering nose to nose (A), nose to tail (B), and nose to body (C). (D-E) Passive social behavior where the mice are in close proximity without directly interacting, either facing the same direction (side by side, D), or facing the opposite direction (side by side reverse, E). (F-H) Individual behavior of the test mouse, showing time sniffing the wall (F), time spent climbing/rearing on the wall (G) and total speed during the trial (H). Social interaction of the stimulus mouse towards the test mouse covering nose to tail (I) and nose to body (J). (K-M) Individual behavior of the stimulus mouse showing time sniffing the wall (K), time spent climbing/rearing on the wall (L), and total speed during the trial (M). (N) Principal component (PC) analysis of 24 behavioral components. PC1-2 shows a significant effect of genotype, with PC1 separating MNK1^KO^ and MNK2^KO^ mice from wild-type mice, and PC2 separating MNK2^KO^ mice from wild-type and MNK1^KO^ mice. Error bars show s.e.m. Significance was determined using a mixed-effect model followed by Tukey’s post-hoc test for A-H, RM two-way ANOVA followed by Tukey’s post-hoc test (trial 1-4) and Kruskal-Wallis test followed by Dunn’s multiple comparison test (trial 5) for I-L, RM two-way ANOVA followed by Tukey’s post-hoc test (trial 1-4) and one-way ANOVA (trial 5) followed by Tukey’s post-hoc test for M, and two-way ANOVA followed by Tukey’s post-hoc test for N.

**Supplementary Figure 3.**
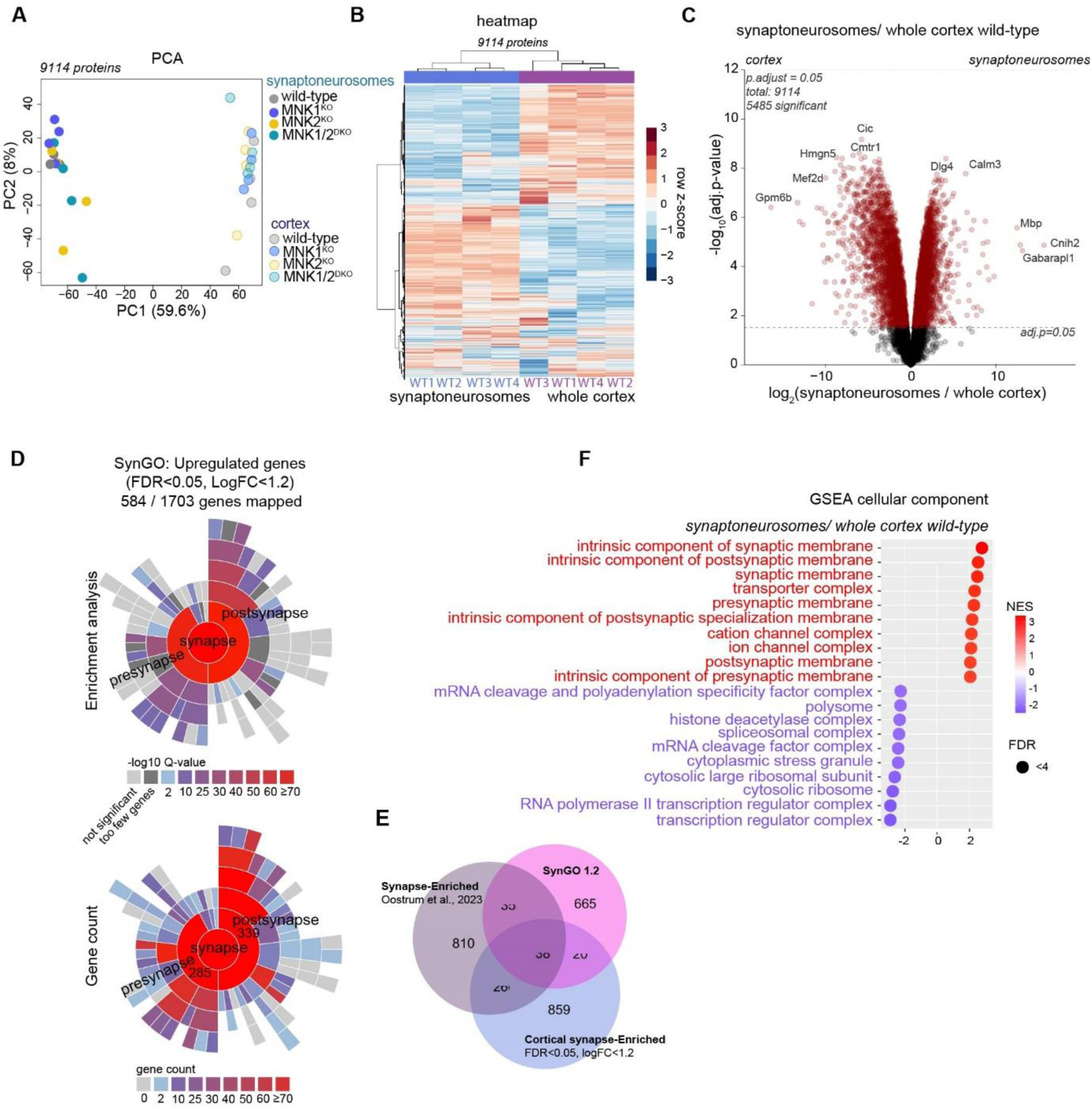
Validation of the synaptoneurosome enrichment. (A) PCA showing a clear separation between the protomes from synaptoneurosomes and cortex in all genotypes. (B) Hierarchical clustering of normalized protein expression of all proteins detected in synaptoneurosomes and corex from wild-type mice. (C) Volcano plot comparing wild-type synaptoneurosomes to cortex. Differentially expressed proteins (P.adjust<0.05) are marked in red. Over half of all identified proteins were significantly altered in the isolated synaptoneurosome fractions. (D) SynGO[43] sunburst plot showing enrichment analysis (top) and gene count (bottom) of upregulated proteins (FDR<0.05, logFC<1.2). We found a significant overrepresentation of pre-and postsynaptic proteins in the synaptoneurosome fraction, with a similar number of pre-and postsynaptic proteins identified. (E) Venn diagram of the overlap between cortical synapse-enriched proteins to proteins identified as synapse-enriched in the SynGO database or the synapse-enriched proteins from Oostrum et al., (2023). We found a good representation of known synaptic proteins enriched in the synaptoneurosome fraction, with around 50% of the enriched proteins present in either dataset. (F) Gene set enrichment analysis (GSEA) of cellular components for wild-type synaptoneurosomes compared to cortex showed enrichment of genes linked to the synapse and decrease of proteins associated with the nucleus and cytosol. Top ten increased (red) and decreased (blue) pathways are shown. NES= normalized enrichment score.

**Supplementary Figure 4.**
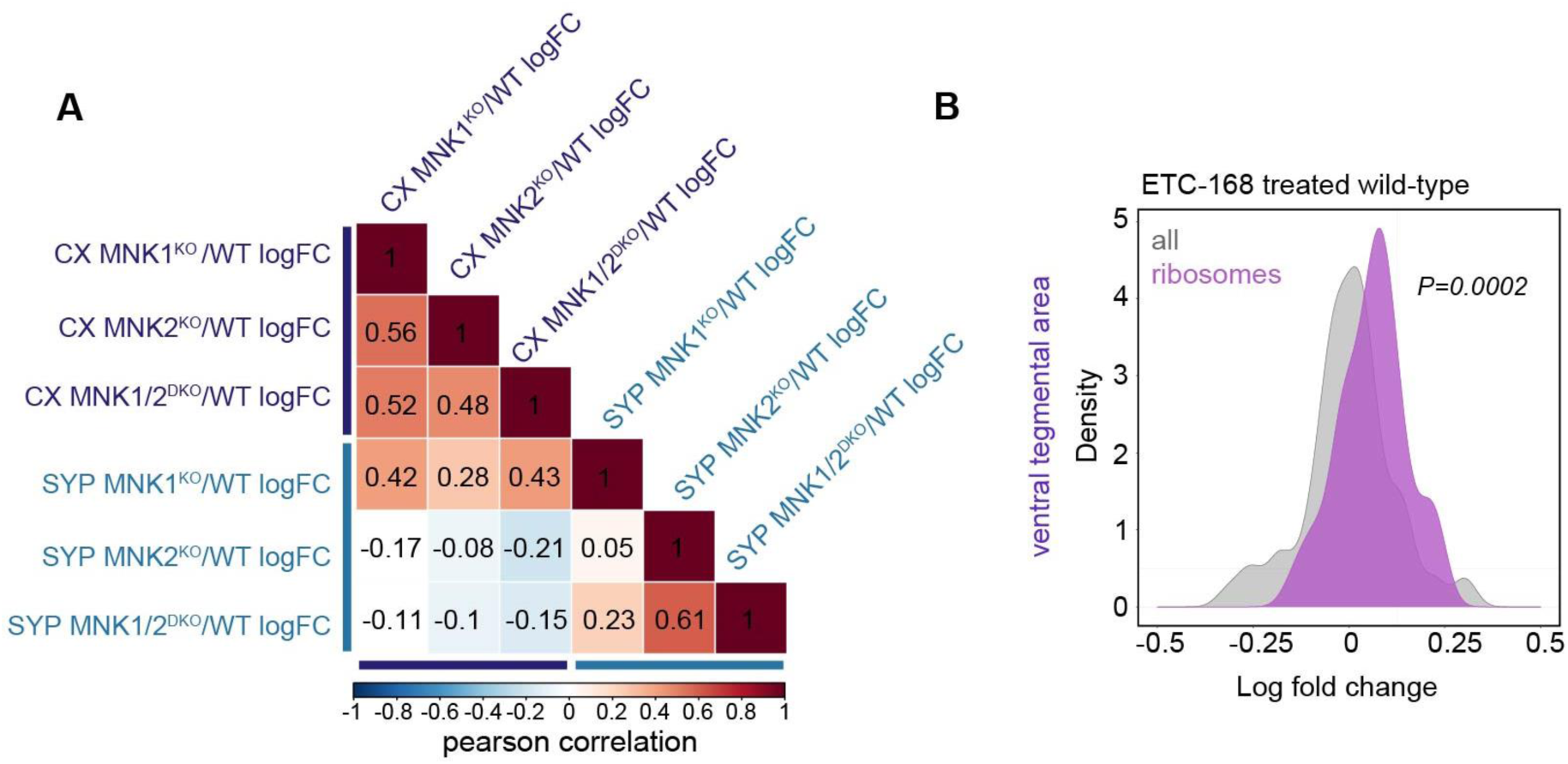
Comparison between the cortical and synaptic proteome shows site-specific effects of MNK2 deletion. (A) Correlation matrix heatmap showing the Pearson correlation of logFC values of the proteome from cortex and synaptoneurosomes in MNK1^KO^, MNK2^KO^, and MNK^DKO^ relative to wild-type. (B) Density plots of logFC ribosomal protein abundance compared to all proteins from wild-type mice treated with ETC-168 compared to vehicle. The proteomic data is from Hörnberg et al., 2020. P-value was calculated using a two-sided Kolmogorov-Smirnov test.

**Supplementary Figure 5.**
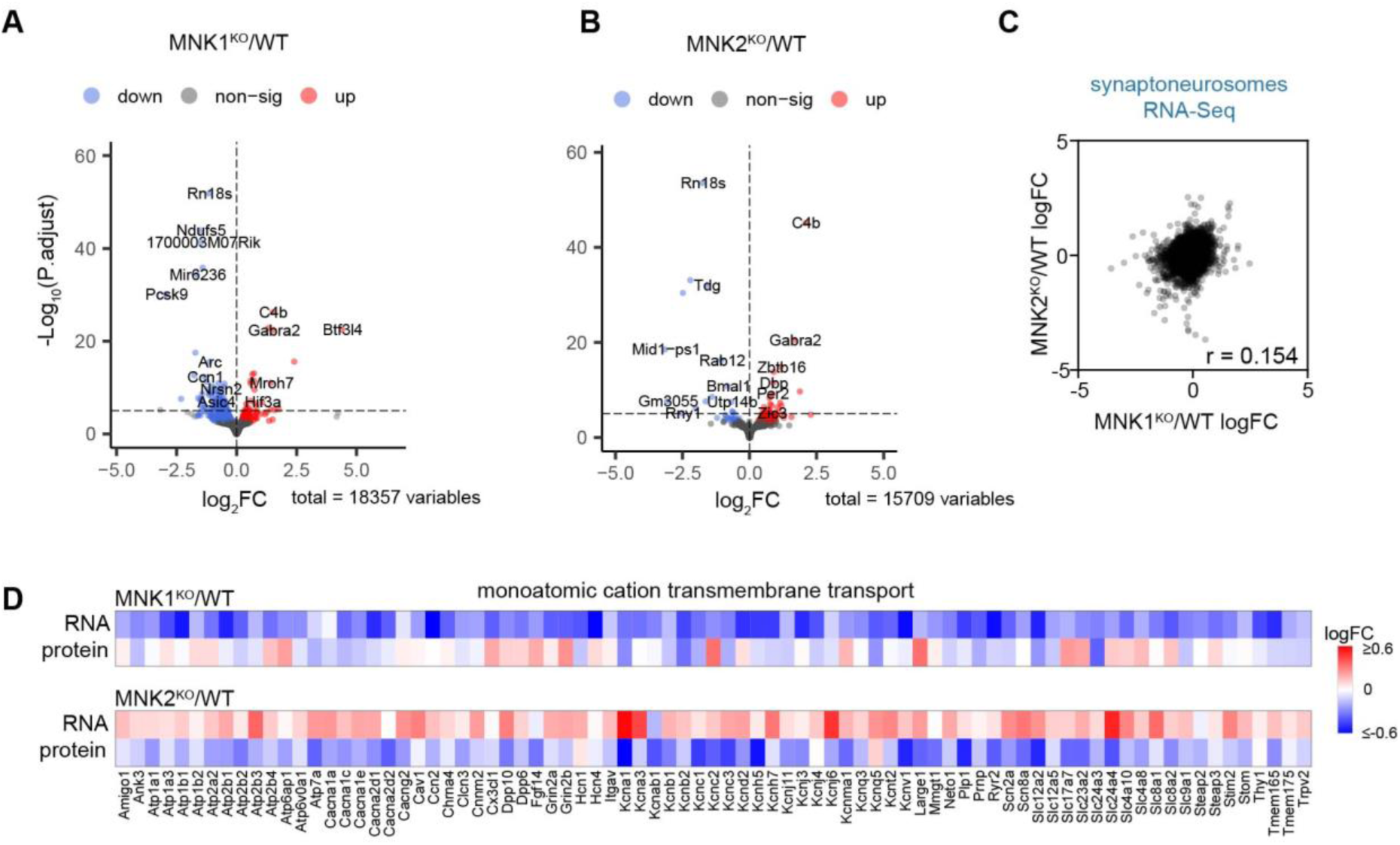
Knockout of MNK1 or MNK2 have different effects on the synaptic transcriptome. (A-B) Volcano plot of mRNA in synaptoneurosomes from (A) MNK1^KO^ and (B) MNK2^KO^ mice. Downregulated genes (P.adjust<0.05) are shown in blue, and upregulated in red. (C) Alterations in mRNA from MNK1^KO^ synaptoneurosomes compared to MNK2^KO^ synaptoneurosomes relative to WT. r was determined by Pearson correlation. (D) Heatmap showing logFC of core enriched proteins and mRNAs in the GO term Monoatomic cation transmembrane transport. Both the mRNA and protein are from isolated synaptoneurosomes from MNK1^KO^ mice (top) and MNK2^KO^ mice (bottom), relative to wild-type mice.

**Supplementary Figure 6.**
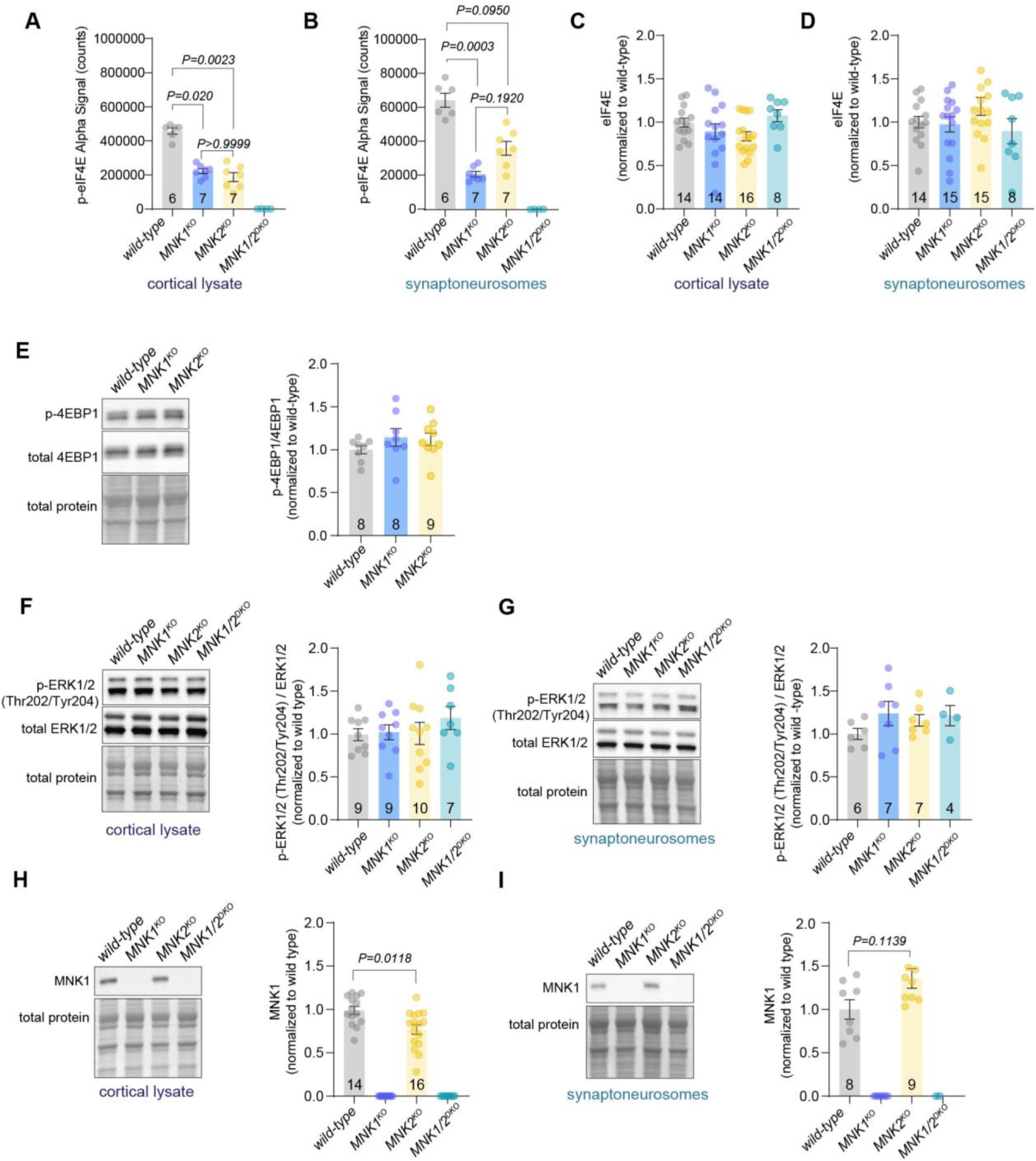
Effect of MNK1 an MNK2 deletion on protein abundance and phosphorylation levels in pathways related to protein synthesis. (A-B) Normalized p-eIF4E AlphaLisa count from (A) cortical lysate and (B) synaptoneurosomes from wild-type, MNK1^KO^, and MNK2^KO^ mice. MNK1/2^DKO^ are included as validation. (C-D) Quantification of eIF4E from wild-type, MNK1^KO^, MNK2^KO^, and MNK1/2^DKO^ mice in (C) cortical lysate and (D) synaptoneurosomes. Images are in Figure 6A-B. (E) Representative image and quantification of p-4EBP1 compared to total 4EBP1 in synaptoneurosomes from wild-type, MNK1^KO^, and MNK2^KO^ mice. (F-G) Representative image and quantification p-ERK1/2 compared to total ERK1/2 in wild-type, MNK1^KO^, MNK2^KO^, and MNK1/2^DKO^ mice in (F) cortical lysate and (G) synaptoneurosomes. (H-I) Representative image and quantification of MNK1 in wild-type and MNK2^KO^ mice in (H) cortical lysate and (I) synaptoneurosomes. MNK1^KO^ and MNK1/2^DKO^ mice are included as validation. Significance was determined by Kruskal-Wallis test followed by Dunn’s multiple comparison test for A-B, one-way ANOVA for C-G, and Mann-Whitney test for H-I.

**Supplementary Figure 7.**
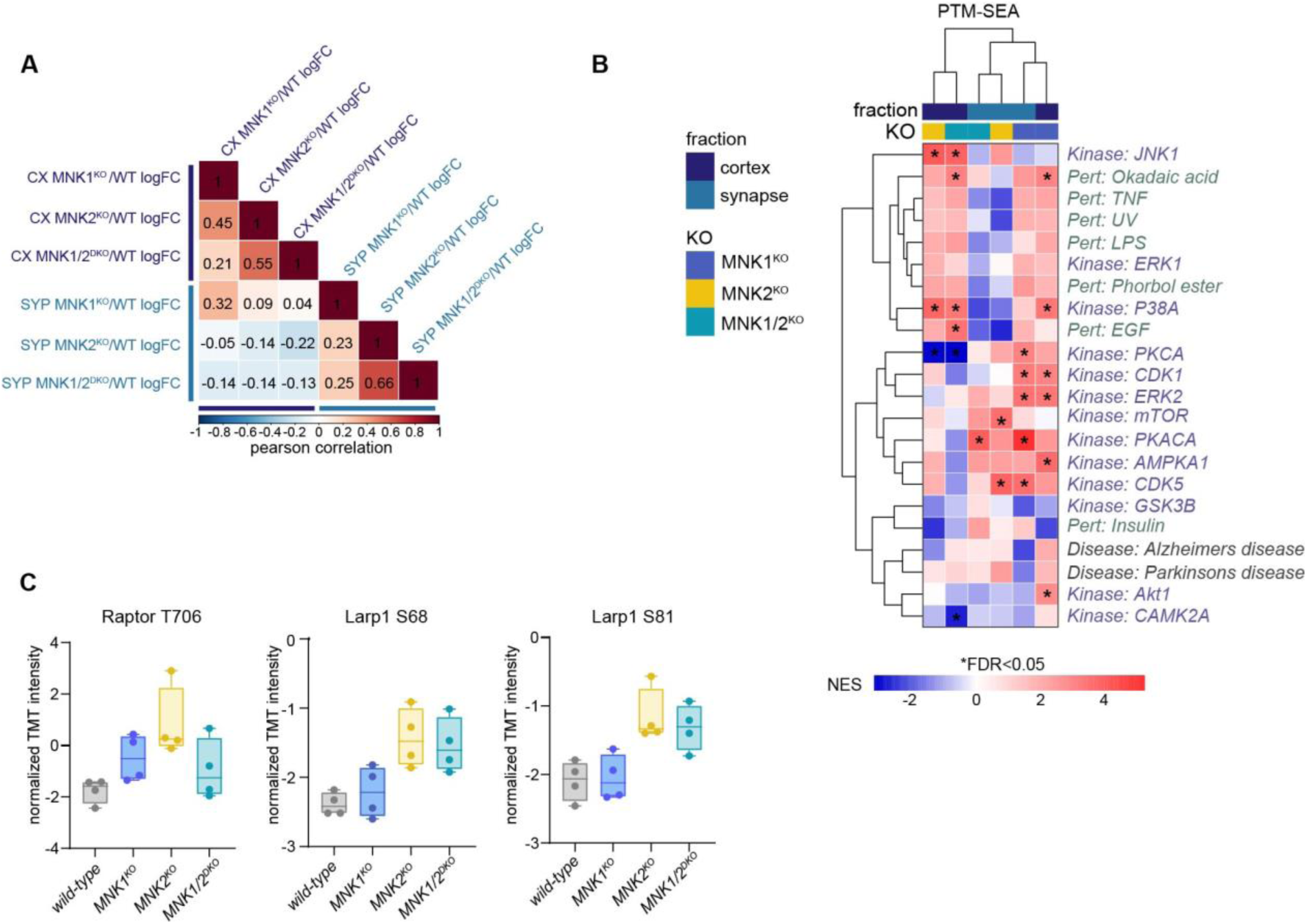
Effect of MNK1 and MNK2 deletion on the cortical and synaptic phosphoproteome. (A) Correlation matrix heatmap showing Pearson correlation of cortical and synaptoneurosomes phosphoproteome logFC values in MNK1^KO^, MNK2^KO^, and MNK1/2^DKO^ relative to wild-type. (B) Heatmap showing hierarchical clustering of phosphosite-specific signature normalized enrichment scores (NES) as determined by PTM-SEA. Significant changes (P.adjust<0.05) are marked with *. (C) Normalized TMT intensities for phosphosites linked to the mTOR pathway increased in synaptoneurosomes from MNK2^KO^ mice compared to MNK1^KO^ mice. Error bars in C show min to max.

