## Supplementary Table 1 for "Distinct roles for MNK1 and MNK2 in social and cognitive behavior through kinase-specific regulation of the synaptic proteome and phosphoproteome"

**PCA behavior, Figure 1O**

| | classical tests | | | | | | | | | | deepOF | | | | | | | | | | | | |
|---|---|---|---|---|---|---|---|---|---|---|---|---|---|---|---|---|---|---|---|---|---|---|---|
| ID | SI trial1 | SR index | NOR DI | NO 1st tria | SO Index | SO S1 | OI trial1 | OH DI | OF distanc | OF time in | nose to no | nose to tai | nose to tai | side by sid | side by sid | nose to bo | nose to bo | climbing te | sniffing tes | speed test | speed stim | climbin sti | sniffing stim |
| Wild type | -1.5003 | 1.644019 | 0.414798 | -1.49547 | 0.433838 | -0.07507 | -0.63106 | -0.47314 | 0.148012 | -0.24563 | -0.68563 | -1.2669 | -0.25903 | -0.40633 | -0.94708 | -1.45626 | -0.39155 | 2.595852 | -0.5554 | 0.452164 | 0.362822 | -0.90663 | -0.81452393 |
| Wild type | -0.8828 | -1.19656 | -0.37482 | -0.24108 | 0.371838 | -1.39918 | -0.47268 | -1.00331 | -0.52533 | -1.07532 | 2.025851 | -1.26409 | 0.918335 | 1.977153 | 1.841524 | -0.1617 | 1.002608 | 2.13361 | 1.597128 | -1.42976 | -0.67511 | -0.29156 | 1.218137488 |
| Wild type | -1.40957 | -0.35491 | 1.866401 | -0.61708 | -1.50908 | 0.088381 | -0.70448 | 0.00969 | 0.115509 | 1.07151 | -1.03117 | -0.59401 | 0.313024 | -0.66225 | -1.17773 | -1.0879 | -0.90729 | 2.591456 | -0.42455 | 0.250057 | 1.251791 | 0.638401 | 0.50314881 |
| Wild type | -0.49735 | -0.47074 | 0.866093 | -0.19707 | 2.351952 | 1.922013 | -0.35415 | -0.12285 | 1.684762 | -0.05895 | -0.98152 | 0.054777 | -0.82443 | 0.01291 | -1.45694 | -0.34877 | -0.49636 | 1.074828 | -1.63071 | 0.15708 | 2.047337 | 1.125574 | -0.86493436 |
| Wild type | 0.549038 | 0.470053 | 0.925229 | 0.315086 | -0.29931 | 0.72142 | -0.30381 | 1.284934 | -1.16899 | 0.285894 | 0.779926 | 0.613176 | -0.69804 | -0.84891 | 1.322996 | 1.27634 | 0.943311 | 0.980313 | -0.41504 | -1.00332 | -0.3894 | -0.43183 | 0.24549548 |
| Wild type | 0.163034 | 0.989006 | -1.57898 | -1.14054 | -0.74648 | -0.45947 | -0.53142 | -1.02698 | 1.40381 | -0.66825 | 1.077804 | 0.006569 | 1.876189 | -0.67975 | 0.629314 | 0.515547 | 0.582015 | 1.323203 | 1.007608 | -0.98735 | -0.71412 | -1.26699 | 0.665582432 |
| Wild type | -0.42367 | 0.807929 | -0.84576 | -0.55133 | -0.56048 | -0.43338 | 0.369578 | 0.125191 | -0.80362 | -1.03125 | -0.16534 | -1.28297 | 0.266462 | -1.28346 | -0.59157 | -0.12005 | -0.60667 | 0.771502 | -0.66246 | -1.4847 | -1.1967 | -1.12006 | -0.4056393 |
| Wild type | -0.91855 | 0.377998 | -0.64785 | -1.2745 | -1.3254 | -1.32039 | -1.52052 | -0.61515 | -0.56004 | -0.64233 | 1.157238 | -0.14006 | -1.15036 | 0.765359 | -0.21525 | 0.746674 | 0.251057 | 1.151758 | -0.27944 | -0.65538 | -0.81547 | -1.12161 | 1.55336619 |
| Wild type | 0.235066 | 0.486745 | 1.745808 | 0.914754 | 0.017665 | 0.594173 | 2.476792 | 1.42505 | 0.223597 | 0.807045 | -0.0303 | -0.65226 | 0.579095 | 0.355595 | -0.48058 | 1.82286 | -0.28675 | 2.877197 | -0.71955 | 0.145178 | 0.147282 | 0.451265 | 0.409329391 |
| Wild type | 0.608973 | -0.58353 | 0.204944 | 2.168047 | -0.92008 | 1.219759 | 1.707958 | -0.79503 | 0.13804 | -0.40638 | -0.81273 | 1.175592 | -1.0905 | 0.861603 | -0.31063 | 0.830939 | -0.79973 | 2.048547 | 0.025077 | -0.28705 | 0.61553 | 0.664693 | -0.04996568 |
| Wild type | -0.8138 | -1.21325 | -1.13961 | 0.879009 | -0.83057 | -0.07427 | 0.109454 | 0.484001 | -0.19562 | -0.28063 | -0.85443 | 0.641296 | 0.678871 | -0.21166 | -0.09038 | -0.27655 | -0.31709 | 3.367354 | -0.09387 | 1.541219 | 1.856048 | 1.756578 | -0.30061756 |
| Wild type | 1.143165 | 0.011797 | -1.12808 | -0.47031 | 0.365638 | 0.279783 | 0.834235 | 0.810623 | -0.83407 | 3.184637 | -0.74124 | -0.78885 | 0.845165 | 0.355595 | 1.973324 | -1.57182 | 3.177276 | 1.164946 | 0.508016 | -1.28577 | -0.65333 | -1.69848 | -1.44185378 |
| Wild type | 0.743139 | -1.08706 | -0.85506 | -0.98162 | 1.387858 | 0.102756 | -0.27758 | -0.92284 | -0.71228 | -0.44787 | -0.15342 | 0.645314 | -1.14371 | -0.51351 | 0.393461 | 0.741859 | -0.24262 | 3.688263 | -1.20249 | -0.32986 | -0.23558 | 0.381669 | -0.95315262 |
| Wild type | -0.8839 | 2.179663 | 0.119404 | 0.535715 | -0.07378 | -1.39918 | -0.4947 | 1.395701 | -0.50676 | -0.34675 | 0.905034 | 0.631253 | -1.01733 | 0.725987 | 0.565148 | 1.160777 | -0.22607 | 3.411314 | 1.680868 | -1.34663 | -1.01682 | -0.4906 | -1.39284363 |
| Wild type | 0.930643 | -0.96693 | 0.542427 | 0.077596 | -0.33961 | -1.39918 | 0.050717 | -1.98601 | -1.41256 | -0.31563 | 1.33795 | -0.98168 | -1.21023 | 1.860495 | -0.23952 | -0.8977 | -0.52945 | 2.635416 | -0.42218 | -1.41503 | -0.94736 | 0.905959 | -0.3356248 |
| Wild type | 1.02137 | -0.60781 | -0.41961 | 0.972167 | 0.726011 | 0.321045 | -1.08208 | 1.130617 | 1.015607 | 0.363677 | -1.00933 | 2.171872 | 1.490386 | -1.39283 | -0.32624 | -0.83992 | -0.31157 | 1.109996 | -0.23186 | -0.32428 | 0.715485 | 1.093095 | -0.14518539 |
| Wild type | 1.935517 | -0.48642 | 0.304666 | 1.106635 | 0.949985 | 1.310802 | 0.823746 | 0.279508 | 1.98992 | -0.19377 | -0.81869 | 1.03097 | 0.426104 | -0.91599 | -0.88985 | -0.33433 | -0.8411 | 0.85063 | 1.818849 | -1.3305 | -0.35241 | 0.310526 | 2.109281257 |
| MNK1KO | -0.08605 | -1.37157 | -0.83298 | -1.88791 | -1.64238 | 0.504196 | -1.94112 | -0.93609 | -2.65047 | -1.48498 | -0.34804 | 0.996824 | 1.337396 | 2.650858 | 5.061948 | 1.459316 | -0.54048 | 1.013283 | 0.158302 | -0.85198 | -0.69064 | -1.47887 | -1.077777 |
| MNK1KO | -1.98638 | -0.76259 | -4.22448 | -1.76474 | -1.1549 | -1.39918 | -1.78379 | -0.88024 | -1.56457 | -0.27285 | -0.43939 | -0.59401 | -1.01068 | 0.884935 | -0.36959 | 0.156818 | -0.86317 | 1.024273 | -3.23892 | -0.66509 | -0.39524 | -1.72941 | -1.32002738 |
| MNK1KO | -0.6821 | -0.17434 | 0.913045 | -1.2052 | -0.30783 | -1.39918 | -1.6254 | -0.01303 | -1.08092 | -0.50232 | 3.19075 | -0.28267 | -0.38541 | 0.839729 | -1.37023 | 1.365421 | 0.50755 | 0.606651 | -2.91538 | -1.53104 | -1.01397 | -1.20667 | 1.036662774 |
| MNK1KO | -1.97978 | -0.96794 | -0.40702 | -1.22898 | -0.38843 | -0.62985 | -2.0072 | -1.83738 | -0.61789 | -1.34497 | 1.971437 | -0.7788 | -0.11934 | 0.715779 | 0.313688 | -0.79177 | 0.383441 | 0.661601 | -2.49191 | -0.58239 | -0.71231 | -1.35823 | -1.33263001 |
| MNK1KO | -0.22902 | -0.72465 | 0.229484 | 0.941395 | -0.67983 | -1.39918 | -1.48276 | -1.93584 | -0.18523 | -0.97939 | 0.033245 | -0.47148 | -0.49184 | 2.391292 | 0.077835 | 0.737044 | 0.775074 | 2.246368 | -0.82423 | -0.50513 | -0.68066 | -1.29482 | -0.1983964 |
| MNK1KO | 0.89985 | -1.39383 | 0.582147 | 0.085015 | 0.567912 | -1.39918 | -1.82469 | -1.93584 | -1.05207 | 0.044764 | 0.8256 | -0.34695 | -0.09939 | 0.027492 | 1.054194 | 1.497837 | 0.422052 | 1.008887 | -0.91939 | -0.48073 | -0.44303 | -1.2608 | -1.04557175 |
| MNK1KO | 1.167084 | -1.24259 | -1.32225 | -1.71003 | -1.11615 | -1.39918 | -1.61701 | -0.689 | -2.66029 | -1.15051 | 0.815671 | 0.086915 | 0.266462 | 0.581622 | 0.095177 | 2.041949 | 0.278637 | 1.732033 | -0.89084 | -0.29444 | -0.66125 | -1.09995 | -0.47145292 |
| MNK1KO | -1.80327 | -1.44946 | -0.56466 | 0.096734 | 0.354013 | -1.39918 | -1.61911 | -1.32141 | -0.8244 | -0.41157 | 0.440345 | -1.13232 | 0.785299 | 1.641759 | 3.513301 | -0.86159 | 0.750252 | 2.710148 | -0.01537 | -1.13054 | -1.33304 | -1.284 | -0.40843988 |
| MNK1KO | 0.569933 | -2.43047 | -0.67191 | -1.24466 | -0.20553 | 0.359379 | -0.58386 | -1.94531 | 0.520839 | 0.228852 | 0.853402 | 0.936565 | -1.2834 | 0.814939 | 1.871006 | 2.311598 | 1.290816 | 2.382644 | 1.359701 | -0.7948 | -1.00973 | -1.16027 | 1.294316362 |
| MNK1KO | 0.894352 | -0.0972 | -1.55205 | -0.4231 | -0.14586 | -1.20911 | 0.860457 | -0.42297 | -1.74748 | 1.473393 | 0.863332 | 0.520779 | 0.106819 | -0.20291 | 4.118539 | 2.925529 | 2.686355 | 1.58037 | -1.20725 | -0.68906 | -0.89831 | -1.44175 | -1.25141311 |
| MNK1KO | 2.119446 | 0.378503 | -0.83298 | -1.79589 | 0.97246 | 1.062697 | -1.27403 | -0.61988 | -1.67898 | -0.82901 | 0.414529 | 0.583046 | 0.632309 | 1.377818 | 2.745047 | 4.028199 | 1.834139 | 0.760512 | -2.68461 | -0.33249 | -0.06084 | -1.34122 | -1.64209437 |
| MNK1KO | 2.130443 | -1.26788 | -0.98126 | -0.62965 | -0.41401 | -0.71769 | -0.6363 | -1.50981 | -0.66661 | 1.017061 | 2.003211 | -0.72256 | 1.091281 | 1.592179 | 4.154958 | 0.888721 | 1.734852 | 0.142871 | -1.54983 | -2.02883 | -1.51419 | -2.18875 | -0.38463284 |
| MNK1KO | 0.990578 | -0.89865 | -2.27742 | -0.46441 | 0.566362 | -1.39918 | -0.25661 | -1.93584 | 0.642628 | 0.335157 | 0.867303 | -0.33891 | 0.652264 | 2.423374 | 1.149576 | -1.38162 | -0.00819 | 0.162653 | -2.41341 | -1.4176 | -1.01139 | -1.42938 | -1.56507835 |
| MNK1KO | -0.73599 | -2.66389 | -0.49992 | -0.9157 | -0.09626 | 0.058565 | -1.04747 | -1.93584 | -1.10048 | 1.027433 | -1.09472 | 1.010884 | 1.237619 | -0.30644 | 2.57336 | -0.66176 | -0.44671 | 0.195623 | -3.12473 | -1.59213 | -1.0301 | -1.58249 | -1.70510748 |
| MNK2KO | -0.3115 | 0.225751 | -0.15539 | -0.42832 | 0.804286 | -0.91202 | -0.02166 | -0.2696 | -0.4062 | -0.33119 | -1.42238 | -0.19028 | 0.239855 | -1.1391 | 0.041417 | -2.09667 | -0.74733 | 1.435301 | -1.87813 | 0.290914 | 2.261186 | 2.081365 | -0.97975661 |
| MNK2KO | 2.087554 | -0.35896 | 0.337245 | 0.92015 | -1.3626 | 0.083589 | 2.244988 | 3.664064 | 1.113343 | 1.13633 | 0.398642 | 0.916479 | 0.406149 | 1.764251 | 0.027543 | -0.01653 | 0.22072 | 1.211104 | -1.68305 | 0.335492 | 0.828777 | -0.97313 | -1.25001282 |
| MNK2KO | 0.309298 | 0.026465 | -0.77358 | 2.172262 | 0.096715 | -0.22148 | -0.23773 | 1.807529 | -0.20982 | 0.853715 | 1.308162 | -0.47148 | -0.21912 | 2.157975 | 2.883783 | 0.900759 | -0.29778 | 1.373757 | -1.61644 | -0.75409 | -0.66992 | -1.54692 | -1.7163098 |
| MNK2KO | -0.0932 | -1.27648 | -0.07409 | 0.848911 | 0.06494 | -0.00639 | -1.04117 | 0.119511 | -0.59985 | 0.16144 | -0.354 | -0.2425 | 2.913864 | 3.845154 | 2.928873 | 1.069289 | -0.56255 | 1.288035 | -3.12473 | -0.60154 | -0.37305 | -1.52526 | -1.8605398 |
| MNK2KO | 0.73929 | 1.057289 | 1.310696 | 0.115871 | -0.07378 | 0.042061 | -0.06571 | -1.09136 | 0.376823 | 0.56073 | 1.228728 | -0.15814 | -1.18362 | 1.071589 | 1.746143 | 2.049172 | 2.143033 | 1.274847 | -2.17788 | -0.7942 | -0.83693 | -1.52217 | -1.74571592 |
| MNK2KO | -0.54904 | -0.30129 | -1.13227 | 0.565728 | -0.58916 | -0.00053 | -0.58596 | -0.49586 | -1.75669 | -0.93531 | 2.960391 | 0.228021 | -0.46856 | 4.330017 | 0.288975 | 1.568861 | -0.62736 | 0.766556 | -3.17588 | -0.79071 | -0.68144 | -2.09054 | -1.76952087 |
| MNK2KO | 0.119045 | 0.408346 | 0.165243 | -0.63976 | 2.058229 | 0.357249 | 2.562801 | 4.360857 | -1.29077 | -0.97939 | -0.80876 | 0.245596 | 0.332979 | -1.09535 | 0.422943 | -0.16098 | 0.984681 | 0.86162 | -3.09618 | -0.47668 | 1.063999 | 0.544063 | -1.56087748 |
| MNK2KO | 0.967483 | 0.787191 | 0.279317 | -0.05637 | 0.243189 | -0.58565 | -0.51673 | -0.33303 | -1.33645 | -0.11858 | 1.292276 | 0.671426 | 1.596815 | 3.190406 | 3.097091 | 4.685466 | 0.984681 | 0.709958 | -2.43006 | -1.34932 | -0.97907 | -1.74642 | -1.24721224 |
| MNK2KO | 2.458711 | -1.76053 | 0.741378 | -0.56777 | -1.07895 | -1.39918 | -0.99397 | -0.60852 | -0.87974 | -0.0771 | -0.79883 | 2.985367 | -1.25014 | -0.47706 | -0.23952 | 6.693384 | -0.50739 | 1.316609 | -2.96296 | -0.25491 | 0.646438 | -0.61742 | -1.33263001 |
| MNK2KO | 0.646364 | -0.76157 | -0.44927 | -0.21874 | -0.848 | -1.39918 | -0.65623 | 2.7855 | -0.77317 | 0.094027 | 0.364883 | 0.564968 | 1.650029 | -0.44352 | 0.556477 | 2.133437 | 0.637175 | 1.345183 | -2.10651 | -0.62441 | -0.48507 | -1.11542 | -1.19120059 |
| MNK2KO | -0.8839 | 0.235867 | 0.812277 | 0.048426 | -0.67673 | 0.042061 | -0.23353 | 1.418423 | -0.5981 | -0.58269 | 0.952695 | -0.87723 | 0.732085 | 2.990627 | 3.563593 | 2.362157 | 1.671418 | 1.202312 | -2.15409 | -1.13378 | -1.00435 | -2.03409 | -1.47966059 |
| MNK2KO | 0.518795 | -0.71049 | 0.046536 | -0.80686 | -1.59433 | -0.41209 | 2.047797 | -1.63004 | -1.13854 | -0.82901 | -0.73329 | 1.466843 | 0.253158 | -0.89411 | 0.433348 | 2.506611 | 0.413778 | 0.734136 | -3.11521 | -0.46683 | 0.914885 | -0.48596 | -1.36063583 |
| MNK2KO | 0.449513 | 0.478652 | 0.883496 | 1.106635 | 0.582637 | 0.791699 | -0.13074 | 3.633769 | -0.28792 | -0.29489 | 0.390699 | 0.550908 | 1.237619 | 0.877644 | 2.280279 | 0.385538 | 0.482728 | 1.51443 | -0.44597 | -1.39593 | -1.16389 | -1.52062 | 1.043660688 |
| MNK2KO | 2.041365 | -0.90421 | -0.91763 | 0.953198 | 0.00604 | -0.21563 | -0.06256 | -0.27812 | -0.72156 | 1.341161 | 0.563468 | 3.045626 | 0.439408 | 0.176232 | -0.77193 | 1.702481 | -0.30881 | 0.890195 | -1.71398 | -0.45562 | -0.34464 | -0.86024 | 0.545157505 |
| MNK2KO | 1.318846 | 0.665799 | 2.263316 | 2.412955 | 1.091034 | 0.56915 | 3.960967 | 0.243532 | 0.042055 | 1.224485 | 1.409441 | -0.54982 | -0.32555 | 1.369069 | 3.879219 | 0.619072 | 1.10879 | 1.430905 | 1.797439 | -1.32154 | -1.47041 | -1.4634 | 0.580164751 |
| MNK2KO | -0.82342 | -0.51323 | 0.597476 | -1.33124 | 0.211414 | -0.71556 | 0.011908 | 2.813902 | 0.741582 | -0.67344 | -1.78182 | 1.272006 | -1.13041 | -0.60538 | -0.4459 | 0.505916 | -1.35685 | 2.116685 | -2.10414 | -0.14449 | 1.209291 | 0.457454 | -0.48965468 |
| MNK2KO | -0.15864 | -0.1799 | 1.23749 | -0.90693 | 0.602787 | 0.709175 | -1.56876 | 3.151884 | 0.863371 | 0.425904 | 0.015373 | 1.16354 | 0.253158 | -0.43331 | 1.853664 | 2.862932 | -0.48532 | 1.844132 | -1.36902 | 0.010509 | -0.16896 | -0.33749 | -0.94474933 |

| classical tests | |
|---|---|
| SI trial1 | social habituation/recognition test, time interacting trial 1 |
| SR index | social habituation/recognition test, recognition index |
| NOR DI | Novel object recognition, discrimination index |
| NO 1st trial | Novel object recognition, total object interaction trial 1 |
| SO Index | social olfaction test, social odor recognition index |
| SO S1 | social olfaction test, social odor interaction trial 1 |
| OI trial1 | Oject habituation/recognition test, time interacing trial 1 |
| OH DI | Oject habituation/recognition test, object recognition index |
| OF distance | Open field, total distance traveled |
| OF time in center | Open field, total time in center |

| deepOF: social habituation/recognition test merged trials |
|---|
| nose to nose |
| nose to tail test |
| nose to tail stim |
| side by side |
| side by side reverse |
| nose to body test |
| nose to body stim |
| climbing test |
| sniffing test |
| speed test |
| speed stim |
| climbin stim |
| sniffing stim |
