## Supplementary Table 2 for "Distinct roles for MNK1 and MNK2 in social and cognitive behavior through kinase-specific regulation of the synaptic proteome and phosphoproteome"

**TMT cortex all proteins**

| protein names | | normalized protein expression | | | | | | | | | | | | | | | | LogFC over WT | | | p-value / WT | | | adjusted p-value / WT | | |
|---|---|---|---|---|---|---|---|---|---|---|---|---|---|---|---|---|---|---|---|---|---|---|---|---|---|---|
| id | GeneName | CX_WT | CX_WT | CX_WT | CX_WT | CX_MNK1K | CX_MNK1K | CX_MNK1K | CX_MNK1K | CX_MNK2K | CX_MNK2K | CX_MNK2K | CX_MNK2K | CX_MNKD | CX_MNKD | CX_MNKD | CX_MNKD | logFC.CX_ | logFC.CX_ | logFC.CX_ | P.Value.CX | P.Value.CX | P.Value.CX | adj.P.Val.C | adj.P.Val.C | adj.P.Val.CX_MNKDKO.over.CX_WT |
| 0610009B2 | Q8R3W2 | 2.55 | -0.8 | -0.06 | -0.94 | -1.55 | -0.86 | -0.45 | -0.88 | -1.65 | 1.4 | -0.36 | 0.18 | -1.23 | -0.75 | -0.27 | -0.09 | -1.1225 | -0.295 | -0.7725 | 0.164084 | 0.750644 | 0.328114 | 0.999971 | 1 | 0.856472 |
| 0610010K1 | D3Z687 | -1.05 | 1.85 | 0.28 | 1.08 | 2.47 | 1.24 | -0.09 | 0.95 | 1.52 | -0.15 | 0.24 | 1.1 | 2.61 | 1.21 | 1.3 | 0.97 | 0.6025 | 0.1375 | 0.9825 | 0.420356 | 0.836167 | 0.159477 | 0.999971 | 1 | 0.799489 |
| 0610012G | A0A0J9YTR | -0.83 | -2.86 | -1.05 | -0.63 | -3.21 | -2.2 | -1.3 | -0.62 | -3.97 | -1.57 | -0.5 | -1.11 | -4.16 | -2.2 | -0.46 | -0.2 | -0.49 | -0.445 | -0.4125 | 0.482883 | 0.594268 | 0.662453 | 0.999971 | 1 | 0.936148 |
| 1110059G1 | Q8K039 | 2.96 | 2.95 | 3.36 | 5.02 | 3 | 2.74 | 3.66 | 3.83 | 2.48 | 2.95 | 3.66 | 2.65 | 0.67 | 2.37 | 2.96 | 2.46 | -0.265 | -0.6375 | -1.4575 | 0.609042 | 0.238347 | 0.045333 | 0.999971 | 1 | 0.710102 |
| 1110065P2 | V9GXF1 | 5.14 | 0.56 | 0.97 | 1.14 | 0.14 | 1.51 | 1.25 | 0.61 | 0.96 | 3.43 | 0.77 | 0.99 | -0.58 | 0.73 | 0.97 | 1.05 | -1.075 | -0.415 | -1.41 | 0.295736 | 0.71061 | 0.188364 | 0.999971 | 1 | 0.809025 |
| 1500009L1 | A0A1W2P6 | -2.17 | -2.24 | -1.78 | -1.82 | -1.8 | -2.38 | -1.48 | -1.66 | -1.98 | -3.37 | -1.16 | -1 | -1.43 | -3.04 | -2.14 | -2.23 | 0.1725 | 0.125 | -0.2075 | 0.502498 | 0.808478 | 0.552571 | 0.999971 | 1 | 0.913595 |
| 1600014C1 | Q8WUR0 | 0.89 | 1.29 | -0.76 | -1.16 | 1.2 | 1.42 | -0.14 | -0.86 | 0.98 | 1.02 | -0.39 | -0.83 | 1.41 | 1.65 | -0.52 | -0.07 | 0.34 | 0.13 | 0.5525 | 0.644493 | 0.852209 | 0.456505 | 0.999971 | 1 | 0.887217 |
| 1700025G | Q8K207 | -2.9 | -1.28 | -0.64 | -1.01 | -1.47 | -1.88 | -0.08 | -1.03 | -0.98 | -3.16 | -0.23 | -0.82 | -0.4 | -2.77 | -0.95 | -1.63 | 0.3425 | 0.16 | 0.02 | 0.556655 | 0.828322 | 0.975382 | 0.999971 | 1 | 0.99925 |
| 1700037H | H7BXB9 | -0.23 | -0.23 | 0 | 0.16 | 0.04 | -0.11 | -0.42 | 0.69 | -0.06 | -0.44 | -0.06 | 0.32 | 0.35 | 0.16 | 0.89 | 0.64 | 0.125 | 0.015 | 0.585 | 0.646839 | 0.945997 | 0.028192 | 0.999971 | 1 | 0.678474 |
| 2210016F1 | G3X8U3 | -0.2 | 1.01 | 0.61 | 0.37 | 0.9 | 1.02 | 0.27 | 0.45 | 0.69 | -0.29 | -0.2 | 0.2 | 0.03 | 0.88 | 0.44 | 0.19 | 0.2125 | -0.3475 | -0.0625 | 0.503902 | 0.315563 | 0.843933 | 0.999971 | 1 | 0.979814 |
| 2310061I0 | B8JJ66 | -0.92 | -0.87 | -1.03 | -0.82 | -1.09 | -0.96 | -0.5 | -0.63 | -1.25 | -0.94 | -1.28 | -1.04 | -1.81 | -0.79 | -1.48 | -1.65 | 0.115 | -0.2175 | -0.5225 | 0.572427 | 0.231181 | 0.066655 | 0.999971 | 1 | 0.733937 |
| 2410002F2 | Q3TE80 | -0.22 | 0.94 | 0.61 | 0.08 | 1.03 | 1.04 | 0.01 | 0.56 | 1.05 | 0.66 | 0.2 | 0.7 | 1.21 | 1.34 | 1.04 | 1.14 | 0.3075 | 0.3 | 0.83 | 0.390862 | 0.355646 | 0.015837 | 0.999971 | 1 | 0.613095 |
| 2410004B1 | Q9CWU4 | 3.74 | 1.4 | 2.25 | 3.99 | 1.88 | 1.89 | 3.16 | 2.39 | 1.57 | 2.7 | 1.61 | 0.8 | 0.36 | 1.16 | 1.56 | 1.51 | -0.515 | -1.175 | -1.6975 | 0.418426 | 0.105394 | 0.021207 | 0.999971 | 1 | 0.640006 |
| 2900026A | A0A1D5RLI | -1.19 | -2.08 | -1.16 | -0.79 | -1.98 | -2.58 | 1.99 | -1.7 | -2.21 | -1.88 | -1.94 | -2.1 | -2.17 | -2.49 | -2.67 | -1.54 | 0.2375 | -0.7275 | -0.9125 | 0.804101 | 0.033248 | 0.030763 | 0.999971 | 1 | 0.678474 |
| 2900026A | A0A1B0GR | 0.83 | -0.28 | -0.83 | -0.71 | 0.09 | -0.18 | -0.33 | -0.47 | -0.28 | 0.56 | -0.52 | -0.94 | -0.79 | -0.48 | -1.19 | -0.9 | 0.025 | -0.0475 | -0.5925 | 0.9478 | 0.918222 | 0.154954 | 0.999971 | 1 | 0.795578 |
| 3110082I1 | Q9CXL3 | 2.69 | 3.06 | 1.89 | 1.91 | 2.09 | 2.13 | 1.68 | 1.4 | 3.07 | 2.87 | 1.69 | 1.34 | 1.57 | 1.96 | 1.82 | 2.45 | -0.5625 | -0.145 | -0.4375 | 0.122134 | 0.764972 | 0.222559 | 0.999971 | 1 | 0.827753 |
| 4931406C | Q91V76 | 0.32 | 0.07 | 0.07 | -0.38 | 0.61 | 0.38 | 0.5 | -0.16 | 0.69 | 0.38 | 0.52 | -0.14 | 0.71 | 0.22 | -0.24 | -0.05 | 0.3125 | 0.3425 | 0.14 | 0.234869 | 0.201803 | 0.609342 | 0.999971 | 1 | 0.92673 |
| 4933402D | Q9D4D3 | -1.65 | -1.69 | -1.35 | -1.77 | -1.22 | -0.84 | -0.96 | -2.22 | -2.61 | -1.26 | -1.38 | -1.16 | -1.16 | -1.21 | -1.52 | -1.18 | 0.305 | 0.0125 | 0.3475 | 0.363032 | 0.971154 | 0.097499 | 0.999971 | 1 | 0.78268 |
| 4933427D1 | A0A0N4SU | 1.56 | 0.69 | -0.77 | -0.27 | 0.58 | 0.99 | 0.26 | -0.35 | 0.96 | 0.89 | -0.47 | -1.16 | 0.61 | -0.58 | -0.89 | -0.59 | 0.0675 | -0.2475 | -0.665 | 0.900838 | 0.712785 | 0.257894 | 0.999971 | 1 | 0.839012 |
| 4933434E2 | H7BX33 | -0.25 | -0.3 | -0.75 | -0.47 | -0.68 | -0.25 | -0.9 | -0.44 | 0.03 | 0.26 | -0.49 | -0.65 | -1.01 | -0.03 | -1.05 | -0.38 | -0.125 | 0.23 | -0.175 | 0.578536 | 0.391959 | 0.546204 | 0.999971 | 1 | 0.911398 |
| 5031439G | B1APX2 | -0.54 | -0.55 | -0.36 | -0.05 | -0.5 | -0.41 | -0.34 | 0 | -0.62 | -0.6 | -0.04 | 0 | -0.26 | -0.4 | -0.23 | 0.24 | 0.0625 | 0.06 | 0.2125 | 0.766713 | 0.801103 | 0.355848 | 0.999971 | 1 | 0.868793 |
| 5730455P1 | Q9CYI0 | 0.81 | 1.17 | 1.22 | 0.33 | 0.66 | 1.15 | 2.22 | -0.03 | 1.1 | 0.8 | 0.57 | 0.15 | 1.32 | 1.37 | 0.04 | -1.03 | 0.1175 | -0.2275 | -0.4575 | 0.806925 | 0.449901 | 0.423582 | 0.999971 | 1 | 0.87919 |
| 6030458C1 | Q8BGC1 | 0.97 | 1.28 | 0.93 | 1.17 | 0.69 | -0.11 | 0.71 | 1.23 | -0.23 | 0.62 | 0.74 | 0.73 | 1.19 | 0.67 | 0.63 | 1.13 | -0.4575 | -0.6225 | -0.1825 | 0.150432 | 0.041848 | 0.409533 | 0.999971 | 1 | 0.875847 |
| 6820408C1 | Q8BJX2 | -0.64 | -0.56 | 0.69 | 0.65 | -0.48 | -0.49 | 0.51 | 0.32 | -0.4 | -0.31 | 0.61 | 0.29 | 0.05 | -0.7 | 0.68 | 0.94 | -0.07 | 0.0125 | 0.2075 | 0.869729 | 0.976157 | 0.669015 | 0.999971 | 1 | 0.938711 |
| 8030462N | Q0VAW6 | -0.39 | 0.54 | 1.57 | 1.89 | 0.87 | 0.21 | 1.41 | 1.72 | 0.61 | -0.59 | 1.62 | 1.85 | 0.38 | -0.3 | 1.57 | 0.79 | 0.15 | -0.03 | -0.2925 | 0.790417 | 0.965419 | 0.624656 | 0.999971 | 1 | 0.930479 |
| A830018L1 | B7ZN57 | -2.22 | -1.98 | -1.56 | -1.65 | -2.14 | -2.96 | -1.41 | -1.24 | -2.11 | -4.92 | -2.31 | -1.79 | -1.92 | -2.91 | -2.02 | -2.03 | -0.085 | -0.93 | -0.3675 | 0.832667 | 0.190246 | 0.225857 | 0.999971 | 1 | 0.828225 |
| Aaas | P58742 | 1.32 | -0.47 | -0.9 | -0.47 | 0.03 | -0.69 | -0.9 | -0.52 | -0.08 | 0.85 | -0.82 | -0.78 | 0.17 | -0.32 | -0.5 | -0.12 | -0.39 | -0.0775 | -0.0625 | 0.438387 | 0.893621 | 0.896336 | 0.999971 | 1 | 0.988571 |
| Aacs | Q9D2R0 | -0.21 | -0.1 | 0.09 | -0.09 | 0.09 | -0.03 | 0.06 | -0.09 | 0.38 | 0.12 | 1.05 | 0.81 | 1.02 | 0.69 | 0.48 | 1.03 | 0.085 | 0.6675 | 0.8825 | 0.619065 | 0.022576 | 0.001934 | 0.999971 | 1 | 0.476373 |
| Aagab | Q8R2R3 | 0.86 | 0.23 | -0.18 | 0.16 | -0.51 | -0.48 | -0.09 | 0.63 | -0.11 | 0.42 | 0.3 | 0.46 | -0.01 | 0.44 | 0.08 | 0.7 | -0.38 | 7.85E-17 | 0.035 | 0.280254 | 1 | 0.902597 | 0.999971 | 1 | 0.990759 |
| Aak1 | Q3UHJ0 | -0.34 | -0.13 | -0.03 | -0.56 | -0.52 | -0.24 | -0.34 | -0.12 | -0.48 | -0.44 | -0.24 | -0.67 | 0.02 | -0.28 | -0.17 | -0.2 | -0.04 | -0.1925 | 0.1075 | 0.842888 | 0.349786 | 0.587344 | 0.999971 | 1 | 0.920758 |
| Aak1_1 | A0A571BEI | 0.95 | -0.49 | -0.62 | -0.04 | -0.67 | -0.74 | -0.39 | 0.15 | -0.39 | -0.3 | -0.57 | -0.24 | -0.58 | -0.63 | -0.15 | 0.01 | -0.3625 | -0.325 | -0.2875 | 0.367561 | 0.372767 | 0.453714 | 0.999971 | 1 | 0.887217 |
| Aak1_2 | A0A571BD | -0.06 | -2.53 | -0.97 | -1.41 | -1.5 | -1.73 | -1.39 | -1.38 | -2.37 | -0.75 | -2.66 | -0.84 | -0.38 | -1.37 | -1.67 | -1.18 | -0.2575 | -0.4125 | 0.0925 | 0.597058 | 0.533261 | 0.863278 | 0.999971 | 1 | 0.983079 |
| Aamdc | D3Z442 | 0.14 | -0.09 | 0.38 | 0.09 | 0.24 | 0.24 | 0.43 | 0.5 | -0.12 | -0.03 | 0.56 | 0.31 | 0 | 0.33 | 0.46 | 0.14 | 0.2225 | 0.05 | 0.1025 | 0.254647 | 0.822812 | 0.610864 | 0.999971 | 1 | 0.92673 |
| Aamp | J3QN89 | 0.82 | 0.84 | 1.17 | 0.54 | 0.73 | 0.84 | 1.18 | 0.91 | 0.68 | 0.93 | 0.91 | 0.66 | 0.46 | 1.06 | 0.74 | 0.45 | 0.0725 | -0.0475 | -0.165 | 0.731467 | 0.813139 | 0.4826 | 0.999971 | 1 | 0.896345 |
| Aar2 | Q9D2V5 | -0.49 | -0.14 | -0.04 | 0.29 | 0.13 | -0.22 | -0.08 | 0.35 | 0.13 | -0.14 | 0.18 | 0.53 | 0.31 | 0 | 0.29 | 0.25 | 0.14 | 0.27 | 0.3075 | 0.559107 | 0.280519 | 0.187134 | 0.999971 | 1 | 0.809025 |
| Aars1 | Q8BGQ7 | 0.41 | 0.43 | 0.64 | 0.64 | 0.23 | 0.23 | 0.59 | 0.51 | 0.46 | 0.37 | 0.46 | 0.76 | 0.12 | 0.33 | 0.36 | 0.53 | -0.14 | -0.0175 | -0.195 | 0.458065 | 0.922129 | 0.30183 | 0.999971 | 1 | 0.847402 |
| Aars2 | Q14CH7 | 0.51 | -0.37 | -0.38 | -0.46 | -0.76 | -0.48 | 0.13 | -0.48 | -0.66 | 0.24 | -0.5 | -0.46 | -0.49 | -0.42 | -0.55 | -0.25 | -0.2225 | -0.17 | -0.2525 | 0.470534 | 0.585386 | 0.348006 | 0.999971 | 1 | 0.86438 |
| Aarsd1 | Q3THG9 | 0.51 | 1.06 | 0.88 | 0.71 | 0.52 | 1.63 | 0.62 | 1.08 | 0.82 | 1.27 | 0.63 | 0.39 | 0.93 | 1.5 | 1.05 | 1.28 | 0.1725 | -0.0125 | 0.4 | 0.557343 | 0.959635 | 0.094427 | 0.999971 | 1 | 0.779538 |
| Aasdhppt | Q9CQF6 | -1.51 | -0.36 | 2.27 | 1.41 | 0.16 | -0.61 | 1.93 | 2.01 | -0.21 | -1.37 | 2.61 | 2.1 | -1.54 | -0.16 | 1.91 | 1.76 | 0.42 | 0.33 | 0.04 | 0.663122 | 0.77275 | 0.96985 | 0.999971 | 1 | 0.998787 |
| Aass | Q99K67 | 0 | -0.65 | -0.76 | -0.94 | -0.61 | -0.93 | -1.24 | -1.77 | -0.86 | -0.35 | -0.92 | -1.21 | -0.68 | -0.93 | -1.23 | -1.05 | -0.55 | -0.2475 | -0.385 | 0.115156 | 0.394893 | 0.163285 | 0.999971 | 1 | 0.800215 |
| Aatf | Q9JKX4 | -0.96 | 1.28 | -2.19 | -2.12 | 0.64 | 0.71 | -2.62 | -1.55 | 1.1 | 0.19 | -1.48 | -2.19 | 1.37 | 1.23 | -3.02 | -1.97 | 0.2925 | 0.4025 | 0.4 | 0.777031 | 0.687011 | 0.745154 | 0.999971 | 1 | 0.961064 |
| Aatk | B1AZF9 | -0.15 | 0.01 | -0.13 | -0.34 | -0.01 | 0.04 | -0.47 | 0.15 | -0.11 | 0.1 | 0.03 | 0.24 | 0.49 | 0.11 | 0.1 | 0.02 | 0.08 | 0.2175 | 0.3325 | 0.700369 | 0.24046 | 0.112505 | 0.999971 | 1 | 0.78268 |
| Abat | P61922 | -1.07 | -1.21 | -1.47 | -0.88 | -1.16 | -1.28 | -1.45 | -0.51 | -1.11 | -1.45 | -1.81 | -1.46 | -0.97 | -0.87 | -1.07 | -1.23 | 0.0575 | -0.3 | 0.1225 | 0.826695 | 0.207747 | 0.550475 | 0.999971 | 1 | 0.913169 |
| Abca1 | P41233 | -1.07 | -0.66 | -0.84 | -1.22 | -0.76 | -0.85 | -1.17 | -1.16 | -0.79 | -1.08 | -0.79 | -0.65 | -0.59 | -1.02 | -0.56 | -0.67 | -0.0375 | 0.12 | 0.2375 | 0.859511 | 0.560094 | 0.281523 | 0.999971 | 1 | 0.844538 |
| Abca2 | A2AJ26 | -0.24 | 1.01 | 0.71 | 0.83 | 1 | 0.84 | 0.73 | 0.64 | 1.29 | 0.13 | 1.14 | 1.07 | 1.18 | 0.92 | 0.93 | 0.8 | 0.225 | 0.33 | 0.38 | 0.458816 | 0.387123 | 0.226321 | 0.999971 | 1 | 0.828236 |
| Abca3 | Q8R420 | -0.79 | 0.03 | -0.67 | -0.7 | -0.18 | -0.33 | -0.73 | -0.37 | -0.09 | -0.51 | 0.07 | -0.16 | 0.69 | 0.02 | -0.6 | -0.44 | 0.13 | 0.36 | 0.45 | 0.605923 | 0.175893 | 0.211014 | 0.999971 | 1 | 0.822884 |
| Abca5 | Q8K448 | -0.3 | 0.16 | -0.9 | -0.72 | -0.1 | 0.65 | -0.96 | -0.99 | -0.19 | 0.12 | -0.88 | -0.87 | 0.09 | 0.29 | -1.29 | -0.62 | 0.09 | -0.015 | 0.0575 | 0.835056 | 0.96485 | 0.888616 | 0.999971 | 1 | 0.987485 |
| Abca6 | Q8K441 | 1.7 | 1.65 | 3.88 | 2.16 | 1.01 | 2.71 | 1.99 | 2.91 | 2.13 | 2.89 | 2.76 | 2.24 | 1.72 | 1.78 | 1.69 | 4.15 | -0.1925 | 0.1575 | -0.0125 | 0.755423 | 0.76098 | 0.986201 | 0.999971 | 1 | 0.99925 |
| Abca7 | E9Q6G4 | 0.8 | 0.78 | 0.33 | 0.43 | -2.63 | -2.93 | -3.44 | -3.85 | 0.15 | 0.05 | 0.29 | 0.12 | 0.49 | -0.19 | -0.13 | 0.35 | -3.7975 | -0.4325 | -0.455 | 9.01E-07 | 0.047032 | 0.086573 | 0.007516 | 1 | 0.760141 |
| Abca8b | Q8K440-2 | -0.19 | -0.26 | -0.43 | -0.23 | -0.61 | -1.06 | -0.52 | -0.21 | -0.68 | -0.49 | -0.51 | -0.49 | -0.12 | -0.56 | 0.11 | 0.43 | -0.3225 | -0.265 | 0.2425 | 0.175462 | 0.134412 | 0.337435 | 0.999971 | 1 | 0.858827 |
| Abca9 | Q8K449 | -0.13 | 0.88 | 1.47 | 1.95 | 1.29 | 0.83 | 1.45 | 0.58 | 0.8 | 0.46 | 0.7 | 0.39 | 1.02 | 1.26 | 1.59 | 0.52 | -0.005 | -0.455 | 0.055 | 0.991312 | 0.310444 | 0.906611 | 0.999971 | 1 | 0.991405 |
| Abcb10 | Q9JI39 | -0.66 | -0.29 | -0.19 | -0.07 | -0.62 | -0.47 | -0.61 | 0.04 | -0.81 | -0.5 | -0.3 | -0.75 | -1.15 | -0.57 | -0.3 | 0.16 | -0.1125 | -0.2875 | -0.1625 | 0.63547 | 0.206335 | 0.601004 | 0.999971 | 1 | 0.92426 |
| Abcb11 | J3QNY6 | -1.31 | -1.46 | -3.78 | -1.38 | -0.9 | -4.23 | -0.42 | -4.11 | -2.92 | -1.94 | -3.33 | -3.39 | -3.49 | -2.89 | -3.2 | -3.21 | -0.4325 | -0.9125 | -1.215 | 0.682566 | 0.172136 | 0.055606 | 0.999971 | 1 | 0.725348 |
| Abcb1a | P21447 | 0.09 | 0.01 | -0.37 | 0.08 | 0.49 | -0.25 | -0.27 | -0.67 | 0.26 | 0.08 | -0.02 | 0.09 | -0.37 | -0.32 | -0.02 | -0.37 | -0.1275 | 0.15 | -0.2225 | 0.651237 | 0.432507 | 0.279893 | 0.999971 | 1 | 0.843812 |
| Abcb6 | Q9DC29 | -0.33 | 0.18 | 0.21 | -0.33 | 0.03 | 0.18 | 0.58 | -0.07 | 0.36 | -0.37 | 0.95 | 0.39 | 0.06 | 0 | 0.42 | -0.21 | 0.2475 | 0.4 | 0.135 | 0.318829 | 0.224048 | 0.571112 | 0.999971 | 1 | 0.91816 |
| Abcb7 | Q61102 | 0.03 | -0.27 | -0.3 | -0.23 | -0.34 | -0.13 | -0.44 | -0.37 | -0.51 | -0.3 | -0.33 | -0.62 | -0.51 | -0.19 | -0.21 | -0.26 | -0.1275 | -0.2475 | -0.1 | 0.484874 | 0.191661 | 0.587497 | 0.999971 | 1 | 0.920758 |
| Abcb8 | Q9CXJ4 | 0.14 | -0.49 | 0 | 0.18 | -0.26 | -0.23 | -0.11 | -0.01 | -0.81 | -0.2 | -0.73 | -0.6 | -0.6 | -0.21 | -0.32 | -0.36 | -0.11 | -0.5425 | -0.33 | 0.608164 | 0.04499 | 0.158757 | 0.999971 | 1 | 0.798217 |
| Abcb9 | Q9JJ59 | -0.58 | -0.09 | -0.57 | -0.46 | -0.07 | -0.41 | -0.74 | -0.65 | -0.38 | -0.62 | -0.35 | -0.55 | 0.08 | -0.21 | -0.62 | -0.49 | -0.0425 | -0.05 | 0.115 | 0.852102 | 0.794908 | 0.621927 | 0.999971 | 1 | 0.929641 |
| Abcc1 | O35379 | -0.54 | -0.14 | -0.6 | -0.24 | -0.42 | -0.22 | -0.8 | -0.21 | -0.38 | -0.56 | -0.84 | -0.6 | -0.42 | -0.41 | -0.61 | -0.87 | -0.0325 | -0.215 | -0.1975 | 0.883099 | 0.299298 | 0.356189 | 0.999971 | 1 | 0.868793 |
| Abcc4 | E9Q236 | -0.5 | -0.91 | -0.27 | -0.04 | -0.55 | -0.98 | 0.36 | -0.38 | -0.82 | -0.75 | -0.45 | 0.3 | -0.6 | -0.7 | -0.12 | 0.08 | 0.0425 | 2.16E-16 | 0.095 | 0.89856 | 1 | 0.735131 | 0.999971 | 1 | 0.958235 |
| Abcc5 | Q9R1X5 | 3.07 | -0.57 | 0.71 | 0.17 | -0.83 | -0.33 | 1.44 | -0.38 | -1.62 | 2.55 | -0.16 | -0.57 | -1.4 | -0.94 | -0.22 | -1.15 | -0.87 | -0.795 | -1.7725 | 0.313642 | 0.462935 | 0.039163 | 0.999971 | 1 | 0.700838 |
| Abcc8 | B2RUS7 | -0.23 | -0.71 | -0.76 | -0.63 | -1.02 | -0.64 | -1.4 | -0.88 | -0.83 | -0.5 | -0.7 | -1.17 | -0.68 | -0.84 | -0.97 | -0.55 | -0.4025 | -0.2175 | -0.1775 | 0.11379 | 0.344494 | 0.399426 | 0.999971 | 1 | 0.871816 |
| Abcd1 | P48410 | -0.22 | -0.41 | 0.02 | -0.8 | -0.37 | -0.85 | -0.52 | 0.34 | -0.37 | 0.15 | -0.67 | -0.36 | -0.75 | -0.79 | -0.58 | -0.23 | 0.0025 | 0.04 | -0.235 | 0.993559 | 0.879146 | 0.352146 | 0.999971 | 1 | 0.867656 |
| Abcd2 | Q61285 | -0.5 | -0.62 | -0.08 | -0.65 | -0.53 | -0.4 | -0.32 | -0.3 | -0.52 | -0.15 | -0.11 | 0.19 | 0.13 | -0.67 | -0.42 | 0 | 0.075 | 0.315 | 0.2225 | 0.708487 | 0.19589 | 0.392021 | 0.999971 | 1 | 0.871816 |
| Abcd3 | P55096 | -0.17 | -1.1 | -0.77 | -0.63 | -0.91 | -0.57 | -0.74 | -0.14 | -0.64 | 0.04 | -0.59 | -0.63 | -0.53 | -0.6 | -0.71 | -0.39 | 0.0775 | 0.2125 | 0.11 | 0.77634 | 0.441824 | 0.64672 | 0.999971 | 1 | 0.933613 |
| Abcd4 | O89016 | -1.31 | -1.64 | 1.47 | -0.45 | -1.39 | -1.67 | 1.15 | 1.67 | -1.3 | -1.42 | 1.75 | 0.7 | -0.81 | -1.3 | 0.83 | 0.06 | 0.4225 | 0.415 | 0.1775 | 0.669826 | 0.660531 | 0.8153 | 0.999971 | 1 | 0.974918 |
| Abce1 | P61222 | 0.86 | 1.08 | 1.48 | 1.38 | 1.04 | 1.1 | 1.13 | 1.57 | 1.21 | 1.09 | 1.22 | 1.07 | 0.88 | 1.22 | 1.47 | 1.51 | 0.01 | -0.0525 | 0.07 | 0.964765 | 0.793502 | 0.768734 | 0.999971 | 1 | 0.962792 |
| Abcf1 | Q6P542 | -0.05 | 1.37 | 0.49 | 0.54 | 1.37 | 1.27 | 0.39 | 0.59 | 1.33 | 0.63 | 0.5 | 0.9 | 1.23 | 1.06 | 0.99 | 0.94 | 0.3175 | 0.2525 | 0.4675 | 0.400925 | 0.4672 | 0.153483 | 0.999971 | 1 | 0.795578 |
| Abcf2 | Q99LE6 | 0.36 | 0.58 | 0.48 | 0.21 | 0.54 | 0.56 | 0.43 | 0.32 | 0.68 | 0.2 | 0.44 | 0.4 | 0.95 | 0.56 | 0.31 | 0.36 | 0.055 | 0.0225 | 0.1375 | 0.758254 | 0.905289 | 0.526366 | 0.999971 | 1 | 0.909459 |
| Abcf3 | Q8K268 | -0.4 | 0.21 | 0.02 | -0.34 | 0.31 | -0.32 | -0.08 | -0.28 | 0.28 | -0.51 | 0.33 | 0.14 | 0.07 | -0.33 | 0.05 | -0.12 | 0.035 | 0.1875 | 0.045 | 0.883287 | 0.482768 | 0.837049 | 0.999971 | 1 | 0.979416 |
| Abcg1 | Q64343 | 1.15 | -0.56 | -0.52 | -0.41 | -0.96 | -0.54 | 0.39 | -1.08 | -1.14 | 0.29 | -0.29 | -0.8 | -1.14 | -1.52 | -1.27 | -0.96 | -0.4625 | -0.4 | -1.1375 | 0.360257 | 0.417878 | 0.020464 | 0.999971 | 1 | 0.635388 |
| Abcg2 | S4R2E1 | -0.73 | -0.44 | -0.01 | -0.38 | -0.64 | -0.69 | -0.2 | -0.39 | -0.6 | -0.62 | 0.18 | -0.15 | -0.65 | -0.51 | -0.31 | -0.31 | -0.09 | 0.0925 | -0.055 | 0.692557 | 0.726056 | 0.799909 | 0.999971 | 1 | 0.970626 |
| Abcg4 | Q91WA9 | 3.13 | -2.18 | -1.67 | -0.48 | -1.73 | -1.61 | -0.91 | -0.37 | -1.73 | 2.38 | -0.94 | -0.65 | -2.14 | -0.39 | -0.4 | -0.53 | -0.855 | 0.065 | -0.565 | 0.44622 | 0.961289 | 0.620389 | 0.999971 | 1 | 0.929305 |
| Abhd10 | Q6PE15-2 | -0.09 | -0.24 | 0.13 | 0.03 | -0.61 | -0.62 | -0.37 | 0.26 | -0.47 | -0.18 | -0.31 | -1.39 | -0.65 | -0.4 | -0.24 | -0.25 | -0.2925 | -0.545 | -0.3425 | 0.259961 | 0.091921 | 0.101297 | 0.999971 | 1 | 0.78268 |
| Abhd11 | Q8K4F5 | -1.3 | -1.4 | -0.61 | -1 | -1.46 | -1.61 | -0.56 | -0.57 | -1.68 | -1.84 | -0.45 | -0.5 | -1.7 | -1.65 | -0.59 | -0.83 | 0.0275 | -0.04 | -0.115 | 0.933875 | 0.919547 | 0.730792 | 0.999971 | 1 | 0.956041 |
| Abhd12 | Q99LR1 | -0.76 | -0.36 | 0.48 | -0.09 | -0.25 | -0.46 | 0.73 | 0.16 | -0.09 | -0.69 | 0.28 | -0.86 | -0.36 | -0.86 | 0.02 | -0.22 | 0.2275 | -0.1575 | -0.1725 | 0.532579 | 0.665375 | 0.59521 | 0.999971 | 1 | 0.92377 |
| Abhd14b | E9QN99 | 1.12 | 0.93 | 0.97 | 1.26 | 0.87 | 0.89 | 0.53 | 1.09 | 0.43 | 0.51 | -0.21 | 0.57 | 0.15 | 0.88 | 0.86 | 0.54 | -0.225 | -0.745 | -0.4625 | 0.274243 | 0.009973 | 0.06815 | 0.999971 | 1 | 0.733937 |
| Abhd16a | Q9Z1Q2 | -0.43 | -0.6 | -0.46 | -0.43 | -0.56 | -0.52 | -0.34 | -0.37 | -0.83 | -0.58 | -0.39 | -0.28 | -0.38 | -0.51 | -0.32 | -0.08 | 0.0325 | -0.04 | 0.1575 | 0.846289 | 0.832837 | 0.39166 | 0.999971 | 1 | 0.871816 |
| Abhd17a | Q99JW1 | 1.55 | -1.42 | -0.78 | -0.5 | -1.19 | -0.41 | 0.04 | -0.66 | -1.76 | -0.23 | -0.83 | -1.34 | -1.71 | -1.72 | -1.04 | -1.05 | -0.2675 | -0.7525 | -1.0925 | 0.671928 | 0.271999 | 0.101843 | 0.999971 | 1 | 0.78268 |
| Abhd17b | Q7M759 | -0.93 | -0.81 | -0.68 | -1.61 | -0.59 | -0.71 | -0.6 | -0.91 | -0.33 | -0.82 | -0.55 | -0.23 | -0.06 | -0.84 | -0.32 | -1.54 | 0.305 | 0.525 | 0.3175 | 0.240178 | 0.074165 | 0.407241 | 0.999971 | 1 | 0.874911 |
| Abhd17c | Q8VCV1 | -0.3 | -0.68 | -0.32 | -0.13 | -0.84 | -1.02 | -0.82 | -0.23 | -0.5 | -0.35 | -0.36 | -0.41 | -0.75 | -0.95 | 0.07 | 0.36 | -0.37 | -0.0475 | 0.04 | 0.148944 | 0.798722 | 0.904901 | 0.999971 | 1 | 0.991091 |
| Abhd3 | Q91ZH7 | -0.12 | -0.57 | -0.02 | -0.05 | -0.1 | -0.01 | 0 | 0.73 | -0.05 | 0.29 | -0.18 | -0.54 | 0.73 | -0.12 | 0.29 | 0.04 | 0.345 | 0.07 | 0.425 | 0.201727 | 0.773714 | 0.120021 | 0.999971 | 1 | 0.78268 |
| Abhd4 | Q3U7M5 | 0.34 | -0.27 | 0.36 | 0.41 | -0.4 | -0.33 | -0.62 | 1.07 | -0.05 | 0.47 | 0.19 | 0.4 | -0.39 | -0.12 | 0.72 | 0.54 | -0.28 | 0.0425 | -0.0225 | 0.488926 | 0.855031 | 0.942803 | 0.999971 | 1 | 0.99692 |
| Abhd5 | Q9DBL9 | 0.21 | -0.83 | -0.83 | -1.13 | -1.01 | -0.88 | -1.13 | -0.37 | -0.93 | -0.38 | -1.36 | -0.91 | -1.33 | -0.97 | -0.47 | -0.66 | -0.2025 | -0.25 | -0.2125 | 0.550337 | 0.48136 | 0.542176 | 0.999971 | 1 | 0.911119 |
| Abhd6 | Q8R2Y0-2 | -2.46 | -2.44 | -1.65 | -1.8 | -2.32 | -2.17 | -1.38 | -1.72 | -2.29 | -2.13 | -1.41 | -1.47 | -2.9 | -2.22 | -1.59 | -1.37 | 0.19 | 0.2625 | 0.0675 | 0.5408 | 0.410999 | 0.862266 | 0.999971 | 1 | 0.982963 |
| Abi1 | B7ZCU3 | -2.12 | -0.31 | -1.45 | -2.64 | -0.28 | -1.91 | -0.51 | -2.17 | -0.45 | -2.7 | -0.67 | -0.62 | -0.8 | -3.13 | -1.97 | -4.61 | 0.4125 | 0.52 | -0.9975 | 0.519129 | 0.44447 | 0.264319 | 0.999971 | 1 | 0.842533 |
| Abi1_1 | B7ZCU4 | 0.39 | -0.34 | -0.02 | -0.82 | -0.48 | -0.5 | 0.23 | -0.64 | -0.18 | 0.18 | 0.05 | -0.1 | 0.06 | -0.51 | -0.54 | -0.69 | -0.15 | 0.185 | -0.2225 | 0.644509 | 0.516369 | 0.479791 | 0.999971 | 1 | 0.894438 |
| Abi2 | A0A087WF | 0.14 | -0.28 | -0.36 | -0.4 | -0.7 | -0.43 | 0.21 | -0.84 | -0.62 | -0.18 | -0.36 | -0.76 | -0.69 | -1.33 | -0.78 | -1.31 | -0.215 | -0.255 | -0.8025 | 0.449544 | 0.26541 | 0.008769 | 0.999971 | 1 | 0.610079 |
| Abi2_1 | A0A087WF | 2.01 | -4.6 | -1.85 | -1.11 | -4.47 | -4.74 | 0.39 | -2.64 | -3.98 | -0.55 | -3.84 | -4.72 | -9.03 | -6.59 | -3.46 | -3.72 | -1.4775 | -1.885 | -4.3125 | 0.366126 | 0.221765 | 0.028547 | 0.999971 | 1 | 0.678474 |
| Abi3 | Q8BYZ1 | 0.33 | 1.5 | 1.1 | 1.13 | 1.14 | 1.2 | 0.52 | 1.12 | 1.21 | 0.81 | 1.32 | 1.41 | 1.27 | 0.9 | 1.07 | 1.68 | -0.02 | 0.1725 | 0.215 | 0.946968 | 0.555898 | 0.487524 | 0.999971 | 1 | 0.897906 |
| Abl1 | P00520-2 | 1.47 | -0.04 | -0.16 | -0.27 | -0.15 | 0.13 | 0.12 | -0.18 | -0.09 | 0.69 | -0.18 | -0.07 | -0.27 | -0.27 | -0.35 | -0.31 | -0.27 | -0.1625 | -0.55 | 0.504735 | 0.708245 | 0.186578 | 0.999971 | 1 | 0.809025 |
| Abl2 | F8VQH0 | 0.82 | -0.37 | 0.08 | -0.16 | -0.43 | -0.25 | 0.18 | -0.19 | -0.4 | 0.28 | -0.06 | -0.13 | -0.17 | -0.45 | -0.21 | -0.32 | -0.265 | -0.17 | -0.38 | 0.385253 | 0.57664 | 0.201199 | 0.999971 | 1 | 0.815758 |
| Ablim1 | E9Q9C0 | -0.27 | -0.78 | -0.65 | -0.81 | -0.84 | -0.27 | -0.43 | -0.86 | -0.76 | -0.28 | -0.91 | -1.14 | -0.38 | -0.76 | -0.68 | -0.84 | 0.0275 | -0.145 | -0.0375 | 0.905104 | 0.561268 | 0.858845 | 0.999971 | 1 | 0.982702 |
| Ablim2 | Q8BL65 | -0.82 | -0.62 | 0.12 | -0.43 | -0.84 | -0.72 | -0.17 | -0.44 | -0.82 | -1.17 | -0.35 | -0.84 | -0.39 | -1.43 | -0.38 | -0.21 | -0.105 | -0.3575 | -0.165 | 0.699129 | 0.22045 | 0.631123 | 0.999971 | 1 | 0.932858 |
| Ablim2_1 | E9Q0W6 | -1.89 | -1.65 | -1.62 | -1.46 | -1.86 | -1.56 | -2.43 | -1.55 | -1.85 | -1.48 | -1.85 | -1.57 | -1.28 | -1.18 | -1.22 | -1.52 | -0.195 | -0.0325 | 0.355 | 0.446453 | 0.864997 | 0.086035 | 0.999971 | 1 | 0.759991 |
| Ablim2_2 | Q8BL65-4 | -1.94 | 1.15 | -2.67 | -3.45 | 0.02 | 0.31 | -2.13 | -0.67 | -0.1 | -1.78 | 0.13 | -1.2 | 0.82 | -0.92 | -0.71 | -1.67 | 1.11 | 0.99 | 1.1075 | 0.29364 | 0.333224 | 0.291568 | 0.999971 | 1 | 0.847402 |
| Ablim3 | Q69ZX8 | -0.26 | -0.5 | -0.76 | -0.76 | -0.7 | -0.24 | -0.73 | -0.03 | -0.59 | 0.11 | -0.93 | -1.04 | 0.04 | -0.67 | -0.17 | -0.67 | 0.145 | -0.0425 | 0.2025 | 0.553458 | 0.885357 | 0.420939 | 0.999971 | 1 | 0.87788 |
| Abr | A0A571BG | 1.84 | -1.44 | -0.34 | -0.05 | -1.23 | -1.35 | 0.38 | 0.92 | 0.11 | 0.92 | 0.12 | 0.33 | -2.13 | -0.53 | 0.77 | 0.34 | -0.3225 | 0.3675 | -0.39 | 0.687717 | 0.573469 | 0.644554 | 0.999971 | 1 | 0.93301 |
| Abr_1 | Q5SSL4 | -0.26 | -0.08 | -0.08 | -0.28 | -0.25 | -0.03 | 0.16 | -0.23 | 0.12 | -0.19 | -0.09 | 0.04 | 0.01 | -0.23 | -0.24 | -0.21 | 0.0875 | 0.145 | 0.0075 | 0.637232 | 0.407032 | 0.965348 | 0.999971 | 1 | 0.998721 |
| Abr_2 | Q5SSL4-4 | -0.49 | -0.92 | -1.13 | -0.42 | 0.06 | -1.96 | -1.27 | -0.39 | -0.47 | -1.44 | -1.02 | -0.28 | -0.33 | -0.59 | -0.59 | -1.57 | -0.15 | -0.0625 | -0.03 | 0.740822 | 0.843764 | 0.926223 | 0.999971 | 1 | 0.99401 |
| Abracl | A0A1W2P7 | -1.11 | 2.66 | 0.88 | 0.2 | 2.46 | 1.2 | 0.26 | 0.35 | 2.26 | -1.01 | 0.63 | -0.36 | 3.37 | 0.99 | 0.73 | -0.16 | 0.41 | -0.2775 | 0.575 | 0.627483 | 0.771428 | 0.558609 | 0.999971 | 1 | 0.915051 |
| Abraxas2 | Z4YJY0 | 0.56 | 0.38 | 0.35 | 0.63 | 0.61 | 0.35 | 0.68 | 0.66 | 0.68 | 0.36 | 0.58 | 0.06 | 0.86 | 0.25 | 0.1 | 0.4 | 0.095 | -0.06 | -0.0775 | 0.601703 | 0.769071 | 0.727565 | 0.999971 | 1 | 0.956041 |
| Abtb2 | Q7TQI7 | -0.01 | -0.07 | -0.3 | -0.74 | -0.09 | -0.33 | -0.63 | -0.59 | -0.01 | -0.32 | -0.65 | -0.91 | -0.06 | -0.45 | 0.12 | -0.49 | -0.13 | -0.1925 | 0.06 | 0.591729 | 0.487201 | 0.811365 | 0.999971 | 1 | 0.973864 |
| Acaa1a | Q921H8 | -0.39 | -1.33 | -0.79 | -0.81 | -1.08 | -1.16 | -0.94 | -0.63 | -1.18 | -0.6 | -0.74 | -0.64 | -0.83 | -0.72 | -0.47 | -0.18 | -0.1225 | 0.04 | 0.28 | 0.629358 | 0.876333 | 0.304189 | 0.999971 | 1 | 0.847402 |
| Acaa2 | Q8BWT1 | -0.95 | -0.62 | -0.77 | -0.74 | -0.4 | -1.06 | -0.96 | -0.47 | -0.49 | -1.01 | -1.6 | -0.96 | -0.4 | -0.93 | -1.04 | -0.85 | 0.0475 | -0.245 | -0.035 | 0.831525 | 0.357538 | 0.86688 | 0.999971 | 1 | 0.983462 |
| Acaca | Q5SWU9 | -0.51 | -0.45 | -0.34 | -0.29 | -0.47 | -0.23 | -0.48 | -0.22 | -0.28 | -0.23 | -0.14 | -0.11 | -0.16 | -0.1 | 0.01 | -0.03 | 0.0475 | 0.2075 | 0.3275 | 0.785937 | 0.223621 | 0.076258 | 0.999971 | 1 | 0.748444 |
| Acad11 | A0A0R4J0I | -0.05 | -0.92 | -1.03 | -1.37 | -0.98 | -0.5 | -0.82 | -0.67 | -0.57 | 0.44 | -0.68 | -0.66 | -0.53 | -0.42 | -1.22 | -0.92 | 0.1 | 0.475 | 0.07 | 0.744159 | 0.229863 | 0.834344 | 0.999971 | 1 | 0.979036 |
| Acad12 | D3Z7X0 | -0.45 | -0.58 | -1.73 | -1.7 | -0.36 | 0.46 | -1.82 | -0.91 | -0.47 | -0.33 | -1.95 | -1.71 | -0.57 | 0.07 | -1.56 | -0.97 | 0.4575 | 0 | 0.3575 | 0.410096 | 1 | 0.443296 | 0.999971 | 1 | 0.883386 |
| Acad8 | D3YTT4 | -0.33 | 0.12 | 0.26 | 0.06 | -0.4 | 0.05 | 0.1 | 0.54 | -0.29 | -0.17 | -0.37 | -0.17 | -0.16 | 0.2 | 0.11 | 0.15 | 0.045 | -0.2775 | 0.0475 | 0.860149 | 0.17707 | 0.817043 | 0.999971 | 1 | 0.974929 |
| Acad9 | Q8JZN5 | -0.25 | -0.46 | -0.53 | -0.4 | -0.59 | -0.44 | -0.39 | -0.27 | -0.49 | -0.41 | -0.42 | -0.38 | -0.23 | -0.61 | -0.48 | -0.31 | -0.0125 | -0.015 | 0.0025 | 0.943042 | 0.926484 | 0.989033 | 0.999971 | 1 | 0.99925 |
| Acadl | A0A0R4J08 | -1.1 | -1.5 | -1.49 | -1.08 | -1.39 | -1.17 | -1.22 | -0.79 | -1.35 | -1.25 | -1.51 | -1.53 | -1.63 | -1.37 | -1.63 | -1.27 | 0.15 | -0.1175 | -0.1825 | 0.496813 | 0.547935 | 0.383453 | 0.999971 | 1 | 0.871816 |
| Acadm | P45952 | -0.86 | -1.09 | -1.02 | -1.15 | -0.87 | -0.8 | -1.13 | -0.74 | -0.63 | -0.61 | -0.96 | -0.73 | -0.73 | -0.78 | -0.93 | -1.04 | 0.145 | 0.2975 | 0.16 | 0.434921 | 0.120295 | 0.380032 | 0.999971 | 1 | 0.871816 |
| Acads | Q07417 | -0.86 | -1.11 | -1.48 | -1.64 | -0.92 | -1.28 | -1.01 | -1.2 | -0.87 | -1.3 | -1.16 | -1.04 | -0.88 | -1.55 | -1.52 | -1.63 | 0.17 | 0.18 | -0.1225 | 0.470897 | 0.447141 | 0.649864 | 0.999971 | 1 | 0.933842 |
| Acadsb | Q9DBL1 | -1.05 | -0.44 | -0.47 | -0.31 | -0.41 | -0.58 | -0.59 | 0.04 | -0.31 | -0.84 | -0.29 | -0.37 | -0.25 | -0.29 | -0.32 | -0.19 | 0.1825 | 0.115 | 0.305 | 0.471005 | 0.63405 | 0.180495 | 0.999971 | 1 | 0.808532 |
| Acadvl | P50544 | -0.86 | -0.57 | -0.29 | -0.55 | -0.66 | -0.7 | -0.48 | -0.28 | -0.48 | -0.84 | -0.41 | -0.32 | -0.19 | -0.78 | -0.31 | -0.2 | 0.0375 | 0.055 | 0.1975 | 0.854991 | 0.793395 | 0.38904 | 0.999971 | 1 | 0.871816 |
| Acan | Q61282 | 1.12 | 0.66 | 0.38 | 0.65 | 0.63 | 0.61 | -0.08 | 0.28 | 0.45 | 0.74 | -0.3 | 0.08 | 0.66 | 0.42 | 0.37 | 0.21 | -0.3425 | -0.46 | -0.2875 | 0.199388 | 0.133881 | 0.218725 | 0.999971 | 1 | 0.827064 |
| Acap2 | D4AFX6 | 0.46 | 0.33 | -0.29 | -0.19 | -0.68 | -1.06 | -0.12 | -0.6 | 0.41 | 0.11 | -0.83 | -0.65 | 0.59 | -1 | -0.33 | 0.04 | -0.6925 | -0.3175 | -0.2525 | 0.034508 | 0.370768 | 0.502473 | 0.999971 | 1 | 0.904573 |
| Acap2_1 | A0A338P6 | -0.46 | -0.57 | -0.04 | -0.34 | -0.67 | -0.32 | -0.07 | -0.25 | -0.44 | -0.4 | 0 | 0 | -0.44 | -0.48 | -0.02 | -0.21 | 0.025 | 0.1425 | 0.065 | 0.907907 | 0.507248 | 0.756406 | 0.999971 | 1 | 0.961116 |
| Acap3 | A2ADA6 | 0.35 | 0.35 | 0.3 | 0.29 | 0.49 | 0.49 | 0.44 | 0.29 | 0.41 | 0.53 | 0.26 | -0.05 | 0.38 | 0.3 | 0.12 | 0.25 | 0.105 | -0.035 | -0.06 | 0.523671 | 0.853108 | 0.7173 | 0.999971 | 1 | 0.952711 |
| Acat1 | Q8QZT1 | -0.74 | -0.78 | -0.69 | -1.1 | -0.72 | -0.55 | 0.31 | -0.75 | -0.43 | -0.79 | -0.4 | 0.08 | -0.72 | -0.72 | -0.68 | -0.85 | 0.4 | 0.4425 | 0.085 | 0.179345 | 0.086149 | 0.639198 | 0.999971 | 1 | 0.93301 |
| Acat2 | Q8CAY6 | -0.44 | -0.73 | -0.77 | -0.82 | -0.62 | -0.86 | -0.52 | -0.72 | -0.64 | -0.69 | -0.83 | -0.2 | -0.24 | -0.68 | -0.29 | -0.24 | 0.01 | 0.1 | 0.3275 | 0.956842 | 0.633365 | 0.124872 | 0.999971 | 1 | 0.78268 |
| Acbd3 | A0A0R4J07 | 0.35 | 0.73 | 1.16 | 1.17 | 0.74 | 0.21 | 0.56 | 1.28 | 0.67 | 0.35 | 1.32 | 1.19 | 0.34 | 0.6 | 1.18 | 1.47 | -0.155 | 0.03 | 0.045 | 0.61301 | 0.92174 | 0.890311 | 0.999971 | 1 | 0.988102 |
| Acbd5 | A0A2I3BRB | 0.13 | -1.24 | -0.83 | -0.87 | -1.05 | -0.96 | -0.73 | -0.42 | -1 | -0.36 | -0.82 | -0.81 | -1.11 | -0.91 | -0.98 | -0.94 | -0.0875 | -0.045 | -0.2825 | 0.788092 | 0.889321 | 0.364589 | 0.999971 | 1 | 0.869526 |
| Acbd6 | Q9D061 | 0.04 | 0.7 | 0.2 | -0.14 | 0.77 | 0.41 | 0.22 | 0.1 | 0.46 | -0.14 | 0.5 | 0.08 | 0.41 | 0.45 | 0.24 | 0.2 | 0.175 | 0.025 | 0.125 | 0.50183 | 0.923138 | 0.589788 | 0.999971 | 1 | 0.922012 |
| Ace | P09470 | -0.43 | 0.87 | 0.92 | -0.02 | 0.61 | -0.14 | 0.8 | -0.31 | 0.51 | -0.75 | -1.2 | 0.24 | 0.34 | -0.29 | 0.26 | -0.35 | -0.095 | -0.635 | -0.345 | 0.817221 | 0.21704 | 0.362741 | 0.999971 | 1 | 0.869526 |
| Ace2 | Q8R0I0 | 0.57 | -0.61 | -1.17 | -0.69 | -0.47 | -0.87 | -0.6 | -0.76 | -0.98 | 0.53 | -0.78 | -0.92 | -0.81 | -0.75 | -0.51 | -0.61 | -0.2 | -0.0625 | -0.195 | 0.590725 | 0.896683 | 0.597481 | 0.999971 | 1 | 0.92377 |
| Ache | P21836 | -0.23 | -0.38 | -1.37 | -0.43 | 0.19 | -0.86 | -1.25 | -0.59 | 0.07 | -0.96 | -1.39 | -0.94 | 0.47 | -0.51 | -0.67 | -1.09 | -0.025 | -0.2025 | 0.1525 | 0.948152 | 0.605511 | 0.705372 | 0.999971 | 1 | 0.950182 |
| Acin1 | Q9JIX8-4 | 1.64 | -0.04 | 0.89 | 0.83 | 0.1 | 0.2 | 0.65 | 0.64 | -0.07 | 0.71 | 0.34 | 0.53 | -0.2 | -0.08 | 0.81 | 1.13 | -0.4325 | -0.4525 | -0.415 | 0.252954 | 0.243449 | 0.365638 | 0.999971 | 1 | 0.869526 |
| Acly | Q91V92 | 1.38 | -0.6 | -0.84 | 0.06 | -0.24 | -0.76 | 0.95 | -0.15 | -0.82 | 0.49 | -1.33 | -1.92 | -2.35 | -0.35 | -1.65 | -1.53 | -0.05 | -0.895 | -1.47 | 0.929156 | 0.194473 | 0.033055 | 0.999971 | 1 | 0.685881 |
| Acly_1 | Q3V117 | -0.19 | -0.43 | -0.38 | -0.3 | -0.2 | -0.57 | 0.02 | -0.39 | -0.04 | -0.38 | -0.13 | 0.03 | 0.1 | -0.42 | -0.37 | -0.47 | 0.04 | 0.195 | 0.035 | 0.839544 | 0.290153 | 0.86153 | 0.999971 | 1 | 0.982963 |
| Aco1 | P28271 | -0.62 | -0.43 | -0.16 | -0.41 | -0.33 | -0.59 | -0.17 | 0.01 | -0.25 | -0.54 | -0.01 | 0.15 | -0.29 | -0.47 | 0.02 | -0.14 | 0.135 | 0.2425 | 0.185 | 0.524181 | 0.28492 | 0.367622 | 0.999971 | 1 | 0.869526 |
| Aco2 | Q99KI0 | -0.62 | -0.28 | -0.51 | -0.44 | -0.54 | -0.78 | -0.66 | -0.5 | -0.72 | -0.83 | -0.68 | -0.38 | -0.17 | -0.49 | -0.59 | -0.29 | -0.1575 | -0.19 | 0.0775 | 0.386044 | 0.320969 | 0.682844 | 0.999971 | 1 | 0.943421 |
| Acot1 | O55137 | 0 | 0.75 | -2.79 | -2.91 | 0.93 | 1.07 | -2.65 | -2.37 | 0.39 | -0.12 | -2.35 | -3.13 | 0.31 | 1.02 | -2.75 | -2.77 | 0.4825 | -0.065 | 0.19 | 0.696288 | 0.95448 | 0.876554 | 0.999971 | 1 | 0.985129 |
| Acot11 | Q8VHQ9 | 0.72 | 0.6 | 0.14 | -0.05 | 0.41 | 0.51 | 0.79 | -0.16 | 0.02 | -0.17 | -0.07 | -0.47 | 1.01 | 0.1 | -0.22 | -0.12 | 0.035 | -0.525 | -0.16 | 0.902086 | 0.05442 | 0.634194 | 0.999971 | 1 | 0.93301 |
| Acot13 | Q9CQR4 | 0.49 | -0.31 | -3.15 | -2.84 | -0.66 | -0.87 | -0.81 | -3.63 | -1.03 | -0.41 | -3.42 | -3.69 | -1.35 | -0.58 | -3.62 | -4.06 | -0.04 | -0.685 | -0.95 | 0.968971 | 0.539322 | 0.401663 | 0.999971 | 1 | 0.871816 |
| Acot2 | Q9QYR9 | -0.96 | -0.53 | -0.32 | -0.64 | -0.54 | -0.59 | -0.6 | -0.14 | -0.82 | -0.96 | -0.62 | -0.43 | -0.72 | -0.49 | -0.42 | -0.31 | 0.145 | -0.095 | 0.1275 | 0.511538 | 0.66499 | 0.548997 | 0.999971 | 1 | 0.911727 |
| Acot7 | Q91V12-2 | 0.3 | -0.06 | 0.51 | 0.15 | -0.42 | 0.37 | 0.81 | 0.37 | -0.15 | 0.02 | 0.59 | -0.63 | -0.58 | 0.45 | 0.07 | 0.11 | 0.0575 | -0.2675 | -0.2125 | 0.84473 | 0.368207 | 0.435723 | 0.999971 | 1 | 0.882419 |
| Acot8 | P58137 | -0.45 | 0.13 | -0.34 | 0.06 | -0.13 | 0.36 | -0.93 | 0.35 | 0.28 | -0.01 | -0.37 | -0.46 | 0.18 | 0.62 | -0.09 | 0.41 | 0.0625 | 0.01 | 0.43 | 0.851489 | 0.967959 | 0.103305 | 0.999971 | 1 | 0.78268 |
| Acot9 | Q9R0X4 | -0.53 | -0.47 | -0.96 | -1.28 | -0.29 | -0.52 | -0.3 | -0.99 | -0.13 | -0.87 | -0.77 | -0.83 | -0.3 | -0.66 | -1.09 | -1.14 | 0.285 | 0.16 | 0.0125 | 0.307897 | 0.564079 | 0.965372 | 0.999971 | 1 | 0.998721 |
| Acox1 | Q9R0H0-2 | -1.2 | -1.41 | -0.89 | -1 | -1.48 | -1.16 | -0.91 | 0.26 | -0.97 | -0.98 | -0.41 | -0.17 | -0.45 | -0.9 | -0.16 | -0.01 | 0.3025 | 0.4925 | 0.745 | 0.436706 | 0.082238 | 0.015782 | 0.999971 | 1 | 0.613095 |
| Acox3 | Q9EPL9 | -0.49 | 0.06 | -0.79 | -0.59 | 0.04 | 0.04 | -0.95 | -0.13 | -0.37 | -0.51 | -0.45 | -0.98 | 0.49 | -0.19 | -0.47 | -0.23 | 0.2025 | -0.125 | 0.3525 | 0.512405 | 0.622923 | 0.241503 | 0.999971 | 1 | 0.835635 |
| Acp1 | Q561M1 | -2.4 | -0.49 | 0.28 | -0.74 | 1.15 | -1.04 | 2.77 | 0.17 | 1.44 | -2.77 | 0.58 | 1.44 | 1.94 | -0.8 | -0.37 | -1.07 | 1.6 | 1.01 | 0.7625 | 0.096546 | 0.34191 | 0.353513 | 0.999971 | 1 | 0.868675 |
| Acp1_1 | Q9D358 | 0.98 | -0.17 | -0.54 | -1.31 | 0.21 | 0.04 | -0.6 | -1.03 | -0.14 | 0.43 | -0.62 | -0.56 | -0.24 | -0.01 | -0.69 | -0.86 | -0.085 | 0.0375 | -0.19 | 0.868724 | 0.939974 | 0.693984 | 0.999971 | 1 | 0.947174 |
| Acp2 | A0A5F8MP | -0.04 | 0.38 | 0.64 | 0.8 | 0.09 | 0.3 | 0.4 | 0.83 | 0.36 | 0.51 | 0.69 | 0.74 | 0.33 | 0.51 | 0.44 | 1.11 | -0.04 | 0.13 | 0.1525 | 0.879002 | 0.584032 | 0.579157 | 0.999971 | 1 | 0.919302 |
| Acp6 | Q8BP40 | 0.02 | 0.22 | 0.14 | 0.29 | 0.25 | 0.01 | -0.08 | -0.14 | 0 | 0.28 | 0.17 | 0.59 | -0.41 | 0.23 | -0.04 | -0.18 | -0.1575 | 0.0925 | -0.2675 | 0.394885 | 0.636851 | 0.209856 | 0.999971 | 1 | 0.82263 |
| Acsbg1 | Q99PU5 | -1.5 | -1.38 | -1.48 | -1.36 | -1.59 | -1.29 | -1.62 | -0.89 | -1.33 | -1.35 | -1.37 | -1.24 | -1.32 | -1.17 | -1.44 | -1.11 | 0.0825 | 0.1075 | 0.17 | 0.706499 | 0.501357 | 0.340178 | 0.999971 | 1 | 0.860511 |
| Acsf2 | Q8VCW8 | -0.11 | -0.01 | -0.75 | -0.86 | 0.27 | -0.03 | -0.81 | -0.75 | 0.63 | -0.2 | -0.77 | -1.07 | 1.42 | -0.16 | -1.08 | -1.19 | 0.1025 | 0.08 | 0.18 | 0.763885 | 0.846293 | 0.759635 | 0.999971 | 1 | 0.961116 |
| Acsf3 | Q3URE1 | -0.17 | -0.44 | -0.4 | -0.77 | -0.6 | -0.5 | -0.32 | -0.48 | -0.54 | -0.18 | -0.05 | 0.07 | -0.14 | -0.54 | -0.46 | -0.58 | -0.03 | 0.27 | 0.015 | 0.879054 | 0.241802 | 0.943114 | 0.999971 | 1 | 0.99692 |
| Acsl1 | D3Z041 | -0.05 | 0.31 | 0.36 | 0.34 | 0.27 | 0.3 | 0.12 | 0.35 | 0.33 | 0.15 | 0.4 | 0.46 | 0.43 | 0.14 | 0.48 | 0.39 | 0.02 | 0.095 | 0.12 | 0.913201 | 0.609907 | 0.534526 | 0.999971 | 1 | 0.910077 |
| Acsl3 | Q9CZW4 | -1.34 | -1.17 | -0.54 | -0.55 | -1.29 | -1.28 | -0.79 | -0.08 | -1.13 | -0.95 | -0.09 | -0.59 | -0.93 | -1.15 | -0.22 | -0.31 | 0.04 | 0.21 | 0.2475 | 0.908226 | 0.508179 | 0.439014 | 0.999971 | 1 | 0.883105 |
| Acsl4 | Q9QUJ7-2 | -1.52 | 0.07 | 0.44 | -0.4 | 0.22 | -0.21 | 0.45 | -0.1 | 0.69 | -1.13 | 0.7 | 0.18 | 0.59 | -0.63 | 0.16 | -0.36 | 0.4425 | 0.4625 | 0.2925 | 0.315151 | 0.417344 | 0.539937 | 0.999971 | 1 | 0.910077 |
| Acsl5 | Q8JZR0 | -0.76 | 0.12 | -0.21 | -0.36 | -0.33 | -0.13 | -0.34 | -0.79 | -0.36 | -0.83 | -0.24 | -0.3 | 0.34 | -0.08 | -0.25 | -0.4 | -0.095 | -0.13 | 0.205 | 0.710677 | 0.608372 | 0.446982 | 0.999971 | 1 | 0.884581 |
| Acsl6 | Q5ICG5 | 2.31 | -0.37 | 0.26 | 0.27 | -0.36 | 0.37 | 1.02 | -0.25 | 0.14 | 1.33 | -0.94 | -0.38 | -0.22 | -0.09 | 0.4 | -0.28 | -0.4225 | -0.58 | -0.665 | 0.491902 | 0.410813 | 0.250712 | 0.999971 | 1 | 0.839012 |
| Acsl6_1 | Q91WC3 | -0.83 | -0.64 | -0.44 | -0.5 | -0.96 | -0.51 | -0.67 | -0.15 | -0.86 | -0.62 | -0.42 | -0.48 | -0.54 | -0.49 | -0.33 | -0.04 | 0.03 | 0.0075 | 0.2525 | 0.895546 | 0.968693 | 0.229777 | 0.999971 | 1 | 0.828687 |
| Acss1 | Q99NB1 | -1.1 | -0.72 | -0.98 | -1.66 | -0.71 | -0.59 | -0.99 | -0.53 | -0.57 | -1.13 | -1.26 | -1.05 | -0.64 | -0.77 | -1.21 | -1.62 | 0.41 | 0.1125 | 0.055 | 0.128607 | 0.675579 | 0.857241 | 0.999971 | 1 | 0.98269 |

| | | | | | | | | | | | | | | | | | | | | | | | | | | |
|---|---|---|---|---|---|---|---|---|---|---|---|---|---|---|---|---|---|---|---|---|---|---|---|---|---|---|
| Acss2 | A2AQN4 | -0.76 | 0.75 | 0.68 | 0.07 | 0.44 | 1 | 0.41 | 0.19 | 0.88 | 0.01 | 0.75 | 0.95 | 1.28 | 0.9 | 0.57 | 0.83 | 0.325 | 0.4625 | 0.71 | 0.399122 | 0.261364 | 0.081725 | 0.999971 | 1 | 0.755254 |
| Actb | P60710 | -0.77 | -0.8 | -1.36 | -1.02 | -0.72 | -0.98 | -0.98 | -1.07 | -0.84 | -0.99 | -0.91 | -1.29 | -0.6 | -0.99 | -1.01 | -0.83 | 0.05 | -0.02 | 0.13 | 0.810037 | 0.92514 | 0.548685 | 0.999971 | 1 | 0.911727 |
| Actbl2 | Q8BFZ3 | -1.16 | -0.13 | -0.6 | -1.05 | 1.75 | -1.02 | 3.95 | -1.28 | 4.35 | -0.43 | 1.19 | 2.65 | 3.93 | -1.56 | -0.97 | -0.57 | 1.585 | 2.675 | 0.9425 | 0.182402 | 0.018776 | 0.417801 | 0.999971 | 1 | 0.877584 |
| Actc1 | P68033 | -0.61 | -1.35 | -1.66 | -0.82 | 0.2 | -0.97 | -1.44 | -1.2 | -0.9 | -0.12 | 0.7 | 1.69 | -1.51 | -0.67 | -1.27 | -0.98 | 0.2575 | 1.4525 | 0.0025 | 0.538722 | 0.027428 | 0.993519 | 0.999971 | 1 | 0.99933 |
| Actg1 | P63260 | -0.19 | -0.6 | -1.14 | -0.64 | -1.01 | -0.69 | -1.37 | -0.61 | -0.81 | -0.75 | -1.25 | -0.85 | -1.03 | -0.38 | -0.82 | -0.86 | -0.2775 | -0.2725 | -0.13 | 0.330567 | 0.293785 | 0.622444 | 0.999971 | 1 | 0.929759 |
| Actl6a | Q9Z2N8 | 0.82 | 1.05 | 0.92 | 0.76 | 0.96 | 0.33 | 0.05 | 0.62 | 0.61 | 0.87 | 0.22 | 0.39 | 0.71 | 1.02 | 0.78 | 0.93 | -0.3975 | -0.365 | -0.0275 | 0.122724 | 0.101983 | 0.877254 | 0.999971 | 1 | 0.985139 |
| Actl6b | Q99MR0 | 1.53 | 1.26 | 1.01 | 1.37 | 1.15 | 1.12 | 0.46 | 1.22 | 1.03 | 1.55 | 0.77 | 0.78 | 0.83 | 1.16 | 0.94 | 1.14 | -0.305 | -0.26 | -0.275 | 0.225971 | 0.29926 | 0.187129 | 0.999971 | 1 | 0.809025 |
| Actn1 | A1BN54 | -0.64 | -0.63 | -0.69 | -0.49 | -0.48 | -0.86 | -0.72 | -0.8 | -0.77 | -0.65 | -0.51 | -0.34 | -0.46 | -0.87 | -0.69 | -0.56 | -0.1025 | 0.045 | -0.0325 | 0.566158 | 0.799733 | 0.856423 | 0.999971 | 1 | 0.98255 |
| Actn1_1 | Q7TPR4 | 0.07 | 1.8 | 2.15 | 0.08 | 1.99 | 1.58 | 4.55 | 0.98 | 2.25 | 0.5 | 2.85 | 4.32 | 4.64 | 1.3 | 1.25 | 1.75 | 1.25 | 1.455 | 1.21 | 0.171553 | 0.122269 | 0.192548 | 0.999971 | 1 | 0.810547 |
| Actn2 | Q9JI91 | -1.2 | -1.11 | -2.24 | -1.97 | -0.99 | -1.74 | -1.98 | -1.65 | -1.13 | -1.42 | -1.68 | -1.55 | -0.88 | -1.25 | -1.73 | -2.04 | 0.04 | 0.185 | 0.155 | 0.907931 | 0.553411 | 0.676394 | 0.999971 | 1 | 0.941944 |
| Actn4 | A0A1L1SV2 | -1.1 | -0.94 | -1.18 | -0.83 | -0.42 | -0.78 | -1.11 | -1.11 | -0.7 | -0.74 | -0.77 | -0.74 | -0.61 | -0.95 | -1.01 | -0.67 | 0.1575 | 0.275 | 0.2025 | 0.488089 | 0.129004 | 0.309483 | 0.999971 | 1 | 0.847402 |
| Actn4_1 | A0A1L1SVJ | -5.98 | -6.19 | -5.25 | -5.14 | -4.3 | -6.61 | -4.52 | -5.44 | -4.45 | -7.53 | -4.95 | -5.39 | -4.98 | -6.1 | -5.33 | -7.07 | 0.4225 | 0.06 | -0.23 | 0.442307 | 0.927827 | 0.643967 | 0.999971 | 1 | 0.93301 |
| Actn4_2 | E9Q2W9 | 0.13 | -1.33 | -0.92 | -1.12 | -1.41 | -1.12 | -1.22 | -1.32 | -1.35 | -0.47 | -1.14 | -1.28 | -1.27 | -1.83 | -1.16 | -0.37 | -0.4575 | -0.25 | -0.3475 | 0.188973 | 0.506194 | 0.417739 | 0.999971 | 1 | 0.877584 |
| Actr10 | Q9QZB7 | -0.72 | 0.05 | 0.19 | -0.09 | 0.09 | 0.3 | 0.22 | 0.24 | 0.27 | -0.27 | 0.51 | 0.48 | 0.59 | 0.26 | 0.37 | 0.21 | 0.355 | 0.39 | 0.5 | 0.162537 | 0.192146 | 0.069738 | 0.999971 | 1 | 0.733937 |
| Actr1a | P61164 | -0.33 | -1.35 | -0.64 | -0.65 | -1.35 | -1.03 | -0.46 | -0.63 | -1.31 | -0.57 | -0.73 | -1.1 | -1.23 | -0.95 | -0.73 | -0.53 | -0.125 | -0.185 | -0.1175 | 0.680362 | 0.522947 | 0.675494 | 0.999971 | 1 | 0.941576 |
| Actr1b | Q8R5C5 | -1.45 | -1.07 | -0.76 | -0.66 | -0.89 | -1.1 | -0.7 | -0.6 | -1 | -1.49 | -0.67 | -0.74 | -0.39 | -0.88 | -0.39 | -0.52 | 0.1625 | 0.01 | 0.44 | 0.506375 | 0.970819 | 0.098641 | 0.999971 | 1 | 0.78268 |
| Actr2 | P61161 | -0.14 | 0.22 | -0.26 | -0.09 | 0.4 | 0.28 | -0.5 | -0.18 | 0.17 | 0.08 | 0.05 | -0.11 | 0.55 | 0.01 | -0.21 | -0.06 | 0.0675 | 0.115 | 0.14 | 0.792972 | 0.537946 | 0.551068 | 0.999971 | 1 | 0.913169 |
| Actr3 | Q99JY9 | -0.52 | -0.52 | -1.03 | -1.13 | -0.46 | -0.43 | -0.84 | -1.12 | -0.54 | -0.68 | -1.2 | -1.41 | -0.62 | -0.57 | -1.15 | -1.18 | 0.0875 | -0.1575 | -0.08 | 0.733849 | 0.574934 | 0.756448 | 0.999971 | 1 | 0.961116 |
| Actr3b | Q641P0 | -0.52 | -0.32 | -0.36 | -0.51 | -0.43 | -0.19 | -0.33 | -0.5 | -0.73 | -0.77 | -0.66 | -0.78 | -0.87 | -0.58 | -0.62 | -0.65 | 0.065 | -0.3075 | -0.2525 | 0.708641 | 0.083271 | 0.170214 | 0.999971 | 1 | 0.802549 |
| Actr6 | A0A0R4J00 | -0.9 | 0.57 | 0.3 | 0.06 | 0.61 | 0.53 | 0.68 | 0.37 | 0.76 | -0.66 | 1.03 | 1.16 | -0.36 | 0.08 | 2.25 | 0.9 | 0.54 | 0.565 | 0.71 | 0.125623 | 0.26959 | 0.256271 | 0.999971 | 1 | 0.839012 |
| Acvr1 | P37172 | -0.38 | 0.22 | 0.51 | 0.64 | 0.04 | 0.19 | -0.12 | 1.09 | 0.2 | -0.16 | 0.49 | -1.51 | 0.52 | 0.2 | 0.86 | 1.05 | 0.0525 | -0.4925 | 0.41 | 0.879918 | 0.307134 | 0.19889 | 0.999971 | 1 | 0.813335 |
| Acvr1b | Q61271 | -0.56 | 0.27 | -0.44 | -0.43 | -0.16 | -0.02 | -0.77 | 0 | 0.34 | -0.16 | -0.48 | -0.9 | 0.64 | 0.19 | 0.14 | -0.3 | 0.4625 | -0.01 | 0.4575 | 0.850191 | 0.975321 | 0.134221 | 0.999971 | 1 | 0.784948 |
| Acvr1c | Q8K348 | -0.39 | -0.14 | -0.04 | 0.07 | -0.12 | -0.93 | -1.11 | 0.32 | -1.46 | -0.78 | -0.49 | -1.17 | -0.57 | -0.44 | 0.38 | 0.14 | -0.335 | -0.85 | 0.0025 | 0.346566 | 0.008956 | 0.992571 | 0.999971 | 1 | 0.99925 |
| Acvr2a | P27038 | -1.62 | -0.28 | -0.49 | -0.16 | -0.09 | -0.45 | -0.71 | -0.17 | 0.12 | -1.78 | -0.57 | -0.55 | 0.31 | -0.67 | -0.3 | 0.1 | 0.2825 | -0.0575 | 0.4975 | 0.435217 | 0.905445 | 0.218178 | 0.999971 | 1 | 0.827064 |
| Acy1 | A0A0R4J05 | 1.06 | 0.6 | 0.35 | 0.65 | 0.37 | 0.58 | -0.11 | 0.75 | 0.49 | 1.15 | 0.21 | 0.15 | 0.44 | 0.69 | 0.5 | 0.39 | -0.2675 | -0.165 | -0.16 | 0.319367 | 0.565962 | 0.45725 | 0.999971 | 1 | 0.88762 |
| Acy3 | Q91XE4 | 0.27 | 1.18 | 1.69 | 1.46 | 0.69 | 0.44 | 1.3 | 1.79 | 0.46 | 0.29 | 1.19 | 0.99 | -0.01 | 0.58 | 1.31 | 1.17 | -0.095 | -0.4175 | -0.3875 | 0.818497 | 0.27417 | 0.361039 | 0.999971 | 1 | 0.869526 |
| Acyp1 | P56376 | -0.56 | 0.56 | 1.85 | 1.19 | 0.2 | 0.66 | 2.15 | 0.73 | 0.16 | -0.46 | 1.03 | 0.3 | 0 | 0.42 | 0.51 | 0.3 | 0.175 | -0.5025 | -0.4525 | 0.772838 | 0.374149 | 0.366568 | 0.999971 | 1 | 0.869526 |
| Acyp2 | P56375 | 1.14 | 0.6 | 0.96 | 1.11 | 0.05 | 0.79 | 1.47 | 0.02 | 0.72 | 0.78 | 0.46 | 0.41 | -0.45 | 0.14 | 0.64 | -0.14 | -0.37 | -0.36 | -0.905 | 0.317028 | 0.106216 | 0.009686 | 0.999971 | 1 | 0.613095 |
| Ada | P03958 | -1.27 | 0.02 | 1.01 | 1.61 | 0.31 | -0.4 | 0.42 | 1.27 | 0.54 | -1.35 | 0.63 | 1.32 | -0.34 | 0.23 | 1.83 | 1.53 | 0.0575 | -0.0575 | 0.47 | 0.929478 | 0.940493 | 0.528127 | 0.999971 | 1 | 0.909725 |
| Adal | Q80SY6 | 1.32 | 0.14 | 0.47 | 0.2 | 0.16 | 0.29 | 0.11 | 0.52 | 0.15 | 0.54 | 0.32 | 0.65 | -0.08 | 0.26 | 0.43 | 0.24 | -0.2625 | -0.1175 | -0.32 | 0.387125 | 0.697759 | 0.303835 | 0.999971 | 1 | 0.847402 |
| Adam10 | O35598 | -0.25 | -0.53 | -0.45 | -0.45 | -0.25 | -0.45 | -0.2 | -0.14 | -0.29 | -0.34 | -0.31 | 0.17 | -0.06 | -0.3 | 0.29 | -0.62 | 0.16 | 0.2275 | 0.2475 | 0.373347 | 0.259354 | 0.311128 | 0.999971 | 1 | 0.847973 |
| Adam11 | Q7TQG7 | 0.32 | -0.68 | -0.92 | -0.81 | -0.88 | -0.3 | -0.87 | -0.63 | -0.95 | -0.25 | -1.11 | -1.23 | -1.04 | -0.9 | -0.84 | -0.67 | -0.1475 | -0.3625 | -0.34 | 0.645204 | 0.319648 | 0.279936 | 0.999971 | 1 | 0.843812 |
| Adam15 | O88839-2 | 0.43 | -0.55 | -1.16 | -0.73 | -0.5 | 0.07 | -0.87 | 0.2 | -0.24 | 0.4 | -1.04 | -1.34 | -0.15 | 0.16 | -0.81 | -0.47 | 0.2275 | -0.0525 | 0.185 | 0.573245 | 0.913598 | 0.630608 | 0.999971 | 1 | 0.932711 |
| Adam17 | Q9Z0F8 | -0.38 | -0.08 | 0.56 | -0.13 | -0.14 | 0.2 | 0.53 | -0.21 | -0.16 | -0.38 | 0.27 | -2.38 | -0.11 | -0.23 | 0.31 | -0.58 | 0.1025 | -0.655 | -0.145 | 0.71404 | 0.270946 | 0.614461 | 0.999971 | 1 | 0.927646 |
| Adam22 | Q9R1V6-14 | -0.52 | -0.81 | -0.61 | -0.61 | -1.01 | -0.47 | -0.39 | -0.49 | -0.88 | -0.6 | -0.74 | -0.86 | -0.58 | -0.44 | -0.48 | -0.52 | 0.0475 | -0.1325 | 0.1325 | 0.818691 | 0.4481 | 0.439861 | 0.999971 | 1 | 0.883105 |
| Adam23 | Q9R1V7-2 | -0.79 | -1.29 | -0.55 | -0.71 | -1.47 | -1.22 | -0.6 | -0.77 | -1.69 | -1.43 | -1.02 | -1.17 | -1.7 | -1.39 | -0.89 | -0.87 | -0.18 | -0.4925 | -0.3775 | 0.516281 | 0.071147 | 0.194167 | 0.999971 | 1 | 0.810645 |
| Adam9 | Q61072 | -0.83 | -0.07 | -0.04 | 0.35 | -0.15 | 0.48 | -0.39 | 0.8 | 0.18 | -0.55 | 0.23 | 0.45 | 0.61 | 0.08 | 0.47 | 0.31 | 0.3325 | 0.225 | 0.515 | 0.368446 | 0.499141 | 0.099017 | 0.999971 | 1 | 0.78268 |
| Adap1 | E9PY16 | -1.01 | -0.15 | -1.18 | -1.08 | -0.06 | -0.21 | -0.89 | -0.89 | -0.07 | -0.6 | -0.7 | -1.02 | 0.08 | -0.26 | -1.07 | -0.75 | 0.3425 | 0.2575 | 0.355 | 0.307152 | 0.419596 | 0.318221 | 0.999971 | 1 | 0.853893 |
| Adar | Q99MU3-5 | 0.44 | 1.45 | 1 | 0.87 | 1.27 | 0.54 | 1.27 | 0.25 | 1.3 | 0.54 | 0.82 | 0.94 | 0.73 | 0.6 | 0.48 | 0.03 | -0.1075 | -0.04 | -0.48 | 0.746157 | 0.884843 | 0.108888 | 0.999971 | 1 | 0.78268 |
| Adarb1 | Q91ZS8-4 | 0.44 | 0.95 | 0.34 | 0.14 | 0.81 | 0.61 | 0.1 | 0.35 | 0.44 | 0.61 | 0.05 | 0.16 | 0.32 | 0.46 | -0.04 | 0.42 | -1.96E-16 | -0.1525 | -0.1775 | 1 | 0.535747 | 0.467203 | 1 | | 0.891731 |
| Adat2 | Q6P6J0 | 1.02 | 1.09 | 0.53 | 0.76 | 1.09 | 0.9 | 0.11 | 0.44 | 0.53 | 0.96 | 0.7 | 0.58 | 0.5 | 0.83 | 0.43 | 0.91 | -0.215 | -0.1575 | -0.1825 | 0.44083 | 0.456606 | 0.416025 | 0.999971 | 1 | 0.876885 |
| Adat3 | Q6PAT0 | 3.11 | 4.15 | 3.34 | 1.17 | 4.49 | 1.17 | 5.66 | 2.32 | 5.74 | 1.04 | 5.06 | 5.44 | 13.99 | 0.03 | 2.31 | 0.69 | 0.4675 | 1.3775 | 1.3125 | 0.66288 | 0.246745 | 0.658417 | 0.999971 | 1 | 0.934638 |
| Adck1 | Q9D0L4 | 0.13 | 0.14 | -0.11 | -0.13 | 0.05 | 0.36 | 0.36 | 0.48 | 0.23 | 0.26 | 0.1 | -0.38 | 0.54 | 0.6 | 0.2 | 0.44 | 0.305 | 0.045 | 0.4375 | 0.130929 | 0.831469 | 0.041294 | 0.999971 | 1 | 0.700838 |
| Adck2 | E9QL72 | 1.78 | 1.51 | 1.95 | 2.49 | 1.16 | 1.83 | 1.19 | 3.09 | 1.37 | 1.96 | 2.48 | 1.43 | 0.9 | 1.57 | 1.85 | 2.05 | -0.115 | -0.1225 | -0.34 | 0.804506 | 0.712122 | 0.312851 | 0.999971 | 1 | 0.849367 |
| Adck5 | E9PUK2 | -0.99 | 0.21 | -0.42 | -0.44 | 0.11 | -0.4 | -0.71 | -0.65 | -0.21 | -1.45 | -0.98 | -0.42 | -0.21 | 0.39 | -0.61 | 0.06 | -0.0025 | -0.355 | 0.3175 | 0.993612 | 0.342443 | 0.342738 | 0.999971 | 1 | 0.861474 |
| Adcy1 | O88444 | -0.52 | -0.03 | 0.07 | -0.02 | -0.41 | 0.14 | -0.48 | 0.06 | -0.47 | -0.14 | -0.27 | -0.67 | -0.04 | -0.15 | -0.42 | -0.21 | -0.0475 | -0.2625 | -0.08 | 0.843264 | 0.249217 | 0.702242 | 0.999971 | 1 | 0.949019 |
| Adcy2 | Q3V1Q3 | -0.71 | -0.75 | -0.77 | -0.99 | -0.51 | -0.4 | -0.53 | -0.87 | -0.7 | -0.69 | -0.81 | -1.58 | -0.25 | -0.56 | -1.22 | -0.77 | 0.2275 | -0.14 | 0.105 | 0.247202 | 0.577033 | 0.669186 | 0.999971 | 1 | 0.938711 |
| Adcy3 | B8JK56 | 0.27 | -0.27 | -0.34 | -0.12 | -0.13 | -0.06 | -0.27 | -0.17 | -0.25 | 0.18 | 0.18 | 0.3 | 0.41 | -0.42 | -0.32 | -0.15 | -0.0425 | 0.2175 | -0.005 | 0.832452 | 0.340343 | 0.984408 | 0.999971 | 1 | 0.99925 |
| Adcy5 | P84309 | -0.7 | -1.46 | -2.5 | -1.82 | -0.97 | -2.71 | -1.7 | -1.71 | -0.8 | -1 | -2.18 | -2.01 | -1.04 | -1.37 | -1.78 | -2.62 | -0.1525 | 0.1225 | -0.0825 | 0.75221 | 0.798537 | 0.861751 | 0.999971 | 1 | 0.982963 |
| Adcy6 | A0A2I3BRC | -1.2 | -2.01 | -1.94 | -1.73 | -1.61 | -1.68 | -1.7 | -1.83 | -1.93 | -1.55 | -1.44 | -1.38 | -1.68 | -1.56 | -1.89 | -1.43 | 0.015 | 0.145 | 0.08 | 0.947515 | 0.562152 | 0.740505 | 0.999971 | 1 | 0.960899 |
| Adcy8 | A0A2I3BQ( | -0.28 | -0.9 | -0.32 | -1.01 | -0.92 | -0.76 | -0.78 | -0.85 | -0.84 | -0.5 | 0.11 | -0.5 | -0.28 | -0.5 | -0.7 | -0.31 | -0.2 | 0.195 | 0.18 | 0.39762 | 0.502037 | 0.469792 | 0.999971 | 1 | 0.892481 |
| Adcy9 | P51830 | -0.63 | -0.86 | -0.35 | -0.53 | -0.88 | -0.86 | -0.53 | -0.7 | -1.13 | -0.8 | -0.74 | -0.8 | -1.04 | -1.22 | -1.03 | -0.81 | -0.15 | -0.275 | -0.4325 | 0.451498 | 0.183823 | 0.052571 | 0.999971 | 1 | 0.724862 |
| Adcyap1 | O70176 | 0.09 | 0.24 | -0.16 | -0.46 | -0.08 | -0.35 | -0.76 | -0.24 | 2.14 | -0.73 | -1.44 | 0.62 | 1.6 | 1.22 | 0.92 | 0.11 | -0.285 | 0.22 | 1.035 | 0.258896 | 0.763759 | 0.014719 | 0.999971 | 1 | 0.613095 |
| Adcyap1r1 | E9Q968 | -1.41 | -0.96 | -1.01 | -1.12 | -0.92 | -0.9 | -1.07 | -1.1 | -1.28 | -1.43 | -0.97 | -1.32 | -1.01 | -0.67 | -0.92 | -0.57 | 0.1275 | -0.125 | 0.3325 | 0.497448 | 0.530329 | 0.126829 | 0.999971 | 1 | 0.783193 |
| Adcyap1r1 | E9Q3E8 | -1.11 | 0.84 | -2.29 | -2.34 | 0.17 | 2.5 | -2 | -1.82 | -0.97 | 0.33 | -2.97 | -1.74 | 0.57 | 1.32 | -2.11 | -2.58 | 0.9375 | -0.1125 | 0.525 | 0.422402 | 0.902182 | 0.63177 | 0.999971 | 1 | 0.932858 |
| Add1 | E9Q1K3 | -0.24 | -1.04 | -0.47 | -0.46 | -1.49 | -0.33 | -0.8 | 0.33 | -1.41 | -0.2 | -1.49 | -0.87 | -1.1 | -0.19 | -0.54 | -0.76 | -0.02 | -0.44 | -0.095 | 0.960205 | 0.217765 | 0.730245 | 0.999971 | 1 | 0.956041 |
| Add1_1 | F8WGR0 | -0.21 | -1.06 | -1.4 | -1.17 | -1.12 | -0.83 | -1.3 | -1.2 | -1.16 | -0.6 | -1.47 | -1.54 | -1.16 | -1.02 | -1.38 | -1.29 | -0.1525 | -0.2325 | -0.2525 | 0.602342 | 0.493284 | 0.387653 | 0.999971 | 1 | 0.871816 |
| Add1_2 | Q9QYC0 | -0.4 | 2.64 | 1.93 | 0.47 | 1.52 | 3.62 | 0.19 | 1.07 | 1.95 | 0.87 | 0.14 | 0.88 | 2.47 | 1.79 | 0.72 | 1.44 | 0.44 | -0.2 | 0.445 | 0.625862 | 0.779148 | 0.532793 | 0.999971 | 1 | 0.910069 |
| Add2 | Q9QYB8 | -0.08 | -0.14 | -0.53 | -0.77 | -0.21 | 0.02 | -0.53 | -0.72 | -0.24 | -0.02 | -0.5 | -0.51 | 0.3 | -0.1 | -0.83 | -0.51 | 0.02 | 0.0625 | 0.095 | 0.937933 | 0.789995 | 0.75471 | 0.999971 | 1 | 0.961116 |
| Add3 | Q9QYB5-2 | 0.59 | -0.59 | -0.41 | -0.15 | -0.67 | -0.92 | -0.55 | -0.49 | -0.64 | 0.54 | -0.52 | -0.31 | -0.37 | -1.05 | -0.75 | 0.19 | -0.5175 | -0.0925 | -0.355 | 0.100343 | 0.799083 | 0.342805 | 0.999971 | 1 | 0.861474 |
| Add3_1 | Q9QYB5 | -0.51 | -2.05 | -0.1 | -1.05 | -1.85 | -2 | -0.39 | -0.69 | -2.36 | -0.81 | -0.03 | 0.97 | -2.43 | -1.78 | -0.94 | 1.42 | -0.305 | 0.37 | -0.005 | 0.574556 | 0.621243 | 0.995276 | 0.999971 | 1 | 1 |
| Adgb | D3YZ77 | -0.82 | -2.31 | -1.97 | -0.9 | -1.65 | -1.31 | -0.34 | -0.87 | -1.92 | -0.38 | -0.51 | 0.18 | -2.48 | -1.47 | -1.47 | -0.12 | 0.4575 | 0.8425 | 0.115 | 0.317007 | 0.145741 | 0.838282 | 0.999971 | 1 | 0.979626 |
| Adgra1 | Q8C4G9 | 0.16 | -1.17 | -1.23 | -0.63 | -1.26 | -0.46 | -0.25 | -0.68 | -1.27 | -0.21 | -1.18 | -2.06 | -0.74 | -0.45 | -1.05 | -0.67 | 0.055 | -0.4625 | -0.01 | 0.88363 | 0.336041 | 0.97665 | 0.999971 | 1 | 0.99925 |
| Adgra2 | J3QMG7 | 3.76 | 1.21 | 1.82 | 1.32 | 0.64 | 3.33 | 3.91 | 0.45 | 0.06 | 4.32 | 5.78 | -1.64 | -1.31 | 3.86 | -0.11 | 0.6 | 0.055 | 0.1025 | -1.2675 | 0.954248 | 0.950314 | 0.275143 | 0.999971 | 1 | 0.843794 |
| Adgrb1 | Q3UHD1 | -0.42 | -1.23 | -0.95 | -1.29 | -1.33 | -0.94 | -0.97 | -1.21 | -1.3 | -0.65 | -0.6 | -0.83 | -0.74 | -1.26 | -1.11 | -0.89 | -0.14 | 0.1275 | -0.0725 | 0.576496 | 0.640966 | 0.914331 | 0.999971 | 1 | 0.992633 |
| Adgrb2 | H7BX74 | 0.19 | -0.36 | -0.16 | -0.23 | -0.59 | -0.09 | 0 | 0.21 | -0.76 | 0.07 | -0.4 | -0.54 | 0.05 | -0.48 | -0.34 | -0.12 | 0.0225 | -0.2675 | -0.0825 | 0.925065 | 0.284449 | 0.701681 | 0.999971 | 1 | 0.949019 |
| Adgrb3 | Q80ZF8 | -0.1 | -0.53 | -0.74 | -0.53 | -0.43 | -0.44 | -0.46 | -0.76 | -0.31 | -0.29 | -0.55 | -0.63 | 0.09 | -0.82 | -1.03 | -0.72 | -0.0475 | 0.03 | -0.145 | 0.819574 | 0.885013 | 0.621618 | 0.999971 | 1 | 0.929641 |
| Adgre1 | Q61549 | 3.25 | 2.24 | 2.22 | 3.24 | 3.04 | 2.01 | 3.29 | 2.39 | 3.52 | 3.16 | 2.24 | 1.82 | 1.81 | 0.59 | 1.22 | -1.3 | -0.055 | -0.0525 | -2.1575 | 0.88976 | 0.909467 | 0.009948 | 0.999971 | 1 | 0.613095 |
| Adgrf5 | G5E8Q8 | -0.83 | -0.33 | 1.92 | 0.99 | 0.61 | -0.48 | 0.28 | 0.68 | 0.13 | -0.57 | 0.58 | 0.84 | 0.29 | 0.15 | 0.69 | 2.34 | -0.165 | -0.1925 | 0.43 | 0.789982 | 0.76334 | 0.557724 | 0.999971 | 1 | 0.915051 |
| Adgrg1 | Q8K209 | 0.07 | -0.61 | -1.38 | -0.58 | -0.63 | -1.15 | -1.54 | -1.17 | -0.82 | -0.12 | -1.07 | -1.04 | -0.51 | -0.65 | -0.55 | -0.59 | -0.4975 | -0.1375 | 0.05 | 0.174585 | 0.704683 | 0.869981 | 0.999971 | 1 | 0.983917 |
| Adgrl1 | H7BX15 | -6.18 | -2.26 | -1.39 | -2.21 | -2.17 | -1.29 | -0.45 | -3.09 | -1.01 | -4.33 | -1.54 | -2.22 | -3.08 | -3.59 | -2.78 | -4.75 | 1.26 | 0.735 | -0.54 | 0.263344 | 0.532604 | 0.604725 | 0.999971 | 1 | 0.924454 |
| Adgrl1_1 | E9Q3V9 | -0.8 | -1 | -0.97 | -0.78 | -0.94 | -1.07 | -0.92 | -0.78 | -0.84 | -0.89 | -0.92 | -0.65 | -0.71 | -1.13 | -0.85 | -0.62 | -0.04 | 0.0625 | 0.06 | 0.816712 | 0.712889 | 0.755112 | 0.999971 | 1 | 0.961116 |
| Adgrl2 | A0A0G2JFV | -0.97 | -1.2 | -1.16 | -1.29 | -0.83 | -0.99 | -0.9 | -1.18 | -0.88 | -1.16 | -1.03 | -1 | -0.64 | -1.17 | -1.15 | -1.03 | 0.18 | 0.1375 | 0.1575 | 0.331863 | 0.431353 | 0.439231 | 0.999971 | 1 | 0.883105 |
| Adgrl3 | D3YTW7 | -0.05 | -0.84 | -1.05 | -0.83 | -0.7 | -0.56 | -0.67 | -0.9 | -0.5 | -0.37 | -0.89 | -0.72 | -0.2 | -0.75 | -0.73 | -0.8 | -0.015 | 0.0725 | 0.0725 | 0.953288 | 0.786491 | 0.794667 | 0.999971 | 1 | 0.970337 |
| Adgrl3_1 | D3Z6H9 | -0.41 | -0.34 | -1.64 | 0.2 | -0.57 | 0.07 | -2.07 | 0.17 | -0.64 | 0.46 | -2.05 | -1.73 | 0.15 | -0.98 | -2.02 | 0.72 | -0.0525 | -0.4425 | 0.015 | 0.929106 | 0.48966 | 0.981751 | 0.999971 | 1 | 0.99925 |
| Adh5 | P28474 | 0.41 | -0.15 | 1.13 | 1.32 | -0.27 | 0.31 | 1.13 | 1.51 | -0.11 | 0.41 | 0.81 | -0.18 | -0.92 | 0.38 | 0.95 | 0.85 | -0.0075 | -0.445 | -0.3625 | 0.987718 | 0.277828 | 0.481593 | 0.999971 | 1 | 0.89485 |
| Adhfe1 | A0A087WF | -1.37 | -1.2 | -0.86 | -0.94 | -2.14 | -1.65 | -0.67 | -1.13 | -1.96 | -1.71 | -1.19 | -0.95 | -1.43 | -2.37 | -1.23 | -1.33 | -0.305 | -0.36 | -0.4975 | 0.376514 | 0.213884 | 0.122277 | 0.999971 | 1 | 0.78268 |
| Adi1 | Q99JT9 | 0.54 | 0.74 | 0.86 | 0.65 | 0.19 | 0.19 | 0.04 | 0.35 | 1.15 | 0.8 | 1.07 | 0.73 | 0.8 | 1 | 1.35 | 1.01 | -0.505 | 0.24 | 0.3425 | 0.017697 | 0.221139 | 0.107729 | 0.999971 | 1 | 0.78268 |
| Adipoq | Q60994 | 0.01 | -0.48 | -1.24 | 2.5 | 4.46 | 1.99 | 0.95 | 2.04 | 0.33 | 1.49 | 1.52 | 2.55 | -0.88 | -0.17 | 0.29 | 0.49 | 2.1625 | 1.275 | -0.265 | 0.051642 | 0.154573 | 0.734414 | 0.999971 | 1 | 0.957849 |
| Adipor1 | Q91VH1 | -1 | -0.44 | -0.03 | -0.15 | -0.42 | -0.35 | -0.22 | 0 | -0.29 | -0.63 | -0.51 | 0.88 | -0.37 | -0.17 | -0.54 | 0.19 | 0.1575 | 0.2675 | 0.1825 | 0.547317 | 0.501693 | 0.523305 | 0.999971 | 1 | 0.907797 |
| Adk | P55264-2 | 0.82 | 1.01 | 0.33 | 0.49 | 0.88 | 0.7 | 0.82 | 0.61 | 0.99 | 0.58 | 0.33 | 0.38 | 0.57 | 0.18 | 0.1 | 0.45 | 0.09 | -0.0925 | -0.3375 | 0.675139 | 0.705352 | 0.166332 | 0.999971 | 1 | 0.800215 |
| Adnp | Q9Z103 | 0.61 | 0.31 | 0.88 | 0.61 | 0.62 | 0.23 | 1.18 | 0.57 | 0.3 | 0.47 | 0.47 | 1.16 | 0.54 | 0.41 | 0.69 | 0.56 | 0.0475 | -0.0025 | -0.0525 | 0.851959 | 0.991999 | 0.787364 | 0.999971 | 1 | 0.96724 |
| Ado | Q6PDY2 | -0.26 | 0.4 | 0.27 | 0.3 | 0.13 | 0.37 | 0.14 | 0.19 | 0.2 | 0.33 | 0.35 | 0.21 | 0.48 | 0.53 | 0.5 | 0.52 | 0.03 | 0.095 | 0.33 | 0.886106 | 0.6436 | 0.133536 | 0.999971 | 1 | 0.784948 |
| Adora1 | Q60612 | 1.56 | 0.6 | 1.41 | 0.9 | 0.3 | 0.9 | 1.79 | -0.48 | 0.36 | 0.08 | 0.08 | 0.03 | 0.12 | -1.22 | -0.19 | -1.09 | -0.49 | -0.98 | -1.7125 | 0.332561 | 0.004198 | 0.001663 | 0.999971 | 0.969702 | 0.473622 |
| Adora2a | Q60613 | 1.92 | -1.09 | -1.57 | -0.62 | -1.99 | -3.48 | -1.01 | -0.6 | -1.5 | -0.48 | -1.41 | -2.05 | -1.21 | -1.84 | -0.67 | -1.87 | -1.43 | -1.02 | -1.0575 | 0.139192 | 0.205258 | 0.181879 | 0.999971 | 1 | 0.808532 |
| Adpgk | Q3UDS7 | -0.92 | 0.11 | 0.37 | 0.42 | 0.26 | 0.17 | 0.29 | 0.76 | 0.08 | 0.18 | 0.4 | 0.8 | 0.46 | 0.55 | 0.76 | 1 | 0.375 | 0.37 | 0.6975 | 0.282786 | 0.301167 | 0.062332 | 0.999971 | 1 | 0.725348 |
| Adprh | P54923 | -0.38 | 0.66 | 0.81 | 0.6 | 0.97 | 0.56 | 1.14 | 1.14 | 1.24 | 0 | 1.25 | 1.07 | 0.74 | 0.45 | 0.89 | 0.81 | 0.53 | 0.4675 | 0.3 | 0.113681 | 0.249707 | 0.327866 | 0.999971 | 1 | 0.856472 |
| Adprm | Q99KS6-2 | -1.32 | -0.81 | -0.68 | -0.87 | -0.64 | -1.16 | -0.4 | -0.3 | -0.74 | -1.75 | -1.2 | -1.81 | -2.54 | 0.09 | -0.42 | -1.35 | 0.295 | -0.455 | -0.135 | 0.275094 | 0.151396 | 0.805325 | 0.999971 | 1 | 0.971701 |
| Adprs | Q8CG72 | 1.05 | 0.62 | 0.45 | 0.37 | 0.21 | 0.81 | 0.54 | 0.29 | 0.34 | 0.57 | 0.32 | 0.35 | 0.16 | 0.62 | 0.27 | -0.12 | -0.16 | -0.2275 | -0.39 | 0.506027 | 0.294372 | 0.139591 | 0.999971 | 1 | 0.784948 |
| Adra1a | P97718 | 0.99 | -1.38 | -0.22 | 0.34 | -1.49 | -0.86 | -0.52 | -0.39 | -1.37 | 0.05 | -0.04 | -0.33 | -1.46 | -1.29 | 0.02 | 0.58 | -0.7475 | -0.355 | -0.47 | 0.172884 | 0.526074 | 0.472245 | 0.999971 | 1 | 0.89314 |
| Adra1b | P97717 | -1.91 | -3.06 | -0.34 | 0.58 | -2.38 | -2.3 | -0.19 | 0.01 | -2.6 | -1.36 | 0.21 | -0.04 | -1.57 | -1.29 | -0.03 | 1.02 | -0.0325 | 0.235 | 0.715 | 0.972056 | 0.80174 | 0.436708 | 0.999971 | 1 | 0.882419 |
| Adra2a | Q01338 | 1.23 | -1.5 | -1.36 | -0.75 | -0.83 | -0.5 | -0.42 | -1.18 | -0.67 | 1.2 | -1.24 | -1.74 | -2.22 | -1.14 | -0.74 | -1.54 | -0.1375 | -0.0175 | -0.1815 | 0.817706 | 0.982822 | 0.225043 | 0.999971 | 1 | 0.828225 |
| Adra2c | Q01337 | -0.89 | -3.24 | -1.03 | -0.34 | -1.75 | -7.15 | -2.37 | 0.41 | -2.9 | -1.2 | -1.93 | -0.55 | -1.36 | -1.71 | -0.95 | -1.85 | -1.34 | -0.27 | -0.0925 | 0.389261 | 0.715806 | 0.879764 | 0.999971 | 1 | 0.985441 |
| Adrbk2 | Q3UYH7 | 0.88 | -0.54 | -0.73 | -0.94 | -0.31 | 0.07 | -0.31 | -0.35 | -0.21 | 0.18 | -0.87 | -0.72 | -0.58 | -0.48 | -0.64 | -0.55 | 0.1075 | -0.0725 | -0.23 | 0.790642 | 0.872031 | 0.566004 | 0.999971 | 1 | 0.917959 |
| Adrm1 | Q9JKV1 | 0.91 | 0.88 | 1.1 | 0.74 | 0.75 | 1.19 | 1.78 | 1.23 | 0.64 | 0.51 | 1.46 | 1.38 | 0.49 | 1.03 | 1.62 | 0.71 | 0.33 | 0.09 | 0.055 | 0.210884 | 0.742588 | 0.841438 | 0.999971 | 1 | 0.979814 |
| Adsl | P54822 | -0.85 | -0.56 | -0.8 | -0.89 | -0.53 | -0.83 | -0.51 | -0.8 | -0.71 | -0.85 | -0.94 | -0.91 | -0.93 | -0.74 | -0.86 | -1.17 | 0.1075 | -0.0775 | -0.15 | 0.565797 | 0.654029 | 0.433556 | 0.999971 | 1 | 0.882409 |
| Adss1 | P28650 | 1.24 | 2.35 | 2.51 | 1.66 | 1.97 | 1.73 | 2.98 | 1.6 | 1.96 | 0.7 | 1.75 | 1.87 | 1.48 | 1.54 | 1.5 | 0.9 | 0.13 | -0.37 | -0.585 | 0.752102 | 0.367701 | 0.106732 | 0.999971 | 1 | 0.78268 |
| Adss2 | P46664 | 0.27 | 0.43 | 0.26 | 0.13 | 0.58 | 0.53 | -0.09 | 0.58 | 0.5 | 0.26 | 0.26 | 0.55 | 0.65 | 0.68 | 0.51 | 0.85 | 0.1275 | 0.12 | 0.4 | 0.56468 | 0.500435 | 0.047396 | 0.999971 | 1 | 0.710102 |
| Afap1 | Q80YS6 | 0.77 | -0.13 | -0.25 | 0.29 | -0.29 | -0.55 | -0.33 | -0.3 | -0.4 | 0.28 | -0.04 | 0.09 | -0.75 | -1.24 | -0.39 | -0.04 | -0.5375 | -0.1875 | -0.775 | 0.065826 | 0.51529 | 0.046793 | 0.999971 | 1 | 0.710102 |
| Afap1l1 | Q8BZI0 | 0.23 | -0.2 | -1.51 | -0.83 | -1.01 | -0.83 | -2.13 | -1.55 | -1.64 | -0.24 | -1.81 | -1.82 | -1.52 | -1.36 | -0.67 | -1.13 | -0.8025 | -0.8 | -0.5925 | 0.102034 | 0.137577 | 0.167516 | 0.999971 | 1 | 0.800215 |
| Afap1l2 | Q5DTU0 | -1.09 | -1.09 | -0.09 | -0.07 | -0.24 | -0.82 | -0.6 | 0.41 | -0.71 | -1.19 | -0.29 | 0.36 | -1.57 | -0.78 | -0.09 | 0.68 | 0.2725 | 0.2725 | 0.145 | 0.482384 | 0.550979 | 0.780642 | 0.999971 | 1 | 0.965859 |
| Afdn | Q9QZQ1 | -0.82 | -1.35 | -0.34 | -0.55 | -1 | -1.05 | 0 | -0.55 | -0.86 | -0.81 | -0.25 | -0.37 | -0.8 | -1.1 | -0.46 | -0.15 | 0.115 | 0.1925 | 0.0875 | 0.726223 | 0.499127 | 0.763063 | 0.999971 | 1 | 0.961892 |
| Afdn_1 | E9PYX7 | -1.05 | -1.28 | -2.04 | -1.67 | -1.79 | -1.89 | -1.16 | -1.43 | -1.65 | -1.5 | -1.08 | -1.42 | -2.54 | -1.65 | -1.37 | -0.72 | -0.0575 | 0.0975 | -0.06 | 0.841719 | 0.71673 | 0.88503 | 0.999971 | 1 | 0.98711 |
| Aff4 | Q9ESC8 | 1.78 | -0.56 | 0.37 | 0.83 | 0.29 | 0.28 | 0.65 | 0.42 | 0.43 | 1.38 | -0.37 | -0.26 | -0.93 | 0.67 | 0.27 | 0.4 | -0.195 | -0.31 | -0.5025 | 0.674642 | 0.596779 | 0.375784 | 0.999971 | 1 | 0.870465 |
| Afg1l | Q3V384 | -0.92 | -0.44 | -0.38 | -0.81 | -0.55 | -0.87 | -0.25 | -0.58 | -0.52 | -1.45 | -0.34 | -0.25 | -0.49 | -0.83 | -0.79 | -0.65 | 0.075 | -0.0025 | -0.0525 | 0.739656 | 0.993577 | 0.800743 | 0.999971 | 1 | 0.971076 |
| Afg3l1 | Q920A7 | 1.6 | -1.11 | -0.79 | -0.73 | -1.89 | -1.25 | -0.16 | -0.62 | -1.77 | 0.24 | -0.1 | -0.41 | -0.7 | -1.08 | -0.38 | -0.3 | -0.7225 | -0.2525 | -0.3575 | 0.291423 | 0.717573 | 0.551481 | 0.999971 | 1 | 0.913379 |
| Afg3l2 | Q8JZQ2 | 0.02 | 0.18 | -0.65 | -0.61 | -0.09 | 0.24 | -0.59 | -0.66 | -0.08 | 0.13 | -0.46 | -0.75 | 0.02 | -0.08 | -0.76 | -0.69 | -0.01 | -0.025 | -0.1125 | 0.974051 | 0.933085 | 0.710836 | 0.999971 | 1 | 0.950695 |
| Afm | O89020 | 0.85 | 2.33 | 0.65 | 3.01 | 3.96 | 1.78 | 2.31 | 1.14 | 1.26 | 0.67 | 1.62 | 1.74 | 1.48 | 1.96 | 1.16 | 1.86 | 0.5875 | -0.3875 | -0.095 | 0.441388 | 0.504034 | 0.863674 | 0.999971 | 1 | 0.983079 |
| Aftph | Q80WT5-2 | 0.4 | 0.34 | 0.44 | 0.41 | 0.34 | 0.42 | 0.97 | 0.14 | 0.13 | 0.59 | 0.4 | 0.48 | 0.43 | 0.43 | 0.08 | 0.19 | 0.07 | 0.0025 | -0.115 | 0.752697 | 0.988682 | 0.519574 | 0.999971 | 1 | 0.906123 |
| Aga | Q64191 | -0.66 | -0.52 | 0.1 | 0.27 | -0.74 | -1.05 | -0.25 | 0.65 | -0.98 | -0.97 | -0.17 | -0.33 | -1.49 | -0.47 | -0.23 | 0.17 | -0.145 | -0.41 | -0.3025 | 0.726839 | 0.214869 | 0.460255 | 0.999971 | 1 | 0.889706 |
| Agap1 | A0A087WF | 1.21 | 0.18 | -0.6 | -0.3 | -0.16 | -0.17 | -1.06 | -0.29 | 0.43 | 1.5 | -0.95 | -0.8 | 0.02 | 0.27 | -0.33 | -0.41 | -0.5425 | -0.0775 | -0.235 | 0.225038 | 0.903383 | 0.567944 | 0.999971 | 1 | 0.91816 |
| Agap1_1 | A0A1D5RLI | 1.22 | -0.63 | 0.13 | -0.32 | -0.45 | -0.42 | 0.15 | 0.53 | -0.02 | 0.86 | 0.01 | -0.25 | -0.27 | -0.13 | 0.23 | -0.01 | -0.2025 | 0.015 | -0.3625 | 0.646332 | 0.972665 | 0.360464 | 0.999971 | 1 | 0.869526 |
| Agap1_2 | E9Q3K3 | 0.23 | -0.23 | -0.81 | -0.37 | -0.25 | -0.5 | -0.76 | -0.27 | -0.14 | 0.23 | -0.55 | -0.72 | 0.02 | -0.31 | -0.35 | -0.43 | 0.15 | -7.85E-17 | 0.20755 | 0.576382 | 1 | 0.915804 | 0.999971 | 1 | 0.992633 |
| Agap2 | Q3UHD9 | -0.09 | -1.15 | -0.68 | -0.47 | -1.2 | -0.67 | 0.03 | -0.6 | -1.21 | -0.56 | -0.78 | -0.99 | -0.82 | -1.03 | -0.56 | -0.45 | -0.0125 | -0.2875 | -0.1175 | 0.970059 | 0.314069 | 0.670572 | 0.999971 | 1 | 0.93891 |
| Agap3 | Q8VHH5 | 0.23 | -0.88 | -1 | -0.87 | -0.72 | -1.06 | -0.79 | -0.99 | -0.64 | -0.08 | -0.73 | -0.62 | -0.95 | -0.88 | -0.99 | -0.98 | -0.26 | 0.1125 | -0.32 | 0.405074 | 0.729513 | 0.300493 | 0.999971 | 1 | 0.847402 |
| Agap3_1 | A0A1D5RN | 1.45 | 0.1 | 0.35 | -0.1 | -0.07 | 0.48 | 0.03 | 0.59 | -0.11 | 0.9 | 0.37 | 0.09 | 0.09 | 0.64 | 0.65 | 0.46 | -0.1925 | -0.1375 | 0.01 | 0.606298 | 0.727627 | 0.977911 | 0.999971 | 1 | 0.99925 |
| Agbl4 | E4W7Y4 | -2.25 | -1.04 | -0.44 | -0.35 | -0.4 | -0.08 | 0.49 | 0.05 | -0.8 | -1.54 | 0.11 | 0.19 | -0.57 | -0.68 | -0.43 | -0.24 | 1.035 | 0.51 | 0.54 | 0.042835 | 0.369364 | 0.225172 | 0.999971 | 1 | 0.828225 |
| Agfg1 | A0A087WN | 0.42 | -0.36 | -0.07 | 0.2 | -0.32 | -0.27 | 0.41 | -0.15 | -0.24 | 0.03 | 0.04 | -0.01 | -0.61 | -0.36 | -0.32 | 0.12 | -0.13 | -0.0925 | -0.34 | 0.620963 | 0.677074 | 0.203456 | 0.999971 | 1 | 0.81617 |
| Agfg2 | Q3U2K8 | 0.35 | -0.12 | 0.5 | 0.39 | 0.09 | 0.09 | 0.85 | 0.15 | 0.46 | -0.2 | 0.32 | 0.25 | 0.14 | 0.04 | 0.33 | 0.05 | 0.015 | -0.0725 | -0.14 | 0.953218 | 0.755121 | 0.504075 | 0.999971 | 1 | 0.905265 |
| Agk | Q9ESW4 | -0.07 | -0.48 | -0.85 | -0.65 | -0.5 | -0.64 | -0.79 | -0.59 | -0.43 | -0.02 | -0.71 | -0.84 | -0.12 | -0.52 | -0.69 | -0.55 | -0.1175 | 0.0125 | 0.0425 | 0.596657 | 0.962384 | 0.859368 | 0.999971 | 1 | 0.982702 |
| Agl | F8VPN4 | -0.77 | -0.54 | -0.65 | -0.61 | -0.42 | -0.24 | -0.33 | -0.41 | -0.44 | -0.81 | -0.64 | -0.45 | -0.15 | -0.24 | -0.68 | -0.44 | 0.2925 | 0.0575 | 0.265 | 0.105996 | 0.745075 | 0.193212 | 0.999971 | 1 | 0.810547 |
| Ago1 | A0A6I8MX: | -0.14 | 0.62 | 0.14 | 0 | 0.27 | -0.03 | -0.18 | -0.11 | 0.29 | -0.04 | 0.07 | 0.47 | 0.34 | 0.47 | -0.08 | -0.21 | -0.1675 | 0.045 | -0.025 | 0.474069 | 0.855773 | 0.92264 | 0.999971 | 1 | 0.993264 |
| Ago2 | Q8CJG0 | 0.9 | 0.88 | 1 | 1.12 | 0.91 | 0.78 | 0.62 | 1.08 | 0.59 | 1.09 | 0.46 | 0.62 | 0.34 | 1.01 | 0.68 | 0.98 | -0.1275 | -0.285 | -0.2225 | 0.49651 | 0.180893 | 0.314682 | 0.999971 | 1 | 0.851546 |
| Ago3 | Q8CJF9 | 1.29 | 1.02 | 0.97 | 0.91 | 0.55 | 1.02 | 0.12 | 1 | 0.18 | 1.1 | 0.56 | 0.64 | 0.39 | 1.12 | 1.11 | 1.33 | -0.375 | -0.4275 | -0.06 | 0.166688 | 0.099905 | 0.810927 | 0.999971 | 1 | 0.973864 |
| Agpat1 | A0A0R4J26 | 0.03 | 0.3 | 0.87 | 0.47 | 0.34 | 0.13 | 1.11 | 1.17 | 1 | 0 | 1.64 | 2.22 | 2.05 | 0.28 | 1.59 | 1.09 | 0.27 | 0.7975 | 0.835 | 0.408891 | 0.120829 | 0.062643 | 0.999971 | 1 | 0.725348 |
| Agpat3 | Q9D517 | -0.58 | -0.47 | -0.19 | 0.03 | -0.6 | -0.15 | -0.36 | 0.73 | -0.4 | -0.15 | 0.02 | -0.29 | -0.2 | -0.04 | 0.33 | 0.22 | 0.2075 | 0.0975 | 0.38 | 0.524903 | 0.646009 | 0.118608 | 0.999971 | 1 | 0.78268 |
| Agpat4 | Q8K4X7 | -0.55 | -0.5 | -0.02 | -0.29 | -0.6 | -0.23 | 0.04 | 0.07 | -0.37 | -0.24 | 0.3 | 0.03 | -0.18 | -0.01 | 0.05 | -0.24 | 0.16 | 0.27 | 0.245 | 0.498289 | 0.255342 | 0.238847 | 0.999971 | 1 | 0.833594 |
| Agpat5 | Q9D1E8 | -0.72 | -1.89 | -0.03 | 0.16 | -1.57 | -2.04 | 0.33 | 0.2 | -1.34 | -1.19 | -0.03 | -0.05 | -1.64 | -1.64 | 0.14 | -0.37 | -0.15 | -0.0325 | -0.2575 | 0.827956 | 0.951989 | 0.665489 | 0.999971 | 1 | 0.93697 |
| Agps | A2AL50 | -0.49 | -0.53 | -0.12 | -0.2 | -0.53 | -0.42 | -0.39 | 0.11 | -0.46 | -0.22 | -0.08 | -0.19 | -0.2 | -0.08 | 0.47 | 0.17 | 0.0275 | 0.0975 | 0.425 | 0.900332 | 0.612376 | 0.084028 | 0.999971 | 1 | 0.755254 |
| Agrn | M0QWP1 | 0.81 | 0.12 | 0.41 | 0.49 | 0.68 | 0.13 | 0.9 | -0.38 | 0.48 | 0.72 | 0.97 | 0.56 | 0.04 | 0.19 | 0.56 | 0.1 | -0.125 | 0.225 | -0.235 | 0.699005 | 0.319848 | 0.312483 | 0.999971 | 1 | 0.849367 |
| Agt | Q3UTR7 | -0.08 | 0.58 | 0.95 | 1.4 | 0.96 | 0.58 | 1.27 | 1.41 | 1.33 | 0.39 | 1.63 | 2.07 | 1.03 | 1.53 | 1.67 | 2.58 | 0.3425 | 0.6425 | 0.99 | 0.348266 | 0.175748 | 0.042749 | 0.999971 | 1 | 0.701725 |
| Agtpbp1 | Q641K1-2 | -0.4 | -0.33 | 0.15 | 0.16 | -0.32 | -0.43 | 0.11 | -0.23 | -0.3 | -0.06 | -0.32 | -0.27 | -0.37 | -0.75 | 0.02 | -0.02 | -0.1125 | -0.1325 | -0.24 | 0.625722 | 0.53157 | 0.347304 | 0.999971 | 1 | 0.864164 |
| Ahctf1 | Q8CJF7 | -0.22 | -1.31 | -0.59 | -0.34 | -1.06 | -1.21 | -1.08 | -0.38 | -1.42 | -0.95 | -0.88 | -0.74 | -1.4 | -0.87 | -0.45 | -0.24 | -0.3175 | -0.3825 | -0.125 | 0.323671 | 0.21623 | 0.720906 | 0.999971 | 1 | 0.954713 |
| Ahcy | P50247 | 0.34 | 1 | 0.38 | 0.25 | 1.05 | 0.91 | 0.3 | 0.28 | 1.03 | 0.03 | 0.13 | 0.26 | 0.81 | 0.57 | 0.21 | 0.26 | 0.1425 | -0.13 | -0.03 | 0.613111 | 0.66021 | 0.904562 | 0.999971 | 1 | 0.991091 |
| Ahcyl1 | Q80SW1 | -1.16 | -1.46 | -0.07 | -0.22 | -1.37 | -1 | -0.37 | 0.28 | -1.07 | -1.17 | -0.22 | -0.21 | -1.16 | -1.07 | 0.05 | -0.04 | 0.1125 | 0.06 | 0.1725 | 0.809616 | 0.884206 | 0.699323 | 0.999971 | 1 | 0.947892 |
| Ahcyl2 | Q68FL4-2 | 1.2 | 2.06 | 2.59 | 0.81 | -0.1 | 1.1 | 0.82 | 0.58 | 1.55 | 1.05 | -3.63 | 1.51 | 0.21 | 1.3 | 0.66 | 1.81 | -1.065 | -1.545 | -0.67 | 0.039388 | 0.21359 | 0.201503 | 0.999971 | 1 | 0.815758 |
| Ahcyl2_1 | F8WGT1 | -1.03 | -0.78 | -0.42 | -0.45 | -0.95 | 0.14 | -0.37 | 0.21 | -0.61 | -0.87 | -0.4 | -0.25 | -0.71 | 0.19 | -0.23 | -0.29 | 0.4275 | 0.1375 | 0.41 | 0.192152 | 0.557364 | 0.14006 | 0.999971 | 1 | 0.784948 |
| Ahdc1 | Q6PAL7 | 0.37 | 0.61 | 0.31 | 0.3 | 0.6 | 0.35 | -0.05 | 0.37 | 0.49 | 0.42 | 0.23 | 0.08 | 0.63 | 0.4 | 0.46 | 0.35 | -0.08 | -0.0925 | 0.0625 | 0.699436 | 0.618956 | 0.726394 | 0.999971 | 1 | 0.956041 |
| Ahi1 | E9QP54 | 0.85 | -1.23 | 0.5 | 0.24 | 0.09 | -0.07 | 0.45 | 1.31 | 0.69 | -0.16 | -1.17 | 0.82 | 0.76 | 1.27 | 1.65 | -0.12 | 0.355 | -0.045 | 0.8 | 0.492604 | 0.939721 | 0.170211 | 0.999971 | 1 | 0.802549 |
| Ahnak | E9Q616 | 0.64 | 0.22 | -0.07 | 0.74 | 1.62 | 0.08 | -0.1 | -0.34 | 0.36 | 1.19 | 0.11 | 0.91 | -0.24 | 0.73 | 0.48 | 0.32 | -0.0675 | 0.26 | -0.06 | 0.881229 | 0.417573 | 0.836745 | 0.999971 | 1 | 0.979416 |
| Ahnak2 | A0A7N9VR | -0.42 | -0.93 | -1.12 | -0.88 | -0.94 | -0.92 | -1.27 | -0.48 | -0.72 | 0.08 | -0.72 | -0.58 | 0.1 | -0.37 | -0.62 | -0.57 | -0.065 | 0.3525 | 0.4725 | 0.793843 | 0.202605 | 0.085948 | 0.999971 | 1 | 0.759991 |
| Ahsa1 | Q8BK64 | -0.1 | 0.61 | 0.59 | 0.66 | 0.87 | 1.02 | 0.33 | 0.85 | 0.14 | 0.81 | 1.1 | 0.56 | 0.34 | 0.37 | 0.4 | 0.385 | 0.0775 | 0.0775 | 0.151234 | 0.191686 | 0.724882 | 0.999971 | 1 | 0.955813 | |
| Ahsa2 | F6S1R2 | 0.41 | 0.34 | 0.78 | -0.48 | 0.4 | 0.8 | -0.21 | -0.6 | 0.42 | 0.38 | -0.4 | -0.47 | 0.32 | 0.9 | -0.41 | -0.02 | -0.165 | -0.28 | -0.065 | 0.675731 | 0.435699 | 0.861706 | 0.999971 | 1 | 0.982963 |
| Ahsg | P29699 | 1.65 | 2.24 | 0.87 | 3.7 | 4.15 | 2.66 | 3.41 | 2.44 | 2.13 | 1.6 | 2.16 | 1.99 | 2.1 | 3.27 | 2.52 | 2.42 | 1.05 | -0.145 | 0.4625 | 0.131714 | 0.797404 | 0.443131 | 0.999971 | 1 | 0.883386 |
| Aida | Q8C4Q6 | -0.51 | -0.51 | -0.63 | -0.71 | -0.34 | -0.68 | -0.93 | -0.84 | -0.65 | -0.69 | -0.96 | -0.44 | -0.23 | -0.49 | -0.85 | -0.49 | -0.1075 | -0.095 | 0.075 | 0.591058 | 0.609304 | 0.705682 | 0.999971 | 1 | 0.950182 |
| Aif1 | O70200 | 0.63 | -0.6 | 0.47 | 0.7 | -0.42 | -0.22 | 0.68 | 0.16 | -0.32 | 0.35 | 0.09 | 0.46 | -0.88 | -0.35 | 0.42 | -0.17 | -0.25 | -0.155 | -0.545 | 0.511138 | 0.65502 | 0.186385 | 0.999971 | 1 | 0.809025 |
| Aifm1 | B1AU25 | -0.32 | -0.36 | -0.47 | -0.55 | -0.66 | -0.53 | -0.21 | -0.41 | -0.36 | -0.72 | -0.55 | -0.33 | -0.35 | -0.37 | -0.55 | -0.52 | -0.0275 | -0.065 | -0.0225 | 0.880875 | 0.716569 | 0.894698 | 0.999971 | 1 | 0.988571 |
| Aifm2 | Q8BUE4 | -0.65 | -1.12 | -1.21 | -1.27 | -0.91 | -0.72 | -1 | -1.31 | -0.74 | -0.53 | -1.1 | -0.46 | -0.78 | -0.25 | -1.38 | -0.91 | 0.0775 | 0.355 | 0.2325 | 0.733361 | 0.155601 | 0.42507 | 0.999971 | 1 | 0.880275 |
| Aifm3 | Q3TY86 | -0.21 | -0.59 | 0.07 | -0.33 | -0.63 | -0.22 | 0.12 | -0.86 | -0.79 | -0.42 | 0.1 | -0.42 | -0.43 | -0.63 | -0.16 | -0.42 | -0.1325 | -0.1175 | -0.145 | 0.631466 | 0.644303 | 0.50653 | 0.999971 | 1 | 0.905775 |
| Aig1 | H7BX91 | 4.89 | -2.98 | 1.46 | -0.22 | -4.81 | 0.7 | 7.19 | -2.92 | -7.56 | 4.09 | -1.39 | -3.24 | -6.44 | -5.39 | -1.97 | -4.04 | -0.7475 | -2.8125 | -5.2475 | 0.785906 | 0.296309 | 0.012175 | 0.999971 | 1 | 0.613095 |
| Aimp1 | P31230 | 0.85 | 1.39 | 1.16 | 1.2 | 1.12 | 1.39 | 1.2 | 1.25 | 1.37 | 1.45 | 1.53 | 1.09 | 1.14 | 1.66 | 1.38 | 1.51 | 0.09 | 0.21 | 0.2725 | 0.640292 | 0.310492 | 0.21491 | 0.999971 | 1 | 0.826893 |
| Aimp2 | Q8R010 | 1.18 | 1.31 | 1.03 | 1.37 | 1.22 | 1.26 | 1.59 | 1.1 | 1.62 | 1.78 | 1.66 | 1.02 | 0.77 | 1.58 | 1.37 | 0.98 | 0.07 | 0.2975 | -0.0475 | 0.718173 | 0.207324 | 0.839763 | 0.999971 | 1 | 0.979728 |
| Aip | O08915 | 0.74 | 0.28 | 0.62 | 0.49 | 0.3 | 0.57 | 0.57 | 0.74 | 0.46 | 0.11 | 0.76 | 0.35 | 0.62 | 0.91 | 0.89 | 0.28 | 0.0125 | -0.1125 | 0.1425 | 0.949077 | 0.598847 | 0.52473 | 0.999971 | 1 | 0.90868 |
| Ajap1 | A2ALI5 | -0.53 | 1.55 | -1.43 | -0.95 | 2.61 | 0.74 | -0.78 | -1.82 | 2.58 | -0.67 | -2.07 | -1.98 | 4.71 | 0.74 | -1.65 | -1.82 | 0.5275 | -0.195 | 0.835 | 0.614175 | 0.863957 | 0.573874 | 0.999971 | 1 | 0.91816 |
| Ajm1 | A2AJA9 | -0.66 | -1.08 | -1.33 | -1.31 | -1.03 | -1.03 | -1.04 | -1.49 | -0.96 | -0.79 | -1.55 | -1.62 | -0.55 | -1.29 | -1.86 | -1.86 | -0.0525 | -0.135 | -0.295 | 0.820119 | 0.626739 | 0.401071 | 0.999971 | 1 | 0.871816 |
| Ak1 | Q9R0Y5 | 0.14 | 0.45 | -0.04 | 0.43 | 0.35 | 0.89 | 0.2 | 0.13 | 0.39 | 0.58 | -0.03 | -0.13 | 0.07 | 0.58 | -0.1 | 0.03 | 0.1475 | -0.0425 | -0.1 | 0.544999 | 0.858069 | 0.665148 | 0.999971 | 1 | 0.93697 |
| Ak2 | Q9WTP6 | 0.55 | -0.38 | -0.22 | 0.17 | -0.51 | -0.49 | -0.54 | 0.12 | -0.24 | 0.34 | -0.08 | 0.27 | -0.25 | -0.11 | 0.03 | 0.09 | -0.385 | 0.0425 | -0.09 | 0.18972 | 0.874266 | 0.720104 | 0.999971 | 1 | 0.954623 |
| Ak3 | Q9WTP7 | -0.14 | -0.67 | -0.8 | -1 | -0.37 | -0.66 | -0.63 | -0.83 | -0.53 | -0.34 | -0.65 | -0.71 | -0.36 | -0.54 | -0.97 | -1.03 | 0.03 | 0.095 | -0.0725 | 0.900381 | 0.686202 | 0.786493 | 0.999971 | 1 | 0.967132 |

| | | | | | | | | | | | | | | | | | | | | | | | | | | |
|---|---|---|---|---|---|---|---|---|---|---|---|---|---|---|---|---|---|---|---|---|---|---|---|---|---|---|
| Ak4 | Q9WUR9 | -0.2 | -0.47 | -0.28 | -0.41 | -0.51 | -0.48 | -0.58 | -0.53 | -0.7 | -0.49 | -0.94 | -0.76 | -0.73 | -0.47 | -0.62 | -0.47 | -0.185 | -0.3825 | -0.2325 | 0.284147 | 0.06049 | 0.207544 | 0.999971 | 1 | 0.820632 |
| Ak5 | Q920P5 | 0.25 | 0.09 | 0.74 | 0.48 | 0.06 | 0.6 | 0.81 | 0.86 | -0.03 | 0.02 | 0.5 | 0.22 | 0.22 | 0.52 | 0.53 | 0.91 | 0.1925 | -0.2125 | 0.155 | 0.460129 | 0.355627 | 0.516331 | 0.999971 | 1 | 0.905815 |
| Akap1 | O08715 | 0.2 | -0.03 | 0.16 | 0.03 | -0.1 | -0.54 | 0.23 | -0.1 | 0.02 | -0.19 | 0.22 | -0.05 | 0.39 | -0.23 | -0.06 | -0.1 | -0.2175 | -0.09 | -0.09 | 0.325024 | 0.613133 | 0.658734 | 0.999971 | 1 | 0.93472 |
| Akap10 | O88845 | -0.74 | -0.52 | -0.5 | -0.62 | -0.1 | -0.81 | -0.12 | -0.48 | -0.2 | -0.75 | 0.08 | 0.3 | -0.18 | -0.93 | -0.48 | -0.56 | 0.2175 | 0.4525 | 0.0575 | 0.3382 | 0.105283 | 0.78774 | 0.999971 | 1 | 0.96724 |
| Akap11 | E9Q777 | -1.87 | -0.51 | 0.43 | 0.07 | 1.03 | -0.22 | -0.2 | -0.24 | 0.54 | -0.97 | -0.01 | -0.13 | -0.77 | -0.76 | -0.13 | -0.25 | 0.5625 | 0.3275 | -0.0075 | 0.317997 | 0.554736 | 0.9879 | 0.999971 | 1 | 0.99925 |
| Akap12 | Q9WTQ5 | 0.06 | -1.01 | -0.63 | -0.53 | -0.7 | -0.89 | -0.34 | -0.61 | -0.65 | -0.13 | -0.77 | -0.49 | -0.69 | -0.49 | -0.18 | -0.45 | -0.1075 | 0.0175 | 0.075 | 0.690557 | 0.949541 | 0.778831 | 0.999971 | 1 | 0.96575 |
| Akap13 | E9Q394-2 | 2.51 | 1.46 | 1.17 | 1.51 | 2.19 | 2.49 | 1.68 | 0.04 | 1.47 | 1.72 | 0.46 | 0.23 | 1.75 | 1.11 | 0.01 | 0.6 | -0.0625 | -0.6925 | -0.795 | 0.912247 | 0.14519 | 0.101241 | 0.999971 | 1 | 0.78268 |
| Akap5 | H3BIV5 | -0.45 | -2.27 | -1.22 | -0.75 | -1.94 | -2.16 | -0.83 | -1.27 | -2.12 | -1.4 | -1.86 | -1.5 | -2.22 | -2.03 | -1.41 | -1.56 | -0.3775 | -0.5475 | -0.6325 | 0.429707 | 0.207047 | 0.158027 | 0.999971 | 1 | 0.797252 |
| Akap6 | E9Q9K8 | 0.69 | 0.45 | 0.38 | 0.39 | 0.75 | 0.2 | 0.43 | 0.62 | 0.52 | 0.64 | 0.78 | 1.01 | 0.42 | 0.31 | 0.42 | 0.38 | 0.0225 | 0.26 | -0.095 | 0.909938 | 0.194262 | 0.582608 | 0.999971 | 1 | 0.920364 |
| Akap7 | Q7TN79 | -0.02 | -1.31 | -0.92 | -0.78 | -1.39 | -0.94 | -0.73 | -0.7 | -1.25 | -0.76 | -0.74 | -1.05 | -0.85 | -1.11 | -0.63 | -0.49 | -0.1825 | -0.1925 | -0.0125 | 0.56811 | 0.530466 | 0.967668 | 0.999971 | 1 | 0.998787 |
| Akap8 | Q9DBR0 | 1.07 | 1.5 | 2.53 | 2.15 | 1.13 | 0.59 | 2.14 | 1.31 | 1.18 | 0.96 | 1.79 | 1.59 | 0.11 | 0.74 | 1.64 | 1.39 | -0.52 | -0.4325 | -0.8425 | 0.248722 | 0.258321 | 0.086418 | 0.999971 | 1 | 0.759991 |
| Akap8l | A0A494B9I | 1.28 | 0.73 | 0.84 | 0.62 | 0.76 | 0.84 | 1.09 | 0.66 | 0.71 | 0.79 | 0.9 | 1.19 | -0.05 | 0.36 | 0.74 | 0.3 | -0.03 | 0.03 | -0.53 | 0.890064 | 0.891259 | 0.056824 | 0.999971 | 1 | 0.725348 |
| Akap9 | E9QQ10 | 1 | 1.17 | 1.13 | 1.65 | 0.97 | 0.48 | 1.16 | 1.49 | 1 | 1.08 | 1.25 | 1.53 | 0.85 | 0.84 | 1.29 | 1.14 | -0.2125 | -0.0225 | -0.2075 | 0.443947 | 0.91961 | 0.365149 | 0.999971 | 1 | 0.869526 |
| Akr1a1 | Q9JII6 | 1.81 | 0.61 | 1.1 | 0.65 | 0.63 | 0.54 | 1.72 | 0.36 | 0.68 | 0.9 | 0.75 | 0.56 | 0.51 | -0.41 | 0.15 | -0.2 | -0.23 | -0.32 | -1.03 | 0.566361 | 0.297039 | 0.013928 | 0.999971 | 1 | 0.613095 |
| Akr1b1 | P45376 | -0.09 | 0.7 | 1.16 | 0.6 | 0.76 | 0.29 | 0.85 | 1.03 | 1.27 | -0.09 | 1.18 | 0.79 | 1.18 | 0.29 | 1.02 | 1.13 | 0.14 | 0.195 | 0.3125 | 0.651488 | 0.618653 | 0.357946 | 0.999971 | 1 | 0.868793 |
| Akr1b10 | G5E895 | -0.98 | 0.74 | 0.97 | -0.22 | 1.23 | 0.55 | 1.24 | -0.08 | 1.83 | -0.97 | 1.27 | 1.82 | 2.74 | 0.16 | 0.73 | 0.25 | 0.6075 | 0.86 | 0.8425 | 0.252085 | 0.258044 | 0.237576 | 0.999971 | 1 | 0.833594 |
| Akr1b7 | P21300 | -0.8 | -0.1 | -0.85 | -1.33 | -0.48 | -0.17 | -0.65 | -0.68 | -0.2 | -0.78 | -0.52 | -1.01 | -0.17 | -0.05 | 0.11 | 0.56 | 0.275 | 0.1425 | 0.8825 | 0.35656 | 0.649231 | 0.017286 | 0.999971 | 1 | 0.617356 |
| Akr1e2 | Q9DCT1 | 0.22 | 0.5 | 0.64 | 0.21 | 0.34 | 0.82 | 1 | 0.89 | 0.67 | 0.09 | 0.83 | 0.92 | 0.26 | 0.48 | 0.73 | 0.5 | 0.37 | 0.235 | 0.1 | 0.123504 | 0.347556 | 0.622216 | 0.999971 | 1 | 0.929759 |
| Akr7a2 | Q8CG76 | 0.58 | 0.58 | 0.48 | 0.16 | 0.06 | 0.33 | 0.26 | 0.04 | 0.16 | 0.36 | 0.26 | 0.25 | -0.11 | 0.18 | 0.17 | -0.07 | -0.2775 | -0.1925 | -0.4075 | 0.169459 | 0.300441 | 0.059223 | 0.999971 | 1 | 0.725348 |
| Akt1 | P31750 | 0.88 | 0.74 | 0.88 | 0.61 | 0.59 | 0.81 | 0.82 | 0.87 | 0.54 | 0.47 | 0.63 | 0.15 | 0.6 | 0.51 | 0.54 | 0.78 | -0.005 | -0.33 | -0.17 | 0.977237 | 0.105602 | 0.346413 | 0.999971 | 1 | 0.864164 |
| Akt1s1 | Q9D1F4 | -0.71 | 1.22 | -0.23 | 0.25 | 1.02 | 1.25 | -0.69 | 0.02 | 1.35 | -0.2 | -0.28 | 0.04 | 2.38 | 1.05 | -0.04 | 0.47 | 0.2675 | 0.095 | 0.8325 | 0.635057 | 0.85519 | 0.193883 | 0.999971 | 1 | 0.810547 |
| Akt2 | Q60823 | 0.22 | 0.19 | -0.13 | 0.05 | -0.19 | 0.21 | -0.19 | 0.95 | -0.14 | 0.07 | 0.1 | -0.02 | -0.03 | 0.14 | -0.01 | 0.5 | 0.1125 | -0.08 | 0.0675 | 0.700716 | 0.648894 | 0.740496 | 0.999971 | 1 | 0.960899 |
| Akt3 | Q9WUA6 | 0.04 | 0.22 | 0.04 | 0.23 | 0.31 | 0.54 | 0.33 | 0.33 | 0.38 | 0.1 | 0.06 | -0.05 | 0.52 | 0.37 | -0.16 | -0.08 | 0.245 | -0.01 | 0.03 | 0.174911 | 0.955404 | 0.891538 | 0.999971 | 1 | 0.988225 |
| Aktip | D3Z632 | -0.36 | 1.4 | 1.2 | 1.05 | 1.14 | 1.31 | 0.61 | 1.45 | 0.93 | 0.45 | 1.17 | 1.23 | 1.12 | 1.5 | 1.17 | 1.6 | 0.305 | 0.1225 | 0.525 | 0.472662 | 0.769507 | 0.212166 | 0.999971 | 1 | 0.823193 |
| Alad | P10518 | 0.41 | 0.84 | 0.33 | 0.68 | 0.85 | 0.72 | 0.02 | 0.14 | 0.71 | 0.46 | 0 | 0.2 | 0.6 | 1.02 | 0.64 | 0.74 | -0.1325 | -0.2225 | 0.185 | 0.614406 | 0.346377 | 0.380848 | 0.999971 | 1 | 0.871816 |
| Alb | P07724 | 1.17 | 3.47 | 0.83 | 3.78 | 5.21 | 3.36 | 3.88 | 1.63 | 2.93 | 0.4 | 2.05 | 1.72 | 2.51 | 2.68 | 1.67 | 2.12 | 1.2075 | -0.5375 | -0.0675 | 0.225185 | 0.525809 | 0.92522 | 0.999971 | 1 | 0.99401 |
| Alcam | E9Q3Q6 | -1.08 | -1.86 | -1.44 | -0.61 | -1.21 | -1.68 | -0.96 | -0.85 | -1.25 | -1.11 | -1.31 | -1.06 | -1.07 | -1.24 | -0.58 | -0.94 | 0.0725 | 0.065 | 0.29 | 0.82318 | 0.819441 | 0.357766 | 0.999971 | 1 | 0.868793 |
| Aldh16a1 | D3Z0B9 | 1.21 | 0.73 | 0.8 | 0.83 | 0.82 | 0.27 | 0.78 | 0.67 | 0.69 | 0.5 | 1.09 | 0.62 | 0.42 | 0.4 | 1.06 | 0.94 | -0.2575 | -0.1675 | -0.1875 | 0.247868 | 0.438711 | 0.439025 | 0.999971 | 1 | 0.883105 |
| Aldh18a1 | Q9Z110 | -0.55 | -0.01 | -0.46 | -0.48 | -0.27 | -0.21 | -0.3 | -0.34 | -0.03 | -0.41 | 0.03 | -0.22 | -0.21 | -0.27 | -0.72 | -0.57 | 0.095 | 0.2175 | -0.0675 | 0.622795 | 0.309129 | 0.758188 | 0.999971 | 1 | 0.961116 |
| Aldh18a1_ | Q9Z110-2 | -0.14 | -1.46 | -2.03 | -0.13 | -0.19 | -1.61 | -0.82 | -2.88 | 0.03 | -3.05 | -1.17 | -0.72 | -2.42 | -1.73 | -0.74 | -0.4 | -0.435 | -0.2875 | -0.3825 | 0.52762 | 0.697762 | 0.533835 | 0.999971 | 1 | 0.910077 |
| Aldh1a1 | P24549 | -1.74 | -1.92 | -1.53 | -1.15 | -1 | -1.84 | -0.65 | -1.28 | -1.57 | -2.1 | -0.93 | -0.58 | -2.17 | -1.85 | -0.97 | -1.12 | 0.3925 | 0.29 | 0.0575 | 0.222575 | 0.437224 | 0.862337 | 0.999971 | 1 | 0.982963 |
| Aldh1a2 | Q62148 | 2.04 | 1.45 | 1.38 | 2.52 | 2.12 | 1.09 | 1.04 | 1.95 | 1.19 | 1.25 | 1.07 | 1.58 | 0.19 | 2.21 | 1.71 | 1.64 | -0.2975 | -0.575 | -0.41 | 0.438023 | 0.080987 | 0.400556 | 0.999971 | 1 | 0.871816 |
| Aldh1a7 | O35945 | -1.29 | -0.49 | 0.22 | 0.71 | -1.22 | -1.48 | 0.28 | 1.24 | -0.34 | -0.05 | 1 | 1 | -1.48 | -0.89 | 0.92 | 1.36 | -0.0825 | 0.615 | 0.19 | 0.906263 | 0.255471 | 0.79633 | 0.999971 | 1 | 0.970435 |
| Aldh1b1 | Q9CZS1 | -0.42 | -0.67 | -0.85 | -0.58 | -0.77 | -0.69 | -0.96 | -0.46 | -0.74 | -0.31 | -0.69 | -0.6 | -1.04 | -0.73 | -0.69 | -0.8 | -0.09 | 0.045 | -0.185 | 0.650351 | 0.814583 | 0.34133 | 0.999971 | 1 | 0.860984 |
| Aldh1l1 | Q8R0Y6 | -0.46 | -0.21 | -0.42 | -0.73 | -0.46 | -0.06 | -0.54 | -0.19 | -0.06 | -0.04 | -0.62 | -0.72 | -0.03 | -0.01 | -0.47 | -0.17 | 0.1425 | 0.095 | 0.285 | 0.498286 | 0.693138 | 0.189404 | 0.999971 | 1 | 0.810547 |
| Aldh1l2 | Q8K009 | -1.19 | -0.07 | -0.06 | -0.36 | 0.18 | -0.21 | -0.15 | -0.14 | 0.04 | -0.93 | -0.61 | -0.04 | 0.1 | -0.58 | -0.66 | -0.7 | 0.34 | 0.035 | -0.04 | 0.26203 | 0.919755 | 0.902556 | 0.999971 | 1 | 0.990759 |
| Aldh2 | P47738 | -0.5 | -0.8 | -1.07 | -1.28 | -0.92 | -1.08 | 0.3 | -1.23 | -1.21 | -1.12 | -1.37 | -0.73 | -1.12 | -0.98 | -1.29 | -1.31 | 0.18 | -0.195 | -0.2625 | 0.634041 | 0.434553 | 0.265736 | 0.999971 | 1 | 0.842677 |
| Aldh3a2 | B1AV77 | -1.74 | -1.39 | -0.89 | -1.37 | -1.19 | -1.35 | -0.86 | -0.76 | -1.41 | -1.71 | -1 | -0.74 | -1.79 | -1.48 | -0.64 | -1.32 | 0.3075 | 0.1325 | 0.04 | 0.23966 | 0.647697 | 0.89607 | 0.999971 | 1 | 0.988571 |
| Aldh3b1 | Q80VQ0 | -1.28 | -1.02 | -0.48 | -0.72 | -1.17 | -1.34 | -1.06 | -0.08 | -1.47 | -1.71 | -1.15 | -0.9 | -1.54 | -1.66 | -0.53 | -0.76 | -0.0375 | -0.4325 | -0.2475 | 0.9099 | 0.133137 | 0.461337 | 0.999971 | 1 | 0.89007 |
| Aldh4a1 | Q8CHT0 | -0.56 | -0.8 | -0.97 | -0.83 | -0.86 | -0.83 | -0.79 | -0.28 | -0.71 | -0.91 | -0.85 | -0.96 | -0.9 | -0.52 | -1.08 | -1.11 | 0.1 | -0.0675 | -0.1125 | 0.638099 | 0.703435 | 0.597071 | 0.999971 | 1 | 0.92377 |
| Aldh5a1 | Q8BWF0 | -0.59 | -1.11 | -0.61 | -1.2 | -0.99 | -1.14 | -0.3 | -0.88 | -0.5 | -1.49 | -0.84 | -0.58 | -0.54 | -1.65 | -0.73 | -1.35 | 0.05 | 0.025 | -0.19 | 0.850698 | 0.930758 | 0.546267 | 0.999971 | 1 | 0.911398 |
| Aldh6a1 | Q9EQ20 | -0.69 | -0.92 | -0.75 | -0.92 | -0.51 | -0.85 | -0.62 | -0.17 | -0.65 | -0.96 | -0.83 | -1.07 | -0.6 | -0.96 | -0.84 | -0.97 | 0.2825 | -0.0575 | -0.0225 | 0.194544 | 0.749766 | 0.901666 | 0.999971 | 1 | 0.990759 |
| Aldh7a1 | Q9DBF1-2 | -0.48 | -0.09 | -0.01 | -0.07 | 0.01 | 0.14 | -0.14 | 0.83 | 0.03 | -0.47 | -0.09 | -0.03 | 0.44 | -0.01 | -0.02 | 0.06 | 0.3725 | 0.0225 | 0.28 | 0.179695 | 0.912671 | 0.199013 | 0.999971 | 1 | 0.813335 |
| Aldh9a1 | Q9JLJ2 | -0.21 | 0.12 | 0.27 | 0.42 | 0.36 | 0.65 | 0.35 | 1.35 | 0.29 | 0.19 | 0.02 | -0.11 | 0.56 | 0.71 | 0.79 | 0.49 | 0.5275 | -0.0525 | 0.4875 | 0.090818 | 0.801651 | 0.039457 | 0.999971 | 1 | 0.700838 |
| Aldoa | A6ZI44 | 0.77 | 0.54 | -1.17 | -0.78 | -0.06 | 0.8 | -0.95 | -0.53 | 0.21 | 1.61 | -0.62 | 1.12 | 0.79 | 0.71 | -1.77 | 0.3 | -0.025 | 0.74 | 0.1675 | 0.964264 | 0.259575 | 0.810276 | 0.999971 | 1 | 0.973864 |
| Aldoa_1 | P05064 | 0.06 | 0.33 | 0.85 | 0.51 | -0.22 | 0.55 | 0.84 | 0.84 | -0.27 | -0.05 | 0.37 | 0.38 | -0.24 | 0.36 | 0.61 | 0.81 | 0.065 | -0.33 | -0.0525 | 0.832115 | 0.217153 | 0.857904 | 0.999971 | 1 | 0.982702 |
| Aldoc | P05063 | -0.15 | -0.55 | -0.26 | -0.42 | -0.7 | -0.43 | -0.59 | -0.12 | -0.59 | -0.58 | -0.46 | -0.24 | -0.73 | -0.32 | -0.38 | 0.04 | -0.115 | -0.1225 | -0.0025 | 0.581378 | 0.514785 | 0.991065 | 0.999971 | 1 | 0.99925 |
| Alg1 | Q921Q3 | -2.02 | -2.13 | -0.45 | 0.24 | -2.01 | -2.21 | -0.33 | -0.18 | -1.61 | -2.06 | -0.54 | -0.6 | -2.4 | -2.19 | -0.31 | -0.36 | -0.0925 | -0.1125 | -0.225 | 0.897457 | 0.859894 | 0.760591 | 0.999971 | 1 | 0.961116 |
| Alg10b | Q3UGP8 | -0.18 | 0.35 | 0.44 | 0.37 | 0.41 | 0.34 | 0.21 | 0.8 | 0.04 | 0.4 | 0.45 | 0.31 | 0.61 | 0.61 | 0.35 | 0.64 | 0.195 | 0.055 | 0.3075 | 0.405475 | 0.797257 | 0.168137 | 0.999971 | 1 | 0.800215 |
| Alg11 | Q3TZM9 | -0.66 | -0.29 | -1.53 | -1.17 | -0.63 | -0.85 | -0.87 | -1.25 | -0.38 | -0.39 | -1.58 | -1.23 | -0.3 | -0.47 | -1.04 | -1.11 | 0.0125 | 0.0175 | 0.1825 | 0.967615 | 0.964424 | 0.592693 | 0.999971 | 1 | 0.923227 |
| Alg13 | Q9D8C3 | -0.58 | -1.3 | -0.39 | -1.21 | -1.38 | -2.22 | -1.18 | -0.72 | -3.84 | -1.2 | -1.55 | -0.4 | 0.32 | -2.55 | -0.6 | -1.51 | -0.505 | -0.8775 | -0.215 | 0.200426 | 0.232684 | 0.720453 | 0.999971 | 1 | 0.954713 |
| Alg2 | Q9DBE8 | -0.02 | 0.35 | 0.13 | -0.01 | 0.23 | 0.12 | -0.03 | 0.14 | 0.41 | 0.12 | 0.25 | -0.08 | 0.83 | 0.64 | 0.45 | 0.44 | 0.0025 | 0.0625 | 0.4775 | 0.988927 | 0.747314 | 0.032877 | 0.999971 | 1 | 0.685881 |
| Alg3 | A0A338P7E | -0.12 | -0.51 | 0.13 | 0.77 | 0.42 | 0.03 | -0.16 | -0.15 | -0.13 | 0.79 | 0 | 0.53 | -1.26 | -0.2 | 1.02 | 1.47 | -0.0325 | 0.23 | 0.19 | 0.915641 | 0.506601 | 0.757414 | 0.999971 | 1 | 0.961116 |
| Alg5 | Q9DB25 | 0.12 | -0.62 | 0.27 | 0.35 | -0.48 | -0.15 | -0.26 | 0.3 | -0.2 | -0.18 | 0.03 | -0.13 | -0.84 | 0.04 | -0.03 | 0.02 | -0.1775 | -0.15 | -0.2325 | 0.542587 | 0.556569 | 0.4645 | 0.999971 | 1 | 0.890905 |
| Alg9 | Q8VDI9 | 2.83 | 0 | 0.32 | 1.28 | -0.22 | 0.84 | 0.93 | -0.04 | -0.19 | 2.5 | -1.24 | -1.39 | -0.72 | -0.12 | -0.63 | -1.69 | -0.73 | -1.1875 | -1.8975 | 0.269885 | 0.250742 | 0.016289 | 0.999971 | 1 | 0.613095 |
| Alk | P97793 | 0.77 | -1.76 | -1.04 | -0.52 | -1.6 | -1.34 | -0.72 | -1.23 | -1.8 | 0.18 | -0.75 | -0.8 | -2.12 | -1.38 | -0.31 | -0.85 | -0.585 | -0.155 | -0.5275 | 0.279277 | 0.800825 | 0.391583 | 0.999971 | 1 | 0.871816 |
| Alkbh3 | Q8K1E6 | -1.72 | 1.14 | 1.28 | -0.76 | 1.19 | 0.37 | 0.96 | -0.61 | 2.51 | -1.24 | -0.02 | -0.64 | -0.9 | -0.79 | -0.6 | -3.12 | 0.4925 | 0.1675 | -1.3375 | 0.518672 | 0.866003 | 0.141119 | 0.999971 | 1 | 0.784967 |
| Alkbh5 | Q3TSG4 | 1.01 | 1.12 | 1.11 | 1.61 | 1.17 | 0.86 | 0.94 | 1.39 | 1.07 | 0.42 | 1.15 | 1.74 | 0.55 | 0.88 | 1.18 | 2.17 | -0.1225 | -0.1175 | -0.0175 | 0.585282 | 0.703205 | 0.96177 | 0.999971 | 1 | 0.998721 |
| Alkbh6 | Q8K2U2 | 0.8 | 0.93 | 1.05 | 0.78 | 0.67 | 1.24 | 0.6 | 0.79 | 0.44 | 0.95 | 1.05 | 1.35 | 0.36 | 1.49 | 0.86 | 1.13 | -0.065 | 0.0575 | 0.07 | 0.755791 | 0.805298 | 0.793765 | 0.999971 | 1 | 0.970141 |
| Alkbh7 | Q9D6Z0 | -1.63 | 1.27 | -2.23 | -1.07 | -1.41 | -1.08 | -1.62 | -0.82 | -1.1 | -1.93 | -3.42 | -1.6 | -0.19 | -0.36 | -1.43 | -0.15 | -0.3175 | -1.0975 | 0.3825 | 0.656359 | 0.207149 | 0.609551 | 0.999971 | 1 | 0.92673 |
| Alkbh8 | Q80Y20-2 | -0.67 | -0.95 | -0.09 | -1.31 | -0.29 | -0.95 | -0.31 | -0.12 | -0.1 | -0.63 | 0.39 | 0.31 | 0.54 | -0.54 | -0.61 | -0.92 | 0.3375 | 0.7475 | 0.3725 | 0.304798 | 0.054277 | 0.356576 | 0.999971 | 1 | 0.868793 |
| Alox5ap | A0A0J9YUC | 0.23 | -0.67 | -1.42 | -0.02 | -1.15 | -1.78 | -1.68 | -0.91 | -0.91 | -0.54 | 0.57 | -0.2 | -1.85 | -0.37 | -0.5 | 0.74 | -0.91 | 0.2 | -0.025 | 0.047972 | 0.663176 | 0.966246 | 0.999971 | 1 | 0.998721 |
| Alpl | P09242 | -0.31 | 0.2 | 0.18 | 0.26 | 0.4 | -0.77 | 0.2 | -1.12 | 0.08 | -0.04 | -0.62 | -0.01 | 0.13 | -0.58 | 0.34 | -0.34 | -0.405 | -0.23 | -0.195 | 0.300389 | 0.347442 | 0.475765 | 0.999971 | 1 | 0.894045 |
| Als2 | Q920R0 | 0.47 | -0.58 | -0.45 | -0.46 | -0.71 | -0.38 | 0.02 | -0.7 | -0.81 | 0.16 | -0.57 | -1.04 | -0.22 | -0.35 | -0.74 | -0.6 | -0.1875 | -0.31 | -0.2225 | 0.543023 | 0.387175 | 0.443057 | 0.999971 | 1 | 0.883386 |
| Alyref | O08583 | -0.34 | 0.66 | 0.59 | -0.14 | 1.05 | 0.49 | 0.68 | 0.05 | 1.25 | -0.25 | 0.56 | 1.02 | 0.91 | -0.14 | 0.3 | -0.45 | 0.375 | 0.4525 | -0.0375 | 0.270744 | 0.275885 | 0.920683 | 0.999971 | 1 | 0.992702 |
| Amacr | O09174 | -1.31 | -1.79 | -0.69 | -0.85 | -1.44 | -1.9 | -0.91 | -0.74 | -1.56 | -1.22 | -0.91 | -0.41 | -1.38 | -1.61 | -0.81 | -0.49 | -0.0875 | 0.135 | 0.0875 | 0.805132 | 0.695427 | 0.803461 | 0.999971 | 1 | 0.971467 |
| Ambra1 | A2AH22-7 | 1.35 | 0.29 | 1.29 | 1.05 | 0.46 | 1.15 | 1.61 | 0.85 | 0.32 | 0.71 | 0.54 | 0.7 | -1.77 | 0.49 | 0.96 | 2.47 | 0.0225 | -0.4275 | -0.4575 | 0.94725 | 0.147615 | 0.579757 | 0.999971 | 1 | 0.91958 |
| Amdhd2 | F6UP77 | -1.17 | -0.23 | -0.51 | -0.53 | 1.04 | -0.18 | 0.3 | -0.77 | 2.17 | -0.48 | -0.16 | -0.24 | 1.87 | -0.03 | -0.89 | -0.61 | 0.7075 | 0.9325 | 0.695 | 0.11215 | 0.145389 | 0.262867 | 0.999971 | 1 | 0.842394 |
| Amer2 | A0A0R4J03 | 1.22 | 0.9 | 0.71 | 0.84 | 0.87 | 1.22 | 0.81 | 0.8 | 1.1 | 1.21 | 0.35 | 0.5 | 1 | 1.24 | 0.55 | 0.68 | 0.0075 | -0.1275 | -0.05 | 0.9705 | 0.627646 | 0.827561 | 0.999971 | 1 | 0.977122 |
| Amer3 | Q6NS69 | 2.26 | 1.13 | 0.74 | 0.52 | 0.7 | 1.62 | 0.34 | 0.36 | 2.21 | 2.11 | -0.31 | -0.97 | 1.22 | -0.85 | 0.65 | 0.6 | -0.4075 | -0.4025 | -0.7575 | 0.385133 | 0.625669 | 0.185604 | 0.999971 | 1 | 0.809025 |
| Amfr | Q9R049-2 | 0.24 | -0.48 | -0.07 | -0.29 | -0.62 | -0.58 | 0.14 | -0.13 | -0.42 | 0.39 | -0.54 | -0.36 | -0.05 | 0.11 | -0.1 | -0.29 | -0.1475 | -0.0825 | 0.0675 | 0.577302 | 0.766063 | 0.759628 | 0.999971 | 1 | 0.961116 |
| Amigo1 | Q80ZD8 | 0.74 | -0.62 | -0.5 | -0.62 | -0.54 | -0.44 | -1.29 | -0.33 | -1.02 | 0.43 | -0.93 | -1.05 | -0.08 | -0.4 | -0.6 | -0.2 | -0.4 | -0.3925 | -0.07 | 0.310513 | 0.402809 | 0.839767 | 0.999971 | 1 | 0.979728 |
| Amigo2 | Q80ZD9 | -0.64 | -1.27 | -1.69 | -0.79 | -1.36 | -0.81 | -1.33 | -0.57 | -0.82 | -0.4 | -2.13 | -0.64 | -1.59 | -0.46 | -1.33 | -0.98 | 0.08 | 0.1 | 0.0075 | 0.798398 | 0.816785 | 0.982347 | 0.999971 | 1 | 0.99925 |
| Amot | Q8VHG2-2 | 0.07 | -0.45 | -0.84 | -0.84 | -0.35 | 0.12 | -0.53 | -0.67 | -0.19 | 0.39 | -0.66 | -0.73 | -0.28 | -0.24 | -0.72 | -1.11 | 0.1575 | 0.2175 | -0.0725 | 0.587944 | 0.52085 | 0.812411 | 0.999971 | 1 | 0.974458 |
| Amotl1 | A0A1Y7VM | -0.48 | -3.9 | -1.01 | -0.57 | -2.75 | -3.67 | -0.9 | -0.93 | -2.69 | -1.66 | -0.71 | -0.41 | -3.31 | -3.36 | -0.87 | -0.5 | -0.5725 | 0.1225 | -0.52 | 0.551198 | 0.887663 | 0.605486 | 0.999971 | 1 | 0.92451 |
| Amotl2 | B8JK87 | 2.26 | -1.06 | 0.62 | 0.44 | -1.49 | -0.31 | 2.27 | -0.9 | -1.62 | 1.3 | -0.81 | 1.27 | -2.66 | -1.4 | -2.38 | -3.76 | -0.6725 | -0.53 | -3.115 | 0.488275 | 0.562216 | 0.002582 | 0.999971 | 1 | 0.560302 |
| Ampd2 | A2AE27 | -0.22 | -0.26 | -0.25 | -0.6 | -0.33 | -0.47 | 0.02 | -0.99 | -0.28 | -0.54 | -0.65 | -0.44 | -0.41 | -0.49 | -0.47 | -0.77 | -0.11 | -0.145 | -0.2025 | 0.666509 | 0.442262 | 0.301488 | 0.999971 | 1 | 0.847402 |
| Ampd3 | A0A1L1SRX | -0.64 | -0.77 | -1.03 | -0.12 | -0.35 | -0.99 | -0.68 | -0.57 | 0.16 | -0.87 | -0.73 | -0.63 | -0.43 | -0.67 | -0.23 | -0.28 | -0.0075 | 0.1225 | 0.2375 | 0.976753 | 0.690029 | 0.347641 | 0.999971 | 1 | 0.864164 |
| Amph | Q7TQF7 | 0 | -0.05 | 0.76 | 1 | -0.29 | 0.37 | 0.81 | 0.74 | 0.05 | -0.03 | 0.81 | -0.34 | -0.45 | -0.22 | 0.95 | 0.4 | -0.02 | -0.305 | -0.2575 | 0.955398 | 0.399515 | 0.521039 | 0.999971 | 1 | 0.906972 |
| Amt | Q8CFA2 | -0.58 | -0.98 | -0.82 | -1.3 | -0.72 | -0.64 | -0.73 | -0.25 | -0.67 | -0.06 | -0.61 | -0.91 | -0.52 | -0.44 | -0.61 | -1.06 | 0.335 | 0.3575 | 0.2625 | 0.166797 | 0.188722 | 0.289213 | 0.999971 | 1 | 0.847402 |
| Amz2 | B1AT59 | 0.83 | 0.26 | 0.11 | 0.33 | 0.13 | 0.14 | 0.84 | -0.1 | -0.25 | -0.28 | 0.01 | 0.04 | -0.58 | -0.7 | 0.18 | -0.94 | -0.13 | -0.5025 | -0.8925 | 0.637398 | 0.044411 | 0.014296 | 0.999971 | 1 | 0.613095 |
| Anapc1 | P53995 | -0.46 | 0.22 | 0.34 | 0.45 | 0.25 | 0.04 | 0.36 | 0.4 | 0.25 | -0.15 | 0.42 | 0.66 | 0.03 | 0.4 | 0.62 | 0.51 | 0.125 | 0.1575 | 0.2525 | 0.617459 | 0.57772 | 0.351927 | 0.999971 | 1 | 0.867656 |
| Anapc13 | Q8R034 | 1.14 | 0.73 | 0.63 | 0.24 | 0.47 | -0.33 | 0.23 | 1.38 | 0.43 | -2.02 | 0.52 | 0.14 | 0.16 | 0.94 | 0.8 | 0.59 | -0.2475 | -0.9175 | -0.0625 | 0.52621 | 0.14004 | 0.817761 | 0.999971 | 1 | 0.974929 |
| Anapc16 | Q9CPV2 | 0.29 | 0.67 | 0.7 | 1.18 | 1.54 | 0.5 | -0.09 | 0.86 | 0.66 | 1.24 | 0.43 | 0.85 | 0.66 | 0.96 | 0.7 | 1.07 | -0.0075 | 0.085 | 0.1375 | 0.983994 | 0.752148 | 0.571882 | 0.999971 | 1 | 0.91816 |
| Anapc2 | Q8BZQ7 | 0.16 | 1.23 | 0.63 | 0.87 | 0.86 | 1.14 | 0.15 | 1.12 | 0.96 | 0.52 | 0.46 | 1.15 | 1.11 | 1.52 | 1.08 | 1.23 | 0.095 | 0.05 | 0.5125 | 0.769403 | 0.863508 | 0.082012 | 0.999971 | 1 | 0.755254 |
| Anapc4 | Q91W96 | -0.17 | 0.67 | 0.13 | 0.13 | 0.34 | 0.41 | -0.42 | 0.17 | 0.26 | 0.25 | -0.13 | 0.08 | 0.4 | 0.75 | 0.1 | 0.3 | -0.065 | -0.075 | 0.1975 | 0.813061 | 0.746725 | 0.438442 | 0.999971 | 1 | 0.882889 |
| Anapc5 | A0A0G2JEC | 0.38 | 0.75 | 0.75 | 0.79 | 0.74 | 0.49 | 0.31 | 0.99 | 0.61 | 0.48 | 0.95 | 1.11 | 0.66 | 1.13 | 1.38 | 1.02 | -0.035 | 0.12 | 0.38 | 0.873821 | 0.583894 | 0.113849 | 0.999971 | 1 | 0.78268 |
| Anapc7 | Q9WVM3 | 0.97 | 1.18 | 0.84 | 0.86 | 0.3 | 0.21 | 0.36 | 0.82 | 1.35 | 0.1 | 0.72 | 1.12 | 1.37 | 0.14 | 0.9 | 0.92 | -0.54 | -0.14 | -0.13 | 0.027387 | 0.636131 | 0.646912 | 0.999971 | 1 | 0.933613 |
| Angel2 | Q8K1C0-3 | 0.15 | -0.66 | -0.55 | -0.87 | -0.98 | -0.62 | -0.72 | -0.74 | -1.23 | -1.01 | -1 | -0.95 | -1.64 | -1.08 | -0.6 | -0.96 | -0.2825 | -0.565 | -0.5875 | 0.290485 | 0.049937 | 0.088385 | 0.999971 | 1 | 0.766448 |
| Ank1 | A0A0R4J1N | -0.67 | -0.6 | -1.36 | -0.5 | -0.45 | -0.88 | -0.87 | -1.31 | -1.03 | -1.11 | -1.54 | -1.41 | -0.67 | -0.79 | -1.51 | -1.42 | -0.095 | -0.49 | -0.315 | 0.733942 | 0.080902 | 0.308721 | 0.999971 | 1 | 0.847402 |
| Ank2 | S4R2D6 | -1.05 | -1.33 | 0.5 | -0.72 | -1 | -0.97 | 0.57 | 0.06 | -0.67 | -1.96 | 0.35 | 0.38 | 0.36 | -2.13 | -0.06 | -1.32 | 0.315 | 0.175 | -0.1375 | 0.547138 | 0.780048 | 0.829268 | 0.999971 | 1 | 0.978248 |
| Ank2_1 | Q8C8R3-7 | 2.56 | -0.98 | -0.71 | -0.76 | -1.15 | -0.08 | 0.71 | -1.85 | -1.19 | 0.97 | -0.41 | -2.07 | -2.21 | -1.13 | -1.48 | -2.02 | -0.62 | -0.7025 | -1.7375 | 0.501768 | 0.470229 | 0.053045 | 0.999971 | 1 | 0.725348 |
| Ank2_2 | Q8C8R3-2 | -0.22 | -0.81 | -1.24 | -1.52 | -0.53 | -0.49 | -1.14 | -1.59 | -0.34 | -0.43 | -0.92 | -0.7 | -0.05 | -0.67 | -1.2 | -1.23 | 0.01 | 0.35 | 0.16 | 0.978662 | 0.28453 | 0.677421 | 0.999971 | 1 | 0.942594 |
| Ank2_3 | Q8C8R3-4 | -0.29 | -1.21 | -0.84 | -0.89 | -1.26 | -0.97 | -0.63 | -1.09 | -1.09 | -0.68 | -1.08 | -0.89 | -0.73 | -1.29 | -0.8 | -0.9 | -0.18 | -0.1275 | -0.1225 | 0.490542 | 0.602492 | 0.632929 | 0.999971 | 1 | 0.93301 |
| Ank3 | S4R2K9 | -0.71 | -1.15 | -0.7 | -0.93 | -1.3 | -1.09 | -0.84 | -0.94 | -1.29 | -0.99 | -1.04 | -0.98 | -1.03 | -1.13 | -0.88 | -0.96 | -0.17 | -0.2025 | -0.1275 | 0.41081 | 0.303838 | 0.505647 | 0.999971 | 1 | 0.905277 |
| Ank3_1 | A0A087WN | 0.13 | -0.86 | -0.7 | -0.43 | -0.88 | -1.15 | -0.77 | -0.45 | -0.85 | -0.33 | -0.94 | -0.98 | -0.69 | -0.95 | -0.94 | -0.76 | -0.3475 | -0.31 | -0.37 | 0.231276 | 0.283925 | 0.169925 | 0.999971 | 1 | 0.802438 |
| Ank3_2 | A0A087WR | -1.53 | -0.4 | -1.18 | -1.42 | -0.69 | -1.64 | -2.5 | -0.63 | -1.22 | -1.35 | -0.83 | -1.85 | -0.86 | -0.92 | -0.93 | -0.94 | -0.2325 | -0.18 | 0.22 | 0.62792 | 0.588324 | 0.430901 | 0.999971 | 1 | 0.881611 |
| Ank3_3 | G5E8K2 | -0.4 | -0.64 | -1.85 | -2.17 | -1.7 | -1.22 | -1.88 | -2.76 | -1.97 | -0.85 | -1.85 | -2.26 | -1.19 | -0.73 | -1.87 | -2.36 | -0.625 | -0.4675 | -0.2725 | 0.23495 | 0.360704 | 0.605958 | 0.999971 | 1 | 0.924649 |
| Ankfn1 | F6X7B3 | -0.71 | -0.32 | 2.27 | 1.54 | 0.54 | -0.02 | 0.18 | 2.08 | 0.18 | -0.79 | 3.31 | 0.41 | 1.02 | -0.15 | 2.8 | 0.88 | 0 | 0.0825 | 0.4425 | 1 | 0.935688 | 0.604578 | 1 | 1 | 0.924454 |
| Ankfy1 | Q810B6 | -0.02 | 0.36 | 0.28 | 0.1 | 0.03 | 0.07 | 0.1 | 0.02 | 0.33 | -0.03 | 0.18 | 0.2 | 0.41 | -0.1 | 0.31 | 0.15 | -0.125 | -0.01 | 0.0125 | 0.480734 | 0.956308 | 0.95015 | 0.999971 | 1 | 0.998038 |
| Ankh | Q9JHZ2 | 0.46 | 1.46 | 1.06 | 0.63 | 0.81 | 1.58 | 0.64 | 1.11 | 0.71 | 0.65 | 1.62 | 1.19 | 1.41 | 1.67 | 0.87 | 1.65 | 0.1325 | 0.14 | 0.4975 | 0.670535 | 0.664404 | 0.12521 | 0.999971 | 1 | 0.78268 |
| Ankhd1 | E9Q1M6 | 1.27 | 0.99 | 0.87 | 1.24 | 0.81 | 0.9 | 0.43 | 0.69 | 0.52 | 0.94 | 0.42 | 0.43 | 0.57 | 0.84 | 0.88 | 0.91 | -0.385 | -0.515 | -0.2925 | 0.080468 | 0.03198 | 0.152944 | 0.999971 | 1 | 0.795578 |
| Ankib1 | Q6ZPS6 | 0.24 | -0.34 | -0.06 | -0.38 | -0.34 | -0.2 | 0.01 | -0.18 | -0.31 | 0.04 | -0.4 | -0.63 | -0.39 | -0.64 | -0.18 | -0.29 | -0.0425 | -0.19 | -0.24 | 0.840293 | 0.424821 | 0.291004 | 0.999971 | 1 | 0.847402 |
| Ankle2 | Q6P1H6 | 0.09 | -0.07 | -0.2 | 0.13 | -0.22 | -0.25 | -0.19 | 0.03 | -0.07 | 0.2 | -0.22 | -0.28 | 0.27 | -0.26 | 0 | -0.06 | -0.145 | -0.08 | 0 | 0.42729 | 0.677687 | 1 | 0.999971 | 1 | 1 |
| Ankmy1 | E0CYY3 | 2.7 | -0.04 | -1.06 | 0.73 | 0.22 | 2.02 | -0.9 | 0.73 | -0.01 | 1.64 | -0.25 | 0.25 | -3.75 | -1 | 0.02 | 0.09 | -0.065 | -0.175 | -1.7425 | 0.942019 | 0.830017 | 0.130692 | 0.999971 | 1 | 0.784948 |
| Ankmy2 | Q3TPE9 | 0.34 | 0.23 | -0.34 | -0.97 | 0.31 | 0.42 | -0.11 | -0.28 | 0.72 | 0.39 | -0.02 | -0.07 | 0.79 | 0.51 | -0.09 | -0.24 | 0.27 | 0.44 | 0.4275 | 0.436327 | 0.228131 | 0.273449 | 0.999971 | 1 | 0.843794 |
| Ankrd13a | Q80UP5 | -0.22 | -0.5 | 0.77 | 0.39 | -0.51 | -0.15 | 0.07 | 0.94 | -0.42 | 0.08 | 0.02 | 0.99 | -0.18 | 0.21 | 0.9 | 1.01 | -0.0225 | 0.0575 | 0.375 | 0.955397 | 0.884664 | 0.348572 | 0.999971 | 1 | 0.865162 |
| Ankrd13c | Q3UX43 | -1.78 | -0.23 | -0.41 | 0.04 | -0.38 | -0.22 | -0.31 | -0.13 | -0.34 | -0.94 | 0.42 | -0.28 | 0.13 | -0.2 | 0.09 | 0.06 | 0.335 | 0.31 | 0.615 | 0.403405 | 0.508143 | 0.146738 | 0.999971 | 1 | 0.793221 |
| Ankrd13d | Q6PD24 | 0.24 | -0.72 | 0.13 | 0.02 | -0.53 | -0.29 | 0.55 | 0.11 | -0.64 | -0.34 | 0.18 | 0.21 | -0.64 | -0.36 | 0.1 | 0.09 | 0.0425 | -0.065 | -0.12 | 0.895175 | 0.831813 | 0.68446 | 0.999971 | 1 | 0.944176 |
| Ankrd17 | A0A0G2JD | 1.57 | 1.81 | 1.68 | 1.9 | 1.57 | 1.62 | 1.58 | 2.02 | 1.42 | 1.5 | 1.87 | 2.07 | 1.47 | 1.82 | 1.84 | 2.09 | -0.0425 | -0.025 | 0.065 | 0.826539 | 0.906937 | 0.749835 | 0.999971 | 1 | 0.961064 |
| Ankrd27 | Q3UMR0-3 | -0.58 | -0.16 | -1.08 | -0.39 | -0.34 | 0.07 | -0.61 | -0.37 | -0.53 | 0.02 | -0.31 | -0.73 | -0.36 | -0.14 | -0.78 | -0.07 | 0.24 | 0.165 | 0.215 | 0.373654 | 0.546464 | 0.437828 | 0.999971 | 1 | 0.882419 |
| Ankrd28 | Q505D1 | -0.26 | -0.14 | 0.03 | 0.05 | -0.41 | -0.22 | 0.22 | -0.16 | -0.12 | 0.15 | 0.03 | 0.05 | -0.25 | -0.33 | -0.12 | -0.26 | -0.0625 | 0.1075 | -0.16 | 0.761217 | 0.538851 | 0.371007 | 0.999971 | 1 | 0.869526 |
| Ankrd29 | D3YTU5 | -0.62 | -0.99 | -0.56 | -0.65 | -1.11 | -1.44 | -1.46 | -0.82 | -1.76 | -1.21 | -1.06 | -1.23 | -1.54 | -1.21 | -0.6 | -0.85 | -0.5025 | -0.61 | -0.345 | 0.047167 | 0.021052 | 0.198035 | 0.999971 | 1 | 0.813335 |
| Ankrd33b | Q3U0L2 | 0.5 | -0.25 | -0.3 | -0.15 | -0.82 | -0.05 | -0.53 | -0.55 | -1.03 | 0.24 | -0.07 | -0.69 | -0.62 | -0.77 | -0.94 | -0.39 | -0.4375 | -0.3375 | -0.63 | 0.127157 | 0.334803 | 0.030214 | 0.999971 | 1 | 0.678474 |
| Ankrd34a | B2RW11 | 1.91 | 0.16 | 0.45 | 0 | -0.29 | -0.02 | 0.55 | -0.21 | -0.04 | 1.53 | 0.68 | 0.72 | -0.24 | -0.29 | 0.16 | 0.19 | -0.6225 | 0.0925 | -0.675 | 0.187844 | 0.854535 | 0.143563 | 0.999971 | 1 | 0.788654 |
| Ankrd34b | Q3UUF8 | 0.25 | 0.11 | 0.89 | 0.2 | 0.34 | 0.2 | 0.24 | 1.14 | 0.05 | 0.54 | 1.35 | 1.5 | 0.51 | 0.18 | 1.43 | 0.83 | 0.1175 | 0.4975 | 0.375 | 0.691374 | 0.20717 | 0.263785 | 0.999971 | 1 | 0.842533 |
| Ankrd40 | Q5SUE8-2 | 0.91 | 0.15 | 1.07 | 0.73 | 0.1 | -0.07 | 0.24 | 0.92 | -0.08 | 0.59 | 0.82 | 1.07 | 0.22 | 0.63 | 0.7 | 1.34 | -0.4175 | -0.115 | 0.0075 | 0.191743 | 0.720153 | 0.980794 | 0.999971 | 1 | 0.99925 |
| Ankrd44 | E9QNQ2 | 0.25 | 0.78 | -0.23 | -0.44 | 0.43 | 0.4 | -0.04 | 0.1 | 0.72 | 0.31 | 0.37 | 0.3 | 1.03 | 0.86 | -0.33 | -0.19 | 0.1325 | 0.335 | 0.2525 | 0.662935 | 0.278155 | 0.553092 | 0.999971 | 1 | 0.913595 |
| Ankrd45 | Q810N6 | 0.48 | 0.95 | 0.55 | 0.73 | 0.09 | 0.92 | 0.28 | 0.94 | 1.08 | 1.22 | 0.95 | 0.97 | 0.75 | 1.13 | 1.13 | 1.45 | -0.12 | 0.3775 | 0.4375 | 0.651444 | 0.069627 | 0.074857 | 0.999971 | 1 | 0.746958 |
| Ankrd46 | Q8BTZ5 | 1.02 | 0.45 | 0.47 | 1.72 | 0.07 | 0.47 | 0.38 | 2.24 | -0.11 | 1.58 | 0.5 | 0.3 | 0.13 | 1.02 | 1.67 | 2.21 | -0.125 | -0.3475 | 0.3425 | 0.813586 | 0.440833 | 0.499314 | 0.999971 | 1 | 0.903717 |
| Ankrd49 | Q8VE42 | -0.49 | 0.83 | 0.73 | -0.46 | 1.17 | 0.44 | 0.64 | 0.58 | 0.81 | -0.09 | 0.61 | 0.42 | 1.38 | -0.19 | 0.3 | -0.12 | 0.555 | 0.285 | 0.19 | 0.171297 | 0.477408 | 0.692608 | 0.999971 | 1 | 0.947174 |
| Ankrd50 | F7BE84 | 0.46 | -0.24 | 0.1 | 0.13 | -0.5 | -0.67 | 0.15 | 0.38 | -0.06 | 0.2 | 0.67 | 0.71 | -0.4 | -0.63 | 0.48 | 0.38 | -0.2725 | 0.2675 | -0.155 | 0.37339 | 0.314824 | 0.626445 | 0.999971 | 1 | 0.930479 |
| Ankrd52 | Q8BTI7 | -1.37 | -0.36 | -0.25 | -0.85 | -0.2 | -0.3 | 1.12 | 4.43 | -0.25 | -1.25 | 1.02 | 0.44 | 0.4 | 0 | -0.59 | 0.68 | 1.97 | 0.6975 | 0.83 | 0.078228 | 0.197246 | 0.04705 | 0.999971 | 1 | 0.710102 |
| Ankrd54 | Q91WK7 | -0.29 | -0.66 | -0.06 | 0.42 | -0.49 | -0.32 | -0.01 | 0.33 | -0.72 | 0.51 | -0.54 | 0.15 | -1.1 | -0.38 | -0.4 | 0.01 | 0.025 | -0.0025 | -0.32 | 0.933146 | 0.994436 | 0.338229 | 0.999971 | 1 | 0.859348 |
| Ankrd63 | A2ARS0 | 0.19 | -0.51 | -1.22 | -0.27 | -0.59 | -1.71 | -1.39 | -0.27 | -0.66 | -0.46 | -1.63 | -0.57 | -0.34 | -0.52 | -0.42 | -1.19 | -0.5375 | -0.3775 | -0.165 | 0.225331 | 0.340988 | 0.637345 | 0.999971 | 1 | 0.93301 |
| Anks1 | A0A3B2WA | -1 | -0.47 | 0.31 | 0.2 | -0.22 | -0.05 | -0.34 | 0.28 | -0.19 | -0.51 | 0.3 | 0.08 | -0.06 | -0.27 | 0.26 | 0.33 | 0.1575 | 0.16 | 0.305 | 0.638331 | 0.647059 | 0.375061 | 0.999971 | 1 | 0.870465 |
| Anks1b | A0A0R4J2A | 0.55 | -0.39 | 0.29 | 0.29 | -0.16 | -0.43 | 0.8 | -0.06 | 0.02 | 0.87 | 1.72 | 2.72 | -0.47 | -0.05 | -0.09 | 0 | -0.1475 | 1.1475 | -0.3375 | 0.658614 | 0.068691 | 0.207751 | 0.999971 | 1 | 0.820739 |
| Anks1b_1 | A0A0R4J2A | -6.14 | -2.21 | -0.63 | -3.43 | -0.59 | -2.29 | 0.25 | -2.39 | 0.5 | -5.26 | -0.27 | -0.1 | 1.27 | -2.14 | -1.81 | -3.39 | 1.8475 | 1.82 | 1.585 | 0.145172 | 0.268611 | 0.261832 | 0.999971 | 1 | 0.842262 |
| Anks1b_2 | S4R286 | -0.37 | -0.82 | -0.43 | -0.8 | -0.7 | -0.58 | -0.61 | -0.5 | -0.79 | -0.83 | -0.41 | -0.41 | -0.07 | -0.79 | -0.53 | -0.59 | 0.0075 | -0.005 | 0.11 | 0.968772 | 0.981132 | 0.636716 | 0.999971 | 1 | 0.93301 |
| Anks1b_3 | S4R1Q0 | -1.97 | -2.27 | -1.76 | -1.83 | -2.33 | -2.04 | -1.98 | -2.1 | -2.03 | -1.73 | -1.36 | -1.24 | -1.65 | -2.26 | -1.37 | -1.1 | -0.155 | 0.3675 | 0.3625 | 0.44003 | 0.154599 | 0.227372 | 0.999971 | 1 | 0.828563 |
| Anks1b_4 | S4R2Q2 | -4.94 | -9.14 | -4.99 | -5.26 | -8.56 | -5.58 | -5.44 | -6.44 | -7.76 | -5.95 | -5.06 | -7.58 | -10.48 | -5.1 | -6.42 | -6.01 | -0.4225 | -0.505 | -0.92 | 0.705022 | 0.644365 | 0.514719 | 0.999971 | 1 | 0.905815 |
| Anks3 | A0A2R8VH | 1.79 | 0.66 | 1.32 | 2.21 | -0.06 | 0.77 | 0.99 | 0.5 | -0.07 | 0.73 | 1.56 | 1.72 | -0.48 | -0.07 | 1.76 | 1.24 | -0.945 | -0.51 | -0.8825 | 0.034984 | 0.317918 | 0.150202 | 0.999971 | 1 | 0.793248 |
| Ankzf1 | J3QM81 | 0.72 | 0.53 | 1.6 | 3.4 | 0.47 | -0.17 | 0.82 | 2.01 | 0.14 | 0.84 | 2.97 | 2.14 | 0.16 | 0.88 | 1.77 | 3.03 | -0.78 | -0.04 | -0.1025 | 0.295992 | 0.961315 | 0.899311 | 0.999971 | 1 | 0.989654 |
| Anln | Q8K298 | -0.21 | -0.33 | -0.8 | -1.13 | -0.48 | -0.35 | -1.61 | -1.91 | -0.62 | -0.32 | -1.38 | -1.32 | -0.79 | -0.37 | -0.9 | -1.78 | -0.47 | -0.2925 | -0.3425 | 0.284919 | 0.393561 | 0.351339 | 0.999971 | 1 | 0.867656 |
| Ano10 | Q8BH79-2 | -1.17 | -0.14 | -0.63 | -0.75 | -0.15 | 0.31 | -0.82 | -0.74 | 0.23 | -0.22 | -0.62 | -0.63 | 0.15 | -0.07 | -0.87 | -0.45 | 0.3225 | 0.3625 | 0.3625 | 0.351854 | 0.248733 | 0.265964 | 0.999971 | 1 | 0.842677 |
| Ano3 | A2AHL1 | -0.82 | -1.1 | 0.89 | 0.07 | -1.02 | -1.23 | 0.17 | 0.3 | -1.29 | -1.14 | 0.49 | 0.34 | -1.56 | -0.8 | 0.25 | 0.17 | -0.205 | -0.16 | -0.245 | 0.711044 | 0.790035 | 0.670154 | 0.999971 | 1 | 0.938809 |
| Ano6 | A0A2I3BPX | -1.4 | -1.34 | -1.3 | -1.1 | -0.69 | -1.53 | -1.19 | -1.31 | -0.78 | -1.78 | -1.04 | -0.57 | -0.55 | -1.23 | -1.11 | -1.42 | 0.105 | 0.2425 | 0.2075 | 0.647495 | 0.404088 | 0.387336 | 0.999971 | 1 | 0.871816 |
| Ano8 | A0A1L1SR9 | -0.03 | -0.08 | -0.12 | -0.27 | -0.61 | -0.1 | -0.36 | -0.28 | -0.41 | 0.1 | -0.4 | -0.44 | -0.17 | -0.02 | -0.14 | -0.13 | -0.2125 | -0.1625 | 0.01 | 0.275338 | 0.41665 | 0.952037 | 0.999971 | 1 | 0.998056 |
| Anp32a | D3Z7M9 | 2.06 | 3.11 | 2.02 | 2.56 | 3.4 | 2.72 | 2.33 | 2.2 | 3.83 | 2.51 | 1.97 | 2.01 | 3.91 | 2.42 | 1.91 | 1.99 | 0.225 | 0.1425 | 0.12 | 0.538829 | 0.763393 | 0.807944 | 0.999971 | 1 | 0.973362 |
| Anp32b | Q9EST5 | 0.46 | 1.29 | 2.71 | 2.05 | 2.03 | 0.62 | 2.27 | 2.15 | 1.12 | 0.51 | 2.47 | 3.05 | 1.42 | 1.14 | 2.9 | 2.15 | 0.14 | 0.16 | 0.275 | 0.805438 | 0.817683 | 0.633909 | 0.999971 | 1 | 0.93301 |
| Anp32e | P97822-2 | 1.34 | 2.93 | 2.97 | 2.62 | 2.73 | 2.64 | 2.47 | 2.92 | 2.52 | 2.06 | 3.11 | 2.98 | 1.72 | 2.55 | 3.23 | 2.51 | 0.225 | 0.2025 | 0.0375 | 0.558126 | 0.638491 | 0.935163 | 0.999971 | 1 | 0.995618 |
| Anpep | P97449 | 0.37 | -0.07 | -0.16 | 0.61 | 0.71 | -0.43 | -0.18 | -1 | 0.39 | 0.12 | -0.25 | 0.41 | -0.73 | 0.35 | 0.8 | 0.24 | -0.4125 | -0.02 | -0.0225 | 0.299888 | 0.938689 | 0.950363 | 0.999971 | 1 | 0.998038 |
| Antkmt | Q501J2 | -0.55 | -0.39 | -0.32 | -1.01 | -0.41 | -1.07 | -0.88 | -0.97 | -0.65 | -0.77 | -1.42 | -0.13 | -0.45 | -0.38 | -0.71 | -0.31 | -0.265 | -0.175 | 0.105 | 0.29379 | 0.577566 | 0.638115 | 0.999971 | 1 | 0.93301 |
| Antxr1 | Q9CZ52 | -0.76 | -0.67 | -1.15 | -2.68 | -0.18 | -0.78 | -0.92 | -2.69 | -0.49 | -1.35 | -1.73 | -0.26 | -0.83 | -1.09 | -1.95 | -2.86 | 0.1725 | 0.3575 | -0.3675 | 0.790322 | 0.512637 | 0.543851 | 0.999971 | 1 | 0.911398 |
| Anxa1 | P10107 | 1.92 | 1.05 | -0.29 | 1.3 | 2.1 | 0.39 | 0.4 | -0.38 | 0.89 | 0.94 | -0.25 | 1.3 | -0.2 | 0.56 | 0.39 | -0.19 | -0.3675 | -0.275 | -0.855 | 0.567521 | 0.607963 | 0.097965 | 0.999971 | 1 | 0.78268 |
| Anxa11 | P97384 | -0.91 | -0.04 | -0.07 | 0.06 | -0.07 | -0.03 | -0.22 | -0.33 | -0.54 | -0.85 | -0.4 | -0.37 | -0.05 | -0.66 | -0.5 | -0.26 | 0.0775 | -0.3 | -0.1275 | 0.765398 | 0.281717 | 0.6492 | 0.999971 | 1 | 0.933842 |
| Anxa2 | P07356 | 1.11 | 0.06 | -0.39 | 0.72 | 1.61 | 0.21 | -0.3 | -0.69 | 0.27 | 0.84 | -0.62 | 0.08 | -0.34 | 0.5 | 0.32 | -0.39 | -0.1675 | -0.2325 | -0.3525 | 0.762876 | 0.589561 | 0.376843 | 0.999971 | 1 | 0.870465 |
| Anxa3 | O35639 | -0.02 | -0.23 | 0.02 | 0.81 | 0.47 | -0.5 | -0.07 | -0.18 | -0.13 | 0.06 | 0.18 | 0.86 | -0.52 | -0.06 | 0.6 | 0.4 | -0.215 | 0.0975 | -0.04 | 0.493887 | 0.758639 | 0.905412 | 0.999971 | 1 | 0.991091 |
| Anxa4 | P97429 | -0.37 | 0.7 | -0.57 | -0.33 | 1.34 | 0.14 | -0.67 | -0.65 | 0.44 | -0.68 | -0.74 | 0.2 | -0.24 | 0.25 | -0.11 | -0.52 | 0.1825 | -0.0525 | -0.0125 | 0.721655 | 0.895161 | 0.969575 | 0.999971 | 1 | 0.998787 |
| Anxa5 | P48036 | -0.89 | -0.55 | -1.01 | -1.03 | -0.23 | -0.32 | -0.99 | -1.43 | -0.81 | -1.01 | -1.28 | -0.8 | -0.7 | -0.26 | -0.95 | -0.98 | 0.1275 | -0.105 | 0.1475 | 0.682952 | 0.61425 | 0.53646 | 0.999971 | 1 | 0.910077 |
| Anxa6 | F8WIT2 | -1.42 | -1.61 | -3.98 | -4.09 | -2.72 | -2.74 | -3.58 | -3.63 | -1.95 | -1.32 | -3.39 | -4.42 | -1.76 | -1.83 | -4 | -3.52 | -0.3925 | 0.005 | -0.0025 | 0.576916 | 0.995615 | 0.997608 | 0.999971 | 1 | 1 |
| Anxa6_1 | P14824 | -1.22 | -1.34 | -1.5 | -1.88 | -1.07 | -0.99 | -1.15 | -1.76 | -1.17 | -0.95 | -1.41 | -1.61 | -1.28 | -1.07 | -1.6 | -1.91 | 0.2425 | 0.2 | 0.02 | 0.351201 | 0.405536 | 0.938282 | 0.999971 | 1 | 0.99575 |
| Anxa7 | Q07076 | -0.43 | -0.5 | -0.45 | -1.02 | -0.47 | -0.09 | -0.33 | -0.77 | -0.46 | -0.55 | -0.48 | -0.73 | -0.74 | -0.72 | -0.46 | -1.27 | 0.185 | 0.045 | -0.1975 | 0.439764 | 0.825634 | 0.437353 | 0.999971 | 1 | 0.882419 |
| Aoc3 | O70423-2 | 0.56 | 0.18 | -1.99 | 0.37 | 5 | 1.14 | -0.13 | -1.32 | 1.02 | 3.84 | 0.75 | 2.97 | -2.35 | 0.65 | -1.68 | -1.35 | 1.3925 | 2.365 | -0.9625 | 0.30938 | 0.022113 | 0.243509 | 0.999971 | 1 | 0.837316 |
| Aox1 | G3X8P9 | -0.64 | 1.15 | 0.69 | 0.8 | 0.26 | 0.94 | 0.72 | 0.88 | 0.3 | 0.13 | 0.63 | 1.84 | 0.47 | 0.82 | 0.55 | 1.06 | 0.2 | 0.225 | 0.225 | 0.621535 | 0.661808 | 0.575197 | 0.999971 | 1 | 0.91816 |
| Ap1b1 | Q5SVG5 | -0.92 | -0.42 | -0.23 | -0.41 | -0.54 | -0.66 | 0.05 | -0.45 | -0.65 | -0.9 | -0.26 | -0.24 | -0.92 | -0.45 | -0.42 | -0.54 | 0.095 | -0.0175 | -0.0875 | 0.699668 | 0.943175 | 0.702241 | 0.999971 | 1 | 0.949019 |
| Ap1g1 | Q8CBB7 | 0 | 0.93 | -0.02 | -0.15 | 0.59 | 0.84 | -0.07 | -0.19 | 0.82 | 0.56 | 0.18 | -0.02 | 0.85 | 0.75 | -0.02 | 0.14 | 0.1025 | 0.195 | 0.24 | 0.768381 | 0.54072 | 0.474328 | 0.999971 | 1 | 0.894044 |
| Ap1m1 | P35585 | -0.49 | 0.26 | -0.23 | -0.14 | 0.26 | 0 | -0.03 | -0.07 | 0.16 | -0.22 | -0.21 | 0.11 | 0.45 | 0.02 | -0.36 | -0.03 | 0.19 | 0.11 | 0.17 | 0.393637 | 0.625713 | 0.50961 | 0.999971 | 1 | 0.905815 |

| | | | | | | | | | | | | | | | | | | | | | | | | | | |
|---|---|---|---|---|---|---|---|---|---|---|---|---|---|---|---|---|---|---|---|---|---|---|---|---|---|---|
| Ap1s1 | P61967 | -0.04 | 0.39 | 0.9 | 0.41 | -0.1 | 0.46 | 0.51 | 0.72 | 0.15 | 0.08 | 0.95 | 0.68 | -0.1 | 0.46 | 1.34 | 0.69 | -0.0175 | 0.05 | 0.1825 | 0.949521 | 0.864964 | 0.60421 | 0.999971 | 1 | 0.924454 |
| Ap1s2 | Q9DB50 | 0.38 | 0.66 | 0.07 | 0.46 | 0.48 | 0.48 | -0.37 | 0.82 | 0.81 | 0.34 | 0.02 | 0.3 | 0.57 | 1.04 | 0.47 | 0.69 | -0.04 | -0.025 | 0.3 | 0.891263 | 0.915747 | 0.195449 | 0.999971 | 1 | 0.811225 |
| Ap2a1 | P17426-2 | -0.74 | -0.29 | -0.61 | -0.76 | -0.52 | -0.38 | -0.57 | -0.73 | -0.57 | -0.61 | -0.71 | -0.97 | -0.18 | -0.46 | -0.46 | -0.65 | 0.05 | -0.115 | 0.1625 | 0.796822 | 0.563051 | 0.43155 | 0.999971 | 1 | 0.881643 |
| Ap2a2 | P17427 | -1.2 | -0.77 | -1.03 | -1.12 | -0.75 | -1.01 | -1.02 | -1.25 | -0.73 | -1.23 | -1.15 | -1.42 | -0.63 | -1.08 | -1 | -1.05 | 0.0225 | -0.1025 | 0.09 | 0.909573 | 0.637049 | 0.654305 | 0.999971 | 1 | 0.934061 |
| Ap2b1 | Q9DBG3 | -0.81 | -0.66 | -0.76 | -0.89 | -0.85 | -0.71 | -0.7 | -0.86 | -0.84 | -1.01 | -0.8 | -0.85 | -0.57 | -0.73 | -0.57 | -0.64 | 1.96E-16 | -0.095 | 0.1525 | 1 | 0.565129 | 0.37125 | 1 | 1 | 0.869526 |
| Ap2m1 | A0A338P6T | 1.39 | 0.7 | 2.77 | 1.05 | -0.62 | 2.15 | 1.44 | 1.83 | -0.44 | -0.1 | 0.73 | -0.18 | -0.91 | 1.29 | 1.67 | 2.03 | -0.2775 | -1.475 | -0.4575 | 0.691687 | 0.013645 | 0.532391 | 0.999971 | 1 | 0.910069 |
| Ap2m1_1 | P84091 | -0.18 | -0.57 | -0.7 | -0.74 | -0.71 | -0.32 | 0.28 | -0.98 | -0.74 | -0.43 | -1.16 | -1.37 | -0.63 | -0.38 | -0.84 | -0.96 | 0.115 | -0.3775 | -0.155 | 0.709528 | 0.179976 | 0.493728 | 0.999971 | 1 | 0.901531 |
| Ap2s1 | P62743 | -2.27 | -0.24 | -0.8 | -3.14 | -0.37 | -0.86 | -0.53 | -2.1 | -0.35 | -2.89 | -0.39 | -0.11 | 0.05 | -1.58 | -1.15 | -3.25 | 0.6475 | 0.6775 | 0.13 | 0.365594 | 0.430501 | 0.879221 | 0.999971 | 1 | 0.985441 |
| Ap3b1 | Q9Z1T1 | 0.86 | 0.02 | -0.06 | 0.01 | 0.28 | 0.28 | -0.3 | 0.09 | 0.2 | 0.57 | -0.2 | -0.08 | -0.49 | 0.12 | -0.15 | -0.3 | -0.12 | -0.085 | -0.4125 | 0.664026 | 0.768715 | 0.155849 | 0.999971 | 1 | 0.796008 |
| Ap3b2 | Q9JME5 | -0.9 | -1.09 | -1.03 | -0.92 | -1.18 | -0.74 | -1.02 | -0.91 | -1 | -0.95 | -0.85 | -0.92 | -1.05 | -0.75 | -0.9 | -0.53 | 0.0225 | 0.055 | 0.1775 | 0.900957 | 0.731923 | 0.36097 | 0.999971 | 1 | 0.869526 |
| Ap3d1 | O54774 | -1.21 | -0.51 | -0.97 | -1.18 | -0.33 | -0.41 | -0.77 | -0.88 | -0.2 | -1.32 | -0.63 | -0.71 | 0.31 | -0.63 | -0.71 | -1.02 | 0.37 | 0.2525 | 0.455 | 0.15188 | 0.397397 | 0.190995 | 0.999971 | 1 | 0.810547 |
| Ap3m1 | Q9JKC8 | -1.36 | -0.65 | -0.5 | -0.52 | -1.01 | -0.78 | -0.71 | -0.28 | -0.84 | -1.03 | -0.63 | -0.78 | -0.89 | -0.58 | -0.42 | -0.23 | 0.0625 | -0.0625 | 0.2275 | 0.818773 | 0.800338 | 0.405811 | 0.999971 | 1 | 0.874911 |
| Ap3m2 | Q8R2R9 | -0.87 | -0.15 | -0.4 | -0.57 | -0.28 | 0.09 | -0.15 | -0.46 | -0.31 | -0.54 | -0.3 | -0.83 | -0.09 | -0.02 | -0.51 | -0.44 | 0.2975 | 0.0025 | 0.2325 | 0.215725 | 0.991345 | 0.331521 | 0.999971 | 1 | 0.857152 |
| Ap3s1 | Q9DCR2 | -1.34 | -0.87 | -0.63 | -0.91 | -0.42 | -1.3 | -0.69 | -0.67 | -0.16 | -1.54 | -0.55 | -0.49 | 0 | -1.18 | -0.69 | -0.32 | 0.1675 | 0.2525 | 0.39 | 0.526333 | 0.453515 | 0.218483 | 0.999971 | 1 | 0.827064 |
| Ap3s2 | Q8BSZ2 | -1.21 | -0.4 | -1.11 | -1.79 | -0.42 | -0.35 | -0.83 | -1.06 | -0.41 | -1.22 | -0.78 | -0.99 | 0.52 | -0.19 | -1.12 | -1.18 | 0.4625 | 0.2775 | 0.635 | 0.185668 | 0.412512 | 0.195641 | 0.999971 | 1 | 0.811225 |
| Ap4b1 | A0A0G2JFY | -1.53 | -0.25 | -0.26 | -0.14 | -0.22 | -0.24 | -0.27 | 0.17 | -0.37 | -1.09 | -0.52 | 0.04 | -0.4 | 0.17 | -0.07 | 0.74 | 0.405 | 0.06 | 0.655 | 0.254852 | 0.877449 | 0.122146 | 0.999971 | 1 | 0.78268 |
| Ap4e1 | Q80V94 | 0.15 | 0.52 | 0.02 | -0.21 | 0.59 | 0.6 | -0.16 | -0.46 | 0.17 | 0.31 | -0.08 | -0.19 | 1.14 | 1.07 | -0.13 | -0.29 | 0.0225 | -0.0675 | 0.3275 | 0.94264 | 0.766837 | 0.415974 | 0.999971 | 1 | 0.876885 |
| Ap4m1 | Q9JKC7 | 1.95 | 0.88 | 0.19 | -0.06 | 0.4 | 0.89 | -0.73 | -0.07 | 0.57 | 1.26 | 0.28 | -0.04 | 0.71 | 0.7 | -0.12 | 0.11 | -0.6175 | -0.2225 | -0.39 | 0.256927 | 0.653466 | 0.411363 | 0.999971 | 1 | 0.875847 |
| Ap4s1 | Q9WVL1 | -1.15 | -1.61 | -1.99 | -0.74 | -0.22 | -1.82 | -2.1 | -0.67 | -0.99 | -0.88 | -1.05 | -0.63 | -0.18 | -0.66 | -1.32 | -1.09 | 0.17 | 0.485 | 0.56 | 0.728804 | 0.128662 | 0.148123 | 0.999971 | 1 | 0.793248 |
| Ap5s1 | A2ANC4 | 0.75 | 2.34 | 2.46 | 2.48 | 1.27 | 0.96 | 1.69 | 2.31 | 1.22 | 0.11 | 2.5 | 2.31 | 2.24 | 1.69 | 2.56 | 3.51 | -0.45 | -0.4725 | 0.4925 | 0.357993 | 0.464179 | 0.360478 | 0.999971 | 1 | 0.869526 |
| Ap5z1 | Q3U829 | 0.65 | 1.09 | -0.04 | -0.13 | 1.61 | 0.44 | -0.18 | -0.02 | 1.21 | 0.97 | -0.06 | 0.19 | 2.16 | 0.37 | -0.19 | 0.07 | 0.07 | 0.185 | 0.21 | 0.880404 | 0.648011 | 0.707136 | 0.999971 | 1 | 0.950182 |
| Apaf1 | G3XA09 | 0.88 | 0.29 | 1.76 | 0.12 | 1.38 | 0.68 | 1.08 | 0.88 | 0.63 | 1.1 | 1.11 | 0.32 | 0.96 | 1.22 | 1.39 | -0.04 | 0.2425 | 0.0275 | 0.12 | 0.532724 | 0.945161 | 0.794042 | 0.999971 | 1 | 0.970141 |
| Apba1 | B2RUJ5 | -0.76 | -0.99 | -0.75 | -0.25 | -0.72 | -0.84 | -0.23 | -0.77 | -0.82 | -0.57 | -0.7 | -0.72 | -0.73 | -0.93 | -0.76 | -0.51 | 0.0475 | -0.015 | -0.045 | 0.843835 | 0.94315 | 0.839449 | 0.999971 | 1 | 0.979728 |
| Apba2 | P98084 | 0.07 | -0.23 | -0.52 | -0.51 | -0.37 | -0.22 | -0.51 | -0.66 | -0.52 | -0.15 | -0.94 | -1.1 | -0.02 | -0.07 | -0.82 | -0.65 | -0.1425 | -0.38 | -0.0925 | 0.513614 | 0.187308 | 0.730041 | 0.999971 | 1 | 0.956041 |
| Apba3 | O88888 | -0.17 | -0.29 | -0.45 | -0.16 | -0.24 | -0.33 | -0.5 | -0.15 | -0.92 | 0.03 | -0.37 | -0.73 | -0.07 | 0.01 | -0.21 | -0.31 | -0.0375 | -0.23 | 0.1225 | 0.834929 | 0.364443 | 0.500932 | 0.999971 | 1 | 0.904237 |
| Apbb1 | Q9QXJ1 | 0.19 | -1.16 | -1.04 | -0.87 | -0.7 | -0.84 | -0.74 | -0.62 | -1 | -0.53 | -0.89 | -1.34 | -0.29 | -0.42 | -0.41 | -0.5 | -0.005 | -0.22 | 0.315 | 0.987357 | 0.530005 | 0.333368 | 0.999971 | 1 | 0.857921 |
| Apbb1ip | Q8R5A3 | 2.21 | 0.06 | 0.47 | -0.01 | 0.4 | 0.35 | -0.2 | 0.12 | 0.57 | 2.21 | 0.37 | 0.11 | -0.91 | -0.12 | 0.72 | 0.78 | -0.515 | 0.1325 | -0.565 | 0.316785 | 0.83691 | 0.358524 | 0.999971 | 1 | 0.86927 |
| Apbb2 | E9PWH3 | 0.55 | -0.5 | 0.25 | 0.55 | -0.25 | -0.33 | 0.84 | 1.28 | -0.22 | -0.33 | 0.27 | -0.18 | 0.17 | -0.38 | 0.9 | 0.78 | 0.1725 | -0.3275 | 0.155 | 0.697977 | 0.278608 | 0.680369 | 0.999971 | 1 | 0.942957 |
| Apc | B2RUG9 | 0.09 | -0.63 | -0.62 | -0.81 | -0.58 | -0.81 | -0.25 | -0.58 | -0.4 | -0.35 | -0.38 | -0.17 | -0.28 | -0.96 | -0.44 | -0.87 | -0.0625 | 0.1675 | -0.145 | 0.806863 | 0.486745 | 0.602203 | 0.999971 | 1 | 0.924453 |
| Apc2 | G5E832 | 1.2 | -0.54 | -0.58 | -0.48 | -0.42 | -0.1 | -0.58 | -0.28 | -0.38 | 0.94 | -0.48 | -0.69 | -0.11 | -0.06 | -0.67 | -0.52 | -0.245 | -0.0525 | -0.24 | 0.565451 | 0.92081 | 0.583725 | 0.999971 | 1 | 0.920364 |
| Apeh | A0A0R4J10 | 1.08 | 0.1 | 0.15 | 0.54 | -0.01 | 0.01 | 0.22 | 0.26 | 0.09 | 0.49 | 0.22 | 0.43 | 0.25 | 0.56 | 0.64 | 0.48 | -0.3475 | -0.16 | 0.015 | 0.204689 | 0.549698 | 0.954701 | 0.999971 | 1 | 0.998056 |
| Apex1 | P28352 | 1.75 | 2.75 | 2.72 | 2.38 | 2.39 | 2.9 | 2.4 | 2.53 | 2.6 | 2.29 | 2.3 | 2.35 | 2.31 | 2.77 | 2.96 | 2.51 | 0.155 | -0.015 | 0.2375 | 0.579866 | 0.954593 | 0.416776 | 0.999971 | 1 | 0.877251 |
| Aph1b | F8WHK7 | -0.45 | -0.27 | -0.2 | -2.05 | -0.13 | -0.11 | 0.04 | -0.18 | 0.11 | -0.37 | -0.82 | -1.11 | -3.23 | 0.06 | -0.51 | -0.29 | 0.6475 | 0.195 | -0.25 | 0.148257 | 0.686248 | 0.751141 | 0.999971 | 1 | 0.961064 |
| Api5 | O35841 | 1.21 | 1.7 | 1.82 | 2.05 | 1.85 | 1.38 | 1.47 | 1.83 | 1.56 | 1.21 | 1.51 | 1.72 | 1.31 | 1.8 | 1.61 | 1.87 | -0.0625 | -0.195 | -0.0475 | 0.798866 | 0.421773 | 0.847645 | 0.999971 | 1 | 0.980635 |
| Apip | Q9WVQ5 | 0.12 | 1.21 | 0.71 | 0.6 | 1.24 | 0.73 | 0.56 | 0.62 | 1.65 | 0.1 | 0.78 | 0.32 | 1.5 | 0.9 | 0.78 | 0.97 | 0.1275 | 0.0525 | 0.3775 | 0.656832 | 0.893885 | 0.211982 | 0.999971 | 1 | 0.823181 |
| Aplf | Q9D842-2 | 3.14 | 1.84 | 2.93 | 3.57 | 1.39 | 1.88 | 3.23 | 2.89 | 3.17 | 3.09 | 2.42 | 2.42 | -1.87 | 0.77 | 3.2 | 1.3 | -0.5225 | -0.095 | -2.02 | 0.330897 | 0.814151 | 0.06781 | 0.999971 | 1 | 0.733937 |
| Aplp1 | Q03157 | 2.93 | 1.1 | 1.04 | 1.37 | 1.06 | 1.31 | 3.1 | 0.83 | 1.16 | 1.89 | 1.01 | 0.85 | 1.05 | 0.76 | 0.98 | 0.61 | -0.035 | -0.3825 | -0.76 | 0.955062 | 0.425283 | 0.106421 | 0.999971 | 1 | 0.78268 |
| Aplp2 | Q60709 | -0.19 | -0.01 | -0.17 | -0.36 | 0.08 | -0.03 | -0.16 | -0.32 | 0.27 | -0.08 | 0.03 | 0.05 | -0.02 | -0.08 | -0.13 | -0.14 | 0.075 | 0.25 | 0.09 | 0.686333 | 0.183035 | 0.602021 | 0.999971 | 1 | 0.924453 |
| Aplp2_1 | Q06335 | -2.02 | -1.44 | -0.25 | 1.05 | -0.8 | -1.28 | -0.36 | 1.41 | -1.97 | -1.55 | 0.31 | 1.47 | -2.35 | -0.84 | 0.27 | 1.41 | 0.4075 | 0.23 | 0.2875 | 0.615882 | 0.808643 | 0.760495 | 0.999971 | 1 | 0.961116 |
| Apmap | Q9D7N9 | -0.49 | -0.55 | -0.69 | -0.54 | -0.59 | -0.42 | -0.67 | -0.46 | -0.64 | -0.83 | -0.55 | -0.44 | -0.49 | -0.43 | -0.48 | -0.44 | 0.0325 | -0.0475 | 0.1075 | 0.847468 | 0.784267 | 0.513414 | 0.999971 | 1 | 0.905815 |
| Apoa1 | Q00623 | 2.07 | 4.13 | 1.44 | 4.79 | 5.06 | 1.45 | 3.51 | -0.29 | 2.62 | 1.46 | 3.26 | 3.15 | 3.93 | 2.41 | 2.54 | 2.59 | -0.675 | -0.485 | -0.24 | 0.595227 | 0.556292 | 0.761959 | 0.999971 | 1 | 0.961309 |
| Apoa2 | P09813 | 4.48 | 2.64 | 0.48 | 3.24 | 3.99 | -0.02 | 1.41 | -2.31 | 1.4 | 2.09 | 1.38 | 1.54 | -0.68 | -0.11 | 1.09 | 1.06 | -1.9425 | -1.1075 | -2.37 | 0.184928 | 0.176739 | 0.019937 | 0.999971 | 1 | 0.635388 |
| Apoa4 | P06728 | 2.83 | 3.86 | 1.55 | 4.98 | 4.55 | 1.59 | 3.66 | 2.37 | 1.1 | 0.33 | 1.62 | 1.06 | 0.63 | 1.53 | 1.67 | 1.73 | -0.2625 | -2.2775 | -1.915 | 0.766267 | 0.010326 | 0.022096 | 0.999971 | 1 | 0.649631 |
| Apob | E9Q1Y3 | 2.09 | 2.41 | 1.4 | 1.52 | 2.62 | -1.09 | 0.81 | 1.51 | 0.29 | 1.83 | 1.58 | 1.96 | 2.06 | -0.54 | 1.85 | 2.29 | -0.8925 | -0.44 | -0.44 | 0.244382 | 0.31955 | 0.495765 | 0.999971 | 1 | 0.901531 |
| Apoc1 | P34928 | -0.74 | 2.52 | 1.11 | 3.34 | 3.48 | -0.48 | 2.9 | -1.28 | 1.47 | -0.1 | 3.6 | 2.58 | 2.31 | 0.29 | 2.2 | 1.22 | -0.4025 | 0.33 | -0.0525 | 0.761228 | 0.758728 | 0.953759 | 0.999971 | 1 | 0.998056 |
| Apoc3 | E9QP56 | -0.6 | 1.72 | 1.35 | 4.09 | 1.9 | -0.28 | 2.46 | -0.89 | 0.94 | 0.12 | 2.79 | 2.24 | 1.84 | -0.28 | 1.91 | 2.29 | -0.8425 | -0.1175 | -0.2 | 0.460342 | 0.908459 | 0.84227 | 0.999971 | 1 | 0.979814 |
| Apod | P51910 | 0.89 | 0.66 | 0.64 | 1.56 | 0.62 | 1.08 | 0.3 | 1.87 | 0.62 | 1.04 | 0.92 | 0.77 | 0.22 | 1.2 | 1.05 | 1.7 | 0.03 | -0.1 | 0.105 | 0.938213 | 0.697864 | 0.774475 | 0.999971 | 1 | 0.963884 |
| Apoe | P08226 | 0.3 | 1.32 | 1.02 | 0.67 | 1.3 | 1.19 | 1.14 | 0.57 | 1.8 | 0.78 | 1.52 | 1.47 | 1.06 | 1.03 | 0.8 | 0.8 | 0.2225 | 0.565 | 0.095 | 0.446436 | 0.098636 | 0.712719 | 0.999971 | 1 | 0.951793 |
| Apoh | Q01339 | 0.72 | 2.24 | 0.57 | 3.31 | 2.89 | 1.5 | 2.03 | 0.88 | 1.63 | 0.91 | 2.04 | 1.99 | 1.8 | 1.6 | 1.15 | 2.33 | 0.115 | -0.0675 | 0.01 | 0.870288 | 0.916328 | 0.987425 | 0.999971 | 1 | 0.99925 |
| Apoo | Q9DCZ4-2 | -1.98 | -2.83 | -0.47 | -0.61 | -2.73 | -2.57 | -0.92 | -0.64 | -2.46 | -2.52 | -0.55 | -0.33 | -2.8 | -2.57 | -0.46 | -0.48 | -0.2425 | 0.0075 | -0.105 | 0.733049 | 0.991955 | 0.891642 | 0.999971 | 1 | 0.988225 |
| Apool | Q78IK4 | -0.49 | -0.71 | -0.1 | -0.44 | -0.35 | -0.45 | 0.66 | -0.15 | -0.47 | -0.6 | -0.48 | -0.18 | -0.68 | -0.76 | -0.43 | -0.72 | 0.3625 | 0.0025 | -0.2125 | 0.235999 | 0.990245 | 0.311277 | 0.999971 | 1 | 0.848125 |
| App | P12023-2 | -1.59 | -2.41 | 0.91 | 0.75 | -1.66 | -2.61 | 1.25 | 0.64 | -2.58 | -0.82 | 1.71 | 0.57 | -2.77 | -2.23 | 2.55 | 2.24 | -0.01 | 0.305 | 0.5325 | 0.992769 | 0.785177 | 0.71647 | 0.999971 | 1 | 0.952576 |
| App_1 | A0A2I3BQ2 | 0.56 | 0.8 | 0.53 | 0.44 | 0.58 | 0.68 | 0.62 | 0.41 | 0.73 | 0.55 | 0.6 | 0.63 | 0.67 | 0.81 | 0.59 | 0.49 | -0.01 | 0.045 | 0.0575 | 0.955155 | 0.791856 | 0.751667 | 0.999971 | 1 | 0.961116 |
| Appbp2 | Q9DAX9 | -0.93 | 0.58 | 2.23 | 1.8 | 0.44 | 0.46 | 2.48 | 1.88 | -0.13 | -0.5 | 1.48 | -1.41 | 1.12 | 0.14 | 1.3 | 0.57 | 0.395 | -1.06 | -0.1375 | 0.618214 | 0.228913 | 0.841128 | 0.999971 | 1 | 0.979814 |
| Appl1 | Q8K3H0 | 0.51 | 0.42 | 0.27 | 0.2 | 0.4 | 0.36 | 0.46 | 0.43 | 0.52 | 0.51 | 0.44 | 0.35 | 0.39 | 0.28 | 0.19 | 0.35 | 0.0625 | 0.105 | -0.0475 | 0.71314 | 0.537395 | 0.784531 | 0.999971 | 1 | 0.966376 |
| Appl2 | Q8K3G9 | -0.79 | -0.46 | -0.88 | -0.96 | -0.43 | -0.59 | -0.89 | -0.93 | -0.47 | -0.81 | -0.73 | -0.77 | -0.1 | -0.6 | -0.7 | -0.55 | 0.0625 | 0.0775 | 0.285 | 0.768568 | 0.689333 | 0.213986 | 0.999971 | 1 | 0.825334 |
| Aprt | P08030 | 0.49 | 0.5 | 0.62 | 0.64 | 0.28 | 0.43 | 0.51 | 0.58 | 0.3 | 0.44 | 0.38 | 0.15 | 0.21 | 0.91 | 0.24 | 0.63 | -0.1125 | -0.245 | -0.065 | 0.513648 | 0.165323 | 0.766219 | 0.999971 | 1 | 0.962472 |
| Aptx | Q7TQC5-4 | -0.34 | 3.06 | 2.36 | 1.3 | 2.46 | 2.3 | 2.03 | 1.84 | 2.22 | -0.14 | 1.8 | 1.72 | 2.64 | 2.29 | 1.87 | 2.03 | 0.5625 | -0.195 | 0.6125 | 0.418065 | 0.812165 | 0.384068 | 0.999971 | 1 | 0.871816 |
| Aqp1 | Q02013 | 0.93 | 2.12 | 1.96 | 1.73 | 0.5 | 1.54 | 1.06 | 1.81 | 0.5 | -0.12 | -1.58 | 0.29 | -0.35 | 1.73 | 0.43 | 1.2 | -0.4575 | -1.9125 | -0.9325 | 0.245375 | 0.004206 | 0.083848 | 0.999971 | 0.969702 | 0.755254 |
| Aqp11 | A0A0X1KG | 1.88 | 1.68 | 3.23 | 2.36 | 1 | 1.98 | 1.87 | 3.25 | 1.05 | 1.03 | 2.56 | 2.11 | 1.16 | 1.95 | 2.38 | 2.65 | -0.2625 | -0.6 | -0.2525 | 0.624175 | 0.235555 | 0.575293 | 0.999971 | 1 | 0.91816 |
| Aqp4 | A0A5F8MC | -2.42 | -2.78 | -0.59 | -1.16 | -1.24 | -1.53 | -1.13 | -2.21 | -0.96 | -1.08 | -0.94 | -1.79 | 1.63 | -1.16 | -0.47 | -0.87 | 0.21 | 0.545 | 1.52 | 0.691074 | 0.306291 | 0.065194 | 0.999971 | 1 | 0.727266 |
| Aqp4_1 | A0A0R4J0Z | -0.95 | -0.7 | -0.42 | -0.75 | -0.99 | -0.49 | -0.79 | -1.67 | -0.52 | -0.07 | -0.59 | -0.58 | -0.36 | -0.18 | -0.08 | -0.25 | -0.28 | 0.265 | 0.4875 | 0.341597 | 0.230855 | 0.029511 | 0.999971 | 1 | 0.678474 |
| Aqr | Q8CFQ3 | 0.93 | 1.08 | 0.34 | 0.45 | 1.16 | 0.55 | 0.05 | 0.24 | 0.87 | 0.93 | 0.25 | 0.52 | 0.69 | 1.06 | -0.2 | 0.55 | -0.2 | -0.0575 | -0.175 | 0.521402 | 0.825968 | 0.590292 | 0.999971 | 1 | 0.922217 |
| Araf | P04627 | 0.8 | 0.41 | 0.51 | 1.23 | 0.28 | 0.86 | 2.05 | 1.3 | 0.55 | 1.03 | 0.69 | 0.04 | 0.95 | 1.05 | 1.03 | 1.22 | 0.385 | -0.16 | 0.325 | 0.346476 | 0.581058 | 0.183402 | 0.999971 | 1 | 0.809025 |
| Araf_1 | Q8CAD1 | 0.5 | 0.36 | 1.15 | 0.6 | 0.1 | 1.52 | 0.45 | 2.01 | 0.45 | 1.69 | 1.12 | -0.58 | 1.26 | 1.7 | -0.37 | 0.76 | 0.3675 | 0.0175 | 0.185 | 0.423406 | 0.971116 | 0.682296 | 0.999971 | 1 | 0.943204 |
| Arap1 | Q4LDD4-2 | 0.71 | 0.76 | -0.02 | -0.06 | 0.43 | 0.5 | -0.5 | -0.28 | 0.52 | 0.49 | -0.19 | -0.03 | 0.73 | 0.64 | -0.06 | -0.24 | -0.31 | -0.15 | -0.08 | 0.366331 | 0.615466 | 0.809416 | 0.999971 | 1 | 0.973762 |
| Arap2 | E9QP44 | -0.35 | -0.49 | -0.3 | -0.44 | -0.4 | -0.24 | 0.03 | -0.35 | -0.4 | -0.07 | 0.05 | 0.09 | 0.04 | -0.24 | -0.09 | 0.25 | 0.155 | 0.3125 | 0.385 | 0.403954 | 0.119166 | 0.063908 | 0.999971 | 1 | 0.725348 |
| Arc | Q9WV31 | -0.85 | -1.04 | -0.53 | -0.25 | -0.76 | -0.43 | -0.31 | -0.38 | -1.63 | -1.12 | -1.19 | -2.02 | -1.6 | -0.77 | -0.9 | -0.72 | 0.1975 | -0.8225 | -0.33 | 0.411977 | 0.017259 | 0.263372 | 0.999971 | 1 | 0.842533 |
| Arcn1 | Q5XJY5 | 1.19 | 1.36 | 1.37 | 1.42 | 1.58 | 1.26 | 1.38 | 1.51 | 1.56 | 1.11 | 1.32 | 1.54 | 1.87 | 1.22 | 1.4 | 1.32 | 0.0975 | 0.0475 | 0.1175 | 0.579029 | 0.796855 | 0.573173 | 0.999971 | 1 | 0.91816 |
| Arel1 | Q8CHG5 | -0.99 | -0.83 | -0.73 | -0.39 | -0.43 | -0.89 | -0.97 | -0.17 | -0.7 | -1.03 | -0.34 | -0.74 | -0.29 | -0.3 | 0.11 | -0.38 | 0.12 | 0.0325 | 0.52 | 0.639239 | 0.886009 | 0.036816 | 0.999971 | 1 | 0.700096 |
| Arf1 | P84078 | -0.29 | 0.35 | 0.63 | 0.67 | 0.23 | 0.15 | -0.11 | 0.63 | 0.05 | -0.1 | 0.58 | 0.79 | -0.13 | 0.24 | 0.79 | 1.11 | -0.115 | -0.01 | 0.1625 | 0.686394 | 0.974313 | 0.644036 | 0.999971 | 1 | 0.93301 |
| Arf2 | Q8BSL7 | 0.38 | 2.98 | 1.37 | 3 | 0.74 | 2.89 | 1.14 | 2.49 | 1.61 | 0.96 | 0.41 | 2.65 | 2.91 | 3.33 | 2.83 | 3 | -0.1175 | -0.525 | 1.085 | 0.87451 | 0.478586 | 0.096513 | 0.999971 | 1 | 0.78268 |
| Arf4 | P61750 | -0.04 | 0.23 | 0.87 | 0.79 | 0.09 | 0.29 | 0.02 | 0.62 | 0.25 | 0.34 | 1 | 1.19 | 0.32 | 0.25 | 0.78 | 0.45 | -0.2075 | 0.2325 | -0.0125 | 0.45801 | 0.478056 | 0.962993 | 0.999971 | 1 | 0.998721 |
| Arf5 | P84084 | -0.99 | -0.81 | -0.47 | -0.27 | -0.64 | -0.57 | -0.43 | -0.03 | -0.54 | -0.99 | 0.08 | -0.25 | -0.43 | -0.37 | -0.05 | 0.65 | 0.2175 | 0.21 | 0.585 | 0.382447 | 0.475748 | 0.081472 | 0.999971 | 1 | 0.755254 |
| Arf6 | P62331 | 0.79 | -0.11 | 0.43 | 0.35 | -0.1 | 0.15 | -0.18 | 0.19 | -0.44 | 0.26 | 0.21 | 0.02 | -0.75 | -0.16 | 0.4 | 0.53 | -0.35 | -0.3525 | -0.36 | 0.16859 | 0.205016 | 0.309497 | 0.999971 | 1 | 0.847402 |
| Arfgap1 | V9GXM1 | -0.95 | -0.89 | -1.34 | -1.38 | -1 | -0.83 | -1.01 | -1.19 | -0.99 | -0.88 | -1.17 | -1.4 | -0.2 | -0.54 | -1.19 | -1.25 | 0.1325 | 0.03 | 0.345 | 0.519475 | 0.888841 | 0.262674 | 0.999971 | 1 | 0.842381 |
| Arfgap2 | Q99K28 | 0.09 | 1.39 | 0.51 | -0.28 | 1.21 | 1.31 | 1.11 | 0.05 | 1.42 | 0.48 | 0.55 | 0.78 | 1.36 | 0.92 | 0.03 | -0.11 | 0.4925 | 0.38 | 0.1225 | 0.277269 | 0.355972 | 0.795121 | 0.999971 | 1 | 0.970361 |
| Arfgap3 | A0A0R4J0T | 0.3 | 0.11 | 0.1 | -0.15 | 0.17 | 0.31 | 0.58 | -0.02 | 0.66 | 0.21 | 0.22 | 0.06 | 0.47 | -0.19 | -0.08 | -0.2 | 0.17 | 0.1975 | -0.09 | 0.423396 | 0.353751 | 0.691027 | 0.999971 | 1 | 0.946786 |
| Arfgef1 | G3X9K3 | -0.24 | 0.09 | 0.09 | 0.18 | -0.03 | 0.14 | 0.16 | 0.19 | 0.27 | 0.1 | 0.24 | 0.1 | 0.08 | 0.26 | 0.45 | 0.25 | 0.085 | 0.1475 | 0.23 | 0.641853 | 0.415737 | 0.24408 | 0.999971 | 1 | 0.837316 |
| Arfgef2 | A2A5R2 | 1.1 | 0 | 0.42 | 0.21 | -0.08 | 0.17 | 0.58 | 0.1 | 0.07 | 0.46 | 0.4 | 0.4 | -0.11 | -0.21 | 0.15 | 0.04 | -0.24 | -0.1 | -0.465 | 0.41503 | 0.713617 | 0.113589 | 0.999971 | 1 | 0.78268 |
| Arfgef3 | Q3UGY8 | -0.06 | 0.47 | 0.28 | 0.08 | 0.3 | 0.28 | 0.29 | 0.16 | 0.45 | 0.05 | 0.28 | 0.4 | 0.6 | 0.35 | 0.13 | 0.21 | 0.065 | 0.1025 | 0.13 | 0.731558 | 0.610656 | 0.53689 | 0.999971 | 1 | 0.910077 |
| Arfip1 | G5E8V9-2 | -0.19 | -0.39 | -0.93 | -0.41 | -0.58 | -0.09 | -0.83 | -0.66 | -0.66 | -0.2 | -0.97 | -1.03 | -0.81 | 0.04 | -0.97 | -0.55 | -0.06 | -0.235 | -0.0925 | 0.81146 | 0.388319 | 0.747185 | 0.999971 | 1 | 0.961064 |
| Arfip2 | A0A1B0GS | 1.66 | 0.3 | 0.41 | 0.51 | -0.03 | 0.79 | 0.09 | 0.66 | 0.2 | 1.22 | 0.35 | 0.16 | 0.31 | 0.39 | 0.23 | 0.91 | -0.3425 | -0.2375 | -0.26 | 0.361634 | 0.544958 | 0.461156 | 0.999971 | 1 | 0.89007 |
| Arfrp1 | Q8BXL7 | 0.09 | 1.01 | 0.81 | 0.96 | -0.03 | 0.78 | 0.58 | 0.76 | 0.74 | 0.2 | 0.53 | 0.91 | 0.55 | 0.49 | 1.14 | 1 | -0.195 | -0.1225 | 0.0775 | 0.514487 | 0.66062 | 0.784504 | 0.999971 | 1 | 0.966376 |
| Arg2 | O08691 | -0.9 | -1.01 | -1.08 | -0.96 | -0.68 | -2.1 | -1.82 | -0.55 | -0.74 | -0.51 | -0.99 | -0.43 | -1.51 | -0.78 | -1.17 | -1.3 | -0.3 | 0.32 | -0.2025 | 0.439462 | 0.122565 | 0.348066 | 0.999971 | 1 | 0.86438 |
| Arglu1 | Q3UL36 | 2.7 | 0.78 | 0.91 | 0.94 | 0.66 | 0.99 | 1.55 | 0.75 | 0.95 | 2.04 | 1.26 | 0.81 | 0.83 | 0.8 | 0.67 | 0.93 | -0.345 | -0.0675 | -0.525 | 0.46648 | 0.891994 | 0.247464 | 0.999971 | 1 | 0.839012 |
| Arhgap1 | A2AH25 | -0.84 | -1.34 | -0.87 | -1.06 | -0.52 | -1 | -0.65 | -1.13 | -0.75 | -0.81 | -0.85 | -0.83 | -1.01 | -1.3 | -1.11 | -0.95 | 0.2025 | 0.2175 | -0.065 | 0.379228 | 0.257112 | 0.744292 | 0.999971 | 1 | 0.961064 |
| Arhgap1_1 | Q5FWK3 | 0.27 | -0.01 | -1.39 | -0.9 | -0.8 | -0.5 | -1.1 | -1.24 | -1.04 | 0.48 | -1.04 | -1.12 | -0.86 | -0.35 | -0.84 | -0.5 | -0.4025 | -0.1725 | -0.13 | 0.329369 | 0.735057 | 0.739899 | 0.999971 | 1 | 0.960899 |
| Arhgap10 | Q6Y5D8-3 | 0.37 | 0.2 | 0.5 | 0.86 | 0.4 | -0.12 | 0.6 | 0.76 | 0.18 | -0.46 | 0.28 | -0.55 | -0.36 | -0.72 | -0.23 | 0.11 | -0.0725 | -0.62 | -0.7825 | 0.781026 | 0.046138 | 0.011582 | 0.999971 | 1 | 0.613095 |
| Arhgap12 | B2RUJ8 | 0.6 | 0.35 | 0.08 | -0.05 | 0.8 | 0.65 | 0.51 | -0.06 | 0.82 | -0.01 | 0.04 | 0.21 | 1.05 | -0.06 | -0.04 | -0.4 | 0.23 | 0.02 | -0.1075 | 0.388014 | 0.938831 | 0.754678 | 0.999971 | 1 | 0.961116 |
| Arhgap17 | Q3UIA2-2 | -0.61 | -1.08 | -0.11 | 0.16 | -1.2 | -0.71 | -0.2 | -0.28 | -1.42 | -0.49 | -0.4 | -0.37 | -1.9 | -1.23 | -0.16 | -0.22 | -0.1875 | -0.26 | -0.4675 | 0.596519 | 0.480749 | 0.333454 | 0.999971 | 1 | 0.857921 |
| Arhgap18 | Q8K0Q5 | -0.76 | 0.03 | -0.94 | 0.5 | 0.25 | -0.07 | -0.15 | -0.86 | 1.45 | -0.51 | -0.34 | 0.42 | -0.22 | 0 | 0.43 | 0.51 | 0.085 | 0.5475 | 0.4725 | 0.829205 | 0.308335 | 0.222991 | 0.999971 | 1 | 0.827753 |
| Arhgap20 | Q6IFT4 | 1.76 | -0.59 | -0.53 | 0.37 | -0.53 | 0.21 | -0.39 | 0.07 | -0.4 | 1 | -0.24 | -0.36 | -0.59 | -0.12 | -0.66 | 0.48 | -0.4125 | -0.2525 | -0.475 | 0.445445 | 0.670498 | 0.40552 | 0.999971 | 1 | 0.874565 |
| Arhgap21 | B7ZCJ1 | -0.24 | -0.5 | -0.83 | -0.27 | -0.49 | -0.6 | -0.66 | -0.81 | -0.47 | -0.31 | -0.67 | -0.84 | -0.12 | -0.65 | -0.77 | -0.12 | -0.18 | -0.1125 | 0.045 | 0.391949 | 0.610978 | 0.856545 | 0.999971 | 1 | 0.98255 |
| Arhgap23 | B1AQY2 | -0.71 | -0.8 | -0.99 | -0.83 | -1 | -0.68 | -0.45 | -1.04 | -0.74 | -0.57 | -0.92 | -1.12 | -0.45 | -0.76 | -1.11 | -0.87 | 0.04 | -0.005 | 0.035 | 0.845634 | 0.979194 | 0.864207 | 0.999971 | 1 | 0.983079 |
| Arhgap25 | Q8BYW1 | -0.44 | 0.39 | 0.04 | -0.25 | -0.05 | 0.28 | 0.02 | -0.01 | -0.34 | -0.5 | 0.46 | 0.09 | -0.27 | -0.4 | -0.53 | -0.12 | 0.125 | -0.0075 | -0.265 | 0.593748 | 0.979517 | 0.280235 | 0.999971 | 1 | 0.844318 |
| Arhgap26 | F6T836 | -0.92 | -0.06 | -0.3 | -0.49 | -0.18 | -0.31 | -0.2 | -0.35 | 0.19 | -0.54 | -0.06 | -0.2 | 0.15 | -0.4 | -0.46 | -0.38 | 0.1825 | 0.29 | 0.17 | 0.429228 | 0.280756 | 0.512917 | 0.999971 | 1 | 0.905815 |
| Arhgap27 | A2AB59 | 1.1 | 0.29 | 0.46 | 0.03 | 0.11 | 0.32 | -0.09 | 0.7 | 0.16 | 0.95 | -0.08 | 0.48 | -0.23 | 0.28 | 0.72 | 0.37 | -0.21 | -0.0925 | -0.185 | 0.481585 | 0.773086 | 0.551531 | 0.999971 | 1 | 0.913379 |
| Arhgap29 | A0A0G2JEY | -0.86 | -1.7 | -1.23 | -0.48 | -1.81 | -1.46 | -1.44 | -0.72 | -1.9 | -0.87 | -1.07 | -0.44 | -1.2 | -1.11 | -1.18 | -0.61 | -0.29 | -0.0025 | 0.0425 | 0.407073 | 0.994849 | 0.88866 | 0.999971 | 1 | 0.987485 |
| Arhgap31 | A6X8Z5 | 0.8 | 0.06 | -0.71 | -0.21 | 0.31 | 0.61 | -0.28 | -0.75 | 0.36 | 0.34 | -0.9 | -0.08 | 0.09 | 0.32 | -0.73 | -0.06 | -0.0125 | -0.055 | -0.08 | 0.975973 | 0.893689 | 0.831472 | 0.999971 | 1 | 0.978898 |
| Arhgap32 | S4R2G6 | 1.85 | 0.4 | 1.24 | 1.71 | 0.84 | 0.84 | 1.09 | 1.76 | 0.37 | 1.53 | 1.3 | 0.02 | 1.42 | 0.66 | 0.42 | 1.31 | -0.1675 | -0.495 | -0.3475 | 0.660239 | 0.297709 | 0.386973 | 0.999971 | 1 | 0.871816 |
| Arhgap32_ | Q811P8 | 0.08 | -0.68 | -0.87 | -0.95 | -0.74 | -0.55 | -0.66 | -1.2 | -0.62 | -0.28 | -0.67 | -0.66 | -0.59 | -0.95 | -1.04 | -0.94 | -0.1825 | 0.0475 | -0.275 | 0.530789 | 0.860641 | 0.32976 | 0.999971 | 1 | 0.856787 |
| Arhgap33 | A0A140LH | -0.81 | -1.59 | -1.4 | -1.32 | -1.94 | -1.35 | -1.19 | -1.08 | -1.49 | -0.54 | -1.05 | -1.13 | -1.24 | -1.04 | -0.93 | -0.74 | -0.11 | 0.2275 | 0.2925 | 0.687407 | 0.415159 | 0.231544 | 0.999971 | 1 | 0.8307 |
| Arhgap35 | Q91YM2 | -0.29 | -0.53 | -0.51 | -0.72 | -0.62 | -0.24 | -0.35 | -0.65 | -0.28 | -0.33 | -0.47 | -0.22 | -0.53 | -0.52 | -0.64 | -0.58 | 0.0475 | 0.1875 | -0.055 | 0.808477 | 0.306861 | 0.756564 | 0.999971 | 1 | 0.961116 |
| Arhgap39 | P59281-2 | 3.43 | -1.3 | 3.22 | 1.25 | 1.03 | 2.38 | 2.88 | 0.82 | -0.53 | 3.59 | 3.11 | -2 | -2.18 | 0.95 | 2.8 | 2.2 | 0.1275 | -0.6075 | -0.7075 | 0.905703 | 0.699482 | 0.61275 | 0.999971 | 1 | 0.927364 |
| Arhgap39_ | H7BX46 | 0.04 | -0.36 | -0.3 | -0.56 | -0.54 | -0.19 | 0.19 | -0.46 | -0.51 | 0.03 | -0.19 | -0.42 | -0.08 | -0.41 | -0.77 | -0.49 | 0.045 | 0.0225 | -0.1425 | 0.851013 | 0.917182 | 0.53789 | 0.999971 | 1 | 0.910077 |
| Arhgap42 | B2RQE8 | 1.22 | 0.49 | -0.32 | -0.36 | 0.13 | 0.57 | -1.21 | -0.97 | 0.49 | 1.29 | -0.71 | 0.2 | -0.06 | 0.41 | -0.84 | -0.35 | -0.6275 | 0.06 | -0.4675 | 0.251874 | 0.907811 | 0.297125 | 0.999971 | 1 | 0.847402 |
| Arhgap44 | Q5SSM3 | 0.72 | -0.06 | -0.09 | 0.33 | -0.16 | 0.42 | -0.12 | 0.15 | -0.16 | 0.64 | 0.08 | -0.1 | -0.41 | -0.06 | 0.16 | 0.32 | -0.1525 | -0.11 | -0.2225 | 0.558206 | 0.694017 | 0.416714 | 0.999971 | 1 | 0.877251 |
| Arhgap5 | E9PYT0 | -0.8 | -0.83 | -0.27 | -0.4 | -0.49 | -0.86 | -0.36 | -0.12 | -0.48 | -0.86 | -0.39 | 0.12 | -0.47 | -0.82 | -0.19 | -0.03 | 0.1175 | 0.1725 | 0.1975 | 0.62862 | 0.52127 | 0.440801 | 0.999971 | 1 | 0.883347 |
| Arhgap6 | G3UZI7 | 0.34 | -1.05 | -0.83 | -0.35 | 0.21 | 0.07 | -1.25 | -0.5 | 0.41 | 0.03 | -1.49 | -1.09 | -0.82 | 0.88 | -0.03 | -1.19 | 0.105 | -0.0625 | 0.1825 | 0.80631 | 0.90192 | 0.722313 | 0.999971 | 1 | 0.955258 |
| Arhgdia | Q99PT1 | 0.05 | -0.96 | -0.03 | -0.19 | -0.75 | -0.74 | -0.02 | -0.05 | -0.89 | -0.37 | -0.14 | -0.39 | -1.4 | -0.6 | -0.17 | 0 | -0.1075 | -0.165 | -0.26 | 0.732157 | 0.573802 | 0.496151 | 0.999971 | 1 | 0.90168 |
| Arhgdib | Q61599 | 0.46 | 1.56 | 0.17 | 1.2 | 1.54 | 1.45 | 0.48 | 1.39 | 1.59 | 1.1 | 0.9 | 1.01 | 1.06 | 1.72 | 1.22 | 0.74 | 0.3675 | 0.3025 | 0.3375 | 0.357824 | 0.397833 | 0.37415 | 0.999971 | 1 | 0.87031 |
| Arhgef1 | E9PUF7 | 0.37 | -0.25 | -0.71 | -0.73 | -0.14 | -0.37 | -0.58 | -0.98 | -0.12 | 0.08 | -0.65 | -0.76 | -0.06 | -0.13 | -0.69 | -0.64 | -0.1875 | -0.0325 | -0.05 | 0.558103 | 0.921072 | 0.872726 | 0.999971 | 1 | 0.984109 |
| Arhgef10 | Q8C033 | -0.55 | -1.21 | -0.01 | -0.35 | -1.03 | -0.54 | -0.62 | -0.09 | -0.69 | -0.5 | -1.11 | -0.66 | -0.83 | -0.5 | -0.07 | -0.57 | -0.04 | -0.21 | 0.0375 | 0.900427 | 0.481262 | 0.902194 | 0.999971 | 1 | 0.990759 |
| Arhgef10l | A2AWP8-3 | 0.19 | -0.35 | -0.38 | -0.25 | -0.19 | -0.01 | -0.1 | -0.22 | -0.28 | 0.11 | -0.22 | -0.32 | 0 | -0.16 | -0.22 | -0.28 | 0.0675 | 0.02 | 0.0325 | 0.735494 | 0.924315 | 0.872544 | 0.999971 | 1 | 0.984109 |
| Arhgef11 | Q68FM7 | -0.76 | -0.87 | -1.2 | -1.27 | -0.92 | -0.66 | -0.66 | -1.04 | -1.03 | -0.9 | -0.82 | -0.81 | -0.59 | -0.79 | -0.99 | -1.12 | 0.205 | 0.135 | 0.1525 | 0.338935 | 0.490827 | 0.488358 | 0.999971 | 1 | 0.898175 |
| Arhgef12 | Q8R4H2 | 0.59 | 0.46 | -1.79 | -1.31 | -0.96 | -0.25 | -1.85 | -1.53 | -0.29 | -0.14 | -2.13 | -1.36 | 1.6 | -0.24 | -2.51 | -1.85 | -0.635 | -0.4675 | -0.2375 | 0.333351 | 0.509187 | 0.809337 | 0.999971 | 1 | 0.973762 |
| Arhgef12_ | F8VQN6 | -0.21 | -0.34 | -0.01 | -0.05 | -0.5 | -0.52 | 0.26 | -0.32 | -0.44 | -0.47 | -0.05 | 0.08 | -0.45 | -0.32 | -0.32 | -0.1 | -0.1175 | -0.0675 | -0.145 | 0.617392 | 0.744567 | 0.435444 | 0.999971 | 1 | 0.882419 |
| Arhgef17 | Q80U35 | -0.43 | 0.54 | 0.31 | -0.57 | 0.23 | 0.51 | 0.74 | -0.36 | 0.04 | -0.43 | 0.52 | 0.23 | 0.11 | 0.17 | -0.3 | -0.61 | 0.3175 | 0.1275 | -0.12 | 0.381235 | 0.70492 | 0.716197 | 0.999971 | 1 | 0.952576 |
| Arhgef18 | A0A5F8MP | 0.02 | 0.05 | -0.24 | -0.22 | 0.06 | 0.03 | -0.33 | -0.17 | 0 | 0.18 | -0.18 | -0.03 | -0.45 | 0.08 | -0.22 | 0.21 | -0.005 | 0.09 | 0.0025 | 0.978816 | 0.618106 | 0.990731 | 0.999971 | 1 | 0.99925 |
| Arhgef19 | A2A832 | 1.12 | 0.06 | -0.39 | 0.7 | -0.7 | -1.12 | 0.69 | 0.96 | -0.25 | 0.97 | 0.2 | 0.98 | -0.65 | -2.43 | -2.73 | 1.16 | -0.415 | 0.1025 | -1.535 | 0.466067 | 0.811087 | 0.102497 | 0.999971 | 1 | 0.78268 |
| Arhgef2 | H3BJU7 | -0.1 | 0.06 | 0.37 | 0.3 | 0.06 | 0.02 | 0.38 | 0.37 | 0.33 | 0.03 | 0.69 | 0.45 | 0.07 | -0.06 | 0.23 | 0.54 | 0.05 | 0.2175 | 0.0375 | 0.804813 | 0.330808 | 0.862356 | 0.999971 | 1 | 0.982963 |
| Arhgef25 | A0A1W2P6 | -1.57 | -1.29 | -0.26 | -0.64 | -0.59 | -1.21 | -0.41 | -0.3 | -1.09 | -1.03 | -0.58 | -0.51 | -0.95 | -1.19 | -0.45 | -0.64 | 0.3125 | 0.1375 | 0.1325 | 0.388265 | 0.680769 | 0.696692 | 0.999971 | 1 | 0.947174 |
| Arhgef26 | D3YYY8 | 0 | -0.27 | 0.12 | -0.25 | -0.53 | -0.1 | -0.08 | 0.03 | -0.75 | 0.18 | -0.69 | -0.54 | -0.6 | 0.14 | -0.23 | 0.13 | -0.07 | -0.35 | -0.04 | 0.737221 | 0.197546 | 0.865398 | 0.999971 | 1 | 0.983318 |
| Arhgef28 | A0A286YD | -0.04 | -0.05 | -0.26 | 0.17 | 0.37 | 0.45 | -0.24 | 0.53 | -0.05 | -0.06 | 0.15 | -0.2 | 0.52 | 0.3 | 0.35 | 0.03 | 0.3225 | 0.005 | 0.345 | 0.189557 | 0.97813 | 0.106756 | 0.999971 | 1 | 0.78268 |
| Arhgef4 | E9PYK8 | 0.18 | 0.69 | 0.43 | 0.52 | 0.93 | 0.68 | 0.2 | 0.73 | 0.02 | 0.32 | 0.52 | 0.7 | 0.37 | 1.08 | 0.57 | 0.46 | 0.18 | -0.065 | 0.165 | 0.436855 | 0.768646 | 0.479871 | 0.999971 | 1 | 0.894438 |
| Arhgef40 | Q3UPH7-3 | -0.43 | -1.02 | -0.42 | -0.23 | -0.59 | -0.4 | -0.5 | -0.5 | -0.84 | -0.28 | -0.59 | -0.23 | -1.24 | -0.78 | -0.2 | -0.52 | 0.0275 | 0.04 | -0.16 | 0.900358 | 0.872381 | 0.585594 | 0.999971 | 1 | 0.920413 |
| Arhgef6 | F6WMJ3 | 0.63 | -1.11 | -0.27 | -0.66 | -1.36 | -0.49 | -0.23 | -0.56 | -1.37 | -0.23 | -0.29 | -0.37 | -1.71 | -0.85 | -0.36 | -0.86 | -0.3075 | -0.2125 | -0.5925 | 0.471011 | 0.62644 | 0.19955 | 0.999971 | 1 | 0.813335 |
| Arhgef7 | Q9ES28 | 0.1 | -0.43 | -0.94 | -0.91 | -0.43 | -0.36 | -0.49 | -1.21 | -0.27 | -0.12 | -0.51 | -0.89 | -0.48 | -0.61 | -1.02 | -0.82 | -0.0775 | 0.0975 | -0.1875 | 0.807278 | 0.748476 | 0.517376 | 0.999971 | 1 | 0.905815 |
| Arhgef7_1 | Q9ES28-5 | 1.32 | 2.27 | 2.94 | 3.27 | 1.57 | 2.63 | 2.66 | 3.02 | 1.33 | 1.75 | 2.34 | 2.83 | 3.05 | 2.73 | 2.53 | 2.52 | 0.02 | -0.3875 | 0.2575 | 0.967658 | 0.450336 | 0.548636 | 0.999971 | 1 | 0.911727 |
| Arhgef9 | B1AXI4 | -0.03 | -1.17 | -1.08 | -0.94 | -0.61 | -0.72 | -0.61 | -0.65 | -0.72 | -0.28 | -0.89 | -0.94 | -0.98 | -0.71 | -0.94 | -0.92 | 0.1575 | 0.0975 | -0.0825 | 0.576532 | 0.751921 | 0.772004 | 0.999971 | 1 | 0.963498 |
| Arid1a | E9QAQ7 | 2.4 | 1.5 | 1.17 | 1.83 | 1.14 | 1.36 | 2.87 | 1.02 | 0.85 | 2.14 | 0.77 | 0.96 | 0.19 | 1.26 | 1.02 | 1.16 | -0.1275 | -0.545 | -0.8175 | 0.786495 | 0.195455 | 0.042861 | 0.999971 | 1 | 0.701725 |
| Arid1b | E9Q6R4 | 2.03 | 2.15 | 1.43 | 2.01 | 1.86 | 1.91 | 1.18 | 1.73 | 1.62 | 1.99 | 1.31 | 1.5 | 1.2 | 2.12 | 1.58 | 1.95 | -0.235 | -0.3 | -0.1925 | 0.37276 | 0.239606 | 0.494339 | 0.999971 | 1 | 0.901531 |
| Arid2 | E9Q7E2 | 1.21 | -0.76 | -0.54 | -0.09 | -0.89 | -0.69 | -0.1 | -0.28 | -0.63 | 0.42 | -0.5 | -0.79 | -0.5 | -0.29 | -0.6 | -0.45 | -0.445 | -0.33 | -0.415 | 0.334527 | 0.501385 | 0.340338 | 0.999971 | 1 | 0.860574 |
| Arid4a | F8VPQ2 | 2.25 | 1.84 | 0.4 | 0.96 | 1.11 | 2.47 | 0.05 | 0.12 | 1.45 | 2.96 | 1.28 | -0.38 | 1.63 | 2.44 | -0.01 | 0.28 | -0.425 | -0.035 | -0.2775 | 0.511412 | 0.961605 | 0.66977 | 0.999971 | 1 | 0.938809 |
| Arid4b | Z4YL44 | 4.86 | 2.87 | 1.69 | 0.96 | 1.92 | 4.48 | 3.43 | -0.45 | 3.2 | 5.25 | 1.36 | -0.86 | -0.05 | 1.74 | -0.31 | -1.49 | -0.25 | -0.3575 | -2.6225 | 0.836983 | 0.797452 | 0.022759 | 0.999971 | 1 | 0.652043 |
| Arih1 | Q9Z1K5 | 0.24 | 1.79 | 1.28 | 0.86 | 1.63 | 1.51 | 1.85 | 0.43 | 1.95 | 0.87 | 1.3 | 0.93 | 1.54 | 0.95 | 0.86 | 0.07 | 0.3125 | 0.22 | -0.1875 | 0.47472 | 0.581199 | 0.659485 | 0.999971 | 1 | 0.935181 |
| Arih2 | Q9Z1K6 | 0.46 | 0.72 | 0.08 | 0.65 | 0.43 | 0.8 | 0.22 | 0.63 | 0.5 | 0.45 | 0.55 | 0.29 | 0.18 | 0.74 | 0.46 | 0.48 | 0.0425 | -0.03 | -0.0125 | 0.852825 | 0.883301 | 0.955714 | 0.999971 | 1 | 0.998056 |
| Arl1 | F8WIB1 | -0.03 | 0.57 | 0.84 | 0.87 | 0.7 | 0.47 | 0.19 | 1.02 | 0.35 | -0.17 | 0.73 | 0.74 | 0.31 | 0.84 | 1.31 | 1.13 | 0.0325 | -0.15 | 0.335 | 0.909513 | 0.625402 | 0.294287 | 0.999971 | 1 | 0.847402 |
| Arl10 | Q9QXJ4 | -1.31 | -0.87 | -1.52 | -1.95 | -1.09 | -1.26 | -1.59 | -1.59 | -0.85 | -1.2 | -1.11 | -0.65 | -0.54 | -0.45 | -0.72 | -1.89 | 0.03 | 0.46 | 0.5125 | 0.912903 | 0.120054 | 0.209482 | 0.999971 | 1 | 0.822138 |
| Arl13b | Q640N2 | -2.41 | -1.12 | -0.72 | -1.16 | -1.55 | -2.31 | -0.86 | 0.2 | -2.47 | -1.79 | -2.11 | -0.48 | -1.68 | -2.7 | -1.36 | -0.9 | 0.2225 | -0.36 | -0.3075 | 0.707376 | 0.499977 | 0.536807 | 0.999971 | 1 | 0.910077 |
| Arl15 | Q8BGR6 | -1.33 | -0.74 | -0.85 | -1.25 | -0.71 | -1.05 | -1.09 | -0.84 | -0.52 | -1.47 | -0.6 | -0.43 | 0.12 | -1.16 | -0.55 | -0.47 | 0.12 | 0.2875 | 0.5275 | 0.583256 | 0.338175 | 0.112731 | 0.999971 | 1 | 0.78268 |
| Arl2 | Q9D0J4 | -0.54 | -0.08 | -0.4 | -0.39 | -0.12 | 0.21 | -0.46 | 0.06 | -0.01 | -0.36 | -0.12 | -0.02 | 0.28 | 0.34 | 0.05 | -0.06 | 0.275 | 0.225 | 0.505 | 0.229416 | 0.253808 | 0.02886 | 0.999971 | 1 | 0.678474 |
| Arl2bp | Q9D385 | -0.42 | 0.93 | 1.48 | 1.31 | 0.75 | 1.07 | 0.78 | 1.81 | 1.09 | -0.38 | 0.94 | 1.59 | 0.74 | 0.55 | 1.64 | 1.6 | 0.2775 | -0.015 | 0.3075 | 0.553823 | 0.978436 | 0.527802 | 0.999971 | 1 | 0.909722 |
| Arl3 | Q9WUL7 | 0.32 | -0.1 | 0.12 | -0.13 | 0.31 | 0.25 | 0.61 | 0.48 | -0.48 | 0.34 | 0.09 | 0.31 | -0.75 | 0.19 | 0.04 | -0.09 | 0.36 | 0.0125 | -0.205 | 0.092715 | 0.959285 | 0.43493 | 0.999971 | 1 | 0.882419 |
| Arl4c | P61208 | -1.56 | -2.23 | 0.47 | -0.76 | -3.05 | -3.56 | 1.2 | 0.06 | -3.57 | -2.79 | 3.88 | -1.02 | -2.99 | -2.09 | -0.18 | -1.03 | -0.3175 | 0.145 | -0.5525 | 0.784022 | 0.926852 | 0.474917 | 0.999971 | 1 | 0.894044 |
| Arl5a | Q80ZU0 | -0.28 | 0.87 | 0.73 | 0.78 | 0.77 | 1.05 | 0 | 0.77 | 0.51 | 0.5 | 0.61 | 0.37 | 0.81 | 1.27 | 0.94 | 0.78 | 0.1225 | -0.0275 | 0.425 | 0.724444 | 0.923432 | 0.182368 | 0.999971 | 1 | 0.808532 |
| Arl6 | O88848 | -0.53 | -0.07 | -0.83 | -0.79 | 0.29 | 0.17 | 0.79 | -0.44 | 0.23 | -0.05 | -0.2 | 0.25 | 0.56 | 0.3 | -0.27 | -0.63 | 0.7575 | 0.6125 | 0.545 | 0.035715 | 0.028273 | 0.119222 | 0.999971 | 1 | 0.78268 |
| Arl6ip1 | Q9JKW0 | 0.44 | -0.64 | -0.11 | 0.37 | 0.07 | 0.22 | 0.47 | 0.65 | 0.71 | 0.76 | 0.83 | 0.14 | 0.13 | -0.18 | 0.29 | -0.04 | 0.3375 | 0.595 | 0.035 | 0.266856 | 0.076549 | 0.902135 | 0.999971 | 1 | 0.990759 |
| Arl6ip4 | Q9JM93 | 1.84 | -0.2 | 0.39 | 0.93 | -0.47 | 0.3 | 0.86 | 0.06 | -0.24 | 1.5 | 0.1 | -0.03 | -0.55 | 0.23 | 0.04 | 0.64 | -0.5525 | -0.4075 | -0.65 | 0.266648 | 0.460147 | 0.187995 | 0.999971 | 1 | 0.809025 |
| Arl6ip5 | Q8R5J9 | -1.3 | -0.62 | -1.8 | -3.09 | -0.15 | -1.35 | -1.23 | -2.94 | -0.19 | -1.41 | -1.13 | -0.84 | 0.01 | -1.62 | -2.51 | -3.32 | 0.285 | 0.81 | -0.1575 | 0.685792 | 0.159735 | 0.843141 | 0.999971 | 1 | 0.979814 |
| Arl6ip6 | Q8BH07 | -3.66 | -4.59 | -3.7 | -5.21 | -3.32 | -4.78 | -3.39 | -4.07 | -3.47 | -3.75 | -3.05 | -1.93 | -3.57 | -5.07 | -4.58 | -4.86 | 0.4 | 1.24 | -0.23 | 0.407288 | 0.035934 | 0.625685 | 0.999971 | 1 | 0.930479 |
| Arl8a | Q8VEH3 | -1.52 | -1.28 | -1.76 | -2.75 | -1.1 | -1.14 | -1.29 | -1.81 | -0.96 | -2.04 | -0.95 | -0.72 | 0.03 | -1.25 | -1.08 | -2.07 | 0.4925 | 0.66 | 0.735 | 0.188386 | 0.140651 | 0.167505 | 0.999971 | 1 | 0.800215 |
| Arl8b | Q9CQW2 | -3.53 | -2.02 | -0.84 | -1.04 | -1.95 | -1.74 | -0.94 | -0.12 | -1.82 | -2.53 | -0.51 | -0.26 | -1.43 | -1.33 | -0.18 | -0.58 | 0.67 | 0.5775 | 0.9775 | 0.333141 | 0.444503 | 0.143423 | 0.999971 | 1 | 0.788395 |
| Armc1 | Q9D7A8 | 0.63 | -0.51 | -0.07 | -0.09 | -0.88 | -0.7 | 0.26 | -0.62 | -1.15 | -0.35 | -0.63 | -0.73 | -0.99 | -0.53 | -0.5 | -0.61 | -0.475 | -0.705 | -0.6475 | 0.189473 | 0.039423 | 0.04246 | 0.999971 | 1 | 0.701725 |
| Armc10 | Q9D0L7 | -2.18 | -0.85 | 0.16 | -0.23 | -0.73 | -0.98 | -0.15 | -0.07 | -0.72 | -1.58 | 0.29 | 0.22 | -0.19 | -0.87 | 0.36 | 0.37 | 0.2925 | 0.3275 | 0.6925 | 0.574167 | 0.600259 | 0.224493 | 0.999971 | 1 | 0.828225 |
| Armc6 | Q8BNU0 | -0.39 | -0.36 | 0.75 | 0.03 | -0.19 | -0.3 | 0.87 | 0.12 | -0.23 | -0.45 | 1.17 | 0.68 | -0.39 | -0.26 | 0.32 | 0.31 | 0.1175 | 0.285 | -0.0125 | 0.747542 | 0.52168 | 0.969361 | 0.999971 | 1 | 0.998787 |
| Armc8 | Q9DBR3-2 | -2.46 | 0.09 | 0.26 | -0.61 | 0.47 | 0.04 | 0.41 | 0.68 | 0.57 | -2.08 | 0.52 | 0.36 | 1.73 | -0.47 | 0.55 | 0.06 | 1.08 | 0.5225 | 1.1475 | 0.090846 | 0.523543 | 0.130964 | 0.999971 | 1 | 0.784948 |

| | | | | | | | | | | | | | | | | | | | | | | | | | | |
|---|---|---|---|---|---|---|---|---|---|---|---|---|---|---|---|---|---|---|---|---|---|---|---|---|---|---|
| Armc8_1 | G3X920 | -0.9 | 0.25 | 1.58 | 0.47 | 0.04 | 0.05 | 0.75 | 0.79 | 0.19 | -0.6 | 0.77 | -0.07 | 0.2 | 0.3 | 0.6 | 0.66 | 0.0575 | -0.2775 | 0.09 | 0.909933 | 0.609431 | 0.852826 | 0.999971 | 1 | 0.98187 |
| Armc9 | Q9D2I5-2 | 0.12 | 0.65 | -0.64 | 0.28 | 0.72 | 1.03 | -0.5 | -0.15 | 0.47 | 0.58 | -0.41 | -0.03 | 1.24 | 0.6 | -0.58 | -0.39 | 0.1725 | 0.05 | 0.115 | 0.686715 | 0.88623 | 0.808266 | 0.999971 | 1 | 0.973379 |
| Armcx2 | Q6A058 | 1.39 | 0.02 | 0.07 | -0.5 | -0.21 | 0.18 | -0.28 | 0.15 | 0.36 | 1.09 | 0.02 | -0.25 | -0.47 | -0.49 | 0.43 | 0.31 | -0.285 | 0.06 | -0.3 | 0.483997 | 0.897495 | 0.505889 | 0.999971 | 1 | 0.905277 |
| Armcx3 | Q8BHS6 | -0.08 | 0.42 | 0.09 | -0.25 | 0.3 | 0.43 | -0.05 | 0 | 0.44 | 0.13 | 0.15 | 0.05 | 0.82 | 0.28 | 0.09 | 0.71 | 0.125 | 0.1475 | 0.43 | 0.581892 | 0.492496 | 0.115675 | 0.999971 | 1 | 0.78268 |
| Armcx4 | E9PWM3 | 0.79 | -0.44 | -0.48 | 0.44 | -0.36 | -0.02 | 0.15 | 0.48 | -0.52 | 0 | 0.57 | 0.33 | -0.78 | -0.86 | -0.03 | 0.23 | -0.015 | 0.0175 | -0.4375 | 0.966325 | 0.96352 | 0.291858 | 0.999971 | 1 | 0.847402 |
| Armh3 | Q6PD19 | -0.69 | 0.62 | 0.03 | -0.09 | 0.25 | 0.05 | -0.04 | -0.02 | 0.52 | -0.34 | 0.03 | 0.44 | 0.79 | -0.1 | 0.09 | 0.35 | 0.0925 | 0.195 | 0.315 | 0.74929 | 0.561591 | 0.353448 | 0.999971 | 1 | 0.868675 |
| Armt1 | A6H630 | 0.1 | 1.46 | 0.77 | 0.48 | 1.7 | 0.91 | 1 | 1.05 | 1.73 | -0.02 | 0.86 | 0.91 | 2.16 | 0.72 | 0.75 | 0.38 | 0.4625 | 0.1675 | 0.3 | 0.193137 | 0.700887 | 0.517953 | 0.999971 | 1 | 0.905815 |
| Arnt2 | Q61324-2 | 0.04 | 1.37 | 0.93 | 1.22 | 1.4 | 1.63 | 0.74 | 0.85 | 1.56 | 1.1 | 0.66 | 0.63 | 1.2 | 1.64 | 0.87 | 1.18 | 0.265 | 0.0975 | 0.3325 | 0.466745 | 0.787421 | 0.335945 | 0.999971 | 1 | 0.858827 |
| Arntl | A0A1B0GS` | 2.53 | 2.37 | 1.82 | 2.73 | 1.71 | 1.41 | 1.6 | 2.24 | 1.31 | 1.74 | 1.7 | 0.66 | -0.41 | 2.09 | 0.73 | 2.38 | -0.6225 | -1.01 | -1.165 | 0.049673 | 0.01119 | 0.085562 | 0.999971 | 1 | 0.759991 |
| Arpc1a | Q9R0Q6 | -1.94 | -0.86 | -1.17 | -1.26 | -0.78 | -0.18 | -0.97 | -1.14 | -0.82 | -1.55 | -1.23 | -1.74 | -1.63 | -0.57 | -1.49 | -1.84 | 0.54 | -0.0275 | -0.075 | 0.111996 | 0.928989 | 0.832549 | 0.999971 | 1 | 0.979036 |
| Arpc1b | Q91Z25 | 0.73 | 0.4 | 0.53 | 0.28 | 0.86 | 0.41 | 0.65 | 0.35 | 0.44 | 0.44 | 0.86 | 0.32 | 0.22 | 0.66 | 0.91 | 0.69 | 0.0825 | 0.03 | 0.135 | 0.689295 | 0.882817 | 0.542385 | 0.999971 | 1 | 0.911119 |
| Arpc2 | Q9CVB6 | -1.17 | 0.03 | -0.99 | -1.63 | -0.3 | -0.18 | -0.86 | -1.19 | -0.32 | -1.27 | -0.69 | -1.18 | -0.28 | -0.62 | -1.16 | -1.24 | 0.3075 | 0.075 | 0.115 | 0.45411 | 0.850531 | 0.774286 | 0.999971 | 1 | 0.963884 |
| Arpc3 | Q9JM76 | -1.06 | -0.55 | 0.4 | -0.28 | -0.5 | -0.09 | 0.61 | 0.17 | -0.48 | -0.58 | 0.38 | 0.65 | -1.3 | -0.38 | 0.04 | -0.5 | 0.42 | 0.365 | -0.1625 | 0.275736 | 0.387445 | 0.683353 | 0.999971 | 1 | 0.943649 |
| Arpc4 | P59999 | -1.86 | -0.49 | -1.03 | -1.14 | -0.52 | -0.91 | -0.87 | -1.59 | -0.73 | -2.27 | -0.75 | -0.8 | -0.13 | -0.56 | -1.72 | -1.08 | 0.1575 | -0.0075 | 0.2575 | 0.656916 | 0.986539 | 0.545553 | 0.999971 | 1 | 0.911398 |
| Arpc5 | Q9CPW4 | 0.08 | 0.16 | -0.09 | 0.22 | 0.24 | 0.55 | 0.19 | 0.3 | 0.14 | 0.18 | 0.08 | -0.11 | 0.26 | 0.1 | -0.08 | 0.33 | 0.2275 | -0.02 | 0.06 | 0.230861 | 0.908049 | 0.747949 | 0.999971 | 1 | 0.961064 |
| Arpc5l | Q9D898 | 0.15 | 0.2 | 1.06 | 1.24 | -0.01 | 0.06 | 0.67 | 0.69 | -0.06 | 0.06 | 0.88 | 1.01 | -0.17 | -0.14 | 0.78 | 0.82 | -0.31 | -0.19 | -0.34 | 0.371441 | 0.621529 | 0.385265 | 0.999971 | 1 | 0.871816 |
| Arpin | Q9D0A3 | 0.17 | 0.59 | 0.04 | 0.35 | 0.61 | 0.46 | -0.19 | 0.18 | 0.42 | 0.23 | 0.06 | 0.23 | 0.54 | 0.92 | 0.18 | 0.35 | -0.0225 | -0.0525 | 0.21 | 0.926323 | 0.789498 | 0.382641 | 0.999971 | 1 | 0.871816 |
| Arpp19 | P56212 | 0.92 | 0.51 | 1.32 | 1.92 | 0.51 | 0.5 | 0.9 | 1.54 | 0.43 | 1.25 | 1.09 | 1.41 | -0.04 | 0.48 | 1.23 | 1.67 | -0.305 | -0.1225 | -0.3325 | 0.424629 | 0.735084 | 0.472732 | 0.999971 | 1 | 0.893755 |
| Arpp21 | Q9DCB4 | 0.35 | -0.89 | 0.14 | 0.4 | -0.66 | -0.75 | 0.73 | -0.03 | -0.25 | 0.13 | -0.41 | -0.6 | -0.3 | -0.59 | 0.22 | -0.69 | -0.1775 | -0.2825 | -0.34 | 0.681807 | 0.411376 | 0.354299 | 0.999971 | 1 | 0.868793 |
| Arpp21_1 | Q9DCB4-2 | 2.69 | -0.7 | 1.32 | 3 | 0.42 | 0.46 | 1.79 | 2.19 | 0.77 | 2.2 | 1.16 | 0.15 | -1.46 | -0.67 | 1.06 | 0.57 | -0.3625 | -0.5075 | -1.7025 | 0.674004 | 0.556407 | 0.089841 | 0.999971 | 1 | 0.771991 |
| Arrb1 | Q8BWG8 | -0.16 | 0.36 | -0.28 | -0.2 | 0.48 | -0.07 | -0.4 | -0.37 | 0.26 | -0.13 | -0.25 | -0.69 | -0.39 | -0.02 | -0.53 | -0.6 | -0.02 | -0.1325 | -0.315 | 0.940892 | 0.618029 | 0.198335 | 0.999971 | 1 | 0.813335 |
| Arrb2 | Q91YI4 | 2.93 | 2.58 | 2.32 | 1.73 | 1.81 | 2.24 | 1.67 | 2.45 | 1.95 | 1.89 | 2.52 | 2.03 | 2.46 | 2.59 | 2.28 | 2.33 | -0.3475 | -0.2925 | 0.025 | 0.287374 | 0.341539 | 0.928569 | 0.999971 | 1 | 0.994278 |
| Arrdc1 | A2AIS9 | 1.21 | 0.28 | -0.42 | -0.68 | 0.13 | 0.23 | -0.32 | 0.09 | 0.1 | 0.76 | -1.53 | -0.64 | 0.03 | 0.26 | -0.82 | -0.63 | -0.065 | -0.425 | -0.3875 | 0.876167 | 0.482389 | 0.413426 | 0.999971 | 1 | 0.876261 |
| Arsa | P50428 | 0.86 | 0.6 | 0.64 | 0.74 | 1.32 | 0.37 | 0.68 | 0.92 | 1.12 | 0.58 | 0.49 | 0.41 | 0.86 | -0.12 | 0.43 | 0.44 | 0.1125 | -0.06 | -0.3075 | 0.642502 | 0.779959 | 0.224973 | 0.999971 | 1 | 0.828225 |
| Arsb | A0A0R4J13 | 0.29 | 0.33 | -0.11 | -0.82 | 0.26 | 0.03 | -0.16 | -0.23 | 0.08 | 0.31 | 0.21 | -0.06 | 0.15 | 0.44 | -0.86 | -0.58 | 0.0525 | 0.2125 | -0.135 | 0.859962 | 0.469942 | 0.72969 | 0.999971 | 1 | 0.956041 |
| Arsg | Q3TYD4 | 0.53 | -1.17 | -2.14 | -1.5 | -1.43 | -1.07 | -1.96 | -0.83 | -0.76 | 0.04 | -1.17 | -1.58 | -0.02 | -0.75 | -2.18 | -1.6 | -0.2525 | 0.2025 | -0.0675 | 0.65864 | 0.741084 | 0.920043 | 0.999971 | 1 | 0.992702 |
| Arsj | Q8BM89 | -0.01 | -2.14 | -1.52 | -2.12 | -2.74 | -0.97 | -0.3 | -1.21 | -3 | 0.16 | -1.07 | -2.05 | -4.65 | -1.68 | -2.34 | -0.7 | 0.1425 | -0.0425 | -0.895 | 0.82703 | 0.955528 | 0.322249 | 0.999971 | 1 | 0.85548 |
| Arvcf | P98203 | -0.59 | -1.38 | -1.06 | -1.09 | -1.41 | -0.96 | -0.84 | -1.09 | -1.24 | -0.79 | -0.96 | -1.19 | -0.99 | -1.17 | -1.01 | -0.95 | -0.045 | -0.015 | 7.85E-17 | 0.850136 | 0.947832 | 1 | 0.999971 | 1 | 1 |
| Arxes2 | C0HK80 | -1.69 | 0.82 | 0.19 | -1.34 | 1.73 | -0.8 | 1.18 | -1.12 | 2.54 | -1.28 | 1.37 | 1.93 | 3.26 | -1.48 | -0.5 | -0.9 | 0.7525 | 1.645 | 0.6 | 0.378322 | 0.105276 | 0.587843 | 0.999971 | 1 | 0.920758 |
| As3mt | Q91WU5 | 0.49 | 0.43 | -0.59 | 0.11 | 0.68 | 0.27 | -0.08 | -0.68 | 0.82 | 0.64 | -0.31 | 0.29 | 0.89 | 0.46 | -0.12 | -0.44 | -0.0625 | 0.25 | 0.0875 | 0.865414 | 0.477031 | 0.815607 | 0.999971 | 1 | 0.974918 |
| Asah1 | Q9WV54 | -0.55 | 0.13 | 0.07 | 0.42 | 0.16 | -0.23 | -0.29 | -0.46 | -0.01 | -0.53 | -0.17 | -0.3 | -0.05 | 0.24 | -0.06 | -0.06 | -0.2225 | -0.27 | 1.96E-17 | 0.409387 | 0.306103 | 1 | 0.999971 | 1 | 1 |
| Asah2 | B7ZN55 | -0.03 | -0.89 | -0.52 | -0.65 | -0.43 | -0.61 | -0.69 | 0.15 | -0.93 | -0.66 | -0.62 | 0.09 | -0.3 | -0.36 | -0.45 | -0.37 | 0.1275 | -0.0075 | 0.1525 | 0.648723 | 0.979565 | 0.504478 | 0.999971 | 1 | 0.905277 |
| Asap1 | H3BJY2 | -3.59 | -0.46 | 1.11 | -0.08 | -1.24 | -1.41 | 0.15 | -0.7 | -1.47 | -3.05 | 0.54 | 0.12 | -2.17 | -2.17 | -0.22 | 0.34 | -0.045 | -0.21 | -0.3 | 0.962132 | 0.856369 | 0.779107 | 0.999971 | 1 | 0.96583 |
| Asap1_1 | E9QMJ1 | 0.02 | -0.03 | -0.27 | -0.27 | -0.28 | 0.03 | -0.1 | -0.68 | -0.29 | -0.08 | -0.24 | -0.44 | -0.31 | -0.22 | -0.64 | -0.14 | -0.12 | -0.125 | -0.19 | 0.585107 | 0.49233 | 0.347232 | 0.999971 | 1 | 0.864164 |
| Asap2 | D3YX85 | 1.88 | -2.44 | 0.92 | 0.54 | -3.06 | -1.32 | 5.27 | -0.06 | -2.68 | 0.2 | 0.75 | -1.46 | -4.67 | -4.72 | -1.14 | -2.63 | -0.0175 | -1.0225 | -3.515 | 0.992191 | 0.362038 | 0.012249 | 0.999971 | 1 | 0.613095 |
| Asap2_1 | E9PX52 | -0.08 | -0.38 | -0.71 | -0.5 | -0.14 | -0.13 | -0.65 | -0.87 | -0.4 | -0.13 | -0.64 | -0.74 | -0.15 | -0.25 | 0.37 | -0.45 | -0.03 | -0.06 | 0.2975 | 0.905894 | 0.791156 | 0.250586 | 0.999971 | 1 | 0.839012 |
| Asb13 | Q8VBX0 | -0.08 | 0.39 | 1.07 | 0.72 | -0.12 | 0.27 | 0.62 | 0.95 | 0.29 | -0.38 | 1.43 | 0.61 | 0.12 | -0.19 | 0.37 | 0.87 | -0.095 | -0.0375 | -0.2325 | 0.776416 | 0.929798 | 0.489388 | 0.999971 | 1 | 0.898346 |
| Asb3 | A0A0N4SV( | 1.29 | -0.79 | -0.2 | -0.75 | -0.18 | -0.34 | -0.11 | -0.31 | -0.48 | -1.12 | 0.06 | 0.87 | 0.91 | -1.76 | 0.34 | -0.87 | -0.1225 | -0.055 | -0.2325 | 0.789527 | 0.925869 | 0.740248 | 0.999971 | 1 | 0.960899 |
| Asb6 | Q91ZU1 | 1.03 | 0.69 | 2.07 | 1.81 | 0.48 | 0.28 | 1.9 | 3.09 | 0.44 | 1.02 | 2.3 | 1.4 | -0.48 | 0.45 | 2.08 | 2.79 | 0.0375 | -0.11 | -0.19 | 0.954979 | 0.817012 | 0.79624 | 0.999971 | 1 | 0.970435 |
| Ascc1 | Q9D8Z1 | -0.39 | 0.51 | 0.28 | -0.13 | 0.38 | -0.37 | -0.45 | 0.13 | 0.45 | -0.21 | 0.22 | 0.3 | 0.32 | 0.35 | 0.01 | -0.09 | -0.145 | 0.1225 | 0.08 | 0.624294 | 0.646944 | 0.755321 | 0.999971 | 1 | 0.961116 |
| Ascc2 | Q91WR3-2 | 0.45 | 1.13 | 0.83 | 1.12 | 0.88 | 1.42 | 1.04 | 0.98 | 0.71 | 0.97 | 1.43 | 0.99 | 0.41 | 1.53 | 1.32 | 1.5 | 0.1975 | 0.1425 | 0.3075 | 0.409377 | 0.566564 | 0.340679 | 0.999971 | 1 | 0.860574 |
| Ascc3 | E9PZJ8 | -0.15 | 0.78 | 0.39 | 0.28 | 0.8 | 0.54 | 0.25 | 0.52 | 0.79 | 0.43 | 0.67 | 0.68 | 1.06 | 0.95 | 0.5 | 0.61 | 0.2025 | 0.3175 | 0.455 | 0.426476 | 0.203806 | 0.105613 | 0.999971 | 1 | 0.78268 |
| Asf1a | Q9CQE6 | 1.1 | 0.84 | 0.77 | 1.04 | 0.89 | 0.93 | 0.53 | 1.14 | 1.07 | 0.8 | 0.29 | 0.17 | 1.37 | 0.9 | 0.88 | 1.38 | -0.065 | -0.355 | 0.195 | 0.750691 | 0.182482 | 0.368391 | 0.999971 | 1 | 0.869526 |
| Ash2l | Q91X20 | 2.01 | 1.73 | 2.35 | 2.73 | 1.65 | 1.05 | 1.81 | 2.5 | 1.25 | 1.77 | 2.16 | 2.31 | 0.9 | 1.65 | 2.16 | 2.6 | -0.4525 | -0.3325 | -0.3775 | 0.229147 | 0.316771 | 0.364166 | 0.999971 | 1 | 0.869526 |
| Asic1 | Q6NXK8 | -0.22 | -0.65 | -0.58 | -0.23 | -0.77 | -0.44 | -0.35 | -0.11 | -0.25 | -0.26 | -0.44 | -0.46 | 0.26 | -0.32 | -0.14 | 0.16 | 0.0025 | 0.0675 | 0.41 | 0.990963 | 0.722142 | 0.089903 | 0.999971 | 1 | 0.771991 |
| Asic2 | Q925H0-2 | -2.08 | -0.16 | -0.35 | -1.09 | 0.26 | -0.18 | -0.13 | -0.71 | 0.04 | -1.95 | 0.14 | 0.33 | 1.37 | -1.29 | -0.72 | -0.97 | 0.73 | 0.56 | 0.5175 | 0.131411 | 0.385515 | 0.450718 | 0.999971 | 1 | 0.886265 |
| Asic2_1 | Q925H0 | 0.13 | 0.48 | -0.34 | 0.5 | 1 | -0.3 | -0.5 | 0.23 | -0.08 | -0.39 | 1.05 | 0.27 | 0.54 | 0.08 | -0.4 | 0.04 | -0.085 | 0.02 | -0.1275 | 0.821118 | 0.955549 | 0.659936 | 0.999971 | 1 | 0.93526 |
| Asic4 | D3YYS9 | 0.73 | -0.46 | -1.13 | 1.02 | -0.55 | -0.74 | 0.13 | 0.86 | -1.04 | 0.05 | -0.41 | 0.11 | -0.23 | -0.48 | 0.15 | 0.26 | -0.115 | -0.3625 | -0.115 | 0.839997 | 0.498933 | 0.816555 | 0.999971 | 1 | 0.974929 |
| Asl | Q91YI0 | -0.44 | 0.26 | 0.09 | 0.07 | 0.28 | 0.07 | -0.11 | 0.18 | 0.07 | -0.49 | -0.15 | 0.17 | -0.08 | 0.03 | -0.03 | 0.08 | 0.11 | -0.095 | 0.005 | 0.616201 | 0.693787 | 0.980822 | 0.999971 | 1 | 0.99925 |
| Asns | Q61024 | 0.65 | 0.95 | 0.56 | 0.91 | 1.01 | 0.38 | 1.06 | 0.92 | 0.75 | 0.54 | 0.79 | 1.21 | 0.52 | 0.78 | 0.7 | 0.68 | 0.075 | 0.055 | -0.0975 | 0.739371 | 0.797432 | 0.600777 | 0.999971 | 1 | 0.92426 |
| Aspa | Q8R3P0 | -1.28 | 1.09 | 1.76 | 0.31 | 1.37 | -0.35 | 0.63 | -0.37 | 2.33 | -1.57 | 0.57 | 0.94 | 3.67 | -0.2 | 1.47 | 0.23 | -0.15 | 0.0975 | 0.8225 | 0.83105 | 0.916937 | 0.407448 | 0.999971 | 1 | 0.874911 |
| Asph | A2AL85 | -0.09 | -0.18 | -0.25 | -0.46 | 0.52 | 0.34 | -0.14 | 0.08 | 0.17 | 0.3 | 0.04 | 0.57 | 0.82 | 0.23 | 0.02 | 0.35 | 0.445 | 0.515 | 0.6 | 0.062337 | 0.025598 | 0.025346 | 0.999971 | 1 | 0.678474 |
| Asph_1 | Q8BSY0-3 | -1.43 | -0.01 | -1 | -0.53 | 0.2 | 0.08 | -1.3 | -0.24 | -0.09 | -0.66 | -0.44 | 0.17 | 0.34 | 0.34 | -0.29 | -0.52 | 0.4275 | 0.4875 | 0.71 | 0.336694 | 0.18857 | 0.080131 | 0.999971 | 1 | 0.755254 |
| Asphd1 | Q2TA57 | 0.57 | 0 | 0.37 | 0.32 | -0.49 | -0.21 | 1.24 | -0.44 | -0.62 | -0.26 | -0.18 | -0.44 | -1.09 | -0.38 | -0.75 | -0.76 | -0.29 | -0.69 | -1.06 | 0.481859 | 0.007916 | 0.001222 | 0.999971 | 1 | 0.412385 |
| Asphd2 | Q80VP9 | -1.32 | -0.9 | -1.88 | -1.59 | -1.01 | -0.92 | -1.39 | -1.32 | -0.74 | -1.58 | -1.59 | -1.25 | -0.66 | -0.92 | -1.32 | -1.67 | 0.2625 | 0.1325 | 0.28 | 0.328794 | 0.657281 | 0.378554 | 0.999971 | 1 | 0.871816 |
| Aspscr1 | Q8VBT9 | -0.51 | 0.38 | 0.28 | -0.07 | 0.31 | -0.29 | 0.67 | -0.02 | 0.03 | -0.52 | 0.75 | 0.73 | 0.99 | -0.37 | -0.01 | 0.06 | 0.1475 | 0.2275 | 0.1475 | 0.623164 | 0.530217 | 0.673253 | 0.999971 | 1 | 0.94048 |
| Asrgl1 | Q8C0M9 | 0.25 | 0.76 | 0.49 | 0.3 | 0.57 | 0.72 | 0.34 | 0.84 | 0.49 | 0.28 | 0.17 | 0.04 | 0.73 | 0.88 | 0.34 | 0.21 | 0.1675 | -0.205 | 0.09 | 0.431896 | 0.324682 | 0.700673 | 0.999971 | 1 | 0.948876 |
| Ass1 | P16460 | 0.57 | 0.42 | 1.02 | 1.3 | 0.42 | 0.14 | 0.62 | 1.64 | 0.3 | 0.39 | 0.75 | 1.21 | 0.13 | 0.61 | 1.67 | 1.17 | -0.1225 | -0.165 | 0.0675 | 0.742877 | 0.582825 | 0.858512 | 0.999971 | 1 | 0.982702 |
| Astn1 | Q61137-2 | -1.11 | -1.54 | -1.14 | -1.06 | -1.3 | -1.38 | -1.14 | -1.13 | -1.39 | -1.33 | -0.85 | -1.09 | -1.32 | -1.49 | -1.42 | -1.13 | -0.025 | 0.0475 | -0.1275 | 0.896312 | 0.822266 | 0.523069 | 0.999971 | 1 | 0.907797 |
| Astn2 | Q80Z10 | 0.39 | -0.28 | -0.16 | -0.06 | -0.19 | 0.03 | -0.19 | -0.07 | -0.13 | -0.08 | -0.13 | -0.77 | 0.28 | -0.57 | -0.67 | -0.21 | -0.0775 | -0.25 | -0.265 | 0.710126 | 0.326405 | 0.352135 | 0.999971 | 1 | 0.867656 |
| Asxl2 | D3Z112 | 1.05 | 2.37 | -2.8 | -2.65 | 1.98 | 2 | 2.42 | -5.02 | 0.97 | 1.24 | 1.54 | -3.8 | 2.81 | 2.01 | -4.35 | -4.72 | 0.8525 | 0.495 | -0.555 | 0.665332 | 0.761333 | 0.794227 | 0.999971 | 1 | 0.970141 |
| Atad1 | Q9D5T0 | 0.21 | 0.31 | 0.02 | 0.15 | 0.19 | 0.97 | 0.59 | 0.5 | 0.43 | 0.7 | 0.2 | 0.09 | 0.92 | 0.82 | 0.24 | 0.29 | 0.39 | 0.1825 | 0.395 | 0.099938 | 0.375393 | 0.109184 | 0.999971 | 1 | 0.78268 |
| Atad2b | E9Q166 | 0.22 | -1.01 | -1.26 | -0.39 | -0.74 | -0.81 | -1.42 | -1.14 | -0.68 | 0.53 | -0.87 | -1.11 | -1.67 | -1.21 | -2.06 | -0.86 | -0.4175 | 0.0775 | -0.84 | 0.262983 | 0.86713 | 0.063614 | 0.999971 | 1 | 0.725348 |
| Atad3 | Q925I1 | -0.01 | -0.79 | -0.63 | -0.34 | -0.71 | -0.96 | -0.2 | -0.65 | -0.57 | -0.39 | -0.45 | -0.67 | -0.78 | -0.83 | -0.96 | -0.97 | -0.1875 | -0.0775 | -0.4425 | 0.473933 | 0.727796 | 0.072504 | 0.999971 | 1 | 0.743312 |
| Atat1 | Q8K341-4 | 0.19 | -1.44 | -0.42 | -0.34 | -0.73 | -1.05 | -0.03 | -0.39 | -0.56 | -0.04 | 0.21 | -0.12 | -0.46 | -1.04 | -0.23 | -0.01 | -0.0475 | 0.375 | 0.0675 | 0.902789 | 0.321591 | 0.862906 | 0.999971 | 1 | 0.982963 |
| Atcay | Q8BHE3 | -3.75 | -6.05 | -2.82 | -3 | -5.17 | -5.55 | -2.31 | -3.06 | -5.31 | -4.81 | -3.04 | -3.18 | -6.22 | -5.26 | -3 | -2.94 | -0.1175 | -0.18 | -0.45 | 0.903408 | 0.831809 | 0.651667 | 0.999971 | 1 | 0.933842 |
| Ate1 | A0A1L1SQ4 | 1.64 | 1.16 | 0.8 | 0.73 | 1.35 | 1.08 | 1.41 | 0.51 | 1.5 | 1.26 | 0.67 | 0.9 | 1.45 | 0.81 | 0.56 | 0.31 | 0.005 | -3.34E-16 | -0.3 | 0.986722 | 1 | 0.365815 | 0.999971 | 1 | 0.869526 |
| Atf2 | P16951 | 1.82 | 2.19 | 2.58 | 2.04 | 1.51 | 1.82 | 2.54 | 2.02 | 1.84 | 1.71 | 2.39 | 2.88 | 1.08 | 1.96 | 1.76 | 1.58 | -0.185 | 0.0475 | -0.5625 | 0.518859 | 0.880436 | 0.061279 | 0.999971 | 1 | 0.725348 |
| Atf7ip | Q7TT18 | 1.48 | 1.86 | 1.19 | 0.76 | 1 | 1.36 | 1.47 | 1.28 | 1.16 | 1.73 | 1.57 | 0.76 | 0.86 | 1.5 | 0.3 | 1.39 | -0.045 | -0.0175 | -0.31 | 0.868435 | 0.956365 | 0.392075 | 0.999971 | 1 | 0.871816 |
| Atg101 | Q9D8Z6 | 0.34 | -0.13 | -0.82 | -0.92 | -0.53 | -0.39 | -0.84 | -0.61 | -0.11 | -0.02 | -0.46 | -0.63 | 0.2 | 0.1 | -0.86 | -0.67 | -0.21 | 0.0775 | 0.075 | 0.511642 | 0.814533 | 0.846114 | 0.999971 | 1 | 0.979924 |
| Atg12 | Q9CQY1 | 0.94 | 2.37 | 1.62 | 1.56 | 1.93 | 2.84 | 1.42 | 1.46 | 2.59 | 2.6 | 1.95 | 1 | 1.85 | 1.92 | 1.24 | 1.5 | 0.29 | 0.4125 | 0.005 | 0.494429 | 0.371416 | 0.987986 | 0.999971 | 1 | 0.99925 |
| Atg13 | Q91YI1-2 | -1.25 | -0.38 | 0.4 | -0.13 | -0.42 | -0.32 | -0.11 | 0.87 | -0.37 | -1.76 | 0.92 | 0.44 | -0.83 | -0.57 | 0.13 | 0.91 | 0.345 | 0.1475 | 0.25 | 0.429779 | 0.812906 | 0.608801 | 0.999971 | 1 | 0.926608 |
| Atg14 | Q8CDJ3 | 2.36 | 1.63 | 1.46 | 1.12 | 1.11 | 1.87 | 1.93 | 1.94 | 1.62 | 2.11 | 1.5 | 1.07 | 1.1 | 1.52 | 1.56 | 0.8 | 0.07 | -0.0675 | -0.3975 | 0.831854 | 0.840621 | 0.23615 | 0.999971 | 1 | 0.832918 |
| Atg16l1 | Q8C0J2 | 0.95 | 0.19 | 0 | 0.07 | 0.11 | -0.44 | 0.07 | -0.58 | -0.25 | 0.34 | -0.02 | 0.05 | -0.76 | -0.29 | -0.8 | -0.26 | -0.5125 | -0.2725 | -0.83 | 0.106658 | 0.325659 | 0.015019 | 0.999971 | 1 | 0.613095 |
| Atg2a | Q6P4T0 | 0 | 0.05 | 0.06 | 0.01 | -0.19 | -0.07 | -0.16 | -0.08 | -0.39 | 0.13 | 0.02 | -0.11 | 0.21 | 0.42 | 0 | 0.3 | -0.155 | -0.1175 | 0.2025 | 0.34403 | 0.524355 | 0.268526 | 0.999971 | 1 | 0.842746 |
| Atg2b | Q80XK6 | 0.01 | 0 | 0.37 | 0.31 | 0 | -0.19 | 0.31 | 0.29 | -0.21 | -0.09 | 0.48 | 0.52 | -0.03 | -0.13 | 0.39 | 0.46 | -0.07 | 0.0025 | 5.89E-17 | 0.736465 | 0.991757 | 1 | 0.999971 | 1 | 1 |
| Atg3 | Q9CPX6 | -0.75 | -0.4 | 0.06 | -0.18 | -0.14 | -0.44 | -0.06 | 0.11 | -0.31 | -0.94 | 0.19 | 0.57 | -0.32 | -0.15 | 0.15 | 0.17 | 0.185 | 0.195 | 0.28 | 0.448313 | 0.591985 | 0.265835 | 0.999971 | 1 | 0.842677 |
| Atg4b | A0A0R4J06 | -0.21 | 0.2 | 0.79 | 1.13 | 0.33 | 0.33 | 0.24 | 1.33 | 0.38 | -0.01 | 0.42 | 0.43 | 0.46 | 0.3 | 0.97 | 1.14 | 0.08 | -0.1725 | 0.24 | 0.833778 | 0.591904 | 0.503341 | 0.999971 | 1 | 0.904824 |
| Atg4c | Q811C2 | 1.06 | 0.17 | 1.22 | 1.25 | 0.28 | 0.69 | -1.22 | -0.89 | 0.04 | 0.71 | 1.27 | 0.89 | -0.07 | -0.06 | 0.98 | 0.47 | -1.21 | -0.1975 | -0.595 | 0.032667 | 0.581415 | 0.118234 | 0.999971 | 1 | 0.78268 |
| Atg5 | Q99J83 | 0.12 | 0.97 | -0.38 | -1.03 | 0.7 | 0.5 | -0.35 | -1.65 | 0.75 | 0.23 | -0.23 | -0.2 | 0.57 | 0.33 | -1.38 | -1.41 | -0.12 | 0.2175 | -0.3925 | 0.846443 | 0.632925 | 0.532211 | 0.999971 | 1 | 0.910069 |
| Atg7 | A0A0A0M( | -0.33 | 0.17 | -0.25 | -0.11 | -0.01 | 0.19 | -0.23 | -0.09 | -0.03 | -0.37 | -0.2 | -0.22 | 0.1 | 0.06 | -0.13 | 0 | 0.095 | -0.075 | 0.1375 | 0.635813 | 0.695629 | 0.475687 | 0.999971 | 1 | 0.894045 |
| Atg9a | Q68FE2 | 0.03 | -0.16 | -0.08 | -0.47 | -0.05 | -0.21 | -0.13 | -0.49 | 0.12 | -0.31 | -0.14 | -0.1 | 0.59 | -0.66 | -0.3 | -0.19 | -0.05 | 0.0625 | 0.03 | 0.803887 | 0.750188 | 0.918986 | 0.999971 | 1 | 0.992633 |
| Atic | Q9CWJ9 | -0.13 | 0.38 | -0.06 | 0.1 | 0.41 | 0.26 | -0.2 | 0.19 | 0.13 | 0.02 | -0.09 | 0.04 | 0.35 | 0.53 | 0.16 | 0.38 | 0.0925 | -0.0475 | 0.2825 | 0.672467 | 0.799418 | 0.177698 | 0.999971 | 1 | 0.808532 |
| Atl1 | Q8BH66 | -0.67 | -0.64 | -0.7 | -1.1 | -0.65 | -0.24 | -0.6 | -0.96 | -0.44 | -0.61 | -0.49 | -0.81 | 0.15 | -0.32 | -0.64 | -0.9 | 0.165 | 0.19 | 0.35 | 0.46891 | 0.341415 | 0.217703 | 0.999971 | 1 | 0.826893 |
| Atl2 | Q6PA06 | 0.2 | 0.26 | 0.67 | 0.84 | 0.38 | 0.42 | 0.36 | 1.13 | 0.33 | 0.11 | 0.74 | 0.91 | 0.68 | 0.6 | 1.02 | 1.03 | 0.08 | 0.03 | 0.34 | 0.762734 | 0.908725 | 0.165879 | 0.999971 | 1 | 0.800215 |
| Atl3 | Q91YH5 | -0.89 | -0.61 | 0.16 | 0.32 | -0.24 | -0.63 | -0.3 | 0.31 | -0.46 | -0.96 | 0.18 | 0.73 | -0.46 | -0.41 | 0.47 | 0.28 | 0.04 | 0.1275 | 0.225 | 0.908094 | 0.774837 | 0.544741 | 0.999971 | 1 | 0.911398 |
| Atox1 | O08997 | 0.25 | 0.48 | 1.14 | 1.7 | 0.43 | 0.19 | 0.86 | 1.27 | 0.38 | 0.23 | 0.76 | 1.38 | 0.66 | 0.6 | 1.32 | 1.84 | -0.205 | -0.205 | 0.2125 | 0.601812 | 0.610721 | 0.614883 | 0.999971 | 1 | 0.927839 |
| Atp11a | E9Q3G7 | 0.1 | -0.64 | -0.43 | -0.27 | -0.6 | -0.67 | -0.29 | -0.71 | -0.49 | -0.33 | -0.61 | -0.43 | 0.02 | -0.69 | -0.28 | -0.38 | -0.2575 | -0.155 | -0.0225 | 0.269173 | 0.471916 | 0.926849 | 0.999971 | 1 | 0.994151 |
| Atp11b | Q6DFW5 | -0.88 | 1 | 1.24 | 0.42 | 0.73 | 0.91 | 0.88 | 1.14 | 0.98 | -0.11 | 0.95 | -0.54 | 1.39 | 0.47 | 1.21 | 0.68 | 0.47 | -0.125 | 0.4925 | 0.31351 | 0.824053 | 0.325264 | 0.999971 | 1 | 0.856472 |
| Atp11c | E9QKK8 | -1.43 | -1.02 | -0.89 | -1.24 | -1.02 | -1.16 | -1.05 | -1.22 | -0.79 | -1.49 | -1.45 | -0.91 | -0.5 | -0.69 | -1.09 | -1.14 | 0.0325 | -0.015 | 0.29 | 0.866163 | 0.951083 | 0.234339 | 0.999971 | 1 | 0.832918 |
| Atp13a1 | Q9EPE9 | -0.18 | 0.32 | 0.27 | 0.09 | 0.3 | 0.33 | -0.03 | 0.17 | 0.39 | 0.22 | 0.06 | 0.03 | 0.76 | 0.55 | 0.21 | 0.43 | 0.0675 | 0.05 | 0.3625 | 0.734941 | 0.799331 | 0.114093 | 0.999971 | 1 | 0.78268 |
| Atp13a2 | E9PYX5 | -0.65 | -0.18 | -2.28 | -2 | -0.58 | -0.02 | -1.99 | -0.9 | -0.25 | -0.02 | -0.94 | -1.19 | -0.4 | 0.08 | -0.77 | -1.55 | 0.405 | 0.6775 | 0.6175 | 0.50515 | 0.229082 | 0.291406 | 0.999971 | 1 | 0.847402 |
| Atp13a3 | Q5XF89 | 1.37 | 0.55 | 0.03 | -0.32 | 0.01 | 1.56 | -0.45 | 1.45 | 0.05 | 0.9 | 0.22 | -0.55 | -0.02 | 1.35 | 1.01 | 0.82 | 0.235 | -0.2525 | 0.3825 | 0.683115 | 0.575244 | 0.397423 | 0.999971 | 1 | 0.871816 |
| Atp13a4 | A0A0R4J0N | -1.07 | -2.12 | -2.06 | -1.13 | -1.54 | -2.15 | -1.79 | -0.39 | -1.88 | -1.55 | -1.36 | -1.22 | -2.22 | -1.35 | -1.02 | -1.88 | 0.1275 | 0.0925 | -0.0225 | 0.775921 | 0.773843 | 0.952667 | 0.999971 | 1 | 0.998056 |
| Atp1a1 | Q8VDN2 | -0.47 | -1.12 | -1.33 | -1.29 | -1.19 | -1.01 | -1.26 | -1.37 | -1.39 | -0.91 | -1.37 | -1.64 | -1.2 | -1.34 | -1.68 | -1.26 | -0.155 | -0.275 | -0.3175 | 0.530058 | 0.32152 | 0.231481 | 0.999971 | 1 | 0.8307 |
| Atp1a2 | Q6PIE5 | -1.03 | -1.55 | -1.01 | -0.98 | -1.59 | -1.23 | -1.25 | -1.11 | -1.46 | -1.21 | -1.33 | -2.24 | -1.44 | -1.36 | -1.05 | -0.97 | -0.1525 | -0.4175 | -0.0625 | 0.488051 | 0.165022 | 0.778649 | 0.999971 | 1 | 0.965656 |
| Atp1a3 | A0A0G2JG) | -0.76 | -1.36 | -1.89 | -1.66 | -1.52 | -1.35 | -1.94 | -1.76 | -1.5 | -1.01 | -2.02 | -2.04 | -1.27 | -1 | -1.79 | -1.58 | -0.225 | -0.225 | 0.0075 | 0.444466 | 0.516159 | 0.980481 | 0.999971 | 1 | 0.99925 |
| Atp1b1 | P14094 | -1.71 | -1.81 | -2.05 | -2.03 | -1.89 | -1.24 | -1.78 | -2.02 | -1.86 | -1.8 | -1.58 | -1.74 | -1.8 | -1.45 | -1.71 | -1.76 | 0.1675 | 0.155 | 0.22 | 0.471649 | 0.393456 | 0.25833 | 0.999971 | 1 | 0.839012 |
| Atp1b2 | P14231 | -2.38 | -2.64 | -1.33 | -1.42 | -2.28 | -2.65 | -1.22 | -1.43 | -2.25 | -2.66 | -1.4 | -1.25 | -2.04 | -2.39 | -1.48 | -1.51 | 0.0475 | 0.0525 | 0.0875 | 0.915312 | 0.906534 | 0.820172 | 0.999971 | 1 | 0.975761 |
| Atp1b3 | P97370 | 0.55 | -0.85 | 0.08 | 0.15 | -0.32 | -0.82 | -0.72 | -0.35 | -0.63 | -0.08 | -0.65 | 0.03 | -0.41 | -0.52 | 0.19 | 0.16 | -0.535 | -0.315 | -0.1275 | 0.125229 | 0.36921 | 0.713166 | 0.999971 | 1 | 0.951793 |
| Atp23 | G3UW46 | 1.05 | 1.44 | -0.37 | -1.13 | 0.9 | 0.5 | -0.25 | 0.08 | -0.21 | 0.45 | 0.21 | -0.08 | 1.35 | 0.01 | 0.15 | -0.51 | 0.06 | -0.155 | 0.0025 | 0.919593 | 0.786277 | 0.996944 | 0.999971 | 1 | 1 |
| Atp2a2 | O55143 | -0.48 | -0.19 | -0.51 | -0.33 | -0.48 | -0.09 | -0.67 | -0.25 | -0.46 | -0.35 | -0.35 | -1.02 | -0.3 | -0.08 | -0.46 | -0.31 | 0.005 | -0.1675 | 0.09 | 0.980339 | 0.450587 | 0.626351 | 0.999971 | 1 | 0.930479 |
| Atp2a3 | B1ATS5 | 1.59 | -0.15 | 0.1 | 0.55 | 2.69 | 0.33 | 0.09 | 0.42 | 0.52 | 2.61 | 0.68 | 1.26 | -1.26 | 0.11 | 0.2 | 0.7 | 0.36 | 0.745 | -0.585 | 0.583734 | 0.208878 | 0.283352 | 0.999971 | 1 | 0.844538 |
| Atp2b1 | G5E829 | -1.24 | -0.6 | 0.23 | 0.15 | -1.19 | -1.02 | -0.1 | -0.01 | -1.21 | -1.32 | -0.19 | -1.93 | -0.76 | -1.19 | -0.28 | -0.17 | -0.215 | -0.7975 | -0.235 | 0.623932 | 0.116969 | 0.561501 | 0.999971 | 1 | 0.916133 |
| Atp2b1_1 | A0A1W2P7 | 0.12 | -1 | -1.7 | -1.37 | -1.57 | -1.33 | -1.43 | -1.61 | -1.28 | -0.7 | -1.6 | -1.53 | -1.77 | -1.18 | -1.43 | -1.46 | -0.4975 | -0.29 | -0.4725 | 0.21902 | 0.49945 | 0.253602 | 0.999971 | 1 | 0.839012 |
| Atp2b2 | F8WHB1 | -0.93 | -1.16 | -1.23 | -1.37 | -1.58 | -1.23 | -1.28 | -1.58 | -1.61 | -1.43 | -1.21 | -1.45 | -1.36 | -1.45 | -1.64 | -1.28 | -0.245 | -0.2525 | -0.26 | 0.229134 | 0.201073 | 0.193526 | 0.999971 | 1 | 0.810547 |
| Atp2b2_1 | S4R1C4 | -1.19 | -0.09 | -0.7 | -0.75 | -0.78 | 0.28 | -1.05 | -0.51 | -0.84 | -0.75 | -0.48 | -0.92 | 0.64 | 0.23 | -0.61 | -0.36 | 0.1675 | -0.065 | 0.6575 | 0.641139 | 0.806506 | 0.091947 | 0.999971 | 1 | 0.773779 |
| Atp2b3 | Q0VF55 | -0.68 | -1.16 | -0.67 | -0.79 | -1.52 | -1.09 | -0.75 | -0.8 | -1.31 | -0.92 | -1.2 | -1.49 | -1.46 | -1.39 | -1.11 | -0.98 | -0.215 | -0.405 | -0.41 | 0.385944 | 0.082603 | 0.079976 | 0.999971 | 1 | 0.755254 |
| Atp2b3_1 | A2ALL9 | -1.42 | -0.75 | -0.76 | -1.09 | -1.21 | -0.25 | -1.25 | -0.89 | -0.79 | -0.69 | -0.81 | -0.83 | -0.17 | -0.39 | -0.64 | -0.4 | 0.105 | 0.225 | 0.605 | 0.719869 | 0.29886 | 0.024228 | 0.999971 | 1 | 0.669136 |
| Atp2b4 | E9Q828 | -2.05 | -1.73 | -1.45 | -1.63 | -1.48 | -1.5 | -0.64 | -1.31 | -1.61 | -1.96 | -1.26 | -1.51 | -0.95 | -1.77 | -1.37 | -1.35 | 0.4825 | 0.13 | 0.355 | 0.091683 | 0.572305 | 0.167573 | 0.999971 | 1 | 0.800215 |
| Atp2c1 | H3BL44 | -0.31 | 0.23 | 0.23 | 0.28 | 0.3 | 0.67 | 0.06 | 0.41 | 0.51 | 0.11 | 0.27 | 0.66 | 0.61 | 0.62 | 0.71 | 0.43 | 0.2525 | 0.28 | 0.485 | 0.284751 | 0.232379 | 0.040397 | 0.999971 | 1 | 0.700838 |
| Atp5c1 | A2AKU9 | 0.35 | -0.4 | -1.09 | -1.05 | -0.13 | 0.14 | -0.18 | -1.45 | -0.02 | -0.17 | -1.17 | -1.49 | -0.42 | -1.04 | -1.41 | -1.83 | 0.1425 | -0.165 | -0.6275 | 0.75687 | 0.724302 | 0.16773 | 0.999971 | 1 | 0.800215 |
| Atp5c1_1 | Q8C2Q8 | 0.08 | -0.57 | -1.79 | -0.75 | -1.42 | -2.03 | -1.34 | -0.98 | -1.37 | -0.96 | -0.75 | -1.53 | -1.83 | -1.3 | -2.24 | -1.28 | -0.685 | -0.395 | -0.905 | 0.130954 | 0.346986 | 0.059637 | 0.999971 | 1 | 0.725348 |
| Atp5f1a | Q03265 | -1.06 | -1.19 | -1.72 | -1.38 | -1.31 | -1.31 | -1.44 | -1.5 | -1.64 | -1.3 | -1.55 | -1.45 | -1.14 | -1.18 | -1.31 | -1.42 | -0.0525 | -0.1475 | 0.075 | 0.798352 | 0.485674 | 0.7209 | 0.999971 | 1 | 0.954713 |
| Atp5f1b | P56480 | -0.52 | -0.87 | -1.18 | -0.43 | -0.88 | -0.79 | -1.3 | -0.68 | -1.16 | -0.67 | -1.36 | -1.19 | -1.06 | -0.55 | -1.06 | -0.33 | -0.1625 | -0.345 | 0 | 0.517184 | 0.194528 | 1 | 0.999971 | 1 | 1 |
| Atp5f1d | Q9D3D9 | -1.32 | 0 | -0.26 | -0.39 | -0.35 | 0.19 | -0.43 | -0.3 | -0.54 | -1.36 | -0.96 | -0.76 | -0.37 | 0.08 | -0.34 | -0.61 | 0.27 | -0.4125 | 0.1825 | 0.410731 | 0.237856 | 0.575861 | 0.999971 | 1 | 0.918194 |
| Atp5f1e | P56382 | 0.86 | 0.98 | 0.65 | 0.13 | -0.17 | 1.05 | 2.25 | 0.34 | -0.02 | 0.96 | -0.05 | 0.29 | 0.99 | 0.45 | 0.63 | 0.4 | 0.2125 | -0.36 | -0.0375 | 0.681032 | 0.259017 | 0.883708 | 0.999971 | 1 | 0.986782 |
| Atp5j2 | F8WHP8 | -0.96 | -0.48 | -0.7 | -0.25 | -0.29 | -0.73 | -1.06 | -0.1 | -0.85 | -0.52 | -0.61 | -0.6 | -0.54 | -0.24 | -0.26 | -0.06 | 0.0525 | -0.0475 | 0.3225 | 0.851228 | 0.823321 | 0.174393 | 0.999971 | 1 | 0.807788 |
| Atp5mc1 | Q9CR84 | 0.46 | -0.12 | 1.74 | 0.94 | -2.24 | 0.48 | 1.98 | 0.64 | -0.6 | -0.69 | 1.95 | -0.77 | -1.37 | -0.21 | -0.01 | -1.13 | -0.54 | -0.7825 | -1.435 | 0.53804 | 0.280981 | 0.014861 | 0.999971 | 1 | 0.613095 |
| Atp5me | Q06185 | -2.22 | -1.62 | -2.15 | -1.78 | -2.16 | -2.05 | -1.58 | -2.91 | -2.74 | -2.55 | -2.57 | -2.64 | -1.71 | -1.39 | -1.65 | -1.46 | -0.2325 | -0.6825 | 0.39 | 0.46714 | 0.008036 | 0.094278 | 0.999971 | 1 | 0.779538 |
| Atp5mg | Q9CPQ8 | -3.33 | -3.5 | -2.79 | -2.59 | -3.43 | -3.48 | -2.38 | -3.12 | -3.75 | -3.57 | -2.95 | -3.14 | -3.65 | -3.12 | -2.18 | -2.27 | -0.05 | -0.3 | 0.2475 | 0.880214 | 0.322639 | 0.53722 | 0.999971 | 1 | 0.910077 |
| Atp5mj | P56379 | -3.87 | -1.17 | -0.94 | -2.13 | -1.12 | -2.32 | 0.1 | -2.04 | -1.39 | -4.48 | -1.28 | -0.74 | -1.02 | -2.41 | -1.34 | -3.2 | 0.6825 | 0.055 | 0.035 | 0.389014 | 0.954741 | 0.962827 | 0.999971 | 1 | 0.998721 |
| Atp5mk | Q78IK2 | -0.63 | -0.69 | -0.47 | -0.11 | -0.37 | -0.46 | -0.78 | 0 | -0.75 | -0.48 | -0.4 | -0.25 | -0.43 | -0.61 | -0.16 | 0.25 | 0.0725 | 0.005 | 0.2375 | 0.762633 | 0.981192 | 0.363185 | 0.999971 | 1 | 0.869526 |
| Atp5pb | Q9CQQ7 | -0.57 | 0.07 | 0.36 | 0.17 | -0.32 | 0.1 | 0.21 | 0.24 | -0.45 | -0.37 | 0.21 | 0.3 | 0.11 | -0.12 | 0.59 | 0.92 | 0.05 | -0.085 | 0.3675 | 0.848763 | 0.77 | 0.260691 | 0.999971 | 1 | 0.841256 |
| Atp5pd | Q9DCX2 | -2.82 | -3.06 | -2.19 | -2.56 | -2.87 | -3.04 | -1.81 | -2.35 | -3.06 | -3.03 | -2.15 | -2.25 | -3.49 | -2.4 | -2.36 | -2.33 | 0.14 | 0.035 | 0.0125 | 0.67547 | 0.910561 | 0.970327 | 0.999971 | 1 | 0.998787 |
| Atp5pf | P97450 | -0.52 | -1.31 | -1.77 | -0.86 | -1.63 | -1.69 | -1.38 | -1.18 | -1.75 | -1.36 | -1.34 | -1.86 | -1.81 | -0.82 | -1.68 | -0.88 | -0.355 | -0.4625 | -0.1825 | 0.260781 | 0.158969 | 0.62102 | 0.999971 | 1 | 0.929388 |
| Atp5po | Q9DB20 | -0.46 | -0.47 | -0.7 | -0.75 | -0.62 | -0.59 | -0.58 | -0.69 | -0.78 | -0.53 | -0.86 | -0.9 | -0.29 | -0.27 | -0.78 | -0.78 | -0.025 | -0.1725 | 0.065 | 0.884126 | 0.356057 | 0.760528 | 0.999971 | 1 | 0.961116 |
| Atp6ap1 | Q9R1Q9 | -1.36 | -1.07 | -1.2 | -1.44 | -1.03 | -1.26 | -1.02 | -1.51 | -1.01 | -1.25 | -0.78 | -1.03 | -1.4 | -1.14 | -1.14 | -1.1 | 0.0625 | 0.25 | 0.0725 | 0.755502 | 0.208964 | 0.694155 | 0.999971 | 1 | 0.947174 |
| Atp6ap2 | Q9CYN9 | -1.11 | -0.94 | -1 | -1.43 | -0.35 | -1.2 | -1.17 | -1.51 | -0.5 | -1.68 | -0.58 | -0.22 | -0.34 | -1.29 | -1 | -1.28 | 0.0625 | 0.375 | 0.1425 | 0.826328 | 0.282773 | 0.598977 | 0.999971 | 1 | 0.924014 |
| Atp6v0a1 | K3W4T3 | -1.14 | -1.3 | -1.59 | -1.62 | -1.47 | -1.29 | -1.33 | -1.64 | -1.5 | -1.39 | -1.33 | -1.72 | -1.49 | -1.25 | -1.65 | -1.46 | -0.02 | -0.0725 | -0.05 | 0.920023 | 0.716234 | 0.803696 | 0.999971 | 1 | 0.971467 |
| Atp6v0a1_ | Q9Z1G4 | 0.37 | 0.85 | 1.08 | 0.54 | 0.88 | -0.15 | -0.37 | 0.67 | -0.72 | -0.37 | -0.07 | 1.03 | -0.03 | -0.64 | -0.09 | 0.51 | -0.4525 | -0.7425 | -0.7725 | 0.206104 | 0.087299 | 0.026079 | 0.999971 | 1 | 0.678474 |
| Atp6v0a2 | P15920 | 1.37 | 0.4 | 0.48 | 0.71 | 0.41 | 0.21 | 0.68 | 0.35 | 0.36 | 1.01 | 0.08 | 0.08 | 0.02 | 0.24 | 0.65 | 0.68 | -0.3275 | -0.3575 | -0.3425 | 0.233978 | 0.273294 | 0.251459 | 0.999971 | 1 | 0.839012 |
| Atp6v0c | E9Q9C5 | -1.96 | -0.06 | -0.44 | -1.01 | -0.27 | -0.4 | -0.7 | -0.27 | -0.62 | -1.7 | 0.01 | -0.81 | -0.82 | -0.1 | -0.49 | -0.5 | 0.4575 | 0.0875 | 0.39 | 0.277119 | 0.862683 | 0.362421 | 0.999971 | 1 | 0.869526 |
| Atp6v0d1 | P51863 | -1.13 | -0.77 | -0.14 | -0.08 | -0.97 | -0.61 | -0.42 | 0.17 | -1.01 | -0.99 | -0.14 | -1.03 | -0.65 | -0.55 | 0.39 | 0.03 | 0.0725 | -0.2625 | 0.335 | 0.832888 | 0.438572 | 0.349535 | 0.999971 | 1 | 0.865435 |
| Atp6v1a | P50516 | -1.16 | -1.12 | -0.3 | -0.59 | -0.97 | -1.23 | -0.18 | -0.12 | -1.1 | -1.67 | -0.07 | 0.03 | -0.68 | -1.2 | -0.17 | -0.18 | 0.1675 | 0.09 | 0.235 | 0.629001 | 0.836833 | 0.475175 | 0.999971 | 1 | 0.894044 |
| Atp6v1b2 | P62814 | -0.88 | -1.33 | -0.94 | -0.93 | -1.36 | -1.45 | -0.81 | -1.11 | -1.59 | -1.43 | -0.89 | -1.02 | -1.43 | -1.19 | -0.98 | -0.75 | -0.1625 | -0.2125 | -0.0675 | 0.468717 | 0.368693 | 0.761898 | 0.999971 | 1 | 0.961309 |
| Atp6v1c1 | Q9Z1G3 | -1.11 | -0.82 | -0.42 | -0.89 | -1.01 | -0.86 | -0.12 | -0.69 | -0.8 | -1.08 | -0.22 | -0.43 | -0.96 | -1 | -0.55 | -0.81 | 0.14 | 0.1775 | -0.02 | 0.598613 | 0.501885 | 0.927794 | 0.999971 | 1 | 0.994229 |
| Atp6v1d | P57746 | -1.16 | -1.04 | -0.9 | -0.86 | -1.13 | -1.02 | -0.93 | -0.83 | -1.04 | -1.36 | -0.68 | -0.6 | -0.94 | -1.18 | -0.71 | -0.64 | 0.0125 | 0.07 | 0.1225 | 0.943639 | 0.756757 | 0.544308 | 0.999971 | 1 | 0.911398 |
| Atp6v1e1 | P50518 | -0.74 | 0.18 | -1.19 | -1.18 | 0.93 | -0.24 | -0.66 | -1.26 | 1.29 | -0.97 | -0.5 | -0.32 | 2.51 | -0.65 | -1.19 | -0.84 | 0.425 | 0.6075 | 0.69 | 0.423019 | 0.2811 | 0.412048 | 0.999971 | 1 | 0.875856 |
| Atp6v1f | Q9D1K2 | -0.35 | -0.07 | -0.86 | -1.14 | 0.08 | -0.04 | -0.62 | -0.96 | 0.21 | -1.36 | -0.11 | -0.24 | 1.47 | -0.08 | -0.88 | -1 | 0.22 | 0.23 | 0.4825 | 0.523419 | 0.570178 | 0.405163 | 0.999971 | 1 | 0.874565 |
| Atp6v1g1 | Q9CR51 | -0.09 | -0.16 | 0.19 | 0.36 | -0.32 | -0.19 | -0.01 | 0.38 | -0.09 | 0.73 | 0.49 | 0.51 | 0.01 | -0.1 | 0.46 | 1 | -0.11 | 0.335 | 0.2675 | 0.637138 | 0.19076 | 0.369435 | 0.999971 | 1 | 0.869526 |
| Atp6v1g2 | Q9WTT4 | 0.1 | -0.82 | -0.83 | -0.87 | -1.17 | -0.45 | -0.49 | -1.03 | -0.9 | -0.26 | -0.9 | -1.52 | -1.47 | -0.71 | -0.89 | -1.49 | -0.18 | -0.29 | -0.535 | 0.559533 | 0.409209 | 0.115321 | 0.999971 | 1 | 0.78268 |
| Atp6v1h | A0A0A6YW | 1.78 | 3.27 | 3.13 | 2.64 | 2.16 | 4.52 | 2.12 | 2.6 | 2.32 | 4.16 | 2.16 | -0.14 | 3.89 | 3.61 | 2.48 | 3.47 | 0.145 | -0.58 | 0.6575 | 0.809668 | 0.501658 | 0.153269 | 0.999971 | 1 | 0.795578 |
| Atp6v1h_1 | Q8BVE3 | -1.09 | -1.23 | -1.16 | -1.01 | -1.15 | -0.71 | -1.02 | -1.17 | -1.19 | -1.05 | -1.04 | -1.01 | -1.17 | -0.63 | -0.94 | -0.79 | 0.11 | 0.05 | 0.24 | 0.56004 | 0.758152 | 0.231023 | 0.999971 | 1 | 0.830179 |
| Atp7a | A2AG68 | -1.03 | -0.11 | -0.27 | -0.34 | 0.35 | -0.11 | -0.7 | 0.04 | 0.08 | -1.26 | -0.59 | -0.1 | -0.36 | -0.01 | -0.21 | -0.35 | 0.3325 | -0.03 | 0.205 | 0.293947 | 0.932599 | 0.418894 | 0.999971 | 1 | 0.87788 |
| Atp8a1 | P70704 | -1.07 | -0.97 | -0.76 | -0.89 | -0.83 | -0.87 | -0.64 | -0.75 | -0.9 | -1.03 | -0.63 | -0.95 | -0.65 | -0.98 | -0.79 | -0.83 | 0.15 | 0.045 | 0.11 | 0.395575 | 0.803044 | 0.540852 | 0.999971 | 1 | 0.910478 |
| Atp8a2 | P98200 | -1.16 | -0.6 | -0.07 | -0.39 | -0.51 | -0.34 | -0.22 | -0.19 | -0.17 | -0.66 | -0.07 | -0.41 | 0.08 | -0.5 | -0.35 | -0.47 | 0.24 | 0.2275 | 0.245 | 0.372695 | 0.42388 | 0.394369 | 0.999971 | 1 | 0.871816 |
| Atp9a | A2AQC3 | -0.52 | -0.54 | -0.39 | -0.16 | -0.45 | 0.02 | 0.16 | 0.05 | -0.19 | -0.25 | 0.24 | -0.08 | 0.15 | 0.18 | 0.12 | 0.06 | 0.3475 | 0.3325 | 0.53 | 0.12621 | 0.117704 | 0.014158 | 0.999971 | 1 | 0.613095 |
| Atp9b | A0A286YC\ | -0.15 | -0.31 | 0.28 | 0.06 | -0.63 | -0.4 | -0.03 | 0.19 | -0.56 | -0.35 | -0.25 | -0.8 | -0.27 | -0.2 | 0.05 | 0.03 | -0.1875 | -0.46 | -0.0675 | 0.462776 | 0.060624 | 0.743904 | 0.999971 | 1 | 0.961064 |
| Atpaf1 | H3BLL2 | 0 | -0.23 | -0.39 | -0.25 | -0.37 | -0.8 | -0.64 | -0.42 | -1.03 | -0.8 | -0.29 | -0.31 | -1.11 | -0.69 | -0.48 | -0.74 | -0.34 | -0.39 | -0.5375 | 0.105345 | 0.122658 | 0.027444 | 0.999971 | 1 | 0.678474 |
| Atpaf2 | Q91YY4 | -1.03 | -0.58 | -0.69 | -0.55 | -0.31 | -0.36 | -0.98 | -0.07 | -0.67 | -0.78 | -0.85 | -0.58 | -0.2 | -0.07 | -0.46 | -0.62 | 0.2825 | -0.0075 | 0.375 | 0.277272 | 0.968212 | 0.107798 | 0.999971 | 1 | 0.78268 |
| Atpif1 | E9PV44 | -0.02 | -2.15 | -1.13 | -1.54 | -1.99 | -2 | 0.03 | -1.64 | -2.28 | -0.68 | -0.85 | -1.86 | -3.81 | -2.53 | -1.93 | -2.77 | -0.19 | -0.2075 | -1.55 | 0.752738 | 0.706006 | 0.018901 | 0.999971 | 1 | 0.635388 |
| Atpsckmt | E0CXY7 | 0.13 | -0.53 | 0.05 | -0.51 | -0.3 | -0.52 | -0.04 | 0.16 | 0.13 | 0.65 | 0.08 | 0.14 | 0.32 | -0.16 | -0.36 | 0.27 | 0.04 | 0.465 | 0.2325 | 0.875893 | 0.088584 | 0.390554 | 0.999971 | 1 | 0.871816 |
| Atraid | G3UXG8 | 0.99 | 2.05 | 2.72 | 1.67 | 1.94 | 2.02 | 4.56 | 1.47 | 1.81 | 2.29 | 1.74 | -1.27 | 0.28 | 1.47 | 0.4 | -1.7 | 0.64 | -0.715 | -1.745 | 0.378765 | 0.385857 | 0.029238 | 0.999971 | 1 | 0.678474 |
| Atrn | Q9WU60 | -0.1 | -0.17 | 0.15 | -0.02 | -0.38 | 0.48 | 0 | 0.49 | -0.27 | -0.03 | 0.12 | 0.08 | 0.22 | 0.17 | 0.32 | -0.14 | 0.1825 | 0.01 | 0.1775 | 0.470694 | 0.956034 | 0.36211 | 0.999971 | 1 | 0.869526 |
| Atrnl1 | Q6A051 | 1.06 | 0.16 | 0.5 | 0.57 | 0.09 | 0.82 | 0.46 | 0.44 | 0.34 | 0.86 | 0.54 | 0.15 | -0.29 | 0.32 | 0.34 | 0.24 | -0.12 | -0.1 | -0.42 | 0.647572 | 0.702582 | 0.135098 | 0.999971 | 1 | 0.784948 |
| Atrx | Q61687 | -0.1 | -0.59 | -0.56 | -0.69 | -0.64 | -0.72 | -0.56 | -0.67 | -0.66 | -0.46 | -0.27 | -0.5 | -0.52 | -0.73 | -0.59 | -0.73 | -0.1625 | 0.0125 | -0.1575 | 0.417973 | 0.951167 | 0.440502 | 0.999971 | 1 | 0.883277 |
| Atxn1 | J3QPR1 | 3.05 | 2.09 | 2.48 | 2.89 | 1.95 | 1.83 | 1.96 | 2.85 | 1.85 | 2.59 | 2.13 | 2.39 | 1.3 | 2.28 | 2.46 | 2.76 | -0.48 | -0.3875 | -0.4275 | 0.160805 | 0.193429 | 0.26859 | 0.999971 | 1 | 0.842746 |

| | | | | | | | | | | | | | | | | | | | | | | | | | | |
|---|---|---|---|---|---|---|---|---|---|---|---|---|---|---|---|---|---|---|---|---|---|---|---|---|---|---|
| Atxn10 | P28658 | -0.15 | 0.61 | 0.44 | 0.53 | 0.76 | 0.61 | 0.23 | 0.72 | 0.68 | 0.16 | 0.34 | 0.59 | 0.77 | 0.98 | 0.9 | 0.99 | 0.2225 | 0.085 | 0.5525 | 0.370415 | 0.723756 | 0.033031 | 0.999971 | 1 | 0.685881 |
| Atxn1l | P0C7T6 | 2.37 | 1.64 | 0.72 | 0.44 | 1.38 | 1.87 | 1.83 | 0.41 | 1.27 | 2.99 | 0.77 | 1.3 | 1.52 | 2.41 | 0.23 | 0.02 | 0.08 | 0.29 | -0.2475 | 0.876406 | 0.631969 | 0.704547 | 0.999971 | 1 | 0.950182 |
| Atxn2 | E9QM77 | 1.13 | 1.79 | 1.23 | 1.47 | 1.58 | 2.07 | 1.46 | 1.52 | 1.55 | 1.57 | 1.05 | 1.29 | 1.41 | 1.79 | 1.45 | 1.41 | 0.2525 | -0.04 | 0.11 | 0.302541 | 0.860572 | 0.617942 | 0.999971 | 1 | 0.928581 |
| Atxn2_1 | F6U2C2 | 0.15 | 1.38 | 2.04 | 2.24 | -0.06 | 0.72 | 1.34 | 2.31 | 0.34 | -0.26 | 1.83 | 1.57 | 0.83 | 1.26 | 1.37 | 2.25 | -0.375 | -0.5825 | -0.025 | 0.552982 | 0.365569 | 0.961344 | 0.999971 | 1 | 0.998721 |
| Atxn2l | Q7TQH0-2 | 1.59 | 1.62 | 1.73 | 1.69 | 1.5 | 1.34 | 1.5 | 1.36 | 1.64 | 1.69 | 1.48 | 1.81 | 1.05 | 1.21 | 1.29 | 1.55 | -0.2325 | -0.0025 | -0.3825 | 0.17971 | 0.988008 | 0.063589 | 0.999971 | 1 | 0.725348 |
| Atxn3 | Q9CVD2 | 2.7 | 0.35 | 0.05 | 0.33 | 0.39 | 0.35 | 0.22 | 0.29 | 0.23 | 0.78 | 0.06 | 0 | -0.38 | -0.48 | 0.12 | -0.68 | -0.545 | -0.59 | -1.2125 | 0.34968 | 0.332463 | 0.064758 | 0.999971 | 1 | 0.726851 |
| Atxn7l3b | Q3UD01 | 1.74 | 1.28 | 0.66 | 0.33 | 1.39 | 1.04 | 0.12 | -0.01 | 1.09 | 1.6 | 0.69 | 1.18 | 1.05 | 1.17 | 0.7 | 0.88 | -0.3675 | 0.1375 | -0.0525 | 0.412092 | 0.702192 | 0.873966 | 0.999971 | 1 | 0.984109 |
| Auh | A0A0R4J02 | 0.13 | 0.03 | -0.52 | -0.53 | -0.48 | -0.16 | -0.79 | -0.92 | -0.34 | -0.36 | -0.67 | -0.73 | -0.07 | -0.06 | -0.92 | -0.47 | -0.365 | -0.3025 | -0.1575 | 0.192683 | 0.220873 | 0.582437 | 0.999971 | 1 | 0.920364 |
| Aup1 | P70295 | -0.75 | -1.38 | -0.42 | -0.19 | -1.05 | -1.24 | -0.85 | -0.36 | -1.68 | -1.16 | -0.62 | -0.77 | -1.99 | -0.87 | -0.57 | -0.19 | -0.19 | -0.3725 | -0.22 | 0.559041 | 0.297755 | 0.619252 | 0.999971 | 1 | 0.929184 |
| Aurkaip1 | Q9DCJ7 | 1.08 | 0.15 | -0.3 | -0.76 | -0.81 | -0.16 | 0.02 | -0.22 | -0.69 | 0.17 | 1.39 | 1.33 | 0.58 | 1 | 1.07 | 0.71 | -0.335 | 0.5075 | 0.7975 | 0.423871 | 0.395865 | 0.069045 | 0.999971 | 1 | 0.733937 |
| Aven | A2AGL5 | 1.14 | 0.7 | -0.42 | -0.49 | -0.85 | 0.52 | -0.23 | -0.89 | 0.29 | 0.94 | 0.15 | 0.63 | 0.45 | 0.15 | -0.21 | -0.33 | -0.595 | 0.27 | -0.2175 | 0.242503 | 0.527585 | 0.608745 | 0.999971 | 1 | 0.926608 |
| Avl9 | Q80U56 | -1.27 | -0.52 | -0.99 | -1.38 | -0.52 | -0.53 | -0.55 | -1.28 | -0.39 | -0.95 | -0.95 | -0.94 | -0.35 | -0.85 | -0.98 | -1.31 | 0.32 | 0.2325 | 0.1675 | 0.274783 | 0.380731 | 0.566495 | 0.999971 | 1 | 0.917959 |
| AW209491 | Q91WD4 | 2.47 | 0.89 | 1.98 | 1.96 | -0.41 | 1.6 | 3.52 | 0.44 | 0.31 | 1.49 | 1.44 | 0.17 | 0.4 | 0.36 | 0.92 | -0.25 | -0.5375 | -0.9725 | -1.4675 | 0.517278 | 0.059352 | 0.00471 | 0.999971 | 1 | 0.582714 |
| AW551984 | Q8BGF0 | -1.22 | -1.55 | -0.81 | -0.88 | -0.44 | -1.42 | -0.96 | 0.06 | -0.16 | -0.84 | -1.13 | 0.51 | 0.63 | 0.18 | 1.11 | -0.48 | 0.425 | 0.71 | 1.475 | 0.250946 | 0.096111 | 0.003141 | 0.999971 | 1 | 0.577271 |
| Axl | Q6PE80 | -1.05 | -0.42 | -1.54 | -1.3 | -0.65 | -0.62 | -0.54 | -1.53 | -0.03 | -0.82 | -1.23 | -1.19 | 0.12 | -0.55 | -0.56 | -0.54 | 0.2425 | 0.26 | 0.695 | 0.473756 | 0.476817 | 0.044049 | 0.999971 | 1 | 0.70308 |
| Azi2 | Q9QYP6 | 2.33 | -0.29 | 0.25 | 0.47 | -0.25 | 0.31 | 0.93 | 0.02 | -0.24 | 1.56 | 0.2 | -0.06 | -0.42 | -0.1 | -0.01 | -0.1 | -0.4375 | -0.325 | -0.8475 | 0.451102 | 0.614571 | 0.138055 | 0.999971 | 1 | 0.784948 |
| B2m | P01887 | 1.55 | 1.04 | 1.1 | 1.63 | 2.5 | 0.14 | 1.93 | 0.93 | 2.63 | 0.47 | 1.63 | 3.04 | 1.8 | 0.24 | 1.17 | 1.17 | 0.045 | 0.6125 | -0.235 | 0.928964 | 0.281456 | 0.507547 | 0.999971 | 1 | 0.905815 |
| B3galnt1 | Q920V1 | -1.75 | -1.27 | -0.44 | -0.22 | -0.75 | -1.35 | -1.06 | 0.34 | -0.95 | -2.05 | -0.35 | -0.46 | -1.57 | -0.82 | 0.31 | 0.1 | 0.215 | -0.0325 | 0.425 | 0.654977 | 0.947338 | 0.423139 | 0.999971 | 1 | 0.87887 |
| B3galt5 | Q9JI67 | 3.11 | 1.63 | 1.42 | 1.73 | 0.84 | 1.11 | 1.09 | 1.75 | 1.5 | 1.17 | 1.7 | 2.38 | -0.11 | 1.27 | 1.64 | 0.83 | -0.775 | -0.285 | -1.065 | 0.085927 | 0.519494 | 0.058578 | 0.999971 | 1 | 0.725348 |
| B3galt6 | Q91Z92 | 0.69 | 0.57 | 0.89 | 0.97 | 0.3 | 0.89 | 0.37 | 0.15 | 0.68 | 0.31 | 0.93 | 0.62 | 0.66 | 1.05 | 0.92 | 0.98 | -0.3525 | -0.145 | 0.1225 | 0.144322 | 0.488014 | 0.528118 | 0.999971 | 1 | 0.909725 |
| B3gat3 | P58158 | 1.7 | 1.02 | 0.85 | 1.03 | 1.21 | 1.13 | 0.71 | 1.29 | 0.97 | 1.31 | 1.2 | 0.72 | 0.07 | 0.71 | 0.88 | 0.76 | -0.065 | -0.1 | -0.545 | 0.798315 | 0.694182 | 0.076991 | 0.999971 | 1 | 0.750479 |
| B3glct | Q8BHT6 | -0.71 | 0.3 | 0.28 | 0.65 | 0.12 | -0.09 | 0.2 | 0.82 | 0.13 | -0.12 | 0.72 | 0.18 | 0.13 | 0.41 | 0.69 | 1.09 | 0.1325 | 0.0975 | 0.45 | 0.703265 | 0.773814 | 0.22215 | 0.999971 | 1 | 0.827753 |
| B4galnt1 | Q09200 | 0.12 | 0.98 | 0.23 | 0.7 | 0.98 | 0.73 | -0.06 | 0.84 | 0.63 | 0.15 | 0.67 | 0.8 | 0.77 | 1.01 | 0.82 | 0.8 | 0.115 | 0.055 | 0.3425 | 0.713828 | 0.836229 | 0.180609 | 0.999971 | 1 | 0.808532 |
| B4galnt4 | Q766D5 | 0.99 | -1.26 | -2.71 | -1.39 | 1.1 | -2.22 | -0.87 | -0.38 | 0.49 | -1.55 | -0.14 | -0.8 | -6.67 | -3.66 | -0.4 | -3.54 | 0.5 | 0.5925 | -2.475 | 0.589856 | 0.466071 | 0.089577 | 0.999971 | 1 | 0.771991 |
| B4galt3 | Q91YY2 | -0.72 | -0.11 | -0.36 | -0.59 | -0.12 | -0.48 | -0.82 | -1.03 | 0.2 | -0.28 | -0.24 | 0.01 | 0.35 | 0.19 | -0.19 | -0.09 | -0.1675 | 0.3675 | 0.51 | 0.528571 | 0.118171 | 0.046512 | 0.999971 | 1 | 0.710102 |
| B4galt7 | D3Z065 | 0.9 | -1.91 | -0.58 | -0.39 | -1.42 | -1.24 | 1.72 | -0.81 | -1.54 | -0.6 | -1.04 | -1.32 | -2.65 | -1.77 | -1.15 | -1.31 | 0.0575 | -0.63 | -1.225 | 0.944967 | 0.28038 | 0.070526 | 0.999971 | 1 | 0.735439 |
| B4gat1 | Q8BWP8 | -0.06 | 1.68 | 1.38 | 1.02 | 1.57 | 1.87 | 1.09 | 1.27 | 1.82 | 0.41 | 1.33 | 0.81 | 1.8 | 1.71 | 1.24 | 1.62 | 0.445 | 0.0875 | 0.5875 | 0.281543 | 0.848932 | 0.153257 | 0.999971 | 1 | 0.795578 |
| B630019K( | Q7TNS5 | -1.89 | -1.92 | -1.52 | -1.44 | -1.51 | -2.09 | -1.64 | -1.75 | -1.68 | -1.95 | -2.1 | -1.94 | -1.39 | -1.53 | -1.64 | -1.65 | -0.055 | -0.225 | 0.14 | 0.802471 | 0.284911 | 0.487305 | 0.999971 | 1 | 0.897906 |
| B9d1 | Q5NCM9 | -1.24 | -0.8 | 0.64 | 0.51 | -0.76 | -1.27 | -0.5 | 1.14 | -1.21 | -3.67 | 0.34 | 0.72 | -0.68 | -0.9 | 1.12 | 0.17 | -0.125 | -0.7325 | 0.15 | 0.844549 | 0.466988 | 0.80301 | 0.999971 | 1 | 0.971467 |
| Baalc | Q8VHV1 | -2.79 | -0.67 | 0.13 | 0.16 | -0.54 | -0.62 | 0.55 | -0.04 | -0.64 | -2.72 | 0.37 | 0.54 | -0.74 | -1.11 | -0.23 | 0.14 | 0.63 | 0.18 | 0.3075 | 0.36195 | 0.844829 | 0.651262 | 0.999971 | 1 | 0.933842 |
| Babam1 | Q3UI43 | 0.77 | 0.77 | 1.04 | 1.3 | 0.55 | 0.7 | 1.46 | 1.17 | 0.79 | 1.06 | 1.23 | 0.6 | 0.97 | 0.89 | 0.8 | 0.92 | -3.93E-17 | -0.05 | -0.075 | 1 | 0.825124 | 0.702341 | 1 | 1 | 0.949019 |
| Babam2 | Q8K3W0 | 0.57 | 0.54 | 0.34 | 0.63 | 0.71 | 0.84 | 0.31 | 0.54 | 0.69 | 0.56 | 0.35 | 0.36 | 0.73 | 0.86 | 0.45 | 0.67 | 0.08 | -0.03 | 0.1575 | 0.681566 | 0.86607 | 0.399684 | 0.999971 | 1 | 0.871816 |
| Bace1 | P56818 | -0.1 | -1.51 | -0.79 | -0.41 | -1.09 | -0.95 | -0.42 | -0.47 | -1.07 | -0.14 | -0.74 | -1.07 | -1.82 | -1.28 | -0.7 | -1.22 | -0.03 | -0.0525 | -0.5525 | 0.93028 | 0.885695 | 0.161137 | 0.999971 | 1 | 0.800215 |
| Bad | Q61337 | 0.25 | -0.33 | 0.28 | 1.12 | -0.2 | 0.1 | 1.25 | 0.64 | 0.04 | 0.54 | -0.31 | -0.41 | -1 | -0.17 | -0.1 | -0.46 | 0.1175 | -0.365 | -0.7625 | 0.777343 | 0.324758 | 0.056837 | 0.999971 | 1 | 0.725348 |
| Bad_1 | Q3U9H3 | 0.73 | -0.19 | -1.98 | -2.04 | -0.75 | -0.36 | -1.09 | -1 | -0.87 | 0.11 | -0.81 | -0.46 | -0.78 | 0.32 | -1.41 | -1.48 | 0.07 | 0.3625 | 0.0325 | 0.912749 | 0.584849 | 0.964181 | 0.999971 | 1 | 0.998721 |
| Bag1 | F6TCF9 | -0.5 | 0.55 | -0.51 | -0.3 | 0.63 | 0.46 | -0.8 | 0.1 | 0.72 | -0.13 | -0.19 | -0.3 | 0.77 | 0.54 | -0.37 | -0.42 | 0.2875 | 0.215 | 0.32 | 0.467906 | 0.532092 | 0.413281 | 0.999971 | 1 | 0.876261 |
| Bag2 | Q91YN9 | 0.35 | -0.34 | -0.97 | -0.87 | -0.5 | -0.19 | -0.11 | -1.3 | -0.46 | 0.14 | -1.14 | -1.23 | -1.14 | -0.78 | -0.85 | -1.66 | -0.0675 | -0.215 | -0.65 | 0.862686 | 0.611104 | 0.095347 | 0.999971 | 1 | 0.78268 |
| Bag3 | Q9JLV1 | 2.65 | 0.67 | 0.42 | 0.72 | 0.69 | 0.97 | 0.51 | 0.66 | 0.7 | 1.82 | 0.26 | 0.07 | 0.17 | 0.64 | 0.76 | 0.15 | -0.4075 | -0.4025 | -0.685 | 0.412993 | 0.504616 | 0.195062 | 0.999971 | 1 | 0.811225 |
| Bag4 | Q8CI61 | 1.55 | 1.06 | 2.14 | 1.9 | 1.72 | 0.55 | 4.86 | 0.53 | 2.92 | 0.83 | 2.66 | 3.07 | 2.4 | -0.24 | 0.83 | 0.71 | 0.2525 | 0.7075 | -0.7375 | 0.787504 | 0.204295 | 0.20132 | 0.999971 | 1 | 0.815758 |
| Bag5 | Q8CI32 | 0.07 | -0.17 | -0.28 | -0.35 | -0.27 | -0.7 | -0.56 | -0.3 | -0.51 | -0.33 | -0.37 | -0.2 | -0.36 | 0.18 | -0.23 | -0.19 | -0.275 | -0.17 | 0.0325 | 0.189246 | 0.362159 | 0.873472 | 0.999971 | 1 | 0.984109 |
| Bag6 | Q3UF95 | -0.21 | 0.37 | 0.17 | 0.14 | 0.4 | 0.04 | 0.11 | 0.09 | 0.32 | -0.19 | -0.04 | 0.15 | 0.5 | 0.2 | 0.07 | -0.11 | 0.0425 | -0.0575 | 0.0475 | 0.833451 | 0.784848 | 0.829704 | 0.999971 | 1 | 0.978384 |
| Baiap2 | Q8BKX1-2 | -1.14 | -1.13 | -1.33 | -1.29 | -0.94 | -1.07 | -0.79 | -1.14 | -0.85 | -1.21 | -1.07 | -0.89 | -0.31 | -1.12 | -1.28 | -1.2 | 0.2375 | 0.2175 | 0.245 | 0.201182 | 0.235608 | 0.35246 | 0.999971 | 1 | 0.867768 |
| Baiap3 | S4R1K2 | 0.45 | -2.26 | -0.14 | -0.3 | -0.28 | -0.99 | -1.01 | 0.17 | 0.34 | -0.05 | -1.73 | 0.52 | -0.23 | 0.43 | 1.64 | -1.14 | 0.035 | 0.3325 | 0.7375 | 0.953259 | 0.641776 | 0.338462 | 0.999971 | 1 | 0.8595 |
| Bak1 | O08734 | 0.73 | 0.33 | -0.32 | -0.5 | 0.87 | 0.91 | -0.72 | -0.37 | 1.23 | 1.21 | -0.66 | -0.09 | 0.54 | 1.25 | -0.94 | 0.31 | 0.1125 | 0.3625 | 0.23 | 0.812542 | 0.487924 | 0.647403 | 0.999971 | 1 | 0.933613 |
| Banf1 | O54962 | -1.12 | -0.76 | -0.79 | -0.81 | -0.56 | -0.86 | 0.28 | -1.22 | -0.55 | -1.31 | -1.6 | -1.12 | -0.21 | -0.48 | -0.55 | -1.24 | 0.28 | -0.275 | 0.25 | 0.406039 | 0.304285 | 0.348365 | 0.999971 | 1 | 0.864885 |
| Basp1 | Q91XV3 | 0.37 | -1.78 | -1.23 | -0.73 | -1.91 | -1.53 | -0.31 | -1.5 | -1.85 | -0.9 | -1.59 | -1.97 | -1.99 | -1.88 | -1.58 | -1.61 | -0.47 | -0.735 | -0.9225 | 0.385525 | 0.154297 | 0.062525 | 0.999971 | 1 | 0.725348 |
| Bax | A0A1B0GT: | -0.01 | 0.81 | -0.01 | 0.51 | 0.77 | 0.46 | -0.08 | 0.24 | 0.7 | 0.4 | 0.55 | 0.24 | 0.96 | 0.7 | 0.29 | 0.48 | 0.0225 | 0.1475 | 0.2825 | 0.93691 | 0.560625 | 0.309908 | 0.999971 | 1 | 0.847402 |
| Baz1b | Q9Z277 | -0.98 | -1.08 | -0.94 | -0.8 | -1.07 | -1.27 | -0.61 | -1.03 | -1.22 | -1.17 | -0.91 | -1.04 | -0.78 | -1.16 | -1.16 | -1.23 | -0.045 | -0.135 | -0.1325 | 0.826157 | 0.441552 | 0.487831 | 0.999971 | 1 | 0.897906 |
| Baz2a | F8VPM0 | 0.83 | -2.31 | -0.72 | -1.02 | -1.95 | -2.42 | -1.05 | -2.35 | -1.75 | 0.36 | -2.41 | -2.93 | -4.11 | -3.24 | -1.17 | -2.67 | -1.1375 | -0.8775 | -1.9925 | 0.108143 | 0.329816 | 0.033103 | 0.999971 | 1 | 0.685881 |
| Bbln | P58686 | -0.1 | 0.51 | -0.07 | -0.53 | -0.14 | 1 | -0.53 | 0.22 | 0.28 | -1 | -0.25 | -0.77 | 0.54 | 0.73 | -1.03 | 0.37 | 0.185 | -0.3875 | 0.2 | 0.625413 | 0.286078 | 0.643873 | 0.999971 | 1 | 0.93301 |
| Bbs1 | Q3V3N7 | -1.06 | -0.2 | -0.08 | -0.03 | -0.07 | -0.55 | -0.45 | -0.21 | -0.39 | -0.8 | -0.4 | -0.07 | -0.65 | 0 | -0.12 | -0.26 | 0.0225 | -0.0725 | 0.085 | 0.936062 | 0.80528 | 0.771359 | 0.999971 | 1 | 0.963498 |
| Bbs2 | Q9CWF6 | 0.37 | 0.23 | 0.34 | 0.1 | 0.42 | 0.34 | -0.02 | 0.21 | 0.53 | 0.69 | -0.62 | 0.46 | 0.25 | 0.52 | 0.49 | 0 | -0.0225 | 0.005 | 0.055 | 0.903596 | 0.98711 | 0.780954 | 0.999971 | 1 | 0.965859 |
| Bbs4 | Q8C1Z7 | 0.89 | 0.18 | -0.19 | -0.72 | 0.22 | -0.19 | 0.91 | -0.12 | 0.18 | 0.63 | 0.25 | 0.59 | 0.61 | 0.73 | -0.14 | -0.19 | 0.165 | 0.3725 | 0.2125 | 0.68302 | 0.304829 | 0.597188 | 0.999971 | 1 | 0.92377 |
| Bbs5 | A2AUC6 | -1.44 | -0.2 | -0.66 | 0.5 | -0.61 | 0.58 | -0.85 | -0.57 | -0.98 | -2.73 | -0.39 | -0.29 | 0.41 | -0.99 | -0.76 | -0.42 | 0.0875 | -0.6475 | 0.01 | 0.855962 | 0.324316 | 0.983224 | 0.999971 | 1 | 0.99925 |
| Bbs7 | E9Q0R0 | 0.39 | 0.31 | 0.74 | 0.69 | 0.38 | 0.12 | 0.37 | 1.02 | -0.06 | 0.16 | 0.37 | 0.72 | -0.3 | 0.48 | 0.86 | 0.84 | -0.06 | -0.235 | -0.0625 | 0.80942 | 0.32525 | 0.835341 | 0.999971 | 1 | 0.979185 |
| Bbs9 | Q811G0-2 | -1.3 | 0.14 | -1.56 | -1.3 | 0.02 | -0.42 | -1.67 | -1.68 | 0.26 | -0.71 | -1.32 | -1.05 | 0.12 | -0.15 | -0.87 | -1.17 | 0.0675 | 0.3 | 0.4875 | 0.899702 | 0.540245 | 0.305123 | 0.999971 | 1 | 0.847402 |
| BC004004 | A0A3Q4L2) | 0.45 | 0.24 | -1.02 | -0.86 | 0.22 | 0.67 | -0.61 | -0.62 | 0.17 | 0.73 | -0.93 | -0.91 | 1.03 | 0.92 | -0.44 | -0.46 | 0.2125 | 0.0625 | 0.56 | 0.646615 | 0.903752 | 0.294242 | 0.999971 | 1 | 0.847402 |
| BC024978 | D3Z7G4 | -1.21 | -0.11 | -0.12 | -0.57 | 0.38 | -0.71 | -0.45 | -0.09 | 0.04 | -0.88 | 0.37 | 0.93 | 0.03 | -0.86 | 0.31 | 0.04 | 0.285 | 0.6175 | 0.3825 | 0.418794 | 0.180422 | 0.299785 | 0.999971 | 1 | 0.847402 |
| Bcam | Q9R069 | 0.6 | 0.68 | 0.19 | 0.66 | 1.81 | 0.21 | -0.03 | 0.13 | 0.37 | 1.56 | 0.69 | 1.18 | -0.23 | 0.64 | 0.67 | 0.14 | -0.0025 | 0.4175 | -0.2275 | 0.995263 | 0.182736 | 0.403242 | 0.999971 | 1 | 0.873348 |
| Bcan | Q61361 | 0.99 | 1.52 | 1.24 | 1.33 | 1.41 | 1.41 | 1.13 | 1.33 | 1.12 | 0.86 | 0.83 | 0.74 | 0.97 | 1.23 | 1.08 | 1.32 | 0.05 | -0.3825 | -0.12 | 0.795769 | 0.076383 | 0.546677 | 0.999971 | 1 | 0.911398 |
| Bcap29 | Q61334 | 0.12 | -0.74 | -0.18 | 0.22 | -0.14 | -0.76 | 0.45 | -0.15 | 0.68 | -0.93 | 0.16 | 0.16 | -0.44 | -1.56 | -0.4 | 0.39 | -0.005 | 0.1625 | -0.3575 | 0.98782 | 0.676204 | 0.416334 | 0.999971 | 1 | 0.877131 |
| Bcap31 | Q61335 | 0.26 | -0.56 | -0.06 | 0.17 | -0.39 | -0.2 | 0.31 | -0.2 | -0.43 | 0.2 | -0.44 | -0.25 | -0.66 | -0.55 | -0.19 | -0.15 | -0.0725 | -0.1825 | -0.34 | 0.781066 | 0.487428 | 0.199895 | 0.999971 | 1 | 0.813335 |
| Bcar1 | Q61140 | 0.34 | -0.36 | -0.68 | -0.61 | -0.38 | -0.45 | -0.9 | -0.76 | -0.45 | -0.02 | -0.74 | -0.83 | -0.61 | -0.25 | -0.8 | -0.6 | -0.295 | -0.1825 | -0.2375 | 0.306814 | 0.550621 | 0.401418 | 0.999971 | 1 | 0.871816 |
| Bcar3 | Q9QZK2 | 0.61 | -0.36 | 0.03 | 0.23 | -0.87 | -0.45 | 0.28 | -0.05 | -0.44 | 0.04 | 0.04 | -0.26 | -0.45 | -0.24 | 0.02 | -0.03 | -0.4 | -0.2825 | -0.3025 | 0.234998 | 0.288916 | 0.255454 | 0.999971 | 1 | 0.839012 |
| Bcas1 | Q80YN3 | -0.56 | -1.16 | -0.75 | -0.71 | -1.47 | -0.86 | -0.29 | -1.13 | -1.15 | -1.19 | -1.48 | -1.24 | -1.05 | -1.05 | -0.83 | -1.31 | -0.1425 | -0.47 | -0.265 | 0.627357 | 0.041611 | 0.232013 | 0.999971 | 1 | 0.831121 |
| Bcas2 | Q9D287 | -0.11 | 0.2 | 0.45 | 0.78 | 0.13 | -0.14 | 0.35 | 0.58 | 0.24 | -0.05 | 0.43 | 0.33 | 0.05 | 0.41 | 0.37 | 0.64 | -0.1 | -0.0925 | 0.0375 | 0.70674 | 0.705291 | 0.881786 | 0.999971 | 1 | 0.986206 |
| Bcas3 | B1AR74 | -0.52 | -0.24 | -0.55 | -0.76 | -0.14 | -0.22 | -0.45 | -0.65 | 0.04 | -0.34 | -0.61 | -0.69 | -0.03 | -0.45 | -0.81 | -0.9 | 0.1525 | 0.1175 | -0.03 | 0.472022 | 0.613431 | 0.905426 | 0.999971 | 1 | 0.991091 |
| Bcas3_1 | Q8CCN5 | 2.82 | -0.26 | -0.75 | -0.78 | -0.79 | -0.16 | -1.23 | -0.46 | -1 | 0.67 | -1.44 | -0.63 | -0.23 | 0.2 | -0.65 | -0.51 | -0.9175 | -0.8575 | -0.555 | 0.2706 | 0.343269 | 0.491057 | 0.999971 | 1 | 0.898873 |
| Bcat1 | Q8CBC8 | 0.42 | 0.11 | -0.55 | -0.48 | 0.28 | 0.4 | -0.43 | -0.25 | 0.3 | 0.19 | -0.45 | -0.09 | 0.09 | 0.72 | -0.33 | -0.51 | 0.125 | 0.1125 | 0.1175 | 0.690861 | 0.705479 | 0.740344 | 0.999971 | 1 | 0.960899 |
| Bcat2 | O35855 | -0.89 | 0.4 | 0.05 | -0.12 | 0.3 | 0.53 | -0.21 | 0.25 | 0.35 | 0.03 | 0.65 | 0.15 | 0.57 | 0.67 | 0.56 | 0.36 | 0.3575 | 0.435 | 0.68 | 0.276533 | 0.183781 | 0.042102 | 0.999971 | 1 | 0.701725 |
| Bccip | Q9CWI3 | 1.65 | 3.05 | 1.98 | 1.78 | 2.5 | 2.9 | 1.82 | 1.64 | 2.98 | 2.14 | 1.83 | 1.12 | 0.68 | 1.62 | 1.42 | 1.72 | 0.1 | -0.0975 | -0.755 | 0.808566 | 0.835476 | 0.075377 | 0.999971 | 1 | 0.746958 |
| Bckdha | Q3U3J1 | 0.7 | -0.5 | -0.61 | -0.51 | -0.78 | -0.67 | -0.53 | -0.39 | -0.7 | -0.04 | -0.59 | -0.6 | -0.21 | -0.36 | -0.73 | -0.5 | -0.3625 | -0.2525 | -0.22 | 0.279951 | 0.466619 | 0.510651 | 0.999971 | 1 | 0.905815 |
| Bckdhb | Q6P3A8 | 1.33 | 1.01 | 1.48 | 1.3 | 0.64 | 0.66 | 0.53 | 1.94 | 0.73 | 1.14 | 1.33 | -0.73 | 1.55 | 1.04 | 1.51 | 1.31 | -0.3375 | -0.6625 | 0.0725 | 0.339359 | 0.166068 | 0.726556 | 0.999971 | 1 | 0.956041 |
| Bckdk | A0A0U1RN | -0.03 | -0.34 | 0.23 | 0.05 | -0.28 | -0.38 | 0.45 | 0.47 | -0.68 | -0.49 | 0.61 | 0.44 | -0.41 | -0.36 | 0.37 | 0.27 | 0.0875 | -0.0075 | -0.01 | 0.75113 | 0.982475 | 0.969394 | 0.999971 | 1 | 0.998787 |
| Bcl10 | Q9Z0H7 | 0.54 | 0.87 | 0.55 | 0.84 | 0.45 | 0.79 | 0.65 | 0.67 | 1.09 | 0.73 | 0.43 | 0.32 | 0.3 | 1.18 | 1.28 | 1 | -0.06 | -0.0575 | 0.24 | 0.747469 | 0.802534 | 0.371573 | 0.999971 | 1 | 0.869526 |
| Bcl11a | Q5STS9 | 0.74 | 1.35 | 1.23 | 1.19 | 1.61 | 1.56 | 1.05 | 1.08 | 1.18 | 0.94 | 0.61 | 0.47 | 1.04 | 1.04 | 0.5 | 1.01 | 0.1975 | -0.3275 | -0.23 | 0.412364 | 0.194746 | 0.328113 | 0.999971 | 1 | 0.856472 |
| Bcl11b | A0A0R4J1E | 2.73 | 0.16 | 0.36 | 0.62 | -0.26 | 0.65 | 2.22 | 0.2 | -0.39 | 1.62 | 0.06 | -0.76 | -0.79 | -0.51 | -0.28 | -0.71 | -0.265 | -0.835 | -1.54 | 0.715299 | 0.26521 | 0.020845 | 0.999971 | 1 | 0.635402 |
| Bcl2 | P10417-2 | -1.06 | -0.56 | -2.87 | -3.7 | -0.56 | -0.35 | -2.03 | -4.82 | -0.55 | -0.58 | -2.38 | -2.67 | -0.87 | 0.24 | -2.87 | -0.19 | 0.1075 | 0.5025 | 1.125 | 0.924097 | 0.555704 | 0.23534 | 0.999971 | 1 | 0.832918 |
| Bcl2l1 | A2AHX9 | -0.28 | 0.12 | -0.08 | 0.43 | 0.39 | 0.3 | -0.27 | 0.47 | 0.03 | -0.14 | 0.16 | 0.48 | 0.34 | 0.72 | 0.57 | 0.67 | 0.175 | 0.085 | 0.5275 | 0.494591 | 0.716817 | 0.036867 | 0.999971 | 1 | 0.700096 |
| Bcl2l13 | P59017 | -0.56 | -1.64 | -1.33 | -0.96 | -1.81 | -1.6 | -0.85 | -1.54 | -1.72 | -1.11 | -1.59 | -1.21 | -1.68 | -1.48 | -1.11 | -0.82 | -0.3275 | -0.285 | -0.15 | 0.315305 | 0.335372 | 0.628984 | 0.999971 | 1 | 0.932176 |
| Bcl7a | Q9CXE2 | 1.5 | 1.84 | 1.84 | 2.03 | 1.98 | 1.79 | 1.41 | 2.13 | 2.01 | 1.57 | 1.95 | 2.01 | 1.77 | 2.02 | 1.67 | 2.11 | 0.025 | 0.0825 | 0.09 | 0.913097 | 0.686841 | 0.663697 | 0.999971 | 1 | 0.936621 |
| Bcl7b | Q921K9 | 0.43 | 1.31 | 0.43 | 0.91 | 1.61 | 1.14 | 0.22 | 1.07 | 0.92 | 0.69 | 0.62 | 0.32 | 1.12 | 1.02 | 0.51 | 1.11 | 0.24 | -0.1325 | 0.17 | 0.501336 | 0.620111 | 0.541555 | 0.999971 | 1 | 0.910988 |
| Bclaf1 | Q8K019-2 | 0.26 | -0.7 | -0.47 | -0.24 | -0.18 | -0.26 | 0.35 | -0.08 | -0.63 | -0.1 | -0.45 | -0.53 | -0.49 | -0.34 | -0.28 | -0.05 | 0.245 | -0.14 | -0.0025 | 0.370238 | 0.590263 | 0.99209 | 0.999971 | 1 | 0.99925 |
| Bcr | Q6PAJ1 | -0.08 | -0.66 | -0.4 | -0.22 | -0.5 | -0.82 | -0.37 | -0.22 | -0.62 | -0.43 | -0.49 | -0.24 | -0.85 | -0.47 | -0.2 | -0.26 | -0.1375 | -0.105 | -0.105 | 0.539587 | 0.604009 | 0.6517 | 0.999971 | 1 | 0.933842 |
| Bcs1l | Q9CZP5 | -1.41 | -1.17 | -0.73 | -0.74 | -0.7 | -1.43 | -1.06 | -0.63 | -0.87 | -2.23 | -0.92 | -0.95 | -1.18 | -1.48 | -0.73 | -0.96 | 0.0575 | -0.23 | -0.075 | 0.830691 | 0.529215 | 0.770972 | 0.999971 | 1 | 0.963498 |
| Bdh1 | Q80XN0 | -1.28 | -0.71 | -0.81 | -0.73 | -0.92 | -0.78 | -0.63 | -0.4 | -0.62 | -0.97 | -0.38 | -0.68 | -0.23 | -0.57 | -0.54 | -0.14 | 0.2 | 0.22 | 0.5125 | 0.372678 | 0.333495 | 0.040946 | 0.999971 | 1 | 0.700838 |
| Bdnf | H9H9S8 | -2.09 | -1.37 | -0.21 | -0.9 | -0.69 | -0.27 | -0.09 | -1.74 | -0.04 | -0.87 | 0.04 | -1.39 | -3.15 | -1.79 | -1.16 | -2.25 | 0.445 | 0.5775 | -0.945 | 0.38601 | 0.257309 | 0.103136 | 0.999971 | 1 | 0.78268 |
| Becn1 | O88597 | 0.59 | -0.53 | 0.45 | 0.44 | -0.57 | -0.18 | 0.61 | 0.22 | -0.44 | -0.34 | 0.25 | 0.19 | -0.55 | -0.76 | 0.34 | 0.21 | -0.2175 | -0.3225 | -0.4275 | 0.544198 | 0.322682 | 0.262184 | 0.999971 | 1 | 0.842262 |
| Begain | F8WIG2 | -0.12 | -1.2 | -0.53 | -0.72 | -1.26 | -1.25 | -0.33 | -1.14 | -1.12 | -0.77 | -0.53 | -0.98 | -1.27 | -1.83 | -1.36 | -1.33 | -0.3525 | -0.2075 | -0.805 | 0.286013 | 0.457278 | 0.016272 | 0.999971 | 1 | 0.613095 |
| Bet1 | O35623 | -0.09 | -0.97 | -1.54 | -1.44 | -2.06 | -1.16 | -1.53 | -1.29 | -1.7 | -0.54 | -2.31 | -1.47 | 0.54 | -1.18 | -1.08 | -1.32 | -0.5 | -0.495 | 0.25 | 0.20368 | 0.302724 | 0.62473 | 0.999971 | 1 | 0.930479 |
| Bex2 | Q9WTZ8 | 1.13 | -0.31 | 3.15 | 2.79 | 0.99 | 0.42 | 4.33 | 1.73 | 1.27 | 1.78 | 2.05 | 1.47 | 0.65 | 0.32 | 1.22 | -0.15 | 0.1775 | -0.0475 | -1.18 | 0.865504 | 0.948876 | 0.149448 | 0.999971 | 1 | 0.793248 |
| Bfar | A0A0R4J04 | -0.27 | -0.31 | -0.62 | -0.88 | -0.41 | -0.36 | -0.44 | -0.48 | -0.24 | -0.35 | -0.2 | -0.2 | 0.26 | -0.02 | 0.18 | -0.07 | 0.0975 | 0.2725 | 0.6075 | 0.631589 | 0.197794 | 0.018003 | 0.999971 | 1 | 0.635388 |
| Bgn | P28653 | 0.56 | 1.59 | 0.56 | -0.1 | 6.66 | 1.97 | 0.37 | -0.31 | 2.94 | 2.58 | 1.62 | 3.78 | 1.56 | 1.88 | 0.67 | -0.11 | 1.52 | 2.0775 | 0.3475 | 0.302556 | 0.003468 | 0.514367 | 0.999971 | 0.957868 | 0.905815 |
| Bhlhe23 | Q8BGW3 | -0.51 | -1.15 | -0.95 | -2.15 | -0.69 | -1.2 | 1.07 | -1.79 | -0.87 | -2.08 | -1.39 | -0.7 | -0.23 | -2.08 | -1.66 | -2.38 | 0.5375 | -0.07 | -0.3975 | 0.412394 | 0.873566 | 0.470875 | 0.999971 | 1 | 0.892857 |
| Bicd1 | G3UXK5 | -0.57 | -0.85 | -0.32 | -0.71 | -0.88 | -0.75 | -0.14 | -0.22 | -0.82 | -0.34 | 0.54 | 0.3 | -0.57 | -0.63 | -0.35 | -0.37 | 0.115 | 0.5325 | 0.1325 | 0.643364 | 0.132086 | 0.505 | 0.999971 | 1 | 0.905277 |
| Bicd2 | Q921C5 | -0.9 | -0.93 | -0.12 | -0.17 | -1.22 | -1.57 | -0.15 | 0.24 | -1.23 | -1.16 | 0.2 | 0.55 | -1.21 | -0.98 | -0.08 | 0.28 | -0.145 | 0.12 | 0.0325 | 0.749654 | 0.801834 | 0.935484 | 0.999971 | 1 | 0.995618 |
| Bicra | F8VPZ9 | 2.07 | 1.35 | 0.56 | 1.6 | 1.51 | 1.71 | 1.56 | 1.21 | 1.07 | 0.93 | 0.51 | -0.64 | 1.58 | 1.71 | 0.25 | -0.29 | 0.1025 | -0.9275 | -0.5825 | 0.757685 | 0.075199 | 0.297512 | 0.999971 | 1 | 0.847402 |
| Bicral | Q8CHH5 | -0.04 | 0.77 | 0.41 | 0.51 | -0.27 | 0.43 | -0.87 | -0.6 | -0.57 | -0.85 | -0.2 | -0.13 | 0.17 | -0.7 | -0.46 | -0.43 | -0.74 | -0.85 | -0.7675 | 0.047168 | 0.009376 | 0.018415 | 0.999971 | 1 | 0.635388 |
| Bid | P70444 | 0.23 | 0.49 | -0.45 | -0.01 | 1.15 | 0.33 | -0.47 | 0.65 | 0.64 | 0.54 | -0.18 | 0 | 0.93 | 1.07 | 0.22 | 0.58 | 0.35 | 0.185 | 0.635 | 0.369291 | 0.532521 | 0.052119 | 0.999971 | 1 | 0.723736 |
| Bin1 | O08539 | -0.93 | -1.23 | -0.35 | -0.74 | -1.24 | -0.95 | -0.48 | -0.56 | -1.07 | -1.17 | -0.47 | -0.32 | -1.14 | -1.04 | -0.37 | -0.7 | 0.005 | 0.055 | 2.16E-16 | 0.985395 | 0.850373 | 1 | 0.999971 | 1 | 1 |
| Bin2 | S4R270 | 1.49 | 0.81 | 0.49 | 1.07 | 1.29 | 1.17 | 1.11 | 0.42 | 1.08 | 0.86 | 0.51 | 0.38 | 0.21 | 0.66 | 0.87 | 0.64 | 0.0325 | -0.2575 | -0.37 | 0.912974 | 0.369107 | 0.197075 | 0.999971 | 1 | 0.812891 |
| Bin3 | Q9JI08 | -0.27 | -0.17 | -0.07 | 0.05 | -0.33 | -0.74 | -0.16 | 0.25 | -0.54 | -0.72 | 0.32 | 0.1 | -0.28 | -0.58 | 0.43 | 0.81 | -0.13 | -0.095 | 0.21 | 0.599991 | 0.730136 | 0.525768 | 0.999971 | 1 | 0.909459 |
| Birc2 | Q62210 | 0.44 | -0.36 | 0.53 | 0.89 | -0.08 | 0.16 | 0.01 | 0.54 | -0.09 | 0.3 | 0.11 | 0.29 | -0.55 | 0.09 | 0.24 | 0.66 | -0.2175 | -0.2225 | -0.265 | 0.480901 | 0.450732 | 0.465027 | 0.999971 | 1 | 0.89116 |
| Birc6 | O88738 | 0.47 | 0.19 | 0.58 | 0.39 | 0.14 | 0.18 | 0.37 | 0.43 | 0.17 | 0.35 | 0.53 | 0.28 | 0.12 | 0.14 | 0.3 | 0.58 | -0.1275 | -0.075 | -0.1225 | 0.492905 | 0.681086 | 0.538145 | 0.999971 | 1 | 0.910077 |
| Bivm | Q8CBX9 | 1.79 | 1.08 | 0.98 | 0.72 | 0.98 | 0.02 | 0.91 | 1.11 | 2.17 | -2.46 | 0.95 | 1.65 | 4.24 | -0.95 | 0.03 | 0.22 | -0.3875 | -0.565 | -0.2575 | 0.266493 | 0.560445 | 0.805451 | 0.999971 | 1 | 0.971701 |
| Bles03 | Q8VD62 | 0.74 | 0.84 | 0.82 | 0.73 | 0.76 | 0.99 | 0.5 | 0.9 | 0.77 | 0.41 | 0.62 | 0.63 | 0.66 | 0.99 | 0.69 | 0.86 | 0.005 | -0.175 | 0.0175 | 0.978325 | 0.312628 | 0.919444 | 0.999971 | 1 | 0.992633 |
| Blmh | Q8R016 | 0.01 | 0.23 | -0.02 | 0.21 | 0.31 | 0.4 | 0.24 | 0.31 | 0.35 | 0.11 | 0.03 | -0.21 | 0.38 | 0.78 | 0.17 | 0.2 | 0.2075 | -0.0375 | 0.275 | 0.240511 | 0.84548 | 0.208639 | 0.999971 | 1 | 0.822138 |
| Bloc1s1 | O55102 | -0.52 | -0.2 | -0.64 | 0.11 | -0.19 | 0.29 | -0.77 | 0.29 | -0.16 | -0.38 | -0.47 | -0.25 | 0.02 | 0.15 | -0.06 | 0.01 | 0.2175 | -0.0025 | 0.3425 | 0.487359 | 0.990993 | 0.145001 | 0.999971 | 1 | 0.79148 |
| Bloc1s2 | Q9CWG9 | 1.88 | 1.03 | 0.81 | 0.86 | 0.65 | 1.26 | 0.57 | 0.61 | 0.65 | 1.59 | 0.17 | 0.7 | 1.15 | 1.07 | 0.54 | 1.05 | -0.3725 | -0.3675 | -0.1925 | 0.242004 | 0.34145 | 0.520874 | 0.999971 | 1 | 0.906972 |
| Bloc1s4 | Q8VED2 | 0.99 | -1.14 | 0.61 | 1.58 | -0.79 | -0.69 | 1.23 | 1.81 | -0.74 | 0.33 | 1.76 | 1.09 | -1.63 | -1.21 | 0.79 | -0.74 | -0.12 | 0.1 | -1.2075 | 0.879945 | 0.889701 | 0.117746 | 0.999971 | 1 | 0.78268 |
| Blvra | Q9CY64 | -0.38 | 0.72 | 0.33 | 0.12 | 0.53 | 0.25 | 0 | -0.07 | 0.52 | -0.29 | 0.45 | 0.53 | 0.61 | 0.45 | 0.24 | 0.46 | -0.02 | 0.105 | 0.2425 | 0.943296 | 0.734018 | 0.370338 | 0.999971 | 1 | 0.869526 |
| Blvrb | Q923D2 | 1.08 | 0 | -0.47 | 0.81 | 0.2 | 0.06 | 0.91 | -0.18 | -0.84 | -0.5 | -0.3 | -0.35 | -0.04 | -0.98 | -0.21 | -0.16 | -0.1075 | -0.8525 | -0.7025 | 0.792273 | 0.043507 | 0.106397 | 0.999971 | 1 | 0.78268 |
| Bmerb1 | Q8R1Y2 | -0.87 | -1.53 | -0.83 | -0.8 | -1.07 | -1.01 | -1.06 | -0.34 | -0.83 | -0.8 | -0.96 | -1.31 | -0.88 | -0.52 | -0.1 | -0.28 | 0.1375 | 0.0325 | 0.5625 | 0.611381 | 0.892389 | 0.058889 | 0.999971 | 1 | 0.725348 |
| Bmp2k | Q91Z96 | 0.18 | -1.22 | -0.94 | -0.52 | -1.63 | -1.72 | -0.84 | -0.82 | -1.9 | -0.76 | -0.83 | -0.58 | -1.9 | -1.11 | -1.29 | -0.76 | -0.6275 | -0.3925 | -0.64 | 0.123455 | 0.349196 | 0.115482 | 0.999971 | 1 | 0.78268 |
| Bmpr1a | Q60607 | -1.6 | 0.04 | -0.95 | -1.15 | 0 | -0.63 | -1.7 | -0.62 | -0.03 | -1.16 | -0.62 | -0.97 | 1.08 | -0.33 | -0.76 | -1.03 | 0.1775 | 0.22 | 0.655 | 0.702135 | 0.592174 | 0.243453 | 0.999971 | 1 | 0.837316 |
| Bmpr1b | P36898 | -0.73 | -2.54 | -0.26 | 0.09 | 0.31 | -0.53 | -1.19 | 0.06 | -1.37 | -0.34 | -1.77 | -0.23 | -0.21 | -1.09 | -0.92 | -0.08 | 0.5225 | -0.0675 | 0.285 | 0.404376 | 0.915677 | 0.627324 | 0.999971 | 1 | 0.931025 |
| Bmpr2 | O35607 | -1.08 | -0.91 | -0.66 | -0.51 | -0.89 | -0.78 | -0.6 | -0.42 | -0.67 | -0.56 | -0.47 | -0.67 | -0.51 | -0.79 | -0.68 | -0.48 | 0.1175 | 0.1975 | 0.175 | 0.583615 | 0.323831 | 0.399817 | 0.999971 | 1 | 0.871816 |
| Bms1 | Q6PGF5 | -1.69 | -1.95 | -1.23 | -1.61 | -1.75 | -2.46 | -1.51 | -1.88 | -1.71 | -1.65 | -0.83 | -1.54 | -1.41 | -1.74 | -1.23 | -1.84 | -0.28 | 0.1875 | 0.065 | 0.314952 | 0.493793 | 0.786525 | 0.999971 | 1 | 0.967132 |
| Bmt2 | A0A0N4SU | 1.52 | -0.04 | 0.85 | 0.76 | 0.56 | 0.62 | 1.68 | 0.86 | 0.55 | 1.16 | 1.11 | 0.74 | 1.07 | 0.96 | 0.65 | 0.35 | 0.1575 | 0.1175 | -0.015 | 0.690316 | 0.735264 | 0.966022 | 0.999971 | 1 | 0.998721 |
| Bnip1 | Q6QD59 | 0.23 | -0.8 | 0.08 | 0.11 | -0.78 | -0.27 | -0.41 | 0.04 | -1.07 | -0.36 | 0.02 | 0.04 | -0.85 | -0.53 | 0.4 | 0.78 | -0.26 | -0.2475 | 0.045 | 0.396833 | 0.481541 | 0.916043 | 0.999971 | 1 | 0.992633 |
| Bnip2 | Q91VL0 | -4.2 | -6.01 | -5.39 | -4.43 | -4.38 | -6.82 | -4.1 | -4.69 | -5.72 | -5.17 | -4.04 | -3.88 | -6.19 | -5.18 | -3.52 | -5.06 | 0.01 | 0.305 | 0.02 | 0.988227 | 0.59239 | 0.974732 | 0.999971 | 1 | 0.99925 |
| Bnip3 | O55003 | 2.54 | 0.25 | -0.06 | 0.48 | -0.56 | -0.5 | 0.81 | -0.62 | -0.62 | 1.13 | 0 | 0.22 | -0.31 | -0.36 | -0.61 | -0.12 | -1.02 | -0.62 | -1.1525 | 0.127493 | 0.341389 | 0.061952 | 0.999971 | 1 | 0.725348 |
| Bnip3l | Q9Z2F7 | -0.6 | -0.11 | -0.55 | -0.66 | 0.44 | 0.8 | -0.39 | -0.78 | 1.2 | 0.57 | -0.92 | -0.95 | 2.2 | 0.32 | -0.65 | -0.1 | 0.4975 | 0.455 | 0.9225 | 0.20515 | 0.388846 | 0.137555 | 0.999971 | 1 | 0.784948 |
| Boc | D3Z763 | 2 | 0.4 | 1.81 | 2.08 | -0.42 | 1.01 | 1.95 | 2.41 | 0.5 | 1.36 | 0.72 | 1.65 | -1.22 | 0.37 | 2.48 | 1.97 | -0.335 | -0.515 | -0.6725 | 0.619177 | 0.271334 | 0.427913 | 0.999971 | 1 | 0.880793 |
| Bod1 | Q5SQY2 | 0.59 | 1.69 | 0.97 | 0.89 | 1.88 | 1.93 | 1.16 | 0.67 | 1.75 | 1.43 | 1.44 | 1.8 | 1.43 | 1.99 | 1.19 | 1.63 | 0.375 | 0.57 | 0.525 | 0.326268 | 0.060751 | 0.105497 | 0.999971 | 1 | 0.78268 |
| Bod1l | E9Q6J5-2 | 4.47 | 2.53 | 2.5 | 2.77 | 2.61 | 3.11 | 2.31 | 2.48 | 2.44 | 3.19 | 2.3 | 2.16 | 1.76 | 1.78 | 2.01 | 1.92 | -0.44 | -0.545 | -1.2 | 0.360237 | 0.283371 | 0.023801 | 0.999971 | 1 | 0.664227 |
| Bok | O35425 | 0.96 | 1.14 | 1.92 | 1.45 | 0.69 | 0.21 | 1.29 | 0.24 | 0.62 | 1 | 0.77 | -0.53 | -0.68 | -0.46 | 1.09 | 0.63 | -0.76 | -0.9025 | -1.2225 | 0.043076 | 0.041998 | 0.021886 | 0.999971 | 1 | 0.645537 |
| Bola1 | Q9D8S9 | -1.04 | -1.51 | -1.32 | -1.63 | -1.21 | -1.64 | -1.26 | -2.05 | -1.44 | -1.38 | -1.04 | -1.4 | -2.02 | -1.76 | -1.49 | -1.4 | -0.165 | 0.06 | -0.2925 | 0.527455 | 0.772312 | 0.223993 | 0.999971 | 1 | 0.828225 |
| Bola2 | Q8BGS2 | 0.29 | 0.44 | 0.44 | 0.13 | 0.38 | 0.72 | 0.04 | 1.08 | 0.25 | 0.41 | 0.48 | 0.27 | 0.1 | 1.26 | 0.95 | 0.62 | 0.23 | 0.0275 | 0.4075 | 0.385733 | 0.873705 | 0.165674 | 0.999971 | 1 | 0.800215 |
| Bola3 | E9Q705 | -1.63 | -3.74 | -0.14 | 0.42 | -3.62 | -3.55 | 0.27 | 0.55 | -2.96 | -2.57 | -0.38 | 0.21 | -4.88 | -2.68 | -0.11 | 0.02 | -0.315 | -0.1525 | -0.64 | 0.81089 | 0.88897 | 0.631796 | 0.999971 | 1 | 0.932858 |
| Bop1 | P97452 | 0.5 | 0.92 | 0.5 | 0.55 | 0.93 | 0.3 | 0.17 | 0.7 | 0.85 | 0.51 | 0.91 | 1.03 | 0.87 | 0.45 | 0.66 | 0.32 | -0.0925 | 0.2075 | -0.0425 | 0.697715 | 0.320499 | 0.839461 | 0.999971 | 1 | 0.979728 |
| Borcs5 | E9Q2W8 | -0.27 | 0.52 | 0.44 | 0.22 | 0.09 | 0.25 | -0.15 | 0.33 | -0.01 | 0.14 | 0.24 | -0.26 | 0.4 | -0.05 | 0.26 | 0.1 | -0.0975 | -0.2 | -0.05 | 0.685356 | 0.412264 | 0.83337 | 0.999971 | 1 | 0.979036 |
| Borcs6 | Q9D6W8 | -0.06 | 0.61 | 0.91 | 0.9 | 0.48 | 0.04 | 0.4 | 1.26 | 0.23 | -0.27 | 0.5 | 0.05 | 0.98 | 0.49 | 0.8 | 1.19 | -0.045 | -0.4625 | 0.275 | 0.894549 | 0.137491 | 0.34901 | 0.999971 | 1 | 0.865162 |
| Borcs7 | Q9CRC6 | -0.35 | -0.09 | -1.59 | -1.14 | 0.09 | 0 | -1.38 | -1.21 | -0.53 | -0.19 | -1.94 | -1.12 | 0.76 | -0.01 | -0.86 | -0.07 | 0.1675 | -0.1525 | 0.7475 | 0.730414 | 0.752988 | 0.124687 | 0.999971 | 1 | 0.78268 |
| Borcs8 | Q9D6Y4 | -1.93 | -0.11 | 0.57 | 0.54 | 0.47 | -0.28 | -0.1 | 0.71 | 0.06 | -0.62 | 0.11 | 0.79 | 0.39 | 0.85 | 0.51 | 0.7 | 0.4325 | 0.3175 | 0.845 | 0.461635 | 0.59982 | 0.149698 | 0.999971 | 1 | 0.793248 |
| Bpgm | P15327 | -0.55 | -1.13 | -0.02 | 0.49 | -0.08 | -1.06 | 1.09 | 0.11 | -0.43 | -1.15 | 0.41 | 0.19 | -0.69 | -1.35 | -0.22 | 0.49 | 0.3175 | 0.0575 | -0.14 | 0.544646 | 0.901286 | 0.773454 | 0.999971 | 1 | 0.963759 |
| Bphl | Q8R164 | -0.82 | -0.25 | 0.48 | -0.15 | -0.24 | -0.51 | 0.34 | 0.36 | -0.23 | -0.89 | 0.58 | 0.85 | -0.11 | -0.67 | 0.3 | -0.1 | 0.1725 | 0.2625 | 0.04 | 0.614092 | 0.563346 | 0.904097 | 0.999971 | 1 | 0.991091 |
| Bpnt1 | Q9Z0S1 | -1.46 | 1.25 | 0.04 | 0.09 | 1.49 | 0.25 | -0.16 | -0.11 | 1.35 | -0.21 | -0.17 | 0.26 | -1.24 | 1.18 | -0.26 | -0.49 | 0.3875 | 0.3275 | -0.1825 | 0.533636 | 0.593659 | 0.789086 | 0.999971 | 1 | 0.96741 |
| Bpnt1_1 | A0A1B0GR | 0.9 | 1.38 | 1.1 | 0.99 | 1.14 | 1.24 | 1.55 | 0.79 | 1.09 | 0.89 | 0.77 | 0.99 | 1.53 | 0.9 | 1.16 | 0.86 | 0.0875 | -0.1575 | 0.02 | 0.701596 | 0.412268 | 0.929731 | 0.999971 | 1 | 0.994744 |
| Bpnt2 | Q80V26 | 0.35 | 1.78 | 1.54 | 0.34 | 2.11 | 1.67 | 1.48 | 1.24 | 1.98 | 1.08 | 1.64 | 1.75 | 2.11 | 2.12 | 1.71 | 0.64 | 0.6225 | 0.61 | 0.6425 | 0.150363 | 0.161594 | 0.205903 | 0.999971 | 1 | 0.817241 |
| Bptf | A2A654 | 1.16 | 0.29 | 0.58 | 0.88 | 0.12 | -0.16 | 0.84 | 0.32 | 0.29 | 0.5 | 0.31 | -0.12 | -1.29 | -0.41 | 0 | -0.14 | -0.4475 | -0.4825 | -1.1875 | 0.152757 | 0.083929 | 0.006851 | 0.999971 | 1 | 0.582714 |
| Braf | P28028-1 | -0.84 | 0.46 | 0.76 | 0.28 | 0.03 | 0.56 | 0.6 | 0.25 | 0.32 | -0.7 | 0.4 | 0.2 | -0.05 | -0.2 | 0.62 | 0.33 | 0.195 | -0.11 | 0.01 | 0.595596 | 0.790076 | 0.979108 | 0.999971 | 1 | 0.99925 |
| Braf_1 | P28028 | -0.29 | -0.54 | -0.71 | -0.42 | -0.62 | -0.86 | -1.44 | -0.24 | -1.14 | -0.22 | -1.36 | -0.62 | -0.29 | -0.74 | -0.61 | -0.14 | -0.3 | -0.345 | 0.045 | 0.302796 | 0.245686 | 0.833555 | 0.999971 | 1 | 0.979036 |
| Brap | Q99MP8 | 1.12 | 1.4 | 1.5 | 1.37 | 1.2 | 1.5 | 1.21 | 1.49 | 1.43 | 1.66 | 1.32 | 1.06 | 0.94 | 1.43 | 1.23 | 1.73 | 0.0025 | 0.02 | -0.015 | 0.989323 | 0.920754 | 0.947084 | 0.999971 | 1 | 0.997651 |
| Brat1 | E9QLK3 | -0.22 | 0.22 | -0.45 | 0.24 | 0.5 | -0.07 | -0.07 | -0.02 | 0.47 | 0.05 | -0.07 | 0.15 | 0.12 | -0.02 | 0.33 | 0.24 | 0.1375 | 0.2025 | 0.22 | 0.582698 | 0.403218 | 0.345439 | 0.999971 | 1 | 0.863502 |
| Brawnin | Q8BTC1 | -0.66 | -1.44 | -0.52 | 0.1 | -1.18 | -1.96 | -0.74 | -1.52 | -1.56 | -1.32 | -1.26 | -0.91 | -2.11 | -1.66 | -1.71 | -0.6 | -0.72 | -0.6325 | -0.89 | 0.093563 | 0.090679 | 0.063521 | 0.999971 | 1 | 0.725348 |
| Brcc3 | P46737 | 0.09 | 1.2 | 0.35 | 0.12 | 0.84 | 1.22 | 0.1 | 0.41 | 1.4 | 1.05 | 0.29 | 0.33 | 1.59 | 1.26 | 0.56 | 0.41 | 0.2025 | 0.3275 | 0.515 | 0.567372 | 0.384249 | 0.190206 | 0.999971 | 1 | 0.810547 |
| Brd1 | G5E8P1 | -0.54 | -1.39 | -1.31 | -3.25 | -1.11 | -1.5 | -1.24 | -2.71 | -1.06 | -1.1 | -1.19 | -1.76 | -1.68 | -2.33 | -1.62 | -3.18 | -0.0175 | 0.345 | -0.58 | 0.977476 | 0.53688 | 0.362367 | 0.999971 | 1 | 0.869526 |
| Brd2 | Q7JJ13 | 0.8 | 0.55 | -0.12 | -0.94 | 0.31 | -0.07 | -0.14 | -0.01 | 0.36 | -0.82 | 0.09 | -0.35 | -1.52 | 0.33 | -0.17 | -0.46 | -0.05 | -0.2525 | -0.5275 | 0.897005 | 0.571275 | 0.317293 | 0.999971 | 1 | 0.853841 |
| Brd3 | Q8K2F0 | 0.9 | -0.76 | 1.07 | 1.31 | -0.38 | -0.62 | 0.54 | 1.24 | -0.15 | -0.2 | 0.98 | 1.74 | -1.63 | -1.03 | 1.46 | 1.29 | -0.435 | -0.0375 | -0.6075 | 0.462313 | 0.950926 | 0.469957 | 0.999971 | 1 | 0.892481 |
| Brd4 | Q3UH70 | 1.17 | 1.66 | 0.32 | 0.76 | 1.65 | 1.5 | 0.4 | 0.53 | 1.71 | 1.44 | 0.45 | 0.44 | 1.75 | 1.51 | 0.34 | 0.58 | 0.0425 | 0.0325 | 0.0675 | 0.917492 | 0.937789 | 0.873783 | 0.999971 | 1 | 0.984109 |
| Brd7 | O88665 | -1.23 | -0.71 | -0.96 | -1.06 | -0.95 | -1.38 | -0.76 | -0.58 | -1.31 | -1.27 | -1.88 | -2.27 | -1.46 | -0.89 | -0.97 | -1.86 | 0.0725 | -0.6925 | -0.305 | 0.760442 | 0.032706 | 0.27819 | 0.999971 | 1 | 0.843812 |
| Brd8 | Q8R3B7-2 | 3.2 | 0.84 | 1.9 | 2.04 | 0.35 | 1.16 | 2.16 | 1.06 | 0.79 | 1.93 | 1.28 | 1.13 | -1.62 | 0.38 | 0.78 | 1.02 | -0.8125 | -0.7125 | -1.855 | 0.171183 | 0.180421 | 0.024834 | 0.999971 | 1 | 0.67615 |
| Brd9 | A0A1Y7VJ6 | 1.77 | 0.03 | 0.32 | -0.4 | -0.38 | -2.11 | 0.31 | 0.08 | 0.24 | 0.8 | -0.56 | 0.25 | -2.18 | 0.78 | 1.08 | 0.37 | -0.955 | -0.2475 | -0.4175 | 0.169707 | 0.629225 | 0.601576 | 0.999971 | 1 | 0.924453 |
| Bri3bp | Q8BXV2 | -1.46 | -0.05 | 0.18 | 0.1 | 0.09 | 0.22 | -0.3 | 0.01 | 0.27 | -2.56 | -0.06 | -2.07 | -0.08 | -0.1 | 0.18 | 0.02 | 0.3125 | -0.7975 | 0.3125 | 0.428253 | 0.293854 | 0.419786 | 0.999971 | 1 | 0.87788 |
| Brinp1 | Q920P3 | -0.18 | -0.16 | -0.15 | -0.47 | 0 | -0.22 | 0.13 | -0.21 | -0.03 | -0.13 | 0.31 | 0.36 | 0.21 | -0.41 | -0.5 | -0.18 | 0.165 | 0.3675 | 0.02 | 0.386494 | 0.091907 | 0.927737 | 0.999971 | 1 | 0.994229 |
| Brinp2 | Q6DFY8 | -1.07 | -0.95 | -1.27 | -1.52 | -0.94 | -0.91 | -0.96 | -1.25 | -0.47 | -0.92 | -1.1 | -0.56 | -0.56 | -1.15 | -1.55 | -1.09 | 0.1875 | 0.44 | 0.115 | 0.36826 | 0.082364 | 0.662386 | 0.999971 | 1 | 0.936148 |
| Brinp3 | Q499E0 | 0.61 | -1.38 | -2.31 | -3.21 | -1.12 | -1.24 | -1.71 | -2.7 | -1.09 | -0.48 | -1.79 | -1.87 | -0.26 | -1.17 | -2.39 | -2.71 | -0.12 | 0.265 | -0.06 | 0.881057 | 0.740166 | 0.946281 | 0.999971 | 1 | 0.997646 |

| | | | | | | | | | | | | | | | | | | | | | | | | | | |
|---|---|---|---|---|---|---|---|---|---|---|---|---|---|---|---|---|---|---|---|---|---|---|---|---|---|---|
| Brix1 | Q9DCA5 | -0.26 | -2.17 | -1.02 | -1.46 | -1.82 | -2.57 | -1.52 | -1.47 | -3.22 | -1.99 | -1.75 | -1.39 | -3.21 | -2.41 | -0.97 | -1.27 | -0.6175 | -0.86 | -0.7375 | 0.1894 | 0.127646 | 0.238393 | 0.999971 | 1 | 0.833594 |
| Brk1 | Q91VR8 | -1.61 | -0.99 | -1.86 | -1.75 | 0.34 | -2.51 | 1.58 | -2.47 | 1.3 | -2.23 | -1.39 | -0.14 | 1 | -2.51 | -1.53 | -1.74 | 0.7875 | 0.9375 | 0.3575 | 0.40925 | 0.215951 | 0.617538 | 0.999971 | 1 | 0.928581 |
| Brms1l | Q3U1T3 | 0.16 | 0.09 | 0.94 | -0.09 | 0.44 | -0.21 | -0.37 | -0.27 | -0.34 | 0.36 | -0.8 | 0.92 | -1.3 | -0.13 | -0.6 | 0.34 | -0.3775 | -0.24 | -0.6975 | 0.230472 | 0.572214 | 0.108743 | 0.999971 | 1 | 0.78268 |
| Brox | Q8K2Q7 | 2.23 | 1.28 | 1.25 | 0.82 | 1.16 | 1.04 | 1.39 | 0.68 | 1.18 | 1.95 | 0.85 | 0.77 | 1.12 | 1.01 | 0.81 | 0.23 | -0.3275 | -0.2075 | -0.6025 | 0.336683 | 0.594788 | 0.114172 | 0.999971 | 1 | 0.78268 |
| Brpf1 | A0A0N4SU | -0.83 | 2.72 | 2.38 | 1.12 | 4.19 | 1.76 | 2.52 | 1.76 | 4.48 | 0.05 | 4.09 | 4.68 | 5.71 | 0.57 | 3.07 | 1.21 | 1.21 | 1.9775 | 1.2925 | 0.194165 | 0.132055 | 0.316365 | 0.999971 | 1 | 0.85361 |
| Brsk1 | Q5RJI5 | -0.44 | -0.7 | -0.12 | -0.26 | -0.72 | -0.63 | -0.07 | -0.09 | -0.49 | -0.45 | 0.02 | -0.16 | -0.31 | -0.72 | -0.09 | 0.09 | 0.0025 | 0.11 | 0.1225 | 0.99182 | 0.614424 | 0.620454 | 0.999971 | 1 | 0.929305 |
| Brsk2 | G3UWV4 | 1.46 | -1.1 | -0.98 | -0.26 | -1.4 | -0.73 | 0.24 | -0.4 | -2.03 | 0.14 | -0.42 | -1.02 | -1.69 | -1 | -0.72 | -1.13 | -0.3525 | -0.6125 | -0.915 | 0.573393 | 0.381897 | 0.136799 | 0.999971 | 1 | 0.784948 |
| Brsk2_1 | Q69Z98 | 0.4 | -0.24 | -0.1 | 0 | -0.37 | 0.04 | -0.18 | -0.06 | -0.14 | 0.01 | -0.15 | -0.19 | -0.25 | -0.27 | -0.14 | 0.32 | -0.1575 | -0.1325 | -0.1 | 0.4653 | 0.510531 | 0.668607 | 0.999971 | 1 | 0.938711 |
| Brwd1 | Q921C3 | 1.92 | -0.27 | -0.05 | 0 | 0.14 | 0.48 | 0.93 | -2.44 | -0.35 | 1.63 | -0.55 | -2.9 | 1.2 | -0.03 | -1.6 | -4.7 | -0.6225 | -0.9425 | -1.6825 | 0.45484 | 0.336545 | 0.192406 | 0.999971 | 1 | 0.810547 |
| Bscl2 | A0A0R4J22 | -0.92 | 0.32 | -0.37 | -0.32 | 0.17 | 0.63 | -0.34 | 0.17 | 0.91 | -0.64 | 0.37 | -0.07 | 1.78 | 0.47 | -0.04 | -0.01 | 0.48 | 0.465 | 0.8725 | 0.162745 | 0.262428 | 0.087823 | 0.999971 | 1 | 0.765196 |
| Bsdc1 | Q80Y55 | 0.99 | -0.06 | -0.25 | -0.48 | -0.64 | -0.12 | -0.04 | -0.47 | -0.66 | 0.16 | -0.61 | -0.44 | -0.23 | -0.28 | -0.5 | -0.24 | -0.3675 | -0.4375 | -0.3625 | 0.307689 | 0.251859 | 0.290229 | 0.999971 | 1 | 0.847402 |
| Bsg | K3W4Q8 | 0.61 | -0.29 | -0.11 | 0.28 | -0.69 | 0.03 | 0.87 | -0.07 | -0.75 | 0.43 | -0.35 | -0.57 | -0.53 | -0.48 | -0.23 | 0.01 | -0.0875 | -0.4325 | -0.43 | 0.812107 | 0.210455 | 0.126138 | 0.999971 | 1 | 0.783071 |
| Bsn | O88737 | -0.42 | -1.38 | -0.93 | -1.01 | -1.45 | -0.95 | -0.76 | -1.27 | -1.45 | -0.88 | -1.15 | -1.3 | -1.01 | -1.29 | -1.19 | -0.94 | -0.1725 | -0.26 | -0.1725 | 0.528436 | 0.323649 | 0.487294 | 0.999971 | 1 | 0.897906 |
| Btaf1 | E9QAE3 | -0.14 | 1.3 | 0.94 | 1.03 | 1.2 | 0.89 | 0.65 | 1.15 | 1.08 | -0.11 | 1.08 | 0.98 | 0.35 | 0.63 | 1 | 0.99 | 0.19 | -0.025 | -0.04 | 0.577783 | 0.951371 | 0.908584 | 0.999971 | 1 | 0.991707 |
| Btbd1 | P58544 | 3.95 | 0.18 | -0.04 | -1.03 | -0.38 | 0.71 | 0.14 | -0.43 | -0.46 | 2.34 | 0.02 | 0.62 | -1.37 | -0.05 | -0.31 | -0.17 | -0.755 | -0.135 | -1.24 | 0.459572 | 0.90433 | 0.2417 | 0.999971 | 1 | 0.835682 |
| Btbd10 | E9Q6L8 | 0.61 | -0.69 | -0.4 | -0.3 | -0.62 | -0.39 | -0.58 | -0.57 | -0.71 | 0.23 | -0.08 | 0.06 | -0.93 | -0.24 | -0.29 | 0.05 | -0.345 | 0.07 | -0.1575 | 0.26153 | 0.838492 | 0.649561 | 0.999971 | 1 | 0.933842 |
| Btbd11 | Q6GQW0 | -0.46 | -0.35 | -0.98 | -0.75 | -0.64 | -0.27 | -1.13 | -0.62 | -0.84 | -0.3 | -1.01 | -1.24 | -0.72 | -0.8 | -1.1 | -0.49 | -0.03 | -0.2125 | -0.1425 | 0.905728 | 0.431823 | 0.539345 | 0.999971 | 1 | 0.910077 |
| Btbd17 | Q9DB72 | -0.45 | -0.26 | -0.45 | -0.57 | -0.55 | -0.25 | -0.5 | -0.71 | -0.57 | -0.29 | -0.7 | -0.86 | -0.43 | -0.37 | -0.96 | -0.39 | -0.07 | -0.1725 | -0.105 | 0.708062 | 0.386166 | 0.61663 | 0.999971 | 1 | 0.928229 |
| Btbd8 | Q3TEG0 | -0.29 | 0.2 | -0.43 | -1.18 | 0.29 | 0.19 | 0.07 | -1.39 | 1.32 | -0.55 | 0.41 | 0.87 | 1.26 | -1.09 | -1.33 | -0.61 | 0.215 | 0.9375 | -0.0175 | 0.640209 | 0.069031 | 0.976586 | 0.999971 | 1 | 0.99925 |
| Btbd9 | Q8C726 | 1.44 | -0.06 | -0.46 | 1.32 | -0.09 | 0.36 | -0.21 | 0.69 | 0.31 | 1.16 | -0.65 | -0.21 | -0.31 | 0.55 | 0.03 | 1.32 | -0.3725 | -0.4075 | -0.1625 | 0.452134 | 0.481309 | 0.76808 | 0.999971 | 1 | 0.962535 |
| Btd | A0A0R4J13 | -1.23 | -0.08 | 1.56 | 0.91 | -0.14 | -0.69 | 0.88 | 0.66 | -0.24 | -1.2 | 2.01 | 1.68 | 0.37 | -0.32 | 1.25 | 1.11 | -0.1125 | 0.2725 | 0.3125 | 0.861175 | 0.758482 | 0.630066 | 0.999971 | 1 | 0.932649 |
| Btf3 | Q64152 | 3.14 | 1.69 | 1.79 | 2.12 | 1.61 | 1.85 | 1.71 | 2.34 | 1.67 | 2.56 | 1.88 | 1.74 | 1.01 | 1.82 | 2.14 | 2.18 | -0.3075 | -0.2225 | -0.3975 | 0.403007 | 0.559006 | 0.344374 | 0.999971 | 1 | 0.862346 |
| Btf3l4 | Q9CQH7 | 0.5 | 2.33 | 2.26 | 2.39 | 2.28 | 2 | 2.05 | 2.31 | 2.27 | 1.35 | 2.48 | 2.63 | 2.28 | 2.02 | 2.21 | 2.84 | 0.29 | 0.3125 | 0.4675 | 0.512728 | 0.538998 | 0.324085 | 0.999971 | 1 | 0.85548 |
| Bub3 | A0A140LH/ | 3.38 | 3.17 | 3.08 | 3.27 | 2.89 | 3.45 | 3.62 | 2.46 | 3.26 | 3.34 | 2.79 | 2.31 | 1.9 | 2.67 | 2.36 | 1.93 | -0.12 | -0.3 | -1.01 | 0.676627 | 0.277373 | 0.001906 | 0.999971 | 1 | 0.476373 |
| Bud13 | Q8R149 | 1.89 | 0.48 | -0.04 | -0.54 | -0.14 | 1.36 | -0.12 | 0.64 | -0.07 | 1.99 | 0.89 | 0.14 | 0.91 | 0.11 | 0.16 | 0.21 | -0.0125 | 0.29 | -0.1 | 0.982795 | 0.652701 | 0.846256 | 0.999971 | 1 | 0.979924 |
| Bud31 | Q6PGH1 | 1.72 | 0.74 | 0.37 | 0.08 | 0.01 | 0.38 | 0.9 | -0.07 | 0.2 | 1.11 | 0.66 | -0.01 | -0.27 | 0.55 | 0.38 | -0.32 | -0.4225 | -0.2375 | -0.6425 | 0.308199 | 0.571201 | 0.137434 | 0.999971 | 1 | 0.784948 |
| Bysl | O54825 | -2.64 | -0.19 | -0.02 | -1.44 | -0.83 | -0.92 | -0.06 | -1.11 | -0.9 | -3.39 | 0.1 | 0.12 | -0.74 | -1.92 | -0.81 | -2.42 | 0.3425 | 0.055 | -0.4 | 0.569154 | 0.952561 | 0.55492 | 0.999971 | 1 | 0.91473 |
| Bzw1 | A0A087WC | 0.97 | 1.54 | 1.94 | 1.83 | 1.28 | 1.39 | 1.6 | 1.58 | 1.61 | 1.45 | 1.72 | 0.58 | 0.8 | 1.15 | 2.1 | 1.82 | -0.1075 | -0.23 | -0.1025 | 0.67556 | 0.499307 | 0.776886 | 0.999971 | 1 | 0.964475 |
| Bzw2 | Q91VK1 | 1.23 | 2.01 | 1.14 | 0.96 | 1.75 | 2.04 | 0.91 | 1.47 | 2.03 | 2.02 | 1.07 | 1.34 | 1.85 | 2.01 | 1.31 | 1.26 | 0.2075 | 0.28 | 0.2725 | 0.537464 | 0.411666 | 0.385041 | 0.999971 | 1 | 0.871816 |
| C030006K1 | Q8VE95 | -0.4 | -0.88 | -0.23 | 0.11 | -1.32 | -0.65 | -0.96 | 0.02 | -1.13 | -0.78 | -0.06 | -1.25 | -1.05 | -1.03 | -0.76 | -0.73 | -0.3775 | -0.455 | -0.5425 | 0.291864 | 0.198447 | 0.055484 | 0.999971 | 1 | 0.725348 |
| C1qa | P98086 | 2.43 | 2.39 | 1.6 | 1.89 | 2.36 | 2.32 | 1.2 | 1.38 | 1.95 | 1.98 | 1.45 | 2.07 | 2 | 2.52 | 1.83 | 1.77 | -0.2625 | -0.215 | -0.0475 | 0.469768 | 0.425685 | 0.865245 | 0.999971 | 1 | 0.983318 |
| C1qb | P14106 | 2.25 | 2.74 | 0.8 | 0.99 | 2.45 | 2.67 | 0.99 | 1.1 | 2.9 | 2.65 | 1.49 | 1.37 | 2.64 | 2.39 | 1.49 | 1.46 | 0.1075 | 0.4075 | 0.3 | 0.855636 | 0.479297 | 0.568601 | 0.999971 | 1 | 0.91816 |
| C1qbp | Q8R5L1 | -1.38 | -0.57 | -1.79 | -1.64 | -0.64 | -0.86 | -1.71 | -1.76 | -0.85 | -1.04 | -1.93 | -2.06 | -0.76 | -0.45 | -1.74 | -2.03 | 0.1025 | -0.125 | 0.1 | 0.788591 | 0.751591 | 0.820522 | 0.999971 | 1 | 0.975761 |
| C1qc | Q02105 | 0.63 | 1.28 | 0.87 | 1.04 | 1.04 | 1.1 | 1.17 | 0.89 | 0.98 | 1.11 | 0.78 | 0.94 | 0.25 | 1.24 | 1 | 1.34 | 0.095 | -0.0025 | 0.0025 | 0.643915 | 0.990205 | 0.993196 | 0.999971 | 1 | 0.99925 |
| C1ql3 | Q9ESN4 | -0.84 | -1.27 | -0.8 | -0.65 | -0.98 | -0.98 | -0.56 | -0.75 | -0.91 | -0.78 | -0.83 | -1.15 | -1.04 | -1.2 | -0.62 | -0.68 | 0.0725 | -0.0275 | 0.005 | 0.736311 | 0.893635 | 0.982752 | 0.999971 | 1 | 0.99925 |
| C1qtnf4 | Q8R066 | -0.78 | -0.92 | -0.71 | -0.81 | -0.45 | -0.99 | -0.57 | -0.48 | -0.57 | -0.9 | -0.59 | -0.18 | -0.23 | -0.67 | -0.52 | -0.62 | 0.1825 | 0.245 | 0.295 | 0.361874 | 0.250138 | 0.134859 | 0.999971 | 1 | 0.784948 |
| C2cd2 | E9Q3C1 | 0.92 | -0.06 | -0.24 | -0.28 | -0.03 | 0.12 | 0.11 | -0.51 | 0.06 | 1.17 | -0.21 | -0.39 | 0.08 | -0.48 | -0.42 | -0.68 | -0.1625 | 0.0725 | -0.46 | 0.614998 | 0.865147 | 0.183065 | 0.999971 | 1 | 0.809025 |
| C2cd2l | Q80X80 | -0.19 | -1.04 | -0.78 | -0.89 | -0.85 | -1.04 | -0.76 | -0.81 | -0.88 | -0.4 | -0.72 | -0.69 | -0.42 | -0.69 | -0.72 | -0.76 | -0.14 | 0.0525 | 0.0775 | 0.551285 | 0.827651 | 0.744114 | 0.999971 | 1 | 0.961064 |
| C2cd4c | Q5HZI2 | 0.62 | -0.25 | -0.19 | -0.08 | -0.56 | 0.5 | -0.23 | -0.19 | -0.23 | 0.45 | -0.25 | -0.55 | -0.07 | 0.1 | -0.09 | 0.14 | -0.145 | -0.17 | -0.005 | 0.638212 | 0.574083 | 0.983522 | 0.999971 | 1 | 0.99925 |
| C2cd5 | Q7TPS5 | 2.27 | -1.72 | -0.95 | -0.94 | -2.55 | -2.04 | -1.53 | -0.79 | 2.46 | 3.22 | -2.34 | -1.2 | -0.84 | -1.83 | -2.96 | -1.86 | -1.3925 | 0.87 | -1.5375 | 0.133503 | 0.552962 | 0.110029 | 0.999971 | 1 | 0.78268 |
| C2cd5_1 | A0A0N4SW | -1.14 | -0.84 | -0.63 | -1.13 | -0.86 | -0.7 | -0.81 | -0.75 | -0.7 | -0.86 | -0.74 | -0.78 | -0.37 | -0.81 | -0.73 | -0.61 | 0.155 | 0.165 | 0.305 | 0.430096 | 0.395247 | 0.168987 | 0.999971 | 1 | 0.80174 |
| C3 | P01027 | 2.19 | 2.96 | 1 | 1.95 | 4.41 | 1.9 | 2.96 | 1.14 | 2.23 | 0.88 | 1.91 | 2.72 | 2.82 | 1.15 | 1.49 | 1.34 | 0.5775 | -0.09 | -0.325 | 0.440565 | 0.862743 | 0.531594 | 0.999971 | 1 | 0.910069 |
| C4b | P01029 | 0.88 | -1.23 | -2.53 | -0.33 | 1.6 | 1.15 | 0.24 | 0.99 | 1.52 | 1.26 | 0.48 | 1.36 | 1.56 | 1.17 | -0.3 | 0.63 | 1.7975 | 1.9575 | 1.5675 | 0.028427 | 0.018642 | 0.060854 | 0.999971 | 1 | 0.725348 |
| C9orf72 | Q6DFW0 | -0.42 | -1.56 | 0.67 | 0.8 | -1.37 | -1.52 | 0.07 | 0.89 | -1.61 | -1.46 | 0.83 | 0.67 | -1.5 | -1.8 | 0.75 | 0.86 | -0.355 | -0.265 | -0.295 | 0.625297 | 0.734472 | 0.716354 | 0.999971 | 1 | 0.952576 |
| Ca1 | P13634 | -0.35 | -0.36 | -1.34 | 1.03 | 1.58 | -1.83 | 0.67 | -0.6 | -1.75 | -2.52 | -1.13 | -1.45 | -0.54 | -1.76 | -1.21 | -0.64 | 0.21 | -1.4575 | -0.7825 | 0.792931 | 0.021433 | 0.1588 | 0.999971 | 1 | 0.798217 |
| Ca10 | P61215 | -0.34 | -1.28 | -2.31 | -1.97 | -1.17 | -1.91 | -2 | -2.13 | -1.04 | -0.99 | -2.47 | -2.52 | -1.84 | -2.02 | -2.45 | -2.43 | -0.3275 | -0.28 | -0.71 | 0.478226 | 0.620983 | 0.129913 | 0.999971 | 1 | 0.784948 |
| Ca11 | O70354 | 2.11 | 0.59 | -0.66 | 0.05 | 1.29 | 0.65 | -0.28 | -0.11 | 1.12 | 1.71 | -0.1 | 0.31 | 1.45 | 0.22 | -0.01 | 0.6 | -0.135 | 0.2375 | 0.0425 | 0.829863 | 0.716666 | 0.944521 | 0.999971 | 1 | 0.997225 |
| Ca14 | Q9WVT6 | -1.2 | -0.99 | -0.68 | -0.78 | -0.94 | -1.51 | -1.43 | -1.86 | -1 | -1.23 | -1.65 | -1.42 | -1.98 | -1.08 | -0.67 | -1.54 | -0.5225 | -0.4125 | -0.405 | 0.061953 | 0.088737 | 0.215777 | 0.999971 | 1 | 0.826893 |
| Ca2 | P00920 | -0.92 | -0.54 | -0.84 | -0.94 | -0.03 | -0.93 | 0.02 | -1.35 | -0.57 | -1.57 | -0.85 | -0.76 | -0.49 | -0.8 | -1.02 | -1.29 | 0.2375 | -0.1275 | -0.09 | 0.498356 | 0.625364 | 0.698415 | 0.999971 | 1 | 0.947781 |
| Ca4 | Q64444 | -0.8 | -0.37 | -0.65 | -0.78 | -0.67 | -0.64 | -1.36 | -1.34 | -0.52 | -0.7 | -0.77 | -0.65 | -0.29 | -0.54 | -1.08 | -0.77 | -0.3525 | -0.01 | -0.02 | 0.184502 | 0.956129 | 0.931475 | 0.999971 | 1 | 0.995069 |
| Ca5b | Q9QZA0 | -0.2 | -0.44 | -0.57 | -1.03 | -0.37 | -0.95 | -1.03 | -0.39 | -0.69 | -0.71 | 0.11 | -1.06 | 0.16 | -0.51 | -0.63 | -0.52 | -0.125 | -0.0275 | 0.185 | 0.643303 | 0.928802 | 0.500347 | 0.999971 | 1 | 0.904223 |
| Ca7 | Q9ERQ8 | 0 | -0.44 | -0.18 | -0.37 | -0.82 | -0.57 | -1.05 | -0.45 | -1.38 | -1.21 | -0.91 | -1.43 | -0.64 | -0.91 | -1.31 | -0.52 | -0.475 | -0.985 | -0.5975 | 0.050267 | 0.001008 | 0.030537 | 0.999971 | 0.540208 | 0.678474 |
| Ca8 | P28651 | -0.01 | 0.59 | 0.55 | 0.5 | -0.15 | 0.56 | 0.11 | 0.55 | 0.03 | 0.06 | 0.53 | 0.55 | 1.15 | 0.84 | 0.88 | 1.22 | -0.14 | -0.115 | 0.615 | 0.580554 | 0.624978 | 0.018399 | 0.999971 | 1 | 0.635388 |
| Caap1 | Q8VDY9 | -0.25 | 0.49 | -7.72 | -5.12 | 0.64 | 0.83 | -6.58 | -3 | 0.06 | -1.05 | -4.19 | -7.38 | 0.85 | 1.62 | -3.69 | -4.67 | 1.1225 | 0.01 | 1.6775 | 0.631854 | 0.99653 | 0.460348 | 0.999971 | 1 | 0.889706 |
| Cab39 | Q06138 | -0.1 | 0.06 | -0.47 | -0.68 | 0.1 | 0.33 | -0.22 | -0.94 | 0.37 | 0.04 | -0.54 | -0.34 | -0.11 | -0.22 | -0.69 | -0.94 | 0.115 | 0.18 | -0.1925 | 0.724106 | 0.522819 | 0.492183 | 0.999971 | 1 | 0.899886 |
| Cab39l | Q9DB16 | 0.16 | 1.13 | 1.22 | 0.98 | 0.92 | 1.06 | 0.78 | 0.69 | 0.51 | 0.03 | -1.14 | 0.72 | 0.52 | 1.09 | 0.92 | 1.11 | -0.01 | -0.8425 | 0.0375 | 0.970822 | 0.090992 | 0.897378 | 0.999971 | 1 | 0.98896 |
| Cabin1 | G3X8Q1 | 1.46 | -0.03 | 0.85 | 0.64 | 0.08 | -0.13 | 0.97 | 0.22 | -0.18 | 1.11 | 0.35 | 0.13 | 0.42 | -0.47 | 0.42 | -0.01 | -0.445 | -0.3775 | -0.64 | 0.256419 | 0.353568 | 0.106736 | 0.999971 | 1 | 0.78268 |
| Cables1 | Q3U148 | -1.22 | -0.94 | -1.25 | -1.1 | -1.12 | -1.02 | -1.45 | -0.89 | -0.8 | -1.41 | -1.84 | -1.3 | -1.07 | -1.14 | -0.94 | -1.07 | 0.0075 | -0.21 | 0.0725 | 0.969823 | 0.411658 | 0.676717 | 0.999971 | 1 | 0.942048 |
| Cables2 | Q8K3M5 | -3.1 | -3.07 | -1.1 | -0.85 | -2.44 | -2.55 | -0.78 | 0.14 | -2.53 | -3.55 | -1.18 | -0.84 | -2.23 | -2.47 | -0.48 | -1.34 | 0.6225 | 0.005 | 0.4 | 0.447371 | 0.994957 | 0.565797 | 0.999971 | 1 | 0.917959 |
| Cabp1 | D3Z1M4 | 1.69 | 0.29 | 1.04 | 1.47 | 0.23 | 0.3 | 2.92 | 0.69 | -0.73 | 0.62 | 1.51 | 1.42 | 0.38 | 0.13 | 0.92 | 1.86 | -0.0875 | -0.4175 | -0.3 | 0.891589 | 0.460966 | 0.520591 | 0.999971 | 1 | 0.906972 |
| Cacfd1 | A6PWX1 | 0.08 | -1.53 | -1.54 | -0.36 | -0.1 | 0.58 | -0.77 | 0 | 0.03 | 0.51 | -1.83 | -0.61 | 0.34 | -0.31 | -0.19 | 0.1 | 0.765 | 0.3625 | 0.8225 | 0.126546 | 0.549861 | 0.075119 | 0.999971 | 1 | 0.746958 |
| Cachd1 | Z4YK18 | 0.06 | -0.04 | -0.25 | -0.2 | 0.21 | 0.5 | -0.68 | 0 | -0.08 | 0.37 | -0.63 | -0.07 | 0.55 | 0.42 | 0.17 | 0.48 | 0.115 | 0.005 | 0.5125 | 0.679849 | 0.983694 | 0.020221 | 0.999971 | 1 | 0.635388 |
| Cacna1a | A0A571BE1 | -0.34 | -0.54 | -0.18 | -0.31 | -0.72 | -0.21 | -0.42 | -0.33 | -0.69 | -0.2 | -0.26 | -0.4 | -0.33 | -0.73 | -0.4 | -0.35 | -0.0775 | -0.045 | -0.11 | 0.691852 | 0.814911 | 0.56491 | 0.999971 | 1 | 0.917442 |
| Cacna1a_1 | A0A571BE7 | -0.44 | -0.78 | -1.64 | -1.56 | -0.54 | 0.76 | -2.04 | -1.86 | 0.19 | 0.18 | -2.48 | -2.14 | -0.04 | 0.13 | -1.66 | -1.46 | 0.185 | 0.0425 | 0.3475 | 0.776085 | 0.952254 | 0.502437 | 0.999971 | 1 | 0.904573 |
| Cacna1b | O55017-2 | -0.49 | -0.83 | -0.78 | -1.02 | -0.89 | -0.86 | -0.38 | -1.17 | -0.74 | -0.75 | -0.62 | -0.68 | -0.15 | -1.18 | -1.05 | -1.16 | -0.045 | 0.0825 | -0.105 | 0.847141 | 0.65325 | 0.71304 | 0.999971 | 1 | 0.951793 |
| Cacna1c | A0A087WF | -1.15 | -1.9 | -1.24 | -1.46 | -1.6 | -1.96 | -1.39 | -1.19 | -1.89 | -1.51 | -1.38 | -1.53 | -1.37 | -1.93 | -1.42 | -1.23 | -0.0975 | -0.14 | -0.05 | 0.707245 | 0.55264 | 0.843865 | 0.999971 | 1 | 0.979814 |
| Cacna1d | A0A589Q4 | 0.28 | -1.19 | -1.07 | -0.29 | -0.96 | -0.01 | -1.06 | -0.58 | -0.75 | -0.46 | -1.2 | -0.74 | -0.7 | -0.91 | -1 | -0.6 | -0.085 | -0.22 | -0.235 | 0.832165 | 0.554229 | 0.510005 | 0.999971 | 1 | 0.905815 |
| Cacna1e | A0A087WS | -0.9 | -0.77 | -0.73 | -0.94 | -0.9 | -0.94 | -0.78 | -0.78 | -0.81 | -0.88 | -0.9 | -0.8 | -0.3 | -1.08 | -1.02 | -0.97 | -0.015 | -0.0125 | -0.0075 | 0.928283 | 0.937852 | 0.973829 | 0.999971 | 1 | 0.99925 |
| Cacna1e_1 | A0A571BF1 | -2.32 | -0.65 | 1.28 | 0.16 | -0.64 | -0.35 | 0.78 | -0.48 | -0.4 | -1.72 | 0.56 | 0.38 | -1.36 | -1.67 | 0.32 | -0.03 | 0.21 | 0.0875 | -0.3025 | 0.777483 | 0.915947 | 0.710024 | 0.999971 | 1 | 0.950695 |
| Cacna1g | Q5SUG2 | 2.22 | -1.76 | -1.49 | -1.74 | -2 | -0.05 | -0.94 | -1.45 | -1.88 | 0.43 | -1.92 | -2.34 | -2.13 | -0.99 | -2.05 | -1.81 | -0.4175 | -0.735 | -1.0525 | 0.66014 | 0.486209 | 0.262456 | 0.999971 | 1 | 0.842262 |
| Cacna1h | E9Q6P9 | -0.66 | -1.25 | -0.44 | -0.32 | -1.11 | -0.97 | -0.04 | 0.1 | -0.68 | -0.39 | -0.83 | -0.58 | -0.37 | -0.87 | 0.05 | -0.49 | 0.1625 | 0.0475 | 0.2475 | 0.656904 | 0.850147 | 0.406882 | 0.999971 | 1 | 0.874911 |
| Cacna1i | E9Q7P2 | 0.62 | -0.43 | 0.14 | 0.15 | -0.92 | -0.24 | -0.07 | -0.02 | -0.62 | 0.32 | -0.08 | -0.31 | -0.5 | -0.73 | -0.5 | -0.17 | -0.4325 | -0.2925 | -0.595 | 0.181108 | 0.342868 | 0.048656 | 0.999971 | 1 | 0.710102 |
| Cacna2d1 | O08532-5 | -0.95 | -1.44 | -1.16 | -1.05 | -1.18 | -1.09 | -1.22 | -1.04 | -1.14 | -1.23 | -1.14 | -1.05 | -0.9 | -1.28 | -1.18 | -1.18 | 0.0175 | 0.01 | 0.015 | 0.924836 | 0.956094 | 0.939079 | 0.999971 | 1 | 0.995869 |
| Cacna2d2 | E9Q683 | -1.42 | -1.58 | -1.08 | -1.2 | -0.94 | -1.57 | -1.29 | -1.03 | -0.67 | -1.91 | -1.45 | -0.86 | -0.86 | -1.44 | -0.76 | -1.47 | 0.1125 | 0.0975 | 0.1875 | 0.615982 | 0.752959 | 0.457111 | 0.999971 | 1 | 0.88754 |
| Cacna2d3 | Q9Z1L5 | -0.44 | -0.62 | -0.83 | -0.89 | -0.47 | -0.62 | -0.85 | -0.38 | -0.36 | -0.43 | -0.77 | -0.94 | 0.13 | -0.4 | -0.76 | -0.7 | 0.115 | 0.07 | 0.2625 | 0.572664 | 0.745728 | 0.317618 | 0.999971 | 1 | 0.853893 |
| Cacnb1 | Q8R3Z5-3 | 0.58 | -1.33 | -1.81 | -0.27 | -1.22 | -1.01 | -1.26 | -0.98 | -2.51 | -0.03 | -0.86 | -2.07 | -2.86 | -0.81 | -2.42 | -1.32 | -0.41 | -0.66 | -1.145 | 0.422646 | 0.364505 | 0.106322 | 0.999971 | 1 | 0.78268 |
| Cacnb1_1 | A2A543 | -1.23 | -1.85 | -1.46 | -1.59 | -1.62 | -1.71 | -1.95 | -1.76 | -1.57 | -1.02 | -1.44 | -1.34 | -1.37 | -1.66 | -1.49 | -1.19 | -0.2275 | 0.19 | 0.105 | 0.279615 | 0.391966 | 0.623968 | 0.999971 | 1 | 0.930479 |
| Cacnb2 | Q8CC27 | -1.68 | -1.23 | -1.69 | -1.41 | -1.25 | -1.4 | -1.64 | -1.38 | -1.31 | -2.06 | -1.84 | -2.33 | -1.32 | -1.48 | -1.64 | -2 | 0.085 | -0.3825 | -0.1075 | 0.669262 | 0.171765 | 0.635236 | 0.999971 | 1 | 0.93301 |
| Cacnb3 | P54285 | -0.9 | -1.02 | -0.79 | -0.9 | -0.99 | -0.86 | -0.72 | -0.94 | -0.98 | -0.95 | -0.68 | -0.8 | -0.79 | -1.11 | -0.9 | -0.85 | 0.025 | 0.05 | -0.01 | 0.883092 | 0.768615 | 0.954095 | 0.999971 | 1 | 0.998056 |
| Cacnb4 | Q8R0S4 | -0.01 | -0.94 | -1.01 | -1.04 | -1.37 | -1.1 | -0.51 | -1.25 | -1.28 | -0.72 | -1.11 | -1.33 | -1.09 | -1.18 | -1.47 | -1.22 | -0.3075 | -0.36 | -0.49 | 0.343675 | 0.240122 | 0.105041 | 0.999971 | 1 | 0.78268 |
| Cacng2 | O88602 | -1.3 | -1.35 | -1.74 | -2.2 | -1.33 | -1.45 | -1.52 | -1.51 | -1.59 | -1.39 | -1.35 | -1.95 | -0.66 | -1.62 | -2.14 | -1.93 | 0.195 | 0.0775 | 0.06 | 0.432855 | 0.772992 | 0.873256 | 0.999971 | 1 | 0.984109 |
| Cacng3 | Q9JJV5 | -1.1 | 0.34 | 0.61 | -1.07 | 0.26 | 0.45 | 2.01 | -0.98 | 0.62 | -0.87 | 0.83 | 0.91 | 1.07 | -0.79 | -0.38 | -0.85 | 0.74 | 0.6775 | 0.0675 | 0.299724 | 0.253455 | 0.907946 | 0.999971 | 1 | 0.991496 |
| Cacng4 | Q9JJV4 | -1.81 | -1.35 | -1.02 | -0.94 | -2.13 | -2.53 | -1.01 | -0.44 | -2.18 | -1.56 | -1.22 | -0.61 | -2.13 | -2.47 | -0.81 | -0.75 | -0.2475 | -0.1125 | -0.26 | 0.613536 | 0.76275 | 0.572687 | 0.999971 | 1 | 0.91816 |
| Cacng7 | P62956 | -1.89 | -1.09 | -1.2 | -1.56 | -0.56 | 0.21 | -0.86 | -1.54 | -0.37 | -0.09 | -2.13 | -0.72 | 0.28 | 0.8 | -0.97 | -1.74 | 0.7475 | 0.6075 | 1.0275 | 0.082022 | 0.208599 | 0.092991 | 0.999971 | 1 | 0.776565 |
| Cacng8 | S4R2F1 | -0.65 | -1.02 | -1.28 | -1.34 | -1.15 | -0.84 | -0.99 | -1.33 | -1.16 | -0.82 | -1.45 | -1.69 | -1.02 | -1.11 | -1.09 | -1.6 | -0.005 | -0.2075 | -0.1325 | 0.982499 | 0.441207 | 0.583686 | 0.999971 | 1 | 0.920364 |
| Cactin | Q9CS00 | -0.67 | -0.14 | 0.19 | -0.82 | -0.34 | -0.68 | -1.51 | 0.03 | -0.42 | -0.94 | -0.57 | -0.29 | -0.35 | 0.19 | -0.72 | 0.33 | -0.265 | -0.195 | 0.2225 | 0.500288 | 0.500404 | 0.51342 | 0.999971 | 1 | 0.905815 |
| Cacul1 | E9PV69 | 0.27 | 0.49 | 2.25 | 2.59 | 0.35 | 0.62 | 1.88 | 0.26 | 0.27 | 0.4 | 1.01 | 1.48 | 0.46 | -0.29 | -0.72 | 1.07 | -0.6225 | -0.61 | -1.27 | 0.34331 | 0.326923 | 0.077701 | 0.999971 | 1 | 0.752571 |
| Cacybp | Q9CXW3 | -1.06 | -0.3 | 1.51 | 1.16 | 0.48 | -0.09 | 1.48 | 1.04 | 0.39 | -1.08 | 1.2 | 1.09 | -0.02 | -0.3 | 1.33 | 1.03 | 0.4 | 0.0725 | 0.1825 | 0.531631 | 0.920661 | 0.781755 | 0.999971 | 1 | 0.965859 |
| Cad | E9QAI5 | -0.43 | -0.37 | -0.1 | -0.01 | -0.16 | -0.46 | -0.1 | 0.38 | -0.15 | -0.32 | 0.13 | 0.13 | 0.15 | -0.03 | 0.3 | 0.24 | 0.1425 | 0.175 | 0.3925 | 0.550522 | 0.397853 | 0.066678 | 0.999971 | 1 | 0.733937 |
| Cadm1 | E0CY16 | -1.38 | -0.87 | -1.56 | -0.74 | -0.8 | -1.93 | -1.3 | -1.44 | -1.33 | -1.21 | -0.79 | -1.35 | -1.75 | -1.13 | -1.81 | -1.65 | -0.23 | -0.0325 | -0.4475 | 0.465307 | 0.899945 | 0.125466 | 0.999971 | 1 | 0.78268 |
| Cadm1_1 | Q8R5M8-4 | -0.71 | -1.42 | 0.56 | 0.41 | -1.3 | -1.14 | 0.08 | 0.43 | -1.3 | -1.23 | 0.08 | 0.41 | -1.09 | -1.17 | 0.48 | 0.5 | -0.1925 | -0.22 | -0.03 | 0.742674 | 0.711681 | 0.960521 | 0.999971 | 1 | 0.998721 |
| Cadm2 | G3UZM4 | -0.62 | -1.02 | -1.21 | -1.25 | -1.37 | -0.53 | -1.46 | -1.66 | -1.24 | -0.67 | -1.76 | -1.82 | -1.15 | -0.82 | -1.6 | -1.41 | -0.23 | -0.3475 | -0.22 | 0.446865 | 0.278729 | 0.390819 | 0.999971 | 1 | 0.871816 |
| Cadm3 | K4DI58 | -0.27 | -2.08 | -1.18 | -0.87 | -1.66 | -0.75 | -0.76 | -1.35 | -1.23 | -0.38 | -1.78 | -2.01 | -1.72 | -0.91 | -1.18 | -1.58 | -0.03 | -0.25 | -0.2475 | 0.942628 | 0.611542 | 0.542911 | 0.999971 | 1 | 0.911251 |
| Cadm4 | Q8R464 | -0.73 | -0.92 | -0.92 | -0.67 | -0.75 | -0.94 | -0.7 | -0.5 | -0.72 | -0.85 | -1.09 | -0.88 | -0.1 | -0.93 | -0.84 | -0.58 | 0.0875 | -0.075 | 0.1975 | 0.637451 | 0.672201 | 0.408532 | 0.999971 | 1 | 0.875014 |
| Cadps | Q80TJ1 | -0.57 | -0.44 | -0.35 | -0.52 | -0.52 | -0.62 | -0.32 | -0.77 | -0.62 | -0.43 | -0.7 | -0.96 | -0.33 | -0.59 | -0.76 | -0.87 | -0.0875 | -0.2075 | -0.1675 | 0.633127 | 0.281503 | 0.395872 | 0.999971 | 1 | 0.871816 |
| Cadps_1 | Q80TJ1-4 | -1.27 | -0.54 | -1.61 | -1.13 | -0.93 | -0.79 | -1.56 | -1.58 | -0.98 | -1.66 | -2.28 | -1.97 | -1.49 | -0.42 | -1.54 | -1.33 | -0.0775 | -0.585 | -0.0575 | 0.802573 | 0.122601 | 0.866215 | 0.999971 | 1 | 0.983462 |
| Cadps_2 | Q80TJ1-2 | 0 | -0.96 | -0.29 | -0.13 | -1.23 | -1.34 | -1.02 | -0.39 | -1.24 | -0.52 | -0.63 | -0.01 | -1.32 | -0.94 | -0.07 | -0.3 | -0.65 | -0.255 | -0.3125 | 0.059437 | 0.447473 | 0.385973 | 0.999971 | 1 | 0.871816 |
| Cadps_3 | A0A286YE( | -2.86 | -1.06 | -1.9 | -2.19 | -1.96 | -2.05 | -2.31 | -1.44 | -1.67 | -1.68 | -1.7 | -2.41 | -4.84 | -0.05 | -1.82 | -2.04 | 0.0625 | 0.1375 | -0.185 | 0.874955 | 0.730572 | 0.845124 | 0.999971 | 1 | 0.979814 |
| Cadps2 | E9Q835 | -0.39 | -0.71 | -0.67 | -1.08 | -0.83 | -0.56 | -0.54 | -1.28 | -0.68 | -0.61 | -0.82 | -1.05 | -0.73 | -0.76 | -1.12 | -1.47 | -0.09 | -0.0775 | -0.3075 | 0.720425 | 0.719497 | 0.243035 | 0.999971 | 1 | 0.837316 |
| Cadps2_1 | Q8BYR5-4 | 1.7 | 1.02 | -0.99 | -0.35 | 1.2 | 0.56 | -0.2 | 0.26 | 0.18 | 0.11 | -0.17 | 0.19 | 2.16 | -0.18 | -0.85 | 1 | 0.11 | -0.2675 | 0.1875 | 0.859424 | 0.642527 | 0.818069 | 0.999971 | 1 | 0.975007 |
| Calb1 | P12658 | 0.62 | 1.02 | 0.65 | 0.5 | 1.13 | 1.15 | 1.01 | 0.77 | 1.31 | 0.68 | 1.17 | 1.57 | 1.42 | 1.14 | 0.86 | 1.01 | 0.3175 | 0.485 | 0.41 | 0.139298 | 0.075162 | 0.081218 | 0.999971 | 1 | 0.755254 |
| Calb2 | Q08331 | -0.06 | -0.64 | 0.23 | 0.24 | 0.34 | -0.14 | 0.85 | 0.54 | 1.42 | 0.65 | 0.29 | 1.1 | 0.91 | 0.06 | 0.92 | -0.13 | 0.455 | 0.9225 | 0.4975 | 0.156014 | 0.018298 | 0.17093 | 0.999971 | 1 | 0.803758 |
| Calcoco1 | Q8CGU1 | 0.81 | -0.19 | -0.72 | -0.52 | -0.49 | -0.45 | -0.65 | -0.82 | -0.31 | 0.15 | -0.84 | -0.92 | -0.44 | -0.55 | -1.17 | -0.98 | -0.4475 | -0.325 | -0.63 | 0.216614 | 0.429808 | 0.116772 | 0.999971 | 1 | 0.78268 |
| Cald1 | Q8VCQ8 | 0.96 | 1.97 | 0.93 | 2.5 | 4.63 | 2.43 | 2.43 | 0.66 | 2.35 | 2.62 | 2.97 | 3.83 | 0.84 | 2.1 | 1.78 | 1.4 | 0.9475 | 1.3525 | -0.06 | 0.260223 | 0.018046 | 0.892848 | 0.999971 | 1 | 0.98828 |
| Cald1_1 | E9QA15 | 1.58 | 0.97 | 0.65 | 2.04 | 4.78 | 2.86 | 1.76 | -0.07 | 2.07 | 3.93 | 3.61 | 5.27 | -0.63 | 1.41 | 0.75 | 0.56 | 1.0225 | 2.41 | -0.7875 | 0.297356 | 0.005495 | 0.134965 | 0.999971 | 1 | 0.784948 |
| Calhm2 | D3Z2E8 | 0.68 | 2.03 | -0.35 | -0.23 | 0.07 | 0.17 | -1.1 | -0.59 | 0.87 | -0.03 | -1.07 | -1.15 | 1.11 | 0.86 | -1.85 | -0.71 | -0.895 | -0.8775 | -0.68 | 0.144254 | 0.209784 | 0.404337 | 0.999971 | 1 | 0.87384 |
| Calhm5 | Q8R100 | -0.8 | 0.15 | 0.28 | 0.58 | 0.27 | -0.31 | -0.95 | 0.28 | -1.53 | 0.2 | -0.69 | -0.72 | -1.64 | 0.19 | -1 | 1.25 | -0.23 | -0.7375 | -0.3525 | 0.567971 | 0.119543 | 0.586762 | 0.999971 | 1 | 0.920758 |
| Calm3 | P0DP28 | -3.36 | -3.53 | -3.47 | -4.12 | -3.58 | -3.36 | -3.14 | -3.73 | -3.73 | -4.16 | -2.85 | -4 | -4.11 | -3.92 | -4.24 | -4.25 | 0.1675 | -0.065 | -0.51 | 0.498733 | 0.846528 | 0.048399 | 0.999971 | 1 | 0.710102 |
| Calr | P14211 | 0.57 | 1.49 | 2.22 | 0.83 | 2.05 | 1.66 | 3.61 | 1.26 | 2.77 | 0.37 | 2.82 | 3.79 | 3.91 | 0.81 | 1.98 | 1.04 | 0.8675 | 1.16 | 0.6575 | 0.159619 | 0.144239 | 0.372788 | 0.999971 | 1 | 0.869526 |
| Calu | Q6XLQ8 | -0.88 | 0.7 | 0.08 | 0.07 | 0.78 | 0.47 | 0.14 | 0.54 | 1.08 | -0.29 | 0.55 | 0.88 | 1.58 | 0.7 | 0.37 | 0.67 | 0.49 | 0.5625 | 0.8375 | 0.181469 | 0.206278 | 0.061948 | 0.999971 | 1 | 0.725348 |
| Calu_1 | O35887 | -1.65 | 1.02 | -0.44 | 0.25 | 2.51 | 0.45 | 0.18 | 0.79 | 3.17 | -1.12 | 0.46 | 0.97 | 4.84 | -0.19 | 0.55 | 0.66 | 1.1875 | 1.075 | 1.67 | 0.116044 | 0.274576 | 0.166549 | 0.999971 | 1 | 0.800215 |
| Caly | Q9DCA7 | 3.44 | -0.68 | 0.35 | 0.81 | -0.27 | -0.13 | 2.29 | 0.1 | 1.25 | 2.63 | -0.42 | -0.1 | -0.33 | -0.89 | -0.31 | -1.22 | -0.4825 | -0.14 | -1.6675 | 0.614212 | 0.889443 | 0.066619 | 0.999971 | 1 | 0.733937 |
| Camk1 | Q91YS8 | -0.45 | -0.24 | -0.67 | -0.72 | -0.28 | -0.46 | -0.46 | -0.33 | -0.42 | -0.13 | 0.02 | -0.13 | -0.27 | -0.03 | 0.14 | -0.25 | 0.1375 | 0.355 | 0.4175 | 0.472515 | 0.10116 | 0.066233 | 0.999971 | 1 | 0.733937 |
| Camk1d | Q8BW96 | -0.39 | -0.42 | 0.18 | 0.07 | -0.09 | -0.18 | -0.05 | 0.16 | -0.15 | -0.85 | -0.27 | -0.2 | 0.29 | -0.39 | 0.1 | 0.07 | 0.1 | -0.2275 | 0.1575 | 0.64637 | 0.374939 | 0.522874 | 0.999971 | 1 | 0.907797 |
| Camk1d_1 | Q8BW96-2 | -3.29 | -1.07 | -1.07 | -1.25 | -0.58 | 0.02 | -1.11 | -1.05 | -1.86 | -2.77 | -2 | -1.6 | -1.57 | 0.49 | -1.06 | -1.41 | 0.99 | -0.3875 | 0.7825 | 0.100528 | 0.488782 | 0.250294 | 0.999971 | 1 | 0.839012 |
| Camk1g | F6YRI7 | -1.24 | -0.87 | -0.94 | -0.23 | -0.45 | -0.97 | -1.24 | -0.35 | -0.97 | -1.04 | -0.99 | -0.6 | -1.02 | -0.92 | -0.83 | -0.12 | 0.0675 | -0.08 | 0.0975 | 0.82568 | 0.756765 | 0.748271 | 0.999971 | 1 | 0.961064 |
| Camk2a | P11798 | -1.73 | -1.36 | -2.6 | -2.74 | -1.05 | -0.25 | -1.99 | -1.68 | -0.91 | -1.88 | -1.97 | -3.53 | -1.16 | -1.69 | -2.85 | -2.47 | 0.865 | 0.035 | 0.065 | 0.096066 | 0.9523 | 0.890798 | 0.999971 | 1 | 0.988184 |
| Camk2a_1 | F8WIS9 | 0.33 | -0.06 | 0.23 | 0.11 | -0.34 | 0.07 | 0.09 | 0.21 | -0.71 | -0.08 | 0.11 | -0.75 | -0.2 | -0.33 | 0.02 | 0.02 | -0.145 | -0.51 | -0.275 | 0.480604 | 0.073974 | 0.171909 | 0.999971 | 1 | 0.804075 |
| Camk2b | P28652 | -0.32 | -0.75 | -0.61 | -0.67 | -0.86 | -0.7 | -0.54 | -0.65 | -0.85 | -0.68 | -0.57 | -0.84 | -0.27 | -0.57 | -0.44 | -0.22 | -0.1 | -0.1475 | 0.2125 | 0.594372 | 0.430256 | 0.283461 | 0.999971 | 1 | 0.844538 |
| Camk2b_1 | Q5SVI1 | 0.55 | -0.29 | -1.78 | -0.88 | -0.99 | -1.12 | -1.79 | -1.08 | -0.73 | -0.43 | -2.73 | -1.57 | 0.22 | 0.61 | -1.36 | -0.99 | -0.645 | -0.765 | 0.22 | 0.20796 | 0.258514 | 0.723858 | 0.999971 | 1 | 0.955292 |
| Camk2d | E9Q1T1 | -0.98 | -1.11 | -0.45 | -0.21 | -0.47 | -0.59 | -0.46 | 0.23 | -0.44 | -0.81 | -0.27 | -0.61 | -0.09 | -0.4 | -0.15 | -0.05 | 0.365 | 0.155 | 0.515 | 0.23514 | 0.560072 | 0.069177 | 0.999971 | 1 | 0.733937 |
| Camk2d_1 | F6RWZ9 | -0.05 | -1.65 | -0.98 | -1.3 | -0.78 | -0.86 | -1.41 | -0.49 | -0.91 | 0.02 | -1.19 | -2.17 | -0.83 | 0.34 | -0.35 | -0.86 | 0.11 | -0.0675 | 0.57 | 0.772433 | 0.897754 | 0.199214 | 0.999971 | 1 | 0.813335 |
| Camk2d_2 | A0A0G2JGS | -0.25 | -3.73 | -1.72 | -1.25 | -3.17 | -1.77 | -0.98 | -1.37 | -2.2 | -5.26 | -3.95 | -3.62 | -3.92 | -0.99 | -2.2 | -1.92 | -0.085 | -2.02 | -0.52 | 0.91373 | 0.043077 | 0.547785 | 0.999971 | 1 | 0.911722 |
| Camk2g | A0A286YCE | -1.42 | -3 | -3.24 | -2.78 | -4.27 | -1.68 | -3.3 | -3.18 | -3.19 | 0.15 | -3.76 | -3.63 | -3.5 | -1.27 | -2.64 | -1.72 | -0.4975 | 0.0025 | 0.3275 | 0.425633 | 0.997812 | 0.580693 | 0.999971 | 1 | 0.920178 |
| Camk2g_1 | A0A286YC\ | -1.12 | -1.03 | -1.47 | -3.56 | -0.92 | -1.08 | -1.77 | -1.79 | -1.88 | -1.66 | -1.28 | -1.24 | -1.37 | -1.26 | -2.47 | -2.96 | 0.405 | 0.28 | -0.22 | 0.493427 | 0.624002 | 0.739767 | 0.999971 | 1 | 0.960899 |
| Camk2g_2 | Q923T9-2 | 4.16 | -1.13 | 0.28 | 0.37 | -1.9 | -1.01 | 2.48 | -1.74 | -1.88 | 1.2 | -2.42 | -2.55 | -3.2 | -2.57 | -1.45 | -3.03 | -1.4625 | -2.3325 | -3.4825 | 0.297237 | 0.097542 | 0.009635 | 0.999971 | 1 | 0.613095 |
| Camk2g_3 | A0A2I3BQF | -0.17 | -0.16 | -0.65 | -1.1 | -0.43 | 0.04 | -0.85 | -0.99 | -0.64 | -0.42 | -1 | -1.05 | -0.15 | -0.32 | -0.78 | -0.59 | -0.0375 | -0.2575 | 0.06 | 0.907779 | 0.374958 | 0.830815 | 0.999971 | 1 | 0.978535 |
| Camk2g_4 | Q923T9-3 | 1.17 | 1.38 | -1.28 | -1.8 | 1.53 | 2.5 | -1.76 | -1.33 | 1.3 | 2.2 | -1.59 | -1.75 | 1.95 | 2.39 | -2.48 | -1.2 | 0.3675 | 0.1725 | 0.2975 | 0.756896 | 0.882033 | 0.817137 | 0.999971 | 1 | 0.974929 |
| Camk2n1 | Q6QWF9 | -0.17 | -0.25 | -0.99 | -0.59 | -0.46 | -1.41 | -0.79 | -0.63 | -0.4 | -1.82 | -0.05 | -0.27 | 0.72 | 0.01 | -0.87 | -0.51 | -0.3225 | -0.135 | 0.3375 | 0.2843 | 0.749225 | 0.385381 | 0.999971 | 1 | 0.871816 |
| Camk4 | Q8BGR3 | -0.49 | 0.88 | 1.14 | 0.75 | 0.84 | 0.14 | 0.85 | 0.9 | 0.97 | -0.79 | 1.18 | 1.18 | 0.8 | 0.2 | 0.84 | 0.46 | 0.1125 | 0.065 | 0.005 | 0.772917 | 0.906489 | 0.989492 | 0.999971 | 1 | 0.99925 |
| Camkk1 | Q8VBY2 | -0.24 | 0.29 | 0.04 | -0.31 | 0.33 | -0.17 | -0.16 | -0.53 | 0.38 | -0.58 | -0.09 | -0.37 | -0.16 | -0.56 | -0.5 | -0.43 | -0.0775 | -0.11 | -0.3575 | 0.758175 | 0.682428 | 0.120488 | 0.999971 | 1 | 0.78268 |
| Camkk2 | Q8C078 | 0.01 | 0.12 | -0.21 | -0.18 | 0.15 | 0.25 | 0.06 | -0.32 | 0.57 | -0.09 | -0.23 | 0.07 | 0.5 | 0.21 | -0.4 | -0.28 | 0.1 | 0.145 | 0.0725 | 0.624689 | 0.529048 | 0.774383 | 0.999971 | 1 | 0.963884 |
| Camkmt | Q3U2J5 | 0.19 | -0.18 | 0.15 | 0.37 | 0.3 | 0.69 | -0.51 | 0.37 | 0.39 | 0.9 | -0.03 | 0.3 | -0.22 | 0.01 | -0.09 | 0.53 | 0.08 | 0.2575 | -0.075 | 0.783867 | 0.317876 | 0.751325 | 0.999971 | 1 | 0.961064 |
| Camkv | Q3UHL1 | -2.1 | -2.18 | -0.74 | -1 | -1.54 | -2.43 | -0.62 | -0.75 | -1.53 | -2.4 | -0.87 | -0.75 | -1.87 | -2.26 | -0.91 | -0.98 | 0.17 | 0.1175 | 1.57E-16 | 0.742449 | 0.812268 | 1 | 0.999971 | 1 | 1 |
| Camlg | P49070 | -0.68 | -0.2 | -1.12 | -0.37 | -0.13 | -0.02 | 0.85 | -0.46 | 0.02 | -0.42 | 0.35 | -1 | -0.2 | -0.6 | -0.76 | -0.69 | 0.6525 | 0.33 | 0.03 | 0.082673 | 0.357625 | 0.908584 | 0.999971 | 1 | 0.991707 |
| Camsap1 | A0A0A0MC | 0.28 | -0.03 | -0.28 | -0.45 | 0.02 | -0.05 | -0.3 | -0.3 | 0.1 | 0.27 | -0.12 | -0.44 | 0.53 | -0.02 | -0.2 | -0.07 | -0.0375 | 0.0725 | 0.18 | 0.865895 | 0.769986 | 0.484141 | 0.999971 | 1 | 0.896352 |
| Camsap2 | A0A0A6YY( | 0.72 | 0.6 | 0.7 | 0.49 | 0.3 | 0.43 | 0.4 | 0.54 | 0.5 | 0.48 | 0.73 | 0.66 | 0.44 | 0.43 | 0.57 | 0.53 | -0.21 | -0.035 | -0.135 | 0.234674 | 0.835407 | 0.427435 | 0.999971 | 1 | 0.880793 |
| Camsap3 | Q80VC9-14 | -0.64 | -1.02 | -1.28 | -1.56 | -1.28 | -1.12 | -1.17 | -1.31 | -0.88 | -0.65 | -1.1 | -1.05 | -0.61 | -0.84 | -1.28 | -1.33 | -0.095 | 0.205 | 0.11 | 0.688154 | 0.416749 | 0.694287 | 0.999971 | 1 | 0.947174 |
| Camsap3_1 | Q80VC9-5 | -1.64 | -0.53 | 1.49 | -0.54 | -2.14 | -0.19 | 0.13 | 1.5 | -0.42 | -2.25 | 1.51 | 3.09 | 2.01 | -0.49 | 1.64 | 1.1 | 0.13 | 0.7875 | 1.37 | 0.883836 | 0.514326 | 0.102726 | 0.999971 | 1 | 0.78268 |
| Camta1 | A2A891-5 | 1.3 | 1.65 | 1.91 | 1.88 | 1.51 | 1.95 | 1.44 | 2.16 | 1.08 | 0.96 | 1.77 | 1.4 | 1.13 | 1.86 | 1.64 | 1.98 | 0.08 | -0.3825 | -0.0325 | 0.750387 | 0.157634 | 0.900263 | 0.999971 | 1 | 0.990583 |
| Camta2 | B0QZH6 | 1.01 | 1.09 | 0.68 | 1.13 | 0.89 | 0.91 | 0.3 | 0.92 | 0.37 | 0.65 | 0.36 | 0.19 | 0.68 | 1.26 | 0.34 | 0.65 | -0.2225 | -0.585 | -0.245 | 0.335156 | 0.014774 | 0.336077 | 0.999971 | 1 | 0.858827 |
| Cand1 | Q6ZQ38 | -0.71 | -0.44 | 0.3 | 0.05 | -0.3 | -0.16 | 0.1 | 0.18 | -0.41 | -0.46 | -0.03 | -0.34 | -0.57 | 0.02 | 0.37 | 0.29 | 0.155 | -0.11 | 0.2275 | 0.573759 | 0.682131 | 0.479217 | 0.999971 | 1 | 0.894438 |
| Cand2 | Q6ZQ73 | -0.77 | 0.08 | 0.1 | 0.16 | 0.46 | -0.13 | -0.45 | 0.34 | 0.48 | 0.11 | 0.02 | 0.27 | 0.68 | 0.6 | -0.18 | 0.32 | 0.1625 | 0.3275 | 0.4625 | 0.60387 | 0.234893 | 0.15273 | 0.999971 | 1 | 0.795578 |
| Cant1 | Q8VCF1-3 | 0.31 | 0.75 | 1.18 | 1.02 | 0.65 | 0.92 | 0.14 | 0.79 | 0.24 | 0.79 | 1.68 | 0.24 | 0.43 | 1.33 | 1.17 | 1.53 | -0.19 | -0.0775 | 0.3 | 0.493428 | 0.837327 | 0.349016 | 0.999971 | 1 | 0.865162 |
| Canx | P35564 | -0.69 | -1.5 | -0.04 | -0.06 | -1.13 | -0.86 | 0.45 | 0.01 | -0.87 | -0.89 | -0.09 | 0.11 | -0.71 | -0.54 | 0.56 | 0.3 | 0.19 | 0.1375 | 0.475 | 0.687762 | 0.739054 | 0.294801 | 0.999971 | 1 | 0.847402 |
| Cap1 | P40124 | -0.34 | -0.17 | 0.15 | -0.08 | 0.01 | -0.17 | 0.53 | -0.18 | 0.3 | -0.37 | 0.2 | 0.3 | 0.22 | -0.47 | -0.07 | 0 | 0.1575 | 0.2175 | 0.03 | 0.503334 | 0.351374 | 0.891939 | 0.999971 | 1 | 0.988225 |
| Cap2 | Q9CYT6 | -0.3 | 0.52 | 0.46 | 0.28 | 0.47 | 0.23 | 0.42 | -0.03 | 0.5 | -0.12 | 0.47 | 0.44 | 0.6 | -0.13 | 0 | 0.33 | 0.0325 | 0.0825 | -0.04 | 0.895569 | 0.751655 | 0.88205 | 0.999971 | 1 | 0.986381 |
| Capg | Q99LB4 | -0.71 | 1.11 | 0.83 | 0.62 | 2.15 | 1.22 | 0.33 | 0.62 | 1.04 | -1.17 | 1.01 | 1.25 | 0.14 | 1.04 | 1.22 | 0.76 | 0.6175 | 0.07 | 0.3275 | 0.258809 | 0.912664 | 0.465136 | 0.999971 | 1 | 0.89116 |
| Capn1 | O35350 | -0.02 | 0.12 | -0.07 | 0.04 | -0.05 | -0.03 | -0.12 | -0.04 | -0.41 | -0.11 | -0.07 | -0.59 | 0.24 | -0.02 | 0.18 | 0.18 | -0.0775 | -0.3125 | 0.1275 | 0.633927 | 0.129521 | 0.458167 | 0.999971 | 1 | 0.888454 |
| Capn10 | Q9ESK3 | 3.02 | 0.7 | -0.07 | 0.55 | 0.27 | 0.41 | 0.22 | 0.11 | 0.46 | 1.87 | -0.51 | -0.18 | -1.28 | -0.12 | -0.24 | 0.07 | -0.7975 | -0.64 | -1.4425 | 0.219568 | 0.419793 | 0.057205 | 0.999971 | 1 | 0.725348 |
| Capn2 | O08529 | -0.21 | -0.26 | -0.6 | -0.47 | -0.31 | -0.04 | -0.77 | -0.14 | -0.39 | -0.29 | -0.81 | -0.61 | -0.08 | -0.07 | -0.29 | -0.45 | 0.07 | -0.14 | 0.1625 | 0.757141 | 0.491641 | 0.411785 | 0.999971 | 1 | 0.875847 |
| Capn5 | O08688 | -0.91 | -0.39 | -0.87 | -0.85 | -0.47 | -0.04 | -0.99 | -0.2 | -0.29 | -0.69 | -0.57 | -0.5 | 0.34 | -0.44 | -0.6 | -0.27 | 0.33 | 0.2425 | 0.5125 | 0.231147 | 0.247741 | 0.077575 | 0.999971 | 1 | 0.752152 |
| Capn7 | Q9R1S8 | 0.16 | 0.42 | 0.01 | -0.08 | -0.04 | 0.17 | -0.39 | -0.42 | 0.32 | 0.11 | 0.2 | -0.09 | -0.32 | 0.16 | -0.4 | -0.29 | -0.2975 | 0.0075 | -0.34 | 0.203233 | 0.969453 | 0.140543 | 0.999971 | 1 | 0.784948 |
| Capns1 | O88456 | 0.37 | -0.17 | -0.69 | -0.21 | -0.15 | -0.26 | -0.63 | 0.03 | -0.38 | -0.09 | -0.29 | -0.43 | -0.01 | 0.25 | -0.29 | -0.18 | -0.0775 | -0.1225 | 0.1175 | 0.778382 | 0.631195 | 0.662337 | 0.999971 | 1 | 0.936148 |
| Caprin1 | Q60865 | 1.06 | 1.96 | 1.45 | 1.42 | 1.95 | 1.8 | 1.39 | 1.7 | 2.3 | 1.48 | 1.74 | 1.9 | 1.89 | 1.72 | 1.45 | 1.62 | 0.2375 | 0.3825 | 0.1975 | 0.351848 | 0.181304 | 0.420393 | 0.999971 | 1 | 0.87788 |
| Capza1 | Q5RKN9 | -0.19 | -0.09 | 0 | -0.42 | -0.1 | 0.12 | -0.47 | 0.09 | -0.02 | -0.4 | -0.18 | -0.41 | 0 | -0.18 | 0.14 | -0.37 | 0.085 | -0.0775 | 0.0725 | 0.689877 | 0.686395 | 0.719623 | 0.999971 | 1 | 0.954401 |
| Capza2 | P47754 | -0.96 | 0.39 | 0.48 | -0.56 | 0.33 | 0.45 | 0.43 | 0.21 | 0.55 | -0.9 | 0.56 | 0.27 | 0.96 | 0.2 | 0.23 | -0.49 | 0.5175 | 0.2825 | 0.3875 | 0.166976 | 0.548496 | 0.385981 | 0.999971 | 1 | 0.871816 |
| Capzb | P47757-2 | -0.81 | -1.45 | -0.87 | -0.85 | -1.41 | -1.24 | -1.08 | -0.72 | -1.63 | -0.84 | -0.71 | -0.93 | -2.06 | -1.38 | -1.01 | -0.8 | -0.1175 | -0.0325 | -0.3175 | 0.630834 | 0.905097 | 0.333605 | 0.999971 | 1 | 0.857921 |

| | | | | | | | | | | | | | | | | | | | | | | | | | | | |
|---|---|---|---|---|---|---|---|---|---|---|---|---|---|---|---|---|---|---|---|---|---|---|---|---|---|---|---|
| Capzb_1 | P47757 | -0.36 | -0.03 | 0.05 | 0.24 | -0.18 | 0.09 | 0.55 | 0.51 | 0.21 | -0.26 | 1.2 | 1.34 | 0.63 | -0.05 | 0.68 | 0.75 | 0.2675 | 0.6475 | 0.5275 | 0.292221 | 0.125656 | 0.061717 | 0.999971 | | 1 | 0.725348 |
| Card19 | Q9D1I2 | 0.27 | -0.3 | 0.28 | -0.13 | -0.26 | -0.38 | -0.92 | -0.11 | -0.82 | 0.41 | 0.2 | 0.03 | -1.41 | -0.4 | 0.79 | 0.39 | -0.4475 | -0.075 | -0.1875 | 0.106014 | 0.809548 | 0.693268 | 0.999971 | | 1 | 0.947174 |
| Carhsp1 | Q9CR86 | -1.08 | -0.85 | -0.96 | -1.35 | -1.2 | -1.69 | -0.64 | -1.24 | -0.68 | -0.75 | -1.35 | -1.13 | -0.21 | -1.23 | -1.01 | -1.65 | -0.1325 | 0.0825 | 0.035 | 0.616502 | 0.718196 | 0.91366 | 0.999971 | | 1 | 0.992621 |
| Carm1 | D3YUP1 | -0.15 | 0.1 | 0.56 | 0.51 | 0 | 0.02 | 0.79 | 0.32 | -0.1 | -0.25 | 0.34 | 0.29 | -0.12 | 0.08 | 0.24 | 0.24 | 0.0275 | -0.185 | -0.145 | 0.918804 | 0.465438 | 0.532209 | 0.999971 | | 1 | 0.910069 |
| Carmil1 | D3Z030 | -1 | -0.99 | -1.06 | -1.02 | -0.94 | -1.06 | -0.89 | -0.89 | -0.66 | -0.78 | -0.53 | -0.73 | -0.75 | -1.01 | -0.87 | -0.8 | 0.0725 | 0.3425 | 0.16 | 0.654548 | 0.058013 | 0.347288 | 0.999971 | | 1 | 0.864164 |
| Carmil2 | S0DHL8 | 0.17 | -0.34 | -0.65 | -0.91 | -0.64 | -0.5 | -0.04 | -0.66 | -0.49 | -0.33 | -0.68 | -0.61 | 0.25 | -0.27 | -0.6 | -0.82 | -0.0275 | -0.095 | 0.0725 | 0.923477 | 0.720198 | 0.825718 | 0.999971 | | 1 | 0.976715 |
| Carmil3 | Q3UFQ8 | 0.16 | -0.6 | -0.69 | -1.28 | -0.37 | -0.7 | -0.33 | -0.51 | 0.04 | 0.21 | -0.96 | -0.42 | -0.54 | -0.5 | -0.2 | -0.74 | 0.125 | 0.32 | 0.1075 | 0.689639 | 0.411869 | 0.736888 | 0.999971 | | 1 | 0.959447 |
| Carnmt1 | Q80UY1 | 4.48 | 1.47 | 2.06 | 2.18 | 1.07 | 1.77 | 2.27 | 1.45 | 1.43 | 3.9 | 1.7 | 1.64 | -0.85 | 0.78 | 0.17 | 0.35 | -0.9075 | -0.38 | -2.435 | 0.183172 | 0.63572 | 0.005726 | 0.999971 | | 1 | 0.582714 |
| Cars1 | Q9ER72 | 0.69 | 1.66 | 1.08 | 0.73 | 1.39 | 1.31 | 1.14 | 0.43 | 1.54 | 1.22 | 1.39 | 1.73 | 1.29 | 1.33 | 0.71 | 0.92 | 0.0275 | 0.43 | 0.0225 | 0.930729 | 0.135348 | 0.93688 | 0.999971 | | 1 | 0.995618 |
| Cars2 | Q8BYM8 | 0.04 | 0.48 | 0.12 | 0.26 | 0.53 | 0.15 | -0.48 | 0.12 | 0.75 | 0.28 | 0.3 | 0.19 | 0.42 | 0.06 | 0.21 | 0.14 | -0.145 | 0.155 | -0.0175 | 0.573637 | 0.461328 | 0.927019 | 0.999971 | | 1 | 0.99417 |
| Cartpt | P56388-2 | -0.69 | -3.02 | -0.88 | -0.42 | -1.52 | -1.96 | -1.43 | -0.65 | -0.47 | -1.22 | -2.52 | -0.2 | -0.29 | -0.86 | 0.32 | -1.27 | -0.1375 | 0.15 | 0.7275 | 0.818399 | 0.834805 | 0.265674 | 0.999971 | | 1 | 0.842677 |
| Casc3 | Q8K3W3 | 2.19 | 1.62 | 1.6 | 1.88 | 1.64 | 1.72 | 1.63 | 1.94 | 1.9 | 2.11 | 1.59 | 1.55 | 0.57 | 1.32 | 1.38 | 1.52 | -0.09 | -0.035 | -0.625 | 0.666612 | 0.877913 | 0.044891 | 0.999971 | | 1 | 0.710102 |
| Casd1 | Q7TN73 | 0.91 | 1.18 | 0.16 | -0.07 | 0.9 | 1.35 | -0.57 | 0.7 | 0.98 | 0.9 | 0.25 | 0.59 | 0.1 | 2.05 | 0.67 | 0.91 | 0.05 | 0.135 | 0.3875 | 0.915856 | 0.690719 | 0.421978 | 0.999971 | | 1 | 0.877992 |
| Cask | B1AUH5 | -3.9 | -0.67 | -2.47 | -3.51 | -1.05 | -1.09 | -2.93 | -3.57 | -0.37 | -3.86 | -2.43 | -3.49 | 0.86 | -1.28 | -2.78 | -3.44 | 0.4775 | 0.1 | 0.9775 | 0.584511 | 0.916875 | 0.371702 | 0.999971 | | 1 | 0.869526 |
| Cask_1 | O70589 | -1.62 | -1.73 | -1.09 | -1.01 | -1.54 | -1.73 | -0.93 | -1.03 | -1.59 | -1.95 | -1.05 | -0.84 | -1.38 | -1.48 | -1.06 | -0.89 | 0.055 | 0.005 | 0.16 | 0.845057 | 0.987325 | 0.535052 | 0.999971 | | 1 | 0.910077 |
| Caskin1 | Q6P9K8 | -0.61 | -1.57 | -0.74 | -0.74 | -1.56 | -1.41 | -0.71 | -0.93 | -1.48 | -0.85 | -0.81 | -1.17 | -1.4 | -1.54 | -1.06 | -0.9 | -0.2375 | -0.1625 | -0.31 | 0.443274 | 0.569955 | 0.287524 | 0.999971 | | 1 | 0.846961 |
| Caskin2 | Q8VHK1 | -1.16 | 0.23 | -0.06 | -0.35 | 0.34 | -0.57 | -0.35 | -0.35 | 0.33 | -2.19 | -0.08 | -0.08 | 0.06 | -0.59 | -0.34 | -0.18 | 0.1025 | -0.17 | 0.0725 | 0.771264 | 0.774193 | 0.825898 | 0.999971 | | 1 | 0.976757 |
| Casp1 | P29452 | -0.76 | -0.99 | 0.04 | 0.62 | -0.79 | -0.35 | -0.43 | 0.66 | -0.08 | -0.84 | 0.04 | 0.73 | 0.57 | -0.4 | -0.93 | 0.3 | 0.045 | 0.235 | 0.1575 | 0.920898 | 0.612917 | 0.738362 | 0.999971 | | 1 | 0.959976 |
| Casp3 | P70677 | -0.6 | 0.21 | 0.09 | -0.03 | 0.35 | 0.31 | 0.07 | 0.12 | 0.92 | 0.49 | 0.41 | -0.38 | 0.89 | 0.71 | 0.2 | 0.65 | 0.295 | 0.4425 | 0.695 | 0.221371 | 0.19722 | 0.022894 | 0.999971 | | 1 | 0.652043 |
| Casp6 | O08738 | 1.9 | 1.6 | 0.81 | 0.34 | 2.35 | 2.55 | 0.27 | 0.64 | 2.54 | 2.48 | -0.15 | 0.05 | 1.1 | 2.6 | 0.62 | 0.66 | 0.29 | 0.0675 | 0.0825 | 0.642922 | 0.927852 | 0.878655 | 0.999971 | | 1 | 0.985441 |
| Casp7 | P97864 | 2.01 | 1.05 | 0.54 | 0.56 | 1.26 | 0.66 | 0.13 | 0.43 | 0.83 | 1.91 | 0.35 | -0.11 | -0.49 | 0.7 | 0.43 | 0.43 | -0.42 | -0.295 | -0.7725 | 0.309907 | 0.569799 | 0.087848 | 0.999971 | | 1 | 0.765196 |
| Casp8 | A0A087WC | -3.1 | -1.42 | 1.31 | 1.89 | -1.23 | -1.59 | 1.18 | 1.69 | -1.15 | -3.03 | 0.82 | 1.57 | -2.26 | -1.72 | 1.8 | 0.66 | 0.3425 | -0.1175 | -0.05 | 0.788561 | 0.932937 | 0.970404 | 0.999971 | | 1 | 0.998787 |
| Casq2 | E9PZ67 | 1.01 | 0.95 | 0.07 | 1.5 | 0.34 | 0.7 | 0.38 | -0.13 | -0.78 | 0.82 | 1.06 | -0.28 | -0.37 | 0.14 | 0.1 | 0.58 | -0.56 | -0.6775 | -0.77 | 0.126237 | 0.194809 | 0.052078 | 0.999971 | | 1 | 0.723736 |
| Cast | Q8C281 | 0.73 | 0.94 | 0.81 | 1.09 | 1.37 | 0.54 | 0.93 | 0.58 | 0.83 | 0.31 | 0.5 | 1.11 | 1.05 | 0.79 | 1.07 | 0.86 | -0.0375 | -0.205 | 0.05 | 0.876132 | 0.382631 | 0.784385 | 0.999971 | | 1 | 0.966376 |
| Castor2 | Q8CAB8 | 1.1 | 0.86 | 0.17 | -0.25 | 0.75 | 1.46 | 0.3 | 0.36 | 0.76 | 1.13 | 0.51 | 0.27 | 0.77 | 1.12 | 0.29 | 0.95 | 0.2475 | 0.1975 | 0.3125 | 0.533085 | 0.580248 | 0.386435 | 0.999971 | | 1 | 0.871816 |
| Cat | P24270 | -0.54 | -0.89 | -0.8 | -0.79 | -0.57 | -0.41 | -0.36 | -0.33 | -0.48 | -0.37 | -0.36 | -0.83 | -0.48 | -0.73 | -0.65 | -0.67 | 0.3375 | 0.245 | 0.1225 | 0.082472 | 0.225048 | 0.495112 | 0.999971 | | 1 | 0.901531 |
| Cav1 | P49817 | -0.1 | -0.51 | 0.32 | 0.33 | 3.71 | 0.71 | 0.84 | -0.56 | 1.09 | 0.64 | 1.52 | 2.84 | -0.87 | -0.15 | 0.76 | -0.3 | 1.165 | 1.5125 | -0.15 | 0.184034 | 0.01167 | 0.694079 | 0.999971 | | 1 | 0.947174 |
| Cavin1 | O54724 | 0.16 | -0.16 | -0.01 | 0.78 | 2.3 | 0.18 | -0.07 | -0.38 | 0.2 | 1.25 | 0.76 | 1.57 | -0.8 | 0.1 | 0.35 | 0.32 | 0.315 | 0.7525 | -0.2 | 0.594489 | 0.060212 | 0.557793 | 0.999971 | | 1 | 0.915051 |
| Cavin2 | Q63918 | 0.6 | -0.27 | -0.26 | 0.99 | 2.16 | 0.01 | 0.83 | -0.78 | 0.28 | 1.05 | 0.17 | 0.89 | -0.78 | -0.06 | 0 | -0.83 | 0.29 | 0.3325 | -0.6825 | 0.65154 | 0.38431 | 0.09696 | 0.999971 | | 1 | 0.78268 |
| Cavin3 | Q91VJ2 | 1.95 | 0.67 | -0.26 | 1.51 | 6.48 | 2.08 | 0.96 | -0.1 | 2.5 | 4.03 | 2.57 | 3.74 | -0.29 | 1.42 | 0.88 | 0.36 | 1.3875 | 2.2425 | -0.375 | 0.320098 | 0.003807 | 0.508595 | 0.999971 | 0.969702 | | 0.905815 |
| Cbarp | Q66L44 | 1.67 | -0.84 | -0.5 | -0.63 | -1.41 | -0.14 | 0.38 | -1.13 | -1.01 | 0.41 | -1.07 | -1.03 | 0.1 | -0.91 | -0.55 | -1.19 | -0.5 | -0.6 | -0.5625 | 0.451904 | 0.35414 | 0.355986 | 0.999971 | | 1 | 0.868793 |
| Cbarp_1 | E9Q6U0 | -0.52 | -4.34 | -3.06 | -3.35 | -4.79 | -2.57 | -2.52 | -1.79 | -1.49 | -1.79 | -2.72 | -1.83 | -5.5 | -5.71 | -2.47 | -3.12 | -0.1 | 0.86 | -1.3825 | 0.91432 | 0.284771 | 0.203879 | 0.999971 | | 1 | 0.81617 |
| Cbfa2t2 | A2AKD9 | 2.89 | 1.54 | 3.5 | 2.25 | 1.98 | 2.9 | 3.28 | 3.32 | 2.39 | 3.19 | 3.44 | 1.17 | 2.62 | 2.15 | 2.61 | 2.62 | 0.325 | 0.0025 | -0.045 | 0.509908 | 0.996712 | 0.913789 | 0.999971 | | 1 | 0.992633 |
| Cbfb | Q08024-2 | -1.23 | 0.27 | 0.2 | 0.67 | 0.71 | 0.29 | -0.18 | 0.71 | 0.17 | 0.01 | 0.18 | -0.1 | 0.42 | 0.68 | 0.82 | 0.67 | 0.405 | 0.0875 | 0.67 | 0.368658 | 0.828283 | 0.125421 | 0.999971 | | 1 | 0.78268 |
| Cbl | P22682 | -0.12 | -0.01 | 0.04 | -0.21 | 0.01 | -0.17 | 0.42 | -0.26 | -0.11 | -0.5 | -0.33 | -0.19 | 0.2 | -0.02 | -0.11 | -0.38 | 0.075 | -0.2075 | -0.0025 | 0.722904 | 0.260882 | 0.989802 | 0.999971 | | 1 | 0.99925 |
| Cblb | B9EKI5 | 0.65 | -0.93 | 0.29 | -0.39 | -0.31 | -0.44 | -0.51 | 0.11 | 0.2 | -0.2 | -0.84 | -0.94 | 0 | 0.05 | -0.47 | 0.09 | -0.1925 | -0.35 | 0.0125 | 0.603327 | 0.416656 | 0.972737 | 0.999971 | | 1 | 0.99925 |
| Cbll1 | E9QM14 | -0.18 | 0.14 | 1.02 | 1.05 | -0.13 | -0.51 | 0.85 | 0.97 | -0.22 | 0.17 | 0.44 | 0.61 | 0.09 | -0.01 | -0.12 | 0.92 | -0.2125 | -0.2575 | -0.2875 | 0.638736 | 0.473327 | 0.45581 | 0.999971 | | 1 | 0.887217 |
| Cbln1 | Q9R171 | 0.7 | -0.48 | -0.99 | -0.85 | -0.53 | 0.06 | -0.01 | -0.45 | 0.13 | 0.87 | -0.52 | -1.92 | -0.59 | 0.63 | -0.62 | -0.76 | 0.1725 | 0.045 | 0.07 | 0.663695 | 0.94434 | 0.881499 | 0.999971 | | 1 | 0.986007 |
| Cbln4 | Q8BME9 | 0.92 | 1.71 | -0.14 | -0.28 | 0.82 | 1.54 | -0.32 | 0.01 | 1 | 0.35 | 0.12 | 0.01 | 1.17 | 2.54 | -0.29 | 0.09 | -0.04 | -0.1825 | 0.325 | 0.944521 | 0.707932 | 0.650867 | 0.999971 | | 1 | 0.933842 |
| Cbr1 | P48758 | 0.5 | -0.11 | 0.42 | 0.11 | 0.14 | 0.48 | 1.15 | 0.26 | 0.41 | 0.02 | 0.09 | 0.29 | 0.04 | 0.24 | 0.36 | -0.23 | 0.2775 | -0.0275 | -0.1275 | 0.337183 | 0.896914 | 0.582907 | 0.999971 | | 1 | 0.920364 |
| Cbr2 | P08074 | -0.14 | 0.48 | -0.47 | -0.21 | 3.11 | 0.76 | -0.52 | -1.97 | 2.12 | 0.82 | 0.53 | 1.94 | -0.2 | 0.38 | 0.09 | -1.13 | 0.43 | 1.4375 | -0.13 | 0.66182 | 0.007871 | 0.728224 | 0.999971 | | 1 | 0.956041 |
| Cbr3 | Q8K354 | 0.99 | 0.39 | 0.41 | 0.2 | 0.4 | 0.11 | 0.07 | 0.37 | 0.27 | 0.85 | 0.23 | 0.12 | 0.25 | 0.48 | 0.66 | 0.16 | -0.26 | -0.13 | -0.11 | 0.275031 | 0.617981 | 0.647342 | 0.999971 | | 1 | 0.933613 |
| Cbr4 | Q91VT4 | -1.14 | -0.91 | -0.02 | -0.48 | -0.57 | -1.28 | -0.08 | -0.09 | -0.47 | -0.96 | 0.02 | 0.12 | -0.46 | -0.87 | -0.07 | -0.57 | 0.1325 | 0.315 | 0.145 | 0.717571 | 0.373895 | 0.636543 | 0.999971 | | 1 | 0.93301 |
| Cbs | Q91WT9-2 | 0.58 | -0.17 | 0.11 | 0.41 | 0 | 0.16 | 0.08 | -0.13 | 0.32 | 0.76 | 0.15 | -0.05 | 0.39 | 0.06 | 0.19 | 0.58 | -0.205 | 0.0625 | 0.0725 | 0.364583 | 0.81057 | 0.759762 | 0.999971 | | 1 | 0.961116 |
| Cbwd1 | Q8VEH6 | -1.36 | -0.17 | -0.9 | -0.65 | -0.11 | -0.36 | -0.64 | -0.46 | -0.08 | -1.17 | -0.56 | -0.28 | -0.29 | -0.17 | -0.83 | -0.43 | 0.3775 | 0.2475 | 0.34 | 0.208088 | 0.473459 | 0.270482 | 0.999971 | | 1 | 0.843794 |
| Cbx1 | P83917 | 0.22 | 0.56 | 0.12 | -0.39 | 0.32 | -0.25 | -0.64 | 0.15 | 0.16 | -0.86 | -0.25 | 0.42 | 1.55 | 0.74 | 0.02 | -0.47 | -0.2325 | -0.26 | 0.3325 | 0.445706 | 0.450013 | 0.470068 | 0.999971 | | 1 | 0.892481 |
| Cbx3 | Q9DCC5 | 0.85 | 0.57 | 0.16 | 0.37 | 0.25 | 0.38 | -0.39 | -0.17 | 0.58 | 0.44 | -0.32 | -0.18 | 0.22 | 0.34 | 0.28 | 0.36 | -0.47 | -0.3575 | -0.1875 | 0.094455 | 0.224788 | 0.373226 | 0.999971 | | 1 | 0.869526 |
| Cbx5 | Q61686 | -0.21 | -0.01 | -0.34 | -0.63 | -0.04 | -0.11 | -0.45 | -0.54 | -0.05 | -0.37 | -0.66 | -0.85 | 0.19 | 0.04 | -0.62 | -0.68 | 0.0125 | -0.185 | 0.03 | 0.955053 | 0.458606 | 0.913439 | 0.999971 | | 1 | 0.992618 |
| Cbx8 | Q9QXV1 | 0.38 | -0.6 | -0.04 | 0.66 | -0.35 | -0.45 | -0.44 | -0.36 | -0.48 | -0.25 | -0.35 | -0.37 | -1.04 | -0.16 | 0.05 | 0.38 | -0.5 | -0.4625 | -0.2925 | 0.109682 | 0.136639 | 0.464146 | 0.999971 | | 1 | 0.890905 |
| Cby1 | Q9D1C2 | 0.01 | -0.12 | -1.21 | -0.03 | 0.27 | 0 | -0.44 | 0.09 | -0.43 | 0.67 | -0.84 | 0.04 | 0.22 | 0.35 | -0.62 | -0.41 | 0.3175 | 0.1975 | 0.2225 | 0.346941 | 0.637063 | 0.54788 | 0.999971 | | 1 | 0.911722 |
| Cc2d1a | Q8K1A6 | -0.29 | -0.25 | -0.27 | -0.06 | 0.03 | -0.2 | -0.27 | -0.18 | 0.04 | -0.11 | -0.13 | 0.12 | 0.5 | -0.02 | -0.02 | 0.19 | 0.0625 | 0.1975 | 0.38 | 0.718768 | 0.259468 | 0.079975 | 0.999971 | | 1 | 0.755254 |
| Cc2d1a_1 | E9PX94 | 1.63 | 1.31 | -0.77 | -0.21 | 0.54 | 1.75 | 0.28 | 0.05 | 1.24 | 1.87 | 0.57 | -0.49 | 2.39 | 0.82 | 0.95 | 0.45 | 0.165 | 0.3075 | 0.6625 | 0.793737 | 0.6622 | 0.326154 | 0.999971 | | 1 | 0.856472 |
| Cc2d1b | Q8BRN9 | 0.88 | -0.51 | -0.35 | 0.07 | -0.5 | -0.52 | -0.25 | -0.13 | -0.46 | -0.06 | -0.1 | -0.19 | -1.04 | -0.71 | -0.37 | -0.23 | -0.3725 | -0.225 | -0.61 | 0.270993 | 0.493922 | 0.112106 | 0.999971 | | 1 | 0.78268 |
| Cc2d2a | Q8CFW7-2 | 2.5 | -1.73 | 5.62 | 2.23 | -1.01 | -0.98 | 9.3 | 1.26 | -0.51 | 0.9 | 3.77 | 2.03 | -3.48 | -2.28 | 2.85 | 1.71 | -0.0125 | -0.6075 | -2.455 | 0.996062 | 0.699707 | 0.218322 | 1 | | 1 | 0.827064 |
| Ccar1 | Q8CH18-3 | 0.86 | 2.22 | 2.34 | 2.48 | 2.1 | 1.62 | 1.93 | 2.24 | 1.84 | 1.1 | 2.46 | 2.5 | 1.79 | 1.81 | 2.16 | 2.47 | -0.0025 | -3.14E-16 | 0.0825 | 0.994796 | 1 | 0.833758 | 0.999971 | | 1 | 0.979036 |
| Ccar2 | Q8VDP4 | 2.83 | 2.42 | 2.17 | 2.28 | 2.01 | 2.32 | 2.11 | 2.02 | 1.93 | 2.39 | 1.91 | 2.05 | 1.83 | 2.4 | 2.15 | 2.34 | -0.31 | -0.355 | -0.245 | 0.166968 | 0.136172 | 0.305527 | 0.999971 | | 1 | 0.847402 |
| Ccdc106 | E9QPN6 | -1.27 | -2.91 | -1.27 | -1.05 | -2.83 | -3.58 | -0.94 | -0.54 | -2.7 | -4.52 | -2.44 | -0.28 | -4.4 | -1.85 | -1.06 | -1.44 | -0.3475 | -0.86 | -0.5625 | 0.651437 | 0.339611 | 0.479323 | 0.999971 | | 1 | 0.894438 |
| Ccdc107 | Q9DCC3 | 2.58 | -0.16 | -0.16 | 1.85 | 0.38 | 0.79 | -0.06 | -0.74 | -0.09 | 1.95 | 1.65 | 0.51 | 0.02 | 2.69 | 0.9 | 1.16 | -0.935 | -0.0225 | 0.165 | 0.204237 | 0.976661 | 0.837698 | 0.999971 | | 1 | 0.979416 |
| Ccdc115 | Q8VE99 | 1.42 | 0.62 | 0.14 | 0.18 | 0.8 | 0.75 | 0.89 | 0.68 | 0.31 | 0.25 | 0.15 | -0.4 | 1.44 | 0.94 | 0.04 | -0.14 | 0.19 | -0.5125 | -0.02 | 0.540078 | 0.15374 | 0.964479 | 0.999971 | | 1 | 0.998721 |
| Ccdc12 | Q8R344 | 2.77 | 1.52 | 2.55 | 2.86 | 1.7 | 2 | 2.95 | 1.51 | 1.55 | 2.15 | 1 | 1.23 | 0.42 | 1.66 | 1.35 | 1.35 | -0.385 | -0.9425 | -1.23 | 0.372365 | 0.034483 | 0.011698 | 0.999971 | | 1 | 0.613095 |
| Ccdc120 | A2AEV7 | -1.6 | -0.67 | -0.07 | -0.9 | -0.98 | -0.49 | -0.74 | -0.57 | -0.95 | -0.35 | -0.5 | -1.06 | 0.05 | -0.69 | -0.54 | -0.26 | 0.115 | 0.095 | 0.45 | 0.730304 | 0.788146 | 0.220799 | 0.999971 | | 1 | 0.827096 |
| Ccdc124 | Q9D8X2 | 2.11 | 1.17 | 1.42 | 1.7 | 0.9 | 1.91 | 1.93 | 1.69 | 1.03 | 1.95 | 1.54 | 1.02 | 0.81 | 1.66 | 1.53 | 1.43 | 0.0075 | -0.215 | -0.2425 | 0.981149 | 0.489874 | 0.411073 | 0.999971 | | 1 | 0.875847 |
| Ccdc127 | E0CYX9 | 0.37 | -0.07 | 0.17 | 0.39 | 0.02 | -0.13 | 0.14 | 0.39 | -0.4 | 0.04 | -0.11 | 0.24 | 0.23 | 0.4 | 0.22 | 0.45 | -0.11 | -0.2725 | 0.11 | 0.597322 | 0.22727 | 0.567434 | 0.999971 | | 1 | 0.91816 |
| Ccdc134 | Q8C7V8 | -0.16 | 0.16 | 0.98 | 0.76 | -0.02 | 0.09 | 0.12 | 0.86 | 0.28 | 0.11 | 1.04 | 0.73 | 0.38 | 0.35 | 1.11 | 1.24 | -0.1725 | 0.105 | 0.335 | 0.604586 | 0.754906 | 0.349438 | 0.999971 | | 1 | 0.865428 |
| Ccdc136 | D3Z165 | -0.2 | -0.36 | -1 | -1.15 | -0.79 | -0.2 | -1.02 | -0.89 | -0.85 | 0.07 | -0.97 | -0.77 | -0.5 | 0.04 | -0.99 | -0.7 | -0.0475 | 0.0475 | 0.14 | 0.875474 | 0.886049 | 0.664953 | 0.999971 | | 1 | 0.93697 |
| Ccdc137 | Q8R0K4 | 0.38 | -0.01 | 0.07 | 0.59 | 0.23 | 0.39 | -0.06 | 0.42 | -0.1 | 0.21 | -0.23 | -0.43 | -0.43 | 0.75 | -0.09 | -0.03 | -0.0125 | -0.395 | -0.2075 | 0.954806 | 0.112046 | 0.489716 | 0.999971 | | 1 | 0.898405 |
| Ccdc141 | E9Q8Q6 | -0.47 | -0.46 | -0.12 | 0.26 | -0.78 | -0.78 | -0.51 | -0.12 | -0.81 | -0.1 | -0.26 | -0.11 | -0.82 | -0.5 | 0.07 | 0.27 | -0.35 | -0.1225 | -0.0475 | 0.19708 | 0.641572 | 0.878595 | 0.999971 | | 1 | 0.985441 |
| Ccdc149 | A0A1D5RM | 2.06 | 0.99 | 0.69 | 0.94 | 0.6 | 0.75 | 1.5 | 0.48 | 0.82 | 1.11 | 0.77 | 0.17 | 0.31 | 0.59 | 0.55 | 0.42 | -0.3375 | -0.4525 | -0.7025 | 0.372636 | 0.224327 | 0.049509 | 0.999971 | | 1 | 0.715605 |
| Ccdc167 | Q9D162 | -0.14 | -3.46 | -0.39 | -0.24 | -2.33 | -4.44 | -0.31 | 0.16 | -4.41 | -1.72 | -0.37 | -0.65 | -2.09 | -2.4 | 0.01 | -0.18 | -0.6725 | -0.73 | -0.1075 | 0.571189 | 0.511033 | 0.90621 | 0.999971 | | 1 | 0.991381 |
| Ccdc177 | Q3UHB8 | 0.17 | 0.02 | 0.19 | 0.1 | -0.48 | -0.13 | 0.26 | -0.32 | -0.38 | -0.03 | -0.21 | -0.26 | -0.52 | -0.54 | -0.2 | -0.48 | -0.2875 | -0.34 | -0.555 | 0.199483 | 0.070703 | 0.010955 | 0.999971 | | 1 | 0.613095 |
| Ccdc184 | Q8BMK5 | -0.48 | 0.57 | -0.98 | -0.52 | 0.54 | 0.82 | -1.68 | -1.52 | 0.45 | 0.52 | -0.73 | -1.24 | 0.92 | 0.3 | -1.01 | -0.33 | -0.1075 | 0.1025 | 0.3225 | 0.872266 | 0.84024 | 0.517182 | 0.999971 | | 1 | 0.905815 |
| Ccdc186 | Q8C9S4 | 1.53 | 1.38 | 1.06 | 1.07 | 1.23 | 1.46 | 1.28 | 0.86 | 1.14 | 1.51 | 0.82 | 0.55 | 0.02 | 0.93 | 0.71 | 0.88 | -0.0525 | -0.255 | -0.625 | 0.809266 | 0.339569 | 0.039137 | 0.999971 | | 1 | 0.700838 |
| Ccdc22 | Q9JIG7 | 0.38 | 0.27 | -0.02 | -0.03 | 0.25 | 0.36 | 0.33 | -0.17 | 0.38 | 0.49 | 0.07 | -0.02 | 0.16 | 0.52 | -0.07 | 0.05 | 0.0425 | 0.08 | 0.015 | 0.840282 | 0.701428 | 0.943929 | 0.999971 | | 1 | 0.99692 |
| Ccdc25 | Q78PG9 | 0.09 | 1.6 | 1.19 | 1.51 | 1.21 | 1.19 | 1.14 | 1.52 | 2.09 | 0.59 | 1.67 | 0.65 | 2.48 | 2.23 | 1.47 | 1.32 | 0.1675 | 0.1525 | 0.7775 | 0.635415 | 0.750093 | 0.094928 | 0.999971 | | 1 | 0.78268 |
| Ccdc28a | Q3UFK1 | -2.66 | -1.48 | -1.96 | -1.93 | -1.21 | -1.36 | -2.02 | -2.08 | -1.51 | -2 | -1.66 | -2.38 | -2.67 | -1.37 | -3.34 | -3.32 | 0.34 | 0.12 | -0.6675 | 0.317609 | 0.703792 | 0.193756 | 0.999971 | | 1 | 0.810547 |
| Ccdc28b | Q8CEG5 | -8.64 | -7.77 | -11.91 | 0.75 | -8.24 | 0.77 | -7.13 | -11.03 | -8.71 | -10.29 | -11.92 | -10.15 | -10.53 | 0.6 | -12.59 | 0.87 | 0.485 | -3.375 | 1.48 | 0.881811 | 0.19617 | 0.70937 | 0.999971 | | 1 | 0.950625 |
| Ccdc32 | Q3UHY7 | 1.69 | 0.76 | -0.09 | 0.01 | 0.68 | 2.01 | -0.3 | -0.13 | 1.96 | 1.31 | -0.04 | 0.16 | 1.9 | 1.22 | -0.55 | -0.24 | -0.0275 | 0.255 | -0.01 | 0.963981 | 0.66129 | 0.987719 | 0.999971 | | 1 | 0.99925 |
| Ccdc33 | A0A1L1SR7 | -0.72 | 0.83 | -0.58 | -0.02 | 0.24 | -0.01 | 0.25 | -1.95 | -0.06 | -0.57 | -1.2 | -1.92 | 1.29 | 0.43 | -1.13 | -1.94 | -0.245 | -0.815 | -0.215 | 0.675294 | 0.129096 | 0.769911 | 0.999971 | | 1 | 0.963498 |
| Ccdc43 | Q9CR29 | 1.81 | 2.29 | 2.35 | 2.71 | 1.99 | 2.61 | 2.08 | 2.55 | 2.25 | 2.55 | 2.47 | 2.04 | 1.75 | 2.05 | 1.83 | 2.6 | 0.0175 | 0.0375 | -0.2325 | 0.947325 | 0.878548 | 0.418836 | 0.999971 | | 1 | 0.87788 |
| Ccdc47 | Q9D024 | -0.51 | -0.35 | -0.3 | -0.66 | -0.02 | -0.69 | -0.21 | -0.49 | 0.31 | -0.37 | 0 | 0.35 | 0.32 | -0.62 | -0.3 | -0.46 | 0.1025 | 0.5275 | 0.19 | 0.636942 | 0.040584 | 0.455074 | 0.999971 | | 1 | 0.887217 |
| Ccdc50 | A0A668KL3 | 0.74 | 1.05 | 0.88 | 1.51 | 0.97 | 0.86 | 0.83 | 1.19 | 1.06 | 1.11 | 0.88 | 0.64 | 0.95 | 0.58 | 1.02 | 0.89 | -0.0825 | -0.1225 | -0.185 | 0.716404 | 0.600824 | 0.433199 | 0.999971 | | 1 | 0.882273 |
| Ccdc51 | Q3URS9-2 | -0.42 | -0.09 | -0.06 | 0.04 | -0.42 | 0.04 | 0.47 | -0.2 | -0.33 | -0.14 | -0.43 | -0.47 | -0.45 | -0.11 | -0.59 | -0.73 | 0.105 | -0.21 | -0.3375 | 0.670065 | 0.281095 | 0.14227 | 0.999971 | | 1 | 0.784967 |
| Ccdc59 | Q32M26 | 3.33 | 0.19 | 0.97 | 2.07 | -0.34 | 0.85 | 1.77 | 0.82 | 0.2 | 1.57 | 1.21 | 0.14 | -1.47 | 0.52 | 0.46 | 0.73 | -0.865 | -0.86 | -1.58 | 0.254529 | 0.241303 | 0.066236 | 0.999971 | | 1 | 0.733937 |
| Ccdc6 | D3YZP9 | 1.43 | -0.58 | -0.02 | 0.21 | -0.71 | 0 | 0.12 | 0.37 | -0.31 | 0.83 | 0.17 | 0.36 | -0.48 | -0.09 | 0.23 | 0.48 | -0.315 | 0.0025 | -0.225 | 0.492849 | 0.995626 | 0.615375 | 0.999971 | | 1 | 0.927967 |
| Ccdc85a | F6TRJ8 | -1.33 | -2.03 | -0.5 | -1.61 | -1.16 | -1.76 | -0.79 | -1.93 | -1.52 | -1.3 | -1.16 | -0.97 | -3.56 | -2.17 | -1.16 | -1.15 | -0.0425 | 0.13 | -0.6425 | 0.915139 | 0.703004 | 0.29935 | 0.999971 | | 1 | 0.847402 |
| Ccdc85a_1 | E0CX32 | 0.31 | -0.16 | -0.35 | -0.38 | -0.31 | 0.3 | 0.05 | -0.68 | -0.15 | 0.24 | -0.34 | -0.68 | 0.01 | -0.24 | -0.36 | -0.48 | -0.015 | -0.0875 | -0.1225 | 0.957466 | 0.744588 | 0.596914 | 0.999971 | | 1 | 0.92377 |
| Ccdc85b | Q6PDY0 | -0.07 | 1.5 | 1.51 | 1.29 | 1.36 | 1.51 | 1.26 | 1.93 | 1.88 | 1.03 | 1.99 | -0.57 | 1.02 | 1.35 | 1.57 | 1.46 | 0.4575 | 0.025 | 0.2925 | 0.260319 | 0.968909 | 0.45367 | 0.999971 | | 1 | 0.887217 |
| Ccdc85c | A0A217FL8 | 0.94 | -0.14 | -0.37 | 0 | -0.59 | -0.23 | -0.17 | -0.37 | -0.76 | 0.32 | -0.4 | 0.32 | 0.16 | -0.36 | -0.29 | -0.42 | -0.4475 | -0.2375 | -0.335 | 0.171354 | 0.53822 | 0.310242 | 0.999971 | | 1 | 0.847402 |
| Ccdc88a | Q5SNZ0-2 | 0.01 | -0.27 | -0.45 | -0.42 | -0.36 | -0.07 | -0.31 | -0.41 | -0.13 | 0.28 | -0.04 | -0.36 | 0.08 | -0.2 | -0.22 | -0.18 | -0.005 | 0.22 | 0.1525 | 0.97936 | 0.318077 | 0.43725 | 0.999971 | | 1 | 0.882419 |
| Ccdc88c | Q6VGS5 | -0.4 | -0.24 | 1.95 | 2.95 | -0.33 | -0.43 | 2.85 | 1.47 | -0.62 | 0.19 | 1.44 | 0.98 | -0.16 | 0.11 | 0.98 | 0.56 | -0.175 | -0.5675 | -0.6925 | 0.863299 | 0.510285 | 0.384463 | 0.999971 | | 1 | 0.871816 |
| Ccdc9 | D3YW42 | 1.33 | 1.03 | 1.41 | 1.1 | 1.07 | 1.13 | 1.32 | 0.9 | 0.67 | 1.46 | 1.32 | 1.36 | 0.45 | 0.97 | 0.8 | 1.15 | -0.1125 | -0.015 | -0.375 | 0.560638 | 0.94893 | 0.115055 | 0.999971 | | 1 | 0.78268 |
| Ccdc90b | Q8C3X2 | -0.17 | -0.71 | -1.75 | -0.62 | -0.74 | -0.99 | -1.2 | -0.84 | -1.09 | -0.4 | -1.39 | -1.21 | -1.24 | -0.71 | -1.26 | -1.04 | -0.13 | -0.21 | -0.25 | 0.706512 | 0.587993 | 0.483849 | 0.999971 | | 1 | 0.896352 |
| Ccdc91 | Q9D8L5 | 0.27 | 0.66 | 0.65 | 0.28 | 0.68 | 0.4 | 0.87 | 0.68 | 0.27 | 0.38 | 0.84 | 1.13 | 0.52 | 0.65 | 0.31 | 0.23 | 0.1925 | 0.19 | -0.0375 | 0.354636 | 0.460474 | 0.853454 | 0.999971 | | 1 | 0.98187 |
| Ccdc92 | Q8VDN4 | 0.34 | -0.53 | 0.19 | 0.02 | -0.57 | -0.73 | 0.6 | -0.03 | -0.8 | -0.1 | 0.04 | 0.16 | -0.49 | -0.52 | 0.03 | 0.07 | -0.1875 | -0.18 | -0.2325 | 0.594629 | 0.547134 | 0.398168 | 0.999971 | | 1 | 0.871816 |
| Ccdc92b | Q5SUE3 | 0.48 | 0.53 | -0.37 | 0.08 | -0.47 | -0.41 | 0.95 | -0.46 | 0.46 | 1.48 | -1.34 | -1.3 | 1.49 | 0.69 | -0.81 | -1.89 | -0.2775 | -0.355 | -0.31 | 0.484059 | 0.59415 | 0.663837 | 0.999971 | | 1 | 0.936621 |
| Ccdc93 | E9QAD4 | 0.95 | -0.21 | -0.4 | 0.67 | -1.26 | 0.27 | 0.47 | 1.46 | -0.89 | 0.65 | -0.05 | 0.1 | 0.68 | 0.87 | -0.38 | 1.31 | -0.0175 | -0.3 | 0.3675 | 0.976498 | 0.495511 | 0.430075 | 0.999971 | | 1 | 0.881611 |
| Ccdc93_1 | Q7TQK5 | 0.35 | 0.4 | 0.25 | 0.12 | 0.24 | 0.47 | 0.21 | 0.13 | 0.28 | 0.21 | 0.19 | 0.19 | 0.34 | 0.13 | 0.14 | 0.15 | -0.0175 | -0.0625 | -0.09 | 0.921505 | 0.702649 | 0.604135 | 0.999971 | | 1 | 0.924454 |
| Ccdc97 | Q9DBT3 | 1.47 | 0.5 | 0.21 | 0.26 | 0.64 | 0.84 | 0.27 | 0.18 | 0.24 | 1.08 | 0.17 | 0.16 | -0.2 | 0.58 | -0.07 | 0.22 | -0.1275 | -0.1975 | -0.4775 | 0.701178 | 0.586958 | 0.182709 | 0.999971 | | 1 | 0.808532 |
| Cck | P09240 | -0.96 | 0.29 | -0.61 | -1.03 | -0.58 | 0.27 | -0.12 | -1.01 | -0.13 | 0.44 | -0.99 | -0.25 | -0.13 | -0.09 | -0.38 | -0.23 | 0.2175 | 0.345 | 0.37 | 0.584243 | 0.403836 | 0.258893 | 0.999971 | | 1 | 0.839012 |
| Cckbr | P56481 | 0.08 | -1.79 | -0.49 | 0.63 | -0.41 | -2.48 | 1.38 | -0.92 | 0.02 | -0.8 | -0.5 | -0.04 | 0.3 | -2.9 | -1.23 | -0.39 | -0.215 | 0.0625 | -0.6625 | 0.800784 | 0.903369 | 0.4045 | 0.999971 | | 1 | 0.87384 |
| Ccm2 | Q8K2Y9-2 | 0.57 | 0.51 | 0.21 | 0.58 | 0.51 | 0.32 | 0.57 | 0.31 | 0.36 | 0.28 | -0.04 | 0.43 | 0.29 | 0.43 | 0.26 | 0.11 | -0.04 | -0.21 | -0.195 | 0.82777 | 0.295512 | 0.30591 | 0.999971 | | 1 | 0.847402 |
| Ccn2 | P29268 | 0.42 | 0.28 | -0.37 | 0.01 | 0.81 | 0.79 | 0.7 | -0.5 | 0.97 | 0.68 | 0.51 | 0.49 | 0.15 | 0.41 | -0.05 | -0.99 | 0.365 | 0.5775 | -0.205 | 0.318291 | 0.035851 | 0.558179 | 0.999971 | | 1 | 0.915051 |
| Ccn3 | Q64299 | 1.49 | 1.74 | 1.54 | 1.89 | 2.15 | 2.51 | 1.76 | 3.37 | 2.69 | 2.16 | 2.57 | 2.02 | 2.74 | 2.97 | 3.17 | 3.09 | 0.7825 | 0.695 | 1.3275 | 0.048786 | 0.012089 | 9.42E-05 | 0.999971 | | 1 | 0.143156 |
| Ccnk | Q3U3M5 | 0.77 | 1.89 | 2.15 | 2.49 | 1.43 | 1.31 | 1.5 | 2.04 | 1.47 | 1.29 | 1.9 | 2.35 | 1.95 | 1.69 | 2.49 | 2.71 | -0.255 | -0.0725 | 0.385 | 0.51816 | 0.862952 | 0.370181 | 0.999971 | | 1 | 0.869526 |
| Ccnl2 | Q9JJA7 | 0.3 | 0.63 | 1.43 | 0.77 | 0.57 | 0.67 | 0.38 | 1.37 | 0.41 | 0.72 | 0.37 | 1.42 | 0.66 | 1.21 | 1.22 | 2.05 | -0.035 | -0.0525 | 0.5025 | 0.913477 | 0.876127 | 0.190549 | 0.999971 | | 1 | 0.810547 |
| Ccnt1 | Q9QWV9 | -0.56 | 0.79 | 3.84 | 0.73 | 0.19 | 0.63 | 3.85 | 2.18 | -0.03 | 0.13 | 3.44 | -2.53 | -0.11 | 0.9 | 2.53 | -0.48 | 0.5125 | -0.9475 | -0.49 | 0.646742 | 0.497409 | 0.635659 | 0.999971 | | 1 | 0.93301 |
| Ccnt2 | Q7TQK0-2 | 5.12 | 1.47 | 3.14 | 3.64 | 1.98 | 2.71 | 2.97 | 2.87 | 1.42 | 3.52 | 2.85 | 2.92 | 0.12 | 0.98 | 3.22 | 3.81 | -0.71 | -0.665 | -1.31 | 0.330912 | 0.410762 | 0.227097 | 0.999971 | | 1 | 0.828236 |
| Ccny | Q8BGU5 | -0.62 | -0.64 | -1.14 | -0.71 | -0.93 | -0.8 | -1.46 | -0.9 | -1.04 | -0.71 | -1.28 | -1.17 | -0.37 | -0.63 | -1.18 | -0.67 | -0.245 | -0.2725 | 0.065 | 0.30326 | 0.229252 | 0.788362 | 0.999971 | | 1 | 0.96741 |
| Ccnyl1 | D3YUJ3 | -1.22 | -2.02 | -1.1 | -0.97 | -1.73 | -2.3 | -0.96 | -0.81 | -1.44 | -1.36 | -0.57 | -0.84 | -1.17 | -1.7 | -1.05 | -0.88 | -0.1225 | 0.275 | 0.1275 | 0.760742 | 0.397327 | 0.675757 | 0.999971 | | 1 | 0.941576 |
| Ccp110 | Q7TSH4 | 1.77 | 0.66 | -0.12 | 0.05 | 0.46 | 0.76 | -0.23 | 0.43 | 0.48 | 1.5 | 0.23 | 0.1 | 0.01 | 0.01 | -0.17 | -0.2 | -0.235 | -0.0125 | -0.6775 | 0.601845 | 0.979895 | 0.126608 | 0.999971 | | 1 | 0.783193 |
| Ccs | A0A494BB! | 1.62 | 1.54 | 0.66 | 0.77 | 1.39 | 0.99 | 0.88 | 0.8 | 1.49 | 1.08 | 0.53 | 0.27 | 1.33 | 0.26 | 0.89 | 0.93 | -0.1325 | -0.305 | -0.295 | 0.653744 | 0.410565 | 0.38772 | 0.999971 | | 1 | 0.871816 |
| Ccsap | Q8QZT2 | 1.91 | 2.71 | 2.2 | 2.3 | 3.14 | 2.45 | 2.05 | 1.91 | 3.25 | 2.06 | 2.22 | 2.04 | 2.51 | 1.82 | 1.72 | 1.81 | 0.1075 | 0.1125 | -0.315 | 0.739785 | 0.73499 | 0.258558 | 0.999971 | | 1 | 0.839012 |
| Ccser1 | Q8C0C4-3 | -1.42 | 0.77 | 0.28 | -0.17 | 0.35 | 0.2 | 0.04 | 0.26 | 0.05 | -1.05 | 0.4 | 0.16 | 1.31 | -0.29 | -0.56 | -0.09 | 0.3475 | 0.025 | 0.2275 | 0.443465 | 0.962171 | 0.693889 | 0.999971 | | 1 | 0.947174 |
| Ccser2 | Q3UHI0 | -0.8 | 0.28 | 0.19 | -0.93 | 0.77 | -0.22 | 0.27 | -0.24 | -0.35 | -0.5 | 0.34 | 0.43 | 1.07 | -0.32 | -0.53 | -1.39 | 0.46 | 0.295 | 0.0225 | 0.250665 | 0.449992 | 0.967529 | 0.999971 | | 1 | 0.998787 |
| Ccsmst1 | Q6RUT7 | 0.02 | 0.57 | 1.3 | 0.69 | 0.77 | 0.85 | 0.4 | 0.77 | 0.43 | 0.56 | 1.08 | 0.92 | 0.61 | 0.77 | 1.22 | 2.15 | 0.0525 | 0.1025 | 0.5425 | 0.857463 | 0.740214 | 0.212082 | 0.999971 | | 1 | 0.823193 |
| Cct2 | P80314 | -0.19 | 0.16 | 0.36 | 0.48 | 0 | 0.03 | 0.38 | 0.34 | 0.03 | -0.44 | 0.21 | 0.52 | -0.13 | 0.22 | 0.54 | 0.69 | -0.015 | -0.1225 | 0.1275 | 0.94581 | 0.648603 | 0.624343 | 0.999971 | | 1 | 0.930479 |
| Cct3 | P80318 | 0.03 | 0.13 | 0.5 | 0.4 | 0.03 | -0.08 | 0.27 | 0.46 | -0.01 | -0.12 | 0.3 | 0.34 | -0.04 | 0.08 | 0.5 | 0.46 | -0.095 | -0.1375 | -0.015 | 0.656925 | 0.512237 | 0.945567 | 0.999971 | | 1 | 0.997427 |
| Cct4 | P80315 | 0.09 | 0.7 | 0.13 | 0.25 | 0.67 | 0.35 | 0.01 | 0.23 | 0.86 | 0.19 | -0.09 | 0.01 | 0.89 | 0.94 | 0.26 | 0.21 | 0.0225 | -0.05 | 0.2825 | 0.922971 | 0.854405 | 0.300997 | 0.999971 | | 1 | 0.847402 |
| Cct5 | P80316 | 0.28 | 0.89 | 0.97 | 0.75 | 0.84 | 0.81 | 0.81 | 0.86 | 0.84 | 0.25 | 0.78 | 1.2 | 0.91 | 0.9 | 1 | 0.89 | 0.1075 | 0.045 | 0.2025 | 0.60683 | 0.866597 | 0.345762 | 0.999971 | | 1 | 0.863794 |
| Cct6a | Q52KG9 | 0.39 | 1.36 | 0.77 | 0.77 | 1.52 | 1.03 | 0.73 | 0.75 | 1.67 | 0.51 | 0.79 | 0.89 | 2.13 | 1.07 | 0.94 | 1.08 | 0.185 | 0.1425 | 0.4825 | 0.521957 | 0.658467 | 0.179483 | 0.999971 | | 1 | 0.808532 |
| Cct7 | P80313 | -0.44 | 0.61 | 0.5 | 0.22 | 0.52 | 0.5 | 0.35 | 0.3 | 0.56 | -0.39 | 0.27 | 0.59 | 0.63 | 0.34 | 0.59 | 0.19 | 0.195 | 0.035 | 0.215 | 0.46674 | 0.914806 | 0.443632 | 0.999971 | | 1 | 0.883386 |
| Cct8 | P42932 | 0.24 | 0.57 | 0.72 | 0.59 | 0.67 | 0.52 | 0.74 | 0.65 | 0.93 | 0.49 | 0.75 | 0.95 | 0.66 | 0.54 | 0.73 | 0.77 | 0.115 | 0.25 | 0.145 | 0.538617 | 0.233852 | 0.445144 | 0.999971 | | 1 | 0.884152 |
| Ccz1 | Q8C1Y8 | -0.42 | -0.86 | 0.28 | -0.19 | -0.67 | -0.83 | -0.05 | 0.17 | -0.71 | -0.66 | 0.11 | -1.09 | -0.84 | -0.74 | 0.5 | 0.1 | -0.0475 | -0.29 | 0.0525 | 0.887372 | 0.406426 | 0.892408 | 0.999971 | | 1 | 0.98828 |
| Cd109 | Q8R422 | -0.11 | -0.41 | -0.71 | -0.82 | -0.11 | -0.05 | -0.86 | -0.21 | -0.29 | 0 | -0.66 | -0.71 | 0.18 | 0.29 | -0.72 | -0.18 | 0.205 | 0.0975 | 0.405 | 0.449842 | 0.703052 | 0.187114 | 0.999971 | | 1 | 0.809025 |
| Cd151 | O35566 | 2.05 | 1.83 | 1.35 | 2.09 | 2.46 | 1.76 | 2.06 | 1.58 | 2.53 | 2.67 | 1.45 | 2.2 | 2.91 | 1.42 | 1.37 | 0.72 | 0.135 | 0.3825 | -0.225 | 0.624896 | 0.254925 | 0.628895 | 0.999971 | | 1 | 0.932176 |
| Cd163 | Q2VLH6 | 1.99 | 0.39 | -0.41 | -0.64 | 1.5 | 0.74 | -0.78 | -0.7 | 0.79 | 2.27 | -0.87 | 0.81 | -0.31 | 0.83 | -0.74 | -0.04 | -0.1425 | 0.4175 | -0.3975 | 0.846504 | 0.601014 | 0.527019 | 0.999971 | | 1 | 0.909459 |
| Cd180 | Q62192 | 0.41 | 0.23 | -0.29 | -0.63 | 0.46 | 0.61 | -0.06 | -0.11 | 0.31 | 0.5 | -0.2 | -0.44 | -0.6 | 0.84 | -0.31 | -0.75 | 0.295 | 0.1125 | -0.135 | 0.348526 | 0.729207 | 0.743621 | 0.999971 | | 1 | 0.961064 |
| Cd200 | Q80VX2 | -1.73 | -1.94 | -0.92 | -0.36 | -0.84 | -1.59 | -0.41 | -0.64 | -0.65 | -2.49 | -0.42 | -0.77 | -0.51 | -1.34 | -0.8 | -0.14 | 0.3675 | 0.155 | 0.54 | 0.395224 | 0.779783 | 0.222831 | 0.999971 | | 1 | 0.827753 |
| Cd276 | Q8VE98 | -2.28 | 0.2 | -1.33 | -2.78 | -1.51 | -1.91 | -2.2 | -1.61 | -0.47 | -1.04 | -3.29 | -0.94 | -1.45 | -2.61 | -2.1 | -1.89 | -0.26 | 0.1125 | -0.465 | 0.673254 | 0.891144 | 0.471578 | 0.999971 | | 1 | 0.892857 |
| Cd2ap | Q9JLQ0 | 0.26 | 0.12 | -0.31 | 0.71 | 0.44 | -0.2 | -0.2 | 0.16 | 0.15 | 0.49 | -0.37 | 0.01 | -0.39 | 0.46 | 0.19 | 0.16 | -0.145 | -0.125 | -0.09 | 0.604207 | 0.664887 | 0.755751 | 0.999971 | | 1 | 0.961116 |
| Cd2bp2 | Q9CWK3 | 2.39 | 2.21 | 2.67 | 2.1 | 1.73 | 1.8 | 1.73 | 1.81 | 2.31 | 2.84 | 1.72 | 1.51 | 1.5 | 2.49 | 1.87 | 1.56 | -0.575 | -0.2475 | -0.4875 | 0.014233 | 0.454915 | 0.105009 | 0.999971 | | 1 | 0.78268 |
| Cd34 | Q64314 | -0.47 | -0.14 | 1.3 | 1.6 | -0.32 | -0.69 | 0.26 | 1.55 | -0.82 | -1.79 | 1.16 | -0.49 | -0.58 | -1.16 | 0.72 | 0.92 | -0.3725 | -1.0575 | -0.5975 | 0.567581 | 0.171508 | 0.373111 | 0.999971 | | 1 | 0.869526 |
| Cd36 | Q08857 | -1.39 | -0.8 | -0.85 | 0.2 | -0.43 | -0.97 | -2.65 | -1.41 | -0.63 | -1.93 | -2.96 | -1.88 | 0.05 | -0.83 | -0.73 | -1.41 | -0.655 | -1.14 | -0.02 | 0.238661 | 0.059578 | 0.962364 | 0.999971 | | 1 | 0.998721 |
| Cd38 | P56528 | -0.27 | 2.35 | 1.55 | 1.42 | 2.61 | 2.38 | 2.31 | 1.91 | 2.42 | 0.41 | 1.31 | 2.41 | 3.61 | 1.36 | 1.85 | 0.62 | 1.04 | 0.375 | 0.5975 | 0.074386 | 0.578716 | 0.439329 | 0.999971 | | 1 | 0.883105 |
| Cd44 | A2APM2 | -1.11 | -1.08 | -1.75 | -1.2 | -0.71 | -0.56 | -1.57 | -1.51 | -0.75 | -0.81 | -1.45 | -0.79 | 0.01 | -0.45 | -1.02 | -0.04 | 0.1975 | 0.335 | 0.91 | 0.530623 | 0.209105 | 0.012657 | 0.999971 | | 1 | 0.613095 |
| Cd47 | A0A2R8VK | -0.77 | 0.69 | 0.41 | 0.12 | 0.24 | 0.91 | 0.3 | 0.39 | 0.42 | -0.11 | -0.19 | -0.76 | 0.9 | 0.96 | 0.01 | 1.24 | 0.3475 | -0.2725 | 0.665 | 0.329764 | 0.484771 | 0.121078 | 0.999971 | | 1 | 0.78268 |
| Cd47_1 | A0A2R8W6 | -1.13 | -0.71 | -1.28 | -0.92 | -0.94 | -0.5 | -1.74 | -1 | -1.1 | -0.77 | -1.36 | -1.44 | -0.48 | -0.61 | -1.24 | -0.74 | -0.035 | -0.1575 | 0.2425 | 0.905655 | 0.500281 | 0.328354 | 0.999971 | | 1 | 0.856472 |
| Cd55 | Q61475 | 0.11 | 0.85 | -0.73 | -0.88 | 0.85 | 0.84 | -1.04 | -0.89 | 1.2 | 0.44 | -0.47 | -0.51 | 1.67 | 1.27 | 0.07 | -0.67 | 0.1025 | 0.3275 | 0.7475 | 0.864725 | 0.541962 | 0.24328 | 0.999971 | | 1 | 0.837316 |
| Cd63 | A0A1W2P8 | 0.12 | -0.02 | 0.55 | -0.66 | -0.35 | 0.71 | -0.22 | -0.2 | 0.16 | 0.1 | -0.12 | 0.17 | 1.4 | 1.99 | 0.97 | -0.53 | -0.0125 | 0.08 | 0.96 | 0.971048 | 0.771721 | 0.106584 | 0.999971 | | 1 | 0.78268 |
| Cd81 | P35762 | 0.21 | -0.43 | 0.05 | -0.66 | -0.8 | -0.3 | -0.3 | -0.05 | -0.91 | -0.27 | -1.13 | -0.64 | -0.62 | -0.64 | -0.04 | -0.43 | -0.155 | -0.53 | -0.225 | 0.57563 | 0.092105 | 0.409903 | 0.999971 | | 1 | 0.875847 |
| Cd82 | P40237 | 0.83 | 0 | -0.15 | -0.84 | 0.03 | 0.51 | -0.96 | -0.49 | 0.39 | 0.03 | -1.38 | -0.87 | -0.34 | -0.18 | -0.71 | -1.33 | -0.1875 | -0.4175 | -0.6 | 0.671766 | 0.409478 | 0.166869 | 0.999971 | | 1 | 0.800215 |
| Cd9 | P40240 | -0.16 | 0.12 | -1.26 | -1.16 | -0.94 | 0.31 | -2.04 | -1.65 | -0.8 | 0.29 | -2.24 | -1.89 | -0.1 | 0.49 | -1.35 | -1.73 | -0.465 | -0.545 | -0.0575 | 0.424117 | 0.386281 | 0.920277 | 0.999971 | | 1 | 0.992702 |
| Cd93 | O89103 | -0.52 | 0.98 | 0.9 | 0.36 | 1.69 | 0.3 | 0.98 | -0.58 | 1.19 | -0.56 | 1.18 | 1.56 | 0.84 | 0.55 | 1.7 | 0.55 | 0.1675 | 0.4125 | 0.48 | 0.759728 | 0.455766 | 0.269306 | 0.999971 | | 1 | 0.843504 |
| Cd99 | Q8VCN6 | 1.49 | 2.21 | 1.06 | 1.51 | 1.8 | 0.98 | 0.4 | 1.14 | 0.89 | 1.92 | 1.02 | 0.92 | 2.83 | 1.78 | 1.4 | 1.71 | -0.4875 | -0.38 | 0.3625 | 0.202997 | 0.279969 | 0.352067 | 0.999971 | | 1 | 0.867656 |
| Cd99l2 | A2AP74 | 0.5 | 0.31 | 0.25 | 0.35 | -0.12 | 0.23 | 0.09 | 0.36 | 0.92 | 1.05 | 0.85 | 1.4 | -0.01 | -0.25 | 0.45 | 1.04 | -0.2125 | 0.7025 | -0.045 | 0.272935 | 0.005301 | 0.880228 | 0.999971 | | 1 | 0.985441 |
| Cda | P56389 | 0.44 | -1.29 | -0.6 | -0.02 | -1.46 | -0.97 | -0.9 | 0.86 | -1.51 | -1.41 | -1.55 | -1.1 | -0.92 | -0.59 | 0.76 | 0.08 | -0.25 | -1.025 | 0.2 | 0.6664 | 0.022368 | 0.685215 | 0.999971 | | 1 | 0.944874 |
| Cdc123 | Q8CII2 | -0.92 | 0.2 | 0.7 | 0.99 | -0.63 | -0.43 | 0.25 | 0.27 | -0.15 | -0.25 | 0.79 | 0.81 | -1.26 | 0.09 | 0.09 | 0.69 | -0.3775 | 0.0575 | -0.34 | 0.411778 | 0.904173 | 0.535659 | 0.999971 | | 1 | 0.910077 |
| Cdc16 | Q8R349 | -0.01 | 1.11 | 0.59 | 0.56 | 0.89 | 0.95 | 0.08 | 0.8 | 1.24 | 1.02 | 0.4 | 0.51 | 1.42 | 1.53 | 0.9 | 1.09 | 0.1175 | 0.23 | 0.6725 | 0.706079 | 0.464678 | 0.040293 | 0.999971 | | 1 | 0.700838 |
| Cdc23 | G3X8W7 | -0.25 | 1.24 | 1.42 | 1.2 | 1.2 | 1.27 | 1.52 | 1.45 | 2.17 | 0.34 | 1.77 | 1.94 | 1.55 | 1.24 | 1.86 | 1.08 | 0.4575 | 0.6525 | 0.53 | 0.248768 | 0.235144 | 0.213216 | 0.999971 | | 1 | 0.824457 |
| Cdc26 | Q99JP4 | -0.33 | -1.17 | -1.72 | -0.58 | -0.72 | -1.43 | -0.93 | -0.8 | -0.84 | -1.01 | -0.56 | -0.58 | -1.68 | -0.95 | -1.52 | -1.67 | -0.02 | 0.2025 | -0.505 | 0.953699 | 0.542636 | 0.174328 | 0.999971 | | 1 | 0.807788 |
| Cdc27 | A2A6Q5 | 0.38 | 0.63 | 0.12 | 0.4 | 0.29 | 0.31 | -0.1 | 0.4 | 0.34 | 0.23 | 0.3 | 0.58 | 0.51 | -0.06 | 0.37 | 0.67 | -0.1575 | -0.02 | -0.01 | 0.451995 | 0.916296 | 0.965025 | 0.999971 | | 1 | 0.998721 |
| Cdc34 | Q8CFI2 | 0.61 | 0.7 | 0.89 | 1.18 | 0.78 | 0.84 | 0.38 | 1.77 | 0.64 | 0.94 | 1.46 | 1.01 | 0.39 | 1.01 | 1.16 | 1.48 | 0.0975 | 0.1675 | 0.165 | 0.762219 | 0.493016 | 0.557016 | 0.999971 | | 1 | 0.915051 |
| Cdc37 | Q61081 | 0.35 | 0.08 | 0.17 | 0.06 | 0.46 | 0.43 | 0.37 | 0.08 | 0.46 | 0.41 | 0.45 | 0.42 | 0.27 | 0.1 | 0.2 | 0.41 | 0.17 | 0.27 | 0.08 | 0.366758 | 0.126277 | 0.653627 | 0.999971 | | 1 | 0.934061 |
| Cdc37l1 | Q9CZP7-2 | 1.12 | 1.39 | 0.5 | 0.84 | 0.93 | 1.3 | 0.68 | 0.87 | 1.22 | 1.25 | 0.95 | 1.02 | 1.07 | 1.33 | 0.48 | 0.48 | -0.0175 | 0.1475 | -0.1225 | 0.945437 | 0.537308 | 0.681552 | 0.999971 | | 1 | 0.943204 |
| Cdc40 | Q9DC48 | -0.44 | 0.21 | 1.01 | 0.96 | -0.02 | -0.16 | 0.4 | 0.97 | 0.12 | -0.27 | 0.83 | 0.78 | 0 | 0 | 0.95 | 0.97 | -0.1375 | -0.07 | 0.045 | 0.736433 | 0.866049 | 0.914627 | 0.999971 | | 1 | 0.992633 |
| Cdc42 | P60766 | -1.01 | -0.81 | -0.01 | -0.16 | -1.06 | -0.59 | -0.1 | -0.33 | -0.75 | -0.87 | -0.23 | -0.23 | -0.85 | -0.48 | 0.09 | 0.5 | -0.0225 | -0.0225 | 0.3125 | 0.944045 | 0.940797 | 0.413518 | 0.999971 | | 1 | 0.876261 |
| Cdc42_1 | P60766-1 | -0.32 | -0.19 | -1.47 | -1.4 | -0.28 | -0.19 | -1.64 | -1.19 | -0.43 | -0.15 | -1.39 | -1.69 | -0.47 | -0.18 | -1.45 | -1.29 | 0.02 | -0.07 | -0.0025 | 0.965299 | 0.882208 | 0.995417 | 0.999971 | | 1 | 1 |
| Cdc42bpa | D3YYN8 | -0.5 | -0.34 | -0.28 | -0.03 | -0.42 | -0.14 | -0.31 | 0.02 | -0.21 | -0.33 | 0.16 | 0.12 | -0.13 | -0.1 | 0 | 0.28 | 0.075 | 0.2225 | 0.3 | 0.705294 | 0.295127 | 0.153832 | 0.999971 | | 1 | 0.795578 |

| | | | | | | | | | | | | | | | | | | | | | | | | | | |
|---|---|---|---|---|---|---|---|---|---|---|---|---|---|---|---|---|---|---|---|---|---|---|---|---|---|---|
| Cdc42bpb | Q7TT50 | -0.09 | 0.12 | 0.07 | -0.22 | 0.15 | 0.2 | 0.25 | -0.14 | 0.42 | 0.1 | 0.31 | 0.23 | 0.34 | -0.01 | -0.16 | -0.37 | 0.145 | 0.295 | -0.02 | 0.446659 | 0.123815 | 0.926187 | 0.999971 | 1 | 0.99401 |
| Cdc42ep1 | A0A0R4J0S | -1.41 | -0.72 | -0.86 | -1.36 | -1.11 | -1.17 | -1.46 | -1.81 | -1.19 | -1.69 | -2.1 | -1.35 | -0.81 | -1.25 | -1.32 | -1.49 | -0.3 | -0.495 | -0.13 | 0.267258 | 0.105255 | 0.611168 | 0.999971 | 1 | 0.92673 |
| Cdc42ep2 | Q8JZX9 | -0.84 | -0.27 | -0.8 | 0.08 | -1.71 | -1.66 | -1.44 | 0.34 | -1.76 | -0.75 | -0.52 | 0.3 | 0.85 | -1.71 | -0.51 | -0.16 | -0.66 | -0.225 | 0.075 | 0.20767 | 0.620166 | 0.887193 | 0.999971 | 1 | 0.987485 |
| Cdc42ep4 | Q9JM96 | 0.15 | -0.39 | -0.44 | -0.34 | -0.23 | -0.67 | 0.2 | -0.5 | -0.2 | -0.13 | -0.26 | 0.08 | -0.19 | -0.83 | -0.71 | -0.35 | -0.045 | 0.1275 | -0.265 | 0.86155 | 0.538875 | 0.28233 | 0.999971 | 1 | 0.844538 |
| Cdc42se2 | Q8BGH7 | -0.16 | -1.15 | -0.69 | -1.11 | -0.79 | -1.15 | -1.49 | -1.31 | -1.11 | -0.56 | -1.26 | -0.91 | -0.71 | -1.05 | -1.17 | -0.58 | -0.4075 | -0.1825 | -0.1 | 0.180253 | 0.530366 | 0.725604 | 0.999971 | 1 | 0.956041 |
| Cdc5l | Q6A068 | 0.09 | 0.16 | -0.06 | -0.37 | 0.42 | -0.15 | 0.27 | -0.65 | 0.37 | 0.05 | -0.42 | -0.53 | 0.16 | 0.29 | -0.53 | -0.58 | 0.0175 | -0.0875 | -0.12 | 0.950473 | 0.738408 | 0.663247 | 0.999971 | 1 | 0.936388 |
| Cdc73 | Q8JZM7 | 0.89 | 0.58 | 0.66 | 0.63 | 0.47 | 0.32 | 0.46 | 0.93 | 0.52 | 0.77 | 0.77 | 0.77 | 0.19 | 0.69 | 0.68 | 0.42 | -0.145 | 0.0175 | -0.195 | 0.484336 | 0.919469 | 0.339644 | 0.999971 | 1 | 0.860218 |
| Cdh10 | P70408 | -0.81 | -1.16 | -0.95 | -0.92 | -1.22 | -0.96 | -0.41 | -0.76 | -1.29 | -0.68 | -0.62 | -0.65 | -1.37 | -0.99 | -1.01 | -0.49 | 0.1225 | 0.15 | -0.005 | 0.590976 | 0.497425 | 0.982838 | 0.999971 | 1 | 0.99925 |
| Cdh11 | P55288 | -1.08 | -1.06 | -1.04 | -1.02 | -1.19 | -1.08 | -1.05 | -0.86 | -1.11 | -0.97 | -0.95 | -0.72 | -1.73 | -1.19 | -1.43 | -1.37 | 0.005 | 0.1125 | -0.38 | 0.976267 | 0.510602 | 0.067118 | 0.999971 | 1 | 0.733937 |
| Cdh12 | Q5RJH3 | -0.96 | -0.36 | -0.74 | -0.45 | -0.89 | -1.02 | -1.1 | -0.75 | -0.88 | -0.87 | -0.7 | -0.24 | -1.09 | -0.68 | -1.32 | -0.53 | -0.3125 | -0.045 | -0.2775 | 0.159809 | 0.848574 | 0.292644 | 0.999971 | 1 | 0.847402 |
| Cdh13 | Q9WTR5 | -0.63 | -1.5 | -1.74 | -1.57 | -1.32 | -1.23 | -1.57 | -1.38 | -1.24 | -0.87 | -1.7 | -1.31 | -1.24 | -1.21 | -1.18 | -1.06 | -0.015 | 0.08 | 0.1875 | 0.956579 | 0.79445 | 0.495504 | 0.999971 | 1 | 0.901531 |
| Cdh18 | E9Q9Q6 | 0.26 | -0.64 | -1.15 | -1.47 | -0.39 | -0.64 | 0.16 | -1.26 | -0.08 | -0.31 | -0.54 | -0.64 | -0.95 | -1.35 | -2.07 | -1.74 | 0.2175 | 0.3575 | -0.7775 | 0.630669 | 0.364377 | 0.095163 | 0.999971 | 1 | 0.78268 |
| Cdh19 | E9Q3A7 | -0.61 | -1.98 | -0.72 | -0.99 | -1.48 | -1.46 | -1.2 | -1.04 | -1.65 | -1.34 | -0.98 | -1.57 | -1.28 | -1.26 | -1.18 | -1.06 | -0.22 | -0.31 | -0.12 | 0.509362 | 0.376714 | 0.707981 | 0.999971 | 1 | 0.950182 |
| Cdh2 | D3YYT0 | -1.02 | -1.21 | -1.2 | -1.15 | -1.07 | -0.9 | -1.18 | -1.13 | -1.03 | -1.2 | -1.32 | -1.08 | -1.23 | -1.06 | -1.54 | -1.13 | 0.075 | -0.0125 | -0.095 | 0.660605 | 0.94045 | 0.613711 | 0.999971 | 1 | 0.927364 |
| Cdh20 | Q9Z0M3 | -0.29 | 0.61 | -1.33 | -1.73 | -0.51 | 0.77 | -0.47 | -1.03 | -0.67 | 0.42 | -0.78 | -1.13 | -1.04 | 0.81 | -1.19 | -1.28 | 0.375 | 0.145 | 0.01 | 0.532934 | 0.801216 | 0.987878 | 0.999971 | 1 | 0.99925 |
| Cdh22 | Q9WTP5 | -3.22 | -0.74 | -0.72 | -0.52 | -0.49 | -1.62 | 0.58 | -0.24 | -0.93 | -3.72 | -0.1 | 0.63 | -0.91 | -1.69 | -0.39 | -1.39 | 0.8575 | 0.27 | 0.205 | 0.247057 | 0.793706 | 0.748954 | 0.999971 | 1 | 0.961064 |
| Cdh4 | P39038 | -1.77 | -1.92 | -0.36 | -0.37 | -1.51 | -1.71 | -0.47 | -0.49 | -1.7 | -1.34 | -0.42 | -0.15 | -2.25 | -1.87 | -0.83 | -1.01 | 0.06 | 0.2025 | -0.385 | 0.904512 | 0.70024 | 0.455533 | 0.999971 | 1 | 0.887217 |
| Cdh5 | P55284 | 0.41 | 0.49 | 0.34 | 1.01 | 1.47 | 0.54 | 0.07 | 0.12 | 0.12 | 0.81 | -0.18 | 0.62 | -0.76 | 0.88 | 0.6 | -0.01 | -0.0125 | -0.22 | -0.385 | 0.971606 | 0.449018 | 0.326969 | 0.999971 | 1 | 0.856472 |
| Cdh6 | P97326 | -1.02 | -1.28 | -2.47 | -2.35 | -1.32 | -1.66 | -2.6 | -2.56 | -1.41 | -1.13 | -2.06 | -2.54 | -1.21 | -1.43 | -2.96 | -2.61 | -0.255 | -0.005 | -0.2725 | 0.581283 | 0.991281 | 0.60597 | 0.999971 | 1 | 0.924649 |
| Cdh7 | Q8BM92 | -2.28 | -2.57 | -1.02 | -1.58 | -2.23 | -2.57 | -0.68 | -1.48 | -2.32 | -2.16 | -0.86 | -0.46 | -1.99 | -2.24 | -1.68 | -1.36 | 0.1225 | 0.4125 | 0.045 | 0.80885 | 0.451177 | 0.906712 | 0.999971 | 1 | 0.991405 |
| Cdh8 | P97291 | -1.13 | -0.72 | -1.08 | -1.83 | -0.2 | -0.66 | -1.34 | -1.08 | -0.02 | -1.04 | -0.86 | -0.64 | -0.08 | -1.06 | -1.81 | -1.66 | 0.37 | 0.55 | 0.0375 | 0.290323 | 0.115701 | 0.930695 | 0.999971 | 1 | 0.994744 |
| Cdh9 | F8WHU6 | -1.12 | -0.97 | -1.32 | -1.01 | -1.15 | -0.74 | -0.3 | -1.08 | -0.39 | -1.01 | -1.19 | -0.71 | -1.14 | -0.88 | -1.05 | -0.91 | 0.2875 | 0.28 | 0.11 | 0.253198 | 0.240407 | 0.545379 | 0.999971 | 1 | 0.911398 |
| Cdip1 | Q9DB75 | 0.8 | -0.45 | 1.25 | 1.27 | -0.49 | 0.21 | 1.35 | 0.69 | 0.11 | 1.02 | -0.12 | -0.94 | 0.29 | -0.15 | 0.41 | 0.23 | -0.2775 | -0.7 | -0.5225 | 0.594763 | 0.208647 | 0.217317 | 0.999971 | 1 | 0.826893 |
| Cdipt | Q8VDP6 | -0.96 | 0.65 | 0.69 | 0.56 | -0.08 | 1.13 | 0.55 | 0.94 | 0.15 | -0.48 | 0.93 | -0.27 | -0.4 | 1.39 | 0.58 | 0.79 | 0.4 | -0.1525 | 0.355 | 0.385961 | 0.748432 | 0.489385 | 0.999971 | 1 | 0.898346 |
| Cdk11b | A2A9P6 | 0.49 | 1.82 | 1.34 | 1.39 | 1.73 | 1.47 | 0.6 | 0.99 | 1.5 | 0.8 | 1.32 | 1.49 | 1.59 | 1.5 | 1.07 | 1.24 | -0.0625 | 0.0175 | 0.09 | 0.863995 | 0.956884 | 0.771329 | 0.999971 | 1 | 0.963498 |
| Cdk13 | Q69ZA1 | 0.02 | -2.15 | -0.77 | -0.85 | -1.75 | -2.95 | -0.08 | -1.22 | -2.61 | -1.35 | -0.19 | -0.52 | -3.31 | -2.14 | -1.07 | -1.81 | -0.5625 | -0.23 | -1.145 | 0.414635 | 0.720847 | 0.080365 | 0.999971 | 1 | 0.755254 |
| Cdk14 | O35495 | -1.18 | 0.21 | 0.09 | -1.19 | 1.02 | 0.6 | 0.83 | -0.89 | 1.22 | -0.93 | 0.37 | 0.69 | 1.47 | -0.24 | -0.62 | -0.73 | 0.9075 | 0.855 | 0.4875 | 0.11753 | 0.148589 | 0.415308 | 0.999971 | 1 | 0.876885 |
| Cdk16 | Q04735 | -0.04 | -0.26 | -0.56 | -0.53 | -0.51 | -0.32 | -0.52 | -0.59 | -0.32 | 0.13 | -0.21 | -0.53 | 0.01 | -0.56 | -0.88 | -0.45 | -0.1375 | 0.115 | -0.1225 | 0.490764 | 0.609144 | 0.626568 | 0.999971 | 1 | 0.930511 |
| Cdk17 | Q8K0D0 | -1.24 | -0.37 | -0.15 | -0.51 | -0.02 | -0.47 | 0.13 | -0.13 | 0.13 | -0.7 | 0.3 | 0.39 | 0.79 | -0.5 | 0.09 | 0 | 0.445 | 0.5975 | 0.6625 | 0.141812 | 0.106516 | 0.08568 | 0.999971 | 1 | 0.759991 |
| Cdk18 | Q04899 | -0.72 | -0.17 | -0.74 | -0.75 | -0.27 | -0.33 | -0.78 | -0.15 | -0.15 | -0.48 | -0.9 | -0.48 | 0.24 | 0.05 | 0.05 | -0.34 | 0.2125 | 0.0925 | 0.595 | 0.374729 | 0.700437 | 0.02633 | 0.999971 | 1 | 0.678474 |
| Cdk19 | Q8BWD8 | 1.6 | 1.82 | 1.72 | 0.94 | 1.27 | 1.32 | 0.33 | 1.52 | 1.26 | 1.49 | 1.59 | 1.77 | 1.32 | 1.86 | 1.11 | 1.24 | -0.41 | 0.0075 | -0.1375 | 0.235539 | 0.976106 | 0.620684 | 0.999971 | 1 | 0.929344 |
| Cdk20 | Q9JHU3 | 1.04 | 0.64 | 0.41 | 0.58 | 0.19 | 0.44 | -0.21 | 0.79 | 0.04 | 0.75 | -0.04 | 0.62 | 0.1 | 0.64 | -0.43 | 0.59 | -0.365 | -0.325 | -0.4425 | 0.197806 | 0.234839 | 0.158521 | 0.999971 | 1 | 0.798217 |
| Cdk2ap1 | F2Z4B3 | 1.99 | 2.71 | 2.49 | 2.06 | 2.08 | 2.68 | 1.72 | 1.94 | 2.01 | 3.62 | 1.43 | -0.52 | 1.39 | 2.35 | 1.16 | 1.87 | -0.2075 | -0.6775 | -0.62 | 0.469988 | 0.4012 | 0.078875 | 0.999971 | 1 | 0.75432 |
| Cdk4 | E9Q9E4 | 0.35 | 1.72 | 0.98 | 1.42 | 1.1 | 0.8 | 0.83 | 1.13 | 0.46 | 0.87 | 0.58 | 1.38 | 0.76 | 1.15 | 1.37 | 1.55 | -0.1525 | -0.295 | 0.09 | 0.629351 | 0.414664 | 0.79155 | 0.999971 | 1 | 0.968737 |
| Cdk5 | P49615 | 0.16 | -0.05 | 0.28 | 0.29 | 0.47 | 0.34 | 0.01 | 0.21 | 0.5 | 0.19 | 0.03 | 0.42 | 0.75 | 0.66 | 0.4 | 0.67 | 0.0875 | 0.115 | 0.45 | 0.649955 | 0.554819 | 0.035167 | 0.999971 | 1 | 0.69205 |
| Cdk5r1 | P61809 | -0.31 | 0.13 | 0.53 | 0.38 | 0.03 | 0.33 | -0.41 | 0.39 | -0.03 | -0.06 | 0.01 | 0.42 | 0.7 | 0.39 | -0.01 | 0.87 | -0.0975 | -0.0975 | 0.305 | 0.724771 | 0.690553 | 0.296221 | 0.999971 | 1 | 0.847402 |
| Cdk5r2 | O35926 | -0.19 | -0.29 | 0.56 | 0.54 | -0.42 | 0.18 | -0.18 | 0.67 | -0.28 | 0.17 | 0.42 | 0.31 | -0.66 | 0 | 0.26 | 0.93 | -0.0925 | 7.85E-17 | -0.0225 | 0.77899 | 1 | 0.953528 | 0.999971 | 1 | 0.998056 |
| Cdk5rap3 | Q99LM2 | 1.14 | -0.29 | 0.19 | 0.39 | -0.02 | 0.2 | -0.17 | 0.4 | -0.18 | 0.56 | 0.44 | -0.07 | -0.08 | 0.32 | 0.24 | 0.43 | -0.255 | -0.17 | -0.13 | 0.439013 | 0.624203 | 0.685743 | 0.999971 | 1 | 0.944874 |
| Cdk7 | A0A286YD( | 2.14 | 2.65 | 1.54 | 1.6 | 2.71 | 2.64 | 1.53 | 1.69 | 2.89 | 2.22 | 1.35 | 1.85 | 2.45 | 2.27 | 1.39 | 1.43 | 0.16 | 0.095 | -0.0975 | 0.681719 | 0.81188 | 0.792217 | 0.999971 | 1 | 0.969163 |
| Cdk9 | Q99J95 | 0.54 | 2.01 | 1.48 | 1.54 | 1.92 | 1.55 | 1.45 | 1.72 | 1.97 | 1.02 | 1.75 | 1.59 | 1.94 | 1.74 | 1.97 | 2.03 | 0.2675 | 0.19 | 0.5275 | 0.420108 | 0.600989 | 0.124166 | 0.999971 | 1 | 0.78268 |
| Cdkal1 | Q91WE6-2 | 1.39 | 1.58 | 0.29 | 0.41 | 1 | 1.01 | 0.15 | 0.35 | 1.02 | 0.75 | 0.61 | -0.24 | 1.65 | 1.42 | 0.06 | 0.18 | -0.29 | -0.3825 | -0.09 | 0.45714 | 0.362492 | 0.854758 | 0.999971 | 1 | 0.982125 |
| Cdkl5 | A0A0G2JG\ | -0.2 | -0.63 | -0.21 | -0.43 | -0.69 | -0.61 | 0.23 | -0.58 | -0.7 | -0.43 | -0.32 | -0.44 | -0.73 | -0.94 | -0.78 | -0.52 | -0.045 | -0.105 | -0.375 | 0.863298 | 0.585843 | 0.083459 | 0.999971 | 1 | 0.755254 |
| Cdkn1b | P46414 | 2.47 | 0.83 | 1.74 | 2.21 | 1.09 | 1.02 | 3.65 | 0.64 | 0.97 | 0.67 | 1.3 | 0.92 | -0.13 | -0.62 | 0.31 | 0.69 | -0.2125 | -0.8475 | -1.75 | 0.763114 | 0.046284 | 0.002969 | 0.999971 | 1 | 0.575711 |
| Cdkn2aip | Q8BI72 | 1.87 | 2.08 | 1.49 | 1.66 | 1.98 | 2.08 | 2.04 | 1.17 | 1.99 | 1.57 | 0.93 | 1.26 | 1.04 | 1.64 | 1.4 | 1.06 | 0.0425 | -0.3375 | -0.49 | 0.875242 | 0.240724 | 0.059199 | 0.999971 | 1 | 0.725348 |
| Cdkn2c | Q60772 | -0.63 | 0.73 | 1.73 | 2.08 | 0.73 | -0.8 | 1.61 | 1.63 | 0.18 | -2.69 | 2.44 | 1.93 | 0.25 | 0.19 | 1.41 | 1.67 | -0.185 | -0.5125 | -0.0975 | 0.805515 | 0.663439 | 0.881382 | 0.999971 | 1 | 0.986007 |
| Cdr2l | A2A6T1 | 0.48 | 1.14 | 0.19 | 0.89 | 0.53 | 0.41 | -0.55 | 0.28 | 0.19 | 0.22 | -0.02 | 0.34 | 0.6 | 0.45 | 0.26 | 0.56 | -0.5075 | -0.4925 | -0.2075 | 0.143733 | 0.075624 | 0.420164 | 0.999971 | 1 | 0.87788 |
| Cds1 | P98191 | 0.6 | 0.23 | -0.4 | -0.54 | 0.22 | 0.32 | 0.18 | -0.09 | 0.13 | 0.73 | -0.3 | -0.27 | 0.29 | 0.73 | -0.55 | 0.01 | 0.185 | 0.1 | 0.1475 | 0.532116 | 0.77734 | 0.690489 | 0.999971 | 1 | 0.946334 |
| Cds2 | Q99L43 | 1.41 | -0.48 | 0.6 | 0.78 | -0.77 | 0.17 | 3.62 | -0.43 | -0.81 | 0.24 | -0.54 | -0.76 | -1.05 | -0.69 | 0.05 | -0.8 | 0.07 | -1.045 | -1.2 | 0.942261 | 0.038539 | 0.020782 | 0.999971 | 1 | 0.635402 |
| Cdv3 | Q4VAA2 | 0.21 | 0.02 | -0.01 | -0.06 | 0.05 | 0.09 | 0.54 | -0.52 | 0.37 | 0.08 | -0.25 | -0.07 | -0.37 | 0.04 | -0.25 | -0.35 | -2.45E-17 | -0.0075 | -0.2725 | 1 | 0.969836 | 0.166687 | 1 | 1 | 0.800215 |
| Cebpz | A0A0R4J04 | 0.75 | -0.88 | 9.2 | 9.49 | -1.13 | -0.04 | 5.06 | 7.96 | -0.15 | 1.67 | 10.44 | 9.42 | 0.15 | 0.27 | 9.89 | 7.51 | -1.6775 | 0.705 | -0.185 | 0.587659 | 0.836226 | 0.954923 | 0.999971 | 1 | 0.998056 |
| Cebpzos | Q8BTE5 | -2.51 | -0.36 | -0.88 | -0.11 | 0.37 | -1 | -1.72 | -0.87 | -0.67 | 0.3 | -1.01 | -0.69 | 0.24 | 0.28 | -1.02 | -1.33 | 0.16 | 0.4475 | 0.5075 | 0.799554 | 0.434608 | 0.424241 | 0.999971 | 1 | 0.879824 |
| Celf1 | A0A0R4J0T | 0.52 | 1.57 | 1.43 | 1.26 | 1.52 | 1.38 | 1.14 | 1.39 | 1.52 | 1.48 | 1.51 | 1.5 | 1.65 | 1.84 | 1.5 | 1.3 | 0.1625 | 0.3075 | 0.3775 | 0.547356 | 0.250861 | 0.196943 | 0.999971 | 1 | 0.812891 |
| Celf2 | V9GX43 | 0.86 | 0.99 | 1.56 | 1.83 | 0.68 | 0.99 | 1.76 | 1.85 | 0.6 | 0.56 | 2.07 | 1.85 | 0.38 | 0.83 | 1.85 | 2.21 | 0.01 | -0.04 | 0.0075 | 0.977807 | 0.927123 | 0.986875 | 0.999971 | 1 | 0.99925 |
| Celf3 | A0A0G2JGI | 0.53 | 1.78 | 1.16 | 1.26 | 1.7 | 1.67 | 1.87 | 0.55 | 1.93 | 1.21 | 0.39 | 0.27 | 2.15 | 1.89 | 0.49 | 0.22 | 0.265 | -0.2325 | 0.005 | 0.494044 | 0.599798 | 0.992169 | 0.999971 | 1 | 0.99925 |
| Celf4 | Q7TSY6-4 | 1.07 | 1.76 | 0.33 | 1.13 | 1.93 | 1.51 | 2.81 | 0.5 | 1.67 | 1.62 | 0.34 | 0.64 | 1.59 | 1.52 | 0.56 | -0.18 | 0.615 | -0.005 | -0.2 | 0.255133 | 0.990614 | 0.677933 | 0.999971 | 1 | 0.942594 |
| Celf5 | D3Z580 | 3.25 | 1.45 | 1.46 | 2.08 | 0.77 | 1.51 | 1.05 | 2.32 | 0.77 | 2.31 | 1.89 | 2.03 | -0.31 | 1.08 | 2.17 | 1.78 | -0.6475 | -0.31 | -0.88 | 0.219079 | 0.543202 | 0.185908 | 0.999971 | 1 | 0.809025 |
| Celf6 | D6RHP1 | 1.59 | 0.41 | 0.9 | 2.05 | 0.49 | 1.22 | 0.58 | 1.67 | 0.21 | 0.45 | 0.22 | 1.79 | 0.25 | 1.56 | 1.68 | 0.18 | -0.2475 | -0.57 | -0.32 | 0.57054 | 0.263464 | 0.531717 | 0.999971 | 1 | 0.910069 |
| Celsr2 | Q9R0M0 | 0.71 | -0.33 | -0.48 | -0.3 | -0.26 | 0.12 | 0.01 | -0.23 | -0.33 | 0.65 | -0.24 | -0.38 | 0.02 | 0.08 | -0.38 | -0.4 | 0.01 | 0.025 | -0.07 | 0.973148 | 0.944257 | 0.82029 | 0.999971 | 1 | 0.975761 |
| Cemip2 | Q5FWI3 | -1.47 | -1.27 | -0.98 | -0.92 | -1.08 | -1.3 | -1.17 | -0.99 | -1.4 | -1.07 | -0.89 | -0.42 | -0.7 | -1.01 | -0.32 | -0.64 | 0.025 | 0.215 | 0.4925 | 0.901239 | 0.421465 | 0.057156 | 0.999971 | 1 | 0.725348 |
| Cend1 | Q9JKC6 | 1.42 | -1.27 | -0.67 | -0.21 | -1.7 | -0.8 | 0.86 | -0.91 | -1.42 | -0.18 | -1.06 | -1.53 | -1.82 | -1.62 | -0.97 | -1.12 | -0.455 | -0.865 | -1.2 | 0.527639 | 0.174121 | 0.057582 | 0.999971 | 1 | 0.725348 |
| Cenpa | A0A0G2JGI | 0.18 | -1.35 | -1.4 | -4.47 | -1.56 | -1.67 | -0.96 | -3.6 | -0.94 | -1.13 | -1.8 | -1.61 | -1.45 | -2.69 | -2.71 | -4.84 | -0.1875 | 0.39 | -1.1625 | 0.85249 | 0.66546 | 0.295204 | 0.999971 | 1 | 0.847402 |
| Cenpv | Q9CXS4 | -0.09 | -1.37 | -1.12 | -1.39 | -1.17 | -1.2 | -0.47 | -1.46 | -0.88 | -0.6 | -1.43 | -1.59 | -1.22 | -1.19 | -1.36 | -1.68 | -0.0825 | -0.1325 | -0.37 | 0.820494 | 0.723545 | 0.276655 | 0.999971 | 1 | 0.843794 |
| Cep112 | Z4YKK6 | 1.01 | 0.37 | 0.93 | 0.94 | 0.02 | 0.82 | -0.18 | 1.41 | -0.21 | 1.23 | 1.28 | 0.72 | -1.38 | 0.16 | 0.61 | 0.22 | -0.295 | -0.0575 | -0.91 | 0.448089 | 0.875318 | 0.063405 | 0.999971 | 1 | 0.725348 |
| Cep128 | Q8BI22 | 0.71 | 0.12 | 2.02 | 0.57 | -1.24 | 0.79 | 2.27 | -0.92 | -0.93 | 1.03 | -0.01 | -1.3 | -0.69 | -0.61 | -0.16 | -0.82 | -0.63 | -1.1575 | -1.425 | 0.448896 | 0.084001 | 0.007051 | 0.999971 | 1 | 0.582714 |
| Cep131 | B1AXI9 | 1.66 | 0.4 | -0.24 | -0.13 | -0.09 | 0.73 | -0.16 | -0.21 | 0.53 | 1.55 | -0.01 | -0.03 | -0.47 | 0.46 | -0.15 | 0.02 | -0.355 | 0.0875 | -0.4575 | 0.446192 | 0.868785 | 0.322657 | 0.999971 | 1 | 0.85548 |
| Cep135 | Q6P5D4 | 0.2 | 0.27 | -0.32 | -0.32 | 0.37 | -0.16 | -0.36 | -1.1 | 0.48 | -0.29 | -0.37 | 0.04 | 0.37 | -0.15 | -0.41 | -0.22 | -0.27 | 0.0075 | -0.06 | 0.436296 | 0.977769 | 0.815748 | 0.999971 | 1 | 0.974918 |
| Cep170 | Q6A065 | -0.47 | -1.2 | -1.2 | -1.46 | -1.19 | -1.26 | -0.82 | -1.58 | -0.98 | -0.87 | -1.18 | -1.12 | -0.87 | -1.27 | -1.64 | -1.41 | -0.13 | 0.045 | -0.215 | 0.643985 | 0.85718 | 0.454728 | 0.999971 | 1 | 0.887217 |
| Cep170_1 | A0A0A6YX! | 0.75 | 1.35 | -1.1 | 0.93 | 1.7 | 1.1 | 0.06 | 1.06 | 1.14 | -0.17 | 2.08 | -0.03 | 0.36 | -1.63 | 2.11 | 0.4 | 0.4975 | 0.2725 | -0.1725 | 0.404991 | 0.694362 | 0.837726 | 0.999971 | 1 | 0.979416 |
| Cep170b | Q80U49 | 0.28 | -0.07 | 0.42 | 0.63 | -0.05 | 0.16 | 1.2 | 0.48 | 0.21 | 0.26 | 0.35 | 0.16 | 0.3 | 0 | 0.21 | 0.24 | 0.1325 | -0.07 | -0.1275 | 0.674797 | 0.731653 | 0.55091 | 0.999971 | 1 | 0.913169 |
| Cep41 | Q99NF3 | 0.81 | 0.92 | 0.49 | 0.78 | 1.03 | 0.57 | 0.17 | 0.84 | 0.99 | 0.93 | 0.24 | 0.51 | 0.11 | 1.12 | 0.56 | 0.54 | -0.0975 | -0.0825 | -0.1675 | 0.686274 | 0.724757 | 0.51436 | 0.999971 | 1 | 0.905815 |
| Cep76 | Q0VEJ0 | 0.79 | 1.66 | 2.44 | 2.42 | 1.02 | 1.54 | 1.47 | 2.21 | 0.97 | 1.49 | 1.09 | 1.32 | 1.89 | 1.58 | 2.32 | 3.38 | -0.2675 | -0.61 | 0.465 | 0.543001 | 0.146483 | 0.376547 | 0.999971 | 1 | 0.870465 |
| Cep97 | Q9CZ62 | 1.64 | 1.41 | 0.65 | 1 | 1.72 | 2.13 | 1.01 | 0.97 | 1.14 | 1.89 | 0.41 | 0.48 | 1.82 | 1.61 | 0.41 | 0.29 | 0.2825 | -0.195 | -0.1425 | 0.429984 | 0.622351 | 0.740768 | 0.999971 | 1 | 0.960899 |
| Cept1 | A0A0A6YXI | -0.42 | 1 | 0.05 | -0.18 | 0.89 | 1.04 | -0.06 | -0.18 | 0.54 | 0.76 | 0.47 | 0.49 | 0.63 | 0.83 | 0.57 | 0.19 | 0.31 | 0.4525 | 0.4425 | 0.467694 | 0.181721 | 0.211272 | 0.999971 | 1 | 0.823077 |
| Cerk | Q8K4Q7 | 1.25 | -1.81 | -1.81 | -1.17 | -2.91 | -2.12 | 1.81 | -1.57 | -2.59 | 0.13 | -3.39 | -2.4 | -2.75 | -1.94 | -2.86 | -1.74 | -0.3125 | -1.1775 | -1.4375 | 0.782204 | 0.235315 | 0.068075 | 0.999971 | 1 | 0.733937 |
| Cers1 | P27545 | -3.21 | -3.44 | -1.52 | -2.54 | -2.19 | -4.28 | -0.34 | -1.61 | -2.15 | -3.4 | -0.09 | 0.1 | -1.94 | -3.12 | -1.43 | -1.87 | 0.5725 | 1.2925 | 0.5875 | 0.497261 | 0.156641 | 0.276457 | 0.999971 | 1 | 0.843794 |
| Cers2 | Q924Z4 | -1.07 | -1.63 | -0.83 | -1.38 | -1.41 | -0.8 | -0.03 | -0.94 | -0.75 | -0.24 | -0.29 | 0.62 | -0.32 | -0.58 | -0.82 | -0.81 | 0.4325 | 1.0625 | 0.595 | 0.217072 | 0.010787 | 0.034712 | 0.999971 | 1 | 0.69205 |
| Cers4 | Q9D6J1 | 1.11 | 0.37 | -0.45 | -0.01 | 0.79 | 0.67 | 0.45 | -0.27 | 0.33 | 1.2 | -0.01 | -0.76 | 0.53 | 0.11 | -1.08 | -0.34 | 0.155 | -0.065 | -0.45 | 0.692624 | 0.894261 | 0.330754 | 0.999971 | 1 | 0.857107 |
| Cers6 | H3BL08 | -1.56 | 0.34 | -0.58 | -4.18 | 0.56 | -0.36 | 0.39 | -2.41 | 1.15 | -1.66 | 1.49 | 1.65 | 1.43 | -0.54 | -0.98 | -4.01 | 1.04 | 2.1525 | 0.47 | 0.34024 | 0.082105 | 0.723283 | 0.999971 | 1 | 0.955258 |
| Cert1 | Q9EQG9 | -0.43 | 1.05 | 0.72 | 0.39 | 1.15 | 1.21 | 0.97 | 0.61 | 1.33 | -0.12 | 1.06 | 0.86 | 0.81 | 0.77 | 0.92 | 0.46 | 0.5525 | 0.35 | 0.3075 | 0.132119 | 0.420119 | 0.366512 | 0.999971 | 1 | 0.869526 |
| Ces1c | P23953 | 3.56 | 2.39 | 0.23 | 3.85 | 4.32 | 2.96 | 2.88 | 1.92 | 2.84 | 1.78 | 2.07 | 2.35 | 1.42 | 2.51 | 1.34 | 1.95 | 0.5125 | -0.2475 | -0.7025 | 0.554848 | 0.74923 | 0.3792 | 0.999971 | 1 | 0.871816 |
| Cetn2 | Q9R1K9 | -1 | -1.23 | -0.79 | -0.76 | -0.94 | -1.03 | -0.67 | -0.87 | -0.77 | -1.31 | -0.97 | -0.47 | -0.76 | -0.98 | -0.56 | -0.84 | 0.0675 | 0.065 | 0.16 | 0.730521 | 0.78534 | 0.431713 | 0.999971 | 1 | 0.881643 |
| Cetn3 | A0A286YDI | -2.77 | -3.38 | -0.25 | -0.69 | -2.36 | -2.47 | -0.18 | -0.57 | -2.55 | -2.76 | -0.44 | -0.27 | -2.48 | -1.89 | -0.68 | -0.35 | 0.3775 | 0.2675 | 0.4225 | 0.665749 | 0.770591 | 0.61083 | 0.999971 | 1 | 0.92673 |
| Cetn4 | Q8K4K1-3 | 1.07 | 0.33 | -0.06 | 0.22 | 0.86 | 1.6 | 0.11 | 1.56 | 0.28 | 0.9 | 0.4 | 0.29 | -1.59 | 0.58 | 0.65 | 0.88 | 0.6425 | 0.0775 | -0.26 | 0.140179 | 0.79112 | 0.65207 | 0.999971 | 1 | 0.933842 |
| Cfap20 | Q8BTU1 | 1.43 | 0.58 | 1.22 | 0.99 | 1 | 1.03 | 1.41 | 1.02 | 1.18 | 1.32 | 1.18 | 1.6 | 0.64 | 0.5 | 0.79 | 0.77 | 0.06 | 0.265 | -0.38 | 0.802645 | 0.28349 | 0.128301 | 0.999971 | 1 | 0.784699 |
| Cfap298 | Q8BL95 | 1.79 | -1.2 | 0.17 | 0.33 | -0.62 | -0.24 | 0.08 | 1.04 | -0.9 | 0.66 | 0.37 | 0.61 | -1.22 | -1.02 | 0.76 | 0.49 | -0.2075 | -0.0875 | -0.52 | 0.747103 | 0.893096 | 0.476309 | 0.999971 | 1 | 0.894333 |
| Cfap300 | Q8CC70 | 1.55 | 0.37 | 2.11 | 0.94 | 0.6 | 0.71 | -0.39 | 2.31 | 0.27 | 0.17 | 1.33 | 0.31 | -0.36 | -0.74 | 1.57 | 1.61 | -0.435 | -0.7225 | -0.7225 | 0.484683 | 0.127301 | 0.291193 | 0.999971 | 1 | 0.847402 |
| Cfap36 | Q8C6E0 | 0.97 | 0.85 | 1.18 | 0.92 | 0.77 | 0.83 | 1.1 | 1.12 | 0.54 | 0.98 | 1.16 | 0.98 | 0.1 | 0.39 | 0.82 | 1.03 | -0.025 | -0.065 | -0.395 | 0.893235 | 0.748185 | 0.140557 | 0.999971 | 1 | 0.784948 |
| Cfap410 | Q8C6G1 | -0.06 | 0.45 | -0.05 | 0.07 | 0.54 | 0.2 | 0.15 | -0.2 | 0.66 | -0.35 | -0.15 | 0.02 | 1.41 | 0.88 | -0.18 | 0.5 | 0.07 | -0.0575 | 0.55 | 0.761924 | 0.830006 | 0.142496 | 0.999971 | 1 | 0.784967 |
| Cfap418 | B1AV75 | 0.36 | 0.37 | 1.4 | 1.17 | 0.25 | 1.43 | 2.72 | 0.86 | 0.77 | -0.54 | 1.15 | 1.04 | -0.8 | -0.44 | 1.2 | 0 | 0.49 | -0.22 | -0.835 | 0.379118 | 0.625204 | 0.108223 | 0.999971 | 1 | 0.78268 |
| Cfap74 | G3UWF6 | 3.51 | 3.46 | 1.74 | 1.96 | 2.49 | 3.63 | 0.88 | 1.35 | 3.08 | 3.84 | 1.09 | 0.61 | 2.55 | 3.13 | 1.33 | 1.73 | -0.58 | -0.5125 | -0.4825 | 0.418 | 0.537494 | 0.408705 | 0.999971 | 1 | 0.875014 |
| Cfdp1 | O88271 | 0.43 | 3.01 | 2.6 | 3.14 | 3.72 | 0.93 | 2.91 | 2.51 | 3.94 | 0.4 | 2.37 | 1.73 | 4.21 | 0.24 | 2.44 | 2.08 | 0.2225 | -0.185 | -0.0525 | 0.774471 | 0.832585 | 0.954598 | 0.999971 | 1 | 0.998056 |
| Cfh | E9Q8I0 | 2.08 | 1.65 | 0.63 | 0.95 | 2.45 | 1.29 | 1.28 | 0.83 | 1.24 | 1.58 | 1.25 | 1.39 | 1.54 | 1.75 | 1.13 | 0.88 | 0.135 | 0.0375 | -0.0025 | 0.763849 | 0.911088 | 0.994637 | 0.999971 | 1 | 1 |
| Cfl1 | P18760 | 0.72 | 0.47 | -0.07 | 0.41 | 0.81 | 0.66 | 0.27 | 0.05 | 0.72 | 0.57 | 0.05 | -0.12 | 1.36 | 0.67 | -0.02 | 0.28 | 0.065 | -0.0775 | 0.19 | 0.804733 | 0.779311 | 0.578028 | 0.999971 | 1 | 0.918596 |
| Cfl2 | P45591 | 0.25 | 0.08 | 0.23 | 0.43 | 0.54 | 1.21 | 0.14 | 0.38 | 1.34 | 0.33 | 0.08 | 0.11 | 1.52 | 0.69 | 0.2 | 0.48 | 0.32 | 0.2175 | 0.475 | 0.243089 | 0.489329 | 0.141853 | 0.999971 | 1 | 0.784967 |
| Cgn | D3YUW7 | 2.83 | 1.16 | 1.85 | 2.15 | 0.62 | 0.8 | 2.15 | 1.45 | 0.84 | 1.71 | 1.09 | 1.12 | -0.35 | 0.25 | 1.57 | 0.62 | -0.7425 | -0.8075 | -1.475 | 0.132821 | 0.059008 | 0.01462 | 0.999971 | 1 | 0.613095 |
| Cgnl1 | B2RSU6 | 0.53 | -0.07 | 0.72 | 0.19 | 0.87 | -0.1 | -0.03 | -0.17 | 0.11 | 0.62 | 0.63 | 1.2 | -0.26 | 0 | 0.86 | 0.35 | -0.2 | 0.2975 | -0.105 | 0.519875 | 0.325332 | 0.732601 | 0.999971 | 1 | 0.956652 |
| Cgref1 | E9QPK1 | 1.21 | 0.73 | 0.55 | 0.67 | 0.75 | 0.77 | 0.36 | 0.9 | 0.66 | 0.17 | 0.58 | 0.51 | -0.14 | 1.11 | 0.89 | 0.93 | -0.095 | -0.31 | -0.0925 | 0.675855 | 0.184412 | 0.773515 | 0.999971 | 1 | 0.963759 |
| Chac2 | Q9CQG1 | 0.53 | 0.33 | 0.9 | 0.14 | 0.13 | -0.02 | 0.31 | 0.27 | 0.11 | 0.6 | 0.87 | 1.02 | -0.67 | 0.04 | 0.28 | 0.37 | -0.3025 | 0.175 | -0.47 | 0.195757 | 0.527813 | 0.139509 | 0.999971 | 1 | 0.784948 |
| Chadl | E9Q7T7 | 0.73 | 0.83 | 1.37 | 0.78 | 0.1 | 0.29 | 0.99 | 1.05 | 0.4 | 0.91 | 0.74 | 0.91 | -0.2 | 0.43 | 0.88 | 0.67 | -0.32 | -0.1875 | -0.4825 | 0.293261 | 0.419576 | 0.123145 | 0.999971 | 1 | 0.78268 |
| Champ1 | Q8K327 | 0.35 | 0.44 | 0.42 | 0.52 | 0.4 | 0.14 | 0.17 | 0.48 | 0.28 | 0.37 | 0.23 | 0.54 | 0.44 | 0.6 | 0.38 | 0.83 | -0.135 | -0.0775 | 0.13 | 0.450536 | 0.644303 | 0.484172 | 0.999971 | 1 | 0.896352 |
| Chat | Q8BQV2 | -0.16 | -0.38 | -1.19 | -0.47 | 0.02 | -0.51 | -1.08 | -0.6 | -0.14 | -0.23 | -1.38 | -1.17 | -0.47 | -0.15 | -0.79 | -1.23 | 0.0075 | -0.18 | -0.11 | 0.981248 | 0.634754 | 0.734078 | 0.999971 | 1 | 0.957685 |
| Chchd1 | Q9CQA6 | 1.09 | 2.23 | 2.12 | 1.87 | 1.6 | 1.48 | 1.06 | 1.67 | 1.93 | 0.85 | 0.53 | 2.59 | 1.76 | 0.97 | 1.79 | 2.09 | -0.375 | -0.3525 | -0.175 | 0.231597 | 0.491209 | 0.616369 | 0.999971 | 1 | 0.928229 |
| Chchd10 | Q7TNL9 | 0.54 | 0.27 | 2.01 | 0.71 | 0.94 | 0.67 | 1.53 | 0.8 | 1.15 | -0.04 | 0.41 | 1.29 | 1.25 | 0.47 | 0.75 | 0.33 | 0.1025 | -0.18 | -0.1825 | 0.802719 | 0.701214 | 0.662253 | 0.999971 | 1 | 0.936148 |
| Chchd2 | Q9D1L0 | 1.03 | 0.42 | 0.36 | 0.37 | 0.38 | 0.89 | 0.42 | 0.5 | 0.57 | 0.98 | 0.84 | 0.1 | 0.3 | 1.19 | 0.33 | -0.08 | 0.0025 | 0.0775 | -0.11 | 0.991509 | 0.774747 | 0.729653 | 0.999971 | 1 | 0.956041 |
| Chchd3 | Q9CRB9 | -0.09 | -0.65 | -0.24 | 0.21 | -0.76 | -0.55 | 0.78 | -0.46 | -0.21 | -0.49 | -0.51 | -0.74 | -0.59 | -1.05 | -0.32 | -0.84 | -0.055 | -0.295 | -0.5075 | 0.884299 | 0.238842 | 0.079572 | 0.999971 | 1 | 0.755254 |
| Chchd3_1 | D3Z0L4 | -2.32 | 5.07 | 5.29 | 2.17 | 9.38 | 4.76 | 6.05 | 2.86 | 11.83 | 2.17 | 6.47 | 7.63 | 17.46 | 2.03 | 3.6 | 0.45 | 3.21 | 4.4725 | 3.3325 | 0.132168 | 0.08694 | 0.390666 | 0.999971 | 1 | 0.871816 |
| Chchd4 | Q8VEA4 | 1.58 | 2.6 | 1.57 | 1.59 | 2.66 | 3.04 | 1.19 | 1.52 | 2.9 | 1.92 | 1.01 | 0.43 | 2.92 | 2.51 | 1.46 | 1.68 | 0.2675 | -0.27 | 0.3075 | 0.578178 | 0.627014 | 0.459321 | 0.999971 | 1 | 0.889191 |
| Chchd5 | Q9CQP3 | 0.68 | 1.61 | -1.15 | -1.88 | 1.28 | 1.13 | -1.24 | -1.55 | 0.83 | 1.04 | -1.45 | -1.56 | 1.71 | 1.03 | -1.17 | -0.87 | 0.09 | -0.1 | 0.36 | 0.927077 | 0.917276 | 0.708102 | 0.999971 | 1 | 0.950182 |
| Chchd6 | Q91VN4 | 0.08 | -0.46 | -0.41 | -0.65 | -0.11 | -0.54 | -0.07 | -0.59 | -0.44 | 0.07 | -0.39 | -0.62 | -0.18 | -0.3 | -0.54 | -0.77 | 0.0325 | 0.015 | -0.0875 | 0.892383 | 0.950702 | 0.71411 | 0.999971 | 1 | 0.952257 |
| Chchd7 | Q8K2Q5 | 0.05 | 1.21 | 1.17 | 0.43 | 1.05 | 0.99 | 2.12 | 1.59 | 0.97 | 0.31 | 1.59 | 2.33 | -0.03 | 0.82 | 1.08 | 0.47 | 0.7225 | 0.585 | -0.13 | 0.081687 | 0.246607 | 0.72175 | 0.999971 | 1 | 0.955137 |
| Chd2 | E9PZM4 | 0.43 | -0.38 | 0.2 | 0.13 | -0.4 | -0.17 | 0.35 | -0.11 | -0.36 | 0.34 | -0.13 | -0.34 | -0.03 | -0.01 | 0.13 | 0.22 | -0.1775 | -0.2175 | -0.0175 | 0.495564 | 0.409488 | 0.937577 | 0.999971 | 1 | 0.995618 |
| Chd3 | B1AR17 | 1.23 | 0.94 | 0.84 | 0.82 | 0.97 | 1.1 | 0.79 | 0.78 | 0.83 | 1.33 | 0.96 | 0.78 | 0.61 | 0.95 | 0.48 | 0.9 | -0.0475 | 0.0175 | -0.2225 | 0.802462 | 0.932134 | 0.29196 | 0.999971 | 1 | 0.847402 |
| Chd4 | E9QAS4 | 1.01 | 1.36 | 1.01 | 1.13 | 1.35 | 1.26 | 0.73 | 0.99 | 1.27 | 0.93 | 1.13 | 1.33 | 1.31 | 1.12 | 1.26 | 1.37 | -0.045 | 0.0375 | 0.1375 | 0.831914 | 0.840592 | 0.451549 | 0.999971 | 1 | 0.886993 |
| Chd5 | E9PYU4 | 0.11 | 0.64 | 1.15 | 1.58 | 0.74 | 1.24 | 0.74 | 0.97 | 2.19 | 0.85 | 0.86 | 1.12 | 1.58 | 1.9 | 0.56 | 1.13 | 0.0525 | 0.385 | 0.4225 | 0.876086 | 0.378735 | 0.318932 | 0.999971 | 1 | 0.853893 |
| Chd6 | A3KFM7-2 | 0.01 | -1.38 | 0.04 | -0.56 | -1.24 | -1.75 | -0.98 | -0.23 | -1.19 | -0.92 | -0.72 | 0 | -1.24 | -1.38 | -0.53 | -1.54 | -0.5775 | -0.235 | -0.7 | 0.205342 | 0.562155 | 0.097316 | 0.999971 | 1 | 0.78268 |
| Chd8 | Q09XV5 | 0.94 | 1.68 | 1.19 | 0.88 | 1.8 | 1.12 | 0.47 | 1.28 | 1.39 | 0.96 | 0.83 | 0.81 | 2.08 | 1.25 | 1.15 | 0.94 | -0.005 | -0.175 | 0.1825 | 0.987849 | 0.492936 | 0.565021 | 0.999971 | 1 | 0.917442 |
| Chd9 | Q8BYH8-2 | 2.03 | 3.01 | 1.59 | 1.38 | 2.07 | 3.29 | 3.12 | -0.56 | 2.69 | 2.74 | -1.34 | -1.3 | 3.2 | 2.67 | -1.93 | -2.22 | -0.0225 | -1.305 | -1.5725 | 0.979099 | 0.252973 | 0.256323 | 0.999971 | 1 | 0.839012 |
| Chdh | Q8BJ64 | 0.14 | -0.3 | 0.24 | -0.12 | -1.31 | -1.45 | -0.19 | -1 | -1.23 | -1.02 | -1.02 | -0.14 | -1.61 | -1.26 | -0.37 | -0.53 | -0.9775 | -0.8425 | -0.9325 | 0.011515 | 0.015284 | 0.016836 | 0.999971 | 1 | 0.616254 |
| Cherp | G5E8I8 | 0.31 | 1.45 | 1.08 | 0.98 | 1.48 | 0.9 | 0.75 | 0.85 | 1.27 | 0.6 | 1.08 | 0.95 | 1.58 | 1.22 | 0.9 | 1.06 | 0.04 | 0.02 | 0.235 | 0.893207 | 0.94456 | 0.428105 | 0.999971 | 1 | 0.880793 |
| Chga | P26339 | 2.12 | 0.89 | 1.1 | 1.52 | 0.8 | 1.49 | 1.44 | 1.43 | 1.13 | 2.2 | 1.06 | 0.92 | 0.89 | 1.03 | 1.22 | 1.56 | -0.1175 | -0.08 | -0.2325 | 0.713716 | 0.835856 | 0.464514 | 0.999971 | 1 | 0.890905 |
| Chgb | P16014 | 0.33 | -0.34 | -0.25 | -0.29 | -0.11 | -0.09 | 0.15 | 0.09 | -0.31 | -0.1 | -0.09 | -0.21 | -0.49 | -0.19 | -0.02 | 0.07 | 0.1475 | -0.04 | -0.02 | 0.500791 | 0.849763 | 0.932292 | 0.999971 | 1 | 0.995069 |
| Chic1 | Q8CBW7 | 0.36 | -0.68 | -0.44 | 0.04 | -0.71 | -0.52 | -0.74 | 0.58 | -1.06 | -0.4 | -0.05 | -0.23 | -0.38 | -0.02 | -0.63 | 0.07 | -0.1675 | -0.255 | -0.06 | 0.658973 | 0.437674 | 0.839208 | 0.999971 | 1 | 0.979728 |
| Chid1 | A0A0R4J24 | 0.39 | 0.75 | 1.19 | 1.26 | 0.42 | 0.55 | 1.05 | 1.18 | 0.34 | 1.06 | 0.78 | 0.78 | -0.01 | 0.59 | 0.8 | 1.15 | -0.0975 | -0.1575 | -0.265 | 0.73556 | 0.563214 | 0.417166 | 0.999971 | 1 | 0.877464 |
| Chka | O54804-2 | -0.22 | 0.84 | 0.34 | 0.11 | 1.1 | 0.15 | -0.11 | 0.27 | 0.53 | -0.56 | -0.21 | 0.17 | 0.35 | -0.28 | -0.41 | 0.1 | 0.085 | -0.285 | -0.3275 | 0.802741 | 0.391044 | 0.28282 | 0.999971 | 1 | 0.844538 |
| Chkb | O55229 | -0.67 | 0.07 | 0.31 | 0.05 | 0.15 | -0.63 | 0.15 | -0.02 | 0.19 | -1.05 | 0.51 | 0.16 | 0.67 | -0.16 | 0.01 | -0.07 | -0.0275 | 0.0125 | 0.1725 | 0.924994 | 0.974323 | 0.562335 | 0.999971 | 1 | 0.916386 |
| Chl1 | P70232-2 | -0.19 | -2.27 | -4.13 | -3.58 | -1.05 | -3.23 | -3.89 | -2.12 | -1.82 | -2.59 | -2.99 | -1.95 | -0.9 | -1.63 | -1.95 | -3.76 | -0.03 | 0.205 | 0.4825 | 0.975068 | 0.805018 | 0.615505 | 0.999971 | 1 | 0.927967 |
| Chl1_1 | P70232 | 0.41 | -0.32 | -0.26 | -0.19 | 0.11 | 0.07 | -0.09 | 0.15 | -0.03 | 0.25 | -0.03 | 0.07 | 0.18 | 0.43 | 0.32 | 0.26 | 0.15 | 0.155 | 0.3875 | 0.50195 | 0.489582 | 0.107006 | 0.999971 | 1 | 0.78268 |
| Chm | A2AD03 | -0.54 | 0.3 | 0.45 | 0.37 | 0.24 | -0.03 | 0.59 | 0.16 | 0.37 | -0.44 | 0.29 | 0.22 | 0.14 | 0.01 | 0.25 | 0.52 | 0.095 | -0.035 | 0.085 | 0.734939 | 0.90792 | 0.75642 | 0.999971 | 1 | 0.961116 |
| Chml | Q9QZD5 | 1.3 | 1.6 | 0.05 | 0.62 | 1.06 | 0.91 | -0.88 | 0.08 | 0.74 | 0.96 | 0.39 | 0.39 | 0.64 | 1.43 | -0.62 | 0.37 | -0.6 | -0.2725 | -0.4375 | 0.268816 | 0.463309 | 0.399318 | 0.999971 | 1 | 0.871816 |
| Chmp1a | Q921W0 | 1.22 | 0.22 | 0.97 | 0.62 | 0.04 | 0.8 | 1.1 | 1 | 0.54 | 0.76 | 1.55 | 0.71 | -0.32 | 0.29 | 1.01 | 1.18 | -0.0225 | 0.1325 | -0.2175 | 0.94468 | 0.676181 | 0.582734 | 0.999971 | 1 | 0.920364 |
| Chmp1b1 | Q99LU0 | 1.13 | 0.8 | 1.1 | 0.13 | 0.98 | 1.52 | 2.09 | 0.47 | 1.31 | 1.13 | 1.91 | 1.66 | 1.56 | 0.98 | 0.84 | 0.03 | 0.475 | 0.7125 | 0.0625 | 0.254697 | 0.038641 | 0.869036 | 0.999971 | 1 | 0.983462 |
| Chmp1b2 | Q9CQD4 | 0.95 | 0.86 | 1.99 | 1.99 | 0.57 | 0.94 | 4.55 | 0.17 | 0.88 | 0.87 | 1.35 | 0.07 | 0.82 | 0.62 | 0.6 | -0.28 | 0.11 | -0.655 | -1.0075 | 0.907245 | 0.124844 | 0.026269 | 0.999971 | 1 | 0.678474 |
| Chmp2a | Q9DB34 | 0.43 | 0.07 | -0.22 | 0.42 | 0.02 | 0.48 | 0.3 | 0.68 | 0.34 | 0.65 | 0.2 | -0.36 | -0.2 | 0.29 | 0.15 | -0.02 | 0.195 | 0.0325 | -0.12 | 0.428456 | 0.906751 | 0.60234 | 0.999971 | 1 | 0.924453 |
| Chmp2b | Q8BJF9 | 0.68 | 0.74 | 0.8 | 0.98 | 0.82 | 1.12 | 1.5 | 1.02 | 0.96 | 0.81 | 0.74 | 0.42 | 0.85 | 0.37 | 0.44 | 0.02 | 0.315 | -0.0675 | -0.38 | 0.156291 | 0.725207 | 0.117612 | 0.999971 | 1 | 0.78268 |
| Chmp3 | Q9CQ10 | -0.38 | -0.26 | -0.49 | -0.55 | -0.1 | -0.33 | -0.2 | -0.27 | -0.35 | -0.41 | -0.13 | -0.16 | -0.43 | -0.34 | -0.46 | -0.6 | 0.195 | 0.1575 | -0.0375 | 0.274616 | 0.37637 | 0.829199 | 0.999971 | 1 | 0.978248 |
| Chmp4b | Q9D8B3 | 0.15 | 0.6 | 0.46 | 0.5 | 0.34 | 0.67 | 0.47 | 0.55 | 0.51 | 0.38 | 0.44 | 0.29 | 1.03 | 0.39 | 0.39 | 0.63 | 0.08 | -0.0225 | 0.1825 | 0.672573 | 0.900354 | 0.421879 | 0.999971 | 1 | 0.877992 |
| Chmp5 | Q9D7S9 | 0.09 | -0.12 | -0.01 | 0.32 | -0.25 | 0.01 | -0.26 | 0.26 | -0.09 | 0.1 | 0.05 | 0.35 | -0.4 | -0.04 | 0.01 | 0.59 | -0.13 | 0.0325 | -0.03 | 0.535628 | 0.865269 | 0.905494 | 0.999971 | 1 | 0.991091 |
| Chmp6 | P0C0A3 | -1.06 | -0.44 | -0.44 | -1.31 | -0.77 | -0.61 | -0.49 | -0.91 | -0.51 | -1.35 | -0.83 | -1.01 | -0.12 | -1.01 | -0.96 | -1.3 | 0.1175 | -0.1125 | -0.035 | 0.655484 | 0.701305 | 0.916796 | 0.999971 | 1 | 0.992633 |
| Chmp7 | Q8R1T1 | -0.02 | -0.02 | -0.21 | -0.23 | -0.05 | 0 | -0.34 | -0.39 | -0.15 | -0.07 | -0.31 | -0.06 | 0.11 | 0.29 | -0.14 | 0.14 | -0.075 | -0.0275 | 0.22 | 0.688447 | 0.870565 | 0.249047 | 0.999971 | 1 | 0.839012 |
| Chn1 | Q91V57 | 0.23 | 0.49 | 0.86 | 0.48 | 0.07 | 0.21 | 0.81 | 0.61 | 0.18 | 0.57 | 0.5 | 0.88 | -0.09 | 0.49 | 0.42 | 0.37 | -0.09 | 0.0175 | -0.2175 | 0.714763 | 0.939069 | 0.349116 | 0.999971 | 1 | 0.865162 |
| Chn2 | Q3V2R3 | 0.42 | -0.35 | -0.84 | -0.2 | -0.29 | -0.69 | -0.5 | -0.96 | -0.21 | 0.18 | -1.05 | -0.6 | -0.39 | -0.19 | -0.91 | -1.08 | -0.3675 | -0.1775 | -0.4 | 0.245704 | 0.625693 | 0.249984 | 0.999971 | 1 | 0.839012 |
| Chordc1 | Q9D1P4 | -0.16 | 0.86 | 1.08 | 0.38 | 0.93 | 1.13 | 0.94 | 0.73 | 0.76 | 0.11 | 0.94 | 0.77 | 1.14 | 0.88 | 0.85 | 0.87 | 0.3925 | 0.105 | 0.395 | 0.208277 | 0.750741 | 0.20306 | 0.999971 | 1 | 0.81617 |
| Chp1 | P61022 | -1.65 | -1.61 | -1.63 | -1.7 | -1.41 | -1.97 | -1.79 | -1.53 | -1.73 | -1.83 | -1.5 | -1.2 | -1.9 | -1.9 | -1.48 | -1.83 | -0.0275 | 0.0825 | -0.13 | 0.886456 | 0.676876 | 0.479981 | 0.999971 | 1 | 0.894438 |
| Chpf | Q6IQX7 | 1.21 | 0.25 | 2.09 | 2.16 | 0.29 | 0.09 | 3.57 | 1.37 | 0.09 | 0.64 | 1.98 | 1.81 | 0.25 | 0.29 | 0.37 | 0.48 | -0.0975 | -0.2975 | -1.08 | 0.905428 | 0.615654 | 0.030612 | 0.999971 | 1 | 0.678474 |
| Chpf2 | Q3UU43 | 0.64 | -0.31 | 2.67 | 2.31 | 0.41 | 0.33 | 2.43 | 2.75 | -0.07 | 1.22 | 0.99 | 1.54 | 0.83 | 0.38 | 2.06 | 1.82 | 0.1525 | -0.4075 | -0.055 | 0.858311 | 0.571319 | 0.939898 | 0.999971 | 1 | 0.995869 |
| Chpt1 | Q8C025 | 2.23 | -0.97 | -1.83 | -1.78 | -0.87 | -0.18 | -0.74 | -0.66 | -0.69 | 2.81 | -0.82 | -0.23 | -1.18 | 1.19 | -1.16 | -1.51 | -0.025 | 0.855 | -0.0775 | 0.97705 | 0.465814 | 0.939516 | 0.999971 | 1 | 0.995869 |
| Chrm1 | P12657 | -0.18 | -0.07 | 0.25 | 0.11 | -0.8 | -0.91 | 0.54 | -0.23 | -1.14 | -0.09 | 0.11 | -0.11 | -0.17 | -0.8 | 0.08 | 0.43 | -0.3775 | -0.335 | -0.1425 | 0.285995 | 0.287673 | 0.624574 | 0.999971 | 1 | 0.930479 |
| Chrm2 | Q9ERZ4 | 0.06 | 0.02 | -1.52 | -1.04 | -0.72 | -0.39 | -1.52 | -1.21 | -0.74 | 0.44 | -2.03 | -2.15 | -1.86 | -1.12 | -2.21 | -1.35 | -0.34 | -0.5 | -1.015 | 0.447399 | 0.458108 | 0.043166 | 0.999971 | 1 | 0.701725 |
| Chrm3 | Q9ERZ3 | -1.97 | -0.45 | 0 | -0.92 | -1.01 | -0.14 | -0.56 | -0.76 | -1.17 | -1.01 | -0.21 | -0.56 | -0.53 | -0.61 | -0.74 | -0.41 | 0.2175 | 0.0975 | 0.2625 | 0.61911 | 0.827956 | 0.525566 | 0.999971 | 1 | 0.909437 |
| Chrm4 | P32211 | 0.69 | -0.35 | -0.14 | 0.47 | -0.02 | -0.71 | -0.65 | -0.08 | -0.29 | 0.19 | -0.1 | -0.13 | -0.44 | -0.62 | -0.29 | -0.09 | -0.5325 | -0.25 | -0.5275 | 0.11436 | 0.38294 | 0.091505 | 0.999971 | 1 | 0.773779 |
| Chrna4 | B7ZBU7 | -0.48 | 0.02 | -0.05 | -0.03 | -0.79 | -0.64 | -0.76 | 0.52 | -0.73 | -0.69 | 0.07 | -0.26 | -0.42 | -0.89 | -0.05 | 0.21 | -0.2825 | -0.2675 | -0.1525 | 0.406474 | 0.298845 | 0.590809 | 0.999971 | 1 | 0.92226 |
| Chrnb2 | A0A0G2JG( | -2.07 | -1.54 | -1.91 | -1.8 | -2.2 | -2.39 | -2.29 | -1.45 | -1.95 | -2.1 | -1.79 | -2.2 | -2.24 | -1.9 | -2.05 | -1.92 | -0.2525 | -0.18 | -0.1975 | 0.35097 | 0.375063 | 0.33196 | 0.999971 | 1 | 0.857152 |
| Chst2 | Q80WV3 | 0.89 | -0.52 | -0.55 | -0.44 | -0.63 | -0.55 | -0.43 | -0.2 | -0.38 | 1.17 | -0.07 | -0.36 | -0.3 | 0.04 | -0.64 | -0.15 | -0.2975 | 0.245 | -0.1075 | 0.410514 | 0.608341 | 0.770445 | 0.999971 | 1 | 0.963498 |
| Chsy3 | Q5DTK1 | 2.41 | 0.34 | 0.94 | 1.91 | 1.91 | -0.11 | 1.03 | 1.46 | 0.87 | 2.9 | 1.57 | -0.2 | -3.66 | 0.81 | 1.03 | 1.01 | -0.3275 | -0.115 | -1.6025 | 0.57719 | 0.874259 | 0.17358 | 0.999971 | 1 | 0.807607 |
| Chtop | E9PW20 | -0.05 | 0.27 | 0.56 | 0.81 | -0.01 | -0.2 | 0.25 | 0.92 | -0.05 | 0.22 | 0.93 | 0.53 | 0.24 | 0.09 | 0.44 | 0.47 | -0.1575 | 0.01 | -0.0875 | 0.61585 | 0.972571 | 0.716057 | 0.999971 | 1 | 0.952576 |
| Chuk | E9Q605 | 0.83 | 0.72 | -0.06 | -0.24 | -0.18 | 0.21 | -0.69 | -0.25 | -0.04 | 0.55 | -0.05 | 0.12 | -0.03 | 0.82 | -0.12 | -0.23 | -0.54 | -0.1675 | -0.2025 | 0.126058 | 0.591506 | 0.572128 | 0.999971 | 1 | 0.91816 |
| Ciao1 | Q99KN2 | 0.7 | 0.22 | -0.54 | -0.21 | 0.09 | 0.35 | 1.35 | -0.6 | 0.41 | 0.2 | -0.95 | -0.22 | -0.36 | 0.08 | -0.8 | -1.15 | 0.255 | -0.1825 | -0.6 | 0.578436 | 0.640354 | 0.130787 | 0.999971 | 1 | 0.784948 |
| Ciao2a | Q9DCL2 | 2.42 | 0.12 | -0.19 | 0.24 | 0.63 | 0.42 | -0.61 | -0.15 | 1.61 | 3.86 | -0.46 | -0.1 | 0.08 | 0.79 | -0.32 | -0.08 | -0.575 | 0.58 | -0.53 | 0.352667 | 0.57917 | 0.379744 | 0.999971 | 1 | 0.871816 |

| | | | | | | | | | | | | | | | | | | | | | | | | | | |
|---|---|---|---|---|---|---|---|---|---|---|---|---|---|---|---|---|---|---|---|---|---|---|---|---|---|---|
| Ciao2b | Q9D187 | -0.04 | 0.49 | 0.08 | 0.45 | 0.51 | 0.02 | -0.48 | -0.39 | 0.35 | -0.14 | 0.53 | 0.55 | -0.15 | 0.21 | 0.19 | 0.67 | -0.33 | 0.0775 | -0.015 | 0.254787 | 0.746214 | 0.951198 | 0.999971 | 1 | 0.998056 |
| Ciao3 | Q7TMW6 | 0.95 | 0.28 | -0.03 | -0.46 | 0.09 | 0.06 | -0.19 | -0.17 | -0.11 | 0.36 | -0.16 | 0.35 | -0.11 | -0.15 | 0.16 | -0.09 | -0.2375 | -0.075 | -0.2325 | 0.452402 | 0.81948 | 0.461712 | 0.999971 | 1 | 0.890095 |
| Ciapin1 | Q8WTY4 | 0.75 | 1.13 | 1.31 | 0.92 | 1.17 | 1.23 | 1.23 | 0.76 | 1.67 | 1.17 | 1.22 | 1.5 | 1.91 | 1.2 | 1 | 1.35 | 0.07 | 0.3625 | 0.3375 | 0.74449 | 0.117752 | 0.210228 | 0.999971 | 1 | 0.822657 |
| Cib1 | A0A0U1RP | -0.01 | -0.86 | -1.23 | -0.31 | -0.17 | -0.3 | -1.33 | -1.52 | -0.58 | 0.04 | -1.62 | -1.06 | -0.13 | -0.32 | -1.82 | -0.57 | -0.2275 | -0.2025 | -0.1075 | 0.590035 | 0.635206 | 0.808037 | 0.999971 | 1 | 0.973362 |
| Cibar1 | Q8BP22 | 0.61 | -0.56 | 0.89 | 0.8 | -0.32 | 0.1 | 0.95 | 0.85 | -0.01 | 0.34 | 1.02 | 0.53 | -1.05 | -0.04 | 0.57 | 0.17 | -0.04 | 0.035 | -0.5225 | 0.925675 | 0.927591 | 0.267322 | 0.999971 | 1 | 0.842677 |
| Cic | Q924A2 | 2.38 | 2.56 | 2.56 | 3 | 2.25 | 1.99 | 1.9 | 1.97 | 2.45 | 2.53 | 1.33 | 1.98 | 1.3 | 2.13 | 2.06 | 2.27 | -0.5975 | -0.5525 | -0.685 | 0.01662 | 0.102604 | 0.031537 | 0.999971 | 1 | 0.678474 |
| Cirbp | P60824 | 2.15 | 0.98 | 1.05 | 0.58 | 0.23 | 0.95 | 1.46 | 0.71 | 0.67 | 0.82 | 0.48 | 0.43 | -0.18 | 0.19 | 0.97 | -0.13 | -0.3525 | -0.59 | -0.9775 | 0.392697 | 0.113722 | 0.038234 | 0.999971 | 1 | 0.700838 |
| Cisd1 | Q91WS0 | -0.64 | 0.58 | 0.04 | 0.55 | 0.91 | 0.74 | 2.03 | 0.31 | 1.66 | 0.46 | 0.41 | 1.63 | 2.16 | 0.69 | 0.24 | -0.8 | 0.865 | 0.9075 | 0.44 | 0.074933 | 0.059837 | 0.483003 | 0.999971 | 1 | 0.896345 |
| Cisd2 | Q9CQB5 | -0.16 | 0.1 | 0.27 | 0.16 | 0.23 | 0.23 | -0.04 | 0.32 | 0.12 | 0.1 | 0.39 | 0.45 | 0.39 | 0.41 | 0.68 | 0.16 | 0.0925 | 0.1725 | 0.3175 | 0.626359 | 0.375882 | 0.138598 | 0.999971 | 1 | 0.784948 |
| Cisd3 | Z4YKM2 | 1.09 | 1.05 | 1.18 | 0.9 | 0.76 | 1.57 | 1.32 | 0.99 | 0.63 | 1.32 | 1.1 | 1.76 | 0.98 | 1.41 | 1.06 | 0.89 | 0.105 | 0.1475 | 0.03 | 0.646131 | 0.57792 | 0.876703 | 0.999971 | 1 | 0.985129 |
| Cit | E9QL53 | -0.62 | -1.24 | -0.88 | -0.85 | -1.21 | -1.11 | -0.67 | -0.79 | -1.16 | -0.74 | -0.74 | -1.06 | -1.16 | -1.08 | -0.84 | -0.73 | -0.0475 | -0.0275 | -0.055 | 0.831535 | 0.897142 | 0.796215 | 0.999971 | 1 | 0.970435 |
| Ciz1 | Q3UUV0 | 0.28 | -1.4 | -1.34 | 0.98 | -2.11 | -1.19 | -0.02 | 1.21 | -0.32 | -0.8 | -0.63 | -0.02 | -0.44 | 0.08 | -0.01 | 1.44 | -0.1575 | -0.0725 | 0.6375 | 0.850841 | 0.898844 | 0.343961 | 0.999971 | 1 | 0.862346 |
| Ckap4 | Q8BMK4 | 0.85 | 0.5 | 0.22 | 0.46 | 0.63 | 0.19 | 0.64 | 0.44 | 1.09 | 0.87 | 1.01 | 0.89 | 0.96 | 0.28 | 0.24 | 0.77 | -0.0325 | 0.4575 | 0.055 | 0.879645 | 0.042121 | 0.826004 | 0.999971 | 1 | 0.976757 |
| Ckap5 | A0A0R4J0K | -0.3 | -0.63 | -0.51 | -0.61 | -0.47 | -0.6 | -0.51 | -0.52 | -0.45 | -0.5 | -0.44 | -0.39 | -0.53 | -0.65 | -0.41 | -0.39 | -0.0125 | 0.0675 | 0.0175 | 0.941948 | 0.688539 | 0.922007 | 0.999971 | 1 | 0.993069 |
| Ckb | Q04447 | -0.25 | 0.29 | 0.41 | 0.37 | -0.12 | -0.19 | -0.07 | 0.33 | -0.25 | -0.75 | -0.21 | -0.07 | -0.31 | -0.06 | 0.26 | 0.32 | -0.2175 | -0.525 | -0.1525 | 0.358291 | 0.055316 | 0.536883 | 0.999971 | 1 | 0.910077 |
| Ckm | A0A0J9YKD | -1.2 | 1.1 | 4.35 | 1.83 | 0.89 | -0.41 | 2.72 | 1.32 | -1.74 | -1.43 | 1.66 | -4.24 | -0.81 | 0.64 | 1.62 | 1.22 | -0.39 | -2.9575 | -0.8525 | 0.738875 | 0.073646 | 0.455803 | 0.999971 | 1 | 0.887217 |
| Ckmt1 | P30275 | -0.25 | -0.62 | 0.23 | 0.13 | -0.88 | -0.53 | 0.04 | 0.16 | -0.78 | -0.56 | 0.04 | 0.01 | -0.55 | -0.33 | 0.13 | 0.36 | -0.175 | -0.195 | 0.03 | 0.582362 | 0.510718 | 0.919044 | 0.999971 | 1 | 0.992633 |
| Ckmt2 | Q6P8J7 | 1.53 | 4.15 | 3.58 | 3.4 | 3.01 | 4.09 | 2.25 | 3.66 | 2.56 | 2.93 | 3.07 | 2.36 | 2.03 | 4.42 | 3.2 | 4.18 | 0.0875 | -0.435 | 0.2925 | 0.889851 | 0.434676 | 0.682339 | 0.999971 | 1 | 0.943204 |
| Clasp1 | A0A087WF | 1.52 | 1.92 | 0.63 | -1.01 | 1.18 | 1.56 | -0.1 | -0.24 | 1.16 | 0.42 | 0.74 | -0.33 | 1.52 | 0.37 | -1.04 | 0.51 | -0.165 | -0.2675 | -0.425 | 0.817592 | 0.686241 | 0.576835 | 0.999971 | 1 | 0.918194 |
| Clasp1_1 | E9QKH0 | -0.13 | 0.29 | 0.48 | 0.57 | -0.79 | 0.37 | -0.24 | -0.45 | -0.43 | -0.22 | 0.01 | 0.77 | -0.5 | 0.88 | 1.42 | 0.61 | -0.58 | -0.27 | 0.3 | 0.078507 | 0.393418 | 0.474218 | 0.999971 | 1 | 0.894044 |
| Clasp1_2 | A0A087WC | 0.02 | 0.12 | 0.24 | -0.33 | 0.18 | 0.17 | 0.85 | -0.45 | 0.47 | -0.14 | 0.15 | 0 | 0.36 | -0.27 | -0.41 | -0.72 | 0.175 | 0.1075 | -0.2725 | 0.56425 | 0.627266 | 0.337211 | 0.999971 | 1 | 0.858827 |
| Clasp2 | Q08EB5 | 1.79 | 3.81 | 2.18 | 2.42 | 3.02 | 3.86 | 2.89 | 1.61 | 4.19 | 4.16 | 2.91 | 2.74 | 5.47 | 3.37 | 1.37 | 2.11 | 0.295 | 0.95 | 0.53 | 0.616216 | 0.108768 | 0.55801 | 0.999971 | 1 | 0.915051 |
| Clasp2_1 | Q8BRT1-5 | -0.32 | 1.8 | 0.36 | -0.07 | 1.01 | 1.49 | -0.15 | -0.03 | 0.97 | 0.16 | 0.15 | 0.31 | 1.29 | 1.3 | 0.04 | 0.27 | 0.1375 | -0.045 | 0.2825 | 0.80883 | 0.925139 | 0.599719 | 0.999971 | 1 | 0.92422 |
| Clasp2_2 | Q08EB6 | -0.43 | -0.64 | -0.76 | -0.84 | -0.62 | -0.68 | -0.6 | -0.74 | -0.2 | -0.35 | -0.55 | -0.55 | 0.02 | -0.78 | -0.73 | -0.65 | 0.0075 | 0.255 | 0.1325 | 0.966231 | 0.197074 | 0.585001 | 0.999971 | 1 | 0.920413 |
| Clasp2_3 | F7DCH5 | -0.6 | -1.57 | -1.35 | -0.26 | -1.42 | -1.26 | -0.65 | 0 | -1.08 | -0.41 | 0.19 | -0.1 | -0.37 | -0.78 | -0.46 | 0.51 | 0.1125 | 0.595 | 0.67 | 0.790645 | 0.158551 | 0.119035 | 0.999971 | 1 | 0.78268 |
| Clasrp | Q8CFC7 | 0.56 | -2 | -1.41 | 1.36 | -1.46 | -2.38 | -1.05 | 0.11 | -1.62 | -0.11 | 0.32 | -0.17 | -1.79 | -1.78 | 0.29 | 0.16 | -0.8225 | -0.0225 | -0.4075 | 0.346554 | 0.977946 | 0.646518 | 0.999971 | 1 | 0.933613 |
| Clcc1 | Q99LI2 | 0.09 | -1 | -0.32 | -0.19 | -0.82 | -0.71 | -0.01 | 0.2 | -0.75 | -0.64 | -0.02 | -0.04 | -0.22 | -1.13 | -0.25 | -0.1 | 0.02 | -0.0075 | -0.07 | 0.952966 | 0.980491 | 0.832676 | 0.999971 | 1 | 0.979036 |
| Clcn2 | A0A338P67 | -0.84 | 0.07 | -0.05 | -0.69 | 0.04 | -0.83 | -0.61 | 0.06 | -0.28 | -0.89 | -0.78 | -0.62 | -0.13 | -0.78 | -0.51 | 0.57 | 0.0425 | -0.265 | 0.165 | 0.895079 | 0.353476 | 0.650609 | 0.999971 | 1 | 0.933842 |
| Clcn3 | Q8K4W8 | -0.57 | -0.45 | -0.75 | -0.55 | -0.31 | -0.8 | -1.45 | -0.68 | -0.62 | -0.28 | -0.73 | -0.12 | -0.49 | -1.2 | -1.73 | -0.13 | -0.23 | 0.1425 | -0.3075 | 0.398109 | 0.494029 | 0.398929 | 0.999971 | 1 | 0.871816 |
| Clcn3_1 | A0A6Q6Q7 | -0.1 | 0 | 0.06 | 0.03 | 0.02 | 0.53 | -0.22 | 0.06 | -0.07 | 0.03 | 0.16 | -0.06 | 0.29 | 0.52 | 0.29 | 0.34 | 0.1 | 0.0175 | 0.3625 | 0.63718 | 0.914228 | 0.054876 | 0.999971 | 1 | 0.725348 |
| Clcn4 | Q3LRV6 | -0.37 | -0.46 | -0.77 | -0.3 | -0.14 | -0.36 | -0.87 | -0.35 | -0.5 | -0.49 | -0.49 | -0.28 | -0.45 | -0.27 | -0.23 | 0.4 | 0.045 | 0.035 | 0.3375 | 0.842623 | 0.849208 | 0.190423 | 0.999971 | 1 | 0.810547 |
| Clcn5 | Q9WVD4 | -1.83 | -0.86 | -0.82 | -0.71 | -0.84 | -1.23 | -1.08 | -0.43 | -1.07 | -1.36 | -1.28 | -1.16 | -1.73 | -1.03 | -1.1 | -0.29 | 0.16 | -0.1625 | 0.0175 | 0.615727 | 0.567639 | 0.963266 | 0.999971 | 1 | 0.998721 |
| Clcn6 | A2A7F6 | -0.65 | -0.33 | -0.05 | -0.39 | -0.42 | -0.18 | -0.47 | -0.1 | -0.28 | -0.48 | -0.35 | -0.28 | -0.14 | -0.01 | 0 | 0.11 | 0.0625 | 0.0075 | 0.345 | 0.762105 | 0.968798 | 0.106012 | 0.999971 | 1 | 0.78268 |
| Clcn7 | F6SUM2 | 1.93 | 0.03 | 0.21 | 0.07 | -0.05 | 0.32 | 0.06 | 0.29 | 0.39 | 0.91 | -0.11 | -0.46 | -0.08 | -0.62 | -0.11 | 0.12 | -0.405 | -0.3775 | -0.7325 | 0.36902 | 0.46552 | 0.134492 | 0.999971 | 1 | 0.784948 |
| Cldn1 | O88551 | -1.67 | -0.36 | 1.43 | 1.32 | -0.81 | -0.71 | 0.55 | 0.45 | 0.3 | -0.91 | 1.2 | 1.12 | -3.53 | -0.06 | 1.61 | 1.51 | -0.31 | 0.2475 | -0.2975 | 0.678137 | 0.758465 | 0.812689 | 0.999971 | 1 | 0.974458 |
| Cldn10 | Q9Z0S6-6 | 0.07 | 0.8 | -0.28 | -0.5 | 0.64 | 0.27 | -0.3 | 0.01 | 1.32 | 0.06 | 0.19 | -0.11 | 1.4 | -0.15 | -0.73 | -0.43 | 0.1325 | 0.3425 | -3.93E-16 | 0.700294 | 0.415163 | 1 | 0.999971 | 1 | 1 |
| Cldn11 | Q60771 | 0.41 | 1.19 | 1.89 | 1.3 | 0.98 | 1.06 | 1.26 | 1.13 | 1.44 | 0.54 | 0.4 | 0.19 | -0.66 | 0.3 | 0.79 | -0.33 | -0.09 | -0.555 | -1.1725 | 0.774657 | 0.183925 | 0.020343 | 0.999971 | 1 | 0.635388 |
| Cldn12 | Q9ET43 | -0.21 | 0.27 | 1.42 | 0.96 | 0.29 | 1.05 | 0.54 | 0.89 | -0.25 | 0.52 | 0.33 | -2.8 | 0.22 | 0.98 | 0.75 | 0.95 | 0.0825 | -1.16 | 0.115 | 0.83032 | 0.157557 | 0.766951 | 0.999971 | 1 | 0.962472 |
| Cldn3 | Q9Z0G9-2 | 4.31 | -4.74 | -1.67 | -1.57 | -4.34 | 1.7 | -1.6 | -2.88 | 3.63 | 7.98 | -6.04 | -3.33 | -5.46 | -2.95 | -3.19 | -3.2 | -0.8625 | 1.4775 | -2.7825 | 0.671251 | 0.657094 | 0.13974 | 0.999971 | 1 | 0.784948 |
| Cldn5 | O54942 | 4.18 | 0.41 | 2.13 | 3.61 | 2.02 | 0.92 | 3.17 | 0.77 | 2.34 | 4.34 | 1.49 | 1.71 | -0.89 | -0.01 | 1.89 | -0.01 | -0.8625 | -0.1125 | -2.3375 | 0.353143 | 0.906418 | 0.029928 | 0.999971 | 1 | 0.678474 |
| Cldnd1 | Q9CQX5 | -1.48 | -0.99 | -1 | -1.32 | -1 | -1.04 | -0.97 | -1.1 | -0.93 | -1.27 | -0.71 | -1.16 | -0.54 | -1.01 | -1.5 | -1.05 | 0.17 | 0.18 | 0.1725 | 0.384582 | 0.415344 | 0.506952 | 0.999971 | 1 | 0.905775 |
| Clec16a | Z4YLG3 | 0.3 | 0.25 | 0.29 | -0.04 | 0.09 | 0.49 | 0.44 | 0.18 | 0.79 | 0.43 | 0.17 | 0.03 | 0.59 | -0.02 | -0.04 | 0.31 | 0.1 | 0.155 | 0.01 | 0.605307 | 0.494767 | 0.963238 | 0.999971 | 1 | 0.998721 |
| Clec2l | P0C7M9 | 0.23 | -0.53 | -0.14 | 0.29 | -0.34 | 0.21 | -0.13 | 0.53 | -0.13 | 0.23 | -0.04 | -0.45 | -0.04 | 0 | 0.21 | 0.34 | 0.105 | -0.06 | 0.165 | 0.712076 | 0.816393 | 0.50174 | 0.999971 | 1 | 0.904573 |
| Clgn | P52194 | 2.02 | 0.5 | 0.56 | 0.83 | 0 | 0.54 | -0.34 | 0.48 | 0.93 | 2.18 | 1.04 | 0.95 | 0.41 | 0.33 | 0.71 | 1.32 | -0.8075 | 0.2975 | -0.285 | 0.066594 | 0.505522 | 0.484803 | 0.999971 | 1 | 0.896611 |
| Clic1 | Q9Z1Q5 | 0.61 | 0.8 | 1.43 | 1.45 | 0.67 | 0.7 | 1 | 0.93 | 0.65 | 0.71 | 1.11 | 1.09 | 0 | 0.93 | 1.12 | 1.33 | -0.2475 | -0.1825 | -0.2275 | 0.346943 | 0.499812 | 0.52885 | 0.999971 | 1 | 0.910069 |
| Clic4 | Q9QYB1 | -0.21 | -0.05 | 0.25 | -0.45 | -0.37 | -0.01 | -0.31 | 0.21 | -0.26 | -0.18 | -0.13 | 0.49 | -0.19 | -0.14 | 0.27 | -0.23 | -0.005 | 0.095 | 0.0425 | 0.983007 | 0.706492 | 0.851793 | 0.999971 | 1 | 0.98187 |
| Clic5 | A0A668KLE | -0.86 | 0.64 | -2.18 | -3.1 | 0.86 | -0.77 | -2.62 | -2.79 | 0.63 | -1.02 | -2.95 | -1.24 | -0.07 | -0.21 | -4.19 | -2.43 | 0.045 | 0.23 | -0.35 | 0.965971 | 0.815432 | 0.758834 | 0.999971 | 1 | 0.961116 |
| Clic6 | Q8BHB9 | 1.69 | 2.82 | 3.54 | 2.72 | 1.58 | 1.96 | 2.65 | 2.9 | 2.16 | 0.64 | -4.59 | 2.79 | 0.61 | 1.66 | 2.43 | 2.44 | -0.42 | -2.4425 | -0.9075 | 0.369828 | 0.138499 | 0.115348 | 0.999971 | 1 | 0.78268 |
| Clint1 | Q5SUH7 | -2.38 | -0.01 | -0.42 | 0 | 0.59 | -0.18 | 1.52 | -1.28 | 1.6 | -0.53 | 0.48 | 0.99 | -1.78 | -1.76 | -1.13 | -2.86 | 0.865 | 1.3375 | -1.18 | 0.261308 | 0.069927 | 0.08138 | 0.999971 | 1 | 0.755254 |
| Clint1_1 | Q5SUH6 | 0.04 | -0.53 | 0.09 | 0.14 | -0.45 | -0.14 | -0.06 | 0.1 | -0.28 | -0.2 | -0.16 | 0.13 | -0.43 | 0 | -0.09 | 0.28 | -0.0725 | -0.0625 | 0.005 | 0.75471 | 0.777068 | 0.98372 | 0.999971 | 1 | 0.99925 |
| Clip1 | D3Z3M7 | 0.44 | 0.69 | -0.07 | -0.27 | -0.59 | -0.13 | -0.56 | -0.02 | -0.27 | 0.8 | 0.67 | 0.74 | 1.17 | -0.3 | -0.84 | -0.45 | -0.5225 | 0.2875 | -0.3025 | 0.090036 | 0.400557 | 0.517861 | 0.999971 | 1 | 0.905815 |
| Clip1_1 | Q922J3 | 0.84 | -0.71 | -0.02 | 0.6 | 0.11 | -0.18 | -0.54 | -0.76 | -0.36 | 0.29 | 0.15 | 1.13 | -0.21 | 0.34 | 0.37 | 1.58 | -0.52 | 0.125 | 0.3425 | 0.197026 | 0.776417 | 0.480878 | 0.999971 | 1 | 0.89485 |
| Clip1_2 | D3Z2Z1 | 0.3 | 0.43 | 0.33 | 0.18 | 0.31 | 0.5 | 0.5 | 0.18 | 0.47 | 0.3 | 0.47 | 0.49 | 0.39 | 0.27 | 0.22 | 0.36 | 0.0625 | 0.1225 | -7.85E-17 | 0.724669 | 0.462347 | 1 | 0.999971 | 1 | 1 |
| Clip2 | Q9Z0H8 | 0.04 | -0.19 | 0 | 0.14 | -0.4 | -0.28 | -0.05 | -0.04 | -0.07 | 0.15 | 0.23 | 0.24 | -0.02 | -0.22 | 0.04 | 0.32 | -0.19 | 0.14 | 0.0325 | 0.319562 | 0.435456 | 0.867688 | 0.999971 | 1 | 0.983462 |
| Clip3 | B9EHT4 | -0.53 | -0.02 | -0.6 | -0.79 | 0.11 | 0.15 | -0.4 | -0.21 | 0.64 | -0.22 | -0.2 | -0.31 | 1.26 | -0.02 | -0.27 | -0.07 | 0.3975 | 0.4625 | 0.71 | 0.127669 | 0.135074 | 0.08529 | 0.999971 | 1 | 0.759325 |
| Clip4 | A0A2R8VH | 0.47 | 0.85 | 0.66 | 0.86 | 0.83 | 0.99 | 0.96 | 1.06 | 0.55 | 0.48 | 0.47 | 0.8 | 1.29 | 1.33 | 0.93 | 0.84 | 0.25 | -0.135 | 0.3875 | 0.191494 | 0.47455 | 0.089683 | 0.999971 | 1 | 0.771991 |
| Clmn | Q8C5W0-4 | 0.41 | -0.3 | -0.7 | -0.76 | -0.47 | -0.13 | -0.62 | -0.43 | -0.46 | 0.06 | -0.73 | -0.66 | -0.51 | -0.37 | -0.51 | -0.61 | -0.075 | -0.11 | -0.1625 | 0.801116 | 0.734785 | 0.575211 | 0.999971 | 1 | 0.91816 |
| Clmp | Q8R373 | 0.29 | -1.18 | -0.21 | 0.18 | -0.77 | -0.87 | 0.21 | -0.39 | -0.69 | 0.15 | -0.56 | -0.62 | -0.84 | -0.72 | -0.52 | -0.56 | -0.225 | -0.2 | -0.43 | 0.573885 | 0.597167 | 0.227075 | 0.999971 | 1 | 0.828236 |
| Cln6 | D6RE34 | -1.46 | -0.2 | -0.33 | 0.06 | 0.52 | 0.05 | -0.97 | 0.24 | -0.06 | -0.74 | -0.41 | 0.2 | 0.57 | -0.05 | 0.48 | -0.18 | 0.4425 | 0.23 | 0.6875 | 0.328742 | 0.549811 | 0.093103 | 0.999971 | 1 | 0.776565 |
| Cln8 | Q9QUK3 | 1.43 | 0.26 | -1.06 | -0.97 | -0.33 | -0.05 | -1.26 | -0.53 | -0.25 | 0.87 | -0.06 | 0 | -0.21 | -0.28 | -0.09 | 0.42 | -0.4575 | 0.225 | 0.045 | 0.443793 | 0.702581 | 0.935852 | 0.999971 | 1 | 0.995618 |
| Clns1a | Q923F1 | 0.25 | 0.58 | 0.19 | 0.5 | 0.79 | 0.31 | 0.05 | 0.56 | 0.28 | 0.33 | 0.43 | 0.62 | 0.72 | 0.77 | 0.6 | 0.42 | 0.0475 | 0.035 | 0.2475 | 0.833492 | 0.850891 | 0.216224 | 0.999971 | 1 | 0.826893 |
| Clock | A0A0J9YU6 | 1.46 | 1.06 | 0.19 | 1.71 | 1.11 | 1.05 | 0.3 | 1.11 | 0.36 | 1.78 | 0.68 | 0.1 | 0.46 | 1.32 | 0.65 | 2.13 | -0.2125 | -0.375 | 0.035 | 0.574298 | 0.430864 | 0.940738 | 0.999971 | 1 | 0.996218 |
| Clpb | Q60649 | -0.19 | -0.38 | -0.16 | -0.16 | -0.44 | -0.21 | -0.4 | 0.17 | -0.58 | -0.38 | -0.11 | 0 | -0.5 | -0.25 | -0.02 | 0.01 | 0.0025 | -0.045 | 0.0325 | 0.990217 | 0.819636 | 0.867222 | 0.999971 | 1 | 0.983462 |
| Clpp | O88696 | -0.41 | -0.94 | -0.54 | -0.09 | -0.74 | -1.06 | -0.85 | -0.13 | -1.17 | -0.75 | -0.85 | -0.6 | -0.79 | -0.72 | -0.22 | -0.23 | -0.2 | -0.3475 | 0.005 | 0.483261 | 0.176457 | 0.984558 | 0.999971 | 1 | 0.99925 |
| Clptm1 | Q8VBZ3 | -0.24 | -0.49 | -0.38 | -0.3 | -0.42 | -0.33 | -0.14 | 0.08 | -0.27 | 0.06 | -0.06 | -0.16 | -0.18 | 0.14 | -0.27 | -0.16 | 0.15 | 0.245 | 0.235 | 0.439051 | 0.177844 | 0.218668 | 0.999971 | 1 | 0.827064 |
| Clptm1l | Q8BXA5 | -1.26 | -0.67 | -0.15 | -0.24 | -0.93 | -0.59 | -0.39 | 0.05 | -0.24 | -0.93 | 0.28 | 0.21 | -0.74 | -0.35 | 0.52 | 0.04 | 0.115 | 0.41 | 0.4475 | 0.725165 | 0.281347 | 0.236739 | 0.999971 | 1 | 0.833594 |
| Clpx | Q9JHS4 | -0.4 | -0.47 | -0.2 | -0.67 | -0.44 | -0.68 | -0.36 | -0.45 | -0.62 | -0.35 | -0.04 | -0.02 | -0.41 | -0.42 | -0.28 | -0.62 | -0.0475 | 0.1775 | 0.0025 | 0.800987 | 0.419546 | 0.989455 | 0.999971 | 1 | 0.99925 |
| Clstn1 | Q9EPL2-2 | 1.92 | 2.6 | 1.94 | 2.19 | 2.69 | 2.95 | 0.94 | 2.27 | 2.69 | 2.72 | 1.73 | 0.51 | 3.22 | 3.06 | 2.08 | 2.28 | 0.05 | -0.25 | 0.4975 | 0.91057 | 0.624091 | 0.151761 | 0.999971 | 1 | 0.795578 |
| Clstn1_1 | Q9EPL2 | 1.47 | 1.73 | 1.41 | 0.9 | 1.82 | 1.85 | 1.52 | 1.3 | 1.89 | 1.18 | 1.82 | 1.59 | 2.3 | 1.37 | 1.4 | 1.24 | 0.245 | 0.2425 | 0.2 | 0.334774 | 0.36057 | 0.518797 | 0.999971 | 1 | 0.905901 |
| Clstn2 | Q9ER65-2 | -1.08 | 0.59 | -0.34 | -0.23 | -0.02 | 0.54 | -0.6 | -0.15 | 0.29 | 0.36 | -0.09 | -0.39 | -0.58 | 0.75 | -1.06 | -0.28 | 0.2075 | 0.3075 | -0.0275 | 0.603796 | 0.418952 | 0.954178 | 0.999971 | 1 | 0.998056 |
| Clstn3 | D3Z601 | 1.29 | 0.51 | 0.56 | 0.71 | 0.47 | 1.13 | 1 | 1.16 | 0.87 | 1.32 | 0.86 | 0.82 | 0.82 | 1 | 0.92 | 0.92 | 0.1725 | 0.2 | 0.1475 | 0.51686 | 0.420408 | 0.517327 | 0.999971 | 1 | 0.905815 |
| Clta | B1AWD8 | -1.47 | -0.21 | -0.94 | -2.74 | -0.31 | -0.84 | -0.73 | -0.73 | 0.1 | -1.3 | 0.37 | 1.07 | 0.69 | -1.04 | -2.22 | -2.27 | 0.6875 | 1.4 | 0.13 | 0.197157 | 0.061207 | 0.869003 | 0.999971 | 1 | 0.983462 |
| Clta_1 | B1AWE0 | -0.69 | -1.08 | -2.34 | -2.82 | -0.65 | -0.8 | 0.1 | -2.85 | 0.02 | -1.4 | -0.76 | -1.81 | -0.49 | -1.61 | -2.46 | -3.03 | 0.6825 | 0.745 | -0.165 | 0.361805 | 0.229543 | 0.808178 | 0.999971 | 1 | 0.973379 |
| Cltb | Q6IRU5-2 | -0.44 | -0.6 | -0.93 | 0.64 | -0.31 | -0.91 | -0.6 | -1.01 | -0.25 | 0.15 | -1 | -1.1 | -0.73 | -0.71 | -1.44 | -1.66 | -0.375 | -0.2175 | -0.8025 | 0.319124 | 0.615815 | 0.071718 | 0.999971 | 1 | 0.739418 |
| Cltb_1 | Q6IRU5 | -0.81 | -1.2 | -2.62 | -2.46 | -1.4 | -1 | -2.25 | -2.37 | -1.6 | -1.37 | -1.99 | -2.49 | -1.15 | -1.08 | -1.98 | -2.15 | 0.0175 | -0.09 | 0.1825 | 0.972988 | 0.851477 | 0.711969 | 0.999971 | 1 | 0.951449 |
| Cltc | Q68FD5 | -1.45 | -1.07 | -0.87 | -1.38 | -0.98 | -0.97 | -0.36 | -1.09 | -1.01 | -1.93 | -0.76 | -0.88 | -0.38 | -1.01 | -0.73 | -1.09 | 0.3425 | 0.0475 | 0.39 | 0.185149 | 0.876265 | 0.134541 | 0.999971 | 1 | 0.784948 |
| Clu | Q06890 | -1.15 | -0.65 | -0.15 | -0.53 | 0 | -0.02 | 0.6 | -0.64 | 0.11 | -0.5 | 0.48 | 0.55 | 0.04 | -0.76 | -0.59 | 0.05 | 0.605 | 0.78 | 0.305 | 0.091403 | 0.035016 | 0.328718 | 0.999971 | 1 | 0.856472 |
| Cluh | A0A0R4J14 | 1.03 | 1.52 | 1.14 | 1.25 | 1.4 | 1.47 | 1.11 | 1.59 | 1.44 | 1.52 | 1.47 | 1.55 | 1.4 | 1.41 | 1.27 | 1.39 | 0.1575 | 0.26 | 0.1325 | 0.444999 | 0.173311 | 0.479884 | 0.999971 | 1 | 0.894438 |
| Clvs1 | Q9D4C9 | -0.05 | 0.29 | 0.2 | 0.37 | 0.3 | 0.37 | -0.11 | 0.22 | 0.52 | 0.41 | 0.24 | 0.38 | 0.7 | 0.47 | 0.25 | 0.49 | -0.0075 | 0.185 | 0.275 | 0.969909 | 0.318664 | 0.180933 | 0.999971 | 1 | 0.808532 |
| Clvs2 | Q8BG92 | -0.14 | 0.26 | 0.27 | 0.48 | 0.25 | 0.06 | 0.08 | 0.18 | 0.12 | 0.09 | 0.45 | 0.47 | 0.32 | 0.31 | 0.39 | 0.6 | -0.075 | 0.065 | 0.1875 | 0.705019 | 0.758701 | 0.367797 | 0.999971 | 1 | 0.869526 |
| Clybl | Q8R4N0 | -0.47 | -0.42 | -0.34 | -0.28 | -0.51 | -0.39 | -0.88 | 0.02 | -0.61 | -0.64 | -0.78 | -1.02 | -0.52 | -0.38 | -0.35 | -0.16 | -0.0625 | -0.385 | 0.025 | 0.785055 | 0.054719 | 0.885944 | 0.999971 | 1 | 0.987222 |
| Cmas | A0A0R4J0E | 1.47 | 2.26 | 1.85 | 2.07 | 2.19 | 2.23 | 2.08 | 2.46 | 2.68 | 1.78 | 2.25 | 2.13 | 2.28 | 2.75 | 2.42 | 2.44 | 0.3275 | 0.2975 | 0.56 | 0.173944 | 0.286807 | 0.037557 | 0.999971 | 1 | 0.700096 |
| Cmbl | Q8R1G2 | -0.28 | 0.83 | 1.98 | 1.11 | 1.3 | 0.82 | 1.94 | 1.17 | 1.73 | -0.16 | 1.84 | 1.36 | 2.08 | 0.93 | 1.87 | 1.23 | 0.3975 | 0.2825 | 0.6175 | 0.421666 | 0.641217 | 0.237359 | 0.999971 | 1 | 0.833594 |
| Cmc1 | Q9CPZ8 | -0.84 | -0.36 | -0.15 | 0.2 | 0.12 | -0.39 | -0.14 | -0.13 | -0.48 | -1.08 | -0.33 | -0.33 | -0.1 | -0.51 | -0.17 | -0.49 | 0.1525 | -0.2675 | -0.03 | 0.565945 | 0.370876 | 0.909651 | 0.999971 | 1 | 0.991822 |
| Cmc2 | Q8K199 | 1.36 | 1.11 | 1.72 | 1.43 | 0.18 | 0.84 | 1.23 | 1.45 | 0.57 | 1.26 | 1.75 | 1.84 | 1.04 | 0.55 | 1.27 | 2.14 | -0.48 | -0.05 | -0.155 | 0.148172 | 0.875416 | 0.659611 | 0.999971 | 1 | 0.935181 |
| Cmc4 | Q61908 | 2.22 | 1.82 | 2.49 | 1.73 | 1.56 | 3.39 | 1.96 | 1.83 | 1.7 | 2.4 | 1.61 | -1.89 | 2.23 | 2.69 | 1.51 | 1.42 | 0.12 | -1.11 | -0.1025 | 0.777075 | 0.230898 | 0.767739 | 0.999971 | 1 | 0.962472 |
| Cmip | Q9D486 | 0.51 | 0.26 | 0.58 | 0.27 | 0.22 | 0.24 | 1.03 | 0.03 | 0.68 | 1.06 | 0.56 | 0.36 | 0.65 | 0.2 | 0.36 | -0.12 | -0.025 | 0.26 | -0.1325 | 0.923334 | 0.242901 | 0.556969 | 0.999971 | 1 | 0.915051 |
| Cmpk1 | Q9DBP5 | -1.05 | -0.17 | -0.35 | -0.4 | 0.09 | -0.58 | 0.08 | -0.58 | 0.14 | -1.04 | -0.04 | 0.41 | -0.2 | -0.53 | -0.5 | -0.45 | 0.245 | 0.36 | 0.0725 | 0.400549 | 0.333772 | 0.763155 | 0.999971 | 1 | 0.961892 |
| Cmpk2 | Q3U5Q7 | -0.34 | -0.3 | -1.38 | -1.38 | -0.5 | -0.27 | -0.94 | -1.7 | -0.65 | -0.43 | -1.41 | -1.81 | -0.49 | -0.35 | -1.45 | -1.48 | -0.0025 | -0.225 | -0.0925 | 0.995209 | 0.5976 | 0.821968 | 0.999971 | 1 | 0.975833 |
| Cmss1 | Q9CZT6 | 1.88 | 0.52 | 0.32 | 1.18 | 1.97 | 0.88 | 0.32 | 1.39 | 0.64 | 0.68 | 0.44 | 1.52 | 1.73 | 1.84 | 1.44 | 1.08 | 0.165 | -0.155 | 0.5475 | 0.72422 | 0.704888 | 0.17382 | 0.999971 | 1 | 0.807788 |
| Cmtm4 | Q8CJ61 | -1.24 | -0.27 | -0.97 | -1.02 | -1.08 | -0.27 | -0.18 | -1.02 | -0.7 | 0.06 | -0.46 | -0.84 | -0.93 | 0.09 | -0.36 | -0.82 | 0.2375 | 0.39 | 0.37 | 0.46562 | 0.211745 | 0.263089 | 0.999971 | 1 | 0.842533 |
| Cmtm5 | Q45TP8 | -0.46 | -0.4 | -0.48 | -0.55 | -0.71 | -0.44 | -1.19 | -0.16 | -1.17 | -0.41 | -0.84 | -0.56 | -0.06 | -0.5 | -0.77 | -0.24 | -0.1525 | -0.2725 | 0.08 | 0.545421 | 0.224766 | 0.704043 | 0.999971 | 1 | 0.950051 |
| Cmtr1 | Q9DBC3 | 2.24 | 2.57 | 2.22 | 2.43 | 2.25 | 2.13 | 1.73 | 1.86 | 2.25 | 1.95 | 2.11 | 2.33 | 1.4 | 1.8 | 2.12 | 2.08 | -0.3725 | -0.205 | -0.515 | 0.092567 | 0.284309 | 0.046103 | 0.999971 | 1 | 0.710102 |
| Cnbp | P53996-3 | 1.61 | 3.69 | 3.52 | 0.99 | 2.38 | 2.78 | 5.34 | 1.33 | 3.17 | 2.6 | 1.25 | -3.35 | 0.92 | 0.63 | 0.94 | 0 | 0.505 | -1.535 | -1.83 | 0.605866 | 0.306997 | 0.018986 | 0.999971 | 1 | 0.635388 |
| Cndp2 | Q9D1A2 | 0.06 | 0.52 | -0.54 | -0.58 | 0.21 | 0.45 | -0.36 | -0.02 | 0.27 | 0.14 | -0.56 | -0.33 | 0.21 | 0.31 | -0.28 | -0.16 | 0.205 | 0.015 | 0.155 | 0.523252 | 0.963409 | 0.615124 | 0.999971 | 1 | 0.927914 |
| Cnep1r1 | D3Z6J3 | -0.3 | -0.15 | -0.66 | 0.1 | 0.43 | 0.62 | -0.4 | 0.01 | -0.45 | -0.38 | -0.22 | -0.59 | 0.53 | 0.35 | 0.04 | 0.43 | 0.4175 | -0.1575 | 0.59 | 0.173777 | 0.477079 | 0.028571 | 0.999971 | 1 | 0.678474 |
| Cnih1 | D6RGU4 | 0.64 | 0.93 | 1.61 | 1.22 | 0.72 | 1.26 | 0.07 | 0.68 | 0.94 | -0.36 | -1.05 | -2.22 | 0.9 | 1.86 | 1.95 | 0.39 | -0.4175 | -1.7725 | 0.175 | 0.215743 | 0.019975 | 0.672197 | 0.999971 | 1 | 0.939777 |
| Cnih2 | A0A494B9 | -7.39 | -16.45 | -11.97 | -12.88 | -14.13 | -14.92 | -13.55 | -13.36 | -21.45 | -6.32 | -8.66 | -12.41 | -21.04 | -10.22 | -8.81 | -11.8 | -1.8175 | -0.0375 | -0.795 | 0.295559 | 0.99117 | 0.78741 | 0.999971 | 1 | 0.96724 |
| Cnih3 | G3XA11 | 1.63 | 0.05 | -0.67 | 0.07 | 0.17 | -0.03 | -0.39 | 0.87 | 0.28 | 0.93 | -1 | -0.78 | 1.11 | 0.31 | -1.6 | -0.02 | -0.115 | -0.4125 | -0.32 | 0.822053 | 0.504024 | 0.638862 | 0.999971 | 1 | 0.93301 |
| Cnih4 | A0A0A6YW | -1.05 | -0.34 | -0.85 | -0.84 | -1.07 | -1.29 | -1.21 | -0.38 | -0.93 | -1.1 | -1.06 | -0.33 | -0.71 | -0.29 | -1.22 | -0.85 | -0.2175 | -0.085 | 0.0025 | 0.436406 | 0.741446 | 0.992502 | 0.999971 | 1 | 0.99925 |
| Cnksr2 | Q80YA9 | -0.82 | -0.69 | -0.6 | -0.8 | -0.74 | -0.66 | -0.81 | -0.68 | -0.34 | -0.4 | -0.08 | -0.08 | 0.07 | -0.41 | -0.18 | 0.13 | 0.005 | 0.5025 | 0.63 | 0.9759 | 0.017961 | 0.010267 | 0.999971 | 1 | 0.613095 |
| Cnn1 | Q08091 | -0.86 | -2.82 | -4.69 | -2.25 | 6.53 | 3.53 | -1.62 | -3.29 | 0.79 | 5.51 | 5.86 | 7.5 | -4.02 | -0.67 | -2.93 | -2.77 | 3.9425 | 7.57 | 0.0575 | 0.091042 | 0.000689 | 0.951538 | 0.999971 | 0.448349 | 0.998056 |
| Cnn2 | Q08093 | 0.82 | 0.97 | 1.66 | 2.35 | 4.55 | 3.19 | 1.45 | 0.94 | 2.12 | 3.52 | 2.73 | 2.69 | 0.6 | 2 | 1.7 | 2.13 | 1.0825 | 1.315 | 0.1575 | 0.202715 | 0.013347 | 0.734551 | 0.999971 | 1 | 0.957874 |
| Cnn3 | Q9DAW9 | 1.35 | 0.38 | 0.5 | 0.38 | 0.79 | 0.9 | 0.15 | -0.84 | 0.81 | 1.6 | 0.45 | 0.94 | 0.48 | 0.6 | 0.53 | 0.83 | -0.4025 | 0.2975 | -0.0425 | 0.368275 | 0.383854 | 0.873878 | 0.999971 | 1 | 0.984109 |
| Cnnm1 | A0A286YDI | -0.35 | -0.55 | -0.96 | -1.02 | -0.58 | -0.79 | -1.24 | -1.11 | -0.44 | -0.39 | -0.67 | -0.61 | 0 | -0.62 | -0.79 | -0.89 | -0.21 | 0.1925 | 0.145 | 0.40876 | 0.386341 | 0.600894 | 0.999971 | 1 | 0.92426 |
| Cnnm2 | Q3TWN3 | -0.53 | -1.04 | -1.8 | -1.67 | -0.91 | -0.9 | -1.92 | -1.65 | -0.73 | -0.53 | -1.77 | -1.31 | -0.42 | -0.8 | -1.83 | -1.48 | -0.085 | 0.175 | 0.1275 | 0.822724 | 0.657228 | 0.758559 | 0.999971 | 1 | 0.961116 |
| Cnnm3 | Q32NY4 | -0.81 | -0.77 | -0.97 | -0.75 | -0.47 | -0.84 | -0.88 | -0.65 | -0.41 | -0.76 | -1.03 | -0.93 | 0.31 | -0.73 | -0.73 | -0.36 | 0.115 | 0.0425 | 0.4475 | 0.533403 | 0.831098 | 0.124807 | 0.999971 | 1 | 0.78268 |
| Cnnm4 | Q69ZF7 | -1.64 | -0.96 | -1.58 | -2.37 | -0.88 | -1.13 | -1.77 | -1.94 | -0.81 | -0.98 | -0.98 | -0.92 | -0.26 | -0.69 | -1.56 | -1.46 | 0.2075 | 0.715 | 0.645 | 0.580184 | 0.038341 | 0.138963 | 0.999971 | 1 | 0.784948 |
| Cnot1 | Q6ZQ08 | 0.67 | 0.89 | 0.61 | 0.62 | 0.82 | 0.96 | 0.32 | 0.61 | 0.75 | 0.84 | 0.61 | 0.51 | 0.74 | 0.99 | 0.77 | 0.91 | -0.02 | -0.02 | 0.155 | 0.922894 | 0.909215 | 0.387873 | 0.999971 | 1 | 0.871816 |
| Cnot10 | Q8BH15-2 | -0.51 | 0.59 | -0.73 | -0.58 | 1.44 | 0.58 | 0.77 | -1.53 | 2.43 | -0.92 | 0.02 | 1.57 | 0.03 | -1.04 | -0.37 | -1.94 | 0.6225 | 1.0825 | -0.5225 | 0.347313 | 0.167602 | 0.302553 | 0.999971 | 1 | 0.847402 |
| Cnot10_1 | Q8BH15 | 0.52 | 1.19 | 1.16 | 1.19 | 1.27 | 1.08 | 1.04 | 1.07 | 1.22 | 0.75 | 1.27 | 1.15 | 1.04 | 0.8 | 1.1 | 1.43 | 0.1 | 0.0825 | 0.0775 | 0.648699 | 0.726931 | 0.749869 | 0.999971 | 1 | 0.961064 |
| Cnot11 | Q9CWN7 | 0.43 | 0.32 | 0.45 | 0.56 | 0.1 | -0.03 | -0.11 | 0.59 | 0.25 | 0.19 | 0.27 | 0.7 | -1.04 | 0.12 | 0.51 | 0.69 | -0.3025 | -0.0875 | -0.37 | 0.179707 | 0.645907 | 0.342745 | 0.999971 | 1 | 0.861474 |
| Cnot2 | Q8C5L3-2 | 0.78 | 0.95 | 1.64 | 0.74 | 0.29 | 1.07 | 2.14 | 0.34 | 0.77 | 0.71 | 0.96 | 1.07 | 0.75 | 0.17 | 0.77 | 0.19 | -0.0675 | -0.15 | -0.5575 | 0.880819 | 0.55366 | 0.075592 | 0.999971 | 1 | 0.746958 |
| Cnot3 | Q8K0V4 | 0.32 | 0.82 | 0.88 | 0.97 | 0.82 | 0.86 | 0.64 | 0.86 | 0.76 | 0.73 | 1.19 | 1.22 | 0.67 | 0.87 | 0.84 | 0.74 | 0.0475 | 0.2275 | 0.0325 | 0.818996 | 0.339864 | 0.875175 | 0.999971 | 1 | 0.984491 |
| Cnot4 | Q05BG1 | 0.24 | 0.92 | 0.75 | 0.93 | 0.33 | 0.32 | 0.72 | 0.72 | 0.1 | -0.32 | 1 | 0.82 | 0.16 | 0.72 | 0.99 | 1.43 | -0.1875 | -0.31 | 0.115 | 0.431614 | 0.379344 | 0.716353 | 0.999971 | 1 | 0.952576 |
| Cnot6l | Q8VEG6-2 | -0.94 | 1.26 | 1.44 | 0.49 | 1.96 | 1.29 | 0.37 | 0.34 | 2.03 | -0.94 | 0.09 | -0.25 | 3.35 | -0.33 | 0.44 | 0.17 | 0.4275 | -0.33 | 0.345 | 0.488851 | 0.664786 | 0.69885 | 0.999971 | 1 | 0.947812 |
| Cnot7 | Q60809 | 0.2 | 0.54 | 1 | 0.82 | 0.49 | 0.55 | 0.65 | 0.43 | 0.39 | 0.62 | 0.26 | 0.84 | 0.35 | 0.91 | 1.25 | 1.3 | -0.11 | -0.1125 | 0.3125 | 0.623974 | 0.647434 | 0.300466 | 0.999971 | 1 | 0.847402 |
| Cnot9 | Q9JKY0 | 0.07 | 0.52 | 0.48 | 0.61 | 0.35 | 0.62 | 0.01 | 0.42 | 0.35 | 0.48 | 0.37 | 0.31 | 0.36 | 1 | 0.4 | 0.97 | -0.07 | -0.0425 | 0.2625 | 0.749865 | 0.821649 | 0.297999 | 0.999971 | 1 | 0.847402 |
| Cnp | P16330 | -1.41 | -1.65 | -0.89 | -1.32 | -2.11 | -1.5 | -1.89 | -1.02 | -1.85 | -1.53 | -1.82 | -1.66 | -1.69 | -1.23 | -0.95 | -1.27 | -0.3125 | -0.3975 | 0.0325 | 0.307713 | 0.094905 | 0.896237 | 0.999971 | 1 | 0.988571 |
| Cnpy2 | Q9QXT0 | 0.72 | 0.32 | 0.91 | 1.09 | 0.12 | 0.47 | 0.89 | 1.24 | 0.4 | 0.65 | 0.69 | 0.41 | 0.56 | 0.73 | 1.12 | 1.17 | -0.08 | -0.2225 | 0.135 | 0.791542 | 0.329838 | 0.593262 | 0.999971 | 1 | 0.923349 |
| Cnpy3 | Q9DAU1 | 0.46 | 1.71 | 1.11 | 1.04 | 0.82 | 1.34 | 0.97 | 1.09 | 1.14 | 1.15 | 1.43 | 0.86 | 0.63 | 1.27 | 1 | 1.06 | -0.025 | 0.065 | -0.09 | 0.931249 | 0.82347 | 0.76316 | 0.999971 | 1 | 0.961892 |
| Cnpy4 | Q8BQ47 | -0.6 | -0.84 | -1.33 | -1.77 | -0.5 | -1.64 | -1.5 | -0.42 | -0.85 | -1.49 | -0.7 | -0.02 | -0.4 | -0.91 | -0.78 | -0.85 | 0.12 | 0.37 | 0.4 | 0.762602 | 0.349595 | 0.198923 | 0.999971 | 1 | 0.813335 |
| Cnr1 | P47746 | -0.69 | -1.46 | -0.79 | -0.77 | -1.46 | -1.43 | -0.91 | -0.87 | -1.58 | -0.71 | -1.21 | -1.3 | -1.78 | -1.2 | -1.14 | -0.53 | -0.24 | -0.2725 | -0.235 | 0.372515 | 0.330395 | 0.464404 | 0.999971 | 1 | 0.890905 |
| Cnrip1 | Q5M8N0 | -0.36 | -0.07 | 0.02 | 0.21 | -0.22 | 0.18 | 0.08 | -0.05 | 0.25 | -0.67 | 0.2 | 0.23 | 0.09 | -0.1 | 0.11 | 0.17 | 0.0475 | 0.0525 | 0.1175 | 0.814969 | 0.846521 | 0.552251 | 0.999971 | 1 | 0.913595 |
| Cnst | Q8CBC4 | 0.4 | 0.37 | 0.38 | 0.67 | 0.44 | 0.02 | 0.91 | 0.51 | 0.4 | 0.08 | 0.27 | 0.48 | 0.45 | 0.2 | 0.05 | 0.3 | 0.015 | -0.1475 | -0.205 | 0.948531 | 0.427584 | 0.285125 | 0.999971 | 1 | 0.845133 |
| Cntfr | O88507 | -0.18 | -1.67 | -0.65 | -0.32 | -1.35 | -1.37 | -0.44 | -0.06 | -1.09 | -0.94 | -0.86 | -0.65 | -1.05 | -1.38 | -0.35 | -0.57 | -0.1 | -0.18 | -0.1325 | 0.821546 | 0.603732 | 0.73671 | 0.999971 | 1 | 0.959447 |
| Cntn1 | P12960 | -0.66 | -1.09 | -0.77 | -0.69 | -0.89 | -1 | -0.46 | -0.59 | -0.8 | -0.88 | -0.74 | -0.56 | -0.66 | -0.86 | -0.69 | -0.48 | 0.0675 | 0.0575 | 0.13 | 0.748908 | 0.75726 | 0.504588 | 0.999971 | 1 | 0.905277 |
| Cntn2 | Q61330 | -1.34 | -0.96 | -0.77 | -0.95 | -1.41 | -0.93 | -1.07 | -0.99 | -1.21 | -1.39 | -1 | -0.82 | -0.75 | -0.94 | -0.66 | -0.69 | -0.095 | -0.1 | 0.245 | 0.65406 | 0.6447 | 0.234911 | 0.999971 | 1 | 0.832918 |
| Cntn3 | Q07409 | -0.91 | -1.16 | -1 | -1.2 | -0.94 | -0.87 | -0.77 | -0.48 | -0.71 | -0.66 | -0.42 | -0.62 | -0.54 | -1 | -0.52 | -1.19 | 0.3025 | 0.465 | 0.255 | 0.137262 | 0.023792 | 0.272353 | 0.999971 | 1 | 0.843794 |
| Cntn4 | Q69Z26 | -0.39 | -1.08 | -1.09 | -0.56 | -0.67 | -0.97 | -1.08 | -0.78 | -1.01 | -0.28 | -0.86 | -1.05 | -0.54 | -1.04 | -0.49 | -0.44 | -0.095 | -0.02 | 0.1525 | 0.688995 | 0.941001 | 0.552411 | 0.999971 | 1 | 0.913595 |
| Cntn5 | P68500 | 0.15 | -1.61 | -1.22 | -1.1 | -0.88 | -0.78 | -0.68 | -1.05 | -0.63 | 0.12 | -0.8 | -0.5 | -1.15 | -1.33 | -0.82 | -1.16 | 0.0975 | 0.4925 | -0.17 | 0.795654 | 0.250714 | 0.658184 | 0.999971 | 1 | 0.934638 |
| Cntn6 | Q9JMB8 | -0.63 | -0.77 | -1.29 | -0.99 | -0.7 | -1.02 | -1.48 | -1 | -0.73 | -0.74 | -0.93 | -0.89 | -0.26 | -0.69 | -1.04 | -0.66 | -0.13 | 0.0975 | 0.2575 | 0.599686 | 0.634216 | 0.310148 | 0.999971 | 1 | 0.847402 |
| Cntnap1 | O54991 | -0.41 | -0.72 | -0.26 | -0.47 | -0.93 | -0.61 | -0.77 | -0.24 | -0.74 | -0.65 | -0.76 | -0.98 | -0.76 | -0.52 | -0.29 | -0.64 | -0.1725 | -0.3175 | -0.0875 | 0.441199 | 0.113778 | 0.661879 | 0.999971 | 1 | 0.936148 |
| Cntnap2 | E9QNF7 | -0.04 | -0.57 | -0.86 | -0.82 | -0.59 | -0.33 | -0.41 | -0.85 | -0.63 | -0.4 | -1.06 | -0.94 | 0.01 | -0.8 | -0.71 | -0.76 | 0.0275 | -0.185 | 0.0075 | 0.912137 | 0.48619 | 0.978936 | 0.999971 | 1 | 0.99925 |
| Cntnap3 | E9PY62 | 0.97 | -1.52 | -0.45 | -0.96 | -0.61 | -0.68 | -0.04 | -0.65 | -0.38 | 0.75 | 0.16 | 0.65 | -1.04 | -1.02 | -0.47 | -0.86 | -0.005 | 0.785 | -0.3575 | 0.992214 | 0.177133 | 0.489154 | 0.999971 | 1 | 0.898346 |
| Cntnap4 | D3YWB9 | -0.53 | -0.66 | -0.54 | -0.09 | -0.69 | -0.67 | -0.54 | -0.27 | -0.8 | -0.39 | -0.56 | -0.73 | -0.76 | -0.61 | -0.43 | -0.02 | -0.0875 | -0.165 | 1.08E-16 | 0.677291 | 0.429968 | 1 | 0.999971 | 1 | 1 |
| Cntnap5a | Q0V8T9 | -0.44 | -0.15 | -0.66 | -0.67 | 0.21 | -0.1 | -0.31 | -0.36 | 0.3 | -0.14 | -0.48 | -0.38 | 0.43 | -0.23 | -0.39 | -0.58 | 0.34 | 0.305 | 0.2875 | 0.150195 | 0.228322 | 0.30461 | 0.999971 | 1 | 0.847402 |
| Cntnap5b | Q0V8T8 | -0.79 | -1.51 | -0.87 | -0.51 | -1.07 | -1.29 | -0.7 | -0.25 | -1.37 | -0.89 | -0.17 | -0.17 | -0.26 | -0.89 | 0.06 | -0.44 | 0.0925 | 0.27 | 0.5375 | 0.769016 | 0.454545 | 0.099841 | 0.999971 | 1 | 0.78268 |
| Cntnap5c | Q0V8T7 | 0.06 | -0.87 | -1.27 | -1.84 | -0.29 | -0.57 | -1.24 | -1.28 | -0.2 | -0.52 | -1.09 | -1.33 | -0.47 | -1.84 | -1.09 | -1.99 | 0.135 | 0.195 | -0.3675 | 0.760432 | 0.665657 | 0.465577 | 0.999971 | 1 | 0.891255 |
| Coa3 | Q9D2R6 | -0.87 | -0.05 | -0.47 | -0.24 | -0.16 | 0.05 | -0.8 | -0.19 | 0.2 | -0.17 | -0.6 | -0.27 | 0.41 | 0.05 | -0.17 | 0.17 | 0.1325 | 0.1975 | 0.5225 | 0.629007 | 0.458733 | 0.057981 | 0.999971 | 1 | 0.725348 |
| COA4 | Q8BT51 | 1.24 | 1.84 | -0.96 | -1.34 | 1.48 | 1.94 | -1.14 | -0.16 | 1.83 | 2.25 | -1.16 | -1.05 | 0.75 | 1.28 | -0.9 | -1.09 | 0.335 | 0.2725 | -0.185 | 0.725787 | 0.801542 | 0.834484 | 0.999971 | 1 | 0.979036 |
| Coa6 | Q8BGD8 | -0.45 | 1.71 | 1.55 | 0.82 | 1.58 | 1.21 | 1.56 | 1.24 | 2.07 | -0.54 | 1.75 | 1.35 | 1.98 | 0.07 | 1.28 | 0.94 | 0.49 | 0.25 | 0.16 | 0.311547 | 0.720183 | 0.782734 | 0.999971 | 1 | 0.965859 |
| Coa7 | Q921H9 | 0.1 | 0.54 | 0.35 | -0.12 | 0.7 | 0.98 | 0.91 | 0.27 | 1.09 | 0.16 | 0.69 | 0.45 | 1.07 | 0.7 | 0.06 | -0.26 | 0.4975 | 0.38 | 0.175 | 0.068579 | 0.17477 | 0.602332 | 0.999971 | 1 | 0.924453 |
| Coa8 | E9Q6C3 | -0.39 | -0.97 | 0.02 | -0.54 | -0.04 | -0.59 | -0.84 | -0.05 | -1.12 | -0.43 | -0.36 | 0.41 | 0.12 | -0.35 | 0.45 | 0.97 | 0.09 | 0.095 | 0.7675 | 0.761383 | 0.794614 | 0.048355 | 0.999971 | 1 | 0.710102 |
| Coasy | Q9DBL7 | -1.87 | -0.76 | -1.28 | -1.75 | -0.6 | -0.68 | -0.47 | -1.44 | -0.21 | -2.02 | -0.82 | -1.22 | -0.48 | -1.12 | -1.57 | -1.67 | 0.6175 | 0.3475 | 0.205 | 0.090648 | 0.430197 | 0.574973 | 0.999971 | 1 | 0.91816 |
| Cobl | G5E8P4 | 0.42 | 0.13 | 0.34 | 0.56 | -0.03 | 0.39 | 0.66 | 0.5 | -0.26 | 0.3 | 0.55 | -0.17 | 0.05 | 0.04 | 0.01 | 0.18 | 0.0175 | -0.2575 | -0.2925 | 0.93595 | 0.303129 | 0.129444 | 0.999971 | 1 | 0.784948 |
| Cobll1 | B1AZ15 | 2.03 | 0.72 | 1.64 | 1.81 | 1.88 | 1.41 | 1.91 | 1.26 | 1.23 | 2.17 | 0.95 | 2.12 | 1.01 | 1.5 | 1.54 | 1.24 | 0.065 | 0.0675 | -0.2275 | 0.844226 | 0.867596 | 0.479875 | 0.999971 | 1 | 0.894438 |
| Coch | Q62507 | -0.36 | 0.73 | 0.4 | 0.16 | 1.16 | 0.39 | 0.28 | 0.13 | 2.06 | 0.22 | 1.06 | 1.17 | 2.54 | 0.89 | 1.55 | 1.63 | 0.2575 | 0.895 | 1.42 | 0.43696 | 0.058401 | 0.005591 | 0.999971 | 1 | 0.582714 |
| Cog1 | Q9Z160 | 1.33 | 1.43 | 1.09 | 1.03 | 1.58 | 1.24 | 1.13 | 1.48 | 1.34 | 1.61 | 1.4 | 1.36 | 0.77 | 1.88 | 1.06 | 1.65 | 0.1375 | 0.2075 | 0.12 | 0.497093 | 0.274156 | 0.677736 | 0.999971 | 1 | 0.942594 |
| Cog2 | Q921L5 | 0.2 | 2.48 | 1.45 | 1.22 | 1.98 | 2.51 | 1.35 | 1.49 | 2.43 | 0.84 | 1.49 | 1.31 | 1.98 | 2.58 | 1.51 | 1.62 | 0.495 | 0.18 | 0.585 | 0.334003 | 0.735915 | 0.252016 | 0.999971 | 1 | 0.839012 |
| Cog3 | E9QL65 | 1.36 | 1.75 | 0.79 | 1.07 | 1.52 | 1.69 | 0.66 | 0.81 | 1.59 | 1.85 | 0.57 | 0.93 | 1.49 | 1.98 | 0.91 | 0.96 | -0.0725 | -0.0075 | 0.0925 | 0.824954 | 0.983 | 0.777062 | 0.999971 | 1 | 0.964475 |
| Cog4 | G3UWH9 | 0.1 | 1.12 | 0.73 | 0.71 | 1.1 | 0.84 | 0.97 | 0.85 | 1.22 | 0.38 | 0.55 | 1 | 1.49 | 1.13 | 0.93 | 0.95 | 0.275 | 0.1225 | 0.46 | 0.284794 | 0.680074 | 0.113661 | 0.999971 | 1 | 0.78268 |
| Cog5 | Q8C0L8 | 0.47 | 0.99 | 0.61 | 0.79 | 1.18 | 0.87 | 0.63 | 1.03 | 0.91 | 0.55 | 0.97 | 1.27 | 0.85 | 0.75 | 0.69 | 1.02 | 0.2125 | 0.21 | 0.1125 | 0.330449 | 0.361114 | 0.570483 | 0.999971 | 1 | 0.91816 |
| Cog6 | A0A0R4J0L | 0.91 | 1.47 | 1.21 | 1.28 | 1.17 | 1.24 | 1.05 | 1.51 | 1.2 | 1.48 | 1.33 | 1.63 | 0.74 | 1.53 | 1.4 | 1.69 | 0.025 | 0.1925 | 0.1225 | 0.903153 | 0.352245 | 0.642396 | 0.999971 | 1 | 0.93301 |
| Cog7 | A0A0R4J0C | -0.07 | 0.75 | 1.01 | 0.62 | 0.41 | 0.75 | 0.69 | 1.17 | 0.37 | 0.06 | 0.67 | 0.87 | 0.69 | 0.87 | 1.15 | 0.93 | 0.1775 | -0.085 | 0.3325 | 0.545499 | 0.776313 | 0.23797 | 0.999971 | 1 | 0.833594 |
| Cog8 | Q9JJA2 | 2.3 | 1.29 | 0.88 | 0.5 | 1.82 | 1.89 | 0.57 | 0.79 | 1.48 | 2.57 | 0.71 | 0.53 | 1.6 | 2.09 | 0.83 | 0.96 | 0.025 | 0.08 | 0.1275 | 0.958547 | 0.885963 | 0.78026 | 0.999971 | 1 | 0.965859 |
| Coil | Q5SU73 | -1.17 | -1.91 | -2.27 | -1.94 | -2.31 | -2.47 | -1.81 | -2.7 | -2.24 | -1.37 | -1.99 | -2.92 | -2.78 | -2.05 | -2.19 | -2.06 | -0.5 | -0.3075 | -0.4475 | 0.128833 | 0.430451 | 0.160172 | 0.999971 | 1 | 0.800144 |
| Col12a1 | E9PX70 | 0.61 | 0.82 | 0.39 | 2.26 | 2.97 | 0.94 | 0.3 | 0.48 | 1.46 | 1.38 | 1.65 | 2.42 | 0.17 | 2.28 | 2.83 | 2.19 | 0.1525 | 0.7075 | 0.8475 | 0.821354 | 0.147686 | 0.217423 | 0.999971 | 1 | 0.826893 |

| | | | | | | | | | | | | | | | | | | | | | | | | | | |
|---|---|---|---|---|---|---|---|---|---|---|---|---|---|---|---|---|---|---|---|---|---|---|---|---|---|---|
| Col18a1 | E9QPX1 | -0.31 | 0.18 | 0.81 | 0.8 | 4.72 | 0.95 | 1.08 | -0.24 | 2 | 1.47 | 2.93 | 4.52 | -0.35 | 0.46 | 1.28 | 0.41 | 1.2575 | 2.36 | 0.08 | 0.224173 | 0.005915 | 0.845077 | 0.999971 | 1 | 0.979814 |
| Col19a1 | F8WHV4 | -0.02 | -0.53 | 0.33 | -0.07 | -0.62 | 0.32 | 0.12 | -0.36 | -0.84 | 0.35 | 0.3 | 0.52 | -0.71 | -0.52 | -0.13 | 0.37 | -0.0625 | 0.155 | -0.175 | 0.829652 | 0.660806 | 0.569235 | 0.999971 | 1 | 0.91816 |
| Col1a1 | P11087 | 4.27 | 0.88 | 3.26 | 3.39 | 5.9 | 2.9 | 5.53 | -0.98 | 2.23 | 5.35 | 2.1 | 4.69 | -3.37 | 1.77 | 2.71 | -1.85 | 0.3875 | 0.6425 | -3.135 | 0.802177 | 0.523547 | 0.053732 | 0.999971 | 1 | 0.725348 |
| Col1a2 | Q01149 | 1.27 | 0.54 | 1.01 | 0.58 | 5.13 | 1.86 | 4.45 | -1.79 | 2.06 | 2.98 | 1.08 | 2.88 | -1.11 | 1.46 | 0.62 | -1.85 | 1.5625 | 1.4 | -1.07 | 0.281547 | 0.011936 | 0.156638 | 0.999971 | 1 | 0.796008 |
| Col25a1 | V9GWX5 | 6.71 | 0.87 | 1.72 | 2.32 | 0.91 | 1.78 | 3.48 | 1.14 | 1.11 | 4.18 | 0.43 | 0.49 | -0.83 | -0.95 | 0.75 | 0.79 | -1.0775 | -1.3525 | -2.965 | 0.40492 | 0.350723 | 0.03764 | 0.999971 | 1 | 0.700096 |
| Col3a1 | P08121 | 3.77 | 1.01 | 3.06 | 3.71 | 6.13 | 2.24 | 3.59 | -0.29 | 2.85 | 5.26 | 6.47 | 8.76 | -2.76 | 1.43 | 3.63 | -1.64 | 0.03 | 2.9475 | -2.7225 | 0.981842 | 0.040002 | 0.081388 | 0.999971 | 1 | 0.755254 |
| Col4a1 | P02463 | 1.16 | 0.55 | 0.33 | 1.25 | 3.3 | 0.9 | 0.26 | -0.01 | 1.56 | 2.21 | 0.95 | 1.83 | 0.73 | 0.37 | 0.6 | 0.03 | 0.29 | 0.815 | -0.39 | 0.68445 | 0.039736 | 0.197381 | 0.999971 | 1 | 0.812891 |
| Col4a2 | P08122 | 1.74 | 0.24 | 1 | 0.9 | 2.52 | 1.13 | 1.46 | 0.28 | 1.05 | 2.21 | 1.05 | 1.77 | -0.2 | 0.8 | 1.26 | -0.01 | 0.3775 | 0.55 | -0.5075 | 0.469418 | 0.195063 | 0.262403 | 0.999971 | 1 | 0.842262 |
| Col4a5 | Q63ZW6 | 2.73 | 4.73 | 0.74 | 0.02 | 3.6 | 4.25 | 5.41 | 0.48 | 4.98 | 4.34 | 0.96 | 6.4 | 5.42 | 4.91 | 2.94 | -0.56 | 1.38 | 2.115 | 1.1225 | 0.31344 | 0.157138 | 0.469417 | 0.999971 | 1 | 0.892481 |
| Col6a1 | Q04857 | 1.04 | 1.26 | 1.02 | 2.54 | 5.07 | 1.48 | 0.85 | -0.01 | 1.76 | 3.17 | 2.54 | 4.29 | -1.09 | 2.4 | 2.24 | 0.45 | 0.3825 | 1.475 | -0.465 | 0.715531 | 0.032992 | 0.56984 | 0.999971 | 1 | 0.91816 |
| Col6a2 | Q02788 | 0.91 | 1.13 | 0.71 | 2.08 | 5.42 | 2.08 | 0.6 | -0.61 | 2.52 | 2.88 | 2.36 | 3.84 | -0.7 | 1.97 | 1.83 | 0.25 | 0.665 | 1.6925 | -0.37 | 0.579653 | 0.003377 | 0.571213 | 0.999971 | 0.957868 | 0.91816 |
| Col6a3 | J3QQ16 | 1.05 | 0.98 | 0.5 | 1.69 | 5.27 | 1.63 | 0.57 | -0.93 | 1.93 | 3.37 | 2.24 | 3.89 | -1.02 | 2.31 | 1.61 | -0.28 | 0.58 | 1.8025 | -0.4 | 0.629237 | 0.005076 | 0.591645 | 0.999971 | 1 | 0.922655 |
| Colec12 | A0A452J8C | 1.08 | 0.29 | 0.59 | 0.87 | 1.15 | 0.17 | 0.59 | 0.91 | 0.32 | 0.6 | 0.47 | 1.13 | -0.53 | 0.7 | 1.02 | 0.89 | -0.0025 | -0.0775 | -0.1875 | 0.993018 | 0.770695 | 0.625863 | 0.999971 | 1 | 0.930479 |
| Colgalt1 | Q8K297 | 0.9 | 0.17 | -0.17 | 0.18 | 0.46 | 0.54 | -0.29 | 0.23 | 0.01 | 1.02 | -0.18 | 0.32 | -0.24 | 0.52 | 0.02 | 0.7 | -0.035 | 0.0225 | -0.02 | 0.907318 | 0.947572 | 0.949662 | 0.999971 | 1 | 0.998038 |
| Commd1 | Q8VI86 | -0.33 | -0.35 | -1.15 | -0.61 | -0.4 | -0.07 | -1.34 | -0.92 | 0.02 | 0.16 | -0.68 | -0.59 | 0.49 | 0.09 | -1.08 | -1.66 | -0.0725 | 0.3375 | 0.07 | 0.829557 | 0.270387 | 0.888102 | 0.999971 | 1 | 0.987485 |
| Commd10 | Q8JZY2 | -0.64 | 0.89 | 0.48 | 0.44 | 0.71 | 0.35 | 0.16 | 0.38 | 0.78 | -0.05 | 0.33 | 0.18 | 1.01 | 0.82 | 0.66 | 1.06 | 0.1075 | 0.0175 | 0.595 | 0.753932 | 0.961431 | 0.105567 | 0.999971 | 1 | 0.78268 |
| Commd2 | Q8BXC6 | -0.27 | -0.37 | -1.44 | -0.88 | -0.27 | -0.27 | -0.6 | -0.89 | -0.47 | -0.08 | -1.44 | -1 | -0.3 | 0.24 | -1.18 | -0.58 | 0.2325 | -0.0075 | 0.285 | 0.463501 | 0.984524 | 0.466866 | 0.999971 | 1 | 0.891477 |
| Commd3 | Q63829 | -0.19 | 0.02 | -0.1 | 0.42 | 0.2 | -0.04 | -0.37 | 0.39 | -0.1 | 0.04 | 0.15 | 0.05 | 0.25 | 0.59 | -0.11 | 0.71 | 0.0075 | -0.0025 | 0.3225 | 0.975377 | 0.989949 | 0.227031 | 0.999971 | 1 | 0.828236 |
| Commd4 | Q9CQ02 | 0.54 | 1.26 | 1.45 | 1.02 | 1.03 | 1.37 | 1.1 | 1.08 | 1.23 | 0.12 | 1.81 | 1.3 | 1.75 | 1.68 | 2.15 | 1.76 | 0.0775 | 0.0475 | 0.7675 | 0.749785 | 0.903346 | 0.013108 | 0.999971 | 1 | 0.613095 |
| Commd5 | A0A0R4J0L | 0.48 | 0.16 | -0.03 | -0.14 | -0.15 | 0.13 | -0.4 | 0.07 | -0.13 | 0.08 | 0.07 | -0.26 | -0.09 | 0.14 | 0.01 | 0.24 | -0.205 | -0.1775 | -0.0425 | 0.370452 | 0.401974 | 0.837773 | 0.999971 | 1 | 0.979416 |
| Commd6 | Q3V4B5 | -0.18 | -0.13 | -0.18 | 0.01 | -0.14 | -0.14 | -0.18 | -0.11 | -0.29 | -0.61 | -0.08 | -0.07 | -0.33 | 0.35 | 0.16 | 0.35 | -0.0225 | -0.1425 | 0.2525 | 0.889533 | 0.467881 | 0.259142 | 0.999971 | 1 | 0.839012 |
| Commd7 | Q8BG94 | 0.56 | 0.46 | 0.51 | 1.01 | 0.55 | 0.99 | 0.04 | 0.86 | 0.72 | 1.28 | 0.46 | 0.56 | 1.1 | 1.08 | 0.56 | 1.29 | -0.025 | 0.12 | 0.3725 | 0.925309 | 0.631793 | 0.141951 | 0.999971 | 1 | 0.784967 |
| Commd8 | Q9CZG3 | 0.12 | 0.4 | -0.35 | -0.18 | 0.6 | 0.33 | -0.48 | -0.13 | 0.2 | 0.26 | -0.31 | -0.03 | 0.85 | 0.6 | 0.09 | 0.11 | 0.0825 | 0.0325 | 0.415 | 0.783572 | 0.892371 | 0.150658 | 0.999971 | 1 | 0.793248 |
| Commd9 | Q8K2Q0 | -1.77 | 0.03 | 0.08 | -0.29 | 0.53 | -0.45 | 0.14 | -0.38 | 0.51 | -1.6 | 0.41 | 0.13 | 0.82 | -0.56 | 0.32 | -0.24 | 0.4475 | 0.35 | 0.5725 | 0.345196 | 0.566196 | 0.2655 | 0.999971 | 1 | 0.842677 |
| Comt | O88587 | -0.67 | 0.5 | -0.75 | -1.18 | 0.52 | 0.51 | -0.48 | -0.78 | 0.49 | -0.49 | -0.43 | -0.63 | 0.7 | 0.72 | -0.36 | -0.77 | 0.4675 | 0.26 | 0.5975 | 0.324767 | 0.54008 | 0.239194 | 0.999971 | 1 | 0.833594 |
| Comtd1 | Q8BIG7 | 1.31 | 0.44 | -0.1 | 0.24 | 0.57 | 0.92 | 0.69 | 0.14 | 0.67 | 1.28 | -0.16 | -0.13 | 0.98 | 0.67 | 0.33 | 0.19 | 0.1075 | -0.0575 | 0.07 | 0.751658 | 0.894666 | 0.839259 | 0.999971 | 1 | 0.979728 |
| Copa | F8WHL2 | 0.79 | 1.27 | 1.88 | 1.69 | 1.26 | 0.92 | 1.76 | 1.89 | 1.39 | 0.97 | 1.87 | 1.87 | 1.31 | 1.2 | 1.91 | 2.06 | 0.05 | 0.1175 | 0.2125 | 0.879427 | 0.718836 | 0.51814 | 0.999971 | 1 | 0.905815 |
| Copb1 | Q9JIF7 | -0.19 | 0.34 | 0.62 | 0.81 | 0.06 | 0.4 | 0.79 | 0.69 | 0.21 | 0.17 | 0.58 | 0.57 | -0.53 | 0.59 | 0.76 | 1.04 | 0.09 | -0.0125 | 0.07 | 0.753338 | 0.962254 | 0.858202 | 0.999971 | 1 | 0.982702 |
| Copb2 | O55029 | 0.75 | 1.57 | 1.31 | 1.04 | 1.53 | 1.5 | 1.32 | 1.21 | 1.68 | 1.07 | 1.3 | 0.96 | 1.3 | 1.52 | 1.39 | 1.14 | 0.2225 | 0.085 | 0.17 | 0.346464 | 0.743713 | 0.470056 | 0.999971 | 1 | 0.892481 |
| Cope | O89079 | 1.17 | 2.23 | 1.75 | 1.6 | 1.47 | 2.59 | 1.16 | 1.8 | 1.68 | 1.96 | 1.77 | 1.73 | 1.48 | 2.49 | 1.64 | 2.22 | 0.0675 | 0.0975 | 0.27 | 0.853971 | 0.69986 | 0.415517 | 0.999971 | 1 | 0.876885 |
| Copg1 | Q9QZE5 | 1.1 | 1.79 | 2.13 | 1.39 | 1.52 | 1.61 | 1.74 | 1.65 | 1.89 | 1.37 | 1.95 | 1.54 | 1.55 | 1.81 | 2.15 | 1.91 | 0.0275 | 0.085 | 0.2525 | 0.91427 | 0.761469 | 0.371357 | 0.999971 | 1 | 0.869526 |
| Copg2 | Q9QXK3-4 | -0.36 | 1.2 | 1.11 | 0.53 | 1.46 | 0.38 | 1.81 | 0.85 | 1.11 | -0.6 | 1.52 | 1.26 | 1.24 | 0.35 | 1.02 | 0.5 | 0.505 | 0.2025 | 0.1575 | 0.279325 | 0.715891 | 0.694089 | 0.999971 | 1 | 0.947174 |
| Coprs | Q9CQ13 | -1.28 | -0.4 | -0.9 | 0.18 | 0.07 | -0.51 | 0.05 | 0.52 | -0.42 | -1.23 | 0.16 | 0.4 | -0.47 | -0.33 | 0.01 | -0.79 | 0.6325 | 0.3275 | 0.205 | 0.113877 | 0.476841 | 0.563536 | 0.999971 | 1 | 0.917087 |
| Cops2 | P61202-2 | -0.46 | 0.33 | -0.25 | -0.81 | 0.68 | -0.03 | 0.1 | -0.54 | 0.75 | -0.62 | 0.12 | 0.33 | 0.87 | -0.24 | -0.24 | -0.78 | 0.35 | 0.4425 | 0.2 | 0.321126 | 0.244595 | 0.622005 | 0.999971 | 1 | 0.929641 |
| Cops3 | O88543 | -0.06 | 0.18 | 1.04 | 0.57 | -0.07 | 0.32 | 0.77 | 0.56 | 0.17 | 0.06 | 0.5 | -0.16 | -0.19 | 0.08 | 0.76 | 0.45 | -0.0375 | -0.29 | -0.1575 | 0.902648 | 0.329066 | 0.625817 | 0.999971 | 1 | 0.930479 |
| Cops4 | O88544 | -0.41 | 0.25 | 0.14 | 0 | 0.22 | 0.12 | 0.05 | -0.13 | 0.26 | -0.23 | 0.11 | 0.11 | -0.15 | 0.05 | 0.01 | -0.16 | 0.07 | 0.0675 | -0.0575 | 0.741399 | 0.758298 | 0.782168 | 0.999971 | 1 | 0.965859 |
| Cops5 | O35864 | 0.03 | 0 | -0.61 | -0.62 | -0.05 | -0.24 | -0.31 | -0.66 | 0.02 | -0.5 | -0.74 | -0.69 | -0.39 | -0.19 | -0.62 | -0.64 | -0.015 | -0.1775 | -0.16 | 0.952124 | 0.515492 | 0.515564 | 0.999971 | 1 | 0.905815 |
| Cops6 | O88545 | -0.41 | -0.19 | -0.88 | -1.06 | 0 | -0.03 | -0.78 | -1.08 | 0.05 | -0.18 | -1.24 | -0.9 | -0.2 | 0.22 | -1.01 | -1.02 | 0.1625 | 0.0675 | 0.1325 | 0.630563 | 0.849604 | 0.714062 | 0.999971 | 1 | 0.952257 |
| Cops7a | Q9CZ04-2 | 0.84 | 0.78 | 0.35 | 0.2 | 0.63 | 0.74 | 0.28 | 0.23 | 0.74 | 0.41 | 0.51 | 0.37 | 0.86 | 0.55 | 0.07 | 0.33 | -0.0725 | -0.035 | -0.09 | 0.759915 | 0.87317 | 0.726215 | 0.999971 | 1 | 0.956041 |
| Cops7b | Q8BV13 | 0.31 | 0.34 | 0.53 | 0.46 | 0.62 | 0.52 | 0.95 | 0.46 | 0.85 | 0.46 | 0.87 | 0.95 | 0.81 | 0 | 0.74 | 0.62 | 0.2275 | 0.3725 | 0.1325 | 0.24865 | 0.072206 | 0.569426 | 0.999971 | 1 | 0.91816 |
| Cops8 | Q8VBV7 | 0.36 | -0.28 | -0.12 | 0.11 | 0.09 | 0.08 | -0.58 | -0.32 | 0.13 | 0.56 | -0.93 | -0.07 | -0.41 | 0.28 | -0.23 | -0.13 | -0.2 | -0.095 | -0.14 | 0.423375 | 0.779863 | 0.558918 | 0.999971 | 1 | 0.915051 |
| Cops9 | Q3U898 | 2.95 | 0.22 | 0.88 | 0.55 | -0.91 | 1.29 | 0.28 | 0.27 | -1.72 | -1.13 | -0.74 | -0.09 | -0.87 | -0.04 | -0.86 | 0.57 | -0.9175 | -2.07 | -1.45 | 0.205721 | 0.010337 | 0.047738 | 0.999971 | 1 | 0.710102 |
| Copz1 | P61924 | -0.78 | 2.03 | 1.07 | 0.32 | 2.9 | 0.35 | 1.37 | 0.64 | 3.3 | -0.5 | 1.58 | 2.38 | 3.53 | -0.23 | 0.8 | 0.3 | 0.655 | 1.03 | 0.44 | 0.388281 | 0.27365 | 0.634316 | 0.999971 | 1 | 0.93301 |
| Copz2 | Q9JHH9 | 0.91 | 1.09 | 1.7 | 1.84 | 0.97 | 1.01 | 1.45 | 1.88 | 0.87 | 0.18 | 1.72 | 1.53 | -0.64 | 1.92 | 2.54 | 2.42 | -0.0575 | -0.31 | 0.175 | 0.855367 | 0.446234 | 0.804247 | 0.999971 | 1 | 0.971467 |
| Coq10a | A0A1W2P7 | 1.25 | 0.62 | -0.43 | -0.23 | 0.62 | 0.6 | -0.67 | 0.08 | 0.75 | 1.13 | -0.47 | -0.25 | 0.71 | 0.73 | -0.27 | -0.2 | -0.145 | -0.0125 | -0.06 | 0.753846 | 0.980418 | 0.893673 | 0.999971 | 1 | 0.98828 |
| Coq2 | F7CDY3 | 0.48 | 1.02 | -1.22 | -1.35 | 0.16 | -0.21 | -2.26 | -0.55 | -1.04 | 0.8 | -1.6 | -1.48 | 0.4 | 1.34 | -2.53 | -0.53 | -0.4475 | -0.5625 | -0.0625 | 0.541007 | 0.455921 | 0.945452 | 0.999971 | 1 | 0.997427 |
| Coq3 | Q8BMS4 | -1.2 | -1.84 | -1.9 | -1.66 | -1.72 | -2.34 | -1.17 | -1.87 | -2.3 | -1.48 | -2.26 | -2.38 | -1.86 | -2.16 | -2.2 | -2.11 | -0.125 | -0.455 | -0.4325 | 0.675905 | 0.128631 | 0.076536 | 0.999971 | 1 | 0.748444 |
| Coq4 | Q8BGB8 | 0.67 | 0 | -0.02 | 0.7 | -0.21 | 0.53 | -0.35 | 0.67 | -0.08 | 0.82 | -0.73 | 0.23 | 0.15 | 0.44 | 0.34 | 1.18 | -0.1775 | -0.2775 | 0.19 | 0.589825 | 0.46044 | 0.542433 | 0.999971 | 1 | 0.911119 |
| Coq5 | Q9CXI0 | -0.35 | -0.54 | -0.37 | -0.43 | -0.44 | -0.5 | -0.61 | -0.14 | -0.73 | -0.62 | 0.13 | -0.75 | -0.53 | -0.21 | -0.29 | -0.25 | 3.93E-17 | -0.07 | 0.1025 | 1 | 0.773419 | 0.55912 | 1 | 1 | 0.915051 |
| Coq6 | Q8R1S0 | -0.57 | 0.27 | -0.18 | -0.64 | -0.14 | -0.21 | -0.23 | -0.4 | -0.69 | -0.78 | -0.49 | -0.91 | -0.42 | -0.18 | -0.61 | -1.47 | 0.035 | -0.4375 | -0.39 | 0.886928 | 0.110218 | 0.277511 | 0.999971 | 1 | 0.843812 |
| Coq7 | P97478 | -0.09 | 0.7 | 1.28 | 1.17 | -0.36 | 0.81 | 0.15 | 0.48 | -0.15 | 0.4 | 0.06 | 0.16 | -0.4 | 0.61 | 0.95 | 1.35 | -0.495 | -0.6475 | -0.1375 | 0.219389 | 0.077394 | 0.76405 | 0.999971 | 1 | 0.962405 |
| Coq8a | Q60936 | -1.32 | -0.78 | -1.28 | -1.43 | -0.98 | -0.74 | -1.39 | -0.82 | -0.93 | -1.49 | -0.81 | -0.93 | -0.45 | -0.33 | -0.89 | -0.92 | 0.22 | 0.1625 | 0.555 | 0.367487 | 0.504458 | 0.044287 | 0.999971 | 1 | 0.704422 |
| Coq8b | Q566J8 | -0.3 | -0.06 | -0.3 | -0.63 | -0.23 | -0.06 | -0.09 | -0.17 | 0.02 | 0.26 | 0.16 | -0.45 | 0.38 | 0.22 | 0.01 | -0.23 | 0.185 | 0.32 | 0.4175 | 0.344482 | 0.189302 | 0.085927 | 0.999971 | 1 | 0.759991 |
| Coq9 | Q8K1Z0 | -0.42 | -0.36 | -0.32 | -0.16 | -0.81 | -0.58 | -0.22 | -0.13 | -0.93 | -0.8 | -0.49 | -0.28 | -0.69 | -0.55 | -0.95 | -0.89 | -0.12 | -0.31 | -0.455 | 0.580058 | 0.15907 | 0.032922 | 0.999971 | 1 | 0.685881 |
| Coro1a | O89053 | 0.36 | 0.65 | 0.1 | 0.37 | 0.71 | 0.62 | 0.64 | 0.09 | 0.8 | 0.46 | 0.23 | -0.04 | 0.51 | 0.39 | -0.32 | -0.16 | 0.145 | -0.0075 | -0.265 | 0.521672 | 0.975099 | 0.318724 | 0.999971 | 1 | 0.853893 |
| Coro1a_1 | A0A0U1RP | -0.02 | 0.76 | 1.42 | 2 | -0.5 | 1.67 | 0.62 | 0.97 | 0.84 | 0.35 | 1 | 0.07 | -0.67 | 1.13 | -0.46 | 1.25 | -0.35 | -0.475 | -0.7275 | 0.546605 | 0.313786 | 0.253351 | 0.999971 | 1 | 0.839012 |
| Coro1b | Q9WUM3 | 0.18 | 0.27 | -0.12 | 0.16 | 0.37 | 0.59 | -0.17 | 0.25 | 0.43 | 0.73 | 0.16 | -0.1 | 0.29 | 0.47 | 0.04 | 0.2 | 0.1375 | 0.1825 | 0.1275 | 0.541537 | 0.438969 | 0.509908 | 0.999971 | 1 | 0.905815 |
| Coro1c | Q9WUM4 | -1.23 | -0.83 | -0.61 | -0.57 | -0.34 | -0.33 | -0.68 | -0.31 | -0.49 | -0.65 | -0.45 | -0.42 | -0.32 | -0.57 | -0.55 | -0.52 | 0.395 | 0.3075 | 0.32 | 0.099167 | 0.164286 | 0.156962 | 0.999971 | 1 | 0.796966 |
| Coro2a | B1AVH5 | 0.37 | 0.07 | -0.41 | -0.03 | -0.04 | 0.5 | -0.46 | -0.31 | -0.16 | 0.15 | -0.42 | -0.56 | 0.07 | 0.38 | -0.36 | -0.05 | -0.0775 | -0.2475 | 0.01 | 0.782775 | 0.33628 | 0.968147 | 0.999971 | 1 | 0.998787 |
| Coro2b | Q8BH44 | -0.33 | -0.35 | -0.73 | -1.18 | -0.79 | -0.89 | -0.95 | -1.05 | -0.79 | -0.68 | -0.97 | -1.33 | -0.6 | -0.63 | -1.26 | -0.98 | -0.2725 | -0.295 | -0.22 | 0.274001 | 0.283183 | 0.429189 | 0.999971 | 1 | 0.881196 |
| Coro6 | Q920M5 | 0.73 | -0.49 | 1.35 | 0.59 | -0.84 | -0.28 | 1.63 | 0.2 | -1.36 | -0.07 | 1.03 | 0.47 | -0.67 | -0.62 | 0.44 | 0.11 | -0.3675 | -0.5275 | -0.73 | 0.541643 | 0.380352 | 0.126989 | 0.999971 | 1 | 0.783603 |
| Coro7 | Q9D2V7 | 0.3 | 0.53 | 0.18 | 0.18 | 0.5 | 0.29 | 0.19 | 0.59 | 0.33 | 0.38 | 0.47 | 0.38 | -0.01 | 0.46 | 0.22 | 0.29 | 0.095 | 0.0925 | -0.0575 | 0.620583 | 0.59166 | 0.766504 | 0.999971 | 1 | 0.962472 |
| Cotl1 | Q9CQI6 | 0.76 | 0.81 | 0.65 | 0.45 | 0.52 | 1.06 | 0.5 | 0.57 | 0.82 | 0.95 | 0.39 | -0.41 | 0.62 | 0.62 | 0.37 | 0.66 | -0.005 | -0.23 | -0.1 | 0.980773 | 0.477742 | 0.585866 | 0.999971 | 1 | 0.920413 |
| Cox10 | Q8CFY5 | 0.13 | -0.76 | -1.66 | -2.7 | -0.69 | -1.74 | -0.84 | -1.1 | -1.35 | -1.43 | -1.68 | -0.98 | 0.06 | -1.34 | -1.61 | -1.95 | 0.155 | -0.1125 | 0.0375 | 0.793982 | 0.84472 | 0.955998 | 0.999971 | 1 | 0.998056 |
| Cox11 | Q6P8I6 | 1.23 | 0.39 | 0.52 | 0.1 | 0.72 | 0.63 | -0.64 | 0.13 | 1.26 | 0.81 | -0.46 | 0.7 | 1.12 | 0.69 | 0.11 | 0.67 | -0.35 | 0.0175 | 0.0875 | 0.368456 | 0.966513 | 0.784787 | 0.999971 | 1 | 0.96643 |
| Cox14 | Q8BH51 | 1.74 | -0.15 | -0.1 | -0.04 | 0.03 | -0.28 | 0.58 | -0.04 | -0.8 | 0.45 | 0.72 | 0.22 | 0.15 | 0.26 | 1.07 | 0.57 | -0.29 | -0.215 | 0.15 | 0.535151 | 0.683965 | 0.750729 | 0.999971 | 1 | 0.961064 |
| Cox15 | Q8BJ03-2 | 0.2 | 0.06 | -0.58 | -0.64 | -0.34 | 0.4 | -0.73 | 0.27 | -0.17 | 0.47 | -0.27 | -0.59 | -0.18 | 0.9 | -0.21 | -0.05 | 0.14 | 0.1 | 0.355 | 0.680486 | 0.750218 | 0.310217 | 0.999971 | 1 | 0.847402 |
| Cox17 | P56394 | 0.77 | 2.3 | 1.58 | 1.85 | 1.78 | 2.56 | 1.41 | 1.75 | 2.52 | 1.09 | 0.78 | 1.18 | 2.7 | 2.83 | 2.15 | 1.61 | 0.25 | -0.2325 | 0.6975 | 0.523706 | 0.623678 | 0.113947 | 0.999971 | 1 | 0.78268 |
| Cox18 | Q8VC74 | 2.2 | -0.64 | -0.9 | 0.54 | -0.95 | 2.13 | 0.27 | -0.17 | -0.42 | 1.76 | -0.3 | -0.52 | -2.81 | -0.85 | -0.31 | -1.17 | 0.02 | -0.17 | -1.585 | 0.98148 | 0.833292 | 0.07416 | 0.999971 | 1 | 0.746069 |
| Cox19 | Q8K0C8 | -0.59 | 0.69 | 0.87 | 0.98 | 0.91 | 0.73 | 0.49 | 1 | 0.45 | 0.52 | 0.78 | 0.53 | 0.31 | 0.98 | 0.46 | 0.59 | 0.295 | 0.0825 | 0.0975 | 0.43358 | 0.820049 | 0.797138 | 0.999971 | 1 | 0.970479 |
| Cox20 | Q9D7J4 | 0.47 | -1.46 | -1.76 | -0.95 | -1.68 | -1.35 | -0.54 | -0.11 | -1.55 | -0.8 | -1.9 | -1.48 | -2.43 | -1.09 | -1.29 | -1.3 | 0.005 | -0.5075 | -0.6025 | 0.992882 | 0.331469 | 0.278374 | 0.999971 | 1 | 0.843812 |
| Cox4i1 | P19783 | -1.33 | -1.39 | -1.06 | -1.17 | -1.64 | -1.44 | -0.8 | -0.92 | -1.84 | -1.9 | -0.94 | -1.3 | -1.9 | -1.41 | -0.99 | -0.94 | 0.0375 | -0.2575 | -0.0725 | 0.87859 | 0.33928 | 0.780598 | 0.999971 | 1 | 0.965859 |
| Cox4i1_1 | M0QWX7 | 2.96 | 3.74 | 1.57 | 5.16 | -0.7 | 3.46 | 1.95 | 0.44 | 3.28 | 4.34 | 0.21 | -1.68 | 4.54 | 1.06 | -0.61 | -1.21 | -2.07 | -1.82 | -2.4125 | 0.07476 | 0.218273 | 0.096651 | 0.999971 | 1 | 0.78268 |
| Cox4i2 | Q91W29 | 1.65 | 1.83 | -3.05 | -2.27 | 3.23 | -0.83 | -2.03 | -2.61 | -1.71 | 0.01 | -2.63 | -3.58 | -2.16 | -1.28 | -2.97 | -3.58 | -0.1 | -1.5175 | -2.0375 | 0.950915 | 0.273716 | 0.1231 | 0.999971 | 1 | 0.78268 |
| Cox5a | P12787 | -0.7 | -0.96 | -1.49 | -0.96 | -0.76 | -1.75 | -1.1 | -1.22 | -0.99 | -2.46 | -1.08 | -0.66 | -0.69 | -1.3 | -1.27 | -1.03 | -0.18 | -0.27 | -0.045 | 0.524681 | 0.517073 | 0.855868 | 0.999971 | 1 | 0.982416 |
| Cox5b-ps | Q9D881 | 0.18 | -1.03 | -1.63 | -1.29 | -1.24 | -1.36 | -1.28 | -1.39 | -1.4 | -0.68 | -1.07 | -1.79 | -1.46 | -1.01 | -1.68 | -1.24 | -0.375 | -0.2925 | -0.405 | 0.338962 | 0.506978 | 0.327615 | 0.999971 | 1 | 0.856472 |
| Cox6a1 | Q9DCW5 | -4.54 | -2.56 | -2.03 | -2.11 | -2.34 | -2.56 | -1.81 | -2.38 | -2.43 | -3.91 | -2.73 | -2.23 | -3.06 | -1.65 | -2.06 | -2.47 | 0.5375 | -0.015 | 0.5 | 0.34983 | 0.981238 | 0.417043 | 0.999971 | 1 | 0.877407 |
| Cox6a2 | P43023 | 0.71 | 0.31 | -3.04 | -1.72 | -1.34 | 0.36 | -2.22 | -3.13 | -1.49 | 0.06 | -2.26 | -1.39 | -2.71 | -0.52 | -1.34 | -2.16 | -0.6475 | -0.335 | -0.7475 | 0.533203 | 0.712383 | 0.414984 | 0.999971 | 1 | 0.876789 |
| Cox6b1 | P56391 | 0.16 | -0.11 | -0.3 | -0.19 | -0.07 | 0.92 | 0.01 | 0.24 | 0.35 | 0.45 | -0.32 | -0.79 | -0.32 | 0.53 | -0.04 | -0.55 | 0.385 | 0.0325 | 0.015 | 0.172662 | 0.917113 | 0.956005 | 0.999971 | 1 | 0.998056 |
| Cox6c | Q9CPQ1 | -2.03 | -2.42 | -1.84 | -2.8 | -2.47 | -2.4 | -1.72 | -2.7 | -2.78 | -1.93 | -1.53 | -1.58 | -2.84 | -2.19 | -2.07 | -2.85 | -0.05 | 0.3175 | -0.215 | 0.870283 | 0.379478 | 0.487648 | 0.999971 | 1 | 0.897906 |
| Cox7a1 | A0A140LIU | -2.68 | -1.25 | -1.2 | -2.07 | -0.93 | -0.7 | -1.82 | -1.65 | -0.47 | -0.65 | -1.39 | -0.42 | 0.28 | -1.14 | -0.32 | -1.07 | 0.525 | 1.0675 | 1.2375 | 0.235492 | 0.025185 | 0.023256 | 0.999971 | 1 | 0.66029 |
| Cox7a2 | P48771 | -2.75 | -2.67 | -1.47 | -2.34 | -2.14 | -2.28 | -1.78 | -3.03 | -2 | -2.08 | -2.34 | -1.68 | -2.11 | -3.37 | -1.9 | -2.33 | 1.86E-16 | 0.2825 | -0.12 | 1 | 0.393593 | 0.773936 | 1 | 1 | 0.963877 |
| Cox7a2l | E9PZS8 | -2.54 | -0.92 | -0.41 | -0.76 | -0.68 | -0.53 | -0.67 | -0.52 | -0.7 | -1.54 | -0.66 | -0.55 | -0.07 | -1.05 | -0.56 | -0.76 | 0.5575 | 0.295 | 0.5475 | 0.232501 | 0.551562 | 0.27343 | 0.999971 | 1 | 0.843794 |
| Cox7c | P17665 | -0.27 | 0.11 | -0.44 | -0.17 | 0.1 | -1.43 | 0.54 | -1.14 | 0.02 | -0.35 | -1.99 | -1.11 | 0.58 | -0.82 | -1.47 | -0.4 | -0.29 | -0.665 | -0.335 | 0.531601 | 0.15281 | 0.436336 | 0.999971 | 1 | 0.882419 |
| Cp | G3X9T8 | -0.04 | 0.85 | 0.03 | 0.99 | 1.69 | 0.2 | 0.8 | -0.2 | 0.74 | -0.42 | 0.2 | 0.92 | 0.17 | 0.39 | -0.05 | -0.14 | 0.165 | -0.0975 | -0.365 | 0.720451 | 0.802436 | 0.247232 | 0.999971 | 1 | 0.839012 |
| Cpd | O89001 | 0.89 | 1.54 | 1.08 | 0.84 | 1.57 | 1.44 | 1.44 | 0.71 | 1.62 | 1.23 | 1.36 | 1.5 | 1.61 | 1.48 | 1.07 | 0.9 | 0.2025 | 0.34 | 0.1775 | 0.462633 | 0.149592 | 0.495677 | 0.999971 | 1 | 0.901531 |
| Cpe | Q00493 | -0.48 | 0.08 | -0.17 | -0.28 | 0.64 | 0.13 | 0.03 | 0.05 | 0.9 | -0.01 | -0.1 | 0 | 1.46 | 0 | -0.12 | -0.08 | 0.425 | 0.41 | 0.5275 | 0.08691 | 0.165127 | 0.195004 | 0.999971 | 1 | 0.811225 |
| Cpeb2 | E9Q969 | 0.63 | 0.78 | 0.1 | 0.44 | 0.67 | 0.78 | -0.14 | 0.51 | 0.96 | 1.08 | 0.35 | 0.65 | 0.66 | 0.97 | 0.5 | 1.05 | -0.0325 | 0.2725 | 0.3075 | 0.904677 | 0.287205 | 0.209323 | 0.999971 | 1 | 0.822138 |
| Cpeb3 | A0A0R4J10 | -0.37 | 0.56 | 0.6 | 0.58 | 0.51 | 0.63 | 0.38 | 0.83 | 0.54 | -0.04 | 0.6 | 0.62 | 0.71 | 0.62 | 0.74 | 0.99 | 0.245 | 0.0875 | 0.4225 | 0.382236 | 0.766769 | 0.144542 | 0.999971 | 1 | 0.790933 |
| Cpeb4 | Q5SU48 | 0.55 | 0.71 | 0.05 | -0.51 | 0.41 | 0.43 | 0.34 | -0.81 | 0.59 | 0.79 | 0.04 | 0 | 1.09 | 1.12 | -0.35 | 0.15 | -0.1075 | 0.155 | 0.3025 | 0.783959 | 0.646888 | 0.488952 | 0.999971 | 1 | 0.898346 |
| Cplx1 | P63040 | 0.56 | -0.59 | -0.48 | -0.49 | -0.56 | 0.37 | -0.5 | -0.48 | -0.67 | 0.11 | -0.9 | -0.87 | -0.65 | -0.09 | -0.53 | -0.24 | -0.0425 | -0.3325 | -0.1275 | 0.901939 | 0.359035 | 0.679389 | 0.999971 | 1 | 0.942957 |
| Cplx2 | P84086 | 0.14 | -0.6 | -0.01 | 0.03 | -0.76 | -0.7 | -0.24 | -0.05 | -0.85 | -0.49 | -0.36 | -0.31 | -0.95 | -0.75 | -0.28 | -0.21 | -0.3275 | -0.3925 | -0.4375 | 0.233343 | 0.125772 | 0.127732 | 0.999971 | 1 | 0.783941 |
| Cplx3 | Q8R1B5 | -0.03 | 0.04 | -0.35 | -0.12 | -0.21 | 0.51 | -0.17 | -0.15 | -0.29 | 0.2 | 0.32 | -0.26 | -0.01 | -0.23 | -0.57 | 0.13 | 0.11 | 0.1075 | -0.055 | 0.634535 | 0.62596 | 0.802576 | 0.999971 | 1 | 0.971467 |
| Cpm | Q80V42 | -0.94 | -0.96 | -0.75 | -1.04 | -0.93 | -1 | -1.08 | -1.04 | -0.94 | -0.62 | -0.97 | -0.9 | -0.76 | -0.62 | -0.38 | -0.87 | -0.09 | 0.065 | 0.265 | 0.595212 | 0.714081 | 0.188129 | 0.999971 | 1 | 0.809025 |
| Cpne1 | Q8C166 | 0.94 | -0.9 | -1.01 | -1.24 | -0.73 | -0.31 | -0.92 | -1.28 | -0.88 | -0.04 | -1.21 | -0.87 | -1.2 | -0.56 | -1.01 | -1.11 | -0.2575 | -0.1975 | -0.4175 | 0.610442 | 0.705527 | 0.401105 | 0.999971 | 1 | 0.871816 |
| Cpne2 | P59108 | -1.08 | -1.86 | -1.37 | -1.25 | -1.58 | -1.71 | -2.12 | -1 | -1.36 | -1.42 | -2.32 | -1.34 | -1.11 | -1.37 | -0.89 | -1.71 | -0.2125 | -0.22 | 0.12 | 0.478867 | 0.468323 | 0.652987 | 0.999971 | 1 | 0.934005 |
| Cpne3 | Q8BT60 | -1.6 | -0.8 | -0.67 | -1.12 | -0.47 | -0.64 | -0.44 | -0.8 | -0.35 | -1.41 | -0.63 | -0.69 | -0.77 | -0.94 | -0.68 | -1.04 | 0.46 | 0.2775 | 0.19 | 0.093767 | 0.383173 | 0.456399 | 0.999971 | 1 | 0.887217 |
| Cpne4 | Q8BLR2 | -0.66 | -0.49 | -0.46 | -0.54 | -0.58 | -0.14 | -0.12 | -0.28 | -0.45 | -0.37 | -0.55 | -0.52 | -0.02 | -0.2 | -0.36 | -0.56 | 0.2575 | 0.065 | 0.2525 | 0.190478 | 0.688331 | 0.208619 | 0.999971 | 1 | 0.822138 |
| Cpne5 | E9Q1V2 | 1.34 | 0.63 | 0.7 | 1.83 | 0.46 | -0.34 | 0.77 | 1.41 | 0.37 | 1.01 | 0.76 | 0.76 | 0.7 | 0.26 | 0.84 | 1.66 | -0.55 | -0.4 | -0.26 | 0.227454 | 0.227342 | 0.512655 | 0.999971 | 1 | 0.905815 |
| Cpne5_1 | Q8JZW4 | 0.03 | 0.07 | -0.83 | -0.47 | 0.21 | -0.28 | -0.98 | -0.26 | -0.06 | 0.09 | -0.97 | -0.55 | 0.54 | 0.17 | -0.34 | -0.48 | -0.0275 | -0.0725 | 0.2725 | 0.932821 | 0.823113 | 0.406398 | 0.999971 | 1 | 0.874911 |
| Cpne6 | Q9Z140 | 1.68 | -0.37 | -0.38 | -0.92 | -0.23 | 0.02 | -0.38 | -0.53 | -0.03 | 1.82 | -0.5 | -0.54 | 0.42 | -0.17 | -0.55 | -0.72 | -0.2825 | 0.185 | -0.2575 | 0.60285 | 0.799083 | 0.655941 | 0.999971 | 1 | 0.934061 |
| Cpne7 | Q0VE82 | -0.47 | -0.61 | 0.06 | -0.28 | -0.82 | -0.64 | -0.44 | 0.6 | -0.44 | -0.28 | -0.18 | -0.29 | 0.25 | -0.24 | 0.34 | 0.04 | 1.96E-16 | 0.0275 | 0.4225 | 1 | 0.89312 | 0.09587 | 1 | 1 | 0.78268 |
| Cpne8 | Q9DC53 | -0.4 | -1.24 | -0.49 | -0.45 | -1.22 | -0.8 | -0.59 | 0.32 | -1.04 | -0.3 | -0.45 | -0.48 | -0.87 | -0.9 | -0.27 | 0.21 | 0.0725 | 0.0775 | 0.1875 | 0.844625 | 0.777205 | 0.575336 | 0.999971 | 1 | 0.91816 |
| Cpne9 | A0A0R4J0J | -0.3 | -0.53 | -0.6 | -1.01 | -1.35 | -0.79 | -1.15 | -1.02 | -1.1 | -0.86 | -0.59 | -0.85 | -0.57 | -0.83 | -0.97 | -0.24 | -0.4675 | -0.24 | -0.0425 | 0.06614 | 0.294054 | 0.863972 | 0.999971 | 1 | 0.983079 |
| Cpox | P36552 | -0.57 | -0.77 | -0.73 | -0.43 | -0.82 | -0.95 | -0.34 | -0.6 | -0.76 | -0.48 | -0.31 | -0.8 | -1.08 | -0.71 | -0.71 | -0.59 | -0.0525 | 0.0375 | -0.1475 | 0.800311 | 0.848754 | 0.458007 | 0.999971 | 1 | 0.888332 |
| Cpped1 | Q8BFS6 | 0.19 | 1.21 | 1.43 | 0.56 | 0.73 | 0.44 | 0.49 | 1.24 | 0.69 | 0.38 | 1.14 | 0.66 | 0.63 | 0.5 | 1.16 | 1.54 | -0.1225 | -0.13 | 0.11 | 0.717262 | 0.692004 | 0.763752 | 0.999971 | 1 | 0.962238 |
| Cpq | Q9WVJ3 | 0.93 | 0.26 | -0.04 | 0.24 | 0.27 | 0.14 | -0.11 | 0.01 | 0.29 | 0.67 | -0.07 | 0.52 | -0.42 | 0.65 | 0.23 | 0.69 | -0.27 | 0.005 | -0.06 | 0.295899 | 0.985569 | 0.855745 | 0.999971 | 1 | 0.982398 |
| Cpsf1 | Q9EPU4 | 1.09 | 1.12 | 1.22 | 1.45 | 1.14 | 0.96 | 0.92 | 1.42 | 0.91 | 0.96 | 1.33 | 1.12 | 0.88 | 1.22 | 1.34 | 1.38 | -0.11 | -0.14 | -0.015 | 0.583854 | 0.46439 | 0.940067 | 0.999971 | 1 | 0.995869 |
| Cpsf2 | O35218 | 0.89 | 2.2 | 1.89 | 1.67 | 1.96 | 2 | 1.45 | 1.66 | 1.8 | 1.37 | 1.89 | 2.13 | 2.43 | 2.16 | 1.99 | 2.15 | 0.105 | 0.135 | 0.52 | 0.737656 | 0.676888 | 0.112897 | 0.999971 | 1 | 0.78268 |
| Cpsf3 | Q9QXK7 | 0.85 | 1.8 | 1.89 | 1.94 | 1.71 | 1.39 | 1.25 | 1.77 | 1.4 | 1.29 | 1.33 | 1.58 | 1.71 | 1.21 | 1.46 | 2 | -0.09 | -0.22 | -0.025 | 0.761974 | 0.440744 | 0.936434 | 0.999971 | 1 | 0.995618 |
| Cpsf4 | E0CXT7 | 0 | 0.35 | 0.45 | 0.19 | 0.9 | -0.57 | 0.62 | 0.27 | 0.64 | -0.33 | 0.71 | 0.95 | 0.22 | -0.68 | 0.22 | 0.34 | 0.0575 | 0.245 | -0.2225 | 0.862606 | 0.430488 | 0.426835 | 0.999971 | 1 | 0.880793 |
| Cpsf6 | H3BJ30 | 2.9 | 2.6 | 2.03 | 3.01 | 2.38 | 2.5 | 2.47 | 2.3 | 2.23 | 3.02 | 2.07 | 1.87 | 1.66 | 2.76 | 2.01 | 2.21 | -0.2225 | -0.3375 | -0.475 | 0.387016 | 0.324942 | 0.163978 | 0.999971 | 1 | 0.800215 |
| Cpsf7 | Q8BTV2-2 | 1.75 | 3.05 | 2.25 | 2.4 | 3.1 | 1.97 | 3.57 | 2.28 | 3.02 | 1.78 | 2.06 | 2.13 | 2.71 | 2.12 | 1.72 | 1.83 | 0.3675 | -0.115 | -0.2675 | 0.402752 | 0.755813 | 0.44501 | 0.999971 | 1 | 0.884152 |
| Cpt1a | P97742 | -0.21 | -0.14 | -1.32 | -1.22 | -0.19 | 0.06 | -1.17 | -0.72 | -0.08 | -0.14 | -1.24 | -1.42 | 0.18 | -0.15 | -1.22 | -1.1 | 0.2175 | 0.0025 | 0.15 | 0.590501 | 0.995548 | 0.735554 | 0.999971 | 1 | 0.958637 |
| Cpt1c | Q8BGD5 | -0.3 | -0.67 | -0.14 | -0.04 | -0.68 | -0.74 | 0.02 | 0.04 | -0.72 | -0.38 | -0.08 | -0.37 | -0.51 | -0.67 | 0.01 | -0.01 | -0.0525 | -0.1 | -0.0075 | 0.847493 | 0.661257 | 0.976092 | 0.999971 | 1 | 0.99925 |
| Cpt2 | P52825 | -1.42 | -1.08 | -0.09 | -0.69 | -0.6 | -1.51 | -0.22 | 0.11 | -0.66 | -1.61 | -0.04 | -0.08 | -0.63 | -1.35 | -0.23 | -0.37 | 0.265 | 0.2225 | 0.175 | 0.539321 | 0.614831 | 0.637233 | 0.999971 | 1 | 0.93301 |
| Cptp | Z4YLU2 | 0.22 | 0.31 | 0.29 | -0.03 | 0.04 | 0.9 | -0.03 | 0.63 | -0.04 | -0.05 | -0.41 | -0.31 | -0.35 | 1.41 | 0.9 | 0.6 | 0.1875 | -0.4 | 0.4425 | 0.481192 | 0.057151 | 0.249269 | 0.999971 | 1 | 0.839012 |
| Cr1l | Q64735-2 | 0.08 | 0.32 | 0.19 | -0.03 | 0.58 | 0.48 | 0.83 | 0.53 | 0.47 | 0.08 | 0.53 | 0.58 | 0.67 | 0.4 | 0.53 | -0.31 | 0.465 | 0.275 | 0.1825 | 0.029732 | 0.18153 | 0.483086 | 0.999971 | 1 | 0.896345 |
| Crabp1 | P62965 | 0.39 | -0.24 | -0.08 | 1.14 | 0.98 | 0.11 | -0.55 | 1.7 | 1.27 | 0.93 | -0.22 | 0.41 | 1.01 | 0.59 | 0.43 | 0.56 | 0.2575 | 0.295 | 0.345 | 0.632767 | 0.494902 | 0.315866 | 0.999971 | 1 | 0.85297 |
| Cracd | E9Q5L4 | 0.33 | -1.03 | -0.04 | -0.3 | -0.58 | -0.69 | -0.05 | 0.01 | -0.5 | 0.21 | 0.09 | 0.32 | -0.65 | -0.6 | -0.23 | 0.14 | -0.0675 | 0.29 | -0.075 | 0.840931 | 0.400842 | 0.824946 | 0.999971 | 1 | 0.976309 |
| Cracdl | F7B0R9 | -0.01 | -0.91 | -1.82 | -1.22 | -0.92 | -1.35 | -1.72 | -2.05 | -1.4 | 0.18 | -2.16 | -1.35 | -0.89 | -0.95 | -0.38 | -0.66 | -0.52 | -0.1925 | 0.27 | 0.241227 | 0.736331 | 0.489337 | 0.999971 | 1 | 0.898346 |
| Cracdl_1 | E9Q3M9 | 0.54 | -0.24 | -0.35 | 0.01 | -0.23 | -0.08 | -0.22 | -0.07 | -0.3 | 0.36 | -0.62 | -0.62 | -0.31 | -0.24 | -0.38 | -0.23 | -0.14 | -0.285 | -0.28 | 0.5588 | 0.368807 | 0.255966 | 0.999971 | 1 | 0.839012 |
| Cradd | O88843 | -1.42 | -0.33 | -1.37 | -1.22 | -0.04 | -0.28 | -1.71 | -0.86 | -0.32 | -1.73 | -1.4 | -0.84 | -0.85 | -0.4 | -0.84 | -0.76 | 0.3625 | 0.0125 | 0.3725 | 0.405173 | 0.974257 | 0.219626 | 0.999971 | 1 | 0.827064 |
| Crat | H7BX88 | -0.07 | -0.13 | -0.3 | -0.52 | -0.11 | -0.06 | -0.07 | -0.35 | -0.07 | -0.05 | -0.21 | 0.13 | 0.12 | -0.21 | -0.42 | -0.5 | 0.1075 | 0.205 | 0.0025 | 0.574338 | 0.290217 | 0.990831 | 0.999971 | 1 | 0.99925 |
| Crbn | Q8C7D2-2 | -0.01 | 0.44 | 0.58 | 0.27 | 0.3 | 0.33 | 0.52 | 0.26 | 0.56 | -0.06 | 0.61 | 0.81 | -0.09 | 0.04 | 0.49 | 0.25 | 0.0325 | 0.16 | -0.1475 | 0.8701 | 0.529955 | 0.513316 | 0.999971 | 1 | 0.905815 |
| Creb1 | Q01147-1 | 1.85 | 1.79 | 1.38 | 2.26 | 1.55 | 1.66 | 1.65 | 2.24 | 1.62 | 1.93 | 2.05 | 2.15 | 1.1 | 1.83 | 2 | 2.59 | -0.045 | 0.1175 | 0.06 | 0.863434 | 0.630414 | 0.864012 | 0.999971 | 1 | 0.983079 |
| Crebbp | A0A0U1RQ | 3.84 | 2.65 | 1.56 | 1.67 | 2.45 | 2.55 | 1.28 | 1.33 | 2 | 2.29 | 1.5 | 1.69 | 1.6 | 2.17 | 1.48 | 1.57 | -0.5275 | -0.56 | -0.725 | 0.373795 | 0.296444 | 0.181345 | 0.999971 | 1 | 0.808532 |
| Creg1 | K4DI63 | 4.3 | 0.69 | 0.94 | 1.14 | 0.63 | 2.81 | 1.69 | 0.94 | 1.15 | 4.33 | 0.19 | -0.01 | -1.9 | -0.17 | 0.08 | 0.1 | -0.25 | -0.3525 | -2.24 | 0.77558 | 0.765276 | 0.02902 | 0.999971 | 1 | 0.678474 |
| Creg2 | Q8BGC9 | 5.04 | 1.29 | 1.3 | 2.09 | 1.14 | 2.2 | 2.45 | 0.68 | 0.73 | 3.37 | 0.73 | 0.06 | -1.94 | -0.51 | -0.23 | 0.01 | -0.8125 | -1.2075 | -3.0975 | 0.368816 | 0.26279 | 0.006754 | 0.999971 | 1 | 0.582714 |
| Creld1 | Q91XD7 | 0.63 | 0.51 | 1.93 | 2.13 | 0.59 | 0.7 | 2.07 | 1.73 | 0.86 | 0.16 | 1.72 | 0.34 | 0.06 | -0.2 | 1.75 | 1.25 | -0.0275 | -0.53 | -0.585 | 0.957736 | 0.315312 | 0.326587 | 0.999971 | 1 | 0.856472 |
| Creld2 | Q9CYA0 | -0.35 | 1.58 | 0.84 | -0.75 | 1.36 | 1.3 | 1.22 | 0.09 | 1.55 | -0.22 | 0.88 | 0.53 | 1.62 | 0.36 | 0.58 | -1.29 | 0.6625 | 0.355 | -0.0125 | 0.259291 | 0.556487 | 0.9863 | 0.999971 | 1 | 0.99925 |
| Crh | Q8CIT0 | 0.34 | -0.86 | -2.07 | -1.33 | -1.31 | -0.85 | -1.44 | -2.44 | -1.31 | -0.09 | -2.11 | -2.56 | -1.57 | -1.14 | -2 | -2.05 | -0.53 | -0.5375 | -0.71 | 0.353692 | 0.434839 | 0.186757 | 0.999971 | 1 | 0.809025 |
| Crhbp | A0A217FL6 | -1.13 | -0.44 | 0.49 | -0.61 | -0.42 | -0.68 | 0.39 | -0.83 | -0.45 | -1.53 | -0.03 | 0.18 | -0.73 | -1.05 | -0.76 | -0.48 | 0.0375 | -0.035 | -0.3325 | 0.927459 | 0.94141 | 0.357038 | 0.999971 | 1 | 0.868793 |
| Crhr1 | P35347 | 0.82 | 1.21 | -1.03 | -1.24 | 0.95 | 0.77 | -1.38 | -1.64 | 0.48 | 0.55 | -0.41 | -0.97 | -1.26 | 0 | -2.13 | -1.09 | -0.265 | -0.0275 | -1.06 | 0.751324 | 0.966876 | 0.15206 | 0.999971 | 1 | 0.795578 |
| Crip1 | A0A0G2JEK | 4.25 | -1.51 | 1.26 | 3.2 | 4.06 | 4.13 | 2.4 | 0.31 | -0.17 | 4.64 | 2.71 | 3.03 | -6.16 | -0.32 | 0.76 | -0.19 | 0.925 | 0.7525 | -3.2775 | 0.507151 | 0.604525 | 0.095129 | 0.999971 | 1 | 0.78268 |
| Crip2 | Q9DCT8 | 0.86 | 1.74 | 0.89 | 0.67 | 1.93 | 1.51 | 0.88 | 0.82 | 1.58 | 1.26 | 0.69 | 1.35 | 1.38 | 1.53 | 1.28 | 0.14 | 0.245 | 0.18 | 0.0425 | 0.490709 | 0.564173 | 0.91181 | 0.999971 | 1 | 0.992551 |
| Cript | O70333 | 2.82 | 0.69 | 1.71 | 2.57 | 0.27 | 2 | 2.68 | 3.37 | 0 | 2.41 | 0.91 | -0.36 | -1.87 | 2.67 | 2.84 | 2.08 | 0.1325 | -1.2075 | -0.5175 | 0.857916 | 0.117738 | 0.634316 | 0.999971 | 1 | 0.93301 |
| Crk | Q64010 | -0.67 | -0.28 | 0.65 | 0.28 | -0.28 | -0.38 | 0.7 | 0.28 | 0.01 | -0.6 | 0.65 | 0.65 | -0.25 | -0.51 | 0.53 | 0.01 | 0.085 | 0.1825 | -0.05 | 0.820406 | 0.651325 | 0.88939 | 0.999971 | 1 | 0.987681 |
| Crk_1 | Q8JZR2 | -0.45 | -0.74 | -0.34 | -1.07 | -0.33 | 0.11 | -0.55 | -0.08 | -0.1 | 0.08 | -0.8 | -1.01 | -1.02 | 0.15 | -0.48 | 0.35 | 0.4375 | 0.1925 | 0.4 | 0.10396 | 0.544579 | 0.267982 | 0.999971 | 1 | 0.842746 |
| Crkl | P47941 | 0.11 | 0.34 | 0.14 | -0.07 | 0.66 | 0.33 | 0.06 | 0.03 | 0.35 | 0.12 | 0.33 | 0.37 | 0.73 | 0.38 | 0 | 0.19 | 0.14 | 0.1625 | 0.195 | 0.521152 | 0.371089 | 0.388829 | 0.999971 | 1 | 0.871816 |
| Crlf3 | Q9Z2L7 | 1.52 | 0.48 | -0.45 | 0.49 | 0.6 | 0.94 | 0.23 | -0.07 | 0.42 | 1.39 | 0.23 | -0.43 | -0.02 | 0.88 | -0.09 | 0.2 | -0.085 | -0.1075 | -0.2675 | 0.844291 | 0.833799 | 0.542278 | 0.999971 | 1 | 0.911119 |
| Crls1 | E9PUW6 | 0.76 | 0.55 | 0.59 | 1.23 | -3.1 | -0.18 | -0.75 | 1.9 | -0.12 | 1.86 | -0.39 | -0.85 | 0.7 | -1.55 | 0.03 | 1.53 | -1.315 | -0.6575 | -0.605 | 0.181363 | 0.266809 | 0.33705 | 0.999971 | 1 | 0.858827 |
| Crmp1 | P97427 | -0.66 | -3.8 | -4.29 | -3.4 | -4.2 | -3.88 | -2.19 | -4.3 | -4.32 | -2.15 | -4.89 | -5.28 | -5.57 | -4.01 | -4.41 | -3.46 | -0.605 | -1.1225 | -1.325 | 0.484048 | 0.263539 | 0.138781 | 0.999971 | 1 | 0.784948 |
| Crmp1_1 | Q6P1J1 | -0.85 | -0.88 | -0.6 | -1.19 | -0.65 | -1.17 | -0.19 | -1.14 | -0.41 | -1.09 | -0.54 | -0.14 | -0.54 | -1.08 | -0.76 | -1 | 0.0925 | 0.335 | 0.035 | 0.739894 | 0.213944 | 0.872532 | 0.999971 | 1 | 0.984109 |
| Crnkl1 | P63154 | -0.17 | 0.61 | 0.51 | 0.54 | 0.66 | 0.53 | 0.16 | 0.47 | 0.42 | 0.13 | 0.6 | 1.05 | -0.32 | 0.78 | 0.84 | 0.84 | 0.0825 | 0.1775 | 0.1625 | 0.734149 | 0.530161 | 0.631568 | 0.999971 | 1 | 0.932858 |
| Crocc | Q8CJ40 | 1.12 | 0.45 | 0.34 | 0.6 | 0.19 | 0.23 | 0.41 | 0.41 | 0.25 | 0.88 | 0.36 | 0.16 | -0.22 | 0.47 | 0.24 | 0.13 | -0.3175 | -0.215 | -0.4725 | 0.180183 | 0.414922 | 0.088286 | 0.999971 | 1 | 0.766425 |
| Crocc_1 | Q8CJ40-2 | -0.03 | -1.04 | 3.81 | 3.19 | 1.54 | -0.21 | 1.86 | 3.83 | -0.21 | 0.97 | 4.77 | 5.79 | -1.63 | 0.97 | 4.2 | 3.68 | 0.2725 | 1.3475 | 0.3225 | 0.832332 | 0.429746 | 0.839684 | 0.999971 | 1 | 0.979728 |
| Crocc2 | F6XLV1 | -1.05 | -1.16 | -0.81 | -0.51 | -0.29 | 0.27 | -0.77 | -0.66 | -0.27 | -0.16 | -1.7 | -1.33 | 1.46 | 0.63 | -0.33 | -0.63 | 0.52 | 0.0175 | 1.165 | 0.098067 | 0.964522 | 0.031845 | 0.999971 | 1 | 0.681307 |
| Crot | Q9DC50 | -0.19 | -1.06 | -0.94 | -0.57 | -0.96 | -0.56 | -0.46 | -0.2 | -0.79 | -0.33 | -0.32 | -0.82 | -0.4 | -0.76 | -0.76 | -0.31 | 0.145 | 0.125 | 0.1325 | 0.595523 | 0.634885 | 0.607062 | 0.999971 | 1 | 0.925193 |
| Crppa | A0A1Y7VM | -0.23 | 0.49 | 0.92 | 0.48 | 0.31 | 0.23 | 0.62 | 1.12 | 0.3 | -0.08 | 0.57 | 1.1 | 0.25 | 0.25 | 1.04 | 0.59 | 0.155 | 0.0575 | 0.1175 | 0.625591 | 0.865813 | 0.704982 | 0.999971 | 1 | 0.950182 |
| Crtac1 | Q8R555 | -1.95 | -0.74 | -0.73 | -0.9 | -0.51 | -1.11 | -0.84 | -0.26 | -0.39 | -1.52 | -0.67 | -0.36 | 0.02 | -1.1 | -0.39 | -0.88 | 0.4 | 0.345 | 0.4925 | 0.26168 | 0.381494 | 0.210584 | 0.999971 | 1 | 0.822657 |
| Crtc1 | Q68ED7 | 0.14 | 0.26 | 0.02 | 0.51 | 0.03 | 0.13 | -0.08 | 0.26 | -0.28 | -0.42 | -0.45 | 0.02 | -0.16 | 0.03 | 0.05 | 0.31 | -0.1475 | -0.515 | -0.175 | 0.451045 | 0.029561 | 0.39466 | 0.999971 | 1 | 0.871816 |
| Crtc2 | Q3U182 | -1.02 | -0.44 | 1.24 | 0.33 | -0.33 | -0.11 | 2.7 | 0.98 | -0.78 | -1.27 | 1.74 | 0.67 | -2.1 | -0.64 | 0.99 | 0.19 | 0.7825 | 0.0625 | -0.4175 | 0.320036 | 0.934819 | 0.577105 | 0.999971 | 1 | 0.918236 |
| Crtc3 | Q91X84 | -1.77 | 0.23 | -1.24 | -2.64 | 0.4 | -0.53 | 1.62 | -1.93 | 0.04 | -1.51 | -0.16 | -0.14 | -0.19 | -1.3 | -1.1 | -3.34 | 1.245 | 0.9125 | -0.1275 | 0.173125 | 0.179245 | 0.874404 | 0.999971 | 1 | 0.984109 |
| Cry2 | Q9R194 | 2.01 | 1.08 | 1.05 | 1.69 | 1.39 | 1.45 | 1.33 | 1.48 | 0.61 | 1.78 | 1.14 | 1.12 | 0.46 | 1.59 | 1.45 | 1.71 | -0.045 | -0.295 | -0.155 | 0.863209 | 0.388509 | 0.670213 | 0.999971 | 1 | 0.938809 |
| Cryab | P23927 | 2.14 | 0.65 | 1.39 | 1.1 | 0.53 | 0.84 | 0.6 | 0.26 | -0.49 | -0.29 | -1.67 | -0.85 | -0.65 | 0.82 | 0.41 | -0.53 | -0.7625 | -2.145 | -1.3075 | 0.044968 | 0.000683 | 0.016304 | 0.999971 | 0.448349 | 0.613095 |
| Crybb1 | E9PYP8 | -0.06 | 0.83 | 0.43 | 1.03 | 0.51 | 0.15 | 0.04 | 1.11 | 1.06 | 0.15 | 0.8 | 1.51 | 0.92 | 0.56 | 0.67 | 1.75 | -0.105 | 0.3225 | 0.4175 | 0.756335 | 0.386023 | 0.258197 | 0.999971 | 1 | 0.839012 |
| Crybg3 | A0A338P6I | 2.12 | -0.74 | -0.51 | -0.02 | -0.1 | -0.88 | -0.24 | -0.58 | -0.58 | 2.1 | 0.11 | 0.45 | -0.23 | 0.95 | 0.68 | 0.11 | -0.6625 | 0.3075 | 0.165 | 0.298419 | 0.69615 | 0.797516 | 0.999971 | 1 | 0.970479 |

| | | | | | | | | | | | | | | | | | | | | | | | | | | |
|---|---|---|---|---|---|---|---|---|---|---|---|---|---|---|---|---|---|---|---|---|---|---|---|---|---|---|
| Cryl1 | Q99KP3 | -0.03 | 1.05 | 0.59 | 0.14 | 0.89 | 0.91 | 1.1 | 0.33 | 1.15 | 0.23 | 0.94 | 0.83 | 0.25 | 0.93 | -0.06 | -0.2 | 0.37 | 0.35 | -0.2075 | 0.240288 | 0.284883 | 0.552043 | 0.999971 | 1 | 0.913595 |
| Crym | O54983 | -0.02 | 0.07 | 0.49 | 0.68 | 0.44 | 0.25 | 0.67 | 1.29 | 0.49 | 0.27 | 0.37 | 0.53 | 0.11 | 0.42 | 1.09 | 0.61 | 0.3575 | 0.11 | 0.2525 | 0.241365 | 0.617424 | 0.379843 | 0.999971 | 1 | 0.871816 |
| Cryz | P47199 | -0.68 | -0.74 | 0.38 | 0.52 | -0.27 | -0.62 | 0.72 | 0.95 | -0.56 | -0.74 | 0.09 | 0.11 | -0.93 | -0.28 | 0.19 | -0.04 | 0.325 | -0.145 | -0.135 | 0.497764 | 0.709082 | 0.734644 | 0.999971 | 1 | 0.957874 |
| Cryzl1 | Q921W4 | -0.57 | 0.99 | 0.25 | -0.8 | 0.77 | 0.93 | 0.2 | -0.23 | 0.97 | -0.53 | 0.32 | -0.01 | 0.93 | 0.47 | 0 | -0.76 | 0.45 | 0.22 | 0.1925 | 0.338601 | 0.649954 | 0.704436 | 0.999971 | 1 | 0.950182 |
| Cryzl2 | Q3UNZ8 | -0.34 | -0.69 | -1.09 | -0.94 | -0.57 | -0.23 | -0.95 | -0.17 | -0.48 | -0.49 | -1.4 | -1.01 | -0.05 | 0.03 | -0.68 | -0.86 | 0.285 | -0.08 | 0.375 | 0.297915 | 0.78142 | 0.217263 | 0.999971 | 1 | 0.826893 |
| Cs | Q9CZU6 | -0.83 | -0.95 | -0.93 | -0.82 | -1.15 | -1.09 | -1.02 | -0.77 | -1.2 | -0.98 | -0.95 | -0.98 | -1.16 | -0.84 | -0.78 | -0.63 | -0.125 | -0.145 | 0.03 | 0.482623 | 0.386553 | 0.87306 | 0.999971 | 1 | 0.984109 |
| Csad | Q9DBE0 | 0.23 | 0.41 | 0.33 | 0.24 | 0.58 | 0.28 | -0.04 | -0.82 | 0.66 | 0.11 | 0.07 | 0.16 | 0.39 | -0.21 | 0.03 | 0.2 | -0.3025 | -0.0525 | -0.2 | 0.341885 | 0.791921 | 0.324002 | 0.999971 | 1 | 0.85548 |
| Csdc2 | Q91YQ3 | 0.07 | 0.24 | 1.78 | 1.01 | 0.27 | -0.16 | 0.71 | 0.43 | 0.3 | 0.7 | 2.07 | 0.57 | 0.63 | 0.42 | 1.61 | 1.24 | -0.4625 | 0.135 | 0.2 | 0.278606 | 0.794439 | 0.658574 | 0.999971 | 1 | 0.934638 |
| Csde1 | Q91W50 | 0.33 | 1.07 | 1.2 | 1.17 | 0.84 | 1.12 | 1.13 | 1.39 | 1.1 | 0.85 | 1.46 | 1.38 | 0.92 | 1.31 | 1.41 | 1.6 | 0.1775 | 0.255 | 0.3675 | 0.498487 | 0.353241 | 0.197928 | 0.999971 | 1 | 0.813335 |
| Cse1l | Q9ERK4 | 0.03 | 0.71 | 1.01 | 0.58 | 0.79 | 0.64 | 0.98 | 0.56 | 0.91 | 0.18 | 0.93 | 1 | 0.67 | 0.48 | 0.84 | 0.7 | 0.16 | 0.1725 | 0.09 | 0.530082 | 0.558072 | 0.717387 | 0.999971 | 1 | 0.952711 |
| Csf1r | P09581 | -0.25 | 0.5 | 0.12 | 0.74 | 0.89 | 0.6 | 0.36 | 0.7 | 0.38 | 0.03 | 0.73 | 0.78 | 0.09 | 1.11 | 0.74 | 0.91 | 0.36 | 0.2025 | 0.435 | 0.197429 | 0.490552 | 0.190347 | 0.999971 | 1 | 0.810547 |
| Csk | P41241 | -0.8 | -0.32 | -0.81 | -0.96 | -0.21 | -0.29 | -0.17 | -0.71 | -0.13 | -0.5 | -0.62 | -0.34 | -0.04 | -0.01 | -0.42 | -0.54 | 0.3775 | 0.325 | 0.47 | 0.122528 | 0.16007 | 0.068194 | 0.999971 | 1 | 0.733937 |
| Csl | Q80X68 | -0.61 | -0.78 | 0.36 | -0.44 | -0.83 | -1.22 | 1.08 | -0.51 | -0.53 | -1.18 | 2.54 | 1.66 | -0.42 | -1.26 | -0.05 | -1.41 | -0.0025 | 0.99 | -0.4175 | 0.996166 | 0.252191 | 0.308569 | 1 | 1 | 0.847402 |
| Csmd1 | Q923L3 | 0.08 | -0.44 | -0.4 | -0.36 | -0.58 | -0.47 | -0.83 | -0.18 | -0.48 | -0.34 | -0.29 | -0.21 | -0.14 | -0.85 | -0.06 | -0.03 | -0.235 | -0.05 | 0.01 | 0.307881 | 0.795747 | 0.968766 | 0.999971 | 1 | 0.998787 |
| Csmd2 | V9GX34 | -0.25 | -0.43 | -0.77 | -0.68 | -0.48 | -0.39 | -0.43 | -0.72 | -0.59 | -0.4 | -0.5 | -1.08 | 0.02 | -0.55 | -0.72 | -0.58 | 0.0275 | -0.11 | 0.075 | 0.889999 | 0.631683 | 0.752283 | 0.999971 | 1 | 0.961116 |
| Csmd3 | E9PWJ0 | -0.01 | -0.15 | -0.8 | -0.35 | -0.22 | -0.47 | -0.59 | -0.57 | -0.22 | 0.04 | -0.88 | -0.99 | 0.12 | -0.03 | -0.48 | -0.65 | -0.135 | -0.185 | 0.0675 | 0.561204 | 0.553524 | 0.803229 | 0.999971 | 1 | 0.971467 |
| Csnk1a1 | E9Q4G7 | -0.24 | 1.23 | 1.17 | 1.23 | 0.65 | 1.97 | 2.29 | -0.04 | 1.44 | 2.69 | -0.29 | 0.04 | 2.7 | -1.74 | 0.05 | -0.3 | 0.37 | 0.1225 | -0.67 | 0.542138 | 0.862084 | 0.459791 | 0.999971 | 1 | 0.889706 |
| Csnk1a1_1 | Q8BK63-2 | -0.6 | -0.21 | -0.1 | -0.12 | -0.12 | 0.01 | -0.37 | 0.02 | -0.22 | -0.37 | 0.26 | 0.2 | 0.39 | 0.02 | 0.17 | 0.37 | 0.1425 | 0.225 | 0.495 | 0.489688 | 0.340546 | 0.035188 | 0.999971 | 1 | 0.69205 |
| Csnk1d | Q9DC28-2 | 1.17 | -1.19 | -0.59 | -0.81 | -1.62 | -1.35 | -0.23 | -1.03 | -0.95 | 1.58 | -0.25 | 0.13 | -1.74 | -0.92 | -0.14 | -0.38 | -0.7025 | 0.4825 | -0.44 | 0.22615 | 0.485041 | 0.455221 | 0.999971 | 1 | 0.887217 |
| Csnk1d_1 | Q9DC28 | -0.51 | -0.54 | -0.32 | -0.37 | -0.58 | -0.66 | -0.24 | -0.19 | -0.54 | -0.85 | -0.18 | -0.89 | -0.33 | -0.56 | -0.21 | -0.04 | 0.0175 | -0.18 | 0.15 | 0.927979 | 0.414303 | 0.438792 | 0.999971 | 1 | 0.883105 |
| Csnk1e | Q9JMK2 | 0.27 | -0.15 | -1.25 | -1.22 | -0.26 | -0.4 | -0.76 | -1.56 | -0.64 | 0.21 | -1.05 | -0.9 | 0.2 | -0.49 | -1.31 | -0.78 | -0.1575 | -0.0075 | -0.0075 | 0.728177 | 0.98665 | 0.987113 | 0.999971 | 1 | 0.99925 |
| Csnk1g1 | A0A0U1RP | -0.89 | -1.45 | -0.83 | -0.81 | -1.03 | -1.91 | -0.31 | -1.01 | -1.04 | -1.24 | -0.79 | -1.21 | -0.89 | -1.77 | -1.15 | -0.47 | -0.07 | -0.075 | -0.075 | 0.843317 | 0.737342 | 0.812648 | 0.999971 | 1 | 0.974458 |
| Csnk1g2 | Q99K78 | 0.83 | -0.07 | -0.54 | -0.28 | -0.33 | 0.39 | -0.87 | -0.65 | -0.44 | 0.84 | -0.7 | -0.79 | -0.15 | -0.75 | -1.06 | -0.37 | -0.35 | -0.2575 | -0.5675 | 0.379789 | 0.573587 | 0.135148 | 0.999971 | 1 | 0.784948 |
| Csnk1g3 | Q8C4X2 | -1.52 | -0.83 | -2.07 | -1.99 | -0.99 | -1.18 | -2.07 | -1.97 | -0.93 | -1.63 | -2.04 | -2.13 | -0.74 | -1.27 | -1.99 | -2.28 | 0.05 | -0.08 | 0.0325 | 0.895452 | 0.833999 | 0.939241 | 0.999971 | 1 | 0.995869 |
| Csnk2a1 | Q60737 | -0.99 | -1.01 | -0.7 | -0.78 | -0.76 | -1.06 | -0.82 | -0.61 | -0.85 | -0.86 | -0.6 | -0.61 | -0.32 | -0.91 | -0.91 | -0.62 | 0.0575 | 0.14 | 0.18 | 0.761783 | 0.441844 | 0.404524 | 0.999971 | 1 | 0.87384 |
| Csnk2a2 | O54833 | -0.98 | -0.43 | -0.46 | -1.05 | -0.07 | -0.6 | -0.29 | -0.73 | -0.12 | -1.37 | -0.01 | 0.21 | 0.39 | -1.06 | -0.31 | -0.94 | 0.3075 | 0.4075 | 0.25 | 0.239792 | 0.299681 | 0.497484 | 0.999971 | 1 | 0.902655 |
| Csnk2b | P67871 | -0.39 | 0.01 | -0.15 | -0.15 | 0.15 | 0.15 | -0.15 | 0.09 | 0.39 | -0.49 | 0.01 | 0.08 | 1.07 | -0.07 | -0.21 | -0.1 | 0.23 | 0.1675 | 0.3425 | 0.231092 | 0.479707 | 0.295011 | 0.999971 | 1 | 0.847402 |
| Cspg4 | Q8VHY0 | -0.87 | -0.88 | -1.38 | -1.01 | -0.21 | -0.89 | -1.29 | -1.22 | -0.4 | -0.42 | -0.97 | -0.54 | -0.39 | -0.57 | -1.04 | -0.8 | 0.1325 | 0.4525 | 0.335 | 0.646175 | 0.065183 | 0.163622 | 0.999971 | 1 | 0.800215 |
| Cspg5 | Q71M36-2 | -2.39 | -3.75 | -3.51 | -2.84 | -2.65 | -3.38 | -2.31 | -2.58 | -3.74 | -1.88 | -3.63 | -3.28 | -4.04 | -2.99 | -3.57 | -1.81 | 0.3925 | -0.01 | 0.02 | 0.309188 | 0.983839 | 0.969828 | 0.999971 | 1 | 0.998787 |
| Cspg5_1 | Q71M36 | -1.05 | 0.36 | -0.58 | -0.65 | -0.31 | 0.17 | -0.76 | -0.43 | 0.22 | -0.08 | -0.42 | -0.52 | 0.71 | 0.12 | -0.4 | 0.04 | 0.1475 | 0.28 | 0.5975 | 0.674152 | 0.419004 | 0.129945 | 0.999971 | 1 | 0.784948 |
| Cspp1 | D3Z378 | 0.08 | 0.78 | 0.84 | 0.16 | -0.29 | -1.07 | 0.33 | 1.28 | -0.64 | 0.57 | 0.35 | 0.71 | -0.62 | 0.89 | 0.83 | 1.43 | -0.4025 | -0.2175 | 0.1675 | 0.426722 | 0.546916 | 0.712645 | 0.999971 | 1 | 0.951793 |
| Csrp1 | P97315 | -0.48 | 1.11 | 0.9 | 0.57 | 1.49 | 1.66 | 0.49 | 0.46 | 0.8 | -0.07 | 1.02 | 0.98 | 0.65 | 0.83 | 1.02 | 0.49 | 0.5 | 0.1575 | 0.2225 | 0.281091 | 0.705513 | 0.543794 | 0.999971 | 1 | 0.911398 |
| Csrp2 | P97314 | 0.05 | -0.62 | 0.61 | 1.68 | 4.8 | 2.74 | 0.82 | -0.02 | 1.49 | 2.55 | 4.55 | 5.77 | -2.53 | 0.67 | 1.47 | 0.74 | 1.655 | 3.16 | -0.3425 | 0.141571 | 0.00982 | 0.707533 | 0.999971 | 1 | 0.950182 |
| Cst3 | P21460 | 0.62 | 0.09 | -0.57 | -1.18 | 0.76 | 0.09 | 0.6 | 0.46 | 0.78 | 0.48 | 0.39 | 0.57 | 0.1 | -0.13 | -0.47 | -0.97 | 0.7375 | 0.815 | -0.1075 | 0.091592 | 0.060602 | 0.803219 | 0.999971 | 1 | 0.971467 |
| Cstb | Q62426 | 1.11 | 0.71 | 1.26 | 1.22 | 0.27 | 0.75 | 0.44 | 0.65 | 0.3 | 1.07 | 0.23 | 0.79 | -0.26 | 0.99 | 0.54 | 1.22 | -0.5475 | -0.4775 | -0.4525 | 0.028576 | 0.092699 | 0.212674 | 0.999971 | 1 | 0.823987 |
| Cstf1 | Q99LC2 | 0.75 | 1.41 | 1.62 | 1.49 | 1.34 | 1.37 | 0.82 | 1.62 | 1.32 | 1.36 | 1.39 | 0.38 | 0.44 | 1.71 | 1.17 | 1.8 | -0.03 | -0.205 | -0.0375 | 0.912851 | 0.520742 | 0.917013 | 0.999971 | 1 | 0.992633 |
| Cstf2 | Q8BIQ5-2 | 0.7 | 1.14 | 1.52 | 1.71 | 0.84 | 0.98 | 1.48 | 1.57 | 0.65 | 0.64 | 1.27 | 1.43 | 0.72 | 0.93 | 1.62 | 1.91 | -0.05 | -0.27 | 0.0275 | 0.866168 | 0.392707 | 0.937829 | 0.999971 | 1 | 0.995618 |
| Cstf2_1 | F6ZKC7 | 1.23 | 3.03 | 1.06 | 1.51 | 2.19 | 2.63 | 0.2 | 2.59 | 3.16 | 1.89 | 0.27 | 2.18 | 3.37 | 3.57 | 1.8 | 3.38 | 0.195 | 0.1675 | 1.3225 | 0.769009 | 0.806611 | 0.040399 | 0.999971 | 1 | 0.700838 |
| Cstf2t | Q8C7E9 | 1.55 | 0.97 | 0.71 | 0.9 | 0.5 | 1.07 | 1.15 | 0.53 | 0.11 | 0.99 | 0.34 | 0.55 | -0.1 | 0.73 | 0.49 | 0.22 | -0.22 | -0.535 | -0.6975 | 0.423543 | 0.079619 | 0.029153 | 0.999971 | 1 | 0.678474 |
| Cstf3 | Q99LI7 | 0.69 | 1.99 | 1.74 | 1.48 | 1.86 | 1.57 | 1.43 | 1.3 | 1.79 | 0.92 | 2.02 | 1.65 | 1.83 | 1.63 | 1.43 | 1.9 | 0.065 | 0.12 | 0.2225 | 0.834504 | 0.739642 | 0.47631 | 0.999971 | 1 | 0.894333 |
| Cstpp1 | Q8BHR8 | 0.21 | 0.16 | -0.09 | 0.16 | 0.39 | 0.58 | -0.6 | 0.23 | 0.25 | 0.34 | -0.44 | -0.16 | 0.8 | 1.04 | 0.17 | 0.59 | 0.04 | -0.1125 | 0.54 | 0.887565 | 0.626298 | 0.043238 | 0.999971 | 1 | 0.701725 |
| Ctbp1 | O88712 | -0.33 | -1.19 | -0.93 | -0.98 | -1.21 | -0.93 | -1.21 | -1.29 | -1.11 | -0.76 | -1.43 | -1.33 | -1.54 | -0.78 | -1.12 | -0.93 | -0.3025 | -0.3 | -0.235 | 0.220932 | 0.265301 | 0.391725 | 0.999971 | 1 | 0.871816 |
| Ctbp2 | Q91YZ2 | 0.38 | 0.26 | 0.63 | 0.66 | 0.06 | 0.15 | 0.48 | 1.01 | 0.09 | 0.39 | 0.17 | -0.01 | -0.41 | 0.21 | 0.44 | 0.09 | -0.0575 | -0.3225 | -0.4 | 0.82474 | 0.11785 | 0.1207 | 0.999971 | 1 | 0.78268 |
| Ctbs | Q8R242 | 0.29 | 1.7 | 1.69 | 1.82 | 2.23 | 1.11 | 1.93 | 1.74 | 1.95 | 0.68 | 1.22 | 1.5 | 2.26 | 0.86 | 1.57 | 1.19 | 0.3775 | -0.0375 | 0.095 | 0.371342 | 0.92997 | 0.830693 | 0.999971 | 1 | 0.978535 |
| Ctc1 | Q5SUQ9-2 | 3.41 | 1.93 | 1.38 | 1.94 | 1.52 | 2.26 | 1.77 | 1.8 | 1.59 | 2.7 | 1.63 | 1.3 | 0.63 | 2.15 | 1.21 | 1.38 | -0.3275 | -0.36 | -0.8225 | 0.459331 | 0.47549 | 0.125997 | 0.999971 | 1 | 0.783071 |
| Ctcf | Q61164 | -0.13 | 1.82 | 1 | -0.17 | 3.23 | 1.1 | 2.11 | 0.15 | 4.14 | 0.67 | 0.37 | 1.23 | 5.27 | 0.61 | 0.33 | -0.22 | 1.0175 | 0.9725 | 0.8675 | 0.191312 | 0.292519 | 0.479424 | 0.999971 | 1 | 0.894438 |
| Ctdnep1 | Q3TP92 | -2.28 | 0.73 | 0.05 | -1.2 | -0.79 | 0.21 | -0.25 | -0.01 | 0.37 | -1.51 | 1 | 1.34 | 2.72 | 0.17 | 0.13 | -0.55 | 0.465 | 0.975 | 1.2925 | 0.472548 | 0.261174 | 0.167526 | 0.999971 | 1 | 0.800215 |
| Ctdp1 | Q7TSG2 | 1.62 | 1.5 | 1.61 | 1.51 | 1.21 | 1.3 | 1.61 | 1.72 | 1.02 | 1.42 | 1.37 | 1.35 | 0.98 | 1.51 | 1.31 | 1.28 | -0.1 | -0.27 | -0.29 | 0.604689 | 0.148728 | 0.146155 | 0.999971 | 1 | 0.791767 |
| Ctdsp1 | P58466 | -0.98 | -0.93 | -0.34 | -0.85 | -1.45 | -1.64 | 0.05 | -0.21 | -1.83 | -1.21 | -0.65 | -0.23 | -0.99 | -0.66 | -0.4 | -0.48 | -0.0375 | -0.205 | 0.1425 | 0.930119 | 0.580219 | 0.546928 | 0.999971 | 1 | 0.911447 |
| Ctdspl | G3UXN0 | -0.54 | -0.67 | -0.64 | 0.03 | -0.6 | -1.25 | -0.89 | 0.25 | -0.64 | -0.72 | 0.05 | 0.08 | -0.11 | -0.65 | -0.51 | 1.17 | -0.1675 | 0.1475 | 0.43 | 0.636665 | 0.606061 | 0.324208 | 0.999971 | 1 | 0.85548 |
| Ctdspl2 | Q8BG15-6 | -1.85 | -1.78 | -1.05 | -1.75 | -1.93 | -2.14 | -1.26 | -1.57 | -1.33 | -1.47 | -0.79 | -1.07 | -1.88 | -1.91 | -0.89 | -1.21 | -0.1175 | 0.4425 | 0.135 | 0.68029 | 0.117379 | 0.672567 | 0.999971 | 1 | 0.940005 |
| Cth | Q8VCN5 | 0.37 | 0.02 | -0.27 | 0.22 | -0.08 | 0.52 | -0.22 | 0.59 | 0.04 | 0.42 | -0.2 | -0.09 | -0.38 | 0.48 | -0.41 | 0.29 | 0.1175 | -0.0425 | -0.09 | 0.662884 | 0.852771 | 0.750615 | 0.999971 | 1 | 0.961064 |
| Ctif | E9Q1U6 | 1.08 | 1.14 | 0.7 | 0.65 | 1.06 | 1.1 | 1.06 | 1.27 | 1.01 | 0.72 | 0.48 | -0.07 | 0.67 | 1 | 0.56 | 0.75 | 0.23 | -0.3575 | -0.1475 | 0.261682 | 0.218572 | 0.487693 | 0.999971 | 1 | 0.897906 |
| Ctif_1 | Q6PEE2 | -1.15 | 3.34 | 2.64 | 0.66 | 3.45 | 2.36 | 2.37 | 1.33 | 3.14 | -1.36 | 1.76 | 2.14 | 5.33 | 1.08 | 1.61 | 1.71 | 1.005 | 0.0475 | 1.06 | 0.322001 | 0.969834 | 0.407255 | 0.999971 | 1 | 0.874911 |
| Ctnna1 | P26231 | -0.1 | -0.34 | -0.89 | -0.78 | -0.27 | -0.39 | -0.95 | -0.37 | -0.28 | -0.32 | -0.92 | -0.81 | -0.57 | -0.4 | -0.83 | -0.84 | 0.0325 | -0.055 | -0.1325 | 0.901671 | 0.83689 | 0.592037 | 0.999971 | 1 | 0.922837 |
| Ctnna2 | Q61301 | -1.12 | -1.42 | -1.07 | -1.15 | -1.43 | -1.14 | -1.11 | -1.02 | -1.24 | -1.01 | -0.86 | -0.95 | -1.51 | -1.26 | -1.35 | -1.14 | 0.015 | 0.175 | -0.125 | 0.936355 | 0.350002 | 0.504879 | 0.999971 | 1 | 0.905277 |
| Ctnna2_1 | Q61301-2 | -0.72 | -0.24 | -1.45 | -1.14 | -1.18 | -1.5 | -1.84 | -0.06 | -1.01 | -0.61 | 0.33 | -0.71 | -1.68 | -0.5 | -1.26 | -0.56 | -0.2575 | 0.3875 | -0.1125 | 0.561857 | 0.320954 | 0.765018 | 0.999971 | 1 | 0.962405 |
| Ctnna3 | Q65CL1 | 1.23 | -0.75 | 1.45 | -0.6 | -1.45 | 0.36 | 0.91 | -1.42 | -0.48 | -0.84 | -0.13 | 0.38 | -1.33 | -1.47 | 0.31 | -0.93 | -0.7325 | -0.6 | -1.1875 | 0.34777 | 0.322466 | 0.093729 | 0.999971 | 1 | 0.776618 |
| Ctnnb1 | Q02248 | -0.84 | -1.15 | -1.06 | -1.18 | -1.14 | -0.76 | -0.66 | -1.13 | -0.73 | -0.75 | -0.8 | -0.96 | -1.23 | -1.12 | -1.22 | -1.21 | 0.135 | 0.2475 | -0.1375 | 0.510383 | 0.177694 | 0.434622 | 0.999971 | 1 | 0.882419 |
| Ctnnbip1 | Q9JJN6 | -0.63 | 0.69 | -0.66 | -0.19 | 0.3 | 0.07 | -0.95 | -0.1 | 0.64 | -0.77 | -0.57 | -0.11 | 1.26 | 0.63 | -0.04 | -0.06 | 0.0275 | -0.005 | 0.645 | 0.944895 | 0.990539 | 0.152976 | 0.999971 | 1 | 0.795578 |
| Ctnnbl1 | Q9CWL8 | 1.07 | 2.13 | 2.53 | 2.38 | 2.43 | 1.8 | 2.3 | 2.09 | 2.06 | 1.56 | 2.77 | 2.83 | 2.02 | 2.02 | 2.66 | 2.31 | 0.1275 | 0.2775 | 0.225 | 0.717043 | 0.519764 | 0.532356 | 0.999971 | 1 | 0.910069 |
| Ctnnd1 | P30999-2 | -1.14 | -1.94 | -2.1 | -1.84 | -1.97 | -2.2 | -0.09 | -1.86 | -0.52 | -2.27 | -0.08 | -0.2 | 0.15 | -3.33 | -2.41 | -1.36 | 0.225 | 0.9875 | 0.0175 | 0.64903 | 0.081617 | 0.980114 | 0.999971 | 1 | 0.99925 |
| Ctnnd1_1 | G3X9V2 | -0.32 | -0.78 | -0.59 | -0.77 | -0.59 | -0.76 | -0.09 | -0.72 | -0.16 | -0.22 | -0.21 | -0.55 | 0.03 | -0.95 | -0.89 | -0.47 | 0.075 | 0.33 | 0.045 | 0.742317 | 0.120076 | 0.868029 | 0.999971 | 1 | 0.983462 |
| Ctnnd2 | E9QKH8 | -1.51 | -1.3 | -0.86 | -1.3 | -1.3 | -1.05 | -0.58 | -1.2 | -0.97 | -1.08 | -1.11 | -1.33 | -2.23 | -1.24 | -2.42 | -2.24 | 0.21 | 0.12 | -0.79 | 0.395611 | 0.563418 | 0.02847 | 0.999971 | 1 | 0.678474 |
| Ctnnd2_1 | A0A2I3BQ\ | -0.8 | -2.17 | -1.76 | -0.76 | -1.32 | -1.77 | -1.26 | -0.42 | -1.27 | -1.8 | -1.79 | -1.71 | -1.48 | -1.51 | -0.84 | -0.68 | 0.18 | -0.27 | 0.245 | 0.674435 | 0.466483 | 0.540618 | 0.999971 | 1 | 0.910252 |
| Ctps1 | P70698 | -0.96 | -0.09 | 0.07 | -0.29 | -0.21 | -0.12 | 0.61 | 0.21 | -0.16 | -0.68 | 0.17 | 0.1 | 0.28 | -0.06 | 0.41 | -0.45 | 0.44 | 0.175 | 0.3625 | 0.168803 | 0.568313 | 0.253787 | 0.999971 | 1 | 0.839012 |
| Ctps2 | P70303 | 0.59 | 0.69 | 0.66 | 0.06 | 0.44 | 0.42 | 0.5 | 0.68 | 0.38 | 0.53 | 0.51 | 0.88 | 1.08 | 0.73 | 1.13 | 0.54 | 0.01 | 0.075 | 0.37 | 0.961999 | 0.736253 | 0.147768 | 0.999971 | 1 | 0.793248 |
| Ctr9 | Q62018 | 0.57 | 0.98 | 0.77 | 1.35 | 0.85 | 0.79 | 0.65 | 1.23 | 0.64 | 0.5 | 0.92 | 1.23 | 0.52 | 1.16 | 1.18 | 1.46 | -0.0375 | -0.095 | 0.1625 | 0.875976 | 0.710987 | 0.561221 | 0.999971 | 1 | 0.91591 |
| Ctsa | G3X8T3 | -0.2 | 0.53 | -0.26 | -0.04 | 0.65 | 0.46 | 0.1 | 0.1 | 0.8 | -0.56 | 0.01 | -0.06 | 0.86 | 0.8 | -0.06 | -0.17 | 0.32 | 0.04 | 0.35 | 0.226918 | 0.904168 | 0.302789 | 0.999971 | 1 | 0.847402 |
| Ctsb | P10605 | 0.44 | 1.07 | 1.11 | 1.17 | 1.04 | 1.23 | 0.88 | 1.58 | 1.22 | 0.51 | 1.21 | 1.11 | 0.82 | 1.18 | 1.11 | 1.39 | 0.235 | 0.065 | 0.1775 | 0.366244 | 0.803758 | 0.467754 | 0.999971 | 1 | 0.89205 |
| Ctsc | P97821 | 0.23 | 0.98 | 2.03 | 0.05 | 1.28 | 0.56 | 0.79 | 1.51 | 0.39 | -0.1 | 1.08 | 1.35 | 0.06 | 1.17 | 2.14 | -0.24 | 0.2125 | -0.1425 | -0.04 | 0.651035 | 0.783508 | 0.950344 | 0.999971 | 1 | 0.998038 |
| Ctsd | P18242 | -0.51 | 0.66 | 0.17 | 0 | 0.41 | 0.62 | 0.06 | 0.39 | 0.71 | -0.09 | 0.45 | 0.39 | 0.68 | 0.96 | 0.45 | 0.32 | 0.29 | 0.285 | 0.5225 | 0.319092 | 0.356181 | 0.100385 | 0.999971 | 1 | 0.78268 |
| Ctsf | Q9R013 | 1.46 | 0.4 | 1.16 | 0.9 | 0.31 | 0.7 | 0.85 | 2.06 | 0.3 | 0.67 | 1.86 | 1.57 | -0.01 | 0.55 | 1.18 | 1.33 | 5.89E-17 | 0.12 | -0.2175 | 1 | 0.770928 | 0.561264 | 1 | 1 | 0.91591 |
| Ctsh | P49935 | 1.1 | -0.06 | 0.11 | 0.34 | -0.17 | 0.96 | -0.24 | 0.51 | 0.2 | 1.82 | 1.04 | -0.56 | -0.74 | 0.54 | 0.46 | 0.52 | -0.1075 | 0.2525 | -0.1775 | 0.773217 | 0.637719 | 0.64984 | 0.999971 | 1 | 0.933842 |
| Ctsl | P06797 | -0.12 | -0.54 | 0.03 | -0.34 | -0.23 | -0.57 | 0.35 | 1.15 | -0.92 | -0.64 | 0.6 | 0.58 | -0.21 | -0.29 | 0.54 | 0.11 | 0.4175 | 0.1475 | 0.28 | 0.291465 | 0.714099 | 0.28628 | 0.999971 | 1 | 0.846029 |
| Ctso | Q8BM88 | 2.63 | 6.87 | 2.19 | 1.23 | 7.06 | 5.13 | 4 | 2.04 | 8.11 | 3.71 | 3.47 | 3.37 | 11.39 | 3.15 | 2.18 | 2.46 | 1.3275 | 1.435 | 1.565 | 0.370435 | 0.356842 | 0.4918 | 0.999971 | 1 | 0.899692 |
| Ctss | F6WR04 | 1.71 | 1.4 | 1.06 | 0.63 | 1.07 | 1.47 | 0.26 | 0.72 | 1 | 1.52 | 0.87 | 0.76 | 0.8 | 1.36 | 0.88 | 1.13 | -0.32 | -0.1625 | -0.1575 | 0.361945 | 0.585457 | 0.577786 | 0.999971 | 1 | 0.918581 |
| Ctsz | Q9WUU7 | -1.77 | 1.7 | 0.96 | -0.18 | 2.17 | 0.91 | 0.83 | 1.22 | 2.88 | -1.05 | 1.14 | 1.3 | 3.98 | 0.63 | 1.49 | -0.06 | 1.105 | 0.89 | 1.3325 | 0.15861 | 0.382255 | 0.220692 | 0.999971 | 1 | 0.827096 |
| Cttn | Q60598 | -1.07 | -2.17 | -0.6 | -0.52 | -2.12 | -1.91 | -0.19 | -0.5 | -1.96 | -1.4 | -0.65 | -0.75 | -2.77 | -2.28 | -0.73 | -0.63 | -0.09 | -0.1 | -0.5125 | 0.87378 | 0.827518 | 0.406814 | 0.999971 | 1 | 0.874911 |
| Cttnbp2 | B9EJA2-5 | -0.3 | -0.72 | -0.24 | -0.14 | -0.46 | -0.57 | -0.4 | -0.28 | -0.5 | -0.33 | -0.18 | -0.27 | -0.52 | -0.63 | -0.37 | 0.02 | -0.0775 | 0.03 | -0.025 | 0.699122 | 0.880043 | 0.913411 | 0.999971 | 1 | 0.992618 |
| Cttnbp2nl | Q99LJ0 | 0.54 | -0.54 | 0.41 | 0.67 | -0.48 | -0.51 | 0.25 | 0.26 | -0.66 | -0.01 | 0.42 | 0.4 | -1.17 | -0.12 | 0.42 | 0.33 | -0.39 | -0.2325 | -0.405 | 0.276341 | 0.528422 | 0.361232 | 0.999971 | 1 | 0.869526 |
| Ctu1 | Q99J10 | 3.15 | 0.57 | 0.61 | 0.42 | 0.35 | 0.98 | 0.15 | 0.85 | 0.04 | 1.82 | 0.1 | 0.2 | -0.7 | 0.39 | 0.64 | 0.3 | -0.605 | -0.6475 | -1.03 | 0.344874 | 0.374841 | 0.141454 | 0.999971 | 1 | 0.784967 |
| Ctu2 | Q3U308 | 0 | 0.67 | -0.64 | -1.2 | 0.25 | 0.23 | -0.81 | -1.19 | 0.25 | -0.11 | -0.1 | -0.44 | 0.45 | 0.24 | -0.78 | -1.35 | -0.0875 | 0.1925 | -0.0675 | 0.862373 | 0.639654 | 0.900598 | 0.999971 | 1 | 0.990593 |
| Cuedc1 | Q8R3V6 | -0.51 | -0.88 | -0.29 | -0.21 | -0.92 | -0.54 | -0.58 | -0.15 | -0.22 | -0.3 | -0.09 | 0.49 | -0.58 | -0.78 | -0.32 | -0.4 | -0.075 | 0.4425 | -0.0475 | 0.761715 | 0.111814 | 0.832174 | 0.999971 | 1 | 0.979036 |
| Cuedc2 | A0A494BB! | 0.25 | 1.44 | 1.59 | 1.47 | 1.15 | 1.28 | 1.1 | 1.36 | 1.09 | 0.79 | 1.46 | 1.75 | 0.84 | 1.45 | 1.59 | 1.79 | 0.035 | 0.085 | 0.23 | 0.913246 | 0.817411 | 0.53413 | 0.999971 | 1 | 0.910077 |
| Cul1 | Q9WTX6 | -0.01 | 0.49 | 0.6 | 0.32 | 0.39 | 0.53 | 0.79 | 0.22 | 0.65 | 0.15 | 0.66 | 0.53 | 0.69 | 0.24 | 0.34 | 0.31 | 0.1325 | 0.1475 | 0.045 | 0.554907 | 0.507469 | 0.834329 | 0.999971 | 1 | 0.979036 |
| Cul2 | Q9D4H8 | 0.11 | -0.03 | -0.2 | -0.58 | -0.01 | 0.01 | 0.31 | -0.55 | 0.15 | -0.16 | -0.15 | -0.53 | -0.24 | -0.13 | -0.54 | -0.67 | 0.115 | 0.0025 | -0.22 | 0.65687 | 0.991543 | 0.357325 | 0.999971 | 1 | 0.868793 |
| Cul3 | Q9JLV5 | -0.82 | -0.36 | -1.22 | -1.4 | -0.25 | -0.22 | -0.46 | -1.41 | 0.15 | -0.67 | -1.01 | -1.08 | -0.33 | -0.54 | -1.31 | -1.5 | 0.365 | 0.2975 | 0.03 | 0.31934 | 0.414382 | 0.933501 | 0.999971 | 1 | 0.995138 |
| Cul4a | Q3TCH7 | -0.37 | 0.4 | 0.8 | 0.84 | 0.23 | 0.19 | 0.73 | 0.51 | 0.24 | -0.09 | 0.7 | 0.73 | 0.08 | 0.21 | 0.57 | 0.76 | -0.0025 | -0.0225 | -0.0125 | 0.993609 | 0.947099 | 0.969163 | 0.999971 | 1 | 0.998787 |
| Cul4b | A2A432 | 1.22 | 0.78 | 0.87 | 0.82 | 0.84 | 0.91 | 0.72 | 0.45 | 0.91 | 1.1 | 0.57 | 0.49 | 0.52 | 1.01 | 0.55 | 0.68 | -0.1925 | -0.155 | -0.2325 | 0.350355 | 0.482393 | 0.276273 | 0.999971 | 1 | 0.843794 |
| Cul5 | Q9D5V5 | -0.2 | 0.13 | 0.52 | 0.66 | 0.16 | 0 | 0.09 | 0.47 | 0.2 | -0.13 | 0.39 | 0.72 | -0.16 | -0.01 | 0.42 | 0.6 | -0.0975 | 0.0175 | -0.065 | 0.695863 | 0.949831 | 0.816705 | 0.999971 | 1 | 0.974929 |
| Cul7 | Q8VE73-3 | 5.16 | 1.53 | 0.15 | 0.31 | 1.29 | 1.8 | 0.56 | 0.62 | 2.12 | 0.84 | -0.42 | 0.01 | 5.74 | 3.13 | 0.59 | 0.03 | -0.72 | -1.15 | 0.585 | 0.506944 | 0.335254 | 0.707844 | 0.999971 | 1 | 0.950182 |
| Cul9 | E9QP09 | -0.32 | -0.31 | -0.54 | -0.96 | -0.13 | -0.32 | -0.67 | -0.59 | -0.23 | -0.6 | -0.51 | -0.71 | -0.27 | -0.57 | -0.58 | -0.71 | 0.105 | 0.02 | 0 | 0.653793 | 0.928342 | 1 | 0.999971 | 1 | 1 |
| Custos | Q3UY34 | 1.56 | 2.21 | 2.12 | 1.9 | 1.78 | 1.54 | 1.29 | 2.37 | 2.24 | 1.43 | 2.07 | 2.18 | 1.74 | 2.57 | 2.27 | 2.13 | -0.2025 | 0.0325 | 0.23 | 0.485136 | 0.899868 | 0.374237 | 0.999971 | 1 | 0.87031 |
| Cuta | D5MCW4 | -1.1 | -0.1 | 0.44 | -0.37 | -0.36 | -0.44 | 0.07 | -0.2 | -0.43 | -1.17 | 0.42 | 0.29 | -0.75 | -0.6 | 0.11 | -0.58 | 0.05 | 0.06 | -0.1725 | 0.88195 | 0.895781 | 0.638134 | 0.999971 | 1 | 0.93301 |
| Cutc | Q9D8X1 | 1.64 | 2.16 | 1.24 | 1.77 | 1.89 | 1.85 | 0.95 | 1.25 | 1.28 | 1.6 | 0.63 | 0.57 | 0.85 | 1.64 | 0.91 | 1.27 | -0.2175 | -0.6825 | -0.535 | 0.482697 | 0.056671 | 0.082847 | 0.999971 | 1 | 0.755254 |
| Cux1 | H3BKT0 | 1.5 | 0.37 | 0.17 | -0.52 | 0.72 | 0.67 | -0.16 | -0.5 | 0.64 | 1.55 | 0.14 | 0.5 | -0.17 | 0.63 | 0.15 | -0.38 | -0.1975 | 0.3275 | -0.3225 | 0.683252 | 0.502779 | 0.476233 | 0.999971 | 1 | 0.894333 |
| Cux1_1 | H3BK24 | 1.84 | 1.43 | 0.54 | 0.88 | 1.22 | 1.13 | 0.44 | 0.49 | 1.05 | 1.33 | 0.36 | 0.58 | 0.42 | 0.83 | 0.51 | 0.31 | -0.3525 | -0.3425 | -0.655 | 0.326258 | 0.34873 | 0.062652 | 0.999971 | 1 | 0.725348 |
| Cwc15 | Q9JHS9 | 1.68 | 1.84 | 0.76 | 0.01 | 1.65 | 1.9 | 0.48 | -0.58 | 1.41 | 1.54 | 0.81 | 0.47 | -0.27 | 0.84 | 0.02 | -0.33 | -0.21 | -0.015 | -1.0075 | 0.74628 | 0.974264 | 0.057881 | 0.999971 | 1 | 0.725348 |
| Cwc22 | B1AYU7 | 0.74 | 0.17 | -0.26 | -0.59 | 0.33 | 0.2 | 0.31 | -0.71 | 0.31 | 0.57 | -0.12 | -0.2 | -0.06 | 0.03 | -0.52 | -1.14 | 0.0175 | 0.125 | -0.4375 | 0.962156 | 0.7119 | 0.268132 | 0.999971 | 1 | 0.842746 |
| Cwc25 | Q9DBF7 | -0.81 | 0.72 | 0.32 | -0.51 | 1.22 | 0.42 | -0.11 | 0 | 0.73 | -1.31 | -0.19 | -0.2 | 0.52 | 0.27 | -0.19 | -0.12 | 0.4525 | -0.1725 | 0.19 | 0.317905 | 0.735827 | 0.619684 | 0.999971 | 1 | 0.929296 |
| Cwc27 | Q3TKY6 | 0.66 | 1.04 | 0.93 | 1.35 | 0.79 | 0.81 | 0.74 | 1.06 | 0.48 | 0.66 | 0.91 | 0.84 | 0.78 | 1.35 | 1 | 1.49 | -0.145 | -0.2725 | 0.16 | 0.496154 | 0.227329 | 0.52129 | 0.999971 | 1 | 0.907001 |
| Cx3cl1 | O35188 | -0.21 | 0.48 | 0.1 | 0.18 | 0.23 | 0.72 | 0.13 | 0.09 | -0.08 | -0.18 | 0.42 | 0.48 | 0.12 | 0.74 | 0.23 | 0.69 | 0.155 | 0.0225 | 0.3075 | 0.518779 | 0.927518 | 0.228506 | 0.999971 | 1 | 0.828687 |
| Cx3cr1 | Q9Z0D9 | 0.06 | 1.31 | 1.34 | 1.31 | 0.96 | 2.15 | 1.3 | 0.57 | 1.99 | 1.23 | 1.28 | 2.63 | 0.74 | 1.85 | 2.58 | 2.09 | 0.24 | 0.7775 | 0.81 | 0.58419 | 0.100711 | 0.111411 | 0.999971 | 1 | 0.78268 |
| Cxadr | P97792 | -0.62 | -2.19 | -2.8 | -2.44 | -1.62 | -1.99 | -1.76 | -2.11 | -1.14 | -1.26 | -1.98 | -2.29 | -0.73 | -1.86 | -2.15 | -1.83 | 0.1425 | 0.345 | 0.37 | 0.757964 | 0.509399 | 0.490981 | 0.999971 | 1 | 0.898873 |
| Cxcl12 | H7BX38 | -1.39 | -1.21 | -1.52 | -3.74 | 0.31 | -1.83 | 0.39 | -4.31 | 0.09 | -1.9 | -1 | -2.18 | -0.63 | -0.88 | -1.84 | -2.3 | 0.605 | 0.7175 | 0.5525 | 0.592223 | 0.329355 | 0.404784 | 0.999971 | 1 | 0.874011 |
| Cxcl14 | Q9WUQ5 | 4.26 | 1.22 | 1 | 1.91 | 0.92 | 2.98 | 1.96 | 0.86 | 1.7 | 3.19 | 0.65 | 0.32 | -0.52 | 0.74 | 0.61 | 0.06 | -0.4175 | -0.6325 | -1.875 | 0.607802 | 0.483916 | 0.027536 | 0.999971 | 1 | 0.678474 |
| Cxxc1 | Q9CWW7 | 4.83 | 4.45 | 3.16 | 3.42 | 3.7 | 2.51 | 1.72 | 3.24 | 3.75 | 4.36 | 2.03 | 2.55 | 0.35 | 1.34 | 2.83 | 2.95 | -1.1725 | -0.7925 | -2.0975 | 0.055517 | 0.218301 | 0.011947 | 0.999971 | 1 | 0.613095 |
| Cyb561 | Q60720 | 1.21 | -0.69 | -1.58 | -1.25 | -0.76 | -0.45 | -0.87 | -0.98 | -1.25 | 0.29 | -1.98 | -0.82 | -0.2 | 0.36 | -1.16 | 0.41 | -0.1875 | -0.3625 | 0.43 | 0.74692 | 0.613536 | 0.517644 | 0.999971 | 1 | 0.905815 |
| Cyb561d2 | A0A0A6YY. | 0.08 | -0.7 | -0.57 | -0.3 | -1.9 | -1.04 | -0.43 | -0.56 | -1.68 | 0.28 | -0.58 | -0.99 | -2.48 | -0.06 | 0.4 | -0.42 | -0.61 | -0.37 | -0.2675 | 0.121566 | 0.392001 | 0.658495 | 0.999971 | 1 | 0.934638 |
| Cyb5a | G5E850 | -1.03 | -0.9 | -1.06 | -1.26 | -0.95 | -0.96 | -0.97 | -1.06 | -0.82 | -1 | -1.17 | -0.93 | -0.93 | -1.11 | -1.06 | -0.46 | 0.0775 | 0.0825 | 0.1725 | 0.652532 | 0.645519 | 0.430081 | 0.999971 | 1 | 0.881611 |
| Cyb5b | Q9CQX2 | -0.13 | -1.42 | -0.98 | -0.79 | -1.15 | -1.21 | -0.94 | -0.68 | -1.51 | -0.82 | -1.2 | -1.19 | -1.71 | -1.44 | -0.7 | -0.78 | -0.165 | -0.35 | -0.3275 | 0.587375 | 0.274134 | 0.371892 | 0.999971 | 1 | 0.869526 |
| Cyb5d2 | Q5SSH8 | -0.66 | -0.41 | 0.7 | 0.81 | 0.02 | -0.95 | 0.05 | 0.96 | 0.3 | -1.35 | 1.06 | 0.49 | -0.1 | -1.95 | 0.8 | 0.72 | -0.09 | 0.015 | -0.2425 | 0.857784 | 0.979639 | 0.71946 | 0.999971 | 1 | 0.954401 |
| Cyb5r1 | Q9DB73 | -0.54 | -0.49 | -2.34 | -1.9 | -0.27 | -0.48 | -1.97 | -2.24 | -0.22 | -0.37 | -1.64 | -1.64 | -0.15 | -0.28 | -1.92 | -1.57 | 0.0775 | 0.35 | 0.3375 | 0.901751 | 0.539667 | 0.574284 | 0.999971 | 1 | 0.91816 |
| Cyb5r3 | Q9DCN2 | -0.47 | -0.74 | -1.3 | -1.2 | -0.54 | -0.76 | -1.18 | -1.27 | -0.64 | -0.46 | -1.22 | -1 | -0.86 | -0.49 | -1.22 | -1.21 | -0.01 | 0.0975 | -0.0175 | 0.971204 | 0.724411 | 0.949777 | 0.999971 | 1 | 0.998038 |
| Cyb5r4 | Q3TDX8 | 0.26 | 0.69 | 0.74 | 1.14 | 0.3 | 0.99 | 0.49 | 1.29 | 0.38 | 0.5 | 1.04 | 0.85 | 0.58 | 0.96 | 0.7 | 1.26 | 0.06 | -0.015 | 0.1675 | 0.840756 | 0.953663 | 0.523561 | 0.999971 | 1 | 0.907797 |
| Cybc1 | Q3TYS2 | 0.14 | -0.12 | -0.76 | -0.7 | 0.14 | 0.21 | -0.2 | -0.12 | -0.61 | 0.23 | 0.27 | -0.09 | -0.28 | 0.34 | -0.52 | -0.87 | 0.3675 | 0.31 | 0.0275 | 0.187074 | 0.3254 | 0.934626 | 0.999971 | 1 | 0.995618 |
| Cyc1 | Q9D0M3 | -0.75 | -1.38 | -1.36 | -1.55 | -1.09 | -1.49 | -0.72 | -1.35 | -1.23 | -0.99 | -1.16 | -0.74 | -0.96 | -1.37 | -1.46 | -1.56 | 0.0975 | 0.23 | -0.0775 | 0.713793 | 0.346617 | 0.755829 | 0.999971 | 1 | 0.961116 |
| Cycs | P62897 | -1.53 | -1.46 | -1.17 | -1.21 | -1.43 | -1.9 | -0.81 | -1.64 | -1.78 | -1.65 | -1.19 | -1.19 | -1.87 | -1.55 | -1.15 | -1.09 | -0.1025 | -0.11 | -0.0725 | 0.704052 | 0.61807 | 0.76134 | 0.999971 | 1 | 0.961116 |
| Cyfip1 | Q7TMB8 | -1.04 | -0.46 | -0.66 | -0.72 | -0.33 | -0.47 | -0.77 | -0.39 | -0.63 | -1.09 | -0.62 | -0.5 | -0.25 | -0.6 | -0.68 | -0.55 | 0.23 | 0.01 | 0.2 | 0.284587 | 0.963467 | 0.34662 | 0.999971 | 1 | 0.864164 |
| Cyfip1_1 | A0A0R4J11 | -0.79 | 0.27 | -1.65 | -2.36 | 0.07 | -0.46 | -1.72 | -1.38 | -0.61 | -0.95 | -1 | -1.51 | -0.17 | -0.28 | -1.81 | -2.09 | 0.26 | 0.115 | 0.045 | 0.68429 | 0.834791 | 0.947676 | 0.999971 | 1 | 0.997651 |
| Cyfip2 | Q5SQX6 | -0.98 | -1.14 | -0.74 | -0.62 | -1.23 | -1.4 | -0.75 | -0.87 | -1.13 | -1.22 | -0.54 | -0.63 | -1.31 | -1.25 | -0.74 | -0.47 | -0.1925 | -0.01 | -0.0725 | 0.411915 | 0.966592 | 0.779886 | 0.999971 | 1 | 0.965847 |
| Cygb | Q9CX80 | 0.75 | 0.48 | 0.86 | 1.19 | 0.82 | 0.64 | 0.96 | 1.75 | 1.06 | 0.75 | 0.92 | 0.99 | 0.82 | 0.98 | 1.69 | 1.05 | 0.2225 | 0.11 | 0.315 | 0.458774 | 0.600989 | 0.251945 | 0.999971 | 1 | 0.839012 |
| Cyhr1 | Q9QXA1 | -0.8 | 1.46 | 1.78 | 1.85 | 0.91 | 1.68 | 1.11 | 1.96 | 0.55 | -0.73 | 2.1 | 1.5 | -0.03 | 1.37 | 1.75 | 2.29 | 0.3425 | -0.2175 | 0.2725 | 0.581012 | 0.785511 | 0.707811 | 0.999971 | 1 | 0.950182 |
| Cyld | Z4YJJ5 | -0.06 | 0.52 | 0.45 | 0.72 | 0.56 | 0.04 | -0.14 | 0.78 | 0.48 | 0.18 | 0.44 | 0.85 | 1.14 | 0.69 | 0.86 | 0.79 | -0.0975 | 0.08 | 0.4625 | 0.733521 | 0.743761 | 0.071282 | 0.999971 | 1 | 0.737413 |
| Cyp20a1 | Q8BKE6 | -0.57 | 0.23 | 0.68 | -0.87 | 0.83 | -0.63 | 1.49 | -0.64 | 1.29 | -0.52 | 0.79 | 1.39 | 1.09 | -0.77 | 0.07 | -0.68 | 0.395 | 0.87 | 0.06 | 0.507123 | 0.124963 | 0.907678 | 0.999971 | 1 | 0.991405 |
| Cyp2d22 | Q9JKY7 | -0.2 | -0.68 | -1.4 | -1.8 | -0.41 | -0.63 | -1.48 | -0.93 | -0.72 | -0.62 | -1.51 | -1.32 | -0.2 | -0.53 | -1.56 | -1.32 | 0.1575 | -0.0225 | 0.1175 | 0.699613 | 0.955382 | 0.795323 | 0.999971 | 1 | 0.970361 |
| Cyp2j6 | O54750 | -2.74 | -0.36 | -0.09 | -0.03 | -0.07 | -0.13 | -0.24 | 0.77 | 0.46 | -1.42 | 0.41 | 0.8 | 0.79 | 0.96 | 0.58 | 0.69 | 0.8875 | 0.8675 | 1.56 | 0.181112 | 0.262959 | 0.026805 | 0.999971 | 1 | 0.678474 |
| Cyp2j9 | Q924D1 | -0.42 | -0.47 | -0.02 | -0.18 | -0.12 | 0.21 | -0.45 | 0.47 | -0.02 | 0.04 | -0.17 | -0.11 | 0.61 | 0.22 | 0.11 | 0.18 | 0.3 | 0.2075 | 0.5525 | 0.254109 | 0.275754 | 0.024413 | 0.999971 | 1 | 0.669611 |
| Cyp2s1 | D3Z7F1 | -1.61 | 0.13 | -0.87 | -0.94 | -0.43 | -1.64 | -1.25 | -0.98 | -0.09 | -1.8 | -1.07 | -0.69 | -0.42 | -0.9 | -1.54 | -2.08 | -0.2525 | -0.09 | -0.4125 | 0.549178 | 0.849679 | 0.397354 | 0.999971 | 1 | 0.871816 |
| Cyp46a1 | Q9WVK8 | 0.01 | -0.09 | -0.31 | -0.27 | -0.09 | 0.2 | -0.45 | -0.01 | -0.03 | 0.17 | -0.06 | 0.19 | 0.56 | 0.43 | 0.06 | 0.27 | 0.0775 | 0.2325 | 0.495 | 0.709677 | 0.209575 | 0.030034 | 0.999971 | 1 | 0.678474 |
| Cyp4f14 | Q9EP75 | -1.39 | -1.59 | -0.99 | -1.17 | -1.35 | -1.37 | -0.19 | -0.75 | -1.46 | -1.14 | -0.05 | -0.26 | -1.54 | -1.23 | -0.64 | -0.85 | 0.37 | 0.5575 | 0.22 | 0.257621 | 0.14449 | 0.411116 | 0.999971 | 1 | 0.875847 |
| Cyp4f16 | Q99N17 | -1.43 | -0.92 | -0.88 | 0.25 | -0.61 | -0.28 | -1.47 | -0.42 | -1.53 | -1.21 | -2.72 | -0.71 | -0.88 | -0.83 | -0.14 | -0.65 | 0.05 | -0.7975 | 0.12 | 0.905339 | 0.14866 | 0.752669 | 0.999971 | 1 | 0.961116 |
| Cyp4x1 | A2A9Y7 | -0.67 | -0.11 | -0.54 | -1.03 | 0.11 | -0.62 | -0.08 | -0.86 | 0.07 | -0.59 | -0.42 | -0.75 | 0.27 | -0.58 | -0.7 | -0.66 | 0.225 | 0.165 | 0.17 | 0.465118 | 0.553522 | 0.581802 | 0.999971 | 1 | 0.920364 |
| Cyp51a1 | Q8K0C4 | -1.39 | -0.91 | -0.61 | -1.06 | -0.8 | -1.07 | -0.52 | -0.91 | -0.7 | -1.27 | -0.55 | -0.65 | -0.05 | -0.43 | -0.15 | -0.43 | 0.1675 | 0.2 | 0.7275 | 0.482003 | 0.438756 | 0.010696 | 0.999971 | 1 | 0.613095 |
| Cyp7b1 | Q60991 | -0.06 | -1.43 | -1.12 | -0.58 | -1.09 | -0.69 | -1.32 | -0.51 | -1.28 | -0.56 | -0.63 | -0.63 | -2.29 | -1.19 | -0.54 | -0.44 | -0.105 | 0.0225 | -0.3175 | 0.763423 | 0.947526 | 0.51895 | 0.999971 | 1 | 0.905901 |
| Cyria | Q8BHZ0 | -0.17 | 0.06 | -0.06 | -0.19 | 0.28 | 0.11 | 0.35 | 0.09 | 0.28 | -0.22 | 0.11 | -0.05 | 0.17 | 0.24 | -0.19 | 0.22 | 0.2975 | 0.12 | 0.2 | 0.114245 | 0.525158 | 0.302625 | 0.999971 | 1 | 0.847402 |
| Cyrib | A0A2I3BQ( | -3.65 | 2.88 | 2.12 | 0.41 | 2.87 | 3.02 | 1.17 | 2.14 | 2.58 | -2.23 | 2.16 | 2.5 | 4.09 | 2.24 | 1.87 | 1.61 | 1.86 | 0.8125 | 2.0125 | 0.190925 | 0.627884 | 0.172014 | 0.999971 | 1 | 0.804075 |
| Cyrib_1 | Q921M7 | 0.48 | 0.21 | 0.11 | 0.36 | 0.04 | 0.31 | 0.14 | 0.12 | 0.16 | 0.32 | -0.01 | -0.32 | -0.28 | 0.01 | -0.15 | -0.05 | -0.1375 | -0.2525 | -0.4075 | 0.450924 | 0.243959 | 0.048393 | 0.999971 | 1 | 0.710102 |
| Cys1 | A0A0R4J21 | 0.77 | -1.62 | -1.45 | -1.18 | -0.88 | -0.53 | 0.86 | -2.43 | -0.71 | 0.39 | -1.37 | -2.58 | -2.82 | -1.87 | -2.7 | -4.61 | 0.125 | -0.1975 | -2.13 | 0.873544 | 0.793878 | 0.015608 | 0.999971 | 1 | 0.613095 |
| Cystm1 | D3YVS7 | -2.49 | -4.29 | -0.64 | -0.64 | -2.41 | -3.58 | -1.02 | -1.4 | -4.5 | -3.48 | -0.81 | -0.74 | -4.39 | -3.09 | -0.14 | -0.45 | -0.0875 | -0.3675 | -0.0025 | 0.925352 | 0.751602 | 0.998341 | 0.999971 | 1 | 1 |
| Cyth1 | Q9QX11-2 | -1.18 | -0.47 | 0.13 | 0.68 | -0.52 | -0.33 | -0.07 | 0.67 | -0.56 | -0.6 | 0.4 | 0.16 | -0.28 | -0.03 | 0.6 | 1.04 | 0.1475 | 0.06 | 0.5425 | 0.742708 | 0.893046 | 0.264721 | 0.999971 | 1 | 0.842533 |
| Cyth2 | A0A1B0GQ | 2.8 | -0.11 | 1.46 | -0.62 | 0.12 | -0.32 | 1.05 | 0.64 | -0.28 | 1.54 | 0.02 | 0.69 | 0.84 | 0.08 | -0.78 | 1.15 | -0.51 | -0.39 | -0.56 | 0.502405 | 0.625194 | 0.490134 | 0.999971 | 1 | 0.898612 |
| Cyth2_1 | Q99KH2 | -0.88 | -1.22 | -1.11 | -0.98 | -1.22 | -1.36 | -1.33 | -0.92 | -1.18 | -1.06 | -0.79 | -1.1 | -1.31 | -1.22 | -1.1 | -0.81 | -0.16 | 0.015 | -0.0625 | 0.412316 | 0.934304 | 0.749662 | 0.999971 | 1 | 0.961064 |
| Cyth3 | O08967 | 1 | 1.46 | 1.11 | 1.18 | 1.16 | 1.31 | 0.43 | 1.3 | 1.01 | 1.34 | 1.1 | 0.4 | 1.95 | 1.28 | 0.83 | 1.3 | -0.1375 | -0.225 | 0.1525 | 0.594666 | 0.377701 | 0.576353 | 0.999971 | 1 | 0.918194 |
| Czib | Q8BHG2 | 1.01 | 1.36 | 1.33 | 0.83 | 1.24 | 1.12 | 1.41 | 0.54 | 1.29 | 0.47 | 0.95 | 0.86 | 0.88 | 0.32 | 0.38 | 0.25 | -0.055 | -0.24 | -0.675 | 0.828791 | 0.334178 | 0.01602 | 0.999971 | 1 | 0.613095 |
| D030056L2 | Q3U2A5 | 4.88 | 1.45 | 2.33 | 2.24 | 1.3 | 1.43 | 2.02 | 2.7 | 1.11 | 2.08 | 2.07 | 1.47 | -0.63 | 0.25 | 0.88 | 1.9 | -0.8625 | -1.0425 | -2.125 | 0.257322 | 0.168276 | 0.028052 | 0.999971 | 1 | 0.678474 |
| D17h6s53e | Q9Z1R4 | 0.84 | -0.1 | -0.88 | -0.55 | 0.01 | 0.56 | -0.52 | -0.62 | -0.08 | 0.59 | -0.95 | -1.47 | 0.38 | 0.38 | -0.82 | -0.45 | 0.03 | -0.305 | 0.045 | 0.945042 | 0.579566 | 0.920623 | 0.999971 | 1 | 0.992702 |
| D2hgdh | E9QN44 | -1.89 | -1.12 | -0.76 | -2.15 | -2.02 | -0.61 | -2.03 | -2.3 | -1.12 | -1.14 | -2.33 | -0.31 | -0.34 | -0.73 | -2.96 | -1.14 | -0.26 | 0.255 | 0.1875 | 0.582058 | 0.607324 | 0.757711 | 0.999971 | 1 | 0.961116 |
| D3Ertd751 | Q8BGN2 | -0.08 | -0.21 | 0.42 | 0.43 | -0.24 | 0.05 | 0.3 | 0.77 | -0.49 | -0.43 | 0.44 | 0.55 | -0.31 | 0.56 | 0.97 | 0.35 | 0.08 | -0.1225 | 0.2525 | 0.779164 | 0.706271 | 0.436307 | 0.999971 | 1 | 0.882419 |
| D430041D | A0A2R8VK | -1.22 | -1.13 | -0.86 | -0.91 | -1.34 | -1.05 | -0.8 | -1.01 | -1.41 | -1.38 | -0.88 | -0.99 | -1.26 | -1.37 | -1.12 | -0.96 | -0.02 | -0.135 | -0.1475 | 0.920077 | 0.521426 | 0.448683 | 0.999971 | 1 | 0.885333 |
| D630045J1 | D3YTS3 | -0.79 | -1.08 | -1.03 | -1.02 | -1.32 | -0.84 | -0.92 | -1.11 | -1.19 | -0.82 | -1.05 | -1.05 | -0.67 | -0.86 | -1 | -0.73 | -0.0675 | -0.0475 | 0.165 | 0.724937 | 0.787999 | 0.369476 | 0.999971 | 1 | 0.869526 |
| D6Wsu163 | Q91YN0 | 0.66 | 0.47 | 0.18 | -0.54 | 0.18 | 0.61 | 1.3 | -0.81 | 0.3 | 0.47 | 0.03 | 0.11 | 0.07 | 0.58 | -0.27 | -0.9 | 0.1275 | 0.035 | -0.3225 | 0.790322 | 0.904382 | 0.419521 | 0.999971 | 1 | 0.87788 |
| D8Ertd738 | Q8R1F0 | 1.86 | 0.74 | 1.69 | -0.28 | 1.76 | -0.57 | 2.46 | -0.22 | 0.52 | 0.46 | 1.64 | 1.22 | 0.6 | 0.94 | 0.24 | -2.39 | -0.145 | -0.0425 | -1.155 | 0.856064 | 0.935816 | 0.18064 | 0.999971 | 1 | 0.808532 |
| Daam1 | Q8BPM0 | -0.79 | -1.91 | -0.76 | -0.87 | -1.87 | -1.41 | -0.62 | -1.06 | -1.54 | -1.05 | -0.8 | -0.71 | -1.97 | -1.93 | -1.1 | -1.05 | -0.1575 | 0.0575 | -0.43 | 0.672987 | 0.862543 | 0.258794 | 0.999971 | 1 | 0.839012 |
| Daam2 | Q80U19 | -0.41 | -0.76 | -1.55 | -1.48 | -0.62 | -0.85 | -1.26 | -1.83 | -0.38 | -0.54 | -1.31 | -1.13 | -0.08 | -1.26 | -1.92 | -1.11 | -0.09 | 0.21 | -0.0425 | 0.809208 | 0.554193 | 0.923734 | 0.999971 | 1 | 0.993382 |
| Dab1 | F2Z465 | -0.24 | 0.5 | -0.04 | 0.45 | 0.1 | 0.27 | -0.45 | -0.01 | 0.39 | 0.13 | 0.24 | 0.24 | 0.63 | 0.23 | -0.31 | 0.24 | -0.19 | 0.0825 | 0.03 | 0.474142 | 0.716982 | 0.915058 | 0.999971 | 1 | 0.992633 |
| Dab2 | E9PX84 | -0.18 | 0.16 | -0.21 | 0.57 | 0.6 | -0.17 | -0.32 | -0.07 | 0.31 | -0.24 | -0.71 | 0.62 | 0.22 | 0.73 | 0.65 | 0.2 | -0.075 | -0.09 | 0.365 | 0.79346 | 0.79287 | 0.177898 | 0.999971 | 1 | 0.808532 |

| | | | | | | | | | | | | | | | | | | | | | | | | | | |
|---|---|---|---|---|---|---|---|---|---|---|---|---|---|---|---|---|---|---|---|---|---|---|---|---|---|---|
| Dab2ip | Q3UHC7 | -1.06 | -1.23 | -0.98 | -0.62 | -1.31 | -1.24 | -1.05 | -0.62 | -1.29 | -1.04 | -0.87 | -1.01 | -0.87 | -1.04 | -1.06 | -0.52 | -0.0825 | -0.08 | 0.1 | 0.72756 | 0.697795 | 0.655108 | 0.999971 | 1 | 0.934061 |
| Dact2 | Q7TN08 | 1.75 | -0.27 | 0.85 | 1.4 | -0.12 | 1.08 | 4.06 | 1.04 | -0.72 | 0.08 | 1.63 | 1.19 | -1.38 | -0.19 | 0.76 | 1.84 | 0.5825 | -0.3875 | -0.675 | 0.518596 | 0.545017 | 0.371279 | 0.999971 | 1 | 0.869526 |
| Dact3 | Q0PHV7 | 1.91 | -0.69 | -0.3 | -0.26 | -0.57 | 0.03 | 0.45 | -0.47 | -0.3 | 1.42 | 0.05 | -0.39 | -0.14 | -0.43 | -0.27 | 0 | -0.305 | 0.03 | -0.375 | 0.602358 | 0.963731 | 0.500669 | 0.999971 | 1 | 0.904223 |
| Dad1 | P61804 | 0.23 | 0.4 | 0.51 | 0.69 | 0.28 | 0.96 | 0.07 | 0.52 | -0.03 | 0.47 | 0.35 | -0.08 | 0.5 | 1.26 | -0.3 | 1.25 | 7.85E-17 | -0.28 | 0.22 | 1 | 0.210132 | 0.558868 | 1 | 1 | 0.915051 |
| Dag1 | Q62165 | -0.44 | -1.28 | -1.21 | -0.86 | -0.74 | -1.31 | -1.02 | -1.28 | -0.36 | -0.38 | -0.71 | -0.7 | -0.37 | -0.98 | -0.78 | -0.56 | -0.14 | 0.41 | 0.275 | 0.590667 | 0.12014 | 0.303548 | 0.999971 | 1 | 0.847402 |
| Dagla | Q6WQJ1 | -1.4 | -0.49 | -1.06 | -1.13 | -0.8 | -0.78 | -1.57 | -1.29 | -0.76 | -1.45 | -1.05 | -1.43 | -0.1 | -1.13 | -1.21 | -0.9 | -0.09 | -0.1525 | 0.185 | 0.753058 | 0.57635 | 0.567515 | 0.999971 | 1 | 0.91816 |
| Daglb | Q91WC9 | -0.73 | 0.37 | 0.8 | 0.9 | 0.4 | 0.51 | 0.55 | 0.85 | 0.42 | 0.07 | 1.01 | 0.42 | 1 | 0.36 | 0.88 | 1.02 | 0.2425 | 0.145 | 0.48 | 0.520887 | 0.719973 | 0.237871 | 0.999971 | 1 | 0.833594 |
| Dalrd3 | Q6PJN8 | 1.15 | -0.04 | 0.2 | 1.2 | 0.07 | 0.21 | 0.14 | 1.01 | -0.51 | 1.37 | -0.07 | 0.15 | -0.85 | 0.64 | 0.1 | 0.8 | -0.27 | -0.3925 | -0.455 | 0.478523 | 0.423789 | 0.337004 | 0.999971 | 1 | 0.858827 |
| Dap | A0A087WF | 3.31 | 0.89 | 0.68 | 1.92 | 0.53 | 1.49 | 4.64 | 0.95 | 4.75 | 0.84 | 1.89 | -0.12 | -0.09 | -1.46 | 0.68 | -0.3 | 0.2025 | 0.14 | -1.9925 | 0.838074 | 0.897597 | 0.015693 | 0.999971 | 1 | 0.613095 |
| Dap3 | G3X9M0 | -1.2 | -0.58 | -0.81 | -0.39 | -0.52 | -0.42 | -0.44 | -0.67 | -0.45 | -0.65 | -0.83 | -0.64 | -0.26 | -0.13 | -0.7 | -0.61 | 0.2325 | 0.1025 | 0.32 | 0.315487 | 0.653803 | 0.222427 | 0.999971 | 1 | 0.827753 |
| Dapk1 | Q80YE7-2 | -0.57 | -0.21 | -0.52 | -0.19 | -0.16 | -0.18 | -0.46 | -0.38 | -0.44 | -0.2 | -0.58 | -0.72 | -0.17 | -0.17 | -0.66 | -0.61 | 0.0775 | -0.1125 | -0.03 | 0.686247 | 0.580414 | 0.88933 | 0.999971 | 1 | 0.987681 |
| Dapk3 | O54784 | 0.12 | 1.04 | -0.64 | -0.77 | 1.15 | 0.68 | -0.36 | -0.96 | 0.56 | 0.37 | -0.78 | -0.3 | 0.41 | 1.36 | -1.38 | -1.11 | 0.19 | 0.025 | -0.1175 | 0.744447 | 0.958825 | 0.866114 | 0.999971 | 1 | 0.983462 |
| Dars1 | Q922B2 | 0.08 | 0.89 | 0.91 | 0.88 | 1.02 | 0.83 | 0.84 | 1.31 | 1.19 | 0.59 | 1.11 | 1.32 | 1.01 | 1.09 | 1.16 | 1.55 | 0.31 | 0.3625 | 0.5125 | 0.247481 | 0.209796 | 0.075575 | 0.999971 | 1 | 0.746958 |
| Dars2 | Q8BIP0 | -0.78 | 0.19 | -1.61 | -1.94 | 0.07 | -0.46 | -1.44 | -2.21 | -0.04 | -0.98 | -1.53 | -1.46 | -0.32 | -0.81 | -2.4 | -2.46 | 0.025 | 0.0325 | -0.4625 | 0.968436 | 0.952195 | 0.489112 | 0.999971 | 1 | 0.898346 |
| Daxx | O35613 | -1.82 | -1.45 | 0.05 | -0.33 | -0.5 | -1.68 | -0.47 | -0.1 | -1.11 | -2.34 | 0.29 | 0.34 | -0.65 | -0.73 | -0.25 | 0.23 | 0.2 | 0.1825 | 0.5375 | 0.700928 | 0.797114 | 0.266278 | 0.999971 | 1 | 0.842677 |
| Dazap1 | Q3UGB5 | 0.97 | 2.01 | 1.53 | 2.09 | 1.87 | 2.21 | 1.35 | 2.36 | 1.95 | 1.56 | 1.99 | 2.14 | 1.37 | 2.35 | 2.19 | 1.48 | 0.2975 | 0.26 | 0.1975 | 0.390409 | 0.389591 | 0.577817 | 0.999971 | 1 | 0.918581 |
| Dbh | Q64237 | -1.43 | -0.47 | 0.01 | -0.7 | -1.24 | -0.29 | -0.98 | 0.21 | -1.14 | -1.02 | -0.42 | -0.89 | -0.22 | -0.74 | -1.31 | -0.3 | 0.0725 | -0.22 | 0.005 | 0.863626 | 0.51819 | 0.98944 | 0.999971 | 1 | 0.99925 |
| Dbi | P31786 | -0.36 | -0.3 | 0.37 | 0.1 | -0.09 | -0.29 | -0.22 | 0.46 | -0.21 | -0.29 | -0.27 | 0.09 | -0.39 | 0.41 | 0.47 | 0.6 | 0.0125 | -0.1225 | 0.32 | 0.962213 | 0.596199 | 0.29347 | 0.999971 | 1 | 0.847402 |
| Dbn1 | Q9QXS6 | -1.04 | -1.1 | -1.36 | -1.52 | -1.26 | -1.02 | -0.88 | -1.52 | -1.01 | -1.23 | -1.11 | -1.27 | -1.33 | -1.45 | -1.34 | -1.57 | 0.085 | 0.1 | -0.1675 | 0.703372 | 0.600241 | 0.39419 | 0.999971 | 1 | 0.871816 |
| Dbn1_1 | A0A0R4J1E | 1.05 | -0.43 | -1.74 | -1.4 | -1.15 | -1.06 | -1.79 | -1.18 | -0.5 | 0.48 | -1.85 | -1.62 | -1.46 | -0.35 | -1.07 | -1.31 | -0.665 | -0.2425 | -0.4175 | 0.278033 | 0.746564 | 0.501637 | 0.999971 | 1 | 0.904573 |
| Dbndd1 | H3BK17 | 3.06 | 0.35 | -0.53 | -0.12 | 0.15 | 0.08 | -0.41 | -0.22 | 1.68 | 1.33 | 0.35 | -0.38 | 0.52 | 0.32 | 0.01 | -0.2 | -0.79 | 0.055 | -0.5275 | 0.301621 | 0.948086 | 0.485617 | 0.999971 | 1 | 0.897386 |
| Dbndd2 | Q9CRD4 | -2.18 | 0.19 | 0.02 | -0.4 | -0.63 | 0.93 | -1.04 | 0.64 | 1.44 | 0.16 | -0.85 | 0.33 | 0.95 | 0.48 | -0.48 | 0.32 | 0.5675 | 0.8625 | 0.91 | 0.397974 | 0.210981 | 0.136376 | 0.999971 | 1 | 0.784948 |
| Dbnl | Q62418 | -0.45 | -0.7 | -0.63 | -0.52 | -0.52 | -0.63 | -0.55 | -0.16 | -0.9 | -0.99 | -0.53 | -0.63 | -0.6 | -0.74 | -0.62 | -0.47 | 0.11 | -0.1875 | -0.0325 | 0.562088 | 0.330245 | 0.850068 | 0.999971 | 1 | 0.98187 |
| Dbr1 | Q923B1 | 0.52 | 2.69 | 2.09 | 1.87 | 2.47 | 2.23 | 1.56 | 1.66 | 2.67 | 0.84 | 1.76 | 1.51 | 0.52 | 1.54 | 1.84 | 1.61 | 0.1875 | -0.0975 | -0.415 | 0.693568 | 0.859094 | 0.420184 | 0.999971 | 1 | 0.87788 |
| Dbt | P53395 | -0.96 | -0.52 | -0.25 | -0.37 | -0.75 | -0.44 | -0.41 | -0.29 | -0.58 | -0.88 | -0.67 | -0.33 | -0.47 | -0.61 | -0.46 | -0.46 | 0.0525 | -0.09 | 0.025 | 0.815351 | 0.694717 | 0.90551 | 0.999971 | 1 | 0.991091 |
| Dcaf1 | Q80TR8-4 | 3.2 | 2.44 | 0.89 | 1.02 | 1.85 | 2.23 | 0.77 | 0.85 | 1.64 | 2.54 | -0.04 | 0.01 | 1.16 | 2.26 | 0.62 | 0.14 | -0.4625 | -0.85 | -0.8425 | 0.455377 | 0.285788 | 0.221167 | 0.999971 | 1 | 0.8275 |
| Dcaf10 | A2AKB9-2 | 0 | 0.87 | 0.42 | 2.07 | 1.11 | 0.58 | 1.14 | 1.68 | 0.68 | 0.44 | 1.41 | 0.89 | 1 | 1.08 | 1.07 | 1.56 | 0.2875 | 0.015 | 0.3375 | 0.542673 | 0.974106 | 0.44989 | 0.999971 | 1 | 0.885596 |
| Dcaf11 | Q91VU6 | -0.32 | 0.07 | 2.54 | 1.13 | -0.1 | -0.14 | 1.33 | 2.05 | 0.14 | 0.15 | 1.7 | 0.73 | 0.17 | -0.03 | 2.5 | 2.48 | -0.07 | -0.175 | 0.425 | 0.926119 | 0.794926 | 0.620072 | 0.999971 | 1 | 0.929296 |
| Dcaf13 | Q6PAC3 | -1.37 | -1.67 | 0.34 | -0.49 | -1.56 | -2.51 | -0.01 | 0.17 | -1.46 | -1.73 | -0.24 | -0.27 | -1.5 | -2.01 | -1.1 | -0.36 | -0.18 | -0.1275 | -0.445 | 0.800063 | 0.817919 | 0.409811 | 0.999971 | 1 | 0.875847 |
| Dcaf5 | Q80T85 | 2.66 | 3.24 | 1.11 | 1.71 | 3.77 | 3.56 | 1.31 | 1.55 | 2.91 | 3.27 | 1.06 | 1.07 | 1.14 | 1.9 | 0.75 | 0.88 | 0.3675 | -0.1025 | -1.0125 | 0.61506 | 0.881839 | 0.070314 | 0.999971 | 1 | 0.735439 |
| Dcaf6 | Q9DC22 | 0.55 | 0.82 | 1.09 | 0.99 | 1.15 | 0.58 | 0.81 | 0.81 | 1.07 | 0.66 | 0.91 | 1.24 | 1.16 | 0.52 | 0.43 | 1.05 | -0.025 | 0.1075 | -0.0725 | 0.907053 | 0.618781 | 0.770789 | 0.999971 | 1 | 0.963498 |
| Dcaf7 | P61963 | 0.16 | 0.13 | -0.11 | -0.25 | 0.06 | 0.25 | -0.39 | 0.13 | 0.35 | 0.16 | -0.11 | -0.04 | 0.27 | 0.47 | 0.23 | 0.08 | 0.03 | 0.1075 | 0.28 | 0.889916 | 0.590349 | 0.171465 | 0.999971 | 1 | 0.804075 |
| Dcaf8 | Q8N7N5 | 0.73 | 0.68 | 1.15 | 0.89 | 0.67 | 0.44 | 0.94 | 0.74 | 0.78 | 1 | 1.12 | 0.55 | 0.91 | 0.84 | 0.99 | 1.14 | -0.165 | 7.85E-17 | 0.1075 | 0.426019 | 1 | 0.577701 | 0.999971 | 1 | 0.918581 |
| Dcakd | Q8BHC4 | -0.63 | -1.08 | -0.61 | -1.11 | -0.62 | -0.9 | -0.6 | -0.44 | -0.47 | -0.51 | -0.41 | -0.22 | -0.11 | -0.71 | -0.48 | -0.28 | 0.2175 | 0.455 | 0.4625 | 0.325468 | 0.048708 | 0.069142 | 0.999971 | 1 | 0.733937 |
| Dcc | P70211-3 | -0.29 | -0.76 | -0.48 | -0.44 | -0.36 | -0.19 | -0.69 | -0.37 | -0.36 | -0.26 | -0.56 | -0.05 | -0.17 | -0.02 | -0.48 | -0.21 | 0.09 | 0.185 | 0.2725 | 0.655771 | 0.364611 | 0.192702 | 0.999971 | 1 | 0.810547 |
| Dchs1 | E9PVD3 | 0.35 | -0.84 | 0.09 | 0.23 | -0.35 | -0.52 | -0.32 | 0.04 | -0.53 | -0.26 | -0.2 | 0.1 | -0.46 | -0.86 | 0 | 0.44 | -0.245 | -0.18 | -0.1775 | 0.427562 | 0.560415 | 0.641047 | 0.999971 | 1 | 0.93301 |
| Dclk1 | Q80VB6 | -2.21 | -1.58 | -0.89 | -1.97 | -2.54 | -0.52 | -2.4 | -2.19 | -0.35 | 0.61 | -0.88 | -1.14 | 3.88 | -0.64 | 0.23 | 0.68 | -0.25 | 1.2225 | 2.7 | 0.62615 | 0.02372 | 0.015926 | 0.999971 | 1 | 0.613095 |
| Dclk1_1 | A0A0G2JG( | -1.85 | 0.44 | -0.51 | -1.96 | -0.48 | 0.18 | -0.46 | -0.87 | 0.1 | -1.41 | 0 | 0.6 | 2.6 | -0.35 | -0.04 | -1.64 | 0.5625 | 0.7925 | 1.1125 | 0.331673 | 0.246953 | 0.25937 | 0.999971 | 1 | 0.839154 |
| Dclk1_2 | Q8CCN4 | 0.23 | 0.29 | -2.09 | -1.41 | -0.78 | -0.52 | -1.63 | -2.05 | -1.48 | -0.94 | -1.45 | -1.16 | 0.01 | -1.3 | -0.64 | -0.73 | -0.5 | -0.5125 | 0.08 | 0.437678 | 0.374432 | 0.893561 | 0.999971 | 1 | 0.98828 |
| Dclk1_3 | H7BX36 | -1.05 | 0.56 | 0.7 | 0.44 | 1.33 | 1.07 | 2.01 | 0.46 | 2.43 | -0.49 | 2.61 | 0.99 | 0.86 | -1.04 | 0.59 | -1.03 | 1.055 | 1.2225 | -0.3175 | 0.053269 | 0.131594 | 0.598185 | 0.999971 | 1 | 0.92377 |
| Dclk1_4 | Q9JLM8 | -0.88 | -0.8 | -0.53 | -0.63 | -0.59 | -0.59 | -0.13 | -0.29 | -0.34 | -0.56 | -0.01 | 0.11 | 0.1 | -0.51 | -0.17 | 0.27 | 0.31 | 0.51 | 0.6325 | 0.144893 | 0.039837 | 0.020397 | 0.999971 | 1 | 0.635388 |
| Dclk2 | Q6PGN3-3 | 0.26 | 0.71 | 0.46 | 0.3 | 0.65 | 0.59 | 0.66 | 0.54 | 0.85 | 0.5 | 0.58 | 0.47 | 0.92 | 0.42 | 0.49 | 0.37 | 0.1775 | 0.1675 | 0.1175 | 0.343175 | 0.395979 | 0.583031 | 0.999971 | 1 | 0.920364 |
| Dcn | P28654 | 1.26 | 0.74 | 0.47 | 0.33 | 3.06 | 0.41 | -0.66 | -0.43 | 0.38 | 1.51 | -0.13 | 1.82 | 0.04 | 1.52 | 0.98 | -0.63 | -0.105 | 0.195 | -0.2225 | 0.894043 | 0.68143 | 0.648836 | 0.999971 | 1 | 0.933842 |
| Dcp1a | Q91YD3 | 0.38 | 0.72 | 0.68 | 0.62 | 0.53 | 0.61 | 0.06 | 0.92 | 0.37 | 0.36 | 0.63 | 0.89 | 0.79 | 0.76 | 0.9 | 0.91 | -0.07 | -0.0375 | 0.24 | 0.762262 | 0.851545 | 0.192069 | 0.999971 | 1 | 0.810547 |
| Dcp1b | B9EIX0 | 0.62 | 1.49 | -0.29 | -0.03 | 0.43 | 0.85 | -0.6 | 0.39 | 1.15 | 0.84 | -0.53 | -0.02 | 0.16 | 1.27 | 0 | -0.67 | -0.18 | -0.0875 | -0.2575 | 0.701913 | 0.86496 | 0.624471 | 0.999971 | 1 | 0.930479 |
| Dcp2 | Q9CYC6 | 2 | 0.87 | 0.53 | 0.87 | 1.04 | 0.97 | 2.17 | 0.58 | 1.11 | 1.59 | 1.12 | 0.83 | 0.76 | 0.25 | -0.24 | 0.49 | 0.1225 | 0.095 | -0.7525 | 0.781689 | 0.787113 | 0.070464 | 0.999971 | 1 | 0.735439 |
| Dcps | Q9DAR7 | 1.82 | 2.27 | 3.01 | 3.66 | 1.87 | 1.79 | 2.83 | 2.45 | 1.5 | 1.79 | 2.21 | 2.46 | 0.87 | 1.96 | 2.41 | 2.68 | -0.455 | -0.7 | -0.71 | 0.322831 | 0.134448 | 0.200081 | 0.999971 | 1 | 0.813352 |
| Dctn1 | E9Q586 | -1.23 | 0.09 | -1.26 | -0.11 | -1.5 | -0.37 | -0.54 | -0.5 | -0.82 | -0.92 | -0.14 | 0.18 | -0.3 | -0.22 | -0.27 | 0.01 | -0.1 | 0.2025 | 0.4325 | 0.812031 | 0.635413 | 0.247363 | 0.999971 | 1 | 0.839012 |
| Dctn1_1 | E9Q3M3 | 0.05 | -0.19 | -0.33 | -0.23 | -0.29 | 0.02 | -0.22 | -0.03 | -0.21 | 0.11 | -0.37 | -0.4 | 0.12 | -0.04 | -0.06 | 0 | 0.045 | -0.0425 | 0.18 | 0.806782 | 0.829667 | 0.321804 | 0.999971 | 1 | 0.85548 |
| Dctn2 | Q99KJ8 | -0.84 | -0.26 | -0.56 | -0.78 | 0.01 | -0.38 | -0.03 | -0.62 | 0.15 | -0.41 | -0.24 | 0.12 | 0.62 | -0.16 | -0.44 | -0.51 | 0.355 | 0.515 | 0.4875 | 0.156032 | 0.046461 | 0.130531 | 0.999971 | 1 | 0.784948 |
| Dctn3 | Q9Z0Y1 | -0.21 | -0.14 | -0.03 | -0.34 | -0.12 | 0.19 | -0.26 | 0.41 | 0.14 | 0.19 | 0.6 | 0.22 | 0.44 | 0.21 | 0.66 | 0.71 | 0.235 | 0.4675 | 0.685 | 0.286886 | 0.032507 | 0.006425 | 0.999971 | 1 | 0.582714 |
| Dctn4 | Q8CBY8-2 | -0.39 | -0.12 | 0.33 | 0.29 | -0.13 | 0.15 | -0.03 | 0.73 | -0.11 | -0.19 | 0.49 | -0.24 | 0.19 | 0.33 | 0.81 | 0.56 | 0.1525 | -0.04 | 0.445 | 0.582995 | 0.878841 | 0.1011 | 0.999971 | 1 | 0.78268 |
| Dctn5 | Q9QZB9 | 0.41 | 0.31 | 0.58 | 1.08 | 0.29 | 0.45 | 0.21 | 1.28 | 0.44 | 0.59 | 0.79 | 0.47 | 0.79 | 0.72 | 1.04 | 1.16 | -0.0375 | -0.0225 | 0.3325 | 0.902409 | 0.920585 | 0.18265 | 0.999971 | 1 | 0.808532 |
| Dctn6 | D3Z6H3 | -1.06 | -0.5 | -0.39 | -0.43 | -0.72 | -0.3 | -0.13 | -0.32 | -0.26 | -0.58 | -0.07 | -0.19 | -0.12 | -0.2 | -0.53 | -0.27 | 0.2275 | 0.32 | 0.315 | 0.347155 | 0.183542 | 0.182203 | 0.999971 | 1 | 0.808532 |
| Dctpp1 | Q9QY93 | 1.42 | 1.01 | 1.07 | 1.56 | 1.2 | 0.87 | 1.16 | 1.81 | 1.64 | 0.32 | 1.43 | 1.8 | 1.82 | 0.55 | 1.67 | 1.67 | -0.005 | 0.0325 | 0.1625 | 0.984721 | 0.926678 | 0.619517 | 0.999971 | 1 | 0.929296 |
| Dcun1d1 | Q9QZ73 | -0.13 | 0.54 | 0.72 | 0.16 | 0.74 | 0.58 | 0.63 | 0.26 | 0.74 | 0 | 0.61 | 0.72 | 0.6 | 0.52 | 0.88 | 0.33 | 0.23 | 0.195 | 0.26 | 0.362817 | 0.484357 | 0.315057 | 0.999971 | 1 | 0.852056 |
| Dcun1d2 | Q8BZJ7 | -0.39 | -0.76 | -0.73 | -0.43 | -0.79 | -0.77 | -0.93 | -0.36 | -1.2 | -1.15 | -0.24 | -0.33 | -0.48 | -0.61 | -0.36 | -0.45 | -0.135 | -0.1525 | 0.1025 | 0.522114 | 0.598346 | 0.582404 | 0.999971 | 1 | 0.920364 |
| Dcun1d3 | Q8K0V2 | 0.48 | -0.54 | -0.21 | -0.13 | -0.63 | -0.24 | -0.19 | -0.39 | -0.48 | 0.36 | -0.32 | -0.56 | -0.66 | -0.57 | -0.39 | -0.47 | -0.2625 | -0.15 | -0.4225 | 0.324597 | 0.624809 | 0.117788 | 0.999971 | 1 | 0.78268 |
| Dcun1d4 | Q8C5X2 | 3.1 | 2.1 | 2.1 | 1.92 | 2.13 | 2.35 | 1.92 | 2.44 | 2.2 | 2.75 | 1.6 | 1.77 | 1.03 | 1.96 | 2.09 | 1.65 | -0.095 | -0.225 | -0.6225 | 0.752604 | 0.539369 | 0.104371 | 0.999971 | 1 | 0.78268 |
| Dcun1d5 | Q9CXV9 | -0.21 | 2.33 | 1.58 | 0.99 | 1.9 | 1.06 | 1.23 | 1.94 | 1.54 | -0.1 | 1.75 | 1.21 | 1.72 | 1.22 | 1.2 | 0.51 | 0.36 | -0.0725 | -0.01 | 0.507291 | 0.907033 | 0.985331 | 0.999971 | 1 | 0.99925 |
| Dcx | O88809 | 1.56 | -0.27 | -0.41 | -0.01 | 0.29 | 0.18 | 0.15 | -0.15 | 1.13 | 1.57 | 0.37 | 1.04 | 0.79 | -0.29 | -0.21 | 0.22 | -0.1 | 0.81 | -0.09 | 0.819468 | 0.122178 | 0.852224 | 0.999971 | 1 | 0.98187 |
| Dcxr | Q91X52 | 0.85 | 0.26 | 0.81 | 0.89 | 0.33 | 0.69 | 0.33 | 0.26 | 0.38 | 0.76 | 0.49 | 0.81 | 0.58 | 0.66 | 0.56 | 1.08 | -0.3 | -0.0925 | 0.0175 | 0.197409 | 0.677094 | 0.939497 | 0.999971 | 1 | 0.995869 |
| Dda1 | Q9D9Z5 | 2.08 | 2.78 | 1.83 | 1.71 | 2.92 | 3.02 | 2.6 | 1.98 | 3.08 | 3.16 | 1.99 | 2.29 | 3.91 | 2.99 | 2.04 | 1.79 | 0.53 | 0.53 | 0.5825 | 0.138442 | 0.173223 | 0.264084 | 0.999971 | 1 | 0.842533 |
| Ddah1 | Q9CWS0 | -0.63 | -0.28 | -1.32 | -1.15 | -0.29 | -0.91 | -0.25 | -1.05 | 0.13 | -0.88 | -1.23 | -1.4 | 0.17 | -1.05 | -1.43 | -1.79 | 0.22 | -2.36E-16 | -0.18 | 0.495851 | 1 | 0.695638 | 0.999971 | 1 | 0.947174 |
| Ddah2 | Q99LD8 | -0.66 | 0.23 | -0.09 | -0.69 | 0.12 | -0.05 | 0.22 | -0.74 | 0.84 | -0.71 | 0 | 0.04 | 0.69 | -0.41 | -0.37 | -0.77 | 0.19 | 0.345 | 0.0875 | 0.551232 | 0.37152 | 0.815745 | 0.999971 | 1 | 0.974918 |
| Ddb1 | Q3U1J4 | -0.11 | 0.69 | 0.88 | 0.67 | 0.49 | 0.43 | 0.81 | 0.5 | 0.55 | -0.08 | 0.77 | 0.46 | 0.04 | 0.56 | 0.66 | 0.7 | 0.025 | -0.1075 | -0.0425 | 0.923118 | 0.715187 | 0.880453 | 0.999971 | 1 | 0.985441 |
| Ddc | O88533 | 0.34 | -0.22 | -0.93 | -0.2 | 0.24 | -0.92 | -0.96 | -0.35 | 0.52 | 0.36 | -0.67 | -0.17 | 0.54 | 0.34 | -0.03 | -0.8 | -0.245 | 0.2625 | 0.265 | 0.515758 | 0.479842 | 0.492703 | 0.999971 | 1 | 0.900198 |
| Ddhd1 | F8WIJ5 | -0.2 | -1.07 | -0.97 | -0.63 | -0.99 | -0.67 | -0.96 | -0.91 | -0.96 | -0.26 | -0.77 | -1.21 | -1.09 | -0.93 | -0.96 | -0.78 | -0.165 | -0.0825 | -0.2225 | 0.500307 | 0.777557 | 0.36638 | 0.999971 | 1 | 0.869526 |
| Ddhd2 | Q80Y98 | -0.12 | -0.04 | 0.15 | -0.05 | -0.2 | 0.09 | -0.04 | -0.07 | -0.25 | -0.03 | -0.1 | -0.43 | -0.41 | -0.1 | -0.19 | -0.3 | -0.04 | -0.1875 | -0.235 | 0.81703 | 0.310291 | 0.20368 | 0.999971 | 1 | 0.81617 |
| Ddi2 | A2ADY9 | -0.05 | 0.29 | 0.43 | 0.36 | 0.16 | 0.37 | -0.03 | 0.56 | -0.06 | 0.12 | -0.03 | 0.33 | 0.41 | 0.61 | 0.57 | 0.61 | 0.0075 | -0.1675 | 0.2925 | 0.972028 | 0.402338 | 0.145544 | 0.999971 | 1 | 0.791767 |
| Ddn | Q80TS7 | 0.14 | 0.03 | -0.52 | -0.42 | -0.28 | -0.47 | -0.53 | -0.16 | -0.28 | -0.29 | -0.45 | -0.62 | -0.23 | -1.29 | -1.03 | -0.8 | -0.1675 | -0.2175 | -0.645 | 0.463776 | 0.33971 | 0.050511 | 0.999971 | 1 | 0.722689 |
| Ddost | O54734 | -0.26 | -0.79 | -0.23 | -0.43 | -0.94 | -0.49 | 0.07 | -0.14 | -0.83 | -0.56 | 0.03 | 0.33 | -0.82 | -0.52 | -0.09 | 0.09 | 0.0525 | 0.17 | 0.0925 | 0.847822 | 0.577124 | 0.727992 | 0.999971 | 1 | 0.956041 |
| Ddr1 | Q03146-2 | -1.15 | -0.6 | -0.04 | -0.47 | -0.33 | -1.69 | -0.39 | -0.08 | -0.06 | -1.68 | 0.09 | -0.03 | 0.01 | -1.77 | -0.03 | -0.19 | -0.0575 | 0.145 | 0.07 | 0.887868 | 0.748787 | 0.877749 | 0.999971 | 1 | 0.985319 |
| Ddrgk1 | Q80WW9 | 2.65 | -0.1 | -0.17 | 0.12 | 0.13 | -0.13 | 1.44 | 0.59 | -0.44 | 1.74 | 0.61 | 0.82 | -0.57 | -0.82 | -0.15 | -0.17 | -0.1175 | 0.0575 | -1.0525 | 0.864338 | 0.937791 | 0.124515 | 0.999971 | 1 | 0.78268 |
| Ddt | O35215 | -0.29 | 0.93 | 0.44 | 0.45 | 0.7 | 0.53 | 0.15 | 0.38 | 0.45 | -0.63 | -0.26 | 0.52 | 0.28 | 0.8 | 0.35 | 0.32 | 0.0575 | -0.3625 | 0.055 | 0.842546 | 0.336492 | 0.85043 | 0.999971 | 1 | 0.98187 |
| Ddx1 | Q91VR5 | 0.13 | 1.87 | 1.99 | 1.61 | 1.58 | 1.46 | 1.8 | 1.58 | 1.72 | 0.89 | 1.96 | 2.12 | 1.34 | 1.58 | 1.73 | 1.68 | 0.205 | 0.2725 | 0.1825 | 0.62333 | 0.57197 | 0.663877 | 0.999971 | 1 | 0.936621 |
| Ddx10 | Q80Y44 | 1.64 | -0.77 | 0.31 | 0.42 | -0.48 | -0.42 | -0.24 | -1.38 | -0.13 | 1.09 | -1.38 | -2.88 | -3.61 | -2.36 | -0.34 | -2.21 | -1.03 | -1.225 | -2.53 | 0.071098 | 0.191155 | 0.008192 | 0.999971 | 1 | 0.602121 |
| Ddx17 | Q3U741 | 1.21 | 2.08 | 1.88 | 1.62 | 2.21 | 1.53 | 1.34 | 1.64 | 2.01 | 0.74 | 1.36 | 1.81 | 1.87 | 1.11 | 1.41 | 1.27 | -0.0175 | -0.2175 | -0.2825 | 0.950201 | 0.522588 | 0.310028 | 0.999971 | 1 | 0.847402 |
| Ddx18 | Q8K363 | 1.48 | -0.94 | -1.48 | -1.53 | -0.94 | -0.97 | -1.36 | -1.89 | -1.13 | -0.13 | -1.19 | -0.88 | -1.44 | -1.02 | -1.18 | -1.42 | -0.6725 | -0.215 | -0.6475 | 0.333235 | 0.754256 | 0.33575 | 0.999971 | 1 | 0.858827 |
| Ddx19a | Q61655 | -0.36 | 0.34 | -0.19 | -0.57 | 0.63 | 0.24 | -0.19 | -0.42 | 0.58 | 0.09 | -0.06 | -0.17 | 0.62 | 0.08 | -0.42 | -0.62 | 0.26 | 0.305 | 0.11 | 0.409648 | 0.280345 | 0.744649 | 0.999971 | 1 | 0.961064 |
| Ddx19b | Q8R3C7 | -0.83 | -0.58 | 0.2 | -0.14 | -0.95 | -1.17 | 0.13 | 0.35 | -0.82 | -1.11 | 0.03 | 0.86 | -1.73 | -1.41 | 0.55 | 0.65 | -0.0725 | 0.0775 | -0.1475 | 0.86326 | 0.86883 | 0.809701 | 0.999971 | 1 | 0.973791 |
| Ddx20 | Q9JJY4 | 0.21 | 1.19 | 1.62 | 1.87 | 1.19 | 0.34 | 1.28 | 1.9 | 0.55 | 0.18 | 1.65 | 1.99 | 1.16 | 0.44 | 1.83 | 2.26 | -0.045 | -0.13 | 0.2 | 0.921278 | 0.804669 | 0.691413 | 0.999971 | 1 | 0.946835 |
| Ddx21 | Q9JIK5 | -0.1 | -0.18 | -0.53 | -0.58 | -0.33 | -0.5 | -0.86 | -0.8 | -0.51 | -0.27 | -0.59 | -1.2 | -0.57 | -0.46 | -0.76 | -0.99 | -0.275 | -0.295 | -0.3475 | 0.230101 | 0.266912 | 0.13397 | 0.999971 | 1 | 0.784948 |
| Ddx23 | D3Z0M9 | 0.97 | 0.47 | 1.09 | 0.49 | 0.56 | 0.25 | 0.77 | 0.56 | 0.28 | 0.47 | 0.4 | 0.46 | -0.07 | 0.52 | 0.69 | 0.7 | -0.22 | -0.3525 | -0.295 | 0.353196 | 0.123506 | 0.28249 | 0.999971 | 1 | 0.844538 |
| Ddx24 | F8WJA0 | 0.81 | 0 | -0.29 | -0.63 | 0.09 | -0.12 | -0.56 | -0.17 | -0.12 | 0.74 | -0.49 | -0.59 | 0 | -0.14 | -0.84 | -0.85 | -0.1625 | -0.0875 | -0.43 | 0.628884 | 0.831732 | 0.264851 | 0.999971 | 1 | 0.842533 |
| Ddx25 | Q9QY15 | 1.16 | 1.3 | 0.68 | 0.82 | 1.14 | 1.25 | 0.63 | 1.3 | 1.29 | 1.48 | 0.73 | 1.07 | 1.42 | 1.75 | 0.97 | 1.09 | 0.09 | 0.1525 | 0.3175 | 0.711334 | 0.537289 | 0.232417 | 0.999971 | 1 | 0.831121 |
| Ddx27 | Q921N6 | -0.59 | -1.23 | -1.52 | -1.06 | -0.89 | -1.63 | -1.99 | -1.75 | -1.39 | -0.76 | -1.17 | -1.57 | -1.42 | -1.38 | -1.61 | -0.87 | -0.465 | -0.1225 | -0.22 | 0.160909 | 0.659728 | 0.425501 | 0.999971 | 1 | 0.880767 |
| Ddx28 | Q9CWT6 | 1.49 | 0.62 | -0.66 | -1.24 | -0.91 | -0.39 | -1.06 | -1.06 | -0.11 | 1.19 | -1.25 | -0.69 | 0.66 | 0.23 | -0.57 | -0.47 | -0.9075 | -0.2675 | -0.09 | 0.145496 | 0.716541 | 0.885017 | 0.999971 | 1 | 0.98711 |
| Ddx39b | Q9Z1N5 | -0.03 | 0.61 | 0.43 | 0.6 | 0.58 | 0.59 | 0.5 | 0.36 | 0.49 | 0.29 | -0.15 | 0.42 | 0.09 | 0.9 | 0.44 | 0.6 | 0.105 | -0.14 | 0.105 | 0.61919 | 0.561155 | 0.679648 | 0.999971 | 1 | 0.942957 |
| Ddx3x | Q62167 | 0.07 | 0.84 | 0.53 | 0.48 | 0.8 | 0.65 | 0.42 | 0.69 | 0.95 | 0.14 | 0.89 | 0.68 | 0.78 | 0.64 | 0.54 | 0.74 | 0.16 | 0.185 | 0.195 | 0.475233 | 0.48849 | 0.376402 | 0.999971 | 1 | 0.870465 |
| Ddx3y | Q62095 | -1.9 | 2.89 | 2.13 | -3.72 | 2.8 | -3.61 | 1.52 | -3.03 | -2.93 | 0.52 | 2.85 | -3.06 | 2.81 | -2.01 | 2.89 | -4.1 | -0.43 | -0.505 | 0.0475 | 0.82961 | 0.791367 | 0.981907 | 0.999971 | 1 | 0.99925 |
| Ddx41 | A0A1S6GW | 0.62 | 1.14 | 0.44 | 0.3 | 0.72 | 0.76 | 0.37 | 0.52 | 1.04 | 0.74 | 0.71 | 0.74 | 1.12 | 1.26 | 0.43 | 0.59 | -0.0325 | 0.1825 | 0.225 | 0.891562 | 0.441937 | 0.440284 | 0.999971 | 1 | 0.883105 |
| Ddx42 | Q810A7 | 2.23 | 2.46 | 2.32 | 2.37 | 2.45 | 2.17 | 1.7 | 2.13 | 2.45 | 2.41 | 2.3 | 2.35 | 1.75 | 1.84 | 2.11 | 2.45 | -0.2325 | 0.0325 | -0.3075 | 0.288711 | 0.840085 | 0.174743 | 0.999971 | 1 | 0.808161 |
| Ddx46 | Q569Z5 | 1.86 | 1.77 | 1.38 | 1.23 | 1.95 | 1.72 | 1.35 | 1.06 | 1.77 | 1.87 | 1.36 | 1.49 | 1.33 | 1.71 | 1.26 | 1.31 | -0.04 | 0.0625 | -0.1575 | 0.881494 | 0.784705 | 0.490687 | 0.999971 | 1 | 0.898612 |
| Ddx47 | F6QKD2 | 1.12 | 0.35 | -0.02 | 0.29 | -0.08 | -0.15 | -0.38 | 0.19 | -0.41 | 0.23 | 0.52 | 0.45 | -0.59 | -0.12 | 0.03 | 0.48 | -0.54 | -0.2375 | -0.485 | 0.083675 | 0.469004 | 0.164904 | 0.999971 | 1 | 0.800215 |
| Ddx5 | Q8BTS0 | 0.28 | 1.28 | 1.38 | 1.02 | 1.38 | 1.02 | 1.09 | 1.4 | 1.18 | 0.24 | 1.5 | 1.24 | 1.2 | 1.11 | 1.52 | 1.14 | 0.2325 | 0.05 | 0.2525 | 0.41869 | 0.8902 | 0.381117 | 0.999971 | 1 | 0.871816 |
| Ddx50 | Q99MJ9 | 0.7 | -0.15 | -0.52 | -0.49 | -0.32 | 0.15 | -0.52 | -0.69 | -0.47 | 0.75 | -0.58 | -0.82 | 0.03 | 0.12 | -0.66 | -0.69 | -0.23 | -0.165 | -0.185 | 0.498381 | 0.701359 | 0.602082 | 0.999971 | 1 | 0.924453 |
| Ddx51 | Q6P9R1 | -0.3 | -1.27 | -1.41 | -1.3 | -1.52 | -1.18 | -1.28 | -1.83 | -1.3 | -0.84 | -1.56 | -2.08 | -1.63 | -1.43 | -1.99 | -1.68 | -0.3825 | -0.375 | -0.6125 | 0.228486 | 0.310557 | 0.063061 | 0.999971 | 1 | 0.725348 |
| Ddx52 | Q8K301 | -3.65 | -0.95 | -1.3 | -5.95 | -0.12 | -2.45 | -0.39 | -4.45 | 0.4 | -3.94 | -0.65 | 0.2 | 0.1 | -3.58 | -3.58 | -4.81 | 1.11 | 1.965 | -0.005 | 0.425421 | 0.177957 | 0.997146 | 0.999971 | 1 | 1 |
| Ddx54 | Q8K4L0 | 0.09 | -0.84 | -0.56 | -1.28 | -0.15 | -1.47 | -0.38 | -0.99 | -0.8 | 0.21 | -1.5 | -1.55 | -0.45 | -0.3 | -1.27 | -1.16 | -0.1 | -0.2625 | -0.1475 | 0.801161 | 0.579888 | 0.689661 | 0.999971 | 1 | 0.945805 |
| Ddx55 | E9Q9T6 | 1.98 | 1.38 | 0.55 | 0.36 | 0.32 | 0.62 | -0.75 | 0.4 | 0.3 | 0.88 | -0.11 | -0.01 | -0.88 | 0.7 | 0.17 | -0.64 | -0.92 | -0.8025 | -1.23 | 0.069214 | 0.08155 | 0.030964 | 0.999971 | 1 | 0.678474 |
| Ddx56 | Q9D0R4 | -0.12 | 0.64 | 0.02 | -0.36 | 0.54 | 0.33 | -0.07 | -0.13 | 0.54 | 0.03 | -0.34 | -0.44 | 1.06 | 0.13 | -1.13 | -0.92 | 0.1225 | -0.0975 | -0.26 | 0.665368 | 0.75504 | 0.612423 | 0.999971 | 1 | 0.927335 |
| Ddx58 | A2AP29 | 2.13 | 2.45 | 0.52 | 0.53 | 2.97 | 1.8 | 0.07 | -0.23 | 2.65 | 1.49 | 0.4 | 0.47 | 3.59 | 0.92 | 0.35 | -0.7 | -0.255 | -0.155 | -0.3675 | 0.755774 | 0.817682 | 0.696472 | 0.999971 | 1 | 0.947174 |
| Ddx6 | P54823 | 0.79 | 1.38 | 1.06 | 1.15 | 1.3 | 1.42 | 0.86 | 1.21 | 1.28 | 1.15 | 1.05 | 1.07 | 1.05 | 1.74 | 1.32 | 1.36 | 0.1025 | 0.0425 | 0.2725 | 0.638937 | 0.825234 | 0.250728 | 0.999971 | 1 | 0.839012 |
| Decr1 | Q9CQ62 | 0.75 | -1.7 | -0.5 | 0.11 | -1.66 | -1.23 | -0.61 | 0.24 | -1.67 | 0.43 | -0.73 | -0.7 | -2.61 | -1.2 | -0.59 | -0.49 | -0.48 | -0.3325 | -0.8875 | 0.436921 | 0.594279 | 0.195474 | 0.999971 | 1 | 0.811225 |
| Decr2 | Q9WV68 | 0.92 | 0.42 | -0.27 | 0.04 | 0.08 | 1.04 | 0 | -0.22 | 0.24 | 1.42 | -0.14 | -0.27 | 0.24 | 0.6 | -0.26 | -0.17 | -0.0525 | 0.035 | -0.175 | 0.886526 | 0.936185 | 0.593931 | 0.999971 | 1 | 0.92342 |
| Def6 | A0A0R4IZX | 3.43 | 0.98 | 3.41 | 1.72 | 0.7 | 0.57 | 2.51 | 1.38 | 2.85 | 2.72 | 3.78 | 2.76 | 1.35 | 0.61 | 1.85 | 1.47 | -1.095 | 0.6425 | -1.065 | 0.137627 | 0.308722 | 0.108649 | 0.999971 | 1 | 0.78268 |
| Def8 | A0A0R4IZV | 1.22 | -1.47 | -0.16 | 1.19 | -1.53 | -1.95 | 1.3 | 0.21 | -1.41 | -0.3 | 0.43 | -1.02 | -1.98 | -1.55 | -0.74 | -0.92 | -0.6875 | -0.77 | -1.4925 | 0.447405 | 0.283134 | 0.041871 | 0.999971 | 1 | 0.701725 |
| Degs1 | O09005 | -1.85 | -0.64 | -0.49 | -0.38 | -1.32 | -1.14 | -0.73 | 0.12 | -0.92 | -1.16 | 0.13 | 0.17 | -0.83 | -0.21 | -0.44 | -0.61 | 0.0725 | 0.395 | 0.3175 | 0.869247 | 0.398684 | 0.386171 | 0.999971 | 1 | 0.871816 |
| Dek | Q7TNV0 | 1.8 | 2.36 | 1.83 | 1.7 | 2.52 | 1.98 | 1.66 | 1.62 | 2.46 | 1.57 | 2.01 | 2.19 | 1.98 | 2.17 | 2.11 | 1.67 | 0.0225 | 0.135 | 0.06 | 0.934364 | 0.60667 | 0.791477 | 0.999971 | 1 | 0.968737 |
| Dennd10 | Q9D8N2 | 0.16 | 0.8 | 0.48 | 1.26 | 0.46 | 0.73 | 0.37 | 1.37 | 0.52 | 0.8 | 0.86 | 0.69 | 0.95 | 0.96 | 1.22 | 1.2 | 0.0575 | 0.0425 | 0.4075 | 0.860035 | 0.872892 | 0.153231 | 0.999971 | 1 | 0.795578 |
| Dennd11 | Q3UHG7 | -0.01 | -0.06 | -0.48 | -0.87 | -0.19 | 0 | -0.27 | -0.82 | -0.67 | -0.12 | -0.63 | -0.34 | 0.13 | 0.22 | -0.44 | -0.97 | 0.035 | -0.085 | 0.09 | 0.901225 | 0.745072 | 0.791323 | 0.999971 | 1 | 0.968737 |
| Dennd1a | Q8K382 | 0.14 | -0.17 | -0.14 | -0.41 | -0.45 | -0.04 | -0.35 | -0.06 | -0.47 | -0.36 | -0.14 | -0.47 | -0.11 | 0.15 | -0.11 | -0.2 | -0.08 | -0.215 | 0.0775 | 0.69902 | 0.28654 | 0.695752 | 0.999971 | 1 | 0.947174 |
| Dennd1b | Q3U1T9-2 | 0.71 | -1.24 | 0.72 | -0.22 | -0.83 | -0.49 | 0.7 | -0.01 | -0.5 | 0.56 | 1.08 | -0.72 | -1 | -0.88 | 0.06 | 0.36 | -0.15 | 0.1125 | -0.3575 | 0.776459 | 0.846897 | 0.507077 | 0.999971 | 1 | 0.905815 |
| Dennd2a | Q8C4S8 | -0.4 | -0.27 | -0.64 | -0.06 | 0.27 | 0.14 | -0.5 | -0.26 | 0.04 | -1.11 | -0.66 | -0.54 | 0.84 | 0.67 | -0.71 | 0.23 | 0.255 | -0.225 | 0.6 | 0.313961 | 0.43074 | 0.122836 | 0.999971 | 1 | 0.78268 |
| Dennd4a | E9Q8V6 | -0.35 | 0.25 | 0.1 | -0.22 | 0.16 | 0.07 | -0.21 | -0.15 | 0.22 | 0.13 | 0.2 | 0.34 | 0.39 | 0.11 | 0.02 | 0.26 | 0.0225 | 0.2775 | 0.25 | 0.91579 | 0.188213 | 0.256666 | 0.999971 | 1 | 0.839012 |
| Dennd4b | A0A0R4J17 | -0.63 | -0.41 | -0.25 | -0.36 | -0.47 | -0.78 | -0.38 | -0.51 | -0.68 | -1.11 | -0.49 | -0.11 | -0.44 | -0.67 | -0.57 | -0.84 | -0.1225 | -0.185 | -0.2175 | 0.518673 | 0.464636 | 0.264851 | 0.999971 | 1 | 0.842533 |
| Dennd4c | A6H8H2-2 | 0.32 | -0.42 | -0.75 | -0.45 | 0.57 | -0.36 | -1.15 | -0.72 | 0.62 | 0.53 | -0.33 | -0.34 | 0.08 | 0.13 | -0.54 | 0.08 | -0.09 | 0.445 | 0.2625 | 0.82635 | 0.217493 | 0.377596 | 0.999971 | 1 | 0.871417 |
| Dennd5a | A0A0R4J1E | -0.02 | -0.3 | -0.42 | -0.35 | -0.43 | -0.19 | -0.37 | -0.03 | -0.26 | 0.07 | -0.35 | -0.04 | -0.24 | -0.26 | -0.24 | -0.19 | 0.0175 | 0.1275 | 0.04 | 0.927257 | 0.510534 | 0.820107 | 0.999971 | 1 | 0.975761 |
| Dennd5b | A2RSQ0 | -0.34 | 0.27 | -0.06 | -0.19 | 0.11 | 0.17 | 0.23 | -0.13 | 0.31 | -0.16 | 0.56 | 0.28 | -0.02 | 0.09 | -0.02 | 0.23 | 0.175 | 0.3275 | 0.15 | 0.404962 | 0.182734 | 0.463466 | 0.999971 | 1 | 0.890905 |
| Dennd6a | Q8BH65 | 0.24 | 0.04 | 0.39 | 0.22 | 0.29 | 0.08 | -0.9 | 0 | -0.13 | 0.24 | -0.05 | -0.23 | -0.31 | 0.53 | -0.14 | 0.46 | -0.355 | -0.265 | -0.0875 | 0.234913 | 0.182522 | 0.728457 | 0.999971 | 1 | 0.956041 |
| Dennd6b | Q9D9V7 | -0.83 | 0.06 | -0.26 | -0.41 | -0.2 | -0.2 | -0.31 | -0.12 | -0.22 | -0.62 | -0.43 | -0.49 | 0.55 | -0.04 | -0.33 | -0.29 | 0.1525 | -0.08 | 0.3325 | 0.509115 | 0.734477 | 0.266548 | 0.999971 | 1 | 0.842677 |
| Denr | Q9CQJ6 | 0.49 | 0.58 | 0.21 | 0.17 | 0.49 | 0.99 | -0.25 | 0.46 | 0.54 | 0.64 | 0.16 | -0.37 | 0.32 | 1.13 | 0.22 | 0.58 | 0.06 | -0.12 | 0.2 | 0.834617 | 0.656841 | 0.440293 | 0.999971 | 1 | 0.883105 |
| Depdc5 | P61460-2 | 1.13 | 0.01 | 0.39 | -0.14 | -0.1 | 0.32 | -0.09 | 0.43 | -0.19 | 0.63 | 0.36 | 0.11 | -0.68 | -0.22 | 0.33 | 0.08 | -0.2075 | -0.12 | -0.47 | 0.519187 | 0.718579 | 0.204185 | 0.999971 | 1 | 0.81617 |
| Deptor | Q570Y9 | -0.86 | 0.14 | 0.51 | 0.75 | -0.13 | -0.44 | 0.05 | 0.38 | 0.05 | -0.17 | 0.66 | 0.2 | 0.06 | -0.14 | 0.65 | 0.52 | -0.17 | 0.05 | 0.1375 | 0.656303 | 0.895923 | 0.722828 | 0.999971 | 1 | 0.955258 |
| Dera | Q91YP3 | 0.33 | -0.28 | 0.08 | 0.37 | -0.42 | -1.76 | 0.75 | 0.26 | -0.51 | 1.03 | 0.16 | 0.31 | -1.19 | 0.09 | -0.41 | 0.01 | -0.4175 | 0.1225 | -0.5 | 0.431996 | 0.723179 | 0.154753 | 0.999971 | 1 | 0.795578 |
| Derl1 | Q99J56 | -1.69 | -0.91 | -0.61 | -0.81 | -0.79 | -0.62 | 2.68 | -0.94 | -0.96 | -0.92 | -0.38 | 0.41 | -0.74 | -0.53 | -0.22 | -0.88 | 1.0875 | 0.5425 | 0.4125 | 0.201398 | 0.182363 | 0.178756 | 0.999971 | 1 | 0.808532 |
| Derl2 | Q3U957 | -0.3 | -0.87 | -2.34 | -0.55 | -1.36 | -0.72 | -1.82 | -0.02 | -1.55 | -1.14 | -2.92 | -0.21 | -0.46 | -1.37 | -0.74 | -1.33 | 0.035 | -0.44 | 0.04 | 0.94943 | 0.511612 | 0.932883 | 0.999971 | 1 | 0.995069 |
| Derpc | P0CG14 | 1.78 | 2.54 | 2.03 | 2.01 | 2.41 | 2.35 | 1.18 | 2.06 | 2.39 | 2.39 | 1.82 | 1.72 | 2.87 | 2.41 | 1.93 | 2.15 | -0.09 | -0.01 | 0.25 | 0.782985 | 0.969525 | 0.376347 | 0.999971 | 1 | 0.870465 |
| Des | P31001 | 0.61 | 0.79 | 0.3 | 0.94 | 2.31 | 0.9 | 0.19 | -0.09 | 0.95 | 1.46 | 0.19 | 1.13 | -0.59 | 0.61 | 0.77 | 0.46 | 0.1675 | 0.2725 | -0.3475 | 0.744113 | 0.386754 | 0.315499 | 0.999971 | 1 | 0.852494 |
| Desi1 | Q9CQT7 | 1.19 | 0.51 | 0.13 | 0.54 | 0.17 | -0.05 | -0.91 | 0.28 | 0.3 | 0.89 | -0.21 | 0.17 | 0.71 | 0.65 | 0.13 | 0.13 | -0.72 | -0.305 | -0.1875 | 0.061518 | 0.352037 | 0.516925 | 0.999971 | 1 | 0.905815 |
| Dffa | O54786-2 | 0.44 | 0.47 | 0.87 | 0.73 | 0.18 | 0.49 | 0.88 | 1.53 | 0.58 | 0.68 | 1.04 | 0.51 | -0.13 | 0.62 | 2 | 1.42 | 0.1425 | 0.075 | 0.35 | 0.650856 | 0.716877 | 0.443372 | 0.999971 | 1 | 0.883386 |
| Dgat1 | Q9Z2A7 | -0.06 | -0.39 | 0.72 | 0.88 | 0.05 | 0.17 | 0.68 | 0.73 | 0.12 | -0.08 | 0.57 | 0.55 | -0.06 | -0.01 | 0.55 | 0.84 | 0.12 | 0.0025 | 0.0425 | 0.729455 | 0.99415 | 0.907584 | 0.999971 | 1 | 0.991405 |
| Dgcr2 | A0A0R4J1N | -0.15 | -0.74 | -0.58 | 0.23 | -0.91 | -0.99 | -0.16 | 0.88 | -0.71 | -0.54 | 0.09 | 0.19 | -0.97 | -0.37 | 0.29 | 0.08 | 0.015 | 0.0675 | 0.0675 | 0.973704 | 0.831099 | 0.846836 | 0.999971 | 1 | 0.980295 |
| Dgcr8 | Q9EQM6 | 0.99 | 2.19 | 1.65 | 0.36 | 1.65 | 1.51 | 0.11 | 1.15 | 1.77 | 0.66 | 1.62 | 0.54 | 1.56 | 2.44 | 1.47 | 2.01 | -0.1925 | -0.15 | 0.5725 | 0.695867 | 0.753616 | 0.207306 | 0.999971 | 1 | 0.820632 |
| Dgkb | Q6NS52-2 | 0.12 | -0.7 | -0.55 | -0.04 | -0.37 | -0.43 | -0.66 | 0.19 | -1.27 | 0.14 | -0.27 | 0.17 | -1.46 | -0.29 | -0.1 | 0 | -0.025 | -0.015 | -0.17 | 0.92928 | 0.968273 | 0.655263 | 0.999971 | 1 | 0.934061 |
| Dgkb_1 | Q6NS52 | -0.7 | 0.22 | 0.02 | 0.33 | 0.3 | 0.09 | -0.2 | 0.76 | 0.34 | -0.34 | 0.05 | 0.27 | 0.5 | 0.26 | 0.69 | 0.23 | 0.27 | 0.1125 | 0.4525 | 0.397077 | 0.697924 | 0.125866 | 0.999971 | 1 | 0.783071 |
| Dgkd | E9PUQ8 | 0.58 | 1.11 | 0.77 | 0.5 | 0.88 | 1.01 | 0.24 | 0.6 | 0.56 | 0.4 | 0.66 | 0.34 | 1.19 | 0.98 | 0.22 | 0.69 | -0.0575 | -0.25 | 0.03 | 0.816172 | 0.241881 | 0.911547 | 0.999971 | 1 | 0.992551 |
| Dgke | Q9R1C6 | -0.8 | -0.14 | -0.27 | -0.14 | -0.34 | 0.12 | 0.09 | 0.03 | -0.12 | -0.62 | -0.22 | -1.05 | -0.13 | -0.06 | -0.45 | -0.5 | 0.3125 | -0.165 | 0.0525 | 0.195078 | 0.557679 | 0.820335 | 0.999971 | 1 | 0.975761 |
| Dgkg | Q91WG7 | -0.01 | 0.52 | 0.86 | 0.22 | 0.25 | 0.74 | 0.5 | 0.84 | 0.25 | -0.26 | 0.12 | 0.21 | 0.26 | 0.51 | 0.96 | 0.5 | 0.185 | -0.3175 | 0.16 | 0.475693 | 0.222464 | 0.545738 | 0.999971 | 1 | 0.911398 |
| Dgkh | A0A2I3BQ4 | 0.36 | 0.79 | 0.09 | 0.14 | 0.62 | 0.94 | -0.01 | 0.22 | 0.71 | 0.85 | 0.26 | 0.33 | 0.77 | 0.8 | 0.05 | 0.17 | 0.0975 | 0.1925 | 0.1025 | 0.728274 | 0.437366 | 0.707642 | 0.999971 | 1 | 0.950182 |
| Dgki | D3YWQ0-2 | -0.11 | 0.17 | 0.8 | 0.85 | -0.35 | -0.38 | 0.3 | 0.74 | -0.25 | -0.31 | 0.33 | -0.64 | -0.5 | -0.27 | 0.16 | 0.29 | -0.35 | -0.645 | -0.5075 | 0.335013 | 0.066417 | 0.125288 | 0.999971 | 1 | 0.78268 |
| Dgkq | Q6P5E8 | -0.75 | -1.09 | -1.13 | -1.05 | -0.82 | -0.6 | -1.41 | -0.95 | -0.9 | -0.72 | -1.44 | -0.95 | -0.89 | -0.73 | -0.62 | -0.98 | 0.06 | 0.0025 | 0.2 | 0.794264 | 0.990849 | 0.30476 | 0.999971 | 1 | 0.847402 |
| Dgkz | Q80UP3 | -5.22 | -13.2 | -7.91 | -8.18 | -11.46 | -8.98 | -6.79 | -9.28 | -12.4 | -6.63 | -5.72 | -9.09 | -17.03 | -10.52 | -8.3 | -8.57 | -0.5 | 0.1675 | -2.4775 | 0.768802 | 0.93298 | 0.303077 | 0.999971 | 1 | 0.847402 |
| Dgkz_1 | A2AHJ7 | 0.89 | 0.82 | 0.69 | 0.57 | 0.65 | 0.88 | 0.72 | 0.8 | 0.48 | 0.43 | 0.29 | 0.05 | 0.44 | 0.65 | 0.44 | 0.33 | 0.02 | -0.43 | -0.2775 | 0.908503 | 0.043051 | 0.147598 | 0.999971 | 1 | 0.793248 |
| Dglucy | E9QMK9 | -0.15 | -0.19 | -0.71 | -0.42 | -0.25 | -0.26 | -1.04 | -0.34 | -0.47 | -0.31 | -1 | -0.77 | -0.15 | -0.28 | -0.87 | -0.44 | -0.105 | -0.27 | -0.0675 | 0.682099 | 0.266822 | 0.776775 | 0.999971 | 1 | 0.964475 |
| Dguok | Q9QX60-2 | 0.54 | -0.4 | -3.02 | -2.46 | -0.81 | -0.75 | -2.49 | -2.39 | -0.85 | 0.49 | -3.04 | -3.05 | -1.44 | -0.22 | -2.99 | -2.65 | -0.275 | -0.2775 | -0.49 | 0.751759 | 0.798693 | 0.605249 | 0.999971 | 1 | 0.924458 |
| Dhcr24 | Q8VCH6 | -1.54 | -0.74 | -0.5 | -1.61 | -0.78 | 0.44 | -0.39 | -0.49 | 0.17 | -0.47 | -0.16 | 0.75 | 0.31 | 1.06 | 0.17 | -0.72 | 0.7925 | 1.17 | 1.3025 | 0.058071 | 0.011577 | 0.014468 | 0.999971 | 1 | 0.613095 |
| Dhcr7 | A0A140LIT | -0.34 | 0.4 | 1.13 | 0.42 | 0.33 | 0.71 | 0.46 | 1.45 | 0.78 | 0.88 | 0.76 | 1.05 | 1.05 | 1.06 | 1.25 | 0.89 | 0.335 | 0.465 | 0.66 | 0.385883 | 0.161286 | 0.060928 | 0.999971 | 1 | 0.725348 |
| Dhdds | A3KGL2 | 1.77 | -1.24 | 0.89 | 1.52 | -1.05 | 0.11 | 1.48 | 0.41 | -1.01 | 1.42 | -0.49 | -0.08 | -1.95 | -0.46 | 0.1 | -0.45 | -0.4975 | -0.775 | -1.425 | 0.524753 | 0.334445 | 0.079407 | 0.999971 | 1 | 0.755254 |
| Dhdh | Q9DBB8 | 1.43 | 0.07 | -0.04 | -0.09 | 0.57 | 1.06 | 0.06 | 0.34 | 0.69 | 1.04 | -0.38 | -0.17 | 0.43 | 1.27 | 0.22 | 0.34 | 0.165 | -0.0475 | 0.2225 | 0.682798 | 0.918962 | 0.594539 | 0.999971 | 1 | 0.92377 |
| Dhfr | P00375 | 0.14 | -0.36 | -1.8 | -1.49 | 0.14 | -0.08 | -1.97 | -0.9 | -0.31 | 0.35 | -1.25 | -1.31 | 0.2 | 0.17 | -1.55 | -1.72 | 0.175 | 0.2475 | 0.1525 | 0.772501 | 0.66078 | 0.810669 | 0.999971 | 1 | 0.973864 |
| Dhodh | O35435 | 0.12 | -0.34 | -0.99 | -1.17 | -0.7 | -0.78 | -1.32 | -1.06 | -0.71 | -0.17 | -1.21 | -1.21 | -1.06 | -0.43 | -1.11 | -1.16 | -0.37 | -0.23 | -0.345 | 0.27838 | 0.544805 | 0.32569 | 0.999971 | 1 | 0.856472 |
| Dhps | Q3TXU5 | -0.16 | 0.67 | 1.15 | 1.22 | 1.01 | 0.88 | 0.83 | 1.55 | 0.82 | 0.37 | 1.06 | 1.55 | 1.63 | 1.18 | 1.55 | 1.77 | 0.3475 | 0.23 | 0.8125 | 0.336682 | 0.556744 | 0.037959 | 0.999971 | 1 | 0.700838 |
| Dhrs1 | Q99L04 | 0.84 | 0.63 | -0.47 | -0.83 | 0.4 | 0.96 | -0.42 | -0.48 | 0.68 | 1.22 | 0.09 | -0.96 | 0.95 | 1.02 | -0.51 | -0.31 | 0.0725 | 0.215 | 0.245 | 0.883901 | 0.7078 | 0.646581 | 0.999971 | 1 | 0.933613 |
| Dhrs11 | Q3U0B3 | -0.19 | 0.34 | -0.77 | -0.33 | 0.2 | -0.49 | -0.73 | -1.09 | -0.23 | -0.7 | -1.32 | -0.35 | -0.4 | -0.26 | -1.18 | -0.84 | -0.29 | -0.4125 | -0.4325 | 0.415877 | 0.236529 | 0.19345 | 0.999971 | 1 | 0.810547 |
| Dhrs13 | Q5SS80 | 0.87 | -0.68 | -0.76 | -0.28 | -0.56 | -0.66 | -0.24 | -0.23 | -0.37 | 0.77 | -0.02 | 0.07 | -0.81 | -0.23 | 0.2 | -0.54 | -0.21 | 0.325 | -0.1325 | 0.581986 | 0.450582 | 0.749127 | 0.999971 | 1 | 0.961064 |
| Dhrs4 | Q99LB2 | -0.36 | -1.05 | -0.68 | -0.74 | -1.06 | -0.91 | -0.93 | -0.57 | -1.3 | -0.74 | -0.95 | -0.93 | -1.18 | -0.88 | -0.94 | -0.97 | -0.16 | -0.2725 | -0.285 | 0.473636 | 0.240987 | 0.195053 | 0.999971 | 1 | 0.811225 |

| | | | | | | | | | | | | | | | | | | | | | | | | | | |
|---|---|---|---|---|---|---|---|---|---|---|---|---|---|---|---|---|---|---|---|---|---|---|---|---|---|---|
| Dhrs7 | A0A1Y7VN | 0.03 | 0.33 | -0.42 | 0.02 | 0.68 | 0.57 | -0.87 | 0.47 | 0.08 | 0.45 | -0.5 | -0.32 | 0.64 | 0.15 | 0.04 | 0.05 | 0.2225 | -0.0625 | 0.23 | 0.562788 | 0.8219 | 0.355478 | 0.999971 | 1 | 0.868793 |
| Dhrs7b | Z4YKT6 | -0.44 | -0.44 | -0.88 | -0.24 | -0.64 | -0.69 | -0.89 | -0.59 | -0.89 | -0.47 | -0.85 | -0.66 | -0.71 | -0.59 | -0.91 | -0.55 | -0.2025 | -0.2175 | -0.19 | 0.336506 | 0.319194 | 0.376133 | 0.999971 | 1 | 0.870465 |
| Dhtkd1 | A2ATU0 | -2.53 | -4.55 | 0.48 | 1.06 | -0.6 | -1.1 | 3.99 | -1.58 | -1.53 | -0.55 | 0.76 | 1.52 | -1.61 | -1.43 | 1.08 | 0.01 | 1.5625 | 1.435 | 0.8975 | 0.350595 | 0.295476 | 0.495896 | 0.999971 | 1 | 0.901531 |
| Dhx15 | O35286 | 0.05 | 1.59 | 1.37 | 1.38 | 1.66 | 1.25 | 1.14 | 1.25 | 1.38 | 0.38 | 1.52 | 1.58 | 1.36 | 1.12 | 1.3 | 1.33 | 0.2275 | 0.1175 | 0.18 | 0.534025 | 0.784137 | 0.611001 | 0.999971 | 1 | 0.92673 |
| Dhx16 | G3X8X0 | 5.49 | 1.26 | 0.1 | 0.16 | 1.36 | 0.47 | -0.63 | -0.44 | 2.55 | 2.07 | -0.72 | 1.09 | 1.44 | 0.93 | -0.03 | 0.03 | -1.5625 | -0.505 | -1.16 | 0.216054 | 0.701073 | 0.338834 | 0.999971 | 1 | 0.86005 |
| Dhx29 | Q6PGC1 | 0.93 | 1 | 1.43 | 1.61 | 0.77 | 1.23 | 1.33 | 1.41 | 1.02 | 1.22 | 1.12 | 0.35 | 0.9 | 1.23 | 1.26 | 1.12 | -0.0575 | -0.315 | -0.115 | 0.816427 | 0.267675 | 0.61287 | 0.999971 | 1 | 0.927364 |
| Dhx30 | A0A0G2JGI | -0.21 | 0.42 | -0.05 | -0.09 | 0.3 | 0.18 | 0.13 | -0.03 | 0.33 | 0.11 | 0.09 | 0.28 | 0.67 | 0.37 | 0.22 | 0.46 | 0.1275 | 0.185 | 0.4125 | 0.542488 | 0.372975 | 0.082397 | 0.999971 | 1 | 0.755254 |
| Dhx32 | D3Z4E3 | 1.75 | 1.62 | 1.46 | 1.7 | 1.21 | 1.4 | 0.82 | 1.84 | 1.4 | 1.6 | 1.23 | 1.85 | 1.75 | 1.98 | 1.56 | 1.81 | -0.315 | -0.1125 | 0.1425 | 0.227606 | 0.578654 | 0.445561 | 0.999971 | 1 | 0.884152 |
| Dhx35 | Q8K1G9 | 1.21 | 0.68 | 0.6 | 1.02 | 1.27 | 0.79 | 0.49 | 1.34 | 0.3 | 1.99 | 0.86 | 0.69 | 1.96 | 0.91 | 0.45 | 1.08 | 0.095 | 0.0825 | 0.2225 | 0.723664 | 0.827247 | 0.522219 | 0.999971 | 1 | 0.90761 |
| Dhx36 | Q8VHK9 | 0.62 | 0.57 | 0.67 | 0.61 | 0.5 | 0.62 | 0.46 | 0.88 | 0.46 | 0.63 | 0.81 | 0.89 | 0.54 | 0.72 | 0.79 | 0.79 | -0.0025 | 0.08 | 0.0925 | 0.988798 | 0.650331 | 0.582378 | 0.999971 | 1 | 0.920364 |
| Dhx38 | Q80X98 | 1.32 | 1.34 | 0.84 | 1.29 | 1.05 | 1.14 | 1.22 | 0.56 | 1.1 | 1.47 | 0.36 | 0.32 | 0.7 | 1.49 | 0.1 | 0.12 | -0.205 | -0.385 | -0.595 | 0.382011 | 0.23732 | 0.111445 | 0.999971 | 1 | 0.78268 |
| Dhx57 | Q6P5D3 | 0.29 | 0.97 | 0.98 | 0.8 | 1.25 | 0.71 | 0.6 | 1.27 | 1.56 | 0.5 | 1.18 | 1.21 | 1.67 | 0.84 | 1.32 | 1.56 | 0.1975 | 0.3525 | 0.5875 | 0.458323 | 0.239094 | 0.051737 | 0.999971 | 1 | 0.723736 |
| Dhx8 | A2A4P0 | 0.63 | 0.15 | 0.97 | 1.09 | -0.01 | -0.94 | 0.18 | 1 | -0.36 | -0.45 | 0.65 | 0.83 | -1.11 | -0.31 | 1.2 | 1.6 | -0.6525 | -0.5425 | -0.365 | 0.151727 | 0.179394 | 0.554606 | 0.999971 | 1 | 0.914543 |
| Dhx9 | E9QNN1 | 0.49 | -0.75 | 0.53 | 1.01 | -0.82 | -0.27 | -0.22 | 0.13 | -0.01 | -0.39 | 0 | 0.19 | -1.18 | -0.56 | -0.36 | -0.4 | -0.615 | -0.3725 | -0.945 | 0.15416 | 0.342922 | 0.041112 | 0.999971 | 1 | 0.700838 |
| Dhx9_1 | A0A087WF | -0.3 | -0.28 | -0.22 | -0.29 | -0.26 | -0.35 | -0.31 | -0.41 | -0.31 | -0.19 | -0.33 | -0.34 | -0.21 | -0.12 | -0.08 | -0.09 | -0.06 | -0.02 | 0.1475 | 0.708478 | 0.898212 | 0.369605 | 0.999971 | 1 | 0.869526 |
| Diablo | Q9JIQ3 | 0.93 | 0.06 | -0.35 | -0.43 | 0.32 | -0.14 | -0.57 | -0.35 | 0.2 | 0.53 | -0.64 | -0.75 | 0.46 | 0.08 | -0.62 | 0.05 | -0.2375 | -0.2175 | -0.06 | 0.511095 | 0.608062 | 0.872166 | 0.999971 | 1 | 0.984109 |
| Diaph1 | E9PV41 | -0.42 | -1.05 | -0.49 | -0.5 | -0.83 | -0.89 | -0.24 | -0.86 | -1.05 | -0.87 | -0.66 | -0.96 | -1.1 | -1.09 | -0.9 | -1.14 | -0.09 | -0.27 | -0.4425 | 0.713225 | 0.226321 | 0.058855 | 0.999971 | 1 | 0.725348 |
| Diaph2 | Q6W4W7 | -0.12 | -0.45 | -0.32 | -0.26 | -0.19 | -0.23 | -0.41 | -0.32 | -0.44 | -0.34 | -0.42 | -0.3 | -0.23 | 0.09 | -0.29 | -0.38 | 0 | -0.0875 | 0.085 | 1 | 0.602439 | 0.657965 | 1 | 1 | 0.934638 |
| Dicer1 | F8VQ54 | 0.78 | -0.17 | -0.4 | -0.17 | 0.3 | -0.42 | 0.69 | -0.09 | 1.16 | 0.46 | -0.03 | 0.44 | 1.1 | 3.61 | 0.18 | 0.27 | 0.11 | 0.4975 | 1.28 | 0.753519 | 0.18275 | 0.119434 | 0.999971 | 1 | 0.78268 |
| Dido1 | Q8C9B9 | 1.8 | 2.23 | 2.06 | 2.65 | 2.35 | 1.68 | 0.9 | 2.31 | 1.93 | 1.32 | 2.06 | 1.6 | 0.85 | 1.18 | 1.69 | 2.45 | -0.375 | -0.4575 | -0.6425 | 0.327868 | 0.110909 | 0.117351 | 0.999971 | 1 | 0.78268 |
| Dimt1 | Q9D0D4 | 1.54 | 0.69 | 0.53 | 1.04 | 0.65 | 1.15 | 0 | 0.23 | 0.34 | 1.94 | -0.12 | 0.69 | 0.73 | 1.44 | 0.16 | 0.57 | -0.4425 | -0.2375 | -0.225 | 0.209409 | 0.611994 | 0.518707 | 0.999971 | 1 | 0.905901 |
| Dip2a | F8WI56 | -0.54 | -0.17 | -0.19 | -0.23 | -0.32 | -0.11 | -0.6 | -0.04 | -0.18 | -0.33 | -0.12 | -0.45 | -0.27 | -0.13 | -0.08 | -0.12 | 0.015 | 0.0125 | 0.1325 | 0.941958 | 0.945482 | 0.465496 | 0.999971 | 1 | 0.891255 |
| Dip2b | B2RQC7 | -1.37 | -1.43 | -0.39 | -0.22 | -1.27 | -1.73 | -1.15 | -0.32 | -1.02 | -0.79 | -0.8 | -0.75 | -1.41 | -1.54 | -0.3 | -1.13 | -0.265 | 0.0125 | -0.2425 | 0.524887 | 0.969248 | 0.552837 | 0.999971 | 1 | 0.913595 |
| Dip2b_1 | Q3UH60 | 0.07 | 0.15 | 0.03 | -0.04 | 0.52 | 0.3 | -0.02 | 0.27 | 0.05 | 0.35 | 0.14 | -0.33 | 0.01 | 0.14 | 0.19 | 0.13 | 0.215 | 0 | 0.065 | 0.270044 | 1 | 0.694274 | 0.999971 | 1 | 0.947174 |
| Dip2c | B2RQ71 | 0.08 | -0.24 | 0.07 | 0.29 | -0.41 | -0.28 | 0.02 | 0.28 | -0.19 | 0.07 | -0.11 | -0.46 | -0.3 | -0.03 | -0.05 | 0.03 | -0.1475 | -0.2225 | -0.1375 | 0.523288 | 0.294536 | 0.487329 | 0.999971 | 1 | 0.897906 |
| Dip2c_1 | B7ZMZ7 | 0.4 | 0.06 | -0.59 | 0.2 | -0.32 | -0.02 | -0.25 | 1 | -0.25 | 0.29 | 0.01 | 0.29 | -0.48 | -0.27 | -0.15 | 0.41 | 0.085 | 0.0675 | -0.14 | 0.815327 | 0.801825 | 0.639115 | 0.999971 | 1 | 0.93301 |
| Dipk1b | Q99ML4 | -0.3 | 0.48 | -0.2 | -0.05 | 0.85 | 0.82 | -0.62 | 0.36 | 0.75 | 0.46 | 0.23 | 0.27 | 1.24 | 1.31 | 0.44 | 0.64 | 0.37 | 0.445 | 0.925 | 0.334499 | 0.091572 | 0.01057 | 0.999971 | 1 | 0.613095 |
| Dipk2a | Q3USZ8 | -1.68 | -0.85 | 3.32 | 2.32 | 0.34 | -1.41 | 2.7 | 3.33 | -0.66 | -1.46 | 3.81 | 4.93 | 0.1 | -0.93 | 3.85 | 3.4 | 0.4625 | 0.8775 | 0.8275 | 0.749349 | 0.625027 | 0.585569 | 0.999971 | 1 | 0.920413 |
| Diras1 | Q91Z61 | -0.31 | 0.11 | -0.1 | 0.02 | -0.02 | 0.49 | -0.36 | 0.68 | 0.35 | 0.03 | 0.18 | 0.07 | 0.69 | 0.77 | 0.39 | 0.81 | 0.2675 | 0.2275 | 0.735 | 0.339896 | 0.236023 | 0.004353 | 0.999971 | 1 | 0.582714 |
| Diras2 | Q5PR73 | 0.33 | 0.37 | 0.31 | 0.7 | 0.03 | 0.77 | 0.14 | 0.67 | 0.28 | 0.7 | 0.31 | 0.21 | -0.11 | 0.34 | 0.29 | 0.88 | -0.025 | -0.0525 | -0.0775 | 0.91697 | 0.79147 | 0.758282 | 0.999971 | 1 | 0.961116 |
| Dis3 | Q9CSH3 | 3.24 | 2.19 | 2.77 | 2.7 | 1.92 | 2.64 | 2.5 | 1.77 | 2 | 2.5 | 2.22 | 1.69 | -0.17 | 1.14 | 2.12 | 2.21 | -0.5175 | -0.6225 | -1.4 | 0.120545 | 0.054687 | 0.029099 | 0.999971 | 1 | 0.678474 |
| Dis3l2 | Q8CI75-2 | -0.01 | 0.52 | -0.16 | 0.02 | 0.51 | 0.46 | -0.13 | 0.02 | 0.44 | 0.31 | 0.26 | 0.23 | 0.68 | 0.72 | 0.43 | 0.49 | 0.1225 | 0.2175 | 0.4875 | 0.621868 | 0.304586 | 0.045136 | 0.999971 | 1 | 0.710102 |
| Disp2 | Q8CIP5 | 1.59 | -2.6 | -0.26 | 0.1 | -4.59 | -1.24 | 0.23 | -0.11 | -5.75 | 1.16 | -0.25 | -0.43 | -8.27 | -0.94 | 0.04 | 0.16 | -1.135 | -1.025 | -1.96 | 0.373322 | 0.517127 | 0.328718 | 0.999971 | 1 | 0.856472 |
| Dixdc1 | Q80Y83-7 | -0.08 | -0.79 | -0.38 | -0.53 | -0.72 | -0.85 | 0.44 | -0.42 | -0.86 | -0.53 | -0.31 | -0.3 | -0.55 | -1.06 | -0.68 | -0.61 | 0.0575 | -0.055 | -0.28 | 0.860068 | 0.812786 | 0.239335 | 0.999971 | 1 | 0.833594 |
| Dkc1 | Q9ESX5 | 0.18 | -0.19 | -1.01 | -0.16 | -0.1 | -0.13 | -1.33 | -1.59 | 0.06 | 0.76 | -1.36 | -1.38 | 0.45 | -0.24 | -1.22 | -0.26 | -0.4925 | -0.185 | -0.0225 | 0.279716 | 0.73525 | 0.955878 | 0.999971 | 1 | 0.998056 |
| Dkk3 | Q9QUN9 | 1.49 | 1.97 | 1.64 | 1.51 | 1.77 | 2.23 | 2.13 | 1.65 | 1.43 | 1.49 | 2.01 | 1.12 | 1.25 | 1.24 | 1.18 | 1.53 | 0.2925 | -0.14 | -0.3525 | 0.208558 | 0.569493 | 0.101589 | 0.999971 | 1 | 0.78268 |
| Dlat | Q8BMF4 | -0.93 | -1.18 | -0.16 | -0.29 | -1.39 | -1.2 | -0.22 | -0.45 | -1.28 | -1.29 | -0.42 | -0.35 | -1.52 | -1.28 | -0.43 | -0.36 | -0.175 | -0.195 | -0.2575 | 0.634223 | 0.58285 | 0.495971 | 0.999971 | 1 | 0.901531 |
| Dlc1 | A0A0R4J17 | 0.4 | -0.05 | -1.25 | -1.21 | -0.19 | 0.89 | -1.4 | -0.74 | -0.2 | -0.01 | -1.02 | -0.8 | -0.23 | 0.43 | -1.26 | -1.03 | 0.1675 | 0.02 | 0.005 | 0.774489 | 0.964703 | 0.992392 | 0.999971 | 1 | 0.99925 |
| Dld | O08749 | -1.16 | -1.75 | -1.27 | -1.22 | -1.46 | -2.19 | -0.83 | -1.25 | -1.2 | -1.6 | -0.96 | -1.11 | -1.67 | -2.06 | -1.37 | -1.6 | -0.0825 | 0.1325 | -0.325 | 0.795361 | 0.565443 | 0.1886 | 0.999971 | 1 | 0.809276 |
| Dlg1 | E9Q9H0 | -1.54 | -1.46 | -1.48 | -1.52 | -1.46 | -1.16 | -1.2 | -1.75 | -1.5 | -1.84 | -1.41 | -1.56 | -1.62 | -1.61 | -1.76 | -1.68 | 0.1075 | -0.0775 | -0.1675 | 0.589926 | 0.657587 | 0.313883 | 0.999971 | 1 | 0.850589 |
| Dlg1_1 | H7BWY4 | -0.01 | -0.87 | -2.31 | -2.32 | 0.11 | -0.25 | -1.97 | -2.11 | -0.21 | -1.23 | -2.6 | -1.92 | 0.25 | -0.98 | -1.55 | -2.43 | 0.3225 | -0.1125 | 0.2 | 0.659864 | 0.871507 | 0.782292 | 0.999971 | 1 | 0.965859 |
| Dlg1_2 | Q811D0-2 | -1.41 | -4.26 | -0.39 | -2.3 | -2.96 | -1.66 | 0.13 | -4.52 | -4.35 | -1.95 | -0.52 | -2.16 | -1.18 | -3.05 | -1.09 | 0.46 | -0.1625 | -0.155 | 0.875 | 0.886788 | 0.879734 | 0.38134 | 0.999971 | 1 | 0.871816 |
| Dlg1_3 | D3Z3B8 | 0.33 | -2.6 | -2.6 | -1.92 | -2.49 | -1.79 | -1.39 | -1.92 | -2.27 | -0.48 | -1.72 | -2.35 | -3.43 | -2.61 | -2.31 | -2.2 | -0.2 | -0.0075 | -0.94 | 0.763152 | 0.991929 | 0.18983 | 0.999971 | 1 | 0.810547 |
| Dlg2 | A0A338P6J | -1.34 | -1.84 | -1.63 | -1.65 | -1.6 | -1.66 | -1.53 | -1.64 | -1.53 | -1.3 | -1.58 | -1.66 | -1.08 | -1.7 | -1.67 | -1.63 | 0.0075 | 0.0975 | 0.095 | 0.967221 | 0.611124 | 0.67163 | 0.999971 | 1 | 0.939561 |
| Dlg2_1 | Q91XM9-6 | -1.73 | -1.01 | -0.74 | -1.48 | -0.67 | -0.73 | -0.8 | -0.53 | -0.31 | -1.01 | -1.11 | -0.25 | -0.01 | 0.39 | 0.54 | 0.71 | 0.5575 | 0.57 | 1.6475 | 0.053798 | 0.102666 | 0.000303 | 0.999971 | 1 | 0.247672 |
| Dlg2_2 | A0A571BEI | 4 | 3.41 | -0.17 | -0.63 | 2.61 | 0.36 | 2.11 | 0.25 | 5.07 | 4.66 | 0.36 | 1.39 | 4.24 | -4.25 | -0.38 | -2.41 | -0.32 | 1.2175 | -2.3525 | 0.788145 | 0.425033 | 0.243806 | 0.999971 | 1 | 0.837316 |
| Dlg3 | P70175 | -1.08 | -0.84 | -1.41 | -1.28 | -0.88 | -0.87 | -1.32 | -1.28 | -0.85 | -0.91 | -1.05 | -1.07 | -0.53 | -1.01 | -1.32 | -1.2 | 0.065 | 0.1825 | 0.1375 | 0.767299 | 0.359032 | 0.577981 | 0.999971 | 1 | 0.918596 |
| Dlg4 | A0A338P6I | -1.6 | -1.56 | -1.43 | -1.39 | -1.48 | -1.46 | -1.34 | -1.52 | -1.73 | -1.67 | -1.36 | -1.55 | -1.25 | -1.88 | -1.63 | -1.37 | 0.045 | -0.0825 | -0.0375 | 0.786998 | 0.638419 | 0.854744 | 0.999971 | 1 | 0.982125 |
| Dlg4_1 | Q62108-3 | 2.61 | 0.21 | 4.65 | 2.15 | 0.8 | 0.23 | -1.78 | 2.53 | 0.25 | 3.09 | 4.64 | 5.62 | 0.76 | 0.13 | 3.79 | 3.03 | -1.96 | 0.995 | -0.4775 | 0.110836 | 0.461267 | 0.673714 | 0.999971 | 1 | 0.940599 |
| Dlg5 | E9Q9I2 | -0.3 | -1.48 | -0.9 | -0.96 | -1.05 | -1.27 | -0.87 | -0.6 | -1.18 | -0.62 | -0.72 | -0.87 | -0.83 | -0.86 | -0.7 | -0.56 | -0.0375 | 0.0625 | 0.1725 | 0.897536 | 0.825626 | 0.529077 | 0.999971 | 1 | 0.910069 |
| Dlgap1 | Q9D415 | -0.37 | -0.51 | -0.89 | -1.25 | -0.68 | 0.08 | -0.97 | -1.06 | -0.92 | -0.32 | -1 | -1.36 | -0.65 | -0.65 | -1.25 | -1.03 | 0.0975 | -0.145 | -0.14 | 0.765531 | 0.631545 | 0.604777 | 0.999971 | 1 | 0.924454 |
| Dlgap1_1 | D3Z508 | -1.07 | -0.3 | -0.31 | -1.54 | -1.31 | -0.95 | -0.95 | -1.85 | -1.84 | -1.22 | -1.08 | -1.72 | -1.97 | -1.88 | -1.37 | -1.12 | -0.46 | -0.66 | -0.78 | 0.225002 | 0.087208 | 0.054266 | 0.999971 | 1 | 0.725348 |
| Dlgap1_2 | Q9D415-3 | -0.66 | -0.65 | -0.28 | -1.08 | -1.03 | -0.88 | -1.6 | -0.01 | -0.98 | -1.23 | -2.14 | -0.01 | 0.35 | 0.23 | -2.22 | -1 | -0.2125 | -0.4225 | 0.0075 | 0.55755 | 0.35201 | 0.989545 | 0.999971 | 1 | 0.99925 |
| Dlgap1_3 | Q3UVU8 | -2.13 | -1.66 | -1.35 | -1.18 | -1.39 | -1.35 | -1.31 | -1.07 | -1.76 | -1.38 | -0.68 | -1.78 | -2.25 | -1.24 | -1.14 | -0.56 | 0.3 | 0.18 | 0.2825 | 0.247845 | 0.588704 | 0.47849 | 0.999971 | 1 | 0.894438 |
| Dlgap1_4 | Q9D415-2 | 0.11 | 1.54 | 2.26 | 1.6 | 2.05 | 1.21 | 0.55 | 1.19 | 1.82 | 0.96 | 0.34 | -0.01 | 3.17 | 1.22 | 1.56 | 1.77 | -0.1275 | -0.6 | 0.5525 | 0.801526 | 0.29914 | 0.347399 | 0.999971 | 1 | 0.864164 |
| Dlgap2 | Q8BJ42 | 0.51 | -0.68 | 0 | -0.88 | -1.54 | -1.09 | -0.85 | 0.49 | -1.56 | -0.99 | -0.46 | -0.23 | -1.73 | -1.7 | -0.62 | 0.62 | -0.485 | -0.5475 | -0.595 | 0.346383 | 0.210007 | 0.325369 | 0.999971 | 1 | 0.856472 |
| Dlgap2_1 | Q0VF59 | -0.25 | -0.83 | -0.71 | -0.67 | -0.79 | -0.59 | -0.5 | -0.81 | -0.79 | -0.19 | -0.49 | -0.77 | -0.34 | -0.79 | -0.9 | -0.56 | -0.0575 | 0.055 | -0.0325 | 0.777483 | 0.808203 | 0.883326 | 0.999971 | 1 | 0.986597 |
| Dlgap3 | B1AS06 | -0.93 | -1.15 | -0.92 | -0.86 | -1.15 | -0.98 | -0.97 | -0.77 | -1.38 | -1 | -0.86 | -0.95 | -0.98 | -1.1 | -1.08 | -0.7 | -0.0025 | -0.0825 | 1.96E-17 | 0.988892 | 0.668105 | 1 | 0.999971 | 1 | 1 |
| Dlgap4 | B7ZNS2 | -0.39 | -0.75 | -0.7 | -0.69 | -0.74 | -0.5 | -0.79 | -0.87 | -0.62 | -0.36 | -0.68 | -0.72 | -0.22 | -0.62 | -0.8 | -0.63 | -0.0925 | 0.0375 | 0.065 | 0.621304 | 0.838006 | 0.750587 | 0.999971 | 1 | 0.961064 |
| Dlgap4_1 | B1AZP2-2 | -0.97 | -1.41 | -1.78 | -1.36 | -1.35 | -1.97 | -0.91 | -1.11 | -1.51 | -1.06 | -1.25 | -2.04 | -1.78 | -1.41 | -1.55 | -1.56 | 0.045 | -0.085 | -0.195 | 0.878276 | 0.764174 | 0.395844 | 0.999971 | 1 | 0.871816 |
| Dlst | Q9D2G2 | -0.11 | -0.5 | -0.87 | -0.52 | -0.38 | -0.18 | -0.93 | -0.84 | -0.27 | -0.13 | -1.17 | -1.06 | -1.02 | -0.03 | -0.98 | -0.75 | -0.0825 | -0.1575 | -0.195 | 0.752445 | 0.616712 | 0.506686 | 0.999971 | 1 | 0.905775 |
| Dmac1 | Q9CQ00 | -1.45 | -3.35 | -1.44 | -1.23 | -2.84 | -2.44 | -0.64 | -0.92 | -3.54 | -1.61 | -1.18 | -1.58 | -2.83 | -2.43 | -1 | -1.04 | 0.1575 | -0.11 | 0.0425 | 0.813952 | 0.868182 | 0.945703 | 0.999971 | 1 | 0.997427 |
| Dmac2 | Q8CA44 | 0.32 | 2.76 | 0.11 | 0.38 | 2.79 | 0.93 | 0.48 | 0.71 | 2.47 | 0.88 | 1 | 0.25 | 4.11 | 1.65 | 0.21 | -0.06 | 0.335 | 0.2575 | 0.585 | 0.651708 | 0.718129 | 0.569615 | 0.999971 | 1 | 0.91816 |
| Dmac2l | Q9CRA7 | -0.29 | -0.66 | -0.2 | -0.13 | -0.61 | -0.63 | -0.42 | 0.06 | 0.09 | -0.33 | -0.4 | -0.5 | -0.33 | -0.21 | -0.14 | 0 | -0.08 | 0.035 | 0.15 | 0.733864 | 0.872083 | 0.456196 | 0.999971 | 1 | 0.887217 |
| Dmap1 | Q9JI44 | -0.6 | -0.29 | 0.78 | 1.09 | 0.33 | -0.38 | 0.41 | 0.42 | -0.11 | -0.18 | 0.79 | 0.67 | -0.2 | 0.41 | 0.65 | 1.14 | -0.05 | 0.0475 | 0.255 | 0.906726 | 0.916447 | 0.585154 | 0.999971 | 1 | 0.920413 |
| Dmd | P11531 | -0.55 | -0.82 | -0.58 | -0.55 | -0.55 | -0.71 | -0.72 | -0.44 | -0.81 | -0.26 | -0.25 | -0.33 | -0.89 | -0.71 | -0.25 | -0.38 | 0.02 | 0.2125 | 0.0675 | 0.909991 | 0.307164 | 0.750318 | 0.999971 | 1 | 0.961064 |
| Dmtn | Q9WV69-2 | -2.74 | -1.5 | -1.26 | -2.86 | -2.02 | -1.32 | -2.99 | -2.45 | -1.89 | -2.55 | -2.07 | -1.81 | -0.16 | -1.8 | -2.41 | -1.57 | -0.105 | 0.01 | 0.605 | 0.834853 | 0.981144 | 0.310767 | 0.999971 | 1 | 0.84774 |
| Dmtn_1 | Q9WV69 | -1.21 | -1.01 | -1.33 | -1.36 | -0.66 | -1.22 | -1.25 | -1.69 | -0.57 | -1.18 | -1.46 | -1.47 | -0.35 | -1.17 | -1.65 | -1.12 | 0.0225 | 0.0575 | 0.155 | 0.928897 | 0.818808 | 0.598702 | 0.999971 | 1 | 0.92377 |
| Dmwd | Q08274 | 0.18 | -0.34 | -0.38 | -0.18 | -0.39 | -0.21 | -0.61 | -0.04 | -0.19 | 0.34 | -0.26 | -0.5 | 0.06 | -0.18 | -0.41 | -0.3 | -0.1325 | 0.0275 | -0.0275 | 0.551756 | 0.910869 | 0.897004 | 0.999971 | 1 | 0.988737 |
| Dmwd_1 | E9Q6J8 | 0.49 | 1 | -0.21 | -0.8 | 1.92 | 0.86 | -0.87 | 0.2 | 2.41 | 0.42 | 0.21 | 0.84 | 2.4 | 0.41 | 1.11 | 0.12 | 0.4075 | 0.85 | 0.89 | 0.529715 | 0.170888 | 0.156353 | 0.999971 | 1 | 0.796008 |
| Dmxl1 | Q6PNC0 | -0.33 | -0.23 | -0.37 | -0.57 | -0.21 | -0.23 | -0.51 | -0.52 | -0.36 | -0.33 | -0.3 | -0.31 | -0.01 | -0.34 | -0.47 | -0.48 | 0.0075 | 0.05 | 0.05 | 0.967556 | 0.762976 | 0.79781 | 0.999971 | 1 | 0.970479 |
| Dmxl2 | Q8BPN8-2 | -0.84 | -0.89 | -1.23 | -1.28 | -0.95 | -0.63 | -1.03 | -1.23 | -0.9 | -0.72 | -1.2 | -1.22 | -0.81 | -0.78 | -1.17 | -1.16 | 0.1 | 0.05 | 0.08 | 0.644535 | 0.813381 | 0.702533 | 0.999971 | 1 | 0.949025 |
| Dnaaf10 | Q8BGF3 | 0.95 | 1.13 | 0.94 | 1.48 | 0.49 | 1.64 | 0.85 | 0.61 | 1.09 | 1.64 | 0.82 | 1.18 | 0.83 | 0.63 | 0.68 | 1.36 | -0.2275 | 0.0575 | -0.25 | 0.451546 | 0.81206 | 0.316831 | 0.999971 | 1 | 0.853713 |
| Dnaaf5 | B9EJR8 | -0.45 | 0.68 | 0.25 | 0.43 | 0.96 | 0.51 | 1.75 | 0.01 | 0.54 | 0.2 | -0.32 | -0.26 | 1.68 | 0.2 | 0.03 | -0.91 | 0.58 | -0.1875 | 0.0225 | 0.189981 | 0.559971 | 0.96684 | 0.999971 | 1 | 0.998787 |
| Dnaaf9 | Q3V0A7 | 0.35 | -0.13 | -0.09 | -0.18 | 0.6 | 0.18 | -0.12 | 0.39 | 0.37 | 0.88 | 0.25 | 0.21 | -0.16 | 0.57 | 0.41 | -0.01 | 0.275 | 0.44 | 0.215 | 0.256587 | 0.084593 | 0.387421 | 0.999971 | 1 | 0.871816 |
| Dnah2 | D3Z667 | 2.35 | 2.25 | 2.9 | 4.85 | -0.28 | 2.72 | 3.27 | 2.39 | 0.92 | 2.05 | 1.67 | -0.7 | -1.1 | 2.71 | 1.68 | 0.34 | -1.0625 | -2.1025 | -2.18 | 0.25202 | 0.023201 | 0.039105 | 0.999971 | 1 | 0.700838 |
| Dnaja1 | P63037 | 0.16 | -0.44 | -0.44 | -0.1 | -0.72 | 0.05 | -0.27 | -0.3 | -0.47 | -0.08 | -0.37 | -0.11 | -0.53 | -0.13 | -0.06 | 0.03 | -0.105 | -0.0525 | 0.0325 | 0.669861 | 0.808918 | 0.887635 | 0.999971 | 1 | 0.987485 |
| Dnaja2 | Q9QYJ0 | -0.35 | -0.53 | -0.31 | -0.62 | -0.32 | -0.56 | 0.26 | -0.32 | 0.14 | -0.67 | 0.26 | 0.25 | 0.3 | -0.93 | -0.1 | -0.43 | 0.2175 | 0.4475 | 0.1625 | 0.353517 | 0.109995 | 0.572108 | 0.999971 | 1 | 0.91816 |
| Dnaja3 | Q99M87-2 | -3.08 | -2.06 | -2.07 | -2.8 | -1.71 | -2.43 | -1.62 | -1.74 | -2.64 | -4.89 | -2.37 | -2.38 | -1.74 | -3.92 | -1.84 | -1.76 | 0.6275 | -0.5675 | 0.1875 | 0.077436 | 0.362979 | 0.732761 | 0.999971 | 1 | 0.956652 |
| Dnaja3_1 | Q99M87 | -0.19 | -0.42 | -1.06 | -0.79 | -0.42 | -0.48 | -1.1 | -0.81 | -0.42 | -0.61 | -0.98 | -1.03 | -0.83 | -0.5 | -0.9 | -1.12 | -0.0875 | -0.145 | -0.2225 | 0.745584 | 0.584766 | 0.397677 | 0.999971 | 1 | 0.871816 |
| Dnaja4 | Q9JMC3 | -0.63 | -0.57 | 0.86 | 0.76 | -0.67 | -0.2 | 0.78 | 0.87 | -0.7 | -0.75 | 0.32 | 0.25 | -0.82 | -0.03 | 0.7 | 0.77 | 0.09 | -0.325 | 0.05 | 0.860754 | 0.49573 | 0.922064 | 0.999971 | 1 | 0.993069 |
| Dnajb1 | Q9QYJ3 | 0.58 | -0.58 | 0.52 | 0.65 | -0.5 | -0.09 | 0.78 | 0.25 | -0.5 | 0.2 | 0.43 | 0.21 | -0.8 | -0.32 | 0.5 | 0.46 | -0.1825 | -0.2075 | -0.3325 | 0.636313 | 0.556581 | 0.426809 | 0.999971 | 1 | 0.880793 |
| Dnajb11 | Q99KV1 | -0.14 | 0.41 | 0.34 | -0.32 | 0.69 | 0.03 | 0.77 | 0.06 | 1.24 | -0.19 | 0.91 | 1.15 | 1.38 | -0.21 | 0.34 | 0.15 | 0.315 | 0.705 | 0.3425 | 0.281373 | 0.081444 | 0.371747 | 0.999971 | 1 | 0.869526 |
| Dnajb12 | Q9QYI4 | 1.26 | 0.52 | 1.1 | 1.47 | 0.17 | 1.03 | 2.37 | 0.4 | 0.49 | 1.08 | 0.56 | 0.14 | 0.18 | 0.39 | 0.49 | -0.58 | -0.095 | -0.52 | -0.9675 | 0.848017 | 0.102204 | 0.013663 | 0.999971 | 1 | 0.613095 |
| Dnajb14 | Q149L6 | -0.49 | 0.59 | -0.38 | -0.25 | 0.92 | 0.6 | -0.23 | -0.15 | 1.02 | 0.02 | -0.19 | -0.01 | 1.1 | 0.12 | -0.52 | -0.32 | 0.4175 | 0.3425 | 0.2275 | 0.27013 | 0.354241 | 0.587182 | 0.999971 | 1 | 0.920758 |
| Dnajb2 | Q9QYI5 | 0.22 | -0.85 | -0.97 | -0.86 | -1.08 | -0.65 | -0.65 | -0.94 | -1.07 | -0.17 | -1.05 | -1.62 | -1.86 | -0.93 | -1.33 | -1.7 | -0.215 | -0.3625 | -0.84 | 0.488684 | 0.369839 | 0.035233 | 0.999971 | 1 | 0.69205 |
| Dnajb4 | Q9D832 | 0.59 | 0.07 | 0.35 | 0.4 | 0.24 | 0.57 | 1.03 | -0.17 | -0.1 | 0.42 | -0.08 | -0.26 | 0.15 | 0.44 | -0.13 | 0.03 | 0.065 | -0.3575 | -0.23 | 0.821739 | 0.133941 | 0.2941 | 0.999971 | 1 | 0.847402 |
| Dnajb5 | A2AG30 | 0.77 | 0.06 | 0.46 | 0.83 | -0.1 | 0.22 | 0.46 | 0.79 | -0.11 | 0.67 | 0.84 | 0.6 | 0.08 | 0.13 | 0.82 | 1.01 | -0.1875 | -0.03 | -0.02 | 0.501397 | 0.916395 | 0.947446 | 0.999971 | 1 | 0.997651 |
| Dnajb6 | O54946 | 0.07 | -0.32 | 0.4 | 0.4 | -0.86 | -0.89 | 0.63 | -0.27 | -1.02 | -0.4 | -0.02 | -0.09 | -1.61 | -0.53 | -0.62 | -0.44 | -0.485 | -0.52 | -0.9375 | 0.224134 | 0.105052 | 0.016836 | 0.999971 | 1 | 0.616254 |
| Dnajb8 | A0A0N4SV | -0.12 | -1.88 | -2.06 | -0.38 | -0.51 | -3.28 | -1.78 | -1.23 | -1.79 | -1.92 | -2.14 | -1.37 | -1.09 | -1.91 | -1.87 | -2.14 | -0.59 | -0.695 | -0.6425 | 0.407571 | 0.181843 | 0.229137 | 0.999971 | 1 | 0.828687 |
| Dnajc1 | Q61712 | -0.34 | -0.28 | -0.52 | -1.36 | -1.25 | -1.08 | -0.2 | -1.62 | -0.11 | 0.35 | -0.65 | -0.49 | -1.71 | -0.71 | -0.57 | 0.05 | -0.4125 | 0.4 | -0.11 | 0.292016 | 0.249892 | 0.793718 | 0.999971 | 1 | 0.970141 |
| Dnajc10 | Q9DC23 | 1.14 | 0.38 | 0.53 | 0.92 | 0.61 | 0.67 | 0.57 | 0.85 | 0.59 | 1.44 | 0.99 | 0.89 | 0.04 | 0.79 | 0.67 | 1.09 | -0.0675 | 0.235 | -0.095 | 0.765362 | 0.38953 | 0.746497 | 0.999971 | 1 | 0.961064 |
| Dnajc11 | Q5U458 | -0.46 | -0.02 | 0.03 | -0.36 | -0.05 | 0.06 | -0.12 | -0.22 | -0.22 | -0.19 | -0.12 | -0.52 | -0.03 | -0.06 | -0.24 | -0.18 | 0.12 | -0.06 | 0.075 | 0.54596 | 0.7673 | 0.702069 | 0.999971 | 1 | 0.949019 |
| Dnajc12 | D3Z0D9 | 0.57 | -0.41 | -0.6 | 0.43 | 0.07 | -0.76 | -0.51 | 0.65 | 0.2 | -0.13 | -0.51 | -0.19 | -1.15 | 0.11 | 0.06 | 0.8 | -0.135 | -0.155 | -0.0425 | 0.74255 | 0.638395 | 0.927672 | 0.999971 | 1 | 0.994229 |
| Dnajc13 | G3X922 | -0.83 | -0.57 | -0.13 | -0.11 | -0.4 | -0.5 | -0.28 | -0.1 | -0.28 | -0.6 | 0.06 | 0.1 | -0.25 | -0.38 | 0.03 | 0.22 | 0.09 | 0.23 | 0.315 | 0.699587 | 0.389743 | 0.22951 | 0.999971 | 1 | 0.828687 |
| Dnajc14 | A0A1W2P6 | 3.02 | -0.82 | 0.27 | 0.17 | -0.51 | -0.06 | 2.65 | -1.38 | -1.6 | 2.28 | -0.74 | -2.37 | -1.57 | -1.02 | -0.86 | -2.99 | -0.485 | -1.2675 | -2.27 | 0.651231 | 0.297105 | 0.025296 | 0.999971 | 1 | 0.678474 |
| Dnajc15 | Q78YY6 | 1.21 | -0.31 | -0.34 | 0.53 | -0.02 | 0.29 | 0.29 | -0.05 | -0.11 | 0.63 | -0.14 | -1.01 | -0.44 | -0.32 | -0.03 | -0.34 | -0.145 | -0.43 | -0.555 | 0.697602 | 0.371987 | 0.160828 | 0.999971 | 1 | 0.800215 |
| Dnajc16 | Q80TN4 | -0.27 | 0.22 | 0.14 | 0.24 | 0.19 | 0.38 | 0.4 | 0.38 | 0.16 | -0.03 | 0.24 | -0.13 | 0.33 | 0.17 | 0.23 | -0.01 | 0.255 | -0.0225 | 0.0975 | 0.209637 | 0.910415 | 0.627014 | 0.999971 | 1 | 0.930921 |
| Dnajc18 | Q9CZJ9 | -0.88 | -0.43 | -1.05 | -0.78 | -0.24 | 0.45 | -1.18 | 0.22 | -0.2 | -0.19 | -0.71 | -0.43 | 0.47 | 0.31 | -0.69 | -0.87 | 0.5975 | 0.4025 | 0.59 | 0.134736 | 0.094608 | 0.126671 | 0.999971 | 1 | 0.783193 |
| Dnajc19 | Q9CQV7 | -1.04 | -0.92 | 0.05 | 0.23 | -0.62 | -0.83 | -0.4 | 0.13 | -0.44 | -0.16 | 0.63 | 0.21 | -0.01 | -0.68 | 0.35 | 0.47 | -0.01 | 0.48 | 0.4525 | 0.978631 | 0.235007 | 0.274807 | 0.999971 | 1 | 0.843794 |
| Dnajc2 | E9Q9H2 | 2.26 | 1.48 | 1.73 | 2.1 | 1.85 | 1.22 | 1.81 | 2.29 | 2.24 | 2.51 | 1.38 | 1.21 | 2.72 | 1.31 | 1.73 | 1.89 | -0.1 | -0.0575 | 0.02 | 0.733341 | 0.871914 | 0.953245 | 0.999971 | 1 | 0.998056 |
| Dnajc24 | Q91ZF0 | -0.48 | -0.06 | 0.28 | 0.38 | 0.02 | 0.06 | -0.2 | 0.33 | -0.06 | -0.23 | -0.11 | 0.17 | -0.59 | 0.29 | 0.36 | 0.6 | 0.0225 | -0.0875 | 0.135 | 0.928352 | 0.717722 | 0.680197 | 0.999971 | 1 | 0.942957 |
| Dnajc25 | A2ALW5 | 0.04 | 0.52 | 1.02 | 0.6 | 0.54 | 0.21 | 0.66 | 1.32 | 0.79 | 0.54 | 0.68 | 1.44 | -0.05 | 0.48 | 1.53 | 1.91 | 0.1375 | 0.3175 | 0.4225 | 0.661088 | 0.29488 | 0.375575 | 0.999971 | 1 | 0.870465 |
| Dnajc27 | Q8CFP6 | 1 | -0.23 | 0.58 | 0.82 | -0.08 | -0.24 | 1.15 | 0.58 | 0.12 | 0.71 | 0.77 | 0.53 | -0.81 | 0.08 | 0.19 | 0.15 | -0.19 | -0.01 | -0.64 | 0.637372 | 0.974444 | 0.098919 | 0.999971 | 1 | 0.78268 |
| Dnajc28 | Q8VCE1 | -1.59 | -0.5 | -0.49 | -0.61 | -0.77 | -0.31 | 0.46 | -1.04 | -1.09 | -0.88 | 0.78 | 0.2 | -0.85 | 0.15 | -0.12 | -0.5 | 0.3825 | 0.55 | 0.4675 | 0.355295 | 0.273872 | 0.194055 | 0.999971 | 1 | 0.810547 |
| Dnajc3 | Q91YW3 | 0.3 | -0.01 | -0.18 | 0.44 | 0.28 | 0.02 | -0.11 | -0.09 | 0.07 | 0.1 | -0.06 | 0.18 | 0.53 | 0.27 | 0.31 | 0.47 | -0.1125 | -0.065 | 0.2575 | 0.603284 | 0.748311 | 0.236153 | 0.999971 | 1 | 0.832918 |
| Dnajc30 | P59041 | 0.43 | -0.28 | -0.09 | -0.13 | -0.36 | -0.34 | -0.15 | -0.34 | -0.28 | 0.11 | -0.12 | -0.33 | -0.26 | -0.15 | -0.51 | 0.12 | -0.28 | -0.1375 | -0.1825 | 0.208397 | 0.541386 | 0.450016 | 0.999971 | 1 | 0.885596 |
| Dnajc5 | P60904 | -0.27 | -3.05 | -1.36 | -1.59 | -3.21 | -2.46 | -0.51 | -2.2 | -2.78 | -1.5 | -1.61 | -1.98 | -4.44 | -3.46 | -1.96 | -2.94 | -0.5275 | -0.4 | -1.6325 | 0.475657 | 0.503074 | 0.041633 | 0.999971 | 1 | 0.701725 |
| Dnajc6 | Q80TZ3-3 | 5.26 | -3.24 | -1.51 | -1.89 | -3.17 | -2.39 | 4.95 | -4.61 | -5.21 | 3.07 | -2.4 | -4.46 | -4.4 | -4.31 | -2 | -4.81 | -0.96 | -1.905 | -3.535 | 0.704945 | 0.43184 | 0.073706 | 0.999971 | 1 | 0.746069 |
| Dnajc6_1 | Q80TZ3-2 | -0.54 | -0.63 | -1.21 | -0.85 | -0.64 | -0.84 | -1.2 | -0.71 | -0.76 | -0.69 | -1.03 | -0.8 | -0.64 | -0.64 | -1.06 | -0.54 | -0.04 | -0.0125 | 0.0875 | 0.862831 | 0.95289 | 0.702984 | 0.999971 | 1 | 0.949466 |
| Dnajc6_2 | Q80TZ3 | -0.58 | -0.56 | 0.26 | -0.17 | -0.42 | -0.14 | 0.21 | -0.15 | -0.37 | -0.41 | -0.34 | 0.02 | -0.09 | -0.08 | 0.16 | 0.12 | 0.1375 | -0.0125 | 0.29 | 0.599675 | 0.959791 | 0.249553 | 0.999971 | 1 | 0.839012 |
| Dnajc6_3 | B1B0B7 | 6.07 | -3.47 | -6.87 | -2.82 | -3.45 | -3.06 | -4.4 | -6.32 | -3.73 | 4.19 | -7.55 | -8.58 | -5.38 | -3.09 | -8.66 | -6.82 | -2.535 | -2.145 | -4.215 | 0.328972 | 0.551058 | 0.138954 | 0.999971 | 1 | 0.784948 |
| Dnajc7 | Q9QYI3 | 0.86 | 0.91 | 1.41 | 1.36 | 1.14 | 1.14 | 1.77 | 1.48 | 1.02 | 1.01 | 1.38 | 1.42 | 0.82 | 0.93 | 1.42 | 0.93 | 0.2475 | 0.0725 | -0.11 | 0.321008 | 0.744815 | 0.640753 | 0.999971 | 1 | 0.93301 |
| Dnajc8 | Q6NZB0 | 1.35 | 2.29 | 2.06 | 2.31 | 2.58 | 1.95 | 2.95 | 1.52 | 2.48 | 1.76 | 1.98 | 2.34 | 1.31 | 1.87 | 1.82 | 1.68 | 0.2475 | 0.1375 | -0.3325 | 0.516887 | 0.636993 | 0.247793 | 0.999971 | 1 | 0.839012 |
| Dnajc9 | Q91WN1 | 2.98 | 2.24 | 3.47 | 3.41 | 1.12 | 4.12 | 3.02 | 3.3 | 1.82 | 3.66 | 2.9 | 1.99 | -0.28 | 2.58 | 2.99 | 2.64 | -0.135 | -0.4325 | -1.0425 | 0.830849 | 0.379243 | 0.1792 | 0.999971 | 1 | 0.808532 |
| Dnal1 | Q05A62-2 | 0.84 | 1.43 | 2.22 | 1.57 | 1.12 | 1.45 | 1.87 | 1.77 | 1.56 | 1.33 | 2.4 | 2.78 | 1.62 | 1.52 | 2.13 | 2.36 | 0.0375 | 0.5025 | 0.3925 | 0.909359 | 0.254292 | 0.271947 | 0.999971 | 1 | 0.843794 |
| Dnal4 | Q8CDT8 | 0.39 | 0.48 | 0.09 | -2.74 | 0.53 | -0.05 | 3.75 | -3.85 | 1.15 | -0.39 | -0.4 | 0.1 | 0.38 | -0.43 | -2.96 | -5.65 | 0.54 | 0.56 | -1.72 | 0.72648 | 0.474849 | 0.236584 | 0.999971 | 1 | 0.833485 |
| Dner | Q8JZM4 | 0.8 | -0.57 | 0.61 | 0.89 | 0.23 | -0.04 | 1.01 | 0.74 | 0.12 | 0.72 | 0.43 | 0.23 | 0.21 | 0.1 | 0.51 | 0.14 | 0.0525 | -0.0575 | -0.1925 | 0.894672 | 0.871812 | 0.582527 | 0.999971 | 1 | 0.920364 |
| Dnlz | Q9D113 | 0 | -0.25 | -1.88 | -1.51 | -0.44 | -0.19 | -0.94 | -2.35 | -0.58 | 0.11 | -2.06 | -2.42 | -0.08 | 0.23 | -1.78 | -1.89 | -0.07 | -0.3275 | 0.03 | 0.908427 | 0.637102 | 0.963526 | 0.999971 | 1 | 0.998721 |
| Dnm1 | P39053-4 | -0.59 | -1.08 | -1.19 | -0.99 | -1.14 | -0.95 | -1.07 | -1.26 | -1.25 | -0.84 | -1.21 | -1.27 | -1.28 | -0.97 | -1.15 | -0.94 | -0.1425 | -0.18 | -0.1225 | 0.487519 | 0.404453 | 0.558113 | 0.999971 | 1 | 0.915051 |
| Dnm1_1 | A0A0J9YUI | -1.68 | -0.35 | 0.41 | -1.3 | -0.68 | -0.4 | 0.27 | -0.37 | -1.09 | -1.62 | 0.5 | -0.13 | -0.21 | -1.16 | 0.72 | -0.95 | 0.435 | 0.145 | 0.33 | 0.374208 | 0.813421 | 0.574181 | 0.999971 | 1 | 0.91816 |
| Dnm1_2 | P39053-6 | -2.21 | -3.25 | -0.72 | -2 | -2.32 | -4.31 | -1.72 | -2.76 | -2.93 | -3.07 | -2.34 | -1.94 | -2.43 | -2.75 | -2.04 | -1.9 | -0.7325 | -0.525 | -0.235 | 0.302141 | 0.34353 | 0.648845 | 0.999971 | 1 | 0.933842 |
| Dnm1_3 | A0A0J9YUS | 0.78 | -0.47 | 0.83 | 1.01 | -0.8 | 0.5 | 0.28 | 0.77 | -0.25 | -0.82 | -0.61 | -0.24 | -1.64 | 0.74 | 0.76 | -0.32 | -0.35 | -1.0175 | -0.6525 | 0.447952 | 0.01906 | 0.296642 | 0.999971 | 1 | 0.847402 |
| Dnm1l | Q8K1M6-3 | 0.1 | 0.33 | 0.54 | 1.04 | 0.33 | 0.85 | 0.43 | 1.07 | 0.91 | 0.49 | 1.03 | 2.01 | 1.18 | 1.52 | 2.27 | 2.65 | 0.1675 | 0.6075 | 1.4025 | 0.55496 | 0.126942 | 0.0049 | 0.999971 | 1 | 0.582714 |
| Dnm1l_1 | E9PUD2 | 0.78 | 0.36 | 0.44 | 0.43 | 0.48 | 1.1 | 0.15 | 0.83 | 1.04 | 0.4 | 0.18 | 0.43 | 1.51 | 1.21 | 0.39 | 1.09 | 0.1375 | 0.01 | 0.5475 | 0.590689 | 0.96648 | 0.071881 | 0.999971 | 1 | 0.739418 |
| Dnm1l_2 | Q8K1M6 | -0.03 | 0.07 | 0.28 | 0.19 | -0.13 | 0.17 | 0.18 | 0.34 | -0.1 | -0.05 | 0.22 | -0.03 | -0.01 | 0.31 | 0.39 | 0.65 | 0.0125 | -0.1175 | 0.2075 | 0.947056 | 0.50912 | 0.32907 | 0.999971 | 1 | 0.856525 |
| Dnm2 | P39054-2 | 0.27 | 0.01 | -0.74 | -0.75 | 0.17 | -0.07 | -1.14 | -0.47 | 0.15 | 0.4 | -0.95 | -0.68 | 0.08 | 0.11 | -0.8 | -0.66 | -0.075 | 0.0325 | -0.015 | 0.841453 | 0.934987 | 0.965722 | 0.999971 | 1 | 0.998721 |
| Dnm3 | E9QLL2 | -0.28 | -0.44 | -0.59 | -0.68 | -0.3 | -0.44 | -0.68 | -0.64 | -0.51 | -0.35 | -0.83 | -0.63 | -0.33 | -0.41 | -0.44 | -0.61 | -0.0175 | -0.0825 | 0.05 | 0.927016 | 0.670618 | 0.784748 | 0.999971 | 1 | 0.96643 |
| Dnmbp | A0A1D5RN | 1.29 | 0.44 | -0.56 | -0.21 | 0.46 | 0.01 | -0.05 | -0.6 | 0.06 | 1.2 | -0.41 | -0.93 | -0.01 | 0.48 | -0.92 | 0.03 | -0.285 | -0.26 | -0.345 | 0.517441 | 0.645729 | 0.469318 | 0.999971 | 1 | 0.892481 |
| Dnmt1 | P13864 | 0.74 | 1.71 | 1.43 | 1.66 | 1.39 | 1.07 | 0.98 | 1.3 | 1 | 1.17 | 1.05 | 1.24 | 0.61 | 1.43 | 1.51 | 1.64 | -0.2 | -0.27 | -0.0875 | 0.457383 | 0.304074 | 0.787793 | 0.999971 | 1 | 0.96724 |
| Dnmt3a | O88508 | 1.31 | -1.97 | -0.71 | -0.63 | -1.45 | -1.68 | -0.45 | -0.71 | -0.92 | 0.6 | -0.46 | -0.92 | -1.8 | -1.74 | -1.03 | -0.94 | -0.5725 | 0.075 | -0.8775 | 0.401772 | 0.914272 | 0.199805 | 0.999971 | 1 | 0.813335 |
| Dnpep | Q3TVK3 | -0.2 | -0.04 | 0.21 | 0.16 | -0.12 | -0.23 | 0.07 | 0.16 | -0.17 | -0.29 | 0.25 | 0.07 | -0.56 | 0.02 | 0.01 | -0.09 | -0.0625 | -0.0675 | -0.1875 | 0.747306 | 0.741436 | 0.393634 | 0.999971 | 1 | 0.871816 |
| Dnph1 | Q80VJ3 | 0.83 | 0.42 | 0.51 | 0.8 | 0.28 | -0.24 | 0.25 | 1.09 | 0.68 | 0.31 | 0.47 | 0.89 | 1.26 | 0.07 | 0.82 | 0.3 | -0.295 | -0.0525 | -0.0275 | 0.341891 | 0.802425 | 0.926031 | 0.999971 | 1 | 0.99401 |
| Dnttip1 | Q99LB0 | 2.27 | -0.28 | -1.48 | -0.06 | 0.03 | -0.49 | -1.25 | -0.47 | -0.5 | 1.55 | -0.38 | -0.77 | -0.81 | -0.58 | 0.43 | -0.78 | -0.6575 | -0.1375 | -0.5475 | 0.388535 | 0.872216 | 0.475174 | 0.999971 | 1 | 0.894044 |
| Dnttip2 | Q8R2M2 | 0.13 | -1.19 | -0.88 | -0.18 | -1 | -1.05 | -1.31 | -0.68 | -1.27 | -0.13 | -1 | -0.62 | -0.72 | -0.78 | -0.85 | -0.27 | -0.48 | -0.225 | -0.125 | 0.171159 | 0.557725 | 0.70721 | 0.999971 | 1 | 0.950182 |
| Doc2a | Q7TNF0 | -1.46 | -1.43 | -1.32 | -1.27 | -1.23 | -1.74 | -1.3 | -1.34 | -0.84 | -1.91 | -0.99 | -1.04 | -0.17 | -1.51 | -0.82 | -1.07 | -0.0325 | 0.175 | 0.4775 | 0.86443 | 0.515357 | 0.131814 | 0.999971 | 1 | 0.784948 |
| Doc2b | P70169 | -0.4 | -0.36 | -1.23 | -1.21 | 0.44 | 0.52 | -0.33 | -1.24 | -0.17 | -0.44 | -0.85 | -0.23 | 0.76 | 0.35 | -0.28 | -0.68 | 0.6475 | 0.3775 | 0.8375 | 0.172805 | 0.224003 | 0.055571 | 0.999971 | 1 | 0.725348 |
| Dock1 | Q8BUR4 | -0.92 | -0.84 | -0.63 | -0.59 | -0.91 | -1.23 | -0.94 | -0.76 | -0.97 | -1.19 | -1.09 | -0.66 | -1.09 | -0.98 | -0.45 | -0.8 | -0.215 | -0.2325 | -0.085 | 0.280485 | 0.255922 | 0.689411 | 0.999971 | 1 | 0.945805 |
| Dock10 | A0A087WC | 0.95 | -0.59 | 0.46 | 0.54 | -0.21 | -0.27 | 0.11 | 0.91 | -0.13 | 0.23 | 0.14 | 0.67 | -0.81 | -0.29 | 0.82 | 0.14 | -0.205 | -0.1125 | -0.375 | 0.616007 | 0.754892 | 0.412309 | 0.999971 | 1 | 0.875881 |
| Dock11 | A2AF47 | 0.23 | -0.18 | -0.24 | -0.52 | 0.05 | -0.06 | -0.2 | -0.37 | 0.07 | 0.52 | -0.28 | 0.15 | -0.28 | -0.24 | -0.4 | -0.37 | 0.0325 | 0.2925 | -0.145 | 0.883605 | 0.262618 | 0.497008 | 0.999971 | 1 | 0.902655 |
| Dock2 | Q8C3J5 | 0.36 | 0.58 | 0.66 | 0.83 | 0.25 | 0.03 | 0.27 | 0.43 | 0.07 | 0.19 | -0.01 | 0.63 | 0.29 | 0.45 | 0.41 | 0.95 | -0.3625 | -0.3875 | -0.0825 | 0.086781 | 0.100203 | 0.708929 | 0.999971 | 1 | 0.950592 |
| Dock3 | F8VPQ1 | -0.76 | -0.32 | -0.15 | -0.1 | -0.28 | -0.61 | -0.11 | -0.28 | -0.3 | -0.43 | -0.17 | -0.01 | -0.15 | -0.41 | -0.07 | 0.09 | 0.0125 | 0.105 | 0.1975 | 0.955516 | 0.630864 | 0.390511 | 0.999971 | 1 | 0.871816 |
| Dock4 | P59764 | -0.73 | -0.32 | -0.15 | -0.58 | -0.31 | -0.29 | -0.33 | -0.27 | -0.18 | -0.48 | -0.14 | -0.14 | 0.09 | -0.22 | -0.14 | -0.27 | 0.145 | 0.21 | 0.31 | 0.462109 | 0.31878 | 0.158929 | 0.999971 | 1 | 0.798217 |
| Dock5 | B2RY04 | -0.71 | -0.28 | -0.44 | -0.13 | -0.51 | -1.26 | -1.19 | -0.81 | -0.8 | -0.88 | -0.99 | -0.49 | -0.26 | -0.39 | -0.55 | -0.42 | -0.5525 | -0.4 | -0.015 | 0.047152 | 0.084535 | 0.939707 | 0.999971 | 1 | 0.995869 |
| Dock6 | A0A1L1SQI | 0.71 | 0.42 | -0.29 | -0.04 | 0.8 | -0.24 | 0.71 | -0.41 | 0.51 | 0.74 | -0.68 | 0.09 | 0.18 | 0.22 | 0.23 | 0.05 | 0.015 | -0.035 | -0.03 | 0.967973 | 0.925263 | 0.906333 | 0.999971 | 1 | 0.991397 |
| Dock7 | A2A9M5 | -0.63 | -0.3 | -0.28 | -0.71 | -0.49 | -0.65 | -0.57 | -0.74 | -0.43 | -0.43 | -0.39 | -0.33 | -0.33 | -0.42 | -0.31 | -0.73 | -0.1325 | 0.085 | 0.0325 | 0.492484 | 0.643408 | 0.873336 | 0.999971 | 1 | 0.984109 |
| Dock9 | E9QMR2 | -0.16 | -0.37 | -0.01 | -0.17 | -0.26 | -0.24 | -0.18 | -0.17 | -0.14 | -0.19 | -0.25 | 0.04 | -0.1 | -0.35 | -0.05 | -0.09 | -0.035 | 0.0425 | 0.03 | 0.837563 | 0.808144 | 0.868037 | 0.999971 | 1 | 0.983462 |
| Dock9_1 | Q8BIK4-2 | 3.18 | -1.29 | -0.18 | -0.2 | -0.7 | 0.43 | 1.49 | -1.36 | -0.3 | 1.92 | -1.15 | -1.63 | -3.33 | -1.49 | -2.18 | -3.03 | -0.4125 | -0.6675 | -2.885 | 0.689965 | 0.555155 | 0.013164 | 0.999971 | 1 | 0.613095 |
| Dohh | D3Z7J7 | 0.3 | 0.41 | -1.01 | 0.14 | 0.1 | 0.03 | -1.01 | -0.22 | 0.35 | 0.3 | -0.42 | 0.06 | 0.81 | 0.5 | -1.43 | -1.21 | -0.235 | 0.1125 | -0.2925 | 0.558049 | 0.757304 | 0.630982 | 0.999971 | 1 | 0.932858 |
| Dok4 | Q99KE3 | -0.13 | 0.05 | -0.21 | 0.2 | -0.5 | -0.25 | -1.02 | -0.59 | -0.45 | 0.54 | -0.13 | -0.41 | -0.34 | 0.23 | -0.6 | -0.16 | -0.5675 | -0.09 | -0.195 | 0.030867 | 0.73572 | 0.414209 | 0.999971 | 1 | 0.876329 |
| Dok6 | Q2MHE5 | 0.54 | 0.44 | 0.5 | 0.27 | 0.65 | 0.63 | 0.07 | 0.32 | 0.14 | 1.35 | 0.3 | 0.21 | 1.36 | 0.63 | 0.15 | 0.39 | -0.02 | 0.0625 | 0.195 | 0.922194 | 0.834599 | 0.497204 | 0.999971 | 1 | 0.902655 |
| Dok7 | A0A0R4J0F | 2.39 | 0.25 | 1.23 | 0.31 | -0.22 | 0.55 | 0.08 | 1.3 | -0.13 | 3.49 | 0.64 | 1.98 | -0.69 | 0.31 | 1.35 | 1.23 | -0.6175 | 0.45 | -0.495 | 0.280553 | 0.598953 | 0.439457 | 0.999971 | 1 | 0.883105 |
| Dolk | Q8R2Y3 | -2.97 | -0.21 | 0.05 | 0.11 | -0.1 | 0.39 | 0.09 | 0.29 | -0.49 | -2.74 | 0.89 | -0.24 | -0.33 | 1 | 0.4 | 1.23 | 0.9225 | 0.11 | 1.33 | 0.197215 | 0.908319 | 0.099856 | 0.999971 | 1 | 0.78268 |
| Dolpp1 | A2AWJ3 | 2.34 | 2.58 | 1.94 | 0.32 | 4.48 | 1.79 | 4.85 | 2.18 | 4.88 | 1.62 | 2.6 | 3.58 | 8.06 | 0.94 | 1.21 | 0.13 | 1.53 | 1.375 | 0.79 | 0.094757 | 0.106336 | 0.642559 | 0.999971 | 1 | 0.93301 |
| Dop1a | H7BWZ9 | -0.28 | 0.77 | 0.65 | 0.53 | 0.8 | 0.59 | 0.63 | 0.26 | 0.67 | 0.01 | 0.73 | 1.1 | 1.54 | 0.61 | 0.79 | 0.26 | 0.1525 | 0.21 | 0.3825 | 0.587668 | 0.527065 | 0.295876 | 0.999971 | 1 | 0.847402 |
| Dop1b | Q3UHQ6 | -0.1 | -0.16 | -0.06 | -0.01 | -0.03 | 0.21 | -0.11 | 0.17 | 0.12 | -0.02 | 0.18 | 0.21 | 0.27 | 0.24 | 0.2 | 0.54 | 0.1425 | 0.205 | 0.395 | 0.419251 | 0.225855 | 0.045931 | 0.999971 | 1 | 0.710102 |
| Dpagt1 | P42867 | -0.07 | -0.25 | 0.68 | 1.77 | 0.08 | 1.03 | 0.61 | 1.92 | -0.43 | -0.88 | 0.83 | 0.77 | -1.03 | 0.62 | 0.88 | 1.9 | 0.3775 | -0.46 | 0.06 | 0.500098 | 0.435548 | 0.930634 | 0.999971 | 1 | 0.994744 |
| Dpcd | Q8BPA8 | -0.36 | 0.1 | 0.27 | 0.22 | 0.17 | 0.11 | 0.33 | 0.75 | 0.22 | 0.09 | 0.37 | 0.58 | -0.25 | 0.38 | 0.59 | 0.95 | 0.2825 | 0.2575 | 0.36 | 0.253714 | 0.259055 | 0.248752 | 0.999971 | 1 | 0.839012 |

| | | | | | | | | | | | | | | | | | | | | | | | | | | |
|---|---|---|---|---|---|---|---|---|---|---|---|---|---|---|---|---|---|---|---|---|---|---|---|---|---|---|
| Dpf1 | D3YYG6 | 2.04 | 2.54 | 1.36 | 1.44 | 2.8 | 1.4 | 1.95 | 1.23 | 2.94 | 1.3 | 1.63 | 1.45 | 1.81 | 0.78 | 0.93 | 1.2 | -6.28E-16 | -0.015 | -0.665 | 1 | 0.972866 | 0.086423 | 1 | 1 | 0.759991 |
| Dpf2 | D3Z5N6 | 1.65 | 2.03 | 2.06 | 2.51 | 1.96 | 1.75 | 1.87 | 2.46 | 1.53 | 0.95 | 1.68 | 2.18 | 1.62 | 1.91 | 2.18 | 1.76 | -0.0525 | -0.4775 | -0.195 | 0.839444 | 0.15248 | 0.433555 | 0.999971 | 1 | 0.882409 |
| Dph1 | Q5NCQ5 | 1.1 | 0.31 | 0.35 | 0.91 | 0.24 | 0.51 | -0.08 | 0.66 | 0.38 | 0.98 | 0.57 | 0.65 | -0.26 | 0.72 | 0.83 | 1.48 | -0.335 | -0.0225 | 0.025 | 0.242267 | 0.930337 | 0.949399 | 0.999971 | 1 | 0.998038 |
| Dph2 | Q9CR25 | -0.02 | 0.52 | 0.12 | 0.65 | 0.24 | -0.04 | 0.31 | 0.22 | 0.05 | 0.23 | 0.61 | 0.13 | -0.44 | 0.11 | 0.31 | 0.67 | -0.135 | -0.0625 | -0.155 | 0.544569 | 0.7902 | 0.599594 | 0.999971 | 1 | 0.92422 |
| Dph5 | Q9CWQ0 | 0.41 | 0.04 | 0.58 | 0.61 | 0.33 | -0.43 | 1.3 | 0.31 | -0.37 | -0.22 | 0.47 | 0.48 | 0.31 | 0.21 | 0.67 | 0.39 | -0.0325 | -0.32 | -0.015 | 0.929453 | 0.264272 | 0.943921 | 0.999971 | 1 | 0.99692 |
| Dph6 | Q9CQ28-2 | 0.53 | 1.19 | 0.41 | 0.5 | 0.75 | 0.52 | -0.12 | 0.67 | 0.39 | 0.84 | 0.37 | -0.02 | 0.13 | 1.11 | 0.56 | -0.32 | -0.2025 | -0.2625 | -0.2875 | 0.4787 | 0.343079 | 0.419335 | 0.999971 | 1 | 0.87788 |
| Dph7 | Q9CYU6 | 1.16 | 1.31 | -0.11 | -0.21 | 1.32 | 1.59 | 0.34 | 0.48 | 0.31 | 0.6 | -0.2 | -0.34 | -0.35 | -0.17 | 0.16 | -0.06 | 0.395 | -0.445 | -0.6425 | 0.414205 | 0.322042 | 0.135592 | 0.999971 | 1 | 0.784948 |
| Dpm1 | O70152 | -0.83 | -1.01 | -1.69 | -1.73 | -0.64 | -1.14 | -1.45 | -1.58 | -0.61 | -0.99 | -1.99 | -1.55 | -0.3 | -0.98 | -1.47 | -1.55 | 0.1125 | 0.03 | 0.24 | 0.72194 | 0.935537 | 0.511399 | 0.999971 | 1 | 0.905815 |
| Dpm3 | Q9D1Q4 | -0.38 | 0.13 | 0.33 | 0.4 | 0.8 | 0.92 | 0 | -0.25 | 0.6 | 1.51 | -0.37 | 0.29 | 1.9 | 0.07 | -0.02 | 0.17 | 0.2475 | 0.3875 | 0.41 | 0.469234 | 0.356786 | 0.384471 | 0.999971 | 1 | 0.871816 |
| Dpp10 | Q6NXK7 | -0.9 | -0.52 | -0.81 | -0.92 | -0.52 | -0.36 | -0.4 | -0.98 | -0.6 | -0.77 | -0.79 | -1.01 | -0.21 | -0.67 | -0.9 | -1.06 | 0.2225 | -0.005 | 0.0775 | 0.318238 | 0.97879 | 0.747595 | 0.999971 | 1 | 0.961064 |
| Dpp10_1 | A0A5F8MP | 0.65 | -1.27 | -0.49 | -0.75 | -1.65 | -0.18 | 0.25 | -0.47 | -1.47 | 0.56 | -0.49 | -1.14 | -0.94 | -1.26 | -1.2 | -0.2 | -0.0475 | -0.17 | -0.435 | 0.92846 | 0.760773 | 0.342498 | 0.999971 | 1 | 0.861474 |
| Dpp3 | Q99KK7 | 0.47 | -0.06 | -0.3 | 0.01 | 0.03 | -0.08 | -0.11 | 0.06 | 0.12 | 0.47 | -0.09 | 0.02 | -0.24 | 0.17 | -0.15 | 0.29 | -0.055 | 0.1 | -0.0125 | 0.797411 | 0.671143 | 0.958248 | 0.999971 | 1 | 0.998622 |
| Dpp4 | P28843 | 0.36 | 0.55 | 0.48 | 1.12 | 0.67 | 0.14 | 0.53 | 0.13 | 0.57 | 0.76 | 0.85 | 0.59 | 0.05 | 0.75 | 0.95 | 0.77 | -0.26 | 0.065 | 0.0025 | 0.307935 | 0.769573 | 0.992811 | 0.999971 | 1 | 0.99925 |
| Dpp6 | E9PWX1 | -1.41 | -1.94 | -2.19 | -2.15 | -1.83 | -1.9 | -2.09 | -2.12 | -1.82 | -1.76 | -2.21 | -2.07 | -1.85 | -1.78 | -2.15 | -2 | -0.0625 | -0.0425 | -0.0225 | 0.786685 | 0.858837 | 0.923407 | 0.999971 | 1 | 0.993265 |
| Dpp6_1 | Q80VM5 | 0.74 | 0.61 | 1.54 | 0.9 | 0.14 | 1.76 | 0.52 | 1.56 | 0.22 | 2.18 | 0.67 | -0.77 | 1.4 | 0.72 | 1.33 | 1.41 | 0.0475 | -0.3725 | 0.2675 | 0.910171 | 0.535799 | 0.353504 | 0.999971 | 1 | 0.868675 |
| Dpp7 | Q9ET22 | -0.1 | 1.18 | 1.13 | 0.93 | 1.03 | 1.08 | 0.97 | 1.01 | 1.21 | 0.34 | 1.34 | 1.54 | 0.82 | 1.16 | 1.35 | 1 | 0.2375 | 0.3225 | 0.2975 | 0.446991 | 0.413127 | 0.368583 | 0.999971 | 1 | 0.869526 |
| Dpp8 | Q80YA7 | 0.32 | 0.41 | -0.13 | 0.1 | 0.42 | 0.18 | 0.69 | 0.05 | 0.23 | -0.06 | -0.56 | -0.18 | 0.09 | 0 | -0.4 | -0.39 | 0.16 | -0.3175 | -0.35 | 0.485242 | 0.201011 | 0.13932 | 0.999971 | 1 | 0.784948 |
| Dpp9 | Q8BVG4 | -0.39 | -0.04 | 0.24 | -0.04 | 0.02 | -0.04 | 0.35 | -0.32 | 0.23 | -0.51 | -0.11 | 0.15 | -0.28 | -0.07 | -0.4 | -0.55 | 0.06 | -0.0025 | -0.2675 | 0.79251 | 0.991703 | 0.230938 | 0.999971 | 1 | 0.830179 |
| Dpy19l1 | A6X919 | -1.3 | -0.24 | -0.9 | -1.65 | 0.1 | -0.41 | -0.17 | -0.77 | 0.11 | -1.12 | -0.1 | 0.22 | 0.5 | -0.18 | -0.45 | -0.95 | 0.71 | 0.8 | 0.7525 | 0.067709 | 0.078999 | 0.09277 | 0.999971 | 1 | 0.776565 |
| Dpy19l3 | Q71B07 | -0.08 | -0.26 | 1 | -1.16 | -0.21 | 0.04 | 1.65 | -0.39 | -0.5 | 0.46 | 0.61 | -1 | 0.17 | 0.15 | -0.21 | -0.98 | 0.3975 | 0.0175 | -0.0925 | 0.505041 | 0.974288 | 0.848233 | 0.999971 | 1 | 0.98119 |
| Dpy19l4 | A2AJQ3-2 | 0.85 | 0.59 | 2.67 | 2.12 | 0.55 | 0.96 | 2.41 | 2.76 | 0.15 | 1.79 | 2.19 | 0.62 | 0.04 | 0.53 | 1.53 | 1.37 | 0.1125 | -0.37 | -0.69 | 0.865865 | 0.563164 | 0.239533 | 0.999971 | 1 | 0.83365 |
| Dpy30 | Q99LT0 | 0.33 | 0.96 | -0.74 | 0.05 | 0.48 | -0.06 | -0.83 | -0.14 | 0.41 | -0.76 | -0.13 | -0.3 | 0.3 | 0.48 | -0.39 | 0.16 | -0.2875 | -0.345 | -0.0125 | 0.499708 | 0.408271 | 0.97409 | 0.999971 | 1 | 0.99925 |
| Dpyd | Q8CHR6 | -0.34 | -0.4 | -0.69 | -0.78 | -0.55 | -1.11 | -0.72 | -0.9 | -1.73 | -0.87 | -1.75 | -1.2 | -0.25 | -0.85 | -0.5 | -1.06 | -0.2675 | -0.835 | -0.1125 | 0.226337 | 0.010504 | 0.644542 | 0.999971 | 1 | 0.93301 |
| Dpys | Q9EQF5 | 0.95 | 0.84 | 0.97 | 2.7 | 1.75 | 0.41 | 0.57 | 2.1 | 1.95 | 1.82 | 2.66 | 1.23 | -0.59 | 2.22 | 2.53 | 2.54 | -0.1575 | 0.55 | 0.31 | 0.77982 | 0.286589 | 0.696717 | 0.999971 | 1 | 0.947174 |
| Dpysl2 | O08553 | -0.21 | -0.09 | -0.35 | -0.06 | -0.32 | -0.42 | -0.54 | -0.16 | -0.42 | -0.57 | -0.58 | -0.28 | -0.07 | 0.01 | -0.13 | 0.11 | -0.1825 | -0.285 | 0.1575 | 0.328473 | 0.130351 | 0.376566 | 0.999971 | 1 | 0.870465 |
| Dpysl3 | E9PWE8 | -0.83 | -1.3 | -1.08 | -0.96 | -1.08 | -1.26 | -0.68 | -0.64 | -0.79 | -0.75 | -0.58 | -0.56 | -0.68 | -0.82 | -0.51 | -0.56 | 0.1275 | 0.3725 | 0.4 | 0.572039 | 0.068845 | 0.060713 | 0.999971 | 1 | 0.725348 |
| Dpysl4 | Q3TMU8 | 0.2 | -0.02 | -0.46 | -0.3 | -0.14 | -0.23 | -0.46 | -0.37 | -0.27 | 0.14 | -0.56 | -0.53 | 0 | 0.25 | -0.52 | -0.2 | -0.155 | -0.16 | 0.0275 | 0.471855 | 0.520549 | 0.911853 | 0.999971 | 1 | 0.992551 |
| Dpysl5 | Q9EQF6 | -1.22 | -1.4 | -0.89 | -0.58 | -1.16 | -1.63 | -0.77 | -0.46 | -0.78 | -1.11 | -0.41 | -0.24 | -1.03 | -1.27 | -0.37 | -0.43 | 0.0175 | 0.3875 | 0.2475 | 0.955631 | 0.190284 | 0.414421 | 0.999971 | 1 | 0.876329 |
| Dr1 | Q91WV0 | 1.09 | 2.27 | 1.56 | 0.48 | 2.08 | 1.62 | 1.16 | 0.88 | 1.91 | 0.87 | 1.72 | 2.12 | 1.82 | 1.42 | 1.44 | 1.18 | 0.085 | 0.305 | 0.115 | 0.845001 | 0.495653 | 0.766315 | 0.999971 | 1 | 0.962472 |
| Drap1 | F6T9K1 | 0.74 | 2.61 | 2.12 | 2.56 | 2.41 | 2.07 | 1.97 | 2.97 | 2.52 | 2.02 | 2.73 | 2.47 | 3.18 | 3.43 | 3.12 | 3.18 | 0.3475 | 0.4275 | 1.22 | 0.457495 | 0.341513 | 0.016677 | 0.999971 | 1 | 0.615351 |
| Drd1 | A0A1Y7VK! | -3.12 | -7.81 | -2.48 | -1.9 | -7.65 | -7.43 | -1.19 | -2.23 | -7.59 | -4.75 | -3.58 | -2.61 | -9.28 | -6.58 | -2.59 | -2.58 | -0.7975 | -0.805 | -1.43 | 0.678842 | 0.604813 | 0.454833 | 0.999971 | 1 | 0.887217 |
| Drg1 | P32233 | 0.82 | 1.69 | 1.19 | 1.15 | 1.65 | 1.64 | 0.93 | 1.1 | 1.65 | 1.26 | 1.38 | 1.49 | 1.58 | 1.69 | 1.2 | 1.35 | 0.1175 | 0.2325 | 0.2425 | 0.670809 | 0.330345 | 0.33248 | 0.999971 | 1 | 0.857921 |
| Drg2 | Q9QXB9 | 0.53 | 1.02 | 0.67 | 0.49 | 1.21 | 0.77 | 0.23 | 0.73 | 1.32 | 0.53 | 0.52 | 0.43 | 2.02 | 0.83 | 0.56 | 0.98 | 0.0575 | 0.0225 | 0.42 | 0.823816 | 0.931346 | 0.236206 | 0.999971 | 1 | 0.832918 |
| Drp2 | Q05AA6 | -0.31 | -0.44 | -0.31 | -0.08 | -0.33 | -0.22 | -0.49 | 0.14 | -0.39 | 0.25 | -0.01 | -0.43 | -0.27 | -0.23 | -0.51 | 0.12 | 0.06 | 0.14 | 0.0625 | 0.771704 | 0.528383 | 0.761174 | 0.999971 | 1 | 0.961116 |
| Dscam | Q9ERC8 | -0.31 | -0.59 | -0.79 | -0.27 | -1.42 | -1.84 | -0.71 | -0.37 | -1.77 | -0.64 | -0.08 | -0.36 | -1.45 | -1.43 | -0.59 | -0.15 | -0.595 | -0.2225 | -0.415 | 0.115692 | 0.560115 | 0.244173 | 0.999971 | 1 | 0.837316 |
| Dscaml1 | E9QPR7 | -0.34 | -0.72 | -0.19 | -0.24 | -0.38 | -0.29 | -0.13 | 0.2 | -0.67 | -0.21 | -0.33 | -0.62 | -0.16 | -0.34 | -0.15 | -0.07 | 0.2225 | -0.085 | 0.1925 | 0.325096 | 0.687202 | 0.339568 | 0.999971 | 1 | 0.860218 |
| Dst | A0A087WF | -1.05 | 2.17 | 0.8 | -0.49 | 1.9 | 1.41 | 0.82 | 0.9 | 2.46 | -0.09 | 0.79 | 2.47 | 4.75 | 0.51 | -0.01 | -0.03 | 0.9 | 1.05 | 0.9475 | 0.212834 | 0.245489 | 0.442093 | 0.999971 | 1 | 0.883386 |
| Dst_1 | S4R1P5 | -0.59 | -0.73 | -0.72 | -0.67 | -0.8 | -0.76 | -0.74 | -0.78 | -0.68 | -0.57 | -0.67 | -0.57 | -0.61 | -0.86 | -0.78 | -0.77 | -0.0925 | 0.055 | -0.0775 | 0.565855 | 0.727621 | 0.643403 | 0.999971 | 1 | 0.93301 |
| Dstn | Q9R0P5 | 0.49 | 1.62 | 1.01 | 1.32 | 1.81 | 1.97 | 1.22 | 1.58 | 1.81 | 1 | 1.45 | 1.67 | 2.22 | 1.65 | 1.47 | 0.93 | 0.535 | 0.3725 | 0.4575 | 0.101437 | 0.242782 | 0.216989 | 0.999971 | 1 | 0.826893 |
| Dstyk | A0A087WF | 2.13 | 1.59 | 0.69 | 0.56 | 1.39 | 1.47 | 0.27 | 0.54 | 1.34 | 1.47 | 0.78 | 1.08 | 2.26 | 1.71 | 0.4 | 0.83 | -0.325 | -0.075 | 0.0575 | 0.477684 | 0.847215 | 0.911992 | 0.999971 | 1 | 0.992583 |
| Dtd1 | Q9DD18 | 2.39 | 0.09 | 1.04 | 1.02 | -0.41 | 0.23 | 0.97 | 0.77 | -0.89 | 0.78 | 0.76 | 0.5 | -1.23 | -0.62 | 0.51 | 0.37 | -0.745 | -0.8475 | -1.3775 | 0.177873 | 0.162686 | 0.038332 | 0.999971 | 1 | 0.700838 |
| Dtd2 | Q8BHA3 | 1.68 | 0.69 | 0.66 | 0.9 | 0.62 | 1.32 | 0.3 | 1.4 | 0.76 | 1.09 | 0.39 | 0.71 | 0.43 | 0.48 | 0.75 | 0.92 | -0.0725 | -0.245 | -0.3375 | 0.836958 | 0.407029 | 0.249334 | 0.999971 | 1 | 0.839012 |
| Dtna | Q8CFR5 | 0.42 | 0.14 | 0.47 | 0.54 | -0.04 | 0.29 | 0.68 | 0.33 | 0.11 | 0.48 | 0.54 | 0.18 | -0.07 | 0.06 | 0.57 | 0.14 | -0.0775 | -0.065 | -0.2175 | 0.72195 | 0.740124 | 0.322781 | 0.999971 | 1 | 0.85548 |
| Dtnb | G3UXK0 | -1.44 | -0.78 | -0.59 | -0.63 | -0.6 | -0.56 | -0.87 | -0.73 | -0.57 | -1.21 | -0.91 | -0.45 | -0.35 | -0.59 | -1.1 | -0.54 | 0.17 | 0.075 | 0.215 | 0.48794 | 0.786936 | 0.4398 | 0.999971 | 1 | 0.883105 |
| Dtnbp1 | A0A1Y7VIX | 2.13 | 0.16 | 1.16 | -0.09 | -0.17 | 0.4 | 1.5 | 1.36 | -0.02 | 2.21 | 0.48 | -0.8 | -0.86 | 0.17 | -0.35 | 0.02 | -0.0675 | -0.3725 | -1.095 | 0.908754 | 0.61719 | 0.059081 | 0.999971 | 1 | 0.725348 |
| Dtx3 | D3YV13 | 2.45 | 1.29 | 1.68 | 1.43 | 1.65 | 1.27 | 1.56 | 1.61 | 2.29 | 0.97 | 1.71 | 1.6 | 2.54 | 0.74 | 1.79 | 1.66 | -0.19 | -0.07 | -0.03 | 0.511778 | 0.847929 | 0.943978 | 0.999971 | 1 | 0.99692 |
| Dtymk | P97930 | -1.66 | 0.04 | -0.62 | -1.34 | 0.16 | -0.25 | 0.03 | -0.48 | 0.2 | -1.44 | -0.58 | -0.51 | 0.49 | -0.74 | -0.78 | -1.33 | 0.76 | 0.3125 | 0.305 | 0.078112 | 0.516039 | 0.547627 | 0.999971 | 1 | 0.911722 |
| Dus2 | Q9D7B1 | 1.8 | -0.35 | 2.05 | 2.65 | 0.23 | 0.65 | 2.49 | 1.85 | 0.61 | 1.31 | 2.22 | 1.22 | -0.69 | 0.36 | 1.28 | 2.42 | -0.2325 | -0.1975 | -0.695 | 0.758335 | 0.768211 | 0.415436 | 0.999971 | 1 | 0.876885 |
| Dus3l | A0A0R4IZY | 1.35 | 1.7 | 1.93 | 1.94 | 1.61 | 1.69 | 1.18 | 1.44 | 1.72 | 1.65 | 0.83 | 1.51 | 0.95 | 2.14 | 2.15 | 1.65 | -0.25 | -0.3025 | -0.0075 | 0.276617 | 0.27312 | 0.98118 | 0.999971 | 1 | 0.99925 |
| Dusp14 | Q9JLY7 | -0.63 | -0.73 | 0.4 | 0.55 | -0.05 | -1.11 | 0.16 | -0.06 | -0.71 | -0.82 | 0.48 | 0.82 | -1.39 | -0.29 | -0.1 | 0.29 | -0.1625 | 0.045 | -0.27 | 0.69874 | 0.927865 | 0.56226 | 0.999971 | 1 | 0.916386 |
| Dusp15 | Q8R4V2 | -0.13 | -0.43 | -0.89 | -0.71 | -0.06 | -0.41 | -1.68 | -0.78 | 0.15 | 0.35 | -2.2 | -1.23 | 0.06 | 0.07 | -0.82 | -1.07 | -0.1925 | -0.1925 | 0.1 | 0.609082 | 0.738534 | 0.767423 | 0.999971 | 1 | 0.962472 |
| Dusp19 | Q99N12 | 0.22 | 0.1 | 1.22 | 1.27 | -0.1 | 0.13 | 0.86 | 0.68 | 0.19 | 0.43 | 0.61 | 0.14 | -0.61 | 0.14 | 0.08 | 0.95 | -0.31 | -0.36 | -0.5625 | 0.417297 | 0.294373 | 0.208216 | 0.999971 | 1 | 0.822138 |
| Dusp23 | Q6NT99 | -0.05 | 0.13 | 0.16 | -0.48 | 0.76 | 0.24 | 0.7 | 0.25 | 0.72 | 0.5 | 0.43 | 0.27 | 0.62 | 0.71 | -0.02 | 0.01 | 0.5475 | 0.54 | 0.39 | 0.043727 | 0.032538 | 0.167464 | 0.999971 | 1 | 0.800215 |
| Dusp26 | Q9D700 | 0.48 | -0.06 | -0.41 | 0.12 | 0.69 | 0.52 | -0.96 | 0.48 | -0.26 | 1.12 | -0.04 | -0.43 | 0.49 | 0.12 | 0.47 | 0.67 | 0.15 | 0.065 | 0.405 | 0.712726 | 0.865129 | 0.129222 | 0.999971 | 1 | 0.784948 |
| Dusp28 | Q8BTR5 | 1.48 | 0.39 | 0.54 | 0.61 | -0.05 | 0.43 | 0.1 | 0.74 | 0.35 | 0.92 | 0.3 | 0.09 | 0.46 | 0.55 | 0.4 | 0.5 | -0.45 | -0.34 | -0.2775 | 0.169329 | 0.287543 | 0.316887 | 0.999971 | 1 | 0.853713 |
| Dusp3 | B1AQF4 | -0.4 | 0.41 | 0.18 | 0.38 | 0.16 | 0.15 | 0.35 | -0.22 | -0.27 | -0.15 | 0.57 | 0.3 | 0.29 | 0.52 | 0.46 | 0.98 | -0.0325 | -0.03 | 0.42 | 0.896647 | 0.915837 | 0.135863 | 0.999971 | 1 | 0.784948 |
| Dusp6 | Q9DBB1 | -1.87 | -1.93 | 1.88 | 1.09 | 0.05 | -0.36 | 0.29 | 2.12 | -0.09 | -1.07 | 1.34 | 1.11 | -1.07 | -0.81 | 1.52 | 1.28 | 0.7325 | 0.53 | 0.4375 | 0.474951 | 0.606948 | 0.684386 | 0.999971 | 1 | 0.944176 |
| Dusp9 | Q7TNL7 | 3.82 | 0.59 | 1.3 | 2.39 | -0.09 | 1.92 | -1.16 | 2 | -1.64 | 2.73 | -0.78 | -1.26 | -3.55 | 1.8 | 2.21 | 2.74 | -1.3575 | -2.2625 | -1.225 | 0.172314 | 0.06618 | 0.405447 | 0.999971 | 1 | 0.874565 |
| Dut | Q9CQ43-2 | -0.69 | -0.72 | -0.28 | 0.03 | -0.67 | -0.66 | 0.2 | -0.36 | -0.59 | -1.22 | -0.9 | -1.53 | -1.7 | -1.27 | -0.8 | -1.51 | 0.0425 | -0.645 | -0.905 | 0.881424 | 0.046133 | 0.009932 | 0.999971 | 1 | 0.613095 |
| Dvl1 | P51141 | 0.06 | 0.16 | 0.3 | 0.09 | -0.15 | 0.2 | 0.46 | 0.13 | 0 | 0.33 | -0.22 | -0.13 | 0.3 | 0.34 | 0.21 | 0.26 | 0.0075 | -0.1575 | 0.125 | 0.969615 | 0.421036 | 0.459201 | 0.999971 | 1 | 0.889191 |
| Dvl2 | Q60838 | 1.97 | 0.65 | 1.85 | 2.49 | 0.7 | 2.25 | 1.68 | 1.43 | 0.23 | 1.75 | 0.82 | 0.94 | -0.36 | 0.92 | 0.8 | 0.84 | -0.225 | -0.805 | -1.19 | 0.63484 | 0.113507 | 0.028522 | 0.999971 | 1 | 0.678474 |
| Dvl3 | E9Q150 | -2.25 | -0.65 | -1.86 | -1.24 | -0.56 | -0.17 | -1.18 | -0.64 | -0.06 | -2.18 | -1.05 | -0.8 | -0.22 | -0.79 | -1.35 | -1.62 | 0.8625 | 0.4775 | 0.505 | 0.051805 | 0.37244 | 0.27189 | 0.999971 | 1 | 0.843794 |
| Dym | Q8CHY3 | 0.53 | 0.14 | 0.81 | 0.94 | 0.63 | 0.13 | 0.86 | 0.99 | 0.15 | 0.43 | 0.96 | 0.92 | -0.36 | 0.81 | 1.26 | 1.51 | 0.0475 | 0.01 | 0.2 | 0.863424 | 0.97137 | 0.641628 | 0.999971 | 1 | 0.93301 |
| Dync1h1 | Q9JHU4 | -0.7 | -0.55 | -0.61 | -0.65 | -0.43 | -0.46 | -0.45 | -0.61 | -0.25 | -0.6 | -0.38 | -0.21 | -0.24 | -0.39 | -0.35 | -0.19 | 0.14 | 0.2675 | 0.335 | 0.401163 | 0.149245 | 0.068466 | 0.999971 | 1 | 0.733937 |
| Dync1i1 | O88485 | 0.26 | -0.16 | -0.01 | 0.15 | -0.18 | 0.18 | 0.75 | 0 | -0.1 | 0.2 | 0.13 | 0.14 | -0.19 | -0.09 | 0.11 | 0.22 | 0.1275 | 0.0325 | -0.0475 | 0.611917 | 0.858572 | 0.807659 | 0.999971 | 1 | 0.973317 |
| Dync1i2 | Q3TPJ8 | -0.58 | -0.28 | -0.72 | -0.98 | -0.35 | -0.1 | -0.6 | -0.81 | -0.25 | -0.17 | -1.05 | -1.2 | -0.05 | -0.09 | -0.87 | -0.84 | 0.175 | -0.0275 | 0.1775 | 0.478259 | 0.928963 | 0.53707 | 0.999971 | 1 | 0.910077 |
| Dync1li1 | Q8R1Q8 | -0.33 | -0.04 | -0.31 | -0.39 | 0.12 | 0.28 | -0.31 | -0.25 | 0.17 | -0.02 | 0.05 | -0.2 | 0.3 | 0.3 | -0.14 | -0.05 | 0.2275 | 0.2675 | 0.37 | 0.299184 | 0.164248 | 0.090632 | 0.999971 | 1 | 0.772703 |
| Dync1li2 | Q6PDL0 | -0.34 | 0.06 | 0.08 | -0.25 | -0.23 | 0.35 | 0.26 | 0.02 | 0.01 | -0.12 | -0.19 | -0.3 | 0.28 | 0.32 | 0.2 | 0.14 | 0.2125 | -0.0375 | 0.3475 | 0.337648 | 0.842282 | 0.089975 | 0.999971 | 1 | 0.771991 |
| Dync2h1 | Q45VK7 | 0.17 | 0.66 | 0.55 | 0.39 | 0.37 | 0.5 | 0.27 | 0.56 | 0.56 | 0.43 | 0.69 | 1.01 | 0.8 | 0.83 | 0.7 | 0.52 | -0.0175 | 0.23 | 0.27 | 0.926991 | 0.290774 | 0.185983 | 0.999971 | 1 | 0.809025 |
| Dync2i1 | Q8C761 | 1.09 | 2.29 | 2.17 | 2.17 | 1.38 | 2.07 | 1.66 | 2 | 1.39 | 1.6 | 2.47 | 1.27 | 2 | 1.61 | 1.33 | 1.57 | -0.1525 | -0.2475 | -0.3025 | 0.640393 | 0.518383 | 0.353616 | 0.999971 | 1 | 0.868693 |
| Dync2i2 | Q5U4F6 | -0.2 | -0.18 | -0.74 | -0.4 | -0.13 | -0.24 | -0.51 | 0.22 | -0.4 | 0.18 | -0.01 | 0.25 | 0.04 | 0.07 | 0.49 | 0.08 | 0.215 | 0.385 | 0.55 | 0.371463 | 0.121788 | 0.029432 | 0.999971 | 1 | 0.678474 |
| Dync2li1 | Q8K0T2 | 1.07 | -1 | 0.44 | 0.87 | 0.35 | -0.77 | -0.37 | 1.14 | -0.3 | -0.32 | -0.13 | 1.08 | 0.74 | 0.44 | 0.61 | 1.6 | -0.2575 | -0.2625 | 0.5025 | 0.656081 | 0.624517 | 0.32589 | 0.999971 | 1 | 0.856472 |
| Dynll1 | P63168 | -0.86 | -0.05 | -0.19 | -1.21 | 0.09 | 0.2 | 0.58 | -0.31 | -0.08 | -0.81 | -0.11 | 0.2 | 0.33 | -0.05 | -0.47 | -0.44 | 0.7175 | 0.3775 | 0.42 | 0.054049 | 0.290303 | 0.228257 | 0.999971 | 1 | 0.828687 |
| Dynll2 | Q9D0M5 | -1.89 | 0.21 | 0.38 | -1.08 | 0.37 | -0.06 | 0.37 | 0.11 | 0.31 | -1.84 | 0.48 | 0.55 | 1.13 | -0.55 | -0.05 | -0.37 | 0.7925 | 0.47 | 0.635 | 0.146313 | 0.517369 | 0.307974 | 0.999971 | 1 | 0.847402 |
| Dynlrb1 | P62627 | -0.88 | -0.56 | -1.13 | -1.12 | -0.49 | -0.53 | -1.02 | -0.97 | -0.53 | -0.91 | -1.05 | -1.02 | -0.35 | -0.25 | -0.84 | -0.79 | 0.17 | 0.045 | 0.365 | 0.469616 | 0.838048 | 0.14921 | 0.999971 | 1 | 0.793248 |
| Dynlt1a | A0A338P7E | -0.37 | -1.83 | 0.15 | 0.32 | -1.38 | -1.91 | -0.1 | 0.17 | -1.3 | -0.84 | 0 | -1.85 | -2.24 | -1.43 | 1.2 | 0.77 | -0.3725 | -0.565 | 0.0075 | 0.561373 | 0.340698 | 0.993101 | 0.999971 | 1 | 0.99925 |
| Dynlt3 | P56387 | -2.43 | -3.11 | -7.23 | -6.81 | -2.74 | -3.14 | -5.56 | -5.83 | -2.2 | -3.15 | -6.97 | -7.25 | -3.15 | -2.77 | -6.54 | -6.31 | 0.5775 | 0.0025 | 0.2025 | 0.660621 | 0.998752 | 0.886224 | 0.999971 | 1 | 0.987414 |
| Dyrk1a | F6U6X3 | -0.6 | -0.81 | -0.12 | -0.2 | -0.65 | -1.2 | 0.29 | -0.32 | -0.95 | -1.31 | 0.07 | 0.33 | -0.59 | -0.87 | -0.27 | -0.38 | -0.0375 | -0.0325 | -0.095 | 0.913807 | 0.936502 | 0.696716 | 0.999971 | 1 | 0.947174 |
| Dyrk1b | Q9Z188-3 | -2.32 | -0.35 | -0.47 | -1.87 | -0.06 | -1.58 | -0.22 | -1.52 | 0.75 | -2.1 | 0.19 | 0.72 | 1.5 | -1.34 | -2.09 | -1.5 | 0.4075 | 1.1425 | 0.395 | 0.493514 | 0.158748 | 0.642882 | 0.999971 | 1 | 0.93301 |
| Dyrk2 | Q5U4C9 | -0.26 | 0.1 | 0.35 | 0.07 | 1.1 | -0.01 | 1.96 | -0.49 | 0.6 | -0.49 | -0.18 | -0.55 | 1.04 | -0.55 | -0.87 | -0.52 | 0.575 | -0.22 | -0.29 | 0.287836 | 0.470585 | 0.499956 | 0.999971 | 1 | 0.903908 |
| Dzank1 | E9Q4Z6 | -0.16 | -0.15 | -0.44 | -0.38 | -0.08 | -0.15 | -0.22 | -0.1 | -0.26 | 0.01 | -0.54 | -0.67 | -0.17 | 0.05 | -0.54 | -0.43 | 0.145 | -0.0825 | 0.01 | 0.408993 | 0.700059 | 0.961343 | 0.999971 | 1 | 0.998721 |
| Dzip3 | E9QNZ2 | 0.71 | 0.58 | 0.88 | 1.17 | 0.72 | 0.74 | 0.91 | 0.99 | 0.63 | 1.07 | 0.69 | 0.83 | 0.88 | 1.09 | 1.02 | 0.86 | 0.005 | -0.03 | 0.1275 | 0.980127 | 0.885458 | 0.527065 | 0.999971 | 1 | 0.909459 |
| E2f3 | Q3TMJ9 | 1.64 | 1.76 | 1.68 | 1.96 | 2.63 | 2.16 | 0.12 | 1.48 | 1.92 | 1.34 | -0.07 | 1.25 | 2.42 | 3.16 | 1.35 | 1.69 | -0.1625 | -0.65 | 0.395 | 0.750301 | 0.137915 | 0.329209 | 0.999971 | 1 | 0.856525 |
| E9Q0B3 | E9Q0B3 | -0.4 | 0.87 | 0.94 | 0.5 | 0.81 | 0.86 | 0.33 | 1.36 | 0.44 | 0.47 | 0.83 | -0.65 | -1.27 | -0.94 | 0.8 | 0.65 | 0.3625 | -0.205 | -0.6675 | 0.332594 | 0.62966 | 0.257164 | 0.999971 | 1 | 0.839012 |
| Ears2 | Q9CXJ1 | -0.62 | -0.73 | -0.59 | -0.49 | -0.55 | -1.13 | -0.53 | -0.61 | -0.94 | -0.89 | -0.79 | -0.54 | -0.58 | -0.64 | -0.64 | -0.67 | -0.0975 | -0.1825 | -0.025 | 0.636681 | 0.317854 | 0.878938 | 0.999971 | 1 | 0.985441 |
| Ebag9 | Q9D0V7 | -0.51 | 0.29 | 0.3 | -0.21 | 0.24 | -0.72 | 0.09 | 0.36 | -0.12 | -0.61 | 0.43 | 1.5 | -0.37 | -0.26 | 1.02 | -0.23 | 0.025 | 0.3325 | 0.0725 | 0.93718 | 0.480825 | 0.845752 | 0.999971 | 1 | 0.979924 |
| Ebna1bp2 | Q9D903 | -0.83 | -1.66 | -1.83 | -2.24 | -1.64 | -1.59 | -2.19 | -1.87 | -1.76 | -1.22 | -2.44 | -2.78 | -1.79 | -1.7 | -2.21 | -1.89 | -0.1825 | -0.41 | -0.2575 | 0.579544 | 0.35702 | 0.429121 | 0.999971 | 1 | 0.881196 |
| Ebp | A2AC29 | 0.42 | 0.75 | 0.5 | 0.1 | -0.15 | 1.21 | -0.18 | 0.73 | -0.21 | 0.41 | 1.08 | -0.79 | -0.08 | 1.7 | -0.4 | 0.23 | -0.04 | -0.32 | -0.08 | 0.911184 | 0.438828 | 0.86008 | 0.999971 | 1 | 0.982702 |
| Ebpl | Q9D0P0 | -0.11 | -0.25 | 0.14 | 0.65 | 0.23 | 0.32 | 0.92 | 0.31 | 0.43 | 0.46 | 0.07 | 0.08 | -0.93 | 0.09 | 0.24 | -0.3 | 0.3375 | 0.1525 | -0.3325 | 0.236148 | 0.547082 | 0.325769 | 0.999971 | 1 | 0.856472 |
| Ecd | Q9CS74 | 1.29 | 1.72 | 0.91 | 1.06 | 0.44 | 1.08 | 0.38 | 0.86 | -0.85 | 2.32 | 0.71 | 0.13 | -2.42 | 0.87 | 0.87 | 0.78 | -0.555 | -0.6675 | -1.22 | 0.061677 | 0.304759 | 0.132028 | 0.999971 | 1 | 0.784948 |
| Ece1 | Q4PZA2-3 | 0.02 | -0.44 | -0.61 | -0.64 | 0.07 | -0.74 | -0.89 | -0.42 | 0.02 | -0.71 | 0 | -0.04 | 0.26 | -0.5 | -0.2 | -0.71 | -0.0775 | 0.235 | 0.13 | 0.780449 | 0.370862 | 0.641012 | 0.999971 | 1 | 0.93301 |
| Ece2 | E9Q1S5 | 3.8 | -0.3 | -0.53 | -0.13 | -2.21 | 1.6 | -0.84 | -0.05 | -1.61 | 2.81 | -0.53 | -0.26 | -4.12 | -1.55 | 0.29 | -0.07 | -1.085 | -0.6075 | -2.0725 | 0.361774 | 0.630748 | 0.132382 | 0.999971 | 1 | 0.784948 |
| Ecel1 | Q9JMI0 | -0.23 | -0.8 | -0.6 | -0.12 | -0.8 | -1.47 | -1.27 | -0.49 | -0.72 | -0.38 | -1 | -0.12 | -0.8 | 0.01 | 0.52 | -0.48 | -0.57 | -0.1175 | 0.25 | 0.072847 | 0.662678 | 0.455838 | 0.999971 | 1 | 0.887217 |
| Ech1 | O35459 | 0.02 | -0.69 | -0.28 | -0.37 | -0.64 | -0.59 | -0.08 | 0.15 | -0.54 | -0.39 | -0.34 | -0.54 | -0.76 | -0.71 | -0.27 | -0.29 | 0.04 | -0.1225 | -0.1775 | 0.879664 | 0.554099 | 0.455317 | 0.999971 | 1 | 0.887217 |
| Echdc1 | Q9D9V3 | -0.78 | 0.32 | -0.2 | -0.01 | 0.02 | 0.31 | -0.37 | -0.36 | 0.03 | -0.37 | -0.29 | -0.36 | 0.29 | 0.38 | -0.04 | 0.1 | 0.0675 | -0.08 | 0.35 | 0.818305 | 0.765559 | 0.216157 | 0.999971 | 1 | 0.826893 |
| Echdc2 | Q3TLP5 | -0.33 | -0.46 | 0.15 | 0.22 | -1.31 | -0.99 | -1.05 | -0.35 | -0.68 | 0.04 | 0.36 | 0.67 | -1.42 | -1 | -0.53 | -0.44 | -0.82 | 0.2025 | -0.7425 | 0.016061 | 0.548318 | 0.030823 | 0.999971 | 1 | 0.678474 |
| Echs1 | Q8BH95 | -1.01 | -1.56 | -1.26 | -1.28 | -1.57 | -1.32 | -1.47 | -1.02 | -1.51 | -1.45 | -1.38 | -1.74 | -1.08 | -1.14 | -1.07 | -1.06 | -0.0675 | -0.2425 | 0.19 | 0.751757 | 0.233672 | 0.323052 | 0.999971 | 1 | 0.85548 |
| Eci1 | P42125 | 0.05 | -0.76 | -0.34 | -0.77 | -0.55 | -0.31 | -0.15 | 0.11 | -0.46 | -0.38 | -0.51 | -0.66 | -0.34 | -0.57 | -0.6 | -0.67 | 0.23 | -0.0475 | -0.09 | 0.391453 | 0.840844 | 0.71035 | 0.999971 | 1 | 0.950695 |
| Eci2 | Q3TCD4 | -0.84 | -0.28 | -0.65 | -0.32 | -0.46 | -0.36 | -0.28 | -0.21 | -0.38 | -0.42 | -0.73 | -0.39 | 0.05 | -0.27 | -0.6 | -0.51 | 0.195 | 0.0425 | 0.19 | 0.346808 | 0.837376 | 0.426787 | 0.999971 | 1 | 0.880793 |
| Eci2_1 | E9PUY9 | -1.65 | -0.73 | -1.15 | -1.25 | 0.47 | -0.84 | -1.59 | -0.68 | 0.18 | -1.21 | -0.92 | -0.44 | -0.39 | -1.24 | -1.11 | -0.37 | 0.535 | 0.5975 | 0.4175 | 0.242807 | 0.116873 | 0.195562 | 0.999971 | 1 | 0.811225 |
| Ecpas | Q6PDI5 | 0.19 | 0.25 | 0.88 | 1 | 0.43 | 0.14 | 0.84 | 0.83 | 0.22 | 0.06 | 0.89 | 0.86 | 0.09 | 0.32 | 0.9 | 0.88 | -0.02 | -0.0725 | -0.0325 | 0.943741 | 0.812821 | 0.913952 | 0.999971 | 1 | 0.992633 |
| Ecsit | J3KMM6 | 2.27 | 1.66 | -0.33 | 1.01 | 0.92 | 2.81 | -0.14 | 1.57 | 2.11 | 1.89 | -0.7 | 0.18 | 0.17 | 2.25 | 0.94 | 1.42 | 0.1375 | -0.2825 | 0.0425 | 0.854331 | 0.722948 | 0.947298 | 0.999971 | 1 | 0.997651 |
| Ecsit_1 | A0A0R4J17 | 0.14 | -0.77 | -0.67 | -0.73 | -0.84 | -0.55 | -0.74 | -0.72 | -0.83 | 0.17 | -0.59 | -0.59 | -0.57 | -0.04 | -0.37 | -0.13 | -0.205 | 0.0475 | 0.23 | 0.424908 | 0.878701 | 0.401873 | 0.999971 | 1 | 0.871904 |
| Edc3 | Q8K2D3 | 0.5 | 0.66 | 0.36 | 0.33 | 0.92 | 0.83 | 0.52 | 0.4 | 1 | 0.54 | -0.03 | 0.01 | 1.12 | 0.88 | 0.39 | 0.04 | 0.205 | -0.0825 | 0.145 | 0.324152 | 0.762439 | 0.598329 | 0.999971 | 1 | 0.92377 |
| Edc4 | G5E896 | 0.65 | 1.03 | 0.83 | 0.9 | 0.73 | 0.67 | 0.86 | 0.83 | 0.69 | 0.74 | 0.92 | 0.72 | 0.61 | 0.76 | 0.8 | 1.05 | -0.08 | -0.085 | -0.0475 | 0.650678 | 0.627403 | 0.802691 | 0.999971 | 1 | 0.971467 |
| Edem3 | A0A087WF | 3.34 | -0.1 | -0.41 | -0.36 | -0.31 | -0.28 | -0.96 | 0.01 | 0.02 | 2.5 | -0.45 | -0.32 | -0.87 | -0.02 | -0.57 | -0.01 | -1.0025 | -0.18 | -0.985 | 0.250376 | 0.861078 | 0.259763 | 0.999971 | 1 | 0.840127 |
| Edf1 | Q9JMG1 | 1.76 | 0.72 | 2.63 | 2.47 | 0.79 | 0.7 | 3.15 | 2.29 | 0.76 | 0.79 | 1.87 | 2.49 | 0.28 | 0.42 | 2.47 | 1.78 | -0.1625 | -0.4175 | -0.6575 | 0.808236 | 0.464063 | 0.30953 | 0.999971 | 1 | 0.847402 |
| Edil3 | Q8C4U8 | 0.59 | 1.08 | 0.25 | 0.13 | 0.79 | 1.01 | -0.12 | 0.96 | 0.72 | 0.57 | 0.31 | 0.17 | 1.08 | 0.82 | 0.44 | 0.33 | 0.1475 | -0.07 | 0.155 | 0.66285 | 0.79248 | 0.593081 | 0.999971 | 1 | 0.923349 |
| Ednrb | P48302 | 0.31 | -0.74 | -0.61 | -1.09 | 0 | -0.35 | -0.5 | -0.72 | -0.73 | 0.25 | -1.16 | -1.49 | -0.7 | 0.24 | -0.28 | -1.17 | 0.14 | -0.25 | 0.055 | 0.675646 | 0.584723 | 0.891914 | 0.999971 | 1 | 0.988225 |
| Edrf1 | D3YZE7 | 4.3 | 2.31 | 0.29 | 0.49 | 2.37 | 1.69 | -0.4 | 0.09 | 2.6 | 3.87 | -0.16 | -0.46 | 0.39 | 1.13 | -0.16 | -0.29 | -0.91 | -0.385 | -1.58 | 0.382315 | 0.761094 | 0.102545 | 0.999971 | 1 | 0.78268 |
| Eea1 | Q8BL66 | 0.63 | 1.06 | 1.04 | 0.91 | 0.99 | 1.14 | 1.24 | 1.06 | 1.03 | 0.46 | 1.18 | 1.26 | 0.9 | 0.89 | 1.08 | 1.09 | 0.1975 | 0.0725 | 0.08 | 0.302389 | 0.760542 | 0.66906 | 0.999971 | 1 | 0.938711 |
| Eed | A0A5F8MP | -0.01 | -0.04 | -0.52 | 0.76 | 0.57 | 0.17 | -0.17 | 0.13 | -0.45 | -0.27 | 0.26 | 0.36 | -0.26 | -0.11 | 0.02 | 0.21 | 0.1275 | -0.0725 | -0.0825 | 0.682427 | 0.826047 | 0.779402 | 0.999971 | 1 | 0.965847 |
| Eef1a1 | P10126 | 0.6 | 0.99 | 0.95 | 0.69 | 1.17 | 1.17 | 1.66 | 0.85 | 1.3 | 0.75 | 0.8 | 1.04 | 1.4 | 1.38 | 0.97 | 0.79 | 0.405 | 0.165 | 0.3275 | 0.106149 | 0.434185 | 0.166727 | 0.999971 | 1 | 0.800215 |
| Eef1a2 | P62631 | -0.28 | 1.33 | 1.08 | -0.01 | 1.52 | 1.07 | 1.89 | -0.03 | 1.51 | -0.21 | 1.61 | 1.94 | 1.83 | 1.11 | 0.65 | 0.34 | 0.5825 | 0.6825 | 0.4525 | 0.288654 | 0.255326 | 0.356719 | 0.999971 | 1 | 0.868793 |
| Eef1akmt1 | A0A0D2X7. | 1.2 | 1.11 | 0.88 | 1.27 | 0.98 | 1.13 | 1.43 | 1.2 | 0.8 | 1.4 | 0.98 | 0.78 | 0.66 | 1.06 | 0.81 | 0.76 | 0.07 | -0.125 | -0.2925 | 0.716237 | 0.5598 | 0.148845 | 0.999971 | 1 | 0.793248 |
| Eef1akmt2 | Q9D853 | -0.78 | 0.47 | 1.27 | 0.23 | 0.83 | 0.27 | 1.36 | 1.24 | 0.84 | -0.46 | 1.33 | 1.4 | 1.42 | 0.03 | 0.74 | 0.17 | 0.6275 | 0.48 | 0.2925 | 0.193985 | 0.399279 | 0.55554 | 0.999971 | 1 | 0.914901 |
| Eef1b | O70251 | 2.16 | 2.41 | 2.19 | 2.37 | 2.38 | 2.72 | 1.97 | 2.5 | 2.59 | 2.76 | 2.58 | 2.7 | 2.34 | 2.63 | 2.64 | 3.07 | 0.11 | 0.375 | 0.3875 | 0.613154 | 0.048273 | 0.095897 | 0.999971 | 1 | 0.78268 |
| Eef1d | Q91VK2 | -1.16 | 0.79 | 0.71 | -0.93 | -0.68 | -0.68 | 0.12 | -0.45 | -1.56 | -1.28 | -0.4 | 1.15 | -0.71 | -0.67 | -0.5 | 1.62 | -0.275 | -0.375 | 0.0825 | 0.594161 | 0.609304 | 0.905441 | 0.999971 | 1 | 0.991091 |
| Eef1d_1 | A0A0R4J1E | 1.45 | 1.53 | 2.18 | 2.01 | 1.49 | 1.81 | 1.81 | 2.13 | 1.66 | 2 | 2.01 | 2.22 | 1.48 | 1.97 | 2.22 | 2.47 | 0.0175 | 0.18 | 0.2425 | 0.94405 | 0.464738 | 0.412057 | 0.999971 | 1 | 0.875856 |
| Eef1d_2 | Q80T06 | 0.32 | 0.97 | 0.78 | 1.15 | 0.54 | 0.02 | 1.02 | 1.17 | 0.34 | 0.55 | 1.32 | 0.98 | -0.5 | 0.88 | 1.34 | 1.43 | -0.1175 | -0.0075 | -0.0175 | 0.712661 | 0.979531 | 0.969051 | 0.999971 | 1 | 0.998787 |
| Eef1e1 | Q9D1M4 | -0.66 | 1.69 | 0.84 | 0.41 | 1.97 | 1.52 | 1.76 | 0.36 | 2.4 | 0.66 | 1.4 | 2.04 | 2.55 | 1.94 | 1.05 | 1.05 | 0.8325 | 1.055 | 1.0775 | 0.160306 | 0.092764 | 0.082093 | 0.999971 | 1 | 0.755254 |
| Eef1g | Q9D8N0 | 0.48 | 0.76 | 1.88 | 1.81 | 1.09 | 0.47 | 1.86 | 2.07 | 0.89 | 0.71 | 2.4 | 2 | 0.54 | 0.95 | 2.23 | 2.18 | 0.14 | 0.2675 | 0.2425 | 0.770027 | 0.602306 | 0.641666 | 0.999971 | 1 | 0.93301 |
| Eef2 | P58252 | -0.6 | 0 | 1.11 | 0.65 | 0.79 | -0.38 | 1.23 | 0.94 | 0.67 | -0.95 | 1.28 | 1.4 | 0.82 | -0.56 | 1.16 | 1.06 | 0.355 | 0.31 | 0.33 | 0.466092 | 0.610089 | 0.520792 | 0.999971 | 1 | 0.906972 |
| Eefsec | Q9JHW4 | 0.66 | 0.65 | 0.41 | 0.58 | 0.53 | 0.76 | 0.27 | 0.58 | 0.59 | 1.01 | 0.52 | 0.77 | 0.67 | 0.99 | 0.44 | 0.96 | -0.04 | 0.1475 | 0.19 | 0.830778 | 0.44018 | 0.357326 | 0.999971 | 1 | 0.868793 |
| Eepd1 | Q3TGW2 | -1.19 | -1.73 | -0.94 | -0.99 | -1.53 | -1.5 | -0.71 | -0.93 | -1.66 | -1.31 | -0.73 | -1.04 | -1.64 | -1.84 | -1.01 | -0.94 | 0.045 | 0.0275 | -0.145 | 0.875283 | 0.922133 | 0.629263 | 0.999971 | 1 | 0.932234 |
| Efcab14 | A2A8T9 | 0.07 | 1.76 | 1.32 | 1.02 | 2.43 | 2.08 | 1.76 | 0.6 | 2.26 | 0.32 | 1.74 | 1.58 | 3.15 | 1.28 | -0.26 | -0.6 | 0.675 | 0.4325 | -0.15 | 0.19672 | 0.404322 | 0.857293 | 0.999971 | 1 | 0.98269 |
| Efemp1 | Q8BPB5 | 1.21 | 1.87 | 0.74 | 2.22 | 1.87 | 1.55 | 0.62 | 1.6 | 0.71 | 2.07 | 0.92 | 2.61 | 0.31 | 0.87 | 1.14 | 0 | -0.1 | 0.0675 | -0.93 | 0.806763 | 0.897228 | 0.043565 | 0.999971 | 1 | 0.701725 |
| Efhd1 | Q9D4J1 | -0.03 | 0.82 | -0.18 | -0.26 | 0.75 | -0.17 | -0.32 | 0.16 | 0.54 | 0.57 | -0.75 | -0.14 | 1.65 | -0.23 | 0.22 | 0.26 | 0.0175 | -0.0325 | 0.3875 | 0.958928 | 0.932968 | 0.398509 | 0.999971 | 1 | 0.871816 |
| Efhd2 | Q8C845 | -0.23 | 0.23 | -0.09 | -0.06 | -0.16 | 0 | -0.05 | -0.25 | -0.37 | 0.05 | -0.1 | -0.31 | -0.6 | -0.19 | -0.44 | 0.25 | -0.0775 | -0.145 | -0.2075 | 0.676291 | 0.463393 | 0.400998 | 0.999971 | 1 | 0.871816 |
| Efl1 | Q8C0D5 | 0 | -0.02 | -0.05 | 0.16 | -0.01 | 0.07 | 0.77 | -0.07 | 0.17 | 0.01 | 0.07 | 0.18 | 0.34 | 0.25 | 0.14 | 0.51 | 0.1675 | 0.085 | 0.2875 | 0.484042 | 0.602991 | 0.130455 | 0.999971 | 1 | 0.784948 |
| Efna3 | O08545 | 0.14 | 0.33 | -0.19 | -0.25 | -0.7 | 0.35 | 0.67 | 0.19 | -1.05 | 1.1 | 0.54 | 0.07 | 0.76 | -0.04 | 0.53 | -0.06 | 0.12 | 0.1575 | 0.29 | 0.712627 | 0.726826 | 0.296602 | 0.999971 | 1 | 0.847402 |
| Efna5 | O08543 | -0.31 | 0.46 | 1.15 | 0.3 | 0.25 | 0.89 | 0.75 | 0.96 | 0.02 | -0.28 | 1.25 | 1.72 | 1.18 | -0.37 | 0.13 | -0.5 | 0.3125 | 0.2775 | -0.29 | 0.36631 | 0.599985 | 0.529544 | 0.999971 | 1 | 0.910069 |
| Efnb1 | P52795 | -0.41 | -0.66 | -1.72 | -1.51 | -0.84 | -0.76 | -1.66 | -1.46 | -0.9 | -0.06 | -1.36 | -1.37 | -0.22 | -0.55 | -1.23 | -1.52 | -0.105 | 0.1525 | 0.195 | 0.780636 | 0.718334 | 0.64148 | 0.999971 | 1 | 0.93301 |
| Efnb2 | P52800 | -0.11 | -0.15 | -1.1 | -0.74 | -0.32 | -0.1 | -0.85 | -1.07 | -0.12 | -0.15 | -1.14 | -1.87 | 0.31 | -0.4 | -0.9 | -0.93 | -0.06 | -0.295 | 0.045 | 0.855213 | 0.523913 | 0.902298 | 0.999971 | 1 | 0.990759 |
| Efnb3 | O35393 | 2.03 | 0.94 | 0.42 | 0.48 | 0.72 | 1.27 | -0.44 | 0.8 | 1.35 | 1.8 | -0.51 | -0.23 | 1.76 | 1.82 | 0.51 | -0.05 | -0.38 | -0.365 | 0.0425 | 0.44083 | 0.563072 | 0.938183 | 0.999971 | 1 | 0.99575 |
| Efr3a | A0A1D5RLI | -0.6 | -0.91 | -0.9 | -1.08 | -1.14 | -0.7 | -1.27 | -1.14 | -1.13 | -0.58 | -1 | -1.12 | -1.09 | -0.89 | -0.94 | -0.95 | -0.19 | -0.085 | -0.095 | 0.377378 | 0.686813 | 0.608486 | 0.999971 | 1 | 0.926452 |
| Efr3b | Q6ZQ18 | -0.68 | -0.94 | -0.67 | -0.68 | -0.86 | -0.45 | -0.68 | -0.69 | -0.91 | -0.81 | -0.67 | -0.57 | -0.35 | -0.32 | -0.56 | -0.63 | 0.0725 | 0.0025 | 0.2775 | 0.692485 | 0.98867 | 0.150646 | 0.999971 | 1 | 0.793248 |
| Efs | A0A2I3BRP | 0.12 | -2.09 | 0.24 | 0.26 | -3.01 | -1.29 | -0.45 | 0.32 | -1.13 | -3.16 | 0.21 | 0.69 | -0.58 | -0.39 | 0.15 | -1.09 | -0.74 | -0.48 | -0.11 | 0.379837 | 0.609569 | 0.84884 | 0.999971 | 1 | 0.981643 |
| Eftud2 | A2AH85 | 0.05 | 0.05 | 0.08 | -0.4 | 0 | 0.08 | 0.71 | -0.48 | 0.08 | -0.09 | -0.04 | -0.21 | 0.04 | 0.19 | -0.5 | -0.46 | 0.1325 | -0.01 | -0.1275 | 0.642909 | 0.958186 | 0.601088 | 0.999971 | 1 | 0.92426 |
| Egfr | Q01279 | -1.21 | -0.6 | -0.75 | -0.33 | -0.17 | -1.03 | -0.43 | -0.44 | -0.05 | -1.31 | -0.45 | -0.43 | 0.61 | -1.46 | -0.58 | -0.49 | 0.205 | 0.1625 | 0.2425 | 0.464682 | 0.61896 | 0.582114 | 0.999971 | 1 | 0.920364 |
| Egln1 | Q91YE3 | 1.21 | 1.51 | 0.94 | 1.17 | 0.64 | 1.52 | 1.52 | 1.3 | 0.92 | 1.24 | 0.87 | 0.45 | 0.17 | 1.88 | 1.06 | 0.77 | 0.0375 | -0.3375 | -0.2375 | 0.885995 | 0.173182 | 0.520539 | 0.999971 | 1 | 0.906972 |
| Egr3 | Q3ZB14 | 1.59 | -0.15 | 0.91 | 0.21 | 0.24 | 0.47 | 0.86 | 0.37 | -0.27 | 0.4 | -0.13 | 0.73 | 0.19 | 0.31 | 0.99 | 1.06 | -0.155 | -0.4575 | -0.0025 | 0.693295 | 0.301015 | 0.99529 | 0.999971 | 1 | 1 |
| Ehbp1 | Q69ZW3 | 0.2 | -0.03 | -0.79 | -0.73 | -0.22 | 0.06 | -0.7 | -0.91 | 0.07 | 0.22 | -0.79 | -1.03 | 0.29 | -0.29 | -0.88 | -0.91 | -0.105 | -0.045 | -0.11 | 0.752029 | 0.906611 | 0.765366 | 0.999971 | 1 | 0.962405 |
| Ehbp1l1 | Q99MS7 | -2.37 | 0.09 | 2.68 | 2.33 | 1.76 | -0.56 | 2.1 | 2.96 | 0.15 | -0.75 | 2.58 | 3.19 | 0.41 | 0.21 | 2.98 | 3.33 | 0.8825 | 0.61 | 1.05 | 0.481083 | 0.651834 | 0.418588 | 0.999971 | 1 | 0.87788 |
| Ehd1 | Q9WVK4 | -0.78 | 0.13 | -0.06 | -0.16 | 0.28 | -0.05 | -0.54 | -0.04 | 0.18 | -0.41 | -0.19 | 0.23 | 0.63 | -0.04 | -0.05 | 0.03 | 0.13 | 0.17 | 0.36 | 0.640116 | 0.53047 | 0.211732 | 0.999971 | 1 | 0.823112 |
| Ehd2 | Q8BH64 | 0.47 | 0.98 | 1.05 | 1.79 | 5.96 | 2.91 | 1.59 | 0.6 | 2.06 | 2.51 | 2.87 | 4.81 | 0.1 | 1.19 | 1.78 | 1.13 | 1.6925 | 1.99 | -0.0225 | 0.138921 | 0.009559 | 0.957271 | 0.999971 | 1 | 0.998578 |
| Ehd3 | Q9QXY6 | -0.49 | 0.38 | -0.79 | -1.15 | 0.53 | 0.26 | -0.45 | -1.05 | 0.45 | -0.75 | -0.66 | -0.47 | 1.43 | 0.04 | -0.98 | -0.94 | 0.335 | 0.155 | 0.4 | 0.467287 | 0.705287 | 0.508601 | 0.999971 | 1 | 0.905815 |
| Ehd4 | Q9EQP2 | -0.34 | -0.16 | -0.43 | -0.41 | 0.42 | -0.1 | -0.43 | -0.28 | -0.09 | -0.16 | -0.3 | 0.3 | -0.43 | -0.22 | -0.25 | -0.38 | 0.2375 | 0.2725 | 0.015 | 0.320862 | 0.191905 | 0.930559 | 0.999971 | 1 | 0.994744 |
| Ehhadh | Q9DBM2 | -0.54 | -0.91 | -1.88 | 0.17 | -1.16 | -0.9 | 0.35 | -1.32 | -1.13 | -0.56 | -1.81 | -1.82 | -0.77 | -1.26 | -1.57 | -1.07 | 0.0325 | -0.54 | -0.3775 | 0.950732 | 0.286897 | 0.394741 | 0.999971 | 1 | 0.871816 |
| Ehmt1 | E9Q5A3 | 0.24 | 0.71 | -0.31 | 0.05 | 0.54 | 1.5 | -0.32 | 0.06 | 1.07 | 1.05 | 0.14 | -0.4 | 0.65 | 1.28 | -0.11 | 0.45 | 0.2725 | 0.2925 | 0.395 | 0.524635 | 0.473039 | 0.278573 | 0.999971 | 1 | 0.843812 |
| Ehmt2 | A2CG76 | 0.82 | 0.57 | -0.1 | -0.03 | 0.28 | 0.44 | 0.13 | -0.22 | 0.16 | 0.12 | -0.77 | -1.45 | 0.25 | 0.64 | -1.11 | -0.2 | -0.1575 | -0.8 | -0.42 | 0.578604 | 0.087696 | 0.328842 | 0.999971 | 1 | 0.856525 |
| Ei24 | A0A0R4J25 | -0.54 | -0.16 | -0.65 | 0.19 | -0.6 | 0.15 | -0.13 | -0.09 | 0.3 | 0.08 | 0.08 | -0.19 | 0.35 | -0.05 | -0.75 | 0.29 | 0.1225 | 0.3575 | 0.25 | 0.649032 | 0.168578 | 0.442693 | 0.999971 | 1 | 0.883386 |
| Eid2 | Q6X7S9 | 2.51 | -0.02 | 0.71 | 0.95 | 0.13 | 0.74 | 1.05 | 0.75 | 0.29 | 1.93 | -0.05 | -0.4 | -0.94 | 0.17 | -0.11 | 0.22 | -0.37 | -0.595 | -1.2025 | 0.485323 | 0.389492 | 0.052869 | 0.999971 | 1 | 0.725348 |
| Eif1 | P48024 | 0.33 | 0.63 | 0.03 | -0.35 | 1.33 | 0.7 | 0.05 | -0.4 | 1.2 | 0.1 | 0.37 | 0.51 | 1.47 | 1.07 | 0.2 | -0.14 | 0.26 | 0.385 | 0.49 | 0.53192 | 0.244726 | 0.250483 | 0.999971 | 1 | 0.839012 |
| Eif1ad | Q3THJ3 | 0.63 | 1.43 | 0.85 | 1.14 | 0.74 | 1 | 0.32 | 1.35 | 0.97 | 1.54 | 0.55 | 0.24 | 1.19 | 0.43 | 1.49 | 1.34 | -0.16 | -0.1875 | 0.1 | 0.584354 | 0.573434 | 0.74085 | 0.999971 | 1 | 0.960899 |
| Eif1ax | Q8BMJ3 | 1.97 | 1.45 | 1.91 | 1.87 | 1.21 | 1.84 | 4.85 | 0.98 | 1.29 | 1.94 | 1.24 | 1.05 | 1.55 | 1.62 | 1.78 | 0.96 | 0.42 | -0.42 | -0.3225 | 0.607141 | 0.122832 | 0.211458 | 0.999971 | 1 | 0.823077 |
| Eif1b | Q9CXU9 | 2.12 | 2.11 | 0.89 | 1.07 | 1.67 | 2.62 | 0.51 | 1.33 | 1.49 | 1.8 | -0.22 | 0.68 | 4.61 | 2.42 | 0.77 | 1.72 | -0.015 | -0.61 | 0.8325 | 0.976349 | 0.259778 | 0.309422 | 0.999971 | 1 | 0.847402 |
| Eif2a | Q8BJW6 | 0.95 | 1.47 | 1.42 | 1.31 | 1.29 | 1.24 | 2.52 | 1.13 | 1.45 | 1.25 | 1.14 | 1.45 | 1.69 | 1.56 | 1.2 | 1.17 | 0.2575 | 0.035 | 0.1175 | 0.459971 | 0.85868 | 0.596322 | 0.999971 | 1 | 0.92377 |

| | | | | | | | | | | | | | | | | | | | | | | | | | | |
|---|---|---|---|---|---|---|---|---|---|---|---|---|---|---|---|---|---|---|---|---|---|---|---|---|---|---|
| Eif2ak2 | Q03963 | 3.27 | -0.63 | 0.12 | -0.51 | 0.97 | 1.07 | -0.08 | -0.43 | 0.96 | 0.93 | -1.59 | -0.21 | -2.17 | 0.32 | -1.48 | -0.77 | -0.18 | -0.54 | -1.5875 | 0.839263 | 0.587522 | 0.121641 | 0.999971 | 1 | 0.78268 |
| Eif2ak3 | Q9Z2B5 | 1.56 | -0.01 | -1.63 | -0.66 | 0.18 | 0.75 | -0.94 | -1.12 | 0.59 | 1.2 | -0.91 | -0.64 | 1.08 | -0.48 | -0.48 | 0.13 | -0.0975 | 0.245 | 0.2475 | 0.893312 | 0.747325 | 0.721728 | 0.999971 | 1 | 0.955137 |
| Eif2ak4 | A2AUM0 | 0.2 | 0.62 | -0.2 | 0.43 | 0.48 | 0.57 | -0.47 | 0.47 | 0.31 | 0.35 | -0.09 | 0.01 | 0.36 | 0.73 | 0.33 | 0.34 | -1.18E-16 | -0.1175 | 0.1775 | 1 | 0.624452 | 0.460762 | 1 | 1 | 0.890033 |
| Eif2b1 | Q99LC8 | 0.15 | -0.33 | -0.16 | -0.22 | -0.29 | -0.66 | -0.33 | -0.32 | -0.15 | -0.26 | -0.28 | -0.34 | -0.75 | -0.43 | -0.44 | -0.21 | -0.26 | -0.1175 | -0.3175 | 0.207705 | 0.521324 | 0.149337 | 0.999971 | 1 | 0.793248 |
| Eif2b2 | Q99LD9 | 1.48 | 0.58 | 0.12 | 0.56 | 0.24 | 0.77 | 0.38 | 0.25 | 0.25 | 1.17 | 0.34 | 0.42 | -0.65 | -0.27 | 0.02 | 0.54 | -0.275 | -0.14 | -0.775 | 0.393171 | 0.689799 | 0.061692 | 0.999971 | 1 | 0.725348 |
| Eif2b3 | B1AUN2 | -0.27 | 0.85 | 1.88 | 1.15 | 0.99 | 0.29 | 1.31 | 1.9 | 0.91 | -0.32 | 1.83 | 1.87 | 0.89 | 0.15 | 1.77 | 1.46 | 0.22 | 0.17 | 0.165 | 0.671365 | 0.785531 | 0.755022 | 0.999971 | 1 | 0.961116 |
| Eif2b4 | D3Z780 | 1.01 | 0.74 | 1.32 | 1.27 | 0.8 | 0.52 | 0.99 | 1.13 | 0.64 | 0.84 | 1.03 | 1.23 | 0.45 | 0.72 | 1.03 | 1.25 | -0.225 | -0.15 | -0.2225 | 0.336116 | 0.506593 | 0.383601 | 0.999971 | 1 | 0.871816 |
| Eif2b5 | Q8CHW4 | -0.16 | 1.76 | -0.31 | -0.14 | 2.08 | 1.42 | 0.07 | -0.07 | 2.31 | 0.48 | -0.08 | -0.11 | 2.47 | 1.41 | -0.5 | -0.28 | 0.5875 | 0.3625 | 0.4875 | 0.378992 | 0.600042 | 0.535054 | 0.999971 | 1 | 0.910077 |
| Eif2d | Q61211 | 0.5 | 0.27 | 0.39 | 0.48 | 0.48 | 0.68 | 0.25 | 0.42 | 0.22 | 0.49 | 0.53 | 0.11 | 0.17 | 0.31 | 1.17 | 0.98 | 0.0475 | -0.0725 | 0.2475 | 0.793118 | 0.693629 | 0.373012 | 0.999971 | 1 | 0.869526 |
| Eif2s1 | Q6ZWX6 | 0.51 | 1.1 | 1.39 | 1.49 | 0.99 | 0.98 | 1.05 | 1.63 | 1.2 | 0.89 | 1.58 | 1.48 | 1.11 | 1.19 | 1.54 | 1.98 | 0.04 | 0.165 | 0.3325 | 0.888016 | 0.563131 | 0.291297 | 0.999971 | 1 | 0.847402 |
| Eif2s2 | Q99L45 | 0.97 | 1.56 | 1.28 | 0.94 | 1.45 | 1.93 | 1.39 | 1.36 | 1.72 | 1.2 | 1.5 | 1.15 | 1.12 | 1.48 | 1.39 | 1.17 | 0.345 | 0.205 | 0.1025 | 0.165373 | 0.388053 | 0.638337 | 0.999971 | 1 | 0.93301 |
| Eif2s3x | Q9Z0N1 | 0.63 | 0.8 | 0.77 | 0.82 | 0.74 | 1.08 | 1.01 | 0.91 | 1.17 | 1.07 | 0.97 | 1.23 | 0.45 | 1.33 | 0.75 | 0.9 | 0.18 | 0.355 | 0.1025 | 0.315077 | 0.056521 | 0.655101 | 0.999971 | 1 | 0.934061 |
| Eif2s3y | Q9Z0N2 | -1.71 | 2.25 | 1.76 | -2.02 | 1.43 | -1 | 1.63 | -1.76 | -1.94 | 0.93 | 1.21 | -2.23 | 0.21 | 0.25 | 1.73 | -1.58 | 0.005 | -0.5775 | 0.0825 | 0.996815 | 0.657677 | 0.943664 | 1 | 1 | 0.99692 |
| Eif3a | P23116 | 1.23 | 0.78 | 1 | 1.14 | 0.88 | 0.74 | 1.36 | 1.29 | 0.93 | 1.11 | 1.23 | 1.17 | 0.58 | 0.94 | 1.1 | 1.16 | 0.03 | 0.0725 | -0.0925 | 0.892945 | 0.695506 | 0.665351 | 0.999971 | 1 | 0.93697 |
| Eif3b | Q8JZQ9 | 0.33 | 1.14 | 0.85 | 1.01 | 1.05 | 1.14 | 0.79 | 1.11 | 1.23 | 0.83 | 1.08 | 1.04 | 1.11 | 1.36 | 1.17 | 1.27 | 0.19 | 0.2125 | 0.395 | 0.421544 | 0.368999 | 0.108995 | 0.999971 | 1 | 0.78268 |
| Eif3c | Q8R1B4 | 0.08 | 1.06 | 1.31 | 0.99 | 0.97 | 0.94 | 1.29 | 1.22 | 1.24 | 0.61 | 1.33 | 1.39 | 1.27 | 1.01 | 1.41 | 1.37 | 0.245 | 0.2825 | 0.405 | 0.413438 | 0.394395 | 0.192772 | 0.999971 | 1 | 0.810547 |
| Eif3d | O70194 | -0.14 | 1.19 | 1.3 | 0.92 | 1.23 | 0.81 | 1.27 | 1.31 | 1.27 | 0.35 | 1.21 | 1.33 | 1.45 | 1.01 | 1.47 | 1.28 | 0.3375 | 0.2225 | 0.485 | 0.340418 | 0.569712 | 0.181126 | 0.999971 | 1 | 0.808532 |
| Eif3e | P60229 | 0.47 | 1.35 | 0.97 | 0.86 | 1.29 | 1.34 | 0.72 | 1.41 | 1.48 | 0.93 | 1.34 | 1.28 | 1.54 | 1.56 | 1.31 | 1.31 | 0.2775 | 0.345 | 0.5175 | 0.306847 | 0.181685 | 0.049258 | 0.999971 | 1 | 0.715024 |
| Eif3f | Q9DCH4 | -0.18 | 0.78 | 0.69 | 0.86 | 0.65 | 0.59 | 0.5 | 0.99 | 0.6 | 0.48 | 0.63 | 0.85 | 0.53 | 0.93 | 0.97 | 1.16 | 0.145 | 0.1025 | 0.36 | 0.606995 | 0.706322 | 0.233167 | 0.999971 | 1 | 0.831526 |
| Eif3g | Q9Z1D1 | 1.29 | 1.58 | 1.18 | 1.47 | 1.59 | 1.9 | 1.24 | 1.43 | 1.84 | 1.83 | 1.27 | 1.25 | 1.55 | 1.9 | 1.12 | 1.39 | 0.16 | 0.1675 | 0.11 | 0.461675 | 0.465664 | 0.629914 | 0.999971 | 1 | 0.932649 |
| Eif3h | Q91WK2 | 0.41 | 1.13 | 1.3 | 0.98 | 0.79 | 1.33 | 2.23 | 1.1 | 0.96 | 0.63 | 1.04 | 1.37 | 0.93 | 0.82 | 1.46 | 1 | 0.4075 | 0.045 | 0.0975 | 0.271597 | 0.865272 | 0.711092 | 0.999971 | 1 | 0.950696 |
| Eif3i | Q9QZD9 | 0.9 | 1.45 | 1.34 | 0.62 | 1.25 | 1.43 | 1.38 | 1.12 | 1.38 | 0.87 | 1.69 | 1.58 | 1.33 | 1.34 | 1.53 | 1.04 | 0.2175 | 0.3025 | 0.2325 | 0.374027 | 0.295916 | 0.360848 | 0.999971 | 1 | 0.869526 |
| Eif3j1 | Q3UGC7 | 0.87 | 1.52 | 1.25 | 1.11 | 1.07 | 2.03 | 1.43 | 1.13 | 1.48 | 1.42 | 1.57 | 1.86 | 1.35 | 1.36 | 1.36 | 1.21 | 0.2275 | 0.395 | 0.1325 | 0.418042 | 0.090266 | 0.513296 | 0.999971 | 1 | 0.905815 |
| Eif3k | Q9DBZ5 | 0.51 | 0.85 | 0.38 | 0.89 | 0.67 | 0.84 | 0.33 | 0.97 | 0.66 | 0.66 | 0.67 | 0.6 | 0.99 | 1.47 | 0.98 | 0.95 | 0.045 | -0.01 | 0.44 | 0.84284 | 0.957822 | 0.073125 | 0.999971 | 1 | 0.745482 |
| Eif3l | Q8QZY1 | 0.23 | 2.39 | 0.87 | 0.79 | 2.64 | 1.81 | 0.75 | 1.07 | 2.93 | 0.5 | 1.17 | 1.4 | 3.78 | 1.53 | 1.16 | 1.26 | 0.4975 | 0.43 | 0.8625 | 0.39528 | 0.502321 | 0.238973 | 0.999971 | 1 | 0.833594 |
| Eif3m | Q99JX4 | -1.1 | 1.44 | 0.79 | 0.75 | 1.17 | 1.28 | 0.8 | 1.07 | 1.4 | -0.05 | 0.95 | 1.23 | 1.24 | 1.52 | 1.06 | 1.26 | 0.61 | 0.4125 | 0.8 | 0.254747 | 0.486904 | 0.145944 | 0.999971 | 1 | 0.791767 |
| Eif4a1 | P60843 | 0.22 | 0.6 | 1.28 | 1.35 | 0.77 | 0.39 | 0.86 | 1.55 | 0.72 | 0.31 | 1.35 | 1.44 | 1.01 | 1.09 | 1.62 | 1.97 | 0.03 | 0.0925 | 0.56 | 0.932894 | 0.80365 | 0.13626 | 0.999971 | 1 | 0.784948 |
| Eif4a2 | P10630 | 0.62 | 1.47 | 1.08 | 0.91 | 1.44 | 1.51 | 1.03 | 1.06 | 1.56 | 1.15 | 1.51 | 1.61 | 1.54 | 1.43 | 1.19 | 1.44 | 0.24 | 0.4375 | 0.38 | 0.343902 | 0.092905 | 0.126837 | 0.999971 | 1 | 0.783193 |
| Eif4a3 | Q91VC3 | -0.03 | 0.45 | 0.43 | 0.48 | 0.25 | 0.33 | 0.06 | 0.39 | 0.06 | -0.04 | 0.37 | 0.13 | 0.01 | 0.44 | 0.33 | 0.48 | -0.075 | -0.2025 | -0.0175 | 0.708043 | 0.329822 | 0.934117 | 0.999971 | 1 | 0.995387 |
| Eif4b | Q8BGD9 | 1.18 | -0.34 | 0.95 | 1.15 | -0.45 | -0.11 | 1.16 | 0.95 | -0.26 | 0.46 | 0.7 | 0.71 | -0.87 | -0.42 | 0.85 | 0.98 | -0.3475 | -0.3325 | -0.6 | 0.489979 | 0.425301 | 0.283809 | 0.999971 | 1 | 0.844538 |
| Eif4e | P63073 | -0.28 | 1.61 | 1.37 | 1.1 | 1.53 | 1.36 | 0.86 | 1.74 | 1.11 | 0.17 | 1.49 | 1.83 | 1.66 | 1.61 | 1.66 | 1.89 | 0.4225 | 0.2 | 0.755 | 0.346148 | 0.698896 | 0.091907 | 0.999971 | 1 | 0.773779 |
| Eif4e2 | D3Z730 | 2.17 | -0.2 | 0.24 | 0.36 | -0.33 | 0.58 | 1.6 | -0.07 | -0.24 | 0.49 | 0.47 | 0.29 | -0.46 | -0.19 | 0.49 | -0.21 | -0.1975 | -0.39 | -0.735 | 0.749263 | 0.453678 | 0.181498 | 0.999971 | 1 | 0.808532 |
| Eif4ebp2 | P70445 | 0.47 | 0.07 | -0.61 | 0.04 | 0.17 | 0.45 | -0.64 | -0.5 | -0.36 | -0.27 | -0.91 | -0.31 | 1.06 | 0.73 | -0.22 | 0 | -0.1225 | -0.455 | 0.4 | 0.719913 | 0.134313 | 0.291056 | 0.999971 | 1 | 0.847402 |
| Eif4enif1 | Q9EST3 | 0.25 | 0.23 | -0.1 | 0.02 | 0.43 | 0.2 | 0.14 | 0.14 | 0.24 | 0.2 | 0.74 | 0.97 | 0.41 | 0.7 | 0.15 | 0.36 | 0.1275 | 0.4375 | 0.305 | 0.493465 | 0.094011 | 0.154012 | 0.999971 | 1 | 0.795578 |
| Eif4g1 | Q6NZJ6 | 0.87 | 1.12 | 1.34 | 1.39 | 1.27 | 1.06 | 1.34 | 1.32 | 1.29 | 1.27 | 1.43 | 1.55 | 0.97 | 1.14 | 1.24 | 1.43 | 0.0675 | 0.205 | 0.015 | 0.731828 | 0.306533 | 0.942159 | 0.999971 | 1 | 0.996889 |
| Eif4g1_1 | D6RCH1 | 1.34 | 1.3 | -0.66 | 0.88 | 1.04 | 0.87 | -1.42 | 0.9 | 0.87 | 1.43 | -0.77 | -2.62 | -0.3 | 1.2 | -0.64 | 0.64 | -0.3675 | -0.9875 | -0.49 | 0.593414 | 0.301392 | 0.407695 | 0.999971 | 1 | 0.874911 |
| Eif4g2 | Q62448 | 0.75 | 0.96 | 1.14 | 0.91 | 1.09 | 0.86 | 1.03 | 0.72 | 1.07 | 1.01 | 1.19 | 1.18 | 0.85 | 0.64 | 0.8 | 1.02 | -0.015 | 0.1725 | -0.1125 | 0.935983 | 0.331881 | 0.548877 | 0.999971 | 1 | 0.911727 |
| Eif4g2_1 | G3XA17 | -1.23 | 0.71 | -0.12 | 0.89 | 0.69 | 0.55 | -0.05 | 0.5 | 0.23 | 0.02 | 0.03 | 0.69 | -0.12 | 0.49 | 0.45 | 1.19 | 0.36 | 0.18 | 0.44 | 0.456711 | 0.706472 | 0.400832 | 0.999971 | 1 | 0.871816 |
| Eif4g3 | Z4YKC4 | 0.67 | 1.8 | 0.9 | -0.34 | 1.1 | 1.26 | 0 | 0.08 | 0.83 | 1.31 | 0.31 | 0.02 | 0.93 | 1.11 | -0.11 | 0.63 | -0.1475 | -0.14 | -0.1175 | 0.772416 | 0.775177 | 0.807097 | 0.999971 | 1 | 0.972962 |
| Eif4g3_1 | A0A0N4SV | 1.14 | 1.73 | 1.47 | 1.66 | 1.56 | 1.87 | 1.37 | 1.56 | 1.79 | 1.62 | 1.72 | 1.51 | 1.44 | 1.7 | 1.5 | 1.91 | 0.09 | 0.16 | 0.1375 | 0.676794 | 0.431104 | 0.530563 | 0.999971 | 1 | 0.910069 |
| Eif4h | Q9WUK2 | 0.59 | 0.1 | -0.98 | -0.71 | 0.15 | 0.23 | -0.32 | -0.88 | 0.47 | 0.54 | -0.54 | -0.92 | -0.51 | -0.24 | -1.23 | -1.53 | 0.045 | 0.1375 | -0.6275 | 0.914649 | 0.775352 | 0.182006 | 0.999971 | 1 | 0.808532 |
| Eif5 | P59325 | 0.06 | 0.42 | 0.54 | 0.44 | 0.53 | 0.56 | 0.7 | 0.18 | 0.43 | -0.05 | 0.56 | 0.16 | 0.9 | 0.47 | 0.74 | 0.57 | 0.1275 | -0.09 | 0.305 | 0.54041 | 0.677607 | 0.15407 | 0.999971 | 1 | 0.795578 |
| Eif5a | P63242 | 0.3 | 0.78 | 1.08 | 0.74 | 1.37 | 0.58 | 1.54 | 0.78 | 1.69 | 0.38 | 1.45 | 1.64 | 1.76 | 0.18 | 0.89 | 0.67 | 0.3425 | 0.565 | 0.15 | 0.259368 | 0.126807 | 0.677918 | 0.999971 | 1 | 0.942594 |
| Eif5a2 | Q8BGY2 | 2.73 | 3.07 | 2.82 | 3.09 | 2.47 | 3.78 | 2.59 | 2.66 | 2.14 | 3.22 | 2.28 | 1.7 | 3.25 | 3.34 | 2.07 | 3.53 | -0.0525 | -0.5925 | 0.12 | 0.869469 | 0.101341 | 0.725172 | 0.999971 | 1 | 0.955918 |
| Eif5b | Q05D44 | 0.67 | 1.26 | 1.12 | 1.05 | 1.41 | 1.18 | 1.03 | 1.14 | 1.6 | 0.78 | 1.28 | 1.53 | 1.27 | 1.09 | 1.17 | 0.97 | 0.165 | 0.2725 | 0.1 | 0.427923 | 0.292805 | 0.620207 | 0.999971 | 1 | 0.929296 |
| Eif6 | O55135 | 0.22 | 1.49 | 0.66 | 0.32 | 1.28 | 1.58 | 0.41 | 0.75 | 1.01 | 0.84 | 0.59 | 0.61 | 1.4 | 1.13 | 0.46 | 0.47 | 0.3325 | 0.09 | 0.1925 | 0.38829 | 0.771842 | 0.600305 | 0.999971 | 1 | 0.92426 |
| Eipr1 | Q8K0G5 | 0.07 | 0.04 | 0.9 | 1.39 | -0.1 | -0.06 | 0.57 | 1.32 | 0.08 | 0.25 | 0.94 | 0.46 | -0.41 | 0.2 | 0.97 | 1.59 | -0.1675 | -0.1675 | -0.0125 | 0.705409 | 0.651587 | 0.980359 | 0.999971 | 1 | 0.99925 |
| Elac1 | Q8VEB6 | 0.25 | 0.24 | 0.02 | -0.24 | -0.06 | 0.14 | -0.06 | 0.47 | -0.06 | -0.12 | -0.65 | 0.06 | 0.11 | 0.08 | -0.27 | -0.73 | 0.055 | -0.26 | -0.27 | 0.799592 | 0.276227 | 0.304158 | 0.999971 | 1 | 0.847402 |
| Elac2 | B1ATP7 | 0.9 | 1.37 | 1.02 | 0.65 | 1.45 | 1.33 | 1.07 | 0.88 | 1.39 | 1.15 | 1.05 | 0.98 | 1.66 | 1.42 | 0.84 | 1.12 | 0.1975 | 0.1575 | 0.275 | 0.407194 | 0.474467 | 0.302579 | 0.999971 | 1 | 0.847402 |
| Elapor1 | A0A0A0M( | 2.83 | 1.3 | 2.51 | 2.74 | 1.27 | 1.83 | 2.83 | 2.25 | 1.24 | 2.14 | 1.65 | 1.84 | -0.03 | 1.42 | 1.76 | 0.83 | -0.3 | -0.6275 | -1.35 | 0.514186 | 0.132531 | 0.021556 | 0.999971 | 1 | 0.642033 |
| Elapor2 | Q3UZV7-2 | 0.3 | 1.48 | 1.44 | 1.45 | 1.57 | 1.41 | 1.08 | 1.61 | 1.51 | 0.29 | 1.56 | 1.51 | 1.03 | 1.53 | 1.98 | 1.52 | 0.25 | 0.05 | 0.3475 | 0.438038 | 0.901766 | 0.327595 | 0.999971 | 1 | 0.856472 |
| Elavl1 | P70372 | -0.07 | -0.24 | 0.12 | 0.15 | -0.06 | -0.34 | -0.16 | 0.08 | -0.31 | -0.38 | -0.17 | -0.17 | -0.43 | -0.08 | 0.09 | 0.21 | -0.11 | -0.2475 | -0.0425 | 0.570462 | 0.18948 | 0.843127 | 0.999971 | 1 | 0.979814 |
| Elavl2 | B1AXZ4 | -0.23 | 1.98 | 0.76 | 0.57 | 1.76 | 1.85 | 1.17 | 0.67 | 2.04 | 0.63 | 1.21 | 1.2 | 2.39 | 1.92 | 0.83 | 0.95 | 0.5925 | 0.5 | 0.7525 | 0.250986 | 0.33534 | 0.192063 | 0.999971 | 1 | 0.810547 |
| Elavl3 | Q60900 | -0.47 | -0.19 | 0.72 | 0.44 | -0.13 | -0.12 | 0.57 | 0.6 | 0.08 | -0.13 | 0.39 | 0.58 | -0.43 | -0.04 | 0.7 | 0.68 | 0.105 | 0.105 | 0.1025 | 0.757571 | 0.742743 | 0.787035 | 0.999971 | 1 | 0.96724 |
| Elavl4 | Q61701-4 | 0.25 | -0.1 | -0.53 | -0.08 | 0.2 | 0.15 | -0.71 | -0.55 | 0.38 | 0.22 | -0.34 | -0.17 | -0.13 | 0.62 | -0.16 | -0.13 | -0.1125 | 0.1375 | 0.165 | 0.703413 | 0.592759 | 0.544057 | 0.999971 | 1 | 0.911398 |
| Elavl4_1 | A2A9R5 | -2.07 | -1.71 | -1.01 | -0.34 | -3.05 | -1.4 | -0.88 | -1.81 | -1.77 | -1.8 | -0.94 | -0.94 | -2.29 | -1.26 | -1.28 | -0.38 | -0.5025 | -0.08 | -0.02 | 0.374148 | 0.852775 | 0.968536 | 0.999971 | 1 | 0.998787 |
| Elfn1 | Q8C8T7 | -0.67 | -1.02 | -0.8 | -0.65 | -0.93 | -0.98 | -0.47 | -0.89 | -0.77 | -1.02 | -0.76 | -0.93 | -0.36 | -0.98 | -0.8 | -1.08 | -0.0325 | -0.085 | -0.02 | 0.872007 | 0.637048 | 0.928633 | 0.999971 | 1 | 0.994278 |
| Elfn2 | Q68FM6 | -0.14 | -1.12 | -0.42 | -0.7 | -1.17 | -0.88 | -0.59 | -0.74 | -1.01 | -0.29 | -0.89 | -0.87 | -0.65 | -1.07 | -0.67 | -0.43 | -0.25 | -0.17 | -0.11 | 0.356919 | 0.546147 | 0.683516 | 0.999971 | 1 | 0.943718 |
| Elmo1 | Q8BPU7 | 0.21 | 0.05 | 0.03 | 0.09 | 0.01 | 0.02 | 0.08 | 0.02 | 0.02 | -0.1 | -0.07 | 0.01 | -0.11 | -0.04 | 0.25 | -0.09 | -0.0625 | -0.13 | -0.0925 | 0.69937 | 0.42228 | 0.604774 | 0.999971 | 1 | 0.924454 |
| Elmo2 | Q8BHL5-2 | -0.32 | -0.85 | -0.2 | -0.45 | -0.53 | -0.77 | 0.23 | -0.78 | -0.49 | -0.52 | -0.55 | -0.57 | -0.77 | -0.85 | -0.6 | -0.93 | -0.0075 | -0.0775 | -0.3325 | 0.979268 | 0.696119 | 0.139402 | 0.999971 | 1 | 0.784948 |
| Elmod1 | E9Q0K9 | -0.8 | -1.07 | -1.71 | -1.18 | -0.9 | -1.32 | -1.33 | -1.13 | -1.19 | -1.2 | -1.15 | -0.8 | -1.17 | -1.06 | -0.9 | -1.41 | 0.02 | 0.105 | 0.055 | 0.935107 | 0.666594 | 0.824902 | 0.999971 | 1 | 0.976309 |
| Elmod2 | A0A1B0GR | -1.26 | -0.39 | 0.32 | 0.01 | -0.04 | -0.75 | 0.09 | 0.34 | 0.15 | -1.17 | 0.16 | -0.63 | 0.35 | -0.93 | 0.12 | -0.07 | 0.24 | -0.0425 | 0.1975 | 0.549384 | 0.923855 | 0.640147 | 0.999971 | 1 | 0.93301 |
| Eln | P54320 | 0.32 | -3.3 | -0.98 | 0.87 | 5 | 1.04 | 0.3 | -0.36 | 3.64 | 5.34 | 4.39 | 7.25 | -5.75 | -0.16 | 0.52 | 1.01 | 2.2675 | 5.9275 | -0.3225 | 0.119669 | 0.000475 | 0.842 | 0.999971 | 0.43267 | 0.979814 |
| Eloa | Q8CB77 | 2.29 | 1.53 | 1.14 | 1.37 | 1.85 | 1.57 | 1.03 | 1.31 | 2 | 1.44 | 1.32 | 1.38 | 1.28 | 1.38 | 1.21 | 1.42 | -0.1425 | -0.0475 | -0.26 | 0.647514 | 0.874792 | 0.35323 | 0.999971 | 1 | 0.868675 |
| Elob | P62869 | -1.08 | -0.7 | -0.64 | -0.8 | -0.6 | -0.82 | -0.52 | -0.61 | -0.33 | -1.17 | -0.54 | -0.54 | -0.67 | -0.73 | -0.62 | -0.71 | 0.1675 | 0.16 | 0.1225 | 0.381877 | 0.506427 | 0.502992 | 0.999971 | 1 | 0.90459 |
| Eloc | A0A087WC | -1.09 | 0.61 | -0.39 | -1.75 | 0.68 | 0 | -0.17 | -1.12 | 0.71 | -1.1 | -0.38 | -0.22 | 1.33 | -0.48 | -0.57 | -1.57 | 0.5025 | 0.4075 | 0.3325 | 0.391657 | 0.486753 | 0.642452 | 0.999971 | 1 | 0.93301 |
| Elovl1 | Q9JLJ5 | 1.14 | -0.33 | -0.74 | -0.22 | 0.24 | 0.33 | -0.44 | -0.2 | 0.21 | 0.95 | -0.88 | -1.06 | 1.22 | 0.22 | -0.07 | -0.06 | 0.02 | -0.1575 | 0.365 | 0.962279 | 0.78469 | 0.450979 | 0.999971 | 1 | 0.886397 |
| Elovl4 | Q9EQC4 | -0.8 | 1.3 | 0.91 | 0.79 | 0.69 | 0.77 | 0.95 | 1.06 | 0.93 | 0.41 | 1.26 | 2.09 | 1.05 | 1.05 | 0.93 | 1.27 | 0.3175 | 0.6225 | 0.525 | 0.479848 | 0.266497 | 0.254105 | 0.999971 | 1 | 0.839012 |
| Elovl5 | D3Z0Z6 | 0.41 | -1.54 | -0.92 | -0.79 | -1.7 | -0.85 | -0.96 | -1.68 | -2.47 | -0.19 | -0.36 | -0.93 | -2.28 | -0.81 | -1.76 | -1.18 | -0.5875 | -0.2775 | -0.7975 | 0.204307 | 0.648566 | 0.127186 | 0.999971 | 1 | 0.783755 |
| Elp1 | Q7TT37 | 0.38 | 0.6 | 0.7 | 0.63 | 0.75 | 0.71 | 0.57 | 1.08 | 0.71 | 0.42 | 0.47 | 0.67 | 0.66 | 1.15 | 0.89 | 0.61 | 0.2 | -0.01 | 0.25 | 0.314854 | 0.954691 | 0.233437 | 0.999971 | 1 | 0.831526 |
| Elp2 | Q91WG4 | 0.28 | 0.86 | 1.5 | 1.01 | 1.01 | 0.96 | 1.29 | 1.57 | 1.05 | 0.57 | 1.14 | 0.52 | 1.07 | 1.07 | 1.49 | 1.17 | 0.295 | -0.0925 | 0.2875 | 0.335098 | 0.761572 | 0.327152 | 0.999971 | 1 | 0.856472 |
| Elp3 | Q9CZX0-2 | 0.56 | 0.57 | 0.56 | 1.1 | 0.39 | 0.74 | 0.31 | 1.49 | 0.56 | 0.54 | 0.42 | 0.44 | 0.83 | 0.79 | 1.09 | 1.12 | 0.035 | -0.2075 | 0.26 | 0.909158 | 0.304912 | 0.237414 | 0.999971 | 1 | 0.833594 |
| Elp4 | Q9ER73 | 1.05 | 0.22 | 0.1 | 0.14 | 0.19 | 0.18 | 0.48 | -0.15 | 0.41 | 0.75 | -0.16 | 0.16 | 0.29 | 0.08 | -0.27 | 0.02 | -0.2025 | -0.0875 | -0.3475 | 0.470839 | 0.773302 | 0.223775 | 0.999971 | 1 | 0.828225 |
| Elp5 | Q99L85 | 0.81 | 0.44 | 0.69 | 0.91 | 0.69 | 0.68 | 0.51 | 0.96 | 0.77 | 1.18 | 0.75 | 1.17 | 0.9 | 1.02 | 0.93 | 1.6 | -0.0025 | 0.255 | 0.4 | 0.989904 | 0.236791 | 0.111233 | 0.999971 | 1 | 0.78268 |
| Elp6 | Q8BK75 | -0.31 | -0.13 | 0.24 | 0.27 | -0.38 | -0.61 | 0.23 | 0.12 | -0.35 | -0.71 | 0.19 | 0.89 | -0.36 | -0.26 | 0.4 | 0.54 | -0.1775 | -0.0125 | 0.0625 | 0.510576 | 0.972758 | 0.825641 | 0.999971 | 1 | 0.976715 |
| Emb | P21995 | -2 | -1.23 | -0.89 | -1.59 | -1.08 | -1.04 | -0.66 | -1.4 | -0.78 | -1.21 | -0.88 | -0.51 | -0.54 | -1.58 | -1.21 | -1.24 | 0.3825 | 0.5825 | 0.285 | 0.214021 | 0.070941 | 0.390584 | 0.999971 | 1 | 0.871816 |
| Emc1 | Q8C7X2 | -0.7 | -0.13 | -0.23 | -0.24 | -0.25 | -0.13 | -0.32 | -0.22 | -0.13 | -0.24 | 0.02 | -0.21 | -0.1 | 0.01 | 0.06 | 0.11 | 0.095 | 0.185 | 0.345 | 0.629306 | 0.35891 | 0.10745 | 0.999971 | 1 | 0.78268 |
| Emc1_1 | Q8C7X2-2 | -2.02 | -0.77 | 0.76 | 0.26 | -0.56 | -1.09 | -0.9 | 1.45 | -0.71 | -0.98 | -0.46 | 1.07 | -0.49 | -0.52 | 0.24 | 0.49 | 0.1675 | 0.1725 | 0.3725 | 0.826537 | 0.805073 | 0.544619 | 0.999971 | 1 | 0.911398 |
| Emc10 | A0A0X1KG | 0.04 | 0.28 | -0.02 | 0.32 | 0.14 | 0.53 | 0.62 | 0.2 | 0.58 | 0.31 | 0.09 | -0.14 | 0.47 | -0.02 | -0.21 | -0.05 | 0.2175 | 0.055 | -0.1075 | 0.298944 | 0.800677 | 0.622357 | 0.999971 | 1 | 0.929759 |
| Emc2 | Q9CRD2 | -0.82 | 0.67 | 0.42 | 0.28 | -0.07 | 0.17 | 0.16 | 0.73 | 0.06 | -0.55 | 0.52 | 0.43 | 0.65 | 0.29 | 0.78 | 0.73 | 0.11 | -0.0225 | 0.475 | 0.761261 | 0.954353 | 0.190752 | 0.999971 | 1 | 0.810547 |
| Emc3 | Q99KI3 | -1.8 | -0.47 | -1.08 | -1 | -0.29 | -0.28 | -1.07 | -0.84 | -0.3 | -0.94 | -0.91 | -0.86 | -0.07 | -0.18 | -0.76 | -0.38 | 0.4675 | 0.335 | 0.74 | 0.18766 | 0.304631 | 0.041609 | 0.999971 | 1 | 0.701725 |
| Emc4 | Q9CZX9 | -0.58 | 0.88 | 1.5 | 0.26 | 0.94 | 0.22 | 1.7 | 0.56 | 1.32 | -0.61 | 1.9 | 1.35 | 2.14 | -0.21 | 1 | 0.21 | 0.34 | 0.475 | 0.27 | 0.506872 | 0.469252 | 0.665428 | 0.999971 | 1 | 0.93697 |
| Emc6 | Q9CQW0 | 0.51 | 0.78 | 0.92 | 1.17 | 0.83 | 0.67 | 0.21 | 0.85 | 0.29 | 0.74 | 0.76 | -0.87 | 1.37 | 0.87 | 1.09 | 1.47 | -0.205 | -0.615 | 0.355 | 0.397541 | 0.142816 | 0.151566 | 0.999971 | 1 | 0.795578 |
| Emc7 | Q9EP72 | -0.12 | -0.11 | 0.27 | 0.23 | -0.33 | -0.14 | -0.25 | 0.02 | -0.36 | -0.21 | -0.08 | -0.19 | -0.05 | 0.28 | -0.09 | 0.56 | -0.2425 | -0.2775 | 0.1075 | 0.231632 | 0.161244 | 0.637475 | 0.999971 | 1 | 0.93301 |
| Emc8 | O70378 | -0.03 | 0.15 | -0.26 | -0.43 | 0.06 | 0.28 | -0.51 | -0.31 | 0.48 | 0.31 | -0.51 | -0.05 | 0.5 | 0.09 | -0.1 | -0.38 | 0.0225 | 0.2 | 0.17 | 0.927618 | 0.467434 | 0.505922 | 0.999971 | 1 | 0.905277 |
| Emd | O08579 | -0.81 | 0.1 | -0.15 | -0.75 | 0.04 | -0.22 | -0.3 | -0.49 | 0.04 | -0.48 | -0.22 | 0.15 | 0.82 | -0.16 | -0.55 | -0.88 | 0.16 | 0.275 | 0.21 | 0.556319 | 0.338465 | 0.612042 | 0.999971 | 1 | 0.927066 |
| Emg1 | O35130 | -0.4 | -0.22 | -1.58 | -0.51 | -0.31 | -0.94 | -0.96 | -1.15 | -0.78 | -0.21 | -1.37 | -0.78 | -0.97 | -0.36 | -0.76 | -0.61 | -0.1625 | -0.1075 | 0.0025 | 0.64495 | 0.775131 | 0.993987 | 0.999971 | 1 | 0.99958 |
| Emilin1 | Q99K41 | -0.02 | -0.29 | -1.67 | 0.29 | 1.81 | -1 | 0.83 | -1.42 | -0.24 | 1.52 | 3.99 | 6.8 | -2.86 | -0.41 | 0.02 | 0.05 | 0.4775 | 3.44 | -0.3775 | 0.548525 | 0.03672 | 0.612021 | 0.999971 | 1 | 0.927066 |
| Eml1 | D3Z4J9 | 0.77 | 0.73 | 0.68 | 0.5 | 0.28 | 0.83 | 0.12 | 0.01 | 0.57 | 1.16 | 0.48 | 0.46 | 0.35 | 0.94 | 0.38 | 0.58 | -0.36 | -0.0025 | -0.1075 | 0.143256 | 0.99084 | 0.601628 | 0.999971 | 1 | 0.924453 |
| Eml2 | Q7TNG5-3 | 0.72 | 0.3 | 0.05 | 0.24 | 0.58 | 0.97 | 0.24 | 0.78 | 0.9 | 0.93 | 0.37 | 0.17 | 1.08 | 1.13 | 0.62 | 0.41 | 0.315 | 0.265 | 0.4825 | 0.215186 | 0.321625 | 0.083556 | 0.999971 | 1 | 0.755254 |
| Eml3 | Q8VC03 | 0.5 | 0.66 | -0.09 | -0.14 | -0.05 | 0.57 | -0.33 | -0.56 | -0.53 | 0.94 | -0.79 | 0.76 | -0.3 | 0.67 | -0.19 | -0.34 | -0.325 | -0.1375 | -0.2725 | 0.324467 | 0.763842 | 0.401008 | 0.999971 | 1 | 0.871816 |
| Eml4 | A7ISP9 | -0.26 | 0.64 | -0.52 | -0.25 | 0.63 | 0.4 | -1.77 | -1.44 | 0.98 | -0.02 | -1.31 | -0.52 | 1.59 | 1.1 | -0.79 | -0.21 | -0.4475 | -0.12 | 0.52 | 0.46902 | 0.812291 | 0.365737 | 0.999971 | 1 | 0.869526 |
| Eml4_1 | F8WJ93 | 0.61 | 0.14 | 0.53 | 0.61 | 0.18 | 0.25 | 0.65 | 0.78 | 0.24 | 0.48 | 0.68 | 0.59 | 0.48 | -0.09 | 0.62 | 0.88 | -0.0075 | 0.025 | 0 | 0.973483 | 0.900702 | 1 | 0.999971 | 1 | 1 |
| Eml5 | Q8BQM8 | 0.07 | 0.1 | -1.19 | -1.24 | -0.3 | 0.01 | -1.09 | -1.24 | -0.12 | 0.04 | -1.24 | -1.74 | -0.21 | -0.44 | -1.37 | -1.25 | -0.09 | -0.2 | -0.2525 | 0.842344 | 0.707153 | 0.575097 | 0.999971 | 1 | 0.91816 |
| Eml6 | Q5SQM0 | -1.09 | -1.67 | -0.17 | -0.25 | -1.95 | 0.39 | -0.66 | -0.27 | -1.69 | -1.2 | -0.47 | -0.59 | -2.32 | 0.1 | -0.56 | -0.51 | 0.1725 | -0.1925 | -0.0275 | 0.758007 | 0.658293 | 0.962123 | 0.999971 | 1 | 0.998721 |
| Emp2 | O88662 | 3.83 | -1.49 | 0.71 | 0.71 | -0.09 | -0.12 | 3.97 | -1.9 | 0.07 | 2.3 | -0.68 | -1.56 | -2.03 | -3.42 | -2.55 | -3.3 | -0.475 | -0.9075 | -3.765 | 0.746693 | 0.467044 | 0.004945 | 0.999971 | 1 | 0.582714 |
| Emsy | Q8BMB0-2 | 3.08 | 1.03 | 0 | 0.26 | 1.34 | 0.76 | -0.23 | 0.06 | 0.38 | 1.93 | -0.07 | 0 | -0.72 | -0.01 | -0.33 | 0.13 | -0.61 | -0.5325 | -1.325 | 0.398213 | 0.49056 | 0.070069 | 0.999971 | 1 | 0.735439 |
| Enah | A0A0A6YW | -0.74 | -1.71 | -2.32 | -2.46 | -0.98 | -1.64 | -1.88 | -2.18 | -1.31 | -1.33 | -1.94 | -2.22 | -0.78 | -1.16 | -2.21 | -1.97 | 0.1375 | 0.1075 | 0.2775 | 0.75508 | 0.802374 | 0.567828 | 0.999971 | 1 | 0.91816 |
| Enc1 | O35709 | 0.33 | 1.61 | 1.96 | 0.16 | 1.09 | 1.29 | 2.53 | 0.84 | 0.62 | 0.54 | 2.38 | 1.83 | 0.51 | 1.38 | 0.22 | -0.08 | 0.4225 | 0.3275 | -0.5075 | 0.442468 | 0.58156 | 0.334041 | 0.999971 | 1 | 0.85856 |
| Endod1 | Q8C522 | 0.37 | -0.13 | -0.62 | -0.45 | -0.15 | -0.14 | -0.75 | -0.47 | -0.29 | 0.45 | -0.78 | -0.36 | -0.05 | 0.27 | -0.64 | -0.62 | -0.17 | -0.0375 | -0.0525 | 0.545388 | 0.910517 | 0.868075 | 0.999971 | 1 | 0.983462 |
| Endog | O08600 | -0.6 | -0.68 | -0.52 | -0.4 | -0.75 | -0.96 | -0.44 | -0.3 | -0.96 | -0.65 | -0.32 | -0.47 | -0.57 | -1.24 | -0.41 | -0.78 | -0.0625 | -0.05 | -0.2 | 0.767217 | 0.804766 | 0.393511 | 0.999971 | 1 | 0.871816 |
| Endou | Q3V188 | 0.38 | -0.96 | 1.79 | 2.18 | -0.77 | -0.3 | 2.27 | 0.44 | -2.24 | -1.01 | 0.33 | -0.19 | -0.1 | 0.13 | -0.16 | -0.72 | -0.4375 | -1.625 | -1.06 | 0.620171 | 0.07509 | 0.140296 | 0.999971 | 1 | 0.784948 |
| Eng | F7A1B4 | 1.43 | -0.05 | 0.44 | 1.26 | 1.37 | 0.75 | 0.93 | -0.77 | 0.57 | 1.38 | 0.36 | 0.56 | -0.85 | 0.25 | 0.82 | 0.28 | -0.2 | -0.0525 | -0.645 | 0.709565 | 0.895146 | 0.187718 | 0.999971 | 1 | 0.809025 |
| Engase | Q8BX80 | -0.01 | 0.63 | 0.6 | 1.89 | 1.08 | 0.36 | 1.36 | 1.23 | 1.08 | -0.54 | 1.29 | 1.32 | 0.9 | 0.2 | 0.68 | 1.16 | 0.23 | 0.01 | -0.0425 | 0.597138 | 0.985546 | 0.920278 | 0.999971 | 1 | 0.992702 |
| Enho | Q8K1D8 | -0.89 | -0.13 | -2.42 | -1.36 | -0.8 | 0.84 | -1.61 | -2.29 | -0.02 | -0.35 | -1.78 | -1.42 | -1.23 | 0.4 | -0.71 | -0.67 | 0.235 | 0.3075 | 0.6475 | 0.753035 | 0.602696 | 0.252269 | 0.999971 | 1 | 0.839012 |
| Eno1 | P17182 | -0.39 | 0.04 | -0.03 | 0.31 | -0.33 | -0.32 | -0.05 | 0.13 | -0.43 | -0.53 | 0.04 | -0.06 | -0.35 | 0.03 | 0.2 | 0.4 | -0.125 | -0.2275 | 0.0875 | 0.579969 | 0.34337 | 0.722514 | 0.999971 | 1 | 0.955258 |
| Eno2 | P17183 | 0.31 | 0.82 | 0.16 | 0.55 | 0.69 | 0.65 | 0 | 0.36 | 0.29 | 0.26 | -0.08 | 0.12 | 0.62 | 1.01 | 0.41 | 0.4 | -0.035 | -0.3125 | 0.15 | 0.886523 | 0.166469 | 0.532938 | 0.999971 | 1 | 0.910069 |
| Eno3 | P21550 | 3.99 | 2.41 | 3.54 | 1.9 | 2.82 | 3.23 | 2.77 | 1.34 | 1.92 | 3.82 | 1.61 | -1.28 | -1.11 | 1.48 | 1.45 | 0.94 | -0.42 | -1.4425 | -2.27 | 0.477546 | 0.190064 | 0.010301 | 0.999971 | 1 | 0.613095 |
| Enoph1 | Q8BGB7 | 0.61 | 0.78 | 0.66 | 0.59 | 0.76 | 0.74 | 0.69 | 0.19 | 0.36 | 0.38 | 0.31 | 0.14 | 0.37 | 0.68 | 0.42 | 0.17 | -0.065 | -0.3625 | -0.25 | 0.746177 | 0.051645 | 0.20234 | 0.999971 | 1 | 0.81617 |
| Enpep | P16406 | 0.64 | -0.08 | -0.2 | 0.46 | 0.18 | -0.08 | 0.17 | -0.75 | -0.14 | 0.45 | -0.29 | -0.05 | -1.2 | -0.12 | -0.07 | -0.26 | -0.325 | -0.2125 | -0.6175 | 0.303336 | 0.448192 | 0.091842 | 0.999971 | 1 | 0.773779 |
| Enpp1 | G3X9S2 | 0.07 | -0.52 | -0.71 | -0.09 | 0.16 | -0.42 | -0.85 | -0.4 | -0.04 | -0.19 | -0.59 | -0.35 | -0.52 | -0.9 | -0.17 | -0.63 | -0.065 | 0.02 | -0.2425 | 0.82159 | 0.934802 | 0.364281 | 0.999971 | 1 | 0.869526 |
| Enpp2 | Q9R1E6-3 | -0.01 | 1.87 | 0.82 | 0.21 | 1.71 | 1.77 | 0.8 | 0.07 | 1.83 | 0.85 | 0.13 | 1.55 | 1.76 | 1.48 | 0.91 | 0.88 | 0.365 | 0.3675 | 0.535 | 0.502994 | 0.491229 | 0.25006 | 0.999971 | 1 | 0.839012 |
| Enpp4 | Q8BTJ4-2 | -0.17 | 0.36 | 1 | 0.95 | 0.35 | -0.06 | 0.67 | 0.88 | 0.55 | 0.01 | 0.71 | 1.38 | 0.44 | 0.15 | 1.31 | 1.15 | -0.075 | 0.1275 | 0.2275 | 0.825625 | 0.739048 | 0.552945 | 0.999971 | 1 | 0.913595 |
| Enpp5 | Q9EQG7 | -0.3 | -0.49 | -1.23 | -0.72 | -0.61 | -0.43 | -1.03 | -0.84 | -0.49 | -0.5 | -1.44 | -1.16 | -1.18 | -0.33 | -1.12 | -0.43 | -0.0425 | -0.2125 | -0.08 | 0.871313 | 0.506294 | 0.794926 | 0.999971 | 1 | 0.970361 |
| Enpp6 | Q8BGN3 | -0.52 | -0.64 | -0.24 | -0.7 | -0.53 | -0.38 | -0.66 | -0.74 | 0 | -0.49 | -0.89 | -0.64 | -0.32 | -0.53 | -0.48 | -0.66 | -0.0525 | 0.02 | 0.0275 | 0.786162 | 0.934225 | 0.885735 | 0.999971 | 1 | 0.98711 |
| Ensa | P60840 | -0.01 | -0.26 | 0.3 | 0.26 | 0.25 | -0.42 | 0.66 | -0.1 | 0.17 | -0.12 | 0.41 | 0.55 | -0.36 | -0.27 | -0.2 | -0.04 | 0.025 | 0.18 | -0.29 | 0.929114 | 0.444119 | 0.178664 | 0.999971 | 1 | 0.808532 |
| Enthd1 | E9Q1Z2 | -2.6 | -0.14 | -0.34 | -0.66 | -0.48 | -0.59 | -0.7 | -0.41 | -0.51 | -1.54 | -0.36 | -0.39 | 0.18 | -0.24 | -0.59 | -0.23 | 0.39 | 0.235 | 0.715 | 0.464515 | 0.68719 | 0.20876 | 0.999971 | 1 | 0.822138 |
| Entpd1 | Q8CDV7 | 0.77 | 1.28 | 0.7 | 1.1 | 2.18 | 1.18 | 0.58 | 0.3 | 1.08 | 1.66 | 1.29 | 1.62 | 0.53 | 1.19 | 1.13 | 1.14 | 0.0975 | 0.45 | 0.035 | 0.814777 | 0.077316 | 0.884571 | 0.999971 | 1 | 0.98711 |
| Entpd2 | O55026 | -1.32 | 0.56 | 0.05 | -0.54 | 0.75 | 0.62 | -0.3 | -0.57 | 0.41 | -0.86 | 0.29 | -0.06 | 1.82 | 0.48 | 0.46 | -0.14 | 0.4375 | 0.2575 | 0.9675 | 0.378161 | 0.583101 | 0.097539 | 0.999971 | 1 | 0.78268 |
| Entpd3 | Q8BFW6 | 0.17 | -0.13 | -0.15 | 0.57 | -0.43 | -0.16 | 0.16 | 0.58 | -0.2 | 0.26 | 0.02 | 0.36 | 0.09 | 0.28 | 0.55 | 0.5 | -0.0775 | -0.005 | 0.24 | 0.78784 | 0.983376 | 0.322312 | 0.999971 | 1 | 0.85548 |
| Entpd5 | Q3TQC7 | 1.97 | -0.13 | 0.71 | 0.92 | -0.18 | 0.23 | 0.71 | 0.48 | 0.26 | 1.28 | 1.04 | 0.42 | 0.12 | 0.41 | 0.18 | 0.88 | -0.5575 | -0.1175 | -0.47 | 0.230161 | 0.801007 | 0.300626 | 0.999971 | 1 | 0.847402 |
| Entpd6 | Q3U0P5 | -0.88 | 0.41 | -0.37 | -0.16 | 0.85 | -0.51 | -0.49 | 0.34 | 0.49 | -1.14 | 0 | -0.47 | 0.89 | 0.76 | 0.34 | 0.43 | 0.2975 | -0.03 | 0.855 | 0.4713 | 0.94263 | 0.019495 | 0.999971 | 1 | 0.635388 |
| Eny2 | A0A2I3BRK | 0.91 | 0.22 | 0.67 | 0.94 | 0.41 | 0.35 | 0.41 | 0.68 | -0.1 | 0.46 | 0.53 | 0.47 | 0.13 | 0.08 | 0.49 | 1.03 | -0.2225 | -0.345 | -0.2525 | 0.333874 | 0.189227 | 0.391919 | 0.999971 | 1 | 0.871816 |
| Eogt | Q8BYW9 | -0.34 | 0.43 | 1.17 | 1.35 | 0.87 | 0.42 | 2.01 | 1.51 | 0.72 | 0.01 | 1.12 | 1.17 | 0.97 | 0.81 | 1.16 | 0.56 | 0.55 | 0.1025 | 0.2225 | 0.274401 | 0.817505 | 0.572926 | 0.999971 | 1 | 0.91816 |
| Ep300 | B2RWS6 | 6.49 | 2.42 | -1.47 | -0.02 | 2.35 | 2.79 | -0.42 | 0.35 | 1.72 | 4.51 | -0.77 | 0.94 | -0.63 | 0.28 | -4.08 | -6.49 | -0.5875 | -0.255 | -4.585 | 0.728238 | 0.889548 | 0.051135 | 0.999971 | 1 | 0.723736 |
| Ep400 | Q8CHI8-3 | 0.77 | 1.39 | 1.23 | 1.35 | 1.41 | 0.76 | 0.25 | 0.96 | 1.33 | 0.88 | 0.59 | 0.49 | 0.39 | 1.24 | 0.95 | 1.17 | -0.34 | -0.3625 | -0.2475 | 0.261295 | 0.184839 | 0.35883 | 0.999971 | 1 | 0.869526 |
| Epb41 | A2A841 | -0.01 | -0.55 | -0.88 | -0.26 | -0.08 | -0.87 | -0.49 | -1.16 | -0.32 | -0.48 | -0.98 | -1.03 | -0.23 | -0.64 | -0.95 | -0.93 | -0.225 | -0.2775 | -0.2625 | 0.469793 | 0.326852 | 0.346858 | 0.999971 | 1 | 0.864164 |
| Epb41l1 | E9PV14 | -0.26 | -0.61 | 0.21 | -0.1 | -0.72 | -0.46 | -0.11 | 0 | -0.6 | -0.34 | -0.21 | 0.1 | -0.12 | -0.46 | 0.07 | 0.25 | -0.1325 | -0.0725 | 0.125 | 0.613535 | 0.772194 | 0.626247 | 0.999971 | 1 | 0.930479 |
| Epb41l1_1 | A2AUK5 | -2.77 | -4.02 | -0.19 | -1.15 | -3.89 | -3.88 | -1.08 | -0.71 | -3.27 | -3.57 | -1.16 | -0.88 | -2.95 | -3.7 | -0.82 | -1.09 | -0.3575 | -0.1875 | -0.1075 | 0.741411 | 0.849642 | 0.913146 | 0.999971 | 1 | 0.992618 |
| Epb41l2 | O70318 | -0.46 | -0.94 | -0.43 | -0.45 | -0.98 | -0.89 | -0.43 | -0.51 | -0.9 | -0.96 | -0.62 | -0.79 | -0.71 | -1.08 | -0.58 | -0.46 | -0.1325 | -0.2475 | -0.1375 | 0.559756 | 0.233997 | 0.544762 | 0.999971 | 1 | 0.911398 |
| Epb41l3 | A0A5F8MP | -1.26 | -2.37 | -0.7 | -1 | -2.38 | -2.3 | -0.87 | -1.12 | -2.06 | -1.9 | -1.12 | -1.09 | -2.96 | -2.37 | -1.05 | -1.14 | -0.335 | -0.21 | -0.5475 | 0.505361 | 0.621699 | 0.3312 | 0.999971 | 1 | 0.857107 |
| Epb41l3_1 | A7YY80 | -1.17 | -1.68 | -1.8 | -1.73 | -2.32 | -2.97 | -1.97 | -0.78 | -0.72 | -2.18 | -1.9 | -0.68 | -1.54 | -1.93 | -1.75 | -2.25 | -0.415 | 0.225 | -0.2725 | 0.370035 | 0.576912 | 0.276288 | 0.999971 | 1 | 0.843794 |
| Epb41l3_2 | Q9WV92-8 | -2.12 | -0.58 | -0.25 | -0.22 | -1.23 | -1.11 | -0.56 | -0.12 | -2.04 | -2.51 | 0.28 | 0.36 | -0.05 | -1.36 | 0.21 | 0.04 | 0.0375 | -0.185 | 0.5025 | 0.937965 | 0.816306 | 0.357867 | 0.999971 | 1 | 0.868793 |
| Epb41l3_3 | D0VYV6 | -0.96 | -1.03 | -0.64 | -0.04 | -1.56 | -0.2 | -0.45 | -0.26 | -1.42 | -0.86 | -0.32 | -0.44 | -0.77 | -0.2 | 0.12 | 0.85 | 0.05 | -0.0925 | 0.6675 | 0.894523 | 0.782058 | 0.115207 | 0.999971 | 1 | 0.78268 |
| Epb41l4b | A2ALK6 | 0.43 | -1.12 | 0.55 | 1.5 | -0.8 | -0.71 | 0.94 | 0.72 | -1.06 | -0.07 | 0.41 | -0.5 | -0.93 | -0.42 | 0.38 | 0.67 | -0.3025 | -0.645 | -0.415 | 0.641377 | 0.280952 | 0.494094 | 0.999971 | 1 | 0.901531 |
| Epb41l5 | Q8BGS1 | -0.18 | -0.96 | -1.31 | -1.34 | -0.36 | -1.07 | -1.09 | -1.41 | -0.35 | -0.62 | -0.99 | -1.41 | -0.24 | -1.05 | -1.12 | -1.14 | -0.035 | 0.105 | 0.06 | 0.919018 | 0.763999 | 0.861054 | 0.999971 | 1 | 0.982963 |
| Epb42 | Q3UV95 | 0.02 | -0.24 | -1.4 | 1.32 | 1.46 | -1.1 | 0.2 | 0.24 | -0.69 | -0.31 | 0.05 | -0.38 | -0.11 | -0.71 | -0.41 | -0.16 | 0.275 | -0.2575 | -0.2725 | 0.692018 | 0.632852 | 0.610272 | 0.999971 | 1 | 0.92673 |
| Epc1 | Q8C9X6 | 4.74 | 1.05 | 1.21 | 2.23 | 1.73 | 1.48 | 3.23 | -0.16 | 0.9 | 2.66 | 0.78 | 0.48 | -4.88 | -0.06 | 0.95 | -3.01 | -0.7375 | -1.1025 | -4.0575 | 0.459511 | 0.235261 | 0.017408 | 0.999971 | 1 | 0.617356 |
| Epdr1 | Q99M71 | 0.74 | 2.26 | 1.3 | 1.22 | 1.83 | 1.84 | 1.38 | 1.88 | 1.73 | 0.6 | 1.43 | 0.88 | 2.25 | 1.33 | 0.81 | 0.82 | 0.3525 | -0.22 | -0.0775 | 0.310785 | 0.579205 | 0.859745 | 0.999971 | 1 | 0.982702 |
| Epg5 | Q80TA9 | -0.06 | -0.44 | -0.32 | -0.43 | -0.31 | -0.35 | -0.66 | -0.54 | -0.48 | -0.14 | -0.54 | -0.29 | -0.16 | -0.05 | -0.55 | -0.31 | -0.1525 | -0.05 | 0.045 | 0.428756 | 0.792064 | 0.821775 | 0.999971 | 1 | 0.975833 |
| Epha10 | F6WVB0 | 0.72 | 0.38 | -1.52 | -1.88 | -0.26 | -0.91 | -1.66 | -1.63 | -0.95 | 0.14 | -0.92 | -1.54 | -0.83 | -0.96 | -0.62 | -1.82 | -0.54 | -0.2425 | -0.4825 | 0.42765 | 0.721649 | 0.461344 | 0.999971 | 1 | 0.89007 |
| Epha3 | B7ZNN5 | 1.75 | -0.74 | -0.48 | -0.27 | -0.18 | -0.45 | -0.89 | 0.21 | -0.46 | 0.88 | -0.41 | -0.67 | -1.04 | -1.18 | -0.67 | -0.09 | -0.3925 | -0.23 | -0.81 | 0.492352 | 0.709097 | 0.178926 | 0.999971 | 1 | 0.808532 |
| Epha4 | Q03137 | -0.36 | -0.61 | -0.8 | -0.82 | -0.67 | -0.07 | -0.35 | -0.57 | -0.51 | -0.23 | -0.55 | -1.03 | -0.27 | -0.63 | -0.66 | -0.59 | 0.2325 | 0.0675 | 0.11 | 0.299423 | 0.77119 | 0.585026 | 0.999971 | 1 | 0.920413 |
| Epha5 | E9PUP9 | -0.45 | -1.71 | -2.7 | -2.66 | -1.07 | -1.05 | -2.23 | -2.16 | -1.16 | -0.37 | -1.97 | -2.19 | -0.91 | -0.85 | -2.17 | -1.9 | 0.2525 | 0.4575 | 0.4225 | 0.659315 | 0.464971 | 0.4699 | 0.999971 | 1 | 0.892481 |
| Epha6 | G1K381 | -1.47 | -1.13 | -0.08 | -0.68 | -0.76 | -0.58 | -0.11 | 0.23 | -1.15 | -1.51 | 0.18 | -0.59 | -0.77 | -1.01 | 0.02 | 1.49 | 0.535 | 0.0725 | 0.7725 | 0.16804 | 0.871515 | 0.209338 | 0.999971 | 1 | 0.822138 |
| Epha7 | Q61772 | 0.92 | 0.11 | -0.37 | -0.46 | -0.14 | 0.46 | 0.08 | -0.6 | 0.15 | 0.7 | -0.21 | -0.44 | 0.19 | -0.15 | -0.46 | -0.35 | -0.1 | -7.85E-17 | -0.2425 | 0.789101 | 1 | 0.486107 | 0.999971 | 1 | 0.897483 |
| Ephb1 | Q8CBF3 | -0.69 | -0.48 | -0.53 | -0.6 | -0.94 | -0.88 | -0.05 | -1.02 | -1.09 | -0.81 | -0.67 | -0.52 | -0.87 | -0.77 | -0.57 | -0.64 | -0.1475 | -0.1975 | -0.1375 | 0.567865 | 0.315654 | 0.434622 | 0.999971 | 1 | 0.882419 |
| Ephb2 | P54763 | -0.14 | -0.28 | -0.55 | -0.68 | -0.37 | 0 | -0.73 | -0.9 | -0.51 | -0.18 | -0.54 | -0.75 | -0.14 | -0.3 | -0.83 | -0.5 | -0.0875 | -0.0825 | -0.03 | 0.736 | 0.701383 | 0.896811 | 0.999971 | 1 | 0.988737 |
| Ephb3 | P54754 | -1.08 | -0.45 | -0.33 | -0.39 | -0.64 | -0.19 | -0.18 | -0.38 | -0.55 | -0.67 | 0.02 | -0.34 | 0.26 | -0.8 | -0.58 | -0.15 | 0.215 | 0.1775 | 0.245 | 0.378557 | 0.4918 | 0.425319 | 0.999971 | 1 | 0.880591 |
| Ephb6 | O08644 | -0.73 | -0.17 | -0.51 | -0.66 | -0.57 | -0.55 | -0.38 | -0.24 | -0.71 | -0.68 | -0.25 | -0.67 | 0.03 | -0.72 | -0.48 | -0.68 | 0.0825 | -0.06 | 0.055 | 0.685464 | 0.776999 | 0.821974 | 0.999971 | 1 | 0.975833 |
| Ephx1 | Q9D379 | -0.16 | -0.46 | 0.07 | 0.03 | -0.06 | -0.37 | -0.34 | 0.45 | -0.4 | -0.43 | -0.41 | 0.22 | -1.19 | -0.6 | -0.17 | -0.23 | 0.05 | -0.125 | -0.4175 | 0.842831 | 0.594532 | 0.161332 | 0.999971 | 1 | 0.800215 |
| Ephx2 | P34914 | -0.06 | 0.49 | 0.14 | 0.28 | 0.18 | 0.42 | -0.23 | 0.85 | 0.1 | 0.21 | -0.05 | 0.34 | 0.28 | 0.53 | 0.32 | 0.19 | 0.0925 | -0.0625 | 0.1175 | 0.734706 | 0.752434 | 0.55656 | 0.999971 | 1 | 0.915051 |
| Ephx4 | Q6IE26 | 1.25 | -0.19 | -0.13 | 0.39 | -0.97 | 0.51 | 0.25 | 1.68 | -0.39 | 0.65 | -0.21 | -0.79 | -0.52 | -0.08 | -1.34 | 0.84 | 0.0375 | -0.515 | -0.605 | 0.948651 | 0.248765 | 0.264309 | 0.999971 | 1 | 0.842533 |
| Epm2a | Q9WUA5 | 0.43 | -0.03 | 0.6 | 0.54 | -0.53 | -0.36 | 0.34 | 1.37 | -0.37 | 0.52 | -0.36 | -0.81 | -0.68 | -0.24 | 0.7 | 0.96 | -0.18 | -0.64 | -0.2 | 0.676533 | 0.069413 | 0.615794 | 0.999971 | 1 | 0.927967 |
| Epm2aip1 | Q8VEH5 | -0.59 | 0.8 | 0.8 | 0.69 | 0.77 | 1.07 | 0.78 | 1.15 | 0.91 | 0.09 | 1 | 0.18 | 1.22 | 1.28 | 1.24 | 1.3 | 0.5175 | 0.12 | 0.835 | 0.161927 | 0.763389 | 0.033426 | 0.999971 | 1 | 0.689248 |
| Epn1 | Q80VP1 | -0.93 | -0.89 | -0.63 | -0.62 | -0.69 | -0.95 | -0.85 | -0.77 | -1.11 | -1.37 | -0.86 | -0.84 | -0.88 | -0.74 | -0.71 | -0.81 | -0.0475 | -0.2775 | -0.0175 | 0.791461 | 0.192713 | 0.921005 | 0.999971 | 1 | 0.992702 |
| Epn1_1 | Q80VP1-2 | -1.13 | -0.61 | -1.37 | -1.4 | -0.46 | -0.24 | -0.83 | -0.07 | -1 | -2.07 | -1.47 | 0.45 | -1.63 | 0.02 | -2.2 | -1.42 | 0.7275 | 0.105 | -0.18 | 0.021798 | 0.841947 | 0.704706 | 0.999971 | 1 | 0.950182 |
| Epn2 | J3QNT7 | 0.82 | -0.35 | 0.08 | -0.07 | -0.59 | -0.18 | -0.17 | 0.28 | -0.38 | -0.02 | 0.27 | 0.41 | -0.39 | -0.46 | 0.42 | 0.4 | -0.285 | -0.05 | -0.1275 | 0.371665 | 0.871654 | 0.711657 | 0.999971 | 1 | 0.951172 |

| | | | | | | | | | | | | | | | | | | | | | | | | | | |
|---|---|---|---|---|---|---|---|---|---|---|---|---|---|---|---|---|---|---|---|---|---|---|---|---|---|---|
| Epn3 | Q91W69 | 0.37 | 0.58 | -0.32 | 0.17 | 0.38 | -0.56 | -0.46 | -0.13 | -0.59 | -0.37 | -0.43 | 1.04 | 1.09 | -0.36 | 0.49 | -0.18 | -0.3925 | -0.2875 | 0.06 | 0.207246 | 0.485709 | 0.872013 | 0.999971 | 1 | 0.984109 |
| Eprs1 | Q8CGC7 | 0.78 | 1.39 | 1.46 | 1.76 | 1.37 | 1.45 | 2.06 | 1.92 | 1.81 | 1.41 | 2.14 | 2.17 | 1.44 | 1.81 | 2.05 | 1.77 | 0.3525 | 0.535 | 0.42 | 0.231821 | 0.08715 | 0.137612 | 0.999971 | 1 | 0.784948 |
| Eps15 | P42567 | -0.8 | -1.48 | -0.48 | -0.73 | -1.24 | -1.62 | -0.62 | -0.54 | -1.28 | -1.47 | -0.69 | -0.46 | -0.91 | -1.42 | -0.27 | -0.23 | -0.1325 | -0.1025 | 0.165 | 0.692125 | 0.750669 | 0.639549 | 0.999971 | 1 | 0.93301 |
| Eps15l1 | Q60902 | -0.47 | -0.81 | -0.4 | -0.25 | -0.75 | -0.86 | -0.54 | -0.37 | -0.74 | -1 | -0.32 | -0.21 | -0.75 | -0.85 | -0.34 | -0.24 | -0.1475 | -0.085 | -0.0625 | 0.490611 | 0.730596 | 0.786238 | 0.999971 | 1 | 0.967132 |
| Eps15l1_1 | Q60902-4 | -0.49 | -0.5 | 0.73 | 0.26 | 0 | 0.38 | -0.56 | 0.56 | -0.12 | -0.23 | -0.17 | -0.71 | -0.29 | 0.56 | 0.82 | 0.45 | 0.095 | -0.3075 | 0.385 | 0.801227 | 0.363728 | 0.3172 | 0.999971 | 1 | 0.853841 |
| Eps8 | Q08509 | -0.18 | -0.72 | 0.01 | -0.85 | -0.56 | -0.73 | -0.08 | -0.19 | -0.71 | -0.55 | 0.01 | 0.28 | -0.21 | -0.84 | 0.11 | -0.32 | 0.045 | 0.1925 | 0.12 | 0.870021 | 0.545422 | 0.687119 | 0.999971 | 1 | 0.945467 |
| Eps8l2 | Q99K30 | -0.5 | 0.86 | 0.99 | 0.68 | 0.79 | 1.21 | -0.22 | 0.24 | 0.32 | -0.16 | -0.15 | 0.37 | -0.39 | 0.89 | 0.34 | 0.49 | -0.0025 | -0.4125 | -0.175 | 0.995428 | 0.273368 | 0.673526 | 0.999971 | 1 | 0.94048 |
| Eral1 | Q9CZU4 | -0.09 | -0.93 | -0.61 | -0.29 | -1.09 | -1.16 | -0.36 | -0.25 | -0.88 | -0.91 | -0.39 | -0.4 | -0.85 | -1.1 | -0.01 | -0.17 | -0.235 | -0.165 | -0.0525 | 0.452055 | 0.525839 | 0.870833 | 0.999971 | 1 | 0.984109 |
| Erap1 | Q9EQH2 | -0.23 | 0.47 | -0.91 | -0.7 | 0.68 | 0.15 | -0.32 | -0.72 | 0.32 | 0.05 | -0.43 | -0.09 | 0.12 | 0.29 | -0.33 | -0.85 | 0.29 | 0.305 | 0.15 | 0.484922 | 0.381016 | 0.697563 | 0.999971 | 1 | 0.947479 |
| Erbb3 | Q61526 | 0.33 | -0.06 | -1.09 | -1.41 | -0.1 | -0.24 | -1.48 | -1.47 | -0.23 | -0.34 | -0.81 | -0.93 | 0.04 | -0.24 | -0.45 | -0.63 | -0.265 | -0.02 | 0.2375 | 0.610813 | 0.962428 | 0.571782 | 0.999971 | 1 | 0.91816 |
| Erbb4 | Q61527-3 | -0.73 | -1.1 | -0.56 | -0.46 | -0.73 | -0.99 | -0.38 | -0.62 | -0.94 | -1.14 | -0.62 | -0.62 | -0.61 | -1.16 | -0.87 | -0.85 | 0.0325 | -0.1175 | -0.16 | 0.886738 | 0.607244 | 0.480096 | 0.999971 | 1 | 0.894438 |
| Erbin | Q80TH2-2 | 0.26 | -1.2 | -0.55 | -0.44 | -1 | -1.21 | -0.42 | -0.73 | -1.13 | -0.22 | -1.21 | -0.85 | -1.17 | -1.32 | -0.91 | -1.07 | -0.3575 | -0.37 | -0.635 | 0.309552 | 0.325118 | 0.070514 | 0.999971 | 1 | 0.735439 |
| Erc1 | V9GWT6 | -0.04 | -0.09 | -0.22 | -0.83 | -0.05 | -0.21 | -1.07 | -0.84 | -0.8 | -0.71 | -0.46 | -0.08 | -0.6 | 0.15 | -0.33 | 0.11 | -0.2475 | -0.2175 | 0.1275 | 0.433784 | 0.418591 | 0.6447 | 0.999971 | 1 | 0.93301 |
| Erc1_1 | V9GXH3 | -0.19 | -0.91 | -0.99 | -0.93 | -0.72 | -0.76 | -0.78 | -0.85 | -0.56 | -0.21 | -0.57 | -0.53 | -0.59 | -0.98 | -0.97 | -0.65 | -0.0225 | 0.2875 | -0.0425 | 0.92182 | 0.248267 | 0.862914 | 0.999971 | 1 | 0.982963 |
| Erc2 | Q3UHT7 | -0.9 | -0.81 | -1.39 | -1.05 | -0.47 | -1.11 | -2.04 | -0.78 | -1.23 | -0.83 | -1.49 | -1.25 | 0.69 | -0.53 | -1.26 | -0.85 | -0.0625 | -0.1625 | 0.55 | 0.86025 | 0.476579 | 0.210212 | 0.999971 | 1 | 0.822657 |
| Erc2_1 | Q6PH08 | -1.12 | -1.59 | -1.06 | -0.94 | -1.62 | -1.45 | -0.9 | -1.21 | -1.51 | -1.22 | -1.14 | -0.98 | -1.41 | -1.66 | -1.35 | -1.05 | -0.1175 | -0.035 | -0.19 | 0.630614 | 0.873993 | 0.416668 | 0.999971 | 1 | 0.877251 |
| Ercc2 | O08811 | 0.6 | 1.33 | 1.67 | 2.38 | 1.46 | 0.85 | 1.31 | 2.09 | 1.1 | 0.64 | 1.38 | 1.57 | 0.62 | 1.72 | 1.63 | 2.45 | -0.0675 | -0.3225 | 0.11 | 0.873988 | 0.433054 | 0.822847 | 0.999971 | 1 | 0.975901 |
| Ercc3 | P49135 | 0.61 | 1.15 | 1.3 | 1.34 | 0.85 | 0.93 | 0.28 | 1.74 | 0.75 | 0.9 | 1.42 | 2.24 | 1.02 | 1.49 | 1.8 | 1.73 | -0.15 | 0.2275 | 0.41 | 0.661335 | 0.539316 | 0.148888 | 0.999971 | 1 | 0.793248 |
| Ercc4 | Q9QZD4 | 1.55 | 1.7 | 1.08 | 1.63 | 1.52 | 1.64 | 1.04 | 1.52 | 1.49 | 1.73 | 1.24 | 1.02 | 1.24 | 1.97 | 1.29 | 1.59 | -0.06 | -0.12 | 0.0325 | 0.79511 | 0.617693 | 0.895846 | 0.999971 | 1 | 0.988571 |
| Erf | P70459 | 0.24 | 0.58 | 1.24 | 0.73 | 0.69 | 0.22 | 0.35 | 1.09 | 0.35 | 0.45 | 1.46 | 0.58 | -0.02 | 0.45 | 0.59 | 1.03 | -0.11 | 0.0125 | -0.185 | 0.710051 | 0.969583 | 0.550871 | 0.999971 | 1 | 0.913169 |
| Erg28 | Q9ERY9 | -4.95 | -3.09 | -3.59 | -3.91 | -1.6 | -3.44 | -3.83 | -3.39 | -2.24 | -4.57 | -3 | -2.59 | -1.87 | -3.13 | -2.78 | -4.35 | 0.82 | 0.785 | 0.8525 | 0.181888 | 0.209483 | 0.174351 | 0.999971 | 1 | 0.807788 |
| Ergic1 | F6SLP4 | -0.36 | 0.16 | 0.2 | 0.26 | 0 | 0.48 | 0.33 | 0.18 | 0.31 | 0.24 | 0.06 | 0.29 | 0.71 | 0.8 | 0.27 | 0.57 | 0.1825 | 0.16 | 0.5225 | 0.416369 | 0.44208 | 0.043007 | 0.999971 | 1 | 0.701725 |
| Ergic2 | Q9CR89 | -0.33 | 1.06 | 0.99 | 0.89 | 0.7 | 1.14 | 1.05 | 0.98 | 0.86 | 0.51 | 0.88 | 0.14 | 1.92 | 1.2 | 0.49 | 0.93 | 0.315 | -0.055 | 0.4825 | 0.365465 | 0.879771 | 0.271988 | 0.999971 | 1 | 0.843794 |
| Ergic3 | Q9CQE7 | 0.04 | 0.42 | 1.17 | 0.06 | 0.71 | 0.52 | 1.7 | 0.54 | 0.75 | -0.1 | 1.8 | 1.99 | 0.76 | 0.22 | 0.89 | 0.32 | 0.445 | 0.6875 | 0.125 | 0.252558 | 0.204768 | 0.69278 | 0.999971 | 1 | 0.947174 |
| Erh | P84089 | 0.37 | 0.2 | 0.14 | -0.2 | 0.05 | 0.56 | 2.21 | -0.85 | 0.04 | 0.48 | -0.34 | -1.29 | -0.05 | 0.45 | -0.21 | -0.12 | 0.365 | -0.405 | -0.11 | 0.544819 | 0.305317 | 0.633843 | 0.999971 | 1 | 0.93301 |
| Eri3 | Q8C460 | 1.44 | 1.45 | 1.08 | 1.15 | 1.14 | 1.7 | 0.84 | 1.2 | 1.33 | 1.56 | 0.88 | 0.64 | 1.86 | 1.57 | 1.28 | 1.38 | -0.06 | -0.1775 | 0.2425 | 0.80033 | 0.491187 | 0.268605 | 0.999971 | 1 | 0.842746 |
| Erlec1 | Q8VEH8 | -0.37 | 0.44 | 0.18 | 0.09 | 0.37 | 0.42 | 0.46 | 0.54 | 0.35 | -0.04 | 0.13 | -0.04 | 0.73 | 0.63 | 0.54 | 0.56 | 0.3625 | 0.015 | 0.53 | 0.123123 | 0.947599 | 0.036553 | 0.999971 | 1 | 0.700096 |
| Erlin1 | Q91X78 | -0.36 | -1.06 | 0.34 | 0.55 | -0.53 | -1.08 | 0.33 | 0.56 | -0.36 | -0.51 | 0.86 | 0.66 | -1.08 | -0.65 | 0.62 | 0.77 | -0.0475 | 0.295 | 0.0475 | 0.922742 | 0.537732 | 0.930015 | 0.999971 | 1 | 0.994744 |
| Erlin2 | Q8BFZ9 | -0.37 | -0.15 | -0.1 | 0 | -0.18 | 0.04 | -0.1 | 0.18 | 0.02 | 0.18 | -0.19 | 0.22 | 0.14 | 0.15 | 0.28 | 0.22 | 0.14 | 0.2125 | 0.3525 | 0.455869 | 0.272854 | 0.069083 | 0.999971 | 1 | 0.733937 |
| Ermn | Q5EBJ4 | -0.29 | -1.08 | -0.9 | -0.94 | -1.54 | -0.94 | -1.69 | -0.92 | -1.46 | -1.02 | -1.82 | -1.33 | -1.16 | -0.97 | -0.89 | -0.98 | -0.47 | -0.605 | -0.1975 | 0.121438 | 0.043578 | 0.392257 | 0.999971 | 1 | 0.871816 |
| Ermp1 | Q3UVK0 | -0.09 | -0.07 | -0.58 | -0.65 | -0.44 | 0.12 | -0.64 | -0.29 | -0.4 | -0.01 | -0.5 | -0.18 | -0.07 | 0.03 | -0.3 | 0.19 | 0.035 | 0.075 | 0.31 | 0.889075 | 0.741671 | 0.195116 | 0.999971 | 1 | 0.811225 |
| Ero1a | Q8R180 | 0.35 | 0.42 | 0.48 | 0.67 | 0.24 | 0.45 | 0.57 | 0.46 | 0.33 | 0.41 | 0.44 | 0.67 | 0.2 | 0.62 | 0.62 | 0.82 | -0.05 | -0.0175 | 0.085 | 0.77962 | 0.920907 | 0.678094 | 0.999971 | 1 | 0.942594 |
| Ero1a_1 | A0A2I3BPN | 0.5 | 2.16 | 1.48 | 3.03 | 1.37 | 1.28 | 0.45 | 1.04 | -0.12 | 1.41 | 1.42 | 1.54 | -0.09 | 2.56 | 2.68 | 2.41 | -0.7575 | -0.73 | 0.0975 | 0.176586 | 0.25132 | 0.89887 | 0.999971 | 1 | 0.989648 |
| Ero1b | Q8R2E9 | -1.33 | 0.24 | 0.08 | -0.09 | 0.06 | -0.36 | 0.08 | -0.73 | -0.48 | -1.06 | -0.25 | 0.42 | 0.45 | -0.27 | -0.61 | 0.23 | 0.0375 | -0.0675 | 0.225 | 0.923417 | 0.879332 | 0.587254 | 0.999971 | 1 | 0.920758 |
| Erp29 | P57759 | -0.6 | -0.01 | 0.19 | 0.25 | 0.08 | 0.16 | 0.13 | 0.37 | 0.2 | -0.51 | 0.58 | 0.65 | 0.64 | 0.4 | 0.71 | 0.65 | 0.2275 | 0.2725 | 0.6425 | 0.351956 | 0.416415 | 0.023662 | 0.999971 | 1 | 0.663563 |
| Erp44 | Q9D1Q6 | 0.43 | 0.87 | 0.53 | 0.68 | 1.17 | 0.77 | 0.65 | 0.85 | 1.48 | 0.69 | 0.76 | 1.14 | 2.01 | 0.77 | 0.89 | 0.81 | 0.2325 | 0.39 | 0.4925 | 0.269723 | 0.128264 | 0.146251 | 0.999971 | 1 | 0.791767 |
| Esam | Q925F2 | 0.49 | 0.2 | -0.32 | 0.39 | 1.18 | -0.06 | -0.43 | -0.4 | 0.45 | 1.03 | -0.19 | 0.14 | -0.49 | 0.83 | 0.49 | 0.05 | -0.1175 | 0.1675 | 0.03 | 0.769674 | 0.601497 | 0.928822 | 0.999971 | 1 | 0.994278 |
| Esd | H3BLJ9 | -0.27 | 1.93 | 1.61 | 0.72 | 1.49 | 1.82 | 1.14 | 1.21 | 1.91 | -0.01 | 1.24 | 0.56 | 1.32 | 1.39 | 1.31 | 0.38 | 0.4175 | -0.0725 | 0.1025 | 0.396633 | 0.902696 | 0.840577 | 0.999971 | 1 | 0.97975 |
| Ess2 | Q3UFM6 | 0.92 | 1.55 | 2.25 | 3.04 | 1.02 | 0.82 | 2.29 | 2.39 | 0.09 | -0.55 | 2.35 | 1.41 | -0.26 | 0.84 | 2.31 | 2.22 | -0.31 | -1.115 | -0.6625 | 0.585595 | 0.14959 | 0.351679 | 0.999971 | 1 | 0.867656 |
| Esyt1 | Q3U7R1 | -0.26 | -0.27 | -0.98 | -0.65 | 0.14 | -0.46 | -1.46 | -1.18 | -0.25 | -0.09 | -0.91 | -0.59 | -0.57 | -0.29 | -0.74 | -0.75 | -0.2 | 0.08 | -0.0475 | 0.606204 | 0.766805 | 0.841843 | 0.999971 | 1 | 0.979814 |
| Esyt2 | Q3TZZ7 | -1.37 | -0.41 | -0.65 | -1.62 | -0.03 | -0.72 | -0.71 | -1.14 | -0.07 | -1.27 | -0.57 | -0.31 | 0.24 | -0.75 | -0.63 | -1.29 | 0.3625 | 0.4575 | 0.405 | 0.327408 | 0.243385 | 0.335408 | 0.999971 | 1 | 0.858827 |
| Etf1 | Q8BWY3 | 0 | 0.39 | 0.8 | -0.02 | 0.53 | 0.14 | 0.86 | 0.16 | 0.68 | 0.12 | 1.02 | 1 | 0.62 | 0.17 | 0.55 | 0.29 | 0.13 | 0.4125 | 0.115 | 0.63917 | 0.186062 | 0.647256 | 0.999971 | 1 | 0.933613 |
| Etfa | Q99LC5 | 1.18 | -0.39 | 0.34 | 0.46 | -0.78 | -0.13 | 0.85 | 0.29 | -0.71 | 0.67 | -0.04 | -0.55 | -1.02 | -0.4 | -0.2 | -0.23 | -0.34 | -0.555 | -0.86 | 0.450405 | 0.213839 | 0.040339 | 0.999971 | 1 | 0.700838 |
| Etfb | Q9DCW4 | -2.04 | -2.27 | -0.06 | -0.11 | -1.76 | -2.57 | -0.15 | 0.15 | -1.72 | -2.47 | -0.15 | -0.15 | -1.63 | -2.46 | -0.13 | 0.07 | 0.0375 | -0.0025 | 0.0825 | 0.962299 | 0.997365 | 0.914533 | 0.999971 | 1 | 0.992633 |
| Etfdh | Q921G7 | -0.12 | -0.35 | -0.39 | -0.28 | -0.56 | -0.43 | -0.72 | -0.09 | -0.43 | -0.13 | -0.58 | -0.47 | -0.34 | -0.31 | -0.43 | -0.23 | -0.165 | -0.1175 | -0.0425 | 0.424479 | 0.524394 | 0.802593 | 0.999971 | 1 | 0.971467 |
| Ethe1 | Q9DCM0 | -0.16 | -1.04 | -0.71 | -0.87 | -0.92 | -0.47 | -0.91 | -0.56 | -0.72 | -0.86 | -0.96 | -1.47 | -1.48 | -0.5 | -0.58 | -0.88 | -0.02 | -0.3075 | -0.165 | 0.936458 | 0.27155 | 0.587427 | 0.999971 | 1 | 0.920758 |
| Etl4 | E9QAU4 | -0.76 | -0.93 | -1.21 | -1.31 | -0.96 | -0.9 | -1.2 | -1.43 | -1.11 | -0.89 | -1.35 | -1.61 | -0.81 | -1.05 | -1.65 | -1.28 | -0.07 | -0.1875 | -0.145 | 0.750273 | 0.430614 | 0.563347 | 0.999971 | 1 | 0.917011 |
| Etl4_1 | D3Z781 | -1.1 | 0.96 | 0.51 | -0.82 | 0.95 | 0.42 | 0.05 | 0.56 | 0.66 | 0.15 | 0.38 | 0.25 | 1.93 | 0.26 | -0.24 | 0.53 | 0.6075 | 0.4725 | 0.7325 | 0.24082 | 0.33911 | 0.258931 | 0.999971 | 1 | 0.839012 |
| Etl4_2 | Q8BHR1 | 4.55 | 0.68 | -0.32 | 3.81 | 0.77 | -1.21 | 4.63 | -0.52 | -1.06 | 0.77 | -1.45 | -1.19 | -0.63 | -3.59 | -2.1 | 0.33 | -1.2625 | -2.9125 | -3.6775 | 0.426902 | 0.030977 | 0.018753 | 0.999971 | 1 | 0.635388 |
| Etnk1 | Q9D4V0 | 0.13 | 0.69 | -0.77 | -0.2 | 0.31 | 0.55 | -1.05 | -0.17 | 0.18 | 0.2 | -0.57 | -0.4 | 0.18 | 0.78 | -0.42 | -0.55 | -0.0525 | -0.11 | 0.035 | 0.904927 | 0.758186 | 0.932228 | 0.999971 | 1 | 0.995069 |
| Etnppl | E9PYH3 | 1.3 | 1.15 | 0.02 | 0.19 | -0.4 | 1.97 | -0.25 | 0.58 | 1.68 | 2.63 | 0.7 | 0.31 | 1.72 | -0.36 | -0.66 | -0.11 | -0.19 | 0.665 | -0.5175 | 0.743586 | 0.259648 | 0.380024 | 0.999971 | 1 | 0.871816 |
| Eva1a | D3Z511 | -2.17 | -1.68 | 0.07 | -0.15 | -0.91 | -1.43 | -0.85 | -0.06 | -0.47 | -1.35 | -0.64 | 0.01 | -0.51 | -0.66 | 0.07 | 0.61 | 0.17 | 0.37 | 0.86 | 0.766074 | 0.52377 | 0.160106 | 0.999971 | 1 | 0.800144 |
| Evi2a | P20934 | 0.73 | 0.53 | -0.9 | -0.49 | -0.08 | 0.34 | -0.49 | -1.77 | -0.04 | -0.35 | -1.55 | -2.72 | -0.22 | -0.02 | -2.36 | -3.2 | -0.4675 | -1.1325 | -1.4175 | 0.407793 | 0.115906 | 0.101403 | 0.999971 | 1 | 0.78268 |
| Evi5 | E9PWR7 | -0.01 | -0.08 | -0.74 | -0.38 | 0.05 | -0.15 | -0.53 | -0.24 | -0.13 | -0.06 | -0.56 | -0.7 | 0.02 | -0.29 | -0.43 | -0.5 | 0.085 | -0.06 | 0.0025 | 0.72272 | 0.813082 | 0.991593 | 0.999971 | 1 | 0.99925 |
| Evi5l | H3BKQ3 | -0.15 | -0.71 | -1.09 | -0.95 | -0.48 | -0.23 | -0.92 | -0.99 | -0.18 | 0.13 | -0.42 | -0.79 | -0.5 | -0.44 | -0.87 | -0.91 | 0.07 | 0.41 | 0.045 | 0.807959 | 0.18671 | 0.864335 | 0.999971 | 1 | 0.983079 |
| Evl | P70429 | -0.52 | -1.49 | -0.76 | -0.9 | -0.82 | -1.45 | -0.64 | -0.71 | -0.97 | -0.84 | -0.6 | -0.36 | -0.84 | -1.19 | -0.42 | -0.6 | 0.0125 | 0.225 | 0.155 | 0.965516 | 0.40723 | 0.584487 | 0.999971 | 1 | 0.920364 |
| Ewsr1 | Q5SUT0 | 1.14 | 2.67 | 2.6 | 2.45 | 2.32 | 1.75 | 2.17 | 2.61 | 2.32 | 1.71 | 2.75 | 2.46 | 2.05 | 2.55 | 2.08 | 2.78 | -0.0025 | 0.095 | 0.15 | 0.994848 | 0.814429 | 0.700561 | 0.999971 | 1 | 0.948864 |
| Exd2 | Q8VEG4 | -0.53 | -0.02 | -0.64 | -0.75 | 0.14 | -0.08 | -0.1 | -0.42 | -0.08 | -0.22 | 0.14 | 0.03 | 0.32 | -0.17 | -0.43 | -0.35 | 0.37 | 0.4525 | 0.3275 | 0.140353 | 0.06533 | 0.225949 | 0.999971 | 1 | 0.828225 |
| Exoc1 | Q5PPR2 | -1.27 | -2.5 | -0.64 | -0.77 | -2.45 | -1.55 | -0.61 | -1.75 | -2.3 | 0.23 | -2.06 | -0.38 | -3.32 | -1.76 | -0.77 | 0.11 | -0.295 | 0.1675 | -0.14 | 0.577429 | 0.807238 | 0.85486 | 0.999971 | 1 | 0.982125 |
| Exoc1_1 | Q6P1Y9 | -0.73 | -0.03 | -0.7 | -0.8 | -0.22 | 0 | -0.51 | -0.8 | 0.09 | -0.23 | -0.57 | -0.8 | 0.15 | -0.03 | -0.64 | -0.79 | 0.1825 | 0.1875 | 0.2375 | 0.503978 | 0.507739 | 0.437097 | 0.999971 | 1 | 0.882419 |
| Exoc2 | Q9D4H1 | 0.12 | 0.11 | 0.15 | 0.14 | -0.01 | 0.09 | 0.32 | 0.11 | 0.07 | 0.03 | 0.1 | 0.21 | -0.05 | 0.1 | 0.04 | 0.14 | -0.0025 | -0.0275 | -0.0725 | 0.988134 | 0.86054 | 0.655799 | 0.999971 | 1 | 0.934061 |
| Exoc3 | Q6KAR6 | -0.83 | 0.24 | -0.31 | -0.17 | 0.16 | 0.33 | -0.41 | -0.11 | 0.38 | -0.16 | -0.15 | -0.05 | 0.5 | 0.16 | -0.01 | 0.2 | 0.26 | 0.2725 | 0.48 | 0.376102 | 0.330775 | 0.099112 | 0.999971 | 1 | 0.78268 |
| Exoc4 | O35382 | 0.24 | 0.56 | 0.45 | 0.22 | 0.55 | 0.48 | 0.66 | -0.04 | 0.58 | 0.31 | 0.56 | -0.07 | 0.67 | 0.27 | 0.18 | 0 | 0.045 | -0.0225 | -0.0875 | 0.837917 | 0.916814 | 0.683016 | 0.999971 | 1 | 0.943421 |
| Exoc5 | Q3TPX4 | 0.37 | 0.39 | 0.26 | -0.31 | 0.18 | 0.74 | 0.11 | -0.07 | 0.58 | 0.62 | 0.42 | 0.35 | 0.32 | 0.19 | 0.1 | 0.38 | 0.0625 | 0.315 | 0.07 | 0.812083 | 0.174209 | 0.752026 | 0.999971 | 1 | 0.961116 |
| Exoc6 | Q3U9D6 | 0.3 | 0.79 | -0.18 | -0.08 | 0.56 | 0.48 | 0.01 | -0.23 | 0.9 | 0.71 | 0.02 | -0.1 | 1.26 | 0.55 | -0.11 | 0.06 | -0.0025 | 0.175 | 0.2325 | 0.993318 | 0.59949 | 0.532176 | 0.999971 | 1 | 0.910069 |
| Exoc6b | A6H5Z3 | -0.38 | 0.09 | 0.23 | 0.17 | -0.21 | 0.08 | -0.01 | 0.15 | -0.11 | -0.28 | 0.09 | 0.33 | -0.05 | 0.09 | 0.1 | 0.08 | -0.025 | -0.02 | 0.0275 | 0.90515 | 0.929911 | 0.891421 | 0.999971 | 1 | 0.988225 |
| Exoc7 | O35250 | -0.41 | -0.54 | -0.36 | -0.42 | -0.78 | -0.7 | -0.45 | -0.5 | -0.76 | -0.64 | -0.13 | -0.15 | -0.89 | -0.69 | -0.45 | -0.36 | -0.175 | 0.0125 | -0.165 | 0.330184 | 0.953113 | 0.401121 | 0.999971 | 1 | 0.871816 |
| Exoc8 | Q6PGF7 | 0.17 | -0.32 | 0.11 | 0.27 | -0.32 | -0.02 | 0.25 | -0.03 | -0.23 | 0.14 | 0.2 | -0.02 | -0.48 | -0.36 | -0.04 | 0.13 | -0.0875 | -0.035 | -0.245 | 0.690788 | 0.866993 | 0.303741 | 0.999971 | 1 | 0.847402 |
| Exog | Q8C163 | -0.02 | 0.44 | -1.15 | -1.15 | 0.22 | 0.36 | -0.36 | -1.46 | 0.24 | -0.41 | -1.15 | -1.1 | -0.15 | 0.42 | -1.38 | -1.52 | 0.16 | -0.135 | -0.1875 | 0.764233 | 0.781617 | 0.743041 | 0.999971 | 1 | 0.961064 |
| Exosc1 | Q9DAA6 | 0.82 | 3.22 | 1.16 | 0.19 | 2.64 | 2.11 | 1.07 | 0.43 | 1.87 | 1.42 | 1.3 | 1.71 | 3.33 | 1.52 | 0.97 | 0.78 | 0.215 | 0.2275 | 0.3025 | 0.772484 | 0.711464 | 0.702553 | 0.999971 | 1 | 0.949025 |
| Exosc10 | P56960 | 1.44 | 1.06 | 0.77 | 0.76 | 1.16 | -0.05 | 0.44 | 0.48 | 0.84 | 1.26 | 0.26 | 0.45 | 0.22 | 0.86 | 0.23 | 0.74 | -0.5 | -0.305 | -0.495 | 0.12595 | 0.301792 | 0.081369 | 0.999971 | 1 | 0.755254 |
| Exosc2 | Q8VBV3 | 0.22 | 1.46 | 0.94 | 1.1 | 1.16 | 1.17 | 0.47 | 0.61 | 1.17 | 0.51 | 0.56 | 0.78 | 1.47 | 1.21 | 1.02 | 0.83 | -0.0775 | -0.175 | 0.2025 | 0.808914 | 0.571466 | 0.508804 | 0.999971 | 1 | 0.905815 |
| Exosc3 | Q7TQK4 | 0.38 | 1.07 | 0.18 | 0.76 | 1.46 | 0.3 | 0.32 | -0.27 | 0.53 | 0.42 | -1.31 | 0.48 | 1.71 | 0.67 | -3.03 | 0.04 | -0.145 | -0.5675 | -0.75 | 0.714895 | 0.238133 | 0.428701 | 0.999971 | 1 | 0.881196 |
| Exosc4 | Q921I9 | -2.4 | -1.02 | -3.07 | -5 | 0.18 | -2.43 | 3.19 | -4.15 | 1.24 | -2.91 | -2.48 | 1.31 | 0.24 | -2.44 | -6.77 | -7.07 | 2.07 | 2.1625 | -1.1375 | 0.216569 | 0.115884 | 0.516243 | 0.999971 | 1 | 0.905815 |
| Exosc5 | Q9CRA8 | -0.39 | 0.31 | 1.53 | 1.55 | 0.7 | -0.55 | 0.57 | 0.9 | -0.07 | 0.4 | 1.47 | 1.19 | -0.79 | -0.08 | 1.47 | 0.78 | -0.345 | -0.0025 | -0.405 | 0.522226 | 0.996368 | 0.522854 | 0.999971 | 1 | 0.907797 |
| Exosc6 | Q8BTW3 | 2.73 | 1.11 | 1.69 | 2.26 | 0.51 | 0.96 | 1.12 | 1.42 | 0.92 | 2.1 | 1.13 | 1.03 | -0.1 | 0.74 | 1.28 | 1.73 | -0.945 | -0.6525 | -1.035 | 0.034009 | 0.148732 | 0.060405 | 0.999971 | 1 | 0.725348 |
| Exosc7 | Q9D0M0 | 2.17 | 2.32 | 1.72 | 2.19 | 2.25 | 1.97 | 1.58 | 1.93 | 2.05 | 2.2 | 1.31 | 0.99 | 1.21 | 2.24 | 1.4 | 1.68 | -0.1675 | -0.4625 | -0.4675 | 0.470703 | 0.173887 | 0.118792 | 0.999971 | 1 | 0.78268 |
| Exosc8 | Q9D753 | 0.68 | 0.84 | 1.16 | 0.61 | 0.96 | 0.53 | 0.76 | 0.36 | 0.46 | 0.84 | 0.88 | 0.99 | 0.4 | 0.54 | 0.35 | 0.51 | -0.17 | -0.03 | -0.3725 | 0.451733 | 0.887892 | 0.082619 | 0.999971 | 1 | 0.755254 |
| Exosc9 | Q9JHI7 | 2.54 | 1.51 | 2.18 | 2.03 | 1.27 | 1.18 | 1.48 | 2.05 | 1.34 | 2.17 | 2.18 | 2.08 | 1.06 | 1.68 | 2.27 | 1.65 | -0.57 | -0.1225 | -0.4 | 0.083143 | 0.685632 | 0.242191 | 0.999971 | 1 | 0.837062 |
| Exph5 | Q0VAV2 | 1.82 | 0 | 0.1 | 1.36 | -0.48 | 0.25 | -0.78 | 0.44 | -1.16 | 1.44 | -0.27 | -0.42 | 0.1 | -0.13 | -0.22 | 1.75 | -0.9625 | -0.9225 | -0.445 | 0.081391 | 0.181637 | 0.461079 | 0.999971 | 1 | 0.89007 |
| Ext2 | P70428 | -0.72 | -0.29 | 1.13 | 1.09 | 0.61 | -0.6 | 0.69 | 1.45 | 0.68 | -0.48 | 1 | 0.72 | -1.31 | -0.31 | 1.46 | 1.15 | 0.235 | 0.1775 | -0.055 | 0.687423 | 0.740285 | 0.939487 | 0.999971 | 1 | 0.995869 |
| Extl1 | Q9JKV7 | -0.73 | -1.63 | 0.92 | 1.02 | 0.5 | -0.07 | 0.94 | 1.23 | -0.48 | 0.17 | 2.03 | 1.93 | -0.77 | 0.69 | 1.24 | 1.47 | 0.755 | 1.0175 | 0.7625 | 0.258511 | 0.234336 | 0.318281 | 0.999971 | 1 | 0.853893 |
| Extl2 | Q3TSR0 | -1.68 | -1.06 | -1.18 | -1.26 | -0.49 | -1.14 | -0.66 | -0.78 | -0.32 | -1.91 | -1.77 | -0.33 | -1.46 | -1.22 | -1.46 | -1.25 | 0.5275 | 0.2125 | -0.0525 | 0.044785 | 0.62695 | 0.798224 | 0.999971 | 1 | 0.970479 |
| Extl3 | Q6P1H4 | 0.36 | 0.96 | 0.82 | 0.75 | 1 | 0.96 | 0.35 | 0.66 | 1.2 | 1.01 | 1.08 | 1.08 | 0.8 | 1.11 | 0.73 | 1.43 | 0.02 | 0.37 | 0.295 | 0.931894 | 0.082815 | 0.238396 | 0.999971 | 1 | 0.833594 |
| Ezr | P26040 | -0.43 | -0.81 | 0.03 | -0.05 | -0.73 | -0.64 | -0.08 | -0.19 | -0.6 | -0.38 | -0.5 | -0.21 | -0.75 | -0.85 | -0.22 | 0.3 | -0.095 | -0.1075 | -0.065 | 0.726389 | 0.656648 | 0.843315 | 0.999971 | 1 | 0.979814 |
| F11r | O88792 | 1.94 | 0.83 | 1.07 | 1.24 | 1.53 | 0.77 | 0.97 | 0.46 | 1.49 | 2.32 | 1.76 | 1.96 | 0.81 | 0.77 | 1.39 | 1.48 | -0.3375 | 0.6125 | -0.1575 | 0.318375 | 0.069474 | 0.613438 | 0.999971 | 1 | 0.927364 |
| F13a1 | Q8BH61 | 2.32 | 1.92 | 0.68 | 1.24 | 2.6 | 1.2 | 0.83 | 0.34 | 1.08 | 2.38 | 0.64 | 0.61 | -0.08 | 0.97 | 0.84 | 0.27 | -0.2975 | -0.3625 | -1.04 | 0.595773 | 0.485945 | 0.03237 | 0.999971 | 1 | 0.683918 |
| F2 | H7BX99 | 1.64 | 2.08 | 0.81 | 3.31 | 4.05 | 1.84 | 2.59 | 1.54 | 1.7 | 1.43 | 2.1 | 2.75 | 1.74 | 2.32 | 1.71 | 2.28 | 0.545 | 0.035 | 0.0525 | 0.439335 | 0.949195 | 0.917555 | 0.999971 | 1 | 0.992633 |
| F3 | P20352 | -1.87 | -0.98 | -0.47 | -0.93 | -0.79 | -0.87 | -0.6 | -0.09 | -0.46 | -1.74 | -0.03 | 0.29 | -0.49 | -1.1 | -0.46 | -0.24 | 0.475 | 0.5775 | 0.49 | 0.184061 | 0.264309 | 0.177251 | 0.999971 | 1 | 0.808532 |
| F8a1 | Q00558 | -0.16 | -0.08 | -1.42 | -0.71 | -0.44 | -0.43 | -1.48 | -1.17 | -0.52 | -0.2 | -1.83 | -1.74 | -0.06 | 0.11 | -1.37 | -1.06 | -0.2875 | -0.48 | -0.0025 | 0.4688 | 0.336704 | 0.995565 | 0.999971 | 1 | 1 |
| Fa2h | Q5MPP0-2 | -0.6 | -0.31 | 0.07 | -0.57 | -0.86 | -0.81 | -0.17 | 0.16 | 0.01 | 0.16 | 0.35 | 0.66 | 0 | -0.35 | 0.14 | 0.15 | -0.0675 | 0.6475 | 0.3375 | 0.822895 | 0.023038 | 0.170762 | 0.999971 | 1 | 0.803758 |
| Faah | O08914 | -0.74 | 0.19 | -0.23 | -0.45 | 0.11 | 0.56 | -0.43 | -0.22 | 0.04 | -0.15 | -0.12 | -0.48 | 0.32 | 0.6 | -0.07 | 0 | 0.3125 | 0.13 | 0.52 | 0.312786 | 0.605854 | 0.080859 | 0.999971 | 1 | 0.755254 |
| Faap100 | A2ACJ2 | 0.38 | -2.89 | -1.53 | -1.14 | -0.48 | -1.24 | -1.3 | -0.26 | 0 | 0.33 | -0.02 | -0.85 | 0.91 | -1 | -0.86 | 0.24 | 0.475 | 1.16 | 1.1175 | 0.47535 | 0.104785 | 0.154016 | 0.999971 | 1 | 0.795578 |
| Fabp3 | P11404 | 0.36 | 0.49 | 0.19 | -0.62 | 0.53 | 0.32 | 1.01 | -0.38 | 1.13 | 0.02 | 0.94 | 1.16 | 1.01 | 0.22 | -0.31 | -0.48 | 0.265 | 0.7075 | 0.005 | 0.480512 | 0.075583 | 0.990019 | 0.999971 | 1 | 0.99925 |
| Fabp5 | Q05816 | -1.35 | -0.66 | -0.23 | -0.82 | -0.06 | -0.55 | -0.15 | 0.14 | 0.23 | -1.46 | 0.12 | 0.48 | 0.1 | -0.9 | -0.02 | -0.51 | 0.61 | 0.6075 | 0.4325 | 0.057975 | 0.215868 | 0.209188 | 0.999971 | 1 | 0.822138 |
| Fabp7 | P51880 | -0.04 | -0.78 | 0.14 | 0.07 | -0.35 | -0.48 | 0.63 | 0.75 | 0.66 | 0.73 | 1.07 | 0.82 | 0.27 | 0.42 | 1 | 0.94 | 0.29 | 0.9725 | 0.81 | 0.444768 | 0.004072 | 0.020566 | 0.999971 | 0.969702 | 0.635388 |
| Fadd | Q61160 | 1.33 | 1.24 | 12.12 | 10.91 | 1.37 | 0.9 | 7.79 | 13.2 | 0.67 | 1.03 | 13.22 | 13.99 | 1.17 | 1.01 | 12.71 | 10.33 | -0.585 | 0.8275 | -0.095 | 0.873325 | 0.844089 | 0.97981 | 0.999971 | 1 | 0.99925 |
| Fads1 | Q920L1 | -1.05 | -0.99 | -0.88 | -1.67 | -1 | -1.12 | -1 | -0.79 | -1.16 | -0.64 | -0.88 | -1.23 | -0.92 | -0.7 | -0.98 | -1.12 | 0.17 | 0.17 | 0.2175 | 0.465136 | 0.501411 | 0.365929 | 0.999971 | 1 | 0.869526 |
| Fads2 | Q9Z0R9 | -0.37 | -1.7 | -3.17 | -3.29 | -0.91 | -0.69 | -2.24 | -3.22 | -0.77 | -0.25 | -2.75 | -3.02 | -0.75 | -0.48 | -2.5 | -2.81 | 0.3675 | 0.435 | 0.4975 | 0.654522 | 0.624928 | 0.547832 | 0.999971 | 1 | 0.911722 |
| Fads3 | Q9JJE7 | 0.21 | -0.55 | -1.48 | -0.34 | -0.61 | -0.96 | -0.91 | -0.7 | -0.92 | -0.21 | -1.02 | -0.43 | -1.03 | -0.52 | -0.81 | -0.23 | -0.255 | -0.105 | -0.1075 | 0.478314 | 0.786666 | 0.777506 | 0.999971 | 1 | 0.964764 |
| Fads6 | Q80UG1 | 0.1 | 1.82 | 0.46 | 0.86 | 1.5 | 0.8 | 0.29 | 1.15 | 0.85 | 0.65 | 2.35 | 1.46 | 2.35 | 1.03 | 1.3 | 1.15 | 0.125 | 0.5175 | 0.6475 | 0.7702 | 0.312773 | 0.175122 | 0.999971 | 1 | 0.808532 |
| Faf1 | P54731 | 0.68 | 0.43 | -0.2 | 0.23 | 0.39 | 0.67 | 0.24 | 0.12 | 0.44 | 0.32 | -0.31 | -0.07 | 0.51 | 0.29 | -0.4 | -0.01 | 0.07 | -0.19 | -0.1875 | 0.778953 | 0.490337 | 0.516409 | 0.999971 | 1 | 0.905815 |
| Faf2 | Q3TDN2 | -0.06 | -0.23 | -0.28 | -0.16 | -0.29 | -0.34 | -0.42 | -0.3 | -0.35 | -0.08 | -0.5 | -0.79 | -0.27 | -0.38 | -0.46 | -0.35 | -0.155 | -0.2475 | -0.1825 | 0.357801 | 0.248004 | 0.289989 | 0.999971 | 1 | 0.847402 |
| Fah | P35505 | 0.62 | 0.85 | 0.36 | 0.27 | 0.5 | 0.58 | -0.24 | 0.19 | 0.24 | 0.41 | 0 | 0.14 | 0.47 | 0.51 | 0.4 | 0.07 | -0.2675 | -0.3275 | -0.1625 | 0.307524 | 0.138995 | 0.455372 | 0.999971 | 1 | 0.887217 |
| Fahd1 | Q8R0F8 | -1.11 | -0.11 | -0.19 | 0.03 | -0.19 | -0.46 | -0.02 | -0.5 | -0.51 | -1.23 | -0.66 | -0.66 | -0.92 | -0.93 | -0.94 | -0.97 | 0.0525 | -0.42 | -0.595 | 0.858063 | 0.197402 | 0.056321 | 0.999971 | 1 | 0.725348 |
| Fahd2a | A0A0R4J09 | -1.14 | -0.74 | -0.33 | -0.73 | -0.79 | -1.01 | -0.79 | -0.27 | -0.9 | -1.5 | -0.64 | -0.59 | -0.63 | -0.75 | -0.21 | -0.41 | 0.02 | -0.1725 | 0.235 | 0.937224 | 0.542856 | 0.338296 | 0.999971 | 1 | 0.859348 |
| Faim | Q9WUD8 | -0.11 | 0.59 | 0.25 | 0.33 | 0.07 | 0.6 | 0.18 | 0.51 | 0.27 | 0.52 | 0.41 | 0.29 | 0.21 | 0.84 | 0.36 | 0.61 | 0.075 | 0.1075 | 0.24 | 0.745468 | 0.603558 | 0.324102 | 0.999971 | 1 | 0.85548 |
| Faim2 | Q8K097-2 | 1.01 | -1.8 | -0.09 | -0.32 | -1.6 | -1.09 | -0.28 | -0.37 | -1.61 | -0.36 | -0.71 | -1.17 | -1.68 | -1.31 | -0.41 | -0.13 | -0.535 | -0.6625 | -0.5825 | 0.383458 | 0.277501 | 0.362797 | 0.999971 | 1 | 0.869526 |
| Fam102a | Q78T81 | 0.91 | 0.92 | 0.51 | 0.04 | 0.31 | 0.56 | -0.01 | 0.02 | 0.33 | 0.43 | 0.29 | 0.16 | -0.23 | 0.47 | -0.18 | -0.6 | -0.375 | -0.2925 | -0.73 | 0.185471 | 0.250285 | 0.04064 | 0.999971 | 1 | 0.700838 |
| Fam107a | A0A5S8E87 | 0.15 | -1.5 | -1.16 | -0.94 | -0.86 | -0.75 | -0.46 | -1.1 | -0.33 | -0.07 | -0.7 | -0.6 | -0.34 | -1.74 | -2.38 | -0.63 | 0.07 | 0.4375 | -0.41 | 0.849681 | 0.259727 | 0.461846 | 0.999971 | 1 | 0.890095 |
| Fam107b | Q3TGF2 | 1.5 | 0.88 | 0.12 | 0.7 | 1.09 | 0.73 | 0.1 | 0.27 | 0.81 | 1.32 | -0.03 | 0.32 | 1.46 | 0.74 | 0.32 | 0.65 | -0.2525 | -0.195 | -0.0075 | 0.482916 | 0.622365 | 0.983515 | 0.999971 | 1 | 0.99925 |
| Fam110b | Q8C739 | 1.27 | -0.01 | 0.26 | 0.51 | -0.18 | 0.39 | -0.12 | 0.47 | 0.37 | 0.86 | 0.28 | 0.58 | 0.28 | 0.12 | 0.35 | 0.78 | -0.3675 | 0.015 | -0.125 | 0.274555 | 0.961247 | 0.691927 | 0.999971 | 1 | 0.947164 |
| Fam114a1 | Q9D281 | 1.25 | -0.13 | 0.61 | 0.9 | 0.31 | 0.19 | 0.89 | 0.34 | -0.09 | 1.58 | 1.05 | 1.18 | -1.01 | -0.09 | 0.71 | 1.37 | -0.225 | 0.2725 | -0.4125 | 0.50251 | 0.538478 | 0.45675 | 0.999971 | 1 | 0.887217 |
| Fam114a2 | Q8VE88-2 | -0.73 | 0.07 | 0.34 | 0.66 | 0.13 | 0.13 | 0.41 | 0.33 | 0.28 | -0.33 | 0.28 | 0.26 | 0.29 | -0.14 | 0.36 | 0.23 | 0.165 | 0.0375 | 0.1 | 0.598138 | 0.909999 | 0.755429 | 0.999971 | 1 | 0.961116 |
| Fam117b | Q3U3E2 | 1.46 | 0.3 | 0.79 | 1.09 | 0.09 | 0.44 | 1.19 | 0.87 | 0.08 | 1.17 | 0.69 | 0.79 | -0.37 | 0.32 | 0.78 | 0.01 | -0.2625 | -0.2275 | -0.725 | 0.448328 | 0.499116 | 0.059458 | 0.999971 | 1 | 0.725348 |
| Fam118a | Q91YN1 | 1.37 | 2.02 | 1.84 | 2.48 | 2.18 | 2.69 | 1.72 | 2.82 | 1.93 | 2.3 | 2.23 | 2.45 | 1.64 | 1.9 | 2.12 | 2.38 | 0.425 | 0.3 | 0.0825 | 0.229515 | 0.286337 | 0.776997 | 0.999971 | 1 | 0.964475 |
| FAM120A | Q6A0A9 | 1.07 | 0.85 | 0.89 | 1.15 | 0.68 | 0.84 | 1.18 | 1.01 | 0.82 | 1.08 | 0.81 | 0.82 | 0.58 | 0.77 | 0.7 | 1.08 | -0.0625 | -0.1075 | -0.2075 | 0.747264 | 0.543682 | 0.300231 | 0.999971 | 1 | 0.847402 |
| Fam120b | Q6RI63 | 0.15 | 0.58 | 1.17 | 0.99 | 0.34 | 0.2 | 0.87 | 0.59 | 0.51 | 0.06 | 0.62 | 0.64 | 0.22 | 0.11 | 0.91 | 0.21 | -0.2225 | -0.265 | -0.36 | 0.442504 | 0.355234 | 0.251727 | 0.999971 | 1 | 0.839012 |
| Fam120c | Q8C3F2 | -0.01 | 0.93 | 0.91 | 0.4 | 0.79 | 0.45 | 0.76 | 0.82 | 0.9 | 0.17 | 0.83 | 1.14 | 0.65 | 0.4 | 0.57 | 0.6 | 0.1475 | 0.2025 | -0.0025 | 0.578704 | 0.519912 | 0.992243 | 0.999971 | 1 | 0.99925 |
| Fam124a | D3Z5V4 | -1.9 | -0.36 | -1.66 | -2.53 | 0.06 | -0.67 | -1.65 | -2.57 | -0.04 | -2.21 | -1.06 | -1.68 | -1.11 | -0.28 | -0.9 | -1.46 | 0.405 | 0.365 | 0.675 | 0.546231 | 0.54645 | 0.187024 | 0.999971 | 1 | 0.809025 |
| Fam126a | Q6P9N1 | -0.94 | -1.7 | -1.88 | -1.6 | -1.48 | -0.93 | -1.77 | -0.98 | -0.96 | -0.52 | -1.67 | -1.27 | -0.56 | -0.96 | -1.44 | -1.23 | 0.24 | 0.425 | 0.4825 | 0.428545 | 0.206857 | 0.12499 | 0.999971 | 1 | 0.78268 |
| Fam126b | Q8C729-2 | -0.41 | -0.73 | -1.15 | -1.08 | -0.89 | -0.63 | -1.36 | -1.28 | -1.06 | -0.38 | -0.84 | -1.32 | -0.26 | -0.76 | -1.32 | -0.6 | -0.1975 | -0.0575 | 0.1075 | 0.461141 | 0.83591 | 0.713301 | 0.999971 | 1 | 0.951793 |
| Fam131a | D3YXU9 | -2.2 | 0.1 | -1.24 | -1.39 | -0.59 | 0.09 | -1.59 | -0.82 | -0.25 | -1.2 | -1.25 | 0.22 | -0.33 | 0.26 | -0.44 | -0.29 | 0.455 | 0.5625 | 0.9825 | 0.410819 | 0.323994 | 0.061996 | 0.999971 | 1 | 0.725348 |
| Fam131b | E9Q8P8 | -0.26 | -0.47 | -0.91 | -0.65 | -0.67 | -0.31 | -0.82 | -0.74 | -0.47 | -0.71 | -0.72 | -0.8 | -0.34 | -0.62 | -1.04 | -0.3 | -0.0625 | -0.1025 | -0.0025 | 0.777849 | 0.619647 | 0.991974 | 0.999971 | 1 | 0.99925 |
| Fam135a | A0A087WF | -0.43 | 0.5 | -1.08 | -1.38 | -0.55 | -0.17 | -1.06 | -0.49 | -0.27 | -0.19 | -0.23 | -0.05 | 1.01 | 0.26 | -1.14 | -0.14 | 0.03 | 0.4125 | 0.595 | 0.944315 | 0.319367 | 0.305377 | 0.999971 | 1 | 0.847402 |
| Fam135b | Q9DAI6 | -0.42 | -0.22 | -0.61 | 0.04 | -0.73 | -0.69 | -0.67 | -0.74 | -0.48 | -0.16 | -0.07 | -1.03 | -0.41 | -0.92 | -0.54 | -0.45 | -0.405 | -0.1325 | -0.2775 | 0.067958 | 0.630945 | 0.236067 | 0.999971 | 1 | 0.832918 |
| Fam136a | Q9CR98 | 0.58 | 0.97 | 1.39 | 1 | 0.49 | 1.38 | 2.8 | 0.8 | 0.71 | 0.39 | 0.89 | 0.26 | 0.75 | 0.6 | 0.83 | -0.04 | 0.3825 | -0.4225 | -0.45 | 0.450961 | 0.114368 | 0.129396 | 0.999971 | 1 | 0.784948 |
| Fam13c | G3X9S1 | 0.19 | -0.2 | -0.24 | -0.2 | 0.01 | 0.03 | -0.47 | -0.22 | -0.3 | 0.63 | -0.87 | -0.43 | 0.3 | 0.06 | -0.13 | 0.15 | -0.05 | -0.13 | 0.2075 | 0.809622 | 0.695021 | 0.308563 | 0.999971 | 1 | 0.847402 |
| Fam151b | D3YUE4 | 0 | 0.97 | 1.96 | 1.32 | 1 | 0.76 | 1.2 | 1.21 | 0.67 | -0.46 | 0.96 | 1.15 | 0.11 | 0.61 | 1.03 | 0.94 | -0.02 | -0.4825 | -0.39 | 0.960413 | 0.354679 | 0.380331 | 0.999971 | 1 | 0.871816 |
| Fam162a | Q9D6U8 | -0.59 | -0.06 | -0.65 | -0.33 | -0.08 | -0.48 | -0.24 | -0.38 | -0.05 | -0.63 | -0.18 | -0.04 | -0.33 | -0.21 | -0.72 | -1.31 | 0.1125 | 0.1825 | -0.235 | 0.596328 | 0.434428 | 0.432095 | 0.999971 | 1 | 0.881643 |
| Fam163b | Q8BUM6 | -1.25 | -0.63 | -1.02 | -1.07 | -0.88 | -0.85 | -1.45 | -0.91 | -1.28 | -1.51 | -1.37 | -0.83 | -0.86 | -1.15 | -0.93 | -0.76 | -0.03 | -0.255 | 0.0675 | 0.896661 | 0.287096 | 0.745976 | 0.999971 | 1 | 0.961064 |
| Fam169a | Q5XG69 | 1.91 | 2.43 | 2.48 | 2.66 | 2.34 | 2.51 | 2.38 | 2.41 | 2.15 | 2.11 | 2.31 | 2.18 | 1.69 | 2.54 | 2.32 | 2.93 | 0.04 | -0.1825 | -3.93E-17 | 0.851318 | 0.399853 | 1 | 0.999971 | 1 | 1 |
| Fam171a1 | A2ATK9 | -0.55 | -0.92 | -1.08 | -0.58 | -0.88 | -0.93 | -1.1 | -0.87 | -1.05 | -0.31 | -0.94 | -0.63 | 0.23 | -0.81 | -0.67 | -0.65 | -0.1625 | 0.05 | 0.3075 | 0.423882 | 0.835695 | 0.297384 | 0.999971 | 1 | 0.847402 |
| Fam171a2 | A2A699 | -1.14 | -1.32 | -1.28 | -1.11 | -1.52 | -1.27 | -1.31 | -0.97 | -1.12 | -0.87 | -1.14 | -1.11 | -0.66 | -1 | -0.97 | -0.36 | -0.055 | 0.1525 | 0.465 | 0.773716 | 0.377942 | 0.051 | 0.999971 | 1 | 0.723736 |
| Fam171b | Q14CH0-2 | -0.23 | -1.65 | -1.17 | -0.91 | -1.23 | -1.23 | -0.65 | -1.1 | -1.31 | -0.74 | -0.85 | -1.1 | -1.08 | -1.26 | -1.08 | -0.84 | -0.0625 | -0.01 | -0.075 | 0.84831 | 0.975306 | 0.81101 | 0.999971 | 1 | 0.973864 |
| Fam172a | E9QMA2 | 0.79 | -0.11 | 0.42 | 0.3 | -0.29 | 0 | -0.12 | 0.6 | -0.87 | 0.1 | 0.05 | 0.41 | -0.76 | -0.18 | 0.52 | 0.68 | -0.3025 | -0.4275 | -0.285 | 0.300408 | 0.218926 | 0.44937 | 0.999971 | 1 | 0.885522 |
| Fam174a | Q9D3L0 | 0.93 | 1.23 | 2.15 | 2.23 | 1.19 | 1.15 | 1.7 | 2.35 | 1.22 | 1.82 | 2.43 | 1.79 | 0.75 | 1.44 | 2.51 | 2.09 | -0.0375 | 0.18 | 0.0625 | 0.927012 | 0.649467 | 0.894617 | 0.999971 | 1 | 0.988571 |
| Fam177a1 | Q8BR63 | 0.39 | -0.31 | 0.55 | 0.32 | -0.68 | 0.17 | 0.66 | 0.78 | -0.73 | 0.15 | -0.45 | -1.29 | -0.58 | -0.59 | -0.23 | -1.03 | -0.005 | -0.8175 | -0.845 | 0.98921 | 0.041296 | 0.011569 | 0.999971 | 1 | 0.613095 |
| Fam184a | E9PW83 | -0.19 | -0.09 | -0.38 | -0.08 | -0.19 | -0.12 | -0.49 | -0.01 | 0.25 | 0.18 | -0.04 | 0.22 | 0.12 | 0.01 | -0.33 | 0.31 | -0.0175 | 0.3375 | 0.2125 | 0.926863 | 0.080086 | 0.31734 | 0.999971 | 1 | 0.853841 |
| Fam184b | Q0KK56 | 0.21 | 0.01 | -3.33 | -0.03 | -0.37 | -0.28 | -2.98 | -1.24 | -0.35 | -1.36 | -3.05 | -1.62 | -0.79 | 0.3 | -2.15 | -0.76 | -0.4325 | -0.81 | -0.065 | 0.648383 | 0.387505 | 0.941419 | 0.999971 | 1 | 0.996526 |
| Fam185a | Q7TPD2 | 0.65 | 0.29 | -0.34 | -0.25 | -0.02 | 0.25 | -0.76 | -0.22 | -0.22 | 0.45 | -0.28 | -0.3 | 0.25 | 0.42 | -0.34 | -0.19 | -0.275 | -0.175 | -0.0525 | 0.398916 | 0.565819 | 0.862312 | 0.999971 | 1 | 0.982963 |
| Fam189a1 | Q6A044-2 | 0.48 | -1.2 | -0.86 | -0.69 | -1.11 | -0.91 | -0.71 | -1.27 | -0.83 | 0.02 | -0.45 | -1.01 | -0.58 | -1.08 | -0.95 | -1.35 | -0.4325 | 3.14E-16 | -0.4225 | 0.264334 | 1 | 0.289001 | 0.999971 | 1 | 0.847402 |
| Fam193a | M0QWZ1 | 1.11 | 1.11 | 1.68 | 1.64 | 1.01 | 1.48 | 2.93 | 0.69 | 1 | 0.95 | 1 | -0.2 | 1.37 | 0.59 | 0.79 | -0.3 | 0.1425 | -0.6975 | -0.7725 | 0.768609 | 0.062755 | 0.062818 | 0.999971 | 1 | 0.725348 |
| Fam20b | Q8VCS3 | -0.25 | 1.06 | 0.2 | 0.15 | 1.27 | 0.62 | 0.18 | 0.4 | 1.39 | -0.36 | 0.79 | 0.72 | 1.68 | 0.9 | 0.84 | 0.13 | 0.3275 | 0.345 | 0.5975 | 0.368101 | 0.434164 | 0.162749 | 0.999971 | 1 | 0.800215 |
| Fam20c | Q5MJS3 | 2.16 | -1.32 | -1.97 | 0.19 | -1.49 | -0.81 | 0.36 | -0.09 | -1.06 | 1.24 | 1.18 | 1.2 | -3.48 | -1.18 | -0.16 | 1.94 | -0.2725 | 0.875 | -0.485 | 0.761969 | 0.378829 | 0.708316 | 0.999971 | 1 | 0.95033 |
| Fam210a | Q8BGY7 | -1.07 | -0.45 | -0.75 | -0.03 | -0.65 | -0.59 | -0.8 | -0.68 | -0.72 | -1.16 | -0.87 | -0.98 | -0.75 | -0.64 | -0.87 | -0.94 | -0.105 | -0.3575 | -0.225 | 0.678913 | 0.194646 | 0.391736 | 0.999971 | 1 | 0.871816 |
| Fam210b | Q9D8B6 | 0.6 | -0.69 | -0.83 | -1.09 | -1.15 | 0.13 | -1.7 | -0.81 | -0.9 | 0.16 | -1.09 | -2.34 | -1.56 | 0.05 | -1.49 | -1.58 | -0.38 | -0.54 | -0.6425 | 0.453854 | 0.368176 | 0.227835 | 0.999971 | 1 | 0.828598 |
| Fam217b | A2AJW5 | -0.24 | 0.06 | 0.41 | 0.12 | 1.48 | 0.5 | -0.02 | 0.18 | -0.53 | 0.23 | -0.86 | -0.45 | 1.56 | 0.43 | -0.49 | 0.54 | 0.4475 | -0.49 | 0.4225 | 0.224551 | 0.106829 | 0.327111 | 0.999971 | 1 | 0.856472 |
| Fam219a | A2ANP1 | 0.37 | 1 | -0.16 | -0.6 | 0.18 | 0.99 | -0.12 | -0.34 | 0.48 | 0.62 | -0.23 | 0.13 | 1.94 | 0.92 | -0.48 | -0.31 | 0.025 | 0.0975 | 0.365 | 0.953266 | 0.798301 | 0.55157 | 0.999971 | 1 | 0.913379 |
| Fam234a | Q8C0Z1 | -0.62 | -0.13 | 0.14 | -0.57 | -0.13 | -0.42 | -0.48 | 0.2 | 0.04 | -0.99 | -0.76 | 0.04 | 0.86 | -1.11 | -0.02 | -0.15 | 0.0875 | -0.1225 | 0.19 | 0.739212 | 0.70697 | 0.653032 | 0.999971 | 1 | 0.934005 |
| Fam234b | Q8BYI8 | 0.48 | 1.22 | 1.57 | 1.07 | 1.01 | 1.35 | 0.67 | 1.35 | 0.76 | 0.88 | 1.21 | 0.99 | 1.17 | 1.52 | 1.45 | 1.72 | 0.01 | -0.125 | 0.38 | 0.972559 | 0.640278 | 0.187893 | 0.999971 | 1 | 0.809025 |
| Fam241b | D3Z7G3 | -0.97 | 0.27 | -0.02 | -0.81 | 0.53 | -0.31 | 1.17 | -0.79 | 0.19 | -1.39 | 0.2 | -0.39 | 1.26 | -0.63 | -0.32 | -1.55 | 0.5325 | 0.035 | 0.0725 | 0.295382 | 0.93824 | 0.903939 | 0.999971 | 1 | 0.991091 |
| Fam3c | G3X9I1 | 0.37 | 1.05 | 1.68 | 1.42 | 1.6 | 0.93 | 1.38 | 2.03 | 1.26 | 0.37 | 1.84 | 1.79 | 1.13 | 1.03 | 2.13 | 2 | 0.355 | 0.185 | 0.4425 | 0.334134 | 0.663037 | 0.273088 | 0.999971 | 1 | 0.843794 |
| Fam50a | Q9WV03 | 1.77 | 1.72 | 2.73 | 2.84 | 1.74 | 1.64 | 1.95 | 2.09 | 1 | 2.01 | 2.98 | 2.09 | 1.12 | 1.65 | 2.72 | 3.45 | -0.41 | -0.245 | -0.03 | 0.221684 | 0.607202 | 0.956922 | 0.999971 | 1 | 0.998513 |
| Fam76b | Q80XP8 | 0.79 | 2.97 | 2.85 | 0.05 | 3.63 | 1.08 | 4.05 | 0.28 | 4.09 | 0.91 | 3.01 | 2.96 | 2.76 | 0.12 | 0.65 | -0.6 | 0.595 | 1.0775 | -0.9325 | 0.576721 | 0.24781 | 0.327013 | 0.999971 | 1 | 0.856472 |
| Fam78b | F6ZXE6 | 0.36 | -1.67 | -1.04 | -0.62 | -0.68 | 0.11 | -1.65 | -0.38 | 1.13 | -1.06 | -1.06 | -1.45 | 0 | -0.94 | -0.26 | -1.19 | 0.0925 | 0.1325 | 0.145 | 0.8592 | 0.841442 | 0.761177 | 0.999971 | 1 | 0.961116 |
| Fam81a | Q3UXZ6 | -2 | -2 | -1.91 | -2.69 | -1.71 | -1.99 | -0.85 | -2.51 | -2.02 | -2.66 | -1.26 | -1.45 | -1.57 | -2.67 | -2.39 | -2.13 | 0.385 | 0.3025 | -0.04 | 0.323354 | 0.405962 | 0.895302 | 0.999971 | 1 | 0.988571 |
| Fam89b | Q05DI7 | 0.15 | 0.31 | 3.8 | -0.2 | 0.63 | -0.02 | 6.96 | 2.32 | 0.29 | 0.49 | 3.91 | 7.66 | 0.72 | 0.7 | 1.33 | -2.9 | 1.4575 | 2.0725 | -1.0525 | 0.38003 | 0.258048 | 0.389824 | 0.999971 | 1 | 0.871816 |

| | | | | | | | | | | | | | | | | | | | | | | | | | | |
|---|---|---|---|---|---|---|---|---|---|---|---|---|---|---|---|---|---|---|---|---|---|---|---|---|---|---|
| Fam8a1 | Q3URQ4 | 2.69 | -0.85 | 0.95 | 1.22 | -0.9 | -0.46 | 1.87 | 0.51 | -1.34 | 0.67 | 0.28 | -0.02 | -2.28 | -1.13 | 0.09 | 0.1 | -0.7475 | -1.105 | -1.8075 | 0.391106 | 0.175197 | 0.053994 | 0.999971 | 1 | 0.725348 |
| Fam91a1 | Q3UVG3 | -0.63 | -0.3 | 0.08 | -0.21 | 0.01 | -0.12 | 0.68 | -0.14 | -0.07 | -0.7 | -0.11 | -0.14 | -0.81 | -0.68 | -0.09 | -0.57 | 0.3725 | 0.01 | -0.2725 | 0.182784 | 0.966638 | 0.284292 | 0.999971 | 1 | 0.844538 |
| Fam98a | Q3TJZ6 | 0.76 | -0.3 | 0.62 | 0.01 | -0.08 | -0.33 | 0.52 | 0.55 | -0.13 | 0.32 | 0.85 | 0.74 | -0.19 | -0.01 | 0.39 | 0.74 | -0.1075 | 0.1725 | -0.04 | 0.746735 | 0.608033 | 0.902572 | 0.999971 | 1 | 0.990759 |
| Fam98b | Q80VD1 | 1.32 | 1.53 | 1.2 | 0.9 | 1.16 | 1.47 | 0.89 | 1.26 | 0.89 | 1.27 | 0.69 | 0.89 | 0.24 | 1.7 | 1.12 | 0.8 | -0.0425 | -0.3025 | -0.2725 | 0.847818 | 0.192143 | 0.420843 | 0.999971 | 1 | 0.87788 |
| Fam98c | E9PYD1 | 2.13 | 0.88 | 1.76 | 1.79 | 0.75 | 0.96 | 1.75 | 1.96 | 0.74 | 1.5 | 1.35 | -0.12 | -0.17 | 0.94 | 1.51 | 1.55 | -0.285 | -0.7725 | -0.6825 | 0.464211 | 0.10097 | 0.157423 | 0.999971 | 1 | 0.797252 |
| Far1 | Q922J9 | -0.55 | -0.94 | -1.54 | -1.21 | -0.45 | -1.09 | -1.73 | -0.97 | -0.58 | -0.42 | -1.27 | -1.12 | 0.6 | -0.42 | -1.27 | -0.38 | -1.18E-16 | 0.2125 | 0.6925 | 1 | 0.486517 | 0.122544 | 1 | 1 | 0.78268 |
| Farp1 | F8VPU2 | -0.18 | -0.42 | -0.76 | -0.85 | -0.37 | -0.32 | -0.65 | -0.77 | 0 | -0.18 | -0.59 | -0.52 | 0.16 | -0.45 | -1.03 | -0.57 | 0.025 | 0.23 | 0.08 | 0.912726 | 0.349626 | 0.789574 | 0.999971 | 1 | 0.96775 |
| Farp2 | Q91VS8 | -0.91 | -0.99 | -1.57 | -1.87 | -0.81 | -1.1 | -1.08 | -2.09 | -0.8 | -0.42 | -1.56 | -0.75 | -0.32 | -0.87 | -1.74 | -1.35 | 0.065 | 0.4525 | 0.265 | 0.855186 | 0.197073 | 0.483984 | 0.999971 | 1 | 0.896352 |
| Fars2 | Q99M01 | 1.08 | -2.32 | -0.42 | -0.28 | -2.44 | -1.89 | 0.16 | -0.3 | -2.51 | 0.48 | -0.3 | -0.33 | -2.75 | -1.8 | -0.43 | -0.23 | -0.6325 | -0.18 | -0.8175 | 0.458484 | 0.833779 | 0.335966 | 0.999971 | 1 | 0.858827 |
| Farsa | Q8C0C7 | 1.34 | 1.71 | 1.83 | 1.53 | 1.61 | 1.6 | 2.11 | 1.68 | 1.68 | 1.83 | 1.95 | 2.06 | 1.51 | 1.76 | 1.76 | 2.04 | 0.1475 | 0.2775 | 0.165 | 0.491458 | 0.176067 | 0.433035 | 0.999971 | 1 | 0.882137 |
| Farsb | Q9WUA2 | 0.68 | 2.28 | 1.85 | 2 | 2.27 | 1.86 | 2.14 | 1.87 | 2.54 | 1.15 | 2.16 | 2.44 | 2.57 | 2.25 | 1.8 | 2.15 | 0.3325 | 0.37 | 0.49 | 0.365531 | 0.418037 | 0.21339 | 0.999971 | 1 | 0.82451 |
| Fasn | P19096 | -0.26 | -0.17 | -0.05 | -0.1 | -0.33 | -0.14 | 0.19 | -0.36 | -0.13 | -0.21 | -0.11 | -0.08 | -0.12 | -0.16 | -0.08 | -0.18 | -0.015 | 0.0125 | 0.01 | 0.938971 | 0.937552 | 0.951228 | 0.999971 | 1 | 0.998056 |
| Fastkd2 | Q922E6 | 0.18 | -0.34 | 0.13 | -0.17 | -0.44 | -0.62 | -0.41 | -0.16 | -0.67 | 0.35 | -0.41 | -0.21 | -1.26 | -0.73 | -0.34 | -0.05 | -0.3575 | -0.185 | -0.545 | 0.112693 | 0.49596 | 0.095814 | 0.999971 | 1 | 0.78268 |
| Fastkd5 | Q7TMV3 | -1.77 | 0.09 | -0.25 | -0.34 | -0.22 | -0.83 | -0.13 | -0.24 | -0.4 | -0.83 | -0.3 | -0.18 | 0.29 | -0.66 | -0.63 | -1 | 0.2125 | 0.14 | 0.0675 | 0.614321 | 0.7364 | 0.88438 | 0.999971 | 1 | 0.98711 |
| Fat1 | F2Z4A3 | -0.23 | -1.09 | -0.77 | -0.79 | -1.16 | -1.19 | -0.21 | -0.95 | -1.31 | -0.63 | -1.02 | -0.94 | -1.23 | -1.17 | -0.7 | -0.56 | -0.1575 | -0.255 | -0.195 | 0.601377 | 0.326214 | 0.471581 | 0.999971 | 1 | 0.892857 |
| Fat3 | E9QK16 | -0.4 | -0.91 | -0.65 | -0.54 | -1.22 | -1.15 | -0.97 | -0.71 | -1.19 | -0.73 | -0.57 | -0.66 | -0.81 | -1.09 | -0.78 | -0.42 | -0.3875 | -0.1625 | -0.15 | 0.090185 | 0.462032 | 0.500825 | 0.999971 | 1 | 0.904223 |
| Fat4 | Q2PZL6 | 0.54 | -0.87 | -1.08 | 0.27 | -0.59 | -0.97 | -0.92 | 0.84 | -0.51 | 0.09 | -1.6 | -1.42 | -0.52 | -0.74 | -0.93 | -0.67 | -0.125 | -0.575 | -0.43 | 0.816997 | 0.290839 | 0.294926 | 0.999971 | 1 | 0.847402 |
| Faxc | Q3UMF9 | -0.84 | -0.65 | 1.07 | 1.07 | -1.17 | -0.79 | 0.5 | 0.56 | -0.83 | 0.18 | 1.33 | 1.68 | 0.23 | -0.47 | 0.47 | 1.38 | -0.3875 | 0.4275 | 0.24 | 0.539791 | 0.549212 | 0.687508 | 0.999971 | 1 | 0.945573 |
| Fbh1 | Q8K2I9 | 1.07 | 2.52 | 2.19 | 1.97 | 2.33 | 2.2 | 1.4 | 1.03 | 2.27 | 0.98 | 0.96 | -0.03 | 1.26 | 1.17 | 1.44 | 1.23 | -0.1975 | -0.8925 | -0.6625 | 0.639019 | 0.116155 | 0.063635 | 0.999971 | 1 | 0.725348 |
| Fbl | P35550 | -1.38 | -1.15 | -2.3 | -2.44 | -0.83 | -1.48 | -1.97 | -2.63 | -0.73 | -1.26 | -2.03 | -2.08 | -0.44 | -1.27 | -2.31 | -2.77 | 0.09 | 0.2925 | 0.12 | 0.847167 | 0.50633 | 0.831941 | 0.999971 | 1 | 0.979036 |
| Fbll1 | Q80WS3 | -0.8 | -1.24 | -1.49 | -1.6 | -0.77 | -1.49 | -0.95 | -1.82 | -0.84 | -0.98 | -1.69 | -1.73 | -0.66 | -1.16 | -1.61 | -1.7 | 0.025 | -0.0275 | -7.85E-17 | 0.934929 | 0.926991 | 1 | 0.999971 | 1 | 1 |
| Fbln1 | Q08879-2 | 2.2 | 1.57 | 0.98 | 1.32 | 1.96 | 1.62 | 0.85 | 0.81 | 1.96 | 2.94 | 1.39 | 1.29 | 1.52 | 1.55 | 1.21 | 1.45 | -0.2075 | 0.3775 | -0.085 | 0.581014 | 0.394739 | 0.764985 | 0.999971 | 1 | 0.962405 |
| Fbln5 | A0A1Y7VJV | 0.19 | -1.7 | -2.45 | 0.53 | 6.13 | 1.44 | 0.95 | -0.95 | 4.02 | 4.91 | 6.09 | 7.93 | -3.03 | 0.57 | 0.01 | -0.22 | 2.75 | 6.595 | 0.19 | 0.090188 | 0.000129 | 0.844351 | 0.999971 | 0.19604 | 0.979814 |
| Fbn1 | Q61554 | 0.74 | 0.29 | 0.49 | 1.54 | 4.96 | 0.43 | 0.7 | 0.06 | 1.3 | 3.14 | 3.69 | 7.02 | -2.73 | 0.7 | 0.85 | 0.16 | 0.7725 | 3.0225 | -1.02 | 0.469745 | 0.021493 | 0.220688 | 0.999971 | 1 | 0.827096 |
| Fbn2 | Q61555 | 3.69 | 3.8 | -0.14 | 0.97 | 5.95 | 2.77 | -0.18 | -0.96 | 2.74 | 4.6 | 0.97 | 3.37 | -3.19 | 5.18 | 1.86 | -0.71 | -0.185 | 0.84 | -1.295 | 0.910331 | 0.459195 | 0.481563 | 0.999971 | 1 | 0.89485 |
| Fbxl15 | Q91W61 | -0.27 | 0.51 | 1.56 | 1.58 | 0.11 | 0.32 | 0.95 | 1.49 | 0.34 | 0.17 | 1.55 | -0.34 | 0.51 | 0.95 | 1.57 | 1.46 | -0.1275 | -0.415 | 0.2775 | 0.80124 | 0.46179 | 0.564226 | 0.999971 | 1 | 0.917087 |
| Fbxl16 | A2RT62 | 0.21 | 1.18 | -0.33 | -0.37 | 0.96 | 0.94 | -0.32 | -0.46 | 0.9 | 0.41 | -0.38 | -0.1 | 0.83 | 0.97 | -0.48 | -0.38 | 0.1075 | 0.035 | 0.0625 | 0.827115 | 0.935725 | 0.898758 | 0.999971 | 1 | 0.989644 |
| Fbxl17 | Q9QZN1 | 1.22 | 1.52 | -1.62 | -1 | 0.82 | 1.57 | -0.89 | 0.34 | 0.72 | 1.29 | -1.17 | -1.33 | -0.28 | 1.4 | -1.53 | -1.45 | 0.43 | -0.1525 | -0.495 | 0.612686 | 0.869161 | 0.598717 | 0.999971 | 1 | 0.92377 |
| Fbxl18 | F8VQK6 | 0.36 | 0.62 | 0.8 | 0.93 | 0.64 | 0.48 | 0.2 | 0.69 | 0.83 | 0.6 | 0.74 | 0.53 | 0.88 | 0.41 | 0.43 | 1.05 | -0.175 | -0.0025 | 0.015 | 0.421827 | 0.989854 | 0.949674 | 0.999971 | 1 | 0.998038 |
| Fbxl2 | Q8BH16 | -0.8 | 1.53 | -0.36 | -2.31 | 0.53 | 1.25 | -0.34 | -0.06 | 0.92 | -1.03 | 0.74 | -0.15 | 1.23 | 0.76 | -1.75 | -1.66 | 0.83 | 0.605 | 0.13 | 0.302397 | 0.469013 | 0.896137 | 0.999971 | 1 | 0.988571 |
| Fbxl20 | Q9CZV8 | 0.32 | 1.56 | 0.48 | 0.34 | 1.27 | 1.67 | -0.12 | 0.49 | 1.58 | 0.96 | 0.18 | -0.56 | 1.47 | 0.91 | 0.37 | 0.53 | 0.1525 | -0.135 | 0.145 | 0.743781 | 0.792726 | 0.698996 | 0.999971 | 1 | 0.947812 |
| Fbxl4 | Q8BH70 | 1.48 | 1.63 | -0.14 | 0.38 | 0.81 | 0.28 | 0.92 | -0.73 | -0.19 | 0.9 | 0.08 | -0.5 | -0.52 | -0.07 | -0.74 | -0.51 | -0.5175 | -0.765 | -1.2975 | 0.339252 | 0.144399 | 0.013559 | 0.999971 | 1 | 0.613095 |
| Fbxo10 | Q7TQF2 | 0.75 | 0.17 | 0.27 | 0.26 | 0.08 | 0.6 | 0.81 | 0.61 | 0.48 | 0.51 | 0.02 | 0.41 | 0.15 | 0.54 | 0.38 | 0.54 | 0.1625 | -0.0075 | 0.04 | 0.500021 | 0.972303 | 0.849818 | 0.999971 | 1 | 0.98187 |
| Fbxo11 | Q7TPD1-3 | 1.99 | 0.5 | 0.78 | 1.37 | 0.27 | 0.84 | 0.48 | 1 | 0.05 | 1.68 | 0.56 | 0.37 | -0.77 | 0.67 | 0.78 | 1.27 | -0.5125 | -0.495 | -0.6725 | 0.180177 | 0.294822 | 0.209463 | 0.999971 | 1 | 0.822138 |
| Fbxo2 | Q80UW2 | -0.12 | 1.01 | -0.79 | -0.9 | 0.4 | 1.05 | -1.19 | -0.59 | 0.45 | -0.18 | -0.97 | -1.23 | 1.16 | 0.9 | -0.65 | -1 | 0.1175 | -0.2825 | 0.3025 | 0.846345 | 0.602844 | 0.636132 | 0.999971 | 1 | 0.93301 |
| Fbxo21 | Q8VDH1 | -0.49 | 0.43 | 0.36 | -0.2 | -0.11 | -0.18 | -0.32 | -0.07 | -0.05 | -0.63 | 0.28 | -0.22 | -0.63 | -0.38 | 0.25 | -0.15 | -0.195 | -0.18 | -0.2525 | 0.45153 | 0.55211 | 0.409259 | 0.999971 | 1 | 0.875789 |
| Fbxo22 | Q78JE5 | 2 | 2.32 | 1.48 | 1.5 | 2.13 | 1.43 | 1.26 | 1.37 | 2.02 | 1.76 | 1.46 | 1.44 | 1.61 | 1.68 | 1.08 | 1.18 | -0.2775 | -0.155 | -0.4375 | 0.358215 | 0.563443 | 0.135476 | 0.999971 | 1 | 0.784948 |
| Fbxo3 | Q9DC63-3 | -0.63 | 0.73 | -0.05 | -0.32 | 0.36 | 0.13 | 0.5 | 0.05 | 0.63 | -0.37 | 0.43 | 0.56 | 0.59 | -0.31 | 0.4 | -0.13 | 0.3275 | 0.38 | 0.205 | 0.311 | 0.310794 | 0.566684 | 0.999971 | 1 | 0.917976 |
| Fbxo30 | Q8BJL1 | -0.14 | -1.1 | -0.22 | -0.22 | -0.29 | -0.71 | -0.17 | 0.26 | -0.68 | -0.16 | -0.37 | -0.6 | -0.46 | -0.46 | -0.09 | 0.03 | 0.1925 | -0.0325 | 0.175 | 0.536874 | 0.90511 | 0.533024 | 0.999971 | 1 | 0.910069 |
| Fbxo31 | Q3TQF0 | 2.3 | -0.07 | -1.07 | -0.7 | -0.16 | 0.59 | -1.41 | -0.88 | -0.38 | 1.56 | -0.48 | -0.79 | -0.63 | 0.18 | -1.16 | -0.53 | -0.58 | -0.1375 | -0.65 | 0.467068 | 0.869546 | 0.383178 | 0.999971 | 1 | 0.871816 |
| Fbxo38 | Q8BMI0 | 1.78 | 0.06 | 0.03 | 0.06 | -0.4 | 0.27 | 0.69 | 0.54 | -0.09 | 1.42 | 0.08 | 0.15 | 0.31 | 0.2 | 0.4 | 0.47 | -0.2075 | -0.0925 | -0.1375 | 0.655847 | 0.857529 | 0.741327 | 0.999971 | 1 | 0.96105 |
| Fbxo4 | Q8CHQ0 | -0.33 | 0.51 | -0.21 | -0.42 | -0.16 | -0.09 | -0.57 | -0.57 | 0.08 | -0.84 | -0.13 | 0.16 | -0.3 | -0.12 | 0.02 | -0.43 | -0.235 | -0.07 | -0.095 | 0.391615 | 0.823857 | 0.714449 | 0.999971 | 1 | 0.952364 |
| Fbxo41 | Q6NS60 | -0.65 | -1.13 | -0.93 | -0.84 | -1.44 | -1.07 | -0.89 | -0.95 | -1.23 | -0.76 | -0.85 | -1.01 | -0.73 | -0.97 | -0.88 | -0.43 | -0.2 | -0.075 | 0.135 | 0.352627 | 0.706239 | 0.521371 | 0.999971 | 1 | 0.907001 |
| Fbxo42 | Q6PDJ6 | 0.6 | 1.14 | 0.87 | 0.46 | 1.28 | 1.14 | 1.09 | 0.49 | 0.95 | 1.41 | 0.78 | -0.23 | 1.39 | 1.02 | 0.48 | 0.31 | 0.2325 | -0.04 | 0.0325 | 0.375049 | 0.913219 | 0.91357 | 0.999971 | 1 | 0.992621 |
| Fbxo44 | A2A7H5 | -0.93 | 1.56 | -0.4 | -0.73 | 0.43 | 1.2 | -0.73 | -0.36 | 1.49 | -0.59 | 0.12 | 0 | 1.53 | 2.37 | 0.05 | -0.18 | 0.26 | 0.38 | 1.0675 | 0.690034 | 0.567841 | 0.180982 | 0.999971 | 1 | 0.808532 |
| Fbxo45 | Q8K3B1 | 0.18 | -1.33 | -1.1 | -1.01 | -0.61 | -0.96 | -1.4 | -1.17 | -0.76 | 0.15 | -1.21 | -1.08 | -1.12 | -0.76 | -1.33 | -0.95 | -0.22 | 0.09 | -0.225 | 0.553029 | 0.835319 | 0.529205 | 0.999971 | 1 | 0.910069 |
| Fbxo6 | Q9QZN4 | -0.07 | -0.1 | 0.09 | -0.21 | -0.53 | 0.52 | -0.14 | 0.33 | -0.08 | -0.63 | 0.16 | -0.55 | 0.18 | 0.71 | 0.34 | 0.34 | 0.1175 | -0.2025 | 0.465 | 0.659911 | 0.394802 | 0.037309 | 0.999971 | 1 | 0.700096 |
| Fbxo7 | Q3U7U3 | 1.31 | 0.26 | 0.32 | 0.73 | 0.03 | -0.28 | 0.54 | 0.22 | 0.22 | 0.55 | 0.63 | 0.24 | -0.23 | 0.15 | 0.4 | 0.59 | -0.5275 | -0.245 | -0.4275 | 0.10991 | 0.390038 | 0.18754 | 0.999971 | 1 | 0.809025 |
| Fbxo9 | Q8BK06-3 | 3.53 | 2.94 | -0.19 | -0.33 | 1.82 | 1.99 | -0.48 | -0.06 | 2.32 | 4.76 | -1.76 | 0.06 | 0.28 | -0.25 | -1.24 | 0.3 | -0.67 | -0.1425 | -1.715 | 0.534825 | 0.926849 | 0.103062 | 0.999971 | 1 | 0.78268 |
| Fbxw11 | Q5SRY7-4 | 1.49 | 0.96 | 1.68 | 1.45 | 0.6 | 0.77 | 1 | 1.54 | 0.85 | 1.15 | 1.39 | 0.81 | 0.42 | 0.11 | 1.39 | 1.37 | -0.4175 | -0.345 | -0.5725 | 0.152829 | 0.170498 | 0.134222 | 0.999971 | 1 | 0.784948 |
| Fbxw2 | Q8BS25 | 1.75 | 1.86 | 0.73 | 0.63 | 1.08 | 2.02 | 2.2 | 0.36 | 0.37 | 0.57 | 1.03 | -0.78 | 1.3 | 1.8 | -0.39 | -0.38 | 0.1725 | -0.945 | -0.66 | 0.730547 | 0.072589 | 0.287421 | 0.999971 | 1 | 0.84693 |
| Fbxw5 | Q9QXW2-2 | 0.1 | -0.01 | 0.13 | -0.19 | -0.63 | -0.73 | 0.72 | 0.09 | 0.11 | -0.2 | 1.68 | 1.94 | 0.08 | 0.14 | 0.08 | -0.09 | -0.145 | 0.875 | 0.045 | 0.673513 | 0.113152 | 0.797631 | 0.999971 | 1 | 0.970479 |
| Fbxw8 | Q8BIA4-4 | -1.36 | -1.54 | -1.03 | -0.85 | -0.9 | -1.82 | -1.62 | -1.15 | -0.62 | -2.07 | -0.92 | -1.38 | -2.23 | -0.73 | 0.05 | -0.8 | -0.1775 | -0.0525 | 0.2675 | 0.528696 | 0.879664 | 0.571177 | 0.999971 | 1 | 0.91816 |
| Fcer1g | P20491 | 0.5 | 0.2 | 0.08 | 0.14 | 0.13 | 0.03 | -0.24 | -0.26 | -0.02 | -0.1 | 0.84 | 0.74 | -0.1 | 0.19 | 0.28 | 0.08 | -0.315 | 0.135 | -0.1175 | 0.135224 | 0.630062 | 0.543798 | 0.999971 | 1 | 0.911398 |
| Fcf1 | E9Q186 | 0.25 | -0.99 | 0.44 | 0.51 | -2.45 | 0.01 | -0.61 | 1.72 | -1.06 | -0.55 | 0.8 | 0.66 | -2.38 | 0.22 | 1.97 | -0.69 | -0.385 | -0.09 | -0.2725 | 0.646226 | 0.866027 | 0.755429 | 0.999971 | 1 | 0.961116 |
| Fcgr1 | P26151 | 1.13 | 1.46 | 1.86 | 1.31 | -0.01 | 0.87 | 0.99 | 1.21 | 0.28 | 0.1 | 1.78 | 1.41 | 0.76 | 1.53 | 1.91 | 2.42 | -0.675 | -0.5475 | 0.215 | 0.056084 | 0.215737 | 0.566827 | 0.999971 | 1 | 0.917976 |
| Fcgr2b | A0A0B4J1G | 1.68 | 0.07 | 0.76 | 1.57 | 0.9 | 0.49 | 0.25 | 1.58 | 1.11 | 0.68 | -0.19 | -0.08 | 0.2 | 1.1 | 0.63 | 0.83 | -0.215 | -0.64 | -0.33 | 0.633138 | 0.188154 | 0.423066 | 0.999971 | 1 | 0.87887 |
| Fcgrt | Q61559 | -2.64 | -2.48 | -0.48 | -0.14 | -2.84 | -3.2 | -0.83 | -0.44 | -3.18 | -2.35 | -0.45 | -0.6 | -5.28 | -2.92 | -0.25 | -0.09 | -0.3925 | -0.21 | -0.7 | 0.648579 | 0.803948 | 0.576462 | 0.999971 | 1 | 0.918194 |
| Fcho1 | Q8K285 | -0.2 | -0.22 | -0.48 | -0.66 | -0.21 | -0.36 | -0.88 | -0.23 | -0.23 | -0.1 | -0.12 | -0.44 | -0.19 | -0.22 | -0.77 | -0.19 | -0.03 | 0.1675 | 0.0475 | 0.89613 | 0.397548 | 0.831644 | 0.999971 | 1 | 0.978898 |
| Fcho2 | Q3UQN2 | -0.12 | -0.16 | -0.24 | -0.74 | 0.21 | -0.11 | 0.1 | -0.32 | 0.53 | -0.2 | -0.34 | 0.16 | 0.26 | -0.48 | -0.37 | -1.11 | 0.285 | 0.3525 | -0.11 | 0.228174 | 0.202912 | 0.73069 | 0.999971 | 1 | 0.956041 |
| Fchsd2 | Q3USJ8 | -0.2 | -0.23 | -0.63 | -0.34 | -0.11 | -0.29 | -0.79 | -0.06 | -0.08 | -0.01 | -0.93 | -0.78 | -0.12 | -0.09 | -0.36 | -0.48 | 0.0375 | -0.1 | 0.0875 | 0.87097 | 0.714645 | 0.659699 | 0.999971 | 1 | 0.935181 |
| Fcsk | Q7TMC8 | -0.45 | -0.06 | 0.37 | 0.15 | -0.3 | -0.31 | 0.24 | 0.44 | -0.43 | -0.32 | 0.47 | 0.31 | -0.74 | -0.44 | 0.43 | 0.24 | 0.015 | 0.005 | -0.13 | 0.956551 | 0.986408 | 0.692898 | 0.999971 | 1 | 0.947174 |
| Fdft1 | P53798 | -0.29 | -0.32 | -0.43 | -0.38 | -0.48 | -0.24 | -0.55 | 0.1 | -0.62 | -0.57 | 0.03 | -0.59 | 0.19 | -0.27 | -0.21 | -0.13 | 0.0625 | -0.0825 | 0.25 | 0.76046 | 0.692126 | 0.196086 | 0.999971 | 1 | 0.812331 |
| Fdps | Q920E5 | 0.06 | 0.7 | -0.89 | -0.75 | 0.15 | 1.13 | -0.16 | -0.33 | 1.08 | 0.49 | -0.15 | -1.63 | 1.43 | 1.62 | 0.05 | -0.39 | 0.4175 | 0.1675 | 0.8975 | 0.378022 | 0.79132 | 0.142368 | 0.999971 | 1 | 0.784967 |
| Fdx1 | P46656 | 0.15 | 1.3 | 1.65 | 0.91 | 0.96 | 2.27 | 2.77 | 0.81 | 1.59 | 0.56 | 1.53 | 1.73 | 1.95 | 2.06 | 1.22 | 0.22 | 0.7 | 0.35 | 0.36 | 0.212999 | 0.393698 | 0.473938 | 0.999971 | 1 | 0.894044 |
| Fdx2 | Q9CPW2 | -0.89 | -0.26 | -0.6 | -0.52 | -0.55 | -1.01 | -1 | -0.54 | -0.64 | -0.79 | -0.64 | -0.59 | 0.23 | -0.57 | -0.82 | -0.49 | -0.2075 | -0.0975 | 0.155 | 0.37032 | 0.618757 | 0.579641 | 0.999971 | 1 | 0.919556 |
| Fdxr | Q61578 | -0.99 | -1.82 | -0.84 | -0.65 | -1.55 | -1.52 | -1.25 | -0.88 | -1.58 | -1.27 | -0.83 | -1.05 | -1.88 | -1.51 | -1.15 | -0.71 | -0.225 | -0.1075 | -0.2375 | 0.470792 | 0.728302 | 0.506863 | 0.999971 | 1 | 0.905775 |
| Fech | P22315 | -0.8 | -0.19 | -0.48 | -1.07 | -0.3 | -0.56 | -0.28 | -0.9 | -0.26 | -1 | -0.44 | -0.38 | -0.17 | -0.68 | -1.23 | -1.18 | 0.125 | 0.115 | -0.18 | 0.635749 | 0.671512 | 0.573906 | 0.999971 | 1 | 0.91816 |
| Fem1aa | Q9Z2G1 | 1.1 | 1.37 | 0.73 | 0.94 | 1.56 | 1.59 | 0.15 | 0.98 | 1.28 | 1.34 | 1.69 | 0.75 | 1.84 | 1.77 | 0.52 | 1.57 | 0.035 | 0.23 | 0.39 | 0.921518 | 0.384823 | 0.262234 | 0.999971 | 1 | 0.842262 |
| Fem1b | Q9Z2G0 | 1.06 | 0.79 | 1.15 | 1.19 | 1.09 | 1.7 | 0.1 | 1.01 | 1.16 | 2.29 | -0.05 | 0.46 | -0.98 | 0.94 | 0.82 | 1.28 | -0.0725 | -0.0825 | -0.5325 | 0.830582 | 0.863887 | 0.285615 | 0.999971 | 1 | 0.845133 |
| Fem1c | Q8CEF1 | -0.08 | 0.82 | 0.76 | 0.6 | 1.1 | -0.58 | 0.12 | 0.6 | 0.91 | -0.06 | -0.3 | 0.42 | -2.65 | 0.59 | -0.04 | -0.93 | -0.215 | -0.2825 | -1.2825 | 0.59003 | 0.410636 | 0.081247 | 0.999971 | 1 | 0.755254 |
| Fer | P70451-2 | 0.37 | 0.2 | 0.32 | 0.41 | 0.55 | 0.21 | -0.16 | 0.59 | 0.56 | 0.51 | 0.63 | 0.65 | 0.7 | 0.52 | 0.36 | 0.72 | -0.0275 | 0.2625 | 0.25 | 0.90171 | 0.128693 | 0.185607 | 0.999971 | 1 | 0.809025 |
| Fermt2 | Q8CIB5 | 0.16 | 0.44 | 0.14 | 0.05 | 0.57 | 0.69 | 0.06 | -0.35 | 0.59 | 0.61 | 0.21 | 0.23 | 0.31 | 0.54 | 0.06 | -0.06 | 0.045 | 0.2125 | 0.015 | 0.868919 | 0.293859 | 0.942975 | 0.999971 | 1 | 0.99692 |
| Fermt2_1 | A6X940 | -0.37 | -0.67 | -0.6 | 0.96 | -1.17 | -0.99 | -0.69 | -0.96 | -0.37 | 0.51 | -0.3 | -0.25 | 0.08 | -0.14 | -1.16 | 1.13 | -0.7825 | 0.0675 | 0.1475 | 0.065744 | 0.870416 | 0.791271 | 0.999971 | 1 | 0.968737 |
| Fermt3 | Q8K1B8 | 0.4 | 0.87 | 0.86 | 0.91 | 1.24 | 0.38 | 1.07 | 0.63 | 0.61 | 0.46 | 1.33 | 1.36 | 0.73 | 0.71 | 1.17 | 1.3 | 0.07 | 0.18 | 0.2175 | 0.78513 | 0.524282 | 0.359892 | 0.999971 | 1 | 0.869526 |
| Fez1 | Q8K0X8 | -0.29 | -0.46 | 0.33 | 0.14 | -0.56 | -0.04 | 0.09 | 0.22 | -0.23 | 0.28 | 0.26 | 0.49 | -0.35 | -0.43 | -0.06 | 0.2 | -0.0025 | 0.27 | -0.09 | 0.992608 | 0.314807 | 0.728365 | 0.999971 | 1 | 0.956041 |
| Fga | E9PV24 | 3.15 | 1.66 | 0.45 | 3.77 | 6.92 | 0.37 | 2.4 | 0.37 | 1.19 | 1.37 | 1.63 | 1.96 | 0.94 | 0.37 | 0.92 | 0.54 | 0.2575 | -0.72 | -1.565 | 0.865458 | 0.317236 | 0.046525 | 0.999971 | 1 | 0.710102 |
| Fgb | Q8K0E8 | 2.56 | 1.66 | 0.37 | 2.84 | 6.11 | 0.21 | 1.51 | -0.12 | 1.05 | 0.93 | 1.31 | 1.65 | 1.14 | 0.37 | 0.63 | 0.65 | 0.07 | -0.6225 | -1.16 | 0.959134 | 0.264164 | 0.054549 | 0.999971 | 1 | 0.725348 |
| Fgd1 | P52734 | -0.38 | 0.18 | -0.37 | -0.42 | -0.24 | 0.56 | -0.71 | 0.25 | 0.05 | 0.02 | -0.24 | -0.54 | -0.23 | -0.17 | -0.48 | 0.07 | 0.2125 | 0.07 | 0.045 | 0.50749 | 0.763463 | 0.841151 | 0.999971 | 1 | 0.979814 |
| Fgd2 | Q8BY35 | -2.04 | 0.49 | 0.7 | 0.61 | 0.62 | 0.38 | 0.06 | 0.33 | 0.82 | -0.87 | 0.85 | 1.23 | 0.75 | 0.75 | 0.48 | 0.51 | 0.4075 | 0.5675 | 0.6825 | 0.510639 | 0.448697 | 0.279165 | 0.999971 | 1 | 0.843812 |
| Fgd4 | Q91ZT5 | -0.48 | -0.41 | -0.53 | -0.46 | -0.16 | -0.12 | -0.57 | -0.48 | -0.13 | -0.31 | -0.64 | -0.38 | -0.04 | -0.49 | -0.64 | -0.46 | 0.1375 | 0.105 | 0.0625 | 0.468366 | 0.563909 | 0.749175 | 0.999971 | 1 | 0.961064 |
| Fgd6 | Q69ZL1 | 0.92 | 0.06 | 0.12 | 0.62 | -0.3 | 0.16 | 0.09 | 0.27 | -0.63 | 0.42 | -0.32 | -0.01 | 0.24 | -0.2 | 0.26 | 0.87 | -0.375 | -0.565 | -0.1375 | 0.177824 | 0.095388 | 0.656937 | 0.999971 | 1 | 0.934351 |
| Fgf1 | P61148 | -0.82 | -1.26 | -1.21 | -0.99 | -1 | -0.68 | -0.87 | -0.54 | -1.04 | -0.94 | -0.93 | -1.1 | -1.06 | -0.57 | -0.84 | -0.73 | 0.2975 | 0.0675 | 0.27 | 0.164369 | 0.709811 | 0.204845 | 0.999971 | 1 | 0.81617 |
| Fgf12 | P61329 | -0.14 | -0.33 | 0.09 | -0.23 | -0.4 | -0.09 | 0.04 | -0.25 | -0.29 | -0.4 | -0.17 | 0.01 | -0.36 | 0.01 | -0.23 | -0.12 | -0.0225 | -0.06 | -0.0225 | 0.907781 | 0.750879 | 0.90527 | 0.999971 | 1 | 0.991091 |
| Fgf13 | P70377 | -1.31 | 0.55 | 0.49 | 0.16 | 0.42 | 1.11 | 0.05 | 0.95 | 0.51 | -1.2 | 1.28 | 0.88 | 0.91 | 0.46 | 0.92 | 1.48 | 0.66 | 0.395 | 0.97 | 0.181606 | 0.540819 | 0.057861 | 0.999971 | 1 | 0.725348 |
| Fgf13_1 | P70377-3 | 0.03 | -0.89 | 0.07 | -0.3 | -0.07 | -0.92 | -1.5 | 0.17 | -0.49 | -0.43 | -1.24 | -1.73 | 0.02 | -0.31 | -1.51 | -0.95 | -0.3075 | -0.7 | -0.415 | 0.472673 | 0.087796 | 0.304291 | 0.999971 | 1 | 0.847402 |
| Fgf14 | O89096 | 1.65 | 1.07 | 0.04 | 0.1 | 0.66 | 1.31 | -0.1 | 0.02 | 0.68 | 0.55 | 0.03 | 0.16 | 1.99 | 1.07 | 0.64 | 0.34 | -0.2425 | -0.36 | 0.295 | 0.611636 | 0.382788 | 0.554393 | 0.999971 | 1 | 0.914357 |
| Fgfr1 | P16092 | -0.13 | -0.26 | 0.4 | 0.37 | -0.44 | -0.62 | -0.21 | 0.67 | -0.7 | -0.32 | 0.42 | -0.51 | -0.32 | -0.24 | 0.79 | 0.38 | -0.245 | -0.3725 | 0.0575 | 0.466756 | 0.241459 | 0.856277 | 0.999971 | 1 | 0.982514 |
| Fgfr1op2 | Q9CRA9-2 | -0.7 | -0.38 | -0.03 | 0.57 | -0.37 | -0.77 | -0.44 | 0.27 | -0.68 | -0.52 | -0.1 | 0.18 | -0.42 | -0.33 | -0.13 | -0.12 | -0.1925 | -0.145 | -0.115 | 0.578667 | 0.665286 | 0.696238 | 0.999971 | 1 | 0.947174 |
| Fgfr2 | E9Q7T0 | 0.18 | 0.1 | 0.05 | -1.61 | -0.01 | 0.14 | 0.12 | -0.81 | 0.73 | -0.23 | 0.54 | 1.07 | 1.65 | -0.82 | -0.07 | -1.63 | 0.18 | 0.8475 | 0.1025 | 0.694048 | 0.1045 | 0.890024 | 0.999971 | 1 | 0.98799 |
| Fgfr3 | Q61563 | -0.04 | 0.5 | -0.8 | 0.17 | 0.52 | 0.34 | -0.93 | -0.94 | 0.51 | 0.19 | -1.04 | -0.6 | 1.42 | 0.03 | -0.74 | -0.58 | -0.21 | -0.1925 | 0.075 | 0.644494 | 0.654076 | 0.885541 | 0.999971 | 1 | 0.98711 |
| Fgg | Q3UER8 | 2.24 | 1.22 | 0.52 | 4.15 | 6.84 | -0.18 | 2.48 | 0.49 | 0.89 | 0.57 | 1.59 | 1.91 | 0.37 | 0.17 | 0.66 | 0.7 | 0.375 | -0.7925 | -1.5575 | 0.810837 | 0.317513 | 0.055978 | 0.999971 | 1 | 0.725348 |
| Fggy | A2AJL3 | -0.33 | 0.34 | 0.51 | 0.99 | 0.62 | -0.08 | 0.94 | 0.75 | 0.53 | -0.63 | 0.31 | 0.21 | 0.85 | 0.7 | 0.28 | 0.32 | 0.18 | -0.2725 | 0.16 | 0.606378 | 0.461385 | 0.611222 | 0.999971 | 1 | 0.92673 |
| Fh | P97807-2 | -0.87 | -1.44 | -0.87 | -0.82 | -1.28 | -1.42 | -0.67 | -1.15 | -1.17 | -1.04 | -0.99 | -0.74 | -2.06 | -1.81 | -1.17 | -1.1 | -0.13 | 0.015 | -0.535 | 0.603291 | 0.944489 | 0.093471 | 0.999971 | 1 | 0.776565 |
| Fhdc1 | Q3ULZ2 | 2.22 | 0.32 | -0.86 | 0.03 | 0.08 | 0.66 | 0.09 | -0.08 | -0.15 | 1.56 | -0.89 | -0.34 | 0.67 | -0.18 | 0.49 | -0.22 | -0.24 | -0.3825 | -0.2375 | 0.694556 | 0.616576 | 0.705502 | 0.999971 | 1 | 0.950182 |
| Fhip1b | Q3U2I3 | 0.08 | 0.8 | 0.98 | 0.93 | 0.43 | 0.47 | 0.56 | 0.74 | 0.38 | 0.53 | 0.73 | 0.69 | 0.5 | 0.71 | 0.72 | 1.01 | -0.1475 | -0.115 | 0.0375 | 0.557251 | 0.647285 | 0.884808 | 0.999971 | 1 | 0.98711 |
| Fhip2a | Q8CDM8 | -0.33 | 0.71 | 1.02 | 1.01 | 0.8 | 0.64 | 1.05 | 1.41 | 0.92 | -0.16 | 1.38 | 1.21 | 0.98 | 0.45 | 1.22 | 1.31 | 0.3725 | 0.235 | 0.3875 | 0.307433 | 0.599775 | 0.30323 | 0.999971 | 1 | 0.847402 |
| Fhip2b | Q80YR2 | -1.38 | 0.65 | 0.13 | -0.01 | 0.81 | 0.32 | -0.08 | 0.06 | 0.4 | -0.94 | 0.39 | 0.15 | 0.47 | 0.45 | 0.34 | 0.01 | 0.43 | 0.1525 | 0.47 | 0.348821 | 0.76125 | 0.284229 | 0.999971 | 1 | 0.844538 |
| Fhit | E9PZ91 | -0.88 | 0 | -0.2 | 0.01 | 0.01 | -0.07 | -0.01 | 0.45 | 0.1 | 0.16 | -0.14 | 0.06 | 0.43 | 0.48 | -0.12 | 0.2 | 0.3625 | 0.3125 | 0.515 | 0.192287 | 0.226815 | 0.083927 | 0.999971 | 1 | 0.755254 |
| Fhl1 | A2AEX8 | 0.63 | 0.88 | 0.38 | 0.5 | 1.41 | 1.64 | 2.12 | 1.01 | 2.42 | 1.2 | 1.04 | 0.29 | 1.61 | 1.26 | 0.94 | 0.11 | 0.9475 | 0.64 | 0.3825 | 0.006811 | 0.163047 | 0.273599 | 0.999971 | 1 | 0.843794 |
| Fhl1_1 | A2AEY2 | 3.52 | 0.75 | 1.24 | 1 | 1.5 | 1.38 | 1.44 | 0.17 | 0.79 | 3.14 | 0.42 | 0.36 | -0.65 | -0.21 | 0.16 | -0.15 | -0.505 | -0.45 | -1.84 | 0.443764 | 0.591134 | 0.013228 | 0.999971 | 1 | 0.613095 |
| Fhl2 | O70433 | 0.27 | 1.3 | 2.31 | 1.81 | 1.03 | 1.45 | 1.58 | 2.14 | 0.82 | 0.45 | 2.01 | -0.4 | 0.43 | 1.02 | 1.69 | 1.84 | 0.1275 | -0.7025 | -0.1775 | 0.782329 | 0.267123 | 0.725782 | 0.999971 | 1 | 0.956041 |
| Fhod1 | Q6P9Q4 | 2.25 | 0.66 | 0.59 | 1.21 | 0.88 | 0.61 | 0.23 | 0.54 | 1.26 | 1.58 | -0.64 | 0.97 | -4.99 | -0.27 | 0.7 | 1.08 | -0.6125 | -0.385 | -2.0475 | 0.142531 | 0.509041 | 0.139961 | 0.999971 | 1 | 0.784948 |
| Fhod3 | Q76LL6-3 | 0.29 | 0.78 | 0.31 | 0.78 | 0.81 | 0.94 | 0.02 | 0.39 | 0.59 | 0.16 | -0.01 | 0.25 | 0.27 | -0.03 | 0.49 | 1.13 | -7.85E-17 | -0.2925 | -0.075 | 1 | 0.216048 | 0.798666 | 1 | 1 | 0.970479 |
| Fibcd1 | A2AV25 | 0.04 | -0.94 | -0.1 | -0.17 | -0.95 | -0.44 | -0.3 | 0.44 | -0.8 | 0.1 | 0 | 0.05 | -0.82 | -0.35 | 0.32 | -0.21 | -0.02 | 0.13 | 0.0275 | 0.954951 | 0.677353 | 0.93224 | 0.999971 | 1 | 0.995069 |
| Fibp | Q8K2D8 | -0.72 | 0.41 | -0.37 | -0.71 | 0.48 | 0.38 | -0.72 | -0.28 | 0.52 | -0.42 | -0.28 | -0.2 | 1.49 | 0.4 | -0.22 | -0.2 | 0.3125 | 0.2525 | 0.715 | 0.415023 | 0.459763 | 0.140292 | 0.999971 | 1 | 0.784948 |
| Fig4 | Q91WF7 | -0.46 | -0.42 | -0.21 | -0.64 | -0.56 | 0.02 | -0.23 | 0.02 | -0.25 | -0.13 | -0.17 | 0.07 | -0.26 | -0.01 | 0.19 | -0.64 | 0.245 | 0.3125 | 0.2525 | 0.266858 | 0.112738 | 0.299827 | 0.999971 | 1 | 0.847402 |
| Filip1 | A0A5F8MP | 1.59 | -1.02 | -1.01 | -0.41 | 0.81 | -1.51 | -1.59 | -1.85 | 0.53 | 1.54 | 0.34 | 0.08 | -1.66 | 0.11 | -1.24 | -0.65 | -0.8225 | 0.835 | -0.6475 | 0.310955 | 0.21187 | 0.340671 | 0.999971 | 1 | 0.860574 |
| Fip1l1 | Q9D824-3 | 1.49 | 1.12 | 1.8 | 1.57 | 1.03 | 0.73 | 1.31 | 1.46 | 0.86 | 1.22 | 1.45 | 1.63 | 0.9 | 0.92 | 1.25 | 1.75 | -0.3625 | -0.205 | -0.29 | 0.163581 | 0.41364 | 0.291744 | 0.999971 | 1 | 0.847402 |
| Fis1 | Q9CQ92 | -1.87 | -1.32 | -1.03 | -0.77 | -1.24 | -1.34 | -0.94 | -1.15 | -1.07 | -1.99 | -0.99 | -1.11 | -1.37 | -1.09 | -0.77 | -1.29 | 0.08 | -0.0425 | 0.1175 | 0.767497 | 0.897937 | 0.681952 | 0.999971 | 1 | 0.943204 |
| Fitm2 | P59266 | 0.55 | 0.56 | 0.02 | -0.51 | 0.06 | 0.99 | -0.88 | 0.17 | 0.11 | 0.82 | 0.18 | -0.68 | 0.26 | 0.88 | -0.68 | 0.51 | -0.07 | -0.0475 | 0.0875 | 0.871819 | 0.90185 | 0.827562 | 0.999971 | 1 | 0.977122 |
| Fkbp15 | Q6P9Q6 | -0.19 | -0.59 | -0.13 | 0.11 | -0.35 | -0.35 | 0.24 | 0.15 | -0.33 | 0.11 | -0.03 | 0.21 | -0.72 | -0.31 | 0.23 | 0.23 | 0.1225 | 0.19 | 0.0575 | 0.619662 | 0.407772 | 0.841024 | 0.999971 | 1 | 0.979814 |
| Fkbp1a | P26883 | 0.47 | -0.28 | -0.21 | -0.1 | -0.52 | -0.37 | 0.2 | -0.08 | -0.52 | 0.17 | -0.04 | -0.15 | -0.91 | -0.2 | -0.13 | -0.18 | -0.1625 | -0.105 | -0.325 | 0.533457 | 0.675403 | 0.251129 | 0.999971 | 1 | 0.839012 |
| Fkbp1b | Q9Z2I2 | -0.9 | -0.15 | -1.44 | -1 | -0.19 | -0.15 | -1.27 | -0.91 | -0.63 | -0.08 | -0.88 | -1.16 | -0.23 | -0.34 | -1.37 | 0.34 | 0.2425 | 0.185 | 0.4725 | 0.520046 | 0.597377 | 0.281476 | 0.999971 | 1 | 0.844538 |
| Fkbp2 | P45878 | -1.16 | -1.63 | -2.13 | -2.73 | -1.37 | -1.71 | -0.89 | -2.8 | -1.14 | -1.66 | -1.95 | -1.88 | -1.88 | -1.77 | -2.07 | -2.66 | 0.22 | 0.255 | -0.1825 | 0.655069 | 0.499495 | 0.631834 | 0.999971 | 1 | 0.932858 |
| Fkbp3 | Q62446 | 1.07 | 2.15 | 1.62 | 1.23 | 2.25 | 1.88 | 1.9 | 1.83 | 2.56 | 1.64 | 2.32 | 2.61 | 2.57 | 2.17 | 1.87 | 2.04 | 0.4475 | 0.765 | 0.645 | 0.131505 | 0.042131 | 0.05252 | 0.999971 | 1 | 0.724862 |
| Fkbp4 | P30416 | -0.17 | 0 | 0.27 | -0.05 | -0.13 | 0.04 | 0.69 | -0.17 | 0.18 | -0.39 | -0.06 | 0.22 | -0.25 | 0.15 | 0.16 | -0.19 | 0.095 | -0.025 | -0.045 | 0.703374 | 0.906491 | 0.823413 | 0.999971 | 1 | 0.976225 |
| Fkbp5 | Q64378 | 0.73 | 1.26 | 1.62 | 1.27 | 1.72 | 1.07 | 1.81 | 1.67 | 1.65 | 1.18 | 2.02 | 1.9 | 1.22 | 0.99 | 1.73 | 1.45 | 0.3475 | 0.4675 | 0.1275 | 0.217838 | 0.118406 | 0.632283 | 0.999971 | 1 | 0.93301 |
| Fkbp8 | O35465-2 | -0.48 | 0.19 | 0.43 | 0.17 | 0.24 | 0.09 | 1.16 | 0.16 | 1.41 | -0.35 | 0.37 | 0.22 | 1.13 | -0.07 | 0.39 | 0.13 | 0.335 | 0.335 | 0.3175 | 0.310667 | 0.410621 | 0.346029 | 0.999971 | 1 | 0.863794 |
| Fkbp9 | Q9Z247 | -0.14 | 0.82 | -0.35 | 0.05 | 1.64 | 0.59 | -0.74 | 0.16 | 1.78 | 0.78 | -0.28 | -0.07 | 2.83 | 0.92 | 0.2 | 0.28 | 0.3175 | 0.4575 | 0.9625 | 0.541201 | 0.370091 | 0.138916 | 0.999971 | 1 | 0.784948 |
| Fkbpl | O35450 | 2.04 | 1.63 | 1.45 | 1.64 | 0.75 | 1.36 | 2.21 | 0.87 | 0.73 | 1.51 | 1.02 | 0.87 | 1.81 | 1.2 | 0.76 | 1.07 | -0.3925 | -0.6575 | -0.48 | 0.277526 | 0.021957 | 0.1053 | 0.999971 | 1 | 0.78268 |
| Flad1 | Q8R123 | -0.22 | 0.39 | 0.39 | 0.07 | 0.38 | 0.3 | 0.35 | 0.15 | 0.36 | -0.21 | 0.05 | -0.27 | 0.32 | 0.09 | 0.08 | -0.08 | 0.1375 | -0.175 | -0.055 | 0.513188 | 0.467464 | 0.799138 | 0.999971 | 1 | 0.970479 |
| Flcn | Q8QZS3 | 4.34 | 1.69 | 0.05 | 0.28 | 1.05 | 1.37 | 0.17 | 0.81 | 0.51 | 3.35 | 0.11 | 0.35 | 1.34 | 1.63 | 0.34 | 1.01 | -0.74 | -0.51 | -0.51 | 0.425306 | 0.649859 | 0.581545 | 0.999971 | 1 | 0.920364 |
| Flii | Q9JJ28 | 0.3 | -0.13 | -0.54 | -0.59 | -0.13 | -0.01 | -0.35 | -0.81 | -0.01 | -0.01 | -0.51 | -0.83 | 0.36 | -0.09 | -0.64 | -0.57 | -0.085 | -0.1 | 0.005 | 0.766405 | 0.737421 | 0.987368 | 0.999971 | 1 | 0.99925 |
| Flna | B7FAU9 | 0.34 | -0.65 | -0.91 | 0.19 | 4.48 | 1.45 | 0.11 | -0.87 | 0.85 | 3.58 | 2.71 | 4.45 | -1.6 | 0.3 | 0.16 | -0.09 | 1.55 | 3.155 | -0.05 | 0.173152 | 0.002435 | 0.919931 | 0.999971 | 0.821787 | 0.992702 |
| Flnb | Q80X90 | 0.82 | 0.57 | -0.03 | 0.59 | 1.41 | 0.45 | -0.02 | -0.31 | 0.59 | 0.91 | -0.04 | 0.51 | 0.19 | 0.85 | 0.43 | 0.32 | -0.105 | 0.005 | -0.04 | 0.792969 | 0.985828 | 0.876206 | 0.999971 | 1 | 0.985129 |
| Flnc | Q8VHX6-2 | 1.97 | -0.12 | 0.59 | 1.3 | 1.91 | 0.93 | 0.12 | 0.5 | -0.16 | 2.72 | 1.71 | 2.37 | -0.1 | -0.02 | 1.02 | 0.75 | -0.07 | 0.725 | -0.5225 | 0.897775 | 0.324165 | 0.305136 | 0.999971 | 1 | 0.847402 |
| Flot1 | O08917 | -0.97 | -1.82 | -1.33 | -1.28 | -1.71 | -1.46 | -0.65 | -1.09 | -1.62 | -0.8 | -1.46 | -1.89 | -2.08 | -1.45 | -1.61 | -1.55 | 0.1225 | -0.0925 | -0.3225 | 0.683186 | 0.757414 | 0.22268 | 0.999971 | 1 | 0.827753 |
| Flot2 | Q60634 | -0.89 | -1.58 | -2.14 | -1.49 | -1.26 | -1.06 | -2.06 | -1.17 | -1.32 | -0.88 | -1.67 | -1.89 | -1.03 | -1.07 | -1.51 | -1.46 | 0.1375 | 0.085 | 0.2575 | 0.686503 | 0.800175 | 0.394798 | 0.999971 | 1 | 0.871816 |
| Flrt1 | Q6RKD8 | -1 | -0.5 | -2.3 | -3.21 | -0.62 | -0.55 | -2.14 | -2.94 | -0.66 | -0.84 | -2.39 | -2.42 | -1.13 | -0.47 | -2.99 | -3.75 | 0.19 | 0.175 | -0.3325 | 0.804366 | 0.805493 | 0.707927 | 0.999971 | 1 | 0.950182 |
| Flrt2 | Q8BLU0 | -0.6 | -0.92 | -1.28 | -1.32 | -0.98 | -0.81 | -1.03 | -1.54 | -1.24 | -0.82 | -1.02 | -0.66 | -1.2 | -0.99 | -1.54 | -0.73 | -0.06 | 0.095 | -0.085 | 0.814917 | 0.694143 | 0.747655 | 0.999971 | 1 | 0.961064 |
| Flrt3 | Q8BGT1 | -0.45 | -0.56 | -0.54 | 0.06 | -0.46 | -0.46 | -0.24 | -0.05 | -0.84 | -0.46 | -0.56 | -0.62 | -0.49 | -0.78 | -0.47 | -0.42 | 0.07 | -0.2475 | -0.1675 | 0.751131 | 0.262115 | 0.444716 | 0.999971 | 1 | 0.884152 |
| Flt1 | P35969 | 1.45 | 0.95 | 1.52 | 1.78 | 0.75 | 1.16 | 1.16 | 0.27 | 0.65 | 1.75 | 1.27 | 0.72 | 0.28 | 0.6 | 0.85 | 1.11 | -0.59 | -0.3275 | -0.715 | 0.065342 | 0.312888 | 0.024466 | 0.999971 | 1 | 0.669611 |
| Flywch1 | Q8CI03 | 0.99 | -0.24 | 0.02 | 0.39 | -0.81 | -1.01 | -0.03 | -0.09 | -1.16 | 0.29 | -0.35 | -0.89 | -1.22 | -0.78 | -0.53 | -0.42 | -0.775 | -0.8175 | -1.0275 | 0.05444 | 0.067351 | 0.010669 | 0.999971 | 1 | 0.613095 |
| Fmn1 | E9Q7P6 | -1.01 | -0.23 | -0.6 | -1.07 | -0.72 | 0.02 | -1.02 | -1.25 | -0.45 | -1.57 | -0.59 | -0.61 | -1.16 | -0.43 | -0.84 | -1.15 | -0.015 | -0.0775 | -0.1675 | 0.964385 | 0.811219 | 0.550716 | 0.999971 | 1 | 0.913169 |
| Fmn2 | Q9JL04 | -0.08 | -0.58 | -1.25 | -1.32 | -0.55 | -0.59 | -0.85 | -1.29 | -0.33 | -0.03 | -0.69 | -0.69 | -0.35 | -0.58 | -1.07 | -1.31 | -0.0125 | 0.3725 | -0.02 | 0.970399 | 0.281119 | 0.955621 | 0.999971 | 1 | 0.998056 |
| Fmnl1 | A2AB60 | 0.21 | -0.31 | -0.42 | -0.47 | -0.69 | -0.33 | -0.49 | -0.81 | -0.79 | -0.35 | -0.85 | -0.9 | -0.56 | -0.58 | -0.8 | -0.69 | -0.3325 | -0.475 | -0.41 | 0.169625 | 0.068593 | 0.082754 | 0.999971 | 1 | 0.755254 |
| Fmnl2 | F8VPR2 | -1.64 | 1.69 | 0.57 | 0.41 | 1.11 | -0.15 | -0.18 | 1.93 | 1.11 | -1.09 | 0.01 | 0.7 | 1.38 | 0.18 | 1.73 | 0.24 | 0.42 | -0.075 | 0.625 | 0.591699 | 0.921862 | 0.396938 | 0.999971 | 1 | 0.871816 |
| Fmnl2_1 | A2APV2-3 | -0.91 | -0.7 | -0.67 | -0.7 | -0.73 | -0.6 | -0.88 | -0.81 | -0.37 | -0.53 | -0.71 | -0.79 | 0.08 | -0.59 | -0.64 | -0.6 | -0.01 | 0.145 | 0.3075 | 0.953682 | 0.43049 | 0.193348 | 0.999971 | 1 | 0.810547 |
| Fmnl3 | Q6ZPF4 | -2.55 | 1.65 | 1.05 | 1.11 | 2.82 | 2.62 | 2.04 | 0.98 | 1.4 | -0.19 | 1.92 | 2.05 | 3.48 | 0.85 | 0.6 | 0.84 | 1.8 | 0.98 | 1.1275 | 0.081906 | 0.333233 | 0.301072 | 0.999971 | 1 | 0.847402 |
| Fmo1 | P50285 | -0.75 | -0.69 | -1.56 | -1.49 | 0.23 | -0.51 | -1.14 | -0.68 | 0.08 | 0.09 | -1.33 | -1.12 | 0.14 | -0.06 | -0.71 | -1.03 | 0.5975 | 0.5525 | 0.7075 | 0.123441 | 0.215177 | 0.071849 | 0.999971 | 1 | 0.739418 |
| Fmo5 | P97872 | 1.23 | -0.03 | 0.09 | 1.86 | 0.27 | 0.27 | 0.5 | 3.45 | 0.16 | 0.71 | 0.1 | 0.04 | -0.38 | 0.01 | -0.12 | 1.48 | 0.335 | -0.535 | -0.54 | 0.680395 | 0.257965 | 0.355084 | 0.999971 | 1 | 0.868793 |
| Fmr1 | P35922-9 | 1.34 | 0.81 | 0.66 | 1.01 | 0.6 | 0.52 | 1.21 | 0.51 | 0.78 | 0.84 | 0.9 | 0.86 | 0.06 | 0.1 | 0.25 | 0.43 | -0.245 | -0.11 | -0.745 | 0.341945 | 0.588353 | 0.007115 | 0.999971 | 1 | 0.582714 |
| Fn1 | B9EHT6 | 0.58 | 0.63 | -0.39 | 1.12 | 3.28 | 0.94 | 1.34 | -0.19 | 1.74 | 1.49 | 1.48 | 2.85 | 0.42 | 0.64 | 0.39 | 0.29 | 0.8575 | 1.405 | -0.05 | 0.248262 | 0.009617 | 0.878026 | 0.999971 | 1 | 0.985441 |
| Fn3k | Q9ER35 | -0.22 | 0.24 | 0.24 | -0.12 | -0.03 | -0.01 | -0.25 | -0.35 | -0.09 | -0.58 | 0.12 | 0.15 | -0.2 | -0.26 | 0.24 | -0.53 | -0.195 | -0.135 | -0.2225 | 0.348652 | 0.574327 | 0.357718 | 0.999971 | 1 | 0.868793 |
| Fn3krp | Q8K274 | 1.96 | 0.4 | -0.57 | -0.53 | -0.28 | 0.77 | 0.48 | -0.53 | -0.43 | 0.97 | -0.78 | -0.79 | -0.49 | 0.44 | -0.63 | -0.98 | -0.205 | -0.5725 | -0.73 | 0.736725 | 0.397377 | 0.249387 | 0.999971 | 1 | 0.839012 |
| Fnbp1 | A2AQ42 | -0.67 | -0.86 | 0.17 | -0.07 | -1.02 | -0.69 | -0.36 | -0.15 | -0.82 | -0.77 | -0.11 | -0.14 | -0.97 | -0.61 | -0.05 | -0.13 | -0.1975 | -0.1025 | -0.0825 | 0.533186 | 0.745048 | 0.800634 | 0.999971 | 1 | 0.971076 |
| Fnbp1l | E9PUI5 | 0.12 | -0.64 | -0.23 | -0.05 | -0.8 | -0.46 | 0.66 | -0.21 | -0.73 | 0.17 | -0.07 | 0.17 | -0.62 | -0.45 | 0.11 | 0.06 | -0.0025 | 0.085 | -0.025 | 0.994239 | 0.763225 | 0.925214 | 0.999971 | 1 | 0.99401 |
| Fndc10 | A2A9Q0 | 0.75 | -0.83 | -0.15 | -0.05 | -1.29 | -0.28 | -0.15 | -0.06 | -1.25 | 0.73 | 0.26 | -0.39 | -1.7 | -0.9 | -0.23 | 0.3 | -0.375 | -0.0925 | -0.5625 | 0.372687 | 0.853454 | 0.279658 | 0.999971 | 1 | 0.843812 |
| Fndc3a | Q8BX90 | 2.3 | 0.46 | 0.67 | -0.05 | 0.26 | 1.42 | 2.09 | -0.5 | 0.35 | 1.77 | -0.27 | -0.6 | -0.62 | -0.08 | -0.39 | -1.26 | -0.0275 | -0.5325 | -1.4325 | 0.968476 | 0.432775 | 0.02182 | 0.999971 | 1 | 0.645537 |
| Fndc3b | A0A0R4J0H | 0.13 | 0.38 | -2.14 | -2.37 | -0.3 | -0.38 | -2.05 | -2.42 | 0.08 | -0.46 | -2.5 | -0.48 | 0.25 | 0.46 | -1.15 | -1.8 | -0.2875 | 0.16 | 0.44 | 0.726658 | 0.847967 | 0.593843 | 0.999971 | 1 | 0.92342 |
| Fndc4 | A0A0J9YUI | -0.53 | -8.71 | -7.73 | -7.45 | -5.74 | -6.82 | -5.79 | -9.38 | -8.76 | -2.63 | -6.57 | -8.59 | -8.16 | -6.93 | -8.29 | -10.63 | -0.8275 | -0.5325 | -2.3975 | 0.651883 | 0.800117 | 0.205561 | 0.999971 | 1 | 0.816688 |
| Fnta | Q61239 | 0.2 | 0.37 | -0.33 | -0.06 | 0.26 | 0.09 | -0.09 | -0.76 | 0.41 | 0.42 | -0.69 | -1.29 | 0.3 | 0.42 | -0.55 | -0.43 | -0.17 | -0.3325 | -0.11 | 0.554282 | 0.445074 | 0.715133 | 0.999971 | 1 | 0.952576 |
| Fntb | Q8K2I1 | -1.85 | 0.08 | -0.14 | -0.22 | -0.11 | -0.23 | 0.09 | -0.12 | 0.22 | -1.42 | 0.28 | 0.53 | 0.5 | -0.05 | 0.02 | 0.21 | 0.44 | 0.435 | 0.7025 | 0.315412 | 0.459999 | 0.133417 | 0.999971 | 1 | 0.784948 |
| Focad | A2AKG8 | -0.37 | -0.33 | -0.32 | -0.64 | -0.06 | -0.27 | -0.35 | -0.4 | -0.15 | -0.58 | -0.01 | 0.22 | -0.18 | -0.15 | 0.14 | -0.16 | 0.145 | 0.285 | 0.3275 | 0.435262 | 0.223368 | 0.101683 | 0.999971 | 1 | 0.78268 |
| Folh1 | O35409 | -1.04 | -1.17 | -1.15 | -1.03 | -0.97 | -1.13 | -1.15 | -1.66 | -1.08 | -1.25 | -1.09 | -1.7 | -0.51 | -0.92 | -0.89 | -1.27 | -0.13 | -0.1825 | 0.2 | 0.534423 | 0.377685 | 0.355346 | 0.999971 | 1 | 0.868793 |
| Folr1 | E9Q543 | 0.58 | 2.57 | 3.65 | 1.1 | 1.61 | 1.55 | 1.98 | 1.87 | 2.37 | 0.04 | -5.94 | 2.8 | 2.1 | 1.6 | 2.89 | 2.71 | -0.2225 | -2.1575 | 0.35 | 0.729646 | 0.2744 | 0.613491 | 0.999971 | 1 | 0.927364 |
| Foxg1 | Q60987 | 1.29 | 1.71 | 0.5 | 0.3 | 1.63 | 0.27 | 1.65 | 3.47 | 0.24 | 1.12 | 4.84 | 2.52 | 0.59 | 2 | 0.71 | 3.71 | 0.805 | 1.23 | 0.8025 | 0.246931 | 0.218166 | 0.283642 | 0.999971 | 1 | 0.844538 |

| | | | | | | | | | | | | | | | | | | | | | | | | | | |
|---|---|---|---|---|---|---|---|---|---|---|---|---|---|---|---|---|---|---|---|---|---|---|---|---|---|---|
| Foxj3 | Q8BUR3-2 | 3.02 | 2.39 | 0.02 | 0.8 | 2.35 | 0.88 | 0.39 | 1.37 | 2.26 | 2.99 | 0.74 | 1.02 | 1.36 | 3.15 | 0.21 | 1.27 | -0.31 | 0.195 | -0.06 | 0.672668 | 0.805047 | 0.942308 | 0.999971 | 1 | 0.996889 |
| Foxk1 | P42128 | 2.33 | 2.29 | 2.71 | 2.88 | 1.98 | 2 | 2.31 | 2.72 | 1.93 | 2.28 | 2.71 | 3.16 | 1.84 | 2.27 | 2.64 | 3.27 | -0.3 | -0.0325 | -0.0475 | 0.253757 | 0.915956 | 0.887027 | 0.999971 | 1 | 0.987485 |
| Foxk2 | Q3UCQ1-2 | 0.26 | 0.88 | 0.99 | 1.33 | 0.71 | 0.67 | 0.75 | 1.39 | 0.4 | -0.06 | 1.15 | 1.06 | -0.22 | 0.74 | 1.12 | 1.91 | 0.015 | -0.2275 | 0.0225 | 0.959013 | 0.527507 | 0.961307 | 0.999971 | 1 | 0.998721 |
| Foxo1 | Q9R1E0 | 0.55 | 1.33 | 1.97 | -1.07 | 1.41 | 0.03 | 1.1 | 0.96 | 0.33 | 1.14 | 1.01 | 1.08 | 2.13 | 0.98 | 1.35 | -1.18 | 0.18 | 0.195 | 0.125 | 0.783196 | 0.755557 | 0.885258 | 0.999971 | 1 | 0.98711 |
| Foxo3 | Q9WVH4 | 2.99 | -0.6 | 0.58 | 0.13 | -0.03 | -0.02 | 0.12 | 1.18 | -0.79 | 2.36 | 0.8 | 1.05 | -1.64 | -0.57 | 0.75 | 0.89 | -0.4625 | 0.08 | -0.9175 | 0.540682 | 0.929968 | 0.312388 | 0.999971 | 1 | 0.849367 |
| Foxp1 | A0A0R4J28 | -0.13 | 1.45 | 2.51 | 2.21 | 1.38 | 0.18 | 1.52 | 2.49 | 1.88 | 1.11 | 2.63 | 2.42 | 0.11 | 0.53 | 2.16 | 2.15 | -0.1175 | 0.5 | -0.2725 | 0.863745 | 0.431477 | 0.706588 | 0.999971 | 1 | 0.950182 |
| Foxred1 | Q3TQB2 | 0.02 | -0.52 | -0.98 | -0.76 | -0.62 | -0.67 | -0.94 | -0.73 | -0.54 | 0.1 | -0.56 | -0.75 | -0.41 | -0.32 | -0.78 | -0.44 | -0.18 | 0.1225 | 0.0725 | 0.48289 | 0.677739 | 0.781703 | 0.999971 | 1 | 0.965859 |
| Fras1 | Q80T14 | -0.11 | -0.36 | -0.9 | -0.22 | -0.95 | 0 | -0.71 | -0.61 | -0.6 | -0.34 | -1.14 | -0.01 | -0.48 | -0.45 | -0.26 | 0.34 | -0.17 | -0.125 | 0.185 | 0.550918 | 0.681049 | 0.508878 | 0.999971 | 1 | 0.905815 |
| Frg1 | P97376 | 1.65 | 1.32 | 0.47 | 0.55 | 1.66 | 1.69 | 0.02 | 0.41 | 1.37 | 2.39 | 0.66 | 1.38 | 1.32 | 2.11 | 0.68 | 1.18 | -0.0525 | 0.4525 | 0.325 | 0.913178 | 0.313472 | 0.422836 | 0.999971 | 1 | 0.87887 |
| Frmd3 | Q8BHD4 | -1.11 | 0.7 | -0.23 | 0.06 | 0.53 | -0.31 | -1.63 | -0.59 | 0.1 | -0.45 | -0.59 | -0.14 | 1.52 | 0.35 | -0.57 | -0.03 | -0.355 | -0.125 | 0.4625 | 0.513515 | 0.749915 | 0.39855 | 0.999971 | 1 | 0.871816 |
| Frmd4a | H3BIZ7 | 0.44 | -0.43 | -0.66 | -0.37 | -0.36 | -0.12 | -0.2 | -0.61 | -0.11 | 0.51 | -0.4 | -0.3 | -0.15 | -0.25 | -0.63 | -0.21 | -0.0675 | 0.18 | -0.055 | 0.808971 | 0.57425 | 0.844196 | 0.999971 | 1 | 0.979814 |
| Frmd4b | Q6ZQ04 | 1.89 | -1.58 | -1.79 | -2.77 | -1.33 | -0.43 | -1.92 | -2.67 | -0.53 | 0.84 | -1.92 | -1.84 | -2.09 | -1.04 | -1.01 | -0.76 | -0.525 | 0.2 | -0.1625 | 0.602805 | 0.85366 | 0.86394 | 0.999971 | 1 | 0.983079 |
| Frmd5 | A2AR56 | 0.46 | 0.47 | -0.34 | -0.41 | -0.43 | -0.17 | -0.53 | -0.17 | -0.2 | 0.83 | -0.74 | -0.77 | 1.06 | 0.13 | -0.47 | 0.03 | -0.37 | -0.265 | 0.1425 | 0.203478 | 0.536308 | 0.713375 | 0.999971 | 1 | 0.951793 |
| Frmd8 | Q3UFK8 | 1.28 | -1.21 | -0.77 | -1.16 | -1.52 | -0.68 | -1.23 | -1.26 | -1.55 | 0.41 | -1.47 | -1.29 | -1.88 | -1.39 | -1.05 | -1.25 | -0.7075 | -0.51 | -0.9275 | 0.230953 | 0.463483 | 0.12809 | 0.999971 | 1 | 0.784371 |
| Frmpd1 | A2AKB4 | 1.13 | 1.35 | 0.29 | 0.78 | 1.44 | 1.2 | 0.21 | 0.41 | 1.28 | 0.81 | 0.02 | -0.39 | 0.32 | 1.21 | -0.61 | -0.44 | -0.0725 | -0.4575 | -0.7675 | 0.843389 | 0.293992 | 0.114041 | 0.999971 | 1 | 0.78268 |
| Frmpd3 | A0A140LIW | 0.09 | -0.64 | -0.66 | -0.58 | -0.74 | -0.28 | -0.19 | -0.65 | -0.96 | -0.36 | -1.06 | -1.21 | -0.39 | -0.73 | -1.04 | -0.81 | -0.0175 | -0.45 | -0.295 | 0.944613 | 0.128913 | 0.261403 | 0.999971 | 1 | 0.842262 |
| Frmpd4 | A0A589MR | 0.58 | 0.42 | -0.04 | 0.11 | 0.5 | 0.61 | -0.3 | 0.31 | 0.51 | 0.56 | 0.08 | 0.24 | 0.28 | 0.35 | 0.2 | 0.39 | 0.0125 | 0.08 | 0.0375 | 0.962722 | 0.718509 | 0.854103 | 0.999971 | 1 | 0.982125 |
| Frrs1l | B1AXV0 | 0.99 | -0.27 | -0.36 | -0.11 | 0.02 | -0.19 | 0.65 | -0.09 | -0.1 | -0.1 | -0.29 | -0.24 | 0.03 | -0.56 | -0.73 | -0.32 | 0.035 | -0.245 | -0.4575 | 0.922019 | 0.450698 | 0.212347 | 0.999971 | 1 | 0.823451 |
| Frs2 | Q8C180 | -0.98 | -1.31 | -1.32 | -1.11 | -1.42 | -1.86 | -1.52 | -0.85 | -1.43 | -1.39 | -1.2 | -1.3 | -0.52 | -1.08 | -1.11 | -1 | -0.2325 | -0.15 | 0.2525 | 0.368633 | 0.401497 | 0.251642 | 0.999971 | 1 | 0.839012 |
| Frs3 | Q91WJ0 | -0.81 | -1.1 | -0.84 | -1.1 | -0.78 | -0.62 | -1.21 | -1.13 | -0.57 | -1.1 | -1.15 | -1.16 | -0.04 | -0.84 | -1.24 | -1.19 | 0.0275 | -0.0325 | 0.135 | 0.89637 | 0.876854 | 0.652565 | 0.999971 | 1 | 0.934005 |
| Fry | A0A5F8MP | -0.24 | -0.03 | 0.14 | 0.14 | 0.04 | 0.01 | 0.1 | 0.29 | 0.04 | 0.17 | 0.26 | 0.38 | 0.28 | 0.17 | 0.24 | 0.56 | 0.1075 | 0.21 | 0.31 | 0.562822 | 0.270591 | 0.132422 | 0.999971 | 1 | 0.784948 |
| Fry_1 | A0A0J9YUF | -0.08 | -0.15 | -0.68 | -0.19 | 0.47 | -0.43 | -0.77 | -0.1 | 0.06 | -0.19 | -0.07 | 0.68 | 0.53 | 0.15 | 0.03 | 0.35 | 0.0675 | 0.395 | 0.54 | 0.824585 | 0.153362 | 0.034534 | 0.999971 | 1 | 0.69205 |
| Fryl | F8VQ05 | 0.16 | -0.11 | 0.03 | 0.41 | -0.01 | 0.01 | -0.18 | 0.36 | -0.23 | -0.2 | 0 | 0.09 | -0.34 | 0.17 | -0.17 | 0.23 | -0.0775 | -0.2075 | -0.15 | 0.712182 | 0.300221 | 0.501859 | 0.999971 | 1 | 0.904573 |
| Fscn1 | Q61553 | -0.77 | -0.6 | -0.92 | -1.63 | -0.51 | -0.3 | 0.22 | -1.45 | -0.19 | -0.81 | -0.6 | -0.53 | -0.9 | -0.51 | -1.2 | -1.5 | 0.47 | 0.4475 | -0.0475 | 0.256568 | 0.131901 | 0.87991 | 0.999971 | 1 | 0.985441 |
| Fsd1 | Q7TPM6 | -0.53 | -0.64 | -0.59 | -0.65 | -0.28 | -0.42 | -0.48 | -0.64 | -0.28 | -0.21 | -0.39 | -0.27 | -0.08 | -0.12 | -0.5 | -0.21 | 0.1475 | 0.315 | 0.375 | 0.399947 | 0.072832 | 0.062431 | 0.999971 | 1 | 0.725348 |
| Fsd1l | E0CXZ1 | -0.74 | -0.52 | 0.08 | 0 | -0.38 | -0.34 | -0.34 | 0.03 | -0.27 | -0.44 | 0.08 | 0.17 | -0.11 | -0.26 | 0.19 | 0.48 | 0.0375 | 0.18 | 0.37 | 0.880471 | 0.503318 | 0.202913 | 0.999971 | 1 | 0.81617 |
| Fstl4 | Q5STE3 | 0.24 | 0.85 | -1.15 | 0.19 | 0.81 | 1.7 | -1 | -0.2 | 0.39 | 0.02 | -0.7 | -0.28 | 0.62 | 0.26 | -0.18 | 0.59 | 0.295 | -0.175 | 0.29 | 0.654981 | 0.700319 | 0.511149 | 0.999971 | 1 | 0.905815 |
| Fth1 | P09528 | 1.36 | 2.01 | 1.18 | 0.44 | 1.34 | 3.23 | 0.48 | 1.05 | 0.52 | 0.71 | 0.99 | -0.21 | 1.08 | 2.42 | 0.88 | 0.61 | 0.2775 | -0.745 | 3.14E-16 | 0.654576 | 0.087607 | 1 | 0.999971 | 1 | 1 |
| Ftl1 | Q9CPX4 | 0.37 | 1.19 | 2.22 | 1.02 | 1.49 | 3.6 | 1.59 | 2.54 | -0.51 | 0.1 | 2.5 | 1.01 | 1.42 | 3.03 | 3.25 | 3.22 | 1.105 | -0.425 | 1.53 | 0.079994 | 0.542698 | 0.018315 | 0.999971 | 1 | 0.635388 |
| Fto | Q8BGW1 | 1.28 | 1.49 | 2.12 | 2.08 | 1.4 | 1.34 | 1.5 | 2.13 | 1.16 | 1.46 | 1.77 | 2.19 | 1.25 | 1.57 | 2.04 | 2.35 | -0.15 | -0.0975 | 0.06 | 0.608136 | 0.753407 | 0.853273 | 0.999971 | 1 | 0.98187 |
| Ftsj1 | F6Q475 | -1.66 | 0.53 | 1.44 | -1.44 | 0.29 | 0.82 | -0.69 | -0.82 | 0.16 | -1.09 | 0.87 | 1.66 | 0.09 | 0.96 | -1.94 | -0.64 | 0.1825 | 0.6825 | -0.1 | 0.812175 | 0.437243 | 0.908829 | 0.999971 | 1 | 0.991707 |
| Ftsj3 | Q9DBE9 | -0.41 | -0.98 | 0.06 | 0.05 | -0.74 | -1.2 | -0.69 | -0.24 | -1.07 | -0.5 | -0.55 | -0.39 | -1.22 | -0.43 | -0.2 | 0.17 | -0.3975 | -0.3075 | -0.1 | 0.231328 | 0.316 | 0.788482 | 0.999971 | 1 | 0.96741 |
| Fubp1 | Q3TUE1 | 2.31 | 2.8 | 2.68 | 2.74 | 2.54 | 2.76 | 2.6 | 2.77 | 2.51 | 2.53 | 2.75 | 2.83 | 2.21 | 2.79 | 2.87 | 2.84 | 0.035 | 0.0225 | 0.045 | 0.854576 | 0.907715 | 0.845216 | 0.999971 | 1 | 0.979814 |
| Fubp3 | A2AJ72 | 1.17 | 1.04 | 1.34 | 1.59 | 0.87 | 0.91 | 1.64 | 1.4 | 1.02 | 1.37 | 1.12 | 1.35 | 0.92 | 1.12 | 1.5 | 1.29 | -0.08 | -0.07 | -0.0775 | 0.749996 | 0.727376 | 0.72271 | 0.999971 | 1 | 0.955258 |
| Fuca1 | Q99LJ1 | -0.51 | 0.43 | 1.29 | 0.06 | 0.36 | 0.01 | 1.24 | 1.33 | -0.08 | 0.3 | 1.08 | 0.11 | 0.12 | 0.57 | 0.08 | 0.64 | 0.4175 | 0.035 | 0.035 | 0.381586 | 0.935316 | 0.928406 | 0.999971 | 1 | 0.994278 |
| Fundc1 | H3BJZ1 | 0.87 | -1.03 | -0.52 | -0.32 | -1.25 | -0.59 | -0.88 | -0.58 | -0.88 | -0.27 | -0.41 | -0.88 | -0.79 | -0.97 | -0.82 | -0.73 | -0.575 | -0.36 | -0.5775 | 0.185776 | 0.393595 | 0.164283 | 0.999971 | 1 | 0.800215 |
| Fundc2 | Q9D6K8 | 0.72 | -0.2 | -0.07 | 0.25 | -0.24 | 0.49 | -0.19 | 0.56 | -0.24 | 0.91 | 0.15 | 0.05 | -0.15 | 0.22 | 0.34 | 0.78 | -0.02 | 0.0425 | 0.1225 | 0.947439 | 0.894617 | 0.676605 | 0.999971 | 1 | 0.942037 |
| Fuom | Q8R2K1 | 0.29 | 0.56 | 0.92 | 0.78 | -0.13 | 0.58 | 0.18 | 0.89 | -0.29 | -0.2 | 0.43 | 0.22 | -0.47 | 0.91 | 1.15 | 1.01 | -0.2575 | -0.5975 | 0.0125 | 0.367212 | 0.036299 | 0.974177 | 0.999971 | 1 | 0.99925 |
| Fus | Q8CFQ9 | 2.11 | 2.43 | 2.73 | 2.38 | 2.19 | 2.33 | 2.81 | 2.34 | 2.1 | 2.5 | 2.21 | 2.21 | 2.06 | 2.37 | 2.37 | 2.34 | 0.005 | -0.1575 | -0.1275 | 0.982326 | 0.447953 | 0.536557 | 0.999971 | 1 | 0.910077 |
| Fut11 | Q8BHC9 | 1.43 | -0.01 | -0.07 | -0.14 | 0.34 | 1.13 | 0.06 | -0.15 | 0.45 | 1.68 | 0.39 | 0.1 | -0.21 | -0.08 | 0.03 | 0.11 | 0.0425 | 0.3525 | -0.34 | 0.923319 | 0.470906 | 0.372848 | 0.999971 | 1 | 0.869526 |
| Fut8 | Q9WTS2 | 1.69 | 0.73 | 1.01 | 1.32 | 0.92 | 0.48 | 0.66 | 1.76 | 0.72 | 1.2 | 1.22 | 1.39 | 0.26 | 0.76 | 1.35 | 1.08 | -0.2325 | -0.055 | -0.325 | 0.50633 | 0.838361 | 0.319179 | 0.999971 | 1 | 0.853893 |
| Fut9 | O88819 | -0.27 | 1.21 | 1.38 | 0.83 | 1.03 | 1.66 | 0.9 | 2.14 | 0.88 | 0.63 | 2.01 | 2.18 | 1.32 | 1.49 | 2.03 | 2.36 | 0.645 | 0.6375 | 1.0125 | 0.169511 | 0.226128 | 0.036945 | 0.999971 | 1 | 0.700096 |
| Fv1 | P70213 | -0.78 | 1.65 | 1 | 0.73 | 1.06 | 2.08 | -0.6 | 1.04 | 0.57 | 0.5 | 1.24 | 0.02 | -1.48 | -0.02 | 0.13 | 1.06 | 0.245 | -0.0675 | -0.7275 | 0.721075 | 0.898666 | 0.292351 | 0.999971 | 1 | 0.847402 |
| Fxn | O35943 | -0.29 | -0.56 | -0.81 | -2.88 | -0.76 | -1.08 | -0.8 | -2.27 | -0.66 | -0.79 | -1.05 | -0.6 | -0.43 | -0.6 | -2.24 | -2.78 | -0.0925 | 0.36 | -0.3775 | 0.882691 | 0.520801 | 0.61815 | 0.999971 | 1 | 0.928581 |
| Fxr1 | Q61584-3 | 0.43 | 0.08 | 1.03 | 1.12 | 0.39 | -0.06 | 1.34 | 1 | 0.19 | 0.26 | 1.2 | 1.08 | 0.21 | 0.29 | 1.26 | 1.25 | 0.0025 | 0.0175 | 0.0875 | 0.994798 | 0.960758 | 0.813852 | 0.999971 | 1 | 0.974826 |
| Fxr2 | Q6P5B5 | 0.5 | 0.46 | 0.16 | 0.44 | 0.51 | 0.15 | 0.16 | 0.52 | 0.39 | 0.46 | 0.37 | 0.42 | 0.46 | 0.54 | 0.51 | 0.46 | -0.055 | 0.02 | 0.1025 | 0.776762 | 0.90528 | 0.55631 | 0.999971 | 1 | 0.915051 |
| Fxyd1 | A0A0U1RP | -0.58 | -1.03 | -0.55 | -0.18 | -1.55 | -0.92 | -1.08 | -0.09 | -1.7 | -0.33 | -1.95 | -1.24 | -1.75 | -0.93 | -0.91 | -0.8 | -0.325 | -0.72 | -0.5125 | 0.359299 | 0.088081 | 0.106591 | 0.999971 | 1 | 0.78268 |
| Fxyd6 | Q9D164-2 | -1.32 | -1.22 | -0.67 | -0.91 | -0.94 | -1.17 | -0.88 | -0.87 | -1.23 | -0.56 | -0.79 | -0.47 | -0.76 | -1.07 | -0.61 | 0.25 | 0.065 | 0.2675 | 0.4825 | 0.760436 | 0.302442 | 0.159233 | 0.999971 | 1 | 0.799146 |
| Fxyd7 | A0A0U1RP | 3.98 | 0.51 | 1.04 | 0.96 | -2.05 | 2.12 | 5.49 | -0.49 | -1.22 | 2.59 | -1.09 | 0.75 | -0.62 | -0.4 | 1.07 | -2.09 | -0.355 | -1.365 | -2.1325 | 0.826722 | 0.226421 | 0.042579 | 0.999971 | 1 | 0.701725 |
| Fyco1 | Q8VDC1 | -0.46 | -0.09 | -0.36 | 0.01 | 0.32 | -1.22 | -0.51 | 0.25 | -0.35 | -1.08 | -0.58 | -0.63 | 0.09 | 0.14 | -0.21 | -0.17 | -0.065 | -0.435 | 0.1875 | 0.859833 | 0.081249 | 0.362846 | 0.999971 | 1 | 0.869526 |
| Fyn | P39688 | -2.06 | -2.18 | -1 | -1.22 | -1.85 | -2.28 | -1.24 | -0.79 | -1.71 | -2.06 | -0.93 | -0.9 | -1.93 | -2.16 | -0.8 | -1.2 | 0.075 | 0.215 | 0.0925 | 0.858091 | 0.591411 | 0.822745 | 0.999971 | 1 | 0.975901 |
| Fyttd1 | Q91Z49 | 0.77 | 1.1 | 0.76 | 1.01 | 1.13 | 0.81 | 1.52 | 0.46 | 0.96 | 1.22 | 0.65 | 0.26 | 0.68 | 0.93 | 0.27 | 0.35 | 0.07 | -0.1375 | -0.3525 | 0.790911 | 0.585126 | 0.135759 | 0.999971 | 1 | 0.784948 |
| Fzd2 | Q9JIP6 | -2.64 | -1.15 | -1.37 | 0.17 | -0.89 | -1.38 | 0.55 | 1.52 | -1.27 | -1.89 | 0.71 | 0.17 | -1.2 | -1.23 | 0.49 | 0.9 | 1.1975 | 0.6775 | 0.9875 | 0.155387 | 0.383005 | 0.196229 | 0.999971 | 1 | 0.812552 |
| Fzd3 | Q61086 | 0.14 | -1.3 | -1.96 | -1.32 | -1.95 | -1.64 | -1.27 | -1.33 | -1.92 | -0.38 | -1.63 | -2.23 | -1.64 | -1.34 | -1.75 | -2.45 | -0.4375 | -0.43 | -0.685 | 0.337114 | 0.44678 | 0.169999 | 0.999971 | 1 | 0.802438 |
| G0s2 | Q61585 | 1.76 | 2.06 | 1.65 | 1.23 | -1.7 | 0.16 | 1.04 | 1.03 | 1.53 | 2.11 | 1.37 | 0.6 | 3.61 | 0.7 | 1.63 | 1.65 | -1.5425 | -0.2725 | 0.2225 | 0.030056 | 0.443941 | 0.703553 | 0.999971 | 1 | 0.949671 |
| G3bp1 | P97855 | -0.44 | 1.17 | 0.93 | 1.01 | 1.17 | 1.07 | 0.71 | 1.24 | 1.1 | 0.42 | 0.77 | 1.07 | 1.01 | 1.4 | 1.23 | 1.16 | 0.38 | 0.1725 | 0.5325 | 0.328444 | 0.65988 | 0.175646 | 0.999971 | 1 | 0.808532 |
| G3bp2 | P97379 | 1.43 | 1.84 | 1.89 | 2.07 | 1.75 | 1.84 | 2.09 | 1.74 | 1.89 | 1.92 | 1.76 | 1.86 | 1.29 | 1.76 | 1.65 | 1.91 | 0.0475 | 0.05 | -0.155 | 0.820406 | 0.798993 | 0.503747 | 0.999971 | 1 | 0.905195 |
| G6pc3 | Q6NSQ9 | 0.36 | 0.76 | -0.63 | -1.27 | 1.86 | 0.06 | -0.58 | -0.87 | 0.9 | -0.11 | -0.97 | -0.17 | 2.11 | 1.03 | -0.43 | -1.08 | 0.3125 | 0.1075 | 0.6025 | 0.654141 | 0.846278 | 0.442412 | 0.999971 | 1 | 0.883386 |
| G6pdx | Q00612 | 0.88 | -0.3 | 0.45 | 0.67 | -0.29 | -0.25 | 0.26 | 0.41 | -0.24 | 0.06 | 0.03 | 0.23 | -0.69 | -0.33 | 0.49 | 0.32 | -0.3925 | -0.405 | -0.4775 | 0.234575 | 0.182249 | 0.216184 | 0.999971 | 1 | 0.826893 |
| Gaa | P70699 | 1.31 | 0.97 | 0.47 | -0.03 | 1.26 | 0.66 | 0.19 | 0.28 | 1.52 | 0.97 | 0.15 | 0.56 | 1.29 | 0.98 | 0.63 | 0.05 | -0.0825 | 0.12 | 0.0575 | 0.82338 | 0.763122 | 0.88039 | 0.999971 | 1 | 0.985441 |
| Gab1 | A0A1B0GS | -0.16 | -0.44 | -0.56 | -0.24 | -0.49 | -0.78 | -0.55 | -0.42 | -0.57 | -0.52 | -0.96 | -0.53 | -0.82 | -0.45 | -0.47 | -0.47 | -0.21 | -0.295 | -0.2025 | 0.283385 | 0.158467 | 0.310498 | 0.999971 | 1 | 0.847402 |
| Gab2 | Q3ZB57 | 0.6 | 0.21 | 0.21 | 0.25 | 0.52 | 0.19 | 0.31 | 0.34 | 0.35 | 0.57 | 0.64 | 0.18 | 0.48 | 0.19 | 0.39 | 0.6 | 0.0225 | 0.1175 | 0.0975 | 0.904248 | 0.555258 | 0.616932 | 0.999971 | 1 | 0.928413 |
| Gabarapl1 | Q8R3R8 | -8.16 | -14 | -6.72 | -7.59 | -8.24 | -10.65 | -2.02 | -8.93 | -7.31 | -6.14 | -5.09 | -5.66 | -8.39 | -10.59 | -7.73 | -8.67 | 1.6575 | 3.0675 | 0.2725 | 0.461459 | 0.072808 | 0.861814 | 0.999971 | 1 | 0.982963 |
| Gabarapl2 | P60521 | -0.2 | -0.46 | -0.69 | -0.83 | -0.32 | -0.24 | -0.71 | -0.52 | -0.55 | -0.35 | -0.67 | -0.5 | 0.15 | -0.24 | -0.31 | -0.28 | 0.0975 | 0.0275 | 0.375 | 0.657187 | 0.89264 | 0.115501 | 0.999971 | 1 | 0.78268 |
| Gabbr1 | Q9WV18-2 | 1.34 | 1.78 | 0.54 | 0.62 | -0.08 | 0.72 | 0.98 | 0.44 | -0.03 | 0.9 | 0.16 | -0.65 | -0.75 | -1.38 | 0.16 | -0.38 | -0.555 | -0.975 | -1.6575 | 0.153311 | 0.041444 | 0.003296 | 0.999971 | 1 | 0.577271 |
| Gabbr1_1 | Q9WV18 | -0.03 | -0.98 | -0.9 | -0.57 | -1.03 | -0.45 | -0.6 | -0.7 | -1.1 | -0.58 | -1.02 | -1.56 | -0.82 | -0.82 | -1.05 | -0.62 | -0.075 | -0.445 | -0.2075 | 0.780142 | 0.165801 | 0.430275 | 0.999971 | 1 | 0.881611 |
| Gabbr2 | Q80T41 | -1.38 | -1.51 | -1.8 | -2.05 | -1.44 | -1.25 | -1.57 | -1.93 | -1.37 | -1.49 | -1.69 | -2.07 | -1.07 | -1.55 | -2.16 | -1.99 | 0.1375 | 0.03 | -0.0075 | 0.570598 | 0.901784 | 0.979822 | 0.999971 | 1 | 0.99925 |
| Gabpa | Q00422 | 1.56 | 2.23 | 1.64 | 2.2 | 2.34 | 2.13 | 1.5 | 1.65 | 1.75 | 2.75 | 1.59 | 2.27 | 0.77 | 1.96 | 2.22 | 2.46 | -0.0025 | 0.1825 | -0.055 | 0.9929 | 0.572078 | 0.890101 | 0.999971 | 1 | 0.98799 |
| Gabra1 | P62812 | -0.43 | -0.42 | -0.79 | -0.6 | -1.02 | -0.2 | -0.4 | -1.11 | -0.67 | -0.59 | -0.66 | -1.43 | -0.61 | -0.48 | -1.48 | -1.18 | -0.1225 | -0.2775 | -0.3775 | 0.643973 | 0.274754 | 0.18805 | 0.999971 | 1 | 0.809025 |
| Gabra2 | P26048 | -2.76 | -3.32 | -4.1 | -4.01 | -0.44 | -0.57 | -0.12 | -0.47 | -0.55 | -0.72 | -0.26 | 0.44 | -0.19 | -0.66 | -0.15 | 0.03 | 3.1475 | 3.275 | 3.305 | 7.59E-06 | 2.30E-05 | 7.53E-06 | 0.023056 | 0.069821 | 0.02288 |
| Gabra3 | B1AVY2 | -1.09 | -0.59 | -1.01 | -0.97 | -1 | -0.75 | -1.04 | -1.38 | -0.41 | -1.5 | -1.41 | -1.71 | -0.63 | -0.97 | -1.51 | -1.01 | -0.1275 | -0.3425 | -0.115 | 0.560163 | 0.291732 | 0.639807 | 0.999971 | 1 | 0.93301 |
| Gabra4 | Q9D6F4 | -0.07 | -0.14 | -0.33 | -0.38 | -0.86 | -1.1 | -0.28 | -1.49 | -0.8 | -1 | -0.75 | -0.87 | -0.31 | -1.16 | -1.02 | -1.25 | -0.7025 | -0.625 | -0.705 | 0.031089 | 0.005535 | 0.020125 | 0.999971 | 1 | 0.635388 |
| Gabra5 | Q8BHJ7 | -0.01 | 0.09 | -1.47 | -2.06 | 0.12 | 0.53 | -1.47 | -1.88 | -0.07 | 0.37 | -0.69 | -1.16 | -0.09 | 0.28 | -2.13 | -2.24 | 0.1875 | 0.475 | -0.1825 | 0.794343 | 0.423555 | 0.812326 | 0.999971 | 1 | 0.974458 |
| Gabrb1 | P50571 | -1.32 | -2.21 | -1.85 | -2.34 | -1.89 | -1.24 | -1.49 | -1.64 | -1.87 | -1.07 | -1.39 | -1.94 | -1.81 | -1.32 | -1.56 | -1.17 | 0.365 | 0.3625 | 0.465 | 0.21564 | 0.263588 | 0.127372 | 0.999971 | 1 | 0.783863 |
| Gabrb2 | P63137-2 | -1.02 | -1.15 | -0.74 | -0.94 | -1 | -0.71 | -0.52 | -0.18 | -1.17 | -1.24 | -0.53 | -1.18 | -0.89 | -0.78 | -0.81 | -0.41 | 0.36 | -0.0675 | 0.24 | 0.143969 | 0.765175 | 0.245667 | 0.999971 | 1 | 0.83858 |
| Gabrb3 | Q8C446 | -0.13 | -0.89 | -1.28 | -1.02 | -1.21 | -0.9 | -1.61 | -1.25 | -1.39 | -0.45 | -1.44 | -1.52 | -1.31 | -1 | -1.46 | -0.91 | -0.4125 | -0.37 | -0.34 | 0.187587 | 0.30283 | 0.261543 | 0.999971 | 1 | 0.842262 |
| Gabrd | P22933 | 0.29 | -1.29 | -0.87 | -0.18 | -1.42 | -1 | -1.22 | -0.58 | -1.49 | -0.62 | -1.1 | -1.71 | -1.88 | -0.87 | -1.44 | -0.97 | -0.5425 | -0.7175 | -0.7775 | 0.179141 | 0.105829 | 0.081188 | 0.999971 | 1 | 0.755254 |
| Gabrg1 | Q9R0Y8 | 0.54 | -0.97 | 0.41 | 0.08 | 1.15 | 1.75 | 0.42 | 0.5 | 1.51 | 1.86 | 0.92 | 1.37 | 2.19 | 1.94 | 0.86 | 1.18 | 0.94 | 1.4 | 1.5275 | 0.055977 | 0.005158 | 0.00671 | 0.999971 | 1 | 0.582714 |
| Gabrg2 | P22723-2 | -0.48 | -0.82 | -0.83 | -1.21 | -0.57 | 0.15 | -0.57 | -0.96 | -0.62 | -0.52 | -0.43 | -0.91 | -0.06 | -0.48 | -1.11 | -0.96 | 0.3475 | 0.215 | 0.1825 | 0.247203 | 0.345289 | 0.538148 | 0.999971 | 1 | 0.910077 |
| Gabrg2_1 | P22723 | -0.26 | -0.75 | -0.26 | -0.35 | -1.59 | -0.63 | -1.09 | -0.99 | -1 | 0.09 | -1.14 | -1.14 | -0.74 | -0.76 | -0.67 | 0.19 | -0.67 | -0.3925 | -0.09 | 0.02597 | 0.242225 | 0.744383 | 0.999971 | 1 | 0.961064 |
| Gabrg3 | P27681 | 1.56 | 4.48 | -1.6 | 0.81 | 6.18 | 3.37 | -1.53 | -0.8 | 6.91 | 0.73 | -1.35 | -1.36 | 8.48 | -1.14 | -1.25 | 0.14 | 0.4925 | -0.08 | 0.245 | 0.800707 | 0.969133 | 0.916553 | 0.999971 | 1 | 0.992633 |
| Gad1 | P48318 | -0.22 | -0.77 | -0.57 | -0.47 | -1.04 | -0.59 | -0.72 | -0.4 | -0.94 | -0.54 | -0.97 | -1.23 | -0.76 | -0.53 | -0.34 | -0.57 | -0.18 | -0.4125 | -0.0425 | 0.423049 | 0.090644 | 0.833011 | 0.999971 | 1 | 0.979036 |
| Gad2 | P48320 | -0.39 | -0.79 | -0.39 | -0.42 | -0.25 | 0.1 | -0.19 | -0.03 | 0.12 | -0.83 | -0.8 | -0.44 | 0.39 | -0.25 | 0.37 | -0.6 | 0.405 | 0.01 | 0.475 | 0.059683 | 0.969568 | 0.115682 | 0.999971 | 1 | 0.78268 |
| Gadd45gip | Q9CR59 | 0.91 | -0.94 | -0.26 | -0.2 | -0.83 | -0.2 | 0.64 | -0.08 | -0.64 | 0.31 | -0.19 | -0.49 | -1.65 | -0.94 | -0.45 | -1.12 | 0.005 | -0.13 | -0.9175 | 0.991252 | 0.755155 | 0.058775 | 0.999971 | 1 | 0.725348 |
| Gak | A0A0R4J0F | -0.19 | -0.38 | -0.26 | -0.29 | -0.53 | -0.11 | -0.1 | -0.29 | -0.41 | -0.05 | -0.25 | -0.45 | -0.26 | -0.39 | -0.28 | -0.32 | 0.0225 | -0.01 | -0.0325 | 0.902197 | 0.954617 | 0.842172 | 0.999971 | 1 | 0.979814 |
| Gal3st3 | P61315 | -0.51 | 0.1 | -0.05 | 0.18 | 0.23 | 0.09 | -0.16 | -0.07 | -0.03 | 0.25 | 0.35 | 0.4 | -0.34 | 0.18 | -0.22 | 0.61 | 0.0925 | 0.3125 | 0.1275 | 0.676137 | 0.182482 | 0.651498 | 0.999971 | 1 | 0.933842 |
| Galc | P54818 | 0.43 | 0.48 | 0.72 | 1.07 | 0.3 | 0.47 | 0.33 | 0.95 | -0.11 | 0.15 | 0.45 | 0.41 | -0.26 | 0.68 | 1.11 | 1.34 | -0.1625 | -0.45 | 0.0425 | 0.506829 | 0.077804 | 0.908901 | 0.999971 | 1 | 0.991707 |
| Gale | Q8R059 | 0.19 | 0.97 | 1.1 | 0.68 | 1.58 | 0.92 | 1.09 | 1.03 | 1.72 | 0.32 | 1.28 | 1.05 | 1.95 | 0.28 | 0.82 | 1.07 | 0.42 | 0.3575 | 0.295 | 0.144352 | 0.321527 | 0.454509 | 0.999971 | 1 | 0.887217 |
| Galk1 | Q9R0N0 | 0.12 | 0.51 | 0.65 | 0.35 | 0.59 | 0.42 | 0.99 | 0.15 | 0.34 | 0 | 0.25 | -0.06 | 0.5 | 0.62 | 0.35 | 0.22 | 0.13 | -0.275 | 0.015 | 0.593365 | 0.195886 | 0.940639 | 0.999971 | 1 | 0.996218 |
| Galk2 | Q68FH4 | 3.02 | 1.7 | 0.82 | 1.8 | 1.47 | 1.57 | 1.23 | 1.18 | 1.74 | 2.37 | 0.91 | 1.54 | 0.54 | 1.25 | 1.46 | 1.56 | -0.4725 | -0.195 | -0.6325 | 0.294521 | 0.700776 | 0.205062 | 0.999971 | 1 | 0.81617 |
| Galm | Q8K157 | 0.27 | 0.73 | 1.03 | 0.98 | 0.66 | 0.18 | 1.07 | 1.05 | 0.26 | 0.14 | 0.48 | 0.7 | 0.73 | 0.18 | 0.89 | 0.59 | -0.0125 | -0.3575 | -0.155 | 0.965027 | 0.165581 | 0.550123 | 0.999971 | 1 | 0.912765 |
| Galns | Q571E4 | 0.01 | 0.22 | 0.48 | 0.58 | -0.08 | -0.12 | 0.42 | 0.68 | -0.19 | -0.21 | 0.07 | -0.09 | -0.14 | 0.42 | -0.07 | 0.15 | -0.0975 | -0.4275 | -0.2325 | 0.706633 | 0.056244 | 0.313144 | 0.999971 | 1 | 0.849656 |
| Galnt1 | O08912 | -0.05 | 0.59 | 0.17 | 0.56 | 0.54 | 0.47 | -0.18 | 0.42 | 0.93 | 0.64 | 0.23 | 0.59 | 0.73 | 1.03 | 0.49 | 0.7 | -0.005 | 0.28 | 0.42 | 0.984258 | 0.264547 | 0.095237 | 0.999971 | 1 | 0.78268 |
| Galnt13 | Q8CF93 | -0.63 | -0.23 | -0.48 | 0.13 | 0.36 | -0.29 | -0.45 | 0.34 | -0.64 | -0.02 | -0.15 | 0.17 | -0.59 | 0.25 | 0.34 | 0.59 | 0.2925 | 0.1425 | 0.45 | 0.315717 | 0.588263 | 0.172766 | 0.999971 | 1 | 0.806242 |
| Galnt14 | Q08EC9 | -1.44 | -1.51 | 0.04 | 1.53 | -1.31 | -2.15 | -1.17 | -0.04 | -1.2 | -1.94 | 0.61 | -1.09 | -1.28 | -1.18 | 0.41 | 1.31 | -0.8225 | -0.56 | 0.16 | 0.293723 | 0.497226 | 0.852316 | 0.999971 | 1 | 0.98187 |
| Galnt16 | Q9JJ61 | 0.7 | 1.34 | 0.96 | 1.03 | 1.35 | 1.35 | 1.46 | 1.49 | 1.67 | 1.2 | 1.24 | 1.61 | 2.26 | 1.64 | 1.43 | 1.16 | 0.405 | 0.4225 | 0.615 | 0.065817 | 0.082037 | 0.054625 | 0.999971 | 1 | 0.725348 |
| Galnt17 | Q7TT15 | 0.52 | 0.77 | 0.55 | 0.7 | 0.7 | 0.9 | 0.16 | 0.89 | 0.61 | 0.84 | 0.76 | 1.14 | 0.3 | 0.96 | 0.94 | 1.64 | 0.0275 | 0.2025 | 0.325 | 0.90257 | 0.300576 | 0.283096 | 0.999971 | 1 | 0.844538 |
| Galnt2 | Q6PB93 | 1.36 | 0.86 | 1.14 | 1.43 | 0.7 | 1.19 | 1.98 | 1.05 | 1.09 | 1.35 | 1.27 | 1.15 | 0.55 | 1.31 | 1.2 | 0.37 | 0.0325 | 0.0175 | -0.34 | 0.915309 | 0.92915 | 0.247711 | 0.999971 | 1 | 0.839012 |
| Galt | Q03249 | -0.19 | 0.28 | 1.07 | 0.92 | 0.19 | 0.04 | 1.87 | 0.74 | 0.62 | 0.15 | 0.73 | 0.02 | 0.01 | 0.62 | 0.44 | 0.21 | 0.19 | -0.14 | -0.2 | 0.689283 | 0.679359 | 0.540415 | 0.999971 | 1 | 0.910077 |
| Gamt | O35969 | -1.57 | -1.73 | -0.54 | -1.02 | -1.66 | -2.11 | -0.69 | -0.66 | -1.35 | -1.9 | -0.68 | 0.15 | -1.72 | -1.76 | -0.23 | -0.63 | -0.065 | 0.27 | 0.13 | 0.879066 | 0.58108 | 0.770746 | 0.999971 | 1 | 0.963498 |
| Gan | Q8CA72 | -0.48 | -0.65 | -0.39 | -0.35 | -0.2 | -0.76 | 0.28 | -0.23 | -0.44 | -0.76 | -0.26 | -0.17 | -0.42 | -0.56 | -0.69 | -0.62 | 0.24 | 0.06 | -0.105 | 0.350804 | 0.764504 | 0.553426 | 0.999971 | 1 | 0.913595 |
| Ganab | Q8BHN3 | -1.65 | -5 | -4.95 | -3.74 | -4 | -6.88 | -3.37 | -2.61 | -3.81 | -2.86 | -2.03 | -2.85 | -4.29 | -4.11 | -2.48 | -3.51 | -0.38 | 0.9475 | 0.2375 | 0.72694 | 0.246546 | 0.765832 | 0.999971 | 1 | 0.962465 |
| Ganab_1 | Q8BHN3-2 | -1.21 | -0.57 | -0.2 | -0.27 | -0.34 | -0.57 | -0.07 | 0.09 | -0.15 | -0.99 | 0.07 | 0.41 | -0.16 | -0.31 | 0.16 | 0.33 | 0.34 | 0.3975 | 0.5675 | 0.253584 | 0.295515 | 0.074241 | 0.999971 | 1 | 0.746069 |
| Ganc | A2AQJ8 | 0.94 | -0.71 | 2.17 | 2.45 | 1.49 | 1.64 | 1.35 | 2.11 | 0.65 | -1.79 | 1.3 | 1.86 | 1.52 | 2.11 | 1.62 | 1.91 | 0.435 | -0.7075 | 0.5775 | 0.520686 | 0.473176 | 0.395744 | 0.999971 | 1 | 0.871816 |
| Gap43 | P06837 | -1.32 | -2.18 | -1.3 | -1.41 | -1.4 | -1.8 | 0.36 | -1.91 | -0.83 | -2.06 | -1.56 | -1.55 | -0.69 | -2.92 | -1.67 | -2.78 | 0.365 | 0.0525 | -0.4625 | 0.492066 | 0.873069 | 0.385985 | 0.999971 | 1 | 0.871816 |
| Gapdh | P16858 | -0.99 | -1.01 | -1.22 | -0.85 | -0.78 | -1.03 | -0.88 | -0.8 | -0.89 | -1.4 | -1.17 | -0.79 | -1.32 | -0.8 | -0.65 | -0.79 | 0.145 | -0.045 | 0.1275 | 0.423076 | 0.827762 | 0.556318 | 0.999971 | 1 | 0.915051 |
| Gapdh_1 | S4R1W8 | -0.54 | 0.66 | -1.76 | 0.29 | -0.03 | 0.9 | -0.48 | -0.29 | -0.14 | -1.15 | 0.58 | -0.57 | -0.65 | 0.43 | -0.22 | -0.87 | 0.3625 | 0.0175 | 0.01 | 0.526656 | 0.976539 | 0.98571 | 0.999971 | 1 | 0.99925 |
| Gapdhs | A0A0R4J0X | -0.09 | 0.37 | -0.55 | -1.31 | -0.52 | -0.09 | -0.95 | -0.65 | -0.78 | 0.12 | -0.95 | -1.06 | 0.16 | 0.05 | -1.08 | -0.53 | -0.1575 | -0.2725 | 0.045 | 0.68383 | 0.526768 | 0.917319 | 0.999971 | 1 | 0.992633 |
| Gapvd1 | Q6PAR5 | -0.45 | -0.19 | -0.15 | -0.43 | -0.36 | -0.23 | -0.16 | -0.12 | -0.14 | -0.11 | 0.23 | 0.29 | 0.23 | -0.07 | 0.09 | -0.16 | 0.0875 | 0.3725 | 0.3275 | 0.623891 | 0.082778 | 0.108574 | 0.999971 | 1 | 0.78268 |
| Gar1 | D3YZ09 | -0.94 | -0.16 | 0.8 | -0.03 | -0.05 | -0.36 | 0.43 | -0.86 | -0.2 | -0.08 | 0.46 | 0.04 | 0.11 | -0.09 | -0.27 | 0.12 | -0.1275 | 0.1375 | 0.05 | 0.763903 | 0.713278 | 0.88957 | 0.999971 | 1 | 0.987761 |
| Garem1 | Q3UFT3 | 0.23 | -0.15 | -0.48 | -0.49 | -0.03 | -0.04 | -0.56 | -0.22 | -0.14 | -0.05 | -0.39 | -0.08 | 0.37 | 0.12 | -0.31 | -0.13 | 0.01 | 0.0575 | 0.235 | 0.967031 | 0.79836 | 0.365411 | 0.999971 | 1 | 0.869526 |
| Garnl3 | Q3V0G7 | -0.39 | 0.42 | -0.3 | -0.11 | 0.3 | 0.27 | -0.41 | 0.04 | 0.29 | 0.08 | 0 | -0.07 | 0.77 | 0.69 | -0.12 | 0.15 | 0.145 | 0.17 | 0.4675 | 0.588183 | 0.470065 | 0.136588 | 0.999971 | 1 | 0.784948 |
| Garre1 | Q8C5X1 | -0.24 | 0.43 | 0.94 | -3.89 | 0.47 | -0.95 | 1.64 | -2.07 | -0.37 | -0.09 | 1.89 | 2.54 | 0.5 | 0.2 | 0.07 | -4.8 | 0.4625 | 1.6825 | -0.3175 | 0.703266 | 0.17675 | 0.830896 | 0.999971 | 1 | 0.978535 |
| Gars1 | Q9CZD3 | 0.51 | 0.69 | 0.78 | 0.34 | 0.69 | 0.66 | 0.74 | 0.32 | 0.91 | 0.61 | 0.93 | 0.97 | 0.36 | 0.67 | 0.61 | 0.36 | 0.0225 | 0.275 | -0.08 | 0.909159 | 0.172484 | 0.680109 | 0.999971 | 1 | 0.942957 |
| Gart | Q64737 | 0.7 | 0.48 | 0.57 | 0.36 | 0.21 | 0.42 | 0.42 | 0.25 | 0.11 | 0.39 | 0.25 | 0.46 | 0.19 | 0.49 | 0.67 | 0.66 | -0.2025 | -0.225 | -0.025 | 0.267497 | 0.229617 | 0.898509 | 0.999971 | 1 | 0.989489 |
| Gas2 | P11862 | -1.36 | -0.64 | -0.18 | -0.11 | -0.37 | -0.15 | -1.09 | -0.82 | -0.96 | -0.98 | -0.4 | -0.39 | -1.37 | -0.17 | -0.49 | -0.71 | -0.035 | -0.11 | -0.1125 | 0.920557 | 0.740259 | 0.763155 | 0.999971 | 1 | 0.961892 |
| Gas2l1 | Q8JZP9 | 1.31 | -1.43 | -0.05 | 0.01 | -1.44 | -0.57 | 0.06 | -0.24 | -1.53 | 0.39 | 0.11 | 0.17 | -1.79 | -1.26 | -0.31 | -0.09 | -0.5075 | -0.175 | -0.8225 | 0.400403 | 0.788524 | 0.211866 | 0.999971 | 1 | 0.823112 |
| Gas6 | Q61592 | -1.74 | 1.11 | -0.8 | -1.31 | 0.84 | 0.83 | -0.79 | -1.07 | -1.24 | 1.52 | -0.26 | 0.07 | -2.95 | -1.73 | -0.98 | -0.6 | 0.6375 | 0.7075 | -0.88 | 0.393976 | 0.370153 | 0.251202 | 0.999971 | 1 | 0.839012 |
| Gas7 | F6QB26 | -1.26 | -0.43 | -0.82 | -1.45 | 0.04 | -0.8 | 0.41 | -1.16 | 0.52 | -1.3 | 0.11 | -0.26 | 0.58 | -1.34 | -1.03 | -1.26 | 0.6125 | 0.7575 | 0.2275 | 0.159644 | 0.10523 | 0.633145 | 0.999971 | 1 | 0.93301 |
| Gatad1 | Q920S3 | -0.09 | 0.31 | 0.44 | 0.77 | -0.12 | -0.39 | 0.16 | 0.67 | -0.05 | -0.43 | 1.39 | 0.83 | 0.5 | -1.02 | 0.9 | 1.07 | -0.2775 | 0.0775 | 0.005 | 0.362803 | 0.856077 | 0.99159 | 0.999971 | 1 | 0.99925 |
| Gatad2a | A0A1I7Q4C | 1.41 | 0.92 | 2.11 | 2.13 | 1.14 | 1.32 | 2.22 | 0.8 | 0.74 | 1.23 | 0.7 | 1.11 | -0.74 | 0.54 | 0.81 | 0.74 | -0.2725 | -0.6975 | -1.305 | 0.504204 | 0.055735 | 0.015252 | 0.999971 | 1 | 0.613095 |
| Gatad2b | A1L3S7 | 2.4 | 1.53 | 1.99 | 2.29 | 1.6 | 1.46 | 1.79 | 1.97 | 1.48 | 2.11 | 2.09 | 1.83 | 0.91 | 1.45 | 1.81 | 2.04 | -0.3475 | -0.175 | -0.5 | 0.190381 | 0.512709 | 0.141812 | 0.999971 | 1 | 0.784967 |
| Gatb | Q99JT1 | -0.09 | -0.06 | -0.24 | -0.29 | -0.4 | -0.24 | -0.14 | -0.29 | -0.66 | -0.02 | -0.2 | -0.18 | -0.24 | -0.27 | -0.51 | -0.43 | -0.0975 | -0.095 | -0.1925 | 0.571975 | 0.639009 | 0.286833 | 0.999971 | 1 | 0.846151 |
| Gatc | Q8CBY0 | 0.86 | -0.2 | -0.69 | 0.05 | -0.43 | -0.23 | -0.22 | -0.15 | -0.5 | -0.21 | -0.18 | -0.33 | -0.62 | -0.25 | -0.55 | -0.65 | -0.2625 | -0.31 | -0.5225 | 0.433417 | 0.361236 | 0.145405 | 0.999971 | 1 | 0.791767 |
| Gatd1 | Q8BFQ8 | -0.41 | 0 | 0.19 | 0.15 | -0.12 | -0.72 | 0.13 | -0.27 | 0.11 | -0.93 | 0.06 | 0.3 | -0.11 | -0.17 | -0.04 | 0.28 | -0.2275 | -0.0975 | 0.0075 | 0.378129 | 0.755561 | 0.972387 | 0.999971 | 1 | 0.99925 |
| Gatd3 | Q9D172 | -0.24 | -0.6 | -0.07 | 0.2 | -0.85 | -0.65 | -0.17 | 0.29 | -0.92 | -0.55 | -0.42 | -0.34 | -1.07 | -0.38 | -0.08 | 0.02 | -0.1675 | -0.38 | -0.2 | 0.592396 | 0.141221 | 0.517413 | 0.999971 | 1 | 0.905815 |
| Gatm | Q9D964 | 0.96 | -0.68 | -0.29 | -0.39 | -0.69 | -0.77 | 0.45 | 0.9 | -0.64 | 0.7 | -0.7 | -0.24 | -0.44 | -0.23 | 0.34 | -0.17 | 0.0725 | -0.12 | -0.025 | 0.88706 | 0.793235 | 0.948164 | 0.999971 | 1 | 0.997756 |
| Gba | P17439 | -0.94 | 0.22 | -0.53 | -0.89 | -0.35 | -0.47 | -0.68 | -0.45 | -0.15 | -0.86 | -0.63 | -0.45 | -0.22 | -0.24 | -0.4 | -0.67 | 0.0475 | 0.0125 | 0.1525 | 0.869334 | 0.967919 | 0.610645 | 0.999971 | 1 | 0.92673 |
| Gba2 | Q69ZF3 | 0.42 | -0.52 | -0.65 | -0.34 | -0.45 | -0.19 | -0.53 | -0.04 | -0.61 | 0.51 | -0.54 | -0.55 | -0.45 | 0.41 | -0.01 | -0.16 | -0.03 | -0.025 | 0.22 | 0.914726 | 0.943607 | 0.479192 | 0.999971 | 1 | 0.894438 |
| Gbe1 | Q9D6Y9 | 0.58 | -0.12 | -0.26 | 0.24 | -0.27 | -0.38 | -0.16 | -0.43 | -0.2 | -0.22 | -0.33 | -0.47 | -0.79 | -0.56 | -0.58 | -0.18 | -0.42 | -0.415 | -0.6375 | 0.099902 | 0.101893 | 0.031456 | 0.999971 | 1 | 0.678474 |
| Gbf1 | Q6DFZ1 | 0.53 | 0.72 | 1 | 1.15 | 0.75 | 0.8 | 0.63 | 1.4 | 0.6 | 0.49 | 0.9 | 1.27 | 0.53 | 1.07 | 1.17 | 1.48 | 0.045 | -0.035 | 0.2125 | 0.856812 | 0.888299 | 0.430156 | 0.999971 | 1 | 0.881611 |
| Gbp11 | A9YVJ5 | 0.61 | 1.76 | 1.41 | 1.65 | 1.15 | 0.66 | 0.69 | 1.15 | 0.61 | 0.82 | 1.23 | 1.69 | 0.97 | 1.85 | 1.05 | 1.49 | -0.445 | -0.27 | -0.0175 | 0.165212 | 0.445102 | 0.957757 | 0.999971 | 1 | 0.998622 |
| Gbp4 | Q61107 | 1.27 | 1.65 | 1.19 | 1.31 | 1.86 | 1.26 | -0.04 | 0.54 | 0.58 | 1.01 | 0.86 | 1.27 | 0.82 | 2.09 | 0.36 | 0.46 | -0.45 | -0.425 | -0.4225 | 0.285889 | 0.078198 | 0.300727 | 0.999971 | 1 | 0.847402 |
| Gbp5 | Q8BMN7 | -1.04 | 2.07 | 0.89 | 0.94 | 2.02 | 2.05 | -0.08 | 0.99 | 2.03 | -0.97 | 0.46 | 0.19 | 2.53 | 2.3 | 1.73 | 0.01 | 0.53 | -0.2875 | 0.9275 | 0.479845 | 0.722815 | 0.251407 | 0.999971 | 1 | 0.839012 |
| Gbp7 | Q91Z40 | 1.08 | 0.86 | 1.17 | 1.45 | 1.14 | 0.63 | 1.2 | 1.38 | 0.72 | 0.87 | 1.33 | 1.69 | 0.82 | 1.31 | 1.27 | 1.26 | -0.0525 | 0.0125 | 0.025 | 0.824407 | 0.963086 | 0.90774 | 0.999971 | 1 | 0.991405 |
| Gbp9 | Q8BTS3 | -3.79 | 5.83 | 0.54 | 0.57 | 8.52 | 3.49 | -0.12 | -0.26 | 7.82 | -2.45 | 0.45 | 0.36 | 12.63 | 1.63 | 0.22 | 0.51 | 2.12 | 0.7575 | 2.96 | 0.409451 | 0.773393 | 0.36054 | 0.999971 | 1 | 0.869526 |
| Gc | P21614 | 0.9 | 3.09 | 1.25 | 3.35 | 4.09 | 1.68 | 3.07 | 0.93 | 1.93 | 0.26 | 2.45 | 2.07 | 1.97 | 1.17 | 2.19 | 2.09 | 0.295 | -0.47 | -0.2925 | 0.72788 | 0.51885 | 0.633397 | 0.999971 | 1 | 0.93301 |
| Gca | Q8VC88 | -0.87 | 0.13 | -0.01 | 0.03 | -0.11 | 0.48 | -0.45 | 0.96 | 0.04 | -0.01 | 0.13 | 0.22 | 0.25 | 0.7 | 0.42 | 0.5 | 0.4 | 0.275 | 0.6475 | 0.303752 | 0.304855 | 0.037601 | 0.999971 | 1 | 0.700096 |
| Gcat | O88986 | -1.09 | -0.3 | -0.58 | -0.9 | -0.67 | -0.91 | -0.95 | -0.94 | -1.16 | -1.37 | -0.81 | -0.61 | -0.78 | -0.74 | -0.84 | -0.64 | -0.15 | -0.27 | -0.0325 | 0.513379 | 0.321627 | 0.883979 | 0.999971 | 1 | 0.986964 |
| Gcc1 | Q9D4H2 | 2.3 | 2.27 | 2.3 | 2.6 | 1.79 | 1.88 | 1.9 | 2.37 | 1.71 | 2.41 | 2.26 | 2.02 | 0.99 | 2.02 | 1.85 | 2.55 | -0.3825 | -0.2675 | -0.515 | 0.090099 | 0.234231 | 0.147672 | 0.999971 | 1 | 0.793248 |
| Gcc2 | E9PWI6 | 0.77 | 0.97 | 1.05 | 0.93 | 0.92 | 0.89 | 0.76 | 0.84 | 0.81 | 1.06 | 0.9 | 1.03 | 0.6 | 1.02 | 0.84 | 0.83 | -0.0775 | 0.02 | -0.1075 | 0.646082 | 0.905837 | 0.55899 | 0.999971 | 1 | 0.915051 |
| Gcdh | Q60759 | -1.11 | -0.53 | -0.79 | -0.77 | -0.2 | -0.69 | 0.04 | -0.4 | -0.03 | -0.81 | -0.89 | -0.29 | 0.48 | -0.63 | -0.57 | -0.86 | 0.4875 | 0.295 | 0.405 | 0.060919 | 0.274841 | 0.23116 | 0.999971 | 1 | 0.830179 |
| Gclc | P97494 | 0 | 0.48 | 0.32 | 0.67 | 0.35 | 0.55 | 0.2 | 0.69 | 0.38 | 0.21 | 0.14 | 0.37 | 0.49 | 0.72 | 0.51 | 0.67 | 0.08 | -0.0925 | 0.23 | 0.718683 | 0.652494 | 0.283761 | 0.999971 | 1 | 0.844538 |
| Gclm | O09172 | 0.26 | 0.72 | -0.02 | 0.36 | 0.59 | 0.6 | 0.01 | 0.55 | 0.67 | 0.55 | 0.02 | 0.12 | 0.57 | 0.87 | 0.05 | 0.31 | 0.1075 | 0.01 | 0.12 | 0.657687 | 0.967738 | 0.643914 | 0.999971 | 1 | 0.93301 |
| Gcn1 | E9PVA8 | 0.83 | 1.23 | 1.2 | 1.12 | 1.15 | 1 | 1.1 | 1.31 | 1.2 | 1.14 | 1.35 | 1.32 | 0.97 | 1.19 | 1.25 | 1.38 | 0.045 | 0.1575 | 0.1025 | 0.807882 | 0.387372 | 0.596429 | 0.999971 | 1 | 0.92377 |
| Gcsh | Q91WK5 | -0.42 | -0.64 | -0.67 | -0.5 | -0.89 | -0.66 | -0.65 | -0.4 | -0.92 | -0.49 | -0.6 | -0.59 | -0.66 | -0.41 | -0.36 | -0.51 | -0.0925 | -0.0925 | 0.0725 | 0.62298 | 0.612032 | 0.681091 | 0.999971 | 1 | 0.943097 |
| Gda | Q9R111 | -0.41 | -0.11 | -3.13 | -0.61 | 0.21 | -0.19 | 0.31 | -0.26 | 0.18 | -0.43 | 0.17 | 0.44 | -0.12 | -0.67 | -0.17 | -0.79 | 1.0825 | 1.155 | 0.6275 | 0.120465 | 0.106784 | 0.348695 | 0.999971 | 1 | 0.865162 |
| Gdap1 | O88741 | -0.87 | -1.36 | -0.7 | -1.12 | -0.93 | -0.77 | -0.44 | -0.75 | -1 | -1.07 | -0.47 | -0.42 | -0.74 | -0.99 | -0.65 | -0.54 | 0.29 | 0.2725 | 0.2825 | 0.21085 | 0.29222 | 0.218766 | 0.999971 | 1 | 0.827064 |
| Gdap1l1 | Q8VE33 | -0.16 | -0.04 | 0.12 | 0.18 | -0.18 | 0.22 | 0.41 | 0.01 | -0.08 | 0.06 | -0.14 | 0.01 | 0.15 | 0.18 | 0.22 | 0.34 | 0.09 | -0.0625 | 0.1975 | 0.661068 | 0.716759 | 0.277565 | 0.999971 | 1 | 0.843812 |
| Gdap2 | Q9DBL2-2 | -0.1 | -0.54 | -0.01 | -0.12 | -0.25 | -0.01 | -0.27 | -0.25 | -0.35 | 0.07 | -0.11 | 0.09 | -0.02 | -0.07 | 0.08 | 0.42 | -0.0025 | 0.1175 | 0.295 | 0.989792 | 0.571471 | 0.187953 | 0.999971 | 1 | 0.809025 |
| Gde1 | Q9JL56 | -0.01 | 0.4 | -0.29 | -0.23 | 0.67 | 0.11 | 0.07 | 0.13 | 0.98 | -0.01 | -0.11 | -0.52 | 1.39 | -0.03 | -0.06 | 0.07 | 0.2775 | 0.1175 | 0.375 | 0.270885 | 0.736833 | 0.328622 | 0.999971 | 1 | 0.856472 |
| Gdi1 | P50396 | -0.06 | 0.27 | 0.31 | 0.06 | -0.09 | 0.2 | 0.05 | 0.18 | -0.06 | -0.1 | 0.18 | 0.04 | -0.15 | 0.17 | 0.26 | 0.27 | -0.06 | -0.13 | -0.0075 | 0.745191 | 0.476253 | 0.969356 | 0.999971 | 1 | 0.998787 |
| Gdi2 | Q61598 | -0.98 | -1.2 | -1.19 | -1.23 | -0.77 | -1.33 | -0.86 | -1.26 | -0.77 | -1.01 | -0.98 | -1.29 | -1.42 | -1.05 | -1.3 | -1.3 | 0.095 | 0.1375 | -0.1175 | 0.645906 | 0.467794 | 0.516504 | 0.999971 | 1 | 0.905815 |
| Gdpd1 | Q9CRY7 | -0.89 | 0.47 | -0.05 | -0.94 | 0.33 | -0.17 | 0.15 | -0.69 | 0.38 | -0.76 | 0.05 | 0.53 | 0.75 | -0.6 | -0.68 | -0.96 | 0.2575 | 0.4025 | -0.02 | 0.516275 | 0.356214 | 0.96663 | 0.999971 | 1 | 0.998787 |
| Gdpd2 | Q9ESM6 | -1.5 | -1.64 | -0.92 | -0.89 | -1.54 | -1.72 | -1.2 | -0.28 | -1.27 | -1.02 | -0.87 | -1.09 | -0.42 | -0.98 | -0.82 | -1.43 | 0.0525 | 0.175 | 0.325 | 0.885458 | 0.474545 | 0.288894 | 0.999971 | 1 | 0.847402 |
| Gdpd5 | A0A1L1STK | -0.39 | -1.28 | -1.76 | -1.56 | -1.14 | -0.95 | -1.57 | -1.65 | -1.2 | -0.3 | -1.43 | -1.9 | -0.82 | -1.2 | -1.52 | -1.27 | -0.08 | 0.04 | 0.045 | 0.815152 | 0.925606 | 0.892836 | 0.999971 | 1 | 0.98828 |
| Gdpgp1 | Q3TLS3 | -0.2 | 0.48 | -0.21 | -0.06 | 0.01 | -0.1 | -0.18 | -0.23 | -0.02 | -0.02 | 0.3 | -0.26 | -0.02 | -0.13 | -0.49 | -0.13 | -0.1275 | -0.0025 | -0.195 | 0.56053 | 0.991427 | 0.408293 | 0.999971 | 1 | 0.875014 |
| Gemin4 | Q6P6L6 | 0.47 | 1.41 | 0.92 | 1.03 | 1.77 | 1.81 | 0.44 | 0.86 | 1.22 | 0.86 | 1.18 | 0.75 | 1.83 | 2.01 | 1.07 | 1.43 | 0.2625 | 0.045 | 0.6275 | 0.493433 | 0.857777 | 0.059281 | 0.999971 | 1 | 0.725348 |

| | | | | | | | | | | | | | | | | | | | | | | | | | | |
|---|---|---|---|---|---|---|---|---|---|---|---|---|---|---|---|---|---|---|---|---|---|---|---|---|---|---|
| Gemin5 | A2AFQ9 | 0.81 | 1.48 | 1.65 | 1.47 | 1.26 | 1.42 | 1.28 | 1.64 | 1.18 | 1.15 | 1.48 | 1.33 | 1.28 | 1.69 | 1.66 | 1.91 | 0.0475 | -0.0675 | 0.2825 | 0.842233 | 0.773081 | 0.283329 | 0.999971 | 1 | 0.844538 |
| Gemin7 | D3Z684 | 1.48 | 1.13 | 1.64 | 1.72 | 0.68 | 1.59 | 0.6 | 1.37 | 1.2 | 1.84 | 0.31 | 1.39 | -0.8 | 1.06 | 1.25 | 0.87 | -0.4325 | -0.3075 | -0.8975 | 0.162527 | 0.380765 | 0.077483 | 0.999971 | 1 | 0.752059 |
| Get1 | Q8K0D7 | -0.69 | -0.29 | 0.28 | 0.53 | -0.31 | 0.55 | 0.39 | 0.35 | 0.09 | 0.23 | 0.44 | -0.61 | -0.37 | 0.02 | 0.34 | -0.05 | 0.2875 | 0.08 | 0.0275 | 0.398525 | 0.819929 | 0.930606 | 0.999971 | 1 | 0.994744 |
| Get3 | O54984 | 0.42 | 0.47 | 0.32 | 0.16 | 0.13 | 0.6 | 0.13 | 0.26 | 0.31 | 0.48 | 0.31 | 0.36 | 0.41 | 0.38 | 0.23 | 0.52 | -0.0625 | 0.0225 | 0.0425 | 0.747844 | 0.893321 | 0.809889 | 0.999971 | 1 | 0.973791 |
| Get4 | Q9D1H7 | 0.53 | 0.99 | 0.71 | 0.64 | 0.89 | 0.84 | 0.55 | 0.37 | 0.9 | 0.54 | 0.4 | -0.01 | 0.98 | 0.81 | 0.39 | 0.37 | -0.055 | -0.26 | -0.08 | 0.792464 | 0.298307 | 0.721868 | 0.999971 | 1 | 0.955155 |
| Gfap | P03995 | -1.49 | -3.33 | -0.49 | -0.77 | -1.97 | -2.94 | -0.57 | -0.9 | -2.37 | -1.96 | -0.96 | -0.73 | -2.37 | -2.17 | -0.28 | -0.9 | -0.075 | 0.015 | 0.09 | 0.920574 | 0.982473 | 0.902432 | 0.999971 | 1 | 0.990759 |
| Gfap_1 | P03995-2 | -1.42 | -2.02 | -2.22 | -3.22 | -1.77 | -2.58 | -2.33 | -2.96 | -2.35 | -1.4 | -1.56 | -3.96 | -1.75 | -2.4 | -2.8 | -1.03 | -0.19 | -0.0975 | 0.225 | 0.656956 | 0.878057 | 0.654725 | 0.999971 | 1 | 0.934061 |
| Gfer | P56213 | 2.39 | 0.64 | 1.06 | 1.09 | 0.77 | 1.59 | 1.68 | 1.21 | 0.74 | 1.4 | 0.82 | 1.05 | 0.93 | 0.97 | 1.09 | 0.64 | 0.0175 | -0.2925 | -0.3875 | 0.966017 | 0.462624 | 0.320976 | 0.999971 | 1 | 0.854682 |
| Gfm1 | Q8K0D5 | -0.64 | -0.29 | -0.11 | -0.17 | 0.25 | -0.33 | 0.36 | -0.18 | -0.16 | -0.07 | 0.08 | 0.42 | -0.27 | -0.35 | -0.01 | -0.1 | 0.3275 | 0.37 | 0.12 | 0.19201 | 0.115087 | 0.553947 | 0.999971 | 1 | 0.913787 |
| Gfm2 | Q8R2Q4-3 | -0.25 | -0.13 | -0.3 | -0.08 | -0.18 | -0.12 | -0.33 | -0.28 | -0.29 | -0.04 | -0.37 | -0.51 | 0.27 | 0.06 | -0.43 | -0.17 | -0.0375 | -0.1125 | 0.1225 | 0.823705 | 0.539628 | 0.5643 | 0.999971 | 1 | 0.917087 |
| Gfod1 | Q3UHD2 | -0.62 | -0.4 | 0.07 | -0.1 | -0.29 | -0.09 | -0.37 | 0 | -0.56 | -0.09 | -0.32 | -0.33 | -0.08 | -0.26 | -0.25 | 0.4 | 0.075 | -0.0625 | 0.215 | 0.733891 | 0.777787 | 0.396384 | 0.999971 | 1 | 0.871816 |
| Gfod2 | Q9CYH5 | -0.09 | -0.68 | -0.51 | -0.35 | -0.38 | -0.7 | -0.5 | -0.16 | -0.02 | 0.43 | -0.71 | -0.51 | -0.54 | -0.63 | -0.6 | -0.28 | -0.0275 | 0.205 | -0.105 | 0.89845 | 0.492293 | 0.610312 | 0.999971 | 1 | 0.92673 |
| Gfpt1 | P47856-2 | -0.13 | 0.49 | 0.41 | -0.06 | 0.72 | 0.19 | 0.41 | 0.18 | 0.87 | -0.07 | 0.75 | 0.92 | 1.28 | 0.31 | 0.36 | 0.35 | 0.1975 | 0.44 | 0.3975 | 0.414671 | 0.157006 | 0.200592 | 0.999971 | 1 | 0.814342 |
| Gfra1 | D6RDD4 | 0.12 | 0.97 | 1.47 | 1.43 | 0.82 | 1.53 | 1.45 | 2.02 | 1.3 | 0.53 | 1.1 | 1.14 | 0.31 | 1.06 | 2.27 | 0.23 | 0.4575 | 0.02 | -0.03 | 0.252947 | 0.954388 | 0.954341 | 0.999971 | 1 | 0.998056 |
| Gfra2 | O08842 | 0.41 | 0.46 | 0.3 | 0.77 | 0.53 | 1.02 | 1.13 | 0.68 | 0.75 | 0.71 | 0.57 | -0.01 | 0.43 | 0.53 | 0.48 | 0.52 | 0.355 | 0.02 | 0.005 | 0.130515 | 0.932191 | 0.978026 | 0.999971 | 1 | 0.99925 |
| Gfus | P23591 | -1.01 | 0.07 | 0.04 | -1.07 | 0.55 | 0.2 | 0.42 | -0.15 | 0.52 | -0.77 | 0.54 | -0.05 | 0.5 | -0.21 | 0.2 | -0.04 | 0.7475 | 0.5525 | 0.605 | 0.055534 | 0.211641 | 0.109073 | 0.999971 | 1 | 0.78268 |
| Gga1 | Q8R0H9 | -0.08 | 0.07 | 0.47 | 0.26 | 0.38 | 0.37 | 0.63 | 0.34 | 0.48 | 0.31 | 0.41 | 0.61 | 0.31 | 0.22 | 0.32 | 0.71 | 0.25 | 0.2725 | 0.21 | 0.226314 | 0.183335 | 0.335538 | 0.999971 | 1 | 0.858827 |
| Gga2 | Q6P5E6 | -0.09 | 0.96 | 0.18 | 0.21 | 0.85 | 0.42 | 0.19 | 0.84 | 0.6 | 0.39 | 0.46 | 0.81 | 1.39 | 1.18 | 0.93 | 0.59 | 0.26 | 0.25 | 0.7075 | 0.380954 | 0.356018 | 0.037383 | 0.999971 | 1 | 0.700096 |
| Gga3 | A2A9W7 | 0.28 | 0.37 | 0.05 | -0.03 | 0.63 | 0.69 | 0.42 | -0.16 | 0.55 | 0.43 | 0.11 | 0.27 | 0.9 | 0.29 | 0.01 | 0.29 | 0.2275 | 0.1725 | 0.205 | 0.366208 | 0.383306 | 0.407873 | 0.999971 | 1 | 0.874911 |
| Ggact | E0CZ04 | -0.44 | 1.37 | 0.88 | 0.33 | 1.17 | 0.54 | 0.12 | 0.94 | 0.54 | -0.31 | -0.11 | 0.11 | 0.69 | 0.61 | 0.28 | 0.51 | 0.1575 | -0.4775 | -0.0125 | 0.713276 | 0.262927 | 0.974036 | 0.999971 | 1 | 0.99925 |
| Ggct | Q9D7X8 | 0.18 | 0.84 | 1.5 | 1.16 | 0.21 | 1.29 | 0.93 | 1.67 | 0.62 | 0.15 | 0.82 | -0.35 | -0.11 | 0.79 | 1.27 | 1.06 | 0.105 | -0.61 | -0.1675 | 0.793374 | 0.128946 | 0.674583 | 0.999971 | 1 | 0.941037 |
| Ggh | Q9Z0L8-2 | 3.22 | 0.94 | 0 | -0.92 | 0.87 | 1.76 | -0.15 | 0.2 | 1.45 | 2.58 | 0.2 | 0.21 | -1.38 | -0.31 | -0.06 | -0.36 | -0.14 | 0.3 | -1.3375 | 0.873572 | 0.752887 | 0.138127 | 0.999971 | 1 | 0.784948 |
| Ggps1 | Q9WTN0 | -0.64 | 0.64 | 0.08 | 0.31 | 0.52 | 0.64 | -0.18 | -0.07 | 0.59 | -0.12 | 0.25 | 0.16 | 0.33 | 0.51 | 0.26 | 0.23 | 0.13 | 0.1225 | 0.235 | 0.701437 | 0.695974 | 0.428046 | 0.999971 | 1 | 0.880793 |
| Ggt1 | Q60928 | -0.89 | -1.35 | -1.51 | -1.59 | -1.66 | -2.01 | -0.9 | -1.64 | -2.09 | -1.72 | -1.35 | -1.25 | -1.87 | -1.56 | -1.87 | -2.03 | -0.2175 | -0.2675 | -0.4975 | 0.463813 | 0.329649 | 0.051862 | 0.999971 | 1 | 0.723736 |
| Ggt5 | Q9Z2A9 | 1.21 | 1.24 | 1.03 | 0.91 | 1.44 | 0.59 | 1.53 | 1.62 | 1.9 | 1.29 | 1.2 | 1.54 | 1.65 | 0.58 | 0.7 | 0.17 | 0.1975 | 0.385 | -0.3225 | 0.470893 | 0.103787 | 0.333981 | 0.999971 | 1 | 0.85856 |
| Ggt7 | Q99JP7 | -0.43 | -0.56 | -0.8 | -1.04 | -0.84 | -0.31 | -0.87 | -0.78 | -0.76 | -0.54 | -0.58 | -0.49 | 0.03 | -0.31 | -0.48 | -0.21 | 0.0075 | 0.115 | 0.465 | 0.973685 | 0.56958 | 0.057906 | 0.999971 | 1 | 0.725348 |
| Ghdc | Q99J23 | 2.45 | -0.49 | -0.3 | 0.17 | -0.58 | -0.2 | 0.09 | 0.18 | -1.05 | 1.32 | -0.73 | -0.73 | -1.59 | -0.69 | -0.07 | -0.57 | -0.585 | -0.755 | -1.1875 | 0.369502 | 0.351047 | 0.107872 | 0.999971 | 1 | 0.78268 |
| Ghitm | Q91VC9 | 0.16 | -0.79 | -0.21 | 0.06 | -0.82 | -1.24 | -0.4 | -0.21 | -0.99 | -0.26 | 0.02 | 0.13 | -0.98 | -0.74 | -0.55 | -0.15 | -0.4725 | -0.08 | -0.41 | 0.160779 | 0.808281 | 0.180345 | 0.999971 | 1 | 0.808532 |
| Gid4 | Q9CPY6 | -0.81 | 0.08 | -1.62 | -1.68 | 0.07 | 0.32 | -1.67 | -1.79 | -0.06 | -0.28 | -2.62 | -1.87 | -0.12 | -0.27 | -2.57 | -1.46 | 0.24 | -0.2 | -0.0975 | 0.704958 | 0.768731 | 0.8794 | 0.999971 | 1 | 0.985441 |
| Gid8 | Q9D7M1 | -0.49 | 0.04 | -0.98 | -0.95 | -0.16 | 0.29 | -0.82 | -0.91 | -0.27 | -0.2 | -0.75 | -0.64 | -0.41 | 0.21 | -0.84 | -0.9 | 0.195 | 0.13 | 0.11 | 0.593452 | 0.652316 | 0.751298 | 0.999971 | 1 | 0.961064 |
| Gigyf1 | Q99MR1 | -0.17 | 0.34 | 0.28 | 0.12 | -0.37 | 0.01 | 0.09 | 0.15 | 0.15 | 0.38 | 0.67 | 0.31 | 0.91 | 0.55 | 0.41 | 0.88 | -0.1725 | 0.235 | 0.545 | 0.425441 | 0.273255 | 0.030455 | 0.999971 | 1 | 0.678474 |
| Gigyf2 | Q6Y7W8 | 1.21 | 1.02 | 0.75 | 0.53 | 1.18 | 1.8 | 2.03 | 0.53 | 1.29 | 1.42 | 1.32 | 0.76 | 1.43 | 1.5 | 0.8 | 0.74 | 0.5075 | 0.32 | 0.24 | 0.18248 | 0.206276 | 0.3864 | 0.999971 | 1 | 0.871816 |
| Gipc1 | Q9Z0G0 | -0.46 | -0.19 | -0.66 | -0.78 | -0.02 | -0.42 | -0.39 | -0.45 | -0.1 | -0.31 | -0.23 | -0.07 | 0.29 | -0.26 | -0.62 | 0.02 | 0.2025 | 0.345 | 0.38 | 0.353438 | 0.107119 | 0.166787 | 0.999971 | 1 | 0.800215 |
| Gipc3 | Q8R5M0 | 0.39 | 0.06 | 1.21 | 1.33 | 0.65 | -0.99 | 2.77 | 0.03 | -0.78 | -0.02 | 0.09 | 0.47 | 0.55 | -0.51 | 0.17 | 1.09 | -0.1325 | -0.8075 | -0.4225 | 0.862822 | 0.065056 | 0.342466 | 0.999971 | 1 | 0.861474 |
| Git1 | Q68FF6 | 0.25 | -0.42 | 0.07 | -0.38 | -0.34 | -0.45 | 0.28 | -0.47 | -0.26 | -0.31 | -0.02 | -0.02 | -0.63 | -0.85 | -0.34 | -0.73 | -0.125 | -0.0325 | -0.5175 | 0.63848 | 0.88372 | 0.051605 | 0.999971 | 1 | 0.723736 |
| Git2 | E9PVA6 | 0.48 | -0.44 | 0.38 | 0.39 | -0.69 | -0.19 | 0.4 | 0.17 | -0.04 | 0.37 | 0.63 | 0.94 | -0.42 | -0.36 | 0.19 | 0.14 | -0.28 | 0.2725 | -0.315 | 0.395234 | 0.383079 | 0.284249 | 0.999971 | 1 | 0.844538 |
| Gja1 | P23242 | -0.79 | -0.88 | -1.06 | -1.21 | -1.11 | -0.41 | -1.15 | -1.13 | -0.99 | -0.74 | -1.48 | -1.65 | -1.5 | -0.51 | -1.2 | -1.39 | 0.035 | -0.23 | -0.165 | 0.882679 | 0.378291 | 0.536324 | 0.999971 | 1 | 0.910077 |
| Gjb1 | P28230 | -0.45 | -1.44 | -4.95 | -3.8 | -1.1 | -0.85 | -5.18 | -5.58 | -1.27 | -2.12 | -6.38 | -6.03 | -1.84 | -1.11 | -4.23 | -4.51 | -0.5175 | -1.29 | -0.2625 | 0.723248 | 0.399233 | 0.826182 | 0.999971 | 1 | 0.976757 |
| Gjb6 | P70689 | -1.03 | -1.18 | -0.18 | -0.15 | -0.82 | -1.09 | -1.21 | 1.52 | -0.86 | -1.23 | -2.6 | 1.48 | -1.36 | -1.03 | 0.45 | 0.88 | 0.235 | -0.1675 | 0.37 | 0.712907 | 0.835409 | 0.516077 | 0.999971 | 1 | 0.905815 |
| Gjc2 | Q8BQU6 | 0.64 | 1.58 | -0.66 | 0.85 | 0.99 | 0.5 | -0.32 | 0.22 | 1.48 | 0.33 | -1.04 | -0.79 | 0.97 | 1.47 | 0.17 | 0.48 | -0.255 | -0.6075 | 0.17 | 0.613419 | 0.380968 | 0.738064 | 0.999971 | 1 | 0.959862 |
| Gjc3 | Q921C1 | -0.84 | -1.58 | -0.98 | -1.21 | -1.77 | -1.11 | -0.88 | -1.46 | -1.83 | -1.03 | -1.55 | -2.42 | -2.12 | -1.6 | -0.71 | -2.23 | -0.1525 | -0.555 | -0.5125 | 0.578594 | 0.120785 | 0.191778 | 0.999971 | 1 | 0.810547 |
| Gjd2 | O54851 | 1.62 | -1.48 | -0.63 | 0.17 | -1.66 | -0.31 | -0.93 | -0.87 | -1.28 | 0.63 | -1.51 | -2.1 | -2.22 | -1.28 | -1.64 | -0.8 | -0.8625 | -0.985 | -1.405 | 0.206459 | 0.239558 | 0.057833 | 0.999971 | 1 | 0.725348 |
| Gk | Q64516-1 | -0.78 | -0.32 | -0.3 | -0.18 | -0.49 | -0.07 | -0.24 | -0.12 | -0.45 | -0.48 | -0.25 | -0.58 | -0.23 | -0.19 | -0.34 | 0.02 | 0.165 | -0.045 | 0.21 | 0.443335 | 0.823377 | 0.323625 | 0.999971 | 1 | 0.85548 |
| Gkap1 | Q9JMB0 | 2.8 | 1.43 | 0.49 | 0.64 | 2.21 | 2.48 | 0.88 | 1.61 | 1.27 | 2.59 | 0.9 | 1.04 | 0.19 | 2.07 | 0.9 | 1.04 | 0.455 | 0.11 | -0.29 | 0.442635 | 0.854924 | 0.63057 | 0.999971 | 1 | 0.932711 |
| Gla | Q8BGZ6 | 0.43 | 0.73 | 0.32 | -0.18 | -0.4 | 1.21 | 0.34 | 0.82 | 0.04 | 0.5 | -0.16 | -0.19 | -0.45 | 0.78 | 0.51 | 0.78 | 0.1675 | -0.2775 | 0.08 | 0.661722 | 0.313173 | 0.816361 | 0.999971 | 1 | 0.974929 |
| Glb1 | P23780 | 0.1 | 0.51 | 0.42 | 0.44 | 0.79 | 0.94 | 0.25 | 0.38 | 0.64 | 0.33 | 0.15 | 0.55 | 0.81 | 1.2 | 0.85 | 0.96 | 0.2225 | 0.05 | 0.5875 | 0.340922 | 0.801014 | 0.013119 | 0.999971 | 1 | 0.613095 |
| Glb1l | A0A087WP | -0.18 | 0.49 | -0.56 | -0.67 | -0.66 | 0.07 | -0.77 | -0.58 | -0.16 | 0.75 | 0.1 | -1.01 | 0.32 | 0.37 | -0.09 | 0.04 | -0.255 | 0.15 | 0.39 | 0.439603 | 0.72562 | 0.208881 | 0.999971 | 1 | 0.822138 |
| Glcci1 | E9QKK4 | 0.1 | 0.63 | 0.65 | 0.37 | 0.11 | -0.49 | 0.23 | 0.57 | 0.57 | 0.49 | 0.38 | 0.27 | -0.03 | 0.33 | 0.12 | 0.41 | -0.3325 | -0.01 | -0.23 | 0.244516 | 0.959911 | 0.296796 | 0.999971 | 1 | 0.847402 |
| Glce | Q9EPS3 | 1.23 | 0.61 | 1.07 | 1.19 | 0.53 | 0.8 | 0.7 | 1.85 | 0.66 | 1.72 | 1.45 | 1.55 | 0.36 | 1.06 | 1.63 | 1.66 | -0.055 | 0.32 | 0.1525 | 0.867635 | 0.281939 | 0.651352 | 0.999971 | 1 | 0.933842 |
| Gldc | Q91W43 | -0.23 | -0.73 | -1.52 | -1.96 | -1.12 | -1.09 | -1.34 | -0.73 | -0.99 | -0.86 | -1.4 | -1.68 | -0.93 | -1.3 | -2.24 | -1.97 | 0.04 | -0.1225 | -0.5 | 0.918597 | 0.766595 | 0.294373 | 0.999971 | 1 | 0.847402 |
| Gle1 | Q8R322 | -0.77 | -0.77 | 0.6 | 0.59 | -0.22 | -0.15 | -0.61 | 0.39 | -0.49 | -0.87 | -0.09 | 0.85 | -0.18 | -0.19 | 0.06 | 0.64 | -0.06 | -0.0625 | 0.17 | 0.886702 | 0.901063 | 0.685478 | 0.999971 | 1 | 0.944874 |
| Glg1 | Q61543 | 1.61 | 2.08 | 1.97 | 1.73 | 1.87 | 2.23 | 1.99 | 1.8 | 2.16 | 1.89 | 1.95 | 1.81 | 1.66 | 1.97 | 2.04 | 1.84 | 0.125 | 0.105 | 0.03 | 0.53818 | 0.586822 | 0.879417 | 0.999971 | 1 | 0.985441 |
| Glipr2 | Q9CYL5 | -1.2 | -0.6 | -0.03 | -0.91 | 0.07 | -1.43 | 0.1 | 0.52 | -1.25 | -1.24 | 0.29 | 0.26 | -0.6 | -1.35 | 0.06 | -0.76 | 0.5 | 0.2 | 0.0225 | 0.296913 | 0.674185 | 0.951744 | 0.999971 | 1 | 0.998056 |
| Glmn | Q8BZM1 | 0.5 | 0.47 | 0.84 | 0.39 | 0 | 0.43 | 0.43 | 0.55 | 0.1 | 0.3 | 0.77 | 0.72 | 0.03 | 0.9 | 0.69 | 0.83 | -0.1975 | -0.0775 | 0.0625 | 0.355726 | 0.733848 | 0.803485 | 0.999971 | 1 | 0.971467 |
| Glo1 | Q9CPU0 | 0.28 | 0.82 | 0.55 | 0.08 | 0.99 | 0.9 | 0.22 | 0.15 | 0.72 | 0.99 | 0.23 | 0.13 | 0.8 | 0.83 | 0.37 | 0.47 | 0.1325 | 0.085 | 0.185 | 0.645412 | 0.758048 | 0.439311 | 0.999971 | 1 | 0.883105 |
| Glod4 | Q9CPV4 | -0.16 | 0.27 | -0.41 | -1.15 | 0.12 | 0.13 | 0.96 | -1.07 | 0.67 | -0.37 | -0.46 | -0.02 | -0.07 | -0.13 | -0.74 | -1.27 | 0.3975 | 0.3175 | -0.19 | 0.414951 | 0.413413 | 0.632282 | 0.999971 | 1 | 0.93301 |
| Glra3 | G5E811 | 1.64 | -2.5 | -0.24 | -0.04 | -2.66 | -0.83 | 0.51 | 0.36 | -2.18 | 0.8 | 0.01 | 0.28 | -2.83 | -2.84 | 0.12 | -0.1 | -0.37 | 0.0125 | -1.1275 | 0.712571 | 0.989671 | 0.301787 | 0.999971 | 1 | 0.847402 |
| Glrb | E9Q3E3 | -0.57 | -0.44 | 0.03 | -0.05 | -0.33 | -0.08 | -0.89 | 0.46 | -0.57 | -0.78 | -0.43 | 0.32 | -0.59 | 0.3 | 0.1 | -0.12 | 0.0475 | -0.1075 | 0.18 | 0.881468 | 0.712724 | 0.500447 | 0.999971 | 1 | 0.904223 |
| Glrx | Q9QUH0 | -0.16 | 0.24 | -0.27 | -0.24 | 0.65 | 0.53 | -0.06 | -0.36 | 0.17 | 0.02 | -0.85 | 0.01 | 0.41 | 0.59 | 0.52 | -0.73 | 0.2975 | -0.055 | 0.305 | 0.307513 | 0.84208 | 0.370862 | 0.999971 | 1 | 0.869526 |
| Glrx2 | Q923X4 | 0.92 | -0.58 | -0.06 | 0.03 | -0.46 | -1 | -0.16 | 0.17 | -0.49 | -0.09 | -0.14 | -0.96 | -0.23 | -0.64 | -0.24 | 0.11 | -0.44 | -0.4975 | -0.3275 | 0.269601 | 0.193119 | 0.353299 | 0.999971 | 1 | 0.868675 |
| Glrx3 | Q9CQM9 | -0.8 | -0.19 | -0.95 | -0.97 | -0.19 | -0.17 | -0.92 | -0.65 | -0.15 | -0.67 | -0.45 | -0.35 | -0.29 | -0.05 | -0.67 | -0.28 | 0.245 | 0.3225 | 0.405 | 0.385662 | 0.205472 | 0.1343 | 0.999971 | 1 | 0.784948 |
| Glrx5 | Q80Y14 | -0.62 | -1.67 | -1.59 | -0.74 | -1.72 | -0.87 | -1.28 | -0.86 | -1.43 | -0.68 | -0.77 | -1.27 | -1.74 | -0.57 | -0.77 | -1.03 | -0.0275 | 0.1175 | 0.1275 | 0.935419 | 0.723136 | 0.728674 | 0.999971 | 1 | 0.956041 |
| Gls | D3Z7P3 | -0.88 | -0.83 | -0.62 | -0.88 | -1.13 | -0.63 | -0.39 | -0.39 | -1.19 | -1.01 | -0.63 | -1.19 | -0.68 | -0.78 | -0.48 | -0.39 | 0.1675 | -0.2025 | 0.22 | 0.465058 | 0.325157 | 0.251769 | 0.999971 | 1 | 0.839012 |
| Gls_1 | D3Z7P3-2 | -0.1 | 0.77 | 0.42 | 0 | 1.07 | 0.13 | 1.31 | 0.33 | 0.71 | 0.25 | 0.22 | 1.84 | 1.28 | 0 | 0.6 | -0.51 | 0.4375 | 0.4825 | 0.07 | 0.22408 | 0.257935 | 0.865837 | 0.999971 | 1 | 0.983454 |
| Gls2 | Q571F8 | -1.25 | -0.76 | -0.71 | -0.69 | -1.51 | -1.24 | -0.93 | -1.16 | -1.96 | -1.77 | -0.51 | -1.28 | -1.56 | -1.18 | -1.36 | -0.98 | -0.3575 | -0.5275 | -0.4175 | 0.133759 | 0.152646 | 0.08997 | 0.999971 | 1 | 0.771991 |
| Gltp | Q9JL62 | 0.28 | -0.57 | -0.79 | -1.62 | -0.81 | 0.01 | -1.31 | -0.91 | -0.18 | 0.38 | -1.13 | -1.07 | -0.16 | -0.06 | -1.01 | -1.65 | -0.08 | 0.175 | -0.045 | 0.858706 | 0.726172 | 0.928698 | 0.999971 | 1 | 0.994278 |
| Glud1 | P26443 | -1.07 | -1.45 | -0.96 | -0.71 | -1.55 | -1.29 | -0.26 | -1.1 | -1.79 | -1.69 | -1.02 | -1.09 | -1.38 | -1.37 | -0.98 | -0.88 | -0.0025 | -0.35 | -0.105 | 0.993772 | 0.216438 | 0.65921 | 0.999971 | 1 | 0.935181 |
| Glul | P15105 | -0.18 | 0.11 | 0.11 | -0.27 | -0.01 | 0.34 | 0.28 | -0.09 | 0.3 | -0.09 | 0.23 | -0.78 | -0.04 | 0.12 | -0.01 | -0.37 | 0.1875 | -0.0275 | -0.0175 | 0.364448 | 0.921664 | 0.930785 | 0.999971 | 1 | 0.994744 |
| Glyctk | A0A0R4J1H | 0.25 | 0.29 | -0.12 | -0.28 | -0.07 | -0.32 | -0.65 | 0.29 | -0.2 | -0.33 | -0.55 | -0.13 | -0.8 | -0.02 | -0.37 | 0.21 | -0.2225 | -0.3375 | -0.28 | 0.40984 | 0.139004 | 0.32799 | 0.999971 | 1 | 0.856472 |
| Glyr1 | D3YYT1 | -0.19 | -0.51 | -0.44 | -0.32 | -0.63 | -0.6 | 1.41 | -1.64 | -0.75 | -0.79 | -1.39 | -1.4 | -0.31 | -0.82 | -0.96 | -1.48 | 9.81E-17 | -0.7175 | -0.5275 | 1 | 0.011698 | 0.077955 | 1 | 1 | 0.754157 |
| Gm11938 | Q9D3H4 | -3.1 | -8.23 | 3.54 | 4.54 | -8.71 | -8.25 | -6.72 | -8.75 | -7.77 | 5.04 | 4.93 | 3.69 | -14.41 | 13.85 | 5.09 | 2.75 | -7.295 | 2.285 | 2.6325 | 0.022251 | 0.55523 | 0.654778 | 0.999971 | 1 | 0.934061 |
| Gm13889 | A0A087WS | 2.52 | 0.76 | 0.58 | -0.13 | 0.96 | 1.51 | 0.07 | 1.15 | 0.71 | 2.9 | 0.39 | 1.13 | -0.52 | 1.28 | 1.68 | 0.09 | -0.01 | 0.35 | -0.3 | 0.986362 | 0.629898 | 0.664959 | 0.999971 | 1 | 0.93697 |
| Gm1673 | Q3UR78 | 0.36 | 1.36 | 1.62 | 1.5 | 1.78 | 0.93 | 0.98 | 1.23 | 0.86 | 0.63 | 0.83 | 1.1 | 1.75 | 1.53 | 1.96 | 1.13 | 0.02 | -0.355 | 0.3825 | 0.95351 | 0.268951 | 0.274031 | 0.999971 | 1 | 0.843794 |
| Gm19410 | A0A140LJC | -0.8 | -1.07 | -0.07 | -0.11 | -1.29 | -0.52 | -0.78 | -0.03 | -0.93 | -0.21 | -0.11 | -0.76 | 0.13 | -0.31 | -0.36 | 0.45 | -0.1425 | 0.01 | 0.49 | 0.689401 | 0.975242 | 0.150422 | 0.999971 | 1 | 0.793248 |
| Gm20390 | E9PZF0 | -0.27 | -0.15 | 0.64 | 0.31 | 0.04 | -0.06 | 0.73 | 0.65 | 0.37 | -0.24 | 0.87 | 0.78 | 0.08 | -0.28 | 0.78 | 0.53 | 0.2075 | 0.3125 | 0.145 | 0.496435 | 0.354439 | 0.65135 | 0.999971 | 1 | 0.933842 |
| Gm20431 | E9PY39 | 0.44 | 0.32 | -0.07 | -0.16 | 0.05 | 0.22 | -0.07 | -0.03 | 0.05 | 0.65 | -0.04 | -0.06 | 0.09 | 0.17 | -0.16 | 0.08 | -0.09 | 0.0175 | -0.0875 | 0.669915 | 0.943943 | 0.6819 | 0.999971 | 1 | 0.943204 |
| Gm20441 | G3UYJ7 | 0.7 | 0.12 | 1.73 | 0.76 | -0.04 | -0.51 | 0.02 | 1.32 | -0.04 | 0.29 | 0.99 | 1.23 | -1.18 | 0.1 | 1.63 | 1.49 | -0.63 | -0.21 | -0.3175 | 0.211388 | 0.622844 | 0.638499 | 0.999971 | 1 | 0.93301 |
| Gm20517 | G3UVU6 | 0.84 | 1.76 | 2.11 | 2.43 | 1.42 | 1.5 | 1.69 | 1.76 | 1.12 | 0.94 | 2.53 | 1.83 | 1.09 | 1.38 | 2.06 | 2.67 | -0.1925 | -0.18 | 0.015 | 0.582347 | 0.702196 | 0.974098 | 0.999971 | 1 | 0.99925 |
| Gm20521 | D3Z5F7 | 2.34 | 1.44 | 1.06 | 1.68 | 0.6 | 1.37 | 1.15 | 0.93 | 0.99 | 1.26 | 1.17 | 0.64 | 0.34 | 1.11 | 1.26 | 1.14 | -0.6175 | -0.615 | -0.6675 | 0.078853 | 0.072023 | 0.07564 | 0.999971 | 1 | 0.746958 |
| Gm20604 | F8WIA3 | -0.42 | 1.19 | -1.14 | 0.04 | -1.09 | -1.07 | -1.03 | -0.02 | 0.36 | -0.38 | 1.46 | 0.04 | 3.82 | -0.64 | -0.52 | 0.06 | -0.72 | 0.4525 | 0.7625 | 0.184059 | 0.441814 | 0.470646 | 0.999971 | 1 | 0.892725 |
| Gm20671 | F6TVX7 | -0.25 | -0.53 | -0.51 | -0.3 | -0.51 | -0.24 | -0.43 | -0.36 | -0.68 | -0.25 | -0.26 | -0.7 | -0.87 | -0.42 | -0.32 | -0.27 | 0.0125 | -0.075 | -0.0725 | 0.943366 | 0.707198 | 0.727994 | 0.999971 | 1 | 0.956041 |
| Gm21985 | Q6P6P5 | 1.66 | 1.32 | 0.59 | 0.86 | -0.45 | 1.65 | 0.64 | 0.25 | 0.68 | 1.58 | -0.22 | -0.39 | -0.88 | 1.13 | 0.11 | 0.55 | -0.585 | -0.695 | -0.88 | 0.230372 | 0.172946 | 0.081077 | 0.999971 | 1 | 0.755254 |
| Gm21992 | F7BGR7 | 0.2 | 0.97 | 0.94 | 0.67 | 0.74 | 1.1 | 0.46 | 0.99 | 0.72 | 0.65 | 0.24 | 0.61 | 0.87 | 0.87 | 0.71 | 0.66 | 0.1275 | -0.14 | 0.0825 | 0.618337 | 0.56169 | 0.717274 | 0.999971 | 1 | 0.952711 |
| Gm2a | Q60648 | 0.42 | 0.62 | 0.88 | 0.81 | 0.64 | 0.82 | 0.55 | 1.68 | 1.12 | 0.83 | 1.16 | 0.88 | 1.22 | 1.18 | 0.97 | 1.33 | 0.24 | 0.315 | 0.4925 | 0.417637 | 0.128687 | 0.029451 | 0.999971 | 1 | 0.678474 |
| Gm3839 | S4R1W1 | -0.61 | 0.91 | 0.8 | 0.8 | 1.48 | 0.75 | 1.07 | 1.58 | 1.19 | -0.32 | 1.64 | 0.41 | 1.33 | 1.81 | 1.71 | 0.01 | 0.745 | 0.255 | 0.74 | 0.085242 | 0.628531 | 0.172238 | 0.999971 | 1 | 0.804599 |
| Gm38393 | A0A087WP | 0.97 | 1.57 | 0.49 | 0.4 | 1.35 | 0.59 | 0.97 | 0.27 | 0.54 | 0.49 | 2.06 | 1.4 | 1.45 | 0.87 | 2.11 | 1.7 | -0.0625 | 0.265 | 0.675 | 0.858183 | 0.54864 | 0.091399 | 0.999971 | 1 | 0.773779 |
| Gm42517 | A0A1Y7VL( | 1.67 | -1.78 | -0.54 | -0.03 | -2.35 | -0.24 | 1.68 | -1.1 | -1.71 | 0.71 | -0.44 | -2.51 | -1.65 | -1.05 | -0.98 | -1.25 | -0.3325 | -0.8175 | -1.0625 | 0.73732 | 0.379084 | 0.136127 | 0.999971 | 1 | 0.784948 |
| Gm42715 | A0A087WP | 1.07 | 0.73 | 1.4 | 1.19 | 0.58 | -0.19 | 1.28 | 0.41 | 0.31 | 0.21 | 1.06 | 0.21 | -0.68 | -0.44 | -0.02 | -1.2 | -0.5775 | -0.65 | -1.6825 | 0.109328 | 0.035859 | 0.000322 | 0.999971 | 1 | 0.247672 |
| Gm43738 | A0A0G2JEA | -1.07 | -0.89 | 0.41 | 0.34 | -0.59 | -0.9 | 0.71 | 0.94 | -0.55 | -1.42 | 0.69 | 0.42 | -0.36 | -0.28 | 0.71 | 0.74 | 0.3425 | 0.0875 | 0.505 | 0.541813 | 0.878554 | 0.293716 | 0.999971 | 1 | 0.847402 |
| Gm44505 | E0CY81 | -0.3 | 1.25 | 4.02 | 2.57 | 0.13 | 2.42 | 2.96 | 3.39 | 0.8 | 0.51 | 3.57 | 3.65 | 0.4 | 1.42 | 3.98 | 2.95 | 0.34 | 0.2475 | 0.3025 | 0.745677 | 0.826411 | 0.780895 | 0.999971 | 1 | 0.965859 |
| Gm45208 | A0A140LJ3 | -0.12 | 1.31 | -0.35 | -0.21 | 0.16 | 0.47 | -0.58 | -0.02 | -0.06 | 0.02 | 0.27 | -0.37 | 0.14 | -0.19 | 0.19 | 0.56 | -0.15 | -0.1925 | 0.0175 | 0.723054 | 0.626786 | 0.964995 | 0.999971 | 1 | 0.998721 |
| Gm45785 | A0A1B0GT( | -1.16 | 0.99 | 0.46 | -0.64 | 1.1 | 0.37 | 0.48 | -0.06 | 0.56 | -1.32 | 0.96 | 0.77 | 0.66 | -0.13 | 0.7 | -0.05 | 0.56 | 0.33 | 0.3825 | 0.286685 | 0.618944 | 0.454124 | 0.999971 | 1 | 0.887217 |
| Gm45808 | A0A1B0GSI | -0.24 | 0.45 | 0.26 | 0.2 | 0.7 | 0.21 | -0.39 | 0.37 | 0.74 | 0.55 | 0.62 | 0.58 | 0.29 | 0.64 | 0.48 | 0.76 | 0.055 | 0.455 | 0.375 | 0.846798 | 0.048792 | 0.118077 | 0.999971 | 1 | 0.78268 |
| Gm45837 | F7D3W5 | -0.26 | -0.91 | -1.03 | -0.94 | -0.81 | -0.86 | -0.74 | -0.84 | -0.52 | -0.5 | -0.71 | -0.83 | -0.42 | -0.8 | -0.87 | -1.07 | -0.0275 | 0.145 | -0.005 | 0.901227 | 0.531795 | 0.984112 | 0.999971 | 1 | 0.99925 |
| Gm49325 | G3UZ60 | -1.67 | 6.55 | 5.96 | 2.75 | 6.3 | 5.77 | 5.61 | 4.58 | 6.55 | 0.1 | 6.07 | 6.6 | 8.39 | 3.83 | 5.42 | 2.84 | 2.1675 | 1.4325 | 1.7225 | 0.223191 | 0.518606 | 0.395914 | 0.999971 | 1 | 0.871816 |
| Gm49416 | Q9D937 | 0.72 | 3.78 | 0.68 | 1.95 | 2.63 | 1.81 | 0.15 | 3.27 | 1.69 | 1.51 | -0.44 | 0.66 | 2.29 | 2.23 | 0.35 | 1.82 | 0.1825 | -0.9275 | -0.11 | 0.837301 | 0.261229 | 0.886787 | 0.999971 | 1 | 0.987485 |
| Gm49486 | E9QAC6 | 0.52 | 1.27 | 0.84 | 0.84 | 0.36 | 0.57 | 1.32 | 0.37 | 0.52 | 0.26 | 0.48 | 0.7 | -0.04 | 0.55 | 0.33 | 0.41 | -0.2125 | -0.3775 | -0.555 | 0.466211 | 0.112798 | 0.039927 | 0.999971 | 1 | 0.700838 |
| Gm49510 | A0A2R8W6 | 4.53 | 0 | -0.47 | -0.58 | -0.05 | -0.03 | -0.59 | -0.51 | -0.23 | 2.35 | -0.64 | -0.58 | -0.01 | 0.39 | -0.65 | -0.68 | -1.165 | -0.645 | -1.1075 | 0.305301 | 0.614215 | 0.335772 | 0.999971 | 1 | 0.858827 |
| Gm49804 | A0A3B2W8 | 0.9 | 1.13 | 1.66 | 1.67 | 1.48 | 0.99 | 1.35 | 1.47 | 1.23 | 1.02 | 1.58 | 1.78 | 0.8 | 1.01 | 1.38 | 1.26 | -0.0175 | 0.0625 | -0.2275 | 0.944572 | 0.819845 | 0.388836 | 0.999971 | 1 | 0.871816 |
| Gm49948 | D3YXH0 | 0.52 | 0.98 | 0.52 | 0.89 | 0.8 | 0.2 | 2 | 0.78 | 0.41 | 0.73 | 0.33 | 0.43 | 0.01 | 0.75 | 0.32 | 0.32 | 0.2175 | -0.2525 | -0.3775 | 0.573245 | 0.231924 | 0.130868 | 0.999971 | 1 | 0.784948 |
| Gm50364 | D6RI20 | -1.35 | -0.49 | -1 | -0.99 | -0.05 | -0.31 | -0.22 | -0.05 | 0.09 | -0.31 | 0.74 | 0.91 | -0.46 | -0.58 | 0.24 | 0.37 | 0.8 | 1.315 | 0.85 | 0.00634 | 0.003438 | 0.020352 | 0.999971 | 0.957868 | 0.635388 |
| Gm55978 | A0A1B0GS | 0.8 | -1.72 | 0.42 | 0.97 | 0.57 | -1.07 | 1.07 | 0.69 | -0.47 | 1.37 | 0.36 | 0.34 | -2.64 | -0.54 | 0.54 | 0.75 | 0.1975 | 0.2825 | -0.59 | 0.780313 | 0.672198 | 0.51379 | 0.999971 | 1 | 0.905815 |
| Gm56451 | A0A338P6I | -1.49 | 0.5 | 1.47 | 0.92 | 0.36 | 0.58 | 0.28 | 1.47 | -0.15 | -1.14 | 1.1 | 1.3 | 0.42 | 0.4 | 1.66 | 1.32 | 0.3225 | -0.0725 | 0.6 | 0.61474 | 0.9259 | 0.3708 | 0.999971 | 1 | 0.869526 |
| Gm7324 | A0A2I3BRL | 1.18 | -0.33 | -0.18 | 0.21 | -0.35 | 0.02 | 0.41 | 0.21 | -0.17 | 0.78 | 0 | -0.29 | -0.56 | -0.3 | -0.08 | 0.02 | -0.1475 | -0.14 | -0.45 | 0.688041 | 0.726674 | 0.227924 | 0.999971 | 1 | 0.828598 |
| Gmds | Q8K0C9 | 0.33 | 0.62 | 0.35 | 0.54 | 0.71 | 0.96 | 0.61 | 0.62 | 1.03 | 0.63 | 0.56 | 0.27 | 0.69 | 0.84 | 0.56 | 0.43 | 0.265 | 0.1625 | 0.17 | 0.172735 | 0.458029 | 0.372918 | 0.999971 | 1 | 0.869526 |
| Gmeb1 | Q9JL60 | 0.06 | -1.26 | 1.91 | 1.96 | 0.02 | -0.52 | 1.15 | 1.28 | -1.79 | -0.14 | 1.63 | 1.5 | -1.73 | 0.35 | 1.37 | 2.02 | -0.185 | -0.3675 | -0.165 | 0.81797 | 0.716124 | 0.870325 | 0.999971 | 1 | 0.984079 |
| Gmeb2 | P58929 | 0.22 | 0.68 | 2.1 | 1.97 | 0.82 | 0.24 | 1.52 | 0.96 | 0.67 | 0.23 | 1.2 | 2.51 | 0.58 | 0.34 | 1.45 | 2.11 | -0.3575 | -0.09 | -0.1225 | 0.479152 | 0.885117 | 0.829668 | 0.999971 | 1 | 0.978384 |
| Gmfb | Q9CQI3 | 0.28 | 0.23 | 0.34 | 0.63 | 0.39 | 0.43 | 0.83 | 0.95 | 0.43 | 0.12 | 0.49 | 0.41 | -0.18 | 0.33 | 0.51 | 0.5 | 0.28 | -0.0075 | -0.08 | 0.213732 | 0.967923 | 0.724512 | 0.999971 | 1 | 0.955602 |
| Gmppa | Q922H4 | 0.45 | -0.14 | -0.29 | -0.43 | -0.28 | 0.16 | -0.51 | -0.66 | -0.16 | 0.39 | -0.77 | -0.46 | -0.06 | 0.13 | -0.34 | -0.24 | -0.22 | -0.1475 | -0.025 | 0.43899 | 0.643152 | 0.919923 | 0.999971 | 1 | 0.992702 |
| Gmppb | Q8BTZ7 | -0.17 | -0.35 | -0.27 | -0.14 | -0.25 | -0.23 | -0.74 | -0.51 | -0.31 | -0.31 | -0.28 | -0.56 | -0.45 | -0.18 | -0.56 | -0.5 | -0.2 | -0.1325 | -0.19 | 0.316959 | 0.440161 | 0.304086 | 0.999971 | 1 | 0.847402 |
| Gmpr | Q9DCZ1 | 0.48 | 0.9 | 0.63 | 0.57 | 0.93 | 1.13 | 0.79 | 1.46 | 0.87 | -0.04 | 0.84 | 1.02 | 1.35 | 1.48 | 1.17 | 1.05 | 0.4325 | 0.0275 | 0.6175 | 0.072579 | 0.919884 | 0.010931 | 0.999971 | 1 | 0.613095 |
| Gmpr2 | Q99L27 | 0.22 | 0.6 | 0.44 | 0.5 | 0.48 | 0.93 | 1.19 | 1.15 | 0.85 | 0.46 | 0.76 | 0.53 | 0.99 | 0.8 | 0.94 | 0.73 | 0.4975 | 0.21 | 0.425 | 0.049614 | 0.279297 | 0.04031 | 0.999971 | 1 | 0.700838 |
| Gmps | Q3THK7 | 0.13 | 1.03 | 0.73 | 0.83 | 0.93 | 0.99 | 0.58 | 0.81 | 0.94 | 0.22 | 0.71 | 0.93 | 0.75 | 0.7 | 1.04 | 0.87 | 0.1475 | 0.02 | 0.16 | 0.550259 | 0.941613 | 0.512284 | 0.999971 | 1 | 0.905815 |
| Gna11 | P21278 | -1.06 | -1.13 | -1.15 | -1.28 | -0.61 | -1.56 | -0.04 | -1.19 | 0.08 | -1.26 | -0.57 | -0.31 | 0.14 | -1.69 | -1.14 | -0.81 | 0.305 | 0.64 | 0.28 | 0.373511 | 0.054772 | 0.462728 | 0.999971 | 1 | 0.890653 |
| Gna12 | P27600 | -0.68 | 0.14 | -0.14 | -1.15 | -1.07 | -0.35 | -0.82 | -0.21 | -0.2 | -0.11 | -0.68 | -0.86 | -0.63 | -0.59 | -0.07 | -0.6 | -0.155 | -0.005 | -0.015 | 0.65501 | 0.988152 | 0.962554 | 0.999971 | 1 | 0.998721 |
| Gna13 | P27601 | -0.62 | -0.63 | -0.83 | -0.97 | -0.5 | -0.56 | -0.66 | -1.01 | -0.09 | -0.71 | -0.79 | -0.72 | -0.04 | -0.59 | -0.59 | -0.48 | 0.08 | 0.185 | 0.3375 | 0.690487 | 0.41604 | 0.132359 | 0.999971 | 1 | 0.784948 |
| Gna14 | P30677 | 3.27 | -0.29 | -1.88 | -0.81 | 0.84 | -0.38 | 2.02 | -1.56 | 2.44 | 2.61 | 0.39 | -0.22 | 1.98 | -0.78 | -2.19 | -1.65 | 0.1575 | 1.2325 | -0.7325 | 0.896035 | 0.315119 | 0.573701 | 0.999971 | 1 | 0.91816 |
| Gnai1 | B2RSH2 | -1.28 | -1.78 | -1.8 | -1.74 | -1.46 | -1.44 | -0.76 | -1.87 | -0.58 | -1 | -1.53 | -1.7 | -0.77 | -1.49 | -1.7 | -1.84 | 0.2675 | 0.4475 | 0.2 | 0.348295 | 0.153567 | 0.486504 | 0.999971 | 1 | 0.897675 |
| Gnai2 | P08752 | -1.35 | -0.73 | -1.11 | -1.3 | -0.5 | -0.71 | -1.23 | -1.24 | -0.39 | -1.5 | -1.26 | -0.93 | -0.01 | -0.84 | -0.9 | -1.46 | 0.2025 | 0.1025 | 0.32 | 0.441008 | 0.724019 | 0.345997 | 0.999971 | 1 | 0.863794 |
| Gnai3 | Q9DC51 | -1.69 | -0.77 | -1.41 | -1.24 | -0.75 | -0.72 | -1.26 | -1.15 | -0.33 | -1.51 | -1.16 | -1.04 | -0.66 | -0.78 | -1.24 | -1.05 | 0.3075 | 0.2675 | 0.345 | 0.256562 | 0.40816 | 0.204374 | 0.999971 | 1 | 0.81617 |
| Gnal | Q8CGK7 | 0.06 | -2.28 | -1.46 | -0.48 | -1.33 | -2.96 | -0.85 | -0.67 | 0.69 | -0.72 | -1.38 | -0.65 | -1.04 | -1.89 | -0.75 | -1.45 | -0.4125 | 0.525 | -0.2425 | 0.540656 | 0.407278 | 0.650032 | 0.999971 | 1 | 0.933842 |
| Gnal_1 | Q66L47 | -0.35 | -1.62 | -2.23 | -1.19 | -0.99 | -1.96 | -0.81 | -2.01 | -1.07 | -1.09 | -0.85 | -1.28 | -1.99 | -1.36 | -2.01 | -1.45 | -0.095 | 0.275 | -0.355 | 0.840438 | 0.486956 | 0.398364 | 0.999971 | 1 | 0.871816 |
| Gnao1 | P18872-2 | -3.8 | 3.25 | 0.98 | -1.85 | 2.67 | 0.5 | 1.83 | -0.54 | 3.95 | -2.47 | 1.71 | 0.6 | 5.78 | -2.01 | 0.57 | -1.93 | 1.47 | 1.3025 | 0.9575 | 0.344197 | 0.482192 | 0.653539 | 0.999971 | 1 | 0.934061 |
| Gnao1_1 | P18872 | -1.32 | -1.33 | -1.32 | -1.68 | -0.99 | -1.21 | 0.06 | -1.44 | -1.05 | -1.62 | -1.08 | -0.74 | -0.7 | -1.39 | -1.52 | -1.7 | 0.5175 | 0.29 | 0.085 | 0.153458 | 0.239016 | 0.744652 | 0.999971 | 1 | 0.961064 |
| Gnaq | P21279 | -1.24 | -0.93 | -0.83 | -1.09 | -1.01 | -0.9 | -0.6 | -0.93 | -0.79 | -1.1 | -0.81 | -0.69 | -0.42 | -1.2 | -0.81 | -0.83 | 0.1625 | 0.175 | 0.2075 | 0.40799 | 0.366367 | 0.368379 | 0.999971 | 1 | 0.869526 |
| Gnas | A0A571BE3 | -0.75 | 2.25 | 0.19 | -0.31 | 2.81 | 0.45 | 4.06 | -1.04 | -1.29 | 0.17 | -1.19 | -1.42 | 4.31 | 1.64 | -0.96 | -0.73 | 1.225 | -1.2775 | 0.72 | 0.31388 | 0.09265 | 0.566548 | 0.999971 | 1 | 0.917959 |
| Gnas_1 | P63094 | -0.96 | -0.76 | -1.36 | -1.97 | -0.86 | -1.1 | -0.54 | -1.96 | -0.73 | -1.03 | -0.99 | -1.16 | -0.48 | -1.08 | -2.08 | -1.98 | 0.1475 | 0.285 | -0.1425 | 0.705268 | 0.342361 | 0.746322 | 0.999971 | 1 | 0.961064 |
| Gnaz | O70443 | -0.56 | -0.37 | -1.32 | -1.31 | -0.23 | -0.39 | -1.19 | -1.45 | 0.25 | -0.45 | -1.25 | -1.2 | 0.75 | -0.71 | -1.58 | -1.25 | 0.075 | 0.2275 | 0.1925 | 0.841666 | 0.585283 | 0.716296 | 0.999971 | 1 | 0.952576 |
| Gnb1 | P62874 | 0 | -1.08 | -1.83 | -1.41 | -1.6 | -1.42 | -1.06 | -1.5 | -1.93 | -1.23 | -1.83 | -2.21 | -1.77 | -1.21 | -1.7 | -1.88 | -0.315 | -0.72 | -0.56 | 0.430287 | 0.114953 | 0.186403 | 0.999971 | 1 | 0.809025 |
| Gnb2 | P62880 | 0.16 | -0.42 | -0.44 | -0.64 | -0.66 | -0.45 | -0.41 | -0.31 | -0.72 | 0.04 | -0.45 | -0.61 | -0.37 | -0.53 | -0.34 | -0.5 | -0.1225 | -0.1 | -0.1 | 0.592589 | 0.703203 | 0.654498 | 0.999971 | 1 | 0.934061 |
| Gnb2_1 | E9PWM7 | -1.37 | 3.2 | 2.79 | 1.38 | 0.76 | 3.56 | 1.25 | 2.55 | 1.99 | 1.22 | 2.03 | -0.67 | 1.3 | 2.87 | 1.89 | 2 | 0.53 | -0.3575 | 0.515 | 0.625756 | 0.742841 | 0.597744 | 0.999971 | 1 | 0.92377 |
| Gnb4 | P29387 | -0.04 | -0.52 | -1.15 | -1.11 | -0.16 | -0.52 | -0.56 | -0.87 | 0.07 | -0.2 | -0.89 | -0.98 | 0.39 | -0.89 | -0.76 | -0.92 | 0.1775 | 0.205 | 0.16 | 0.567539 | 0.573497 | 0.685969 | 0.999971 | 1 | 0.94497 |
| Gnb5 | P62881-2 | 0.19 | -0.19 | 0.22 | -0.29 | -0.46 | 0.03 | 0.39 | -0.24 | -0.4 | -0.81 | 0.05 | -0.13 | -0.29 | -0.26 | -0.27 | -0.33 | -0.0525 | -0.305 | -0.27 | 0.834913 | 0.247081 | 0.189972 | 0.999971 | 1 | 0.810547 |
| Gne | Q3UW64 | 0.78 | 1.58 | 1.84 | 1.31 | 1.72 | 1.32 | 0.63 | 1.87 | 1.42 | 1.01 | 1 | 1.38 | 1.81 | 1.66 | 1.59 | 1.69 | 0.0075 | -0.175 | 0.31 | 0.982953 | 0.523769 | 0.247405 | 0.999971 | 1 | 0.839012 |
| Gng10 | Q9CXP8 | -1.23 | -2.65 | -1.46 | -1.81 | -2.07 | -2.29 | -2.11 | -2.52 | -2.19 | -1.73 | -2.7 | -3.05 | -3.26 | -2.39 | -2.68 | -2.18 | -0.46 | -0.63 | -0.84 | 0.184419 | 0.14777 | 0.050005 | 0.999971 | 1 | 0.717713 |
| Gng11 | P61953 | -0.75 | -0.25 | 0.4 | 0.54 | -0.42 | -0.08 | -0.88 | -0.48 | -0.97 | 0.1 | 0.06 | 0.33 | -0.7 | 0 | 0.3 | -0.1 | -0.45 | -0.105 | -0.11 | 0.206066 | 0.792906 | 0.759224 | 0.999971 | 1 | 0.961116 |
| Gng12 | Q9DAS9 | -0.74 | -1.78 | -1.48 | -1.68 | -1.79 | -1.79 | -1.58 | -1.94 | -1.64 | -1.33 | -1.74 | -2.09 | -1.87 | -1.59 | -1.89 | -1.82 | -0.355 | -0.28 | -0.3725 | 0.206138 | 0.351447 | 0.186265 | 0.999971 | 1 | 0.809025 |
| Gng13 | Q9JMF3 | -1.29 | -1.95 | -1.6 | -1.59 | -1.82 | -1.6 | -2.08 | -2.09 | -1.99 | -1.18 | -1.54 | -2.34 | -1.78 | -1.57 | -2.12 | -1.21 | -0.29 | -0.155 | -0.0625 | 0.212808 | 0.603726 | 0.809177 | 0.999971 | 1 | 0.973762 |
| Gng2 | P63213 | -0.85 | -1.77 | 0.83 | -0.09 | -1.56 | -1.37 | -0.57 | -1.07 | -2.05 | -0.1 | -0.82 | -3.45 | -0.97 | -1.76 | -0.93 | -0.08 | -0.6725 | -1.135 | -0.465 | 0.238166 | 0.195891 | 0.442864 | 0.999971 | 1 | 0.883386 |
| Gng3 | P63216 | -1.24 | 0.48 | -1.4 | 0.13 | -0.27 | -0.19 | -1.11 | -0.87 | -1.04 | -1.45 | -0.72 | -0.95 | 1.87 | 1.04 | 0.38 | 1.25 | -0.1025 | -0.5325 | 1.6425 | 0.83404 | 0.274119 | 0.011534 | 0.999971 | 1 | 0.613095 |
| Gng4 | P50153 | 0.12 | -1.3 | -0.18 | 0.04 | -0.8 | -1.11 | 0.03 | -0.01 | -1.01 | -0.04 | 0.01 | -0.65 | -0.58 | -0.92 | 0.04 | -0.55 | -0.1425 | -0.0925 | -0.1725 | 0.731617 | 0.815188 | 0.646384 | 0.999971 | 1 | 0.933613 |
| Gng5 | Q80SZ7 | -1.37 | -1.72 | -1.5 | -2.41 | -1.22 | -1.15 | -1.33 | -2.09 | -1.54 | -0.89 | -2.45 | -3.22 | -1.68 | -1.88 | -2.4 | -2.79 | 0.3025 | -0.275 | -0.4375 | 0.356812 | 0.600277 | 0.219384 | 0.999971 | 1 | 0.827064 |
| Gng7 | Q61016 | -0.67 | -2.34 | -1.51 | -1.45 | -1.9 | -2.46 | -1.93 | -1.83 | -2.01 | -1.35 | -1.66 | -1.68 | -1.85 | -2.04 | -1.23 | -2.22 | -0.5375 | -0.1825 | -0.3425 | 0.1624 | 0.613649 | 0.38907 | 0.999971 | 1 | 0.871816 |
| Gnl1 | P36916 | -1.02 | 0.34 | 0.17 | 0.06 | 0.13 | 0.13 | -0.14 | 0.32 | 0.24 | -1.05 | -0.08 | 0.14 | 0.42 | 0.04 | 0.33 | -0.23 | 0.2225 | -0.075 | 0.2525 | 0.497264 | 0.854078 | 0.463798 | 0.999971 | 1 | 0.890905 |
| Gnl3 | Q8CI11 | 0.31 | 0 | 0.11 | 0.13 | -0.22 | -0.01 | -0.38 | 0.15 | -0.23 | 0.44 | 0.06 | 0.2 | 0.44 | 0.34 | -0.03 | 0.55 | -0.2525 | -0.02 | 0.1875 | 0.217768 | 0.92198 | 0.361974 | 0.999971 | 1 | 0.869526 |
| Gnl3l | Q6PGG6 | 0.46 | 0.77 | -0.13 | 0.26 | 0.91 | 0.71 | -0.4 | -0.01 | 0.86 | 0.34 | -0.1 | 0.54 | 0.79 | 0.37 | -0.14 | 0.23 | -0.0375 | 0.07 | -0.0275 | 0.915272 | 0.807619 | 0.922809 | 0.999971 | 1 | 0.993264 |
| Gnpat | P98192 | -0.64 | -0.17 | -0.55 | -0.29 | -0.24 | 0.2 | -0.76 | 0.24 | 0.03 | -0.19 | -0.54 | -0.33 | 0.52 | 0.31 | 0.24 | -0.17 | 0.2725 | 0.155 | 0.6375 | 0.336016 | 0.46646 | 0.01814 | 0.999971 | 1 | 0.635388 |
| Gnpda1 | O88958 | -0.79 | 0.58 | -0.06 | -0.56 | 0.89 | 0.52 | 0.61 | -0.53 | 0.85 | -0.71 | 0.38 | 0.89 | 1.22 | 0.39 | -0.28 | -0.71 | 0.58 | 0.56 | 0.3625 | 0.184351 | 0.237065 | 0.461155 | 0.999971 | 1 | 0.89007 |
| Gnpda2 | Q9CRC9 | -0.12 | 0.47 | 0.11 | 0.1 | 0.3 | 0.66 | -0.05 | 0.7 | 0.06 | -0.21 | -0.14 | -0.07 | 0.21 | 0.66 | 0.37 | 0.27 | 0.2625 | -0.23 | 0.2375 | 0.299067 | 0.25438 | 0.275219 | 0.999971 | 1 | 0.843794 |
| Gnpnat1 | Q9JK38 | -0.39 | 0.52 | 0.38 | 0.26 | 0.4 | 0 | -0.51 | 0.28 | -0.19 | -0.27 | -0.31 | 0.24 | -0.33 | 0.73 | 1 | 0.58 | -0.15 | -0.325 | 0.3025 | 0.614083 | 0.231648 | 0.394203 | 0.999971 | 1 | 0.871816 |
| Gnptg | A0A0R4J0H | -0.48 | 0.05 | -0.76 | 0 | 0.74 | -0.97 | -1.21 | 0.61 | -0.22 | -0.36 | -0.56 | 0.37 | 0.47 | 0.24 | 0.88 | -0.12 | 0.09 | 0.105 | 0.665 | 0.859241 | 0.718279 | 0.048522 | 0.999971 | 1 | 0.710102 |
| Gns | Q8BFR4 | -0.3 | 0.61 | 0.45 | -0.22 | 0.52 | 0.24 | 0.12 | 0.81 | 0.51 | -0.2 | 0.65 | 0.34 | 1.39 | 0.58 | 0.48 | 0.51 | 0.2875 | 0.19 | 0.605 | 0.334662 | 0.53562 | 0.086124 | 0.999971 | 1 | 0.759991 |
| Golga1 | Q9CW79-2 | 1.22 | 1.45 | 1.15 | 1.43 | 1.84 | 1.59 | 0.71 | 1.58 | 1.27 | 1.57 | 0.98 | 1.23 | 1.18 | 1.86 | 1.16 | 1.72 | 0.1175 | -0.05 | 0.1675 | 0.671379 | 0.800531 | 0.479626 | 0.999971 | 1 | 0.894438 |
| Golga2 | E9PUQ5 | 0.7 | 1.49 | 1.62 | 1.71 | 1.65 | 1.42 | 1.34 | 1.94 | 1.17 | 1.1 | 1.5 | 1.48 | 2.2 | 1.52 | 1.52 | 1.79 | 0.2075 | -0.0675 | 0.3775 | 0.469037 | 0.803313 | 0.219438 | 0.999971 | 1 | 0.827064 |
| Golga3 | E9QP99 | -0.06 | 0.49 | 0.31 | 0.16 | 0.42 | 0.27 | 0.25 | 0.27 | 0.5 | 0.23 | 0.37 | 0.51 | 0.44 | 0.45 | 0.31 | 0.29 | 0.0775 | 0.1775 | 0.1475 | 0.685 | 0.36966 | 0.448799 | 0.999971 | 1 | 0.885351 |
| Golga4 | Q91VW5 | 1.02 | 1.37 | 1.56 | 1.55 | 1.58 | 1.15 | 1.18 | 1.78 | 1.82 | 1.24 | 1.95 | 2.18 | 1.73 | 1.55 | 1.85 | 2.08 | 0.0475 | 0.4225 | 0.4275 | 0.839698 | 0.129568 | 0.074123 | 0.999971 | 1 | 0.746069 |
| Golga5 | Q9QYE6 | 1.44 | 0.84 | 1.51 | 1.44 | 0.61 | 0.59 | 1.85 | 1.24 | 0.65 | 1.01 | 1.18 | 1.4 | 0.12 | 0.98 | 1.17 | 1.16 | -0.235 | -0.2475 | -0.45 | 0.490395 | 0.334234 | 0.162914 | 0.999971 | 1 | 0.800215 |
| Golga7 | Q91W53 | -0.86 | -0.74 | -0.99 | -1.44 | -0.84 | -0.79 | -1.22 | -1.43 | -0.77 | -0.77 | -1.03 | -1.07 | -0.65 | -0.71 | -1.38 | -1.5 | -0.0625 | 0.0975 | -0.0525 | 0.799977 | 0.653526 | 0.853229 | 0.999971 | 1 | 0.98187 |

| | | | | | | | | | | | | | | | | | | | | | | | | | | |
|---|---|---|---|---|---|---|---|---|---|---|---|---|---|---|---|---|---|---|---|---|---|---|---|---|---|---|
| GOLGA7B | Q9D428 | -0.63 | -0.63 | -1 | -0.91 | -1 | -0.44 | -0.95 | -1.35 | -0.69 | -0.63 | -0.75 | -0.88 | -0.21 | -0.78 | -1.17 | -1.13 | -0.1425 | 0.055 | -0.03 | 0.560019 | 0.761227 | 0.909609 | 0.999971 | 1 | 0.991822 |
| Golgb1 | E9PVZ8 | 1.05 | 1.61 | 1.46 | 1.27 | 1.47 | 1.48 | 1.2 | 1.4 | 1.48 | 1.42 | 1.74 | 1.69 | 1.37 | 1.65 | 1.36 | 1.4 | 0.04 | 0.235 | 0.0975 | 0.839719 | 0.256261 | 0.627151 | 0.999971 | 1 | 0.930921 |
| Golim4 | D3YVW2 | 1.67 | 0.86 | 0.37 | -0.1 | 1.41 | 1.03 | 0.13 | 0.79 | 1 | 0.94 | 0.95 | 0.63 | 0.52 | 0.77 | 1 | 0.85 | 0.14 | 0.18 | 0.085 | 0.74936 | 0.633945 | 0.822519 | 0.999971 | 1 | 0.975901 |
| Golm2 | E9PZJ6 | 0.59 | 0.59 | 1.59 | 1.24 | 0.61 | 0.38 | 1.35 | 1.91 | 0.67 | 0.53 | 1.68 | 1.63 | 0.59 | 0.38 | 1.46 | 1.63 | 0.06 | 0.125 | 0.0125 | 0.883298 | 0.743636 | 0.974027 | 0.999971 | 1 | 0.99925 |
| Golph3 | Q9CRA5 | 1.88 | 0.65 | 1.88 | 1.57 | 0.66 | 1.21 | 1.9 | 1.87 | 0.64 | 1.08 | 2.21 | 1.15 | 0.14 | 0.57 | 1.54 | 1.31 | -0.085 | -0.225 | -0.605 | 0.830699 | 0.595839 | 0.170014 | 0.999971 | 1 | 0.802438 |
| Golph3l | H3BJ07 | -0.65 | 0.8 | 1.54 | 1.15 | 0.96 | 0.29 | 1.07 | 1.69 | 0.38 | -0.26 | 0.56 | 1.33 | 0.18 | 1.1 | 1.17 | 0.88 | 0.2925 | -0.2075 | 0.1225 | 0.57355 | 0.699853 | 0.80353 | 0.999971 | 1 | 0.971467 |
| Golt1b | Q9CR60 | -1.11 | 0.41 | -0.11 | -0.63 | 0.35 | 0.6 | -0.82 | -0.17 | 0.06 | 0.51 | -0.02 | -0.54 | -0.05 | 0.52 | -0.25 | -0.01 | 0.35 | 0.3625 | 0.4125 | 0.423785 | 0.352751 | 0.269291 | 0.999971 | 1 | 0.843504 |
| Gopc | Q8BH60 | 0.73 | 1 | 0.54 | 0.72 | 0.86 | 0.96 | 0.56 | 0.84 | 0.88 | 1.06 | 0.69 | 0.9 | 1.18 | 1.24 | 0.54 | 1.1 | 0.0575 | 0.135 | 0.2675 | 0.765808 | 0.475852 | 0.258349 | 0.999971 | 1 | 0.839012 |
| Gorab | Q8BRM2 | 0.53 | 0.68 | 0.46 | 1.15 | 0.29 | 0.48 | -0.04 | 1.12 | 0.08 | 0.34 | 0.58 | 0.17 | 0.29 | 0.4 | 0.62 | 1.53 | -0.2425 | -0.4125 | 0.005 | 0.425163 | 0.096704 | 0.987663 | 0.999971 | 1 | 0.99925 |
| Gorasp1 | Q91X51 | 0.23 | 0.28 | -0.42 | 0.24 | 0.05 | -0.25 | -0.62 | -0.37 | -0.37 | -0.02 | -0.88 | 0.23 | -0.41 | 0.22 | 0.49 | 0.48 | -0.38 | -0.3425 | 0.1125 | 0.150154 | 0.272929 | 0.693579 | 0.999971 | 1 | 0.947174 |
| Gorasp2 | A2ATI9 | 1.64 | 1.26 | 0.51 | 0.54 | 1.14 | 1.18 | 0.53 | 0.86 | 0.97 | 1.47 | 1.21 | 1.5 | 0.89 | 1.45 | 1.21 | 0.97 | -0.06 | 0.3 | 0.1425 | 0.850688 | 0.345738 | 0.648891 | 0.999971 | 1 | 0.933842 |
| Gosr1 | O88630 | 0.72 | 0.82 | 0.83 | 0.91 | 1.14 | 1.14 | 2.12 | 0.91 | 1.2 | 0.81 | 1.15 | 0.38 | 1.21 | 1.13 | 1.06 | 0.92 | 0.5075 | 0.065 | 0.26 | 0.103044 | 0.77682 | 0.151904 | 0.999971 | 1 | 0.795578 |
| Gosr2 | A2A9I0 | -0.64 | -1.24 | -2.3 | -0.95 | -1.19 | -1.98 | -1.81 | -0.39 | -0.6 | -1.09 | -0.85 | -1.26 | -1.05 | -1.16 | -1.06 | -1.62 | -0.06 | 0.3325 | 0.06 | 0.899438 | 0.388033 | 0.872431 | 0.999971 | 1 | 0.984109 |
| Got1 | P05201 | -0.65 | 0.02 | -0.69 | -0.68 | -0.24 | -0.43 | -0.38 | -0.61 | -0.09 | -0.73 | -0.5 | -0.61 | -0.48 | -0.34 | -0.62 | -0.61 | 0.085 | 0.0175 | -0.0125 | 0.710846 | 0.943849 | 0.955972 | 0.999971 | 1 | 0.998056 |
| Got2 | P05202 | -1.65 | -0.86 | -0.12 | 0.46 | -0.81 | -1.01 | -0.32 | -0.25 | -0.87 | -1.4 | -0.59 | -1.18 | -0.57 | -0.6 | 0.25 | 0.14 | -0.055 | -0.4675 | 0.3475 | 0.905137 | 0.32554 | 0.47289 | 0.999971 | 1 | 0.893778 |
| Gpaa1 | Q9WTK3 | -0.45 | -0.18 | 0.08 | -0.13 | -0.18 | 0.03 | -0.5 | 0.19 | -0.14 | 0.16 | 0.16 | 0.02 | 0.43 | 0.47 | 0.13 | 0.55 | 0.055 | 0.22 | 0.565 | 0.807348 | 0.268151 | 0.01925 | 0.999971 | 1 | 0.635388 |
| Gpalpp1 | Q69ZC8 | 0.7 | 1.82 | 0.91 | 3.19 | 1.92 | 2.1 | 2.33 | 2.3 | 2.03 | 0.92 | 1.52 | 1.26 | 0.78 | 2.39 | 1.62 | 1.66 | 0.5075 | -0.2225 | -0.0425 | 0.350812 | 0.69491 | 0.943379 | 0.999971 | 1 | 0.99692 |
| Gpam | A0A494BA | -1.24 | 0.13 | -0.66 | -0.13 | -0.46 | -0.79 | -0.47 | 0.69 | -1.2 | -0.85 | -0.36 | -0.06 | -0.85 | -0.63 | 0.66 | 0.13 | 0.2175 | -0.1425 | 0.3025 | 0.608347 | 0.710124 | 0.494087 | 0.999971 | 1 | 0.901531 |
| Gpat3 | Q8C0N2 | 0.14 | -0.06 | -0.06 | 0.68 | 0.2 | -0.72 | -0.55 | 0.21 | 0 | -0.57 | 0.25 | -0.61 | -0.64 | -0.39 | -1.3 | -0.12 | -0.39 | -0.4075 | -0.7875 | 0.225277 | 0.180358 | 0.030853 | 0.999971 | 1 | 0.678474 |
| Gpat4 | Q8K2C8 | 0.67 | 0.18 | -0.11 | -0.05 | -0.53 | 0.2 | 0.45 | 0.2 | -0.3 | 0.06 | -0.01 | -0.05 | -1.17 | -0.24 | -0.08 | 0.03 | -0.0925 | -0.2475 | -0.5375 | 0.748993 | 0.296889 | 0.127539 | 0.999971 | 1 | 0.783863 |
| Gpatch2 | M0QWJ1 | 1.06 | -0.41 | 0.65 | 0.34 | -0.8 | -0.06 | 0.45 | 0.79 | 0.64 | 1.56 | -0.16 | 0.45 | -1.02 | -0.2 | 1.14 | -0.03 | -0.315 | 0.2125 | -0.4375 | 0.478552 | 0.635843 | 0.395706 | 0.999971 | 1 | 0.871816 |
| Gpatch2l | Q6PE65 | 1.1 | 0.05 | 1.19 | -0.64 | 0.21 | 0.18 | -0.08 | 0.36 | -0.12 | 1.35 | -0.61 | 1.48 | -1.13 | 0.45 | 0.3 | 0.96 | -0.2575 | 0.1 | -0.28 | 0.548516 | 0.873084 | 0.628672 | 0.999971 | 1 | 0.932176 |
| Gpatch8 | A2A6A1 | 1.98 | 0.37 | 0.58 | 0.82 | 0.36 | 0.24 | 0.38 | 0.82 | 0.37 | 1.45 | 0.76 | 0.11 | 0.31 | 0.04 | 0.73 | 0.55 | -0.4875 | -0.265 | -0.53 | 0.20994 | 0.549352 | 0.184292 | 0.999971 | 1 | 0.809025 |
| Gpbp1 | Q6NXH3 | 2.28 | 0.36 | -0.07 | -0.59 | -1.5 | -0.53 | 0.91 | 1.03 | 0.54 | 0.94 | 0.26 | 1.43 | 0.3 | 0.76 | 0.23 | -0.07 | -0.5175 | 0.2975 | -0.19 | 0.514354 | 0.632935 | 0.749482 | 0.999971 | 1 | 0.961064 |
| Gpc1 | Q9QZF2 | -0.22 | -0.92 | -0.22 | 0.26 | -0.7 | -0.51 | 0.03 | 0.1 | -0.43 | -0.37 | -0.78 | -1.21 | -1.14 | -0.57 | -0.18 | -0.49 | 0.005 | -0.4225 | -0.32 | 0.987381 | 0.201592 | 0.329135 | 0.999971 | 1 | 0.856525 |
| Gpc4 | P51655 | -1.54 | -1.58 | -1.19 | -1.12 | -0.76 | -1.79 | -1.06 | -1.36 | -1.07 | -1.88 | -1.13 | -0.94 | -1.48 | -1.74 | -1.18 | -1.3 | 0.115 | 0.1025 | -0.0675 | 0.670637 | 0.697963 | 0.75597 | 0.999971 | 1 | 0.961116 |
| Gpc5 | Q8CAL5 | 0.93 | -0.41 | -0.26 | -0.07 | -0.3 | -0.13 | -0.26 | -0.45 | 0.08 | 0.63 | 0.12 | 0.34 | 0.24 | -0.46 | 0.05 | 0.1 | -0.3325 | 0.245 | -0.065 | 0.304658 | 0.461708 | 0.847054 | 0.999971 | 1 | 0.980295 |
| Gpc6 | Q8R3X6 | -1.01 | -0.87 | -0.88 | -1.17 | -0.47 | -0.7 | -0.72 | -1.5 | -0.33 | -1.36 | -0.61 | -0.64 | 0.19 | -0.64 | -0.77 | -0.85 | 0.135 | 0.2475 | 0.465 | 0.605266 | 0.34431 | 0.112316 | 0.999971 | 1 | 0.78268 |
| Gpcpd1 | Q8C0L9 | 0.54 | 0.84 | 0.87 | 0.3 | 0.78 | 0.83 | 0.66 | 0.94 | 0.56 | 0.75 | 0.89 | 0.79 | 0.3 | 0.73 | 0.27 | 0.73 | 0.165 | 0.11 | -0.13 | 0.424982 | 0.59095 | 0.570664 | 0.999971 | 1 | 0.91816 |
| Gpd1 | P13707 | 0.51 | 0.61 | 0.61 | 0.58 | 0.28 | 0.65 | 0.39 | 0.34 | 0.14 | 0.27 | 0.3 | -0.27 | -0.44 | 0.35 | 0.11 | 0.33 | -0.1625 | -0.4675 | -0.49 | 0.361192 | 0.036145 | 0.055133 | 0.999971 | 1 | 0.725348 |
| Gpd1l | Q3ULJ0 | 0.4 | 0.74 | 0.57 | -0.01 | 0.48 | 0.95 | 0.61 | 0.36 | 0.66 | 0.54 | 0.73 | 0.57 | 0.65 | 0.79 | 0.23 | 0.2 | 0.175 | 0.2 | 0.0425 | 0.470169 | 0.357905 | 0.864217 | 0.999971 | 1 | 0.983079 |
| Gpd2 | Q64521 | -0.89 | -0.46 | -0.58 | -0.65 | -0.66 | -0.62 | -0.56 | -0.81 | -0.84 | -1.01 | -0.55 | -0.68 | -0.86 | -0.89 | -0.91 | -0.74 | -0.0175 | -0.125 | -0.205 | 0.923262 | 0.523177 | 0.271658 | 0.999971 | 1 | 0.843794 |
| Gphn | A0JNY3 | 0.1 | -0.69 | -0.58 | -0.45 | -0.45 | -0.38 | -0.65 | -0.56 | -0.52 | 0.03 | -0.91 | -1.22 | -0.6 | -0.41 | -0.52 | -0.63 | -0.105 | -0.25 | -0.135 | 0.643686 | 0.446501 | 0.551595 | 0.999971 | 1 | 0.913379 |
| Gpi | P06745 | -0.52 | -0.32 | -0.87 | -0.45 | -0.38 | -0.35 | 0.23 | -0.85 | -0.43 | -0.72 | -0.98 | -1.3 | -0.33 | -0.13 | -0.98 | -1.03 | 0.2025 | -0.3175 | -0.0775 | 0.460279 | 0.219966 | 0.778104 | 0.999971 | 1 | 0.965243 |
| Gpkow | Q56A08 | 1.27 | 2.47 | 1.8 | 1.75 | 2.24 | 1.72 | 1.6 | 1.62 | 2.46 | 1.47 | 1.87 | 2.11 | 1.98 | 1.74 | 2.3 | 2.34 | -0.0275 | 0.155 | 0.2675 | 0.926441 | 0.634023 | 0.37662 | 0.999971 | 1 | 0.870465 |
| Gpld1 | Q8VCU2 | 2.39 | 1.17 | 0.58 | 0.89 | 0.35 | 0.65 | 0.49 | 0.17 | 0.36 | 1.38 | 0.09 | 0.7 | -0.1 | 1.14 | 0.47 | 0.39 | -0.8425 | -0.625 | -0.7825 | 0.056643 | 0.194379 | 0.105629 | 0.999971 | 1 | 0.78268 |
| Gpm6a | P35802 | 0.19 | 1.05 | -0.28 | -0.05 | 0 | 0.39 | -0.56 | -0.1 | -0.19 | 0.71 | -0.28 | -0.55 | 0.19 | 0.01 | -0.4 | 0.15 | -0.295 | -0.305 | -0.24 | 0.402678 | 0.436891 | 0.464027 | 0.999971 | 1 | 0.890905 |
| Gpm6b | A2AEG6 | -0.67 | -0.66 | -0.43 | -0.36 | -0.62 | -0.26 | -0.67 | -0.55 | -0.33 | -0.24 | -0.48 | -0.41 | -0.35 | -0.52 | -0.73 | -0.25 | 0.005 | 0.165 | 0.0675 | 0.978914 | 0.355704 | 0.730438 | 0.999971 | 1 | 0.956041 |
| Gpm6b_1 | P35803-8 | 2.78 | 3.89 | 4.66 | 3.37 | 3.39 | 2.51 | 3.8 | 2.29 | 4.61 | 7.91 | 3.32 | 6.3 | 4.72 | 1.78 | 3.96 | 2.71 | -0.6775 | 1.86 | -0.3825 | 0.196237 | 0.082236 | 0.584205 | 0.999971 | 1 | 0.920364 |
| Gpn1 | Q8VCE2 | 2.28 | 0.62 | -0.08 | 0.08 | 1.5 | 0.62 | 0.1 | 0.06 | 0.13 | 1.75 | -0.7 | -0.63 | 0.87 | 0.31 | -0.28 | -0.35 | -0.155 | -0.5875 | -0.5875 | 0.789823 | 0.419547 | 0.310507 | 0.999971 | 1 | 0.847402 |
| Gpr151 | Q7TSN6 | 0.37 | 0.78 | -0.85 | -1.19 | 0.71 | -1.97 | -0.59 | -0.68 | -0.57 | -0.41 | 0.06 | 0.29 | -0.88 | -1.09 | 0.13 | -0.03 | -0.41 | 0.065 | -0.245 | 0.536636 | 0.892388 | 0.63959 | 0.999971 | 1 | 0.93301 |
| Gpr155 | A2AWR2 | 1.46 | -0.03 | 0.19 | 0.69 | -0.53 | 0.13 | 0.7 | -0.07 | -0.43 | 0.29 | -1.14 | -0.65 | -0.79 | -0.04 | -0.87 | -1.57 | -0.52 | -1.06 | -1.395 | 0.214581 | 0.031837 | 0.009854 | 0.999971 | 1 | 0.613095 |
| Gpr158 | Q8C419 | -0.13 | -0.71 | -0.75 | -0.73 | -0.88 | -0.5 | -0.75 | -0.91 | -0.7 | -0.35 | -0.96 | -1.3 | -0.7 | -0.72 | -1.07 | -1.09 | -0.18 | -0.2475 | -0.315 | 0.423357 | 0.369698 | 0.186969 | 0.999971 | 1 | 0.809025 |
| Gpr162 | Q3UN16 | 0.14 | -0.14 | -0.11 | -0.42 | -0.67 | 0.26 | -0.67 | -0.6 | -0.39 | 0.62 | -0.31 | -0.78 | 0.37 | -0.28 | -0.4 | 0.15 | -0.2875 | -0.0825 | 0.0925 | 0.307441 | 0.796665 | 0.706849 | 0.999971 | 1 | 0.950182 |
| Gpr17 | Q6NS65 | 0.34 | -0.82 | -0.06 | 0.3 | -0.5 | -0.48 | 1.55 | -0.45 | -0.32 | 0.33 | -0.3 | -0.24 | 0.01 | -0.36 | -0.18 | -0.44 | 0.09 | -0.0725 | -0.1825 | 0.864828 | 0.817442 | 0.543428 | 0.999971 | 1 | 0.911398 |
| Gpr176 | Q80WT4 | -1.55 | -0.75 | -1.4 | -1.23 | -0.83 | -0.76 | -1.1 | -1.18 | -1.07 | -1.35 | -1.1 | -0.48 | -0.48 | -0.6 | -1.25 | -0.82 | 0.265 | 0.2325 | 0.445 | 0.27915 | 0.400757 | 0.120098 | 0.999971 | 1 | 0.78268 |
| Gpr180 | Q8BPS4 | -0.09 | -0.09 | 1.83 | 0.72 | -0.26 | -0.53 | 0.82 | 1.55 | 0.26 | 0.19 | 0.6 | 0.87 | 0.17 | 0.22 | 1.47 | 0.53 | -0.1975 | -0.1125 | 0.005 | 0.744534 | 0.804162 | 0.992108 | 0.999971 | 1 | 0.99925 |
| Gpr22 | G3X9C3 | 0.02 | 0.2 | -0.16 | 0.06 | 0.38 | -0.44 | 0.17 | -0.83 | 0.12 | 0.17 | 0.64 | 0.06 | 1.21 | -0.31 | 0.64 | 0.26 | -0.21 | 0.2175 | 0.42 | 0.485812 | 0.300848 | 0.223016 | 0.999971 | 1 | 0.827753 |
| Gpr37 | Q9QY42 | 0.62 | 0.8 | -0.12 | 0.17 | -0.1 | 0.71 | -0.97 | 0.16 | 0.14 | 0.3 | -0.55 | -0.82 | 0.42 | 0.86 | 0.01 | -0.31 | -0.4175 | -0.6 | -0.1225 | 0.302229 | 0.103725 | 0.710913 | 0.999971 | 1 | 0.950695 |
| Gpr37l1 | Q99JG2 | 0.56 | 1.04 | -3.2 | -3.3 | 1 | 0.39 | -2.57 | -3.6 | 0.84 | 0.44 | -3.62 | -3.64 | 1.47 | 0.18 | -3.46 | -2.87 | 0.03 | -0.27 | 0.055 | 0.983325 | 0.859181 | 0.97037 | 0.999971 | 1 | 0.998787 |
| Gpr68 | Q8BFQ3 | -2.1 | -1.12 | 0.65 | 0.35 | -1.47 | -1.4 | -0.9 | 1.08 | -2.92 | -1.05 | 0.32 | 0.55 | -1.72 | -0.54 | 0.26 | 1.3 | -0.1175 | -0.22 | 0.38 | 0.881683 | 0.811835 | 0.643122 | 0.999971 | 1 | 0.93301 |
| Gpr88 | Q9EPB7 | 0.08 | 0.9 | -1.49 | -1.96 | -0.08 | -0.64 | -1.69 | -0.18 | 0.02 | -3.39 | -1.36 | -1.01 | 0.99 | 1.23 | -1.08 | -1.09 | -0.03 | -0.8175 | 0.63 | 0.965274 | 0.366431 | 0.454789 | 0.999971 | 1 | 0.887217 |
| Gpr89 | Q8BS95 | -0.22 | 0.87 | 0.21 | 0.43 | 0.62 | -0.32 | -0.27 | 0.53 | 0.24 | -0.21 | 0.27 | 0.53 | 0.9 | 0.36 | 0.76 | 0.54 | -0.1825 | -0.115 | 0.3175 | 0.590704 | 0.688986 | 0.266229 | 0.999971 | 1 | 0.842677 |
| Gprasp1 | Q5U4C1 | 0.42 | 0.29 | -0.04 | 0.25 | 0.51 | 0.8 | -0.11 | 0.69 | 0.57 | 0.8 | 0.08 | 0.43 | 0.51 | 0.86 | 0.79 | 0.58 | 0.2425 | 0.24 | 0.455 | 0.349117 | 0.291886 | 0.040426 | 0.999971 | 1 | 0.700838 |
| Gprasp2 | Q8BUY8 | 1.63 | 0.44 | 0.98 | 1.16 | 0.55 | 1.12 | 1.32 | 1.79 | 0.73 | 1.21 | 0.94 | 1.03 | 0.28 | 1.07 | 1.75 | 1.18 | 0.1425 | -0.075 | 0.0175 | 0.684646 | 0.788807 | 0.963007 | 0.999971 | 1 | 0.998721 |
| Gprc5b | Q923Z0 | -0.21 | -0.88 | -1.24 | -1.43 | -1.05 | -0.66 | -1.43 | -1.11 | -0.94 | -0.14 | -1.89 | -1.25 | -1.13 | -0.66 | -1.3 | -1.01 | -0.1225 | -0.115 | -0.085 | 0.698683 | 0.788882 | 0.782662 | 0.999971 | 1 | 0.965859 |
| Gprin1 | Q3UNH4 | 0.25 | -1.45 | -1.03 | -1 | -1.52 | -0.97 | -0.56 | -1.11 | -1.32 | -0.05 | -0.97 | -1.34 | -0.96 | -1.36 | -1.37 | -0.94 | -0.2325 | -0.1125 | -0.35 | 0.563713 | 0.802147 | 0.36271 | 0.999971 | 1 | 0.869526 |
| Gprin2 | A0A2I3BRN | 0.45 | -2.17 | -1.22 | -1.48 | -1.37 | -1.44 | 0.28 | -1.22 | -0.77 | -1.04 | -1.27 | -0.9 | -1.37 | -1.68 | -0.9 | -2.15 | 0.1675 | 0.11 | -0.42 | 0.789683 | 0.834086 | 0.464213 | 0.999971 | 1 | 0.890905 |
| Gprin3 | Q8BWS5 | -0.17 | -1.47 | -1.41 | -1.03 | -1.11 | -1.47 | -1.05 | -1.07 | -0.95 | -0.27 | -1.27 | -0.7 | -0.86 | -1.1 | -0.91 | -1.09 | -0.155 | 0.2225 | 0.03 | 0.628431 | 0.539439 | 0.923213 | 0.999971 | 1 | 0.993265 |
| Gps1 | A0A140LJB | 0.98 | 1.18 | 1.17 | 0.5 | 1.32 | 1.39 | 0.84 | 1.32 | 0.68 | 0.91 | 0.96 | 0.61 | 0.09 | 1.23 | 0.69 | -0.11 | 0.26 | -0.1675 | -0.4825 | 0.290613 | 0.455848 | 0.180584 | 0.999971 | 1 | 0.808532 |
| Gps1_1 | G3UXW9 | -0.28 | 0.39 | 0.83 | 0.31 | 0.27 | 0.2 | 1.01 | 0.12 | 0.42 | -0.01 | 0.33 | -0.42 | 0.39 | 0.38 | 0.28 | -0.02 | 0.0875 | -0.2325 | -0.055 | 0.779299 | 0.451981 | 0.837677 | 0.999971 | 1 | 0.979416 |
| Gps2 | Q921N8 | 1.05 | 1.57 | 2.38 | 1.86 | 1.02 | 1.3 | 2.54 | 2.4 | 1.37 | 2.15 | 3.36 | 3.95 | -0.23 | 1.25 | 2.25 | 1.45 | 0.1 | 0.9925 | -0.535 | 0.822376 | 0.121748 | 0.336825 | 0.999971 | 1 | 0.858827 |
| Gpsm1 | Q6IR34 | 0.45 | 0.09 | -0.43 | -0.28 | -0.01 | 0.11 | -0.32 | -0.21 | 0.46 | 0.41 | -0.12 | 0.01 | 0.56 | 0.27 | 0.2 | 0 | -0.065 | 0.2325 | 0.3 | 0.79378 | 0.389892 | 0.258789 | 0.999971 | 1 | 0.839012 |
| Gpt | Q8QZR5 | -0.18 | 0.38 | 0.18 | 0.22 | 0.36 | 0.67 | -0.25 | -1.17 | 0.59 | 0.03 | 0.28 | -0.1 | 0.75 | 0.28 | 0.1 | 0.41 | -0.2475 | 0.05 | 0.235 | 0.543462 | 0.826864 | 0.30929 | 0.999971 | 1 | 0.847402 |
| Gpt2 | Q8BGT5 | -0.34 | -1.13 | -0.7 | -0.45 | -1.31 | -0.79 | -0.33 | 0.01 | -1.16 | -0.26 | -0.31 | -0.27 | -1.35 | -0.79 | -0.67 | -0.33 | 0.05 | 0.155 | -0.13 | 0.880967 | 0.598129 | 0.653904 | 0.999971 | 1 | 0.934061 |
| Gpx1 | P11352 | -0.73 | -0.15 | 0.16 | 0.76 | 0.4 | -0.24 | 0.04 | 0.34 | -0.03 | -0.5 | -0.07 | 0.54 | -0.13 | 0.09 | 0.33 | 0.48 | 0.125 | -0.025 | 0.1825 | 0.714297 | 0.945656 | 0.590858 | 0.999971 | 1 | 0.92226 |
| Gpx3 | P46412 | -0.41 | 0.13 | -0.88 | 0.84 | 1.6 | -0.17 | -0.47 | 0.66 | 0.09 | -0.35 | 0.15 | 1.95 | 0.14 | -0.24 | -0.15 | 0.14 | 0.485 | 0.54 | 0.0525 | 0.385119 | 0.363466 | 0.887649 | 0.999971 | 1 | 0.987485 |
| Gpx4 | O70325-2 | 0.95 | 0.41 | 1.1 | 0.53 | 0.31 | 0.46 | 0.69 | 0.71 | 0.92 | 1.2 | 1.17 | 0.67 | -0.06 | 0.66 | 0.81 | 0.63 | -0.205 | 0.2425 | -0.2375 | 0.382454 | 0.322269 | 0.395666 | 0.999971 | 1 | 0.871816 |
| Gpx7 | Q99LJ6 | -0.18 | -0.4 | -0.27 | -0.84 | 0.53 | 0.06 | -0.63 | 0.76 | 0.43 | 0.7 | 0.96 | -0.01 | 0.63 | 0.46 | 0.75 | -0.23 | 0.6025 | 0.9425 | 0.825 | 0.101309 | 0.006825 | 0.015423 | 0.999971 | 1 | 0.613095 |
| Gpx8 | Q9D7B7 | -3.47 | -0.33 | -1.31 | -2.07 | -0.55 | -0.87 | -1.71 | -2.7 | -0.11 | -2.75 | -2.79 | -1.4 | -0.83 | -0.84 | -0.81 | -2.49 | 0.3375 | 0.0325 | 0.5525 | 0.649165 | 0.968786 | 0.443176 | 0.999971 | 1 | 0.883386 |
| Gramd1a | F8WJ74 | -1.28 | -1.15 | -0.66 | -0.64 | -1.24 | -1.2 | -0.86 | 0.22 | -1.22 | -0.73 | -0.19 | -0.15 | -0.24 | -0.77 | -0.12 | 0.3 | 0.1625 | 0.36 | 0.725 | 0.660372 | 0.260992 | 0.031223 | 0.999971 | 1 | 0.678474 |
| Gramd1b | A0A1L1SQ | -0.38 | -0.02 | -0.72 | -0.87 | -0.11 | -0.22 | 0.3 | -0.83 | 0.14 | -0.36 | -0.16 | -0.07 | 0.35 | -0.3 | -0.73 | -1.02 | 0.2825 | 0.385 | 0.0725 | 0.36908 | 0.14148 | 0.835042 | 0.999971 | 1 | 0.979036 |
| Gramd3 | Q9D2W5 | -2.02 | -2.28 | 0.63 | 0.2 | -1.18 | -2.42 | 0.32 | 1.3 | -0.52 | -2.4 | 2.07 | 1.04 | -2.9 | -4.14 | 1.27 | 1.48 | 0.3725 | 0.915 | -0.205 | 0.707628 | 0.415888 | 0.886586 | 0.999971 | 1 | 0.987485 |
| Gramd4 | Q8CB44 | -0.24 | -0.73 | 0.06 | 0.13 | -0.74 | -0.66 | -0.33 | 0.26 | -0.66 | -0.57 | 0.06 | -0.07 | -0.69 | -0.44 | 0.22 | 0.51 | -0.1725 | -0.115 | 0.095 | 0.57655 | 0.682028 | 0.779548 | 0.999971 | 1 | 0.965847 |
| Grb10 | Q5SUW3 | 0.15 | -0.52 | 0.65 | 0.8 | -0.04 | -0.16 | 0.52 | 0.61 | -0.01 | -0.04 | 0.03 | -0.02 | -0.42 | 0.09 | 0.5 | 0.09 | -0.0375 | -0.28 | -0.205 | 0.914482 | 0.369701 | 0.559345 | 0.999971 | 1 | 0.915051 |
| Grb14 | Q9JLM9 | 2.55 | -0.21 | 0.83 | 0.43 | -0.34 | 0.68 | 1.28 | -0.08 | 0.45 | 2.48 | 1.15 | 0.33 | -1.84 | -0.78 | -0.2 | 0.84 | -0.515 | 0.2025 | -1.395 | 0.424552 | 0.772768 | 0.084894 | 0.999971 | 1 | 0.758256 |
| Grb2 | Q60631 | 0.44 | -0.35 | -0.48 | -0.27 | -0.3 | -0.09 | -0.23 | -0.19 | -0.02 | 0.2 | -0.5 | -0.4 | -0.18 | 0.06 | -0.44 | -0.18 | -0.0375 | -0.015 | -0.02 | 0.877032 | 0.956937 | 0.93773 | 0.999971 | 1 | 0.995618 |
| Grcc10 | O35127 | -0.36 | 0.91 | -0.89 | -0.15 | 0.51 | 0.74 | -0.64 | 0.1 | 0.62 | -0.24 | -0.45 | -0.55 | 0.91 | 0.39 | -0.68 | -0.66 | 0.3 | -0.0325 | 0.1125 | 0.514054 | 0.940713 | 0.824592 | 0.999971 | 1 | 0.976309 |
| Grhpr | Q91Z53 | -0.81 | -0.6 | -0.53 | -0.41 | -0.75 | -0.71 | -0.8 | -0.71 | -1.26 | -1.63 | -1.01 | -0.83 | -1.1 | -0.57 | -0.73 | -0.72 | -0.155 | -0.595 | -0.1925 | 0.384403 | 0.027013 | 0.349062 | 0.999971 | 1 | 0.865162 |
| Gria1 | Q7TNB5 | -0.38 | -0.63 | -0.88 | -1.07 | -0.8 | -0.48 | -1.08 | -0.9 | -0.81 | -0.3 | -0.89 | -1.09 | -0.36 | -0.57 | -1.03 | -0.86 | -0.075 | -0.0325 | 0.035 | 0.747562 | 0.89684 | 0.88569 | 0.999971 | 1 | 0.98711 |
| Gria2 | G5E8H1 | -1.65 | -0.99 | -1.21 | -1.48 | -1.95 | 0.01 | -2.69 | -0.96 | -2.13 | -1.48 | -1.12 | -1.07 | -1.48 | -1.41 | -2.13 | -1.32 | -0.065 | -0.1175 | -0.2525 | 0.906854 | 0.690615 | 0.344299 | 0.999971 | 1 | 0.862346 |
| Gria2_1 | P23819-4 | -2.13 | -3.22 | -0.95 | -0.86 | -2.56 | -2.96 | -0.85 | -0.92 | -2.86 | -2.69 | -0.86 | -1.03 | -2.9 | -2.84 | -0.99 | -0.64 | -0.0325 | -0.07 | -0.0525 | 0.963244 | 0.920333 | 0.943252 | 0.999971 | 1 | 0.99692 |
| Gria3 | B0QZW1 | -1.14 | -0.79 | -1.1 | -0.79 | -0.72 | -0.43 | -0.91 | -1.04 | -1 | -1.14 | -0.87 | -1.35 | -0.3 | -0.97 | -0.88 | -0.43 | 0.18 | -0.135 | 0.31 | 0.406449 | 0.498143 | 0.200261 | 0.999971 | 1 | 0.813722 |
| Gria3_1 | Q9Z2W9 | -2.28 | -1.05 | -1 | -1.34 | -0.65 | -0.92 | -1.19 | -1.36 | -0.9 | -1.66 | -0.75 | -1.09 | -1.16 | -1.46 | -1.15 | -1.09 | 0.3875 | 0.3175 | 0.2025 | 0.264109 | 0.377904 | 0.523346 | 0.999971 | 1 | 0.907797 |
| Gria4 | G5E863 | -1.06 | -1.1 | -1.01 | -1.63 | -1.43 | -0.91 | -1.15 | -1.06 | -1.34 | -1.01 | -1.33 | -1.46 | -1.06 | -0.71 | -1.09 | -0.97 | 0.0625 | -0.085 | 0.2425 | 0.779783 | 0.695517 | 0.279088 | 0.999971 | 1 | 0.843812 |
| Grid1 | Q61627 | -1.16 | -1.24 | -0.88 | -1.04 | -1.11 | -1.12 | -0.62 | -1.04 | -0.96 | -0.89 | -0.8 | -0.81 | -0.66 | -1.01 | -1.11 | -0.92 | 0.1075 | 0.215 | 0.155 | 0.594514 | 0.229205 | 0.427363 | 0.999971 | 1 | 0.880793 |
| Grid2 | Q61625 | -1.14 | -0.37 | -0.95 | -0.78 | -0.72 | -0.58 | -0.99 | -0.62 | -0.6 | -1.06 | -0.67 | -0.56 | -0.14 | -0.46 | -0.41 | -0.06 | 0.0825 | 0.0875 | 0.5425 | 0.717891 | 0.708887 | 0.040232 | 0.999971 | 1 | 0.700838 |
| Grik1 | E9Q2Y1 | 0.59 | 1.71 | 2.29 | 2.16 | 0.77 | 1.64 | 1.21 | 3.04 | 0.8 | 0.79 | 1.04 | -2.93 | 2.25 | 1.69 | 4.14 | 2.88 | -0.0225 | -1.7625 | 1.0525 | 0.96861 | 0.084763 | 0.105422 | 0.999971 | 1 | 0.78268 |
| Grik2 | P39087 | -1.26 | -1.03 | -1.88 | -1.5 | -0.38 | -1.14 | -1.66 | -1.35 | -0.61 | -1.19 | -1.48 | -1.71 | 0.24 | -0.89 | -1.29 | -1.24 | 0.285 | 0.17 | 0.6225 | 0.394748 | 0.580794 | 0.133506 | 0.999971 | 1 | 0.784948 |
| Grik3 | B1AS29 | -0.38 | -0.87 | -0.94 | -0.73 | -0.64 | -0.48 | -0.87 | -0.33 | -0.74 | -0.3 | -0.53 | -0.79 | -0.25 | -0.72 | -0.67 | -0.32 | 0.15 | 0.14 | 0.24 | 0.493919 | 0.51527 | 0.289197 | 0.999971 | 1 | 0.847402 |
| Grik5 | Q61626 | 0.05 | -0.81 | -1.16 | -0.95 | -0.87 | -0.82 | -1.06 | -1.06 | -0.86 | 0.09 | -1.4 | -1.28 | -0.46 | -0.71 | -1.23 | -1.05 | -0.235 | -0.145 | -0.145 | 0.420822 | 0.724742 | 0.651961 | 0.999971 | 1 | 0.933842 |
| Grin1 | A2AI14 | -0.63 | -0.08 | -1.61 | -0.43 | -0.08 | -0.39 | -1.06 | -0.76 | -0.94 | -0.97 | -0.81 | -0.83 | -0.2 | -0.42 | 0.24 | -1.58 | 0.115 | -0.2 | 0.1975 | 0.761405 | 0.548138 | 0.678837 | 0.999971 | 1 | 0.942697 |
| Grin1_1 | A2AI21 | -1.21 | -1.12 | -1.08 | -1.11 | -1.22 | -0.87 | -0.93 | -1.04 | -1.06 | -1.11 | -0.84 | -1.48 | -0.88 | -1.07 | -1.54 | -1.14 | 0.115 | 0.0075 | -0.0275 | 0.510021 | 0.969136 | 0.890949 | 0.999971 | 1 | 0.988184 |
| Grin2a | P35436 | -1.15 | -1.49 | -0.79 | -1 | -1.6 | -1.14 | -0.72 | -1.25 | -1.73 | -1.27 | -0.58 | -0.97 | -1.55 | -1.68 | -1.32 | -1.02 | -0.07 | -0.03 | -0.285 | 0.786396 | 0.918563 | 0.254636 | 0.999971 | 1 | 0.839012 |
| Grin2b | G3X9V4 | -0.98 | -1.19 | -1.04 | -1.09 | -0.95 | -1 | -0.71 | -1 | -1.06 | -1.15 | -1.11 | -1.09 | -0.46 | -1.21 | -1.14 | -1.02 | 0.16 | -0.0275 | 0.1175 | 0.365827 | 0.862174 | 0.597371 | 0.999971 | 1 | 0.92377 |
| Grin2d | Q03391 | 0.36 | -0.62 | 0.37 | 1.96 | -0.79 | -0.85 | 0.04 | 1.86 | -1.16 | -0.88 | 0.21 | -0.14 | -0.44 | -0.39 | 1.21 | 1.97 | -0.4525 | -1.01 | 0.07 | 0.547994 | 0.105573 | 0.923079 | 0.999971 | 1 | 0.993264 |
| Grin3a | A2AIR5-2 | -0.2 | -1.57 | -1.23 | -1.01 | -1.13 | -0.53 | -0.96 | -0.84 | -0.73 | -0.04 | -1.42 | -0.97 | -1.1 | -0.69 | -1.02 | -0.95 | 0.1375 | 0.2125 | 0.0625 | 0.668685 | 0.592539 | 0.840473 | 0.999971 | 1 | 0.97975 |
| Grip1 | D3YZL8 | -1.09 | -1.88 | -1.26 | -1.54 | -1.9 | -1.52 | -1.27 | -1.25 | -1.57 | -1.38 | -0.93 | -1.35 | -1.36 | -1.42 | -1.18 | -1.1 | -0.0425 | 0.135 | 0.1775 | 0.867755 | 0.58715 | 0.445142 | 0.999971 | 1 | 0.884152 |
| Grip2 | G3XA20 | -0.65 | -1.2 | -1.44 | -1.18 | -1.55 | -1.08 | -1.73 | -1.16 | -1.28 | -0.74 | -1.43 | -1.26 | -1.42 | -1.55 | -1.48 | -1.26 | -0.2625 | -0.06 | -0.31 | 0.316381 | 0.810709 | 0.185953 | 0.999971 | 1 | 0.809025 |
| Gripap1 | A2AEW8 | 0.04 | 0.26 | 0.04 | -0.21 | 0.27 | -0.05 | 0.1 | -0.11 | 0.34 | -0.25 | 0.15 | 0.04 | 0.4 | -0.24 | -0.12 | -0.19 | 0.02 | 0.0375 | -0.07 | 0.917349 | 0.855397 | 0.751806 | 0.999971 | 1 | 0.961116 |
| Gripap1_1 | Q8VD04 | 0.39 | 0.42 | 0.49 | -0.2 | 0.53 | 0.33 | 0.13 | 0.88 | -0.18 | 0.56 | -0.01 | 0.3 | 0.12 | 0.83 | 0.6 | 0.66 | 0.1925 | -0.1075 | 0.2775 | 0.453828 | 0.672888 | 0.283567 | 0.999971 | 1 | 0.844538 |
| Gripap1_2 | A2AEW6 | -0.55 | 0.79 | 2 | 0.09 | 0.91 | 0.59 | 1.04 | 1.22 | 1.74 | -0.45 | 0.84 | 2.36 | 1.98 | 0.03 | 1.32 | 1.01 | 0.3575 | 0.54 | 0.5025 | 0.496792 | 0.474318 | 0.426886 | 0.999971 | 1 | 0.880793 |
| Grk2 | Q99MK8 | -0.28 | 0.29 | 0.32 | 0.17 | 0.01 | 0.23 | 0.34 | 0.41 | 0.16 | -0.05 | 0.54 | 0.06 | 0.3 | -0.01 | 0.29 | 0.52 | 0.1225 | 0.0525 | 0.15 | 0.568369 | 0.816536 | 0.502805 | 0.999971 | 1 | 0.904573 |
| Grk5 | Q8VEB1 | 0.85 | 0.42 | -0.2 | -0.05 | -0.35 | 0.37 | -0.42 | -0.03 | -0.36 | 0.72 | -0.62 | -0.6 | -0.32 | 0.4 | 0.28 | 0.44 | -0.3625 | -0.47 | -0.055 | 0.254567 | 0.240722 | 0.856632 | 0.999971 | 1 | 0.98255 |
| Grk6 | O70293-2 | 0.34 | -0.09 | -0.64 | -0.55 | -0.12 | -0.25 | -0.9 | -0.31 | -0.11 | 0.38 | -0.7 | -0.49 | 0.12 | 0.03 | -0.52 | -0.37 | -0.16 | 0.005 | 0.05 | 0.59057 | 0.987811 | 0.861937 | 0.999971 | 1 | 0.982963 |
| Grm1 | P97772 | -1.36 | -0.82 | -0.72 | -0.57 | -0.76 | -0.54 | -1.07 | -0.53 | -0.52 | -0.66 | -0.7 | -0.45 | -0.04 | -0.37 | -0.37 | -0.4 | 0.1425 | 0.285 | 0.5725 | 0.563383 | 0.219954 | 0.032742 | 0.999971 | 1 | 0.685881 |
| Grm2 | Q14BI2 | -0.81 | -0.65 | -0.73 | -0.65 | -0.98 | -0.96 | -1.03 | -0.77 | -1.08 | -1.09 | -0.87 | -1.01 | -0.77 | -1.03 | -1.23 | -0.72 | -0.225 | -0.3025 | -0.2275 | 0.202815 | 0.090221 | 0.254965 | 0.999971 | 1 | 0.839012 |
| Grm3 | Q9QYS2 | -1.45 | -1.81 | -1.13 | -0.93 | -1.72 | -1.88 | -0.77 | -1.39 | -1.84 | -2.01 | -1.14 | -1.72 | -1.27 | -2.03 | -1.58 | -1.22 | -0.11 | -0.3475 | -0.195 | 0.72808 | 0.239692 | 0.496792 | 0.999971 | 1 | 0.902655 |
| Grm4 | A0A338P7 | -0.71 | -0.59 | -0.81 | -0.7 | -0.82 | -0.75 | -1.27 | -0.61 | -0.83 | -1.34 | -1.02 | -0.87 | -0.35 | -0.49 | -0.93 | -0.71 | -0.16 | -0.3125 | 0.0825 | 0.441881 | 0.123593 | 0.676819 | 0.999971 | 1 | 0.942048 |
| Grm5 | Q3UVX5 | -0.49 | -0.64 | -0.58 | -0.51 | -0.78 | -0.82 | -0.62 | -0.64 | -0.66 | -0.44 | -0.64 | -0.74 | -0.48 | -0.71 | -0.5 | -0.52 | -0.16 | -0.065 | 0.0025 | 0.346684 | 0.695499 | 0.988042 | 0.999971 | 1 | 0.99925 |
| Grm7 | Q68ED2 | -0.67 | -1.27 | -1.61 | -1.25 | -1.43 | -1.17 | -1.33 | -1.29 | -1.45 | -0.96 | -1.58 | -1.62 | -0.83 | -0.94 | -1.44 | -1.34 | -0.105 | -0.2025 | 0.0625 | 0.659028 | 0.454787 | 0.814976 | 0.999971 | 1 | 0.974918 |
| Grm8 | Q05BD6 | -0.63 | -1.09 | -0.85 | -0.88 | -1.12 | -0.95 | -0.68 | -1.02 | -1.09 | -0.68 | -1.06 | -1.06 | -0.74 | -0.85 | -0.87 | -1.21 | -0.08 | -0.11 | -0.055 | 0.683561 | 0.57423 | 0.782521 | 0.999971 | 1 | 0.965859 |
| Grm8_1 | P47743 | 0.96 | 1.47 | -0.69 | -0.58 | 1.65 | 1.62 | -0.54 | 0.1 | 1.64 | 2.55 | -0.76 | 0.57 | 4.75 | 1.64 | 1.79 | 0.33 | 0.4175 | 0.71 | 1.8375 | 0.55503 | 0.392334 | 0.084619 | 0.999971 | 1 | 0.758256 |
| Grn | P28798 | 0.63 | 2.79 | 2.48 | 3.8 | 3.03 | 2.51 | 2.24 | 4.12 | 2.92 | 0.9 | 3.73 | 2.87 | 3.47 | 2.79 | 3.25 | 3.73 | 0.55 | 0.18 | 0.885 | 0.443768 | 0.823961 | 0.183876 | 0.999971 | 1 | 0.809025 |
| Grp | Q8R1I2 | -0.76 | -2.28 | -0.63 | -0.52 | -1.34 | -0.87 | -1.11 | 0.57 | -1.58 | -1.82 | -0.16 | -0.47 | -2 | -1.25 | -0.25 | 0.27 | 0.36 | 0.04 | 0.24 | 0.516643 | 0.940649 | 0.689657 | 0.999971 | 1 | 0.945805 |
| Grpel1 | Q99LP6 | 1.83 | 0.3 | -0.05 | 0.09 | -0.24 | 0.56 | 0.71 | -0.46 | -0.79 | 0.26 | -0.44 | -0.66 | 0.39 | -0.32 | 0.07 | -0.22 | -0.4 | -0.95 | -0.5625 | 0.419719 | 0.067125 | 0.220602 | 0.999971 | 1 | 0.827096 |
| Grpel2 | O88396 | -2.01 | 0.61 | -0.91 | -0.22 | 0.38 | 1.05 | -1.24 | -0.82 | -0.49 | -0.56 | -0.4 | -0.85 | 2.5 | 0.67 | 0.57 | 0.45 | 0.475 | 0.0575 | 1.68 | 0.498851 | 0.912291 | 0.031475 | 0.999971 | 1 | 0.678474 |
| Grsf1 | E9Q179 | -0.79 | -0.76 | -0.05 | 0.13 | -0.87 | -0.85 | -0.2 | 0.34 | -0.9 | -0.66 | 0.05 | 0.24 | -1.21 | -0.82 | 0.29 | 0.38 | -0.0275 | 0.05 | 0.0275 | 0.939936 | 0.88823 | 0.949908 | 0.999971 | 1 | 0.998038 |
| Grtp1 | Q9D3N8-4 | -0.44 | -0.77 | -0.47 | -0.66 | -0.84 | -1.01 | -0.74 | -1.07 | -1.12 | -0.52 | -1.42 | -1.14 | -0.36 | -0.48 | -1.1 | -0.36 | -0.33 | -0.465 | 0.01 | 0.099672 | 0.076502 | 0.965577 | 0.999971 | 1 | 0.998721 |
| Grwd1 | Q810D6 | -0.11 | 0.31 | 0.89 | 0.52 | 0.27 | 0.02 | 0.59 | 0.6 | -0.05 | -0.49 | 1.02 | 1.03 | 0.72 | 0.01 | 1.11 | 0.46 | -0.0325 | -0.025 | 0.1725 | 0.904233 | 0.952128 | 0.587797 | 0.999971 | 1 | 0.920758 |
| Gsdme | Q9Z2D3 | -0.68 | -0.68 | -0.13 | 0 | -0.23 | -0.79 | -0.38 | 0.15 | -0.36 | -1.13 | -0.08 | 0.2 | -0.24 | -0.55 | 0.36 | 0 | 0.06 | 0.03 | 0.265 | 0.830401 | 0.928711 | 0.3558 | 0.999971 | 1 | 0.868793 |
| Gse1 | Q3U3C9-4 | 4.39 | 2.15 | 1.2 | 1.18 | 0.88 | 4.34 | 0.92 | 1.05 | 2.48 | 4.83 | 1.28 | 1.06 | 0.28 | 1.7 | 1 | 1.42 | -0.4325 | 0.1825 | -1.13 | 0.670745 | 0.859467 | 0.151129 | 0.999971 | 1 | 0.7948 |
| Gsg1l | D3Z7H4 | -1.06 | -0.93 | -1.64 | -1.17 | -0.87 | -0.68 | -1.84 | -1.29 | -1.1 | -1.36 | -1.48 | -2.11 | -1.11 | -1.13 | -1.54 | -1.43 | 0.03 | -0.3125 | -0.1025 | 0.92194 | 0.28016 | 0.655434 | 0.999971 | 1 | 0.934061 |
| Gsk3a | Q2NL51 | -0.85 | 0.59 | -0.07 | -0.46 | 0.64 | 0.5 | -0.16 | -0.41 | 0.53 | -0.91 | -0.33 | -0.27 | 0.91 | 0.21 | 0 | -0.23 | 0.34 | -0.0475 | 0.42 | 0.38633 | 0.907182 | 0.286972 | 0.999971 | 1 | 0.846151 |
| Gsk3b | E9QAQ5 | -0.8 | 0.3 | 0.49 | 0.1 | 0.32 | 0.12 | 0.62 | 0.17 | 0.3 | -0.34 | 0.4 | 0.6 | 0.38 | 0.19 | 0.27 | 0.23 | 0.285 | 0.2175 | 0.245 | 0.372035 | 0.533752 | 0.42061 | 0.999971 | 1 | 0.87788 |
| Gskip | A0A1Y7VN | 0.55 | 0.61 | 0.68 | 0.67 | 0.12 | 0.86 | 0.31 | 1.34 | 0.44 | 0.28 | 0.09 | 0.14 | 0.68 | 0.77 | 0.93 | 1.62 | 0.03 | -0.39 | 0.3725 | 0.91751 | 0.044425 | 0.155554 | 0.999971 | 1 | 0.795578 |
| Gsn | P13020-2 | 0.86 | -0.14 | 0.36 | 0.45 | 0.16 | -0.05 | 1.04 | -0.1 | 0.4 | 0.74 | 0.19 | 0.54 | -0.22 | -0.08 | 0.29 | 0.21 | -0.12 | 0.085 | -0.3325 | 0.719918 | 0.742566 | 0.224991 | 0.999971 | 1 | 0.828225 |
| Gsn_1 | P13020 | -0.09 | -0.78 | 0.25 | 0.49 | 0.65 | 0.14 | 0.07 | 0.19 | -0.26 | 0.07 | -0.16 | 0.98 | -0.36 | 0.63 | 0.07 | 0.55 | 0.295 | 0.19 | 0.255 | 0.355178 | 0.621327 | 0.477245 | 0.999971 | 1 | 0.894438 |
| Gspt1 | Q8R050-2 | -0.15 | 0.47 | 0.39 | 0.04 | 0.9 | 0.23 | 0.57 | 0.22 | 1.04 | -0.09 | 0.64 | 0.96 | 1.19 | 0.01 | 0.42 | 0.08 | 0.2925 | 0.45 | 0.2375 | 0.256012 | 0.162921 | 0.454972 | 0.999971 | 1 | 0.887217 |
| Gspt2 | Q149F3 | -1.46 | -0.34 | -0.68 | -1.24 | -0.18 | -0.17 | -0.5 | -0.45 | -0.06 | -1.3 | -0.26 | -0.35 | 0.42 | 0.12 | -0.53 | -0.48 | 0.605 | 0.4375 | 0.8125 | 0.058387 | 0.252736 | 0.03931 | 0.999971 | 1 | 0.700838 |
| Gsr | P47791-2 | 1.02 | 0.19 | 0.19 | 0.29 | 0.74 | 0.51 | 0.63 | 0.4 | 0.62 | 0.58 | 0.31 | 0.68 | 0.61 | 0.5 | 0.58 | 0 | 0.1475 | 0.125 | -1.18E-16 | 0.550091 | 0.612034 | 1 | 0.999971 | 1 | 1 |
| Gss | P51855 | 0.2 | 0.78 | -0.17 | -0.33 | 0.57 | 0.7 | -0.09 | -0.06 | 0.9 | 0.5 | 0.08 | 0.13 | 0.93 | 0.47 | -0.35 | -0.53 | 0.16 | 0.2825 | 0.01 | 0.622631 | 0.380782 | 0.980269 | 0.999971 | 1 | 0.99925 |
| Gsta4 | P24472 | 0.58 | 0.6 | 0.63 | 0.52 | 0.6 | 0.52 | 0.58 | 0.7 | 0.43 | 0.06 | 0.46 | 0.94 | 0.39 | 1.13 | 0.35 | 0.72 | 0.0175 | -0.11 | 0.065 | 0.913594 | 0.621943 | 0.772673 | 0.999971 | 1 | 0.963573 |
| Gsta4_1 | A0A1L1SS6 | 0.39 | 1.66 | -1.33 | -0.63 | 0.18 | 1 | -1.73 | -1.35 | 1.6 | 0.97 | -1.95 | -2.16 | -0.51 | 2.21 | -2.16 | 0.51 | -0.4975 | -0.4075 | -0.01 | 0.548231 | 0.699417 | 0.992043 | 0.999971 | 1 | 0.99925 |
| Gstk1 | Q9DCM2 | -2.32 | -0.72 | -1.47 | -1.55 | -0.63 | -0.72 | -1.07 | -1.07 | -0.93 | -2.3 | -1.49 | -1.76 | 0.04 | -1.09 | -1.43 | -1.37 | 0.6425 | -0.105 | 0.5525 | 0.087909 | 0.799885 | 0.236033 | 0.999971 | 1 | 0.832918 |
| Gstm1 | P10649 | -1.01 | -1.37 | -0.96 | -1.01 | -1.15 | -1.06 | -0.77 | -0.4 | -1.4 | -1.85 | -1.57 | -1.33 | -2.03 | -0.7 | -1 | -1.18 | 0.2425 | -0.45 | -0.14 | 0.309768 | 0.049692 | 0.650662 | 0.999971 | 1 | 0.933842 |
| Gstm2 | P15626 | 2.28 | 1.05 | 0.95 | 1.06 | 3.1 | 0.89 | 1.33 | -0.11 | 1.82 | 0.9 | 0.82 | 1.54 | 1.31 | 1.06 | 1.26 | 0.44 | -0.0325 | -0.065 | -0.3175 | 0.961358 | 0.866023 | 0.394816 | 0.999971 | 1 | 0.871816 |
| Gstm4 | Q8R5I6 | 0.88 | 0.85 | 0.8 | 0.97 | 1.67 | 1.01 | 0.71 | 1.51 | 1.12 | 0.88 | 0.88 | 0.5 | 1.38 | 1.54 | 1 | 0.93 | 0.35 | -0.03 | 0.3375 | 0.18822 | 0.87617 | 0.128469 | 0.999971 | 1 | 0.784699 |
| Gstm5 | P48774 | 0.16 | 0.55 | 1.07 | 0.92 | 0.47 | 0.93 | 0.96 | 0.86 | 0.22 | 0.27 | 0.42 | 0 | 0.1 | 0.64 | 0.61 | 1.05 | 0.13 | -0.4475 | -0.075 | 0.616229 | 0.098664 | 0.798269 | 0.999971 | 1 | 0.970479 |
| Gstm7 | Q80W21 | 0.75 | 0.32 | 1.01 | 0.89 | 0.27 | 0.52 | 1.2 | 0.92 | 0.41 | 0.39 | 0.96 | 1.03 | -0.22 | 0.51 | 1.33 | 1.15 | -0.015 | -0.045 | -0.05 | 0.956236 | 0.858964 | 0.892906 | 0.999971 | 1 | 0.98828 |
| Gsto1 | O09131 | 0.16 | 0.62 | 0.98 | 0.97 | 0.38 | 0.21 | 1.1 | 1.18 | 1.02 | 0.75 | 1.16 | 0.02 | 0.47 | 0.46 | 0.64 | 0.78 | 0.035 | 0.055 | -0.095 | 0.911937 | 0.863791 | 0.694578 | 0.999971 | 1 | 0.947174 |
| Gstp1 | P19157 | -0.68 | -0.26 | -0.65 | -0.69 | 0.08 | -0.04 | -0.78 | -0.32 | -0.08 | -1.26 | -1.23 | -0.6 | -0.6 | -0.03 | -0.44 | -0.72 | 0.305 | -0.2225 | 0.1225 | 0.236882 | 0.474404 | 0.58899 | 0.999971 | 1 | 0.921663 |
| Gstt1 | Q64471 | -1.6 | -1.74 | 0.61 | 0.69 | -0.92 | -0.85 | 0.46 | 0.25 | -2.34 | -1.36 | 0.35 | 0.65 | -1.75 | -0.52 | 1.19 | 0.17 | 0.245 | -0.165 | 0.2825 | 0.722821 | 0.851428 | 0.731007 | 0.999971 | 1 | 0.956041 |
| Gstt2 | A0A1W2P7 | -1.3 | 1.15 | 0.89 | 1.24 | 1.85 | 0.12 | 0.27 | 1.24 | 1.15 | 0.82 | 0.53 | -0.18 | 1.1 | 1.59 | 1.92 | 2.51 | 0.375 | 0.085 | 1.285 | 0.574036 | 0.88928 | 0.061666 | 0.999971 | 1 | 0.725348 |
| Gstz1 | Q9WVL0 | -0.04 | -0.28 | -2 | -1.89 | -0.17 | 0.26 | -1.38 | -1.86 | -0.44 | -0.22 | -2.27 | -2.3 | -0.48 | 0.02 | -2.07 | -1.77 | 0.265 | -0.255 | -0.0225 | 0.685746 | 0.715938 | 0.972599 | 0.999971 | 1 | 0.99925 |
| Gtdc1 | G8JL42 | -1.11 | 0.83 | -0.66 | -0.5 | 0.69 | 0.76 | -0.89 | -0.42 | 0.84 | 0 | -0.32 | -0.13 | 1.18 | 1.15 | -0.67 | -0.37 | 0.395 | 0.4575 | 0.6825 | 0.470037 | 0.334475 | 0.265388 | 0.999971 | 1 | 0.842677 |
| Gtf2a1 | Q99PM3 | 1.71 | 2.32 | 2.25 | 2.97 | 1.89 | 1.42 | 1.72 | 2.52 | 1.7 | 0.81 | 2.77 | 2.63 | 1.34 | 2.27 | 2.05 | 2.36 | -0.425 | -0.335 | -0.3075 | 0.235633 | 0.499463 | 0.381251 | 0.999971 | 1 | 0.871816 |
| Gtf2a2 | D3Z793 | 0.88 | 2.14 | 2.42 | 2.92 | 1.3 | 0.85 | 1.79 | 2.54 | 1.5 | 0.95 | 2.19 | 2.08 | 1.75 | 2.08 | 2.07 | 2.36 | -0.47 | -0.41 | -0.025 | 0.379076 | 0.409161 | 0.9534 | 0.999971 | 1 | 0.998056 |
| Gtf2b | P62915 | 3.41 | 2.89 | 1.7 | 2.22 | 1.79 | 2.02 | 1.42 | 1.94 | 1.42 | 2.17 | 1.85 | 2.02 | 2.11 | 2.44 | 1.66 | 2.75 | -0.7625 | -0.69 | -0.315 | 0.073111 | 0.106915 | 0.460374 | 0.999971 | 1 | 0.889706 |
| Gtf2e1 | Q9D0D5 | 0.53 | 1.43 | 1.86 | 2.17 | 1.78 | 1.34 | 1.57 | 2.33 | 1.61 | 0.57 | 2.39 | 2.45 | 0.76 | 1.25 | 2.2 | 2.81 | 0.2575 | 0.2575 | 0.2575 | 0.52136 | 0.62577 | 0.634184 | 0.999971 | 1 | 0.93301 |
| Gtf2e2 | Q9D902 | 0.86 | 1.79 | 1.55 | 2.21 | 1.71 | 1.32 | 1.28 | 1.95 | 1.21 | 1.3 | 1.38 | 1.54 | 1.21 | 1.99 | 1.53 | 1.98 | -0.0375 | -0.245 | 0.075 | 0.908273 | 0.421886 | 0.824482 | 0.999971 | 1 | 0.976309 |
| Gtf2f1 | Q3THK3 | 1.6 | 1.73 | 2.14 | 2.38 | 1.45 | 1.06 | 1.63 | 2.21 | 1.16 | 1.11 | 2.04 | 2.61 | 0.92 | 1.35 | 2.31 | 2.9 | -0.375 | -0.2325 | -0.0925 | 0.240719 | 0.555418 | 0.839034 | 0.999971 | 1 | 0.979728 |
| Gtf2f2 | Q8R0A0 | 1.2 | 2.28 | 2.7 | 2.46 | 2.73 | 1.88 | 2.42 | 2.58 | 2.67 | 1.53 | 3.07 | 3.21 | 2.66 | 2.04 | 2.57 | 2.64 | 0.2425 | 0.46 | 0.3175 | 0.515882 | 0.344639 | 0.383445 | 0.999971 | 1 | 0.871816 |
| Gtf2h1 | G3X8R4 | 1.54 | 2.72 | 0.22 | 0.81 | 2.68 | 1.99 | 2.2 | 0.33 | 2.86 | 2.06 | 0.6 | 0.22 | 3.8 | 2.94 | -0.76 | 1.31 | 0.4775 | 0.1125 | 0.5 | 0.483901 | 0.879831 | 0.625472 | 0.999971 | 1 | 0.930479 |
| Gtf2h4 | O70422 | 0.85 | 1.81 | 0.8 | 1.24 | 1.85 | 1.49 | 0.9 | 1.57 | 1.52 | 1.51 | 1.6 | 1.51 | 2.19 | 1.8 | 1.12 | 1.62 | 0.2775 | 0.36 | 0.5075 | 0.384479 | 0.18572 | 0.143643 | 0.999971 | 1 | 0.788654 |
| Gtf2h5 | Q8K2X8 | -0.33 | 0.42 | 0.5 | 0.93 | 0.56 | -0.12 | -0.08 | 1.34 | 0.29 | -0.58 | 0.36 | 1.88 | -0.55 | 0.04 | -0.3 | 0.06 | 0.045 | 0.1075 | -0.5675 | 0.912591 | 0.839907 | 0.091799 | 0.999971 | 1 | 0.773779 |
| Gtf2i | Q3UHU8 | -0.17 | -0.09 | 0.49 | 0.43 | 0.09 | -0.5 | 0.06 | 0.15 | 0.08 | -0.25 | 0.18 | 0.52 | -0.16 | -0.06 | 0.38 | 0.42 | -0.215 | -0.0325 | -0.02 | 0.408029 | 0.899148 | 0.937219 | 0.999971 | 1 | 0.995618 |
| Gtf3c1 | A0A0U1RP | -0.03 | -0.29 | 0.18 | -0.03 | -0.22 | -0.32 | 0.33 | -0.35 | -0.36 | -0.1 | -0.12 | 0.06 | -0.17 | -0.23 | -0.06 | -0.25 | -0.0975 | -0.0875 | -0.135 | 0.667915 | 0.648128 | 0.46681 | 0.999971 | 1 | 0.891477 |

| | | | | | | | | | | | | | | | | | | | | | | | | | | |
|---|---|---|---|---|---|---|---|---|---|---|---|---|---|---|---|---|---|---|---|---|---|---|---|---|---|---|
| Gtf3c2 | D3Z7I0 | 0.92 | -0.32 | -0.33 | -0.12 | -0.21 | -0.23 | -0.03 | 0.03 | -0.27 | 1.33 | -0.53 | 0.19 | 0.49 | 0.34 | 0.14 | 0.54 | -0.1475 | 0.1425 | 0.34 | 0.636475 | 0.764966 | 0.297857 | 0.999971 | 1 | 0.847402 |
| Gtf3c3 | Q3TMP1 | -0.23 | -0.81 | -0.64 | -1.02 | -0.64 | -0.88 | -0.87 | -0.59 | -0.55 | -0.37 | -1.07 | -0.65 | -0.78 | -0.51 | -0.72 | -0.37 | -0.07 | 0.015 | 0.08 | 0.756017 | 0.95205 | 0.729785 | 0.999971 | 1 | 0.956041 |
| Gtf3c4 | Q8BMQ2 | -1.61 | -1.43 | -0.21 | -0.7 | -1.8 | -2.01 | 0.03 | -0.39 | -1.65 | -1.78 | -0.05 | -0.21 | -2.57 | -1.03 | -0.43 | -0.7 | -0.055 | 0.065 | -0.195 | 0.920735 | 0.900938 | 0.715857 | 0.999971 | 1 | 0.952576 |
| Gtf3c5 | Q8R2T8 | 0.33 | 0.02 | -0.34 | -0.25 | 0.53 | 0.06 | 0.12 | -0.03 | 0.71 | 0.62 | -0.51 | -0.58 | 0.59 | -0.07 | 0.7 | -0.62 | 0.23 | 0.12 | -0.14 | 0.335541 | 0.746646 | 0.675957 | 0.999971 | 1 | 0.941711 |
| Gtpbp1 | O08582 | 2.11 | 0.12 | 0.03 | 0.21 | -0.03 | 0.59 | 0.39 | 0.66 | -0.03 | 2.27 | -0.42 | 0.13 | -0.3 | 0.28 | 0.11 | 0.02 | -0.215 | -0.13 | -0.59 | 0.659697 | 0.855363 | 0.238806 | 0.999971 | 1 | 0.833594 |
| Gtpbp10 | Q8K013 | -1.12 | -0.3 | 0.98 | -0.38 | -0.51 | -0.51 | -1.08 | -1.04 | -0.5 | -0.65 | -0.13 | 0.35 | -0.37 | -0.63 | -1.13 | 1.12 | -0.58 | -0.0275 | -0.0475 | 0.207377 | 0.952322 | 0.936393 | 0.999971 | 1 | 0.995618 |
| Gtpbp2 | Q3UJK4 | 0.43 | -0.34 | 0.19 | 0.51 | -0.81 | -0.5 | 0.11 | 0.26 | -0.85 | -0.14 | 0.25 | 0.1 | -0.58 | -0.1 | 0.75 | 0.38 | -0.4325 | -0.3575 | -0.085 | 0.198683 | 0.272497 | 0.804625 | 0.999971 | 1 | 0.971467 |
| Gtpbp3 | E9PWV7 | 0.09 | -0.21 | -0.15 | -0.04 | 0.33 | -0.35 | -0.75 | -0.66 | -0.1 | -0.26 | -1.07 | -0.9 | 0.77 | -0.49 | -0.98 | -0.01 | -0.28 | -0.505 | -0.1 | 0.31812 | 0.08458 | 0.786436 | 0.999971 | 1 | 0.967132 |
| Gtpbp4 | Q99ME9 | -0.05 | 0.34 | 0.08 | 0.33 | 0.45 | 0.5 | -0.17 | 0.49 | 0.17 | 0.04 | 0.26 | 0.28 | 0.94 | 0.68 | 0.05 | 0.19 | 0.1425 | 0.0125 | 0.29 | 0.536854 | 0.94494 | 0.274372 | 0.999971 | 1 | 0.843794 |
| Gtpbp6 | Q3U6U5 | 0.06 | -1.01 | 1.08 | 0.34 | -0.37 | -0.44 | -0.3 | 0.29 | -0.44 | -0.17 | -0.39 | -0.78 | -0.84 | -0.41 | 0.16 | 0.39 | -0.3225 | -0.5625 | -0.2925 | 0.468149 | 0.211565 | 0.546802 | 0.999971 | 1 | 0.911403 |
| Gtpbp8 | F6XXP5 | -0.38 | -0.95 | -0.86 | -1.05 | -1.16 | -1.27 | -1.28 | -1.53 | -1.1 | -1.03 | -0.58 | -1.05 | -0.98 | -1.03 | -0.69 | -0.76 | -0.5 | -0.13 | -0.055 | 0.042181 | 0.57174 | 0.800095 | 0.999971 | 1 | 0.970723 |
| Gucy1a1 | Q9ERL9 | -0.44 | 0.5 | -0.11 | -0.1 | 0.76 | 0.15 | 0.21 | 0.34 | 0.49 | -0.1 | -0.18 | 0.23 | 1.09 | 0.62 | -0.39 | -0.09 | 0.4025 | 0.1475 | 0.54 | 0.150221 | 0.585323 | 0.116473 | 0.999971 | 1 | 0.78268 |
| Gucy1a2 | F8VQK3 | -0.44 | -0.58 | -0.66 | -0.86 | -0.52 | -0.45 | -0.51 | -0.87 | -0.56 | -0.64 | -1.04 | -1 | -0.17 | -0.22 | -0.67 | -0.85 | 0.0475 | -0.175 | 0.1575 | 0.806345 | 0.397906 | 0.497194 | 0.999971 | 1 | 0.902655 |
| Gucy1b1 | O54865 | -0.86 | -1.02 | -0.63 | -0.48 | -0.72 | -0.97 | -0.4 | -0.35 | -0.96 | -1.16 | -0.5 | -0.42 | -0.68 | -0.73 | -0.36 | -0.42 | 0.1375 | -0.0125 | 0.2 | 0.550068 | 0.959052 | 0.344951 | 0.999971 | 1 | 0.862993 |
| Guf1 | Q8C3X4 | -0.02 | 0.07 | -0.77 | -0.5 | -0.36 | -0.3 | -1.03 | -1.01 | -0.36 | -0.42 | -0.81 | -0.74 | -0.39 | -0.55 | -0.95 | -0.77 | -0.37 | -0.2775 | -0.36 | 0.227455 | 0.290166 | 0.18812 | 0.999971 | 1 | 0.809025 |
| Guk1 | Q64520-2 | -0.27 | -0.12 | -0.29 | -0.01 | -0.41 | 0.06 | 0.05 | -0.02 | -0.34 | -0.23 | -0.57 | -0.48 | -0.42 | 0.11 | -0.17 | -0.12 | 0.0925 | -0.2325 | 0.0225 | 0.63484 | 0.209834 | 0.907182 | 0.999971 | 1 | 0.991405 |
| Gulp1 | E0CXC5 | -0.42 | 0.2 | -0.61 | -0.58 | 0.45 | -0.22 | -0.78 | -1.1 | 0.49 | -0.37 | -1.04 | -0.76 | -0.17 | -0.63 | -0.24 | 0.4 | -0.06 | -0.0675 | 0.1925 | 0.873239 | 0.855975 | 0.5193 | 0.999971 | 1 | 0.906123 |
| Gusb | P12265 | -1.86 | -0.19 | -0.47 | -0.47 | 0.43 | -1.04 | -0.56 | 0.11 | 0.42 | -1.63 | -0.37 | 0.03 | 0.53 | -0.55 | -0.42 | -0.17 | 0.4825 | 0.36 | 0.595 | 0.317668 | 0.509894 | 0.184887 | 0.999971 | 1 | 0.809025 |
| Gvin1 | L7N451 | 0.56 | 1.35 | 0.44 | 1.13 | 3.11 | 0.66 | 0.3 | 0.42 | 1.46 | 0.78 | 1.53 | 1.68 | 2.04 | 2.74 | 0.2 | 1.29 | 0.2525 | 0.4925 | 0.6975 | 0.693298 | 0.131614 | 0.218721 | 0.999971 | 1 | 0.827064 |
| Gyg1 | Q9R062 | -1.33 | -0.8 | 0.02 | -1.02 | -0.25 | -0.63 | -0.77 | -0.43 | 0.83 | -0.38 | -0.06 | 0.31 | 1.21 | -0.36 | 0.64 | 0.05 | 0.2625 | 0.9575 | 1.1675 | 0.413086 | 0.030095 | 0.02153 | 0.999971 | 1 | 0.642033 |
| Gypa | P14220 | -1.57 | -2.64 | -2.59 | 0.36 | 0.57 | -2.9 | -1.66 | -0.64 | -1.71 | -3.98 | -1.17 | -2.33 | -2.9 | -1.95 | -1.17 | -1.02 | 0.4525 | -0.6875 | -0.15 | 0.62165 | 0.421942 | 0.840127 | 0.999971 | 1 | 0.97975 |
| Gys1 | A0A1B0GT! | -0.63 | -0.66 | -0.67 | -0.93 | -0.89 | -0.63 | -0.48 | -0.84 | -0.68 | -1.02 | -1.36 | -0.92 | -0.97 | -0.79 | -0.93 | -1.1 | 0.0125 | -0.2725 | -0.225 | 0.946873 | 0.209233 | 0.227312 | 0.999971 | 1 | 0.828563 |
| H1-0 | P10922 | -2.63 | -2.87 | -1.91 | -2.19 | -2.1 | -3.01 | -0.89 | -2.75 | -2.26 | -2.97 | -1.61 | -2.49 | -2.76 | -3.34 | -2.98 | -3.23 | 0.2125 | 0.0675 | -0.6775 | 0.661948 | 0.846935 | 0.03197 | 0.999971 | 1 | 0.681356 |
| H1-1 | P43275 | 0.25 | -1.45 | -1.38 | -0.58 | -1.23 | -1.04 | -0.25 | -1.33 | -1.28 | 0.04 | -1.42 | -1.83 | -1.72 | -0.68 | -1.73 | -1.26 | -0.1725 | -0.3325 | -0.5575 | 0.696573 | 0.532804 | 0.229273 | 0.999971 | 1 | 0.828687 |
| H1-2 | P15864 | -2.61 | -4.38 | -1.83 | -1.89 | -3.05 | -4.17 | -0.09 | -1.6 | -3.95 | -3.74 | -1.56 | -2.01 | -5.36 | -4.58 | -2.08 | -2.89 | 0.45 | -0.1375 | -1.05 | 0.638713 | 0.857846 | 0.24303 | 0.999971 | 1 | 0.837316 |
| H13 | A3KGR9 | -0.2 | 0.64 | 0.15 | -0.12 | 0.46 | 0.89 | -0.1 | 0.17 | 0.72 | -0.02 | 0.49 | 0.29 | 0.88 | 1 | 0.37 | 0.45 | 0.2375 | 0.2525 | 0.5575 | 0.428813 | 0.354545 | 0.062156 | 0.999971 | 1 | 0.725348 |
| H1-3 | P43277 | -0.66 | -1.53 | -1.28 | -0.84 | -1.95 | -1.42 | -0.29 | -1.84 | -1.99 | -0.74 | -1.13 | -2.12 | -1.4 | -1.41 | -1.3 | -1.29 | -0.2975 | -0.4175 | -0.2725 | 0.472829 | 0.285485 | 0.268617 | 0.999971 | 1 | 0.842746 |
| H1-4 | P43274 | -0.14 | -1.7 | -1.75 | -1.79 | -2.03 | -1.44 | -0.76 | -2.32 | -2.47 | -0.82 | -1.98 | -3.37 | -2.44 | -1.36 | -2.26 | -2.35 | -0.2925 | -0.815 | -0.7575 | 0.556139 | 0.205388 | 0.116888 | 0.999971 | 1 | 0.78268 |
| H1-5 | P43276 | -0.73 | -1.28 | -1.52 | -0.82 | -1.18 | -1.24 | -1.25 | -1.52 | -1.3 | -0.64 | -1.25 | -2.05 | -1.65 | -1.07 | -1.52 | -1.81 | -0.21 | -0.2225 | -0.425 | 0.386296 | 0.519009 | 0.138433 | 0.999971 | 1 | 0.784948 |
| H1f10 | Q80ZM5 | 3.09 | 0.03 | 0.25 | 1.37 | 0.08 | 0.09 | 1.09 | 1.15 | 0.08 | 1.98 | 0.07 | -0.2 | -1.71 | -0.43 | 0.17 | -0.02 | -0.5825 | -0.7025 | -1.6825 | 0.407151 | 0.379705 | 0.045822 | 0.999971 | 1 | 0.710102 |
| H2ac21 | Q64522 | -2.02 | 3.87 | 3.19 | 0.3 | 6.81 | 3.53 | 3.76 | 2.39 | 9.5 | -2.64 | 5.66 | 6.49 | 15.82 | 1.91 | -0.57 | -2.3 | 2.7875 | 3.4175 | 2.38 | 0.084653 | 0.213215 | 0.539796 | 0.999971 | 1 | 0.910077 |
| H2ax | P27661 | 0.22 | -2.91 | -2.06 | -2.42 | -3.18 | -2.14 | -1.47 | -3.3 | -3.44 | -1.45 | -2.73 | -2.83 | -4.73 | -3.16 | -2.58 | -3.69 | -0.73 | -0.82 | -1.7475 | 0.336773 | 0.282264 | 0.041989 | 0.999971 | 1 | 0.701725 |
| H2az2 | Q3THW5 | -1.1 | -1.93 | -2.14 | -1.36 | -1.89 | -2.77 | -2.59 | -1.91 | -2.28 | -1.46 | -1.94 | -2.54 | -1.77 | -1.96 | -1.85 | -1.61 | -0.6575 | -0.4225 | -0.165 | 0.073917 | 0.227735 | 0.548903 | 0.999971 | 1 | 0.911727 |
| H2bc15 | P10853 | -1.39 | -1.84 | -2.27 | -2.49 | -1.69 | -1.99 | -1.04 | -2.95 | -1.93 | -1.85 | -2.61 | -2.44 | -1.48 | -2.01 | -2.78 | -2.99 | 0.08 | -0.21 | -0.3175 | 0.855328 | 0.505647 | 0.443049 | 0.999971 | 1 | 0.883386 |
| H2bu2 | Q9D2U9 | -3.08 | -4.94 | -2.12 | -2.54 | -4.28 | -4.83 | -1.74 | -2.31 | -4.49 | -4.48 | -2.63 | -2.7 | -4.83 | -4.3 | -2.33 | -3.26 | -0.12 | -0.405 | -0.51 | 0.890391 | 0.586945 | 0.503675 | 0.999971 | 1 | 0.905195 |
| H2-D1 | P01899 | 0.64 | 0.18 | 0.23 | 0.9 | 1.74 | -0.15 | -0.25 | -0.26 | 0.14 | 0.19 | 0.44 | 1.4 | -0.28 | 1.07 | 0.65 | 0.73 | -0.2175 | 0.055 | 0.055 | 0.654634 | 0.870388 | 0.86953 | 0.999971 | 1 | 0.983849 |
| H2-K1 | A0A494BA! | 1.71 | 0.21 | 0.59 | 1.1 | 0.57 | 0.29 | -0.35 | -0.17 | -0.52 | -1.21 | 0.01 | 1.06 | 0.56 | 1.66 | 0.58 | 0.74 | -0.8175 | -1.0675 | -0.0175 | 0.05355 | 0.073299 | 0.96505 | 0.999971 | 1 | 0.998721 |
| H2-Q10 | P01898 | 2.39 | 3.23 | 1.18 | 3.29 | 3.01 | 0.72 | 2.06 | -0.08 | 0.64 | 0.32 | 1.41 | 1.64 | 0.21 | 1.97 | 1.5 | 1.13 | -1.095 | -1.52 | -1.32 | 0.175099 | 0.019393 | 0.042297 | 0.999971 | 1 | 0.701725 |
| H3c14 | A0A1W2P7 | 0.1 | -1.65 | -1.8 | -1.06 | -1.5 | -1.57 | -2.08 | -1.68 | -1.63 | -0.16 | -1.85 | -1.87 | -1.51 | -1.16 | -1.45 | -1.06 | -0.605 | -0.275 | -0.1925 | 0.180121 | 0.619287 | 0.650571 | 0.999971 | 1 | 0.933842 |
| H3f3a | F8WI35 | 3.56 | -1.3 | -0.31 | 0.24 | -0.79 | -0.05 | 0.93 | -1.52 | -1.31 | 1.84 | -2.54 | -2.48 | -2.31 | -2.06 | -1.49 | -2.71 | -0.905 | -1.67 | -2.69 | 0.398336 | 0.22629 | 0.020532 | 0.999971 | 1 | 0.635388 |
| H4f16 | P62806 | -2.91 | -4.37 | -3.46 | -2.51 | -4.15 | -4.83 | -3.18 | -3.5 | -4.15 | -4.22 | -3.33 | -3.21 | -4.82 | -4.37 | -2.92 | -2.84 | -0.6025 | -0.415 | -0.425 | 0.251297 | 0.37297 | 0.47821 | 0.999971 | 1 | 0.894438 |
| H6pd | Q8CFX1 | 0.42 | 0.24 | 0.35 | 0.32 | 0.6 | 0.68 | 0.38 | 0.25 | 0.4 | 0.82 | 0.54 | 0.63 | 1 | 0.27 | 0.01 | 0.47 | 0.145 | 0.265 | 0.105 | 0.434259 | 0.154513 | 0.669194 | 0.999971 | 1 | 0.938711 |
| Habp4 | E9QKB2 | 0.55 | 1.65 | 1.14 | 0.8 | 1.83 | 1.45 | 1.6 | 1.15 | 2.52 | 0.86 | 1.73 | 1.86 | 1.97 | 0.88 | 1.03 | 1.24 | 0.4725 | 0.7075 | 0.245 | 0.129335 | 0.104179 | 0.474372 | 0.999971 | 1 | 0.894044 |
| Hacd2 | Q9D3B1 | 2.43 | -1.4 | 1.17 | 0.82 | -1.49 | -0.57 | 3.27 | -0.24 | -1.03 | 0.99 | 0.22 | -1.06 | -1.95 | -1.94 | -0.06 | -0.47 | -0.5125 | -0.975 | -1.86 | 0.66226 | 0.27023 | 0.051254 | 0.999971 | 1 | 0.723736 |
| Hacd3 | Q8K2C9 | -0.8 | -0.76 | -1.68 | -1.52 | -0.85 | -0.76 | -1.3 | -1.38 | -0.82 | -0.69 | -1.24 | -1.42 | -0.63 | -0.56 | -1.43 | -1.2 | 0.1175 | 0.1475 | 0.235 | 0.692144 | 0.627168 | 0.473449 | 0.999971 | 1 | 0.894044 |
| Hace1 | Q3U0D9 | -0.49 | -0.13 | -0.31 | -0.34 | 0.07 | -0.42 | -0.17 | -0.65 | -0.22 | -0.63 | -0.36 | -0.14 | -0.22 | -0.37 | -0.48 | -0.28 | 0.025 | -0.02 | -0.02 | 0.908504 | 0.916638 | 0.910194 | 0.999971 | 1 | 0.991822 |
| Hacl1 | D3YZ54 | 0.02 | -2.66 | -1.14 | -0.97 | -1.41 | -1.86 | -1.37 | -0.32 | -1.4 | -1.37 | -2.01 | -1.23 | -2.11 | -1.6 | -1.42 | -1.14 | -0.0525 | -0.315 | -0.38 | 0.928673 | 0.5613 | 0.49066 | 0.999971 | 1 | 0.898612 |
| Hadh | Q61425 | -1.73 | -4.18 | -1.54 | -1.71 | -3.78 | -3.63 | -1.28 | -1.35 | -3.72 | -2.76 | -1.65 | -1.97 | -4.89 | -3.81 | -1.78 | -1.95 | -0.22 | -0.235 | -0.8175 | 0.793431 | 0.741718 | 0.365288 | 0.999971 | 1 | 0.869526 |
| Hadha | Q8BMS1 | -0.94 | -1.44 | -0.78 | -0.8 | -1.45 | -1.48 | -1.05 | -0.59 | -1.48 | -1.02 | -1.08 | -0.84 | -1.47 | -1.16 | -0.89 | -0.7 | -0.1525 | -0.115 | -0.065 | 0.584148 | 0.629314 | 0.798592 | 0.999971 | 1 | 0.970479 |
| Hadhb | Q99JY0 | -1.27 | -1.67 | -1.29 | -1.09 | -1.57 | -1.81 | -1.46 | -0.88 | -1.65 | -1.13 | -1.32 | -1.11 | -1.77 | -1.47 | -1.37 | -1.22 | -0.1 | 0.0275 | -0.1275 | 0.698088 | 0.899101 | 0.558937 | 0.999971 | 1 | 0.915051 |
| Hagh | G5E8T9 | -0.3 | -0.01 | -0.45 | -1.14 | -0.05 | -0.28 | 0.76 | -0.83 | 0.73 | -0.68 | -0.47 | 0.4 | -0.01 | -0.68 | -0.78 | -1.38 | 0.375 | 0.47 | -0.2375 | 0.350967 | 0.257314 | 0.515891 | 0.999971 | 1 | 0.905815 |
| Haghl | Q9DB32 | -0.82 | 0.9 | 1.06 | 0.96 | 1.2 | 1.49 | 1.05 | 1.59 | 1.41 | -0.59 | 1.57 | 1.43 | 1.57 | 1.12 | 1.4 | 1.58 | 0.8075 | 0.43 | 0.8925 | 0.093557 | 0.498616 | 0.067129 | 0.999971 | 1 | 0.733937 |
| Hap1 | O35668 | 1.09 | -1.34 | 0.93 | 0.3 | 0.34 | -0.17 | 0.69 | 1.36 | 0.94 | 0.5 | -0.79 | 1.52 | 1.19 | 1.32 | 2.02 | -0.33 | 0.31 | 0.2975 | 0.805 | 0.5997 | 0.661317 | 0.252383 | 0.999971 | 1 | 0.839012 |
| Hapln1 | Q9QUP5 | -0.7 | -0.22 | -0.32 | -0.11 | -0.36 | -0.04 | -0.29 | 0.07 | -0.17 | -0.7 | -0.48 | -0.62 | -0.64 | -0.34 | -0.22 | -0.13 | 0.1825 | -0.155 | 0.005 | 0.400617 | 0.479983 | 0.981582 | 0.999971 | 1 | 0.99925 |
| Hapln2 | Q9ESM3 | -0.04 | 0.32 | -2.03 | -0.87 | -0.18 | -0.5 | -2.92 | -0.89 | -0.51 | -0.72 | -1.2 | -1.12 | -1.39 | -0.1 | -0.45 | -0.81 | -0.4675 | -0.2325 | -0.0325 | 0.525905 | 0.649487 | 0.952273 | 0.999971 | 1 | 0.998056 |
| Hapln3 | Q80WM5 | 0.73 | 1.05 | 0.83 | -0.22 | 0.76 | 1.76 | 0.48 | -0.21 | 1.52 | 0.9 | -0.16 | 0.15 | 1.63 | 1.71 | 0.22 | -0.43 | 0.1 | 0.005 | 0.185 | 0.829253 | 0.991019 | 0.737912 | 0.999971 | 1 | 0.959801 |
| Hapln4 | Q80WM4 | 0.04 | 0.56 | 0.23 | 0.44 | 0.1 | 0.41 | -0.12 | 0.13 | 0.19 | -0.13 | 0.1 | -0.17 | 0.13 | 0.15 | -0.42 | -0.16 | -0.1875 | -0.32 | -0.3925 | 0.381513 | 0.134505 | 0.103655 | 0.999971 | 1 | 0.78268 |
| Hars1 | Q61035 | 0.9 | 0.88 | 0.78 | 0.47 | 1.11 | 1.45 | 1.14 | 1 | 1.27 | 1.07 | 1.25 | 1.25 | 1.55 | 1.23 | 1.24 | 1.09 | 0.4175 | 0.4525 | 0.52 | 0.060413 | 0.031866 | 0.026538 | 0.999971 | 1 | 0.678474 |
| Hars2 | Q99KK9 | -0.34 | 0.3 | 0.15 | -0.01 | -0.07 | 0.24 | -0.44 | -0.13 | -0.04 | -0.19 | 0.25 | -0.16 | 0.4 | 0.23 | 0.1 | 0.25 | -0.125 | -0.06 | 0.22 | 0.593489 | 0.779536 | 0.300386 | 0.999971 | 1 | 0.847402 |
| Hax1 | O35387 | 0.85 | -0.92 | -0.59 | -0.64 | -0.39 | -0.7 | 0.37 | -0.79 | -0.98 | -0.03 | -1.26 | -1.05 | -0.91 | -0.92 | -1.19 | -2.03 | -0.0525 | -0.505 | -0.9375 | 0.906759 | 0.28445 | 0.062779 | 0.999971 | 1 | 0.725348 |
| Hba-a1 | Q91VB8 | -1.98 | -1.68 | -3.89 | -0.34 | -0.01 | -3.1 | 0.63 | -3.13 | -2.53 | -3.61 | -2.49 | -3.45 | -2.5 | -2.88 | -3.7 | -3.38 | 0.57 | -1.0475 | -1.1425 | 0.60737 | 0.169868 | 0.131942 | 0.999971 | 1 | 0.784948 |
| Hbb-b1 | P02088 | -0.92 | -1.32 | -3.47 | 0.43 | 0.6 | -3.08 | -0.27 | -2.18 | -1.91 | -2.37 | -1.87 | -3.06 | -1.66 | -2.38 | -3.75 | -2.57 | 0.0875 | -0.9825 | -1.27 | 0.933092 | 0.226178 | 0.1498 | 0.999971 | 1 | 0.793248 |
| Hbb-bs | A8DUK4 | -0.64 | -1.67 | -4.98 | 0.26 | 0.22 | -3.23 | -0.54 | -3.14 | -2.5 | -2.56 | -3.54 | -3.86 | -2.03 | -2.5 | -4.12 | -2.67 | 0.085 | -1.3575 | -1.0725 | 0.947225 | 0.227276 | 0.341809 | 0.999971 | 1 | 0.861042 |
| Hbs1l | A3TGM7 | 0.03 | 0.53 | 0.58 | 0.43 | 0.48 | 0.5 | 0.37 | 0.78 | 0.66 | 0.47 | 1.1 | 0.89 | 0.67 | 0.63 | 0.75 | 0.81 | 0.14 | 0.3875 | 0.3225 | 0.502276 | 0.110795 | 0.125249 | 0.999971 | 1 | 0.78268 |
| Hccs | P53702 | -0.34 | 0.57 | 0.56 | -1.28 | 1.23 | 0.58 | 1.61 | -0.28 | 1.6 | -0.33 | 1.78 | 2.35 | 1.96 | 0.39 | 0.14 | -0.75 | 0.9075 | 1.4725 | 0.5575 | 0.128964 | 0.051602 | 0.401969 | 0.999971 | 1 | 0.871904 |
| Hcfc1 | Q61191 | 1.3 | 1.25 | 1.34 | 1.43 | 1.21 | 1.11 | 1.03 | 1.17 | 1.04 | 0.96 | 1.01 | 1.14 | 0.64 | 1.14 | 1.11 | 1.38 | -0.2 | -0.2925 | -0.2625 | 0.242252 | 0.09469 | 0.233473 | 0.999971 | 1 | 0.831526 |
| Hcls1 | P49710 | 0.13 | 1.64 | 1.09 | 0.52 | 1.36 | 1.64 | 0.08 | 0.45 | 1.6 | 0.8 | 0.09 | 0.5 | 1.76 | 1.53 | 0.8 | 0.42 | 0.0375 | -0.0975 | 0.2825 | 0.935397 | 0.822754 | 0.516761 | 0.999971 | 1 | 0.905815 |
| Hcn1 | O88704 | -0.29 | -0.69 | -0.4 | -0.4 | -0.62 | -0.53 | -0.69 | -0.71 | -0.98 | -0.46 | -0.72 | -1.23 | -0.76 | -0.61 | -0.56 | -0.52 | -0.1925 | -0.4025 | -0.1675 | 0.294376 | 0.100785 | 0.365556 | 0.999971 | 1 | 0.869526 |
| Hcn2 | O88703 | -0.66 | -1.2 | -1.13 | -1.48 | -1.61 | -1.77 | -0.52 | -1.69 | -1.81 | -1.14 | -1.83 | -1.81 | -1.56 | -1.69 | -1.81 | -1.71 | -0.28 | -0.53 | -0.575 | 0.415786 | 0.068767 | 0.027308 | 0.999971 | 1 | 0.678474 |
| Hcn3 | O88705 | 0.42 | -1.39 | -1.99 | -1.52 | -1.55 | -1.3 | -1.55 | -1.45 | -1.16 | -0.43 | -1.59 | -1.06 | -0.99 | -1.43 | -1.62 | -1.01 | -0.3425 | 0.06 | -0.1425 | 0.494354 | 0.911095 | 0.780849 | 0.999971 | 1 | 0.965859 |
| Hcn4 | B2RY58 | -0.26 | -0.53 | -0.8 | -0.47 | -0.95 | -0.42 | -1.05 | -0.64 | -0.72 | -0.2 | -0.83 | -0.4 | -0.63 | -0.6 | -0.74 | -0.37 | -0.25 | -0.0225 | -0.07 | 0.281414 | 0.91922 | 0.723782 | 0.999971 | 1 | 0.955292 |
| Hdac1 | O09106 | 2.26 | 2.08 | 2.12 | 1.17 | 1.27 | 2.28 | 1.92 | 1.19 | 2.68 | 1.73 | 1.43 | 1.26 | 2.83 | 0.9 | 1.49 | 0.33 | -0.2425 | -0.1325 | -0.52 | 0.499295 | 0.733652 | 0.352454 | 0.999971 | 1 | 0.867768 |
| Hdac10 | Q6P3E7 | 0.75 | 0.15 | 0.01 | 0.14 | 0.12 | -0.31 | -0.11 | 0.12 | -0.44 | 0.33 | 0.01 | 0.59 | -0.08 | 0.67 | -0.2 | 0.36 | -0.3075 | -0.14 | -0.075 | 0.207791 | 0.629373 | 0.787294 | 0.999971 | 1 | 0.96724 |
| Hdac11 | Q91WA3 | -1.38 | -0.76 | -1.21 | -1.69 | -0.71 | -0.83 | -1.12 | -1.08 | -0.86 | -1.26 | -1.15 | -1.08 | -0.33 | -0.9 | -1.21 | -1.58 | 0.325 | 0.1725 | 0.255 | 0.210475 | 0.481185 | 0.445756 | 0.999971 | 1 | 0.884152 |
| Hdac2 | P70288 | 1.61 | 1.72 | 1.61 | 1.42 | 1.66 | 1.73 | 1.44 | 1.58 | 1.84 | 1.64 | 1.56 | 1.72 | 1.43 | 1.47 | 1.55 | 1.34 | 0.0125 | 0.1 | -0.1425 | 0.942911 | 0.561508 | 0.414328 | 0.999971 | 1 | 0.876329 |
| Hdac3 | O88895 | 1.14 | 1.12 | 2.31 | 0.03 | 1.5 | 0.72 | 1.73 | 1.44 | 2.02 | 0.37 | 1.94 | 1.58 | 1.69 | 0.03 | 1.69 | 0.58 | 0.1975 | 0.3275 | -0.1525 | 0.681361 | 0.559516 | 0.790134 | 0.999971 | 1 | 0.968174 |
| Hdac4 | Q6NZM9 | -0.34 | -0.26 | 0.08 | -0.17 | -0.2 | -0.33 | 0.5 | -0.15 | -0.21 | -0.28 | 0.17 | 0.33 | 0.16 | -0.18 | -0.26 | 0.12 | 0.1275 | 0.175 | 0.1325 | 0.597585 | 0.426846 | 0.511843 | 0.999971 | 1 | 0.905815 |
| Hdac5 | Q9Z2V6 | 0.34 | 0.43 | 0.66 | 0.86 | 0.09 | 0.52 | 0.87 | 0.49 | 0.39 | 0.64 | 1.1 | 0.45 | -0.04 | 0.26 | 0.58 | 0.94 | -0.08 | 0.0725 | -0.1375 | 0.732964 | 0.755844 | 0.604922 | 0.999971 | 1 | 0.924454 |
| Hdac6 | Q9Z2V5 | 0.75 | 0.4 | 0.26 | 0.38 | 0.46 | 0.41 | 0.59 | 0.34 | 0.54 | 0.86 | 0.34 | -0.05 | 0.74 | 0.31 | 0.36 | 0.27 | 0.0025 | -0.025 | -0.0275 | 0.989366 | 0.918818 | 0.893725 | 0.999971 | 1 | 0.98828 |
| Hdac8 | Q8VH37 | -0.63 | -0.55 | -0.16 | -0.64 | -0.26 | 1.05 | -1.1 | -0.43 | -0.29 | -0.54 | -2.21 | -0.51 | -0.53 | 0.91 | -0.26 | -0.28 | 0.31 | -0.3925 | 0.455 | 0.484709 | 0.377898 | 0.203088 | 0.999971 | 1 | 0.81617 |
| Hdac9 | A0A1B0GS! | 0.23 | -0.16 | 0.06 | -0.16 | -0.31 | -0.47 | 0.71 | -1.03 | -1.1 | 1.23 | -0.56 | -1.03 | -1.05 | -0.21 | -1.2 | -1.93 | -0.2675 | -0.3575 | -1.09 | 0.470432 | 0.493008 | 0.013344 | 0.999971 | 1 | 0.613095 |
| Hddc2 | Q3SXD3 | -0.94 | -0.83 | -0.76 | -0.67 | 0.19 | -1.15 | 0.1 | -0.23 | -0.3 | -0.93 | -0.18 | 0.42 | 0.36 | -1.02 | -0.55 | -0.47 | 0.5275 | 0.5525 | 0.38 | 0.121119 | 0.086303 | 0.228402 | 0.999971 | 1 | 0.828687 |
| Hddc3 | Q9D114 | 1.06 | 0.82 | 0.45 | 0.44 | 0.88 | -0.01 | 0.29 | 0.66 | 0.57 | 0.51 | -0.45 | -0.34 | 0.31 | 0.72 | -0.02 | 0.02 | -0.2375 | -0.62 | -0.435 | 0.386952 | 0.07517 | 0.114897 | 0.999971 | 1 | 0.78268 |
| Hdgf | P51859 | 2.05 | 3.08 | 2.66 | 2.72 | 2.5 | 2.96 | 2.19 | 2.61 | 2.44 | 2.18 | 2.25 | 2.36 | 2.33 | 2.87 | 2.51 | 2.71 | -0.0625 | -0.32 | -0.0225 | 0.824232 | 0.218788 | 0.932496 | 0.999971 | 1 | 0.995069 |
| Hdgfl2 | Q3UMU9 | 2.55 | 1.49 | 0.89 | 1.15 | 1.46 | 1.55 | 1.65 | 0.72 | 1.93 | 1.6 | 0.68 | 0.68 | 0.56 | 0.64 | 0.35 | 0.88 | -0.175 | -0.2975 | -0.9125 | 0.66534 | 0.520095 | 0.033907 | 0.999971 | 1 | 0.689803 |
| Hdgfl3 | Q9JMG7 | 2.01 | 2.1 | 2.36 | 2.22 | 2.31 | 1.97 | 1.78 | 2.28 | 1.96 | 2.01 | 2.29 | 2.37 | 1.48 | 1.78 | 2.18 | 2.23 | -0.0875 | -0.015 | -0.255 | 0.668957 | 0.936754 | 0.287064 | 0.999971 | 1 | 0.846151 |
| Hdhd2 | Q3UGR5 | 0.03 | 0.02 | 0.31 | 0.21 | -0.31 | 0.3 | 0.22 | -0.07 | 0.17 | 0.08 | 0.03 | 0.05 | -0.51 | 0.22 | 0 | 0.02 | -0.1075 | -0.06 | -0.21 | 0.60869 | 0.720459 | 0.347662 | 0.999971 | 1 | 0.864164 |
| Hdhd3 | Q9CYW4 | 0.11 | -0.05 | -0.56 | -1.03 | 0.08 | -0.64 | -0.36 | -1 | 0.01 | 0.12 | -0.4 | -0.39 | 0.11 | -0.53 | -1.34 | -1.3 | -0.0975 | 0.2175 | -0.3825 | 0.775328 | 0.474629 | 0.36545 | 0.999971 | 1 | 0.869526 |
| Hdhd5 | Q91WM2 | -0.63 | -0.67 | -1.2 | -1.12 | -1.13 | -0.83 | -1.24 | -1.45 | -0.98 | -0.85 | -1.36 | -1.79 | -1.35 | -0.97 | -1.86 | -1.43 | -0.2575 | -0.34 | -0.4975 | 0.286274 | 0.236112 | 0.082362 | 0.999971 | 1 | 0.755254 |
| Hdlbp | Q8VDJ3 | 0.83 | 1.37 | 1.08 | 1.06 | 1.44 | 1.26 | 0.84 | 1.02 | 1.51 | 1.15 | 1.03 | 1.42 | 1.58 | 1.23 | 1.18 | 1.38 | 0.055 | 0.1925 | 0.2575 | 0.800673 | 0.363877 | 0.221978 | 0.999971 | 1 | 0.827753 |
| Heatr1 | G3X9B1 | -0.72 | -1.01 | -0.72 | -0.61 | -1.25 | -1.19 | -1.12 | -0.66 | -1.15 | -0.85 | -0.9 | -0.9 | -1.4 | -0.73 | -0.79 | -0.6 | -0.29 | -0.185 | -0.115 | 0.190722 | 0.32037 | 0.625651 | 0.999971 | 1 | 0.930479 |
| Heatr3 | Q8BQM4 | 0.5 | 0.56 | 0.64 | 0.48 | 0.52 | 1.55 | 1.7 | 0.36 | -0.09 | 1.57 | 0.12 | 0.05 | 0.81 | 1.77 | -0.19 | 0.2275 | -0.345 | -0.13 | 0.576748 | 0.436249 | 0.771884 | 0.999971 | 1 | 0.963498 |
| Heatr5a | Q5PRF0 | -0.38 | -1.51 | -1.6 | -0.88 | -1.88 | -0.77 | -0.77 | -0.44 | -1.17 | -0.65 | 0.21 | 0.24 | -0.62 | -0.17 | -1.28 | -1.04 | 0.1275 | 0.75 | 0.315 | 0.753836 | 0.105914 | 0.400807 | 0.999971 | 1 | 0.871816 |
| Heatr5b | Q8C547 | 0.14 | -0.04 | 0.18 | 0.14 | 0.2 | -0.23 | 0.25 | 0.04 | 0.05 | 0 | 0.21 | 0.1 | -0.32 | -0.23 | 0.07 | 0.02 | -0.04 | -0.015 | -0.22 | 0.831714 | 0.926859 | 0.249548 | 0.999971 | 1 | 0.839012 |
| Heatr6 | Q6P1G0 | 1.85 | -0.56 | 0.13 | 0.48 | 0.18 | -0.31 | 0 | 0.4 | -0.06 | 0.57 | -0.27 | 0.29 | -0.81 | -0.1 | -0.13 | 0.54 | -0.4075 | -0.3425 | -0.6 | 0.41586 | 0.501556 | 0.277629 | 0.999971 | 1 | 0.843812 |
| Hebp1 | A0A140T8J | -2.34 | -4.39 | -2.8 | -2.91 | 0.35 | 0.52 | 0.54 | 0.65 | -0.12 | 0.5 | 0.27 | 0.34 | 0.06 | 0.1 | 0.65 | 0.53 | 3.625 | 3.3575 | 3.445 | 1.82E-05 | 4.44E-05 | 3.65E-05 | 0.041372 | 0.095451 | 0.083255 |
| Hebp2 | Q9WU63 | -0.33 | 0.49 | 0.09 | -0.03 | 0.85 | -0.22 | 1.45 | -0.65 | 1.46 | -0.37 | -0.19 | 0.98 | 0.69 | 0.65 | -0.2 | -0.29 | 0.3025 | 0.415 | 0.1575 | 0.529928 | 0.368161 | 0.622963 | 0.999971 | 1 | 0.929759 |
| Heca | Q3V1N5 | 0.51 | 1.09 | 0.99 | 0.92 | 0.98 | 0.72 | -0.01 | 1.05 | 0.93 | 0.72 | 0.11 | 1.08 | 0.31 | 1 | 1.15 | 1.4 | -0.1925 | -0.1675 | 0.0875 | 0.507495 | 0.53536 | 0.756504 | 0.999971 | 1 | 0.961116 |
| Hectd1 | F8WIE5 | 0.63 | 0.84 | 0.88 | 0.38 | 0.83 | 0.72 | 0.94 | 0.66 | 0.94 | 0.6 | 0.86 | 0.98 | 0.82 | 0.59 | 0.73 | 0.44 | 0.105 | 0.1625 | -0.0375 | 0.591341 | 0.421281 | 0.851803 | 0.999971 | 1 | 0.98187 |
| Hectd2 | B2RPZ5 | 0.39 | 0.16 | 2.65 | 1.2 | -0.3 | 0.46 | 1.82 | 1.64 | 0.47 | 0.67 | 2.19 | 1.09 | 1.53 | 0.41 | 1.67 | 1.43 | -0.195 | 0.005 | 0.16 | 0.775457 | 0.993599 | 0.782657 | 0.999971 | 1 | 0.965859 |
| Hectd3 | Q3U487 | 0.45 | 0.46 | 0.3 | 0.23 | 0.18 | 0.38 | -0.07 | 0.49 | 0.3 | 0.41 | 0.23 | 0.4 | 0.74 | 0.53 | 0.39 | 0.54 | -0.115 | -0.025 | 0.19 | 0.5625 | 0.879528 | 0.298874 | 0.999971 | 1 | 0.847402 |
| Hectd4 | E9Q2E4 | -0.49 | -0.31 | -0.26 | -0.41 | -0.29 | -0.24 | -0.03 | -0.18 | -0.16 | -0.09 | -0.06 | -0.04 | -0.09 | -0.22 | -0.05 | 0.1 | 0.1675 | 0.245 | 0.315 | 0.340213 | 0.154386 | 0.096895 | 0.999971 | 1 | 0.78268 |
| Hecw1 | Q8K4P8 | 0.56 | 0.05 | 0.27 | 0.3 | -0.27 | 0.33 | -0.02 | 0.25 | -0.5 | 0.31 | 0.27 | 0 | 0.56 | 0.06 | -0.03 | 0.61 | -0.2225 | -0.275 | 0.005 | 0.320968 | 0.275232 | 0.982885 | 0.999971 | 1 | 0.99925 |
| Hecw2 | Q6I6G8 | -0.12 | -0.48 | -0.19 | -0.42 | -0.68 | -0.46 | -0.24 | -0.28 | -0.59 | -0.28 | -0.12 | 0.04 | -0.44 | -0.75 | -0.33 | -0.22 | -0.1125 | 0.065 | -0.1325 | 0.568351 | 0.755276 | 0.516649 | 0.999971 | 1 | 0.905815 |
| Helz | Q6DFV5-2 | 1.19 | 0.64 | 0.22 | 1.01 | 0.6 | 0.78 | 0.36 | 0.09 | 0.77 | 0.96 | 0.49 | 0.18 | 0.68 | 0.91 | 0.57 | 0.75 | -0.3075 | -0.165 | -0.0375 | 0.285679 | 0.567695 | 0.882367 | 0.999971 | 1 | 0.986576 |
| Hepacam | Q640R3 | -2.01 | -2.55 | -0.05 | -0.22 | -1.96 | -2.28 | -0.65 | 0.04 | -2.61 | -2.13 | -0.83 | -0.45 | -2.33 | -2.03 | -0.26 | -0.13 | -0.005 | -0.2975 | 0.02 | 0.994683 | 0.688308 | 0.979256 | 0.999971 | 1 | 0.99925 |
| Heph | Q9Z0Z4 | -0.12 | -1.55 | -0.72 | -0.52 | -1.37 | -0.71 | -0.28 | -0.06 | -0.76 | -0.42 | -1.12 | -0.63 | -0.97 | -0.68 | -0.9 | -1.06 | 0.1225 | -0.005 | -0.175 | 0.759168 | 0.988021 | 0.583119 | 0.999971 | 1 | 0.920364 |
| Herc1 | E9PZP8 | 0.32 | 0.19 | 0.06 | -0.26 | 0.14 | 0.25 | -0.28 | 0.02 | -0.04 | 0.55 | 0.07 | 0.18 | 0.28 | 0.42 | -0.02 | 0.22 | -0.045 | 0.1125 | 0.1475 | 0.834763 | 0.610789 | 0.484448 | 0.999971 | 1 | 0.896402 |
| Herc2 | Q4U2R1 | 0.03 | -0.2 | 0.17 | 0.29 | -0.45 | -0.33 | -0.03 | 0.06 | -0.43 | -0.07 | 0.2 | 0.13 | -0.06 | -0.15 | 0.14 | 0.38 | -0.26 | -0.115 | 0.005 | 0.237146 | 0.600512 | 0.980966 | 0.999971 | 1 | 0.99925 |
| Herc3 | A6H6S0 | -0.6 | -0.07 | 0.78 | 0.06 | -0.33 | -0.37 | 0.38 | 0.42 | -0.32 | -0.45 | 1.1 | 0.8 | -0.46 | -0.06 | 0.61 | 0.65 | -0.0175 | 0.24 | 0.1425 | 0.960193 | 0.600919 | 0.708087 | 0.999971 | 1 | 0.950182 |
| Herc4 | Q6PAV2-2 | 0.21 | 0.22 | -0.37 | 0.01 | 0.26 | 0.37 | -0.18 | -0.02 | 0.24 | 0.71 | -0.04 | -0.04 | 0.06 | 0.52 | -0.13 | 0.37 | 0.09 | 0.2 | 0.1875 | 0.692656 | 0.432555 | 0.437216 | 0.999971 | 1 | 0.882419 |
| Herpud1 | Q8C4N0 | 1.58 | 0.93 | 0.98 | 1.61 | -0.07 | 0.07 | 1.18 | 1.44 | 0.49 | -0.1 | 0.49 | 0.75 | 0.67 | -0.6075 | -0.4 | -1.0675 | 0.154509 | 0.172081 | 0.01299 | 0.999971 | 1 | 0.613095 | | | |
| Herpud2 | Q9JJC9 | -0.62 | -0.92 | -0.9 | -1.66 | 0.53 | 0.72 | -0.6 | 0.05 | 0.2 | 1.25 | 0.49 | -0.02 | 1.04 | 0.58 | -0.12 | 0.13 | 1.2 | 1.505 | 1.4325 | 0.008457 | 0.002078 | 0.002191 | 0.999971 | 0.788955 | 0.508206 |
| Hexa | P29416 | -0.2 | 0.34 | -0.33 | 0.05 | 0.5 | 0.35 | -0.05 | 0.61 | 0.33 | 0.34 | 0.45 | 0.32 | 0.82 | 1.02 | 0.27 | 0.41 | 0.3875 | 0.395 | 0.665 | 0.132407 | 0.07815 | 0.026784 | 0.999971 | 1 | 0.678474 |
| Hexb | P20060 | 0.17 | 0.2 | 0.19 | 0.31 | 0.25 | 0.58 | 0.53 | 1.05 | 0.12 | 0.58 | 0.88 | 0.51 | -0.12 | 0.61 | 0.84 | 1.21 | 0.385 | 0.305 | 0.4175 | 0.10141 | 0.167143 | 0.179348 | 0.999971 | 1 | 0.808532 |
| Hexdc | F6UV46 | -0.37 | -0.46 | 0.99 | -0.06 | 3.13 | -1.61 | 1.5 | 0.56 | 5.47 | 0.25 | 1.14 | 1.25 | 5.21 | -1.55 | 0.68 | -0.12 | 0.87 | 2.0025 | 1.03 | 0.363366 | 0.094254 | 0.447046 | 0.999971 | 1 | 0.884581 |
| Hexim1 | Q8R409 | 1.75 | 1.79 | 2.12 | 2.31 | 1.72 | 1.64 | 1.53 | 2.32 | 1.34 | 1.74 | 2.08 | 2.44 | 2.13 | 2.13 | 2.19 | 2.67 | -0.19 | -0.0925 | 0.2875 | 0.454919 | 0.744968 | 0.228097 | 0.999971 | 1 | 0.828687 |
| Hgh1 | Q8C3I8 | -0.26 | 0.33 | 0.54 | -0.02 | 0.45 | 0.34 | 0.41 | 0.23 | 0.51 | -0.07 | 0.58 | 0.39 | -1.06 | 0.85 | 0.43 | 0.07 | 0.21 | 0.205 | -0.075 | 0.363656 | 0.42931 | 0.859607 | 0.999971 | 1 | 0.982702 |
| Hgs | Q3UMA3 | 0.15 | 0.05 | 0.61 | 0.35 | -0.3 | 0.1 | 0.99 | 0.37 | -0.28 | 0.04 | 0.56 | 0.32 | -0.08 | -0.16 | 0.2 | 0.32 | 1.08E-16 | -0.13 | -0.22 | 1 | 0.600795 | 0.322417 | 1 | 1 | 0.85548 |
| Hhatl | Q9D1G3 | 0.42 | 0.49 | 1.62 | 1.82 | 0.78 | 0.55 | 0.16 | -0.09 | 0.3 | 0.38 | -0.04 | 1.6 | 0.12 | 0.68 | 1.1 | 0.75 | -0.7375 | -0.5275 | -0.425 | 0.091182 | 0.289501 | 0.30633 | 0.999971 | 1 | 0.847402 |
| Hibadh | Q99L13 | -1.16 | -1.21 | -2.19 | -2.32 | -0.81 | -1.52 | -2.08 | -2.33 | -1.29 | -1.51 | -2.3 | -2.5 | -1.22 | -1.25 | -2.36 | -2.65 | 0.035 | -0.18 | -0.15 | 0.93545 | 0.66153 | 0.742051 | 0.999971 | 1 | 0.961064 |
| Hibch | Q8QZS1 | 0.4 | -0.75 | -0.79 | -0.7 | -0.92 | -0.66 | -0.86 | -0.38 | -0.84 | -0.36 | -0.75 | -0.94 | -0.78 | -0.49 | -0.74 | -0.73 | -0.245 | -0.2625 | -0.225 | 0.445437 | 0.416753 | 0.464946 | 0.999971 | 1 | 0.89116 |
| Hid1 | Q8R1F6 | 0.76 | 0.95 | 0.99 | 0.79 | 0.87 | 0.83 | 0.94 | 0.79 | 0.92 | 1.27 | 0.48 | 0.36 | 0.82 | 1.08 | 0.67 | 0.66 | -0.015 | -0.115 | -0.065 | 0.928336 | 0.640439 | 0.727943 | 0.999971 | 1 | 0.956041 |
| Hif1an | Q8BLR9 | 0.65 | 0.97 | 0.47 | 0.7 | 1.31 | 1.4 | -0.06 | 1.17 | 1.14 | 0.96 | 0.76 | 0.86 | 1.13 | 1.06 | 0.99 | 0.93 | 0.2575 | 0.2325 | 0.33 | 0.469551 | 0.245201 | 0.102865 | 0.999971 | 1 | 0.78268 |
| Higd1a | A0A1L1SR( | 0.13 | 0.05 | 0.13 | 0.57 | 0.15 | 0.19 | 0.37 | 0.17 | -0.93 | 0.26 | 0.04 | 0.23 | -1.36 | 0.22 | 0.08 | 0.84 | 0 | -0.32 | -0.275 | 1 | 0.316528 | 0.546699 | 1 | 1 | 0.911398 |
| Hikeshi | Q9DD02 | -0.05 | 1.1 | 0.68 | 0.79 | 0.58 | 1.14 | -0.02 | 0.63 | 0.71 | 0.17 | 1.38 | 0.89 | 1.06 | 1.16 | 0.81 | 1.25 | -0.0475 | 0.1575 | 0.44 | 0.887941 | 0.649166 | 0.140306 | 0.999971 | 1 | 0.784948 |
| Hint1 | P70349 | -0.55 | 0.39 | 0.19 | 0.23 | 0.16 | 0.07 | 0.09 | 0.2 | 0.31 | -0.19 | -0.19 | -0.22 | 0.2 | 0.23 | -0.02 | -0.2 | 0.065 | -0.1375 | -0.1075 | 0.789316 | 0.606584 | 0.961305 | 0.999971 | 1 | 0.998721 |
| Hint2 | Q9D0S9 | -1.41 | -0.54 | -0.66 | -0.72 | -0.71 | -1.17 | -0.93 | -0.89 | -1.05 | -1.14 | -0.98 | -0.81 | -0.99 | -0.64 | -0.85 | -0.98 | -0.0925 | -0.1625 | -0.0325 | 0.709043 | 0.502179 | 0.894027 | 0.999971 | 1 | 0.988375 |
| Hint3 | Q9CPS6 | -1.07 | -0.54 | -0.27 | -0.2 | -0.29 | 0.73 | -0.16 | 0.46 | -0.56 | -0.93 | -0.79 | -0.19 | -0.47 | 0.81 | 0.41 | -0.32 | 0.705 | -0.0975 | 0.6275 | 0.050039 | 0.720967 | 0.104751 | 0.999971 | 1 | 0.78268 |
| Hip1 | Q8VD75 | -0.38 | -0.3 | -0.25 | -0.01 | -0.28 | -0.3 | -0.22 | -0.32 | -0.09 | -0.03 | -0.13 | -0.02 | 0.11 | -0.1 | -0.23 | 0.09 | -0.045 | 0.1675 | 0.2025 | 0.794539 | 0.337136 | 0.293938 | 0.999971 | 1 | 0.847402 |
| Hip1r | Q9JKY5 | -0.39 | -0.2 | 0.03 | -0.28 | -0.12 | -0.32 | 0.1 | -0.24 | 0.07 | -0.34 | 0.22 | 0 | 0.32 | -0.42 | -0.31 | -0.14 | 0.065 | 0.1975 | 0.0725 | 0.736338 | 0.339532 | 0.749661 | 0.999971 | 1 | 0.961064 |
| Hirip3 | Q8BLH7 | 0.73 | 1.32 | 2.6 | 2.37 | 1.67 | 1.27 | 1.94 | 2.35 | 0.77 | 0.96 | 1.26 | 1.94 | 0.14 | 1.32 | 1.8 | 1.51 | -0.5625 | -0.5225 | -0.5625 | 0.909805 | 0.291026 | 0.30315 | 0.999971 | 1 | 0.847402 |
| Hist1h2ap | C0HKE9 | -0.12 | 0.83 | 0.43 | -2.32 | 2.76 | -0.99 | 1.5 | -2.24 | 0.48 | -0.33 | 2.42 | 5.99 | 8.65 | -1.46 | 2.56 | 2.23 | 0.5525 | 2.435 | 3.29 | 0.644456 | 0.111145 | 0.119899 | 0.999971 | 1 | 0.78268 |
| Hist2h2aa2 | Q6GSS7 | -0.99 | -2.37 | -1.05 | -1.2 | -1.71 | -2.51 | -1.12 | -1.4 | -1.92 | -1.79 | -1.13 | -1 | -2.16 | -2.03 | -1.36 | -1.35 | -0.2825 | -0.0575 | -0.3225 | 0.50666 | 0.881325 | 0.403668 | 0.999971 | 1 | 0.873348 |
| Hivep2 | Q3UHF7 | 0.41 | 0.05 | 0.42 | 0.6 | 0.03 | 0.45 | 0.43 | 0.41 | 0.05 | 0.61 | 0.44 | 0.56 | -0.27 | 0.34 | 0.46 | 0.43 | -0.04 | 0.045 | -0.13 | 0.846215 | 0.83419 | 0.591817 | 0.999971 | 1 | 0.922655 |
| Hk1 | G3UVV4 | -0.05 | -0.34 | -0.4 | -0.29 | -0.5 | -0.13 | 0.18 | -0.43 | -0.67 | -0.4 | -0.41 | -0.82 | -0.48 | -0.45 | -0.45 | -0.42 | 0.05 | -0.305 | -0.18 | 0.819058 | 0.135379 | 0.309739 | 0.999971 | 1 | 0.847402 |
| Hk2 | O08528 | 0.37 | 0.12 | -0.32 | 0.11 | -0.07 | -0.23 | -0.78 | -0.11 | -0.24 | 0.48 | -0.36 | -0.18 | -0.7 | 0.06 | -0.24 | -0.03 | -0.3675 | -0.145 | -0.2975 | 0.162406 | 0.578936 | 0.254377 | 0.999971 | 1 | 0.839012 |
| Hkdc1 | Q91W97 | 0.02 | -0.45 | -0.68 | -0.86 | -0.82 | -0.33 | -0.68 | -0.02 | -0.9 | -0.8 | -0.29 | -0.11 | -0.63 | -0.58 | -0.62 | -0.17 | 0.03 | -0.0325 | -0.0075 | 0.914033 | 0.908676 | 0.975977 | 0.999971 | 1 | 0.99925 |
| Hlcs | A0A2I3BP( | 0.79 | -0.32 | -0.67 | -0.46 | 0 | 0.25 | -0.39 | -0.19 | 0.19 | 0.55 | 0.12 | 0.08 | 0.45 | 0.23 | -0.48 | -0.31 | 0.0825 | 0.4 | 0.175 | 0.812733 | 0.258795 | 0.718667 | 0.999971 | 1 | 0.953825 |
| Hltf | Q6PCN7 | 2.19 | 2.36 | 2.11 | 1.39 | 2.4 | 2.45 | 1.76 | 1.7 | 2.25 | 2.53 | 1.76 | 1.79 | 2.42 | 2.91 | 2.74 | 3.03 | 0.065 | 0.07 | 0.7625 | 0.829443 | 0.811659 | 0.019213 | 0.999971 | 1 | 0.635388 |
| Hmbs | P22907 | 0.26 | 1.01 | 0.08 | 0.43 | 1.39 | 0.66 | 0.21 | -0.05 | 0.81 | 0.41 | -0.17 | 0.38 | 0.92 | 0.86 | 0.09 | 0.08 | 0.1075 | -0.0875 | 0.0425 | 0.768286 | 0.766557 | 0.891928 | 0.999971 | 1 | 0.988225 |
| Hmcn1 | D3YXG0-2 | 0.55 | -0.21 | 0.11 | 0.52 | 0.44 | 0.8 | 0.18 | 0.3 | -0.19 | 0.85 | 0.6 | 0.09 | 0.11 | -0.01 | -0.55 | 0.1875 | 0.1225 | -0.3325 | 0.464656 | 0.683631 | 0.226022 | 0.999971 | 1 | 0.828225 | |
| Hmg20a | Q9DC33 | 0.32 | -0.2 | -0.57 | -0.46 | -0.19 | -0.6 | -0.45 | -0.62 | -0.22 | 1.37 | -0.73 | -0.62 | -0.43 | -0.48 | -0.78 | -0.57 | -0.2375 | 0.1775 | -0.3375 | 0.35437 | 0.718028 | 0.190852 | 0.999971 | 1 | 0.810547 |
| Hmga1 | A0A338P6! | -1.31 | -4.31 | -5.47 | -2.66 | -1.42 | -4.83 | -3.69 | -2.62 | -3.78 | -4.18 | -4.01 | -4.65 | -4.27 | -5.79 | -4.46 | -4.25 | 0.2975 | -0.7175 | -1.255 | 0.775626 | 0.404467 | 0.180136 | 0.999971 | 1 | 0.808532 |
| Hmga1_1 | Q3TE85 | 2.71 | 0.26 | 1.46 | 2.94 | 0.65 | 0.34 | 2.1 | 1.86 | 0.26 | 1.4 | 0.83 | 0.44 | -0.68 | 0.09 | 0.91 | 1.5 | -0.605 | -1.11 | -1.3875 | 0.388365 | 0.09823 | 0.076652 | 0.999971 | 1 | 0.74875 |
| Hmga2 | Q6NSP9 | 1.41 | 0.06 | 0.2 | 0.74 | 1.09 | 0.42 | -0.65 | 0.84 | 0.12 | 1.59 | -0.83 | -0.03 | -1 | 0.79 | 0.46 | 0.03 | -0.1775 | -0.39 | -0.5325 | 0.700592 | 0.481383 | 0.269604 | 0.999971 | 1 | 0.843592 |
| Hmgb1 | A0A0J9YUI | 1.17 | 2.73 | 1.06 | 1.67 | 3.1 | 1.29 | 1.48 | 1.4 | 2.19 | 0.81 | 1.9 | 1.99 | 3.07 | 2.78 | 2.09 | 1.5 | 0.16 | 0.065 | 0.7025 | 0.762823 | 0.888061 | 0.171846 | 0.999971 | 1 | 0.804075 |
| Hmgb2 | P30681 | 0.16 | 1.07 | 0.44 | 1.4 | 1.65 | 0.07 | 0.8 | 1.67 | 1.43 | -0.32 | 1.07 | 1.29 | 0.81 | 0.92 | 2.17 | 0.91 | 0.28 | 0.1 | 0.435 | 0.536867 | 0.829084 | 0.305332 | 0.999971 | 1 | 0.847402 |
| Hmgb3 | A2AP78 | 1.67 | 2.27 | 1.58 | 2.57 | 2.19 | 1.23 | 1.3 | 2.38 | 2.05 | 1.53 | 1.92 | 1.89 | 2.45 | 2.46 | 2.67 | 1.79 | -0.2475 | -0.175 | 0.32 | 0.508702 | 0.533572 | 0.318661 | 0.999971 | 1 | 0.853893 |
| Hmgcl | P38060 | -0.27 | -1.02 | -1.15 | -1.13 | -1.04 | -0.9 | -0.98 | -1.07 | -0.97 | -0.7 | -1.42 | -1.15 | -0.93 | -0.75 | -1.09 | -1.15 | -0.105 | -0.1675 | -0.0875 | 0.668085 | 0.546171 | 0.731804 | 0.999971 | 1 | 0.956087 |
| Hmgcll1 | D3YVU9 | 0.72 | -0.53 | 0.53 | 1.41 | 0.05 | 0.21 | 0.25 | 1.65 | -0.42 | 0.38 | -0.08 | 1.6 | -0.17 | 0.3 | 1.54 | 0.99 | 0.0075 | -0.1625 | -0.1325 | 0.988183 | 0.768567 | 0.795325 | 0.999971 | 1 | 0.970361 |
| Hmgcr | F2Z470 | -0.25 | -0.2 | 0.06 | -0.12 | -0.77 | -0.9 | -0.62 | 0.64 | -0.59 | -0.51 | -0.25 | -1.18 | 0.44 | 0.54 | 0.21 | -0.29 | -0.285 | -0.505 | 0.3525 | 0.430112 | 0.059765 | 0.157555 | 0.999971 | 1 | 0.797252 |
| Hmgcs1 | Q8JZK9 | -0.06 | 0.39 | 0.6 | -0.12 | 0.31 | 0.57 | 0.49 | 1.5 | 0.46 | 0.39 | 0.34 | 0.15 | 0.75 | 0.86 | 1.03 | 0.36 | 0.515 | 0.1325 | 0.5475 | 0.134834 | 0.559411 | 0.05484 | 0.999971 | 1 | 0.725348 |
| Hmgcs2 | P54869 | -1.17 | -1.38 | -1.02 | -0.78 | 0.02 | -0.5 | -0.58 | 0.58 | -0.04 | -0.07 | -0.54 | -0.06 | -0.94 | -0.67 | -0.03 | -0.28 | 0.9675 | 0.91 | 0.6075 | 0.010617 | 0.002392 | 0.042664 | 0.999971 | 0.821787 | 0.701725 |
| Hmgn1 | A0A3B2W3 | 3.67 | 2.23 | 2.82 | 2.92 | 1.01 | 2.73 | 5.39 | 1.08 | 1.69 | 1.18 | 1.13 | 0.74 | 0.21 | 0.35 | 1.57 | -0.76 | -0.3575 | -1.725 | -2.5675 | 0.70868 | 0.000878 | 0.000866 | 0.999971 | 0.499846 | 0.365812 |
| Hmgn2 | A3KGL9 | 5.21 | 1.77 | 2.29 | 3.51 | 1.15 | 2.25 | 6.83 | 0.53 | 1.89 | 3.07 | -1.92 | -0.51 | -0.19 | -1.43 | 0.35 | -0.78 | -0.505 | -2.5625 | -3.7075 | 0.725387 | 0.061993 | 0.001024 | 0.999971 | 1 | 0.387721 |
| Hmgn3 | A0A087WS | 3.12 | -1.53 | 0.81 | 1.08 | 2.61 | 0.38 | 1.98 | 1.1 | 1.79 | 1.72 | 2.69 | -0.77 | 1.81 | 0.89 | 1.06 | 0.1 | 0.6475 | 0.4875 | 0.095 | 0.503835 | 0.65455 | 0.916702 | 0.999971 | 1 | 0.992633 |
| Hmgn5 | Q9JL35 | 3.22 | 3.04 | 3.16 | 3.7 | 3.18 | 3.32 | 3.14 | 2.66 | 2.94 | 3.19 | 2.76 | 3.06 | 1.5 | 2.85 | 3.2 | 2.65 | -0.205 | -0.2925 | -0.73 | 0.399029 | 0.196873 | 0.083343 | 0.999971 | 1 | 0.755254 |

| | | | | | | | | | | | | | | | | | | | | | | | | | | |
|---|---|---|---|---|---|---|---|---|---|---|---|---|---|---|---|---|---|---|---|---|---|---|---|---|---|---|
| Hmox1 | P14901 | 0.33 | -0.33 | 0.42 | 0.04 | 0.25 | -0.09 | -0.04 | 0.1 | -0.32 | -0.22 | 0.58 | 0.83 | -0.47 | 0.02 | -0.17 | 0.95 | -0.06 | 0.1025 | -0.0325 | 0.790799 | 0.757716 | 0.925034 | 0.999971 | 1 | 0.99401 |
| Hmox2 | O70252 | -0.3 | -0.93 | -0.32 | -0.42 | -0.76 | -1.22 | -0.02 | -0.42 | -0.86 | -1.31 | -0.13 | -0.14 | -0.92 | -1.42 | -0.3 | -0.24 | -0.1125 | -0.1175 | -0.2275 | 0.710741 | 0.718811 | 0.482858 | 0.999971 | 1 | 0.896345 |
| Hnmt | Q91VF2 | 0.71 | 0.78 | 0.56 | -0.15 | 0.52 | 0.75 | 1.73 | 0.22 | 0.93 | -0.07 | 0.26 | -0.03 | 0.64 | -0.09 | -0.28 | -0.58 | 0.33 | -0.2025 | -0.5525 | 0.392262 | 0.52804 | 0.126003 | 0.999971 | 1 | 0.783071 |
| Hnrnpa0 | Q9CX86 | 0.34 | 0.97 | 1.27 | 1.1 | 0.82 | 0.52 | 1.27 | 1.21 | 0.98 | 0.45 | 1.14 | 0.92 | 0.28 | 0.4 | 1.22 | 1.13 | 0.035 | -0.0475 | -0.1625 | 0.901493 | 0.859995 | 0.613734 | 0.999971 | 1 | 0.927364 |
| Hnrnpa1 | Q5EBP8 | -0.58 | -1.3 | 0.17 | 0.49 | -0.93 | -1.09 | 0.12 | 0.73 | -0.6 | -0.69 | 0.58 | 0.64 | -1.08 | -1.06 | 0.85 | 1.11 | 0.0125 | 0.2875 | 0.26 | 0.981608 | 0.571416 | 0.690959 | 0.999971 | 1 | 0.946786 |
| Hnrnpa2b1 | O88569 | -0.38 | -0.11 | -0.87 | -1.04 | -0.19 | -0.44 | 0.06 | -1.13 | -0.24 | -0.32 | -0.85 | -1.01 | -0.53 | -0.36 | -1.06 | -1.19 | 0.175 | -0.005 | -0.185 | 0.601919 | 0.986529 | 0.545505 | 0.999971 | 1 | 0.911398 |
| Hnrnpa3 | A2AL12 | 1.93 | 0.4 | 0.86 | 0.59 | -0.08 | 0.36 | 1.23 | 0.61 | -0.19 | 0.92 | 0.58 | 0.32 | 0.03 | 0.39 | 0.81 | 0.39 | -0.415 | -0.5375 | -0.54 | 0.332088 | 0.19981 | 0.166848 | 0.999971 | 1 | 0.800215 |
| Hnrnpa3_1 | Q8BG05-2 | -0.27 | -2.97 | -3.01 | -2.18 | -1.43 | -4.02 | -1.49 | -3.61 | -2.67 | -1.58 | -2.32 | -3.84 | -2.61 | -2.53 | -2.55 | -3.46 | -0.53 | -0.495 | -0.68 | 0.533512 | 0.499632 | 0.289751 | 0.999971 | 1 | 0.847402 |
| Hnrnpab | Q99020 | 0.76 | -0.21 | 0.95 | 1.01 | -0.71 | 0.02 | 1.59 | 0.38 | -0.84 | -0.11 | 0.72 | 0.92 | -1.48 | -0.61 | 1.12 | 0.9 | -0.3075 | -0.455 | -0.645 | 0.554933 | 0.341016 | 0.31547 | 0.999971 | 1 | 0.852494 |
| Hnrnpc | Q9Z204-2 | -0.46 | 0.07 | -0.03 | -0.04 | 0.12 | -0.32 | -0.73 | -0.45 | -0.73 | -0.05 | -0.88 | 0.16 | 0.23 | -0.13 | 0.14 | -0.01 | -0.23 | -0.26 | 0.1725 | 0.357521 | 0.381937 | 0.400801 | 0.999971 | 1 | 0.871816 |
| Hnrnpc_1 | Q9Z204 | -1.89 | -2.32 | 0.07 | -0.02 | -1.97 | -2.05 | 1.14 | -0.12 | -1.76 | -2.02 | -0.12 | 0 | -2.07 | -1.92 | -0.02 | -0.08 | 0.29 | 0.065 | 0.0175 | 0.744243 | 0.930152 | 0.981516 | 0.999971 | 1 | 0.99925 |
| Hnrnpd | Q60668-3 | -1.82 | 0.15 | -0.95 | -1.54 | 0.03 | -0.05 | 0.09 | -1.36 | 0.14 | -1.76 | -0.77 | -0.74 | -0.14 | -0.1 | -0.74 | -1.28 | 0.7175 | 0.2575 | 0.475 | 0.187505 | 0.635751 | 0.339292 | 0.999971 | 1 | 0.860218 |
| Hnrnpdl | Q9Z130 | 0.25 | 0.71 | -1.13 | -0.59 | 0.37 | 0.18 | -1.19 | -0.75 | 0.39 | 0.63 | -0.95 | -0.8 | 0.3 | 0.71 | -1.19 | -0.56 | -0.1575 | 0.0075 | 0.005 | 0.759761 | 0.988804 | 0.992699 | 0.999971 | 1 | 0.99925 |
| Hnrnpdl_1 | D3YTQ3 | -0.57 | 0.15 | -0.03 | -0.02 | 0 | 0.15 | 0.3 | -0.39 | 0.05 | -0.65 | 0.16 | 0.02 | 0.28 | -0.07 | -0.07 | 0.05 | 0.1325 | 0.0125 | 0.165 | 0.592421 | 0.961955 | 0.462873 | 0.999971 | 1 | 0.890653 |
| Hnrnpf | Q9Z2X1 | 0.66 | 1.78 | 1.01 | 0.25 | 1.62 | 1.04 | 1 | -0.07 | 1.64 | 1.18 | 1.15 | 1.53 | 1.42 | 1.18 | 0.4 | -0.28 | -0.0275 | 0.45 | -0.245 | 0.951187 | 0.212095 | 0.606617 | 0.999971 | 1 | 0.925135 |
| Hnrnph1 | Q8C2Q7 | 0.39 | 1.07 | 0.94 | 1.16 | 0.96 | 0.93 | 0.62 | 1.3 | 0.97 | 0.86 | 0.9 | 1.13 | 1.2 | 1.61 | 1.2 | 1.39 | 0.0625 | 0.075 | 0.46 | 0.802562 | 0.736186 | 0.076732 | 0.999971 | 1 | 0.74875 |
| Hnrnph2 | P70333 | 0.8 | 1.29 | 1.03 | 0.84 | 1.46 | 1.08 | 1.53 | 0.8 | 1.35 | 1 | 0.99 | 1.4 | 1.46 | 1.29 | 1.05 | 0.74 | 0.2275 | 0.195 | 0.145 | 0.351827 | 0.357001 | 0.534487 | 0.999971 | 1 | 0.910077 |
| Hnrnph3 | D3Z3N4 | -0.24 | 0.26 | -0.11 | 0.31 | 0.34 | -0.08 | 0.09 | 0.2 | 0.3 | 0.09 | -0.38 | 0.15 | 0.27 | 0.3 | -0.16 | -0.15 | 0.0825 | -0.015 | 0.01 | 0.697787 | 0.94871 | 0.964817 | 0.999971 | 1 | 0.998721 |
| Hnrnpk | B2M1R6 | 0.85 | 1.18 | 1.1 | 1.41 | 1.21 | 0.78 | 1.16 | 1.32 | 1.03 | 0.95 | 1.25 | 1.42 | 0.89 | 1.35 | 1.18 | 1.33 | -0.0175 | 0.0275 | 0.0525 | 0.934501 | 0.893917 | 0.802074 | 0.999971 | 1 | 0.971467 |
| Hnrnpl | G5E924 | 0.07 | -0.15 | 0.28 | 0.56 | -0.13 | -0.24 | 0.85 | 0.35 | -0.19 | 0.23 | 0.33 | 0.3 | -0.4 | 0.13 | 0.44 | 0.41 | 0.0175 | -0.0225 | -0.045 | 0.953473 | 0.921773 | 0.866505 | 0.999971 | 1 | 0.983462 |
| Hnrnpll | Q921F4 | 1.52 | 1.75 | 1.71 | 1.95 | 1.82 | 1.65 | 1.73 | 1.86 | 2 | 1.61 | 1.8 | 1.43 | 1.62 | 1.68 | 1.83 | 1.92 | 0.0325 | -0.0225 | 0.03 | 0.856352 | 0.911551 | 0.871521 | 0.999971 | 1 | 0.984109 |
| Hnrnpm | Q9D0E1-2 | 0.93 | 0.8 | 0.62 | 0.74 | 0.85 | 0.72 | 0.6 | 0.62 | 0.76 | 0.83 | 0.48 | 0.47 | 0.25 | 0.76 | 0.52 | 0.57 | -0.075 | -0.1375 | -0.2475 | 0.667915 | 0.458922 | 0.217218 | 0.999971 | 1 | 0.826893 |
| Hnrnpr | Q8VHM5 | -0.03 | 0.6 | 0.31 | 0.21 | 0.65 | 0.38 | 0.73 | 0.22 | 0.93 | -0.27 | 0.33 | 0.06 | 0.85 | 0.24 | 0.29 | -0.31 | 0.2225 | -0.01 | -0.005 | 0.326336 | 0.972914 | 0.985988 | 0.999971 | 1 | 0.99925 |
| Hnrnpu | Q8VEK3 | -0.36 | -0.27 | 0.25 | 0.3 | -0.2 | -0.08 | 0.33 | 0.34 | -0.05 | -0.4 | -0.02 | -0.05 | -0.33 | 0.03 | 0.58 | 0.27 | 0.1175 | -0.11 | 0.1575 | 0.639395 | 0.633652 | 0.571345 | 0.999971 | 1 | 0.91816 |
| Hnrnpul1 | Q8VDM6 | 1.62 | 1.8 | 2.15 | 1.58 | 1.61 | 1.63 | 1.77 | 1.77 | 1.57 | 1.58 | 1.82 | 1.89 | 1.36 | 1.52 | 1.61 | 1.52 | -0.0925 | -0.0725 | -0.285 | 0.64148 | 0.723392 | 0.178056 | 0.999971 | 1 | 0.808532 |
| Hnrnpul2 | Q00PI9 | 0.04 | 0.12 | 0.23 | 0.17 | 0.09 | -0.2 | -0.04 | -0.05 | 0.09 | 0.09 | 0.1 | -0.14 | -0.27 | -0.23 | -0.07 | -0.14 | -0.19 | -0.105 | -0.3175 | 0.277976 | 0.529167 | 0.082946 | 0.999971 | 1 | 0.755254 |
| Homer1 | Q9Z2Y3 | -0.71 | -1.02 | -1.3 | -1.22 | -1.12 | -1.14 | -1.39 | -1.4 | -1.3 | -0.88 | -1.02 | -1.15 | -1.28 | -1.26 | -1.41 | -1.16 | -0.2 | -0.025 | -0.215 | 0.344489 | 0.904024 | 0.299472 | 0.999971 | 1 | 0.847402 |
| Homer1_1 | Q9Z2Y3-5 | -0.23 | 1.52 | 2.02 | 1.19 | 1.18 | 0.95 | 1.21 | 2.34 | 0.8 | 0.06 | 2.26 | 1.94 | 2.06 | 0.5 | 2.37 | 2.44 | 0.295 | 0.14 | 0.7175 | 0.580971 | 0.827369 | 0.255759 | 0.999971 | 1 | 0.839012 |
| Homer2 | Q9QWW1- | -0.56 | -1.08 | -0.31 | -0.4 | -1.18 | -0.66 | -0.2 | -0.89 | -0.8 | -0.46 | -0.53 | -0.86 | -0.72 | -1.22 | -0.88 | -0.77 | -0.145 | -0.075 | -0.31 | 0.611413 | 0.747347 | 0.217527 | 0.999971 | 1 | 0.826893 |
| Homer3 | Q99JP6-2 | 0.82 | 0.48 | 0.37 | -0.18 | -0.22 | 0.76 | -0.21 | 0.29 | 0.17 | 0.1 | -0.28 | -0.35 | 0.53 | 0.08 | -0.24 | 0.29 | -0.2175 | -0.4625 | -0.2075 | 0.497456 | 0.108809 | 0.46632 | 0.999971 | 1 | 0.891431 |
| Hook1 | Q8BIL5 | 0.89 | 0.66 | 0.73 | 0.54 | 1.24 | 0.84 | 1.1 | 1.17 | 0.48 | 0.8 | 0.98 | 1.08 | 0.54 | 0.69 | 0.69 | 0.93 | 0.3825 | 0.13 | 0.0075 | 0.065245 | 0.525736 | 0.967493 | 0.999971 | 1 | 0.998787 |
| Hook2 | A0A1B0GS( | 1.11 | 1.57 | 2.25 | 1.48 | 1.46 | 1.85 | 2.48 | 1.47 | 2.15 | 1.5 | 1.64 | 0.61 | -1.57 | 0.56 | 1.24 | 1.63 | 0.2125 | -0.1275 | -1.1375 | 0.530874 | 0.741164 | 0.122552 | 0.999971 | 1 | 0.78268 |
| Hook3 | Q8BUK6 | 0.71 | 0.72 | 0.61 | 0.45 | 0.72 | 0.76 | 0.85 | 0.49 | 0.66 | 0.47 | 0.72 | 0.55 | 0.37 | 0.48 | 0.3 | 0.53 | 0.0825 | -0.0225 | -0.2025 | 0.646994 | 0.894416 | 0.260509 | 0.999971 | 1 | 0.841256 |
| Hopx | Q8R1H0 | 0.9 | 1.04 | 1.64 | 1.07 | 0.56 | 0 | 1.76 | 2.73 | 0.84 | -0.35 | 2.36 | 1.72 | 0.68 | -0.05 | 2.14 | 1.26 | 0.1 | -0.02 | -0.155 | 0.862865 | 0.971584 | 0.737063 | 0.999971 | 1 | 0.959519 |
| Hp1bp3 | Q3TEA8-2 | 0.67 | -0.41 | -0.3 | -0.23 | -1.4 | -1.47 | -0.92 | -1.73 | -1.34 | -0.84 | -1.37 | -1.02 | -1.89 | -1.35 | -1.8 | -1.96 | -1.3125 | -1.075 | -1.6825 | 0.00207 | 0.004891 | 0.000326 | 0.999971 | 1 | 0.247672 |
| Hpca | P84075 | 0.19 | 1.18 | 0.35 | 0.38 | 0.81 | 1.44 | -0.2 | 1.05 | 0.82 | 0.64 | -0.14 | -0.27 | 1.35 | 1.57 | 0.43 | 0.74 | 0.25 | -0.2625 | 0.4975 | 0.533313 | 0.457373 | 0.170898 | 0.999971 | 1 | 0.803758 |
| Hpcal1 | P62748 | -1.91 | -0.92 | -0.17 | -0.33 | -0.74 | -0.87 | 0.6 | -0.05 | -1.08 | -2.15 | -0.1 | 0.09 | -1.2 | -1.02 | -0.43 | -0.68 | 0.5675 | 0.0225 | 1.18E-16 | 0.259795 | 0.969827 | 1 | 0.999971 | 1 | 1 |
| Hpcal4 | Q8BGZ1 | -1.64 | -1.74 | -1.87 | -1.88 | -1.02 | -1.25 | -1.15 | -1.56 | -1.25 | -1.62 | -1.32 | -1.32 | -1.15 | -1.23 | -1.28 | -1.52 | 0.5375 | 0.405 | 0.4875 | 0.019982 | 0.044645 | 0.022358 | 0.999971 | 1 | 0.652043 |
| Hpgds | Q9JHF7 | 0.83 | 2.63 | 0.46 | -0.28 | 2.48 | 2.25 | 0.37 | -0.44 | 2.55 | 1.17 | 0.65 | -0.07 | 2.97 | 2.15 | 0.14 | 0.14 | 0.255 | 0.165 | 0.44 | 0.763624 | 0.826542 | 0.607786 | 0.999971 | 1 | 0.925695 |
| Hprt1 | P00493 | -0.8 | 2.11 | 1.55 | 0.69 | 2.04 | 1.57 | 1.35 | 1.51 | 2.12 | -0.5 | 1.52 | 1.64 | 2.4 | 1.41 | 1.53 | 0.87 | 0.73 | 0.3075 | 0.665 | 0.239593 | 0.693846 | 0.317403 | 0.999971 | 1 | 0.853841 |
| Hpx | Q91X72 | 3.72 | 2.32 | 0.7 | 2.46 | 3.53 | 0.62 | 2.67 | 1.56 | 0.86 | 0.2 | 1.87 | 2.69 | 2.7 | 4.09 | 2.48 | 2.33 | -0.205 | -0.895 | 0.6 | 0.797327 | 0.253001 | 0.382494 | 0.999971 | 1 | 0.871816 |
| Hras | Q61411 | -1.97 | -0.74 | -1.16 | -1.13 | -0.82 | -0.75 | -1.22 | -0.94 | -0.93 | -1.99 | -1.23 | -0.77 | -0.49 | -0.91 | -0.87 | -0.94 | 0.3175 | 0.02 | 0.4475 | 0.290528 | 0.956248 | 0.150356 | 0.999971 | 1 | 0.793248 |
| Hrg | Q9ESB3 | 2.83 | 3.48 | 1.24 | 3.61 | 4.37 | 2.1 | 4.39 | 0.47 | 2.48 | 0.62 | 1.49 | 2.24 | 2.14 | 0.52 | -0.39 | 0.97 | 0.0425 | -1.0825 | -1.98 | 0.965365 | 0.112981 | 0.017155 | 0.999971 | 1 | 0.617356 |
| Hrh1 | P70174 | -1.1 | -1.65 | -1.2 | -1.13 | -1.22 | -1.39 | -1.05 | -1.04 | -0.84 | -0.97 | -0.97 | -1.33 | -0.17 | -0.96 | -1.68 | -0.45 | 0.095 | 0.2425 | 0.455 | 0.646792 | 0.270242 | 0.215897 | 0.999971 | 1 | 0.826893 |
| Hrh3 | E9Q7T5 | 1.1 | -1.88 | -0.79 | -0.18 | -0.94 | -1 | -0.03 | -0.23 | -1.57 | -0.72 | 0.07 | -0.51 | 0.51 | -0.56 | -0.21 | -0.22 | -0.1125 | -0.245 | 0.3175 | 0.853486 | 0.706038 | 0.601914 | 0.999971 | 1 | 0.924453 |
| Hs1bp3 | Q3TC93 | 0.55 | 0.21 | 0.12 | 0.53 | 0.16 | 0.21 | 0.92 | -0.16 | -0.01 | 0.6 | -0.36 | -0.34 | -0.25 | 0.17 | -0.39 | -0.24 | -0.07 | -0.38 | -0.53 | 0.796801 | 0.180742 | 0.032368 | 0.999971 | 1 | 0.683918 |
| Hs2st1 | Q8R3H7 | -1.29 | 1.25 | 2.38 | 0.95 | 1.75 | 1.89 | 2.43 | 1.31 | 1.4 | 1.1 | 1.72 | 1.48 | 1.89 | 1.03 | 1.93 | 0.85 | 1.0225 | 0.6025 | 0.6025 | 0.182095 | 0.405348 | 0.424273 | 0.999971 | 1 | 0.879824 |
| Hs3st4 | D3YVV6 | -1.05 | -1.69 | -2.09 | -1.23 | -2.82 | -2.02 | -1.32 | -2.93 | -2.77 | -1.02 | -2.04 | -1.73 | -3.67 | -2.26 | -1.92 | -0.78 | -0.7575 | -0.375 | -0.6425 | 0.099126 | 0.374769 | 0.289708 | 0.999971 | 1 | 0.847402 |
| Hs6st1 | Q9QYK5 | 0.48 | 0.2 | 1.57 | 2.2 | 0.7 | 0.49 | 2.21 | 2.51 | 0.59 | 0.79 | 2.12 | 2.05 | 0.74 | 1.14 | 2.15 | 1.52 | 0.365 | 0.275 | 0.275 | 0.567445 | 0.631702 | 0.595983 | 0.999971 | 1 | 0.92377 |
| Hsbp1 | Q9CQZ1 | -0.08 | -1.66 | -1.84 | -1.12 | -2.09 | -2.18 | 0.04 | -1.53 | -2.09 | -2.93 | -1.05 | -1.43 | -2.99 | -1.94 | -1.65 | -2 | -0.265 | -0.7 | -0.97 | 0.65624 | 0.209105 | 0.060937 | 0.999971 | 1 | 0.725348 |
| Hscb | Q8K3A0 | -1.34 | -0.06 | -0.35 | -0.93 | -0.07 | -0.36 | -0.95 | -0.5 | -0.1 | -1.35 | -0.38 | -0.37 | 0.04 | -0.88 | -1.08 | -1.15 | 0.2 | 0.12 | -0.0975 | 0.557846 | 0.755021 | 0.799833 | 0.999971 | 1 | 0.970626 |
| Hsd11b1 | P50172 | 0.17 | -0.23 | 0.13 | 0.11 | -0.62 | 0.62 | 0.38 | -0.16 | -0.85 | 0.44 | 0.02 | -0.7 | -1.35 | -1.27 | -0.34 | -0.45 | 0.01 | -0.3175 | -0.8975 | 0.973468 | 0.334791 | 0.012845 | 0.999971 | 1 | 0.613095 |
| Hsd17b10 | Q99N15 | -0.71 | -0.83 | -0.8 | -0.89 | -0.81 | -0.96 | -1.12 | -0.9 | -0.9 | -0.87 | -1.15 | -0.9 | -0.85 | -0.75 | -0.84 | -0.85 | -0.14 | -0.1475 | -0.015 | 0.4195 | 0.386607 | 0.926303 | 0.999971 | 1 | 0.99401 |
| Hsd17b11 | Q9EQ06 | -0.41 | -0.5 | -0.86 | -0.78 | -0.23 | -0.24 | -0.9 | 0.13 | -0.22 | -0.2 | -0.98 | -0.93 | -0.41 | 0.03 | -0.59 | -0.75 | 0.3275 | 0.055 | 0.2075 | 0.233184 | 0.833716 | 0.389938 | 0.999971 | 1 | 0.871816 |
| Hsd17b12 | O70503 | -1.12 | -0.73 | 0.36 | 0.16 | -0.46 | -0.53 | -0.08 | 0.09 | -0.67 | -1.02 | -0.27 | -0.02 | -0.58 | -0.4 | 0.29 | 0.08 | 0.0875 | -0.1625 | 0.18 | 0.814119 | 0.685516 | 0.647991 | 0.999971 | 1 | 0.933842 |
| Hsd17b4 | P51660 | -0.16 | -1.4 | -0.83 | -0.78 | -0.91 | -0.52 | -0.3 | -0.43 | -0.11 | 0.04 | -0.65 | -0.78 | -0.21 | -0.4 | -0.57 | -0.35 | 0.2525 | 0.4175 | 0.41 | 0.402737 | 0.219362 | 0.168536 | 0.999971 | 1 | 0.801027 |
| Hsd17b7 | O88736 | -0.3 | -0.78 | -0.62 | -0.59 | -0.73 | -0.75 | -0.66 | -0.19 | -0.66 | -0.73 | -0.4 | -0.32 | -0.24 | -0.51 | -0.56 | -0.39 | -0.01 | 0.045 | 0.1475 | 0.962653 | 0.819251 | 0.447489 | 0.999971 | 1 | 0.885073 |
| Hsd17b8 | P50171 | 0.1 | 0.04 | 0.38 | 0.06 | 0.11 | 0.01 | 0.09 | 0.57 | -0.03 | -0.04 | 0.39 | 0.32 | 0.22 | -0.03 | 0.38 | 0.07 | 0.05 | 0.015 | 0.015 | 0.806909 | 0.938801 | 0.936789 | 0.999971 | 1 | 0.995618 |
| Hsdl1 | Q8BTX9 | -1.07 | -0.82 | -0.5 | -0.47 | -0.59 | -0.8 | -0.41 | -0.07 | -0.66 | -0.99 | -0.29 | 0.1 | -0.46 | -0.56 | 0.03 | -0.39 | 0.2475 | 0.255 | 0.37 | 0.321108 | 0.384999 | 0.135974 | 0.999971 | 1 | 0.784948 |
| Hsdl2 | Q2TPA8 | -0.08 | -0.62 | -1.27 | -0.99 | -0.53 | -0.45 | -1.21 | -0.42 | -0.38 | -0.3 | -1.18 | -0.77 | 0.15 | -0.14 | -0.87 | -1.15 | 0.0875 | 0.0825 | 0.2375 | 0.784923 | 0.801023 | 0.541303 | 0.999971 | 1 | 0.910733 |
| Hsf1 | A0A2R8W( | 1.97 | 2.4 | 1.64 | 5.94 | 2.86 | 2.86 | 4.98 | 2.47 | 1.43 | 1.69 | 2.32 | 0.39 | 1.6 | 2.06 | 0.55 | 3.17 | 0.305 | -1.53 | -1.1425 | 0.766087 | 0.140479 | 0.277599 | 0.999971 | 1 | 0.843812 |
| Hsp90aa1 | P07901 | -0.33 | 0.88 | 0.32 | -0.04 | 0.89 | 0.83 | 0.53 | -0.06 | 1.08 | 0.62 | 0.4 | 0.9 | 1.43 | 0.73 | 0.48 | 0.33 | 0.34 | 0.5425 | 0.535 | 0.326998 | 0.104432 | 0.153607 | 0.999971 | 1 | 0.795578 |
| Hsp90ab1 | P11499 | 0.05 | 0.63 | 0.31 | 0.38 | 0.98 | 0.6 | 0.73 | 0.55 | 1.1 | 0.46 | 0.54 | 0.88 | 1.56 | 0.46 | 0.48 | 0.76 | 0.3725 | 0.4025 | 0.4725 | 0.098362 | 0.104452 | 0.135712 | 0.999971 | 1 | 0.784948 |
| Hsp90b1 | P08113 | 0.25 | -0.15 | -0.39 | 0.16 | -0.07 | 0.06 | -0.26 | -0.03 | -0.05 | 0.47 | -0.24 | 0.14 | 0.21 | 0.52 | 0.23 | 0.56 | -0.0425 | 0.1125 | 0.4125 | 0.840587 | 0.642248 | 0.086949 | 0.999971 | 1 | 0.761242 |
| Hspa12a | Q8K0U4 | -0.46 | -0.02 | -0.2 | -0.52 | -0.11 | -0.14 | -0.35 | -0.33 | -0.07 | -0.66 | -0.42 | -0.24 | -0.12 | -0.3 | -0.46 | -0.33 | 0.0675 | -0.0475 | -0.0025 | 0.729763 | 0.825341 | 0.989892 | 0.999971 | 1 | 0.99925 |
| Hspa12a_1 | A0A494B9: | -1.61 | -1.08 | -1.01 | -0.3 | -0.13 | -2.12 | -1.22 | -0.18 | -1.1 | -1.89 | -1.13 | -0.96 | -1.82 | -0.58 | -0.91 | -0.48 | 0.0875 | -0.27 | 0.0525 | 0.862439 | 0.433457 | 0.893211 | 0.999971 | 1 | 0.98828 |
| Hspa12b | Q9CZJ2 | 0.62 | 0.14 | 0.03 | 0.22 | 0.48 | -0.55 | 0.19 | -0.32 | 0.24 | 0.39 | 0.28 | 0.46 | -0.34 | -0.01 | 0.41 | 0.34 | -0.3025 | 0.09 | -0.1525 | 0.299971 | 0.646826 | 0.540167 | 0.999971 | 1 | 0.910077 |
| Hspa13 | Q8BM72 | 0.05 | -0.31 | -0.91 | -1.37 | 0.07 | -0.2 | -0.23 | -0.97 | -0.06 | -0.05 | -0.49 | 0.09 | 0.78 | 0.09 | -0.73 | -0.77 | 0.3025 | 0.5075 | 0.4775 | 0.427058 | 0.157572 | 0.310949 | 0.999971 | 1 | 0.84774 |
| Hspa14 | Q99M31 | 0.98 | 1.37 | 1.08 | 1.12 | 1.37 | 1.23 | 1.07 | 1.4 | 1.05 | 1.12 | 1.38 | 0.99 | 1.3 | 1.4 | 1.28 | 1.33 | 0.13 | -0.0025 | 0.19 | 0.488501 | 0.989233 | 0.294168 | 0.999971 | 1 | 0.847402 |
| Hspa1a | Q61696 | 0.12 | 0.48 | 0.29 | 0.54 | -0.41 | 0.59 | -0.18 | -0.45 | 0.31 | -0.21 | -0.4 | -0.71 | -0.91 | 0.24 | -0.81 | -0.37 | -0.47 | -0.61 | -0.82 | 0.116317 | 0.03934 | 0.019043 | 0.999971 | 1 | 0.635388 |
| Hspa2 | P17156 | -0.13 | -1.26 | -0.43 | -0.9 | -1.26 | -1.11 | -0.11 | -0.83 | -1.42 | 0.17 | -0.91 | -1.08 | -2.33 | -1.36 | -1.13 | -1.03 | -0.1475 | -0.13 | -0.7825 | 0.675494 | 0.749549 | 0.063486 | 0.999971 | 1 | 0.725348 |
| Hspa4 | Q3U2G2 | 0.03 | -0.02 | 0.34 | -0.03 | -0.1 | 0 | 0.55 | -0.08 | 0.16 | -0.01 | 0.37 | 0.28 | -0.35 | -0.15 | -0.07 | -0.17 | 0.0125 | 0.12 | -0.265 | 0.954797 | 0.523491 | 0.171887 | 0.999971 | 1 | 0.804075 |
| Hspa4l | P48722 | 0.67 | 0.18 | -0.06 | 0.03 | -0.21 | 0.51 | 0.22 | -0.22 | 0.08 | 0.42 | -0.19 | -0.45 | -0.18 | 0.27 | -0.14 | -0.16 | -0.13 | -0.24 | -0.2575 | 0.623467 | 0.379215 | 0.285693 | 0.999971 | 1 | 0.845133 |
| Hspa5 | P20029 | -0.68 | -1.19 | -0.09 | -0.39 | -0.9 | -1.01 | 0.18 | -0.36 | -0.83 | -0.95 | -0.03 | 0.16 | -0.95 | -1.08 | -0.11 | -0.29 | 0.065 | 0.175 | -0.02 | 0.853996 | 0.626327 | 0.952214 | 0.999971 | 1 | 0.998056 |
| Hspa8 | P63017 | -0.32 | -0.45 | -0.5 | -0.62 | -0.17 | -0.48 | -0.26 | -0.67 | -0.15 | -0.39 | -0.3 | -0.08 | 0.32 | -0.36 | -0.5 | -0.37 | 0.0775 | 0.2425 | 0.245 | 0.689201 | 0.186599 | 0.308403 | 0.999971 | 1 | 0.847402 |
| Hspa9 | P38647 | -1.08 | -0.98 | -0.9 | -0.94 | -1.1 | -0.9 | -0.9 | -0.97 | -0.98 | -1.17 | -0.98 | -1.03 | -1.18 | -1.06 | -0.94 | -1.01 | 0.0075 | -0.065 | -0.0725 | 0.963834 | 0.688166 | 0.665913 | 0.999971 | 1 | 0.937173 |
| Hspb1 | P14602 | 2.58 | 1.29 | 1.61 | 2.82 | 3.6 | 2.32 | 4.08 | 0.3 | 1.57 | 3.16 | 1.94 | 3 | -0.66 | 2.1 | 1.66 | 1.34 | 0.5 | 0.3425 | -0.965 | 0.549665 | 0.500644 | 0.162523 | 0.999971 | 1 | 0.800215 |
| Hspb11 | Q9D6H2 | -1.44 | -0.66 | 0.29 | -0.28 | -0.68 | -0.57 | -0.17 | 0.76 | -1.19 | -1.24 | 0.5 | 0.22 | -1.37 | -0.48 | 0.98 | 0.29 | 0.3575 | 0.095 | 0.3775 | 0.442843 | 0.860563 | 0.513888 | 0.999971 | 1 | 0.905815 |
| Hspb8 | Q9JK92 | -1.96 | 0.44 | 0.38 | -0.75 | 0.78 | -0.08 | 0.37 | -0.06 | 0.71 | -1.84 | 0.64 | 0.79 | -0.6 | -0.85 | 0.25 | -0.1 | 0.725 | 0.5475 | 0.1475 | 0.211912 | 0.485668 | 0.795065 | 0.999971 | 1 | 0.970361 |
| Hspbp1 | Q99P31 | 0.96 | 0.96 | 0.78 | 0.95 | 0.82 | 0.76 | 0.82 | 0.94 | 0.8 | 0.85 | 0.92 | 0.29 | 0.23 | 0.66 | 0.75 | 0.59 | -0.0775 | -0.1975 | -0.355 | 0.640119 | 0.342556 | 0.089664 | 0.999971 | 1 | 0.771991 |
| Hspd1 | P63038 | -0.58 | -1.08 | -0.58 | -0.62 | -1.17 | -1.06 | -0.9 | -0.75 | -1.4 | -1.12 | -0.96 | -0.65 | -0.99 | -0.87 | -0.61 | -0.57 | -0.255 | -0.3175 | -0.045 | 0.236587 | 0.195445 | 0.830635 | 0.999971 | 1 | 0.978535 |
| Hspe1 | Q64433 | -3.35 | -4.38 | -1.97 | -2.35 | -4.34 | -4.08 | -1.5 | -2.34 | -4.54 | -3.6 | -2.1 | -2.21 | -5.47 | -4.13 | -2.33 | -2.27 | -0.0525 | -0.1 | -0.5375 | 0.946601 | 0.890136 | 0.53025 | 0.999971 | 1 | 0.910069 |
| Hspg2 | E9PZ16 | 0.8 | -0.24 | -0.11 | 0.61 | 2.45 | 0.03 | 0.55 | -0.51 | 0.47 | 1.92 | 1.21 | 2.26 | -0.92 | -0.1 | 0.32 | -0.01 | 0.365 | 1.2 | -0.4425 | 0.566593 | 0.024013 | 0.240599 | 0.999971 | 1 | 0.834089 |
| Hsph1 | Q61699 | -0.41 | -0.5 | 0.14 | -0.23 | -0.45 | -0.34 | 0.25 | -0.27 | -0.37 | -0.48 | 0.07 | -0.31 | -0.4 | -0.27 | -0.07 | 0.09 | 0.0475 | -0.0225 | 0.0875 | 0.844392 | 0.919834 | 0.694578 | 0.999971 | 1 | 0.947174 |
| Htatip2 | Q3U816 | -1.88 | -0.78 | -0.91 | -0.89 | -1.04 | -0.66 | -1.06 | -0.65 | -0.72 | -0.8 | -0.56 | -0.29 | -0.62 | -0.38 | -0.41 | -0.1 | 0.2625 | 0.5225 | 0.7375 | 0.379883 | 0.099957 | 0.029871 | 0.999971 | 1 | 0.678474 |
| Htatsf1 | Q8BGC0 | 1.94 | 2.9 | 2.8 | 2.05 | 3.01 | 2.72 | 2.76 | 2.76 | 2.9 | 1.55 | 3.06 | 3.03 | 2.48 | 2.42 | 2.63 | 1.81 | 0.39 | 0.2125 | -0.0875 | 0.180082 | 0.61472 | 0.779827 | 0.999971 | 1 | 0.965847 |
| Htr1a | Q64264 | 0.07 | -1.05 | -0.43 | -0.37 | -0.85 | -1.51 | -0.01 | -0.25 | -1.17 | -0.29 | 0.26 | -0.29 | -0.38 | -1.34 | 0.07 | -0.37 | -0.21 | 0.0725 | -0.06 | 0.59368 | 0.842796 | 0.870023 | 0.999971 | 1 | 0.983917 |
| Htr1b | P28334 | -0.63 | -0.97 | 0.24 | 0.63 | 2.39 | -1.16 | -0.15 | 0.19 | -3.39 | -3.37 | 0.35 | 0.27 | 3.98 | 0.74 | 0 | 0.21 | 0.5 | -1.3525 | 1.415 | 0.511659 | 0.205448 | 0.141286 | 0.999971 | 1 | 0.784967 |
| Htr1f | Q02284 | -1.02 | -1.24 | -1.34 | -0.9 | -1.69 | -1.42 | -1.93 | -0.78 | -1.3 | -1.33 | -0.43 | -0.26 | -0.55 | -2.05 | -1.42 | -0.92 | -0.33 | 0.295 | -0.11 | 0.260934 | 0.347622 | 0.745428 | 0.999971 | 1 | 0.961064 |
| Htr2a | P35363 | 0.45 | -2.12 | -1.07 | -0.57 | -1.61 | -1.21 | -0.93 | -1.63 | -2.14 | -0.48 | -1.15 | -1.03 | -1.69 | -1.96 | -1.22 | -0.82 | -0.5175 | -0.3725 | -0.595 | 0.331659 | 0.529401 | 0.291369 | 0.999971 | 1 | 0.847402 |
| Htr5a | P30966 | 0.59 | 0.12 | 0.25 | 0.01 | 0.7 | -0.13 | -0.4 | 1.1 | -0.91 | 0.52 | 0.25 | 0.87 | 1.28 | 0.3 | -0.31 | 1.32 | 0.075 | -0.06 | 0.405 | 0.836264 | 0.877868 | 0.324847 | 0.999971 | 1 | 0.856456 |
| Htra1 | Q9R118 | 0.01 | -0.48 | -0.29 | -0.38 | 0.23 | -0.12 | 0.06 | -0.41 | -0.1 | 0.02 | 0.45 | 0.92 | -0.34 | -0.25 | 0.08 | 0.04 | 0.225 | 0.6075 | 0.1675 | 0.317154 | 0.049146 | 0.421905 | 0.999971 | 1 | 0.877992 |
| Htra2 | Q9JIY5 | -1.52 | -0.64 | -0.7 | -0.94 | -0.68 | -1.05 | -0.8 | -1.04 | -0.63 | -1.39 | -0.5 | -0.36 | -1.09 | -0.9 | -0.45 | -0.51 | 0.0575 | 0.23 | 0.2125 | 0.81748 | 0.465334 | 0.443421 | 0.999971 | 1 | 0.883386 |
| Htt | G3X9H5 | -0.35 | -0.2 | 0.1 | -0.08 | -0.28 | -0.19 | -0.1 | -0.08 | -0.2 | -0.22 | -0.04 | -0.24 | -0.2 | -0.21 | -0.15 | 0.07 | -0.03 | -0.0425 | 0.01 | 0.869088 | 0.812269 | 0.957387 | 0.999971 | 1 | 0.998584 |
| Huwe1 | A2AFQ0 | 0.57 | 0.71 | 0.53 | 0.57 | 0.78 | 0.76 | 0.7 | 0.34 | 0.74 | 0.67 | 0.36 | 0.32 | 0.53 | 0.69 | 0.49 | 0.53 | 0.05 | -0.0725 | -0.035 | 0.786315 | 0.692593 | 0.832958 | 0.999971 | 1 | 0.979036 |
| Hyi | Q8R1F5 | -2.76 | -4.76 | -5.3 | -5.45 | 2.73 | 2.02 | 2.36 | 3.24 | -4.74 | -2.95 | -3.75 | -5.46 | 3.1 | 2.37 | 3.27 | 3 | 7.155 | 0.3425 | 7.5025 | 1.65E-06 | 0.650892 | 9.28E-07 | 0.007516 | 1 | 0.004371 |
| Hykk | Q5U5V2 | 0.44 | -0.12 | 0.11 | 0.8 | 0.02 | 0.52 | -0.98 | 0.79 | -0.3 | 0.26 | -0.23 | -0.32 | -0.15 | 0.47 | 0.94 | 1.38 | -0.22 | -0.455 | 0.3525 | 0.599947 | 0.111408 | 0.357429 | 0.999971 | 1 | 0.868793 |
| Hyou1 | Q9JKR6 | -0.17 | 0.1 | 0.08 | -0.05 | 0.05 | 0.08 | 0.12 | 0.14 | 0.12 | -0.13 | 0.1 | -0.27 | 0.37 | 0.24 | 0.75 | 0.18 | 0.1075 | -0.035 | 0.395 | 0.524803 | 0.84782 | 0.076196 | 0.999971 | 1 | 0.748444 |
| Hypk | Q9CR41 | 1.15 | 2.12 | 0.92 | -0.16 | 2.18 | 1.89 | 1.23 | 0.64 | 2.73 | 1.37 | 1.17 | 1.95 | 2.5 | 1.84 | 0.64 | -0.19 | 0.4775 | 0.7975 | 0.19 | 0.383447 | 0.165882 | 0.783565 | 0.999971 | 1 | 0.966376 |
| Iah1 | Q9DB29 | 0.26 | 0.61 | 1.57 | 0.28 | 0.99 | 0.1 | 0.69 | 0.91 | 0.66 | -0.1 | 1.62 | 0.42 | 0.39 | 0.4 | 1 | 0.32 | -0.0075 | -0.03 | -0.1525 | 0.983304 | 0.946418 | 0.657825 | 0.999971 | 1 | 0.934638 |
| Iap | P03975 | -0.25 | 0.24 | 1.27 | -0.17 | 0.55 | 1.09 | 0.17 | 0.29 | 0.09 | -0.32 | 0.1 | -0.54 | 0.74 | 0.31 | -0.49 | -0.73 | 0.2525 | -0.44 | -0.315 | 0.521115 | 0.256954 | 0.498637 | 0.999971 | 1 | 0.903494 |
| Iars1 | Q8BU30 | 0.72 | 1.61 | 1.23 | 1.09 | 1.4 | 1.29 | 1.13 | 1.02 | 1.6 | 1.24 | 1.58 | 1.84 | 1.34 | 1.39 | 1.27 | 1.65 | 0.0475 | 0.4025 | 0.25 | 0.840994 | 0.131694 | 0.307214 | 0.999971 | 1 | 0.847402 |
| Iars2 | Q8BIJ6 | -0.64 | -0.24 | -0.36 | -0.54 | -0.56 | -0.66 | -0.55 | -0.75 | -0.5 | -0.69 | -0.47 | -0.41 | -0.74 | -0.68 | -0.75 | -0.75 | -0.185 | -0.0725 | -0.285 | 0.319162 | 0.688284 | 0.133327 | 0.999971 | 1 | 0.784948 |
| Iba57 | Q8CAK1 | -0.27 | -0.47 | 0.01 | -0.04 | -0.69 | -0.59 | -0.5 | -0.29 | -0.62 | -0.34 | -0.59 | 0.04 | -0.3 | -0.2 | 0.27 | 0.03 | -0.325 | -0.185 | 0.1425 | 0.129089 | 0.422031 | 0.514982 | 0.999971 | 1 | 0.905815 |
| Ica1 | D3Z118 | 0.16 | 0.47 | -0.09 | 0 | 0.52 | 0.72 | -0.4 | -0.14 | 0.66 | 0.48 | -0.07 | 0.27 | 0.97 | 0.61 | -0.2 | 0.46 | 0.04 | 0.2 | 0.325 | 0.894154 | 0.400843 | 0.275994 | 0.999971 | 1 | 0.843794 |
| Ica1l | A0A087WS | -0.44 | 0.24 | -0.25 | 0.13 | 0.57 | 0.34 | -0.19 | 0.6 | 0.37 | 0.32 | 0.07 | -0.54 | 1.14 | 0.78 | 0.9 | 0.51 | 0.41 | 0.135 | 0.9125 | 0.146778 | 0.629154 | 0.004122 | 0.999971 | 1 | 0.582714 |
| Icam2 | G5E8M9 | -0.15 | 0.25 | -0.19 | 0.36 | 0.99 | -0.66 | -0.56 | -0.43 | 0.38 | 0.53 | 0.09 | 0.6 | 0.59 | 0.27 | 0.98 | 0.61 | -0.2325 | 0.3325 | 0.545 | 0.559742 | 0.156424 | 0.0438 | 0.999971 | 1 | 0.702458 |
| Icam5 | Q60625 | -0.37 | 0.14 | -0.79 | -0.92 | -0.01 | 0.32 | -0.77 | -0.6 | 0.1 | -0.25 | -0.77 | -0.83 | 0.52 | 0.01 | -0.94 | -0.76 | 0.22 | 0.0475 | 0.1925 | 0.527531 | 0.884148 | 0.631416 | 0.999971 | 1 | 0.932858 |
| Icmt | Q3U4N2 | -1.46 | -1.3 | -1.75 | 0.62 | -0.28 | -1.58 | -0.75 | 1.25 | -1.35 | -2.02 | 0.2 | 0.53 | -2.4 | -2.01 | -0.39 | 0.64 | 0.6325 | 0.3125 | -0.0675 | 0.392825 | 0.674107 | 0.93284 | 0.999971 | 1 | 0.995069 |
| Ide | Q8CGB9 | -0.08 | 0.2 | 0 | -0.3 | 0.44 | 0.3 | -0.2 | -0.22 | 0.48 | 0.14 | 0.35 | 0.53 | 0.94 | 0.42 | 0.07 | 0.41 | 0.125 | 0.42 | 0.505 | 0.596894 | 0.054888 | 0.061141 | 0.999971 | 1 | 0.725348 |
| Idh1 | O88844 | -1.12 | -1.56 | 0.17 | 0.1 | -1.13 | -1.53 | 0.34 | 0.34 | -1 | -1.54 | 0.15 | -0.34 | -1.15 | -1.24 | 0.43 | 0.21 | 0.1075 | -0.08 | 0.165 | 0.857197 | 0.879943 | 0.771885 | 0.999971 | 1 | 0.963498 |
| Idh2 | P54071 | -0.66 | -0.57 | -0.54 | -0.91 | -0.42 | -0.89 | -0.73 | -0.82 | -0.53 | -1.34 | -0.97 | -1.04 | -0.31 | -0.94 | -0.83 | -0.91 | -0.045 | -0.3 | -0.0775 | 0.818122 | 0.205088 | 0.721685 | 0.999971 | 1 | 0.955137 |
| Idh3a | Q9D6R2 | -0.29 | -0.31 | -0.45 | -0.35 | -0.29 | -0.71 | 0.06 | -0.6 | -0.35 | -0.78 | -0.38 | -0.45 | -0.01 | -0.8 | -0.7 | -0.49 | -0.035 | -0.14 | -0.15 | 0.873751 | 0.441397 | 0.506054 | 0.999971 | 1 | 0.905277 |
| Idh3b | Q91VA7 | -0.57 | -0.16 | -0.61 | -0.61 | -0.43 | -0.48 | -0.77 | -0.79 | -0.5 | -0.45 | -0.7 | -1 | -0.1 | -0.47 | -0.69 | -0.88 | -0.13 | -0.175 | -0.0475 | 0.524052 | 0.417728 | 0.840511 | 0.999971 | 1 | 0.97975 |
| Idh3g | P70404 | -0.14 | -0.3 | -0.11 | -0.37 | -0.76 | -0.54 | -0.02 | -0.47 | -0.94 | -0.17 | -0.58 | -1.1 | -0.98 | -0.53 | -0.7 | -0.49 | -0.2175 | -0.4675 | -0.445 | 0.325658 | 0.082852 | 0.043933 | 0.999971 | 1 | 0.702458 |
| Idi1 | P58044 | -1.22 | -0.4 | -1.2 | -1.78 | -0.39 | -0.34 | -0.72 | -1.29 | 0.13 | -1 | -0.89 | -0.82 | 0.31 | 0.13 | -0.72 | -1.48 | 0.465 | 0.505 | 0.71 | 0.208398 | 0.200569 | 0.155939 | 0.999971 | 1 | 0.796008 |
| Ids | Q08890 | -0.61 | -0.04 | -0.22 | -1.65 | 0.43 | -0.84 | 0.53 | -0.84 | 0.33 | -1 | 0.53 | 0.76 | 0.27 | -0.47 | -0.55 | -1.3 | 0.45 | 0.785 | 0.1175 | 0.368195 | 0.142049 | 0.795733 | 0.999971 | 1 | 0.970435 |
| Iffo1 | D6RH90 | 0.05 | 0.04 | -0.2 | 0.65 | 0.15 | 0.22 | -0.93 | 1.08 | -0.01 | -0.21 | -0.31 | -0.11 | 0.88 | 0.22 | -0.07 | 1.12 | -0.005 | -0.295 | 0.4025 | 0.990618 | 0.218257 | 0.244635 | 0.999971 | 1 | 0.837316 |
| Ifi47 | A0A1L1SQ( | 2.25 | 1.27 | 1.71 | 2.57 | 2.08 | -0.02 | 0 | -0.03 | 0.43 | 1.28 | 0.58 | 1.84 | 0.17 | 1.95 | 2.39 | 1.07 | -1.4425 | -0.9175 | -0.555 | 0.026011 | 0.051571 | 0.307783 | 0.999971 | 1 | 0.847402 |
| Ifngr1 | P15261 | -0.34 | -0.25 | 0.54 | -0.11 | -0.38 | -0.08 | -0.5 | 0.31 | -0.07 | -0.28 | 0.53 | 0.01 | 0.04 | -0.38 | 0.37 | 0.47 | -0.1225 | 0.0875 | 0.165 | 0.666704 | 0.753444 | 0.572014 | 0.999971 | 1 | 0.91816 |
| Ifngr2 | Q63953 | -3.05 | -2.82 | 1.19 | 1.07 | -2.7 | -1.74 | 0.23 | 2.36 | -2.1 | -2.95 | 2.01 | 1.74 | -2.86 | -2.03 | 1.88 | 2.13 | 0.44 | 0.5775 | 0.6825 | 0.760296 | 0.711041 | 0.66185 | 0.999971 | 1 | 0.936148 |
| Ift122 | E9Q9G8 | 0.54 | 0.6 | 0.02 | 0.15 | 0.52 | 0.41 | -0.49 | 0.45 | 0.23 | 0.65 | -0.17 | 0.25 | 0.47 | 0.76 | 0.22 | 0.56 | -0.105 | -0.0875 | 0.175 | 0.718132 | 0.724139 | 0.443096 | 0.999971 | 1 | 0.883386 |
| Ift140 | E9PY46 | -0.06 | 0.42 | 0.29 | 0.02 | 0.3 | 0.15 | 0.01 | 0.51 | 0.02 | 0.38 | 0.16 | 0.22 | -0.17 | 0.79 | 0.48 | 0.41 | 0.075 | 0.0275 | 0.21 | 0.718836 | 0.887046 | 0.421162 | 0.999971 | 1 | 0.87788 |
| Ift172 | Q6VH22 | 0.81 | 0.98 | 1.35 | 1.22 | 0.19 | 1.61 | 0.91 | 1.29 | 0.65 | 1.72 | 1.08 | 0.89 | 0.46 | 1.2 | 1.3 | 1.79 | -0.09 | -0.005 | 0.0975 | 0.784263 | 0.98548 | 0.751325 | 0.999971 | 1 | 0.961064 |
| Ift22 | Q9DAI2 | 0.48 | 1.04 | 0.51 | 0.57 | 1.25 | 1.38 | 1.64 | 1.19 | 2.65 | 1.27 | 1.15 | 1.31 | 3.25 | 2.08 | 1.25 | 1.51 | 0.715 | 0.945 | 1.3725 | 0.008128 | 0.028628 | 0.011557 | 0.999971 | 1 | 0.613095 |
| Ift27 | Q9D0P8 | 2.36 | 0.66 | 2.22 | 2.03 | 0.48 | 0.75 | 0.63 | 2.52 | 0.1 | 1.77 | 1.62 | 2.34 | -1.36 | 0.45 | 2.48 | 2.4 | -0.7225 | -0.36 | -0.825 | 0.223559 | 0.532234 | 0.366896 | 0.999971 | 1 | 0.869526 |
| Ift46 | A0A1L1SR1 | -0.21 | -0.74 | 0.2 | 0.7 | -0.83 | -0.32 | -0.48 | 0.51 | -0.4 | 0.31 | 0.1 | 0.38 | -0.65 | 0.15 | 0.22 | 1.34 | -0.2675 | 0.11 | 0.2775 | 0.50805 | 0.752076 | 0.5644 | 0.999971 | 1 | 0.917087 |
| Ift52 | Q62559 | 0.49 | 0.48 | 0.31 | 0.09 | -0.04 | 0.36 | -0.28 | -0.07 | 0.17 | 0.88 | 0.21 | 0.22 | -0.05 | 0.78 | 0.01 | 0.47 | -0.35 | 0.0275 | -0.04 | 0.126015 | 0.904655 | 0.871854 | 0.999971 | 1 | 0.984109 |
| Ift57 | F7B458 | -0.21 | 0.23 | 1.02 | 0.04 | 0.33 | -0.1 | 1.32 | 0.13 | 0.58 | -0.05 | 0.54 | 0.72 | -0.12 | -0.24 | -0.03 | -0.53 | 0.15 | 0.1775 | -0.5 | 0.7039 | 0.580037 | 0.119799 | 0.999971 | 1 | 0.78268 |
| Ift74 | Q8BKE9 | 1.7 | 0.2 | -0.03 | 0.49 | -0.06 | 0.46 | -0.42 | -0.08 | 0.06 | 1.38 | -0.1 | 0.86 | -0.28 | 0.37 | 0.04 | 0.41 | -0.615 | -0.04 | -0.455 | 0.155422 | 0.934175 | 0.273372 | 0.999971 | 1 | 0.843794 |
| Ift80 | Q8K057-2 | -0.69 | -0.9 | 0.1 | 0.25 | -0.78 | -0.77 | -0.37 | 0.43 | -1.2 | -0.83 | -0.35 | -0.46 | -1.71 | -0.21 | 0.07 | 0.27 | -0.0625 | -0.4 | -0.085 | 0.87161 | 0.259317 | 0.863573 | 0.999971 | 1 | 0.983079 |
| Ift81 | O35594 | 0.79 | 0.66 | 0.65 | 0.87 | 0.37 | 0.5 | 1.49 | 0.63 | 0.06 | 0.75 | 0.52 | 0.6 | 0.13 | 0.74 | 0.35 | 0.56 | 0.005 | -0.26 | -0.2975 | 0.985517 | 0.229602 | 0.163913 | 0.999971 | 1 | 0.800215 |
| Ift88 | G3X9Z7 | 1.83 | 0.79 | 1 | 0.77 | 0.42 | 1.06 | 0.64 | 0.96 | 0.66 | 1.41 | 0.82 | 0.45 | 0.49 | 0.96 | 0.68 | 0.61 | -0.3275 | -0.2625 | -0.4125 | 0.289601 | 0.427325 | 0.170883 | 0.999971 | 1 | 0.803758 |
| Igbp1 | Q61249 | 0.1 | -0.04 | 0.52 | 0.25 | -0.06 | -0.21 | 1.07 | 0.05 | 0.12 | -0.19 | 0.31 | 0.5 | -0.37 | -0.35 | 0.26 | -0.03 | 0.005 | -0.0225 | -0.33 | 0.987418 | 0.920856 | 0.176517 | 0.999971 | 1 | 0.808532 |
| Igbp1b | A0A0B4J1F | 0.41 | -1.19 | -0.33 | 0.54 | 0.44 | 0.76 | 1.46 | 0.98 | 1.37 | 1.06 | 1.46 | 1.19 | -1.47 | -0.44 | 0.57 | -0.46 | 1.0525 | 1.4125 | -0.3075 | 0.033973 | 0.005807 | 0.566924 | 0.999971 | 1 | 0.917976 |
| Igdcc4 | Q9EQS9 | -0.46 | -1.14 | -1.73 | -0.83 | -1.11 | -1.49 | -1.41 | -1.85 | -1.68 | -0.7 | -1.74 | -2.29 | -1.2 | -1.2 | -1.99 | -1.5 | -0.425 | -0.5625 | -0.4325 | 0.197708 | 0.191437 | 0.209549 | 0.999971 | 1 | 0.822138 |
| Igf1r | E9QNX9 | -0.41 | -0.86 | -0.99 | -1.14 | -0.9 | -1.18 | -1.03 | -0.83 | -1.12 | -0.52 | -0.98 | -1.07 | -1.1 | -1.39 | -1.14 | -0.62 | -0.135 | -0.0725 | -0.2125 | 0.542939 | 0.762436 | 0.41012 | 0.999971 | 1 | 0.875847 |
| Igf2bp1 | O88477 | 5.6 | -0.38 | -0.1 | 0.47 | 1.16 | 0.5 | -0.25 | 0.68 | -0.24 | 2.19 | -0.92 | 0.27 | 0.1 | 0.76 | 0.63 | 1.15 | -0.875 | -1.0725 | -0.7375 | 0.500121 | 0.450445 | 0.565409 | 0.999971 | 1 | 0.917745 |
| Igf2r | Q07113 | 0.95 | 0.89 | 0.77 | 0.85 | 0.9 | 1.16 | 0.98 | 1.04 | 0.95 | 1.16 | 0.47 | 0.57 | 0.98 | 1.24 | 1.06 | 0.77 | 0.155 | -0.0775 | 0.1475 | 0.365966 | 0.714994 | 0.427197 | 0.999971 | 1 | 0.880793 |
| Igfbp4 | P47879 | 1.83 | -0.68 | 0.19 | 0.6 | -0.52 | 0.11 | 0.39 | 0.74 | -1.1 | 1.07 | 0.07 | -0.21 | -1.41 | -1.05 | 1.52 | 1.36 | -0.305 | -0.5275 | -0.63 | 0.574997 | 0.411396 | 0.413386 | 0.999971 | 1 | 0.876261 |
| Igfbp5 | Q07079 | 0.64 | -0.52 | 1.93 | 0.99 | -0.4 | 0.01 | 1.55 | 2.56 | -0.28 | 1.03 | 2.2 | 1.14 | 0.18 | 0.2 | 2.39 | 1.96 | 0.17 | 0.2625 | 0.4225 | 0.824798 | 0.689756 | 0.54864 | 0.999971 | 1 | 0.911727 |
| Igfbp6 | P47880 | 1.53 | 1.22 | 0.35 | 0.4 | 0.85 | 2.22 | -0.37 | 0.48 | 0.98 | 1.77 | 0.03 | -0.54 | 1.44 | 1.7 | 0.25 | 1 | -0.08 | -0.315 | -0.2225 | 0.887105 | 0.567508 | 0.593317 | 0.999971 | 1 | 0.923349 |
| Igfbp7 | E9Q5D9 | 0.18 | -0.36 | -0.07 | 0.67 | 1.07 | -0.3 | -0.05 | 0.18 | 0.12 | 1.59 | -0.42 | 1.43 | 0.11 | 0.35 | 0.7 | 1.11 | 0.12 | 0.575 | 0.4625 | 0.739908 | 0.270609 | 0.16516 | 0.999971 | 1 | 0.800215 |
| Ighg1 | A0A075B5I | 5.63 | -1.24 | -1.83 | -2.33 | -0.13 | 6.03 | -0.85 | 6.54 | 3.69 | 1.3 | 1.99 | -1.92 | -3.08 | 0.42 | -1.1 | -2.12 | 2.84 | 1.2075 | -1.5275 | 0.255076 | 0.544162 | 0.402538 | 0.999971 | 1 | 0.872468 |
| Ighg2b | A0A075B5I | 4.38 | 2.18 | 2.3 | 3.78 | 1.43 | 0.47 | 2.65 | 0.95 | 0.94 | 0.46 | 0.24 | 2.44 | 0.7 | 3.53 | 1.21 | 0.68 | -1.785 | -2.14 | -1.63 | 0.021738 | 0.010888 | 0.063361 | 0.999971 | 1 | 0.725348 |
| Ighg2c | F6TQW2 | 1.46 | 0.83 | -1.26 | 0.68 | 3.09 | 2.56 | 0.64 | 2.32 | -0.15 | -1.41 | -0.16 | 3 | -1.3 | 3.93 | 1.54 | 0.05 | 1.725 | -0.1075 | 0.6275 | 0.036526 | 0.914109 | 0.581135 | 0.999971 | 1 | 0.920364 |
| Ighm | A0A075B5I | 3.22 | 2.19 | 0.48 | 1.28 | 3.23 | 1.59 | 0.48 | 0.57 | 1.63 | 1.23 | 0.09 | 0.54 | 1.24 | 2.99 | 0.36 | 0.26 | -0.325 | -0.92 | -0.58 | 0.679131 | 0.167118 | 0.463957 | 0.999971 | 1 | 0.890905 |
| Ighmbp2 | D3Z5P5 | 1.8 | 0.94 | 2.36 | 2.81 | 0.92 | 0.73 | 4.73 | -0.86 | 0.79 | 0.76 | 0.46 | -0.65 | 0.96 | 1.04 | 0.94 | -1.76 | -0.5975 | -1.6375 | -1.6825 | 0.594842 | 0.008638 | 0.041448 | 0.999971 | 1 | 0.701725 |
| Ighv6-7 | A0A0A6YW | 0.78 | 0.61 | -1.04 | -0.21 | -0.31 | 0.29 | -2.21 | -0.03 | -0.06 | 0.91 | -0.83 | -0.61 | 0.42 | 0.3 | -0.22 | -0.51 | -0.6 | -0.1825 | -0.0375 | 0.358464 | 0.730791 | 0.932782 | 0.999971 | 1 | 0.995069 |
| Igkc | A0A5H1ZRI | 4.09 | 1.27 | 1.05 | 2.2 | 2.6 | 4.26 | 4.4 | 5.68 | 4.35 | 0.49 | 4.94 | 4.98 | 0.05 | 2.44 | 1.41 | -1.04 | 2.0825 | 1.5375 | -1.4375 | 0.033054 | 0.203501 | 0.145606 | 0.999971 | 1 | 0.791767 |
| Iglon5 | Q8HW98 | -1.68 | -1.04 | -0.83 | -0.68 | -0.85 | -1.37 | -1.3 | -0.82 | -1.23 | -1.43 | -0.82 | -1.02 | -0.41 | -0.94 | -1 | -0.48 | -0.0275 | -0.0675 | 0.35 | 0.921549 | 0.805032 | 0.236925 | 0.999971 | 1 | 0.833594 |
| Igsf1 | Q7TQA1-5 | -0.02 | -1.76 | 0 | -0.88 | -0.4 | -0.8 | -0.82 | -0.18 | 0.23 | -0.14 | -1.99 | 0.79 | 0.26 | 0.76 | 1.71 | -0.2 | 0.115 | 0.3875 | 1.2975 | 0.786168 | 0.566115 | 0.037379 | 0.999971 | 1 | 0.700096 |
| Igsf11 | D3Z6S7 | 0.22 | -0.67 | -0.69 | -0.29 | 0.54 | 0.54 | -0.99 | -0.05 | -0.35 | 0.67 | -1.08 | -0.53 | 0.06 | 0.53 | -0.79 | -0.56 | 0.3675 | 0.035 | 0.1675 | 0.370969 | 0.931075 | 0.643736 | 0.999971 | 1 | 0.93301 |
| Igsf21 | Q7TNR6 | -0.91 | -1.08 | -1.08 | -1.21 | -0.82 | -1.07 | -1.06 | -1.12 | -0.69 | -0.8 | -0.86 | -0.66 | -0.02 | -0.84 | -0.85 | -0.63 | 0.0525 | 0.3175 | 0.485 | 0.765721 | 0.084766 | 0.06834 | 0.999971 | 1 | 0.733937 |
| Igsf3 | Q6ZQA6 | -0.75 | -0.92 | -0.59 | -0.41 | -0.69 | -0.12 | -0.26 | -0.34 | -0.6 | -0.29 | -0.76 | -0.7 | -0.31 | -0.5 | -0.43 | -0.27 | 0.315 | 0.08 | 0.29 | 0.162941 | 0.69422 | 0.153013 | 0.999971 | 1 | 0.795578 |
| Igsf8 | G3UYZ1 | 0.16 | -0.36 | 0.86 | 0.65 | -0.59 | -0.41 | 0.29 | 1.17 | -0.81 | -0.33 | 0.22 | 0.12 | -0.93 | -0.36 | 0.48 | 1.01 | -0.2125 | -0.5275 | -0.2775 | 0.641227 | 0.161344 | 0.564078 | 0.999971 | 1 | 0.917087 |
| Igsf9b | A0A1L1SR8 | 0.89 | -0.71 | -0.53 | -0.68 | -1.16 | -0.43 | -0.96 | -0.45 | -1.53 | -0.19 | -0.51 | -1.04 | -0.68 | -0.89 | -0.62 | -0.35 | -0.4925 | -0.56 | -0.3775 | 0.245255 | 0.240365 | 0.341806 | 0.999971 | 1 | 0.861042 |
| Igtp | Q9DCE9 | -0.36 | 0.35 | -0.21 | -0.06 | 1.07 | 0.58 | -0.66 | -0.66 | 0.01 | 0.33 | -0.7 | 0.01 | 0.79 | 1.12 | -0.47 | 0.17 | 0.1525 | -0.0175 | 0.4725 | 0.728973 | 0.950132 | 0.225661 | 0.999971 | 1 | 0.828225 |

| | | | | | | | | | | | | | | | | | | | | | | | | | | |
|---|---|---|---|---|---|---|---|---|---|---|---|---|---|---|---|---|---|---|---|---|---|---|---|---|---|---|
| Ik | Q9Z1M8 | -0.55 | -0.37 | 1.12 | 1.23 | -0.21 | -0.29 | 0.67 | 1 | -0.32 | 0 | 1.01 | 1.07 | -0.75 | -0.22 | 1.24 | 1.17 | -0.065 | 0.0825 | 0.0025 | 0.901897 | 0.879897 | 0.996819 | 0.999971 | 1 | 1 |
| Ikbkb | A0A0R4J0T | -0.08 | 0.59 | -0.38 | -0.46 | 0.36 | 0.7 | -0.4 | -0.29 | 0.48 | 0.11 | -0.01 | 0.13 | 0.46 | 0.27 | -0.28 | 0.01 | 0.175 | 0.26 | 0.1975 | 0.618509 | 0.359378 | 0.512494 | 0.999971 | 1 | 0.905815 |
| Ikbkg | E9Q2Y3 | -0.46 | 0.35 | -0.43 | 0.01 | 0.37 | -0.08 | -0.56 | -0.09 | 0.65 | -0.32 | 0.42 | 0.16 | -0.48 | -0.21 | -0.56 | 0.17 | 0.0425 | 0.36 | -0.1375 | 0.881307 | 0.240766 | 0.617007 | 0.999971 | 1 | 0.928413 |
| Il16 | O54824-2 | 0.8 | 0.21 | 3.9 | 0.73 | 0.21 | 0.63 | 9.02 | 2.23 | -0.01 | 1.19 | 5.58 | 6.46 | -0.14 | 0.7 | 2.53 | 1.29 | 1.6125 | 1.895 | -0.315 | 0.418891 | 0.258794 | 0.728695 | 0.999971 | 1 | 0.956041 |
| Il17ra | Q60943 | -0.94 | -0.68 | 0.02 | 0.14 | -0.95 | -0.53 | 0.31 | 0.18 | -1.2 | -1.19 | -0.42 | -1.12 | -0.2 | -0.66 | -1.3 | -1.34 | 0.1175 | -0.6175 | -0.51 | 0.759613 | 0.084453 | 0.191449 | 0.999971 | 1 | 0.810547 |
| Il18 | P70380 | -0.67 | -0.26 | -0.97 | -1.27 | -0.09 | -0.73 | -1.26 | -1.03 | -0.2 | -0.58 | -1.03 | -0.64 | 0 | -0.05 | -0.66 | -1.48 | 0.015 | 0.18 | 0.245 | 0.963889 | 0.534468 | 0.536049 | 0.999971 | 1 | 0.910077 |
| Il19 | A0A087WS | -5.74 | 3.26 | -1.69 | -1.2 | -0.92 | 0.12 | -2.02 | -7.56 | -4.93 | -4.46 | -1.36 | 1.36 | 5.18 | 0.14 | 0.43 | 2.84 | -1.2525 | -1.005 | 3.49 | 0.576162 | 0.634082 | 0.098963 | 0.999971 | 1 | 0.78268 |
| Il1rap | Q61730 | -0.19 | -1.02 | -0.9 | -0.24 | -0.81 | -0.63 | -0.41 | -0.21 | -0.97 | -0.58 | -0.61 | -0.6 | -0.7 | -0.83 | -0.43 | -0.14 | 0.0725 | -0.1025 | 0.0625 | 0.78978 | 0.692688 | 0.82418 | 0.999971 | 1 | 0.976309 |
| Il1rapl1 | B1ASU0 | -1.23 | -0.7 | -1.95 | -1.7 | -1.06 | -0.85 | -1.89 | -1.79 | -1.08 | -1.16 | -1.73 | -2.06 | -0.75 | -0.77 | -1.8 | -1.59 | -0.0025 | -0.1125 | 0.1675 | 0.994573 | 0.751589 | 0.657898 | 0.999971 | 1 | 0.934638 |
| Il1rapl2 | Q9ERS6 | -0.04 | 0.41 | 0.71 | 0.29 | -0.01 | 0.65 | -0.08 | 0.56 | -0.05 | 0.56 | 0.51 | -0.14 | 0.53 | 0.91 | 0.4 | 1 | -0.0625 | -0.1225 | 0.3675 | 0.814129 | 0.640977 | 0.157718 | 0.999971 | 1 | 0.797252 |
| Il6st | V9GX00 | 0.48 | -0.54 | -0.31 | -0.18 | -1.21 | -0.88 | -0.4 | -0.95 | -0.96 | -0.02 | -0.72 | -0.29 | -0.45 | -0.68 | -0.43 | -0.32 | -0.7225 | -0.36 | -0.3325 | 0.031646 | 0.262899 | 0.217837 | 0.999971 | 1 | 0.826893 |
| Ildr2 | A0A0A6YV' | -0.25 | -0.95 | -1.05 | -1.09 | -0.89 | -0.67 | -0.86 | -1.08 | -0.87 | -0.52 | -0.99 | -1.1 | -0.63 | -1.15 | -1.4 | -1.14 | -0.04 | -0.035 | -0.245 | 0.870149 | 0.891548 | 0.381946 | 0.999971 | 1 | 0.871816 |
| Ilf2 | Q9CXY6 | 0.65 | 0.89 | 0.62 | 0.53 | 0.86 | 0.93 | 0.4 | 0.52 | 0.72 | 0.65 | 0.4 | 0.47 | 0.35 | 0.48 | 0.29 | 0.59 | 0.005 | -0.1125 | -0.245 | 0.980481 | 0.535748 | 0.199309 | 0.999971 | 1 | 0.813335 |
| Ilf3 | Q9Z1X4-3 | 0.72 | 0.53 | 0.18 | 0.4 | 0.45 | 0.6 | 0.03 | 0.53 | 0.72 | 0.77 | 0.48 | 0.33 | 0.62 | 0.73 | 0.71 | 0.66 | -0.055 | 0.1175 | 0.2225 | 0.800019 | 0.568429 | 0.254848 | 0.999971 | 1 | 0.839012 |
| Ilk | O55222 | 0.03 | 0.5 | 0.44 | 0.43 | 1.32 | 0.66 | 0.65 | 0.32 | 0.83 | 0.54 | 0.8 | 0.73 | 0.46 | 0.43 | 0.31 | 0.17 | 0.3875 | 0.375 | -0.0075 | 0.160576 | 0.073722 | 0.968917 | 0.999971 | 1 | 0.998787 |
| Ilkap | Q8R0F6 | 3.22 | 2.6 | 1.57 | 1.81 | 2.7 | 2.33 | 2.25 | 1.41 | 2.77 | 3.24 | 1.63 | 1.27 | 2.52 | 2.37 | 1.48 | 1.17 | -0.1275 | -0.0725 | -0.415 | 0.77159 | 0.895644 | 0.387816 | 0.999971 | 1 | 0.871816 |
| Ilrun | A0A3B2W3 | 0.72 | 2.92 | 1.95 | 2.8 | 2.51 | 2.24 | 1.42 | 2.91 | 2 | 1.35 | 2.12 | 2.43 | 2.78 | 2.66 | 2.61 | 2.65 | 0.1725 | -0.1225 | 0.5775 | 0.753705 | 0.81264 | 0.244432 | 0.999971 | 1 | 0.837316 |
| Ilvbl | Q8BU33 | -0.29 | -0.3 | -1.09 | -1.11 | -0.35 | -0.61 | -1.25 | -0.65 | -0.55 | -0.58 | -0.88 | -1.08 | -0.21 | -0.37 | -0.71 | -0.89 | -0.0175 | -0.075 | 0.1525 | 0.954448 | 0.788465 | 0.603761 | 0.999971 | 1 | 0.924454 |
| Immp1l | Q9CQU8 | 0.73 | 0.92 | -1.05 | 0.25 | 0.4 | 0.15 | -0.8 | 0.26 | 0.48 | 0.79 | -0.08 | -0.83 | 0.96 | 0.03 | -0.71 | 0.5 | -0.21 | -0.1225 | -0.0175 | 0.666138 | 0.816423 | 0.973435 | 0.999971 | 1 | 0.99925 |
| Immp2l | Q8BPT6 | -1.62 | -0.35 | -0.55 | -0.75 | -0.6 | -0.47 | -0.59 | -0.37 | -0.71 | -1.97 | -1.16 | -1.32 | -0.95 | 0.08 | 0 | -0.23 | 0.31 | -0.4725 | 0.5425 | 0.308928 | 0.225294 | 0.155291 | 0.999971 | 1 | 0.795578 |
| Immt | E9Q800 | 0.01 | 0.97 | -2.23 | -1.71 | 1.3 | 1.32 | -1.76 | -2.26 | 1.51 | -0.05 | -1.27 | -1.94 | 2.05 | 0.76 | -2.43 | -2 | 0.39 | 0.3025 | 0.335 | 0.719911 | 0.752226 | 0.774844 | 0.999971 | 1 | 0.963884 |
| Immt_1 | Q8CAQ8-3 | -0.73 | -1.23 | 0.44 | 0.39 | -0.05 | 0.19 | 0.02 | -0.14 | -0.14 | 0.36 | -0.32 | 0.57 | -0.92 | -1.18 | -0.52 | -0.81 | 0.2875 | 0.4 | -0.575 | 0.481352 | 0.374766 | 0.191012 | 0.999971 | 1 | 0.810547 |
| Immt_2 | Q8CAQ8-2 | -0.3 | -0.37 | -0.44 | -0.71 | -0.1 | -0.44 | -0.03 | -0.51 | -0.1 | -0.72 | -0.18 | -0.03 | 0.3 | -0.72 | -0.52 | -0.55 | 0.185 | 0.1975 | 0.0825 | 0.377144 | 0.382673 | 0.757734 | 0.999971 | 1 | 0.961116 |
| Immt_3 | Q8CAQ8 | 1.2 | 0.73 | 1.36 | 1.04 | 0.39 | 1.96 | 0.38 | 1.37 | 1.18 | 1.97 | 0.44 | 1.08 | 0.68 | 1.42 | 0.42 | 2.1 | -0.0575 | 0.085 | 0.0725 | 0.88394 | 0.801792 | 0.851793 | 0.999971 | 1 | 0.98187 |
| Immt_4 | E9PVS5 | -4.67 | -2.66 | -6.8 | -0.63 | 1.91 | -1.96 | -6.2 | 1.51 | -6.61 | -5.89 | -6.91 | -6.93 | 2.65 | 0.39 | -5.91 | -1.77 | 2.505 | -2.895 | 2.53 | 0.238883 | 0.037947 | 0.226167 | 0.999971 | 1 | 0.828225 |
| Imp3 | Q921Y2 | -1.29 | -1.57 | -1.3 | -1.67 | -1.86 | -1.12 | -1.5 | -1.35 | -1.98 | -0.78 | -1.32 | -0.37 | -0.66 | -2.26 | -1.69 | -1.3 | 3.93E-16 | 0.345 | -0.02 | 1 | 0.344069 | 0.953812 | 1 | 1 | 0.998056 |
| Impa1 | Q924B0 | -0.16 | 0.38 | 0.41 | 0.58 | 0.17 | 0.34 | 0.1 | 0.44 | 0.26 | 0.12 | 0.43 | 0.4 | -0.12 | 0.5 | 0.41 | 0.45 | -0.04 | 0 | 0.0075 | 0.856575 | 1 | 0.975697 | 0.999971 | 1 | 0.99925 |
| Impa2 | Q91UZ5 | 3.15 | 1.01 | 1.28 | 0.87 | 0.22 | 1.43 | 2.08 | 0.98 | 1.06 | 3.07 | 1.93 | 1.66 | 1.27 | 0.5 | 0.74 | 0.07 | -0.4 | 0.3525 | -0.9325 | 0.511413 | 0.573786 | 0.112305 | 0.999971 | 1 | 0.78268 |
| Impact | O55091 | 0.24 | 0.45 | 0.56 | 0.28 | 0.72 | 0.28 | 1.06 | 0.24 | 0.82 | -0.07 | 0.17 | 0.62 | 0.38 | 0.23 | 0.59 | 0.2 | 0.1925 | 0.0025 | -0.0325 | 0.432876 | 0.991847 | 0.862518 | 0.999971 | 1 | 0.982963 |
| Impdh1 | F7DEU6 | 0.46 | 1.22 | 0.74 | 0.59 | 1.03 | 0.74 | 0.56 | 1.27 | 1.04 | 0.96 | 0.93 | 1.28 | 1.13 | 1.09 | 1.03 | 0.56 | 0.1475 | 0.3 | 0.2 | 0.565599 | 0.201423 | 0.42169 | 0.999971 | 1 | 0.877992 |
| Impdh2 | P24547 | -0.67 | 0.31 | 0.36 | 0.11 | 0.22 | -0.32 | 0.13 | 0.25 | 0.46 | -0.26 | 0.5 | 0.47 | 0.79 | 0.19 | 0.41 | 0.03 | 0.0425 | 0.265 | 0.3275 | 0.881899 | 0.398622 | 0.290825 | 0.999971 | 1 | 0.847402 |
| Ina | P46660 | -0.61 | -0.86 | -1.12 | -1.34 | -1 | -0.61 | -1.57 | -1.42 | -0.95 | -0.72 | -1.4 | -1.25 | -1 | -0.68 | -1.25 | -1.8 | -0.1675 | -0.0975 | -0.2 | 0.55749 | 0.693848 | 0.504097 | 0.999971 | 1 | 0.905265 |
| Inafm2 | A0A1L1STD | -0.32 | 0.55 | 0.11 | 0.18 | 0.71 | 0.55 | 0.14 | 0.17 | 0.64 | 0.2 | -0.57 | -0.5 | 1.15 | 0.92 | -0.17 | -0.03 | 0.2625 | -0.1875 | 0.3375 | 0.315838 | 0.582172 | 0.370257 | 0.999971 | 1 | 0.869526 |
| Inf2 | E9QLA5 | 0.23 | 0.62 | 0.04 | 0.01 | 0.47 | -0.07 | -0.15 | 0.12 | 0.44 | -0.03 | -0.16 | 0.13 | 0.3 | 0.28 | 0.06 | -0.36 | -0.1325 | -0.13 | -0.155 | 0.573341 | 0.571125 | 0.52519 | 0.999971 | 1 | 0.909237 |
| Ing2 | Q9ESK4 | 0.09 | 0.23 | 2.43 | 0.77 | 1.23 | -0.84 | 2.45 | 0.61 | 1.68 | -0.48 | 2.33 | 1.82 | -1.87 | 0.04 | 1.24 | 0.33 | -0.0175 | 0.4575 | -0.945 | 0.982128 | 0.543874 | 0.23618 | 0.999971 | 1 | 0.832918 |
| Inip | Q3TXT3 | 1.7 | 1.95 | 1.79 | 1.68 | 1.6 | 1.25 | 1.83 | 1.45 | 1.28 | 1.22 | 1.26 | 0.78 | 0.7 | 0.74 | 2.25 | 1.02 | -0.2475 | -0.645 | -0.6025 | 0.230823 | 0.008414 | 0.122842 | 0.999971 | 1 | 0.78268 |
| Inpp1 | P49442 | -1.07 | -0.61 | -0.6 | -1.81 | -0.64 | -0.86 | -0.26 | -1.43 | -0.6 | -1.48 | -0.19 | -0.46 | -0.86 | -0.94 | -0.75 | -1.61 | 0.225 | 0.34 | -0.0175 | 0.54202 | 0.387948 | 0.959071 | 0.999971 | 1 | 0.998721 |
| Inpp4a | F6V2U0 | -0.34 | 0.2 | 0.04 | 0.08 | 0.05 | 0.08 | -0.03 | -0.1 | 0.19 | -0.1 | -0.06 | 0.02 | 0.09 | 0.12 | -0.06 | 0.1 | 0.005 | 0.0175 | 0.0675 | 0.979044 | 0.927572 | 0.725066 | 0.999971 | 1 | 0.955916 |
| Inpp4b | E9PVM1 | -0.3 | -1.03 | -0.4 | -0.08 | -0.65 | -0.99 | -0.56 | -0.17 | -0.94 | -0.56 | -0.41 | -0.16 | -1.25 | -0.53 | -0.57 | -0.21 | -0.14 | -0.065 | -0.1875 | 0.619366 | 0.814271 | 0.544615 | 0.999971 | 1 | 0.911398 |
| Inpp5a | Q7TNC9 | -0.75 | -0.49 | -0.29 | -0.25 | -0.51 | -0.29 | -0.45 | -0.03 | -0.37 | -0.42 | -0.29 | -0.71 | 0.01 | -0.15 | 0.02 | -0.08 | 0.125 | -0.0025 | 0.395 | 0.552889 | 0.990019 | 0.063284 | 0.999971 | 1 | 0.725348 |
| Inpp5b | Q8K337 | 0.16 | -0.22 | 0.22 | -0.1 | -0.23 | -0.52 | -0.03 | -0.13 | -0.09 | -0.13 | 0.1 | 0.08 | -0.3 | -0.41 | -0.13 | -0.22 | -0.2425 | -0.025 | -0.28 | 0.253584 | 0.892868 | 0.16504 | 0.999971 | 1 | 0.800215 |
| Inpp5d | Q9ES52-3 | 0.5 | 0.81 | 0.34 | 0.43 | 1.07 | 0.37 | 0.58 | 0.83 | 0.45 | 0.73 | 1.6 | 0.77 | -0.24 | 0.54 | 0.78 | 1.11 | 0.1925 | 0.3675 | 0.0275 | 0.400848 | 0.214294 | 0.929428 | 0.999971 | 1 | 0.994693 |
| Inpp5e | Q9JII1 | -0.34 | -0.57 | -1.3 | -0.8 | -0.65 | -0.57 | -0.22 | -1.13 | -0.86 | -0.31 | -0.97 | -0.82 | -1.16 | -0.47 | -1.23 | -0.85 | 0.11 | 0.0125 | -0.175 | 0.705265 | 0.96305 | 0.541687 | 0.999971 | 1 | 0.911042 |
| Inpp5f | Q8CDA1 | -0.07 | 0.02 | 0.06 | 0.03 | -0.28 | -0.07 | -0.41 | -0.14 | 0.22 | 0.03 | -0.36 | -0.19 | 0.44 | 0.03 | 0.13 | 0.15 | -0.235 | -0.085 | 0.1775 | 0.194828 | 0.657644 | 0.330645 | 0.999971 | 1 | 0.857107 |
| Inpp5j | P59644 | 0.67 | -0.19 | 0.32 | 0.58 | -0.57 | 0.09 | 0.22 | 0.29 | -0.12 | 0.59 | 0.17 | 0.02 | 0.06 | -0.01 | 0.25 | 0.52 | -0.3375 | -0.18 | -0.14 | 0.260457 | 0.504348 | 0.584373 | 0.999971 | 1 | 0.920364 |
| Inpp5k | Q8C5L6 | -1.25 | -0.96 | -0.11 | -0.43 | -0.63 | -0.76 | -0.86 | 0.04 | -1.2 | -0.74 | -0.21 | -0.73 | -0.09 | 0.07 | 0.03 | 0.12 | 0.135 | -0.0325 | 0.72 | 0.681302 | 0.920719 | 0.027353 | 0.999971 | 1 | 0.678474 |
| Inppl1 | Q6P549 | 0.62 | -0.01 | 0.2 | 0 | -0.15 | 0.06 | 0.04 | 0.01 | -0.16 | 0.66 | 0.25 | 0.07 | 0.05 | 0.2 | 0.13 | 0.6 | -0.2125 | 0.0025 | 0.0425 | 0.320325 | 0.992085 | 0.853629 | 0.999971 | 1 | 0.981948 |
| Insc | A0A0U1RN | -1.06 | -1.37 | -1.01 | -0.95 | -1.51 | 0.16 | -1.96 | -0.45 | -1.45 | -0.13 | -1.76 | -0.63 | -2.67 | -0.11 | -1.02 | -1.08 | 0.1575 | 0.105 | -0.1225 | 0.733597 | 0.779045 | 0.807172 | 0.999971 | 1 | 0.972962 |
| Insr | P15208 | -0.2 | -0.13 | -0.54 | -0.69 | -0.38 | -0.22 | -0.91 | -0.41 | -0.27 | -0.36 | -0.51 | -0.36 | 0.11 | -0.48 | -0.49 | -0.42 | -0.09 | 0.015 | 0.07 | 0.703607 | 0.939605 | 0.76517 | 0.999971 | 1 | 0.962405 |
| Insyn1 | Q8CD60 | -1.84 | -1.06 | -1.93 | -1.08 | -1.66 | -1.74 | -1.4 | -1.36 | -1.77 | -0.67 | -1.64 | -1.07 | -1.37 | -1.43 | -0.34 | -1 | -0.0625 | 0.19 | 0.4425 | 0.818673 | 0.58365 | 0.216664 | 0.999971 | 1 | 0.826893 |
| Insyn2a | E9PZW7 | 1.17 | -0.61 | -0.38 | 0.04 | -0.36 | -0.05 | -0.88 | -0.3 | -0.24 | 0.44 | -0.59 | -0.46 | 0.07 | 0.31 | 0.01 | -0.65 | -0.4525 | -0.2675 | -0.12 | 0.287879 | 0.541211 | 0.776939 | 0.999971 | 1 | 0.964475 |
| Ints1 | K3W4P2 | -0.09 | 0.47 | 0.34 | -0.66 | 1.08 | 0.61 | 1.68 | -0.43 | 0.67 | -0.12 | 0.36 | 0.51 | 0.08 | 0.3 | -0.28 | -1.38 | 0.72 | 0.34 | -0.335 | 0.157942 | 0.291224 | 0.442107 | 0.999971 | 1 | 0.883386 |
| Ints10 | E9Q360 | 0.72 | 1.02 | 0.85 | 0.98 | 0.9 | 1.11 | 0.34 | 1 | 0.72 | 1.37 | 0.77 | 0.66 | 0.65 | 1.17 | 0.83 | 1.19 | -0.055 | -0.0125 | 0.0675 | 0.807163 | 0.954501 | 0.742065 | 0.999971 | 1 | 0.961064 |
| Ints11 | Q9CWS4 | -0.1 | 0.72 | 0.75 | 0.79 | 1.01 | 0.71 | 1.02 | 0.93 | 0.89 | 0.55 | 1.14 | 0.96 | 1.07 | 0.9 | 1.01 | 0.72 | 0.3775 | 0.345 | 0.385 | 0.160144 | 0.216915 | 0.155515 | 0.999971 | 1 | 0.795578 |
| Ints12 | Q9D168 | 0.87 | 0.71 | 1.3 | 1.11 | 0.94 | 0.35 | 1.26 | 1.41 | 0.72 | 0.95 | 0.86 | 0.52 | 0.79 | 1.06 | 1.01 | 0.77 | -0.0075 | -0.235 | -0.09 | 0.978821 | 0.276756 | 0.662605 | 0.999971 | 1 | 0.936148 |
| IntS13 | Q8QZV7 | 0.84 | 1.57 | 1.77 | 2.04 | 1.94 | 1.4 | 1.13 | 1.98 | 1.72 | 1.19 | 1.36 | 1.82 | 1.57 | 1.84 | 1.45 | 2.45 | 0.0575 | -0.0325 | 0.2725 | 0.861684 | 0.914643 | 0.429164 | 0.999971 | 1 | 0.881196 |
| Ints14 | Q8R3P6 | 1.87 | 3.08 | 1.65 | 1.39 | 2.7 | 1.51 | 2.39 | 0.98 | 3.95 | 0.93 | 1.49 | 0.98 | 3.3 | 1.76 | 1.32 | 0.99 | -0.1025 | -0.16 | -0.155 | 0.838936 | 0.827314 | 0.789575 | 0.999971 | 1 | 0.96775 |
| Ints15 | Q8BGA7 | -0.6 | 0.58 | -1.23 | -0.53 | -0.14 | -0.31 | -0.24 | -0.4 | -0.12 | -0.16 | -0.77 | 0.33 | -0.26 | -0.11 | -0.76 | -0.86 | 0.1725 | 0.265 | -0.0525 | 0.642137 | 0.530847 | 0.895814 | 0.999971 | 1 | 0.988571 |
| Ints2 | Q80UK8 | -0.01 | 0.52 | -0.47 | 0.29 | 0.54 | 0.08 | -0.25 | -0.15 | -0.01 | 0.08 | -0.12 | -0.08 | 0.14 | 0.76 | 0.11 | -0.14 | -0.0275 | -0.115 | 0.135 | 0.924122 | 0.641861 | 0.651398 | 0.999971 | 1 | 0.933842 |
| Ints3 | A0A0G2JFJ | 0.57 | 1.05 | 0.61 | 1.15 | 1.1 | 0.99 | 0.51 | 1.04 | 1.09 | 1 | 1.25 | 1.11 | 0.5 | 1.17 | 0.79 | 0.97 | 0.065 | 0.2675 | 0.0125 | 0.783234 | 0.217236 | 0.958381 | 0.999971 | 1 | 0.998622 |
| Ints4 | Q8CIM8 | 1.18 | 1.86 | 1.07 | 1.23 | 1.73 | 0.95 | 1.05 | 1.82 | 1.14 | 1.44 | 1.27 | 1.09 | 2.37 | 1.68 | 0.65 | 1.41 | 0.0525 | -0.1 | 0.1925 | 0.859589 | 0.665002 | 0.618564 | 0.999971 | 1 | 0.928581 |
| Ints5 | Q8CHT3 | 0.58 | 0.76 | 0.76 | 0.72 | 0.78 | 0.25 | 0.15 | 0.89 | 0.55 | 1.01 | 0.62 | 0.6 | 0.38 | 0.53 | 0.37 | 0.85 | -0.1875 | -0.01 | -0.1725 | 0.422344 | 0.956368 | 0.374058 | 0.999971 | 1 | 0.87031 |
| Ints6 | Q6PCM2-2 | 0.05 | 0.36 | -0.48 | -0.05 | 0.98 | 0.34 | -0.15 | -0.42 | 0.79 | -0.14 | -0.66 | -0.12 | 0.9 | -0.06 | -1.05 | -0.78 | 0.2175 | -0.0025 | -0.2175 | 0.535898 | 0.994185 | 0.625257 | 0.999971 | 1 | 0.930479 |
| Ints7 | A0A0R4J0E | 0.74 | 0.86 | 0.21 | 0.19 | 0.91 | 0.81 | -0.24 | 0.26 | 0.47 | 0.8 | -0.17 | 0.09 | 1.18 | 0.75 | -0.14 | 0.36 | -0.065 | -0.2025 | 0.0375 | 0.839456 | 0.487213 | 0.909745 | 0.999971 | 1 | 0.991822 |
| Ints9 | A0A0R4J0J | -0.12 | -0.02 | -0.42 | -0.65 | 0.71 | 0.72 | -0.27 | 0.04 | 0.16 | 0.33 | -0.07 | -0.13 | 0.27 | 1.56 | -0.11 | 0.4 | 0.6025 | 0.375 | 0.8325 | 0.06792 | 0.114994 | 0.050835 | 0.999971 | 1 | 0.723736 |
| Intu | Q059U7-2 | 1.43 | 0.57 | 1.4 | 1.53 | -0.9 | 0.38 | 1.78 | 1.44 | 0.08 | 2.27 | 1.1 | 1.36 | 0.04 | 0.2 | 0.45 | 1.41 | -0.5575 | -0.03 | -0.7075 | 0.355676 | 0.949183 | 0.082352 | 0.999971 | 1 | 0.755254 |
| Ip6k1 | Q6PD10 | -0.49 | 0.35 | 0.49 | 0.64 | 0.71 | 0.54 | -0.15 | 0.82 | 0.52 | 0.17 | 0.56 | 0.44 | 0.42 | 0.45 | 0.54 | 0.46 | 0.2325 | 0.175 | 0.22 | 0.490311 | 0.538502 | 0.429342 | 0.999971 | 1 | 0.881311 |
| Ipcef1 | Q5DU31 | 0.35 | 0.52 | 0.92 | 0.8 | 0.24 | 0.9 | 1.11 | 0.85 | 0.33 | 0.08 | 1 | 0.72 | 0.29 | -0.03 | 0.64 | 0.53 | 0.1275 | -0.115 | -0.29 | 0.61789 | 0.662898 | 0.23584 | 0.999971 | 1 | 0.832918 |
| Ipo11 | Q8K2V6 | 1.29 | 0.58 | 0.63 | 0.65 | 0.87 | 0.68 | 0.8 | 0.93 | 0.56 | 1.01 | 0.63 | 0.6 | 0.38 | 0.55 | 0.62 | 0.64 | 0.0325 | -0.0875 | -0.24 | 0.882671 | 0.707031 | 0.296097 | 0.999971 | 1 | 0.847402 |
| Ipo13 | Q8K0C1 | 1.91 | 0.24 | 1.12 | 1.51 | 0.25 | 0.49 | 2.5 | 0.38 | 0.18 | 0.74 | 1.15 | -0.01 | -0.04 | 0.07 | 0.31 | -0.83 | -0.29 | -0.68 | -1.3175 | 0.623468 | 0.134782 | 0.010679 | 0.999971 | 1 | 0.613095 |
| Ipo4 | Q8VI75 | 0.43 | 0.38 | 0.5 | 0.15 | -0.03 | 0.09 | 0.27 | -0.02 | 0.28 | 0.59 | 0.37 | -0.21 | -0.17 | 0.37 | 0.39 | 0.43 | -0.2875 | -0.1075 | -0.11 | 0.138305 | 0.632858 | 0.606057 | 0.999971 | 1 | 0.924649 |
| Ipo5 | Q8BKC5 | -0.57 | 0.03 | 0.1 | -0.11 | 0.11 | -0.08 | 0.56 | -0.11 | 0.12 | -0.74 | 0 | -0.1 | 0.11 | -0.07 | -0.09 | -0.25 | 0.2575 | -0.0425 | 0.0625 | 0.310142 | 0.872115 | 0.772258 | 0.999971 | 1 | 0.963498 |
| Ipo7 | Q9EPL8 | -0.63 | 0.25 | 0.22 | 0.1 | 0.23 | 0.12 | 0.23 | 0.14 | 0.15 | -0.23 | 0.44 | 0.31 | 0.1 | 0.4 | 0.51 | 0.35 | 0.195 | 0.1825 | 0.355 | 0.42831 | 0.505956 | 0.183926 | 0.999971 | 1 | 0.809025 |
| Ipo8 | Q7TMY7 | -0.16 | 1.19 | 0.8 | 0.11 | 0.77 | 0.79 | 0.05 | 0.97 | 1.02 | 0.5 | 0.43 | 0.65 | 0.5 | 1.01 | 0.43 | 0.55 | 0.16 | 0.165 | 0.1375 | 0.659695 | 0.624306 | 0.683085 | 0.999971 | 1 | 0.943421 |
| Ipo9 | E9QKZ2 | 0.3 | 0.56 | 0.6 | 0.41 | 0.62 | 0.73 | 0.86 | 0.56 | 0.9 | 0.48 | 1.03 | 0.94 | 1.06 | 0.7 | 0.69 | 1.05 | 0.225 | 0.37 | 0.4075 | 0.226632 | 0.087252 | 0.058744 | 0.999971 | 1 | 0.725348 |
| Iqcb1 | Q8BP00-2 | -0.04 | -0.31 | 0.59 | 0.29 | -0.26 | -0.65 | 1.48 | -0.13 | -0.28 | -0.03 | 0.67 | 1.13 | -0.39 | -0.36 | 0.12 | -0.06 | -0.0225 | 0.24 | -0.305 | 0.962147 | 0.519029 | 0.254203 | 0.999971 | 1 | 0.839012 |
| Iqgap1 | Q9JKF1 | 0.59 | 0.85 | 0.91 | 1.14 | 1.83 | 0.75 | 0.67 | 0.75 | 1.03 | 1.14 | 0.8 | 1.39 | 0.73 | 0.93 | 1.28 | 1.03 | 0.1275 | 0.2175 | 0.12 | 0.678197 | 0.319983 | 0.573927 | 0.999971 | 1 | 0.91816 |
| Iqgap2 | Q3UQ44 | 0.46 | 0.71 | 0.88 | 0.61 | 0.38 | 0.87 | 0.35 | 0.5 | 0.2 | 0.25 | 0.59 | 0.32 | 0.2 | 0.38 | 0.43 | 0.72 | -0.14 | -0.325 | -0.2325 | 0.497899 | 0.110897 | 0.262277 | 0.999971 | 1 | 0.842262 |
| Iqsec1 | E9PUA3 | -0.91 | -0.49 | -0.47 | -0.64 | -1.05 | -0.98 | -1.3 | -0.42 | -0.83 | -0.43 | -0.73 | -0.61 | -0.14 | -0.33 | -0.26 | -0.35 | -0.31 | -0.0225 | 0.3575 | 0.223224 | 0.906938 | 0.081054 | 0.999971 | 1 | 0.755254 |
| Iqsec1_1 | A0A1D5RN | -0.23 | -0.7 | -0.29 | -0.47 | -0.94 | -0.56 | -0.28 | -0.61 | -0.92 | -0.39 | -0.14 | -0.33 | -0.65 | -0.86 | -0.4 | -0.43 | -0.175 | -0.0225 | -0.1625 | 0.429381 | 0.92263 | 0.437717 | 0.999971 | 1 | 0.882419 |
| Iqsec2 | A4GZ26 | -0.68 | -1.23 | -1.34 | -1.33 | -1.49 | -0.85 | -0.87 | -1.74 | -1.53 | -0.75 | -1.4 | -1.97 | -1.19 | -1.23 | -1.89 | -1.75 | -0.0925 | -0.2675 | -0.37 | 0.747349 | 0.389288 | 0.181296 | 0.999971 | 1 | 0.808532 |
| Iqsec3 | Q3TES0 | 0.31 | -0.72 | -0.57 | -0.58 | -0.59 | -0.43 | 0.58 | -0.8 | -0.5 | -0.17 | -0.61 | -0.54 | -0.49 | -0.6 | -0.51 | -0.8 | 0.08 | -0.065 | -0.21 | 0.830724 | 0.811425 | 0.440669 | 0.999971 | 1 | 0.883277 |
| Irak1 | B1AUW9 | 1.66 | 0.46 | 0.86 | -0.34 | 0.21 | 0.5 | -0.12 | 0.71 | 2.24 | 0.71 | 0.41 | 0.6 | 0.19 | 0.18 | 1.09 | 0.83 | -0.335 | 0.33 | -0.0875 | 0.442841 | 0.550846 | 0.845091 | 0.999971 | 1 | 0.979814 |
| Ireb2 | Q811J3 | 0.89 | 0.87 | 1 | 0.94 | 0.88 | 0.49 | 1.69 | 0.46 | 1.33 | 0.65 | 0.44 | 0.44 | 0.7 | -0.45 | -0.37 | -0.33 | -0.045 | -0.21 | -1.0375 | 0.879629 | 0.396302 | 0.005496 | 0.999971 | 1 | 0.582714 |
| Irf2bp1 | Q8R3Y8-2 | 3.3 | 2.23 | 3.13 | 2.83 | 2.19 | 2.16 | 2.62 | 2.41 | 3.28 | 2.65 | 2.8 | 1.79 | 1.89 | 2.37 | 2.36 | 2.39 | -0.5275 | -0.2425 | -0.62 | 0.082505 | 0.525534 | 0.051085 | 0.999971 | 1 | 0.723736 |
| Irf2bp2 | E9Q1P8 | -0.89 | 0.81 | 0.88 | 1.23 | 0.05 | -0.86 | 1.51 | 1.13 | -0.83 | -1.08 | 1.04 | 0.89 | -0.24 | -0.11 | 1.43 | 1.13 | -0.05 | -0.5025 | 0.045 | 0.938679 | 0.459454 | 0.938511 | 0.999971 | 1 | 0.995869 |
| Irf2bpl | Q8K3X4 | 1.76 | 2.19 | 2.3 | 2.62 | 2.16 | 1.9 | 2.23 | 2.49 | 1.23 | 1.96 | 1.87 | 1.58 | 1.22 | 2.28 | 1.94 | 1.93 | -0.0225 | -0.5575 | -0.375 | 0.926857 | 0.059211 | 0.226044 | 0.999971 | 1 | 0.828225 |
| Irf3 | P70671 | 1.78 | 1.39 | 1.77 | 1.6 | 0.76 | 1.38 | 1.52 | 1.43 | 0.98 | 1.6 | 1.31 | 0.92 | 0.6 | 1.26 | 1.26 | 0.96 | -0.3625 | -0.4325 | -0.615 | 0.145206 | 0.078271 | 0.021371 | 0.999971 | 1 | 0.640714 |
| Irgm1 | Q60766-2 | -1.69 | -0.44 | -1.04 | -0.96 | -0.14 | -1.57 | -1.35 | -1.12 | -0.54 | -2.13 | -1.02 | -0.48 | -0.08 | -1.52 | -1.35 | -0.93 | -0.0125 | -0.01 | 0.0625 | 0.974412 | 0.981669 | 0.874226 | 0.999971 | 1 | 0.984109 |
| Irgq | Q8VIM9 | 0.61 | 0.45 | 0.57 | 0.9 | 0.59 | 0.44 | 0.99 | 0.66 | 0.6 | 0.37 | 0.84 | 0.66 | 0.57 | 0.59 | 0.62 | 1.08 | 0.0375 | -0.015 | 0.0825 | 0.854707 | 0.938384 | 0.69297 | 0.999971 | 1 | 0.947174 |
| Irs1 | Q543V3 | -1.41 | -0.45 | -0.3 | -0.78 | -0.23 | -0.64 | -0.52 | -0.71 | 0.13 | -1.82 | 0.99 | 0.46 | 0.43 | -1.02 | -0.62 | -0.17 | 0.21 | 0.675 | 0.39 | 0.464592 | 0.282984 | 0.323753 | 0.999971 | 1 | 0.85548 |
| Irs2 | P81122 | 0.72 | 0.9 | 1.38 | 1.3 | 0.55 | 1.18 | 0.63 | 1.55 | 0.63 | 0.91 | 1.64 | 1.28 | 0.67 | 1.37 | 1.51 | 2.07 | -0.0975 | 0.04 | 0.33 | 0.741481 | 0.887844 | 0.330093 | 0.999971 | 1 | 0.856869 |
| Isca1 | Q9D924 | -0.12 | -0.35 | -0.12 | -0.37 | -0.18 | -0.16 | -0.14 | -0.16 | -0.22 | -0.2 | -0.03 | 0.05 | 0.17 | -0.31 | 0.18 | 0.26 | 0.08 | 0.14 | 0.315 | 0.636507 | 0.4308 | 0.148178 | 0.999971 | 1 | 0.793248 |
| Isca2 | A0A1Y7VK: | -0.14 | -0.22 | -0.29 | -0.25 | -0.37 | -0.26 | -0.42 | -0.28 | -0.59 | -0.23 | -0.35 | -0.49 | -0.18 | -0.01 | -0.52 | -0.23 | -0.1075 | -0.19 | -0.01 | 0.513705 | 0.281439 | 0.956865 | 0.999971 | 1 | 0.998513 |
| Iscu | Q9D7P6 | 0.15 | 0.26 | 0 | 0.12 | 0.29 | 0.35 | 0.08 | 0.24 | 0.16 | 0.09 | 0.21 | -0.03 | 0.57 | 0.61 | 0.16 | 0.07 | 0.1075 | -0.025 | 0.22 | 0.534789 | 0.880381 | 0.298226 | 0.999971 | 1 | 0.847402 |
| Isg15 | Q64339 | -0.1 | 0.13 | -0.89 | 1.58 | 1.46 | -0.86 | -0.91 | -0.3 | 0.37 | -0.67 | -0.93 | -0.81 | -0.91 | 1.04 | -1.25 | 0.4 | -0.3325 | -0.69 | -0.36 | 0.629657 | 0.230979 | 0.59791 | 0.999971 | 1 | 0.92377 |
| Isg20l2 | A0A0R4J0R | 1.66 | -0.83 | -0.38 | 0.33 | -0.81 | -0.62 | -0.53 | -0.53 | -1.21 | 0.5 | -0.99 | -0.87 | -2.21 | -0.94 | -0.61 | -0.51 | -0.8175 | -0.8375 | -1.2625 | 0.133562 | 0.19497 | 0.064964 | 0.999971 | 1 | 0.727266 |
| Islr2 | E9Q9N7 | -0.96 | -0.92 | -0.33 | -0.54 | -0.79 | -1.21 | -0.82 | 0.12 | -0.74 | -0.7 | -0.4 | -0.5 | -0.62 | -1.01 | -0.71 | -0.01 | 0.0125 | 0.1025 | 0.1 | 0.968986 | 0.636468 | 0.718678 | 0.999971 | 1 | 0.953825 |
| Isoc1 | Q91V64 | -0.26 | -1.36 | -0.25 | -0.02 | -0.9 | -0.84 | -0.18 | -0.03 | -0.62 | -0.51 | -0.38 | 0.37 | -0.92 | -0.32 | 0.42 | 0.38 | -0.015 | 0.1875 | 0.3625 | 0.967161 | 0.610696 | 0.395518 | 0.999971 | 1 | 0.871816 |
| Isoc2a | P85094 | -0.22 | -0.08 | 0.78 | -0.77 | 0.03 | -0.7 | 0.55 | 0.16 | 0.06 | -0.83 | 0.53 | 0.6 | 0.32 | -0.52 | 0.07 | -0.93 | 0.0825 | 0.1625 | -0.1925 | 0.83504 | 0.70989 | 0.639469 | 0.999971 | 1 | 0.93301 |
| Isoc2b | Q9DCC7 | -0.59 | -0.28 | -1.23 | -0.43 | -0.76 | -1.31 | -1.54 | 0.24 | -1.18 | -0.57 | -1.04 | -0.47 | -1.72 | 0 | -1.45 | -0.38 | -0.21 | -0.1825 | -0.255 | 0.622794 | 0.526151 | 0.564032 | 0.999971 | 1 | 0.917087 |
| Ist1 | Q9CX00 | -0.66 | -0.81 | 0.29 | 0.11 | -0.71 | -0.34 | 0.12 | 0.43 | -0.76 | -0.42 | 0.31 | -0.16 | -0.66 | 0 | 0.86 | 1.06 | 0.1425 | 0.01 | 0.5825 | 0.695197 | 0.977132 | 0.221368 | 0.999971 | 1 | 0.8275 |
| Isy1 | Q69ZQ2 | 0.52 | 0.75 | 0.94 | 1.14 | 0.91 | 0.86 | 0.53 | 0.63 | 0.37 | 0.47 | 1.15 | 1.68 | 0.04 | 0.56 | 0.76 | 1.33 | -0.105 | 0.08 | -0.165 | 0.620689 | 0.810404 | 0.59182 | 0.999971 | 1 | 0.922655 |
| Isyna1 | Q9JHU9 | 0.3 | 0.16 | 0.65 | 0.57 | 0.43 | 0.2 | 0.84 | 0.65 | 0.03 | 0.09 | 0.61 | 1.17 | -0.32 | 0.73 | 1.45 | 0.71 | 0.11 | 0.055 | 0.2225 | 0.622822 | 0.853436 | 0.552879 | 0.999971 | 1 | 0.913595 |
| Itch | Q8C863 | -1.29 | -0.78 | -0.58 | -0.75 | -0.49 | -1.01 | -0.33 | -0.28 | -0.51 | -1.8 | -0.61 | -0.4 | -0.21 | -0.95 | -0.44 | -0.38 | 0.3225 | 0.02 | 0.355 | 0.22382 | 0.954844 | 0.179706 | 0.999971 | 1 | 0.808532 |
| Itfg1 | Q99KW9 | -2.11 | -0.7 | -2.05 | -1.85 | -0.55 | -0.23 | -1.65 | -1.33 | -0.53 | -1.61 | -1.2 | -1.4 | 0.14 | 0.13 | -1.37 | -1.17 | 0.7375 | 0.4925 | 1.11 | 0.121093 | 0.228115 | 0.046395 | 0.999971 | 1 | 0.710102 |
| Itfg2 | Q91WI7-2 | 1.23 | 1.34 | 0.38 | 0.27 | 1.17 | 0.88 | 0.72 | 0.31 | 1.47 | 1.16 | 0.48 | 0.4 | 1.5 | 0.28 | 0.45 | 0.13 | -0.035 | 0.0725 | -0.215 | 0.915655 | 0.845174 | 0.593758 | 0.999971 | 1 | 0.92342 |
| Itga1 | Q3V3R4 | 1.32 | 0.59 | 0.52 | 0.53 | 1.62 | 0.76 | 0.86 | -0.51 | 0.89 | 1.57 | 1.25 | 1.57 | 0.15 | 0.23 | 0.72 | -0.04 | -0.0575 | 0.58 | -0.475 | 0.898783 | 0.056949 | 0.108101 | 0.999971 | 1 | 0.78268 |
| Itga3 | Q62470 | 0.03 | -0.96 | -0.84 | -0.28 | -0.18 | -0.78 | -1 | -0.56 | -0.72 | 0.22 | -0.49 | -0.15 | -0.78 | -0.89 | -0.04 | -0.23 | -0.1175 | 0.2275 | 0.0275 | 0.696647 | 0.476047 | 0.930678 | 0.999971 | 1 | 0.994744 |
| Itga4 | Q792F9 | -0.55 | -1.11 | -2.67 | -3.22 | 1.97 | 0.27 | -2.14 | -2.96 | -2.01 | -0.07 | -2.79 | -2.52 | 1.36 | 0.77 | -2.43 | -2.45 | 1.1725 | 0.04 | 1.2 | 0.322341 | 0.959938 | 0.279575 | 0.999971 | 1 | 0.843812 |
| Itga5 | P11688 | -0.31 | -0.78 | -1.46 | -1.45 | 0.13 | -1.61 | -1.87 | -1.33 | -0.53 | 0.14 | -0.78 | -0.61 | -1.21 | -0.65 | -1.2 | -1.35 | -0.17 | 0.555 | -0.1025 | 0.729322 | 0.130991 | 0.75078 | 0.999971 | 1 | 0.961064 |
| Itga6 | Q61739-2 | -0.18 | 0.29 | -0.21 | 0.1 | 0.03 | 0.01 | -0.35 | -0.81 | 0.27 | 0.17 | -0.14 | -0.12 | 0.6 | 0.36 | 0.31 | 0.09 | -0.28 | 0.045 | 0.34 | 0.290206 | 0.827627 | 0.132545 | 0.999971 | 1 | 0.784948 |
| Itga7 | G3X9Q1 | 0.05 | 0.08 | 0.24 | 0.49 | 1.28 | -0.14 | 0.19 | -0.07 | 0.34 | 0.68 | 0.48 | 0.64 | -0.4 | 0.33 | 0.31 | 0.44 | 0.1 | 0.32 | -0.045 | 0.769505 | 0.118809 | 0.855536 | 0.999971 | 1 | 0.982398 |
| Itgam | E9Q604 | 0.25 | 1.43 | -0.01 | -0.16 | 1.03 | 0.86 | -0.53 | -0.22 | 0.52 | 0.33 | 0.1 | -0.02 | 0.59 | 1.28 | 0.42 | 0.14 | -0.0925 | -0.145 | 0.23 | 0.850857 | 0.696227 | 0.581998 | 0.999971 | 1 | 0.920364 |
| Itgav | P43406 | -1.02 | -1.1 | -0.58 | -0.7 | -1.12 | -1.08 | -0.61 | -0.98 | -0.98 | -0.97 | -0.61 | -0.68 | -0.49 | -0.94 | -0.5 | -0.68 | -0.0975 | 0.04 | 0.1975 | 0.654409 | 0.846466 | 0.365816 | 0.999971 | 1 | 0.869526 |
| Itgb1 | P09055 | 0.27 | 0.13 | 0.66 | 0.94 | 1.01 | 0.2 | 0.56 | 0.3 | 0.63 | 0.98 | 1.42 | 1.61 | 0.43 | 0.48 | 1.27 | 0.96 | 0.0175 | 0.66 | 0.285 | 0.949319 | 0.049939 | 0.334301 | 0.999971 | 1 | 0.858827 |
| Itgb1bp1 | O35671 | 1.19 | 1.6 | 0.78 | -0.03 | 0.75 | 0.45 | 0.02 | 0.41 | 0.54 | 0.97 | -0.56 | 0.02 | 0.32 | 0.99 | 0.76 | 0.23 | -0.4775 | -0.6425 | -0.31 | 0.216309 | 0.180179 | 0.422925 | 0.999971 | 1 | 0.87887 |
| Itgb2 | Q542I8 | -0.13 | 0.95 | 0.49 | 0.29 | 1 | 0.85 | 0.72 | 0.04 | 0.84 | 0.11 | 0.07 | 0.27 | 0.67 | 0.71 | 0.14 | -0.42 | 0.2525 | -0.0775 | -0.125 | 0.42849 | 0.793215 | 0.717139 | 0.999971 | 1 | 0.952711 |
| Itgb3 | O54890 | 0.64 | -0.25 | -0.84 | 0.17 | 0.62 | 0.27 | 0.24 | -0.81 | 0.35 | 0.79 | 0.14 | 0.18 | 0.02 | -0.6 | -0.1 | -0.4 | 0.15 | 0.435 | -0.2 | 0.720678 | 0.226958 | 0.562101 | 0.999971 | 1 | 0.916386 |
| Itgb4 | A2A864 | 0.26 | 0.71 | -0.11 | -0.98 | -0.49 | -0.85 | -0.76 | -0.19 | 0.15 | 0.68 | -0.25 | -0.05 | -1.49 | -0.07 | 1.23 | -0.76 | -0.5425 | 0.1625 | -0.2425 | 0.173036 | 0.68165 | 0.697126 | 0.999971 | 1 | 0.94731 |
| Itgb5 | Q6PE70 | -1.89 | 0.6 | 1.33 | 0.37 | 0.55 | 0.4 | 0.88 | 0.84 | 0.58 | -1.59 | 1.05 | 1.74 | 0.84 | -0.15 | 1.35 | 0.06 | 0.565 | 0.3425 | 0.4225 | 0.388178 | 0.704987 | 0.552256 | 0.999971 | 1 | 0.913595 |
| Itgb8 | Q0VBD0 | -0.98 | -0.47 | 0.26 | -0.04 | -0.59 | -0.1 | -0.01 | -0.04 | -0.46 | -0.71 | -0.39 | -0.46 | 0.42 | -0.22 | 0.14 | -0.01 | 0.1225 | -0.1975 | 0.39 | 0.691918 | 0.501027 | 0.226669 | 0.999971 | 1 | 0.828236 |
| Itih1 | F8WJ05 | 0.17 | 0.99 | -0.79 | 0.37 | 1.22 | 0.4 | -0.11 | -0.5 | 0.31 | 0.32 | 0.23 | 0.63 | 1.13 | 0.95 | 0.29 | 0.43 | 0.0675 | 0.1875 | 0.515 | 0.88964 | 0.613973 | 0.222317 | 0.999971 | 1 | 0.827753 |
| Itih2 | Q61703 | 0.3 | 1.82 | 0.11 | 1.58 | 2.8 | 1.27 | 0.9 | 1.47 | 1.6 | -0.31 | 0.41 | 1.34 | 1.79 | 1.03 | 0.57 | 0.75 | 0.6575 | -0.1925 | 0.0825 | 0.253047 | 0.736214 | 0.863032 | 0.999971 | 1 | 0.982963 |
| Itih3 | Q61704 | 2.4 | 1.56 | 0.13 | 0.26 | 2.13 | 1.36 | -0.68 | -0.22 | 1.41 | 1 | -0.45 | 0.66 | 0.44 | 2.26 | 0.98 | -0.33 | -0.44 | -0.4325 | -0.25 | 0.57081 | 0.490299 | 0.720749 | 0.999971 | 1 | 0.954713 |
| Itih4 | E9Q5L2 | 2.54 | 2.5 | 1.56 | 3.74 | 3.2 | 0.82 | 3.13 | 0.61 | 1.35 | 0.13 | 2.06 | 1.74 | 2.46 | 1.07 | 0.57 | 0.78 | -0.645 | -1.265 | -1.365 | 0.402652 | 0.049743 | 0.037067 | 0.999971 | 1 | 0.700096 |
| Itih5 | Q8BJD1 | -0.43 | -0.02 | 0.4 | 0.32 | 1.08 | -0.01 | 0.1 | 0.26 | 0.16 | 0.16 | 0.55 | 1.03 | -0.1 | 0.12 | 1.51 | 0.94 | 0.29 | 0.4075 | 0.55 | 0.368998 | 0.185199 | 0.192883 | 0.999971 | 1 | 0.810547 |
| Itm2a | Q61500 | -0.04 | -0.02 | -0.28 | -0.34 | 0.41 | 0.26 | -0.49 | -0.6 | -0.31 | 0.38 | -0.4 | -0.21 | -0.3 | 0.09 | -0.64 | -0.16 | 0.065 | 0.035 | -0.0825 | 0.818874 | 0.878748 | 0.707032 | 0.999971 | 1 | 0.950182 |
| Itm2b | O89051 | -0.65 | 0.46 | -1.35 | -1.61 | 0.26 | 0.72 | -1.45 | -1.07 | 0.36 | -0.55 | -0.69 | -0.65 | 0.58 | 0.51 | -0.93 | -1.01 | 0.4025 | 0.405 | 0.575 | 0.52929 | 0.418583 | 0.338228 | 0.999971 | 1 | 0.859348 |
| Itm2c | Q91VK4 | -0.48 | 0.24 | -0.92 | -0.28 | -0.07 | 0.01 | -1.47 | -0.66 | 0.15 | -0.68 | -1.14 | -0.74 | 0.29 | 0.21 | -0.64 | -0.18 | -0.1875 | -0.2425 | 0.28 | 0.64085 | 0.500535 | 0.396966 | 0.999971 | 1 | 0.871816 |
| Itpa | Q9D892 | -0.19 | 0.8 | 1.13 | 1.32 | 0.91 | 0.92 | 0.74 | 1.18 | 0.94 | 0.57 | 0.88 | 0.65 | 0.77 | 0.92 | 0.93 | 0.94 | 0.1725 | -0.005 | 0.125 | 0.617954 | 0.988365 | 0.711339 | 0.999971 | 1 | 0.950887 |
| Itpk1 | Q8BYN3 | 0.3 | 0.16 | 0.02 | -0.21 | 0.53 | 0.58 | 0.08 | 0.37 | 0.54 | 0.15 | 0.23 | 0.69 | -0.03 | 0.14 | 0.34 | 0.07 | 0.3225 | 0.335 | 0.0625 | 0.147635 | 0.141622 | 0.751026 | 0.999971 | 1 | 0.961064 |
| Itpka | Q8R071 | -0.33 | -0.55 | 0.58 | 0.44 | -0.8 | -0.98 | 0.05 | -0.01 | -0.89 | -0.84 | 0.54 | -0.08 | -2.09 | -1.5 | -0.09 | 0.22 | -0.47 | -0.3525 | -0.9 | 0.22901 | 0.410509 | 0.140557 | 0.999971 | 1 | 0.784948 |
| Itpkc | Q7TS72 | 1.54 | -0.54 | 0.67 | 1.54 | 0.32 | 0.69 | 1.09 | -0.07 | -0.89 | 1.3 | -0.88 | 0.04 | -0.62 | 1.12 | 0.36 | 0.21 | -0.295 | -0.91 | -0.535 | 0.566708 | 0.187314 | 0.351347 | 0.999971 | 1 | 0.867656 |
| Itpr1 | P11881-4 | -0.57 | 0.14 | -2.3 | -0.64 | -1.58 | -0.45 | -0.43 | 0.39 | -1.96 | -1.64 | -1.8 | -0.15 | -0.86 | -1.6 | -0.47 | -0.05 | 0.325 | -0.545 | 0.0975 | 0.590378 | 0.382549 | 0.862499 | 0.999971 | 1 | 0.982963 |
| Itpr1_1 | P11881-2 | -0.03 | -0.04 | -0.3 | -0.03 | -0.21 | -0.45 | -0.03 | -0.33 | -0.2 | -0.11 | 0.02 | -0.17 | -0.44 | -0.56 | -0.28 | -0.24 | -0.155 | -0.015 | -0.28 | 0.411239 | 0.929268 | 0.14619 | 0.999971 | 1 | 0.791767 |
| Itpr2 | Q9Z329-3 | -0.47 | -0.61 | -0.72 | -0.72 | -0.3 | -0.63 | -0.52 | -0.72 | -0.53 | -0.35 | -0.01 | -0.19 | 0.13 | -0.66 | -0.5 | -0.32 | 0.0875 | 0.36 | 0.2925 | 0.635054 | 0.083776 | 0.212742 | 0.999971 | 1 | 0.823987 |
| Itpr3 | P70227 | -0.86 | -3.76 | 0.24 | 0.67 | -1.19 | 1.1 | 0.87 | -0.76 | -2.74 | -0.83 | 0.44 | -0.02 | -6.37 | 0.38 | -0.03 | -0.99 | 0.9325 | 0.14 | -0.825 | 0.375173 | 0.898167 | 0.618748 | 0.999971 | 1 | 0.928581 |
| Itprid2 | A2AQD6 | 0.99 | -0.99 | -1.17 | -0.74 | -1.09 | -0.8 | -1.32 | -0.74 | -3.15 | 0 | -2.04 | -2.69 | 0.32 | -1.8 | -1.61 | -1.03 | -0.51 | -1.4925 | -0.5525 | 0.303542 | 0.081329 | 0.39047 | 0.999971 | 1 | 0.871816 |
| Itsn1 | E9Q0N0 | 0.24 | -0.36 | 0 | 0.01 | -0.44 | -0.33 | -0.3 | -0.05 | -0.44 | -0.03 | -0.18 | -0.28 | -0.03 | -0.02 | -0.08 | 0.1 | -0.2525 | -0.205 | 0.02 | 0.236095 | 0.326219 | 0.917973 | 0.999971 | 1 | 0.992633 |
| Itsn2 | E9QNG1 | -0.08 | -0.21 | -0.71 | -0.77 | -0.34 | -0.38 | -0.57 | -0.89 | -0.42 | -0.2 | -0.67 | -0.82 | -0.25 | -0.35 | -1.07 | -0.81 | -0.1025 | -0.085 | -0.1775 | 0.677042 | 0.732612 | 0.527296 | 0.999971 | 1 | 0.909612 |
| Ivd | Q9JHI5 | -0.95 | -1.3 | -0.88 | -1.29 | -1.17 | -1.17 | -0.66 | -0.82 | -0.61 | -1.05 | -0.46 | -0.14 | -0.83 | -1.31 | -1.01 | -0.99 | 0.15 | 0.54 | 0.07 | 0.493401 | 0.052827 | 0.733274 | 0.999971 | 1 | 0.957047 |
| Ivns1abp | Q920Q8-4 | 0.72 | 0.37 | 1.54 | 1.59 | -0.25 | 1.48 | 1.71 | 3.06 | 0.16 | 0.06 | 1.44 | 1.4 | 1.93 | 2.14 | 3.14 | 2.57 | 0.445 | -0.29 | 1.39 | 0.514279 | 0.530243 | 0.005911 | 0.999971 | 1 | 0.582714 |
| Iws1 | A0A1D5RL' | 1.05 | 0.98 | 1.25 | 1.32 | 1 | 1.37 | 0.38 | 0.96 | 1.27 | 1.27 | 0.23 | 0.65 | 0.31 | 1.24 | 1.46 | 0.59 | -0.2225 | -0.295 | -0.25 | 0.38164 | 0.309179 | 0.403806 | 0.999971 | 1 | 0.873348 |
| Jagn1 | Q5XKN4 | 1.22 | -0.71 | -1.1 | 0.67 | -0.83 | -0.59 | 0.43 | -0.21 | -0.54 | -0.23 | 0.52 | -0.81 | -0.62 | -0.58 | -1.56 | -1.58 | -0.32 | -0.285 | -1.105 | 0.574397 | 0.62117 | 0.078637 | 0.999971 | 1 | 0.754157 |
| Jak1 | B1ASP2 | -0.42 | 0.2 | 0.14 | 0.36 | 0.11 | 0.09 | -0.27 | 0.16 | 0.19 | -0.1 | -0.01 | 0.2 | 0.62 | 0 | 0.06 | 0.42 | -0.0475 | -1.96E-17 | 0.205 | 0.838407 | 1 | 0.426174 | 0.999971 | 1 | 0.880793 |
| Jak2 | Q62120 | -0.32 | -0.67 | -1.02 | -0.93 | -0.48 | -0.41 | -0.57 | -0.9 | -1.11 | -0.26 | -0.69 | -0.76 | -0.41 | -0.39 | -0.7 | -0.77 | 0.145 | 0.03 | 0.1675 | 0.531755 | 0.907155 | 0.466203 | 0.999971 | 1 | 0.891431 |
| Jakmip1 | D3Z4U5 | -0.51 | -0.09 | -0.12 | 0.13 | -0.12 | -0.2 | -0.31 | 0.15 | 0.15 | -0.27 | 0.08 | 0.08 | -0.06 | -0.03 | 0.11 | 0.3 | 0.0275 | 0.1575 | 0.2275 | 0.897455 | 0.460673 | 0.292851 | 0.999971 | 1 | 0.847402 |
| Jakmip2 | D3YXK0 | -0.45 | -0.38 | -1.46 | -1.38 | -0.58 | -0.47 | -1.17 | -1.18 | -0.5 | -0.32 | -1.06 | -0.98 | -0.07 | -0.52 | -1.19 | -1.3 | 0.0675 | 0.2025 | 0.1475 | 0.843852 | 0.554558 | 0.709518 | 0.999971 | 1 | 0.950683 |
| Jakmip3 | Q5DTN8 | -0.92 | -1.59 | -0.63 | -0.61 | -1.26 | -1.61 | -0.91 | -0.53 | -1.38 | -0.98 | -0.62 | -0.33 | -1.26 | -1.19 | -0.6 | -0.4 | -0.14 | 0.11 | 0.075 | 0.669136 | 0.734325 | 0.813099 | 0.999971 | 1 | 0.974559 |
| Jam2 | Q9JI59 | -1.62 | -1.4 | -0.88 | -1.08 | -1.04 | -1.38 | -0.76 | -0.17 | -1.2 | -1.53 | -1.16 | -0.36 | -1.26 | -1.36 | -0.52 | -0.88 | 0.4075 | 0.1825 | 0.24 | 0.21049 | 0.552818 | 0.388731 | 0.999971 | 1 | 0.871816 |
| Jam3 | Q9D8B7 | -0.72 | -0.68 | 0.42 | 0.37 | -0.41 | -0.43 | -0.14 | -0.19 | -0.24 | -0.72 | -0.28 | -0.19 | -0.28 | -1.14 | 0.06 | -0.08 | -0.14 | -0.205 | -0.2075 | 0.668668 | 0.54678 | 0.605145 | 0.999971 | 1 | 0.924454 |
| Jcad | Q5DTX6 | -0.86 | -0.74 | -0.79 | -0.9 | -0.64 | -1.02 | -0.7 | -1.01 | -0.86 | -1.01 | -0.66 | -0.75 | -0.75 | -1.12 | -0.84 | -1.03 | -0.02 | 0.0025 | -0.1125 | 0.912649 | 0.988214 | 0.52988 | 0.999971 | 1 | 0.910069 |
| Jmjd6 | G3UVU9 | 0.09 | 1.78 | 0.64 | 0.15 | 1.3 | 1.41 | 0.08 | 0.55 | 1.13 | 0.09 | -0.16 | 0.15 | 1.35 | 1.13 | 0.91 | 0.11 | 0.17 | -0.3625 | 0.21 | 0.71824 | 0.434675 | 0.641173 | 0.999971 | 1 | 0.93301 |
| Jmy | A0A0R4J0V | 0.09 | 0.08 | -0.34 | -0.29 | 0.38 | 0.39 | 0.02 | 0.06 | 0.87 | 0.07 | -0.3 | -0.23 | 0.55 | 0.19 | -0.2 | -0.2 | 0.3275 | 0.2175 | 0.2 | 0.138873 | 0.475003 | 0.424044 | 0.999971 | 1 | 0.879749 |
| Josd1 | Q9DBJ6 | 0.23 | -0.35 | 0.01 | -0.16 | -0.28 | -0.39 | -0.46 | -0.56 | -0.41 | -0.28 | 0.29 | 0.71 | 0.49 | 0.38 | 0.96 | -0.03 | -0.355 | 0.145 | 0.5175 | 0.099291 | 0.627034 | 0.074491 | 0.999971 | 1 | 0.746069 |
| Josd2 | D3YTZ7 | 0.18 | 0 | 0.05 | -0.22 | -0.95 | 0.83 | -0.52 | -0.17 | 0.26 | 0.44 | -0.43 | -0.11 | 0.39 | 0.24 | 0.25 | 0.68 | -0.205 | 0.0375 | 0.3875 | 0.588312 | 0.876293 | 0.07405 | 0.999971 | 1 | 0.746069 |
| Jph1 | Q9ET80 | 0.91 | 0.02 | -0.52 | -0.89 | -0.64 | 0.68 | -0.68 | -0.68 | -0.54 | 0.7 | -0.95 | -1.8 | -0.58 | 0.17 | -0.93 | -0.28 | -0.21 | -0.5275 | -0.285 | 0.66358 | 0.388423 | 0.513668 | 0.999971 | 1 | 0.905815 |

| | | | | | | | | | | | | | | | | | | | | | | | | | | |
|---|---|---|---|---|---|---|---|---|---|---|---|---|---|---|---|---|---|---|---|---|---|---|---|---|---|---|
| Jph3 | Q9ET77 | 0.43 | -0.8 | -0.31 | -0.48 | -0.89 | -0.14 | -0.34 | -0.26 | -0.78 | -0.04 | -0.22 | -0.71 | -0.33 | -0.6 | -0.36 | -0.43 | -0.1175 | -0.1475 | -0.14 | 0.708461 | 0.646391 | 0.622899 | 0.999971 | 1 | 0.929759 |
| Jph4 | Q80WT0 | -0.92 | 0.3 | -0.1 | -0.22 | 0.09 | 0 | 0.79 | 0.21 | -0.03 | -0.28 | -0.34 | -0.33 | 0.68 | 0.07 | -0.43 | -0.17 | 0.5075 | -0.01 | 0.2725 | 0.132691 | 0.971358 | 0.436596 | 0.999971 | 1 | 0.882419 |
| Jpt1 | P97825 | 1.39 | 0.18 | 0.42 | 0.28 | 0.18 | 0.81 | 0.7 | 0.81 | 0.02 | 0.67 | 0.55 | 0.37 | 0.73 | 0.79 | 0.87 | 0.86 | 0.0575 | -0.165 | 0.245 | 0.856983 | 0.603787 | 0.411653 | 0.999971 | 1 | 0.875847 |
| Jpt2 | Q6PGH2 | 2.52 | 0.25 | 0.16 | 0.6 | 0.4 | 0.36 | 0.92 | -0.13 | 0.8 | 2.22 | 0.1 | 0.49 | 1.52 | 0.69 | 0.77 | 0.56 | -0.495 | 0.02 | 0.0025 | 0.376197 | 0.975745 | 0.996353 | 0.999971 | 1 | 1 |
| Jtb | D3Z4D1 | 2.12 | 1.95 | 1.37 | 1.32 | 1.94 | 3.02 | 0.54 | 1.14 | 2.44 | 3.06 | 0.31 | 1.12 | 2.44 | 3.21 | 1.51 | 1.8 | -0.03 | 0.0425 | 0.55 | 0.954689 | 0.943622 | 0.201445 | 0.999971 | 1 | 0.815758 |
| Junb | P09450 | 3.26 | 4.48 | 2.09 | 2.08 | 3.86 | 4.02 | 2.36 | 1.51 | 1.94 | 3.92 | 0.01 | 0.32 | 1.59 | 2.04 | 0.96 | 0.49 | -0.04 | -1.43 | -1.7075 | 0.957436 | 0.161273 | 0.019528 | 0.999971 | 1 | 0.635388 |
| Jund | P15066 | 1.44 | 1.96 | 1.83 | 1.9 | 1.34 | 2.48 | 1.27 | 1.5 | 1.75 | 1.87 | 1.91 | 1.07 | 1.04 | 2.2 | 0.94 | 2.88 | -0.135 | -0.1325 | -0.0175 | 0.664948 | 0.603381 | 0.969233 | 0.999971 | 1 | 0.998787 |
| Jup | Q02257 | -0.71 | -0.77 | -1.16 | -1.29 | -0.76 | -1.02 | -1.03 | -1.34 | -1 | -0.8 | -0.92 | -1.06 | -1.15 | -1.02 | -1.5 | -1.53 | -0.055 | 0.0375 | -0.3175 | 0.807968 | 0.8543 | 0.19148 | 0.999971 | 1 | 0.810547 |
| Kalrn | D3Z534 | -3.05 | -1.64 | -2.41 | -2.15 | -0.59 | -3.21 | -1.6 | -3 | -0.42 | -4.01 | -2.67 | -1.06 | 0.69 | -2.16 | -3.8 | -1.81 | 0.2125 | 0.2725 | 0.5425 | 0.73325 | 0.727527 | 0.538806 | 0.999971 | 1 | 0.910077 |
| Kalrn_1 | A2CG49 | -0.51 | -0.4 | -0.38 | -0.58 | -0.56 | -0.46 | 0.12 | -0.87 | -0.29 | -0.41 | -0.2 | -0.29 | -0.18 | -0.77 | -0.91 | -0.71 | 0.025 | 0.17 | -0.175 | 0.918089 | 0.311524 | 0.424518 | 0.999971 | 1 | 0.879931 |
| Kalrn_2 | A2CG49-9 | -1.99 | -0.47 | 0.72 | 0.18 | 0.5 | -1.57 | 2.47 | -1.07 | 1.32 | -1.43 | 1.73 | 2.12 | 1.71 | -1.53 | 1.84 | 0.52 | 0.4725 | 1.325 | 1.025 | 0.62711 | 0.16569 | 0.260735 | 0.999971 | 1 | 0.841256 |
| Kank1 | E9Q238 | 4.72 | 0.85 | 0.96 | 2.22 | 0.69 | 1.21 | 1.77 | 0.85 | 0 | 2.89 | 0.33 | -0.71 | -0.82 | 0.19 | 0.61 | 0.73 | -1.0575 | -1.56 | -2.01 | 0.226548 | 0.170855 | 0.042953 | 0.999971 | 1 | 0.701725 |
| Kank2 | Q8BX02 | 1.01 | 0.34 | 0.15 | 0.59 | 2.11 | 0.23 | 0.5 | -0.11 | 0.7 | 1.78 | 0.43 | 1.5 | -0.35 | 0.81 | 0.46 | -0.25 | 0.16 | 0.58 | -0.355 | 0.743914 | 0.135986 | 0.30494 | 0.999971 | 1 | 0.847402 |
| Kank3 | Q9Z1P7 | 0.43 | 0.74 | -0.27 | 0.96 | 0.66 | 0.52 | -0.07 | 0.19 | 0.87 | 0.39 | 0.28 | 0.29 | 0.66 | 0.85 | 0.39 | 0.56 | -0.14 | -0.0075 | 0.15 | 0.66075 | 0.980528 | 0.614274 | 0.999971 | 1 | 0.927517 |
| Kank4 | Q6P9J5 | 2.51 | 0.35 | -0.64 | 0.5 | 0.21 | 0.18 | -0.25 | -0.46 | 0.69 | 2.43 | -0.47 | -0.96 | -0.38 | 0.63 | -0.38 | -0.16 | -0.76 | -0.2575 | -0.7525 | 0.240179 | 0.775864 | 0.25841 | 0.999971 | 1 | 0.839012 |
| Kars | Q8R2P8 | 0.57 | 1.36 | 1.19 | 1.31 | 1.29 | 1.21 | 0.96 | 1.48 | 1.59 | 0.91 | 1.42 | 1.59 | 1.38 | 1.59 | 1.62 | 1.68 | 0.1275 | 0.27 | 0.46 | 0.602904 | 0.320736 | 0.073832 | 0.999971 | 1 | 0.746069 |
| Kars1 | Q99MN1 | 1.57 | 1.33 | 0.57 | 1.7 | 0.43 | 0.96 | 0.33 | 1.87 | 1.24 | 1.5 | 1.06 | 0.62 | 1.17 | 1.69 | 1.43 | 1.78 | -0.395 | -0.1875 | 0.225 | 0.351065 | 0.557144 | 0.457386 | 0.999971 | 1 | 0.887696 |
| Kat2a | Q6P3Z8 | 0.45 | 0.09 | 0.02 | -0.04 | -0.37 | -0.25 | -0.13 | -0.55 | -0.79 | 0.56 | -0.34 | -0.32 | 0.05 | -0.15 | 0.09 | 0.08 | -0.455 | -0.3525 | -0.1125 | 0.045871 | 0.271198 | 0.560227 | 0.999971 | 1 | 0.915694 |
| Kat5 | A0A494BAI | 5.53 | 2.21 | 0.48 | 0.27 | 1.76 | 2.74 | -0.41 | 1.02 | 2.45 | 4.01 | -0.12 | -0.03 | -1.38 | 0.94 | 0.12 | -0.36 | -0.845 | -0.545 | -2.2925 | 0.498249 | 0.700367 | 0.076166 | 0.999971 | 1 | 0.748444 |
| Kat7 | Q5SVQ0-3 | 3.38 | 0.84 | 0.89 | 1.09 | 0.14 | 1.43 | 1.72 | -0.22 | 0.22 | 4.49 | -0.42 | -0.49 | 0.55 | 1.18 | -0.44 | -0.16 | -0.7825 | -0.6 | -1.2675 | 0.281573 | 0.619419 | 0.07743 | 0.999971 | 1 | 0.752059 |
| Katnal1 | Q8K0T4 | 0.42 | -0.1 | -0.03 | 0.21 | 0.14 | 0.15 | -0.14 | 0.26 | -0.03 | 0.29 | -0.08 | -0.02 | 0.31 | 0.32 | 0.17 | 0.58 | -0.0225 | -0.085 | 0.22 | 0.911444 | 0.671558 | 0.295383 | 0.999971 | 1 | 0.847402 |
| Katnb1 | Q8BG40 | 0.05 | 0.19 | -0.83 | -0.76 | 0.13 | 0.08 | -0.18 | -0.59 | 0.24 | 0.01 | -0.39 | -0.68 | 0.54 | 0.29 | -0.51 | -0.63 | 0.1975 | 0.1325 | 0.26 | 0.536756 | 0.692706 | 0.500366 | 0.999971 | 1 | 0.904223 |
| Kazn | G3UY74 | -0.09 | -0.66 | -0.11 | 0.15 | -0.34 | -0.93 | -0.21 | -0.51 | 0.17 | 0.17 | -0.34 | -0.53 | -1.17 | -0.51 | 0.41 | 0.43 | -0.32 | 0.045 | -0.0325 | 0.233644 | 0.866008 | 0.936046 | 0.999971 | 1 | 0.995618 |
| Kazn_1 | Q69ZS8-2 | -0.13 | 0.09 | -0.45 | 0.03 | 0.02 | -0.01 | -0.2 | -0.14 | 0.28 | 0.04 | -0.24 | -0.14 | 0.13 | 0.16 | -0.24 | -0.55 | 0.0325 | 0.1 | -0.01 | 0.867423 | 0.638807 | 0.966828 | 0.999971 | 1 | 0.998787 |
| Kbtbd11 | Q8BNW9 | -0.1 | 0.08 | -0.26 | -0.04 | 0.03 | -0.26 | -0.34 | -0.08 | -0.4 | -0.29 | -0.64 | -0.35 | -0.47 | 0.03 | -0.29 | -0.46 | -0.0825 | -0.34 | -0.2175 | 0.65553 | 0.083678 | 0.288613 | 0.999971 | 1 | 0.847402 |
| Kbtbd2 | G3X9X1 | 0.5 | 0.87 | 1.19 | 1.16 | 0.73 | 0.87 | 0.96 | 1.01 | 1.02 | 0.55 | 1.28 | 0.15 | 0.48 | 0.88 | 1.24 | 1.13 | -0.0375 | -0.18 | 0.0025 | 0.863049 | 0.557773 | 0.992262 | 0.999971 | 1 | 0.99925 |
| Kbtbd3 | Q8BHI4 | -1.04 | -0.4 | -1.09 | -1.02 | -0.59 | -0.32 | -0.9 | -0.4 | -0.57 | -0.94 | -0.95 | -1.12 | 0 | -0.38 | -0.69 | -0.44 | 0.335 | -0.0075 | 0.51 | 0.187321 | 0.974352 | 0.064646 | 0.999971 | 1 | 0.726597 |
| Kbtbd4 | Q8R179-2 | 0.86 | -0.84 | 0.57 | 0.29 | -0.89 | -0.43 | -0.3 | 0.29 | -0.63 | 0.05 | 1.06 | 0.93 | 0.24 | -0.12 | 0.24 | 0.68 | -0.5525 | 0.1325 | 0.04 | 0.211844 | 0.793793 | 0.918468 | 0.999971 | 1 | 0.992633 |
| Kbtbd7 | G5E8C2 | -0.16 | 0.14 | -0.87 | -0.72 | 0.44 | -0.05 | -0.56 | -0.07 | -0.03 | -0.64 | -0.43 | -0.17 | 0.21 | -0.12 | -0.56 | -0.58 | 0.3425 | 0.085 | 0.14 | 0.294857 | 0.766118 | 0.652968 | 0.999971 | 1 | 0.934005 |
| Kcmf1 | Q80UY2 | 0.67 | -0.24 | 0.28 | 0.72 | -0.94 | -0.4 | 2.34 | 1.02 | -0.84 | 0.38 | 0.27 | -0.26 | -2.21 | -1.06 | -0.49 | -1.14 | 0.1475 | -0.47 | -1.5825 | 0.832423 | 0.203833 | 0.003525 | 0.999971 | 1 | 0.577271 |
| Kcna1 | P16388 | -0.7 | -0.46 | -0.4 | -0.54 | -0.78 | -0.62 | -1.13 | -0.7 | -1.26 | -0.83 | -0.91 | -0.73 | -0.98 | -0.5 | -0.85 | -1.11 | -0.2825 | -0.4075 | -0.335 | 0.169517 | 0.059293 | 0.126302 | 0.999971 | 1 | 0.783071 |
| Kcna2 | P63141 | -0.85 | -1.65 | -1.41 | -1.21 | -1.81 | -1.22 | -1.82 | -1.49 | -1.85 | -1.25 | -1.99 | -1.63 | -2.44 | -1.54 | -1.53 | -1.06 | -0.305 | -0.4 | -0.3625 | 0.242439 | 0.145672 | 0.293842 | 0.999971 | 1 | 0.847402 |
| Kcna3 | P16390 | -1.45 | -1.06 | -1.17 | -0.91 | -0.85 | -1.19 | -1.63 | -1.25 | -0.85 | -0.85 | -1.45 | -1.05 | -0.58 | -0.89 | -0.21 | -1.21 | -0.0825 | 0.0975 | 0.425 | 0.72386 | 0.661315 | 0.135016 | 0.999971 | 1 | 0.784948 |
| Kcna4 | Q61423 | -0.74 | -0.83 | -3.55 | -2.71 | -0.94 | -0.74 | -2.58 | -2.71 | -0.55 | -1.01 | -2.96 | -3.12 | -0.52 | -1.2 | -2.81 | -3.72 | 0.215 | 0.0475 | -0.105 | 0.784366 | 0.956281 | 0.907755 | 0.999971 | 1 | 0.991405 |
| Kcna6 | Q61923 | -0.89 | -1.07 | -1.27 | -1.18 | -1.11 | -0.76 | -1.42 | -1.23 | -0.86 | -0.7 | -1.31 | -1.28 | -0.91 | -0.76 | -1.04 | -0.93 | -0.0275 | 0.065 | 0.1925 | 0.896274 | 0.763755 | 0.301743 | 0.999971 | 1 | 0.847402 |
| Kcnab1 | P63143 | -1.01 | -0.36 | -0.7 | -0.22 | -0.19 | -0.54 | -0.72 | -0.22 | -0.58 | -1.27 | -0.53 | -0.33 | -0.14 | -0.63 | -0.37 | -0.32 | 0.155 | -0.105 | 0.2075 | 0.536118 | 0.712227 | 0.394969 | 0.999971 | 1 | 0.871816 |
| Kcnab2 | P62482 | -0.88 | -0.6 | -1.1 | -1.55 | -0.8 | -0.53 | -1.48 | -1.28 | -0.99 | -1 | -1.45 | -1.36 | -0.62 | -0.96 | -1.21 | -1.5 | 0.01 | -0.1675 | -0.04 | 0.973663 | 0.519102 | 0.889361 | 0.999971 | 1 | 0.987681 |
| Kcnab3 | Q8VD73 | 0.37 | 0.33 | 0.05 | -1.37 | 0.13 | 0.35 | -0.45 | -1.49 | -0.6 | 0.15 | -0.22 | -0.71 | -0.51 | -0.13 | -1.13 | -1.01 | -0.21 | -0.19 | -0.54 | 0.695997 | 0.661477 | 0.244613 | 0.999971 | 1 | 0.837316 |
| Kcnb1 | Q03717 | -0.51 | -0.87 | -0.42 | -0.62 | -1.13 | -0.72 | 0.09 | -1.01 | -0.97 | -0.95 | -0.86 | -1.08 | -0.31 | -1.17 | -0.98 | -0.79 | -0.0875 | -0.36 | -0.2075 | 0.770806 | 0.071591 | 0.400651 | 0.999971 | 1 | 0.871816 |
| Kcnb2 | A6H8H5 | -0.76 | -0.69 | -0.07 | -0.65 | -1.34 | -0.89 | -0.64 | -0.59 | -0.88 | -0.65 | -0.17 | -0.78 | 0.03 | -0.82 | -0.55 | -0.39 | -0.3225 | -0.0775 | 0.11 | 0.232264 | 0.756687 | 0.676169 | 0.999971 | 1 | 0.941862 |
| Kcnc1 | P15388-3 | -0.37 | -0.73 | 0.12 | -0.03 | -1.11 | -0.08 | -0.07 | -0.44 | -1.2 | -0.46 | -0.4 | -0.84 | -0.79 | -0.65 | -0.35 | -0.22 | -0.1725 | -0.4725 | -0.25 | 0.584808 | 0.118495 | 0.343067 | 0.999971 | 1 | 0.861474 |
| Kcnc2 | Q14B80 | -1.14 | -0.47 | -0.37 | -0.9 | -1.02 | 0.16 | -0.79 | -0.03 | -1.01 | -0.8 | -0.61 | -0.76 | 0.19 | -0.2 | -0.51 | -0.11 | 0.3 | -0.075 | 0.5625 | 0.383685 | 0.747864 | 0.052903 | 0.999971 | 1 | 0.725348 |
| Kcnc3 | E9QKI2 | -0.52 | -0.29 | -0.68 | -0.47 | -0.09 | 0.42 | -1.42 | -1.28 | -1.09 | -0.66 | -0.99 | -0.93 | 0.84 | 0.08 | -1.45 | -0.57 | -0.1025 | -0.4275 | 0.215 | 0.812581 | 0.045149 | 0.643621 | 0.999971 | 1 | 0.93301 |
| Kcnc4 | A0A571BG | -0.04 | -0.64 | -1.22 | -0.82 | -0.73 | 0.06 | -0.74 | -0.47 | -0.57 | 0.53 | -0.96 | -1.44 | -0.1 | -0.24 | -0.72 | 0.03 | 0.21 | 0.07 | 0.4225 | 0.507751 | 0.878324 | 0.187334 | 0.999971 | 1 | 0.809025 |
| Kcnd1 | Q03719 | -2.64 | -1.57 | -1.28 | -1.18 | -0.78 | -2.53 | -1.96 | -0.93 | -0.8 | -2.59 | -1.45 | -1 | -0.04 | -1.99 | -0.81 | -0.97 | 0.1175 | 0.2075 | 0.715 | 0.813319 | 0.671845 | 0.166975 | 0.999971 | 1 | 0.800215 |
| Kcnd2 | Q9Z0V2 | -1.01 | -0.83 | -0.82 | -0.54 | -0.85 | -1.06 | -0.87 | -0.56 | -0.75 | -0.8 | -0.58 | -0.84 | -0.45 | -0.87 | -0.98 | -0.66 | -0.035 | 0.0575 | 0.06 | 0.86105 | 0.752753 | 0.771815 | 0.999971 | 1 | 0.963498 |
| Kcnd3 | Q9Z0V1 | -0.76 | -1.28 | -1.83 | -1.8 | -0.9 | -1.05 | -2.02 | -1.37 | -1.01 | -1.09 | -2.43 | -1.5 | -0.83 | -1.42 | -1.77 | -1.59 | 0.0825 | -0.09 | 0.015 | 0.813074 | 0.820356 | 0.963272 | 0.999971 | 1 | 0.998721 |
| Kcne5 | Q9QZ26 | -4.37 | -3.25 | -2.43 | -2.23 | -2.39 | -3.04 | -1.65 | -1.46 | -1.51 | -2.8 | -2.06 | -1.85 | -1.6 | -1.58 | -0.66 | -2.41 | 0.935 | 1.015 | 1.5075 | 0.120111 | 0.077252 | 0.022565 | 0.999971 | 1 | 0.652043 |
| Kcnf1 | E9PYC0 | -0.3 | -1.43 | -0.19 | -0.66 | -2.36 | -1.41 | -0.47 | -0.35 | -2.25 | -1.13 | -0.31 | -1.59 | -0.19 | -2.37 | -0.79 | -0.57 | -0.5025 | -0.675 | -0.335 | 0.334186 | 0.171609 | 0.520093 | 0.999971 | 1 | 0.906681 |
| Kcnh1 | A0A1L1M1 | 0.49 | -0.23 | 0.18 | -0.36 | -0.24 | -0.11 | -0.3 | -0.07 | -0.46 | 0.58 | -0.14 | -0.02 | 0.16 | 0.09 | -0.06 | -0.09 | -0.2 | -0.03 | 0.005 | 0.407294 | 0.920145 | 0.983229 | 0.999971 | 1 | 0.99925 |
| Kcnh2 | Q53Z09 | 1.69 | -1.04 | -0.25 | -0.69 | -1.37 | -1.66 | -0.15 | -0.46 | -2.03 | 0.24 | -0.66 | 0.29 | -1.97 | -1.22 | -0.34 | -0.34 | -0.8375 | -0.4675 | -0.895 | 0.214583 | 0.532639 | 0.197695 | 0.999971 | 1 | 0.813335 |
| Kcnh3 | Q9WVJ0 | -0.69 | 0.26 | -0.69 | -0.75 | -0.08 | 0.25 | -0.97 | -0.63 | -0.55 | 0.12 | -0.87 | -0.86 | 1.04 | -0.69 | -0.93 | -0.33 | 0.11 | -0.0725 | 0.24 | 0.758704 | 0.828283 | 0.612371 | 0.999971 | 1 | 0.927335 |
| Kcnh5 | Q920E3 | -0.66 | 0.22 | -0.56 | -0.64 | -0.34 | -0.36 | 0.01 | -1.23 | 0.18 | -0.52 | -0.24 | -0.37 | 0.86 | -0.14 | -1.23 | -1.82 | -0.07 | 0.1725 | -0.1725 | 0.834875 | 0.535329 | 0.765574 | 0.999971 | 1 | 0.962405 |
| Kcnh7 | Q9ER47 | -0.69 | -0.95 | -0.31 | -0.2 | -0.99 | -0.74 | -0.67 | -0.29 | -1.26 | -0.78 | -0.08 | -0.38 | -0.88 | -1.25 | -0.54 | -0.02 | -0.135 | -0.0875 | -0.135 | 0.59569 | 0.779833 | 0.671941 | 0.999971 | 1 | 0.939563 |
| Kcnip1 | Q3YAB5 | -2.44 | -2.12 | -0.02 | -0.25 | -2.05 | -1.57 | -0.93 | -0.22 | -1.54 | -1.74 | -0.78 | -0.35 | -1.58 | -1.81 | 0.02 | 0.37 | 0.015 | 0.105 | 0.4575 | 0.982128 | 0.870356 | 0.547309 | 0.999971 | 1 | 0.911722 |
| Kcnip2 | Q3YAB3 | -1.41 | -0.99 | -0.71 | -0.67 | -1.11 | -0.83 | -1.25 | -0.51 | -0.76 | -1.05 | -0.72 | -0.55 | -0.11 | -0.51 | -0.88 | -0.89 | 0.02 | 0.175 | 0.3475 | 0.938601 | 0.460442 | 0.221782 | 0.999971 | 1 | 0.827753 |
| Kcnip3 | Q9QXT8 | -2 | 0.02 | -0.66 | -0.42 | -0.05 | 0.05 | -0.8 | -0.64 | -0.09 | -0.43 | -0.01 | 0.07 | 0.46 | -0.24 | -0.51 | -0.22 | 0.405 | 0.65 | 0.6375 | 0.383078 | 0.153577 | 0.183411 | 0.999971 | 1 | 0.809025 |
| Kcnip4 | Q6PHZ8 | -0.73 | 0.07 | -0.22 | 0.26 | -0.16 | -0.12 | -0.62 | -0.07 | 0.17 | -0.09 | -0.16 | -0.75 | 0.75 | 0.09 | -0.37 | -0.05 | -0.0875 | -0.0525 | 0.26 | 0.746188 | 0.860302 | 0.428006 | 0.999971 | 1 | 0.880793 |
| Kcnj10 | Q9JM63 | -0.31 | -1.81 | -1.94 | -2.2 | -1.76 | -0.57 | -1.78 | -2.93 | -1.52 | -1.01 | -2.89 | -2.53 | -1.98 | 0.08 | -1.84 | -2.45 | -0.195 | -0.4225 | 0.0175 | 0.740618 | 0.460062 | 0.97813 | 0.999971 | 1 | 0.99925 |
| Kcnj11 | Q61743 | -0.46 | -0.5 | -0.56 | -1 | -0.69 | -0.46 | -1.02 | -0.79 | -1.17 | -0.14 | -0.45 | -0.57 | -1.37 | -0.64 | -0.92 | -0.32 | -0.11 | 0.0475 | -0.1825 | 0.614217 | 0.859283 | 0.510766 | 0.999971 | 1 | 0.905815 |
| Kcnj12 | B1ATI1 | -0.47 | -0.38 | -0.8 | -1.03 | -0.15 | -0.43 | -0.74 | -1.51 | 0.37 | 0.15 | -0.07 | -0.68 | 0.54 | -0.76 | -1.41 | -0.9 | -0.0375 | 0.6125 | 0.0375 | 0.909222 | 0.056073 | 0.928719 | 0.999971 | 1 | 0.994278 |
| Kcnj16 | Q9Z307 | 0.78 | -1.05 | -1.45 | -0.61 | -0.68 | -0.87 | 0.01 | -2.45 | 0.16 | 0.38 | -1.58 | -0.51 | -0.26 | -0.5 | -1.19 | -0.37 | -0.415 | 0.195 | 0.0025 | 0.525322 | 0.746397 | 0.995941 | 0.999971 | 1 | 1 |
| Kcnj3 | P63250 | -0.82 | -0.65 | -0.54 | -1.17 | -1.11 | -0.61 | -1.02 | -1 | -1.03 | -0.89 | -0.75 | -1.04 | -0.52 | -0.92 | -1.37 | -0.53 | -0.14 | -0.1325 | -0.04 | 0.530136 | 0.521521 | 0.880319 | 0.999971 | 1 | 0.985441 |
| Kcnj4 | Q8R435 | -1.62 | -1.97 | -1.08 | -0.75 | -1.88 | -2.31 | -1.29 | -0.85 | -1.86 | -1.78 | -0.89 | -1.2 | -1.51 | -2.02 | -0.8 | -0.67 | -0.2275 | -0.0775 | 0.105 | 0.575114 | 0.825843 | 0.793098 | 0.999971 | 1 | 0.969851 |
| Kcnj6 | Q0VB45 | -0.56 | -0.4 | -0.35 | -0.39 | -0.58 | -0.35 | -0.07 | -0.74 | -0.65 | -0.68 | -0.71 | -1.17 | -0.64 | -0.93 | -1.51 | -0.67 | -0.01 | -0.3775 | -0.5125 | 0.961284 | 0.075839 | 0.057964 | 0.999971 | 1 | 0.725348 |
| Kcnj9 | P48543 | 2.22 | -0.07 | -0.1 | 0.8 | -0.34 | 0.87 | 0.39 | -0.13 | 0.03 | 1.55 | -0.9 | -0.94 | -1.64 | -0.53 | -1.06 | -0.61 | -0.515 | -0.7775 | -1.6725 | 0.367974 | 0.299815 | 0.013758 | 0.999971 | 1 | 0.613095 |
| Kcnk1 | O08581 | -2.06 | -2.97 | -1.76 | -1.52 | -2.56 | -3.01 | -1.52 | -0.7 | -2.61 | -2.33 | -1.52 | -1.9 | -1.41 | -2.36 | -0.77 | -0.77 | 0.13 | -0.0125 | 0.75 | 0.81612 | 0.974005 | 0.130935 | 0.999971 | 1 | 0.784948 |
| Kcnk10 | Q8BUW1 | 0.8 | -0.99 | -0.64 | 0.33 | -0.22 | 0.29 | -1.25 | 0.53 | -1.01 | 0.32 | -1.09 | -0.58 | 0.1 | 0.14 | -1.19 | -0.2 | -0.0375 | -0.465 | -0.1625 | 0.943393 | 0.357231 | 0.737597 | 0.999971 | 1 | 0.959801 |
| Kcnk2 | Q6P6P9 | -0.26 | -1.07 | -0.27 | -0.81 | -1.9 | -0.93 | -1.03 | -0.52 | -1.41 | 0.48 | -0.29 | -0.67 | -0.25 | -0.48 | 0.02 | 0.01 | -0.4925 | 0.13 | 0.4275 | 0.181036 | 0.757878 | 0.126122 | 0.999971 | 1 | 0.783071 |
| Kcnk4 | O88454 | 1.14 | -0.87 | 1.52 | 0.78 | -1.19 | -0.04 | 0.37 | -0.65 | -0.97 | 0.35 | -1.76 | -1.68 | -0.67 | -0.49 | 0.69 | -0.96 | -1.02 | -1.6575 | -1 | 0.103359 | 0.030832 | 0.114914 | 0.999971 | 1 | 0.78268 |
| Kcnma1 | E3VRY9 | -1.55 | -0.69 | -0.58 | -0.75 | -0.74 | -0.72 | -0.76 | -0.85 | -0.83 | -1.11 | -0.81 | -0.55 | -0.77 | -0.93 | -1.08 | -0.68 | 0.125 | 0.0675 | 0.0275 | 0.621108 | 0.801476 | 0.916457 | 0.999971 | 1 | 0.992633 |
| Kcnmb4 | Q9JIN6 | -0.79 | 1.64 | -0.12 | -0.09 | 0.75 | 1.2 | -0.31 | -0.26 | 0.78 | -0.07 | 0.03 | -0.29 | 1.48 | 0.45 | -0.68 | -0.31 | 0.185 | -0.0475 | 0.075 | 0.752491 | 0.928171 | 0.906879 | 0.999971 | 1 | 0.991405 |
| Kcnn2 | P58390 | -1.76 | -1.24 | -1.63 | -1.69 | -0.96 | -1.46 | -0.92 | -2.02 | -0.66 | -1.57 | -1.01 | -0.8 | -0.29 | -1.62 | -2.29 | -1.95 | 0.24 | 0.57 | 0.0425 | 0.422565 | 0.04926 | 0.921037 | 0.999971 | 1 | 0.992702 |
| Kcnn3 | P58391 | 1.24 | -0.42 | -2.37 | -1.04 | -0.65 | -1.22 | -0.98 | -1.7 | -0.77 | 0.39 | -2.19 | -0.92 | -1.24 | -1.27 | -1.16 | -1.33 | -0.49 | -0.225 | -0.6025 | 0.49346 | 0.786176 | 0.386758 | 0.999971 | 1 | 0.871816 |
| Kcnq2 | B7ZBW2 | 0.38 | -0.49 | -0.61 | -0.58 | -1.04 | -0.66 | -0.76 | -0.5 | -1.16 | -0.34 | -0.48 | -0.78 | -0.48 | 0.12 | -0.55 | -0.64 | -0.415 | -0.365 | -0.0625 | 0.161195 | 0.250265 | 0.83575 | 0.999971 | 1 | 0.979185 |
| Kcnq2_1 | B7ZBW1 | -0.52 | -2.02 | -0.99 | -1.4 | -1.58 | -0.9 | -2.32 | -1.31 | -1.3 | -0.36 | -1.13 | -1.39 | -1.2 | -1.06 | -1.88 | -1.71 | -0.295 | 0.1875 | -0.23 | 0.483267 | 0.625933 | 0.533509 | 0.999971 | 1 | 0.910069 |
| Kcnq3 | Q8K3F6 | 0.1 | -0.47 | -0.47 | -0.3 | -0.51 | -0.31 | -0.79 | -0.42 | -0.65 | -0.06 | -0.33 | -0.37 | -0.02 | 0.01 | -0.15 | -0.08 | -0.2225 | -0.0675 | 0.225 | 0.317488 | 0.760263 | 0.276509 | 0.999971 | 1 | 0.843794 |
| Kcnq5 | G3UYG5 | 0.45 | -0.89 | -0.07 | -0.2 | -0.99 | -0.24 | 0.8 | -1.39 | -1.44 | -0.19 | -0.2 | -0.99 | -1.12 | -1.21 | -1.12 | -1.28 | -0.2775 | -0.5275 | -1.005 | 0.591583 | 0.207264 | 0.006995 | 0.999971 | 1 | 0.582714 |
| Kcns2 | O35174 | -0.98 | -0.3 | -0.17 | -0.16 | -0.94 | -1.21 | -0.46 | 0.47 | -2.23 | -0.19 | 0.07 | -0.69 | -1.24 | -0.8 | -1.02 | 0.22 | -0.1325 | -0.3575 | -0.3075 | 0.74055 | 0.491143 | 0.412379 | 0.999971 | 1 | 0.875881 |
| Kcnt1 | A2AHB9 | -0.78 | -0.78 | -0.45 | -0.64 | -0.77 | -0.79 | -1.06 | -0.45 | -0.88 | -0.8 | -0.79 | -0.86 | -0.53 | -0.46 | -0.58 | -0.39 | -0.105 | -0.17 | 0.1725 | 0.60774 | 0.327786 | 0.339528 | 0.999971 | 1 | 0.860218 |
| Kcnt2 | D3Z592 | -0.75 | -0.73 | -0.44 | -0.53 | -1.25 | -0.56 | -0.93 | -0.73 | -1.2 | -0.67 | -0.48 | -0.99 | -1.06 | -0.84 | -0.75 | -0.73 | -0.255 | -0.2225 | -0.2325 | 0.253217 | 0.324383 | 0.226185 | 0.999971 | 1 | 0.828225 |
| Kcnv1 | Q8BZN2 | -1.7 | -0.45 | -1.54 | -0.52 | 0.14 | 0.02 | -1.08 | -1.19 | -0.08 | -1.24 | -1.28 | -1.58 | 0.21 | -0.25 | -0.62 | -0.15 | 0.525 | 0.0075 | 0.85 | 0.264602 | 0.986423 | 0.04123 | 0.999971 | 1 | 0.700838 |
| Kctd1 | A0A3Q4EG | -0.96 | 1.05 | 1.37 | 1.54 | 1.39 | 0.71 | 1.29 | 1.39 | 1.21 | -0.38 | 1.66 | 1.68 | 1.65 | 0.45 | 1.2 | 1.48 | 0.445 | 0.2925 | 0.445 | 0.429253 | 0.672521 | 0.453268 | 0.999971 | 1 | 0.887217 |
| Kctd10 | Q922M3 | -0.53 | -0.18 | -0.04 | 0.07 | -0.14 | -0.19 | 0.58 | -0.06 | -0.82 | -0.49 | 2.53 | 1.16 | -0.33 | -0.04 | -0.33 | -0.01 | 0.2175 | 0.765 | -0.0075 | 0.39455 | 0.301814 | 0.971427 | 0.999971 | 1 | 0.99925 |
| Kctd12 | A0A0R4J2E | 0.04 | -0.23 | -0.29 | -0.25 | -0.41 | -0.1 | -0.07 | -0.25 | -0.25 | -0.02 | -0.56 | -0.56 | -0.19 | 0.03 | -0.18 | -0.21 | -0.025 | -0.165 | 0.045 | 0.891475 | 0.42506 | 0.800338 | 0.999971 | 1 | 0.970888 |
| Kctd12b | Q8C7J6 | 0.08 | -0.08 | -0.59 | 0.23 | 0.31 | 0.13 | -0.37 | -0.27 | 0.08 | 0.77 | -1.12 | -1.16 | -0.67 | 0.49 | -0.61 | -0.73 | 0.04 | -0.2675 | -0.29 | 0.878991 | 0.575973 | 0.402426 | 0.999971 | 1 | 0.872434 |
| Kctd13 | Q8BGV7 | -0.09 | 0.15 | 0.35 | 0.81 | 0.25 | -0.08 | 0.18 | 0.96 | 0.48 | 0.02 | 0.41 | 0.25 | 0.26 | 0.09 | -0.18 | 0.83 | 0.0225 | -0.015 | -0.055 | 0.94039 | 0.951051 | 0.852966 | 0.999971 | 1 | 0.98187 |
| Kctd16 | Q5DTY9 | -1.55 | -1.94 | -1.96 | -1.98 | -2.12 | -1.59 | -1.58 | -2.43 | -2 | -1.85 | -1.64 | -1.88 | -1.83 | -1.87 | -2.02 | -1.67 | -0.0725 | 0.015 | 0.01 | 0.778597 | 0.936845 | 0.958467 | 0.999971 | 1 | 0.998622 |
| Kctd17 | E0CYQ0 | -1.25 | 0.2 | 0.97 | 0.02 | 0.03 | 0.3 | 0.84 | 0.42 | 0.22 | -0.8 | 0.34 | 1.14 | 0.42 | -0.17 | 0.75 | -0.07 | 0.4125 | 0.24 | 0.2475 | 0.38012 | 0.670494 | 0.604953 | 0.999971 | 1 | 0.924454 |
| Kctd2 | Q4VBE7 | -1.83 | -0.65 | -1.02 | -1.49 | -0.51 | -0.8 | -0.85 | -1.13 | -0.32 | -2.25 | -1.01 | -1 | -0.52 | -2.06 | -0.97 | -1.42 | 0.425 | 0.1025 | 0.005 | 0.177997 | 0.820339 | 0.99004 | 0.999971 | 1 | 0.99925 |
| Kctd21 | B2RTJ2 | 0.2 | 0.63 | -0.07 | -0.31 | 0.98 | 0.78 | 0.67 | -0.11 | 0.35 | 0.19 | 0.56 | 0.45 | 0.57 | 0.13 | -0.45 | -0.48 | 0.4675 | 0.275 | -0.17 | 0.16405 | 0.278418 | 0.602937 | 0.999971 | 1 | 0.924454 |
| Kctd3 | Q8BFX3 | 0.02 | -0.27 | 0.67 | 0.43 | -0.44 | -0.1 | 0.49 | 0.53 | -0.03 | -0.46 | 0.84 | 0.89 | -0.6 | -0.95 | 0.86 | 1.05 | -0.0925 | 0.0975 | -0.1225 | 0.771559 | 0.797773 | 0.809545 | 0.999971 | 1 | 0.973762 |
| Kctd4 | Q9D7X1 | -1.16 | -1.81 | -0.72 | -0.99 | -1.78 | -1.01 | -1.26 | -0.41 | -1.66 | -1.45 | -1.25 | -0.91 | -1.37 | -0.79 | -0.44 | -0.67 | 0.055 | -0.1475 | 0.3525 | 0.878035 | 0.615526 | 0.274459 | 0.999971 | 1 | 0.843794 |
| Kctd5 | A0A0R4J00 | -2.55 | 0.63 | -0.63 | -0.88 | 0.62 | 0.39 | -0.48 | -0.47 | 0.79 | -0.09 | 0.54 | -0.01 | 0.92 | 0.69 | -0.14 | -0.47 | 0.8725 | 1.165 | 1.1075 | 0.201154 | 0.091992 | 0.123243 | 0.999971 | 1 | 0.78268 |
| Kctd6 | Q8BNL5 | -0.38 | 1.17 | 0.43 | 0.41 | 0.51 | 0.66 | -0.2 | 0.9 | 0.42 | 0 | 0.1 | 0.27 | 1.04 | 0.41 | 0.21 | 0.83 | 0.06 | -0.21 | 0.215 | 0.874842 | 0.528411 | 0.555522 | 0.999971 | 1 | 0.914901 |
| Kctd7 | F6QXA3 | -0.85 | 3 | 2.76 | 1.22 | 3.17 | 1.93 | 2.62 | 1.58 | 2.82 | -1.47 | 3.36 | 3.93 | 5.92 | 0.69 | 3.38 | 0.85 | 0.7925 | 0.6275 | 1.1775 | 0.366558 | 0.645674 | 0.394078 | 0.999971 | 1 | 0.871816 |
| Kctd8 | Q50H33 | -0.92 | -1.09 | -0.31 | -0.26 | -1.05 | -0.86 | -0.57 | 0.45 | -1.06 | -0.62 | -0.18 | -0.2 | 0.2 | -0.31 | 0.16 | -0.08 | 0.1375 | 0.13 | 0.6375 | 0.719016 | 0.669418 | 0.036728 | 0.999971 | 1 | 0.700096 |
| Kdelr1 | Q99JH8 | 1.76 | 0.79 | 3.95 | -4.87 | -0.99 | 0.02 | 6.95 | -3.23 | -0.82 | 1.03 | 4.61 | 3.12 | -3.28 | 0.88 | -0.15 | -5.48 | 0.28 | 1.5775 | -2.415 | 0.912551 | 0.434845 | 0.269627 | 0.999971 | 1 | 0.843592 |
| Kdm1a | A3KG93 | 0.73 | 2.13 | 1.27 | 1.1 | 2.62 | 1.4 | 1.88 | 1.01 | 2.27 | 0.98 | 1.49 | 1.9 | 2.24 | 1.74 | 1.26 | 1.26 | 0.42 | 0.3525 | 0.3175 | 0.343066 | 0.378492 | 0.398592 | 0.999971 | 1 | 0.871816 |
| Kdm2a | F6YRW4 | -0.24 | 0.66 | 3.1 | 2.29 | -0.32 | 0.78 | 1.34 | 2.25 | -0.15 | 0.07 | 1.53 | 1.19 | 0.45 | 0.22 | 1.88 | 3.36 | -0.44 | -0.7925 | 0.025 | 0.600621 | 0.325798 | 0.978776 | 0.999971 | 1 | 0.99925 |
| Kdm3b | Q6ZPY7 | 1.33 | 1.35 | 2.1 | 2.12 | 0.63 | 1.59 | 1.17 | 2.07 | 0.9 | 1.5 | 1.63 | 1.37 | 1.38 | 0.8 | 2.24 | 1.83 | -0.36 | -0.375 | -0.1625 | 0.341143 | 0.211705 | 0.662357 | 0.999971 | 1 | 0.936148 |
| Kdm4a | A2A8L9 | 3.03 | -1.31 | 2.51 | 2.25 | 0.51 | 1.31 | 3.61 | 3.14 | 2.23 | 2.93 | 1.87 | -1.75 | 0.93 | 0.96 | 0.6 | 0.81 | 0.5225 | -0.3 | -0.795 | 0.636341 | 0.81615 | 0.383322 | 0.999971 | 1 | 0.871816 |
| Kdm5a | Q3UXZ9 | -0.3 | -0.04 | -0.19 | 1.44 | -0.61 | 1.98 | 0.01 | 0.68 | -0.73 | -0.02 | 0.93 | 0.18 | -0.42 | 0.98 | 0.5 | 1.38 | 0.2875 | -0.1375 | 0.3825 | 0.648012 | 0.781864 | 0.469382 | 0.999971 | 1 | 0.892481 |
| Kdm5b | Q80Y84-2 | -4.41 | -5.62 | 1.03 | 0.52 | -5.82 | -5.54 | 0.37 | 1.11 | -5.67 | -5.53 | 1.86 | 1.63 | -7.25 | -5.73 | 0.94 | 0.68 | -0.35 | 0.1925 | -0.72 | 0.874806 | 0.936462 | 0.765315 | 0.999971 | 1 | 0.962405 |
| Kdsr | Q6GV12 | -0.22 | 0.71 | 1.26 | 0.91 | 0.34 | 1.24 | 0.51 | 1.44 | 0.32 | 0.47 | 1.03 | -0.27 | 0.06 | 1.23 | 1.51 | 1.39 | 0.2175 | -0.2775 | 0.3825 | 0.587355 | 0.491204 | 0.389908 | 0.999971 | 1 | 0.871816 |
| Keap1 | Q9Z2X8 | 0.17 | -0.6 | 0.07 | -0.13 | -0.32 | -0.13 | 0.07 | 0.19 | -0.55 | -0.03 | 0.02 | -0.2 | -0.73 | -0.23 | 0.14 | 0.03 | 0.075 | -0.0675 | -0.075 | 0.753369 | 0.781463 | 0.786221 | 0.999971 | 1 | 0.967132 |
| Khdc4 | Q3TCX3-2 | 3.19 | 2.69 | 1.66 | 1.2 | 2.45 | 3.68 | 1.97 | 1.49 | 2.63 | 2.92 | 1.82 | 1.28 | -0.18 | 1.76 | 2.09 | 1.65 | 0.2125 | -0.0225 | -0.855 | 0.72336 | 0.967146 | 0.195316 | 0.999971 | 1 | 0.811225 |
| Khdrbs1 | Q60749 | 1.78 | 1.66 | 1.86 | 1.92 | 1.75 | 1.25 | 1.28 | 2 | 1.47 | 1.57 | 1.71 | 2.09 | 1.3 | 1.58 | 1.9 | 2.07 | -0.235 | -0.095 | -0.0925 | 0.321502 | 0.637461 | 0.67975 | 0.999971 | 1 | 0.942957 |
| Khdrbs2 | Q9WU01 | -0.2 | 1.07 | 1.05 | 0.55 | 1.35 | 0.43 | 0.76 | 0.74 | 1.01 | -0.48 | 0.28 | 1.13 | 1.1 | 0.24 | 1.14 | 0.35 | 0.2025 | -0.1325 | 0.09 | 0.564982 | 0.768671 | 0.808707 | 0.999971 | 1 | 0.973762 |
| Khdrbs3 | Q9R226 | 0.78 | -0.1 | 0.13 | 0.32 | -0.21 | -0.01 | -0.2 | 0.02 | -0.68 | 0.04 | -0.07 | -0.04 | -0.56 | -0.01 | -0.04 | 0.85 | -0.3825 | -0.47 | -0.2225 | 0.127336 | 0.107255 | 0.521818 | 0.999971 | 1 | 0.907417 |
| Khk | A0A0J9YTU | 0.51 | -0.35 | -1.55 | -0.79 | 0.38 | -0.16 | -0.87 | -0.2 | -0.27 | 0.16 | -1.12 | -0.34 | 0.33 | -0.01 | -0.11 | -1.04 | 0.3325 | 0.1525 | 0.3375 | 0.483489 | 0.748326 | 0.492768 | 0.999971 | 1 | 0.900198 |
| Khnyn | Q80U38 | -1.07 | -1.18 | -0.76 | -0.74 | -0.65 | -0.67 | -0.76 | 0.16 | -0.91 | -1.63 | -0.02 | -0.89 | -1.12 | -0.37 | -0.89 | -0.35 | 0.4575 | 0.075 | 0.255 | 0.109711 | 0.827079 | 0.323095 | 0.999971 | 1 | 0.85548 |
| Khsrp | Q3U0V1 | 2.11 | 3 | 2.83 | 2.84 | 2.85 | 2.66 | 2.58 | 2.5 | 2.8 | 2.43 | 2.71 | 2.65 | 2.53 | 2.77 | 2.56 | 2.55 | -0.0475 | -0.0475 | -0.0925 | 0.845381 | 0.844941 | 0.701284 | 0.999971 | 1 | 0.949019 |
| Kiaa0100 | Q5SYL3 | 0.86 | -0.14 | -0.16 | 0.3 | -0.36 | 0 | -0.3 | 0.32 | -0.52 | 0.65 | -0.1 | -0.3 | -0.45 | -0.02 | 0.03 | 0.17 | -0.3 | -0.2825 | -0.2825 | 0.325278 | 0.421751 | 0.340182 | 0.999971 | 1 | 0.860511 |
| Kiaa0319 | Q5SZV5 | 1.86 | -1.75 | 1.35 | 0.09 | -3.64 | -0.51 | 2.16 | -0.28 | -3.2 | 2.27 | 1.47 | 1.48 | -5.24 | -1.41 | 0.25 | 0.07 | -0.955 | 0.1175 | -1.97 | 0.460965 | 0.929483 | 0.166649 | 0.999971 | 1 | 0.800215 |
| Kiaa0319l | Q8K135 | 0.61 | 1.38 | 0.73 | 0.96 | 1.58 | 0.79 | 0.44 | 0.83 | 1.94 | 1.06 | 0.93 | 0.99 | 2.5 | 0.81 | 0.9 | 1.42 | -0.01 | 0.31 | 0.4875 | 0.97353 | 0.316747 | 0.248566 | 0.999971 | 1 | 0.839012 |
| Kiaa0513 | Q8R0A7 | -0.85 | 0.01 | -0.86 | -1.95 | -0.09 | -0.28 | -0.39 | -2 | 0.04 | -0.94 | -1.02 | -1.25 | 0.44 | -0.64 | -1.47 | -2.11 | 0.2225 | 0.12 | -0.0325 | 0.685945 | 0.795606 | 0.958268 | 0.999971 | 1 | 0.998622 |
| Kiaa1107 | Q80TK0 | 0.55 | -0.09 | 0.15 | 0.19 | -0.18 | 0.31 | 0.41 | 0.05 | -0.06 | 0.45 | 0.19 | 0.08 | -0.48 | 0.1 | -0.2 | 0.17 | -0.0525 | -0.035 | -0.3025 | 0.817283 | 0.870287 | 0.220374 | 0.999971 | 1 | 0.827096 |
| Kiaa1109 | A2AAE1-6 | -0.5 | -0.64 | -0.35 | -0.45 | -0.71 | -0.29 | -0.33 | -0.43 | -0.5 | -0.3 | -0.52 | -0.49 | -2.09 | -0.52 | -1.8 | -1.68 | 0.045 | 0.0325 | -1.0375 | 0.808058 | 0.846467 | 0.014345 | 0.999971 | 1 | 0.613095 |
| Kiaa1522 | A2A7S8-4 | 0.1 | -1.58 | -0.5 | -1.12 | -2.08 | -0.79 | -0.25 | -0.81 | -2.13 | -0.39 | -0.75 | -1.38 | -1.7 | -1.51 | -0.5 | -0.75 | -0.2075 | -0.3875 | -0.34 | 0.676505 | 0.440864 | 0.448376 | 0.999971 | 1 | 0.885289 |
| Kidins220 | E9Q9B7 | -0.71 | -0.82 | -1.5 | -1.49 | -0.82 | -0.68 | -1.51 | -1.1 | -0.75 | -0.83 | -1.41 | -1.11 | -0.43 | -0.67 | -1.34 | -1.25 | 0.1025 | 0.105 | 0.2075 | 0.725742 | 0.70395 | 0.511324 | 0.999971 | 1 | 0.905815 |
| Kif11 | Q6P9P6 | -1.5 | -0.94 | -1.57 | -1.71 | -1.52 | -0.91 | -0.92 | -1.62 | -1.68 | -1.47 | -2.36 | -2.4 | -1.61 | -0.69 | -1.76 | -0.58 | 0.1875 | -0.5475 | 0.27 | 0.497618 | 0.094426 | 0.442271 | 0.999971 | 1 | 0.883386 |
| Kif13a | Q9EQW7 | -2.17 | 0.18 | -2.64 | -3.31 | -0.7 | -2.97 | -1.68 | -1.48 | -1.79 | -2.19 | -3.1 | -1.51 | -3.77 | -0.69 | -1.24 | -2.53 | 0.2775 | -0.1625 | -0.0725 | 0.730055 | 0.829989 | 0.936996 | 0.999971 | 1 | 0.995618 |
| Kif13b | A0A286YC\ | 0.54 | 0.27 | -0.65 | 0.08 | 0.37 | 0.07 | -0.7 | -0.39 | -0.06 | 0.5 | -0.62 | -0.32 | 0.24 | 0.34 | -0.24 | -0.17 | -0.2225 | -0.185 | -0.0175 | 0.522601 | 0.593684 | 0.953725 | 0.999971 | 1 | 0.998056 |
| Kif1a | E9Q9G6 | -0.95 | 0.24 | 0.64 | 0.38 | -0.01 | 0.25 | -0.01 | 0.23 | -0.3 | -0.48 | 0.5 | 0.4 | 0.85 | 0.11 | 0.6 | 0.88 | 0.0375 | -0.0475 | 0.5325 | 0.91527 | 0.907817 | 0.186999 | 0.999971 | 1 | 0.809025 |
| Kif1a_1 | Q6TA13 | -0.41 | -0.22 | -0.15 | -0.22 | -0.22 | -0.31 | 0.12 | -0.02 | -0.25 | -0.42 | -0.02 | 0.12 | 0.03 | -0.18 | -0.06 | -0.08 | 0.1425 | 0.1075 | 0.1775 | 0.448833 | 0.579211 | 0.309696 | 0.999971 | 1 | 0.847402 |
| Kif1b | A0A5F8MP | -0.28 | -0.05 | -0.8 | -0.52 | 0.06 | -0.26 | -0.78 | -0.46 | 0.14 | -0.18 | -0.47 | -0.47 | 0.1 | -0.35 | -0.66 | -0.5 | 0.0525 | 0.1675 | 0.06 | 0.840918 | 0.499833 | 0.814635 | 0.999971 | 1 | 0.974918 |
| Kif1c | O35071 | 2.1 | -0.11 | -0.16 | -0.16 | 0.01 | 0.14 | -0.57 | 0.25 | -0.47 | 1.4 | 0.59 | 0.3 | -0.93 | 0.23 | -0.59 | -0.17 | -0.46 | 0.0375 | -0.7825 | 0.406399 | 0.952015 | 0.189735 | 0.999971 | 1 | 0.810547 |
| Kif21a | F8WGN6 | 3.02 | 0.42 | 0.25 | -0.75 | -0.61 | 1.27 | 1.24 | -0.36 | -1.88 | 2.52 | -1.67 | -0.15 | -2.18 | -0.21 | -2.07 | 0.47 | -0.35 | -1.03 | -1.7325 | 0.682184 | 0.385965 | 0.091306 | 0.999971 | 1 | 0.773779 |
| Kif21a_1 | Q9QXL2-2 | -0.25 | -1.35 | -1.33 | -0.9 | -0.86 | -0.85 | -0.39 | -0.92 | -0.55 | -0.54 | -1.42 | -0.42 | -0.42 | -0.28 | -2.37 | -0.49 | 0.2025 | 0.225 | 0.0675 | 0.498771 | 0.516022 | 0.896035 | 0.999971 | 1 | 0.988571 |
| Kif21a_2 | Q9QXL2-3 | 0.49 | 0.23 | 1.44 | 0.44 | 0.9 | 0.61 | 0.55 | 0.51 | 0.66 | 0.41 | -1.64 | 1.77 | 0.27 | 1.5 | 0.3 | 0.45 | -0.0075 | -0.35 | -0.02 | 0.979618 | 0.615678 | 0.958421 | 0.999971 | 1 | 0.998622 |
| Kif21a_3 | E9Q0J5 | 0.25 | -0.04 | -0.34 | -0.15 | -0.23 | 0.17 | -0.13 | -0.25 | -0.1 | 0.14 | -0.52 | -0.48 | -0.16 | -0.04 | -0.24 | -0.31 | -0.04 | -0.17 | -0.1175 | 0.847922 | 0.473976 | 0.556433 | 0.999971 | 1 | 0.915051 |
| Kif21b | Q9QXL1 | -0.77 | -0.76 | -0.85 | -0.81 | -0.66 | -0.67 | -0.49 | -0.84 | -0.39 | -0.03 | -0.08 | -0.46 | 0.26 | -0.58 | -0.73 | -0.43 | 0.1325 | 0.5575 | 0.4275 | 0.443118 | 0.01262 | 0.113274 | 0.999971 | 1 | 0.78268 |
| Kif2a | P28740-2 | 0.68 | -1.47 | -1.71 | -0.1 | -0.8 | -1.28 | -0.91 | -0.68 | -0.74 | 0.77 | 0.05 | -1.7 | -1.27 | -0.63 | -0.51 | 0.31 | -0.2675 | 0.245 | 0.125 | 0.620159 | 0.729587 | 0.834447 | 0.999971 | 1 | 0.979036 |
| Kif2a_1 | E0CZ72 | -0.09 | 0.13 | -0.17 | -0.21 | 0.14 | -0.13 | -0.02 | -0.1 | 0.43 | 0.23 | 0.08 | -0.06 | 0.66 | -0.32 | -0.02 | 0.22 | 0.0575 | 0.255 | 0.22 | 0.747656 | 0.204071 | 0.388773 | 0.999971 | 1 | 0.871816 |
| Kif3a | Q3TET1 | 1.89 | 0.37 | 0.47 | 0.8 | 0.16 | 0.6 | 1.14 | 0.4 | 0.33 | 1.34 | 0.36 | -0.02 | -0.25 | 0.32 | 0.27 | 0.36 | -0.3075 | -0.38 | -0.7075 | 0.438478 | 0.389002 | 0.08086 | 0.999971 | 1 | 0.755254 |
| Kif3b | Q61771 | 1.06 | 0.82 | 1 | 0.95 | 0.65 | 0.87 | 0.98 | 0.85 | 0.87 | 0.93 | 1.16 | 1.05 | 0.28 | 0.43 | 0.76 | 1.12 | -0.12 | 0.045 | -0.31 | 0.495753 | 0.79025 | 0.203643 | 0.999971 | 1 | 0.81617 |
| Kif3c | O35066 | 1.87 | 0.69 | 0.05 | 0.32 | 0.32 | 1.23 | 0.47 | 0.72 | 0.5 | 1.39 | 0.46 | 0.26 | 0.59 | 0.86 | 0.53 | 0.67 | -0.0475 | -0.08 | -0.07 | 0.910784 | 0.857897 | 0.858266 | 0.999971 | 1 | 0.982702 |
| Kif5a | P33175 | -0.1 | -0.45 | -0.75 | -0.56 | -0.39 | -0.55 | -0.73 | -0.33 | -0.4 | -0.2 | -0.74 | -0.72 | -0.57 | -0.38 | -0.48 | -0.61 | -0.035 | -0.05 | -0.045 | 0.869042 | 0.825124 | 0.824627 | 0.999971 | 1 | 0.976309 |
| Kif5b | Q61768 | -0.04 | -0.75 | -0.18 | -0.29 | -0.53 | -0.46 | -0.07 | -0.46 | -0.3 | -0.08 | -0.47 | -0.32 | -0.58 | -0.75 | -0.32 | -0.51 | -0.065 | 0.0225 | -0.225 | 0.774195 | 0.917068 | 0.324637 | 0.999971 | 1 | 0.856365 |
| Kif5c | P28738 | -0.14 | -0.58 | -0.61 | -0.5 | -0.67 | -0.31 | -0.41 | -0.54 | -0.51 | -0.38 | -0.58 | -0.66 | -0.13 | -0.5 | -0.6 | -0.5 | -0.025 | -0.075 | 0.025 | 0.898297 | 0.69046 | 0.902996 | 0.999971 | 1 | 0.990937 |
| Kifap3 | P70188-2 | 0.1 | 0.22 | 0.15 | 0.36 | 0.25 | 0.07 | 0 | 0.43 | 0.03 | 0.27 | 0.44 | 0.44 | 0.11 | 0.24 | 0.28 | 0.57 | -0.02 | 0.0875 | 0.0925 | 0.913782 | 0.632728 | 0.620952 | 0.999971 | 1 | 0.929388 |
| Kifbp | Q6ZPU9 | -0.49 | -0.43 | 0.99 | 0.88 | -0.18 | -0.44 | 0.75 | 0.87 | -0.49 | -0.34 | 0.94 | 0.61 | -0.98 | -0.1 | 1.06 | 0.74 | 0.0125 | -0.0575 | -0.0575 | 0.979376 | 0.908157 | 0.918245 | 0.999971 | 1 | 0.992633 |
| Kifc2 | A0A2R8W6 | 2.57 | 0.49 | -0.37 | 0.22 | 0.18 | 0.72 | -0.17 | -0.24 | 0.13 | 1.4 | -0.77 | -0.71 | -0.2 | 0.2 | -0.5 | 0.33 | -0.605 | -0.715 | -0.77 | 0.339668 | 0.347725 | 0.22767 | 0.999971 | 1 | 0.828598 |
| Kifc3 | O35231 | 0.08 | -0.37 | -0.83 | -1 | -0.8 | -0.31 | -0.9 | -0.97 | -0.5 | -0.4 | -0.72 | -0.93 | -1.06 | -0.51 | -1.07 | -0.5 | -0.215 | -0.1075 | -0.255 | 0.473182 | 0.705313 | 0.406597 | 0.999971 | 1 | 0.874911 |
| Kirrel2 | Q7TSU7-2 | 1.82 | 1.62 | -1.98 | -0.85 | 2.19 | 2.81 | -2.14 | -2.75 | 2.49 | 2.71 | -0.68 | -0.63 | 2.64 | 2.1 | -1.06 | -0.52 | -0.125 | 0.82 | 0.6375 | 0.934395 | 0.496263 | 0.589496 | 0.999971 | 1 | 0.921872 |
| Kirrel3 | Q8BR86-2 | -0.75 | -1 | -1.61 | -1.56 | -0.81 | -1.1 | -1.02 | -1.36 | -1.21 | -0.86 | -2.03 | -1.69 | -1.17 | -0.95 | -1.74 | -1.64 | 0.1575 | -0.2175 | -0.145 | 0.552559 | 0.51796 | 0.624648 | 0.999971 | 1 | 0.930479 |
| Kit | P05532 | -0.43 | -0.9 | -1.1 | -1.01 | -1.07 | -0.23 | -1.39 | -0.62 | -1.05 | -0.35 | -1.45 | -1.32 | -0.47 | -0.5 | -0.8 | -0.61 | 0.0325 | -0.1825 | 0.265 | 0.914329 | 0.541958 | 0.238117 | 0.999971 | 1 | 0.833594 |
| Kitlg | P20826-2 | 0.01 | -0.27 | 0.6 | 0.52 | -0.39 | -1.18 | -0.16 | 1.07 | -1.49 | 0.16 | 0.62 | 1.75 | 0.08 | -1.43 | 1.14 | 1.26 | -0.38 | 0.045 | 0.0475 | 0.432972 | 0.944171 | 0.93691 | 0.999971 | 1 | 0.995618 |
| Kl | O35082-2 | 1.51 | 0.98 | 2.12 | 2.65 | 0.14 | 0.4 | 0.5 | 2.16 | 0.3 | 0.78 | 1.3 | 1.13 | -0.85 | 0.28 | 1.59 | 1.65 | -1.015 | -0.9375 | -1.1475 | 0.086858 | 0.044072 | 0.09862 | 0.999971 | 1 | 0.78268 |
| Klc1 | Q8CD76 | -0.3 | 0.1 | -0.03 | -0.09 | 0.09 | 0.03 | -0.1 | 0.08 | 0.26 | -0.08 | 0.02 | 0.08 | 0.74 | -0.01 | 0.02 | 0.02 | 0.105 | 0.15 | 0.2725 | 0.557212 | 0.416156 | 0.267347 | 0.999971 | 1 | 0.842677 |
| Klc2 | D3YXZ3 | -0.77 | -0.66 | -0.52 | -1.04 | -0.85 | -0.4 | -0.81 | -0.58 | -0.88 | -0.83 | -0.45 | -0.86 | -0.01 | -0.46 | -0.52 | -0.56 | 0.0875 | -0.0075 | 0.36 | 0.672261 | 0.97038 | 0.122683 | 0.999971 | 1 | 0.78268 |
| Klc4 | Q9DBS5 | 0.15 | -0.82 | -0.58 | -0.16 | -0.69 | -0.52 | -0.52 | -0.37 | -0.32 | -0.15 | -0.55 | -0.39 | -1.22 | -0.72 | -0.53 | -0.32 | -0.1725 | -5.89E-17 | -0.345 | 0.50023 | 1 | 0.266808 | 0.999971 | 1 | 0.842677 |
| Klhdc10 | Q6PAR0 | 3.9 | 1.6 | 1.81 | 1.56 | -0.04 | 2.03 | 1.53 | 1.63 | 1.02 | 0.8 | 1.35 | 1.09 | 1.31 | 0.76 | 1.43 | 2.54 | -0.93 | -1.1525 | -0.7075 | 0.181278 | 0.055536 | 0.269536 | 0.999971 | 1 | 0.843592 |
| Klhdc2 | Q4G5Y1 | -0.62 | 0 | 0.61 | 1.19 | 0.42 | -0.08 | 0.68 | 0.34 | -0.12 | 0.21 | 0.76 | 0.6 | -0.28 | 0.59 | 1 | 1.25 | 0.045 | 0.0675 | 0.345 | 0.910736 | 0.871287 | 0.478965 | 0.999971 | 1 | 0.894438 |
| Klhdc3 | A0A3B2W( | -0.12 | 1.02 | 0.68 | 0.74 | 0.67 | 1.13 | 1.75 | 1.32 | 0.65 | 0.94 | 1.89 | 1.39 | 1.14 | 1.06 | 1.4 | 1.04 | 0.6375 | 0.6375 | 0.58 | 0.080894 | 0.102398 | 0.061086 | 0.999971 | 1 | 0.725348 |
| Klhdc4 | G3UZG5 | 0.82 | 0.02 | 1.17 | 0.97 | 0.71 | 0.38 | 0.52 | 1.54 | -0.06 | -0.1 | 0.79 | 0.1 | -0.23 | 0.86 | 1.06 | 1.25 | 0.0425 | -0.5625 | -0.01 | 0.904518 | 0.11247 | 0.979989 | 0.999971 | 1 | 0.99925 |
| Klhdc7a | A2APT9 | 0.39 | -0.77 | -0.97 | -0.56 | -0.63 | 0 | -1.16 | -0.89 | -0.71 | 0.26 | -1.56 | -0.37 | -0.92 | -0.73 | -1.39 | -0.38 | -0.1925 | -0.1175 | -0.3775 | 0.61219 | 0.796799 | 0.309529 | 0.999971 | 1 | 0.847402 |

| | | | | | | | | | | | | | | | | | | | | | | | | | | |
|---|---|---|---|---|---|---|---|---|---|---|---|---|---|---|---|---|---|---|---|---|---|---|---|---|---|---|
| Klhdc8b | Q9D2D9 | -1.01 | -1.29 | -0.58 | 0.05 | 0.13 | -0.81 | -0.84 | -0.52 | -0.71 | -1.3 | -0.59 | -0.65 | -0.79 | -1.3 | -0.42 | -0.55 | 0.1975 | -0.105 | -0.0575 | 0.586154 | 0.753106 | 0.868026 | 0.999971 | 1 | 0.983462 |
| Klhdc9 | Q3USL1 | 0.18 | 0.16 | -1.58 | -1.81 | -0.57 | -0.37 | -0.65 | 0.06 | -0.35 | -0.9 | -1.95 | 0.41 | -0.61 | 0.35 | -1.34 | -2.02 | 0.38 | 0.065 | -0.1425 | 0.471872 | 0.922578 | 0.832359 | 0.999971 | 1 | 0.979036 |
| Klhl11 | Q8CE33 | -0.46 | 0.45 | 1.21 | 0.98 | 0.06 | 0.22 | 0.77 | 0.61 | 0.15 | -0.36 | 0.52 | 0.47 | -0.03 | 0.11 | 0.43 | 0.63 | -0.13 | -0.35 | -0.26 | 0.739398 | 0.397383 | 0.506077 | 0.999971 | 1 | 0.905277 |
| Klhl13 | Q80TF4-5 | -2.11 | 1.02 | -0.35 | -1.28 | 0.9 | -0.04 | -0.46 | -0.43 | 0.14 | -2.49 | -2.89 | -0.44 | 2.6 | -0.56 | 0.09 | -1.24 | 0.6725 | -0.74 | 0.9025 | 0.331344 | 0.422943 | 0.359867 | 0.999971 | 1 | 0.869526 |
| Klhl2 | Q8JZP3 | -3.88 | -0.31 | -0.41 | -1.79 | -0.29 | -1.45 | -0.4 | -0.37 | -1.26 | -3.79 | 0.12 | 0.49 | -0.35 | -2.83 | -0.59 | -1.6 | 0.97 | 0.4875 | 0.255 | 0.238761 | 0.671937 | 0.777659 | 0.999971 | 1 | 0.964822 |
| Klhl20 | Q8VCK5 | 4.41 | -0.57 | 1.29 | 1.21 | -1.28 | -0.22 | 1.83 | 0.97 | -0.65 | 2.7 | 1.08 | 0.25 | -1.7 | -1.74 | 0.32 | 0.46 | -1.26 | -0.74 | -2.25 | 0.271227 | 0.516643 | 0.061374 | 0.999971 | 1 | 0.725348 |
| Klhl22 | Q99JN2 | 0.35 | 0.12 | -0.26 | -0.37 | 0.07 | 0.31 | -0.3 | -0.39 | -0.09 | 0.39 | 0.18 | -0.01 | 0.16 | 0.28 | -0.31 | -0.17 | -0.0375 | 0.1575 | 0.03 | 0.884334 | 0.503609 | 0.903284 | 0.999971 | 1 | 0.991035 |
| Klhl26 | Q8BGY4 | -0.61 | -2.42 | -1.99 | -1.66 | -2.18 | -1.27 | -1.02 | -1.59 | -1.85 | -0.87 | -1.34 | -1.13 | -2.62 | -1.9 | -0.95 | -1.25 | 0.155 | 0.3725 | -0.01 | 0.721883 | 0.383733 | 0.983947 | 0.999971 | 1 | 0.99925 |
| Klhl3 | E0CZ16 | -1.29 | 0.1 | 0.02 | -0.06 | 0.27 | 0.26 | 0.04 | 0.28 | 0.09 | -0.28 | 0.31 | 0.28 | 0.23 | 0.15 | 0.07 | 0.53 | 0.52 | 0.4075 | 0.5525 | 0.144293 | 0.263357 | 0.132292 | 0.999971 | 1 | 0.784948 |
| Klhl36 | Q8R124 | 0.62 | 0.13 | -0.36 | 1.37 | 0.5 | 0.83 | -0.64 | 1.44 | -0.54 | 0.98 | -0.16 | 0.08 | 1.37 | 0.67 | 0.49 | 1.82 | 0.0925 | -0.35 | 0.6475 | 0.860676 | 0.455434 | 0.177162 | 0.999971 | 1 | 0.808532 |
| Klhl4 | Q8BLJ6 | 0.69 | -0.38 | -0.77 | -0.31 | 0.99 | 0.92 | -0.42 | -0.41 | 1.16 | 0.88 | 0.26 | 0.14 | 0.75 | 0.59 | -0.11 | 0.55 | 0.4625 | 0.8025 | 0.6375 | 0.338743 | 0.061823 | 0.10202 | 0.999971 | 1 | 0.78268 |
| Klhl41 | A2AUC9 | 0.65 | -3.09 | -0.55 | -0.85 | -0.76 | -0.4 | -1.04 | -0.11 | 0.48 | -2.91 | -0.23 | -1.13 | 0.2 | 0.18 | -0.03 | 0.41 | 0.3825 | 0.0125 | 1.15 | 0.601835 | 0.989634 | 0.132637 | 0.999971 | 1 | 0.784948 |
| Klhl42 | Q8BFQ9 | 1.29 | 1.88 | 0.04 | -1.08 | 1.38 | 1.16 | -0.18 | -1.16 | 1.64 | 0.79 | 0.07 | -0.41 | 2.68 | 1.29 | -1.17 | 0.09 | -0.2325 | -0.01 | 0.19 | 0.771507 | 0.988974 | 0.840646 | 0.999971 | 1 | 0.97975 |
| Klhl5 | E9Q1L2 | -0.69 | -0.54 | -1.86 | -1.2 | -0.95 | -0.52 | -0.86 | -1.97 | -1.64 | -0.12 | -1.91 | -1.32 | -2.74 | -0.31 | -2.04 | -1.19 | -0.0025 | -0.175 | -0.4975 | 0.995139 | 0.707385 | 0.382223 | 0.999971 | 1 | 0.871816 |
| Klhl8 | A2AH60 | -0.93 | 0.79 | -0.6 | -0.59 | 0.72 | 0.89 | -0.36 | -0.91 | 0.74 | -0.72 | -1.64 | 0.03 | -0.25 | 0.6 | -2.52 | -1.14 | 0.4175 | -0.065 | -0.495 | 0.440459 | 0.911671 | 0.483524 | 0.999971 | 1 | 0.896352 |
| Klhl9 | Q6ZPT1 | 0.18 | 1.27 | 1.19 | 1.04 | 1.34 | 0.5 | 1.22 | 1.15 | 1.99 | 0.23 | 1.58 | 1.64 | 1.99 | 0.03 | 1.6 | 1.47 | 0.1325 | 0.44 | 0.3525 | 0.677321 | 0.328615 | 0.456395 | 0.999971 | 1 | 0.887217 |
| Kmo | Q91WN4-2 | -1.51 | -1.54 | 0.46 | -0.13 | 0.34 | 0.75 | -0.06 | 0.31 | 0.74 | 0.13 | 0.08 | 0.08 | 0.94 | -0.15 | -0.12 | 1.2 | 1.015 | 0.9375 | 1.1475 | 0.064703 | 0.084585 | 0.067961 | 0.999971 | 1 | 0.733937 |
| Kmt2a | P55200-2 | -1.82 | -0.74 | -0.03 | 0.24 | -0.86 | -0.22 | 0.27 | -0.43 | -0.71 | -1.47 | -0.25 | -1.68 | -0.19 | -1.06 | -1.2 | 0.28 | 0.2775 | -0.44 | 0.045 | 0.567407 | 0.412923 | 0.932945 | 0.999971 | 1 | 0.995069 |
| Kndc1 | Q0KK55 | 0.67 | 0.38 | 0.9 | 0.82 | 0.19 | 0.58 | 0.62 | 1.08 | 0.32 | 0.96 | 0.39 | 0.27 | 0.32 | 1.09 | 0.65 | 0.7 | -0.075 | -0.2075 | -0.0025 | 0.760252 | 0.381197 | 0.991416 | 0.999971 | 1 | 0.99925 |
| Kng1 | O08677-2 | 3.26 | 1.56 | 0.43 | 3.11 | 3.25 | 1.28 | 3 | 1.18 | 1.01 | 0.74 | 0.65 | 0.95 | 1.06 | 1.42 | 0.91 | 0.76 | 0.0875 | -1.2525 | -1.0525 | 0.910829 | 0.070818 | 0.1207 | 0.999971 | 1 | 0.78268 |
| Kng2 | Q6S9I3 | -1.03 | -1.09 | -1.24 | -1.14 | -1.19 | -0.3 | -1.74 | -1.1 | -0.97 | -0.73 | -1.61 | -1.29 | -0.48 | -0.32 | -0.89 | -0.71 | 0.0425 | -0.025 | 0.525 | 0.889618 | 0.913808 | 0.023832 | 0.999971 | 1 | 0.664227 |
| Kpna1 | Q60960 | 2.17 | 2.08 | -0.1 | 0.46 | 1.44 | 2.6 | 0 | 0.43 | 1.81 | 2.69 | 0 | -0.55 | 1.32 | 2.52 | -0.19 | -0.12 | -0.035 | -0.165 | -0.27 | 0.961962 | 0.847652 | 0.729642 | 0.999971 | 1 | 0.956041 |
| Kpna2 | P52293 | -0.41 | 0.52 | -0.54 | -0.25 | -0.33 | 0.09 | -0.37 | 0.08 | -0.1 | -0.76 | 2.33 | -0.44 | -0.15 | 0.25 | 0.11 | 0.02 | 0.0375 | 0.4275 | 0.2275 | 0.89457 | 0.532614 | 0.411717 | 0.999971 | 1 | 0.875847 |
| Kpna3 | O35344 | 1.54 | -0.3 | 0.67 | 1.02 | -0.21 | 0.29 | 1.02 | 0.76 | 0.01 | 1.11 | 0.64 | 0.33 | -0.37 | 0.11 | 0.46 | 0.56 | -0.2675 | -0.21 | -0.5425 | 0.551688 | 0.627398 | 0.217686 | 0.999971 | 1 | 0.826893 |
| Kpna4 | O35343 | -0.16 | 0.61 | -0.09 | 0 | 0.48 | 0.36 | -0.34 | 0.3 | 0.16 | 0.07 | -0.11 | -0.32 | 0.4 | 0.49 | -0.02 | -0.1 | 0.11 | -0.14 | 0.1025 | 0.688141 | 0.559182 | 0.690311 | 0.999971 | 1 | 0.946255 |
| Kpna6 | O35345 | 0.56 | 0.4 | 0.43 | 0.5 | 0.14 | 0.89 | 0.65 | 0.55 | 0.14 | 0.56 | 0.57 | 0.24 | 0.28 | 0.81 | 0.56 | 0.72 | 0.085 | -0.095 | 0.12 | 0.687752 | 0.607766 | 0.533026 | 0.999971 | 1 | 0.910069 |
| Kpnb1 | P70168 | 0.52 | -0.05 | -0.1 | -0.14 | 0.03 | 0.26 | 0.03 | -0.24 | 0.01 | 0.04 | -0.44 | -0.07 | 0 | 0.22 | 0.14 | -0.18 | -0.0375 | -0.1725 | -0.0125 | 0.868278 | 0.455529 | 0.955073 | 0.999971 | 1 | 0.998056 |
| Kptn | G3X8R1 | 0.68 | -0.29 | -0.2 | 0.1 | -0.31 | -0.35 | 0.23 | -0.67 | 0.06 | 0.45 | 0.05 | 0.07 | -0.13 | 0.37 | -0.65 | -0.51 | -0.3475 | 0.085 | -0.3025 | 0.260919 | 0.745242 | 0.356377 | 0.999971 | 1 | 0.868793 |
| Kras | P32883-2 | -1.57 | -1.62 | -1.11 | -0.83 | -1.28 | -1.41 | -1.14 | -0.78 | -1.57 | -1.38 | -0.92 | -0.96 | -1.68 | -1.43 | -0.94 | -0.69 | 0.13 | 0.075 | 0.0975 | 0.616354 | 0.778871 | 0.748075 | 0.999971 | 1 | 0.961064 |
| Kri1 | V9GXJ9 | -2.73 | 0.78 | 0.69 | 0.68 | 0.82 | 0.47 | 0.04 | 0.8 | 0.79 | -0.43 | 0.96 | 0.81 | 1.18 | 1.08 | 0.45 | 0.78 | 0.6775 | 0.6775 | 1.0175 | 0.402354 | 0.424602 | 0.219697 | 0.999971 | 1 | 0.827064 |
| Krit1 | A0A0G2JG( | 0.14 | -0.12 | 0.14 | 0.73 | 0 | -0.19 | -0.11 | 0.75 | 0.08 | 0.05 | 0.23 | 0.38 | -0.27 | 0.02 | 0.31 | 0.71 | -0.11 | -0.0375 | -0.03 | 0.707539 | 0.870624 | 0.917266 | 0.999971 | 1 | 0.992633 |
| Krr1 | Q8BGA5 | 1.65 | 0.35 | -0.39 | -0.86 | -0.3 | 0.28 | -0.13 | -0.04 | -0.15 | 1.28 | 0.7 | 0.02 | -0.98 | -0.51 | -0.64 | -0.59 | -0.235 | 0.275 | -0.8675 | 0.651809 | 0.641463 | 0.11997 | 0.999971 | 1 | 0.78268 |
| Krt12 | Z4YJD9 | -1.53 | -0.13 | 0.62 | 0.36 | 0.38 | -1.64 | 0.48 | 0.68 | -0.35 | -1.02 | 0.68 | 0.89 | -0.05 | -0.28 | 0.7 | 1.14 | 0.145 | 0.22 | 0.5475 | 0.82518 | 0.715702 | 0.322203 | 0.999971 | 1 | 0.85548 |
| Krt33a | Q8K0Y2 | -0.73 | -1.15 | 0.08 | 1.64 | -1.28 | -0.68 | -2.34 | -2.23 | -1.42 | 1.39 | 1.12 | -0.29 | -1.73 | 6.48 | 0.55 | -0.08 | -1.5925 | 0.24 | 1.345 | 0.037565 | 0.767961 | 0.431814 | 0.999971 | 1 | 0.881643 |
| Krt76 | Q3UV17 | 2.07 | 2.01 | -0.54 | -1.1 | 4.31 | 1.62 | 0.11 | 2.1 | -1.49 | -0.7 | -0.77 | -4.03 | -2.49 | 5.39 | -5.11 | -2.41 | 1.425 | -2.3575 | -1.765 | 0.20654 | 0.044854 | 0.419005 | 0.999971 | 1 | 0.87788 |
| Krt81 | Q9ERE2 | -5.3 | -8.22 | 2.46 | 3.3 | -8.55 | -8.41 | -4.1 | -5.16 | -7.45 | 5.3 | 5.3 | 3.41 | -11.56 | 13.07 | 4.35 | 1.23 | -4.615 | 3.58 | 3.7125 | 0.115682 | 0.349508 | 0.478715 | 0.999971 | 1 | 0.894438 |
| Krtap3-1 | A2A591 | -2.9 | -4.55 | 1.13 | 2.55 | -4.19 | -5.63 | -3.8 | -7.9 | -3.95 | 1.39 | 4.63 | 2.7 | -12.55 | 11.16 | 3.17 | 1.86 | -4.4375 | 2.135 | 1.8525 | 0.025898 | 0.347705 | 0.688466 | 0.999971 | 1 | 0.945753 |
| Krtcap2 | A0A0A0MC | -0.85 | 0.12 | 0.88 | -1 | -0.15 | 0.16 | 0.74 | -0.23 | -0.14 | -2.03 | 0.43 | 0.19 | -0.89 | 0.19 | -0.1 | 0.04 | 0.3425 | -0.175 | 0.0225 | 0.464663 | 0.788288 | 0.961766 | 0.999971 | 1 | 0.998721 |
| Ksr1 | Q61097-2 | 0.09 | 0.71 | 0.42 | 0.27 | 0.25 | 0.44 | 0.35 | 0.53 | 0.1 | -0.05 | 0.17 | 0.12 | 0.56 | -0.04 | 0.43 | 0.23 | 0.02 | -0.2875 | -0.0775 | 0.920845 | 0.167829 | 0.732753 | 0.999971 | 1 | 0.956652 |
| Ksr2 | M0QW59 | 0.04 | -0.26 | 0.17 | -0.46 | 0.22 | -0.1 | -0.03 | -0.12 | 0.07 | 0.21 | 0.35 | 0.45 | 0.38 | -0.33 | 0.01 | -0.23 | 0.12 | 0.3975 | 0.085 | 0.574848 | 0.086987 | 0.728754 | 0.999971 | 1 | 0.956041 |
| Kti12 | Q9D1R2 | 0.87 | 0.62 | 0.03 | 0.05 | 0.79 | 0.79 | -0.38 | -0.08 | 0.99 | 0.29 | -0.14 | -0.37 | -0.31 | 0.63 | -0.36 | -0.28 | -0.1125 | -0.2 | -0.4725 | 0.754046 | 0.579314 | 0.164864 | 0.999971 | 1 | 0.800215 |
| Ktn1 | A0A087WC | 0.17 | 0.13 | -0.07 | -0.13 | 0.1 | 0.18 | 0.17 | 0 | 0.36 | 0.2 | 0.24 | 0.2 | 0.11 | 0.14 | -0.11 | 0.11 | 0.0875 | 0.225 | 0.0375 | 0.616202 | 0.20635 | 0.832912 | 0.999971 | 1 | 0.979036 |
| Ktn1_1 | A0A087WF | 0.34 | -0.06 | -1.37 | -0.83 | -0.4 | -0.11 | -0.7 | -0.74 | -0.41 | -0.95 | -0.43 | -0.14 | 0.2 | -0.24 | -1 | 0.43 | -0.0075 | -0.0025 | 0.3275 | 0.984777 | 0.995028 | 0.487321 | 0.999971 | 1 | 0.897906 |
| Kxd1 | E9Q4P0 | -1.14 | 0.35 | -0.1 | 1.05 | 0.36 | -0.12 | 0.38 | 0.66 | 0.5 | -0.03 | 0.65 | 0.24 | -0.04 | 0.08 | 0.01 | -1.28 | 0.28 | 0.3 | -0.3475 | 0.543593 | 0.514714 | 0.509751 | 0.999971 | 1 | 0.905815 |
| Kyat1 | Q8BTY1 | -0.36 | 0.33 | 0.65 | 0.32 | 0.32 | 0.28 | -0.01 | 0.79 | 0.14 | 0.05 | -0.02 | 0.27 | 0.26 | 0.27 | 0.09 | 0.57 | 0.11 | -0.125 | 0.0625 | 0.699168 | 0.616983 | 0.809827 | 0.999971 | 1 | 0.973791 |
| Kyat3 | Q71RI9-2 | -0.54 | 0.33 | 0.27 | -0.16 | -0.03 | 0.08 | -0.3 | 0.6 | 0.04 | -0.47 | -0.04 | 0.34 | -0.03 | -0.08 | 0.38 | 0.06 | 0.1125 | -0.0075 | 0.1075 | 0.698539 | 0.978465 | 0.674542 | 0.999971 | 1 | 0.941037 |
| L1cam | Q6PGJ3 | -1.05 | -1.7 | -1.75 | -2.06 | -1.46 | -1.51 | -1.22 | -1.99 | -1.06 | -1 | -1.29 | -1.13 | -1.02 | -1.4 | -1.36 | -1.64 | 0.095 | 0.52 | 0.285 | 0.736338 | 0.061537 | 0.30429 | 0.999971 | 1 | 0.847402 |
| L2hgdh | Q91YP0 | -0.15 | 0.23 | -0.44 | -0.74 | 0.32 | 0.14 | -0.81 | -0.77 | -0.09 | -0.21 | -0.96 | -0.99 | 0.87 | 0.08 | -1.02 | -0.64 | -0.005 | -0.2875 | 0.0975 | 0.988733 | 0.378539 | 0.824517 | 0.999971 | 1 | 0.976309 |
| L3hypdh | Q9CXA2 | 0.16 | 0.69 | 0.52 | 1.06 | 1.74 | 0.29 | 0.91 | 0.43 | 2.72 | 0.57 | 0.15 | 0.02 | 1.76 | 0.41 | 0.12 | 0.25 | 0.235 | 0.2575 | 0.0275 | 0.526234 | 0.670642 | 0.945772 | 0.999971 | 1 | 0.997427 |
| Lactb | Q9EP89 | -1.43 | -0.35 | -0.51 | -1.23 | -0.49 | -0.82 | -0.43 | -1.05 | -0.66 | -1.5 | -0.5 | -0.24 | -0.77 | -1 | -0.84 | -1.22 | 0.1825 | 0.155 | -0.0775 | 0.557523 | 0.676419 | 0.792585 | 0.999971 | 1 | 0.969483 |
| Lactb2 | Q99KR3 | 0.11 | 1.47 | 0.94 | 0.99 | 1.32 | 1.4 | 0.93 | 1.02 | 1 | 0.25 | 0.73 | 0.66 | 1.37 | 1.35 | 0.8 | 0.86 | 0.29 | -0.2175 | 0.2175 | 0.361415 | 0.506487 | 0.506836 | 0.999971 | 1 | 0.905775 |
| Lama1 | P19137 | 0.72 | 0.78 | 0.73 | 1.24 | 2.11 | 0.56 | 0.85 | 0.01 | 1.25 | 0.83 | 0.7 | 0.46 | -0.08 | 1.16 | 1.82 | 0.55 | 0.015 | -0.0575 | -0.005 | 0.972499 | 0.809503 | 0.990188 | 0.999971 | 1 | 0.99925 |
| Lama2 | A0A1W2P7 | -0.12 | -1.87 | -0.05 | 0.67 | -0.18 | -0.26 | -0.99 | -0.48 | -0.05 | 0.45 | -0.12 | 0.49 | -0.87 | -0.04 | 0.18 | 0.41 | -0.135 | 0.535 | 0.2625 | 0.797375 | 0.321027 | 0.640239 | 0.999971 | 1 | 0.93301 |
| Lama2_1 | Q60675 | 0.27 | 0.24 | 0.21 | 0.62 | 1.07 | 0.21 | 0.38 | -0.05 | 0.55 | 0.55 | 0.2 | 0.26 | 0.05 | 0.34 | 0.54 | -0.06 | 0.0675 | 0.055 | -0.1175 | 0.806381 | 0.775884 | 0.58803 | 0.999971 | 1 | 0.920843 |
| Lama4 | P97927 | 0.18 | -0.03 | 0.04 | 0.77 | 1.31 | 0.03 | 0.43 | -0.18 | 0.4 | 0.87 | 1.04 | 1.2 | -0.26 | 0.03 | 0.62 | 0.11 | 0.1575 | 0.6375 | -0.115 | 0.668442 | 0.039586 | 0.678322 | 0.999971 | 1 | 0.942697 |
| Lama5 | Q61001 | 1.43 | 0.75 | 0.58 | 1.14 | 3.25 | 0.75 | 0.5 | -0.17 | 1.46 | 2.89 | 2.14 | 3.49 | -0.15 | 0.66 | 0.84 | 0.49 | 0.1075 | 1.52 | -0.515 | 0.87757 | 0.008431 | 0.111554 | 0.999971 | 1 | 0.78268 |
| Lamb1 | E9QN70 | 0.3 | -0.19 | 0.02 | 0.3 | 0.82 | -0.2 | 0.05 | -0.3 | 0.8 | 0.93 | 0.48 | 0.72 | 0.43 | 0.11 | 0.52 | 0.63 | -0.015 | 0.625 | 0.315 | 0.958902 | 0.012751 | 0.164467 | 0.999971 | 1 | 0.800215 |
| Lamb2 | Q61292 | 1.1 | 0.32 | 0.42 | 1.21 | 2.55 | 0.38 | 0.77 | -0.18 | 1.06 | 1.84 | 1.07 | 1.78 | -0.25 | 0.35 | 0.97 | 0.33 | 0.1175 | 0.675 | -0.4125 | 0.839119 | 0.05847 | 0.240566 | 0.999971 | 1 | 0.834089 |
| Lamc1 | F8VQJ3 | 0.68 | 0.61 | 0.42 | 0.94 | 2.28 | 0.52 | 0.69 | -0.16 | 1.31 | 1.5 | 1.23 | 1.62 | 0.45 | 0.68 | 0.9 | 0.32 | 0.17 | 0.7525 | -0.075 | 0.7293 | 0.003982 | 0.728337 | 0.999971 | 0.969702 | 0.956041 |
| Lamp1 | P11438 | 0.73 | 0.89 | 0.47 | 0.08 | 0.78 | 1.12 | 0.2 | 1.37 | 0.84 | 0.98 | 0.43 | 0.3 | 0.96 | 0.92 | 0.82 | 0.73 | 0.325 | 0.095 | 0.315 | 0.3144 | 0.7168 | 0.186732 | 0.999971 | 1 | 0.809025 |
| Lamp2 | P17047-3 | 0.43 | 0.66 | -0.09 | -0.15 | 0.62 | 0.54 | -0.64 | 0.41 | 0.69 | 0.41 | -0.46 | -0.28 | 0.85 | 1.2 | 0.15 | 0.28 | 0.02 | -0.1225 | 0.4075 | 0.954203 | 0.716119 | 0.223042 | 0.999971 | 1 | 0.827753 |
| Lamp5 | Q9D387 | -1.3 | 0.53 | 1.68 | 0.28 | 0.99 | 1.35 | 1.46 | 1.72 | 1.32 | 0.09 | 1.54 | 2.5 | 2.75 | 0.56 | 1.91 | 0.6 | 1.0825 | 1.065 | 1.1575 | 0.088415 | 0.163232 | 0.14164 | 0.999971 | 1 | 0.784967 |
| Lamtor1 | Q9CQ22 | -1.73 | -1.3 | -0.47 | -0.58 | -1.24 | -1.58 | -0.53 | -0.31 | -1.23 | -1.99 | -0.79 | -0.32 | -1.29 | -1.24 | -0.69 | -0.61 | 0.105 | -0.0625 | 0.0625 | 0.794707 | 0.886858 | 0.856241 | 0.999971 | 1 | 0.982514 |
| Lamtor2 | Q9JHS3 | -0.2 | 0.75 | 0.42 | 0.06 | 0.6 | 1.01 | 0.11 | 0.5 | 0.89 | 0.63 | 0.42 | -0.36 | 1.28 | 1.22 | 0.67 | 0.33 | 0.2975 | 0.1375 | 0.6175 | 0.319578 | 0.684833 | 0.075365 | 0.999971 | 1 | 0.746958 |
| Lamtor3 | O88653 | -1.15 | -1.03 | 0.24 | 0.49 | -1.77 | -1.57 | 0.32 | 0.21 | -1.5 | -1.9 | -0.13 | 0.24 | -1.7 | -1.16 | 0.34 | 0.92 | -0.34 | -0.46 | -0.0375 | 0.597211 | 0.461259 | 0.955876 | 0.999971 | 1 | 0.998056 |
| Lamtor4 | Q8CF66 | 0.32 | 1.97 | 0.27 | 1.99 | 0.32 | 3.34 | -0.51 | 0.07 | 4.07 | 3.52 | 1.43 | 1.16 | 4.81 | 3.59 | 0.27 | 1.23 | -0.3325 | 1.4075 | 1.3375 | 0.708557 | 0.10489 | 0.216028 | 0.999971 | 1 | 0.826893 |
| Lamtor5 | Q9D1L9 | 0.84 | 1.82 | -0.13 | 0.4 | 1.75 | 1.13 | 0.24 | 0.7 | 1.85 | 0.7 | 0.87 | 0.79 | 2.99 | 1.35 | 0.52 | 0.59 | 0.2225 | 0.32 | 0.63 | 0.649313 | 0.495412 | 0.341275 | 0.999971 | 1 | 0.860984 |
| Lancl1 | O89112 | 0.96 | 1.32 | 1.23 | 1.06 | 1.2 | 1.32 | 0.61 | 1.09 | 1.2 | 1.35 | 0.49 | -0.02 | 1.71 | 1.71 | 1.04 | 0.93 | -0.0875 | -0.3875 | 0.205 | 0.69158 | 0.258012 | 0.426854 | 0.999971 | 1 | 0.880793 |
| Lancl2 | F6RJV6 | -0.65 | -0.54 | -0.7 | -0.79 | -0.57 | -0.43 | -0.95 | -0.85 | -0.56 | -0.71 | -0.81 | -0.82 | -0.41 | -0.35 | -0.86 | -0.39 | -0.03 | -0.055 | 0.1675 | 0.877421 | 0.744025 | 0.400189 | 0.999971 | 1 | 0.871816 |
| Lancl3 | Q8CD19 | 1.16 | 1.28 | 1.49 | 1.56 | 1.4 | 1.43 | 1.19 | 1.67 | 1.31 | 1.8 | 0.94 | 0.46 | 1.52 | 1.55 | 1.9 | 1.24 | 0.05 | -0.245 | 0.18 | 0.799231 | 0.430176 | 0.408542 | 0.999971 | 1 | 0.875014 |
| Lap3 | Q9CPY7 | -0.33 | 0.05 | -0.34 | -0.42 | 0.01 | -0.05 | -0.18 | -0.04 | -0.05 | -0.34 | -0.43 | -0.27 | 0.13 | 0.01 | -0.24 | -0.37 | 0.195 | -0.0125 | 0.1425 | 0.308253 | 0.948175 | 0.499718 | 0.999971 | 1 | 0.903837 |
| Laptm4a | E9QLR3 | -1.15 | -0.25 | -0.68 | -0.82 | -1.3 | -0.6 | -1.43 | -0.05 | -1.52 | -3.71 | -1.06 | 0.55 | -2 | -0.97 | 0.58 | -0.05 | -0.12 | -0.71 | 0.115 | 0.741112 | 0.392961 | 0.833086 | 0.999971 | 1 | 0.979036 |
| Large1 | Q9Z1M7 | 2.6 | 1.9 | 1.39 | 1.13 | 1.68 | 2 | 2.44 | 0.22 | 1.56 | 2.68 | 1.03 | 0.37 | 0.46 | 1.57 | 0.74 | -0.13 | -0.17 | -0.345 | -1.095 | 0.750725 | 0.529361 | 0.03591 | 0.999971 | 1 | 0.699952 |
| Larp1 | Z4YJT3 | 0.38 | 0.35 | 0.21 | 0.33 | 0.41 | 0.22 | 0.62 | 0.4 | 0.59 | 0.37 | 0.31 | 0.35 | 0.78 | 0.06 | 0.34 | 0.34 | 0.095 | 0.0875 | 0.0625 | 0.591136 | 0.600252 | 0.763373 | 0.999971 | 1 | 0.961895 |
| Larp1b | F6U5V1 | 3.69 | 0.23 | 2.6 | 2.92 | 1.19 | 0.54 | 4.94 | 1.29 | 1.07 | 2.68 | 1.99 | 0.26 | -2.58 | -0.84 | 0.55 | -0.15 | -0.37 | -0.86 | -3.115 | 0.73915 | 0.313682 | 0.00697 | 0.999971 | 1 | 0.582714 |
| Larp4 | E9Q066 | 1.92 | 1.99 | 1.63 | 1.58 | 1.46 | 1.89 | 1.56 | 1.67 | 1.7 | 2.22 | 1.38 | 1.05 | 1.71 | 1.85 | 0.85 | 1.36 | -0.135 | -0.1925 | -0.3375 | 0.501034 | 0.501635 | 0.226976 | 0.999971 | 1 | 0.828236 |
| Larp4b | Q6A0A2 | 1.62 | 1.05 | 1.15 | 1.14 | 1.15 | 1.21 | 1.45 | 1.58 | 1.05 | 1.19 | 1.52 | 1.15 | 1.35 | 1.09 | 1 | 1.55 | 0.1075 | -0.0125 | 0.0075 | 0.615355 | 0.952587 | 0.973122 | 0.999971 | 1 | 0.99925 |
| Larp6 | Q8BN59 | 0.3 | 0.48 | 0.1 | 0.21 | 0.46 | 0.71 | -0.06 | 0.85 | 0.57 | 0.75 | 0.61 | 0.58 | 0.95 | 0.55 | 0.64 | 0.97 | 0.2175 | 0.355 | 0.505 | 0.386993 | 0.065317 | 0.028345 | 0.999971 | 1 | 0.678474 |
| Larp7 | Q05CL8 | 3.66 | 0.67 | 0.87 | 1.44 | 0.83 | 1.49 | 0.62 | 1.1 | 0.78 | 2.85 | 0.38 | 0.68 | -1.38 | 0.15 | 0.77 | 0.78 | -0.65 | -0.4875 | -1.58 | 0.328652 | 0.548883 | 0.066252 | 0.999971 | 1 | 0.733937 |
| Lars1 | Q8BMJ2 | 0.44 | 1.36 | 1.31 | 1.01 | 1.22 | 1.01 | 1.18 | 1.05 | 1.38 | 0.76 | 1.48 | 1.66 | 0.95 | 1.06 | 1.3 | 1.15 | 0.085 | 0.29 | 0.085 | 0.731146 | 0.34177 | 0.736446 | 0.999971 | 1 | 0.959447 |
| Lars2 | Q8VDC0 | -0.54 | -0.53 | -0.42 | -0.44 | -0.68 | -0.77 | -0.23 | -0.28 | -0.58 | -0.89 | -0.62 | -0.44 | -0.53 | -0.69 | -0.42 | -0.59 | -0.0075 | -0.15 | -0.075 | 0.969975 | 0.403755 | 0.655616 | 0.999971 | 1 | 0.934061 |
| Las1l | A2BE28-2 | 1.41 | 1.63 | 1.36 | 1.46 | 1.77 | 1.03 | 1.39 | 1.2 | 1.57 | 1.35 | 0.97 | 1.31 | 1.5 | 0.98 | 1.11 | 1.14 | -0.1175 | -0.165 | -0.2825 | 0.589389 | 0.406961 | 0.167052 | 0.999971 | 1 | 0.800215 |
| Lasp1 | A2A6H1 | -1.83 | 2.05 | 1.93 | -0.53 | 1.81 | 1.16 | 3.12 | -0.74 | 2.83 | -0.4 | 2.17 | 1.37 | 3.01 | -0.61 | -1.28 | -0.49 | 0.9325 | 1.0875 | -0.2475 | 0.410387 | 0.320407 | 0.83778 | 0.999971 | 1 | 0.979416 |
| Lasp1_1 | A2A6G9 | 3.31 | 0.1 | 0.35 | 0.44 | 0.15 | 1.55 | 5.36 | 1.37 | -1.45 | 2.12 | 0.09 | -2.53 | 0.98 | -2.66 | -0.42 | -0.55 | 1.0575 | -1.4925 | -1.7125 | 0.391655 | 0.209881 | 0.100124 | 0.999971 | 1 | 0.78268 |
| Lasp1_2 | Q61792 | 0.29 | 0.28 | 1.06 | 0.63 | 0.27 | 0.8 | 1.65 | 0.87 | 0.29 | 0.53 | 0.98 | -0.46 | -0.21 | 0.27 | 0.81 | 0.5 | 0.3325 | -0.23 | -0.2225 | 0.336488 | 0.513491 | 0.456872 | 0.999971 | 1 | 0.887264 |
| Lbh | Q9CX60 | 0.05 | -0.55 | -0.27 | 0.02 | -0.33 | -0.75 | -0.77 | -0.35 | -0.74 | -0.57 | -0.25 | -0.14 | -0.64 | -0.29 | 0.4 | 0.07 | -0.3625 | -0.2375 | 0.0725 | 0.13542 | 0.320884 | 0.79665 | 0.999971 | 1 | 0.970479 |
| Lbhd2 | G3UW55 | 0.4 | 0.34 | 0.34 | 0.43 | 1.04 | -0.61 | -0.04 | 0.55 | 1.33 | 0.08 | -0.1 | 0.89 | 1.58 | 1.06 | 1.07 | -0.11 | -0.1425 | 0.1725 | 0.5225 | 0.686672 | 0.610204 | 0.163856 | 0.999971 | 1 | 0.800215 |
| Lbr | Q3U9G9 | -0.24 | -0.96 | -0.6 | -0.24 | -0.21 | -0.77 | -0.58 | -0.58 | -0.67 | -0.24 | -0.61 | -0.68 | -1.15 | -0.73 | -0.26 | -0.74 | -0.025 | -0.04 | -0.21 | 0.917252 | 0.864507 | 0.445701 | 0.999971 | 1 | 0.884152 |
| Lcat | P16301 | 1.25 | 1.14 | 0.76 | 1.92 | 1.13 | 0.63 | 0.96 | 1.49 | 1.16 | 1.36 | 1.33 | 0.62 | 1.63 | 1.92 | 1.12 | 0.94 | -0.215 | -0.15 | 0.135 | 0.489615 | 0.622889 | 0.684422 | 0.999971 | 1 | 0.944176 |
| Lclat1 | Q3UN02 | -0.64 | 0.19 | -0.17 | 0.12 | -0.06 | 0.07 | -0.38 | -0.12 | 0.18 | -0.63 | 0.01 | -0.22 | 0.41 | 0.28 | 0.26 | -0.08 | 0.0025 | -0.04 | 0.3425 | 0.991764 | 0.883818 | 0.188019 | 0.999971 | 1 | 0.809025 |
| Lcmt1 | A2RTH5 | -0.51 | 1.21 | 0.72 | 0.66 | 1.25 | 0.9 | 0.49 | 0.73 | 1.48 | -0.01 | 0.51 | 0.75 | 1.55 | 0.89 | 0.61 | 0.63 | 0.3225 | 0.1625 | 0.4 | 0.410127 | 0.719431 | 0.339149 | 0.999971 | 1 | 0.860218 |
| Lcp1 | Q61233 | 0.54 | 1.07 | 1.17 | 0.65 | 0.62 | 0.92 | 0.73 | 0.42 | 0.68 | 0.44 | 0.91 | 1.11 | 0.7 | 0.74 | 0.92 | 0.55 | -0.185 | -0.0725 | -0.13 | 0.423256 | 0.764297 | 0.555618 | 0.999971 | 1 | 0.914901 |
| Ldah | A0A0F6AIX | -1.68 | -2.65 | -1.07 | -0.93 | -2.92 | -2.11 | -0.4 | 0.04 | -2.74 | -2.8 | -0.75 | -0.23 | -3.82 | -2.77 | -0.04 | -0.72 | 0.235 | -0.0475 | -0.255 | 0.745854 | 0.946113 | 0.76825 | 0.999971 | 1 | 0.962535 |
| Ldb1 | P70662 | 1.56 | 2.6 | 1.94 | 1.48 | 2.79 | 2.24 | 2.02 | 1.88 | 2.58 | 1.72 | 1.41 | 1.98 | 2.23 | 2.35 | 1.67 | 1.61 | 0.3375 | 0.0275 | 0.07 | 0.314557 | 0.93736 | 0.827427 | 0.999971 | 1 | 0.977122 |
| Ldha | P06151 | -1.05 | -0.3 | -0.55 | 0.04 | -0.4 | -0.52 | 0.05 | -0.46 | -0.63 | -0.91 | -0.02 | -0.21 | -0.36 | -0.38 | -0.25 | 0.53 | 0.1325 | 0.0225 | 0.35 | 0.63728 | 0.942068 | 0.289554 | 0.999971 | 1 | 0.847402 |
| Ldhb | A0A6I8MX | 0.08 | -0.04 | -0.34 | 0.12 | -0.21 | -0.14 | -0.01 | 0.03 | -0.33 | -0.13 | -0.06 | -0.42 | -0.42 | 0.11 | -0.15 | 0.07 | -0.0375 | -0.19 | -0.0525 | 0.841804 | 0.338666 | 0.803808 | 0.999971 | 1 | 0.971467 |
| Ldhd | Q7TNG8 | -0.8 | -0.18 | -0.7 | -0.52 | 0.07 | -0.45 | -0.44 | -0.55 | -0.26 | -0.45 | -0.23 | -0.12 | 0.42 | -0.04 | 0.02 | -0.82 | 0.2075 | 0.285 | 0.445 | 0.382401 | 0.185622 | 0.165338 | 0.999971 | 1 | 0.800215 |
| Ldlr | A0A1L1SRE | 0.17 | -1.03 | -0.45 | -0.53 | -1.22 | -1.04 | -0.63 | -0.88 | -1.1 | -0.27 | -0.53 | -0.36 | -0.82 | -0.65 | -0.54 | -0.56 | -0.4825 | -0.105 | -0.1825 | 0.121422 | 0.737563 | 0.509601 | 0.999971 | 1 | 0.905815 |
| Ldlrap1 | Q8C142 | 0.19 | -0.3 | 0.08 | -0.9 | -1.19 | -0.13 | 0.48 | -0.11 | -0.13 | 0.37 | 0.67 | 0.25 | -3.57 | -1.4 | -0.18 | -1.13 | -0.005 | 0.5225 | -1.3375 | 0.990162 | 0.112455 | 0.078309 | 0.999971 | 1 | 0.754157 |
| Lemd2 | Q6DVA0 | -0.48 | -0.09 | -0.27 | 0.21 | 0.93 | -0.5 | -0.21 | 0.25 | 3 | -0.29 | -0.02 | 0.05 | 1.48 | 0.23 | -0.05 | -0.53 | 0.275 | 0.8425 | 0.44 | 0.428247 | 0.259326 | 0.32024 | 0.999971 | 1 | 0.853962 |
| Lemd3 | E9QP59 | 0.19 | -0.31 | -0.36 | -0.34 | -0.11 | -0.45 | -0.09 | -0.37 | -0.17 | 0.09 | -0.22 | -0.52 | 0.05 | -0.49 | -0.35 | -0.57 | -0.05 | 0 | -0.135 | 0.812476 | 1 | 0.560974 | 0.999971 | 1 | 0.91591 |
| Leng1 | Q9DB98 | 5.03 | -0.16 | 4.07 | 5.9 | 0.78 | 2.25 | 5.41 | 3.24 | 0.43 | 2.98 | 0.65 | 1.57 | -3.13 | -1.11 | 2.67 | 2.41 | -0.79 | -2.3025 | -3.5 | 0.593737 | 0.106434 | 0.068763 | 0.999971 | 1 | 0.733937 |
| Leo1 | Q5XJE5 | 1.66 | 1.12 | 0.45 | 0.71 | 0.6 | 0.91 | 0.11 | 0.39 | 0.64 | 1.51 | 0.33 | 0.97 | 2.3 | 1.09 | 0.34 | 0.98 | -0.4825 | -0.1225 | 0.1925 | 0.153204 | 0.732504 | 0.675026 | 0.999971 | 1 | 0.941133 |
| Leprotl1 | Q9CQ74 | -0.51 | 1.25 | 3.15 | 0.39 | 0.85 | 1.82 | 2.23 | 0.82 | 1.56 | 0.61 | 1.7 | 0.41 | 1.97 | 1.47 | 1.25 | 0.65 | 0.36 | -2.36E-16 | 0.265 | 0.642579 | 1 | 0.723518 | 0.999971 | 1 | 0.955258 |
| Letm1 | Q9Z2I0 | 0.12 | 0.47 | -0.78 | -0.78 | 0.54 | 0.38 | -0.75 | -0.78 | 0.16 | 0.06 | -0.81 | -1.43 | 0.77 | 0.06 | -0.9 | -0.91 | 0.09 | -0.2625 | -0.0025 | 0.840834 | 0.575051 | 0.995861 | 0.999971 | 1 | 1 |
| Letmd1 | Q924L1 | -0.45 | 0.01 | -0.28 | -0.26 | -0.14 | -0.15 | -0.43 | -0.46 | 0.01 | 0.09 | -0.26 | -0.82 | 0.14 | -0.34 | -0.2 | -0.21 | -0.05 | -3.93E-17 | 0.0925 | 0.795974 | 1 | 0.645148 | 0.999971 | 1 | 0.93301 |
| Lgals1 | P16045 | 0.83 | 1.47 | 1.62 | 1.63 | 1.76 | 0.96 | 1.06 | 0.03 | 1.11 | 0.93 | 0.94 | 1.26 | 0.18 | 0.19 | 0.48 | 0.59 | -0.435 | -0.3275 | -1.0275 | 0.278698 | 0.190435 | 0.00241 | 0.999971 | 1 | 0.535772 |
| Lgals3bp | Q07797 | -0.14 | 0.34 | -0.14 | -0.5 | -0.06 | 0.21 | -0.93 | -0.49 | 0.37 | 0.73 | -0.26 | -0.17 | -0.01 | 0.18 | -0.46 | -0.35 | -0.2075 | 0.2775 | -0.05 | 0.507486 | 0.364589 | 0.844175 | 0.999971 | 1 | 0.979814 |
| Lgals8 | Q9JL15 | -1.99 | -1.42 | 0.2 | -0.6 | -1.13 | -1.23 | -0.23 | 0.42 | -1.78 | -1.88 | 0.21 | -0.19 | -1.75 | -1.43 | 0.07 | 0.54 | 0.41 | 0.0425 | 0.31 | 0.476054 | 0.948392 | 0.644579 | 0.999971 | 1 | 0.93301 |
| Lgals9 | B1AQR8 | 0.12 | 0.83 | 0.59 | 0.24 | 0.58 | 0 | -0.13 | -0.47 | 0.39 | -0.35 | -0.17 | 0.77 | 0.45 | 0.56 | 0.45 | -0.31 | -0.45 | -0.285 | -0.1575 | 0.141664 | 0.368683 | 0.571998 | 0.999971 | 1 | 0.91816 |
| Lgalsl | Q8VED9 | -0.62 | -0.3 | 0.74 | 0.96 | -0.31 | -0.44 | 0.2 | 0.15 | -0.47 | -0.63 | 0.26 | 0.57 | -0.59 | -0.12 | 0.43 | 0.89 | -0.295 | -0.2625 | -0.0425 | 0.468013 | 0.56659 | 0.927989 | 0.999971 | 1 | 0.994278 |
| Lgi1 | Q9JIA1 | -0.69 | 0.03 | -0.36 | -0.64 | -0.19 | 0.23 | -0.43 | -0.55 | -0.01 | -0.68 | -0.81 | -0.8 | 0.09 | 0 | -0.55 | -1.08 | 0.18 | -0.16 | 0.03 | 0.496987 | 0.557432 | 0.925386 | 0.999971 | 1 | 0.99401 |
| Lgi2 | Q8K4Z0-2 | -1.69 | -0.89 | -1.17 | -1.8 | -1.27 | -1.18 | -1.29 | -1.27 | -1.25 | -1.78 | -1.3 | -1.14 | -0.53 | -0.95 | -1.38 | -1.53 | 0.135 | 0.02 | 0.29 | 0.586865 | 0.941758 | 0.370761 | 0.999971 | 1 | 0.869526 |
| Lgi3 | Q8K406 | -0.24 | 0.08 | -0.35 | -0.86 | -0.27 | 0.14 | -0.57 | -0.39 | 0.27 | -0.07 | -0.33 | -0.59 | 1.23 | 0.06 | -0.32 | -0.42 | 0.07 | 0.1625 | 0.48 | 0.793455 | 0.567624 | 0.25731 | 0.999971 | 1 | 0.839012 |
| Lgi4 | Q8K1S1 | -0.23 | -0.07 | 0.44 | 0.64 | 0.56 | -0.1 | 0.4 | 0.69 | -1.75 | -1.42 | 0.09 | 0.24 | 0.53 | -0.32 | 1.04 | 0.26 | 0.1925 | -0.905 | 0.1825 | 0.502385 | 0.104958 | 0.600647 | 0.999971 | 1 | 0.92426 |
| Lgmn | O89017 | -0.25 | 1.04 | 1.2 | 1.37 | 1.13 | 0.94 | 2.24 | 1.22 | 1.37 | 0.34 | 0.47 | 0.89 | 1.91 | 0.54 | 1.29 | 0.51 | 0.5425 | -0.0725 | 0.2225 | 0.240893 | 0.861679 | 0.635909 | 0.999971 | 1 | 0.93301 |
| Lhfpl2 | Q8BGA2 | 0.12 | 0.09 | 0.15 | 0.85 | -0.43 | 0.4 | 0.33 | -0.36 | -0.59 | -0.44 | 0.34 | -0.14 | -1.69 | -0.64 | 0.6 | -0.61 | -0.3175 | -0.51 | -0.8875 | 0.299715 | 0.10289 | 0.086359 | 0.999971 | 1 | 0.759991 |
| Lhfpl3 | Q9CTN8 | 0.43 | -0.47 | -1.27 | -0.33 | -0.93 | 0.42 | -1.35 | -0.14 | 0.57 | -0.5 | -0.93 | -1.76 | 0.47 | 0.57 | -1.03 | -0.49 | -0.09 | -0.245 | 0.29 | 0.854268 | 0.658537 | 0.553683 | 0.999971 | 1 | 0.913787 |
| Lhfpl4 | Q5U4E0 | 0.24 | 0.83 | 1.15 | 0.17 | 0.52 | 1.57 | 0.51 | 0.73 | 0.64 | 1.11 | 0.82 | 0.6 | 1.23 | 0.84 | 0.95 | 1.41 | 0.235 | 0.195 | 0.51 | 0.496544 | 0.489461 | 0.100285 | 0.999971 | 1 | 0.78268 |
| Lhpp | Q9D7I5 | -1.07 | 0.33 | 0.52 | -0.55 | 0.47 | 0.71 | 0.27 | 0.2 | 0.95 | -0.97 | 0.34 | 0.77 | 0.31 | -0.13 | 0.62 | -0.66 | 0.605 | 0.465 | 0.2275 | 0.135984 | 0.391709 | 0.60768 | 0.999971 | 1 | 0.925689 |
| Lias | Q99M04 | 0.06 | 0.09 | -0.07 | -0.26 | -0.06 | 0.18 | -0.29 | -0.08 | 0.27 | 0.46 | 0.09 | -0.19 | 1.05 | 0.26 | 0.15 | 0.27 | -0.0175 | 0.2025 | 0.4775 | 0.927003 | 0.343146 | 0.083958 | 0.999971 | 1 | 0.755254 |
| Lifr | P42703 | 0.43 | -0.45 | -0.56 | -0.9 | 0.05 | -0.16 | 0.27 | -0.09 | -0.55 | 0.56 | 0.1 | 0.28 | -0.72 | 0.09 | -0.52 | -0.39 | 0.3875 | 0.4675 | -0.015 | 0.225745 | 0.216712 | 0.963859 | 0.999971 | 1 | 0.998721 |
| Lig3 | Q3UC82 | 1.89 | 2.58 | 2.02 | 2.07 | 2.68 | 2.45 | 1.43 | 2.02 | 2.89 | 2.91 | 1.81 | 2.12 | 1.95 | 2.28 | 2.01 | 2.34 | 0.005 | 0.2925 | 0.005 | 0.987397 | 0.370049 | 0.982093 | 0.999971 | 1 | 0.99925 |
| Lima1 | Q9ERG0 | 0.55 | -0.1 | 0.01 | 0.13 | 0.36 | 0.33 | 0.05 | -0.28 | 0.32 | 1.07 | -0.05 | 0.39 | 0.15 | 0.25 | 0.1 | 0.36 | -0.0325 | 0.285 | 0.0675 | 0.891852 | 0.332471 | 0.745175 | 0.999971 | 1 | 0.961064 |
| Limch1 | Q3UH68 | 0.5 | -0.16 | 0.23 | 0.15 | -0.32 | -0.1 | 0.15 | 0.34 | 0.01 | 0.21 | -0.24 | -0.17 | -0.12 | -0.17 | 0.05 | -0.14 | -0.1625 | -0.2275 | -0.275 | 0.493345 | 0.302729 | 0.197093 | 0.999971 | 1 | 0.812891 |
| Limd1 | Q9QXD8 | 1.42 | 1.5 | 0.62 | 1.22 | 1.85 | 1.33 | 2.04 | -0.21 | 1.16 | 2.43 | 0.89 | 1.37 | 0.86 | 1.08 | 0.9 | 0.64 | 0.0625 | 0.2725 | -0.32 | 0.901787 | 0.478822 | 0.218991 | 0.999971 | 1 | 0.827064 |
| Limd2 | Q8BGB5 | 1.83 | 0.2 | 0.85 | 0.44 | 0.1 | 0.96 | 3.04 | 0.4 | 0.27 | 0.72 | 0.61 | 2.8 | 0.72 | 0.61 | 1.59 | -0.39 | 0.295 | 0.27 | -0.1975 | 0.666964 | 0.665248 | 0.695583 | 0.999971 | 1 | 0.947174 |
| Limk1 | P53668 | 0.22 | -0.63 | -0.5 | -0.2 | -0.43 | -0.28 | -0.68 | -0.05 | -0.46 | -0.35 | -1.16 | -0.8 | -0.36 | -0.13 | -0.48 | -0.17 | -0.0825 | -0.415 | -0.0075 | 0.747555 | 0.160804 | 0.975015 | 0.999971 | 1 | 0.99925 |
| Limk2 | Q5NC03 | 0.8 | 0.15 | -0.89 | -0.15 | 0.39 | 0.33 | -0.1 | -0.55 | 0 | 0.82 | -0.27 | -0.09 | 0.36 | 0.41 | -0.38 | -0.25 | 0.04 | 0.1375 | 0.0575 | 0.919388 | 0.736246 | 0.882874 | 0.999971 | 1 | 0.986576 |
| Lims1 | A0A5F8MP | 0.62 | 1.8 | 1.34 | 1.44 | 1.88 | 1.91 | 1.58 | 1.66 | 1.31 | 1.92 | 1.63 | 0.24 | 0.98 | 1.85 | 1.11 | 0.81 | 0.4575 | -0.025 | -0.1125 | 0.12537 | 0.952588 | 0.738182 | 0.999971 | 1 | 0.959879 |
| Lin7a | Q8JZS0 | -0.36 | -0.98 | -1.16 | -1.43 | -1.04 | -0.85 | -1.22 | -1.38 | -1.18 | -0.47 | -1.7 | -1.95 | -1.22 | -0.87 | -1.8 | -1.18 | -0.14 | -0.3425 | -0.285 | 0.610143 | 0.384292 | 0.363937 | 0.999971 | 1 | 0.869526 |
| Lin7b | O88951 | 0.02 | -0.96 | -0.36 | -0.33 | -0.86 | -0.72 | 0.13 | -0.41 | -1.15 | -0.11 | -0.51 | -0.56 | -0.62 | -0.71 | -0.63 | -0.1 | -0.0575 | -0.175 | -0.1075 | 0.850838 | 0.566448 | 0.689401 | 0.999971 | 1 | 0.945805 |
| Lin7c | O88952 | 0.91 | -0.25 | -0.44 | -0.25 | -0.43 | -0.27 | -0.28 | -0.75 | -0.44 | 0.54 | -1.06 | -1.85 | -0.51 | -0.32 | -0.86 | -0.71 | -0.425 | -0.695 | -0.5925 | 0.21754 | 0.22567 | 0.100686 | 0.999971 | 1 | 0.78268 |
| Lin9 | A0A0A6YW | -0.91 | 2.34 | 3.06 | 1.22 | 2.69 | 1.75 | 1.75 | 2.97 | 2.34 | -0.01 | 2.75 | 3.38 | 1.2 | 1.6 | 3.13 | 2.05 | 0.8625 | 0.6875 | 0.5675 | 0.312697 | 0.507508 | 0.515909 | 0.999971 | 1 | 0.905815 |
| Lingo1 | A9DA50 | -0.4 | -0.17 | -1.07 | -1.06 | -0.43 | -0.02 | -1.12 | -1 | -0.39 | -0.21 | -0.85 | -1.13 | 0.52 | -0.06 | -0.98 | -0.73 | 0.0325 | 0.03 | 0.3625 | 0.923936 | 0.924059 | 0.368927 | 0.999971 | 1 | 0.869526 |
| Lingo2 | Q3URE9 | -1.9 | -0.87 | -1.15 | -1.46 | -0.18 | -0.68 | -1.93 | -1.21 | -0.19 | -1.56 | -1.23 | -1 | 0.92 | -0.76 | -1.57 | -0.8 | 0.345 | 0.35 | 0.7925 | 0.41339 | 0.342227 | 0.157921 | 0.999971 | 1 | 0.797252 |
| Lingo3 | Q6GQU6 | 0.15 | -1.12 | -0.58 | -0.21 | -0.84 | -1.68 | -0.46 | -0.43 | -0.98 | -0.56 | -1.01 | -0.64 | -1.16 | -1.1 | -0.36 | -0.9 | -0.4125 | -0.3575 | -0.44 | 0.298176 | 0.255685 | 0.202118 | 0.999971 | 1 | 0.81617 |
| Lipa | Q9Z0M5 | -0.51 | 0.52 | 0.44 | 0.15 | 1.29 | 2.46 | 0.07 | 0.94 | 1.21 | 0.5 | 0.62 | 0.13 | 1.83 | 1.75 | 0.49 | 0.99 | 1.04 | 0.465 | 1.115 | 0.066114 | 0.176724 | 0.016511 | 0.999971 | 1 | 0.613095 |
| Lipe | E9Q4M2 | -0.54 | -0.81 | -0.08 | -0.43 | -0.9 | -0.4 | -0.87 | -0.44 | -1.27 | -0.28 | -0.56 | -0.2 | -0.59 | -0.36 | 0.05 | -0.36 | -0.1875 | -0.1125 | 0.15 | 0.436859 | 0.704442 | 0.531249 | 0.999971 | 1 | 0.910069 |
| Lipt1 | Q8VCM4 | 1.57 | 2.67 | -1.24 | -0.29 | 0.87 | 1.33 | -0.83 | -0.02 | 0.99 | 0.2 | -0.21 | -0.09 | 0.85 | 1.11 | -1.16 | -0.22 | -0.34 | -0.455 | -0.5325 | 0.70651 | 0.589087 | 0.566224 | 0.999971 | 1 | 0.917959 |
| Lipt2 | Q9D009 | 1.02 | -0.2 | -0.06 | 0.22 | -0.47 | -0.61 | -0.57 | 0.48 | -0.79 | 0.98 | -0.34 | -0.69 | -1.62 | 0.01 | -0.36 | 0.35 | -0.5375 | -0.455 | -0.65 | 0.166945 | 0.338171 | 0.195367 | 0.999971 | 1 | 0.811225 |
| Lix1 | Q6P566 | -0.28 | -1.1 | -1.17 | -0.29 | -0.79 | -0.97 | -1.36 | 1.13 | -1.46 | -1.04 | -1.42 | 0.35 | -1.07 | -0.63 | 1.64 | -1.18 | 0.2125 | -0.1825 | 0.4 | 0.703943 | 0.693594 | 0.537532 | 0.999971 | 1 | 0.910077 |
| Llgl1 | A0A0R4J0S | -0.88 | -0.16 | -0.7 | -1.02 | -0.28 | -0.45 | -0.67 | -1.05 | 0.07 | -0.9 | -0.96 | -0.92 | 0.26 | -0.83 | -0.98 | -1.19 | 0.0775 | 0.0125 | 0.005 | 0.774768 | 0.968378 | 0.989067 | 0.999971 | 1 | 0.99925 |
| Lman1 | Q9D0F3 | -0.71 | 0.24 | 1.46 | 1.09 | 0.1 | 0.18 | 1.61 | 1.36 | 0.02 | -0.13 | 1.54 | 1.32 | -0.39 | 0.31 | 1.08 | 1.06 | 0.2925 | 0.1675 | -0.005 | 0.610235 | 0.778756 | 0.992733 | 0.999971 | 1 | 0.99925 |
| Lman2 | Q9DBH5 | -0.64 | 1.26 | 0.19 | 0.03 | 1.4 | 1.05 | 0.21 | 0.74 | 1.52 | -0.41 | 0.59 | 0.91 | 2.03 | 1.35 | 0.83 | 0.8 | 0.64 | 0.4425 | 1.0425 | 0.170853 | 0.407421 | 0.04596 | 0.999971 | 1 | 0.710102 |
| Lman2l | D3Z4P2 | 0.79 | 1.31 | 0.7 | 0.88 | 1.3 | 1.58 | 0.38 | 1.52 | 1.52 | 1.56 | 1.08 | 0.93 | 2.03 | 1.9 | 1.43 | 1.59 | 0.275 | 0.3525 | 0.8175 | 0.390954 | 0.164897 | 0.006172 | 0.999971 | 1 | 0.582714 |
| Lmbr1 | J3QM78 | 0.95 | 1.16 | 0.25 | 0.74 | 4.92 | 3.72 | 0.39 | 0.64 | -2.05 | 0.05 | 0.32 | -0.38 | 4.09 | 2.21 | 0.36 | 0.53 | 1.6425 | -1.29 | 1.0225 | 0.133924 | 0.034614 | 0.224398 | 0.999971 | 1 | 0.828225 |
| Lmbrd1 | Q8K0B2-3 | -0.28 | 0.26 | 0.95 | 1 | -0.14 | -0.54 | 1.93 | 1.56 | -0.41 | -0.6 | 0.48 | 0.07 | -0.27 | -0.37 | 0.36 | 0.75 | 0.22 | -0.5975 | -0.365 | 0.724473 | 0.140894 | 0.361206 | 0.999971 | 1 | 0.869526 |
| Lmbrd2 | Q8C561 | -0.06 | -0.73 | -0.58 | -0.64 | -0.74 | -0.36 | 1.7 | -1.01 | -0.57 | -0.91 | -0.84 | -1.98 | -0.73 | -1.04 | -0.84 | -1.3 | 0.4 | -0.5725 | -0.475 | 0.496009 | 0.120984 | 0.067663 | 0.999971 | 1 | 0.733937 |
| Lmcd1 | Q8VEE1 | 0.44 | 0.68 | 0.77 | 0.47 | 0.95 | 1.18 | 0.69 | -0.02 | 0.75 | 0.96 | 1.36 | 1.24 | 0.56 | 0.98 | 0.62 | 0.78 | 0.11 | 0.4875 | 0.145 | 0.701118 | 0.040918 | 0.455921 | 0.999971 | 1 | 0.887217 |
| Lmf1 | Z4YJR1 | -0.78 | 0.77 | 0.53 | 0.59 | 0.59 | 0.8 | -0.2 | 0.31 | 0.56 | -0.3 | 0.3 | -0.03 | 1.24 | 0.68 | 0.27 | 0.99 | 0.0975 | -0.145 | 0.5175 | 0.806879 | 0.709711 | 0.213962 | 0.999971 | 1 | 0.825334 |
| Lmf2 | Q8C3X8 | -1.07 | -1.58 | -1.3 | -1.21 | -1.63 | -0.97 | -1.54 | -1.14 | -1.76 | -1.3 | -1.62 | -1.02 | -2.6 | -1.23 | -0.93 | -1.11 | -0.03 | -0.135 | -0.1775 | 0.895942 | 0.564175 | 0.645558 | 0.999971 | 1 | 0.93301 |
| Lmln | Q8BMN4 | -0.26 | -0.07 | 0.01 | 0.09 | -0.04 | -0.07 | -0.58 | 0.34 | 0.09 | -0.2 | -0.34 | 0.01 | 0.16 | 0.13 | -0.01 | 0.3 | -0.03 | -0.0525 | 0.2025 | 0.89935 | 0.779819 | 0.277155 | 0.999971 | 1 | 0.843812 |
| Lmna | P48678 | -0.33 | -0.96 | -0.67 | -0.48 | -0.76 | -1.17 | -0.47 | -1.11 | -1.22 | -0.54 | -0.4 | -0.59 | -1.72 | -1.38 | -1.07 | -0.82 | -0.2675 | -0.0775 | -0.6375 | 0.288574 | 0.758904 | 0.035043 | 0.999971 | 1 | 0.69205 |
| Lmnb1 | P14733 | -0.6 | -1.04 | -0.9 | -0.77 | -0.71 | -1.01 | -0.78 | -1.09 | -0.63 | -0.64 | -0.86 | -1.06 | -0.78 | -0.83 | -1 | -0.93 | -0.07 | 0.03 | -0.0575 | 0.719086 | 0.878138 | 0.753763 | 0.999971 | 1 | 0.961116 |
| Lmnb2 | P21619 | -0.84 | -1.58 | -1.03 | -0.8 | -1.73 | -1.71 | -0.94 | -1.31 | -1.57 | -1.08 | -1.17 | -1.35 | -1.95 | -1.37 | -1.08 | -1.13 | -0.36 | -0.23 | -0.32 | 0.214918 | 0.350916 | 0.276707 | 0.999971 | 1 | 0.843794 |
| Lmo4 | P61969 | -0.9 | 0.25 | 0.16 | 0.78 | 0.26 | -0.11 | -0.58 | 0.29 | 0.05 | 0.49 | 0.52 | 0.65 | 0.91 | 0.67 | -0.09 | 1.4 | -0.1075 | 0.355 | 0.65 | 0.783119 | 0.345373 | 0.167121 | 0.999971 | 1 | 0.800215 |
| Lmo7 | E9PYI7 | -0.38 | -1.01 | -0.98 | -0.88 | -1.18 | -1.07 | -1.12 | -1.22 | -1.11 | -0.48 | -1.01 | -1.07 | -0.75 | -1.07 | -0.97 | -1 | -0.335 | -0.105 | -0.135 | 0.128793 | 0.661743 | 0.529768 | 0.999971 | 1 | 0.910069 |

| | | | | | | | | | | | | | | | | | | | | | | | | | | | |
|---|---|---|---|---|---|---|---|---|---|---|---|---|---|---|---|---|---|---|---|---|---|---|---|---|---|---|---|
| Lmtk2 | Q3TYD6 | -1.29 | -0.85 | -0.58 | -0.61 | -0.85 | -0.7 | -0.59 | -0.56 | -1.04 | -1.12 | -0.48 | -0.43 | 0.08 | -0.86 | -0.63 | -0.16 | 0.1575 | 0.065 | 0.44 | 0.481321 | 0.806032 | 0.148259 | 0.999971 | | 1 | 0.793248 |
| Lmtk3 | A0A3B2WC | 0.33 | -0.41 | -0.45 | -0.51 | -0.57 | -0.3 | -0.26 | -0.35 | -0.5 | 0.16 | 0.01 | -0.46 | 0.15 | -0.21 | -0.29 | -0.01 | -0.11 | 0.0625 | 0.17 | 0.650545 | 0.819952 | 0.502428 | 0.999971 | | 1 | 0.904573 |
| Lmtk3_1 | A0A1B0GS | -0.36 | -0.42 | -0.24 | -0.01 | -0.25 | -0.23 | 0.36 | -0.04 | -0.52 | -0.32 | -0.41 | -0.37 | 0.01 | -0.09 | -0.33 | -0.2 | 0.2175 | -0.1475 | 0.105 | 0.326353 | 0.412533 | 0.579296 | 0.999971 | | 1 | 0.919302 |
| Lnpep | Q8C129 | -0.14 | -0.21 | 0.23 | 0.33 | -0.07 | -0.05 | 0.02 | 0.51 | -0.29 | 0.12 | 0.32 | 0.24 | -0.26 | -0.09 | 0.39 | 0.73 | 0.05 | 0.045 | 0.14 | 0.827806 | 0.842927 | 0.618444 | 0.999971 | | 1 | 0.928581 |
| Lnpk | Q7TQ95 | 0.01 | 0.15 | -0.54 | -0.22 | 0.13 | 0.6 | -0.6 | -0.23 | 0.14 | 0.42 | -0.18 | -0.21 | 0.63 | 0.52 | -0.19 | 0.23 | 0.125 | 0.1925 | 0.4475 | 0.682442 | 0.433724 | 0.113739 | 0.999971 | | 1 | 0.78268 |
| Lonp1 | Q8CGK3 | -0.35 | -0.84 | -0.75 | -0.73 | -0.9 | -1.19 | -0.69 | -1.06 | -0.62 | -0.92 | -0.84 | -0.66 | -1.29 | -1.48 | -1.23 | -0.81 | -0.2925 | -0.0925 | -0.535 | 0.180353 | 0.6303 | 0.036773 | 0.999971 | | 1 | 0.700096 |
| Lonp2 | Q9DBN5 | -0.58 | -0.62 | -1.11 | -0.96 | -0.69 | -0.32 | -1.19 | -0.51 | -0.93 | 0.16 | -0.8 | -0.53 | -1.02 | -0.16 | -1.09 | -0.5 | 0.14 | 0.2925 | 0.125 | 0.584093 | 0.323193 | 0.650701 | 0.999971 | | 1 | 0.933842 |
| Lonrf2 | F6ZE64 | 1.74 | 2.41 | 2.27 | 2.66 | 1.53 | 1.75 | 4.36 | 2.48 | 2.2 | 1.16 | 1.92 | 0.16 | 1.16 | 2.17 | 2.21 | 1.52 | 0.26 | -0.91 | -0.505 | 0.672299 | 0.077972 | 0.144871 | 0.999971 | | 1 | 0.79148 |
| Lpar1 | P61793-2 | 2.27 | 0.75 | 0.98 | 0.82 | 0 | 0.56 | 0.6 | 1.19 | 0.71 | 1.72 | 0.31 | 0.36 | 0.26 | 0.18 | 1.03 | 0.65 | -0.6175 | -0.43 | -0.675 | 0.15926 | 0.359193 | 0.112968 | 0.999971 | | 1 | 0.78268 |
| Lpcat1 | Q3TFD2 | 0.08 | 0.82 | 0.29 | -0.13 | 0.45 | 0.42 | 0.32 | 0.45 | 0.41 | 0.27 | 0.62 | 0.6 | 1.6 | 0.43 | 0.62 | 0.39 | 0.145 | 0.21 | 0.495 | 0.548763 | 0.405594 | 0.17694 | 0.999971 | | 1 | 0.808532 |
| Lpcat2 | Q8BYI6 | 0.75 | -1.95 | 0.13 | 0.32 | 0.3 | 0.35 | -0.04 | 0.14 | -0.07 | 0.41 | -0.02 | 0.22 | -0.1 | 0.07 | 0.81 | 0.59 | 0.375 | 0.3225 | 0.53 | 0.506987 | 0.571218 | 0.376406 | 0.999971 | | 1 | 0.870465 |
| Lpcat3 | Q91V01 | 0.08 | 0.4 | -0.01 | -0.8 | -0.19 | 0.27 | -0.2 | 0.18 | -0.01 | 0.37 | 0.32 | 0.22 | -0.09 | 0.57 | 0.23 | 0.23 | 0.0975 | 0.3075 | 0.3175 | 0.740298 | 0.292237 | 0.302918 | 0.999971 | | 1 | 0.847402 |
| Lpcat4 | Q6NVG1 | -0.83 | 0.02 | -1.14 | -0.94 | 0.37 | -0.16 | -0.82 | -0.83 | 0.17 | -0.71 | -0.57 | -0.44 | 0.54 | -0.4 | -0.89 | -0.53 | 0.3625 | 0.335 | 0.4025 | 0.345563 | 0.313799 | 0.310314 | 0.999971 | | 1 | 0.847402 |
| Lpgat1 | E9QL80 | -0.98 | -1.03 | -0.65 | -0.56 | -0.78 | -0.6 | -0.05 | -0.6 | -0.14 | -1.22 | -0.39 | -0.28 | -0.44 | -1.28 | -0.53 | -0.24 | 0.2975 | 0.2975 | 0.1825 | 0.223559 | 0.308827 | 0.511379 | 0.999971 | | 1 | 0.905815 |
| Lpin1 | A0A1Y7VL | 1.69 | 0.7 | -0.3 | 0.35 | 0.75 | 0.12 | 0.5 | 1.2 | -0.03 | 1.37 | 0.03 | -0.17 | 0.67 | 0.75 | -0.3 | -0.38 | 0.0325 | -0.31 | -0.425 | 0.9417 | 0.548076 | 0.387206 | 0.999971 | | 1 | 0.871816 |
| Lpin2 | E9PWN0 | -0.2 | 1.24 | 0.09 | 0.12 | 1.52 | 0.65 | -0.2 | -0.03 | 1.49 | 0.25 | 0.07 | -0.04 | 2.62 | 0.89 | -0.04 | 0.73 | 0.1725 | 0.13 | 0.7375 | 0.714492 | 0.772381 | 0.232401 | 0.999971 | | 1 | 0.831121 |
| Lpp | Q8BFW7 | 0.54 | -0.12 | -0.14 | 0.06 | 2.32 | 0.09 | -0.52 | 0.06 | 0.24 | 1.54 | 0.47 | 1.85 | -0.26 | -0.1 | 0.17 | 0.2 | 0.4025 | 0.94 | -0.0825 | 0.50116 | 0.043786 | 0.72216 | 0.999971 | | 1 | 0.955258 |
| Lratd1 | A0A0R4J01 | -0.3 | 2.6 | 2.67 | 0.01 | 4.24 | 2.58 | 5.83 | 1.72 | 4.4 | 0.02 | 5.27 | 7.12 | 6.1 | 1.13 | 3.48 | 3.57 | 2.3475 | 2.9575 | 2.325 | 0.054742 | 0.079777 | 0.070442 | 0.999971 | | 1 | 0.735439 |
| Lratd2 | D3YXJ5 | 0.46 | 0.61 | -1.17 | -0.7 | 0.26 | -0.15 | -0.98 | -0.95 | -0.75 | 0.99 | -0.82 | -0.5 | 0.18 | 0.12 | -0.45 | -0.82 | -0.255 | -0.07 | -0.0425 | 0.608728 | 0.900832 | 0.927253 | 0.999971 | | 1 | 0.99417 |
| Lrba | E9Q3Y4 | 0.34 | 0.55 | 0.69 | 0.74 | 0.38 | -0.01 | 0.53 | 0.47 | 0.37 | 0.42 | 0.28 | 0.53 | -0.02 | 0.39 | 0.52 | 0.58 | -0.2375 | -0.18 | -0.2125 | 0.265349 | 0.326249 | 0.33095 | 0.999971 | | 1 | 0.857107 |
| Lrch1 | P62046 | -0.12 | 0.26 | -0.25 | -0.15 | 0.18 | 0.06 | -0.93 | -0.34 | 0.11 | -0.21 | -0.46 | -0.26 | 1.01 | 0.37 | -0.45 | -0.23 | -0.1925 | -0.14 | 0.24 | 0.507849 | 0.509578 | 0.490307 | 0.999971 | | 1 | 0.898612 |
| Lrch2 | F6VQ19 | -0.7 | -0.69 | 0.24 | 0.17 | -0.87 | -0.69 | -0.3 | 0.3 | -0.67 | -0.57 | 0.16 | 0.45 | -0.27 | -0.15 | 0.51 | 0.43 | -0.145 | 0.0875 | 0.375 | 0.687192 | 0.812222 | 0.271105 | 0.999971 | | 1 | 0.843794 |
| Lrch3 | B2RXP1 | 0.18 | 0.51 | -0.28 | 0.87 | 0.05 | -0.07 | -0.19 | -0.4 | -0.01 | -0.21 | 0.8 | 1.19 | 0.19 | 0.12 | 0.58 | 0.99 | -0.4725 | 0.1225 | 0.15 | 0.117129 | 0.757328 | 0.640802 | 0.999971 | | 1 | 0.93301 |
| Lrfn1 | Q2WF71 | -1.09 | 0.13 | -0.96 | -0.76 | 0.46 | -0.54 | -0.95 | -0.77 | 0.58 | -0.74 | -0.88 | -0.85 | 1.24 | -0.88 | -0.85 | -0.48 | 0.22 | 0.1975 | 0.4275 | 0.585117 | 0.643816 | 0.42804 | 0.999971 | | 1 | 0.880793 |
| Lrfn2 | Q80TG9 | -0.84 | -0.93 | -1.52 | -1.12 | -1.06 | -1.22 | -1.19 | -0.7 | -1.24 | -1.06 | -1.47 | -1.23 | -1.02 | -1.29 | -1.77 | -1.19 | 0.06 | -0.1475 | -0.215 | 0.79467 | 0.500134 | 0.399288 | 0.999971 | | 1 | 0.871816 |
| Lrfn3 | Q8BLY3 | -1.33 | -1.32 | -1.09 | -1.12 | -1.11 | -1.29 | -0.87 | -1.08 | -0.96 | -1.37 | -1.04 | -1.03 | -0.89 | -1.27 | -0.94 | -1.07 | 0.1275 | 0.115 | 0.1725 | 0.491418 | 0.531595 | 0.358359 | 0.999971 | | 1 | 0.869102 |
| Lrfn4 | Q80XU8 | -0.29 | -0.94 | -0.91 | -0.51 | -1.19 | -0.89 | -0.99 | -0.59 | -1.29 | -0.5 | -1.1 | -1.26 | -0.7 | -1.01 | -0.67 | -0.3 | -0.2525 | -0.375 | -0.0075 | 0.301246 | 0.17858 | 0.975627 | 0.999971 | | 1 | 0.99925 |
| Lrfn5 | Q8BXA0 | -1.25 | -1.23 | -1.66 | -1.54 | -1.05 | -1.31 | -1.17 | -1.5 | -0.81 | -1.09 | -1.37 | -1.3 | -0.76 | -1.33 | -1.87 | -1.17 | 0.1625 | 0.2775 | 0.1375 | 0.428392 | 0.21094 | 0.616378 | 0.999971 | | 1 | 0.928229 |
| Lrif1 | G5E8M8 | -0.32 | 0.38 | 0.04 | -0.28 | 0.55 | -1.2 | -0.55 | -0.71 | 0.03 | -0.46 | 0.33 | 0.45 | 1.31 | -0.16 | 0.07 | -0.41 | -0.4325 | 0.1325 | 0.2475 | 0.282296 | 0.633054 | 0.538786 | 0.999971 | | 1 | 0.910077 |
| Lrig1 | P70193 | 0.37 | -0.79 | -0.8 | -0.78 | -1.2 | -0.97 | -1.13 | -0.85 | -0.78 | -0.34 | -1 | -1.12 | -1.06 | -0.45 | -0.67 | -0.9 | -0.5375 | -0.31 | -0.27 | 0.107048 | 0.366891 | 0.411057 | 0.999971 | | 1 | 0.875847 |
| Lrp1 | A0A0R4J0 | -0.46 | -0.31 | -0.19 | -0.34 | -0.14 | -0.18 | 0.05 | -0.2 | 0.06 | -0.31 | -0.16 | 0 | 0.33 | -0.27 | -0.09 | -0.14 | 0.2075 | 0.2225 | 0.2825 | 0.246245 | 0.228245 | 0.182749 | 0.999971 | | 1 | 0.808532 |
| Lrp11 | Q8CB67 | 0.27 | 0.57 | 1.22 | 1.41 | 0.51 | 0.87 | 1.09 | 1.32 | 1.21 | 0.59 | 1.24 | 1.56 | 0.97 | 1.23 | 1.08 | 1.26 | 0.08 | 0.2825 | 0.2675 | 0.803265 | 0.408113 | 0.367623 | 0.999971 | | 1 | 0.869526 |
| Lrp1b | A2API5 | -0.04 | -0.3 | -0.71 | -0.72 | -0.4 | -0.22 | -0.17 | -0.56 | -0.2 | 0.12 | -0.36 | -0.47 | 0.38 | -0.23 | -0.56 | -0.62 | 0.105 | 0.215 | 0.185 | 0.646665 | 0.381852 | 0.534129 | 0.999971 | | 1 | 0.910077 |
| Lrp4 | Q8VI56-2 | 1.04 | -0.89 | -0.71 | -0.6 | -0.85 | -0.46 | -0.27 | -0.84 | -1.14 | 0.43 | -0.73 | -1.1 | -1.28 | -1.13 | -0.92 | -0.92 | -0.315 | -0.345 | -0.7725 | 0.482702 | 0.524595 | 0.100942 | 0.999971 | | 1 | 0.78268 |
| Lrp6 | O88572 | 0.37 | -0.05 | -0.71 | -0.82 | 0.11 | -0.2 | -0.82 | -0.9 | -0.29 | 0.2 | -0.62 | -0.84 | -0.36 | -0.03 | -1.19 | -0.45 | -0.15 | -0.085 | -0.205 | 0.680707 | 0.81041 | 0.57682 | 0.999971 | | 1 | 0.918194 |
| Lrp8 | B1AXJ4 | 0.57 | 0.01 | -0.53 | -0.22 | 0.28 | -0.1 | -0.31 | -0.41 | -0.08 | 0.57 | -0.37 | -0.48 | -0.38 | -0.16 | -0.52 | -0.21 | -0.0925 | -0.0475 | -0.275 | 0.750111 | 0.885493 | 0.319006 | 0.999971 | | 1 | 0.853893 |
| Lrpap1 | P55302 | -0.55 | -1.13 | -1.06 | -0.87 | -0.92 | -1.08 | -0.57 | -0.64 | -0.58 | -0.6 | -0.72 | -0.86 | -0.67 | -0.67 | -0.72 | -1.09 | 0.1 | 0.2125 | 0.115 | 0.651483 | 0.302067 | 0.593673 | 0.999971 | | 1 | 0.92342 |
| Lrpprc | Q6PB66 | -0.7 | -0.44 | -0.13 | -0.14 | -0.52 | -0.51 | -0.12 | 0.07 | -0.22 | -0.93 | -0.09 | 0.14 | 0.09 | -0.35 | 0.17 | 0.34 | 0.0825 | 0.0775 | 0.415 | 0.726736 | 0.783422 | 0.106259 | 0.999971 | | 1 | 0.78268 |
| Lrrc1 | Q80VQ1 | -0.08 | -0.82 | -1.04 | -0.63 | -0.42 | -1.56 | -1.38 | -0.83 | -0.76 | -0.41 | -1.51 | -0.63 | -1.8 | -1.26 | -0.42 | -0.7 | -0.405 | -0.185 | -0.4025 | 0.240989 | 0.563778 | 0.282084 | 0.999971 | | 1 | 0.844538 |
| Lrrc10b | G3XA50 | 1.39 | -0.19 | 0.24 | 0.09 | -0.94 | -0.58 | -0.51 | 0.4 | -1.03 | -0.34 | -0.7 | -0.42 | -1.17 | -0.38 | 0.32 | -0.49 | -0.79 | -1.005 | -0.8125 | 0.090608 | 0.022892 | 0.090273 | 0.999971 | | 1 | 0.771991 |
| Lrrc14 | Q8VC16 | -0.98 | -0.49 | -0.25 | -0.19 | -0.62 | -0.74 | -0.4 | 0.19 | -0.57 | -0.55 | -0.39 | 0.02 | -0.86 | -0.55 | 0.18 | 0.29 | 0.085 | 0.105 | 0.2425 | 0.767257 | 0.677227 | 0.471912 | 0.999971 | | 1 | 0.892857 |
| Lrrc20 | Q8CI70 | 0.62 | 0.78 | 0.22 | 0.28 | 0.11 | 0.55 | -0.06 | 0.23 | 0.75 | 0.33 | 0 | 0.4 | -0.31 | 0.94 | 0.21 | 0.28 | -0.2675 | -0.105 | -0.195 | 0.256288 | 0.658472 | 0.519525 | 0.999971 | | 1 | 0.906123 |
| Lrrc24 | Q8BHA1 | 0.23 | -0.15 | 0.55 | 0.72 | -0.14 | -0.03 | 0.63 | 0.38 | -0.21 | 0.04 | 0.87 | 0.33 | -0.03 | 0.15 | 0.53 | 0.92 | -0.1275 | -0.08 | 0.055 | 0.648816 | 0.794377 | 0.852488 | 0.999971 | | 1 | 0.98187 |
| Lrrc4 | Q99PH1 | -0.64 | 0.69 | -1.45 | -0.96 | -0.71 | -0.22 | -1.57 | -0.99 | -1.34 | -0.73 | -1.51 | -1.27 | 0.52 | -1.03 | -0.8 | -0.58 | -0.2825 | -0.6225 | 0.1175 | 0.574451 | 0.198658 | 0.824233 | 0.999971 | | 1 | 0.976309 |
| Lrrc40 | A0A0R4J0 | 0.07 | -0.4 | -0.03 | 0.05 | -0.46 | -0.28 | -0.03 | 0.5 | -0.37 | 0.04 | -0.15 | -0.01 | -0.24 | 0.03 | 0.21 | 0.26 | 0.01 | -0.045 | 0.1425 | 0.969226 | 0.82037 | 0.50226 | 0.999971 | | 1 | 0.904573 |
| Lrrc47 | Q505F5 | 0.01 | 0.46 | 0.16 | 0.2 | 0.56 | 0.58 | 0.37 | 0.28 | 0.78 | 0.55 | 0.23 | 0.36 | 0.94 | 0.65 | 0.37 | 0.35 | 0.24 | 0.2725 | 0.37 | 0.223484 | 0.202842 | 0.113156 | 0.999971 | | 1 | 0.78268 |
| Lrrc49 | D3YWN5 | 0.17 | -0.09 | -0.06 | -0.11 | -0.03 | -0.05 | -0.27 | -0.23 | 0 | 0.17 | 0 | -0.06 | 0.22 | -0.3 | -0.22 | 0.19 | -0.1225 | 0.05 | -0.005 | 0.490557 | 0.767445 | 0.980559 | 0.999971 | | 1 | 0.99925 |
| Lrrc4b | P0C192 | -1.55 | -0.5 | -1.28 | -1.48 | -0.45 | -0.76 | -1.17 | -1.2 | 0.06 | -1.37 | -1.01 | -0.85 | 0.42 | -1.03 | -1.13 | -1.07 | 0.3075 | 0.41 | 0.5 | 0.330052 | 0.292867 | 0.256437 | 0.999971 | | 1 | 0.839012 |
| Lrrc4c | Q8C031 | -1.69 | -1.28 | -1.1 | -0.94 | -1.21 | -1.77 | -1.17 | -1.07 | -1.37 | -1.84 | -1.24 | -0.98 | -1.24 | -1.98 | -0.85 | -1.15 | -0.0525 | -0.105 | -0.0525 | 0.835378 | 0.69041 | 0.860563 | 0.999971 | | 1 | 0.982963 |
| Lrrc57 | Q8JZX5 | -0.59 | -0.99 | -0.39 | -1.03 | -1.11 | -0.98 | -0.08 | -0.75 | -0.79 | -0.51 | -0.54 | -0.63 | -0.94 | -1 | -0.76 | -0.9 | 0.02 | 0.1325 | -0.15 | 0.944808 | 0.53811 | 0.488252 | 0.999971 | | 1 | 0.898175 |
| Lrrc58 | Q3UGP9 | -0.05 | 0.22 | 0.94 | 0.76 | -0.47 | 0.02 | 0.15 | 1.15 | -0.46 | -0.38 | -0.03 | -0.81 | 0.25 | -0.34 | 0.44 | 0.58 | -0.255 | -0.8875 | -0.235 | 0.522075 | 0.013315 | 0.459106 | 0.999971 | | 1 | 0.889191 |
| Lrrc59 | Q922Q8 | 0.25 | 1.12 | 1.44 | 1.06 | 0.63 | 1.07 | 1.24 | 1.28 | 0.8 | 0.34 | 1.55 | 1.39 | 1.03 | 1.21 | 1.23 | 1.36 | 0.0875 | 0.0525 | 0.24 | 0.772111 | 0.886065 | 0.399839 | 0.999971 | | 1 | 0.871816 |
| Lrrc7 | A0A571BE | 2.03 | -2.15 | -0.6 | -0.52 | -2.35 | -0.36 | -0.26 | -1.05 | -2.41 | 0.56 | -0.99 | -0.96 | -2.87 | -2.35 | -1.17 | -0.66 | -0.695 | -0.64 | -1.4525 | 0.441297 | 0.506786 | 0.133874 | 0.999971 | | 1 | 0.784948 |
| Lrrc7_1 | A0A0G2JD | -1.09 | -1.8 | -1.56 | -1.39 | -1.78 | -1.84 | -1.63 | -1.36 | -1.8 | -1.36 | -1.45 | -1.51 | -1.61 | -1.67 | -1.48 | -1.25 | -0.1925 | -0.07 | -0.0425 | 0.401822 | 0.749176 | 0.847244 | 0.999971 | | 1 | 0.980295 |
| Lrrc73 | B2RWC4 | -0.8 | -0.79 | -2.07 | -2.08 | -0.98 | -0.94 | -1.7 | -2.44 | -1.01 | -1.12 | -1.79 | -2 | -0.84 | -1.06 | -1.92 | -1.85 | -0.08 | -0.045 | 0.0175 | 0.866798 | 0.914909 | 0.967868 | 0.999971 | | 1 | 0.998787 |
| Lrrc75a | Q7TSF4 | -1.52 | -0.71 | -1.64 | -1.84 | -0.92 | -1.1 | -1.44 | -2.54 | -1.19 | -1.38 | -1.48 | -1.91 | -1.16 | -1.54 | -2.26 | -2.09 | -0.0725 | -0.0625 | -0.335 | 0.862063 | 0.834516 | 0.35049 | 0.999971 | | 1 | 0.866844 |
| Lrrc8a | Q80WG5 | -0.71 | -1.03 | -1.11 | -1.35 | -0.84 | -1.05 | -1.11 | -1.14 | -1 | -0.87 | -0.94 | -1.01 | -0.68 | -0.81 | -0.96 | -0.77 | 0.015 | 0.095 | 0.245 | 0.941263 | 0.626979 | 0.245539 | 0.999971 | | 1 | 0.838458 |
| Lrrc8b | Q5DU41 | -1.32 | -0.89 | -0.91 | -1.02 | -0.85 | -0.73 | -1.14 | -0.97 | -0.67 | -1.32 | -0.9 | -0.92 | -0.38 | -1.09 | -0.77 | -0.51 | 0.1125 | 0.0825 | 0.3475 | 0.567585 | 0.698963 | 0.150991 | 0.999971 | | 1 | 0.794534 |
| Lrrc8c | Q8R502 | -0.63 | -0.6 | -0.61 | -0.96 | -0.69 | -0.57 | -0.49 | -1.12 | -0.84 | -0.68 | -0.67 | -0.96 | -0.99 | -0.76 | -0.97 | -1.1 | -0.0175 | -0.0875 | -0.255 | 0.93457 | 0.631836 | 0.193244 | 0.999971 | | 1 | 0.810547 |
| Lrrc8d | Q8BGR2 | -0.83 | -1.11 | -0.58 | -0.72 | -1.45 | -1.03 | -0.81 | -0.23 | -1.37 | -0.87 | -0.65 | -0.61 | -1.27 | -0.79 | -0.28 | -0.25 | -0.07 | -0.065 | 0.1625 | 0.809241 | 0.785531 | 0.568172 | 0.999971 | | 1 | 0.91816 |
| Lrrc9 | Q8CDN9 | -0.6 | -2.05 | 3.27 | 3.12 | -1.94 | -1.12 | 0.81 | 3.33 | -1.75 | -1.34 | 2.67 | 1.38 | -1.48 | -0.65 | 2.25 | 2.89 | -0.665 | -0.695 | -0.1825 | 0.674305 | 0.651213 | 0.904455 | 0.999971 | | 1 | 0.991091 |
| Lrrfip1 | A0A087WN | 0.17 | -0.34 | -0.35 | 0.4 | -0.27 | 0.01 | -0.2 | 0.14 | -0.34 | 0.25 | 0.7 | 1.07 | 0.57 | 0.55 | -0.04 | 0.64 | -0.05 | 0.45 | 0.46 | 0.836365 | 0.221722 | 0.112001 | 0.999971 | | 1 | 0.78268 |
| Lrrfip1_1 | Q3UZ39-2 | -0.04 | -0.76 | 1.18 | 0.01 | -0.97 | -0.67 | 1.06 | -0.02 | -0.37 | -0.04 | 1.22 | 0.14 | -0.59 | -0.44 | 0.47 | 0.35 | -0.2475 | 0.14 | -0.15 | 0.656289 | 0.776921 | 0.741963 | 0.999971 | | 1 | 0.961064 |
| Lrrfip1_2 | Q3UZ39 | -0.58 | -1.71 | 1.68 | 1.89 | -1.29 | -1.22 | 3.26 | 0.09 | -1.31 | -0.98 | 0.74 | -1.32 | -0.81 | -1.6 | 0.53 | -1.19 | -0.11 | -1.0375 | -1.0875 | 0.928457 | 0.271178 | 0.241668 | 0.999971 | | 1 | 0.835682 |
| Lrrfip2 | A0A1L1SSC | 0.29 | 0.48 | -0.28 | -0.03 | 0.26 | 0.52 | 0.35 | -0.83 | 0.19 | 0.15 | -0.63 | -0.7 | 0.95 | 0.43 | -0.49 | -0.41 | -0.04 | -0.3625 | 0.005 | 0.907599 | 0.249418 | 0.989316 | 0.999971 | | 1 | 0.99925 |
| Lrrk2 | Q5S006 | -0.43 | 0.57 | -0.08 | 0.38 | 0.37 | 0.07 | -0.02 | -0.01 | -0.1 | -0.18 | 0.48 | 0.2 | -0.09 | 0.49 | 0.02 | 0.33 | -0.0075 | -0.01 | 0.0775 | 0.977401 | 0.971909 | 0.781796 | 0.999971 | | 1 | 0.965859 |
| Lrrn1 | Q61809 | -2.01 | -2.15 | -0.57 | -0.71 | -2.1 | -2.29 | -0.31 | -0.44 | -2.01 | -1.76 | -0.18 | -0.06 | -2.37 | -2.34 | -0.4 | -0.05 | 0.075 | 0.3575 | 0.07 | 0.902542 | 0.559596 | 0.917636 | 0.999971 | | 1 | 0.992633 |
| Lrrtm1 | Q8K377 | -1.74 | -1.39 | -1.64 | -1.78 | -1.43 | -1.07 | -1.61 | -1.83 | -1.3 | -1.4 | -1.41 | -1.38 | -0.65 | -1.48 | -1.75 | -1.06 | 0.1525 | 0.265 | 0.4025 | 0.502347 | 0.150612 | 0.168126 | 0.999971 | | 1 | 0.800215 |
| Lrrtm2 | Q8BGA3 | -1.32 | -0.67 | -1.01 | -0.9 | -0.86 | -0.72 | -1.17 | -1.06 | -0.93 | -0.88 | -0.96 | -1.05 | -0.66 | -0.77 | -1.23 | -0.88 | 0.0225 | 0.02 | 0.09 | 0.916734 | 0.918757 | 0.690328 | 0.999971 | | 1 | 0.946255 |
| Lrrtm3 | Q8BZ81 | -0.8 | -0.48 | -1.06 | -0.73 | -0.55 | -0.79 | -0.87 | -0.94 | -0.46 | -0.95 | -0.53 | -0.77 | -0.23 | -1.05 | -0.64 | -0.74 | -0.02 | 0.09 | 0.1025 | 0.921303 | 0.670234 | 0.671463 | 0.999971 | | 1 | 0.939561 |
| Lrrtm4 | A0A0R4J1T | -2.49 | -1.29 | -0.97 | -1.64 | -0.97 | -1 | -1.2 | -1.24 | -1.01 | -1.73 | -0.68 | -0.67 | -0.62 | -1.68 | -1.39 | -1.33 | 0.495 | 0.575 | 0.3425 | 0.162484 | 0.170963 | 0.383251 | 0.999971 | | 1 | 0.871816 |
| Lrsam1 | Q80ZI6 | -0.26 | 0 | -0.53 | -0.9 | -0.05 | 0.37 | -0.83 | -0.29 | 0.12 | -0.03 | -0.47 | -0.48 | -0.1 | 0.03 | -0.5 | -0.37 | 0.2225 | 0.2075 | 0.1875 | 0.490849 | 0.442692 | 0.468189 | 0.999971 | | 1 | 0.89236 |
| Lrtm2 | Q8BGX3 | 0.15 | -1.87 | -1.35 | -1.2 | -1.59 | -1.32 | -1.27 | -1.06 | -1.6 | -0.6 | -1.59 | -1.24 | -1.78 | -1.36 | -0.71 | -0.68 | -0.2425 | -0.19 | -0.065 | 0.568156 | 0.681496 | 0.890738 | 0.999971 | | 1 | 0.988184 |
| Lsamp | Q3TYE5 | -1.33 | -1.01 | -1.11 | -1.2 | -1.01 | -1.21 | -1.1 | -1 | -1.18 | -1.57 | -1.13 | -0.9 | -1.11 | -1.24 | -0.87 | -0.81 | 0.0825 | -0.0325 | 0.155 | 0.6356 | 0.873925 | 0.424484 | 0.999971 | | 1 | 0.879931 |
| Lsg1 | Q3UM18-2 | 1.27 | 0.67 | 0.22 | 0.42 | 0.43 | 1.48 | -0.34 | 0.55 | 1.61 | 1.44 | -0.19 | 0.53 | -0.41 | 1.3 | 0.24 | 0.19 | -0.115 | 0.2025 | -0.315 | 0.782041 | 0.654413 | 0.44356 | 0.999971 | | 1 | 0.883386 |
| Lsm1 | Q8VC85 | -0.61 | 0.6 | -1.13 | -1.82 | 0.15 | -0.59 | -0.64 | -1.22 | -0.43 | -0.76 | -1.22 | -0.54 | -0.07 | -0.86 | -1.5 | -0.76 | 0.165 | 0.0025 | -0.0575 | 0.758902 | 0.996029 | 0.915534 | 0.999971 | | 1 | 0.992633 |
| Lsm11 | Q8BUV6 | -0.09 | -0.37 | -0.47 | -0.86 | -0.44 | -0.05 | -0.37 | -0.31 | -0.51 | -0.13 | -0.23 | -0.76 | 0.12 | -0.31 | -0.97 | -0.14 | 0.155 | 0.04 | 0.1225 | 0.492708 | 0.869049 | 0.677601 | 0.999971 | | 1 | 0.942594 |
| Lsm12 | Q9D0R8 | 1.04 | 1.4 | 1.58 | 1.75 | 1.19 | 1.27 | 1.22 | 1.87 | 1.12 | 0.94 | 1.69 | 2.25 | 0.91 | 1.21 | 1.82 | 1.79 | -0.055 | 0.0575 | -0.01 | 0.825884 | 0.862491 | 0.971978 | 0.999971 | | 1 | 0.99925 |
| Lsm14a | Q8K2F8 | 1.62 | 0.84 | 0.53 | 0.41 | 1.32 | 0.66 | 0 | 0.37 | 0.76 | 1.08 | 0.04 | 0.77 | 0.96 | 0.92 | 0.5 | 0.54 | -0.2625 | -0.1875 | -0.12 | 0.492061 | 0.590322 | 0.695707 | 0.999971 | | 1 | 0.947174 |
| Lsm14b | Q8BM41 | 1.29 | 1.25 | 0.33 | 0.8 | 0.9 | 2.06 | 0.64 | 1.48 | 0.76 | 1.39 | 1.32 | 0.91 | -0.16 | 0.97 | 0.85 | 1.13 | 0.3525 | 0.1775 | -0.22 | 0.360455 | 0.538129 | 0.545926 | 0.999971 | | 1 | 0.911398 |
| Lsm14b_1 | Q8CGC4 | 1.85 | 1.6 | 0.96 | 1.14 | 1.61 | 1.86 | 1.14 | 1.28 | 1.25 | 0.69 | 1.09 | 1.34 | 1.12 | 1.45 | 1.45 | 1.11 | 0.085 | -0.295 | -0.105 | 0.759979 | 0.288997 | 0.680247 | 0.999971 | | 1 | 0.942957 |
| Lsm2 | O35900 | 1.71 | 1.76 | 0.16 | -0.69 | 1.53 | 1.38 | 0.23 | -0.57 | 1.93 | 0.91 | 0.21 | 0.19 | 1.65 | 1.22 | -0.21 | -0.49 | -0.0925 | 0.075 | -0.1925 | 0.895734 | 0.910125 | 0.79036 | 0.999971 | | 1 | 0.968321 |
| Lsm3 | P62311 | 0.49 | 1.31 | 0.45 | 0.83 | 1.28 | 0.67 | -0.25 | 0.57 | 1.07 | 0.78 | 0.25 | 0.09 | 1.19 | 1.52 | 0.15 | 0.48 | -0.2025 | -0.2225 | 0.065 | 0.579935 | 0.476949 | 0.858224 | 0.999971 | | 1 | 0.982702 |
| Lsm4 | Q9CY46 | 1.16 | 1.19 | 0.95 | 0.77 | 1.54 | 1.16 | 0.71 | 0.68 | 1 | 1.41 | 0.69 | 0.93 | 0.71 | 1.38 | 1 | 1.14 | 0.005 | -0.01 | 0.04 | 0.984234 | 0.963668 | 0.854145 | 0.999971 | | 1 | 0.982125 |
| Lsm5 | P62322 | 0.62 | 0.47 | 0.7 | 0.82 | -0.6 | 0.3 | 0.26 | 0.36 | -0.06 | 0.23 | 0.69 | 0.13 | 0.53 | 0.27 | 0.86 | 1.11 | -0.5725 | -0.405 | 0.04 | 0.052948 | 0.089942 | 0.864713 | 0.999971 | | 1 | 0.983195 |
| Lsm6 | P62313 | -0.11 | 0.55 | 1.02 | 0.53 | 0.71 | -0.01 | 0.73 | 0.44 | 1.16 | 0.23 | 1.2 | 1.46 | 0.92 | 0.23 | 0.99 | 1.26 | -0.03 | 0.515 | 0.3525 | 0.919773 | 0.166211 | 0.290133 | 0.999971 | | 1 | 0.847402 |
| Lsm7 | Q9CQQ8 | 2.47 | 0.53 | 0.08 | 0.18 | 0.61 | 0.06 | -0.39 | -0.37 | 0.41 | 0.54 | -0.55 | 0.05 | 0.33 | 0.26 | 0.15 | 0.24 | -0.8375 | -0.7025 | -0.57 | 0.158487 | 0.233542 | 0.289403 | 0.999971 | | 1 | 0.847402 |
| Lsm8 | Q6ZWM4 | -0.36 | 1.43 | 1.07 | 1.1 | 0.75 | 0.41 | 0.6 | 0.34 | 1.41 | -0.06 | 2.83 | 0.87 | 2.96 | 0.67 | 0.54 | 0.69 | -0.285 | 0.4525 | 0.405 | 0.474376 | 0.49935 | 0.53302 | 0.999971 | | 1 | 0.910069 |
| Lsp1 | A2A6J4 | 0.33 | 5.25 | 2.51 | 2.11 | 4.65 | 3.11 | 1.05 | 1.27 | 4.19 | 0.19 | 1.65 | 2.41 | 8.51 | 1.51 | 2.03 | 0.98 | -0.03 | -0.44 | 0.7075 | 0.979639 | 0.709684 | 0.696098 | 0.999971 | | 1 | 0.947174 |
| Lsr | Q99KG5-3 | 3.75 | -0.63 | -0.78 | -0.12 | 0.55 | -0.36 | -1.86 | -1.09 | 1.64 | 4.18 | -1.75 | -1.22 | -3.42 | -1.42 | -1.26 | 0.04 | -1.245 | 0.1575 | -2.07 | 0.259639 | 0.919428 | 0.099301 | 0.999971 | | 1 | 0.78268 |
| Lss | Q8BLN5 | -0.92 | -0.71 | -0.76 | -1.37 | -0.84 | -1.41 | -0.41 | -0.92 | -0.33 | -0.41 | -0.3 | -0.17 | 0.19 | -0.77 | -0.53 | -0.94 | 0.045 | 0.6375 | 0.4275 | 0.868782 | 0.012613 | 0.178451 | 0.999971 | | 1 | 0.808532 |
| Lta4h | P24527 | 0.46 | 1.08 | 1.23 | 1.28 | 1.07 | 0.76 | 0.89 | 1.22 | -0.27 | 0.1 | 0.51 | 0.61 | 0.76 | 0.73 | 1.13 | 1.26 | -0.0275 | -0.775 | -0.0425 | 0.910458 | 0.023558 | 0.86852 | 0.999971 | | 1 | 0.983462 |
| Ltbp4 | E9QPS5 | 1.07 | -3.3 | -4.13 | -0.2 | 7.05 | 2.26 | 0.94 | -1.62 | 4.68 | 7.05 | 7.63 | 9.53 | -5.59 | 0.62 | -0.91 | -0.7 | 3.7975 | 8.8625 | -0.005 | 0.077714 | 0.000192 | 0.997549 | 0.999971 | 0.218167 | | 1 |
| Ltn1 | Q6A009 | 0.53 | 1.05 | 1.4 | 1.06 | 0.82 | 0.91 | 1.55 | 2.4 | 1.02 | 0.93 | 1.24 | 1.26 | 1.09 | 1.03 | 1.13 | 1.12 | 0.41 | 0.1025 | 0.0825 | 0.309087 | 0.659622 | 0.713158 | 0.999971 | | 1 | 0.951793 |
| Luc7l | Q9CYI4 | 1.07 | 1.47 | 1.88 | 1.42 | 1.74 | 1.65 | 2.28 | 2.08 | 1.85 | 1.48 | 2.18 | 2.24 | 2.18 | 1.75 | 2.02 | 1.81 | 0.4775 | 0.4775 | 0.48 | 0.082707 | 0.095868 | 0.063622 | 0.999971 | | 1 | 0.725348 |
| Luc7l2 | Q7TNC4-2 | 2.06 | 3.11 | 2.52 | 2.56 | 3.25 | 3.05 | 1.96 | 2.29 | 4.09 | 2.05 | 2.06 | 1.8 | 4.01 | 2.47 | 2.09 | 2.51 | 0.075 | -0.0625 | 0.2075 | 0.836877 | 0.90655 | 0.644825 | 0.999971 | | 1 | 0.93301 |
| Luc7l3 | Q5SUF2 | 1.42 | 1.19 | 2.09 | 1.98 | 1.77 | 1.7 | 1.97 | 1.97 | 1.88 | 2.04 | 1.99 | 1.99 | 1.56 | 1.48 | 1.9 | 2.22 | 0.1825 | 0.305 | 0.12 | 0.478995 | 0.237207 | 0.679042 | 0.999971 | | 1 | 0.942839 |
| Lum | P51885 | 0.54 | 1.32 | 1.15 | 3.44 | 1.85 | 0.93 | 1.54 | 1.98 | 0.47 | 0.51 | 1.48 | 3.06 | -0.13 | 1.12 | 1.47 | 0.55 | -0.0375 | -0.2325 | -0.86 | 0.951222 | 0.769741 | 0.212413 | 0.999971 | | 1 | 0.823451 |
| Lurap1 | Q9D6I9 | 0.5 | -0.43 | 0.81 | 0.91 | -0.5 | -0.48 | 0.11 | 0.8 | -0.61 | 0.14 | 0.49 | 0.42 | -0.47 | -0.02 | 0.85 | 1.2 | -0.465 | -0.3375 | -0.0575 | 0.27654 | 0.388444 | 0.900588 | 0.999971 | | 1 | 0.990593 |
| Lurap1l | Q8K2P1 | 0.82 | 1.17 | 1.47 | 0.79 | 1.03 | 1.15 | 1.69 | 1.73 | 0.36 | 1.12 | 2.47 | 0.37 | 1.17 | 1.22 | 0.61 | -1.23 | 0.3375 | 0.0175 | -0.62 | 0.222296 | 0.971333 | 0.276714 | 0.999971 | | 1 | 0.843794 |
| Luzp1 | Q8R4U7 | 0.47 | 0.66 | 0.26 | 0.21 | 0.34 | 0.6 | 0.47 | -0.05 | 0.31 | 0.3 | 0.2 | 0.02 | 0.63 | 0.37 | -0.1 | 0.03 | -0.06 | -0.1925 | -0.1675 | 0.784353 | 0.319496 | 0.478265 | 0.999971 | | 1 | 0.894438 |
| Luzp2 | Q8BGY3 | 1.18 | 0.61 | -0.53 | -0.35 | 1.18 | 0.75 | -0.85 | -0.45 | 1.44 | 1.94 | -0.53 | -0.52 | 2.31 | 0.97 | -0.31 | -0.06 | -0.07 | 0.355 | 0.5 | 0.903087 | 0.612063 | 0.452605 | 0.999971 | | 1 | 0.887217 |
| Lxn | P70202 | 0.7 | 0.74 | 0.27 | 0.15 | 0.54 | 1.07 | -0.08 | 0.29 | 0.3 | 0.95 | 0.13 | 0.16 | 0.86 | 1.03 | 0.4 | 0.59 | -0.01 | -0.08 | 0.255 | 0.97286 | 0.76189 | 0.302037 | 0.999971 | | 1 | 0.847402 |
| Ly6a | P05533 | 1.38 | 2.78 | 2.09 | 2.27 | 2.59 | 3.48 | 0.87 | 2.31 | 2 | 1.7 | 1.98 | 1.38 | 2.18 | 3.71 | 2.31 | 2.1 | 0.1825 | -0.365 | 0.445 | 0.746394 | 0.27867 | 0.336715 | 0.999971 | | 1 | 0.858827 |
| Ly6c1 | A0A087WF | -3.4 | 1.59 | -0.15 | 1.05 | 2.03 | 0.88 | 0.43 | 0.03 | 1.82 | -1.41 | 0.54 | 1.83 | 1.12 | 1.15 | 0.44 | -0.09 | 1.07 | 0.9225 | 0.8825 | 0.330599 | 0.454836 | 0.40327 | 0.999971 | | 1 | 0.873348 |
| Ly6e | A0A087WC | 0.64 | 2.59 | 1.74 | 1.46 | 2.82 | 4.86 | 0.86 | 2.69 | 3.38 | 2.04 | 1.21 | 0.53 | 3.29 | 4.55 | 2.82 | 2.41 | 1.2 | 0.1825 | 1.66 | 0.167385 | 0.785292 | 0.015648 | 0.999971 | | 1 | 0.613095 |
| Ly6g6e | Q8K1T6-2 | -5.48 | -7.3 | 2.98 | 3.12 | -9.31 | -7.9 | 1.69 | 1.98 | -6.8 | -6.87 | 1.59 | 2.36 | -9.39 | -7.57 | 0.98 | 1.72 | -1.715 | -0.76 | -1.895 | 0.636645 | 0.819905 | 0.592356 | 0.999971 | | 1 | 0.923175 |
| Ly6h | Q8K356 | -0.85 | 2.37 | 2.04 | 0.92 | 2.28 | 2.89 | 2.09 | 1.79 | 2.74 | -0.52 | 1.92 | 2.01 | 3.42 | 1.66 | 2.46 | 0.2 | 1.1425 | 0.4175 | 0.815 | 0.124685 | 0.650098 | 0.373054 | 0.999971 | | 1 | 0.869526 |
| Ly75 | A2AW86 | 0.57 | 0.45 | -0.01 | 2.63 | 3.17 | 2.23 | -0.07 | 0.61 | -2.84 | -1.24 | -0.27 | -0.7 | 1.77 | 1.68 | 0.22 | 0.16 | 0.575 | -2.1725 | 0.0475 | 0.50287 | 0.015834 | 0.943247 | 0.999971 | | 1 | 0.99692 |
| Lyar | Q08288 | 0.87 | 0.92 | 0.77 | 0.5 | 0.47 | 0.62 | 1.19 | 0.05 | 0.86 | 1.24 | 0.86 | 0.49 | 0.77 | 0.96 | 0.25 | 0.25 | -0.1825 | 0.0975 | -0.2075 | 0.507925 | 0.659322 | 0.395376 | 0.999971 | | 1 | 0.871816 |
| Lyn | P25911-2 | -0.39 | -0.04 | 0.29 | 0.03 | 0.13 | -0.07 | -0.09 | -0.03 | 0.11 | 0.1 | 0.53 | 0.38 | 0.31 | -0.26 | 0.5 | 0.03 | 0.0125 | 0.3075 | 0.1725 | 0.951077 | 0.181769 | 0.491676 | 0.999971 | | 1 | 0.899647 |
| Lynx1 | P0DP60 | 1.29 | 1.2 | 0.82 | 0.77 | 0.95 | 1.92 | -0.17 | 1.29 | 1.09 | 1.46 | 0.25 | -0.06 | 2.22 | 2.32 | 1.18 | 1.24 | -0.0225 | -0.335 | 0.72 | 0.958401 | 0.374735 | 0.055468 | 0.999971 | | 1 | 0.725348 |
| Lypd1 | Q8BLC3 | -0.41 | -0.08 | -0.34 | -0.5 | -0.25 | -0.28 | -0.7 | 0.11 | -0.4 | -0.69 | -0.34 | -0.86 | -0.55 | 0.14 | -0.72 | -0.59 | 0.0525 | -0.24 | -0.0975 | 0.817947 | 0.257186 | 0.691629 | 0.999971 | | 1 | 0.946899 |
| Lypd6 | Q8BPP5 | 2.6 | -0.53 | -0.16 | -0.47 | -0.96 | 0.18 | 1.36 | -0.71 | -0.98 | 1.2 | -0.93 | -0.68 | -3.52 | -3.99 | -1.51 | -1.66 | -0.3925 | -0.7075 | -3.03 | 0.635152 | 0.401909 | 0.007264 | 0.999971 | | 1 | 0.582714 |
| Lypla1 | P97823 | -0.07 | 0.7 | 0.08 | -0.04 | 0.79 | 0.94 | 0.74 | -0.54 | 0.43 | 0.22 | -0.2 | -0.63 | 0.99 | 0.83 | -0.33 | -0.68 | 0.315 | -0.2125 | 0.035 | 0.41096 | 0.489298 | 0.934986 | 0.999971 | | 1 | 0.995618 |
| Lypla2 | Q9WTL7 | 0.7 | -0.14 | 0.28 | 0.93 | -0.1 | -0.13 | -0.05 | 0.75 | -0.42 | -0.07 | 0.38 | 0.13 | -0.12 | 0.19 | 0.64 | 0.84 | -0.325 | -0.4375 | -0.055 | 0.323536 | 0.168071 | 0.864746 | 0.999971 | | 1 | 0.983195 |
| Lyplal1 | E9QLB2 | 0.34 | 0.35 | 0.56 | -0.18 | 0 | 0.2 | 0.04 | 0.77 | 0.26 | -0.14 | 0.26 | -0.16 | 0.2 | 0.27 | 0.25 | -0.34 | -0.015 | -0.2125 | -0.1725 | 0.954082 | 0.370549 | 0.488692 | 0.999971 | | 1 | 0.898346 |
| Lyrm1 | Q9CQB7 | 2.67 | 2.24 | -0.01 | -0.07 | 0.79 | 2.15 | -0.54 | 0.31 | 0.77 | 3.22 | 0.47 | 0.17 | 1.94 | 1.97 | 0.08 | 0.49 | -0.53 | -0.05 | -0.0875 | 0.524996 | 0.956029 | 0.911551 | 0.999971 | | 1 | 0.992551 |
| Lyrm2 | Q8R033 | 2.35 | -1.59 | -0.97 | -0.31 | -0.66 | -1.26 | -0.02 | -0.11 | -1.31 | 0.75 | -0.55 | -1.18 | -1.6 | -0.39 | -1.39 | -0.91 | -0.3825 | -0.4425 | -0.9425 | 0.642616 | 0.621379 | 0.266913 | 0.999971 | | 1 | 0.842677 |
| Lyrm4 | Q8K215 | 1.17 | 1.34 | 1.01 | 0.67 | 2.11 | 1.3 | 1.22 | 0.54 | 2.02 | 1.07 | 1.27 | 1.45 | 3.3 | 1.13 | 0.44 | 0.47 | 0.245 | 0.405 | 0.2875 | 0.485731 | 0.156503 | 0.648868 | 0.999971 | | 1 | 0.933842 |
| Lyrm7 | Q9DA03 | 0.29 | 0.19 | -0.89 | -0.02 | 0.43 | 0.55 | -0.75 | -0.59 | 0.2 | -0.34 | -0.38 | -0.3 | 1 | 0.87 | -0.72 | 0 | 0.0175 | -0.0975 | 0.395 | 0.965952 | 0.751128 | 0.395572 | 0.999971 | | 1 | 0.871816 |
| Lyrm9 | E9PX24 | 0.52 | -0.7 | -0.07 | 0.05 | -0.39 | -0.12 | -0.76 | 0.46 | -0.66 | 0.24 | -0.14 | -0.43 | -0.19 | -0.05 | 0.29 | -0.25 | -0.1525 | -0.1975 | 1.47E-17 | 0.66673 | 0.540984 | 1 | 0.999971 | | 1 | 1 |
| Lysmd1 | Q9D0E3 | -1.41 | -0.79 | -0.31 | -1.47 | -0.75 | -0.93 | -0.06 | -0.68 | -0.88 | -1.41 | 0.38 | 0.77 | -0.04 | -0.78 | -0.07 | -1.05 | 0.39 | 0.71 | 0.51 | 0.259751 | 0.211167 | 0.187041 | 0.999971 | | 1 | 0.809025 |
| Lysmd2 | G3X915 | 0.79 | 1.03 | 1.31 | 1.1 | 0.87 | 1.56 | 1.43 | 0.65 | 0.8 | 0.95 | 0.54 | 0.59 | 1.77 | 1.15 | 0.92 | 0.48 | 0.07 | -0.3375 | 0.0225 | 0.791763 | 0.116112 | 0.939953 | 0.999971 | | 1 | 0.995869 |
| Lyst | G5E8Q0 | -0.17 | -0.54 | -1.02 | -0.92 | -0.72 | -0.59 | -0.92 | -0.86 | -0.79 | -0.6 | -0.99 | -1.03 | -0.96 | -0.64 | 0.54 | -1.23 | -0.11 | -0.19 | 0.09 | 0.64886 | 0.447169 | 0.82829 | 0.999971 | | 1 | 0.977728 |
| Lzic | Q8K3C3 | -0.85 | -0.96 | -0.35 | 0.21 | -0.72 | -1.46 | -0.63 | -0.07 | -0.88 | -0.49 | -0.36 | 0.11 | -0.87 | -0.9 | 0.13 | 0.34 | -0.2325 | 0.0825 | 0.1625 | 0.542896 | 0.805416 | 0.688509 | 0.999971 | | 1 | 0.945753 |
| Lztfl1 | Q9JHQ5 | -0.24 | 0.7 | 0.75 | 0.61 | 0.47 | 0.52 | 0.9 | 0.51 | 0.52 | 0.14 | 0.35 | 0.66 | 0.78 | 0.68 | 0.47 | 0.45 | 0.145 | -0.0375 | 0.14 | 0.597561 | 0.89147 | 0.604205 | 0.999971 | | 1 | 0.924454 |
| Lztr1 | Q9CQ33 | -0.52 | -0.67 | -0.35 | -0.64 | -0.64 | -0.04 | -0.46 | -0.47 | -0.61 | -0.19 | -0.33 | -0.46 | -0.8 | -0.3 | -0.68 | -0.3 | 0.1425 | 0.1475 | 0.025 | 0.48864 | 0.430341 | 0.902592 | 0.999971 | | 1 | 0.990759 |
| Lzts1 | P60853 | -1.97 | -2.36 | -1.06 | -1.08 | -1.27 | -2.35 | -0.55 | -0.66 | -1.49 | -1.91 | -0.67 | -0.31 | -0.79 | -2.23 | -0.69 | -0.62 | 0.41 | 0.5225 | 0.535 | 0.41103 | 0.27572 | 0.274204 | 0.999971 | | 1 | 0.843794 |
| Lzts3 | A2AHG0 | -0.08 | -0.75 | -1.38 | -0.93 | -0.82 | -0.46 | -0.62 | -0.71 | -1.06 | -0.46 | -1.29 | -1.54 | -0.74 | -0.47 | -1.48 | -1.29 | 0.1325 | -0.3025 | -0.21 | 0.651289 | 0.39714 | 0.554826 | 0.999971 | | 1 | 0.91473 |
| M6pr | P24668 | 0.69 | 1.14 | 0.83 | 0.87 | 0.91 | 1.25 | 1.83 | 0.57 | 1.46 | 1.13 | 0.82 | 0.75 | 1.52 | 0.8 | 0.66 | 0.33 | 0.2575 | 0.1575 | -0.055 | 0.392993 | 0.490856 | 0.84603 | 0.999971 | | 1 | 0.979924 |
| Macf1 | A0A0A0M | 0.55 | 1.25 | 0.53 | 0.63 | 1.03 | 0.98 | 0.67 | -0.36 | 0.67 | 0.53 | 0.08 | 0.38 | 0.95 | 1.72 | -0.08 | -0.1 | -0.16 | -0.325 | -0.1175 | 0.656327 | 0.202617 | 0.792087 | 0.999971 | | 1 | 0.969134 |
| Macf1_1 | F6Q750 | -1.58 | -1.46 | -0.99 | -1.49 | -2.32 | -1.16 | -1.07 | -0.71 | -2.58 | -0.7 | -2.1 | -0.77 | -0.57 | -1.87 | -0.7 | -0.92 | 0.065 | -0.1575 | 0.365 | 0.858043 | 0.734045 | 0.278 | 0.999971 | | 1 | 0.843812 |
| Macf1_2 | F7ACR9 | -2.28 | -2.55 | -1.26 | -0.83 | -2.04 | -2.97 | -1.26 | -0.92 | -2.42 | -1.6 | -1.19 | -1.24 | -2.46 | -2.07 | -1.01 | -0.91 | -0.0675 | 0.1175 | 0.1175 | 0.904236 | 0.801687 | 0.821315 | 0.999971 | | 1 | 0.975833 |
| Macf1_3 | E9PVY8 | -0.51 | -0.65 | -0.81 | -0.89 | -0.61 | -0.46 | -0.61 | -0.8 | -0.53 | -0.45 | -0.73 | -0.86 | -0.22 | -0.56 | -0.74 | -0.74 | 0.095 | 0.0725 | 0.15 | 0.607943 | 0.702095 | 0.470654 | 0.999971 | | 1 | 0.892725 |
| Macf1_4 | Q9QXZ0-3 | -1.43 | -0.49 | -0.82 | -0.99 | -0.23 | -0.61 | -1.03 | -0.96 | -0.83 | -0.06 | -1.89 | -1.45 | 0.28 | -1.09 | -0.22 | -0.56 | 0.225 | -0.125 | 0.535 | 0.434777 | 0.76708 | 0.146712 | 0.999971 | | 1 | 0.793221 |
| Maco1 | Q7TQE6 | 0.24 | 0.43 | 0.74 | 0.4 | 0.68 | 0.55 | 0.7 | 0.51 | 0.59 | 0.22 | 0.61 | 0.69 | 0.79 | 0.28 | 0.3 | 0.12 | 0.1575 | 0.075 | -0.08 | 0.407393 | 0.709843 | 0.719745 | 0.999971 | | 1 | 0.954424 |
| Macrod1 | Q922B1 | 0.75 | 0.41 | 0.51 | 0.29 | 0.21 | 1.16 | 0.47 | 0.86 | 0.52 | 0.87 | -0.02 | -0.21 | 0.33 | 1.06 | 0.13 | -0.1 | 0.185 | -0.2 | -0.135 | 0.477188 | 0.481938 | 0.636644 | 0.999971 | | 1 | 0.93301 |
| Macrod2 | Q3UYG8 | 0.74 | 0.31 | 1.79 | 1.48 | 0.3 | 0.52 | 2.01 | 2.16 | 0.64 | 0.23 | 2.15 | 1.44 | 0.78 | -0.17 | 2 | 1.18 | 0.1675 | 0.035 | -0.1325 | 0.758922 | 0.944888 | 0.799554 | 0.999971 | | 1 | 0.970592 |
| Macroh2a1 | Q9QZQ8 | -0.79 | -1.59 | -1.49 | -1.01 | -1.55 | -1.7 | -1.71 | -1.3 | -1.53 | -1.02 | -1.62 | -2 | -1.33 | -1.22 | -1.27 | -1.51 | -0.345 | -0.3225 | -0.1125 | 0.182087 | 0.282138 | 0.636819 | 0.999971 | | 1 | 0.93301 |
| Macroh2a2 | Q8CCK0 | -0.62 | -0.97 | -1.51 | -1.29 | -1.07 | -1.34 | -1.69 | -1.41 | -1.04 | -0.64 | -1.42 | -2.05 | -0.5 | -0.82 | -1.76 | -1.57 | -0.28 | -0.19 | -0.065 | 0.292304 | 0.591377 | 0.8533 | 0.999971 | | 1 | 0.98187 |
| Mad1l1 | Q9WTX8 | 1.17 | 0.68 | 0.84 | 1.13 | 0.85 | 0.66 | 1.41 | 0.74 | 0.81 | 1.28 | 0.99 | 0.48 | 0.29 | 0.66 | 0.47 | 0.56 | -0.04 | -0.065 | -0.46 | 0.867264 | 0.783724 | 0.044467 | 0.999971 | | 1 | 0.706051 |
| Mad2l1 | Q9Z1B5 | -0.12 | -0.27 | 0.63 | 0.56 | 0.22 | -0.29 | 0.03 | 0.35 | -0.08 | -0.56 | 0.29 | 0.04 | -1.08 | -0.78 | 0.85 | -0.2 | -0.1225 | -0.2775 | -0.5025 | 0.667075 | 0.366086 | 0.286184 | 0.999971 | | 1 | 0.84602 |
| Mad2l1bp | Q9DCX1 | 2.25 | 1.49 | 0.97 | 1.86 | 1.13 | 1.53 | 1.72 | 0.96 | 0.73 | 2.03 | 1.27 | 0.14 | -0.67 | 0.76 | 0.26 | -0.3 | -0.3075 | -0.6 | -1.63 | 0.356534 | 0.211962 | 0.002765 | 0.999971 | | 1 | 0.572658 |
| Madd | A2AGQ8 | 3.95 | -1.38 | 0.75 | 0.46 | -3.02 | 1.06 | 4.18 | -0.07 | -7.53 | 0.22 | -2.97 | -3.06 | -4.51 | -3.05 | -0.81 | -1 | -0.4075 | -4.28 | -3.2875 | 0.804145 | 0.033703 | 0.026774 | 0.999971 | | 1 | 0.678474 |
| Madd_1 | E9QN47 | -0.24 | -0.66 | -0.51 | -0.64 | -0.64 | -0.27 | -0.49 | -0.69 | -0.55 | -0.32 | -0.69 | -0.59 | -0.52 | -0.37 | -0.6 | -0.48 | -0.01 | -0.025 | 0.02 | 0.959364 | 0.894172 | 0.913291 | 0.999971 | | 1 | 0.992618 |
| Madd_2 | A2AGR0 | -2.53 | -2.66 | -2.86 | -2.59 | -2.06 | -2.98 | 1.16 | -5.01 | -0.39 | -2.91 | -1.86 | -1.46 | -2.82 | -3.03 | -3.38 | -3.5 | 0.4375 | 1.005 | -0.5225 | 0.703319 | 0.067077 | 0.037858 | 0.999971 | | 1 | 0.700838 |
| Maea | Q4VC33 | 0.53 | 1.04 | 0.44 | 0.34 | 0.57 | 1.26 | 0.29 | 1.12 | 0.77 | 0.9 | 0.86 | 0.48 | 0.59 | 1.21 | 0.41 | 0.56 | 0.2225 | 0.165 | 0.105 | 0.448878 | 0.464283 | 0.687652 | 0.999971 | | 1 | 0.945573 |
| Mafg | O54790 | 2.35 | 0.89 | 0.26 | 0.79 | 0.52 | 0.51 | 0.37 | 0.81 | 0.16 | 1.76 | 0.45 | 0.49 | 1.21 | 1.03 | 0.36 | 0.68 | -0.52 | -0.3575 | -0.2525 | 0.248088 | 0.505796 | 0.583231 | 0.999971 | | 1 | 0.920364 |
| Mag | P20917-2 | -2.1 | -6.85 | -6.01 | -4.99 | -4.69 | -6.55 | -5.47 | -5.7 | -5.1 | -4.76 | -4.29 | -4.2 | -6.65 | -5.7 | -5.68 | -6.97 | -0.615 | 0.4 | -1.2625 | 0.536812 | 0.675347 | 0.215865 | 0.999971 | | 1 | 0.826893 |
| Mag_1 | P20917 | 0.11 | -0.88 | -1.52 | -1.46 | -1.5 | -0.86 | -1.83 | -1.7 | -1.13 | -0.55 | -2.43 | -2.14 | -1.31 | -0.77 | -1.33 | -1.88 | -0.535 | -0.625 | -0.385 | 0.217328 | 0.262175 | 0.369984 | 0.999971 | | 1 | 0.869526 |
| Maged1 | Q9QYH6 | 0.29 | 0.81 | 0.14 | 0.42 | 0.5 | 0.52 | 0.82 | 0.56 | 0.65 | 0.45 | 0.47 | 0.5 | 0.76 | 0.48 | 0.23 | 0.5 | 0.185 | 0.1025 | 0.0775 | 0.392002 | 0.615037 | 0.728928 | 0.999971 | | 1 | 0.956041 |
| Maged2 | Q9ER67 | 0.53 | -0.35 | -0.1 | 0.12 | -0.08 | -0.17 | 0.06 | -0.18 | -0.24 | 0.17 | -0.15 | 0.17 | 0 | 0.11 | 0.98 | 0.38 | -0.1425 | -0.0625 | 0.3175 | 0.543019 | 0.797756 | 0.302475 | 0.999971 | | 1 | 0.847402 |
| Magee1 | Q6PCZ4 | 0.34 | 0.61 | 0.27 | 0.39 | 0.55 | 0.72 | 0.34 | 1.03 | 0.76 | 0.13 | 0.3 | 0.31 | 1.53 | 0.83 | 0.77 | 1.14 | 0.2575 | -0.0275 | 0.665 | 0.244889 | 0.892773 | 0.01646 | 0.999971 | | 1 | 0.613095 |
| Magee2 | Q52KG3 | -2 | -0.07 | 1.33 | 0.91 | -0.2 | -0.6 | 3.75 | -0.49 | -0.65 | -0.63 | 2.29 | -0.8 | 0.13 | 0.25 | -0.37 | -1.69 | 0.5725 | 0.01 | -0.4625 | 0.618612 | 0.991569 | 0.556512 | 0.999971 | | 1 | 0.915051 |
| Magi1 | A0A0N4SU | -0.62 | -0.46 | -1.45 | -1.33 | -0.67 | -0.65 | -1.55 | -1.46 | -0.59 | -0.58 | -1.06 | -1.01 | -0.26 | -0.47 | -1.28 | -0.9 | -0.1175 | 0.155 | 0.2375 | 0.73349 | 0.597466 | 0.485464 | 0.999971 | | 1 | 0.897286 |
| Magi2 | Q9WVQ1 | -0.29 | 0.55 | -0.69 | -0.66 | 0.7 | -0.79 | -0.39 | -1.21 | 0.08 | 0.38 | -1.49 | 0.04 | 2.4 | -0.56 | -1.73 | -0.51 | -0.15 | 0.025 | 0.1725 | 0.749271 | 0.958252 | 0.835829 | 0.999971 | | 1 | 0.979185 |
| Magi2_1 | A0A0G2JE | -0.42 | -0.88 | -0.9 | -1.16 | -0.88 | -0.91 | -0.94 | -1.23 | -0.94 | -0.78 | -0.72 | -0.73 | -0.91 | -1.3 | -1.21 | -1.15 | -0.15 | 0.0475 | -0.3025 | 0.498515 | 0.820477 | 0.192631 | 0.999971 | | 1 | 0.810547 |

| | | | | | | | | | | | | | | | | | | | | | | | | | | |
|---|---|---|---|---|---|---|---|---|---|---|---|---|---|---|---|---|---|---|---|---|---|---|---|---|---|---|
| Magi3 | G5E8T6 | -0.34 | -0.43 | -1.05 | -1.11 | -0.35 | -0.38 | -0.89 | -0.83 | -0.35 | -0.45 | -0.84 | -0.79 | 0.05 | -0.68 | -1.19 | -1.36 | 0.12 | 0.125 | -0.0625 | 0.655853 | 0.630473 | 0.864427 | 0.999971 | 1 | 0.983079 |
| Magoh | P61327 | -0.26 | 1.29 | 1.55 | 0.58 | 1.02 | 1.29 | 1.93 | 0.98 | 1.39 | 0.11 | 1.98 | 1.49 | 1.2 | 1.1 | 1.23 | 0.96 | 0.515 | 0.4525 | 0.3325 | 0.255629 | 0.40136 | 0.408664 | 0.999971 | 1 | 0.875014 |
| Magt1 | A2ADH1 | -0.37 | 0.45 | -0.66 | -1.5 | 0.54 | 1 | 0.56 | -1.27 | 0.84 | 1.08 | 0.56 | -0.36 | 0.71 | 1.02 | -1.14 | -1.58 | 0.7275 | 1.05 | 0.2725 | 0.237303 | 0.05289 | 0.695622 | 0.999971 | 1 | 0.947174 |
| Maip1 | Q8BHE8 | -1.31 | -0.57 | 0.27 | 0.25 | -0.42 | -0.16 | 0.24 | 0.34 | -0.17 | -1.06 | -0.02 | 0.3 | 0.03 | -0.47 | 0.62 | 0.37 | 0.34 | 0.1025 | 0.4775 | 0.404112 | 0.819867 | 0.276728 | 0.999971 | 1 | 0.843794 |
| Mal2 | Q8BI08 | -0.71 | -0.69 | -1 | -0.71 | -0.49 | -0.42 | -1.11 | -0.52 | -0.86 | -0.35 | -0.89 | -0.77 | -0.59 | -0.23 | -0.8 | -0.36 | 0.1425 | 0.06 | 0.2825 | 0.522297 | 0.764023 | 0.188224 | 0.999971 | 1 | 0.809025 |
| Malsu1 | Q9CWV0 | -0.36 | -2.01 | -0.71 | -0.86 | -1.21 | -0.18 | -1.33 | -0.32 | -0.76 | -1.8 | -1.54 | -1.33 | 0.4 | 0.15 | -0.84 | -0.14 | 0.225 | -0.3725 | 0.8775 | 0.609869 | 0.367107 | 0.064323 | 0.999971 | 1 | 0.726597 |
| Man1a1 | P45700 | 1.02 | 1.16 | 1 | 1.36 | 1.15 | 0.96 | 1.1 | 1.14 | 1.29 | 0.98 | 1.13 | 1.28 | 0.5 | 1.33 | 1.46 | 1.15 | -0.0475 | 0.035 | -0.025 | 0.788701 | 0.846843 | 0.921898 | 0.999971 | 1 | 0.993069 |
| Man1a2 | P39098-2 | -0.49 | 1.07 | 1.12 | 0.95 | 1.04 | 1.1 | 0.91 | 1.2 | 1.17 | -0.15 | 1.26 | 1.94 | 1.15 | 0.99 | 1.59 | 1.67 | 0.4 | 0.3925 | 0.6875 | 0.305003 | 0.473788 | 0.114675 | 0.999971 | 1 | 0.78268 |
| Man1b1 | A2AJ15 | 0.62 | 0.88 | 0.56 | 0.85 | 0.68 | 0.98 | -0.21 | 1.03 | 0.91 | 1.38 | 0.64 | 1.04 | 0.84 | 1.25 | 0.69 | 0.9 | -0.1075 | 0.265 | 0.1925 | 0.725326 | 0.240809 | 0.352477 | 0.999971 | 1 | 0.867768 |
| Man2a1 | P27046 | 1.26 | -0.27 | -1.91 | -0.52 | 0.49 | 0.6 | -1.39 | -1.02 | 0.27 | 0.7 | -3.82 | -4.15 | -0.59 | 0.43 | -1.96 | -1.44 | 0.03 | -1.39 | -0.53 | 0.967833 | 0.300182 | 0.487439 | 0.999971 | 1 | 0.897906 |
| Man2a2 | Q8BRK9 | 0.71 | 1.01 | 0.79 | 1.02 | 0.96 | 0.98 | 0.43 | 1.19 | 0.87 | 1.46 | 0.49 | 0.98 | 0.9 | 1.18 | 1.1 | 1.08 | 0.0075 | 0.0675 | 0.1825 | 0.973065 | 0.781743 | 0.323837 | 0.999971 | 1 | 0.85548 |
| Man2b1 | O09159 | 0.25 | -0.03 | -0.16 | -0.12 | -0.01 | -0.2 | -0.02 | 1.1 | 0.07 | 0.13 | 0.31 | 0.17 | 0.46 | 0.13 | 0.45 | 0.28 | 0.2325 | 0.185 | 0.345 | 0.467588 | 0.316069 | 0.096784 | 0.999971 | 1 | 0.78268 |
| Man2b2 | O54782 | 1.07 | 0.39 | -0.06 | 0.22 | 0.97 | 0.68 | -0.22 | 0.57 | 0.96 | 0.93 | -0.52 | -0.44 | 0.82 | 1.03 | -0.16 | 0.39 | 0.095 | -0.1725 | 0.115 | 0.783325 | 0.702233 | 0.743583 | 0.999971 | 1 | 0.961064 |
| Man2c1 | F8WIE1 | 1.08 | 0.44 | 0.61 | 0.58 | 0.25 | 0.16 | 0.37 | 0.08 | -0.19 | 0.38 | 0.21 | 0.11 | -0.31 | 0.2 | 0.29 | 0.47 | -0.4625 | -0.55 | -0.515 | 0.048119 | 0.033549 | 0.063043 | 0.999971 | 1 | 0.725348 |
| Manba | A0A0R4J09 | 0.08 | 0.57 | 0.85 | 0.91 | -0.02 | 0.66 | 0.67 | 1.05 | 0.27 | 0.03 | 0.41 | 0.14 | -0.37 | 0.63 | 0.64 | 0.27 | -0.0125 | -0.39 | -0.31 | 0.966786 | 0.127922 | 0.330807 | 0.999971 | 1 | 0.857107 |
| Manbal | A2BGN7 | 2.38 | 0.53 | 0.34 | 0.58 | 0.64 | 0.18 | 2.09 | 0.34 | 1.49 | 2.28 | 1.12 | 0.95 | 1.01 | 0.27 | -0.17 | 0.17 | -0.145 | 0.5025 | -0.6375 | 0.806165 | 0.348138 | 0.223556 | 0.999971 | 1 | 0.828225 |
| Manea | Q6NXH2 | -0.02 | 4.03 | 0.6 | 0.17 | 3.86 | 2.27 | 1.91 | -0.35 | 4.12 | 1.56 | 0.74 | 0.8 | 2.43 | 1.55 | -0.29 | -0.54 | 0.7275 | 0.61 | -0.4075 | 0.53068 | 0.58695 | 0.703273 | 0.999971 | 1 | 0.949575 |
| Maneal | Q6P1J0 | 2.61 | 1.32 | 0.49 | 0.83 | 1.01 | 1.53 | 1.1 | 1.04 | 1.16 | 3.06 | 0.74 | 0.61 | -0.82 | 1.17 | 0.51 | 1 | -0.1425 | 0.08 | -0.8475 | 0.752464 | 0.904829 | 0.177152 | 0.999971 | 1 | 0.808532 |
| Manf | Q80ZP8 | -1.59 | 0.06 | 0.19 | 0.2 | -0.01 | 0.03 | 0.27 | 0.27 | 0.33 | -0.49 | 0.49 | 0.87 | 0.4 | 0.31 | 0.57 | 0.48 | 0.425 | 0.585 | 0.725 | 0.325846 | 0.25021 | 0.110465 | 0.999971 | 1 | 0.78268 |
| Maoa | Q64133 | -0.59 | -0.09 | -0.23 | -0.57 | 0.29 | -0.18 | -0.17 | -0.04 | 0.48 | -0.32 | 0.16 | 0.29 | 0.64 | -0.18 | -0.1 | -0.26 | 0.345 | 0.5225 | 0.395 | 0.133425 | 0.054946 | 0.160839 | 0.999971 | 1 | 0.800215 |
| Maob | Q8BW75 | -0.64 | -0.69 | -0.63 | -0.42 | -0.36 | -0.36 | -0.64 | 0 | -0.61 | -0.7 | -0.72 | -0.97 | -0.61 | -0.61 | -0.52 | -0.43 | 0.255 | -0.155 | 0.0525 | 0.226436 | 0.387412 | 0.758276 | 0.999971 | 1 | 0.961116 |
| Map1a | A2ARP8 | 0.78 | 0.98 | 1.24 | 1.17 | 0.85 | 1.04 | 1.59 | 0.95 | 0.93 | 0.64 | 1.22 | 1.18 | 0.59 | 0.62 | 0.92 | 0.92 | 0.065 | -0.05 | -0.28 | 0.779665 | 0.814969 | 0.182618 | 0.999971 | 1 | 0.808532 |
| Map1b | P14873 | 0.44 | 0.28 | 0.23 | 0.09 | 0.06 | 0.42 | 0.47 | -0.08 | 0.27 | 0.57 | 0.08 | 0.32 | 0.1 | 0.29 | 0.15 | 0.09 | -0.0425 | 0.05 | -0.1025 | 0.836816 | 0.790367 | 0.560829 | 0.999971 | 1 | 0.91591 |
| Map1lc3a | Q91VR7 | -0.78 | -0.23 | -0.12 | 0.27 | -0.21 | 0.24 | -0.11 | 0 | -0.1 | -0.09 | -0.56 | 0.15 | -0.53 | 0.35 | -0.08 | 0.7 | 0.195 | 0.065 | 0.325 | 0.46153 | 0.815036 | 0.352955 | 0.999971 | 1 | 0.868675 |
| Map1lc3b | Q9CQV6 | 1.71 | 1.01 | 0.46 | 0.69 | -0.04 | 0.02 | -0.01 | 0.39 | 0.27 | 0.77 | 0.25 | 0.66 | 0.47 | 0.6 | 0.48 | 0.76 | -0.8775 | -0.48 | -0.39 | 0.015703 | 0.145783 | 0.204665 | 0.999971 | 1 | 0.81617 |
| Map1s | Q8C052 | 1.16 | 0.39 | 0.41 | 0.39 | 0.43 | 0.35 | 0.84 | 0.43 | 0.74 | 0.76 | 0.3 | -0.13 | 0.2 | -0.07 | 0.26 | 0.25 | -0.075 | -0.17 | -0.4275 | 0.763744 | 0.566938 | 0.102084 | 0.999971 | 1 | 0.78268 |
| Map2 | G3UZJ2 | 0.31 | 2.33 | 1.02 | 0.72 | 1.69 | 1.83 | 0.53 | 1.12 | 1.46 | 0.68 | 1.01 | 1.41 | 1.53 | 2.2 | 1.15 | 1.78 | 0.1975 | 0.045 | 0.57 | 0.688389 | 0.919739 | 0.235008 | 0.999971 | 1 | 0.832918 |
| Map2_1 | A0A668KL( | 0.7 | 1.08 | 1.2 | 1.11 | 1.07 | 1.32 | 1.43 | 1.08 | 0.99 | 0.51 | 1.19 | 1.44 | 1.01 | 1.18 | 1.26 | 1.48 | 0.2025 | 0.01 | 0.21 | 0.326352 | 0.968206 | 0.317911 | 0.999971 | 1 | 0.853893 |
| Map2k1 | P31938 | -0.48 | -0.87 | -1.53 | -1.16 | -0.52 | -0.8 | -1.62 | -1.27 | -0.92 | -0.78 | -1.78 | -1.46 | -1.1 | -0.55 | -1.2 | -1.34 | -0.0425 | -0.225 | -0.0375 | 0.897393 | 0.492632 | 0.898206 | 0.999971 | 1 | 0.989395 |
| Map2k2 | Q91YS7 | 0.02 | 0.13 | 0.4 | 0 | 1.08 | -0.29 | 0.32 | 0.01 | 2.3 | -0.19 | 0.29 | 0.39 | 1.8 | -0.44 | 0.29 | -0.01 | 0.1425 | 0.56 | 0.2725 | 0.650391 | 0.295577 | 0.561141 | 0.999971 | 1 | 0.91591 |
| Map2k3 | O09110 | -1.21 | -2.9 | -1.88 | -0.86 | -3.31 | -3.22 | -1.47 | -1.05 | -1.49 | -1.78 | -0.57 | -0.61 | -3.12 | -3.11 | -1.33 | -0.91 | -0.55 | 0.6 | -0.405 | 0.419785 | 0.255651 | 0.548214 | 0.999971 | 1 | 0.911727 |
| Map2k4 | P47809 | 0.43 | 0.28 | 0.36 | 0.36 | 0.46 | 0.58 | 0.84 | -0.04 | 0.64 | 0.14 | 0.09 | 0.05 | 0.51 | 0.43 | 0.12 | -0.21 | 0.1025 | -0.1275 | -0.145 | 0.65322 | 0.522744 | 0.505857 | 0.999971 | 1 | 0.905277 |
| Map2k5 | Q9WVS7 | -0.07 | 0.32 | -0.03 | 1.12 | 0.25 | 0.24 | -0.28 | 0.71 | -0.14 | 0.19 | -0.55 | 0.27 | 0.04 | 0.36 | 0.23 | 1.03 | -0.105 | -0.3925 | 0.08 | 0.757151 | 0.256264 | 0.817162 | 0.999971 | 1 | 0.974929 |
| Map2k6 | P70236 | -0.89 | 0.46 | 1.06 | 0.68 | 0.18 | 0.44 | 1.31 | 0.9 | 0.55 | -0.42 | 1.4 | 1.24 | 0.41 | 0.31 | 0.83 | 0.96 | 0.38 | 0.365 | 0.3 | 0.41893 | 0.510911 | 0.490499 | 0.999971 | 1 | 0.898612 |
| Map2k7 | Q8CE90-2 | -0.44 | -0.44 | 0.19 | 0.12 | 0.05 | -0.88 | 0.05 | -0.31 | 0.07 | -0.79 | -0.01 | 0.01 | -0.02 | -0.67 | -0.1 | 0.45 | -0.13 | -0.0375 | 0.0575 | 0.656427 | 0.893761 | 0.84667 | 0.999971 | 1 | 0.980253 |
| Map3k10 | D3YXM8 | 0.07 | 0.17 | -0.13 | 0.08 | -0.17 | -0.1 | -0.36 | 0.11 | -0.01 | 0.31 | 0 | -0.6 | 0.23 | -0.01 | 0.06 | -0.06 | -0.1775 | -0.1225 | 0.0075 | 0.35461 | 0.602876 | 0.965979 | 0.999971 | 1 | 0.998721 |
| Map3k11 | Q80XI6 | 0.19 | -0.99 | -0.75 | -0.87 | -1.09 | -1.17 | -1.15 | -0.39 | -1.31 | 0.17 | -0.8 | -1.28 | -0.57 | -0.85 | -0.74 | -0.18 | -0.345 | -0.2 | 0.02 | 0.30862 | 0.633904 | 0.948882 | 0.999971 | 1 | 0.998038 |
| Map3k12 | Q60700 | 1.41 | -0.53 | -0.36 | -0.72 | -0.93 | 0.15 | -0.25 | -0.8 | 0.09 | 1.89 | -0.47 | -0.38 | -1.02 | -1.68 | -0.35 | -0.62 | -0.4075 | 0.3325 | -0.8675 | 0.433567 | 0.623537 | 0.127111 | 0.999971 | 1 | 0.783755 |
| Map3k13 | Q1HKZ5 | -0.49 | -1.34 | -1.19 | -1.2 | -2.9 | -2.89 | -0.4 | -1.73 | -1.65 | -1.36 | -1.11 | -1.77 | -1.51 | -1.77 | -1.61 | -0.97 | -0.925 | -0.4175 | -0.41 | 0.132321 | 0.13877 | 0.162886 | 0.999971 | 1 | 0.800215 |
| Map3k2 | G5E8L8 | 0.52 | 0.1 | 1.3 | 1.22 | 1.19 | 1.09 | 1.35 | 2.12 | 1.61 | 0.66 | 1.41 | 0.62 | -0.18 | -0.09 | 0.87 | 0.82 | 0.6525 | 0.29 | -0.43 | 0.098505 | 0.445166 | 0.286395 | 0.999971 | 1 | 0.846095 |
| Map3k20 | Q9ESL4-2 | 0.81 | 1.53 | -0.02 | 0.75 | 1.43 | 1.56 | -0.8 | -0.17 | 0.03 | 0.07 | -0.42 | -0.05 | 1.76 | 1.32 | 0.63 | -0.41 | -0.2625 | -0.86 | 0.0575 | 0.667183 | 0.028508 | 0.912882 | 0.999971 | 1 | 0.992618 |
| Map3k4 | O08648 | -0.16 | 0.21 | 0.33 | 0.04 | 0.16 | 0.05 | 0 | 0 | 0.11 | -0.32 | 0.07 | 0.06 | 0.12 | 0.38 | -0.04 | -0.09 | -0.0525 | -0.125 | -0.0125 | 0.777378 | 0.536654 | 0.951421 | 0.999971 | 1 | 0.998056 |
| Map3k5 | E9PWG9 | -0.8 | -0.06 | 0.15 | -0.1 | -0.35 | -0.22 | -0.09 | -0.52 | -0.61 | -0.2 | 0.11 | -0.22 | -0.22 | -0.04 | -0.36 | 0.14 | -0.0925 | -0.0275 | 0.0825 | 0.715258 | 0.919165 | 0.75012 | 0.999971 | 1 | 0.961064 |
| Map3k7 | Q62073 | 0.68 | -1.3 | -0.5 | -0.02 | -1.56 | -0.76 | -0.46 | -0.14 | -1.56 | -0.13 | -0.21 | -0.38 | -1.24 | -1.48 | -0.37 | -0.05 | -0.445 | -0.285 | -0.5 | 0.366002 | 0.570402 | 0.331783 | 0.999971 | 1 | 0.857152 |
| Map3k9 | A0A1Y7VK) | -2.17 | -0.8 | -0.48 | -0.68 | -1.17 | -0.85 | -0.95 | -0.36 | -1.19 | -1.64 | -0.3 | -0.62 | -0.27 | -1.07 | -0.43 | 0.22 | 0.2 | 0.095 | 0.645 | 0.621298 | 0.835688 | 0.16912 | 0.999971 | 1 | 0.80174 |
| Map4 | E9PZ43 | 1.02 | 0.66 | 0.62 | 0.52 | 0.24 | 1 | 1.08 | 0.58 | 0.55 | 0.95 | 0.34 | 0.57 | 0.39 | 0.79 | 0.5 | 0.69 | 0.02 | -0.1025 | -0.1125 | 0.936471 | 0.631628 | 0.578431 | 0.999971 | 1 | 0.919075 |
| Map4_1 | P27546-2 | 1.24 | 0.61 | 0.9 | 0.89 | 0.51 | 0.9 | 1 | 0.93 | 0.77 | 0.79 | 0.73 | 0.77 | 0.55 | 0.67 | 0.72 | 0.81 | -0.075 | -0.145 | -0.2225 | 0.729289 | 0.454711 | 0.282503 | 0.999971 | 1 | 0.844538 |
| Map4_2 | P27546 | 0.39 | 2.05 | 1.59 | 0.43 | 2.27 | 1.73 | 0.99 | 1.35 | 2.18 | 1.8 | 0.96 | 1.95 | 2.88 | 1.88 | 2.3 | 2.22 | 0.47 | 0.6075 | 1.205 | 0.328289 | 0.214016 | 0.021805 | 0.999971 | 1 | 0.645537 |
| Map4k2 | Q61161 | -1 | -0.19 | -0.87 | -0.54 | -0.41 | -0.45 | -0.51 | -0.43 | -0.24 | -1.17 | -0.46 | -0.56 | -0.18 | -0.64 | -0.5 | 0.07 | 0.2 | 0.0425 | 0.3375 | 0.383199 | 0.880399 | 0.222966 | 0.999971 | 1 | 0.827753 |
| Map4k3 | A0A3Q4EG | 0.39 | -0.22 | -0.35 | -0.83 | -0.31 | -0.17 | -0.88 | -0.84 | -0.67 | 0.33 | -0.52 | -0.67 | -0.52 | -0.26 | -0.67 | -0.49 | -0.2975 | -0.13 | -0.2325 | 0.355781 | 0.705814 | 0.417368 | 0.999971 | 1 | 0.877483 |
| Map4k4 | A0A0A6YW | 0.33 | -0.67 | -0.42 | -0.34 | -0.35 | -0.62 | -0.37 | -0.14 | -0.49 | 0.45 | -0.16 | -0.22 | -0.48 | -0.58 | -0.37 | -0.1 | -0.095 | 0.17 | -0.1075 | 0.714632 | 0.573618 | 0.681967 | 0.999971 | 1 | 0.943204 |
| Map4k4_1 | B2RUE8 | -0.27 | 0.13 | 0.04 | -0.07 | -0.18 | -0.48 | 0 | 0.7 | -1.11 | -1.27 | 0.05 | 0.26 | 0.23 | -0.53 | 0.54 | 0.58 | 0.0525 | -0.475 | 0.2475 | 0.851377 | 0.240384 | 0.396462 | 0.999971 | 1 | 0.871816 |
| Map4k5 | E9PX30 | -1.3 | 0.29 | 0.39 | 0.05 | 0.35 | 0.01 | 0.17 | 0.01 | 0.57 | -0.57 | 0.39 | 0.25 | 0.95 | -0.21 | 0.45 | 0.39 | 0.2775 | 0.3025 | 0.5375 | 0.478233 | 0.498579 | 0.235986 | 0.999971 | 1 | 0.832918 |
| Map6 | Q7TSJ2 | 0.65 | -0.26 | 0.14 | 0.31 | -0.16 | -0.12 | 0.28 | -0.09 | -0.25 | 0.05 | -0.26 | -0.22 | -0.19 | -0.17 | -0.13 | -0.12 | -0.2325 | -0.38 | -0.3625 | 0.355451 | 0.133498 | 0.140366 | 0.999971 | 1 | 0.784948 |
| Map6d1 | Q14BB9 | 2.34 | 1.99 | 2.1 | 2.16 | 1.37 | 1.66 | 2.6 | 1.6 | 1.29 | 1.06 | 1.4 | 0.96 | -0.22 | 0.69 | 0.48 | 0.49 | -0.34 | -0.97 | -1.7875 | 0.26319 | 0.000662 | 5.47E-05 | 0.999971 | 0.448349 | 0.099736 |
| Map7 | E9QMU3 | 0.81 | 0.74 | 0.8 | 0.81 | 1.19 | 0.41 | 0.93 | 1.08 | 1.28 | 0.51 | 0.81 | 0.84 | 1.82 | 0.51 | 0.36 | 0.6 | 0.1125 | 0.07 | 0.0325 | 0.609067 | 0.73669 | 0.922718 | 0.999971 | 1 | 0.993264 |
| Map7_1 | A0A1L1SUI | 0.51 | 1.43 | 1.01 | 0.26 | 1.64 | 0.67 | -0.41 | 0.94 | 1.16 | 1.02 | 0.76 | 0.54 | 2.3 | 1.85 | 0.78 | 0.72 | -0.0925 | 0.0675 | 0.61 | 0.842828 | 0.823242 | 0.193938 | 0.999971 | 1 | 0.810547 |
| Map7d1 | A2AJI0 | 1.22 | 0.51 | 0.34 | 0.58 | 0.45 | 0.51 | 0.73 | 0.38 | 0.45 | 1.04 | 0.09 | 0.39 | 0.55 | 0.44 | 0.21 | 0.55 | -0.145 | -0.17 | -0.225 | 0.549473 | 0.559113 | 0.363782 | 0.999971 | 1 | 0.869526 |
| Map7d1_1 | A8Y5P4 | 1.39 | 1.38 | 2.1 | 2.22 | 1.41 | 1.53 | 1.55 | 1.82 | 1.11 | 1.84 | 1.16 | 1.46 | 0.48 | 1.78 | 2.18 | 1.95 | -0.195 | -0.38 | -0.175 | 0.465943 | 0.214269 | 0.678826 | 0.999971 | 1 | 0.942697 |
| Map7d2 | A2AG50 | 0.74 | 0.72 | 0.62 | 0.68 | 0.71 | 0.94 | 0.81 | 0.61 | 0.86 | 0.8 | 0.5 | 0.63 | 0.5 | 0.87 | 0.67 | 0.74 | 0.0775 | 0.0075 | 0.005 | 0.650374 | 0.96487 | 0.976917 | 0.999971 | 1 | 0.99925 |
| Map9 | Q3TRR0 | 0.26 | 0.36 | 0.58 | 0.75 | 0.69 | 0.11 | 0.9 | 0.24 | 0.35 | 0.34 | 0.64 | 0.66 | 0.35 | 0.17 | 0.18 | 0.39 | -0.0025 | 0.01 | -0.215 | 0.991881 | 0.959471 | 0.278926 | 0.999971 | 1 | 0.843812 |
| Mapk1 | P63085 | -0.31 | -0.29 | -0.14 | -0.23 | 0.28 | -0.37 | -0.08 | -0.05 | 0.39 | -0.49 | -0.18 | 0.1 | 0.54 | -0.29 | 0.01 | -0.03 | 0.1875 | 0.1975 | 0.3 | 0.356658 | 0.397527 | 0.199549 | 0.999971 | 1 | 0.813335 |
| Mapk10 | Q3TQZ7 | -0.96 | -0.62 | -0.26 | -0.25 | -0.29 | -0.66 | 1 | -0.23 | 0.12 | -0.92 | -0.2 | -0.15 | 0 | -0.78 | -0.09 | -0.29 | 0.4775 | 0.235 | 0.2325 | 0.234373 | 0.426961 | 0.391278 | 0.999971 | 1 | 0.871816 |
| Mapk11 | Q9WUI1 | -0.13 | 0.78 | 0.15 | 0.97 | 0.39 | 0.75 | 0.13 | 1.11 | 0.65 | 0.65 | 1.28 | 0.97 | 0.54 | 0.94 | 0.9 | 1.39 | 0.1525 | 0.445 | 0.5 | 0.649701 | 0.170768 | 0.140492 | 0.999971 | 1 | 0.784948 |
| Mapk14 | P47811 | 0.43 | 0.64 | -0.01 | 0.52 | 0.7 | 0.34 | 0.04 | 0.69 | 0.31 | 0.76 | 0.67 | 0.2 | 0.85 | 0.85 | 0.56 | 1.52 | 0.0475 | 0.09 | 0.55 | 0.845299 | 0.697886 | 0.066564 | 0.999971 | 1 | 0.733937 |
| Mapk1ip1l | A0A2I3BRS | 0.35 | 0.51 | 0.49 | 0.93 | 1.19 | -0.47 | -0.69 | 0.56 | 0.49 | -0.89 | 0.77 | 1.34 | 0.33 | 1.04 | 0.28 | 1.11 | -0.4225 | -0.1425 | 0.12 | 0.342837 | 0.757427 | 0.6633 | 0.999971 | 1 | 0.936388 |
| Mapk3 | Q63844 | -0.09 | -0.34 | -0.69 | -0.7 | 0 | -0.17 | -0.6 | -0.13 | 0.12 | -0.02 | -0.44 | -0.32 | 0.42 | 0.42 | -0.02 | -0.22 | 0.23 | 0.29 | 0.605 | 0.337649 | 0.23065 | 0.035217 | 0.999971 | 1 | 0.69205 |
| Mapk4 | Q6P5G0 | -0.18 | -0.07 | -0.43 | -0.34 | 0.47 | -0.17 | -0.15 | -0.09 | -0.14 | -0.12 | -0.58 | 0.03 | -0.43 | 0.05 | -0.83 | 0.03 | 0.27 | 0.0525 | -0.04 | 0.235662 | 0.797561 | 0.87414 | 0.999971 | 1 | 0.984109 |
| Mapk7 | Q9WVS8-4 | 0.77 | 1.09 | 1.12 | 1.17 | 0.8 | 1.66 | 1.85 | 1 | 0.91 | 1.02 | 0.84 | 1.37 | -0.94 | -0.25 | 1.49 | 1.09 | 0.29 | -0.0025 | -0.69 | 0.320581 | 0.990077 | 0.215951 | 0.999971 | 1 | 0.826893 |
| Mapk8 | Q91Y86 | -0.64 | -0.56 | -0.84 | -0.55 | -0.56 | -0.62 | -0.62 | -0.88 | -0.64 | -0.41 | -0.88 | -0.76 | -0.87 | -0.47 | -0.86 | -0.65 | -0.0225 | -0.025 | -0.065 | 0.899789 | 0.893113 | 0.730124 | 0.999971 | 1 | 0.956041 |
| Mapk8ip1 | Q9WVI9-3 | -0.03 | -0.53 | -0.6 | -0.59 | -0.57 | -0.69 | -0.36 | -0.19 | -0.61 | -0.01 | -0.2 | -0.33 | 0.2 | -0.42 | -0.36 | -0.23 | -0.015 | 0.15 | 0.235 | 0.945595 | 0.508889 | 0.327614 | 0.999971 | 1 | 0.856472 |
| Mapk8ip2 | Q9ERE9 | -0.41 | -0.37 | -0.1 | -0.42 | -0.52 | -0.58 | -0.78 | 0.3 | -0.12 | 0.04 | 0.07 | 0.18 | 0.36 | 0.2 | 0.5 | 0.77 | -0.07 | 0.3675 | 0.7825 | 0.79529 | 0.061739 | 0.003622 | 0.999971 | 1 | 0.579062 |
| Mapk8ip3 | J3QNR6 | -0.13 | -0.03 | -0.26 | -0.21 | -0.08 | -0.26 | -0.12 | -0.44 | -0.3 | -0.23 | -0.28 | -0.16 | 0.26 | -0.27 | -0.47 | -0.35 | -0.0675 | -0.085 | -0.05 | 0.705109 | 0.600951 | 0.81729 | 0.999971 | 1 | 0.974929 |
| Mapk8ip3_ | Q9ESN9-5 | -1.54 | -1.72 | -2.57 | -2.04 | -1.05 | -1.65 | -2.23 | -0.01 | -1.39 | -0.2 | -1.74 | -0.99 | -0.63 | -1.09 | -0.28 | -0.67 | 0.7325 | 0.8875 | 1.3 | 0.159012 | 0.044845 | 0.001637 | 0.999971 | 1 | 0.473622 |
| Mapk9 | Q9WTU6-4 | 2.45 | 2.24 | 3.47 | 4.39 | -1.09 | 0.02 | 3.97 | -0.66 | 2.84 | 3.63 | 2.72 | 1.35 | 0.63 | 2.45 | 2.24 | 2.91 | -2.5775 | -0.5025 | -1.08 | 0.04442 | 0.432446 | 0.118259 | 0.999971 | 1 | 0.78268 |
| Mapk9_1 | Q9WTU6-3 | 0.65 | 1.22 | 0.81 | 0.75 | -0.83 | -0.99 | 1.7 | -0.86 | 0.86 | 0.57 | 1.53 | 1.85 | 1.27 | 0.91 | 1 | 0.98 | -1.1025 | 0.345 | 0.1825 | 0.094901 | 0.300032 | 0.38272 | 0.999971 | 1 | 0.871816 |
| Mapkap1 | Q8BKH7-2 | -1.29 | 0.38 | -0.01 | -0.3 | 0 | 0.1 | -0.28 | 0.13 | 0.15 | -1.09 | -0.25 | -0.21 | 0.73 | -0.06 | -0.02 | -0.14 | 0.2925 | -0.045 | 0.4325 | 0.42534 | 0.91488 | 0.289342 | 0.999971 | 1 | 0.847402 |
| Mapkapk2 | P49138 | 0.52 | 0.16 | 0.15 | 0.56 | -0.06 | 0.36 | 0.75 | 0.43 | 0.1 | 0.26 | 0.62 | 0.28 | -0.17 | 0.3 | 0.71 | 0.84 | 0.0225 | -0.0325 | 0.0725 | 0.923832 | 0.874405 | 0.790661 | 0.999971 | 1 | 0.968447 |
| Mapkbp1 | A0A0A0M( | 0.71 | 0.43 | -0.3 | 0.19 | 0.64 | 0.88 | -0.53 | 0.2 | 0.69 | 1.28 | 0.2 | 0.1 | 0.71 | 0.97 | -0.15 | 0.3 | 0.04 | 0.31 | 0.2 | 0.912932 | 0.375076 | 0.544481 | 0.999971 | 1 | 0.911398 |
| Mapre1 | Q61166 | -0.08 | -0.96 | -0.77 | -1.04 | -0.38 | -0.88 | -0.57 | -0.86 | 0.07 | 0.32 | -0.16 | 0.07 | 0.2 | -0.61 | -0.49 | -0.21 | 0.04 | 0.7875 | 0.435 | 0.881773 | 0.013866 | 0.164391 | 0.999971 | 1 | 0.800215 |
| Mapre2 | Q8R001 | -1.75 | -2.16 | 0.11 | -1.17 | -1.19 | -2.6 | 0.63 | -0.97 | -0.66 | -2.39 | 0.37 | 0.89 | -1.29 | -2.95 | -0.56 | -1.42 | 0.21 | 0.795 | -0.3125 | 0.778089 | 0.330624 | 0.628258 | 0.999971 | 1 | 0.931957 |
| Mapre2_1 | E9Q6X0 | -0.45 | -0.68 | -0.17 | -0.28 | -1.43 | -0.47 | -1.92 | -0.82 | -1.27 | 0.28 | -0.91 | -1 | -2.3 | -1.39 | -0.01 | -0.66 | -0.765 | -0.33 | -0.695 | 0.046599 | 0.363925 | 0.165978 | 0.999971 | 1 | 0.800215 |
| Mapre3 | D3Z6G3 | 0.7 | 0.31 | 1.23 | 0.6 | 0.05 | 0.34 | 1.01 | 0.74 | 0.05 | 0.3 | 0.68 | 0.57 | 0.16 | -0.07 | 0.83 | 1.76 | -0.175 | -0.31 | -0.04 | 0.55831 | 0.252812 | 0.92569 | 0.999971 | 1 | 0.99401 |
| Mapre3_1 | Q6PER3 | -1.08 | 1.09 | 0.41 | 0 | 1.02 | 0.99 | 0.83 | 0.19 | 1.21 | -0.45 | 1.06 | 1.33 | 1.46 | 0.99 | 0.53 | 0.54 | 0.6525 | 0.6825 | 0.775 | 0.182215 | 0.249505 | 0.128071 | 0.999971 | 1 | 0.784371 |
| Mapt | P10637-5 | 0.42 | -1.52 | -2.36 | -2.3 | -2.33 | -1.68 | -1.11 | -2.24 | -2.15 | -0.33 | -2.33 | -2.71 | -1.72 | -2.11 | -2.63 | -1.42 | -0.4 | -0.44 | -0.53 | 0.538197 | 0.566134 | 0.414855 | 0.999971 | 1 | 0.876789 |
| Mapt_1 | P10637-4 | -1.29 | -2.07 | -2.28 | -2.21 | -1.29 | -2.35 | -2.11 | -1.72 | -0.89 | -0.91 | -1.4 | -0.6 | -0.42 | -1.05 | -0.71 | -0.48 | 0.095 | 1.0125 | 1.2975 | 0.771152 | 0.007 | 0.001393 | 0.999971 | 1 | 0.437254 |
| Mapt_2 | A2A5Y6 | -0.53 | -0.36 | -0.09 | -0.45 | -0.46 | -0.34 | 0.2 | -0.45 | -0.28 | -0.42 | -0.11 | 0.05 | 0.06 | -0.37 | -0.27 | -0.11 | 0.095 | 0.1675 | 0.185 | 0.67376 | 0.403708 | 0.359469 | 0.999971 | 1 | 0.869526 |
| Mapt_3 | P10637-2 | 0.13 | -0.01 | -0.83 | -1.17 | -0.89 | -1.63 | -1.16 | -0.32 | -3.37 | -1.06 | -1.69 | -2.5 | -1.9 | -1.76 | -2.34 | -0.97 | -0.53 | -1.685 | -1.2725 | 0.206541 | 0.012796 | 0.011642 | 0.999971 | 1 | 0.613095 |
| Marchf1 | D3YVY0 | -2.93 | -3.48 | -0.44 | -0.32 | -4.29 | -3.3 | -1.04 | -1.54 | -3.02 | -3.37 | -1.57 | -1.58 | -6.13 | -4.47 | 0.05 | -0.3 | -0.75 | -0.5925 | -0.92 | 0.459934 | 0.495671 | 0.555461 | 0.999971 | 1 | 0.914901 |
| Marchf5 | Q3KNM2 | -0.05 | 0.3 | 0.27 | 0.54 | 0.32 | 0.36 | -0.07 | 0.85 | 0.6 | -0.27 | 0.15 | -0.73 | 0.51 | 0.54 | 0.77 | 0.29 | 0.1 | -0.3275 | 0.2625 | 0.691934 | 0.311262 | 0.229388 | 0.999971 | 1 | 0.828687 |
| Marchf6 | Q6ZQ89 | -0.71 | 0.47 | -1.49 | -0.41 | 0.24 | 0.42 | 0.05 | -1.39 | 0.06 | -0.11 | 0.85 | 0.92 | 0.13 | 0.67 | 0.64 | -0.79 | 0.365 | 0.965 | 0.6975 | 0.499452 | 0.05994 | 0.181119 | 0.999971 | 1 | 0.808532 |
| Marchf7 | Q9WV66 | 1.88 | -0.36 | 1.44 | 1.54 | 0.22 | 0.65 | 0.86 | 0.45 | 0.52 | 1.67 | 0.92 | 0.22 | -0.8 | 0.16 | 0.94 | 1.47 | -0.58 | -0.2925 | -0.6825 | 0.251781 | 0.596865 | 0.303801 | 0.999971 | 1 | 0.847402 |
| Marcks | P26645 | 0.4 | -0.05 | -0.4 | -0.04 | 0.19 | 0.51 | 0.2 | -0.41 | 0.67 | 0.69 | -0.13 | -0.12 | 1.38 | 0.58 | -0.21 | -0.08 | 0.145 | 0.3 | 0.44 | 0.59613 | 0.322262 | 0.271934 | 0.999971 | 1 | 0.843794 |
| Marcksl1 | P28667 | -0.85 | -2.5 | -1.39 | -1.41 | -2.24 | -1.48 | 0.41 | -1.54 | -1.02 | -0.14 | -1.15 | -1.22 | -0.5 | -1.86 | -1.07 | -0.9 | 0.325 | 0.655 | 0.455 | 0.594739 | 0.136038 | 0.301062 | 0.999971 | 1 | 0.847402 |
| Mark1 | Q8VHJ5 | 0.6 | -0.27 | -0.23 | -0.05 | -0.33 | -0.29 | 0.77 | -0.36 | -0.1 | 0.04 | -0.12 | 0.02 | -0.22 | -0.73 | -0.37 | -0.38 | -0.065 | -0.0525 | -0.4375 | 0.84738 | 0.824948 | 0.113705 | 0.999971 | 1 | 0.78268 |
| Mark2 | E9QMP6 | -0.13 | -0.12 | 0.21 | 0.01 | -0.35 | -0.12 | 0.06 | -0.1 | -0.25 | 0.29 | 0.16 | -0.41 | -0.12 | -0.37 | -0.24 | 0.03 | -0.12 | -0.045 | -0.1675 | 0.525318 | 0.839876 | 0.383393 | 0.999971 | 1 | 0.871816 |
| Mark3 | Q03141 | -0.06 | 0.74 | 0.17 | -0.6 | -0.02 | -0.29 | 0.32 | -0.29 | 0.23 | -0.2 | 0.92 | 0.54 | -0.28 | -1.46 | -0.46 | -0.85 | -0.1325 | 0.31 | -0.825 | 0.676641 | 0.396484 | 0.049948 | 0.999971 | 1 | 0.717713 |
| Mark3_1 | A0A1Y7VN: | 0.38 | -0.56 | -1.07 | -1.16 | -0.7 | -0.24 | 0.61 | -1.59 | -0.56 | 0.05 | -1 | -1.09 | -0.74 | -0.3 | -1.63 | -1.77 | 0.1225 | -0.0475 | -0.5075 | 0.818419 | 0.909453 | 0.29453 | 0.999971 | 1 | 0.847402 |
| Mark4 | Q8CIP4-2 | 0.44 | -0.86 | -0.52 | -0.51 | -0.82 | -0.49 | -0.27 | -0.33 | -0.78 | 0.03 | -0.57 | -0.46 | -0.53 | -0.66 | -0.46 | -0.42 | -0.115 | -0.0825 | -0.155 | 0.712061 | 0.801495 | 0.602498 | 0.999971 | 1 | 0.924453 |
| Mars1 | E9QB02 | 0.85 | 1.85 | 1.25 | 1.49 | 1.9 | 1.51 | 1.16 | 1.58 | 2.22 | 1.35 | 1.81 | 1.73 | 2.48 | 1.89 | 1.42 | 2.02 | 0.1775 | 0.4175 | 0.5925 | 0.525148 | 0.171194 | 0.082243 | 0.999971 | 1 | 0.755254 |
| Mars2 | Q499X9 | -1.67 | -0.82 | -0.73 | -0.88 | -0.98 | -1.03 | -0.59 | -0.76 | -0.84 | -1.03 | -0.26 | -0.21 | -0.64 | -0.97 | -0.7 | -0.81 | 0.185 | 0.44 | 0.245 | 0.487024 | 0.174579 | 0.350391 | 0.999971 | 1 | 0.866844 |
| Mas1 | P30554 | -4.54 | -0.73 | -1.13 | -1.57 | -1.51 | -1.66 | -1.2 | -1.1 | -1.97 | -2.54 | -0.07 | -2.05 | -0.53 | -2.09 | 0.13 | -1.46 | 0.625 | 0.335 | 1.005 | 0.43591 | 0.717094 | 0.27815 | 0.999971 | 1 | 0.843812 |
| Masp1 | A0A2R8VH | 3.73 | 1.85 | 1.97 | 2.53 | 2.01 | 1.29 | 2.37 | 0.9 | 3.01 | 3.28 | 2.4 | 2.09 | -0.48 | 1.71 | 0.65 | 0.84 | -0.8775 | 0.175 | -1.84 | 0.109164 | 0.714474 | 0.010106 | 0.999971 | 1 | 0.613095 |
| Mast1 | Q9R1L5 | -0.17 | 0.21 | -0.19 | -0.61 | 0.61 | 0.15 | 0.34 | -0.56 | 0.58 | -0.2 | 0.11 | 0.22 | 0.37 | -0.58 | -0.36 | -0.69 | 0.325 | 0.3675 | -0.125 | 0.305756 | 0.175434 | 0.678779 | 0.999971 | 1 | 0.942697 |
| Mast2 | B1AST8 | 1.54 | -0.3 | -1.94 | -1.07 | 0.66 | 1.01 | -2.11 | -2.15 | 0.55 | 1.48 | -2.29 | -1.27 | 1.5 | 0.57 | -0.81 | -2.07 | -0.205 | 0.06 | 0.24 | 0.839447 | 0.952887 | 0.8036 | 0.999971 | 1 | 0.971467 |
| Mast3 | A0A1D5RN | -0.02 | -0.05 | -0.24 | -0.26 | -0.3 | -0.33 | -0.15 | -0.57 | -0.3 | -0.03 | -0.15 | -0.07 | -0.12 | -0.19 | -0.55 | -0.34 | -0.195 | 0.005 | -0.1575 | 0.301757 | 0.976592 | 0.408239 | 0.999971 | 1 | 0.875014 |
| Mast4 | A0A0A6YX( | 0.23 | -0.05 | -0.37 | -0.11 | -0.09 | -0.18 | 0.11 | -0.27 | -0.34 | 0.11 | -0.55 | -0.9 | -0.5 | -0.24 | -0.12 | -0.67 | -0.0325 | -0.345 | -0.3075 | 0.872916 | 0.214028 | 0.18614 | 0.999971 | 1 | 0.809025 |
| Mat2a | Q3THS6 | 0.62 | 1.21 | 0.73 | 0.52 | 1.57 | 0.67 | 0.84 | 0.9 | 1.57 | 0.42 | 0.74 | 1.09 | 1.26 | 0.66 | 0.43 | 0.39 | 0.225 | 0.185 | -0.085 | 0.41322 | 0.539485 | 0.754958 | 0.999971 | 1 | 0.961116 |
| Mat2b | Q99LB6-2 | 0.8 | 0.5 | -0.22 | -0.36 | 0.42 | 0.66 | -0.08 | -0.12 | 0.31 | 0.57 | -0.66 | -0.76 | 0.43 | 0.93 | -0.29 | -0.28 | 0.04 | -0.315 | 0.0175 | 0.905262 | 0.457736 | 0.964364 | 0.999971 | 1 | 0.998721 |
| Matk | A0A0R4J1P | 0.79 | -0.19 | 0.79 | 0.68 | 0.36 | 0.25 | 1.02 | 1.35 | -0.13 | 0.67 | 1.26 | 0.9 | -0.13 | 0.29 | 1.15 | 0.92 | 0.2275 | 0.1575 | 0.04 | 0.519382 | 0.670291 | 0.913163 | 0.999971 | 1 | 0.992618 |
| Matn2 | A0A0A0M( | -0.63 | -0.67 | -0.15 | -1.05 | 0.14 | -0.69 | -0.88 | -0.12 | -0.58 | -0.33 | -0.48 | -0.9 | 0.35 | 0.04 | -0.16 | -0.31 | 0.2375 | 0.0525 | 0.605 | 0.447976 | 0.831968 | 0.041112 | 0.999971 | 1 | 0.700838 |
| Matn4 | F2Z468 | -0.27 | -0.11 | -0.23 | -0.28 | 0.44 | 0.12 | -0.05 | -0.36 | 0.77 | 0.73 | 0.92 | 0.8 | 0.46 | 0.21 | 0.45 | 0.46 | 0.26 | 1.0275 | 0.6175 | 0.250693 | 0.000152 | 0.005313 | 0.999971 | 0.197262 | 0.582714 |
| Matr3 | A0A494B9( | 0.48 | 3.1 | 2.87 | 2.7 | 1.34 | 3.58 | 0.13 | 1.21 | 0.74 | 2.45 | 1.28 | 0.94 | 1.06 | 3.36 | 2.93 | 2.63 | -0.7225 | -0.935 | 0.2075 | 0.404119 | 0.178887 | 0.771365 | 0.999971 | 1 | 0.963498 |
| Matr3_1 | Q8K310 | 1.17 | 1.12 | 1.17 | 1.04 | 0.55 | 1.15 | 1.51 | 0.86 | 0.91 | 1.14 | 0.89 | 0.16 | 0.53 | 0.93 | 0.97 | 0.68 | -0.1075 | -0.35 | -0.3475 | 0.655807 | 0.17544 | 0.08635 | 0.999971 | 1 | 0.759991 |
| Mau2 | Z4YNA3 | -1.27 | -0.43 | -0.51 | -0.52 | -0.36 | -0.03 | -0.62 | -0.89 | -0.48 | -0.67 | -0.72 | -0.68 | -0.35 | 0 | -0.32 | -0.16 | 0.2075 | 0.045 | 0.475 | 0.470952 | 0.84887 | 0.07866 | 0.999971 | 1 | 0.754157 |
| Mavs | Q8VCF0 | 1.32 | -0.17 | -0.56 | -0.33 | -0.66 | -0.11 | -0.13 | -0.53 | -0.63 | 0.59 | -0.81 | -1 | -0.94 | -0.87 | -0.71 | -0.84 | -0.4225 | -0.5275 | -0.905 | 0.333488 | 0.322883 | 0.051077 | 0.999971 | 1 | 0.723736 |
| Max | P28574 | 2.64 | 2.98 | 2.21 | 1.44 | 3.27 | 2.6 | 1.86 | 2.25 | 2.59 | 1.45 | 2.45 | 2.41 | 3.13 | 2.26 | 1.96 | 1.43 | 0.1775 | -0.0925 | -0.1225 | 0.676457 | 0.819241 | 0.78892 | 0.999971 | 1 | 0.96741 |
| Mb21d2 | D3Z742 | -0.54 | -0.42 | -0.63 | -0.47 | -0.7 | -0.38 | -0.58 | -0.75 | -0.34 | -0.18 | -0.24 | -0.31 | -0.19 | -0.33 | -0.72 | -0.32 | -0.0875 | 0.2475 | 0.125 | 0.623542 | 0.150284 | 0.518173 | 0.999971 | 1 | 0.905815 |
| Mbd1 | A0A0R4J15 | 0.43 | -0.71 | 0.34 | 0.3 | -0.51 | -0.1 | 0.52 | -0.29 | -0.2 | 0.54 | 0.83 | -0.43 | -0.73 | -0.64 | -0.11 | -0.39 | -0.185 | 0.095 | -0.5575 | 0.592893 | 0.806179 | 0.098183 | 0.999971 | 1 | 0.78268 |
| Mbd2 | Q9Z2E1 | 0.51 | 1.61 | 0.51 | 0.63 | 1.53 | 1.34 | 0.09 | 0.76 | 1.23 | 1.04 | 0.36 | 0.96 | 1.71 | 1.5 | 0.4 | 0.83 | 0.115 | 0.0825 | 0.295 | 0.774627 | 0.800548 | 0.454866 | 0.999971 | 1 | 0.887217 |
| Mbd3 | Q9Z2D8 | 2.35 | 2.02 | 1.73 | 1.13 | 1.74 | 1.5 | 0.98 | 0.55 | 1.18 | 1.67 | 1.68 | 2.25 | 0.4 | 1.8 | 1.03 | 1.27 | -0.615 | -0.1125 | -0.6825 | 0.116767 | 0.738695 | 0.097762 | 0.999971 | 1 | 0.78268 |
| Mblac1 | Q8BWY4 | 0.92 | 0.53 | -0.24 | 0.55 | 0.47 | 0.45 | 0.46 | 0.74 | 0.55 | 0.62 | 0.77 | 0.19 | 0.88 | 0.77 | 0.78 | 0.59 | 0.09 | 0.0925 | 0.315 | 0.741442 | 0.746401 | 0.26363 | 0.999971 | 1 | 0.842533 |
| Mblac2 | Q8BL86 | -0.77 | -0.81 | -0.68 | -1.02 | -0.56 | -0.7 | -0.36 | -0.8 | -0.15 | -0.61 | -0.37 | -0.29 | -0.27 | -0.75 | -0.72 | -1.01 | 0.215 | 0.465 | 0.1325 | 0.272037 | 0.031946 | 0.545371 | 0.999971 | 1 | 0.911398 |
| Mbnl1 | A0A0A6YXI | -2.21 | -3.96 | 3.22 | 4.58 | 1.17 | -3.59 | 5.23 | 2.64 | 10.44 | -1.15 | 1.79 | 1.39 | 5.46 | -2.67 | 2.88 | 1.19 | 0.955 | 2.71 | 1.3075 | 0.697813 | 0.363383 | 0.584467 | 0.999971 | 1 | 0.920364 |
| Mbnl2 | Q8C181-4 | 1.61 | 1.83 | 2.08 | 2.02 | 1.48 | 1.27 | 3.03 | 1.5 | 1.71 | 1.38 | 1.44 | 1.93 | 0.8 | 1.8 | 1.6 | 1.78 | -0.065 | -0.27 | -0.39 | 0.871324 | 0.223027 | 0.18239 | 0.999971 | 1 | 0.808532 |
| Mboat2 | A0A217FL6 | -0.33 | -0.5 | 0.07 | 0.09 | 1.05 | 0.93 | 0.44 | 0.46 | 0.55 | 0.16 | 0.3 | 0.61 | 0.8 | 0.59 | 0.19 | 0.16 | 0.8875 | 0.5725 | 0.6025 | 0.005455 | 0.027673 | 0.034277 | 0.999971 | 1 | 0.69205 |
| Mboat7 | Q8CHK3 | 2.71 | -0.06 | 0.69 | -0.33 | -0.14 | 0.22 | 2.17 | 0.28 | -0.77 | 0.59 | 0.72 | -0.15 | -1.1 | -0.98 | 0.33 | -0.79 | -0.12 | -0.655 | -1.3875 | 0.877001 | 0.362364 | 0.070826 | 0.999971 | 1 | 0.736129 |
| Mbp | P04370-11 | -8.11 | -11.18 | -6.5 | -8.79 | -10.34 | -8.72 | -6.49 | -9.26 | -9.89 | -5.58 | -6.05 | -7.88 | -13.94 | -9.79 | -7.37 | -8.69 | -0.0575 | 1.295 | -1.3025 | 0.959237 | 0.310861 | 0.403791 | 0.999971 | 1 | 0.873348 |
| Mbp_1 | P04370-4 | -1.31 | -0.88 | -1.2 | -1.43 | -0.69 | -1.28 | -0.91 | -1.29 | -0.36 | -2.37 | -1.95 | -1.74 | 0.42 | -1.68 | -1.27 | -1.76 | 0.1625 | -0.4 | 0.1325 | 0.482199 | 0.36254 | 0.785737 | 0.999971 | 1 | 0.967132 |
| Mbp_2 | P04370 | -1.62 | -0.94 | -1.09 | -1.1 | -0.76 | -0.93 | -1.35 | -1.09 | -0.86 | -1.23 | -1.22 | -1.39 | -0.02 | -0.41 | -1.04 | -1 | 0.155 | 0.0125 | 0.57 | 0.508984 | 0.955519 | 0.082281 | 0.999971 | 1 | 0.755254 |
| Mbp_3 | P04370-10 | 2.38 | -3.18 | -0.98 | -0.89 | -4.59 | -5.3 | 0.48 | -0.87 | -7.89 | -1.13 | -1.35 | -0.94 | -6.38 | -4.78 | -1.31 | -1.6 | -1.9025 | -2.16 | -2.85 | 0.255389 | 0.2551 | 0.084012 | 0.999971 | 1 | 0.755254 |
| Mbtps2 | Q8CHX6 | 2.75 | 2.52 | 2.58 | 2.48 | 1.9 | 2.81 | 2.17 | 1.88 | 2.45 | 2.6 | 1.76 | 1.83 | 1.58 | 2.25 | 2.44 | 3.2 | -0.3925 | -0.4225 | -0.215 | 0.145434 | 0.11607 | 0.528028 | 0.999971 | 1 | 0.909725 |
| Mcam | Q8R2Y2 | -0.53 | -1.11 | -1.2 | -1.45 | -0.47 | -1.08 | -0.55 | -1.41 | -0.55 | -0.83 | -0.55 | 0.12 | -0.65 | -1.1 | -1.19 | -1.41 | 0.195 | 0.62 | -0.015 | 0.525349 | 0.058362 | 0.955889 | 0.999971 | 1 | 0.998056 |
| Mcat | Q8R3F5 | 0.13 | -0.36 | -0.89 | -0.74 | -0.32 | -0.41 | -0.86 | -0.75 | -0.26 | -0.03 | -1.03 | -0.72 | -0.56 | -0.43 | -0.89 | -0.73 | -0.12 | -0.045 | -0.1875 | 0.667831 | 0.888603 | 0.492203 | 0.999971 | 1 | 0.899886 |
| Mcc | G3UW40 | 0.51 | -0.43 | -0.13 | -0.39 | -0.56 | -0.56 | 0.27 | 0.1 | -0.34 | 0.23 | 0.2 | 0.08 | -0.81 | -0.64 | -0.54 | -0.38 | -0.0775 | 0.1525 | -0.4825 | 0.804049 | 0.57731 | 0.090282 | 0.999971 | 1 | 0.771991 |
| Mccc1 | Q99MR8 | -0.46 | -0.33 | -0.55 | -0.57 | -0.51 | -0.24 | -0.62 | -0.39 | -0.38 | -0.37 | -0.59 | -0.42 | -0.33 | -0.21 | -0.53 | -0.38 | 0.0375 | 0.0375 | 0.115 | 0.83394 | 0.821679 | 0.515404 | 0.999971 | 1 | 0.905815 |
| Mccc2 | Q3ULD5 | -0.67 | -0.29 | 0.02 | -0.47 | -0.41 | -0.59 | -0.05 | -0.37 | -0.35 | -0.64 | -0.06 | -0.18 | -0.13 | -0.45 | -0.54 | -0.45 | -0.0025 | 0.045 | -0.04 | 0.991138 | 0.844103 | 0.854515 | 0.999971 | 1 | 0.982125 |
| Mcee | Q9D1I5 | -1.25 | -1.94 | -3.18 | -2.5 | -2.2 | -1.95 | -2.12 | -2.73 | -2.51 | -1.68 | -4.24 | -3.49 | -2.69 | -1.72 | -3.08 | -2.78 | -0.0325 | -0.7625 | -0.35 | 0.938317 | 0.249686 | 0.465752 | 0.999971 | 1 | 0.891404 |
| Mcf2l | E9PXE2 | -0.72 | -0.61 | -0.41 | -0.21 | -0.35 | -0.71 | -0.52 | -0.28 | -0.43 | -0.68 | -0.2 | -0.39 | 0.1 | -0.15 | -0.18 | -0.02 | 0.0225 | 0.0625 | 0.425 | 0.911882 | 0.757368 | 0.052959 | 0.999971 | 1 | 0.725348 |
| Mcfd2 | Q8K5B2 | -0.63 | 2.54 | 1.12 | 0.79 | 3.8 | 2.14 | 0.6 | 1.2 | 3.12 | -1.15 | 1.8 | 1.95 | 4.44 | 1.66 | 1.69 | 1.2 | 0.98 | 0.475 | 1.2925 | 0.269922 | 0.638541 | 0.168573 | 0.999971 | 1 | 0.801027 |
| Mchr1 | Q8JZL2 | -0.03 | -0.67 | 0.46 | 1.75 | -0.7 | -0.84 | 1.08 | 0.35 | -0.36 | -0.49 | -0.18 | -1.44 | -1.55 | -1.96 | -0.49 | -3.22 | -0.405 | -0.995 | -2.1825 | 0.521049 | 0.093763 | 0.011144 | 0.999971 | 1 | 0.613095 |
| Mcl1 | P97287 | 0.15 | 0.52 | 0.17 | -0.12 | 0.68 | 0.32 | -0.89 | 0.26 | -0.6 | 0.98 | 0.06 | -0.36 | -0.03 | 0.2 | 0.08 | 0.05 | -0.0875 | -0.16 | -0.105 | 0.806429 | 0.660957 | 0.601598 | 0.999971 | 1 | 0.924453 |
| Mcm3ap | Q9WUU9 | -1.05 | -0.74 | -1.63 | -1.08 | -0.72 | -1.08 | -1.08 | -1.35 | -0.9 | -0.88 | -0.9 | -1.54 | -1.13 | -0.92 | -1.73 | -1.64 | 0.0675 | 0.07 | -0.23 | 0.78947 | 0.792149 | 0.427501 | 0.999971 | 1 | 0.880793 |
| Mcm4 | P49717 | 1.05 | 0.36 | 1.36 | 1.97 | 1.01 | -1.54 | 2.63 | 1.91 | 0.56 | -0.01 | 1.88 | 0.95 | 1.69 | 0.91 | 1.48 | 0.7 | -0.1825 | -0.34 | 0.01 | 0.833614 | 0.490189 | 0.979646 | 0.999971 | 1 | 0.99925 |
| Mcmbp | H9KV04 | 0.63 | 2.07 | 1.65 | 1.81 | 2.09 | 1.4 | 1.71 | 2.33 | 2.32 | 0.95 | 1.97 | 1.38 | 2.97 | 1.68 | 1.76 | 1.92 | 0.3425 | 0.115 | 0.5425 | 0.361744 | 0.783381 | 0.212985 | 0.999971 | 1 | 0.823987 |
| Mcoln1 | Q99J21 | 1.57 | 1.13 | 1.92 | 1.39 | 1.01 | 0.39 | 4.65 | 1.4 | 1.79 | 0.52 | 2.38 | 2.15 | 1.92 | 0.41 | 1.57 | 1.4 | 0.36 | 0.2075 | -0.1775 | 0.678876 | 0.627294 | 0.621163 | 0.999971 | 1 | 0.929449 |
| Mcrip1 | Q3UGS4 | 1.86 | 1.78 | 1.74 | 1.97 | 1.5 | 1.47 | 1.61 | 1.54 | 1.27 | 1.74 | 1.67 | 1.56 | 1.2 | 1.73 | 1.18 | 1.71 | -0.3075 | -0.2775 | -0.3825 | 0.089524 | 0.156198 | 0.098214 | 0.999971 | 1 | 0.78268 |
| Mcrip2 | A0A494BB( | 3.21 | 2.66 | 0.91 | -0.54 | 2.86 | 2.84 | -1.12 | 0.55 | 3.49 | 2.81 | -0.33 | -0.2 | 3.59 | 2.71 | 1.08 | 0.69 | -0.2775 | -0.1175 | 0.4575 | 0.809126 | 0.920362 | 0.641967 | 0.999971 | 1 | 0.93301 |
| Mcrs1 | Q3TJY1 | -1.12 | -0.49 | 0.32 | 0.5 | 0.12 | -1.51 | 0.3 | 0.64 | -0.39 | -1.23 | 0.36 | 0.47 | -1.69 | -0.37 | 0.76 | 0.58 | 0.085 | 1.57E-16 | 0.0175 | 0.878819 | 1 | 0.97731 | 0.999971 | 1 | 0.99925 |
| Mctp1 | E9PV86 | -0.4 | 0 | -0.78 | -0.4 | -0.15 | -0.39 | -0.87 | -0.2 | 0.03 | -0.03 | -0.54 | -0.7 | 0.15 | -0.41 | -0.69 | -0.59 | -0.0075 | 0.085 | 0.01 | 0.976473 | 0.746806 | 0.970132 | 0.999971 | 1 | 0.998787 |
| Mcts1 | Q9DB27 | -1.2 | -0.15 | 0.36 | -0.08 | 0.1 | -0.48 | 0.85 | 0.24 | 0.51 | -0.98 | 0.62 | 0.87 | 0.69 | -0.19 | 0.35 | -0.2 | 0.445 | 0.5225 | 0.43 | 0.292566 | 0.309305 | 0.278758 | 0.999971 | 1 | 0.843812 |
| Mcu | Q3UMR5 | -0.88 | -0.36 | -0.95 | -0.55 | -0.42 | -0.59 | -1.2 | -0.73 | -0.57 | -0.61 | -0.86 | -0.91 | -0.53 | -0.44 | -0.92 | -0.64 | -0.05 | -0.0525 | 0.0525 | 0.839629 | 0.802894 | 0.810935 | 0.999971 | 1 | 0.973864 |
| Mcub | Q810S1 | -1.99 | -1.67 | -4.59 | -3.06 | -1.42 | -2.17 | -3.93 | -3.9 | -1.24 | -2.46 | -3.26 | -3.49 | -2.96 | -1.62 | -2.5 | -2.17 | -0.0275 | 0.215 | 0.515 | 0.973167 | 0.775946 | 0.437481 | 0.999971 | 1 | 0.882419 |
| Mcur1 | Q9CXD6 | 0.12 | 0.32 | -1.38 | -0.68 | -0.34 | 0.12 | -1.28 | -1.08 | -0.08 | 0.34 | -1.38 | -1.49 | -0.52 | 0.12 | -1.51 | -1.56 | -0.24 | -0.2475 | -0.4625 | 0.615003 | 0.658528 | 0.386256 | 0.999971 | 1 | 0.871816 |
| Mdc1 | Q5PSV9 | 3.2 | 4.9 | 1.04 | 2.16 | 6.9 | 3.06 | 0.44 | 1.02 | 7.21 | 1.5 | 1.85 | -1.95 | 10.64 | 2.73 | 2.31 | 1.6 | 0.03 | -0.6725 | 1.495 | 0.98386 | 0.715431 | 0.464835 | 0.999971 | 1 | 0.89116 |
| Mdga1 | D3Z499 | -1.01 | -1.11 | -0.73 | -0.5 | -0.8 | -0.77 | -0.12 | -1.05 | -0.3 | -0.45 | -0.39 | -0.34 | -0.93 | -1.42 | -1.06 | -1.38 | 0.1525 | 0.4675 | -0.36 | 0.566858 | 0.039724 | 0.136338 | 0.999971 | 1 | 0.784948 |
| Mdga2 | Q3UV20 | -0.34 | -0.5 | -1.25 | -0.9 | -0.41 | -0.48 | -0.26 | -1.11 | -0.27 | -0.29 | -1.1 | -1.06 | -0.32 | -0.71 | -1.05 | -1.06 | 0.1825 | 0.0675 | -0.0375 | 0.533133 | 0.829135 | 0.894962 | 0.999971 | 1 | 0.988571 |
| Mdh1 | P14152 | -0.26 | -0.63 | -0.42 | -0.1 | -1.05 | -0.87 | -0.29 | -0.58 | -0.95 | -0.99 | -0.12 | -0.44 | -1.51 | -0.65 | -0.22 | -0.14 | -0.345 | -0.2725 | -0.2775 | 0.169092 | 0.310273 | 0.414325 | 0.999971 | 1 | 0.876329 |

| | | | | | | | | | | | | | | | | | | | | | | | | | | |
|---|---|---|---|---|---|---|---|---|---|---|---|---|---|---|---|---|---|---|---|---|---|---|---|---|---|---|
| Mdh2 | P08249 | 0.41 | 0.66 | 0.22 | 0.2 | 0.37 | 0.85 | -0.21 | 0.29 | 0.08 | 0.64 | 0.09 | -0.52 | 0.59 | 0.77 | 0.11 | 0.29 | -0.0475 | -0.3 | 0.0675 | 0.857092 | 0.293166 | 0.764308 | 0.999971 | 1 | 0.962405 |
| Mdk | P12025 | 2.76 | 1.35 | 1.52 | 1.64 | 1.21 | 2.42 | 1.92 | 3.45 | 1.29 | 2.27 | 1.63 | 2.03 | 1.53 | 1.46 | 2.27 | 0.75 | 0.4325 | -0.0125 | -0.315 | 0.419264 | 0.973356 | 0.463922 | 0.999971 | 1 | 0.890905 |
| Mdm1 | E9Q0B1 | 1.71 | 0.89 | 1.36 | 1.49 | 0.19 | 0.36 | 0.95 | 1.8 | -0.57 | 1.89 | 1.28 | 1.55 | 0.53 | 0.97 | 1.71 | 1.48 | -0.5375 | -0.325 | -0.19 | 0.188796 | 0.547278 | 0.555659 | 0.999971 | 1 | 0.914901 |
| Mdm4 | A0A087WF | 2.09 | 0.23 | 0.99 | 1.05 | 0.14 | 1.01 | 1.03 | 1.01 | 0.18 | 1.57 | 0.03 | -0.5 | 0.81 | 1.3 | 0.14 | 0.87 | -0.2925 | -0.77 | -0.31 | 0.490165 | 0.178915 | 0.475076 | 0.999971 | 1 | 0.894044 |
| Mdn1 | J3QMC5 | 2.37 | 0.64 | 1.22 | 1.29 | 0.69 | 0.54 | 0.24 | 1.1 | 0.31 | 0.88 | 0.91 | 1.27 | 0.11 | 0.18 | 1.35 | 0.86 | -0.7375 | -0.5375 | -0.755 | 0.083838 | 0.197787 | 0.113491 | 0.999971 | 1 | 0.78268 |
| Mdp1 | Q9D967 | 0.27 | -0.03 | -0.16 | -0.06 | 0.03 | -0.07 | 2.52 | -0.52 | -0.26 | -0.23 | -0.6 | -0.37 | -0.67 | -0.05 | -0.46 | -0.41 | 0.485 | -0.37 | -0.4025 | 0.449553 | 0.0763 | 0.082486 | 0.999971 | 1 | 0.755254 |
| Me1 | P06801 | -0.61 | -0.36 | 0.2 | -0.03 | -0.37 | -0.34 | 0.09 | -0.23 | -0.5 | -0.35 | 0.13 | 0.35 | -0.67 | -0.29 | 0.07 | 0.2 | -0.0125 | 0.1075 | 0.0275 | 0.958426 | 0.70349 | 0.921936 | 0.999971 | 1 | 0.993069 |
| Me2 | Q99KE1 | -0.01 | -0.23 | -0.27 | -0.3 | -0.7 | -0.92 | -0.74 | -0.3 | -0.6 | -0.47 | -0.78 | -0.49 | -0.49 | -0.36 | -0.44 | -0.69 | -0.4625 | -0.3825 | -0.2925 | 0.045201 | 0.053206 | 0.128545 | 0.999971 | 1 | 0.784699 |
| Me3 | Q8BMF3 | 0.18 | 0.11 | 0.09 | 0.23 | -0.26 | -0.63 | -0.04 | -0.19 | -0.44 | -0.43 | -0.51 | -0.36 | -0.84 | -0.63 | -0.57 | -0.49 | -0.4325 | -0.5875 | -0.785 | 0.048911 | 0.004676 | 0.001439 | 0.999971 | 1 | 0.437254 |
| Mea1 | Q64327 | -1.64 | -3.18 | 0.2 | -1.19 | -3.21 | -2.42 | 0.46 | 0.07 | -2.53 | -2.35 | 0.84 | 1.05 | -3.72 | -1.87 | 0.64 | 0.22 | 0.1775 | 0.705 | 0.27 | 0.861792 | 0.518482 | 0.80506 | 0.999971 | 1 | 0.971701 |
| Meaf6 | V9GX42 | -0.05 | 0.03 | 1 | 0.87 | 0.12 | 0 | 0.56 | 1.03 | -0.21 | -0.12 | 0.39 | 0.73 | 0.06 | 0.65 | 1.08 | 1.24 | -0.035 | -0.265 | 0.295 | 0.921097 | 0.453295 | 0.434318 | 0.999971 | 1 | 0.882419 |
| Meak7 | Q8K0P3 | -0.02 | -0.68 | -0.92 | -1.24 | -0.43 | -0.76 | -0.91 | -0.85 | -0.43 | -0.54 | -1 | -0.56 | 0.3 | -0.38 | -1.11 | -1.01 | -0.0225 | 0.0825 | 0.165 | 0.938357 | 0.780959 | 0.680517 | 0.999971 | 1 | 0.943018 |
| Mecp2 | Q9Z2D6-2 | -0.6 | -1.36 | -0.04 | -0.08 | -1.08 | -1.3 | -0.23 | -0.38 | -1.24 | -0.84 | -0.75 | -0.76 | -1.56 | -0.96 | -0.23 | -0.45 | -0.2275 | -0.3775 | -0.28 | 0.561317 | 0.268878 | 0.498112 | 0.999971 | 1 | 0.902911 |
| Mecr | Q9DCS3 | -0.47 | -0.15 | 0.01 | -0.13 | -0.24 | -0.51 | -0.49 | 0.01 | -0.5 | -0.38 | -0.16 | -0.05 | -0.24 | -0.59 | -0.31 | -0.14 | -0.1225 | -0.0875 | -0.135 | 0.562954 | 0.661228 | 0.504872 | 0.999971 | 1 | 0.905277 |
| Med1 | Q925J9 | 3.48 | 1.38 | 1.74 | 1.19 | 2.09 | 2.16 | 1.6 | 1.11 | 2.1 | 2.71 | 1.17 | 1.3 | 1.36 | 1.24 | 0.71 | 1.36 | -0.2075 | -0.1275 | -0.78 | 0.697622 | 0.827475 | 0.149706 | 0.999971 | 1 | 0.793248 |
| Med12 | A2AGH8 | 3.11 | 1.93 | 1.7 | 2.79 | 1.68 | 1.67 | 2.6 | 2.15 | 2.03 | 3.18 | 1.38 | 1.54 | 0.29 | 1.62 | 1.65 | 1.13 | -0.3575 | -0.35 | -1.21 | 0.368868 | 0.48438 | 0.020818 | 0.999971 | 1 | 0.635402 |
| Med13l | A0A0J9YUA | 0.6 | -3.25 | 1.89 | 1.8 | 1.15 | -1.13 | -0.41 | 0.3 | 0.5 | 1.2 | -0.03 | 1.97 | -0.78 | -0.13 | -0.15 | 0.95 | -0.2825 | 0.65 | -0.2875 | 0.807295 | 0.575249 | 0.797992 | 0.999971 | 1 | 0.970479 |
| Med14 | A2ABV5 | 0.43 | 3.24 | -1.06 | 0 | 1.59 | 1.69 | -1.16 | 2.02 | 1.56 | 1.35 | 0.74 | 0.76 | 2.43 | 1.01 | 1.74 | -0.13 | 0.3825 | 0.45 | 0.61 | 0.716224 | 0.599279 | 0.528278 | 0.999971 | 1 | 0.909812 |
| Med15 | A0A338P73 | 1.36 | -0.73 | 1.69 | 0.87 | -0.2 | 0.72 | 1.09 | 2.4 | -0.15 | 0.85 | 2.04 | 1.45 | -0.41 | 0.12 | 1.3 | 1.45 | 0.205 | 0.25 | -0.1825 | 0.766059 | 0.70151 | 0.775516 | 0.999971 | 1 | 0.96393 |
| Med16 | E9QP84 | 1.41 | 4.57 | 1.96 | 1.99 | 4.23 | 4.05 | 1.13 | 1.43 | 4.54 | 0.98 | 1.9 | 1.71 | 6.06 | 3.94 | 1.95 | 2.45 | 0.2275 | -0.2 | 1.1175 | 0.8153 | 0.832872 | 0.298368 | 0.999971 | 1 | 0.847402 |
| Med17 | Q8VCD5 | 1.37 | 2.59 | 1.27 | 2.03 | 2.02 | 1.73 | 1.27 | 1.72 | 1.77 | 2.26 | 1.76 | 1.68 | 1.57 | 2.55 | 1.57 | 1.91 | -0.13 | 0.0525 | 0.085 | 0.704236 | 0.874996 | 0.820353 | 0.999971 | 1 | 0.975761 |
| Med19 | Q8C1S0 | -1.43 | 1.22 | -0.21 | 1.44 | 1.97 | 1.64 | 0.8 | 0.33 | 1.25 | 0.99 | 0.06 | 0.79 | 1.7 | 1.18 | 0.42 | 0.78 | 0.93 | 0.5175 | 0.765 | 0.203731 | 0.438047 | 0.26385 | 0.999971 | 1 | 0.842533 |
| Med22 | A2ALA4 | 2.75 | 0.88 | 0.85 | 0.27 | 0.9 | 1.34 | 0.23 | 0.7 | 0.76 | 2.45 | -0.05 | 0.1 | 0.51 | 1.53 | 0.7 | 0.04 | -0.395 | -0.3725 | -0.4925 | 0.471111 | 0.605265 | 0.398997 | 0.999971 | 1 | 0.871816 |
| Med23 | F8WJB0 | -0.52 | 1.1 | 2.02 | 1.57 | 1.51 | -0.71 | 1.51 | 1.72 | 2 | -1.87 | 2.25 | 1.91 | 1.72 | -0.43 | 1.85 | 1.65 | -0.035 | 0.03 | 0.155 | 0.961196 | 0.976381 | 0.82552 | 0.999971 | 1 | 0.976715 |
| Med24 | A6PW47 | 1.55 | 2.28 | 0.53 | -1.33 | 1.88 | 2.51 | 0.58 | 0.01 | 2.18 | 2.45 | 0.91 | 1.31 | 2.34 | 2.34 | 0.63 | -0.04 | 0.4875 | 0.955 | 0.56 | 0.578647 | 0.242014 | 0.532904 | 0.999971 | 1 | 0.910069 |
| Med28 | Q920D3 | 2.62 | 4.73 | 1.55 | 1.99 | 4.75 | 4.91 | 0.86 | 1.7 | 4.81 | 4.41 | 0.73 | 0.25 | 7.97 | 4.83 | 1.32 | 1.65 | 0.3325 | -0.1725 | 1.22 | 0.76643 | 0.889573 | 0.430319 | 0.999971 | 1 | 0.881611 |
| Med29 | Q9DB91 | 0.35 | 1.61 | -0.12 | 0.78 | 2.12 | 2.34 | -0.84 | 0.64 | 1.77 | 1.32 | 0.42 | 0.78 | 4.05 | 2.17 | 0.67 | 1.3 | 0.41 | 0.4175 | 1.3925 | 0.584525 | 0.361926 | 0.087904 | 0.999971 | 1 | 0.765196 |
| Med30 | Q9CQI9 | 1.01 | 1.38 | 0.95 | 1.47 | 0.62 | 1.76 | 0.28 | 1.22 | 0.01 | 2.42 | 0.65 | 0.61 | -0.5 | 2.87 | 1.16 | 1.99 | -0.2325 | -0.28 | 0.1775 | 0.508144 | 0.578707 | 0.788923 | 0.999971 | 1 | 0.96741 |
| Med4 | Q9CQA5-2 | 1.26 | 1.56 | 0.73 | 0.93 | 1.41 | 1.02 | 0.44 | 0.85 | 1.32 | 1.71 | 1.05 | 0.67 | 0.57 | 1.62 | 0.46 | 1.14 | -0.19 | 0.0675 | -0.1725 | 0.510815 | 0.819014 | 0.600167 | 0.999971 | 1 | 0.92426 |
| Med6 | Q921D4 | 2.1 | 2.53 | 2.74 | 2.96 | 2.04 | 2 | 1.5 | 2.05 | 1.91 | 1.61 | 2.5 | 3.17 | 2.32 | 2.13 | 2.72 | 3.12 | -0.685 | -0.285 | -0.01 | 0.022614 | 0.458486 | 0.973089 | 0.999971 | 1 | 0.99925 |
| Med8 | A2A9C6 | 1.6 | -0.12 | 1.23 | 2.97 | -0.67 | 0.48 | 1.57 | 1.96 | -2.27 | -0.19 | 2.25 | 1.79 | -1.8 | -1.48 | 0.96 | 2.63 | -0.585 | -1.025 | -1.3425 | 0.46088 | 0.359392 | 0.240579 | 0.999971 | 1 | 0.834089 |
| Med9 | Q8VCS6 | -1.62 | 0.67 | 0.03 | -2.3 | 0.74 | -0.95 | 0.41 | -1.37 | 0.34 | -1.85 | 0.03 | 0.45 | 0.21 | -1.3 | -1.13 | -2.21 | 0.5125 | 0.5475 | -0.3025 | 0.514105 | 0.496882 | 0.695631 | 0.999971 | 1 | 0.947174 |
| Mef2c | Q8CFN5 | 2.82 | 2.39 | 2.06 | 2.26 | 1.75 | 2.07 | 2.04 | 1.98 | 1.74 | 2.79 | 2.19 | 2.05 | 1.09 | 2.1 | 1.73 | 1.94 | -0.4225 | -0.19 | -0.6675 | 0.081885 | 0.510662 | 0.042552 | 0.999971 | 1 | 0.701725 |
| Mef2d | E9QKT0 | 3.34 | 2.54 | 3.82 | 4.09 | 1.73 | 2.23 | 3.4 | 3.31 | 1.67 | 3.08 | 3.02 | 2.84 | 0.98 | 2 | 3.29 | 3.54 | -0.78 | -0.795 | -0.995 | 0.141173 | 0.103824 | 0.138178 | 0.999971 | 1 | 0.784948 |
| Megf10 | A0A140T8S | 1.93 | 1.25 | 0.39 | 1.02 | 0.62 | 1.34 | 0.85 | 1.35 | 1.22 | 1.5 | 0.12 | 0.27 | 1.05 | 0.79 | 0.13 | 0.22 | -0.1075 | -0.37 | -0.6 | 0.764447 | 0.412009 | 0.137406 | 0.999971 | 1 | 0.784948 |
| Megf6 | B1ASH4 | 0.58 | 0.05 | -0.89 | 1.77 | 0.52 | 0.72 | 0.92 | -0.36 | 0.7 | 1.97 | 1.14 | 0.13 | -0.07 | 0.96 | -1.83 | -2.05 | 0.0725 | 0.6075 | -1.125 | 0.898729 | 0.341192 | 0.192856 | 0.999971 | 1 | 0.810547 |
| Megf8 | P60882 | 0.45 | 0.18 | 0 | 0.37 | 0.02 | -0.02 | 0.13 | 0.44 | -0.1 | 0.65 | 0.18 | 0.04 | -0.25 | -0.16 | 0.19 | 0.1 | -0.1075 | -0.0575 | -0.28 | 0.597036 | 0.8009 | 0.190399 | 0.999971 | 1 | 0.810547 |
| Megf9 | Q8BH27 | -1.12 | -1.58 | -0.37 | -0.8 | -1.78 | -1.35 | 0.76 | -0.91 | -1.68 | -1.3 | -0.06 | -0.28 | -1.35 | -2.02 | -0.91 | -1 | 0.1475 | 0.1375 | -0.3525 | 0.792814 | 0.756115 | 0.331953 | 0.999971 | 1 | 0.857152 |
| Meis2 | Q3TYM2 | 1.86 | 2.42 | 0.72 | 1.72 | 1.77 | 1.37 | 0.54 | 1.2 | 1.95 | 2.74 | 0.29 | 0.83 | 1.47 | 1.61 | 1.08 | 0.67 | -0.46 | -0.2275 | -0.4725 | 0.284896 | 0.706123 | 0.252085 | 0.999971 | 1 | 0.839012 |
| Memo1 | Q91VH6 | -1.09 | 0.15 | -0.32 | -0.61 | 0 | 0.31 | -0.33 | -0.28 | 0.05 | -0.83 | -0.17 | -0.12 | -0.07 | 0.08 | -0.01 | -0.08 | 0.3925 | 0.2 | 0.4475 | 0.22041 | 0.542146 | 0.135564 | 0.999971 | 1 | 0.784948 |
| Men1 | F8WHD9 | 0.14 | 0.36 | 0.8 | 0.94 | 0.34 | 0.08 | 0.35 | 0.81 | 0.11 | 0.27 | 0.62 | 0.44 | -0.31 | 0.34 | 0.93 | 0.99 | -0.165 | -0.2 | -0.0725 | 0.534838 | 0.422901 | 0.836377 | 0.999971 | 1 | 0.979416 |
| Mepce | Q8K3A9 | 1.95 | 2.89 | 2.57 | 2.54 | 2.93 | 2.39 | 2.23 | 2.65 | 2.93 | 2.71 | 2.18 | 2.23 | 3.31 | 2.7 | 2.77 | 2.4 | 0.0625 | 0.025 | 0.3075 | 0.816165 | 0.929269 | 0.299342 | 0.999971 | 1 | 0.847402 |
| Mertk | Q60805 | 0.16 | -0.32 | -0.22 | -0.02 | -0.72 | -0.58 | -0.13 | -0.33 | 0.15 | -0.65 | -0.29 | 0.12 | 0.43 | -1.39 | -0.75 | -0.51 | -0.34 | -0.0675 | -0.455 | 0.141317 | 0.783534 | 0.250157 | 0.999971 | 1 | 0.839012 |
| Mesd | Q9ERE7 | -0.79 | 2.16 | 0.96 | 1.08 | 2.36 | 2.11 | 1.04 | 1.41 | 2.76 | 0.15 | 1.49 | 1.64 | 3.55 | 1.53 | 1.6 | 1.46 | 0.8775 | 0.6575 | 1.1825 | 0.182009 | 0.383467 | 0.126233 | 0.999971 | 1 | 0.783071 |
| Met | F8VQL0 | 0.49 | 1.15 | -2.17 | -2.76 | 1.08 | 1.71 | -1.97 | -1.38 | 2.2 | 1.02 | -1.76 | -1.58 | 2.46 | 0.92 | -2.11 | -1.62 | 0.6825 | 0.7925 | 0.735 | 0.565242 | 0.524597 | 0.571503 | 0.999971 | 1 | 0.91816 |
| Metap1 | Q8BP48 | 1.4 | 1.07 | 1.32 | 1.15 | 0.9 | 1.47 | 1.87 | 1.67 | 0.83 | 1.48 | 1.3 | 0.7 | 0.44 | 1.2 | 1.23 | 1.32 | 0.2425 | -0.1575 | -0.1875 | 0.345709 | 0.506505 | 0.456588 | 0.999971 | 1 | 0.887217 |
| Metap1d | Q9CPW9-2 | 5.77 | -0.58 | -1.62 | 0.51 | -0.46 | 0.58 | -1.22 | 0.04 | -0.29 | 4.73 | -1.31 | -1.72 | -1.26 | 0.83 | -1.39 | 0.48 | -1.285 | -0.6675 | -1.355 | 0.399959 | 0.736237 | 0.391053 | 0.999971 | 1 | 0.871816 |
| Metap2 | Q3UI33 | 0.63 | 0.51 | 1.56 | 1.59 | 0.42 | 0.51 | 1.27 | 1.65 | 0.69 | 0.66 | 0.9 | 1.35 | 0.79 | 0.63 | 1.37 | 1.46 | -0.11 | -0.1725 | -0.01 | 0.782815 | 0.604829 | 0.977286 | 0.999971 | 1 | 0.99925 |
| Metap2_1 | M0QW48 | 1.25 | 2.64 | 2.19 | 2.46 | 2.25 | 1.63 | 1.45 | 2.36 | 1.71 | 2.45 | 1.97 | 2.06 | 3.69 | 2.06 | 1.93 | 2.49 | -0.2125 | -0.0875 | 0.4075 | 0.570149 | 0.797855 | 0.399438 | 0.999971 | 1 | 0.871816 |
| Mettl1 | Q9Z120 | 0.94 | 1.7 | -0.82 | 0.03 | 3.4 | 1.29 | -0.66 | -0.46 | 2.2 | 2.25 | -2.35 | -0.48 | 1.52 | 1.38 | 0.33 | 0.19 | 0.43 | -0.0575 | 0.3925 | 0.660384 | 0.958887 | 0.512858 | 0.999971 | 1 | 0.905815 |
| Mettl13 | Q91YR5 | 0.38 | 0.38 | 0.02 | 0.11 | 0.38 | -0.12 | 0.43 | 0.04 | 0.4 | 0.2 | -0.22 | 0.33 | -0.01 | 0.04 | 0 | -0.01 | -0.04 | -0.045 | -0.2175 | 0.850079 | 0.832116 | 0.24088 | 0.999971 | 1 | 0.834478 |
| Mettl14 | Q3UIK4-2 | 3.96 | 2.51 | 3.23 | 3.5 | 3.37 | 3.04 | 2.81 | 3 | 3.78 | 3.1 | 2.01 | 2.11 | 4.22 | 1.76 | 2.69 | 2.22 | -0.245 | -0.55 | -0.5775 | 0.459096 | 0.27593 | 0.320897 | 0.999971 | 1 | 0.854682 |
| Mettl15 | Q9DCL4 | -1.02 | -0.45 | 0.28 | -0.8 | -0.72 | -0.38 | -0.67 | -0.09 | -0.67 | -1.03 | -0.8 | -0.3 | 0.3 | -0.35 | -1.18 | -0.15 | 0.0325 | -0.2025 | 0.1525 | 0.919397 | 0.536205 | 0.705791 | 0.999971 | 1 | 0.950182 |
| Mettl16 | Q5SW15 | 0.58 | 1.64 | 1.41 | 1.65 | 1.48 | 1.32 | -0.16 | 0.89 | 1.38 | 0.6 | 1.31 | 0.97 | 1.62 | 1.73 | 1.18 | 1.33 | -0.4375 | -0.255 | 0.145 | 0.316883 | 0.424975 | 0.624394 | 0.999971 | 1 | 0.930479 |
| Mettl17 | E9Q7K9 | -0.42 | -0.04 | -0.07 | -0.22 | -0.04 | 0 | -0.64 | 0.26 | -0.05 | -0.27 | 0.05 | 0.52 | 0.3 | 0.11 | 0.06 | 0.5 | 0.0825 | 0.25 | 0.43 | 0.733145 | 0.284675 | 0.052251 | 0.999971 | 1 | 0.723736 |
| Mettl2 | Q8BMK1 | 0.81 | 0.46 | 0.96 | -0.27 | 0.5 | 0.87 | 1.42 | 0.63 | 0.67 | 1.04 | 0.9 | 0.82 | 0.01 | 0.12 | 0.77 | 0.06 | 0.365 | 0.3675 | -0.25 | 0.296921 | 0.232685 | 0.453261 | 0.999971 | 1 | 0.887217 |
| Mettl21A | Q9CQL0 | -0.87 | 1.54 | 2.05 | 0.85 | 2.17 | 0.97 | 1.84 | 1.66 | 2.45 | 0.31 | 1.92 | 2.19 | 2.1 | 0.48 | 1.07 | 0.65 | 0.7675 | 0.825 | 0.1825 | 0.238774 | 0.273711 | 0.784291 | 0.999971 | 1 | 0.966376 |
| Mettl22 | Q8R1C6 | 1.53 | -0.14 | 0.11 | 0.53 | 0.15 | 0.02 | -2.08 | 1.17 | -1.41 | 1.14 | 0.51 | -0.09 | -1.11 | 0.46 | 0.18 | 0.88 | -0.6925 | -0.47 | -0.405 | 0.336559 | 0.443728 | 0.446369 | 0.999971 | 1 | 0.884467 |
| Mettl26 | Q9DCS2 | -0.16 | -0.93 | 0.35 | -0.17 | -0.55 | -0.57 | 0.89 | 0.19 | -0.58 | -0.17 | 0 | 0.07 | 0.08 | 0.26 | 0.27 | 0.47 | 0.2175 | 0.0575 | 0.4975 | 0.602909 | 0.851376 | 0.112144 | 0.999971 | 1 | 0.78268 |
| Mettl3 | A0A0R4J04 | 1.35 | 1.55 | 1.61 | 2.39 | 1.59 | 1.28 | 1.31 | 2.09 | 1.36 | 0.68 | 1.75 | 2.08 | 1.39 | 1.56 | 1.59 | 2.77 | -0.1575 | -0.2575 | 0.1025 | 0.605985 | 0.490263 | 0.786848 | 0.999971 | 1 | 0.96724 |
| Mettl7a1 | Q8C6B0 | -0.93 | -0.15 | 2.16 | 2.33 | -0.06 | -0.57 | 0.77 | 2.15 | -0.4 | -0.43 | 1.39 | 0.76 | -0.75 | -0.32 | 1.64 | 1.84 | -0.28 | -0.5225 | -0.25 | 0.757712 | 0.540717 | 0.791503 | 0.999971 | 1 | 0.968737 |
| Mettl9 | Q9EPL4 | -1.99 | -1.94 | -2.32 | -1.64 | -2.15 | -2.16 | -1.58 | -2.04 | -1.92 | -2.49 | -2.04 | -1.42 | -1.98 | -1.97 | -1.69 | -2.05 | -0.01 | 0.005 | 0.05 | 0.965608 | 0.985526 | 0.813042 | 0.999971 | 1 | 0.974559 |
| Mex3c | Q05A36 | 0.84 | 0.35 | 0.96 | 1.06 | 0.6 | 1.63 | 0.54 | 0.75 | 1.17 | 1.1 | 0.86 | 1.51 | 1.14 | 1.01 | 0.95 | 2.01 | 0.0775 | 0.3575 | 0.475 | 0.799845 | 0.158863 | 0.141977 | 0.999971 | 1 | 0.784967 |
| Mfap1b | C0HKD9 | 0.76 | -1.07 | -0.21 | 0.73 | -0.81 | -0.89 | 0.55 | -0.05 | -0.68 | 0.04 | -0.01 | -0.58 | -1.65 | -0.68 | -0.39 | -0.51 | -0.3525 | -0.36 | -0.86 | 0.497202 | 0.430887 | 0.104892 | 0.999971 | 1 | 0.78268 |
| Mfap3l | Q9D3X9-2 | -0.83 | -0.61 | -1.47 | -1.03 | -0.12 | -1.42 | -1.54 | -0.82 | -1.56 | -0.08 | -1.67 | -0.79 | -0.02 | -0.18 | -1.07 | -1 | 0.01 | -0.04 | 0.4175 | 0.978044 | 0.919699 | 0.225277 | 0.999971 | 1 | 0.828225 |
| Mff | Q6PCP5 | -0.65 | -0.38 | -0.44 | -0.84 | -0.34 | -0.47 | -0.23 | -0.65 | 0.06 | -0.38 | -0.44 | -0.15 | 0.22 | -0.58 | -0.68 | -0.5 | 0.155 | 0.35 | 0.1925 | 0.442413 | 0.115406 | 0.460014 | 0.999971 | 1 | 0.889706 |
| Mff_1 | Q6PCP5-2 | 0.74 | -0.53 | -0.83 | -0.2 | -1.09 | 1.72 | -1.66 | -0.8 | 1.19 | 1.02 | -0.71 | -0.43 | 1.03 | 1.57 | -0.95 | -0.48 | -0.2525 | 0.4725 | 0.4975 | 0.733682 | 0.400533 | 0.436971 | 0.999971 | 1 | 0.882419 |
| Mfge8 | P21956-2 | -0.69 | -0.01 | -0.64 | -0.52 | 1.11 | 0.4 | -0.17 | -0.95 | 0.29 | 0.26 | 0.22 | 0.58 | 0.07 | -0.07 | -0.25 | -0.58 | 0.5625 | 0.8025 | 0.2575 | 0.219567 | 0.005005 | 0.303816 | 0.999971 | 1 | 0.847402 |
| Mfhas1 | Q3V1N1 | 0.2 | -0.06 | -0.32 | -0.19 | -0.97 | -1.18 | -0.38 | -0.14 | -0.47 | 0.73 | -0.46 | -1.09 | -0.89 | -1.12 | -0.28 | -0.4 | -0.575 | -0.23 | -0.58 | 0.067883 | 0.55229 | 0.045722 | 0.999971 | 1 | 0.710102 |
| Mfn1 | Q811U4 | -0.77 | -0.49 | -0.94 | -1.13 | -0.42 | 0.14 | -0.49 | -0.55 | -0.38 | -0.63 | -0.19 | 0.15 | -0.25 | -0.14 | -0.5 | -0.61 | 0.5025 | 0.57 | 0.4575 | 0.062641 | 0.040709 | 0.062494 | 0.999971 | 1 | 0.725348 |
| Mfn2 | Q80U63 | -0.74 | -0.16 | -0.35 | -0.3 | -0.09 | -0.44 | -0.47 | -0.47 | -0.21 | -0.35 | -0.2 | -0.12 | 0.07 | -0.31 | -0.5 | -0.27 | 0.02 | 0.1675 | 0.135 | 0.923146 | 0.395032 | 0.540275 | 0.999971 | 1 | 0.910077 |
| Mfsd1 | Q9DC37 | 0.16 | -1.88 | -1.21 | -1.22 | -0.88 | -0.34 | 1.74 | -1.63 | -1.45 | -0.78 | -1.49 | -1.92 | -1.98 | -2.24 | -1.74 | -2.69 | 0.76 | -0.3725 | -1.125 | 0.329458 | 0.427156 | 0.031098 | 0.999971 | 1 | 0.678474 |
| Mfsd10 | Q9D2V8 | -0.51 | -0.05 | 1.78 | 1.54 | 0.38 | 0.06 | 1.7 | 0.91 | -0.08 | 0.13 | 2.04 | 2.06 | -0.58 | 0.25 | 0.69 | 0.22 | 0.0725 | 0.3475 | -0.545 | 0.905825 | 0.641578 | 0.356864 | 0.999971 | 1 | 0.868793 |
| Mfsd11 | Q8BJ51 | 3.57 | -0.17 | 1.11 | 2.27 | 0.39 | 0.75 | 2.97 | 3 | -0.35 | 3.45 | -0.07 | 0.87 | -3.22 | -2.57 | 0.65 | -0.72 | 0.0825 | -0.72 | -3.16 | 0.930717 | 0.501935 | 0.014902 | 0.999971 | 1 | 0.613095 |
| Mfsd2a | Q9DA75 | 0.27 | 0.7 | -0.78 | -0.43 | -0.86 | 1.16 | -0.8 | -0.77 | -0.65 | -0.05 | -0.57 | 0.22 | -1.19 | 0.81 | -0.89 | 0.79 | -0.2575 | -0.2025 | -0.06 | 0.639978 | 0.598166 | 0.917276 | 0.999971 | 1 | 0.992633 |
| Mfsd3 | A0A0R4J00 | 0.19 | -0.82 | -0.34 | -0.59 | -0.99 | -1.47 | -0.8 | 0 | -1.19 | 0.44 | -0.19 | 0.43 | -1.4 | -1.05 | 0.22 | -0.31 | -0.425 | 0.2625 | -0.245 | 0.26291 | 0.536068 | 0.549781 | 0.999971 | 1 | 0.912364 |
| Mfsd4a | Q6PDC8 | 0.44 | 0.12 | 0.2 | 0.32 | -0.41 | 0.86 | -0.25 | 0.14 | 0.01 | 0.87 | 0 | -0.25 | 0.55 | 0.36 | 0.18 | 0.96 | -0.185 | -0.1125 | 0.2425 | 0.54176 | 0.680373 | 0.29525 | 0.999971 | 1 | 0.847402 |
| Mfsd6 | F6U3S2 | -0.98 | -0.18 | -1.27 | -0.53 | -0.58 | -0.64 | -1.36 | -1.95 | 0.01 | -0.78 | -0.73 | -1.88 | -0.72 | 0.27 | -1.08 | 0.52 | -0.3925 | -0.105 | 0.4875 | 0.326979 | 0.808899 | 0.274488 | 0.999971 | 1 | 0.843794 |
| Mfsd6_1 | D3Z183 | -0.47 | -1.08 | -1.42 | -1.23 | -0.78 | -1.18 | -1.19 | -1.53 | -0.82 | -1.49 | -1.26 | -1.09 | -0.62 | -1.08 | -1.06 | -1.06 | -0.12 | -0.115 | 0.095 | 0.663102 | 0.669224 | 0.714469 | 0.999971 | 1 | 0.952364 |
| Mfsd9 | Q8C0T7 | 0.92 | -1.24 | 1.17 | 0.11 | -0.32 | 0.03 | 1.59 | -0.03 | -0.01 | 1.08 | 0.19 | 1.01 | -0.96 | -1.62 | -0.26 | -0.01 | 0.0775 | 0.3275 | -0.9525 | 0.902055 | 0.564463 | 0.137456 | 0.999971 | 1 | 0.784948 |
| Mgat1 | J3JS24 | 1.46 | 1.79 | 1.78 | 3.11 | 2.41 | 1.71 | 1.65 | 4.2 | 1.7 | 2.32 | 3.86 | 2.27 | 0.56 | 3.1 | 1.91 | 3.63 | 0.4575 | 0.5025 | 0.265 | 0.477792 | 0.369295 | 0.706517 | 0.999971 | 1 | 0.950182 |
| Mgat3 | Q5RKT9 | -0.4 | -0.02 | 0.19 | 0.41 | 0.76 | 0.23 | -0.23 | 0.59 | -0.25 | 0.01 | 0.54 | 0.78 | -1.54 | 0.24 | 0.6 | -0.01 | 0.2925 | 0.225 | -0.2225 | 0.328163 | 0.460976 | 0.637529 | 0.999971 | 1 | 0.93301 |
| Mgat5 | Q8R4G6 | 0.36 | 1.29 | 0.52 | 0.22 | 1.6 | 0.3 | 0.11 | 0.47 | 0.54 | -0.07 | 0.65 | 0.81 | 1.87 | 1.29 | 0.36 | 0.36 | 0.0225 | -0.115 | 0.3725 | 0.954427 | 0.712453 | 0.385965 | 0.999971 | 1 | 0.871816 |
| Mgll | E9Q3B9 | -2.34 | -2.45 | -1.47 | -1.24 | -2.33 | -2.4 | -1.42 | -1.53 | -2.45 | -2.9 | -1.24 | -1.27 | -2.82 | -2.46 | -1.36 | -1.11 | -0.045 | -0.09 | -0.0625 | 0.906775 | 0.852828 | 0.896482 | 0.999971 | 1 | 0.988571 |
| Mgrn1 | Q9D074 | -0.92 | -0.29 | -0.32 | -0.96 | -0.59 | -0.65 | -0.32 | -0.31 | -0.52 | -0.84 | -0.2 | -0.47 | 0.39 | -0.28 | -0.22 | -1.03 | 0.155 | 0.115 | 0.3375 | 0.519703 | 0.648694 | 0.336294 | 0.999971 | 1 | 0.858827 |
| Mgst1 | D3YU60 | -2.26 | -0.91 | -1.2 | 0.31 | 0.18 | -0.78 | -0.44 | 1.27 | 0.39 | -1.57 | -1.41 | 1.06 | 0.38 | -0.32 | 0.43 | 0.13 | 1.0725 | 0.6325 | 1.17 | 0.116608 | 0.41658 | 0.046185 | 0.999971 | 1 | 0.710102 |
| Mgst3 | Q9CPU4 | -1.38 | -2.18 | -2.65 | -2.49 | -1.77 | -1.52 | -1.41 | -2.54 | -2.53 | -1.65 | -3.14 | -3.62 | -2.57 | -1.23 | -2.41 | -2.71 | 0.365 | -0.56 | -0.055 | 0.336624 | 0.260319 | 0.8956 | 0.999971 | 1 | 0.988571 |
| Mia2 | E9PY90 | 0.02 | -0.38 | -0.28 | 0.22 | -0.24 | -0.31 | -0.26 | 0.05 | -0.23 | -0.03 | -0.1 | 0.11 | -0.4 | -0.12 | -0.12 | 0.43 | -0.085 | 0.0425 | 0.0525 | 0.687317 | 0.835738 | 0.834026 | 0.999971 | 1 | 0.979036 |
| Mia3 | A0A0R4J2D | 1.1 | 0.43 | 0.63 | 0.79 | 0.39 | 0.78 | 0.79 | 0.55 | 0.49 | 0.89 | 0.53 | 0.43 | 0.57 | 0.42 | 0.49 | 0.62 | -0.11 | -0.1525 | -0.2125 | 0.614918 | 0.489392 | 0.313525 | 0.999971 | 1 | 0.850436 |
| Mib1 | Q80SY4 | -0.26 | -0.3 | 0.3 | -0.25 | -0.39 | -0.22 | 0.47 | 0.26 | -0.14 | -0.45 | 0.43 | 0.02 | -0.29 | -0.34 | 0.03 | -0.37 | 0.1575 | 0.0925 | -0.115 | 0.559397 | 0.718366 | 0.599118 | 0.999971 | 1 | 0.924075 |
| Mib2 | Q8R516-2 | 0.11 | 0.07 | 0.05 | -0.43 | -0.07 | 0.2 | 0.33 | -0.32 | -0.08 | 0.18 | 0.01 | -0.38 | 0.62 | 0.1 | -0.27 | -0.33 | 0.085 | -0.0175 | 0.08 | 0.713203 | 0.935366 | 0.769208 | 0.999971 | 1 | 0.963122 |
| Mical1 | Q8VDP3-3 | -0.09 | 0.14 | 0.24 | -0.25 | 0.15 | -0.24 | 0.18 | -0.46 | -0.14 | -0.08 | 0.27 | 0.66 | 0.2 | -0.04 | 0.09 | 0.06 | -0.1025 | 0.1675 | 0.0675 | 0.656596 | 0.497904 | 0.723374 | 0.999971 | 1 | 0.955258 |
| Mical2 | Q8BML1-2 | 0.32 | -0.31 | 0.24 | -0.12 | -0.03 | 0.1 | 0.4 | -0.41 | -0.06 | 0.23 | 0.45 | 0.39 | -0.53 | -0.35 | 0.05 | 0.16 | -0.0175 | 0.22 | -0.2 | 0.944502 | 0.342306 | 0.432083 | 0.999971 | 1 | 0.881643 |
| Mical3 | A0A5F8MP | 0.23 | -0.32 | -0.3 | -0.37 | -0.59 | -0.35 | -0.53 | -0.55 | -0.5 | -0.08 | -0.39 | -0.71 | -0.4 | -0.58 | -0.46 | -0.49 | -0.315 | -0.23 | -0.2925 | 0.150038 | 0.328514 | 0.174007 | 0.999971 | 1 | 0.807788 |
| Micall1 | Q8BGT6 | -0.75 | 0.48 | -0.43 | -0.35 | 0.31 | 0.77 | -0.35 | 0.57 | 0.2 | 0 | -0.76 | -0.05 | 0.95 | 0.67 | -0.4 | -0.16 | 0.5875 | 0.11 | 0.5275 | 0.121686 | 0.742332 | 0.208894 | 0.999971 | 1 | 0.822138 |
| Micos10 | Q7TNS2 | -1.32 | 0.77 | 0.44 | 0.52 | -0.1 | 1.34 | 0.49 | 0.38 | 0.14 | -0.62 | -0.02 | -0.64 | -0.99 | 1.31 | 0.02 | -1.42 | 0.425 | -0.3875 | -0.3725 | 0.424324 | 0.434078 | 0.597707 | 0.999971 | 1 | 0.92377 |
| Micos13 | Q8R404 | -1.52 | -1.19 | -1.05 | -0.9 | -1.23 | -1.14 | -1.33 | -0.58 | -1.43 | -1.72 | -0.91 | -0.87 | -1.56 | -0.81 | -0.7 | -0.62 | 0.095 | -0.0675 | 0.2425 | 0.698515 | 0.799143 | 0.38466 | 0.999971 | 1 | 0.871816 |
| Micu1 | Q8VCX5 | -0.32 | -0.72 | -0.03 | 0.31 | -0.69 | -0.71 | 0.4 | -0.02 | -0.37 | -0.29 | 0.17 | -0.49 | -1.11 | -1.02 | -0.19 | -0.38 | -0.065 | -0.055 | -0.485 | 0.849943 | 0.84271 | 0.155223 | 0.999971 | 1 | 0.795578 |
| Micu2 | Q8CD10 | -1.19 | -0.47 | -1.18 | -1.33 | -1.24 | -1.78 | -0.44 | -0.75 | -0.57 | -1.09 | -0.91 | -0.77 | -0.49 | 0.29 | -1.06 | -1.64 | -0.01 | 0.2075 | 0.3175 | 0.976931 | 0.414367 | 0.468287 | 0.999971 | 1 | 0.89236 |
| Micu3 | Q9CTY5 | -0.86 | -0.56 | -0.33 | -0.44 | -0.14 | -1.47 | -0.4 | -0.53 | -0.17 | -0.84 | -0.13 | -0.21 | 0.01 | -0.88 | -0.65 | -0.41 | -0.0875 | 0.21 | 0.065 | 0.781608 | 0.384136 | 0.795488 | 0.999971 | 1 | 0.970368 |
| Mid1ip1 | Q9CQ20 | -0.07 | 0.05 | -0.38 | -0.73 | -0.36 | -0.28 | -0.4 | -0.69 | -0.27 | -0.14 | -0.52 | -0.26 | 0.28 | -0.11 | -0.83 | -0.54 | -0.15 | -0.015 | -0.0175 | 0.524075 | 0.947509 | 0.954445 | 0.999971 | 1 | 0.998056 |
| Mid2 | B1AVF4 | -1.23 | -0.55 | 0.04 | 0.58 | -0.4 | -0.93 | 0.84 | 0.15 | -0.66 | -0.78 | 0.52 | 0 | -0.43 | -0.34 | 0.51 | 0.03 | 0.205 | 0.06 | 0.2325 | 0.684965 | 0.896934 | 0.584594 | 0.999971 | 1 | 0.920364 |
| Mief1 | Q8BGV8 | -0.13 | -0.21 | 0.28 | 0.05 | -0.19 | -0.16 | -0.08 | 0.11 | -0.37 | -0.19 | -0.07 | -0.53 | -0.59 | -0.11 | 0.01 | -0.24 | -0.0775 | -0.2875 | -0.23 | 0.688823 | 0.177557 | 0.303972 | 0.999971 | 1 | 0.847402 |
| Mien1 | Q9CQ86 | 0.16 | 0.67 | 0.07 | 0.12 | 1.05 | 0.71 | 0.81 | 0.19 | 0.89 | -0.06 | -0.08 | -0.01 | 1.7 | 0.5 | 0.1 | 0.45 | 0.435 | -0.07 | 0.4325 | 0.114671 | 0.806722 | 0.257145 | 0.999971 | 1 | 0.839012 |
| Mif | P34884 | -2.11 | -1.02 | -0.56 | -0.94 | -0.17 | -0.97 | -0.13 | -1.87 | -0.62 | -2.55 | -1.35 | -0.4 | -0.43 | -1.44 | -1.32 | -1.39 | 0.3725 | -0.0725 | 0.0125 | 0.455385 | 0.893877 | 0.974673 | 0.999971 | 1 | 0.99925 |
| Mif4gd | A2A9W1 | -4.96 | -4.3 | -2.43 | -3.54 | -4.19 | -4.86 | -2.75 | -3.39 | -3.49 | -6.03 | -2.85 | -2.82 | -3.27 | -4.35 | -3.45 | -3.73 | 0.01 | 0.01 | 0.1075 | 0.987645 | 0.990534 | 0.843832 | 0.999971 | 1 | 0.979814 |
| Miga1 | Q4QQM5 | -0.19 | 0.08 | -0.52 | -0.54 | -0.15 | 0.52 | -0.54 | -0.31 | 0.11 | 0 | -0.7 | -0.58 | -0.11 | 0.39 | -0.22 | -0.4 | 0.1725 | -5.89E-17 | 0.2075 | 0.549385 | 1 | 0.420344 | 0.999971 | 1 | 0.87788 |
| Miga2 | Q8BK03 | -0.39 | -0.3 | -0.11 | -0.05 | -0.1 | -0.59 | -0.53 | 0.26 | 0 | -0.22 | 0.12 | -0.27 | 0.4 | -0.06 | 0 | 0.26 | -0.0275 | 0.12 | 0.3625 | 0.91053 | 0.525139 | 0.092805 | 0.999971 | 1 | 0.776565 |
| Minar2 | Q8C4X7-2 | 0.67 | 0.67 | 0.42 | 0.04 | 0.96 | 0.9 | 0.11 | -0.33 | 2.23 | 0.68 | 0 | 0.53 | -0.54 | -0.33 | -0.37 | 0.17 | -0.04 | 0.41 | -0.7175 | 0.90705 | 0.394532 | 0.015795 | 0.999971 | 1 | 0.613095 |
| Mindy1 | Q76LS9-2 | 0.57 | 0.44 | -0.24 | -0.32 | -0.19 | 0.61 | -0.58 | 0 | 0.3 | 0.69 | -0.7 | -0.48 | -0.02 | 0.45 | -0.35 | -0.18 | -0.1525 | -0.16 | -0.1375 | 0.650918 | 0.678833 | 0.644891 | 0.999971 | 1 | 0.93301 |
| Mindy2 | A0A1L1STF | 0.27 | -0.06 | -0.04 | -0.16 | -0.25 | 0.03 | 0.12 | -0.49 | 0.35 | 0.07 | -0.27 | -0.9 | 0.17 | -0.06 | -0.54 | -0.45 | -0.15 | -0.19 | -0.2225 | 0.490478 | 0.523476 | 0.346389 | 0.999971 | 1 | 0.864164 |
| Mindy3 | Q9CV28-2 | 1.69 | 2.46 | 2.46 | 1.33 | 1.64 | 1.76 | 3 | 1.52 | 0.95 | 1.28 | 3.19 | 3.6 | 1.42 | 1.6 | 2.39 | 2.12 | -0.005 | 0.27 | -0.1025 | 0.990541 | 0.684453 | 0.773524 | 0.999971 | 1 | 0.963759 |
| Mindy4 | A0A0N4SV | 2.21 | 1.14 | -3.18 | -2.9 | 0.53 | 1.94 | -3.6 | -2.39 | 1.01 | 1.35 | -2.8 | -1.87 | 1.73 | 0.72 | -2.78 | -3.38 | -0.1975 | 0.105 | -0.245 | 0.90552 | 0.9456 | 0.882648 | 0.999971 | 1 | 0.986576 |
| Mink1 | J3QM71 | 4.82 | -0.53 | 3.39 | 0.45 | 1.27 | -1.86 | 6.35 | -0.17 | 5.76 | 1.28 | 5.82 | 7.23 | 2.9 | -3.3 | 1.41 | -1.69 | -0.635 | 2.99 | -2.2025 | 0.740817 | 0.090101 | 0.21046 | 0.999971 | 1 | 0.822657 |
| Mink1_1 | Q9JM52 | -0.14 | -0.41 | -0.02 | -0.24 | -0.43 | -0.13 | 0.09 | 0.06 | -0.14 | -0.03 | -0.16 | -0.27 | -0.22 | -0.59 | -0.17 | -0.23 | 0.1 | 0.0525 | -0.1 | 0.622805 | 0.764026 | 0.607097 | 0.999971 | 1 | 0.925193 |
| Minpp1 | Q9Z2L6 | -0.09 | 0.74 | 0.46 | 0.36 | 0.76 | 0.27 | 0.12 | 0.1 | 0.58 | 0.19 | 0.67 | 0.85 | 0.5 | 0.26 | 0.32 | 0.38 | -0.055 | 0.205 | -0.0025 | 0.83011 | 0.418227 | 0.991048 | 0.999971 | 1 | 0.99925 |
| Mios | Q8VE19 | -0.92 | 0.92 | 0.6 | 0.33 | 0.72 | 0.62 | 0.8 | 0.29 | 0.92 | -0.11 | 0.81 | 0.27 | 0.45 | 0.44 | 0.21 | -0.1 | 0.375 | 0.24 | 0.0175 | 0.360345 | 0.591178 | 0.965449 | 0.999971 | 1 | 0.998721 |
| Mipep | A6H611 | -0.22 | 0.29 | -0.21 | -0.57 | 0.33 | -0.05 | 0.05 | -0.51 | 0.67 | -0.84 | -0.41 | -0.57 | 1.18 | -0.31 | -0.76 | -1.04 | 0.1325 | -0.11 | -0.055 | 0.623939 | 0.764185 | 0.910362 | 0.999971 | 1 | 0.991836 |
| Mipol1 | E9Q8Z7 | 5.58 | 0.36 | 0.72 | 1.19 | -0.16 | 1.31 | 3.44 | 1.15 | -0.43 | 4.68 | 1.7 | -0.45 | -1.69 | 0.92 | 0.55 | 0.07 | -0.5275 | -0.5875 | -2 | 0.678655 | 0.702832 | 0.122867 | 0.999971 | 1 | 0.78268 |
| Mix23 | Q8R3Q6 | 1.87 | 1.92 | -0.33 | -0.52 | 1.65 | 1.24 | -0.04 | -0.21 | 2.04 | 1.26 | -0.2 | -0.39 | 1.42 | 1.22 | -0.71 | -0.79 | -0.075 | -0.0575 | -0.45 | 0.918612 | 0.943026 | 0.581469 | 0.999971 | 1 | 0.920364 |
| Mkln1 | O89050 | -0.68 | -0.21 | -0.1 | -0.09 | 0 | 0.04 | 0.51 | 0.09 | -0.25 | -0.8 | -0.52 | -0.29 | -0.31 | -0.08 | -0.45 | -0.71 | 0.43 | -0.195 | -0.1175 | 0.081858 | 0.397902 | 0.612891 | 0.999971 | 1 | 0.927364 |
| Mkrn2 | Q9ERV1 | 0.06 | 1.35 | 0.62 | 0.57 | 1.15 | 1.43 | 0.79 | 1.11 | 1.28 | 0.37 | 1.33 | 0.97 | 1.25 | 1.35 | 0.96 | 0.97 | 0.47 | 0.3375 | 0.4825 | 0.147333 | 0.336862 | 0.128147 | 0.999971 | 1 | 0.784371 |
| Mlc1 | E9QP87 | 0.13 | -0.06 | -1.02 | -0.8 | -0.45 | 0.19 | -0.73 | -0.38 | -0.12 | 0.21 | -0.75 | -0.78 | -0.04 | 0.09 | -0.7 | -0.75 | 0.095 | 0.0775 | 0.0875 | 0.778153 | 0.83045 | 0.802413 | 0.999971 | 1 | 0.971467 |
| Mlec | Q6ZQI3 | -0.47 | -0.31 | -0.3 | -0.92 | -0.16 | -0.18 | 0.27 | -0.6 | 0.28 | -0.13 | -0.22 | -0.48 | -0.26 | -0.03 | -0.29 | -0.55 | 0.3325 | 0.3625 | 0.2175 | 0.214386 | 0.161696 | 0.34322 | 0.999971 | 1 | 0.8615 |
| Mlf2 | Q99KX1 | -0.06 | 0.9 | 0.6 | 0.3 | 0.74 | 0.41 | 0.68 | 0.14 | 0.31 | -0.54 | 0.43 | 0.54 | -0.16 | 0.08 | 0.25 | -0.29 | 0.0575 | -0.25 | -0.465 | 0.82995 | 0.445589 | 0.103151 | 0.999971 | 1 | 0.78268 |
| Mlip | V9GXA0 | -1.03 | -0.92 | -1.05 | 0.19 | -0.92 | -1.72 | -0.95 | 0 | -1.31 | -1.79 | -0.83 | -0.25 | -0.71 | -1.35 | 0.12 | 0.27 | -0.195 | -0.3425 | 0.285 | 0.655904 | 0.427184 | 0.534022 | 0.999971 | 1 | 0.910077 |
| Mlip_1 | V9GWW6 | 0.97 | -0.61 | 0.27 | 0.44 | -0.75 | -0.45 | 1.11 | -0.77 | -0.97 | -0.08 | -0.55 | -1.18 | -1.23 | -1.14 | -0.87 | -0.96 | -0.4825 | -0.9625 | -1.3175 | 0.359627 | 0.035016 | 0.003547 | 0.999971 | 1 | 0.577271 |
| Mllt11 | P97783 | 2.16 | 1.67 | -0.37 | 0.32 | 0.53 | 0.83 | 4.12 | 0.12 | 0.23 | 0.25 | -0.95 | 0.41 | -0.04 | 0.97 | 0.19 | 0.3 | 0.455 | -0.96 | -0.59 | 0.641822 | 0.142359 | 0.318428 | 0.999971 | 1 | 0.853893 |
| Mllt6 | B1AR09 | 2.96 | 2.58 | 1.72 | 1.39 | 2.12 | 3.13 | 2.07 | 1.19 | 1.9 | 3.17 | 1.46 | 0.42 | 1.81 | 2.72 | 1.38 | -0.14 | -0.035 | -0.425 | -0.72 | 0.944131 | 0.49894 | 0.27637 | 0.999971 | 1 | 0.843794 |
| Mlst8 | Q9DCJ1 | 0.43 | -0.58 | 0.69 | -1.11 | -0.2 | -1.19 | 1.16 | -0.34 | -0.26 | -0.49 | 0.9 | 0.65 | -0.21 | -1 | 0.05 | -0.98 | 1.57E-16 | 0.3425 | -0.3925 | 1 | 0.503295 | 0.412077 | 1 | 1 | 0.875856 |
| Mlycd | Q99J39-2 | -0.59 | -0.26 | -0.64 | -0.57 | -0.32 | -0.06 | -0.86 | -0.45 | -0.37 | -0.31 | -0.75 | -0.67 | -0.31 | -0.07 | -0.55 | -0.65 | 0.0925 | -0.01 | 0.12 | 0.685393 | 0.959212 | 0.568723 | 0.999971 | 1 | 0.91816 |
| Mmaa | Q8C7H1 | -0.76 | -0.4 | -1.02 | -1.01 | -0.46 | 0 | -0.72 | -0.45 | -0.23 | -0.63 | -0.43 | -0.67 | 0.12 | 0 | -0.83 | -0.76 | 0.39 | 0.3075 | 0.43 | 0.132144 | 0.183856 | 0.17359 | 0.999971 | 1 | 0.807607 |
| Mmab | Q9D273 | -0.64 | -0.32 | -0.62 | -1.05 | 0.18 | -0.15 | -0.03 | -0.64 | 0.05 | -0.6 | -0.54 | -0.57 | 0.86 | -0.47 | -0.77 | -1.26 | 0.4975 | 0.2425 | 0.2475 | 0.077869 | 0.334888 | 0.585094 | 0.999971 | 1 | 0.920413 |
| Mme | Q61391 | -1.14 | -1.03 | -0.93 | -0.79 | -0.57 | -0.96 | -0.75 | -0.42 | -0.44 | -0.87 | -0.64 | -0.51 | -0.14 | -0.94 | -0.58 | -0.79 | 0.2975 | 0.3575 | 0.36 | 0.158666 | 0.081675 | 0.142108 | 0.999971 | 1 | 0.784967 |
| Mmgt1 | Q8K273 | -0.39 | 0.4 | 1.02 | -0.49 | 1 | 0.28 | 1.49 | 0.04 | 0.72 | -0.75 | 1.43 | 1.56 | 1.3 | -0.3 | 0.66 | 0.06 | 0.5675 | 0.605 | 0.295 | 0.234322 | 0.317915 | 0.533141 | 0.999971 | 1 | 0.910069 |
| Mmp17 | Q9R0S3 | -0.39 | -0.14 | -1.11 | -1.42 | -0.16 | 0.02 | -1.08 | -0.8 | -0.42 | -0.01 | -0.42 | -0.62 | -0.28 | -0.17 | -0.73 | -0.49 | 0.26 | 0.3975 | 0.3475 | 0.502458 | 0.243453 | 0.302108 | 0.999971 | 1 | 0.847402 |
| Mms19 | Q9D071 | -0.08 | -0.04 | -0.23 | -0.31 | 0 | -0.45 | 0.31 | -0.41 | 0.2 | -0.17 | -0.06 | 0.23 | 0.06 | -0.12 | -0.1 | 0.19 | 0.0275 | 0.215 | 0.1725 | 0.904606 | 0.261868 | 0.347974 | 0.999971 | 1 | 0.86438 |
| Mmtag2 | Q99LX5 | 0.48 | 0.65 | 0.46 | -0.02 | -0.12 | 0.2 | 0.06 | 0.6 | 0.86 | 0.75 | 1.07 | 1.25 | -0.22 | 0.54 | 1.13 | 1.19 | -0.2075 | 0.59 | 0.2675 | 0.401168 | 0.024573 | 0.455316 | 0.999971 | 1 | 0.887217 |
| Mmut | P16332 | -0.73 | -0.97 | -0.65 | -0.9 | -0.82 | -1.14 | -0.86 | -0.37 | -1.17 | -0.85 | -0.78 | -0.82 | -0.88 | -0.82 | -0.8 | -0.68 | 0.015 | -0.0925 | 0.0175 | 0.94553 | 0.617301 | 0.919963 | 0.999971 | 1 | 0.992702 |
| Mnat1 | P51949 | 1.52 | 2.08 | 1.91 | 2.31 | 1.49 | 2.07 | 1.59 | 2.13 | 1.58 | 1.96 | 1.44 | 0.67 | 1.03 | 2.89 | 1.72 | 2.69 | -0.135 | -0.5425 | 0.1275 | 0.603211 | 0.117344 | 0.771784 | 0.999971 | 1 | 0.963498 |
| Moap1 | Q9ERH6 | -1.1 | -1.18 | -1.58 | -1.89 | -1.48 | -1.19 | -1.49 | -0.94 | -1.78 | -1.33 | -2.3 | -1.49 | -0.13 | -0.19 | -1.6 | -1.11 | 0.1625 | -0.2875 | 0.68 | 0.525103 | 0.33814 | 0.10783 | 0.999971 | 1 | 0.78268 |
| Mob1b | Q3UDM0 | 0.76 | 0.77 | -0.95 | -0.63 | 0.87 | 0.6 | -1.06 | -0.6 | 0.54 | 0.88 | -0.89 | -0.45 | 0.03 | 0.72 | -1 | -0.64 | -0.035 | 0.0325 | -0.21 | 0.952883 | 0.954189 | 0.701033 | 0.999971 | 1 | 0.949019 |
| Mob2 | A0A1B0GR | -0.65 | -0.71 | -0.37 | -1.78 | -0.08 | -0.65 | -0.38 | -1.74 | -0.26 | -0.39 | -0.14 | -0.49 | -0.62 | -1.07 | -1.43 | -2.23 | 0.165 | 0.5575 | -0.46 | 0.713188 | 0.109826 | 0.306353 | 0.999971 | 1 | 0.847402 |
| Mob4 | E0CXA9 | -0.07 | 0.88 | 0.45 | 0.17 | 0.44 | 0.49 | 0.37 | 0.58 | 0.92 | 0.32 | 0.47 | -0.32 | 1.12 | 0.7 | 0.61 | 0.22 | 0.1125 | -0.01 | 0.305 | 0.642574 | 0.975615 | 0.306099 | 0.999971 | 1 | 0.847402 |
| Mobp | Q9D2P8 | -0.39 | -4.45 | -1.29 | -1.87 | -3.57 | -2.86 | -1.79 | -2.25 | -2.73 | -1.44 | -2.71 | -2.63 | -4.07 | -2.79 | -1.92 | -3.26 | -0.6175 | -0.3775 | -1.01 | 0.476449 | 0.653685 | 0.269235 | 0.999971 | 1 | 0.843504 |
| Mocs1 | Q5RKZ7 | -1.44 | -1.05 | 0.11 | -0.85 | -1.23 | -0.81 | -1.47 | -0.23 | -1.16 | -0.91 | 0.11 | 0.12 | -0.12 | -1.11 | 0.48 | 0.55 | -0.1275 | 0.3475 | 0.7575 | 0.754521 | 0.442503 | 0.136063 | 0.999971 | 1 | 0.784948 |
| Mocs2 | F6ZGI7 | 0.08 | 1.06 | 0.03 | 0.38 | 1.09 | 0.62 | 0.24 | 0.94 | 0.98 | 0.13 | 0.43 | 0.53 | 0.98 | 1.55 | 0.68 | 0.63 | 0.335 | 0.13 | 0.5725 | 0.29401 | 0.668645 | 0.101502 | 0.999971 | 1 | 0.78268 |
| Mocs2_1 | E9PVP5 | -1.32 | -1.14 | -2.46 | -1.2 | 0.35 | -2.29 | -1.15 | -0.45 | -0.49 | -1.49 | -0.47 | -0.37 | -0.31 | -2.12 | -0.81 | -2.24 | 0.645 | 0.825 | 0.16 | 0.2878 | 0.061532 | 0.762481 | 0.999971 | 1 | 0.961701 |
| Mocs3 | A2BDX3 | 0.26 | 0.23 | 0.13 | 0.18 | 0.05 | 0.32 | -0.32 | 0.33 | 0.15 | 0.33 | 0.03 | 0.01 | 0.13 | 0.38 | 0.03 | 0.34 | -0.105 | -0.07 | 0.02 | 0.615662 | 0.677767 | 0.909272 | 0.999971 | 1 | 0.991822 |
| Mog | Q61885 | -1.21 | -0.73 | 0.17 | -1.11 | -1.21 | -0.94 | -0.3 | -0.78 | -0.72 | -1.4 | -0.86 | -0.46 | -0.58 | -1.49 | -0.37 | -1.22 | -0.0875 | -0.14 | -0.195 | 0.807638 | 0.700523 | 0.622951 | 0.999971 | 1 | 0.929759 |
| Mogs | Q80UM7 | -0.61 | -0.34 | -0.06 | 0.11 | -0.27 | -0.32 | 0.09 | 0.17 | -0.01 | -0.32 | 0.33 | 0.14 | 0.35 | 0.38 | 0.35 | 0.62 | 0.1425 | 0.26 | 0.65 | 0.550163 | 0.295033 | 0.014528 | 0.999971 | 1 | 0.613095 |
| Mon1a | A0A0R4J0D | -1.64 | -0.19 | -0.43 | -0.73 | 0.07 | -0.61 | -0.51 | -0.66 | 0.44 | -1.31 | -0.18 | -0.29 | 0.45 | -1.01 | -0.42 | -0.62 | 0.32 | 0.4125 | 0.3475 | 0.375509 | 0.375193 | 0.418117 | 0.999971 | 1 | 0.877685 |
| Mon1b | Q8BMQ8 | -0.04 | -0.4 | 0.13 | 0.19 | -0.38 | -0.11 | 0.73 | 0.08 | -0.43 | -0.1 | 0.22 | 0.46 | -0.34 | -0.22 | 0.08 | 0.04 | 0.11 | 0.0675 | -0.08 | 0.700271 | 0.793288 | 0.710979 | 0.999971 | 1 | 0.950695 |
| Mon2 | Q80TL7 | 0.43 | 0.59 | 0.8 | 0.67 | 0.51 | 0.63 | 0.53 | 0.63 | 0.62 | 0.75 | 0.72 | 0.13 | 0.7 | 0.82 | 0.82 | 0.82 | -0.0475 | -0.0675 | 0.1675 | 0.784011 | 0.748729 | 0.347557 | 0.999971 | 1 | 0.864164 |
| Mon2_1 | B9EKJ3 | 1.24 | -1.59 | -0.41 | -0.29 | 0.21 | 0.1 | 1.22 | 0.79 | -0.7 | -0.45 | -1.66 | -0.87 | -0.55 | -0.35 | -0.11 | -0.41 | 0.8425 | -0.6575 | -0.0925 | 0.172598 | 0.279256 | 0.864367 | 0.999971 | 1 | 0.983079 |
| Mon2_2 | Q80TL7-2 | -0.46 | 0.48 | 0.28 | 0.29 | 0.06 | -0.87 | -0.45 | -0.51 | -0.64 | -1.74 | -0.73 | -0.1 | 0.14 | 0.04 | -0.26 | -1.58 | -0.59 | -0.95 | -0.5625 | 0.070358 | 0.034371 | 0.208653 | 0.999971 | 1 | 0.822138 |
| Morc2a | Q69ZX6 | 2.12 | 2.27 | 1.86 | 2.26 | 2.36 | 2.67 | 1.41 | 1.74 | 2.64 | 2.32 | 1.85 | 1.81 | 2.49 | 2.2 | 1.56 | 2.19 | -0.0825 | 0.0275 | -0.0175 | 0.789042 | 0.911459 | 0.943688 | 0.999971 | 1 | 0.99692 |
| Morf4l1 | P60762-2 | -2.82 | -1.36 | 1.17 | 0.99 | -2.72 | -3.9 | 1.22 | 0.86 | -1.43 | -2.6 | 0.48 | 1.24 | -3.52 | -3.56 | 0.68 | 0.17 | -0.63 | -0.0725 | -1.0525 | 0.659097 | 0.95045 | 0.43821 | 0.999971 | 1 | 0.882619 |

| | | | | | | | | | | | | | | | | | | | | | | | | | | |
|---|---|---|---|---|---|---|---|---|---|---|---|---|---|---|---|---|---|---|---|---|---|---|---|---|---|---|
| Morf4l2 | Q9R0Q4 | 0.76 | 1.72 | 1.42 | 2.62 | 1.77 | 2.18 | 2.42 | 2.01 | 1.88 | 1.52 | 1.55 | 1.16 | 1.31 | 1.61 | 1.47 | 1.18 | 0.465 | -0.1025 | -0.2375 | 0.255761 | 0.796376 | 0.540399 | 0.999971 | 1 | 0.910077 |
| Morn4 | Q6PGF2 | 0.37 | 0.12 | 0.59 | 1.02 | -0.24 | 0.33 | 1 | 0.48 | -0.08 | 0.29 | 0.57 | -1.48 | -0.19 | 0.21 | 0.64 | 0.69 | -0.1325 | -0.7 | -0.1875 | 0.680682 | 0.157135 | 0.526739 | 0.999971 | 1 | 0.909459 |
| Mospd1 | Q8VEL0 | -0.02 | -0.37 | -0.2 | -0.07 | -0.88 | -0.88 | -0.06 | 0.65 | -0.67 | -0.35 | -0.38 | -0.51 | -1.44 | -0.57 | 0.05 | 0.48 | -0.1275 | -0.3125 | -0.205 | 0.72868 | 0.10912 | 0.615508 | 0.999971 | 1 | 0.927967 |
| Mospd2 | B1AU74 | 0.21 | 0.09 | 0.31 | -0.1 | 0.1 | 0.36 | 0.48 | 0.02 | 0.33 | 0.76 | 0.7 | 0.75 | 0.37 | 0.34 | -0.29 | -0.12 | 0.1125 | 0.5075 | -0.0525 | 0.574854 | 0.026371 | 0.817869 | 0.999971 | 1 | 0.974929 |
| Mov10 | D3Z3E8 | 2.17 | 0.67 | -0.59 | -0.95 | 0.95 | 0.45 | -0.75 | -0.52 | 1.94 | 1.49 | -1.99 | -0.36 | 1.16 | -0.26 | 0.02 | -0.79 | -0.2925 | -0.055 | -0.2925 | 0.690839 | 0.95737 | 0.693123 | 0.999971 | 1 | 0.947174 |
| Moxd1 | Q9CXI3 | -0.66 | -1.61 | 0.31 | 0.02 | -1.1 | -1.34 | -0.09 | 0.33 | -1.16 | -0.91 | -0.14 | 0.02 | -1.6 | -0.9 | 0.17 | 0.44 | -0.065 | -0.0625 | 0.0125 | 0.903721 | 0.896654 | 0.982913 | 0.999971 | 1 | 0.99925 |
| Mpc1 | D3YWY6 | -3 | -0.13 | 0.03 | -0.51 | -0.25 | -0.03 | 0.43 | -0.29 | 0.31 | -1.22 | 0.74 | 0.14 | 0.28 | -0.47 | -0.72 | -0.37 | 0.8675 | 0.895 | 0.5825 | 0.210687 | 0.251786 | 0.396049 | 0.999971 | 1 | 0.871816 |
| Mpc2 | Q9D023 | -1.4 | -0.28 | -1.13 | -1.53 | -0.33 | 0.1 | -1.06 | -1.38 | -0.41 | -0.84 | -1.55 | -2.2 | -0.58 | 0.18 | -1.58 | -1.56 | 0.4175 | -0.165 | 0.2 | 0.330598 | 0.718221 | 0.675634 | 0.999971 | 1 | 0.941576 |
| Mpdu1 | Q8R0J2 | 1.17 | 0.79 | 0.61 | 0.41 | 0.12 | 0.99 | -0.01 | 0.84 | 0.99 | 1.72 | 0.5 | 0.08 | 1.26 | 1.26 | 0.6 | 1.54 | -0.26 | 0.0775 | 0.42 | 0.404765 | 0.836634 | 0.152518 | 0.999971 | 1 | 0.795578 |
| Mpdz | D3YUX2 | 0.08 | -0.5 | -0.73 | -1.08 | -0.95 | -0.64 | -1.08 | -1.47 | -0.36 | -0.22 | -0.78 | -0.48 | -0.32 | -0.67 | -0.84 | -0.85 | -0.4775 | 0.0975 | -0.1125 | 0.143656 | 0.732087 | 0.696217 | 0.999971 | 1 | 0.947174 |
| Mphosph6 | Q9D1Q1 | 0.58 | 1.55 | 0.47 | 0.94 | 1.59 | 1.24 | 0.62 | 1.04 | 2.27 | 3.01 | 1 | 1.06 | 6.17 | 0.9 | 1.4 | 0.51 | 0.2375 | 0.95 | 1.36 | 0.463752 | 0.08924 | 0.273576 | 0.999971 | 1 | 0.843794 |
| Mphosph8 | Q3TYA6 | 2.06 | 2.95 | 3.66 | 2.41 | 3.42 | 3.02 | 1.07 | 3.34 | 3.53 | 3.28 | 3.34 | 4.59 | 5.61 | 2.92 | 3.62 | 3.57 | -0.0575 | 0.915 | 1.16 | 0.923274 | 0.063186 | 0.088298 | 0.999971 | 1 | 0.766425 |
| Mpi | Q924M7 | -0.05 | 0.89 | 0.51 | 0.25 | 0.62 | 1.18 | 0.26 | 0.51 | 0.78 | 0.56 | 0.2 | -0.09 | 0.76 | 0.75 | 0.28 | 0.54 | 0.2425 | -0.0375 | 0.1825 | 0.41292 | 0.896506 | 0.481587 | 0.999971 | 1 | 0.89485 |
| Mpp1 | P70290 | 0.4 | -1.02 | -1.04 | -0.73 | -1.14 | -1.08 | -0.33 | -1.05 | -1.19 | -0.65 | -1.44 | -1.11 | -1.63 | -1.29 | -1.31 | -1.66 | -0.3025 | -0.5 | -0.875 | 0.430836 | 0.19773 | 0.031691 | 0.999971 | 1 | 0.679598 |
| Mpp2 | Q9WV34 | -0.09 | -1.1 | -1.05 | -0.93 | -1.4 | -0.77 | -0.39 | -1.47 | -1.06 | -0.5 | -1.25 | -1.46 | -0.96 | -1.36 | -1.62 | -1.18 | -0.215 | -0.275 | -0.4875 | 0.538703 | 0.395759 | 0.11806 | 0.999971 | 1 | 0.78268 |
| Mpp3 | Q6XE40 | -0.49 | -0.19 | -0.59 | -0.69 | -0.49 | -0.07 | -0.81 | -0.78 | -0.31 | -0.47 | -0.78 | -1.09 | 0.06 | -0.54 | -0.56 | -0.43 | -0.0475 | -0.1725 | 0.1225 | 0.841262 | 0.471579 | 0.587259 | 0.999971 | 1 | 0.920758 |
| Mpp7 | G5E8S8 | -0.23 | -0.05 | -2.42 | -1.83 | -0.4 | 0 | -1.8 | -1.51 | -0.34 | 0.24 | -1.88 | -1.54 | 0.13 | -0.48 | -1.39 | -0.7 | 0.205 | 0.2525 | 0.5225 | 0.75675 | 0.719378 | 0.400133 | 0.999971 | 1 | 0.871816 |
| Mpped1 | E9PYE1 | -1.22 | 0.54 | 0.78 | 1 | 1.16 | 0.48 | 0.89 | 1.17 | 0.51 | 0.68 | 0.95 | 0.58 | -1.48 | 1.86 | 0.64 | 1.77 | 0.65 | 0.405 | 0.4225 | 0.210661 | 0.411386 | 0.614894 | 0.999971 | 1 | 0.927839 |
| Mpped2 | Q9CZJ0 | 0.21 | 1.01 | 0.62 | 0.55 | 1.05 | 0.92 | 0.36 | 0.92 | 1.38 | 0.02 | 0.61 | 1.04 | 0.74 | 0.66 | 0.64 | 0.1 | 0.215 | 0.165 | -0.0625 | 0.403546 | 0.623831 | 0.80221 | 0.999971 | 1 | 0.971467 |
| Mprip | Q5SWZ5 | -0.36 | -0.56 | -0.69 | -0.77 | -0.62 | -0.42 | -0.6 | -0.65 | -0.52 | -0.18 | -0.42 | -0.37 | -0.13 | -0.49 | -0.79 | -0.59 | 0.0225 | 0.2225 | 0.095 | 0.901053 | 0.244223 | 0.65844 | 0.999971 | 1 | 0.934638 |
| Mprip_1 | P97434 | 0.74 | -0.4 | 0.05 | -0.38 | -0.34 | -0.55 | -0.35 | -0.06 | -0.21 | -0.24 | 0.67 | 0.16 | 0.14 | 0.17 | 0.09 | 1.19 | -0.3275 | 0.0925 | 0.395 | 0.284463 | 0.784592 | 0.297506 | 0.999971 | 1 | 0.847402 |
| Mpst | Q99J99 | 0.24 | -0.18 | -1.1 | -1.42 | -0.08 | -0.49 | -0.58 | -0.95 | -0.19 | -0.38 | -0.8 | -0.95 | 0.51 | -0.54 | -1.2 | -1.26 | 0.09 | 0.035 | -0.0075 | 0.824977 | 0.931511 | 0.988538 | 0.999971 | 1 | 0.99925 |
| Mpv17 | D3YWI4 | -0.84 | -0.14 | -0.29 | -0.79 | 0.08 | -1.05 | 0.66 | -0.62 | -0.06 | -0.89 | -0.92 | -0.17 | 0.77 | -1.37 | -0.8 | -1.54 | 0.2825 | 0.005 | -0.22 | 0.484894 | 0.986577 | 0.670039 | 0.999971 | 1 | 0.938809 |
| Mpv17l | D3Z6M3 | -0.22 | 3.4 | 2.6 | 1.64 | 2.5 | 3.37 | 1.88 | 3.04 | 3.11 | 0.72 | 2.79 | 2.8 | 3.06 | 3.06 | 3.11 | 3.32 | 0.8425 | 0.5 | 1.2825 | 0.285297 | 0.565349 | 0.097525 | 0.999971 | 1 | 0.78268 |
| Mpv17l2 | Q8VIK2 | -0.28 | -0.45 | 0.84 | 0.73 | -0.15 | -0.24 | 0.69 | 1.24 | -0.2 | 0.02 | 0.32 | 0.6 | 0.79 | 0.1 | 0.17 | 1.35 | 0.175 | -0.025 | 0.3925 | 0.702264 | 0.945712 | 0.365157 | 0.999971 | 1 | 0.869526 |
| Mpzl1 | A0A0A6YW | -1.94 | -1.68 | 0.14 | -0.8 | -1.95 | -0.94 | -0.04 | 0.47 | -0.93 | -0.63 | 1.03 | 0.68 | -0.7 | -1.52 | 0.5 | -0.11 | 0.455 | 1.1075 | 0.6125 | 0.487021 | 0.100707 | 0.310461 | 0.999971 | 1 | 0.847402 |
| Mras | O08989 | -0.95 | -0.91 | -0.78 | -0.66 | -0.94 | -0.88 | -1.06 | -0.8 | -1.25 | -1.28 | -0.73 | -0.75 | -0.88 | -0.57 | -0.95 | -0.34 | -0.095 | -0.1775 | 0.14 | 0.587735 | 0.410544 | 0.508076 | 0.999971 | 1 | 0.905815 |
| Mrc1 | Q61830 | 0.54 | 0.98 | 1.32 | 1.94 | 1.79 | 0.28 | 0.96 | 1.12 | 1.13 | 0.66 | 1.47 | 0.78 | 1.32 | 0.87 | 2.36 | 1.8 | -0.1575 | -0.185 | 0.3925 | 0.700346 | 0.592716 | 0.356519 | 0.999971 | 1 | 0.868793 |
| Mrc2 | A2AAA9 | 0.3 | 1.1 | 0.75 | 1.24 | 1.75 | 0.14 | -0.51 | 0.53 | 0.49 | 0.18 | 0.96 | -0.01 | -0.45 | 1.34 | 1.14 | 0.96 | -0.37 | -0.4425 | -0.1 | 0.451718 | 0.171039 | 0.817379 | 0.999971 | 1 | 0.974929 |
| Mre11 | Q61216-2 | 2.83 | 2.92 | 1.88 | 2.43 | 2.86 | 3.39 | 1.84 | 1.68 | 2.71 | 3.2 | 1.35 | 1.26 | 1.61 | 2.09 | 1.97 | 1.43 | -0.0725 | -0.385 | -0.74 | 0.870583 | 0.453059 | 0.030874 | 0.999971 | 1 | 0.678474 |
| Mri1 | Q9CQT1 | 0.5 | 0.35 | 0.25 | 0.39 | 0.07 | 0.36 | 0.61 | 0.05 | 0.29 | 0.37 | 0.24 | 0.19 | 0 | 0.09 | 0.13 | 0.27 | -0.1 | -0.1 | -0.25 | 0.620359 | 0.54349 | 0.169008 | 0.999971 | 1 | 0.80174 |
| Mrm1 | Q99J25 | -0.23 | -0.19 | -1.12 | -0.6 | -0.5 | -0.43 | -1.11 | -1.48 | -0.45 | 0.07 | -0.84 | -0.86 | -0.47 | -0.12 | -1.12 | -0.41 | -0.345 | 0.015 | 0.005 | 0.312022 | 0.961507 | 0.987049 | 0.999971 | 1 | 0.99925 |
| Mrm3 | Q5ND52 | -1.96 | 0.7 | 0.11 | -0.1 | 0.32 | -0.21 | -0.65 | -0.46 | 0.75 | -1.6 | -0.43 | 0.05 | 0.96 | -0.27 | -0.04 | 0.27 | 0.0625 | 0.005 | 0.5425 | 0.911251 | 0.994215 | 0.362627 | 0.999971 | 1 | 0.869526 |
| Mroh1 | E0CZ22 | -0.05 | 0.32 | 0.55 | -0.02 | 0.49 | 0.18 | -0.13 | 0.33 | -0.03 | -0.03 | 0.77 | 1.05 | 1 | 0.66 | 0.78 | 1.02 | 0.0175 | 0.24 | 0.665 | 0.939835 | 0.454436 | 0.012475 | 0.999971 | 1 | 0.613095 |
| Mrpl1 | Q99N96 | -0.58 | -0.55 | -1.97 | -1.75 | -0.38 | -0.53 | -1.4 | -1.47 | -0.22 | -0.92 | -1.74 | -1.27 | 0.12 | -0.39 | -1.59 | -1.35 | 0.2675 | 0.175 | 0.41 | 0.551037 | 0.707199 | 0.430826 | 0.999971 | 1 | 0.881611 |
| Mrpl10 | Q3TBW2 | 0.26 | -0.95 | -1.72 | -1.08 | -0.88 | -0.92 | -1.42 | -1.2 | -0.71 | -0.44 | -1.29 | -1.11 | -0.43 | -0.71 | -1.24 | -0.98 | -0.2325 | -0.015 | 0.0325 | 0.575271 | 0.972277 | 0.939055 | 0.999971 | 1 | 0.995869 |
| Mrpl11 | Q9CQF0 | -0.35 | 0.19 | -1.07 | -1.58 | 0.01 | -0.06 | -0.93 | -1.05 | 0.53 | -0.15 | -0.76 | -0.49 | 0.76 | -0.17 | -0.94 | -1.21 | 0.195 | 0.485 | 0.3125 | 0.666748 | 0.299573 | 0.568784 | 0.999971 | 1 | 0.91816 |
| Mrpl12 | Q9DB15 | -1.4 | -0.71 | -1.29 | -1.55 | -0.25 | -1.08 | -1.35 | -1.11 | -0.18 | -1.34 | -1.16 | -0.97 | 0.26 | -1.14 | -1.35 | -0.98 | 0.29 | 0.325 | 0.435 | 0.359157 | 0.320758 | 0.283086 | 0.999971 | 1 | 0.844538 |
| Mrpl13 | Q9D1P0 | -0.28 | -0.16 | -0.8 | -0.97 | -0.12 | -0.52 | -0.35 | -0.41 | -0.17 | -0.34 | -0.47 | -0.3 | 0.23 | -0.25 | -0.71 | -0.47 | 0.2025 | 0.2325 | 0.2525 | 0.416859 | 0.341289 | 0.398151 | 0.999971 | 1 | 0.871816 |
| Mrpl14 | Q9D1I6 | -1.15 | -0.34 | -0.57 | -0.59 | -0.52 | -0.69 | -0.04 | -0.69 | -0.72 | -1.12 | -0.35 | 0.11 | -1.03 | -0.71 | -0.3 | 0.08 | 0.1775 | 0.1425 | 0.1725 | 0.494011 | 0.6543 | 0.57454 | 0.999971 | 1 | 0.91816 |
| Mrpl15 | Q9CPR5 | -0.54 | -0.95 | -0.17 | -0.5 | -0.76 | -1.01 | -0.87 | -0.23 | -0.99 | -0.69 | -0.44 | -0.62 | -0.78 | -0.61 | -0.18 | -0.23 | -0.1775 | -0.145 | 0.09 | 0.497284 | 0.535125 | 0.716294 | 0.999971 | 1 | 0.952576 |
| Mrpl16 | Q99N93 | -0.92 | -0.48 | -0.96 | -0.88 | -0.81 | -0.69 | -0.8 | -0.38 | -0.6 | -0.96 | -0.87 | -0.35 | -0.63 | -0.37 | -0.61 | -0.7 | 0.14 | 0.115 | 0.2325 | 0.49944 | 0.601719 | 0.254136 | 0.999971 | 1 | 0.839012 |
| Mrpl17 | Q9D8P4 | -0.51 | -0.93 | -0.98 | -1.05 | -0.7 | -1.03 | -0.82 | -0.86 | -0.52 | -0.79 | -0.87 | -0.91 | -0.09 | -0.69 | -1.04 | -0.6 | 0.015 | 0.095 | 0.2625 | 0.939809 | 0.640288 | 0.32124 | 0.999971 | 1 | 0.855075 |
| Mrpl18 | Q9CQL5 | -0.16 | -0.12 | -0.05 | -0.06 | 0.04 | 0.18 | -0.03 | 0.42 | 0.28 | 0.38 | 0.21 | -0.22 | 0.7 | 0.61 | 0.4 | 0.64 | 0.25 | 0.26 | 0.685 | 0.190069 | 0.202062 | 0.00286 | 0.999971 | 1 | 0.575374 |
| Mrpl19 | Q9D338 | -1.02 | -0.7 | -1.13 | -0.88 | -0.75 | -0.93 | -1.2 | -0.83 | -0.74 | -0.85 | -1.48 | -0.76 | -0.61 | -0.37 | -0.73 | 0.03 | 0.005 | -0.025 | 0.5125 | 0.979671 | 0.914358 | 0.049882 | 0.999971 | 1 | 0.717713 |
| Mrpl2 | Q9D773 | 0.13 | 0.74 | 0.29 | 0.24 | 0.12 | 0.39 | 0.61 | 0.66 | 0.17 | 0.5 | 0.56 | 0.2 | 0.59 | 0.24 | 0.23 | 0.42 | 0.095 | 0.0075 | 0.02 | 0.672663 | 0.971835 | 0.92425 | 0.999971 | 1 | 0.993702 |
| Mrpl20 | Q9CQL4 | -0.81 | -0.81 | -1.09 | -1.86 | -0.07 | -0.84 | -1.27 | -1.17 | -0.33 | -1.74 | -0.97 | -0.66 | 0.3 | -0.94 | -0.97 | -1.62 | 0.305 | 0.2175 | 0.335 | 0.405979 | 0.568822 | 0.458541 | 0.999971 | 1 | 0.888801 |
| Mrpl21 | Q9D1N9 | -1 | -0.64 | -2.59 | -2.89 | -0.84 | -0.96 | -2.32 | -2.72 | -0.81 | -1.07 | -2.27 | -2.79 | -0.51 | -0.64 | -2.33 | -2.27 | 0.07 | 0.045 | 0.3425 | 0.916388 | 0.946471 | 0.617819 | 0.999971 | 1 | 0.928581 |
| Mrpl22 | Q8BU88 | 0.35 | -0.35 | -1.92 | -1.54 | -0.14 | -0.33 | -1.56 | -1.31 | -0.52 | 0.15 | -1.82 | -1.45 | 0.01 | -0.33 | -1.53 | -1.19 | 0.03 | -0.045 | 0.105 | 0.958599 | 0.943098 | 0.85714 | 0.999971 | 1 | 0.98269 |
| Mrpl23 | O35972 | -0.73 | -0.8 | -0.12 | -0.04 | -0.33 | -0.26 | -0.15 | 0.24 | -0.64 | -0.64 | -0.58 | -0.03 | -0.03 | -0.01 | 0.14 | 0.33 | 0.2975 | -0.05 | 0.53 | 0.270091 | 0.85145 | 0.056093 | 0.999971 | 1 | 0.725348 |
| Mrpl24 | Q9CQ06 | -0.14 | -1.06 | -1.13 | -0.83 | -0.88 | -0.94 | -1.02 | -0.68 | -1.06 | -0.5 | -1.06 | -1.08 | -1.08 | -0.67 | -0.86 | -0.68 | -0.09 | -0.135 | -0.0325 | 0.730486 | 0.632661 | 0.903086 | 0.999971 | 1 | 0.990937 |
| Mrpl27 | Q99N92 | -0.7 | -0.99 | -0.26 | -0.8 | -0.83 | -0.71 | -1.05 | -0.3 | -0.66 | -0.52 | -0.54 | 0.04 | -0.41 | -0.53 | -1.22 | -0.26 | -0.035 | 0.2675 | 0.0825 | 0.888202 | 0.295484 | 0.768082 | 0.999971 | 1 | 0.962535 |
| Mrpl28 | Q9D1B9 | -3.08 | -4.7 | -1.75 | -1.42 | -4.67 | -4.81 | -1.17 | -1.37 | -4.12 | -3.35 | -1.4 | -1.49 | -5.6 | -4.1 | -1.27 | -1.37 | -0.2675 | 0.1475 | -0.3475 | 0.810121 | 0.871156 | 0.764524 | 0.999971 | 1 | 0.962405 |
| Mrpl3 | Q99N95 | -0.71 | -0.08 | -0.62 | -0.24 | 0.04 | 0.1 | -0.67 | 0.03 | 0.35 | -0.1 | -0.39 | -0.36 | 0.8 | -0.3 | -0.43 | -0.16 | 0.2875 | 0.2875 | 0.39 | 0.285737 | 0.273203 | 0.243705 | 0.999971 | 1 | 0.837316 |
| Mrpl30 | Q9D7N6 | 1.65 | 0.41 | -0.72 | -0.58 | 0.42 | 0.92 | 0.13 | -0.3 | 1.15 | 1.65 | -0.79 | -0.39 | 0.39 | 0.32 | -0.43 | 0 | 0.1025 | 0.215 | -0.12 | 0.853483 | 0.768955 | 0.822353 | 0.999971 | 1 | 0.975901 |
| Mrpl32 | Q9DCI9 | -1.01 | -0.76 | 0.38 | 0.56 | -0.99 | -0.69 | 0.2 | 1.28 | -0.65 | -0.87 | 0.46 | -1.04 | -0.37 | -0.06 | 1.12 | 0.76 | 0.1575 | -0.3175 | 0.57 | 0.789768 | 0.519296 | 0.263858 | 0.999971 | 1 | 0.842533 |
| Mrpl33 | Q9CQP0 | -0.65 | 0.25 | -0.37 | -0.18 | -0.08 | -0.26 | -0.38 | -0.16 | -0.64 | -0.51 | 0.37 | 0.25 | 0.52 | 0.11 | 0.11 | 1.01 | 0.0175 | 0.105 | 0.675 | 0.940613 | 0.744395 | 0.045563 | 0.999971 | 1 | 0.710102 |
| Mrpl34 | Q99N91 | 0.83 | -0.04 | -0.07 | 0.35 | -0.49 | -0.44 | -0.94 | -0.44 | -0.94 | 0.72 | -0.57 | -1.16 | -0.31 | 0.28 | -1.08 | 0.92 | -0.845 | -0.755 | -0.315 | 0.0104 | 0.116891 | 0.487734 | 0.999971 | 1 | 0.897906 |
| Mrpl35 | Q9CQL6 | -1.24 | 0.73 | 3.03 | -1.09 | 1.54 | 0.39 | 3.15 | 1.46 | 1.19 | -1.24 | 3.38 | 4.25 | 1.84 | 0.4 | 1.44 | -0.63 | 1.2775 | 1.5375 | 0.405 | 0.232791 | 0.293675 | 0.692252 | 0.999971 | 1 | 0.947174 |
| Mrpl37 | Q921S7 | -0.38 | -0.44 | -0.57 | -0.67 | -0.2 | -0.44 | -0.55 | -0.33 | -0.16 | -0.28 | -0.77 | -0.49 | 0.33 | -0.21 | -0.48 | -0.3 | 0.135 | 0.09 | 0.35 | 0.458838 | 0.656551 | 0.148965 | 0.999971 | 1 | 0.793248 |
| Mrpl38 | Q8K2M0 | 0.83 | -0.03 | 0.16 | 0.53 | -0.03 | 0.02 | 0.58 | 0.35 | 0.09 | 0.31 | -0.05 | -0.01 | 0.16 | -0.3 | 0.3 | 0.58 | -0.1425 | -0.2875 | -0.1875 | 0.589966 | 0.248466 | 0.510624 | 0.999971 | 1 | 0.905815 |
| Mrpl39 | Q9JKF7 | -1.13 | -1.68 | -0.81 | -0.97 | -1.29 | -1.42 | -0.28 | -0.71 | -0.81 | -1.48 | -0.36 | -0.43 | -1.27 | -1.29 | -0.63 | -0.5 | 0.2225 | 0.3775 | 0.225 | 0.500416 | 0.257954 | 0.450237 | 0.999971 | 1 | 0.885703 |
| Mrpl4 | Q9DCU6 | 3.28 | -0.64 | -0.11 | -0.29 | -0.9 | -0.61 | 0.22 | -0.07 | -1.34 | 0.97 | -0.63 | -0.06 | -0.94 | -0.82 | -0.12 | 0.26 | -0.9 | -0.825 | -0.965 | 0.30535 | 0.387469 | 0.279063 | 0.999971 | 1 | 0.843812 |
| Mrpl40 | D3Z7C0 | 2.36 | 0.68 | -0.91 | -0.22 | 0.71 | 0.69 | -0.41 | -0.3 | 3.35 | 1.82 | -0.87 | -0.32 | 0.7 | 0.31 | -0.43 | -0.41 | -0.305 | 0.5175 | -0.435 | 0.662863 | 0.634512 | 0.532322 | 0.999971 | 1 | 0.910069 |
| Mrpl41 | Q9CQN7 | -1.37 | -0.44 | -0.66 | -0.74 | -0.46 | -0.67 | -0.09 | -0.57 | -0.69 | -0.95 | -0.44 | -0.51 | 0 | -0.44 | -0.57 | -0.22 | 0.355 | 0.155 | 0.495 | 0.195109 | 0.545731 | 0.083943 | 0.999971 | 1 | 0.755254 |
| Mrpl42 | Q9CPV3 | -0.33 | -0.94 | 1.48 | 0.41 | -1.2 | -0.79 | 0.54 | 0.48 | -1.12 | 0.07 | -0.56 | 0.69 | 0.02 | -0.77 | 1.04 | 0.91 | -0.3975 | -0.385 | 0.145 | 0.527607 | 0.524865 | 0.812911 | 0.999971 | 1 | 0.974559 |
| Mrpl43 | Q5RL20 | 0.4 | -0.81 | -0.72 | -0.51 | -1.08 | -0.71 | 0.16 | -0.49 | -1.17 | -0.1 | -0.88 | -0.82 | -0.9 | -1.04 | -0.57 | -0.82 | -0.12 | -0.3325 | -0.4225 | 0.74564 | 0.357802 | 0.186608 | 0.999971 | 1 | 0.809025 |
| Mrpl44 | Q9CY73 | -0.07 | -0.25 | -1.16 | -1.34 | -0.25 | -0.13 | -0.61 | -1.27 | -0.18 | -0.29 | -1.44 | -1.71 | -0.09 | -0.38 | -1.45 | -1.45 | 0.14 | -0.2 | -0.1375 | 0.721961 | 0.673988 | 0.760143 | 0.999971 | 1 | 0.961116 |
| Mrpl45 | Q9D0Q7 | -1.17 | 0.25 | -0.43 | -0.58 | 0.65 | -0.2 | -0.24 | -0.4 | 0.86 | -0.92 | -0.36 | 0.08 | 1.64 | -0.08 | -0.52 | -0.1 | 0.435 | 0.3975 | 0.7175 | 0.25319 | 0.386764 | 0.190736 | 0.999971 | 1 | 0.810547 |
| Mrpl46 | Q9EQI8 | -0.45 | 0.09 | -0.06 | 0.05 | 0.09 | 0 | -0.41 | 0.4 | 0.08 | -0.35 | -0.33 | 0.07 | 0.64 | 0.1 | -0.01 | 0.6 | 0.1125 | -0.04 | 0.425 | 0.64152 | 0.852616 | 0.105448 | 0.999971 | 1 | 0.78268 |
| Mrpl47 | Q8K2Y7 | -0.24 | -0.23 | 0 | 0.47 | 0.07 | 0.12 | 0.15 | 0.68 | -0.24 | -0.39 | 0.17 | 0.34 | -0.47 | 0.2 | 0.37 | 0.97 | 0.255 | -0.03 | 0.2675 | 0.318547 | 0.908092 | 0.436659 | 0.999971 | 1 | 0.882419 |
| Mrpl48 | A0A140LH | -0.12 | 0.13 | -1.05 | -0.62 | -0.48 | -0.12 | -1.23 | -0.89 | -0.1 | -0.44 | -1.09 | -0.92 | -0.29 | -0.3 | -1.08 | -0.4 | -0.265 | -0.2225 | -0.1025 | 0.457226 | 0.521075 | 0.753211 | 0.999971 | 1 | 0.961116 |
| Mrpl49 | Q9CQ40 | -1.87 | 0.2 | -0.67 | -0.22 | 0.29 | -0.1 | -0.68 | 0.13 | 0.37 | -1.13 | -0.76 | -0.59 | 1.2 | 0.49 | -0.37 | -0.12 | 0.55 | 0.1125 | 0.94 | 0.253923 | 0.825805 | 0.100801 | 0.999971 | 1 | 0.78268 |
| Mrpl49_1 | A0A494B9 | -1.45 | 0.48 | -1.3 | -1.93 | -1.15 | -1.82 | -1.6 | -0.69 | -0.98 | -1.17 | -1.49 | -0.81 | -0.02 | -1.45 | -0.69 | -1.15 | -0.265 | -0.0625 | 0.2225 | 0.624258 | 0.902217 | 0.693616 | 0.999971 | 1 | 0.947174 |
| Mrpl50 | Q8VDT9 | -0.02 | 0.22 | 0.17 | 0.49 | 0.04 | -0.01 | -0.23 | 0.58 | 0.14 | -0.15 | 0.12 | 0.58 | 0.79 | 0.05 | 0.41 | 0.86 | -0.12 | -0.0425 | 0.3125 | 0.614057 | 0.848857 | 0.224848 | 0.999971 | 1 | 0.828225 |
| Mrpl51 | Q9CPY1 | -1.15 | -0.07 | 0.35 | 1.2 | -0.45 | -0.25 | -0.53 | 1.24 | -0.25 | -0.49 | 0.28 | 0.84 | 0.01 | -0.35 | 0.89 | 1.16 | -0.08 | 0.0125 | 0.345 | 0.891476 | 0.98114 | 0.539633 | 0.999971 | 1 | 0.910077 |
| Mrpl52 | Q9D0Y8 | -1.56 | -0.75 | 1.2 | -1.1 | -1.49 | -1.35 | 5.64 | -0.19 | -0.68 | -2.13 | 1.46 | 2.69 | -0.1 | -0.83 | -0.06 | -1.22 | 1.205 | 0.8875 | 3.93E-17 | 0.455676 | 0.432447 | 1 | 0.999971 | 1 | 1 |
| Mrpl53 | Q9D1H8 | 0.37 | -0.03 | 0.79 | 0.36 | -0.04 | -0.18 | 0.83 | 1.1 | 0.43 | -0.68 | 0.69 | 0.13 | 0.64 | -0.32 | 0.83 | 0.36 | 0.055 | -0.23 | 0.005 | 0.875587 | 0.502088 | 0.987068 | 0.999971 | 1 | 0.99925 |
| Mrpl54 | Q9CPW3 | 1.29 | 0.41 | -2.13 | -1.86 | 0.13 | 1.23 | -2.02 | -1.16 | 0.84 | 1.26 | -2.28 | -1.66 | 1.64 | 1.16 | -1.67 | -1.81 | 0.1175 | 0.1125 | 0.4025 | 0.905069 | 0.918223 | 0.717083 | 0.999971 | 1 | 0.952711 |
| Mrpl55 | Q9CZ83 | 0.99 | -0.12 | 0.87 | 0.89 | -0.3 | 0.35 | 1.63 | 0.62 | -0.2 | 0.56 | 0.25 | 0.16 | 0.27 | 0.19 | 0.45 | 0.43 | -0.0825 | -0.465 | -0.3225 | 0.854255 | 0.158046 | 0.272709 | 0.999971 | 1 | 0.843794 |
| Mrpl57 | Q9CQF8 | -0.24 | -1.27 | 0 | 0.18 | -1.2 | -1.24 | -0.05 | 0.16 | -1.06 | -0.78 | 0.16 | 0.19 | -1.02 | -0.92 | 0.23 | 0.78 | -0.25 | -0.04 | 0.1 | 0.590472 | 0.926261 | 0.843813 | 0.999971 | 1 | 0.979814 |
| Mrpl58 | Q8R035 | 3.05 | -0.07 | 0.08 | 1.39 | -0.14 | -0.06 | -0.09 | 0.8 | -0.07 | 1.12 | 0.19 | -1 | -0.42 | -0.37 | 1.32 | 2.51 | -0.985 | -1.0525 | -0.3525 | 0.175734 | 0.193529 | 0.69916 | 0.999971 | 1 | 0.947812 |
| Mrpl9 | Q99N94 | -0.52 | -0.81 | -1.07 | -0.48 | -0.26 | -0.74 | -1.17 | -0.89 | -0.64 | -0.78 | -0.81 | -0.63 | -0.23 | -0.7 | -0.73 | -0.63 | -0.045 | 0.005 | 0.1475 | 0.862073 | 0.979987 | 0.514163 | 0.999971 | 1 | 0.905815 |
| Mrps10 | E9QJS0 | 0.04 | 0.18 | 0.06 | 0.3 | 0.13 | 0.05 | -0.33 | 0.21 | 0.14 | -0.31 | -0.36 | 0.63 | 0.51 | 0.45 | 0.82 | 1.17 | -0.13 | -0.12 | 0.5925 | 0.512088 | 0.64652 | 0.023511 | 0.999971 | 1 | 0.663406 |
| Mrps11 | Q3U8Y1 | -0.35 | -0.86 | -1.3 | -1.28 | -0.52 | -0.92 | 0.37 | -1.26 | -0.85 | -1.22 | -0.63 | -0.53 | -0.87 | -0.45 | -1.76 | -1.2 | 0.365 | 0.14 | -0.1225 | 0.371484 | 0.623985 | 0.727254 | 0.999971 | 1 | 0.956041 |
| Mrps12 | O35680 | 0.63 | 0.34 | -0.79 | -0.44 | 0.66 | 0.23 | -0.97 | -0.76 | 1.05 | -0.38 | -0.08 | -0.15 | -0.84 | -1.13 | -0.78 | 0.02 | -0.145 | 0.175 | -0.6175 | 0.761554 | 0.689145 | 0.145267 | 0.999971 | 1 | 0.791767 |
| Mrps14 | D3Z2K2 | -0.26 | -0.63 | 0.29 | 0.89 | -0.63 | -1.16 | 0.58 | 0.71 | -0.02 | -0.99 | 0.45 | 0.67 | -0.68 | -0.68 | 0.14 | 0.93 | -0.1975 | -0.045 | -0.145 | 0.70618 | 0.922925 | 0.760827 | 0.999971 | 1 | 0.961116 |
| Mrps15 | Q9DC71 | -0.37 | -0.33 | -1.07 | -1.07 | -0.21 | -0.38 | -0.81 | -1.01 | -0.02 | -0.49 | -1.02 | -0.55 | -0.87 | -0.63 | -1.04 | -1.45 | 0.1075 | 0.19 | -0.2875 | 0.712118 | 0.530594 | 0.327297 | 0.999971 | 1 | 0.856472 |
| Mrps16 | Q9CPX7 | 0.89 | -0.66 | -1.58 | -1.41 | -0.59 | -0.52 | -1.38 | -1.19 | -1.22 | -0.22 | -1.88 | -1.12 | -0.87 | -0.45 | -1.08 | -1.1 | -0.23 | -0.42 | -0.185 | 0.679044 | 0.493266 | 0.731568 | 0.999971 | 1 | 0.956053 |
| Mrps17 | Q9CQE3 | 0.3 | -0.35 | 0.86 | 1.02 | -0.3 | -0.61 | 0.53 | 0.61 | 0.89 | -0.53 | 0.31 | 0.57 | 1.39 | -0.16 | 0.66 | 0.98 | -0.4 | -0.1475 | 0.26 | 0.345364 | 0.722636 | 0.547894 | 0.999971 | 1 | 0.911722 |
| Mrps18a | Q99N85 | 0.69 | -0.07 | -0.29 | -0.07 | -0.05 | -0.11 | -0.87 | -0.08 | 0.02 | 0.6 | -0.3 | -0.64 | 0.06 | 0.49 | 0.24 | 0.26 | -0.3425 | -0.145 | 0.1975 | 0.272062 | 0.668764 | 0.450974 | 0.999971 | 1 | 0.886397 |
| Mrps18b | Q99N84 | -1.8 | -1.67 | 0.17 | 0.14 | -1.09 | -1.03 | 0.8 | 0.29 | -0.91 | -1.55 | 0.37 | 0.95 | -0.98 | -1.31 | 0.53 | -0.6 | 0.5325 | 0.505 | 0.2 | 0.426272 | 0.489262 | 0.74675 | 0.999971 | 1 | 0.961064 |
| Mrps18c | Q8R2L5 | -0.57 | 0.04 | -1.61 | -1.22 | 0.62 | -0.89 | -0.96 | -2.12 | -0.71 | -0.77 | -0.21 | -0.63 | -0.89 | 0.28 | 0.26 | -0.81 | 0.0025 | 0.26 | 0.55 | 0.996725 | 0.492608 | 0.248542 | 1 | 1 | 0.839012 |
| Mrps2 | Q924T2 | -1.56 | -0.68 | -0.87 | -1.68 | -0.8 | -1.49 | -0.26 | -1.42 | -0.81 | -2.19 | -1.24 | -0.5 | -0.39 | -2.16 | -1.5 | -1.85 | 0.205 | 0.0125 | -0.2775 | 0.582593 | 0.976343 | 0.527398 | 0.999971 | 1 | 0.909612 |
| Mrps21 | P58059 | -0.07 | 0.16 | 0.94 | 0.93 | -0.22 | 0.05 | 0.97 | 1.83 | 0.4 | -0.44 | 0.61 | 0.61 | 0.48 | 0.28 | 1.31 | 0.61 | 0.1675 | -0.195 | 0.18 | 0.736236 | 0.585509 | 0.600531 | 0.999971 | 1 | 0.92426 |
| Mrps22 | Q9CXW2 | -0.79 | -0.97 | -2.08 | -2.13 | -0.91 | -0.99 | -2.05 | -2.02 | -0.72 | -0.73 | -2.19 | -1.61 | -0.43 | -0.91 | -1.88 | -2.14 | 1.18E-16 | 0.18 | 0.1525 | 1 | 0.705012 | 0.76039 | 1 | 1 | 0.961116 |
| Mrps23 | Q8VE22 | -0.57 | -0.23 | 0.23 | -0.27 | -0.24 | -0.23 | -0.33 | -0.62 | 0.04 | -0.76 | -0.09 | 0.73 | 0.23 | 0.03 | 0.31 | 0 | -0.145 | 0.19 | 0.3525 | 0.529343 | 0.582792 | 0.142096 | 0.999971 | 1 | 0.784967 |
| Mrps24 | Q9CQV5 | 0.27 | 0.43 | 0.29 | -0.5 | 1.88 | 0.2 | -0.41 | -0.19 | 2.11 | -0.01 | -0.49 | -0.09 | 3.02 | -0.27 | -0.28 | -0.25 | 0.2475 | 0.2575 | 0.4325 | 0.634398 | 0.655236 | 0.575226 | 0.999971 | 1 | 0.91816 |
| Mrps25 | Q9D125 | -1.33 | -1.62 | -2.3 | -1.92 | -1.35 | -1.78 | -1.77 | -1.75 | -1.36 | -1.89 | -2 | -1.53 | -1.41 | -1.3 | -1.59 | -1.64 | 0.13 | 0.0975 | 0.3075 | 0.615615 | 0.721741 | 0.240248 | 0.999971 | 1 | 0.834057 |
| Mrps26 | Q80ZS3 | 0.49 | -0.47 | -0.72 | -1.25 | -0.65 | -0.89 | 0.56 | -1.33 | -0.05 | -0.78 | -0.37 | -0.04 | 0.35 | -1.05 | -1.04 | -1.57 | -0.09 | 0.1775 | -0.34 | 0.858277 | 0.650033 | 0.510022 | 0.999971 | 1 | 0.905815 |
| Mrps27 | Q8BK72 | -1.12 | -1 | -1.25 | -1.06 | -0.68 | -1.06 | -1.33 | -1.16 | -0.75 | -1.61 | -1.14 | -1 | -0.34 | -1.11 | -0.75 | -0.52 | 0.05 | -0.0175 | 0.4275 | 0.805881 | 0.938161 | 0.07841 | 0.999971 | 1 | 0.754157 |
| Mrps28 | Q9CY16 | -2.27 | -2.1 | -0.82 | -1.25 | -1.47 | -2.15 | 0.08 | -1.23 | -1.58 | -2.69 | -0.35 | 0.21 | -1.45 | -2.26 | -1.13 | -1.81 | 0.4175 | 0.5075 | -0.0525 | 0.442948 | 0.456189 | 0.896474 | 0.999971 | 1 | 0.988571 |
| Mrps30 | Q9D0G0 | -0.6 | 0.22 | -1.14 | -1.33 | 0.13 | 0.19 | -1.15 | -1.01 | 0.05 | -0.25 | -0.99 | -0.7 | 0.43 | 0.23 | -0.88 | -0.74 | 0.2525 | 0.24 | 0.4725 | 0.591579 | 0.552698 | 0.311576 | 0.999971 | 1 | 0.848685 |
| Mrps31 | Q61733 | -0.98 | -0.93 | -1.36 | -0.87 | -1.04 | -1.32 | -1.06 | -1.1 | -1.03 | -1.17 | -1.54 | -0.98 | -0.43 | -1.13 | -0.94 | -0.8 | -0.095 | -0.145 | 0.21 | 0.624514 | 0.501602 | 0.365176 | 0.999971 | 1 | 0.869526 |
| Mrps34 | Q9JIK9 | -0.74 | -0.7 | -0.82 | -0.99 | -0.24 | -0.68 | -0.86 | -0.75 | -0.32 | -0.56 | -1.02 | -0.45 | -0.1 | -0.57 | -0.67 | -0.15 | 0.18 | 0.225 | 0.44 | 0.39021 | 0.302729 | 0.06157 | 0.999971 | 1 | 0.725348 |
| Mrps35 | Q8BJZ4 | 0.79 | -0.8 | -0.19 | 0.01 | -0.82 | -0.49 | 0.46 | 0.03 | -0.27 | -0.22 | -0.4 | -0.25 | -1.34 | -0.93 | -0.36 | -0.47 | -0.1575 | -0.2375 | -0.7275 | 0.703024 | 0.478018 | 0.085531 | 0.999971 | 1 | 0.759991 |
| Mrps36 | Q9CQX8 | -0.54 | -1.09 | -1.16 | -1.03 | -0.67 | -1.27 | -0.25 | -1.11 | -0.72 | -0.49 | -1.02 | -0.75 | -1.27 | -1.22 | -1.47 | -1.12 | 0.13 | 0.21 | -0.315 | 0.64857 | 0.350609 | 0.159816 | 0.999971 | 1 | 0.800134 |
| Mrps5 | Q99N87 | -0.09 | -1.1 | -0.57 | -0.59 | -0.65 | -0.94 | -0.19 | -0.55 | -0.58 | -0.67 | -0.87 | -1.15 | -1.08 | -0.44 | -0.55 | -0.08 | 0.005 | -0.23 | 0.05 | 0.985487 | 0.391703 | 0.868058 | 0.999971 | 1 | 0.983462 |
| Mrps6 | P58064 | -0.73 | -0.37 | -0.52 | -0.77 | 0.28 | -0.35 | -0.01 | -0.53 | 0.21 | -0.91 | -0.57 | -0.01 | 0.62 | -0.26 | -0.88 | -0.59 | 0.445 | 0.2775 | 0.32 | 0.087598 | 0.343989 | 0.354365 | 0.999971 | 1 | 0.868793 |
| Mrps7 | Q80X85 | 0.9 | -0.08 | -0.44 | -0.68 | 0.84 | -0.46 | -0.53 | -0.4 | 0.31 | -0.02 | -1.12 | -1.43 | 0.9 | -0.2 | -0.56 | -0.12 | -0.0625 | -0.49 | 0.08 | 0.88904 | 0.347684 | 0.856186 | 0.999971 | 1 | 0.982514 |
| Mrps9 | Q9D7N3 | -0.75 | -0.05 | -0.78 | -0.64 | -0.37 | -0.31 | -0.51 | -0.65 | -0.19 | -0.66 | -0.73 | -0.32 | 0.18 | -0.05 | -0.53 | -0.07 | 0.095 | 0.08 | 0.4375 | 0.676836 | 0.743285 | 0.111962 | 0.999971 | 1 | 0.78268 |
| Mrrf | Q9D6S7 | -0.17 | -0.99 | 0.23 | -0.13 | -1.18 | -1.03 | -0.16 | 0.21 | -1.25 | -0.53 | 0.3 | 0.01 | -1.26 | -0.72 | 0.15 | 0.23 | -0.275 | -0.1025 | -0.135 | 0.502046 | 0.802024 | 0.748237 | 0.999971 | 1 | 0.961064 |
| Mrs2 | Q5NCE8 | -1.2 | 0.13 | -0.42 | -0.44 | 0.32 | 0.3 | -0.12 | -0.47 | -0.07 | -0.23 | 0.08 | -0.07 | 0.18 | 0 | -0.23 | -0.36 | 0.49 | 0.41 | 0.38 | 0.164085 | 0.183127 | 0.234661 | 0.999971 | 1 | 0.832918 |
| Mrtfa | A0A5F8MC | 0.83 | -0.67 | 0.89 | 1.11 | -0.33 | -0.35 | 0.72 | 0.8 | -0.22 | 0.34 | 1.29 | 0.56 | -0.64 | 0.24 | 1.07 | 0.87 | -0.33 | -0.0475 | -0.155 | 0.498851 | 0.921224 | 0.765532 | 0.999971 | 1 | 0.962405 |
| Mrtfb | P59759 | 0.34 | 0.1 | 0.28 | 0.64 | 0.11 | -0.11 | 0.48 | 0.34 | -0.09 | 0.39 | 0.71 | 0.43 | -0.14 | 0.1 | 0.39 | 0.69 | -0.135 | 0.02 | -0.08 | 0.538847 | 0.931742 | 0.743345 | 0.999971 | 1 | 0.961064 |
| Mrto4 | Q9D0I8 | 0.93 | 0.81 | 1.38 | 0.93 | 0.7 | 0.85 | 1.04 | 1.19 | 0.52 | 0.64 | 1.05 | 0.9 | 0.84 | 0.9 | 1.35 | 0.66 | -0.0675 | -0.235 | -0.075 | 0.752362 | 0.294775 | 0.746479 | 0.999971 | 1 | 0.961064 |
| Msh2 | P43247 | 1.24 | 2.61 | 2.11 | 3.15 | 2.59 | 2.05 | 1.09 | 2.48 | 2.12 | 1.66 | 1.88 | 1.98 | 1.95 | 2.38 | 2.27 | 2.72 | -0.225 | -0.3675 | 0.0525 | 0.64949 | 0.370397 | 0.899267 | 0.999971 | 1 | 0.989654 |
| Msh6 | P54276 | 2.37 | 2.83 | 2.36 | 1.89 | 2.61 | 2.96 | 2.3 | 1.76 | 2.37 | 2.71 | 2.71 | 2.04 | 1.88 | 2.78 | 1.55 | 1.65 | 0.045 | 0.095 | -0.3975 | 0.888351 | 0.724309 | 0.25827 | 0.999971 | 1 | 0.839012 |
| Msi1 | F8WJA5 | 1.12 | 0.79 | 0 | -0.43 | 1.1 | 0.3 | 0.96 | -0.81 | 1.11 | 0.88 | -0.36 | 0 | 1.6 | 0.46 | 0.08 | -0.64 | 0.0175 | 0.0375 | 0.005 | 0.97308 | 0.936112 | 0.992624 | 0.999971 | 1 | 0.99925 |
| Msi2 | Q920Q6 | -0.85 | 0.07 | 0.19 | -0.07 | -0.01 | -0.14 | -0.05 | -0.02 | -0.06 | -0.57 | -0.3 | -0.11 | -0.27 | 0.07 | -0.08 | 0.05 | 0.11 | -0.095 | 0.1075 | 0.673464 | 0.732047 | 0.689688 | 0.999971 | 1 | 0.945805 |
| Msl1 | Q6PDM1-2 | 1.21 | -0.13 | 0.38 | 0.66 | -0.09 | -0.07 | 0.2 | 0.38 | -0.4 | 0.5 | 0.33 | -0.34 | -0.9 | 0.18 | 0.57 | 0.22 | -0.425 | -0.5075 | -0.5125 | 0.190471 | 0.177347 | 0.22693 | 0.999971 | 1 | 0.828236 |
| Msl3 | Q9WVG9-2 | -0.16 | 0.51 | 0.28 | -0.6 | 0.81 | 0.96 | 0.1 | -0.24 | -0.27 | 0.68 | 0.06 | 0.31 | 0.56 | 0.56 | 0.26 | -0.35 | 0.4 | 0.1875 | 0.25 | 0.291778 | 0.560749 | 0.45241 | 0.999971 | 1 | 0.887217 |
| Msn | P26041 | 0.62 | 0.69 | 0.59 | -0.28 | 1.16 | 0.89 | 0.94 | -0.42 | 0.8 | 0.54 | 0.53 | 1.05 | 0.12 | 0.24 | 0.55 | -0.65 | 0.2375 | 0.325 | -0.34 | 0.562407 | 0.258255 | 0.331865 | 0.999971 | 1 | 0.857152 |
| Msra | Q9D6Y7-2 | 0.14 | 0.63 | 0.02 | -0.18 | 0.47 | -0.13 | 0.34 | -0.38 | 0.53 | -0.02 | -0.39 | -0.26 | 1.19 | 0.09 | -0.32 | -0.48 | -0.0775 | -0.1875 | -0.0325 | 0.781521 | 0.509381 | 0.934717 | 0.999971 | 1 | 0.995618 |
| Msrb2 | Q78J03 | -0.21 | -0.33 | 0.11 | 0.14 | -0.08 | -0.31 | 0.2 | -0.33 | -0.49 | -0.25 | 0.22 | -0.46 | -0.58 | -0.25 | -0.65 | -0.23 | -0.0575 | -0.1725 | -0.355 | 0.790719 | 0.468624 | 0.119575 | 0.999971 | 1 | 0.78268 |
| Msto1 | E9PUB7 | 0.27 | 0.21 | 0.28 | 0.25 | -0.15 | 0.22 | 0.25 | 0.61 | 0.45 | 0.29 | 0.28 | 0.48 | -0.26 | 0.69 | 0.72 | 0.09 | -0.02 | 0.1225 | 0.0575 | 0.923362 | 0.451745 | 0.826885 | 0.999971 | 1 | 0.976804 |
| Mt1 | P02802 | 0.59 | 1.46 | 1.44 | 0.52 | 1.96 | 1.19 | 3.01 | 2.58 | 3.11 | 0.49 | 1.4 | 0.71 | 0.93 | -0.38 | 0.22 | 0.03 | 1.1825 | 0.425 | -0.8025 | 0.024586 | 0.481467 | 0.053269 | 0.999971 | 1 | 0.725348 |
| Mt2 | P02798 | -0.16 | 2.25 | 2.28 | 0.1 | 2.52 | 2.05 | 3.82 | 1.77 | 3.94 | -0.13 | 2.64 | 1.53 | 2.27 | -0.11 | 0.6 | -0.32 | 1.4225 | 0.8775 | -0.5075 | 0.076869 | 0.38182 | 0.529801 | 0.999971 | 1 | 0.910069 |
| Mt3 | P28184 | 1.59 | 1.23 | 0.83 | 0.53 | 0.8 | 1.61 | 2.17 | 0.72 | 1.7 | 0.88 | -0.19 | -0.62 | -0.46 | -0.73 | -0.34 | -0.97 | 0.28 | -0.6025 | -1.67 | 0.48817 | 0.274979 | 0.000273 | 0.999971 | 1 | 0.247672 |
| Mta1 | Q8K4B0 | 0.71 | 1.01 | 1.04 | 1.06 | 1.15 | 0.72 | 0.85 | 0.98 | 1.07 | 0.41 | 1 | 1.23 | 1.03 | 0.81 | 0.71 | 0.74 | -0.03 | -0.0275 | -0.1325 | 0.874788 | 0.905376 | 0.479322 | 0.999971 | 1 | 0.894438 |
| Mta2 | Q9R190 | 1.55 | 1.76 | 1.8 | 2.28 | 1.71 | 1.19 | 1.4 | 1.64 | 1.52 | 1.44 | 1.62 | 1.74 | 1.32 | 1.44 | 1.74 | 2 | -0.3625 | -0.2675 | -0.2225 | 0.143866 | 0.229071 | 0.379775 | 0.999971 | 1 | 0.871816 |
| Mta3 | Q3UII8 | 0.75 | 1.4 | 1.38 | 1 | 1.02 | 1.13 | 1.57 | 1.56 | 0.94 | 0.51 | 1.51 | 1.18 | 1.28 | 1.66 | 1.22 | 1.39 | 0.1875 | -0.0975 | 0.255 | 0.449384 | 0.726244 | 0.27733 | 0.999971 | 1 | 0.843812 |
| Mtap | Q9CQ65 | 0.32 | 0.57 | 0.47 | 0.14 | 0.09 | -0.24 | 0.07 | -0.52 | 1.36 | 0.35 | 0.65 | 1.23 | 1.18 | 0.41 | 0.14 | -0.06 | -0.525 | 0.5225 | 0.0425 | 0.036656 | 0.083101 | 0.886387 | 0.999971 | 1 | 0.987475 |
| Mtarc2 | Q922Q1 | -0.3 | -0.44 | -0.16 | -0.22 | -0.15 | -0.42 | -0.28 | 0.07 | -0.41 | -0.32 | -0.22 | -0.43 | -0.69 | -0.5 | -0.25 | 0.02 | 0.085 | -0.065 | -0.075 | 0.654605 | 0.69845 | 0.727136 | 0.999971 | 1 | 0.956041 |
| Mtatp6 | P00848 | -1.55 | 1.4 | 2.03 | 0.12 | 1.26 | 2.66 | 2.37 | 1.5 | 1.48 | 0.51 | 1.6 | 2.01 | 2.12 | 1.77 | 1.54 | 0.38 | 1.4475 | 0.9 | 0.9525 | 0.088343 | 0.262749 | 0.247043 | 0.999971 | 1 | 0.839012 |
| Mtatp8 | P03930 | -1.4 | -1.07 | -2.02 | -1.9 | -0.7 | -1.19 | -1.89 | -2.09 | -1.32 | -1.25 | -1.99 | -1.46 | -0.19 | -1.38 | -1.43 | -1.93 | 0.13 | 0.0925 | 0.365 | 0.730519 | 0.749187 | 0.384956 | 0.999971 | 1 | 0.871816 |
| Mtch1 | Q791T5-2 | -0.65 | -0.66 | -0.97 | -0.7 | -0.71 | -0.52 | -0.91 | -0.37 | -0.65 | -0.5 | -0.86 | -1.34 | -0.36 | -0.21 | -0.13 | -0.44 | 0.1175 | -0.0925 | 0.46 | 0.558361 | 0.691477 | 0.030212 | 0.999971 | 1 | 0.678474 |
| Mtch2 | Q791V5 | -1.62 | -0.55 | -0.67 | -0.73 | -0.66 | -1.35 | -0.93 | -0.57 | -0.31 | -1.68 | -0.69 | -0.53 | -0.51 | -1.25 | -0.72 | -0.72 | 0.015 | 0.09 | 0.0925 | 0.961047 | 0.811694 | 0.758929 | 0.999971 | 1 | 0.961116 |
| Mtcl1 | Q3UHU5-3 | 1.29 | 1.17 | 1.14 | 1.51 | 1.06 | 1.17 | 1.31 | 1.17 | 0.98 | 1.17 | 1.05 | 0.91 | 0.82 | 1.08 | 0.77 | 0.97 | -0.1 | -0.25 | -0.3675 | 0.579474 | 0.181251 | 0.073 | 0.999971 | 1 | 0.745263 |
| Mtcl1_1 | H3BL83 | -0.33 | 0.9 | -0.93 | -0.48 | 0.88 | 0.2 | 1.24 | 0.16 | -0.06 | -0.87 | 1.15 | 2.29 | -1.28 | -0.72 | 1.12 | -0.37 | 0.83 | 0.8375 | -0.1025 | 0.088987 | 0.265401 | 0.862173 | 0.999971 | 1 | 0.982963 |
| Mtco1 | P00397 | -0.74 | -1.5 | -1.3 | -1.22 | -1.57 | -2.15 | -0.36 | -1.6 | -1.96 | -1.81 | -1.41 | -1.88 | -1.9 | -1.17 | -1.37 | -1.08 | -0.23 | -0.575 | -0.19 | 0.562693 | 0.034838 | 0.481762 | 0.999971 | 1 | 0.894981 |
| Mtco2 | P00405 | -0.11 | 0.01 | 0.19 | 0.01 | -0.54 | -0.1 | 0.09 | 0.02 | -0.59 | -0.43 | -0.04 | -0.27 | -0.69 | -0.09 | -0.05 | 0.28 | -0.1575 | -0.3575 | -0.1625 | 0.454486 | 0.090559 | 0.509687 | 0.999971 | 1 | 0.905815 |
| mt-Co3 | P00416 | 0.56 | -5.97 | -3.81 | -3.58 | -6.41 | -5.02 | -1.76 | -3.76 | -5.99 | -2.17 | -4.67 | -5.02 | -7.83 | -5.29 | -4.63 | -4.26 | -1.0375 | -1.2625 | -2.3025 | 0.49292 | 0.385122 | 0.128774 | 0.999971 | 1 | 0.784948 |
| Mt-Cyb | P00158 | -1.27 | 1.57 | 0.49 | 0.36 | 1.03 | 1.19 | 0.37 | 1.02 | 1.19 | 0.4 | 0.72 | 0.39 | 0.46 | 1.18 | 0.65 | 0.66 | 0.615 | 0.3875 | 0.45 | 0.290855 | 0.50028 | 0.428014 | 0.999971 | 1 | 0.880793 |
| Mtdh | Q80WJ7 | 0.28 | 0 | 0.3 | 0.39 | 0.43 | -0.02 | 0.57 | 0.4 | 0.28 | 0.3 | 0.54 | 0.49 | 0.55 | 0.03 | 0.44 | 0.64 | 0.1025 | 0.16 | 0.1725 | 0.621265 | 0.383822 | 0.421441 | 0.999971 | 1 | 0.87788 |
| Mtdh_1 | F6ZSG0 | 0.35 | -0.17 | -0.19 | -1.24 | -0.59 | 1.37 | -0.4 | -0.49 | -0.71 | 0.35 | -1.07 | -0.23 | 0.36 | 0.94 | -0.75 | -0.49 | 0.285 | -0.1025 | 0.3275 | 0.593212 | 0.811863 | 0.499365 | 0.999971 | 1 | 0.903717 |
| Mtdh_2 | F6ZQL0 | -0.14 | 0.02 | -0.24 | 1.02 | 0.93 | -0.25 | 0.43 | -0.27 | 0.48 | -0.18 | 0.71 | 1.12 | 0.35 | 0.24 | 1.15 | 1.46 | 0.045 | 0.3675 | 0.635 | 0.908803 | 0.351625 | 0.138535 | 0.999971 | 1 | 0.784948 |
| Mterf2 | Q8BKY8 | 0.89 | -0.37 | -0.52 | -0.16 | 0.72 | 0.05 | -1 | -0.2 | 0.09 | 0.76 | -1.08 | -0.69 | -0.11 | 0.38 | -0.16 | -0.19 | -0.0675 | -0.19 | 0.02 | 0.880257 | 0.696556 | 0.953396 | 0.999971 | 1 | 0.998056 |
| Mtfmt | Q9D799 | -0.97 | -0.54 | -0.51 | -2.3 | -0.64 | -0.83 | -0.07 | -1.62 | -1.17 | -1.61 | -0.76 | -0.67 | -1.37 | -0.78 | -1.15 | -2.54 | 0.29 | 0.0275 | -0.38 | 0.558386 | 0.950743 | 0.474896 | 0.999971 | 1 | 0.894044 |
| Mtfp1 | Q9CRB8 | -0.2 | -0.35 | -1.28 | -1.15 | -0.77 | -0.82 | -1.42 | -1.2 | -0.74 | -0.73 | -1.23 | -1.28 | -0.2 | 0 | -1.16 | -0.83 | -0.3075 | -0.25 | 0.1975 | 0.346949 | 0.43718 | 0.600067 | 0.999971 | 1 | 0.92426 |
| Mtfr1l | Q9CWE0 | -0.96 | -1.22 | -0.96 | -1.06 | -1.17 | -1.01 | -1.33 | -0.79 | -1.32 | -0.92 | -1.34 | -1.01 | -1.51 | -1 | -1.14 | -1.25 | -0.025 | -0.0975 | -0.175 | 0.897471 | 0.605763 | 0.37256 | 0.999971 | 1 | 0.869526 |
| Mtg1 | Q8R2R6 | -0.73 | -0.13 | 0.04 | -0.04 | 0.03 | 0 | -0.23 | -0.3 | 0.08 | 0 | -0.21 | -0.14 | 0.02 | 0.02 | -0.29 | 0.5 | 0.09 | 0.1475 | 0.2775 | 0.697926 | 0.517162 | 0.306396 | 0.999971 | 1 | 0.847402 |
| Mthfd1 | Q922D8 | -0.08 | -0.19 | -0.24 | -0.13 | -0.3 | -0.27 | -0.33 | 0 | -0.3 | -0.29 | -0.04 | -0.14 | -0.12 | -0.09 | -0.12 | -0.01 | -0.065 | -0.0325 | 0.075 | 0.708095 | 0.844055 | 0.645368 | 0.999971 | 1 | 0.93301 |
| Mthfd1l | Q3V3R1 | -0.74 | -0.41 | -0.58 | -0.7 | -0.39 | -0.52 | -0.63 | -0.61 | -0.46 | -1.08 | -0.53 | -0.56 | -0.24 | -0.75 | -0.81 | -0.59 | 0.07 | -0.05 | 0.01 | 0.692461 | 0.810418 | 0.960881 | 0.999971 | 1 | 0.998721 |
| Mthfd2l | D3YZG8 | 0.31 | 0.1 | -0.07 | -0.29 | -0.01 | -0.34 | -0.55 | 0.14 | 0.33 | 0.4 | 0.73 | 0.28 | 0.2 | 0.09 | 0.52 | 0.02 | -0.2025 | 0.4225 | 0.195 | 0.40157 | 0.070285 | 0.378363 | 0.999971 | 1 | 0.871727 |
| Mthfsl | L7N466 | -0.71 | -0.12 | 0.44 | 0.65 | -0.29 | -1.86 | 0.47 | -0.63 | -0.7 | -1.14 | -0.77 | -0.75 | -0.75 | -1.32 | -0.49 | 0.12 | -0.6425 | -0.905 | -0.675 | 0.244171 | 0.019662 | 0.125109 | 0.999971 | 1 | 0.78268 |
| Mtif2 | Q91YJ5 | -0.66 | -0.73 | 0.48 | 0.67 | -0.77 | -1.02 | -0.22 | 0.61 | -1.09 | -1.23 | 0.28 | 0.17 | -0.88 | -0.84 | 0.32 | 0.94 | -0.29 | -0.4075 | -0.055 | 0.550348 | 0.430772 | 0.918172 | 0.999971 | 1 | 0.992633 |
| Mtif3 | Q9CZD5 | -0.04 | 0.95 | 0.45 | -0.54 | 1.77 | 0.99 | 2.29 | 0.46 | 2.64 | 0.05 | 1.59 | 2.74 | 4 | 0.01 | 0.64 | 0.19 | 1.1725 | 1.55 | 1.005 | 0.035408 | 0.03705 | 0.276629 | 0.999971 | 1 | 0.843794 |

| | | | | | | | | | | | | | | | | | | | | | | | | | | |
|---|---|---|---|---|---|---|---|---|---|---|---|---|---|---|---|---|---|---|---|---|---|---|---|---|---|---|
| Mtm1 | Q9Z2C5 | -0.01 | -0.58 | -0.32 | -0.26 | -0.44 | -0.12 | -0.72 | -0.66 | -0.44 | -0.35 | -0.68 | -0.5 | -0.68 | -0.02 | -0.38 | -0.26 | -0.1925 | -0.2 | -0.0425 | 0.396171 | 0.318377 | 0.849089 | 0.999971 | 1 | 0.98178 |
| Mtmr1 | F6T9H9 | -0.74 | -2.48 | -0.19 | 0.89 | -1.39 | -1.93 | -0.65 | 0.96 | -1.22 | -0.44 | 1 | 0.95 | 2.1 | -1.16 | 1.14 | 0.81 | -0.1225 | 0.7025 | 1.3525 | 0.884759 | 0.39369 | 0.15066 | 0.999971 | 1 | 0.793248 |
| Mtmr1_1 | Q9Z2C4 | 0.37 | -0.69 | -0.98 | -0.96 | -0.72 | -0.31 | -0.75 | -1.36 | -0.98 | -0.13 | -1 | -1.03 | -1.15 | -0.89 | -1.38 | -1.15 | -0.22 | -0.22 | -0.5775 | 0.559193 | 0.561127 | 0.110537 | 0.999971 | 1 | 0.78268 |
| Mtmr10 | Q7TPM9 | -0.13 | 0.33 | -0.68 | -1.02 | 0.01 | 0.79 | -0.8 | -0.66 | 0.01 | 0.14 | -1 | -0.99 | -0.21 | 0.39 | -0.88 | -0.77 | 0.21 | -0.085 | 0.0075 | 0.637275 | 0.836085 | 0.985019 | 0.999971 | 1 | 0.99925 |
| Mtmr12 | Q80TA6 | 0.61 | 0.34 | 0.3 | 0.4 | 0.2 | 0.54 | 0.19 | 0.11 | 0.03 | 0.25 | -0.02 | 0.07 | 0.51 | 0.18 | -0.08 | -0.07 | -0.1525 | -0.33 | -0.2775 | 0.425425 | 0.082638 | 0.204987 | 0.999971 | 1 | 0.81617 |
| Mtmr14 | Q8VEL2 | 1.67 | 0.17 | -0.21 | 0.41 | 0.21 | 1.13 | 0.09 | -0.54 | 0.09 | 1.37 | -0.38 | -0.67 | 0.56 | 0.85 | -0.21 | -0.32 | -0.2875 | -0.4075 | -0.29 | 0.564963 | 0.474008 | 0.538717 | 0.999971 | 1 | 0.910077 |
| Mtmr2 | Q9Z2D1 | -0.43 | -0.23 | -0.26 | -0.24 | -0.39 | -0.35 | -0.79 | -0.08 | -0.51 | -0.27 | -0.64 | -0.24 | 0.05 | -0.34 | -0.25 | -0.24 | -0.1125 | -0.125 | 0.095 | 0.589406 | 0.492478 | 0.598474 | 0.999971 | 1 | 0.92377 |
| Mtmr3 | B1ATD5 | -0.07 | -0.44 | 0.11 | -0.08 | -0.44 | -0.29 | 0.62 | 0.18 | -0.52 | -0.34 | 0.41 | 0.46 | -0.52 | -0.4 | 0.23 | 0.05 | 0.1375 | 0.1225 | -0.04 | 0.627412 | 0.672916 | 0.869919 | 0.999971 | 1 | 0.983917 |
| Mtmr6 | Q8VE11 | -0.63 | -0.14 | -0.13 | -0.45 | -0.13 | -0.54 | -0.18 | -0.28 | -0.05 | -0.55 | 0.07 | 0.49 | -0.2 | -0.5 | -0.59 | -0.21 | 0.055 | 0.3275 | -0.0375 | 0.789954 | 0.238674 | 0.858237 | 0.999971 | 1 | 0.982702 |
| Mtmr7 | Q9Z2C9 | -0.56 | -0.01 | 0.08 | -0.33 | 0.15 | -0.17 | -0.12 | 0.09 | 0.02 | -0.57 | 0.09 | 0.3 | 0.55 | -0.06 | 0.1 | 0.18 | 0.1925 | 0.165 | 0.3975 | 0.380559 | 0.529721 | 0.115604 | 0.999971 | 1 | 0.78268 |
| Mtmr9 | Q9Z2D0 | -0.37 | -0.17 | 0.03 | -0.01 | -0.08 | -0.22 | 0.32 | 0.16 | -0.24 | -0.29 | 0.21 | -0.62 | -0.08 | -0.08 | 0.04 | 0.07 | 0.175 | -0.105 | 0.1175 | 0.403801 | 0.64841 | 0.518866 | 0.999971 | 1 | 0.905901 |
| Mtnd1 | P03888 | -1.28 | 0.2 | 0.32 | 0.2 | 0.17 | -0.07 | 0.2 | 0.47 | 0 | -0.7 | 0.18 | -0.13 | -0.36 | 0.54 | 0.52 | 0.36 | 0.3325 | -0.0225 | 0.405 | 0.394995 | 0.955891 | 0.342962 | 0.999971 | 1 | 0.861474 |
| mt-Nd2 | P03893 | 1.24 | 0 | 1.11 | 1.38 | -1.11 | 0.27 | 3.08 | 0.86 | -1.44 | 0.22 | 0.18 | -1.1 | -0.07 | 0.08 | 0.76 | 0.51 | -0.1575 | -1.4675 | -0.6125 | 0.849867 | 0.015702 | 0.117232 | 0.999971 | 1 | 0.78268 |
| mt-Nd3 | P03899 | 1.43 | 1.11 | 1.44 | 1.16 | 0.89 | 0.84 | 0.71 | 1.01 | 0.57 | 0.89 | 0.22 | 1.07 | 1.17 | 1.03 | 0.85 | 2.06 | -0.4225 | -0.5975 | -0.0075 | 0.043077 | 0.031287 | 0.979629 | 0.999971 | 1 | 0.99925 |
| Mtnd4 | P03911 | -0.75 | -0.7 | -1.63 | -1.8 | -1.19 | -1.16 | -0.58 | -2.11 | -1.11 | -1.84 | -1.61 | -2.04 | -2.03 | -0.9 | -2 | -2.64 | -0.04 | -0.43 | -0.6725 | 0.921746 | 0.234779 | 0.150143 | 0.999971 | 1 | 0.793248 |
| Mtnd4l | P03903 | -1.35 | 0.4 | -0.77 | -1.9 | 1.08 | 0.07 | 0.55 | -1.81 | 3.35 | -1.36 | 0.77 | 0.91 | 1.02 | -2.87 | -1.7 | -2.8 | 0.8775 | 1.8225 | -0.6825 | 0.243533 | 0.08877 | 0.468324 | 0.999971 | 1 | 0.89236 |
| Mtnd5 | P03921 | -0.33 | -0.21 | -2.03 | -1.28 | -0.56 | -0.18 | -0.69 | -1.78 | -0.37 | -0.67 | -2.65 | -3.02 | -0.77 | -0.24 | -2.64 | -1.13 | 0.16 | -0.715 | -0.2325 | 0.753434 | 0.339166 | 0.704958 | 0.999971 | 1 | 0.950182 |
| Mto1 | G5E889 | 0.76 | -0.09 | -0.05 | 0.19 | -0.81 | -0.23 | 0.22 | 0.05 | -0.67 | 0.28 | -0.53 | -0.33 | -0.74 | -0.42 | -0.23 | -0.11 | -0.395 | -0.515 | -0.5775 | 0.217773 | 0.109222 | 0.051885 | 0.999971 | 1 | 0.723736 |
| Mtor | Q9JLN9 | -0.2 | -0.27 | -0.37 | -0.65 | -0.07 | -0.42 | -0.21 | -0.67 | -0.08 | -0.25 | -0.35 | -0.24 | -0.48 | -0.5 | -0.52 | -0.51 | 0.03 | 0.1425 | -0.13 | 0.887769 | 0.443288 | 0.476791 | 0.999971 | 1 | 0.894438 |
| Mtpap | Q9D0D3 | -0.52 | -0.68 | -0.17 | -0.63 | -0.72 | -0.72 | -0.22 | -0.48 | -0.59 | -0.88 | -0.46 | -0.68 | -0.83 | -0.94 | -0.59 | -0.78 | -0.035 | -0.1525 | -0.285 | 0.869882 | 0.451666 | 0.17415 | 0.999971 | 1 | 0.807788 |
| Mtpn | P62774 | -0.65 | 1.49 | -0.52 | -0.76 | 1.37 | 0.99 | -0.18 | -0.56 | 1.46 | -1.26 | -0.3 | -0.63 | 1.47 | 0.44 | -0.9 | -0.76 | 0.515 | -0.0725 | 0.1725 | 0.430882 | 0.919652 | 0.805486 | 0.999971 | 1 | 0.971701 |
| Mtr | A6H5Y3 | 0.82 | 0.66 | 0.14 | 0.17 | 0.99 | 0.86 | -0.01 | 0.21 | 0.5 | 0.43 | 0.53 | 0.2 | 0.49 | 0.74 | 0.22 | 0.35 | 0.065 | -0.0325 | 0.0025 | 0.831387 | 0.88523 | 0.991651 | 0.999971 | 1 | 0.99925 |
| Mtres1 | Q9CQF4 | -0.62 | 0.22 | -3.25 | -2.63 | -0.14 | -0.13 | -2.86 | -2.97 | 0.15 | -0.33 | -2.44 | -2.97 | 0.59 | 0.05 | -3.03 | -3.18 | 0.045 | 0.1725 | 0.1775 | 0.964866 | 0.864352 | 0.87716 | 0.999971 | 1 | 0.985139 |
| Mtrex | Q9CZU3 | 0.84 | 1.31 | 1.41 | 1.77 | 1.07 | 0.5 | 0.91 | 1.2 | 1.06 | 1.07 | 1.5 | 1.58 | 0.7 | 1.37 | 1.49 | 1.69 | -0.4125 | -0.03 | -0.02 | 0.146544 | 0.907531 | 0.94639 | 0.999971 | 1 | 0.997646 |
| Mtrf1l | Q8BJU9 | -1.27 | -0.13 | -1.76 | -0.31 | 0.7 | -0.07 | -1.5 | -0.82 | 0.18 | -0.58 | -2.05 | -0.8 | 0.77 | 0.58 | -1.11 | -1.24 | 0.445 | 0.055 | 0.6175 | 0.437254 | 0.9214 | 0.322235 | 0.999971 | 1 | 0.85548 |
| Mtrfr | Q80VP5 | 0.48 | 0.02 | 0.09 | -0.6 | -0.47 | 0.45 | 0.45 | 0.18 | -0.33 | 0.61 | 0.53 | 0.65 | 0.23 | 0.07 | -0.27 | -0.55 | 0.155 | 0.3675 | -0.1275 | 0.625042 | 0.27428 | 0.6667 | 0.999971 | 1 | 0.937557 |
| Mtrr | Q8C1A3 | 1.32 | 0.77 | 2.87 | 1.88 | 0.61 | 1.01 | 3.54 | 1.83 | 0.81 | 1.38 | 2.58 | 1.61 | 0.17 | 0.94 | 2.17 | 2.83 | 0.0375 | -0.115 | -0.1825 | 0.957971 | 0.830604 | 0.788098 | 0.999971 | 1 | 0.96724 |
| Mtss1 | G3X9H7 | 0.22 | -0.26 | -0.38 | 0.29 | -0.74 | -0.88 | -0.93 | -0.24 | -0.44 | 0.44 | -0.91 | 0.15 | 0.46 | 0.12 | -0.1 | 0.08 | -0.665 | -0.1575 | 0.1725 | 0.027031 | 0.646764 | 0.47744 | 0.999971 | 1 | 0.894438 |
| Mtss2 | F7D291 | 0.96 | -0.32 | 0.31 | -0.14 | -0.55 | 0.01 | 2.94 | -0.95 | 0.57 | 1.34 | -0.77 | -0.78 | 0.43 | 0.6 | -0.06 | -0.22 | 0.16 | -0.1125 | -0.015 | 0.847338 | 0.838197 | 0.965005 | 0.999971 | 1 | 0.998721 |
| Mtss2_1 | Q6P9S0 | -0.22 | -0.85 | -0.19 | -0.53 | -0.85 | -1.33 | 0.62 | -1.08 | -0.38 | -0.97 | -0.76 | -0.58 | -0.27 | -1.22 | -0.41 | -0.53 | -0.2125 | -0.225 | -0.16 | 0.62938 | 0.347529 | 0.569195 | 0.999971 | 1 | 0.91816 |
| Mturn | A0A087WC | -1.07 | 0.73 | -1.28 | 0.33 | 0.69 | 0.45 | 0.05 | 0.04 | 0.16 | -0.41 | -0.7 | -0.66 | 0.26 | -0.35 | -1.18 | -0.41 | 0.63 | -0.08 | -0.0975 | 0.218822 | 0.873273 | 0.855711 | 0.999971 | 1 | 0.982398 |
| Mtus2 | Q3UHD3 | -0.22 | -0.89 | -0.27 | 0.04 | -0.78 | -1.07 | -0.25 | -0.43 | -1.05 | -0.45 | -0.56 | -0.58 | -1.36 | -1.27 | -0.86 | -0.43 | -0.2975 | -0.325 | -0.645 | 0.308771 | 0.231638 | 0.056137 | 0.999971 | 1 | 0.725348 |
| Mtx1 | Q8R5C0 | -0.44 | -1.36 | -0.71 | -0.56 | -1.33 | -1.45 | -1.1 | -0.8 | -1.32 | -0.57 | -1.07 | -1.73 | -1.42 | -0.87 | -0.87 | -0.45 | -0.4025 | -0.405 | -0.135 | 0.16073 | 0.226511 | 0.650384 | 0.999971 | 1 | 0.933842 |
| Mtx2 | O88441 | -1.21 | 0.5 | -0.51 | -1.28 | 0.93 | -0.17 | 0.02 | -0.82 | 1.34 | -1.2 | -0.66 | -0.79 | 2.21 | -0.52 | -0.89 | -1.23 | 0.615 | 0.2975 | 0.5175 | 0.245955 | 0.644983 | 0.522407 | 0.999971 | 1 | 0.907762 |
| Mtx3 | D3YTP3 | -0.67 | -0.12 | -0.42 | -0.67 | -0.28 | -0.11 | -0.59 | -0.33 | -0.37 | -0.35 | -0.83 | -1.49 | -0.02 | -0.02 | -0.32 | -0.79 | 0.1425 | -0.29 | 0.1825 | 0.508612 | 0.35318 | 0.475546 | 0.999971 | 1 | 0.894045 |
| Muc5b | E9Q5I3 | -1.93 | -0.49 | -0.29 | -0.34 | -1.23 | -0.25 | -0.37 | -0.67 | -1.2 | -0.61 | -0.73 | -1.04 | -0.63 | -0.82 | -0.78 | -0.09 | 0.1325 | -0.1325 | 0.1825 | 0.755397 | 0.739722 | 0.655653 | 0.999971 | 1 | 0.934061 |
| Mug1 | P28665 | 1.88 | 2.99 | 0.94 | 3.07 | 4.18 | 1.05 | 3.56 | -0.29 | 0.25 | 1.91 | 2.47 | 1.22 | 2 | 0.69 | 2.24 | 0.96 | -0.095 | -0.7575 | -0.7475 | 0.927157 | 0.253825 | 0.219528 | 0.999971 | 1 | 0.827064 |
| Mul1 | Q8VCM5 | -0.29 | 0.09 | -0.68 | -0.54 | 0.38 | -0.34 | -0.25 | -0.54 | 0.06 | -0.31 | -0.35 | -0.11 | 0.41 | -0.16 | -0.32 | -0.31 | 0.1675 | 0.1775 | 0.26 | 0.549822 | 0.446814 | 0.338308 | 0.999971 | 1 | 0.859348 |
| Mvb12a | Q78HU3 | -0.3 | -0.34 | 0.86 | 0.24 | 0.01 | -1.27 | -0.22 | -0.04 | -0.8 | -0.33 | 0.22 | 0.14 | -1.38 | 0.23 | 0.02 | 0.54 | -0.495 | -0.3075 | -0.2625 | 0.230592 | 0.402292 | 0.584328 | 0.999971 | 1 | 0.920364 |
| Mvb12b | Q6KAU4 | 0.5 | -0.31 | -0.16 | -0.16 | -0.47 | -0.07 | 0.15 | 0.29 | -0.5 | 0.01 | 0.09 | -0.23 | -0.4 | -0.48 | -0.09 | 0.24 | 0.0075 | -0.125 | -0.15 | 0.977507 | 0.619777 | 0.576372 | 0.999971 | 1 | 0.918194 |
| Mvd | Q99JF5 | 0.57 | 0.38 | 0.18 | 0.37 | 0.5 | 0.76 | 0.58 | 0.43 | 0.87 | 0.52 | 0.54 | 0.63 | 0.75 | 0.38 | 0.26 | 0.76 | 0.1925 | 0.265 | 0.1625 | 0.307261 | 0.171221 | 0.437918 | 0.999971 | 1 | 0.882419 |
| Mvk | Q9R008 | -1.51 | -0.64 | -0.42 | -1.23 | -0.64 | -0.41 | -0.66 | -0.36 | -0.32 | -0.83 | -0.38 | -0.19 | -0.19 | -0.41 | 0.09 | -0.33 | 0.4325 | 0.52 | 0.74 | 0.148532 | 0.107942 | 0.028843 | 0.999971 | 1 | 0.678474 |
| Mvp | Q9EQK5 | -0.41 | -0.54 | 0 | -0.57 | -0.43 | -0.73 | -0.6 | -0.41 | -0.74 | 0.01 | -0.81 | -0.54 | -0.87 | -0.15 | -0.05 | -0.41 | -0.1625 | -0.14 | 0.01 | 0.437446 | 0.582193 | 0.968429 | 0.999971 | 1 | 0.998787 |
| Mx1 | A0A338P7: | 0.14 | -0.26 | -0.51 | -0.28 | -0.66 | 0.13 | -0.78 | -0.36 | -0.82 | 0.08 | -0.12 | -0.87 | -1.05 | -0.29 | -0.67 | 0.09 | -0.19 | -0.205 | -0.2525 | 0.47987 | 0.483568 | 0.397538 | 0.999971 | 1 | 0.871816 |
| Mxra7 | Q9D1T2 | 0.68 | 0.29 | 0.63 | 1.03 | -0.32 | 0.13 | 0.51 | 0.5 | -0.21 | 0.6 | 0.5 | 0.73 | -0.37 | -0.35 | 0.07 | 0.01 | -0.4525 | -0.2525 | -0.8175 | 0.118258 | 0.370602 | 0.00597 | 0.999971 | 1 | 0.582714 |
| Myadm | O35682 | -0.53 | -1.07 | -2.11 | -1.48 | -1.45 | -0.64 | -1.3 | -1.66 | -1.19 | -0.32 | -1.73 | -2.22 | -2.11 | -0.95 | -2.2 | -3.23 | 0.035 | -0.0675 | -0.825 | 0.92744 | 0.890634 | 0.145513 | 0.999971 | 1 | 0.791767 |
| Mybbp1a | Q7TPV4 | -1.12 | -1.25 | -0.22 | -0.72 | -1.21 | -1.53 | 0.51 | -1.01 | -1.28 | -1.04 | -0.46 | -0.65 | -0.75 | -0.99 | -0.73 | -1.06 | 0.0175 | -0.03 | -0.055 | 0.97054 | 0.921188 | 0.83722 | 0.999971 | 1 | 0.979416 |
| Mycbp | Q9EQS3 | 0.49 | 0.72 | 1.61 | 1.77 | 1.23 | 1.1 | 1.14 | 1.7 | 0.52 | 0.81 | 1.11 | 1.11 | 1.03 | 1.09 | 1.37 | 1.73 | 0.145 | -0.26 | 0.1575 | 0.67378 | 0.457279 | 0.655186 | 0.999971 | 1 | 0.934061 |
| Mycbp2 | Q7TPH6 | -0.02 | -0.05 | 0.05 | -0.24 | -0.12 | -0.14 | 0.14 | -0.12 | 0.09 | 0.02 | 0.02 | -0.01 | 0.11 | -0.36 | -0.2 | -0.2 | 0.005 | 0.095 | -0.0975 | 0.977302 | 0.5642 | 0.606085 | 0.999971 | 1 | 0.924649 |
| Mydgf | Q9CPT4 | -1.4 | 0.15 | -0.98 | -1 | -0.37 | 0.08 | -0.17 | -0.4 | 0.24 | -0.79 | -0.53 | -0.87 | 0.33 | 0.2 | -0.57 | -0.78 | 0.5925 | 0.32 | 0.6025 | 0.114264 | 0.43394 | 0.169346 | 0.999971 | 1 | 0.801824 |
| Myef2 | Q8C854 | 0.93 | 0.3 | 0.89 | 0.64 | 0.33 | 0.24 | 1.23 | 0.42 | 0.37 | 0.63 | 0.85 | 0.49 | -0.28 | 0.24 | 0.35 | 0.05 | -0.135 | -0.105 | -0.6 | 0.636704 | 0.633743 | 0.029649 | 0.999971 | 1 | 0.678474 |
| Myf6 | P15375 | -4.42 | -0.97 | 0.76 | -2.3 | -0.12 | -0.18 | 0.86 | -1.18 | 2.18 | -3.12 | -0.57 | 1.45 | 3.46 | -1.8 | -0.14 | -1.76 | 1.5775 | 1.7175 | 1.6725 | 0.156017 | 0.25347 | 0.271706 | 0.999971 | 1 | 0.843794 |
| Myg1 | Q9JK81 | -0.32 | 0.32 | 0.55 | 0.75 | 0.7 | -0.15 | 0.52 | 1.22 | 0.43 | -0.44 | 0.6 | 0.99 | 0.5 | 0.17 | 0.84 | 1.03 | 0.2475 | 0.07 | 0.31 | 0.495058 | 0.850126 | 0.327029 | 0.999971 | 1 | 0.856472 |
| Myh10 | Q5SV64 | -0.14 | -0.53 | -0.37 | -0.1 | -0.34 | -0.53 | -0.28 | -0.45 | -0.2 | 0.04 | -0.16 | -0.01 | -0.3 | -0.4 | -0.16 | 0.04 | -0.115 | 0.2025 | 0.08 | 0.541998 | 0.288126 | 0.689181 | 0.999971 | 1 | 0.945805 |
| Myh10_1 | Q61879 | 1.26 | 0.64 | -0.08 | 0.41 | 0.45 | 0.8 | 1.88 | 0.1 | 0.69 | 0.97 | 3.5 | 2.95 | 1.37 | 0.12 | -0.81 | 2.52 | 0.25 | 1.47 | 0.2425 | 0.57876 | 0.058191 | 0.731349 | 0.999971 | 1 | 0.956041 |
| Myh11 | A0A338P6I | 0.74 | -1.01 | -1.54 | 1.08 | 7.59 | 0.48 | 0.41 | -1.05 | 2.12 | 5.91 | 5.61 | 7.97 | -2.65 | 0.11 | -0.02 | 0.06 | 2.04 | 5.585 | -0.4425 | 0.2776 | 0.001519 | 0.600473 | 0.999971 | 0.629148 | 0.92426 |
| Myh14 | Q6URW6-2 | -0.23 | -0.87 | -0.58 | -0.56 | -0.86 | -0.55 | -0.58 | -0.46 | -0.95 | -0.58 | -0.81 | -0.8 | -0.75 | -0.69 | -0.63 | -0.66 | -0.0525 | -0.225 | -0.1225 | 0.801362 | 0.284894 | 0.536989 | 0.999971 | 1 | 0.910077 |
| Myh9 | Q8VDD5 | 0.4 | 0.19 | -0.3 | 0.13 | 0.75 | 0.31 | -0.29 | -0.15 | 0.5 | 0.8 | 0.02 | 0.31 | 0.2 | 0.3 | 0.07 | 0.02 | 0.05 | 0.3025 | 0.0425 | 0.863068 | 0.241225 | 0.840444 | 0.999971 | 1 | 0.97975 |
| Myl1 | A0A087WF | -2.1 | -0.06 | -0.99 | -1.62 | 0.94 | -0.57 | -0.34 | -1.91 | 0.35 | -0.54 | 0.7 | 0.92 | 0.22 | -0.92 | -1.03 | -1.68 | 0.7225 | 1.55 | 0.34 | 0.292473 | 0.013333 | 0.537309 | 0.999971 | 1 | 0.910077 |
| Myl12a | Q6ZWQ9 | 0.91 | -0.72 | -4.5 | -4.41 | -1.29 | -1.08 | -3.74 | -4.46 | -2.17 | -1.95 | -3.79 | -4.05 | -3.71 | -0.37 | -3.75 | -4.97 | -0.4625 | -0.81 | -1.02 | 0.745333 | 0.539599 | 0.499088 | 0.999971 | 1 | 0.903717 |
| Myl12b | Q3THE2 | -0.63 | -0.22 | -0.05 | 0.2 | 0.3 | -0.26 | -0.1 | -0.15 | -0.2 | 0.22 | 0.24 | 0.69 | -0.36 | -0.2 | 0.11 | 0.43 | 0.1225 | 0.4125 | 0.17 | 0.617828 | 0.152392 | 0.530624 | 0.999971 | 1 | 0.910069 |
| Myl4 | A2A6Q8 | 0.47 | 0.38 | -0.09 | 0.43 | 0.53 | 0.72 | 0.21 | -0.29 | 0.36 | 0.26 | 0.3 | 0.76 | 0.11 | -0.1 | -0.57 | 0.35 | -0.005 | 0.1225 | -0.35 | 0.985435 | 0.575129 | 0.200498 | 0.999971 | 1 | 0.814323 |
| Myl6 | Q60605-2 | 0.01 | 1.41 | -0.62 | -0.85 | 2.32 | 1.08 | 0.18 | -0.92 | 1.2 | 0.41 | 0.39 | 0.89 | 1.5 | 0.85 | -0.36 | -0.95 | 0.6775 | 0.735 | 0.2725 | 0.388837 | 0.172268 | 0.691478 | 0.999971 | 1 | 0.946835 |
| Myl6b | Q8CI43 | 0.94 | 0.47 | 0.03 | -0.14 | 0.46 | 0.5 | -0.35 | -0.14 | 0.25 | 1.19 | -0.04 | -0.19 | 0.79 | 0.53 | 0.14 | 0.38 | -0.2075 | -0.0225 | 0.135 | 0.526639 | 0.952759 | 0.643784 | 0.999971 | 1 | 0.93301 |
| Myl9 | Q9CQ19 | 0.75 | 0.2 | -1.12 | 0.79 | 6.02 | 0.85 | -0.34 | -1.03 | 1.31 | 4.89 | 2.83 | 6.11 | -1.54 | 0.57 | -0.06 | -0.05 | 1.22 | 3.63 | -0.425 | 0.415664 | 0.006775 | 0.470253 | 0.999971 | 1 | 0.892522 |
| Mylk | B1B1A8 | -2.13 | 0.46 | -0.3 | 1.29 | 3.19 | 0.69 | 0.94 | -0.44 | 1.17 | 0.14 | 4 | 6.54 | 0.33 | -0.77 | 0.45 | -0.06 | 1.265 | 3.1325 | 0.1575 | 0.201964 | 0.055438 | 0.823482 | 0.999971 | 1 | 0.976225 |
| Myo16 | F8WGF8 | -0.25 | -0.17 | -0.41 | -0.2 | -0.13 | -0.29 | -0.94 | -0.54 | -0.55 | 0.27 | -0.76 | -0.6 | 0.02 | -0.14 | -0.67 | -0.05 | -0.2175 | -0.1525 | 0.0475 | 0.34726 | 0.560085 | 0.824744 | 0.999971 | 1 | 0.976309 |
| Myo18a | B2RRE2 | -0.77 | -1.33 | -1.69 | -1.42 | -1.35 | -1.41 | -1.7 | -1.88 | -1.34 | -1.1 | -1.57 | -1.54 | -1.3 | -1.5 | -1.72 | -1.42 | -0.2825 | -0.085 | -0.1825 | 0.285722 | 0.73272 | 0.462183 | 0.999971 | 1 | 0.890369 |
| Myo1b | E9QNH6 | 0.14 | -0.75 | -0.18 | -0.08 | -0.55 | -0.68 | 0.31 | -0.49 | -0.45 | 0.01 | -0.06 | -0.26 | -0.61 | -0.61 | -0.37 | -0.43 | -0.135 | 0.0275 | -0.2875 | 0.656188 | 0.910368 | 0.242338 | 0.999971 | 1 | 0.83725 |
| Myo1c | Q9WTI7-3 | -0.56 | -0.75 | -0.64 | -0.08 | 0.45 | -0.81 | -0.22 | -0.57 | -0.71 | -0.19 | -0.04 | 0.47 | -1.16 | -0.64 | -0.47 | -0.36 | 0.22 | 0.39 | -0.15 | 0.491028 | 0.207128 | 0.562561 | 0.999971 | 1 | 0.916551 |
| Myo1d | Q5SYD0 | -1.24 | -0.84 | -0.04 | -0.41 | -1.17 | -1.15 | -0.99 | -0.59 | -0.8 | -1.12 | -0.97 | -0.39 | -1.1 | -1.16 | -0.39 | -0.82 | -0.3425 | -0.1875 | -0.235 | 0.273139 | 0.548368 | 0.466572 | 0.999971 | 1 | 0.891477 |
| Myo1e | E9Q634 | -0.85 | -0.73 | -0.64 | -1.04 | -0.5 | -0.69 | -0.48 | -1.52 | -0.64 | -0.7 | -0.58 | -0.07 | -0.65 | -0.39 | -1.08 | -1.35 | 0.0175 | 0.3175 | -0.0525 | 0.949554 | 0.161854 | 0.838269 | 0.999971 | 1 | 0.979626 |
| Myo1f | Q8CG29 | 2.53 | 0.59 | 0.92 | 1.02 | 0.55 | 0.59 | 0.6 | 0.61 | 0.61 | 2.08 | 0.03 | 0.17 | -0.59 | 1.09 | 0.91 | -0.59 | -0.6775 | -0.5425 | -1.06 | 0.126125 | 0.36695 | 0.094383 | 0.999971 | 1 | 0.779538 |
| Myo5a | D3YZ62 | -1.5 | -1.72 | -1.18 | -1.2 | -1.74 | -1.54 | -0.94 | -1.18 | -1.63 | -1.5 | -0.96 | -1.21 | -2.1 | -1.81 | -1.23 | -1.12 | 0.05 | 0.075 | -0.165 | 0.841053 | 0.747194 | 0.563133 | 0.999971 | 1 | 0.917003 |
| Myo5b | G5E8G6 | -0.15 | -0.38 | 0.36 | -0.43 | -0.45 | 0.14 | -0.59 | 0.82 | 0.3 | 0.4 | 0.36 | 0.96 | 0 | 0.46 | 0.8 | 0.48 | 0.13 | 0.655 | 0.585 | 0.718841 | 0.030448 | 0.052028 | 0.999971 | 1 | 0.723736 |
| Myo6 | E9Q3L1 | -0.25 | -0.9 | -0.63 | -0.96 | -1.06 | -0.68 | -0.32 | -0.81 | -0.55 | -0.48 | -0.76 | -0.79 | -0.77 | -1.06 | -0.68 | -0.78 | -0.0325 | 0.04 | -0.1375 | 0.896913 | 0.855549 | 0.543783 | 0.999971 | 1 | 0.911398 |
| Myo9a | Q8C170 | -0.04 | -0.07 | -0.12 | -0.3 | -0.14 | 0.15 | -0.23 | -0.28 | -0.1 | -0.07 | -0.02 | 0.01 | 0.22 | 0.19 | -0.29 | 0.2 | 0.0075 | 0.0875 | 0.2125 | 0.967653 | 0.5932 | 0.299165 | 0.999971 | 1 | 0.847402 |
| Myo9b | E9QKV6 | 0.76 | -0.15 | -0.37 | -0.4 | -0.57 | -0.37 | -0.12 | -0.55 | -0.86 | 0.64 | -0.16 | -0.03 | 0.04 | -0.38 | -0.32 | -1.06 | -0.3625 | -0.0625 | -0.39 | 0.246836 | 0.87416 | 0.283614 | 0.999971 | 1 | 0.844538 |
| Myoc | O70624 | 0.69 | 1.4 | -0.1 | 0.48 | 0.92 | 1.63 | 0.2 | 0.65 | 0.56 | 1.91 | 0.9 | -0.36 | 1.78 | 1.39 | 0.09 | 0.55 | 0.2325 | 0.135 | 0.335 | 0.574128 | 0.795938 | 0.476457 | 0.999971 | 1 | 0.894424 |
| Myof | A0A286YDI | -0.11 | -0.13 | 0.15 | 0.55 | 1.32 | 0.07 | 0.19 | -0.42 | -0.04 | 0.45 | -0.39 | 1.28 | -0.03 | 0.22 | 0.16 | 0.08 | 0.175 | 0.21 | -0.0075 | 0.651492 | 0.58566 | 0.972199 | 0.999971 | 1 | 0.99925 |
| Myorg | Q69ZQ1 | -1.26 | -2.09 | -0.7 | 0.11 | -1.43 | -1.93 | -0.81 | 0.61 | -1.58 | -1.66 | -0.32 | -0.59 | -1.37 | -1.99 | -0.31 | 0.61 | 0.095 | -0.0525 | 0.22 | 0.88419 | 0.921488 | 0.743279 | 0.999971 | 1 | 0.961064 |
| Mypop | Q8R4U1 | -1.47 | -1.29 | -0.83 | -0.54 | -1.72 | -1.14 | 0.92 | -0.59 | -1.54 | -1.43 | -0.46 | -1.67 | -2.29 | -1.81 | -1.04 | -2.48 | 0.4 | -0.2425 | -0.8725 | 0.477998 | 0.487037 | 0.041813 | 0.999971 | 1 | 0.701725 |
| Myrip | Q8K3I4 | 0.5 | -0.74 | -1.21 | -1.14 | -0.74 | -0.41 | -0.8 | -0.71 | -0.91 | 0.19 | -0.66 | -0.88 | -0.76 | -0.88 | -0.86 | -0.58 | -0.0175 | 0.0825 | -0.1225 | 0.964133 | 0.853316 | 0.752468 | 0.999971 | 1 | 0.961116 |
| Myt1l | P97500-2 | 0.15 | 1.22 | 1.52 | 1.56 | 1.33 | 1.49 | 1.46 | 1.62 | 1.6 | -0.44 | 0.71 | -0.2 | 0.03 | 0.18 | 1.07 | 0.67 | 0.3625 | -0.695 | -0.625 | 0.293421 | 0.211449 | 0.136663 | 0.999971 | 1 | 0.784948 |
| Mzt1 | Q8BUR9 | -1.77 | 0.75 | -0.66 | -0.2 | 1.18 | 0.39 | -1.08 | -0.32 | 0.35 | -1.16 | -0.72 | 0.25 | 1.08 | 0.06 | -0.3 | -0.66 | 0.5125 | 0.15 | 0.515 | 0.436633 | 0.799183 | 0.392885 | 0.999971 | 1 | 0.871816 |
| Mzt2 | Q9CQ25 | -0.43 | -1.11 | 0.48 | 0.33 | -1.45 | -1.08 | -0.59 | 0.55 | -1.52 | -0.79 | 0.08 | 0.48 | -1.65 | -1.16 | -0.91 | 0.23 | -0.46 | -0.255 | -0.69 | 0.391833 | 0.636642 | 0.193945 | 0.999971 | 1 | 0.810547 |
| N4bp1 | Q6A037 | 0.13 | 0.3 | 0.63 | 0.46 | 0.14 | -0.34 | 0.68 | 0.65 | -0.01 | -0.1 | 0.61 | 0.91 | 0 | 0.16 | 1.24 | 0.42 | -0.0975 | -0.0275 | 0.075 | 0.728284 | 0.921803 | 0.805149 | 0.999971 | 1 | 0.971701 |
| N4bp2l2 | Q8JZS6 | 0.21 | -2 | -1.86 | -2.43 | -0.4 | -1.43 | -1.82 | -1.62 | -1.42 | -0.33 | -1.81 | -1.42 | 0.8 | -0.96 | -1.93 | -2.37 | 0.2025 | 0.275 | 0.405 | 0.740248 | 0.656009 | 0.625636 | 0.999971 | 1 | 0.930479 |
| N4bp3 | Q8C7U1 | 0.84 | -0.94 | -1.12 | -0.6 | -0.9 | -1.08 | -0.7 | -0.79 | -1.42 | 0.18 | -0.93 | -1.07 | -1.36 | -1.22 | -0.74 | -1.15 | -0.4125 | -0.355 | -0.6625 | 0.348033 | 0.502973 | 0.156686 | 0.999971 | 1 | 0.796008 |
| N6amt1 | Q6SKR2 | -0.77 | 0.64 | 0.06 | 0.06 | 0.83 | 0.31 | -0.22 | 0.02 | 0.64 | -0.41 | -0.16 | -0.08 | 0.16 | 0.7 | 0.08 | -0.03 | 0.2375 | -2.36E-16 | 0.23 | 0.513709 | 1 | 0.494447 | 0.999971 | 1 | 0.901531 |
| Naa10 | Q3V4D5 | 3.8 | -0.07 | 1.53 | 3.13 | 1.63 | 2.01 | 2.08 | 2.86 | 1.03 | 2.89 | 2.41 | 0.83 | -1.18 | 0.35 | 2.46 | 2.56 | 0.0475 | -0.3075 | -1.05 | 0.953232 | 0.734514 | 0.358332 | 0.999971 | 1 | 0.869102 |
| Naa10_1 | B1AUY7 | 0.26 | 0.41 | -0.02 | 0.09 | 0.06 | -0.02 | 0.82 | -0.16 | 0.07 | 0.48 | 0.05 | 0.29 | -0.33 | 0.05 | -0.09 | 0.06 | -0.01 | 0.0375 | -0.2625 | 0.969512 | 0.848073 | 0.201324 | 0.999971 | 1 | 0.815758 |
| Naa12 | Q9CQX6 | 1.52 | 1.74 | 0.97 | -0.7 | 1.94 | 1.37 | 1.18 | -0.65 | 1.6 | 1.65 | 0.86 | 1.45 | 0.85 | 1.09 | 0.24 | -0.06 | 0.0775 | 0.5075 | -0.3525 | 0.913048 | 0.35812 | 0.535623 | 0.999971 | 1 | 0.910077 |
| Naa15 | G3X8Y3 | 0.34 | 1.35 | 1.9 | 1.65 | 1.17 | 1.22 | 2.04 | 1.75 | 1.22 | 0.82 | 1.94 | 1.8 | 1.04 | 1.37 | 1.71 | 1.55 | 0.235 | 0.135 | 0.1075 | 0.547487 | 0.742924 | 0.767375 | 0.999971 | 1 | 0.962472 |
| Naa20 | P61600 | 1.19 | 1.13 | 0.65 | 0.6 | 1.09 | 0.3 | 0.5 | 0.8 | 0.87 | 1.02 | 0.86 | 1.05 | 1.21 | 1.17 | 0.79 | 0.82 | -0.22 | 0.0575 | 0.105 | 0.402328 | 0.784262 | 0.650118 | 0.999971 | 1 | 0.933842 |
| Naa25 | Q8BWZ3 | 1.19 | 1.52 | 1.61 | 1.46 | 1.67 | 1.37 | 1.81 | 1.49 | 1.78 | 1.25 | 1.64 | 1.86 | 1.76 | 1.21 | 1.67 | 1.7 | 0.14 | 0.1875 | 0.14 | 0.479753 | 0.382381 | 0.507223 | 0.999971 | 1 | 0.905815 |
| Naa30 | E9QMB7 | 0.36 | 0.92 | 0.62 | 1.06 | 0.64 | 0.62 | 0.53 | 0.89 | 0.76 | 0.87 | 0.66 | 0.84 | 0.44 | 0.88 | 0.65 | 1.53 | -0.07 | 0.0425 | 0.135 | 0.749769 | 0.839633 | 0.648374 | 0.999971 | 1 | 0.933842 |
| Naa35 | Q6PHQ8 | 0.09 | 0.76 | 0.61 | 0.76 | 0.91 | 0.73 | 0.49 | 0.74 | 0.8 | 0.31 | 0.53 | 0.81 | 0.2 | 0.53 | 0.88 | 0.8 | 0.1625 | 0.0575 | 0.0475 | 0.47305 | 0.805105 | 0.849323 | 0.999971 | 1 | 0.981827 |
| Naa38 | Q9D2U5 | 0.6 | 0.67 | 1.36 | 0.91 | 1 | 0.89 | 1.34 | 1.63 | 1.12 | 1.18 | 1.9 | 1.75 | 1.01 | 0.99 | 1.65 | 1.86 | 0.33 | 0.6025 | 0.4925 | 0.229923 | 0.054289 | 0.119465 | 0.999971 | 1 | 0.78268 |
| Naa40 | A0A494B9I | 0.78 | 1.85 | 0.7 | 0.75 | 1.28 | 1.43 | -0.09 | 0.93 | 1.17 | 1.12 | -0.36 | -0.21 | 0.92 | 1.64 | 0.55 | 0.69 | -0.1325 | -0.59 | -0.07 | 0.751693 | 0.228574 | 0.845972 | 0.999971 | 1 | 0.979924 |
| Naa50 | Q6PGB6-2 | 0.86 | 1.59 | 2.05 | 2.01 | 1.64 | 1.56 | 1.34 | 1.84 | 1.35 | 1.14 | 2.05 | 1.85 | 1.32 | 1.66 | 1.89 | 2 | -0.0325 | -0.03 | 0.09 | 0.91438 | 0.930387 | 0.777505 | 0.999971 | 1 | 0.964764 |
| Naaa | Q9D7V9 | -0.88 | 0.26 | 0.09 | -0.35 | 0.44 | 0.2 | -0.2 | 0.2 | 0 | -0.52 | 0.01 | -0.33 | -0.15 | -0.03 | -0.1 | 0.41 | 0.38 | 0.01 | 0.2525 | 0.221842 | 0.972948 | 0.403521 | 0.999971 | 1 | 0.873348 |
| Nab2 | Q3TYF1 | 4.27 | 2.66 | 1.19 | 1.48 | 1.77 | 2.28 | 1.14 | 2.15 | 1.71 | 3.46 | 1.61 | 1.64 | 1.06 | 1.64 | 1.78 | 2.24 | -0.565 | -0.295 | -0.72 | 0.411919 | 0.697327 | 0.300402 | 0.999971 | 1 | 0.847402 |
| Nabp2 | E9Q199 | 0.71 | 1.9 | 0.19 | 0.12 | 1.33 | 1.03 | 0.09 | -0.07 | 1.27 | 0.64 | -1 | -1.41 | 0.54 | 1.48 | -0.39 | -0.08 | -0.135 | -0.855 | -0.3425 | 0.78656 | 0.23898 | 0.528707 | 0.999971 | 1 | 0.910069 |
| Naca | Q60817 | 0.37 | 1.33 | 1.14 | 1.21 | 1.27 | 1.15 | 0.96 | 1.36 | 1.27 | 0.37 | 1.28 | 1.41 | 1.22 | 1.32 | 1.19 | 1.57 | 0.1725 | 0.07 | 0.3125 | 0.509875 | 0.829217 | 0.24736 | 0.999971 | 1 | 0.839012 |
| Nacad | Q5SWP3 | 1.68 | 1.07 | 1.18 | 1.27 | 0.47 | 0.86 | 1.22 | 1.16 | 1.16 | 1.5 | 1.18 | 0.79 | 0.84 | 0.79 | 0.96 | 0.73 | -0.3725 | -0.1425 | -0.47 | 0.156563 | 0.542398 | 0.041163 | 0.999971 | 1 | 0.700838 |
| Nacc1 | Q7TSZ8 | 1.54 | 1.43 | 0.92 | 1.5 | 1.12 | 1.31 | 0.8 | 1.26 | 0.73 | 1.66 | 1.2 | 1.18 | 1.15 | 1.66 | 1.3 | 1.91 | -0.225 | -0.155 | 0.1575 | 0.331791 | 0.555245 | 0.537273 | 0.999971 | 1 | 0.910077 |
| Nacc2 | Q9DCM7 | 2.85 | 1.07 | 2.23 | 0.68 | 0.69 | 0.83 | 0.56 | 0.43 | 0.75 | 2.02 | 0.66 | 1.61 | -0.03 | 1 | 1.1 | 1.58 | -1.08 | -0.4475 | -0.795 | 0.046459 | 0.430208 | 0.178621 | 0.999971 | 1 | 0.808532 |
| Nadk | P58058 | -0.76 | 1.04 | 0.13 | 0.33 | 0.3 | 0.31 | -0.26 | 0.41 | 0.16 | 0.64 | 0.23 | 0.19 | 1.37 | 1.18 | 0.12 | 0.38 | 0.005 | 0.12 | 0.5775 | 0.989638 | 0.749881 | 0.221455 | 0.999971 | 1 | 0.8275 |
| Nadk2 | Q8C5H8-2 | -1.22 | -0.82 | -0.73 | -0.88 | -0.78 | -1.11 | -0.84 | -1.04 | -0.37 | -1.42 | -0.68 | -0.69 | -0.6 | -1.38 | -1.15 | -1.04 | -0.03 | 0.1225 | -0.13 | 0.877886 | 0.647391 | 0.580168 | 0.999971 | 1 | 0.91994 |
| Nae1 | E0CZE0 | -0.1 | 1.43 | 1.43 | 0.85 | 1.72 | 1.02 | 1.38 | 1.04 | 1.31 | -0.11 | 1.5 | 1.26 | 1.4 | 0.85 | 0.83 | 1.21 | 0.3875 | 0.0875 | 0.17 | 0.325752 | 0.856416 | 0.65252 | 0.999971 | 1 | 0.934005 |
| Naf1 | E9QJT2 | -0.45 | 0.92 | 1.48 | 1.24 | 0.66 | -0.12 | 0.51 | 0.73 | -0.36 | -0.21 | 0.45 | 0.57 | 0.92 | -0.22 | 1.2 | 2 | -0.3525 | -0.685 | 0.1775 | 0.436383 | 0.163003 | 0.759095 | 0.999971 | 1 | 0.961116 |
| Naga | Q9QWR8 | -0.87 | 0.46 | 0.41 | 0.14 | 0.6 | 0.63 | 0.21 | 1.28 | 0.81 | -0.39 | 0.32 | 0.45 | 0.87 | 0.46 | 0.77 | 0.37 | 0.645 | 0.2625 | 0.5825 | 0.108382 | 0.501358 | 0.106687 | 0.999971 | 1 | 0.78268 |
| Nagk | Q9QZ08 | 1.05 | 0.4 | 0.25 | 0.16 | 0.39 | 0.16 | 0.02 | 0.11 | 0.12 | 0.69 | 0.28 | 0.03 | 0.62 | 0.68 | -0.01 | 0.51 | -0.295 | -0.185 | -0.015 | 0.249806 | 0.495087 | 0.956307 | 0.999971 | 1 | 0.998257 |
| Naglu | O88325 | 1.24 | 0.58 | 1.11 | 1.95 | 0.18 | 0.37 | 0.47 | 2.02 | 0.31 | 0.93 | 0.81 | 0.86 | 0.15 | 0.77 | 1.17 | 2 | -0.46 | -0.4925 | -0.1975 | 0.346141 | 0.147791 | 0.662941 | 0.999971 | 1 | 0.936388 |
| Nalcn | E9QLE4 | 2.33 | -0.78 | -0.43 | -0.16 | -0.63 | -0.22 | -0.81 | -0.31 | -0.96 | 0.62 | -0.24 | -0.4 | -0.69 | -0.6 | 0.1 | -0.03 | -0.7325 | -0.485 | -0.545 | 0.280491 | 0.500103 | 0.423525 | 0.999971 | 1 | 0.87919 |
| Nampt | Q99KQ4 | 0.71 | 1.79 | 1.78 | 1.5 | 1.68 | 1.65 | 1.5 | 1.24 | 1.98 | 1.52 | 1.81 | 1.74 | 1.76 | 1.58 | 1.58 | 1.83 | 0.0725 | 0.3175 | 0.2425 | 0.800312 | 0.281319 | 0.394795 | 0.999971 | 1 | 0.871816 |
| Nanp | Q9CPT3 | 0.53 | 0.67 | 2.33 | 2.38 | 0.99 | 0.84 | 1.9 | 2.73 | 0.9 | 0.58 | 2.14 | 0.87 | 0.55 | 1.03 | 2.57 | 2.58 | 0.1375 | -0.355 | 0.205 | 0.822189 | 0.535657 | 0.757597 | 0.999971 | 1 | 0.961116 |
| Nans | Q99J77 | 0.6 | 0.46 | 0.7 | 0.59 | 0.83 | 0.78 | 0.94 | 1.07 | 1.2 | 0.76 | 1.03 | 1.28 | 1.36 | 0.94 | 1.22 | 0.81 | 0.3175 | 0.48 | 0.495 | 0.092139 | 0.029591 | 0.031279 | 0.999971 | 1 | 0.678474 |
| Nap1l1 | Q3TF41 | -2.31 | -2.56 | 0.08 | -0.04 | -2.08 | -2.75 | 0.01 | -0.28 | -2.26 | -2.98 | -0.03 | -0.13 | -3.29 | -2.31 | -0.17 | -0.23 | -0.0675 | -0.1425 | -0.2925 | 0.938642 | 0.878126 | 0.756814 | 0.999971 | 1 | 0.961116 |
| Nap1l4 | B7ZNL2 | -0.57 | -0.07 | -0.71 | -0.87 | 0.14 | 0.18 | 0.95 | -0.74 | 0.23 | -0.22 | -0.57 | -0.34 | 0.39 | 0.2 | -0.59 | -0.99 | 0.6875 | 0.33 | 0.3075 | 0.093327 | 0.229065 | 0.404369 | 0.999971 | 1 | 0.87384 |
| Nap1l5 | Q9JJF0 | -2.67 | -0.44 | -0.58 | -0.44 | 0.13 | -2.04 | 0.44 | -0.96 | 0.31 | -2.21 | -1.15 | 0.5 | -1.53 | -1.9 | -1.18 | -2.87 | 0.425 | 0.395 | -0.8375 | 0.553201 | 0.607699 | 0.187198 | 0.999971 | 1 | 0.809025 |
| Napa | Q9DB05 | -1.14 | -0.46 | -0.35 | -1.03 | -0.92 | -0.87 | -0.89 | -0.95 | -0.7 | -1.38 | -0.53 | -0.78 | -0.59 | -0.71 | -0.46 | -0.69 | -0.1625 | -0.1025 | 0.1325 | 0.495178 | 0.71927 | 0.584796 | 0.999971 | 1 | 0.920413 |
| Napb | P28663 | -0.36 | -0.36 | -1.13 | -1.31 | -0.25 | 0.33 | -0.82 | -1.12 | 0.17 | 0.26 | -1.13 | -1.37 | 0.4 | 0.25 | -0.82 | -0.91 | 0.325 | 0.2725 | 0.52 | 0.414997 | 0.561345 | 0.223726 | 0.999971 | 1 | 0.828225 |
| Napepld | Q8BH82 | -0.35 | -0.01 | -1.04 | -1.42 | -0.18 | 0.35 | -0.63 | -1.16 | 0.15 | -0.15 | -0.98 | -1.74 | -0.09 | -0.49 | -1.2 | -1.6 | 0.3 | 0.025 | -0.14 | 0.490629 | 0.959877 | 0.751517 | 0.999971 | 1 | 0.961116 |
| Napg | Q9CWZ7 | -1.06 | -0.95 | -0.99 | -1.29 | -0.73 | -0.77 | -0.42 | -1.29 | -0.73 | -1.4 | -0.79 | -0.59 | -0.7 | -1.2 | -0.97 | -1.17 | 0.27 | 0.195 | 0.0625 | 0.263885 | 0.406154 | 0.75365 | 0.999971 | 1 | 0.961116 |
| Naprt | Q8CC86 | 0.49 | 0.72 | 0.74 | 0.94 | 0.46 | 0.49 | 0.52 | -0.28 | 0.71 | 0.16 | 0.51 | -0.11 | 0.24 | -0.23 | 0.09 | -0.3 | -0.425 | -0.405 | -0.7725 | 0.109167 | 0.113931 | 0.004949 | 0.999971 | 1 | 0.582714 |
| NARS1 | Q8BP47 | -0.26 | 0.34 | 0.54 | 0.19 | 0.4 | 0.28 | 0.76 | 0.33 | 0.6 | -0.29 | 0.57 | 0.61 | 0.47 | 0.17 | 0.51 | 0.49 | 0.24 | 0.17 | 0.2075 | 0.324495 | 0.561214 | 0.374783 | 0.999971 | 1 | 0.870465 |
| Nars2 | Q8BGV0 | -0.51 | -0.12 | 0.05 | 0.3 | 0.01 | -0.23 | 0.1 | 0.47 | 0.11 | -0.49 | 0.06 | 0.47 | 0.21 | 0.04 | -0.11 | 0.74 | 0.1575 | 0.1075 | 0.29 | 0.535148 | 0.69857 | 0.301113 | 0.999971 | 1 | 0.847402 |
| Nasp | B1AU75 | 1.43 | 2.62 | 2.62 | 2.71 | 2.19 | 2.19 | 2.44 | 2.09 | 2.2 | 1.97 | 2.7 | 2.86 | 1.88 | 2.23 | 2.14 | 2.7 | -0.1175 | 0.0875 | -0.1075 | 0.712295 | 0.80894 | 0.756577 | 0.999971 | 1 | 0.961116 |
| Nat1 | A0A1D5RL( | -0.51 | -0.39 | -0.3 | -0.24 | -0.09 | -0.15 | -0.16 | -0.2 | -0.59 | -0.62 | -0.37 | 0.34 | -0.57 | -0.35 | 0.05 | -0.32 | 0.21 | 0.05 | 0.0625 | 0.227298 | 0.844562 | 0.755743 | 0.999971 | 1 | 0.961116 |
| Nat10 | Q8K224 | -0.45 | -0.06 | -0.23 | -0.49 | -0.19 | -0.63 | -0.22 | -0.34 | -0.6 | -0.64 | -0.19 | 0.18 | 0.21 | -0.47 | -0.24 | -0.15 | -0.0375 | -0.005 | 0.145 | 0.851384 | 0.983728 | 0.513218 | 0.999971 | 1 | 0.905815 |
| Nat14 | Q8BVG8 | -0.67 | -1.12 | -0.52 | -0.49 | -0.44 | -1 | -0.82 | -0.56 | -0.78 | -1.16 | -0.74 | -0.38 | -0.82 | -0.69 | -0.37 | -0.78 | -0.005 | -0.065 | 0.035 | 0.982672 | 0.790314 | 0.874512 | 0.999971 | 1 | 0.984109 |
| Nat2 | P50295 | 0.38 | -0.23 | -0.95 | -0.38 | -0.44 | -0.33 | -0.87 | -1.02 | -0.79 | -0.13 | -0.59 | -1.5 | -0.39 | -0.27 | 0.25 | -0.47 | -0.37 | -0.4575 | 0.075 | 0.268094 | 0.250581 | 0.815123 | 0.999971 | 1 | 0.974918 |
| Nat8f1 | E9PUL7 | -2.72 | -0.56 | -1.29 | -2.87 | -0.81 | -0.67 | -1.57 | -0.86 | -0.79 | -2.93 | -0.99 | -0.88 | -0.14 | -1.88 | -1.5 | -2.43 | 0.8825 | 0.4625 | 0.3725 | 0.13385 | 0.509359 | 0.584037 | 0.999971 | 1 | 0.920364 |
| Nat8l | A0A0R4J0F | 0.25 | -0.17 | 0.23 | -0.23 | -0.51 | 0.32 | 0.48 | -0.02 | -0.44 | -0.81 | 0.12 | 0.27 | 0.48 | -0.42 | 0.26 | -0.41 | 0.0475 | -0.235 | -0.0425 | 0.861338 | 0.428836 | 0.87941 | 0.999971 | 1 | 0.985441 |
| Natd1 | Q9DBW3 | 0.08 | 1.14 | -0.73 | -1.57 | 0.82 | 0.46 | -0.62 | -1.43 | 1.15 | 1.2 | -0.87 | -0.27 | 0.23 | 1.05 | -1.12 | -1.72 | 0.0775 | 0.5725 | -0.12 | 0.911652 | 0.42683 | 0.876233 | 0.999971 | 1 | 0.985129 |
| Nav1 | Q8CH77 | -0.27 | -0.72 | -1.16 | -1.05 | -0.58 | -0.43 | -0.64 | -0.91 | -0.32 | -0.01 | -0.68 | -0.65 | -0.24 | -0.5 | -0.89 | -0.8 | 0.16 | 0.385 | 0.1925 | 0.528708 | 0.180995 | 0.481287 | 0.999971 | 1 | 0.89485 |
| Nav2 | V9GXT3 | 1.06 | 0.28 | 0.5 | 0.96 | 0.08 | 0.81 | 0.07 | 1.02 | -0.35 | 1.19 | 0.78 | 0.83 | -0.16 | 0.19 | 1 | 0.53 | -0.205 | -0.0875 | -0.31 | 0.517303 | 0.813708 | 0.337489 | 0.999971 | 1 | 0.858827 |
| Nav3 | A0A668KM | -0.36 | -0.65 | -0.58 | -0.54 | -0.77 | -0.51 | -0.12 | -0.76 | -0.24 | -0.05 | -0.22 | -0.33 | -0.18 | -0.76 | -1.02 | -0.45 | -0.0075 | 0.3225 | -0.07 | 0.97188 | 0.085222 | 0.76262 | 0.999971 | 1 | 0.961744 |
| Naxd | J3QMM7 | -0.84 | -0.55 | -0.56 | -0.33 | -0.53 | -0.55 | -0.51 | -0.42 | -0.42 | -0.61 | -0.42 | -0.51 | -0.5 | -0.3 | -0.61 | -0.21 | 0.0675 | 0.08 | 0.165 | 0.713527 | 0.663138 | 0.416689 | 0.999971 | 1 | 0.877251 |
| Naxd_1 | Q9CZ42-2 | -1.45 | -1.02 | 0.93 | 1.9 | -0.3 | -0.47 | -0.41 | 1.38 | -0.3 | -1.18 | 0.17 | 0 | -0.99 | -2.31 | 1.49 | 2.24 | -0.04 | -0.4175 | 0.0175 | 0.960944 | 0.590888 | 0.988163 | 0.999971 | 1 | 0.99925 |
| Naxe | Q8K4Z3 | -0.84 | -0.14 | 0.19 | 0.31 | -0.32 | -0.27 | -0.09 | -0.16 | -0.24 | -0.49 | -0.34 | -0.44 | -0.68 | -0.36 | -0.06 | -0.3 | -0.09 | -0.2575 | -0.23 | 0.747689 | 0.367607 | 0.448095 | 0.999971 | 1 | 0.885289 |
| Nbas | E9Q411 | -0.79 | -0.49 | -1.06 | -0.9 | -0.26 | -0.75 | -1.2 | -1.05 | -0.51 | -0.81 | -0.98 | -1.03 | -0.26 | -0.41 | -1.17 | -1.02 | -0.005 | -0.0225 | 0.095 | 0.984769 | 0.915745 | 0.728541 | 0.999971 | 1 | 0.956041 |
| Nbdy | A0A0N4SU | 1.62 | 0.3 | 0.77 | 1.09 | 0.6 | 0.51 | 1.18 | 1.46 | 0.51 | 1.03 | 0.92 | 0.47 | 0.24 | 0.69 | 0.65 | 0.57 | -0.0075 | -0.2125 | -0.4075 | 0.983013 | 0.505423 | 0.202525 | 0.999971 | 1 | 0.81617 |
| Nbea | Q9EPN1-4 | 0.07 | 0.12 | 0.19 | -0.01 | 0.1 | 0.37 | 0.62 | 0.04 | 0.23 | 0.18 | 0.38 | 0.12 | 0.21 | 0.28 | 0.24 | 0.2 | 0.19 | 0.135 | 0.14 | 0.35261 | 0.421172 | 0.399275 | 0.999971 | 1 | 0.871816 |
| Nbeal1 | E9PYP2 | 0 | -0.12 | -0.35 | -0.79 | -0.07 | -0.48 | -1.01 | -0.53 | -0.09 | -1.12 | -0.94 | 0.43 | 0.14 | -0.34 | -0.07 | 0.05 | -0.2075 | -0.115 | 0.26 | 0.460135 | 0.767881 | 0.291109 | 0.999971 | 1 | 0.847402 |
| Nbr1 | Q05BC8 | 1.31 | -0.38 | -0.44 | -0.61 | -0.56 | 0.02 | -0.76 | -0.11 | -0.34 | 1.76 | 0.12 | 0.05 | -0.35 | -0.01 | 0.13 | 0.61 | -0.3225 | 0.4275 | 0.125 | 0.484953 | 0.478645 | 0.786494 | 0.999971 | 1 | 0.967132 |
| Ncald | Q91X97 | -2.15 | -0.21 | -0.24 | -0.49 | -0.44 | -0.43 | -0.15 | -0.4 | -0.38 | -1.82 | 0.03 | -0.09 | -0.46 | -0.98 | -0.7 | -0.5 | 0.4175 | 0.2075 | 0.1125 | 0.35649 | 0.721323 | 0.802813 | 0.999971 | 1 | 0.971467 |
| Ncam1 | E9QB01 | -5.71 | -1.8 | -1.38 | -2.19 | -1.27 | -0.91 | -1.49 | -1.74 | -0.32 | -2.6 | -0.5 | -0.93 | 0.21 | -1.28 | -0.95 | -1.61 | 1.4175 | 1.6825 | 1.8625 | 0.142852 | 0.122813 | 0.078692 | 0.999971 | 1 | 0.754157 |
| Ncam1_1 | A0A0A6YY! | -0.77 | -1.61 | -1.55 | -1.53 | -1.59 | -1.43 | -1.23 | -1.5 | -1.31 | -1.18 | -1.44 | -1.6 | -1.32 | -1.55 | -1.54 | -1.45 | -0.0725 | -0.0175 | -0.1 | 0.766791 | 0.943223 | 0.678142 | 0.999971 | 1 | 0.942594 |
| Ncam1_2 | A0A0A6YY4 | 2.52 | 3 | 0.92 | -1.02 | 5.17 | 1.79 | 1.3 | 0.1 | 7.42 | 0.16 | 2.99 | 2.79 | 11.3 | 0.81 | 0.25 | -0.32 | 0.735 | 1.985 | 1.655 | 0.56262 | 0.227581 | 0.526275 | 0.999971 | 1 | 0.909459 |
| Ncam1_3 | P13595 | -1.17 | -3.8 | -4.82 | -4.25 | -3.43 | -3.9 | -1.06 | -5.19 | -3.44 | -1.82 | -4.29 | -4.87 | -3.07 | -3.92 | -4.53 | -5.71 | 0.115 | -0.095 | -0.7975 | 0.912957 | 0.919526 | 0.375863 | 0.999971 | 1 | 0.870465 |
| Ncam2 | O35136 | -1.09 | -1.71 | -1.67 | -1.58 | -1.39 | -1.06 | -1.29 | -1.31 | -1.31 | -1.06 | -1.31 | -1.24 | -0.99 | -1.16 | -1.09 | -1.27 | 0.25 | 0.2825 | 0.385 | 0.253851 | 0.192012 | 0.09161 | 0.999971 | 1 | 0.773779 |
| Ncan | A0A0R4IZX | 1.3 | 1.56 | 1.07 | 1.51 | 1.61 | 1.43 | 1.34 | 1.35 | 1.65 | 1.36 | 1.49 | 1.12 | 1.38 | 1.45 | 1.14 | 1.44 | 0.0725 | 0.045 | -0.0075 | 0.707913 | 0.827935 | 0.969482 | 0.999971 | 1 | 0.998787 |
| Ncbp1 | Q3UYV9 | -0.14 | 1.98 | 1.75 | 1.19 | 1.98 | 1.75 | 1.6 | 1.78 | 2.33 | 0.47 | 1.76 | 1.79 | 1.87 | 1.54 | 2.08 | 1.39 | 0.5825 | 0.3925 | 0.525 | 0.21888 | 0.496482 | 0.279618 | 0.999971 | 1 | 0.843812 |
| Ncbp2 | Q9CQ49 | 1.7 | 1.61 | 1.74 | 2.25 | 1.5 | 1.54 | 2.41 | 1.68 | 1.17 | 1.24 | 1.4 | 1.43 | 0.25 | 1.24 | 1.34 | 1.79 | -0.0425 | -0.515 | -0.67 | 0.876938 | 0.032194 | 0.082669 | 0.999971 | 1 | 0.755254 |
| Ncbp3 | Q8BZR9 | 1.11 | -0.44 | 0.34 | 0.95 | -0.1 | 0.29 | 0.16 | 0.44 | 0.31 | 1.01 | -0.25 | 0 | -0.32 | 1.01 | 0.16 | 0.35 | -0.2925 | -0.2225 | -0.19 | 0.426391 | 0.601761 | 0.655695 | 0.999971 | 1 | 0.934061 |

| | | | | | | | | | | | | | | | | | | | | | | | | | | |
|---|---|---|---|---|---|---|---|---|---|---|---|---|---|---|---|---|---|---|---|---|---|---|---|---|---|---|
| Ncdn | Q9Z0E0-2 | 1.96 | 0.79 | 1.16 | 3.17 | 2.02 | 0.57 | 1.48 | 2.4 | -0.68 | 1.5 | 1.03 | 0.13 | 2.14 | 0.13 | -0.14 | 2.15 | -0.1525 | -1.275 | -0.7 | 0.800048 | 0.078249 | 0.354979 | 0.999971 | 1 | 0.868793 |
| Ncdn_1 | Q9Z0E0 | 0.38 | 0.87 | 0.89 | 0.65 | 0.74 | 0.9 | 0.85 | 1.03 | 0.68 | 0.37 | 0.89 | 0.48 | 0.85 | 0.78 | 1.3 | 0.86 | 0.1825 | -0.0925 | 0.25 | 0.362831 | 0.66289 | 0.265324 | 0.999971 | 1 | 0.842677 |
| Nceh1 | Q8BLF1 | -1.57 | -0.19 | 0.2 | -1.05 | -0.1 | -0.15 | 0.17 | -0.43 | 0.25 | -1.34 | 0.22 | 0.35 | 0.44 | -0.25 | 0.29 | -0.58 | 0.525 | 0.5225 | 0.6275 | 0.213253 | 0.337047 | 0.178806 | 0.999971 | 1 | 0.808532 |
| Ncf1 | Q09014 | 2.12 | 1.13 | 1.77 | 1.85 | 0.76 | 0.97 | 1.5 | 1.58 | 1.19 | 2.67 | 1.34 | 1.7 | -0.49 | 0.24 | 1.47 | 0.89 | -0.515 | 0.0075 | -1.19 | 0.11195 | 0.984253 | 0.023929 | 0.999971 | 1 | 0.664905 |
| Ncf2 | O70145 | 4.55 | 0.99 | 1.47 | 1.19 | 0.63 | -0.06 | 0.75 | 1.38 | 0.14 | 2.67 | 1.49 | 0.77 | -0.13 | 0.21 | 0.08 | 1.32 | -1.375 | -0.7825 | -1.68 | 0.112678 | 0.395768 | 0.063744 | 0.999971 | 1 | 0.725348 |
| Nck1 | Q99M51 | -0.87 | 0.09 | -0.27 | -0.28 | 0.05 | 0.16 | -0.59 | -0.16 | 0.27 | -0.41 | -0.12 | -0.01 | 0.24 | 0.15 | 0.02 | 0.19 | 0.1975 | 0.265 | 0.4825 | 0.480373 | 0.329283 | 0.069355 | 0.999971 | 1 | 0.733937 |
| Nck2 | O55033 | -0.72 | -0.58 | -0.44 | -0.23 | -0.76 | -0.79 | -0.43 | -0.45 | -0.88 | -0.87 | -0.23 | -0.51 | -0.65 | -0.75 | -0.6 | 0.02 | -0.115 | -0.13 | -0.0025 | 0.569813 | 0.56824 | 0.991615 | 0.999971 | 1 | 0.99925 |
| Nckap1 | A2AS98 | -0.84 | -0.42 | -0.47 | -0.65 | -0.6 | -0.66 | -0.52 | -0.52 | -0.42 | -0.73 | -0.24 | -0.34 | -0.39 | -0.63 | -0.41 | -0.35 | 0.02 | 0.1625 | 0.15 | 0.911544 | 0.420541 | 0.430557 | 0.999971 | 1 | 0.881611 |
| Nckap1l | Q8K1X4 | 1 | 1.72 | 1.38 | 1.46 | 0.86 | 1.84 | 1.15 | 1.49 | 1.05 | 1.62 | 1.38 | 1.25 | 0.84 | 1.77 | 1.43 | 1.57 | -0.055 | -0.065 | 0.0125 | 0.842064 | 0.775331 | 0.963003 | 0.999971 | 1 | 0.998721 |
| Nckap5l | Q6GQX2 | -0.44 | -0.76 | -0.17 | 2.66 | -0.6 | -1.17 | 1.2 | -0.18 | -1.17 | -0.79 | 1.77 | 1.7 | -0.17 | -0.5 | 2.31 | 3.17 | -0.51 | 0.055 | 0.88 | 0.547783 | 0.956182 | 0.421214 | 0.999971 | 1 | 0.87788 |
| Nckipsd | Q9ESJ4 | 0.26 | 0.14 | 0.5 | 0 | -0.12 | 0.26 | 0.7 | -0.1 | -0.11 | -0.08 | 0.13 | -0.07 | 0.02 | -0.12 | -0.1 | -0.24 | -0.04 | -0.2575 | -0.335 | 0.872073 | 0.18884 | 0.103279 | 0.999971 | 1 | 0.78268 |
| Ncl | P09405 | 2.67 | 2.34 | 1.5 | 2.02 | 2.07 | 1.57 | 1.15 | 1.23 | 2.04 | 2.25 | 1.24 | 1.48 | 1.08 | 1.66 | 1.08 | 1.2 | -0.6275 | -0.38 | -0.8775 | 0.081036 | 0.28091 | 0.01493 | 0.999971 | 1 | 0.613095 |
| Ncln | Q8VCM8 | -0.61 | 0.6 | -0.09 | 0.04 | 0.73 | 0.41 | 0.04 | 0.43 | 0.92 | -0.12 | 0.26 | 0.23 | 1.27 | 0.53 | 0.32 | 0.89 | 0.4175 | 0.3375 | 0.7675 | 0.182076 | 0.320451 | 0.040396 | 0.999971 | 1 | 0.700838 |
| Ncoa1 | P70365-4 | 0.89 | 1.4 | 1.2 | 0.39 | 1.22 | 0.89 | 1.44 | 0.89 | 0.77 | 0.54 | 0.5 | 0.7 | 1.59 | 1.38 | 0.12 | -0.42 | 0.14 | -0.3425 | -0.3025 | 0.613477 | 0.199798 | 0.546561 | 0.999971 | 1 | 0.911398 |
| Ncoa2 | Q61026 | 4.46 | 2.61 | 3.48 | 3.62 | 3.06 | 3.19 | 4.21 | 3.39 | 2.42 | 3.21 | 2.73 | 3.33 | 2.33 | 2.78 | 2.82 | 0.3 | -0.08 | -0.62 | -1.485 | 0.853314 | 0.160129 | 0.043526 | 0.999971 | 1 | 0.701725 |
| Ncoa5 | B7ZC24 | 1.82 | -3.05 | 0.71 | 0.51 | -4.62 | -2.42 | 1.23 | 0.7 | -4.14 | 0.64 | 0.55 | -1.71 | -7.31 | 1.34 | -0.22 | -0.96 | -1.275 | -1.1625 | -1.785 | 0.415785 | 0.410644 | 0.366475 | 0.999971 | 1 | 0.869526 |
| Ncoa5_1 | Q91W39 | 0.15 | -0.68 | -0.1 | -0.14 | -0.54 | -0.59 | -0.01 | -0.02 | -0.58 | -0.28 | 0.05 | -0.13 | -0.65 | -0.63 | -0.25 | -0.05 | -0.0975 | -0.0425 | -0.2025 | 0.708444 | 0.863312 | 0.435736 | 0.999971 | 1 | 0.882419 |
| Ncoa7 | A0A1L1SSS | -0.3 | -0.27 | -0.3 | -0.59 | 0.07 | -0.16 | -0.24 | -0.4 | -0.09 | -0.25 | 0.06 | 0.06 | 0.06 | -0.27 | -0.35 | -0.4 | 0.1825 | 0.31 | 0.125 | 0.35094 | 0.110518 | 0.523256 | 0.999971 | 1 | 0.907797 |
| Ncor1 | Q5RIM6 | 1.34 | 1.54 | 1.45 | 1.53 | 1.14 | 1.48 | 1.63 | 1.59 | 1.24 | 1.6 | 1.35 | 0.93 | 0.46 | 0.98 | 0.95 | 0.82 | -0.005 | -0.185 | -0.6625 | 0.978876 | 0.367047 | 0.00729 | 0.999971 | 1 | 0.582714 |
| Ncor2 | F8VQL9 | 1.46 | 1.43 | 0.98 | 1.26 | 1.43 | 1.62 | 0.95 | 0.98 | 1.39 | 1.59 | 0.69 | 1.02 | 0.47 | 1.39 | 0.59 | 0.57 | -0.0375 | -0.11 | -0.5275 | 0.873119 | 0.664713 | 0.071228 | 0.999971 | 1 | 0.737413 |
| Ncs1 | Q8BNY6 | -0.72 | -0.01 | -0.39 | -1.14 | -0.44 | 0.69 | -0.8 | -0.69 | -0.35 | -0.34 | -0.6 | -0.86 | -0.29 | 0.78 | -0.81 | -0.47 | 0.255 | 0.0275 | 0.3675 | 0.528 | 0.922575 | 0.371603 | 0.999971 | 1 | 0.869526 |
| Ncstn | P57716 | 0.11 | -0.42 | -0.07 | -0.49 | -0.71 | 0.02 | -0.46 | -0.14 | -0.81 | 0.04 | -0.42 | -0.67 | -0.72 | 0.28 | -0.56 | 0 | -0.105 | -0.2475 | -0.0325 | 0.671463 | 0.349735 | 0.910091 | 0.999971 | 1 | 0.991822 |
| Ndc1 | Q8VCB1 | -0.47 | -0.84 | 0.13 | 0.09 | -0.45 | -0.57 | 0.77 | 0.01 | -0.54 | -0.86 | -0.1 | -0.27 | -0.28 | -0.47 | -0.32 | -0.6 | 0.2125 | -0.17 | -0.145 | 0.57069 | 0.568821 | 0.589408 | 0.999971 | 1 | 0.921872 |
| Nde1 | Q9CZA6-3 | -0.04 | -0.08 | -0.02 | -0.24 | -0.47 | 0.12 | -0.52 | 0.38 | -0.17 | 0.41 | -0.01 | 0.13 | -0.02 | 0.08 | -0.4 | -0.26 | -0.0275 | 0.185 | -0.055 | 0.913588 | 0.349506 | 0.772276 | 0.999971 | 1 | 0.963498 |
| Ndel1 | Q9ERR1 | -0.01 | -0.35 | -0.63 | -0.42 | -0.42 | -0.34 | -0.9 | -0.46 | -0.48 | -0.07 | -0.46 | -0.51 | -0.38 | -0.58 | -0.51 | -0.36 | -0.1775 | -0.0275 | -0.105 | 0.433476 | 0.896364 | 0.601092 | 0.999971 | 1 | 0.92426 |
| Ndfip1 | A0A494BA/ | -0.35 | 2.64 | 1.69 | 0.86 | 2.73 | 2.05 | 1 | 1.67 | 2.78 | -0.16 | 2.13 | 2 | 2.72 | 1.22 | 1.59 | 1.19 | 0.6525 | 0.4775 | 0.47 | 0.338091 | 0.561567 | 0.484121 | 0.999971 | 1 | 0.896352 |
| Ndfip2 | F6RT43 | 1.19 | -1.34 | 0.26 | 0.31 | -1.88 | -1.02 | 0.56 | 0.14 | -1.29 | 0.23 | -0.13 | -0.16 | -1.67 | -2.01 | -0.05 | -0.87 | -0.655 | -0.4425 | -1.255 | 0.356604 | 0.447645 | 0.070915 | 0.999971 | 1 | 0.736129 |
| Ndn | P25233 | 0.13 | -0.54 | 0.61 | -0.82 | -1.13 | -1.47 | 0.84 | 0.03 | -1.38 | -0.53 | 0.28 | 0.6 | -0.56 | -1.27 | 0.42 | 0.21 | -0.2775 | -0.1025 | -0.145 | 0.628716 | 0.840894 | 0.758677 | 0.999971 | 1 | 0.961116 |
| Ndor1 | A2AI05 | -0.85 | 0.28 | -1.68 | -0.64 | 0.04 | 0.06 | -1.87 | -0.78 | 0.38 | -0.47 | -1.03 | -1.08 | 0.5 | 0.58 | -0.97 | -1.27 | 0.085 | 0.1725 | 0.4325 | 0.878762 | 0.726404 | 0.46073 | 0.999971 | 1 | 0.890033 |
| Ndrg1 | Q62433 | -0.9 | -0.97 | -1.33 | -1.7 | -0.82 | -1.22 | -1.39 | -1.62 | -0.79 | -1.49 | -1.77 | -1.22 | -1.14 | -1.19 | -1.17 | -1.66 | -0.0375 | -0.0925 | -0.065 | 0.889241 | 0.749732 | 0.795544 | 0.999971 | 1 | 0.970368 |
| Ndrg2 | Q9QYG0 | -1.76 | -1.12 | -0.45 | -0.79 | -0.66 | -0.72 | -0.35 | 0.23 | -0.6 | -1.69 | -0.46 | 0.08 | -0.67 | -0.97 | -0.79 | -0.68 | 0.655 | 0.3625 | 0.2525 | 0.087006 | 0.418606 | 0.406318 | 0.999971 | 1 | 0.874911 |
| Ndrg2_1 | A0A2I3BQ( | -1.12 | 0.21 | 1.48 | -0.16 | 0.54 | -0.97 | 0.6 | 2.41 | -0.44 | -1.04 | 0.62 | 1.4 | -0.74 | -0.85 | 1.15 | 0.9 | 0.5425 | 0.0325 | 0.0125 | 0.496792 | 0.962729 | 0.985396 | 0.999971 | 1 | 0.99925 |
| Ndrg2_2 | Q9QYG0-2 | -1.89 | -1.29 | -0.37 | -0.29 | -0.51 | -2.04 | 3.6 | 0.57 | -0.48 | -1.32 | 1.9 | 3.02 | -0.65 | -1.49 | 1.47 | -0.63 | 1.365 | 1.74 | 0.635 | 0.240944 | 0.101678 | 0.356349 | 0.999971 | 1 | 0.868793 |
| Ndrg3 | Q9QYF9 | -1.64 | -0.04 | -0.3 | -0.41 | 0.07 | -0.28 | -0.38 | -0.17 | 0.25 | -0.84 | -0.12 | 0.09 | 0.45 | -0.06 | -0.12 | -0.06 | 0.4075 | 0.4425 | 0.65 | 0.27562 | 0.297599 | 0.106418 | 0.999971 | 1 | 0.78268 |
| Ndrg4 | Q8BTG7-3 | -1.98 | -1.06 | -1.06 | -1.08 | -0.4 | -0.61 | -0.78 | -0.83 | -0.58 | -1.5 | -0.96 | -0.44 | 0.13 | -0.2 | -0.41 | -0.27 | 0.64 | 0.425 | 1.1075 | 0.038225 | 0.218083 | 0.002904 | 0.999971 | 1 | 0.575374 |
| Ndufa1 | O35683 | -1.8 | -0.53 | -1.63 | -1.54 | -0.51 | -1.57 | -1.65 | -1.62 | -1.13 | -2.29 | -1.93 | -1.19 | -1.51 | -0.94 | -0.97 | -1.59 | 0.0375 | -0.26 | 0.1225 | 0.922052 | 0.509464 | 0.714534 | 0.999971 | 1 | 0.952364 |
| Ndufa10 | Q99LC3 | -0.91 | -0.98 | 0.4 | 0.6 | -1.08 | -0.78 | 0.46 | 0.41 | -0.87 | -1.07 | 0.24 | -0.72 | -1.18 | -1 | 0.57 | 0.19 | -0.025 | -0.3825 | -0.1325 | 0.962559 | 0.432161 | 0.811252 | 0.999971 | 1 | 0.973864 |
| Ndufa11 | G5E814 | 0.42 | 1.47 | 1.2 | 0.87 | 0.16 | 2.67 | 1.94 | 1.49 | 0.23 | 0.96 | 0.58 | -1.32 | 0.02 | 2.53 | 0.74 | 0.66 | 0.575 | -0.8775 | -0.0025 | 0.293819 | 0.113473 | 0.996294 | 0.999971 | 1 | 1 |
| Ndufa12 | Q7TMF3 | -1.01 | -0.3 | -0.29 | -0.97 | 0.15 | -0.63 | 0.71 | -0.73 | 0.58 | -1.16 | 0.56 | 1.09 | 1.2 | -0.65 | -0.41 | -0.59 | 0.5175 | 0.91 | 0.53 | 0.198008 | 0.09392 | 0.262682 | 0.999971 | 1 | 0.842381 |
| Ndufa13 | Q9ERS2 | -0.28 | -0.72 | -0.99 | -0.92 | -0.86 | -0.95 | -1.2 | -1.04 | -1.22 | -0.74 | -1.09 | -1.12 | -0.94 | -0.64 | -1.05 | -0.88 | -0.285 | -0.315 | -0.15 | 0.216211 | 0.189897 | 0.508965 | 0.999971 | 1 | 0.905815 |
| Ndufa2 | Q9CQ75 | -0.62 | -0.28 | -0.93 | -0.87 | -0.55 | -0.49 | -0.89 | -0.98 | -0.29 | -1.41 | -1.44 | -0.94 | -0.65 | -1.03 | -0.97 | -1.05 | -0.0525 | -0.345 | -0.25 | 0.81935 | 0.284931 | 0.273448 | 0.999971 | 1 | 0.843794 |
| Ndufa3 | Q9CQ91 | -3.14 | -1.4 | 0.57 | -0.02 | -1.2 | -1.32 | 0.52 | 0.59 | -1.23 | -2.85 | 0.97 | 0.94 | -0.9 | -1.61 | 0.75 | 0.43 | 0.645 | 0.455 | 0.665 | 0.468297 | 0.683787 | 0.463188 | 0.999971 | 1 | 0.890823 |
| Ndufa4 | Q62425 | -0.33 | -0.56 | -0.08 | 0.21 | -0.78 | -0.86 | -0.59 | -0.4 | -0.94 | -0.1 | -0.42 | -0.46 | -0.22 | -0.45 | -0.37 | 0.23 | -0.4675 | -0.29 | -0.0125 | 0.069874 | 0.283649 | 0.960461 | 0.999971 | 1 | 0.998721 |
| Ndufa4l2 | Q4FZG9 | 0.39 | -0.49 | -0.91 | -0.25 | 0.08 | -0.14 | -1.09 | -2.43 | -0.32 | 0.91 | -1.06 | -1.87 | -0.48 | 0.04 | -1.65 | -1.57 | -0.58 | -0.27 | -0.6 | 0.330372 | 0.652789 | 0.21881 | 0.999971 | 1 | 0.827064 |
| Ndufa5 | Q9CPP6 | 0.08 | -0.26 | -0.67 | -0.65 | -0.66 | -0.38 | -0.71 | -0.58 | -0.72 | 0.04 | -0.68 | -0.66 | -0.58 | 0.16 | -0.66 | -0.68 | -0.2075 | -0.13 | -0.065 | 0.379437 | 0.634914 | 0.819022 | 0.999971 | 1 | 0.975633 |
| Ndufa6 | Q9CQZ5 | -1.55 | -2.52 | -1.15 | -1.26 | -1.81 | -2.42 | -0.29 | -1.51 | -1.31 | -2 | -0.94 | -0.66 | -2.17 | -2.25 | -1.43 | -1.31 | 0.1125 | 0.3925 | -0.17 | 0.823857 | 0.348266 | 0.658465 | 0.999971 | 1 | 0.934638 |
| Ndufa7 | Q9Z1P6 | -0.23 | -0.62 | -0.61 | -0.74 | -0.55 | -0.95 | -0.67 | -0.89 | -0.45 | -0.88 | -0.71 | -0.93 | -0.4 | -0.89 | -1.04 | -0.85 | -0.215 | -0.1925 | -0.245 | 0.302582 | 0.359996 | 0.285042 | 0.999971 | 1 | 0.845133 |
| Ndufa8 | Q9DCJ5 | 0.47 | 0.23 | 1.64 | 1.39 | -0.32 | 0.94 | 1.26 | 1.33 | -0.06 | 0.92 | 0.94 | 0.93 | -0.46 | 0.72 | 1.32 | 1.34 | -0.13 | -0.25 | -0.2025 | 0.786634 | 0.541823 | 0.689611 | 0.999971 | 1 | 0.945805 |
| Ndufa9 | A0A0R3P9( | -1.88 | -1.21 | -0.07 | -0.2 | -1.18 | -1.55 | -0.05 | -0.36 | -1.33 | -2.22 | -0.27 | -0.16 | -1.41 | -1.53 | -0.17 | 0.03 | 0.055 | -0.155 | 0.07 | 0.914434 | 0.79511 | 0.897831 | 0.999971 | 1 | 0.989101 |
| Ndufab1 | F6ZFT1 | -1.88 | -4.1 | -1.54 | -1.53 | -4.28 | -3.96 | -1.5 | -1.62 | -4.13 | -3.25 | -1.56 | -1.71 | -5.28 | -3.66 | -1.57 | -1.42 | -0.5775 | -0.4 | -0.72 | 0.509902 | 0.616375 | 0.474481 | 0.999971 | 1 | 0.894044 |
| Ndufaf1 | A0A0R4J08 | 0.86 | -0.05 | -0.88 | -0.89 | -0.06 | -0.11 | -0.8 | -0.52 | -0.04 | 0.56 | -0.74 | -0.34 | 0.09 | -0.12 | -0.58 | -0.32 | -0.1325 | 0.1 | 0.0075 | 0.756929 | 0.830716 | 0.985684 | 0.999971 | 1 | 0.99925 |
| Ndufaf2 | Q59J78 | -0.59 | -0.76 | -0.77 | -0.49 | -0.18 | -0.67 | 0.04 | -0.51 | -0.03 | -0.54 | -0.02 | -0.2 | 0.44 | -0.58 | -0.47 | -0.39 | 0.3225 | 0.455 | 0.4025 | 0.165383 | 0.042727 | 0.154801 | 0.999971 | 1 | 0.795578 |
| Ndufaf3 | Q9JKL4 | 3.51 | -0.36 | -0.35 | 0.23 | -0.4 | -0.31 | 0.18 | -0.22 | -0.34 | 2.05 | 0.15 | 0.33 | -0.43 | -0.09 | -0.06 | -0.27 | -0.945 | -0.21 | -0.97 | 0.278174 | 0.826474 | 0.265277 | 0.999971 | 1 | 0.842677 |
| Ndufaf4 | Q9D1H6 | -0.96 | -0.58 | -0.58 | -0.34 | -0.58 | -0.66 | -0.6 | -0.35 | -0.77 | -0.78 | -0.25 | -0.34 | -0.77 | -0.16 | -0.28 | -0.35 | 0.0675 | 0.08 | 0.225 | 0.738719 | 0.724796 | 0.333232 | 0.999971 | 1 | 0.857921 |
| Ndufaf5 | A2APY7 | 1.34 | -0.13 | 0.1 | -0.08 | -0.26 | -0.23 | 0.28 | 0.62 | -0.07 | 0.52 | 0.41 | -0.06 | 0.77 | 0.18 | 0.66 | 0.36 | -0.205 | -0.1075 | 0.185 | 0.602781 | 0.771805 | 0.614086 | 0.999971 | 1 | 0.927387 |
| Ndufaf6 | A2AIL4 | 1.07 | -1.22 | -0.95 | -0.25 | -0.72 | -0.81 | -1.67 | -0.12 | -1.01 | -0.08 | -0.71 | -0.59 | -0.96 | -0.33 | -0.64 | -0.17 | -0.4925 | -0.26 | -0.1875 | 0.384775 | 0.612123 | 0.709771 | 0.999971 | 1 | 0.950695 |
| Ndufaf7 | Q9CWG8 | -0.28 | -0.43 | -0.39 | -0.26 | -0.48 | -0.51 | -0.25 | 0.17 | -0.53 | -0.5 | -0.37 | -0.42 | 0.04 | -0.55 | -0.45 | -0.11 | 0.0725 | -0.115 | 0.0725 | 0.732735 | 0.479488 | 0.721318 | 0.999971 | 1 | 0.954981 |
| Ndufaf8 | A0A087WC | 2.51 | -0.04 | 1.8 | 1.75 | -0.03 | 0.7 | 2.47 | 0.31 | 1.08 | 3.95 | 0.54 | 3.05 | 1.1 | -1.24 | 0.51 | 0.58 | -0.6425 | 0.65 | -1.2675 | 0.372256 | 0.465743 | 0.088153 | 0.999971 | 1 | 0.766425 |
| Ndufb1 | P0DN34 | -1.14 | -0.39 | -0.64 | -0.52 | -0.38 | -0.59 | -0.89 | -1.1 | -0.72 | -0.8 | -0.58 | -0.98 | -0.04 | -0.64 | -0.68 | -0.19 | -0.0675 | -0.0975 | 0.285 | 0.791047 | 0.663417 | 0.282391 | 0.999971 | 1 | 0.844538 |
| Ndufb10 | Q9DCS9 | -0.71 | -1.4 | 0.11 | 0.52 | -1.47 | -1.59 | -0.13 | 0.35 | -1.62 | -1.38 | -0.06 | -0.05 | -1.88 | -1.23 | -0.06 | -0.2 | -0.34 | -0.4075 | -0.4725 | 0.568198 | 0.469005 | 0.408437 | 0.999971 | 1 | 0.875014 |
| Ndufb11 | O09111 | -0.71 | -0.59 | -0.98 | -0.99 | -0.82 | -0.69 | -1.15 | -0.78 | -0.93 | -0.95 | -1.08 | -0.86 | -0.67 | -0.4 | -0.91 | -0.91 | -0.0425 | -0.1375 | 0.095 | 0.831728 | 0.454644 | 0.651869 | 0.999971 | 1 | 0.933842 |
| Ndufb2 | Q9CPU2 | 0.31 | -0.36 | -0.46 | 0.12 | -0.34 | -0.39 | -0.9 | -0.45 | -0.56 | 0.06 | -0.3 | -0.6 | -0.08 | -0.28 | -0.43 | 0.19 | -0.4225 | -0.2525 | -0.0525 | 0.121397 | 0.346896 | 0.837215 | 0.999971 | 1 | 0.979416 |
| Ndufb3 | Q9CQZ6 | -0.69 | -0.77 | -0.39 | -1.65 | -1.06 | -0.83 | -0.66 | -1.25 | -1.37 | -1.12 | -0.67 | -0.7 | -1.11 | -1.17 | -0.82 | -1.84 | -0.075 | -0.09 | -0.36 | 0.806862 | 0.779618 | 0.309447 | 0.999971 | 1 | 0.847402 |
| Ndufb4 | Q9CQC7 | -2.21 | -1.91 | -2.11 | -2.5 | -2.01 | -2.62 | -1.68 | -2.49 | -1.95 | -2.44 | -1.88 | -1.54 | -1.83 | -2.32 | -2.24 | -2.45 | -0.0175 | 0.23 | -0.0275 | 0.948136 | 0.367377 | 0.902313 | 0.999971 | 1 | 0.990759 |
| Ndufb5 | Q9CQH3 | 0.3 | 0.07 | 0.35 | 0.28 | 0.3 | 1.37 | -0.27 | -0.05 | 0.6 | 0.88 | -1 | -0.43 | 0.19 | 1.61 | 0.1 | -0.17 | 0.0875 | -0.2375 | 0.1825 | 0.808117 | 0.577171 | 0.641444 | 0.999971 | 1 | 0.93301 |
| Ndufb6 | Q3UIU2 | 0.22 | -0.37 | -0.17 | -0.47 | -0.38 | -0.52 | -0.43 | -0.56 | -0.58 | -0.43 | -0.77 | -0.36 | -0.54 | -0.5 | -0.44 | -0.62 | -0.275 | -0.3375 | -0.3275 | 0.210757 | 0.149107 | 0.143968 | 0.999971 | 1 | 0.78996 |
| Ndufb7 | Q9CR61 | -0.02 | 1.03 | 0.15 | 0.56 | 0.6 | 0.74 | -0.2 | 0.22 | 0.62 | 0.27 | -0.3 | 0.07 | 1.51 | 1 | 0.15 | 0.5 | -0.09 | -0.265 | 0.36 | 0.777395 | 0.400107 | 0.341702 | 0.999971 | 1 | 0.861042 |
| Ndufb8 | Q9D6J5 | -0.21 | -0.71 | -0.71 | -0.46 | -0.82 | -0.99 | -0.92 | -0.58 | -1 | -0.89 | -0.57 | -0.5 | -0.97 | -0.7 | -0.61 | -0.46 | -0.305 | -0.2175 | -0.1625 | 0.16081 | 0.324863 | 0.449922 | 0.999971 | 1 | 0.885596 |
| Ndufb9 | Q9CQJ8 | -1.13 | -1.85 | -0.26 | -0.41 | -1.76 | -1.99 | -0.34 | -0.03 | -1.92 | -1.9 | -0.17 | -0.73 | -2.03 | -1.73 | -0.08 | -0.41 | -0.1175 | -0.2675 | -0.15 | 0.834875 | 0.615005 | 0.787375 | 0.999971 | 1 | 0.96724 |
| Ndufc2 | Q9CQ54 | -0.94 | -0.62 | -1 | -1.27 | -0.71 | -1 | -0.7 | -1.24 | -0.89 | -1.08 | -0.94 | -0.92 | -0.85 | -0.99 | -1.28 | -1.59 | 0.045 | 1.57E-16 | -0.22 | 0.842317 | 1 | 0.377683 | 0.999971 | 1 | 0.871417 |
| Ndufs1 | Q91VD9 | -0.59 | 0.2 | -0.25 | -0.38 | -0.02 | -0.01 | -0.23 | -0.33 | -0.16 | -0.85 | -0.39 | -0.4 | 0.35 | -0.05 | -0.24 | -0.46 | 0.1075 | -0.195 | 0.155 | 0.635641 | 0.439364 | 0.55926 | 0.999971 | 1 | 0.915051 |
| Ndufs1_1 | A0A087WS | 0.28 | 0.78 | 1.87 | 0.46 | -0.23 | 1.39 | 1.33 | 1.6 | -0.26 | 1.08 | 1.62 | -0.18 | -0.06 | 1.11 | 0.84 | 2.27 | 0.175 | -0.2825 | 0.1925 | 0.732388 | 0.605647 | 0.727231 | 0.999971 | 1 | 0.956041 |
| Ndufs2 | Q91WD5 | -1.05 | -0.55 | -0.65 | -0.76 | -0.7 | -0.88 | -0.81 | -0.85 | -0.8 | -1.24 | -0.75 | -0.61 | -0.56 | -0.7 | -0.69 | -0.75 | -0.0575 | -0.0975 | 0.0775 | 0.758126 | 0.654059 | 0.680301 | 0.999971 | 1 | 0.942957 |
| Ndufs2_1 | A0A0A6YXI | 14.58 | 0.94 | -0.39 | -0.7 | 0.82 | 0.58 | -0.66 | -0.07 | 0.54 | 9.55 | -0.74 | -1.8 | 1.88 | 0.51 | -1.24 | -1.78 | -3.44 | -1.72 | -3.765 | 0.306465 | 0.668987 | 0.276263 | 0.999971 | 1 | 0.843794 |
| Ndufs3 | Q9DCT2 | -1.27 | -1.27 | -0.52 | -0.54 | -1.05 | -1.74 | -0.7 | -0.68 | -1.13 | -1.78 | -0.62 | -0.47 | -1.24 | -1.62 | -0.53 | -0.34 | -0.1425 | -0.1 | -0.0325 | 0.664646 | 0.779897 | 0.928096 | 0.999971 | 1 | 0.994278 |
| Ndufs4 | Q9CXZ1 | 0.17 | 0.11 | -0.03 | 0.3 | -0.31 | 0.32 | 0.35 | -0.18 | 0.15 | 0.35 | -0.16 | -0.25 | -0.16 | -0.1 | -0.06 | -0.32 | -0.0925 | -0.115 | -0.2975 | 0.681331 | 0.577907 | 0.117866 | 0.999971 | 1 | 0.78268 |
| Ndufs5 | Q99LY9 | 0.5 | 0.61 | 1.08 | 0.77 | 1.21 | 0.69 | 1.07 | 0.65 | 1.26 | 0.32 | 0.9 | 0.86 | 1.85 | 0.32 | 1.04 | 0.45 | 0.165 | 0.095 | 0.175 | 0.474564 | 0.710352 | 0.63144 | 0.999971 | 1 | 0.932858 |
| Ndufs6 | P52503 | -0.19 | -0.44 | -0.28 | -0.89 | -0.68 | 0.16 | -0.32 | -0.56 | -0.03 | 0.31 | -1.13 | -1.21 | 0.38 | -0.09 | -0.4 | -0.69 | 0.1 | -0.065 | 0.25 | 0.705849 | 0.87034 | 0.398264 | 0.999971 | 1 | 0.871816 |
| Ndufs7 | Q9DC70 | -3.17 | -0.26 | -1.83 | -4.08 | 0.21 | -1.73 | -0.95 | -3.7 | 0.62 | -3.14 | -1.35 | -1.11 | 0.96 | -2.98 | -3.12 | -4.27 | 0.7925 | 1.09 | -0.0175 | 0.454708 | 0.300815 | 0.988877 | 0.999971 | 1 | 0.99925 |
| Ndufs8 | Q8K3J1 | 0.39 | -0.01 | 0.03 | -0.01 | -0.37 | 0.26 | 0.51 | -0.34 | -0.14 | -0.19 | -0.04 | -0.58 | -0.62 | -0.27 | -0.2 | -0.69 | -0.085 | -0.3375 | -0.545 | 0.746731 | 0.125049 | 0.026717 | 0.999971 | 1 | 0.678474 |
| Ndufv1 | D3YUM1 | -1.09 | -0.44 | -0.54 | -0.65 | -0.27 | -0.7 | -0.36 | -0.71 | -0.14 | -1.14 | -0.3 | -0.25 | 0.11 | -0.74 | -0.6 | -0.81 | 0.17 | 0.2225 | 0.17 | 0.456232 | 0.441299 | 0.5393 | 0.999971 | 1 | 0.910077 |
| Ndufv2 | Q9D6J6 | -0.64 | -0.17 | -0.06 | -0.11 | -0.29 | -0.54 | -0.22 | 0.24 | -0.42 | -0.8 | -0.05 | 0.37 | -0.22 | -0.4 | 0.17 | -0.01 | 0.0425 | 0.02 | 0.13 | 0.860689 | 0.945635 | 0.566284 | 0.999971 | 1 | 0.917959 |
| Ndufv3 | Q3U422 | -0.56 | -1.52 | -0.95 | -0.11 | -1.44 | -2.08 | -0.58 | -0.95 | -1.05 | -1.15 | -1.11 | -1.05 | -1.42 | -1.61 | -1.09 | -1.23 | -0.4775 | -0.305 | -0.5525 | 0.272101 | 0.333071 | 0.113757 | 0.999971 | 1 | 0.78268 |
| Ndufv3_1 | Q8BK30 | -2.74 | -5.48 | -6.49 | -6.91 | -6.09 | -5.7 | -6.15 | -6.67 | -6.01 | -4.36 | -5.77 | -7.35 | -8.42 | -5.26 | -7.61 | -7.04 | -0.7475 | -0.4675 | -1.6775 | 0.394674 | 0.644323 | 0.130217 | 0.999971 | 1 | 0.784948 |
| Nebl | Q9DC07 | 0.43 | 0.39 | 0.56 | 0.52 | 0.04 | 0.22 | 0.59 | 0.39 | 0.03 | 0.34 | 0.07 | -0.22 | -0.56 | -0.12 | -0.16 | -0.04 | -0.165 | -0.42 | -0.695 | 0.39727 | 0.047758 | 0.005214 | 0.999971 | 1 | 0.582714 |
| Necab1 | Q8BG18 | -0.22 | 0.03 | -1.08 | -0.71 | 0.56 | 0.03 | -0.42 | 0.07 | 0.26 | -0.14 | 0.02 | -0.3 | 0.28 | 0.27 | -0.48 | -0.45 | 0.555 | 0.455 | 0.4 | 0.11102 | 0.139905 | 0.243136 | 0.999971 | 1 | 0.837316 |
| Necab2 | Q91ZP9 | 0.04 | -0.21 | -0.02 | 0.01 | 0.16 | 0.04 | -0.15 | 0.61 | 0.63 | 0.27 | -0.38 | 0.37 | 0.65 | 0.51 | 0.6 | 0.18 | 0.21 | 0.2675 | 0.53 | 0.346214 | 0.297991 | 0.019572 | 0.999971 | 1 | 0.635388 |
| Necab3 | Q9D6J4 | 0.78 | 0.02 | -0.37 | 0.37 | 0.08 | 0.24 | -0.52 | 0.3 | 0.13 | 1.41 | 0.08 | -0.34 | 0.17 | -0.02 | -0.45 | 0.33 | -0.175 | 0.12 | -0.1925 | 0.579472 | 0.779508 | 0.53292 | 0.999971 | 1 | 0.910069 |
| Necap1 | Q9CR95 | -0.53 | -0.71 | -1.04 | -1.05 | -0.75 | -0.45 | -0.15 | -1.16 | -0.71 | -0.83 | -1.04 | -0.91 | -0.74 | -0.56 | -1.05 | -1.12 | 0.205 | -0.04 | -0.035 | 0.455423 | 0.841175 | 0.876576 | 0.999971 | 1 | 0.985129 |
| Necap2 | Q9D1J1 | 0.06 | 0.13 | -0.12 | 0.02 | 0.11 | 0.18 | -0.15 | -0.23 | 0.25 | 0.06 | 0 | 0.28 | 0.44 | 0.5 | 0.1 | 0.57 | -0.045 | 0.125 | 0.38 | 0.808031 | 0.471275 | 0.0693 | 0.999971 | 1 | 0.733937 |
| Nectin1 | Q9JKF6 | -0.66 | -1.47 | -2.98 | -2.32 | -1.2 | -1.31 | -2.54 | -2.47 | -1.16 | -0.82 | -2.55 | -2.44 | -1.11 | -1.28 | -2.61 | -2.97 | -0.0225 | 0.115 | -0.135 | 0.968418 | 0.851482 | 0.829614 | 0.999971 | 1 | 0.978384 |
| Nectin2 | P32507-2 | 1.1 | 0.88 | -0.13 | 0.32 | 0.52 | 0.73 | -0.45 | 0.23 | 1.06 | 1.69 | -0.69 | 0.15 | 1.66 | 1.28 | 0.45 | 0.67 | -0.285 | 0.01 | 0.4725 | 0.446379 | 0.985369 | 0.234505 | 0.999971 | 1 | 0.832918 |
| Nectin3 | Q9JLB9 | -0.71 | -1.77 | -1.48 | -1.52 | -1.99 | -1.36 | -1.63 | -1.31 | -1.93 | -1.32 | -1.16 | -1.15 | -1.2 | -1.36 | -1.5 | -0.56 | -0.2025 | -0.02 | 0.215 | 0.490039 | 0.947017 | 0.499432 | 0.999971 | 1 | 0.903717 |
| Nedd4 | P46935 | 0.56 | -0.14 | 0 | 0.24 | 0.14 | 0.02 | -0.07 | 0.39 | 0.08 | 0.33 | -0.31 | -0.07 | -0.56 | -0.05 | 0.02 | -0.15 | -0.045 | -0.1575 | -0.35 | 0.840992 | 0.509856 | 0.163983 | 0.999971 | 1 | 0.800215 |
| Nedd4l | A0A494B9! | -0.93 | -0.49 | -0.73 | -0.77 | -0.52 | -0.77 | -0.61 | -0.9 | -0.83 | -0.98 | -0.63 | -0.59 | -0.5 | -0.79 | -0.9 | -0.69 | 0.03 | -0.0275 | 0.01 | 0.875007 | 0.884981 | 0.958353 | 0.999971 | 1 | 0.998622 |
| Nedd8 | P29595 | 1.91 | 0.3 | -0.06 | 0.36 | 0.31 | 0.82 | 0.28 | 0.5 | 0.21 | 1.54 | -0.06 | -0.94 | -0.21 | 0.64 | -0.25 | 0.08 | -0.15 | -0.44 | -0.5625 | 0.727729 | 0.483074 | 0.2361 | 0.999971 | 1 | 0.832918 |
| Nefh | P19246 | 0.51 | -1.33 | -1.17 | -0.88 | -1.34 | -0.92 | -0.73 | -1.3 | -1.17 | -0.01 | -1.69 | -1.1 | -1.11 | -1.21 | -0.62 | -1.55 | -0.355 | -0.275 | -0.405 | 0.409677 | 0.592788 | 0.3653 | 0.999971 | 1 | 0.869526 |
| Nefl | P08551 | -1.37 | -1.62 | -1.25 | -1.63 | -1.71 | -0.81 | -1.24 | -1.36 | -1.35 | -1.28 | -1.55 | -1.22 | -1.55 | -1.16 | -0.94 | -1.25 | 0.1875 | 0.1175 | 0.2425 | 0.443969 | 0.530097 | 0.26587 | 0.999971 | 1 | 0.842677 |
| Nefm | A0A0R4J03 | -1.16 | -1.26 | -0.91 | -0.99 | -1.08 | -0.93 | -1.01 | -0.65 | -0.68 | -1.04 | -0.79 | -0.72 | -0.11 | -0.59 | -0.34 | -0.57 | 0.1625 | 0.2725 | 0.6775 | 0.403425 | 0.160954 | 0.007456 | 0.999971 | 1 | 0.582714 |
| Negr1 | D3Z4T6 | -1.38 | -0.95 | -0.85 | -1.39 | -0.71 | -0.87 | -0.6 | -0.98 | -0.8 | -1.19 | -1.04 | -0.51 | -0.27 | -0.98 | -0.99 | -1.53 | 0.3525 | 0.2575 | 0.2 | 0.124967 | 0.293762 | 0.514381 | 0.999971 | 1 | 0.905815 |
| Nek1 | B2RXQ4 | 0.26 | 0.21 | 0.93 | 0.92 | 0.02 | -0.21 | 0.04 | 1.2 | 0.06 | 0.16 | 1.41 | 1.49 | -0.42 | 0.26 | 0.77 | 0.77 | -0.3175 | 0.2 | -0.235 | 0.394825 | 0.632531 | 0.4981 | 0.999971 | 1 | 0.902911 |
| Nek4 | A0A2I3BPN | -0.22 | -0.13 | -0.28 | -0.26 | -0.08 | 0.13 | -0.42 | 0.39 | -0.35 | -0.05 | -0.49 | -0.05 | 0.02 | 0.04 | -0.14 | 0.01 | 0.2275 | -0.0125 | 0.205 | 0.314048 | 0.945645 | 0.232632 | 0.999971 | 1 | 0.831128 |
| Nek6 | Q9ES70 | -1.68 | -2.46 | -2.11 | -1.65 | -2.14 | -2.49 | -1.86 | -2.24 | -2.29 | -2.43 | -2.05 | -2.97 | -2.98 | -1.85 | -2.33 | -2.3 | -0.2075 | -0.46 | -0.39 | 0.428798 | 0.134896 | 0.227459 | 0.999971 | 1 | 0.828563 |
| Nek7 | Q9ES74 | -0.54 | -0.09 | -0.01 | -0.17 | -0.19 | 0.32 | -0.39 | -0.3 | -0.28 | -0.25 | -0.22 | -0.27 | -0.34 | 0.2 | 0.01 | 0.15 | 0.0625 | -0.0525 | 0.2075 | 0.789151 | 0.776572 | 0.35058 | 0.999971 | 1 | 0.866844 |
| Nek9 | Q8K1R7 | 0.95 | 1.04 | 0.37 | 0.24 | 1.07 | 1.23 | 0.52 | 0.62 | 1.88 | 1 | 0.34 | 0.61 | 0.82 | 0.74 | 0.31 | -0.03 | 0.21 | 0.3075 | -0.19 | 0.461121 | 0.426689 | 0.523147 | 0.999971 | 1 | 0.907797 |
| Nelfa | Q8BG30 | 1.28 | 1.52 | 1.35 | 1.11 | 1.46 | 0.99 | 1.17 | 0.96 | 1.55 | 1.12 | 1.16 | 1.25 | 0.86 | 1.46 | 0.84 | 1.28 | -0.17 | -0.045 | -0.205 | 0.406575 | 0.813352 | 0.365458 | 0.999971 | 1 | 0.869526 |
| Nelfb | A0A5H1ZRI | 1.85 | 1.01 | 0.99 | 0.67 | 1.17 | 1.11 | 0.82 | 0.7 | 1.2 | 1.26 | 1.14 | 0.95 | 0.82 | 0.67 | 1.03 | 1.15 | -0.18 | 0.0075 | -0.2125 | 0.53756 | 0.978338 | 0.467125 | 0.999971 | 1 | 0.891731 |
| Nelfcd | Q3TW27 | 0.18 | 0.61 | 2.12 | 2.45 | 0.74 | 0.56 | 2.89 | 1.28 | 0.25 | 0.12 | 1.3 | 0.69 | 1.14 | 1.14 | 1.5 | 0.82 | 0.0275 | -0.75 | -0.19 | 0.968402 | 0.210028 | 0.720737 | 0.999971 | 1 | 0.954713 |
| Nelfe | P19426 | -0.64 | 1.08 | 1.02 | 1.11 | 1.63 | 0.43 | -0.02 | 0.59 | 0.26 | -1.77 | -0.16 | -0.13 | 0.1 | 1.35 | 0.51 | 1.41 | 0.015 | -1.0925 | 0.2 | 0.976543 | 0.083771 | 0.688902 | 0.999971 | 1 | 0.945805 |
| Nell2 | A0A2R8VH | 0.47 | 1.95 | 1.75 | 1.07 | 1.94 | 1.27 | 2.09 | 1.68 | 2.09 | 0.65 | 1.98 | 1.87 | 2.09 | 1.1 | 1.59 | 1.5 | 0.435 | 0.3375 | 0.26 | 0.259547 | 0.459637 | 0.500818 | 0.999971 | 1 | 0.904223 |
| Nemf | Q8CCP0 | 0.92 | 0.81 | 0.37 | 0.66 | 0.71 | 0.41 | 0.34 | 0.52 | 0.67 | 0.66 | 0.36 | 0.33 | 0.55 | 0.69 | 0.75 | 0.51 | -0.195 | -0.185 | -0.065 | 0.345782 | 0.373029 | 0.740242 | 0.999971 | 1 | 0.960899 |
| Nenf | Q9CQ45 | 0.69 | 1.08 | -0.64 | -0.06 | 0.43 | 1.11 | -0.63 | 0.01 | 0.34 | 0.76 | -0.89 | -0.72 | 0.77 | 1.01 | -0.12 | -0.41 | -0.0375 | -0.395 | 0.045 | 0.939213 | 0.452232 | 0.925243 | 0.999971 | 1 | 0.99401 |
| Neo1 | E9QK04 | -0.03 | -0.59 | -0.74 | -0.58 | -0.58 | -0.76 | -0.67 | -0.49 | -0.1 | 0.03 | -0.51 | -0.46 | 0.01 | -0.56 | -0.71 | -0.35 | -0.14 | 0.225 | 0.0825 | 0.518498 | 0.354819 | 0.741715 | 0.999971 | 1 | 0.961064 |
| Nes | Q6P5H2-2 | 2.54 | 1.78 | 1.34 | 2.35 | 1.55 | 1.32 | 1.46 | 1.13 | 1.3 | 1.95 | 1.08 | 1.19 | 0.71 | 1.6 | 2.26 | 1.83 | -0.6375 | -0.6225 | -0.4025 | 0.057043 | 0.090109 | 0.336674 | 0.999971 | 1 | 0.858827 |
| Neto1 | Q8R4I7 | 0.45 | -0.04 | 0 | 0.04 | -0.15 | 0.63 | 0.02 | 0.3 | -0.1 | 0.6 | -0.39 | -0.31 | 0.28 | 0.15 | -0.25 | -0.34 | 0.0875 | -0.1625 | -0.1525 | 0.714695 | 0.551406 | 0.510243 | 0.999971 | 1 | 0.905815 |
| Neto2 | B2RXQ6 | -1.4 | -1.06 | -1.09 | -0.72 | -0.9 | -1.06 | -0.91 | -0.75 | -0.96 | -1.06 | -0.92 | -0.5 | -0.39 | -0.98 | -0.65 | -0.7 | 0.1625 | 0.2075 | 0.3875 | 0.438882 | 0.366824 | 0.113306 | 0.999971 | 1 | 0.78268 |
| Neu1 | O35657 | 0.52 | 2.71 | 0.92 | -0.17 | 2.8 | 1.7 | 0.07 | 1.93 | 3.46 | 0.77 | 2.52 | -0.08 | 4.18 | 1.23 | 0.42 | 0.33 | 0.63 | 0.6725 | 0.545 | 0.413371 | 0.468747 | 0.580186 | 0.999971 | 1 | 0.91994 |
| Neu2 | E9Q0P5 | 4.11 | 0.58 | 1.12 | 2.14 | -0.6 | 0.06 | 1.27 | 0.73 | -0.66 | 2.1 | 1.3 | 0.32 | -6.21 | -1.57 | 0.69 | 0.17 | -1.6225 | -1.2225 | -3.7175 | 0.065581 | 0.191345 | 0.038142 | 0.999971 | 1 | 0.700838 |
| Neu3 | Q9JMH7 | -2.95 | -0.37 | -1.78 | -1.43 | -0.22 | -0.48 | -1.6 | -1.24 | -0.19 | -1.51 | -2.06 | -1.65 | 0.15 | -0.08 | -1.16 | -1.3 | 0.7475 | 0.28 | 1.035 | 0.211461 | 0.649484 | 0.108105 | 0.999971 | 1 | 0.78268 |
| Neu4 | Q8BZL1 | -0.57 | 0.4 | -0.94 | 0 | 0.14 | -0.43 | -0.2 | -1 | 0.73 | -0.12 | -0.91 | -0.71 | 1.03 | -0.29 | -1.49 | -1.25 | -0.095 | 0.025 | -0.2225 | 0.797783 | 0.9553 | 0.707089 | 0.999971 | 1 | 0.950182 |
| Neurl1 | Q923S6 | -1.33 | 2.5 | -0.97 | -0.19 | 3.98 | 1.31 | -0.85 | 0.34 | 4 | -0.59 | 0.01 | -0.17 | 6.28 | 0.08 | -0.57 | 0.16 | 1.1925 | 0.81 | 1.485 | 0.332312 | 0.51642 | 0.370706 | 0.999971 | 1 | 0.869526 |
| Neurl4 | H3BK72 | 0.29 | -0.42 | -0.27 | -0.27 | -0.32 | -0.44 | -0.5 | -0.44 | -0.49 | 0.21 | -0.11 | -0.54 | -0.45 | -0.05 | -0.32 | -0.22 | -0.2575 | -0.065 | -0.0925 | 0.242253 | 0.801269 | 0.677977 | 0.999971 | 1 | 0.942594 |
| Nexn | A0A0G2JE) | 0.51 | -0.74 | -1.07 | 0.18 | -0.25 | -1.76 | -1.67 | 0.37 | -0.61 | 0.49 | -0.81 | 0.27 | -0.78 | -0.05 | 0.03 | -0.74 | -0.5475 | 0.115 | -0.105 | 0.36604 | 0.803727 | 0.798912 | 0.999971 | 1 | 0.970479 |
| Nf1 | Q04690-2 | -0.8 | 0 | -0.11 | -0.24 | 0.11 | 0.04 | -0.14 | -0.18 | 0.34 | -0.53 | -0.09 | -0.18 | 0.52 | -0.08 | -0.01 | 0.06 | 0.245 | 0.1725 | 0.41 | 0.301448 | 0.527752 | 0.130039 | 0.999971 | 1 | 0.784948 |
| Nf1_1 | Q5SYH9 | 1.17 | -0.45 | -0.31 | -0.24 | -0.67 | 0.15 | -0.62 | -0.55 | -0.35 | 0.89 | -0.93 | -0.17 | -0.74 | -0.03 | -0.38 | -0.72 | -0.465 | -0.1825 | -0.51 | 0.269019 | 0.715747 | 0.220732 | 0.999971 | 1 | 0.827096 |
| Nf2 | P46662-2 | -0.08 | 0.27 | -1.42 | -1.02 | 0.4 | 0 | -1.25 | -0.89 | 0.26 | 0 | -1.22 | -1.4 | 0.47 | -0.59 | -1.28 | -0.92 | 0.1275 | -0.0275 | -0.0175 | 0.802295 | 0.958914 | 0.972414 | 0.999971 | 1 | 0.99925 |
| Nfasc | A0A087WF | -0.45 | -0.45 | -0.6 | -0.76 | -0.74 | -0.35 | -0.69 | -0.76 | -0.74 | -0.68 | -0.95 | -0.75 | -0.4 | -0.54 | -0.54 | -0.81 | -0.07 | -0.215 | -0.0075 | 0.712608 | 0.236967 | 0.96785 | 0.999971 | 1 | 0.998787 |
| Nfasc_1 | Q810U3 | -0.61 | -1.97 | -3.52 | -3.8 | -2.15 | -2.33 | -3.54 | -4.27 | -1.83 | -1.82 | -3.89 | -4.59 | -2.62 | -1.73 | -3.99 | -3.99 | -0.5975 | -0.5575 | -0.6075 | 0.46502 | 0.550996 | 0.471957 | 0.999971 | 1 | 0.892857 |
| Nfasc_2 | E9Q171 | 0.51 | 1.29 | -0.8 | -1.59 | -0.72 | 2.01 | -1.72 | -1.39 | 0.45 | 1.13 | -0.89 | -1.33 | -0.59 | 1.15 | -2.13 | -2.57 | -0.3075 | -0.0125 | -0.8875 | 0.747192 | 0.987232 | 0.362763 | 0.999971 | 1 | 0.869526 |
| Nfat5 | A0A1U6UR | -0.52 | 0.03 | 0.1 | 0.04 | 0.06 | -0.02 | 0.2 | -0.09 | 0.11 | -0.03 | -0.17 | 0.32 | -0.18 | 0.17 | 0.2 | -0.12 | 0.125 | 0.145 | 0.105 | 0.553931 | 0.514295 | 0.635139 | 0.999971 | 1 | 0.93301 |
| Nfia | Q02780-3 | 0.98 | 2.12 | 1.81 | 2.45 | 1.99 | 1.71 | 1.77 | 2.15 | 1.51 | 1.64 | 1.56 | 1.81 | 0.64 | 1.4 | 1.71 | 2.57 | 0.065 | -0.21 | -0.26 | 0.843963 | 0.520546 | 0.587415 | 0.999971 | 1 | 0.920758 |
| Nfib | A2BG76 | -1.22 | 4.3 | 3.54 | 2.36 | 3.03 | 3.85 | 2.08 | 2.87 | 3.22 | 1.02 | 2.62 | 2.48 | 2.81 | 3.99 | 2.42 | 3.36 | 0.7125 | 0.09 | 0.9 | 0.534452 | 0.938757 | 0.434734 | 0.999971 | 1 | 0.882419 |
| Nfic | P70255-3 | 1.56 | 0.19 | 0.78 | 0.51 | 0.61 | 1.27 | -0.29 | 0.42 | 0.38 | 0.44 | -0.12 | -0.49 | 1.27 | 1.07 | -1.06 | 1.45 | -0.2575 | -0.7075 | -0.0775 | 0.537342 | 0.075417 | 0.896973 | 0.999971 | 1 | 0.988737 |
| Nfix | P70257-1 | 0.72 | 3.33 | 2.63 | 2.11 | 2.39 | 3.04 | 1.67 | 2.21 | 2.37 | 1.31 | 2.15 | 1.89 | 2.52 | 2.82 | 1.67 | 2.75 | 0.13 | -0.2675 | 0.2425 | 0.819148 | 0.630279 | 0.667941 | 0.999971 | 1 | 0.938711 |
| Nfkb1 | P25799-4 | -0.45 | 0.65 | -0.51 | -0.11 | 0.25 | 0.12 | -0.65 | -0.07 | -0.21 | -0.6 | -0.75 | -0.04 | 0.29 | 0.38 | -0.02 | 0.01 | 0.0175 | -0.295 | 0.27 | 0.957843 | 0.363709 | 0.372681 | 0.999971 | 1 | 0.869526 |
| Nfrkb | Q6PIJ4 | 3.07 | 0.18 | -0.5 | -0.11 | 0.09 | 0.92 | 0.13 | -0.21 | 0.61 | 2 | -1.11 | -0.95 | -1.12 | -0.19 | 0.92 | 0.15 | -0.4275 | -0.5225 | -0.72 | 0.57972 | 0.598892 | 0.393818 | 0.999971 | 1 | 0.871816 |
| Nfs1 | Q9Z1J3 | 0.37 | 0 | 0.12 | 0.18 | -0.27 | 0.18 | -0.27 | 0.28 | -0.19 | 0.31 | 0.18 | -0.1 | 0.04 | 0.1 | 0.12 | 0.26 | -0.1875 | -0.1175 | -0.0375 | 0.39004 | 0.553991 | 0.831576 | 0.999971 | 1 | 0.978898 |
| Nfu1 | Q9QZ23 | -0.46 | -0.37 | -0.27 | -0.63 | -0.07 | -0.36 | -0.08 | -0.48 | -0.05 | -0.15 | 0.06 | 0.54 | 0.31 | 0.05 | -0.47 | -0.01 | 0.185 | 0.5325 | 0.4025 | 0.350062 | 0.032972 | 0.098029 | 0.999971 | 1 | 0.78268 |
| Nfx1 | B1AY10 | 0.28 | 0.34 | 0.2 | 0.14 | 0.09 | 0.4 | 0.06 | 0.92 | -0.01 | -0.07 | 0.65 | 0.5 | 0.09 | 0.34 | 0.74 | 0.6 | 0.1275 | 0.0275 | 0.2025 | 0.595213 | 0.90228 | 0.338151 | 0.999971 | 1 | 0.859348 |
| Nfxl1 | E9Q8I7 | 0.9 | 0.75 | 0.31 | 0.85 | 1.03 | 0.99 | 0.12 | 0.84 | 1.1 | 0.93 | 0.21 | -0.75 | 0.61 | 1.26 | 0.15 | 0.5 | 0.0425 | -0.33 | -0.0725 | 0.875102 | 0.441008 | 0.798109 | 0.999971 | 1 | 0.970479 |
| Nfyc | P70353 | 0.87 | 2.17 | 2.02 | 2.5 | 1.5 | 0.76 | 1 | 2.2 | 1.57 | 1.19 | 2.15 | 2.13 | 2.28 | 1.12 | 3.19 | 2.21 | -0.525 | -0.13 | 0.31 | 0.259639 | 0.749194 | 0.548936 | 0.999971 | 1 | 0.911727 |
| Ngb | D3YW07 | -2.87 | -1.1 | -1.16 | -3.36 | -0.39 | -0.04 | -0.36 | -0.87 | -1.99 | -3.86 | -2.04 | 0.46 | 1.24 | -0.36 | -0.44 | -0.12 | 1.7075 | 0.265 | 2.2025 | 0.012804 | 0.781591 | 0.007138 | 0.999971 | 1 | 0.582714 |
| Ngef | Q8CHT1-2 | -0.77 | -0.84 | -1.55 | -1.18 | -1.19 | -1.23 | -1.37 | -1.14 | -1.31 | -1.17 | -1.63 | -1.51 | -1.2 | -1.08 | -1.47 | -1.25 | -0.1475 | -0.32 | -0.165 | 0.519127 | 0.201179 | 0.486561 | 0.999971 | 1 | 0.897675 |
| Ngly1 | Q9JI78 | 0.07 | 0.29 | 0.18 | -0.01 | 0.3 | 0.11 | -0.11 | 0.6 | 0.16 | 0.15 | 0.2 | 0.08 | 0.03 | 0.22 | 0.09 | -0.07 | 0.0925 | 0.015 | -0.065 | 0.665268 | 0.927337 | 0.712576 | 0.999971 | 1 | 0.951793 |
| Ngp | O08692 | 3.56 | 1 | -0.68 | 1.75 | -1.77 | -2.55 | 1.84 | -0.34 | -2.11 | -1.85 | -1.18 | -2.09 | -4.19 | -1.31 | -2.01 | -1.71 | -2.1125 | -3.215 | -3.7125 | 0.096037 | 0.003578 | 0.004153 | 0.999971 | 0.959217 | 0.582714 |
| Nhlrc2 | Q8BZW8 | 0.22 | 0.65 | 1.46 | 1.3 | 0.61 | 0.5 | 1.22 | 0.76 | 0.36 | 0.32 | 0.71 | 1.02 | 0.66 | 0.29 | 0.6 | 1.03 | -0.135 | -0.305 | -0.2625 | 0.682569 | 0.368817 | 0.4308 | 0.999971 | 1 | 0.881611 |
| Nhlrc3 | Q8CCH2 | -0.44 | 0.57 | -0.25 | -0.47 | 0.34 | -0.03 | -0.37 | 0.16 | 0.09 | -1.26 | 0.01 | -0.65 | 0.68 | 0.3 | 0.15 | -0.52 | 0.1725 | -0.305 | 0.3 | 0.564928 | 0.437341 | 0.395813 | 0.999971 | 1 | 0.871816 |
| Nhp2 | Q9CRB2 | 0.15 | -0.21 | 0.14 | -0.16 | -0.62 | -0.65 | -0.27 | -0.07 | -0.67 | -0.33 | -0.35 | -0.9 | -1.04 | -0.47 | -0.27 | -0.74 | -0.3825 | -0.5425 | -0.61 | 0.104263 | 0.029217 | 0.025343 | 0.999971 | 1 | 0.678474 |
| Nhsl1 | A0A140LIZ | 0.67 | -0.07 | -0.06 | 0.17 | 0.1 | 0.44 | 0.43 | 0.13 | -0.05 | 0.64 | -0.79 | -0.37 | -0.07 | 0.17 | -0.46 | -0.57 | 0.0975 | -0.32 | -0.41 | 0.676881 | 0.363951 | 0.149075 | 0.999971 | 1 | 0.793248 |
| Nhsl2 | B1AXH1 | 0.34 | -0.05 | -0.21 | -0.25 | -0.02 | -0.11 | -0.49 | 0.17 | -0.22 | 0.29 | -0.44 | -0.05 | 0.13 | 0.22 | -0.1 | 0.01 | -0.07 | -0.0625 | 0.1075 | 0.762172 | 0.791772 | 0.604914 | 0.999971 | 1 | 0.924454 |
| Niban1 | E9PYV4 | -0.56 | 1.16 | 0.32 | 0.34 | 1.08 | 0.58 | -0.21 | -0.11 | 0.5 | -0.19 | 0.19 | 0.36 | 1.26 | 0.19 | 0.3 | 0.23 | 0.02 | -0.1 | 0.18 | 0.963526 | 0.787724 | 0.66529 | 0.999971 | 1 | 0.93697 |
| Niban2 | Q8R1F1 | 0.2 | 0.29 | 0.22 | 0.73 | 0.7 | 0.34 | 0.03 | -0.12 | 0.68 | 0.41 | -0.01 | 0.11 | 0.36 | 0.15 | 0.25 | 0.64 | -0.1225 | -0.0625 | -0.01 | 0.62471 | 0.788604 | 0.962549 | 0.999971 | 1 | 0.998721 |
| Nicn1 | Q9CQM0 | -0.68 | -0.6 | 0.79 | 0.34 | -0.01 | -0.48 | -0.66 | 0.08 | -0.56 | -0.7 | 1.59 | 1.04 | -0.23 | 0.14 | 1.46 | 1.77 | -0.23 | 0.38 | 0.8225 | 0.556123 | 0.544503 | 0.166096 | 0.999971 | 1 | 0.800215 |
| Nid1 | P10493 | 0.95 | 0.42 | -0.06 | 0.57 | 2.06 | 0.63 | 0.43 | -0.31 | 0.98 | 1.85 | 1.18 | 1.39 | 0.01 | 0.75 | 0.9 | 0.25 | 0.2325 | 0.88 | 0.0075 | 0.642981 | 0.013763 | 0.980264 | 0.999971 | 1 | 0.99925 |
| Nid2 | O88322 | 0.88 | -0.04 | -0.24 | 0.41 | 0.9 | 0.01 | 0.15 | -0.84 | 0.18 | 0.77 | -0.09 | 0.25 | -0.96 | -0.05 | 0.02 | -0.34 | -0.1975 | 0.025 | -0.585 | 0.635059 | 0.936007 | 0.107188 | 0.999971 | 1 | 0.78268 |
| Nif3l1 | Q9EQ80 | 0.66 | 0.14 | 0.25 | -0.07 | -0.04 | 0.26 | 0.94 | -0.12 | 0.1 | 0.62 | -0.17 | -0.29 | -0.49 | 0.06 | -0.04 | -0.24 | 0.015 | -0.18 | -0.4225 | 0.959486 | 0.512227 | 0.09714 | 0.999971 | 1 | 0.78268 |
| Nifk | Q91VE6-2 | 1.81 | -0.47 | 0.32 | 0 | -0.33 | 0.02 | 0.28 | 0.29 | -0.48 | 0.61 | 0.16 | 0.09 | -0.23 | -0.06 | 0.26 | -0.33 | -0.35 | -0.32 | -0.505 | 0.471361 | 0.53007 | 0.304379 | 0.999971 | 1 | 0.847402 |
| Nin | A0A1Y7VN | 2.24 | 3.06 | 2.04 | 1.33 | 1.93 | 2.43 | 1.38 | 2.12 | 3.09 | 2.68 | 0.81 | 0.65 | 2.48 | 2.58 | 2.21 | 2.49 | -0.2025 | -0.36 | 0.2725 | 0.615228 | 0.588239 | 0.453696 | 0.999971 | 1 | 0.887217 |

| | | | | | | | | | | | | | | | | | | | | | | | | | | |
|---|---|---|---|---|---|---|---|---|---|---|---|---|---|---|---|---|---|---|---|---|---|---|---|---|---|---|
| Nip7 | E9Q109 | 1.66 | -0.18 | 0.53 | 0.05 | -0.45 | 0.08 | 0.13 | 0.45 | -0.25 | 1.58 | -0.04 | 0.33 | -1.27 | 0.07 | 0.04 | -0.14 | -0.4625 | -0.11 | -0.84 | 0.293808 | 0.837492 | 0.109852 | 0.999971 | 1 | 0.78268 |
| Nipa1 | Q8BHK1 | 1.13 | 1.25 | 1.35 | 1.29 | -0.03 | 1.53 | 0.73 | 1.08 | 1.06 | 0.94 | 1.07 | -0.2 | 0.12 | 1.06 | 0.68 | 1 | -0.4275 | -0.5375 | -0.54 | 0.218935 | 0.114724 | 0.054819 | 0.999971 | 1 | 0.725348 |
| Nipal3 | Q8BGN5 | 0.61 | -0.49 | -0.77 | -0.39 | -1.29 | 0.48 | -0.4 | -0.26 | -0.41 | 0.11 | -0.18 | -0.7 | -0.1 | -0.04 | -0.17 | -0.13 | -0.1075 | -0.035 | 0.15 | 0.808647 | 0.918536 | 0.629196 | 0.999971 | 1 | 0.932234 |
| Nipbl | Q6KCD5 | 0.75 | -0.67 | -0.39 | 0.09 | -0.4 | -0.41 | -0.22 | -0.09 | -0.61 | 0.3 | -0.7 | -1.56 | -0.49 | -0.66 | -0.44 | -0.56 | -0.225 | -0.5875 | -0.4825 | 0.490638 | 0.224871 | 0.155123 | 0.999971 | 1 | 0.795578 |
| Nipsnap1 | O55125 | -2.16 | -0.48 | 0.39 | -0.5 | -0.12 | -0.74 | 0.64 | -0.08 | -0.01 | -2.21 | 0.42 | -0.38 | 0.23 | -1.48 | 0.1 | -0.93 | 0.6125 | 0.1425 | 0.1675 | 0.285672 | 0.842084 | 0.785505 | 0.999971 | 1 | 0.967053 |
| Nipsnap2 | O55126 | -2.22 | -0.63 | -0.45 | -0.55 | -0.28 | -0.88 | -0.64 | -0.77 | -0.63 | -2.17 | -1.15 | -0.96 | -0.74 | -1.16 | -0.8 | -1.4 | 0.32 | -0.265 | -0.0625 | 0.451687 | 0.598679 | 0.883182 | 0.999971 | 1 | 0.986597 |
| Nipsnap3b | Q9CQE1 | 0.63 | 0.58 | -0.07 | 0.02 | 0.92 | 0.45 | -0.28 | 0 | 0.95 | 0.33 | -0.02 | 0.41 | 1.2 | 0.75 | 0.21 | 0.06 | -0.0175 | 0.1275 | 0.265 | 0.956621 | 0.65549 | 0.419068 | 0.999971 | 1 | 0.87788 |
| Nisch | Q80TM9 | -0.1 | 0.34 | 0.4 | 0.37 | 0.37 | 0.29 | 0.75 | 0.58 | 0.29 | 0.27 | 0.59 | 0.75 | 0.27 | 0.42 | 0.56 | 0.6 | 0.245 | 0.2225 | 0.21 | 0.25958 | 0.309373 | 0.308595 | 0.999971 | 1 | 0.847402 |
| Nit1 | Q8VDK1-2 | -0.56 | 0.03 | -0.04 | -0.31 | 0.02 | -0.3 | -0.32 | -0.4 | 0.03 | -0.45 | -0.6 | -0.41 | 0.5 | -0.07 | -0.16 | -0.16 | -0.03 | -0.1375 | 0.2475 | 0.887694 | 0.549966 | 0.32027 | 0.999971 | 1 | 0.853962 |
| Nit2 | Q9JHW2 | -0.38 | -0.07 | 0.01 | 0.24 | -0.09 | -0.06 | -0.12 | 0.24 | 0.02 | -0.32 | -0.07 | 0.1 | 0.47 | 0.07 | 0.13 | -0.07 | 0.0425 | -0.0175 | 0.2 | 0.836792 | 0.932336 | 0.371021 | 0.999971 | 1 | 0.869526 |
| Nkain4 | Q9JMG4 | -0.79 | -1.89 | -1.03 | 2.33 | -1.08 | -0.64 | -0.49 | -0.07 | -1.88 | -0.38 | 4.38 | 5.9 | -2.97 | -0.97 | 0.49 | 0.82 | -0.225 | 2.35 | -0.3125 | 0.790919 | 0.226366 | 0.78172 | 0.999971 | 1 | 0.965859 |
| Nkiras1 | Q8CEC5 | 1.06 | 0.93 | 0.66 | 0.96 | 0.64 | 0.83 | 0.7 | 0.96 | 0.77 | 0.65 | 0.97 | 0.16 | 0.7 | 0.76 | 0.61 | 0.69 | -0.12 | -0.265 | -0.2125 | 0.520722 | 0.264538 | 0.24834 | 0.999971 | 1 | 0.839012 |
| Nkiras2 | Q9CR56 | -0.4 | -0.95 | -1.06 | -1.21 | -0.5 | -0.82 | -1.12 | -0.72 | -0.56 | 0.03 | -0.63 | -1.21 | -0.76 | -0.56 | -0.83 | -0.49 | 0.115 | 0.3125 | 0.245 | 0.644178 | 0.332045 | 0.306719 | 0.999971 | 1 | 0.847402 |
| Nkrf | A0A668KL6 | -0.22 | 0.22 | -0.32 | -0.1 | -0.23 | -0.12 | -0.33 | -0.07 | -0.2 | 0.1 | -0.04 | -0.35 | 0.15 | 0.21 | -0.17 | 0.25 | -0.0825 | -0.0175 | 0.215 | 0.67249 | 0.931426 | 0.31207 | 0.999971 | 1 | 0.849271 |
| Nlgn1 | E9QK34 | -1.33 | -0.71 | -1.47 | -1.18 | -0.89 | -0.6 | -1.56 | -1.15 | -0.6 | -1.06 | -1.11 | -1.33 | -0.78 | -0.66 | -1.08 | -0.8 | 0.1225 | 0.1475 | 0.3425 | 0.661477 | 0.560308 | 0.158322 | 0.999971 | 1 | 0.798092 |
| Nlgn2 | Q69ZK9 | -0.89 | -1.17 | -0.77 | -0.46 | -1.1 | -0.96 | -0.95 | -0.58 | -1.09 | -0.74 | -0.6 | -0.3 | -0.59 | -0.9 | -0.53 | -0.37 | -0.075 | 0.14 | 0.225 | 0.739719 | 0.57465 | 0.333016 | 0.999971 | 1 | 0.857921 |
| Nlgn3 | A2AGI2 | -1.28 | -1.26 | -1.3 | -1.05 | -1.12 | -1.17 | -1.47 | -1.05 | -1.01 | -1.1 | -1.23 | -1.31 | -0.74 | -1.04 | -1.3 | -0.94 | 0.02 | 0.06 | 0.2175 | 0.91323 | 0.727076 | 0.281018 | 0.999971 | 1 | 0.844538 |
| Nlgn4l | B0F2B4 | 1.83 | -1.82 | -0.71 | -0.48 | -1.95 | -1.45 | -0.4 | -1.01 | -1.44 | 0.09 | -0.76 | -1.25 | -1.89 | -1.64 | -1.13 | -0.73 | -0.9075 | -0.545 | -1.0525 | 0.246899 | 0.480427 | 0.174773 | 0.999971 | 1 | 0.808161 |
| Nlk | O54949 | -0.56 | 0.03 | 0.65 | -0.98 | -0.21 | 0.27 | 0.1 | -0.45 | -0.12 | -0.41 | -0.26 | -0.86 | -0.09 | -0.44 | -0.87 | -0.71 | 0.1425 | -0.1975 | -0.3125 | 0.706078 | 0.603597 | 0.421267 | 0.999971 | 1 | 0.87788 |
| Nln | Q91YP2 | -0.24 | -0.56 | -0.39 | -0.44 | -0.29 | -0.81 | 0.03 | -0.14 | 0 | -0.33 | -0.19 | -0.29 | -0.02 | -0.68 | -0.35 | -0.33 | 0.105 | 0.205 | 0.0625 | 0.650846 | 0.262628 | 0.761384 | 0.999971 | 1 | 0.961116 |
| Nlrx1 | Q3TL44 | -0.47 | 0.13 | -0.23 | -0.08 | -0.36 | -0.15 | -0.53 | -0.33 | -0.48 | -0.29 | -0.32 | -0.2 | -0.14 | -0.41 | -0.27 | -0.14 | -0.18 | -0.16 | -0.0775 | 0.387705 | 0.423253 | 0.700523 | 0.999971 | 1 | 0.948864 |
| Nmd3 | Q99L48 | 0.48 | 1.8 | 0.87 | 1.14 | 1.38 | 1.92 | 0.86 | 0.97 | 1.58 | 0.69 | 1.01 | 0.91 | 0.06 | 0.63 | 0.93 | 0.82 | 0.21 | -0.025 | -0.4625 | 0.562045 | 0.940341 | 0.193254 | 0.999971 | 1 | 0.810547 |
| Nme1 | P15532 | -0.38 | 0.73 | -0.66 | 0.38 | 0.27 | 0.31 | 0.02 | -0.23 | 0.13 | -0.07 | -0.62 | -0.87 | 0.06 | 0.97 | -0.62 | -0.37 | 0.075 | -0.375 | -0.0075 | 0.826807 | 0.343618 | 0.986637 | 0.999971 | 1 | 0.99925 |
| Nme3 | Q9WV85 | 0.86 | 0.09 | -0.23 | -0.15 | -0.04 | 0.37 | -0.1 | 0.2 | 0.51 | 0.75 | -0.15 | 0.18 | 1.13 | 0.89 | 0.4 | 0.55 | -0.035 | 0.18 | 0.6 | 0.902217 | 0.575796 | 0.076433 | 0.999971 | 1 | 0.748444 |
| Nme7 | Q3UMG6 | 0.17 | 0.27 | -0.03 | 0.69 | 0.12 | 0.54 | 0.02 | 0.59 | 0.28 | 0.32 | 0.27 | 0.25 | 0.28 | 0.53 | 0.51 | 0.71 | 0.0425 | 0.005 | 0.2325 | 0.860372 | 0.980377 | 0.307014 | 0.999971 | 1 | 0.847402 |
| Nmnat1 | Q9EPA7 | -0.07 | 0.9 | 1.94 | 1.65 | 0.54 | -0.11 | 0.44 | 1.58 | 0.52 | 0.01 | 2.09 | 2.36 | -0.66 | 0.39 | 2.44 | 1.89 | -0.4925 | 0.14 | -0.09 | 0.361828 | 0.833933 | 0.905089 | 0.999971 | 1 | 0.991091 |
| Nmnat2 | Q8BNJ3 | 1.12 | -0.31 | -0.07 | 0.11 | -0.33 | 0.04 | 0.47 | 0.91 | -0.35 | 0.68 | 0.33 | 0.79 | -0.54 | 0.37 | 0.18 | 0.14 | 0.06 | 0.15 | -0.175 | 0.879437 | 0.702259 | 0.632322 | 0.999971 | 1 | 0.93301 |
| Nmnat3 | Q99JR6 | -0.34 | -0.41 | -0.88 | 0.16 | -0.46 | -0.78 | -0.96 | -0.15 | -0.89 | -0.58 | -1.12 | -0.65 | -1.25 | -0.36 | -0.3 | -0.55 | -0.22 | -0.4425 | -0.2475 | 0.455362 | 0.122598 | 0.434265 | 0.999971 | 1 | 0.882419 |
| Nmral1 | D3YU12 | 0.35 | 0.4 | 0.87 | 0.48 | -0.02 | 0.34 | 0.06 | 0.08 | 0.6 | 0.69 | 0.51 | -0.3 | 0.52 | 0.47 | 0.48 | -0.09 | -0.41 | -0.15 | -0.18 | 0.06606 | 0.586305 | 0.437636 | 0.999971 | 1 | 0.882419 |
| Nmt1 | O70310 | 2.39 | 2.19 | 1.48 | 1.75 | 2.13 | 2.46 | 1.87 | 1.62 | 2.34 | 2.75 | 1.45 | 1.68 | 1.62 | 2.26 | 1.47 | 1.59 | 0.0675 | 0.1025 | -0.2175 | 0.814092 | 0.773687 | 0.455481 | 0.999971 | 1 | 0.887217 |
| Nmt2 | O70311 | 0.78 | 0.63 | 0.55 | -0.01 | 0.45 | 0.69 | 1.14 | 0.32 | 0.59 | 0.3 | 0.71 | 0.61 | 0.33 | 0.19 | 0.54 | -0.09 | 0.1625 | 0.065 | -0.245 | 0.550289 | 0.777121 | 0.335934 | 0.999971 | 1 | 0.858827 |
| Nnt | Q61941 | -4.99 | -6.87 | -5.39 | -5.09 | -6.74 | 0.71 | 0.02 | 0.37 | 0.22 | 0.39 | 0.01 | 3.74 | -2.73 | 0.71 | 3.74 | 3.66 | 4.175 | 6.675 | 6.93 | 0.028476 | 5.24E-05 | 0.000908 | 0.999971 | 0.095451 | 0.365812 |
| Noa1 | Q9JJG9 | 1.78 | -0.71 | -0.29 | -0.5 | -0.41 | -0.33 | -0.64 | -0.15 | -0.47 | 0.96 | -0.1 | -0.18 | -1.99 | -0.86 | -0.71 | 0.16 | -0.4525 | -0.0175 | -0.92 | 0.410542 | 0.976783 | 0.187129 | 0.999971 | 1 | 0.809025 |
| Nob1 | Q8BW10 | -1.72 | -2.84 | -5.49 | -4.57 | -1.92 | -2.82 | -4.41 | -2.44 | -2.55 | -2.56 | -2.65 | -3.28 | -2.21 | -1.64 | -2.81 | -3.26 | 0.7575 | 0.895 | 1.175 | 0.408435 | 0.271736 | 0.178977 | 0.999971 | 1 | 0.808532 |
| Noc2l | J3QK52 | -0.6 | -0.82 | 0.21 | -0.63 | -0.83 | -1.1 | -0.27 | -0.56 | -0.19 | -0.45 | -0.36 | -0.85 | -0.52 | -1.06 | -0.2 | -1.18 | -0.23 | -0.0025 | -0.28 | 0.449122 | 0.99291 | 0.400324 | 0.999971 | 1 | 0.871816 |
| Noc3l | Q8VI84 | -0.36 | -1.12 | -1.41 | -0.52 | -0.55 | 0.25 | -1.26 | 0.83 | -0.33 | 0.29 | 0.33 | -0.25 | 0.43 | -0.16 | -1.06 | -0.68 | 0.67 | 0.8625 | 0.485 | 0.189756 | 0.019777 | 0.236472 | 0.999971 | 1 | 0.833415 |
| Noc4l | Q8BHY2 | -1.07 | -1.72 | -2.97 | -2.83 | -1.37 | -1.81 | -2.22 | -3.09 | -1.25 | -1.31 | -3.83 | -3.65 | -1.7 | -0.97 | -3.73 | -2.62 | 0.025 | -0.3625 | -0.1075 | 0.962877 | 0.637856 | 0.874423 | 0.999971 | 1 | 0.984109 |
| Noct | O35710 | -0.79 | -0.13 | 0.27 | 0.57 | 0.02 | -0.88 | -0.5 | -0.59 | -0.14 | -1.09 | -0.1 | -0.69 | -0.46 | 0.03 | 0.1 | -0.27 | -0.4675 | -0.485 | -0.13 | 0.197428 | 0.209932 | 0.689188 | 0.999971 | 1 | 0.945805 |
| Nol10 | Q5RJG1 | -0.23 | -0.84 | 0.34 | 0.64 | -1.34 | -0.79 | 0.76 | 0.35 | -1.41 | -0.15 | -0.34 | 0.64 | -1.61 | -0.27 | 0.29 | 0.42 | -0.2325 | -0.2925 | -0.27 | 0.66853 | 0.560685 | 0.609419 | 0.999971 | 1 | 0.92673 |
| Nol11 | Q8BJW5-2 | 2.69 | -0.77 | -0.34 | -0.4 | -0.13 | 0.9 | -0.47 | 0.35 | -0.75 | 1.78 | -0.58 | -0.61 | -2.37 | 0.46 | -0.79 | -0.61 | -0.1325 | -0.335 | -1.1225 | 0.863591 | 0.713193 | 0.2288 | 0.999971 | 1 | 0.828687 |
| Nol3 | Q9D1X0 | 1.42 | 1.41 | 2.1 | 2.17 | 0.74 | 1.52 | 2.41 | 1.84 | 1.07 | 1.6 | 1.61 | -0.3 | 0.74 | 1.84 | 1.69 | 2.03 | -0.1475 | -0.78 | -0.2 | 0.706075 | 0.121411 | 0.570694 | 0.999971 | 1 | 0.91816 |
| Nol4 | F7BIN0 | 3.31 | 2.34 | 2.17 | 2.36 | 1.17 | 2.54 | 1.52 | 1.96 | 1.37 | 3.22 | 1.72 | 1.74 | 0.24 | 1.74 | 1.84 | 1.87 | -0.7475 | -0.5325 | -1.1225 | 0.075287 | 0.263206 | 0.030818 | 0.999971 | 1 | 0.678474 |
| Nol6 | Q8R5K4-2 | -0.58 | -0.38 | 0.2 | -0.21 | -0.39 | -0.67 | -0.06 | -0.2 | -0.95 | -0.98 | -0.18 | -0.41 | -0.67 | -0.87 | -0.11 | 0.01 | -0.0875 | -0.3875 | -0.1675 | 0.71971 | 0.183155 | 0.560131 | 0.999971 | 1 | 0.915694 |
| Nol7 | Q9D7Z3-2 | -1.29 | -0.25 | -0.87 | -0.98 | -1.04 | -1.36 | -1.05 | -0.66 | -2.45 | -0.78 | -3.22 | -2.98 | -2.22 | -1.46 | -1.11 | -0.65 | -0.18 | -1.51 | -0.5125 | 0.521433 | 0.021199 | 0.203852 | 0.999971 | 1 | 0.81617 |
| Nol8 | E9QKD1 | 0.64 | -1.09 | -0.56 | -0.41 | -0.36 | 0.15 | -0.66 | -0.35 | 0.14 | 0.24 | -0.56 | -0.4 | -1.43 | -0.9 | -0.07 | -0.74 | 0.05 | 0.21 | -0.43 | 0.896453 | 0.599001 | 0.335001 | 0.999971 | 1 | 0.858827 |
| Nol9 | Q3TZX8-2 | 1.26 | 0.63 | 1.08 | 1.37 | 0.51 | 0.71 | 0.76 | 1.13 | 0.61 | 1.2 | 1.04 | 1.47 | 0.45 | 0.69 | 0.83 | 1.78 | -0.3075 | -0.005 | -0.1475 | 0.222526 | 0.98483 | 0.659745 | 0.999971 | 1 | 0.935181 |
| Nolc1 | A0A286YD/ | 1.47 | -0.06 | -0.18 | -0.43 | -0.45 | -0.5 | 0.66 | -0.76 | -0.19 | -0.01 | -0.06 | -0.04 | -0.53 | -0.5 | -0.88 | -1.15 | -0.4625 | -0.275 | -0.965 | 0.362283 | 0.510413 | 0.049544 | 0.999971 | 1 | 0.715605 |
| Nomo1 | Q6GQT9 | -0.56 | 0.13 | 0.01 | -0.09 | 0.08 | 0.12 | 0.23 | 0.06 | 0.18 | 0.13 | 0.19 | -0.08 | 0.4 | 0.51 | 0.03 | 0.25 | 0.25 | 0.2325 | 0.425 | 0.248665 | 0.285863 | 0.086163 | 0.999971 | 1 | 0.759991 |
| Nono | Q99K48 | 1.34 | 2.02 | 2.13 | 2.38 | 2.02 | 1.59 | 1.92 | 2.03 | 1.84 | 1.72 | 2.05 | 2.1 | 1.31 | 1.84 | 2.1 | 2.42 | -0.0775 | -0.04 | -0.05 | 0.771135 | 0.878087 | 0.87764 | 0.999971 | 1 | 0.985318 |
| Nop10 | Q9CQS2 | 0.11 | -1.47 | -1.04 | -1.38 | -1.51 | -1.64 | -0.86 | -1.25 | -1.35 | -0.42 | -1.23 | -1.43 | -1.53 | -1.18 | -1.43 | -1.37 | -0.37 | -0.1625 | -0.4325 | 0.351704 | 0.694504 | 0.25248 | 0.999971 | 1 | 0.839012 |
| Nop14 | Q8R3N1 | -1.31 | 0.08 | -1.43 | -1.2 | 0.53 | -0.45 | -1.35 | -2.13 | 0.49 | -0.53 | -1.48 | -1 | 1.46 | 0.16 | -0.84 | -1.11 | 0.115 | 0.335 | 0.8825 | 0.851202 | 0.5165 | 0.179304 | 0.999971 | 1 | 0.808532 |
| Nop16 | Q9CPT5 | 1.58 | 1.38 | 0.75 | 0.63 | 1.37 | 1.22 | 1.44 | 0.84 | 1.7 | 1.52 | 0.63 | 0.24 | 1.8 | 1.69 | 0.73 | 1.25 | 0.1325 | -0.0625 | 0.2825 | 0.641651 | 0.87671 | 0.409663 | 0.999971 | 1 | 0.875847 |
| Nop2 | E9QN31 | -1.42 | -0.64 | -0.59 | -1.47 | -0.22 | -1.64 | -0.77 | -1.83 | -0.28 | -1.56 | -0.3 | -0.63 | 0.08 | -1.37 | -1.39 | -1.5 | -0.085 | 0.3375 | -0.015 | 0.840908 | 0.377383 | 0.971715 | 0.999971 | 1 | 0.99925 |
| Nop53 | Q8BK35 | 0.45 | -0.72 | -0.26 | 0.35 | -0.5 | -0.44 | -0.47 | 0.3 | -0.86 | -0.17 | -0.35 | -0.38 | -1.13 | -0.38 | -0.16 | -0.1 | -0.2325 | -0.395 | -0.3975 | 0.491916 | 0.230634 | 0.281405 | 0.999971 | 1 | 0.844538 |
| Nop56 | Q9D6Z1 | -0.74 | -1.24 | -0.68 | -0.78 | -1.1 | -1.37 | -0.86 | -0.86 | -1.12 | -0.56 | -0.78 | -0.91 | -1.09 | -1.14 | -0.7 | -0.6 | -0.1875 | 0.0175 | -0.0225 | 0.40487 | 0.935518 | 0.921191 | 0.999971 | 1 | 0.992702 |
| Nop58 | Q6DFW4 | -0.25 | -0.43 | -0.32 | -0.94 | -0.74 | -0.65 | -0.27 | -0.96 | -0.47 | -0.41 | -0.53 | -0.8 | -0.4 | -0.77 | -0.76 | -1.03 | -0.17 | -0.0675 | -0.255 | 0.491018 | 0.758644 | 0.299583 | 0.999971 | 1 | 0.847402 |
| Nop9 | Q8BMC4 | -1.4 | 0.51 | 0.61 | 0.73 | -0.06 | 0.24 | -0.04 | 0.02 | -0.29 | -0.25 | -0.2 | 0.43 | -1.53 | 0.75 | 0.93 | 0.83 | -0.0725 | -0.19 | 0.1325 | 0.878927 | 0.703746 | 0.851048 | 0.999971 | 1 | 0.98187 |
| Nopchap1 | Q9CX66 | 1.26 | 1.75 | 1 | 1.58 | 0.47 | 0.94 | 1.25 | 1.71 | 0.82 | 1.26 | 0.75 | 0.9 | -0.04 | 1.22 | 0.93 | 1.98 | -0.305 | -0.465 | -0.375 | 0.344349 | 0.074234 | 0.390054 | 0.999971 | 1 | 0.871816 |
| Nos1 | F8WGF2 | -0.06 | -0.78 | -0.53 | -0.71 | -0.44 | -0.45 | -0.43 | -0.53 | -0.22 | -0.2 | -0.88 | -0.51 | -0.05 | -0.11 | -0.42 | -0.85 | 0.0575 | 0.0675 | 0.1625 | 0.787029 | 0.789116 | 0.546 | 0.999971 | 1 | 0.911398 |
| Nos1ap | Q9D3A8 | -0.12 | -0.66 | 0.5 | -0.42 | -0.11 | -0.34 | 0.41 | -0.18 | -0.5 | -0.42 | 0.34 | 0.21 | -0.35 | -0.62 | -0.06 | -0.08 | 0.12 | 0.0825 | -0.1025 | 0.695375 | 0.802318 | 0.728462 | 0.999971 | 1 | 0.956041 |
| Nos3 | P70313 | -0.21 | 1.93 | -0.19 | -0.28 | 4.56 | 1.25 | -0.03 | -1.77 | 3.15 | -0.3 | -0.03 | 1.15 | 5.38 | 0.57 | 1.08 | 0.84 | 0.69 | 0.68 | 1.655 | 0.592984 | 0.437645 | 0.166924 | 0.999971 | 1 | 0.800215 |
| Nosip | A0A1B0GT( | 2.48 | 1.52 | 3.63 | 4.33 | 1.22 | 1.41 | 4.38 | 1.74 | 1.17 | 1.88 | 1.97 | 0.91 | 0.25 | 1.57 | 1.06 | -0.01 | -0.8025 | -1.5075 | -2.2725 | 0.365384 | 0.035447 | 0.006861 | 0.999971 | 1 | 0.582714 |
| Nostrin | Q6WKZ7 | 1 | 1.43 | -0.22 | 1.5 | 0.71 | 1.08 | 0.68 | 0.6 | 0.77 | 1.41 | 0.87 | 2.25 | 0.43 | 1.67 | -0.13 | 0.24 | -0.16 | 0.3975 | -0.375 | 0.686544 | 0.424864 | 0.474372 | 0.999971 | 1 | 0.894044 |
| Notch1 | Q01705-4 | -0.53 | -1.73 | 0.79 | 0.96 | -1.1 | -0.83 | 0.18 | 1.74 | -0.85 | -0.41 | 0.54 | 0.5 | -0.33 | -0.85 | 1.32 | 0.88 | 0.125 | 0.0725 | 0.3825 | 0.877001 | 0.911811 | 0.603308 | 0.999971 | 1 | 0.924454 |
| Notch2 | O35516 | 2.1 | -1.04 | -0.77 | -1.08 | -0.27 | 0.29 | 0.26 | -1.08 | -0.43 | 1.42 | -1.94 | -1.86 | -2.59 | -0.37 | -1.49 | -2.86 | -0.0025 | -0.505 | -1.63 | 0.997338 | 0.612602 | 0.084944 | 1 | 1 | 0.758256 |
| Nova1 | Q9JKN6 | 1.78 | 1.62 | 1.9 | 1.78 | 1.38 | 1.9 | 1.46 | 1.67 | 1.36 | 1.76 | 1.26 | 2.03 | 0.88 | 1.99 | 2.14 | 1.78 | -0.1675 | -0.1675 | -0.0725 | 0.397601 | 0.464618 | 0.807966 | 0.999971 | 1 | 0.973362 |
| Nova2 | A0A1W2P8 | 1.14 | 1.69 | 1.57 | 1.81 | 1.53 | 1.42 | 1.4 | 1.65 | 1.42 | 1.36 | 1.59 | 1.64 | 1.41 | 2.05 | 1.76 | 1.73 | -0.0525 | -0.05 | 0.185 | 0.801448 | 0.810445 | 0.43626 | 0.999971 | 1 | 0.882419 |
| Npc1 | O35604 | 0.4 | 0.69 | -0.19 | -0.36 | 0.63 | 0.44 | -0.01 | 0.48 | 0.62 | 0.39 | -0.32 | 0.17 | 1.09 | 0.68 | -0.41 | -0.3 | 0.25 | 0.08 | 0.13 | 0.403747 | 0.802563 | 0.757948 | 0.999971 | 1 | 0.961116 |
| Npc1l1 | Z4YJC9 | 0.89 | 1.77 | -0.06 | 0.43 | 0.23 | 0.7 | -0.01 | 0.36 | 0.46 | 0.95 | -0.02 | 0.03 | 1.67 | 1.45 | -0.18 | 0.69 | -0.4375 | -0.4025 | 0.15 | 0.288959 | 0.358866 | 0.776777 | 0.999971 | 1 | 0.964475 |
| Npc2 | Q9Z0J0 | -0.84 | 0.64 | 1.41 | 1.08 | 0.45 | 0.53 | 0.45 | 2.15 | 0.94 | 0.36 | 0.84 | 1.61 | 0.9 | 0.73 | 1.8 | 1.56 | 0.3225 | 0.365 | 0.675 | 0.589652 | 0.488528 | 0.214287 | 0.999971 | 1 | 0.826146 |
| Npdc1 | Q64322 | 1.09 | -0.27 | 0.25 | 0.16 | -0.16 | 0.54 | 0.33 | 0.36 | 0.32 | 0.75 | -0.28 | -0.14 | -0.73 | 0.12 | 0.36 | -0.31 | -0.04 | -0.145 | -0.4475 | 0.901336 | 0.688427 | 0.239136 | 0.999971 | 1 | 0.833594 |
| Npepl1 | Q6NSR8 | 0.08 | -0.05 | 0.29 | 0.28 | 0.16 | 0.16 | 0.3 | 0.35 | -0.35 | -0.04 | 0.35 | 0.58 | -0.37 | 0.19 | 0.58 | 0.9 | 0.0925 | -0.015 | 0.175 | 0.6055 | 0.951813 | 0.558431 | 0.999971 | 1 | 0.915051 |
| Npepps | Q11011 | -0.5 | 0.25 | -0.07 | -0.15 | 0.33 | 0.35 | 0.09 | -0.17 | 0.34 | -0.19 | -0.02 | 0.26 | 0.57 | 0.22 | 0.11 | 0.21 | 0.2675 | 0.215 | 0.395 | 0.268927 | 0.365276 | 0.107258 | 0.999971 | 1 | 0.78268 |
| Nphp1 | A2APS1 | 0.13 | -0.09 | -0.88 | -0.14 | -0.27 | 0.32 | -1.67 | -0.44 | -0.04 | 0.42 | -0.99 | -0.57 | 1.34 | 0.25 | -0.85 | -0.01 | -0.27 | -0.05 | 0.4275 | 0.547407 | 0.891852 | 0.374576 | 0.999971 | 1 | 0.870318 |
| Nphp4 | P59240 | 1.04 | -0.19 | -0.52 | -0.42 | 0.04 | 0.16 | -0.52 | -0.31 | 0.02 | 0.49 | -0.22 | -0.02 | -0.54 | -0.22 | -0.62 | -0.18 | -0.135 | 0.09 | -0.3675 | 0.722909 | 0.812186 | 0.331964 | 0.999971 | 1 | 0.857152 |
| Npl | Q9DCJ9 | -0.25 | 0.72 | 0.91 | 0.5 | 0.6 | 0.55 | 0.22 | 0.98 | 0.72 | 0.02 | 0.53 | 1.33 | 0.97 | 1.03 | 1.12 | 1.27 | 0.1175 | 0.18 | 0.6275 | 0.700974 | 0.62189 | 0.047956 | 0.999971 | 1 | 0.710102 |
| Nploc4 | P60670 | -0.17 | 0.03 | 0.71 | 0.89 | 0.15 | 0.41 | 0.38 | 1.1 | 0.31 | 0.24 | 1.01 | 1.04 | 0.19 | 0.37 | 1.14 | 1.38 | 0.145 | 0.285 | 0.405 | 0.660581 | 0.404291 | 0.297233 | 0.999971 | 1 | 0.847402 |
| Npm1 | Q61937 | 0.81 | 1.87 | 1.52 | 1.03 | 1.63 | 1.31 | 1.24 | 1.36 | 1.8 | 0.76 | 1.51 | 1.3 | 1.23 | 1.14 | 1.27 | 0.76 | 0.0775 | 0.035 | -0.2075 | 0.776255 | 0.914193 | 0.468406 | 0.999971 | 1 | 0.89236 |
| Npm3 | Q9CPP0 | 0.66 | 0.14 | -0.16 | 1.04 | 0.32 | 0.69 | -0.02 | 0.3 | -0.35 | 0.59 | -0.36 | 0.16 | -0.41 | 0.1 | -0.32 | -0.74 | -0.0975 | -0.41 | -0.7625 | 0.753203 | 0.255999 | 0.039076 | 0.999971 | 1 | 0.700838 |
| Npr2 | B1AWI7 | -0.89 | -1.13 | -1.3 | -1.21 | -1.49 | -1.38 | -1.22 | -0.81 | -1.25 | -0.63 | -1.78 | -1.42 | -1.49 | -1.36 | -1.3 | -1.24 | -0.0925 | -0.1375 | -0.215 | 0.673002 | 0.616525 | 0.255524 | 0.999971 | 1 | 0.839012 |
| Nprl2 | Q9WUE4 | 0.29 | 0.85 | 0.15 | -0.07 | 0.44 | 0.7 | -0.2 | 0.32 | 0.9 | -0.56 | -0.14 | 0.19 | 0.56 | 0.89 | 0.28 | -0.31 | 0.01 | -0.2075 | 0.05 | 0.972061 | 0.565691 | 0.877042 | 0.999971 | 1 | 0.985139 |
| Nprl3 | A7M7S2 | 2.2 | -0.22 | 0.27 | 0.21 | -0.38 | -0.41 | 0.71 | 0.18 | -0.59 | 0.34 | -0.77 | -0.84 | -1.23 | -0.48 | -0.64 | -0.63 | -0.59 | -1.08 | -1.36 | 0.30138 | 0.08041 | 0.026951 | 0.999971 | 1 | 0.678474 |
| Nptn | P97300-3 | 1.18 | 0.4 | 0.21 | 0.32 | -0.41 | 0.93 | -0.1 | 0.01 | -0.07 | 1.02 | 0.29 | -0.12 | 0.63 | 0.47 | -0.2 | 0.57 | -0.42 | -0.2475 | -0.16 | 0.25651 | 0.473629 | 0.598527 | 0.999971 | 1 | 0.92377 |
| Nptn_1 | P97300 | 0.41 | -0.66 | -1.04 | -0.3 | -1.31 | -0.5 | 0.64 | -1.3 | -1.09 | -0.15 | -1.39 | -1.69 | -1.15 | -1 | -1.48 | -1.36 | -0.22 | -0.6825 | -0.85 | 0.669082 | 0.140124 | 0.027356 | 0.999971 | 1 | 0.678474 |
| Nptx1 | Q62443 | -0.32 | -0.27 | -0.27 | -0.54 | -0.33 | 0.02 | -0.07 | -0.34 | -0.33 | -0.21 | -0.45 | -0.95 | -0.33 | -0.5 | -0.85 | -0.57 | 0.17 | -0.135 | -0.2125 | 0.369595 | 0.538572 | 0.286556 | 0.999971 | 1 | 0.846151 |
| Nptx2 | O70340 | -1.14 | -1.64 | 0.03 | 0.48 | -0.8 | -1.52 | 0.32 | 0 | -1.12 | -1.02 | -0.08 | -0.76 | -0.5 | -1.48 | -0.01 | -0.83 | 0.0675 | -0.1775 | -0.1375 | 0.908667 | 0.727989 | 0.798254 | 0.999971 | 1 | 0.970479 |
| Nptxr | E9PZM8 | -0.35 | -0.57 | -0.57 | -0.53 | -0.23 | 0.09 | -0.39 | -0.24 | -0.24 | -0.2 | -0.19 | -0.56 | -0.38 | -0.11 | -0.3 | -0.46 | 0.3125 | 0.2075 | 0.1925 | 0.119638 | 0.261245 | 0.292875 | 0.999971 | 1 | 0.847402 |
| Npy | P57774 | -0.53 | -2.02 | -1.03 | -1.79 | -1.62 | -0.08 | -1.14 | -1.55 | -1.86 | -1.26 | -1.84 | -2.79 | -1.61 | -1.7 | -0.78 | 0.56 | 0.245 | -0.595 | 0.46 | 0.599035 | 0.201177 | 0.431889 | 0.999971 | 1 | 0.881643 |
| Npy1r | Q04573 | 2.21 | 1.27 | -1.19 | -0.6 | 1.05 | 2.68 | -0.24 | -1.01 | 1.28 | 2.69 | -0.27 | -0.62 | 1.92 | 0.94 | -0.8 | -0.78 | 0.1975 | 0.3475 | -0.1025 | 0.84496 | 0.726138 | 0.912263 | 0.999971 | 1 | 0.992618 |
| Nqo1 | Q64669 | 0.48 | 0.98 | 0.18 | 0.6 | 0.87 | 0.89 | 0.04 | 0.43 | 0.33 | 0.58 | -0.74 | 0.2 | 0.56 | 0.86 | 0.19 | 0.37 | -0.0025 | -0.4675 | -0.065 | 0.992813 | 0.182833 | 0.793874 | 0.999971 | 1 | 0.970141 |
| Nqo2 | Q9JI75 | -0.73 | 0.24 | -0.41 | -0.85 | 0.23 | 0.27 | 0.32 | -0.41 | -0.12 | -0.14 | 0.02 | 0.19 | 0.49 | -0.01 | -0.9 | -0.82 | 0.54 | 0.425 | 0.1275 | 0.104211 | 0.145461 | 0.748598 | 0.999971 | 1 | 0.961064 |
| Nr1d2 | A0A286YD! | 0.06 | 0.3 | -0.46 | -0.2 | -0.17 | -1 | 0.99 | -0.17 | -0.39 | 0.02 | 2.06 | -0.79 | -0.6 | 0.08 | -1.51 | -0.28 | -0.0125 | 0.3 | -0.5025 | 0.976122 | 0.619991 | 0.195641 | 0.999971 | 1 | 0.811225 |
| Nr2c2 | G3X9W4 | 0.06 | 0.28 | 1.2 | 0.87 | 0.45 | -0.2 | 1.05 | 0.6 | -0.21 | 0.68 | 1.35 | 1.92 | 0.65 | 0.24 | 0.97 | 0.91 | -0.1275 | 0.3325 | 0.09 | 0.723785 | 0.505532 | 0.77552 | 0.999971 | 1 | 0.96393 |
| Nr2c2ap | Q3TV70 | -0.71 | 0.28 | -1.15 | -1.48 | 0.15 | -0.32 | -0.57 | -2.45 | -0.58 | -1.57 | -0.99 | -0.22 | 0.54 | -0.02 | -1.18 | -1.72 | -0.0325 | -0.075 | 0.17 | 0.958482 | 0.868213 | 0.773479 | 0.999971 | 1 | 0.963759 |
| Nr2f1 | Q32NY6 | 2.12 | 1.36 | 1.6 | 1.36 | 1.43 | 1.66 | 1.43 | 1.25 | 1.4 | 2 | 1.36 | 1.04 | 0.73 | 1.75 | 0.98 | 1.23 | -0.1675 | -0.16 | -0.4375 | 0.48034 | 0.574141 | 0.161522 | 0.999971 | 1 | 0.800215 |
| Nr3c1 | P06537-5 | 2.29 | 2.65 | 2.36 | 2.01 | 2.13 | 2.26 | 2.09 | 1.85 | 1.9 | 1.92 | 1.45 | 1.29 | 1.56 | 1.53 | 1.93 | 1.25 | -0.245 | -0.6875 | -0.76 | 0.258868 | 0.017047 | 0.008878 | 0.999971 | 1 | 0.613003 |
| Nr4a3 | F6R7A2 | 1.85 | 2.93 | 1.81 | 1.67 | 2 | 2.52 | 1.65 | 0.98 | 2.05 | 2.16 | 1.76 | 1.36 | 2.14 | 2.15 | 1.58 | 2.28 | -0.2775 | -0.2325 | -0.0275 | 0.507413 | 0.497948 | 0.933556 | 0.999971 | 1 | 0.995138 |
| Nras | A0A0G2JGI | 1.61 | -1 | -2.15 | -3.66 | -0.82 | 0.28 | -2.1 | -3.09 | -0.54 | -0.01 | -1.95 | -1.78 | -2.04 | -0.98 | -1.82 | -4.38 | -0.1325 | 0.23 | -1.005 | 0.910964 | 0.832168 | 0.406071 | 0.999971 | 1 | 0.874911 |
| Nrbf2 | Q8VCQ3 | 0.02 | 1.03 | 0.16 | 0.18 | 1.03 | 1.13 | 0.04 | 0.51 | 0.96 | 0.28 | -0.07 | 0.57 | 1.45 | 1.02 | 0.11 | 0.76 | 0.33 | 0.0875 | 0.4875 | 0.342954 | 0.784398 | 0.194517 | 0.999971 | 1 | 0.811225 |
| Nrbp1 | D3YUV1 | 0.51 | 0.82 | 0.81 | 0.66 | 0.43 | 0.78 | 0.59 | 0.18 | 0.6 | 0.59 | 0.37 | -0.44 | 0.87 | 0.57 | 0.6 | 0.44 | -0.205 | -0.42 | -0.08 | 0.326276 | 0.150266 | 0.671317 | 0.999971 | 1 | 0.939555 |
| Nrbp2 | Q91V36 | -0.32 | -0.29 | 0.19 | -0.4 | -0.13 | -0.04 | 0.07 | -0.23 | 0.09 | 0.05 | 0.05 | 0.18 | 0.62 | -0.13 | 0.09 | -0.32 | 0.1225 | 0.2975 | 0.27 | 0.551488 | 0.152813 | 0.323453 | 0.999971 | 1 | 0.85548 |
| Nrcam | Q810U4-2 | -1.73 | 0.2 | -0.69 | -1.77 | -0.39 | -0.19 | -0.65 | -1.23 | -0.24 | -3.05 | -0.55 | -0.24 | -0.29 | -0.79 | -1.17 | -1.77 | 0.3825 | -0.0225 | -0.0075 | 0.439229 | 0.97607 | 0.988537 | 0.999971 | 1 | 0.99925 |
| Nrcam_1 | Q810U4 | 1.35 | 1.5 | 0.33 | 1.16 | 0.43 | 2 | 0.13 | 0.46 | 1.59 | 2.23 | 0.53 | 0.94 | 3.25 | 2 | 1.44 | 1.6 | -0.33 | 0.2375 | 0.9875 | 0.483147 | 0.585412 | 0.055215 | 0.999971 | 1 | 0.725348 |
| Nrcam_2 | Q810U4-3 | 0.21 | 0.76 | 0.8 | 0.54 | 0.43 | 0.79 | 0.7 | 0.68 | 0.66 | 0.49 | 0.65 | 1.1 | 0.99 | 0.54 | 0.89 | 0.91 | 0.0725 | 0.1475 | 0.255 | 0.727938 | 0.518467 | 0.256479 | 0.999971 | 1 | 0.839012 |
| Nrcam_3 | A0A1W2P7 | 0.04 | -0.17 | -1.07 | -1.6 | 0.5 | 0.11 | 2.64 | -0.51 | 5.1 | -0.64 | -0.71 | 0.08 | 0.38 | -0.76 | -0.67 | -1.18 | 1.385 | 1.6575 | 0.1425 | 0.077514 | 0.221763 | 0.764397 | 0.999971 | 1 | 0.962405 |
| Nrd1 | A2A9Q2 | 0.26 | 0.94 | 0.79 | 0.42 | 0.55 | 0.25 | 0.45 | 0.35 | 1.27 | 0.61 | 1.26 | 1.45 | 0.98 | 0.6 | 0.57 | 0.58 | -0.2025 | 0.545 | 0.08 | 0.362386 | 0.064314 | 0.725648 | 0.999971 | 1 | 0.956041 |
| Nrf1 | Q99K73 | 0.5 | 1.02 | 0.85 | 1.34 | 1.06 | 0.57 | 0.59 | 1.37 | 0.43 | 0.85 | 0.97 | 1.36 | 0.22 | 1.05 | 1.22 | 1.72 | -0.03 | -0.025 | 0.125 | 0.913883 | 0.927571 | 0.723159 | 0.999971 | 1 | 0.955258 |
| Nrg3 | G3V023 | -1.2 | -1.89 | -0.43 | 0.2 | -1.48 | -1.6 | 0.36 | -0.06 | -0.97 | -1.01 | 0.24 | -0.58 | -1.63 | -1.22 | -0.2 | -0.17 | 0.135 | 0.25 | 0.025 | 0.825746 | 0.621144 | 0.962982 | 0.999971 | 1 | 0.998721 |
| Nrgn | P60761 | 1.46 | 0.27 | 1.13 | 1.26 | 0.89 | 0.3 | 1.23 | 0.57 | 0.47 | 1.04 | 1 | 0.8 | -0.56 | 0.03 | -0.13 | 0.61 | -0.2825 | -0.2025 | -1.0425 | 0.401784 | 0.505818 | 0.015063 | 0.999971 | 1 | 0.613095 |
| Nrip2 | Q9JHR9-2 | 1.19 | -0.09 | 0.38 | 0.77 | 0.67 | 0.67 | 1.25 | 0.35 | 0.47 | 0.85 | 0.07 | -0.08 | 0.52 | 0.34 | 0.4 | -0.34 | 0.1725 | -0.235 | -0.3325 | 0.604439 | 0.496211 | 0.333521 | 0.999971 | 1 | 0.857921 |
| Nrip3 | Q9JJR9 | 0.1 | 1.45 | 0.93 | 1.19 | 1.18 | 1.5 | 0.72 | 1.82 | 1.15 | 0.55 | 1.33 | 1.28 | 1.28 | 1.52 | 1.19 | 1.42 | 0.3875 | 0.16 | 0.435 | 0.304187 | 0.639842 | 0.182209 | 0.999971 | 1 | 0.808532 |
| Nrm | Q8VC65 | -1.43 | -0.07 | -0.15 | -0.53 | -0.3 | -0.48 | -0.34 | -0.37 | -0.46 | -1.29 | 0.21 | -0.11 | -0.6 | -0.3 | 0.37 | 0.15 | 0.1725 | 0.1325 | 0.45 | 0.589414 | 0.756361 | 0.244323 | 0.999971 | 1 | 0.837316 |
| Nrn1 | A0A286YC\ | -0.39 | 1.06 | -0.13 | -0.64 | 0.47 | 1.28 | -0.55 | -0.55 | 0.04 | -0.51 | -0.72 | -1.14 | 0.7 | 0.91 | -0.8 | -0.07 | 0.1875 | -0.5575 | 0.21 | 0.726938 | 0.21383 | 0.677792 | 0.999971 | 1 | 0.942594 |
| Nrp1 | P97333 | -0.38 | -0.01 | -0.43 | -0.7 | -0.46 | -0.12 | -1.09 | -0.14 | -0.2 | -0.17 | -0.77 | -0.76 | 0.03 | -0.1 | -0.37 | -0.46 | -0.0725 | -0.095 | 0.155 | 0.797246 | 0.701308 | 0.495969 | 0.999971 | 1 | 0.901531 |
| Nrp2 | O35375-5 | -0.68 | -0.67 | -0.56 | -0.74 | 0.03 | -0.41 | -0.32 | -0.67 | 0.1 | -0.75 | -0.91 | -0.42 | 0.07 | -0.5 | -0.5 | -1.2 | 0.32 | 0.1675 | 0.13 | 0.143434 | 0.510629 | 0.64346 | 0.999971 | 1 | 0.93301 |
| Nrsn1 | P97799 | 2.48 | -2.23 | 0.83 | -1.02 | -1.05 | -0.44 | 2.2 | -0.29 | -0.12 | 1.13 | 1.09 | 1.07 | -0.8 | -1.09 | 0.63 | -1.09 | 0.09 | 0.7775 | -0.6025 | 0.935927 | 0.431913 | 0.549232 | 0.999971 | 1 | 0.911952 |
| Nrxn1 | Q9CS84-4 | -1.13 | -0.86 | -0.95 | -0.79 | -0.91 | -0.85 | -1.04 | -0.89 | -0.83 | -0.9 | -0.81 | -0.96 | -0.94 | -0.92 | -0.81 | -0.6 | 0.01 | 0.0575 | 0.115 | 0.953927 | 0.733688 | 0.535322 | 0.999971 | 1 | 0.910077 |
| Nrxn1_1 | H3BK41 | -0.55 | -0.25 | -0.56 | 0.16 | -0.4 | -1.19 | -1.1 | -0.39 | -0.65 | 0.18 | -0.97 | -0.25 | -0.65 | -0.73 | -0.25 | 0.24 | -0.47 | -0.1225 | -0.0475 | 0.129488 | 0.690838 | 0.870553 | 0.999971 | 1 | 0.984079 |
| Nrxn2 | E9PUM9 | -0.53 | -0.68 | -0.76 | -0.68 | -0.68 | -0.48 | -0.76 | -0.54 | -0.4 | -0.62 | -0.66 | -0.5 | -0.14 | -0.51 | -0.56 | -0.49 | 0.0475 | 0.1175 | 0.2375 | 0.782331 | 0.487326 | 0.217547 | 0.999971 | 1 | 0.826893 |
| Nrxn3 | Q6P9K9 | -1.32 | -0.83 | -1 | -0.85 | -0.76 | -0.56 | -0.89 | -0.74 | -0.8 | -1.19 | -0.91 | -0.77 | -0.55 | -0.69 | -0.63 | -0.65 | 0.2625 | 0.0825 | 0.37 | 0.200725 | 0.682838 | 0.075647 | 0.999971 | 1 | 0.746958 |
| Nsd1 | E9QAE4 | 1.62 | -0.73 | 0.7 | 0.99 | -0.28 | 0.02 | 2.75 | -0.17 | 0.05 | 1.13 | -0.04 | -0.5 | -0.73 | 0.24 | 0.12 | 0.44 | -0.065 | -0.485 | -0.6275 | 0.934424 | 0.395694 | 0.245853 | 0.999971 | 1 | 0.838611 |
| Nsd2 | D3Z3E4 | 2.14 | 0.44 | 1.31 | 1.72 | 0.56 | 0.38 | 1.71 | 1.25 | 0.02 | 1.25 | 0.57 | 0.93 | -0.36 | 0.22 | 0.91 | 0.48 | -0.4275 | -0.71 | -1.09 | 0.35225 | 0.12319 | 0.029157 | 0.999971 | 1 | 0.678474 |
| Nsd3 | D3Z197 | 0.48 | 0.07 | -0.45 | -0.31 | -0.1 | 0.17 | -0.83 | 0.3 | -0.8 | 0.14 | -0.39 | -0.3 | -1.26 | -0.05 | -0.11 | 0.46 | -0.0625 | -0.285 | -0.1875 | 0.848692 | 0.345379 | 0.642229 | 0.999971 | 1 | 0.93301 |
| Nsdhl | Q9R1J0 | -0.59 | -0.73 | -0.36 | -0.38 | -1.02 | -0.96 | -0.07 | 0.01 | -0.82 | -0.57 | -0.02 | -0.62 | -0.18 | -0.35 | 0.53 | 0.56 | 0.005 | 0.0075 | 0.655 | 0.986691 | 0.973804 | 0.036895 | 0.999971 | 1 | 0.700096 |
| Nsf | P46460 | -0.61 | -0.82 | -0.19 | -0.66 | -0.96 | -0.72 | -0.1 | -0.51 | -1.01 | -0.94 | -0.2 | -0.13 | -0.82 | -0.94 | -0.3 | -0.16 | -0.0025 | 7.85E-17 | 0.015 | 0.992109 | 1 | 0.953734 | 0.999971 | 1 | 0.998056 |
| Nsfl1c | Q9CZ44 | 0.85 | 0.35 | 0.52 | -0.58 | 0.78 | 0.18 | 1 | -0.15 | 1.07 | -0.22 | 0.56 | 0.88 | 1.6 | -0.38 | 0.19 | -0.79 | 0.1675 | 0.2875 | -0.13 | 0.668441 | 0.479854 | 0.815414 | 0.999971 | 1 | 0.974918 |
| Nsg1 | Q62092 | -1.4 | -2.34 | -1.38 | -0.75 | -1.26 | -2.36 | -0.88 | -0.27 | -1.17 | -1.82 | -0.43 | -0.71 | -1.67 | -2.13 | -0.3 | -0.45 | 0.275 | 0.435 | 0.33 | 0.590774 | 0.320608 | 0.527563 | 0.999971 | 1 | 0.909612 |
| Nsg2 | P47759 | -1.97 | -0.4 | 1 | 0.14 | 0.12 | -0.21 | 1.36 | -0.05 | 0.03 | -1.69 | 0.97 | 0.78 | -0.3 | -1.16 | 0.39 | -0.64 | 0.6125 | 0.33 | -0.12 | 0.360812 | 0.676745 | 0.851452 | 0.999971 | 1 | 0.98187 |
| Nsmaf | O35242 | -0.02 | -0.71 | -0.3 | -1.39 | 0.57 | -0.05 | -0.92 | -0.86 | -0.6 | -0.04 | 0.08 | -0.89 | 0.18 | 0.67 | -1.63 | 0.62 | 0.29 | 0.2425 | 0.565 | 0.512269 | 0.513524 | 0.333242 | 0.999971 | 1 | 0.857921 |
| Nsmce3 | Q9CPR8 | 1.14 | 1.34 | -1.91 | 1.44 | 3.24 | 0.37 | 0.77 | 0.38 | 3.64 | 0.96 | -1.28 | -0.39 | 5.66 | 1.64 | -0.58 | 0.56 | 0.6875 | 0.23 | 1.3175 | 0.475104 | 0.848344 | 0.360236 | 0.999971 | 1 | 0.869526 |
| Nsmf | Q99NF2-2 | -0.67 | 0.91 | -1.46 | -1.37 | 0.7 | 1.37 | 0.24 | -0.9 | 2.2 | 1.61 | 0.66 | -0.87 | 3.23 | 1.54 | -1.39 | -0.4 | 1 | 1.5475 | 1.3925 | 0.155176 | 0.075526 | 0.20361 | 0.999971 | 1 | 0.81617 |
| Nsmf_1 | Q99NF2-3 | -0.66 | -0.51 | -0.24 | -0.34 | -0.64 | -0.14 | -0.21 | -0.36 | -0.37 | -0.41 | -0.1 | -0.19 | -0.09 | -0.54 | -0.15 | 0.3 | 0.1 | 0.17 | 0.3175 | 0.622103 | 0.369989 | 0.195952 | 0.999971 | 1 | 0.812143 |
| Nsrp1 | Q5NCR9 | 2.49 | 4.59 | 3.59 | 1.64 | 4.34 | 5.39 | 4.09 | 2.66 | 4.89 | 2.87 | 4.65 | 4.35 | 4.02 | 2.51 | 2.74 | 0.58 | 1.0425 | 1.1125 | -0.615 | 0.198177 | 0.146639 | 0.481057 | 0.999971 | 1 | 0.89485 |
| Nsun2 | Q1HFZ0 | 1.31 | 1.75 | 1.54 | 1.14 | 1.59 | 1.36 | 0.94 | 1.2 | 1.64 | 1.35 | 1.33 | 1.55 | 1.31 | 1.87 | 1.41 | 1.32 | -0.1625 | 0.0325 | 0.0425 | 0.484905 | 0.873399 | 0.852263 | 0.999971 | 1 | 0.98187 |
| Nsun4 | Q9CZ57 | -0.11 | 0.08 | -0.49 | -0.46 | 0.07 | 0.14 | -0.6 | 0.56 | 0.35 | -0.22 | -0.68 | -0.18 | 0.77 | 0.53 | 0.12 | -0.69 | 0.2875 | 0.0625 | 0.4275 | 0.334032 | 0.817 | 0.23595 | 0.999971 | 1 | 0.832918 |
| Nt5c | Q9JM14 | 0.69 | 1.99 | 2.31 | 2.03 | 1.67 | 2.66 | 1.16 | 2.62 | 1.75 | 1.29 | 1.7 | 1.84 | 2.82 | 2.81 | 2.73 | 2.54 | 0.2725 | -0.11 | 0.97 | 0.574546 | 0.767408 | 0.023357 | 0.999971 | 1 | 0.661101 |
| Nt5c1a | A3KFX0 | -0.4 | 0.14 | -1.01 | -0.39 | 1.2 | 0.27 | -0.48 | -0.54 | 0.95 | -0.07 | -0.49 | -0.43 | 1.47 | 0.05 | -0.7 | -0.65 | 0.5275 | 0.405 | 0.4575 | 0.252154 | 0.317231 | 0.386579 | 0.999971 | 1 | 0.871816 |
| Nt5c2 | A0A494B9) | -0.17 | 0.53 | 0.45 | 0.32 | 0.4 | 0.15 | 0.02 | 0.88 | 0.22 | 0.13 | 0.08 | 0.36 | 0.08 | 0.47 | 0.43 | 0.15 | 0.08 | -0.085 | 3.93E-17 | 0.764691 | 0.691221 | 1 | 0.999971 | 1 | 1 |
| Nt5c3a | Q9D020-1 | 0.3 | 0.69 | 0.48 | 0.66 | 0.72 | 0.5 | 0.83 | 0.46 | 0.49 | 0.39 | 0.51 | -0.2 | 0.48 | 0.35 | 0.47 | 0.28 | 0.095 | -0.235 | -0.1375 | 0.622989 | 0.316272 | 0.455935 | 0.999971 | 1 | 0.887217 |
| Nt5c3b | Q3UFY7 | -0.23 | 0.59 | 0.9 | 0.7 | 0.55 | 0.13 | 0.59 | 0.7 | 0.46 | -0.2 | 0.6 | 0.93 | 0.45 | 0.46 | 0.7 | 1.02 | 0.0025 | -0.0425 | 0.1675 | 0.993113 | 0.900476 | 0.571947 | 0.999971 | 1 | 0.91816 |
| Nt5dc2 | A0A2I3BRL | -0.86 | -1.12 | -1.04 | -1.14 | -0.09 | -1.15 | -1.33 | -0.64 | -0.45 | -0.97 | -0.5 | -0.48 | -0.49 | -0.8 | -0.09 | -0.44 | 0.2375 | 0.44 | 0.585 | 0.430797 | 0.048401 | 0.020124 | 0.999971 | 1 | 0.635388 |
| Nt5dc3 | Q3UHB1 | -0.32 | -0.51 | -0.46 | -0.42 | -0.76 | -0.52 | -0.91 | -0.45 | -0.98 | -0.61 | -0.91 | -0.64 | -0.72 | -0.17 | -0.67 | -0.17 | -0.2325 | -0.3575 | -0.005 | 0.231641 | 0.070038 | 0.98093 | 0.999971 | 1 | 0.99925 |
| Nt5e | Q61503 | -0.81 | -0.88 | -1.35 | -0.39 | -0.56 | -1.56 | -1.79 | -0.36 | -0.82 | -0.76 | -1.82 | -1.16 | -0.81 | -0.85 | -0.32 | -0.96 | -0.21 | -0.2825 | 0.1225 | 0.593706 | 0.382214 | 0.645587 | 0.999971 | 1 | 0.93301 |
| Nt5m | Q8VCE6 | -0.37 | -0.07 | -0.54 | -0.06 | -0.2 | -0.29 | -0.84 | -0.23 | 0.15 | -0.71 | -0.81 | -0.98 | 0.95 | -0.18 | -0.3 | -0.25 | -0.13 | -0.3275 | 0.315 | 0.575061 | 0.275587 | 0.343854 | 0.999971 | 1 | 0.862346 |
| Ntan1 | Q64311 | -0.45 | -0.9 | 2.12 | 2.68 | -0.24 | -0.44 | 1.62 | 3.42 | -0.94 | -1.1 | 2.77 | 1.22 | -1.44 | 0.21 | 1.76 | 1.18 | 0.2275 | -0.375 | -0.435 | 0.841171 | 0.746598 | 0.670715 | 0.999971 | 1 | 0.93891 |
| Ntaq1 | Q80WB5 | 0.04 | 0.16 | 0.82 | 0.59 | 0.03 | -0.04 | 1.51 | 0.06 | 2.4 | 0.51 | 1.29 | 0.78 | 0.66 | 0.46 | -0.29 | -0.08 | -0.0125 | 0.8425 | -0.215 | 0.974918 | 0.078445 | 0.477462 | 0.999971 | 1 | 0.894438 |
| Ntm | D3Z396 | -1.29 | -1.71 | -1.53 | -1.07 | -1.59 | -1.56 | -1.71 | -1.21 | -1.75 | -1.29 | -1.4 | -1.15 | -1.7 | -1.35 | -1.42 | -0.95 | -0.1175 | 0.0025 | 0.045 | 0.59637 | 0.991184 | 0.852167 | 0.999971 | 1 | 0.98187 |
| Ntmt1 | Q8R2U4 | -0.5 | 0.77 | -0.07 | -0.06 | 0.66 | -0.14 | 0.04 | -0.03 | 0.54 | -0.05 | 0.21 | 0.23 | 0.74 | 0.06 | 0.01 | 0.27 | 0.0975 | 0.1975 | 0.235 | 0.762604 | 0.514972 | 0.466095 | 0.999971 | 1 | 0.891431 |
| Ntng1 | Q8R4G0-5 | -1.67 | -0.94 | -0.89 | -0.68 | 0.61 | -1.51 | -0.87 | -0.88 | 1.65 | -1.87 | -0.67 | -0.55 | 1.41 | -1.34 | -0.47 | -0.92 | 0.3825 | 0.685 | 0.715 | 0.421126 | 0.338886 | 0.24508 | 0.999971 | 1 | 0.837316 |
| Ntng2 | J3QNA0 | -0.37 | -0.14 | 0.06 | 0.06 | -0.48 | 1.12 | -0.22 | -0.01 | -0.7 | -0.52 | 0.02 | -0.18 | -0.45 | 0.33 | -0.1 | 0.02 | 0.2 | -0.2475 | 0.0475 | 0.580348 | 0.294705 | 0.836458 | 0.999971 | 1 | 0.979416 |
| Ntpcr | Q9CQA9 | 0.38 | 0.37 | 1.88 | 1.12 | 0.13 | -0.32 | 1.92 | 1.76 | -0.04 | 0 | 2.33 | 3.17 | 0.46 | -0.3 | 1.88 | 1.41 | -0.065 | 0.4275 | -0.075 | 0.915333 | 0.59978 | 0.892824 | 0.999971 | 1 | 0.98828 |
| Ntrk2 | P15209 | -0.43 | -0.44 | -0.13 | 0.05 | -0.23 | -0.15 | -0.18 | -0.05 | -0.4 | -0.36 | -0.3 | -0.51 | -0.13 | -0.39 | -0.29 | 0.12 | 0.085 | -0.155 | 0.065 | 0.658944 | 0.423142 | 0.760798 | 0.999971 | 1 | 0.961116 |
| Ntrk3 | Q6VNS1-3 | -2.14 | -1.1 | -1.03 | -1.03 | -0.62 | -1.68 | -0.55 | -0.93 | -0.99 | -1.77 | -1.69 | 0.11 | -0.08 | -1.64 | -0.32 | -1.15 | 0.38 | 0.24 | 0.5275 | 0.313443 | 0.619721 | 0.239649 | 0.999971 | 1 | 0.83365 |
| Ntrk3_1 | Q6VNS1 | -1.29 | -1.35 | -1.29 | -1.17 | -1.06 | -0.97 | -1.03 | -0.94 | -0.51 | -0.89 | -1.09 | -1.29 | -0.69 | -0.92 | -1.02 | -0.8 | 0.275 | 0.33 | 0.4175 | 0.116576 | 0.150063 | 0.03643 | 0.999971 | 1 | 0.700096 |
| Nts | Q9D3P9 | 0.63 | -2.1 | -0.77 | -0.71 | 0.83 | -0.31 | -0.76 | -0.22 | 1.27 | 0.38 | -1.61 | 0.24 | 2.03 | 1.51 | 1.32 | -0.9 | 0.6225 | 0.8075 | 1.7275 | 0.310442 | 0.296 | 0.04859 | 0.999971 | 1 | 0.710102 |
| Ntsr2 | P70310 | -0.68 | -2.42 | -0.88 | -0.83 | -2.74 | -2.37 | 0.21 | -1.75 | -1.78 | -2.17 | -0.87 | -2.09 | -1.47 | -1.56 | -1.62 | -2.43 | -0.46 | -0.525 | -0.5675 | 0.515617 | 0.285085 | 0.21773 | 0.999971 | 1 | 0.826893 |
| Nuak1 | Q641K5 | 2.67 | 0.18 | 0.7 | -0.62 | 0.37 | 0.46 | 1.18 | -0.26 | 0.64 | 1.2 | 0.09 | -0.5 | 0.7 | -0.79 | -0.49 | -1.56 | -0.295 | -0.375 | -1.2675 | 0.669353 | 0.604224 | 0.122857 | 0.999971 | 1 | 0.78268 |
| Nub1 | A0A0G2JG( | 0.42 | 0.5 | 0.77 | 0.89 | 0.45 | 0.65 | 0.71 | 0.92 | 0.73 | 0.72 | 0.81 | 0.78 | 0.44 | 0.82 | 0.84 | 0.9 | 0.0375 | 0.115 | 0.105 | 0.853429 | 0.532695 | 0.613969 | 0.999971 | 1 | 0.927364 |

| | | | | | | | | | | | | | | | | | | | | | | | | | | |
|---|---|---|---|---|---|---|---|---|---|---|---|---|---|---|---|---|---|---|---|---|---|---|---|---|---|---|
| Nubp1 | Q9R060 | 1.1 | 0.57 | 0.45 | 0.28 | 0.83 | 1.16 | -0.31 | 0.12 | 1.06 | 1.12 | 0.16 | 0.52 | 0.47 | 1.12 | 0.68 | 0.02 | -0.15 | 0.115 | -0.0275 | 0.683984 | 0.70035 | 0.926608 | 0.999971 | 1 | 0.99401 |
| Nubp2 | A0A3Q4L2 | -0.02 | -0.19 | -0.11 | 0.05 | 0.61 | -0.9 | -0.43 | -0.74 | -0.61 | 0.49 | 0.11 | 0.07 | -0.01 | -1.11 | -0.04 | 0.3 | -0.2975 | 0.0825 | -0.1475 | 0.391995 | 0.749129 | 0.643189 | 0.999971 | 1 | 0.93301 |
| Nubpl | Q9CWD8 | -0.51 | -0.18 | -0.57 | -0.44 | -0.96 | -0.71 | -0.7 | 0.1 | -1.26 | -0.55 | -0.66 | -0.63 | -0.75 | -0.6 | 0.04 | -0.13 | -0.1425 | -0.35 | 0.065 | 0.595961 | 0.143456 | 0.786872 | 0.999971 | 1 | 0.96724 |
| Nucb1 | Q02819 | 0.45 | 2.85 | 1.65 | 1.47 | 2.54 | 2.66 | 1.65 | 1.53 | 2.42 | 1.08 | 1.61 | 1.64 | 3 | 2.46 | 1.72 | 1.7 | 0.49 | 0.0825 | 0.615 | 0.365673 | 0.874716 | 0.271566 | 0.999971 | 1 | 0.843794 |
| Nucb2 | P81117 | 0.85 | 0.73 | 1.08 | 1.22 | 0.72 | 0.87 | 0.87 | 1.3 | 1.1 | 0.79 | 1.18 | 1.28 | 0.81 | 0.63 | 0.95 | 1.62 | -0.03 | 0.1175 | 0.0325 | 0.888708 | 0.567925 | 0.902322 | 0.999971 | 1 | 0.990759 |
| Nucks1 | A0A087WF | 2.87 | 3.03 | 2.83 | 3.74 | 2.41 | 2.08 | 2.68 | 2.77 | 2.29 | 2.73 | 2.47 | 2.04 | 1.74 | 2.87 | 2.28 | 2.55 | -0.6325 | -0.735 | -0.7575 | 0.046221 | 0.023667 | 0.040585 | 0.999971 | 1 | 0.700838 |
| Nudc | O35685 | -0.62 | 0.34 | 0.44 | 0.15 | 0.2 | 0.57 | 0.63 | 0.54 | 0.34 | -0.45 | 0.51 | 0.72 | 0.76 | 0.46 | 0.61 | 0.45 | 0.4075 | 0.2025 | 0.4925 | 0.164827 | 0.561883 | 0.096356 | 0.999971 | 1 | 0.78268 |
| Nudcd1 | Q6PIP5 | -1.41 | -0.56 | 0.08 | 0.43 | -0.83 | -0.91 | -0.07 | 0.08 | -0.86 | -0.92 | 0.02 | 0.22 | -1.6 | -1.05 | 0.17 | -0.05 | -0.0675 | -0.02 | -0.2675 | 0.880388 | 0.965968 | 0.620244 | 0.999971 | 1 | 0.929296 |
| Nudcd2 | Q9CQ48 | 0.42 | 1.03 | 0.86 | 0.19 | 1.84 | -0.06 | 0.74 | 0.07 | 2.4 | -0.7 | 0.91 | 0.64 | 2.94 | -0.41 | -0.1 | 0.47 | 0.0225 | 0.1875 | 0.1 | 0.959808 | 0.758783 | 0.887961 | 0.999971 | 1 | 0.987485 |
| Nudcd3 | Q8R1N4 | -0.37 | 0.21 | -0.16 | 0.07 | 0.6 | 0.18 | 0.56 | 0.09 | 0.8 | -0.18 | 0.36 | 0.25 | 0.64 | 0.05 | -0.01 | 0.34 | 0.42 | 0.37 | 0.3175 | 0.087692 | 0.179327 | 0.19731 | 0.999971 | 1 | 0.812891 |
| Nudt11 | P0C028 | -0.24 | -0.27 | -0.25 | -0.17 | -0.1 | -0.4 | -0.1 | 0.16 | -0.14 | -0.29 | 0.16 | 0.53 | -0.46 | -0.13 | 0.46 | -0.19 | 0.1225 | 0.2975 | 0.1525 | 0.517742 | 0.20291 | 0.517322 | 0.999971 | 1 | 0.905815 |
| Nudt12 | Q9DCN1 | 0.85 | -0.38 | 0.36 | -0.02 | -0.28 | 0.52 | 0.69 | 0.44 | -0.2 | 1.45 | 0.68 | 0.34 | -0.69 | -0.45 | 0.34 | -0.66 | 0.14 | 0.365 | -0.5675 | 0.678922 | 0.389379 | 0.133236 | 0.999971 | 1 | 0.784948 |
| Nudt13 | Q8JZU0 | -2.47 | -1.31 | 0.62 | 0.78 | -1.91 | -1.65 | 3.76 | 0.54 | -2.92 | -0.7 | -1.18 | -0.91 | -1.93 | -1.97 | -0.5 | -2.23 | 0.78 | -0.8325 | -1.0625 | 0.568883 | 0.338057 | 0.201657 | 0.999971 | 1 | 0.815758 |
| Nudt14 | Q9D142 | -0.78 | -0.29 | 0.97 | 0.45 | 2.39 | -0.44 | 0.55 | 1.59 | 0.53 | -0.09 | 0.2 | 0.6 | -0.74 | -0.3 | 0.44 | 0.77 | 0.935 | 0.2225 | -0.045 | 0.180916 | 0.583055 | 0.925741 | 0.999971 | 1 | 0.99401 |
| Nudt15 | Q8BG93 | 1.39 | 0.74 | 1.14 | 0.84 | 0.36 | 0.71 | 0.88 | 0.32 | 0.58 | 1.55 | 0.59 | 0.2 | -1.17 | 0.95 | 0.39 | 0.36 | -0.46 | -0.2975 | -0.895 | 0.0777 | 0.370955 | 0.073348 | 0.999971 | 1 | 0.746069 |
| Nudt16 | Q6P3D0 | -0.36 | 0.73 | 0.09 | -0.08 | 0.82 | 0.83 | 0.09 | 0.32 | 0.84 | 0.1 | -0.01 | 0.32 | 0.84 | 1.04 | 0.62 | -0.15 | 0.42 | 0.2175 | 0.4925 | 0.189841 | 0.481804 | 0.177263 | 0.999971 | 1 | 0.808532 |
| Nudt16l1 | Q8VHN8 | -0.01 | 0.94 | 1.1 | 0.75 | 1.28 | 0.56 | 0.68 | 0.68 | 1.16 | 0.61 | 0.78 | 1.38 | 1.22 | 1.18 | 1.32 | 1.28 | 0.105 | 0.2875 | 0.555 | 0.728978 | 0.361966 | 0.064377 | 0.999971 | 1 | 0.726597 |
| Nudt17 | Q9CWD3-2 | 1.56 | 0.59 | 0.55 | 0.52 | -0.19 | 0.75 | 0.91 | -0.36 | 0.18 | 1.16 | -0.12 | -0.37 | -0.32 | 0.31 | -0.64 | -0.68 | -0.5275 | -0.5925 | -1.1375 | 0.201892 | 0.166081 | 0.008109 | 0.999971 | 1 | 0.602121 |
| Nudt18 | Q3U2V3 | -0.97 | 0.05 | 0.26 | 0.24 | 0.73 | -0.46 | 0.16 | 0.11 | -0.43 | -0.86 | 0.1 | 0.54 | 0.24 | -0.24 | -0.14 | 0.15 | 0.24 | -0.0575 | 0.1075 | 0.520503 | 0.886962 | 0.735649 | 0.999971 | 1 | 0.958637 |
| Nudt19 | P11930 | -1.24 | -0.55 | -0.08 | -1.64 | -0.59 | -1.16 | -0.28 | -1.11 | -0.64 | -1.95 | -0.16 | -1.27 | -0.18 | -1.14 | -0.54 | -1.76 | 0.0925 | -0.1275 | -0.0275 | 0.813225 | 0.793888 | 0.952398 | 0.999971 | 1 | 0.998056 |
| Nudt2 | P56380 | -0.13 | 1.08 | 0.48 | -0.31 | 0.97 | 1.13 | 0.81 | 0.95 | 0.86 | -0.02 | 0.66 | 0.97 | 1.3 | 1.08 | 0.67 | 0.05 | 0.685 | 0.3375 | 0.495 | 0.059455 | 0.37903 | 0.237453 | 0.999971 | 1 | 0.833594 |
| Nudt21 | Q9CQF3 | 0.53 | 2.36 | 2.59 | 2.39 | 2.39 | 1.96 | 2.34 | 2.32 | 2.41 | 1.36 | 2.62 | 2.6 | 1.45 | 2.09 | 2.38 | 2.55 | 0.285 | 0.28 | 0.15 | 0.540292 | 0.597607 | 0.764948 | 0.999971 | 1 | 0.962405 |
| Nudt3 | I1E4X7 | -0.4 | 1.63 | 1.19 | 0.08 | 2.97 | 0.13 | 3.67 | -0.27 | 3.77 | -0.96 | 2.22 | 2.03 | 1.04 | -1.88 | -0.84 | -1.49 | 1 | 1.14 | -1.4175 | 0.322712 | 0.266079 | 0.077281 | 0.999971 | 1 | 0.752059 |
| Nudt4 | Q8R2U6 | -0.04 | 0.2 | -0.57 | 0.33 | -0.22 | 0 | -0.8 | -0.23 | 0.34 | -0.24 | -0.65 | -0.42 | -0.17 | 0.36 | -0.8 | 0.22 | -0.2925 | -0.2225 | -0.0775 | 0.308674 | 0.463148 | 0.813202 | 0.999971 | 1 | 0.974559 |
| Nudt5 | Q9JKX6 | 0.57 | 0.72 | 1.5 | 0.82 | 0.72 | 0.9 | 0.89 | 0.83 | 0.81 | 0.87 | 0 | 1.12 | 0.56 | 0.96 | 0.92 | 0.8 | -0.0675 | -0.2025 | -0.0925 | 0.780347 | 0.531627 | 0.714813 | 0.999971 | 1 | 0.952456 |
| Nudt7 | Q99P30-3 | -0.33 | -0.93 | -1.15 | -0.82 | -0.38 | -0.29 | -0.91 | -0.26 | -0.31 | -0.19 | -1.38 | -1.14 | -0.35 | -0.64 | -1.18 | -1.06 | 0.3475 | 0.0525 | -1.03E-16 | 0.197523 | 0.877277 | 1 | 0.999971 | 1 | 1 |
| Nudt8 | Q9CR24 | -0.35 | 0.03 | -0.17 | -0.59 | -0.33 | 0.07 | -0.62 | -0.16 | -0.43 | -0.4 | -0.78 | -1.25 | -0.36 | 0.08 | -0.23 | -0.29 | 0.01 | -0.445 | 0.07 | 0.965702 | 0.114142 | 0.743323 | 0.999971 | 1 | 0.961064 |
| Nudt9 | Q8BVU5 | 0.93 | 0.16 | 1.25 | 0.87 | 0.11 | 0.19 | 0.98 | 1.12 | 0.38 | 0.12 | 1.62 | 1.41 | -0.29 | -0.6 | 0.7 | 0.4 | -0.2025 | 0.08 | -0.75 | 0.559745 | 0.847628 | 0.067983 | 0.999971 | 1 | 0.733937 |
| Nufip2 | Q5F2E7 | 1.03 | 1.53 | 1.62 | 1.76 | 1.51 | 1.48 | 1.47 | 1.7 | 1.51 | 1.56 | 1.53 | 1.56 | 1.7 | 1.48 | 1.64 | 1.81 | 0.055 | 0.055 | 0.1725 | 0.79829 | 0.791276 | 0.439004 | 0.999971 | 1 | 0.883105 |
| Numa1 | E9Q7G0 | 0.29 | -0.39 | -0.34 | -0.37 | -0.28 | -0.54 | -0.17 | -0.72 | -0.43 | 0.07 | -0.71 | -0.83 | -0.41 | -0.49 | -0.84 | -0.93 | -0.225 | -0.2725 | -0.465 | 0.359708 | 0.335681 | 0.080954 | 0.999971 | 1 | 0.755254 |
| Numb | Q9QZS3-4 | 0.52 | -0.58 | -0.76 | -0.63 | -0.44 | 0.23 | -0.05 | -1.46 | -0.28 | 0.34 | -0.51 | -1.09 | -0.68 | -0.51 | -1.01 | -1.23 | -0.0675 | -0.0225 | -0.495 | 0.879551 | 0.955256 | 0.16628 | 0.999971 | 1 | 0.800215 |
| Numbl | O08919 | -0.51 | -0.24 | -0.62 | -0.9 | -0.08 | -0.2 | -0.4 | -0.6 | -0.2 | -0.52 | -0.68 | -0.83 | 0.39 | -0.02 | -0.8 | -0.72 | 0.2475 | 0.01 | 0.28 | 0.280719 | 0.964998 | 0.392045 | 0.999971 | 1 | 0.871816 |
| Nup107 | Q8BH74 | -0.88 | -1.17 | -0.27 | -0.41 | -1.1 | -1.28 | -0.37 | -0.34 | -1.05 | -1.12 | -0.32 | -0.37 | -1.21 | -1.1 | -0.2 | 0.12 | -0.09 | -0.0325 | 0.085 | 0.78035 | 0.915093 | 0.821935 | 0.999971 | 1 | 0.975833 |
| Nup133 | Q8R0G9 | -0.08 | -0.22 | -0.55 | -0.38 | -0.19 | -0.13 | -0.79 | -0.51 | -0.37 | -0.01 | -0.46 | -0.59 | 0.13 | 0.01 | -0.5 | -0.32 | -0.0975 | -0.05 | 0.1375 | 0.665905 | 0.81039 | 0.539155 | 0.999971 | 1 | 0.910077 |
| Nup153 | E9Q3G8 | -0.11 | -0.24 | -0.6 | -0.2 | -0.26 | -0.11 | -0.69 | -0.26 | -0.31 | -0.26 | -0.3 | -0.5 | 0.01 | -0.03 | -0.09 | 0.03 | -0.0425 | -0.055 | 0.2675 | 0.842131 | 0.767735 | 0.173152 | 0.999971 | 1 | 0.806803 |
| Nup155 | A0A2R8VH | -1.13 | -0.86 | -0.88 | -0.78 | -1.05 | -1.18 | -0.69 | -0.92 | -1.14 | -1.16 | -0.91 | -0.79 | -1.06 | -0.7 | -0.69 | -0.84 | -0.0475 | -0.0875 | 0.09 | 0.806037 | 0.637536 | 0.632356 | 0.999971 | 1 | 0.93301 |
| Nup160 | Q9Z0W3 | -1.71 | -1.16 | -0.85 | -1.28 | -1.24 | -1.32 | -0.88 | -1.18 | -1.59 | -1.65 | -0.89 | -0.73 | -1.25 | -1.13 | -1.26 | -1.07 | 0.095 | 0.035 | 0.0725 | 0.689429 | 0.907913 | 0.748541 | 0.999971 | 1 | 0.961064 |
| Nup188 | Q6ZQH8 | 0.73 | -0.06 | -0.33 | -0.26 | 0.33 | -0.01 | -0.23 | -0.16 | -0.08 | 0.34 | -0.34 | -0.53 | 0.21 | 0.02 | -0.39 | -0.6 | -0.0375 | -0.1725 | -0.21 | 0.895831 | 0.583471 | 0.505933 | 0.999971 | 1 | 0.905277 |
| Nup205 | B9EJ54 | -0.8 | -0.12 | 0.04 | -0.3 | -0.01 | -0.31 | -0.25 | -0.22 | 0.13 | -0.85 | 0 | -0.18 | 0.53 | -0.29 | -0.33 | -0.3 | 0.0975 | 0.07 | 0.1975 | 0.674321 | 0.811662 | 0.501767 | 0.999971 | 1 | 0.904573 |
| Nup210 | Q9QY81 | -0.39 | -0.25 | -0.6 | -0.57 | -0.37 | -0.19 | -1.01 | -0.65 | -0.39 | -0.17 | -0.41 | -0.63 | -0.22 | -0.13 | -0.42 | -0.09 | -0.1025 | 0.0525 | 0.2375 | 0.661649 | 0.780526 | 0.220247 | 0.999971 | 1 | 0.827096 |
| Nup214 | Q80U93 | -0.56 | -0.29 | -0.43 | -0.38 | -0.34 | -0.63 | -0.35 | -0.45 | -0.27 | -0.71 | -0.32 | -0.41 | -0.02 | -0.31 | -0.55 | -0.48 | -0.0275 | -0.0125 | 0.075 | 0.874801 | 0.94536 | 0.701392 | 0.999971 | 1 | 0.949019 |
| Nup35 | Q8R4R6 | 0.47 | -1.22 | -0.77 | -0.94 | -1.17 | -1.14 | -0.97 | -1.23 | -1.43 | -0.27 | -0.99 | -1.18 | -1.65 | -1.11 | -1.11 | -1.11 | -0.5125 | -0.3525 | -0.63 | 0.186423 | 0.417918 | 0.127509 | 0.999971 | 1 | 0.783863 |
| Nup37 | Q9CWU9 | 1.2 | 0.05 | 0.65 | 0.74 | -0.23 | 0.43 | 0.97 | 0.26 | -0.07 | 0.57 | 0.36 | -0.63 | -0.31 | -0.24 | -0.07 | -0.46 | -0.3025 | -0.6025 | -0.93 | 0.383269 | 0.112335 | 0.007073 | 0.999971 | 1 | 0.582714 |
| Nup43 | P59235 | 1.22 | 0.32 | -0.73 | -0.96 | 0.01 | 0.16 | -0.95 | -1.23 | -0.2 | 0.67 | -0.68 | -1.12 | 0.35 | 0.5 | -0.99 | -0.37 | -0.465 | -0.295 | -0.09 | 0.415632 | 0.61482 | 0.872399 | 0.999971 | 1 | 0.984109 |
| Nup50 | Q9JIH2 | 0.3 | -0.43 | -0.11 | -0.42 | -0.63 | -0.39 | 0.09 | -0.2 | -0.38 | -0.21 | -0.42 | -0.4 | -1.11 | -0.21 | -0.08 | -0.36 | -0.1175 | -0.1875 | -0.275 | 0.647005 | 0.402934 | 0.367518 | 0.999971 | 1 | 0.869526 |
| Nup54 | Q8BTS4 | -0.92 | -0.93 | -0.96 | -0.68 | -0.72 | -1.33 | -0.58 | -0.9 | -0.94 | -1.15 | -1.28 | -0.97 | -0.59 | -0.9 | -0.89 | -0.75 | -0.01 | -0.2125 | 0.09 | 0.963634 | 0.250992 | 0.617913 | 0.999971 | 1 | 0.928581 |
| Nup58 | Q8R332-4 | -0.5 | -2.37 | -1.5 | -1.26 | -1.84 | -2.16 | -1.27 | -1.49 | -2.13 | -1.02 | -1.27 | -1.12 | -2.29 | -1.82 | -1.34 | -1.12 | -0.2825 | 0.0225 | -0.235 | 0.497776 | 0.958852 | 0.595431 | 0.999971 | 1 | 0.92377 |
| Nup62 | Q63850 | -0.06 | -0.01 | -1.21 | -1.28 | -0.04 | -0.44 | -0.77 | -1.97 | -0.28 | -0.05 | -1.15 | -1.35 | 0.3 | 0.63 | -1.38 | -1.71 | -0.165 | -0.0675 | 0.1 | 0.743675 | 0.879911 | 0.872592 | 0.999971 | 1 | 0.984109 |
| Nup85 | Q8R480 | -0.31 | 0.04 | -1.27 | -1.41 | -0.19 | -0.53 | -0.7 | -1.64 | -0.13 | -0.35 | -1.19 | -1.38 | -0.44 | -0.34 | -1.75 | -1.65 | -0.0275 | -0.025 | -0.3075 | 0.950597 | 0.955095 | 0.530841 | 0.999971 | 1 | 0.910069 |
| Nup88 | Q8CEC0-3 | -0.82 | -0.76 | -0.95 | -1.15 | -0.63 | -1 | -0.92 | -1.24 | -0.85 | -0.85 | -0.74 | -1.16 | -1.06 | -0.83 | -1.25 | -1.3 | -0.0275 | 0.02 | -0.19 | 0.893784 | 0.915374 | 0.350933 | 0.999971 | 1 | 0.867482 |
| Nup93 | Q8BJ71 | 0.11 | -0.34 | -0.18 | -0.33 | -0.35 | -0.52 | 0.1 | -0.62 | -0.3 | -0.37 | -0.39 | -0.25 | 0.08 | -0.66 | -0.57 | -0.57 | -0.1625 | -0.1425 | -0.245 | 0.484699 | 0.438912 | 0.315327 | 0.999971 | 1 | 0.852494 |
| Nup98 | A0A1B0GS | -0.69 | -0.63 | 0.01 | -0.35 | -0.47 | -0.7 | -0.23 | -0.32 | -0.34 | -0.78 | -0.29 | -0.65 | -0.12 | -0.81 | -0.17 | -0.27 | -0.015 | -0.1 | 0.0725 | 0.947676 | 0.66917 | 0.774757 | 0.999971 | 1 | 0.963884 |
| Nupl2 | E9QL43 | 0.68 | 0.39 | 2.36 | 0.92 | 0.22 | -0.33 | 1.51 | 2.29 | 0.28 | 0.41 | 3.79 | 2.02 | 1.72 | -0.49 | 1.79 | 0.96 | -0.165 | 0.5375 | -0.0925 | 0.80581 | 0.528227 | 0.882519 | 0.999971 | 1 | 0.986576 |
| Nutf2 | P61971 | 0.24 | 0.87 | 0.44 | 0.41 | 0.56 | 0.54 | 0.33 | 0.56 | 0.37 | -0.31 | 0.47 | 0.82 | 0.06 | 0.79 | 0.47 | 0.23 | 0.0075 | -0.1525 | -0.1025 | 0.970354 | 0.594835 | 0.671853 | 0.999971 | 1 | 0.939563 |
| Nvl | Q9DBY8 | 3.54 | -0.05 | 0.33 | 0.48 | 1.34 | 0.98 | -0.21 | 0.73 | 1.01 | 2.56 | -0.06 | 0.29 | -0.74 | -0.7 | 0.69 | 1.04 | -0.365 | -0.125 | -1.0025 | 0.650831 | 0.890985 | 0.258901 | 0.999971 | 1 | 0.839012 |
| Nwd1 | E0CX99 | 0.33 | -0.14 | 0.06 | -0.23 | -0.28 | 0.3 | -0.47 | 0 | -0.22 | 0.43 | 0.26 | -1.12 | 0.05 | 0.02 | -0.24 | -0.34 | -0.1175 | -0.1675 | -0.1325 | 0.62821 | 0.644329 | 0.531474 | 0.999971 | 1 | 0.910069 |
| Nwd2 | Q6P5U7 | -0.72 | -0.5 | -0.42 | -0.79 | -0.7 | -0.29 | -0.58 | -0.43 | -0.64 | -0.66 | -0.38 | -0.52 | -0.06 | -0.45 | -0.57 | -0.52 | 0.1075 | 0.0575 | 0.2075 | 0.577611 | 0.750487 | 0.320998 | 0.999971 | 1 | 0.854682 |
| Nxf1 | Q99JX7 | 0.52 | 0.51 | 0.31 | 0.3 | 0.37 | 0.44 | 0.12 | -0.07 | 0.16 | 0.7 | 0.17 | 0.16 | 0.42 | 0.81 | 0.15 | 0.5 | -0.195 | -0.1125 | 0.06 | 0.331017 | 0.578112 | 0.769726 | 0.999971 | 1 | 0.963498 |
| Nxn | P97346 | -0.4 | -0.19 | 0.18 | -0.5 | -0.16 | -0.23 | 0.09 | -0.16 | 0.19 | -0.62 | 0.2 | 0.7 | -0.17 | -0.13 | 0.39 | -0.24 | 0.1125 | 0.345 | 0.19 | 0.601741 | 0.290452 | 0.439639 | 0.999971 | 1 | 0.883105 |
| Nxph1 | Q61200 | -1.28 | -0.79 | -1.41 | -2.13 | -1.09 | -0.27 | -0.74 | -1.67 | -0.47 | -1.01 | -1.28 | -0.63 | -1.09 | -0.77 | -1.56 | -1.56 | 0.46 | 0.555 | 0.1575 | 0.255734 | 0.121053 | 0.641274 | 0.999971 | 1 | 0.93301 |
| Nxph3 | Q91VX5 | 0.2 | -0.01 | -0.37 | -0.79 | 0.01 | 0.98 | 0.22 | 1.16 | 0.03 | 0.27 | 0.89 | 0.41 | -0.31 | 0.56 | 0.37 | 0.39 | 0.835 | 0.6425 | 0.495 | 0.038091 | 0.052627 | 0.125379 | 0.999971 | 1 | 0.78268 |
| Nxt1 | Q9QZV9 | 0.82 | 0.86 | 0.1 | -0.38 | 0.32 | 0.76 | 0.93 | -0.48 | 1.15 | 1.11 | 0.03 | -1.46 | 0.96 | 0.48 | 0.21 | -0.47 | 0.0325 | -0.1425 | -0.055 | 0.937235 | 0.819767 | 0.89176 | 0.999971 | 1 | 0.988225 |
| Nyap1 | E9PXJ0 | 0.41 | -0.57 | -0.19 | -0.22 | -0.34 | -0.36 | 0.46 | 0.01 | -0.75 | 0.05 | -0.08 | -0.44 | -0.63 | -0.46 | -0.49 | -0.32 | 0.085 | -0.1625 | -0.3325 | 0.770777 | 0.571783 | 0.197102 | 0.999971 | 1 | 0.812891 |
| Nyap2 | A0A0R4J1X | 1.19 | -0.43 | 0.2 | 0.14 | -0.97 | -0.05 | 0.95 | -0.22 | -0.97 | 0.42 | -0.16 | -0.29 | -0.53 | -0.59 | -0.35 | -0.37 | -0.3475 | -0.525 | -0.735 | 0.478378 | 0.232372 | 0.054932 | 0.999971 | 1 | 0.725348 |
| Oard1 | Q8R5F3 | 1.73 | 2.13 | 2.81 | 2.61 | 3.25 | 0.01 | 2.6 | 2 | 2.39 | 1.1 | 2.2 | 1.77 | 2.31 | 1.84 | 0.97 | 0.75 | -0.355 | -0.455 | -0.8525 | 0.599617 | 0.234667 | 0.067666 | 0.999971 | 1 | 0.733937 |
| Oat | P29758 | -1.32 | -1.43 | -1.82 | -1.5 | -1.01 | -1.31 | -1.62 | -1.22 | -1.25 | -1.56 | -1.76 | -1.54 | -0.84 | -0.93 | -1.42 | -1.45 | 0.2275 | -0.01 | 0.3575 | 0.303532 | 0.960514 | 0.147974 | 0.999971 | 1 | 0.793248 |
| Obp1a | Q9D3H2 | 2.52 | 6.68 | 5.09 | 6.25 | -8.6 | 2.33 | -4.27 | 6.47 | 4.06 | 0.73 | 2.13 | -2.91 | 9.09 | 7.06 | -0.87 | -14.93 | -6.1525 | -4.1325 | -5.0475 | 0.071949 | 0.025163 | 0.316424 | 0.999971 | 1 | 0.85361 |
| Obscn | H7BX05 | -11.19 | 0.48 | 0.26 | -0.49 | 0.25 | 0.22 | 0.55 | -0.39 | -0.2 | -5.06 | 0.43 | -0.6 | 0.23 | -0.53 | -1.44 | -1.04 | 2.8925 | 1.3775 | 2.04 | 0.266051 | 0.619098 | 0.427167 | 0.999971 | 1 | 0.880793 |
| Ociad1 | Q9CRD0 | -0.74 | -0.91 | -0.38 | -0.64 | -0.61 | -0.69 | -0.21 | -0.31 | -0.84 | -1.02 | -0.31 | -0.35 | -0.97 | -0.92 | -0.42 | -0.9 | 0.2125 | 0.0375 | -0.135 | 0.326979 | 0.875559 | 0.537904 | 0.999971 | 1 | 0.910077 |
| Ociad1_1 | Q9CRD0-3 | -1.81 | -2.56 | -2.35 | -2.19 | -1.76 | -1.98 | -1.63 | -2.72 | -1.39 | -1.82 | -1.78 | -2.39 | -1.97 | -3.91 | -1.79 | -3.48 | 0.205 | 0.3825 | -0.56 | 0.499031 | 0.189072 | 0.293206 | 0.999971 | 1 | 0.847402 |
| Ociad2 | Q9D8W7 | 0.64 | 1.57 | 1.37 | 1 | 1.74 | 1.53 | 1.7 | 1.08 | 2.43 | 0.7 | 1.78 | 2.07 | 2.43 | 1.16 | 1.1 | 1.68 | 0.3675 | 0.6 | 0.4475 | 0.201456 | 0.166209 | 0.237069 | 0.999971 | 1 | 0.833594 |
| Ocln | Q61146 | -1.01 | -0.89 | 0.35 | 0.32 | -0.73 | -1.3 | 0.29 | 0.24 | -1.07 | -0.37 | 0.22 | -0.24 | -1.31 | -1.11 | 0.4 | 0.77 | -0.0675 | -0.0575 | -0.005 | 0.892133 | 0.894458 | 0.99322 | 0.999971 | 1 | 0.99925 |
| Ocrl | Q6NVF0 | -0.59 | -0.34 | -0.91 | -1.23 | -0.24 | -0.25 | -0.66 | -1.22 | -0.17 | -0.57 | -0.78 | -0.83 | -0.03 | -0.39 | -0.93 | -1.44 | 0.175 | 0.18 | 0.07 | 0.572433 | 0.502117 | 0.844725 | 0.999971 | 1 | 0.979814 |
| Odr4 | Q4PJX1 | -0.75 | -0.46 | 0.09 | -0.12 | -0.41 | -0.79 | -0.15 | 0.09 | -0.23 | -0.78 | 0.08 | 0.08 | -0.15 | -1.03 | -0.34 | 0.01 | -0.005 | 0.0975 | -0.0675 | 0.985728 | 0.734411 | 0.823748 | 0.999971 | 1 | 0.976286 |
| Oga | Q9EQQ9 | 0.26 | 0.46 | -0.17 | 0.08 | 0.57 | 0.69 | 0.63 | 0.25 | 0.82 | 0.49 | 0.18 | 0.01 | 0.64 | 0.98 | 0.1 | 0.25 | 0.3775 | 0.2175 | 0.335 | 0.104854 | 0.393581 | 0.223501 | 0.999971 | 1 | 0.828225 |
| Ogdh | Q60597-3 | -1.28 | -0.88 | -1.27 | -1.35 | -1.01 | -1.13 | -1.13 | -1.47 | -0.87 | -0.96 | -1.32 | -1.32 | -1.05 | -1.08 | -1.27 | -1.51 | 0.01 | 0.0775 | -0.0325 | 0.960497 | 0.709708 | 0.874285 | 0.999971 | 1 | 0.984109 |
| Ogdh_1 | Q60597 | -0.99 | -1.42 | -1.59 | -1.4 | -1.62 | -1.04 | -1.51 | -1.79 | -0.77 | -0.32 | -1.74 | -1.55 | -1.43 | -1.32 | -1.55 | -0.87 | -0.14 | 0.255 | 0.0575 | 0.561065 | 0.47364 | 0.805274 | 0.999971 | 1 | 0.971701 |
| Ogdhl | B2RXT3 | -0.93 | -0.7 | -0.76 | -0.7 | -0.8 | -1.2 | -0.77 | -0.98 | -0.77 | -1.25 | -0.81 | -0.66 | -0.8 | -1.11 | -1.04 | -0.88 | -0.165 | -0.1 | -0.185 | 0.384495 | 0.613852 | 0.308864 | 0.999971 | 1 | 0.847402 |
| Ogfod1 | Q3U0K8-2 | 1.59 | 2.61 | 2.18 | 1.84 | 2.56 | 3.08 | 1.56 | 2.17 | 2.44 | 1.89 | 1.98 | 1.88 | 2.93 | 2.7 | 2.11 | 1.96 | 0.2875 | -0.0075 | 0.37 | 0.452708 | 0.978147 | 0.269787 | 0.999971 | 1 | 0.843794 |
| Ogfod3 | Q9D136 | 1.76 | -1.5 | -0.11 | 0.27 | -0.23 | -0.4 | 0.5 | 0.62 | -0.2 | 1.12 | -0.71 | -0.54 | -0.88 | -0.2 | -0.15 | 0.07 | 0.0175 | -0.1875 | -0.395 | 0.97853 | 0.79375 | 0.540299 | 0.999971 | 1 | 0.910077 |
| Ogfr | Q99PG2 | 0.11 | -0.11 | 0.01 | 0.07 | 0.23 | 0.01 | 0.01 | 0.22 | 0.61 | -0.05 | 0.25 | 0.44 | -0.03 | -0.33 | 0.19 | 0.21 | 0.0975 | 0.2925 | -0.01 | 0.572388 | 0.171981 | 0.959451 | 0.999971 | 1 | 0.998721 |
| Ogfrl1 | Q8VE52 | 0.85 | -0.7 | -0.16 | 0.23 | -0.44 | -0.04 | 0.6 | -0.18 | -0.88 | 0.1 | -0.43 | -0.2 | -0.47 | -0.77 | -0.28 | -0.63 | -0.07 | -0.4075 | -0.5925 | 0.853985 | 0.293835 | 0.109212 | 0.999971 | 1 | 0.78268 |
| Ogt | Q8CGY8-2 | -0.58 | 0.31 | -1.13 | -0.77 | 0.29 | 0.19 | -1.76 | -0.28 | -0.11 | 0.63 | -0.59 | -0.73 | -0.28 | -0.63 | -0.11 | -0.57 | 0.1525 | 0.3425 | 0.145 | 0.769849 | 0.417106 | 0.661973 | 0.999971 | 1 | 0.936148 |
| Ogt_1 | Q8CGY8 | -0.68 | -0.29 | -0.54 | -0.37 | -0.45 | -0.34 | -0.63 | -0.62 | -0.37 | -0.46 | -0.37 | -0.36 | -0.07 | -0.21 | -0.35 | 0.14 | -0.04 | 0.08 | 0.3475 | 0.828827 | 0.644137 | 0.106124 | 0.999971 | 1 | 0.78268 |
| Ola1 | Q9CZ30 | -0.04 | 1.8 | 1.42 | 0.87 | 1.75 | 2.38 | 1.67 | 1.34 | 1.99 | 0.91 | 1.75 | 2.19 | 1.77 | 1.58 | 1.59 | 1.08 | 0.7725 | 0.6975 | 0.4925 | 0.099613 | 0.155288 | 0.246213 | 0.999971 | 1 | 0.839012 |
| Olfm1 | Q8R357 | -0.57 | -0.34 | -0.8 | -0.67 | -0.3 | -0.07 | -0.91 | -0.59 | -0.44 | -0.22 | -0.54 | -0.81 | 0.03 | -0.32 | -0.71 | -0.22 | 0.1275 | 0.0925 | 0.29 | 0.597062 | 0.654719 | 0.217797 | 0.999971 | 1 | 0.826893 |
| Olfm2 | A0A1L1SV4 | -0.64 | -1.01 | -1.05 | -0.63 | -1.29 | -0.69 | -0.55 | -0.84 | -1.25 | -0.84 | -0.82 | -0.96 | -0.8 | -1.08 | -0.97 | -0.25 | -0.01 | -0.135 | 0.0575 | 0.965778 | 0.511124 | 0.816117 | 0.999971 | 1 | 0.974929 |
| Olfm3 | P63056 | -1.93 | -0.58 | -1.33 | -1.1 | -0.81 | -0.92 | -1.68 | -0.32 | -1.07 | -1.84 | -1.41 | -1.58 | -0.36 | -0.66 | -1.33 | -0.54 | 0.3025 | -0.24 | 0.5125 | 0.437194 | 0.465768 | 0.164474 | 0.999971 | 1 | 0.800215 |
| Olfml3 | A0A0R4J08 | 2.15 | 1.33 | 3.39 | 3.95 | 0.66 | 1.83 | 5.79 | 2.47 | 0.94 | 1.88 | 1.94 | 1.65 | -0.8 | 1.06 | 2.24 | 0.86 | -0.0175 | -1.1025 | -1.865 | 0.987431 | 0.087018 | 0.037561 | 0.999971 | 1 | 0.700096 |
| Olig1 | Q9JKN5 | 0.63 | 0.05 | 0.96 | 0.76 | 0.02 | 0.29 | -0.04 | 0.57 | 0.15 | 0.71 | -0.37 | 0.37 | -0.27 | 0.5 | 0.63 | 0.4 | -0.39 | -0.385 | -0.285 | 0.162823 | 0.228528 | 0.342653 | 0.999971 | 1 | 0.861474 |
| Oma1 | Q9D8H7 | -1.28 | -0.22 | -0.35 | -0.42 | 0.23 | -0.37 | -0.4 | 0.24 | -0.11 | -0.08 | 0.19 | -1.02 | 0.69 | -0.26 | -0.15 | 0.13 | 0.4925 | 0.3125 | 0.67 | 0.134807 | 0.384725 | 0.064003 | 0.999971 | 1 | 0.725525 |
| Omg | G3XA53 | -0.31 | -0.14 | -0.85 | -0.75 | -0.28 | -0.28 | -0.63 | -0.97 | -0.4 | -0.81 | -1.19 | -0.91 | 0.37 | -0.24 | -0.73 | -0.66 | -0.0275 | -0.315 | 0.1975 | 0.915984 | 0.243535 | 0.529927 | 0.999971 | 1 | 0.910069 |
| Omp | Q64288 | 2.41 | -3.51 | -5.75 | -5.79 | -3.06 | -1.29 | -3.65 | -5.35 | 3.06 | 6.26 | -6.14 | -5.46 | -4.23 | -3.12 | -4.42 | -5.79 | -0.1775 | 2.59 | -1.23 | 0.92399 | 0.433478 | 0.495438 | 0.999971 | 1 | 0.901531 |
| Opa1 | P58281-2 | -0.47 | -0.35 | -1.27 | -1.18 | -0.43 | -0.5 | -1.17 | -1.27 | -0.4 | -0.51 | -1.26 | -1.85 | -0.25 | -0.38 | -1.44 | -1.26 | -0.025 | -0.1875 | -0.015 | 0.93846 | 0.639734 | 0.967901 | 0.999971 | 1 | 0.998787 |
| Opa3 | Q505D7 | -0.72 | -0.84 | -1.26 | -1.66 | -1.1 | -1.02 | -1.35 | -1.6 | -1.49 | -1.15 | -1.21 | -1.45 | -1.53 | -1.22 | -1.5 | -1.19 | -0.1475 | -0.205 | -0.24 | 0.588311 | 0.428942 | 0.363687 | 0.999971 | 1 | 0.869526 |
| Opalin | Q7M750 | 1.46 | -0.81 | -0.35 | -0.43 | -1.62 | -0.66 | -0.66 | -0.39 | -1.24 | -0.15 | -0.91 | -0.71 | -1.3 | -1.1 | -0.13 | -0.75 | -0.8 | -0.72 | -0.7875 | 0.156802 | 0.188055 | 0.159727 | 0.999971 | 1 | 0.800134 |
| Opcml | Q6DFY2 | -1.07 | -0.99 | -1.23 | -1.39 | -1.14 | -0.73 | -1.33 | -1.41 | -1.11 | -0.82 | -1.14 | -1.51 | -0.99 | -1.03 | -1.74 | -1.25 | 0.0175 | 0.025 | -0.0825 | 0.936504 | 0.906152 | 0.722745 | 0.999971 | 1 | 0.955258 |
| Opcml_1 | G5E8G3 | -2.55 | -1.28 | -3.96 | -3.17 | -1.25 | -1.78 | -3.48 | -3.6 | -1.5 | -2.47 | -3.23 | -3.18 | -0.85 | -1.36 | -3.47 | -4.29 | 0.2125 | 0.145 | 0.2475 | 0.774324 | 0.819764 | 0.782845 | 0.999971 | 1 | 0.965866 |
| Ophn1 | Q99J31 | -0.21 | -0.25 | -0.48 | -0.63 | -0.28 | 0.01 | -0.39 | -0.78 | -0.27 | -0.05 | -0.12 | -0.27 | -0.14 | -0.19 | -0.83 | -0.69 | 0.0325 | 0.215 | -0.07 | 0.887328 | 0.257976 | 0.76742 | 0.999971 | 1 | 0.962472 |
| Oplah | Q8K010 | -0.09 | -0.37 | 0.08 | -0.1 | -0.27 | -0.18 | 0.15 | -0.03 | -0.33 | -0.32 | -0.3 | -0.13 | -0.58 | -0.17 | -0.35 | -0.26 | 0.0375 | -0.15 | -0.22 | 0.846908 | 0.409741 | 0.273491 | 0.999971 | 1 | 0.843794 |
| Oprk1 | A0A5F8MP | 1.31 | -0.99 | 3.37 | 1.63 | -0.56 | 0.26 | 3.02 | 1.83 | -0.73 | 1.32 | 3.72 | 3.6 | -0.92 | -0.85 | 3.65 | 1.43 | -0.1925 | 0.6475 | -0.5025 | 0.857184 | 0.604754 | 0.689798 | 0.999971 | 1 | 0.945813 |
| Oprl1 | B0R0C0 | -0.17 | -0.7 | -0.75 | -0.62 | -0.22 | 0.44 | -1.52 | -0.34 | 0.07 | 1.23 | -1.19 | -0.9 | -1.29 | -0.28 | -2.03 | -1.48 | 0.15 | 0.3625 | -0.71 | 0.714286 | 0.49373 | 0.086843 | 0.999971 | 1 | 0.761242 |
| Oprm1 | P42866-14 | -0.66 | -3.07 | 0.33 | 0.38 | -2.91 | -1.8 | 0.76 | 0.66 | -2.12 | -0.71 | -0.37 | -1.37 | -5.44 | -1.84 | 0.94 | -0.41 | -0.0675 | -0.3875 | -0.9325 | 0.950389 | 0.634544 | 0.515346 | 0.999971 | 1 | 0.905815 |
| Optn | Q8K3K8 | 0.23 | -0.59 | -0.82 | -0.66 | -0.34 | -0.43 | -0.39 | -0.18 | -0.43 | 0.11 | -0.58 | -0.71 | -0.52 | 0.06 | -0.46 | 0.05 | 0.125 | 0.0575 | 0.2425 | 0.637076 | 0.849435 | 0.419177 | 0.999971 | 1 | 0.87788 |
| Orai1 | Q8BWG9 | -3.85 | -0.8 | -0.58 | -2.13 | -0.22 | -1.1 | 1.13 | -1.87 | -0.74 | -4.17 | -0.52 | -1.11 | -0.06 | -2.05 | -1.18 | -2.31 | 1.325 | 0.205 | 0.44 | 0.160441 | 0.841149 | 0.592926 | 0.999971 | 1 | 0.923349 |
| Orai2 | Q8BH10 | -0.23 | -0.03 | -1 | -0.57 | -1.39 | -0.73 | -1.58 | -0.97 | -0.96 | -0.21 | -1.83 | -1.87 | -0.79 | -1.13 | -1.49 | -0.61 | -0.71 | -0.76 | -0.5475 | 0.037944 | 0.102835 | 0.09336 | 0.999971 | 1 | 0.776565 |
| Ormdl3 | Q9CPZ6 | 0.95 | 0.49 | 0.53 | 0.66 | -0.08 | -0.61 | 0.08 | 0.66 | 0.56 | 0.95 | 0.08 | 0.15 | -0.34 | 0.49 | 0.39 | 0.56 | -0.645 | -0.2225 | -0.3825 | 0.051298 | 0.388033 | 0.163213 | 0.999971 | 1 | 0.800215 |
| Os9 | Q8K2C7 | 0.66 | -0.09 | 0.28 | 0.51 | 0.02 | 0.22 | 0.25 | 0.9 | 0.06 | 0.57 | 0.95 | 0.32 | 0.35 | 0.33 | 1.01 | 1.05 | 0.0075 | 0.135 | 0.345 | 0.977855 | 0.617535 | 0.231601 | 0.999971 | 1 | 0.8307 |
| Osbp | Q3B7Z2 | -0.19 | -0.64 | -0.84 | -0.82 | -0.59 | -0.45 | -0.01 | -0.86 | -0.15 | 0.05 | -0.71 | -0.82 | -0.6 | -0.69 | -0.77 | -0.82 | 0.145 | 0.215 | -0.0975 | 0.576896 | 0.443624 | 0.644624 | 0.999971 | 1 | 0.93301 |
| Osbp2 | Q8K0C7 | -0.45 | -0.09 | 0.81 | 0.75 | -0.73 | -0.19 | -1.5 | -0.4 | -0.9 | -0.56 | -1.56 | -0.69 | -0.24 | 0.07 | -0.93 | -0.35 | -0.96 | -1.1825 | -0.6175 | 0.039255 | 0.011003 | 0.119171 | 0.999971 | 1 | 0.78268 |
| Osbp2_1 | Q5QNQ6 | 0.35 | 0.29 | -0.14 | 0.05 | 0.23 | 0.29 | 0.27 | 0.12 | 0.26 | 0.05 | -0.03 | 0.01 | 0.2 | -0.04 | -0.45 | -0.33 | 0.09 | -0.065 | -0.2925 | 0.634851 | 0.734307 | 0.217645 | 0.999971 | 1 | 0.826893 |
| Osbpl10 | S4R1M9 | -0.29 | -0.02 | -0.39 | -0.41 | -0.07 | -0.06 | -0.24 | 0.03 | 0.09 | -0.08 | -0.39 | -0.45 | 0.04 | 0.02 | -0.17 | -0.18 | 0.1925 | 0.07 | 0.205 | 0.306388 | 0.734451 | 0.281674 | 0.999971 | 1 | 0.844538 |
| Osbpl11 | G5E8A0 | 0.77 | 0.21 | -0.09 | -0.23 | 0.48 | 0.3 | 0.32 | -0.26 | 0.56 | 0.38 | -0.01 | -0.05 | -0.17 | 0.2 | -0.13 | -0.46 | 0.045 | 0.055 | -0.305 | 0.875444 | 0.844816 | 0.287803 | 0.999971 | 1 | 0.847258 |
| Osbpl1a | Q91XL9 | -0.52 | 0.68 | 0.3 | -0.26 | 0.71 | 1.17 | 0.35 | 0.09 | 0.88 | -0.09 | 0.09 | 0.21 | 0.56 | 0.73 | 0.1 | -0.17 | 0.53 | 0.2225 | 0.255 | 0.157141 | 0.518364 | 0.458844 | 0.999971 | 1 | 0.889191 |
| Osbpl2 | Q8BX94 | -1.24 | -1.41 | -0.05 | -0.78 | -1.33 | -1.53 | -0.33 | -0.1 | -1.26 | -0.53 | 0.12 | 0.08 | -1.38 | -1.16 | -0.01 | -0.14 | 0.0475 | 0.4725 | 0.1975 | 0.914134 | 0.280992 | 0.653167 | 0.999971 | 1 | 0.934005 |
| Osbpl3 | F8WH20 | 0.67 | 0.39 | -0.07 | -0.25 | -0.11 | 0.84 | -0.46 | -0.1 | 0.36 | 0.62 | 0.19 | -0.3 | 0.22 | 0.29 | -0.44 | -0.34 | -0.1425 | 0.0325 | -0.2525 | 0.680407 | 0.912284 | 0.399962 | 0.999971 | 1 | 0.871816 |
| Osbpl5 | Q9ER64 | -0.05 | -1.09 | -0.51 | 0.27 | -0.38 | -0.47 | -0.22 | -0.08 | -0.95 | -0.62 | 0.55 | 0.94 | -0.66 | -0.64 | -0.13 | 0.29 | 0.0575 | 0.325 | 0.06 | 0.853913 | 0.524262 | 0.869081 | 0.999971 | 1 | 0.983462 |
| Osbpl6 | F8WIZ7 | -0.21 | 0.18 | 0.31 | 0.09 | 0.02 | 0.27 | 0.53 | 0.08 | 0.19 | 0.14 | 0.41 | 0.33 | 0.44 | 0.12 | -0.19 | -0.06 | 0.1325 | 0.175 | -0.015 | 0.532658 | 0.367277 | 0.945698 | 0.999971 | 1 | 0.997427 |
| Osbpl7 | F8VQM4 | 0.64 | -0.53 | -0.24 | -0.05 | 1.25 | -0.24 | -0.44 | 0.06 | -0.02 | -0.02 | -0.35 | -0.01 | -1.07 | -0.82 | 0.07 | -0.45 | 0.2025 | -0.055 | -0.5225 | 0.638214 | 0.84288 | 0.157529 | 0.999971 | 1 | 0.797252 |
| Osbpl8 | B9EJ86 | -0.51 | -0.79 | -0.71 | -0.32 | -0.41 | -0.71 | 0.19 | -0.53 | -0.39 | -0.35 | -0.56 | -0.45 | -0.32 | -0.24 | -0.17 | -0.38 | 0.2175 | 0.145 | 0.305 | 0.394295 | 0.43729 | 0.129825 | 0.999971 | 1 | 0.784948 |
| Osbpl9 | A0A0A6YXI | -0.09 | 0.22 | -1.39 | -0.46 | 1.2 | 0.68 | -1.41 | -1.78 | 1.03 | 0.52 | -0.03 | -2.08 | -0.84 | 0.72 | -1.25 | -2.05 | 0.1025 | 0.29 | -0.425 | 0.889789 | 0.678826 | 0.498216 | 0.999971 | 1 | 0.902911 |
| Osbpl9_1 | E9PXZ2 | -1.01 | -1.13 | -0.11 | -0.39 | -0.74 | -0.93 | -0.06 | -0.15 | -0.57 | -0.82 | 0.02 | 0.08 | -0.77 | -0.69 | 0.06 | -0.42 | 0.19 | 0.3375 | 0.205 | 0.563974 | 0.320547 | 0.518031 | 0.999971 | 1 | 0.905815 |
| Oscp1 | A0ZV96 | -0.47 | 0.19 | 0.07 | -0.37 | 0.6 | -0.14 | 0.68 | -0.42 | 0.67 | -0.88 | -0.1 | 0.3 | 0.49 | -0.37 | -0.16 | -0.6 | 0.325 | 0.1425 | -0.015 | 0.323119 | 0.694201 | 0.95951 | 0.999971 | 1 | 0.998721 |
| Osgep | A0A0R4J1Y | 0.63 | 1.93 | 1.44 | 1.29 | 1.65 | 1.36 | 1.25 | 1.41 | 1.85 | 1 | 0.86 | 0.74 | 1.47 | 2 | 1.04 | 0.85 | 0.095 | -0.21 | 0.0175 | 0.745694 | 0.563062 | 0.961411 | 0.999971 | 1 | 0.998721 |
| Ostc | Q78XF5 | 1.18 | 0.26 | 2.33 | 0.4 | 0.82 | 1.61 | 1.94 | 1.96 | 1.68 | 2.21 | 2.01 | 0.32 | 0.46 | 1.21 | 1.2 | 0.99 | 0.54 | 0.5125 | -0.0775 | 0.300209 | 0.392885 | 0.869831 | 0.999971 | 1 | 0.983917 |
| Ostf1 | Q62422 | 0.82 | 0.67 | 0.35 | 0.22 | 0.53 | 0.66 | 0.42 | 0.26 | 0.22 | 0.54 | 0.05 | 0.41 | 0.37 | 1.07 | 0.3 | 0.54 | -0.0475 | -0.21 | 0.055 | 0.822758 | 0.347623 | 0.826697 | 0.999971 | 1 | 0.976804 |
| Ostm1 | Q8BGT0 | -0.89 | 1.37 | 1.3 | 1.16 | 1.45 | 2 | 1.07 | 1.33 | 1.46 | -0.8 | 1.22 | 1.59 | 1.34 | 1.27 | 1.24 | 1.26 | 0.7275 | 0.1325 | 0.5425 | 0.195013 | 0.852041 | 0.298791 | 0.999971 | 1 | 0.847402 |
| Otof | A0A0R4J1K | 5.12 | 0.65 | 0.09 | 3.28 | -1.21 | 2.2 | -1.25 | 2.41 | -0.71 | 6.43 | -1.9 | -0.49 | -1.73 | 5.56 | 1.7 | 3.59 | -1.7475 | -1.4525 | -0.005 | 0.226556 | 0.471197 | 0.997684 | 0.999971 | 1 | 1 |
| Otub1 | Q7TQI3 | 0.15 | 0.64 | 0.99 | 0.57 | 0.47 | 0.45 | 0.83 | 0.81 | 0.31 | -0.17 | 0.59 | 0.05 | 0.05 | 0.58 | 0.82 | 0.47 | 0.0525 | -0.3925 | -0.1075 | 0.82413 | 0.156668 | 0.680741 | 0.999971 | 1 | 0.943097 |
| Otub2 | Q9CQX0 | -0.03 | 0.39 | 0.63 | 0.84 | 0.54 | 0.47 | 0.15 | 0.94 | 0.1 | -0.23 | 0.51 | 0.8 | -0.42 | 1.01 | 0.93 | 1.26 | 0.0675 | -0.1625 | 0.2375 | 0.801076 | 0.592098 | 0.560067 | 0.999971 | 1 | 0.915694 |
| Otud4 | A0A0R4J26 | 1.32 | 0.84 | 1.05 | 1.17 | 0.75 | 0.81 | 0.96 | 1.64 | 0.71 | 1.15 | 0.99 | 1.22 | 0.52 | 0.97 | 1.31 | 1.47 | -0.055 | -0.0775 | -0.0275 | 0.828937 | 0.703941 | 0.915202 | 0.999971 | 1 | 0.992633 |
| Otud5 | A2AES3 | -0.52 | 0.13 | 0.44 | 1.02 | 0.79 | 0.99 | -1.67 | 0.73 | 0.98 | -0.07 | -0.14 | 0.65 | 2.29 | 0.32 | 1.05 | 1.46 | -0.0575 | 0.0875 | 1.0125 | 0.928506 | 0.828664 | 0.062905 | 0.999971 | 1 | 0.725348 |
| Otud6b | Q8K2H2 | -0.37 | 0.57 | 0.83 | 0.32 | 0.98 | 0.02 | 1.42 | 0.59 | 1.03 | -0.44 | 1.3 | 1.67 | 1.34 | 0.1 | 0.31 | -0.06 | 0.415 | 0.5525 | 0.085 | 0.291179 | 0.281079 | 0.828236 | 0.999971 | 1 | 0.977728 |
| Otud7a | Q8R554 | -0.48 | 0.2 | -0.18 | -0.19 | 0.03 | 0.01 | 0.24 | -0.32 | -0.02 | -0.21 | 0.03 | -0.46 | 0.14 | 0.02 | -0.45 | -0.45 | 0.1525 | -0.0025 | -0.0225 | 0.50019 | 0.990893 | 0.925705 | 0.999971 | 1 | 0.99401 |
| Otud7b | B2RUR8 | -0.14 | -0.8 | -0.73 | -0.34 | -0.99 | -0.79 | -0.95 | -0.53 | -0.6 | -0.56 | -1.05 | -0.76 | -1.39 | -1.01 | -1.06 | -0.5 | -0.3125 | -0.24 | -0.4875 | 0.194041 | 0.309672 | 0.092619 | 0.999971 | 1 | 0.776565 |
| Otulin | Q3UCV8 | 1.21 | 1.28 | 1.43 | 1.4 | 1.23 | 1.32 | 1.08 | 1.15 | 1.42 | 1.11 | 0.73 | 1.11 | 1.22 | 0.75 | 0.5 | 1.15 | -0.135 | -0.2375 | -0.425 | 0.433657 | 0.259329 | 0.082165 | 0.999971 | 1 | 0.755254 |
| Otulinl | A0A2I3BPS | -0.66 | 0.09 | -0.2 | 0.58 | -0.13 | 0.14 | -0.8 | 0.46 | 0.19 | 0.04 | -0.15 | -0.01 | 0.08 | 0.35 | 0.22 | 0.72 | -0.035 | 0.065 | 0.39 | 0.92324 | 0.818177 | 0.219047 | 0.999971 | 1 | 0.827064 |
| Ovos | Q3UU35 | -0.97 | -0.15 | 0.11 | 0.59 | -0.72 | -0.3 | -0.55 | -0.65 | -0.58 | -0.44 | -0.06 | -0.86 | -0.65 | -0.01 | -0.37 | 0.28 | -0.45 | -0.38 | -0.0825 | 0.204022 | 0.305378 | 0.825314 | 0.999971 | 1 | 0.976618 |
| Oxa1l | Q8BGA9 | 0.76 | -0.02 | 0.13 | 0.49 | 0.69 | 0.33 | -0.36 | 1.18 | 1.02 | 0.65 | 0.5 | -0.63 | 0.26 | 0.37 | 0.4 | 0.74 | 0.12 | 0.045 | 0.1025 | 0.739548 | 0.90649 | 0.66925 | 0.999971 | 1 | 0.938711 |
| Oxct1 | Q9D0K2 | -1.11 | -0.18 | -0.25 | -0.35 | -0.15 | -0.55 | -0.03 | -0.09 | 0 | -0.86 | 0.15 | -0.33 | 0.44 | -0.85 | -0.23 | 0.14 | 0.2675 | 0.2125 | 0.3475 | 0.329202 | 0.504287 | 0.330916 | 0.999971 | 1 | 0.857107 |
| Oxld1 | Q9CR10 | -0.92 | -1.18 | -0.94 | -1.31 | -0.81 | -1.55 | -0.78 | -0.95 | -1.31 | -1.61 | -0.69 | -0.92 | -1.04 | -1.03 | -0.91 | -0.41 | 0.065 | -0.045 | 0.24 | 0.784442 | 0.857516 | 0.293853 | 0.999971 | 1 | 0.847402 |
| Oxnad1 | Q8VE38-2 | -0.7 | 0.36 | 0.37 | 0.22 | 0.55 | 0.21 | -0.09 | 0.41 | 0.79 | -0.27 | -0.15 | 0.27 | 1.5 | 0 | 0.15 | 0.07 | 0.2075 | 0.0975 | 0.3675 | 0.495153 | 0.778478 | 0.390909 | 0.999971 | 1 | 0.871816 |
| Oxr1 | Q4KMM3 | -0.95 | -0.63 | -0.5 | -0.49 | -0.62 | -0.48 | 0.3 | -0.89 | -0.74 | -0.99 | -0.38 | -0.44 | -0.67 | -0.8 | -0.7 | -0.52 | 0.22 | 0.005 | -0.03 | 0.454394 | 0.981749 | 0.874809 | 0.999971 | 1 | 0.984322 |
| Oxr1_1 | E9Q0A7 | -0.97 | 0.91 | -0.64 | -0.99 | 0.08 | 0.55 | -1.18 | -0.15 | 0.01 | -0.53 | -1.3 | -0.84 | -0.63 | 0.48 | -0.32 | 0.29 | 0.2475 | -0.2425 | 0.3775 | 0.645326 | 0.625271 | 0.444567 | 0.999971 | 1 | 0.88409 |
| Oxsm | Q9D404 | -0.25 | -0.47 | -0.65 | -0.58 | -0.43 | -0.5 | -0.65 | -0.67 | -0.56 | -0.99 | -0.97 | -0.67 | -0.83 | -0.89 | -0.78 | -1.44 | -0.075 | -0.31 | -0.4975 | 0.681196 | 0.139884 | 0.047655 | 0.999971 | 1 | 0.710102 |
| Oxsr1 | Q6P9R2 | -0.07 | 0.24 | 0.07 | -0.08 | 0.23 | 0.5 | -0.35 | -0.1 | 0.12 | 0.17 | -0.03 | -0.45 | 0.59 | 0.54 | 0.03 | 0.25 | 0.03 | -0.0875 | 0.3125 | 0.898651 | 0.674858 | 0.154681 | 0.999971 | 1 | 0.795578 |
| P10404 | P10404 | -0.42 | -0.1 | -1.46 | -1.52 | -0.02 | -0.08 | -0.93 | -1.24 | 0.37 | -0.15 | -1.05 | -1 | -0.3 | 0.21 | -1.93 | -0.74 | 0.3075 | 0.4175 | 0.185 | 0.495429 | 0.384108 | 0.731193 | 0.999971 | 1 | 0.956041 |
| P2ry12 | Q9CPV9 | 0.63 | 1.62 | 1.41 | 1.18 | 1.43 | 2.3 | 1.04 | 1.27 | 1.46 | 1.11 | 1.3 | 1.23 | 1.11 | 1.47 | 1.4 | 1.53 | 0.3 | 0.065 | 0.1675 | 0.393189 | 0.795722 | 0.521129 | 0.999971 | 1 | 0.906972 |
| P33monox | Q9DBN4 | 1.03 | 0.26 | 0.04 | -0.21 | 0.16 | 0.6 | 0.11 | -0.36 | 0.58 | 1.31 | -0.31 | -0.42 | 0.17 | 0.02 | -0.49 | -0.31 | -0.1525 | 0.01 | -0.4325 | 0.646856 | 0.982577 | 0.190171 | 0.999971 | 1 | 0.810547 |
| P4ha1 | Q60715-2 | 0.79 | 0.66 | 0.09 | 0.09 | 0.85 | 0.49 | 0.43 | 0.06 | 0.46 | 1.08 | 0.05 | 0.41 | 0.9 | 0.73 | 0.25 | 0.3 | 0.05 | 0.0925 | 0.1375 | 0.85124 | 0.752558 | 0.60848 | 0.999971 | 1 | 0.926452 |
| P4hb | P09103 | -0.55 | -0.61 | -0.24 | -0.29 | -0.77 | -0.67 | -0.14 | -0.24 | -0.45 | -0.57 | -0.36 | -0.13 | -0.53 | -0.63 | -0.4 | -0.21 | -0.0325 | 0.045 | -0.02 | 0.884127 | 0.814366 | 0.917917 | 0.999971 | 1 | 0.992633 |
| P4htm | Q8BG58 | -0.23 | 0.48 | -0.07 | 0.01 | 0.47 | 0.59 | -0.19 | 0.45 | 0.45 | 0.6 | 0.03 | 0.06 | 1.19 | 1.12 | 0.11 | 1 | 0.2825 | 0.2375 | 0.8075 | 0.289422 | 0.334644 | 0.024193 | 0.999971 | 1 | 0.669136 |
| Pa2g4 | P50580 | 0.8 | 1.71 | 1.51 | 1.28 | 1.62 | 1.47 | 1.58 | 1.46 | 1.71 | 1.27 | 1.63 | 1.8 | 1.87 | 1.89 | 1.41 | 1.59 | 0.2075 | 0.2775 | 0.365 | 0.388286 | 0.288659 | 0.175798 | 0.999971 | 1 | 0.808532 |
| Paat | Q9D2Q3 | -2.32 | 0.49 | -1.24 | -1.78 | 0.25 | -0.56 | -1.96 | -1.64 | -0.3 | -1.38 | -0.99 | -1.18 | 0.58 | 0.83 | -2.02 | -0.59 | 0.235 | 0.25 | 0.9125 | 0.743075 | 0.677853 | 0.273576 | 0.999971 | 1 | 0.843794 |
| Pabpc1 | P29341 | 0.68 | 0.06 | 0.51 | 0.48 | 0.03 | 0.27 | 0.76 | 0.34 | 0.09 | 0.44 | 0.33 | 0.57 | -0.37 | 0.45 | 0.53 | 0.19 | -0.0825 | -0.075 | -0.2325 | 0.727336 | 0.724147 | 0.390533 | 0.999971 | 1 | 0.871816 |
| Pabpc4 | Q6PHQ9 | 0.24 | 1.13 | 0.33 | 0.38 | 0.96 | 1.11 | 0.61 | 0.38 | 1.34 | 0.63 | 0.67 | 0.85 | 1.06 | 0.68 | 0.51 | 0.6 | 0.245 | 0.3525 | 0.1925 | 0.391377 | 0.225192 | 0.469374 | 0.999971 | 1 | 0.892481 |
| Pabpn1 | G3UY42 | -0.46 | -0.78 | -0.49 | 1.65 | -1.24 | -0.65 | -1.93 | 1.4 | -1.65 | 0.42 | -1.29 | -3.33 | -0.57 | -1.23 | 0.2 | 2.33 | -0.585 | -1.4425 | 0.2025 | 0.480944 | 0.120948 | 0.813504 | 0.999971 | 1 | 0.97473 |
| Pacs1 | Q8K212 | -1.55 | -1.06 | -0.79 | -0.77 | -0.88 | -0.9 | -0.39 | -0.67 | -0.74 | -1.45 | -0.63 | -0.83 | -0.5 | -0.92 | -0.78 | -0.55 | 0.3325 | 0.13 | 0.355 | 0.200292 | 0.637852 | 0.164717 | 0.999971 | 1 | 0.800215 |

| | | | | | | | | | | | | | | | | | | | | | | | | | | |
|---|---|---|---|---|---|---|---|---|---|---|---|---|---|---|---|---|---|---|---|---|---|---|---|---|---|---|
| Pacs2 | A0A1Y7VLZ | 0.2 | -0.6 | 0.48 | -0.08 | -0.43 | -0.23 | 0.44 | 0.27 | 0.04 | -0.03 | 0.45 | -0.44 | -0.05 | -0.57 | 0.15 | -0.47 | 0.0125 | 0.005 | -0.235 | 0.968075 | 0.986734 | 0.43725 | 0.999971 | 1 | 0.882419 |
| Pacsin1 | Q61644 | -1.01 | -0.84 | -0.74 | -0.8 | -1.22 | -1.06 | -0.27 | -0.86 | -0.97 | -1.27 | -0.62 | -0.89 | -1.46 | -1.19 | -0.88 | -0.95 | -0.005 | -0.09 | -0.2725 | 0.983741 | 0.65386 | 0.199698 | 0.999971 | 1 | 0.813335 |
| Pacsin2 | A0A2R8W6 | -1.23 | -1.15 | -0.76 | -1.86 | -1.49 | -0.97 | -1.3 | -2.33 | -0.77 | -3.42 | -0.3 | -0.47 | -0.46 | -1.46 | -1.88 | -0.73 | -0.2725 | 0.01 | 0.1175 | 0.45642 | 0.988569 | 0.760305 | 0.999971 | 1 | 0.961116 |
| Pacsin2_1 | Q9WVE8 | 0.98 | -0.7 | 0.07 | -0.03 | -0.99 | -0.54 | -0.06 | -0.39 | -0.82 | 0.15 | -0.2 | -0.24 | -0.92 | -0.5 | -0.09 | 0.08 | -0.575 | -0.3575 | -0.4375 | 0.15733 | 0.366097 | 0.285337 | 0.999971 | 1 | 0.845133 |
| Pacsin3 | Q99JB8 | -0.91 | -0.35 | 0.24 | -0.1 | -0.38 | -0.38 | -0.74 | 0.16 | -0.64 | -0.58 | -0.43 | -0.5 | -0.44 | -0.16 | 0.16 | 0.2 | -0.055 | -0.2575 | 0.22 | 0.859298 | 0.346278 | 0.46418 | 0.999971 | 1 | 0.890905 |
| Padi2 | Q08642 | -0.87 | -1.01 | -0.3 | -0.65 | -1.07 | -0.61 | -0.46 | -0.79 | -0.84 | -0.8 | -0.68 | -0.32 | -0.95 | -0.76 | -0.45 | -0.61 | -0.025 | 0.0475 | 0.015 | 0.915967 | 0.836395 | 0.947523 | 0.999971 | 1 | 0.997651 |
| Paf1 | Q8K2T8 | 1.23 | 0.7 | 0.87 | 0.99 | 0.91 | 0.81 | 0.73 | 0.95 | 0.84 | 1.53 | 0.91 | 1.23 | 0.71 | 1.17 | 0.35 | 0.25 | -0.0975 | 0.18 | -0.3275 | 0.609922 | 0.440939 | 0.229459 | 0.999971 | 1 | 0.828687 |
| Pafah1b1 | P63005 | -0.32 | -0.23 | -0.31 | -0.27 | -0.19 | 0.04 | -0.31 | -0.13 | 0.12 | 0 | -0.18 | -0.07 | 0.37 | -0.01 | -0.16 | 0.07 | 0.135 | 0.25 | 0.35 | 0.436044 | 0.15208 | 0.087414 | 0.999971 | 1 | 0.763848 |
| Pafah1b2 | A0A1L1SVK | -0.94 | 0.35 | 0.15 | -0.09 | 0.61 | 0.38 | 0.42 | -0.23 | 0.51 | 0.03 | -0.01 | -0.03 | -0.86 | -0.07 | -0.22 | -0.76 | 0.4275 | 0.2575 | -0.345 | 0.22336 | 0.422767 | 0.32699 | 0.999971 | 1 | 0.856472 |
| Pafah1b3 | Q61205 | -0.31 | -0.55 | 0.18 | -0.33 | -0.27 | -0.52 | 0.86 | -0.61 | 0.47 | -0.09 | 0.39 | 0.79 | -0.6 | -0.23 | 0.05 | 0.16 | 0.1175 | 0.6425 | 0.0975 | 0.746783 | 0.033522 | 0.703747 | 0.999971 | 1 | 0.949793 |
| Pafah2 | E9QNW6 | -1.44 | -1.32 | -0.75 | -1.65 | -1.12 | -1.3 | -1 | -1.24 | -1.11 | -1.34 | -0.84 | -0.52 | -1.55 | -0.87 | -0.14 | -1.12 | 0.125 | 0.3375 | 0.37 | 0.601643 | 0.242733 | 0.303649 | 0.999971 | 1 | 0.847402 |
| Pag1 | Q3U1F9 | -1.63 | -1.44 | -1.26 | -1.5 | -1.42 | -0.86 | -1.37 | -1.47 | -1.18 | -1.06 | -1.04 | -1.06 | -0.76 | -1.27 | -1.33 | -1.33 | 0.1775 | 0.3725 | 0.285 | 0.409796 | 0.052164 | 0.197775 | 0.999971 | 1 | 0.813335 |
| Paics | Q9DCL9 | -0.08 | -0.05 | -0.3 | -0.41 | -0.23 | -0.43 | -0.38 | -0.19 | -0.13 | -0.38 | -0.15 | -0.7 | -0.5 | -0.43 | -0.53 | -0.66 | -0.0975 | -0.13 | -0.32 | 0.594038 | 0.534934 | 0.102772 | 0.999971 | 1 | 0.78268 |
| Paip1 | F6Y616 | 0.21 | -0.26 | 0.18 | -0.13 | -0.31 | -0.3 | 0.53 | -0.13 | -0.53 | -0.14 | 0.42 | -0.03 | -0.53 | -0.25 | 0.01 | -0.1 | -0.0525 | -0.07 | -0.2175 | 0.837147 | 0.781529 | 0.323582 | 0.999971 | 1 | 0.85548 |
| Paip2b | Q91W45 | 1.88 | 0.47 | 0.03 | 0.9 | 1.21 | 1.66 | 1.29 | -0.03 | 1 | 1.23 | 0.31 | -1.11 | 0.96 | 1.2 | -0.98 | -0.29 | 0.2125 | -0.4625 | -0.5975 | 0.672488 | 0.452066 | 0.32993 | 0.999971 | 1 | 0.856787 |
| Pak1 | G5E884 | -0.78 | -0.16 | 0.17 | -0.02 | -0.21 | -0.16 | 0.04 | -0.29 | -0.21 | -0.54 | -0.13 | -0.23 | -0.16 | -0.15 | -0.07 | -0.04 | 0.0425 | -0.08 | 0.0925 | 0.863235 | 0.749682 | 0.702128 | 0.999971 | 1 | 0.949019 |
| Pak1ip1 | Q9DCE5 | 0.91 | 1.09 | 2.98 | 2.76 | 1.36 | 1.34 | 2.74 | 2.52 | 1.03 | 1.14 | 2.7 | 2.55 | 0.03 | 1.51 | 2.54 | 2.33 | 0.055 | -0.08 | -0.3325 | 0.926962 | 0.900843 | 0.642347 | 0.999971 | 1 | 0.93301 |
| Pak2 | Q8CIN4 | -0.35 | -0.51 | -0.64 | -0.72 | -0.47 | -0.54 | -0.8 | -0.9 | -0.58 | -0.5 | -0.57 | -0.53 | -0.63 | -0.14 | -0.28 | -0.41 | -0.1225 | 0.01 | 0.19 | 0.532745 | 0.952842 | 0.344354 | 0.999971 | 1 | 0.862346 |
| Pak3 | Q61036 | 0.95 | 0.53 | -0.32 | 0.38 | 0.23 | 0.71 | 0.06 | 0.54 | 0.49 | 0.41 | 0.01 | 0.06 | 1.02 | 1.08 | 0.34 | 0.61 | -3.93E-17 | -0.1425 | 0.3775 | 1 | 0.635418 | 0.257333 | 1 | 1 | 0.839012 |
| Pak4 | Q8BTW9 | 0.17 | 0.03 | 0.35 | 1.23 | 1.47 | -0.83 | 1.75 | -0.26 | -0.32 | 0.7 | 0.31 | -0.12 | -0.4 | 0.02 | -0.09 | -0.97 | 0.0875 | -0.3025 | -0.805 | 0.889105 | 0.394814 | 0.042018 | 0.999971 | 1 | 0.701725 |
| Pak5 | Q8C015 | -0.08 | -0.3 | -0.6 | -0.79 | -0.7 | -0.68 | -0.69 | -0.98 | -0.53 | -0.53 | -0.62 | -0.61 | -0.09 | -0.77 | -1.25 | -0.86 | -0.32 | -0.13 | -0.3 | 0.167675 | 0.535702 | 0.32833 | 0.999971 | 1 | 0.856472 |
| Pak6 | Q3ULB5 | 0.25 | -1 | -0.61 | -0.43 | -1.18 | -0.29 | -0.34 | -0.02 | -1.09 | 0.02 | -0.17 | -0.12 | 0.09 | -0.67 | -0.38 | 0.34 | -0.01 | 0.1075 | 0.2925 | 0.977481 | 0.763322 | 0.403796 | 0.999971 | 1 | 0.873348 |
| Pakap | F7AA26 | -0.47 | -0.94 | -0.88 | -0.85 | -1.06 | -1.06 | -0.81 | -0.84 | -0.97 | 0.19 | -0.76 | -0.86 | -0.92 | -1 | -1.03 | -1.03 | -0.1575 | 0.185 | -0.21 | 0.421238 | 0.536781 | 0.273309 | 0.999971 | 1 | 0.843794 |
| Pald1 | A0A0R4J00 | 0.05 | 0.74 | 0.17 | 0.83 | 0.31 | 0.21 | 0.04 | 0.78 | 0.69 | 1.1 | 0.33 | 0.46 | 0.77 | 0.78 | 0.74 | 0.93 | -0.1125 | 0.1975 | 0.3575 | 0.68011 | 0.480048 | 0.157105 | 0.999971 | 1 | 0.797246 |
| Palld | Q9ET54-3 | -0.51 | -0.35 | -0.38 | -0.73 | 4.97 | 1.31 | -0.51 | -1.09 | 1.11 | 3.86 | 2.11 | 4.08 | -0.85 | 0.61 | -0.63 | -0.5 | 1.6625 | 3.2825 | 0.15 | 0.195025 | 0.000822 | 0.656518 | 0.999971 | 0.499699 | 0.934061 |
| Palm | Q9Z0P4-2 | -1.26 | -2.27 | -2.73 | -2.16 | -1.57 | -1.02 | -2.05 | -2.17 | -1.98 | -0.92 | -1.78 | -1.83 | -1 | -1.4 | -2.32 | -1.5 | 0.4025 | 0.4775 | 0.55 | 0.314905 | 0.227658 | 0.189853 | 0.999971 | 1 | 0.810547 |
| Palm_1 | Q9Z0P4 | -0.82 | -1.66 | -1.78 | -1.56 | -1.88 | -1.58 | -1.48 | -1.55 | -1.63 | -1.34 | -1.57 | -2.13 | -1.39 | -1.85 | -1.88 | -1.86 | -0.1675 | -0.2125 | -0.29 | 0.521311 | 0.463681 | 0.295363 | 0.999971 | 1 | 0.847402 |
| Palm2 | Q8BR92 | -0.83 | -1.21 | -1.03 | -0.97 | -1.41 | -1.39 | -0.61 | -1.45 | -1.46 | -0.92 | -1.41 | -1.08 | -1.25 | -1.37 | -1.47 | -1.21 | -0.205 | -0.2075 | -0.315 | 0.414947 | 0.322257 | 0.106807 | 0.999971 | 1 | 0.78268 |
| Palm3 | A2TJV2 | -0.74 | -2.24 | -1.58 | -1.38 | -1.37 | -1.22 | -1.53 | -0.85 | -1.35 | -0.63 | -1.13 | -1.22 | -1.22 | -1.49 | -0.99 | -0.73 | 0.2425 | 0.4025 | 0.3775 | 0.48006 | 0.259268 | 0.290637 | 0.999971 | 1 | 0.847402 |
| Palmd | A0A0R4J0J | 0.1 | -0.35 | 0.33 | 0.37 | -0.37 | 0 | 0.58 | 0.31 | -0.33 | -0.13 | 0.37 | 0 | -0.7 | -0.36 | 0.23 | 0.55 | 0.0175 | -0.135 | -0.1825 | 0.949906 | 0.589369 | 0.580981 | 0.999971 | 1 | 0.920364 |
| Pals1 | Q9JLB2 | -0.71 | -0.8 | -0.6 | -0.67 | -1.08 | -0.8 | -1.25 | -1.3 | -0.62 | -0.49 | -0.27 | -0.43 | -0.47 | -0.65 | -0.55 | -0.64 | -0.4125 | 0.2425 | 0.1175 | 0.053189 | 0.177327 | 0.485139 | 0.999971 | 1 | 0.896868 |
| Pals2 | Q9JLB0 | -1.03 | -1.41 | -0.56 | -1.45 | -1.63 | -2.2 | -1.68 | -0.78 | -1.21 | -1.34 | -1.94 | -1.64 | -1.71 | -0.16 | -1.23 | -0.17 | -0.46 | -0.42 | 0.295 | 0.214066 | 0.15708 | 0.487868 | 0.999971 | 1 | 0.897906 |
| Pals2_1 | Q9JLB0-2 | -1.38 | -1.41 | -0.79 | -0.83 | -1.26 | -1.44 | -0.99 | -0.7 | -1.48 | -1.55 | -0.84 | -0.73 | -1.32 | -1.13 | -0.83 | -0.66 | 0.005 | -0.0475 | 0.1175 | 0.984534 | 0.867267 | 0.643637 | 0.999971 | 1 | 0.93301 |
| Pam | P97467 | -0.22 | -0.41 | -0.18 | -0.5 | -0.55 | -0.62 | -0.26 | -0.38 | -0.15 | -0.48 | -0.74 | -0.25 | -0.04 | -0.26 | 0.08 | -0.43 | -0.125 | -0.0775 | 0.165 | 0.504432 | 0.703853 | 0.414375 | 0.999971 | 1 | 0.876329 |
| Pam16 | Q9CQV1 | 0.74 | -0.89 | -0.66 | 0.29 | -0.82 | -0.82 | -0.54 | -0.17 | -1.14 | 0.18 | -0.28 | -0.8 | -1.15 | -1.01 | -0.75 | 0.13 | -0.4575 | -0.38 | -0.565 | 0.268569 | 0.413361 | 0.232831 | 0.999971 | 1 | 0.831513 |
| Pan2 | Q8BGF7-2 | 0.28 | 0.09 | 0.54 | 0.5 | -0.39 | -0.91 | 1.15 | -0.04 | -0.52 | -0.5 | 0.38 | 0.09 | -0.78 | -0.46 | 0.63 | 0.3 | -0.4 | -0.49 | -0.43 | 0.359674 | 0.092706 | 0.227901 | 0.999971 | 1 | 0.828598 |
| Pan3 | H3BKF3 | 0.64 | 0.01 | 0.13 | 0.08 | 0.21 | -0.48 | -0.46 | -0.05 | 0.18 | 0.52 | -0.13 | -0.16 | 0.5 | 0.31 | -0.19 | 0.42 | -0.41 | -0.1125 | 0.045 | 0.126573 | 0.645497 | 0.853358 | 0.999971 | 1 | 0.98187 |
| Pank1 | Q8K4K6 | -0.89 | 2.27 | 1.86 | 0.86 | 1.07 | 1.05 | 0.19 | 1.57 | 1.61 | 0.55 | 2.08 | 3.34 | 3.35 | 0.22 | 2.01 | 0.99 | -0.055 | 0.87 | 0.6175 | 0.936159 | 0.306742 | 0.487207 | 0.999971 | 1 | 0.897906 |
| Pank2 | Q7M753 | -0.1 | 0.02 | -0.64 | -0.85 | -0.37 | -0.38 | -0.38 | -0.68 | -0.29 | -0.28 | -0.74 | -0.58 | 0.34 | -0.33 | -0.94 | -0.79 | -0.06 | -0.08 | -0.0375 | 0.810815 | 0.758549 | 0.914717 | 0.999971 | 1 | 0.992633 |
| Pank4 | Q80YV4-2 | -0.55 | 0.79 | 0.17 | 0.04 | 0.91 | 0.66 | -0.12 | -0.31 | 0.8 | -0.02 | -0.17 | 0.19 | 0.83 | 0.6 | -0.09 | -0.25 | 0.1725 | 0.0875 | 0.16 | 0.658232 | 0.799112 | 0.666471 | 0.999971 | 1 | 0.937524 |
| Panx1 | Q9JIP4 | -0.89 | -1.67 | -1.38 | -1.36 | -1.08 | -1.18 | -1.69 | -1.21 | -1.52 | -1.05 | -1.22 | -0.92 | 0.17 | -1.15 | -1.43 | -1.34 | 0.035 | 0.1475 | 0.3875 | 0.885255 | 0.540123 | 0.336352 | 0.999971 | 1 | 0.858827 |
| Panx2 | Q6IMP4-2 | -0.28 | -0.01 | -0.43 | -0.43 | -0.44 | -0.34 | -0.89 | 0.04 | -0.58 | -0.28 | -0.52 | -0.39 | 0.53 | -0.28 | -0.33 | -0.56 | -0.12 | -0.155 | 0.1275 | 0.627093 | 0.413254 | 0.645655 | 0.999971 | 1 | 0.93301 |
| Paox | Q8C0L6 | 0.18 | 0.84 | 0.91 | 0.83 | 1.01 | 1.04 | 0.69 | 1.18 | 0.86 | 0.58 | 0.8 | 0.74 | 1.62 | 1.17 | 0.76 | 1.37 | 0.29 | 0.055 | 0.54 | 0.237739 | 0.803751 | 0.071637 | 0.999971 | 1 | 0.739418 |
| Papola | Q61183 | 0.67 | 0.81 | 2.37 | 2.01 | 0.99 | 0.62 | 1.83 | 1.93 | 0.61 | 0.65 | 1.88 | 1.74 | 0.13 | 1.15 | 1.96 | 2.07 | -0.1225 | -0.245 | -0.1375 | 0.804491 | 0.631916 | 0.80886 | 0.999971 | 1 | 0.973762 |
| Papss1 | Q6NZM8 | 1 | 1.96 | 1.73 | 1.47 | 1.96 | 1.55 | 1.59 | 1.92 | 1.82 | 1.02 | 1.95 | 2.16 | 2.11 | 1.95 | 2.19 | 1.72 | 0.215 | 0.1975 | 0.4525 | 0.41247 | 0.54647 | 0.105002 | 0.999971 | 1 | 0.78268 |
| Papss2 | A0A494B92 | -0.1 | -0.1 | 0.18 | 0.38 | 0.24 | 0.05 | 0.44 | 0.82 | 0.04 | 0.05 | 0.57 | 0.43 | 0.09 | 0.12 | 0.69 | 0.44 | 0.2975 | 0.1825 | 0.245 | 0.229445 | 0.414726 | 0.294472 | 0.999971 | 1 | 0.847402 |
| Paqr4 | Q9JJE4 | -2.64 | -1.12 | -0.68 | -0.14 | -0.96 | -2.15 | -0.61 | -0.03 | -1.22 | -2.67 | -0.23 | -0.73 | -0.79 | -2.05 | -0.02 | -0.62 | 0.2075 | -0.0675 | 0.275 | 0.744499 | 0.921445 | 0.661668 | 0.999971 | 1 | 0.936148 |
| Paqr7 | Q80ZE4 | 0.81 | -0.17 | -2.91 | -4.19 | -0.23 | 0.22 | -1.48 | -2.1 | -0.31 | 0.56 | 0.81 | 0.56 | -0.35 | 0.53 | -1.12 | -3.37 | 0.7175 | 2.02 | 0.5375 | 0.534226 | 0.086355 | 0.674713 | 0.999971 | 1 | 0.941037 |
| Paqr9 | Q6TCG2 | 4.64 | -0.27 | 1.63 | 1.54 | 0.23 | 1.7 | 2.7 | 0.83 | -0.08 | 1.85 | 0.94 | -0.98 | -3.17 | -1.83 | -0.22 | 0.2 | -0.52 | -1.4525 | -3.14 | 0.614708 | 0.197038 | 0.021129 | 0.999971 | 1 | 0.639761 |
| Pard3 | Q99NH2 | -0.56 | -0.45 | -0.17 | 0.18 | -0.38 | -0.89 | -0.18 | -0.13 | -0.54 | -0.05 | -0.25 | -0.45 | -0.81 | -0.42 | -0.1 | -0.11 | -0.145 | -0.0725 | -0.11 | 0.583447 | 0.755235 | 0.673314 | 0.999971 | 1 | 0.94048 |
| Pard6a | Q9Z101-3 | -1.11 | 0.15 | -0.05 | -0.24 | -0.12 | 0.16 | -0.52 | 0.02 | 0.08 | -0.7 | 0.28 | 0.08 | 0.45 | 0.06 | -0.11 | 0.35 | 0.1975 | 0.2475 | 0.5 | 0.537666 | 0.482237 | 0.135342 | 0.999971 | 1 | 0.784948 |
| Pard6b | Q9JK83 | -0.66 | -0.54 | -0.89 | -0.43 | -0.9 | -0.5 | -1.11 | -0.12 | -1.42 | -0.13 | -1.4 | -0.64 | -0.85 | 0.21 | -0.98 | -0.93 | -0.0275 | -0.2675 | -0.0075 | 0.916592 | 0.424479 | 0.980509 | 0.999971 | 1 | 0.99925 |
| Parg | G3X8U8 | 0.14 | 0.61 | 0.56 | 0.82 | 1.19 | 0.02 | 0.54 | 1.03 | 0.85 | 0.1 | 1.05 | 1.12 | 1.07 | 0.2 | 0.8 | 0.89 | 0.1625 | 0.2475 | 0.2075 | 0.598691 | 0.397157 | 0.434252 | 0.999971 | 1 | 0.882419 |
| Park7 | Q99LX0 | 0.35 | 0.25 | 0.07 | 0.42 | 0.23 | 0.37 | 0.6 | 0.18 | 0.24 | 0.09 | 0.24 | 0.01 | 0.42 | 0.59 | 0.53 | 0.58 | 0.0725 | -0.1275 | 0.2575 | 0.702604 | 0.471133 | 0.164977 | 0.999971 | 1 | 0.800215 |
| Parl | A0A338P6U | -0.07 | -0.87 | -1.15 | -1.04 | -0.95 | -0.63 | -0.6 | -0.63 | -1.25 | -0.63 | -0.19 | -0.98 | -0.85 | -1.13 | -1.15 | -1.34 | 0.08 | 0.02 | -0.335 | 0.771706 | 0.952105 | 0.25186 | 0.999971 | 1 | 0.839012 |
| Parm1 | Q923D3 | 4.8 | -1.71 | 1.18 | 1.78 | -2.06 | 0.11 | 2.05 | 0.08 | -1.96 | 2.04 | -0.87 | -1.53 | -3.66 | -1.9 | -0.45 | -1.44 | -1.4675 | -2.0925 | -3.375 | 0.309491 | 0.171394 | 0.029909 | 0.999971 | 1 | 0.678474 |
| Parn | Q8VDG3 | 1.07 | 1.58 | 1.3 | 1.56 | 1.39 | 0.26 | 0.68 | 1.27 | 0.79 | 0.87 | 1.24 | 1.31 | 1.35 | 1.29 | 1.41 | 1.31 | -0.4775 | -0.325 | -0.0375 | 0.136674 | 0.163013 | 0.844581 | 0.999971 | 1 | 0.979814 |
| Parp1 | Q921K2 | 1.43 | 1.8 | 2.15 | 2.14 | 1.8 | 1.32 | 1.74 | 1.48 | 1.73 | 1.54 | 1.97 | 1.76 | 0.87 | 1.5 | 1.36 | 1.99 | -0.295 | -0.13 | -0.45 | 0.235734 | 0.572305 | 0.155447 | 0.999971 | 1 | 0.795578 |
| Parp12 | Q8BZ20 | -0.1 | 0.05 | -0.2 | -0.46 | 0.74 | 0.27 | -0.54 | -0.14 | -0.29 | 0.42 | -0.49 | -0.79 | 0.02 | 0.53 | -0.43 | -0.66 | 0.26 | -0.11 | 0.0425 | 0.400085 | 0.705092 | 0.885356 | 0.999971 | 1 | 0.98711 |
| Pars2 | Q8CFI5 | -0.48 | -1.03 | -0.81 | -0.69 | -0.58 | -0.34 | -0.73 | -0.22 | -1.18 | -0.34 | -1.49 | -1.08 | -0.5 | 0.05 | -0.21 | -0.01 | 0.285 | -0.27 | 0.585 | 0.201494 | 0.352471 | 0.022838 | 0.999971 | 1 | 0.652043 |
| Parva | Q9EPC1 | 0.35 | 0.83 | 0.92 | 0.66 | 1.57 | 1.15 | 0.9 | 0.74 | 0.99 | 1.32 | 1.47 | 1.82 | 0.6 | 0.86 | 1.04 | 1.3 | 0.4 | 0.71 | 0.26 | 0.134299 | 0.016004 | 0.279507 | 0.999971 | 1 | 0.843812 |
| Parvb | Q9ES46 | -0.04 | 0.22 | 0.11 | 0.42 | 0.72 | 0.18 | 0.16 | 0.37 | 0.59 | 0.11 | -0.13 | 0.3 | 0.55 | 0.32 | 0.4 | 0.09 | 0.18 | 0.04 | 0.1625 | 0.404635 | 0.856098 | 0.420583 | 0.999971 | 1 | 0.87788 |
| Patl1 | Q3TC46 | 3.02 | 0.79 | 0.16 | 0.32 | 0.04 | 0.31 | 0.05 | 0.06 | -0.1 | 1.6 | 0 | 0.05 | -0.18 | 1.15 | -0.03 | 0.17 | -0.9575 | -0.685 | -0.795 | 0.141212 | 0.345962 | 0.248682 | 0.999971 | 1 | 0.839012 |
| Patz1 | Q9JMG9 | 0.86 | 3.6 | 0.9 | 1.48 | 3.56 | 3.45 | 1.39 | 1.37 | 3.8 | 1.17 | 1.12 | 0.87 | 4.36 | 2.64 | 1.44 | 1.57 | 0.7325 | 0.03 | 0.7925 | 0.371857 | 0.971922 | 0.357694 | 0.999971 | 1 | 0.868793 |
| Pawr | Q925B0-2 | 1.57 | 0.31 | 0.02 | 0.99 | 2.73 | 1.28 | 0.66 | -0.75 | 0.74 | 1.99 | 0.5 | 1.28 | -0.84 | 0.63 | 0.23 | 0.07 | 0.2575 | 0.405 | -0.7 | 0.722444 | 0.381971 | 0.13854 | 0.999971 | 1 | 0.784948 |
| Paxbp1 | P58501 | -0.67 | 0.52 | -0.88 | -0.77 | 0.65 | 0.45 | -0.1 | -1.1 | 0.24 | -0.1 | -1.41 | -1.13 | 0.01 | 0.35 | -0.92 | -0.95 | 0.425 | -0.15 | 0.0725 | 0.382411 | 0.755563 | 0.868433 | 0.999971 | 1 | 0.983462 |
| Paxx | A2AJ30 | -1.14 | 0.72 | -0.21 | -0.86 | -0.08 | -0.2 | -0.6 | 0.86 | 0.37 | 0.08 | 0.26 | -1.37 | -0.97 | -0.54 | 0.83 | -1 | 0.3675 | 0.2075 | -0.0475 | 0.451903 | 0.699882 | 0.930955 | 0.999971 | 1 | 0.994809 |
| Pbdc1 | Q9D0B6 | -0.86 | -0.65 | 0.31 | 0.19 | -0.67 | -0.26 | 0.17 | -0.06 | -0.28 | -0.41 | 0.6 | 0.63 | -1.67 | -0.32 | 0.34 | 0.5 | 0.0475 | 0.3875 | -0.035 | 0.888959 | 0.334857 | 0.947379 | 0.999971 | 1 | 0.997651 |
| Pbrm1 | D3Z1N4 | -1.45 | -1.44 | -0.95 | -1.53 | -1.29 | -1.43 | -0.98 | -1.36 | -1.3 | -1.46 | -0.85 | -0.82 | -0.53 | -1.08 | -1.1 | -1.3 | 0.0775 | 0.235 | 0.34 | 0.717532 | 0.339253 | 0.186631 | 0.999971 | 1 | 0.809025 |
| Pbx1 | P41778-2 | 0.63 | 2 | 1.45 | 1.84 | 1.89 | 1.82 | 1.54 | 1.81 | 1.63 | 1.14 | 1.14 | 1.91 | 1.95 | 1.67 | 1.38 | 1.77 | 0.285 | -0.025 | 0.2125 | 0.381644 | 0.943597 | 0.523171 | 0.999971 | 1 | 0.907797 |
| Pbxip1 | Q3TVI8 | 0.61 | -0.17 | -0.3 | 0 | -0.82 | -0.1 | -0.7 | 1.95 | -0.15 | 0.48 | 0.41 | 0.15 | 0.85 | 0.03 | -0.08 | 0.25 | 0.0475 | 0.1875 | 0.2275 | 0.938175 | 0.487461 | 0.454037 | 0.999971 | 1 | 0.887217 |
| Pcbd1 | P61458 | 0.13 | -0.31 | -0.24 | 0.22 | 0.37 | 0.37 | -0.34 | 0.78 | 0.46 | 0.05 | -0.03 | 0.53 | 1.58 | 1.02 | 1.08 | 0.9 | 0.345 | 0.3025 | 1.195 | 0.241514 | 0.209419 | 0.000708 | 0.999971 | 1 | 0.365812 |
| Pcbd2 | A0A0R4J1R | -0.22 | 0.03 | 0.08 | 0.23 | 0.29 | 0.09 | 0.18 | 0.74 | -0.08 | 0.15 | -0.52 | 0.14 | 0.1 | 0.13 | -0.12 | 0.24 | 0.295 | -0.1075 | 0.0575 | 0.197984 | 0.630372 | 0.762077 | 0.999971 | 1 | 0.961324 |
| Pcbp1 | P60335 | 0.54 | 0.54 | 0.6 | 0.55 | 0.38 | 0.16 | 0.3 | 0.4 | 0.29 | 0.25 | 0.1 | -0.55 | 0.26 | 0.36 | 0.39 | 0.46 | -0.2475 | -0.535 | -0.19 | 0.157454 | 0.042831 | 0.260066 | 0.999971 | 1 | 0.840512 |
| Pcbp2 | Q61990-2 | -2.07 | -0.51 | -0.22 | -1.15 | 0.12 | 0.03 | -0.21 | -0.68 | 0.12 | -3.88 | 0.23 | 0.74 | -0.18 | -0.11 | 0.45 | -1.59 | 0.8025 | 0.29 | 0.63 | 0.085283 | 0.778397 | 0.270049 | 0.999971 | 1 | 0.843794 |
| Pcbp2_1 | B2M1R7 | 0.34 | 1.76 | 1.88 | 0.27 | 1.59 | 1.39 | 2.1 | 0.48 | 1.88 | 0.45 | 1.7 | 1.77 | 1.68 | 1.08 | 0.83 | 0.14 | 0.3275 | 0.3875 | -0.13 | 0.527158 | 0.45768 | 0.796299 | 0.999971 | 1 | 0.970435 |
| Pcbp3 | P57722 | 0.85 | 1.6 | 1.37 | 1.29 | 1.5 | 1.31 | 1.69 | 1.27 | 1.55 | 1.1 | 1.25 | 1.16 | 1.57 | 1.41 | 1.35 | 1.23 | 0.165 | -0.0125 | 0.1125 | 0.470494 | 0.955496 | 0.608894 | 0.999971 | 1 | 0.926608 |
| Pcbp4 | A0A0R4J04 | 1.8 | 1.23 | 2.29 | 1.71 | 0.96 | 1.44 | 2.08 | 2.27 | 1.55 | 1.49 | 1.82 | 1.53 | 0.14 | 1.5 | 2.3 | 1.92 | -0.07 | -0.16 | -0.2925 | 0.8465 | 0.533322 | 0.549405 | 0.999971 | 1 | 0.912072 |
| Pcca | Q91ZA3 | -0.89 | -0.85 | -0.47 | -0.64 | -1.05 | -1.17 | -0.4 | -0.69 | -1.1 | -1.22 | -0.66 | -0.38 | -0.88 | -1.15 | -0.73 | -0.55 | -0.115 | -0.1275 | -0.115 | 0.627729 | 0.606497 | 0.589059 | 0.999971 | 1 | 0.921663 |
| Pccb | Q99MN9 | -0.68 | -0.08 | -0.65 | -1.25 | -0.22 | -0.64 | -0.37 | -1.04 | -0.18 | -0.78 | -0.48 | -0.31 | 0.03 | -0.64 | -1.23 | -1.51 | 0.0975 | 0.2275 | -0.1725 | 0.750235 | 0.432545 | 0.667277 | 0.999971 | 1 | 0.938223 |
| Pcdh1 | E0CXD4 | 0.26 | -1.6 | -0.3 | 0.01 | -0.8 | -0.76 | -0.59 | 0.05 | -1.03 | -0.06 | -0.54 | -0.78 | -1.56 | -0.82 | -0.08 | -0.51 | -0.1175 | -0.195 | -0.335 | 0.786138 | 0.657574 | 0.494095 | 0.999971 | 1 | 0.901531 |
| Pcdh1_1 | F7BJK1 | -0.94 | -1.23 | -0.94 | -1.41 | -1.38 | -1.05 | -0.8 | -1.2 | -1.19 | -0.98 | -0.63 | -0.9 | -0.54 | -1.31 | -1.11 | -0.87 | 0.0225 | 0.205 | 0.1725 | 0.916785 | 0.342958 | 0.474004 | 0.999971 | 1 | 0.894044 |
| Pcdh1_2 | Q8CFX3 | -1.36 | -1.42 | -1.28 | -0.62 | -1.35 | -1.31 | -1.09 | -0.97 | -1.35 | -1.56 | -1.15 | -0.7 | -0.45 | -0.7 | -0.46 | -0.82 | -0.01 | -0.02 | 0.5625 | 0.966655 | 0.942271 | 0.041265 | 0.999971 | 1 | 0.700838 |
| Pcdh10 | Q925I8 | -2.35 | -3.66 | -1.18 | -0.07 | -3.31 | -3.38 | -1.13 | -0.37 | -3.33 | -2.62 | -0.81 | -0.58 | -3.83 | -3.33 | -0.37 | 0.36 | -0.2325 | -0.02 | 0.0225 | 0.810694 | 0.982707 | 0.984497 | 0.999971 | 1 | 0.99925 |
| Pcdh11x | F6ZNL5 | -0.19 | -0.52 | 0.68 | -0.01 | -0.76 | -0.65 | -0.09 | -0.03 | -1.04 | -0.01 | 0.18 | 0.51 | -1.55 | 0.04 | 0.39 | 0.01 | -0.3725 | -0.08 | -0.2675 | 0.260257 | 0.842234 | 0.57213 | 0.999971 | 1 | 0.91816 |
| Pcdh15 | Q99PJ1 | 0.03 | -4.64 | -2.81 | -1.49 | -2.1 | -5.95 | 1.68 | -1.83 | -4.99 | -2.94 | -1.74 | -1.67 | -1.99 | -5.9 | -2.84 | -1.3 | 0.1775 | -0.6075 | -0.78 | 0.913533 | 0.593506 | 0.540981 | 0.999971 | 1 | 0.910526 |
| Pcdh17 | E9PXF0 | -0.88 | -0.85 | -0.83 | -0.84 | -0.6 | -1.05 | -0.59 | -0.48 | -0.66 | -0.85 | -0.33 | 0.08 | -0.06 | -0.81 | -0.35 | -0.44 | 0.17 | 0.41 | 0.435 | 0.387135 | 0.110179 | 0.06205 | 0.999971 | 1 | 0.725348 |
| Pcdh19 | Q80TF3 | -0.57 | -1.22 | -1.07 | -1.15 | -1.27 | -0.9 | -1.09 | -1.13 | -1.83 | -1.23 | -2.46 | -2.43 | -0.78 | -0.81 | -0.95 | -0.85 | -0.095 | -0.985 | 0.155 | 0.658586 | 0.014068 | 0.460099 | 0.999971 | 1 | 0.889706 |
| Pcdh7 | E9Q2S0 | -0.43 | -1.06 | -1.15 | -1.11 | -1.11 | -0.85 | -0.88 | -1.21 | -0.99 | -0.5 | -1.32 | -1.26 | -0.56 | -0.88 | -1.38 | -1.13 | -0.075 | -0.08 | -0.05 | 0.744616 | 0.767703 | 0.850985 | 0.999971 | 1 | 0.98187 |
| Pcdh8 | Q7TSK3 | 0.36 | -0.42 | 0.27 | 0.01 | -0.06 | -0.06 | 0.19 | 0.63 | -0.25 | 0.22 | 0.24 | -0.04 | 0.22 | -0.21 | 0.61 | 0.24 | 0.12 | -0.0125 | 0.16 | 0.64831 | 0.958464 | 0.549663 | 0.999971 | 1 | 0.912334 |
| Pcdh9 | A0A0A6YY( | -0.4 | -1.44 | -1.14 | -1.2 | -1.62 | -0.66 | -1.06 | -1.11 | -1.46 | -0.76 | -1.3 | -1.32 | -1.52 | -1.27 | -1.31 | -1.3 | -0.0675 | -0.165 | -0.305 | 0.825109 | 0.566176 | 0.255087 | 0.999971 | 1 | 0.839012 |
| Pcdh9_1 | A0A0A6YY3 | -1 | -2 | -1.62 | -2.16 | -2.15 | -2.14 | -1.72 | -1.28 | -1.1 | -0.69 | -0.64 | -0.79 | -1.29 | -2.02 | -2.17 | -1.92 | -0.1275 | 0.89 | -0.155 | 0.700118 | 0.012658 | 0.634701 | 0.999971 | 1 | 0.93301 |
| Pcdha12 | Q91Y18 | -0.74 | -1.43 | -0.24 | 0.43 | -2.06 | -2.02 | -0.56 | 0.41 | -2.19 | -0.93 | -0.24 | -0.1 | -2.56 | -1.41 | 0.06 | -0.12 | -0.5625 | -0.37 | -0.5125 | 0.398289 | 0.521348 | 0.446203 | 0.999971 | 1 | 0.884449 |
| Pcdhac2 | Q91Y09 | 0.07 | -0.86 | -0.97 | -0.66 | -1.11 | -0.92 | -0.78 | -0.76 | -0.71 | -0.21 | -0.9 | -0.7 | -0.85 | -1.04 | -1.07 | -1.03 | -0.2875 | -0.025 | -0.3925 | 0.299073 | 0.930762 | 0.160325 | 0.999971 | 1 | 0.800215 |
| Pcdhb12 | Q91Y07 | -0.65 | -1.73 | -0.62 | 1.63 | -0.47 | -0.93 | -1.75 | 1.08 | -0.96 | -0.41 | -1.62 | -0.67 | -1.95 | -0.87 | -0.17 | 0.2 | -0.175 | -0.5725 | -0.355 | 0.832742 | 0.413602 | 0.645116 | 0.999971 | 1 | 0.93301 |
| Pcdhb14 | Q6PB90 | -0.54 | -0.83 | -1.17 | -1.04 | -1.1 | -0.55 | -1.16 | -1 | -0.78 | -0.49 | -1.53 | -0.87 | -1.12 | -0.75 | -1.06 | -0.71 | -0.0575 | -0.0225 | -0.015 | 0.804372 | 0.934756 | 0.945298 | 0.999971 | 1 | 0.997427 |
| Pcdhb17 | Q91VD8 | -0.83 | -1.11 | -2.97 | -1.4 | -1 | -1.82 | -1.35 | -1.6 | -1.02 | -0.29 | -0.76 | -3.06 | -1 | -1.15 | -0.75 | -2.34 | 0.135 | 0.295 | 0.2675 | 0.776714 | 0.677578 | 0.627436 | 0.999971 | 1 | 0.931041 |
| Pcdhb18 | Q91Y02 | 0.35 | -1.83 | -1.27 | -0.76 | -1.29 | -1.23 | -0.99 | -1.04 | -2.33 | 0.24 | -1.5 | -1.26 | -1.25 | -0.92 | -1.43 | -0.15 | -0.26 | -0.335 | -0.06 | 0.560067 | 0.607602 | 0.905229 | 0.999971 | 1 | 0.991091 |
| Pcdhb20 | Q91XZ9 | -0.53 | -0.09 | -0.84 | -1.98 | -0.76 | -1.14 | -0.84 | -1.75 | -1.67 | -0.16 | -0.63 | -0.43 | -0.05 | -0.27 | -2.1 | -1.82 | -0.2625 | 0.1375 | -0.2 | 0.550967 | 0.778158 | 0.741508 | 0.999971 | 1 | 0.96105 |
| Pcdhb21 | Q91V48 | -0.51 | -0.44 | -0.16 | 0.3 | 0.09 | -1.16 | -0.89 | 0.02 | -0.93 | -0.93 | -0.23 | -0.36 | -0.39 | -0.87 | -0.3 | -0.1 | -0.2825 | -0.41 | -0.2125 | 0.438454 | 0.163448 | 0.435008 | 0.999971 | 1 | 0.882419 |
| Pcdhb9 | E9Q5G2 | 0.11 | -0.17 | -1.08 | -0.9 | -0.03 | -1.61 | -0.94 | -0.73 | -0.71 | 0.02 | -1.26 | -0.53 | -0.27 | -0.49 | -0.99 | -1.11 | -0.3175 | -0.11 | -0.205 | 0.448394 | 0.770623 | 0.555941 | 0.999971 | 1 | 0.914924 |
| Pcdhga10 | Q91XY9 | -1.91 | -0.45 | -0.33 | -0.53 | -0.82 | 0.8 | -1.11 | 0.18 | -0.76 | -0.79 | -0.62 | 0.38 | -0.65 | -0.59 | -1.47 | -0.55 | 0.5675 | 0.3575 | -0.01 | 0.30205 | 0.424099 | 0.980557 | 0.999971 | 1 | 0.99925 |
| Pcdhga11 | Q91XY8 | -0.23 | -0.45 | -1.01 | -0.84 | -0.46 | -0.97 | -1.1 | -0.84 | -0.89 | -0.58 | -1.05 | -0.97 | -0.12 | -0.73 | -1.48 | -1.4 | -0.21 | -0.24 | -0.3 | 0.41465 | 0.326321 | 0.411658 | 0.999971 | 1 | 0.875847 |
| Pcdhga12 | Q91XY7 | -0.76 | -1.18 | -0.73 | -0.83 | -1.03 | -1.08 | -0.71 | -0.68 | -1.36 | -1.01 | -0.62 | -0.64 | -1.29 | -1.28 | -0.74 | -0.95 | 1.57E-16 | -0.0325 | -0.19 | 1 | 0.890425 | 0.391333 | 1 | 1 | 0.871816 |
| Pcdhga3 | Q91XY5 | -0.78 | -1 | -1.22 | -1.03 | -0.61 | -1.27 | -1.18 | -0.93 | -1.27 | -0.56 | -1.13 | -1.04 | -0.63 | -0.88 | -0.89 | -0.76 | 0.01 | 0.0075 | 0.2175 | 0.963415 | 0.972704 | 0.256742 | 0.999971 | 1 | 0.839012 |
| Pcdhga5 | Q91XY3 | -0.77 | -0.93 | 0.36 | 1.07 | -0.64 | -1.46 | 0.25 | 0.38 | -1.08 | -0.56 | -0.25 | 0.04 | -1.22 | -0.3 | -0.16 | 1.17 | -0.3 | -0.395 | -0.06 | 0.610841 | 0.435019 | 0.923338 | 0.999971 | 1 | 0.993265 |
| Pcdhga6 | Q91XY2 | -0.16 | -0.62 | -0.54 | -0.23 | -0.71 | -1 | -0.65 | -0.8 | -1.55 | -0.11 | -0.45 | -0.3 | -0.82 | -0.51 | -0.62 | -0.71 | -0.4025 | -0.215 | -0.2775 | 0.067212 | 0.531166 | 0.179069 | 0.999971 | 1 | 0.808532 |
| Pcdhga7 | Q6DD96 | -0.51 | -2.81 | -1.09 | -0.43 | -2.78 | -1.37 | -1.05 | -1.65 | -3.45 | -1.11 | -0.45 | -0.86 | -3.42 | -2.29 | -0.09 | -0.54 | -0.5025 | -0.2575 | -0.375 | 0.419133 | 0.745031 | 0.662314 | 0.999971 | 1 | 0.936148 |
| Pcdhga8 | Q91XY0 | -0.31 | -0.71 | -1.32 | -0.81 | -1.39 | -0.83 | -1.45 | -0.98 | -0.82 | -0.2 | -1.09 | -0.83 | -0.68 | -1.19 | -1.56 | -1.26 | -0.375 | 0.0525 | -0.385 | 0.195829 | 0.856945 | 0.206238 | 0.999971 | 1 | 0.817241 |
| Pcdhgb1 | Q91XX8 | -0.19 | -0.52 | 1.23 | -0.22 | -0.61 | -0.52 | 0.39 | -0.88 | -0.74 | -0.8 | -0.12 | 0.42 | -0.1 | -0.16 | 0.73 | 1.48 | -0.48 | -0.385 | 0.4125 | 0.302228 | 0.410163 | 0.428728 | 0.999971 | 1 | 0.881196 |
| Pcdhgb2 | Q91XX7 | -0.34 | -1.13 | -0.34 | 0.43 | -0.62 | -0.64 | -1.47 | 1.09 | -1.81 | -0.81 | -0.11 | -0.04 | 0.36 | -0.61 | -0.05 | 0.59 | -0.065 | -0.3475 | 0.4175 | 0.909496 | 0.481223 | 0.308102 | 0.999971 | 1 | 0.847402 |
| Pcdhgb5 | Q91XX5 | -0.77 | -0.77 | -1.17 | -0.8 | -1.42 | -1.32 | -1.96 | -0.73 | -1.2 | -0.73 | -1.16 | -0.82 | -0.14 | -0.62 | -1.12 | -0.88 | -0.48 | -0.1 | 0.1875 | 0.118314 | 0.626655 | 0.472766 | 0.999971 | 1 | 0.893755 |
| Pcdhgb6 | Q91XX4 | -0.48 | -1.19 | -0.54 | -0.59 | -0.88 | -0.93 | -0.74 | -0.8 | -0.8 | -0.75 | -0.55 | 0.11 | -1.12 | -1.47 | -0.85 | -0.35 | -0.1375 | 0.2025 | -0.2475 | 0.529944 | 0.477264 | 0.415643 | 0.999971 | 1 | 0.876885 |
| Pcdhgc3 | Q91XX1 | 0.72 | -0.94 | -0.7 | -0.51 | -0.71 | -0.36 | -0.8 | -0.58 | -0.45 | 0.74 | -1.08 | -1.01 | -0.29 | -0.72 | -0.56 | -0.83 | -0.255 | -0.0925 | -0.2425 | 0.497131 | 0.858856 | 0.524093 | 0.999971 | 1 | 0.908269 |
| Pcdhgc5 | Q91XW9 | 0.49 | 0.01 | -0.12 | -0.14 | -0.41 | -0.05 | -0.25 | -0.56 | -0.46 | 0.03 | -0.18 | -0.5 | -0.28 | -0.38 | -0.55 | -0.16 | -0.3775 | -0.3375 | -0.4025 | 0.119717 | 0.165436 | 0.089802 | 0.999971 | 1 | 0.771991 |
| Pcf11 | G3X9Z4 | 1.92 | 2.29 | 1.9 | 2.09 | 1.83 | 2.14 | 1.39 | 1.83 | 1.62 | 2.22 | 1.32 | 1.22 | 1.43 | 2.4 | 2.39 | 2.6 | -0.2525 | -0.455 | 0.155 | 0.272788 | 0.111171 | 0.59659 | 0.999971 | 1 | 0.92377 |
| Pcgf2 | P23798 | 0.76 | 0.45 | 0.82 | 0.74 | 0.26 | 0.47 | -0.08 | 1.01 | 0.09 | 0.67 | 0.64 | 0.71 | 0.05 | 0.55 | 0.97 | 0.9 | -0.2775 | -0.165 | -0.075 | 0.309544 | 0.44651 | 0.767564 | 0.999971 | 1 | 0.962472 |
| Pcid2 | Q8BFV2 | 1.74 | 1.2 | 0.21 | 0.42 | 0.77 | 1.47 | 0.71 | 0.16 | 0.81 | 1.21 | 0.2 | -0.21 | -0.14 | 0.33 | 0.11 | -0.45 | -0.115 | -0.39 | -0.93 | 0.784998 | 0.394005 | 0.034469 | 0.999971 | 1 | 0.69205 |
| Pcif1 | P59114 | 3.06 | 3.72 | 3.16 | 3.13 | 3.01 | 2.91 | 2.6 | 3.08 | 3.27 | 3.02 | 2.78 | 2.03 | 1.96 | 3.24 | 3.1 | 2.87 | -0.3675 | -0.4925 | -0.475 | 0.130461 | 0.141249 | 0.170909 | 0.999971 | 1 | 0.803758 |
| Pck2 | A0A0R4J0G | -0.35 | -0.66 | -0.37 | -0.51 | -0.89 | -0.64 | -0.44 | -0.56 | -0.62 | -0.57 | -0.63 | -0.47 | -0.62 | -0.12 | -0.1 | -0.17 | -0.16 | -0.1 | 0.22 | 0.405686 | 0.556613 | 0.29119 | 0.999971 | 1 | 0.847402 |
| Pclo | Q9QYX7 | -0.11 | -1.22 | -1.07 | -0.98 | -1.32 | -1 | -1.02 | -1.39 | -1.27 | -0.48 | -1.28 | -1.42 | -1.12 | -1.19 | -1.34 | -1.19 | -0.3375 | -0.2675 | -0.365 | 0.253282 | 0.424439 | 0.204206 | 0.999971 | 1 | 0.81617 |
| Pcm1 | Q9R0L6-2 | 1.09 | 0.34 | 0.46 | 0.72 | 0.47 | 0.26 | 0.74 | 0.49 | 0.26 | 0.77 | 0.64 | 0.47 | 0.62 | 0.65 | 0.58 | 0.39 | -0.1625 | -0.1175 | -0.0925 | 0.48729 | 0.61648 | 0.676434 | 0.999971 | 1 | 0.941944 |
| Pcmt1 | F7D432 | -1.12 | -1.34 | -0.78 | -1.31 | -1.2 | -1.35 | 0.24 | -1.56 | -1.23 | -1.44 | -0.84 | -0.88 | -1.72 | -1.56 | -1.64 | -1.63 | 0.17 | 0.04 | -0.5 | 0.678645 | 0.861219 | 0.029218 | 0.999971 | 1 | 0.678474 |
| Pcmtd1 | P59913 | 0.16 | 0.43 | 0.93 | -1.19 | 0.29 | -0.14 | 1.43 | -0.44 | 0.42 | -0.12 | 1.42 | 1.61 | 0.38 | -0.18 | -0.2 | -1.47 | 0.2025 | 0.75 | -0.45 | 0.718677 | 0.206726 | 0.423235 | 0.999971 | 1 | 0.87887 |
| Pcmtd2 | Q8BHD8 | 0.61 | 0.34 | 0.67 | 1.13 | 0.38 | -0.2 | 0.15 | 1.3 | 0.33 | 0.99 | 1.03 | 0.94 | 0.18 | 0.89 | 0.74 | 1.57 | -0.28 | 0.135 | 0.1575 | 0.436094 | 0.60084 | 0.635085 | 0.999971 | 1 | 0.93301 |
| Pcna | P17918 | 0.09 | 1.97 | 1.68 | 1.32 | 2.24 | 1.27 | 1.13 | 1.45 | 1.98 | -0.53 | 1.3 | 1.06 | 2.21 | 1.42 | 1.7 | 1.28 | 0.2575 | -0.3125 | 0.3875 | 0.572713 | 0.615033 | 0.385564 | 0.999971 | 1 | 0.871816 |
| Pcnp | Q6P8I4 | 1.89 | 2.44 | 2.32 | 2.98 | 2.32 | 2.22 | 2.14 | 2.39 | 2.6 | 2.11 | 2.62 | 2.73 | 2.36 | 2.85 | 2.69 | 3.23 | -0.14 | 0.1075 | 0.375 | 0.587832 | 0.700281 | 0.229335 | 0.999971 | 1 | 0.828687 |
| Pcnt | A0A1W2P7 | 0.3 | 0.07 | 0.21 | 0.23 | 0.13 | -0.09 | 0.03 | 0.26 | -0.01 | 0.03 | 0.11 | -0.01 | 0.44 | 0.02 | 0.25 | 0.49 | -0.12 | -0.1725 | 0.0975 | 0.498486 | 0.298451 | 0.606826 | 0.999971 | 1 | 0.925161 |
| Pcnx | A0A1Y7VKI | -0.1 | -1.25 | -0.45 | 0.19 | -1.37 | -0.91 | 0.1 | -0.3 | -1.05 | -0.39 | -0.76 | -1.13 | -1.37 | -0.99 | -0.48 | -0.95 | -0.2175 | -0.43 | -0.545 | 0.612054 | 0.237488 | 0.150283 | 0.999971 | 1 | 0.793248 |
| Pcnx2 | Q5DU28 | 0.45 | 0 | 0.41 | -0.31 | -0.26 | -0.2 | 0.62 | -0.43 | 0.2 | 0.49 | 1.1 | 0.54 | 0.63 | -0.75 | -0.32 | -0.71 | -0.205 | 0.445 | -0.425 | 0.504516 | 0.134086 | 0.257349 | 0.999971 | 1 | 0.839012 |
| Pcnx3 | Q8VI59 | -0.58 | -0.98 | 0.05 | -0.04 | -0.56 | -0.64 | -0.36 | 0.06 | -0.9 | -0.25 | -0.16 | 0.34 | -0.54 | -0.51 | 0.24 | 0.18 | 0.0125 | 0.145 | 0.23 | 0.966417 | 0.676589 | 0.483857 | 0.999971 | 1 | 0.896352 |
| Pcolce | D3YWH0 | 0.86 | 0.38 | 0.37 | 0.37 | 0.75 | -0.26 | 2.34 | -0.89 | -0.01 | 0.29 | 0.23 | 1.12 | -1.36 | 0.08 | -0.08 | -1.04 | -0.01 | -0.0875 | -1.095 | 0.987691 | 0.760541 | 0.014368 | 0.999971 | 1 | 0.613095 |
| Pcp4l1 | Q6W8Q3 | 2.6 | 1.35 | 0.76 | 2.02 | 1.78 | 0.35 | 1.19 | 1.64 | 2.02 | 2.66 | 0.8 | 1.51 | 1.42 | 0.75 | 1.93 | 1.17 | -0.4425 | 0.065 | -0.365 | 0.366948 | 0.900633 | 0.419127 | 0.999971 | 1 | 0.87788 |
| Pcsk1 | P63239 | -0.08 | -1.59 | -0.84 | -1.18 | -0.72 | -0.11 | -0.05 | -0.6 | -0.36 | -0.07 | -0.79 | -0.92 | -0.68 | -0.34 | -0.27 | -0.7 | 0.5525 | 0.3875 | 0.425 | 0.145506 | 0.305417 | 0.228676 | 0.999971 | 1 | 0.828687 |
| Pcsk1n | Q9QXV0 | -0.07 | -1.29 | -1.03 | -0.58 | -0.74 | -0.93 | -0.83 | -0.34 | -1.15 | -1.62 | -1.3 | -0.72 | -0.79 | -0.15 | 0.14 | -0.5 | 0.0325 | -0.455 | 0.4175 | 0.914918 | 0.187863 | 0.233257 | 0.999971 | 1 | 0.831526 |
| Pcsk2 | P21661 | -0.12 | -0.03 | -0.1 | -0.4 | -0.11 | -0.1 | 0.92 | 0.23 | -0.15 | -0.13 | 0.14 | 0.47 | 0.64 | 0.18 | 0.32 | -0.02 | 0.3975 | 0.245 | 0.4425 | 0.170284 | 0.266109 | 0.061918 | 0.999971 | 1 | 0.725348 |
| Pcx | G5E8R3 | -1.62 | -1.28 | -0.67 | -1.09 | -1.38 | -1.53 | -0.84 | -1.04 | -1.33 | -1.58 | -0.88 | -0.84 | -1.27 | -1.2 | -1.02 | -1.12 | -0.0325 | 0.0075 | 0.0125 | 0.904686 | 0.978688 | 0.958311 | 0.999971 | 1 | 0.998622 |
| Pcyox1 | Q9CQF9 | -0.84 | -0.73 | -0.87 | -0.86 | -0.9 | -0.64 | -0.82 | -0.48 | -0.76 | -0.9 | -0.61 | -0.38 | -0.55 | -0.51 | -0.44 | -0.39 | 0.115 | 0.1625 | 0.3525 | 0.52617 | 0.387349 | 0.055166 | 0.999971 | 1 | 0.725348 |
| Pcyox1l | Q8C7K6 | -0.6 | -0.18 | -0.28 | -1.35 | -0.16 | -0.22 | -0.5 | -0.49 | -0.16 | -0.3 | 0.26 | 0.14 | 0.16 | 0.01 | -0.04 | -0.32 | 0.26 | 0.5875 | 0.555 | 0.382468 | 0.078781 | 0.086473 | 0.999971 | 1 | 0.759991 |
| Pcyt1a | P49586 | 1.72 | 2.27 | 1.81 | 2.11 | 2.16 | 1.8 | 1.88 | 1.73 | 2.13 | 1.54 | 1.78 | 2.51 | 1.59 | 1.9 | 1.47 | 1.98 | -0.085 | 0.0125 | -0.2425 | 0.686731 | 0.962546 | 0.289906 | 0.999971 | 1 | 0.847402 |
| Pcyt1b | Q811Q9 | 0.07 | 0.28 | 0.49 | -0.2 | 0.92 | -0.29 | 0.76 | -0.48 | 0.94 | 0.13 | 1.03 | 0.84 | 0.25 | -0.61 | -0.38 | -0.5 | 0.0675 | 0.575 | -0.47 | 0.856805 | 0.058767 | 0.103261 | 0.999971 | 1 | 0.78268 |
| Pcyt2 | A2ABY3 | -0.41 | 0.23 | 0.07 | -0.11 | 0.01 | 0.4 | -0.2 | 0.06 | 0.06 | -0.33 | -0.2 | -0.09 | 0.01 | 0.19 | 0.03 | -0.04 | 0.1225 | -0.085 | 0.1025 | 0.590312 | 0.684217 | 0.61614 | 0.999971 | 1 | 0.928229 |
| Pdap1 | Q3UHX2 | 0.56 | -0.16 | 0.36 | 0.04 | 0.07 | 0.34 | 0.89 | 0.17 | 0.2 | 0.42 | 0.68 | 0.51 | 0.39 | 0.03 | 0.32 | 0.37 | 0.1675 | 0.2525 | 0.0775 | 0.531916 | 0.282195 | 0.73004 | 0.999971 | 1 | 0.956041 |
| Pdcd10 | Q8VE70 | 0.89 | 1.4 | 0.66 | 0.59 | 0.81 | 1.37 | 0.15 | 0.55 | 0.76 | 1.07 | 0.55 | 0.61 | 0.55 | 1.4 | 0.67 | 0.58 | -0.165 | -0.1375 | -0.085 | 0.605911 | 0.57798 | 0.76704 | 0.999971 | 1 | 0.962472 |
| Pdcd11 | Q6NS46 | -1.12 | -1.75 | -1.4 | -1.02 | -1.96 | -2.09 | -1.69 | -1.73 | -1.94 | -1.4 | -1.46 | -1.2 | -1.78 | -1.53 | -1.52 | -1.83 | -0.545 | -0.1775 | -0.3425 | 0.038119 | 0.487034 | 0.153491 | 0.999971 | 1 | 0.795578 |
| Pdcd2 | A2T4N6 | -1.82 | -1.37 | -2.82 | 0.03 | -0.38 | -1.53 | -1.22 | -0.21 | -1.03 | -1.66 | -1.27 | -0.78 | -2.16 | -0.64 | -1.64 | -1.24 | 0.66 | 0.31 | 0.075 | 0.297847 | 0.590821 | 0.902788 | 0.999971 | 1 | 0.990848 |
| Pdcd4 | Q61823 | 0.12 | 0.3 | 0.26 | 0.54 | 0.29 | 0.04 | 0.49 | 0.23 | 0.72 | 0.26 | 0.85 | 0.96 | 0.94 | 0.82 | 0.88 | 1.07 | -0.0425 | 0.3925 | 0.6225 | 0.825292 | 0.099868 | 0.007194 | 0.999971 | 1 | 0.582714 |
| Pdcd5 | P56812 | -0.37 | 0.18 | 0.7 | -0.03 | 0.81 | -0.29 | 1.38 | 0 | 1.17 | -0.28 | 0.81 | 1.56 | 0.93 | -0.11 | 0.45 | -0.19 | 0.355 | 0.695 | 0.15 | 0.407025 | 0.1343 | 0.662304 | 0.999971 | 1 | 0.936148 |
| Pdcd6 | P12815-2 | -0.91 | 0.02 | 0.36 | 0.38 | 0.07 | 0.11 | 0.95 | 0.15 | -0.09 | -0.63 | 0.08 | 0.01 | -0.24 | -0.18 | -0.06 | -0.59 | 0.3575 | -0.12 | -0.23 | 0.334304 | 0.724417 | 0.485112 | 0.999971 | 1 | 0.896868 |
| Pdcd6ip | Q9WU78 | -0.13 | -0.44 | 0.07 | -0.13 | -0.32 | -0.43 | 0.18 | -0.25 | -0.44 | -0.56 | 0.03 | -0.19 | -0.53 | -0.65 | -0.19 | -0.11 | -0.0475 | -0.1325 | -0.2125 | 0.826319 | 0.538735 | 0.337929 | 0.999971 | 1 | 0.859348 |
| Pdcd6ip_1 | Q9WU78-3 | -3.54 | -0.54 | -0.5 | -0.84 | -2.22 | 0.1 | -1.36 | -1.84 | -1.2 | -1.53 | -1.61 | -0.05 | -3.33 | -0.79 | -0.95 | -1.62 | 0.025 | 0.2575 | -0.3175 | 0.975063 | 0.728876 | 0.706368 | 0.999971 | 1 | 0.950182 |
| Pdcl | Q9DBX2 | 1.31 | 0.42 | 0.54 | 1.02 | 0.3 | 0.59 | 0.12 | 0.56 | 0.5 | 1.18 | 0.7 | 0.96 | -0.03 | 0.77 | 0.56 | 0.87 | -0.43 | 0.0125 | -0.28 | 0.124571 | 0.963367 | 0.361502 | 0.999971 | 1 | 0.869526 |
| Pdcl3 | Q8BVF2 | 0.1 | -0.13 | 0.92 | 1.28 | 0.54 | 0.26 | 0.77 | 1.28 | 0.2 | 0.11 | 1.14 | 1.11 | 0.25 | 0.36 | 1.25 | 1.2 | 0.17 | 0.0975 | 0.2225 | 0.658621 | 0.814605 | 0.588457 | 0.999971 | 1 | 0.921285 |
| Pde10a | Q8CA95-3 | 0.05 | -0.53 | -1.4 | -0.54 | -0.31 | -1.58 | -1.82 | 0.04 | -0.39 | -0.4 | -1.44 | -0.78 | 0.09 | -0.14 | -0.24 | -1.31 | -0.3125 | -0.1475 | 0.205 | 0.542522 | 0.695635 | 0.620574 | 0.999971 | 1 | 0.929332 |
| Pde11a | P0C1Q2 | -1.09 | -0.88 | -1.73 | -1.58 | -1.23 | -0.84 | -2.11 | -1.43 | -1.33 | -1.24 | -1.27 | -1.8 | -1.21 | -0.99 | -1.28 | -1.15 | -0.0825 | -0.09 | 0.1625 | 0.803731 | 0.73086 | 0.508047 | 0.999971 | 1 | 0.905815 |
| Pde12 | Q3TIU4 | -0.25 | 0.24 | 0.12 | 0 | 0.09 | 0.01 | -0.01 | 0 | 0.03 | -0.11 | -0.05 | 0.14 | -0.2 | 0.24 | 0.02 | 0.03 | -0.005 | -0.025 | -0.005 | 0.978147 | 0.892267 | 0.979902 | 0.999971 | 1 | 0.99925 |
| Pde1a | Q61481 | -0.41 | -0.24 | -0.08 | 0.04 | -0.34 | 0.19 | 0.15 | 0.07 | -0.12 | -0.08 | 0.27 | 0.29 | -0.8 | -0.06 | 0.31 | 0.59 | 0.19 | 0.2625 | 0.1825 | 0.372913 | 0.212705 | 0.571582 | 0.999971 | 1 | 0.91816 |
| Pde1b | Q01065 | 1.18 | 0.62 | 0.05 | 1 | 0.29 | -0.3 | 0.01 | 1.28 | 0.63 | 1.15 | 0.12 | 0.87 | 0.51 | 1.17 | 1.5 | 0.84 | -0.3925 | -0.02 | 0.2925 | 0.344659 | 0.951737 | 0.38475 | 0.999971 | 1 | 0.871816 |

| | | | | | | | | | | | | | | | | | | | | | | | | | | | |
|---|---|---|---|---|---|---|---|---|---|---|---|---|---|---|---|---|---|---|---|---|---|---|---|---|---|---|---|
| Pde1c | Q8CDV2 | 0.54 | -0.11 | 0.89 | 1.25 | 0.11 | -0.47 | 0.58 | 2.94 | 0.51 | 0.43 | 0.64 | 2.07 | 0.45 | 0.59 | 3.01 | 2 | 0.1475 | 0.27 | 0.87 | 0.838564 | 0.557785 | 0.182112 | 0.999971 | | 1 | 0.808532 |
| Pde2a | Q5PR72 | -1.42 | -2 | -0.96 | 0.03 | -1.54 | -0.77 | 0.45 | -0.35 | -0.84 | -0.55 | -0.79 | -1.01 | -2.52 | -1.99 | -0.42 | -1.63 | 0.535 | 0.29 | -0.5525 | 0.342834 | 0.494375 | 0.344126 | 0.999971 | | 1 | 0.862346 |
| Pde3b | E9QLQ3 | -0.13 | -0.17 | 0.15 | 0.96 | 0.34 | -1.57 | 1.27 | -0.21 | 0.62 | -0.08 | 0.41 | 1.36 | 1.7 | 0.18 | -0.3 | -0.08 | -0.245 | 0.375 | 0.1725 | 0.680639 | 0.342847 | 0.724071 | 0.999971 | | 1 | 0.955315 |
| Pde4a | O89084 | -0.42 | -0.79 | -1.03 | -0.55 | -1.09 | -1.13 | -0.91 | -0.73 | -0.98 | -0.29 | -1.15 | -0.92 | -0.77 | -0.91 | -0.59 | -0.56 | -0.2675 | -0.1375 | -0.01 | 0.228311 | 0.592796 | 0.961929 | 0.999971 | | 1 | 0.998721 |
| Pde4b | Q8VBU5 | -0.89 | -0.7 | -0.82 | -0.87 | -0.88 | -0.93 | -0.78 | -0.96 | -0.66 | -0.66 | -0.72 | -0.73 | -1.01 | -1.21 | -0.69 | -0.58 | -0.0675 | 0.1275 | -0.0525 | 0.683326 | 0.428652 | 0.798896 | 0.999971 | | 1 | 0.970479 |
| Pde4d | E9PVU7 | -0.13 | -0.17 | -0.67 | -0.68 | 0.75 | -0.12 | -0.69 | -0.67 | 1.01 | -0.02 | -0.59 | -0.5 | 1.89 | -0.54 | -0.58 | -0.47 | 0.23 | 0.3875 | 0.4875 | 0.529013 | 0.327129 | 0.404609 | 0.999971 | | 1 | 0.87384 |
| Pde4dip | Q80YT7-2 | 1.59 | 1.88 | 1.36 | 1.63 | 1.57 | 1.56 | 1.3 | 1.79 | 1.35 | 2.08 | 1.68 | 1.65 | 1.17 | 1.85 | 1.47 | 1.52 | -0.06 | 0.075 | -0.1125 | 0.7674 | 0.737047 | 0.612016 | 0.999971 | | 1 | 0.927066 |
| Pde5a | A0A0G2JF6 | 1.74 | 0.05 | -0.23 | 0.12 | 0.42 | 1.08 | -0.26 | 1.19 | 0.62 | 1.12 | 0.52 | -0.96 | 0.26 | 0.25 | 0.49 | 0.18 | 0.1875 | -0.095 | -0.125 | 0.717148 | 0.870284 | 0.770545 | 0.999971 | | 1 | 0.963498 |
| Pde6d | O55057 | -1.58 | 0.11 | -0.19 | -0.83 | 0.68 | -0.68 | -0.33 | -0.42 | 1.36 | -1.5 | -0.4 | 0.37 | 1.6 | -1.18 | -0.54 | -0.94 | 0.435 | 0.58 | 0.3575 | 0.346607 | 0.383509 | 0.596338 | 0.999971 | | 1 | 0.92377 |
| Pde7a | P70453 | -0.39 | -0.37 | 1.37 | 1.89 | 0.28 | -0.55 | 1.12 | 1.47 | -0.07 | -0.99 | 1.43 | 0.95 | -1.22 | -0.55 | 1.26 | 1.76 | -0.045 | -0.295 | -0.3125 | 0.946636 | 0.685937 | 0.707869 | 0.999971 | | 1 | 0.950182 |
| Pde7b | E9PW23 | 0.37 | 0.7 | 0.04 | 1.66 | 0.86 | 0.45 | -0.27 | 0.87 | 0.45 | 0.66 | -0.25 | 0.86 | 0.06 | 1.16 | -0.4 | -0.23 | -0.215 | -0.2625 | -0.545 | 0.609395 | 0.523441 | 0.259138 | 0.999971 | | 1 | 0.839012 |
| Pde8a | O88502 | -2.92 | -2.64 | 1.87 | 0.84 | -3.74 | -2.61 | 0.4 | 0.91 | -3.48 | -2.97 | 0.65 | 0.31 | -4.95 | -2.91 | 1.08 | 0.58 | -0.5475 | -0.66 | -0.8375 | 0.710848 | 0.65078 | 0.618335 | 0.999971 | | 1 | 0.928581 |
| Pde8b | E9Q8J6 | 0.79 | -1.92 | -1.77 | -0.62 | -1.92 | -1.89 | 1.28 | -1.09 | -1.74 | -0.75 | -1.56 | -1.43 | -3.12 | -1.52 | -1.94 | -1.38 | -0.025 | -0.49 | -1.11 | 0.97726 | 0.430005 | 0.126288 | 0.999971 | | 1 | 0.783071 |
| Pde8b_1 | E9PX38 | -0.96 | -1.65 | 0.7 | -0.14 | -1.47 | -1.98 | 0.15 | 0.34 | -1.91 | -0.55 | 1.08 | 0.84 | -1.68 | -1.41 | 0.58 | 0.49 | -0.2275 | 0.3775 | 0.0075 | 0.744646 | 0.629791 | 0.991607 | 0.999971 | | 1 | 0.99925 |
| Pde9a | A0A494BAI | 0.01 | -0.33 | -1.45 | -0.8 | -0.7 | -0.72 | -1.8 | -1.59 | -0.35 | -0.72 | -1.7 | -0.79 | 0.53 | -0.41 | -1.06 | -0.82 | -0.56 | -0.2475 | 0.2025 | 0.192688 | 0.548451 | 0.650571 | 0.999971 | | 1 | 0.933842 |
| Pdf | S4R2K0 | 0.02 | -0.96 | -0.55 | -0.71 | -0.76 | -0.67 | -0.96 | -0.16 | -1.01 | 0.56 | -0.47 | -0.62 | -0.17 | -0.39 | -0.45 | 0.03 | -0.0875 | 0.165 | 0.305 | 0.757694 | 0.666677 | 0.258972 | 0.999971 | | 1 | 0.839012 |
| Pdgfra | P26618 | -0.26 | -0.86 | -0.19 | 0.37 | -0.18 | -1.19 | -0.08 | 0.28 | -0.7 | -0.08 | 0.11 | 0.43 | -0.68 | -0.47 | -0.18 | 0.18 | -0.0575 | 0.175 | -0.0525 | 0.881992 | 0.611642 | 0.868466 | 0.999971 | | 1 | 0.983462 |
| Pdgfrb | E9QN12 | -0.24 | 0.88 | -0.62 | -0.79 | 0.97 | 0.79 | -1.07 | -0.94 | 0.91 | 0.48 | -0.64 | -0.52 | 0.95 | 0.21 | -0.84 | -0.7 | 0.13 | 0.25 | 0.0975 | 0.829721 | 0.617818 | 0.851413 | 0.999971 | | 1 | 0.98187 |
| Pdha1 | P35486 | -1.33 | -0.83 | -0.69 | -0.67 | -0.87 | -0.93 | -0.59 | -0.68 | -1.04 | -1.47 | -0.57 | -0.86 | -1.27 | -1.02 | -0.77 | -0.89 | 0.1125 | -0.105 | -0.1075 | 0.609162 | 0.691629 | 0.639266 | 0.999971 | | 1 | 0.93301 |
| Pdhb | Q9D051 | -1.09 | -0.69 | -0.87 | -0.72 | -0.69 | -0.92 | -0.95 | -0.78 | -0.99 | -1.21 | -0.94 | -0.86 | -0.83 | -0.69 | -0.92 | -0.71 | 0.0075 | -0.1575 | 0.055 | 0.967445 | 0.404344 | 0.764165 | 0.999971 | | 1 | 0.962405 |
| Pdhx | Q8BKZ9 | -0.28 | -0.9 | -0.28 | -0.14 | -1.27 | -0.93 | -0.58 | -0.18 | -1.16 | -0.55 | -0.18 | -0.21 | -1.2 | -0.86 | -0.48 | -0.17 | -0.34 | -0.125 | -0.2775 | 0.272064 | 0.671847 | 0.356149 | 0.999971 | | 1 | 0.868793 |
| Pdia3 | P27773 | -0.77 | -0.43 | -0.17 | -0.08 | -0.35 | -0.37 | 0.04 | 0.05 | -0.12 | -0.57 | 0.14 | 0.25 | -0.13 | -0.37 | 0.15 | 0.34 | 0.205 | 0.2875 | 0.36 | 0.386455 | 0.287052 | 0.173001 | 0.999971 | | 1 | 0.806514 |
| Pdia4 | A0A0R4J0Z | -0.26 | -0.21 | 0.05 | 0.2 | 0.1 | 0.03 | 0.24 | 0.18 | 0.4 | -0.04 | 0.54 | 0.52 | 0.57 | 0.09 | 0.24 | 0.45 | 0.1925 | 0.41 | 0.3925 | 0.320252 | 0.085317 | 0.084699 | 0.999971 | | 1 | 0.758256 |
| Pdia5 | Q921X9 | 0.61 | 0.46 | -1.3 | 0.26 | 1.07 | -0.17 | -0.47 | -0.09 | 1.01 | 1.03 | -0.61 | 0.02 | 1.23 | 0.59 | 0.75 | 1.23 | 0.0775 | 0.355 | 0.9425 | 0.880167 | 0.524453 | 0.059218 | 0.999971 | | 1 | 0.725348 |
| Pdia6 | Q3TML0 | 1.91 | 0.7 | 0.55 | 0.88 | 0.24 | 0.85 | 1.08 | 0.71 | 0.49 | 1.23 | 0.55 | 0.55 | 0.5 | 0.66 | 0.61 | 0.77 | -0.29 | -0.305 | -0.375 | 0.415362 | 0.392312 | 0.255505 | 0.999971 | | 1 | 0.839012 |
| Pdk1 | Q8BFP9 | -1.16 | -0.39 | -0.03 | -0.1 | -0.5 | -0.32 | 0.38 | -0.73 | -0.86 | -1.22 | -0.19 | -0.42 | -0.49 | -0.94 | -0.47 | -0.68 | 0.1275 | -0.2525 | -0.225 | 0.714145 | 0.467272 | 0.449703 | 0.999971 | | 1 | 0.885596 |
| Pdk2 | Q9JK42 | -0.77 | -0.5 | -0.56 | -0.72 | -0.38 | -0.11 | -0.22 | -0.25 | -0.21 | -0.69 | -0.35 | -0.41 | 0.29 | -0.18 | -0.33 | -0.33 | 0.3975 | 0.2225 | 0.5 | 0.044581 | 0.250044 | 0.039413 | 0.999971 | | 1 | 0.700838 |
| Pdk3 | Q922H2 | -0.51 | -0.29 | -0.49 | -1.04 | -0.45 | -0.47 | -0.46 | -0.95 | -0.48 | -0.8 | -0.93 | -0.55 | -0.33 | -0.91 | -0.84 | -1.37 | 9.81E-17 | -0.1075 | -0.28 | 1 | 0.63941 | 0.33486 | 1 | | 1 | 0.858827 |
| Pdk4 | O70571 | -1.62 | 2.41 | -0.99 | -0.78 | -0.75 | 0.3 | -0.9 | -0.58 | -0.65 | -0.45 | -0.39 | -2.37 | 1.27 | -1.47 | -1.46 | -0.97 | -0.2375 | -0.72 | -0.4125 | 0.77885 | 0.441137 | 0.680077 | 0.999971 | | 1 | 0.942957 |
| Pdlim1 | O70400 | -1.06 | -0.42 | -1.27 | -1.69 | 1.59 | 0.07 | -0.8 | -1.25 | -0.05 | 0.54 | 0.16 | 0.52 | -0.3 | -0.05 | -0.72 | -0.58 | 1.0125 | 1.4025 | 0.6975 | 0.12857 | 0.00143 | 0.047836 | 0.999971 | 0.620535 | 0.710102 | |
| Pdlim3 | O70209 | -0.7 | -0.68 | -0.97 | -0.89 | 3.67 | -0.07 | -0.36 | -1.75 | 0.19 | 0.37 | 0.92 | 3.44 | -1.08 | -0.82 | -0.43 | -1.81 | 1.1825 | 2.04 | -0.225 | 0.271285 | 0.015418 | 0.469984 | 0.999971 | | 1 | 0.892481 |
| Pdlim4 | P70271 | -0.03 | -0.22 | 0.98 | 1.52 | 0.24 | -0.49 | 0.86 | 2.15 | -0.7 | 0.19 | 1.46 | 1.58 | -0.67 | 0.23 | 1.28 | 1.68 | 0.1275 | 0.07 | 0.0675 | 0.840276 | 0.910898 | 0.912386 | 0.999971 | | 1 | 0.992618 |
| Pdlim5 | Q8CI51 | 0.47 | 0.18 | 0.17 | 0.03 | 1.07 | 0.52 | 0.68 | 0.19 | 0.49 | 0.86 | 0.49 | 1.14 | 0.04 | 0.46 | 0.29 | 0.31 | 0.4025 | 0.5325 | 0.0625 | 0.118382 | 0.03789 | 0.746551 | 0.999971 | | 1 | 0.961064 |
| Pdlim7 | Q3TJD7 | -0.79 | -0.69 | -1.4 | -1.03 | 4.84 | 0.5 | -0.01 | -2 | 0.92 | 3.16 | 3.63 | 5.98 | -1.67 | 0.38 | -0.94 | -1.1 | 1.81 | 4.4 | 0.145 | 0.183623 | 0.001298 | 0.740048 | 0.999971 | 0.620535 | 0.960899 | |
| Pdp1 | A2AJP9 | -1.18 | -0.68 | -0.82 | -0.85 | -0.8 | -0.94 | -0.95 | -0.87 | -1.31 | -1.14 | -0.89 | -0.75 | -1.04 | -0.7 | -1.07 | -0.86 | -0.0075 | -0.14 | -0.035 | 0.967577 | 0.511201 | 0.859596 | 0.999971 | | 1 | 0.982702 |
| Pdp2 | Q504M2 | -0.37 | -1.14 | 0 | -0.75 | -0.9 | -1 | 0.09 | 0.21 | -1.08 | -0.65 | -0.42 | -0.01 | -0.57 | -0.61 | 0.18 | 0.42 | 0.165 | 0.025 | 0.42 | 0.671288 | 0.939695 | 0.253216 | 0.999971 | | 1 | 0.839012 |
| Pdpk1 | Q9Z2A0 | -0.25 | -0.01 | -0.5 | -0.5 | -0.04 | 0.05 | -0.52 | -0.35 | 0.03 | -0.28 | -0.85 | -0.61 | 0.05 | 0.15 | -0.45 | -0.68 | 0.1 | -0.1125 | 0.0825 | 0.652467 | 0.655319 | 0.748453 | 0.999971 | | 1 | 0.961064 |
| Pdpr | Q7TSQ8 | -0.65 | -0.44 | -0.9 | -1.05 | -0.24 | -0.48 | -0.46 | -0.57 | -0.67 | -0.84 | -0.89 | -0.92 | -0.33 | -0.45 | -1.23 | -1.25 | 0.3225 | -0.07 | -0.055 | 0.143146 | 0.726742 | 0.851016 | 0.999971 | | 1 | 0.98187 |
| Pds5a | Q6A026 | 1.32 | 1.8 | 1.88 | 2.5 | 1.8 | 1.41 | 1.63 | 1.95 | 1.53 | 1.44 | 1.42 | 2.06 | 1.08 | 1.78 | 2.22 | 2.48 | -0.1775 | -0.2625 | 0.015 | 0.535053 | 0.384366 | 0.968327 | 0.999971 | | 1 | 0.998787 |
| Pds5b | F8WHU5 | -0.32 | 0.85 | 0.6 | 0.68 | 1.03 | 0.47 | 0.57 | 0.44 | 0.73 | -0.21 | 0.57 | 0.76 | 0.75 | 0.56 | 0.58 | 0.56 | 0.175 | 0.01 | 0.16 | 0.567897 | 0.97675 | 0.574244 | 0.999971 | | 1 | 0.91816 |
| Pdxdc1 | A0A0R4J03 | 0.78 | 1.78 | 1.46 | 0.97 | 1.65 | 1.44 | 1.38 | 1.15 | 1.58 | 0.91 | 1.48 | 1.85 | 1.4 | 1.49 | 1.38 | 1.26 | 0.1575 | 0.2075 | 0.135 | 0.562919 | 0.505228 | 0.603759 | 0.999971 | | 1 | 0.924454 |
| Pdxk | Q8K183 | 0.59 | -0.36 | 0.13 | 0.36 | -0.28 | -0.24 | 0.29 | 0.03 | -0.48 | -0.09 | -0.64 | -0.29 | -0.86 | 0.45 | 0.19 | 0.23 | -0.23 | -0.555 | -0.1775 | 0.394463 | 0.056268 | 0.615602 | 0.999971 | | 1 | 0.927967 |
| Pdxp | P60487 | 0.12 | 1.49 | 0.6 | 0.61 | 1.19 | 1.37 | 0.35 | 0.96 | 0.91 | 0.6 | 0.6 | 0.46 | 1.62 | 1.33 | 0.93 | 1.1 | 0.2625 | -0.0625 | 0.54 | 0.46605 | 0.838518 | 0.122856 | 0.999971 | | 1 | 0.78268 |
| Pdyn | O35852 | -0.08 | -0.79 | -1.59 | -0.65 | 0.26 | -0.51 | -0.93 | -0.39 | 0.16 | -0.39 | -1.46 | -0.42 | 0.54 | -0.15 | 0.2 | -0.76 | 0.385 | 0.25 | 0.735 | 0.328585 | 0.569428 | 0.09342 | 0.999971 | | 1 | 0.776565 |
| Pdzd11 | Q9CZG9 | -1.08 | -1.31 | -1.61 | -1.45 | -1.17 | -0.89 | -1.35 | -1.13 | -1.13 | -0.43 | -1.52 | -1.63 | -0.57 | -0.88 | -1.32 | -1.11 | 0.2275 | 0.185 | 0.3925 | 0.279581 | 0.543154 | 0.119692 | 0.999971 | | 1 | 0.78268 |
| Pdzd2 | E9Q1M1 | 0.04 | -0.6 | -0.13 | -0.16 | -0.48 | -0.55 | -0.12 | -0.21 | -0.02 | -0.32 | 0 | -0.2 | -0.55 | -0.57 | 0.08 | -0.3 | -0.1275 | 0.0775 | -0.1225 | 0.560921 | 0.707442 | 0.609941 | 0.999971 | | 1 | 0.92673 |
| Pdzd4 | A2AFG3 | 1.07 | -0.46 | -1.34 | -1.63 | -0.62 | -0.41 | -0.84 | -0.98 | -0.67 | 1.14 | -1.17 | -1.39 | 0.06 | -0.33 | -1.43 | -2.44 | -0.1225 | 0.0675 | -0.445 | 0.829159 | 0.928944 | 0.555933 | 0.999971 | | 1 | 0.914924 |
| Pdzd8 | B9EJ80 | -0.3 | -1.22 | -0.78 | -0.6 | -1.21 | -1.14 | -0.66 | -0.52 | -1.01 | -0.55 | -0.37 | -0.11 | -1.22 | -1.1 | -0.6 | -0.36 | -0.1575 | 0.215 | -0.095 | 0.570737 | 0.455381 | 0.745668 | 0.999971 | | 1 | 0.961064 |
| Pdzrn3 | Q69ZS0-2 | -0.57 | 0.02 | -0.41 | -0.63 | 0.05 | -0.27 | -0.55 | -0.84 | -0.25 | -0.46 | -0.17 | -0.04 | 0.51 | -0.16 | -0.5 | -0.19 | -0.005 | 0.1675 | 0.3125 | 0.984802 | 0.44357 | 0.275632 | 0.999971 | | 1 | 0.843794 |
| Pea15 | Q62048 | -1.32 | -1 | -0.68 | -0.86 | -1.01 | -0.51 | -0.59 | -0.4 | -0.89 | -1.17 | -0.66 | -1.07 | -1.51 | -0.42 | -0.55 | -0.51 | 0.3375 | 0.0175 | 0.2175 | 0.165082 | 0.935833 | 0.473579 | 0.999971 | | 1 | 0.894044 |
| Peak1 | Q69Z38 | -0.59 | -0.86 | -0.12 | -0.27 | -0.9 | -0.67 | -0.49 | 0 | -1.18 | -0.67 | 0 | -0.23 | -0.95 | -1.24 | -0.29 | -0.16 | -0.055 | -0.06 | -0.2 | 0.8395 | 0.847682 | 0.527534 | 0.999971 | | 1 | 0.909612 |
| Pebp1 | P70296 | -0.7 | -0.29 | 0.37 | 0.35 | -0.22 | -0.47 | 0.33 | 0.68 | -0.02 | -0.85 | 0.51 | 0.71 | -0.4 | -0.24 | 0.61 | 0.36 | 0.1475 | 0.155 | 0.15 | 0.683124 | 0.709639 | 0.669145 | 0.999971 | | 1 | 0.938711 |
| Pecam1 | B1ARB3 | 1.62 | 0.42 | -0.41 | 0.06 | 2.08 | 0.64 | -0.45 | -0.36 | 0.69 | 1.52 | 0.23 | 0.81 | 0.02 | 0.03 | -0.03 | -0.27 | 0.055 | 0.39 | -0.485 | 0.933801 | 0.422851 | 0.264634 | 0.999971 | | 1 | 0.842533 |
| Pecr | Q3UXD9 | -0.05 | 0.54 | 0.35 | 0.33 | 0.17 | 0.1 | -0.72 | 0.86 | -0.04 | 0.22 | -0.4 | -0.37 | 0.11 | 0.06 | -0.92 | -0.27 | -0.19 | -0.44 | -0.5475 | 0.581847 | 0.080983 | 0.079664 | 0.999971 | | 1 | 0.755254 |
| PEDS1 | Q99LQ7 | -5.45 | -6.8 | -8.3 | -6.72 | -4.28 | -8.5 | -6.68 | -6.16 | -6.79 | -6.04 | -5.21 | -5.5 | -4.68 | -5.87 | -5.6 | -7.2 | 0.4125 | 0.9325 | 0.98 | 0.660496 | 0.159547 | 0.189791 | 0.999971 | | 1 | 0.810547 |
| Pef1 | Q8BFY6 | -0.02 | -0.42 | -0.08 | 0.3 | -0.49 | 0.5 | -0.16 | 0.82 | -1.02 | 0.03 | -0.24 | -0.44 | -0.53 | 0.83 | 0.79 | 0.42 | 0.2225 | -0.3625 | 0.4325 | 0.508715 | 0.218929 | 0.231003 | 0.999971 | | 1 | 0.830179 |
| Peg3 | A0A5F8MP | 6.04 | -0.49 | 1.22 | 2.29 | 0.27 | 1.25 | 2.43 | 2.05 | 0.45 | 5.13 | 0.65 | 0.41 | -1.46 | 0.51 | 1.61 | 0.91 | -0.765 | -0.605 | -1.8725 | 0.559612 | 0.708041 | 0.19245 | 0.999971 | | 1 | 0.810547 |
| Peli1 | Q8C669 | 0.16 | 0.3 | 0.13 | 0.81 | -1.25 | -0.21 | 0.14 | 0.93 | 1.2 | -0.35 | 0.29 | 0.69 | 0.28 | 0.71 | 0.56 | 0.94 | -0.4475 | 0.1075 | 0.2725 | 0.333598 | 0.763097 | 0.278471 | 0.999971 | | 1 | 0.843812 |
| Pelo | Q80X73 | 0.13 | 0.12 | 0.91 | -0.51 | 0.81 | 0.42 | 3.33 | -0.68 | 1.43 | -0.45 | 1.81 | 1.33 | 1.03 | -1.79 | -0.07 | -1.1 | 0.8075 | 0.8675 | -0.645 | 0.329928 | 0.134442 | 0.314234 | 0.999971 | | 1 | 0.851094 |
| Pelp1 | Q9DBD5 | -0.31 | 1.42 | 1.81 | 1.55 | 1.6 | 0.66 | 1.07 | 1.85 | 1.24 | 0.15 | 1.62 | 1.47 | 0.86 | 1.42 | 1.98 | 1.82 | 0.1775 | 0.0025 | 0.4025 | 0.728634 | 0.996318 | 0.433434 | 0.999971 | | 1 | 0.882409 |
| Penk | P22005 | -0.13 | -1.2 | -0.75 | 0.87 | 0.72 | -1.91 | -0.77 | 1.39 | 1.01 | -1.83 | -0.86 | 1.23 | 0.52 | 0.46 | 2.09 | -0.5 | 0.16 | 0.19 | 0.945 | 0.837442 | 0.808549 | 0.162947 | 0.999971 | | 1 | 0.800215 |
| Pepd | Q11136 | 0.38 | 0.25 | 0.98 | 0.66 | 0.34 | 0.62 | 0.56 | 0.54 | 0.53 | 0.18 | 0.71 | 0.63 | 0.2 | 0.57 | 0.89 | 1.01 | -0.0525 | -0.055 | 0.1 | 0.809933 | 0.81367 | 0.707222 | 0.999971 | | 1 | 0.950182 |
| Pes1 | Q5SQ20 | -0.02 | 0.18 | -0.11 | 0.24 | -0.35 | -0.65 | -0.45 | -0.13 | -0.57 | 0.18 | -0.46 | -0.04 | -1.1 | -0.05 | -0.08 | 0.2 | -0.4675 | -0.295 | -0.33 | 0.038957 | 0.22092 | 0.297945 | 0.999971 | | 1 | 0.847402 |
| Pex1 | Q5BL07 | -0.55 | -0.38 | -0.38 | -0.39 | -0.55 | -0.48 | -0.68 | -0.48 | -0.52 | -0.06 | -0.66 | -0.24 | -0.45 | -0.28 | -0.65 | -0.15 | -0.1225 | 0.055 | 0.0425 | 0.467962 | 0.780691 | 0.820856 | 0.999971 | | 1 | 0.975833 |
| Pex10 | B1AUE5 | -0.13 | -0.74 | -1.34 | -0.74 | -0.67 | -0.47 | -1.27 | -0.7 | -0.91 | 0.05 | -1.13 | -1.34 | -0.23 | -0.27 | -1.2 | -0.92 | -0.04 | -0.095 | 0.0825 | 0.896298 | 0.803295 | 0.80946 | 0.999971 | | 1 | 0.973762 |
| Pex11b | Q9Z210 | -0.49 | 0.01 | 0.42 | 0.01 | 0.3 | 0 | 0.88 | 0.37 | 0.65 | 0.12 | 0.94 | 0.4 | 0.59 | -0.33 | 0.22 | -0.12 | 0.4 | 0.54 | 0.1025 | 0.173809 | 0.074194 | 0.722568 | 0.999971 | | 1 | 0.955258 |
| Pex12 | Q8VC48 | -1.75 | -0.66 | -0.34 | -1.47 | -0.66 | -0.35 | -0.98 | -0.03 | -1.02 | -1.76 | -0.62 | -0.17 | 0.26 | -0.44 | -0.08 | -0.85 | 0.55 | 0.1625 | 0.7775 | 0.169195 | 0.716855 | 0.074574 | 0.999971 | | 1 | 0.746069 |
| Pex13 | Q9D0K1 | -0.58 | -0.61 | -0.5 | -0.55 | -0.76 | -1.3 | -0.76 | 0.17 | -1.19 | -0.09 | -0.94 | -0.68 | -0.76 | -0.58 | -0.61 | -0.51 | -0.1025 | -0.165 | -0.055 | 0.742346 | 0.529022 | 0.740138 | 0.999971 | | 1 | 0.960899 |
| Pex14 | Q9R0A0 | 0.4 | -0.61 | 0.13 | 0.52 | -0.69 | -0.14 | 0.35 | 0.14 | -0.3 | 0.16 | -0.47 | -0.73 | 0.01 | -0.35 | -0.29 | -0.84 | -0.195 | -0.445 | -0.4775 | 0.565964 | 0.185432 | 0.153744 | 0.999971 | | 1 | 0.795578 |
| Pex16 | Q91XC9 | 1.28 | -0.2 | -0.58 | -0.68 | 0.03 | -0.08 | -0.21 | -0.31 | 0.35 | 1.02 | -0.63 | -0.84 | 0.35 | 0.38 | -0.58 | -0.35 | -0.0975 | 0.02 | -0.005 | 0.822226 | 0.972369 | 0.991692 | 0.999971 | | 1 | 0.99925 |
| Pex19 | Q8VCI5 | -1 | 0.41 | -0.01 | -0.2 | -0.01 | -0.03 | 0.17 | 0.29 | 0.38 | -0.41 | 0.11 | 0.11 | 0.55 | -0.04 | 0 | 0.34 | 0.305 | 0.2475 | 0.4125 | 0.339328 | 0.468684 | 0.229864 | 0.999971 | | 1 | 0.828687 |
| Pex2 | Q91YZ5 | -0.01 | -0.29 | -0.24 | 0.57 | 0.26 | -0.44 | -0.72 | 0.71 | -0.5 | 0.64 | -0.35 | 0.11 | -0.57 | 0.01 | 0.08 | 0.49 | -0.055 | -0.0325 | -0.005 | 0.881671 | 0.920138 | 0.986801 | 0.999971 | | 1 | 0.99925 |
| Pex26 | Q8BGI5-2 | -0.23 | -0.18 | -0.39 | -0.42 | -0.46 | -0.45 | -0.19 | 0.09 | 0.18 | 0.36 | 0.85 | 0.39 | -0.22 | 0.31 | -0.08 | 0.27 | 0.0525 | 0.75 | 0.375 | 0.793823 | 0.005145 | 0.089891 | 0.999971 | | 1 | 0.771991 |
| Pex3 | Q9QXY9 | 0.36 | -0.42 | -0.39 | -0.33 | -1.02 | -0.87 | -1.14 | -0.88 | -0.75 | -0.02 | -0.47 | -0.33 | -0.69 | -0.4 | -0.28 | -0.23 | -0.7825 | -0.1975 | -0.205 | 0.008058 | 0.457146 | 0.410927 | 0.999971 | | 1 | 0.875847 |
| Pex5 | O09012-2 | -1.37 | -1.55 | -3.97 | -3.04 | -1.66 | -3.13 | -2.04 | -2.24 | -1.53 | -2.3 | -2.43 | -1.94 | -1.19 | -2.7 | -2.41 | -1.98 | 0.215 | 0.4325 | 0.4125 | 0.734294 | 0.477603 | 0.523972 | 0.999971 | | 1 | 0.908231 |
| Pex5_1 | O09012 | -0.43 | -0.09 | 0.52 | 0.24 | -0.1 | -0.25 | 0.08 | 0.62 | 0.21 | -0.12 | 0.66 | 0 | 0.5 | -0.28 | 0.87 | 0.86 | 0.0275 | 0.1275 | 0.4275 | 0.9247 | 0.652267 | 0.226372 | 0.999971 | | 1 | 0.828236 |
| Pex5l | F8SLQ1 | -1.45 | -1.33 | -0.52 | -0.53 | -1.14 | -0.92 | -0.75 | -0.5 | -1.17 | -1.19 | -0.66 | -0.55 | -0.92 | -1 | -0.52 | -0.28 | 0.13 | 0.065 | 0.2775 | 0.661164 | 0.832478 | 0.380791 | 0.999971 | | 1 | 0.871816 |
| Pex5l_1 | D3YYH0 | -0.42 | 0.09 | 0.48 | 0.02 | -0.4 | 0.01 | -0.73 | 0.72 | -0.29 | -0.06 | 0 | -1.01 | 0.82 | 0.12 | -0.26 | 1.03 | -0.1425 | -0.3825 | 0.385 | 0.689438 | 0.228193 | 0.285744 | 0.999971 | | 1 | 0.845133 |
| Pex6 | Q99LC9 | -1.01 | -0.41 | -0.56 | -0.61 | -0.16 | -0.64 | -0.8 | -0.88 | -0.09 | -0.29 | -0.76 | -0.39 | 0.04 | -0.61 | -1.02 | -0.43 | 0.0275 | 0.265 | 0.1425 | 0.90839 | 0.26166 | 0.604701 | 0.999971 | | 1 | 0.924454 |
| Pex7 | P97865 | -0.15 | 0.76 | -1.51 | -0.34 | 0.24 | 0.35 | 1.34 | -0.55 | 0.97 | -0.21 | 1.13 | 0.47 | -0.32 | -0.49 | -1.03 | -1.05 | 0.655 | 0.9 | -0.4125 | 0.258235 | 0.109009 | 0.39096 | 0.999971 | | 1 | 0.871816 |
| Pfas | Q5SUR0 | 1.28 | 1.56 | 1.35 | 1.56 | 1.52 | 1.25 | 1.22 | 1.68 | 1.07 | 1.08 | 0.98 | 1.21 | 0.96 | 1.28 | 1.3 | 1.29 | -0.02 | -0.3525 | -0.23 | 0.918174 | 0.064695 | 0.233445 | 0.999971 | | 1 | 0.831526 |
| Pfdn1 | Q9CQF7 | -1.07 | 0.58 | 0.49 | -0.04 | 0.77 | 0.35 | 0.85 | 0.25 | 0.86 | 0 | 0.89 | 0.64 | 1.42 | 0.59 | 0.59 | -0.16 | 0.565 | 0.6075 | 0.62 | 0.173082 | 0.166002 | 0.206209 | 0.999971 | | 1 | 0.817241 |
| Pfdn2 | O70591 | 0.66 | 0.38 | 0.63 | 0.89 | 0.37 | 0.74 | 0.32 | 0.9 | 0.35 | 0.65 | 0.42 | 0.6 | -0.09 | 0.5 | 0.75 | 0.69 | -0.0575 | -0.135 | -0.1775 | 0.793824 | 0.4818 | 0.480813 | 0.999971 | | 1 | 0.89485 |
| Pfdn4 | E9Q6U4 | -1.84 | -2.58 | -0.26 | -0.11 | -2.01 | -2 | -0.47 | -0.04 | -2.11 | -1.94 | -0.22 | 0.02 | -2.75 | -1.98 | -0.09 | 0.26 | 0.0675 | 0.135 | 0.0575 | 0.924789 | 0.856599 | 0.946006 | 0.999971 | | 1 | 0.997559 |
| Pfdn5 | Q9WU28 | 0.42 | 0.28 | -0.14 | 0.14 | 0.42 | 0.17 | -0.2 | 0.11 | 0.47 | 0.05 | -0.07 | 0.03 | 0.81 | 0.62 | -0.03 | 0.3 | -0.05 | -0.055 | 0.25 | 0.819674 | 0.796919 | 0.329967 | 0.999971 | | 1 | 0.856787 |
| Pfdn6 | Q03958 | 0.8 | 0.64 | 0.76 | 0.92 | 0.64 | 0.77 | 0.36 | 1 | 0.77 | 0.67 | 0.44 | 0.3 | 0.59 | 0.68 | 0.64 | 0.95 | -0.0875 | -0.235 | -0.065 | 0.666528 | 0.228992 | 0.718698 | 0.999971 | | 1 | 0.953825 |
| Pfkfb2 | A0A087WF | -0.67 | 0.47 | 0.03 | -0.06 | 0.24 | 0.15 | -0.07 | -0.06 | 0.38 | -0.11 | 0.12 | 0.22 | 0.75 | 0.21 | 0.48 | 0.02 | 0.1225 | 0.21 | 0.4225 | 0.648758 | 0.449468 | 0.176238 | 0.999971 | | 1 | 0.808532 |
| Pfkfb2_1 | B2Z893 | -0.21 | 2.5 | 0.63 | 1.07 | 2.76 | 2.85 | -0.16 | 0.67 | 3 | 2.19 | -0.54 | -0.06 | 2.11 | 3.62 | 1.39 | 2.24 | 0.5325 | 0.15 | 1.3425 | 0.534079 | 0.871285 | 0.07031 | 0.999971 | | 1 | 0.735439 |
| Pfkfb4 | Q6DTY7-5 | 0.33 | -0.93 | -1.22 | -4.13 | 0.09 | -0.57 | 2.9 | -2.56 | 0.32 | 0 | -0.99 | -0.57 | -0.37 | -1.31 | -3.34 | -5.05 | 1.4525 | 1.1775 | -1.03 | 0.283212 | 0.208778 | 0.420627 | 0.999971 | | 1 | 0.87788 |
| Pfkl | P12382 | -0.5 | -0.66 | -0.87 | -0.98 | -0.63 | -0.61 | -0.39 | -1.26 | -0.91 | -0.97 | -1.01 | -1.09 | -1.19 | -0.65 | -1.03 | -1.14 | 0.03 | -0.2425 | -0.25 | 0.902766 | 0.206955 | 0.258408 | 0.999971 | | 1 | 0.839012 |
| Pfkm | P47857 | -0.8 | -0.33 | -0.22 | -0.41 | -0.31 | -0.27 | 0.17 | -0.2 | -0.36 | -0.88 | -0.24 | -0.26 | -0.69 | -0.47 | -0.34 | -0.22 | 0.2875 | 0.005 | 0.01 | 0.203199 | 0.982719 | 0.962313 | 0.999971 | | 1 | 0.998721 |
| Pfkp | Q8C605 | -0.57 | -0.94 | -0.38 | -0.48 | -0.84 | -0.85 | 0.32 | -0.39 | -1.06 | -0.84 | -0.23 | -0.55 | -1.39 | -0.94 | -0.4 | -0.39 | 0.1525 | -0.0775 | -0.1875 | 0.621827 | 0.75263 | 0.515063 | 0.999971 | | 1 | 0.905815 |
| Pfkp_1 | D3YUA3 | 1.81 | -4.82 | -0.34 | 0.65 | -4.06 | 0.7 | -1.46 | -3.19 | -0.39 | -0.28 | -2.21 | -3.73 | -0.37 | -3.12 | -2.65 | -2.38 | -1.3275 | -0.9775 | -1.455 | 0.412066 | 0.516822 | 0.312788 | 0.999971 | | 1 | 0.849367 |
| Pfn2 | Q9JJV2 | -0.99 | 0.34 | -0.73 | -0.44 | 0.07 | 0.42 | -0.14 | -0.34 | 0.15 | -0.24 | -0.08 | 0.07 | 0.06 | 0.42 | -0.08 | -0.54 | 0.4575 | 0.43 | 0.42 | 0.189126 | 0.184538 | 0.244285 | 0.999971 | | 1 | 0.837316 |
| Pgam1 | Q9DBJ1 | -2.59 | -1.87 | -0.87 | -0.76 | -0.72 | -2.44 | -0.16 | -0.9 | -0.85 | -2.88 | -0.77 | 0.42 | -0.94 | -2.02 | -0.57 | -1.45 | 0.4675 | 0.5025 | 0.2775 | 0.441288 | 0.500854 | 0.581315 | 0.999971 | | 1 | 0.920364 |
| Pgam2 | O70250 | 0.78 | 0.09 | 1.4 | 1.09 | 0.9 | 0.94 | 1.44 | 2.26 | 0.85 | 0.24 | 1.39 | 2.13 | 1.33 | 0.43 | 1.33 | 1.61 | 0.545 | 0.3125 | 0.335 | 0.199734 | 0.503177 | 0.376701 | 0.999971 | | 1 | 0.870465 |
| Pgam5 | Q8BX10-2 | -1.13 | -2.14 | -1.26 | -0.99 | -1.12 | -2.19 | -1.46 | -0.78 | -1.8 | -0.78 | -0.73 | -1.32 | -2.33 | -1.02 | -1.2 | -0.69 | -0.0075 | 0.2225 | 0.07 | 0.984359 | 0.53523 | 0.867221 | 0.999971 | | 1 | 0.983462 |
| Pgam5_1 | Q8BX10 | 0.34 | -0.08 | -0.09 | -0.39 | -0.18 | 0.05 | -0.08 | 0.04 | 0.04 | 0.22 | 0.13 | -0.06 | 0.19 | -0.14 | -0.43 | 0.06 | 0.0125 | 0.1375 | -0.025 | 0.952524 | 0.515744 | 0.915962 | 0.999971 | | 1 | 0.992633 |
| Pgap1 | Q3UUQ7 | -0.01 | 0.43 | 0.4 | 0.29 | 0.5 | 0.46 | 0.5 | 1 | 0.92 | 0.49 | 0.78 | 0.59 | 0.97 | 0.9 | 0.75 | 0.79 | 0.3375 | 0.4175 | 0.575 | 0.138243 | 0.058229 | 0.01221 | 0.999971 | | 1 | 0.613095 |
| Pgap3 | A2A559-2 | -0.02 | -0.64 | -0.38 | 0.37 | 0.01 | -0.87 | 0.32 | 0.52 | -0.16 | 0.22 | 1.39 | 0.92 | 1.47 | 0.29 | -0.08 | 0.94 | 0.1625 | 0.76 | 0.8225 | 0.659384 | 0.081808 | 0.061655 | 0.999971 | | 1 | 0.725348 |
| Pgbd5 | D3YZI9 | -0.16 | -0.45 | -0.1 | -0.22 | -0.53 | -0.54 | 0 | 0.01 | -0.5 | -0.4 | 0.09 | -0.13 | -0.66 | -0.28 | -0.25 | -0.07 | -0.0325 | -0.0025 | -0.0825 | 0.881746 | 0.990229 | 0.685561 | 0.999971 | | 1 | 0.944874 |
| Pgd | Q9DCD0 | 0.42 | 0.26 | 0.8 | 0.77 | 0.33 | 0.51 | 0.45 | 1.29 | 0.47 | 0.56 | 0.76 | 1.14 | 0.83 | 0.65 | 1.47 | 1.44 | 0.0825 | 0.17 | 0.535 | 0.763247 | 0.472481 | 0.073022 | 0.999971 | | 1 | 0.745263 |
| Pgghg | Q8BP56 | -1.48 | 0.63 | -0.38 | 0.08 | 1 | 0.49 | 0.09 | -0.38 | 0.53 | -0.95 | -0.33 | -0.21 | 0.71 | 0.27 | -0.16 | 0.33 | 0.5875 | 0.0475 | 0.575 | 0.25605 | 0.924833 | 0.22582 | 0.999971 | | 1 | 0.828225 |
| Pggt1b | Q8BUY9 | -0.2 | 0.59 | 0.64 | 0.26 | 0.67 | 0.96 | 0.49 | 0.86 | 0.75 | -0.01 | 0.48 | 0.51 | 0.14 | 0.84 | 0.41 | 0.08 | 0.4225 | 0.11 | 0.045 | 0.116063 | 0.68376 | 0.870956 | 0.999971 | | 1 | 0.984109 |
| Pgk1 | P09411 | -0.13 | 0.58 | 0.08 | 0 | 0.23 | 0.36 | 0.03 | -0.12 | 0.19 | -0.44 | 0.07 | 0.16 | 0.5 | 0.18 | 0.28 | 0.26 | -0.0075 | -0.1375 | 0.1725 | 0.973684 | 0.574836 | 0.434821 | 0.999971 | | 1 | 0.882419 |
| Pgls | Q9CQ60 | 0.39 | 0.13 | -0.41 | -0.17 | 0.23 | 0.15 | -0.52 | -0.06 | 0.33 | 0.25 | -0.1 | -0.3 | 0.19 | 0.3 | -0.12 | 0.11 | -0.035 | 0.06 | 0.135 | 0.894503 | 0.812846 | 0.566283 | 0.999971 | | 1 | 0.917959 |
| Pgm1 | Q9D0F9 | -0.2 | 0.02 | 0.64 | 0.61 | -0.95 | -0.58 | -0.17 | -0.52 | 0.35 | -0.11 | 0.45 | 0.5 | 0 | 0.33 | 0.55 | 0.27 | -0.8225 | 0.03 | 0.02 | 0.015712 | 0.91169 | 0.939308 | 0.999971 | | 1 | 0.995869 |
| Pgm2 | Q7TSV4 | 1.55 | 1.51 | 1.34 | 1.14 | 1.27 | 1.23 | 1.19 | 1.06 | 1.05 | 1.3 | 0.72 | 0.69 | 0.29 | 0.79 | 0.8 | 0.63 | -0.1975 | -0.445 | -0.7575 | 0.292611 | 0.065073 | 0.004972 | 0.999971 | | 1 | 0.582714 |
| Pgm2l1 | Q8CAA7 | -0.08 | 1.07 | 0.42 | 0.14 | 0.69 | 1 | 0.66 | 0.26 | 0.91 | 0.52 | 1.03 | -0.03 | 0.78 | 0.69 | 0.26 | 0.64 | 0.265 | 0.22 | 0.205 | 0.388344 | 0.524399 | 0.482932 | 0.999971 | | 1 | 0.896345 |
| Pgm3 | Q9CYR6 | 0.14 | 0.26 | 0.28 | 0.14 | 0.02 | 0.46 | 0.6 | 0.07 | 0.09 | -0.12 | 0.09 | 0.32 | -0.28 | 0.24 | 0.19 | -0.13 | 0.0825 | -0.11 | -0.2 | 0.686514 | 0.534712 | 0.319982 | 0.999971 | | 1 | 0.853962 |
| Pgm5 | Q8BZF8 | 1.48 | -0.71 | 1.46 | 1.94 | 3.73 | 0.61 | 1.82 | 1.14 | 0.24 | -0.2 | 1.57 | 3.49 | 1.49 | 1.07 | 1.99 | 1.52 | 0.7825 | 0.2325 | 0.475 | 0.348392 | 0.800455 | 0.415889 | 0.999971 | | 1 | 0.876885 |
| Pgp | Q8CHP8 | 0.57 | 1.17 | -0.18 | -0.34 | 1.02 | 1.14 | 0.03 | -0.25 | 1.3 | 0.36 | -0.53 | -0.36 | 1.23 | 0.69 | -0.08 | -0.49 | 0.18 | -0.1125 | 0.0325 | 0.698132 | 0.824261 | 0.946527 | 0.999971 | | 1 | 0.997646 |
| Pgpep1 | Q9ESW8 | -1.82 | 0.16 | 0.89 | 0.59 | 1.65 | -0.43 | 0.25 | 2.56 | 2.56 | -2.89 | -0.22 | 0.74 | 1.88 | -1.54 | 1.9 | 1.8 | 1.0525 | 0.0925 | 1.055 | 0.219246 | 0.936183 | 0.278001 | 0.999971 | | 1 | 0.843812 |
| Pgrmc1 | O55022 | -0.95 | -0.73 | -1.04 | -0.57 | -0.48 | -0.66 | -1.11 | -0.34 | -0.49 | -0.73 | -1.05 | -0.53 | -0.34 | -0.34 | -0.32 | -0.3 | 0.175 | 0.1225 | 0.4975 | 0.462215 | 0.566792 | 0.022436 | 0.999971 | | 1 | 0.652043 |
| Pgrmc2 | Q80UU9 | -1.37 | -2.09 | -0.37 | 0.08 | -1.88 | -2.38 | -0.44 | 0.16 | -2.02 | -1.82 | 0.08 | 0.67 | -2.44 | -1.51 | 0.24 | 0.03 | -0.1975 | 0.165 | 0.0175 | 0.777265 | 0.827269 | 0.980793 | 0.999971 | | 1 | 0.99925 |
| Pgs1 | Q8BHF7 | -1.18 | -0.68 | -0.55 | -0.6 | -0.51 | -0.3 | -0.65 | -0.43 | -0.79 | -0.94 | -0.6 | -0.48 | -0.58 | -0.56 | -0.55 | -0.11 | 0.28 | 0.05 | 0.3025 | 0.208515 | 0.819112 | 0.202938 | 0.999971 | | 1 | 0.81617 |
| Phactr1 | Q2M3X8 | -0.35 | -0.14 | 0.05 | 0.13 | -0.07 | -0.4 | 0.19 | -0.04 | -0.07 | -0.34 | 0.26 | 0.25 | -0.15 | -0.4 | 0.24 | -0.11 | -0.0025 | 0.1025 | -0.0275 | 0.990558 | 0.643033 | 0.899018 | 0.999971 | | 1 | 0.989654 |
| Phactr2 | F7D4H5 | 1.04 | -0.49 | -0.25 | -0.5 | -0.31 | -0.11 | -0.13 | -0.65 | -0.46 | 0.25 | -0.66 | -0.37 | -0.93 | -0.23 | -0.77 | -0.91 | -0.25 | -0.26 | -0.66 | 0.511251 | 0.520746 | 0.116418 | 0.999971 | | 1 | 0.78268 |
| Phactr3 | Q8BYK5 | -0.07 | 0.44 | 0.56 | 0.87 | 0.24 | 0.14 | 1.36 | 0.33 | 0.65 | 0.06 | 0.78 | 0.79 | 0.41 | 0.07 | 0.39 | 0.07 | 0.0675 | 0.12 | -0.215 | 0.842965 | 0.665844 | 0.395664 | 0.999971 | | 1 | 0.871816 |
| Phactr4 | Q501J7-2 | 0 | -1.27 | 0.37 | 0.21 | -0.52 | -0.82 | 0.41 | -0.18 | -1.2 | -0.62 | -0.28 | -0.06 | -2.27 | -1.07 | -0.13 | -0.35 | -0.105 | -0.3675 | -0.7825 | 0.808039 | 0.398897 | 0.186727 | 0.999971 | | 1 | 0.809025 |
| Phaf1 | Q922R1 | -0.93 | -0.56 | -0.63 | -0.93 | -0.69 | -0.43 | -0.71 | -0.62 | -0.44 | -0.83 | -0.87 | -0.82 | -0.61 | -1.13 | -0.67 | -0.71 | 0.15 | 0.0225 | -0.0175 | 0.431578 | 0.908795 | 0.932684 | 0.999971 | | 1 | 0.995069 |
| Phax | Q9JJT9 | -0.03 | 2.14 | 2.06 | 1.68 | 2.39 | 1.42 | 1.82 | 2.3 | 2.04 | 0.81 | 2.19 | 2.58 | 2.56 | 1.82 | 2.56 | 2.1 | 0.52 | 0.4425 | 0.7975 | 0.325636 | 0.456645 | 0.138064 | 0.999971 | | 1 | 0.784948 |
| Phb1 | P67778 | 0.49 | -0.42 | 0.22 | 0.38 | -0.23 | 0.35 | -0.19 | 0.2 | -0.16 | 0.47 | -0.27 | -0.22 | -0.34 | 0.65 | 0.3 | 0.17 | -0.135 | -0.2125 | 0.0275 | 0.617968 | 0.457134 | 0.926585 | 0.999971 | | 1 | 0.99401 |
| Phb2 | O35129 | 0.62 | -0.7 | -0.45 | -0.14 | -0.89 | -0.56 | -0.05 | -0.19 | -0.9 | 0.05 | -0.25 | -0.59 | -0.85 | -0.71 | -0.44 | -0.14 | -0.255 | -0.255 | -0.3675 | 0.460401 | 0.470075 | 0.279279 | 0.999971 | | 1 | 0.843812 |
| Phc1 | Q3V116 | -1.46 | -0.14 | -3.08 | -0.78 | -0.35 | -1.89 | -0.62 | -1.41 | -1.41 | -4.01 | -2.17 | -0.14 | -2.09 | 0.07 | -0.94 | -1.92 | 0.2975 | -0.5675 | 0.145 | 0.652534 | 0.543941 | 0.842117 | 0.999971 | | 1 | 0.979814 |
| Phc2 | Q9QWH1-2 | 0.63 | 0 | 0.8 | 0.51 | -0.41 | 0.42 | -0.07 | 0.73 | -0.16 | 0.45 | 0.01 | 0.17 | -1.22 | 0.08 | 0.99 | 0.85 | -0.3175 | -0.3675 | -0.31 | 0.322254 | 0.157875 | 0.537383 | 0.999971 | | 1 | 0.910077 |
| Phf10 | K4DI61 | -1.12 | -1.11 | -0.87 | -0.53 | -2.58 | -2.54 | -0.65 | -1.13 | -0.81 | -2.74 | -0.24 | -0.26 | -1.41 | -1.33 | -1.24 | -0.81 | -0.8175 | -0.105 | -0.29 | 0.113499 | 0.851243 | 0.230301 | 0.999971 | | 1 | 0.829302 |
| Phf14 | Q9D4H9-2 | -0.48 | -0.09 | -0.19 | -0.18 | -0.13 | -0.1 | -0.31 | -0.13 | -0.05 | -0.04 | -0.39 | -0.74 | 0.08 | 0.18 | -0.38 | -0.5 | 0.0675 | -0.07 | 0.08 | 0.706234 | 0.755764 | 0.726465 | 0.999971 | | 1 | 0.956041 |
| Phf2 | Q9WTU0 | 2.05 | -0.97 | 0.35 | 0.98 | -0.88 | -0.41 | 0.84 | -0.05 | -0.48 | 0.79 | -0.55 | -0.86 | -2.13 | -1.12 | -0.14 | -0.48 | -0.7275 | -0.8775 | -1.57 | 0.287508 | 0.209624 | 0.047412 | 0.999971 | | 1 | 0.710102 |
| Phf20l1 | A0A2R8VH | 0.74 | 1.89 | 1.87 | 1.73 | 1.47 | 1.64 | 1.74 | 0.95 | 1.06 | 0.42 | 1.94 | 1.6 | 1.82 | 1.62 | 0.76 | 0.66 | -0.1075 | -0.3025 | -0.3425 | 0.742545 | 0.469429 | 0.388876 | 0.999971 | | 1 | 0.871816 |
| Phf24 | Q80TL4 | -0.31 | -0.2 | -0.83 | -0.6 | -0.37 | -0.06 | -0.54 | -0.61 | -0.37 | -0.37 | -0.69 | -1.23 | -0.49 | -0.6 | -1.02 | -0.81 | 0.09 | -0.18 | -0.245 | 0.693525 | 0.505751 | 0.294869 | 0.999971 | | 1 | 0.847402 |
| Phf24_1 | Q80TL4-2 | -1.48 | -3.04 | -4.21 | -7.4 | -3.02 | -3.42 | -5.12 | -7.88 | -2.23 | -2.17 | -6.21 | -7.04 | -5.32 | -2.5 | -6.89 | -8.07 | -0.8275 | -0.38 | -1.6625 | 0.579675 | 0.813021 | 0.297782 | 0.999971 | | 1 | 0.847402 |
| Phf3 | B2RQG2 | 1.46 | 0.55 | 0.55 | 0.76 | 0.18 | 0.31 | 0.63 | 0.51 | 0.18 | 1.32 | 0.53 | 0.24 | -0.26 | 0.61 | 0.28 | 0.56 | -0.4225 | -0.2625 | -0.5325 | 0.132202 | 0.444101 | 0.1056 | 0.999971 | | 1 | 0.78268 |
| Phf5a | P83870 | -0.64 | 1.38 | 2.41 | 1.32 | 1.37 | 1.54 | 3.09 | 1.81 | 1.64 | 0.41 | 2.34 | 2.96 | 1.6 | 1.36 | 2.35 | 2.26 | 0.835 | 0.72 | 0.775 | 0.237379 | 0.357104 | 0.233591 | 0.999971 | | 1 | 0.831621 |
| Phf6 | Q9D4J7 | 2.26 | 2.28 | 2.31 | 1.48 | 2 | 1.93 | 2.26 | 1.99 | 1.65 | 1.83 | 2.46 | 2.98 | 1.86 | 2.48 | 2.42 | 1.79 | -0.0375 | 0.1475 | 0.055 | 0.878095 | 0.680986 | 0.847115 | 0.999971 | | 1 | 0.980295 |
| Phf8 | Q80TJ7 | 2.65 | 0.44 | 2.12 | 4.06 | 1.67 | 1.36 | 3.06 | 3.17 | 2.02 | 2.11 | 2.29 | 1.78 | -0.84 | 1.11 | 2.35 | 1.76 | -0.0025 | -0.2675 | -1.2225 | 0.997479 | 0.698494 | 0.203711 | 1 | | 1 | 0.81617 |
| Phgdh | Q61753 | -0.94 | -0.2 | -0.5 | -0.88 | -0.07 | 0.01 | -0.29 | -0.04 | 0.08 | -0.42 | -0.19 | -0.25 | -0.04 | -0.03 | -0.31 | -0.45 | 0.5325 | 0.435 | 0.4225 | 0.040065 | 0.091355 | 0.102122 | 0.999971 | | 1 | 0.78268 |
| Phip | F8VQ93 | -1.12 | -1.19 | -0.98 | -1.68 | -1.11 | -1.24 | -0.9 | -1.68 | -1.17 | -1.22 | -1.05 | -1.18 | -0.8 | -1.2 | -1.09 | -2.05 | 0.01 | 0.0875 | -0.0425 | 0.968259 | 0.672339 | 0.892176 | 0.999971 | | 1 | 0.98828 |
| Phka1 | A2AI89 | -1.39 | -1.27 | -0.66 | -0.57 | -1.03 | -0.89 | -0.64 | -0.44 | -1.08 | -1.06 | -0.52 | -0.31 | -0.94 | -0.95 | -0.46 | -0.07 | 0.2225 | 0.23 | 0.3675 | 0.414076 | 0.442705 | 0.247962 | 0.999971 | | 1 | 0.839012 |
| Phka2 | A2AHQ7 | -0.39 | 0.09 | -0.12 | -0.06 | 0.04 | -0.27 | -0.71 | -0.39 | -0.47 | -0.39 | -0.06 | -0.4 | -0.44 | 0.07 | -0.19 | -0.09 | -0.2125 | -0.21 | -0.0425 | 0.358184 | 0.29592 | 0.834523 | 0.999971 | | 1 | 0.979036 |
| Phkb | E0CX43 | -0.58 | 0.35 | -1.37 | -1.3 | -0.58 | -0.71 | -1.49 | -0.76 | -0.64 | -1.32 | -0.7 | -1.08 | -0.16 | -0.76 | -1.22 | -0.28 | -0.16 | -0.21 | 0.12 | 0.707959 | 0.612772 | 0.786061 | 0.999971 | | 1 | 0.967132 |
| Phkb_1 | Q7TSH2 | -0.8 | -0.77 | -0.05 | -0.41 | -0.55 | -0.47 | -0.36 | -0.08 | -0.69 | -0.94 | -0.44 | -0.27 | -0.66 | -0.67 | -0.09 | 0.1 | 0.1425 | -0.0775 | 0.1775 | 0.553228 | 0.760282 | 0.53186 | 0.999971 | | 1 | 0.910069 |
| Phkg1 | P07934 | -1.55 | -0.61 | -0.94 | -1.05 | -0.33 | -0.25 | -1.03 | -0.69 | -0.27 | -0.9 | -0.82 | -0.5 | -0.01 | -0.12 | -1.01 | -0.44 | 0.4625 | 0.415 | 0.6425 | 0.125624 | 0.141918 | 0.060348 | 0.999971 | | 1 | 0.725348 |
| Phkg2 | Q9DB30 | -1.62 | -0.81 | -0.22 | -0.06 | -0.04 | -0.57 | -0.94 | 0.09 | 0.07 | -0.85 | -0.51 | -0.49 | 0.05 | -0.39 | -0.34 | 0 | 0.3125 | 0.2325 | 0.5075 | 0.449781 | 0.552114 | 0.185641 | 0.999971 | | 1 | 0.809025 |
| Phlda1 | Q62392 | 0.13 | 0.1 | 0.51 | 0.77 | 0.07 | 0.54 | 0.04 | 1 | -0.01 | 0.59 | 0.67 | 0.45 | -0.04 | 0.6 | 0.87 | 1.18 | 0.035 | 0.0475 | 0.275 | 0.903836 | 0.848044 | 0.387455 | 0.999971 | | 1 | 0.871816 |
| Phlda3 | Q9WV95 | -2.82 | -0.15 | 0.16 | -0.93 | -0.59 | -0.16 | 0.03 | 0.11 | -0.39 | -1.76 | 0.12 | -0.16 | -0.71 | -1.07 | -0.55 | -0.43 | 0.7825 | 0.3875 | 0.245 | 0.231417 | 0.591819 | 0.695189 | 0.999971 | | 1 | 0.947174 |
| Phldb1 | Q6PDH0-2 | -0.19 | -0.21 | -0.57 | -0.34 | -0.44 | -0.62 | -0.81 | -0.41 | -0.54 | -0.17 | -0.96 | -0.85 | -0.56 | -0.61 | -0.64 | -0.67 | -0.2425 | -0.3025 | -0.2925 | 0.228445 | 0.214183 | 0.124229 | 0.999971 | | 1 | 0.78268 |
| Phldb1_1 | E9PWW4 | 0.94 | -0.51 | 0.73 | 1.94 | -0.82 | -0.04 | 2.73 | 0.94 | -0.7 | 1.11 | 1.21 | -0.35 | -0.35 | -0.83 | 0.2 | -0.47 | -0.0725 | -0.4575 | -1.1375 | 0.929679 | 0.483547 | 0.048508 | 0.999971 | | 1 | 0.710102 |
| Phldb2 | F6QU68 | 0.97 | 1.36 | 1.07 | 1.29 | 2.04 | 0.94 | 1.1 | 0.4 | 0.9 | 1.27 | 0.76 | 1.65 | 1.73 | -0.09 | 0.48 | 1.29 | -0.0525 | -0.0275 | -0.32 | 0.879931 | 0.911243 | 0.432204 | 0.999971 | | 1 | 0.881643 |
| Phlpp1 | Q8CHE4 | 0.36 | -0.21 | 0.47 | 0.42 | -0.01 | -0.18 | -0.03 | 0.54 | -0.67 | -0.24 | 0.72 | 0.1 | -0.82 | -0.05 | 0.34 | 0.23 | -0.18 | -0.2825 | -0.335 | 0.479928 | 0.40493 | 0.298292 | 0.999971 | | 1 | 0.847402 |
| Phlpp2 | J3QM82 | 0.44 | 0.39 | -0.23 | -0.09 | 0.33 | 0.46 | 0.1 | 0.14 | 0.41 | 0.73 | -0.15 | -0.3 | 0.86 | 0.53 | 0.19 | 0.56 | 0.13 | 0.045 | 0.4075 | 0.571783 | 0.881167 | 0.126502 | 0.999971 | | 1 | 0.783193 |
| Phospho1 | Q8R2H9 | 1.27 | 1.85 | 1.16 | 0.98 | 1.85 | 1.3 | 1.43 | 0.49 | 1.85 | 1.12 | 0.98 | 1.28 | 2.22 | 1.01 | 0.53 | 0.5 | -0.0475 | -0.0075 | -0.25 | 0.888202 | 0.978732 | 0.557346 | 0.999971 | | 1 | 0.915051 |
| Phospho2 | Q9D9M5 | 1.24 | 1.62 | 0.26 | 0.55 | 1.03 | 1.05 | -0.26 | 0.42 | 0.29 | 1.96 | 0.08 | -0.01 | 0.35 | 1.5 | 0.29 | 0.51 | -0.3575 | -0.3375 | -0.255 | 0.401598 | 0.520822 | 0.531457 | 0.999971 | | 1 | 0.910069 |
| Phpt1 | Q9DAK9 | 0.29 | 0.08 | 0.46 | 0.41 | 0.6 | 1.42 | 0.74 | 0.91 | 1.19 | 0.87 | 0.45 | 0.78 | 0.99 | 1.04 | 1 | 0.7 | 0.6075 | 0.5125 | 0.6225 | 0.027096 | 0.039778 | 0.00849 | 0.999971 | | 1 | 0.604493 |
| Phrf1 | A6H619-2 | 2.17 | -0.23 | -0.43 | 0.48 | -0.01 | 0.65 | -0.13 | 0.72 | -0.3 | 2.43 | -0.21 | -1.29 | -1.35 | 0.5 | -0.85 | -0.37 | -0.19 | -0.34 | -1.015 | 0.742379 | 0.704666 | 0.142541 | 0.999971 | | 1 | 0.784967 |
| Phyh | O35386 | 0.11 | -1.43 | -1.02 | -0.53 | -1.53 | -1.47 | -1 | -0.55 | -0.9 | 0.11 | -0.64 | -0.21 | -1.67 | -1.32 | -0.75 | -0.8 | -0.42 | 0.3075 | -0.4175 | 0.294692 | 0.434551 | 0.29383 | 0.999971 | | 1 | 0.847402 |
| Phyhd1 | A2AQZ2 | 0.47 | 0.79 | 0.62 | 0.04 | 0.27 | 0.18 | 0.22 | 1.24 | -0.18 | -0.61 | -0.11 | -0.05 | -0.26 | 0.23 | 0.54 | 0.61 | -0.0025 | -0.7175 | -0.2 | 0.993501 | 0.01361 | 0.471867 | 0.999971 | | 1 | 0.892857 |
| Phyhip | Q8K0S0 | 0.65 | 1.5 | 1.42 | 1.22 | 1.53 | 1.77 | 1.89 | 1.79 | 2.49 | 1.21 | 1.8 | 0.01 | 2.13 | 1.66 | 1.33 | 1.46 | 0.5475 | 0.18 | 0.4475 | 0.044955 | 0.729672 | 0.133177 | 0.999971 | | 1 | 0.784948 |

| | | | | | | | | | | | | | | | | | | | | | | | | | | |
|---|---|---|---|---|---|---|---|---|---|---|---|---|---|---|---|---|---|---|---|---|---|---|---|---|---|---|
| Phyhipl | Q8BGT8 | -0.32 | 1.16 | 0.83 | 0.2 | 1.97 | 0.77 | 1.5 | 0.79 | 2.46 | -0.12 | 1.19 | 1.41 | 3.34 | 0.41 | 0.63 | 0.5 | 0.79 | 0.7675 | 0.7525 | 0.085778 | 0.204793 | 0.303147 | 0.999971 | 1 | 0.847402 |
| Pi4k2a | Q2TBE6 | -0.72 | -0.15 | 0 | -0.5 | 0.03 | -0.05 | 0.16 | 0.07 | 0.3 | -0.46 | 0.33 | 0.64 | 0.84 | 0.07 | 0.13 | 0.22 | 0.395 | 0.545 | 0.6575 | 0.0946 | 0.091982 | 0.032379 | 0.999971 | 1 | 0.683918 |
| Pi4ka | E9Q3L2 | -0.65 | -0.6 | -0.94 | -1.15 | -0.62 | -0.47 | -0.76 | -1.04 | -0.6 | -0.65 | -0.81 | -0.93 | -0.06 | -0.58 | -0.73 | -0.91 | 0.1125 | 0.0875 | 0.265 | 0.61264 | 0.666084 | 0.308295 | 0.999971 | 1 | 0.847402 |
| Pi4kb | Q8BKC8-2 | -0.39 | 0.46 | 0.81 | 0.66 | -0.12 | 0.08 | 0.35 | 0.8 | 0.54 | 0.3 | 0.42 | 0.97 | 0.93 | 0.17 | 0.57 | 1.11 | -0.1075 | 0.1725 | 0.31 | 0.746977 | 0.581128 | 0.370077 | 0.999971 | 1 | 0.869526 |
| Pianp | A0A0N4SV | 0.55 | -3.39 | -0.94 | -0.6 | -2.42 | -3.88 | 0.3 | -1.09 | -3.08 | -1.5 | -0.69 | -1.77 | -3.52 | -3.14 | -1.41 | -2.28 | -0.6775 | -0.665 | -1.4925 | 0.537411 | 0.453358 | 0.108628 | 0.999971 | 1 | 0.78268 |
| Pias4 | Q9JM05 | 1.3 | 1.26 | 2.48 | 2.99 | 1.74 | 0.68 | 2.82 | 3.27 | 0.58 | 1.92 | 2.8 | 1.59 | 0.9 | 1.75 | 1.26 | 2.29 | 0.12 | -0.285 | -0.4575 | 0.854857 | 0.625602 | 0.363931 | 0.999971 | 1 | 0.869526 |
| Picalm | Q7M6Y3-6 | -0.62 | -1.4 | -1.2 | -1.17 | 0.17 | -1.61 | -1.06 | -0.91 | -1.69 | -2.12 | -0.94 | 0.2 | -0.45 | -1.58 | -1.08 | -1.07 | 0.245 | -0.04 | 0.0525 | 0.536217 | 0.936005 | 0.859222 | 0.999971 | 1 | 0.982702 |
| Picalm_1 | A0A1L1SUI | -2.97 | 1.82 | -3.86 | -3.2 | -2.09 | -1.2 | -3.89 | -3.82 | -0.03 | -3.21 | -3.75 | -3.95 | -2.16 | 0.72 | -2.96 | -2.99 | -0.6975 | -0.6825 | 0.205 | 0.594296 | 0.633973 | 0.883259 | 0.999971 | 1 | 0.986597 |
| Picalm_2 | Q7M6Y3-5 | -0.22 | -0.18 | -0.33 | -0.14 | -0.12 | -0.11 | -0.28 | -0.42 | -0.18 | -0.29 | -0.32 | 0 | -0.18 | -0.07 | -0.27 | -0.06 | -0.015 | 0.02 | 0.0725 | 0.930948 | 0.905917 | 0.666341 | 0.999971 | 1 | 0.937485 |
| Pick1 | E9PUZ5 | 1.3 | -0.71 | -0.32 | -0.44 | -0.9 | -0.38 | 0.24 | -0.72 | -0.74 | 0.13 | -0.6 | -0.4 | -0.73 | -0.55 | -0.88 | -0.71 | -0.3975 | -0.36 | -0.675 | 0.420115 | 0.445447 | 0.146841 | 0.999971 | 1 | 0.793248 |
| Pid1 | Q3UBG2-2 | -0.94 | -0.51 | -1.11 | 0.43 | -0.05 | -1.43 | -0.48 | 0.01 | -0.36 | -0.81 | 0.32 | -1.63 | -0.15 | -0.36 | -0.07 | -1.45 | 0.045 | -0.0875 | 0.025 | 0.920233 | 0.86069 | 0.955002 | 0.999971 | 1 | 0.998056 |
| Pigb | Q9JJQ0 | 0.62 | -1.01 | -1.21 | 0.92 | -1.89 | -1 | 1.49 | 0.94 | -1.47 | 0.58 | -0.37 | 0.87 | -2.68 | -1.58 | -1.45 | 1.05 | 0.055 | 0.0725 | -0.995 | 0.949326 | 0.916624 | 0.266677 | 0.999971 | 1 | 0.842677 |
| Pigg | D3Z3Y1 | -1.62 | 0.33 | 0.76 | 0.21 | 0.37 | 0.52 | 0.02 | 1.33 | 0.16 | -0.98 | 0.88 | 1.04 | 0.54 | 0.62 | 0.07 | 1.38 | 0.64 | 0.355 | 0.7325 | 0.260854 | 0.582139 | 0.202475 | 0.999971 | 1 | 0.81617 |
| Pigk | Q9CXY9 | -0.5 | -0.66 | -0.77 | -0.71 | -0.89 | -0.65 | 0.29 | -0.98 | -0.89 | -0.51 | -0.67 | -0.86 | -1 | -0.65 | -0.69 | -0.87 | 0.1025 | -0.0725 | -0.1425 | 0.736482 | 0.686658 | 0.437571 | 0.999971 | 1 | 0.882419 |
| Pign | A0A087WN | -0.72 | -0.25 | -0.83 | -1.45 | -1.65 | -0.46 | -1.79 | -1.66 | -0.61 | 0.22 | -0.28 | -0.62 | -0.56 | -1.19 | -2.37 | -2.24 | -0.5775 | 0.49 | -0.7775 | 0.157294 | 0.1501 | 0.123094 | 0.999971 | 1 | 0.78268 |
| Pigo | A2AG36 | 2.38 | -0.23 | 1.18 | 0.62 | 0.28 | 0.25 | 1.25 | 1.15 | 0.18 | 1.46 | 1.19 | 1.41 | 0.03 | -0.21 | 1.32 | 1 | -0.255 | 0.0725 | -0.4525 | 0.650608 | 0.899488 | 0.460837 | 0.999971 | 1 | 0.890033 |
| Pigp | B8JK67 | -0.63 | -0.08 | -0.55 | 0.29 | 0.4 | 0.16 | -0.66 | 0.78 | -0.19 | 0.32 | -0.15 | 0.18 | -0.04 | 0.41 | 0.11 | -0.15 | 0.4125 | 0.2825 | 0.325 | 0.274062 | 0.306589 | 0.245113 | 0.999971 | 1 | 0.837316 |
| Pigs | Q6PD26 | -1.2 | 0.35 | -0.43 | -0.58 | 0.55 | 0.17 | -0.6 | -0.59 | 0.58 | -0.64 | -0.14 | -0.53 | 1.43 | 0.23 | -0.51 | -0.11 | 0.3475 | 0.2825 | 0.725 | 0.404513 | 0.490289 | 0.164105 | 0.999971 | 1 | 0.800215 |
| Pigt | Q8BXQ2 | -1.11 | -0.24 | 0.04 | 0.13 | -0.14 | -0.01 | 0.22 | 0.14 | -0.2 | -0.45 | 0.12 | 0.06 | 0 | 0.31 | 0.23 | 0.19 | 0.3475 | 0.1775 | 0.4775 | 0.267805 | 0.576385 | 0.137732 | 0.999971 | 1 | 0.784948 |
| Pigu | Q3TAA8 | 0.32 | 0.2 | -0.58 | -0.04 | 0.6 | 0.94 | -0.88 | 0.34 | 0.68 | 0.27 | 0.05 | -0.7 | 0.26 | 1.1 | 0.31 | 0.45 | 0.275 | 0.1 | 0.555 | 0.518327 | 0.773218 | 0.083398 | 0.999971 | 1 | 0.755254 |
| Pih1d1 | Q9CQJ2 | 0.82 | 0.7 | 0.81 | 0.56 | 0.8 | 0.9 | 0.26 | 0.75 | 0.74 | 0.16 | 0.33 | 0.9 | 0.99 | 0.87 | 0.85 | -0.03 | -0.045 | -0.19 | -0.0525 | 0.828338 | 0.402845 | 0.843135 | 0.999971 | 1 | 0.979814 |
| Pik3ap1 | Q9EQ32 | -1.71 | 0.64 | 0.62 | -0.18 | 0.31 | 0.34 | 0.97 | 0.26 | -0.78 | -0.32 | -0.72 | -1.03 | -2.47 | 1.24 | 0.04 | 0.3 | 0.6275 | -0.555 | -0.065 | 0.256614 | 0.310104 | 0.940143 | 0.999971 | 1 | 0.995869 |
| Pik3c2a | F8VPL2 | -0.39 | -0.32 | -0.53 | -0.68 | -0.12 | -0.21 | -0.73 | -0.7 | -0.25 | -0.47 | -0.58 | 0.02 | -0.37 | -0.06 | -0.46 | -0.41 | 0.04 | 0.16 | 0.155 | 0.856783 | 0.442763 | 0.422879 | 0.999971 | 1 | 0.87887 |
| Pik3c3 | E9QLS6 | -0.69 | -0.63 | -0.01 | -0.15 | -0.4 | -0.4 | -0.06 | -0.2 | -0.32 | -0.63 | 0.07 | 0.09 | -0.37 | -0.3 | -0.07 | 0.01 | 0.105 | 0.1725 | 0.1875 | 0.647575 | 0.516817 | 0.426495 | 0.999971 | 1 | 0.880793 |
| Pik3ca | P42337 | -0.16 | 1.04 | 0.99 | 0.64 | 0.68 | 1.27 | 0.51 | 1.25 | 0.78 | 0.3 | 0.79 | 0.44 | 0.82 | 0.96 | 0.76 | 0.95 | 0.3 | -0.05 | 0.245 | 0.383627 | 0.871181 | 0.412833 | 0.999971 | 1 | 0.876032 |
| Pik3cb | Q8BTI9 | 0.41 | 0.55 | -0.08 | -0.3 | 0.8 | 0.06 | 0.31 | -0.82 | 1.2 | 0.05 | 0.13 | 0.67 | 1.27 | 0.09 | -0.42 | -0.48 | -0.0575 | 0.3675 | -0.03 | 0.879765 | 0.286637 | 0.944165 | 0.999971 | 1 | 0.997001 |
| Pik3cd | B0QZL5 | -0.63 | 0.27 | -0.15 | -0.02 | 0.49 | -0.26 | -0.37 | -0.55 | 0.43 | -0.25 | 1.17 | 0.77 | 0.91 | -0.11 | 0.08 | 0.06 | -0.04 | 0.6625 | 0.3675 | 0.895001 | 0.085073 | 0.246564 | 0.999971 | 1 | 0.839012 |
| Pik3ip1 | Z4YN57 | 2.33 | 0.76 | 0.07 | 0.34 | 0.76 | 1.16 | 0.28 | 0.6 | 0.84 | 1.29 | 0.42 | 0.23 | 1.88 | 1.02 | 0.91 | 1.26 | -0.175 | -0.18 | 0.3925 | 0.726214 | 0.728909 | 0.44926 | 0.999971 | 1 | 0.885522 |
| Pik3r1 | P26450 | -1.26 | -0.12 | -0.3 | -0.84 | 0.06 | -0.23 | -0.17 | -0.47 | 0.21 | -0.91 | -0.02 | 0.16 | 0.08 | -0.59 | -0.47 | -0.25 | 0.4275 | 0.49 | 0.3225 | 0.16891 | 0.197421 | 0.307488 | 0.999971 | 1 | 0.847402 |
| Pik3r2 | O08908 | -0.01 | 0.19 | -0.11 | -0.1 | 0.15 | -0.12 | -0.23 | -0.25 | 0.22 | -0.19 | -0.34 | -0.16 | 0.06 | 0.08 | -0.1 | 0.08 | -0.105 | -0.11 | 0.0375 | 0.576898 | 0.576671 | 0.828661 | 0.999971 | 1 | 0.977963 |
| Pik3r4 | Q8VD65 | -0.23 | -0.11 | 0.52 | -0.29 | -0.3 | -0.25 | 0.71 | 0.06 | -0.38 | -0.4 | 0.65 | 0.12 | -0.17 | -0.27 | 0.44 | -0.06 | 0.0825 | 0.025 | 0.0125 | 0.786852 | 0.9365 | 0.962475 | 0.999971 | 1 | 0.998721 |
| Pikfyve | Q9Z1T6 | -0.65 | -0.73 | -0.08 | -0.37 | -0.6 | -0.85 | -0.29 | -0.29 | -0.68 | -0.68 | -0.09 | 0.03 | -0.72 | -0.77 | -0.29 | -0.27 | -0.05 | 0.1025 | -0.055 | 0.831923 | 0.695831 | 0.81568 | 0.999971 | 1 | 0.974918 |
| Pin1 | Q9QUR7 | -0.85 | -0.72 | -0.9 | -0.76 | -0.35 | -0.72 | -0.36 | -0.89 | -0.77 | -1.01 | -0.71 | -0.85 | -0.98 | -0.59 | -0.84 | -1.03 | 0.2275 | -0.0275 | -0.0525 | 0.271915 | 0.869185 | 0.774893 | 0.999971 | 1 | 0.963884 |
| Pin4 | Q9CWW6 | 0.91 | 0.85 | 0.49 | 0.47 | 0.17 | -0.26 | 0.29 | 1.18 | 0.61 | -0.33 | 0.84 | 0.85 | 0.44 | 0.3 | 0.46 | 0.46 | -0.335 | -0.1875 | -0.265 | 0.317243 | 0.547079 | 0.188321 | 0.999971 | 1 | 0.809025 |
| Pip4k2a | O70172 | 0.68 | 0.18 | -0.36 | -0.16 | -0.31 | 0.31 | -0.31 | -0.33 | -0.48 | 0.69 | -0.42 | -0.42 | -0.67 | 0.12 | -0.55 | -0.12 | -0.245 | -0.2425 | -0.39 | 0.405809 | 0.501277 | 0.217807 | 0.999971 | 1 | 0.826893 |
| Pip4k2b | Q80XI4 | -0.7 | 0.51 | 0.22 | 0.19 | 0.54 | -0.12 | 0.68 | 0.16 | 0.7 | -0.39 | 0.46 | 0.24 | 1.18 | 0.2 | 0.09 | 0.35 | 0.26 | 0.1975 | 0.4 | 0.426486 | 0.571762 | 0.276366 | 0.999971 | 1 | 0.843794 |
| Pip4k2c | Q91XU3 | 0.24 | 0.18 | 0.63 | 0.18 | 0.05 | 0.37 | 1.07 | 0.07 | 0.07 | -0.25 | 0.13 | -0.62 | 0.21 | -0.58 | 0.22 | 0.15 | 0.0825 | -0.475 | -0.3075 | 0.766991 | 0.07047 | 0.24027 | 0.999971 | 1 | 0.834057 |
| Pip4p1 | Q3TWL2 | -0.68 | 0.69 | 1.44 | 0.3 | 0.47 | 0.43 | 1.72 | 0.59 | 0.44 | 0.15 | 0.6 | -0.98 | 0.76 | 0.38 | 0.31 | -0.84 | 0.365 | -0.385 | -0.285 | 0.471235 | 0.471379 | 0.585762 | 0.999971 | 1 | 0.920413 |
| Pip4p2 | Q9CZX7 | -0.56 | 1.02 | 0.84 | -0.48 | 0.71 | 1.19 | 0.46 | 0.26 | 0.92 | -0.12 | 0.53 | 0.19 | 1.55 | 0.26 | 0.42 | -0.33 | 0.45 | 0.175 | 0.27 | 0.320127 | 0.697932 | 0.613902 | 0.999971 | 1 | 0.927364 |
| Pip5k1a | P70182-3 | 0.42 | -1.17 | -0.37 | -0.2 | -1.34 | -0.77 | -0.24 | -0.46 | -1.34 | -0.51 | -0.72 | -0.54 | -2.39 | -1.47 | -0.44 | -0.52 | -0.3725 | -0.4475 | -0.875 | 0.351421 | 0.246411 | 0.120721 | 0.999971 | 1 | 0.78268 |
| Pip5k1b | P70181 | 1.06 | -0.95 | -0.04 | 0.03 | -0.14 | 0.87 | -0.33 | -0.14 | 0.2 | 0.88 | -0.33 | -0.21 | 0.75 | 0.2 | -0.33 | 0.07 | 0.04 | 0.11 | 0.1475 | 0.930815 | 0.812542 | 0.738717 | 0.999971 | 1 | 0.960301 |
| Pip5k1c | F8WHW6 | -0.83 | -0.46 | -0.94 | -1.23 | -0.7 | -0.39 | -0.9 | -1.38 | -0.74 | -0.88 | -1.27 | -1.44 | -0.37 | -0.65 | -1.3 | -1.22 | 0.0225 | -0.2175 | -0.02 | 0.935412 | 0.400001 | 0.944677 | 0.999971 | 1 | 0.997225 |
| Pir | A2AIH8 | 0.46 | 0.2 | 0.08 | 0.05 | 1.13 | 0.42 | 0.23 | 0 | 0.81 | 0.5 | 0.19 | 0.33 | 1.28 | 0.04 | 0.13 | 0.43 | 0.2475 | 0.26 | 0.2725 | 0.38361 | 0.23575 | 0.383049 | 0.999971 | 1 | 0.871816 |
| Pithd1 | Q8BWR2 | 0.68 | 0.61 | 0.85 | 0.99 | 0.71 | 0.38 | 1.08 | 0.27 | 0.66 | 0.05 | 0.35 | 0.14 | 0.45 | 0.35 | 0.5 | 0.23 | -0.1725 | -0.4825 | -0.4 | 0.47209 | 0.042967 | 0.052215 | 0.999971 | 1 | 0.723736 |
| Pitpna | P53810 | -0.46 | -0.16 | 0.34 | -0.25 | -0.09 | 0.04 | 0.47 | 0.05 | 0.2 | -0.43 | 0.42 | 0.4 | -0.44 | -0.06 | 0.2 | -0.5 | 0.25 | 0.28 | -0.0675 | 0.314732 | 0.326116 | 0.795975 | 0.999971 | 1 | 0.970435 |
| Pitpnb | Q8JZZ5 | -1.18 | -0.36 | 0.1 | -0.4 | -0.4 | -0.33 | 0.22 | -0.02 | -0.15 | -0.69 | 0.33 | 0.03 | -0.56 | 0.03 | 0.13 | -0.09 | 0.3275 | 0.34 | 0.3375 | 0.303116 | 0.329174 | 0.29548 | 0.999971 | 1 | 0.847402 |
| Pitpnc1 | Q8K4R4-2 | 1.58 | 0.85 | -0.04 | -0.17 | 0.53 | 0.6 | 1.38 | 0.34 | 0.74 | 1.7 | 0.61 | 1.16 | 1.09 | -0.35 | -1.02 | -1.68 | 0.1575 | 0.4975 | -1.045 | 0.722381 | 0.286955 | 0.137318 | 0.999971 | 1 | 0.784948 |
| Pitpnc1_1 | Q8K4R4 | -0.32 | -0.83 | 0.19 | 0.17 | -1.15 | -0.57 | -0.25 | 0.44 | -0.95 | -0.19 | 0.11 | 0.1 | -0.92 | -0.4 | 0.45 | 0.64 | -0.185 | -0.035 | 0.14 | 0.63995 | 0.918614 | 0.737741 | 0.999971 | 1 | 0.959801 |
| Pitpnm1 | O35954 | 0 | -0.41 | -0.15 | -0.62 | 0.02 | -0.08 | -0.05 | -0.19 | -0.15 | -0.14 | -0.01 | 0.09 | 0.26 | -0.23 | -0.32 | -0.24 | 0.22 | 0.2425 | 0.1625 | 0.291389 | 0.248508 | 0.486161 | 0.999971 | 1 | 0.897483 |
| Pitpnm2 | E9PYJ7 | -0.4 | 0.12 | -0.09 | -0.13 | 0 | -0.15 | 0 | -0.26 | -0.08 | -0.57 | -0.22 | -0.27 | 0.03 | -0.24 | -0.24 | -0.03 | 0.0225 | -0.16 | 0.005 | 0.906085 | 0.435046 | 0.979357 | 0.999971 | 1 | 0.99925 |
| Pitpnm3 | Q3UHE1 | -0.71 | -0.36 | -0.43 | -0.69 | -0.36 | -0.69 | -0.66 | -0.72 | -0.17 | -0.88 | -0.74 | -0.43 | -0.06 | -0.82 | -0.57 | -0.14 | -0.06 | -0.0075 | 0.15 | 0.752474 | 0.97301 | 0.530594 | 0.999971 | 1 | 0.910069 |
| Pitrm1 | Q8K411 | -0.71 | -0.47 | -0.56 | -0.52 | -0.32 | -0.61 | -0.41 | -0.52 | 0.05 | -0.59 | -0.53 | -0.58 | -0.05 | -0.57 | -0.58 | -0.49 | 0.1 | 0.1525 | 0.1425 | 0.565405 | 0.474275 | 0.477578 | 0.999971 | 1 | 0.894438 |
| Pja1 | O55176 | 2.2 | 0.81 | 1.63 | 1.19 | 0.82 | 0.78 | 2.1 | 1.68 | 1.11 | 1.46 | 2.3 | 1.97 | 0.92 | 0.53 | 2.19 | 0.84 | -0.1125 | 0.2525 | -0.3375 | 0.78885 | 0.517321 | 0.456304 | 0.999971 | 1 | 0.887217 |
| Pja2 | Q80U04-2 | 2.35 | 1.19 | 1.63 | 1.74 | 0.74 | 1.59 | 1.94 | 1.13 | 1 | 2.01 | 1.34 | 0.63 | 0.63 | 1.12 | 1.34 | 0.94 | -0.3775 | -0.4825 | -0.72 | 0.295842 | 0.212839 | 0.03445 | 0.999971 | 1 | 0.69205 |
| Pkd2 | O35245-5 | 0.48 | 0.33 | -0.31 | -0.39 | 0 | 0.73 | -0.46 | -0.15 | 0.74 | 0.13 | -0.2 | -0.22 | -0.59 | 0.72 | 0.02 | 0.36 | 0.0025 | 0.085 | 0.1 | 0.994011 | 0.7889 | 0.775523 | 0.999971 | 1 | 0.96393 |
| Pkia | P63248 | 2.03 | -0.42 | -0.94 | 0.39 | -0.55 | -0.38 | 0.9 | 0.06 | -0.89 | -1.27 | 0.08 | 0.08 | -0.74 | -0.59 | -0.07 | -0.2 | -0.2575 | -0.765 | -0.665 | 0.69651 | 0.271791 | 0.292294 | 0.999971 | 1 | 0.847402 |
| Pkib | Q04758-2 | 1.64 | 1.09 | 0.78 | 1.32 | 1.17 | 1.01 | 0.39 | 0.95 | 0.84 | 1.32 | 1.17 | 0.85 | 0.99 | 0.74 | 1.45 | 1.4 | -0.3275 | -0.1625 | -0.0625 | 0.242706 | 0.513073 | 0.816579 | 0.999971 | 1 | 0.974929 |
| Pkig | O70139 | 1.44 | 1.49 | 0.3 | 1.16 | 0.5 | 0.8 | 0.16 | 1.11 | 0.84 | 0.02 | 0.48 | 0.68 | 0.96 | 1.33 | 0.54 | 1.11 | -0.455 | -0.5925 | -0.1125 | 0.202622 | 0.098327 | 0.72908 | 0.999971 | 1 | 0.956041 |
| Pkm | P52480 | -0.66 | -0.39 | -0.24 | -0.43 | -0.22 | -0.39 | -0.36 | -0.61 | -0.55 | -1.01 | -0.66 | -0.29 | -0.51 | -0.01 | -0.21 | -0.54 | 0.035 | -0.1975 | 0.1125 | 0.852376 | 0.371809 | 0.590526 | 0.999971 | 1 | 0.922217 |
| Pkm_1 | P52480-2 | -0.35 | -0.87 | -0.62 | -0.3 | -0.93 | -0.87 | -0.35 | -0.25 | -1.4 | -1.34 | -0.91 | -0.91 | -1.59 | -0.65 | -0.55 | -0.44 | -0.065 | -0.605 | -0.2725 | 0.793419 | 0.023678 | 0.380624 | 0.999971 | 1 | 0.871816 |
| Pkn1 | P70268 | 0.59 | 0.32 | -0.44 | 0.07 | 0.31 | 0.26 | -0.33 | -0.49 | 0.46 | 0.79 | -0.01 | 0.2 | 0.26 | 0.25 | -0.29 | 0.15 | -0.1975 | 0.225 | -0.0425 | 0.523119 | 0.445593 | 0.876503 | 0.999971 | 1 | 0.985129 |
| Pkn2 | G3UXH4 | -0.02 | 0.22 | -0.12 | 0.18 | 0.06 | 0.05 | 0.02 | 0.22 | -0.34 | -0.02 | 0.1 | -0.09 | 0.06 | 0.41 | 0.12 | 0.68 | 0.0225 | -0.1525 | 0.2525 | 0.898516 | 0.424856 | 0.256222 | 0.999971 | 1 | 0.839012 |
| Pkp2 | Q9CQ73 | -0.55 | -0.88 | -1.06 | -1.45 | -0.59 | -0.96 | -0.23 | -1.06 | -0.87 | -0.76 | -1.25 | -1.39 | -0.37 | -1.38 | -1.53 | -1.78 | 0.275 | -0.0825 | -0.28 | 0.341871 | 0.752804 | 0.439081 | 0.999971 | 1 | 0.883105 |
| Pkp4 | Q68FH0 | -0.75 | -1.07 | -1.16 | -1.31 | -1.24 | -1.07 | -0.96 | -1.41 | -1.13 | -1.11 | -1.06 | -1.14 | -0.55 | -1.2 | -1.15 | -1.23 | -0.0975 | -0.0375 | 0.04 | 0.639458 | 0.839928 | 0.865471 | 0.999971 | 1 | 0.983318 |
| Pla2g15 | Q8VEB4 | -0.01 | 0.79 | 0.2 | 0.6 | 1.34 | 0.03 | 0.45 | 0.88 | 1.84 | -0.32 | 0.3 | 0 | 1.99 | 0.18 | -0.08 | 0.35 | 0.28 | 0.06 | 0.215 | 0.411144 | 0.900367 | 0.649391 | 0.999971 | 1 | 0.933842 |
| Pla2g4e | Q50L42 | 0.04 | 0.48 | -0.51 | -0.4 | 0.99 | 0.25 | -1.04 | -0.16 | 0.69 | 0.3 | -0.39 | -0.87 | 1.4 | 0.49 | -0.51 | -0.17 | 0.1075 | 0.03 | 0.4 | 0.811497 | 0.939952 | 0.384565 | 0.999971 | 1 | 0.871816 |
| Pla2g6 | P97819-2 | -0.49 | 0.49 | 0.11 | -0.21 | 0.44 | 0.11 | -0.05 | 0.07 | 0.59 | -0.09 | 0.21 | 0.12 | 0.9 | -0.04 | -0.07 | 0.26 | 0.1675 | 0.2325 | 0.2875 | 0.523293 | 0.40213 | 0.371008 | 0.999971 | 1 | 0.869526 |
| Pla2g7 | Q60963 | -0.41 | -0.17 | 0.22 | 0.16 | -0.5 | -0.16 | -0.26 | 1.01 | 0.07 | 0.31 | 0.56 | 0.64 | -0.08 | 0.46 | 0.3 | 0.75 | 0.0725 | 0.445 | 0.4075 | 0.84003 | 0.080927 | 0.135608 | 0.999971 | 1 | 0.784948 |
| Plaa | P27612 | 0.06 | 0.44 | 0.25 | -0.03 | 0.34 | 0.47 | 0.22 | 0.16 | 0.57 | 0.32 | 0.51 | 0.09 | 0.56 | 0.48 | 0.01 | 0.22 | 0.1175 | 0.1925 | 0.1375 | 0.543287 | 0.354209 | 0.52361 | 0.999971 | 1 | 0.907797 |
| Plaat3 | Q8R3U1-2 | -0.65 | -0.45 | 2.21 | 1.54 | -0.02 | -0.25 | 0.93 | 2.8 | -0.37 | -0.01 | 0.28 | 0.59 | -0.04 | 0.13 | 2.75 | 1.86 | 0.2025 | -0.54 | 0.5125 | 0.82035 | 0.434139 | 0.565569 | 0.999971 | 1 | 0.917842 |
| Plat | P11214 | -0.6 | 0.12 | -0.11 | 0.3 | 0.51 | -0.13 | -0.97 | -0.02 | 0.66 | -0.67 | 0.3 | 0.27 | 0.61 | 0.07 | 0.57 | 0.66 | -0.08 | 0.2125 | 0.55 | 0.821936 | 0.537583 | 0.060981 | 0.999971 | 1 | 0.725348 |
| Plbd2 | Q3TCN2 | -0.93 | -0.05 | 0.79 | 0.11 | 0.1 | -0.09 | 0.69 | 0.59 | 0.15 | -0.88 | 0.4 | 0.7 | 0.28 | -0.07 | 0.47 | 0.16 | 0.3425 | 0.1125 | 0.23 | 0.385498 | 0.808209 | 0.530985 | 0.999971 | 1 | 0.910069 |
| Plcb1 | Q9Z1B3 | 0.12 | -0.08 | 0.02 | 0.03 | -0.38 | -0.18 | -0.1 | -0.3 | -0.49 | -0.14 | -0.18 | -0.06 | -0.75 | -0.36 | -0.25 | -0.11 | -0.2625 | -0.24 | -0.39 | 0.14798 | 0.199554 | 0.079847 | 0.999971 | 1 | 0.755254 |
| Plcb1_1 | Q9Z1B3-2 | 0.76 | -1.5 | -1.91 | -1.18 | -1.72 | -2.07 | -1.25 | -1.45 | -2.22 | -1.63 | -2.59 | -1.3 | -4.09 | -1.91 | -1.78 | -0.41 | -0.665 | -0.9775 | -1.09 | 0.259055 | 0.133135 | 0.22879 | 0.999971 | 1 | 0.828687 |
| Plcb3 | P51432 | 0.29 | 0.33 | -0.58 | 0.17 | 0.04 | 0.26 | -0.85 | -0.5 | 0.07 | 0.36 | -0.63 | -0.61 | 0.4 | 0 | -0.51 | -0.34 | -0.315 | -0.255 | -0.165 | 0.352782 | 0.444169 | 0.587875 | 0.999971 | 1 | 0.920758 |
| Plcb4 | Q91UZ1 | 0.34 | 0.73 | 0.83 | 0.7 | 0.45 | 1.02 | 0.12 | 0.49 | 0.57 | 0.93 | 0.36 | 0.42 | 0.15 | 0.69 | 0.24 | 0.8 | -0.13 | -0.08 | -0.18 | 0.59854 | 0.706946 | 0.445403 | 0.999971 | 1 | 0.884152 |
| Plcd1 | Q8R3B1 | -0.45 | 0.07 | -0.36 | -0.52 | -0.14 | 0.07 | -0.72 | -0.24 | -0.15 | -0.64 | -0.53 | -0.2 | 0.09 | -0.14 | -0.37 | -0.23 | 0.0575 | -0.065 | 0.1525 | 0.813854 | 0.767956 | 0.481245 | 0.999971 | 1 | 0.89485 |
| Plcd3 | Q8K2J0 | -1.33 | -0.95 | 0.63 | 0.22 | -1.13 | -1.45 | -0.21 | -0.47 | -2.07 | -1.71 | -0.02 | -1.13 | -1.37 | -1.23 | -0.18 | -0.32 | -0.4575 | -0.875 | -0.4175 | 0.377861 | 0.166335 | 0.427423 | 0.999971 | 1 | 0.880793 |
| Plcd4 | Q8K3R3-2 | 0.26 | 1.48 | 0.03 | -0.28 | 0.01 | 1.1 | -1.45 | 1.22 | 0.73 | 0.81 | -0.1 | -0.29 | 1.39 | 0.68 | 0.76 | 0.61 | -0.1525 | -0.085 | 0.4875 | 0.817783 | 0.850226 | 0.249804 | 0.999971 | 1 | 0.839012 |
| Plcg1 | A2A4A6 | 0.47 | 0.38 | -0.17 | -0.11 | 0.23 | 0.22 | -0.12 | -0.4 | 0.35 | 1.24 | -0.14 | 0.17 | 0.64 | 0.22 | -0.21 | -0.26 | -0.16 | 0.2625 | -0.045 | 0.528616 | 0.443326 | 0.873442 | 0.999971 | 1 | 0.984109 |
| Plcg2 | Q8CIH5 | 0.23 | 1.09 | 0.45 | 0.39 | 1.24 | 1.04 | 0.09 | -0.12 | 0.66 | 0.71 | 0.48 | 0.79 | 0.42 | 0.94 | 0.46 | 0.89 | 0.0225 | 0.12 | 0.1375 | 0.952135 | 0.61001 | 0.59818 | 0.999971 | 1 | 0.92377 |
| Plch2 | H3BIW6 | -0.26 | -0.33 | -0.32 | -0.11 | -0.21 | -0.41 | -0.5 | -0.4 | -0.48 | -0.53 | -0.57 | -0.48 | -0.29 | -0.35 | -0.26 | -0.29 | -0.125 | -0.26 | -0.0425 | 0.472782 | 0.131552 | 0.796345 | 0.999971 | 1 | 0.970435 |
| Plch2_1 | A2AP18-4 | 1.79 | -1 | -1.52 | -0.42 | -0.52 | -0.26 | -0.96 | -1.18 | 0.07 | 1.88 | -1.41 | -2.67 | 0.29 | -0.31 | -0.29 | -1.37 | -0.4425 | -0.245 | -0.1325 | 0.523342 | 0.82312 | 0.855658 | 0.999971 | 1 | 0.982398 |
| Plcl1 | Q3USB7 | -0.45 | -0.23 | -0.78 | -1.22 | -0.47 | -0.24 | -1.05 | -1.09 | -0.48 | -0.18 | -0.94 | -1.34 | -0.45 | -0.47 | -0.95 | -1.12 | -0.0425 | -0.065 | -0.0775 | 0.890173 | 0.844984 | 0.787881 | 0.999971 | 1 | 0.96724 |
| Plcl2 | Q8K394 | -0.25 | -0.32 | 0.28 | 0.27 | -0.33 | -0.52 | 0 | 0.2 | -0.68 | -0.64 | 0.39 | 0.1 | -0.64 | -0.51 | 0.04 | 0.34 | -0.1575 | -0.2025 | -0.1875 | 0.540934 | 0.526796 | 0.527826 | 0.999971 | 1 | 0.909722 |
| Plcxd2 | B2RXA1 | 0.19 | 0.35 | -0.2 | 0.04 | 0.02 | 0.86 | -0.2 | 0.93 | 0.22 | 0.49 | 0.15 | 0.48 | 0.01 | 0.94 | 0.02 | 0.66 | 0.3075 | 0.24 | 0.3125 | 0.342088 | 0.249655 | 0.279064 | 0.999971 | 1 | 0.843812 |
| Plcxd3 | G3X9A7 | -1.31 | -0.13 | -1.04 | -1.36 | 0.45 | -0.48 | -0.55 | -1.35 | 0.14 | -1.08 | -1 | -0.51 | 0.36 | -0.43 | -0.57 | -1.17 | 0.4775 | 0.3475 | 0.5075 | 0.29322 | 0.379805 | 0.232377 | 0.999971 | 1 | 0.831121 |
| Pld1 | Q6NVF2 | -0.03 | -1.21 | -0.72 | -0.43 | -0.77 | -1.06 | -0.55 | -0.75 | -0.59 | -0.01 | -1.53 | -1.35 | -0.91 | -0.9 | -0.82 | -0.99 | -0.185 | -0.2725 | -0.3075 | 0.519477 | 0.512853 | 0.273941 | 0.999971 | 1 | 0.843794 |
| Pld2 | Q6NV49 | -1.02 | -0.69 | -0.57 | -1.13 | -0.32 | -0.64 | -0.34 | -1.04 | -0.26 | -1.22 | -0.8 | -0.9 | 0.22 | -1.07 | -0.77 | -0.88 | 0.2675 | 0.0575 | 0.2275 | 0.291163 | 0.825667 | 0.484077 | 0.999971 | 1 | 0.896352 |
| Pld3 | O35405 | -1.08 | -1.48 | -0.13 | -0.12 | -0.96 | -1.44 | -0.42 | -0.27 | -1.27 | -1.24 | 0.19 | 0.17 | -1.7 | -0.95 | -0.09 | 0.42 | -0.07 | 0.165 | 0.1225 | 0.865545 | 0.742633 | 0.819225 | 0.999971 | 1 | 0.975699 |
| Plec | Q9QXS1-6 | -0.24 | -0.9 | -0.89 | -0.79 | -0.98 | -0.78 | -0.82 | -1.07 | -0.94 | -0.54 | -0.78 | -0.88 | -0.72 | -0.88 | -0.93 | -0.8 | -0.2075 | -0.08 | -0.1275 | 0.351732 | 0.717411 | 0.553789 | 0.999971 | 1 | 0.913787 |
| Plek | Q9JHK5 | 2.76 | 1.17 | -0.29 | -0.22 | 1.54 | 0.48 | -0.1 | -0.44 | 0.33 | 2.33 | -1.06 | -0.16 | -0.63 | 1.07 | -0.03 | -0.34 | -0.485 | -0.495 | -0.8375 | 0.525931 | 0.591468 | 0.270171 | 0.999971 | 1 | 0.843794 |
| Plek2 | Q9WV52 | 1.23 | 1.6 | 1.18 | 1.15 | 0.6 | 1.54 | 0.72 | 1.06 | 0.92 | 1.71 | 1.34 | 0.76 | 1.61 | 1.89 | 0.56 | 1.45 | -0.31 | -0.1075 | 0.0875 | 0.250796 | 0.680607 | 0.779022 | 0.999971 | 1 | 0.96583 |
| Plekha1 | Q8BUL6 | -0.6 | 0.6 | -0.38 | -0.67 | 0.34 | 0.39 | -0.81 | -0.81 | 0.69 | 0.16 | -0.58 | -0.58 | 0.8 | 0.36 | -0.32 | -0.24 | 0.04 | 0.185 | 0.4125 | 0.924953 | 0.652178 | 0.296997 | 0.999971 | 1 | 0.847402 |
| Plekha2 | Q9ERS5 | 0.29 | 0.74 | 0.06 | 0.12 | 0.4 | -0.01 | -0.21 | -0.05 | -0.39 | 0.3 | -0.11 | -0.02 | -0.55 | 0.16 | 0.54 | 0.14 | -0.27 | -0.3575 | -0.23 | 0.27077 | 0.161535 | 0.432628 | 0.999971 | 1 | 0.882096 |
| Plekha5 | E9Q6H8 | -0.24 | -0.59 | -1.04 | -0.96 | -0.57 | -0.44 | -0.69 | -0.77 | -0.46 | -0.14 | -0.92 | -1 | 0.15 | -0.3 | -1.02 | -0.8 | 0.09 | 0.0775 | 0.215 | 0.701213 | 0.786389 | 0.510295 | 0.999971 | 1 | 0.905815 |
| Plekha5_1 | A0A0N4SW | 0.93 | -0.7 | -1.9 | -1.18 | -1.07 | -0.97 | -1.8 | -2.44 | -1.48 | 0.24 | -2.77 | -1.82 | -3.46 | -0.29 | 0.2 | -3.02 | -0.8575 | -0.745 | -0.93 | 0.196291 | 0.356679 | 0.360592 | 0.999971 | 1 | 0.869526 |
| Plekha6 | Q7TQG1 | 0.72 | -0.54 | -0.35 | -0.35 | -0.84 | -0.19 | -0.11 | -0.47 | -0.91 | 0.27 | -0.53 | -0.8 | -0.65 | -0.79 | -0.78 | -0.61 | -0.2725 | -0.3625 | -0.5775 | 0.418451 | 0.35198 | 0.080985 | 0.999971 | 1 | 0.755254 |
| Plekha7 | A0A0U1RP | -0.57 | -1.39 | -1.31 | -0.87 | -1.51 | -1.42 | -1.19 | -1.18 | -1.45 | -1.15 | -1.17 | -1.46 | -1.46 | -1.61 | -1.07 | -0.82 | -0.29 | -0.2725 | -0.205 | 0.249567 | 0.275352 | 0.471273 | 0.999971 | 1 | 0.892857 |
| Plekhb1 | Q9QYE9-4 | -1.07 | 0.3 | -0.04 | -0.07 | -0.02 | 0.77 | -0.47 | 0.38 | 0.77 | -0.08 | 0.14 | 0.25 | 0.87 | 0.73 | 0.62 | 0.79 | 0.385 | 0.49 | 0.9725 | 0.329183 | 0.176941 | 0.010942 | 0.999971 | 1 | 0.613095 |
| Plekhd1 | B2RPU2 | 0.7 | -0.93 | 0.13 | -0.82 | -0.49 | 1 | -0.56 | 0.17 | -0.35 | 0.89 | -0.31 | -0.35 | -0.58 | 0.39 | -0.58 | -0.71 | 0.26 | 0.2 | -0.14 | 0.602034 | 0.669596 | 0.751094 | 0.999971 | 1 | 0.961064 |
| Plekhf2 | Q91WB4 | 0.28 | -0.01 | -0.39 | 0.31 | -0.12 | -0.31 | -0.58 | -0.01 | -0.14 | 0.51 | 0.16 | -0.24 | -0.31 | -0.11 | -0.24 | 0.26 | -0.3025 | 0.025 | -0.1475 | 0.225913 | 0.92251 | 0.54333 | 0.999971 | 1 | 0.911398 |
| Plekhg1 | Q3UX37 | 0.75 | -0.62 | 0.36 | 0.33 | -0.87 | 0.4 | 1.07 | 0.2 | 0.1 | 0.98 | -0.02 | -0.88 | -1.27 | -0.52 | -1 | -1.25 | -0.005 | -0.16 | -1.215 | 0.991398 | 0.723872 | 0.005645 | 0.999971 | 1 | 0.582714 |
| Plekhg3 | Q4VAC9-2 | -0.82 | -0.87 | -1.24 | -0.12 | -1.04 | -1.56 | -0.15 | -0.29 | -1.83 | -0.75 | -0.69 | -1.34 | -1.86 | -0.44 | -1.41 | -1.62 | 0.0025 | -0.39 | -0.57 | 0.994865 | 0.283435 | 0.156661 | 0.999971 | 1 | 0.796008 |
| Plekhg5 | B1AS67 | 0.54 | 0.75 | 0.67 | 0.44 | 0.46 | 0.6 | 0.13 | 0.55 | 0.19 | 0.26 | 0.71 | 0.38 | 0.33 | 0.08 | 0.16 | 0.79 | -0.165 | -0.215 | -0.26 | 0.399801 | 0.283132 | 0.254007 | 0.999971 | 1 | 0.839012 |
| Plekhh2 | Q8C115-2 | -0.45 | -2.06 | -0.44 | -1.09 | -1.51 | -0.54 | -0.92 | -1.88 | -1.32 | -1.21 | -0.59 | -0.18 | -1.08 | -1.29 | -1.4 | -0.32 | -0.2025 | 0.185 | -0.0125 | 0.657532 | 0.676275 | 0.976718 | 0.999971 | 1 | 0.99925 |
| Plekhm2 | A2ADE0 | -0.63 | -0.31 | -0.18 | -0.12 | -0.25 | -0.22 | 0.7 | -0.12 | 0.2 | -0.32 | 0.27 | 0.02 | 0 | -0.64 | -0.56 | -0.44 | 0.3375 | 0.3525 | -0.1 | 0.234567 | 0.130744 | 0.657972 | 0.999971 | 1 | 0.934638 |
| Plekho1 | F6XQM2 | 1.38 | -0.23 | -1.68 | -1.89 | -0.07 | 0.73 | -0.93 | -1.24 | -0.49 | 0.12 | -1.16 | -0.56 | -0.89 | -0.76 | -1.83 | -1.87 | 0.2275 | 0.0825 | -0.7325 | 0.773556 | 0.909728 | 0.3332 | 0.999971 | 1 | 0.857921 |
| Plekho2 | Q8K124 | -0.14 | -1.23 | 0.07 | 0.38 | -1.21 | -1.61 | -0.6 | 0.39 | -1.61 | -0.72 | -0.02 | 0.34 | -1.59 | -1.44 | 0.08 | 0.44 | -0.5275 | -0.2725 | -0.3975 | 0.321496 | 0.599642 | 0.494531 | 0.999971 | 1 | 0.901531 |
| Plg | P20918 | 1.96 | 2.05 | 0.54 | 3.77 | 3.3 | 2.29 | 3.29 | 1.13 | 1.64 | 1.13 | 1.65 | 1.72 | 1.29 | 2.02 | 0.66 | 2.02 | 0.4225 | -0.545 | -0.5825 | 0.578923 | 0.389174 | 0.395025 | 0.999971 | 1 | 0.871816 |
| Plgrkt | Q9D3P8 | 1.01 | 0.09 | 0.15 | 0.28 | 0.23 | 0.03 | -0.2 | 0.14 | 0.25 | 0.64 | 0.47 | 0.12 | -1.29 | -1.42 | -0.19 | -0.26 | -0.3325 | -0.0125 | -1.1725 | 0.217488 | 0.962015 | 0.012251 | 0.999971 | 1 | 0.613095 |
| Plin3 | Q9DBG5 | 1.04 | 1.2 | -0.87 | -0.87 | 1.21 | 1.31 | -0.44 | -0.91 | 0.97 | 0.95 | -1.5 | -1.4 | 0.67 | 1.21 | -1.07 | -1.3 | 0.1675 | -0.37 | -0.2475 | 0.818301 | 0.652293 | 0.74717 | 0.999971 | 1 | 0.961064 |
| Pllp | Q9DCU2 | 2.44 | -0.37 | 0.02 | -0.2 | -1.01 | 0.3 | 0.05 | -0.28 | -0.98 | 1.25 | -0.96 | -1.4 | -1.58 | -0.73 | -1.47 | -1.62 | -0.7075 | -0.995 | -1.8225 | 0.294829 | 0.23843 | 0.01714 | 0.999971 | 1 | 0.617356 |
| Plod1 | Q9R0E2 | -1.15 | -1.09 | 0.16 | -0.02 | -0.47 | -1.04 | -0.02 | 0.32 | -0.95 | -1.3 | -0.24 | -0.34 | 0.02 | -0.52 | -0.01 | -0.13 | 0.2225 | -0.1825 | 0.365 | 0.60673 | 0.657107 | 0.324859 | 0.999971 | 1 | 0.856456 |
| Plod3 | Q9R0E1 | 1.35 | 0.45 | 0.72 | 1.18 | 0.61 | 0.78 | 0.79 | 0.41 | 0.82 | 1.14 | 0.08 | 0.5 | -0.57 | 0.11 | 1.27 | -0.26 | -0.2775 | -0.29 | -0.7875 | 0.288063 | 0.363547 | 0.093602 | 0.999971 | 1 | 0.776618 |
| Plp1 | P60202 | 0.12 | 1.76 | 1.29 | 1.46 | 1.36 | 2.04 | -0.39 | 1.3 | 1.18 | 0.51 | -0.13 | 0.77 | 0.75 | 2.19 | 1.68 | 1.09 | -0.08 | -0.575 | 0.27 | 0.889515 | 0.203436 | 0.553915 | 0.999971 | 1 | 0.913787 |
| Plp2 | Q9R1Q7 | 2.03 | 0.41 | 2.86 | 3.31 | 2.18 | -0.03 | 4.28 | 1.56 | 2.58 | -0.38 | 1.14 | 1.27 | 3.91 | -1.73 | 0.62 | -0.26 | -0.155 | -1 | -1.5175 | 0.874165 | 0.23063 | 0.229381 | 0.999971 | 1 | 0.828687 |
| Plpbp | A0A1B0GR | -0.52 | -0.17 | 0.17 | 0.42 | -0.21 | -0.28 | -0.19 | 0.21 | -0.36 | -0.53 | 0.04 | 0.06 | -0.38 | -0.32 | 0.27 | 0.48 | -0.0925 | -0.1725 | 0.0375 | 0.720041 | 0.5275 | 0.90177 | 0.999971 | 1 | 0.990759 |
| Plpp1 | Q61469-2 | 0.16 | -1.02 | -1.15 | -0.34 | -0.03 | 0.24 | 0.99 | -0.42 | -0.4 | -0.21 | -0.84 | -1.34 | 0.03 | -0.3 | -1.26 | -1.37 | 0.7825 | -0.11 | -0.1375 | 0.081399 | 0.77416 | 0.753872 | 0.999971 | 1 | 0.961116 |
| Plpp3 | Q99JY8 | -0.64 | -0.07 | -0.14 | 0 | -0.46 | -0.29 | -0.75 | 0.33 | -0.47 | -0.7 | -0.72 | 0.05 | -0.29 | 0.1 | 0.11 | -0.14 | -0.08 | -0.2475 | 0.1575 | 0.778786 | 0.344425 | 0.480567 | 0.999971 | 1 | 0.89485 |
| Plpp4 | Q0VBU9 | -0.28 | -0.23 | 0.31 | 0.05 | -0.81 | 0.25 | -0.71 | -0.21 | -1.1 | 0.06 | -1.27 | -1.42 | -0.78 | -0.61 | 0.01 | -0.41 | -0.3325 | -0.895 | -0.41 | 0.272224 | 0.031642 | 0.125351 | 0.999971 | 1 | 0.78268 |
| Plpp6 | Q9D4F2 | 0.97 | 0.84 | 0.7 | 1.12 | 0.57 | 0.66 | 1.09 | 0.83 | 0.64 | 0.89 | 0.45 | 0.13 | 0.12 | 0.22 | 0.62 | 0.32 | -0.12 | -0.38 | -0.5875 | 0.556453 | 0.115384 | 0.015668 | 0.999971 | 1 | 0.613095 |
| Plpp7 | Q91WB2 | -0.08 | -1.12 | 0.58 | 0.64 | -1.43 | -0.77 | 0.46 | 0.39 | -0.47 | -0.64 | 0.85 | -0.46 | -0.28 | -0.33 | 0.12 | 0.62 | -0.3425 | -0.185 | 0.0275 | 0.54824 | 0.711835 | 0.949951 | 0.999971 | 1 | 0.998038 |
| Plppr1 | Q8BFZ2-2 | -0.69 | -2.67 | -0.01 | 0.68 | -1.43 | -2.64 | -0.45 | 0.03 | -1.29 | -0.75 | 0.35 | 0.36 | -1.21 | -2.08 | 0.47 | 0.9 | -0.45 | 0.34 | 0.1925 | 0.593523 | 0.653867 | 0.831106 | 0.999971 | 1 | 0.978643 |
| Plppr2 | A0A087WC | -0.14 | -1.15 | -0.77 | -0.67 | -1.55 | -0.6 | -0.74 | -0.45 | -0.9 | -0.27 | -0.62 | -0.66 | -0.57 | -0.58 | -0.81 | -0.57 | -0.1525 | 0.07 | 0.05 | 0.639082 | 0.791976 | 0.839568 | 0.999971 | 1 | 0.979728 |
| Plppr3 | A0A0R3P9I | -0.32 | -1.08 | -0.9 | -1.34 | -0.93 | -1.1 | 0 | -0.89 | -0.32 | 0.19 | -0.67 | -0.73 | -0.25 | -0.85 | -1.15 | -0.8 | 0.18 | 0.5275 | 0.1475 | 0.586878 | 0.114063 | 0.621471 | 0.999971 | 1 | 0.929641 |
| Plppr4 | A0A0G2JG | -0.62 | -1.67 | -1.08 | -1.19 | -1.46 | -1.26 | -0.72 | -1.09 | -1.37 | -0.43 | -0.86 | -1.01 | -1.25 | -1.45 | -1.34 | -1.05 | 0.0075 | 0.2225 | -0.1325 | 0.978722 | 0.463306 | 0.609061 | 0.999971 | 1 | 0.926708 |
| Plppr5 | A0A668KM | 0.54 | -1.47 | -0.35 | -0.54 | -0.4 | -0.52 | 0.26 | -0.96 | -0.98 | 0.93 | -0.36 | -0.76 | -2.04 | -1.25 | -1.36 | -1.05 | 0.05 | 0.1625 | -0.97 | 0.912084 | 0.767472 | 0.051204 | 0.999971 | 1 | 0.723736 |
| Plrg1 | Q922V4 | 0.45 | -0.55 | 0.13 | 0.09 | -0.85 | -0.94 | 0.97 | -0.28 | -0.67 | 0 | 0.09 | -0.41 | -1.37 | -0.9 | -0.24 | -0.2 | -0.305 | -0.2775 | -0.7075 | 0.509558 | 0.3465 | 0.067172 | 0.999971 | 1 | 0.733937 |
| Pls1 | Q3V0K9 | -0.15 | 0.23 | 0.33 | 0.35 | -0.29 | 0.09 | -0.42 | -0.39 | -0.51 | -0.28 | -0.43 | -0.38 | -0.5 | 0.24 | 0.59 | 0.08 | -0.4425 | -0.59 | -0.0875 | 0.063777 | 0.011769 | 0.750007 | 0.999971 | 1 | 0.961064 |
| Pls3 | B1AX58 | 0.01 | -0.23 | 0.2 | -0.2 | -0.08 | -0.16 | 0.25 | -0.29 | -0.07 | -0.51 | 0.47 | 0.62 | -0.43 | -0.48 | -0.03 | -0.1 | -0.015 | 0.1825 | -0.205 | 0.941947 | 0.532395 | 0.334431 | 0.999971 | 1 | 0.858827 |
| Plscr3 | Q9JIZ9 | -1.07 | -4.27 | -1.63 | -2.03 | -4.92 | -3.25 | -1.87 | -2.02 | -4.27 | -2.14 | -1.94 | -2.68 | -4.38 | -3.35 | -2.56 | -2.61 | -0.765 | -0.5075 | -0.975 | 0.40072 | 0.528002 | 0.210573 | 0.999971 | 1 | 0.822657 |
| Pltp | P55065 | 1.2 | 1.76 | 2.08 | 2.07 | 1.39 | 1.73 | 1.53 | 2.52 | 1.42 | 1.66 | 2.66 | 1.8 | 1.13 | 2.11 | 2.22 | 2.19 | 0.015 | 0.1075 | 0.135 | 0.963297 | 0.750322 | 0.68622 | 0.999971 | 1 | 0.945172 |
| Plxdc1 | Q91ZV7 | 0.49 | 0.14 | -0.94 | -1.04 | -0.2 | -0.06 | -1.7 | -0.37 | -0.6 | 0.2 | -0.95 | -1.4 | 0.41 | 0.43 | -0.5 | -0.21 | -0.245 | -0.35 | 0.37 | 0.625985 | 0.471923 | 0.394857 | 0.999971 | 1 | 0.871816 |
| Plxdc2 | Q9DC11 | -1.04 | -0.15 | -1.24 | -1.28 | 0.33 | 0.04 | -1.03 | -1.26 | 0.6 | -0.63 | -0.97 | -0.76 | -0.05 | -0.21 | -0.55 | -1.03 | 0.4475 | 0.4875 | 0.4675 | 0.328046 | 0.264958 | 0.192095 | 0.999971 | 1 | 0.810547 |
| Plxna1 | P70206 | -0.48 | -0.83 | -0.98 | -0.98 | -0.87 | -0.62 | -0.97 | -0.68 | -0.64 | -0.33 | -0.72 | -0.65 | -0.03 | -0.61 | -0.77 | -0.5 | 0.0325 | 0.2325 | 0.34 | 0.871465 | 0.263852 | 0.172037 | 0.999971 | 1 | 0.804075 |
| Plxna2 | P70207 | -0.61 | -0.32 | -0.72 | -0.91 | -0.5 | -0.22 | -0.89 | -0.83 | -0.55 | -0.35 | -0.55 | -1 | 0.13 | -0.39 | -0.68 | -0.73 | 0.03 | 0.0275 | 0.2225 | 0.898088 | 0.901964 | 0.398703 | 0.999971 | 1 | 0.871816 |
| Plxna3 | P70208 | -0.5 | -1.28 | -1.32 | -1.15 | -0.87 | -1.5 | -0.78 | -1.23 | -0.78 | -0.31 | -0.64 | -0.59 | -0.48 | -0.95 | -0.92 | -0.79 | -0.0325 | 0.4825 | 0.2775 | 0.904784 | 0.074311 | 0.281759 | 0.999971 | 1 | 0.844538 |
| Plxna4 | Q80UG2 | -0.62 | -0.45 | -0.73 | -1.02 | -0.62 | -0.38 | -0.62 | -0.82 | -0.56 | -0.62 | -0.57 | -0.69 | -0.14 | -0.71 | -0.75 | -0.81 | 0.095 | 0.095 | 0.1025 | 0.644165 | 0.614893 | 0.662259 | 0.999971 | 1 | 0.936148 |
| Plxnb1 | Q8CJH3 | -0.6 | -0.48 | -0.71 | -0.93 | -0.86 | -0.58 | -1.02 | -0.94 | -0.61 | -0.61 | -0.83 | -0.58 | 0.02 | -0.41 | -0.69 | -0.62 | -0.17 | 0.0225 | 0.255 | 0.397496 | 0.901422 | 0.278936 | 0.999971 | 1 | 0.843812 |
| Plxnb2 | B2RXS4 | -0.2 | -0.48 | -0.47 | -0.49 | -0.47 | -0.62 | -0.47 | -0.39 | -0.25 | -0.29 | -0.59 | -0.6 | -0.01 | -0.52 | -0.29 | -0.61 | -0.0775 | -0.0225 | 0.0525 | 0.656833 | 0.902874 | 0.799045 | 0.999971 | 1 | 0.970479 |
| Plxnb3 | Q9QY40 | -1.79 | -0.53 | -0.32 | -0.86 | -1.05 | -1.05 | -1.22 | -1 | -0.9 | -1.33 | -0.65 | -0.84 | -0.57 | -0.29 | -0.16 | -0.54 | -0.205 | -0.055 | 0.485 | 0.536538 | 0.874818 | 0.175383 | 0.999971 | 1 | 0.808532 |
| Plxnc1 | Q9QZC2 | -0.07 | -1.12 | -0.73 | -0.47 | -0.65 | -1.02 | -0.5 | -0.01 | -0.49 | -0.63 | -0.88 | -0.13 | -0.38 | -0.92 | -0.35 | -0.64 | 0.0525 | 0.065 | 0.025 | 0.865275 | 0.818827 | 0.927693 | 0.999971 | 1 | 0.994229 |
| Plxnd1 | Q3UH93 | -1.13 | -1.24 | -1.18 | -1.15 | -0.76 | -0.95 | -1.01 | -0.78 | -0.64 | -0.93 | -0.82 | -0.89 | -0.14 | -0.9 | -1.04 | -0.95 | 0.3 | 0.355 | 0.4175 | 0.099983 | 0.055346 | 0.111716 | 0.999971 | 1 | 0.78268 |
| Pm20d1 | Q8C165 | 0.11 | -0.31 | -0.68 | -0.04 | 0.54 | 0.05 | -0.49 | -0.03 | 0.15 | -0.26 | -0.4 | -0.35 | -0.72 | -0.05 | -0.22 | -0.45 | 0.2475 | 0.015 | -0.13 | 0.397484 | 0.95068 | 0.610651 | 0.999971 | 1 | 0.92673 |
| Pmch | P56942 | -0.35 | 0.3 | 0.4 | 1.08 | 0.85 | 0.82 | 0.08 | 0.73 | 0.29 | -0.55 | -0.29 | 0.96 | 1.06 | 1.14 | 1.72 | 0.45 | 0.2625 | -0.255 | 0.735 | 0.448961 | 0.551012 | 0.079388 | 0.999971 | 1 | 0.755254 |
| Pml | D3Z3A6 | 0.91 | 0.23 | 0.46 | 1.41 | 0.57 | -0.68 | -0.28 | -0.15 | -0.19 | 0.33 | 0.3 | 0.18 | -1.08 | -0.29 | 0.54 | 0.63 | -0.8875 | -0.5975 | -0.8025 | 0.033678 | 0.06984 | 0.102055 | 0.999971 | 1 | 0.78268 |
| Pmm1 | O35621 | -0.95 | -0.19 | -0.14 | 0.09 | -0.1 | -0.17 | -0.27 | 0.19 | -0.24 | -0.58 | 0.18 | 0.13 | -0.17 | -0.45 | 0.17 | 0.37 | 0.21 | 0.17 | 0.2775 | 0.439936 | 0.569554 | 0.365968 | 0.999971 | 1 | 0.869526 |
| Pmm2 | Q9Z2M7 | -0.48 | -0.2 | 0.25 | -0.03 | 0.09 | -0.39 | 0.39 | -0.29 | -0.14 | -0.55 | 0.82 | 1.33 | -0.15 | -0.15 | 0.34 | 0.16 | 0.065 | 0.48 | 0.165 | 0.802402 | 0.285621 | 0.484761 | 0.999971 | 1 | 0.896611 |
| Pmpca | Q9DC61 | -0.3 | -0.81 | -0.53 | -0.48 | -0.68 | -0.41 | -0.4 | -0.58 | -0.44 | -0.18 | -0.53 | -0.41 | -0.26 | -0.05 | -0.47 | -0.58 | 0.0125 | 0.14 | 0.19 | 0.947931 | 0.469541 | 0.376531 | 0.999971 | 1 | 0.870465 |
| Pmpcb | Q9CXT8 | -0.26 | -0.19 | -0.09 | -0.43 | -0.04 | 0.13 | -0.49 | -0.05 | 0.1 | -0.22 | -0.04 | 0.23 | 0.62 | -0.3 | 0.23 | 0.02 | 0.13 | 0.26 | 0.385 | 0.530552 | 0.185906 | 0.134795 | 0.999971 | 1 | 0.784948 |
| Pmvk | D3YXP6 | 1.68 | 0.9 | 1.59 | 1.11 | 0.48 | 0.89 | 1.69 | 2.21 | 0.84 | 0.7 | 2.01 | 0.56 | 0.54 | 0.57 | 1.87 | 2.02 | -0.0025 | -0.2925 | -0.07 | 0.995144 | 0.439062 | 0.868164 | 0.999971 | 1 | 0.983462 |
| Pnck | Q9QYK9 | -0.06 | 0.05 | 0.34 | -0.34 | 0.07 | -0.3 | -0.06 | -0.03 | 0.76 | -0.07 | -0.07 | 0.11 | 0.33 | 0.33 | 1.04 | 0.08 | -0.0775 | 0.185 | 0.4475 | 0.714214 | 0.487627 | 0.12475 | 0.999971 | 1 | 0.78268 |
| Pnisr | A2AJT4 | 0.2 | -0.73 | 1.57 | 1.1 | -0.92 | -0.62 | 2.23 | -0.07 | -1.1 | -0.15 | 0.45 | -0.18 | -1.04 | -0.68 | -0.14 | -1.57 | -0.38 | -0.78 | -1.3925 | 0.631654 | 0.18418 | 0.02911 | 0.999971 | 1 | 0.678474 |
| Pnkd | Q69ZP3 | 0.03 | -0.23 | -0.3 | -0.02 | -0.18 | 0 | -0.27 | 0.36 | -0.32 | -0.3 | -0.1 | 0.13 | -0.53 | 0.17 | 0.08 | 0.69 | 0.1075 | -0.0175 | 0.2325 | 0.612457 | 0.927312 | 0.414419 | 0.999971 | 1 | 0.876329 |
| Pnkd_1 | A0A0R4J1J | 0.04 | -0.97 | -1.44 | -0.49 | -0.35 | -0.8 | 0.58 | -1.94 | -0.79 | -0.17 | -0.11 | -0.37 | -1.39 | -1.68 | -2.82 | -3.56 | 0.0875 | 0.355 | -1.6475 | 0.876109 | 0.322092 | 0.01429 | 0.999971 | 1 | 0.613095 |
| Pnkp | G5E8N7 | 1.79 | 2.39 | 1.65 | 1.51 | 2.12 | 1.46 | 1.04 | 0.88 | 1.73 | 1.91 | 1.21 | 0.94 | 1.74 | 1.67 | 1.3 | 1.62 | -0.46 | -0.3875 | -0.2525 | 0.1938 | 0.223318 | 0.321438 | 0.999971 | 1 | 0.855353 |
| Pnma2 | Q8BHK0 | 0.31 | -0.48 | 0.32 | 0.85 | -0.78 | -0.34 | 0.88 | 0.27 | -0.47 | -0.29 | 0.24 | -0.85 | 0.35 | 0.28 | 0.01 | 0.02 | -0.2425 | -0.5925 | -0.085 | 0.575327 | 0.117871 | 0.775659 | 0.999971 | 1 | 0.96393 |

| | | | | | | | | | | | | | | | | | | | | | | | | | | |
|---|---|---|---|---|---|---|---|---|---|---|---|---|---|---|---|---|---|---|---|---|---|---|---|---|---|---|
| Pnma8a | Q80VM8 | -0.15 | -0.89 | -1.74 | -1.41 | -0.86 | -1.02 | -1.33 | -1.84 | -1.15 | -0.25 | -1.67 | -1.14 | -1.05 | -0.72 | -1.35 | -1.28 | -0.215 | -0.005 | -0.0525 | 0.586165 | 0.990742 | 0.885697 | 0.999971 | 1 | 0.98711 |
| Pnma8b | G3X9N3 | 1.78 | 0.77 | 0.94 | 1.19 | 1.05 | 0.9 | 1.61 | 1.39 | 0.94 | 1.18 | 1.05 | 1 | 0.48 | 0.68 | 1.14 | 1.11 | 0.0675 | -0.1275 | -0.3175 | 0.813822 | 0.615016 | 0.286957 | 0.999971 | 1 | 0.846151 |
| Pnn | O35691 | 2.62 | -0.67 | -0.15 | 0.43 | -0.73 | -0.45 | 0.48 | -0.12 | -0.56 | 1.56 | -0.05 | -0.32 | -1.29 | -0.67 | -0.33 | -0.13 | -0.7625 | -0.4 | -1.1625 | 0.289435 | 0.613203 | 0.121433 | 0.999971 | 1 | 0.78268 |
| Pno1 | Q9CPS7 | 0.16 | -1.82 | -1.13 | -0.56 | -0.81 | -2.76 | -0.62 | -1.24 | -1.81 | -1.01 | -0.9 | -1.53 | -2.35 | -2.6 | -1.38 | -3.16 | -0.52 | -0.475 | -1.535 | 0.386436 | 0.303196 | 0.015369 | 0.999971 | 1 | 0.613095 |
| Pnoc | Q64387 | -0.49 | -0.21 | -0.67 | -1.71 | 0.3 | 0.22 | -1.56 | -1.49 | -0.08 | 0.21 | -0.73 | -0.35 | -0.08 | 1.01 | -0.06 | -1.1 | 0.1375 | 0.5325 | 0.7125 | 0.806264 | 0.177854 | 0.180516 | 0.999971 | 1 | 0.808532 |
| Pnp | Q543K9 | 0.54 | 0.4 | 0.47 | 0.23 | 0.41 | 0.43 | 0.69 | 0.68 | 0.07 | 0.35 | 0.66 | 0.48 | 0.05 | 0.16 | 0.44 | 0.6 | 0.1425 | -0.02 | -0.0975 | 0.437419 | 0.919015 | 0.630157 | 0.999971 | 1 | 0.932649 |
| Pnpla2 | Q8BJ56-3 | -3.76 | -5.49 | 1.29 | 2.44 | -3.52 | -5.49 | 3.56 | 1.18 | -4.07 | -4.61 | 0.86 | 1.19 | -3.8 | -5.91 | 0.33 | 1.25 | 0.3125 | -0.2775 | -0.6525 | 0.900529 | 0.899634 | 0.773899 | 0.999971 | 1 | 0.963877 |
| Pnpla6 | A0A140LHU | -0.18 | -0.37 | -0.19 | -0.2 | -0.34 | -0.18 | -0.24 | -0.32 | -0.4 | -0.15 | -0.35 | -0.16 | -0.43 | 0.01 | 0.02 | -0.12 | -0.035 | -0.03 | 0.105 | 0.831929 | 0.857964 | 0.577298 | 0.999971 | 1 | 0.918236 |
| Pnpla7 | A2AJ88-2 | -2.44 | -0.96 | 0.68 | 0.05 | -0.66 | -0.8 | 0.12 | 0.65 | -1.54 | -2.06 | 0.24 | 0.97 | -1.4 | -1.3 | 0.78 | 0.91 | 0.495 | 0.07 | 0.415 | 0.478753 | 0.937399 | 0.621999 | 0.999971 | 1 | 0.929641 |
| Pnpla8 | Q8K1N1 | -0.67 | -0.91 | -0.54 | -0.68 | -0.75 | -0.79 | -0.32 | -0.35 | -0.71 | -0.26 | -0.66 | -0.54 | -0.22 | -0.46 | -0.44 | -0.15 | 0.1475 | 0.1575 | 0.3825 | 0.474816 | 0.414761 | 0.063772 | 0.999971 | 1 | 0.725348 |
| Pnpo | Q91XF0 | -2.22 | -2.71 | -1.95 | -1.26 | -2.01 | -2.49 | -1.21 | -1.19 | -2.64 | -2.51 | -1.49 | -1.79 | -2.75 | -2.11 | -1.59 | -1.35 | 0.31 | -0.0725 | 0.085 | 0.46441 | 0.854309 | 0.836956 | 0.999971 | 1 | 0.979416 |
| Pnpt1 | Q8K1R3 | -0.5 | -1.08 | -0.68 | -0.8 | -1.07 | -1.02 | -0.78 | -0.77 | -0.95 | -0.37 | -0.52 | -0.5 | -0.76 | -0.78 | -0.65 | -0.79 | -0.145 | 0.18 | 0.02 | 0.478985 | 0.417088 | 0.917172 | 0.999971 | 1 | 0.992633 |
| Podxl | Q9R0M4 | -1.44 | 1.33 | 1.2 | 0.83 | 2.57 | 1.22 | 1.04 | 1.03 | 2.18 | -0.9 | 1.79 | 1.77 | 2.71 | 1.34 | 1.81 | 0.95 | 0.985 | 0.73 | 1.2225 | 0.170376 | 0.410106 | 0.100319 | 0.999971 | 1 | 0.78268 |
| Podxl2 | G3X9D3 | -0.5 | -0.74 | -1.43 | -1.19 | -0.69 | -0.34 | -0.89 | -1.37 | -0.29 | -0.39 | -0.89 | -0.94 | 0.83 | 0.12 | -0.51 | -0.67 | 0.1425 | 0.3375 | 0.9075 | 0.644659 | 0.252015 | 0.041114 | 0.999971 | 1 | 0.700838 |
| Pofut1 | A2AMC3 | -1.85 | -1.24 | -1.03 | -1.32 | -0.32 | -1.23 | -0.66 | -1.03 | -0.89 | -1.52 | -1.52 | -1.08 | -0.92 | -0.24 | -0.73 | -1.23 | 0.55 | 0.1075 | 0.58 | 0.077311 | 0.678784 | 0.068177 | 0.999971 | 1 | 0.733937 |
| Pofut2 | Q8VHI3 | 0.21 | -0.02 | -1.02 | -0.21 | -0.4 | -0.13 | -1 | -0.53 | 1.29 | 1.23 | 1.21 | 1.18 | 2.28 | 1.41 | 0.89 | 1.14 | -0.255 | 1.4875 | 1.69 | 0.440035 | 0.000568 | 0.001925 | 0.999971 | 0.448349 | 0.476373 |
| Poglut1 | Q8BYB9 | -0.1 | 0.59 | 1.42 | 1.26 | 0.54 | 1.03 | 0.77 | 1.48 | 0.91 | 0.99 | 1.42 | 1.69 | 1.34 | 1.1 | 1.37 | 1.39 | 0.1625 | 0.46 | 0.5075 | 0.675491 | 0.246772 | 0.170999 | 0.999971 | 1 | 0.803758 |
| Poglut3 | G5E897 | -1.98 | -0.59 | -0.08 | 0.5 | -0.45 | -0.8 | -0.3 | -0.07 | -0.81 | -1.11 | 0.3 | 0.27 | -0.89 | -0.53 | 0.59 | 0.98 | 0.1325 | 0.2 | 0.575 | 0.795301 | 0.735777 | 0.373437 | 0.999971 | 1 | 0.869657 |
| Pogz | D3YUW8 | 0.8 | -0.04 | -0.21 | -0.28 | -0.04 | -0.2 | -0.17 | -0.33 | -0.04 | 0.41 | -0.68 | -0.45 | -0.84 | -0.08 | -0.23 | -0.3 | -0.2525 | -0.2575 | -0.43 | 0.368732 | 0.458995 | 0.184667 | 0.999971 | 1 | 0.809025 |
| Polb | Q8K409 | 1.83 | 2.03 | 1.72 | 2.06 | 1.91 | 1.89 | 2.01 | 2.05 | 1.41 | 2.16 | 1.78 | 1.68 | 1.34 | 2.17 | 1.72 | 2.34 | 0.055 | -0.1525 | -0.0175 | 0.754446 | 0.489189 | 0.946932 | 0.999971 | 1 | 0.997651 |
| Pold2 | A8Y5G7 | 0.24 | -0.32 | 0.43 | 0.95 | -0.41 | -0.71 | 1.1 | 1.64 | -0.08 | -0.44 | 1.76 | 2.01 | -0.31 | -0.19 | 0.92 | 1.65 | 0.08 | 0.4875 | 0.1925 | 0.889332 | 0.441123 | 0.700476 | 0.999971 | 1 | 0.948864 |
| Poldip2 | Q91VA6 | 0.09 | -0.17 | -0.39 | -0.6 | -0.21 | -0.22 | 0.09 | -0.72 | -0.33 | -0.37 | -0.53 | -0.63 | -0.39 | -0.45 | -0.51 | -0.57 | 0.0025 | -0.1975 | -0.2125 | 0.99205 | 0.359969 | 0.319498 | 0.999971 | 1 | 0.853962 |
| Poldip3 | Q8BG81 | 1.32 | 0.47 | 0.74 | 1.1 | 0.7 | 0.53 | 0.61 | 1.39 | 0.35 | 1.13 | 0.85 | 0.94 | 0.32 | 0.68 | 1.05 | 1.02 | -0.1 | -0.09 | -0.14 | 0.727424 | 0.739023 | 0.611307 | 0.999971 | 1 | 0.92673 |
| Polg | Q75WC0 | -1.16 | -0.64 | 2.73 | 2.39 | -1.43 | -2.02 | 2.49 | 2.6 | -1.4 | -2.16 | 2.38 | 1.38 | -2.86 | -1.7 | 1.38 | 3.05 | -0.42 | -0.78 | -0.8625 | 0.767045 | 0.560213 | 0.569862 | 0.999971 | 1 | 0.91816 |
| Poll | Q9QXE2 | 4.96 | 2.17 | 3.94 | 3.4 | 2.79 | 2.98 | 4.71 | 0.87 | 2.13 | 3.4 | -0.31 | 0.35 | -0.89 | 0.26 | 2.89 | -0.69 | -0.78 | -2.225 | -3.225 | 0.383646 | 0.037336 | 0.007166 | 0.999971 | 1 | 0.582714 |
| Polr1c | P52432 | 3.53 | 0.64 | 0.13 | 0.34 | 0.39 | 0.15 | 0.02 | 0.91 | -0.1 | 1.87 | 0.37 | 0.42 | 1.04 | 0.23 | 0.67 | 1 | -0.7925 | -0.52 | -0.425 | 0.300222 | 0.529497 | 0.569037 | 0.999971 | 1 | 0.91816 |
| Polr1d | P97304 | 3.11 | 0.71 | 2.2 | 1.89 | -0.46 | 0.46 | 1.98 | 1.48 | 0.06 | 1.98 | 0.95 | -1.51 | -1.79 | 0.14 | 1.73 | 0.41 | -1.1125 | -1.6075 | -1.855 | 0.12214 | 0.07284 | 0.041224 | 0.999971 | 1 | 0.700838 |
| Polr1h | Q791N7 | -0.53 | 0.33 | -0.67 | -1.16 | -0.05 | -0.28 | -1.29 | 0.06 | -0.16 | -0.37 | 0.11 | 0.47 | 0.2 | -0.02 | -0.44 | -0.78 | 0.1175 | 0.52 | 0.2475 | 0.777529 | 0.165974 | 0.508133 | 0.999971 | 1 | 0.905815 |
| Polr2a | A0A0R4J0\ | 0.66 | 0.24 | 0.16 | 0.22 | 0.07 | -0.04 | 0.11 | 0.02 | 0 | 0.44 | 0.12 | 0.03 | 0.08 | 0.14 | 0.06 | 0.39 | -0.28 | -0.1725 | -0.1525 | 0.162296 | 0.406785 | 0.450086 | 0.999971 | 1 | 0.885596 |
| Polr2b | Q8CFI7 | 0.46 | 0.6 | 0.92 | 0.86 | 0.67 | 0.42 | 0.82 | 0.81 | 0.74 | 0.63 | 0.89 | 0.68 | 0.4 | 0.22 | 0.88 | 0.92 | -0.03 | 0.025 | -0.105 | 0.881152 | 0.893543 | 0.662617 | 0.999971 | 1 | 0.936148 |
| Polr2c | P97760 | 0.37 | 0.11 | 0.2 | 0.42 | 1.02 | -0.51 | 0.31 | 0.65 | -0.41 | 0.9 | 0.22 | 0.69 | 1.5 | 0.63 | 0.24 | 0.66 | 0.0925 | 0.075 | 0.4825 | 0.781376 | 0.806161 | 0.121356 | 0.999971 | 1 | 0.78268 |
| Polr2d | Q9D7M8 | 0.55 | 0.97 | 0.25 | 0.5 | 0.63 | 1 | 0.03 | 0.15 | 0.2 | 0.97 | 0.05 | 0.2 | -0.82 | 1.49 | 0.49 | 0.72 | -0.115 | -0.2125 | -0.0975 | 0.686048 | 0.444213 | 0.835864 | 0.999971 | 1 | 0.979185 |
| Polr2e | Q80UW8 | 0.36 | 0.37 | 0.69 | 0.71 | 0.33 | -0.31 | 0.67 | 0.59 | 0.08 | 0.48 | 0.76 | 0.8 | -0.07 | 0.04 | 0.68 | 0.52 | -0.2125 | -0.0025 | -0.24 | 0.429765 | 0.991251 | 0.331219 | 0.999971 | 1 | 0.857107 |
| Polr2g | P62488 | 0.4 | 0.46 | 0.14 | -0.08 | 0.67 | 0.3 | -0.26 | -0.26 | 0.88 | 0.24 | -0.35 | 0.31 | 1.45 | 0.7 | -0.07 | 0.02 | -0.1175 | 0.04 | 0.295 | 0.672305 | 0.890645 | 0.428123 | 0.999971 | 1 | 0.880793 |
| Polr2h | Q923G2 | 0.81 | 0.9 | 0.76 | 0.87 | 1.48 | 2.22 | 0.63 | 0.94 | 1.02 | 1.01 | 1.08 | 0.01 | 0.62 | 0.61 | 0.86 | 0.64 | 0.4825 | -0.055 | -0.1525 | 0.185723 | 0.841503 | 0.376 | 0.999971 | 1 | 0.870465 |
| Polr2j | D3YZ20 | -0.01 | -0.26 | 1.13 | 1.11 | -0.39 | -0.3 | 0.97 | 1.41 | -0.55 | 0.12 | 0.64 | 0.36 | -0.41 | 0.35 | 1.22 | -0.06 | -0.07 | -0.35 | -0.2175 | 0.895989 | 0.418033 | 0.648173 | 0.999971 | 1 | 0.933842 |
| Polr2l | P62876 | -1.9 | -0.43 | 0.26 | -0.91 | -0.08 | -0.9 | -0.13 | -0.63 | 0.08 | -3.1 | 0.43 | 0.72 | -0.89 | -1.07 | 0.01 | -0.7 | 0.31 | 0.2775 | 0.0825 | 0.508817 | 0.757884 | 0.862889 | 0.999971 | 1 | 0.982963 |
| Polr2m | Q6P6I6 | 1.56 | -0.63 | -0.37 | -0.19 | 0.07 | 0.1 | 0.83 | -0.24 | 0 | 1.18 | -0.1 | -0.43 | -1.65 | -0.74 | -0.47 | -0.61 | 0.0975 | 0.07 | -0.96 | 0.847246 | 0.90078 | 0.093366 | 0.999971 | 1 | 0.776565 |
| Polr2m_1 | A0A3B2WE | 0.42 | 1.17 | 0.76 | 0.62 | 1.72 | 1.7 | 1.33 | -0.03 | 1.66 | 0.8 | 0.74 | 1.71 | 2.25 | 0.4 | 0.54 | 0.8 | 0.4375 | 0.485 | 0.255 | 0.312619 | 0.147098 | 0.557244 | 0.999971 | 1 | 0.915051 |
| Polr3a | B2RXC6 | -0.45 | 1.19 | 1.45 | 1.24 | 1.37 | 0.7 | 1.1 | 1.76 | 0.51 | -0.62 | 1.57 | 1.51 | 0.29 | 0.71 | 1.22 | 1.52 | 0.375 | -0.115 | 0.0775 | 0.425183 | 0.852928 | 0.872284 | 0.999971 | 1 | 0.984109 |
| Polr3b | P59470 | 0.18 | 0.6 | 1.53 | 1.4 | 0.62 | 0.16 | 1.12 | 1.44 | 0.35 | 0.58 | 1.41 | 1.43 | -0.02 | 0.44 | 1.36 | 1.75 | -0.0925 | 0.015 | -0.045 | 0.820726 | 0.970671 | 0.925956 | 0.999971 | 1 | 0.99401 |
| Polr3c | Q9D483 | 1.38 | 2.4 | 1.71 | 1.25 | 2.74 | 2.71 | 4.39 | 1.92 | 6.46 | 0.47 | 2.52 | 2.27 | 1.91 | 3.38 | -0.83 | 2.2 | 1.255 | 1.245 | -0.02 | 0.040625 | 0.297642 | 0.980822 | 0.999971 | 1 | 0.99925 |
| Polr3d | Q91WD1 | -2.46 | 3.48 | 0.79 | 0.37 | 4.03 | 2.64 | 0.79 | 0.82 | 3.28 | -0.92 | 0.67 | 1.27 | 4.59 | 2.83 | 0.29 | 0.49 | 1.525 | 0.53 | 1.505 | 0.254296 | 0.692824 | 0.302645 | 0.999971 | 1 | 0.847402 |
| Polr3e | Q9CZT4-2 | 1.24 | 0.46 | 1.3 | 1.36 | 0.95 | 0.02 | 0.75 | 1.67 | 1.01 | 0.71 | 1.35 | 1.69 | 1.19 | 0.34 | 1.29 | 1.46 | -0.2425 | 0.1 | -0.02 | 0.533187 | 0.743846 | 0.951316 | 0.999971 | 1 | 0.998056 |
| Polr3f | Q9CZ23 | -0.28 | 1.21 | 1.03 | 1.64 | 1.63 | 0.81 | 1.05 | 0.81 | 1.1 | 0.51 | 1.35 | 1.3 | 1.44 | 1 | 1.07 | 1.15 | 0.175 | 0.165 | 0.265 | 0.686107 | 0.70385 | 0.519068 | 0.999971 | 1 | 0.905934 |
| Polrmt | Q8BKF1 | -0.77 | -1.01 | -0.38 | -0.13 | -1.15 | -0.68 | -0.05 | -0.71 | -1.34 | -1.79 | -0.72 | -0.9 | -1.49 | -0.57 | -0.75 | -0.57 | -0.075 | -0.615 | -0.2725 | 0.806606 | 0.076705 | 0.379378 | 0.999971 | 1 | 0.871816 |
| Pom121 | Q8K3Z9 | 1.45 | -1.04 | 0.09 | 0.69 | -0.76 | -0.39 | 0.2 | -0.06 | -0.74 | 0.53 | -1 | -0.79 | -0.73 | -0.76 | -0.14 | 0.01 | -0.55 | -0.7975 | -0.7025 | 0.307408 | 0.193083 | 0.199779 | 0.999971 | 1 | 0.813335 |
| Pomgnt2 | Q8BW41-2 | -0.36 | -0.82 | -0.73 | -0.9 | -0.16 | -0.78 | -0.43 | -0.51 | -0.4 | -0.92 | -0.89 | -0.48 | 0.15 | -0.09 | -0.46 | -0.87 | 0.2325 | 0.03 | 0.385 | 0.304583 | 0.891794 | 0.181345 | 0.999971 | 1 | 0.808532 |
| Pomk | Q3TUA9 | -0.79 | -0.4 | -0.98 | -0.21 | 0.02 | -0.44 | -0.62 | 0.02 | -0.24 | -0.95 | -0.15 | -0.09 | -0.44 | -0.37 | -0.09 | 0.22 | 0.34 | 0.2375 | 0.425 | 0.216891 | 0.407066 | 0.125444 | 0.999971 | 1 | 0.78268 |
| Pomp | Q9CQT5 | -0.67 | 0.9 | 1.09 | 1.45 | 0.24 | -0.16 | 0.28 | 1.59 | 0.02 | -1.51 | 0.75 | 0.56 | -0.2 | 0.55 | 3.4 | 1.69 | -0.205 | -0.7375 | 0.6675 | 0.712237 | 0.264359 | 0.42496 | 0.999971 | 1 | 0.880275 |
| Pomt2 | Q8BGQ4-2 | -2.15 | -2.18 | -4.47 | -3.71 | -2.06 | -2.41 | -3.83 | -2.8 | -1.42 | -1.93 | -1.8 | -3.76 | -1.89 | -2.42 | -3.47 | -5.2 | 0.3525 | 0.9 | -0.1175 | 0.578893 | 0.225275 | 0.888163 | 0.999971 | 1 | 0.987485 |
| Pon2 | Q62086 | -1.06 | -0.85 | -0.86 | -1.35 | -0.97 | -1.02 | -0.86 | -0.85 | -0.65 | -1.06 | -0.54 | -0.18 | -0.52 | -0.89 | -0.71 | -0.79 | 0.105 | 0.4225 | 0.3025 | 0.585493 | 0.111 | 0.156538 | 0.999971 | 1 | 0.796008 |
| Pon3 | Q62087 | 0.22 | -1.13 | -0.56 | -1.43 | -0.24 | -1.39 | -0.34 | -1.82 | -0.23 | -0.27 | -2.03 | 0.03 | -0.97 | -1.07 | -1.26 | -1.34 | -0.2225 | 0.1 | -0.435 | 0.654509 | 0.855947 | 0.251675 | 0.999971 | 1 | 0.839012 |
| Pop7 | Q9DCH2 | 0.53 | 0.83 | -0.27 | 0.18 | 0.5 | 0.45 | -0.19 | 0.09 | 0.96 | 0.9 | -0.26 | 0.08 | 1.32 | 0.89 | 0 | 0.69 | -0.105 | 0.1025 | 0.4075 | 0.724025 | 0.783809 | 0.270725 | 0.999971 | 1 | 0.843794 |
| Por | P37040 | -0.92 | -1.28 | -1.33 | -1.23 | -1.38 | -1.43 | -1.03 | -1.43 | -1.31 | -0.97 | -1.32 | -1.11 | -1.17 | -1.13 | -1.26 | -1.21 | -0.1275 | 0.0125 | -0.0025 | 0.519708 | 0.94704 | 0.988832 | 0.999971 | 1 | 0.99925 |
| Porcn | Q9JJJ7-4 | -0.54 | 1 | 1.18 | 0.46 | -0.25 | 1.42 | 0.59 | 1.33 | -0.19 | 0.38 | 1.45 | -1.76 | 0.11 | 1.83 | 0.24 | 0.55 | 0.2475 | -0.555 | 0.1575 | 0.627719 | 0.438555 | 0.758468 | 0.999971 | 1 | 0.961116 |
| Postn | Q62009-5 | 1.37 | 0.64 | 0.35 | 0.99 | 6.06 | 1.32 | 0.49 | -0.19 | 1.75 | 3.3 | 0.98 | 3.54 | -1.24 | 2.21 | 0.32 | -0.04 | 1.0825 | 1.555 | -0.525 | 0.404347 | 0.027895 | 0.447196 | 0.999971 | 1 | 0.884685 |
| Pou3f1 | P21952 | 2.75 | 2.53 | 1.82 | 1.99 | 2.09 | 2.25 | 1.11 | 1.97 | 1.44 | 2.07 | 2.03 | 1.48 | 0.71 | 2.15 | 1.39 | 2.33 | -0.4175 | -0.5175 | -0.6275 | 0.2327 | 0.101224 | 0.153821 | 0.999971 | 1 | 0.795578 |
| Pou3f3 | P31361 | 2.82 | 1.63 | 1.49 | 4.16 | 0.91 | 1.15 | 2.53 | 3.2 | 0.85 | 2.2 | 4.36 | 3.04 | 3.57 | 2.55 | 2.25 | 3.78 | -0.5775 | 0.0875 | 0.5125 | 0.447494 | 0.919802 | 0.445249 | 0.999971 | 1 | 0.884152 |
| Pou6f1 | Q5U4D4 | -0.78 | -0.86 | 0.51 | 0.43 | 0.16 | -0.08 | -0.33 | 1.03 | -0.44 | -0.49 | 0.6 | 1 | -0.81 | 0.81 | 0.15 | 0.38 | 0.37 | 0.3425 | 0.3075 | 0.417496 | 0.493564 | 0.521156 | 0.999971 | 1 | 0.906972 |
| Pou6f1_1 | H3BKJ6 | -0.23 | -1.97 | -0.45 | 0.2 | -1.8 | -0.04 | -1.04 | -1.66 | -2.5 | -0.25 | -1.07 | -1.74 | -1.5 | 0.63 | -1.86 | 0.45 | -0.5225 | -0.7775 | 0.0425 | 0.368998 | 0.228818 | 0.95311 | 0.999971 | 1 | 0.998056 |
| Ppa1 | Q9D819 | -0.4 | -0.08 | 0.37 | 0.7 | 0.23 | 0.23 | 0.35 | 0.75 | 0.43 | -0.12 | 0.59 | 0.88 | 0.16 | 0.52 | 0.72 | 0.99 | 0.2425 | 0.2975 | 0.45 | 0.405649 | 0.367921 | 0.16684 | 0.999971 | 1 | 0.800215 |
| Ppa2 | Q91VM9 | 0.71 | 0.09 | 0.13 | -0.18 | -0.19 | -0.45 | 0.28 | 0.39 | -0.24 | -0.11 | 0.15 | 0.25 | -0.1 | -0.08 | -0.14 | 0.08 | -0.18 | -0.175 | -0.2475 | 0.533272 | 0.483824 | 0.301306 | 0.999971 | 1 | 0.847402 |
| Ppan | Q91YU8 | -0.19 | -1.39 | -1.6 | -0.79 | -1.53 | -1.68 | -0.5 | -1.61 | -1.81 | -0.9 | -1.83 | -1.62 | -2.18 | -1.57 | -1.66 | -2.25 | -0.3375 | -0.5475 | -0.9225 | 0.412178 | 0.16897 | 0.027564 | 0.999971 | 1 | 0.678474 |
| Ppat | Q8CIH9 | 0.31 | 0.59 | 0.11 | 0.05 | 0.71 | 0.58 | -0.1 | 0.27 | 0.7 | 0.5 | 0.28 | 0.52 | 0.82 | 0.75 | 0.37 | 0.41 | 0.1 | 0.235 | 0.3225 | 0.686787 | 0.262451 | 0.16004 | 0.999971 | 1 | 0.800144 |
| Ppbp | Q9EQI5 | 0.84 | 1.11 | -1.87 | 1.67 | 2.8 | 0.47 | 0.83 | -2.53 | -0.12 | 0.36 | -0.68 | 0.31 | -1.04 | 0.91 | 1.1 | 1.16 | -0.045 | -0.47 | 0.095 | 0.970132 | 0.53452 | 0.911105 | 0.999971 | 1 | 0.992329 |
| Ppcdc | Q8BZB2 | -0.45 | 0.55 | -0.84 | -0.73 | 0.35 | -0.14 | -0.44 | -0.54 | -0.09 | -1.07 | -0.79 | -0.6 | -0.54 | -1.05 | -0.66 | -0.75 | 0.175 | -0.27 | -0.3825 | 0.633221 | 0.470257 | 0.270267 | 0.999971 | 1 | 0.843794 |
| Ppcs | Q8VDG5 | -0.08 | -0.07 | -0.15 | -0.2 | 0.17 | 0.09 | -0.17 | 0.06 | -0.12 | -0.22 | -0.2 | -0.04 | 0.21 | 0.74 | 0.04 | 0.37 | 0.1625 | -0.02 | 0.465 | 0.355686 | 0.899946 | 0.047831 | 0.999971 | 1 | 0.710102 |
| Ppdpf | Q9CR37 | 0.56 | -1.24 | -0.82 | -0.31 | -0.7 | -1.32 | -1.29 | 0.34 | -1.51 | -0.62 | -0.45 | -1.09 | -2.58 | -1.97 | -0.11 | 0.79 | -0.29 | -0.465 | -0.515 | 0.571219 | 0.29808 | 0.519497 | 0.999971 | 1 | 0.906123 |
| Ppfia1 | B2RXW8 | -0.55 | -0.2 | -0.54 | -0.72 | -0.01 | -0.47 | -0.54 | -0.84 | 0.01 | -0.68 | -0.57 | -0.57 | 0.08 | -0.77 | -0.61 | -1.24 | 0.0375 | 0.05 | -0.1325 | 0.874441 | 0.825877 | 0.663079 | 0.999971 | 1 | 0.936388 |
| Ppfia2 | B8QI34 | 0.32 | -0.66 | -0.53 | -0.45 | -0.66 | -0.36 | -0.15 | -0.5 | -0.25 | -0.04 | -0.49 | -0.53 | -0.09 | -0.61 | -0.51 | -0.42 | -0.0875 | 0.0025 | -0.0775 | 0.743415 | 0.992533 | 0.773444 | 0.999971 | 1 | 0.963759 |
| Ppfia3 | P60469 | -0.77 | -0.87 | -0.77 | -0.83 | -0.93 | -1 | -0.61 | -0.8 | -0.66 | -0.9 | -0.66 | -0.78 | -0.77 | -1.15 | -0.94 | -0.75 | -0.025 | 0.06 | -0.0925 | 0.886432 | 0.712415 | 0.607253 | 0.999971 | 1 | 0.925193 |
| Ppfia4 | B8QI36 | 0.52 | -0.13 | 0.25 | 0.1 | -0.19 | 0.13 | 0.09 | 0.2 | 0.22 | 0.19 | 0.22 | 0.37 | 0.5 | -0.45 | 0.05 | 0.12 | -0.1275 | 0.065 | -0.13 | 0.549629 | 0.742822 | 0.621778 | 0.999971 | 1 | 0.929641 |
| Ppfibp1 | Q8C8U0 | 0.25 | -0.25 | -0.5 | -0.16 | -0.06 | -0.54 | -0.64 | -0.56 | 0.1 | 0.15 | -0.65 | -0.02 | -1.41 | -0.52 | -0.14 | -0.19 | -0.285 | 0.06 | -0.4 | 0.250464 | 0.819405 | 0.248474 | 0.999971 | 1 | 0.839012 |
| Ppfibp2 | G3X957 | 1 | 2.95 | -0.29 | -0.31 | 2.53 | 1.96 | 7.11 | -1 | 2.45 | 0.38 | -1.35 | -0.33 | 2.24 | 0.53 | 0.44 | -2.4 | 1.8125 | -0.55 | -0.635 | 0.284039 | 0.585311 | 0.566513 | 0.999971 | 1 | 0.917959 |
| Pphln1 | G3UWD4 | -3.42 | -5.05 | -0.99 | -0.33 | -3.89 | -5.64 | -0.72 | -0.17 | -4.54 | -4.05 | -0.32 | -0.36 | -4.68 | -4.86 | -0.12 | 0.01 | -0.1575 | 0.13 | 0.035 | 0.916648 | 0.926769 | 0.981959 | 0.999971 | 1 | 0.99925 |
| Ppia | P17742 | -1.85 | -0.69 | -1.07 | -1.49 | -0.56 | -0.22 | -1.09 | -1.11 | -1.18 | -1.92 | -0.99 | -1.18 | 0.19 | -0.96 | -0.99 | -1.06 | 0.53 | -0.0425 | 0.57 | 0.136017 | 0.896071 | 0.157931 | 0.999971 | 1 | 0.797252 |
| Ppib | P24369 | -1.84 | -0.94 | -0.3 | -0.78 | -0.49 | -0.93 | -0.35 | -0.17 | -0.28 | -1.4 | -0.15 | 0.24 | -0.23 | -1.16 | -0.35 | -0.31 | 0.48 | 0.5675 | 0.4525 | 0.197378 | 0.228468 | 0.249078 | 0.999971 | 1 | 0.839012 |
| Ppic | P30412 | -0.39 | 0.13 | 0.03 | 0.66 | 1.05 | 0.23 | -0.21 | -0.09 | 0.51 | -0.3 | 0.66 | 0.72 | 1.34 | 0.05 | 1.09 | -0.02 | 0.1375 | 0.29 | 0.5075 | 0.695797 | 0.378855 | 0.220886 | 0.999971 | 1 | 0.827096 |
| Ppid | Q9CR16 | -0.74 | -0.28 | -0.17 | -0.76 | 0.11 | -0.64 | 0.01 | -0.96 | 0.08 | -0.7 | 0.39 | 0.55 | -0.25 | -0.72 | -0.64 | -0.44 | 0.1175 | 0.5675 | -0.025 | 0.7021 | 0.103152 | 0.912217 | 0.999971 | 1 | 0.992618 |
| Ppie | Q9QZH3 | 0.21 | -3.23 | 4.48 | 4.33 | -0.69 | -4.41 | 5.13 | 2.36 | -1.4 | -1.92 | 2.95 | 1.1 | 0.36 | -2.69 | 1.83 | -0.25 | -0.85 | -1.265 | -1.635 | 0.72785 | 0.51788 | 0.384474 | 0.999971 | 1 | 0.871816 |
| Ppif | Q99KR7 | -0.06 | 0.33 | -0.6 | -1.02 | 0.4 | 0.07 | -0.44 | -0.44 | 0.82 | -0.15 | -0.11 | -0.33 | 1.72 | -0.12 | -0.12 | -0.31 | 0.235 | 0.395 | 0.63 | 0.512225 | 0.315081 | 0.246102 | 0.999971 | 1 | 0.839012 |
| Ppig | A2AR01 | 1.58 | -0.21 | -1.15 | -0.97 | -0.05 | -0.28 | -1.17 | -0.97 | -0.51 | 2.26 | -0.76 | -0.19 | 0.53 | 0.26 | -1.23 | -1.21 | -0.43 | 0.3875 | -0.225 | 0.49261 | 0.648148 | 0.750593 | 0.999971 | 1 | 0.961064 |
| Ppig_1 | A2AR02 | 0.98 | 1.18 | 0.54 | 0.37 | 0.49 | 0.4 | 0.51 | 0.4 | 0.73 | 1.61 | 0.18 | -0.01 | 0.87 | 0.71 | 0.23 | -0.36 | -0.3175 | -0.14 | -0.405 | 0.189292 | 0.722695 | 0.244864 | 0.999971 | 1 | 0.837316 |
| Ppih | Q9D868-2 | -0.92 | 0.51 | 0.48 | 0.52 | 0.7 | -0.1 | 0.21 | 0.21 | 0.56 | -1.15 | 0.69 | 0.44 | 0.74 | -0.01 | 0.06 | 0.92 | 0.1075 | -0.0125 | 0.28 | 0.776946 | 0.980775 | 0.49814 | 0.999971 | 1 | 0.902911 |
| Ppil1 | Q9D0W5 | -1.4 | 0.61 | 0.99 | -0.26 | 1.25 | -0.02 | 1.17 | -0.09 | 1.2 | -0.89 | 1.17 | 0.4 | 0.86 | -0.51 | 0.38 | -0.99 | 0.5925 | 0.485 | -0.05 | 0.328621 | 0.469011 | 0.935345 | 0.999971 | 1 | 0.995618 |
| Ppil2 | Q9D787 | 1.24 | 1.2 | 0.6 | 0.88 | 1.19 | 1.05 | 1 | 1.11 | 1.07 | 0.94 | 0.58 | 0.98 | 0.66 | 1.09 | 0.55 | 0.79 | 0.1075 | -0.0875 | -0.2075 | 0.607567 | 0.696377 | 0.377348 | 0.999971 | 1 | 0.871112 |
| Ppil3 | Q9D6L8 | -0.76 | 0.39 | 1.07 | 0.94 | 0.3 | 0.02 | 0.57 | 0.85 | 0.19 | -0.68 | 1.04 | 0.93 | 0.08 | 0.07 | 0.97 | 0.97 | 0.025 | -0.04 | 0.1125 | 0.953436 | 0.940198 | 0.806994 | 0.999971 | 1 | 0.972962 |
| Ppil4 | Q9CXG3 | 2.15 | 2.17 | 2.47 | 2.92 | 2.2 | 1.73 | 2.29 | 2.25 | 2 | 1.83 | 2.59 | 2.41 | 1.83 | 1.57 | 2.1 | 2.47 | -0.31 | -0.22 | -0.435 | 0.235489 | 0.423723 | 0.146131 | 0.999971 | 1 | 0.791767 |
| Ppip5k1 | A2ARP1 | 0.53 | -0.27 | 0.16 | 0.42 | -0.21 | -0.09 | 0.45 | 0.28 | -0.54 | -0.05 | 0.12 | 0.07 | -0.9 | -0.17 | 0.24 | 0.06 | -0.1025 | -0.31 | -0.4025 | 0.693842 | 0.246528 | 0.219334 | 0.999971 | 1 | 0.827064 |
| Ppip5k2 | Q6ZQB6-3 | 0.19 | 0.41 | -0.22 | -1.81 | 0.51 | 0.11 | 0.37 | -1.28 | 1 | 0.34 | 0.26 | 0.9 | 1.08 | -0.11 | -1.21 | -1.93 | 0.285 | 0.9825 | -0.185 | 0.632355 | 0.076397 | 0.804456 | 0.999971 | 1 | 0.971467 |
| Ppm1a | P49443 | 0.01 | 0.2 | 0.39 | 0.03 | 0.22 | -0.17 | 0.23 | 0.09 | 0.44 | 0.13 | 0.32 | 0.44 | 0.14 | -0.02 | 0.1 | 0.12 | -0.065 | 0.175 | -0.0725 | 0.736914 | 0.352421 | 0.685709 | 0.999971 | 1 | 0.944874 |
| Ppm1b | P36993 | -1.33 | -0.24 | 0.47 | 0.22 | 0 | -0.66 | 0.04 | 1.01 | 0.2 | -1.89 | 0.93 | 1.17 | -0.19 | -0.34 | 0.18 | -0.1 | 0.3175 | 0.3225 | 0.1075 | 0.521108 | 0.65943 | 0.786454 | 0.999971 | 1 | 0.967132 |
| Ppm1b_1 | Q99NF7 | 0.12 | 0.22 | -0.03 | 0.17 | 0.3 | 0.31 | 0.03 | 0.51 | 0.44 | 0.3 | 0.24 | 0.29 | 0.41 | 0.21 | 0.31 | 0.49 | 0.1675 | 0.1975 | 0.235 | 0.37696 | 0.249076 | 0.197231 | 0.999971 | 1 | 0.812891 |
| Ppm1e | Q80TL0 | 0.44 | 0.82 | 0.39 | 0.2 | 0.78 | 1.18 | 0.94 | 0.33 | 0.76 | 0.91 | 0.53 | -0.16 | 0.52 | 0.79 | 0.13 | 0.11 | 0.345 | 0.0475 | -0.075 | 0.190197 | 0.866702 | 0.757093 | 0.999971 | 1 | 0.961116 |
| Ppm1f | Q8CGA0 | -0.09 | 0.2 | 0 | 0.02 | 0.34 | 0.53 | -0.28 | 0.06 | -0.04 | 0 | -0.22 | -0.21 | 0.54 | 0.52 | 0.14 | -0.08 | 0.13 | -0.15 | 0.2475 | 0.569578 | 0.386814 | 0.263375 | 0.999971 | 1 | 0.842533 |
| Ppm1g | Q61074 | 2.9 | 2.94 | 2.92 | 2.92 | 2.91 | 2.62 | 3.57 | 2.65 | 2.81 | 3.24 | 2.73 | 2.77 | 1.89 | 2.37 | 2.62 | 1.58 | 0.0175 | -0.0325 | -0.805 | 0.943952 | 0.860706 | 0.011934 | 0.999971 | 1 | 0.613095 |
| Ppm1h | Q3UYC0 | -0.35 | -0.29 | -1.24 | -1.07 | -0.26 | -0.12 | -1.13 | -1.29 | -0.24 | -0.17 | -1.13 | -1.12 | -0.12 | -0.31 | -1.1 | -0.82 | 0.0375 | 0.0725 | 0.15 | 0.919796 | 0.837748 | 0.65315 | 0.999971 | 1 | 0.934005 |
| Ppm1k | Q8BXN7 | 0.52 | 0.5 | 1.23 | 0.88 | 0.09 | 0.54 | 0.39 | 1.2 | 0.1 | 0.56 | 0.34 | -0.64 | 0.58 | 0.66 | 1.28 | 1.44 | -0.2275 | -0.6925 | 0.2075 | 0.455485 | 0.052883 | 0.480724 | 0.999971 | 1 | 0.89485 |
| Ppm1l | Q8BHN0 | -0.26 | -0.25 | -0.08 | 0.6 | -0.08 | -0.59 | -0.03 | 0.09 | -0.28 | 0.06 | 0.08 | 0.03 | -0.42 | -0.38 | 0.14 | 0.53 | -0.155 | -0.03 | -0.035 | 0.571618 | 0.903392 | 0.910204 | 0.999971 | 1 | 0.991822 |
| Ppm1m | A0A8I4Q8Y | 5.46 | -0.38 | 3.47 | 3.29 | -1.13 | -0.19 | 3.55 | 3.63 | -1.26 | 1.84 | 5.44 | 5.64 | -4.5 | -3.04 | 4.05 | 1.92 | -1.495 | -0.045 | -3.3525 | 0.345091 | 0.980292 | 0.135197 | 0.999971 | 1 | 0.784948 |
| Ppme1 | Q8BVQ5 | 0.19 | 0.97 | 1.01 | 1.08 | 0.76 | 0.96 | 1.06 | 0.59 | 0.73 | 0.52 | 0.96 | 0.98 | 0.42 | 0.79 | 0.84 | 0.97 | 0.03 | -0.015 | -0.0575 | 0.907349 | 0.953555 | 0.826971 | 0.999971 | 1 | 0.976804 |
| Ppox | P51175 | 0.26 | -1.25 | -0.5 | -0.35 | -1.11 | -1.24 | -0.62 | -0.35 | -1.55 | -0.18 | -0.74 | -0.67 | -1.34 | -1.04 | -0.21 | -0.39 | -0.37 | -0.325 | -0.285 | 0.324039 | 0.428804 | 0.475487 | 0.999971 | 1 | 0.894045 |
| Ppp1ca | P62137 | 0.12 | 0.75 | 1.76 | 0.87 | 0.8 | 0.55 | 1.36 | 1.71 | 1.16 | 0.2 | 1.56 | -0.28 | 1.78 | 0.72 | 1.88 | 1.11 | 0.23 | -0.215 | 0.4975 | 0.576289 | 0.671497 | 0.25094 | 0.999971 | 1 | 0.839012 |
| Ppp1cb | P62141 | -0.17 | 0.52 | 1.28 | 0.65 | 0.46 | 0.65 | 0.8 | 0.94 | 0.5 | -0.08 | 1.08 | 1.09 | 1.04 | 0.42 | 1.28 | 0.85 | 0.1425 | 0.0775 | 0.3275 | 0.655261 | 0.8434 | 0.354797 | 0.999971 | 1 | 0.868793 |
| Ppp1cc | P63087-2 | -3 | -4.67 | -1.31 | -0.88 | -3.55 | -4.77 | -1.24 | -0.63 | -3.84 | -3.92 | -0.52 | -0.51 | -3.92 | -4.12 | -0.46 | -0.51 | -0.0825 | 0.2675 | 0.2125 | 0.943122 | 0.818722 | 0.858538 | 0.999971 | 1 | 0.982702 |
| Ppp1r10 | Q80W00 | 0.87 | 1.55 | 1.92 | 2.14 | 1.7 | 1.09 | 2.09 | 1.74 | 1.42 | 0.82 | 1.94 | 1.42 | 1.17 | 1.26 | 1.31 | 1 | 0.035 | -0.22 | -0.435 | 0.918606 | 0.538199 | 0.167632 | 0.999971 | 1 | 0.800215 |
| Ppp1r11 | G3UZ30 | -0.86 | -0.05 | -0.89 | -0.61 | -0.12 | -1.27 | -0.59 | -1.08 | 0.11 | -1.29 | -0.76 | -0.49 | -0.27 | -0.63 | -1.36 | -0.97 | -0.1625 | -0.005 | -0.205 | 0.61874 | 0.988438 | 0.513468 | 0.999971 | 1 | 0.905815 |
| Ppp1r12a | Q9DBR7 | 0.34 | -0.04 | 0.22 | 0.2 | 0.4 | 0.29 | 0.82 | 0.1 | 0.24 | 0.45 | 0.59 | 0.64 | -0.16 | -0.09 | 0.19 | 0.29 | 0.2225 | 0.3 | -0.1225 | 0.320937 | 0.133086 | 0.538112 | 0.999971 | 1 | 0.910077 |
| Ppp1r12a_ | A0A1W2P7 | 5.12 | -0.37 | 1.23 | 2.19 | 0.55 | 1.36 | 3.04 | 0.66 | 0.22 | 3.27 | -0.68 | -1.39 | -2.61 | -1.2 | -1.21 | -1.91 | -0.64 | -1.6875 | -3.775 | 0.57998 | 0.242126 | 0.006388 | 0.999971 | 1 | 0.582714 |
| Ppp1r12b | Q8BG95 | -0.02 | -0.3 | 0.3 | 0.18 | -0.09 | -0.01 | 1.25 | 0.29 | -0.2 | -0.09 | 0.4 | 0.27 | 0.15 | 0.05 | 0.26 | 0.56 | 0.32 | 0.055 | 0.215 | 0.349546 | 0.810651 | 0.337409 | 0.999971 | 1 | 0.858827 |
| Ppp1r12c | F6XWD4 | 2.02 | 0.56 | -0.4 | -0.1 | 1.01 | -0.13 | -0.09 | 0.25 | 0.07 | 1.25 | 0.41 | 0.3 | 0 | 0.54 | 0.09 | 0.47 | -0.26 | -0.0125 | -0.245 | 0.639018 | 0.981899 | 0.635908 | 0.999971 | 1 | 0.93301 |
| Ppp1r12c_ | Q3UMT1 | 0.8 | 0.56 | 0.34 | 0.54 | 0.61 | 0.73 | 0.42 | 0.64 | 0.59 | 0.78 | 0.69 | 0.58 | 0.49 | 0.6 | 0.3 | 0.16 | 0.04 | 0.1 | -0.1725 | 0.829919 | 0.579063 | 0.393168 | 0.999971 | 1 | 0.871816 |
| Ppp1r13b | Q62415 | 0.87 | -0.36 | -0.59 | -0.25 | -0.5 | -0.08 | -0.45 | -0.33 | -0.25 | -0.08 | -0.36 | -0.29 | -0.39 | -0.45 | -0.57 | -0.61 | -0.2575 | -0.1625 | -0.4225 | 0.451329 | 0.624619 | 0.221538 | 0.999971 | 1 | 0.8275 |
| Ppp1r14a | Q91VC7 | 0.84 | 0.35 | 0.47 | -0.01 | 0.51 | 0.97 | -0.06 | -0.45 | 0.59 | 1.18 | -0.26 | -0.19 | 0.81 | 1.13 | 0.1 | 0.03 | -0.17 | -0.0825 | 0.105 | 0.630871 | 0.825291 | 0.746121 | 0.999971 | 1 | 0.961064 |
| Ppp1r14b | Q62084 | 1.28 | 0.69 | 1.52 | 1.63 | 0.38 | 0.78 | 0.93 | 2.14 | 0.55 | 0.82 | 1.5 | -1.29 | 0.28 | 0.87 | 1.95 | 1.63 | -0.2225 | -0.885 | -0.0975 | 0.592153 | 0.154324 | 0.812658 | 0.999971 | 1 | 0.974458 |
| Ppp1r14c | Q8R4S0 | 0.67 | 2.11 | -0.23 | 0.83 | 2.09 | 2.41 | 3.56 | -1.05 | 2.86 | 1.66 | 0.17 | -0.63 | 3.34 | 2.28 | -0.89 | -0.35 | 0.9075 | 0.17 | 0.25 | 0.366324 | 0.83704 | 0.804233 | 0.999971 | 1 | 0.971467 |
| Ppp1r16a | Q923M0 | -0.41 | 0.43 | -0.36 | -0.29 | 1.02 | 0.26 | -0.53 | -0.03 | 0.83 | -0.29 | -0.2 | 0.07 | 1.55 | 0.06 | -0.24 | -0.16 | 0.3375 | 0.26 | 0.46 | 0.37147 | 0.429597 | 0.310006 | 0.999971 | 1 | 0.847402 |
| Ppp1r16b | Q8VHQ3 | -2.06 | -2.47 | -1.75 | -1.62 | -2.74 | -2.56 | -1.74 | -1.98 | -2.64 | -1.97 | -2.04 | -2.25 | -2.91 | -3 | -2.34 | -2.17 | -0.28 | -0.25 | -0.63 | 0.375199 | 0.353621 | 0.055387 | 0.999971 | 1 | 0.725348 |
| Ppp1r18 | Q8BQ30-2 | 0.46 | -0.32 | 0.84 | 0.48 | 0.15 | -0.44 | 0.87 | 0.2 | 0.31 | 0.09 | 0.41 | 0.66 | 0.32 | -0.21 | 0.47 | 0.7 | -0.17 | 0.0025 | -0.045 | 0.634147 | 0.992975 | 0.886664 | 0.999971 | 1 | 0.987485 |
| Ppp1r1a | Q9ERT9 | 0.62 | 0.06 | 0.02 | 0.14 | -0.11 | 0.03 | 0.24 | 0.13 | -0.13 | 0.69 | 0.04 | 0.16 | 0.13 | 0.33 | 0.01 | 0.22 | -0.1375 | -0.02 | -0.0375 | 0.51576 | 0.936317 | 0.856881 | 0.999971 | 1 | 0.982642 |
| Ppp1r1b | Q60829 | 1.28 | 0.77 | 0.42 | 1.58 | 1.77 | -0.82 | 1.31 | 2.35 | 1.09 | 1.02 | -0.11 | 1.56 | 1.32 | 1.23 | 1.92 | 0.29 | 0.14 | -0.1225 | 0.1775 | 0.834178 | 0.769764 | 0.663512 | 0.999971 | 1 | 0.936542 |
| Ppp1r2 | D3Z3A0 | 0.77 | 0.68 | 0.18 | 0.66 | 1.75 | 1.08 | 0.4 | 0.47 | 1.81 | 0.89 | -0.32 | 0.05 | 1.6 | 1.21 | 0.37 | -0.12 | 0.3525 | 0.035 | 0.1925 | 0.312905 | 0.939647 | 0.629422 | 0.999971 | 1 | 0.932318 |
| Ppp1r21 | Q3TDD9 | -0.4 | -0.13 | -0.51 | -0.36 | -0.28 | -0.3 | -0.34 | -0.52 | -0.11 | -0.15 | -0.45 | -0.51 | -0.48 | -0.13 | -0.61 | -0.29 | -0.01 | 0.045 | -0.0275 | 0.95522 | 0.813645 | 0.888206 | 0.999971 | 1 | 0.987485 |
| Ppp1r37 | Q8BKR5 | 1.1 | 2.08 | 1.46 | 1.29 | 1.96 | 1.65 | 0.95 | 1.09 | 1.83 | 1.28 | 1.45 | 1.37 | 2.41 | 1.53 | 1.14 | 1.44 | -0.07 | -1.57E-16 | 0.1475 | 0.827065 | 1 | 0.668288 | 0.999971 | 1 | 0.938711 |
| Ppp1r3d | A2AJW4 | -0.01 | -0.44 | -0.43 | -0.51 | -0.41 | 0.1 | -0.13 | -0.66 | 0.31 | 0.22 | -0.35 | -0.14 | -0.52 | -0.14 | -0.45 | -0.58 | 0.0725 | 0.3575 | -0.075 | 0.75889 | 0.14347 | 0.715864 | 0.999971 | 1 | 0.952576 |
| Ppp1r3f | Q9JIG4-2 | -0.55 | -0.04 | -0.01 | -0.1 | -0.49 | -0.4 | 0.4 | -0.48 | -0.56 | -0.51 | -0.18 | -0.59 | -0.49 | -0.35 | -0.64 | -0.84 | -0.0675 | -0.285 | -0.405 | 0.80216 | 0.191808 | 0.084931 | 0.999971 | 1 | 0.758256 |
| Ppp1r3g | Q9CW07 | -0.21 | 0.37 | 0.21 | 0.45 | 0.48 | 0.66 | 1.21 | -0.62 | 0.73 | 0.17 | 0.55 | 0.12 | 0.09 | 0 | -0.61 | -0.1 | 0.2275 | 0.1875 | -0.36 | 0.567078 | 0.441382 | 0.167859 | 0.999971 | 1 | 0.800215 |
| Ppp1r7 | Q3UM45 | 0.67 | 0.27 | -0.1 | 0.34 | 0.22 | 0.05 | -0.03 | 0.27 | 0.14 | 0.45 | -0.04 | -0.05 | -0.08 | 0.33 | 0.04 | 0.22 | -0.1675 | -0.17 | -0.1675 | 0.45029 | 0.468917 | 0.463201 | 0.999971 | 1 | 0.890823 |
| Ppp1r8 | A2ADR8 | 0.89 | 1.16 | 1.63 | 1.72 | 1.45 | 0.5 | 1.29 | 1.12 | 0.97 | 1.25 | 1.47 | 1.33 | 1.58 | 1.13 | 1.55 | 1.7 | -0.26 | -0.095 | 0.14 | 0.39018 | 0.70389 | 0.590174 | 0.999971 | 1 | 0.922217 |
| Ppp1r9a | H3BJD6 | -0.24 | -1.14 | -1.14 | -1.06 | -1.06 | -0.93 | -0.89 | -1.2 | -1.21 | -0.67 | -1.28 | -1.38 | -0.82 | -1.17 | -1.47 | -1.56 | -0.125 | -0.24 | -0.36 | 0.626935 | 0.40737 | 0.233308 | 0.999971 | 1 | 0.831526 |
| Ppp1r9b | Q6R891 | -0.23 | -0.76 | -0.06 | 0.09 | -1.12 | -0.73 | -0.29 | 0 | -0.85 | -0.2 | -0.27 | 0.07 | -0.48 | -0.8 | -0.35 | 0.28 | -0.295 | -0.0725 | -0.0975 | 0.357588 | 0.797011 | 0.746901 | 0.999971 | 1 | 0.961064 |
| Ppp2ca | P63330 | 0.31 | 1.57 | 0.69 | 0.63 | 1.18 | 1.2 | 0.74 | 0.78 | 1.75 | 0.99 | 0.72 | 0.1 | 1.8 | 1.48 | 0.48 | 0.21 | 0.175 | 0.09 | 0.1925 | 0.568736 | 0.828486 | 0.664912 | 0.999971 | 1 | 0.93697 |
| Ppp2cb | P62715 | 0.01 | 0.01 | 0.62 | 0.44 | 0.13 | -0.08 | 0.67 | 0.64 | 0.16 | -0.27 | 0.68 | 0.03 | 0.15 | -0.09 | 0.81 | 0.33 | 0.07 | -0.12 | 0.03 | 0.791322 | 0.657702 | 0.910435 | 0.999971 | 1 | 0.991836 |
| Ppp2r1a | Q76MZ3 | -0.24 | 0.46 | -0.26 | -0.09 | 0.13 | 0.03 | -0.36 | -0.09 | 0.05 | 0.04 | -0.3 | -0.42 | 0.33 | 0.27 | -0.22 | 0.01 | -0.04 | -0.125 | 0.13 | 0.86458 | 0.601966 | 0.595595 | 0.999971 | 1 | 0.92377 |
| Ppp2r1a_1 | G3UWL2 | -1.98 | 2.99 | 3.68 | 0.85 | 1.11 | 4.53 | 1.15 | 2.17 | 1.52 | -0.11 | 2.63 | 1.91 | 0.75 | 3.31 | -0.14 | 3.18 | 0.855 | 0.1025 | 0.39 | 0.526775 | 0.934729 | 0.776151 | 0.999971 | 1 | 0.964396 |
| Ppp2r1b | H3BK50 | -0.41 | 1.67 | 3.18 | 1.99 | 0.16 | 3.09 | 1.54 | 1.48 | 0.75 | 1.62 | 1.89 | 0.68 | 1.92 | 2.55 | 1.98 | 3.05 | -0.04 | -0.3725 | 0.7675 | 0.962775 | 0.6138 | 0.300683 | 0.999971 | 1 | 0.847402 |
| Ppp2r1b_1 | Q3TTF6 | -0.86 | -0.5 | -0.7 | -0.4 | -1.07 | -0.25 | -0.6 | -0.27 | -1.23 | -0.32 | -0.36 | -0.59 | -1.26 | -0.74 | -0.28 | 0.15 | 0.0675 | -0.01 | 0.0825 | 0.784856 | 0.968919 | 0.798184 | 0.999971 | 1 | 0.970479 |
| Ppp2r2a | Q6P1F6 | 0.1 | 0.54 | -0.01 | 0.18 | 0.36 | 0.28 | 0.23 | -0.02 | 0.58 | 0.12 | 0.3 | 0.59 | 0.35 | 0.03 | 0.04 | -0.13 | 0.01 | 0.195 | -0.13 | 0.960396 | 0.367562 | 0.537595 | 0.999971 | 1 | 0.910077 |
| Ppp2r2b | A0A668KL/ | -0.76 | 0.39 | -0.48 | -0.54 | -0.37 | 0.73 | -0.54 | -0.4 | -0.01 | -0.76 | -0.31 | 0.09 | 0.7 | 0.33 | -0.45 | -0.06 | 0.2025 | 0.1 | 0.4775 | 0.592456 | 0.754426 | 0.195017 | 0.999971 | 1 | 0.811225 |
| Ppp2r2c | Q8BG02 | -0.01 | 0.39 | 0.97 | 0.55 | -0.13 | 0.47 | 1.15 | 0.98 | 0.17 | -0.39 | 1.16 | 1.25 | 0.62 | 0.58 | 1.5 | 0.46 | 0.1425 | 0.0725 | 0.315 | 0.682419 | 0.86409 | 0.334837 | 0.999971 | 1 | 0.858827 |
| Ppp2r2d | Q925E7 | -1.26 | -2.61 | -1.26 | -1.25 | -1.86 | -2.3 | -1.34 | -1.31 | -1.99 | -1.43 | -1.52 | -1.53 | -2.52 | -2 | -1.49 | -1.52 | -0.1075 | -0.0225 | -0.2875 | 0.78578 | 0.949298 | 0.47793 | 0.999971 | 1 | 0.894438 |
| Ppp2r5a | Q6PD03 | -0.57 | -0.14 | 0 | 0 | -0.1 | -0.43 | -0.09 | -0.11 | 0.04 | -0.45 | 0.09 | 0.05 | 0.18 | -0.65 | 0 | 0.1 | -0.005 | 0.11 | 0.085 | 0.980923 | 0.625922 | 0.742323 | 0.999971 | 1 | 0.961064 |
| Ppp2r5b | Q6PD28 | -0.32 | 0.53 | 0.3 | -1.08 | 0.32 | -0.01 | 1.37 | -0.97 | 0.83 | -0.35 | 1.07 | 1.29 | 1.02 | 0.15 | -0.59 | -1.12 | 0.32 | 0.8525 | 0.0075 | 0.566451 | 0.103858 | 0.988949 | 0.999971 | 1 | 0.99925 |
| Ppp2r5c | Q60996-3 | -1.2 | -0.47 | -0.48 | -0.62 | -0.37 | -0.86 | -0.3 | -0.68 | -0.23 | -1.32 | -0.13 | 0.11 | 0.08 | -1.05 | -0.5 | -0.52 | 0.14 | 0.3 | 0.195 | 0.573672 | 0.407112 | 0.518205 | 0.999971 | 1 | 0.905815 |
| Ppp2r5c_1 | A0A1Y7VIR | -0.49 | -1.62 | -0.15 | -0.43 | -1.74 | -0.9 | 1.39 | -0.54 | -1.4 | -1.14 | -0.25 | -0.85 | -2.67 | -2.58 | -1.4 | -1.91 | 0.225 | -0.2375 | -1.4675 | 0.736886 | 0.549157 | 0.00661 | 0.999971 | 1 | 0.582714 |
| Ppp2r5d | Q91V89 | -1.1 | -0.25 | -1.04 | -0.7 | -0.09 | -0.96 | -0.58 | -0.9 | 0.05 | -0.99 | -0.84 | -0.76 | 0.62 | -0.71 | -0.71 | -0.6 | 0.14 | 0.1375 | 0.4225 | 0.63197 | 0.658069 | 0.270022 | 0.999971 | 1 | 0.843794 |
| Ppp2r5e | Q61151 | -1.2 | -0.3 | -0.33 | -0.38 | -0.24 | -0.69 | -0.59 | -0.19 | -0.48 | -1.26 | -0.05 | -0.33 | 0.02 | -0.54 | -0.09 | -0.11 | 0.125 | 0.0225 | 0.3725 | 0.644558 | 0.946461 | 0.190593 | 0.999971 | 1 | 0.810547 |
| Ppp3ca | P63328 | -0.77 | 0.46 | -0.25 | -0.07 | 0.28 | 0.25 | -0.5 | -0.39 | -0.06 | -0.62 | -0.49 | -0.39 | 0.42 | 0.07 | -0.36 | -0.07 | 0.0675 | -0.2325 | 0.1725 | 0.836565 | 0.433024 | 0.578551 | 0.999971 | 1 | 0.919106 |
| Ppp3ca_1 | P63328-2 | 0.53 | 0.48 | 0.92 | 0.43 | -0.24 | 0.79 | 0.02 | 0.6 | -0.04 | 0.53 | 0.45 | -0.06 | -0.63 | 0.07 | 0.34 | 0.3 | -0.2975 | -0.37 | -0.57 | 0.305754 | 0.132391 | 0.060622 | 0.999971 | 1 | 0.725348 |
| Ppp3cb | E0CZ78 | -0.28 | 0.06 | 0.31 | -0.4 | -0.38 | -0.22 | -0.33 | -0.4 | -0.18 | -0.15 | -0.1 | -0.08 | 0.31 | -0.16 | -0.18 | -0.03 | -0.255 | -0.05 | 0.0625 | 0.2534 | 0.811954 | 0.789895 | 0.999971 | 1 | 0.968012 |
| Ppp3cc | A0A2I3BPC | -0.12 | -0.35 | -0.38 | -0.29 | -0.6 | -0.56 | -0.54 | -0.07 | -0.43 | -0.32 | 0.35 | -0.51 | -0.26 | -0.59 | -0.64 | -0.24 | -0.1575 | 0.0575 | -0.1475 | 0.434429 | 0.808179 | 0.445313 | 0.999971 | 1 | 0.884152 |
| Ppp3r1 | Q63810-2 | -2.46 | 0.47 | -0.33 | -0.59 | 0.21 | -0.13 | -0.41 | -0.67 | 0.04 | -1.9 | -0.08 | -0.58 | 0.41 | -0.52 | -0.58 | -1.01 | 0.4775 | 0.0975 | 0.3025 | 0.428594 | 0.888275 | 0.631781 | 0.999971 | 1 | 0.932858 |
| Ppp4c | P97470 | 0.84 | 1.44 | 0.34 | 0.1 | 1.42 | 1.15 | 0.18 | 0.45 | 1.39 | 1.11 | 0.34 | 0.46 | 1.42 | 1.32 | 0.67 | 0.43 | 0.12 | 0.145 | 0.28 | 0.763043 | 0.702072 | 0.459425 | 0.999971 | 1 | 0.889191 |
| Ppp4r1 | E9PX53 | 1.41 | 0.53 | 0.82 | 0.65 | 0.63 | 0.58 | 0.79 | 0.89 | 0.35 | 1.25 | 0.73 | 0.16 | -0.05 | 0.78 | 0.76 | 0.99 | -0.13 | -0.23 | -0.2325 | 0.591679 | 0.469943 | 0.457594 | 0.999971 | 1 | 0.88791 |
| Ppp4r2 | A0A0R4J0L | 2.68 | 2.41 | 2.98 | 2.73 | 1.92 | 2.41 | 2.24 | 2.85 | 2.03 | 2.75 | 2.47 | 1.78 | 1.42 | 2.22 | 2.8 | 2.43 | -0.345 | -0.4425 | -0.4825 | 0.195643 | 0.12361 | 0.154832 | 0.999971 | 1 | 0.795578 |
| Ppp4r3a | E9Q481 | 1.18 | 2.61 | 2.61 | 2.65 | 2.58 | 2.24 | 1.87 | 2.7 | 2.37 | 1.94 | 2.53 | 2.53 | 2.14 | 2.52 | 2.35 | 2.85 | 0.085 | 0.08 | 0.2025 | 0.827573 | 0.831247 | 0.595599 | 0.999971 | 1 | 0.92377 |
| Ppp4r4 | Q8C0Y0 | 0.03 | 0.26 | 0.56 | 0.35 | 0.31 | 0.19 | 0.25 | 0.61 | 0.21 | -0.26 | 0.4 | 0.2 | 0.13 | 0.38 | 0.62 | 0.59 | 0.04 | -0.1625 | 0.13 | 0.842521 | 0.465598 | 0.539881 | 0.999971 | 1 | 0.910077 |

| | | | | | | | | | | | | | | | | | | | | | | | | | | |
|---|---|---|---|---|---|---|---|---|---|---|---|---|---|---|---|---|---|---|---|---|---|---|---|---|---|---|
| Ppp5c | Q60676 | -0.78 | 0.72 | 0.77 | 0.51 | 0.6 | 0.85 | 0.6 | 0.71 | -0.29 | -0.64 | -0.06 | -1.41 | -0.55 | -0.32 | -0.06 | -0.12 | 0.385 | -0.905 | -0.5675 | 0.302963 | 0.068122 | 0.152857 | 0.999971 | 1 | 0.795578 |
| Ppp6c | Q9CQR6 | -0.57 | -1.21 | -0.1 | -0.45 | -1.12 | -1.03 | -0.23 | -0.32 | -0.62 | -0.96 | -0.06 | -0.2 | -1.42 | -1.13 | -0.23 | -0.29 | -0.0925 | 0.1225 | -0.185 | 0.778259 | 0.696627 | 0.618068 | 0.999971 | 1 | 0.928581 |
| Ppp6r1 | Q7TSI3 | 0.02 | 0.53 | 0.34 | 0.43 | 1.07 | 0.35 | 0.78 | 0.56 | 0.74 | -0.22 | 0.4 | 0.52 | 0.59 | 0.23 | 0.38 | 0.77 | 0.36 | 0.03 | 0.1625 | 0.141378 | 0.906845 | 0.451393 | 0.999971 | 1 | 0.886993 |
| Ppp6r2 | Q8R3Q2 | -0.33 | 0.03 | 0.21 | 0.02 | 0 | 0.02 | 0.09 | 0.26 | -0.07 | -0.45 | 0.06 | -0.03 | 0.19 | 0.09 | 0.25 | 0.11 | 0.11 | -0.105 | 0.1775 | 0.571231 | 0.615527 | 0.359865 | 0.999971 | 1 | 0.869526 |
| Ppp6r3 | G5E8R4 | 0.06 | 0.55 | 0.17 | 0.09 | 0.38 | 0.27 | -0.02 | 0.12 | 0.32 | 0.03 | 0.3 | 0.52 | 0.6 | 0.24 | 0.15 | 0.35 | -0.03 | 0.075 | 0.1175 | 0.881147 | 0.71259 | 0.56997 | 0.999971 | 1 | 0.91816 |
| Ppp6r3_1 | A0A286YC( | -0.05 | 1.01 | 0.58 | 1.62 | -0.44 | 0.09 | 0.63 | 1.23 | 0.15 | -1.72 | 0.38 | 1.84 | 1.31 | -1.06 | 1.22 | 1.43 | -0.4125 | -0.6275 | -0.065 | 0.389974 | 0.404328 | 0.917895 | 0.999971 | 1 | 0.992633 |
| Ppt1 | O88531 | -0.52 | 0.11 | -0.31 | -0.35 | 1.08 | 1.39 | 0.26 | 0.63 | 0.25 | -0.61 | -0.46 | -0.23 | 1.67 | 1.24 | 0.65 | 1.2 | 1.1075 | 0.005 | 1.4575 | 0.00425 | 0.98431 | 0.000456 | 0.999971 | 1 | 0.312038 |
| Pptc7 | Q6NVE9 | -1.43 | 0.25 | 0.78 | 0.37 | 0.53 | 0.18 | 0.82 | 1.19 | 0.71 | -0.84 | 0.94 | 1.39 | 0.9 | 0.03 | 1.02 | 0.49 | 0.6875 | 0.5575 | 0.6175 | 0.188166 | 0.386663 | 0.23624 | 0.999971 | 1 | 0.832918 |
| Ppwd1 | Q8CEC6 | 0.94 | 1.3 | -0.44 | -0.7 | 1.02 | 0.81 | -0.33 | -0.3 | 0.89 | 0.63 | -0.48 | -0.35 | -0.01 | 1.16 | -0.38 | -0.53 | 0.025 | -0.1025 | -0.215 | 0.964414 | 0.854119 | 0.709236 | 0.999971 | 1 | 0.950625 |
| Pqbp1 | Q91VJ5 | 1.53 | 1.97 | 1.38 | 1.59 | 2 | 1.04 | 0.69 | 1.51 | 1.47 | 1.59 | 1.23 | 1.32 | 1.65 | 1.7 | 1.43 | 1.89 | -0.3075 | -0.215 | 0.05 | 0.34231 | 0.301719 | 0.811415 | 0.999971 | 1 | 0.973864 |
| Pradc1 | A0A0N4SV. | -4.3 | -0.8 | -1.87 | -1.77 | 0.87 | -1.18 | 7.39 | -11.01 | 2.68 | -4.26 | -0.53 | 0.54 | 2.74 | -1.95 | -3.52 | -1.2 | 1.2025 | 1.7925 | 1.2025 | 0.726202 | 0.239672 | 0.3863 | 0.999971 | 1 | 0.871816 |
| Praf2 | Q9JIG8 | 0.26 | -0.56 | -0.19 | -0.24 | -0.27 | 0.39 | -0.36 | 0.3 | -0.12 | 0.65 | 0.04 | 0.33 | 0.46 | 0.29 | 0.07 | 0 | 0.1975 | 0.4075 | 0.3875 | 0.477184 | 0.144361 | 0.126204 | 0.999971 | 1 | 0.783071 |
| Prag1 | Q571I4 | 0.77 | -0.46 | -0.18 | -0.48 | -0.59 | -0.02 | -0.38 | -0.61 | -0.76 | 0.34 | -0.38 | -0.79 | -0.44 | -0.49 | -0.82 | -0.67 | -0.3125 | -0.31 | -0.5175 | 0.34931 | 0.425405 | 0.124284 | 0.999971 | 1 | 0.78268 |
| Prcc | Q9EQC8 | 3.48 | 2.3 | 2.89 | 3.24 | 1.83 | 2.07 | 3.16 | 2.48 | 1.74 | 2.45 | 2.35 | 1.96 | 0.64 | 1.9 | 2.33 | 2.67 | -0.5925 | -0.8525 | -1.0925 | 0.140753 | 0.021484 | 0.045493 | 0.999971 | 1 | 0.710102 |
| Prcp | Q7TMR0 | 0.23 | 1.59 | 1.09 | 1 | 0.69 | 0.68 | 0.68 | 1.15 | 0.18 | 0.68 | 0.85 | 0.4 | 0.63 | 0.45 | 0.28 | 0.79 | -0.1775 | -0.45 | -0.44 | 0.569916 | 0.18356 | 0.178133 | 0.999971 | 1 | 0.808532 |
| Prdm2 | A2A7B5 | 4.13 | -0.33 | -1.6 | -0.73 | 0.27 | -0.06 | -1.01 | -1.18 | -0.38 | 3.8 | -1.69 | -1.19 | -0.2 | 1.06 | -1.75 | -0.58 | -0.8625 | -0.2325 | -0.735 | 0.472588 | 0.884367 | 0.560877 | 0.999971 | 1 | 0.91591 |
| Prdx1 | P35700 | -0.16 | -0.15 | 0.24 | 0.44 | -0.05 | -0.49 | 0.21 | 0.97 | -0.07 | -0.44 | 0.49 | 0.7 | 0.07 | -0.07 | 0.67 | 1.1 | 0.0675 | 0.0775 | 0.35 | 0.841518 | 0.800036 | 0.282945 | 0.999971 | 1 | 0.844538 |
| Prdx2 | Q61171 | 0.19 | 0.34 | 0.48 | 0.52 | 0.5 | 0.68 | 0.34 | 0.48 | 0.28 | 0.33 | 0.24 | 0.41 | -0.01 | 0.92 | 0.56 | 0.65 | 0.1175 | -0.0675 | 0.1475 | 0.520338 | 0.691468 | 0.546631 | 0.999971 | 1 | 0.911398 |
| Prdx3 | P20108 | -0.75 | -0.49 | -0.22 | -0.42 | -0.6 | -0.38 | -0.25 | 0.08 | -0.81 | -0.83 | 0 | 0.39 | -0.28 | -0.42 | 0.12 | 0.01 | 0.1825 | 0.1575 | 0.3275 | 0.420441 | 0.6287 | 0.153066 | 0.999971 | 1 | 0.795578 |
| Prdx4 | O08807 | -0.35 | -0.65 | -0.55 | -0.11 | -0.26 | -0.24 | -0.4 | 0 | -0.33 | -0.02 | -0.19 | -0.17 | -0.1 | 0.09 | 0.2 | 0.26 | 0.19 | 0.2375 | 0.5275 | 0.359319 | 0.240507 | 0.02621 | 0.999971 | 1 | 0.678474 |
| Prdx5 | P99029-2 | -1 | -0.23 | 0.2 | -0.22 | -0.21 | -0.66 | 0.73 | -0.02 | -0.54 | -1.11 | 0.09 | 0.67 | 0.16 | -0.51 | -0.06 | -0.32 | 0.2725 | 0.09 | 0.13 | 0.469685 | 0.836187 | 0.665743 | 0.999971 | 1 | 0.937078 |
| Prdx6 | Q6GT24 | -1.37 | -1.96 | -0.07 | -0.05 | -1.6 | -1.93 | -0.1 | -0.03 | -1.79 | -2.32 | -0.21 | 0.14 | -2.56 | -1.86 | -0.21 | 0.03 | -0.0525 | -0.1825 | -0.2875 | 0.93318 | 0.793629 | 0.689199 | 0.999971 | 1 | 0.945805 |
| Preb | Q9WUQ2 | 0.32 | -0.16 | -0.02 | -0.07 | -0.19 | -0.01 | 0.09 | 0.12 | -0.41 | 0.17 | 0.08 | -0.09 | -0.37 | 0.04 | -0.03 | 0.26 | -0.015 | -0.08 | -0.0425 | 0.93761 | 0.705809 | 0.843673 | 0.999971 | 1 | 0.979814 |
| Prelid3a | A0A494B9I | -0.24 | -0.64 | -0.41 | 0.16 | -0.64 | -0.65 | -0.09 | 0.19 | -0.66 | -0.49 | -0.3 | -0.7 | -0.11 | -0.13 | -0.34 | -0.02 | -0.015 | -0.255 | 0.1325 | 0.957652 | 0.281021 | 0.558527 | 0.999971 | 1 | 0.915051 |
| Prelp | Q9JK53 | 0.81 | 1.79 | 0.12 | 0.7 | 5 | 1.41 | 0.28 | -0.91 | 1.46 | 2.17 | 0.63 | 2.29 | -1.06 | 2.01 | 0.8 | -0.76 | 0.59 | 0.7825 | -0.6075 | 0.618189 | 0.132296 | 0.407779 | 0.999971 | 1 | 0.874911 |
| Prep | Q9QUR6 | -0.01 | 0.21 | 0.07 | 0.08 | 0.26 | 0.3 | 0 | 0.41 | 0.48 | 0.16 | 0.25 | 0.18 | 0.45 | 0.37 | 0.26 | 0.47 | 0.155 | 0.18 | 0.3 | 0.39624 | 0.307166 | 0.101063 | 0.999971 | 1 | 0.78268 |
| Prepl | Q8C167-2 | -0.05 | 1.44 | 0.96 | 0.49 | 1.41 | 1.54 | 0.82 | 1.07 | 1.32 | -0.11 | 0.76 | 0.86 | 2.31 | 1.74 | 1.43 | 1.13 | 0.5 | -0.0025 | 0.9425 | 0.17951 | 0.995204 | 0.037055 | 0.999971 | 1 | 0.700096 |
| Prex1 | Q69ZK0 | -0.46 | -0.15 | 0.19 | -0.11 | 0 | -0.5 | 0.45 | -0.21 | 0.16 | -0.23 | 0.41 | 0.56 | 0.17 | -0.41 | 0.08 | 0.05 | 0.0675 | 0.3575 | 0.105 | 0.797287 | 0.170569 | 0.645372 | 0.999971 | 1 | 0.93301 |
| Prex2 | Q3LAC4 | -1.02 | -0.42 | -0.69 | -1.16 | -0.4 | -0.51 | -0.5 | -0.08 | -0.21 | -0.65 | -0.47 | -0.53 | 0.36 | -0.58 | -1.29 | -0.9 | 0.45 | 0.3575 | 0.22 | 0.078749 | 0.142177 | 0.5637 | 0.999971 | 1 | 0.917087 |
| Prickle1 | Q3U5C7 | 0.1 | -1.49 | -1.65 | -1.95 | -1.71 | -1.02 | -1.57 | -1.33 | -1.57 | -0.4 | -1.47 | -1.91 | -1.26 | -1.11 | -1.84 | -1.32 | -0.16 | -0.09 | -0.135 | 0.724889 | 0.863295 | 0.767703 | 0.999971 | 1 | 0.962472 |
| Prickle2 | A7YQ68 | -0.68 | -1 | -0.88 | -0.97 | -1.16 | -0.81 | -0.9 | -0.59 | -1.12 | -0.43 | -0.71 | -0.97 | -0.69 | -0.82 | -0.75 | -0.62 | 0.0175 | 0.075 | 0.1625 | 0.929568 | 0.725281 | 0.362152 | 0.999971 | 1 | 0.869526 |
| Prkaa1 | Q3TUQ7 | -1.08 | -0.41 | -0.53 | -0.65 | -0.07 | -0.66 | -0.91 | -0.58 | -0.15 | -0.86 | -0.61 | -0.65 | 0.16 | -0.55 | -0.84 | -0.73 | 0.1125 | 0.1 | 0.1775 | 0.660303 | 0.677634 | 0.535351 | 0.999971 | 1 | 0.910077 |
| Prkaa2 | Q8BRK8 | -0.64 | -0.44 | -0.48 | -0.9 | -0.4 | -0.39 | -0.12 | -0.71 | -0.29 | -0.47 | -0.28 | -0.42 | -0.28 | -0.58 | -0.52 | -0.37 | 0.21 | 0.25 | 0.1775 | 0.33152 | 0.195609 | 0.367011 | 0.999971 | 1 | 0.869526 |
| Prkab1 | Q9R078 | -0.47 | -0.35 | -0.47 | -0.72 | -0.27 | -0.52 | -0.53 | -0.49 | -0.47 | -1.1 | -0.48 | -0.19 | -0.07 | -0.54 | -0.71 | -0.83 | 0.05 | -0.0575 | -0.035 | 0.78055 | 0.809503 | 0.876715 | 0.999971 | 1 | 0.985129 |
| Prkab2 | Q6PAM0 | -2.29 | -2.45 | -1.36 | -1.44 | -1.49 | -2.78 | 0 | -1.3 | -1.24 | -2.34 | -0.95 | -0.59 | -1.33 | -2.53 | -1.4 | -1.18 | 0.4925 | 0.605 | 0.275 | 0.406314 | 0.197737 | 0.499347 | 0.999971 | 1 | 0.903717 |
| Prkaca | P05132 | 0.28 | -1.49 | -0.28 | -0.29 | -1.61 | -0.58 | 1.32 | -0.62 | -0.98 | -0.48 | -0.47 | -0.62 | -1.5 | -1.6 | -0.28 | -1.13 | 0.0725 | -0.1925 | -0.6825 | 0.911213 | 0.614069 | 0.155484 | 0.999971 | 1 | 0.795578 |
| Prkacb | P68181 | -0.09 | -0.26 | -0.61 | -0.19 | 0.2 | -0.72 | -0.46 | -0.36 | 0.39 | 0.36 | -0.54 | 0.02 | 0.5 | -0.47 | 0.1 | 0.17 | -0.0475 | 0.345 | 0.3625 | 0.850246 | 0.213495 | 0.182228 | 0.999971 | 1 | 0.808532 |
| Prkag1 | O54950 | -0.11 | 0.36 | 0.22 | -0.13 | 0.15 | 0.39 | -0.05 | 0.06 | 0.01 | -0.02 | 0.47 | 0.49 | 0.23 | 0.22 | 0.1 | 0.43 | 0.0525 | 0.1525 | 0.16 | 0.799669 | 0.501841 | 0.431499 | 0.999971 | 1 | 0.881643 |
| Prkag2 | Q91WG5 | -0.44 | -0.28 | -0.23 | -0.81 | -0.46 | -0.34 | -0.51 | -0.56 | -0.39 | -0.46 | -0.51 | -0.25 | -0.48 | -0.32 | -0.47 | -0.59 | -0.0275 | 0.0375 | -0.025 | 0.88992 | 0.849784 | 0.90102 | 0.999971 | 1 | 0.990759 |
| Prkar1a | Q9DBC7 | -1.28 | -1.92 | -0.1 | -0.03 | -1.76 | -1.87 | -0.27 | -0.22 | -1.66 | -1.79 | -0.08 | -0.59 | -2.6 | -2.04 | -0.01 | -0.01 | -0.1975 | -0.1975 | -0.3325 | 0.739101 | 0.730789 | 0.6549 | 0.999971 | 1 | 0.934061 |
| Prkar1b | P12849 | -0.03 | 1.18 | 1.44 | 0.36 | 0.66 | 0.78 | 1.15 | 0.2 | 1.04 | 1.48 | 1.1 | -0.65 | 0.58 | 0.23 | 0.56 | 0.65 | -0.04 | 0.005 | -0.2325 | 0.916477 | 0.992632 | 0.513193 | 0.999971 | 1 | 0.905815 |
| Prkar2a | Q8K1M3 | -0.87 | -0.05 | -0.65 | -1.2 | -0.08 | -0.04 | -0.57 | -1.11 | -0.01 | -0.99 | -0.78 | -0.63 | 0.16 | -0.41 | -0.65 | -1.43 | 0.2425 | 0.09 | 0.11 | 0.487127 | 0.780347 | 0.779947 | 0.999971 | 1 | 0.965847 |
| Prkar2b | P31324 | -0.44 | -0.18 | 0.15 | 0.05 | -0.15 | -0.53 | -0.17 | 0.22 | -0.18 | -0.38 | -0.05 | -0.07 | 0.3 | -0.26 | 0.37 | 0.25 | -0.0525 | -0.065 | 0.27 | 0.824318 | 0.749295 | 0.264007 | 0.999971 | 1 | 0.842533 |
| Prkca | Q4VA93 | 1.36 | 0.24 | 1.41 | 1.23 | -0.1 | 0.7 | 1.06 | 1.03 | -0.12 | 0.59 | 0.73 | -2.24 | -0.05 | -0.44 | 0.89 | 0.62 | -0.3875 | -1.32 | -0.805 | 0.31513 | 0.077404 | 0.067599 | 0.999971 | 1 | 0.733937 |
| Prkcb | P68404-2 | -0.28 | -0.03 | -0.76 | -1.06 | -0.39 | 0.03 | -0.52 | -1.01 | -0.48 | -0.21 | -0.72 | -1.13 | -0.45 | 0.07 | -0.97 | -1.4 | 0.06 | -0.1025 | -0.155 | 0.850546 | 0.74018 | 0.685471 | 0.999971 | 1 | 0.944874 |
| Prkcb_1 | P68404 | 0.12 | -1.22 | -0.87 | -1.62 | -1.83 | 0.54 | -0.33 | -1.13 | -2.14 | 0.05 | -1.41 | -2.32 | -1.82 | -1.53 | -1.72 | -2.75 | 0.21 | -0.5575 | -1.0575 | 0.717391 | 0.366078 | 0.035942 | 0.999971 | 1 | 0.699952 |
| Prkcd | P28867 | -0.68 | -0.08 | 0.09 | -0.41 | -0.25 | 0.34 | -0.26 | 0.24 | -0.27 | -0.39 | 0.11 | -0.12 | -0.34 | 0.04 | 0.1 | 0.05 | 0.2875 | 0.1025 | 0.2325 | 0.282459 | 0.664519 | 0.337378 | 0.999971 | 1 | 0.858827 |
| Prkce | P16054 | 0.4 | 0.62 | 0.55 | 0.31 | 0.55 | 0.81 | 0.37 | 0.6 | 0.71 | 0.42 | 0.49 | 0.68 | 0.78 | 0.53 | 0.64 | 0.49 | 0.1125 | 0.105 | 0.14 | 0.549816 | 0.555419 | 0.441113 | 0.999971 | 1 | 0.883386 |
| Prkcg | P63318 | -0.93 | -0.91 | 0.21 | -0.28 | -0.81 | -0.87 | -0.14 | 0.14 | -0.6 | -0.92 | 0.03 | -1.18 | -0.71 | -1.13 | -0.03 | 0.05 | 0.0575 | -0.19 | 0.0225 | 0.873617 | 0.608872 | 0.952832 | 0.999971 | 1 | 0.998056 |
| Prkch | P23298 | -0.8 | 0.51 | 0.82 | 0.36 | 1.78 | -0.66 | -0.23 | 0.74 | 0.54 | -0.22 | 0.57 | 0.89 | 2.65 | -0.09 | 0.73 | 0.83 | 0.185 | 0.2225 | 0.8075 | 0.755153 | 0.587438 | 0.213779 | 0.999971 | 1 | 0.825334 |
| Prkci | Q62074 | 0.06 | 0.4 | 0.96 | 0.53 | 0.32 | 0.16 | 1.2 | 1.16 | 0.39 | -0.64 | 1.14 | 1.03 | 0.21 | -0.25 | 0.67 | 0.44 | 0.2225 | -0.0075 | -0.22 | 0.505761 | 0.985944 | 0.448113 | 0.999971 | 1 | 0.885289 |
| Prkcq | Q02111 | 1.08 | -0.15 | 0.24 | 0.02 | 0.17 | 0.28 | 0.38 | -0.4 | 0.38 | 1 | -0.24 | -0.9 | -0.03 | 0.34 | 0.35 | -0.12 | -0.19 | -0.2375 | -0.1625 | 0.562158 | 0.609366 | 0.598619 | 0.999971 | 1 | 0.92377 |
| Prkcsh | O08795-2 | 0.32 | -0.14 | 0.3 | 0.25 | -0.15 | 0.41 | 0.05 | 0.44 | 0.01 | 0.44 | 0.21 | 0.57 | -0.29 | 0.97 | 0.76 | 0.4 | 0.005 | 0.125 | 0.2775 | 0.982032 | 0.557242 | 0.373975 | 0.999971 | 1 | 0.87031 |
| Prkcz | Q02956-2 | 0.22 | 1.78 | 2.21 | 1.79 | 1.61 | 2.14 | 1.46 | 2.22 | 1.63 | 1.04 | 2.41 | 2.67 | 2.44 | 1.74 | 2.51 | 2.4 | 0.3575 | 0.4375 | 0.7725 | 0.433868 | 0.420719 | 0.110719 | 0.999971 | 1 | 0.78268 |
| Prkd1 | Q62101 | -2.04 | -0.17 | 0.22 | -0.31 | -1.47 | -1.35 | 0.29 | -0.04 | 0.79 | -1.44 | 0.42 | 0.15 | 1.26 | -0.43 | -0.34 | -0.86 | -0.0675 | 0.555 | 0.4825 | 0.912225 | 0.397522 | 0.446791 | 0.999971 | 1 | 0.884581 |
| Prkdc | P97313 | 3.41 | 0.64 | 2.26 | 1.57 | -0.52 | 1.69 | 2.57 | 0.63 | -0.49 | 2.25 | 0.44 | -0.09 | -1.69 | -0.81 | 0.28 | -0.49 | -0.8775 | -1.4425 | -2.6475 | 0.287862 | 0.086009 | 0.002761 | 0.999971 | 1 | 0.572658 |
| Prkg1 | P0C605-2 | -0.47 | 0 | -0.28 | -0.28 | 1.22 | 0.97 | -0.43 | -0.22 | 0.18 | 0.37 | -0.24 | 0.24 | 0.07 | 0.38 | -0.54 | -0.26 | 0.6425 | 0.395 | 0.17 | 0.14043 | 0.087544 | 0.503013 | 0.999971 | 1 | 0.90459 |
| Prkg2 | E9QPH0 | -1.1 | -1.05 | -1.06 | -1.32 | -0.45 | -0.95 | -0.8 | -1.22 | -0.65 | -1.28 | -1.17 | -0.93 | -0.54 | -1.08 | -1.45 | -1.58 | 0.2775 | 0.125 | -0.03 | 0.224101 | 0.544599 | 0.90951 | 0.999971 | 1 | 0.991822 |
| Prkn | B8YP90 | -2.91 | -1.1 | -0.79 | -1.99 | -1.18 | -1.08 | 0.51 | -0.81 | -1.32 | -4.8 | -1.21 | 0.36 | -0.85 | -3.67 | -1.25 | -3.12 | 1.0575 | -0.045 | -0.525 | 0.088845 | 0.966344 | 0.494449 | 0.999971 | 1 | 0.901531 |
| Prkra | Q9WTX2 | 1.28 | 1.19 | 1.02 | 0.99 | 1.18 | 1 | 0.77 | 1.92 | 1.37 | 1.28 | 1.06 | 1.57 | 1.58 | 1.44 | 1.62 | 1.99 | 0.0975 | 0.2 | 0.5375 | 0.724073 | 0.305956 | 0.022294 | 0.999971 | 1 | 0.652043 |
| Prmt1 | Q9JIF0-2 | 0.29 | 0.36 | 0.47 | 0.49 | 0.33 | 0.23 | 0.52 | 0.58 | 0.53 | 0.12 | 0.75 | 0.46 | 0.08 | 0.43 | 0.55 | 0.25 | 0.0125 | 0.0625 | -0.075 | 0.943691 | 0.750005 | 0.688405 | 0.999971 | 1 | 0.945753 |
| Prmt2 | M0QW88 | 2.64 | 2.45 | 0.09 | -1.29 | 3.99 | 3.03 | 2.06 | 0.18 | 3.53 | 1.63 | 1.24 | 1.56 | 5.49 | 2.53 | 0.53 | -0.89 | 1.3425 | 1.0175 | 0.9425 | 0.247918 | 0.312589 | 0.531783 | 0.999971 | 1 | 0.910069 |
| Prmt3 | Q922H1 | -0.62 | 0.24 | 0.91 | 0.35 | 0.53 | -0.17 | 0.33 | -0.11 | 0.3 | -0.48 | 0.56 | 1.03 | 0.06 | 0.41 | 0.45 | 0.48 | -0.075 | 0.1325 | 0.13 | 0.831367 | 0.755528 | 0.695441 | 0.999971 | 1 | 0.947174 |
| Prmt5 | A0A0R4J04 | 0.52 | -0.28 | -0.44 | -0.58 | 0.07 | -0.27 | -0.12 | -0.28 | 0.24 | 0.47 | -0.45 | -0.75 | -0.26 | -0.4 | -0.4 | -0.71 | 0.045 | 0.0725 | -0.2475 | 0.870574 | 0.843681 | 0.387798 | 0.999971 | 1 | 0.871816 |
| Prmt7 | Q922X9 | 0.58 | 0.23 | 0.8 | 0.39 | 0.42 | 0 | 0.42 | 0.75 | 0.29 | 0.82 | 0.72 | 0.17 | 0.23 | 0.05 | 0.51 | 0.18 | -0.1025 | 7.85E-17 | -0.2575 | 0.661778 | 1 | 0.238345 | 0.999971 | 1 | 0.833594 |
| Prmt8 | Q6PAK3 | -0.77 | -0.64 | -1.27 | -1.89 | -0.37 | -0.42 | -1.25 | -1.41 | -0.36 | -0.92 | -0.75 | -0.78 | -0.46 | -0.63 | -1.28 | -1.38 | 0.28 | 0.44 | 0.205 | 0.467611 | 0.183063 | 0.570571 | 0.999971 | 1 | 0.91816 |
| Prmt9 | F8WIU7 | 0.14 | 0.76 | -0.19 | 0.11 | 0.81 | 0.85 | -0.54 | 0.37 | 0.57 | 0.34 | -0.07 | -0.13 | 0.25 | 1.36 | -0.16 | 0.18 | 0.1675 | -0.0275 | 0.2025 | 0.651306 | 0.920567 | 0.591407 | 0.999971 | 1 | 0.922643 |
| Prnp | P04925 | -0.14 | -0.27 | 0.59 | 0.12 | 0.13 | 0.46 | 0.23 | 0.53 | 0.31 | 0.01 | 0.11 | 0.32 | -0.04 | -0.08 | 0.8 | 0.33 | 0.2625 | 0.1125 | 0.1775 | 0.296656 | 0.636421 | 0.546544 | 0.999971 | 1 | 0.911398 |
| Prodh | Q9WU79 | -0.87 | -0.51 | -1.07 | -1.02 | -0.56 | -0.59 | -1.04 | -1.03 | -0.71 | -0.78 | -1.42 | -1.69 | -0.67 | -0.85 | -1.49 | -1.43 | 0.0625 | -0.2825 | -0.2425 | 0.781158 | 0.334824 | 0.370464 | 0.999971 | 1 | 0.869526 |
| Prom1 | Q8CDK8 | -1.1 | -1.08 | -2.17 | -2.31 | -1.1 | -1.13 | -2.22 | -2.12 | -1.06 | -1.2 | -2.33 | -2.22 | -0.89 | -1.32 | -2.03 | -2.38 | 0.0225 | -0.0375 | 0.01 | 0.957971 | 0.932718 | 0.98211 | 0.999971 | 1 | 0.99925 |
| Prorp | V9GXP4 | -0.97 | -0.83 | -0.58 | -1.09 | -0.47 | -0.84 | -1.21 | -0.68 | -0.7 | -1.21 | -1.22 | -0.23 | -0.56 | -0.96 | -0.72 | -0.44 | 0.0675 | 0.0275 | 0.1975 | 0.768858 | 0.920685 | 0.357867 | 0.999971 | 1 | 0.868793 |
| Prorsd1 | Q9D820 | -0.97 | -1.68 | -0.41 | -3.85 | -2.68 | -3.71 | 2.64 | -6.23 | -2.73 | -1.04 | 6.16 | -6.46 | -3.33 | -2.57 | -7.5 | -5.41 | -0.7675 | 0.71 | -2.975 | 0.668046 | 0.772418 | 0.032417 | 0.999971 | 1 | 0.683918 |
| Prpf18 | Q8BM39 | 2.11 | 1.96 | 1.35 | 1.95 | 1.88 | 1.96 | 0.34 | 1.73 | 1.17 | 2.01 | 1.66 | 1.49 | 0.94 | 1.57 | 1 | 1.73 | -0.365 | -0.26 | -0.5325 | 0.37226 | 0.337021 | 0.082335 | 0.999971 | 1 | 0.755254 |
| Prpf19 | Q99KP6-2 | 1.22 | 0.93 | 0.85 | 1.19 | 0.52 | 0.81 | 0.92 | 0.88 | 0.76 | 1.24 | 0.85 | 0.61 | 0.8 | 1.31 | 1.03 | 1.2 | -0.265 | -0.1825 | 0.0375 | 0.194101 | 0.395211 | 0.852955 | 0.999971 | 1 | 0.98187 |
| Prpf3 | Q922U1 | 2.29 | 0.62 | 1.21 | 1.02 | 0.67 | 0.53 | 0.59 | 0.71 | 0.47 | 1.32 | 0.61 | 0.6 | -0.01 | 0.45 | 0.79 | 0.67 | -0.66 | -0.535 | -0.81 | 0.088178 | 0.194504 | 0.060502 | 0.999971 | 1 | 0.725348 |
| Prpf31 | Q8CCF0 | 2.4 | 1.38 | 1.68 | 1.44 | 1.05 | 1.12 | 1.38 | 0.83 | 1.19 | 1.81 | 1.08 | 1.1 | 0.29 | 0.82 | 1.02 | 1.02 | -0.63 | -0.43 | -0.9375 | 0.045758 | 0.175526 | 0.011674 | 0.999971 | 1 | 0.613095 |
| Prpf38a | Q4FK66 | -1.63 | -0.63 | -0.26 | -0.36 | 0.01 | -0.89 | -0.75 | -0.51 | -0.34 | -0.79 | -1.13 | -0.52 | -0.86 | -0.22 | 0.59 | -0.24 | 0.185 | 0.025 | 0.5375 | 0.611226 | 0.943188 | 0.21397 | 0.999971 | 1 | 0.825334 |
| Prpf38b | Q80SY5 | 0.97 | 1.47 | 1.23 | 0.89 | 1.48 | 1.51 | 0.75 | 0.88 | 1.89 | 1.22 | 0.7 | 1.2 | 1.39 | 1.7 | 1.07 | 0.26 | 0.015 | 0.1125 | -0.035 | 0.954133 | 0.697857 | 0.916785 | 0.999971 | 1 | 0.992633 |
| Prpf39 | E9QJV4 | -0.35 | 1.3 | 2.86 | 2.69 | 0.88 | 0.95 | 1.9 | 3.37 | 0.41 | -0.4 | 2.1 | 3.05 | 1.57 | 1.29 | 2.63 | 3.51 | 0.15 | -0.335 | 0.625 | 0.859383 | 0.731459 | 0.449465 | 0.999971 | 1 | 0.885522 |
| Prpf4 | Q9DAW6 | 0.9 | 0.4 | 1.21 | 0.79 | 0.24 | 0.26 | 0.64 | 0.6 | 0.06 | 0.72 | 0.69 | 0.43 | 0.15 | 0.61 | 1.13 | 0.82 | -0.39 | -0.35 | -0.1475 | 0.123012 | 0.188279 | 0.602508 | 0.999971 | 1 | 0.924453 |
| Prpf40a | A0A1B0GR | 0.01 | 0.18 | 0.82 | 0.57 | 0.44 | 0.16 | 0.55 | 0.65 | 0.43 | 0.41 | 0.88 | 1.03 | 0.68 | 0.28 | 0.83 | 0.77 | 0.055 | 0.2925 | 0.245 | 0.821227 | 0.282944 | 0.341408 | 0.999971 | 1 | 0.860984 |
| Prpf40b | D3Z4N6 | 2.86 | 2.5 | 2.72 | 2.74 | 2.75 | 3.19 | 3.27 | 1.05 | 1.74 | 2.64 | 2.03 | 1 | 1.2 | 2.42 | 0.94 | 2.01 | -0.14 | -0.8525 | -1.0625 | 0.773828 | 0.033364 | 0.013216 | 0.999971 | 1 | 0.613095 |
| Prpf4b | Q61136 | 0.08 | -0.04 | 0.97 | 0.72 | 0.03 | -0.23 | 0.49 | 0.48 | -0.21 | 0.22 | 0.89 | 0.84 | 0.22 | 0.33 | 0.9 | 1.17 | -0.24 | 0.0025 | 0.2225 | 0.443828 | 0.994343 | 0.509892 | 0.999971 | 1 | 0.905815 |
| Prpf6 | Q91YR7 | 0.97 | 0.72 | 1.28 | 0.61 | 0.78 | 0.97 | 0.96 | 0.32 | 0.76 | 0.55 | 0.74 | 0.35 | 0.52 | 0.79 | 0.37 | 0.41 | -0.1375 | -0.295 | -0.3725 | 0.576113 | 0.200047 | 0.118788 | 0.999971 | 1 | 0.78268 |
| Prpf8 | Q99PV0 | 0.04 | 0.32 | 0.39 | 0.16 | 0.32 | 0.1 | 0.21 | -0.08 | 0.24 | 0.02 | 0.08 | 0.21 | 0.29 | 0.31 | 0.07 | 0.11 | -0.09 | -0.09 | -0.0325 | 0.632291 | 0.607621 | 0.856928 | 0.999971 | 1 | 0.982642 |
| Prph | G5E846 | 0.16 | 0.17 | -0.2 | -0.67 | -0.39 | -0.26 | -0.48 | -0.45 | 0 | 0.77 | -0.22 | -0.58 | -0.25 | -0.17 | -0.54 | -0.7 | -0.26 | 0.1275 | -0.28 | 0.290932 | 0.711107 | 0.295333 | 0.999971 | 1 | 0.847402 |
| Prps1l1 | Q8C5R8 | 4.41 | 1.68 | 0.14 | 1.74 | 0.41 | 4.66 | -0.27 | 1.88 | -0.88 | 5.95 | -1.17 | -1.79 | 0.7 | 5.77 | 1.1 | 2.87 | -0.3225 | -1.465 | 0.6175 | 0.796626 | 0.42583 | 0.6354 | 0.999971 | 1 | 0.93301 |
| Prps1l3 | G3UXL2 | -0.17 | 0.62 | 0.74 | 0.63 | 0.65 | 1.24 | 0.82 | 0.86 | 0.7 | 0.17 | 0.9 | 0.33 | 1.39 | 1.07 | 0.67 | 0.98 | 0.4375 | 0.07 | 0.5725 | 0.1261 | 0.804114 | 0.063065 | 0.999971 | 1 | 0.725348 |
| Prps2 | Q9CS42 | -0.27 | -0.08 | -0.25 | -0.42 | -0.19 | 0.09 | 0.02 | -0.03 | -0.29 | -0.45 | -0.05 | -0.61 | -0.19 | 0.44 | -0.03 | 0.21 | 0.2275 | -0.095 | 0.3625 | 0.218014 | 0.629068 | 0.108469 | 0.999971 | 1 | 0.78268 |
| Prpsap1 | B1AT82 | -0.58 | -0.37 | -0.15 | -0.04 | -0.37 | 0.12 | -0.01 | 0.25 | -0.47 | 0.38 | -0.06 | -0.14 | 0.28 | -0.01 | 0.26 | 0.04 | 0.2825 | 0.2125 | 0.4275 | 0.224617 | 0.389997 | 0.057805 | 0.999971 | 1 | 0.725348 |
| Prpsap2 | Q8R574 | -1 | -0.47 | -0.14 | 0.4 | -0.37 | -0.82 | -0.17 | 0.14 | -0.7 | -1.3 | -0.33 | -0.14 | -0.59 | -0.55 | 0.06 | 0.33 | -0.0025 | -0.315 | 0.115 | 0.99429 | 0.41331 | 0.751928 | 0.999971 | 1 | 0.961116 |
| Prr12 | E9PYL2 | 2.62 | 0.9 | 2.02 | 2.25 | 1.57 | 1.48 | 1.63 | 2.13 | 1.14 | 2.2 | 1.71 | 2.31 | -0.08 | 1.21 | 1.5 | 2.36 | -0.245 | -0.1075 | -0.7 | 0.52764 | 0.804048 | 0.243341 | 0.999971 | 1 | 0.837316 |
| Prr14 | Q7TPN9 | 0.09 | -1.06 | -0.71 | -0.63 | -0.09 | -0.87 | -1.08 | -0.82 | -0.74 | -1.76 | -0.87 | -1.05 | -0.64 | -0.28 | -1.14 | -0.46 | -0.1375 | -0.5275 | -0.0525 | 0.67324 | 0.137525 | 0.865583 | 0.999971 | 1 | 0.983318 |
| Prr18 | Q6PAN7 | 3.4 | -0.35 | 1.25 | 0.96 | 0.42 | 0.85 | 1.4 | 0.56 | 0.2 | 2.49 | 0.67 | -0.18 | -0.62 | -0.28 | 0.41 | 0.23 | -0.5075 | -0.52 | -1.38 | 0.491764 | 0.558952 | 0.087355 | 0.999971 | 1 | 0.763848 |
| Prr3 | Q811B5 | -1.84 | 0.55 | 1.23 | 0.44 | 1.2 | 0.58 | 0.37 | 0.48 | 1.38 | 0.01 | -0.22 | 1.91 | 0.25 | 0.08 | 0.94 | 1.4 | 0.5625 | 0.675 | 0.5725 | 0.383533 | 0.389715 | 0.401368 | 0.999971 | 1 | 0.871816 |
| Prr36 | E9PV26 | 1.39 | -0.92 | -0.07 | 0.16 | -0.77 | -0.23 | 0.92 | -0.51 | -0.38 | 0.79 | -0.21 | -0.86 | -0.89 | -0.8 | -0.33 | -0.36 | -0.2875 | -0.305 | -0.735 | 0.607396 | 0.579342 | 0.142058 | 0.999971 | 1 | 0.784967 |
| Prr7 | Q3V0I2 | -0.81 | -1.37 | -2.45 | -2.38 | -1.39 | -1.35 | -2.26 | -2.5 | -1.54 | -0.95 | -2.07 | -2.36 | -1.2 | -1.61 | -2.68 | -2.4 | -0.1225 | 0.0225 | -0.22 | 0.792282 | 0.962112 | 0.655264 | 0.999971 | 1 | 0.934061 |
| Prrc1 | Q3UPH1 | -0.65 | 0.57 | 0.57 | 0.26 | 0.75 | 0.89 | 0.69 | 0.69 | 0.75 | 0.64 | 1.08 | 0.73 | 0.06 | 0.51 | 0.42 | 0.64 | 0.5675 | 0.6125 | 0.22 | 0.086119 | 0.07442 | 0.494905 | 0.999971 | 1 | 0.901531 |
| Prrc2a | Q7TSC1 | 0.86 | 0.18 | 0.79 | 0.64 | 0.33 | 0.55 | 0.71 | 1.17 | 0.33 | 0.78 | 0.83 | 0.99 | 0.32 | 0.32 | 0.74 | 0.87 | 0.0725 | 0.115 | -0.055 | 0.779566 | 0.632533 | 0.8202 | 0.999971 | 1 | 0.975761 |
| Prrc2b | F8WHT3 | 1.42 | 0.94 | 1.6 | 1.7 | 1.2 | 1.27 | 1.88 | 1.88 | 1.21 | 1.51 | 1.47 | 1.79 | 0.6 | 1.19 | 1.71 | 1.44 | 0.1425 | 0.08 | -0.18 | 0.601314 | 0.736974 | 0.55255 | 0.999971 | 1 | 0.913595 |
| Prrc2c | A0A0A0M( | 1.5 | 1.32 | 1.2 | 1.38 | 1.52 | 1.46 | 1.46 | 1.29 | 1.61 | 1.52 | 1.12 | 1.16 | 1.27 | 1.22 | 1.23 | 1.3 | 0.0825 | 0.0025 | -0.095 | 0.632786 | 0.989807 | 0.574164 | 0.999971 | 1 | 0.91816 |
| Prrt1 | O35449 | -0.04 | -0.29 | -1.15 | -1 | -0.44 | -0.62 | -1.19 | -1.2 | -0.36 | -0.09 | -1.19 | -1.27 | 0.16 | -0.72 | -1.37 | -0.76 | -0.2425 | -0.1075 | -0.0525 | 0.471516 | 0.780915 | 0.894863 | 0.999971 | 1 | 0.988571 |
| Prrt2 | E9PUL5 | 1.18 | 0.07 | -1.14 | -0.56 | -0.42 | -0.41 | -0.71 | -0.97 | -0.28 | -0.59 | -1.1 | -1.03 | -0.21 | -1.79 | -1.47 | -1.19 | -0.515 | -0.6375 | -1.0525 | 0.298637 | 0.221476 | 0.084802 | 0.999971 | 1 | 0.758256 |
| Prrt3 | Q6PE13 | 0.75 | 0.24 | -0.01 | -0.01 | -0.31 | -0.03 | -0.18 | -0.39 | -0.08 | 0.21 | -0.05 | -0.41 | -0.1 | 0 | -0.42 | -0.05 | -0.47 | -0.325 | -0.385 | 0.069211 | 0.210394 | 0.131898 | 0.999971 | 1 | 0.784948 |
| Prune1 | Q8BIW1 | -0.27 | 0.22 | 0.13 | 0.06 | 0.1 | -0.11 | -0.03 | 0.06 | 0.17 | -0.08 | 0.29 | -0.09 | 0.14 | -0.1 | -0.07 | 0.2 | -0.03 | 0.0375 | 0.0075 | 0.872629 | 0.849599 | 0.969247 | 0.999971 | 1 | 0.998787 |
| Prune2 | Q52KR3 | 0.07 | -1.34 | -0.66 | -0.54 | -1.04 | -1.08 | -0.95 | -0.44 | -1.38 | -0.34 | -0.77 | -0.95 | -2.34 | -1.07 | -0.46 | -0.51 | -0.26 | -0.2425 | -0.4775 | 0.432977 | 0.498885 | 0.341749 | 0.999971 | 1 | 0.861042 |
| Prxl2a | Q3U125 | -0.96 | -1.3 | -1.33 | -1.33 | -1.2 | -0.81 | -1.33 | -0.49 | -1.31 | -0.69 | -1.55 | -1.5 | -1.61 | -1 | -1.34 | -1.21 | 0.2725 | -0.0325 | -0.06 | 0.27901 | 0.894507 | 0.7743 | 0.999971 | 1 | 0.963884 |
| Prxl2b | Q9DB60 | -0.4 | 0.38 | -0.46 | -1.27 | -0.12 | 1.42 | -0.64 | -0.05 | 0.47 | 0.56 | -0.39 | -0.7 | -0.19 | 1.62 | -0.14 | -0.91 | 0.59 | 0.4225 | 0.5325 | 0.269437 | 0.345595 | 0.371744 | 0.999971 | 1 | 0.869526 |
| Psap | E9PZ00 | 1.69 | 3.08 | 2.33 | 2.5 | 2.24 | 2.23 | 2.1 | 2.78 | 2.62 | 1.4 | 3.03 | 2.92 | 2.55 | 2.35 | 2.76 | 2.95 | -0.0625 | 0.0925 | 0.2525 | 0.847179 | 0.83532 | 0.435347 | 0.999971 | 1 | 0.882419 |
| Psat1 | Q99K85 | -0.95 | -0.62 | -0.03 | -0.19 | -0.24 | -0.37 | -0.08 | -0.13 | -0.37 | -0.86 | -0.08 | -0.14 | -0.35 | -0.51 | 0.18 | 0.19 | 0.2425 | 0.085 | 0.325 | 0.34075 | 0.766677 | 0.278426 | 0.999971 | 1 | 0.843812 |
| Psd | Q5DTT2 | -0.3 | -0.47 | -0.05 | -0.48 | -0.42 | -0.4 | -0.25 | -0.06 | -0.52 | -0.42 | 0.11 | -0.02 | -0.94 | -0.9 | -0.4 | -0.26 | 0.0425 | 0.1125 | -0.3 | 0.826757 | 0.614823 | 0.224732 | 0.999971 | 1 | 0.828225 |
| Psd_1 | Q5DTT2-2 | -1.82 | -2.02 | -0.89 | -1.62 | -2.8 | -2.71 | -2.41 | -1.52 | -1.71 | -1.98 | -1.96 | -1.3 | -1.3 | -2.16 | -1.08 | -0.62 | -0.7725 | -0.15 | 0.2975 | 0.062755 | 0.619916 | 0.454155 | 0.999971 | 1 | 0.887217 |
| Psd2 | Q6P1I6-2 | -0.9 | -0.59 | -0.41 | -0.69 | -0.6 | -0.37 | -0.55 | -0.63 | -0.55 | -0.43 | -0.67 | -0.51 | -0.15 | -0.31 | -0.42 | -0.34 | 0.11 | 0.1075 | 0.3425 | 0.561443 | 0.560108 | 0.095663 | 0.999971 | 1 | 0.78268 |
| Psd3 | E9PUC5 | -0.4 | -0.02 | -1.03 | -0.88 | 0 | -0.21 | -0.3 | -1.58 | 0.18 | -0.39 | -0.07 | 0 | 0.28 | -0.29 | -0.96 | 0.06 | 0.06 | 0.5125 | 0.355 | 0.882475 | 0.090162 | 0.325661 | 0.999971 | 1 | 0.856472 |
| Psd3_1 | F6Z9E6 | -0.26 | -0.1 | -0.3 | -0.73 | 0.13 | 0.27 | 0.33 | -0.68 | 0.04 | -0.08 | 0.13 | 0.07 | -0.14 | -0.11 | -0.49 | -0.6 | 0.36 | 0.3875 | 0.0125 | 0.226592 | 0.076433 | 0.955613 | 0.999971 | 1 | 0.998056 |
| Psen1 | P49769 | -0.32 | -1.36 | -0.46 | -0.03 | -1.34 | -0.64 | -1.12 | -0.23 | -1.09 | 0.25 | -1.08 | -0.81 | -1.24 | -0.37 | -0.41 | 0.03 | -0.29 | -0.14 | 0.045 | 0.439534 | 0.732759 | 0.905459 | 0.999971 | 1 | 0.991091 |
| Psip1 | A2BI12 | 0.06 | 2.17 | 1.7 | 2.56 | 4.18 | 2.14 | 1.46 | 2.35 | 2.2 | 2.44 | 1.68 | 2.05 | 2.47 | 2.32 | 1.62 | 1.86 | 0.91 | 0.47 | 0.445 | 0.228516 | 0.386715 | 0.418138 | 0.999971 | 1 | 0.877685 |
| Psip1_1 | Q99JF8 | 1.48 | 1.73 | 1.79 | 2.35 | 2.28 | 1.94 | 2.62 | 2.37 | 1.72 | 1.38 | 1.39 | 1.54 | 0.25 | 0.92 | 1.36 | 0.78 | 0.465 | -0.33 | -1.01 | 0.096848 | 0.181934 | 0.008276 | 0.999971 | 1 | 0.602121 |
| Psma1 | Q9R1P4 | 0.27 | 1.14 | 1 | 0.35 | 1.97 | 0.69 | 1.52 | 0.63 | 2.38 | 0.12 | 1.38 | 1.33 | 2.8 | 0.21 | 0.84 | 0.31 | 0.5125 | 0.6125 | 0.35 | 0.201323 | 0.223755 | 0.555673 | 0.999971 | 1 | 0.914901 |
| Psma2 | P49722 | -0.24 | 0.18 | 0.19 | -0.06 | 0.35 | -0.05 | 0.09 | 0.11 | -0.01 | -0.29 | 0.4 | 0.56 | 0.03 | 0.38 | 0.01 | -0.11 | 0.1075 | 0.1475 | 0.06 | 0.584963 | 0.553821 | 0.7691 | 0.999971 | 1 | 0.963119 |
| Psma3 | O70435 | 0.35 | 0.38 | 0.15 | 0.19 | 0.5 | 0.28 | 0.16 | 0.2 | 0.72 | 0.18 | 0.28 | 0.16 | 0.38 | -0.01 | 0 | 0.18 | 0.0175 | 0.0675 | -0.13 | 0.921402 | 0.733892 | 0.48603 | 0.999971 | 1 | 0.897483 |
| Psma4 | Q9R1P0 | 1.26 | 1.37 | 1.28 | 1.38 | 0.92 | 1.45 | 2.02 | 1.72 | 1.16 | 1.33 | 1.91 | 1.36 | 0.93 | 1.33 | 1.61 | 1.57 | 0.205 | 0.1175 | 0.0375 | 0.437601 | 0.581555 | 0.858613 | 0.999971 | 1 | 0.982702 |
| Psma5 | Q9Z2U1 | -0.99 | 0.33 | 0.49 | 0.46 | 0.45 | 0.3 | 0.15 | 0.44 | 0.2 | -0.48 | 0.36 | 0.76 | 0.05 | 0.4 | 0.47 | 0.42 | 0.2625 | 0.1375 | 0.2625 | 0.466935 | 0.743233 | 0.473598 | 0.999971 | 1 | 0.894044 |
| Psma6 | Q9QUM9 | 0.76 | 2.36 | 1.23 | 1.74 | 1.77 | 2.04 | 1.01 | 1.82 | 1.89 | 1.69 | 1.08 | 0.2 | 2.13 | 2.21 | 1.72 | 1.79 | 0.1375 | -0.3075 | 0.44 | 0.727371 | 0.526069 | 0.238676 | 0.999971 | 1 | 0.833594 |
| Psma7 | Q9Z2U0 | 0.55 | 0.05 | -1.46 | -1.27 | -0.14 | 0.1 | -0.18 | -1.61 | -0.28 | 0.04 | -1.65 | -1.8 | -0.44 | -0.12 | -1.39 | -1.55 | 0.075 | -0.39 | -0.3425 | 0.896115 | 0.536026 | 0.542647 | 0.999971 | 1 | 0.911199 |
| Psmb1 | O09061 | 0.33 | 0.47 | 0.08 | 0 | 0.38 | 0.7 | 0.1 | 0.59 | 0.5 | 0.61 | 0.08 | 0.28 | 0.13 | 0.54 | 0.47 | 0.13 | 0.2225 | 0.1475 | 0.0975 | 0.320133 | 0.485263 | 0.640812 | 0.999971 | 1 | 0.93301 |
| Psmb10 | O35955 | 1.12 | 1.21 | 0.85 | 1.29 | 0.94 | 0.33 | 0.12 | 0.24 | 0.15 | 0.44 | 1.11 | 0.83 | 0.15 | 0.66 | 0.71 | 0.86 | -0.71 | -0.485 | -0.5225 | 0.014906 | 0.08503 | 0.041821 | 0.999971 | 1 | 0.701725 |
| Psmb2 | Q9R1P3 | -1.02 | 0.02 | 1.99 | 0.07 | -0.22 | -0.64 | 1.45 | 0.16 | -0.22 | 3.11 | 0.35 | 1.36 | -0.51 | -0.52 | 0.63 | 0.45 | -0.0775 | 0.885 | -0.2525 | 0.911612 | 0.32346 | 0.692496 | 0.999971 | 1 | 0.947174 |
| Psmb3 | Q9R1P1 | -0.33 | 0.72 | 0.32 | 0.31 | 0.56 | 0.48 | 0.13 | 0.37 | 0.48 | -0.29 | 0.45 | 0.52 | 0.56 | 0.61 | 0.64 | 0.59 | 0.13 | 0.035 | 0.345 | 0.619344 | 0.906758 | 0.187501 | 0.999971 | 1 | 0.809025 |
| Psmb4 | P99026 | -0.81 | -0.08 | -0.18 | 0.03 | 0.36 | -0.47 | -0.1 | -0.04 | 0.66 | -0.83 | -0.11 | 0.29 | 0.46 | -0.37 | -0.09 | -0.03 | 0.1975 | 0.2625 | 0.2525 | 0.474698 | 0.474994 | 0.368745 | 0.999971 | 1 | 0.869526 |
| Psmb5 | O55234 | -0.38 | 0.32 | 0.29 | 0.48 | 0.19 | 0.08 | 0.28 | 0.38 | 0.47 | -0.23 | 0.49 | 0.39 | 0.03 | 0.12 | 0.29 | 0.23 | 0.055 | 0.1025 | -0.01 | 0.815709 | 0.708192 | 0.966063 | 0.999971 | 1 | 0.998721 |
| Psmb6 | Q60692 | 0.38 | 1.1 | 1.19 | 0.95 | 0.94 | 1.03 | 1 | 1.09 | 1.1 | 0.86 | 1.27 | 0.95 | 0.75 | 1.14 | 1.09 | 1.06 | 0.11 | 0.14 | 0.105 | 0.627581 | 0.555907 | 0.659778 | 0.999971 | 1 | 0.935181 |
| Psmb7 | P70195 | -0.09 | 0.55 | 1.65 | 1.36 | 0.18 | 0.63 | 1.1 | 1.52 | 0.67 | 0.4 | 1.57 | 1.49 | 0.13 | 0.34 | 1.89 | 1.72 | -0.01 | 0.165 | 0.1525 | 0.982591 | 0.721905 | 0.783863 | 0.999971 | 1 | 0.966376 |
| Psmb8 | P28063 | 1.45 | 0.42 | 0.38 | 0.96 | 0.99 | 0.18 | 0.21 | 0.17 | 0.35 | 0.73 | 0.33 | 0.72 | 0.1 | 1.19 | 0.57 | -0.07 | -0.415 | -0.27 | -0.355 | 0.22201 | 0.360943 | 0.349141 | 0.999971 | 1 | 0.865162 |
| Psmc1 | P62192 | 0.38 | 0.65 | -0.04 | 0.25 | 0.6 | 0.7 | 0.18 | 0.13 | 0.62 | 0.71 | 0.2 | 0.22 | 0.67 | 0.74 | 0.29 | 0.39 | 0.0925 | 0.1275 | 0.2125 | 0.698184 | 0.583118 | 0.353239 | 0.999971 | 1 | 0.868675 |
| Psmc2 | P46471 | 0.02 | 0.27 | 0.04 | 0.26 | 0.31 | 0.19 | -0.06 | 0.46 | 0.24 | 0.24 | 0.13 | 0.3 | 0.45 | 0.57 | 0.28 | 0.3 | 0.0775 | 0.08 | 0.2525 | 0.689801 | 0.633877 | 0.181887 | 0.999971 | 1 | 0.808532 |
| Psmc3 | A0A087WF | -0.41 | -0.12 | -1.95 | 0.13 | 0.25 | -0.11 | -0.86 | 0.74 | 0.09 | -0.1 | -0.23 | -0.46 | 0.53 | 0.58 | 0.24 | -0.07 | 0.5925 | 0.4125 | 0.9075 | 0.281803 | 0.374874 | 0.075523 | 0.999971 | 1 | 0.746958 |
| Psmc3_1 | O88685 | 0.5 | 0.32 | 0.54 | 0.33 | 0.21 | 0.3 | 1.02 | 0.37 | 0.55 | 0.43 | 0.78 | 0.65 | 0.34 | 0.41 | 0.65 | 0.77 | 0.0525 | 0.18 | 0.12 | 0.820161 | 0.314932 | 0.526881 | 0.999971 | 1 | 0.909459 |
| Psmc4 | P54775 | 0.12 | 0.3 | 0.27 | -0.04 | 0.39 | 0.17 | 1.75 | 0.14 | 0.34 | 0.2 | 0.54 | 0.6 | 0.25 | 0.33 | -0.02 | -0.06 | 0.45 | 0.2575 | -0.0375 | 0.253024 | 0.190375 | 0.844969 | 0.999971 | 1 | 0.979814 |
| Psmc5 | P62196 | 0.37 | 0.81 | 0.45 | 0.63 | 1.04 | 0.55 | 0.52 | 0.53 | 1.42 | 0.53 | 0.55 | 0.64 | 2.04 | 0.66 | 0.63 | 0.95 | 0.095 | 0.22 | 0.505 | 0.653598 | 0.402645 | 0.165427 | 0.999971 | 1 | 0.800215 |
| Psmc6 | P62334 | -0.29 | 0.54 | 0.39 | 0.11 | 0.52 | 0.27 | 0.54 | 0.35 | 0.75 | -0.06 | 0.76 | 0.65 | 0.38 | 0.01 | 0.5 | 0.38 | 0.2325 | 0.3375 | 0.13 | 0.328946 | 0.250658 | 0.595988 | 0.999971 | 1 | 0.92377 |
| Psmd1 | Q3TXS7 | 0.37 | 0.5 | 0.48 | 0.38 | 0.41 | 0.45 | 0.34 | 0.46 | 0.56 | 0.4 | 0.44 | 0.43 | 0.44 | 0.5 | 0.6 | 0.69 | -0.0175 | 0.025 | 0.125 | 0.913499 | 0.874634 | 0.462405 | 0.999971 | 1 | 0.890421 |
| Psmd10 | Q9Z2X2 | -0.31 | -0.19 | 0.49 | 0.59 | 0.18 | -0.3 | -0.15 | 0.66 | 0.08 | -0.1 | 0.32 | -0.07 | 0.17 | -0.21 | 0.1 | 0.08 | -0.0475 | -0.0875 | -0.11 | 0.880692 | 0.744633 | 0.681305 | 0.999971 | 1 | 0.943107 |
| Psmd11 | Q8BG32 | -0.35 | -0.91 | 0.2 | 0.25 | -0.64 | -0.96 | 0.45 | 0.37 | -0.69 | -0.45 | 0.59 | 0.5 | -0.92 | -0.43 | 0.34 | 0.45 | 0.0075 | 0.19 | 0.0625 | 0.985874 | 0.642043 | 0.877487 | 0.999971 | 1 | 0.985267 |
| Psmd12 | Q9D8W5 | -1.07 | -0.38 | 0.02 | -0.18 | -0.14 | -0.61 | 0.18 | 0.02 | -0.09 | -1 | 0.25 | 0.33 | -0.48 | -0.55 | 0.13 | 0.07 | 0.265 | 0.275 | 0.195 | 0.388344 | 0.469947 | 0.526832 | 0.999971 | 1 | 0.909459 |
| Psmd13 | Q9WVJ2 | -0.14 | 0.97 | 0.27 | -0.11 | 0.72 | 0.94 | 0.33 | 0.11 | 0.92 | 0.5 | 0.3 | -0.17 | 0.97 | 0.83 | 0.44 | 0.32 | 0.2775 | 0.14 | 0.3925 | 0.3975 | 0.681779 | 0.223724 | 0.999971 | 1 | 0.828225 |
| Psmd14 | O35593 | -0.66 | 0.08 | -0.5 | -0.24 | 0.22 | 0.19 | -0.05 | -0.31 | 0.28 | -0.27 | -0.23 | 0.02 | 0.25 | 0.12 | -0.32 | -0.23 | 0.3425 | 0.28 | 0.285 | 0.173768 | 0.257364 | 0.260943 | 0.999971 | 1 | 0.841256 |
| Psmd2 | Q8VDM4 | -0.59 | 0.36 | 0.18 | 0.2 | 0.44 | 0.2 | 0.32 | 0.37 | 0.6 | -0.27 | 0.53 | 0.28 | 0.53 | 0.41 | 0.42 | 0.48 | 0.295 | 0.2475 | 0.4225 | 0.253217 | 0.416481 | 0.11279 | 0.999971 | 1 | 0.78268 |
| Psmd3 | P14685 | 0.29 | -0.39 | 0.08 | -0.03 | -0.41 | -0.32 | 0.7 | 0.07 | -0.23 | -0.21 | 0.36 | 0.3 | -0.78 | -0.37 | 0.1 | -0.03 | 0.0225 | 0.0675 | -0.2575 | 0.939897 | 0.781147 | 0.344658 | 0.999971 | 1 | 0.862646 |
| Psmd4 | O35226 | -2.29 | -2.94 | -2.58 | -1.76 | -1.8 | -4.07 | -2.22 | -1.96 | -1.59 | -3.46 | -2.27 | -2.02 | -2.95 | -3.73 | -1.92 | -2.3 | -0.12 | 0.0575 | -0.3325 | 0.822977 | 0.897205 | 0.45931 | 0.999971 | 1 | 0.889191 |
| Psmd4_1 | O35226-2 | -0.07 | 0.88 | 0.78 | 0.37 | 1.01 | 0.98 | 1.03 | 1.01 | 1.8 | 0.35 | 1.02 | 1.11 | 1.35 | 1.04 | 0.94 | 0.72 | 0.5175 | 0.58 | 0.5225 | 0.061121 | 0.133697 | 0.082094 | 0.999971 | 1 | 0.755254 |
| Psmd5 | Q8BJY1 | 0.34 | 0.29 | 0.37 | 0.54 | 0.27 | 0.43 | 0.37 | 0.62 | 0.24 | 0.37 | 0.4 | 0.34 | 0.31 | 0.67 | 0.73 | 0.72 | 0.0375 | -0.0475 | 0.2225 | 0.831378 | 0.770875 | 0.252236 | 0.999971 | 1 | 0.839012 |
| Psmd6 | Q99JI4 | 0.5 | 0 | 0.42 | 0.59 | 0.22 | 0.28 | 0.51 | 0.66 | 0.3 | 0.49 | 0.83 | 0.6 | 0.13 | 0.2 | 0.62 | 0.87 | 0.04 | 0.1775 | 0.0775 | 0.851581 | 0.417916 | 0.755572 | 0.999971 | 1 | 0.961116 |
| Psmd7 | P26516 | -0.74 | -0.48 | 0.07 | 0.21 | -0.62 | -0.28 | -0.04 | -0.14 | -0.49 | -0.57 | -0.21 | -0.06 | -0.71 | -0.22 | -0.27 | 0.25 | -0.035 | -0.0975 | -0.0025 | 0.898754 | 0.720419 | 0.99347 | 0.999971 | 1 | 0.99933 |
| Psmd8 | Q9CX56 | 0.93 | 0.63 | 1.67 | 1.36 | 0.84 | 1 | 1.55 | 1.72 | 0.77 | 1 | 1.91 | 1.82 | 0.77 | 0.85 | 2.04 | 1.99 | 0.13 | 0.2275 | 0.265 | 0.681948 | 0.53192 | 0.512909 | 0.999971 | 1 | 0.905815 |
| Psmd9 | Q9CR00 | 0.13 | 0.85 | 0.43 | 0.51 | 0.94 | 0.23 | 0.88 | 0.03 | 1.03 | 0.71 | 0.57 | 0.58 | 0.46 | 0.04 | 0.11 | 0.15 | 0.04 | 0.2425 | -0.29 | 0.888736 | 0.291856 | 0.209453 | 0.999971 | 1 | 0.822138 |
| Psme1 | G3UXZ5 | 0.31 | 1.16 | 0.83 | 0.92 | 1.26 | 0.91 | 0.66 | 0.48 | 1.37 | 0.53 | 0.4 | 1.04 | 1.21 | 1.35 | 1.17 | 0.95 | 0.0225 | 0.03 | 0.365 | 0.932746 | 0.919393 | 0.145208 | 0.999971 | 1 | 0.791767 |
| Psme2 | P97372 | 0.71 | 1.56 | 1.12 | 0.91 | 1.42 | 1.46 | 0.97 | 0.96 | 1.74 | 1.5 | 0.91 | 1.42 | 1.57 | 1.48 | 1.48 | 0.95 | 0.1275 | 0.3175 | 0.295 | 0.618268 | 0.258795 | 0.268453 | 0.999971 | 1 | 0.842746 |

| | | | | | | | | | | | | | | | | | | | | | | | | | | |
|---|---|---|---|---|---|---|---|---|---|---|---|---|---|---|---|---|---|---|---|---|---|---|---|---|---|---|
| Psme3 | P61290 | 0.96 | 2.51 | 2.11 | 2.46 | 2.33 | 2.11 | 1.86 | 1.95 | 2.32 | 1.52 | 2.28 | 2.49 | 2 | 2.09 | 2.24 | 2.47 | 0.0525 | 0.1425 | 0.19 | 0.885692 | 0.724363 | 0.606485 | 0.999971 | 1 | 0.925106 |
| Psme3ip1 | Q91WE2 | 1.1 | 2.27 | 1.94 | 2.22 | 2.07 | 1.91 | 1.54 | 1.94 | 2.03 | 1.62 | 2.17 | 2.12 | 1.73 | 2.01 | 2.06 | 2.11 | -0.0175 | 0.1025 | 0.095 | 0.953682 | 0.737064 | 0.747818 | 0.999971 | 1 | 0.961064 |
| Psme4 | Q5SSW2 | -0.88 | 0.26 | -0.43 | -0.6 | -0.05 | -0.08 | -0.5 | -0.46 | 0.29 | -0.2 | -0.16 | -0.06 | 0.04 | -0.22 | -0.31 | -0.29 | 0.14 | 0.38 | 0.2175 | 0.624976 | 0.199325 | 0.43615 | 0.999971 | 1 | 0.882419 |
| Psmf1 | Q8BHL8 | -0.49 | 1.03 | 0.46 | 0.7 | 0.9 | 0.42 | 0.3 | 0.46 | 0.66 | 0.26 | 0.31 | 0.59 | 0.81 | 0.72 | 0.57 | 0.68 | 0.095 | 0.03 | 0.27 | 0.784288 | 0.929304 | 0.422978 | 0.999971 | 1 | 0.87887 |
| Psmg1 | Q9JK23 | 0.88 | 1 | -1.35 | -1.12 | 1.07 | 0.99 | -1.07 | -0.83 | 0.62 | 0.86 | -0.89 | -0.98 | 0.12 | 1.09 | -0.85 | -1.25 | 0.1875 | 0.05 | -0.075 | 0.807015 | 0.944835 | 0.919192 | 0.999971 | 1 | 0.992633 |
| Psmg2 | Q9EST4-3 | 0.66 | 0.49 | 0.78 | 0.99 | 0.6 | 0.68 | 0.35 | 1.12 | 0.44 | 0.49 | 0.2 | 0.43 | -0.09 | 0.88 | 1.03 | 0.67 | -0.0425 | -0.34 | -0.1075 | 0.853337 | 0.097357 | 0.706641 | 0.999971 | 1 | 0.950182 |
| Psmg3 | Q9CZH3 | -0.78 | 1.99 | 0.39 | 0.04 | 1.23 | 0.38 | -0.51 | -0.47 | 1.31 | -0.37 | 0.31 | -0.09 | 2.44 | 1.46 | -0.49 | -0.53 | -0.2525 | -0.12 | 0.31 | 0.697252 | 0.848764 | 0.71414 | 0.999971 | 1 | 0.952257 |
| Pspc1 | Q8R326 | 1.12 | 2.24 | 2.44 | 2.3 | 2.53 | 1.66 | 2.09 | 2.36 | 2.14 | 1.3 | 2.26 | 2.3 | 2.44 | 2.15 | 2.29 | 2.27 | 0.135 | -0.025 | 0.2625 | 0.702345 | 0.946631 | 0.414059 | 0.999971 | 1 | 0.876329 |
| Psph | Q99LS3 | -0.3 | 0.21 | -0.33 | 0.08 | -0.01 | -0.31 | -0.48 | 0.04 | -0.35 | -0.87 | 0.09 | -0.17 | -0.5 | -0.14 | -0.31 | -0.33 | -0.105 | -0.24 | -0.235 | 0.642194 | 0.375062 | 0.275669 | 0.999971 | 1 | 0.843794 |
| Psrc1 | A2AEF3 | 0.81 | 0.51 | 0.23 | 0.96 | 1.21 | 0.59 | 1.72 | 1.33 | 1.49 | 0.09 | -0.23 | 0.01 | -0.78 | -0.81 | -0.48 | -0.93 | 0.585 | -0.2875 | -1.3775 | 0.073864 | 0.483076 | 0.000219 | 0.999971 | 1 | 0.247672 |
| Ptar1 | A0A494B9\ | -0.26 | 0.27 | 0.56 | 0.96 | 0.09 | 0.15 | 0.57 | 0.77 | -0.02 | -0.01 | 0.44 | 1.18 | 0.29 | 0.14 | 0.48 | 0.97 | 0.0125 | 0.015 | 0.0875 | 0.967801 | 0.96769 | 0.783212 | 0.999971 | 1 | 0.966188 |
| Ptbp1 | Q8BGJ5 | 0.53 | 0.26 | 1.13 | 0.97 | 0.5 | 0.08 | 1.85 | 0.44 | 0.31 | 0.35 | 0.85 | 0.83 | -0.04 | 0.33 | 0.91 | 0.02 | -0.005 | -0.1375 | -0.4175 | 0.990378 | 0.609339 | 0.192224 | 0.999971 | 1 | 0.810547 |
| Ptbp2 | Q91Z31 | 2.03 | 2.07 | 2.06 | 2.4 | 1.77 | 1.82 | 2.24 | 2.2 | 1.64 | 1.66 | 2.09 | 1.62 | 1.52 | 1.67 | 2.03 | 1.78 | -0.1325 | -0.3875 | -0.39 | 0.523407 | 0.077 | 0.0765 | 0.999971 | 1 | 0.748444 |
| Ptbp2_1 | A0A0G2JF\ | 1.12 | 2.2 | 0.73 | 1.71 | 2.73 | -1.65 | 1.32 | 1.58 | 1.44 | 1.7 | 2.83 | 2.01 | 1.63 | 1.75 | -0.35 | 2.97 | -0.445 | 0.555 | 0.06 | 0.617251 | 0.210178 | 0.930472 | 0.999971 | 1 | 0.994744 |
| Ptbp3 | G8JL74 | 2.38 | 1.37 | 1.13 | 0.95 | 1.22 | 1.89 | 1.08 | 1 | 1.58 | 2.39 | 0.92 | 0.86 | 0.87 | 1.01 | 1.04 | 1.17 | -0.16 | -0.02 | -0.435 | 0.664807 | 0.964623 | 0.205323 | 0.999971 | 1 | 0.816688 |
| Ptcd1 | Q8C2E4 | -0.32 | 0.26 | -0.03 | -0.49 | 0.16 | -0.47 | -0.38 | -0.74 | 0.14 | -0.22 | -0.53 | -0.3 | 0.02 | -0.18 | -0.55 | -0.35 | -0.2125 | -0.0825 | -0.12 | 0.439537 | 0.736079 | 0.618196 | 0.999971 | 1 | 0.928581 |
| Ptcd2 | Q8R3K3 | 0.2 | 0.62 | -0.3 | -0.2 | 0.37 | 0.41 | -0.47 | -0.12 | 0.29 | 0.26 | -0.56 | -0.62 | 0.77 | 0.82 | -0.25 | 0.04 | -0.0325 | -0.2375 | 0.265 | 0.914871 | 0.473986 | 0.440574 | 0.999971 | 1 | 0.883277 |
| Ptcd3 | Q14C51 | -0.12 | 0 | -0.55 | -0.57 | -0.01 | -0.16 | 0.32 | -0.59 | -0.01 | -0.67 | -0.7 | -0.34 | 0.29 | -0.27 | -0.28 | -0.32 | 0.2 | -0.12 | 0.165 | 0.452426 | 0.627851 | 0.497818 | 0.999971 | 1 | 0.902907 |
| Ptdss1 | Q99LH2 | -1.48 | 1.36 | -2.5 | -2.47 | 1 | 0.73 | -1.6 | -2.04 | 1.01 | -0.69 | -2.27 | -1.87 | 0.41 | 1.17 | -1.92 | -2.06 | 0.795 | 0.3175 | 0.6725 | 0.463944 | 0.762863 | 0.541931 | 0.999971 | 1 | 0.911119 |
| Ptdss2 | Q9Z1X2 | -1.6 | 0.02 | 0.29 | 0 | 0.36 | 0.17 | 0.29 | 0.98 | 0.49 | -0.88 | 0.88 | 0.25 | 0.36 | 0.52 | 0.72 | 0.71 | 0.7725 | 0.5075 | 0.9 | 0.106649 | 0.35251 | 0.056187 | 0.999971 | 1 | 0.725348 |
| Pten | O08586 | -0.52 | 0.16 | 0.21 | -0.08 | 0.22 | 0.52 | 0.26 | 0.14 | 0.16 | -0.24 | 0.39 | 0.05 | -0.02 | 0.15 | 0.22 | 0.02 | 0.3425 | 0.1475 | 0.15 | 0.155422 | 0.54558 | 0.502079 | 0.999971 | 1 | 0.904573 |
| Pter | A2AUR3 | 0.18 | 0.13 | -0.43 | -0.04 | 0.12 | -0.12 | -0.5 | -0.14 | -0.45 | 0.22 | -0.4 | 0.11 | -0.41 | 0.05 | -0.32 | 0.06 | -0.12 | -0.09 | -0.115 | 0.600924 | 0.717072 | 0.613607 | 0.999971 | 1 | 0.927364 |
| Ptgds | O09114 | 2.15 | 3.3 | 3.12 | 3.31 | 3.36 | 3.17 | 2.94 | 3.11 | 3.34 | 3.09 | 3.21 | -1.13 | 1.73 | 3.6 | 3.27 | 3.37 | 0.175 | -0.8425 | 0.0225 | 0.561945 | 0.413175 | 0.962185 | 0.999971 | 1 | 0.998721 |
| Ptges2 | Q8BWM0 | -0.84 | -1.04 | -0.27 | -0.1 | -1.07 | -1.09 | -0.06 | -0.28 | -0.92 | -0.67 | -0.33 | -0.39 | -1.33 | -1.22 | -0.44 | -0.58 | -0.0625 | -0.015 | -0.33 | 0.855757 | 0.956855 | 0.317498 | 0.999971 | 1 | 0.853845 |
| Ptges3 | Q9R0Q7 | -0.3 | 1.38 | 1.84 | 1.41 | 1.58 | 1.51 | 2.46 | 1.88 | 2.12 | -0.09 | 2.66 | 2.54 | 1.68 | 1.61 | 2.45 | 1.51 | 0.775 | 0.725 | 0.73 | 0.135741 | 0.33179 | 0.15745 | 0.999971 | 1 | 0.797252 |
| Ptgfrn | Q9WV91 | -0.64 | -0.85 | -0.66 | -0.36 | -0.79 | -0.6 | -0.23 | -0.37 | -0.65 | -0.49 | -0.2 | -0.51 | -0.02 | -0.6 | -0.46 | 0.06 | 0.13 | 0.165 | 0.3725 | 0.540687 | 0.40836 | 0.132134 | 0.999971 | 1 | 0.784948 |
| Ptgis | Q8BXC0 | 0.53 | 1.56 | -1.35 | -0.43 | 4.66 | 1.11 | -1.29 | -1.09 | 1.24 | 2.94 | 1.82 | 2.92 | 1.97 | 1.12 | 0.24 | -1.61 | 0.77 | 2.1525 | 0.3525 | 0.570867 | 0.011788 | 0.69227 | 0.999971 | 1 | 0.947174 |
| Ptgr1 | Q91YR9 | -0.17 | 0.78 | 0.26 | 0.23 | 0.3 | 0.03 | 0.02 | 0.06 | 0.25 | -1.13 | -0.23 | 0.99 | -0.13 | 0.31 | -0.06 | -1.33 | -0.1725 | -0.305 | -0.5775 | 0.477167 | 0.509709 | 0.164514 | 0.999971 | 1 | 0.800215 |
| Ptgr2 | Q8VDQ1 | -0.23 | -0.2 | -0.16 | 0.06 | -0.25 | -0.45 | -0.46 | 0.58 | -0.26 | -0.42 | -0.21 | -0.11 | -0.49 | -0.21 | 0.06 | -0.03 | -0.0125 | -0.1175 | -0.035 | 0.963464 | 0.502661 | 0.860162 | 0.999971 | 1 | 0.982702 |
| Ptgr3 | Q8BGC4 | -0.01 | -0.47 | -0.49 | -0.27 | -0.57 | -0.47 | -0.8 | -0.49 | -0.6 | -0.22 | -0.66 | -0.47 | -0.37 | -0.24 | -0.63 | -0.27 | -0.2725 | -0.1775 | -0.0675 | 0.188794 | 0.388317 | 0.737869 | 0.999971 | 1 | 0.959801 |
| Ptgs1 | P22437 | 0.09 | 0.53 | -1.26 | -1.18 | 0.64 | 1.05 | -1.27 | -1.49 | 0.61 | 0.4 | -0.97 | -1.05 | 0.26 | 1.33 | -0.4 | -0.77 | 0.1875 | 0.2025 | 0.56 | 0.793008 | 0.727595 | 0.35532 | 0.999971 | 1 | 0.868793 |
| Ptgs2 | Q05769 | 0.75 | -0.39 | -0.5 | -0.54 | -1.34 | -1 | -0.69 | 0.19 | -1.37 | -0.02 | -0.76 | -1.55 | -1.59 | -0.63 | -0.76 | -0.45 | -0.54 | -0.755 | -0.6875 | 0.226158 | 0.112796 | 0.101358 | 0.999971 | 1 | 0.78268 |
| Ptk2 | A0A5F8MP | 0.17 | 0.45 | 0.03 | 0.02 | 0.24 | 0.4 | 0.06 | -0.08 | 0.31 | 0.11 | 0.17 | 0.3 | 0.06 | 0 | -0.17 | 0.05 | -0.0125 | 0.055 | -0.1825 | 0.950556 | 0.7625 | 0.340995 | 0.999971 | 1 | 0.860894 |
| Ptk2b | Q9QVP9 | 0.36 | 0.79 | 0.68 | 0.62 | 0.87 | 0.78 | 0.49 | 0.79 | 0.66 | 0.35 | 0.61 | 0.31 | 0.83 | 0.78 | 0.68 | 0.76 | 0.12 | -0.13 | 0.15 | 0.533426 | 0.498846 | 0.411146 | 0.999971 | 1 | 0.875847 |
| Ptk7 | Q8BKG3 | 0.56 | -0.2 | -0.54 | -0.51 | -0.38 | -0.58 | -0.5 | -0.6 | -0.84 | 0.18 | -0.62 | -0.44 | -0.56 | -0.34 | -0.42 | -0.46 | -0.3425 | -0.2575 | -0.2725 | 0.237126 | 0.449849 | 0.339525 | 0.999971 | 1 | 0.860218 |
| Ptma | P26350 | 3.84 | 2.97 | 3.08 | 3.63 | 3.15 | 3.21 | 3.33 | 2.69 | 3.26 | 4.17 | 2.7 | 2.35 | 1.82 | 2.52 | 2.78 | 2.29 | -0.285 | -0.26 | -1.0275 | 0.30952 | 0.546679 | 0.007544 | 0.999971 | 1 | 0.582714 |
| Ptms | Q9D0J8 | 5.08 | 2.18 | 3.57 | 5.26 | 2.54 | 2.35 | 4.01 | 3.46 | 2.5 | 4.07 | 3.04 | 3.39 | -0.49 | 0.29 | 3.31 | 2.3 | -0.9325 | -0.7725 | -2.67 | 0.228561 | 0.300311 | 0.025936 | 0.999971 | 1 | 0.678474 |
| Ptn | P63089 | 1.21 | -0.11 | 1.32 | 1.2 | 0.47 | 0.7 | 2.51 | 1.48 | 3.11 | 0.69 | 1.48 | 0.36 | 0.88 | 0.01 | 1.77 | 0.77 | 0.385 | 0.505 | -0.0475 | 0.472334 | 0.439289 | 0.918393 | 0.999971 | 1 | 0.992633 |
| Ptov1 | Q91VU8 | 1.55 | 2.41 | 1.13 | 0.86 | 2.32 | 2.37 | 1.4 | 0.72 | 2.64 | 1.49 | 1.19 | 1.04 | 1.61 | 1.44 | 0.89 | 0.8 | 0.215 | 0.1025 | -0.3025 | 0.659056 | 0.825874 | 0.435222 | 0.999971 | 1 | 0.882419 |
| Ptp4a1 | Q63739 | 0.78 | -0.83 | -0.32 | 0 | -0.71 | -1.37 | -0.08 | -0.79 | -1.19 | -0.18 | -0.56 | -0.76 | -1.72 | -0.74 | -0.74 | -0.61 | -0.645 | -0.58 | -0.86 | 0.140381 | 0.156997 | 0.059358 | 0.999971 | 1 | 0.725348 |
| Ptp4a2 | O70274 | -0.08 | 0.07 | 0.95 | 0.67 | 0.35 | 0.18 | 1.06 | 0.61 | 0.49 | -0.12 | 0.23 | 0.08 | 0.71 | 0.15 | 0.9 | 0.69 | 0.1475 | -0.2325 | 0.21 | 0.640683 | 0.427561 | 0.491996 | 0.999971 | 1 | 0.89987 |
| Ptp4a3 | Q9D658 | 0.28 | -0.98 | -1.37 | -1.27 | -1.14 | -0.71 | -1.66 | -1.38 | -1.64 | -0.24 | -1.62 | -2.02 | -0.73 | -0.15 | -1.13 | -0.97 | -0.3875 | -0.545 | 0.09 | 0.357066 | 0.299685 | 0.828811 | 0.999971 | 1 | 0.977963 |
| Ptpa | P58389 | -0.29 | -0.27 | -0.09 | -0.25 | 0.15 | -0.29 | 0.95 | -0.29 | 0.6 | -0.57 | 0.08 | 0.3 | 0.29 | -0.04 | -0.25 | -0.49 | 0.355 | 0.3275 | 0.1025 | 0.260534 | 0.245415 | 0.640157 | 0.999971 | 1 | 0.93301 |
| Ptpdc1 | A0A5F8MP | -0.51 | -0.47 | 0.13 | 0.37 | -0.33 | -0.71 | 0.09 | 0.55 | -0.24 | -0.34 | -0.11 | 0.1 | -0.38 | -0.51 | 0.34 | 0.55 | 0.02 | -0.0275 | 0.12 | 0.953619 | 0.915855 | 0.724258 | 0.999971 | 1 | 0.955404 |
| Ptpmt1 | Q66GT5 | 0.03 | -0.69 | 0.67 | 0.72 | -0.96 | -0.69 | -0.23 | 0.39 | -1.27 | -0.32 | -0.57 | 0.02 | -1.12 | -0.47 | 0.49 | 0.75 | -0.555 | -0.7175 | -0.27 | 0.209167 | 0.107762 | 0.596032 | 0.999971 | 1 | 0.92377 |
| Ptpn1 | P35821 | -0.22 | -0.07 | -0.12 | -0.03 | -0.19 | 0.23 | -0.51 | -0.07 | -0.21 | 0.02 | 0.05 | -0.27 | 0.49 | 0.27 | -0.16 | 0.21 | -0.025 | 0.0075 | 0.3125 | 0.90494 | 0.965226 | 0.145026 | 0.999971 | 1 | 0.79148 |
| Ptpn11 | P35235-1 | 0.25 | 0.37 | 0.47 | 0.3 | 0.39 | -0.17 | 0.6 | 0.08 | 0.69 | -0.2 | 0.48 | 0.5 | 0.21 | -0.26 | -0.02 | -0.19 | -0.1225 | 0.02 | -0.4125 | 0.581001 | 0.931805 | 0.051406 | 0.999971 | 1 | 0.723736 |
| Ptpn11_1 | P35235 | -0.89 | -0.44 | 1.39 | 0.4 | -1.49 | 0.83 | 5.02 | -1.93 | -1.54 | -1.16 | 0.35 | -1 | -3.06 | -3.37 | -0.92 | -3.09 | 0.4925 | -0.9525 | -2.725 | 0.739596 | 0.137325 | 0.003291 | 0.999971 | 1 | 0.577271 |
| Ptpn12 | P35831 | 1.08 | -0.25 | 0.52 | 0.62 | -0.34 | 0.24 | 0.25 | 0.83 | -0.28 | 0.54 | 0.56 | 0.07 | -1.18 | 0.04 | 0.53 | 0.68 | -0.2475 | -0.27 | -0.475 | 0.494034 | 0.43426 | 0.328136 | 0.999971 | 1 | 0.856472 |
| Ptpn13 | G5E8B1 | 0.11 | -0.93 | -1.38 | -0.82 | -1.37 | -1.2 | -1.57 | -0.75 | -0.72 | -0.63 | -1.21 | -1.36 | -1.05 | -1.68 | -1.12 | -1.18 | -0.4675 | -0.225 | -0.5025 | 0.20618 | 0.529706 | 0.166005 | 0.999971 | 1 | 0.800215 |
| Ptpn2 | Q06180 | 0 | 0.53 | 0.48 | 0.28 | 0.3 | 0.14 | 0.13 | -0.04 | 0.66 | -0.18 | 0.58 | 0.04 | -0.39 | -0.76 | -0.02 | 0.13 | -0.19 | -0.0475 | -0.5825 | 0.351352 | 0.854723 | 0.047213 | 0.999971 | 1 | 0.710102 |
| Ptpn23 | Q6PB44 | 0.06 | 0.55 | 0.6 | 0.52 | 0.53 | 0.52 | 0.26 | 0.23 | 0.39 | 0.06 | 0.44 | 0.63 | 1.04 | 0.39 | 0.3 | 0.47 | -0.0475 | -0.0525 | 0.1175 | 0.816157 | 0.80776 | 0.629019 | 0.999971 | 1 | 0.932176 |
| Ptpn3 | A2ALK8 | -0.33 | -0.21 | 0.36 | 0.03 | -0.58 | -0.03 | -0.15 | 0.02 | -0.16 | -0.09 | 0.37 | 0.07 | -0.3 | -0.48 | -0.04 | 0.43 | -0.1475 | 0.085 | -0.06 | 0.540022 | 0.710702 | 0.824104 | 0.999971 | 1 | 0.976309 |
| Ptpn4 | Q9WU22 | -0.87 | -0.59 | -1.48 | -1.83 | -0.8 | -0.6 | -1.41 | -1.56 | -1.01 | -0.81 | -1.5 | -1.52 | -0.39 | -0.53 | -1.65 | -1.43 | 0.1 | -0.0175 | 0.1925 | 0.779935 | 0.957952 | 0.63685 | 0.999971 | 1 | 0.93301 |
| Ptpn5 | P54830 | 0.21 | 0.64 | 0.19 | 0.54 | 0.58 | 0.16 | 0.21 | 0.88 | 0.74 | -0.17 | 0.3 | 0.58 | 1.35 | 0.61 | 1.05 | 0.54 | 0.0625 | -0.0325 | 0.4925 | 0.793101 | 0.898144 | 0.076468 | 0.999971 | 1 | 0.748444 |
| Ptpn6 | P29351 | 2.33 | 1.23 | 1.32 | 1.67 | 0.73 | 0.21 | 0.83 | 0.73 | 0.78 | 1.34 | 1.1 | 1.32 | 1.04 | 0.85 | 1 | 0.85 | -1.0125 | -0.5025 | -0.7025 | 0.007345 | 0.112574 | 0.028369 | 0.999971 | 1 | 0.678474 |
| Ptpn9 | O35239 | -0.92 | -0.93 | -1.13 | -1.37 | -0.95 | -0.7 | -0.96 | -1.32 | -0.9 | -0.72 | -1.25 | -1.08 | -0.7 | -0.74 | -1.04 | -1.19 | 0.105 | 0.1 | 0.17 | 0.625559 | 0.62865 | 0.427799 | 0.999971 | 1 | 0.880793 |
| Ptpra | Q91V35 | -0.39 | -1.08 | -0.9 | -0.72 | -1.03 | -1.1 | -1.12 | -0.66 | -1.25 | -0.6 | -0.81 | -0.93 | -0.73 | -0.95 | -0.97 | -0.72 | -0.205 | -0.125 | -0.07 | 0.371561 | 0.595536 | 0.742194 | 0.999971 | 1 | 0.961064 |
| Ptprc | S4R1M0 | 1.28 | 1 | 1.78 | 2.26 | 1.38 | 0.65 | 0.82 | 1.52 | 0.95 | 1.41 | 0.83 | 0.99 | 0.16 | 1.18 | 1.31 | 1.12 | -0.4875 | -0.535 | -0.6375 | 0.181304 | 0.11217 | 0.11445 | 0.999971 | 1 | 0.78268 |
| Ptprd | Q64487-5 | -1.38 | -1.72 | -1.35 | -1.23 | -1.64 | -1.6 | -1.39 | -1.34 | -1.59 | -1.42 | -1.3 | -1.29 | -1.42 | -1.52 | -1.4 | -1.06 | -0.0725 | 0.02 | 0.07 | 0.70853 | 0.91565 | 0.730461 | 0.999971 | 1 | 0.956041 |
| Ptprd_1 | Q64487-1 | 0.99 | -0.12 | -5.29 | -5.08 | -0.68 | 0.12 | -4.17 | -4.33 | -0.48 | 0.14 | -2.17 | -3.65 | -0.23 | 1.11 | -4.65 | -2.45 | 0.11 | 0.835 | 0.82 | 0.950543 | 0.615397 | 0.656663 | 0.999971 | 1 | 0.934108 |
| Ptprd_2 | G3UY75 | 0.91 | -0.19 | 0.16 | 0.32 | -0.9 | 0.58 | -0.08 | 0.3 | -0.64 | -0.08 | -0.19 | -0.86 | -0.94 | 1.43 | 0.41 | 0.74 | -0.325 | -0.7425 | 0.11 | 0.40364 | 0.034155 | 0.828712 | 0.999971 | 1 | 0.977963 |
| Ptpre | A0A1B0GR | -1.38 | -1.32 | 0.55 | 0 | -1.37 | -1.59 | -0.52 | -0.28 | -1.72 | -1.48 | 0.34 | 0.19 | -1.1 | -1.24 | -0.16 | 0.74 | -0.4025 | -0.13 | 0.0975 | 0.457405 | 0.844324 | 0.872801 | 0.999971 | 1 | 0.984109 |
| Ptprf | F6Y3V0 | -1.51 | -1.19 | -2.01 | -3.05 | -1.41 | -0.86 | -1.2 | -1 | -1.34 | -1.25 | -1.75 | -2.97 | -0.29 | -0.5 | -1.18 | -1.61 | 0.8225 | 0.1125 | 1.045 | 0.068118 | 0.830848 | 0.052025 | 0.999971 | 1 | 0.723736 |
| Ptprf_1 | A0A1Y7VN | -0.28 | 0.61 | -4.2 | -5.17 | 0.24 | 0.05 | -3.65 | -5.22 | 0.9 | 0 | -4.16 | -4.94 | 2.57 | -0.52 | -3.69 | -3.64 | 0.115 | 0.21 | 0.94 | 0.947373 | 0.908203 | 0.610031 | 0.999971 | 1 | 0.92673 |
| Ptprf_2 | A2A8L5 | -1.17 | -1.05 | -1.66 | -1.4 | -0.81 | -1.18 | -1.53 | -1.36 | -0.81 | -0.93 | -1.27 | -1.18 | -0.46 | -1.08 | -1.26 | -1.53 | 0.1 | 0.2725 | 0.2375 | 0.676147 | 0.226884 | 0.406681 | 0.999971 | 1 | 0.874911 |
| Ptprg | Q05909 | -0.01 | -1.15 | -1.22 | -1.14 | -0.86 | -1.22 | -0.82 | -1.14 | -1.09 | -0.83 | -1.18 | -1.06 | -1.5 | -1.63 | -1.19 | -1.2 | -0.13 | -0.16 | -0.5 | 0.678098 | 0.603622 | 0.139237 | 0.999971 | 1 | 0.784948 |
| Ptprj | A2AWF8 | -1.15 | -0.82 | -1.06 | -1.46 | -1.01 | -1.27 | -0.73 | -1.75 | -0.92 | -1.26 | -0.68 | -0.85 | -0.1 | -1.31 | -1.27 | -1.37 | -0.0675 | 0.195 | 0.11 | 0.804437 | 0.386924 | 0.740967 | 0.999971 | 1 | 0.960899 |
| Ptprk | A0A1W2P7 | 0.37 | -0.63 | -0.63 | -0.38 | -0.61 | -0.29 | -0.43 | -0.21 | -0.57 | -0.28 | -0.52 | -0.49 | -0.48 | -0.4 | -0.61 | -0.11 | -0.0675 | -0.1475 | -0.0825 | 0.803614 | 0.580874 | 0.766076 | 0.999971 | 1 | 0.962472 |
| Ptprm | Q68FM4 | -0.36 | 0.24 | -0.11 | 0.24 | -0.49 | -3.14 | -1.22 | 0.3 | -0.66 | 0.22 | -1.37 | 0.47 | 1.7 | 0.41 | 0.39 | 0.21 | -1.14 | -0.3375 | 0.675 | 0.120545 | 0.434865 | 0.091611 | 0.999971 | 1 | 0.773779 |
| Ptprn | A6MDD2 | 0.01 | -1.38 | -0.37 | -0.35 | -0.85 | -0.94 | -0.74 | -0.28 | -0.63 | -0.55 | -0.91 | -0.08 | -0.76 | -0.26 | 0.6 | -0.18 | -0.18 | -0.02 | 0.3725 | 0.590406 | 0.953217 | 0.357291 | 0.999971 | 1 | 0.868793 |
| Ptprn2 | P80560 | -0.46 | -1.39 | -0.8 | -0.76 | -0.98 | -0.8 | -0.57 | -0.53 | -0.71 | -0.52 | -0.73 | -0.65 | -0.53 | -0.35 | -0.41 | -0.66 | 0.1325 | 0.2 | 0.365 | 0.598044 | 0.402067 | 0.154099 | 0.999971 | 1 | 0.795578 |
| Ptpro | E9Q4I1 | -1.84 | -1.29 | -1.01 | -1 | -1.19 | -2.12 | -1.59 | -0.91 | -1.78 | -1.82 | -0.2 | -0.72 | -1.97 | -1.16 | -0.48 | -0.86 | -0.1675 | 0.155 | 0.1675 | 0.612166 | 0.715981 | 0.647136 | 0.999971 | 1 | 0.933613 |
| Ptprr | Q62132 | -1.12 | -1.38 | -1.14 | -0.94 | -1.07 | -1.12 | -0.71 | -1.19 | -1.25 | -0.78 | -1.01 | -0.59 | -0.95 | -1.03 | -1.44 | -0.92 | 0.1225 | 0.2375 | 0.06 | 0.542754 | 0.283278 | 0.770879 | 0.999971 | 1 | 0.963498 |
| Ptprs | B0V2N1-2 | -1.36 | -1.24 | -1.2 | -1.25 | -1.28 | -1.24 | -1.4 | -1.15 | -1.22 | -1.36 | -1.21 | -1.12 | -0.9 | -1.32 | -1.22 | -1.16 | -0.005 | 0.035 | 0.1125 | 0.975934 | 0.828589 | 0.53342 | 0.999971 | 1 | 0.910069 |
| Ptprt | Q99M80-3 | -0.9 | -1.25 | -1.8 | -1.78 | -1.22 | -1.28 | -1.61 | -1.71 | -1.17 | -1 | -1.45 | -1.41 | -1 | -1.19 | -1.47 | -1.77 | -0.0225 | 0.175 | 0.075 | 0.93334 | 0.511441 | 0.795273 | 0.999971 | 1 | 0.970361 |
| Ptpru | B1AUH1 | 0 | -0.18 | -0.28 | -0.57 | -0.69 | 0.14 | -0.82 | -0.37 | -0.73 | 0.08 | -0.48 | -0.66 | -0.98 | 0.29 | -0.25 | -0.63 | -0.1775 | -0.19 | -0.135 | 0.509632 | 0.448345 | 0.659368 | 0.999971 | 1 | 0.935181 |
| Ptprz1 | B9EKR1 | 1.28 | 1.61 | 1.47 | 1.57 | 1.45 | 1.61 | 1.42 | 1.34 | 1.51 | 1.4 | 1.27 | 1.41 | 1.43 | 1.44 | 1.46 | 1.49 | -0.0275 | -0.085 | -0.0275 | 0.876237 | 0.622426 | 0.871939 | 0.999971 | 1 | 0.984109 |
| Ptrh1 | Q8BW00 | 0.31 | -0.11 | 0.02 | -0.36 | -0.68 | -0.07 | -0.12 | 0.1 | 0.01 | 0.35 | 0.55 | 0.44 | 1.03 | 0.37 | 0.06 | 1.26 | -0.1575 | 0.3725 | 0.715 | 0.530231 | 0.119926 | 0.047309 | 0.999971 | 1 | 0.710102 |
| Ptrh2 | Q8R2Y8 | -0.84 | -0.22 | -0.02 | -0.33 | -0.31 | 0.45 | 0.08 | -0.14 | 0.3 | -0.21 | 0.21 | 0.02 | 0.44 | 0.71 | 0.04 | -0.2 | 0.3725 | 0.4325 | 0.6 | 0.179622 | 0.097703 | 0.05913 | 0.999971 | 1 | 0.725348 |
| Ptrhd1 | D3Z4S3 | 0.04 | 0.62 | 1.13 | 0.74 | 0.99 | 0.73 | 0.82 | 0.11 | 1.14 | 0.03 | 0.98 | 0.59 | 1.36 | 0.36 | 0.94 | 0.97 | 0.03 | 0.0525 | 0.275 | 0.921176 | 0.874637 | 0.387758 | 0.999971 | 1 | 0.871816 |
| Pts | Q9R1Z7 | 1.3 | 0.16 | 1.76 | -0.02 | 0.66 | 0.46 | 2.19 | 1.47 | 0.69 | 0.59 | 2.07 | 2.96 | 0.66 | -0.38 | 1.25 | 0.79 | 0.395 | 0.7775 | -0.22 | 0.471955 | 0.254741 | 0.669342 | 0.999971 | 1 | 0.938711 |
| Pttg1ip | Q8R143 | -5.5 | -9.18 | 1.25 | 1.33 | -6.71 | -7.66 | 1.39 | 1.4 | -8.56 | -4.39 | 1 | 0.57 | -8.38 | -6.16 | 0.9 | 0.13 | 0.13 | 0.18 | -0.3525 | 0.9673 | 0.953192 | 0.908498 | 0.999971 | 1 | 0.991707 |
| Puf60 | Q3UEB3-2 | 0.45 | 1.24 | 1.41 | 1.33 | 1.31 | 1.01 | 0.81 | 1.35 | 1.1 | 0.97 | 1.27 | 1.17 | 0.81 | 1.52 | 1.5 | 1.51 | 0.0125 | 0.02 | 0.2275 | 0.963578 | 0.937318 | 0.446946 | 0.999971 | 1 | 0.884581 |
| Pum1 | Q80U78-3 | 0.1 | 1.12 | 0.89 | 0.49 | 0.97 | 0.79 | 0.92 | 0.68 | 0.9 | 0.29 | 1.08 | 1.36 | 0.91 | 0.63 | 0.93 | 1.11 | 0.19 | 0.2575 | 0.245 | 0.46906 | 0.430931 | 0.370856 | 0.999971 | 1 | 0.869526 |
| Pum1_1 | Q80U78-2 | -0.98 | -0.6 | 0.47 | 1.29 | -0.51 | -1 | -0.37 | 0.59 | -0.05 | -0.99 | -0.18 | 0.85 | -0.38 | -0.67 | -0.35 | 1.33 | -0.3675 | -0.1375 | -0.0625 | 0.519059 | 0.815091 | 0.920544 | 0.999971 | 1 | 0.992702 |
| Pum2 | Q3UR91 | 0.59 | 1.92 | 0.96 | 0.98 | 1.03 | 0.96 | 0.63 | 1.33 | 1.36 | 1.21 | 0.74 | 1.03 | 1.92 | 1.82 | 1.25 | 1.55 | -0.125 | -0.0275 | 0.5225 | 0.697273 | 0.930658 | 0.132721 | 0.999971 | 1 | 0.784948 |
| Pum3 | Q8BKS9 | -0.41 | -0.81 | -0.96 | -1.08 | -0.31 | -0.64 | -0.31 | -0.97 | -0.86 | -0.29 | -0.93 | -1.12 | -0.95 | -0.65 | -0.89 | -1.24 | 0.2575 | 0.015 | -0.1175 | 0.309152 | 0.952957 | 0.610826 | 0.999971 | 1 | 0.92673 |
| Pura | P42669 | 0.47 | 1.53 | 1.01 | 0.82 | 1.37 | 1.14 | 1.02 | 0.72 | 0.98 | 0.34 | 1.06 | 0.95 | 0.88 | 1.17 | 0.69 | 0.6 | 0.105 | -0.125 | -0.1225 | 0.704802 | 0.665726 | 0.655858 | 0.999971 | 1 | 0.934061 |
| Purb | O35295 | 0.08 | 0.94 | 1.18 | 0.94 | 1.05 | 0.8 | 1.21 | 1.43 | 1.24 | 0.3 | 2.06 | 1.84 | 0.85 | 1.16 | 1.71 | 1.43 | 0.3375 | 0.575 | 0.5025 | 0.260389 | 0.21107 | 0.13177 | 0.999971 | 1 | 0.784948 |
| Purg | Q8R4E6 | 0.48 | 1.64 | 1.7 | 1.46 | 1.45 | 1.95 | 1.5 | 2.03 | 1.76 | 1.19 | 2.3 | 1.49 | 2.13 | 1.83 | 1.97 | 2.21 | 0.4125 | 0.365 | 0.715 | 0.222982 | 0.326848 | 0.040881 | 0.999971 | 1 | 0.700838 |
| Pus7 | Q91VU7-2 | 1.34 | 1.69 | 1.67 | 1.86 | 1.77 | 0.83 | 1.22 | 1.78 | 1.13 | 1.36 | 1.63 | 2.03 | 0.85 | 1.87 | 1.55 | 1.9 | -0.24 | -0.1025 | -0.0975 | 0.390227 | 0.681094 | 0.730741 | 0.999971 | 1 | 0.956041 |
| Pusl1 | A2ADA5 | 0.29 | -0.32 | -0.98 | -0.14 | -0.47 | -0.42 | 0.17 | -0.2 | -0.69 | 0.11 | -1.08 | -0.61 | -0.47 | -0.3 | -1.15 | -0.36 | 0.0575 | -0.28 | -0.2825 | 0.851646 | 0.438844 | 0.4013 | 0.999971 | 1 | 0.871816 |
| Pvalb | P32848 | 0.76 | 1.35 | 1.99 | 1.36 | -0.45 | 2.15 | 1.36 | 1.14 | -0.89 | -0.45 | 0.72 | -0.63 | -1.2 | 0.7 | 0.22 | 1.45 | -0.315 | -1.6775 | -1.0725 | 0.57114 | 0.003039 | 0.083694 | 0.999971 | 0.923185 | 0.755254 |
| Pvr | Q8K094 | 1.43 | 0.54 | 0.63 | -0.33 | 0.22 | 0.53 | 0.8 | -0.33 | -0.1 | 0.79 | 0.58 | -0.87 | -0.37 | 0.03 | -0.24 | -0.42 | -0.2625 | -0.4675 | -0.8175 | 0.529427 | 0.349189 | 0.048696 | 0.999971 | 1 | 0.710102 |
| Pwp1 | Q99LL5 | 0.4 | -0.12 | -0.35 | -0.19 | 0.1 | -0.54 | -0.56 | -0.45 | -0.37 | 0.07 | -1.05 | -0.64 | -1.25 | -0.05 | -0.14 | -0.34 | -0.2975 | -0.4325 | -0.38 | 0.256945 | 0.167882 | 0.256921 | 0.999971 | 1 | 0.839012 |
| Pwp2 | Q8BU03 | 3.98 | -1.9 | -1.8 | -0.85 | -2.34 | -0.58 | -0.37 | -2.15 | -1.86 | 1.41 | -2.07 | -2.67 | -3.48 | -1.86 | -2.31 | -3.09 | -1.2175 | -1.155 | -2.5425 | 0.36796 | 0.446941 | 0.074144 | 0.999971 | 1 | 0.746069 |
| Pwwp3a | Q6DID5 | 3.17 | 2.74 | 2.41 | 2.1 | 2.55 | 2.16 | 1.32 | 0.73 | 2.04 | 3.18 | 1.31 | 1.61 | 1.31 | 2.99 | 1.21 | 1.78 | -0.915 | -0.57 | -0.7825 | 0.064304 | 0.220949 | 0.104106 | 0.999971 | 1 | 0.78268 |
| Pxdn | Q3UQ28 | -1.04 | 0.14 | 0.15 | 0.44 | 0.45 | -0.38 | -0.17 | 0.22 | 0.27 | -0.45 | 0.74 | 0.92 | 0.64 | 0.06 | 0.58 | 0.91 | 0.1075 | 0.4475 | 0.625 | 0.770102 | 0.30947 | 0.113755 | 0.999971 | 1 | 0.78268 |
| Pxk | A0A2C9F2E | -0.1 | 0.48 | 0.76 | 0.64 | 0.05 | 0.5 | 0.7 | 0.84 | 0.61 | 0.64 | 1.16 | 0.73 | -0.38 | 0.85 | 0.6 | 1.22 | 0.0775 | 0.34 | 0.1275 | 0.777943 | 0.202926 | 0.737231 | 0.999971 | 1 | 0.9596 |
| Pxmp2 | D6REH1 | -0.06 | -1.45 | -0.12 | -0.22 | -1.06 | -1 | -0.8 | 1.21 | -1.45 | -0.02 | -0.78 | -0.37 | -0.55 | -0.81 | -0.73 | -0.07 | 0.05 | -0.1925 | -0.0775 | 0.931466 | 0.654948 | 0.830457 | 0.999971 | 1 | 0.978535 |
| Pxmp4 | Q9JJW0 | 0.87 | 0.3 | 3.57 | 1.41 | 0.43 | -0.05 | 0.13 | 2.06 | 0.75 | 1.57 | 1.42 | 1.61 | 1.29 | 0.43 | 2.16 | 1.7 | -0.895 | -0.2 | -0.1425 | 0.266921 | 0.767839 | 0.844443 | 0.999971 | 1 | 0.979814 |
| Pxn | F8VQ28 | 1.25 | 1.96 | 1.6 | 1.86 | 2.11 | 1.74 | 2.68 | 0.83 | 1.91 | 1.45 | 0.92 | 1.36 | 1.1 | 1.32 | 2.01 | 0.86 | 0.1725 | -0.2575 | -0.345 | 0.668277 | 0.359655 | 0.27214 | 0.999971 | 1 | 0.843794 |
| Pycard | Q9EPB4 | -0.8 | 0.66 | 1.22 | 1.51 | 0.78 | 0.8 | 0.51 | 1 | 0.75 | -0.04 | 0.97 | 1.43 | 0.46 | 0.56 | 1.08 | 1.69 | 0.125 | 0.13 | 0.3 | 0.797503 | 0.81421 | 0.582083 | 0.999971 | 1 | 0.920364 |
| Pycr1 | Q922W5 | 0.75 | -0.87 | -1.23 | 0.45 | -1.3 | -1.31 | -0.48 | -0.95 | -1.18 | -0.42 | -0.9 | -1.54 | -2.48 | -1.64 | -0.46 | -0.13 | -0.785 | -0.785 | -0.9525 | 0.133665 | 0.145449 | 0.174427 | 0.999971 | 1 | 0.807788 |
| Pycr2 | Q922Q4 | -0.49 | 0.01 | -0.17 | -0.53 | -0.08 | -0.22 | 0.17 | -0.32 | 0.26 | -0.51 | 0.02 | -0.04 | 1.07 | -0.09 | -0.41 | -0.3 | 0.1825 | 0.2275 | 0.3625 | 0.405673 | 0.352169 | 0.325144 | 0.999971 | 1 | 0.856472 |
| Pycr3 | Q9DCC4 | 1.58 | 0.36 | 0.75 | 1.33 | 0.21 | 0.8 | 1.22 | 0.78 | 0.39 | 1.18 | 1.06 | 0.77 | -0.01 | 0.74 | 1.07 | 0.63 | -0.2525 | -0.155 | -0.3975 | 0.466896 | 0.638041 | 0.275997 | 0.999971 | 1 | 0.843794 |
| Pygb | Q8CI94 | -0.69 | -0.56 | -0.47 | -0.89 | -0.59 | -0.33 | -0.43 | -0.7 | -0.61 | -0.76 | -0.74 | -0.54 | -0.68 | -0.47 | -0.61 | -0.84 | 0.14 | -0.01 | 0.0025 | 0.46857 | 0.955322 | 0.989442 | 0.999971 | 1 | 0.99925 |
| Pygl | Q9ET01 | 0.26 | -0.2 | -0.18 | -0.06 | 0.31 | 0.23 | -0.37 | 0.99 | 0.14 | 0.4 | -0.25 | 0.21 | 0.42 | 0.18 | 0.52 | 0.51 | 0.335 | 0.17 | 0.4525 | 0.288579 | 0.439498 | 0.043233 | 0.999971 | 1 | 0.701725 |
| Pygm | Q9WUB3 | -0.61 | -0.26 | -0.47 | -0.23 | -0.64 | -0.27 | -0.93 | -0.71 | -0.84 | -1.09 | -1.14 | -0.76 | -0.59 | -0.43 | -0.87 | -0.71 | -0.245 | -0.565 | -0.2575 | 0.267801 | 0.015182 | 0.207426 | 0.999971 | 1 | 0.820632 |
| Pygo2 | Q3V113 | 1.78 | 1.92 | 0.29 | 0.66 | 1.93 | 2.11 | 0.63 | 0.33 | 2.11 | 1.51 | 0.14 | 0.35 | 0.11 | 2.71 | 0.52 | 0.6 | 0.0875 | -0.135 | -0.1775 | 0.874867 | 0.813431 | 0.784142 | 0.999971 | 1 | 0.966376 |
| Pym1 | Q8CHP5-2 | 1.05 | 1.42 | 1.58 | -0.17 | 1.41 | 1.18 | 1.65 | 0.77 | 1.63 | 0.8 | 1.99 | 2.15 | 1.56 | 1.45 | 1.47 | 0.51 | 0.2825 | 0.6725 | 0.2775 | 0.502628 | 0.174904 | 0.53324 | 0.999971 | 1 | 0.910069 |
| Pyroxd1 | Q3TMV7 | 0.62 | 1.83 | 0.65 | 0.13 | 1.24 | -0.2 | 0.59 | 0.58 | 0.51 | -0.65 | 0.72 | 0.56 | -2.63 | 0.35 | 0.26 | 0.16 | -0.255 | -0.5225 | -1.2725 | 0.564979 | 0.26604 | 0.108844 | 0.999971 | 1 | 0.78268 |
| Pyroxd2 | A0A494BA! | 4.81 | 0.83 | -1 | -2.06 | -0.02 | -0.05 | 4.5 | -1.03 | -2.85 | 1.03 | -1.5 | -1.81 | 2.37 | -1.46 | -1.8 | -2.26 | 0.205 | -1.9275 | -1.4325 | 0.905572 | 0.231128 | 0.393014 | 0.999971 | 1 | 0.871816 |
| Pyurf | Q9D1C3 | 3.08 | -0.99 | -1.01 | -0.56 | 0.16 | -0.33 | -0.78 | -0.53 | -0.32 | -0.23 | -0.41 | -0.56 | -1.07 | -1.81 | -0.71 | -0.8 | -0.5 | -0.51 | -1.2275 | 0.582746 | 0.571894 | 0.202159 | 0.999971 | 1 | 0.81617 |
| Pzp | Q61838 | 2.68 | 2.88 | 1.08 | 3.72 | 4.32 | 1.13 | 3.32 | 1.52 | 1.28 | 0.83 | 2.2 | 2.26 | 2.64 | 1.89 | 2.08 | 1.62 | -0.0175 | -0.9475 | -0.5325 | 0.983299 | 0.141479 | 0.342529 | 0.999971 | 1 | 0.861474 |
| Q0VG49 | Q0VG49 | -1.38 | -2.79 | -4.85 | -1.22 | -1.87 | -2.45 | -4.26 | -1.71 | -3.5 | -2.12 | -2.1 | -4.09 | -4.61 | -1.52 | -1.63 | -2.17 | -0.0125 | -0.3925 | 0.0775 | 0.989097 | 0.65836 | 0.937644 | 0.999971 | 1 | 0.995618 |
| Q3TQI7 | Q3TQI7 | 2.52 | 2.34 | 2.81 | 2.57 | 1.54 | 1.93 | 2.15 | 3.05 | 1.57 | 1.99 | 2.34 | 2.28 | 2.02 | 2.59 | 2.01 | 2.79 | -0.3925 | -0.515 | -0.2075 | 0.257388 | 0.05196 | 0.416323 | 0.999971 | 1 | 0.877131 |
| Q3TTL0 | Q3TTL0 | 0.2 | 1.2 | 0.57 | 0.93 | 0.32 | 0.7 | 0.57 | 1.16 | 1.01 | 0.9 | 0.82 | 1.73 | 0.47 | 1.7 | 1.06 | 0.95 | -0.0375 | 0.39 | 0.32 | 0.897486 | 0.225385 | 0.349704 | 0.999971 | 1 | 0.865616 |
| Q5EBG8 | Q5EBG8 | 0.44 | 0.32 | -0.54 | 0.17 | 0.84 | 0.8 | -1.02 | -0.12 | 0.56 | 0.74 | -0.36 | 0.2 | 0.91 | 1.85 | 0.04 | 0.59 | 0.0275 | 0.1875 | 0.75 | 0.952414 | 0.570107 | 0.09968 | 0.999971 | 1 | 0.78268 |
| Q80X32 | Q80X32 | 0.51 | 0.28 | 1 | 1.13 | 0.52 | -0.05 | 0.71 | 0.54 | 0.37 | 0.54 | -0.18 | 0.34 | 0.17 | 0.6 | 0.36 | 0.41 | -0.3 | -0.4625 | -0.345 | 0.295026 | 0.11568 | 0.189238 | 0.999971 | 1 | 0.810547 |
| Q8BGK9 | Q8BGK9 | -0.27 | -2.32 | -2.36 | -1.62 | -1.51 | -1.08 | -2.84 | -1.51 | -1.66 | -1.82 | -2.55 | -1.88 | -1.95 | -2.06 | -1.83 | -2.02 | -0.0925 | -0.335 | -0.3225 | 0.870549 | 0.501048 | 0.489691 | 0.999971 | 1 | 0.898405 |
| Q8BH50 | Q8BH50 | 0.47 | 0.08 | 0.25 | 0.24 | -0.25 | -0.27 | 0.25 | 0.41 | -0.59 | 0.85 | 1.2 | 0.16 | -0.07 | 0.6 | -0.72 | -0.1 | -0.225 | 0.145 | -0.3325 | 0.340321 | 0.710342 | 0.274251 | 0.999971 | 1 | 0.843794 |
| Q8BN57 | Q8BN57 | 0.58 | -0.24 | -0.11 | 0.66 | -0.21 | 0.37 | -0.13 | 0.61 | 0.23 | 0.76 | 0.43 | -0.28 | 0.47 | -0.31 | -0.16 | 0.16 | -0.0625 | 0.0625 | -0.1825 | 0.840061 | 0.845047 | 0.545968 | 0.999971 | 1 | 0.911398 |
| Q8BP99 | Q8BP99 | 0.36 | 0.07 | -0.43 | 0.23 | -0.07 | 0.3 | -0.29 | -0.09 | 0.23 | 0.45 | -0.44 | -0.04 | 1.12 | 0.13 | 0.2 | -0.13 | -0.095 | -0.0075 | 0.2725 | 0.697327 | 0.978167 | 0.410938 | 0.999971 | 1 | 0.875847 |
| Q8C3W1 | Q8C3W1 | -0.16 | -0.38 | -0.32 | -0.21 | -0.13 | -0.12 | -0.64 | 0.15 | -0.36 | -0.11 | -0.53 | -0.21 | 0.29 | 0.31 | -0.09 | 0.06 | 0.0825 | -0.035 | 0.41 | 0.705807 | 0.844403 | 0.049511 | 0.999971 | 1 | 0.715605 |
| Q8C5K5 | Q8C5K5 | -1.5 | 1.19 | -0.6 | -2.1 | 1.63 | 1.05 | 0.17 | -1.65 | 1.32 | -0.94 | 0.05 | 0.3 | 1.48 | 0.69 | -0.9 | -1.74 | 1.0525 | 0.935 | 0.635 | 0.264202 | 0.247222 | 0.495445 | 0.999971 | 1 | 0.901531 |
| Q91WE4 | Q91WE4 | 3.12 | -0.74 | 0.48 | 1.32 | -1.11 | 0.43 | 3.11 | -0.65 | -1.39 | 1.57 | -0.89 | -0.66 | -0.82 | -0.78 | -0.36 | -0.68 | -0.6 | -1.3875 | -1.705 | 0.591279 | 0.165115 | 0.043579 | 0.999971 | 1 | 0.701725 |
| Q922M7 | Q922M7 | -0.03 | 1.08 | 0.86 | 0.85 | 1.29 | 0.39 | 0.72 | 0.17 | 0.95 | 0 | 0.55 | 0.57 | 0.11 | 1.09 | 0.59 | 0.49 | -0.0475 | -0.1725 | -0.12 | 0.889481 | 0.589167 | 0.709321 | 0.999971 | 1 | 0.950625 |
| Q99M08 | Q99M08 | 0.18 | 0.46 | 0.14 | 0.68 | 0.34 | 0.34 | 0.76 | 0.81 | 0.09 | 0.59 | -0.71 | -0.02 | -0.11 | -0.75 | -0.48 | 0.62 | 0.1975 | -0.3775 | -0.545 | 0.386624 | 0.233751 | 0.120522 | 0.999971 | 1 | 0.78268 |
| Q9CPS8 | Q9CPS8 | 1.06 | 1.67 | -0.32 | 0.15 | 0.83 | 1.13 | -1.11 | -0.74 | 0.16 | 1.27 | -1.2 | -0.19 | -0.16 | 1.13 | -1.09 | -0.62 | -0.6125 | -0.63 | -0.825 | 0.358055 | 0.325138 | 0.191598 | 0.999971 | 1 | 0.810547 |
| Q9CQ90 | Q9CQ90 | 3.92 | 1.78 | 3.94 | 3.46 | 1.68 | 4.61 | 3.43 | 4.38 | 1.37 | 2.54 | 1.7 | 1.9 | 0.74 | -0.08 | 2.7 | 0.76 | 0.25 | -1.3975 | -2.245 | 0.74133 | 0.02501 | 0.010808 | 0.999971 | 1 | 0.613095 |
| Q9CRC3 | Q9CRC3 | -1.3 | -0.27 | -1.85 | -1.03 | -0.37 | -0.52 | -1.02 | -1.02 | -0.68 | -1.43 | -1.38 | -1.36 | -0.53 | -0.41 | -1.12 | -0.69 | 0.38 | -0.1 | 0.425 | 0.307494 | 0.784138 | 0.252721 | 0.999971 | 1 | 0.839012 |
| Q9CYS6 | Q9CYS6 | -0.68 | -0.56 | -1.34 | -1.68 | -1.03 | -0.68 | -1.44 | 0.69 | -0.84 | -0.89 | -1.63 | -0.54 | -0.4 | 1.46 | -1.15 | -1.27 | 0.45 | 0.09 | 0.725 | 0.375068 | 0.796233 | 0.264009 | 0.999971 | 1 | 0.842533 |
| Q9D7E4 | Q9D7E4 | -0.77 | -0.74 | -1.16 | -1.21 | -0.09 | -0.52 | -1.42 | -0.56 | -0.66 | -0.1 | -1.11 | -1.11 | -1.17 | -0.54 | -1.04 | -1.56 | 0.3225 | 0.225 | -0.1075 | 0.313924 | 0.435594 | 0.687758 | 0.999971 | 1 | 0.945577 |
| Q9D9H8 | Q9D9H8 | 0.22 | 0.1 | -0.17 | 0.24 | -0.35 | -0.21 | 0.45 | -0.04 | -0.16 | 0.01 | -0.23 | -0.13 | -0.5 | -0.14 | -0.63 | -0.13 | -0.135 | -0.225 | -0.4475 | 0.56783 | 0.230996 | 0.058021 | 0.999971 | 1 | 0.725348 |
| Qars1 | Q8BML9 | 0.29 | 1.28 | 0.95 | 0.83 | 1.54 | 1.02 | 0.94 | 1.46 | 1.5 | 0.64 | 1.42 | 1.43 | 1.59 | 1.01 | 1.49 | 1.4 | 0.4025 | 0.41 | 0.535 | 0.166351 | 0.192152 | 0.069089 | 0.999971 | 1 | 0.733937 |
| Qdpr | Q8BVI4 | -0.33 | 0.17 | 0.35 | 0.02 | -0.14 | -0.12 | 0.14 | -0.07 | -0.08 | -0.46 | -0.12 | -0.11 | 0 | 0.25 | 0.45 | -0.02 | -0.1 | -0.245 | 0.1175 | 0.634612 | 0.270428 | 0.603704 | 0.999971 | 1 | 0.924454 |
| Qki | Q9QYS9-8 | 1.11 | 0.37 | 0.22 | 0.06 | 0.41 | 0.61 | -0.24 | 0.39 | 0.6 | 0.79 | -0.04 | 0.27 | 0.82 | 0.86 | 0.44 | 0.29 | -0.1475 | -0.035 | 0.1625 | 0.6293 | 0.907841 | 0.573001 | 0.999971 | 1 | 0.91816 |
| Qpct | A0A3B2W7 | -0.82 | -0.3 | -1.06 | 0.17 | 0.07 | -0.67 | -0.77 | 0.09 | -0.77 | -0.44 | -0.72 | -0.22 | -0.25 | 0.22 | 2.18 | -0.2 | 0.1825 | -0.035 | 0.99 | 0.607267 | 0.909543 | 0.116804 | 0.999971 | 1 | 0.78268 |
| Qpctl | Q8BH73 | 0.71 | -0.04 | 0.69 | -0.21 | -0.39 | -0.01 | -0.16 | -0.18 | -0.2 | -0.02 | 0.87 | 0.48 | -0.13 | 0.98 | 1.5 | 1.13 | -0.4725 | -0.005 | 0.5825 | 0.109262 | 0.988214 | 0.176559 | 0.999971 | 1 | 0.808532 |
| Qrich1 | G3X8R5 | 1.13 | 2.25 | 1.97 | 1.78 | 2.54 | 2.23 | 1.52 | 1.56 | 2.11 | 1.5 | 1.25 | 2.19 | 2.02 | 2.2 | 2.18 | 2.12 | 0.18 | -0.02 | 0.3475 | 0.601933 | 0.951641 | 0.210421 | 0.999971 | 1 | 0.822657 |
| Qrsl1 | Q9CZN8 | -0.07 | -0.02 | -0.2 | 0 | -0.38 | 0.09 | -0.36 | 0.06 | -0.52 | 0.08 | -0.06 | -0.22 | -0.1 | 0.06 | -0.12 | -0.29 | -0.075 | -0.1075 | -0.04 | 0.70456 | 0.583596 | 0.818397 | 0.999971 | 1 | 0.97527 |
| Qsox1 | Q8BND5-3 | 0.38 | 1.69 | 0.15 | 0.76 | 2.12 | 0.84 | 0.91 | 0.5 | 1.36 | 0.48 | 0.04 | 0.46 | 2.09 | 0.79 | 1.31 | 1.08 | 0.3475 | -0.16 | 0.5725 | 0.456576 | 0.702569 | 0.193921 | 0.999971 | 1 | 0.810547 |
| Qsox2 | Q3TMX7 | 0.92 | 1.84 | 1.61 | 1.71 | 1.93 | 1.68 | 0.67 | 2.37 | 1.59 | 1.1 | 1.2 | 1.76 | 1.16 | 1.49 | 2.23 | 2.38 | 0.1425 | -0.1075 | 0.295 | 0.72038 | 0.696521 | 0.411298 | 0.999971 | 1 | 0.875847 |
| Qtrt1 | Q9JMA2 | 0.8 | 1.15 | 0.5 | 0.73 | 0.64 | 0.43 | -0.21 | 0.4 | 0.04 | 0.76 | 0.6 | 0.28 | 0.33 | 0.33 | 0.37 | 0.93 | -0.48 | -0.375 | -0.305 | 0.085219 | 0.144899 | 0.21693 | 0.999971 | 1 | 0.826893 |
| Qtrt2 | A0A5F8MP | 0.35 | 1.46 | 1.14 | 0.81 | 0.9 | 0.99 | 0.4 | 1.04 | 0.96 | 0.37 | 0.41 | 0.9 | 0.78 | 2.43 | 0.93 | 0.97 | -0.1075 | -0.28 | 0.3375 | 0.712429 | 0.353557 | 0.440202 | 0.999971 | 1 | 0.883105 |
| R3hcc1 | E9PUE1 | 1.48 | 0.23 | 0.92 | 0.96 | 0.25 | 0.5 | 0.78 | 1.56 | 0.68 | 1.06 | 1.5 | 1.56 | 0.02 | 0.49 | 1.79 | 1.51 | -0.125 | 0.3025 | 0.055 | 0.73681 | 0.369283 | 0.904762 | 0.999971 | 1 | 0.991091 |
| R3hdm1 | E9Q9Q2 | 0.09 | 0.67 | -0.26 | -0.35 | 0.56 | 0.54 | -0.18 | -0.09 | 0.54 | 0.43 | -0.37 | -0.15 | 0.56 | 0.36 | -0.03 | -0.63 | 0.17 | 0.075 | 0.0275 | 0.586564 | 0.815501 | 0.936488 | 0.999971 | 1 | 0.995618 |
| R3hdm2 | Q80TM6-2 | 0.9 | -0.49 | -0.36 | 0.02 | -0.29 | -0.16 | 0.02 | -0.59 | -0.42 | 0.16 | -0.11 | -0.26 | -0.51 | -0.53 | -0.53 | -0.18 | -0.2725 | -0.175 | -0.455 | 0.426598 | 0.603869 | 0.187049 | 0.999971 | 1 | 0.809025 |
| Rab10 | P61027 | -3.11 | -1.05 | -1.11 | -2.65 | -1.01 | -1.83 | -0.59 | -2.33 | -0.45 | -3.65 | -0.52 | -0.81 | -0.88 | -2.9 | -1.85 | -2.58 | 0.54 | 0.6225 | -0.0725 | 0.379814 | 0.467021 | 0.908486 | 0.999971 | 1 | 0.991707 |
| Rab11b | P46638 | 0.19 | 0.64 | 0.17 | -0.19 | 0.81 | 0.99 | 0.04 | 0.06 | 0.81 | 0.23 | 0.51 | 0.51 | 1.52 | 0.47 | 0.31 | 0.44 | 0.2725 | 0.3125 | 0.4825 | 0.385603 | 0.211992 | 0.167127 | 0.999971 | 1 | 0.800215 |
| Rab11fip2 | G3XA57-2 | -0.76 | -1.07 | -0.71 | -0.75 | -1.04 | -0.89 | -0.9 | -0.66 | -1.16 | -0.79 | -0.6 | -0.67 | -0.74 | -0.77 | -0.71 | -0.29 | -0.05 | 0.0175 | 0.195 | 0.788469 | 0.930905 | 0.343057 | 0.999971 | 1 | 0.861474 |
| Rab11fip3 | Q8CHD8-3 | 1.07 | 0.84 | -0.33 | 0.09 | 0.38 | 0.69 | -0.08 | 0.6 | -0.01 | 1.19 | 0.14 | -0.16 | 0.11 | 0.81 | 0.8 | 0.41 | -0.02 | -0.1275 | 0.115 | 0.955572 | 0.764356 | 0.749265 | 0.999971 | 1 | 0.961064 |
| Rab11fip4 | Q8BQP8 | -0.67 | -0.92 | 0.33 | -0.03 | -0.68 | -2.09 | -0.53 | 1.09 | -0.58 | -0.65 | 0.33 | -0.26 | -2.04 | -0.94 | -0.56 | -0.2 | -0.23 | 0.0325 | -0.6125 | 0.722411 | 0.927274 | 0.206146 | 0.999971 | 1 | 0.817241 |
| Rab11fip5 | Q8R361 | -3.04 | -11.17 | -8.07 | -10.13 | -9.07 | -8.86 | -6.95 | -8.18 | -8.75 | -4.91 | -6.57 | -8.46 | -10.3 | -8.57 | -7.45 | -8.98 | -0.1625 | 0.93 | -0.7225 | 0.921669 | 0.607773 | 0.669467 | 0.999971 | 1 | 0.938711 |
| Rab11fip5_ | A0A0N4SW | -0.47 | -0.41 | -0.38 | -0.51 | -0.72 | -0.45 | -0.6 | -0.35 | -0.53 | -0.42 | -0.25 | -0.12 | -0.07 | -0.65 | -0.3 | -0.12 | -0.0875 | 0.1125 | 0.1575 | 0.617785 | 0.523495 | 0.432309 | 0.999971 | 1 | 0.881643 |
| Rab12 | P35283 | 0.24 | 2.27 | 1.9 | 0.64 | -0.94 | -0.26 | -0.91 | -0.91 | -0.71 | -0.73 | -0.94 | -1.37 | -1 | -0.45 | -0.85 | -0.92 | -2.0175 | -2.2 | -2.0675 | 0.002316 | 0.001407 | 0.001794 | 0.999971 | 0.620535 | 0.476373 |

| | | | | | | | | | | | | | | | | | | | | | | | | | | |
|---|---|---|---|---|---|---|---|---|---|---|---|---|---|---|---|---|---|---|---|---|---|---|---|---|---|---|
| Rab13 | Q9DD03 | -4.43 | -4.77 | -0.83 | -1.5 | -2.36 | -5.57 | -0.21 | -1.54 | -0.95 | -5.5 | -1 | -0.13 | -2.34 | -8.02 | -1.27 | -1.83 | 0.4625 | 0.9875 | -0.4825 | 0.732038 | 0.490258 | 0.770356 | 0.999971 | 1 | 0.963498 |
| Rab14 | Q91V41 | -1.08 | -1.06 | -0.75 | -1.01 | -0.83 | -0.87 | -0.8 | -0.74 | -0.99 | -1.21 | -0.68 | -0.44 | -1.18 | -0.82 | -0.62 | -0.91 | 0.165 | 0.145 | 0.0925 | 0.350668 | 0.522728 | 0.644463 | 0.999971 | 1 | 0.93301 |
| Rab15 | Q3TYH2 | 0.21 | -0.55 | -0.62 | -0.66 | -0.78 | -0.12 | -0.79 | -0.24 | -0.75 | -0.15 | -0.79 | -0.73 | -0.09 | -0.15 | -0.43 | -0.01 | -0.0775 | -0.2 | 0.235 | 0.785942 | 0.470803 | 0.365171 | 0.999971 | 1 | 0.869526 |
| Rab18 | P35293 | 1 | -0.04 | -0.69 | -0.82 | -0.04 | -0.02 | 0 | -0.37 | -0.47 | 0.21 | -0.55 | -0.59 | -0.02 | 0 | -0.64 | -1.01 | 0.03 | -0.2125 | -0.28 | 0.940933 | 0.625586 | 0.54222 | 0.999971 | 1 | 0.911119 |
| Rab1a | Q5SW88 | -0.87 | -0.71 | -0.65 | -0.62 | -0.66 | -0.6 | -0.54 | -0.63 | -0.53 | -0.81 | -0.42 | -0.42 | -0.92 | -0.94 | -0.6 | -0.72 | 0.105 | 0.1675 | -0.0825 | 0.530951 | 0.363753 | 0.648322 | 0.999971 | 1 | 0.933842 |
| Rab1A | P62821 | -0.95 | -0.09 | -0.61 | 0.08 | -0.53 | -0.15 | -0.74 | -0.18 | -0.36 | -0.41 | -0.43 | -0.48 | -0.19 | -0.14 | -0.42 | 0.22 | -0.0075 | -0.0275 | 0.26 | 0.979248 | 0.915646 | 0.374207 | 0.999971 | 1 | 0.87031 |
| Rab1b | Q9D1G1 | 0.54 | -0.48 | -0.54 | -0.85 | -0.84 | 0.14 | 0.11 | -0.87 | -0.55 | -0.24 | -0.86 | -0.94 | -0.5 | -1.02 | -0.85 | -0.96 | -0.0325 | -0.315 | -0.5 | 0.934563 | 0.364446 | 0.150104 | 0.999971 | 1 | 0.793248 |
| Rab21 | P35282 | 1.82 | 0.22 | 2.98 | -0.4 | -0.27 | -0.14 | 2.07 | -0.67 | -0.33 | 2.54 | -0.86 | -2.03 | -0.16 | -0.3 | -0.67 | -1.17 | -0.9075 | -1.325 | -1.73 | 0.318499 | 0.25241 | 0.038072 | 0.999971 | 1 | 0.700838 |
| Rab22a | P35285 | 0.93 | 0.05 | -0.11 | 0.06 | 0.13 | 0.74 | -0.38 | 0.07 | 0.23 | 1.05 | -0.08 | -0.01 | 0.57 | 0.61 | 0.33 | 0.49 | -0.0925 | 0.065 | 0.2675 | 0.778894 | 0.850849 | 0.32841 | 0.999971 | 1 | 0.856472 |
| Rab23 | Q9D4I9 | -0.42 | -0.26 | -0.97 | -1.12 | -0.23 | 0.06 | -1.28 | -0.9 | -0.35 | -0.23 | -0.81 | -1.07 | 0.28 | -0.16 | -1.1 | -0.91 | 0.105 | 0.0775 | 0.22 | 0.771616 | 0.793328 | 0.559246 | 0.999971 | 1 | 0.915051 |
| Rab24 | P35290 | 0.44 | 0.4 | 0.56 | -0.02 | -0.12 | 0.82 | 0.22 | 0.49 | 0.35 | 0.84 | 0.6 | 0.09 | 0.68 | 1.33 | 0.92 | 0.78 | 0.0075 | 0.125 | 0.5825 | 0.976947 | 0.600345 | 0.03007 | 0.999971 | 1 | 0.678474 |
| Rab26 | Q504M8 | -1.73 | -1.45 | -0.36 | -1.21 | -0.61 | -1.26 | 0.38 | -0.77 | -0.81 | -1.93 | -0.15 | -0.92 | -0.51 | -1.47 | -0.68 | -1.18 | 0.6225 | 0.235 | 0.2275 | 0.171391 | 0.600549 | 0.533619 | 0.999971 | 1 | 0.910069 |
| Rab27a | Q9ERI2 | 0.09 | 0.07 | -0.64 | 0.47 | 0.49 | 0.61 | -0.34 | -0.45 | 0.69 | 0.22 | -1.43 | -1.15 | 1.62 | 0.99 | -0.69 | -1.14 | 0.08 | -0.415 | 0.1975 | 0.820531 | 0.437502 | 0.756942 | 0.999971 | 1 | 0.961116 |
| Rab27b | Q99P58 | -0.41 | -1.14 | -1.57 | -1.01 | -0.86 | -0.95 | -1.4 | -1.37 | -1.23 | -1.02 | -1.69 | -1.26 | -1.11 | -0.92 | -1.36 | -1.21 | -0.1125 | -0.2675 | -0.1175 | 0.698561 | 0.366629 | 0.670611 | 0.999971 | 1 | 0.93891 |
| Rab28 | Q99KL7 | 1.37 | 0.39 | 0.5 | 1.52 | 0.39 | 0.58 | 0.05 | 1.08 | 0.21 | 0.97 | 0.1 | 0.42 | -0.17 | 0.65 | 0.72 | 1.18 | -0.42 | -0.52 | -0.35 | 0.255325 | 0.157602 | 0.380231 | 0.999971 | 1 | 0.871816 |
| Rab29 | E9QLQ7 | -0.02 | -0.06 | -0.62 | 0.44 | 0.03 | 0.12 | -0.24 | -0.16 | -0.41 | 0.07 | 0.12 | 0.16 | -0.98 | -0.04 | -0.1 | 1.07 | 0.0025 | 0.05 | 0.0525 | 0.992234 | 0.853979 | 0.906102 | 0.999971 | 1 | 0.991381 |
| Rab2a | P53994 | -1.19 | 0.04 | -0.2 | -0.85 | 0 | 0.39 | 0.51 | 0 | 0.07 | -1.03 | 0.51 | 0.8 | 1.11 | 0.23 | 0.33 | -0.19 | 0.775 | 0.6375 | 0.92 | 0.034674 | 0.192038 | 0.036405 | 0.999971 | 1 | 0.700096 |
| Rab2b | P59279 | -1.47 | -1.75 | -1.16 | -1.62 | -1.51 | -1.44 | -0.8 | -1.04 | -1.02 | -1.62 | -1.51 | -1.44 | -0.59 | -1.54 | -1.39 | -1.49 | 0.3025 | 0.1025 | 0.2475 | 0.232546 | 0.646141 | 0.379308 | 0.999971 | 1 | 0.871816 |
| Rab30 | Q923S9 | -0.33 | -0.97 | -0.94 | -0.53 | -1.18 | -0.72 | -1.3 | -0.41 | -0.9 | -0.32 | -0.92 | -0.9 | -0.41 | -0.39 | -0.53 | -0.39 | -0.21 | -0.0675 | 0.2625 | 0.454671 | 0.781975 | 0.234956 | 0.999971 | 1 | 0.832918 |
| Rab31 | Q3TXV4 | 0.2 | -1.36 | -0.23 | -0.34 | -1.65 | -1.44 | -0.3 | -0.05 | -1.47 | -0.58 | -0.23 | -0.25 | -1.74 | -1.51 | -0.46 | -0.2 | -0.4275 | -0.2 | -0.545 | 0.387473 | 0.635133 | 0.266094 | 0.999971 | 1 | 0.842677 |
| Rab33a | P97950 | -0.11 | 0.45 | -1.26 | -1.27 | 0.31 | 0.64 | -1.04 | -0.88 | 0.34 | -0.17 | -1.03 | -1.12 | 0.79 | 0.08 | -1.07 | -0.83 | 0.305 | 0.0525 | 0.29 | 0.584026 | 0.918841 | 0.605743 | 0.999971 | 1 | 0.924649 |
| Rab33b | O35963 | 1.26 | -0.32 | -0.81 | -1.21 | 0.26 | -0.41 | -0.33 | -0.96 | 0.05 | 0.1 | -0.76 | -0.34 | 0.6 | -0.3 | -0.85 | -1.54 | -0.09 | 0.0325 | -0.2525 | 0.869485 | 0.951479 | 0.695164 | 0.999971 | 1 | 0.947174 |
| Rab34 | Q0PD20 | 0.72 | 0.43 | 0.36 | 0.44 | 0.43 | 0.66 | -0.02 | 0.62 | 0.34 | 0.49 | 0.29 | 0.2 | 0.56 | 0.43 | 0.52 | 0.53 | -0.065 | -0.1575 | 0.0225 | 0.767329 | 0.382916 | 0.897067 | 0.999971 | 1 | 0.988737 |
| Rab35 | Q6PHN9 | -0.86 | -1.01 | -1.74 | -1.76 | -1.12 | -0.8 | -0.98 | -1.84 | -0.47 | -1.02 | -1.44 | -1.7 | -1.11 | -1.07 | -1.8 | -2.02 | 0.1575 | 0.185 | -0.1575 | 0.633906 | 0.601997 | 0.641792 | 0.999971 | 1 | 0.93301 |
| Rab36 | Q8CAM5 | 1.69 | -1.04 | -0.83 | -0.83 | -1.16 | -1.1 | -1.63 | -0.95 | -1.07 | 0.82 | -1.55 | -1.53 | -1.44 | -0.56 | -0.75 | -0.6 | -0.9575 | -0.58 | -0.585 | 0.141112 | 0.463334 | 0.358772 | 0.999971 | 1 | 0.869526 |
| Rab37 | Q8BQX0 | 0.97 | -0.41 | -0.78 | -0.19 | -0.15 | 0.07 | -0.05 | -0.48 | -0.31 | 0.97 | 0.15 | -0.54 | -0.57 | -0.24 | -1.07 | 0.3 | -0.05 | 0.17 | -0.2925 | 0.895665 | 0.719675 | 0.518131 | 0.999971 | 1 | 0.905815 |
| Rab37_1 | Q544E8 | 2.83 | 2.69 | 0.19 | 0.45 | -1.82 | 1.1 | -0.7 | 0.31 | -0.92 | 0.3 | -1.09 | -3.38 | -2.28 | 1.31 | -0.72 | -0.06 | -1.8175 | -2.8125 | -1.9775 | 0.057554 | 0.014682 | 0.056633 | 0.999971 | 1 | 0.725348 |
| Rab39b | Q8BHC1 | -0.32 | -0.92 | -1.36 | -0.89 | -0.78 | -0.28 | -1.19 | -0.93 | -0.53 | -0.22 | -1.04 | -1.12 | -0.2 | -0.07 | -1.03 | -0.56 | 0.0775 | 0.145 | 0.4075 | 0.793827 | 0.639092 | 0.209796 | 0.999971 | 1 | 0.82263 |
| Rab3a | P63011 | -1.92 | -0.84 | -0.8 | -1.7 | -0.83 | -0.4 | -1.11 | -1.26 | -0.94 | -2.28 | -0.93 | -1.64 | -0.93 | -1.31 | -1.14 | -1.72 | 0.415 | -0.1325 | 0.04 | 0.244442 | 0.749457 | 0.904485 | 0.999971 | 1 | 0.991091 |
| Rab3b | Q9CZT8 | -0.63 | 0.16 | 0.24 | 0.4 | -1.48 | -1.06 | -0.62 | -1.73 | 0.78 | 0.08 | 0.16 | -0.3 | -1.49 | -1.67 | -1.08 | -1.63 | -1.265 | 0.1375 | -1.51 | 0.004185 | 0.670768 | 0.000498 | 0.999971 | 1 | 0.312038 |
| Rab3c | P62823 | -0.71 | -0.77 | -0.98 | -1.67 | -0.36 | -0.39 | -0.22 | -1 | -0.36 | -1.31 | -0.68 | -0.97 | -0.16 | -0.87 | -0.81 | -1.55 | 0.54 | 0.2025 | 0.185 | 0.091099 | 0.512679 | 0.60331 | 0.999971 | 1 | 0.924454 |
| Rab3d | P35276 | -1.5 | -1.29 | -1.21 | -1.79 | -1.63 | -0.87 | -1.19 | -1.1 | -1.28 | -1 | -0.5 | -0.84 | -2.01 | -0.94 | -1.48 | -0.87 | 0.25 | 0.5425 | 0.1225 | 0.310228 | 0.046373 | 0.688001 | 0.999971 | 1 | 0.945753 |
| Rab3gap1 | Q80UJ7 | -0.56 | -1.31 | -0.27 | -0.44 | -0.66 | -1.3 | -0.89 | -0.16 | -0.86 | -1.08 | -0.43 | -0.38 | -0.46 | -0.39 | -0.25 | 0.19 | -0.1075 | -0.0425 | 0.4175 | 0.745011 | 0.88525 | 0.169356 | 0.999971 | 1 | 0.801824 |
| Rab3gap1_ | A0A1D5RL( | -0.02 | 0.19 | -1.11 | -1.08 | 0.27 | 0.5 | -0.7 | -1.09 | 0.46 | 0.25 | -1.06 | -1.06 | 0.47 | 0.31 | -0.92 | -0.98 | 0.25 | 0.1525 | 0.225 | 0.603534 | 0.760342 | 0.64353 | 0.999971 | 1 | 0.93301 |
| Rab3gap2 | A0A0A6YW | -0.27 | -0.17 | 0.21 | 0.21 | -0.03 | -0.02 | -0.02 | 0.22 | -0.11 | -0.2 | 0.1 | 0.31 | -0.06 | 0.21 | 0.41 | 0.58 | 0.0425 | 0.03 | 0.29 | 0.830913 | 0.888403 | 0.22315 | 0.999971 | 1 | 0.827753 |
| Rab3il1 | D3YZ86 | 0.41 | 0.41 | 2.28 | 0.27 | 0.21 | -0.21 | 2.01 | 0.72 | 0.62 | -0.03 | 3.17 | 1.77 | 0.94 | 0.79 | 0.4 | 0.99 | -0.16 | 0.54 | -0.0625 | 0.796291 | 0.490094 | 0.893437 | 0.999971 | 1 | 0.98828 |
| Rab3ip | Q68EF0 | 0.47 | -0.08 | 2.03 | 1.53 | -0.51 | 0.54 | 2.02 | 1.07 | 0.06 | 0.74 | 1.81 | 1.02 | -0.86 | 1.44 | 1.22 | 1.34 | -0.2075 | -0.08 | -0.2025 | 0.749615 | 0.885655 | 0.760925 | 0.999971 | 1 | 0.961116 |
| Rab40c | E9PXR4 | 0.66 | 2.33 | -0.3 | -0.53 | 0.95 | 2.41 | 0.16 | -0.55 | 0.73 | 1.88 | -0.36 | 0.02 | 0.02 | 2.06 | -0.23 | -0.75 | 0.2025 | 0.0275 | -0.265 | 0.804047 | 0.970331 | 0.742878 | 0.999971 | 1 | 0.961064 |
| Rab43 | Q8CG50 | -0.36 | -0.27 | 0.09 | -0.07 | -0.29 | -0.16 | 0.38 | 0.22 | 0.15 | -0.33 | 0.31 | 0.46 | -0.16 | -1.56 | 0.71 | -0.05 | 0.19 | 0.3 | -0.1125 | 0.411786 | 0.219124 | 0.804653 | 0.999971 | 1 | 0.971467 |
| Rab4a | P56371 | -2.9 | -0.51 | -0.58 | -0.59 | 0.07 | -0.45 | -0.46 | 0.15 | -0.03 | -1.75 | -0.15 | -0.05 | 0.29 | -0.51 | 0.11 | 0.29 | 0.9725 | 0.65 | 1.19 | 0.108578 | 0.336804 | 0.060682 | 0.999971 | 1 | 0.725348 |
| Rab4b | Q91ZR1 | -1.79 | 0.27 | -0.5 | -1.11 | 0.2 | 0.23 | -0.48 | -0.34 | -0.03 | -1.62 | -0.5 | -0.57 | 0.3 | 0.31 | -0.18 | -0.56 | 0.685 | 0.1025 | 0.75 | 0.15111 | 0.841779 | 0.127599 | 0.999971 | 1 | 0.783863 |
| Rab5a | Q9CQD1 | -0.57 | -1.48 | -0.77 | -1.21 | -1.05 | -1.17 | -0.19 | -1.25 | -0.83 | -1.25 | -0.5 | -0.38 | -1.06 | -1.25 | -0.89 | -1.4 | 0.0925 | 0.2675 | -0.1425 | 0.774322 | 0.373976 | 0.585781 | 0.999971 | 1 | 0.920413 |
| Rab5b | P61021 | -0.41 | -0.77 | -1.06 | -1.07 | -0.77 | -0.58 | -0.91 | -0.91 | -0.78 | -1 | -0.98 | -0.88 | -0.87 | -0.75 | -0.98 | -1.04 | 0.035 | -0.0825 | -0.0825 | 0.872888 | 0.69575 | 0.703394 | 0.999971 | 1 | 0.949598 |
| Rab5c | P35278 | 0.14 | -1.31 | -1.17 | -1.36 | -1.11 | -1.03 | -1.03 | -1.17 | -1.17 | -1.04 | -1.03 | -1.02 | -1 | -1.15 | -1.28 | -1.38 | -0.16 | -0.14 | -0.2775 | 0.651268 | 0.692399 | 0.447623 | 0.999971 | 1 | 0.885146 |
| Rab5if | Q9CQT9 | -1.74 | -1.15 | 1.1 | 0.98 | -1.12 | -1.76 | 0.22 | 0.74 | -1.58 | -1.13 | 0.78 | 0.83 | 0.21 | -0.74 | 1.25 | 1.88 | -0.2775 | -0.0725 | 0.8525 | 0.740033 | 0.933447 | 0.32199 | 0.999971 | 1 | 0.85548 |
| Rab6a | P35279 | -0.84 | -0.93 | -0.66 | -0.55 | -0.87 | -0.84 | -0.89 | -0.61 | -0.78 | -0.88 | -0.4 | -0.48 | -0.84 | -0.72 | -0.63 | -0.1 | -0.0575 | 0.11 | 0.1725 | 0.75382 | 0.582977 | 0.452847 | 0.999971 | 1 | 0.887217 |
| Rab6a_1 | P35279-2 | -0.72 | 0.96 | -0.96 | -0.92 | -0.19 | -0.05 | -1.2 | -0.25 | 0.24 | -0.15 | -1.21 | -0.12 | 1.39 | 0.37 | 0.42 | -1.26 | -0.0125 | 0.1 | 0.64 | 0.979692 | 0.846508 | 0.339495 | 0.999971 | 1 | 0.860218 |
| Rab6b | P61294 | 0.55 | -0.75 | -1.49 | -1.53 | -0.67 | -0.49 | -1.27 | -1.39 | -0.75 | -0.08 | -1.45 | -1.22 | 0.02 | -0.35 | -1.02 | -0.78 | -0.15 | -0.07 | 0.2725 | 0.762444 | 0.894959 | 0.588616 | 0.999971 | 1 | 0.921285 |
| Rab7a | P51150 | -1.21 | -0.91 | -0.98 | -0.95 | -0.78 | -1.02 | -1.01 | -0.91 | -0.9 | -0.95 | -0.55 | -1.02 | -1.1 | -0.88 | -0.7 | -0.68 | 0.0825 | 0.1575 | 0.1725 | 0.638351 | 0.411939 | 0.372628 | 0.999971 | 1 | 0.869526 |
| Rab8a | P55258 | -0.52 | -1.32 | -0.57 | -0.86 | -0.8 | -0.78 | -0.22 | -0.97 | -1.07 | -0.74 | -0.4 | -0.4 | -1.65 | -1.16 | -0.76 | -0.93 | 0.125 | 0.165 | -0.3075 | 0.640818 | 0.536479 | 0.292292 | 0.999971 | 1 | 0.847402 |
| Rab8b | P61028 | 0.28 | 0.51 | -0.62 | -0.81 | -0.02 | 1.32 | 1.02 | -0.91 | 0.51 | 0.85 | -0.86 | -1.05 | -0.15 | 0.74 | -1.23 | -1.29 | 0.5125 | 0.0225 | -0.3225 | 0.36794 | 0.9665 | 0.553341 | 0.999971 | 1 | 0.913595 |
| Rab9a | Q9R0M6 | -0.35 | -0.25 | -0.67 | -0.38 | -0.25 | -0.27 | -0.84 | -0.11 | -0.16 | -0.09 | -0.62 | -0.08 | 0.36 | 0.11 | 0.17 | -0.03 | 0.045 | 0.175 | 0.565 | 0.841907 | 0.406258 | 0.014799 | 0.999971 | 1 | 0.613095 |
| Rab9b | Q8BHH2 | -0.66 | -0.02 | -0.16 | -0.4 | 0.41 | 0.16 | -0.27 | 0.29 | 0.52 | -0.6 | -0.01 | 0.65 | 1.42 | 0.37 | 0.72 | -0.11 | 0.4575 | 0.45 | 0.91 | 0.08144 | 0.183616 | 0.026158 | 0.999971 | 1 | 0.678474 |
| Rabac1 | Q9Z0S9 | 1.01 | 1.99 | 3.39 | 2.43 | 1.42 | 1.5 | 3.02 | 3.11 | 1.87 | 2.42 | 2.37 | -1.55 | 1.93 | 2.1 | 3.56 | 3.05 | 0.0575 | -0.9275 | 0.455 | 0.925701 | 0.34936 | 0.438094 | 0.999971 | 1 | 0.88258 |
| Rabep1 | O35551 | 0.74 | 0.53 | 0.09 | 0.21 | 0.37 | 0.65 | 0.89 | 0 | 0.48 | 0.51 | 0.16 | 0.22 | 0.32 | 0.41 | -0.04 | 0.07 | 0.085 | -0.05 | -0.2025 | 0.748015 | 0.817076 | 0.378051 | 0.999971 | 1 | 0.871417 |
| Rabep2 | Q91WG2 | 0.65 | 0.23 | -0.29 | 0.11 | 0.03 | 0.03 | 0.1 | -0.78 | -0.08 | 0.22 | -0.77 | 0.13 | 0.11 | 0.01 | -0.74 | -0.86 | -0.33 | -0.3 | -0.545 | 0.280567 | 0.335819 | 0.115399 | 0.999971 | 1 | 0.78268 |
| Rabepk | B0R0S4 | 0 | 0.18 | 0.75 | 1.4 | 0.2 | 0.4 | 0.3 | 1.29 | 0.47 | 1.93 | 0.87 | 0.86 | -0.1 | 0.69 | 0.91 | 1.48 | -0.035 | 0.45 | 0.1625 | 0.928008 | 0.303702 | 0.706851 | 0.999971 | 1 | 0.950182 |
| Rabgap1 | A2AWA9 | 0.46 | 0.13 | -0.07 | -0.04 | 0.23 | 0.38 | 0.05 | -0.02 | 0.37 | 0.25 | 0.26 | 0.14 | 0.31 | 0.04 | 0.2 | 0.17 | 0.04 | 0.135 | 0.06 | 0.84564 | 0.486372 | 0.760449 | 0.999971 | 1 | 0.961116 |
| Rabgap1l | A6H6A9 | -0.1 | 0.37 | 0.03 | 0.3 | 0.15 | 0.25 | -0.09 | 0.26 | -0.02 | 0.2 | 0.1 | 0.31 | 0.49 | 0.3 | 0.39 | 0.44 | -0.0075 | -0.0025 | 0.255 | 0.969725 | 0.989584 | 0.199309 | 0.999971 | 1 | 0.813335 |
| Rabgap1l_ | A6H6A9-3 | 0.74 | -0.17 | -0.45 | 0.35 | 0.01 | 0.3 | 0.3 | 0.34 | 0.35 | 1.09 | 0.48 | 0.04 | 0.27 | 0.05 | 0.23 | 0.82 | 0.12 | 0.3725 | 0.225 | 0.678935 | 0.291964 | 0.484494 | 0.999971 | 1 | 0.896402 |
| Rabgef1 | Q9JM13 | 0.23 | 2.09 | 1.53 | 0.35 | 1.68 | 1.32 | 1.46 | 0.9 | 1.47 | 0.99 | 2.2 | 2.64 | 2.06 | 1.12 | 1.32 | 0.91 | 0.29 | 0.775 | 0.3025 | 0.526858 | 0.176966 | 0.536098 | 0.999971 | 1 | 0.910077 |
| Rabggta | Q9JHK4 | 0.62 | 1.01 | 1.36 | 1.25 | 0.89 | 0.99 | 1.22 | 1.24 | 0.83 | 0.79 | 1.16 | 0.63 | 0.64 | 0.86 | 1.26 | 1.19 | 0.025 | -0.2075 | -0.0725 | 0.911745 | 0.382597 | 0.770789 | 0.999971 | 1 | 0.963498 |
| Rabggtb | Q3TVF4 | 0.87 | 1.56 | 1.34 | 1.47 | 1.01 | 2 | 0.71 | 1.37 | 1.16 | 1.2 | 1.03 | 1.02 | 1.06 | 1.49 | 1.32 | 1.42 | -0.0375 | -0.2075 | 0.0125 | 0.906487 | 0.332682 | 0.955291 | 0.999971 | 1 | 0.998056 |
| Rabif | Q91X96 | -1.16 | 0.07 | 0.42 | 0.2 | 0.02 | 0.15 | 0.16 | 0.5 | 0.16 | -0.52 | 0.26 | 0.23 | -0.16 | 0.21 | 0.87 | 0.42 | 0.325 | 0.15 | 0.4525 | 0.3787 | 0.698863 | 0.27356 | 0.999971 | 1 | 0.843794 |
| Rabl2 | E9Q9D5 | 1.55 | 1.25 | 0.17 | -2.27 | 1.25 | 1.69 | 0.49 | -1.6 | 1.27 | 1.68 | 0.88 | 1.12 | 0.74 | 0.97 | -0.99 | -1.98 | 0.2825 | 1.0625 | -0.49 | 0.780116 | 0.206819 | 0.626336 | 0.999971 | 1 | 0.930479 |
| Rabl3 | Q9D4V7 | -1.49 | -0.12 | 0.19 | -0.82 | -0.21 | -0.43 | 0.28 | -0.21 | -0.29 | -1.83 | 0.5 | 0.54 | -0.78 | -1.1 | 0.05 | -0.69 | 0.4175 | 0.29 | -0.07 | 0.298463 | 0.638412 | 0.868901 | 0.999971 | 1 | 0.983462 |
| Rabl6 | Q5U3K5 | -0.13 | -0.82 | -1.09 | -1.04 | -0.87 | -0.54 | -1.15 | -0.84 | -0.76 | -0.58 | -0.89 | -1.02 | -0.24 | -0.56 | -0.94 | -0.83 | -0.08 | -0.0425 | 0.1275 | 0.769337 | 0.870833 | 0.656528 | 0.999971 | 1 | 0.934061 |
| Rac1 | Q3TLP8 | -1.28 | -1.9 | -1.12 | -1.18 | -1.61 | -1.63 | -1.65 | -1.15 | -1.72 | -1.27 | -1.15 | -1.3 | -1.87 | -1.42 | -1.04 | -1.16 | -0.14 | 0.01 | -0.0025 | 0.572848 | 0.967545 | 0.992746 | 0.999971 | 1 | 0.99925 |
| Rac2 | Q05144 | 0.08 | 0.99 | 0.13 | 0.82 | -0.22 | 0.18 | -0.14 | 0.15 | -0.47 | -0.4 | 0.16 | 1.01 | -0.51 | 0.5 | 0.48 | 1.28 | -0.5125 | -0.43 | -0.0675 | 0.087406 | 0.296336 | 0.870514 | 0.999971 | 1 | 0.984079 |
| Rac3 | P60764 | -1.66 | -1.78 | -8.98 | -9.83 | -1.44 | -1.68 | -8.51 | -9.39 | -1.47 | -0.75 | -6.9 | -10.14 | -0.49 | -1.03 | -9.3 | -12.99 | 0.3075 | 0.7475 | -0.39 | 0.910133 | 0.790702 | 0.907751 | 0.999971 | 1 | 0.991405 |
| Rack1 | P68040 | 0.34 | 1.91 | 1.12 | 0.59 | 2.02 | 1.7 | 1.32 | 1.11 | 2.43 | 0.64 | 1.63 | 1.66 | 2.27 | 1.6 | 1.2 | 0.81 | 0.5475 | 0.6 | 0.48 | 0.180619 | 0.22657 | 0.292336 | 0.999971 | 1 | 0.847402 |
| Rad21 | Q61550 | 0.53 | 0.21 | 0.27 | -0.02 | 0.56 | 0.29 | -0.27 | -0.26 | 0.21 | 0.28 | -0.11 | 0.28 | 0.21 | 0.18 | 0.25 | 0.3 | -0.1675 | -0.0825 | -0.0125 | 0.523312 | 0.681324 | 0.946811 | 0.999971 | 1 | 0.997651 |
| Rad23a | Q8CAP3 | 0.93 | 0.46 | 0.8 | 1.08 | 0.78 | 0.97 | 1.33 | 0.65 | 0.76 | 0.99 | 1.07 | 1.2 | 0.4 | 0.87 | 0.79 | 0.65 | 0.115 | 0.1875 | -0.14 | 0.625597 | 0.380428 | 0.521093 | 0.999971 | 1 | 0.906972 |
| Rad23b | P54728 | -0.81 | -0.75 | -0.5 | -0.11 | -0.67 | -1.65 | -0.24 | -0.39 | -1.28 | -1.63 | 0.02 | 0.12 | -1.14 | -0.64 | 0.14 | 0.29 | -0.195 | -0.15 | 0.205 | 0.579319 | 0.737963 | 0.576086 | 0.999971 | 1 | 0.918194 |
| Rad50 | Q5SV02 | 1.26 | 0.62 | 1.92 | 2.12 | 0.56 | 0.5 | 1.19 | 2.51 | -0.14 | 0.79 | 2.16 | 2.42 | 0.1 | 0.93 | 2.31 | 2.64 | -0.29 | -0.1725 | 0.015 | 0.58897 | 0.785085 | 0.980817 | 0.999971 | 1 | 0.99925 |
| Radil | D3YWJ9 | 0.18 | -0.52 | -0.17 | -0.19 | 0.01 | -0.73 | -0.04 | 0.32 | 0.24 | -0.41 | -0.1 | -0.05 | 0.36 | -0.33 | 0.32 | -0.17 | 0.065 | 0.095 | 0.22 | 0.816109 | 0.681306 | 0.394306 | 0.999971 | 1 | 0.871816 |
| Rae1 | Q8C570 | -0.33 | 0.06 | 0 | -0.84 | 0.41 | -0.36 | 0.07 | -0.84 | 0.46 | -0.42 | 0.16 | 0.4 | 0.77 | -0.54 | -0.43 | -1.15 | 0.0975 | 0.4275 | -0.06 | 0.772924 | 0.174258 | 0.88843 | 0.999971 | 1 | 0.987485 |
| Raf1 | Q99N57 | 0.4 | 0.13 | 0.19 | -0.14 | -0.29 | -0.3 | 0.35 | -0.07 | 0.26 | 0.14 | -0.24 | -0.23 | 0.28 | -0.43 | 0.15 | -0.18 | -0.2225 | -0.1625 | -0.19 | 0.342024 | 0.454869 | 0.423766 | 0.999971 | 1 | 0.879372 |
| Rai1 | Q61818 | 0.05 | -0.34 | -0.32 | -0.44 | -0.19 | -0.39 | -0.05 | -0.53 | -0.1 | 0.08 | -0.33 | -0.24 | 0.45 | -0.39 | -0.97 | -0.95 | -0.0275 | 0.115 | -0.2025 | 0.893335 | 0.561716 | 0.562153 | 0.999971 | 1 | 0.916386 |
| Rai14 | A0A2I3BPY | -0.77 | -0.53 | -0.39 | 0.25 | -0.41 | -0.35 | -0.25 | 0.32 | -0.66 | -0.62 | -0.48 | 0.13 | -0.52 | -0.34 | 0.06 | 0.24 | 0.1875 | -0.0475 | 0.22 | 0.519449 | 0.871705 | 0.458325 | 0.999971 | 1 | 0.888571 |
| Rala | P63321 | -0.04 | -0.4 | -0.08 | 0.19 | -0.77 | 0.21 | -0.53 | -0.05 | -0.31 | 0.45 | -0.16 | -0.34 | -0.3 | -0.2 | -0.17 | 0.64 | -0.2025 | -0.0075 | 0.075 | 0.464043 | 0.975801 | 0.781175 | 0.999971 | 1 | 0.965859 |
| Ralb | Q9JIW9 | 0.09 | -1.39 | -0.42 | -0.88 | -1.41 | -0.69 | -0.94 | -0.68 | -0.93 | -0.04 | -0.47 | -0.47 | -0.61 | -0.83 | -0.78 | -0.62 | -0.28 | 0.1725 | -0.06 | 0.435628 | 0.631358 | 0.852775 | 0.999971 | 1 | 0.98187 |
| Ralbp1 | Q62172 | -0.31 | -0.32 | -0.44 | -0.38 | -0.44 | -0.23 | -0.6 | -0.62 | -0.24 | -0.28 | -0.2 | -0.59 | -0.13 | -0.25 | -0.66 | -0.62 | -0.11 | 0.035 | -0.0525 | 0.540009 | 0.840332 | 0.790676 | 0.999971 | 1 | 0.968447 |
| Ralgapa1 | A0A1W2P8 | 0.35 | 0.05 | 0.23 | 0.26 | -0.24 | 0 | 0.19 | 0.03 | -0.16 | 0.23 | 0.21 | 0.35 | 0.06 | -0.01 | 0.13 | 0.38 | -0.2275 | -0.065 | -0.0825 | 0.235432 | 0.732063 | 0.653509 | 0.999971 | 1 | 0.934061 |
| Ralgapa2 | A0A0A0M( | -0.62 | -0.04 | 0.27 | 0.03 | -0.29 | 0.04 | 0.18 | -0.09 | -0.63 | -0.27 | 0.51 | 0.55 | -1.27 | -0.16 | 0.16 | 0.19 | 0.05 | 0.13 | -0.18 | 0.837825 | 0.706173 | 0.636641 | 0.999971 | 1 | 0.93301 |
| Ralgapb | A2ACC6 | -0.14 | -0.04 | -0.32 | -0.54 | 0.2 | 0.04 | -0.05 | -0.28 | 0.47 | 0.07 | -0.02 | -0.23 | 0.5 | 0.11 | -0.21 | -0.26 | 0.2375 | 0.3325 | 0.295 | 0.262639 | 0.160575 | 0.240066 | 0.999971 | 1 | 0.834057 |
| Ralgapb_1 | Q8BQZ4 | -1.72 | 0.86 | 1.77 | 0.27 | 0.3 | 1.2 | 0.94 | 0.91 | -0.51 | -0.86 | -0.01 | 0.67 | 0.52 | 0.11 | 1.1 | 1.88 | 0.5425 | -0.4725 | 0.6075 | 0.440234 | 0.525731 | 0.427851 | 0.999971 | 1 | 0.880793 |
| Ralgps1 | A2AR50-3 | 0.3 | -0.21 | 0.21 | 0.23 | 0.04 | -0.09 | 0.25 | 0 | 0.51 | 0.24 | 0.15 | -0.31 | 0.76 | 0 | 0.14 | 0.41 | -0.0825 | 0.015 | 0.195 | 0.676933 | 0.94961 | 0.422138 | 0.999971 | 1 | 0.877992 |
| Ralgps2 | A0A087WC | 0.35 | 0.7 | 0.11 | -0.14 | 0.9 | 0.96 | 0.1 | 0.08 | 1.12 | 0.88 | 0.09 | 0.1 | 0.86 | 0.47 | -0.04 | 0.24 | 0.255 | 0.2925 | 0.1275 | 0.416435 | 0.375049 | 0.648571 | 0.999971 | 1 | 0.933842 |
| Raly | Q64012-2 | 0.4 | -0.27 | -0.48 | -0.25 | -0.41 | -0.18 | -0.41 | -0.59 | -0.31 | 0.01 | -0.61 | -0.87 | -0.59 | -0.15 | -0.78 | -0.48 | -0.2475 | -0.295 | -0.35 | 0.318679 | 0.311785 | 0.197079 | 0.999971 | 1 | 0.812891 |
| Ralyl | Q8BTF8 | 0.32 | -0.38 | 0.25 | 0.18 | -0.07 | 0.18 | 1.07 | 0.11 | 0.45 | 0.09 | 0.18 | 0.32 | -0.36 | -0.53 | -0.17 | 0.1 | 0.23 | 0.1675 | -0.3325 | 0.460997 | 0.452986 | 0.193107 | 0.999971 | 1 | 0.810547 |
| Ralyl_1 | A0A0A6YX( | -1.75 | 0.1 | 2.13 | 1.46 | 0.59 | -0.25 | 2.54 | -0.39 | 0.7 | -1.47 | -0.16 | -1.26 | -0.18 | 0.32 | 0.76 | -0.22 | 0.1375 | -1.0325 | -0.315 | 0.887602 | 0.268163 | 0.694379 | 0.999971 | 1 | 0.947174 |
| Ramac | Q9CQY2 | 1.68 | 3.19 | 2.61 | 3.11 | 2.8 | 2.78 | 2.97 | 2.78 | 2.96 | 2.06 | 2.83 | 2.1 | 2.48 | 2.48 | 2.33 | 2.79 | 0.185 | -0.16 | -0.1275 | 0.594912 | 0.692212 | 0.719574 | 0.999971 | 1 | 0.954401 |
| Ran | P62827 | -1.69 | -0.68 | -0.63 | -1.94 | -0.19 | -0.68 | 0 | -1.42 | 0.49 | -1.75 | -0.4 | -0.65 | 0.4 | -0.96 | -0.74 | -1.72 | 0.6625 | 0.6575 | 0.48 | 0.155473 | 0.236015 | 0.362212 | 0.999971 | 1 | 0.869526 |
| Ranbp1 | P34022 | -0.9 | 1.11 | 0.43 | 0.39 | 1.19 | 0.6 | -0.02 | 0.87 | 1.27 | -0.44 | 0.43 | 0.55 | 1.53 | 1.14 | 0.59 | 0.24 | 0.4025 | 0.195 | 0.6175 | 0.392697 | 0.702543 | 0.215477 | 0.999971 | 1 | 0.826893 |
| Ranbp10 | Q6VN19 | 0.05 | 0.67 | -0.92 | -1.19 | 0.95 | 0.28 | -1.13 | -0.83 | 0.72 | -0.13 | -0.45 | -0.77 | 0.71 | 0.43 | -0.9 | -0.81 | 0.165 | 0.19 | 0.205 | 0.780771 | 0.705235 | 0.710869 | 0.999971 | 1 | 0.950695 |
| Ranbp17 | Q99NF8 | 0.88 | 1.22 | 0.59 | 1.93 | 0.37 | 1.47 | 1.04 | 1.52 | 1.65 | 1.02 | 1.25 | 0.19 | 1.13 | 0.79 | 0.91 | 1.23 | -0.055 | -0.1275 | -0.14 | 0.884567 | 0.752895 | 0.654629 | 0.999971 | 1 | 0.934061 |
| Ranbp2 | Q9ERU9 | 0.27 | -0.15 | -0.3 | -0.19 | -0.24 | -0.07 | -0.21 | -0.36 | -0.26 | 0.1 | -0.24 | -0.63 | -0.15 | -0.26 | -0.46 | -0.5 | -0.1275 | -0.165 | -0.25 | 0.525003 | 0.47937 | 0.243241 | 0.999971 | 1 | 0.837316 |
| Ranbp3 | A0A3B2W( | 1.95 | 1.93 | 2.42 | 2.17 | 2.11 | 1.55 | 2.63 | 1.79 | 2.07 | 1.85 | 2.25 | 2.3 | 1.7 | 1.92 | 1.85 | 1.69 | -0.0975 | -1.57E-16 | -0.3275 | 0.725353 | 1 | 0.117493 | 0.999971 | 1 | 0.78268 |
| Ranbp6 | Q8BIV3 | 0.42 | 0.27 | -0.22 | -0.14 | 0.51 | 0.6 | 0.2 | 0.33 | 0.49 | 0.4 | 0.02 | -0.09 | 1.27 | 0.76 | 0.31 | 0.17 | 0.3275 | 0.1225 | 0.545 | 0.165454 | 0.612842 | 0.098273 | 0.999971 | 1 | 0.78268 |
| Ranbp9 | E9Q5D6 | -0.36 | 0.54 | 0.1 | 0.02 | 0.28 | 0.35 | 0 | 0.14 | 0.23 | 0.01 | 0.08 | 0 | 0.6 | 0.27 | -0.1 | -0.1 | 0.1175 | 0.005 | 0.0925 | 0.62007 | 0.9825 | 0.732743 | 0.999971 | 1 | 0.956652 |
| Rangap1 | P46061 | -0.29 | 0.09 | 0.52 | 0.5 | 0.56 | 0.07 | 0.33 | 0.54 | 0.17 | -0.04 | 0.59 | 0.68 | 0.55 | 0.28 | 0.49 | 0.68 | 0.17 | 0.145 | 0.295 | 0.504158 | 0.599322 | 0.24345 | 0.999971 | 1 | 0.837316 |
| Rap1a | P62835 | -1.75 | -0.03 | 0 | -0.7 | -0.34 | 0.15 | -0.56 | -0.21 | -0.11 | -1.68 | -0.08 | -0.26 | 0.39 | -0.25 | -0.26 | -0.1 | 0.38 | 0.0875 | 0.565 | 0.370807 | 0.866759 | 0.19802 | 0.999971 | 1 | 0.813335 |
| Rap1b | Q99JI6 | -1.29 | -0.38 | -1.04 | -1.3 | -0.43 | -0.43 | -1.15 | -1.03 | -0.48 | -1.54 | -0.87 | -0.89 | -0.27 | -0.84 | -0.96 | -0.89 | 0.2425 | 0.0575 | 0.2625 | 0.424954 | 0.853834 | 0.365996 | 0.999971 | 1 | 0.869526 |
| Rap1gap | A2ALS5 | 1.05 | 0.06 | 0.11 | 0.26 | -0.01 | -0.42 | 0.31 | 0.3 | 0.41 | 0.97 | 0.2 | 0.38 | 0.12 | 0.34 | 0.48 | 0.45 | -0.325 | 0.12 | -0.0225 | 0.290632 | 0.684418 | 0.932627 | 0.999971 | 1 | 0.995069 |
| Rap1gap_1 | A2ALS7 | 1.55 | 0.74 | -1.31 | 0.17 | -0.09 | 0.43 | -0.69 | -0.09 | 0.66 | 0.94 | 0.16 | -0.34 | 1.27 | 0.44 | -0.23 | -0.16 | -0.3975 | 0.0675 | 0.0425 | 0.505539 | 0.91192 | 0.946496 | 0.999971 | 1 | 0.997646 |
| Rap1gap2 | A0A571BEI | 0.06 | 0.43 | -0.06 | 0.1 | 0.12 | 0.25 | 0.43 | -0.14 | 0.27 | 0.11 | 0.05 | -0.11 | 0.39 | -0.17 | -0.07 | 0.11 | 0.0325 | -0.0525 | -0.0675 | 0.876935 | 0.784221 | 0.750628 | 0.999971 | 1 | 0.961064 |
| Rap1gds1 | E9Q912 | -0.27 | 0.51 | 0.21 | 0.2 | 0.23 | 0.44 | 0.37 | 0.39 | 0.48 | 0.17 | 0.4 | 0.28 | 0.79 | 0.4 | 0.28 | 0.52 | 0.195 | 0.17 | 0.335 | 0.375196 | 0.441862 | 0.174009 | 0.999971 | 1 | 0.807788 |
| Rap1gds1_ | E9Q912-3 | -0.72 | 0.64 | -0.24 | -0.25 | -0.61 | -0.2 | 0.17 | 0.23 | -0.19 | -1.07 | -0.39 | 0.16 | 0.46 | 0.34 | -0.1 | 0.22 | 0.04 | -0.23 | 0.3725 | 0.90635 | 0.541933 | 0.253934 | 0.999971 | 1 | 0.839012 |
| Rap1gds1_ | E9Q912-2 | -0.09 | 0.23 | -0.21 | 0.13 | 0.04 | 0.15 | -0.62 | 0.31 | -0.1 | 0.32 | 0.08 | 0.17 | 0.08 | 0.19 | 0.15 | 0.32 | -0.045 | 0.1025 | 0.17 | 0.859296 | 0.597827 | 0.372074 | 0.999971 | 1 | 0.869526 |
| Rap2a | Q80ZJ1 | -0.44 | -1.51 | -1.8 | -1.49 | -1.34 | -1.37 | -0.96 | -1.85 | -1.79 | -0.86 | -1.93 | -2.12 | -1.54 | -1.2 | -1.84 | -1.6 | -0.07 | -0.365 | -0.235 | 0.839208 | 0.366012 | 0.479415 | 0.999971 | 1 | 0.894438 |
| Rap2b | P61226 | 0.18 | -1.93 | -1.17 | -1.49 | -1.73 | -1.35 | -1.22 | -1.63 | -0.84 | 0.04 | -1.02 | -1.38 | -1.66 | -1.42 | -1.37 | -1.12 | -0.38 | 0.3025 | -0.29 | 0.400657 | 0.555548 | 0.516707 | 0.999971 | 1 | 0.905815 |
| Rap2c | Q8BU31 | 1.18 | -1.17 | -0.05 | 0.06 | -0.87 | -0.33 | 0.27 | -0.2 | -0.48 | 0.15 | -0.41 | -0.14 | -1.34 | -1.13 | -0.09 | -0.59 | -0.2875 | -0.225 | -0.7925 | 0.565691 | 0.63435 | 0.150218 | 0.999971 | 1 | 0.793248 |
| Rapgef1 | A1L338 | 0.96 | 0.43 | 0.83 | 0.6 | 0 | 0.22 | 0.73 | 0.74 | 0.01 | 0.63 | 1.04 | 1.03 | 0.03 | 0.19 | 0.66 | 0.44 | -0.2825 | -0.0275 | -0.375 | 0.273948 | 0.922431 | 0.121772 | 0.999971 | 1 | 0.78268 |
| Rapgef2 | A0A0A6YW | -0.12 | -0.26 | -0.78 | -0.74 | -0.4 | -0.05 | -0.76 | -0.78 | -0.49 | -0.16 | -0.85 | -0.96 | -0.52 | -0.15 | -0.71 | -0.61 | -0.0225 | -0.14 | -0.0225 | 0.931712 | 0.601821 | 0.925426 | 0.999971 | 1 | 0.99401 |
| Rapgef3 | H3BK39 | -0.4 | -0.51 | -0.46 | -1.1 | -1.07 | -1.53 | -0.76 | -1.15 | -0.53 | -1.1 | -1.39 | -1.02 | -0.35 | -1.03 | -1.06 | -0.48 | -0.51 | -0.3925 | -0.1125 | 0.070163 | 0.160086 | 0.67475 | 0.999971 | 1 | 0.941037 |
| Rapgef4 | Q9EQZ6 | -0.64 | -0.98 | -0.63 | -0.73 | -1.1 | -0.74 | -0.41 | -0.86 | -1.1 | -0.81 | -0.92 | -1.14 | -0.86 | -0.99 | -0.96 | -0.84 | -0.0325 | -0.2475 | -0.1675 | 0.878892 | 0.197291 | 0.35394 | 0.999971 | 1 | 0.868793 |
| Rapgef5 | Q8C0Q9 | -1.54 | -1.2 | -0.08 | -0.71 | -1.11 | -0.52 | -0.16 | -0.37 | -0.91 | -1.37 | -0.41 | 0.09 | -1.2 | -0.64 | -0.32 | -0.17 | 0.3425 | 0.2325 | 0.3 | 0.362359 | 0.587325 | 0.436213 | 0.999971 | 1 | 0.882419 |
| Rapgef6 | Q5NCJ4 | 0.33 | -0.64 | -0.54 | -0.61 | -1.31 | -0.91 | -0.8 | -0.23 | -1.1 | -0.39 | -0.76 | -0.28 | -1.3 | -0.24 | -0.36 | -0.02 | -0.4475 | -0.2675 | -0.115 | 0.190089 | 0.390537 | 0.748837 | 0.999971 | 1 | 0.961064 |
| Rapgefl1 | Q68EF8 | -0.13 | 0.43 | 0.3 | 0.25 | 0.43 | 0.68 | 0.91 | 0.25 | 0.48 | 0.21 | 0.46 | 0.22 | -0.07 | 0.3 | 0.06 | 0.01 | 0.355 | 0.13 | -0.1375 | 0.144579 | 0.51466 | 0.502038 | 0.999971 | 1 | 0.904573 |
| Raph1 | F2Z3U3 | 0.16 | -0.54 | -0.67 | -0.69 | -0.55 | -0.16 | -0.08 | -0.7 | -0.63 | -0.01 | -0.49 | -0.87 | -0.49 | -0.63 | -0.61 | -0.76 | 0.0625 | -0.065 | -0.1875 | 0.81711 | 0.818837 | 0.445613 | 0.999971 | 1 | 0.884152 |
| Rars1 | Q9D0I9 | -0.05 | 1.64 | 0.57 | 0.23 | 1.83 | 0.9 | 0.73 | 0.11 | 2.13 | 0.33 | 1.02 | 0.68 | 1.69 | 0.53 | 0.35 | 0.23 | 0.295 | 0.4425 | 0.1025 | 0.541561 | 0.387851 | 0.826578 | 0.999971 | 1 | 0.976804 |
| Rars2 | Q3U186 | -0.27 | -0.36 | -0.83 | -0.34 | -0.33 | -0.43 | -0.51 | -0.18 | -0.27 | -0.73 | -0.82 | -0.58 | 0.77 | -0.04 | -0.4 | -0.43 | 0.0875 | -0.15 | 0.425 | 0.667448 | 0.496968 | 0.197294 | 0.999971 | 1 | 0.812891 |
| Rasa1 | E9PYG6 | -0.16 | -0.28 | -0.12 | -0.08 | -0.05 | -0.25 | -0.3 | -0.11 | -0.13 | 0.01 | -0.31 | 0.1 | -0.31 | -0.04 | -0.13 | -0.02 | -0.0175 | 0.0775 | 0.035 | 0.917559 | 0.661269 | 0.838876 | 0.999971 | 1 | 0.979728 |
| Rasa2 | P58069 | -0.25 | 0.05 | -0.39 | -1.19 | 0.05 | 0.17 | -0.48 | -0.5 | -0.31 | -0.05 | -0.65 | -0.77 | 0.25 | 0.12 | -0.83 | -0.91 | 0.255 | 0 | 0.1025 | 0.433537 | 1 | 0.792758 | 0.999971 | 1 | 0.969564 |
| Rasa3 | Q60790 | -0.73 | -0.66 | -1.04 | -1.24 | -0.71 | -0.73 | -1.25 | -1.13 | -0.61 | -0.75 | -1.29 | -1.37 | -0.14 | -0.59 | -1.14 | -1.11 | -0.0375 | -0.0875 | 0.1725 | 0.870978 | 0.733938 | 0.552496 | 0.999971 | 1 | 0.913595 |
| Rasal1 | Q9Z268 | -0.58 | -0.5 | -0.49 | -0.85 | -0.5 | -0.53 | -0.26 | -0.49 | -0.27 | -0.84 | -0.28 | -0.54 | -0.3 | -0.62 | -0.51 | -0.58 | 0.16 | 0.1225 | 0.1025 | 0.389053 | 0.558268 | 0.582474 | 0.999971 | 1 | 0.920364 |
| Rasal2 | D3YUS5 | -1.15 | -0.65 | -0.74 | -0.84 | -1.02 | -0.73 | -0.52 | -1.14 | -0.54 | -0.96 | -0.34 | -0.42 | -0.38 | -0.99 | -0.87 | -0.55 | -0.0075 | 0.28 | 0.1475 | 0.972894 | 0.220037 | 0.511528 | 0.999971 | 1 | 0.905815 |
| Rasd2 | P63032 | 0.58 | 0.18 | 0.01 | 0.55 | -0.68 | -1.09 | -0.5 | 1.26 | -0.07 | -0.28 | -0.08 | 0.46 | 0.49 | 0.1 | 0.72 | 1.18 | -0.5825 | -0.3225 | 0.2925 | 0.261461 | 0.204127 | 0.313899 | 0.999971 | 1 | 0.850589 |
| Rasgef1a | A6H650 | -0.08 | -0.6 | -0.5 | -0.56 | -0.26 | 0.05 | -0.34 | 0.01 | -0.36 | 0.09 | -0.59 | -0.64 | -0.26 | -0.1 | -0.51 | -0.07 | 0.3 | 0.06 | 0.2 | 0.172122 | 0.800341 | 0.351839 | 0.999971 | 1 | 0.867656 |
| Rasgef1c | B1ATE5 | 2.03 | 0.81 | -0.04 | 1.13 | 0.94 | 2.22 | -0.17 | 1.38 | 1.34 | 2.62 | 1.79 | 1.58 | 0.73 | 1.79 | 2.09 | 1.4 | 0.11 | 0.85 | 0.52 | 0.854201 | 0.103043 | 0.298697 | 0.999971 | 1 | 0.847402 |
| Rasgrf1 | P27671 | 0.49 | -0.04 | 0.95 | 0.55 | 0.24 | 0.15 | 0.98 | 0.76 | 0.25 | 0.12 | 0.55 | 0 | -0.44 | -0.26 | 0.53 | 0.33 | 0.045 | -0.2575 | -0.4475 | 0.878983 | 0.332018 | 0.177785 | 0.999971 | 1 | 0.808532 |
| Rasgrf2 | P70392 | -0.41 | -0.27 | 0.46 | -0.13 | -0.2 | -0.17 | 0.41 | -0.11 | -0.25 | -0.36 | 0.31 | 0.09 | -0.05 | -0.16 | -0.12 | -0.06 | 0.07 | 0.035 | -0.01 | 0.789505 | 0.894968 | 0.965489 | 0.999971 | 1 | 0.998721 |
| Rasgrp1 | Q9Z1S3 | -0.43 | -0.19 | -0.23 | -0.22 | -0.46 | -0.41 | -0.54 | -0.09 | -0.97 | -0.35 | -0.09 | -0.39 | 0.05 | -0.23 | -0.33 | -0.21 | -0.1075 | -0.1825 | 0.0875 | 0.565655 | 0.4344 | 0.627943 | 0.999971 | 1 | 0.931641 |
| Rasgrp2 | Q9QUG9 | 0.78 | 0.07 | -0.18 | 0.95 | 0.36 | -0.79 | 0.3 | 0.93 | 0.47 | 0.58 | 0.21 | 0.55 | 0.32 | 0.49 | 0.78 | 0.03 | -0.205 | 0.0475 | -3.93E-17 | 0.632953 | 0.871892 | 1 | 0.999971 | 1 | 1 |
| Rasip1 | Q3U0S6 | 0.74 | 1.49 | 0.38 | 0.35 | 2.05 | 1.03 | -0.04 | -0.27 | 1.12 | 1.18 | -0.03 | -0.12 | -0.11 | 1.53 | 0.08 | 0.15 | -0.0475 | -0.2025 | -0.3275 | 0.930832 | 0.632748 | 0.459343 | 0.999971 | 1 | 0.889191 |
| Rassf2 | Q8BMS9 | 0.11 | -0.52 | -0.47 | -0.47 | -0.32 | -0.2 | -0.08 | -0.02 | -0.14 | -0.1 | -0.57 | -0.59 | -0.77 | -0.6 | -0.44 | -0.65 | 0.1825 | -0.0125 | -0.2775 | 0.400745 | 0.957282 | 0.215985 | 0.999971 | 1 | 0.826893 |
| Rassf5 | Q5EBH1 | 1.57 | 0.47 | 0.54 | 1.13 | 0.41 | 1.3 | 2.01 | 1.93 | 0.81 | 1.05 | 0.84 | 0.44 | -0.29 | 0.49 | 0.92 | 0.42 | 0.485 | -0.1425 | -0.5425 | 0.274759 | 0.634464 | 0.152454 | 0.999971 | 1 | 0.795578 |
| Rassf8 | Q8CJ96 | 0.83 | -1.12 | -1.86 | -1.06 | 1.13 | -0.68 | -1.55 | -1.26 | -0.46 | 0.54 | -1.27 | -0.96 | 0.33 | 0.3 | -0.84 | -1.74 | 0.2125 | 0.265 | 0.315 | 0.776839 | 0.679064 | 0.649277 | 0.999971 | 1 | 0.933842 |
| Raver1 | Q9CW46 | -0.16 | 0.24 | 1.8 | 2.46 | 0.07 | -0.53 | 1.48 | 2.07 | -0.08 | 0.19 | 1.47 | 1.37 | -0.94 | 0.38 | 1.54 | 1.86 | -0.3125 | -0.3475 | -0.375 | 0.690037 | 0.609169 | 0.641698 | 0.999971 | 1 | 0.93301 |
| Rb1cc1 | Q9ESK9 | -0.05 | -0.05 | -0.09 | -0.01 | 0.21 | 0.2 | 0.13 | -0.05 | 0.44 | 0.37 | -0.13 | -0.25 | 0.46 | -0.06 | -0.16 | -0.2 | 0.1725 | 0.1575 | 0.06 | 0.313839 | 0.474969 | 0.77274 | 0.999971 | 1 | 0.963573 |
| Rbbp4 | Q60972 | 0.56 | 1.6 | 1.41 | 1.29 | 1.66 | 1.47 | 1.41 | 1.33 | 1.8 | 1.03 | 1.59 | 1.8 | 1.43 | 1.63 | 1.29 | 1.61 | 0.2525 | 0.34 | 0.275 | 0.346656 | 0.273827 | 0.312838 | 0.999971 | 1 | 0.849367 |
| Rbbp5 | Q8BX09-2 | 1.45 | 0.31 | 1.9 | 1.38 | 0.44 | -0.6 | 0.11 | 0.54 | 0.69 | 0.56 | 2.55 | 2.39 | 1.33 | 0.1 | 0.18 | 0.04 | -1.1375 | 0.2875 | -0.8475 | 0.019283 | 0.623193 | 0.076396 | 0.999971 | 1 | 0.748444 |
| Rbbp5_1 | A0A0R4J2E | 1.61 | 1.95 | 1.42 | 1.65 | 1.91 | 1.88 | 1.1 | 1.55 | 1.81 | 1.87 | 1.19 | 1.29 | 1.36 | 2.27 | 1.39 | 1.61 | -0.0475 | -0.1175 | -2.36E-16 | 0.847707 | 0.623414 | 1 | 0.999971 | 1 | 1 |
| Rbbp6 | P97868-2 | 1.32 | 1.13 | 0.72 | 0.82 | 1.16 | 1.01 | 0.67 | 0.64 | 0.86 | 1.14 | 0.47 | 0.75 | 0.7 | 0.87 | 0.68 | 0.79 | -0.1275 | -0.1925 | -0.2375 | 0.579084 | 0.413651 | 0.260014 | 0.999971 | 1 | 0.840512 |
| Rbbp7 | Q60973 | 1.12 | 1.31 | 1.04 | 1.27 | 1.48 | 0.73 | 1.06 | 1.18 | 1.4 | 0.81 | 1.24 | 1.99 | 1.51 | 1.24 | 1.38 | 1.04 | -0.0725 | 0.175 | 0.1075 | 0.736398 | 0.521998 | 0.572874 | 0.999971 | 1 | 0.91816 |
| Rbbp9 | O88851 | 0.33 | 0.49 | 0.55 | 0.48 | 0.64 | 0.42 | 0.95 | 0.46 | 0.37 | 0.46 | 0.16 | 0.29 | 0.47 | 0.63 | 0.26 | 0.33 | 0.155 | -0.1425 | -0.04 | 0.431791 | 0.405683 | 0.822196 | 0.999971 | 1 | 0.975901 |
| Rbck1 | Q9WUB0 | 0.44 | 0.23 | -0.3 | 0.12 | 0.39 | 0.72 | -0.3 | 0.11 | 0.2 | 0.57 | 0.07 | 0.23 | 1.28 | 0.97 | 0.29 | 0.44 | 0.1075 | 0.145 | 0.6225 | 0.703431 | 0.526009 | 0.057026 | 0.999971 | 1 | 0.725348 |
| Rbfa | Q6P3B9 | -0.34 | 0.42 | 0.05 | -0.5 | -0.36 | -0.76 | 0.25 | -0.39 | -0.47 | -0.48 | 0.18 | 0.39 | -0.49 | -0.03 | -0.33 | -1.32 | -0.2225 | -0.0025 | -0.45 | 0.467724 | 0.993539 | 0.210189 | 0.999971 | 1 | 0.822657 |
| Rbfox1 | A0A2R8VK | 0.54 | 1.05 | 0.55 | 0.79 | 1.02 | 1.06 | 0.34 | 0.89 | 0.89 | 0.56 | 0.54 | 1.28 | 0.74 | 0.94 | 0.8 | 1.23 | 0.095 | 0.085 | 0.195 | 0.692038 | 0.725317 | 0.370863 | 0.999971 | 1 | 0.869526 |
| Rbfox1_1 | Q9JJ43-6 | 2.82 | 3.78 | 2.82 | 2.56 | 4.03 | 2.69 | 1.07 | 2.42 | 2.52 | 3.17 | 2.28 | 2.49 | 1.96 | 3.8 | 1.94 | 3.14 | -0.4425 | -0.38 | -0.285 | 0.471451 | 0.267879 | 0.568177 | 0.999971 | 1 | 0.91816 |
| Rbfox2 | Q8BP71-7 | -0.57 | -0.06 | 0.77 | 0.58 | -0.29 | -0.03 | -0.34 | 0.49 | -0.3 | -0.77 | -1.09 | -0.16 | -0.78 | -0.58 | -0.07 | 0.1 | -0.2225 | -0.76 | -0.5125 | 0.533763 | 0.063311 | 0.181273 | 0.999971 | 1 | 0.808532 |
| Rbfox3 | Q8BIF2-2 | -0.02 | 1.59 | 1.44 | 1.1 | 1.7 | 1.34 | 1.41 | 1.37 | 1.57 | -0.41 | 1.74 | 1.84 | 1.84 | 0.92 | 1.57 | 1.51 | 0.4275 | 0.1575 | 0.4325 | 0.258577 | 0.791173 | 0.291374 | 0.999971 | 1 | 0.847402 |
| Rbks | Q8R1Q9 | 2.91 | 1.66 | 2.04 | 1.68 | 1.41 | 2.01 | 2.65 | 2.53 | 1.84 | 2.99 | 0.63 | -0.51 | -1.08 | 0.56 | 0.32 | -1.5 | 0.0775 | -0.835 | -2.4975 | 0.842767 | 0.274786 | 0.001329 | 0.999971 | 1 | 0.432702 |
| Rbl2 | A0A1B0GR | 2.4 | 0.84 | 1.5 | 1.9 | 0.65 | 0.79 | 1.57 | 1.39 | -0.05 | 1.62 | 0.8 | 0.11 | -0.61 | 0.44 | 1.39 | 1.18 | -0.56 | -1.04 | -1.06 | 0.169567 | 0.052217 | 0.066716 | 0.999971 | 1 | 0.733937 |
| Rbm10 | Q99KG3-3 | 0.07 | 1.18 | 1.5 | 1.51 | 1.49 | 1.23 | 1.75 | 1.44 | 1.54 | 0.61 | 1.19 | 1.29 | 1.82 | 1.42 | 1.27 | 1.36 | 0.4125 | 0.0925 | 0.4025 | 0.258127 | 0.808019 | 0.274796 | 0.999971 | 1 | 0.843794 |
| Rbm12 | Q8R4X3 | 1.38 | 1.93 | 2.15 | 2.23 | 1.88 | 1.62 | 1.61 | 1.83 | 1.62 | 1.79 | 1.69 | 2.3 | 1.32 | 2.26 | 1.66 | 1.7 | -0.1875 | -0.0725 | -0.1875 | 0.438868 | 0.784985 | 0.518624 | 0.999971 | 1 | 0.905901 |
| Rbm12b2 | Q66JV4 | 0.02 | 0.08 | 0.86 | 0.86 | -0.09 | -0.22 | 0.99 | 0.81 | -0.5 | 0.08 | 1.18 | 0.25 | -0.88 | 0.01 | 0.64 | 0.38 | -0.0825 | -0.2025 | -0.4175 | 0.825659 | 0.617085 | 0.301712 | 0.999971 | 1 | 0.847402 |

| | | | | | | | | | | | | | | | | | | | | | | | | | | |
|---|---|---|---|---|---|---|---|---|---|---|---|---|---|---|---|---|---|---|---|---|---|---|---|---|---|---|
| Rbm14 | Q8C2Q3 | 1.27 | 1.01 | 1.47 | 1.5 | 1.14 | 1.17 | 1.15 | 1.47 | 1.23 | 1.32 | 1.09 | 1.02 | 0.59 | 1.57 | 1.34 | 1.46 | -0.08 | -0.1475 | -0.0725 | 0.687513 | 0.44992 | 0.78807 | 0.999971 | 1 | 0.96724 |
| Rbm15 | Q0VBL3-2 | 0.32 | -0.05 | 0.68 | 0.57 | 0.2 | -0.3 | 0.45 | 0.73 | -0.01 | -0.19 | 0.83 | 0.72 | 0.1 | -0.37 | 0.54 | 0.87 | -0.11 | -0.0425 | -0.095 | 0.701074 | 0.89029 | 0.765487 | 0.999971 | 1 | 0.962405 |
| Rbm15b | Q6PHZ5 | 1.34 | 0.61 | 0.24 | 0.08 | -0.32 | 0.56 | -0.19 | 0.15 | 0.26 | 0.9 | 0.6 | 0.24 | 0.58 | -0.02 | 0.36 | 0.28 | -0.5175 | -0.0675 | -0.2675 | 0.152982 | 0.83423 | 0.401533 | 0.999971 | 1 | 0.871816 |
| Rbm17 | Q8JZX4 | 1.26 | 1.59 | 1.91 | 1.25 | 1.58 | 1.63 | 1.18 | 1.25 | 1.37 | 1.34 | 1.73 | 1.15 | 1.3 | 1.46 | 1.26 | 1.06 | -0.0925 | -0.105 | -0.2325 | 0.69051 | 0.65306 | 0.309372 | 0.999971 | 1 | 0.847402 |
| Rbm19 | Q8R3C6 | -0.59 | -1.69 | -0.29 | -0.16 | -1.42 | -2.17 | 2.59 | -1.21 | -1.9 | -1.25 | -0.87 | -1.03 | -1.97 | -2.22 | -0.85 | -2.19 | 0.13 | -0.58 | -1.125 | 0.89667 | 0.170568 | 0.031251 | 0.999971 | 1 | 0.678474 |
| Rbm22 | Q8BHS3 | 1.34 | 0.61 | 0.56 | 0.86 | 0.56 | 0.64 | 0.6 | 0.5 | 0.49 | 1.15 | 0.26 | -0.08 | 0.06 | 0.79 | 0.35 | 0.33 | -0.2675 | -0.3875 | -0.46 | 0.24973 | 0.242561 | 0.102687 | 0.999971 | 1 | 0.78268 |
| Rbm25 | B2RY56 | 1.75 | 0.55 | 1.72 | 2.09 | 1.17 | 1.22 | 1.88 | 1.63 | 0.94 | 1.39 | 1.72 | 1.6 | 0.23 | 0.62 | 1.63 | 1.85 | -0.0525 | -0.115 | -0.445 | 0.886023 | 0.755002 | 0.366895 | 0.999971 | 1 | 0.869526 |
| Rbm26 | E9Q640 | 1.7 | 2.15 | 2.41 | 2 | 1.87 | 1.95 | 2.07 | 2.16 | 1.98 | 1.81 | 2.58 | 2.81 | 1.67 | 2.02 | 2.12 | 2.23 | -0.0525 | 0.23 | -0.055 | 0.803899 | 0.437819 | 0.811446 | 0.999971 | 1 | 0.973864 |
| Rbm27 | Q5SFM8 | 0.98 | 2.56 | 1.84 | 1.07 | 2.14 | 2.61 | 1.22 | 1.3 | 2.22 | 0.45 | 1.48 | 1.69 | -0.22 | 1.22 | 1.04 | 0.67 | 0.205 | -0.1525 | -0.935 | 0.662377 | 0.755811 | 0.068004 | 0.999971 | 1 | 0.733937 |
| Rbm28 | Q8CGC6 | 0.58 | 0.09 | 1.02 | 0.81 | -0.06 | 0.25 | 0.49 | 1.2 | 0.43 | 0.24 | 0.8 | 1.2 | -0.03 | 0.01 | 0.8 | 1.02 | -0.155 | 0.0425 | -0.175 | 0.643295 | 0.887008 | 0.603446 | 0.999971 | 1 | 0.924454 |
| Rbm3 | O89086 | 2.14 | 0.69 | 1.53 | 0.8 | 1.5 | 1.24 | 1.36 | 0.54 | 1.34 | 1.9 | 0.12 | 0.09 | 0.72 | 1.49 | 0.64 | -0.55 | -0.13 | -0.4275 | -0.715 | 0.736596 | 0.424073 | 0.178945 | 0.999971 | 1 | 0.808532 |
| Rbm34 | A0A1D5RLI | 0.76 | 0.19 | -0.56 | -0.63 | 0.23 | -0.2 | -0.66 | -0.12 | 0.45 | 0.01 | -0.43 | -0.27 | 1.02 | 0.38 | -0.99 | -0.18 | -0.1275 | -1.47E-16 | 0.1175 | 0.729058 | 1 | 0.814742 | 0.999971 | 1 | 0.974918 |
| Rbm39 | Q8VH51-2 | 0.25 | 0.65 | 0.73 | 0.44 | 0.94 | 0.73 | 0.57 | 0.8 | 0.99 | 0.41 | 0.68 | 0.92 | 1.07 | 1.23 | 1.1 | 0.7 | 0.2425 | 0.2325 | 0.5075 | 0.23464 | 0.294268 | 0.035895 | 0.999971 | 1 | 0.699952 |
| Rbm42 | D3YYW8 | 0.61 | 1.41 | 1.22 | 1.13 | 1.05 | 0.57 | 1.58 | 0.48 | 0.89 | 0.79 | 0.84 | 1.01 | 0.52 | 0.49 | 0.34 | 0.99 | -0.1725 | -0.21 | -0.5075 | 0.581786 | 0.350161 | 0.068618 | 0.999971 | 1 | 0.733937 |
| Rbm45 | Q8BHN5 | 0.82 | 0.71 | 1.67 | 0.97 | 0.87 | 0.46 | 1.17 | 1.47 | 0.25 | 1.11 | 0.36 | 1.75 | -0.23 | 1.16 | 1.49 | 0.69 | -0.05 | -0.175 | -0.265 | 0.871928 | 0.65995 | 0.524637 | 0.999971 | 1 | 0.90868 |
| Rbm4b | Q8VE92 | -3.14 | -1.9 | -0.45 | -1.76 | -2.61 | -4.13 | -1.08 | -1.59 | -2.82 | -5.18 | -1.18 | -2.45 | -2.05 | -0.46 | 0.46 | -1.17 | -0.54 | -1.095 | 1.0075 | 0.495347 | 0.24474 | 0.171291 | 0.999971 | 1 | 0.804075 |
| Rbm5 | Q91YE7-2 | 0.7 | 0.74 | 1.04 | 1.37 | 0.69 | 0.39 | 2 | 0.93 | 0.46 | 0.8 | 0.3 | 0.42 | 0.59 | 0.88 | 0.63 | 0.67 | 0.04 | -0.4675 | -0.27 | 0.914216 | 0.064616 | 0.232194 | 0.999971 | 1 | 0.831121 |
| Rbm6 | S4R1W5 | 2.36 | 1.26 | 1.37 | 1.56 | 1.17 | 1.02 | 1.42 | 2.17 | 1.68 | 1.58 | 1.57 | 1.73 | 0.89 | 0.75 | 1.49 | 1.43 | -0.1925 | 0.0025 | -0.4975 | 0.585622 | 0.992594 | 0.141527 | 0.999971 | 1 | 0.784967 |
| Rbm7 | Q9CQT2 | 3.7 | 1.25 | 1.45 | 2.11 | 1.18 | 1.48 | 1.33 | 1.21 | 1.79 | 3.42 | 1.11 | 1.16 | -0.23 | -0.09 | 0.58 | 0.55 | -0.8275 | -0.2575 | -1.925 | 0.136139 | 0.715562 | 0.006412 | 0.999971 | 1 | 0.582714 |
| Rbm8a | Q9CWZ3 | 0.83 | 1.2 | 0.79 | 0.64 | 2 | 0.97 | 0.92 | 0.89 | 2.3 | 1.18 | 0.87 | 1.1 | 2.58 | 1.27 | 0.38 | 0.96 | 0.33 | 0.4975 | 0.4325 | 0.290921 | 0.166384 | 0.351499 | 0.999971 | 1 | 0.867656 |
| Rbms3 | G3UZJ9 | 2.56 | 1.82 | 1.56 | 1.03 | 1.5 | 1.63 | 1 | 1.72 | 1.14 | 2.28 | 1.58 | 1.17 | 1.47 | 1.98 | 1.11 | 1.24 | -0.28 | -0.2 | -0.2925 | 0.432547 | 0.617622 | 0.429178 | 0.999971 | 1 | 0.881196 |
| Rbmx | Q9WV02 | 0.55 | 0.19 | -0.14 | 0.25 | -0.36 | -0.34 | 0.92 | -0.32 | 0.1 | -0.49 | 0.19 | -0.15 | -0.74 | -1.11 | -0.36 | -0.31 | -0.2375 | -0.3 | -0.8425 | 0.492764 | 0.2299 | 0.009509 | 0.999971 | 1 | 0.613095 |
| Rbmx2 | Q8R0F5 | 1.25 | 0.51 | 1.59 | 0.11 | -0.93 | -0.76 | 3.04 | 1.28 | 0.97 | 1.38 | 2.38 | 3.09 | 0.1 | 1.1 | 1.1 | 0.66 | -0.2075 | 1.09 | -0.125 | 0.816469 | 0.071652 | 0.752915 | 0.999971 | 1 | 0.961116 |
| Rbp1 | Q00915 | -0.26 | 0.69 | 0.01 | 0.04 | 1.26 | 0.14 | 0.4 | 0.64 | 0.85 | 0.12 | -0.7 | 0.07 | 0.28 | 0.43 | 0.23 | -0.69 | 0.49 | -0.035 | -0.0575 | 0.147466 | 0.923726 | 0.859952 | 0.999971 | 1 | 0.982702 |
| Rbsn | Q80Y56 | -0.14 | -0.15 | 0.02 | 0.02 | 0.07 | -0.16 | 0.25 | -0.18 | -0.22 | -0.18 | -0.12 | -0.26 | 0.3 | 0.04 | -0.1 | -0.31 | 0.0575 | -0.1325 | 0.045 | 0.756803 | 0.418322 | 0.820263 | 0.999971 | 1 | 0.975761 |
| Rbx1 | P62878 | 0.52 | -0.77 | -0.5 | -0.65 | -0.67 | -0.6 | -0.16 | -0.35 | -0.87 | 0.51 | -0.93 | -1.47 | -1.47 | -0.27 | -0.45 | -1.05 | -0.095 | -0.34 | -0.46 | 0.766841 | 0.486868 | 0.256521 | 0.999971 | 1 | 0.839012 |
| Rc3h1 | Q4VGL6 | 1.2 | -0.31 | -0.2 | -0.05 | 0.2 | 0.4 | 0.38 | -0.07 | -0.5 | 0.78 | -0.62 | -0.28 | 0 | 0.44 | -0.59 | 0.06 | 0.0675 | -0.315 | -0.1825 | 0.850763 | 0.488023 | 0.645239 | 0.999971 | 1 | 0.93301 |
| Rc3h2 | A2AVP4 | 0.98 | 0.89 | 2.07 | 2.4 | 0.7 | 1.14 | 2.62 | 2.4 | 0.81 | 0.87 | 2.18 | 1.68 | 1.15 | 1.33 | 2.08 | 1.53 | 0.13 | -0.2 | -0.0625 | 0.815105 | 0.674203 | 0.879248 | 0.999971 | 1 | 0.985441 |
| Rcan1 | Q9JHG6 | 0.76 | 1 | 0.92 | 1.12 | 0.47 | 0.85 | 0.12 | 1.34 | 0.27 | 0.72 | 1.08 | 0.91 | 0.17 | 0.83 | 1.44 | 1.78 | -0.255 | -0.205 | 0.105 | 0.383129 | 0.377857 | 0.768251 | 0.999971 | 1 | 0.962535 |
| Rcan2 | A0A2R8VKI | 0.62 | -0.17 | 0.62 | -1.51 | -0.74 | -0.69 | 2.02 | -0.09 | -1.12 | -0.98 | 2.03 | 1.33 | -2.84 | -0.61 | -0.47 | -1.13 | 0.235 | 0.425 | -1.1525 | 0.751177 | 0.621176 | 0.11401 | 0.999971 | 1 | 0.78268 |
| Rcan3 | Q9JKK0 | -0.85 | 0.06 | -0.48 | 0.52 | -0.21 | -0.45 | 0.35 | 0.74 | -0.11 | -0.4 | -0.48 | 1.02 | 0.29 | 0.03 | 0.39 | 0.66 | 0.295 | 0.195 | 0.53 | 0.454729 | 0.655542 | 0.133437 | 0.999971 | 1 | 0.784948 |
| Rcbtb1 | A0A0R4J02 | -0.54 | -0.06 | -1.42 | -0.51 | 0.27 | -0.6 | -1.26 | -1.09 | 0.33 | -1.38 | -0.81 | -0.76 | 0.05 | -0.4 | -1.09 | -0.4 | -0.0375 | -0.0225 | 0.1725 | 0.929183 | 0.958495 | 0.634977 | 0.999971 | 1 | 0.93301 |
| Rcbtb2 | E9Q3B1 | -0.02 | 0.98 | 0.2 | 0.52 | 0.8 | 0.89 | -0.43 | 0.33 | 0.12 | 0.63 | -0.03 | -0.05 | 1.47 | 1.24 | 0.39 | 0.78 | -0.0225 | -0.2525 | 0.55 | 0.95045 | 0.382413 | 0.118437 | 0.999971 | 1 | 0.78268 |
| Rcc1 | Q8VE37 | -0.33 | 0.26 | 0.12 | 0.39 | 0.28 | 0.38 | 0.09 | 0.42 | 0.16 | 0.07 | 0.13 | 0.03 | -0.32 | 0.14 | 0.57 | 0.37 | 0.1825 | -0.0125 | 0.08 | 0.412515 | 0.951941 | 0.765954 | 0.999971 | 1 | 0.962472 |
| Rcc2 | Q8BK67 | 2.52 | 2.6 | 2.79 | 3.03 | 2.1 | 2.34 | 3.18 | 2.18 | 1.91 | 2.44 | 1.98 | 1.66 | 1.02 | 2.47 | 2.23 | 1.53 | -0.285 | -0.7375 | -0.9225 | 0.333401 | 0.010894 | 0.024322 | 0.999971 | 1 | 0.669611 |
| Rchy1 | G3UWD8 | 1.11 | 1.6 | 1.22 | 1.55 | 1.43 | 1.66 | 1.01 | 2.05 | 1.95 | 1.48 | 1.45 | 0.64 | 1.92 | 1.73 | 1.42 | 1.59 | 0.1675 | 0.01 | 0.295 | 0.53749 | 0.973733 | 0.186918 | 0.999971 | 1 | 0.809025 |
| Rcl1 | Q9JJT0 | -0.87 | -0.45 | -0.67 | -0.63 | -0.36 | -0.92 | -1.04 | -0.9 | -0.73 | -0.79 | -1.27 | -1.3 | 0.05 | -0.72 | -0.86 | -1.16 | -0.15 | -0.3675 | -0.0175 | 0.499073 | 0.11791 | 0.951148 | 0.999971 | 1 | 0.998056 |
| Rcn1 | Q05186 | -0.8 | -0.25 | -0.73 | -0.51 | 0.26 | -0.71 | 0.01 | 0 | 0.99 | -0.15 | 0.33 | 1.02 | 1 | 0.3 | 0.08 | -0.14 | 0.4625 | 1.12 | 0.8825 | 0.107542 | 0.005799 | 0.013113 | 0.999971 | 1 | 0.613095 |
| Rcn2 | Q8BP92 | -1.07 | 0.08 | 0.13 | -0.67 | 0.45 | -0.27 | 1.24 | 0.22 | 1.1 | -1.16 | 0.9 | 1.41 | 1.36 | -0.63 | 0.07 | -0.94 | 0.7925 | 0.945 | 0.3475 | 0.079893 | 0.141986 | 0.526869 | 0.999971 | 1 | 0.909459 |
| Rcor1 | A0A140T8F | 2.1 | 1.47 | 1.34 | 0.97 | 1.06 | 1.18 | 0.74 | 0.98 | 1.03 | 1.67 | 0.98 | 1.22 | 0.52 | 1.57 | 1.07 | 1.15 | -0.48 | -0.245 | -0.3925 | 0.104404 | 0.412342 | 0.242587 | 0.999971 | 1 | 0.837316 |
| Rcsd1 | Q3UZA1-2 | 2.43 | 0.61 | 1.27 | 1.71 | 1.35 | 0.87 | 2.27 | 0.38 | 1.02 | 1.64 | 0.65 | 0.1 | -0.55 | 0.68 | 0.34 | -0.75 | -0.2875 | -0.6525 | -1.575 | 0.57883 | 0.189052 | 0.009233 | 0.999971 | 1 | 0.613095 |
| Rdh11 | Q9R1R8 | -1.57 | -1.16 | -0.69 | -1.74 | -0.78 | -0.94 | -0.92 | -1.33 | -0.64 | -1.24 | -0.12 | 0.17 | -0.22 | -1.46 | -0.27 | -0.83 | 0.2975 | 0.8325 | 0.595 | 0.301075 | 0.051492 | 0.129171 | 0.999971 | 1 | 0.784948 |
| Rdh13 | Q8CEE7 | 0.12 | -0.23 | -0.44 | -0.42 | -0.23 | -0.19 | -0.67 | -0.47 | -0.18 | 0.19 | -0.27 | -0.62 | -0.13 | -0.75 | -0.79 | -0.42 | -0.1475 | 0.0225 | -0.28 | 0.502665 | 0.925514 | 0.256958 | 0.999971 | 1 | 0.839012 |
| Rdh14 | Q9ERI6 | 0.07 | -0.77 | -0.24 | -0.48 | -0.44 | -0.52 | 0.03 | -0.25 | -0.2 | 0.17 | -0.09 | -0.51 | -0.08 | -0.8 | -0.16 | -0.23 | 0.06 | 0.1975 | 0.0375 | 0.807483 | 0.441657 | 0.887247 | 0.999971 | 1 | 0.987485 |
| Rdx | P26043 | -0.83 | -1.05 | 0.57 | 0.49 | -0.7 | -0.65 | 1.09 | -0.07 | -0.54 | -1.05 | -0.13 | -0.22 | -0.59 | -0.85 | 0.27 | -0.07 | 0.1225 | -0.28 | -0.105 | 0.823162 | 0.536671 | 0.821336 | 0.999971 | 1 | 0.975833 |
| Recql5 | Q8VID5 | 3.2 | 0.74 | 2.78 | 1.8 | 3.6 | 1.6 | 15.05 | -0.86 | 12.93 | 2.68 | 3.59 | 6.41 | 4.8 | 0.97 | 0.32 | 3.58 | 2.7175 | 4.2725 | 0.2875 | 0.397818 | 0.070758 | 0.78771 | 0.999971 | 1 | 0.96724 |
| Reep1 | Q8BGH4 | -0.31 | -1.29 | -0.83 | -0.85 | -1.21 | -1.43 | -0.93 | -0.61 | -0.93 | -0.47 | -0.44 | -0.59 | -0.93 | -0.91 | -0.36 | -0.55 | -0.225 | 0.2125 | 0.1325 | 0.43395 | 0.412245 | 0.620982 | 0.999971 | 1 | 0.929388 |
| Reep2 | Q8VCD6 | -0.09 | -1.27 | -0.14 | -0.59 | -1.45 | -1.08 | 0.02 | -0.66 | -1.37 | -0.97 | -0.36 | -0.61 | -1.58 | -1.23 | -0.41 | -0.42 | -0.27 | -0.305 | -0.3875 | 0.50421 | 0.388891 | 0.331407 | 0.999971 | 1 | 0.857152 |
| Reep3 | A0A1W2P8 | 0.78 | -0.59 | -0.26 | -0.14 | 0.71 | -0.11 | -0.01 | -0.16 | -0.4 | 1.44 | -0.55 | -0.83 | -0.24 | -0.71 | -0.65 | -0.82 | 0.16 | -0.0325 | -0.5525 | 0.649815 | 0.952714 | 0.113526 | 0.999971 | 1 | 0.78268 |
| Reep5 | G3X8R0 | -1.34 | -3.29 | 1.09 | 0.57 | -2.56 | -2.55 | 1.39 | 1.24 | -1.98 | -1.64 | 1.13 | 1.12 | -3.01 | -2.87 | 1.14 | 0.57 | 0.1225 | 0.4 | -0.3 | 0.926497 | 0.734061 | 0.820483 | 0.999971 | 1 | 0.975761 |
| Reep6 | Q9JM62 | -0.54 | 0.96 | -0.42 | -0.77 | 1.43 | 1.11 | -1.02 | -0.38 | 1.04 | -0.07 | -0.86 | -0.12 | 2.03 | 0.97 | -0.81 | -0.81 | 0.4775 | 0.19 | 0.5375 | 0.463365 | 0.712712 | 0.466003 | 0.999971 | 1 | 0.891431 |
| Rel | A4QPD3 | 0.87 | 0 | 1.46 | 1.51 | 0.32 | 0.53 | -0.23 | 1.41 | 0.64 | 0.5 | 1.57 | 0.72 | -0.28 | -0.28 | 2.14 | 3 | -0.4525 | -0.1025 | 0.185 | 0.337451 | 0.801745 | 0.821565 | 0.999971 | 1 | 0.975833 |
| Rela | A0A494BA | 0.07 | 0.25 | 0.23 | 0.25 | 0.3 | -0.08 | -0.17 | 0.31 | -0.19 | 0.06 | 0.18 | 0.09 | -0.31 | 0.41 | -0.08 | 0.13 | -0.11 | -0.165 | -0.1625 | 0.576627 | 0.351056 | 0.448375 | 0.999971 | 1 | 0.885289 |
| Relch | E9QM90 | -0.21 | 0.01 | -0.13 | -0.32 | -0.08 | -0.12 | -0.09 | -0.41 | -0.14 | -0.06 | -0.2 | -0.16 | -0.06 | -0.3 | -0.43 | -0.26 | -0.0125 | 0.0225 | -0.1 | 0.945127 | 0.892443 | 0.585437 | 0.999971 | 1 | 0.920413 |
| Relch_1 | A0A087WS | 0.52 | -0.4 | -2.45 | 0.07 | -1.46 | -0.75 | -2.17 | -1.14 | -0.91 | 0.2 | -1.64 | -2.09 | -2.07 | -0.63 | -1.54 | -0.81 | -0.815 | -0.545 | -0.6975 | 0.233759 | 0.473302 | 0.31183 | 0.999971 | 1 | 0.848872 |
| Rell1 | Q8K2J7-2 | 2.66 | 1.6 | -1.36 | -1.28 | 2.29 | 2.26 | -1.51 | -0.28 | 1.84 | 3 | -0.13 | -0.72 | 3.11 | 1.34 | -1.1 | -0.13 | 0.285 | 0.5925 | 0.4 | 0.817904 | 0.622898 | 0.743755 | 0.999971 | 1 | 0.961064 |
| Rell2 | Q8BRJ3 | 0.69 | -0.87 | -0.04 | 0.28 | -0.73 | -0.43 | -0.18 | -0.4 | -0.68 | 0.21 | -0.58 | -0.85 | -1.09 | -0.63 | -0.72 | -0.41 | -0.45 | -0.49 | -0.7275 | 0.21373 | 0.230995 | 0.065638 | 0.999971 | 1 | 0.731328 |
| Reln | Q60841-3 | -0.6 | 0.11 | 0.25 | 0.28 | 0.59 | -0.16 | 0.27 | 0.43 | 0.14 | -0.44 | 0.12 | 0.17 | 0.28 | 0.23 | 0.41 | 0.11 | 0.2725 | -0.0125 | 0.2475 | 0.340984 | 0.963132 | 0.329414 | 0.999971 | 1 | 0.856787 |
| Rem2 | E9Q4D5 | 0.71 | 0.18 | -0.37 | 0.14 | 0.18 | -0.79 | -1.3 | 0.23 | 0.61 | 0.77 | -0.41 | 0.16 | 1.23 | 0.37 | 0.68 | 0.41 | -0.585 | 0.1175 | 0.5075 | 0.181873 | 0.73094 | 0.122577 | 0.999971 | 1 | 0.78268 |
| Renbp | D6RHA2 | 1.07 | 1.64 | 1.95 | 1.95 | 1.23 | 1.17 | 1.2 | 1.83 | 0.71 | 1.29 | 1.64 | 1.67 | 1.36 | 1.98 | 2.87 | 3.21 | -0.295 | -0.325 | 0.7025 | 0.303213 | 0.309428 | 0.139128 | 0.999971 | 1 | 0.784948 |
| Reps1 | O54916 | -0.07 | 0.55 | 0.58 | 0.11 | 0.4 | 0.3 | 0.15 | 0.1 | 0.76 | 0.04 | -0.02 | 0.14 | 1.64 | 0.14 | 0.26 | 0.15 | -0.055 | -0.0625 | 0.255 | 0.802599 | 0.811895 | 0.513491 | 0.999971 | 1 | 0.905815 |
| Reps2 | B9EI38 | 0.74 | 0.54 | 0.29 | 0.47 | 0.39 | 0.22 | 0.29 | 0.3 | 0.13 | 0.72 | -0.02 | 0.06 | 0.25 | 0.38 | 0.31 | 0.47 | -0.21 | -0.2875 | -0.1575 | 0.260252 | 0.229482 | 0.397685 | 0.999971 | 1 | 0.871816 |
| Rer1 | Q9CQU3 | -0.02 | 0.41 | 0.77 | 0.83 | 0.19 | 0.56 | 0.57 | 1.38 | 0.59 | 1.09 | 0.82 | 1.13 | 0.24 | 1.36 | 1.3 | 1.85 | 0.1775 | 0.41 | 0.69 | 0.582883 | 0.13634 | 0.096194 | 0.999971 | 1 | 0.78268 |
| Retreg1 | Q8VE91 | 0.43 | -0.54 | -1.28 | -0.09 | -0.59 | -0.6 | -1.31 | -0.62 | -0.93 | 0.33 | -1.34 | -0.89 | -0.56 | -0.71 | -0.36 | -0.51 | -0.41 | -0.3375 | -0.165 | 0.305834 | 0.486998 | 0.649741 | 0.999971 | 1 | 0.933842 |
| Retreg2 | Q6NS82-3 | 0.81 | -0.09 | -0.78 | -0.2 | -0.3 | -0.15 | -0.69 | -0.31 | -0.46 | 0.33 | -0.77 | -0.92 | -0.32 | -0.21 | -0.02 | -0.16 | -0.2975 | -0.39 | -0.1125 | 0.397013 | 0.355988 | 0.736591 | 0.999971 | 1 | 0.959447 |
| Retreg3 | Q9CQV4 | 2.4 | 0.98 | 0.09 | -1.3 | -0.33 | 1.49 | 0.19 | -1.17 | 0.33 | 2.26 | -0.53 | -0.12 | 0.01 | 0.99 | -0.91 | -1.89 | -0.4975 | -0.0575 | -0.9925 | 0.565125 | 0.948703 | 0.282074 | 0.999971 | 1 | 0.844538 |
| Retsat | Q64FW2 | 0.58 | -2.15 | 0.27 | 2.79 | -1.46 | -1.79 | 1.58 | -0.12 | -2.84 | 0.65 | 1.56 | 1.19 | -1.67 | -1.89 | 1.95 | 0.56 | -0.82 | -0.2325 | -0.635 | 0.474282 | 0.855805 | 0.604832 | 0.999971 | 1 | 0.924454 |
| Rexo2 | Q9D8S4 | -0.03 | 0.26 | 0.09 | -0.5 | 0.5 | -0.09 | 0.15 | 0.2 | -0.27 | -0.62 | 0.68 | 0.42 | 0.27 | 0.46 | 0.31 | -0.07 | 0.235 | 0.0975 | 0.2875 | 0.335691 | 0.773882 | 0.240126 | 0.999971 | 1 | 0.834057 |
| Rexo4 | A2ALB1 | 1.02 | 0.63 | 0.78 | 0.88 | 0.46 | 0.83 | -0.04 | 0.56 | 0.7 | 1.06 | 0.87 | 0.87 | 0.68 | 1.46 | 0.98 | 1.01 | -0.375 | 0.0475 | 0.205 | 0.136882 | 0.79324 | 0.370806 | 0.999971 | 1 | 0.869526 |
| Rfc2 | Q9WUK4 | 1.36 | 1.72 | 1.54 | 1.5 | 0.9 | 2.5 | 0.89 | 0.77 | 1.27 | 2.3 | 0.77 | 0.75 | 0.28 | 1.68 | 0.8 | 2.04 | -0.265 | -0.2575 | -0.33 | 0.513643 | 0.483306 | 0.411083 | 0.999971 | 1 | 0.875847 |
| Rfc3 | Q8R323 | -1.39 | -0.17 | -0.07 | -0.4 | -1.67 | -1.27 | -0.38 | -0.02 | -0.72 | -1.78 | -0.07 | -0.21 | -1.92 | -1.34 | -0.12 | -0.62 | -0.3275 | -0.1875 | -0.4925 | 0.480016 | 0.686111 | 0.306802 | 0.999971 | 1 | 0.847402 |
| Rfc4 | D6RGM1 | 0.25 | -0.85 | 1.51 | 0.8 | -1.41 | -1.09 | 0.72 | 0.93 | -0.73 | -0.98 | 0.53 | 0.68 | -0.42 | -0.63 | 0.92 | 0.91 | -0.64 | -0.5525 | -0.2325 | 0.376886 | 0.37024 | 0.696858 | 0.999971 | 1 | 0.947174 |
| Rfc5 | Q9D0F6 | 0.76 | -0.51 | 2.11 | 1.73 | 0.25 | -0.15 | 2.47 | -0.48 | 0.63 | 0.74 | -0.25 | -1.1 | -0.28 | -0.66 | 0.9 | -1.16 | -0.5 | -1.0175 | -1.3225 | 0.534671 | 0.150886 | 0.072952 | 0.999971 | 1 | 0.745263 |
| Rfesd | Q8K2P6 | 1.91 | 1.32 | 0.5 | 0.87 | 0.89 | 1.22 | 0.49 | 0.29 | 0.86 | 1.47 | 0.17 | 0.58 | 0.43 | 0.71 | 0.27 | 0.04 | -0.4275 | -0.38 | -0.7875 | 0.253423 | 0.346872 | 0.040199 | 0.999971 | 1 | 0.700838 |
| Rfk | Q8CFV9 | -1.98 | 0.01 | -0.31 | -0.79 | 1.35 | -0.31 | 0.4 | -0.61 | 1.61 | -1.41 | 0.4 | 1.04 | 1.93 | -1.36 | -0.4 | -0.79 | 0.975 | 1.1775 | 0.6125 | 0.111854 | 0.127324 | 0.429816 | 0.999971 | 1 | 0.881611 |
| Rft1 | A0A2I3BRR | 1.66 | 0.58 | 1.97 | 1.94 | 0.1 | 0.83 | 3.04 | 2.06 | 1.08 | 1.93 | 0.73 | -2.31 | 0.47 | 0.25 | 2.08 | -0.87 | -0.03 | -1.18 | -1.055 | 0.96375 | 0.205135 | 0.120744 | 0.999971 | 1 | 0.78268 |
| Rftn1 | Q6A0D4 | -1 | -2.35 | -2.37 | -2.59 | -1.83 | -1.95 | -2.24 | -2.33 | -1.66 | -0.97 | -2.22 | -1.74 | -1.67 | -1.7 | -1.78 | -1.75 | -0.01 | 0.43 | 0.3525 | 0.978443 | 0.324579 | 0.33742 | 0.999971 | 1 | 0.858827 |
| Rftn2 | Q8CHX7 | 0.15 | -0.61 | -0.51 | -0.33 | -0.47 | -0.8 | 0.3 | -1.13 | -0.17 | -0.67 | -1.31 | -0.99 | 0.17 | -1.23 | -1.36 | -1.04 | -0.2 | -0.46 | -0.54 | 0.565488 | 0.15464 | 0.177174 | 0.999971 | 1 | 0.808532 |
| Rfx3 | G3UWP0 | 2.54 | 2.07 | 0.45 | 1.17 | 2.25 | 2.21 | -0.11 | 0.5 | 2.24 | 2.81 | 1.51 | -0.03 | 1.95 | 2.66 | 0.44 | 0.77 | -0.345 | 0.075 | -0.1025 | 0.61824 | 0.91484 | 0.871384 | 0.999971 | 1 | 0.984109 |
| Rfxap | F7DBT7 | 0.66 | 0.62 | 1.12 | 0.02 | 0.82 | 0.2 | 0.1 | 1.1 | 0.47 | -0.98 | 0.63 | 0.63 | 0.23 | -0.03 | 1.47 | 2.12 | -0.05 | -0.4175 | 0.3425 | 0.879472 | 0.34348 | 0.512343 | 0.999971 | 1 | 0.905815 |
| Rgcc | Q9DBX1 | 0.01 | 0.9 | 0.23 | 0.71 | 1.51 | 0.87 | -0.41 | 0.62 | 0.45 | -0.74 | 0.2 | -0.33 | -0.43 | 1.24 | 0.4 | 1.4 | 0.185 | -0.5675 | 0.19 | 0.665134 | 0.118075 | 0.669581 | 0.999971 | 1 | 0.938711 |
| Rgl1 | E9Q8M9 | -0.11 | -0.29 | -1.17 | -0.44 | -0.39 | -0.46 | -0.73 | -0.39 | -0.53 | -0.26 | -1.46 | -1 | -0.38 | -0.3 | -1.34 | -0.92 | 0.01 | -0.31 | -0.2325 | 0.97007 | 0.382921 | 0.49464 | 0.999971 | 1 | 0.901531 |
| Rgl2 | Q61193 | -0.2 | -0.47 | -0.33 | 0.34 | -0.16 | -0.91 | -0.28 | -0.01 | -0.57 | -0.13 | -0.49 | -0.34 | -0.39 | -0.58 | -0.24 | 0.02 | -0.175 | -0.2175 | -0.1325 | 0.537564 | 0.366122 | 0.595661 | 0.999971 | 1 | 0.92377 |
| Rgma | Q8C4C4 | 0.97 | -0.63 | 0.43 | 0.2 | -0.93 | 0.24 | 1.23 | -0.44 | -0.5 | 0.84 | -0.15 | -0.48 | -0.62 | -0.75 | -0.16 | 0.15 | -0.2175 | -0.315 | -0.5875 | 0.682293 | 0.474608 | 0.147983 | 0.999971 | 1 | 0.793248 |
| Rgmb | Q7TQ33 | -0.47 | -0.52 | -1.23 | -0.23 | -0.8 | -0.84 | -0.54 | -0.24 | 0.34 | -2.06 | -0.75 | -0.75 | -1.99 | 0.56 | -1.44 | -2 | 0.0075 | -0.1925 | -0.605 | 0.978133 | 0.701066 | 0.319617 | 0.999971 | 1 | 0.853962 |
| Rgp1 | Q8BHT7 | -0.92 | -2.33 | 0.16 | 0.98 | -1.3 | -3.38 | -2.09 | -0.28 | -1.53 | -1.38 | -1.25 | 0.16 | -1.8 | -1.31 | -0.46 | 0.04 | -1.235 | -0.4725 | -0.355 | 0.179453 | 0.528245 | 0.636377 | 0.999971 | 1 | 0.93301 |
| Rgs10 | Q9CQE5 | 0.3 | 0.92 | 0.44 | 0.72 | 1.2 | 0.93 | 0.64 | 0.55 | 1.08 | 0.56 | 0.4 | 0.77 | 1.58 | 0.65 | 0.94 | 0.91 | 0.235 | 0.1075 | 0.425 | 0.335431 | 0.649137 | 0.134478 | 0.999971 | 1 | 0.784948 |
| Rgs12 | E9Q652 | -0.29 | -1.45 | -1.04 | -0.85 | -1.09 | -1.51 | -0.3 | -1.14 | -1.13 | -1.1 | -0.88 | -1.38 | -1.08 | -2.07 | -1.65 | -1.41 | -0.1025 | -0.215 | -0.645 | 0.767053 | 0.445717 | 0.071017 | 0.999971 | 1 | 0.73635 |
| Rgs14 | P97492 | -0.52 | -0.05 | 0 | -0.27 | -0.16 | -0.6 | 0.01 | -0.48 | -0.03 | -0.87 | -0.1 | 0.17 | 0.35 | -0.6 | -0.2 | -0.56 | -0.0975 | 0.0025 | -0.0425 | 0.666436 | 0.992703 | 0.875004 | 0.999971 | 1 | 0.98442 |
| Rgs17 | G5E8E0 | -1.92 | -1.61 | -1.43 | -1.63 | -1.67 | -1.29 | -1.52 | -1.52 | -1.51 | -2.06 | -1.55 | -1.72 | -0.73 | -1.15 | -1.51 | -1.52 | 0.1475 | -0.0625 | 0.42 | 0.452063 | 0.764916 | 0.112684 | 0.999971 | 1 | 0.78268 |
| Rgs19 | Q9CX84 | -0.67 | -1.21 | 0.06 | 0.01 | -1.08 | -1.24 | -0.39 | -0.55 | -1.41 | -0.87 | -0.33 | -0.42 | -0.84 | -1.11 | 0.24 | 0.21 | -0.3625 | -0.305 | 0.0775 | 0.324552 | 0.429219 | 0.859386 | 0.999971 | 1 | 0.982702 |
| Rgs20 | Q9QZB1 | 0.07 | -0.78 | -1.2 | -1.33 | -0.49 | -1.04 | -0.61 | -1.8 | -0.48 | -0.53 | -1.78 | -1.18 | -0.93 | -1.71 | -1.79 | -1.96 | -0.175 | -0.1825 | -0.7875 | 0.672337 | 0.665063 | 0.06281 | 0.999971 | 1 | 0.725348 |
| Rgs3 | Q9DC04-4 | 0.43 | -0.63 | -0.11 | 0.2 | -0.97 | -1.28 | -0.04 | 0.25 | -0.6 | -0.28 | -0.01 | 0.27 | -1.16 | -0.86 | 0 | 0.35 | -0.4825 | -0.1275 | -0.39 | 0.25887 | 0.674539 | 0.347695 | 0.999971 | 1 | 0.864164 |
| Rgs4 | O08899 | -1.54 | -0.47 | 1.15 | 1.5 | 0.48 | -1.35 | 0.95 | 1.39 | -1.18 | -1.34 | 1.33 | 0.24 | -1.55 | -0.54 | 0.5 | 0 | 0.2075 | -0.3975 | -0.5575 | 0.803992 | 0.645283 | 0.466699 | 0.999971 | 1 | 0.891477 |
| Rgs6 | A0A087WS | -0.31 | -0.51 | -0.7 | -1.19 | -0.69 | -0.55 | -0.98 | -1.29 | -0.83 | -0.86 | -0.78 | -0.92 | -0.54 | -0.76 | -1.7 | -0.99 | -0.2 | -0.17 | -0.32 | 0.465055 | 0.461856 | 0.328604 | 0.999971 | 1 | 0.856472 |
| Rgs6_1 | A0A0J9YTS | -1.42 | 0.89 | -0.72 | 0.07 | -0.23 | 0.52 | -1.23 | -1.36 | -0.36 | -0.26 | -0.09 | -1.29 | 1.59 | 0.74 | -1.97 | -0.34 | -0.28 | -0.205 | 0.3 | 0.647181 | 0.697606 | 0.716343 | 0.999971 | 1 | 0.952576 |
| Rgs7 | A0A0A6YW | -1.08 | -0.67 | -1.07 | -1.1 | -1.01 | -0.69 | -1.3 | -1.16 | -0.73 | -0.83 | -1.3 | -1.11 | -0.65 | -0.82 | -1.47 | -1.25 | -0.06 | -0.0125 | -0.0675 | 0.779908 | 0.952903 | 0.784306 | 0.999971 | 1 | 0.966376 |
| Rgs7_1 | O54829 | -0.23 | -1.29 | -0.82 | -0.36 | -0.86 | -0.27 | -1.44 | -0.08 | -1.81 | -0.59 | -0.65 | -1.32 | 0.37 | -1.33 | -0.9 | -0.3 | 0.0125 | -0.4175 | 0.135 | 0.973565 | 0.274463 | 0.748085 | 0.999971 | 1 | 0.961064 |
| Rgs7bp | Q8BQP9 | -1.11 | -0.69 | -0.73 | -0.39 | -1.04 | -0.89 | -0.21 | -0.29 | -0.76 | -1.22 | 0.21 | -0.21 | -0.38 | -1.33 | -0.51 | 0.36 | 0.1225 | 0.235 | 0.265 | 0.656403 | 0.497875 | 0.4768 | 0.999971 | 1 | 0.894438 |
| Rgs8 | D3Z289 | -1.11 | -0.82 | -1.12 | -0.16 | -0.87 | -1.88 | -1.3 | -0.42 | -1.09 | -1.29 | -1.05 | -0.54 | -0.66 | -0.98 | -0.64 | -1.26 | -0.315 | -0.19 | -0.0825 | 0.407285 | 0.514252 | 0.771757 | 0.999971 | 1 | 0.963498 |
| Rgs9 | O54828 | -0.82 | -1.75 | -0.76 | -0.12 | -1.82 | -3.56 | -0.95 | 0.24 | -1.9 | -1.25 | -0.89 | -0.33 | -1.46 | -1.36 | -0.03 | -0.68 | -0.66 | -0.23 | -0.02 | 0.407263 | 0.607084 | 0.96414 | 0.999971 | 1 | 0.998721 |
| Rhbdd2 | Q8VEK2 | 1.17 | 0.77 | 0.88 | 1.16 | 0.32 | 1.06 | 1.28 | 0.28 | 0.42 | 0.61 | 0.35 | 0.53 | -0.15 | 0.26 | 0.45 | 0.46 | -0.26 | -0.5175 | -0.74 | 0.376785 | 0.019229 | 0.007965 | 0.999971 | 1 | 0.602121 |
| Rhbdf1 | Q6PIX5 | 0.54 | 0.19 | -0.26 | -0.64 | 0.41 | -0.08 | -0.28 | -0.98 | -0.04 | 0.49 | 0.02 | -0.37 | 0.26 | 0.24 | -0.58 | -0.52 | -0.19 | 0.0675 | -0.1075 | 0.614218 | 0.8308 | 0.754437 | 0.999971 | 1 | 0.961116 |
| Rhbdl3 | P58873 | 1.47 | -1.54 | -0.63 | -0.27 | -0.94 | -1.18 | -0.21 | -0.35 | -1.33 | 0.82 | -0.38 | -1.41 | -1.63 | -0.9 | 0.12 | -0.8 | -0.4275 | -0.3325 | -0.56 | 0.490936 | 0.65579 | 0.405439 | 0.999971 | 1 | 0.874565 |
| Rheb | Q921J2 | -0.87 | -1.11 | -2.99 | -3.5 | -0.66 | -1.07 | -2.16 | -3.15 | -0.73 | -1.53 | -2.49 | -2.96 | -0.76 | -1.63 | -3.51 | -3.28 | 0.3575 | 0.19 | -0.1775 | 0.648424 | 0.800407 | 0.832696 | 0.999971 | 1 | 0.979036 |
| Rhoa | Q9QUI0 | -0.5 | -0.08 | 0.47 | -0.6 | -0.16 | -0.52 | 0.84 | 0.29 | -0.17 | -0.74 | 1.23 | 1.48 | 0.72 | -0.22 | 0.59 | -0.49 | 0.29 | 0.6275 | 0.3275 | 0.441496 | 0.268865 | 0.39141 | 0.999971 | 1 | 0.871816 |
| Rhob | P62746 | -2.55 | -4.29 | -0.48 | -0.64 | -3.38 | -4.32 | -0.79 | -0.22 | -4.21 | -5.23 | -0.53 | -0.07 | -4.88 | -3.89 | -0.22 | -0.38 | -0.1875 | -0.52 | -0.3525 | 0.874741 | 0.713267 | 0.791362 | 0.999971 | 1 | 0.968737 |
| Rhobtb2 | Q91V93 | -0.88 | -1.15 | -0.84 | -1.15 | -1.15 | -1.23 | -1.29 | -1.23 | -0.59 | -0.59 | -1.15 | -1.18 | 0.25 | -0.6 | -0.72 | -0.68 | -0.22 | 0.1275 | 0.5675 | 0.229634 | 0.573295 | 0.058475 | 0.999971 | 1 | 0.725348 |
| Rhoc | Q62159 | -2 | -0.95 | 0.19 | 0.1 | -0.21 | -2.07 | 0.16 | 0.25 | -1.2 | -2.21 | -0.15 | 1.24 | -1.23 | -0.61 | 1.18 | 0.67 | 0.1975 | 0.085 | 0.6675 | 0.771049 | 0.916763 | 0.34507 | 0.999971 | 1 | 0.863054 |
| Rhof | Q8BYP3 | -0.4 | -0.77 | -0.36 | -0.51 | -0.72 | -0.55 | -0.51 | 0.16 | -0.46 | -0.01 | 0.22 | -0.44 | -0.15 | -0.23 | -0.22 | 0.22 | 0.105 | 0.3375 | 0.415 | 0.669631 | 0.163147 | 0.063871 | 0.999971 | 1 | 0.725348 |
| Rhog | P84096 | 0.13 | -0.3 | 0.48 | 0.34 | -0.79 | 0.12 | -0.38 | 0.3 | -0.45 | 0.13 | -0.05 | -1.19 | -0.61 | -0.2 | 0.24 | -0.03 | -0.35 | -0.5525 | -0.3125 | 0.27123 | 0.127692 | 0.261509 | 0.999971 | 1 | 0.842262 |
| Rhoq | Q8R527 | 1.13 | -0.56 | -3.55 | -3.39 | -0.01 | 0.25 | -3.21 | -6.2 | 0.23 | 1.41 | -1.49 | -1.49 | 0.34 | 0.1 | -4.42 | -2.65 | -0.7 | 1.2575 | -0.065 | 0.678577 | 0.310957 | 0.963783 | 0.999971 | 1 | 0.998721 |
| Rhot1 | Q8BG51-3 | -1.37 | -0.88 | 0.5 | 0.08 | -0.89 | -0.87 | 0.52 | 0.29 | -0.71 | -1.28 | 0.23 | -0.03 | -0.78 | -0.92 | 0.21 | -0.08 | 0.18 | -0.03 | 0.025 | 0.732415 | 0.953033 | 0.957973 | 0.999971 | 1 | 0.998622 |
| Rhot2 | Q8JZN7 | -1.23 | -0.8 | -0.38 | -0.07 | -0.73 | -1.08 | -0.05 | -0.21 | -0.84 | -0.91 | -0.43 | -1.27 | -0.49 | -0.52 | 0.11 | -0.22 | 0.1025 | -0.2425 | 0.34 | 0.764918 | 0.44257 | 0.27618 | 0.999971 | 1 | 0.843794 |
| Rhou | Q9EQT3 | -1.69 | -0.5 | -0.9 | -1.32 | 0.4 | -1.38 | 0.14 | -0.24 | -0.58 | -1.7 | -0.62 | 0.76 | 1 | -1.03 | -1.06 | -0.12 | 0.8325 | 0.5675 | 0.8 | 0.086821 | 0.296254 | 0.143119 | 0.999971 | 1 | 0.787195 |
| Rhpn2 | Q8BWR8 | 0.84 | 1.06 | -0.81 | -0.27 | 0.47 | 1.1 | -0.43 | -0.19 | 0.77 | 1.19 | -0.04 | 0 | 0.97 | 0.9 | 0.36 | 0.53 | 0.0325 | 0.275 | 0.485 | 0.950174 | 0.586734 | 0.290366 | 0.999971 | 1 | 0.847402 |
| Ric1 | E9QPA1 | 1.5 | 1.51 | 0.05 | 0.29 | 1.54 | 1.09 | 0.32 | 0.54 | 1.32 | 1.77 | 0.58 | 0.38 | 1.77 | 1.63 | 0.72 | 0.57 | 0.035 | 0.175 | 0.335 | 0.937571 | 0.712752 | 0.477217 | 0.999971 | 1 | 0.894438 |
| Ric3 | D3YTZ9 | -0.32 | -0.22 | -0.16 | -1.58 | -0.58 | -0.88 | 0.86 | -0.7 | -0.72 | -0.92 | -0.46 | -0.69 | -0.37 | -1.36 | -1.25 | -1.96 | 0.245 | -0.1275 | -0.665 | 0.617238 | 0.713504 | 0.159869 | 0.999971 | 1 | 0.800134 |
| Ric8a | Q3TIR3 | -0.45 | -0.27 | 0.21 | 0.1 | -0.18 | -0.31 | -0.43 | 0.54 | -0.26 | -0.16 | 0.39 | 0.47 | -0.22 | -0.13 | 0.39 | 0.57 | 0.0075 | 0.2125 | 0.255 | 0.978752 | 0.427877 | 0.355132 | 0.999971 | 1 | 0.868793 |
| Ric8b | Q80XE1-1 | -0.44 | 1.13 | -0.28 | 0.52 | 1.1 | 0.6 | 0.02 | 0.56 | 1.61 | 0.32 | 0.16 | 0.37 | 1.65 | 0.62 | 0.68 | 0.54 | 0.3375 | 0.3825 | 0.64 | 0.416234 | 0.420525 | 0.15855 | 0.999971 | 1 | 0.798217 |
| Rictor | Q6QI06 | -0.59 | -1.08 | -0.02 | -0.17 | -0.8 | -0.95 | -0.23 | 0.03 | -0.78 | -0.68 | 0.09 | 0.23 | -0.55 | -0.54 | 0.14 | 0.29 | -0.0225 | 0.18 | 0.3 | 0.945782 | 0.606375 | 0.369744 | 0.999971 | 1 | 0.869526 |
| Rida | P52760 | -1.27 | -0.06 | 0.32 | -0.13 | 0.06 | 0.22 | -0.42 | 1.47 | -0.02 | -0.8 | -0.6 | -0.06 | -0.4 | 0.18 | -0.33 | -0.19 | 0.6175 | -0.085 | 0.1 | 0.229816 | 0.823741 | 0.780853 | 0.999971 | 1 | 0.965859 |
| Rif1 | Q6PR54-2 | 0.72 | 0.12 | 0 | 0.89 | -0.53 | -0.2 | 0.58 | 0.54 | -0.37 | -0.11 | -0.02 | -0.43 | 0.32 | 0.07 | 0.46 | -0.31 | -0.335 | -0.665 | -0.2975 | 0.347111 | 0.029894 | 0.319154 | 0.999971 | 1 | 0.853893 |
| Rilpl1 | Q9JJC6 | -0.22 | -1.18 | -0.8 | 0.1 | 1.31 | 1.55 | 1.39 | 1.52 | 1.19 | 0.87 | 1.51 | 1.29 | 1.39 | 1.23 | 1.12 | 1.35 | 1.9675 | 1.74 | 1.7975 | 0.000124 | 0.000457 | 0.000244 | 0.226166 | 0.43267 | 0.247672 |
| Rimbp2 | A0A0G2JFE | -1.75 | -1 | -1.45 | -2.13 | -1.16 | -1.31 | -1.6 | -1.66 | -1.27 | -1.25 | -2.3 | -2.93 | -0.78 | -1.25 | -2.61 | -1.55 | 0.15 | -0.355 | 0.035 | 0.597132 | 0.437687 | 0.935261 | 0.999971 | 1 | 0.995618 |
| Rimbp2_1 | A0A571BD | -0.72 | -0.96 | -0.99 | -1.07 | -1.21 | -0.79 | -1.06 | -0.79 | -1.23 | -0.75 | -0.89 | -1.14 | -0.97 | -0.92 | -0.97 | -0.83 | -0.0275 | -0.0675 | 0.0125 | 0.886689 | 0.727339 | 0.942433 | 0.999971 | 1 | 0.996905 |
| Rimoc1 | Q8BR90 | -0.14 | 1.07 | 1.27 | 0.7 | 1.1 | 0.63 | 1.27 | 1.03 | 1.17 | 0.47 | 1.64 | 0.5 | 1.29 | 1.31 | 1.07 | 1.01 | 0.2825 | 0.22 | 0.445 | 0.411581 | 0.588133 | 0.191799 | 0.999971 | 1 | 0.810547 |
| Rims1 | F6TZK4 | 3.63 | 4.5 | 2.98 | 2.31 | 5.66 | 2.79 | 5.91 | 1.24 | 8.13 | 2.26 | 4.06 | 5.21 | 0.36 | 1.91 | 2.15 | 0.6 | 0.545 | 1.56 | -2.1 | 0.620211 | 0.209199 | 0.006304 | 0.999971 | 1 | 0.582714 |
| Rims1_1 | E9PWP6 | -1.22 | -1.56 | -1.18 | -1.08 | -1.73 | -1.37 | -1.32 | -1.05 | -1.67 | -1.24 | -1.25 | -1.4 | -1.52 | -1.42 | -1.11 | -0.99 | -0.1075 | -0.13 | 2.65E-16 | 0.625738 | 0.518738 | 1 | 0.999971 | 1 | 1 |
| Rims2 | Q0VF51 | -0.17 | -0.65 | -0.75 | -0.55 | -0.87 | -0.53 | -0.42 | -0.6 | -0.79 | -0.36 | -0.88 | -1.04 | -0.56 | -0.51 | -0.67 | -0.79 | -0.075 | -0.2375 | -0.1025 | 0.72141 | 0.314343 | 0.611651 | 0.999971 | 1 | 0.927066 |
| Rims3 | Q80U57 | -0.43 | 2.55 | 0.34 | 0.18 | 0.54 | 0.66 | -0.33 | 0.4 | -0.07 | 0.01 | -0.01 | -0.18 | 1.06 | 0.38 | -0.15 | 0.45 | -0.3425 | -0.7225 | -0.225 | 0.58749 | 0.248188 | 0.723738 | 0.999971 | 1 | 0.955292 |
| Rims4 | P60191 | -0.12 | 0.54 | 0.07 | -0.07 | 0.2 | 0.68 | -0.39 | 0.35 | 0.27 | 0.29 | 0.34 | 0.26 | 0.73 | 0.8 | -0.11 | 0.6 | 0.105 | 0.185 | 0.4 | 0.712201 | 0.373748 | 0.171527 | 0.999971 | 1 | 0.804075 |
| Rin1 | A0A2C9F2 | -0.61 | -0.92 | 0.52 | 0.59 | -0.84 | -0.65 | 0.32 | 0.33 | -0.75 | -0.55 | 0.48 | 0.28 | -1.5 | -0.37 | 0.62 | 0.53 | -0.105 | -0.03 | -0.075 | 0.821083 | 0.948131 | 0.896146 | 0.999971 | 1 | 0.988571 |
| Ring1 | O35730 | 3.53 | 0.8 | -0.1 | 0.29 | 1.61 | 1.97 | -0.43 | 0.33 | -0.4 | 2.02 | -0.79 | -0.19 | 1.32 | 0.51 | -0.53 | -0.29 | -0.26 | -0.97 | -0.8775 | 0.770181 | 0.314198 | 0.305569 | 0.999971 | 1 | 0.847402 |
| Rint1 | Q8BZ36-2 | -0.79 | 0.4 | -1.02 | -0.55 | 0.17 | 0.33 | -1.14 | -1.22 | 0.36 | -0.21 | -0.89 | -0.52 | 0.64 | 0.28 | -0.91 | -0.81 | 0.025 | 0.175 | 0.29 | 0.958645 | 0.657971 | 0.538729 | 0.999971 | 1 | 0.910077 |
| Riok3 | A0A3Q4EG | -0.21 | 0.1 | 0.73 | 0.86 | 1.07 | 0.06 | 2.18 | 0.73 | 0.3 | 0.36 | 1.14 | 0.48 | 1.75 | 0.94 | 0.72 | 0.8 | 0.64 | 0.2 | 0.6825 | 0.201898 | 0.538372 | 0.074034 | 0.999971 | 1 | 0.746069 |
| Riox1 | Q9JJF3 | -0.82 | -2.05 | -0.84 | -0.95 | -1.97 | -1.04 | 1.64 | -1.48 | -2.02 | -1.3 | -0.86 | -1.32 | -1.73 | -1.24 | -0.8 | -0.99 | 0.4525 | -0.21 | -0.025 | 0.562196 | 0.574503 | 0.943392 | 0.999971 | 1 | 0.99692 |
| Ripk1 | F7D1J2 | -0.12 | 0.55 | 0.39 | 0.16 | -0.06 | -0.98 | 0.2 | 0.51 | -0.09 | 0.1 | 0.46 | 0.92 | -0.9 | 0.15 | 0.59 | 0.55 | -0.3275 | 0.1025 | -0.1475 | 0.357905 | 0.715145 | 0.688819 | 0.999971 | 1 | 0.945805 |
| Ripor1 | Q68FE6 | 0.07 | 0.02 | 0.3 | 0.35 | -0.01 | -0.14 | -0.04 | 0.31 | 0 | 0.25 | 0.48 | 0.56 | -0.28 | -0.17 | 0.43 | 0.8 | -0.155 | 0.1375 | 0.01 | 0.42933 | 0.502438 | 0.97179 | 0.999971 | 1 | 0.99925 |
| Ripor2 | A6PW28 | 0.23 | 0.07 | -0.32 | -0.41 | 0.27 | 0.18 | -0.43 | 0.28 | 0.17 | 0.14 | 0.07 | 0.1 | -0.37 | 0.64 | 0.05 | 0.11 | 0.1825 | 0.2275 | 0.215 | 0.480072 | 0.284277 | 0.443595 | 0.999971 | 1 | 0.883386 |
| Ripor2_1 | A0A5F8MP | 0.25 | 0.38 | 0.25 | -1.89 | -0.4 | -0.4 | 0.15 | -2.04 | 0.1 | 0.63 | 0.32 | -1.35 | -0.07 | 0.38 | -1.43 | -0.14 | -0.42 | 0.1775 | -0.0625 | 0.527004 | 0.782109 | 0.918666 | 0.999971 | 1 | 0.992633 |
| Rit1 | P70426 | -1.75 | -0.49 | -0.92 | -0.81 | -0.97 | -0.81 | -1.34 | -0.42 | -0.95 | -1.45 | -1.49 | -1.15 | -1.54 | -0.47 | -0.81 | -1.01 | 0.1075 | -0.2675 | 0.035 | 0.744327 | 0.389721 | 0.919246 | 0.999971 | 1 | 0.992633 |
| Rit2 | P70425 | 0.04 | 0.95 | 0.47 | 0.12 | 0.91 | 0.73 | 0.81 | 0.02 | 0.99 | 0.47 | 1.43 | 1.96 | 0.43 | 0.56 | 0.12 | 0.08 | 0.2225 | 0.8175 | -0.0975 | 0.463471 | 0.050959 | 0.710503 | 0.999971 | 1 | 0.950695 |
| Rlbp1 | Q9Z275 | 2.17 | 0.93 | -1.84 | -1.22 | 0.81 | 0.1 | -1.39 | -1.91 | 0.68 | 1.63 | -1.22 | -1.95 | 0.25 | 0.59 | -1.82 | -1.45 | -0.6075 | -0.225 | -0.6175 | 0.550081 | 0.84067 | 0.538425 | 0.999971 | 1 | 0.910077 |
| Rlf | E9Q532 | 1.04 | 0.33 | -0.54 | 0.71 | 0.33 | 1.09 | -0.81 | -0.68 | 0.47 | 1.12 | 0.11 | -1.05 | 0.37 | 1.06 | 0.4 | 0.7 | -0.4025 | -0.2225 | 0.2475 | 0.446934 | 0.674377 | 0.505907 | 0.999971 | 1 | 0.905277 |
| Rlim | Q9WTV7 | -0.63 | 1.87 | 3.04 | 3.44 | 2.18 | 1.36 | 3.78 | 1.53 | 3.9 | -0.06 | 1.89 | 3.37 | -1.82 | -1.35 | 2.66 | -1.47 | 0.2825 | 0.345 | -2.425 | 0.76776 | 0.763265 | 0.078262 | 0.999971 | 1 | 0.754157 |
| Rln3 | Q8CHK2 | -0.96 | 0.18 | 0.83 | -0.81 | -0.6 | -0.78 | 0.07 | 0.32 | 0.22 | -2.5 | 0 | 0.6 | -1.41 | 1.21 | -0.36 | 0.02 | -0.0575 | -0.23 | 0.055 | 0.901756 | 0.758281 | 0.92995 | 0.999971 | 1 | 0.994744 |
| Rmc1 | Q8VC42 | -0.8 | -0.23 | 0.2 | -0.13 | 0.15 | -0.43 | 0.25 | 0.1 | 0.08 | -0.72 | 0.53 | 0.73 | 0.35 | -0.35 | 0.17 | -0.03 | 0.2575 | 0.395 | 0.275 | 0.361682 | 0.304769 | 0.331119 | 0.999971 | 1 | 0.857107 |
| Rmdn1 | Q9DCV4 | -0.87 | -0.55 | -0.82 | -0.93 | -0.87 | -0.34 | -0.77 | -0.75 | -0.78 | -0.89 | -0.92 | -0.71 | -1.06 | -0.31 | -0.86 | -0.74 | 0.11 | -0.0325 | 0.05 | 0.587899 | 0.852669 | 0.822491 | 0.999971 | 1 | 0.975901 |
| Rmdn2 | Q8BSE0 | -1.25 | -0.3 | 0.03 | -1.56 | -0.61 | -0.37 | 0.17 | -0.59 | -1.02 | -0.97 | -0.09 | 0.11 | -0.35 | -0.16 | -0.73 | -1.28 | 0.42 | 0.2775 | 0.14 | 0.310144 | 0.54279 | 0.744134 | 0.999971 | 1 | 0.961064 |
| Rmdn3 | Q3UJU9 | -0.88 | -0.3 | -0.65 | -0.58 | -0.44 | -0.27 | -0.3 | -0.63 | -0.31 | -0.66 | -0.45 | -0.21 | -0.2 | -0.03 | -0.47 | -0.41 | 0.1925 | 0.195 | 0.325 | 0.352955 | 0.353296 | 0.145885 | 0.999971 | 1 | 0.791767 |
| Rmnd1 | Q8CI78 | -0.57 | -0.24 | -0.83 | -0.4 | -0.26 | -0.28 | -1.16 | -0.38 | -0.5 | -0.49 | -0.59 | -0.38 | -0.19 | -0.03 | -0.44 | -0.09 | -0.01 | 0.02 | 0.3225 | 0.970312 | 0.917283 | 0.147556 | 0.999971 | 1 | 0.793248 |
| Rmnd5a | Q80YQ8 | 0.3 | 0.41 | 0.56 | 0.59 | 0.45 | 0.44 | 1.46 | 0.88 | 0.3 | 0.31 | 0.75 | 0.57 | 0.4 | 0.45 | 0.79 | 0.77 | 0.3425 | 0.0175 | 0.1375 | 0.224776 | 0.926553 | 0.478446 | 0.999971 | 1 | 0.894438 |
| Rnase4 | Q8C7E4 | 1.66 | 1.38 | 1.4 | 2.02 | 2.26 | 2.18 | 3.17 | 3.24 | 2.15 | 2.07 | 2.96 | 1.26 | 1.27 | 2.53 | 2.38 | 3.47 | 1.0975 | 0.495 | 0.7975 | 0.00765 | 0.202052 | 0.102056 | 0.999971 | 1 | 0.78268 |
| Rnaseh2a | Q05C51 | 1.66 | 3.07 | 0.59 | -0.87 | 2.77 | 1.88 | 0.61 | -1.05 | 2.57 | 2.5 | 0.25 | 0.82 | 1.56 | 2.38 | -0.61 | -0.66 | -0.06 | 0.4225 | -0.445 | 0.954283 | 0.647782 | 0.662393 | 0.999971 | 1 | 0.936148 |
| Rnaseh2b | Q80ZV0 | 3.07 | 1.83 | 1.67 | 1.89 | 1.99 | 2.21 | 2.84 | 1.73 | 2.05 | 2.41 | 1.63 | 1.54 | 0.66 | 1.73 | 1.22 | 1.28 | 0.0775 | -0.2075 | -0.8925 | 0.840355 | 0.577431 | 0.03978 | 0.999971 | 1 | 0.700838 |
| Rnasek | H3BLR9 | 2.67 | 3.2 | 3.16 | 1.66 | 2.36 | 4.42 | 2.31 | 1.81 | 3.43 | 5.21 | 2.99 | 1.52 | 3.98 | 3.03 | 2.62 | 2.68 | 0.0525 | 0.615 | 0.405 | 0.932396 | 0.428452 | 0.378054 | 0.999971 | 1 | 0.871417 |
| Rnasel | S4R1J5 | -0.36 | 0.02 | 0.3 | 0.86 | -0.06 | 0.65 | 0.39 | 1.18 | -0.83 | 0.13 | 0.8 | -0.25 | 0.74 | 0.48 | -0.09 | 0.84 | 0.335 | -0.2425 | 0.2875 | 0.360077 | 0.556934 | 0.395232 | 0.999971 | 1 | 0.871816 |

| | | | | | | | | | | | | | | | | | | | | | | | | | | |
|---|---|---|---|---|---|---|---|---|---|---|---|---|---|---|---|---|---|---|---|---|---|---|---|---|---|---|
| Rnaset2b | A0A338P7E | -3.4 | -3.35 | -3.92 | -4.59 | -3.33 | -3.5 | -3.09 | -3.84 | -2.82 | -3.69 | -3.23 | -3.93 | -4.52 | -3.21 | -3.53 | -4.41 | 0.375 | 0.3975 | -0.1025 | 0.271856 | 0.298383 | 0.804257 | 0.999971 | 1 | 0.971467 |
| Rnd2 | Q9QYM5 | -1.46 | -1.38 | -1.42 | -1.59 | -1.33 | -1.46 | -1.64 | -1.43 | -0.96 | -1.22 | -1.1 | -0.86 | -0.5 | -0.67 | -1.25 | -0.99 | -0.0025 | 0.4275 | 0.61 | 0.988347 | 0.033132 | 0.019929 | 0.999971 | 1 | 0.635388 |
| Rnd3 | E9Q8D7 | -2.45 | -0.91 | -2.42 | -1.84 | -1.29 | -1.6 | -1.24 | -1.39 | -2.29 | -2.17 | -2.36 | -1.61 | -1.61 | -0.28 | -0.47 | -1.69 | 0.525 | -0.2025 | 0.8925 | 0.16975 | 0.601425 | 0.091478 | 0.999971 | 1 | 0.773779 |
| Rnf10 | F7AN16 | -0.65 | 1.31 | 1.41 | 1.32 | 2.17 | 1.05 | 3.23 | 0.68 | 3.27 | -1.87 | 0.58 | 0.77 | 0.73 | -3.88 | -0.03 | -0.38 | 0.935 | -0.16 | -1.7375 | 0.198811 | 0.878207 | 0.115092 | 0.999971 | 1 | 0.78268 |
| Rnf11 | Q9QYK7 | -0.43 | -1 | -1.21 | -0.63 | -1.41 | -1.05 | -1.43 | -1.33 | -1.2 | -0.44 | -0.71 | -0.93 | -1.46 | -1 | -0.74 | -1.04 | -0.4875 | -0.0025 | -0.2425 | 0.061901 | 0.992334 | 0.357298 | 0.999971 | 1 | 0.868793 |
| Rnf112 | Q96DY5-2 | 0.24 | 1.24 | -0.57 | -0.59 | 1.53 | 0.75 | -0.55 | -0.46 | 1.99 | 0.36 | -0.44 | -0.46 | 3.77 | 0.45 | -0.59 | -0.25 | 0.2375 | 0.2825 | 0.765 | 0.695052 | 0.668693 | 0.440226 | 0.999971 | 1 | 0.883105 |
| Rnf113a2 | Q14B01 | 2.15 | 2.58 | 2.54 | 2.68 | 2.44 | 2.39 | 1.93 | 3.06 | 2.53 | 2.6 | 3.48 | 2.98 | 2.27 | 3.01 | 2.92 | 3.27 | -0.0325 | 0.41 | 0.38 | 0.906333 | 0.149728 | 0.175622 | 0.999971 | 1 | 0.808532 |
| Rnf114 | Q9ET26 | 1.38 | 0.75 | 0.97 | -0.06 | 0.89 | 1.39 | 0.18 | 1.45 | 0.71 | 1.59 | 0.2 | -1.39 | -0.18 | 1.46 | 1.15 | 0.42 | 0.2175 | -0.4825 | -0.0475 | 0.592302 | 0.455661 | 0.915712 | 0.999971 | 1 | 0.992633 |
| Rnf115 | Q9D0C1 | 2.09 | 0.29 | -1.74 | -1.34 | -0.05 | 1.74 | -0.8 | 1.14 | 0.29 | 2.1 | -1.38 | -1.35 | 0.72 | 0.6 | -1.69 | -1.99 | 0.6825 | 0.09 | -0.415 | 0.471623 | 0.933479 | 0.68361 | 0.999971 | 1 | 0.943718 |
| Rnf121 | Q8R1Z9 | 1.56 | 2.2 | -0.47 | 0.39 | 1.38 | 1.95 | 0.12 | 0.59 | 1.54 | 2.02 | 1.75 | 1.3 | 0.75 | 2.13 | 1.54 | 1.21 | 0.09 | 0.7325 | 0.4875 | 0.890597 | 0.218684 | 0.429572 | 0.999971 | 1 | 0.881563 |
| Rnf123 | Q5XPI3 | -0.18 | -0.16 | -0.04 | 0.16 | -0.06 | -0.15 | 0.05 | 0.07 | -0.22 | 0.03 | -0.09 | -0.12 | -0.22 | -0.2 | -0.13 | 0.14 | 0.0325 | -0.045 | -0.0475 | 0.854159 | 0.795555 | 0.799278 | 0.999971 | 1 | 0.970507 |
| Rnf126 | Q91YL2 | 0.07 | 0.11 | 0.83 | -0.47 | -0.1 | -0.12 | 0.78 | -0.15 | -0.22 | -0.11 | 0.94 | -0.78 | -0.58 | -0.36 | -0.25 | -0.94 | -0.0325 | -0.1775 | -0.6675 | 0.924843 | 0.677486 | 0.057649 | 0.999971 | 1 | 0.725348 |
| Rnf13 | Q8C4F9 | -0.1 | 1.86 | 0.96 | 0.47 | 1.78 | 1.81 | 0.61 | 1.32 | 1.67 | 0.22 | 1.17 | 0.22 | 2.86 | 2.04 | 0.56 | 0.48 | 0.5825 | 0.0225 | 0.6875 | 0.233779 | 0.964886 | 0.30609 | 0.999971 | 1 | 0.847402 |
| Rnf130 | Q5SVR5 | -0.73 | 0.54 | 0.86 | -0.11 | 0.51 | 0.08 | -0.67 | 1.15 | -0.4 | 0.14 | -0.04 | 0.49 | 0.98 | 0.71 | 0.95 | 0.86 | 0.1275 | -0.0925 | 0.735 | 0.792479 | 0.81032 | 0.062786 | 0.999971 | 1 | 0.725348 |
| Rnf14 | Q9JI90 | 0.4 | 0.93 | 1.59 | 1.43 | 0.67 | 1.26 | 2 | 1.56 | 0.78 | 0.86 | 1.72 | 0.81 | 0.17 | 1.23 | 1.68 | 1.25 | 0.285 | -0.045 | -0.005 | 0.454814 | 0.896584 | 0.990048 | 0.999971 | 1 | 0.99925 |
| Rnf141 | H3BJB4 | 0.11 | 0.32 | 0.12 | -0.5 | 0.41 | -0.04 | 0.71 | -0.51 | 1.24 | -0.15 | 1.06 | 0.88 | 2.39 | -0.19 | 0.45 | -0.25 | 0.13 | 0.745 | 0.5875 | 0.68752 | 0.059906 | 0.331989 | 0.999971 | 1 | 0.857152 |
| Rnf146 | Q9CZW6 | -0.21 | 1.54 | 0.85 | 0.35 | 2.5 | 1.94 | 2.63 | 0.71 | 2.91 | 0.93 | 2.4 | 2.43 | 2.58 | 0.71 | 1.44 | 0.77 | 1.3125 | 1.535 | 0.7425 | 0.033055 | 0.016576 | 0.184122 | 0.999971 | 1 | 0.809025 |
| Rnf149 | Q3U2C5 | 0.9 | -0.24 | 0.85 | 0.16 | -0.43 | -0.58 | -0.11 | -0.28 | -0.54 | -0.05 | 0.22 | 0.47 | -0.58 | -0.06 | 0.19 | -0.11 | -0.7675 | -0.3925 | -0.5575 | 0.029769 | 0.274914 | 0.111056 | 0.999971 | 1 | 0.78268 |
| Rnf150 | Q5DTZ6 | 0.37 | -0.26 | -2.45 | -1.63 | 0.16 | -0.18 | -1.33 | -1.65 | 1.17 | 0.24 | -1.38 | -2.2 | 2.32 | -0.1 | -1.9 | -1.38 | 0.2425 | 0.45 | 0.7275 | 0.730129 | 0.618923 | 0.480971 | 0.999971 | 1 | 0.89485 |
| Rnf157 | E2AAP0 | 0.43 | -0.06 | 1.58 | -0.12 | -0.5 | 0.15 | 2.37 | 0.39 | -0.24 | -0.36 | 1 | 1.42 | -0.43 | -0.83 | 0.7 | -0.54 | 0.145 | -0.0025 | -0.7325 | 0.827344 | 0.996364 | 0.155066 | 0.999971 | 1 | 0.795578 |
| Rnf168 | E9PYW4 | 3.88 | 2.26 | 3.07 | 2.17 | 1.35 | 3.61 | 3.26 | 1.95 | 1.08 | 2.09 | 2.74 | 1.83 | -0.21 | 1.18 | 1.25 | 1.86 | -0.3025 | -0.91 | -1.825 | 0.621373 | 0.09216 | 0.008381 | 0.999971 | 1 | 0.602121 |
| Rnf170 | A0A180GR | 2.23 | 0.14 | 0.64 | 1.44 | 0.26 | 0.33 | 0.49 | 1.06 | 0.59 | 1.59 | 0.07 | -0.23 | -0.52 | -0.77 | 0.01 | -0.38 | -0.5775 | -0.6075 | -1.5275 | 0.231756 | 0.295721 | 0.00839 | 0.999971 | 1 | 0.602121 |
| Rnf181 | Q9CY62 | 0.86 | 0.81 | -11.31 | -12.03 | 1.14 | 1.48 | -6.85 | -9.95 | 0.68 | 0.47 | -13.01 | -14.47 | 1.11 | 1.28 | -11.65 | -13.27 | 1.8725 | -1.165 | -0.215 | 0.647456 | 0.811791 | 0.963729 | 0.999971 | 1 | 0.998721 |
| Rnf185 | Q91YT2 | 2.02 | -0.03 | 1.4 | 1.34 | 0.01 | 0.79 | 2.66 | 0.91 | 0.37 | 1.85 | 0.3 | -0.47 | -0.62 | 0.23 | 0.35 | -0.51 | -0.09 | -0.67 | -1.32 | 0.888464 | 0.279336 | 0.018914 | 0.999971 | 1 | 0.635388 |
| Rnf2 | Q9CQJ4 | 1.57 | 0.68 | 0.95 | 1.2 | 0 | 0.39 | 0.92 | 0.67 | 0.84 | 0.64 | 0.54 | 0.73 | 1.06 | 0.31 | 0.84 | 0.68 | -0.605 | -0.4125 | -0.3775 | 0.059852 | 0.104565 | 0.182613 | 0.999971 | 1 | 0.808532 |
| Rnf20 | Q5DTM8 | 1.18 | 1.96 | 2.36 | 2.21 | 1.99 | 1.44 | 1.87 | 2.36 | 1.59 | 1.33 | 2.26 | 2.47 | 1.43 | 1.99 | 2.64 | 2.28 | -0.0125 | -0.015 | 0.1575 | 0.96925 | 0.967349 | 0.662363 | 0.999971 | 1 | 0.936148 |
| Rnf213 | A0A8I4P9C | 1.63 | 0.18 | 0.41 | -0.27 | -0.22 | 0.11 | -0.17 | -0.54 | 0.39 | 0.93 | 0.02 | 0.52 | -2.22 | 0.6 | -0.09 | -0.14 | -0.6925 | -0.0225 | -0.95 | 0.116355 | 0.957748 | 0.17787 | 0.999971 | 1 | 0.808532 |
| Rnf214 | Q8BFU3 | -0.17 | 0.78 | 0.84 | 0.43 | 0.76 | 0.66 | 0.86 | 0.76 | 0.91 | -0.11 | 0.74 | 0.54 | 0.77 | 0.5 | 0.72 | 0.83 | 0.29 | 0.05 | 0.235 | 0.280456 | 0.876854 | 0.386028 | 0.999971 | 1 | 0.871816 |
| Rnf214_1 | Q8BFU3-2 | -2.39 | 1.11 | -0.03 | -0.17 | 0.72 | 0.62 | 0.83 | 1.17 | 0.73 | -1.66 | 0.88 | 0.96 | 1.23 | 0.51 | 0.59 | -0.96 | 1.205 | 0.5975 | 0.7125 | 0.100237 | 0.499132 | 0.372938 | 0.999971 | 1 | 0.869526 |
| Rnf216 | P58283-2 | 2.41 | 0.84 | 1.42 | 1.59 | 1.15 | 1.27 | 1.34 | 1.55 | 1.11 | 2.23 | 1.34 | 1.28 | 0.02 | 0.73 | 0.84 | 1.13 | -0.2375 | -0.075 | -0.885 | 0.482488 | 0.849184 | 0.045003 | 0.999971 | 1 | 0.710102 |
| Rnf220 | Q9D4L6 | 1.87 | 2.07 | 2.32 | 2.85 | 1.72 | 2.04 | 2.01 | 2.56 | 2.11 | 2.32 | 2.44 | 2.64 | 2.15 | 2.34 | 2.23 | 3.03 | -0.195 | 0.1 | 0.16 | 0.503117 | 0.702179 | 0.598093 | 0.999971 | 1 | 0.92377 |
| Rnf25 | E9Q446 | 0.47 | 0.04 | 0.59 | 0.99 | 0.11 | 0.34 | 0.72 | 0.51 | 0.09 | 0.53 | 0.34 | 0.33 | -0.41 | 0.09 | 0.25 | 0.92 | -0.1025 | -0.2 | -0.31 | 0.692878 | 0.421238 | 0.367727 | 0.999971 | 1 | 0.869526 |
| Rnf26 | A0A1L1SQ0 | 0.14 | -0.55 | -0.62 | -0.62 | -0.45 | -0.46 | -0.06 | -0.73 | -0.45 | 0.2 | -0.08 | -0.24 | 0.75 | -0.97 | -0.78 | -0.85 | -0.0125 | 0.27 | -0.05 | 0.961005 | 0.304108 | 0.905957 | 0.999971 | 1 | 0.991343 |
| Rnf31 | Q924T7-2 | 0.31 | -0.38 | -0.64 | -0.23 | -0.12 | -0.33 | -0.57 | -0.05 | -0.29 | -0.02 | -0.12 | -0.47 | -0.1 | 0.08 | -0.18 | -0.14 | -0.0325 | 0.01 | 0.15 | 0.899289 | 0.967995 | 0.538994 | 0.999971 | 1 | 0.910077 |
| Rnf34 | Q99KR6 | 0.42 | 0 | 0.52 | 0.39 | 0.34 | 0.14 | 0.12 | 0.88 | 0.58 | 0.74 | 0.05 | 0.39 | 1.15 | 0.55 | 0.85 | 0.63 | 0.0375 | 0.1075 | 0.4625 | 0.876999 | 0.634953 | 0.061599 | 0.999971 | 1 | 0.725348 |
| Rnf40 | Q3U319 | 1.49 | 1.02 | 2.03 | 2.75 | 1.46 | 0.97 | 1.71 | 1.68 | 1.4 | 0.89 | 2.3 | 2.13 | 0.21 | 0.37 | 2.19 | 2.14 | -0.3675 | -0.1425 | -0.595 | 0.361295 | 0.759912 | 0.336094 | 0.999971 | 1 | 0.858827 |
| Rnf41 | Q8BH75 | 3.08 | -1.19 | -0.43 | 0.11 | -0.07 | 0.19 | 0.39 | -1.25 | 0.24 | 1.61 | -1.52 | -2.76 | -1.98 | -1.26 | -1.67 | -3.38 | -0.5775 | -1 | -2.465 | 0.525323 | 0.414944 | 0.025388 | 0.999971 | 1 | 0.678474 |
| Rnf5 | O35445 | 4.98 | 1.8 | -4.34 | -3.48 | 2.58 | 3.86 | -2.77 | -3.88 | 2.11 | 4.88 | -3.11 | -3.78 | 1.29 | 2.16 | -3.08 | -5.16 | 0.2075 | 0.285 | -0.9375 | 0.936002 | 0.916089 | 0.70801 | 0.999971 | 1 | 0.950182 |
| Rnf7 | Q9WTZ1 | 1.82 | 2.36 | -1.9 | -1.41 | 1.51 | 2.13 | -1.18 | -1.88 | 1.49 | 2.15 | -1.5 | -0.58 | 1.18 | 2.55 | -1.56 | -2.16 | -0.0725 | 0.1725 | -0.215 | 0.95566 | 0.88955 | 0.876864 | 0.999971 | 1 | 0.985139 |
| Rnft2 | Q3UF64 | 1.65 | 1.32 | 2.28 | 1.61 | 1.11 | 1.1 | 1.25 | 2.4 | 1.21 | 1.58 | 2.58 | 1.37 | 1.24 | 1.55 | 1.61 | 2.68 | -0.25 | -0.03 | 0.055 | 0.497365 | 0.933537 | 0.880025 | 0.999971 | 1 | 0.985441 |
| Rngtt | Q9DCC1 | 2.12 | 2.15 | 2.66 | 3.01 | 2.01 | 1.86 | 2.68 | 2.34 | 2.07 | 2.13 | 1.85 | 1.27 | 0.99 | 2.2 | 2.09 | 0.95 | -0.2625 | -0.655 | -0.9275 | 0.380737 | 0.052773 | 0.038003 | 0.999971 | 1 | 0.700838 |
| Rnh1 | Q91VI7 | 0.23 | 0.54 | 1.08 | 0.72 | 0.34 | 0.48 | 0.53 | 0.88 | 0.32 | 0.34 | 0.66 | -0.19 | 0.17 | 0.56 | 1 | 0.66 | -0.085 | -0.36 | -0.045 | 0.726978 | 0.202557 | 0.866425 | 0.999971 | 1 | 0.983462 |
| Rnmt | Q9D0L8 | 1.03 | 2.91 | 2.85 | 2.47 | 2.8 | 2.15 | 2.44 | 2.7 | 2.62 | 1.34 | 2.83 | 2.92 | 2.26 | 2.22 | 2.54 | 2.6 | 0.2075 | 0.1125 | 0.085 | 0.636387 | 0.832269 | 0.842239 | 0.999971 | 1 | 0.979814 |
| Rnpep | Q8VCT3 | 0.44 | 0.3 | 0.18 | 0.21 | 0.28 | 0.25 | 0.06 | -0.11 | 0.44 | 0.21 | 0.13 | 0.38 | 0.13 | 0.59 | 0.36 | 0.32 | -0.1625 | 0.0075 | 0.0675 | 0.385732 | 0.965429 | 0.716381 | 0.999971 | 1 | 0.952576 |
| Rnps1 | Q99M28 | 0.47 | 0.62 | 0.38 | -0.09 | 0.75 | 0.6 | 0.73 | -0.03 | 0.86 | -0.17 | 0.41 | 0.37 | 0.75 | 0.11 | -0.54 | -0.53 | 0.1675 | 0.0225 | -0.3975 | 0.527409 | 0.935006 | 0.262403 | 0.999971 | 1 | 0.842262 |
| RO60 | O08848 | 0.56 | 0.44 | 0.57 | 0 | 0.67 | 0.47 | 0.72 | 0.31 | 0.39 | -0.03 | 0.44 | 0.42 | 0.76 | 0.41 | 0.51 | -0.29 | 0.15 | -0.0875 | -0.045 | 0.48704 | 0.688103 | 0.871695 | 0.999971 | 1 | 0.984109 |
| Robo1 | G5E843 | 0.11 | -1.53 | -1.7 | -1.24 | -0.65 | -1.32 | -1.04 | -1.12 | -0.78 | -0.78 | -1.4 | -1.15 | -0.61 | -1.27 | -1.03 | -1.41 | 0.0575 | 0.0625 | 0.01 | 0.889212 | 0.880945 | 0.981167 | 0.999971 | 1 | 0.99925 |
| Robo2 | A0A2I3BR7 | -0.9 | -1.21 | -1.25 | -0.91 | -0.94 | -0.96 | -1.21 | -0.71 | -0.89 | -0.88 | -1.15 | -0.89 | -0.53 | -0.91 | -0.72 | -0.72 | 0.1125 | 0.115 | 0.3475 | 0.574397 | 0.534967 | 0.09535 | 0.999971 | 1 | 0.78268 |
| Rock1 | P70335-2 | 0.58 | -0.12 | 0.1 | 0.03 | 0.32 | 0.03 | -0.02 | -0.14 | -0.02 | 0.83 | -0.07 | 0.5 | -0.31 | -0.07 | 0.35 | -0.01 | -0.1 | 0.1625 | -0.1575 | 0.654793 | 0.563268 | 0.513989 | 0.999971 | 1 | 0.905815 |
| Rock2 | F8VPK5 | -0.32 | -0.38 | -0.06 | -0.39 | -0.6 | -0.42 | 0.3 | -0.68 | -0.48 | -0.66 | 0.03 | 0.07 | -0.99 | -0.68 | -0.5 | -0.66 | -0.0625 | 0.0275 | -0.42 | 0.810124 | 0.905779 | 0.054568 | 0.999971 | 1 | 0.725348 |
| Rogdi | Q3TDK6 | -0.5 | -0.62 | -1.34 | -1.43 | -0.55 | -0.63 | -1.2 | -1.32 | -0.69 | -0.46 | -1.36 | -1.81 | -0.46 | -0.78 | -1.51 | -1.26 | 0.0475 | -0.1075 | -0.03 | 0.879622 | 0.777031 | 0.928645 | 0.999971 | 1 | 0.994278 |
| Romo1 | P60603 | -3.64 | 0.5 | -0.49 | -1.37 | -0.08 | 0.32 | -0.3 | -0.71 | 0.38 | -2.99 | 1.37 | -0.32 | 1.34 | 0.11 | -0.74 | -1.52 | 1.0575 | 0.66 | 1.0475 | 0.217017 | 0.462873 | 0.292867 | 0.999971 | 1 | 0.847402 |
| Rp2 | Q9EPK2 | -0.71 | -1.17 | -0.3 | -0.74 | -1.12 | -0.98 | -0.32 | -0.66 | -0.96 | -0.63 | -0.4 | -0.4 | -1.04 | -1.03 | -0.26 | -0.78 | -0.04 | 0.1325 | -0.0475 | 0.882497 | 0.596371 | 0.86219 | 0.999971 | 1 | 0.982963 |
| Rpa1 | Q5SWN2 | 0.97 | 2.32 | 0.91 | 0.54 | 1.88 | 1.84 | 0.52 | 0.73 | 1.92 | 1.63 | 0.99 | 0.93 | 1.57 | 2.17 | 0.96 | 0.81 | 0.0575 | 0.1825 | 0.1925 | 0.906967 | 0.676358 | 0.681595 | 0.999971 | 1 | 0.943204 |
| Rpa2 | Q62193 | 2.78 | 1.16 | 1.52 | 1.72 | 1.44 | 0.86 | 0.82 | 1.4 | 0.78 | 2.02 | 1.02 | -0.32 | 2.13 | 1.54 | 2.08 | 1.89 | -0.665 | -0.92 | 0.115 | 0.102904 | 0.121027 | 0.75265 | 0.999971 | 1 | 0.961116 |
| Rpa3 | Q9CQ71 | 0.24 | 1.15 | 1.34 | 0.83 | 1.28 | 0.6 | 0.94 | 1.43 | 0.88 | 0.56 | 1.24 | 1.56 | 0.84 | 1.19 | 1.67 | 1.11 | 0.1725 | 0.17 | 0.3125 | 0.580245 | 0.603353 | 0.319449 | 0.999971 | 1 | 0.853962 |
| Rpap1 | Q80TE0 | -0.02 | -0.33 | 0.24 | 0.21 | -0.36 | -0.61 | 0.09 | 0.06 | -0.56 | 0 | 0.24 | 0.25 | -0.68 | -0.34 | 0.07 | 0.23 | -0.23 | -0.0425 | -0.205 | 0.360362 | 0.867532 | 0.448689 | 0.999971 | 1 | 0.885333 |
| Rpap3 | Q9D706 | -0.55 | -0.35 | 2.05 | 2.09 | -0.24 | -0.59 | 1.24 | 1.31 | -0.25 | -0.53 | 2.18 | 1.64 | -0.7 | -0.45 | 1.53 | 1.88 | -0.38 | -0.05 | -0.245 | 0.632972 | 0.955447 | 0.781998 | 0.999971 | 1 | 0.965859 |
| Rpe | B2KGF0 | 1.42 | 1.41 | 1.33 | 1.55 | 1.11 | 0.96 | 1.54 | 1.1 | 1.49 | 1.98 | 0.38 | -0.41 | -0.07 | 0.85 | 1.11 | 0.51 | -0.25 | -0.5675 | -0.8275 | 0.222715 | 0.278387 | 0.014367 | 0.999971 | 1 | 0.613095 |
| Rpf2 | G3X926 | 0.38 | -1.2 | -0.51 | -0.59 | -1.22 | -1.25 | -0.39 | -0.72 | -1 | -0.33 | -1.03 | -1.31 | -1.61 | -1.02 | -1.42 | -0.81 | -0.415 | -0.4625 | -0.735 | 0.284079 | 0.240274 | 0.070045 | 0.999971 | 1 | 0.735439 |
| Rpgr | Q9R0X5-2 | 0.12 | -0.11 | 0.41 | 0.21 | 0.31 | 0.16 | 0.07 | 0.52 | 0.41 | -0.09 | 0.34 | 0.77 | 0.21 | 0.44 | 0.67 | 0.45 | 0.1075 | 0.2 | 0.285 | 0.598082 | 0.41042 | 0.181225 | 0.999971 | 1 | 0.808532 |
| Rpgrip1l | Q8CG73 | -0.68 | -1.43 | 0.19 | 0.61 | -0.96 | -1.26 | -0.04 | 0.69 | -1.34 | -0.99 | -0.04 | 0.31 | -0.98 | -1.01 | 0.03 | 0.69 | -0.1875 | -0.01 | 0.01 | 0.910969 | 0.735148 | 0.985862 | 0.999971 | 1 | 0.99925 |
| Rph3a | P47708 | -0.73 | -1.75 | -1.66 | -1.81 | -1.5 | -1.55 | -0.7 | -1.99 | -1.29 | -1.33 | -1.57 | -1.85 | -1.51 | -1.91 | -1.92 | -2.22 | 0.0525 | -0.0225 | -0.4025 | 0.884286 | 0.939062 | 0.205018 | 0.999971 | 1 | 0.81617 |
| Rpia | P47968 | -0.04 | -0.3 | -0.11 | 0.43 | 0.28 | -0.23 | 0.08 | -0.1 | -0.32 | -0.29 | -0.39 | -0.58 | -1.08 | -0.27 | -0.56 | -0.81 | 0.0125 | -0.39 | -0.675 | 0.956456 | 0.094543 | 0.026182 | 0.999971 | 1 | 0.678474 |
| Rpl10 | I7HLV2 | 0.16 | 1.46 | 2.16 | 1.75 | 1.98 | 1.17 | 2.59 | 2.28 | 2.05 | 0.79 | 2.74 | 2.86 | 2.04 | 1.35 | 2.16 | 1.55 | 0.6225 | 0.7275 | 0.3925 | 0.226267 | 0.238397 | 0.389658 | 0.999971 | 1 | 0.871816 |
| Rpl10a | A0A3B2WE | -0.5 | 0.77 | -0.55 | -1.08 | 0.85 | 0.6 | -0.07 | -0.69 | 1 | 0.56 | 0.23 | 0.78 | 0.78 | 0.75 | 0.04 | -1.05 | 0.5125 | 0.9825 | 0.47 | 0.308336 | 0.03705 | 0.391637 | 0.999971 | 1 | 0.871816 |
| Rpl11 | Q9CXW4 | 0.96 | 1.42 | 0.65 | 0.36 | 1.34 | 1.44 | 0.83 | 0.85 | 1.23 | 1.42 | 1.16 | 0.77 | 1.2 | 1.74 | 1.3 | 1.23 | 0.2675 | 0.2975 | 0.52 | 0.36757 | 0.302704 | 0.087509 | 0.999971 | 1 | 0.763943 |
| Rpl12 | P35979 | 0.72 | 2.96 | 2.04 | 1.63 | 3.4 | 2.2 | 1.46 | 1.94 | 3.55 | 0.93 | 1.93 | 1.49 | 4.81 | 2.11 | 2.05 | 1.95 | 0.4125 | 0.1375 | 0.8925 | 0.475545 | 0.836585 | 0.255706 | 0.999971 | 1 | 0.839012 |
| Rpl13 | P47963 | 2.61 | 1.65 | 0.92 | 0.96 | 1.95 | 1.94 | 1.01 | 0.94 | 1.53 | 2.72 | 1.13 | 1.15 | 1.69 | 1.02 | 0.85 | 1.15 | -0.075 | 0.0975 | -0.3575 | 0.868913 | 0.847412 | 0.39881 | 0.999971 | 1 | 0.871816 |
| Rpl13a | P19253 | 0.96 | 1.52 | 1.58 | 1.59 | 1.45 | 1.42 | 1.37 | 1.68 | 1.75 | 1.37 | 1.66 | 1.58 | 1.33 | 1.6 | 1.71 | 1.88 | 0.0675 | 0.1775 | 0.2175 | 0.753145 | 0.419014 | 0.356477 | 0.999971 | 1 | 0.868793 |
| Rpl14 | Q9CR57 | 0.09 | 2.26 | 2.21 | 1.78 | 2.16 | 2.38 | 1.73 | 2.34 | 2.07 | 1.15 | 2.49 | 2.09 | 1.87 | 2.59 | 2.27 | 2.55 | 0.5675 | 0.365 | 0.735 | 0.268095 | 0.504759 | 0.165535 | 0.999971 | 1 | 0.800215 |
| Rpl15 | Q9CZM2 | -0.8 | -0.28 | 0.61 | 0.71 | 0.35 | -0.68 | 1.25 | 0.82 | 0.95 | -1.43 | 0.76 | 0.68 | -0.43 | 0.01 | 0.42 | 0.81 | 0.375 | 0.18 | 0.1425 | 0.468939 | 0.768878 | 0.73916 | 0.999971 | 1 | 0.960671 |
| Rpl17 | Q6ZWZ7 | 0.44 | 0.51 | 0.4 | 0.22 | 0.71 | 0.42 | 0.72 | 0.52 | 0.63 | 0.27 | 0.42 | 0.67 | 0.27 | 0.66 | 0.43 | 0.57 | 0.2 | 0.105 | 0.09 | 0.278578 | 0.567474 | 0.624392 | 0.999971 | 1 | 0.930479 |
| Rpl18 | P35980 | 1.8 | 1.79 | 1.27 | 1.59 | 1.72 | 1.69 | 1.26 | 1.21 | 1.66 | 1.6 | 1.64 | 1.65 | 1.63 | 1.61 | 1.38 | 2.1 | -0.1425 | 0.025 | 0.0675 | 0.531348 | 0.894249 | 0.77233 | 0.999971 | 1 | 0.963498 |
| Rpl18a | A0A1D5RLY | 1.44 | 1.36 | 1.39 | 1.58 | 1.44 | 1.76 | 1.08 | 1.71 | 1.78 | 1.8 | 1.38 | 1.71 | 0.87 | 1.3 | 1.71 | 1.9 | 0.055 | 0.225 | 0.0025 | 0.795825 | 0.233479 | 0.992289 | 0.999971 | 1 | 0.99925 |
| Rpl19 | A2A547 | 0.72 | 1.93 | 1.54 | 1.33 | 2.31 | 1.71 | 1.68 | 1.34 | 2.32 | 0.96 | 1.42 | 1.79 | 3.16 | 1.44 | 1.27 | 1.18 | 0.38 | 0.2425 | 0.3825 | 0.260035 | 0.5198 | 0.447964 | 0.999971 | 1 | 0.885289 |
| Rpl21 | Q9CQM8 | 1.33 | 0.46 | 1.55 | 2.14 | 0.83 | 0.07 | 2.23 | 1.73 | 1.16 | 0.45 | 1.39 | 1.38 | 0.18 | -0.13 | 1.52 | 1.39 | -0.155 | -0.275 | -0.63 | 0.776227 | 0.493925 | 0.233232 | 0.999971 | 1 | 0.831526 |
| Rpl22 | P67984 | 0.38 | 1.59 | 0.74 | 1.46 | 1.13 | 0.94 | 0.46 | 0.95 | 0.76 | 0.6 | 0.95 | 1.19 | 1.39 | 1.89 | 0.95 | 1.77 | -0.1725 | -0.1675 | 0.4575 | 0.597272 | 0.601508 | 0.216656 | 0.999971 | 1 | 0.826893 |
| Rpl22l1 | Q9D7S7-2 | 1 | 1.26 | 1.82 | 1.54 | 1.29 | 1.37 | 1.94 | 2.81 | 0.92 | 0.93 | 2.06 | 2.02 | 1.76 | 2.46 | 2.6 | 2.36 | 0.4475 | 0.0775 | 0.89 | 0.256791 | 0.829299 | 0.009667 | 0.999971 | 1 | 0.613095 |
| Rpl23 | P62830 | 1.51 | 1.17 | 1.34 | 1.14 | 1.11 | 0.89 | 0.9 | 1.29 | 1 | 0.81 | 1.46 | 1.14 | 1.12 | 0.84 | 1.42 | 1.7 | -0.2425 | -0.1875 | -0.02 | 0.229474 | 0.380732 | 0.933243 | 0.999971 | 1 | 0.995138 |
| Rpl23a | P62751 | 0.88 | 1.43 | 1.05 | 1.18 | 1.2 | 1.31 | 0.86 | 1.32 | 1.32 | 1.35 | 1.36 | 1.06 | 1.5 | 1.7 | 1.11 | 1.59 | 0.0375 | 0.1375 | 0.34 | 0.857816 | 0.485452 | 0.146295 | 0.999971 | 1 | 0.791767 |
| Rpl24 | Q8BP67 | 1.13 | 1.48 | 2.17 | 2.33 | 1.47 | 1.99 | 3.42 | 1.82 | 1.72 | 0.93 | 2.1 | 1.31 | 0.73 | 1.49 | 1.73 | 1.17 | 0.3975 | -0.2625 | -0.4975 | 0.416089 | 0.484414 | 0.181094 | 0.999971 | 1 | 0.808532 |
| Rpl26 | P61255 | 0.22 | 1.55 | 1.07 | 0.18 | 1.51 | 1.3 | 1.57 | 0.67 | 1.74 | 0.27 | 1.81 | 1.85 | 1.55 | 1.14 | 1.14 | 0.21 | 0.5075 | 0.6625 | 0.255 | 0.204 | 0.188885 | 0.545806 | 0.999971 | 1 | 0.911398 |
| Rpl27 | P61358 | 0.24 | 0.89 | 0.99 | 0.9 | 0.94 | 0.91 | 0.87 | 0.89 | 0.95 | 0.33 | 1.19 | 1.12 | 0.76 | 0.65 | 1 | 1.27 | 0.1475 | 0.1425 | 0.165 | 0.505437 | 0.609174 | 0.514633 | 0.999971 | 1 | 0.905815 |
| Rpl27a | P14115 | -0.35 | 2.06 | 2.55 | 1.39 | 1.75 | 2.21 | 1.73 | 2.53 | 2.28 | 0.67 | 2.72 | 2.19 | 2.24 | 2.32 | 2.34 | 1.54 | 0.6425 | 0.5525 | 0.6975 | 0.30355 | 0.44191 | 0.2673 | 0.999971 | 1 | 0.842677 |
| Rpl28 | P41105 | 0.14 | 1.33 | 1.09 | 0.61 | 1.42 | 0.74 | 1.12 | 1.03 | 1.54 | 0.17 | 1.6 | 0.75 | 0.98 | 0.32 | 1.12 | 1.2 | 0.285 | 0.2225 | 0.1125 | 0.363064 | 0.592427 | 0.734759 | 0.999971 | 1 | 0.957888 |
| Rpl29 | P47915 | 1.4 | 0.55 | 0.82 | 1.13 | 1.25 | 0.78 | 2.24 | 0.82 | 0.94 | 1.25 | 0.93 | 0.7 | 0.55 | 0.27 | 0.89 | 0.43 | 0.2975 | -0.02 | -0.44 | 0.435057 | 0.934776 | 0.110285 | 0.999971 | 1 | 0.78268 |
| Rpl3 | P27659 | -0.06 | 2.02 | 1.82 | 1.39 | 1.99 | 1.79 | 1.9 | 1.59 | 2.35 | 0.56 | 2.06 | 1.84 | 2.12 | 1.15 | 1.63 | 1.22 | 0.525 | 0.41 | 0.2375 | 0.260551 | 0.475464 | 0.626412 | 0.999971 | 1 | 0.930479 |
| Rpl30 | P62889 | 1.96 | 1.05 | 1.12 | 1.79 | 0.84 | 1.25 | 1.4 | 1.07 | 1.13 | 1.84 | 1.07 | 0.92 | 0.12 | 1.08 | 0.98 | 1.03 | -0.34 | -0.24 | -0.6775 | 0.240177 | 0.450644 | 0.063527 | 0.999971 | 1 | 0.725348 |
| Rpl31 | P62900 | 3.2 | 2.04 | 1.71 | 1.36 | 1.6 | 2.01 | 2.61 | 1.14 | 1.92 | 1.96 | 1.36 | 1.47 | 1.9 | 1.14 | 1.16 | 1.28 | -0.2375 | -0.4 | -0.7075 | 0.617758 | 0.341454 | 0.116709 | 0.999971 | 1 | 0.78268 |
| Rpl32 | P62911 | 1.09 | 1.47 | 1.12 | 1.56 | 1.35 | 1.4 | 1.65 | 1.63 | 1.76 | 1.53 | 1 | 1.59 | 1.67 | 1.51 | 1.78 | 0.2275 | 0.2175 | 0.375 | 0.243849 | 0.352176 | 0.068273 | 0.999971 | 1 | 0.733937 | |
| Rpl34 | Q9D1R9 | 0.12 | 1.89 | 1.66 | 0.7 | 2 | 1.77 | 2.26 | 1.08 | 2.62 | 0.25 | 2.26 | 2.27 | 1.4 | 1.15 | 1.55 | 0.91 | 0.685 | 0.7575 | 0.16 | 0.157821 | 0.244503 | 0.701761 | 0.999971 | 1 | 0.949019 |
| Rpl35 | Q6ZWV7 | 0.72 | 0.65 | 1.02 | 1.03 | 0.23 | 0.77 | 3.03 | 0.44 | 0.33 | 0.73 | 0.3 | 0.67 | 0.81 | 0.52 | 0.86 | 0.97 | 0.2625 | -0.3475 | -0.065 | 0.661972 | 0.112625 | 0.744202 | 0.999971 | 1 | 0.961064 |
| Rpl35a | O55142 | 0.74 | 3.08 | 2.72 | 2.13 | 3.02 | 2.82 | 3.37 | 2.57 | 3.49 | 0.89 | 3.2 | 2.62 | 3.32 | 1.82 | 2.49 | 2.67 | 0.7775 | 0.3825 | 0.4075 | 0.14749 | 0.591448 | 0.467489 | 0.999971 | 1 | 0.891731 |
| Rpl36 | Q6ZWZ4 | 1.23 | 1.41 | 0.79 | 1.21 | 1.32 | 1.34 | 0.86 | 1.6 | 1.33 | 1.59 | 1.56 | 1.16 | 1.23 | 1.7 | 1.41 | 1.83 | 0.12 | 0.25 | 0.3825 | 0.614657 | 0.256654 | 0.121754 | 0.999971 | 1 | 0.78268 |
| Rpl36a | P83882 | 0.91 | 2.17 | 1.08 | 1.1 | 2.53 | 2.16 | 1.51 | 1.67 | 3.71 | 1.68 | 1.64 | 0.68 | 3.52 | 1.76 | 1.09 | 0.67 | 0.6525 | 0.6125 | 0.445 | 0.098392 | 0.351575 | 0.486098 | 0.999971 | 1 | 0.897483 |
| Rpl37 | Q9D823 | 1.95 | -1.58 | 1.48 | 0.74 | -2.35 | -1.22 | 4.24 | -0.89 | -2.17 | 0.64 | 1.82 | 1.79 | -2.56 | -2.5 | 2.12 | 0.56 | -0.7025 | -0.1275 | -1.2425 | 0.635375 | 0.907361 | 0.333106 | 0.999971 | 1 | 0.857921 |
| Rpl37a | P61514 | 1.35 | 1.78 | 2.58 | -0.3 | 1.07 | 2 | 3.16 | 1.43 | 1.75 | 1.32 | 2.66 | 0.12 | 0.62 | 1.47 | 1.89 | -0.29 | 0.5625 | 0.11 | -0.43 | 0.422236 | 0.880245 | 0.544613 | 0.999971 | 1 | 0.911398 |
| Rpl38 | Q9JJI8 | -0.85 | -1.49 | -0.97 | -0.95 | -1.57 | -1.44 | -1.2 | -0.8 | -1.83 | -1.08 | -0.49 | -0.89 | -2.49 | -0.89 | -0.73 | -0.24 | -0.1875 | -0.0075 | -0.0225 | 0.460767 | 0.981163 | 0.962181 | 0.999971 | 1 | 0.998721 |
| Rpl39 | P62892 | 0.21 | 1.36 | 1.33 | 1.83 | 1.32 | 0.91 | 0.95 | 1.72 | 1.59 | 0.88 | 1.82 | 1.09 | 1.42 | 1.4 | 1.44 | 1.4 | 0.0425 | 0.1625 | 0.2325 | 0.910605 | 0.679416 | 0.501199 | 0.999971 | 1 | 0.904273 |
| Rpl4 | Q9D8E6 | 0.35 | 1.63 | 1.27 | 1.13 | 1.72 | 1.62 | 1.32 | 1.48 | 1.84 | 1.04 | 1.43 | 1.39 | 1.49 | 1.93 | 1.45 | 1.1 | 0.44 | 0.33 | 0.3975 | 0.159749 | 0.314688 | 0.23685 | 0.999971 | 1 | 0.833594 |
| Rpl5 | P47962 | 0.53 | 1.18 | 1.25 | 1.59 | 1.24 | 0.99 | 1 | 1.57 | 1.36 | 0.84 | 1.4 | 1.57 | 1.41 | 1.41 | 1.68 | 1.94 | 0.0625 | 0.1875 | 0.4725 | 0.821376 | 0.528457 | 0.111823 | 0.999971 | 1 | 0.78268 |
| Rpl6 | P47911 | 1.23 | 1.83 | 1.68 | 1.08 | 2.14 | 1.94 | 2.13 | 1.38 | 2.04 | 1.9 | 1.97 | 2.08 | 1.97 | 1.44 | 1.54 | 1.63 | 0.4425 | 0.5425 | 0.19 | 0.129615 | 0.035181 | 0.444446 | 0.999971 | 1 | 0.88409 |
| Rpl7 | P14148 | -0.22 | 0.48 | 1.01 | 0.2 | 0.78 | -0.17 | 1.28 | 0.69 | 0.86 | -0.29 | 1.78 | 1.88 | 1.21 | 0.33 | 1.07 | 0.8 | 0.2775 | 0.69 | 0.485 | 0.474505 | 0.211848 | 0.160446 | 0.999971 | 1 | 0.800215 |
| Rpl7a | P12970 | 0.14 | 1.13 | 1.18 | 1.22 | 1.63 | 0.57 | 1.03 | 1.56 | 1.22 | 0.91 | 1.74 | 1.54 | 1.92 | 1.23 | 1.55 | 1.97 | 0.28 | 0.435 | 0.75 | 0.435485 | 0.196702 | 0.039404 | 0.999971 | 1 | 0.700838 |
| Rpl8 | P62918 | 1.34 | 1.65 | 1.54 | 1.42 | 1.17 | 1.79 | 2.31 | 0.94 | 1.01 | 1.22 | 1.23 | 0.98 | 1.17 | 1.26 | 0.87 | 0.47 | 0.065 | -0.3775 | -0.545 | 0.838817 | 0.055011 | 0.039236 | 0.999971 | 1 | 0.700838 |
| Rpl9 | P51410 | -0.84 | 1.57 | 1.79 | 0.59 | 1.59 | 1.46 | 1.55 | 1.3 | 1.82 | -0.22 | 2.08 | 2.4 | 1.74 | 0.65 | 2.03 | 1.66 | 0.6975 | 0.7425 | 0.7425 | 0.228364 | 0.344423 | 0.244938 | 0.999971 | 1 | 0.837316 |
| Rplp0 | P14869 | 0.69 | 1.79 | 1.62 | 1.67 | 1.54 | 1.67 | 1.44 | 1.83 | 1.67 | 1.47 | 2.23 | 2.27 | 1.86 | 2.02 | 1.93 | 2.27 | 0.1775 | 0.4675 | 0.5775 | 0.53343 | 0.17399 | 0.067825 | 0.999971 | 1 | 0.733937 |
| Rplp1 | P47955 | 0.28 | 1.29 | 2.08 | 2.09 | 1.57 | 1.66 | 1.72 | 2.58 | 1.1 | 0.85 | 2.01 | 2.39 | 1.03 | 2.18 | 2.6 | 2.73 | 0.4475 | 0.1525 | 0.7 | 0.341383 | 0.770766 | 0.211093 | 0.999971 | 1 | 0.822884 |
| Rplp2 | P99027 | 0.5 | 0.88 | 0.89 | 0.88 | 1 | 0.22 | 0.58 | 1.02 | 0.83 | 0.47 | 0.96 | 1.38 | 1.05 | 0.84 | 0.97 | 1.24 | -0.0825 | 0.1225 | 0.2375 | 0.736178 | 0.61345 | 0.239022 | 0.999971 | 1 | 0.833594 |
| Rpn1 | Q91YQ5 | -0.61 | -1.22 | -1.73 | -1.49 | -1.12 | -0.7 | -0.8 | -1.26 | -0.79 | -0.63 | -1.39 | -1.3 | -1.34 | -0.36 | -1.32 | -1.43 | 0.2925 | 0.235 | 0.15 | 0.32359 | 0.455926 | 0.665149 | 0.999971 | 1 | 0.93697 |
| Rpn2 | A2ACG7 | 0.89 | -0.13 | -0.24 | -0.09 | -0.02 | 0.02 | -0.05 | -0.03 | -0.13 | 0.55 | -0.12 | -0.15 | 0.05 | 0.26 | -0.1275 | 0.015 | -0.0025 | -0.03 | 0.649481 | 0.961181 | 0.918068 | 0.999971 | 1 | 0.992633 | |
| Rpp14 | J3QMX0 | -1.41 | 0.58 | -0.52 | 0.16 | 0.02 | -0.16 | -0.45 | -0.4 | -0.18 | -2.42 | 0.36 | -0.38 | 0.46 | 0.41 | -0.04 | 0.28 | 0.05 | -0.3575 | 0.575 | 0.906123 | 0.602737 | 0.199987 | 0.999971 | 1 | 0.813335 |
| Rpp25 | Q91WE3 | 4.48 | 2.97 | 3.03 | 2.83 | 2.7 | 2.94 | 3.31 | 2.45 | 3.37 | 3.76 | 2.63 | 1.5 | -0.22 | 1.41 | 1.12 | 0.38 | -0.4775 | -0.5125 | -2.655 | 0.260084 | 0.387542 | 0.000514 | 0.999971 | 1 | 0.312038 |
| Rpp30 | O88796 | 1.43 | 1.8 | 2.09 | 1.38 | 1.75 | 1.86 | 1.22 | 1.04 | 1.83 | 1.62 | 0.95 | 1.96 | 1.64 | 1.78 | 1.42 | 1.11 | -0.2075 | -0.085 | -0.1875 | 0.460934 | 0.770202 | 0.461531 | 0.999971 | 1 | 0.890095 |
| Rprd1a | Q8VDS4 | 0.57 | 1.81 | 2.13 | 2.49 | 1.55 | 1.51 | 1.27 | 2.18 | 1.26 | 0.79 | 2.13 | 2.46 | 1.52 | 1.6 | 2.41 | 2.8 | -0.1225 | -0.09 | 0.3325 | 0.778122 | 0.864221 | 0.499454 | 0.999971 | 1 | 0.903717 |
| Rprd1b | A0A0R4J19 | 1.22 | 1.68 | 1.63 | 1.79 | 1.58 | 0.97 | 0.98 | 1.01 | 1.06 | 0.93 | 0.86 | 1.44 | 1.51 | 1.66 | 1.4 | 1.7 | -0.445 | -0.5075 | -0.0125 | 0.081116 | 0.043846 | 0.950403 | 0.999971 | 1 | 0.998038 |
| Rprd2 | A0A0G2JFV | 1.51 | 1.39 | 0.84 | 1.02 | 0.76 | 1.11 | 0.66 | 0.81 | 0.95 | 1.38 | 0.75 | 0.75 | 0.65 | 1.06 | 0.64 | 0.94 | -0.355 | -0.2325 | -0.3675 | 0.14084 | 0.353759 | 0.134489 | 0.999971 | 1 | 0.784948 |
| Rps10 | P63325 | -1.64 | -3.41 | -0.98 | -0.86 | -1.93 | -2.98 | -0.94 | -1.06 | -3.14 | -1.81 | -0.48 | -0.98 | -3.25 | -2.49 | -0.68 | -1.16 | -0.005 | 0.12 | -0.1725 | 0.994148 | 0.872875 | 0.818908 | 0.999971 | 1 | 0.975625 |
| Rps11 | P62281 | 0.13 | 1.39 | 0.62 | 0.73 | 1.22 | 1.04 | 0.44 | 1.02 | 1.24 | 0.77 | 1.31 | 1.41 | 1.3 | 1.24 | 1.32 | 1.24 | 0.2125 | 0.465 | 0.5575 | 0.503902 | 0.150529 | 0.070762 | 0.999971 | 1 | 0.736129 |
| Rps12 | Q6ZWZ6 | 0.98 | 2.25 | 1.52 | 1.49 | 1.8 | 2.46 | 1.24 | 1.89 | 2.27 | 2.02 | 1.58 | 1.4 | 1.95 | 2.86 | 1.74 | 2.1 | 0.2875 | 0.2575 | 0.6025 | 0.425724 | 0.44106 | 0.114148 | 0.999971 | 1 | 0.78268 |
| Rps13 | P62301 | 0.75 | 1.4 | 1 | 1.34 | 1.33 | 1.23 | 0.72 | 1.36 | 1.38 | 1.2 | 1.37 | 1.56 | 1.23 | 1.95 | 1.54 | 1.99 | 0.0375 | 0.255 | 0.555 | 0.877816 | 0.251879 | 0.057984 | 0.999971 | 1 | 0.725348 |
| Rps14 | P62264 | 0.49 | 1.4 | 0.98 | 1.03 | 1.39 | 1.25 | 0.58 | 1.59 | 1.27 | 0.69 | 1.27 | 1.51 | 1.67 | 1.3 | 1.31 | 1.54 | 0.2275 | 0.21 | 0.48 | 0.451673 | 0.448829 | 0.072719 | 0.999971 | 1 | 0.744673 |
| Rps15 | D3YTQ9 | 0.62 | 1.21 | 1.55 | 1.74 | 1.23 | 1.45 | 1.49 | 1.53 | 1.24 | 1.35 | 0.83 | 1.33 | 1.04 | 1.47 | 1.28 | 1.57 | 0.145 | -0.0925 | 0.06 | 0.598089 | 0.747137 | 0.833981 | 0.999971 | 1 | 0.979036 |
| Rps15a | P62245 | 0.24 | 1.12 | 0.5 | 0.73 | 1.18 | 0.68 | 1.05 | 0.44 | 1.29 | -0.24 | 1.16 | 0.92 | 1.38 | 0.45 | 0.77 | 1.03 | 0.19 | 0.135 | 0.26 | 0.489525 | 0.725057 | 0.374422 | 0.999971 | 1 | 0.87031 |
| Rps16 | P14131 | 1.57 | 1.43 | 0.56 | 0.38 | 1.67 | 1.43 | 0.8 | 0.39 | 1.56 | 1.7 | 0.59 | 0.65 | 1.42 | 1.46 | 0.62 | 0.67 | 0.0875 | 0.14 | 0.0575 | 0.827246 | 0.728724 | 0.87604 | 0.999971 | 1 | 0.985129 |
| Rps17 | P63276 | 0.04 | 1.03 | 1.13 | 0.4 | 1.23 | 0.8 | 1.02 | 1.03 | 1.09 | -0.32 | 1.68 | 1.82 | 1.27 | 0.55 | 1.34 | 1.21 | 0.37 | 0.4175 | 0.4425 | 0.219028 | 0.426118 | 0.191025 | 0.999971 | 1 | 0.810547 |
| Rps18-ps5 | A0A1Y7VKY | 1.08 | 1.53 | 1.85 | 1.32 | 1.48 | 1.37 | 1.53 | 1.61 | 1.78 | 1.54 | 2.12 | 1.98 | 1.57 | 1.67 | 1.86 | 1.8 | 0.0525 | 0.41 | 0.28 | 0.808848 | 0.112016 | 0.22546 | 0.999971 | 1 | 0.828225 |
| Rps19 | Q9CZX8 | 0.55 | 0.64 | 1.41 | 1.79 | 0.59 | 0.53 | 1.34 | 1.78 | 0.78 | 0.81 | 1.63 | 1.76 | 0.1 | 0.77 | 1.63 | 1.72 | -0.0375 | 0.1475 | -0.0425 | 0.926467 | 0.702431 | 0.925953 | 0.999971 | 1 | 0.99401 |
| Rps2 | P25444 | 0.78 | 1.64 | 1.35 | 1.43 | 1.48 | 1.65 | 1.3 | 1.81 | 1.86 | 1.26 | 1.75 | 1.78 | 1.59 | 1.87 | 1.9 | 1.91 | 0.26 | 0.3625 | 0.5175 | 0.303618 | 0.176882 | 0.051865 | 0.999971 | 1 | 0.723736 |
| Rps20 | P60867 | -1.39 | 2.07 | 2.72 | 0.84 | 2.1 | 1.4 | 4.11 | 1.4 | 2.22 | -1.51 | 2.45 | -0.96 | 2.22 | 0.7 | 2.53 | 0.32 | 1.1925 | -0.51 | 0.3825 | 0.247879 | 0.680506 | 0.687415 | 0.999971 | 1 | 0.945573 |
| Rps21 | Q9CQR2 | 2 | 0.82 | 1.09 | 1.5 | 0.54 | 0.88 | 1.58 | 1.18 | 0.89 | 1.52 | 0.92 | 0.55 | 0.83 | 0.94 | 0.9 | 0.82 | -0.3075 | -0.3825 | -0.48 | 0.3726 | 0.260899 | 0.109776 | 0.999971 | 1 | 0.78268 |
| Rps23 | P62267 | 0.63 | 3.01 | 2.11 | 1.12 | 3.34 | 2.41 | 3.08 | 1.83 | 3.68 | 1.25 | 3.08 | 2.98 | 4.79 | 1.72 | 1.44 | 1.36 | 0.9475 | 1.03 | 0.61 | 0.126989 | 0.155033 | 0.493759 | 0.999971 | 1 | 0.901531 |
| Rps24 | P62849-2 | -0.59 | 0.47 | 0.89 | 1.15 | 0.05 | 0.18 | 0.77 | 1.24 | -0.04 | -0.9 | 1.12 | 1.11 | -0.42 | 0.9 | 1.36 | 1.64 | 0.08 | -0.1575 | 0.39 | 0.856961 | 0.783701 | 0.483566 | 0.999971 | 1 | 0.896352 |
| Rps25 | A0A1L1SQA | 1.08 | 1.4 | 1.33 | 1.48 | 1.45 | 1.42 | 1.75 | 1.31 | 1.63 | 1.6 | 1.87 | 2.27 | 1.54 | 1.17 | 1.24 | 1.69 | 0.16 | 0.52 | 0.0875 | 0.415666 | 0.038976 | 0.671767 | 0.999971 | 1 | 0.939563 |
| Rps26 | P62855 | 1.79 | 3.15 | 4.01 | 4.53 | 2.26 | 3.59 | 1.62 | 5.03 | 2.9 | 3.12 | 3.3 | -1.46 | 2.46 | 3.03 | 4.19 | 4.35 | -0.245 | -1.405 | 0.1375 | 0.777027 | 0.2453 | 0.840395 | 0.999971 | 1 | 0.97975 |
| Rps27 | A0A0G2JD\ | -0.79 | 2.02 | 2.9 | 1.75 | 2 | 1.44 | 2.9 | 2.83 | 2.1 | -0.01 | 3.46 | 3.46 | 3.02 | 1.16 | 3.18 | 2.21 | 0.8225 | 0.7825 | 0.9225 | 0.306781 | 0.452765 | 0.280923 | 0.999971 | 1 | 0.844538 |
| Rps27a | P62983 | -0.14 | -0.4 | -1.25 | -0.68 | -1.13 | -0.37 | -1.08 | -1.13 | -1.05 | -0.76 | -1.28 | -1.95 | -1.89 | -0.38 | -0.99 | -0.13 | -0.52 | -0.6425 | -0.23 | 0.328409 | 0.088866 | 0.598519 | 0.999971 | 1 | 0.92377 |
| Rps28 | P62858 | 2.4 | 2.05 | 1.67 | 1.84 | 1.66 | 1.65 | 1.73 | 1.8 | 1.45 | 1.97 | 1.62 | 1.61 | 1.66 | 1.86 | 1.71 | 1.75 | -0.28 | -0.3275 | -0.245 | 0.206875 | 0.175487 | 0.267748 | 0.999971 | 1 | 0.842677 |
| Rps29 | P62274 | -0.05 | 3.99 | 2.37 | 2.47 | 4.64 | 4.12 | 2.49 | 2.88 | 4.57 | 2.38 | 2.42 | 1.56 | 5.01 | 4 | 3.56 | 2.2 | 1.3375 | 0.5375 | 1.4975 | 0.152252 | 0.574737 | 0.127856 | 0.999971 | 1 | 0.784171 |
| Rps3 | P62908 | 0.62 | 1.56 | 1.44 | 1.31 | 1.35 | 1.64 | 1.19 | 1.51 | 1.62 | 1.24 | 1.68 | 1.59 | 1.56 | 1.7 | 1.68 | 2.06 | 0.19 | 0.3 | 0.5175 | 0.465512 | 0.260198 | 0.073505 | 0.999971 | 1 | 0.746069 |
| Rps3a | P97351 | 0.64 | 0.96 | 1.23 | 1.54 | 0.78 | 0.94 | 1.65 | 1.52 | 0.97 | 0.98 | 1.45 | 1.48 | 0.6 | 1.16 | 1.62 | 1.72 | 0.13 | 0.1275 | 0.1825 | 0.662708 | 0.626184 | 0.57457 | 0.999971 | 1 | 0.91816 |
| Rps4x | P62702 | 0.85 | 1.72 | 0.89 | 0.77 | 1.75 | 1.42 | 1.26 | 0.83 | 1.97 | 1.21 | 1.13 | 1.38 | 2.05 | 1.83 | 1.12 | 0.71 | 0.2575 | 0.365 | 0.37 | 0.402627 | 0.243825 | 0.333281 | 0.999971 | 1 | 0.857921 |
| Rps5 | Q91V55 | 0.77 | 1.61 | 0.99 | 0.8 | 1.68 | 1.65 | 1.71 | 0.85 | 1.98 | 1.03 | 1.13 | 1.08 | 1.8 | 1.51 | 0.91 | 0.51 | 0.43 | 0.2625 | 0.14 | 0.170125 | 0.39956 | 0.687066 | 0.999971 | 1 | 0.945467 |
| Rps6 | P62754 | 1.64 | 2.21 | 2.43 | 2.1 | 2.11 | 2.63 | 3.51 | 2.05 | 2.41 | 1.58 | 2.85 | 2.48 | 1.62 | 1.29 | 1.84 | 1 | 0.48 | 0.235 | -0.6575 | 0.212123 | 0.466916 | 0.034846 | 0.999971 | 1 | 0.69205 |
| Rps6ka1 | E9PWV3 | 0.02 | 0.68 | 0.92 | 0.46 | 0.11 | 0.42 | 0.55 | 0.94 | -0.13 | 0.39 | 0.5 | 0.38 | 0.07 | 0.61 | 0.92 | 1.01 | -0.015 | -0.235 | 0.1325 | 0.956406 | 0.376717 | 0.6564 | 0.999971 | 1 | 0.934061 |
| Rps6ka2 | Q9WUT3 | -0.35 | 0.03 | 0.31 | 0.49 | 0.94 | 0.9 | 0.29 | 1.29 | 0.88 | 0.15 | 0.07 | 0.9 | 1.94 | 1.33 | 1.79 | 0.49 | 0.735 | 0.38 | 1.2675 | 0.029596 | 0.224862 | 0.00686 | 0.999971 | 1 | 0.582714 |
| Rps6ka3 | B1AXN9 | 0.17 | 0.18 | -0.4 | 0.22 | 0.7 | -0.14 | -0.02 | -0.24 | 0.42 | 0.08 | 0.27 | 0.06 | 0.45 | 0.23 | 0 | 0.2 | 0.0325 | 0.165 | 0.1775 | 0.906302 | 0.449624 | 0.427693 | 0.999971 | 1 | 0.880793 |
| Rps6ka4 | Q9Z2B9 | 3.5 | 2.4 | 1.55 | 1.78 | 2 | 1.88 | 1.02 | 1.42 | 1.75 | 2.47 | 1.73 | 1.61 | 1.01 | 1.77 | 1.58 | 1.76 | -0.7275 | -0.4175 | -0.7775 | 0.139759 | 0.366566 | 0.107622 | 0.999971 | 1 | 0.78268 |
| Rps6ka5 | Q8C050 | 1.32 | 2.71 | 2.27 | 2.18 | 2.65 | 1.71 | 1.94 | 2.33 | 2.44 | 1.68 | 1.99 | 2.74 | 2.22 | 2.64 | 2.69 | 1.96 | 0.0375 | 0.0925 | 0.2575 | 0.914857 | 0.799946 | 0.452874 | 0.999971 | 1 | 0.887217 |
| Rps6kb1 | Q8BSK8-2 | -0.69 | 1.47 | 1.33 | -0.4 | 1.98 | 0.64 | 1.48 | -0.11 | 1.87 | 0.12 | 2.03 | 2.29 | 2.41 | -0.07 | 0.5 | -0.14 | 0.57 | 1.15 | 0.2475 | 0.399088 | 0.120243 | 0.739213 | 0.999971 | 1 | 0.960671 |
| Rps6kc1 | E9QMX4 | 0.96 | 0.37 | 0.24 | 0.43 | 0.54 | 0.4 | 0.25 | 0.32 | 0.12 | 0.55 | 0.28 | 0.11 | 0.01 | 0.15 | 0.11 | 0.21 | -0.1225 | -0.235 | -0.38 | 0.574799 | 0.313786 | 0.10215 | 0.999971 | 1 | 0.78268 |
| Rps7 | P62082 | 0.73 | 1.35 | 0.55 | 0.81 | 1.76 | 1.16 | 0.38 | 0.5 | 1.59 | 0.84 | 0.53 | 0.65 | 2.39 | 1.5 | 0.6 | 0.92 | 0.09 | 0.0425 | 0.4925 | 0.800329 | 0.887656 | 0.248678 | 0.999971 | 1 | 0.839012 |
| Rps8 | P62242 | 0.36 | 1.71 | 1.81 | 1.36 | 1.81 | 1.82 | 1.1 | 1.79 | 1.78 | 0.89 | 1.95 | 1.46 | 1.84 | 2.03 | 1.98 | 1.72 | 0.32 | 0.21 | 0.5825 | 0.391514 | 0.593312 | 0.111053 | 0.999971 | 1 | 0.78268 |
| Rps9 | Q6ZWN5 | 0.17 | 0.85 | 0.29 | 0.43 | 1.13 | 0.29 | 0.41 | 0.6 | 1.21 | 0.4 | 1.17 | 0.97 | 1.27 | 0.6 | 0.5 | 0.63 | 0.1725 | 0.5025 | 0.315 | 0.513356 | 0.080362 | 0.238694 | 0.999971 | 1 | 0.833594 |
| Rpsa | P14206 | -0.3 | 0.49 | 0.3 | 0.24 | 0.71 | -0.12 | 0.65 | 0.32 | 0.54 | -0.68 | 0.47 | 0.82 | 0.86 | 0.42 | 0.67 | 0.54 | 0.2075 | 0.105 | 0.44 | 0.454794 | 0.772841 | 0.084787 | 0.999971 | 1 | 0.758256 |
| Rptor | A2ACM0 | -0.59 | 0.14 | 0.07 | 0.07 | 0.22 | -0.14 | -0.1 | -0.16 | 0.11 | -0.42 | 0.11 | 0.15 | -0.2 | -0.28 | -0.22 | -0.08 | 0.0325 | 0.065 | -0.1175 | 0.888049 | 0.792184 | 0.59794 | 0.999971 | 1 | 0.92377 |
| Rpusd3 | Q14AI6 | -0.63 | 0.32 | 0.35 | 0.14 | 0.09 | -0.19 | 0.13 | -0.21 | -0.31 | -0.69 | 0.19 | 0.06 | 0.86 | 0.37 | 0.03 | 0.19 | -0.09 | -0.2325 | 0.3175 | 0.736869 | 0.458581 | 0.306839 | 0.999971 | 1 | 0.847402 |

| | | | | | | | | | | | | | | | | | | | | | | | | | |
|---|---|---|---|---|---|---|---|---|---|---|---|---|---|---|---|---|---|---|---|---|---|---|---|---|---|
| Rpusd4 | Q9CWX4 | -0.27 | 0.2 | -0.46 | -0.02 | -0.06 | -0.87 | 0.36 | -0.03 | 0.03 | -0.59 | -1.08 | -0.27 | 0.01 | -1.12 | 0.16 | -0.06 | -0.0125 | -0.34 | -0.115 | 0.967082 | 0.25841 | 0.72605 | 0.999971 | 1 | 0.956041 |
| Rraga | Q80X95 | 0.92 | 0.29 | 0.2 | 0.57 | -0.88 | -0.92 | 0.83 | -0.27 | -0.79 | 1.06 | 1.71 | -0.8 | -1.66 | -0.07 | -0.57 | 0.95 | -0.805 | -0.2 | -0.8325 | 0.080401 | 0.742949 | 0.138516 | 0.999971 | 1 | 0.784948 |
| Rragb | Q6NTA4 | 0.98 | 1.08 | 0.44 | 0.57 | 0.49 | 1.57 | 0.62 | 0.79 | 0.79 | 0.46 | 0.56 | 0.61 | 0.46 | 1.74 | 1.01 | 0.54 | 0.1 | -0.1625 | 0.17 | 0.736978 | 0.455301 | 0.611039 | 0.999971 | 1 | 0.92673 |
| Rragc | Q99K70 | 0.99 | 0.64 | -0.09 | -0.1 | 0.46 | 0.91 | 0.19 | -0.05 | 0.73 | 0.5 | -0.04 | -0.41 | 0.31 | 1.03 | -0.08 | -0.08 | 0.0175 | -0.165 | -0.065 | 0.958694 | 0.653702 | 0.859731 | 0.999971 | 1 | 0.982702 |
| Rragd | B1AWT2 | 1.21 | 0.56 | 1.62 | 2.25 | 0.6 | 1.15 | 2.14 | 1.92 | 0.52 | 0.82 | 1.54 | 1.04 | -0.42 | 0.5 | 1.31 | 0.17 | 0.0425 | -0.43 | -1.02 | 0.927675 | 0.296973 | 0.056 | 0.999971 | 1 | 0.725348 |
| Rras | P10833 | -1.5 | 0.33 | -0.91 | -0.99 | 1.01 | 0.4 | -1.11 | -0.95 | 0.47 | -0.83 | -0.82 | -0.41 | 0.91 | -0.34 | -1.05 | -1.06 | 0.605 | 0.37 | 0.3825 | 0.32061 | 0.43405 | 0.497462 | 0.999971 | 1 | 0.902655 |
| Rras2 | P62071 | -1.26 | -0.93 | -0.81 | -0.89 | -0.38 | -0.72 | -1.14 | -0.71 | -0.43 | -0.88 | -0.57 | -0.27 | 0.08 | -1.06 | -0.67 | -0.81 | 0.235 | 0.435 | 0.3575 | 0.312177 | 0.064031 | 0.224059 | 0.999971 | 1 | 0.828225 |
| Rrbp1 | A2AVJ7 | 0.79 | -0.5 | -0.07 | -0.12 | -0.11 | 0.1 | 0.88 | -0.01 | 0.21 | 0.84 | 0.14 | 0.09 | 0.04 | 0.12 | 0.15 | 0.16 | 0.19 | 0.295 | 0.0925 | 0.588283 | 0.37499 | 0.748143 | 0.999971 | 1 | 0.961064 |
| Rrm1 | P07742 | -0.42 | 0.36 | 0.02 | -0.06 | 0.65 | -0.53 | -0.12 | -0.97 | 0.39 | 0.2 | -0.51 | -0.56 | 0.14 | 0.4 | 0.23 | -0.26 | -0.2175 | -0.095 | 0.1525 | 0.559176 | 0.751132 | 0.536968 | 0.999971 | 1 | 0.910077 |
| Rrm2b | Q6PEE3 | 1.09 | 1.75 | 0.61 | 0.04 | 1.79 | 1.56 | 0.4 | 0.77 | 1.69 | 0.99 | 0.87 | 0.79 | 0.77 | 1.58 | 0.69 | 0.34 | 0.2575 | 0.2125 | -0.0275 | 0.576904 | 0.59815 | 0.948318 | 0.999971 | 1 | 0.997766 |
| Rrp1 | P56183-2 | 3.34 | 0.79 | 1.1 | 1.78 | 1.66 | 1.76 | 3.63 | 0.85 | 1.1 | 2.09 | 0.67 | -1.14 | -0.47 | 0.72 | 0.42 | -0.93 | 0.2225 | -1.0725 | -1.8175 | 0.763175 | 0.202723 | 0.016548 | 0.999971 | 1 | 0.613095 |
| Rrp12 | Q6P5B0 | 0.54 | 0.34 | 0.11 | -0.12 | 0.56 | 0.02 | -0.26 | -0.21 | 0.33 | 0.21 | -0.1 | -0.29 | 0.12 | -0.13 | -0.23 | -0.62 | -0.19 | -0.18 | -0.4325 | 0.470941 | 0.450115 | 0.101643 | 0.999971 | 1 | 0.78268 |
| Rrp15 | Q9CYX7 | -1.16 | -0.28 | -0.25 | -0.16 | -0.14 | -1.53 | -0.38 | 0.01 | -1.08 | -0.71 | -0.11 | -0.28 | -0.69 | -1.11 | 0.12 | -0.56 | -0.0475 | -0.0825 | -0.0975 | 0.90577 | 0.797672 | 0.776439 | 0.999971 | 1 | 0.964475 |
| Rrp9 | Q91WM3 | 0.6 | 0.01 | -0.29 | -0.22 | 0.08 | -0.11 | -0.83 | -0.77 | -0.42 | 0.83 | -0.14 | -0.27 | -0.62 | -0.3 | -0.83 | -0.46 | -0.4325 | -0.025 | -0.5775 | 0.188277 | 0.941859 | 0.04819 | 0.999971 | 1 | 0.710102 |
| Rrs1 | Q9CYH6 | -0.73 | -0.93 | -0.34 | -0.08 | -0.44 | -0.87 | -0.63 | -0.49 | -1.14 | -0.4 | 0.05 | -0.73 | -0.93 | -0.75 | -0.71 | -0.72 | -0.0875 | -0.035 | -0.2575 | 0.721257 | 0.912454 | 0.289103 | 0.999971 | 1 | 0.847402 |
| Rsf1 | E9PWW9 | -0.37 | -1.47 | -1.49 | -1.12 | -1.18 | -1.25 | -1.81 | -1.13 | -1.43 | -0.85 | -1.65 | -1.5 | -1.31 | -0.92 | -1.3 | -1.02 | -0.23 | -0.245 | -0.025 | 0.466057 | 0.448269 | 0.931669 | 0.999971 | 1 | 0.995069 |
| Rsl1d1 | Q8BVY0 | 2.15 | 0.39 | 2.3 | 1.82 | -0.38 | 0.9 | 2.19 | 2.14 | -0.6 | 0.89 | 2.68 | 2.63 | -0.26 | 0.39 | 2.78 | 1.84 | -0.4525 | -0.265 | -0.4775 | 0.510153 | 0.745081 | 0.521909 | 0.999971 | 1 | 0.907417 |
| Rsl24d1 | E9PY85 | -0.89 | 0.72 | -0.25 | -1.18 | 0.11 | 0.6 | 0.59 | -2.17 | 0.41 | -0.09 | -0.91 | -1.5 | 0.62 | 0.43 | -1.64 | -0.26 | 0.1825 | -0.1225 | 0.1875 | 0.796684 | 0.824506 | 0.757273 | 0.999971 | 1 | 0.961116 |
| Rspry1 | Q8BVR6 | -0.7 | -1.02 | -0.69 | -0.65 | -1.07 | -0.62 | -1.1 | -0.43 | -1.14 | -0.53 | -0.56 | -0.83 | -0.81 | -0.78 | -0.47 | -0.48 | -0.04 | 1.57E-16 | 0.13 | 0.860079 | 1 | 0.505182 | 0.999971 | 1 | 0.905277 |
| Rsrc1 | Q3TGY1 | -0.56 | 0 | 0.66 | -0.21 | 0.68 | -0.35 | 0.01 | -0.01 | 0.02 | 0.09 | 0.37 | 0.41 | 0.03 | -0.58 | -0.25 | 0.6 | 0.11 | 0.25 | -0.0225 | 0.742087 | 0.393785 | 0.94904 | 0.999971 | 1 | 0.998038 |
| Rsrc2 | S4R2L4 | 1.39 | 1.76 | 2.76 | 2.16 | 1.83 | 1.67 | 2.73 | 2.39 | 1.77 | 1.29 | 2.57 | 1.69 | 1.32 | 1.61 | 1.77 | 1.4 | 0.1375 | -0.1875 | -0.4925 | 0.711925 | 0.627374 | 0.144293 | 0.999971 | 1 | 0.790933 |
| Rsu1 | Q9D031 | 0.81 | 0.39 | -0.06 | -0.07 | 0.92 | 0.45 | 0.56 | -0.11 | 0.63 | 0.88 | 0.46 | 0.86 | 0.4 | 0.45 | 0.16 | -0.13 | 0.1875 | 0.44 | -0.0475 | 0.54438 | 0.113262 | 0.859944 | 0.999971 | 1 | 0.982702 |
| RtcA | Q9D7H3 | 2.74 | 1.75 | 1.59 | 1.68 | 0.87 | 1.76 | 1.94 | 1.35 | 1.09 | 1.86 | 1.31 | 1.25 | 1.09 | 1.18 | 1.7 | 1.33 | -0.46 | -0.5625 | -0.615 | 0.213422 | 0.105297 | 0.071784 | 0.999971 | 1 | 0.739418 |
| Rtcb | Q99LF4 | 0.27 | 1.52 | 1.09 | 0.12 | 1.75 | 0.72 | 1.27 | 0.34 | 1.78 | 0.21 | 1.43 | 1.82 | 1.52 | 0.36 | 0.36 | 0.09 | 0.27 | 0.56 | -0.1675 | 0.534534 | 0.255057 | 0.70197 | 0.999971 | 1 | 0.949019 |
| Rtf1 | A2AQ19 | 1.38 | 2 | 1.74 | 1.79 | 1.77 | 1.83 | 1.21 | 1.94 | 1.84 | 1.67 | 1.87 | 1.87 | 1.58 | 1.99 | 1.96 | 2.25 | -0.04 | 0.085 | 0.2175 | 0.8678 | 0.664459 | 0.354218 | 0.999971 | 1 | 0.868793 |
| Rtkn | A0A1D5RL | 0.18 | -1.06 | 0.07 | -0.12 | -1.13 | 0.68 | -0.67 | -0.12 | -0.49 | -0.51 | -0.72 | -0.47 | 0.23 | -0.66 | 0.03 | 0.39 | -0.0775 | -0.315 | 0.23 | 0.863665 | 0.305092 | 0.52547 | 0.999971 | 1 | 0.909437 |
| Rtkn_1 | Q8C6B2-2 | -0.04 | -0.07 | 0.52 | 0.7 | -0.58 | -0.3 | 0.6 | 0.88 | 0.03 | -0.3 | 1.04 | 1.1 | -0.25 | -0.36 | 0.99 | 1.05 | -0.1275 | 0.19 | 0.08 | 0.74161 | 0.628333 | 0.845512 | 0.999971 | 1 | 0.979908 |
| Rtkn_2 | D3Z4N1 | 2.68 | -0.37 | 0.58 | 1.4 | -0.1 | 0.76 | 1.17 | 0.55 | 0.12 | 2.27 | 0.99 | -0.14 | -0.8 | -0.54 | -0.54 | -0.51 | -0.4775 | -0.2625 | -1.67 | 0.459453 | 0.732363 | 0.019426 | 0.999971 | 1 | 0.635388 |
| Rtl8b | Q9D1F0 | 0.22 | 0.07 | 1.05 | 0.54 | 0.2 | -0.47 | 1.14 | 0.56 | 0.51 | 0.28 | 0.9 | 2.04 | -0.62 | -0.01 | 0.91 | 1.23 | -0.1125 | 0.4625 | -0.0925 | 0.770789 | 0.292998 | 0.836509 | 0.999971 | 1 | 0.979416 |
| Rtn1 | Q8K0T0 | -0.25 | -0.84 | -0.18 | 0.12 | -0.72 | -0.78 | -0.23 | 0.08 | -0.6 | -0.59 | -0.2 | -0.22 | -0.45 | -0.76 | 0.05 | 0.14 | -0.125 | -0.115 | 0.0325 | 0.67494 | 0.652243 | 0.914047 | 0.999971 | 1 | 0.992633 |
| Rtn1_1 | Q7M6W1 | 0.87 | -0.1 | 0.41 | 0.15 | 0.01 | 1.11 | 0.8 | 1.31 | 0.8 | 0.38 | 0.85 | 0.76 | -0.32 | 0.76 | 0.86 | 0.07 | 0.475 | 0.365 | 0.01 | 0.194921 | 0.179235 | 0.976911 | 0.999971 | 1 | 0.99925 |
| Rtn2 | O70622 | 0.18 | -0.66 | -0.84 | -1.2 | -1 | -0.84 | -0.94 | -0.79 | -0.77 | 0.13 | -1.21 | -1.95 | -0.94 | -0.51 | -1.03 | -1.2 | -0.2625 | -0.32 | -0.29 | 0.397434 | 0.51829 | 0.387515 | 0.999971 | 1 | 0.871816 |
| Rtn3 | Q9ES97-2 | -0.21 | -0.65 | -0.1 | -0.06 | -0.86 | -0.74 | -0.28 | -0.29 | -0.8 | -0.4 | -0.01 | -0.1 | -0.84 | -0.73 | -0.25 | 0.19 | -0.2875 | -0.0725 | -0.1525 | 0.244804 | 0.77193 | 0.597635 | 0.999971 | 1 | 0.92377 |
| Rtn3_1 | Q9ES97-3 | -1.09 | -0.65 | -0.85 | -0.15 | -1.38 | 0.13 | 0.29 | -0.72 | -1.41 | -0.69 | -1.42 | -1.04 | -2.67 | 0.99 | -0.68 | -0.52 | 0.265 | -0.455 | -0.035 | 0.528402 | 0.130447 | 0.960282 | 0.999971 | 1 | 0.998721 |
| Rtn4 | Q99P72-1 | -4.59 | -0.66 | -1.38 | -2.81 | -0.2 | -1.41 | -0.91 | -2.31 | -0.28 | -5.25 | -1.13 | -0.5 | -0.37 | -1.55 | -1.57 | -2.59 | 1.1525 | 0.57 | 0.84 | 0.208922 | 0.662995 | 0.351754 | 0.999971 | 1 | 0.867656 |
| Rtn4_1 | Q99P72 | -0.47 | -0.66 | -0.85 | -0.78 | -0.68 | -0.66 | -0.65 | -0.57 | -0.51 | -0.56 | -0.65 | -0.23 | -0.15 | -0.5 | -0.51 | -0.35 | 0.05 | 0.2025 | 0.3125 | 0.774361 | 0.296146 | 0.12462 | 0.999971 | 1 | 0.78268 |
| Rtn4ip1 | A0A0R4J0S | -0.59 | -0.99 | -1.3 | -0.97 | -0.92 | -1.31 | -1.64 | -0.48 | -1.29 | -1.01 | -1.61 | -1.26 | -1.06 | -1.09 | -1.28 | -1.16 | -0.125 | -0.33 | -0.185 | 0.676655 | 0.171201 | 0.383049 | 0.999971 | 1 | 0.871816 |
| Rtn4r | Q99PI8 | -0.41 | -0.26 | -1.65 | -1.32 | -0.33 | 0.06 | -1.47 | -1.02 | -0.37 | -0.19 | -1.22 | -1.35 | 0.33 | -0.39 | -1.48 | -0.89 | 0.22 | 0.1275 | 0.3025 | 0.629298 | 0.765494 | 0.532156 | 0.999971 | 1 | 0.910069 |
| Rtn4rl1 | Q8K0S5 | -1.36 | 0.35 | 0.58 | 0.11 | 1.01 | 0.04 | 0.28 | -0.08 | 0.7 | -1.7 | -0.28 | 0.47 | 1.43 | -0.37 | 0.24 | 0.04 | 0.3925 | -0.1225 | 0.415 | 0.411189 | 0.847396 | 0.448599 | 0.999971 | 1 | 0.885333 |
| Rtn4rl2 | A2ATS6 | -0.93 | -0.53 | -0.84 | -0.83 | -0.49 | -0.29 | -0.28 | -0.5 | -0.61 | -0.6 | -0.38 | -0.42 | -0.53 | -0.62 | -0.57 | -0.57 | 0.3925 | 0.28 | 0.21 | 0.055902 | 0.143937 | 0.251729 | 0.999971 | 1 | 0.839012 |
| RTRAF | Q9CQE8 | 2.17 | 1.87 | 1.68 | 1.79 | 2.12 | 1.71 | 1.74 | 1.65 | 3.07 | 2.04 | 1.74 | 2.01 | 2.12 | 1.49 | 1.45 | 1.78 | -0.0725 | 0.3375 | -0.1675 | 0.723622 | 0.29958 | 0.469076 | 0.999971 | 1 | 0.892481 |
| Rubcn | D3Z7B1 | -0.16 | 0.8 | 0.16 | -0.74 | 0.71 | 0.23 | 0.58 | -0.25 | 0.52 | -1.22 | 0.68 | 0.36 | -0.46 | -0.35 | -0.59 | -1.43 | 0.3025 | 0.07 | -0.7225 | 0.427408 | 0.89011 | 0.090996 | 0.999971 | 1 | 0.773779 |
| Rufy1 | Q8BIJ7 | 0.31 | 1.38 | 1.1 | 0.82 | 1.34 | 1.2 | 0.97 | 1.07 | 1.37 | 0.48 | 1.34 | 1.01 | 0.88 | 0.99 | 1.08 | 1.19 | 0.2425 | 0.1475 | 0.1325 | 0.369743 | 0.638417 | 0.614669 | 0.999971 | 1 | 0.927806 |
| Rufy2 | Q8R4C2 | 0.66 | 0.03 | -0.44 | -0.23 | 0.15 | 0.51 | 0.02 | 0.11 | 0.33 | 0.82 | 0.07 | -0.28 | 0.91 | 0.57 | 0 | 0.24 | 0.1925 | 0.23 | 0.425 | 0.494561 | 0.493666 | 0.199982 | 0.999971 | 1 | 0.813335 |
| Rufy2_1 | D3Z4D2 | -1.32 | -2.73 | -1.63 | -1.05 | -1.41 | -1.03 | -1.64 | -0.27 | -0.53 | -0.12 | 0.82 | -1.55 | -1.47 | -0.84 | 1.05 | -0.83 | 0.595 | 1.3375 | 1.16 | 0.205594 | 0.040288 | 0.080877 | 0.999971 | 1 | 0.755254 |
| Rufy3 | Q9D394-3 | -0.29 | -2.07 | -2.82 | -1.37 | -2.06 | -1.78 | -2.2 | -2.53 | -1.28 | -0.84 | -1.14 | -2.67 | -3.67 | -0.69 | -2 | -1.67 | -0.505 | 0.155 | -0.37 | 0.342516 | 0.802076 | 0.620065 | 0.999971 | 1 | 0.929296 |
| Rufy3_1 | Q9D394-4 | 0.43 | -0.05 | -0.76 | -0.31 | -0.6 | 0.08 | 1.39 | -1.29 | -0.17 | 1.38 | -0.96 | -0.74 | 0.49 | -0.26 | -0.57 | -0.98 | 0.0675 | 0.05 | -0.1575 | 0.905784 | 0.926152 | 0.683072 | 0.999971 | 1 | 0.943421 |
| Rufy3_2 | Q9D394 | -0.38 | -0.24 | -0.34 | -0.52 | -0.14 | -0.21 | -0.18 | -0.59 | 0.26 | -0.3 | -0.02 | -0.03 | 0.24 | -0.34 | -0.14 | -0.5 | 0.09 | 0.3475 | 0.185 | 0.635047 | 0.094944 | 0.403765 | 0.999971 | 1 | 0.873348 |
| Rundc1 | Q0VDN7 | 1.97 | 1.57 | 1.05 | 1.18 | 2.25 | 1.37 | 0.48 | 1.3 | 1.35 | 1.61 | 1.45 | 1.18 | 2.21 | 2.01 | 1.01 | 1.54 | -0.0925 | -0.045 | 0.25 | 0.816688 | 0.857994 | 0.46511 | 0.999971 | 1 | 0.89116 |
| Rundc3a | O08576-4 | -0.22 | -0.08 | -0.38 | 0.15 | -0.03 | 0.56 | -1.29 | -0.73 | 0.29 | 0.13 | 0.17 | -0.6 | -0.27 | 0.76 | -1.03 | -0.43 | -0.24 | 0.13 | -0.11 | 0.553024 | 0.612603 | 0.770648 | 0.999971 | 1 | 0.963498 |
| Rundc3a_1 | O08576 | -0.81 | -0.24 | -0.5 | -0.64 | -0.14 | -0.54 | -0.76 | -0.21 | -0.04 | -0.51 | -0.17 | -0.17 | 0.67 | -0.17 | -0.23 | -0.07 | 0.135 | 0.325 | 0.5975 | 0.557394 | 0.141535 | 0.046858 | 0.999971 | 1 | 0.710102 |
| Rundc3b | D3YUQ5 | -0.19 | -0.4 | -0.41 | -0.54 | -0.35 | -0.32 | -0.72 | -0.31 | -0.29 | 0.07 | -0.07 | -0.16 | 0.13 | 0.08 | 0 | 0.36 | -0.04 | 0.2725 | 0.5275 | 0.833279 | 0.152814 | 0.017052 | 0.999971 | 1 | 0.617356 |
| Rusc1 | Q8BG26 | 0.44 | -1.02 | 0.41 | 1.28 | -1.59 | -0.22 | -0.01 | 0.96 | -0.86 | -0.07 | 1.11 | -0.59 | -0.13 | -0.65 | 0.8 | 1.68 | -0.4925 | -0.38 | 0.1475 | 0.452475 | 0.527253 | 0.817533 | 0.999971 | 1 | 0.974929 |
| Rusc2 | G8JL85 | -0.78 | -1.27 | -1.02 | -0.86 | -1.1 | -0.16 | -1.15 | -0.7 | -1.16 | -0.49 | -0.63 | -0.45 | -1.24 | -0.3 | -0.44 | -0.92 | 0.205 | 0.3 | 0.2575 | 0.458311 | 0.215937 | 0.344177 | 0.999971 | 1 | 0.862346 |
| Rusf1 | Q91W34 | -0.32 | 0.31 | -1.69 | -2.38 | -0.95 | -0.55 | -1.15 | -1.18 | -0.36 | 0.11 | -1.22 | -0.32 | -1.63 | -0.88 | -2.32 | -1.78 | 0.0625 | 0.5725 | -0.6325 | 0.913991 | 0.36776 | 0.325184 | 0.999971 | 1 | 0.856472 |
| Rusf1_1 | A0A0R4J0E | 0.43 | -0.4 | -0.4 | -0.16 | -0.66 | -0.55 | -0.61 | -0.38 | -0.58 | 0.13 | -0.76 | -0.78 | -0.97 | 0 | -0.44 | -0.29 | -0.4175 | -0.365 | -0.2925 | 0.107363 | 0.241394 | 0.333358 | 0.999971 | 1 | 0.857921 |
| Ruvbl1 | P60122 | 0.2 | 0.25 | 0.41 | -0.04 | 0.22 | 0.16 | 0.16 | -0.12 | 0.25 | 0.32 | 0.09 | -0.01 | 0.13 | 0.42 | 0.09 | -0.18 | -0.1 | -0.0425 | -0.09 | 0.599509 | 0.818949 | 0.666232 | 0.999971 | 1 | 0.937477 |
| Ruvbl2 | Q9WTM5 | -0.65 | -0.25 | 0.21 | -0.07 | -0.1 | -0.27 | -0.09 | -0.02 | -0.29 | -0.58 | 0.12 | 0.4 | -0.23 | -0.28 | 0.21 | 0.02 | 0.07 | 0.1025 | 0.12 | 0.758654 | 0.726381 | 0.626105 | 0.999971 | 1 | 0.930479 |
| Rwdd1 | Q9CQK7 | 0.87 | 0.32 | 0.01 | 0.19 | 0.63 | 0.52 | -0.12 | 0.33 | 0.47 | 0.72 | -0.05 | 0.07 | 0.72 | 0.38 | 0.23 | 0.44 | -0.0075 | -0.045 | 0.095 | 0.977684 | 0.869317 | 0.6978 | 0.999971 | 1 | 0.94766 |
| Rwdd2a | Q14BN8 | -0.11 | 0.42 | 0.55 | -0.73 | 1.23 | 0.63 | 3.33 | -0.18 | 1.95 | -0.9 | 0.97 | 1.87 | 2.21 | -0.2 | 0.87 | 0.04 | 1.22 | 0.94 | 0.6975 | 0.120164 | 0.179623 | 0.238831 | 0.999971 | 1 | 0.833594 |
| Rwdd4 | Q9CPR1 | 1.71 | 0.14 | 1.14 | 0.61 | 0.45 | 0.06 | 1.54 | 0.33 | 0.86 | 0.34 | 0.84 | 0.42 | 0.24 | -0.96 | 0.9 | 0.23 | -0.305 | -0.285 | -0.7975 | 0.496016 | 0.433974 | 0.123878 | 0.999971 | 1 | 0.78268 |
| Rxrb | Q8VCR0 | -0.28 | 1.02 | 1.2 | 1.71 | 0.93 | 0.36 | 1.67 | 1.96 | 0.58 | -0.12 | 2.04 | 2.82 | 0.24 | 1.22 | 2.15 | 2.49 | 0.3175 | 0.4175 | 0.6125 | 0.541901 | 0.566032 | 0.324154 | 0.999971 | 1 | 0.85548 |
| Ryr2 | E9Q401 | -0.25 | -0.11 | -0.39 | -0.48 | -0.34 | -0.18 | -0.48 | -0.68 | -0.48 | -0.45 | -0.57 | -0.73 | -0.24 | -0.41 | -0.81 | -0.58 | -0.1125 | -0.25 | -0.2025 | 0.568891 | 0.183197 | 0.33303 | 0.999971 | 1 | 0.857921 |
| Ryr2_1 | A0A1Y7VK( | -1.56 | -0.78 | -3.46 | -3.09 | -0.75 | -0.75 | -2.78 | -3.29 | -1.05 | -3.4 | -2.95 | -3.64 | -1.17 | -1.3 | -4.33 | -2.67 | 0.33 | -0.5375 | -0.145 | 0.690471 | 0.497933 | 0.867835 | 0.999971 | 1 | 0.983462 |
| Ryr2_2 | A0A1Y7VJF | -0.18 | 0.04 | 0.21 | 0.55 | -0.12 | -0.26 | 0.51 | 0.41 | -1.21 | -0.19 | -0.35 | -0.27 | 0.45 | -0.34 | -0.57 | 0.46 | -0.02 | -0.66 | -0.155 | 0.940046 | 0.047764 | 0.623002 | 0.999971 | 1 | 0.929759 |
| Ryr3 | A0A140LJK | -0.16 | -0.51 | -0.34 | -0.37 | -0.65 | -1.01 | -0.78 | -0.19 | -0.55 | -0.2 | -0.26 | -0.16 | 0.04 | -0.37 | -0.07 | -0.18 | -0.3125 | 0.0525 | 0.2 | 0.191857 | 0.774227 | 0.299051 | 0.999971 | 1 | 0.847402 |
| S100a1 | Q91V77 | 0.37 | 1.5 | -0.53 | -0.17 | 0.88 | 1.32 | 0.01 | 0.14 | 0.66 | 0.27 | -0.79 | -0.42 | 1.18 | 1.19 | -0.29 | 0.22 | 0.295 | -0.3625 | 0.2825 | 0.560007 | 0.48568 | 0.597795 | 0.999971 | 1 | 0.92377 |
| S100a10 | P08207 | 0.07 | -0.26 | -1.48 | -0.36 | 0.73 | -1 | -1.7 | -1.7 | -0.66 | -0.17 | -1.76 | -1.59 | -0.83 | 0.1 | -1.04 | -1.49 | -0.41 | -0.5375 | -0.3075 | 0.504521 | 0.275955 | 0.497579 | 0.999971 | 1 | 0.902655 |
| S100a11 | P50543 | 0.29 | 0.74 | 0.99 | 1.23 | 2.71 | 1.62 | 0.81 | 0.03 | 1.06 | 0.64 | 0.65 | 2.55 | -0.93 | 1.14 | 1.34 | 0.34 | 0.48 | 0.4125 | -0.34 | 0.399356 | 0.386086 | 0.512222 | 0.999971 | 1 | 0.905815 |
| S100a13 | A0A0A0M( | -1.2 | -1.45 | -1.45 | -0.68 | -1.54 | -1.28 | -1.09 | 0.06 | -1.44 | -1.57 | -0.46 | -0.54 | -1.9 | -1.28 | -0.24 | -0.4 | 0.2325 | 0.1925 | 0.24 | 0.547557 | 0.574912 | 0.561927 | 0.999971 | 1 | 0.916386 |
| S100a16 | Q9D708 | 0.27 | 1.02 | 0.6 | 0.65 | 0.55 | 0.81 | 0.52 | 1.06 | 0.99 | 0.38 | 0.46 | 0.69 | 0.39 | 1.14 | 0.74 | 0.58 | 0.1 | -0.005 | 0.0775 | 0.671228 | 0.983166 | 0.757197 | 0.999971 | 1 | 0.961116 |
| S100a4 | A0A0G2JGI | -0.32 | 0.96 | 0.55 | -0.01 | 1.39 | 0.86 | 0.81 | -0.29 | 1.25 | -0.25 | 1.01 | 1.08 | 0.9 | 0.22 | 0.25 | -0.64 | 0.3975 | 0.4775 | -0.1125 | 0.366229 | 0.279298 | 0.782426 | 0.999971 | 1 | 0.965859 |
| S100a6 | P14069 | 3.91 | 0.58 | 1.11 | -0.17 | 0.96 | -0.23 | 0.97 | 0.37 | 1.06 | 3.09 | 1.53 | 1.9 | 0.73 | 0.1 | 0.8 | 1.25 | -0.84 | 0.5375 | -0.6375 | 0.330762 | 0.551489 | 0.449345 | 0.999971 | 1 | 0.885522 |
| S100a8 | P27005 | 8.28 | 2.91 | 0.23 | 1.59 | -0.89 | -1.05 | 1.59 | -0.02 | -2.5 | 0.23 | -0.61 | -0.84 | -4.52 | -1.1 | -0.16 | -1.01 | -3.345 | -4.1825 | -4.95 | 0.06883 | 0.030808 | 0.020362 | 0.999971 | 1 | 0.635388 |
| S100a9 | P31725 | 4.7 | 1.43 | 1.28 | 3.53 | 0.49 | -1.33 | 3.73 | 1.38 | -1.54 | -0.62 | 1.99 | -0.33 | -3.57 | -0.69 | 0.36 | -0.77 | -1.6675 | -2.86 | -3.9025 | 0.185849 | 0.018745 | 0.004915 | 0.999971 | 1 | 0.582714 |
| S100b | P50114 | -0.77 | -0.11 | 0.46 | 0.83 | -0.64 | -0.15 | -0.22 | 1.34 | -0.77 | -1.16 | -0.79 | 0.65 | -1.21 | -0.21 | 0.7 | 1.01 | -0.02 | -0.62 | -0.03 | 0.968935 | 0.231555 | 0.957257 | 0.999971 | 1 | 0.998578 |
| S1pr1 | O08530 | -0.17 | -0.3 | -0.91 | -0.41 | -0.54 | 0.05 | 0.1 | -0.8 | -0.64 | -0.07 | -0.91 | -1.02 | 0.07 | -0.28 | -1.29 | -0.75 | 0.15 | -0.2125 | -0.115 | 0.604291 | 0.456758 | 0.732204 | 0.999971 | 1 | 0.956473 |
| S1pr5 | Q91X56 | 3.05 | -1.23 | 0.36 | 0.24 | -1.38 | -0.93 | 0.18 | -0.58 | -1.49 | 0.85 | -0.83 | -0.98 | -2.51 | -1.61 | -2.17 | -2.31 | -1.2825 | -1.2175 | -2.755 | 0.157862 | 0.210482 | 0.008178 | 0.999971 | 1 | 0.602121 |
| Saal1 | Q9D2C2-2 | -0.06 | 1.41 | 0.94 | 0.88 | 0.86 | 0.85 | 0.66 | 0.97 | 0.88 | 0.88 | 0.32 | 0.92 | 0.69 | 1.08 | 0.55 | 1.6 | 0.0425 | -0.0425 | 0.1875 | 0.893442 | 0.899596 | 0.620117 | 0.999971 | 1 | 0.929296 |
| Sacm1l | Q9EP69 | -1.1 | -0.64 | -0.79 | -0.92 | -0.22 | -0.46 | -0.43 | -0.81 | -0.08 | -0.95 | -0.74 | -0.9 | 0.15 | -0.6 | -0.72 | -1.02 | 0.3825 | 0.195 | 0.315 | 0.093508 | 0.442525 | 0.282305 | 0.999971 | 1 | 0.844538 |
| Sacs | E9QNY8 | -0.55 | -0.09 | -0.12 | -0.3 | -0.18 | 0.01 | -0.45 | -0.42 | -0.15 | -0.28 | -0.17 | -0.06 | -0.29 | 0.16 | -0.32 | -0.2 | 0.005 | 0.1 | 0.1025 | 0.980561 | 0.58844 | 0.622733 | 0.999971 | 1 | 0.929759 |
| Sae1 | Q9R1T2 | 2.15 | 3.16 | 2.79 | 2.47 | 2.63 | 2.92 | 2.42 | 2.45 | 2.93 | 2.42 | 2.7 | 2.53 | 2.22 | 2.71 | 2.33 | 2.1 | -0.0375 | 0.0025 | -0.3025 | 0.887811 | 0.992422 | 0.284391 | 0.999971 | 1 | 0.844554 |
| Safb | S4R1M2 | 0.3 | -0.1 | 0.74 | 0.61 | -0.17 | 0.11 | 1.54 | 0.6 | 0.04 | 0.09 | 0.5 | 0.44 | -0.25 | -0.08 | 0.65 | 0.54 | 0.1325 | -0.12 | -0.1725 | 0.741448 | 0.630329 | 0.569307 | 0.999971 | 1 | 0.91816 |
| Safb2 | Q80YR5 | 0.34 | 0.26 | -0.25 | -0.27 | 0.32 | 0.5 | -0.4 | -0.08 | 0.03 | 0.42 | -0.74 | -0.62 | 1.17 | 0.94 | -0.09 | 0.31 | 0.065 | -0.2475 | 0.5625 | 0.814079 | 0.44791 | 0.116664 | 0.999971 | 1 | 0.78268 |
| Sall2 | A0A2I3BQ\ | 2.33 | 1.35 | -0.11 | 1.07 | 1.28 | 0.02 | 0.56 | 0.98 | 0.68 | 0.33 | -0.06 | 0.07 | 0.76 | 1.15 | 0.68 | 0.13 | -0.45 | -0.905 | -0.48 | 0.403382 | 0.094113 | 0.354806 | 0.999971 | 1 | 0.868793 |
| Samd12 | Q0VE29 | 0.64 | 0.1 | -1.46 | -1.79 | -0.15 | 0.49 | -1.31 | -0.92 | -0.18 | 0.33 | -1.2 | -2.31 | -0.36 | 0.14 | -3.29 | -1.72 | 0.155 | -0.2125 | -0.68 | 0.811029 | 0.778187 | 0.442545 | 0.999971 | 1 | 0.883386 |
| Samd14 | Q8K070 | 3.04 | -1.44 | -1.88 | -1.47 | -1.23 | -0.24 | -1.35 | -1.39 | -1.44 | 1.64 | -1.82 | -1.58 | -1.9 | -1.18 | -1.39 | -1.42 | -0.615 | -0.3625 | -1.035 | 0.56713 | 0.776111 | 0.337076 | 0.999971 | 1 | 0.858827 |
| Samd4a | Q8CBY1-3 | 0.36 | -0.57 | 0.73 | -1.56 | -1.18 | -0.15 | 0.84 | -3.87 | -0.97 | 0.53 | 1.25 | -0.33 | -1.17 | 0.77 | -1.27 | -4.49 | -0.83 | 0.38 | -1.28 | 0.42192 | 0.560011 | 0.253025 | 0.999971 | 1 | 0.839012 |
| Samd4b | G5E8A7 | -0.48 | -0.39 | -0.73 | -0.28 | -0.31 | -0.32 | -0.6 | 0.02 | -0.53 | -0.14 | -0.04 | -0.11 | -0.16 | 0.07 | -0.23 | -0.02 | 0.1675 | 0.265 | 0.385 | 0.433061 | 0.207526 | 0.066639 | 0.999971 | 1 | 0.733937 |
| Samd5 | A0A0R4J18 | 4 | -0.71 | 0.68 | 0.62 | -0.4 | -0.16 | 2.01 | 1.04 | -0.2 | 2.32 | 0.32 | 1.53 | -2.15 | -1.07 | 0.28 | -0.41 | -0.525 | -0.155 | -1.985 | 0.610869 | 0.881212 | 0.075915 | 0.999971 | 1 | 0.748444 |
| Samd8 | Q9DA37 | 0.82 | -1.02 | -0.37 | -0.16 | -0.88 | -0.85 | -0.36 | -0.1 | -0.19 | 1.05 | 0.05 | 0.1 | 0.13 | -0.58 | 0.2 | 0.75 | -0.365 | 0.435 | 0.3075 | 0.380229 | 0.340022 | 0.492722 | 0.999971 | 1 | 0.900198 |
| Samd9l | E9PX59 | -0.54 | -6.3 | 1.48 | 1.12 | -8.63 | -4.33 | 3.49 | 1.68 | -5.5 | -3.15 | 1.46 | 2.94 | -10.95 | -5.55 | 2.84 | 3.18 | -0.8875 | -0.0025 | -1.56 | 0.762149 | 0.999159 | 0.650706 | 0.999971 | 1 | 0.933842 |
| Samhd1 | E0CXZ5 | 0.35 | 0.76 | -0.12 | 1.23 | -0.18 | -1 | -0.14 | -1.28 | 0.19 | -0.08 | 1.69 | 1.01 | 0.4 | -0.18 | -0.38 | 1.03 | -1.205 | 0.1475 | -0.3375 | 0.012558 | 0.751547 | 0.418673 | 0.999971 | 1 | 0.87788 |
| Samhd1_1 | Q60710 | 0.97 | 1.17 | 1.53 | 1.8 | 1.57 | 1.07 | 1.22 | 1.36 | 1.1 | 0.67 | 1.45 | 1.86 | 1.12 | 1.13 | 1.63 | 1.49 | -0.0625 | -0.0975 | -0.025 | 0.798108 | 0.758402 | 0.921208 | 0.999971 | 1 | 0.992702 |
| Samm50 | Q8BGH2 | -0.65 | 0.32 | 0.46 | -0.01 | -0.02 | 0.45 | 0.27 | 0.19 | 0.11 | -0.13 | 0.18 | 0.28 | 0.24 | 0.38 | 0.14 | 0.21 | 0.1925 | 0.08 | 0.2125 | 0.498838 | 0.773282 | 0.442431 | 0.999971 | 1 | 0.883386 |
| Sanbr | M0QWF3 | -1.22 | -0.27 | -0.22 | -0.64 | -0.28 | -0.7 | 0.2 | -0.37 | -0.33 | -0.57 | -0.34 | -0.25 | -0.18 | -0.37 | -0.51 | -0.51 | 0.3 | 0.215 | 0.195 | 0.336765 | 0.420631 | 0.469838 | 0.999971 | 1 | 0.892481 |
| Sap130 | A0A0R4J06 | 4.77 | 0.57 | 0.96 | 1.19 | 0.7 | 0.45 | 1.35 | 0.64 | 0.22 | 2.8 | 0.32 | 0.61 | -1.77 | 0.81 | 0.01 | 1.33 | -1.0875 | -0.885 | -1.7775 | 0.241635 | 0.401785 | 0.12076 | 0.999971 | 1 | 0.78268 |
| Sap18 | E9Q317 | 1.49 | -0.91 | 0.07 | 0.23 | -0.37 | -0.89 | 0.41 | 0.11 | -0.62 | 0.4 | 0.16 | 0.02 | -0.7 | -0.44 | 0.08 | -0.06 | -0.405 | -0.23 | -0.5 | 0.448226 | 0.648581 | 0.320399 | 0.999971 | 1 | 0.853962 |
| Sap30bp | Q02614 | 1.69 | 2.21 | 1.75 | 2.04 | 2.26 | 1.05 | 0.94 | 1.75 | 1.85 | 2.04 | 1.63 | 1.67 | 2.68 | 1.73 | 2.09 | 1.93 | -0.4225 | -0.125 | 0.185 | 0.224937 | 0.545638 | 0.486453 | 0.999971 | 1 | 0.897675 |
| Sap30l | Q5SQF8 | 1.72 | 1.34 | 1.4 | 1.64 | 1 | 1.61 | 1.32 | 1.5 | 1.21 | 1.83 | 1.68 | 1.47 | 1.05 | 1.36 | 1.41 | 2.43 | -0.1675 | 0.0225 | 0.0375 | 0.436672 | 0.914496 | 0.905911 | 0.999971 | 1 | 0.991343 |
| Sar1a | Q99JZ4 | 2.14 | 0.49 | 0.52 | 1.08 | 0.39 | 0.77 | 0.96 | 0.39 | 0.74 | 1.37 | 0.54 | 0.84 | -0.03 | 0.31 | 0.73 | 0.28 | -0.43 | -0.185 | -0.735 | 0.291681 | 0.650377 | 0.092749 | 0.999971 | 1 | 0.776565 |
| Sar1b | Q9CQC9 | -0.71 | -1.59 | -0.03 | -0.66 | -1.41 | -1.44 | -0.14 | -0.29 | -1.33 | -1.44 | -0.14 | 0.17 | -2.2 | -0.95 | -0.01 | -0.19 | -0.0725 | 0.0625 | -0.09 | 0.870985 | 0.897657 | 0.868899 | 0.999971 | 1 | 0.983462 |
| Saraf | A0A0R4J0D | 0.65 | 0.78 | 0.55 | 0.4 | 0.8 | 1.65 | 1.15 | 0.62 | 1.33 | 0.48 | 0.12 | 0.46 | 1.42 | 0.53 | 0.55 | 0 | 0.46 | 0.0025 | 0.03 | 0.107332 | 0.992946 | 0.923008 | 0.999971 | 1 | 0.993264 |
| Sardh | Q99LB7 | -0.39 | -0.61 | -0.59 | -0.79 | -1.26 | -0.85 | -1.17 | -0.58 | -1.22 | -0.76 | -0.74 | -0.68 | -1.02 | -0.68 | -0.66 | -0.43 | -0.37 | -0.255 | -0.1025 | 0.119391 | 0.226962 | 0.61668 | 0.999971 | 1 | 0.928229 |
| Sarm1 | Q6PDS3 | -1.02 | -1.05 | -0.61 | -0.62 | -0.91 | -0.73 | -0.67 | -0.41 | -0.75 | -0.82 | -0.18 | -0.4 | -0.47 | -0.76 | -0.27 | 0.04 | 0.145 | 0.2875 | 0.46 | 0.495972 | 0.230407 | 0.083119 | 0.999971 | 1 | 0.755254 |
| Sarnp | Q9D1J3 | 1.75 | 1.04 | 1.73 | 1.63 | 1.21 | 1.06 | 0.85 | 1.46 | 1.24 | 1.32 | 0.8 | 1.28 | 0.51 | 0.76 | 1.05 | 0.93 | -0.3925 | -0.3775 | -0.725 | 0.132508 | 0.139606 | 0.013432 | 0.999971 | 1 | 0.613095 |
| Sars | Q8C483 | 0.36 | 0.78 | 1.08 | 0.91 | 0.69 | 0.69 | 0.88 | 0.96 | 0.75 | 0.46 | 1.13 | 0.79 | 0.61 | 0.75 | 1.07 | 1.06 | 0.0225 | -1.18E-16 | 0.09 | 0.916677 | 1 | 0.697378 | 0.999971 | 1 | 0.94737 |
| Sars2 | Q9JJL8 | -0.65 | -0.99 | 0.15 | -0.24 | -0.71 | -1.44 | 0.41 | -0.6 | -0.67 | -0.84 | -0.02 | 0.63 | -0.99 | -1.17 | -0.36 | -0.81 | -0.1525 | 0.2075 | -0.4 | 0.722958 | 0.606364 | 0.217076 | 0.999971 | 1 | 0.826893 |
| Sart1 | Q9Z315 | 0.24 | 1.04 | 1.19 | 0.96 | 0.67 | 0.52 | 0.61 | 0.64 | 0.55 | -0.16 | 0.66 | 0.26 | 0.45 | 0.4 | 0.67 | 0.58 | -0.2475 | -0.53 | -0.3325 | 0.325575 | 0.095489 | 0.204864 | 0.999971 | 1 | 0.81617 |
| Sart3 | Q9JLI8 | 1.9 | 2.25 | 2.46 | 2.49 | 1.98 | 2.15 | 1.93 | 2 | 1.94 | 1.99 | 2.19 | 2.15 | 1.52 | 2.01 | 2.23 | 2.27 | -0.26 | -0.2075 | -0.2675 | 0.21802 | 0.317963 | 0.298635 | 0.999971 | 1 | 0.847402 |
| Sash1 | F8VQK5 | -0.38 | -0.27 | -0.26 | -0.19 | -0.47 | -0.68 | -0.25 | 0.02 | -0.53 | -0.26 | -0.05 | 0.01 | -0.32 | -0.75 | -0.09 | -0.19 | -0.07 | 0.0675 | -0.0625 | 0.736512 | 0.723214 | 0.761594 | 0.999971 | 1 | 0.961247 |
| Sat2 | Q6P8J2 | 1.16 | 0.72 | 0.78 | 0.18 | 0.91 | 1.24 | 1.08 | 1.58 | 0.51 | 1.16 | 0.73 | 1.23 | 1.05 | 1.64 | 1.12 | 0.62 | 0.4925 | 0.1975 | 0.3975 | 0.092743 | 0.486541 | 0.207502 | 0.999971 | 1 | 0.820632 |
| Satb1 | E9PVB7 | 1.41 | 2.33 | 2.22 | 1.78 | 2.16 | 2.77 | 2.32 | 1.82 | 1.67 | 1.68 | 2.14 | 1.81 | 1.42 | 2.34 | 1.63 | 2.18 | 0.3325 | -0.11 | -0.0425 | 0.282237 | 0.673612 | 0.891035 | 0.999971 | 1 | 0.988184 |
| Satb2 | H3BKH3 | 1.37 | 2.07 | 3.61 | 3.92 | 2.13 | 2.08 | 4.21 | 2.3 | 1.53 | 1.45 | 2.7 | 2.06 | 1.1 | 1.91 | 1.72 | 1.96 | -0.0625 | -0.8075 | -1.07 | 0.930701 | 0.214472 | 0.096157 | 0.999971 | 1 | 0.78268 |
| Saysd1 | Q8K190 | -1.31 | -0.8 | -0.42 | 0.22 | -0.91 | -0.11 | 0.11 | 0.72 | -0.8 | 0.18 | -1.44 | 1.73 | 0.4 | -1.33 | -0.84 | -0.21 | 0.53 | 0.495 | 0.0825 | 0.247289 | 0.481892 | 0.858934 | 0.999971 | 1 | 0.982702 |
| Sbds | P70122 | 1.05 | 1.07 | 0.92 | 0.8 | 0.86 | 1.3 | 0.73 | 0.82 | 0.93 | 1.18 | 1.13 | 1.29 | 0.32 | 1.16 | 0.72 | 0.68 | -0.0325 | 0.1725 | -0.24 | 0.870926 | 0.339314 | 0.302326 | 0.999971 | 1 | 0.847402 |
| Sbf1 | Q6ZPE2 | -0.55 | -0.31 | -1 | -1.03 | -0.49 | -0.35 | -1.12 | -1.08 | -0.54 | -0.72 | -1.02 | -1.04 | -0.26 | -0.24 | -0.87 | -0.72 | -0.0375 | -0.1075 | 0.2 | 0.893609 | 0.659447 | 0.452469 | 0.999971 | 1 | 0.887217 |
| Sbf2 | E9Q0D4 | -0.52 | -0.64 | -0.55 | -0.5 | -0.53 | -0.69 | -0.68 | -0.52 | -0.29 | -0.64 | -0.52 | -0.25 | -0.68 | -0.96 | -0.67 | -0.51 | -0.0525 | 0.1275 | -0.1525 | 0.749594 | 0.474323 | 0.405892 | 0.999971 | 1 | 0.874911 |
| Sbno1 | F7CVW0 | 1.31 | 2.05 | 1.62 | 1.94 | 1.49 | 1.82 | 0.9 | 1.58 | 1.27 | 1.3 | 1.27 | 1.57 | 1.47 | 1.79 | 1.71 | 1.87 | -0.2825 | -0.3775 | -0.02 | 0.318441 | 0.116197 | 0.930114 | 0.999971 | 1 | 0.994744 |
| Sc5d | O88822 | 0.61 | 0.15 | -0.52 | -0.77 | 0.08 | -0.01 | 0 | -0.48 | 0.46 | 1.2 | -0.02 | 0.08 | 1.24 | -0.04 | -0.56 | -0.34 | 0.03 | 0.5625 | 0.2075 | 0.92896 | 0.185685 | 0.665242 | 0.999971 | 1 | 0.93697 |
| Scaf1 | Q5U4C3 | 0.15 | -0.32 | 0.57 | 0.55 | -0.12 | -0.62 | 0.74 | -0.07 | -0.41 | -0.05 | 0.03 | 0.22 | -0.35 | -0.39 | 0.04 | -0.01 | -0.255 | -0.29 | -0.415 | 0.467783 | 0.293498 | 0.138748 | 0.999971 | 1 | 0.784948 |
| Scaf11 | A0A2I3BQl | 2.42 | -0.2 | 0.07 | 0.89 | -0.52 | 0 | 0.69 | 0.24 | -0.14 | 1.11 | -0.8 | -0.55 | -1.4 | -0.58 | -0.2 | -0.75 | -0.6925 | -0.89 | -1.5275 | 0.256189 | 0.202454 | 0.026941 | 0.999971 | 1 | 0.678474 |
| Scaf4 | Q7TSH6 | 1.82 | 2.27 | 3.17 | 1.92 | 2.16 | 2.04 | 5.39 | 2.72 | 2.44 | 2.25 | 4.28 | 4.64 | 0.85 | 2.43 | 2.71 | 1.72 | 0.7825 | 1.1075 | -0.3675 | 0.317539 | 0.106722 | 0.453329 | 0.999971 | 1 | 0.887217 |
| Scaf8 | Q6DID3 | 3.86 | 2.91 | 1.75 | 1.78 | 2.84 | 2.6 | 1.84 | 2.08 | 2.6 | 3.44 | 2.12 | 2.33 | 3.17 | 2.88 | 1.85 | 1.96 | -0.235 | 0.0475 | -0.11 | 0.649357 | 0.92982 | 0.843062 | 0.999971 | 1 | 0.979814 |
| Scai | Q8C8N2 | 0.02 | -0.29 | -0.48 | -0.61 | -0.46 | -0.03 | -0.66 | -0.88 | -0.49 | -0.22 | -0.4 | -0.7 | -0.16 | -0.56 | -0.84 | -0.33 | -0.1675 | -0.1125 | -0.1325 | 0.513565 | 0.601313 | 0.578834 | 0.999971 | 1 | 0.919236 |
| Scai_1 | B0R0E3 | -2.08 | -8.25 | -6.25 | -6.29 | -6.5 | -6.09 | -3.25 | -6.43 | -8.55 | -3.47 | -5.21 | -8.8 | -12.27 | -6.53 | -7.91 | -6.03 | 0.15 | -0.79 | -2.4675 | 0.911037 | 0.632554 | 0.172859 | 0.999971 | 1 | 0.80626 |
| Scamp1 | Q8K021 | -0.18 | -1.16 | -0.66 | -0.73 | -0.9 | -0.68 | -0.47 | -0.69 | -0.8 | -0.35 | -0.78 | -0.46 | -0.91 | -0.66 | -0.59 | -0.87 | -0.0025 | 0.085 | -0.075 | 0.991947 | 0.738956 | 0.761024 | 0.999971 | 1 | 0.961116 |
| Scamp2 | Q9ERN0 | 1.01 | -1.94 | -1.15 | 0.03 | -2.07 | -1.44 | -0.66 | -0.36 | -2.49 | -0.08 | -0.8 | -0.22 | -3.36 | -3.04 | -0.44 | -0.41 | -0.62 | -0.385 | -1.3 | 0.376013 | 0.620791 | 0.184254 | 0.999971 | 1 | 0.809025 |
| Scamp3 | E9Q855 | -1.93 | 1.3 | 0.41 | -0.39 | 1.52 | 0.96 | -0.35 | 0.24 | 2.22 | -0.5 | 0.07 | 0.19 | 3.3 | -0.13 | 0.74 | 1.09 | 0.745 | 0.6475 | 1.4025 | 0.3163 | 0.438523 | 0.144666 | 0.999971 | 1 | 0.790933 |
| Scamp3_1 | Q3UXS0 | -0.24 | -0.79 | -0.41 | -0.27 | -0.7 | -0.31 | -0.18 | -0.38 | -0.7 | -0.4 | -0.48 | -0.57 | -0.83 | -0.74 | -0.63 | -0.48 | 0.035 | -0.11 | -0.2425 | 0.870692 | 0.58069 | 0.253194 | 0.999971 | 1 | 0.839012 |
| Scamp4 | Q9JKV5 | 2.48 | -0.29 | 0.29 | -0.31 | -1.08 | 1.24 | 1.16 | -0.22 | -0.66 | 1.62 | -0.11 | -0.11 | -1.2 | -1.77 | -0.46 | -0.6 | -0.2675 | -0.3575 | -1.55 | 0.732377 | 0.635398 | 0.04077 | 0.999971 | 1 | 0.700838 |
| Scamp5 | Q9JKD3 | -2.69 | -1.45 | -0.73 | -1.15 | -0.91 | -1.88 | -0.41 | -0.74 | -0.68 | -2.72 | -0.34 | -0.25 | -0.28 | -2.07 | -0.83 | -0.78 | 0.52 | 0.5075 | 0.515 | 0.304052 | 0.447242 | 0.340484 | 0.999971 | 1 | 0.860574 |
| Scap | Q6GQT6 | 0.78 | 0.02 | -0.01 | -0.25 | -0.41 | -0.34 | 0.2 | 0.18 | -0.12 | 0.09 | -0.12 | 0.02 | -0.29 | -0.15 | -0.7 | 0.14 | -0.2275 | -0.1675 | -0.385 | 0.439577 | 0.514353 | 0.21341 | 0.999971 | 1 | 0.82451 |
| Scaper | B2RXS3 | 0.75 | 0.77 | 0.94 | 1.21 | 1.04 | 0.83 | 1.13 | 1.38 | 1.09 | 0.97 | 1.24 | 1.15 | 0.69 | 1.08 | 1.21 | 1.62 | 0.1775 | 0.195 | 0.2325 | 0.403392 | 0.310202 | 0.362224 | 0.999971 | 1 | 0.869526 |
| Scara3 | Q8C850 | -0.94 | -1.58 | -0.11 | 0.05 | -1.33 | -1.48 | 0.59 | 0.46 | -0.69 | 0.21 | 0.38 | -0.11 | -0.93 | -0.67 | 0.01 | 0.88 | 0.205 | 0.5925 | 0.4675 | 0.73955 | 0.187242 | 0.374617 | 0.999971 | 1 | 0.870318 |
| Scarb2 | O35114 | 0.42 | 0.4 | 0.29 | 0.19 | 0.25 | 0.87 | 0.65 | -0.06 | 0.61 | 1.35 | -0.12 | -0.27 | 0.04 | 0.32 | -0.15 | -0.46 | 0.1025 | 0.0675 | -0.3875 | 0.676299 | 0.853947 | 0.103364 | 0.999971 | 1 | 0.78268 |
| Sccpdh | Q8R127 | -0.2 | -0.43 | 0.06 | 0.05 | -0.27 | -0.16 | -0.17 | 0.55 | -0.1 | -0.1 | -0.23 | -0.02 | -0.11 | -0.17 | 0.61 | 0.01 | 0.1175 | 0.0175 | 0.215 | 0.640481 | 0.92581 | 0.390779 | 0.999971 | 1 | 0.871816 |
| Scd2 | A0A494BA\ | -0.09 | 2.98 | 1.07 | 0.95 | 2.01 | 2.45 | 0.42 | 1.44 | 1.7 | 0.33 | 0.78 | 0.51 | 2.44 | 1.9 | 0.93 | 1.37 | 0.3525 | -0.3975 | 0.4325 | 0.617379 | 0.543566 | 0.513122 | 0.999971 | 1 | 0.905815 |
| Scfd1 | Q8BRF7 | -0.76 | -0.07 | -0.56 | -1.51 | 0.19 | -0.59 | -0.09 | -1.37 | 0.35 | -0.93 | -0.11 | 0.13 | 0.26 | -1.04 | -0.96 | -1.64 | 0.26 | 0.585 | -0.12 | 0.549041 | 0.16311 | 0.7972 | 0.999971 | 1 | 0.970479 |
| Scfd2 | Q8BTY8 | -1.23 | -0.5 | -1.07 | -1.15 | -0.26 | -0.55 | -0.59 | -1.06 | -0.43 | -1.29 | -0.43 | -0.11 | -0.14 | -0.24 | -0.9 | -0.94 | 0.3725 | 0.4225 | 0.4325 | 0.174007 | 0.193687 | 0.153466 | 0.999971 | 1 | 0.795578 |
| Scg2 | Q03517 | -0.56 | -1.42 | -0.96 | -0.94 | -0.78 | -0.91 | -0.84 | -0.53 | -0.61 | -1.08 | -1 | -0.42 | -0.35 | -0.08 | 0.44 | -0.21 | 0.205 | 0.1925 | 0.92 | 0.386983 | 0.463476 | 0.007135 | 0.999971 | 1 | 0.582714 |
| Scg3 | P47867 | -1.68 | -2.18 | -1.3 | -1.93 | -1.73 | -1.7 | -0.17 | -1.84 | -1.49 | -1.78 | -0.6 | -0.91 | -1.93 | -1.99 | -1.11 | -1.66 | 0.4125 | 0.5775 | 0.1 | 0.33691 | 0.1055 | 0.729304 | 0.999971 | 1 | 0.956041 |
| Scg5 | P12961 | -1.44 | -0.84 | -1.25 | -0.8 | -0.17 | -0.7 | -1.06 | -0.23 | -0.25 | -2 | -0.71 | -0.61 | -0.11 | -0.01 | -0.13 | -0.37 | 0.5425 | 0.19 | 0.9275 | 0.07776 | 0.634046 | 0.00223 | 0.999971 | 1 | 0.508206 |
| Scgn | Q91WD9 | 8.78 | -4.78 | -2.98 | -1.9 | -2.2 | 2.54 | -1.8 | -2.4 | 2.78 | 6.8 | -4.99 | -3.5 | -8.47 | -3.6 | -2.2 | -2.05 | -0.745 | 0.4925 | -3.86 | 0.796878 | 0.892893 | 0.221855 | 0.999971 | 1 | 0.827753 |
| Schip1 | A0A0A6YX( | 0.33 | 1.2 | 0.26 | 0.32 | 2.02 | 0.93 | -0.65 | -0.43 | 0.72 | 0.38 | 0.67 | 0.52 | 2.24 | 0.23 | 1.4 | 0.28 | -0.06 | 0.045 | 0.51 | 0.920941 | 0.862261 | 0.318527 | 0.999971 | 1 | 0.853893 |
| Scly | A0A0R4J06 | 1.03 | 2.83 | 2.23 | 1.04 | 2.06 | 2.47 | 2.44 | 1.64 | 2.2 | 1.7 | 2.52 | 2.41 | 2.14 | 2.66 | 1.52 | 0.58 | 0.37 | 0.425 | -0.0575 | 0.428016 | 0.363476 | 0.92103 | 0.999971 | 1 | 0.992702 |
| Scn1a | A2APX8-3 | -0.83 | -0.63 | -0.59 | -1.12 | -0.87 | -0.59 | -1.17 | -1.38 | -1.15 | -1.12 | -1.13 | -1.19 | -0.6 | -0.98 | -1.37 | -1.23 | -0.21 | -0.355 | -0.2525 | 0.396895 | 0.08471 | 0.310307 | 0.999971 | 1 | 0.847402 |
| Scn1b | P97952 | -0.58 | 0.69 | 0.51 | 0.02 | 0.49 | 0.43 | 0.29 | -0.05 | 0.67 | -0.05 | 0.35 | -0.33 | 1.61 | 0.49 | 0.33 | 0.6 | 0.13 | -1.18E-16 | 0.5975 | 0.679403 | 1 | 0.152143 | 0.999971 | 1 | 0.795578 |
| Scn2a | A0A5H1ZRI | -0.8 | -0.82 | -0.66 | -0.71 | -0.89 | -0.86 | -0.95 | -0.76 | -0.7 | -0.95 | -0.8 | -0.77 | -0.72 | -0.96 | -0.71 | -0.57 | -0.1175 | -0.0575 | 0.0075 | 0.479954 | 0.724956 | 0.965961 | 0.999971 | 1 | 0.998721 |
| Scn2b | Q56A07 | 1.27 | -2.98 | -0.43 | 0.21 | -3.05 | -1.92 | -0.03 | -0.63 | -2.78 | -0.52 | -0.72 | -0.41 | -4.62 | -3.11 | -0.58 | -0.91 | -0.925 | -0.625 | -1.8225 | 0.369584 | 0.518989 | 0.145856 | 0.999971 | 1 | 0.791767 |
| Scn3a | A2ASI5 | -1.51 | -1.97 | -1.47 | -1.79 | -2.28 | -2.16 | -1.32 | -1.04 | -2.2 | -1.68 | -1.47 | -1.13 | -2.2 | -1.79 | -1.15 | -1.54 | -0.015 | 0.065 | 0.015 | 0.963541 | 0.810596 | 0.955718 | 0.999971 | 1 | 0.998056 |
| Scn3b | H3BJR6 | -1.45 | -1.46 | -2.31 | -1.6 | -1.45 | -1.74 | -2.02 | -1.36 | -0.96 | -1.16 | -1.98 | -1.8 | -0.69 | -1.48 | -1.97 | -1.9 | 0.0625 | 0.23 | 0.195 | 0.818373 | 0.480663 | 0.583151 | 0.999971 | 1 | 0.920364 |
| Scn4b | Q7M729 | 3.89 | -0.55 | -0.83 | -0.11 | -0.98 | -0.39 | -1.83 | 0.6 | -0.5 | 1.59 | -1.09 | -0.77 | -1.5 | -0.57 | 0.58 | -0.51 | -1.25 | -0.7925 | -1.1 | 0.267811 | 0.490036 | 0.313955 | 0.999971 | 1 | 0.850589 |
| Scn7a | B1AYL1 | -0.46 | -1.13 | -1.45 | -1.37 | -0.67 | -0.81 | -1.09 | -1.36 | -0.69 | -0.07 | -1.07 | -0.72 | 0.16 | -0.68 | -0.88 | -0.54 | 0.12 | 0.465 | 0.6175 | 0.67553 | 0.160162 | 0.081689 | 0.999971 | 1 | 0.755254 |
| Scn8a | A0A0J9YU\ | -0.97 | -0.64 | -0.77 | -1.08 | -0.88 | -0.96 | -0.86 | -0.81 | -1.43 | -1.25 | -0.9 | -0.46 | -1.13 | -1.22 | -1.04 | -1.34 | -0.0125 | -0.145 | -0.3175 | 0.944924 | 0.57745 | 0.118961 | 0.999971 | 1 | 0.78268 |
| Scnm1 | A0A0R4J26 | 2.93 | 2.46 | 1.87 | 1.8 | 2.92 | 2.57 | 1.61 | 1.71 | 2.38 | 2.64 | 1.77 | 1.61 | 3.06 | 3.1 | 1.97 | 2.31 | -0.0625 | -0.165 | 0.345 | 0.875642 | 0.64385 | 0.369737 | 0.999971 | 1 | 0.869526 |
| Sco1 | Q5SUD5 | -2.2 | -3.08 | -2.03 | -2.3 | -2.67 | -2.69 | -1.82 | -1.99 | -2.83 | -2.74 | -1.9 | -2.26 | -3.89 | -2.7 | -2.04 | -1.76 | 0.11 | -0.03 | -0.195 | 0.73587 | 0.925249 | 0.69254 | 0.999971 | 1 | 0.947174 |
| Sco2 | Q8VCL2 | -1.11 | -0.55 | 0.57 | 0.12 | -0.2 | -0.5 | -0.12 | 0.07 | -0.45 | -0.7 | 0.26 | -0.45 | 0.28 | -0.24 | 0.7 | 1.04 | 0.055 | -0.0925 | 0.6875 | 0.883466 | 0.819568 | 0.141402 | 0.999971 | 1 | 0.784967 |

| | | | | | | | | | | | | | | | | | | | | | | | | | | |
|---|---|---|---|---|---|---|---|---|---|---|---|---|---|---|---|---|---|---|---|---|---|---|---|---|---|---|
| Scoc | Q78YZ6-2 | -3.56 | -5.55 | -1.5 | -1.1 | -4.83 | -4.85 | -1.21 | -1.09 | -4.75 | -4.61 | -1.49 | -0.78 | -7.37 | -4.53 | -1.29 | -1.32 | -0.0675 | 0.02 | -0.7 | 0.958969 | 0.98775 | 0.660237 | 0.999971 | 1 | 0.935541 |
| Scp2 | P32020 | -0.93 | -0.84 | -0.92 | -1.11 | -0.84 | -0.53 | -0.85 | -0.48 | -0.56 | -0.41 | -0.91 | -0.95 | -0.12 | -0.34 | -0.54 | -0.29 | 0.275 | 0.2425 | 0.6275 | 0.164132 | 0.242488 | 0.006999 | 0.999971 | 1 | 0.582714 |
| Scpep1 | Q920A5 | 0.59 | 0.65 | 0.84 | 0.4 | 0.9 | 0.97 | 0.83 | 1.12 | 0.77 | 1.04 | 0.91 | 0.85 | 0.61 | 0.7 | 0.37 | 0.61 | 0.335 | 0.2725 | -0.0475 | 0.095759 | 0.15491 | 0.799764 | 0.999971 | 1 | 0.970626 |
| Scrg1 | O88745 | -0.46 | 0.97 | 1.12 | 1.24 | 1.12 | 0.98 | 0.38 | 0.86 | 0.42 | 0.12 | 0.92 | 1.22 | 1.19 | 0.96 | 1.12 | 1.36 | 0.1175 | -0.0475 | 0.44 | 0.773413 | 0.91422 | 0.276896 | 0.999971 | 1 | 0.843812 |
| Scrib | Q80U72 | -0.7 | -1.68 | -1.43 | -1.29 | -1.46 | -1.91 | -1.45 | -1.24 | -1.66 | -1.38 | -1.37 | -1.04 | -1.39 | -1.65 | -1.44 | -1.21 | -0.24 | -0.0875 | -0.1475 | 0.38578 | 0.740665 | 0.56499 | 0.999971 | 1 | 0.917442 |
| Scrn1 | Q9CZC8 | 0.3 | 0.15 | 0.31 | 0.06 | 0.23 | 0.04 | 0.47 | -0.12 | 0.48 | 0.32 | 0.13 | -0.28 | -0.17 | 0.14 | 0.01 | -0.28 | -0.05 | -0.0425 | -0.28 | 0.802291 | 0.844783 | 0.156443 | 0.999971 | 1 | 0.796008 |
| Scrn2 | Q8VCA8 | -1.02 | 0.87 | 1.81 | 1.02 | 0.3 | 0.91 | 2.22 | 0.63 | -0.02 | -1.02 | 0.21 | -1.24 | -0.51 | -0.37 | 0.98 | -0.29 | 0.345 | -1.1875 | -0.7175 | 0.606036 | 0.091114 | 0.273332 | 0.999971 | 1 | 0.843794 |
| Scrn3 | Q3TMH2 | -0.7 | -0.04 | 0.13 | -0.01 | 0.02 | 0.02 | 0.08 | 0.3 | -0.16 | -1.34 | -0.41 | 0.06 | -0.82 | -0.59 | -0.13 | -0.39 | 0.26 | -0.3075 | -0.3275 | 0.282267 | 0.394126 | 0.229761 | 0.999971 | 1 | 0.828687 |
| Scrt1 | Q99M85 | 0.33 | 1.11 | 0.24 | 0.38 | -0.08 | 0.2 | 1.12 | 0.47 | 0.29 | 0.2 | 0.33 | 0.48 | -1.7 | 0.47 | 0.72 | 0.89 | -0.0875 | -0.19 | -0.42 | 0.78869 | 0.435801 | 0.478255 | 0.999971 | 1 | 0.894438 |
| Scube1 | A2A442 | 1.33 | 0.46 | -0.23 | -0.08 | 0.53 | 0.59 | 0.25 | -0.25 | 0.32 | 0.87 | -0.28 | 0.57 | 0.44 | -0.1 | 0.57 | -0.01 | -0.09 | 0 | -0.145 | 0.815759 | 1 | 0.701704 | 0.999971 | 1 | 0.949019 |
| Scyl1 | R4H4V1 | -0.02 | -0.34 | 0.01 | -0.32 | -0.43 | 0.1 | 0.64 | 0.04 | -0.28 | -0.35 | 0.28 | 0.43 | -0.05 | -0.01 | 0.4 | -0.27 | 0.255 | 0.1875 | 0.185 | 0.342499 | 0.45275 | 0.40199 | 0.999971 | 1 | 0.871904 |
| Scyl2 | Q8CFE4 | 0.02 | -0.53 | -0.36 | -0.78 | -0.55 | -0.42 | -0.14 | -0.59 | -0.02 | 0.39 | -0.11 | 0.13 | -0.24 | -0.39 | -0.21 | -0.58 | -0.0125 | 0.51 | 0.0575 | 0.957151 | 0.053645 | 0.801297 | 0.999971 | 1 | 0.971405 |
| Scyl3 | Q9DBQ7-3 | -0.13 | -0.08 | 0.78 | 0.7 | 0.55 | -0.89 | 0.41 | 0.68 | -0.12 | -0.12 | 0.62 | 0.59 | 0.05 | -0.34 | 0.72 | 0.95 | -0.13 | -0.075 | 0.0275 | 0.755112 | 0.816516 | 0.941332 | 0.999971 | 1 | 0.996526 |
| Sdad1 | A0A0J9YV2 | 0.92 | 0.56 | 0.63 | 0.35 | 0 | 0.8 | 0.19 | 0.82 | 0.04 | 0.78 | -0.16 | 1.59 | 0.16 | 0.56 | 1.29 | 1.37 | -0.1625 | -0.0525 | 0.23 | 0.541095 | 0.895023 | 0.476899 | 0.999971 | 1 | 0.894438 |
| Sdc2 | P43407 | -1.33 | -0.7 | 0.38 | 1 | -0.43 | -0.81 | 0.79 | -0.18 | 0.08 | -0.26 | 0.23 | 0.79 | 0.4 | -0.85 | 0.55 | -0.21 | 0.005 | 0.3725 | 0.135 | 0.993026 | 0.485953 | 0.810946 | 0.999971 | 1 | 0.973864 |
| Sdc3 | Q64519 | 0.72 | -1.74 | -0.15 | -0.01 | -1.37 | -0.65 | 0.21 | -0.41 | -0.43 | 0.83 | -0.12 | 0.24 | -0.44 | -1.14 | -0.19 | 0.2 | -0.26 | 0.425 | -0.0975 | 0.645213 | 0.44033 | 0.857596 | 0.999971 | 1 | 0.982702 |
| Sdc4 | O35988 | 0.15 | -1.06 | -1.17 | -0.58 | -1.06 | -0.56 | -1.32 | -0.29 | -1.31 | 0.49 | -1.31 | -1.14 | -1.35 | -0.98 | -1.37 | -0.42 | -0.1425 | -0.1525 | -0.365 | 0.700679 | 0.758551 | 0.331127 | 0.999971 | 1 | 0.857107 |
| Sdcbp | Q3TMX0 | -1.06 | 0.52 | -0.79 | -0.86 | 0.92 | 1.17 | -0.8 | -0.64 | 0.73 | -0.75 | -0.6 | -0.24 | 1.07 | 0.92 | -0.94 | -0.51 | 0.71 | 0.3325 | 0.6825 | 0.236606 | 0.47764 | 0.250431 | 0.999971 | 1 | 0.839012 |
| Sde2 | Q8K1J5 | 2.24 | 1.38 | 1.55 | 1.89 | 1.54 | 1.21 | 1.55 | 1.59 | 0.91 | 2.2 | 1.25 | 0.82 | 0.14 | 1.17 | 1.31 | 1.17 | -0.2925 | -0.47 | -0.8175 | 0.24616 | 0.214569 | 0.033634 | 0.999971 | 1 | 0.689803 |
| Sdf2 | A0A0R4IZV | -1.42 | 0.37 | 0.7 | -0.64 | 1.39 | 0.47 | 2.77 | 0.04 | 3.18 | -0.9 | 1.59 | 1.9 | 2.6 | -1.13 | 0.47 | -0.59 | 1.415 | 1.69 | 0.585 | 0.069266 | 0.083649 | 0.501112 | 0.999971 | 1 | 0.904273 |
| Sdf2l1 | Q9ESP1 | -0.3 | -0.13 | 1.07 | -0.01 | -0.1 | 0.42 | 2.65 | -0.19 | 0.24 | -0.81 | 0.85 | 0.44 | -0.26 | -1.09 | 0.41 | -1.17 | 0.5375 | 0.0225 | -0.685 | 0.427937 | 0.959551 | 0.15896 | 0.999971 | 1 | 0.798217 |
| Sdf4 | Q61112 | 0.62 | 1.46 | 2.27 | 1.87 | 1.31 | 1.13 | 2.19 | 2.06 | 1.53 | 1.13 | 2.57 | 2.55 | 1.54 | 1.05 | 1.92 | 2.1 | 0.1175 | 0.39 | 0.0975 | 0.779272 | 0.421134 | 0.809462 | 0.999971 | 1 | 0.973762 |
| Sdha | Q8K2B3 | -0.46 | -0.12 | -0.2 | -0.6 | -0.41 | -0.23 | -0.24 | -0.18 | -0.28 | -0.5 | -0.32 | -1.84 | 0.1 | -0.24 | -0.17 | -0.57 | 0.08 | -0.39 | 0.125 | 0.675091 | 0.315171 | 0.575985 | 0.999971 | 1 | 0.918194 |
| Sdhaf1 | Q3U276 | 1.4 | -1.5 | -0.18 | -0.32 | -0.52 | -1.33 | -0.55 | 0.37 | -0.24 | 0.54 | -1.12 | -0.2 | -1.07 | 0.54 | 0.1 | 0.1 | -0.3575 | -0.105 | 0.0675 | 0.572094 | 0.867091 | 0.914239 | 0.999971 | 1 | 0.992633 |
| Sdhaf2 | A0A494B9I | 2.03 | -0.69 | 0.09 | -0.01 | -0.42 | -0.64 | -0.02 | -0.25 | -0.96 | 0.6 | -0.49 | -0.25 | -0.34 | -0.22 | -0.01 | 0.19 | -0.6875 | -0.63 | -0.45 | 0.232403 | 0.320021 | 0.421486 | 0.999971 | 1 | 0.87788 |
| Sdhaf3 | Q8BQU3 | 1.09 | -0.54 | 0.09 | -0.37 | -0.38 | 0.52 | 0.88 | 0.51 | -0.25 | 0.61 | -0.47 | -1.64 | 0.74 | 0.82 | 0.87 | 0.65 | 0.315 | -0.505 | 0.7025 | 0.469634 | 0.367309 | 0.078637 | 0.999971 | 1 | 0.754157 |
| Sdhaf4 | Q8BTE0 | 0.09 | -0.21 | 0.36 | 0.05 | -0.25 | -0.27 | -0.42 | 1.07 | -0.87 | 0 | 0.14 | -0.14 | 0.19 | 0.18 | 0.42 | 0.99 | -0.04 | -0.29 | 0.3725 | 0.9111 | 0.300521 | 0.163364 | 0.999971 | 1 | 0.800215 |
| Sdhb | Q9CQA3 | -2.01 | -2.07 | -1.61 | -1.74 | -1.7 | -2.47 | -0.86 | -1.43 | -1.8 | -2.3 | -1.16 | -1.29 | -1.87 | -2.1 | -1.45 | -1.8 | 0.2425 | 0.22 | 0.0525 | 0.490244 | 0.459428 | 0.810737 | 0.999971 | 1 | 0.973864 |
| Sdhc | Q9CZB0 | 0.31 | 2.27 | 1.26 | -0.29 | 2.1 | 2.03 | 1.94 | 0.46 | 2.47 | 0.13 | 2.02 | 1.65 | 3.34 | 1.92 | 1.21 | -0.09 | 0.745 | 0.68 | 0.7075 | 0.250948 | 0.337569 | 0.397077 | 0.999971 | 1 | 0.871816 |
| Sdhd | Q9CXV1 | 0.05 | -1.39 | -0.26 | -0.26 | -2.04 | -0.98 | -0.11 | 0.12 | -1.4 | -0.61 | 0.25 | -0.09 | -3.06 | -1.18 | 0.24 | 0.41 | -0.2875 | 0.0025 | -0.4325 | 0.595482 | 0.995576 | 0.581939 | 0.999971 | 1 | 0.920364 |
| Sdk1 | Q3UH53 | 4.88 | -0.59 | -2.14 | -1.68 | -0.01 | -0.3 | -1.92 | -2.65 | -0.9 | 4.33 | -2.36 | -1.47 | -0.59 | -0.53 | -2.29 | -2.38 | -1.3375 | -0.2175 | -1.565 | 0.396126 | 0.912146 | 0.314886 | 0.999971 | 1 | 0.851847 |
| Sdk2 | Q6V4S5 | -1.13 | -1.51 | -1.13 | -1.09 | -1.53 | -1.34 | -1.19 | -0.66 | -1.25 | -0.99 | -0.71 | -0.91 | -1.15 | -1.5 | -0.94 | -0.56 | 0.035 | 0.25 | 0.1775 | 0.885292 | 0.23564 | 0.48291 | 0.999971 | 1 | 0.896345 |
| Sdr39u1 | Q5M8N4 | -1.76 | -0.82 | -1.31 | -1.29 | -0.69 | -1.1 | -0.77 | -1.16 | -0.44 | -1.85 | -1.11 | -0.53 | 0.01 | -1.39 | -1.09 | -1.51 | 0.365 | 0.3125 | 0.3 | 0.172091 | 0.405409 | 0.44134 | 0.999971 | 1 | 0.883386 |
| Sdsl | Q8R238 | 0.53 | -0.07 | -0.38 | -0.08 | -0.05 | 0.45 | -0.44 | 0.04 | -0.13 | 0.23 | -0.72 | -0.34 | -0.74 | 0.35 | -0.63 | -0.21 | 3.93E-17 | -0.24 | -0.3075 | 1 | 0.4129 | 0.344408 | 1 | 1 | 0.862346 |
| Sec11a | D3YWT0 | 0.52 | 0.09 | 0.23 | 0.57 | -0.01 | 0.47 | -0.01 | 0.42 | -0.07 | 0.81 | 0.52 | 0.73 | -0.41 | 0.67 | 0.37 | 0.88 | -0.135 | 0.145 | 0.025 | 0.542637 | 0.571503 | 0.935867 | 0.999971 | 1 | 0.995618 |
| Sec11c | D3Z5H3 | -0.01 | -0.06 | -0.94 | -0.36 | -0.78 | -0.02 | -1.14 | -0.71 | -0.24 | -0.15 | -0.74 | -0.41 | 0.13 | 0.43 | -0.62 | -0.4 | -0.32 | -0.0425 | 0.2275 | 0.330335 | 0.874181 | 0.488246 | 0.999971 | 1 | 0.898175 |
| Sec13 | Q9D1M0 | 0.95 | 1.05 | 0.14 | -0.01 | 1.08 | 0.9 | 0.76 | 0.09 | 1.23 | 0.38 | 0.77 | 0.16 | 0.79 | 0.46 | 0.35 | -0.18 | 0.175 | 0.1025 | -0.1775 | 0.612663 | 0.771968 | 0.601164 | 0.999971 | 1 | 0.92426 |
| Sec14l1 | A8Y5H7 | 0.25 | 0.38 | -0.05 | -0.39 | 0.18 | 0.08 | -0.62 | -0.3 | 0.05 | 0.5 | -0.02 | -0.62 | -0.47 | 0.25 | -0.36 | -0.13 | -0.2125 | -0.07 | -0.225 | 0.440041 | 0.81308 | 0.395111 | 0.999971 | 1 | 0.871816 |
| Sec14l2 | Q99J08 | -0.2 | 0.02 | 0.53 | 0.63 | 0.06 | 0.34 | 0.33 | 0.61 | -0.03 | -0.38 | 0.06 | -0.11 | -0.52 | -0.2 | 0.17 | 0.34 | 0.09 | -0.36 | -0.2975 | 0.72467 | 0.171256 | 0.319712 | 0.999971 | 1 | 0.853962 |
| Sec16a | E9QAT4-2 | 1.06 | 1.29 | 1.33 | 1.23 | 1.13 | 1.1 | 1.25 | 1.34 | 1.06 | 1.14 | 1.24 | 1.37 | 1.35 | 1.37 | 1.12 | 1.45 | -0.0225 | -0.025 | 0.095 | 0.895975 | 0.884284 | 0.594839 | 0.999971 | 1 | 0.92377 |
| Sec22b | O08547 | -0.01 | -0.19 | -0.22 | -0.1 | -0.57 | -0.37 | 0.18 | -0.01 | -0.45 | -0.3 | -0.09 | -0.33 | -0.3 | -0.4 | -0.22 | -0.05 | -0.0625 | -0.1625 | -0.1125 | 0.77684 | 0.356096 | 0.52609 | 0.999971 | 1 | 0.909459 |
| Sec23a | E9Q1S3 | 0.29 | 0.79 | 0.54 | 0.33 | 0.78 | 0.68 | 0.54 | 0.58 | 1.06 | 0.48 | 0.67 | 0.6 | 0.52 | 0.88 | 0.49 | 0.49 | 0.1575 | 0.215 | 0.1075 | 0.421471 | 0.329633 | 0.602477 | 0.999971 | 1 | 0.924453 |
| Sec23b | Q9D662 | 0.5 | 1.19 | 0.59 | 0.47 | 0.99 | 0.92 | 0.45 | 0.82 | 0.92 | 0.68 | 0.05 | 0.55 | 1.01 | 1 | 0.82 | 0.49 | 0.1075 | -0.1375 | 0.1425 | 0.656338 | 0.610758 | 0.558998 | 0.999971 | 1 | 0.915051 |
| Sec23ip | G3X928 | 0.85 | 1.63 | 1.1 | 1.46 | 1.69 | 1.25 | 1.5 | 1.17 | 1.95 | 1 | 1.17 | 1.28 | 1.72 | 1.31 | 1.28 | 1.18 | 0.1425 | 0.09 | 0.1125 | 0.561835 | 0.752579 | 0.646683 | 0.999971 | 1 | 0.933613 |
| Sec24a | A2AA71 | 0.38 | 0.41 | 0.45 | 0.31 | 0.57 | 0.13 | -0.07 | 0.59 | 0.31 | 0.65 | 0.18 | 0.61 | 0.73 | 0.48 | 0.44 | 0.7 | -0.0825 | 0.05 | 0.2 | 0.70127 | 0.78713 | 0.2646 | 0.999971 | 1 | 0.842533 |
| Sec24b | Q80ZX0 | 0.19 | 0.14 | 0.27 | 0.41 | 0.28 | 0.16 | 0.79 | 0.22 | 0.07 | 0.09 | 0.53 | 0.56 | -0.25 | 0.21 | 0.26 | 0.6 | 0.11 | 0.06 | -0.0475 | 0.599934 | 0.764836 | 0.833478 | 0.999971 | 1 | 0.979036 |
| Sec24c | A0A286YDI | -0.25 | -0.54 | -1.66 | -2.35 | -1.02 | -1.31 | -1.59 | -1.21 | -0.97 | -0.29 | -0.92 | -0.79 | -0.71 | -1.46 | -1.82 | -1.39 | -0.0825 | 0.4575 | -0.145 | 0.860641 | 0.353961 | 0.773303 | 0.999971 | 1 | 0.963759 |
| Sec24c_1 | G3X972 | 0.37 | 0.83 | 0.58 | 0.61 | 0.93 | 0.67 | 1.12 | 0.62 | 0.78 | 0.64 | 0.68 | 0.73 | 0.77 | 0.69 | 0.58 | 0.75 | 0.2375 | 0.11 | 0.1 | 0.264235 | 0.53612 | 0.585365 | 0.999971 | 1 | 0.920413 |
| Sec24d | Q6NXL1 | 2.34 | 1.29 | 0.01 | 0.72 | 1.15 | 1.3 | 0.21 | 0.59 | 1.26 | 1.8 | 0.4 | 0.26 | 0.73 | 1.6 | 0.19 | 0.76 | -0.2775 | -0.16 | -0.27 | 0.590446 | 0.777261 | 0.611799 | 0.999971 | 1 | 0.927066 |
| Sec31a | Q3UPL0 | 0.87 | 1.07 | 0.78 | 0.79 | 0.95 | 1.25 | 0.82 | 0.75 | 1.23 | 1.19 | 0.96 | 0.87 | 1.1 | 1.02 | 0.95 | 1.24 | 0.065 | 0.185 | 0.2 | 0.737561 | 0.321749 | 0.275826 | 0.999971 | 1 | 0.843794 |
| Sec61a1 | P61620 | 0.98 | 0.95 | 0.57 | 0.79 | 0.93 | 1.29 | 0.3 | 0.88 | 1.08 | 1.27 | 0.8 | 0.74 | 1.03 | 1.56 | 0.9 | 1.41 | 0.0275 | 0.15 | 0.4025 | 0.913125 | 0.471779 | 0.099847 | 0.999971 | 1 | 0.78268 |
| Sec61a2 | Q9JLR1 | -0.17 | 0.01 | 1.27 | 1.32 | 0.24 | 0.43 | 0.93 | 1.18 | 0.52 | 0.63 | 1.64 | 1.83 | 0.25 | 0.52 | 1.78 | 1.92 | 0.0875 | 0.5475 | 0.51 | 0.838512 | 0.279975 | 0.356773 | 0.999971 | 1 | 0.868793 |
| Sec61b | Q9CQS8 | -0.47 | -0.29 | 0.42 | 0.48 | 0.3 | 0.38 | 1.77 | 0.07 | 0.2 | -0.21 | 0.31 | 0.77 | 0.26 | 0.72 | 0.66 | 0.93 | 0.595 | 0.2325 | 0.6075 | 0.190572 | 0.471439 | 0.063187 | 0.999971 | 1 | 0.725348 |
| Sec61g | P60060 | 3.61 | 1.15 | 0.16 | -0.94 | -0.41 | 0.22 | 1.05 | -1.02 | 0.14 | 3.11 | 0.49 | 0.71 | 0.39 | -1.53 | -0.21 | -0.36 | -1.035 | 0.1175 | -1.4225 | 0.294001 | 0.911847 | 0.156197 | 0.999971 | 1 | 0.796008 |
| Sec62 | Q8BU14 | 0.88 | -0.79 | 0.34 | 0.51 | -0.75 | -0.33 | 0.69 | 0.34 | -0.49 | 0.83 | -0.48 | -1.6 | -1.43 | -0.61 | -0.18 | -0.27 | -0.2475 | -0.67 | -0.8575 | 0.588868 | 0.255729 | 0.074855 | 0.999971 | 1 | 0.746958 |
| Sec63 | Q8VHE0 | -0.69 | -0.29 | -0.25 | -0.25 | -0.36 | -0.34 | -0.56 | -0.4 | -0.25 | -0.46 | 0.18 | 0.03 | -0.11 | -0.3 | -0.13 | 0.12 | -0.045 | 0.245 | 0.265 | 0.810839 | 0.281701 | 0.204408 | 0.999971 | 1 | 0.81617 |
| Secisbp2l | Q6A098 | 0.06 | -0.3 | 0.71 | 0.54 | -0.42 | -0.26 | 0.26 | 0.65 | -0.09 | -0.36 | 0.61 | 1.12 | -0.46 | 0.01 | 0.4 | 1.14 | -0.195 | 0.0675 | 0.02 | 0.563255 | 0.86319 | 0.959397 | 0.999971 | 1 | 0.998721 |
| Seh1l | Q8R2U0-2 | 1.37 | 0.64 | 0.79 | 0.86 | 0.34 | 1.1 | 0.93 | 0.56 | 0.52 | 1.18 | 0.42 | 0.1 | -0.3 | 0.45 | 0.28 | 0.66 | -0.1825 | -0.36 | -0.6425 | 0.486476 | 0.232433 | 0.042851 | 0.999971 | 1 | 0.701725 |
| Sel1l | Q9Z2G6 | -0.01 | 0.61 | 0.23 | -0.42 | 0.83 | 0.9 | 0.86 | -0.37 | 0.71 | -0.17 | 0.28 | 0.17 | 0.49 | 0.39 | -0.05 | -0.48 | 0.4525 | 0.145 | -0.015 | 0.237162 | 0.622046 | 0.96195 | 0.999971 | 1 | 0.998721 |
| Selenbp1 | P17563 | -0.66 | 0.05 | 0.29 | 0.51 | -0.06 | -0.28 | 0.04 | 0.51 | 0.05 | -0.86 | -0.07 | 0.33 | -0.15 | -0.1 | 0.22 | 0.4 | 0.005 | -0.185 | 0.045 | 0.987079 | 0.603152 | 0.879247 | 0.999971 | 1 | 0.985441 |
| Selenof | A0A0R4J0K | -0.42 | -0.58 | -0.19 | -0.14 | -0.1 | -0.47 | -0.27 | -0.09 | -0.06 | -0.44 | -0.02 | 0.12 | 0.13 | -0.38 | 0.09 | 0.37 | 0.1 | 0.2325 | 0.385 | 0.614115 | 0.278129 | 0.118118 | 0.999971 | 1 | 0.78268 |
| Selenoh | A0A087WP | 0.33 | 2.21 | 3.58 | 2.6 | 1.81 | 2.03 | 1.56 | 4.39 | 2.01 | 1.24 | 4.21 | -3.66 | 1.95 | 1.74 | 4.96 | 1.54 | 0.2675 | -1.23 | 0.3675 | 0.752515 | 0.450649 | 0.698733 | 0.999971 | 1 | 0.947812 |
| Selenoi | D6RE20 | -0.75 | -0.34 | -1.36 | -0.26 | -1.28 | -0.72 | -0.95 | -0.76 | -1.05 | -0.44 | -0.55 | -0.26 | -0.94 | 0.11 | -0.72 | 0.37 | -0.25 | 0.1025 | 0.3825 | 0.403435 | 0.740163 | 0.339689 | 0.999971 | 1 | 0.860218 |
| Selenom | Q8VHC3 | -0.43 | 0.01 | 0.49 | 0.41 | 0.21 | 0.12 | -0.21 | 0.41 | 0.33 | -0.15 | 0.03 | 0.86 | 0.26 | -0.02 | 0.61 | 0.89 | 0.0125 | 0.1475 | 0.315 | 0.962742 | 0.636556 | 0.308745 | 0.999971 | 1 | 0.847402 |
| Selenon | D3Z2R5 | 0.57 | 0.18 | -0.54 | -0.9 | 0.58 | 0.45 | -0.16 | -0.64 | 1.05 | 1.89 | 0.61 | 0.71 | -0.12 | 1.2 | 0.17 | -0.71 | 0.23 | 1.2375 | 0.3075 | 0.58337 | 0.016031 | 0.531128 | 0.999971 | 1 | 0.910069 |
| Selenoo | Q9DBC0 | 0.51 | 0.47 | -0.35 | -0.01 | 0.06 | 0.35 | -0.5 | -0.58 | -0.05 | 0.33 | -0.61 | -0.63 | 0.4 | 0.28 | -0.59 | -0.31 | -0.3225 | -0.395 | -0.21 | 0.312356 | 0.229733 | 0.513288 | 0.999971 | 1 | 0.905815 |
| Selenop | P70274 | 0.44 | 1.57 | 1.29 | 1.27 | 1.84 | 1.57 | 1.39 | 0.82 | 1.45 | 0.78 | 2.22 | 2.21 | 1.29 | 1.44 | 1.09 | 2.56 | 0.2625 | 0.5225 | 0.4525 | 0.429631 | 0.219151 | 0.269127 | 0.999971 | 1 | 0.843504 |
| Selenos | Q9BCZ4 | 0.44 | 2.39 | 1.01 | 0.91 | 2.22 | 2.72 | 0.87 | 0.83 | 2.2 | 0.82 | 0.67 | 0.88 | 2.81 | 1.8 | 1.08 | 1.03 | 0.4725 | -0.045 | 0.4925 | 0.425884 | 0.929805 | 0.377018 | 0.999971 | 1 | 0.870571 |
| Selenot | P62342 | -0.1 | -0.39 | -0.92 | -0.99 | 0.25 | -0.1 | -0.44 | -1.13 | -0.4 | -0.11 | -1.1 | -0.9 | -0.11 | 0.03 | -1.01 | -1.09 | 0.245 | -0.0275 | 0.055 | 0.497124 | 0.930351 | 0.877264 | 0.999971 | 1 | 0.985139 |
| Selenow | P63300 | 3.01 | 0.62 | 2.02 | 2.42 | 1.23 | 1.64 | 2.08 | 0.4 | 1.96 | 2.48 | 1.81 | 1.13 | -1.54 | 0.03 | 0.91 | 0.88 | -0.68 | -0.1725 | -1.9475 | 0.251669 | 0.748353 | 0.019585 | 0.999971 | 1 | 0.635388 |
| Selplg | Q3TA56 | -1.1 | -0.9 | -0.3 | -0.48 | -0.32 | -1.92 | -0.77 | 0.11 | -1.45 | -1.96 | 0.23 | 0.23 | -0.91 | -0.4 | -0.21 | -0.19 | -0.03 | -0.0425 | 0.2675 | 0.946297 | 0.938555 | 0.335091 | 0.999971 | 1 | 0.858827 |
| Sema4a | Q62178 | -0.82 | -0.95 | -0.82 | -0.67 | -0.81 | -0.55 | -0.77 | -0.23 | -0.86 | -0.53 | -0.31 | -0.45 | -0.56 | -0.67 | 0.03 | -0.53 | 0.225 | 0.2775 | 0.3825 | 0.281833 | 0.171179 | 0.102926 | 0.999971 | 1 | 0.78268 |
| Sema4b | Q62179 | 0.05 | -0.72 | -1.1 | -0.44 | -0.46 | -0.59 | -0.93 | -0.79 | -0.5 | 0.09 | -1.18 | -0.88 | -0.29 | -0.35 | -0.89 | -0.77 | -0.14 | -0.065 | -0.0225 | 0.618629 | 0.85583 | 0.939252 | 0.999971 | 1 | 0.995869 |
| Sema4c | Q64151 | -1.09 | -1.07 | -0.36 | -0.46 | -0.44 | -0.47 | -0.49 | -0.16 | -0.45 | -1.03 | -0.77 | -0.71 | -0.7 | -0.67 | -0.61 | -1.24 | 0.355 | 0.005 | -0.06 | 0.166398 | 0.984196 | 0.821286 | 0.999971 | 1 | 0.975833 |
| Sema4d | O09126 | -1.15 | -0.66 | 0.03 | -0.03 | -0.62 | -0.56 | -0.33 | 0.32 | -0.36 | -0.79 | -0.34 | -0.14 | -0.06 | -0.45 | 0.38 | 0.34 | 0.155 | 0.045 | 0.505 | 0.657236 | 0.886313 | 0.161601 | 0.999971 | 1 | 0.800215 |
| Sema4f | Q9Z123 | 0.92 | 0.37 | -2.07 | -1.78 | 0.63 | 0.15 | -2.41 | -1.87 | 0.23 | 1.05 | -2.42 | -1.84 | 0.37 | 0.3 | -1.74 | -1.58 | -0.235 | -0.105 | -0.0225 | 0.804015 | 0.916671 | 0.978901 | 0.999971 | 1 | 0.99925 |
| Sema4g | Q9WUH7 | -0.18 | -0.19 | -1.21 | -1.16 | -1.13 | -0.01 | -1.78 | -0.33 | -0.79 | -0.56 | -0.85 | -1.17 | -0.59 | -0.11 | -0.52 | 0.09 | -0.1275 | -0.1575 | 0.4025 | 0.782474 | 0.622259 | 0.244711 | 0.999971 | 1 | 0.837316 |
| Sema5a | Q3UPZ0 | -1.47 | -0.4 | 0.94 | -0.15 | -0.26 | -0.23 | 1.46 | -0.32 | -0.06 | -2.49 | 0.09 | 0.15 | -1.1 | -2.1 | 0.45 | -1.1 | 0.4325 | -0.3075 | -0.6925 | 0.477424 | 0.67658 | 0.307286 | 0.999971 | 1 | 0.847402 |
| Sema5b | Q60519-2 | 1.32 | 0 | 0.72 | 0.48 | 0.17 | 0.43 | 0.72 | 0.17 | -0.05 | 1.02 | -0.19 | -0.04 | -1.12 | -0.84 | -0.08 | -1.11 | -0.2575 | -0.445 | -1.4175 | 0.414803 | 0.260428 | 0.003427 | 0.999971 | 1 | 0.577271 |
| Sema6a | K3W4S5 | -0.09 | -1.18 | -1.21 | -1.11 | -0.66 | -0.89 | -1.27 | -0.58 | -1.47 | -0.74 | -1.13 | -1.26 | -1.89 | -1.07 | -0.68 | -1.02 | 0.0475 | -0.2525 | -0.2675 | 0.880035 | 0.430258 | 0.469221 | 0.999971 | 1 | 0.892481 |
| Sema6b | O54951 | -0.19 | -0.25 | -1.67 | -1.62 | -0.37 | 0 | -1.9 | -1.38 | -0.2 | 0.1 | -1.78 | -1.91 | 0.32 | 0.27 | -1.62 | -1.37 | 0.02 | -0.015 | 0.3325 | 0.971121 | 0.980361 | 0.586038 | 0.999971 | 1 | 0.920413 |
| Sema6d | A2AW73 | -0.2 | -0.28 | -0.2 | -0.02 | -0.47 | -0.2 | -0.24 | -0.18 | -0.21 | -0.17 | -0.57 | -0.42 | -0.43 | -0.31 | -0.17 | -0.68 | -0.0975 | -0.1675 | -0.2225 | 0.57895 | 0.36485 | 0.260665 | 0.999971 | 1 | 0.841256 |
| Sema7a | Q9QUR8 | -0.43 | -0.94 | -0.15 | -0.16 | -1.14 | -0.98 | -0.63 | -0.14 | -1.02 | -0.25 | -0.52 | -0.29 | -0.86 | -0.92 | -0.17 | 0.02 | -0.3025 | -0.1 | -0.0625 | 0.324246 | 0.714833 | 0.839541 | 0.999971 | 1 | 0.979728 |
| Senp2 | A0A338P6/ | -1.29 | -0.98 | -2.81 | -0.65 | -0.47 | 0.61 | -1.77 | -1.28 | 0.08 | -0.62 | -2.2 | -1.43 | 1.39 | -0.39 | -1.78 | -0.37 | 0.705 | 0.39 | 1.145 | 0.288565 | 0.539679 | 0.142461 | 0.999971 | 1 | 0.784967 |
| Senp3 | Q9EP97 | 3.19 | -0.06 | 0.08 | 1.23 | -0.56 | 0.14 | -0.44 | 1.28 | -1.24 | 2.47 | -0.37 | -0.27 | -0.51 | 0.81 | 0.2 | 0.23 | -1.005 | -0.9625 | -0.9275 | 0.216274 | 0.344402 | 0.220519 | 0.999971 | 1 | 0.827096 |
| Senp7 | Q8BUH8 | 0.6 | -0.24 | 1.17 | -0.28 | 0.12 | -0.31 | 2 | 0.65 | 0.37 | 0.09 | 2.72 | 3.61 | 0.7 | -0.31 | 1.18 | 0.73 | 0.3025 | 1.385 | 0.2625 | 0.592841 | 0.128967 | 0.558328 | 0.999971 | 1 | 0.915051 |
| Senp8 | Q9D2Z4 | 1.94 | 0.21 | -0.49 | -0.39 | 0.61 | 1.05 | -0.7 | -0.29 | 0.17 | 2.76 | -0.72 | -0.95 | -0.42 | 0.56 | -0.66 | -0.96 | -0.15 | -0.0025 | -0.6875 | 0.811519 | 0.997826 | 0.267207 | 0.999971 | 1 | 0.842677 |
| Sephs1 | Q8BH69 | 0.29 | 0.89 | 1.24 | 1.55 | 0.69 | 0.62 | 0.93 | 1.22 | 0.53 | 0.68 | 1.02 | 0.98 | 0.28 | 0.71 | 1.33 | 1.44 | -0.1275 | -0.19 | -0.0525 | 0.680334 | 0.533963 | 0.887936 | 0.999971 | 1 | 0.987485 |
| Sephs2 | P97364 | 0.26 | 0.36 | -0.34 | -0.26 | 0.93 | 0.21 | -0.26 | -0.46 | 0.21 | -0.22 | -0.87 | -0.33 | -0.57 | 0.45 | -0.83 | -0.22 | 0.1 | -0.3075 | -0.2975 | 0.775497 | 0.311432 | 0.37838 | 0.999971 | 1 | 0.871727 |
| Sepsecs | Q6P6M7 | 1.24 | 1.11 | 1.13 | 0.98 | 0.72 | 1.71 | 0.75 | 0.68 | 0.96 | 1.79 | 0.57 | 0.13 | 1.02 | 1.15 | 0.57 | 0.82 | -0.15 | -0.2525 | -0.225 | 0.585391 | 0.478083 | 0.274334 | 0.999971 | 1 | 0.843794 |
| Septin10 | A0A0R4J23 | -0.7 | 0.28 | 0.06 | 0.09 | 0.25 | 0.08 | -0.65 | 0.47 | 0.16 | -0.59 | -0.26 | 0.07 | 0.36 | -0.12 | 0.32 | -0.18 | 0.105 | -0.0875 | 0.1625 | 0.747678 | 0.76118 | 0.560692 | 0.999971 | 1 | 0.91591 |
| Septin11 | A0A0J9YUL | -0.64 | -0.59 | -0.4 | -0.26 | -0.73 | -0.33 | -0.71 | -0.42 | -0.74 | -0.8 | -0.42 | -0.48 | -0.54 | -0.56 | -0.31 | -0.15 | -0.075 | -0.1375 | 0.0825 | 0.702841 | 0.476441 | 0.674141 | 0.999971 | 1 | 0.941037 |
| Septin2 | E9Q3V6 | 2.76 | 2.33 | 2.59 | 2.95 | 0.64 | 4.31 | 2.02 | 2.01 | 1.82 | 3.91 | 1.96 | 0.96 | 1.11 | 3.05 | 1.51 | 2.83 | -0.4125 | -0.495 | -0.5325 | 0.558379 | 0.407978 | 0.27125 | 0.999971 | 1 | 0.843794 |
| Septin2_1 | P42208 | -0.49 | -1.32 | -1.25 | -1.15 | -1.23 | -0.88 | -1.21 | -1.16 | -1.2 | -0.62 | -1.22 | -1.64 | -1.64 | -1.39 | -1.31 | -1.37 | -0.0675 | -0.1175 | -0.375 | 0.778896 | 0.689713 | 0.142351 | 0.999971 | 1 | 0.784967 |
| Septin3 | Q9Z1S5-2 | -2.41 | -1.04 | -0.99 | -1.57 | -0.95 | -1.13 | -0.58 | -1.19 | -0.61 | -1.96 | -0.83 | -0.71 | -0.45 | -1.35 | -1.11 | -1.22 | 0.54 | 0.475 | 0.47 | 0.149666 | 0.288496 | 0.230432 | 0.999971 | 1 | 0.829447 |
| Septin3_1 | A0A5F8MP | 0.01 | -0.06 | -0.71 | -0.33 | -0.33 | -0.31 | -0.62 | -0.01 | -0.66 | -0.3 | -0.55 | -0.16 | -0.61 | -0.1 | -0.41 | 0.16 | -0.045 | -0.145 | 0.0325 | 0.850506 | 0.538066 | 0.900428 | 0.999971 | 1 | 0.990593 |
| Septin3_2 | A0A2R8VH | -2.75 | 0.02 | -1.56 | -1.32 | -0.5 | -0.88 | -0.45 | -0.19 | 0.36 | -2.36 | -0.96 | 0.14 | -0.13 | -0.87 | -1.11 | -1.08 | 0.8975 | 0.6975 | 0.605 | 0.1229 | 0.373208 | 0.298523 | 0.999971 | 1 | 0.847402 |
| Septin4 | A0A5F8MP | -0.54 | 0.25 | 0.3 | -0.5 | -0.21 | 0.14 | -0.39 | -0.48 | -0.19 | -0.56 | -0.23 | -0.06 | 0.18 | -0.1 | -0.21 | -0.66 | -0.1125 | -0.1375 | -0.075 | 0.691343 | 0.614139 | 0.801792 | 0.999971 | 1 | 0.971467 |
| Septin5 | Q9Z2Q6 | -1.25 | -1.14 | -0.99 | -1.54 | -1.01 | -1.17 | -0.51 | -1.39 | -0.66 | -1.48 | -0.77 | -1.09 | -0.92 | -1.54 | -1.19 | -1.56 | 0.21 | 0.23 | -0.0725 | 0.408248 | 0.361334 | 0.754052 | 0.999971 | 1 | 0.961116 |
| Septin5_1 | A0A338P7! | 2.72 | 1.83 | 2.23 | 2.67 | 0.95 | 4.11 | 2.17 | 1.3 | 1.92 | 3.45 | 1.48 | 0.23 | 0.59 | 1.85 | 1.47 | 1.46 | -0.23 | -0.5925 | -1.02 | 0.731371 | 0.364853 | 0.013699 | 0.999971 | 1 | 0.613095 |
| Septin5_2 | A0A338P7( | 0.28 | -0.16 | -0.97 | -1.22 | -0.73 | -0.14 | -1.4 | -0.75 | -0.86 | 0.27 | -0.44 | -0.5 | -0.43 | -0.48 | -1.29 | 0.02 | -0.2375 | 0.135 | -0.0275 | 0.56826 | 0.738709 | 0.947844 | 0.999971 | 1 | 0.997651 |
| Septin6 | Q9R1T4-2 | -0.45 | 0.02 | 0.29 | 0.23 | -0.43 | 0.3 | -0.01 | 0.3 | -0.56 | -0.58 | -0.26 | -0.39 | -0.17 | 0 | 0.31 | 0.19 | 0.0175 | -0.47 | 0.06 | 0.946918 | 0.060506 | 0.798489 | 0.999971 | 1 | 0.970479 |
| Septin7 | O55131 | -1.25 | -1.18 | -0.92 | -1.06 | -1.18 | -1.24 | -0.28 | -1.24 | -1.18 | -1.33 | -0.8 | -1.25 | -1.29 | -1.4 | -1.19 | -1.29 | 0.1175 | -0.0375 | -0.19 | 0.661352 | 0.847703 | 0.291411 | 0.999971 | 1 | 0.847402 |
| Septin7_1 | E9Q1G8 | -2.75 | -0.1 | 0.2 | -1.08 | -0.16 | -0.51 | 0.9 | -0.8 | -0.03 | -2.82 | 0.79 | -0.25 | 0.83 | -0.83 | -0.23 | -0.84 | 0.79 | 0.355 | 0.665 | 0.269553 | 0.701686 | 0.35432 | 0.999971 | 1 | 0.868793 |
| Septin8 | B7ZC46 | -1.59 | 0.06 | -2.07 | -2.01 | -0.43 | -0.09 | -1.46 | -1.76 | -0.78 | -0.59 | -1.95 | -0.33 | 0.04 | 0.1 | -2.28 | -1.42 | 0.4675 | 0.49 | 0.5125 | 0.432732 | 0.397364 | 0.467484 | 0.999971 | 1 | 0.891731 |
| Septin8_1 | B1AQZ0 | -0.84 | -1.58 | -2.92 | -2.69 | -1.79 | -1.42 | -2.63 | -3.03 | -1.85 | -1.09 | -2.83 | -3.22 | -1.96 | -1.38 | -2.71 | -2.62 | -0.21 | -0.24 | -0.16 | 0.709145 | 0.702891 | 0.764583 | 0.999971 | 1 | 0.962405 |
| Septin9 | Q80UG5-3 | -0.28 | -0.4 | -0.73 | -0.93 | -0.59 | -0.68 | -0.22 | -1.16 | -0.63 | -0.37 | -0.64 | -0.88 | -0.41 | -1.04 | -1.27 | -0.78 | -0.0775 | -0.045 | -0.29 | 0.770706 | 0.839143 | 0.284091 | 0.999971 | 1 | 0.844538 |
| Serac1 | Q3U213 | 0.44 | 0.13 | 0.39 | 0.45 | 0.15 | 0.13 | 0.34 | 0.42 | 0.22 | 0.24 | 0.6 | 0.37 | 0.42 | 0.06 | 0.37 | 0.74 | -0.0925 | 0.005 | 0.045 | 0.612233 | 0.978236 | 0.830423 | 0.999971 | 1 | 0.978535 |
| Serbp1 | Q9CY58 | 1.83 | 1.97 | 1.74 | 2.09 | 2.06 | 1.93 | 1.94 | 2.35 | 2.15 | 2 | 1.85 | 1.79 | 1.99 | 2.12 | 1.91 | 1.89 | 0.1625 | 0.04 | 0.07 | 0.404932 | 0.825772 | 0.694316 | 0.999971 | 1 | 0.947174 |
| Serbp1_1 | Q9CY58-2 | -0.72 | 5.91 | 5.74 | 3.42 | 5.72 | 5.34 | 3.88 | 4.63 | 5.51 | 0.81 | 5.31 | 5.22 | 6.7 | 5.01 | 4.96 | 4.94 | 1.305 | 0.625 | 1.815 | 0.368566 | 0.715692 | 0.22408 | 0.999971 | 1 | 0.828225 |
| Serf2 | P84102 | 0.55 | 2.41 | 2.08 | 2.57 | 2.65 | 2.6 | 2.49 | 2.9 | 1.99 | 1.44 | 2.15 | 2.34 | 0.69 | 2.71 | 2.05 | 2.32 | 0.7575 | 0.0775 | 0.04 | 0.11426 | 0.869028 | 0.94533 | 0.999971 | 1 | 0.997427 |
| Sergef | Q80YD6 | -0.28 | 0.73 | 1.04 | -0.18 | 1.08 | 0.64 | 0.21 | 0.36 | 0.49 | -0.09 | -0.24 | 0.19 | 0.88 | -0.02 | 0.54 | 0.38 | 0.245 | -0.24 | 0.1175 | 0.512194 | 0.508433 | 0.750156 | 0.999971 | 1 | 0.961064 |
| Serhl | Q9EPB5 | -3.33 | -2.63 | -0.25 | -0.64 | -2.04 | -2.78 | -0.33 | -0.21 | -2.01 | -2.36 | -0.83 | -0.27 | -2.3 | -2.41 | -0.02 | 0.13 | 0.3725 | 0.345 | 0.5625 | 0.673677 | 0.671889 | 0.54396 | 0.999971 | 1 | 0.911398 |
| Serinc1 | Q9QZI8 | 0.51 | 0.33 | 0.29 | 0.4 | 0.64 | 0.81 | 0.19 | 0.5 | 0.84 | 0.69 | 0.13 | 0.35 | 1.09 | 1.11 | 0.72 | 0.9 | 0.1525 | 0.12 | 0.5725 | 0.451256 | 0.576375 | 0.011191 | 0.999971 | 1 | 0.613095 |
| Serinc3 | Q9QZI9 | 0.22 | 1.57 | 1.39 | 0.28 | 2.07 | 1.04 | 3.51 | -1.2 | 2.22 | 0.02 | 0.39 | -0.24 | 3.16 | 0.73 | -0.45 | -2.34 | 0.49 | -0.2675 | -0.59 | 0.60531 | 0.660741 | 0.586368 | 0.999971 | 1 | 0.920454 |
| Serinc5 | Q8BHJ6 | 1.3 | 1.28 | 3.39 | 2.34 | 0.96 | 1.01 | 4.37 | 0.87 | 1.22 | 0.89 | 1.56 | 0.43 | 1.71 | 0.43 | 1.53 | -0.17 | -0.275 | -1.0525 | -1.2025 | 0.75745 | 0.068663 | 0.077421 | 0.999971 | 1 | 0.752059 |
| Serpina1b | P22599 | 1.78 | 3.46 | 0.95 | 4.01 | 4.9 | 2.67 | 3.29 | 2.1 | 2.85 | 1.07 | 2.61 | 3.1 | 2.38 | 3.05 | 2.44 | 3.47 | 0.69 | -0.1425 | 0.285 | 0.419893 | 0.852823 | 0.680238 | 0.999971 | 1 | 0.942957 |
| Serpina1c | A0A0R4J0X | 0.75 | 2.86 | 0.84 | 3.28 | 4.8 | 1.92 | 4.19 | 1.36 | 2.23 | 0.62 | 2.87 | 2.31 | 2.54 | 1.8 | 1.81 | 1.63 | 1.135 | 0.075 | 0.0125 | 0.254643 | 0.919708 | 0.984182 | 0.999971 | 1 | 0.99925 |
| Serpina1d | Q00897 | 1.72 | 2 | 0.4 | 3.29 | 4.1 | 2.2 | 2.58 | 1.65 | 2.43 | 1.17 | 2.11 | 2.53 | 1.97 | 1.98 | 1.54 | 2.63 | 0.78 | 0.2075 | 0.1775 | 0.291961 | 0.735725 | 0.760361 | 0.999971 | 1 | 0.961116 |
| Serpina1e | Q00898 | 1.58 | -0.16 | -2.18 | -2.51 | 3.94 | -0.88 | 1.06 | 0.09 | -1.49 | 2.44 | 4.31 | 1.52 | 7.38 | -0.15 | 2.07 | -0.33 | 1.87 | 2.5125 | 3.06 | 0.161196 | 0.095992 | 0.11705 | 0.999971 | 1 | 0.78268 |
| Serpina3b | Q8BYY9 | 1.75 | 3.07 | 3.52 | 2.52 | 2.81 | 4.29 | 2.17 | 2.87 | 2.55 | 2.62 | 2.85 | 2.74 | 3.51 | 3.02 | 2.97 | 3.54 | 0.32 | -0.025 | 0.545 | 0.557819 | 0.946826 | 0.191071 | 0.999971 | 1 | 0.810547 |
| Serpina3k | A0A0R4J0I: | -0.09 | 1.69 | -0.27 | 0.54 | 4.29 | 1.23 | 3.08 | -0.71 | 1.29 | 1.87 | 4.26 | 3.59 | 4.72 | 1.12 | 2.55 | 3.2 | 1.505 | 2.285 | 2.43 | 0.176776 | 0.013665 | 0.012109 | 0.999971 | 1 | 0.613095 |
| Serpina3m | Q03734 | 2.45 | 2.62 | 1.32 | 3.24 | 3.14 | 2.06 | 2.52 | 1.42 | 2.54 | 1.22 | 2.16 | 1.63 | 3 | 2.25 | 1.42 | 2.95 | -0.1225 | -0.52 | -0.0025 | 0.807008 | 0.280744 | 0.996053 | 0.999971 | 1 | 1 |
| Serpina3n | G3X8T9 | 1.24 | 0.9 | 1.69 | 2.23 | 0.73 | 1.04 | 1.34 | 0.77 | -0.02 | 0.52 | 0.67 | 1.75 | 0.65 | 0.41 | 2.25 | 2.15 | -0.545 | -0.785 | -0.15 | 0.118036 | 0.104687 | 0.774072 | 0.999971 | 1 | 0.963884 |
| Serpina6 | Q06770 | 0.53 | 2.81 | 1.27 | 5.86 | 3.65 | 2.55 | 3.45 | 3.55 | 1.69 | -0.81 | -0.02 | -0.75 | 0.39 | 1.61 | 0.66 | 0.44 | 0.6825 | -2.59 | -1.8425 | 0.53039 | 0.052811 | 0.116403 | 0.999971 | 1 | 0.78268 |
| Serpinb1a | Q9D154 | 0.27 | 1.09 | 1.02 | 0.29 | 0.77 | 0.67 | 0.51 | 0.38 | -0.18 | -0.45 | -0.17 | 0.04 | -0.57 | 0.34 | 0.77 | -0.66 | -0.085 | -0.8575 | -0.6975 | 0.746715 | 0.009948 | 0.107353 | 0.999971 | 1 | 0.78268 |
| Serpinb1b | Q8VHP7 | 0.53 | -0.15 | 0.76 | 0.52 | 0.1 | 1.49 | 0.14 | 0.81 | 0.15 | 0.6 | -0.35 | 0.67 | 0.15 | 0.82 | 0.82 | -0.08 | 0.22 | -0.1475 | 0.0125 | 0.556997 | 0.637558 | 0.967768 | 0.999971 | 1 | 0.998787 |
| Serpinb6 | Q60854 | 0.33 | -0.6 | -0.03 | 0.21 | -0.11 | -0.39 | -0.06 | 0.19 | -0.62 | -0.06 | -0.19 | 0.27 | -1.32 | -0.38 | 0.08 | -0.1 | -0.07 | -0.1275 | -0.4075 | 0.789005 | 0.659773 | 0.281365 | 0.999971 | 1 | 0.844538 |
| Serpinb6a | F8WIV2 | 3.95 | 2.03 | 3.28 | 3.26 | 1.14 | 2.94 | 3.34 | 3.95 | 2.07 | 3.6 | 1.35 | 1.46 | -1.5 | 0.83 | 1.97 | 0.74 | -0.2875 | -1.01 | -2.62 | 0.663393 | 0.120515 | 0.006576 | 0.999971 | 1 | 0.582714 |
| Serpinb8 | O08800 | 1.2 | 1.68 | 1.1 | 1.46 | 1.06 | 1.54 | 0.77 | 1.01 | 0.53 | 0.86 | 0.7 | 1.11 | 0.4 | 0.86 | 0.54 | 1.17 | -0.265 | -0.56 | -0.6175 | 0.287086 | 0.029867 | 0.031564 | 0.999971 | 1 | 0.678474 |
| Serpinb9 | O08797 | 0.23 | 1.03 | 0.36 | 0.83 | 1.23 | 0.92 | 0.26 | 0.94 | 0.72 | -0.05 | 0.03 | 0.25 | 0.84 | 0.68 | 0.38 | 0.64 | 0.225 | -0.375 | 0.0225 | 0.447147 | 0.193552 | 0.926484 | 0.999971 | 1 | 0.99401 |
| Serpinc1 | P32261 | -0.28 | 2.18 | 0.54 | 2.22 | 3.46 | 1.96 | 2 | 0.87 | 2.14 | -0.18 | 1.05 | 1.85 | 1.96 | 1.5 | 0.94 | 1.57 | 0.9075 | 0.05 | 0.3275 | 0.238359 | 0.945686 | 0.587704 | 0.999971 | 1 | 0.920758 |
| Serpine2 | Q07235 | 1.13 | 0.82 | 0.6 | 0.91 | 1.29 | 0.98 | 0.62 | 1.23 | 1.47 | 1.49 | 0.87 | 1 | 1 | 1.01 | 0.87 | 0.78 | 0.165 | 0.3425 | 0.05 | 0.474617 | 0.160768 | 0.793585 | 0.999971 | 1 | 0.970141 |
| Serpinf1 | P97298 | -0.59 | 1.8 | 1.53 | 1.51 | 1.42 | 0.85 | 0.82 | 1.1 | 1.33 | -0.92 | 0.9 | 1.58 | -0.06 | 1.11 | 1.86 | 1.9 | -0.015 | -0.34 | 0.14 | 0.977284 | 0.638737 | 0.830591 | 0.999971 | 1 | 0.978535 |
| Serpinf2 | Q61247 | 0.77 | 4.11 | 2.03 | 2.85 | 5.49 | 2.32 | 4.21 | 3.2 | 3.54 | 0.27 | 4.06 | 4.36 | 4.87 | 2.52 | 3.23 | 1.9 | 1.365 | 0.6175 | 0.69 | 0.145531 | 0.562854 | 0.427646 | 0.999971 | 1 | 0.880793 |
| Serping1 | A2ATR8 | 2.27 | 3.14 | 0.65 | 3.79 | 3.75 | 2.13 | 2.86 | 1.87 | 1.68 | 2.41 | 2.17 | 2.53 | 2.37 | 2.58 | 1.51 | 2.83 | 0.19 | -0.265 | -0.14 | 0.792574 | 0.682296 | 0.834434 | 0.999971 | 1 | 0.979036 |
| Serpinh1 | P19324 | 0.06 | 0 | -0.73 | -0.33 | 0.74 | -0.01 | -0.01 | -0.61 | 0.17 | 0.52 | -0.34 | 0.32 | 0.72 | 0.51 | -0.04 | -0.57 | 0.2775 | 0.4175 | 0.405 | 0.410075 | 0.154184 | 0.252535 | 0.999971 | 1 | 0.839012 |
| Serpini1 | O35684 | 0.09 | 0.2 | 0.38 | 0.59 | 1.01 | 0.39 | 0.91 | 0.79 | 1.14 | 0.26 | 1.38 | 1.19 | 2.1 | 0.61 | 1.07 | 1.7 | 0.46 | 0.6775 | 1.055 | 0.061006 | 0.039096 | 0.012908 | 0.999971 | 1 | 0.613095 |
| Sesn3 | Q9CYP7 | 0.35 | 0.35 | 0.74 | 0.77 | 0.41 | -0.24 | 2.28 | 0.81 | 0.49 | 0.28 | 1.64 | 1.9 | 0.07 | 0.36 | 2.51 | 2.71 | 0.2625 | 0.525 | 0.86 | 0.606722 | 0.216299 | 0.203003 | 0.999971 | 1 | 0.81617 |
| Sestd1 | Q80UK0 | -0.81 | -0.49 | -0.41 | -0.53 | -0.47 | -0.26 | -0.04 | -0.49 | 0.01 | -0.56 | -0.01 | -0.15 | -0.31 | -0.65 | -0.51 | -0.38 | 0.245 | 0.3825 | 0.0975 | 0.234821 | 0.092112 | 0.604818 | 0.999971 | 1 | 0.924454 |
| Set | A2BE93 | 2.27 | 3.4 | 2 | 2.56 | 3.07 | 2.56 | 1.82 | 2.24 | 3.44 | 2.68 | 2.03 | 2.51 | 2.51 | 3.05 | 1.99 | 2.33 | -0.135 | 0.1075 | -0.0875 | 0.727565 | 0.790201 | 0.811238 | 0.999971 | 1 | 0.973864 |
| Set_1 | Q9EQU5-2 | 6.26 | 1.94 | 4.47 | 5.1 | 2.61 | 4 | 5.26 | 2.69 | 3.19 | 6.2 | 3.16 | 3.34 | -1.63 | 0.91 | 2.65 | 1.19 | -0.8025 | -0.47 | -3.6625 | 0.425708 | 0.658334 | 0.010067 | 0.999971 | 1 | 0.613095 |
| Setd2 | E9Q5F9 | 0.69 | 0.62 | 0.34 | 0.49 | 0.73 | 0.14 | 0.29 | 0.54 | 0.91 | 1.05 | 0.95 | 0.79 | 0.68 | 0.9 | 0.3 | 0.99 | -0.11 | 0.39 | 0.1825 | 0.595744 | 0.048404 | 0.412397 | 0.999971 | 1 | 0.875881 |
| Setd3 | Q91WC0 | 0.3 | 1.59 | 1.84 | 2.17 | 1.38 | 1.52 | 1.97 | 1.98 | 1.41 | 0.78 | 1.94 | 1.89 | 1.17 | 1.73 | 1.86 | 2.07 | 0.2375 | 0.03 | 0.2325 | 0.571016 | 0.948038 | 0.590453 | 0.999971 | 1 | 0.922217 |
| Setd7 | Q8VHL1 | 0 | 0.9 | 1 | 0.94 | 0.93 | 0.96 | 0.45 | 1.22 | 0.65 | -0.05 | 0.67 | 0.18 | 0.63 | 0.65 | 0.74 | 1.3 | 0.18 | -0.3475 | 0.12 | 0.547758 | 0.270585 | 0.686559 | 0.999971 | 1 | 0.945467 |
| Setdb1 | G5E8N3 | 1.4 | -1.28 | 0.7 | 1.17 | -1.36 | -0.93 | 0.48 | 0.59 | -2.36 | 0.95 | 0.82 | -0.13 | -2.79 | -1.21 | 0.83 | 1.08 | -0.8025 | -0.6775 | -1.02 | 0.275338 | 0.45087 | 0.314384 | 0.999971 | 1 | 0.851245 |
| Sez6 | Q7TSK2 | 0.19 | 0.94 | 0.65 | 0.65 | 0.98 | 0.76 | 0.59 | 0.74 | 1.48 | 0.59 | 0.38 | 0.74 | 1.33 | 0.89 | 0.95 | 0.7 | 0.16 | 0.19 | 0.36 | 0.472153 | 0.52372 | 0.156464 | 0.999971 | 1 | 0.796008 |
| Sez6l | A0A1D5RN | -0.15 | 0.97 | 0.46 | 0.24 | 1 | 0.58 | 1.07 | 0.51 | 1.05 | -0.01 | 1.09 | 0.89 | 1.34 | 0.58 | 0.2 | 0.65 | 0.41 | 0.375 | 0.3125 | 0.177245 | 0.292626 | 0.359847 | 0.999971 | 1 | 0.869526 |
| Sez6l2 | Q4V9Z5-2 | 0.27 | 0.06 | 0.09 | 0.01 | -0.07 | 0.05 | -0.05 | 0.13 | 0.24 | 0.32 | 0.37 | 0.29 | 0.58 | 0.3 | 0.26 | 0.28 | -0.0925 | 0.1975 | 0.2475 | 0.588038 | 0.243945 | 0.187973 | 0.999971 | 1 | 0.809025 |
| Sez6l2_1 | Q4V9Z5 | 3.97 | -1.4 | 1.29 | 2.01 | -1.41 | 0.85 | 2.94 | 0.08 | -0.06 | 1.74 | 0.27 | -1.21 | -2.02 | -1.01 | -0.59 | -1.74 | -0.8525 | -1.2825 | -2.8075 | 0.508503 | 0.276827 | 0.022633 | 0.999971 | 1 | 0.652043 |
| Sf1 | F6SRV1 | 2.56 | 0.36 | 1.77 | 2.73 | -0.44 | 0.85 | 3.44 | 0.9 | -0.64 | 1.54 | 1.14 | 0.6 | -1.86 | -0.66 | 0.18 | 1.47 | -0.6675 | -1.195 | -2.0725 | 0.452622 | 0.096587 | 0.027163 | 0.999971 | 1 | 0.678474 |
| Sf1_1 | E9Q4Q2 | 1.39 | 2.7 | 3.01 | 2.55 | 2.71 | 2.3 | 2.35 | 2.71 | 2.34 | 1.87 | 2.68 | 2.6 | 1.79 | 2.41 | 2.7 | 2.53 | 0.105 | -0.04 | -0.055 | 0.772063 | 0.917099 | 0.887954 | 0.999971 | 1 | 0.987485 |
| Sf3a1 | Q8K4Z5 | 0.2 | 0.58 | 0.77 | 1.08 | 0.75 | 0.26 | 0.4 | 0.67 | 0.69 | 0.23 | 0.6 | 0.83 | 0.81 | 0.49 | 0.87 | 1.16 | -0.1375 | -0.07 | 0.175 | 0.580113 | 0.779835 | 0.49984 | 0.999971 | 1 | 0.903877 |
| Sf3a2 | G3UVU2 | -0.11 | 0.18 | 0.69 | 0.56 | 0.37 | -0.38 | 0.9 | 0.25 | 0.09 | -0.53 | 1.02 | 0.58 | -0.12 | -0.17 | 0.87 | 0.28 | -0.045 | -0.04 | -0.115 | 0.888649 | 0.913663 | 0.710072 | 0.999971 | 1 | 0.950695 |
| Sf3a3 | Q9D554 | 0.78 | 1.04 | 0.76 | 0.47 | 1.09 | 0.77 | 0.88 | 0.37 | 1.04 | 1.03 | 0.91 | 0.62 | 0.74 | 0.86 | 0.5 | 0.02 | 0.015 | 0.1375 | -0.2325 | 0.947799 | 0.504569 | 0.362224 | 0.999971 | 1 | 0.869526 |
| Sf3b1 | G5E866 | 0.19 | 0.67 | 0.67 | 0.42 | 0.62 | 0.57 | 0.62 | 0.65 | 0.77 | 0.58 | 0.79 | 0.66 | 0.3 | 0.41 | 0.69 | 0.44 | 0.1275 | 0.2125 | -0.0275 | 0.501302 | 0.277502 | 0.890984 | 0.999971 | 1 | 0.988184 |
| Sf3b2 | Q3UJB0 | 1.01 | 0.63 | 0.94 | 1.03 | 0.44 | 0.76 | 0.62 | 0.88 | 0.55 | 1.01 | 0.74 | 0.69 | 0.16 | 0.72 | 0.75 | 1.14 | -0.2275 | -0.155 | -0.21 | 0.262318 | 0.430633 | 0.411667 | 0.999971 | 1 | 0.875847 |
| Sf3b3 | Q921M3 | -0.1 | 0.86 | 0.95 | 0.75 | 0.48 | 0.77 | 0.67 | 0.93 | 0.58 | 0.23 | 0.88 | 0.37 | 0.27 | 1.16 | 0.65 | 0.78 | 0.0975 | -0.1 | 0.1 | 0.724991 | 0.731522 | 0.747382 | 0.999971 | 1 | 0.961064 |

| | | | | | | | | | | | | | | | | | | | | | | | | | | |
|---|---|---|---|---|---|---|---|---|---|---|---|---|---|---|---|---|---|---|---|---|---|---|---|---|---|---|
| Sf3b4 | Q8QZY9 | -1.45 | 0.98 | -0.57 | -1.14 | 1.39 | 0.41 | -1.08 | -0.85 | 1.57 | -1.79 | -0.81 | -0.08 | 1.66 | 0 | -1.01 | -0.72 | 0.5125 | 0.2675 | 0.5275 | 0.478914 | 0.740581 | 0.475646 | 0.999971 | 1 | 0.894045 |
| Sf3b5 | Q923D4 | -0.65 | -1.79 | -0.71 | -1.11 | -2.38 | -2.37 | 0.16 | -1.48 | -1.99 | -1.65 | -1.29 | 0.6 | -3.46 | -1.37 | -0.85 | -1.68 | -0.4525 | -0.0175 | -0.775 | 0.45512 | 0.976087 | 0.199199 | 0.999971 | 1 | 0.813335 |
| Sf3b6 | P59708 | 0.4 | 1.15 | 0.3 | 0.52 | 0.72 | 1.15 | 0.16 | 0.41 | 0.88 | 0.88 | 0.3 | 0.43 | 1 | 1.03 | 0.66 | 0.65 | 0.0175 | 0.03 | 0.2425 | 0.952839 | 0.909397 | 0.341095 | 0.999971 | 1 | 0.860909 |
| Sfn | O70456 | -0.29 | 0.45 | 0.27 | 0.52 | -0.17 | 0.41 | 0.39 | 0.51 | 0.05 | 0.28 | 0.56 | -0.21 | -0.36 | 0.44 | -0.11 | 0.81 | 0.0475 | -0.0675 | -0.0425 | 0.8562 | 0.799603 | 0.895527 | 0.999971 | 1 | 0.988571 |
| Sfpq | Q8VIJ6 | 2.09 | 1.83 | 1.92 | 2.01 | 1.83 | 1.87 | 1.88 | 1.91 | 1.82 | 1.87 | 1.8 | 1.94 | 1.28 | 1.8 | 1.84 | 1.82 | -0.09 | -0.105 | -0.2775 | 0.588308 | 0.523921 | 0.195547 | 0.999971 | 1 | 0.811225 |
| Sfr1 | Q8BP27 | 1.17 | 1.56 | 3.42 | 3.43 | 1.95 | 1.03 | 2.99 | 3.2 | 1.61 | 1.11 | 3 | 3.42 | 1.1 | 0.91 | 3.28 | 3.52 | -0.1025 | -0.11 | -0.1925 | 0.8846 | 0.881654 | 0.815599 | 0.999971 | 1 | 0.974918 |
| Sfrp1 | Q8C4U3 | -0.65 | -0.03 | -0.74 | -0.86 | 0.01 | -0.29 | -0.74 | -0.21 | -0.45 | -1.52 | -1.32 | -0.1 | -0.08 | -0.38 | -0.43 | -0.67 | 0.2625 | -0.2775 | 0.18 | 0.334973 | 0.467537 | 0.478282 | 0.999971 | 1 | 0.894438 |
| Sfswap | Q3USH5 | 2.17 | 1.82 | 2.35 | 2.87 | 2.68 | 2.21 | 3.31 | 2 | 2.36 | 2.46 | 1.12 | 1.56 | 3.45 | 3.27 | 2.25 | 1.32 | 0.2475 | -0.4275 | 0.27 | 0.492848 | 0.27554 | 0.593511 | 0.999971 | 1 | 0.923397 |
| Sft2d2 | A0A0A6YX. | -4.76 | -3.21 | -0.36 | -0.61 | -2.96 | -1.95 | -0.95 | -0.72 | -1.59 | -3.32 | -0.28 | -1.05 | -3.79 | -3.11 | -0.67 | -1.12 | 0.59 | 0.675 | 0.0625 | 0.577349 | 0.548566 | 0.95699 | 0.999971 | 1 | 0.998513 |
| Sfxn1 | Q99JR1 | 0.84 | -0.25 | -0.24 | 0.21 | -0.43 | -0.19 | 0.07 | -0.12 | -0.48 | 0.2 | -0.5 | -0.49 | -0.82 | -0.13 | -0.52 | -0.4 | -0.3075 | -0.4575 | -0.6075 | 0.303204 | 0.169279 | 0.070889 | 0.999971 | 1 | 0.736129 |
| Sfxn2 | Q925N2 | 1.54 | 0.85 | 0.91 | 1.84 | 1.84 | 1.24 | 2.45 | 0.19 | 1.02 | 0.9 | 0.51 | 0.23 | -0.31 | 0.19 | 0.76 | -0.33 | 0.145 | -0.62 | -1.2075 | 0.771984 | 0.070511 | 0.006935 | 0.999971 | 1 | 0.582714 |
| Sfxn3 | Q91V61 | 0.05 | -0.36 | -0.28 | -0.19 | -0.14 | -0.52 | -0.34 | -0.04 | -0.48 | -0.32 | 0.04 | -0.19 | -0.16 | -0.29 | 0.06 | 0.35 | -0.065 | -0.0425 | 0.185 | 0.743851 | 0.829331 | 0.398663 | 0.999971 | 1 | 0.871816 |
| Sfxn4 | Q925N1 | 0.37 | 3.76 | -0.46 | -0.63 | 4.86 | 2.47 | 0.45 | -0.03 | 5.59 | 0.32 | 0.75 | 1.13 | 8.86 | 2.45 | 0.37 | -0.82 | 1.1775 | 1.1875 | 1.955 | 0.391212 | 0.41504 | 0.367106 | 0.999971 | 1 | 0.869526 |
| Sfxn5 | Q925N0 | -0.54 | -0.69 | -1.39 | -1.16 | -0.91 | -0.75 | -1.2 | -0.64 | -0.56 | -0.83 | -1.4 | -1.24 | -0.36 | -0.74 | -1.16 | -1.15 | 0.07 | -0.0625 | 0.0925 | 0.786 | 0.827969 | 0.749249 | 0.999971 | 1 | 0.961064 |
| Sgcd | P82347 | 1.62 | -1.63 | -0.95 | 0.01 | -0.53 | -0.96 | -0.18 | -0.47 | -0.93 | 1.49 | -0.48 | -0.2 | -1.37 | -0.93 | -0.57 | -1.15 | -0.2975 | 0.2075 | -0.7675 | 0.651744 | 0.79487 | 0.26277 | 0.999971 | 1 | 0.842381 |
| Sgce | F6WKE4 | 0.18 | -0.13 | -0.15 | 0.12 | -0.04 | -0.71 | -0.58 | 0.16 | -0.58 | -0.31 | 0.29 | 0.21 | -0.34 | 0.32 | 0.42 | 0.49 | -0.2975 | -0.1025 | 0.2175 | 0.257697 | 0.683931 | 0.379357 | 0.999971 | 1 | 0.871816 |
| Sgcz | Q8BX51 | -1.38 | -1.93 | -1.24 | -0.85 | -1.99 | -2.28 | -1.05 | -0.89 | -1.95 | -1.96 | -1.04 | -1.9 | -0.8 | -1.13 | -1.2 | -0.99 | -0.2025 | -0.3625 | 0.32 | 0.608587 | 0.273846 | 0.244146 | 0.999971 | 1 | 0.837316 |
| Sgip1 | Q8VD37 | -0.96 | -0.15 | -0.31 | -0.79 | -0.68 | -2.01 | -1.17 | -0.86 | -1.03 | -2.51 | -0.99 | -0.09 | -0.3 | -1.02 | -0.19 | -0.44 | -0.6275 | -0.6025 | 0.065 | 0.09746 | 0.248397 | 0.817896 | 0.999971 | 1 | 0.974929 |
| Sgip1_1 | Q8VD37-3 | -1.81 | -1.51 | -1.55 | -1.03 | -2.27 | -2.58 | -2.58 | -0.69 | -2.37 | -1.76 | -0.19 | -1.43 | 0.31 | -3.17 | -1.64 | -1.6 | -0.555 | 0.0375 | -0.05 | 0.239129 | 0.934722 | 0.939895 | 0.999971 | 1 | 0.995869 |
| Sgip1_2 | Q8VD37-8 | -0.37 | -0.92 | -0.18 | -0.9 | -0.84 | -1.1 | -0.15 | -0.62 | -0.9 | -0.76 | -0.12 | 0.22 | -0.49 | -1.18 | -0.61 | -0.92 | -0.085 | 0.2025 | -0.2075 | 0.768044 | 0.537747 | 0.44293 | 0.999971 | 1 | 0.883386 |
| Sgk2 | Q9QZS5-2 | 1.91 | 3.69 | 5.13 | 3.04 | 2.3 | 2.08 | 4.18 | 3.81 | 2.05 | 2.99 | 5.75 | 8.35 | 5.99 | 0.15 | 5.78 | 2.12 | -0.35 | 1.3425 | 0.0675 | 0.650878 | 0.353408 | 0.961599 | 0.999971 | 1 | 0.998721 |
| Sgpl1 | Q8R0X7 | 0.2 | 0.24 | 0.13 | 0.14 | 0.26 | 0.41 | 0.68 | 0.19 | 0.65 | 0.77 | 0.62 | 0.55 | 0.52 | 0.33 | 0.28 | 0.71 | 0.2075 | 0.47 | 0.2825 | 0.278868 | 0.015607 | 0.14461 | 0.999971 | 1 | 0.790933 |
| Sgpp1 | Q9JI99 | 1.56 | -0.06 | -0.13 | 0.3 | 0.4 | 0.83 | 0.06 | -0.35 | 0.39 | 1.39 | -0.4 | -0.61 | 0.19 | 0.69 | -0.54 | -0.84 | -0.1825 | -0.225 | -0.5425 | 0.678667 | 0.685774 | 0.283639 | 0.999971 | 1 | 0.844538 |
| Sgsh | Q9EQ08 | -0.31 | 0.06 | 0.72 | 1.29 | 0.01 | -0.21 | 0.04 | 2.81 | -0.48 | -0.78 | -0.31 | 0.1 | -0.34 | 0.01 | 1.73 | 1.74 | 0.2225 | -0.8075 | 0.345 | 0.758566 | 0.062039 | 0.569236 | 0.999971 | 1 | 0.91816 |
| Sgsm1 | B2RQR5 | -0.49 | -0.94 | -0.62 | -0.48 | -0.74 | -0.74 | -0.83 | -0.41 | -0.53 | -0.38 | -0.44 | -0.42 | -0.13 | -0.48 | 0 | -0.18 | -0.0475 | 0.19 | 0.435 | 0.812405 | 0.311523 | 0.057938 | 0.999971 | 1 | 0.725348 |
| Sgsm2 | Q80U12 | 1.08 | -0.19 | -0.07 | -0.28 | -0.65 | -0.17 | -0.48 | 0.26 | -0.64 | 0.52 | -0.35 | -0.02 | -0.92 | -0.03 | 0.26 | 0.41 | -0.395 | -0.2575 | -0.205 | 0.296024 | 0.513412 | 0.623448 | 0.999971 | 1 | 0.930272 |
| Sgsm3 | Q8VCZ6 | 1.47 | 1.78 | 0.56 | 1.32 | 1.39 | 1.6 | 0.85 | 1.09 | 1.67 | 1.96 | 1.01 | 0.74 | 1.43 | 2.27 | 1.09 | 1.58 | -0.05 | 0.0625 | 0.31 | 0.872576 | 0.866823 | 0.390314 | 0.999971 | 1 | 0.871816 |
| Sgta | Q8BJU0 | 0.29 | 0.4 | 0.56 | 0.29 | 0.72 | 0.03 | 1.44 | 0.14 | 1.56 | -0.37 | 0.99 | 1.36 | 1.56 | -1.03 | 0.36 | -0.19 | 0.1975 | 0.5 | -0.21 | 0.552583 | 0.251889 | 0.680113 | 0.999971 | 1 | 0.942957 |
| Sgtb | Q8VD33 | 0.13 | -0.12 | -0.16 | 0.15 | -0.03 | 0.22 | -0.38 | 0.09 | -0.27 | -0.12 | -0.41 | -0.07 | -0.34 | 0.55 | 0.1 | 0.19 | -0.025 | -0.2175 | 0.125 | 0.903339 | 0.251256 | 0.599261 | 0.999971 | 1 | 0.924139 |
| Sh2b1 | Q91ZM2-2 | 3.65 | 0.27 | 1.87 | 0.67 | -0.36 | 1 | 3.3 | -0.52 | 1.77 | 1.48 | 0.94 | 1.47 | -2.06 | -1.14 | -0.47 | -2.81 | -0.76 | -0.2 | -3.235 | 0.470988 | 0.777847 | 0.003485 | 0.999971 | 1 | 0.577271 |
| Sh2b1_1 | Q91ZM2-4 | 0.41 | 0.4 | 0.47 | 0.44 | 0.33 | 0.24 | 0.11 | 0.26 | 0.3 | 0.38 | 0.12 | 0.09 | 0.5 | 0.39 | 0.25 | 0.74 | -0.195 | -0.2075 | 0.04 | 0.24932 | 0.230284 | 0.826469 | 0.999971 | 1 | 0.976804 |
| Sh2b2 | A0A0G2JEI | -0.18 | -0.27 | -1.32 | -1.22 | -0.53 | -1.18 | -0.04 | -2.12 | -0.73 | -0.14 | -0.95 | -1.35 | -1.46 | 0.14 | -1.14 | -0.94 | -0.22 | -0.045 | -0.1025 | 0.662936 | 0.906003 | 0.813986 | 0.999971 | 1 | 0.974858 |
| Sh2d3c | Q9QZS8-2 | 0.22 | -0.06 | -0.81 | -0.83 | -0.04 | 0.12 | -1.1 | -0.72 | 0.02 | 0.21 | -1.08 | -1 | 0.34 | 0.58 | -0.98 | -0.45 | -0.065 | -0.0925 | 0.2425 | 0.863421 | 0.821393 | 0.571195 | 0.999971 | 1 | 0.91816 |
| Sh2d5 | Q8JZW5 | -0.34 | -0.06 | 0.83 | 0.52 | -0.05 | -0.67 | 0.54 | 0.68 | 0.13 | -0.23 | 0.68 | 0.47 | 0.03 | -0.23 | 0.75 | 0.33 | -0.1125 | 0.025 | -0.0175 | 0.774048 | 0.939861 | 0.958628 | 0.999971 | 1 | 0.998622 |
| Sh3bgrl | Q9JJU8 | 0.17 | 0.03 | 0.71 | 0.52 | 0.33 | -0.37 | 0.85 | 0.55 | 0.16 | 0.06 | 1.02 | 1.24 | -0.17 | -0.26 | 0.83 | 0.65 | -0.0175 | 0.2625 | -0.095 | 0.954734 | 0.441421 | 0.768349 | 0.999971 | 1 | 0.962535 |
| Sh3bgrl2 | Q8BG73 | 0.02 | 0.76 | 0.3 | 0.44 | 0.35 | 0.46 | 0.5 | -0.22 | 0.41 | -0.87 | -0.09 | -0.12 | 0.43 | 0.14 | -0.07 | -0.23 | -0.1075 | -0.5475 | -0.3125 | 0.672978 | 0.105118 | 0.218919 | 0.999971 | 1 | 0.827064 |
| Sh3bgrl3 | Q91VW3 | -0.77 | 0.8 | 0.35 | 0.11 | 0.3 | 0.64 | 1.14 | 0 | 0.15 | -0.7 | 0.77 | 0.49 | 0.11 | 0.18 | 0.62 | -0.01 | 0.3975 | 0.055 | 0.1025 | 0.32711 | 0.899247 | 0.77115 | 0.999971 | 1 | 0.963498 |
| Sh3bp1 | P55194 | 1.47 | -0.18 | 0.05 | 0.27 | -0.32 | -0.02 | 1.02 | -0.79 | -1.27 | 0.14 | 0.05 | -0.9 | -1.13 | -0.44 | -1.14 | -1.27 | -0.43 | -0.8975 | -1.3975 | 0.393731 | 0.086518 | 0.006313 | 0.999971 | 1 | 0.582714 |
| Sh3bp4 | Q921I6 | -0.45 | -0.5 | -0.11 | -0.04 | -0.11 | -0.65 | -0.32 | 0.16 | -0.51 | -0.12 | 0.02 | -0.03 | -0.23 | -0.16 | -0.02 | 0.38 | 0.045 | 0.115 | 0.2675 | 0.851226 | 0.591585 | 0.249522 | 0.999971 | 1 | 0.839012 |
| Sh3bp5 | Q9Z131 | -0.83 | 0.97 | -0.01 | -0.4 | 1.19 | 0.24 | -0.28 | -0.05 | 1.24 | -1.1 | -0.09 | 0.21 | 1.53 | 0.07 | -0.33 | -0.16 | 0.3425 | 0.1325 | 0.345 | 0.471886 | 0.815615 | 0.519818 | 0.999971 | 1 | 0.906375 |
| Sh3bp5l | Q99LH9 | 2.37 | -0.49 | 0.32 | 1.39 | -0.49 | -0.01 | 1.94 | 0.47 | 0 | 1.69 | 0.51 | -0.25 | -0.69 | -0.58 | -0.07 | -0.13 | -0.42 | -0.41 | -1.265 | 0.571405 | 0.556657 | 0.056621 | 0.999971 | 1 | 0.725348 |
| Sh3d19 | A0A571BEI | 1.72 | -0.29 | -0.81 | -0.53 | -0.37 | 0.94 | 0.15 | -0.5 | -0.22 | 1.07 | 0.05 | -0.38 | -1.25 | -0.17 | -0.82 | -0.21 | 0.0325 | 0.1075 | -0.635 | 0.957018 | 0.859377 | 0.28956 | 0.999971 | 1 | 0.847402 |
| Sh3gl1 | Q62419 | -0.39 | 0.18 | -0.34 | -0.13 | -0.49 | 0.13 | -0.67 | -0.29 | -0.24 | -0.29 | -0.21 | -0.59 | 0.17 | -0.29 | -0.69 | -0.08 | -0.16 | -0.1625 | -0.0525 | 0.519403 | 0.43807 | 0.834748 | 0.999971 | 1 | 0.979036 |
| Sh3gl2 | A2ALV3 | -1.12 | -1.39 | -0.84 | -0.84 | -1.81 | -1.62 | -0.37 | -1.37 | -1.74 | -1.62 | -0.54 | -1.26 | -2.18 | -1.56 | -1.2 | -0.91 | -0.245 | -0.2425 | -0.415 | 0.480697 | 0.436962 | 0.203319 | 0.999971 | 1 | 0.81617 |
| Sh3gl3 | A0A0R4J0E | -0.45 | -0.86 | -0.76 | -0.66 | -0.92 | -0.99 | -1.13 | -0.54 | -0.95 | -0.67 | -1.14 | -1.2 | -0.71 | -0.6 | -0.76 | -0.7 | -0.2125 | -0.3075 | -0.01 | 0.318657 | 0.151941 | 0.955085 | 0.999971 | 1 | 0.998056 |
| Sh3glb1 | Q9JK48-2 | -0.38 | -0.71 | -0.39 | -0.35 | -0.53 | -0.18 | -0.43 | 0.05 | -0.54 | -0.16 | 0.05 | 0.27 | -0.26 | -0.37 | 0.01 | 0.46 | 0.185 | 0.3625 | 0.4175 | 0.384013 | 0.138756 | 0.106704 | 0.999971 | 1 | 0.78268 |
| Sh3glb2 | A2AWI9 | -0.55 | -0.87 | -2.8 | -2.28 | -1.1 | -0.91 | -1.95 | -2.57 | -0.46 | -0.81 | -2.16 | -2.15 | -0.43 | -0.88 | -2.52 | -2.97 | -0.0075 | 0.23 | -0.075 | 0.99013 | 0.720751 | 0.91944 | 0.999971 | 1 | 0.992633 |
| Sh3glb2_1 | A2AWI7 | 0.61 | -0.22 | 1.37 | 1.33 | -0.39 | -0.24 | 0.45 | 1.18 | -0.27 | 0.84 | 1.02 | 0.08 | -0.06 | -0.12 | 0.87 | 1.83 | -0.5225 | -0.355 | -0.1425 | 0.295871 | 0.443919 | 0.794285 | 0.999971 | 1 | 0.970141 |
| Sh3glb2_2 | Q8R3V5-1 | -0.43 | 0.34 | -0.01 | -0.19 | 0.1 | 0.11 | 0.39 | -0.14 | 0.17 | -0.55 | 0.42 | -0.36 | 0.91 | 0.05 | 0.16 | 0.08 | 0.1875 | -0.0075 | 0.3725 | 0.427816 | 0.979281 | 0.203289 | 0.999971 | 1 | 0.81617 |
| Sh3kbp1 | B0R0Y8 | 0.31 | -0.72 | -0.29 | -0.07 | -1.43 | -0.25 | -0.4 | -0.24 | -0.5 | -0.23 | -0.36 | -0.5 | 0.07 | -0.31 | 0.01 | 0.37 | -0.3875 | -0.205 | 0.2275 | 0.286736 | 0.422007 | 0.416931 | 0.999971 | 1 | 0.877375 |
| Sh3kbp1_1 | Q8R550 | -0.44 | -1.11 | -1.46 | -1.37 | -1.02 | -1.07 | -1.21 | -1.41 | -1.08 | -0.4 | -1.56 | -1.04 | -0.57 | -0.63 | -1.31 | -1.08 | -0.0825 | 0.075 | 0.1975 | 0.75786 | 0.820557 | 0.516484 | 0.999971 | 1 | 0.905815 |
| Sh3pxd2a | O89032-2 | -0.22 | -0.7 | -0.16 | -0.27 | -0.74 | -0.49 | -0.18 | -0.07 | -0.76 | -0.49 | -0.2 | -0.2 | -0.57 | -0.82 | -0.39 | -0.57 | -0.0325 | -0.075 | -0.25 | 0.888691 | 0.735862 | 0.244953 | 0.999971 | 1 | 0.837316 |
| Sh3pxd2b | A0A5F8MP | -0.06 | -0.26 | -0.24 | -0.41 | -0.28 | -0.12 | -0.08 | -0.17 | -0.01 | 0.33 | 0.12 | 0.35 | -0.15 | -0.05 | -0.53 | 0.16 | 0.08 | 0.44 | 0.1 | 0.645763 | 0.03659 | 0.638789 | 0.999971 | 1 | 0.93301 |
| Sh3rf1 | E9QQ33 | 1.27 | 0.19 | -0.4 | -0.04 | 0.78 | 0.16 | -0.59 | -0.27 | 0.69 | 0.8 | -0.49 | -0.39 | 0.49 | 0.05 | -0.38 | 0.29 | -0.235 | -0.1025 | -0.1425 | 0.595205 | 0.826244 | 0.715775 | 0.999971 | 1 | 0.952576 |
| Sh3rf2 | Q8BZT2 | -0.42 | 1.07 | -0.49 | 0.02 | -0.57 | -0.56 | -0.52 | 0.67 | -0.27 | -1.46 | -0.96 | -0.65 | -3.36 | -0.27 | -1.72 | 0.23 | -0.29 | -0.88 | -1.325 | 0.518575 | 0.061172 | 0.123111 | 0.999971 | 1 | 0.78268 |
| Sh3rf3 | Q8C120 | -0.11 | -0.18 | 0.49 | 0.47 | -0.08 | 0.36 | -0.24 | 0.84 | -0.42 | -0.02 | 0.71 | 0.44 | 0.31 | 0.33 | 0.52 | 0.74 | 0.0525 | 0.01 | 0.3075 | 0.864772 | 0.974437 | 0.222385 | 0.999971 | 1 | 0.827753 |
| Sh3yl1 | O08641-3 | 0.36 | 0.76 | 0.08 | -0.2 | 1.11 | -0.31 | -0.29 | 0.13 | 0.25 | 0.54 | -0.17 | -0.08 | 0.62 | -0.16 | -0.17 | -0.36 | -0.09 | -0.115 | -0.2675 | 0.811628 | 0.679094 | 0.392721 | 0.999971 | 1 | 0.871816 |
| Shank1 | D3YZU5 | -0.73 | -1.55 | -1.22 | -1.33 | -1.54 | -1.26 | -1.02 | -1.6 | -1.84 | -1.22 | -1.43 | -1.59 | -1.52 | -1.78 | -1.89 | -1.54 | -0.1475 | -0.3125 | -0.475 | 0.555907 | 0.223839 | 0.067557 | 0.999971 | 1 | 0.733937 |
| Shank1_1 | D3YZU4 | -1.98 | -0.8 | -2.2 | -2.43 | -1.58 | -0.97 | -2.82 | -2.15 | -1.78 | -2.18 | -2.15 | -2.24 | -1.19 | -1.22 | -2.79 | -2.25 | -0.0275 | -0.235 | -0.01 | 0.955874 | 0.52743 | 0.983979 | 0.999971 | 1 | 0.99925 |
| Shank2 | Q80Z38-2 | -1.23 | -1.64 | -1.3 | -1.32 | -1.76 | -1.63 | -1.3 | -1.66 | -1.6 | -1.21 | -1.36 | -1.54 | -1.33 | -1.72 | -1.68 | -1.16 | -0.215 | -0.055 | -0.1 | 0.290637 | 0.771964 | 0.641183 | 0.999971 | 1 | 0.93301 |
| Shank2_1 | D3Z5K8 | -1.55 | 4.69 | 1.77 | 0.2 | 4.87 | 4.05 | 0.9 | 0.72 | 4.68 | 0.36 | 2.41 | 2.31 | 7.85 | 2.05 | 1.63 | 0.2 | 1.3575 | 1.1625 | 1.655 | 0.379092 | 0.423182 | 0.394198 | 0.999971 | 1 | 0.871816 |
| Shank2_2 | A0A1C7ZM | -3.43 | -3.71 | -1.76 | -2.37 | -3.23 | -3.88 | -2.54 | -2.3 | -4.43 | -3.2 | -2.02 | -2.13 | -5.84 | -4.25 | -2.35 | -1.6 | -0.17 | -0.1275 | -0.6925 | 0.750293 | 0.846915 | 0.471997 | 0.999971 | 1 | 0.892857 |
| Shank3 | Q4ACU6 | -0.42 | -1.56 | -1.28 | -1.18 | -1.43 | -1.2 | -1.3 | -1.2 | -1.58 | -0.96 | -1.7 | -1.47 | -1.28 | -1.45 | -1.37 | -1.28 | -0.1725 | -0.3175 | -0.235 | 0.526644 | 0.306622 | 0.391392 | 0.999971 | 1 | 0.871816 |
| Shank3_1 | Q4ACU6-9 | -0.16 | 0.04 | -2.08 | -1.38 | -0.91 | -0.41 | -2.12 | -1.4 | -1.16 | -0.16 | -1.68 | -1.84 | -1.01 | -0.8 | -1.47 | -1.61 | -0.315 | -0.315 | -0.3275 | 0.583711 | 0.59053 | 0.518335 | 0.999971 | 1 | 0.905869 |
| Sharpin | Q91WA6 | -1.28 | -0.66 | -0.47 | 0.32 | -0.15 | -0.38 | -0.44 | 0.72 | -0.28 | -0.86 | 0.78 | 0.28 | 0.18 | 0.22 | 0.29 | 0.22 | 0.46 | 0.5025 | 0.75 | 0.274382 | 0.287283 | 0.04653 | 0.999971 | 1 | 0.710102 |
| Shb | Q6PD21 | 0.7 | -0.16 | 0.47 | 0.58 | -0.36 | -0.35 | 0.17 | 0.51 | -0.45 | 0.48 | 0.78 | 0.75 | -0.29 | 0.19 | 0.24 | 0.23 | -0.405 | -0.0075 | -0.305 | 0.194331 | 0.982484 | 0.253562 | 0.999971 | 1 | 0.839012 |
| Shc2 | Q8BMC3 | 0.12 | 0.82 | 0.07 | -0.54 | 0.48 | 0.63 | -0.72 | -0.04 | 0.18 | 1.21 | 0.48 | 0.52 | 1.68 | 1.23 | 0.73 | 0.38 | -0.03 | 0.48 | 0.8875 | 0.939455 | 0.191546 | 0.043559 | 0.999971 | 1 | 0.701725 |
| Shc3 | A0A5F8MP | -0.68 | 0.73 | -0.21 | -0.05 | 1 | 0.42 | -0.29 | 0.16 | 0.65 | -0.33 | 0.08 | -0.15 | 1.69 | 0.61 | 0.07 | 0.15 | 0.375 | 0.115 | 0.6825 | 0.341523 | 0.746672 | 0.152539 | 0.999971 | 1 | 0.795578 |
| Shd | O88834-2 | 1.43 | -0.12 | 0.64 | 0.68 | -0.57 | -0.35 | 0.65 | 0.45 | -0.05 | 0.57 | 0.38 | 0.04 | -0.1 | 0.12 | 0.35 | 0.63 | -0.6125 | -0.4225 | -0.4075 | 0.163487 | 0.239628 | 0.260917 | 0.999971 | 1 | 0.841256 |
| Shf | A2AQ87 | 2.44 | -0.64 | 0.06 | -0.06 | -0.22 | 0.34 | 0.54 | 0.27 | -0.55 | 0.76 | 0 | -0.07 | -1.15 | 0.07 | -0.16 | 0.05 | -0.2175 | -0.415 | -0.7475 | 0.732922 | 0.53917 | 0.282812 | 0.999971 | 1 | 0.844538 |
| Shfl | Q8CAK3 | 1.18 | 1.5 | 1.05 | 0.99 | 1.53 | 1.61 | 0.57 | 0.96 | 1.6 | 1.38 | 1.09 | 1.01 | 1.59 | 1.57 | 0.82 | 1.4 | -0.0125 | 0.09 | 0.165 | 0.964966 | 0.681738 | 0.505462 | 0.999971 | 1 | 0.905277 |
| Shisa4 | Q8CA71 | -1.05 | -0.87 | -0.16 | -0.34 | -0.66 | -1.23 | 0.41 | -0.24 | -1.54 | -0.77 | 0.01 | 0.6 | -3.34 | 0.14 | -0.51 | -0.09 | 0.175 | 0.18 | -0.345 | 0.654706 | 0.707896 | 0.648917 | 0.999971 | 1 | 0.933842 |
| Shisa6 | Q3UH99-2 | -0.92 | -1 | -1.42 | -1.71 | -1.32 | -0.72 | -1.16 | -1.59 | -1.29 | -1.19 | -1.74 | -2.17 | -0.83 | -1.43 | -2.09 | -1.89 | 0.065 | -0.335 | -0.2975 | 0.814297 | 0.280779 | 0.384809 | 0.999971 | 1 | 0.871816 |
| Shisa7 | Q8C3Q5-2 | -0.56 | -1.6 | -0.75 | -0.75 | -1.81 | -1.31 | 0.87 | -0.94 | -1.69 | -0.99 | -0.88 | -1.41 | -1.55 | -2.06 | -1.05 | -0.99 | 0.1175 | -0.3275 | -0.4975 | 0.838139 | 0.299548 | 0.167389 | 0.999971 | 1 | 0.800215 |
| Shisa9 | Q9CZN4-3 | -1.26 | -1 | -0.58 | -0.99 | -0.93 | -1.05 | -0.87 | -0.55 | -0.74 | -1.19 | -0.75 | -0.56 | -0.15 | -1.32 | -0.74 | -0.7 | 0.1075 | 0.1475 | 0.23 | 0.627725 | 0.525415 | 0.436113 | 0.999971 | 1 | 0.882419 |
| Shmt1 | G3UZ26 | 0.92 | 0.34 | 0.07 | 0.11 | -0.47 | -0.05 | -0.52 | -0.1 | 0.11 | 0.97 | -0.49 | 0.2 | -0.46 | 0.51 | -0.19 | -0.6 | -0.645 | -0.1625 | -0.545 | 0.030747 | 0.645898 | 0.11447 | 0.999971 | 1 | 0.78268 |
| Shmt2 | Q9CZN7-2 | -0.47 | -1.06 | -0.58 | -0.61 | -1.4 | -1.36 | -0.61 | -0.63 | -1.35 | -0.56 | -0.43 | -1.15 | -1.9 | -1.11 | -0.82 | -0.64 | -0.32 | -0.1925 | -0.4375 | 0.260407 | 0.489538 | 0.185514 | 0.999971 | 1 | 0.809025 |
| Shoc2 | O88520 | -0.4 | 0.04 | 0.32 | -0.57 | 0.18 | 0.4 | 0.85 | -0.24 | 0.28 | -0.64 | 0.31 | 0.35 | -0.24 | -0.56 | 0.11 | -0.03 | 0.45 | 0.2275 | -0.0275 | 0.171311 | 0.478795 | 0.91889 | 0.999971 | 1 | 0.992633 |
| Shpk | Q9D5J6 | 0.12 | 0.35 | -0.66 | 0.01 | 0.24 | 0.07 | -0.14 | 0.54 | 0.24 | 0.34 | -0.39 | -0.57 | 0.26 | 0.43 | 0.12 | 0.17 | 0.2225 | -0.05 | 0.29 | 0.430219 | 0.874335 | 0.272084 | 0.999971 | 1 | 0.843794 |
| Shprh | Q7TPQ3-2 | -0.36 | 0.65 | 1.98 | 1.64 | 0.74 | 0.74 | 1.65 | 1.98 | -0.03 | 0.13 | 2.51 | 2.06 | -0.4 | 0.36 | 1.79 | 2.31 | 0.3 | 0.19 | 0.0375 | 0.597901 | 0.802982 | 0.959523 | 0.999971 | 1 | 0.998721 |
| Shq1 | Q7TMX5-2 | -0.82 | 1.34 | 0.68 | 0.18 | -1.35 | -0.1 | 0.43 | 0.15 | -1.18 | -0.38 | 0.64 | 0.97 | -1.2 | -0.73 | -0.51 | -0.11 | -0.5625 | -0.3325 | -0.9825 | 0.322951 | 0.590917 | 0.064655 | 0.999971 | 1 | 0.726597 |
| Shroom1 | D3Z3D5 | 0.07 | 1.08 | -0.23 | 0.04 | 1.09 | -0.23 | 0.56 | 2.4 | 1.46 | -1.03 | 1.46 | 0.81 | -0.54 | -0.3 | 0.82 | 0.28 | 0.715 | 0.435 | -0.175 | 0.231234 | 0.476431 | 0.664581 | 0.999971 | 1 | 0.93697 |
| Shroom2 | A7TU71 | -0.48 | 0.22 | 0.24 | 0.18 | 0.47 | -0.02 | 0.22 | 0.48 | 0.39 | -0.36 | 0.45 | 0.4 | 0.93 | -0.07 | 0.7 | 0.44 | 0.2475 | 0.18 | 0.46 | 0.321531 | 0.519321 | 0.13827 | 0.999971 | 1 | 0.784948 |
| Shroom4 | Q1W617-2 | 3.13 | -2.35 | 0.03 | 0.6 | 0.96 | 0.13 | -0.86 | 0.21 | -0.1 | 1.29 | -0.95 | 0.56 | 2.8 | -2.02 | 1.01 | 1.66 | -0.2425 | -0.1525 | 0.51 | 0.818473 | 0.889325 | 0.707654 | 0.999971 | 1 | 0.950182 |
| Shtn1 | Q8K2Q9-2 | -0.18 | -0.73 | -0.81 | -0.1 | -0.47 | -0.68 | -0.67 | -0.37 | -0.11 | 0.24 | -0.27 | -0.2 | -0.08 | -0.36 | -0.17 | 0.18 | -0.0925 | 0.37 | 0.3475 | 0.694151 | 0.155204 | 0.181603 | 0.999971 | 1 | 0.808532 |
| Siae | A0A1N9M4 | 1.25 | 2.61 | 1.38 | 1.17 | 1.38 | 2.24 | 1.15 | 2.7 | 1.58 | 1.47 | 0.86 | 0.17 | 2.03 | 2.31 | 0.75 | 0.82 | 0.265 | -0.5825 | -0.125 | 0.571728 | 0.211374 | 0.799149 | 0.999971 | 1 | 0.970479 |
| Sidt1 | E9Q247 | -0.91 | -0.92 | -0.59 | -1.04 | -0.83 | -0.21 | -0.8 | -0.68 | -0.76 | -0.14 | -0.22 | -0.25 | -0.05 | -0.5 | -0.26 | 0.25 | 0.235 | 0.5225 | 0.725 | 0.297572 | 0.035554 | 0.010418 | 0.999971 | 1 | 0.613095 |
| Sidt2 | E0CXC7 | 2.47 | -0.4 | -1.54 | -1.33 | 0.9 | -1.53 | -1.17 | -0.98 | -1.03 | -0.33 | -0.29 | 0.03 | -1.87 | -0.04 | -1.04 | -1.02 | -0.495 | -0.205 | -0.7925 | 0.607768 | 0.81132 | 0.38622 | 0.999971 | 1 | 0.871816 |
| Sigmar1 | O55242 | 1.33 | -0.67 | -1.14 | -2.78 | 0.08 | 0.43 | -0.96 | -1.62 | -0.2 | 2.13 | -0.48 | -0.83 | 0.19 | -0.21 | -1.85 | -2.07 | 0.2975 | 0.97 | -0.17 | 0.732316 | 0.332401 | 0.852496 | 0.999971 | 1 | 0.98187 |
| Sik2 | H3BJC9 | -0.68 | -0.53 | -0.52 | -0.99 | -0.03 | -0.8 | 0.12 | -1.49 | -0.25 | -1.08 | -0.24 | -0.16 | -1.04 | -0.84 | -1.21 | -1.95 | 0.13 | 0.2475 | -0.58 | 0.729946 | 0.359717 | 0.065178 | 0.999971 | 1 | 0.727266 |
| Sik3 | Q6P4S6 | -0.47 | 0.84 | 0.36 | 0.29 | 1.27 | -0.28 | 0.43 | 0.05 | 1.89 | -0.21 | 0.25 | 0.58 | 1.78 | -0.52 | 0.15 | 0.1 | 0.1125 | 0.3725 | 0.1225 | 0.783715 | 0.455846 | 0.813687 | 0.999971 | 1 | 0.974756 |
| Sike1 | Q9CPR7 | 0.47 | 0.62 | 0.07 | 0.34 | 0.42 | 0.13 | 0.53 | 1.06 | 0.42 | 0.51 | 0.78 | 0.9 | 0.76 | 0.56 | 0.82 | 0.43 | 0.16 | 0.2775 | 0.2675 | 0.531986 | 0.207388 | 0.211559 | 0.999971 | 1 | 0.823077 |
| Sil1 | A0A494B9I | -1.67 | -0.17 | 0.58 | 0.22 | -0.38 | -0.29 | -0.89 | 0.11 | -0.55 | -0.42 | 0.39 | 0.91 | 0.34 | 0.51 | 0.78 | 0.74 | -0.1025 | 0.3425 | 0.8525 | 0.836521 | 0.542829 | 0.098522 | 0.999971 | 1 | 0.78268 |
| Sin3a | Q60520-1 | 0.77 | 1.79 | 0.88 | 0.68 | 1.53 | 1.23 | 1.21 | 0.5 | 2.09 | 0.63 | 1.08 | 1.34 | 2.84 | 0.62 | 1.1 | 0.54 | 0.0875 | 0.255 | 0.245 | 0.794093 | 0.513496 | 0.656118 | 0.999971 | 1 | 0.934061 |
| Sin3b | Q62141-1 | -0.36 | 0.79 | -0.03 | 0.09 | 0.62 | 0.33 | -0.13 | -0.29 | 0.22 | -0.1 | 0.01 | -0.13 | 0.18 | 0.69 | -0.21 | 0.18 | 0.01 | -0.1225 | 0.0875 | 0.975158 | 0.653924 | 0.778274 | 0.999971 | 1 | 0.965323 |
| Sipa1 | P46062 | 1.55 | -0.88 | -0.03 | 0.62 | -0.22 | -1.66 | 0.17 | 0.74 | -0.37 | -0.12 | 0.86 | 0.82 | -1.13 | -3.64 | 1.02 | 1.66 | -0.5575 | -0.0175 | -0.8375 | 0.406496 | 0.974923 | 0.47759 | 0.999971 | 1 | 0.894438 |
| Sipa1l1 | Q8C0T5 | -0.92 | -0.99 | -0.96 | -0.84 | -1.01 | -0.7 | -0.87 | -0.73 | -0.96 | -0.96 | -0.95 | -1.01 | -0.35 | -1.04 | -1.07 | -0.96 | 0.1 | -0.0425 | 0.0725 | 0.562645 | 0.785171 | 0.741012 | 0.999971 | 1 | 0.960899 |
| Sipa1l2 | A0A1D5RL\ | 0.32 | -0.35 | 0.01 | -0.23 | -0.47 | -0.14 | 0.29 | 0.01 | -0.35 | 0.26 | 0.26 | 0.16 | -0.45 | -0.09 | -0.51 | -0.37 | -0.015 | 0.145 | -0.2925 | 0.951388 | 0.547787 | 0.205803 | 0.999971 | 1 | 0.817241 |
| Sipa1l3 | G3X9J0 | -0.31 | -0.34 | -0.2 | -0.46 | -0.47 | -0.22 | -0.49 | -0.31 | -0.77 | -0.63 | -0.45 | -0.36 | -0.44 | -0.58 | -0.58 | -0.3 | -0.045 | -0.225 | -0.1475 | 0.795192 | 0.230048 | 0.407889 | 0.999971 | 1 | 0.874911 |
| Sirpa | P97797-2 | -0.96 | -1.16 | -1.65 | -1.79 | -0.74 | -1.04 | -0.68 | -1.68 | -0.92 | -1.27 | -1.56 | -1.39 | -0.54 | -1.42 | -1.98 | -1.54 | 0.355 | 0.105 | 0.02 | 0.2672 | 0.688075 | 0.954906 | 0.999971 | 1 | 0.998056 |
| Sirpa_1 | P97797 | -0.55 | -1.6 | -1.46 | -1.28 | -0.39 | -1.26 | -1.85 | -1.41 | -0.52 | -1.22 | -2.23 | -1.52 | 0.49 | -0.74 | -2.47 | -1.74 | -0.005 | -0.15 | 0.1075 | 0.989288 | 0.712716 | 0.862929 | 0.999971 | 1 | 0.982963 |
| Sirt2 | Q8VDQ8 | -1.66 | -1.41 | -1.33 | -1.83 | -1.71 | -1.24 | -2.03 | -1.6 | -1.43 | -1.84 | -2.03 | -2.01 | -1.46 | -1.47 | -1.29 | -1.81 | -0.0875 | -0.27 | 0.05 | 0.710315 | 0.241385 | 0.812006 | 0.999971 | 1 | 0.974279 |
| Sirt3 | Q8R104 | -0.14 | 1.01 | 1.64 | 1.24 | 0.01 | 0.6 | 0.64 | 1.43 | 0.13 | -0.27 | 0.45 | -1.71 | -0.35 | 0.69 | 0.68 | 0.39 | -0.2675 | -1.2875 | -0.585 | 0.556957 | 0.04564 | 0.19655 | 0.999971 | 1 | 0.812891 |
| Sirt4 | Q8R216 | -2.47 | -2.28 | 0.01 | -0.64 | -1.98 | -1.32 | -0.26 | 0.31 | -2.62 | -1.95 | -0.93 | -1.86 | -1.86 | -1.98 | 0.25 | -0.55 | 0.5325 | -0.495 | 0.31 | 0.467226 | 0.448892 | 0.674399 | 0.999971 | 1 | 0.941037 |
| Sirt5 | A0A1Y7VJY | 0.65 | 0.5 | 0.22 | 0.14 | 0.05 | 0.5 | 0.23 | 0.31 | 0 | 0.77 | 0.11 | -0.69 | -0.12 | -0.12 | -0.16 | -0.39 | -0.105 | -0.33 | -0.575 | 0.611741 | 0.321656 | 0.01632 | 0.999971 | 1 | 0.613095 |
| Skiv2l | Q6NZR5 | 0.34 | 0.3 | -0.43 | -0.27 | 0.29 | 0.36 | -0.19 | -0.19 | 0.19 | 0.43 | -0.23 | -0.1 | 0.17 | 0.57 | -0.26 | -0.09 | 0.0825 | 0.0875 | 0.1125 | 0.757923 | 0.742228 | 0.691376 | 0.999971 | 1 | 0.946835 |
| Skp1 | Q9WTX5 | -1.77 | -0.11 | -0.88 | -2.3 | 0.06 | -0.3 | -0.09 | -1.92 | 0.31 | -1.73 | -0.46 | -0.7 | 0.16 | -0.69 | -1.49 | -2.72 | 0.7025 | 0.62 | 0.08 | 0.26604 | 0.30881 | 0.909785 | 0.999971 | 1 | 0.991822 |
| Skp2 | A0A087WC | 0.63 | -1.04 | 0.37 | 0.07 | -0.74 | -0.86 | 0.6 | 0.13 | -1.5 | 0.44 | -0.41 | -0.1 | -1.1 | -1.33 | -0.55 | -1.48 | -0.225 | -0.4 | -1.1225 | 0.636794 | 0.442218 | 0.020003 | 0.999971 | 1 | 0.635388 |
| Slain1 | Q68FF7 | 1.47 | 2.35 | 0.61 | 0.24 | 1.37 | 2.62 | 0.41 | 0.47 | 2.3 | 2.86 | -0.36 | -0.23 | 0.52 | 1.88 | 0.31 | 0.06 | 0.05 | -0.025 | -0.475 | 0.937199 | 0.97699 | 0.415017 | 0.999971 | 1 | 0.876789 |
| Slain2 | Q8CI08-2 | 0.38 | 0.34 | 1.06 | -0.29 | 0.29 | 0.26 | 1.87 | -0.22 | 0.04 | 0.23 | 0.49 | 0.64 | 0.05 | -0.33 | 0.24 | -0.67 | 0.1775 | -0.0225 | -0.55 | 0.72018 | 0.942243 | 0.132264 | 0.999971 | 1 | 0.784948 |
| Slc10a4 | Q3UEZ8 | 0.22 | -0.67 | 1.88 | 1.06 | -0.83 | 0.3 | 0.85 | 1.86 | -0.42 | 0.32 | 1.8 | 1.02 | -0.66 | 0.07 | 1.82 | 2.49 | -0.0775 | 0.0575 | 0.3075 | 0.912799 | 0.930733 | 0.710192 | 0.999971 | 1 | 0.950695 |
| Slc11a2 | P49282-2 | -1.24 | -2.59 | 0.35 | 0.67 | -2.05 | -2.8 | -0.1 | 1.15 | -1.69 | -2.15 | 1.5 | 0.84 | -2.85 | -2.72 | 0.4 | 0.15 | -0.2475 | 0.3275 | -0.5525 | 0.813487 | 0.757771 | 0.597788 | 0.999971 | 1 | 0.92377 |
| Slc11a2_1 | P49282 | 0.74 | -0.37 | -0.38 | 0.61 | -0.48 | -2.05 | 0.08 | -0.4 | -0.63 | 0.11 | -0.55 | -0.18 | -0.22 | -0.63 | -1.11 | -1.09 | -0.8625 | -0.4625 | -0.9125 | 0.119336 | 0.202831 | 0.031092 | 0.999971 | 1 | 0.678474 |
| Slc12a2 | E9QM38 | -0.85 | -1.02 | -0.84 | -0.99 | -1.16 | -1.31 | -1.46 | -1.03 | -0.87 | -1.06 | -1.78 | -1.08 | -0.89 | -0.99 | -0.94 | -1.07 | -0.315 | -0.2725 | -0.0475 | 0.109564 | 0.268514 | 0.775332 | 0.999971 | 1 | 0.96393 |
| Slc12a4 | Q9JIS8 | 0.25 | 0.25 | -0.34 | -0.59 | 0.15 | 0.08 | -0.49 | -0.56 | 0.16 | 0.4 | -0.49 | -0.2 | -0.45 | -0.19 | -0.31 | -0.7 | -0.0975 | 0.075 | -0.305 | 0.740037 | 0.801224 | 0.263887 | 0.999971 | 1 | 0.842533 |
| Slc12a5 | Q91V14 | -0.75 | -1.24 | -1.79 | -1.41 | -1.37 | -1.17 | -1.55 | -1.79 | -1.42 | -1.06 | -1.71 | -2.05 | -1.11 | -1.18 | -1.87 | -1.46 | -0.1725 | -0.2625 | -0.1075 | 0.529998 | 0.402721 | 0.710987 | 0.999971 | 1 | 0.950695 |
| Slc12a5_1 | Q91V14-2 | -0.75 | -0.34 | -2.27 | -2.18 | -1 | -1.25 | -2.63 | -1.84 | -0.73 | -0.9 | -2.25 | -2.19 | -0.83 | -0.72 | -2.51 | -1.69 | -0.295 | -0.1325 | -0.0525 | 0.601571 | 0.821681 | 0.92918 | 0.999971 | 1 | 0.994545 |
| Slc12a6 | Q924N4 | -1.43 | -1.77 | -1.27 | -1.59 | -1.69 | -1.8 | -1.28 | -1.98 | -1.45 | -1.32 | -1.51 | -1.73 | -1.55 | -1.73 | -1.9 | -2.02 | -0.1725 | 0.0125 | -0.285 | 0.449803 | 0.948794 | 0.186856 | 0.999971 | 1 | 0.809025 |
| Slc12a9 | Q99MR3 | 0.02 | 0.57 | -0.35 | -0.38 | 0.54 | 0.05 | -0.33 | 0.19 | 0.28 | 0.18 | -0.43 | -0.56 | 1.07 | 0.02 | -0.42 | -0.27 | 0.1475 | -0.0975 | 0.135 | 0.619683 | 0.754228 | 0.72841 | 0.999971 | 1 | 0.956041 |
| Slc13a3 | A2A4U9 | 0.13 | -1 | -0.38 | 0.13 | -0.68 | -0.81 | -0.93 | -0.58 | -0.86 | 0.04 | -0.32 | -0.32 | -0.83 | -0.41 | 0.44 | -0.1 | -0.47 | -0.085 | 0.055 | 0.132521 | 0.794769 | 0.88143 | 0.999971 | 1 | 0.986007 |
| Slc13a4 | Q8BZ82 | 1.75 | 0.88 | 2 | 2.66 | 2.05 | 0.61 | 2.91 | 0.27 | 1.49 | 1.18 | 0.08 | 2.6 | -1.72 | 1.13 | 1.76 | 0.05 | -0.3625 | -0.485 | -1.5175 | 0.582028 | 0.416416 | 0.073374 | 0.999971 | 1 | 0.746069 |
| Slc13a5 | Q5NBV0 | -0.59 | -0.08 | -0.99 | -0.99 | -0.62 | 0.57 | -1.31 | -1.13 | -0.42 | -0.35 | -1.01 | -1.53 | -0.29 | -0.01 | -1.06 | -0.55 | 0.04 | -0.165 | 0.185 | 0.928582 | 0.635697 | 0.560947 | 0.999971 | 1 | 0.91591 |
| Slc14a1 | Q8VHL0 | 0.3 | -0.2 | 0.02 | -0.3 | -0.21 | 0.79 | 0.13 | 0.18 | 0.04 | 0.83 | -0.32 | -0.51 | 0.06 | 0.45 | 0.13 | -0.24 | 0.2675 | 0.055 | 0.145 | 0.331127 | 0.865631 | 0.536827 | 0.999971 | 1 | 0.910077 |
| Slc15a2 | Q9JM03 | -0.75 | -0.81 | -1.82 | -1.4 | 0.07 | -0.21 | -0.33 | -1.98 | 0.85 | -0.68 | -0.38 | -0.71 | 0.97 | -0.75 | 0.08 | 0.04 | 0.5825 | 0.965 | 1.28 | 0.254942 | 0.046741 | 0.012332 | 0.999971 | 1 | 0.613095 |
| Slc15a4 | Q91W98 | 1.4 | -0.28 | 0.73 | 1.02 | 0.19 | 0.73 | 0.38 | 0.91 | -0.78 | 0.89 | 0.81 | -0.23 | 0.19 | 0.1 | 0.32 | 1.41 | -0.165 | -0.545 | -0.2125 | 0.666474 | 0.297884 | 0.634665 | 0.999971 | 1 | 0.93301 |
| Slc16a1 | P53986 | 0.5 | -1.04 | 0.25 | 0.08 | -0.45 | 0 | 0.19 | -0.02 | -0.23 | 0.92 | -0.55 | -0.76 | -0.68 | -0.47 | -0.12 | -0.12 | -0.0175 | -0.1025 | -0.295 | 0.960961 | 0.828937 | 0.421221 | 0.999971 | 1 | 0.87788 |
| Slc16a10 | Q3U9N9 | -1.51 | -0.83 | -0.29 | -1.33 | -0.68 | -2.57 | -1.21 | -1.66 | -2.01 | 0.31 | -1.8 | -0.3 | -1.31 | -2.14 | -0.97 | -1.59 | -0.54 | 0.04 | -0.5125 | 0.254429 | 0.944917 | 0.180363 | 0.999971 | 1 | 0.808532 |
| Slc16a2 | O70324 | 0.16 | 0.13 | -0.7 | -0.43 | -0.25 | -0.41 | -1.19 | 0.32 | -0.23 | 0.4 | -0.81 | -0.76 | 0.35 | -0.38 | -0.85 | -0.6 | -0.1725 | -0.14 | -0.16 | 0.639816 | 0.688551 | 0.633675 | 0.999971 | 1 | 0.93301 |
| Slc16a3 | P57787 | -0.04 | -0.35 | 1.68 | -0.55 | 0.02 | -0.39 | 2.1 | 1.65 | 0.07 | -0.02 | 2.07 | 3.12 | 0.51 | -0.81 | 0.36 | 0.89 | 0.66 | 1.125 | 0.0525 | 0.368365 | 0.201769 | 0.92724 | 0.999971 | 1 | 0.99417 |
| Slc16a7 | O70451 | -1.41 | -1.24 | -1.39 | -1 | -1.98 | -1.03 | -0.3 | -1.36 | -1.59 | -1.32 | -0.35 | -1.47 | -1.44 | -0.8 | -1.12 | -1.03 | 0.0925 | 0.0775 | 0.1625 | 0.794563 | 0.799351 | 0.452636 | 0.999971 | 1 | 0.887217 |
| Slc17a5 | Q8BN82 | 1.2 | -0.05 | 0.46 | -0.1 | 0.77 | 1.51 | 0.16 | 2.62 | 0.95 | 0.93 | 1.22 | 0.08 | 0.69 | 1.04 | 1.15 | 1.71 | 0.8875 | 0.4175 | 0.77 | 0.138926 | 0.28681 | 0.057942 | 0.999971 | 1 | 0.725348 |
| Slc17a6 | A0A0R4J0A | -0.51 | -0.7 | -1.11 | -1 | -0.61 | -0.39 | -1.29 | -1.29 | -0.42 | -0.56 | -1.17 | -1.1 | 0.03 | -0.4 | -1.33 | -0.99 | -0.065 | 0.0175 | 0.1575 | 0.819072 | 0.94558 | 0.637416 | 0.999971 | 1 | 0.93301 |
| Slc17a7 | Q3TXX4 | -0.23 | -0.91 | -0.73 | -0.97 | -0.9 | -0.11 | -0.25 | -1.28 | -0.51 | 0.04 | -0.46 | -0.75 | -0.34 | -0.82 | -0.98 | -0.82 | 0.075 | 0.29 | -0.03 | 0.816863 | 0.279336 | 0.903546 | 0.999971 | 1 | 0.991091 |
| Slc17a8 | Q8BFU8 | -2.93 | -0.98 | -0.27 | 0.5 | -1.09 | -2.8 | -0.58 | 0.34 | -3.45 | -3.47 | -1.57 | -1.28 | 1.27 | -1.83 | 0.45 | 0.71 | -0.1125 | -1.5225 | 1.07 | 0.898658 | 0.101633 | 0.253284 | 0.999971 | 1 | 0.839012 |
| Slc18a2 | Q8BRU6 | 2 | -1.45 | -2.08 | -1.03 | -1.62 | -1.52 | -1.1 | -1.34 | -1.84 | 0.2 | -2.32 | -2.43 | -1.95 | -1.91 | -1.23 | -2.57 | -0.755 | -0.9575 | -1.275 | 0.368916 | 0.344069 | 0.159321 | 0.999971 | 1 | 0.799149 |
| Slc18a3 | Q3TYJ1 | -1.49 | -1.42 | -1.05 | -0.62 | -0.86 | -1.4 | -1.19 | -1.08 | -1.55 | -1.4 | -1.34 | -1.41 | -1.58 | -0.78 | -0.99 | -1.38 | 0.0125 | -0.28 | -0.0375 | 0.960867 | 0.255943 | 0.895153 | 0.999971 | 1 | 0.988571 |
| Slc19a1 | P41438-2 | -1.58 | -1.16 | 0.4 | 0.42 | -0.75 | -1.8 | -0.37 | 0.37 | -0.73 | -1.18 | -0.1 | 1.48 | -0.38 | -1.33 | 1.72 | 0.91 | -0.1575 | 0.3475 | 0.71 | 0.801914 | 0.626787 | 0.368267 | 0.999971 | 1 | 0.869526 |
| Slc19a2 | Q9EQN9 | 0.71 | 2.54 | -1.64 | -1.77 | 1.41 | 2.39 | -1.5 | -1.3 | 0.88 | 0.39 | -0.73 | 0.01 | 2.85 | 3.38 | -0.64 | -0.87 | 0.29 | 0.1775 | 1.22 | 0.818093 | 0.855839 | 0.379538 | 0.999971 | 1 | 0.871816 |
| Slc1a1 | P51906 | -0.57 | -0.27 | -0.09 | -0.1 | -0.58 | -0.68 | -0.3 | 0.05 | -0.54 | -0.24 | -0.54 | -0.27 | -0.22 | -0.52 | 0.07 | 0.41 | -0.12 | -0.14 | 0.1925 | 0.610844 | 0.481426 | 0.45771 | 0.999971 | 1 | 0.887945 |
| Slc1a2 | P43006 | -1.24 | -1.53 | -0.66 | -0.54 | -1.24 | -1.49 | -0.87 | -0.85 | -1.58 | -1.94 | -0.94 | -1.66 | -1.31 | -1.98 | -1.06 | -0.86 | -0.12 | -0.5375 | -0.31 | 0.682974 | 0.119081 | 0.369811 | 0.999971 | 1 | 0.869526 |
| Slc1a2_1 | A2APL5 | 1.27 | -0.1 | -0.73 | -1.23 | 1.03 | -0.25 | -1.39 | -1.93 | -0.73 | 0.2 | -1.61 | 0.52 | -0.36 | 2.2 | -0.95 | -0.1 | -0.4375 | -0.2075 | 0.395 | 0.571143 | 0.754058 | 0.619542 | 0.999971 | 1 | 0.929296 |
| Slc1a3 | P56564 | 0.1 | 0.44 | -0.37 | -0.28 | 0.26 | 0.57 | -1.13 | 0.09 | 0.04 | 0.24 | -1.34 | -1.68 | 0.62 | 0.55 | -0.38 | -0.34 | -0.025 | -0.6575 | 0.14 | 0.949915 | 0.197973 | 0.673429 | 0.999971 | 1 | 0.94048 |
| Slc1a4 | O35874 | -0.01 | -0.62 | -0.63 | -0.51 | -1.09 | -0.71 | -0.84 | -0.86 | -0.98 | -0.04 | -1.06 | -1.08 | -1.09 | -0.9 | 0.26 | -0.61 | -0.4325 | -0.3475 | -0.1425 | 0.069641 | 0.263528 | 0.669372 | 0.999971 | 1 | 0.938711 |
| Slc1a5 | Q9ESU7 | -0.87 | 2.78 | -1.44 | -0.05 | 1.22 | -0.16 | -1.02 | 0.61 | 0.74 | 1.02 | 1.28 | 0.46 | 0.63 | -1.26 | -0.83 | -0.04 | 0.0575 | 0.77 | -0.48 | 0.951279 | 0.382078 | 0.604209 | 0.999971 | 1 | 0.924454 |
| Slc1a6 | O35544 | 0.45 | -0.37 | -1.82 | -1.69 | -0.75 | -0.63 | -1.1 | -1.16 | -0.97 | 0.16 | -2.07 | -1.61 | -0.4 | -0.26 | -1.34 | -1.56 | -0.0525 | -0.265 | -0.0325 | 0.919158 | 0.691366 | 0.955479 | 0.999971 | 1 | 0.998056 |
| Slc20a1 | Q61609 | -0.12 | -1.13 | -0.97 | -0.99 | -1.03 | -0.32 | -0.97 | -1.27 | -1.16 | -0.33 | -0.76 | -0.98 | -0.99 | -0.29 | -1.07 | -0.58 | -0.095 | -0.005 | 0.07 | 0.760968 | 0.986663 | 0.816994 | 0.999971 | 1 | 0.974929 |
| Slc20a2 | Q80UP8 | 0.31 | -0.22 | -0.61 | 0.09 | 0.16 | -0.01 | -0.3 | -0.61 | -0.14 | 0.05 | -1.08 | -0.8 | -0.14 | -0.26 | -0.68 | -1.19 | -0.0825 | -0.385 | -0.46 | 0.766991 | 0.265426 | 0.169162 | 0.999971 | 1 | 0.80174 |
| Slc22a17 | A0A2I3BQ( | 1.16 | 0.77 | 0.35 | 0.38 | -0.32 | 1.28 | -0.01 | 0.42 | 0.8 | 1.34 | 0.8 | 0.92 | 1.49 | 1.39 | 0.53 | 1.39 | -0.3225 | 0.3 | 0.535 | 0.408315 | 0.256651 | 0.10478 | 0.999971 | 1 | 0.78268 |
| Slc22a23 | Q3UHH2 | -0.55 | -1.51 | -0.52 | -0.33 | -1.49 | -1.28 | -1.05 | -0.54 | -1.43 | -0.47 | -0.7 | -0.72 | -1.79 | -1.29 | -0.59 | -0.17 | -0.3625 | -0.1025 | -0.2325 | 0.293199 | 0.760026 | 0.587155 | 0.999971 | 1 | 0.920758 |
| Slc22a4 | Q9Z306 | -0.68 | -1.3 | -0.96 | -1.36 | -0.53 | -0.89 | -1.5 | -0.98 | -1.56 | -2.01 | -2.43 | -1.36 | -1.12 | -0.93 | 0.09 | -0.56 | 0.1 | -0.765 | 0.445 | 0.715302 | 0.027865 | 0.181739 | 0.999971 | 1 | 0.808532 |
| Slc22a5 | Q9Z0E8 | 0.42 | 0.51 | -1.56 | -1.14 | -0.64 | -0.98 | -2.24 | -1.24 | -0.7 | 0.23 | -1.04 | -1.07 | -0.77 | 0.65 | -1.04 | -1.16 | -0.8325 | -0.2025 | -0.1375 | 0.175184 | 0.720471 | 0.823668 | 0.999971 | 1 | 0.976286 |
| Slc23a2 | Q9EPR4 | 0.48 | -0.14 | -0.9 | -0.61 | -0.48 | 0.33 | -0.56 | -1.64 | -0.32 | -0.09 | -0.64 | -0.76 | -0.23 | -0.66 | -1.41 | -0.88 | -0.295 | -0.16 | -0.5025 | 0.535413 | 0.635456 | 0.205072 | 0.999971 | 1 | 0.81617 |
| Slc24a2 | Q8BUN9-2 | 0.03 | 1.44 | 0.6 | 0.35 | 0.74 | 1.77 | 0.08 | 0.47 | 0.7 | 0.58 | 0.35 | -0.13 | 1.37 | 0.98 | 0.02 | 0.58 | 0.16 | -0.23 | 0.1325 | 0.718736 | 0.513714 | 0.741075 | 0.999971 | 1 | 0.960899 |
| Slc24a2_1 | Q8BUN9 | 0.9 | -0.77 | -1.33 | -8.35 | -0.65 | -1.44 | 0.68 | -10.08 | -0.27 | -1.9 | 0.92 | 1.78 | 1.15 | -4.45 | -5.65 | -11.04 | -0.485 | 2.52 | -2.61 | 0.862966 | 0.219594 | 0.372464 | 0.999971 | 1 | 0.869526 |
| Slc24a3 | Q8VD29 | -4.18 | -2.39 | -0.95 | -0.98 | -0.89 | -4.22 | -1.4 | -1.2 | -0.99 | -5.13 | -0.94 | -0.52 | -0.44 | -5.14 | -0.63 | -1.17 | 0.1975 | 0.23 | 0.28 | 0.83847 | 0.846435 | 0.815431 | 0.999971 | 1 | 0.974918 |
| Slc24a4 | E9PUL6 | -2.31 | -1.17 | -0.81 | -0.83 | -0.81 | -1.16 | -1.16 | -0.46 | -1.08 | -1.73 | -0.99 | -0.57 | -0.55 | -0.89 | -0.41 | -0.86 | 0.3825 | 0.1875 | 0.6025 | 0.325616 | 0.647744 | 0.124668 | 0.999971 | 1 | 0.78268 |
| Slc25a1 | Q8JZU2 | 0.09 | 0.15 | 0.39 | 0.08 | 0.49 | 0.74 | 0.9 | 0.74 | 1.18 | 1.11 | 0.56 | 0.65 | 1.1 | 0.89 | 0.87 | 0.86 | 0.54 | 0.6975 | 0.7525 | 0.015979 | 0.010058 | 0.002151 | 0.999971 | 1 | 0.508206 |
| Slc25a10 | Q9QZD8 | -1.25 | -0.63 | -0.08 | -0.91 | -0.06 | -1.14 | 1.09 | -0.12 | 0.18 | -1.02 | 0.42 | 1.18 | 0.49 | -1.2 | -0.32 | -0.68 | 0.66 | 0.9075 | 0.29 | 0.194847 | 0.089112 | 0.488409 | 0.999971 | 1 | 0.898175 |

| | | | | | | | | | | | | | | | | | | | | | | | | | | |
|---|---|---|---|---|---|---|---|---|---|---|---|---|---|---|---|---|---|---|---|---|---|---|---|---|---|---|
| Slc25a11 | Q9CR62 | 0.03 | -0.06 | -0.01 | -0.46 | -0.31 | -0.18 | -0.12 | -0.16 | -0.27 | -0.04 | -0.09 | -0.26 | -0.55 | -0.18 | -0.02 | 0 | -0.0675 | -0.04 | -0.0625 | 0.721739 | 0.832985 | 0.773915 | 0.999971 | 1 | 0.963877 |
| Slc25a12 | Q8BH59 | -0.48 | -0.57 | -0.14 | -0.16 | -0.49 | -0.84 | -0.23 | -0.4 | -0.84 | -0.61 | -0.3 | -0.16 | -1.07 | -0.7 | -0.34 | -0.33 | -0.1525 | -0.14 | -0.2725 | 0.485995 | 0.539725 | 0.275425 | 0.999971 | 1 | 0.843794 |
| Slc25a13 | Q9QXX4 | -0.44 | -0.45 | -0.01 | -0.24 | -0.76 | -0.41 | -0.55 | 0.1 | -0.32 | -0.59 | -0.77 | 0.56 | -0.71 | -0.09 | 0.96 | -0.19 | -0.12 | 0.005 | 0.2775 | 0.622427 | 0.987345 | 0.446028 | 0.999971 | 1 | 0.884449 |
| Slc25a14 | A0A0A0MC | 0.39 | 0.74 | 0.83 | 0.65 | 0.81 | 1.09 | 0.35 | 0.9 | 0.87 | 0.9 | 0.92 | 0.94 | 0.39 | 1.12 | 0.89 | 0.93 | 0.135 | 0.255 | 0.18 | 0.55196 | 0.169259 | 0.431624 | 0.999971 | 1 | 0.881643 |
| Slc25a15 | Q9WVD5 | -0.75 | -0.3 | -1.71 | -1.12 | -0.27 | -0.56 | -1.4 | -1.98 | -0.59 | -0.6 | -1.43 | -1.34 | -0.55 | -0.16 | -0.86 | -0.9 | -0.0825 | -0.02 | 0.3525 | 0.857763 | 0.956414 | 0.316568 | 0.999971 | 1 | 0.85361 |
| Slc25a16 | D3YVC9 | -0.19 | 0.68 | 0.33 | 0.1 | 0.27 | 0.72 | 0.04 | 0.67 | 0.43 | 0.03 | 0.16 | 0 | 0.74 | 0.27 | 0.31 | 0.46 | 0.195 | -0.075 | 0.215 | 0.471304 | 0.754233 | 0.389369 | 0.999971 | 1 | 0.871816 |
| Slc25a17 | O70579 | -2.12 | -0.87 | -0.7 | -0.9 | -0.69 | -0.76 | -1.27 | -0.46 | -0.26 | -1.03 | -0.86 | -0.27 | -0.35 | -1.1 | -1.12 | -0.61 | 0.3525 | 0.5425 | 0.3525 | 0.341305 | 0.170304 | 0.351427 | 0.999971 | 1 | 0.867656 |
| Slc25a18 | Q9DB41 | -0.55 | -1.01 | -0.44 | -0.57 | -1.64 | -0.8 | -0.15 | -1.3 | -1.29 | -0.75 | -0.97 | -1.22 | -1.01 | -0.94 | -0.56 | -0.79 | -0.33 | -0.415 | -0.1825 | 0.349192 | 0.084134 | 0.398043 | 0.999971 | 1 | 0.871816 |
| Slc25a19 | Q9DAM5 | 0.27 | 0.26 | 0.55 | 0.25 | 0.26 | -0.04 | 0.24 | -0.08 | 0.39 | 0.51 | 0.58 | 0.46 | 0.28 | -0.31 | -0.2 | -0.93 | -0.2375 | 0.1525 | -0.6225 | 0.224447 | 0.379026 | 0.047874 | 0.999971 | 1 | 0.710102 |
| Slc25a20 | Q9Z2Z6 | -0.2 | -1.06 | 0.02 | -0.11 | -1.04 | -1.01 | -0.28 | 0.32 | -1.37 | -0.79 | 0.16 | -0.3 | -1.52 | -0.55 | 0.32 | 0.04 | -0.165 | -0.2375 | -0.09 | 0.674454 | 0.550777 | 0.840528 | 0.999971 | 1 | 0.97975 |
| Slc25a21 | B6CI26 | -0.56 | -0.37 | 0.08 | -0.56 | -0.1 | -0.15 | -0.53 | 1.15 | -0.22 | -0.58 | 0.64 | -0.71 | -0.2 | -0.15 | 0.46 | 0.19 | 0.445 | 0.135 | 0.4275 | 0.26181 | 0.689752 | 0.110371 | 0.999971 | 1 | 0.78268 |
| Slc25a22 | Q9D6M3 | -0.81 | 0.33 | -0.24 | -0.43 | 0.35 | 0.52 | -0.26 | -0.05 | 0.42 | -0.25 | 0.05 | -0.09 | 0.57 | 0.28 | 0.01 | -0.06 | 0.4275 | 0.32 | 0.4875 | 0.184173 | 0.284722 | 0.120042 | 0.999971 | 1 | 0.78268 |
| Slc25a23 | Q6GQS1 | -0.56 | -1.03 | -0.41 | -0.55 | -1.01 | -0.94 | -0.23 | -0.28 | -1.2 | -0.66 | -0.25 | -0.63 | -1.39 | -0.89 | -0.5 | -0.37 | 0.0225 | -0.0475 | -0.15 | 0.933163 | 0.854751 | 0.597409 | 0.999971 | 1 | 0.92377 |
| Slc25a24 | Q8BMD8 | 0.42 | -0.52 | -1.21 | -0.41 | 0.29 | -0.38 | -0.88 | -1.17 | 0.12 | 0.87 | -0.88 | -0.28 | -0.54 | -0.27 | -1.17 | -0.11 | -0.105 | 0.3875 | -0.0925 | 0.80984 | 0.415345 | 0.813558 | 0.999971 | 1 | 0.97473 |
| Slc25a25 | A2ASZ8-2 | -1.49 | 0.21 | -0.63 | 0.17 | -0.62 | 0.43 | -0.28 | -0.9 | -0.4 | -0.35 | -1 | 0.22 | -0.38 | 0.59 | 0.03 | -1.03 | 0.0925 | 0.0525 | 0.2375 | 0.841218 | 0.906235 | 0.63041 | 0.999971 | 1 | 0.932711 |
| Slc25a25_1 | A2ASZ8-4 | 0.18 | -0.46 | 0 | 0.14 | -0.49 | -0.2 | 0.36 | -0.23 | -0.38 | -0.26 | -0.17 | -0.03 | -0.44 | -0.31 | -0.08 | -0.19 | -0.105 | -0.175 | -0.22 | 0.683516 | 0.415496 | 0.320143 | 0.999971 | 1 | 0.853962 |
| Slc25a26 | Q5U680 | -1.01 | 0.28 | -0.55 | -0.06 | -0.41 | -0.32 | -1.09 | 0.12 | -0.52 | -0.41 | -0.38 | -0.68 | -0.71 | 0.94 | 0.01 | 0.34 | -0.09 | -0.1625 | 0.48 | 0.806212 | 0.589142 | 0.273879 | 0.999971 | 1 | 0.843794 |
| Slc25a27 | Q9D6D0 | -0.39 | -0.52 | -0.13 | -0.28 | -0.07 | -0.45 | 0.22 | -0.37 | -0.47 | -0.85 | -0.5 | -0.55 | -0.78 | -0.6 | -0.34 | -0.73 | 0.1625 | -0.2625 | -0.2825 | 0.464538 | 0.182416 | 0.169759 | 0.999971 | 1 | 0.802438 |
| Slc25a28 | A0A494BA( | 3.57 | 0.64 | 1.17 | -2.36 | -0.58 | -0.29 | 1.42 | -2.14 | 1.62 | 0.39 | 1.4 | 0.2 | 0.06 | -1.89 | 0.01 | -1.75 | -1.1525 | 0.1475 | -1.6475 | 0.372501 | 0.896831 | 0.188567 | 0.999971 | 1 | 0.809276 |
| Slc25a29 | Q8BL03 | -0.68 | -0.14 | -0.52 | -0.23 | -0.27 | 0.06 | -0.63 | -0.2 | -0.27 | 0.02 | -0.3 | -0.71 | 0.16 | 0.05 | -0.27 | -0.16 | 0.1325 | 0.0775 | 0.3375 | 0.565858 | 0.737404 | 0.135431 | 0.999971 | 1 | 0.784948 |
| Slc25a3 | Q8VEM8 | -0.55 | 0.02 | -0.4 | -0.03 | -0.35 | -0.39 | -0.73 | -0.37 | -0.65 | -0.11 | -0.56 | -0.58 | -0.89 | 0.16 | -0.47 | 0.02 | -0.22 | -0.235 | -0.055 | 0.318846 | 0.310296 | 0.849684 | 0.999971 | 1 | 0.98187 |
| Slc25a31 | Q3V132 | -0.47 | -1.21 | -0.58 | -0.45 | -1.37 | -1.3 | -0.49 | -0.76 | -1.39 | -1.26 | -0.63 | -0.42 | -1.25 | -1.21 | -0.66 | -0.35 | -0.3025 | -0.2475 | -0.19 | 0.312613 | 0.424005 | 0.523622 | 0.999971 | 1 | 0.907797 |
| Slc25a32 | Q8BMG8 | 0.28 | -0.09 | -0.5 | 0.26 | -0.79 | 0.28 | -0.64 | 0.66 | -1.41 | 0.64 | 0.2 | -0.21 | -0.71 | 0.7 | -0.19 | -0.02 | -0.11 | -0.1825 | -0.0425 | 0.774022 | 0.68632 | 0.900841 | 0.999971 | 1 | 0.99074 |
| Slc25a33 | Q3TZX3 | -1.5 | -0.26 | 0.36 | 0.09 | 0.04 | -0.24 | 0.31 | 0.51 | -0.4 | -1.08 | 1.16 | 0.74 | 0.1 | -0.14 | 0.11 | -0.27 | 0.4825 | 0.4325 | 0.2775 | 0.268222 | 0.480762 | 0.497259 | 0.999971 | 1 | 0.902655 |
| Slc25a35 | Q5SWT3 | 0.46 | -0.04 | -0.6 | -0.7 | -0.1 | -0.08 | -0.89 | 0 | -0.45 | 0.75 | -0.99 | -0.19 | -0.3 | 0.33 | -1.07 | -0.64 | -0.0475 | -1.57E-16 | -0.2 | 0.88808 | 1 | 0.606708 | 0.999971 | 1 | 0.925135 |
| Slc25a36 | Q922G0 | -0.19 | -0.27 | -1.24 | -1.78 | -0.77 | -0.15 | -0.25 | -1.14 | -0.72 | 0.49 | -0.49 | -1.39 | -1.15 | -0.12 | -1.07 | -1.62 | 0.2925 | 0.3425 | -0.12 | 0.498053 | 0.507349 | 0.797059 | 0.999971 | 1 | 0.970479 |
| Slc25a4 | P48962 | -1.03 | -0.37 | -0.42 | -0.53 | -0.42 | -0.31 | -0.54 | -0.57 | -0.53 | -0.57 | -0.47 | -0.62 | -0.33 | -0.18 | -0.38 | -0.39 | 0.1275 | 0.04 | 0.2675 | 0.551391 | 0.845106 | 0.224768 | 0.999971 | 1 | 0.828225 |
| Slc25a40 | Q8BGP6 | -1.06 | -0.24 | -0.05 | 0.1 | -0.1 | -0.29 | -0.42 | 0.08 | -0.53 | -0.99 | 0.02 | 0.26 | -0.66 | -0.19 | 0.09 | 0.28 | 0.13 | 0.0025 | 0.1925 | 0.657843 | 0.994617 | 0.563241 | 0.999971 | 1 | 0.917003 |
| Slc25a42 | Q8R0Y8 | -0.41 | -0.37 | -0.59 | 0.13 | -0.79 | -0.36 | -0.55 | -0.02 | -0.63 | -0.38 | -0.55 | -0.46 | -1.08 | -0.99 | -0.19 | -0.28 | -0.12 | -0.195 | -0.325 | 0.635179 | 0.365543 | 0.281807 | 0.999971 | 1 | 0.844538 |
| Slc25a44 | Q3TMC4 | 0.1 | -0.53 | 0 | 0.28 | -0.09 | -0.42 | -0.13 | 0.15 | -0.49 | -0.11 | -0.27 | -0.77 | -1.42 | -0.51 | -0.29 | -0.27 | -0.085 | -0.3725 | -0.585 | 0.725753 | 0.16295 | 0.09858 | 0.999971 | 1 | 0.78268 |
| Slc25a46 | Q9CQS4 | -0.35 | 0.51 | 0.22 | 0.09 | 0.36 | 0.62 | 0.03 | -0.05 | 0.69 | 0.28 | 0.08 | -0.4 | 0.86 | 0.35 | -0.02 | 0.29 | 0.1225 | 0.045 | 0.2525 | 0.639452 | 0.879562 | 0.368784 | 0.999971 | 1 | 0.869526 |
| Slc25a5 | P51881 | -0.18 | 0.86 | 0.95 | 0.93 | 0.46 | 1.14 | 0.43 | 1.11 | 0.63 | 0.62 | 0.78 | 0.79 | 0.57 | 1.06 | 1.38 | 0.88 | 0.145 | 0.065 | 0.3325 | 0.666674 | 0.822151 | 0.319701 | 0.999971 | 1 | 0.853962 |
| Slc25a51 | Q5HZI9 | 0.04 | -0.26 | -0.49 | -0.36 | -0.15 | -0.32 | -0.55 | -0.47 | -0.54 | -0.13 | -0.5 | -0.43 | -0.81 | -0.49 | -0.35 | -0.41 | -0.105 | -0.1325 | -0.2475 | 0.603783 | 0.512649 | 0.250514 | 0.999971 | 1 | 0.839012 |
| Slc25a53 | A2AF34 | -0.55 | -0.28 | 0.54 | 0.39 | -0.07 | -0.21 | 0.04 | 0.59 | -0.42 | -0.84 | 0.11 | 0.57 | -0.91 | 0.15 | 0.16 | 0.45 | 0.0625 | -0.17 | -0.0625 | 0.844089 | 0.663768 | 0.870598 | 0.999971 | 1 | 0.984079 |
| Slc26a2 | E9PZ06 | -1.5 | -0.27 | -1.45 | 0.04 | 0.52 | -0.74 | -1.16 | -0.93 | -0.26 | -1.49 | -0.36 | 0.1 | -0.99 | -1.39 | -1.16 | 0.46 | 0.2175 | 0.2925 | 0.025 | 0.669131 | 0.555267 | 0.962532 | 0.999971 | 1 | 0.998721 |
| Slc26a6 | Q8CIW6-3 | -1.46 | -1.13 | 0.01 | -0.52 | -0.07 | -0.81 | 1.24 | -0.54 | 0.67 | -1.56 | -0.54 | -0.81 | 1.64 | -1.06 | -1 | -1.08 | 0.73 | 0.215 | 0.4 | 0.182189 | 0.683752 | 0.558822 | 0.999971 | 1 | 0.915051 |
| Slc27a1 | Q60714 | -0.6 | -0.69 | -0.52 | -1.13 | -0.76 | -0.67 | -0.29 | -1.02 | -0.55 | -0.46 | -0.21 | -0.2 | -0.32 | -0.52 | -0.77 | -0.87 | 0.05 | 0.38 | 0.115 | 0.833304 | 0.097158 | 0.61308 | 0.999971 | 1 | 0.927364 |
| Slc27a2 | O35488 | 0.21 | -0.68 | 0.13 | 0.57 | -0.48 | -0.7 | -0.22 | 0.71 | -0.39 | 0.31 | -0.13 | 0.52 | -0.78 | -0.32 | 0.29 | 0.2 | -0.23 | 0.02 | -0.21 | 0.559928 | 0.952099 | 0.55896 | 0.999971 | 1 | 0.915051 |
| Slc27a3 | G3X8Y7 | 1.37 | 0.11 | -0.06 | -0.09 | 0.98 | 0.36 | 0.4 | -0.31 | 0.49 | 1.69 | -0.12 | 0.22 | 2.22 | 0.8 | -0.56 | -0.25 | 0.025 | 0.2375 | 0.22 | 0.951939 | 0.630737 | 0.736638 | 0.999971 | 1 | 0.959447 |
| Slc27a4 | Q91VE0 | -0.96 | -0.68 | -0.38 | -0.42 | -0.59 | -0.55 | -0.38 | -0.48 | -0.45 | -0.84 | -0.07 | -0.17 | -0.18 | -0.76 | -0.31 | -0.11 | 0.11 | 0.2275 | 0.27 | 0.585752 | 0.367258 | 0.268488 | 0.999971 | 1 | 0.842746 |
| Slc29a1 | Q9JIM1-2 | 0.09 | 0.35 | 0.23 | -0.53 | 1.01 | 0.58 | 0.26 | 0.24 | 1.71 | -0.49 | 0.43 | 0.8 | 2.36 | 0.08 | 0.54 | 0.01 | 0.4875 | 0.5775 | 0.7125 | 0.110063 | 0.235687 | 0.208405 | 0.999971 | 1 | 0.822138 |
| Slc29a2 | A0A494BA( | -0.23 | -0.95 | -0.34 | -0.3 | -1.61 | -0.47 | -0.33 | -0.75 | -0.51 | -0.05 | -0.34 | -0.56 | 0.33 | -0.98 | -0.84 | -0.11 | -0.335 | 0.09 | 0.055 | 0.32574 | 0.702823 | 0.873945 | 0.999971 | 1 | 0.984109 |
| Slc2a1 | P17809 | -1.11 | -2.59 | -2.19 | -2.3 | -2.28 | -2.86 | -0.09 | -3.15 | -2.23 | -1.99 | -2 | -1.67 | -3.36 | -2.84 | -2.32 | -2.46 | -0.0475 | 0.075 | -0.6975 | 0.945158 | 0.825518 | 0.097352 | 0.999971 | 1 | 0.78268 |
| Slc2a13 | Q3UHK1 | -0.29 | 0.11 | 0.59 | 0.43 | 0.19 | 0.08 | 0.47 | 0.24 | 0.26 | -0.1 | 0.4 | 0.66 | 0.53 | -0.12 | 0.33 | 0.33 | 0.035 | 0.095 | 0.0575 | 0.885168 | 0.72461 | 0.826169 | 0.999971 | 1 | 0.976757 |
| Slc2a3 | P32037 | -1.08 | -1.25 | -1.4 | -1.46 | -0.73 | -0.89 | -0.61 | -1.62 | -0.88 | -0.84 | -1.2 | -1.59 | -0.63 | -1.32 | -1.6 | -1.88 | 0.335 | 0.17 | -0.06 | 0.225296 | 0.465361 | 0.837738 | 0.999971 | 1 | 0.979416 |
| Slc2a6 | Q3UDF0 | 0.11 | 3.75 | -0.47 | -0.97 | 1.97 | 2.03 | -0.72 | -0.68 | 1.08 | 2.21 | -0.53 | -0.45 | 2.64 | 1.98 | -0.15 | 0.38 | 0.045 | -0.0275 | 0.6075 | 0.969493 | 0.980524 | 0.590701 | 0.999971 | 1 | 0.92226 |
| Slc30a1 | Q60738 | 0.04 | -1.26 | -0.37 | -0.28 | -1.03 | -0.94 | -0.06 | -0.46 | -1 | -0.79 | -0.51 | -0.61 | -1.32 | -0.94 | -0.37 | -0.81 | -0.155 | -0.26 | -0.3925 | 0.660887 | 0.403736 | 0.264622 | 0.999971 | 1 | 0.842533 |
| Slc30a10 | Q3UVU3 | 0.46 | -0.89 | -0.54 | 0.05 | -0.75 | -0.68 | -0.05 | -0.24 | -0.66 | -0.33 | -1.14 | -0.67 | -0.17 | -1.09 | -0.79 | -0.9 | -0.2 | -0.47 | -0.5075 | 0.561844 | 0.191251 | 0.176812 | 0.999971 | 1 | 0.808532 |
| Slc30a3 | P97441 | -0.59 | -0.19 | -1.11 | -0.8 | -0.45 | -0.24 | -1.14 | -0.92 | -0.8 | -0.77 | -0.98 | -1.18 | -0.28 | -0.27 | -1.17 | -1.02 | -0.015 | -0.26 | -0.0125 | 0.959237 | 0.301998 | 0.968046 | 0.999971 | 1 | 0.998787 |
| Slc30a6 | J3QMX8 | 0.51 | -0.06 | 0.03 | 0.17 | -0.22 | -0.86 | 0.01 | 0.28 | -0.55 | 1.56 | 0.54 | 1.88 | -1.72 | 0.31 | -0.35 | 0.58 | -0.36 | 0.695 | -0.4575 | 0.230008 | 0.206741 | 0.364611 | 0.999971 | 1 | 0.869526 |
| Slc30a7 | Q9JKN1 | 0.32 | 4.78 | 0.42 | 1.28 | 4.75 | 5.3 | 0.14 | 1.62 | 5.3 | 0.13 | 2.63 | 1.36 | 6.18 | 3.19 | 2.75 | 0.75 | 1.2525 | 0.655 | 1.5175 | 0.394706 | 0.633128 | 0.283481 | 0.999971 | 1 | 0.844538 |
| Slc30a9 | Q5IRJ6 | -0.94 | -0.62 | -1.05 | -0.84 | -0.71 | -0.82 | -1.12 | -0.76 | -0.73 | -0.53 | -0.9 | -1.15 | -0.83 | -0.75 | -0.86 | -0.72 | 0.01 | 0.035 | 0.0725 | 0.958778 | 0.866382 | 0.686921 | 0.999971 | 1 | 0.945467 |
| Slc31a1 | Q8K211 | -0.24 | -0.62 | 0.41 | 0.24 | -0.66 | -0.94 | -0.49 | -0.52 | -0.94 | -0.54 | -0.08 | 0.23 | -0.9 | -0.63 | -0.88 | 0.94 | -0.6 | -0.28 | -0.315 | 0.052309 | 0.42376 | 0.505373 | 0.999971 | 1 | 0.905277 |
| Slc32a1 | O35633-2 | -0.27 | 0.36 | -0.03 | -0.29 | 0.32 | 0.54 | -0.3 | -0.17 | 0.58 | 0.2 | -0.12 | -0.14 | 1.57 | 0.82 | 0.21 | -0.04 | 0.155 | 0.1875 | 0.6975 | 0.568667 | 0.46375 | 0.091921 | 0.999971 | 1 | 0.773779 |
| Slc33a1 | Q99J27 | 0.27 | 1.33 | 1.34 | 0.71 | 1.45 | 1.81 | 1.02 | 1.14 | 1.2 | 0.38 | 0.54 | 0.88 | 1.22 | 0.97 | 0.93 | 0.23 | 0.4425 | -0.1625 | -0.075 | 0.186929 | 0.613694 | 0.822728 | 0.999971 | 1 | 0.975901 |
| Slc35a2 | A0A140LHI | -0.18 | 0.62 | -0.74 | -0.06 | -1.39 | -1.12 | -0.73 | 0.3 | -0.45 | 0.69 | 3.4 | -0.62 | 2.48 | 0.01 | 0.46 | -0.56 | -0.645 | 0.845 | 0.6875 | 0.165505 | 0.34749 | 0.307855 | 0.999971 | 1 | 0.847402 |
| Slc35a3 | E9Q3V2 | 3.33 | -0.5 | 0.76 | 0.49 | 0.3 | 1.67 | 1.46 | 0.62 | 1.75 | 3.63 | 0.61 | -1.36 | -1.38 | -0.33 | 0.88 | -0.96 | -0.0075 | 0.1375 | -1.4675 | 0.992424 | 0.907809 | 0.113856 | 0.999971 | 1 | 0.78268 |
| Slc35a4 | A0A087WC | -0.63 | 0 | -0.26 | 0.11 | -0.67 | 1.03 | -0.08 | 0.51 | -0.55 | 0.19 | -0.16 | -0.27 | -0.09 | 1.28 | 0.2 | 0.78 | 0.3925 | -0.0025 | 0.7375 | 0.325091 | 0.992022 | 0.056568 | 0.999971 | 1 | 0.725348 |
| Slc35a5 | M0QWE5 | 2.31 | 0.3 | 2.43 | 2.18 | 1.34 | 1.96 | 2.36 | 1.2 | 1.39 | 2.12 | 1.7 | 0.46 | -2.77 | 0.08 | 0.97 | 0.21 | -0.09 | -0.3875 | -2.1825 | 0.864583 | 0.500331 | 0.030797 | 0.999971 | 1 | 0.678474 |
| Slc35b2 | Q91ZN5-2 | -0.33 | 1.38 | 2.06 | 1.94 | 0.43 | 0.37 | 1.32 | 2.59 | 0.66 | 0.51 | 3.39 | 2.39 | 0.57 | 1.23 | 2.95 | 2.81 | -0.085 | 0.475 | 0.6275 | 0.901137 | 0.558514 | 0.399243 | 0.999971 | 1 | 0.871816 |
| Slc35d3 | Q8BGF8 | 0.54 | -2.58 | -2.88 | -0.71 | -0.85 | -2.85 | -1.98 | -1.19 | -1.38 | -0.44 | -2.68 | -0.9 | -1.8 | -0.99 | -0.66 | -2.37 | -0.31 | 0.0575 | -0.0475 | 0.708827 | 0.945997 | 0.952956 | 0.999971 | 1 | 0.998056 |
| Slc35f1 | Q8BGK5 | 0.05 | -0.53 | 1.25 | -0.09 | -0.99 | -1.34 | 9.19 | 0.21 | -0.98 | 0.1 | 1.06 | 1.46 | -0.92 | -0.84 | -0.2 | -2.24 | 1.5975 | 0.24 | -1.22 | 0.480492 | 0.694685 | 0.043859 | 0.999971 | 1 | 0.702458 |
| Slc35f3 | Q1LZI2 | 2.58 | 1.55 | 2.39 | 2.39 | 1.04 | 1.73 | 1.78 | 2.53 | 1.2 | 2.45 | 2.47 | 2.03 | 0.3 | 1.83 | 2.6 | 3.41 | -0.4575 | -0.19 | -0.1925 | 0.237099 | 0.606616 | 0.763359 | 0.999971 | 1 | 0.961895 |
| Slc35f6 | A0A0G2JE( | 1.71 | 0.68 | 1.24 | 1.52 | 0.52 | 1.92 | 1.61 | 2.48 | 0.84 | 2.14 | 1.98 | 0.37 | 0.33 | 1.95 | 1.91 | 2.19 | 0.345 | 0.045 | 0.3075 | 0.442892 | 0.921678 | 0.50275 | 0.999971 | 1 | 0.904573 |
| Slc35g2 | D3YVE8 | -2.38 | -2.37 | -1.69 | 0.26 | -0.05 | -0.55 | -1.58 | 2.56 | -1.85 | -2.56 | -2.09 | -1.91 | 0.32 | -0.54 | -2.05 | -0.04 | 1.64 | -0.5575 | 0.9675 | 0.116092 | 0.358517 | 0.209945 | 0.999971 | 1 | 0.82263 |
| Slc36a1 | Q8K4D3 | 0.02 | 0.94 | 0.57 | 0.47 | 0.75 | 0.92 | 0.24 | 0.92 | 1 | 0.68 | 1.69 | 0.78 | 0.87 | 1.14 | 0.54 | 1.92 | 0.2075 | 0.5375 | 0.6175 | 0.447332 | 0.103545 | 0.101607 | 0.999971 | 1 | 0.78268 |
| Slc36a4 | D3Z0L1 | 0.45 | 1.45 | 0.84 | 1.07 | 1.14 | 0.83 | 0.73 | 1.37 | 0.5 | 0.86 | 1.28 | 1.09 | -0.36 | 1.03 | 1.15 | 0.81 | 0.065 | -0.02 | -0.295 | 0.812266 | 0.943388 | 0.456683 | 0.999971 | 1 | 0.887217 |
| Slc37a4 | A0A1L1SUI | -0.49 | -0.44 | -0.07 | -1.04 | -0.36 | -0.31 | -0.62 | 0.63 | -0.01 | -0.01 | 0.63 | 0.53 | 0.16 | 0.17 | 0.19 | 0.34 | 0.345 | 0.795 | 0.725 | 0.319882 | 0.018002 | 0.013372 | 0.999971 | 1 | 0.613095 |
| Slc38a1 | Q8K2P7 | 0.03 | 0.59 | 0.79 | 0.63 | 0.31 | 0.7 | 0.17 | -0.42 | 0.47 | 0.52 | 0.24 | 0.11 | 0.33 | 0.7 | 0.36 | 0.74 | -0.32 | -0.175 | 0.0225 | 0.294949 | 0.450659 | 0.923581 | 0.999971 | 1 | 0.993334 |
| Slc38a10 | J3QNE8 | 0.78 | -0.49 | 2.62 | 2.69 | 0.05 | -1.03 | 1.09 | 2.94 | 0.38 | 0.25 | 1.7 | 2.23 | 0.45 | -1.09 | 2.06 | 3.23 | -0.6375 | -0.26 | -0.2375 | 0.536228 | 0.752993 | 0.826907 | 0.999971 | 1 | 0.976804 |
| Slc38a2 | Q8CFE6 | -0.49 | 0.85 | 1.19 | 1.25 | 0.2 | 0.3 | 0.84 | -1.68 | 0.03 | -0.26 | 0.82 | 0.96 | -0.93 | -0.21 | 0.69 | 1.69 | -0.785 | -0.3125 | -0.39 | 0.228222 | 0.513463 | 0.543558 | 0.999971 | 1 | 0.911398 |
| Slc38a3 | Q9DCP2 | -0.59 | -0.47 | 0.03 | -0.79 | -0.58 | -0.85 | 1.34 | -0.58 | -0.07 | -1.11 | -1.06 | -0.44 | 0.43 | -1.45 | -0.79 | -1.25 | 0.2875 | -0.215 | -0.31 | 0.56658 | 0.494861 | 0.479489 | 0.999971 | 1 | 0.894438 |
| Slc38a7 | A0A1D5RL) | 0.12 | 0.62 | 0.5 | 0.41 | 0.26 | 0.45 | -0.06 | 0.93 | 0.5 | 0.91 | 0.6 | 0.93 | 0.94 | 1.5 | 1.22 | 0.86 | -0.0175 | 0.3225 | 0.7175 | 0.945759 | 0.139544 | 0.010129 | 0.999971 | 1 | 0.613095 |
| Slc39a10 | Q6P5F6 | -0.81 | -1.37 | -0.32 | -0.37 | -1.17 | -1.16 | -0.19 | -0.4 | -1.1 | -1.25 | -0.11 | -0.18 | -0.53 | -1.47 | -0.24 | -0.26 | -0.0125 | 0.0575 | 0.0925 | 0.971244 | 0.877929 | 0.801941 | 0.999971 | 1 | 0.971467 |
| Slc39a11 | Q8BWY7 | 2.54 | -0.96 | 1.03 | 0.6 | -1.04 | -0.46 | 0.83 | 1.01 | -0.59 | 1.16 | 0.84 | -0.36 | -3.56 | -2.18 | 0.78 | -0.13 | -0.7175 | -0.54 | -2.075 | 0.373456 | 0.48492 | 0.083302 | 0.999971 | 1 | 0.755254 |
| Slc39a12 | Q5FWH7 | -0.07 | -1.21 | -0.67 | -0.45 | -1.16 | -0.41 | -0.29 | -1.85 | -0.73 | -0.22 | -0.75 | -1.09 | -1.07 | -0.83 | -0.78 | -0.72 | -0.3275 | -0.0975 | -0.25 | 0.435782 | 0.749271 | 0.367029 | 0.999971 | 1 | 0.869526 |
| Slc39a14 | Q75N73-3 | 0.84 | -0.86 | -0.03 | 0.88 | -0.73 | 0.55 | 1.48 | -0.19 | -1.16 | -0.94 | 0.23 | -1.16 | -1.38 | -1.08 | -0.39 | -0.67 | 0.07 | -0.965 | -1.0875 | 0.903653 | 0.078592 | 0.033806 | 0.999971 | 1 | 0.689803 |
| Slc39a3 | A0A452J89 | 4.52 | -2.45 | 1.63 | 1.51 | -3.08 | -2.48 | 1.14 | 2.13 | -3.57 | 1.71 | 1.9 | 1.86 | -3.73 | -3.13 | 1.7 | 2.22 | -1.875 | -0.8275 | -2.0375 | 0.289461 | 0.639263 | 0.294724 | 0.999971 | 1 | 0.847402 |
| Slc39a6 | Q8C145 | -0.65 | -1.17 | -0.46 | -0.04 | -0.68 | -0.45 | 0.2 | -0.14 | -0.41 | -0.5 | -0.11 | -0.05 | 0.07 | -0.83 | 0.16 | 0.25 | 0.3125 | 0.3125 | 0.4925 | 0.325048 | 0.273921 | 0.172571 | 0.999971 | 1 | 0.805746 |
| Slc39a7 | Q31125 | -1.61 | 0.43 | 1.26 | 0.42 | 0.4 | -0.51 | 3.13 | -0.76 | -0.36 | -1.63 | -0.47 | -0.38 | 1.77 | 0.87 | 0.34 | -0.46 | 0.44 | -0.835 | 0.505 | 0.64999 | 0.205522 | 0.475569 | 0.999971 | 1 | 0.894045 |
| Slc3a2 | P10852-2 | -0.92 | -0.94 | -1.4 | -1.58 | -1.05 | -1.04 | -0.62 | -2.12 | -0.81 | -1.25 | -1.32 | -1.49 | -0.75 | -1.2 | -1.75 | -1.93 | 0.0025 | -0.0075 | -0.1975 | 0.99435 | 0.975771 | 0.540286 | 0.999971 | 1 | 0.910077 |
| Slc3a2_1 | P10852 | 4.83 | 0.04 | 0.67 | 2.75 | -0.33 | 0.45 | 0.89 | 1.45 | -0.71 | 3.3 | 0.58 | -0.16 | -0.58 | 1.22 | 1.13 | 1.9 | -1.4575 | -1.32 | -1.155 | 0.179366 | 0.30951 | 0.300096 | 0.999971 | 1 | 0.847402 |
| Slc43a2 | Q8CGA3 | 2.68 | -0.83 | 0.3 | 0.77 | -1.47 | 0.13 | 2.12 | 0.88 | -1.37 | 1.61 | -0.64 | -1.01 | -2.47 | -1.68 | -0.67 | -2.31 | -0.315 | -1.0825 | -2.5125 | 0.737247 | 0.246703 | 0.008623 | 0.999971 | 1 | 0.609224 |
| Slc44a1 | A2AMH5 | -0.45 | 0.09 | 0.71 | 0 | -0.47 | 0.4 | -0.09 | 0.5 | -0.05 | -0.59 | -0.21 | -0.43 | -0.3 | -0.16 | 0.1 | -0.16 | -0.0025 | -0.4075 | -0.2175 | 0.993916 | 0.171633 | 0.432974 | 0.999971 | 1 | 0.882137 |
| Slc44a2 | A0A1L1SU4 | 1.13 | -2 | -1.06 | -0.74 | -1.9 | -1.07 | -0.52 | -1.48 | -2.12 | -0.21 | -2.33 | -1.77 | -2.53 | -1.04 | -1.12 | -1.11 | -0.575 | -0.94 | -0.7825 | 0.389013 | 0.224557 | 0.267613 | 0.999971 | 1 | 0.842677 |
| Slc44a2_1 | Q8BY89-2 | -0.13 | -1.69 | -0.84 | -0.98 | -1.54 | -1.51 | -1.23 | -0.68 | -1.68 | -1.35 | -0.94 | -1.03 | -1.61 | -1.33 | -0.99 | -0.82 | -0.33 | -0.34 | -0.2775 | 0.379018 | 0.350451 | 0.446128 | 0.999971 | 1 | 0.884449 |
| Slc45a1 | Q8BIV7 | 0.31 | -1.98 | 0.28 | 0.73 | -1.89 | -0.98 | 0.53 | 0.1 | -1.09 | -0.11 | 0.72 | -0.39 | -2.29 | -1.75 | -0.18 | -0.13 | -0.395 | -0.0525 | -0.9225 | 0.596366 | 0.936191 | 0.235589 | 0.999971 | 1 | 0.832918 |
| Slc45a4 | F7BEQ1 | 0.45 | -1.2 | -0.24 | -0.5 | -0.71 | -0.01 | -0.56 | -0.01 | 0.11 | 0.31 | -0.98 | -1.32 | 0.66 | -1.68 | -1.18 | -0.25 | 0.05 | -0.0975 | -0.24 | 0.89373 | 0.842695 | 0.674444 | 0.999971 | 1 | 0.941037 |
| Slc47a1 | D6RGQ3 | 3.81 | -0.22 | -0.64 | 1.12 | -0.23 | 0.99 | 0.21 | -0.89 | -0.75 | 2.38 | -1.25 | -0.34 | -3.32 | 0.35 | -0.81 | -0.11 | -0.9975 | -1.0075 | -1.99 | 0.315455 | 0.394648 | 0.112032 | 0.999971 | 1 | 0.78268 |
| Slc49a4 | D6RI31 | -1.75 | -1.22 | -0.33 | -0.34 | -2.06 | -2.52 | -0.28 | 0.21 | -1.62 | -1.73 | -0.12 | -0.02 | -2.34 | -1.52 | -0.14 | 0.02 | -0.2525 | 0.0375 | -0.085 | 0.71132 | 0.944356 | 0.888672 | 0.999971 | 1 | 0.987485 |
| Slc4a1 | P04919-2 | -0.6 | -1.39 | -5.48 | -3.18 | 0.41 | -2.12 | -2.62 | -5.04 | -1.53 | -1.71 | -3.78 | -5.08 | -0.96 | -1.86 | -5.24 | -4.43 | 0.32 | -0.3625 | -0.46 | 0.816761 | 0.769577 | 0.728655 | 0.999971 | 1 | 0.956041 |
| Slc4a10 | Q5DTL9 | -0.47 | -1.01 | -0.58 | -1.12 | -1.33 | -0.8 | -0.08 | -0.9 | -1.22 | -1.1 | -0.68 | -0.64 | -0.68 | -1.4 | -1.16 | -0.88 | 0.0175 | -0.115 | -0.235 | 0.954879 | 0.640144 | 0.363335 | 0.999971 | 1 | 0.869526 |
| Slc4a10_1 | Q5DTL9-2 | -1.57 | -2.06 | -1.69 | -1.71 | -2.16 | -1.59 | -2.03 | -1.47 | -2.57 | -0.98 | -1.32 | -1.51 | -1.27 | -1.68 | -1.69 | -1 | -0.055 | 0.1625 | 0.3475 | 0.814022 | 0.646557 | 0.164169 | 0.999971 | 1 | 0.800215 |
| Slc4a1ap | E9Q585 | 0.71 | 1.05 | 2.13 | 1.57 | 1.22 | 1.24 | 2.34 | 1.89 | 0.54 | 1.51 | 1.97 | 2.13 | 0.02 | 1.06 | 2.08 | 1.5 | 0.3075 | 0.1725 | -0.2 | 0.444321 | 0.700488 | 0.688451 | 0.999971 | 1 | 0.945753 |
| Slc4a2 | A0A0R4J1K | 0.99 | 0.99 | 1.54 | -0.24 | -0.2 | 0.06 | 0.84 | -0.22 | 0.26 | 0.45 | -1.64 | 1.52 | -1.2 | 0.34 | 0.22 | 0.02 | -0.7 | -0.6725 | -0.975 | 0.12777 | 0.343262 | 0.069568 | 0.999971 | 1 | 0.733937 |
| Slc4a3 | P16283 | 0.55 | -0.39 | -0.95 | -0.61 | -0.55 | -0.34 | -0.76 | -1.05 | -0.49 | 0.19 | -1.06 | -0.87 | -0.67 | -0.38 | -0.8 | -0.78 | -0.325 | -0.2075 | -0.3075 | 0.36425 | 0.611296 | 0.369904 | 0.999971 | 1 | 0.869526 |
| Slc4a4 | O88343 | -2.39 | -1.58 | -1.26 | -0.59 | -1.35 | -2.04 | -1.04 | 0.04 | -1.43 | -2.72 | -1.44 | -0.94 | -1.78 | -1.75 | -0.97 | -0.6 | 0.3575 | -0.1775 | 0.18 | 0.503494 | 0.722047 | 0.688459 | 0.999971 | 1 | 0.945753 |
| Slc4a4_1 | A7E1Z5 | 2.12 | -0.64 | -0.42 | -0.14 | -1.62 | 0.44 | 2.77 | -1.08 | -0.84 | 0.56 | -1.97 | -3.17 | -2.18 | -1.09 | -1.75 | -1.81 | -0.1025 | -1.585 | -1.9375 | 0.921756 | 0.11252 | 0.011565 | 0.999971 | 1 | 0.613095 |
| Slc4a4_2 | A0A571BE! | 0.85 | -0.3 | 0.19 | -0.11 | -0.92 | -0.37 | -0.2 | 1.24 | 0.44 | 0.14 | 0.57 | -0.44 | -0.98 | -0.4 | -0.04 | 0.35 | -0.22 | 0.02 | -0.425 | 0.653782 | 0.952388 | 0.267416 | 0.999971 | 1 | 0.842677 |
| Slc4a4_3 | E9Q8N8 | -0.46 | -1.64 | -1.91 | -1.77 | -1.49 | -1.08 | -1.66 | -2 | -1.43 | -0.93 | -2.08 | -1.93 | -1.17 | -0.86 | -1.92 | -1.99 | -0.1125 | -0.1475 | -0.04 | 0.76279 | 0.716473 | 0.922936 | 0.999971 | 1 | 0.993264 |
| Slc4a7 | F8VQC9 | 0.32 | -0.63 | -0.9 | -1.27 | -0.75 | -0.61 | -0.99 | -0.72 | -0.37 | 0.74 | -0.69 | -0.85 | -0.17 | -0.52 | -1.2 | -0.57 | -0.1475 | 0.3275 | 0.005 | 0.669788 | 0.486597 | 0.989686 | 0.999971 | 1 | 0.99925 |
| Slc4a8 | Q8JZR6 | -1.87 | -1.84 | -2.24 | -1.97 | -1.54 | -1.67 | -1.75 | -1.88 | -1.57 | -1.56 | -2.21 | -2.18 | -1.5 | -1.82 | -2.38 | -2.26 | 0.27 | 0.1 | -0.01 | 0.172649 | 0.672867 | 0.968173 | 0.999971 | 1 | 0.998787 |
| Slc52a2 | Q9D8F3-2 | 1.71 | -0.21 | 0.7 | 0.38 | -0.88 | -0.01 | 0.28 | 0.25 | -0.95 | 1.78 | -0.08 | 0.15 | 0.33 | -0.51 | 0.34 | 0.75 | -0.735 | -0.42 | -0.4175 | 0.132728 | 0.515565 | 0.368126 | 0.999971 | 1 | 0.869526 |
| Slc5a3 | Q9JKZ2 | 0.29 | -0.5 | -0.83 | -0.4 | 0.17 | -0.2 | -0.94 | -1.28 | -0.47 | 0.2 | -1.45 | -0.83 | 0.11 | 0.04 | -0.07 | -0.44 | -0.2025 | -0.2775 | 0.27 | 0.606981 | 0.494287 | 0.349936 | 0.999971 | 1 | 0.865956 |
| Slc5a6 | Q5U4D8 | 2.51 | 1.97 | 0.46 | 0.23 | 2.28 | 7.56 | 0.79 | 1.74 | 0.77 | 1.61 | -0.92 | -1.54 | -3.39 | -1.81 | -0.84 | 6.8 | 1.8 | -1.3125 | -1.1025 | 0.231148 | 0.141498 | 0.59651 | 0.999971 | 1 | 0.92377 |
| Slc5a7 | Q8BGY9 | 0.35 | -0.4 | -1.05 | -0.09 | -0.19 | -0.81 | -0.74 | -0.19 | -0.64 | -0.43 | -1.28 | -0.9 | -1 | -0.82 | -0.76 | -1.09 | -0.185 | -0.515 | -0.62 | 0.586064 | 0.158594 | 0.072065 | 0.999971 | 1 | 0.739635 |
| Slc66a2 | Q80XM9-2 | -1.27 | 3.05 | 1.36 | 0.07 | 2.62 | 2.51 | 1.17 | -0.2 | 2.39 | 0.23 | 1.93 | 1.88 | 2.52 | 2.01 | -0.74 | -0.1 | 0.7225 | 0.805 | 0.12 | 0.481949 | 0.398822 | 0.911845 | 0.999971 | 1 | 0.992551 |
| Slc6a1 | P31648 | -0.33 | -0.44 | 0.09 | -0.58 | -0.66 | -0.19 | 0.35 | -0.57 | -0.33 | -0.21 | -0.25 | -0.59 | -0.11 | -0.27 | -0.63 | -0.79 | 0.0475 | -0.03 | -0.135 | 0.867567 | 0.887768 | 0.584412 | 0.999971 | 1 | 0.920364 |
| Slc6a11 | P31650 | 0.21 | -0.55 | -1.41 | -1.3 | -0.65 | -0.36 | -1.53 | -1.29 | 0.02 | 0 | -1.55 | -1.04 | 0.09 | -0.19 | -0.93 | -1.52 | -0.195 | 0.12 | 0.125 | 0.657618 | 0.812343 | 0.79801 | 0.999971 | 1 | 0.970479 |
| Slc6a12 | E9PXG1 | 2.95 | 0.06 | 0.5 | 2.19 | 3.79 | -0.54 | 3.14 | 0.51 | 3.05 | 4.28 | -2.7 | 3.57 | -1.77 | 2.66 | 0.88 | 0.57 | 0.3 | 0.625 | -0.84 | 0.786323 | 0.689331 | 0.418469 | 0.999971 | 1 | 0.87788 |
| Slc6a13 | D3YWZ5 | 2.66 | 0.16 | 0.91 | 1.99 | 2.69 | 0.3 | 0.93 | 0.78 | 1.08 | 2.23 | 1.14 | 1.04 | -0.75 | 1.71 | 2.53 | 0.76 | -0.255 | -0.0575 | -0.3675 | 0.712636 | 0.920411 | 0.650727 | 0.999971 | 1 | 0.933842 |
| Slc6a15 | Q8BG16 | 0.11 | 0.74 | 0.95 | 0.25 | 0.58 | 0.5 | -0.05 | 0.14 | 0.31 | 0.17 | -0.22 | 0.56 | 1.08 | 0.86 | 0.71 | 1.14 | -0.22 | -0.3075 | 0.435 | 0.421448 | 0.278701 | 0.110069 | 0.999971 | 1 | 0.78268 |
| Slc6a17 | A0A0R4J08 | 0.5 | 0.09 | 0.27 | 0.21 | -0.11 | 0.55 | -0.32 | 0.2 | -0.03 | 0.79 | 0.06 | 0.28 | 0.61 | 0.38 | 0.39 | 0.78 | -0.1875 | 0.0075 | 0.2725 | 0.443969 | 0.974543 | 0.184017 | 0.999971 | 1 | 0.809025 |
| Slc6a20a | E9Q0H0 | -2.06 | -2 | 0.37 | 1.1 | -1.46 | -1.83 | -0.04 | 0.42 | -1.44 | 0.07 | 0.53 | 0.95 | -3.44 | 4.33 | 0.53 | 1.1 | -0.08 | 0.675 | 1.2775 | 0.927083 | 0.44637 | 0.429658 | 0.999971 | 1 | 0.881563 |
| Slc6a3 | Q61327 | -0.34 | 0.42 | -1.27 | -0.1 | 0.49 | -1.16 | -1.24 | 0.4 | 0.76 | -0.22 | -0.41 | -0.2 | 0.88 | 0.47 | 0.11 | -0.94 | -0.055 | 0.305 | 0.4525 | 0.919503 | 0.474234 | 0.367716 | 0.999971 | 1 | 0.869526 |
| Slc6a4 | Q60857 | 1.31 | -0.56 | 0.75 | 0.2 | 0.69 | 0.32 | -0.03 | 0.99 | 0.02 | 0.65 | 0.38 | -0.02 | -0.78 | 0.56 | 1.01 | -0.05 | 0.0675 | -0.1675 | -0.24 | 0.87551 | 0.684244 | 0.643398 | 0.999971 | 1 | 0.93301 |
| Slc6a6 | O35316 | 0.82 | -0.84 | -0.72 | -0.25 | -1.04 | -0.89 | -0.49 | -0.76 | -1.28 | 0.17 | -1.3 | -1.58 | -0.88 | -0.56 | -1.1 | -0.27 | -0.5475 | -0.75 | -0.455 | 0.175696 | 0.166021 | 0.275176 | 0.999971 | 1 | 0.843794 |
| Slc6a7 | Q6PGE7 | -0.93 | -1.36 | -1.17 | -1.54 | -1.3 | -0.96 | -0.95 | -0.93 | -0.88 | -0.63 | -1.03 | -1.1 | 0.15 | -0.66 | -0.55 | -0.49 | 0.215 | 0.34 | 0.8625 | 0.318779 | 0.135988 | 0.007363 | 0.999971 | 1 | 0.582714 |
| Slc6a8 | E9Q151 | -1.05 | -0.34 | -1.24 | -1.28 | -0.27 | -0.62 | -1.32 | -1.89 | -0.9 | -1.08 | -0.73 | -1.15 | -0.12 | -0.3 | -0.83 | -1.54 | -0.0475 | 0.0125 | 0.28 | 0.906149 | 0.961565 | 0.462404 | 0.999971 | 1 | 0.890421 |
| Slc6a9 | P28571-1 | 0.38 | -0.53 | 0.21 | 0.06 | -0.26 | -0.48 | 0.52 | -0.77 | 1.11 | 0.38 | 0.11 | 0.43 | 0.23 | -0.05 | 0.89 | 0.02 | -0.2775 | 0.4775 | 0.2425 | 0.419605 | 0.136658 | 0.428723 | 0.999971 | 1 | 0.881196 |
| Slc7a1 | E9Q3N1 | -0.51 | 0.7 | -1.33 | -0.39 | 0.33 | -0.47 | -1.13 | -1.1 | -0.16 | 0.6 | -0.94 | -0.78 | -0.03 | 0.1 | -1.11 | -0.53 | -0.21 | 0.0625 | -0.01 | 0.67613 | 0.901734 | 0.982919 | 0.999971 | 1 | 0.99925 |
| Slc7a10 | P63115 | -2.23 | 0.69 | 0.31 | 0.2 | 0.17 | 0.87 | -0.02 | -0.2 | 0.65 | -1.83 | 0.33 | 0.2 | 1.08 | 0.32 | 0.42 | 0.03 | 0.4625 | 0.095 | 0.72 | 0.4773 | 0.904216 | 0.277516 | 0.999971 | 1 | 0.843812 |
| Slc7a11 | Q9WTR6 | -0.43 | -0.58 | -1.1 | -0.8 | -0.17 | -0.34 | -0.61 | -1.15 | -0.67 | -0.04 | -1.21 | -1.19 | -0.47 | -0.65 | -0.9 | -0.8 | 0.16 | -0.05 | 0.0225 | 0.565789 | 0.873859 | 0.918068 | 0.999971 | 1 | 0.992633 |
| Slc7a14 | Q8BXR1 | -0.39 | -1.13 | -1.83 | -1.73 | -1 | -0.75 | -1.49 | -1.6 | -1.17 | -0.92 | -2.2 | -2.21 | -1.24 | -0.89 | -1.94 | -1.7 | 0.06 | -0.355 | -0.1725 | 0.872918 | 0.436007 | 0.660564 | 0.999971 | 1 | 0.935713 |
| Slc7a2 | P18581-2 | -0.63 | -0.34 | -0.8 | -0.63 | 0.04 | -0.08 | -0.39 | -1.61 | -0.36 | -0.58 | -0.38 | -0.32 | 0.32 | -0.5 | -0.14 | -0.62 | 0.09 | 0.19 | 0.365 | 0.811242 | 0.311277 | 0.180365 | 0.999971 | 1 | 0.808532 |
| Slc7a3 | P70423 | -2.25 | -0.46 | -0.4 | -0.54 | -0.25 | -1.06 | -0.09 | 0.09 | -0.27 | -2.43 | -0.11 | 0.11 | -0.24 | -1.17 | 0.85 | -0.21 | 0.585 | 0.2375 | 0.72 | 0.241809 | 0.725663 | 0.219643 | 0.999971 | 1 | 0.827064 |
| Slc7a4 | A0A338P6\ | -1.15 | -1.87 | -0.13 | -0.78 | -1.08 | -1.7 | -0.39 | -0.64 | -0.93 | -0.65 | 0.34 | 0.46 | -1.21 | -0.7 | 0.27 | 0.8 | 0.03 | 0.7875 | 0.7725 | 0.945152 | 0.12306 | 0.175355 | 0.999971 | 1 | 0.808532 |
| Slc7a5 | Q9Z127 | 0.72 | -0.11 | -0.15 | 0.17 | -0.41 | -0.16 | -0.32 | -0.18 | 0.27 | 0.79 | -0.38 | -0.88 | -0.72 | -0.2 | -0.53 | 0.09 | -0.425 | -0.2075 | -0.4975 | 0.105388 | 0.606706 | 0.106968 | 0.999971 | 1 | 0.78268 |
| Slc7a6 | Q8BGK6 | -0.27 | -0.19 | -0.22 | -0.47 | -0.61 | -0.39 | -0.72 | -1.04 | -0.16 | 0.61 | -0.71 | -0.23 | 0.48 | 0.08 | -0.98 | -0.4 | -0.4025 | 0.165 | 0.0825 | 0.075213 | 0.573918 | 0.798684 | 0.999971 | 1 | 0.970479 |
| Slc7a6os | Q7TPE5 | 1.07 | 0.33 | 0.43 | 1.28 | 0.04 | 0.01 | 0.89 | 1.01 | 0.26 | 0.38 | 1.24 | 2.8 | -0.39 | 0.72 | 1.7 | 0.71 | -0.29 | 0.3925 | -0.0925 | 0.416633 | 0.504596 | 0.839726 | 0.999971 | 1 | 0.979728 |
| Slc7a8 | Q9QXW9 | -0.03 | 0.71 | 0.58 | -0.26 | 0.63 | 0.94 | 0.36 | -0.44 | 0.39 | 0.74 | 0.52 | -0.43 | 0.37 | 0.43 | -0.19 | -0.01 | 0.1225 | 0.055 | -0.1 | 0.738952 | 0.872395 | 0.731349 | 0.999971 | 1 | 0.956041 |
| Slc8a1 | Q68FL0 | -1.07 | -1.76 | -1.42 | -1.61 | -1.7 | -1.33 | -0.79 | -1.82 | -1.55 | -1.14 | -1.57 | -1.57 | -1.74 | -1.55 | -1.6 | -1.62 | 0.055 | 0.0075 | -0.1625 | 0.848478 | 0.973035 | 0.442223 | 0.999971 | 1 | 0.883386 |
| Slc8a1_1 | G3X9J1 | -0.17 | -0.96 | -1.67 | -3.04 | -1.89 | -0.97 | -3.77 | -2.66 | -2.04 | -2.74 | -3.8 | -2.67 | -1.84 | -1.35 | -0.72 | -2.11 | -0.8625 | -1.3525 | -0.045 | 0.277522 | 0.06319 | 0.942222 | 0.999971 | 1 | 0.996889 |
| Slc8a2 | Q8K596 | -2.01 | -1.44 | -2.49 | -2.47 | -1.25 | -1.27 | -2.35 | -2.57 | -1.63 | -1.68 | -2.11 | -2.65 | -1.02 | -1.58 | -2.49 | -2.61 | 0.2425 | 0.085 | 0.1775 | 0.555876 | 0.8022 | 0.680271 | 0.999971 | 1 | 0.942957 |
| Slc8a3 | S4R2P9-2 | -1.33 | -1.81 | -1.07 | -0.58 | -1.92 | -1.86 | -0.97 | -0.82 | -2.26 | -1.06 | -0.85 | -0.9 | -2.84 | -1.22 | -0.77 | -0.91 | -0.195 | -0.07 | -0.2375 | 0.604967 | 0.862181 | 0.639059 | 0.999971 | 1 | 0.93301 |
| Slc9a1 | Q61165 | -0.82 | -1.24 | -0.88 | -1.14 | -1.43 | -1.55 | -1.2 | -1.33 | -1.35 | -1.13 | -1.48 | -1.23 | -1.52 | -1.41 | -1.22 | -1.33 | -0.3575 | -0.2775 | -0.35 | 0.088922 | 0.167188 | 0.09139 | 0.999971 | 1 | 0.773779 |
| Slc9a3r1 | P70441 | -0.87 | -1.3 | -1.03 | -0.93 | -1.13 | -1.25 | -1.12 | -1.55 | -1.22 | -0.99 | -0.97 | -1.25 | -0.61 | -1.25 | -1.33 | -0.65 | -0.23 | -0.075 | 0.0725 | 0.264148 | 0.688015 | 0.767681 | 0.999971 | 1 | 0.962472 |
| Slc9a3r2 | A0A0R4IZX | 1.11 | 0.06 | 0.21 | 1.02 | -0.04 | 0.3 | 0.83 | 0.2 | 0.27 | 1.14 | 0.34 | 0.23 | -0.13 | 0.06 | 0.41 | 0.25 | -0.2775 | -0.105 | -0.4525 | 0.406371 | 0.759584 | 0.161055 | 0.999971 | 1 | 0.800215 |
| Slc9a5 | B2RXE2 | -1.77 | 0.27 | -1.03 | -0.33 | -0.44 | -0.45 | -0.03 | 0.31 | -0.13 | -0.45 | -1.23 | -0.01 | -0.87 | -0.2 | -0.42 | -0.36 | 0.5625 | 0.26 | 0.2525 | 0.230329 | 0.595143 | 0.567729 | 0.999971 | 1 | 0.91816 |
| Slc9a6 | D3Z0Q9 | 1.09 | -0.32 | 0.07 | 0.21 | -0.96 | -0.2 | -0.39 | -0.2 | -1.08 | 1.14 | -0.34 | -0.34 | -1.32 | 0.72 | -0.02 | 0.4 | -0.7 | -0.4175 | -0.3175 | 0.067537 | 0.425482 | 0.530032 | 0.999971 | 1 | 0.910069 |
| Slc9a6_1 | A1L3P4 | -1.55 | -0.94 | -0.74 | -1.24 | -0.72 | -0.32 | -0.78 | -1.01 | -0.59 | -1.33 | -0.55 | -0.51 | 0.22 | -0.38 | -0.53 | -0.6 | 0.41 | 0.3725 | 0.795 | 0.13308 | 0.204081 | 0.016976 | 0.999971 | 1 | 0.617356 |
| Slc9a7 | Q8BLV3 | -0.18 | -0.63 | -0.46 | -0.17 | -0.71 | 0.25 | 0.52 | -0.5 | -0.51 | 0.13 | -0.34 | -0.51 | -0.5 | -0.48 | -0.63 | -1.19 | 0.25 | 0.0525 | -0.34 | 0.43966 | 0.816227 | 0.175156 | 0.999971 | 1 | 0.808532 |
| Slc9a9 | Q8BZ00 | -0.41 | -0.13 | 0.15 | 0.4 | -0.04 | -0.23 | -0.02 | 1.09 | -0.37 | -0.56 | 0.99 | 1.51 | 0.29 | -0.08 | 0.06 | 1.13 | 0.1975 | 0.39 | 0.3475 | 0.568615 | 0.44409 | 0.300147 | 0.999971 | 1 | 0.847402 |
| Slc9b2 | Q5BKR2 | 0.86 | 1.04 | -0.46 | -1.88 | -0.24 | -1.77 | 0.2 | -0.54 | 0.35 | 5.11 | -0.25 | -1.65 | 0.7 | -1.81 | 0.25 | -2.82 | -0.4775 | 1 | -0.81 | 0.513129 | 0.49503 | 0.410228 | 0.999971 | 1 | 0.875847 |
| Slco1a4 | Q9EP96 | 0.11 | -0.79 | -1.13 | -0.38 | 0.72 | -0.26 | 0.12 | -2.58 | 0.14 | -0.37 | -1.97 | -2.49 | -2.4 | -0.78 | -1.27 | -2.33 | 0.0475 | -0.625 | -1.1475 | 0.945545 | 0.331993 | 0.030098 | 0.999971 | 1 | 0.678474 |
| Slco1a6 | Q99J94 | -2.11 | 1.61 | 0.18 | 0.5 | 2.13 | 0.64 | 0.1 | -0.78 | 1.03 | -0.29 | -0.23 | 1 | 2.53 | 1.67 | 0.28 | 0.63 | 0.4775 | 0.3325 | 1.2325 | 0.593431 | 0.672184 | 0.166936 | 0.999971 | 1 | 0.800215 |
| Slco1c1 | A0A0N4SU | -0.83 | -0.39 | -0.31 | -0.14 | -0.41 | -0.88 | -0.64 | -1.48 | -0.22 | -0.41 | -0.64 | -0.53 | 0.28 | -1.32 | -1.36 | 0.09 | -0.435 | -0.0325 | -0.16 | 0.154285 | 0.8801 | 0.716783 | 0.999971 | 1 | 0.952711 |
| Slco2b1 | Q3V1K7 | 1.24 | -0.91 | 0.33 | 0.63 | -1.15 | -1.24 | 0.67 | 0.85 | -1.06 | 0.59 | 1.1 | 1.2 | -1.52 | -1.37 | 1.13 | 0.77 | -0.54 | 0.135 | -0.57 | 0.420343 | 0.831137 | 0.454426 | 0.999971 | 1 | 0.887217 |
| Slco3a1 | Q8R3L5-2 | -0.06 | -2.09 | -1.14 | -0.53 | -1.72 | -0.68 | -0.79 | -0.81 | -1.34 | 0 | -0.88 | -1.46 | -2.55 | -1.61 | -1.57 | -1.19 | -0.045 | 0.035 | -0.775 | 0.923249 | 0.945318 | 0.139609 | 0.999971 | 1 | 0.784948 |
| Slirp | Q9D8T7 | -0.22 | -0.14 | 0.91 | -0.38 | 0.52 | -0.16 | 3.28 | 0.12 | 1.18 | -0.59 | 1.65 | 1.77 | 1.51 | -0.08 | -0.28 | -1.5 | 0.8975 | 0.96 | -0.13 | 0.25715 | 0.120374 | 0.834959 | 0.999971 | 1 | 0.979036 |
| Slit1 | Q80TR4 | -0.27 | -0.54 | -0.45 | -0.4 | -0.54 | -0.41 | -0.7 | -0.13 | -0.64 | -0.4 | -0.41 | -0.3 | -0.72 | -0.34 | -0.37 | -0.11 | -0.03 | -0.0225 | 0.03 | 0.877952 | 0.896228 | 0.879929 | 0.999971 | 1 | 0.985441 |
| Slit2 | A0A140T81 | -0.69 | -0.58 | -0.66 | -0.65 | -0.57 | -0.34 | -0.63 | -0.53 | -0.14 | -0.61 | -0.4 | -0.39 | 0.21 | -0.57 | -0.4 | -0.28 | 0.1275 | 0.26 | 0.385 | 0.453629 | 0.164602 | 0.102659 | 0.999971 | 1 | 0.78268 |
| Slit3 | Q9WVB4 | -0.69 | -0.46 | 0.98 | 0.92 | -0.27 | -0.24 | 1.24 | 1.02 | -0.58 | -0.49 | 1.36 | 1.54 | -0.15 | -0.24 | 1.54 | 0.61 | 0.25 | 0.27 | 0.2525 | 0.651011 | 0.684678 | 0.652069 | 0.999971 | 1 | 0.933842 |
| Slitrk1 | Q810C1 | 0.56 | 0.51 | -0.04 | 0.01 | 0.36 | 0.58 | -0.2 | -0.13 | 0.47 | 0.36 | -0.11 | -0.1 | 0.77 | -0.04 | -0.32 | 0.34 | -0.1075 | -0.105 | -0.0725 | 0.689198 | 0.672394 | 0.806646 | 0.999971 | 1 | 0.972713 |
| Slitrk2 | Q810C0 | 1.21 | -0.52 | -0.73 | -0.15 | -0.76 | -0.25 | -0.9 | -0.38 | -0.54 | 0.09 | -0.22 | 0.11 | 0.09 | -0.23 | -0.5 | -0.37 | -0.525 | -0.0925 | -0.205 | 0.248161 | 0.832553 | 0.635584 | 0.999971 | 1 | 0.93301 |
| Slitrk3 | Q810B9 | -1.15 | -1.14 | -0.91 | -0.77 | -0.92 | -0.88 | -1.24 | -0.4 | -1.15 | -0.76 | -0.7 | -0.48 | -1.86 | -0.92 | -0.82 | 0.01 | 0.1325 | 0.22 | 0.095 | 0.572875 | 0.315734 | 0.803127 | 0.999971 | 1 | 0.971467 |
| Slitrk4 | Q810B8 | 0 | -1.02 | -1.21 | -1.02 | -0.76 | -0.96 | -1.23 | -0.65 | -0.94 | -0.47 | -1.27 | -1.15 | -0.44 | -1.3 | -1.32 | -0.63 | -0.0875 | -0.145 | -0.11 | 0.77689 | 0.658483 | 0.754348 | 0.999971 | 1 | 0.961116 |
| Slitrk5 | Q810B7 | -1.38 | -0.89 | -0.99 | -1.29 | -0.61 | -1.07 | -0.7 | -1.02 | -0.76 | -1.13 | -0.5 | -0.53 | -0.37 | -1.06 | -1 | -0.8 | 0.2875 | 0.4075 | 0.33 | 0.199368 | 0.097357 | 0.180516 | 0.999971 | 1 | 0.808532 |

| | | | | | | | | | | | | | | | | | | | | | | | | | | |
|---|---|---|---|---|---|---|---|---|---|---|---|---|---|---|---|---|---|---|---|---|---|---|---|---|---|---|
| Slk | O54988-2 | 0.07 | -0.17 | -0.1 | -0.07 | -0.06 | -0.34 | 0.21 | -0.18 | 0.11 | -0.27 | -0.23 | -0.08 | -0.35 | -0.54 | -0.34 | -0.5 | -0.025 | -0.05 | -0.365 | 0.896348 | 0.776928 | 0.056114 | 0.999971 | 1 | 0.725348 |
| Slmap | F6WY34 | -1.28 | -0.18 | -0.36 | -0.26 | -0.55 | -0.59 | -0.67 | -0.23 | -0.43 | 0.1 | 1.04 | -1.7 | 0.84 | 0.8 | -0.35 | 0.13 | 0.01 | 0.2725 | 0.875 | 0.972142 | 0.638577 | 0.040991 | 0.999971 | 1 | 0.700838 |
| Slmap_1 | D3Z7V3 | 1.12 | 0.29 | 0.09 | 0.4 | 0.96 | 0.46 | 0.67 | 0.19 | 0.71 | 1.22 | 0.54 | 0.76 | 0.62 | 0.21 | 0.14 | -0.15 | 0.095 | 0.3325 | -0.27 | 0.743251 | 0.256211 | 0.361211 | 0.999971 | 1 | 0.869526 |
| Sltm | Q8CH25-2 | 0.74 | 0.41 | 0.62 | 0.07 | 0.57 | 0.21 | 1.13 | -0.05 | 0.29 | 0.26 | 0.4 | 0.13 | 0.1 | 0.12 | 0.24 | -0.2 | 0.005 | -0.19 | -0.395 | 0.986791 | 0.368723 | 0.099197 | 0.999971 | 1 | 0.78268 |
| Slu7 | Q8BHJ9 | 3.2 | 0.08 | 2.35 | 0.3 | -0.45 | 0.68 | 3.95 | -0.92 | -0.03 | 1.7 | 0.55 | -0.4 | -1.59 | -0.38 | -0.01 | -0.81 | -0.6675 | -1.0275 | -2.18 | 0.578354 | 0.225651 | 0.017344 | 0.999971 | 1 | 0.617356 |
| Smad2 | Q62432-2 | 0.98 | 0.6 | -0.51 | -0.35 | 0.75 | 0.71 | -0.06 | -0.17 | 0.53 | 1.24 | -0.01 | -0.55 | 0.31 | 0.57 | 0.07 | -0.24 | 0.1275 | 0.1225 | -0.0025 | 0.759264 | 0.803404 | 0.994837 | 0.999971 | 1 | 1 |
| Smad3 | Q8BUN5 | 1.69 | 0.13 | 1.46 | 1.4 | 0.95 | 0.86 | 0.75 | 1.25 | 0.58 | 1.61 | 1.06 | 0.63 | 0.5 | 1.38 | 0.94 | 2 | -0.2175 | -0.2 | 0.035 | 0.549364 | 0.62551 | 0.937686 | 0.999971 | 1 | 0.995618 |
| Smad4 | P97471 | 0.87 | 0.6 | 1.03 | 0.96 | 0.34 | 1.08 | 1.18 | 0.36 | 0.22 | 0.87 | 0.65 | 0.34 | -0.5 | 0.79 | 0.15 | 0.36 | -0.125 | -0.345 | -0.665 | 0.640307 | 0.139075 | 0.047818 | 0.999971 | 1 | 0.710102 |
| Smad5 | P97454 | 2.64 | 0.03 | 0.2 | 0.12 | -0.01 | 0.88 | 0.14 | 0.2 | -0.45 | 4.1 | 0.16 | 0.21 | -1.12 | 1.26 | 0.05 | 0.07 | -0.445 | 0.2575 | -0.6825 | 0.468039 | 0.814036 | 0.356338 | 0.999971 | 1 | 0.868793 |
| Smap | Q9R0P4 | 2.14 | 2.57 | 3.76 | 3.32 | 2.78 | 2.74 | 2.36 | 2.75 | 2.54 | 2.78 | 3 | 2.98 | 1.69 | 2.7 | 3.21 | 3.52 | -0.29 | -0.1225 | -0.1675 | 0.437934 | 0.740862 | 0.739675 | 0.999971 | 1 | 0.960899 |
| Smap1 | Q91VZ6 | -0.13 | -0.1 | -0.34 | -0.35 | -0.06 | 0.45 | -0.21 | -0.12 | 0.09 | 0.13 | -0.21 | -0.3 | 0.15 | 0.1 | -0.08 | -0.18 | 0.245 | 0.1575 | 0.2275 | 0.265896 | 0.414273 | 0.230039 | 0.999971 | 1 | 0.828687 |
| Smap2 | Q7TN29 | 0.34 | 0.29 | 0.8 | 0.9 | 0.05 | -0.02 | 1 | 0.12 | 0.23 | -0.25 | 0.39 | 0.68 | -0.36 | -0.25 | 0.46 | 0.08 | -0.295 | -0.32 | -0.6 | 0.332288 | 0.252051 | 0.046255 | 0.999971 | 1 | 0.710102 |
| Smarca1 | Q6PGB8 | -0.14 | -1.9 | -0.69 | -0.47 | -1.32 | -2.05 | -0.98 | -0.02 | -1.46 | -0.02 | -0.39 | -0.55 | -0.97 | -1.64 | 0.1 | 0.1 | -0.2925 | 0.195 | 0.1975 | 0.581187 | 0.673914 | 0.71075 | 0.999971 | 1 | 0.950695 |
| Smarca2 | Q6DIC0 | 0.51 | 1.43 | 1.12 | 1.52 | 1.38 | 1.32 | 0.62 | 1.4 | 1 | 0.84 | 1.03 | 1.37 | 1.26 | 1.67 | 1.28 | 1.49 | 0.035 | -0.085 | 0.28 | 0.907948 | 0.754601 | 0.312819 | 0.999971 | 1 | 0.849367 |
| Smarca4 | A0A0R4J17 | 0.68 | 0.96 | 1.34 | 1.53 | 0.83 | 0.65 | 1.23 | 1.34 | 0.71 | 0.89 | 1.21 | 1.41 | 0.06 | 0.65 | 1.27 | 1.31 | -0.115 | -0.0725 | -0.305 | 0.671691 | 0.785608 | 0.390505 | 0.999971 | 1 | 0.871816 |
| Smarca5 | Q91ZW3 | -0.15 | -0.14 | -0.76 | -0.83 | -0.07 | -0.28 | -0.86 | -0.87 | -0.23 | -0.27 | -0.86 | -0.9 | -0.02 | -0.28 | -0.62 | -0.97 | -0.05 | -0.095 | -0.0025 | 0.862819 | 0.732458 | 0.99315 | 0.999971 | 1 | 0.99925 |
| Smarcal1 | Q8BJL0 | 1.94 | 1.41 | 1.29 | 1.93 | 1.48 | 1.75 | 1.27 | 1.85 | 1.44 | 1.5 | 1.21 | 1.35 | 0.41 | 1.44 | 1.66 | 1.85 | -0.055 | -0.2675 | -0.3025 | 0.822842 | 0.247262 | 0.406397 | 0.999971 | 1 | 0.874911 |
| Smarcb1 | Q9Z0H3 | 0.63 | 1.01 | 1.17 | 1.49 | 0.89 | 0.79 | 0.68 | 1.35 | 1.01 | 1.21 | 0.54 | 0.76 | 0.42 | 1.45 | 1.09 | 1.42 | -0.1475 | -0.195 | 0.02 | 0.568938 | 0.451613 | 0.947846 | 0.999971 | 1 | 0.997651 |
| Smarcc1 | Q3UNN4 | 0.81 | 1.1 | 1.01 | 0.74 | 0.87 | 0.37 | 0.58 | 0.45 | 0.65 | 0.53 | 0.73 | 1.21 | 0.53 | 1.5 | 0.48 | 0.8 | -0.3475 | -0.135 | -0.0875 | 0.106691 | 0.534359 | 0.746306 | 0.999971 | 1 | 0.961064 |
| Smarcc2 | Q3UID0 | 1.69 | 1.84 | 2.47 | 2.38 | 1.94 | 1.7 | 3.01 | 1.88 | 1.95 | 1.75 | 2.14 | 2.42 | 1.43 | 1.77 | 2.19 | 1.75 | 0.0375 | -0.03 | -0.31 | 0.91429 | 0.908658 | 0.267922 | 0.999971 | 1 | 0.842746 |
| Smarcd1 | Q61466 | 0.98 | 1.14 | 1.42 | 1.2 | 1.06 | 1.08 | 0.85 | 1.21 | 1.03 | 0.73 | 1.15 | 1.43 | 0.67 | 0.96 | 1.24 | 1.44 | -0.135 | -0.1 | -0.1075 | 0.47843 | 0.643508 | 0.642152 | 0.999971 | 1 | 0.93301 |
| Smarcd3 | Q6P9Z1 | 1.25 | 1.54 | 0.88 | 1.48 | 1.54 | 1.04 | 0.85 | 1.25 | 1.13 | 0.79 | 1.02 | 1.38 | 1.44 | 1.6 | 1.75 | 1.64 | -0.1175 | -0.2075 | 0.32 | 0.629566 | 0.376437 | 0.15789 | 0.999971 | 1 | 0.797252 |
| Smarce1 | O54941 | 1.75 | 1.92 | 1.89 | 2.31 | 2.09 | 1.88 | 1.58 | 1.86 | 1.83 | 1.71 | 1.41 | 2.11 | 1.55 | 1.74 | 2.14 | 1.82 | -0.115 | -0.2025 | -0.155 | 0.58695 | 0.380854 | 0.483714 | 0.999971 | 1 | 0.896352 |
| Smc1a | Q9CU62 | 0.84 | -0.28 | 0.06 | 0.33 | -0.27 | -0.23 | -0.13 | -0.15 | -0.14 | 0.39 | 0.03 | -0.01 | -0.89 | -0.36 | 0.13 | -0.02 | -0.4325 | -0.17 | -0.5225 | 0.125086 | 0.544289 | 0.137791 | 0.999971 | 1 | 0.784948 |
| Smc1b | Q920F6 | 0.61 | 0.65 | -0.82 | -0.33 | 0.43 | 0.98 | -0.11 | -0.86 | 0.9 | 1.15 | -1.02 | -1.49 | 0.26 | 1.01 | -1.53 | -1.82 | 0.0825 | -0.1425 | -0.5475 | 0.867723 | 0.836704 | 0.444131 | 0.999971 | 1 | 0.883802 |
| Smc2 | Q8CG48 | 0.01 | 0.54 | -3.61 | -1.52 | 0.7 | -0.13 | -0.31 | -2.41 | -0.43 | 0.06 | -1.65 | -1.68 | 1.09 | 0 | -1.48 | -1.93 | 0.6075 | 0.22 | 0.565 | 0.554745 | 0.812488 | 0.588591 | 0.999971 | 1 | 0.921285 |
| Smc3 | Q9CW03 | 0.6 | 0.42 | 0.89 | 0.83 | 0.43 | 0.31 | 0.74 | 0.54 | 0.45 | 0.6 | 0.72 | 0.57 | 0.11 | 0.37 | 0.72 | 0.36 | -0.18 | -0.1 | -0.295 | 0.379299 | 0.594943 | 0.192217 | 0.999971 | 1 | 0.810547 |
| Smc5 | A0A286YD( | -1.35 | -1.77 | -0.45 | 4.01 | 0.04 | -3 | -0.73 | 4.16 | -0.94 | -0.89 | -1.07 | -1.4 | 1.41 | -1.04 | 1.81 | 7.93 | 0.0075 | -1.185 | 2.4175 | 0.996613 | 0.335589 | 0.259326 | 1 | 1 | 0.839154 |
| Smchd1 | Q6P5D8 | 0.14 | -0.59 | -0.22 | -0.11 | -0.59 | -0.75 | -0.42 | -0.31 | -0.4 | -0.44 | -0.58 | -0.24 | -0.36 | -0.49 | -0.4 | -0.18 | -0.3225 | -0.22 | -0.1625 | 0.171597 | 0.315227 | 0.455311 | 0.999971 | 1 | 0.887217 |
| Smcr8 | Q3UMB5 | -0.46 | -0.38 | -0.68 | -1.05 | -0.61 | -0.42 | -0.74 | -0.86 | -0.67 | -0.42 | -0.91 | -0.87 | -0.34 | -0.42 | -0.84 | -1.04 | -0.015 | -0.075 | -0.0175 | 0.945745 | 0.739522 | 0.944705 | 0.999971 | 1 | 0.997225 |
| Smdt1 | Q9DB10 | -0.81 | 1.81 | 1.77 | 0.67 | 1.05 | 1.33 | 1.44 | 0.78 | 1.48 | -0.31 | 1.43 | 1.87 | 1.29 | 0.24 | 0.85 | 0.56 | 0.29 | 0.2575 | -0.125 | 0.619219 | 0.719007 | 0.834765 | 0.999971 | 1 | 0.979036 |
| Smg1 | Q8BKX6 | -0.08 | 0.1 | -0.04 | -0.05 | 0.28 | 0.24 | 0.25 | -0.12 | 0.14 | -0.04 | 0.13 | -0.21 | -0.53 | -0.23 | -0.17 | -0.03 | 0.18 | 0.0225 | -0.2225 | 0.333277 | 0.896362 | 0.251063 | 0.999971 | 1 | 0.839012 |
| Smg5 | Q6ZPY2 | -0.47 | 0.39 | 0.92 | 0.17 | 0.25 | 0.54 | 0.05 | 0.67 | 0.1 | -0.53 | 1.63 | -0.97 | 0.19 | 0.88 | 0.24 | 0.27 | 0.125 | -0.195 | 0.1425 | 0.698634 | 0.739836 | 0.667826 | 0.999971 | 1 | 0.938705 |
| Smg6 | P61406 | 0.41 | 0.3 | 0.22 | 0.41 | 0.87 | 0.46 | 0.23 | 0.41 | 0.34 | 0.13 | 0.29 | 0.45 | 0.74 | 0.43 | 0.21 | 0.57 | 0.1575 | -0.0325 | 0.1525 | 0.440701 | 0.847092 | 0.431163 | 0.999971 | 1 | 0.881611 |
| Smg7 | Q5RJH6-3 | -0.19 | 0.55 | -0.42 | -0.34 | 0.71 | 0.64 | -0.45 | -0.11 | 0.53 | -0.13 | -0.15 | -0.22 | 1.12 | 0.55 | -0.61 | -0.09 | 0.2975 | 0.1075 | 0.3425 | 0.409947 | 0.714253 | 0.419665 | 0.999971 | 1 | 0.87788 |
| Smg8 | Q8VE18 | 0.23 | 0.28 | 0.22 | 0.22 | 0.14 | -0.1 | 0.22 | -0.08 | -0.01 | 0.33 | 0.27 | 0.1 | -0.32 | -0.04 | -0.53 | -0.19 | -0.1925 | -0.065 | -0.5075 | 0.280605 | 0.699649 | 0.020097 | 0.999971 | 1 | 0.635388 |
| Smg9 | Q9DB90 | 0.76 | 1.05 | 0.71 | 0.96 | 0.67 | 0.91 | 0.95 | 0.77 | 0.69 | 0.97 | 0.75 | 0.09 | 1.42 | 0.82 | 0.71 | 0.63 | -0.045 | -0.245 | 0.025 | 0.803852 | 0.315777 | 0.91475 | 0.999971 | 1 | 0.992633 |
| Smim12 | Q78RX3 | -1.01 | -0.66 | 0.66 | 0.61 | -0.5 | -0.09 | 0.73 | 0.55 | -0.23 | -0.77 | 0.66 | 0.2 | 0.36 | -0.95 | 0.52 | 0.05 | 0.2725 | 0.065 | 0.095 | 0.574194 | 0.894922 | 0.85031 | 0.999971 | 1 | 0.98187 |
| Smim13 | E9Q942 | 3.02 | -0.69 | 0.19 | 0.73 | -0.82 | 0.86 | 1.26 | -0.15 | -1 | 2.03 | -0.45 | -1.57 | -2.65 | -0.84 | -1.17 | -1.06 | -0.525 | -1.06 | -2.2425 | 0.530696 | 0.308594 | 0.019988 | 0.999971 | 1 | 0.635388 |
| Smim18 | J3QNP2 | -2.33 | -1.06 | -0.99 | -1.01 | -1.38 | -0.57 | -0.89 | -1.02 | -0.28 | -0.78 | -0.78 | -0.75 | -1.19 | -1.26 | -1.07 | -0.99 | 0.3825 | 0.7 | 0.22 | 0.303286 | 0.06916 | 0.513833 | 0.999971 | 1 | 0.905815 |
| Smim20 | D3Z7Q2 | -1.44 | -0.71 | -1.12 | -1.4 | -1.03 | -1.08 | -1.68 | -0.8 | -0.3 | -1.28 | -0.99 | -0.31 | -0.44 | -0.75 | -0.15 | -0.95 | 0.02 | 0.4475 | 0.595 | 0.941118 | 0.167549 | 0.048401 | 0.999971 | 1 | 0.710102 |
| Smim26 | Q3V460 | -3.28 | -0.48 | 0.32 | -1.41 | -0.86 | -0.37 | 2.04 | -1.51 | -0.05 | -2.56 | -0.15 | -0.17 | 0.01 | -1.83 | -0.74 | -3.04 | 1.0375 | 0.48 | -0.1875 | 0.304481 | 0.592301 | 0.83756 | 0.999971 | 1 | 0.979416 |
| Smim4 | A0A0R3P9( | 0.16 | 1.4 | -0.02 | 0.02 | 1.39 | 0.3 | -0.31 | 0.01 | -0.05 | 0.53 | -0.46 | -0.09 | -0.34 | 0.54 | -1.18 | 0.21 | -0.0425 | -0.4075 | -0.5825 | 0.927585 | 0.303017 | 0.238721 | 0.999971 | 1 | 0.833594 |
| Smim7 | F8WIU9 | 2.52 | 1.32 | 3.24 | 3.01 | 1.57 | 2.06 | 2.7 | 2.45 | 1.18 | 2.57 | 1.87 | 1.45 | -0.82 | 1.62 | 2.24 | 2.31 | -0.3275 | -0.755 | -1.185 | 0.484831 | 0.149629 | 0.148782 | 0.999971 | 1 | 0.793248 |
| Smim8 | Q9CQQ0 | 0.3 | 2.34 | 0.79 | 0.12 | 1.3 | 1.97 | 0.5 | 0.62 | 1.11 | 0.93 | 0.38 | 0.32 | 1.8 | 1.4 | 0.31 | 0.39 | 0.21 | -0.2025 | 0.0875 | 0.707646 | 0.689238 | 0.878791 | 0.999971 | 1 | 0.985441 |
| Smn1 | P97801 | -0.6 | 1.2 | 1.32 | 0.68 | 0.75 | -0.1 | 0.63 | 1.05 | 0.81 | -1.08 | 1.47 | -0.19 | -0.19 | 0.37 | 0.61 | 0.96 | -0.0675 | -0.3975 | -0.2125 | 0.885546 | 0.545367 | 0.652338 | 0.999971 | 1 | 0.934005 |
| Smndc1 | Q8BGT7 | 0.74 | 1.13 | 0.49 | 0.42 | 1.4 | 0.78 | 0.19 | 0.45 | 1.11 | 1.06 | 0.62 | -0.14 | 0.34 | 1.23 | 0.21 | 0.33 | 0.01 | -0.0325 | -0.1675 | 0.97435 | 0.921474 | 0.574943 | 0.999971 | 1 | 0.91816 |
| Smoc1 | D3Z3A3 | 2.24 | 0.85 | -0.59 | 0.35 | 1.58 | 0.75 | -0.49 | 0.3 | 1.54 | 2.19 | -0.15 | -0.04 | 2.84 | 1.3 | 0.54 | -0.55 | -0.1775 | 0.1725 | 0.32 | 0.789216 | 0.818603 | 0.701314 | 0.999971 | 1 | 0.949019 |
| Smpd1 | Q04519 | 0.09 | 0.59 | 0.46 | 0.23 | 0.51 | 0.93 | 1.01 | 0.97 | 0.55 | 0.72 | 0.45 | 0.4 | 0.39 | 0.97 | 0.49 | 1.07 | 0.5125 | 0.1875 | 0.3875 | 0.035824 | 0.343297 | 0.130704 | 0.999971 | 1 | 0.784948 |
| Smpd2 | O70572 | -1.11 | -0.38 | 0.1 | -0.1 | -0.14 | -0.27 | -0.15 | 0.49 | -0.25 | -0.75 | 0.08 | 0.82 | 0.11 | -0.43 | 0.23 | -0.1 | 0.355 | 0.3475 | 0.325 | 0.282032 | 0.4007 | 0.30752 | 0.999971 | 1 | 0.847402 |
| Smpd3 | Q9JJY3 | 0.09 | -1.38 | -1.16 | -0.94 | -1.29 | -1.35 | -1.02 | -1.24 | -1.11 | -0.23 | -1.23 | -1.9 | -1.29 | -1.25 | -1.68 | -1.82 | -0.3775 | -0.27 | -0.6625 | 0.273177 | 0.549342 | 0.085314 | 0.999971 | 1 | 0.759325 |
| Smpd4 | Q6ZPR5 | -0.69 | -0.83 | -0.24 | -0.32 | -0.67 | -0.97 | -0.23 | -0.46 | -0.89 | -0.43 | -0.55 | -0.69 | -0.79 | -0.93 | -0.35 | -0.16 | -0.0625 | -0.12 | -0.0375 | 0.797502 | 0.582064 | 0.883691 | 0.999971 | 1 | 0.986782 |
| Smpdl3a | P70158 | -1.14 | -0.69 | -0.12 | -0.05 | -0.54 | -0.62 | 0.18 | 0.26 | -0.51 | -1.43 | 0.33 | 0.34 | -0.64 | -0.76 | 0.06 | 0.02 | 0.32 | 0.1825 | 0.17 | 0.36197 | 0.694834 | 0.613491 | 0.999971 | 1 | 0.927364 |
| Smpdl3b | P58242 | -0.49 | -1.95 | -0.78 | -0.65 | -1.2 | -1.67 | -1.06 | -0.56 | -1.13 | -1.2 | -1.2 | -0.93 | -1.39 | -1.17 | -0.4 | -1.26 | -0.155 | -0.1475 | -0.0875 | 0.690638 | 0.662625 | 0.820925 | 0.999971 | 1 | 0.975833 |
| Sms | P97355 | 1.2 | 1.74 | 2.38 | 2.09 | 1.49 | 2.33 | 1.71 | 2.05 | 1.83 | 1.92 | 2.33 | 2.05 | 1.7 | 1.69 | 2.49 | 2.36 | 0.0425 | 0.18 | 0.2075 | 0.893354 | 0.536233 | 0.535209 | 0.999971 | 1 | 0.910077 |
| Smu1 | Q3UKJ7 | 0.5 | 1.05 | 0.56 | 0.05 | 0.98 | 1.01 | 0.69 | 0.26 | 1.02 | 0.53 | 0.5 | 0.43 | 0.86 | 0.84 | 0.36 | 0.08 | 0.195 | 0.08 | -0.005 | 0.496594 | 0.763021 | 0.986312 | 0.999971 | 1 | 0.99925 |
| Smug1 | Q6P5C5 | -0.55 | 0.42 | 0.6 | 0.66 | 0.21 | 0.32 | 0.19 | 0.63 | -0.32 | -0.21 | 0.25 | 0.93 | -0.61 | 0.13 | 0.54 | 0.57 | 0.055 | -0.12 | -0.125 | 0.857469 | 0.756521 | 0.743371 | 0.999971 | 1 | 0.961064 |
| Smurf2 | A2A5Z6 | 0.16 | 0.2 | -0.06 | -0.19 | 0.38 | 0.07 | 0.36 | 0.16 | 0.58 | 0.28 | -0.42 | -0.2 | 0.14 | -0.1 | -0.25 | -0.32 | 0.215 | 0.0325 | -0.16 | 0.272138 | 0.902196 | 0.428747 | 0.999971 | 1 | 0.881196 |
| Smyd2 | Q8R5A0 | -0.37 | 0.39 | -0.52 | -0.17 | -0.52 | 0.28 | -1.12 | -0.46 | -0.51 | -0.1 | -1.35 | -2.07 | -0.05 | -0.17 | -0.33 | 0.37 | -0.2875 | -0.84 | 0.1225 | 0.413364 | 0.092304 | 0.651351 | 0.999971 | 1 | 0.933842 |
| Smyd3 | Q9CWR2 | 0.06 | 0.21 | -0.06 | 0.19 | 1.1 | 0.05 | -0.2 | 0.1 | 0.81 | -0.02 | -0.52 | -0.46 | 0.08 | 0.28 | 0.14 | 0.03 | 0.1625 | -0.1475 | 0.0325 | 0.592892 | 0.64268 | 0.851416 | 0.999971 | 1 | 0.98187 |
| Smyd5 | Q3TYX3 | 0.36 | -0.22 | 0.76 | 0.81 | -0.02 | 0.08 | 2.55 | 0.82 | 0.6 | -0.17 | 0.66 | 0.62 | -0.26 | -0.3 | 0.78 | 0.53 | 0.43 | 0 | -0.24 | 0.469519 | 1 | 0.506673 | 0.999971 | 1 | 0.905775 |
| Snap23 | Q9D3L3 | -0.1 | -0.05 | -0.94 | -0.94 | -0.05 | -0.33 | -0.95 | -1.29 | -0.14 | -0.26 | -0.81 | -0.48 | -0.62 | -0.01 | -0.9 | -0.53 | -0.1475 | 0.085 | -0.0075 | 0.688962 | 0.775932 | 0.981031 | 0.999971 | 1 | 0.99925 |
| Snap25 | P60879 | -1.26 | -0.88 | -1.62 | -1.27 | -1.08 | -1.39 | -1.14 | -1.83 | -1.2 | -2 | -1.11 | -1.28 | -0.26 | -1.49 | -1.39 | -1.03 | -0.1025 | -0.14 | 0.215 | 0.687681 | 0.608237 | 0.507479 | 0.999971 | 1 | 0.905815 |
| Snap25_1 | P60879-2 | -0.75 | -1.84 | -2.8 | -2.57 | -1.7 | -1.67 | -1.88 | -2.82 | -1.02 | -0.5 | -1.81 | -2.17 | -1.44 | -1.64 | -2.54 | -1.67 | -0.0275 | 0.615 | 0.1675 | 0.955782 | 0.281928 | 0.731279 | 0.999971 | 1 | 0.956041 |
| Snap29 | Q9ERB0 | -0.35 | -0.95 | -1.62 | -1.26 | -0.63 | -0.56 | -0.62 | -1.34 | -0.91 | -0.56 | -1.71 | -1.64 | -0.71 | -0.7 | -1.94 | -0.91 | 0.2575 | -0.16 | -0.02 | 0.438884 | 0.672715 | 0.958601 | 0.999971 | 1 | 0.998622 |
| Snap47 | Q8R570 | -0.18 | 0.03 | 0.13 | 0.13 | 0.01 | 0.56 | 0.28 | 0.51 | 0.21 | 0.23 | 0.56 | 0.35 | 0.41 | 0.55 | 0.53 | 0.9 | 0.3125 | 0.31 | 0.57 | 0.147952 | 0.112909 | 0.015506 | 0.999971 | 1 | 0.613095 |
| Snap91 | Q61548-3 | -0.82 | -0.57 | -1 | -0.97 | -0.71 | -0.51 | -0.92 | -0.95 | -0.65 | -0.9 | -0.8 | -0.69 | -0.34 | -0.52 | -0.78 | -0.38 | 0.0675 | 0.08 | 0.335 | 0.736589 | 0.66265 | 0.120455 | 0.999971 | 1 | 0.78268 |
| Snapin | Q9Z266 | 0.53 | 0.58 | 0.62 | 0.43 | 0.49 | 0.65 | 0.35 | 2.29 | 0.59 | 0.96 | 1.42 | 0.34 | 1.15 | 0.99 | 1.49 | 1.5 | 0.405 | 0.2875 | 0.7425 | 0.358126 | 0.286299 | 0.004237 | 0.999971 | 1 | 0.582714 |
| Snca | O55042 | -0.85 | -1.31 | -0.62 | -0.34 | -1.7 | -1.24 | -0.26 | -0.4 | -2.19 | -1.59 | -0.5 | -0.76 | -1.37 | -1.71 | -0.56 | -0.42 | -0.12 | -0.48 | -0.235 | 0.755735 | 0.269018 | 0.524276 | 0.999971 | 1 | 0.908412 |
| Sncb | Q91ZZ3 | -1.12 | -0.7 | -1.71 | -2.04 | -1.03 | -1.07 | -1.09 | -2.69 | -1.88 | -1.65 | -1.92 | -1.81 | -0.53 | -1.25 | -0.97 | -1.51 | -0.0775 | -0.4225 | 0.3275 | 0.869345 | 0.196114 | 0.372157 | 0.999971 | 1 | 0.869526 |
| Sncg | Q9Z0F7 | 0.54 | -0.47 | -0.04 | 0.26 | -0.34 | -0.89 | -0.21 | 0.13 | -0.36 | 0.07 | -0.34 | 0.28 | 0.27 | 0.51 | 0.57 | 0.34 | -0.4 | -0.16 | 0.35 | 0.216572 | 0.572931 | 0.192137 | 0.999971 | 1 | 0.810547 |
| Snd1 | Q78PY7 | 0.11 | 0.34 | 0.35 | 0.13 | 0.41 | 0.25 | 0.55 | 0.43 | 0.56 | 0.12 | 0.55 | 0.51 | 0.57 | 0.38 | 0.52 | 0.52 | 0.1775 | 0.2025 | 0.265 | 0.325787 | 0.297438 | 0.147985 | 0.999971 | 1 | 0.793248 |
| Snf8 | Q9CZ28 | -0.3 | -0.38 | 0.12 | -0.13 | -0.37 | -0.12 | -0.57 | 0.33 | -0.47 | -0.3 | 0.17 | 0.36 | -0.41 | 0.09 | 0.31 | 0.67 | -0.01 | 0.1125 | 0.3375 | 0.968172 | 0.654177 | 0.234033 | 0.999971 | 1 | 0.832869 |
| Snip1 | Q8BIZ6 | 3.52 | 0.57 | 0.5 | 0.95 | 1.26 | 2.08 | 1.85 | 1.14 | 1.18 | 1.04 | 0.71 | 0.85 | 0.36 | -1.13 | -0.04 | -0.04 | 0.1975 | -0.44 | -1.5975 | 0.772459 | 0.512069 | 0.048427 | 0.999971 | 1 | 0.710102 |
| Snn | P61807 | -0.15 | 0.4 | -1.91 | -1.61 | -0.32 | 0.46 | -1.64 | -1.35 | 0.77 | 1.01 | -0.24 | -0.67 | 1.35 | -0.37 | -1.49 | -1.47 | 0.105 | 1.035 | 0.3225 | 0.875256 | 0.127709 | 0.682423 | 0.999971 | 1 | 0.943204 |
| Snph | Q80U23-2 | -2.16 | -3.88 | -0.4 | -0.37 | -3.91 | -3.34 | -0.43 | -0.52 | -3.58 | -2.65 | -0.72 | -0.66 | -5.06 | -3.24 | -0.38 | -0.14 | -0.3475 | -0.2 | -0.5025 | 0.753761 | 0.84096 | 0.697935 | 0.999971 | 1 | 0.947682 |
| Snrk | Q8VDU5 | -0.78 | -1.58 | -1.13 | -1.13 | -1.34 | -1.6 | -0.96 | -1.33 | -1.55 | -1.11 | -1.05 | -1.11 | -1.46 | -1.25 | -1.04 | -0.8 | -0.1525 | -0.05 | 0.0175 | 0.533226 | 0.830807 | 0.943443 | 0.999971 | 1 | 0.99692 |
| Snrnp200 | Q6P4T2 | 0.36 | 0.4 | 0.48 | 0.18 | 0.49 | 0.31 | 0.12 | -0.06 | 0.66 | 0.47 | 0.12 | 0.13 | 0.18 | 0.14 | 0.11 | 0.16 | -0.14 | -0.01 | -0.2075 | 0.482031 | 0.960173 | 0.236055 | 0.999971 | 1 | 0.832918 |
| Snrnp27 | Q8K194 | 0.77 | 0.03 | 0.32 | -0.47 | -1.11 | 0.64 | -0.18 | 0.27 | -1.33 | 0.36 | 0.55 | 0.82 | -0.57 | -0.82 | 1.16 | 0.13 | -0.2575 | -0.0625 | -0.1875 | 0.555797 | 0.902855 | 0.696922 | 0.999971 | 1 | 0.947174 |
| Snrnp40 | Q6PE01 | -0.06 | 0.09 | 0.51 | -1.05 | 0.17 | -0.24 | 0.41 | -0.05 | 0.03 | -0.48 | 0.97 | 0.7 | 0.13 | -0.5 | 0.29 | -1.07 | 0.2 | 0.4325 | -0.16 | 0.573433 | 0.339665 | 0.710753 | 0.999971 | 1 | 0.950695 |
| Snrnp70 | Q62376 | 1.12 | 1.11 | 0.47 | 0.37 | 1.83 | 1.22 | 0.32 | 0.58 | 1.7 | 1.26 | 0.46 | 0.5 | 1.9 | 0.95 | 0.43 | 0.5 | 0.22 | 0.2125 | 0.1775 | 0.566489 | 0.554792 | 0.643287 | 0.999971 | 1 | 0.93301 |
| Snrpa | Q62189 | 0.08 | 1.27 | 1.32 | 1.02 | 1 | 1.21 | 2 | 0.82 | 1.41 | -0.08 | 1.5 | 1.33 | 0.84 | 0.49 | 1.06 | 0.3 | 0.335 | 0.1175 | -0.25 | 0.383507 | 0.792646 | 0.461775 | 0.999971 | 1 | 0.890095 |
| Snrpa1 | P57784 | 0.96 | 0.87 | 0.67 | 0.49 | 0.97 | 0.95 | 0.41 | 0.7 | 0.9 | 1.24 | 0.68 | 0.67 | 0.66 | 1.35 | 0.42 | 0.74 | 0.01 | 0.125 | 0.045 | 0.962887 | 0.56294 | 0.858279 | 0.999971 | 1 | 0.982702 |
| Snrpb | P27048 | 0.97 | 1.82 | 0.88 | 0.5 | 1.98 | 1.59 | 0.84 | 0.46 | 2.28 | 1.14 | 0.83 | 1.21 | 2.7 | 1.54 | 1 | 0.89 | 0.175 | 0.3225 | 0.49 | 0.678953 | 0.43275 | 0.310017 | 0.999971 | 1 | 0.847402 |
| Snrpb2 | Q9CQI7 | 1.51 | 0.35 | 0.33 | 0.67 | 0.47 | 0.63 | 1.38 | -0.07 | 0.73 | 1.05 | -0.07 | -0.33 | -0.72 | 0.06 | -0.33 | -0.22 | -0.1125 | -0.37 | -1.0175 | 0.773807 | 0.376466 | 0.01114 | 0.999971 | 1 | 0.613095 |
| Snrpc | Q62241 | 0.39 | 0.71 | 1.16 | 0.85 | 1.14 | 0.4 | 1.11 | 0.34 | 0.7 | 1.08 | 0.84 | 0.51 | 0.01 | 0.78 | 0.55 | 0.36 | -0.03 | 0.005 | -0.3525 | 0.915934 | 0.982883 | 0.190133 | 0.999971 | 1 | 0.810547 |
| Snrpd1 | P62315 | 2.12 | 2.24 | 0.73 | 0.87 | 1.58 | 2.58 | 0 | 0.71 | 1.72 | 2.4 | 0.29 | -0.03 | 1.9 | 2.63 | 0.62 | 1.31 | -0.2725 | -0.395 | 0.125 | 0.664004 | 0.543184 | 0.817035 | 0.999971 | 1 | 0.974929 |
| Snrpd2 | P62317 | 0.27 | 0.73 | 0.13 | -0.07 | 1 | 0.57 | -0.26 | 0.02 | 1.1 | 0.21 | -0.4 | 0.16 | 1.2 | 0.65 | 0.11 | 0.1 | 0.0675 | 0.0025 | 0.25 | 0.837244 | 0.994264 | 0.437882 | 0.999971 | 1 | 0.882419 |
| Snrpd3 | P62320 | 0.63 | 1.37 | 0.9 | 1.03 | 0.87 | 0.97 | 0.24 | 0.79 | 0.76 | 0.85 | 0.39 | 0.87 | 0.93 | 1.17 | 0.98 | 1.17 | -0.265 | -0.265 | 0.08 | 0.307677 | 0.261604 | 0.710283 | 0.999971 | 1 | 0.950695 |
| Snrpe | P62305 | -0.63 | 0.7 | 2.74 | 1.24 | 0.46 | 0.66 | 1.81 | 2.5 | 0.29 | 0.28 | 2.87 | 2.3 | 0.06 | 0.95 | 2.96 | 1.94 | 0.345 | 0.4225 | 0.465 | 0.65308 | 0.631368 | 0.583818 | 0.999971 | 1 | 0.920364 |
| Snrpf | P62307 | 0.51 | 1.74 | 1.73 | 0.91 | 1.53 | 1.52 | 1.07 | 0.77 | 2.05 | 0.82 | 0.06 | 0.82 | 2.21 | 1.21 | 1.42 | 0.28 | -7.85E-17 | -0.285 | 0.0575 | 1 | 0.557201 | 0.90245 | 1 | 1 | 0.990759 |
| Snrpg | P62309 | 1.52 | 0.48 | 0.78 | 0.93 | 0.22 | -0.3 | 0.57 | 0.85 | 0.21 | -0.04 | 0.67 | 0.72 | -0.03 | 0.23 | 0.71 | 0.93 | -0.5925 | -0.5375 | -0.4675 | 0.099749 | 0.095797 | 0.16229 | 0.999971 | 1 | 0.800215 |
| Snta1 | A2AKD7 | -0.11 | -0.23 | -0.36 | -0.11 | 0.03 | -0.12 | -0.07 | -0.36 | 0.16 | 0.15 | -0.11 | -0.46 | 0.13 | -0.19 | 0.05 | 0.13 | 0.0725 | 0.1375 | 0.2325 | 0.689038 | 0.510864 | 0.215354 | 0.999971 | 1 | 0.826893 |
| Sntb1 | Q99L88 | 0.4 | -0.13 | 0.37 | 0.09 | 0.32 | 0.16 | -0.08 | 0.27 | 0.36 | 0.07 | 0.2 | 0.57 | -0.21 | -0.11 | 0.61 | 0.13 | -0.015 | 0.1175 | -0.0775 | 0.942115 | 0.581285 | 0.757412 | 0.999971 | 1 | 0.961116 |
| Sntb2 | Q61235 | 0.18 | -0.07 | -0.31 | -0.36 | 0.1 | -0.24 | -0.78 | -0.84 | -0.18 | -0.1 | -0.39 | 0.07 | -0.23 | 0 | -0.23 | -0.22 | -0.3 | -0.01 | -0.03 | 0.291031 | 0.961224 | 0.879442 | 0.999971 | 1 | 0.985441 |
| Snu13 | Q9D0T1 | 2.18 | 0.27 | 0.45 | 0.19 | -0.09 | 0.17 | 4.45 | -1.38 | 0.03 | 0.81 | -1.49 | -0.77 | 0.15 | -1.44 | -0.44 | -1.37 | 0.015 | -1.1275 | -1.5475 | 0.99003 | 0.100381 | 0.020753 | 0.999971 | 1 | 0.635402 |
| Snupn | Q80W37 | -0.5 | 0.2 | 0.82 | 0.41 | 0.31 | 0.31 | 0.61 | 0.76 | 0.41 | 0.16 | 0.16 | 0.75 | 0.66 | 0.98 | 0.48 | 1.16 | 0.265 | 0.1375 | 0.5875 | 0.395208 | 0.662168 | 0.092976 | 0.999971 | 1 | 0.776565 |
| Snw1 | A0A0B4J1E | 0.22 | 0.16 | 0.55 | 0.51 | 0.26 | -0.08 | 0.18 | 0.35 | 0.13 | 0.4 | 0.4 | 0.75 | -0.1 | 0.22 | 0.62 | 0.91 | -0.1825 | 0.06 | 0.0525 | 0.365791 | 0.774095 | 0.843107 | 0.999971 | 1 | 0.979814 |
| Snx1 | Q6NZD2 | -1 | -0.03 | -0.53 | -0.6 | 0.12 | -0.03 | 0.08 | -0.69 | -0.14 | -0.86 | -0.6 | -0.5 | 0.27 | -0.03 | -0.6 | -0.52 | 0.41 | 0.015 | 0.32 | 0.178092 | 0.955228 | 0.297536 | 0.999971 | 1 | 0.847402 |
| Snx10 | D3YW62 | 1.27 | 0.62 | 1.09 | 2.32 | 0.76 | 0.24 | 0.52 | 3.96 | 1.2 | 1.26 | 1.64 | 1.59 | 1.34 | 1.45 | 1.51 | 2.1 | 0.045 | 0.0975 | 0.275 | 0.957418 | 0.7904 | 0.480069 | 0.999971 | 1 | 0.894438 |
| Snx11 | A2A880 | -0.55 | -2.34 | 1.04 | 1.29 | -2.18 | -2.6 | -0.22 | 1.38 | -2.74 | -0.99 | 1.74 | 1.23 | -2.53 | -2.51 | 1.21 | 1.66 | -0.765 | -0.05 | -0.4025 | 0.496307 | 0.966579 | 0.750514 | 0.999971 | 1 | 0.961064 |
| Snx12 | Q6ZWQ5 | 0.89 | 0.56 | 0.94 | 0.83 | 0.56 | 0.46 | 1.28 | 0.62 | 0.57 | 0.69 | 0.54 | 0.37 | 0.42 | 0.56 | 0.3 | 0.45 | -0.075 | -0.2625 | -0.3725 | 0.753442 | 0.169123 | 0.064594 | 0.999971 | 1 | 0.726597 |
| Snx13 | Q6PHS6 | -0.91 | -0.3 | -0.34 | -0.58 | -0.08 | -0.56 | -0.3 | -0.49 | -0.18 | -0.65 | 0.12 | 0.14 | 0.06 | -0.53 | -0.48 | -0.25 | 0.175 | 0.39 | 0.2325 | 0.435664 | 0.152502 | 0.330898 | 0.999971 | 1 | 0.857107 |
| Snx14 | Q8BHY8 | -0.85 | -0.34 | -0.56 | -1.03 | -0.37 | -0.62 | -0.6 | -0.74 | -0.36 | -0.51 | -0.26 | -0.22 | 0.22 | -0.49 | -0.57 | -0.56 | 0.1125 | 0.3575 | 0.345 | 0.606991 | 0.118489 | 0.216925 | 0.999971 | 1 | 0.826893 |
| Snx15 | Q91WE1 | -0.09 | 0 | 0.36 | 1.04 | -0.44 | -0.29 | 0.39 | 0.65 | -0.38 | -0.33 | 0.65 | 0.06 | -0.35 | -0.04 | 0.24 | 0.34 | -0.25 | -0.3275 | -0.28 | 0.492001 | 0.355632 | 0.373211 | 0.999971 | 1 | 0.869526 |
| Snx16 | Q8C080 | -0.8 | 0.18 | -0.39 | -0.26 | -0.12 | 0.29 | -0.52 | -0.2 | 0.29 | -0.14 | -0.15 | -0.22 | 0.55 | 0.51 | -0.19 | 0.1 | 0.18 | 0.2625 | 0.56 | 0.522277 | 0.320466 | 0.074565 | 0.999971 | 1 | 0.746069 |
| Snx17 | Q8BVL3 | 0.38 | 1.23 | 0.91 | 0.55 | 1.04 | 1.16 | 0.67 | 0.49 | 0.8 | 0.79 | 0.53 | 0.61 | 0.79 | 1.79 | 1.1 | 0.91 | 0.0725 | -0.085 | 0.38 | 0.785926 | 0.717487 | 0.229028 | 0.999971 | 1 | 0.828687 |
| Snx18 | Q8C788 | -0.3 | -0.78 | -0.2 | -0.21 | -0.66 | -0.76 | -0.47 | -0.19 | -0.66 | -0.15 | 0.02 | 0.17 | -0.88 | -0.52 | -0.07 | -0.13 | -0.1475 | 0.2175 | -0.0275 | 0.519714 | 0.398484 | 0.91523 | 0.999971 | 1 | 0.992633 |
| Snx19 | Q6P4T1 | -0.48 | -0.63 | -0.32 | 0 | -0.47 | -0.54 | -0.24 | -0.2 | -0.37 | -0.34 | -0.14 | -0.04 | -0.19 | -0.26 | -0.28 | -0.09 | -0.005 | 0.135 | 0.1525 | 0.980959 | 0.517305 | 0.454838 | 0.999971 | 1 | 0.887217 |
| Snx2 | Q9CWK8 | -0.33 | -0.71 | -0.88 | -0.81 | -0.76 | -0.62 | -0.59 | -0.89 | -0.48 | -0.4 | -0.81 | -0.77 | -0.99 | -0.84 | -1.09 | -0.76 | -0.0325 | 0.0675 | -0.2375 | 0.870546 | 0.74599 | 0.257592 | 0.999971 | 1 | 0.839012 |
| Snx21 | G3UYX2 | -0.1 | 0.21 | 1.92 | 2.17 | -0.23 | 0.08 | 0.47 | 1.9 | -0.58 | 0.09 | 1.65 | 2.27 | -0.5 | -0.2 | 2.69 | 2.35 | -0.495 | -0.1925 | 0.035 | 0.471211 | 0.809422 | 0.969292 | 0.999971 | 1 | 0.998787 |
| Snx24 | Q9CRB0 | -2.06 | -0.44 | -0.38 | -1.19 | -0.76 | -1.12 | 0.57 | -1.24 | -0.48 | -1.5 | -0.58 | -0.13 | -0.9 | -0.62 | -1.15 | -1.17 | 0.38 | 0.345 | 0.0575 | 0.477816 | 0.461416 | 0.884831 | 0.999971 | 1 | 0.98711 |
| Snx25 | E9Q1K0 | -0.46 | -0.04 | -0.01 | -0.23 | -0.08 | -0.07 | -0.1 | -0.21 | -0.28 | -0.5 | 0.13 | -0.36 | 0.21 | 0 | 0.21 | 0.04 | 0.07 | -0.0675 | 0.3 | 0.703747 | 0.753419 | 0.137714 | 0.999971 | 1 | 0.784948 |
| Snx27 | Q3UHD6-2 | -1.05 | -0.84 | -0.84 | -0.72 | -0.83 | -1.17 | -0.93 | -0.65 | -1.17 | -1.08 | -0.75 | -0.36 | -0.95 | -0.97 | -0.37 | -0.65 | -0.0325 | 0.0225 | 0.1275 | 0.866209 | 0.922288 | 0.547403 | 0.999971 | 1 | 0.911722 |
| Snx29 | Q9D3S3 | 1.3 | 0.04 | -0.59 | 0.43 | 0.03 | 0.33 | 0.11 | -0.13 | -0.46 | 1.15 | 0.23 | 0.57 | 0.3 | 0.35 | -0.28 | 0.26 | -0.21 | 0.0775 | -0.1375 | 0.593695 | 0.873066 | 0.734021 | 0.999971 | 1 | 0.957685 |
| Snx3 | Q78ZM0 | -0.38 | -1.08 | 0.04 | 0.04 | -0.41 | -1.66 | 0.78 | -0.13 | -0.7 | -0.69 | -0.04 | 0.34 | -0.81 | -0.95 | -0.31 | 0.22 | -0.01 | 0.0725 | -0.1175 | 0.984778 | 0.840317 | 0.748437 | 0.999971 | 1 | 0.961064 |
| Snx30 | Q8CE50 | -0.58 | -0.29 | -0.27 | -0.19 | -0.36 | -0.47 | -0.29 | -0.14 | -0.35 | -0.46 | 0.13 | 0.39 | -0.17 | -0.34 | -0.29 | 0.09 | 0.0175 | 0.26 | 0.155 | 0.924136 | 0.305585 | 0.432221 | 0.999971 | 1 | 0.881643 |
| Snx32 | Q80ZJ7 | 0.82 | 0.6 | 0.97 | 0.55 | 0.46 | 0.28 | 0.79 | 0.88 | 0.54 | 0.67 | 0.17 | 0.04 | 0.5 | 0.36 | 0.46 | 0.34 | -0.1325 | -0.38 | -0.32 | 0.545705 | 0.11068 | 0.106931 | 0.999971 | 1 | 0.78268 |
| Snx33 | Q4VAA7 | 2.04 | -0.38 | -0.91 | -0.06 | -0.46 | 0.25 | -1.14 | -0.39 | -0.85 | 1.54 | -0.54 | -0.64 | -0.14 | 0.61 | -1.42 | -0.28 | -0.6075 | -0.295 | -0.48 | 0.356341 | 0.704087 | 0.497557 | 0.999971 | 1 | 0.902655 |
| Snx4 | Q91YJ2 | -0.75 | -0.91 | -0.52 | -0.64 | -0.77 | -0.63 | -0.68 | -0.47 | -0.64 | -0.71 | -0.43 | -0.43 | -0.62 | -0.6 | -0.38 | -0.22 | 0.0675 | 0.1525 | 0.25 | 0.710659 | 0.408921 | 0.216236 | 0.999971 | 1 | 0.826893 |
| Snx5 | Q9D8U8 | 0.17 | 0.09 | -0.03 | -0.12 | 0 | 0.06 | 0.14 | -0.17 | -0.17 | -0.08 | -0.23 | -0.45 | -0.47 | -0.12 | -0.33 | -0.29 | -0.02 | -0.26 | -0.33 | 0.909562 | 0.167091 | 0.09214 | 0.999971 | 1 | 0.77469 |
| Snx6 | Q6P8X1 | 0.21 | 0.03 | -0.04 | 0.14 | -0.27 | 0.1 | -0.37 | 0.4 | -0.12 | 0.22 | 0.21 | 0.44 | -0.06 | 0.18 | 0.4 | 0.7 | -0.12 | 0.1025 | 0.22 | 0.59797 | 0.592565 | 0.32653 | 0.999971 | 1 | 0.856472 |
| Snx7 | F8WI30 | -1.32 | -0.23 | -0.56 | -0.83 | -0.15 | -0.51 | -0.54 | -0.76 | 0.07 | -1.17 | -0.16 | -0.34 | 0.47 | -0.79 | -0.7 | -0.74 | 0.245 | 0.335 | 0.295 | 0.390035 | 0.350623 | 0.434898 | 0.999971 | 1 | 0.882419 |
| Snx9 | Q91VH2 | 2.2 | 1.21 | 0.76 | -0.19 | -0.04 | 0.83 | -0.73 | 0.07 | 0.05 | 1.15 | 0.34 | 0.15 | 0.05 | 1.12 | 0.12 | 0.55 | -0.9625 | -0.5725 | -0.535 | 0.103669 | 0.285047 | 0.31279 | 0.999971 | 1 | 0.849367 |
| Soat1 | Q61263 | -1.56 | -0.26 | -0.71 | -1.2 | 0.59 | -0.49 | -0.57 | -1.55 | 0.36 | -1.05 | -0.93 | -0.8 | 0.47 | -0.72 | -0.76 | -1.39 | 0.4275 | 0.3275 | 0.3325 | 0.388879 | 0.435695 | 0.469224 | 0.999971 | 1 | 0.892481 |
| Sobp | Q0P5V2 | 1.42 | 3.78 | 0.91 | 1.23 | 3.26 | 3.18 | 0.65 | 0.72 | 2.68 | 1.53 | 0.23 | 0.86 | 3.39 | 2.6 | 1.01 | 0.33 | 0.1175 | -0.51 | -0.0025 | 0.893792 | 0.508765 | 0.997689 | 0.999971 | 1 | 1 |
| Sod1 | P08228 | 2.54 | 0.24 | -0.33 | 0.43 | -0.09 | 0.16 | 3.94 | -0.9 | -0.01 | 0.93 | -0.44 | 0.27 | 0.11 | 0.25 | 0.18 | -0.27 | 0.0575 | -0.5325 | -0.6525 | 0.95866 | 0.409289 | 0.281948 | 0.999971 | 1 | 0.844538 |
| Sod2 | P09671 | -0.84 | -0.5 | -0.62 | -0.86 | -0.16 | -0.48 | -0.36 | -0.82 | -0.23 | -0.8 | -0.42 | -0.26 | 0.11 | -0.58 | -0.86 | -0.89 | 0.25 | 0.2775 | 0.15 | 0.258335 | 0.201908 | 0.580328 | 0.999971 | 1 | 0.91995 |
| Sod3 | O09164 | 1.71 | 0.96 | 1.22 | 1.87 | 0.94 | 0.57 | 1.73 | 1.32 | 1.21 | 1.55 | 1.68 | 1.96 | 0.58 | 1.06 | 1.22 | 1.69 | -0.3 | 0.16 | -0.3025 | 0.370735 | 0.568003 | 0.351293 | 0.999971 | 1 | 0.867656 |
| Soga1 | A2ACV6 | -0.1 | -0.02 | 0.3 | 0.23 | 0.22 | -0.12 | 0.5 | 0.43 | 0.54 | 0.27 | 0.63 | 0.63 | 0.76 | -0.07 | 0.74 | 0.46 | 0.155 | 0.415 | 0.37 | 0.479291 | 0.053656 | 0.158892 | 0.999971 | 1 | 0.798217 |
| Soga3 | Q6NZL0 | -0.38 | -0.86 | -0.73 | -0.56 | -0.91 | -0.53 | -0.72 | -0.67 | -0.63 | -0.44 | -0.49 | -0.43 | -0.64 | -0.91 | -0.69 | -0.45 | -0.075 | 0.135 | -0.04 | 0.700182 | 0.467263 | 0.841646 | 0.999971 | 1 | 0.979814 |
| Son | H9KV00 | 0.25 | -0.12 | -0.07 | 0.03 | -0.2 | -0.25 | -0.26 | -0.36 | -0.2 | -0.01 | -0.29 | -0.1 | -0.19 | -0.12 | -0.22 | -0.1 | -0.29 | -0.1725 | -0.18 | 0.124198 | 0.343793 | 0.318462 | 0.999971 | 1 | 0.853893 |
| Son_1 | Q9QX47-2 | -0.23 | -1.74 | -1.26 | -1.48 | -0.71 | -1.14 | -1.8 | -1.2 | -1.31 | -0.13 | -1.84 | -0.93 | -0.07 | -0.79 | -1.21 | -1.38 | -0.035 | 0.125 | 0.315 | 0.927427 | 0.785885 | 0.459931 | 0.999971 | 1 | 0.889706 |
| Sorbs1 | A0A286YCI | -0.54 | -0.45 | -0.5 | -0.52 | -0.53 | -0.72 | -0.28 | -0.57 | -0.81 | -0.84 | -0.53 | -0.42 | -0.08 | -0.78 | -0.55 | -0.43 | -0.0225 | -0.1475 | 0.0425 | 0.898682 | 0.418664 | 0.83485 | 0.999971 | 1 | 0.979036 |
| Sorbs1_1 | A0A668KL( | -0.73 | 0.77 | -0.91 | -1.24 | 1.8 | -1.41 | -1.83 | -1.08 | 1.84 | -0.24 | -1.85 | -1.79 | 2.39 | -2.61 | -1.64 | -0.3 | -0.1025 | 0.0175 | -0.0125 | 0.902837 | 0.98411 | 0.990468 | 0.999971 | 1 | 0.99925 |
| Sorbs1_2 | A0A5F8MP | -0.73 | -1.22 | -0.82 | -0.15 | -1.06 | -1.2 | 0.31 | -0.62 | -1.4 | -1.14 | -1.08 | -1.71 | -1.2 | -1.37 | -0.77 | -1.51 | 0.0875 | -0.6025 | -0.4825 | 0.822457 | 0.055113 | 0.119624 | 0.999971 | 1 | 0.78268 |
| Sorbs2 | Q3UTJ2-2 | -0.84 | -0.88 | -0.71 | -1.01 | -0.34 | -0.84 | -0.69 | -1.31 | -0.82 | -0.43 | -0.32 | -0.06 | -0.67 | -1.21 | -1.18 | -0.96 | 0.065 | 0.4525 | -0.145 | 0.788578 | 0.05966 | 0.473883 | 0.999971 | 1 | 0.894044 |
| Sorbs2_1 | D3Z080 | -1.23 | -3.5 | -0.54 | -1.37 | -2.38 | -2.08 | -0.38 | -2.76 | -2.47 | -0.45 | -2.84 | -0.76 | -2.11 | -1.55 | -2.16 | -1.86 | -0.24 | 0.03 | -0.26 | 0.748129 | 0.969822 | 0.665563 | 0.999971 | 1 | 0.93697 |
| Sorbs2_2 | B7ZWM6 | -0.77 | -2.37 | 0.22 | 0.2 | -0.6 | -2.45 | -0.97 | -0.26 | -1.65 | 0.42 | -0.44 | 2.6 | -1.63 | -1.59 | -0.83 | -0.11 | -0.39 | 0.9125 | -0.36 | 0.581714 | 0.361747 | 0.57938 | 0.999971 | 1 | 0.919302 |
| Sorbs3 | Q8K0M3 | 0.25 | 0.71 | 0.58 | 0.7 | 1.31 | 0.55 | 0.22 | 0.31 | 1.15 | 0.79 | 0.28 | 1.03 | 1.41 | 0.6 | 0.71 | 0.83 | 0.0375 | 0.2525 | 0.3275 | 0.894846 | 0.322392 | 0.199792 | 0.999971 | 1 | 0.813335 |
| Sorcs1 | Q9JLC4-2 | -1.71 | -1.16 | -0.31 | -0.3 | -0.76 | -1.86 | 0.34 | -0.31 | -0.61 | -2.3 | 0.29 | -0.32 | -0.13 | -1.48 | -0.62 | -0.39 | 0.2225 | 0.135 | 0.215 | 0.677078 | 0.821928 | 0.618354 | 0.999971 | 1 | 0.928581 |
| Sorcs2 | Q9EPR5 | -0.56 | -0.07 | -0.56 | -0.46 | -0.18 | -0.24 | -0.48 | -0.42 | -0.08 | -0.52 | -0.73 | -0.4 | 0.36 | 0.1 | -0.27 | -0.52 | 0.0825 | -0.02 | 0.33 | 0.677231 | 0.927415 | 0.215911 | 0.999971 | 1 | 0.826893 |
| Sorcs3 | Q8VI51 | -0.86 | -0.34 | 0.09 | -0.67 | -0.07 | -0.5 | 0.48 | -0.83 | -0.03 | -0.47 | 0.19 | -0.48 | 0.04 | -0.55 | -0.45 | -0.82 | 0.215 | 0.2475 | -7.85E-17 | 0.539104 | 0.388548 | 1 | 0.999971 | 1 | 1 |
| Sord | Q64442 | 0.89 | 0.55 | 0.29 | 0.13 | 0.68 | 0.49 | 0.37 | 0.2 | 0.48 | 0.17 | 0.44 | 0.53 | 0.55 | 0.2 | 0.18 | -0.18 | -0.03 | -0.06 | -0.2775 | 0.89686 | 0.788349 | 0.286643 | 0.999971 | 1 | 0.846151 |
| Sorl1 | O88307 | 0.67 | 1.16 | 0.69 | 0.64 | 0.99 | 1.11 | 0.62 | 0.54 | 0.77 | 0.61 | 0.97 | 0.9 | 0.89 | 1.11 | 0.74 | 0.94 | 0.025 | 0.0225 | 0.13 | 0.911943 | 0.910502 | 0.526236 | 0.999971 | 1 | 0.909459 |
| Sort1 | Q6PHU5 | 0.12 | 0.65 | 0.98 | 0.27 | 0.99 | 0.79 | 1.3 | 0.59 | 0.98 | -0.11 | 1.04 | 0.88 | 1.65 | 0.11 | 0.82 | 0.48 | 0.4125 | 0.1925 | 0.26 | 0.147716 | 0.565658 | 0.489622 | 0.999971 | 1 | 0.898405 |
| Sos1 | Q62245 | 0.03 | -0.11 | 0.14 | -0.17 | -0.09 | -0.22 | 0.21 | -0.05 | -0.05 | -0.17 | 0.41 | 0.15 | 0.1 | -0.17 | 0.13 | -0.23 | -0.01 | 0.1125 | -0.015 | 0.957057 | 0.575466 | 0.936114 | 0.999971 | 1 | 0.995618 |
| Sos2 | S4R2C4 | 0.58 | 1 | 0.35 | 0.65 | 0.8 | 0.89 | 0.37 | 0.78 | 0.71 | 0.97 | 0.39 | 0.08 | 1.26 | 0.31 | 0.32 | -0.09 | 0.065 | -0.1075 | -0.195 | 0.768981 | 0.678183 | 0.546031 | 0.999971 | 1 | 0.911398 |
| Sowaha | Q8BLS7 | -0.48 | -0.69 | -0.57 | -0.14 | -0.34 | -0.52 | -0.78 | -0.07 | -0.71 | -0.64 | -0.43 | -0.34 | -0.9 | -0.47 | -0.65 | -0.12 | 0.0425 | -0.06 | -0.065 | 0.852843 | 0.764735 | 0.78443 | 0.999971 | 1 | 0.966376 |
| Sowahb | Q8BZW2 | -0.37 | 0.42 | -0.28 | 0.32 | 0.51 | 0.31 | -0.31 | 0.33 | -0.58 | -0.61 | 0.59 | -0.09 | 0.53 | -0.19 | 0.13 | 0.45 | 0.1875 | -0.195 | 0.2075 | 0.514702 | 0.572264 | 0.462709 | 0.999971 | 1 | 0.890653 |
| Sowahc | Q8C0J6 | 0.3 | -0.01 | -0.07 | 0.05 | 0.3 | -0.02 | 0.34 | -0.01 | 0.16 | 0.09 | 0.03 | -0.27 | 0.55 | -0.61 | -0.21 | -0.06 | 0.085 | -0.065 | -0.15 | 0.659714 | 0.73052 | 0.585938 | 0.999971 | 1 | 0.920413 |
| Sox1 | P53783 | 1.69 | 0.63 | -0.28 | 1.31 | 0.55 | 1.01 | 1.51 | 1.73 | 0.72 | 1.19 | -1.11 | 0.2 | 0.38 | 1.37 | 0.17 | -0.14 | 0.3625 | -0.5875 | -0.3925 | 0.450717 | 0.344356 | 0.443196 | 0.999971 | 1 | 0.883386 |
| Sox13 | D3Z7I3 | 4.94 | 0.82 | 2.6 | 3.43 | 2.68 | 2.3 | 3.62 | 2.29 | 1.95 | 4.15 | 1.77 | 1.3 | -2.58 | 0.6 | 2.36 | -0.28 | -0.225 | -0.655 | -2.9225 | 0.784113 | 0.501258 | 0.034619 | 0.999971 | 1 | 0.69205 |
| Sox5 | B2KFM4 | 3.02 | 1.93 | 2.18 | 2.3 | 1.97 | 2.4 | 2.39 | 1.56 | 1.37 | 2.8 | 2.05 | 1.49 | 1.15 | 2.32 | 1.96 | 0.1 | -0.2775 | -0.43 | -0.975 | 0.385273 | 0.284353 | 0.081041 | 0.999971 | 1 | 0.755254 |
| Sox9 | Q04887 | 3.11 | 1.25 | 1.61 | 1.83 | 0.43 | 1.33 | 4.54 | 0.15 | 0.3 | 2.03 | -0.69 | -0.72 | 1.33 | 0.45 | 0.87 | -0.25 | -0.3375 | -1.72 | -1.35 | 0.728491 | 0.033282 | 0.021075 | 0.999971 | 1 | 0.639761 |
| Sp3 | O70494-3 | 3.19 | 2.24 | 2.24 | 2.43 | 1.97 | 2.92 | 1.68 | 1.94 | 2.6 | 3.5 | 1.27 | 1.55 | 2.2 | 2.73 | 1.75 | 2.5 | -0.3975 | -0.295 | -0.23 | 0.271499 | 0.573267 | 0.471422 | 0.999971 | 1 | 0.892857 |
| Spacdr | D3YZZ4 | -0.22 | -0.98 | -1.04 | -0.6 | -0.93 | -0.16 | -0.96 | 0.84 | -0.7 | 0.03 | -0.55 | -0.13 | -0.57 | -0.85 | -0.36 | -0.13 | 0.4075 | 0.3725 | 0.2325 | 0.363828 | 0.195945 | 0.392843 | 0.999971 | 1 | 0.871816 |
| Spag1 | Q80ZX8-2 | -0.75 | -1.1 | -1.09 | -0.5 | -0.86 | -1.24 | -1.07 | -1.02 | -1.18 | -0.72 | -1.39 | -1.54 | -1.12 | -0.92 | -0.8 | -0.57 | -0.1875 | -0.3475 | 0.0075 | 0.389785 | 0.194846 | 0.973533 | 0.999971 | 1 | 0.99925 |

| | | | | | | | | | | | | | | | | | | | | | | | | | | |
|---|---|---|---|---|---|---|---|---|---|---|---|---|---|---|---|---|---|---|---|---|---|---|---|---|---|---|
| Spag7 | Q7TNE3 | 1.66 | 2.9 | 2.67 | 2.72 | 3.22 | 2.06 | 2 | 2.44 | 2.46 | 1.55 | 2.31 | 2.55 | 3.13 | 3.12 | 2.64 | 2.75 | -0.0575 | -0.27 | 0.4225 | 0.880396 | 0.453772 | 0.199598 | 0.999971 | 1 | 0.813335 |
| Spag9 | Q58A65-2 | 0.32 | 1.08 | 1.01 | 0.87 | 0.87 | 0.84 | 1.02 | 0.66 | 1.12 | 0.42 | 0.95 | 1.21 | 1.08 | 0.9 | 0.66 | 0.79 | 0.0275 | 0.105 | 0.0375 | 0.90358 | 0.694275 | 0.871524 | 0.999971 | 1 | 0.984109 |
| Sparc | Q5NCU4 | -0.12 | -0.05 | 1.5 | 0.47 | 0.8 | -0.06 | 2.4 | 1.6 | 0.97 | -0.26 | 2.6 | 3.61 | 1.46 | 0.82 | 2.71 | 1.5 | 0.735 | 1.28 | 1.1725 | 0.234309 | 0.15612 | 0.04256 | 0.999971 | 1 | 0.701725 |
| Sparcl1 | P70663 | 0.18 | 1.14 | 1.55 | 0.57 | 2.15 | 1.09 | 2.48 | 0.96 | 2.56 | -0.06 | 1.83 | 1.99 | 3.01 | 0.26 | 1.4 | 0.87 | 0.81 | 0.72 | 0.525 | 0.102514 | 0.244331 | 0.396438 | 0.999971 | 1 | 0.871816 |
| Spart | Q8R1X6 | 0.03 | -0.2 | -0.23 | -0.53 | -0.37 | -0.15 | -0.36 | -0.34 | -0.33 | -0.4 | -0.73 | -0.55 | -0.26 | -0.08 | -0.27 | -0.61 | -0.0725 | -0.27 | -0.0725 | 0.706449 | 0.198719 | 0.731566 | 0.999971 | 1 | 0.956053 |
| Spast | A0A286YE2 | 0.55 | -0.32 | -0.15 | -0.25 | -0.07 | -0.48 | 0.6 | -0.59 | -0.12 | 0.49 | -0.22 | -0.22 | -0.61 | -0.41 | -0.64 | -0.42 | -0.0925 | 0.025 | -0.4775 | 0.782077 | 0.928346 | 0.075252 | 0.999971 | 1 | 0.746958 |
| Spata1 | Q9D5R4 | 1.87 | 4.1 | -1.08 | -1.59 | 4.86 | 3.85 | -0.95 | -0.77 | 6.13 | 3.61 | -1.24 | -2.57 | 9.27 | 3.45 | -1.5 | -1.87 | 0.9225 | 0.6575 | 1.5125 | 0.608452 | 0.762436 | 0.563241 | 0.999971 | 1 | 0.917003 |
| Spata2 | Q8K004 | 0.8 | -0.07 | 0.11 | -0.06 | -0.43 | -0.4 | 0.03 | 0.04 | -0.87 | 0.14 | 0.08 | 0.14 | -1.19 | -1.18 | -0.03 | 0.12 | -0.385 | -0.3225 | -0.765 | 0.170863 | 0.332603 | 0.079964 | 0.999971 | 1 | 0.755254 |
| Spata2l | Q8BNN1 | -0.03 | 0.06 | -0.3 | 0.5 | 0.07 | -0.42 | -0.46 | 0.27 | -0.06 | -0.2 | -0.27 | -0.2 | 0.01 | 0.17 | 0.29 | 0.63 | -0.1925 | -0.24 | 0.2175 | 0.476649 | 0.28232 | 0.383367 | 0.999971 | 1 | 0.871816 |
| Spata5 | A0A0G2JFY | 0.27 | -0.23 | -0.07 | -0.05 | -0.27 | -0.27 | 0.01 | 0.03 | -0.32 | -0.05 | -0.12 | -0.01 | -0.29 | 0.03 | -0.04 | -0.22 | -0.105 | -0.105 | -0.11 | 0.594999 | 0.580827 | 0.573934 | 0.999971 | 1 | 0.91816 |
| Spats2 | Q8K1N4 | 1.46 | 0.66 | 1.04 | 1.15 | 0.85 | 1.12 | 1.1 | 1.26 | 0.69 | 1.13 | 1.2 | 1.11 | 0.57 | 0.91 | 1.07 | 1.54 | 0.005 | -0.045 | -0.055 | 0.982341 | 0.84787 | 0.84307 | 0.999971 | 1 | 0.979814 |
| Spats2l | E9PVQ3 | 1.3 | 1.07 | 1.47 | 1.32 | 1.29 | 1.82 | 1.64 | 1.71 | 1.59 | 1.4 | 1.33 | 1.32 | 1.98 | 1.49 | 1.21 | 1.63 | 0.325 | 0.12 | 0.2875 | 0.130365 | 0.505634 | 0.219348 | 0.999971 | 1 | 0.827064 |
| Spcs1 | A0A2I3BRV | -0.8 | -0.89 | -0.58 | 0.08 | -0.88 | -0.82 | -0.49 | -0.09 | -0.95 | -0.41 | -0.78 | -0.29 | -0.63 | -0.27 | 0.26 | 0.9 | -0.0225 | -0.06 | 0.6125 | 0.939089 | 0.831404 | 0.138081 | 0.999971 | 1 | 0.784948 |
| Spcs2 | A0A140LH( | -2.34 | -2.3 | -2.91 | -3.68 | -2.08 | -2.31 | -2.29 | -3.62 | -1.96 | -1.7 | -2.27 | -2.98 | -2.48 | -2.04 | -3.16 | -4.05 | 0.2325 | 0.58 | -0.125 | 0.606821 | 0.178396 | 0.804571 | 0.999971 | 1 | 0.971467 |
| Spcs3 | Q6ZWQ7 | -0.39 | -0.04 | 0.07 | 0.51 | -0.12 | -0.27 | -0.22 | 0.5 | -0.37 | -0.19 | 0.57 | 0.48 | 0.3 | -0.07 | 0.34 | 0.83 | -0.065 | 0.085 | 0.3125 | 0.81322 | 0.781488 | 0.279601 | 0.999971 | 1 | 0.843812 |
| Specc1 | A0A0J9YUF | 0.21 | -0.61 | -1.38 | -1.17 | -1.05 | -0.96 | -1.97 | -0.73 | -0.33 | -0.27 | -0.93 | -1.14 | -0.45 | -0.37 | -2.21 | -0.18 | -0.44 | 0.07 | -0.065 | 0.314579 | 0.860787 | 0.904937 | 0.999971 | 1 | 0.991091 |
| Specc1_1 | A0A0J9YTU | 0.29 | 0.14 | 0.28 | 0.23 | 0.16 | 0.05 | 0.09 | 0.33 | 0.12 | 0.28 | 0.16 | 0.32 | 0.27 | 0.04 | 0.33 | 0.36 | -0.0775 | -0.015 | 0.015 | 0.647445 | 0.925772 | 0.930649 | 0.999971 | 1 | 0.994744 |
| Specc1l | Q2KN98 | -0.47 | -0.32 | -0.08 | -0.1 | -0.4 | -0.67 | -0.46 | -0.24 | -0.45 | -0.09 | 0.27 | 0.24 | -0.04 | -0.19 | -0.1 | 0.3 | -0.2 | 0.235 | 0.235 | 0.315714 | 0.318888 | 0.260807 | 0.999971 | 1 | 0.841256 |
| Speg | E9QQ25 | 0.27 | -0.54 | -0.46 | -0.12 | -0.54 | -0.39 | -0.32 | -0.27 | -0.52 | 0.3 | -0.01 | -0.54 | -0.51 | -0.39 | -0.25 | -0.11 | -0.1675 | 0.02 | -0.1025 | 0.475639 | 0.944498 | 0.668955 | 0.999971 | 1 | 0.938711 |
| Spen | A2ADB1 | 1.97 | 0.67 | -0.51 | -0.44 | -0.11 | 1.06 | -0.45 | -0.94 | 0.25 | 1.71 | -0.99 | -1.11 | 0.18 | 0.5 | -1.18 | -1.01 | -0.5325 | -0.4575 | -0.8 | 0.424732 | 0.568795 | 0.24095 | 0.999971 | 1 | 0.834478 |
| Spg11 | Q3UHA3 | 3.94 | 1.2 | -0.82 | -0.76 | 1.24 | 1.3 | 0.1 | -0.53 | 1.41 | 3.33 | -1.37 | -0.67 | 0.75 | 1.56 | -0.94 | -1.71 | -0.3625 | -0.215 | -0.975 | 0.73637 | 0.876253 | 0.425774 | 0.999971 | 1 | 0.880793 |
| Spg21 | Q9CQC8 | -0.15 | 0.09 | -0.36 | 0.09 | 0.08 | 0.42 | -0.2 | 0.24 | -0.03 | 0.23 | -0.3 | -0.22 | 0.22 | 0.37 | -0.1 | 0.47 | 0.2175 | 0.0025 | 0.3225 | 0.329457 | 0.990402 | 0.157398 | 0.999971 | 1 | 0.797252 |
| Spg7 | D3YZN4 | -0.26 | -0.35 | -0.35 | -0.15 | -0.45 | -0.44 | -0.37 | -0.43 | -0.44 | -0.07 | -0.08 | -0.19 | -0.69 | -0.51 | -0.42 | -0.22 | -0.145 | 0.0825 | -0.1825 | 0.384429 | 0.640382 | 0.335486 | 0.999971 | 1 | 0.858827 |
| Sphk2 | Q9JIA7 | 0.25 | 0.06 | -0.89 | -1.09 | -0.07 | -0.04 | -0.73 | -1.12 | -0.02 | 0.25 | -0.86 | -0.88 | -0.05 | 0.03 | -0.98 | -1.02 | -0.0725 | 0.04 | -0.0875 | 0.858424 | 0.924333 | 0.834708 | 0.999971 | 1 | 0.979036 |
| Sphkap | E9PUC4 | 0.19 | 0.09 | 0.41 | 0.11 | 0.02 | 0.4 | 0.54 | 0.25 | 0.13 | 0.59 | 0.54 | 0.34 | 0.62 | 0.09 | 0.38 | 0.54 | 0.1025 | 0.2 | 0.2075 | 0.602744 | 0.308006 | 0.311796 | 0.999971 | 1 | 0.848872 |
| Spin1 | Q61142-2 | 1.32 | 1.21 | 1.32 | 1.95 | 1.33 | 1.08 | 0.99 | 1.76 | 1.67 | 1.15 | 1.19 | 1.75 | 0.91 | 1.33 | 1.32 | 1.86 | -0.16 | -0.01 | -0.095 | 0.547562 | 0.968606 | 0.731047 | 0.999971 | 1 | 0.956041 |
| Spire1 | A0A5H1ZRI | 0.11 | -0.34 | -0.25 | -0.55 | -0.5 | -0.22 | 0.14 | -0.95 | -1.43 | -0.83 | -1.94 | -2.19 | -0.41 | -0.72 | -1.5 | -0.96 | -0.125 | -1.34 | -0.64 | 0.659905 | 0.002924 | 0.047168 | 0.999971 | 0.918989 | 0.710102 |
| Spire2 | Q8K1S6 | -0.47 | -0.11 | -0.01 | 0.09 | -0.22 | 0.04 | -0.18 | -0.31 | -0.3 | 0.47 | 0.04 | -0.01 | -0.34 | 0.75 | 0 | 0.3 | -0.0425 | 0.175 | 0.3025 | 0.832321 | 0.461166 | 0.29388 | 0.999971 | 1 | 0.847402 |
| Spns1 | D3YTR6 | 0.45 | -0.12 | -0.15 | 0.18 | -0.49 | -0.67 | -0.34 | 0.48 | -0.52 | -0.04 | -0.16 | 0.06 | -0.18 | 0.67 | 0.04 | 0.12 | -0.345 | -0.255 | 0.0725 | 0.267457 | 0.278198 | 0.777053 | 0.999971 | 1 | 0.964475 |
| Spns2 | Q91VM4 | 0.59 | -0.17 | -0.25 | -0.32 | 0.24 | 0.66 | 0.13 | -0.35 | -0.46 | 1.42 | 0.05 | 0.36 | 0.1 | -0.64 | 0.55 | -0.47 | 0.2075 | 0.38 | -0.0775 | 0.499993 | 0.384794 | 0.820679 | 0.999971 | 1 | 0.975821 |
| Spock1 | Q6DIC9 | 0.72 | 1.13 | 0.63 | 0.46 | 1.39 | 1.3 | 1.01 | 0.89 | 1.32 | 1.12 | 0.91 | 0.94 | 0.66 | 0.85 | 0.7 | 0.12 | 0.4125 | 0.3375 | -0.1525 | 0.094554 | 0.142364 | 0.538162 | 0.999971 | 1 | 0.910077 |
| Spock2 | Q9ER58 | 0.8 | 0.81 | 0.2 | 0.62 | 0.97 | 1.42 | 0.31 | 1.47 | 0.92 | 1.08 | 0.84 | 0.53 | 1.11 | 1.45 | 1.67 | 0.98 | 0.435 | 0.235 | 0.695 | 0.184046 | 0.307007 | 0.018138 | 0.999971 | 1 | 0.635388 |
| Spock3 | Q8BKV0 | -0.07 | 0.02 | -0.52 | 0.69 | 0.18 | -0.11 | -1.17 | 0.85 | 1.02 | -0.29 | 0 | -0.27 | 2.15 | 0.62 | 0.59 | 0.42 | -0.0925 | 0.085 | 0.915 | 0.839942 | 0.824726 | 0.066915 | 0.999971 | 1 | 0.733937 |
| Spon1 | Q8VCC9 | 0.25 | 1.12 | 0.75 | 0.45 | 1.17 | 1.11 | 0.99 | 0.82 | 1.31 | 0.5 | 1.29 | 1.21 | 0.84 | 0.73 | 1 | 0.46 | 0.38 | 0.435 | 0.115 | 0.137133 | 0.151517 | 0.647189 | 0.999971 | 1 | 0.933613 |
| Spout1 | Q3UHX9+C | 1.42 | -0.38 | 1.9 | 0.92 | -0.06 | 0.09 | 2.92 | 1.06 | 0.93 | 1.15 | 3.18 | 1.4 | -1.1 | -1.31 | 1.41 | -0.26 | 0.0375 | 0.7 | -1.28 | 0.960565 | 0.297749 | 0.100596 | 0.999971 | 1 | 0.78268 |
| Sppl2a | Q9JJF9 | 1.27 | -0.44 | 0.05 | 1.15 | -0.35 | 0.13 | 0.86 | -0.04 | -0.14 | 1.09 | 0.49 | 0.02 | -0.06 | -0.52 | 0.44 | 0.34 | -0.3575 | -0.1425 | -0.4575 | 0.444687 | 0.762312 | 0.318803 | 0.999971 | 1 | 0.853893 |
| Sppl2b | Q3TD49 | 0.12 | -0.99 | -0.83 | -0.89 | -0.61 | -0.33 | -0.1 | -1.29 | -1.13 | -0.11 | -1.01 | -1.65 | -0.16 | -0.2 | -1.78 | -1.67 | 0.065 | -0.3275 | -0.305 | 0.85542 | 0.416636 | 0.530943 | 0.999971 | 1 | 0.910069 |
| Spr | Q91XH5 | -0.05 | 0.24 | 0.28 | 0.31 | 0.1 | 0.4 | -0.08 | 0.23 | -0.12 | -0.07 | 0.05 | 0.16 | 0.03 | 0.32 | 0.55 | 0.57 | -0.0325 | -0.19 | 0.1725 | 0.867178 | 0.302183 | 0.411488 | 0.999971 | 1 | 0.875847 |
| Spred1 | Q924S8 | -1.17 | -1.02 | -0.74 | -0.96 | -1.23 | -1.34 | -1.1 | -0.64 | -1.64 | -0.94 | -0.87 | -1.06 | -1.01 | -1.25 | -0.84 | -0.63 | -0.105 | -0.155 | 0.04 | 0.636919 | 0.508143 | 0.849174 | 0.999971 | 1 | 0.98178 |
| Spred2 | Q924S7 | -0.93 | -0.52 | -0.31 | -1.01 | -0.49 | -0.58 | 0.18 | -0.73 | -0.66 | -1.12 | -0.33 | 0.08 | -0.49 | -0.94 | -0.62 | -0.81 | 0.2875 | 0.185 | -0.0225 | 0.315463 | 0.553311 | 0.922727 | 0.999971 | 1 | 0.993264 |
| Spred3 | A0A0R4J0K | -0.05 | -1.59 | -0.95 | -0.65 | -1.57 | -1.34 | -0.89 | -0.86 | -1.36 | -0.37 | -1.13 | -0.87 | -1.08 | -1.02 | -0.92 | -0.95 | -0.355 | -0.1225 | -0.1825 | 0.333503 | 0.743333 | 0.576807 | 0.999971 | 1 | 0.918194 |
| Sprn | A0A1B0GSI | -0.79 | 0.19 | -0.79 | -1.3 | 0.35 | -0.6 | -1.12 | -1.12 | -0.12 | -0.92 | -1.66 | -1.48 | -1.51 | 0.25 | -1.29 | -1.11 | 0.05 | -0.3725 | -0.2425 | 0.909259 | 0.407734 | 0.609898 | 0.999971 | 1 | 0.92673 |
| Spry2 | Q9QXV8 | -0.38 | -1.43 | -2.86 | -3.03 | -0.92 | -1.12 | -0.38 | -4.33 | -1.03 | -0.41 | -1.82 | -1.76 | -0.99 | -1.67 | -2.28 | -3.57 | 0.2375 | 0.67 | -0.2025 | 0.807793 | 0.317672 | 0.787985 | 0.999971 | 1 | 0.96724 |
| Spryd3 | E9Q9B3 | -0.4 | 1.28 | 0.07 | 0.04 | 1.21 | 0.45 | 1.14 | -0.49 | 1.34 | 0.14 | 0.23 | 0.19 | 1.87 | 0.26 | -0.34 | -1.12 | 0.33 | 0.2275 | -0.08 | 0.511367 | 0.605611 | 0.90381 | 0.999971 | 1 | 0.991091 |
| Spryd4 | Q91WK1 | -0.45 | -0.37 | -0.09 | -0.3 | -0.16 | -0.28 | -0.32 | -0.25 | -0.19 | -0.08 | -0.05 | -0.04 | 0.09 | -0.32 | -0.3 | -0.26 | 0.05 | 0.2125 | 0.105 | 0.773279 | 0.231883 | 0.586244 | 0.999971 | 1 | 0.920418 |
| Spryd7 | Q3TFQ1 | 0.08 | 1.36 | 0.57 | -0.53 | 0.68 | 1.56 | 0.23 | 0.22 | 0.86 | 0.57 | 0.38 | 0.83 | -0.21 | 1.43 | 0.58 | 0.28 | 0.3025 | 0.29 | 0.15 | 0.52831 | 0.473737 | 0.760341 | 0.999971 | 1 | 0.961116 |
| Spta1 | P08032 | -0.58 | -0.63 | -2.33 | -0.06 | 0.69 | -1.5 | -0.06 | -1.16 | -0.74 | -1.24 | -0.74 | -1.17 | -0.65 | -1.27 | -2.03 | -1.33 | 0.3925 | -0.0725 | -0.42 | 0.544439 | 0.879688 | 0.43294 | 0.999971 | 1 | 0.882137 |
| Sptan1 | A3KGU9 | -1 | -1.49 | -1.01 | -1.03 | -1.48 | -1.22 | -0.43 | -1.11 | -1.39 | -1.19 | -0.94 | -1.15 | -1.35 | -1.5 | -1.08 | -1.09 | 0.0725 | -0.035 | -0.1225 | 0.790179 | 0.86279 | 0.562715 | 0.999971 | 1 | 0.916637 |
| Sptan1_1 | E9Q447 | -1.09 | -1.63 | -2.82 | -2.67 | -1.33 | -0.86 | -3.02 | -2.9 | -0.92 | -0.8 | -2.75 | -2.54 | -0.97 | -0.96 | -3.13 | -1.48 | 0.025 | 0.3 | 0.4175 | 0.968078 | 0.624626 | 0.494995 | 0.999971 | 1 | 0.901531 |
| Sptan1_2 | A3KGU7 | -0.45 | -0.95 | -0.66 | -0.31 | -1.15 | -0.96 | -1 | -0.28 | -0.93 | -1.37 | -1.05 | -0.14 | -0.62 | -0.53 | -0.06 | -0.3 | -0.255 | -0.28 | 0.215 | 0.342542 | 0.36727 | 0.357738 | 0.999971 | 1 | 0.868793 |
| Sptan1_3 | P16546 | 3.51 | 3.04 | 0.37 | 1.64 | 2.99 | 4.12 | 0.3 | -0.73 | 2.31 | 4.42 | -0.51 | 0.17 | -0.93 | 2.69 | 0.51 | -0.55 | -0.47 | -0.5425 | -1.71 | 0.693628 | 0.649247 | 0.104258 | 0.999971 | 1 | 0.78268 |
| Sptb | Q3UGX2 | -1.03 | -0.93 | -0.93 | -0.68 | -0.64 | -1.41 | -0.87 | -0.97 | -1.09 | -1.5 | -1.03 | -0.93 | -0.9 | -1.28 | -1.17 | -1.16 | -0.08 | -0.245 | -0.235 | 0.718297 | 0.24021 | 0.224459 | 0.999971 | 1 | 0.828225 |
| Sptbn1 | Q62261 | -0.87 | -1.08 | -1.03 | -1.2 | -1.05 | -1.07 | -0.82 | -1.11 | -0.96 | -1.03 | -1.05 | -1.09 | -0.61 | -1.17 | -1.06 | -1.09 | 0.0325 | 0.0125 | 0.0625 | 0.854571 | 0.939863 | 0.757214 | 0.999971 | 1 | 0.961116 |
| Sptbn2 | Q68FG2 | -0.98 | -0.69 | -1.67 | -2 | -0.79 | -0.56 | -1.35 | -1.79 | -0.96 | -1.1 | -1.35 | -1.88 | -0.41 | -0.87 | -1.94 | -1.64 | 0.2125 | 0.0125 | 0.12 | 0.592044 | 0.971995 | 0.783975 | 0.999971 | 1 | 0.966376 |
| Sptbn4 | E9PX29 | -0.54 | -0.65 | -0.87 | -0.83 | -0.68 | -0.63 | -1.11 | -0.78 | -0.62 | -0.6 | -0.98 | -1.08 | -0.35 | -0.75 | -0.83 | -0.87 | -0.0775 | -0.0975 | 0.0225 | 0.692583 | 0.627101 | 0.910808 | 0.999971 | 1 | 0.992125 |
| Sptlc1 | O35704 | 1.11 | -0.32 | -0.33 | -0.37 | -0.31 | -0.19 | -0.87 | -0.12 | 0.17 | 0.41 | -0.15 | -0.16 | 0.19 | -0.37 | -0.1 | 0.08 | -0.395 | 0.045 | -0.0725 | 0.320426 | 0.904681 | 0.845474 | 0.999971 | 1 | 0.979908 |
| Sptlc2 | P97363 | 0.18 | -0.34 | -0.81 | -0.86 | -0.56 | -0.55 | -0.92 | -0.44 | -0.44 | -0.15 | -0.42 | -0.64 | 0 | -0.07 | -0.1 | -0.67 | -0.16 | 0.045 | 0.2475 | 0.57086 | 0.871227 | 0.415383 | 0.999971 | 1 | 0.876885 |
| Sqle | P52019 | -1.32 | -0.26 | 0.76 | 0.66 | -0.56 | -0.89 | 0.63 | 0.29 | -0.57 | -0.76 | 0.82 | -0.66 | 0.92 | 0.23 | 0.5 | 1.24 | -0.0925 | -0.2525 | 0.7625 | 0.866846 | 0.655981 | 0.150402 | 0.999971 | 1 | 0.793248 |
| Sqor | Q9R112 | 1.46 | 0.62 | 0.14 | -0.22 | 0.74 | 0.2 | -0.26 | -0.03 | 0.85 | 0.8 | 0.17 | 0.55 | 1.35 | 0.58 | -0.13 | -0.31 | -0.3375 | 0.0925 | -0.1275 | 0.411517 | 0.808804 | 0.794294 | 0.999971 | 1 | 0.970141 |
| Sqstm1 | Q64337-2 | 1.51 | -0.05 | -0.07 | 0.11 | -0.41 | 0.62 | 1 | 2.26 | -0.1 | 1.26 | 1.94 | 1.36 | -0.07 | 0.61 | 0.48 | 1.42 | 0.4925 | 0.74 | 0.235 | 0.428639 | 0.18935 | 0.611312 | 0.999971 | 1 | 0.92673 |
| Sqstm1_1 | Q64337 | 0.34 | 0.52 | 0.24 | -0.52 | 0.41 | 0.45 | 0.73 | 0.01 | 0.36 | 0.09 | 0.32 | 0.37 | 0.99 | 0.85 | 0.24 | -0.34 | 0.255 | 0.14 | 0.29 | 0.383806 | 0.593305 | 0.442815 | 0.999971 | 1 | 0.883386 |
| Sra1 | Q80VJ2 | 2.65 | 1.11 | 0.49 | 0.69 | 0.47 | 1.33 | 0.79 | 0.33 | 0.71 | 2.06 | 0.28 | 0.33 | -0.63 | 0.68 | 0.08 | 0.67 | -0.505 | -0.39 | -1.035 | 0.325165 | 0.514179 | 0.07982 | 0.999971 | 1 | 0.755254 |
| Srbd1 | F8WGW3 | -0.39 | -0.88 | 0.34 | 0.54 | -0.62 | -0.71 | 0.84 | 0.83 | -1.41 | -1.06 | 0.57 | 0.29 | -1.83 | -0.85 | 0.58 | 0.67 | 0.1825 | -0.305 | -0.26 | 0.717976 | 0.579113 | 0.678753 | 0.999971 | 1 | 0.942697 |
| Src | P05480-1 | -1.43 | -1.07 | -1.27 | -1.4 | -0.62 | -1.39 | -1.3 | -1.04 | -0.62 | -1.4 | -1.22 | -1.08 | -0.16 | -1.22 | -1 | -1.08 | 0.205 | 0.2125 | 0.4275 | 0.382676 | 0.355686 | 0.14271 | 0.999971 | 1 | 0.785422 |
| Srcin1 | A0A571BD( | -0.77 | -1.28 | -1.11 | -1.13 | -1.33 | -1.16 | -0.97 | -1.14 | -1.25 | -0.95 | -1.01 | -1.2 | -1.03 | -1.48 | -1.39 | -1.05 | -0.0775 | -0.03 | -0.165 | 0.69077 | 0.875075 | 0.440037 | 0.999971 | 1 | 0.883105 |
| Srcin1_1 | B1AQX7 | 2.9 | -0.76 | -1.29 | -1.14 | -1.05 | 0.06 | -0.71 | -1.34 | -0.59 | 1.37 | -0.19 | -1.26 | -1.29 | -1.41 | -1.68 | -0.83 | -0.6875 | -0.095 | -1.23 | 0.467419 | 0.926218 | 0.198599 | 0.999971 | 1 | 0.813335 |
| Srcin1_2 | B1AQX9 | -1.4 | -0.34 | -0.96 | -0.6 | -1.23 | -1.12 | -0.8 | -0.67 | -0.71 | -0.66 | -0.7 | -0.34 | -0.23 | -0.89 | -0.82 | -0.23 | -0.13 | 0.2225 | 0.2825 | 0.644739 | 0.412081 | 0.361086 | 0.999971 | 1 | 0.869526 |
| Srd5a3 | Q9WUP4 | 0.01 | 0.68 | -0.68 | 0.58 | 0.52 | 1.23 | -0.7 | 0.02 | 0.33 | 0.7 | -0.69 | -0.97 | 0.28 | 1.57 | -0.31 | 1.02 | 0.12 | -0.305 | 0.4925 | 0.802028 | 0.526189 | 0.32246 | 0.999971 | 1 | 0.85548 |
| Srebf2 | Q3U1N2 | -0.91 | -2.18 | 2.12 | 1.68 | -1.04 | -1.26 | 1.83 | 2.08 | -1.08 | -2.22 | 1.53 | 0.88 | -2.31 | -1.62 | 1.77 | 1.38 | 0.225 | -0.4 | -0.3725 | 0.853183 | 0.74096 | 0.774694 | 0.999971 | 1 | 0.963884 |
| Srek1 | Q8BZX4-2 | 1.69 | 0.71 | 1.01 | 1.71 | 1.51 | 1.32 | 1.35 | 0.98 | 1.23 | 2.18 | 0.62 | 0.82 | -0.25 | 0.69 | 0.85 | 0.67 | 0.01 | -0.0675 | -0.79 | 0.972157 | 0.868732 | 0.0468 | 0.999971 | 1 | 0.710102 |
| Srgap1 | Q91Z69 | 0.87 | -0.65 | -0.29 | 0.02 | -0.4 | -0.43 | -0.25 | 0.19 | -0.58 | 0.28 | -0.3 | -0.25 | -0.63 | -0.54 | 0.03 | -0.37 | -0.21 | -0.2 | -0.365 | 0.551084 | 0.584025 | 0.313085 | 0.999971 | 1 | 0.849656 |
| Srgap2 | Q91Z67 | -0.01 | -0.12 | 0.1 | -0.02 | -0.01 | -0.26 | 0.47 | -0.09 | 0.33 | 0.14 | 0.29 | 0.15 | 0.44 | -0.27 | 0.01 | 0.01 | 0.04 | 0.24 | 0.06 | 0.850354 | 0.167034 | 0.772469 | 0.999971 | 1 | 0.963498 |
| Srgap3 | E9QN14 | -3.14 | 1.12 | 3.56 | -1.32 | 1.14 | -1.05 | 4.65 | 0.79 | 1.31 | -2.02 | 2.82 | -7.8 | 2.19 | -2.6 | 0.92 | -1.64 | 1.3275 | -1.4775 | -0.3375 | 0.434176 | 0.553207 | 0.835546 | 0.999971 | 1 | 0.979185 |
| Srgap3_1 | F8VPQ4 | -0.32 | -0.36 | -0.2 | -0.3 | -0.29 | 0.03 | -0.06 | -0.12 | -0.47 | -0.34 | -0.22 | -0.2 | -0.08 | -0.26 | -0.21 | 0.04 | 0.185 | -0.0125 | 0.1675 | 0.293395 | 0.939549 | 0.340589 | 0.999971 | 1 | 0.860574 |
| Sri | Q6P069-2 | -0.68 | 0.69 | 0.31 | -0.03 | 0.81 | 0.19 | 0.19 | 0.12 | 0.9 | -0.13 | 0.26 | -0.52 | 1.11 | 0.04 | -0.02 | -0.01 | 0.255 | 0.055 | 0.2075 | 0.449626 | 0.891172 | 0.594914 | 0.999971 | 1 | 0.92377 |
| Srm | Q64674 | -0.88 | 0.07 | -0.52 | 0.1 | -0.05 | -0.11 | -0.52 | -0.27 | -0.03 | -0.4 | -0.68 | -0.04 | -0.24 | 0.22 | -0.49 | -0.29 | 0.07 | 0.02 | 0.1075 | 0.800683 | 0.945805 | 0.714763 | 0.999971 | 1 | 0.952456 |
| Srp14 | P16254 | 1.06 | 1.19 | 0.77 | 0.92 | 1.12 | 1.25 | 0.36 | 0.67 | 1.26 | 1.17 | 0.53 | 0.76 | 0.83 | 1.34 | 1.04 | 0.57 | -0.135 | -0.055 | -0.04 | 0.594365 | 0.810902 | 0.860215 | 0.999971 | 1 | 0.982702 |
| Srp19 | Q9D104 | 0.12 | 1.92 | 1.58 | 1.49 | 1.56 | 1.86 | 3.14 | 1.3 | 1.51 | 0.49 | 1.18 | 0.47 | 0.92 | 1.79 | 1.33 | 0.79 | 0.6875 | -0.365 | -0.07 | 0.21257 | 0.42341 | 0.871238 | 0.999971 | 1 | 0.984109 |
| Srp54 | P14576 | 0.92 | 1.24 | 0.9 | 0.76 | 1.15 | 1.34 | 0.53 | 0.92 | 1.17 | 1.14 | 0.88 | 0.53 | 0.26 | 0.99 | 0.82 | 0.77 | 0.03 | -0.025 | -0.245 | 0.898929 | 0.909544 | 0.297969 | 0.999971 | 1 | 0.847402 |
| Srp68 | Q8BMA6 | 2.13 | 1.06 | 0.95 | 1 | 1.36 | 1.39 | 1.03 | 1.1 | 1.09 | 1.99 | 1.29 | 1.04 | 0.46 | 1.38 | 1.04 | 1.41 | -0.065 | 0.0675 | -0.2125 | 0.830716 | 0.847681 | 0.550899 | 0.999971 | 1 | 0.913169 |
| Srp72 | F8VQC1 | 1.01 | 1.48 | 1.77 | 1.71 | 1.69 | 1.49 | 1.83 | 1.74 | 1.86 | 1.25 | 1.72 | 1.53 | 1.54 | 1.4 | 1.64 | 1.64 | 0.195 | 0.0975 | 0.0625 | 0.40011 | 0.691987 | 0.78069 | 0.999971 | 1 | 0.965859 |
| Srp9 | P49962 | 1.52 | 2.33 | 3.25 | 1.83 | 3.16 | 2.1 | 4.39 | 2.16 | 3.78 | 1.58 | 3.66 | 3.86 | 4.13 | 2.13 | 2.58 | 1.29 | 0.72 | 0.9875 | 0.3 | 0.249299 | 0.133117 | 0.64229 | 0.999971 | 1 | 0.93301 |
| Srpk1 | O70551 | -0.02 | 1.62 | 0.8 | 0.78 | 1.92 | 0.6 | 1.83 | 0.24 | 2.08 | 0.26 | 2.17 | 2.45 | 2.3 | 0.74 | 0.89 | 0.9 | 0.3525 | 0.945 | 0.4125 | 0.489252 | 0.116527 | 0.385799 | 0.999971 | 1 | 0.871816 |
| Srpk2 | A0A0R4J12 | 0.39 | 0.86 | 1.09 | 0.63 | 1.03 | 0.77 | 1.38 | 0.77 | 1.82 | 0.6 | 1.46 | 1.2 | 1.64 | 0.44 | 0.95 | 1 | 0.245 | 0.5275 | 0.265 | 0.323669 | 0.110504 | 0.385111 | 0.999971 | 1 | 0.871816 |
| Srpra | Q9DBG7 | 0.2 | 0.41 | -0.12 | 0.06 | -0.17 | 0.14 | -0.16 | 0.14 | 0.25 | 0.49 | -0.06 | 0.15 | 0.5 | 0.21 | 0.06 | 0.52 | -0.15 | 0.07 | 0.185 | 0.461512 | 0.737021 | 0.389863 | 0.999971 | 1 | 0.871816 |
| Srprb | P47758 | -0.01 | -0.02 | 0.2 | 0.35 | 0.23 | -0.09 | -0.2 | 1.1 | -0.07 | 0.47 | 0.37 | 0.5 | 0.17 | 0.06 | 0.7 | 1.1 | 0.13 | 0.1875 | 0.3775 | 0.678436 | 0.377943 | 0.194388 | 0.999971 | 1 | 0.811195 |
| Srr | Q9QZX7-2 | -0.35 | -0.56 | -0.53 | -0.63 | -0.63 | -0.72 | -0.91 | -0.45 | -0.96 | -0.59 | -0.4 | -0.58 | -1.37 | -0.43 | -0.54 | -0.29 | -0.16 | -0.115 | -0.14 | 0.398159 | 0.553133 | 0.607153 | 0.999971 | 1 | 0.925193 |
| Srrm1 | E9QKA4 | 0.83 | 0.22 | 0.92 | 0.89 | 0.27 | 0.26 | 1.48 | 0.58 | 0.26 | 0.85 | 0.7 | 0.18 | 0.44 | 0.72 | 0.58 | 1.07 | -0.0675 | -0.2175 | -0.0125 | 0.838372 | 0.406316 | 0.959316 | 0.999971 | 1 | 0.998721 |
| Srrm2 | Q8BTI8 | 0.25 | -0.47 | 0 | -0.04 | -0.4 | -0.51 | 0.01 | -0.14 | -0.53 | 0.17 | -0.08 | -0.05 | -0.61 | -0.26 | -0.04 | 0.26 | -0.195 | -0.0575 | -0.0975 | 0.405146 | 0.810674 | 0.709314 | 0.999971 | 1 | 0.950625 |
| Srrm3 | D3Z3C2 | 1.83 | 1.32 | 1.35 | -0.08 | 1.71 | 0.95 | 1.76 | 0.27 | -0.24 | 0.66 | 0.37 | 0.46 | -0.93 | 0.74 | 1.01 | 1.56 | 0.0675 | -0.7925 | -0.51 | 0.893141 | 0.094178 | 0.417786 | 0.999971 | 1 | 0.877584 |
| Srrt | Q99MR6-3 | 1.88 | 2.15 | 2.53 | 2.4 | 1.84 | 1.82 | 1.99 | 2.18 | 2.18 | 1.87 | 2.28 | 2.4 | 1.39 | 1.88 | 2.37 | 2.36 | -0.2825 | -0.0575 | -0.24 | 0.208913 | 0.796086 | 0.413408 | 0.999971 | 1 | 0.876261 |
| Srsf1 | H7BX95 | 0.48 | 0.33 | 0.03 | -0.01 | 0.41 | 0.22 | 0.11 | 0.08 | 0.47 | 0.18 | 0.1 | 0.39 | 0.56 | 0.16 | -0.09 | -0.02 | -0.0025 | 0.0775 | -0.055 | 0.989969 | 0.699763 | 0.809502 | 0.999971 | 1 | 0.973762 |
| Srsf10 | Q3TFP0 | -0.82 | -1.29 | -0.39 | -0.3 | -1.27 | -1.37 | -0.6 | -0.79 | -1.16 | -1.19 | -0.55 | -0.46 | -0.98 | -1.27 | -0.39 | -0.41 | -0.3075 | -0.14 | -0.0625 | 0.322139 | 0.64809 | 0.844243 | 0.999971 | 1 | 0.979814 |
| Srsf11 | Q3UIX4 | 0.54 | 1.01 | 1.5 | 1.24 | 1.29 | 0.83 | 1.01 | 1.4 | 1.22 | 0.56 | 1.37 | 1.52 | 0.27 | 1.12 | 1.52 | 1.39 | 0.06 | 0.095 | 0.0025 | 0.820087 | 0.752838 | 0.994187 | 0.999971 | 1 | 0.999672 |
| Srsf12 | A3KG57 | 0.6 | 0.32 | -0.23 | 0.29 | 0.24 | 0.19 | -0.71 | 0.73 | 0.21 | 0.56 | 0.45 | 0.35 | 1.12 | 0.62 | 0.68 | 0.76 | -0.1325 | 0.1475 | 0.55 | 0.69964 | 0.518675 | 0.044599 | 0.999971 | 1 | 0.706921 |
| Srsf2 | Q62093 | 0.12 | 1.4 | 0.45 | 0.09 | 1.5 | 0.73 | 0.6 | 0.21 | 1.2 | 0.19 | 0.56 | 0.7 | 1.53 | 0.45 | -0.18 | 0.18 | 0.245 | 0.1475 | -0.02 | 0.536196 | 0.68492 | 0.964553 | 0.999971 | 1 | 0.998721 |
| Srsf3 | P84104-2 | -1.07 | 0.05 | 0.5 | 0.68 | 0.14 | 0.42 | 0.45 | 0.75 | 0.44 | -0.15 | 0.37 | -0.12 | -0.16 | 0.58 | 0.7 | 0.59 | 0.4 | 0.095 | 0.3875 | 0.325686 | 0.814633 | 0.366476 | 0.999971 | 1 | 0.869526 |
| Srsf4 | Q8VE97 | 0.85 | 0.4 | -0.01 | 0.19 | 0.2 | 0.35 | 0.09 | -0.15 | 0.2 | 0.86 | 0.04 | -0.09 | 0.48 | 0.32 | -0.05 | 0.21 | -0.235 | -0.105 | -0.1175 | 0.347306 | 0.718795 | 0.635114 | 0.999971 | 1 | 0.93301 |
| Srsf5 | O35326 | -0.54 | -0.33 | 0.22 | 0.03 | -0.35 | -0.96 | 0.54 | 0 | -0.47 | -0.72 | -0.01 | 0.06 | -0.42 | -0.57 | -0.05 | -0.21 | -0.0375 | -0.13 | -0.1575 | 0.915015 | 0.633589 | 0.516997 | 0.999971 | 1 | 0.905815 |
| Srsf6 | Q3TWW8 | 1.79 | -0.26 | 0.26 | 0.76 | 0.17 | -0.15 | 0.82 | -0.05 | -0.09 | 0.81 | 0.31 | 0.09 | -0.08 | -0.23 | -0.26 | 0.26 | -0.44 | -0.3575 | -0.715 | 0.349186 | 0.436509 | 0.122868 | 0.999971 | 1 | 0.78268 |
| Srsf7 | A0A3Q4L3! | -0.78 | -0.59 | 0.59 | 0.3 | -0.76 | -0.5 | 0.55 | 0.31 | -0.92 | -0.73 | 0.25 | 0.08 | -1.08 | -0.71 | 0.43 | 0.36 | 0.02 | -0.21 | -0.13 | 0.963105 | 0.620329 | 0.783717 | 0.999971 | 1 | 0.966376 |
| Srsf9 | Q9D0B0 | -0.42 | 0.24 | 0.52 | -0.15 | 0.48 | 0.35 | 0.71 | 0.26 | 0.28 | -0.42 | 0.63 | 1.21 | -0.16 | -0.27 | 0.36 | -0.45 | 0.4025 | 0.3775 | -0.1775 | 0.140986 | 0.340917 | 0.538362 | 0.999971 | 1 | 0.910077 |
| Srxn1 | A2AQU8 | -0.86 | 0.47 | 0.36 | 0.58 | -0.51 | 0.92 | 0.01 | 1.02 | 0.04 | 0.35 | 1.22 | 0.66 | -0.37 | 1.13 | 1.37 | 0.99 | 0.2225 | 0.43 | 0.6425 | 0.634873 | 0.301068 | 0.204407 | 0.999971 | 1 | 0.81617 |
| Ss18 | A0A3Q4EI( | 1.82 | 1.47 | 0.06 | 1.18 | 0.73 | 1.06 | 0.06 | 1.12 | 1.41 | 1.88 | 0.67 | 1.17 | -0.15 | 1.9 | 0.97 | 1.17 | -0.39 | 0.15 | -0.16 | 0.37298 | 0.72952 | 0.761725 | 0.999971 | 1 | 0.96128 |
| Ss18l1 | Q8BW22 | 0.57 | 2.07 | 2.64 | 2.23 | 2.29 | 2.59 | 1.63 | 2.3 | 1.76 | 0.72 | 2.04 | 2.18 | 2.14 | 2.05 | 2.7 | 2.38 | 0.325 | -0.2025 | 0.44 | 0.489343 | 0.697896 | 0.338868 | 0.999971 | 1 | 0.86005 |
| Ssb | P32067 | 0.66 | 1.08 | 0.77 | 0.94 | 1.11 | 0.79 | 0.76 | 0.66 | 0.88 | 0.46 | 0.65 | 0.91 | 0.69 | 0.98 | 0.89 | 0.7 | -0.0325 | -0.1375 | -0.0475 | 0.868333 | 0.490692 | 0.800813 | 0.999971 | 1 | 0.971076 |
| Ssbp1 | Q8R2K3 | -1.26 | -0.29 | -0.72 | -0.5 | -0.58 | -0.18 | -0.88 | -0.32 | -0.44 | -0.62 | -0.72 | -0.75 | 0.58 | 0.15 | -0.25 | -0.02 | 0.2025 | 0.06 | 0.8075 | 0.470424 | 0.808243 | 0.018981 | 0.999971 | 1 | 0.635388 |
| Ssbp2 | A0A338P6( | -0.03 | 3.54 | 1.33 | 1.88 | 3.54 | 2.71 | 2.95 | 1.21 | 3.69 | 0.06 | 0.55 | 0.76 | 2.78 | 2.09 | 0.85 | 1.1 | 0.9225 | -0.415 | 0.025 | 0.266826 | 0.677509 | 0.974341 | 0.999971 | 1 | 0.99925 |
| Ssbp3 | B1AS34 | 4.34 | 2.9 | 2.89 | 2.4 | 3.27 | 4.3 | 3.48 | 2.03 | 3.48 | 4.27 | 2.58 | 1.84 | 3.14 | 3.12 | 1.69 | 1.04 | 0.1375 | -0.09 | -0.885 | 0.811373 | 0.88424 | 0.174393 | 0.999971 | 1 | 0.807788 |
| Ssh1 | Q76I79-2 | -1.12 | -0.92 | -1.01 | -0.29 | -1.23 | -1.14 | -0.71 | -0.22 | -0.89 | -1.3 | -0.44 | -0.04 | -0.53 | -0.26 | -0.19 | 0.33 | 0.01 | 0.1675 | 0.6725 | 0.973739 | 0.614045 | 0.035506 | 0.999971 | 1 | 0.69591 |
| Ssh2 | Q5SW75 | 0.09 | 0.12 | -0.21 | 0 | 0.03 | 0.04 | -0.29 | 0.03 | 0.04 | 0.22 | -0.28 | 0.01 | 0.28 | 0.04 | -0.05 | 0.33 | -0.0475 | -0.0025 | 0.15 | 0.796504 | 0.98947 | 0.435078 | 0.999971 | 1 | 0.882419 |
| Ssh3 | A0A498WG | -0.65 | 1.28 | 0.21 | 0.28 | 1.54 | 1.77 | 0.37 | 0.5 | 1.72 | 0.32 | 0.44 | 0.66 | 2.41 | 1.68 | 0.63 | 1.43 | 0.765 | 0.505 | 1.2575 | 0.147378 | 0.304636 | 0.031373 | 0.999971 | 1 | 0.678474 |
| Ssna1 | Q9JJ94 | -0.07 | 1.38 | 0.34 | 0.61 | 0.66 | 1.04 | -0.15 | 0.92 | -0.26 | 1.21 | 0.03 | 0.91 | 0.4 | 0.02 | 0.97 | 0.84 | 0.0525 | -0.0925 | -0.0075 | 0.892873 | 0.833259 | 0.983608 | 0.999971 | 1 | 0.99925 |
| Sspn | Q62147 | 1.85 | -0.35 | -0.16 | 0.34 | -0.39 | -0.36 | -0.57 | 0.15 | 0 | 2.01 | -0.52 | 0.06 | -0.44 | -0.15 | 0.23 | 0.07 | -0.7125 | -0.0325 | -0.4925 | 0.167193 | 0.961879 | 0.323586 | 0.999971 | 1 | 0.85548 |
| Ssr1 | Q9CY50 | -0.51 | -0.01 | 0.32 | 0.14 | -0.12 | 0.26 | -0.06 | 0.26 | 0.04 | -0.65 | 0.1 | 0.81 | 0.21 | -0.02 | 0.45 | 0.72 | 0.1 | 0.09 | 0.355 | 0.676623 | 0.793365 | 0.198869 | 0.999971 | 1 | 0.813335 |
| Ssr3 | Q9DCF9-2 | 1.25 | 1.07 | 1.01 | 0.18 | 0.87 | 1.17 | 1.19 | 0.51 | 0.63 | 0.88 | 0.5 | 0.63 | 0.96 | 0.14 | 0.65 | 0.31 | 0.0575 | -0.2175 | -0.3625 | 0.846094 | 0.427832 | 0.256635 | 0.999971 | 1 | 0.839012 |
| Ssr4 | Q9D8L3 | -2.4 | 0.56 | -1.17 | -1.66 | 0.25 | -0.8 | 1.2 | -2.6 | 1.08 | -2.69 | -0.67 | 0.32 | 1.05 | -1.8 | -1.85 | -2.17 | 0.68 | 0.6775 | -0.025 | 0.466806 | 0.472409 | 0.977295 | 0.999971 | 1 | 0.99925 |
| Ssrp1 | Q08943-2 | -0.95 | 0.07 | -0.38 | -0.77 | 0.2 | -0.28 | -0.41 | -1.25 | 0.24 | -0.47 | -0.58 | 0.01 | 0.66 | -0.1 | -0.43 | -0.9 | 0.0725 | 0.3075 | 0.315 | 0.843255 | 0.32753 | 0.422118 | 0.999971 | 1 | 0.877992 |
| Sst | P60041 | 0.48 | -0.7 | 0.12 | 0.19 | -0.1 | -0.54 | 1.08 | -0.53 | -0.14 | -0.18 | -0.03 | -0.24 | -1.13 | -1.09 | -0.22 | -0.64 | -0.045 | -0.17 | -0.7925 | 0.917023 | 0.538708 | 0.038121 | 0.999971 | 1 | 0.700838 |
| Sstr1 | P30873 | 3.03 | 0.19 | -0.21 | 1.06 | 0.26 | 1.31 | 0.48 | 1.15 | 0.26 | 2.13 | 0.49 | 0.58 | 3.3 | 0.83 | 0.7 | 0.84 | -0.2175 | -0.1525 | 0.4 | 0.75366 | 0.841015 | 0.643305 | 0.999971 | 1 | 0.93301 |
| Sstr2 | P30875 | -0.46 | -0.06 | -0.08 | -1.52 | -0.36 | 0.15 | 0.25 | -0.81 | -0.7 | -0.3 | -0.11 | -0.78 | -0.16 | -0.07 | -0.7 | -1.42 | 0.3375 | 0.0575 | -0.0575 | 0.411195 | 0.875838 | 0.895207 | 0.999971 | 1 | 0.988571 |
| Ssu72 | Q9CY97-2 | 2.72 | 1.63 | 1.14 | 1.8 | 0.8 | 1.21 | 1.37 | 1.29 | 0.74 | 2.78 | 0.35 | 0.34 | 0 | 1.96 | 0.86 | 1.54 | -0.655 | -0.77 | -0.7325 | 0.086995 | 0.231449 | 0.169717 | 0.999971 | 1 | 0.802438 |
| St13 | Q99L47 | 0.31 | 0.23 | 0.82 | 0.53 | 0.2 | 0.25 | 0.65 | 0.88 | 0.28 | -0.09 | 0.67 | 0.72 | 0.63 | -0.27 | 1.05 | 0.93 | 0.0225 | -0.0775 | 0.1125 | 0.925793 | 0.761258 | 0.731701 | 0.999971 | 1 | 0.956087 |
| St3gal5 | O88829 | 1.3 | 0.07 | 1.05 | 1.55 | 0.1 | 0.5 | 1.7 | 0.93 | 0 | 0.34 | 0.51 | 0.06 | 0.31 | 0.29 | 0.54 | 0.64 | -0.185 | -0.765 | -0.5475 | 0.677848 | 0.049047 | 0.128461 | 0.999971 | 1 | 0.784699 |
| St6galnac5 | Q6GTI0 | 4.25 | 0.56 | 1.44 | 1.07 | -0.29 | 0.7 | 2.19 | 0.4 | -0.15 | 2.51 | 0.07 | -0.59 | -0.93 | -0.14 | -0.67 | -1.16 | -1.08 | -1.37 | -2.555 | 0.237976 | 0.183586 | 0.008754 | 0.999971 | 1 | 0.610079 |
| St7 | Q99M96-9 | 0.73 | 0.54 | -0.34 | -0.42 | -0.04 | -0.02 | -0.54 | -0.26 | -0.68 | 0.79 | -0.53 | -0.11 | -0.47 | -0.05 | -0.45 | -0.2 | -0.3425 | -0.26 | -0.42 | 0.302814 | 0.542162 | 0.207638 | 0.999971 | 1 | 0.820648 |
| St8sia3 | Q64689 | -0.27 | 0.41 | 0.29 | 0.62 | 0.28 | 0.17 | 0.91 | 0.63 | 0.18 | -0.26 | 0.62 | 0.82 | 0.72 | 0.46 | 1.02 | 0.78 | 0.235 | 0.0775 | 0.4825 | 0.39891 | 0.803376 | 0.081195 | 0.999971 | 1 | 0.755254 |
| St8sia5 | P70126-3 | 1.72 | 0.57 | 0.81 | 1.06 | 0.69 | 1.14 | 1.24 | 0.43 | 0.38 | 1.84 | 1.26 | 0.32 | -0.25 | 0.48 | 0.22 | 0.59 | -0.165 | -0.09 | -0.78 | 0.604196 | 0.830693 | 0.033113 | 0.999971 | 1 | 0.685881 |
| Stab1 | G3X973 | 1.37 | 0.91 | 0.21 | 1.39 | 0.89 | 0.34 | 0.55 | -0.36 | 0.49 | 1.24 | 0.16 | 0.62 | 0.31 | 0.84 | 0.16 | 0.58 | -0.615 | -0.3425 | -0.4975 | 0.124191 | 0.342785 | 0.144341 | 0.999971 | 1 | 0.790933 |
| Stac2 | Q8R1B0 | -0.2 | 0.49 | 3.02 | 0.84 | 0.24 | 0.82 | 1.52 | 2.85 | 0.26 | 0.41 | 2.52 | 2.78 | 0.87 | 0.12 | 2.7 | 1.67 | 0.32 | 0.455 | 0.3025 | 0.691428 | 0.604572 | 0.706148 | 0.999971 | 1 | 0.950182 |
| Stag1 | F8WHU7 | -0.39 | 0.17 | -0.89 | -0.75 | 0.05 | -0.79 | -0.53 | -1.17 | 0.17 | -0.42 | -0.94 | -1.08 | -0.4 | -0.13 | -0.76 | -0.48 | -0.145 | -0.1025 | 0.0225 | 0.674773 | 0.776795 | 0.936878 | 0.999971 | 1 | 0.995618 |
| Stag2 | A2AFF6 | -0.27 | 0.08 | 0.26 | 0.4 | 0.21 | -0.51 | 0.03 | 0.18 | 0.19 | -0.2 | 0.16 | 0.29 | -0.17 | -0.19 | 0.18 | 0.15 | -0.14 | -0.0075 | -0.125 | 0.577926 | 0.972825 | 0.574421 | 0.999971 | 1 | 0.91816 |
| Stam | Q3UGN9 | 0.01 | -0.05 | 0.05 | -0.13 | -0.05 | -0.06 | -0.34 | -0.06 | -0.04 | -0.02 | -0.21 | 0.14 | 0.42 | 0.08 | -0.1 | 0.05 | -0.0975 | -0.0025 | 0.1425 | 0.574352 | 0.988166 | 0.455617 | 0.999971 | 1 | 0.887217 |
| Stam2 | O88811 | 0.19 | -0.59 | -0.3 | 0.22 | -0.5 | -1.32 | -0.75 | 0.1 | -0.71 | -0.62 | -0.26 | -0.13 | -0.26 | -0.17 | 0 | 0.12 | -0.4975 | -0.31 | 0.0425 | 0.177639 | 0.256151 | 0.862546 | 0.999971 | 1 | 0.982963 |
| Stambp | Q9CQ26 | 0.63 | 0.58 | -0.03 | -0.15 | 0.84 | 0.91 | -0.35 | 0 | 0.73 | 0.83 | -0.22 | -0.02 | 0.72 | 1.11 | 0.27 | 0.17 | 0.0925 | 0.0725 | 0.31 | 0.798657 | 0.82685 | 0.323209 | 0.999971 | 1 | 0.85548 |
| Stambpl1 | Q76N33 | -0.23 | -0.1 | -0.51 | -1 | -0.39 | 0.06 | -0.92 | -0.89 | -0.39 | -0.47 | -1.01 | -0.71 | -1.16 | -0.3 | -0.72 | -0.62 | -0.075 | -0.185 | -0.24 | 0.809659 | 0.488846 | 0.405322 | 0.999971 | 1 | 0.874565 |
| Stard10 | Q9JMD3 | 0.7 | 0.26 | 0.27 | 0.55 | 0.35 | 0.07 | 0.06 | 0.45 | 0 | -0.24 | -0.59 | 0.27 | -0.19 | 0.49 | 0.35 | -0.17 | -0.2125 | -0.585 | -0.325 | 0.310038 | 0.036457 | 0.198831 | 0.999971 | 1 | 0.813335 |
| Stard3 | Q61542 | -0.37 | -0.16 | -0.06 | -0.54 | -0.69 | -0.55 | 0.32 | 0 | -0.48 | 1.08 | 0.15 | 0.05 | -0.45 | -0.23 | -0.06 | -0.79 | 0.0525 | 0.4825 | -0.1 | 0.849324 | 0.17882 | 0.665101 | 0.999971 | 1 | 0.93697 |
| Stard3nl | A0A1Y7VJA | 1.69 | 0.7 | 0.14 | 0.16 | 0.58 | 0.93 | 0.07 | 0.64 | 0.9 | 1.67 | 0.63 | 0.68 | 0.79 | 1.62 | 0.56 | 1.02 | -0.1175 | 0.2975 | 0.325 | 0.762907 | 0.480428 | 0.435515 | 0.999971 | 1 | 0.882419 |
| Stard5 | D3YU00 | -1.92 | -0.98 | 0.09 | 0.41 | -0.71 | 0 | -0.05 | 0.42 | -0.02 | -2.18 | 0.46 | -0.19 | -0.53 | -1 | 0.18 | -0.03 | 0.515 | 0.1175 | 0.255 | 0.348137 | 0.869692 | 0.642565 | 0.999971 | 1 | 0.93301 |
| Stard7 | Q8R1R3 | -0.34 | 1.19 | 0.42 | 0.01 | 1.58 | 0.89 | 1.21 | 0.38 | 1.83 | 0.08 | 1.02 | 1.24 | 2.67 | 0.34 | 0.18 | -0.03 | 0.695 | 0.7225 | 0.47 | 0.108001 | 0.143834 | 0.474676 | 0.999971 | 1 | 0.894044 |
| Stat1 | Q8C3V4 | 0.04 | 0.78 | 0.56 | 0.73 | 0.34 | -0.09 | -0.41 | 0.43 | 0.34 | 0.32 | 0.32 | 0.71 | 0.16 | 0.59 | 0.37 | 0.45 | -0.46 | -0.105 | -0.135 | 0.12211 | 0.649498 | 0.561776 | 0.999971 | 1 | 0.916386 |
| Stat2 | Q9QXJ2 | 0.1 | 0.78 | 0.29 | 0.4 | 0.58 | -0.1 | 0.42 | -0.06 | 0.79 | -0.1 | 0.56 | 0.54 | 0.77 | 1.18 | -0.09 | 0.1 | -0.1825 | 0.055 | 0.0975 | 0.473228 | 0.832747 | 0.76694 | 0.999971 | 1 | 0.962472 |
| Stat3 | B7ZC18 | 0.63 | 0.71 | -0.14 | 0.24 | 0.94 | 0.73 | -0.31 | 0.04 | 0.81 | 0.78 | -0.32 | 0.07 | 0.64 | 0.86 | -0.05 | 0.26 | -0.01 | -0.025 | 0.0675 | 0.976943 | 0.940776 | 0.817796 | 0.999971 | 1 | 0.974929 |
| Stat5b | P42232 | -0.08 | -0.16 | 0.24 | 0.07 | 0.27 | -0.11 | 0.26 | 0.39 | 0.14 | -0.51 | 0.33 | 1.15 | 0.44 | 0.16 | 0.56 | 0.49 | 0.185 | 0.26 | 0.395 | 0.364398 | 0.462058 | 0.065008 | 0.999971 | 1 | 0.727266 |
| Stat6 | P52633 | -1.98 | 0.06 | -1.03 | -0.89 | 0.07 | -1.1 | -0.89 | -1.05 | -1.11 | -1.91 | -0.99 | -0.41 | -1.01 | -0.43 | -0.06 | -0.75 | 0.2175 | -0.145 | 0.3975 | 0.643145 | 0.765523 | 0.376877 | 0.999971 | 1 | 0.870465 |
| Stau1 | A2A5S3 | 2.87 | 1.48 | 1.68 | 1.54 | 1.23 | 1.54 | 1.55 | 1.41 | 1.24 | 1.83 | 1.59 | 1.49 | 0.78 | 1.34 | 1.25 | 1.63 | -0.46 | -0.355 | -0.6425 | 0.192942 | 0.319484 | 0.105149 | 0.999971 | 1 | 0.78268 |
| Stau2 | Q8CJ67-2 | 0.77 | 0.99 | 0.77 | 0.85 | 0.81 | 0.97 | 0.35 | 1.13 | 0.88 | 0.88 | 0.96 | 1.1 | 0.7 | 1.13 | 0.83 | 1.36 | -0.03 | 0.11 | 0.16 | 0.89148 | 0.512354 | 0.452899 | 0.999971 | 1 | 0.887217 |
| Steap2 | D3YUM6 | -2.1 | -1.76 | -0.71 | -1.19 | -0.52 | -1.61 | -0.43 | -0.14 | -0.66 | -3.14 | -1.11 | 1.43 | -1.96 | -1.21 | 0.5 | -1.34 | 0.765 | 0.57 | 0.4375 | 0.097165 | 0.526689 | 0.443271 | 0.999971 | 1 | 0.883386 |
| Steap3 | E9QN92 | -0.59 | -0.33 | -0.65 | -0.58 | 0.05 | 0.14 | -0.62 | -1.3 | -0.09 | -0.41 | -0.25 | 0.06 | 0.19 | -0.17 | -0.04 | -0.2 | 0.105 | 0.365 | 0.4825 | 0.757404 | 0.078762 | 0.027503 | 0.999971 | 1 | 0.678474 |
| Steep1 | Q8VDP2 | 3.69 | 0.86 | 0.96 | 1.39 | 0.86 | 2.35 | 0.81 | 1 | 1.32 | 3.17 | 0.5 | 0.31 | -0.12 | 0.18 | 0.68 | 1.09 | -0.47 | -0.4 | -1.2675 | 0.499829 | 0.636781 | 0.078789 | 0.999971 | 1 | 0.754291 |
| Stim1 | A0A1B0GR. | 0.1 | -0.63 | -0.49 | -0.31 | -0.5 | -0.47 | -0.65 | -0.47 | -0.36 | 0.45 | -0.65 | -0.33 | -0.18 | -0.32 | -0.54 | -0.27 | -0.19 | 0.11 | 0.005 | 0.382975 | 0.708728 | 0.981897 | 0.999971 | 1 | 0.99925 |
| Stim2 | P83093 | -0.18 | -0.21 | 0.35 | -0.36 | -0.29 | -0.01 | -0.3 | -0.26 | -0.33 | -0.5 | -0.01 | 0.02 | 0.05 | -0.15 | 0.17 | 0.11 | -0.115 | -0.105 | 0.145 | 0.597521 | 0.654544 | 0.509367 | 0.999971 | 1 | 0.905815 |
| Sting1 | Q3TBT3-3 | 0.26 | 0.7 | -0.15 | -0.31 | 0.66 | 0.01 | -1.34 | -0.3 | -0.51 | 0.09 | -1.02 | -0.74 | 0.36 | 0.02 | -0.55 | 0.28 | -0.3675 | -0.67 | -0.0975 | 0.419043 | 0.066681 | 0.754673 | 0.999971 | 1 | 0.961116 |
| Stip1 | Q60864 | 0.33 | 0.85 | 0.79 | 0.45 | 0.85 | 0.67 | 0.75 | 0.16 | 1.17 | 0.15 | 0.71 | 0.92 | 0.85 | 0.14 | 0.5 | 0.31 | 0.0025 | 0.1325 | -0.155 | 0.991498 | 0.625413 | 0.514481 | 0.999971 | 1 | 0.905815 |
| Stk11 | Q9WTK7-2 | 0.24 | -1.53 | 0.08 | -1.45 | -2.22 | -1.35 | -0.42 | -0.74 | -2.56 | -0.08 | -0.3 | -0.72 | -3.2 | -0.49 | -0.64 | -1.38 | -0.5175 | -0.25 | -0.7625 | 0.374219 | 0.711644 | 0.298994 | 0.999971 | 1 | 0.847402 |
| Stk11ip | Q3TAA7 | 0.8 | 1.01 | 1.49 | 1.41 | 0.41 | 0.92 | 1.03 | 1.25 | 0.31 | 0.38 | 0.69 | 1.19 | 0.44 | 0.92 | 0.96 | 1.7 | -0.275 | -0.535 | -0.1725 | 0.312436 | 0.078578 | 0.584333 | 0.999971 | 1 | 0.920364 |
| Stk16 | A0A087WC | 0.5 | -0.05 | -0.47 | -0.52 | 0.37 | 0.03 | -0.51 | -0.19 | 0.38 | 0.76 | 0.34 | 0.46 | 1.09 | 0.53 | 0 | -0.19 | 0.06 | 0.62 | 0.4925 | 0.84487 | 0.045737 | 0.198975 | 0.999971 | 1 | 0.813335 |

| | | | | | | | | | | | | | | | | | | | | | | | | | | |
|---|---|---|---|---|---|---|---|---|---|---|---|---|---|---|---|---|---|---|---|---|---|---|---|---|---|---|
| Stk24 | Q99KH8 | -0.58 | 0.02 | -0.42 | -1.16 | 0.31 | -0.53 | 0.12 | -1.24 | 0.42 | -1.06 | -0.4 | 0 | 0.36 | -0.84 | -1.02 | -1.32 | 0.2 | 0.275 | -0.17 | 0.625726 | 0.481167 | 0.686895 | 0.999971 | 1 | 0.945467 |
| Stk25 | Q9Z2W1 | 1.5 | 0.75 | 0.24 | 0.19 | 0.39 | 0.66 | 0.86 | 0 | 0.42 | 0.66 | 0.17 | 0.09 | 0.32 | 0.42 | 0.05 | 0.02 | -0.1925 | -0.335 | -0.4675 | 0.585747 | 0.324452 | 0.171779 | 0.999971 | 1 | 0.804075 |
| Stk26 | A2AD84 | 2.13 | 0.19 | 0.12 | 0.97 | 0.15 | -0.18 | 0.01 | 0.88 | -0.41 | 1.41 | 0.1 | -0.44 | 0.29 | 0.5 | 0.27 | 0.21 | -0.6375 | -0.6875 | -0.535 | 0.211176 | 0.259944 | 0.248519 | 0.999971 | 1 | 0.839012 |
| Stk3 | A0A2I3BQE | -0.02 | 0.39 | 1 | 0.46 | 0.37 | 0.62 | 0.65 | 0.72 | 0.54 | 0.49 | 0.65 | 1.05 | 0.17 | 0.65 | 0.46 | 0.63 | 0.1325 | 0.225 | 0.02 | 0.59985 | 0.405488 | 0.938887 | 0.999971 | 1 | 0.995869 |
| Stk32c | Q8QZV4 | -0.73 | -0.32 | -0.66 | -1.05 | -0.89 | -0.27 | -0.51 | -0.56 | -0.5 | -0.46 | -0.44 | -0.47 | -0.62 | -0.41 | -0.6 | -0.74 | 0.1325 | 0.2225 | 0.0975 | 0.57323 | 0.288262 | 0.650055 | 0.999971 | 1 | 0.933842 |
| Stk38 | Q91VJ4 | -0.34 | 0.63 | -0.32 | 0.22 | 0.45 | 0.33 | -0.17 | -0.07 | 1.02 | 0.82 | 0.29 | 0.36 | 1.01 | 0.19 | -0.16 | 0.75 | 0.0875 | 0.575 | 0.4 | 0.763108 | 0.083037 | 0.269322 | 0.999971 | 1 | 0.843504 |
| Stk38l | Q7TSE6 | -0.33 | 0.22 | 1.14 | 0.51 | 0.42 | -0.17 | 1.06 | 0.13 | -0.04 | -0.61 | 0.75 | -0.56 | -0.57 | -0.33 | 0.83 | -0.07 | -0.025 | -0.5 | -0.42 | 0.948522 | 0.252283 | 0.323297 | 0.999971 | 1 | 0.85548 |
| Stk39 | Q9Z1W9 | 0.17 | 1.09 | 0.52 | 0.19 | 0.41 | 0.78 | 0.19 | 0.2 | 0.47 | 0.82 | -0.11 | -0.78 | 0.45 | 0.74 | 0.43 | 0.29 | -0.0975 | -0.3925 | -0.015 | 0.72204 | 0.334593 | 0.953828 | 0.999971 | 1 | 0.998056 |
| Stk4 | Q9JI11 | 0.67 | 0.59 | -0.08 | 0.2 | 0.62 | 0.49 | -0.37 | 0.21 | 0.86 | 0.26 | 0.15 | 0.47 | 0.94 | 0.52 | 0.19 | 0.48 | -0.1075 | 0.09 | 0.1875 | 0.713695 | 0.727335 | 0.47494 | 0.999971 | 1 | 0.894044 |
| Stmn1 | P54227 | 1.56 | 0.99 | 0.13 | -0.41 | 0.21 | 1.25 | 0.75 | -0.17 | 0.93 | 1.58 | -0.34 | 0.26 | 0.26 | -0.08 | 0.21 | -0.15 | -0.0575 | 0.04 | -0.5075 | 0.908106 | 0.942753 | 0.254193 | 0.999971 | 1 | 0.839012 |
| Stmn2 | P55821 | -1.31 | -0.07 | -0.23 | -0.34 | -0.76 | -0.75 | 0.18 | -0.03 | -0.42 | -0.71 | -0.11 | 0.01 | 0.41 | -0.08 | 0.12 | 0.2 | 0.1475 | 0.18 | 0.65 | 0.684565 | 0.58173 | 0.058187 | 0.999971 | 1 | 0.725348 |
| Stmn3 | O70166 | 0.1 | -1 | -0.66 | -0.49 | -0.65 | -0.25 | 0.06 | -0.49 | 0.71 | 0.36 | -0.15 | -0.37 | 0 | 0.02 | -0.26 | -0.37 | 0.18 | 0.65 | 0.36 | 0.538194 | 0.078748 | 0.204761 | 0.999971 | 1 | 0.81617 |
| Stmn4 | Q05DI3 | -2.57 | -0.83 | 1 | -0.18 | -1.18 | -1.35 | 0.55 | 1.15 | -0.35 | -2.21 | 0.97 | 0.87 | -0.69 | -0.64 | 0.69 | 0.9 | 0.4375 | 0.465 | 0.71 | 0.617584 | 0.625206 | 0.370056 | 0.999971 | 1 | 0.869526 |
| Stn1 | Q3UXU8 | 0.37 | 1.62 | 2.11 | 2.39 | 1.33 | 1.64 | 1.24 | 2.25 | 1.38 | 0.96 | 1.56 | 2.34 | 1.31 | 1.78 | 1.92 | 2.4 | -0.0075 | -0.0625 | 0.23 | 0.987216 | 0.899697 | 0.62537 | 0.999971 | 1 | 0.930479 |
| Stom | P54116 | -1.27 | -1.46 | -0.05 | 0.96 | 0.06 | -1.51 | 0.84 | -0.16 | -1.47 | -1.01 | 0.02 | 0.38 | -2.37 | -1.58 | -0.18 | 0.4 | 0.2625 | -0.065 | -0.4775 | 0.699203 | 0.920347 | 0.53776 | 0.999971 | 1 | 0.910077 |
| Stoml2 | Q99JB2 | -0.37 | -0.87 | -0.46 | -0.5 | -1.09 | -0.88 | -0.47 | -0.41 | -1.12 | -0.81 | -0.23 | -0.41 | -1.2 | -0.61 | -0.44 | -0.19 | -0.1625 | -0.0925 | -0.06 | 0.492259 | 0.715408 | 0.820336 | 0.999971 | 1 | 0.975761 |
| Ston2 | Q8BZ60 | 0 | -0.73 | -0.19 | -0.67 | -0.45 | -0.26 | -0.34 | -0.1 | -0.34 | 0.02 | -0.3 | -0.39 | 0.41 | -0.38 | -0.2 | -0.41 | 0.11 | 0.145 | 0.2525 | 0.636333 | 0.54108 | 0.376336 | 0.999971 | 1 | 0.870465 |
| Stox2 | Q499E5 | -0.08 | 0.09 | 0.29 | 0.74 | 0.17 | -0.16 | 0.19 | 1.9 | 0.04 | -0.04 | 0.2 | 0.6 | 0.81 | 0.12 | 0.74 | 0.53 | 0.265 | -0.06 | 0.29 | 0.572503 | 0.811968 | 0.281759 | 0.999971 | 1 | 0.844538 |
| Strada | Q3UUJ4 | -0.87 | -1.38 | -0.23 | -0.57 | -1.02 | -1.35 | -0.38 | -0.77 | -1.14 | -0.88 | -0.28 | -0.21 | -1.61 | -1.16 | -0.5 | -0.02 | -0.1175 | 0.135 | -0.06 | 0.714679 | 0.685651 | 0.883152 | 0.999971 | 1 | 0.986597 |
| Stradb | Q8K4T3 | 0.76 | 0.09 | -0.85 | -0.18 | -0.46 | 0.24 | -0.16 | -0.48 | -0.27 | 1.13 | -0.6 | -0.32 | 0.53 | 0.16 | -0.59 | -1.21 | -0.17 | 0.03 | -0.2325 | 0.642061 | 0.950023 | 0.62879 | 0.999971 | 1 | 0.932176 |
| Strap | Q9Z1Z2 | -0.44 | 0.42 | 0.67 | -0.65 | 0.49 | 0.51 | 0.88 | -0.13 | 0.67 | -0.31 | 1.23 | 1.39 | 0.59 | 0.5 | 0.48 | -0.17 | 0.4375 | 0.745 | 0.35 | 0.258468 | 0.139865 | 0.342748 | 0.999971 | 1 | 0.861474 |
| Strbp | Q91WM1 | 0.11 | 1.32 | 1.63 | 1.38 | 1.25 | 1.31 | 1.93 | 1.14 | 1.29 | 0.48 | 1.75 | 1.21 | 1.41 | 1.04 | 1.2 | 1.32 | 0.2975 | 0.0725 | 0.1325 | 0.432293 | 0.859753 | 0.702198 | 0.999971 | 1 | 0.949019 |
| Strip1 | Q8C079 | -1.19 | -0.6 | -0.65 | -0.89 | -0.6 | -0.77 | -0.67 | -0.73 | -0.75 | -1.33 | -0.52 | -0.51 | -0.55 | -0.74 | -0.82 | -0.78 | 0.14 | 0.055 | 0.11 | 0.48806 | 0.83097 | 0.592644 | 0.999971 | 1 | 0.923227 |
| Strip2 | Q8C9H6 | 0.4 | -0.05 | -0.45 | 0.58 | -0.02 | -0.4 | 0 | 0.14 | -0.14 | 0.17 | -0.14 | 0.12 | -0.11 | -0.32 | 0.47 | -0.22 | -0.19 | -0.1175 | -0.165 | 0.496918 | 0.65974 | 0.586197 | 0.999971 | 1 | 0.920418 |
| Strn | O55106 | -0.91 | -0.01 | 0.08 | -0.23 | 0.23 | -0.61 | 0.11 | 0.02 | 0.31 | -0.89 | 0.27 | 0.75 | 0.55 | -0.33 | 0.09 | -0.04 | 0.205 | 0.3775 | 0.335 | 0.500613 | 0.35704 | 0.279925 | 0.999971 | 1 | 0.843812 |
| Strn3 | Q9ERG2 | -0.49 | -0.29 | 0.05 | -0.06 | -0.6 | -0.49 | -0.05 | -0.2 | -0.53 | -0.57 | -0.26 | 0.15 | -0.73 | -0.68 | -0.38 | -0.08 | -0.1375 | -0.105 | -0.27 | 0.535478 | 0.659068 | 0.261672 | 0.999971 | 1 | 0.842262 |
| Strn3_1 | B2RQS1 | -0.12 | 0.49 | -0.46 | -1.03 | 0.14 | 0.44 | -0.61 | -0.45 | 0.12 | -0.76 | -0.22 | -0.39 | 0.54 | 0.57 | -0.57 | -0.3 | 0.16 | -0.0325 | 0.34 | 0.680543 | 0.92779 | 0.417674 | 0.999971 | 1 | 0.877584 |
| Strn4 | P58404-2 | -0.53 | -0.17 | -0.55 | -0.45 | -0.14 | -0.32 | -0.72 | -0.42 | -0.07 | -0.36 | -0.47 | -0.2 | -0.01 | -0.39 | -0.59 | -0.05 | 0.025 | 0.15 | 0.165 | 0.902634 | 0.433425 | 0.447949 | 0.999971 | 1 | 0.885289 |
| Sts | P50427 | 1.47 | -0.64 | -0.42 | -0.63 | -0.52 | -0.86 | -0.85 | -0.71 | -0.6 | 1.47 | 0.71 | -0.62 | 0.16 | -0.3 | -0.25 | -0.37 | -0.68 | 0.295 | -0.135 | 0.181922 | 0.657511 | 0.782586 | 0.999971 | 1 | 0.965859 |
| Stt3a | P46978 | -0.21 | 0.85 | 0.63 | 0.4 | 1.54 | 1.03 | 1.23 | 0.76 | 1.87 | 0.38 | 1.14 | 1.32 | 2.26 | 0.49 | 0.87 | 0.63 | 0.7225 | 0.76 | 0.645 | 0.033325 | 0.06777 | 0.168121 | 0.999971 | 1 | 0.800215 |
| Stt3b | A0A0R4J0D | 0.43 | -0.4 | -0.23 | 0.51 | -0.02 | -0.05 | 0.64 | 0.1 | -0.07 | 0.22 | 0.01 | -0.24 | -0.19 | 0 | -0.33 | -0.2 | 0.09 | -0.0975 | -0.2575 | 0.75815 | 0.716437 | 0.340829 | 0.999971 | 1 | 0.860715 |
| Stub1 | Q9WUD1 | -0.21 | 0.11 | 0.01 | 0.25 | 0.3 | 0.28 | 0.35 | 0.54 | 0.34 | 0.1 | 0.27 | -0.05 | 0.84 | 0.18 | 0.54 | 0.41 | 0.3275 | 0.125 | 0.4525 | 0.104974 | 0.518539 | 0.061506 | 0.999971 | 1 | 0.725348 |
| Stum | A0A1B0GS( | -1.33 | -1.64 | -1.22 | -1.15 | -1.34 | -1.31 | -1.21 | -0.94 | -1.53 | -1.35 | -1.05 | -1.55 | -1.29 | -1.71 | -1.34 | -1.27 | 0.135 | -0.035 | -0.0675 | 0.504937 | 0.865738 | 0.742764 | 0.999971 | 1 | 0.961064 |
| Stx12 | Q9ER00 | -1 | -0.89 | -1.29 | -1.4 | -0.66 | -0.84 | -0.59 | -1.36 | -0.57 | -0.93 | -0.98 | -0.81 | -0.36 | -0.83 | -1.42 | -1.65 | 0.2825 | 0.3225 | 0.08 | 0.262928 | 0.137914 | 0.801973 | 0.999971 | 1 | 0.971467 |
| Stx16 | Q8BVI5 | 0.1 | 0.07 | -0.19 | -0.48 | -0.04 | 0.05 | -0.06 | -0.13 | 0.21 | -0.4 | -0.28 | -0.07 | 0.2 | -0.05 | -0.15 | -0.28 | 0.08 | -0.01 | 0.055 | 0.689046 | 0.964801 | 0.799561 | 0.999971 | 1 | 0.970592 |
| Stx17 | Q9D0I4 | 0.19 | 0.06 | -0.08 | -0.18 | 0.24 | 0.33 | -0.29 | 0.11 | 0.67 | 0.26 | -0.6 | 0 | 1.29 | 0.55 | 0.09 | 1.07 | 0.1 | 0.085 | 0.7525 | 0.635587 | 0.769152 | 0.028402 | 0.999971 | 1 | 0.678474 |
| Stx18 | Q8VDS8-3 | 0.51 | 0.15 | -0.05 | 0.43 | 0.04 | 0.12 | 0.94 | 0.35 | 0.16 | 0.29 | 0.07 | 0.56 | 0.31 | 0.08 | 0.26 | -0.09 | 0.1025 | 0.01 | -0.12 | 0.69785 | 0.96242 | 0.570998 | 0.999971 | 1 | 0.91816 |
| Stx1a | O35526 | -1.09 | 0.2 | -0.5 | -0.97 | 0.08 | -0.42 | -0.56 | -0.78 | 0.04 | -1.15 | -0.31 | -0.18 | 0.98 | -0.73 | -0.83 | -0.68 | 0.17 | 0.19 | 0.275 | 0.620369 | 0.618591 | 0.575493 | 0.999971 | 1 | 0.918194 |
| Stx1b | P61264 | -0.68 | -1.33 | -1.89 | -1.31 | -1.39 | -1.23 | -1.63 | -2.01 | -1.22 | -0.85 | -1.91 | -2.02 | -1.23 | -1.48 | -2.05 | -2.41 | -0.2625 | -0.1975 | -0.49 | 0.401413 | 0.590777 | 0.194018 | 0.999971 | 1 | 0.810547 |
| Stx2 | Q3TJ55 | -0.11 | -0.37 | -0.06 | -0.33 | -0.29 | -0.54 | -0.05 | 0.18 | -0.07 | -0.27 | -0.2 | 0.06 | -0.08 | -0.59 | -0.1 | -0.14 | 0.0425 | 0.0975 | -0.01 | 0.845675 | 0.589548 | 0.960489 | 0.999971 | 1 | 0.998721 |
| Stx3 | A0A1B0GR | 0.37 | 0.21 | -0.69 | 0.06 | 2.25 | -0.34 | -0.42 | -0.76 | 0.36 | 0.36 | -0.51 | 0.38 | 0.58 | 0.19 | 0.29 | -0.42 | 0.195 | 0.16 | 0.1725 | 0.769673 | 0.62189 | 0.591124 | 0.999971 | 1 | 0.922461 |
| Stx4 | P70452 | -0.15 | 1.72 | -1.16 | -0.79 | -0.19 | 0.06 | -1.13 | -1.05 | -0.05 | 0.37 | -0.84 | -1.07 | 0.08 | -0.32 | -0.61 | -0.95 | -0.4825 | -0.3025 | -0.355 | 0.459823 | 0.648553 | 0.568666 | 0.999971 | 1 | 0.91816 |
| Stx5 | Q8K1E0 | -0.7 | 0.31 | 0.3 | 0.51 | 0.48 | -0.24 | -0.1 | 0.34 | 0.14 | -0.04 | 0.28 | 0.33 | 0.34 | 0.12 | 0.42 | 0.51 | 0.015 | 0.0725 | 0.2425 | 0.962993 | 0.805623 | 0.421506 | 0.999971 | 1 | 0.87788 |
| Stx6 | Q9JKK1-2 | -1.18 | -0.56 | -0.56 | -0.32 | -0.71 | -0.92 | -1.1 | -0.56 | -0.68 | -0.86 | -0.42 | -0.36 | -0.49 | -0.58 | -0.31 | -0.17 | -0.1675 | 0.075 | 0.2675 | 0.504527 | 0.760342 | 0.28143 | 0.999971 | 1 | 0.844538 |
| Stx7 | Q8BH40 | 0.25 | -0.45 | -0.6 | -1.21 | -0.52 | -0.35 | -0.54 | -0.68 | 0.16 | -0.08 | -0.32 | -0.87 | 0.22 | -0.72 | -0.22 | -0.79 | -0.02 | 0.225 | 0.125 | 0.948835 | 0.5402 | 0.736902 | 0.999971 | 1 | 0.959447 |
| Stx8 | O88983 | 0.41 | 0.71 | 0.53 | 0.56 | 0.63 | 1.12 | 0.34 | 1.2 | 0.63 | 0.81 | 0.55 | 0.62 | 0.78 | 1.11 | 1.01 | 1.36 | 0.27 | 0.1 | 0.5125 | 0.285772 | 0.558547 | 0.027019 | 0.999971 | 1 | 0.678474 |
| Stxbp1 | O08599 | -0.96 | -0.78 | -1.34 | -1.38 | -1.11 | -0.88 | -1.41 | -1.46 | -0.98 | -0.94 | -1.74 | -1.98 | -1.04 | -1.02 | -1.64 | -1.46 | -0.1 | -0.295 | -0.175 | 0.671904 | 0.350759 | 0.480586 | 0.999971 | 1 | 0.89485 |
| Stxbp1_1 | O08599-2 | -1.45 | -2.75 | -3.14 | -3.41 | -2.53 | -3.25 | -2.73 | -3.41 | -2.82 | -2.94 | -3.17 | -3.68 | -3.67 | -2.58 | -2.65 | -3.6 | -0.2925 | -0.465 | -0.4375 | 0.522526 | 0.314096 | 0.382112 | 0.999971 | 1 | 0.871816 |
| Stxbp2 | Q64324 | 0.1 | 0.33 | -3.22 | -2.25 | 0.54 | 0 | -3 | -1.87 | 0.52 | 0.22 | -1.39 | -2.79 | 1.04 | 0.56 | -1.56 | -1.88 | 0.1775 | 0.4 | 0.8 | 0.868155 | 0.703202 | 0.44213 | 0.999971 | 1 | 0.883386 |
| Stxbp3 | Q60770 | -0.45 | -1.32 | -0.7 | -1.08 | -1.46 | -1.27 | -1.04 | -1.3 | -1.49 | -1.17 | -1.14 | -0.76 | -1.66 | -1.35 | -0.68 | -1.05 | -0.38 | -0.2525 | -0.2975 | 0.144511 | 0.352816 | 0.329082 | 0.999971 | 1 | 0.856525 |
| Stxbp4 | Q9WV89-4 | -1 | -0.36 | -0.53 | -0.1 | -1.26 | -1.01 | -0.37 | -0.6 | -1.14 | -0.39 | -0.37 | 0.02 | -3.01 | -0.73 | -1.48 | 0.03 | -0.3125 | 0.0275 | -0.8 | 0.294386 | 0.929657 | 0.216525 | 0.999971 | 1 | 0.826893 |
| Stxbp5 | D3Z079 | -0.19 | -0.28 | 0.09 | -0.53 | -0.54 | 0.13 | -0.29 | -0.21 | -0.3 | 0.03 | -0.24 | -0.42 | 0.11 | -0.07 | -0.16 | -0.13 | 1.57E-16 | -0.005 | 0.165 | 1 | 0.980769 | 0.420012 | 1 | 1 | 0.87788 |
| Stxbp5l | Q5DQR4 | -1.18 | -0.24 | -0.15 | -0.59 | -0.63 | -0.17 | -0.34 | -0.56 | -0.69 | -0.98 | -0.73 | -0.89 | -0.29 | -0.69 | -0.64 | -0.57 | 0.115 | -0.2825 | -0.0075 | 0.675905 | 0.299914 | 0.977831 | 0.999971 | 1 | 0.99925 |
| Stxbp5l_1 | Q5DQR4-3 | -0.3 | -1.77 | -0.77 | -0.65 | -0.18 | -0.06 | -1.07 | -0.16 | -0.5 | -2.21 | -1.06 | -0.15 | -2.35 | 0.24 | -0.12 | -0.55 | 0.505 | -0.1075 | 0.1775 | 0.206314 | 0.833606 | 0.767509 | 0.999971 | 1 | 0.962472 |
| Stxbp6 | Q3TYA4 | -0.31 | 0.02 | 0.69 | -0.18 | 0.27 | 0.5 | 0.5 | 0.37 | 0.25 | -0.29 | 0.77 | 0.45 | -0.09 | -0.24 | 0.48 | -0.43 | 0.355 | 0.24 | -0.125 | 0.187409 | 0.456751 | 0.682513 | 0.999971 | 1 | 0.943204 |
| Sub1 | P11031 | 0.68 | 1.98 | 1.33 | 1.86 | 2.14 | 2.18 | 1.03 | 2.46 | 2.02 | 0.87 | 1.79 | 2.17 | 2.33 | 2.6 | 2.37 | 2.64 | 0.49 | 0.25 | 1.0225 | 0.253556 | 0.535595 | 0.009262 | 0.999971 | 1 | 0.613095 |
| Sucla2 | Q9Z2I9 | -0.74 | -0.16 | -0.93 | -1.55 | -0.29 | -0.83 | -0.51 | -1.55 | -0.27 | -1.14 | -0.62 | -0.35 | 0.03 | -0.81 | -1.43 | -1.84 | 0.05 | 0.25 | -0.1675 | 0.895968 | 0.473143 | 0.720099 | 0.999971 | 1 | 0.954623 |
| Suclg1 | Q9WUM5 | -1.38 | 0.13 | -0.44 | -0.68 | 0.02 | -0.44 | 0.23 | -0.9 | 0.4 | -0.92 | -0.86 | -0.48 | 0.77 | -0.46 | -0.62 | -0.81 | 0.32 | 0.1275 | 0.3125 | 0.41655 | 0.75962 | 0.490717 | 0.999971 | 1 | 0.898612 |
| Suclg2 | Q9Z2I8 | -0.99 | -1.06 | -0.85 | -0.68 | -1.17 | -1.18 | -0.98 | -0.86 | -1.29 | -1.22 | -1.16 | -1.15 | -1.03 | -1.18 | -1.01 | -0.79 | -0.1525 | -0.31 | -0.1075 | 0.421459 | 0.099061 | 0.570278 | 0.999971 | 1 | 0.91816 |
| Suco | Q8C341 | -1.55 | -0.27 | 0.14 | -0.49 | -0.67 | -1.55 | 0.21 | -0.52 | -0.82 | -1.48 | -0.33 | -0.2 | 0.43 | -1.34 | -1 | -0.62 | -0.09 | -0.165 | -0.09 | 0.849912 | 0.707171 | 0.854239 | 0.999971 | 1 | 0.982125 |
| Suds3 | A0A0R4J24 | -0.36 | 1.1 | 0.62 | 1.15 | 1.02 | 0.96 | 0.06 | 0.62 | 0.44 | 0.73 | 0.38 | 0.53 | 0.56 | 0.74 | 0.22 | 0.99 | 0.0375 | -0.1075 | -9.81E-17 | 0.924489 | 0.760522 | 1 | 0.999971 | 1 | 1 |
| Sugct | Q7TNE1 | 0.75 | -1.17 | -0.54 | -0.95 | -1.18 | -0.69 | -0.53 | -0.89 | -1.06 | 0.11 | -1.36 | -1.78 | -1.77 | -1.51 | -1.52 | -2.34 | -0.345 | -0.545 | -1.3075 | 0.4282 | 0.332561 | 0.015691 | 0.999971 | 1 | 0.613095 |
| Sugp1 | Q8CH02 | 3.13 | 1.3 | 1.93 | 1.56 | 1.19 | 1.01 | 2.06 | 1.62 | 1.16 | 1.73 | 1.28 | 1.72 | 0.13 | 0.39 | 0.96 | 1.32 | -0.51 | -0.5075 | -1.28 | 0.265315 | 0.237983 | 0.019587 | 0.999971 | 1 | 0.635388 |
| Sugp2 | Q8CH09 | 0.06 | 0.13 | 0.45 | 0.06 | 0.22 | 0.11 | 0.23 | 0.28 | 0.26 | 0.27 | 0.27 | 0.47 | 0.27 | 0.21 | 0.23 | 0.21 | 0.035 | 0.1425 | 0.055 | 0.84542 | 0.434748 | 0.757649 | 0.999971 | 1 | 0.961116 |
| Sugt1 | Q9CX34 | 0.1 | 0.7 | 0.3 | 0.5 | 0.81 | 0.71 | 0.29 | 0.34 | 0.86 | 0.15 | 0.44 | 0.2 | 0.46 | 0.43 | 0.23 | 0.3 | 0.1375 | 0.0125 | -0.045 | 0.544817 | 0.958117 | 0.821908 | 0.999971 | 1 | 0.975833 |
| Sulf2 | Q8CFG0 | -0.24 | 0.5 | -0.08 | -1.18 | 0.41 | 0.37 | 0.31 | -0.65 | 0.28 | 0.36 | 0.44 | 0.66 | 0.98 | 0.43 | -0.73 | -1.16 | 0.36 | 0.685 | 0.13 | 0.391051 | 0.078576 | 0.815927 | 0.999971 | 1 | 0.974929 |
| Sult1a1 | Q91W19 | 0.48 | -1.1 | 0.68 | 1.34 | 1.15 | 0.37 | 0.53 | 2.17 | 1.77 | 1.28 | 2.19 | 2.62 | 0.68 | -0.22 | -0.07 | 2.76 | 0.705 | 1.615 | 0.4375 | 0.259566 | 0.015595 | 0.575584 | 0.999971 | 1 | 0.918194 |
| Sult4a1 | P63046 | -0.53 | -0.03 | 0.33 | -0.36 | -0.32 | -0.57 | 0.56 | -0.31 | -0.32 | -0.19 | 0.23 | 0.25 | -0.39 | -0.35 | 0.05 | -0.16 | -0.0125 | 0.14 | -0.065 | 0.968361 | 0.594166 | 0.791802 | 0.999971 | 1 | 0.968915 |
| Sumf2 | Q8BPG6 | 0.54 | 0.39 | 0.6 | 0.77 | 0.04 | 0.11 | 0.01 | 1.04 | 0.06 | 0.87 | 0.36 | 0.82 | -0.39 | 0.39 | 0.5 | 1.38 | -0.275 | -0.0475 | -0.105 | 0.333895 | 0.842757 | 0.772335 | 0.999971 | 1 | 0.963498 |
| Sumo1 | P63166 | 1.43 | -0.71 | 0.48 | 0.5 | 0.13 | -0.29 | 0.9 | 0.51 | 0.06 | 0.49 | 0.09 | -0.09 | -1.33 | -0.19 | 0.32 | -0.17 | -0.1125 | -0.2875 | -0.7675 | 0.812179 | 0.511164 | 0.164562 | 0.999971 | 1 | 0.800215 |
| Sumo2 | H7BWX9 | -0.33 | 1.29 | 1.58 | 0.87 | 1.38 | 0.8 | 1.75 | 1.15 | 1.02 | -0.16 | 1.38 | 1.75 | 0.81 | 1.08 | 0.52 | 0.53 | 0.4175 | 0.145 | -0.1175 | 0.353659 | 0.790519 | 0.779517 | 0.999971 | 1 | 0.965847 |
| Sun1 | Q9D666-3 | -0.26 | 0.02 | -0.27 | -0.63 | 0.01 | -0.18 | -0.17 | -0.51 | 0.03 | -0.17 | -0.15 | -0.57 | 0.63 | -0.32 | -0.43 | -0.2 | 0.0725 | 0.07 | 0.205 | 0.739325 | 0.753833 | 0.484141 | 0.999971 | 1 | 0.896352 |
| Sun2 | Q8BJS4-3 | -0.78 | -0.74 | -1 | -0.41 | 0.19 | -0.69 | -1.01 | -0.64 | -0.67 | -0.2 | -0.4 | -0.04 | -0.31 | -0.51 | -0.56 | -0.11 | 0.195 | 0.405 | 0.36 | 0.512385 | 0.095513 | 0.114671 | 0.999971 | 1 | 0.78268 |
| Suox | Q8R086 | -0.14 | -0.85 | -0.59 | -0.49 | -0.55 | -0.89 | -0.49 | -0.42 | -0.68 | -0.53 | -0.7 | -0.95 | -0.52 | -0.2 | -0.84 | -0.7 | -0.07 | -0.1975 | -0.0475 | 0.753622 | 0.369286 | 0.841194 | 0.999971 | 1 | 0.979814 |
| Supt16 | G3X956 | 0.87 | 0.64 | 0 | 0.05 | 0.35 | 0.45 | 0.11 | -0.43 | 0.38 | 1 | -0.51 | -0.3 | 0.34 | 0.44 | -0.11 | -0.31 | -0.27 | -0.2475 | -0.3 | 0.380202 | 0.531048 | 0.31999 | 0.999971 | 1 | 0.853962 |
| Supt20 | A0A0G2JEY | -2.47 | 1.95 | 0.34 | -0.05 | 1.85 | -0.02 | -0.01 | 2.01 | 1.31 | -2.21 | 0.14 | 0.43 | 1.36 | -0.43 | 0.87 | -0.59 | 1.015 | -0.025 | 0.36 | 0.305172 | 0.981199 | 0.698024 | 0.999971 | 1 | 0.947682 |
| Supt4h1b | Q9Z199 | 0 | 0.54 | 0.8 | 0.04 | 1.55 | 0.7 | 0.73 | 0.01 | 1.54 | 0.09 | 0.02 | 1.29 | 1.48 | 1.3 | 0.32 | 0.31 | 0.4025 | 0.39 | 0.5075 | 0.282698 | 0.365781 | 0.183482 | 0.999971 | 1 | 0.809025 |
| Supt5h | O55201 | 0.58 | 0.91 | 1.02 | 1.2 | 0.88 | 0.87 | 0.68 | 1.24 | 0.85 | 0.7 | 0.98 | 1.03 | 0.46 | 1.01 | 1.08 | 1.35 | -0.01 | -0.0375 | 0.0475 | 0.963615 | 0.852694 | 0.851956 | 0.999971 | 1 | 0.98187 |
| Supt6h | Q62383 | 0.26 | 1.21 | 0.48 | 0.41 | 1.21 | 1.12 | 0.07 | 0.51 | 1.04 | 0.87 | 0.49 | 0.54 | 0.95 | 1.26 | 0.28 | 0.72 | 0.1375 | 0.145 | 0.2125 | 0.686132 | 0.591584 | 0.490273 | 0.999971 | 1 | 0.898612 |
| Supv3l1 | Q80YD1 | -0.71 | -0.6 | -0.27 | -0.54 | -0.72 | -0.6 | -0.46 | -0.35 | -0.81 | -0.87 | -0.49 | -0.52 | -0.72 | -0.6 | -0.29 | -0.33 | -0.0025 | -0.1425 | 0.045 | 0.989521 | 0.469151 | 0.821936 | 0.999971 | 1 | 0.975833 |
| Surf1 | A0A0A6YVI | 0.42 | -0.67 | -0.44 | -0.02 | -0.61 | -0.41 | -0.98 | -0.15 | -0.6 | 0.36 | -0.42 | -0.66 | -0.87 | -0.38 | -0.34 | -0.34 | -0.36 | -0.1525 | -0.305 | 0.255928 | 0.650298 | 0.303595 | 0.999971 | 1 | 0.847402 |
| Surf2 | P09926 | 2.49 | 1.34 | 2.65 | 3.06 | 1.87 | 2.33 | 2.45 | 1.88 | 2.03 | 3.19 | 2.93 | 1.83 | -1.3 | 0.37 | 1.02 | 3.46 | -0.2525 | 0.11 | -1.4975 | 0.515409 | 0.813628 | 0.138973 | 0.999971 | 1 | 0.784948 |
| Surf4 | Q64310 | -0.58 | 1.01 | 0.84 | 0.72 | 0.76 | 0.82 | 0.59 | 1.27 | 0.94 | 0.01 | 1.02 | 0.71 | 1.01 | 0.73 | 1.46 | 1.04 | 0.3625 | 0.1725 | 0.5625 | 0.351103 | 0.676091 | 0.16517 | 0.999971 | 1 | 0.800215 |
| Surf6 | P70279 | 0.13 | -1.02 | -0.68 | -1.27 | -0.88 | -0.88 | -0.74 | -0.71 | -0.77 | -0.61 | -0.55 | -0.46 | -0.86 | -0.57 | -0.92 | -0.99 | -0.0925 | 0.1125 | -0.125 | 0.767731 | 0.721841 | 0.698303 | 0.999971 | 1 | 0.947781 |
| Susd2 | Q9DBX3 | 0.32 | -1.77 | -0.89 | -0.17 | -0.82 | -0.84 | -0.4 | -0.56 | -0.65 | -0.44 | -0.93 | -0.46 | -0.66 | -0.65 | -0.25 | -0.6 | -0.0275 | 0.0075 | 0.0875 | 0.950088 | 0.986473 | 0.842063 | 0.999971 | 1 | 0.979814 |
| Susd5 | G3UW60 | 1.75 | 0.38 | 0.19 | 1.48 | 1.21 | 1.81 | 0.82 | 0.89 | 1.31 | 2.35 | 1.51 | 1.43 | 1.54 | 1.05 | -0.14 | 1.31 | 0.2325 | 0.7 | -0.01 | 0.588456 | 0.132736 | 0.984072 | 0.999971 | 1 | 0.99925 |
| Sv2a | Q9JIS5 | -0.96 | -1.4 | 0.95 | 0.09 | -1.51 | -1.3 | 0.65 | 0.34 | -1.18 | -1.33 | -0.01 | -1.34 | -1.23 | -1.33 | 0.39 | 0.51 | -0.125 | -0.635 | -0.085 | 0.856735 | 0.282774 | 0.897683 | 0.999971 | 1 | 0.989101 |
| Sv2b | Q8BG39 | -0.68 | 0.23 | 0.61 | 0.71 | 0.18 | 0.24 | 0.27 | 0.74 | 0.23 | -0.19 | 0.61 | 0.16 | 0.85 | 0.12 | 1 | 0.78 | 0.14 | -0.015 | 0.47 | 0.680566 | 0.965823 | 0.217428 | 0.999971 | 1 | 0.826893 |
| Sv2c | Q69ZS6 | 0.63 | -1.61 | 0.88 | 1.3 | -0.64 | -1.02 | 0.62 | 1.65 | -0.77 | -0.13 | 0 | -1.58 | -1.58 | -0.43 | 2.32 | 0.84 | -0.1475 | -0.92 | -0.0125 | 0.854011 | 0.199056 | 0.989495 | 0.999971 | 1 | 0.99925 |
| Svbp | A2A7P9 | 1.37 | 0.48 | 0.94 | 0.72 | 0.38 | 1.45 | 1.73 | 1.05 | 0.9 | 1.52 | 1.5 | 1.01 | -0.33 | 0.33 | 0.67 | 0.65 | 0.275 | 0.355 | -0.5475 | 0.432644 | 0.207355 | 0.102215 | 0.999971 | 1 | 0.78268 |
| Svil | Q8K4L2 | 0.21 | -0.02 | -0.57 | -0.35 | 1.02 | 0.15 | -0.41 | -0.74 | 0.45 | 0.25 | -0.43 | -0.06 | 0.08 | -0.08 | -0.65 | -0.42 | 0.1875 | 0.235 | -0.085 | 0.643508 | 0.402172 | 0.746397 | 0.999971 | 1 | 0.961064 |
| Svip | Q3UZP4 | -0.58 | -1.18 | -0.47 | -0.62 | -1.19 | -1 | -0.89 | -0.8 | -1.65 | -1.13 | -0.46 | -0.7 | -1.43 | -1.33 | -0.54 | -0.2 | -0.2575 | -0.2725 | -0.1625 | 0.265243 | 0.390988 | 0.632677 | 0.999971 | 1 | 0.93301 |
| Svop | Q8BFT9 | -0.2 | -0.41 | -0.26 | -0.16 | -0.39 | 0.12 | -0.24 | -0.12 | -0.26 | 0.09 | -0.23 | -0.63 | 0.3 | 0.16 | 0.08 | 0.02 | 0.1 | -3.93E-17 | 0.3975 | 0.599943 | 1 | 0.044243 | 0.999971 | 1 | 0.704422 |
| Swap70 | Q6A028 | 0.84 | 0.51 | 0.08 | 0.34 | 0.75 | 0.64 | 0.62 | -0.4 | 0.54 | 0.97 | -0.01 | -0.18 | 0.54 | 0.85 | -0.07 | -0.26 | -0.04 | -0.1125 | -0.1775 | 0.899212 | 0.718883 | 0.57073 | 0.999971 | 1 | 0.91816 |
| Swi5 | A0A0A0MC | 1.2 | 1.56 | 2.17 | 2.33 | 1.36 | 1.11 | 1.39 | 2.34 | 1.74 | 1.51 | 2.45 | 2.02 | -0.18 | 1.46 | 2.19 | 2.59 | -0.265 | 0.115 | -0.3 | 0.477231 | 0.729144 | 0.624121 | 0.999971 | 1 | 0.930479 |
| Syap1 | Q9D5V6 | 0.21 | 0.56 | 0.08 | 0 | 0.48 | 0.71 | -0.06 | -0.19 | 0.81 | 0.48 | 0.05 | 0.14 | 0.52 | 0.56 | 0.05 | -0.05 | 0.0225 | 0.1575 | 0.0575 | 0.933021 | 0.52067 | 0.80768 | 0.999971 | 1 | 0.973317 |
| Sybu | Q8BHS8-2 | 1.25 | 0.33 | 0.73 | 0.46 | -0.15 | -0.15 | 0.67 | 0.2 | 0.04 | 0.93 | 1.12 | 0.26 | -0.82 | 0.31 | 0.1 | 0.2 | -0.55 | -0.105 | -0.745 | 0.087038 | 0.750296 | 0.047359 | 0.999971 | 1 | 0.710102 |
| Syf2 | Q9D198 | 0.78 | 1.14 | 0.72 | 1.38 | 0.91 | 0.93 | -0.08 | 1.38 | 0.53 | 0.65 | 0.93 | 0.5 | 0.62 | 0.89 | 1.02 | 1.26 | -0.22 | -0.3525 | -0.0575 | 0.524093 | 0.14038 | 0.810157 | 0.999971 | 1 | 0.973855 |
| Syk | E9PWE9 | 1.84 | 0.18 | 1.16 | 2.71 | 0.4 | 0 | 3.62 | 2.35 | -0.15 | 0.95 | 0.29 | 1.43 | -0.32 | 0.8 | 1.81 | -0.34 | 0.12 | -0.8425 | -0.985 | 0.893595 | 0.176622 | 0.168959 | 0.999971 | 1 | 0.80174 |
| Sympk | Q80X82 | 1.56 | 0.87 | 0.49 | 0.56 | 0.87 | 0.98 | 0.43 | 0.33 | 0.65 | 1.13 | 0.56 | 0.81 | 0.03 | 1 | 0.35 | 0.71 | -0.2175 | -0.0825 | -0.3475 | 0.475873 | 0.773777 | 0.300651 | 0.999971 | 1 | 0.847402 |
| Syn1 | O88935 | -0.79 | -1.28 | -0.33 | -1.06 | -1.19 | -1.09 | -0.63 | -1.29 | -1.27 | -1.14 | -1.44 | -0.9 | -0.92 | -1.1 | -0.87 | -1.41 | -0.185 | -0.3225 | -0.21 | 0.499627 | 0.230543 | 0.431229 | 0.999971 | 1 | 0.881611 |
| Syn1_1 | O88935-1 | -0.91 | -1.06 | 1.65 | -0.75 | -0.04 | -0.15 | -0.28 | -0.51 | 0.27 | -0.18 | -1.48 | 0.82 | 1.68 | 0.35 | 0.44 | -0.51 | 0.0225 | 0.125 | 0.7575 | 0.969732 | 0.864733 | 0.302172 | 0.999971 | 1 | 0.847402 |
| Syn2 | Q64332 | -1.22 | -1.42 | -0.83 | -1.21 | -1.44 | -1.19 | -0.69 | -1.07 | -1.39 | -1.31 | -1.16 | -0.98 | -1.21 | -1.02 | -0.86 | -1 | 0.0725 | -0.04 | 0.1475 | 0.757688 | 0.844325 | 0.470528 | 0.999971 | 1 | 0.892725 |
| Syn2_1 | Q64332-2 | 1.22 | 1.64 | 2.21 | 1.78 | 1.14 | 1.66 | 1.27 | 2.38 | 1.4 | 2.09 | 1.58 | -2.17 | 2.91 | 1.95 | 1.91 | 2.2 | -0.1 | -0.9875 | 0.53 | 0.769996 | 0.289221 | 0.118316 | 0.999971 | 1 | 0.78268 |
| Syn3 | Q8JZP2 | -0.23 | -0.85 | -0.61 | -0.65 | -1.04 | -0.98 | -0.71 | -0.89 | -1.19 | -0.99 | -1.1 | -0.79 | -1.01 | -1.2 | -0.74 | -0.93 | -0.32 | -0.4325 | -0.385 | 0.142152 | 0.060629 | 0.095955 | 0.999971 | 1 | 0.78268 |
| Syncrip | A0A0R4J25 | 0.16 | 0.63 | 0.49 | -0.22 | 1.51 | 2.03 | -0.72 | 0.92 | 2.04 | 0.75 | 0.16 | 0.22 | 2.08 | 1.18 | 0.74 | 0.39 | 0.67 | 0.5275 | 0.8325 | 0.261023 | 0.259112 | 0.060559 | 0.999971 | 1 | 0.725348 |
| Syncrip_1 | Q7TMK9 | -1.4 | 0.52 | 0.08 | -0.16 | 0.65 | 0.4 | 0.47 | 0.17 | 0.83 | -0.95 | 0.62 | 0.58 | 0.41 | 0.03 | 0.26 | -0.32 | 0.6625 | 0.51 | 0.335 | 0.12793 | 0.356024 | 0.43301 | 0.999971 | 1 | 0.882137 |
| Syndig1 | A2ANU3 | -0.37 | 0.01 | 0.12 | -0.66 | -0.1 | -0.16 | -0.38 | -0.12 | -0.22 | -0.88 | 0.54 | 0.08 | 0.5 | -0.44 | 0.03 | -0.62 | 0.035 | 0.105 | 0.0925 | 0.87846 | 0.76001 | 0.768423 | 0.999971 | 1 | 0.962535 |
| Syne1 | Q6ZWR6-4 | -0.32 | -0.7 | -0.57 | -0.65 | -0.7 | -0.77 | -0.67 | -0.8 | -0.69 | -0.53 | -0.57 | -0.7 | -0.69 | -0.84 | -0.77 | -0.78 | -0.175 | -0.0625 | -0.21 | 0.332167 | 0.720162 | 0.252477 | 0.999971 | 1 | 0.839012 |
| Syne2 | Q6ZWQ0 | 0.97 | 0 | 0.35 | -0.3 | 0.28 | -0.34 | 0.18 | -0.42 | -0.27 | 0.58 | 0.05 | 0.19 | -0.09 | -0.19 | -0.49 | -0.43 | -0.33 | -0.1175 | -0.555 | 0.326144 | 0.718669 | 0.090438 | 0.999971 | 1 | 0.771991 |
| Syngap1 | F6SEU4 | -1.65 | -1.78 | -1.56 | -1.93 | -1.88 | -1.83 | -1.48 | -1.75 | -1.82 | -1.93 | -1.57 | -1.83 | -1.53 | -1.82 | -1.8 | -1.68 | -0.005 | -0.0575 | 0.0225 | 0.978853 | 0.751816 | 0.901723 | 0.999971 | 1 | 0.990759 |
| Syngap1_1 | A0A0J9YUN | -2.93 | -2.07 | -4.37 | -3.79 | -2.2 | -2.67 | -4.63 | -3.84 | -2.52 | -2.06 | -3.34 | -4.15 | -2.13 | -2.09 | -3.92 | -3.94 | -0.045 | 0.2725 | 0.27 | 0.946904 | 0.664295 | 0.68406 | 0.999971 | 1 | 0.944176 |
| Syngap1_2 | J3QQ18 | -1.12 | -0.86 | -1.5 | -1.6 | -0.94 | -1.02 | -1.71 | -1.53 | -1.28 | -0.86 | -1.03 | -1.55 | -0.7 | -1.12 | -1.65 | -1.1 | -0.03 | 0.09 | 0.1275 | 0.91251 | 0.72252 | 0.647158 | 0.999971 | 1 | 0.933613 |
| Syngap1_3 | A0A0J9YVH | 0.61 | -1.25 | -1.33 | -1.08 | -1.42 | -1.5 | -1.41 | -0.61 | -1.38 | -0.53 | -0.8 | -1.46 | -1.31 | -1.12 | -1.16 | -0.66 | -0.4725 | -0.28 | -0.3 | 0.330943 | 0.563335 | 0.512741 | 0.999971 | 1 | 0.905815 |
| Syngr1 | O55100-2 | -0.63 | -0.33 | 0.08 | 0.05 | -1.19 | 0.39 | 0.2 | -0.1 | -1.29 | 0.01 | 0.5 | -1.38 | -1.21 | 0.23 | -1.06 | -0.59 | 0.0325 | -0.3325 | -0.45 | 0.931429 | 0.484621 | 0.229973 | 0.999971 | 1 | 0.828687 |
| Syngr2 | H3BJY1 | 0.44 | 1.92 | 0.55 | 0.79 | 1.66 | 2.62 | 0.2 | 1.29 | 1.49 | 1.58 | 0.11 | 0.83 | 0.54 | 2.34 | 0.68 | 0.7 | 0.5175 | 0.0775 | 0.14 | 0.363238 | 0.864295 | 0.782246 | 0.999971 | 1 | 0.965859 |
| Syngr3 | Q8R191 | 0.74 | 0.76 | 1.51 | 1.28 | -0.04 | 1.69 | 1.32 | 1.38 | 0.25 | 1.39 | 1.38 | -0.24 | -0.13 | 1.08 | 1.06 | 1.18 | 0.015 | -0.3775 | -0.275 | 0.970725 | 0.391025 | 0.449919 | 0.999971 | 1 | 0.885596 |
| Synj1 | F7BQW7 | -0.28 | -0.08 | -0.44 | -0.93 | -0.27 | -0.28 | -0.24 | -1.12 | -0.27 | -0.53 | -0.61 | -0.98 | -0.52 | -0.44 | -0.96 | -1.09 | -0.045 | -0.165 | -0.32 | 0.877472 | 0.524893 | 0.246337 | 0.999971 | 1 | 0.839012 |
| Synj1_1 | A0A338P6( | 2.05 | -2.99 | -0.7 | -0.71 | -1.03 | -0.38 | -0.98 | -1.84 | -1.41 | 0.72 | -3.36 | -2.13 | -5.46 | 0.38 | -1.08 | -2.07 | -0.47 | -0.9575 | -1.47 | 0.62627 | 0.433805 | 0.319893 | 0.999971 | 1 | 0.853962 |
| Synj1_2 | F6XIX0 | 0.43 | 0.81 | 0.31 | 0.43 | 2.04 | 0.73 | 0.56 | 0.28 | 1.2 | 0.3 | 0.49 | 0.33 | 0.58 | 0.9 | 0.28 | 0.79 | 0.4075 | 0.085 | 0.1425 | 0.310545 | 0.743541 | 0.521626 | 0.999971 | 1 | 0.907272 |
| Synj2 | D3YZB2 | -0.06 | -0.46 | 0.13 | 0.07 | -0.98 | -0.52 | -0.65 | 0.03 | -1.23 | 0.36 | -0.34 | 0.35 | -1.28 | -0.54 | -0.07 | 0.17 | -0.45 | -0.135 | -0.35 | 0.120821 | 0.726501 | 0.321786 | 0.999971 | 1 | 0.85548 |
| Synj2bp | Q9D6K5 | 1.57 | 0.05 | 0.14 | -0.02 | -0.4 | -0.02 | 0.38 | -0.52 | -0.5 | 0.63 | -0.35 | -0.46 | -0.96 | -0.39 | -0.4 | -0.37 | -0.575 | -0.605 | -0.965 | 0.184854 | 0.19293 | 0.033565 | 0.999971 | 1 | 0.689803 |
| Synm | Q70IV5-2 | 1.17 | 0.58 | 0.76 | 1.32 | 1.31 | 1.07 | 0.78 | 0.79 | 0.6 | 1.66 | 1.32 | 1.82 | 0.05 | 0.54 | 0.8 | 0.53 | 0.03 | 0.3925 | -0.4775 | 0.902374 | 0.243752 | 0.091599 | 0.999971 | 1 | 0.773779 |
| Synpo | Q8CC35-3 | -0.81 | -1.8 | -1.3 | -1.18 | -1.65 | -1.63 | -1.23 | -1.39 | -1.86 | -1.35 | -1.55 | -1.7 | -1.88 | -1.68 | -1.55 | -1.25 | -0.2025 | -0.3425 | -0.3175 | 0.433868 | 0.202475 | 0.251171 | 0.999971 | 1 | 0.839012 |
| Synpo2 | D3YVV9 | 0.43 | -0.9 | -0.2 | -0.09 | 0.87 | -0.36 | -0.34 | 0.11 | -0.04 | 1.51 | 0.92 | 1.02 | -1.65 | -0.83 | -0.16 | -0.1 | 0.26 | 1.0425 | -0.495 | 0.502605 | 0.02889 | 0.267666 | 0.999971 | 1 | 0.842677 |
| Synpr | Q8BGN8-2 | 0.21 | -0.17 | -0.01 | -0.27 | -0.92 | 0.17 | -0.59 | -0.09 | -0.7 | 0.04 | -0.34 | -0.5 | -0.32 | -0.3 | -0.17 | 0.28 | -0.2975 | -0.315 | -0.0675 | 0.306085 | 0.187075 | 0.75929 | 0.999971 | 1 | 0.961116 |
| Synrg | Q5SV85-2 | 0 | 0.67 | 0.58 | 0.57 | 0.57 | 0.1 | 0.88 | 0.28 | 0.22 | -0.12 | 0.46 | 0.4 | 0.34 | 0.19 | 0.36 | 0.28 | 0.0025 | -0.215 | -0.1625 | 0.992175 | 0.370661 | 0.446752 | 0.999971 | 1 | 0.884581 |
| Syp | Q62277 | 1.05 | -0.38 | 1 | -1.21 | -0.28 | 0.22 | 5.77 | -0.83 | -0.38 | 0.17 | 1.03 | 2.44 | 0.43 | -1.19 | 0.24 | -1.9 | 1.105 | 0.7 | -0.72 | 0.45168 | 0.362847 | 0.327363 | 0.999971 | 1 | 0.856472 |
| Sypl1 | O09117-2 | 0.39 | 0.72 | 1.6 | 1.48 | 0.46 | 0.82 | 0.35 | 2.03 | 0.2 | 0.57 | 0.37 | 0.87 | -0.07 | 1.06 | 1.39 | 1.38 | -0.1325 | -0.545 | -0.1075 | 0.770652 | 0.122766 | 0.802361 | 0.999971 | 1 | 0.971467 |
| Sys1 | Q78S06 | 0.84 | 1.18 | 2.09 | 2.37 | 0.19 | 0.93 | 1.37 | 2.71 | 0.54 | 1.32 | 1.69 | 1.96 | 1.42 | 2.09 | 1.2 | 2.49 | -0.32 | -0.2425 | 0.18 | 0.588567 | 0.592646 | 0.685796 | 0.999971 | 1 | 0.944874 |
| Syt1 | P46096 | -0.88 | -1.26 | -1.23 | -1.31 | -1.18 | -1.38 | -1.25 | -1.36 | -1.42 | -1.25 | -1.24 | -1.04 | -1.08 | -0.93 | -1.21 | -0.89 | -0.1225 | -0.0675 | 0.1425 | 0.509781 | 0.720935 | 0.462158 | 0.999971 | 1 | 0.890369 |
| Syt10 | Q9R0N4 | -1.15 | -0.96 | -1.1 | -1.05 | 0 | -0.67 | -1.15 | 0.38 | -0.29 | -1.22 | -1.2 | -0.02 | 0.86 | -0.58 | -0.24 | -1.25 | 0.705 | 0.3825 | 0.7625 | 0.063863 | 0.24522 | 0.0973 | 0.999971 | 1 | 0.78268 |
| Syt11 | Q9R0N3 | -0.63 | -0.12 | -0.51 | -0.73 | -0.26 | 0.01 | -0.86 | -0.19 | -0.16 | 0.11 | -0.7 | -0.17 | 0.63 | 0.24 | 0.04 | 0.41 | 0.1725 | 0.2675 | 0.8275 | 0.505183 | 0.290295 | 0.004974 | 0.999971 | 1 | 0.582714 |
| Syt12 | Q920N7 | -0.75 | -1.37 | -1.23 | -0.99 | -1.5 | -0.98 | -1.38 | -1 | -1.47 | -1.07 | -0.98 | -1.23 | -1.07 | -1.13 | -0.95 | -0.75 | -0.13 | -0.1025 | 0.11 | 0.573368 | 0.637718 | 0.605088 | 0.999971 | 1 | 0.924454 |
| Syt13 | Q9EQT6 | -0.19 | -1.19 | -0.53 | -0.01 | -0.71 | -0.04 | 0.14 | 0.16 | -0.4 | 0.02 | 0.11 | -0.29 | 0.21 | 0.12 | -0.28 | -0.03 | 0.3675 | 0.34 | 0.485 | 0.28267 | 0.269234 | 0.124809 | 0.999971 | 1 | 0.78268 |
| Syt16 | Q7TN83-2 | 0.21 | -0.65 | 0.09 | 0.51 | 0.55 | -0.06 | -0.39 | 0.42 | 0.11 | 0.5 | 0.81 | 0.52 | 0.15 | 0.23 | 0.49 | 0.78 | 0.09 | 0.445 | 0.3725 | 0.784263 | 0.15676 | 0.228369 | 0.999971 | 1 | 0.828687 |
| Syt17 | Q920M7 | 0.31 | -1.39 | -1.5 | -1.19 | -1.12 | -1.06 | -0.59 | -1.07 | -1.3 | -0.15 | -1.16 | -1.4 | -0.84 | -1.07 | -0.88 | -0.71 | -0.0175 | -0.06 | 0.0675 | 0.96655 | 0.900213 | 0.868999 | 0.999971 | 1 | 0.983462 |
| Syt2 | A0A0R4J2C | -0.7 | -0.73 | -0.84 | -0.62 | -1.36 | -0.23 | -1.52 | -0.77 | -1.1 | -0.96 | -1.33 | -1.6 | -0.56 | -0.34 | -1.01 | -0.75 | -0.2475 | -0.525 | 0.0575 | 0.425979 | 0.026413 | 0.779375 | 0.999971 | 1 | 0.965847 |
| Syt3 | G3X9Y1 | -0.31 | -0.41 | -0.96 | -1.27 | -0.58 | -0.54 | -0.33 | -1.52 | -0.6 | -0.66 | -0.87 | -1.39 | 0.31 | -0.56 | -1.23 | -1.44 | -0.005 | -0.1425 | 0.0075 | 0.988411 | 0.634942 | 0.986102 | 0.999971 | 1 | 0.99925 |
| Syt4 | P40749 | -0.61 | -0.02 | -0.74 | -0.79 | -0.03 | 0.03 | -1.1 | -0.44 | -0.14 | -0.86 | -0.76 | -1.47 | 0.66 | 0.26 | -0.21 | -0.16 | 0.155 | -0.2675 | 0.6775 | 0.627911 | 0.42029 | 0.038693 | 0.999971 | 1 | 0.700838 |
| Syt5 | Q9R0N5 | -0.26 | -0.4 | -0.76 | -0.6 | -0.19 | -0.4 | -0.73 | -0.72 | 0.13 | -0.07 | -0.66 | -0.58 | 0.45 | -0.3 | -0.49 | -0.43 | -0.005 | 0.21 | 0.3125 | 0.981607 | 0.407734 | 0.258825 | 0.999971 | 1 | 0.839012 |
| Syt6 | Q8C8S6 | 0.23 | -2.65 | -0.81 | -0.95 | -2.39 | -2.24 | -0.07 | -0.89 | -1.68 | -0.18 | -0.84 | 0.3 | -2.87 | -1.79 | -0.61 | -1.76 | -0.3525 | 0.445 | -0.7125 | 0.633723 | 0.512085 | 0.312718 | 0.999971 | 1 | 0.849367 |
| Syt7 | A0A2C9F2( | -2 | -0.59 | -1.73 | -2.07 | -1.72 | -2.11 | -1.53 | -1.55 | -1.34 | -2.57 | -1.66 | -0.66 | -0.07 | -1.37 | -1.66 | -1.48 | -0.13 | 0.04 | 0.4525 | 0.718998 | 0.934901 | 0.347031 | 0.999971 | 1 | 0.864164 |
| Syt7_1 | Q9R0N7-4 | -1.26 | -0.41 | -0.01 | -1.85 | -1.09 | -0.87 | -0.83 | -1.14 | -0.78 | -1.42 | 0.43 | 0.3 | -0.55 | -0.05 | -0.69 | -1.35 | -0.1 | 0.515 | 0.2225 | 0.804037 | 0.369972 | 0.632767 | 0.999971 | 1 | 0.93301 |
| Syt7_2 | A0A498WG | -0.89 | -1.15 | -0.89 | -0.8 | -1.13 | -0.94 | -0.87 | -0.76 | -0.97 | -0.96 | -0.86 | -0.95 | -0.88 | -1.24 | -0.87 | -1.01 | 0.0075 | -0.0025 | -0.0675 | 0.967328 | 0.988116 | 0.71877 | 0.999971 | 1 | 0.953825 |
| Syt9 | Q9R0N9 | 1.62 | 1.2 | -2.74 | -3.27 | -0.87 | -1.13 | -1.33 | -2.65 | -2.21 | -0.57 | -1.47 | -1.02 | -1.52 | -1.53 | -4.47 | -3.38 | -0.6975 | -0.52 | -1.9275 | 0.562455 | 0.663904 | 0.166691 | 0.999971 | 1 | 0.800215 |
| Sytl2 | A0A5F8MP | -1.12 | 0.18 | 0.82 | 0.47 | -0.29 | -0.21 | 0.31 | 0.62 | -0.38 | -0.88 | 0.77 | -0.88 | -0.31 | -0.28 | 0.54 | 1.22 | 0.02 | -0.43 | 0.205 | 0.964252 | 0.428481 | 0.693537 | 0.999971 | 1 | 0.947174 |
| Sytl5 | Q80T23-2 | -0.44 | -0.87 | -0.82 | -0.67 | -0.65 | -1.24 | -1.4 | -0.81 | -1.1 | -1.26 | -1.34 | -0.8 | 0.14 | -1.1 | -0.47 | -0.76 | -0.325 | -0.425 | 0.1525 | 0.19186 | 0.063054 | 0.604232 | 0.999971 | 1 | 0.924454 |
| Syvn1 | E9QQ99 | -1.4 | 0.71 | 1.01 | 0.11 | 0.62 | 1.54 | 1.6 | 0.73 | 0.81 | -0.79 | 1.23 | 0.62 | 1.4 | 0.64 | 0.77 | 0.21 | 1.015 | 0.36 | 0.6475 | 0.091056 | 0.573424 | 0.253073 | 0.999971 | 1 | 0.839012 |
| Szrd1 | A2AA85 | 3.04 | 1.1 | 0.92 | 1.36 | 0.88 | 1.52 | 1.29 | 0.71 | 0.92 | 2.4 | 0.71 | 0.05 | -0.11 | 0.62 | 0.71 | 0.13 | -0.505 | -0.585 | -1.2675 | 0.312696 | 0.369567 | 0.027488 | 0.999971 | 1 | 0.678474 |
| Szt2 | A2A9C3 | 1.99 | 1.85 | 0.79 | 0.67 | 0.82 | -0.02 | -0.24 | -0.55 | 1.91 | 1.37 | 0.52 | 0.44 | 0.7 | -0.6 | -0.57 | -0.46 | -1.3225 | -0.265 | -1.5575 | 0.012448 | 0.571531 | 0.006174 | 0.999971 | 1 | 0.582714 |
| Tab1 | Q8CF89 | -0.53 | -0.04 | 0.18 | -0.23 | -0.23 | 0.08 | -0.01 | 0.44 | 0.04 | -0.35 | 0.05 | 0.22 | -0.02 | -0.12 | 0.11 | 0.17 | 0.225 | 0.145 | 0.19 | 0.358212 | 0.530801 | 0.384285 | 0.999971 | 1 | 0.871816 |
| Tab2 | D3Z564 | -1.16 | 0.18 | -0.37 | -0.34 | 0.12 | 0.02 | 0.41 | -0.11 | 0.31 | -0.68 | 0.51 | -0.27 | -0.59 | -0.19 | -0.35 | -0.33 | 0.5325 | 0.39 | 0.0575 | 0.107682 | 0.314996 | 0.847208 | 0.999971 | 1 | 0.980295 |
| Tab3 | Q571K4 | -0.46 | -0.03 | -0.7 | -0.98 | 0.12 | 0.45 | -0.54 | -0.66 | 0.04 | -0.15 | -0.26 | -0.86 | -0.26 | -0.22 | -0.66 | -0.57 | 0.385 | 0.235 | 0.115 | 0.264427 | 0.427305 | 0.654569 | 0.999971 | 1 | 0.934061 |
| Tacc1 | F8VQ95 | -0.24 | 0.28 | -0.49 | -0.04 | 0.29 | 0.43 | -0.35 | -0.23 | 0.24 | 0.19 | -0.53 | -0.19 | 0.37 | 0.16 | -0.3 | -0.01 | 0.1575 | 0.05 | 0.1775 | 0.563928 | 0.849334 | 0.47779 | 0.999971 | 1 | 0.894438 |
| Tacc2 | E9QL08 | 0.26 | 0.42 | 0.34 | -0.42 | 0.22 | 0.44 | -0.24 | 0.53 | 0.74 | 0.78 | -0.14 | 0.09 | 0.11 | 0.63 | 0.3 | 0.57 | 0.0875 | 0.2175 | 0.2525 | 0.751302 | 0.484932 | 0.33501 | 0.999971 | 1 | 0.858827 |
| Tacc2_1 | E9Q8T1 | -0.15 | -0.52 | 0.16 | 0.04 | -0.74 | -0.54 | -0.36 | -0.2 | -0.98 | -0.26 | -0.24 | 0.11 | -0.68 | -0.19 | -0.19 | 0.22 | -0.3425 | -0.225 | -0.0925 | 0.158362 | 0.438627 | 0.723489 | 0.999971 | 1 | 0.955258 |
| Taco1 | Q8K0Z7 | 0.29 | -0.33 | -0.85 | -0.82 | -0.86 | -0.3 | -0.31 | -0.8 | -0.4 | 0.23 | -1.09 | -1.25 | -0.23 | -0.16 | -1.26 | -0.93 | -0.14 | -0.2 | -0.2175 | 0.655348 | 0.630218 | 0.559432 | 0.999971 | 1 | 0.915051 |

| | | | | | | | | | | | | | | | | | | | | | | | | | | | |
|---|---|---|---|---|---|---|---|---|---|---|---|---|---|---|---|---|---|---|---|---|---|---|---|---|---|---|---|
| Tacr1 | P30548 | -1.64 | -2.64 | -1.95 | -1.67 | -2.25 | -2.74 | -2.67 | -1.1 | -1.37 | -1.19 | -1.77 | -1.11 | -0.14 | -2.45 | -0.5 | -1.04 | -0.215 | 0.615 | 0.9425 | 0.612453 | 0.057151 | 0.095642 | 0.999971 | | 1 | 0.78268 |
| Tada1 | Q99LM9 | 1.73 | 1.07 | 0.91 | 1.19 | 1.28 | 2.15 | 0.73 | 1.28 | 0.55 | -0.43 | 0.85 | 1.1 | -0.44 | 0.74 | 1.07 | 1.63 | 0.135 | -0.7075 | -0.475 | 0.692221 | 0.082846 | 0.301643 | 0.999971 | | 1 | 0.847402 |
| Taf1 | S4R1B9 | -1.57 | 4.26 | 0.24 | 0.84 | 4.01 | 4.36 | 0.67 | 0.88 | 2.02 | -0.67 | -0.33 | 1.26 | 8.2 | 1.45 | -0.8 | -0.78 | 1.5375 | -0.3725 | 1.075 | 0.286242 | 0.76301 | 0.622702 | 0.999971 | | 1 | 0.929759 |
| Taf10 | Q8K0H5 | 0.98 | 0.15 | 0.86 | 0.82 | 1.14 | 0.56 | 0.08 | 0.98 | 0.84 | 0.93 | 0.23 | 0.69 | 0.52 | 1.36 | 0.83 | 1.19 | -0.0125 | -0.03 | 0.2725 | 0.967602 | 0.909355 | 0.345542 | 0.999971 | | 1 | 0.863523 |
| Taf11 | Q99JX1 | 1.56 | 1.06 | 1.13 | 1.87 | 1.77 | 1.33 | 1.94 | 0.67 | 2.52 | 0.46 | 1.63 | 0.5 | 1.64 | 0.49 | -0.16 | 1.08 | 0.0225 | -0.1275 | -0.6425 | 0.946918 | 0.796858 | 0.144608 | 0.999971 | | 1 | 0.790933 |
| Taf15 | Q8BQ46 | 0.54 | -1.7 | 1.42 | 1.34 | -0.77 | -0.75 | 3.74 | -0.29 | 0.01 | -0.34 | 0.51 | -0.23 | -2.15 | -1.75 | -0.05 | -1.15 | 0.0825 | -0.4125 | -1.675 | 0.943561 | 0.551175 | 0.055339 | 0.999971 | | 1 | 0.725348 |
| Taf2 | B9EJX5 | -1.07 | 2.02 | 0.53 | 0.22 | 0.44 | -0.86 | -0.33 | -0.53 | -0.8 | -0.79 | -0.66 | -0.07 | 0.02 | 1.33 | -0.35 | 0.64 | -0.745 | -1.005 | -0.015 | 0.255628 | 0.123503 | 0.982019 | 0.999971 | | 1 | 0.99925 |
| Taf5 | F8VPY2 | 0.25 | -1.3 | -1.03 | -1.59 | -0.54 | -0.68 | -0.47 | -1.63 | -0.88 | -1.24 | -1.31 | -0.97 | -2.15 | -1.29 | -1.28 | -1.75 | 0.0875 | -0.1825 | -0.7 | 0.848049 | 0.651068 | 0.131853 | 0.999971 | | 1 | 0.784948 |
| Taf5l | Q91WQ5 | 1.89 | 2.17 | 1.79 | 1.24 | 2.74 | 1.97 | 2.78 | 1.17 | 3.14 | 0.97 | 1.8 | 1.36 | 2.17 | 0.64 | 1.75 | 0.72 | 0.3925 | 0.045 | -0.4525 | 0.348325 | 0.924987 | 0.284258 | 0.999971 | | 1 | 0.844538 |
| Taf6 | D3Z0T0 | -1.08 | -2.24 | 1.3 | 2.55 | -1.89 | -2.61 | 1.67 | -0.63 | -1.94 | -2.04 | -0.3 | -2.31 | -3.01 | -2.17 | -1.42 | -1.72 | -0.9975 | -1.78 | -2.2125 | 0.44341 | 0.121501 | 0.055381 | 0.999971 | | 1 | 0.725348 |
| Taf6l | K3W4L1 | -0.59 | -0.53 | 0.4 | 1.32 | -0.66 | -0.8 | 0.42 | 0.5 | -0.67 | -0.75 | 0.87 | 0.45 | -0.64 | -0.6 | 0.39 | 0.87 | -0.285 | -0.175 | -0.145 | 0.589555 | 0.754884 | 0.788622 | 0.999971 | | 1 | 0.96741 |
| Taf8 | Q9EQH4-5 | 2.66 | 0.37 | 0.58 | 1.14 | 0.84 | 0.31 | -1.19 | 1.94 | 2.44 | 1.76 | 1.75 | 1.93 | 2.03 | 1.01 | 1.68 | 1.17 | -0.7125 | 0.7825 | 0.285 | 0.354017 | 0.147518 | 0.590135 | 0.999971 | | 1 | 0.922217 |
| Taf9 | Q8VI33 | 1.39 | 1.86 | 1.38 | 1.58 | 1.14 | 1.42 | 0.96 | 1.29 | 1.25 | 1.38 | 1.21 | 1.09 | 0.68 | 1.28 | 0.75 | 1.42 | -0.35 | -0.32 | -0.52 | 0.11409 | 0.11868 | 0.060894 | 0.999971 | | 1 | 0.725348 |
| Tafa1 | Q7TPG8 | 0.63 | 1.43 | 1.32 | 1.06 | 1.22 | 2.7 | 1.1 | 1.9 | 1.61 | 1.44 | 1.54 | 1.01 | 1.41 | 2.26 | 1.64 | 1.07 | 0.62 | 0.29 | 0.485 | 0.140344 | 0.263245 | 0.147008 | 0.999971 | | 1 | 0.793248 |
| Tafa5 | Q91WE9-2 | 2.91 | -0.95 | -0.75 | 0.56 | -1.45 | -0.01 | 0.2 | 0.12 | -1.45 | 1.33 | -1.68 | -2.4 | -3.24 | -0.76 | -1.72 | -1.78 | -0.7275 | -1.4925 | -2.3175 | 0.412121 | 0.192587 | 0.030741 | 0.999971 | | 1 | 0.678474 |
| Tafazzin | J3JS92 | -2.13 | -1.71 | -0.56 | -0.91 | -1.78 | -2.92 | -0.56 | -1.06 | -1.62 | -2.38 | -0.02 | 0.26 | -2.83 | -2.4 | -1.3 | -0.68 | -0.2525 | 0.3875 | -0.475 | 0.661017 | 0.563971 | 0.40738 | 0.999971 | | 1 | 0.874911 |
| Tagln | P37804 | 0.31 | -0.92 | -1.44 | 0.75 | 7.91 | 5.5 | 0.21 | -1.49 | 1.6 | 4.38 | 5.33 | 7.42 | -1.99 | 1.15 | -0.33 | -0.56 | 3.3575 | 5.0075 | -0.1075 | 0.120485 | 0.002235 | 0.884825 | 0.999971 | 0.814943 | | 0.98711 |
| Tagln2 | Q9WVA4 | -0.44 | 1.64 | 1.33 | 1.16 | 3.82 | 2.91 | 1.36 | 0.33 | 2.38 | 0.78 | 2.02 | 2.32 | 1.17 | 2.34 | 1.58 | 1.09 | 1.1825 | 0.9525 | 0.6225 | 0.171637 | 0.111644 | 0.239359 | 0.999971 | | 1 | 0.833594 |
| Tagln3 | Q9R1Q8 | -0.94 | -0.31 | -0.25 | -0.5 | -0.29 | -0.14 | -0.3 | -0.26 | -0.32 | -0.91 | -0.07 | 0.08 | -0.09 | -0.28 | 0.07 | -0.28 | 0.2525 | 0.195 | 0.355 | 0.249923 | 0.49472 | 0.135648 | 0.999971 | | 1 | 0.784948 |
| Taldo1 | A0A1B0GR | 1.13 | 1.89 | 2.04 | 1.78 | 1.72 | 1.67 | 1.64 | 1.73 | 1.83 | 1.32 | 1.88 | 1.93 | 1.47 | 1.35 | 1.77 | 1.84 | -0.02 | 0.03 | -0.1025 | 0.932499 | 0.909834 | 0.692089 | 0.999971 | | 1 | 0.947174 |
| Tamalin | Q9JJA9 | 1.58 | 0.76 | -0.48 | 0.12 | 0.73 | 1.09 | -0.22 | -0.15 | 0.46 | 1.35 | -0.14 | -0.57 | 0.53 | 0.89 | 0.11 | 0.36 | -0.1325 | -0.22 | -0.0225 | 0.794162 | 0.695381 | 0.959528 | 0.999971 | | 1 | 0.998721 |
| Tamm41 | G5E881 | 1.22 | -0.05 | -0.14 | -0.06 | -0.43 | -0.18 | -0.56 | -0.05 | 0.03 | 0.16 | -0.62 | -0.64 | -0.04 | -0.04 | -0.46 | -0.32 | -0.5475 | -0.51 | -0.4575 | 0.136303 | 0.199326 | 0.201787 | 0.999971 | | 1 | 0.815923 |
| Tanc1 | E9QAF9 | 1.62 | 1.07 | 0.71 | 0.39 | 0.73 | 0.99 | 0.23 | 0.49 | 0.76 | 0.98 | 0.15 | 0.07 | 0.77 | 1.06 | 0.34 | 0.78 | -0.3375 | -0.4575 | -0.21 | 0.298433 | 0.204432 | 0.502538 | 0.999971 | | 1 | 0.904573 |
| Tanc2 | A2A690-2 | -0.84 | -0.66 | -0.55 | -0.92 | -0.82 | -0.3 | -0.16 | -0.69 | -0.59 | -0.5 | -0.39 | -0.54 | -0.36 | -0.43 | -0.71 | -0.87 | 0.25 | 0.2375 | 0.15 | 0.275668 | 0.195286 | 0.467314 | 0.999971 | | 1 | 0.891731 |
| Tanc2_1 | F6VCI2 | 2.27 | 3.24 | 2.23 | -0.44 | 3.02 | 0.9 | 5.73 | 1.09 | 3.25 | 0.02 | 5.68 | 2.72 | 4.73 | -2.06 | 1.17 | -0.45 | 0.86 | 1.0925 | -0.9775 | 0.487015 | 0.395621 | 0.510985 | 0.999971 | | 1 | 0.905815 |
| Tanc2_2 | A0A140LJD | 0.4 | 0.02 | -1.38 | -0.55 | -0.38 | 0.04 | -1.29 | -0.6 | 0.01 | 0.81 | -1.56 | -1.42 | 0.72 | 0.52 | -1.22 | -0.58 | -0.18 | -0.1625 | 0.2375 | 0.68876 | 0.797509 | 0.668967 | 0.999971 | | 1 | 0.938711 |
| Tanc2_3 | E9Q2K2 | -0.13 | 0.9 | -3 | -3.66 | 0.11 | -1.69 | -1.79 | -3.02 | 0.55 | 0.52 | -2.96 | -3.17 | 0.21 | -0.54 | -2.5 | -3.11 | -0.125 | 0.2075 | -0.0125 | 0.912248 | 0.87823 | 0.991716 | 0.999971 | | 1 | 0.99925 |
| Tango2 | Q8BTN3 | -0.23 | 0.4 | 0.42 | 0.67 | 0.25 | 0.7 | 0.79 | 0.99 | 0.24 | -0.02 | 0.91 | -0.09 | 0.41 | 0.82 | 0.77 | 0.83 | 0.3675 | -0.055 | 0.3925 | 0.193161 | 0.85655 | 0.138122 | 0.999971 | | 1 | 0.784948 |
| Tango6 | Q8C3S2-2 | -0.14 | -0.38 | -0.34 | -0.37 | -0.55 | -0.67 | -0.89 | -0.54 | -0.51 | -0.3 | -0.36 | -0.67 | -0.95 | -0.21 | -0.32 | 0.19 | -0.355 | -0.1525 | -0.015 | 0.07396 | 0.397692 | 0.95483 | 0.999971 | | 1 | 0.998056 |
| Tank | A2AR78 | 0.98 | -0.2 | 0.52 | 1.45 | 0.29 | 0.05 | 0.5 | 1.34 | -0.27 | 0.7 | 0.86 | 0.99 | -0.85 | 0.04 | 0.64 | 1.6 | -0.1425 | -0.1175 | -0.33 | 0.738348 | 0.784925 | 0.567672 | 0.999971 | | 1 | 0.91816 |
| Taok1 | Q5F2E8 | -0.13 | -0.21 | 0.08 | 0.1 | -0.24 | -0.18 | 0.36 | 0.19 | -0.14 | -0.09 | 0.49 | 0.61 | 0.13 | -0.38 | 0.19 | 0.06 | 0.0725 | 0.2575 | 0.04 | 0.733859 | 0.298028 | 0.845785 | 0.999971 | | 1 | 0.979924 |
| Taok2 | Q6ZQ29-2 | -0.3 | -0.85 | -1.52 | -1.47 | -0.79 | -1.05 | -1.1 | -1.48 | -0.73 | -0.27 | -1.01 | -0.88 | -1.05 | -0.84 | -1.6 | -1.49 | -0.07 | 0.3125 | -0.21 | 0.82849 | 0.356179 | 0.538817 | 0.999971 | | 1 | 0.910077 |
| Taok3 | Q8BYC6 | 0.4 | 1.22 | 1.6 | 1.58 | 0.82 | 1.09 | 1.39 | 1.56 | 0.91 | 0.39 | 1.55 | 1.4 | 1.06 | 1.19 | 1.03 | 1.62 | 0.015 | -0.1375 | 0.025 | 0.963198 | 0.713374 | 0.936909 | 0.999971 | | 1 | 0.995618 |
| Tapbp | Q3TCU5 | 1.43 | 1.26 | -0.75 | -0.76 | 2.42 | 1.34 | -0.74 | 0.13 | 1.09 | 1.15 | 0.32 | 0.92 | 1.53 | 1.75 | -1.05 | -0.8 | 0.4925 | 0.575 | 0.0625 | 0.554831 | 0.339634 | 0.942129 | 0.999971 | | 1 | 0.996889 |
| Tapbpl | Q8VD31 | 1.88 | -0.87 | -0.18 | 0.33 | -0.16 | -0.55 | -0.18 | -1.28 | -0.63 | 0.67 | -0.47 | -1.37 | -1.83 | -1.05 | -0.41 | -1.22 | -0.8325 | -0.74 | -1.4175 | 0.179198 | 0.278589 | 0.039283 | 0.999971 | | 1 | 0.700838 |
| Tapt1 | Q4VBD2 | -0.59 | -0.67 | -0.39 | -0.33 | -0.53 | -0.53 | -0.58 | 0.02 | -0.62 | -0.28 | -0.21 | -0.11 | -0.06 | -0.47 | -0.13 | 0.17 | 0.09 | 0.19 | 0.3725 | 0.673134 | 0.34261 | 0.100791 | 0.999971 | | 1 | 0.78268 |
| Tarbp2 | D3Z282 | 1.4 | -0.07 | 1.43 | 1.81 | -0.1 | 0.18 | 0.65 | 1.56 | -0.44 | 0.26 | 1.25 | 1.39 | 0.11 | -0.22 | 1.44 | 1.45 | -0.57 | -0.5275 | -0.4475 | 0.281446 | 0.353502 | 0.428893 | 0.999971 | | 1 | 0.881196 |
| Tardbp | Q921F2 | -0.52 | 1.82 | 1.54 | 1.79 | 1.81 | 1.72 | 1.43 | 1.7 | 1.91 | 0.05 | 1.5 | 1.37 | 1.82 | 1.5 | 1.7 | 1.64 | 0.5075 | 0.05 | 0.5075 | 0.34654 | 0.936993 | 0.346003 | 0.999971 | | 1 | 0.863794 |
| Tars1 | Q9D0R2 | 0.43 | 1.08 | 0.78 | 0.91 | 0.97 | 1.07 | 1.07 | 0.74 | 1.14 | 0.88 | 0.98 | 1.04 | 0.88 | 1.17 | 0.95 | 1.03 | 0.1625 | 0.21 | 0.2075 | 0.445578 | 0.311662 | 0.328072 | 0.999971 | | 1 | 0.856472 |
| Tars2 | Q3UQ84 | -0.58 | -0.57 | -0.44 | -0.74 | -0.32 | -0.09 | 0.08 | -0.23 | -0.19 | -0.36 | 0.02 | 0.18 | 0.3 | -0.12 | -0.57 | -0.46 | 0.4425 | 0.495 | 0.37 | 0.035894 | 0.028622 | 0.148158 | 0.999971 | | 1 | 0.793248 |
| Tars3 | Q8BLY2 | 1.17 | 1.11 | 1.15 | 1.17 | 1.05 | 1.27 | 1.48 | 1.03 | 1.34 | 1.21 | 1.62 | 1 | 0.84 | 1.08 | 1.13 | 1.1 | 0.0575 | 0.1425 | -0.1125 | 0.753297 | 0.463404 | 0.509689 | 0.999971 | | 1 | 0.905815 |
| Tasor | Q69ZR9-2 | -0.34 | 0.4 | 0.12 | -0.05 | 0.45 | -0.19 | -0.22 | -0.28 | -0.11 | -0.32 | 0.16 | 0.18 | 0 | 0.22 | -0.18 | -0.29 | -0.0925 | -0.055 | -0.095 | 0.718704 | 0.811497 | 0.680942 | 0.999971 | | 1 | 0.943097 |
| Tasor2 | Q5DTT3 | 0.6 | 0.3 | -1.35 | -1.21 | 2.73 | 1.78 | -2.21 | -2.4 | 0.77 | 2.45 | -1.35 | -1.79 | -0.05 | 1 | -1.99 | -1.37 | 0.39 | 0.435 | -0.1875 | 0.758111 | 0.663318 | 0.804391 | 0.999971 | | 1 | 0.971467 |
| Tatdn1 | Q6P8M1-2 | -0.15 | 0.27 | 0.35 | 0.38 | 0.02 | 0.12 | -0.21 | 0.43 | 0.09 | 0.42 | 0.18 | 0.29 | 0.21 | 0.49 | 0.32 | 0.51 | -0.1225 | 0.0325 | 0.17 | 0.586043 | 0.869336 | 0.407646 | 0.999971 | | 1 | 0.874911 |
| Tax1bp1 | Q3UKC1 | -0.2 | -0.5 | -0.94 | -0.79 | -0.49 | -0.36 | -0.76 | -0.62 | -0.54 | -0.11 | -1.13 | -0.98 | -0.25 | -0.48 | -0.81 | -0.7 | 0.05 | -0.0825 | 0.0475 | 0.824598 | 0.778248 | 0.842771 | 0.999971 | | 1 | 0.979814 |
| Tax1bp3 | Q9DBG9 | 0.41 | -0.04 | -0.44 | 0.15 | 0.18 | -0.58 | 0.45 | -0.34 | -0.12 | -0.66 | 0.11 | 0.25 | -0.24 | -1.43 | -0.47 | -1.09 | -0.0925 | -0.125 | -0.8275 | 0.760731 | 0.659167 | 0.03084 | 0.999971 | | 1 | 0.678474 |
| Tbc1d1 | Q60949 | -0.93 | -0.29 | -0.2 | -0.62 | 0.15 | -1.01 | -0.67 | -0.82 | -0.61 | -0.94 | -0.31 | -0.43 | -0.82 | -1.01 | -0.84 | -0.53 | -0.0775 | -0.0625 | -0.29 | 0.80285 | 0.798143 | 0.232155 | 0.999971 | | 1 | 0.831121 |
| Tbc1d10a | Q5SPX8 | -0.39 | 0.41 | 0.73 | -0.25 | 0.44 | 0.26 | 0.72 | -0.09 | 0.44 | -0.35 | 0.48 | 0.65 | 0.73 | -0.03 | 0.27 | -0.16 | 0.2075 | 0.18 | 0.0775 | 0.519958 | 0.602843 | 0.815307 | 0.999971 | | 1 | 0.974918 |
| Tbc1d10b | Q8BHL3 | -0.68 | -0.43 | -0.33 | -0.74 | -0.42 | -0.43 | -0.44 | -0.79 | -0.39 | -0.78 | -0.64 | -0.82 | -0.17 | -0.53 | -0.77 | -0.8 | 0.025 | -0.1125 | -0.0225 | 0.898191 | 0.568352 | 0.918642 | 0.999971 | | 1 | 0.992633 |
| Tbc1d12 | A0A494BA: | -0.67 | 0.18 | -0.09 | -0.18 | 0.07 | -0.27 | -0.24 | -0.1 | -0.49 | -0.84 | -0.27 | -0.11 | -0.1 | -0.78 | 0.12 | -0.29 | 0.055 | -0.2375 | -0.0725 | 0.812019 | 0.371772 | 0.794634 | 0.999971 | | 1 | 0.970337 |
| Tbc1d13 | Q8R3D1 | -0.89 | 0.79 | 0.54 | 0.09 | 0.8 | 0.59 | 0.35 | 0.46 | 0.9 | -0.25 | 0.69 | 0.64 | 1.36 | 0.7 | 0.63 | 0.72 | 0.4175 | 0.3625 | 0.72 | 0.278223 | 0.405783 | 0.093733 | 0.999971 | | 1 | 0.776618 |
| Tbc1d14 | E9PYB1 | 0.01 | 0.65 | 0.22 | 0.14 | 0.73 | 0.23 | -0.1 | 0.32 | 0.58 | 0 | 0.32 | 0.06 | 0.01 | 0.94 | 0.8 | 0.84 | 0.04 | -0.015 | 0.3925 | 0.872125 | 0.947527 | 0.176534 | 0.999971 | | 1 | 0.808532 |
| Tbc1d15 | Q9CXF4 | 0.39 | 0.77 | 0.62 | 0.85 | 0.61 | 0.44 | 0.34 | 0.94 | 0.47 | 0.51 | 0.72 | 0.51 | 0.63 | 0.88 | 0.82 | 0.68 | -0.075 | -0.105 | 0.095 | 0.726564 | 0.571227 | 0.615702 | 0.999971 | | 1 | 0.927967 |
| Tbc1d17 | Q8BYH7 | 0.65 | 0.33 | 0.85 | 0.01 | 0.43 | 0.65 | 1.01 | -0.04 | 0.54 | 0.05 | 0.65 | 0.44 | 0.77 | 0.31 | 0.4 | 0.03 | 0.0525 | -0.04 | -0.0825 | 0.85936 | 0.873427 | 0.753933 | 0.999971 | | 1 | 0.961116 |
| Tbc1d19 | Q8VDV7 | -0.61 | -1.9 | -1.38 | -1.3 | -0.91 | -1.37 | -1.37 | -0.93 | -1.15 | -1.17 | -1.3 | -0.93 | -0.79 | -1.02 | -0.54 | -1.06 | 0.1525 | 0.16 | 0.445 | 0.616801 | 0.581146 | 0.16341 | 0.999971 | | 1 | 0.800215 |
| Tbc1d20 | Q9D9I4 | -1.56 | -1.72 | -0.67 | -0.61 | -1.37 | -1.39 | -0.86 | -0.39 | -1.48 | -1.67 | -0.88 | -0.13 | -1.42 | -1.36 | -0.45 | 0.11 | 0.1375 | 0.1 | 0.36 | 0.707799 | 0.815995 | 0.426361 | 0.999971 | | 1 | 0.880793 |
| Tbc1d22a | Q8R5A6-2 | 0.32 | 1.14 | 0.52 | 0.43 | 0.65 | 1.07 | 0.38 | 0.65 | 0.7 | 0.48 | 0.73 | 0.5 | 0.58 | 0.82 | 0.63 | 0.42 | 0.085 | -1.18E-16 | 0.01 | 0.741664 | 1 | 0.966285 | 0.999971 | | 1 | 0.998721 |
| Tbc1d22b | Q80VE5 | -1.38 | -0.26 | 0.33 | -0.46 | -0.23 | -0.55 | 0.54 | 0.02 | -0.26 | -0.83 | 0.68 | 0.85 | -0.14 | -0.66 | 0.13 | -0.23 | 0.3875 | 0.5525 | 0.2175 | 0.349827 | 0.283467 | 0.569015 | 0.999971 | | 1 | 0.91816 |
| Tbc1d23 | A0A2I3BRD | 0.11 | 0.79 | 1.08 | 1.01 | 0.42 | 0.81 | 0.41 | 1.15 | 0.79 | 0.92 | 0.63 | 0.36 | 1.33 | 1.34 | 1.28 | 1.58 | -0.05 | -0.0725 | 0.635 | 0.864931 | 0.788454 | 0.033307 | 0.999971 | | 1 | 0.688337 |
| Tbc1d24 | Q3UUG6-2 | -0.81 | -0.5 | -0.6 | -0.69 | -0.44 | -0.61 | -0.66 | -0.63 | -0.57 | -1.07 | -0.66 | -0.42 | -0.38 | -0.37 | -0.46 | -0.37 | 0.065 | -0.03 | 0.255 | 0.707383 | 0.883252 | 0.156641 | 0.999971 | | 1 | 0.796008 |
| Tbc1d25 | A0A6I8MX: | -0.83 | -0.36 | 0.25 | 0.72 | -0.42 | -0.28 | 0.28 | 0.89 | -0.87 | -0.52 | 0.77 | 0.19 | 0.23 | 0.05 | 0.94 | 0.71 | 0.1725 | -0.0525 | 0.5375 | 0.688027 | 0.910796 | 0.185803 | 0.999971 | | 1 | 0.809025 |
| Tbc1d30 | Q69ZT9 | 0.71 | 1.06 | 0.59 | 1.16 | 2.05 | 0.64 | 0.92 | 0.98 | 1.31 | 0.47 | 1.25 | 0.88 | 2.91 | 0.36 | 0.63 | 1.11 | 0.2675 | 0.0975 | 0.3725 | 0.43464 | 0.707802 | 0.498996 | 0.999971 | | 1 | 0.903717 |
| Tbc1d32 | Q3URV1 | -0.04 | -0.17 | 0.05 | 0.64 | 0.03 | -0.04 | -0.42 | 0.47 | 0.05 | 0.06 | -0.2 | 0.52 | -0.35 | -0.01 | 0.43 | 0.54 | -0.11 | -0.0125 | 0.0325 | 0.688819 | 0.961119 | 0.909614 | 0.999971 | | 1 | 0.991822 |
| Tbc1d4 | E9Q6Q8 | 0.86 | 0.96 | 0.1 | 0.24 | 0.94 | 0.99 | -0.64 | 0.25 | 0.3 | 0.87 | -0.2 | 0.09 | 0.56 | 1.43 | 0.86 | 0.41 | -0.155 | -0.275 | 0.275 | 0.71046 | 0.394833 | 0.395501 | 0.999971 | | 1 | 0.871816 |
| Tbc1d5 | A0A286YDI | 1.11 | -0.27 | 0.67 | 0.73 | -0.31 | -0.26 | 0.56 | 0.25 | -0.24 | 0.26 | 0.36 | 0.34 | -0.32 | -0.55 | 0.18 | 0.24 | -0.5 | -0.38 | -0.6725 | 0.181123 | 0.262671 | 0.079225 | 0.999971 | | 1 | 0.755254 |
| Tbc1d7 | Q9D0K0 | -0.16 | 0.38 | -0.23 | -0.24 | 0.15 | -0.21 | -0.61 | -0.07 | 0.43 | 0 | 0.06 | 0.33 | 0.82 | 0.53 | 0.06 | 0.08 | -0.1225 | 0.2675 | 0.435 | 0.622805 | 0.246398 | 0.122439 | 0.999971 | | 1 | 0.78268 |
| Tbc1d8 | Q9Z1A9 | -1.17 | 0.37 | -1.29 | -0.1 | -0.76 | -1.16 | -1.26 | 0.04 | -0.47 | -0.9 | -0.19 | -0.22 | -1.06 | -0.38 | -0.65 | -0.61 | -0.2375 | 0.1025 | -0.1275 | 0.61511 | 0.806195 | 0.756529 | 0.999971 | | 1 | 0.961116 |
| Tbc1d8b | A3KGB4 | -0.54 | 0.18 | -0.01 | -0.41 | -0.31 | -0.19 | -0.45 | -0.5 | -0.18 | -0.12 | -0.54 | -0.76 | -0.96 | 0.15 | -0.36 | -0.11 | -0.1675 | -0.205 | -0.125 | 0.461767 | 0.424384 | 0.678499 | 0.999971 | | 1 | 0.942697 |
| Tbc1d9 | Q3UYK3 | -2.44 | -3.53 | -1.76 | -1.21 | -2.99 | -3.42 | -2.18 | -1.03 | -3.37 | -2.87 | -1.32 | -1.79 | -3.48 | -3.25 | -1.84 | -1.22 | -0.17 | -0.1025 | -0.2125 | 0.796199 | 0.870667 | 0.752369 | 0.999971 | | 1 | 0.961116 |
| Tbc1d9b | Q5SVR0-2 | 0.47 | -0.35 | -0.29 | -1.03 | -0.01 | -0.03 | 0.12 | -0.71 | 0.1 | 0.3 | -0.07 | 0.22 | -0.13 | -0.21 | -0.33 | -0.58 | 0.1425 | 0.4375 | -0.0125 | 0.687107 | 0.194055 | 0.969181 | 0.999971 | | 1 | 0.998787 |
| Tbca | A0A1Y7VJ4 | 1.84 | 1.83 | 0.91 | 0.6 | 2.05 | 1.77 | 0.65 | 0.59 | 1.7 | 2.04 | 0.62 | 0.47 | 2.05 | 1.82 | 0.79 | 1.12 | -0.03 | -0.0875 | 0.15 | 0.94812 | 0.852888 | 0.717408 | 0.999971 | | 1 | 0.952711 |
| Tbcb | Q9D1E6 | 0.1 | 0.22 | 0.82 | 0.74 | -0.25 | 0.31 | 1.44 | 0.48 | -0.05 | -0.16 | 0.46 | 0.36 | -0.13 | 0.44 | 0.33 | 0.54 | 0.025 | -0.3175 | -0.175 | 0.947682 | 0.240064 | 0.504825 | 0.999971 | | 1 | 0.905277 |
| Tbcc | A0A0R4J0N | 1.47 | 0.9 | 0.83 | 0.67 | 0.69 | 0.98 | 0.69 | 1 | 1.05 | 1.13 | 0.89 | 0.19 | 0.9 | 0.71 | 0.85 | 0.69 | -0.1275 | -0.1525 | -0.18 | 0.585377 | 0.598955 | 0.430833 | 0.999971 | | 1 | 0.881611 |
| Tbcd | Q8BYA0 | 0.2 | 0.26 | 0.4 | 0.45 | 0.58 | 0.28 | 0.33 | 0.81 | 0.87 | 0.13 | 0.44 | 0.56 | 1.17 | 0.22 | 0.89 | 1.05 | 0.1725 | 0.1725 | 0.505 | 0.391303 | 0.421255 | 0.068823 | 0.999971 | | 1 | 0.733937 |
| Tbce | Q8CIV8 | -1.04 | -0.98 | 0.93 | 0.55 | -1.17 | -1.25 | 0.73 | 0.86 | -1.12 | -1.02 | 0.75 | 0.37 | -1.23 | -0.97 | 0.53 | 0.66 | -0.0725 | -0.12 | -0.1175 | 0.917277 | 0.851183 | 0.855664 | 0.999971 | | 1 | 0.982398 |
| Tbcel | Q8C5W3-2 | 0.73 | 0.62 | -0.02 | 0.04 | 0.52 | 0.44 | 0.99 | 0.16 | 1 | 0.66 | 0.12 | 0.78 | 0.83 | 0.47 | -0.44 | 0.32 | 0.185 | 0.2975 | -0.0475 | 0.508439 | 0.308178 | 0.885581 | 0.999971 | | 1 | 0.98711 |
| Tbck | E9Q1W7 | 0.19 | 0.38 | 0.2 | 0.03 | 0.59 | 0.29 | -0.21 | -0.06 | 0.65 | 0.1 | -0.13 | -0.03 | 1.11 | -0.09 | -0.18 | 0.04 | -0.0475 | -0.0525 | 0.02 | 0.837036 | 0.816089 | 0.949085 | 0.999971 | | 1 | 0.998038 |
| Tbk1 | Q9WUN2 | -0.57 | -0.42 | 0.11 | -0.24 | -0.21 | -0.23 | 0.19 | -0.07 | -0.19 | -0.46 | 0.11 | 0.23 | -0.52 | -0.53 | 0.14 | -0.13 | 0.2 | 0.2025 | 0.02 | 0.374209 | 0.413215 | 0.935794 | 0.999971 | | 1 | 0.995618 |
| Tbkbp1 | Z4YKN2 | 0.38 | -0.28 | -0.63 | -0.64 | -0.24 | -0.09 | -0.88 | -0.08 | -0.38 | 0.2 | -0.46 | -1.2 | -0.78 | -0.33 | -0.99 | -0.31 | -0.03 | -0.1675 | -0.31 | 0.922914 | 0.647813 | 0.318881 | 0.999971 | | 1 | 0.853893 |
| Tbl1x | Q9QXE7 | 0.9 | 0.7 | 0.21 | 0.43 | 1.28 | -0.38 | -0.43 | 0.45 | 0.37 | 1.31 | -0.23 | 0.09 | 1.94 | 0.92 | 0.18 | 0.67 | -0.33 | -0.175 | 0.3675 | 0.43018 | 0.626358 | 0.354253 | 0.999971 | | 1 | 0.868793 |
| Tbl1xr1 | Q8BHJ5 | 0.96 | 1.27 | 1.65 | 1.43 | 1.27 | 1.17 | 1.34 | 1.36 | 1.3 | 1.11 | 1.39 | 1.68 | 0.75 | 1.34 | 1.18 | 1.45 | -0.0425 | 0.0425 | -0.1475 | 0.836123 | 0.850307 | 0.548764 | 0.999971 | | 1 | 0.911727 |
| Tbl2 | Q9R099 | -0.47 | -0.57 | 0.51 | 0.34 | -0.49 | -0.57 | 0.24 | 0.87 | -0.44 | -0.16 | 0.47 | 0.44 | -0.42 | -0.19 | 0.8 | 0.82 | 0.06 | 0.125 | 0.3 | 0.885005 | 0.721927 | 0.468517 | 0.999971 | | 1 | 0.892385 |
| Tbl3 | Q8C4J7 | -0.69 | 0.01 | 0.15 | -0.28 | -0.54 | -0.28 | -0.34 | -0.56 | -0.09 | -0.25 | -0.39 | -0.14 | 0.29 | -0.16 | -0.04 | -0.58 | -0.2275 | -0.015 | 0.08 | 0.344689 | 0.948285 | 0.772403 | 0.999971 | | 1 | 0.963498 |
| Tbpl1 | P62340 | 1.06 | 2.03 | 1.66 | 2.28 | 1.79 | 1.58 | 1.28 | 1.89 | 1.61 | 1.33 | 1.93 | 2.07 | 1.19 | 2.39 | 2.2 | 2.29 | -0.1225 | -0.0225 | 0.26 | 0.688545 | 0.943126 | 0.492359 | 0.999971 | | 1 | 0.899991 |
| Tbr1 | Q64336 | 2.4 | 1.55 | 1.38 | 1.3 | 1.3 | 1.29 | 1.34 | 1.21 | 1.11 | 1.47 | 1.62 | 1.03 | 0.36 | 0.9 | 0.89 | 1.65 | -0.3725 | -0.35 | -0.7075 | 0.195113 | 0.259384 | 0.075541 | 0.999971 | | 1 | 0.746958 |
| Tbrg4 | Q91YM4 | 0.06 | -0.03 | -0.66 | -0.48 | -0.6 | -0.49 | -0.78 | -0.49 | 0.02 | -0.23 | -0.85 | -0.91 | 0.3 | -0.37 | -0.65 | -0.16 | -0.3125 | -0.215 | 0.0575 | 0.191534 | 0.475573 | 0.837971 | 0.999971 | | 1 | 0.979514 |
| Tcaf1 | Q8BNE1 | 0.33 | 0.56 | 1.1 | 1.03 | 0.24 | 0.33 | 0.84 | 1.05 | 0.45 | 0.59 | 1.36 | 0.8 | -0.03 | 0.97 | 1.43 | 1.62 | -0.14 | 0.045 | 0.2425 | 0.624332 | 0.874612 | 0.544809 | 0.999971 | | 1 | 0.911398 |
| TCAIM | Q66JZ4 | -1.7 | -0.43 | -0.65 | -0.72 | -0.09 | -0.18 | -0.8 | -0.23 | 0.13 | -0.87 | -1.25 | -0.47 | -0.21 | -0.9 | -0.02 | -0.65 | 0.55 | 0.26 | 0.43 | 0.118023 | 0.513148 | 0.230766 | 0.999971 | | 1 | 0.830179 |
| Tcea1 | P10711 | 1.3 | 1.9 | 1.93 | 1.89 | 1.73 | 1.54 | 2.97 | 1.28 | 1.43 | 1.32 | 1.26 | 0.94 | 0.98 | 1.3 | 1.23 | 1.12 | 0.125 | -0.5175 | -0.5975 | 0.748215 | 0.043131 | 0.020322 | 0.999971 | | 1 | 0.635388 |
| Tceal1 | Q921P9 | -0.27 | 2.04 | 1.08 | 0.96 | 2.02 | 1.89 | 0.54 | 1.44 | 2.58 | 1.65 | 0.95 | 1.93 | 2.15 | 3.06 | 1.89 | 1.31 | 0.52 | 0.825 | 1.15 | 0.343785 | 0.151936 | 0.062217 | 0.999971 | | 1 | 0.725348 |
| Tceal3 | Q8R0A5 | 3.49 | 5.22 | 2.12 | 2.64 | 3.13 | 4.88 | 1.6 | 0.36 | 3.86 | 4.23 | 0.93 | 0.79 | 2.99 | 3.75 | 0.16 | -0.15 | -0.875 | -0.915 | -1.68 | 0.418354 | 0.385369 | 0.142313 | 0.999971 | | 1 | 0.784967 |
| Tceal5 | Q8CCT4 | 3.65 | 2.54 | 3.92 | 3.83 | 2.14 | 2.8 | 5.45 | 2.88 | 2.47 | 2.87 | 3.57 | 2.94 | 1.47 | 1.77 | 2.46 | 1.74 | -0.1675 | -0.5225 | -1.625 | 0.815971 | 0.192923 | 0.001875 | 0.999971 | | 1 | 0.476373 |
| Tcerg1 | Q8CGF7-2 | 1.75 | 2.57 | 1.85 | 1.61 | 2.71 | 2.21 | 2.07 | 1.51 | 2.43 | 2.19 | 1.98 | 2.16 | 2.17 | 2.44 | 1.57 | 1.78 | 0.18 | 0.245 | 0.045 | 0.58502 | 0.352146 | 0.880346 | 0.999971 | | 1 | 0.985441 |
| Tcf20 | Q9EPQ8 | -0.63 | -0.88 | -0.45 | 0.07 | -0.97 | -1.01 | -0.62 | -0.24 | -1.09 | -0.79 | -0.7 | -0.95 | -1.11 | -0.81 | -0.43 | -0.25 | -0.2375 | -0.41 | -0.1775 | 0.412082 | 0.123266 | 0.546081 | 0.999971 | | 1 | 0.911398 |
| Tcf25 | B2ZAC8 | 0.91 | 1.6 | 1.27 | 1.16 | 2.06 | 1.85 | 1.32 | 1.41 | 1.84 | 1.22 | 1.71 | 2.06 | 2.71 | 1.7 | 1.56 | 1.74 | 0.425 | 0.4725 | 0.6925 | 0.120728 | 0.089048 | 0.047436 | 0.999971 | | 1 | 0.710102 |
| Tcf4 | A0A0J9YTV | 0.9 | 1.29 | 1.73 | 1.71 | 1.02 | 1.15 | 1.67 | 1.6 | 1.06 | 0.57 | 1.71 | 1.32 | 0.39 | 1.18 | 1.32 | 1.28 | -0.0475 | -0.2425 | -0.365 | 0.861905 | 0.44737 | 0.24849 | 0.999971 | | 1 | 0.839012 |
| Tcirg1 | Q9JHF5 | 1.04 | -0.68 | -0.21 | -1.08 | 0.44 | -0.19 | -0.68 | 0.06 | -0.91 | 0.6 | -0.73 | -1.47 | -0.09 | -0.61 | -0.36 | -0.45 | 0.14 | -0.395 | -0.145 | 0.771244 | 0.504864 | 0.745277 | 0.999971 | | 1 | 0.961064 |
| Tcn2 | O88968 | -1.74 | -0.09 | 0.26 | 0.98 | 0.37 | -1.06 | -0.53 | 1.12 | -0.1 | -0.87 | 0.17 | 0.83 | -0.09 | -0.49 | 1.03 | 1.24 | 0.1225 | 0.155 | 0.57 | 0.856998 | 0.802196 | 0.390507 | 0.999971 | | 1 | 0.871816 |
| Tcof1 | H3BL37 | 1.96 | 1.21 | 1.56 | 2.19 | 0.92 | 0.86 | 1.6 | 1.22 | 1.4 | 1.72 | 1.52 | 1.2 | 0.63 | 0.88 | 1.06 | 1.12 | -0.58 | -0.27 | -0.8075 | 0.069974 | 0.319682 | 0.013215 | 0.999971 | | 1 | 0.613095 |
| Tcp1 | P11983 | -0.26 | 0.63 | 0.95 | 0.24 | 0.51 | 0.5 | 0.78 | 0.44 | 0.74 | -0.18 | 0.82 | 0.9 | 0.82 | 0.39 | 1.08 | 0.49 | 0.1675 | 0.18 | 0.305 | 0.560738 | 0.615169 | 0.339934 | 0.999971 | | 1 | 0.860511 |
| Tcp11l1 | Q8BTG3 | 0.4 | 0.06 | -0.68 | -0.43 | -0.24 | 0.27 | -0.75 | -0.17 | 0.28 | 0.52 | -0.56 | -0.07 | 0.16 | 0.27 | -0.33 | 0.35 | -0.06 | 0.205 | 0.275 | 0.852053 | 0.54507 | 0.366231 | 0.999971 | | 1 | 0.869526 |
| Tctn3 | Q8R2Q6 | 2.85 | 0.13 | 0.13 | 1.46 | -0.42 | 0.56 | 0.25 | 0.67 | -1.07 | 1.82 | -0.81 | 0.32 | -0.65 | -0.13 | 0.1 | 0.32 | -0.8775 | -1.0775 | -1.2325 | 0.1884 | 0.219442 | 0.074299 | 0.999971 | | 1 | 0.746069 |
| Tdg | P56581-2 | 0.92 | 1.85 | 2.51 | 2.69 | 1.67 | 1.59 | 2.49 | 3 | 1.71 | 1.76 | 2.95 | 3.07 | 1.08 | 2.21 | 2.72 | 2.96 | 0.195 | 0.38 | 0.25 | 0.690349 | 0.460898 | 0.641722 | 0.999971 | | 1 | 0.93301 |
| Tdp1 | B8JJC1 | -0.65 | 0.5 | -1.15 | -0.82 | -0.13 | -0.39 | -1.68 | -1.43 | -0.53 | -0.68 | -0.83 | 0.15 | -0.68 | -0.62 | -1.56 | -0.14 | -0.3775 | 0.0575 | -0.22 | 0.445813 | 0.88606 | 0.618636 | 0.999971 | | 1 | 0.928581 |
| Tdp2 | Q9JJX7 | 1.71 | 1.88 | -0.77 | -0.16 | 1.13 | 0.94 | -0.89 | -1.1 | 0.44 | 1.65 | -0.81 | -0.59 | 0.1 | 0.86 | -0.54 | -0.19 | -0.645 | -0.4925 | -0.6075 | 0.427835 | 0.537064 | 0.370623 | 0.999971 | | 1 | 0.869526 |
| Tdrd1 | Q99MV1 | 2.06 | 2.11 | 2.13 | 2.41 | 1.27 | 2.69 | 0.91 | 1.65 | 0.81 | 2.61 | 1.57 | 0.37 | 0.61 | 2.82 | 1.59 | 3.19 | -0.5475 | -0.8375 | -0.125 | 0.173076 | 0.100479 | 0.819699 | 0.999971 | | 1 | 0.975761 |
| Tdrd3 | Q91W18-2 | 0.93 | 0.91 | 1.71 | 1.03 | 0.76 | 0.83 | 1.99 | 1.14 | 0.44 | 0.98 | 0.91 | 0.68 | 0.2 | 0.38 | 0.78 | 0.2 | 0.035 | -0.3925 | -0.755 | 0.917353 | 0.144776 | 0.016246 | 0.999971 | | 1 | 0.613095 |
| Tdrd7 | A0A0A0MC | -0.16 | 1.8 | 0.68 | 0.24 | 2.27 | 0.87 | 0.62 | 0.98 | 2.13 | -0.01 | 0.86 | 1.14 | 2.7 | 0.62 | 0.85 | 0.77 | 0.545 | 0.39 | 0.595 | 0.308963 | 0.494965 | 0.329803 | 0.999971 | | 1 | 0.856787 |
| Tdrkh | A0A0G2JFE | -0.49 | -0.71 | -1 | -1.01 | -0.4 | -0.57 | -0.7 | -0.6 | -0.34 | -0.36 | -0.48 | -0.67 | -0.07 | -0.52 | -0.82 | -0.66 | 0.235 | 0.34 | 0.285 | 0.256752 | 0.116542 | 0.251792 | 0.999971 | | 1 | 0.839012 |
| Tdrp | Q8C5P7 | 0.67 | 1.51 | -0.83 | -0.95 | 1.57 | 1.67 | -0.64 | -0.28 | 1.22 | 0.86 | -0.46 | -0.42 | 1.24 | 1.9 | -0.16 | -0.91 | 0.48 | 0.2 | 0.4175 | 0.535104 | 0.766731 | 0.599541 | 0.999971 | | 1 | 0.92422 |
| Tecpr1 | Q80VP0 | 0.17 | -0.27 | -0.88 | -0.8 | -0.21 | 0.25 | -0.93 | -0.57 | 0.31 | 0.67 | -1.01 | -1.06 | -0.18 | 0.2 | -0.77 | -0.72 | 0.08 | 0.1725 | 0.0775 | 0.817621 | 0.718565 | 0.817726 | 0.999971 | | 1 | 0.974929 |
| Tecpr2 | Q3UH45 | 0.16 | 0.66 | 0.44 | 0.4 | 0.4 | 0.79 | 0.12 | 0.4 | 0.73 | 0.67 | 0.27 | 0.16 | 0.98 | 0.37 | 0.34 | 0.81 | 0.0125 | 0.0425 | 0.21 | 0.954082 | 0.845036 | 0.37155 | 0.999971 | | 1 | 0.869526 |
| Tecr | Q9CY27 | -1.41 | -0.2 | -1.61 | -2.76 | -0.56 | -0.86 | -0.88 | -2.37 | -0.28 | -1.31 | -0.73 | -0.67 | 0.07 | -0.46 | -1.64 | -2.5 | 0.3275 | 0.7475 | 0.3625 | 0.591504 | 0.179119 | 0.610756 | 0.999971 | | 1 | 0.92673 |
| Tef | Q9JLC6 | 5.13 | 0.97 | -1.15 | -0.25 | -0.28 | 2.54 | -1 | -0.17 | -1.13 | 4.71 | -1.73 | -1.35 | -0.85 | 2.04 | -0.28 | -0.39 | -0.9025 | -1.05 | -1.045 | 0.52724 | 0.572781 | 0.450647 | 0.999971 | | 1 | 0.886265 |
| Tefm | Q5SSK3 | -1.25 | -1.42 | -1.31 | -1.06 | -1.23 | -1.65 | -0.86 | -1.11 | -1.09 | -1.56 | -0.87 | -1.41 | -0.51 | -1.06 | -1.27 | -1.28 | 0.0475 | 0.0275 | 0.23 | 0.83161 | 0.898571 | 0.337538 | 0.999971 | | 1 | 0.858827 |
| Telo2 | Q9DC40 | 0.76 | -0.11 | 0.06 | 0.1 | 0.38 | 0.26 | -0.19 | 0.44 | 0.1 | 0.58 | 0.19 | 0.14 | -0.59 | 0.63 | 0.33 | 0.44 | 0.02 | 0.05 | 1.57E-16 | 0.938968 | 0.840119 | 1 | 0.999971 | | 1 | 1 |
| Ten1 | Q9D7K2 | -3.33 | -0.31 | -0.18 | 0.6 | 0.58 | -1.07 | 0.37 | 0.43 | 0.79 | -2.8 | 1.23 | 1.28 | -2.39 | 1.38 | 0.47 | 0.38 | 0.8825 | 0.93 | 0.765 | 0.313876 | 0.436317 | 0.47765 | 0.999971 | | 1 | 0.894438 |
| Tenm1 | Q9WTS4 | -0.25 | -1.02 | -1 | -0.88 | -0.79 | -0.4 | -0.44 | -0.92 | -0.68 | -0.09 | -0.8 | -0.92 | -1.07 | -1.03 | -0.88 | -1.21 | 0.15 | 0.165 | -0.26 | 0.553231 | 0.552322 | 0.279974 | 0.999971 | | 1 | 0.843812 |
| Tenm2 | A0A0A0MC | -1.39 | -1.09 | -1.03 | -1.1 | -1.08 | -0.92 | -1.17 | -0.6 | -1.13 | -1.1 | -0.79 | -0.86 | -0.54 | -0.62 | -0.71 | -0.51 | 0.21 | 0.1825 | 0.5575 | 0.31836 | 0.33633 | 0.011213 | 0.999971 | | 1 | 0.613095 |
| Tenm3 | G3X907 | -0.47 | -0.8 | -0.85 | -0.73 | -0.75 | -1.14 | -0.81 | -0.85 | -0.66 | -0.45 | -0.87 | -0.49 | -0.73 | -0.82 | -0.77 | -0.49 | -0.175 | 0.095 | 0.01 | 0.367139 | 0.618963 | 0.956984 | 0.999971 | | 1 | 0.998513 |
| Tenm4 | Q3UHK6-4 | -0.79 | -1.31 | -1.15 | -1.25 | -1.28 | -1.09 | -0.74 | -1.29 | -1.11 | 0.21 | -1 | -0.65 | -1.26 | -1.48 | -1 | -1.21 | 0.025 | 0.4875 | -0.1125 | 0.908807 | 0.156245 | 0.589405 | 0.999971 | | 1 | 0.921872 |
| Tent4b | F8WI26 | 1.62 | 0.84 | 1.11 | 0.88 | 1.04 | 0.66 | 0.46 | 1.25 | 0.51 | 0.87 | 1.4 | 1.48 | -0.53 | 0.82 | 1.48 | 1.64 | -0.26 | -0.0475 | -0.26 | 0.351816 | 0.873732 | 0.597976 | 0.999971 | | 1 | 0.92377 |
| Tepsin | Q3U3N6 | 0.64 | 0.08 | -0.03 | 0.89 | -0.21 | -0.22 | -0.39 | 0.75 | 0.54 | 0.1 | -0.73 | 0.9 | -0.4 | -0.18 | 0.33 | -0.77 | -0.4125 | -0.1925 | -0.65 | 0.241858 | 0.63095 | 0.069366 | 0.999971 | | 1 | 0.733937 |
| Terf2 | D3YZ08 | 1.78 | 0.88 | 0.2 | 1.16 | 0.86 | 0.86 | 0.91 | 0.02 | 0.78 | 1.66 | 0 | 0.04 | 0.12 | 0.76 | 0.25 | -0.65 | -0.3425 | -0.385 | -0.885 | 0.376664 | 0.428772 | 0.059209 | 0.999971 | | 1 | 0.725348 |
| Terf2ip | Q91VL8 | 0.86 | 1.23 | -0.43 | -0.77 | 1.29 | 1.5 | -0.15 | 0.16 | 0.97 | 1.42 | 0.18 | 1.02 | 1.42 | 1.51 | -0.04 | -0.36 | 0.4775 | 0.675 | 0.41 | 0.41996 | 0.210234 | 0.517681 | 0.999971 | | 1 | 0.905815 |
| Tes | Q3ULB1 | 0.22 | 0.46 | 1.25 | 1.52 | 1.54 | 0.19 | 4.25 | 1.21 | 0.54 | 1.28 | 2.91 | 4.53 | -0.42 | 0.34 | 1.36 | 0.92 | 0.935 | 1.4525 | -0.3125 | 0.275457 | 0.115392 | 0.505808 | 0.999971 | | 1 | 0.905277 |
| Tesc | Q9JKL5 | -1.2 | 0.29 | -0.93 | -1.67 | -0.04 | -0.36 | -1.31 | -1.58 | -0.1 | -0.84 | -1.05 | -0.47 | 0.16 | -0.25 | -1.15 | -2.16 | 0.055 | 0.2625 | 0.0275 | 0.91498 | 0.556556 | 0.963777 | 0.999971 | | 1 | 0.998721 |
| Tesk1 | O70146 | -0.07 | -0.2 | 0.47 | 0.66 | -0.2 | -0.27 | 0.71 | 1.49 | -0.54 | -0.5 | 1.24 | 1.02 | -0.64 | -0.37 | 1.2 | 1.17 | 0.2175 | 0.09 | 0.125 | 0.622541 | 0.85347 | 0.801068 | 0.999971 | | 1 | 0.971257 |
| Tex10 | Q3URQ0 | -0.46 | 0.25 | 0.76 | 1.04 | 0.32 | -0.06 | 0.52 | 0.64 | -0.17 | 0.34 | 1 | 1.02 | 0.19 | 0.32 | 0.99 | 1.15 | -0.0425 | 0.15 | 0.265 | 0.90478 | 0.719308 | 0.504098 | 0.999971 | | 1 | 0.905265 |
| Tex2 | Q6ZPJ0 | -0.64 | -0.42 | -0.37 | -0.62 | -0.88 | -0.47 | -0.48 | -0.72 | -0.52 | -0.22 | -0.63 | -0.25 | 0.03 | -0.53 | -0.5 | -0.77 | -0.125 | 0.1075 | 0.07 | 0.51403 | 0.568642 | 0.755644 | 0.999971 | | 1 | 0.961116 |
| Tex264 | E9Q137 | 0.12 | -0.35 | -0.13 | 0.01 | -0.29 | 0.34 | 0.05 | 0.08 | 0.03 | 0.29 | -0.11 | 0.13 | 0.17 | 0.23 | 0.05 | 0.15 | 0.1325 | 0.1725 | 0.2375 | 0.538144 | 0.380087 | 0.218145 | 0.999971 | | 1 | 0.827064 |
| Tf | Q921I1 | 1.06 | 1.78 | 0.35 | 1.92 | 2.42 | 1.36 | 1.57 | 0.99 | 2.1 | 0.98 | 1.32 | 1.13 | 1.89 | 2.74 | 1.48 | 1.83 | 0.3075 | 0.105 | 0.7075 | 0.494606 | 0.802064 | 0.124515 | 0.999971 | | 1 | 0.78268 |
| Tfam | A0A0R4J1N | 0.47 | -0.06 | -0.92 | -0.71 | -0.16 | 0.09 | -1 | -1.07 | 0.27 | 0.4 | -0.66 | -0.89 | -0.33 | 0.01 | -0.85 | -0.89 | -0.23 | 0.085 | -0.21 | 0.578852 | 0.843436 | 0.576479 | 0.999971 | | 1 | 0.918194 |
| Tfb1m | Q8JZM0 | -0.84 | -0.49 | -1.26 | -0.99 | 0.14 | -0.56 | -0.28 | -0.58 | 0.39 | -0.79 | -1.03 | 0.15 | -0.05 | -0.39 | -1.41 | -0.61 | 0.575 | 0.575 | 0.28 | 0.048773 | 0.148743 | 0.406898 | 0.999971 | | 1 | 0.874911 |
| Tfcp2 | A0A2R8VKI | -0.25 | 0.95 | 0.17 | -0.54 | 0.73 | 0.73 | -0.01 | 0.47 | 0.51 | -0.53 | 0.3 | -0.28 | 0.76 | 0.71 | -0.05 | 0.14 | 0.3975 | -0.0825 | 0.3075 | 0.285609 | 0.832522 | 0.417358 | 0.999971 | | 1 | 0.877483 |
| Tfdp1 | Q08639 | -0.8 | 2.97 | 0.99 | 1.11 | 2.82 | 2.85 | 0.26 | 1.22 | 2.59 | -1.02 | -0.14 | 1.11 | 3.22 | 3.23 | -0.08 | 1.08 | 0.72 | -0.4325 | 0.795 | 0.429181 | 0.663382 | 0.4379 | 0.999971 | | 1 | 0.882419 |
| Tfe3 | Q64092 | 2.06 | 0.59 | 0.45 | 0.35 | 0.26 | 1.17 | 0.18 | -0.43 | 0.49 | 1.75 | -0.1 | 0.03 | -0.01 | 0.08 | -0.09 | 0.15 | -0.5675 | -0.32 | -0.83 | 0.259831 | 0.557065 | 0.058731 | 0.999971 | | 1 | 0.725348 |
| Tfeb | Q3UKG7 | 1.05 | -0.47 | 0.75 | -0.17 | 0.08 | 0.2 | 1.19 | -1.23 | 0.29 | 0.34 | -1.02 | 0.31 | -0.6 | -0.28 | 0.24 | -0.75 | -0.23 | -0.31 | -0.6375 | 0.684543 | 0.508714 | 0.14208 | 0.999971 | | 1 | 0.784967 |
| Tfg | Q9Z1A1 | -0.29 | -0.12 | -0.23 | -0.21 | -0.11 | -0.49 | -0.23 | -0.1 | -0.1 | -0.67 | -0.15 | 0.01 | 0.03 | -0.18 | -0.13 | -0.21 | -0.02 | -0.015 | 0.09 | 0.910711 | 0.941662 | 0.593379 | 0.999971 | | 1 | 0.923349 |
| Tfip11 | Q9ERA6 | 1.7 | 0.88 | 1.17 | 1.21 | 0.92 | 0.65 | 0.83 | 1.1 | 0.51 | 0.93 | 0.95 | 1.01 | -0.14 | 0.21 | 1.06 | 1.29 | -0.365 | -0.39 | -0.635 | 0.14103 | 0.126754 | 0.113493 | 0.999971 | | 1 | 0.78268 |
| Tfrc | Q62351 | 0.11 | 0.31 | -0.12 | 0.3 | -0.17 | -0.25 | -0.01 | -0.32 | 0.25 | 0.29 | -0.25 | -0.47 | -0.79 | -0.61 | -0.98 | -0.67 | -0.3375 | -0.195 | -0.9125 | 0.101826 | 0.429642 | 0.001166 | 0.999971 | | 1 | 0.408862 |
| Tgfb1i1 | Q62219-2 | 1.58 | 1.42 | 0.1 | 1.16 | 3.61 | 1.42 | 0.08 | 0.2 | 2.51 | 4.24 | 1.33 | 2.56 | 1.94 | 8.02 | 0.93 | 0.55 | 0.2625 | 1.595 | 1.795 | 0.741642 | 0.030063 | 0.272851 | 0.999971 | | 1 | 0.843794 |
| Tgfbi | Q3UXJ2 | 1.22 | 4.83 | 4.29 | 2.77 | 6.86 | 4.07 | 2.41 | 2.67 | 5.04 | -2.2 | 3.63 | 6.17 | 4.08 | 2.23 | 3.37 | 3.77 | 0.725 | -0.1175 | 0.085 | 0.536431 | 0.948222 | 0.917024 | 0.999971 | | 1 | 0.992633 |
| Tgfbrap1 | Q3UR70 | 0.2 | 0.39 | 0.3 | 0.14 | 0.18 | 0.44 | 0.06 | 0.49 | 0.22 | 0.31 | 0.35 | 0.36 | 0.06 | 0.54 | 0.56 | 0.49 | 0.035 | 0.0525 | 0.155 | 0.851745 | 0.747357 | 0.434782 | 0.999971 | | 1 | 0.882419 |
| Tgm2 | P21981 | 1.5 | 1.91 | 1.25 | 2.02 | 5.06 | 2.19 | 1.25 | 0.98 | 2.29 | 2.91 | 2.19 | 3.03 | -0.54 | 2.6 | 1.75 | 0.47 | 0.7 | 0.935 | -0.6 | 0.42045 | 0.01002 | 0.368334 | 0.999971 | | 1 | 0.869526 |
| Tgoln1 | Q62313 | 1.94 | 0.7 | 2.77 | 2.61 | 1.08 | 1.06 | 2.24 | 2.61 | 1.13 | 1.69 | 2.35 | 1.57 | 0.41 | 1.61 | 2.49 | 2.43 | -0.2575 | -0.32 | -0.27 | 0.650431 | 0.525391 | 0.663115 | 0.999971 | | 1 | 0.936388 |
| Tgs1 | Q923W1 | 0.35 | 0.88 | 0.76 | 0.95 | 1.36 | -0.24 | 0.34 | 0.56 | 1.5 | -1.47 | 0.47 | 0.71 | 1.74 | -1.29 | 0.28 | 0.51 | -0.23 | -0.4325 | -0.425 | 0.517297 | 0.472293 | 0.472975 | 0.999971 | | 1 | 0.893778 |
| Th | P24529 | -0.11 | -1.62 | -2.46 | -0.47 | -0.92 | -2.32 | -1.1 | 0.1 | -0.65 | -0.54 | -1.19 | -1.36 | -0.97 | -0.8 | -0.51 | -2 | 0.105 | 0.23 | 0.095 | 0.874258 | 0.667294 | 0.86905 | 0.999971 | | 1 | 0.983462 |
| Tha1 | Q6XPS7 | 0.01 | -0.68 | -1.3 | -0.83 | -0.45 | -0.87 | -1.15 | -0.97 | -1.14 | -0.44 | -0.99 | -1.26 | -0.21 | -0.11 | -0.93 | -0.57 | -0.16 | -0.2575 | 0.245 | 0.611981 | 0.438288 | 0.464246 | 0.999971 | | 1 | 0.890905 |
| Thada | A8C756-3 | 0.98 | 0.83 | -0.87 | -0.67 | 0.69 | 0.34 | -0.7 | -0.46 | 0.66 | 0.97 | -0.28 | -0.62 | 0.55 | 0.35 | -0.77 | -0.18 | -0.1 | 0.115 | -0.08 | 0.85295 | 0.839356 | 0.879066 | 0.999971 | | 1 | 0.985441 |
| Thbd | P15306 | 0.86 | 0.64 | 0.67 | 1.02 | 1.36 | 0.69 | 0.22 | 1.58 | 0.56 | 0.84 | 0.25 | 0.02 | 1.46 | 0.81 | 1.33 | 0.15 | 0.165 | -0.38 | 0.14 | 0.613888 | 0.130433 | 0.658533 | 0.999971 | | 1 | 0.934638 |
| Thbs1 | Q80YQ1 | -0.14 | -0.97 | -1.91 | 0.84 | 2.43 | -1.14 | 0.27 | -1.98 | -1.36 | -0.91 | -0.42 | -0.04 | -1.61 | -0.81 | 0.81 | -0.19 | 0.44 | -0.1375 | 0.095 | 0.663561 | 0.818741 | 0.892597 | 0.999971 | | 1 | 0.98828 |
| Thbs4 | Q9Z1T2 | 2.38 | 2.9 | 1.45 | 0.73 | 3 | 3.78 | 1.47 | 1.85 | 3.45 | 3.13 | 2.52 | 2.29 | 3.23 | 2.02 | 1.23 | 1.86 | 0.66 | 0.9825 | 0.22 | 0.324879 | 0.083082 | 0.707761 | 0.999971 | | 1 | 0.950182 |
| Them4 | Q3UUI3 | -0.87 | -0.6 | 0.24 | 0.2 | -0.6 | -0.54 | -0.14 | 0.22 | -0.64 | -1.02 | -0.02 | 0.03 | -1.11 | -0.41 | 0.15 | 0.04 | -0.0075 | -0.155 | -0.075 | 0.982241 | 0.675429 | 0.846278 | 0.999971 | | 1 | 0.979924 |
| Them6 | Q80ZW2 | -0.58 | 0.17 | -0.31 | -0.16 | 0.11 | 0.03 | -0.25 | -0.08 | 0.03 | -0.32 | -0.32 | -0.24 | -0.04 | 0.28 | -0.26 | -0.3 | 0.1725 | 0.0075 | 0.14 | 0.439531 | 0.972585 | 0.56236 | 0.999971 | | 1 | 0.916386 |
| Thg1l | Q9CQT0 | 0.62 | 1.05 | 0.32 | 0.55 | 0.81 | 0.31 | 0.38 | -0.01 | 0.71 | 0.25 | 0.53 | 0.17 | 0.92 | 0.64 | 0.17 | 0.1 | -0.2625 | -0.22 | -0.1775 | 0.315816 | 0.35482 | 0.513988 | 0.999971 | | 1 | 0.905815 |
| Thnsl1 | Q8BH55 | -0.73 | -0.51 | -0.23 | -0.69 | 0.14 | -0.67 | 0.31 | -0.26 | 0.38 | -0.69 | -0.23 | 0.05 | 0.15 | -0.71 | -0.52 | -0.61 | 0.42 | 0.4175 | 0.1175 | 0.142162 | 0.149614 | 0.644344 | 0.999971 | | 1 | 0.93301 |
| Thnsl2 | Q80W22 | -0.46 | 0.58 | 0.74 | 0.42 | 0.38 | 0.58 | -0.07 | 1.58 | 0.24 | -0.2 | 0.4 | 0.7 | 1.15 | 1.17 | 1.07 | 0.65 | 0.2975 | -0.035 | 0.69 | 0.482325 | 0.914743 | 0.045701 | 0.999971 | | 1 | 0.710102 |

| | | | | | | | | | | | | | | | | | | | | | | | | | |
|---|---|---|---|---|---|---|---|---|---|---|---|---|---|---|---|---|---|---|---|---|---|---|---|---|---|
| Thoc1 | Q8R3N6 | 1.52 | 1.2 | 1.3 | 1.31 | 0.76 | 0.55 | 1.76 | 0.57 | 0.92 | 1.13 | 1.09 | 0.46 | 0.68 | 0.81 | 0.46 | 0.17 | -0.4225 | -0.4325 | -0.8025 | 0.186469 | 0.068204 | 0.003826 | 0.999971 | 1 | 0.582714 |
| Thoc2 | B1AZI6 | 0.28 | -0.02 | 0.96 | 0.75 | -0.46 | -0.38 | 1.12 | 0.42 | -0.13 | -0.28 | 0.77 | 0.45 | -1.11 | -0.68 | 0.4 | -0.12 | -0.3175 | -0.29 | -0.87 | 0.449185 | 0.391221 | 0.046568 | 0.999971 | 1 | 0.710102 |
| Thoc3 | Q8VE80 | -0.6 | 1.04 | 0.98 | 0.12 | 0.8 | 0.64 | 0.98 | 1.21 | 0.79 | -0.36 | 1.12 | 1.2 | 0.49 | 0.71 | 0.96 | 0.36 | 0.5225 | 0.3025 | 0.245 | 0.205147 | 0.545592 | 0.539694 | 0.999971 | 1 | 0.910077 |
| Thoc5 | A0A0R4J0J | -0.14 | 0.67 | 1.31 | 1.08 | 0.3 | 0.26 | 0.49 | 1.07 | 0.63 | -0.14 | 1.01 | 1.17 | -0.09 | 0.03 | 1.09 | 1.26 | -0.2 | -0.0625 | -0.1575 | 0.582336 | 0.879543 | 0.724876 | 0.999971 | 1 | 0.955813 |
| Thoc6 | D3Z132 | 0.44 | -0.02 | 0.32 | 0.96 | 0.07 | -0.6 | 0.21 | 0.32 | 0.04 | 0.53 | 0.52 | 0.81 | -0.63 | 0.57 | 0.54 | 0.77 | -0.425 | 0.05 | -0.1125 | 0.178916 | 0.855524 | 0.760182 | 0.999971 | 1 | 0.961116 |
| Thoc7 | A0A286YD | 2.07 | 0.61 | 0.85 | 1.22 | 0.14 | 0.39 | 1.34 | 0.03 | 0.23 | 1.44 | -0.16 | -0.23 | -1.17 | -0.29 | -1.07 | -0.33 | -0.7125 | -0.8675 | -1.9025 | 0.11347 | 0.093146 | 0.000838 | 0.999971 | 1 | 0.365812 |
| Thop1 | A0A0R4IZY | 0.24 | 1.19 | 1.06 | 0.71 | 1.18 | 0.99 | 1.02 | 1.25 | 1.09 | 0.27 | 1.33 | 1.2 | 1.1 | 1 | 1.06 | 1.12 | 0.31 | 0.1725 | 0.27 | 0.234707 | 0.59442 | 0.287811 | 0.999971 | 1 | 0.847258 |
| Thra | A0A0R4J1F | 0.63 | 0.58 | 0.67 | 0.93 | 0.23 | 1.08 | 0.58 | 1.55 | 0.87 | 1.17 | 0.93 | 0.84 | 0.46 | 1.17 | 1 | 0.95 | 0.1575 | 0.25 | 0.1925 | 0.608893 | 0.188813 | 0.387566 | 0.999971 | 1 | 0.871816 |
| Thrap3 | Q569Z6 | 0.48 | 0.54 | 0.27 | 0.26 | 0.05 | -0.39 | 0.18 | -0.18 | -0.13 | 0.31 | 0.34 | -0.01 | 0.24 | -0.4 | -0.06 | -0.16 | -0.4725 | -0.26 | -0.4825 | 0.041124 | 0.204808 | 0.040875 | 0.999971 | 1 | 0.700838 |
| Thsd7a | E9PWD2 | 0 | -0.26 | -0.37 | -0.27 | -0.18 | -0.13 | 0.15 | -0.08 | 0.15 | 0.21 | -0.15 | 0.19 | 0.65 | 0.18 | -0.11 | -0.41 | 0.165 | 0.325 | 0.3025 | 0.378295 | 0.104177 | 0.267409 | 0.999971 | 1 | 0.842677 |
| Thsd7b | Q6P4U0 | 0.16 | 0.31 | -1.15 | -1.44 | 0.64 | 0.21 | -1.4 | -1.06 | 0.13 | -0.89 | -2.33 | -0.65 | 1.07 | 0.43 | -0.63 | -0.67 | 0.1275 | -0.405 | 0.58 | 0.833131 | 0.520996 | 0.320446 | 0.999971 | 1 | 0.853962 |
| Thtpa | Q8JZL3 | 0.25 | 0.54 | 0.85 | 1.21 | 0.34 | 0.27 | 0.54 | 1.25 | 0.23 | 0.38 | 0.51 | 0.54 | -0.14 | 0.67 | 0.71 | 0.98 | -0.1125 | -0.2975 | -0.1575 | 0.717312 | 0.245887 | 0.625453 | 0.999971 | 1 | 0.930479 |
| Thumpd1 | Q99J36 | 1.5 | 2.79 | 2.47 | 2.81 | 2.53 | 2.47 | 1.88 | 2.79 | 2.48 | 2.07 | 2.46 | 1.48 | 2.21 | 2.81 | 2.74 | 2.74 | 0.025 | -0.27 | 0.2325 | 0.943809 | 0.478155 | 0.495073 | 0.999971 | 1 | 0.901531 |
| Thumpd3 | P97770 | 0.36 | 1.24 | 1.41 | 0.76 | 1.52 | 1.27 | 1.2 | 1.53 | 1.67 | 0.47 | 1.1 | 1.49 | 1.85 | 0.24 | 1.51 | 1.12 | 0.4375 | 0.24 | 0.2375 | 0.133618 | 0.499519 | 0.559261 | 0.999971 | 1 | 0.915051 |
| Thy1 | A0A1L1SU | -0.37 | -0.31 | -1.06 | -0.65 | -0.45 | -0.21 | -1.32 | -0.54 | -0.46 | -0.22 | -0.33 | -0.99 | -0.01 | -0.11 | -0.86 | -0.63 | -0.0325 | 0.0975 | 0.195 | 0.914395 | 0.711009 | 0.495247 | 0.999971 | 1 | 0.901531 |
| Thyn1 | Q91YJ3 | 3.05 | 2.74 | 3.15 | 3.3 | 2.71 | 2.75 | 2.67 | 3.2 | 2.81 | 2.48 | 2.74 | 2.91 | 2.35 | 2.36 | 3.18 | 3.15 | -0.2275 | -0.325 | -0.3 | 0.309914 | 0.134502 | 0.298374 | 0.999971 | 1 | 0.847402 |
| Tia1 | P52912 | 3.5 | 0.71 | 1.93 | 2.09 | -0.33 | 1.44 | 1.59 | 2.15 | 1.37 | 2.67 | 1.63 | 2.18 | -1.45 | 1.37 | 1.8 | 1.82 | -0.845 | -0.095 | -1.1725 | 0.251946 | 0.871795 | 0.201446 | 0.999971 | 1 | 0.815758 |
| Tia1_1 | Q8CII5 | 0.97 | 0.03 | 1.34 | 1.58 | -0.27 | 0.5 | 2.19 | 1.07 | -0.21 | 0.28 | 0.49 | 0.38 | -0.64 | -0.37 | 0.81 | 0.72 | -0.1075 | -0.745 | -0.85 | 0.849859 | 0.067607 | 0.098773 | 0.999971 | 1 | 0.78268 |
| Tial1 | Q921W2 | 0.18 | 1.33 | 1.42 | 1.56 | 1.28 | 0.88 | 1.92 | 1.42 | 1.17 | 0.14 | 1.66 | 1.89 | 1.51 | 1.01 | 1.7 | 1.5 | 0.2525 | 0.0925 | 0.3075 | 0.502606 | 0.844229 | 0.385227 | 0.999971 | 1 | 0.871816 |
| Tiam1 | G3UWG2 | 0.74 | -0.57 | 0.15 | 0.32 | -0.24 | -0.12 | 0.33 | 0 | -0.35 | 0.5 | -0.13 | 0 | -0.38 | -0.13 | -0.02 | -0.39 | -0.1675 | -0.155 | -0.39 | 0.587101 | 0.637634 | 0.213004 | 0.999971 | 1 | 0.823987 |
| Tiam2 | Q6ZPF3-3 | 0.82 | 0.36 | 1.22 | 2.21 | -0.14 | -0.05 | 1.12 | 1.82 | -0.45 | -0.26 | 1.8 | 0.36 | -0.04 | -0.88 | 1.04 | 0.6 | -0.465 | -0.79 | -0.9725 | 0.4197 | 0.204996 | 0.093922 | 0.999971 | 1 | 0.777481 |
| Tiam2_1 | Q6ZPF3-2 | -0.61 | -0.37 | -0.99 | -0.4 | -0.2 | -0.3 | -0.82 | -0.61 | -0.25 | -0.38 | -0.63 | -0.76 | -0.45 | -0.12 | -0.44 | -0.21 | 0.11 | 0.0875 | 0.2875 | 0.643533 | 0.695761 | 0.20213 | 0.999971 | 1 | 0.81617 |
| Tigar | Q8BZA9 | -1.46 | -0.72 | 0.65 | 0.49 | -0.63 | -1.15 | 0.41 | 0.68 | -0.85 | -0.82 | 0.41 | 0.35 | -0.66 | -0.7 | 0.41 | 0.41 | 0.0875 | 0.0325 | 0.125 | 0.884845 | 0.954154 | 0.819292 | 0.999971 | 1 | 0.975699 |
| Timm10 | P62073 | -0.06 | 0.1 | -0.32 | 0.06 | -0.23 | -0.03 | -0.04 | -0.11 | -0.02 | -0.14 | -0.37 | -0.57 | 0.1 | 0.05 | -0.27 | -0.04 | -0.0475 | -0.22 | 0.015 | 0.794194 | 0.298976 | 0.937697 | 0.999971 | 1 | 0.995618 |
| Timm10b | Q9WV96 | -0.33 | -0.01 | 1.03 | 1.14 | 0.2 | -0.61 | 0.79 | 1.41 | 0.4 | -0.75 | 0.96 | 0.28 | 0.32 | -0.19 | 1.28 | 1.03 | -0.01 | -0.235 | 0.1525 | 0.984729 | 0.626704 | 0.744448 | 0.999971 | 1 | 0.961064 |
| Timm13 | P62075 | 0.5 | 0.54 | 0.47 | 0.83 | 0.99 | 0.91 | 0.5 | 0.6 | 0.91 | 0.55 | 0.55 | 0.17 | 1.33 | 0.92 | 0.29 | 0.18 | 0.165 | -0.04 | 0.095 | 0.422102 | 0.852764 | 0.74767 | 0.999971 | 1 | 0.961064 |
| Timm17a | Q9Z0V8 | -0.63 | -0.78 | -0.39 | -0.37 | -0.39 | -1.41 | -0.01 | 0.04 | -1.05 | -1.09 | -0.68 | -0.48 | -0.39 | -0.49 | -0.85 | 0.13 | 0.1 | -0.2825 | 0.1425 | 0.772658 | 0.218479 | 0.576401 | 0.999971 | 1 | 0.918194 |
| Timm17b | Q9Z0V7 | 1.28 | 0.08 | 0.32 | -0.15 | -0.22 | 0.1 | 0.57 | 0.07 | -0.03 | 0.37 | 0.41 | 0.44 | -0.25 | 0.37 | 0.25 | -0.37 | -0.2525 | -0.085 | -0.3825 | 0.475269 | 0.797916 | 0.299462 | 0.999971 | 1 | 0.847402 |
| Timm21 | Q8CCM6 | -0.38 | -0.4 | -0.66 | -0.31 | -0.19 | 0.08 | -1.24 | -0.6 | -0.61 | -0.33 | -0.53 | -0.57 | -0.21 | -0.01 | -0.73 | -0.39 | -0.05 | -0.0725 | 0.1025 | 0.86943 | 0.681216 | 0.639576 | 0.999971 | 1 | 0.93301 |
| Timm22 | Q9CQ85 | -0.54 | -0.66 | 1.09 | 0.94 | -0.76 | -0.89 | 0.59 | 2.05 | -0.63 | -1.11 | 1.02 | -0.17 | -0.95 | -0.68 | 1.42 | 0.48 | 0.04 | -0.43 | -0.14 | 0.95738 | 0.480339 | 0.83066 | 0.999971 | 1 | 0.978535 |
| Timm23 | A0A2I3BS4 | 1.36 | 0.02 | 0.01 | 0.59 | -0.26 | 0.43 | 0.09 | 0.36 | -0.31 | 0.96 | -0.21 | 0.28 | 0.03 | 0.31 | 0.11 | 0.43 | -0.34 | -0.315 | -0.275 | 0.343357 | 0.45181 | 0.41594 | 0.999971 | 1 | 0.876885 |
| Timm29 | Q8BGX2 | -1 | 0.46 | 0.3 | 0.44 | 0.63 | 0.35 | 0.11 | 0.25 | 0.67 | -0.59 | 0.07 | 0.06 | 0.7 | 0.68 | 0.41 | 0.52 | 0.285 | 0.0025 | 0.5275 | 0.436751 | 0.995194 | 0.160222 | 0.999971 | 1 | 0.800144 |
| Timm44 | O35857 | 0.06 | -0.07 | -0.18 | 0.02 | -0.35 | 0.03 | 0.21 | -0.17 | -0.11 | 0.15 | 0.4 | 0.04 | 0.28 | -0.33 | -0.05 | 0.01 | -0.0275 | 0.1625 | 0.02 | 0.887878 | 0.39239 | 0.919342 | 0.999971 | 1 | 0.992633 |
| Timm50 | Q9D880 | 0.82 | -1.01 | -0.27 | -0.19 | -1.27 | -0.8 | 0.23 | -0.18 | -1 | 0.31 | -0.24 | -0.21 | -1.65 | -0.88 | -0.3 | -0.21 | -0.3425 | -0.1225 | -0.5975 | 0.471266 | 0.779936 | 0.224327 | 0.999971 | 1 | 0.828225 |
| Timm8a1 | Q9WVA2 | -1.85 | -0.32 | 0.02 | 0.61 | -0.58 | -0.8 | -0.09 | 0.16 | -0.67 | -2.32 | 0.22 | -0.24 | -0.42 | -0.56 | 0.41 | 0.62 | 0.0575 | -0.3675 | 0.3975 | 0.912706 | 0.599431 | 0.479399 | 0.999971 | 1 | 0.894438 |
| Timm8b | P62077 | 0.38 | 1.14 | 1.35 | 0.1 | 0.87 | 1.01 | 2.1 | 0.58 | 1.25 | -0.23 | 1.59 | -0.06 | 1.15 | -0.07 | 1.43 | -0.36 | 0.3975 | -0.105 | -0.205 | 0.360068 | 0.836618 | 0.680792 | 0.999971 | 1 | 0.943097 |
| Timm9 | Q9WV98 | -0.09 | 0.11 | 0.01 | 0.14 | 0.13 | 0.44 | -0.34 | 0.28 | -0.04 | -0.33 | -0.16 | -0.38 | 0.05 | 0.86 | 0.76 | 0.04 | 0.085 | -0.27 | 0.385 | 0.700323 | 0.146504 | 0.156141 | 0.999971 | 1 | 0.796008 |
| Timmdc1 | Q8BUY5 | 1.32 | -0.17 | -0.33 | 0.12 | -0.55 | -0.02 | -0.05 | 0.41 | -0.03 | 0.11 | -0.21 | -0.59 | -0.69 | -0.28 | -0.36 | 0.15 | -0.2875 | -0.415 | -0.53 | 0.481809 | 0.301728 | 0.202855 | 0.999971 | 1 | 0.81617 |
| Timp2 | P25785 | -0.38 | -0.67 | -1.04 | -1.26 | 0.14 | -0.53 | -0.59 | -0.41 | 0.44 | -0.53 | -0.7 | -0.58 | 0.22 | -0.04 | -0.41 | -1.22 | 0.49 | 0.495 | 0.475 | 0.101294 | 0.156738 | 0.211051 | 0.999971 | 1 | 0.822884 |
| Tinagl1 | Q99JR5 | 0.46 | -0.36 | 0.22 | 1.7 | 4.53 | 0.2 | 1.28 | 0.44 | 0.84 | 2.57 | 2.79 | 4.01 | -2.36 | 0.55 | 1.47 | 0.45 | 1.1075 | 2.0475 | -0.4775 | 0.272991 | 0.017755 | 0.573276 | 0.999971 | 1 | 0.91816 |
| Tiprl | Q8BH58 | 0.08 | -1.2 | 0.12 | -0.49 | -0.56 | -1.18 | 0.13 | 0.12 | -0.7 | -1.13 | 0.08 | 0.21 | -0.97 | -1.23 | -0.1 | -0.28 | 7.85E-17 | -0.0125 | -0.2725 | 1 | 0.976379 | 0.495736 | 1 | 1 | 0.901531 |
| Tjp1 | B9EHJ3 | -0.02 | -0.85 | -0.56 | -0.56 | -0.87 | -0.8 | -0.52 | -0.84 | -0.7 | -0.17 | -0.75 | -0.66 | -0.92 | -0.72 | -0.6 | -0.72 | -0.26 | -0.0725 | -0.2425 | 0.274814 | 0.769156 | 0.300568 | 0.999971 | 1 | 0.847402 |
| Tjp2 | A0A3B2WC | -0.64 | -1.11 | -0.72 | -0.85 | -1.2 | -1.36 | -0.73 | -1.06 | -0.85 | -0.9 | -1.01 | -1.09 | -0.95 | -1.35 | -0.81 | -1.24 | -0.2575 | -0.1325 | -0.2575 | 0.251841 | 0.477892 | 0.244916 | 0.999971 | 1 | 0.837316 |
| Tk2 | Q9R088 | -0.38 | -0.24 | -0.33 | -0.41 | -0.45 | -0.45 | -0.45 | 0.13 | -0.32 | -0.13 | 0.12 | 0.13 | -0.15 | -0.06 | 0.06 | 0.42 | 0.035 | 0.29 | 0.4075 | 0.864054 | 0.139758 | 0.061822 | 0.999971 | 1 | 0.725348 |
| Tkfc | Q8VC30 | -0.48 | 0.45 | -0.19 | -0.14 | 0 | -0.39 | 0.32 | 0 | -0.05 | -0.67 | -0.14 | -0.32 | 0.35 | -0.18 | -0.41 | -0.16 | 0.0725 | -0.205 | -0.01 | 0.784418 | 0.438768 | 0.970636 | 0.999971 | 1 | 0.998914 |
| Tkt | P40142 | -0.75 | -0.59 | 0.48 | 0.47 | -0.46 | -0.78 | 0.4 | 0.15 | -0.44 | -0.92 | 0.26 | 0.02 | -0.67 | -0.51 | 0.24 | 0.51 | -0.075 | -0.1725 | -0.01 | 0.854305 | 0.671104 | 0.980846 | 0.999971 | 1 | 0.99925 |
| Tlcd1 | Q99JT6 | -0.7 | -0.16 | -0.95 | -1.75 | -0.97 | 0.19 | -1.3 | -0.91 | -0.7 | -0.33 | -0.82 | -0.68 | 0.16 | 0.68 | -1.45 | -0.96 | 0.1425 | 0.2575 | 0.4975 | 0.744623 | 0.460158 | 0.373473 | 0.999971 | 1 | 0.869657 |
| Tlcd3b | Q7TNV1 | -1.32 | -1.08 | -0.51 | -0.81 | -1.42 | -1.47 | 0.87 | -0.49 | -1.39 | -1.77 | -0.11 | -1.02 | -0.94 | -0.91 | -0.57 | -0.76 | 0.3025 | -0.1425 | 0.135 | 0.571513 | 0.71055 | 0.564338 | 0.999971 | 1 | 0.917087 |
| Tlcd4 | Q8CGF5 | 0.08 | 0.26 | 0.72 | 0.58 | 0.08 | -0.12 | 0.5 | 0.35 | 0.06 | -0.74 | 0.94 | 0.56 | 0.2 | 0.39 | 0.59 | 1.16 | -0.2075 | -0.205 | 0.175 | 0.389986 | 0.591437 | 0.526289 | 0.999971 | 1 | 0.909459 |
| Tle1 | Q5SQA2 | 1.17 | 3.27 | 2.25 | 1.61 | 2.48 | 1.22 | 0.85 | 1.68 | 0.68 | 1.82 | 1.83 | 1.74 | 1.54 | 0.06 | 2.3 | 2.21 | -0.5175 | -0.5575 | -0.5475 | 0.342661 | 0.281324 | 0.394343 | 0.999971 | 1 | 0.871816 |
| Tle3 | Q08122-4 | -0.14 | 1.64 | 1.87 | 2.62 | 1.5 | 1.78 | 2 | 2.27 | 1.3 | 0.33 | 2.63 | 2.75 | 1.03 | 1.79 | 1.99 | 2.08 | 0.39 | 0.255 | 0.225 | 0.489949 | 0.732935 | 0.698512 | 0.999971 | 1 | 0.947781 |
| Tle4 | F6ZBY9 | 0.83 | 1.37 | 1.77 | 1.59 | 1.57 | 1.44 | 1.71 | 1.83 | 1.23 | 1.21 | 1.95 | 1.33 | 0.74 | 1.76 | 1.39 | 1.78 | 0.2475 | 0.04 | 0.0275 | 0.334218 | 0.887245 | 0.931487 | 0.999971 | 1 | 0.995069 |
| Tle5 | P63002 | 2.17 | 0.89 | 2.8 | 2.51 | 0.53 | 1 | 2.89 | 2.6 | 0.82 | 1.46 | 1.98 | 0.44 | -0.64 | 0.38 | 1.76 | 0.95 | -0.3375 | -0.9175 | -1.48 | 0.607442 | 0.096806 | 0.033884 | 0.999971 | 1 | 0.689803 |
| Tln1 | P26039 | 0.88 | 0.55 | 0.09 | 0.45 | 1.64 | 0.76 | 0.38 | -0.16 | 0.89 | 1.56 | 0.47 | 0.96 | -0.02 | 0.6 | 0.54 | 0.32 | 0.1625 | 0.4775 | -0.1325 | 0.681925 | 0.12515 | 0.592657 | 0.999971 | 1 | 0.923227 |
| Tln2 | E9PUM4 | 0.05 | -0.48 | -0.31 | -0.41 | -0.44 | -0.5 | -0.14 | -0.49 | -0.38 | -0.2 | -0.27 | -0.49 | -0.55 | -0.63 | -0.53 | -0.55 | -0.105 | -0.0475 | -0.2775 | 0.605916 | 0.805772 | 0.167869 | 0.999971 | 1 | 0.800215 |
| Tln2_1 | A0A1L1SRI | 0.78 | 1.89 | -0.56 | -0.68 | 1 | 3.13 | -1.38 | -0.95 | 1.39 | 2.04 | -0.63 | -0.73 | 1.84 | 2.39 | -0.76 | -0.16 | 0.0925 | 0.16 | 0.47 | 0.930898 | 0.849114 | 0.594044 | 0.999971 | 1 | 0.923438 |
| Tlnrd1 | Q9ERE8 | 2.34 | 0.56 | 1.97 | 1.85 | 0.04 | 0.84 | 1.96 | 1.83 | 0.18 | 1.23 | 0.97 | 0.83 | -1.84 | -0.12 | 1.37 | 1.17 | -0.5125 | -0.8775 | -1.535 | 0.361029 | 0.065273 | 0.067814 | 0.999971 | 1 | 0.733937 |
| Tm2d1 | Q99MB3-2 | -1.52 | 1.2 | 0.83 | -0.75 | 0.18 | 1.4 | -0.34 | 0.45 | 0.15 | -1.04 | -0.73 | 0.54 | 0.13 | 0.85 | 0.53 | 1.51 | 0.4825 | -0.21 | 0.815 | 0.478682 | 0.756991 | 0.226841 | 0.999971 | 1 | 0.828236 |
| Tm7sf2 | Q71KT5 | 0.23 | 0.17 | 0.81 | 0.52 | -0.41 | 0.21 | 0.7 | 0.87 | 0.17 | 0.54 | 0.62 | 0.71 | 0.28 | 0.59 | 0.72 | 0.53 | -0.09 | 0.0775 | 0.0975 | 0.781345 | 0.732527 | 0.65824 | 0.999971 | 1 | 0.934638 |
| Tm9sf1 | D6RGM8 | 0.92 | 0.62 | 1.32 | 0.93 | 0.17 | 0.4 | 0.25 | 1.7 | 0.08 | 0.26 | 0.85 | 1.48 | -0.54 | 0.56 | 2.08 | 0.96 | -0.3175 | -0.28 | -0.1825 | 0.406848 | 0.424106 | 0.725386 | 0.999971 | 1 | 0.956041 |
| Tm9sf2 | P58021 | -0.78 | 2.17 | 1.35 | 1.37 | 2.03 | 2.13 | 0.89 | 1.38 | 1.86 | 0.26 | 1.25 | 1.44 | 1.89 | 2.41 | 1.4 | 1.81 | 0.58 | 0.175 | 0.85 | 0.37105 | 0.789578 | 0.185885 | 0.999971 | 1 | 0.809025 |
| Tm9sf3 | A0A498WG | 1.21 | 2.48 | 2.28 | 2.41 | 2.45 | 2.37 | 2.86 | 2.56 | 2.69 | 2.14 | 3.05 | 3.56 | 1.61 | 1.95 | 2.38 | 1.87 | 0.465 | 0.765 | -0.1425 | 0.169913 | 0.085998 | 0.673339 | 0.999971 | 1 | 0.94048 |
| Tm9sf4 | Q8BH24 | 0.88 | 1.47 | 1.32 | 1.21 | 1.23 | 1.62 | 1.01 | 1.67 | 1.14 | 1.19 | 1.39 | 1.43 | 1.27 | 1.77 | 1.61 | 1.71 | 0.1625 | 0.0675 | 0.37 | 0.497857 | 0.73496 | 0.113423 | 0.999971 | 1 | 0.78268 |
| Tma16 | Q9CR02 | 1.33 | 1.3 | 1.3 | 2.36 | 0.88 | 1.2 | 1.28 | 1.7 | 1.54 | 1.78 | 2.03 | 2.29 | -0.35 | 1.8 | 2.19 | 2.73 | -0.3075 | 0.3375 | 0.02 | 0.343335 | 0.296345 | 0.97573 | 0.999971 | 1 | 0.99925 |
| Tma7 | Q8K003 | 1.49 | 0.91 | 1.94 | 1.66 | 1.11 | 2.03 | 3.02 | 1.62 | 0.43 | 1.66 | 1.46 | 1.36 | -0.02 | 0.35 | 2.13 | 1.23 | 0.445 | -0.2725 | -0.5775 | 0.319632 | 0.437413 | 0.258443 | 0.999971 | 1 | 0.839012 |
| Tmbim1 | D3Z0A5 | -0.14 | -0.31 | -1.22 | -1.21 | 0.19 | -0.85 | -1.19 | 0.11 | -0.42 | -0.88 | -0.81 | -1.14 | -1.75 | -1.76 | -1.69 | -1.18 | 0.285 | -0.0925 | -0.875 | 0.508358 | 0.776028 | 0.022478 | 0.999971 | 1 | 0.652043 |
| Tmcc1 | F8WJ98 | -0.38 | -0.31 | -1.07 | -0.88 | -0.2 | 0.04 | -0.87 | -1.03 | -0.16 | -0.14 | -0.39 | -0.66 | 0.14 | -0.44 | -1.18 | -0.77 | 0.145 | 0.3225 | 0.0975 | 0.652586 | 0.217059 | 0.770973 | 0.999971 | 1 | 0.963498 |
| Tmcc2 | Q80W04 | -0.19 | -0.58 | 0.33 | -0.59 | -0.52 | -0.81 | -0.56 | 0 | -0.46 | -0.37 | -0.1 | -0.16 | -0.28 | -0.47 | 0.33 | 0.14 | -0.215 | -0.015 | 0.1875 | 0.462733 | 0.953262 | 0.530469 | 0.999971 | 1 | 0.910069 |
| Tmcc3 | Q8R310-2 | -0.67 | -1.06 | -0.32 | 0.05 | -1 | -0.89 | -0.98 | 0.06 | -0.95 | -0.75 | -0.59 | 0.15 | -1.44 | -0.76 | -0.4 | -0.1 | -0.2025 | -0.035 | -0.175 | 0.560018 | 0.916823 | 0.6335 | 0.999971 | 1 | 0.93301 |
| Tmco1 | A0A0A6YV | 0.01 | -0.68 | -0.94 | -0.89 | -1.16 | -0.77 | -0.53 | -0.55 | -1.17 | -1.36 | -0.69 | -0.84 | -1.14 | -0.65 | -1.02 | -0.78 | -0.1275 | -0.39 | -0.2725 | 0.649681 | 0.188854 | 0.322389 | 0.999971 | 1 | 0.85548 |
| Tmco6 | Q8BQX5 | 1.2 | 0.49 | -0.05 | 0.05 | 0.43 | 0.59 | -0.64 | 0.36 | 1.18 | 0.47 | 0.1 | -0.06 | 1.6 | -0.05 | 0.71 | 0.21 | -0.2375 | -3.53E-16 | 0.195 | 0.540485 | 1 | 0.656514 | 0.999971 | 1 | 0.934061 |
| Tmed10 | Q9D1D4 | 0.64 | 0.8 | 0.12 | 0.05 | 0.92 | 0.76 | 0.71 | -0.1 | 0.61 | 0.76 | 0.41 | 0.64 | 0.45 | 1.14 | 0.31 | 0.17 | 0.17 | 0.2025 | 0.115 | 0.577747 | 0.396637 | 0.698402 | 0.999971 | 1 | 0.947781 |
| Tmed2 | F6V6T4 | 1.73 | 0.46 | 1.83 | 2.06 | 0.85 | 1.05 | 2.3 | 1.67 | 1.23 | 1.14 | 1.61 | 1.79 | 0.73 | 0.39 | 1.17 | 0.06 | -0.0525 | -0.0775 | -0.9325 | 0.908328 | 0.838072 | 0.046695 | 0.999971 | 1 | 0.710102 |
| Tmed3 | Q78IS1 | 1.65 | 1.42 | 0.9 | 1.21 | 1.64 | 1.6 | 0.66 | 1.1 | 1.89 | 1.77 | 0.73 | 1.65 | 0.76 | 2.01 | 1.53 | 1.99 | -0.045 | 0.215 | 0.2775 | 0.877643 | 0.497732 | 0.413761 | 0.999971 | 1 | 0.876329 |
| Tmed4 | Q8R1V4 | 2.3 | 1.31 | 1.44 | 1.93 | 1.24 | 1.46 | 2.01 | 1.42 | 1.21 | 1.65 | 1.02 | 1.04 | 0.83 | 1.27 | 1.05 | 1.24 | -0.2125 | -0.515 | -0.6475 | 0.47507 | 0.097247 | 0.037504 | 0.999971 | 1 | 0.700096 |
| Tmed5 | Q9CXE7 | -2.02 | -0.96 | -0.84 | -0.71 | 0.55 | -0.36 | -0.16 | -0.08 | 0.35 | -1.3 | 0.14 | 0.57 | 0.74 | -1.33 | -0.3 | 0.11 | 1.12 | 1.0725 | 0.9375 | 0.010683 | 0.0513 | 0.083716 | 0.999971 | 1 | 0.755254 |
| Tmed7 | E9Q7G1 | -0.68 | 1.04 | 1.33 | 0.37 | 0.62 | 0.92 | 1.33 | 0.9 | 0.56 | -0.4 | 1.47 | 1.54 | -0.38 | 0.63 | 1.11 | 0.33 | 0.4275 | 0.2775 | -0.0925 | 0.346547 | 0.638194 | 0.854726 | 0.999971 | 1 | 0.982125 |
| Tmed8 | Q3UHI4 | 0.13 | 1.31 | 1.05 | 0.82 | 1.41 | 1.28 | 0.65 | 0.86 | 1.33 | 0.9 | 1.1 | 1.14 | 1.71 | 1.19 | 1.13 | 1.02 | 0.2225 | 0.29 | 0.435 | 0.484068 | 0.318289 | 0.176847 | 0.999971 | 1 | 0.808532 |
| Tmed9 | Q99KF1 | 0.57 | 1.16 | 0.91 | 0.97 | 1.13 | 1.29 | 1.61 | 0.83 | 1.19 | 0.78 | 0.68 | 1 | 1.27 | 1.27 | 0.93 | 1.01 | 0.3125 | 0.01 | 0.2175 | 0.210867 | 0.962462 | 0.306819 | 0.999971 | 1 | 0.847402 |
| Tmeff1 | A0A0A0MC | 0.12 | -1.12 | -1.19 | -1.91 | -1.02 | -0.57 | -0.51 | -1.43 | -0.38 | 0.39 | -1.22 | -1.22 | -0.02 | -0.7 | -1.15 | -1.59 | 0.1425 | 0.4175 | 0.16 | 0.749208 | 0.439519 | 0.749564 | 0.999971 | 1 | 0.961064 |
| Tmeff2 | Q9QYM9 | 0.48 | -0.81 | 0.41 | -0.29 | -0.17 | -0.2 | 1.62 | -0.72 | 0.55 | 0.44 | 0.07 | 1.33 | 0.77 | -1.15 | -0.35 | -1.08 | 0.185 | 0.65 | -0.4 | 0.736327 | 0.123299 | 0.434481 | 0.999971 | 1 | 0.882419 |
| Tmem106b | Q80X71 | 1.83 | 1.71 | 1.64 | 2.43 | 1.32 | 1.72 | 1.71 | 2.37 | 1.54 | 2.39 | 1.9 | 1.68 | 2.16 | 1.94 | 2.16 | 2.59 | -0.1225 | -0.025 | 0.31 | 0.677454 | 0.927516 | 0.240986 | 0.999971 | 1 | 0.834478 |
| Tmem109 | Q3UBX0 | -1.15 | -0.3 | -0.1 | -0.28 | -0.42 | -0.5 | 0.1 | -0.03 | -0.58 | -1.34 | 0.19 | -0.25 | -0.56 | -0.86 | -0.27 | -0.58 | 0.245 | -0.0375 | -0.11 | 0.406578 | 0.922435 | 0.695974 | 0.999971 | 1 | 0.947174 |
| Tmem11 | Q8BK08 | 1.02 | -0.35 | 0.36 | 0.94 | -0.71 | -0.32 | 0.81 | -0.2 | -0.52 | 0.44 | -0.19 | -0.89 | -1.41 | -0.6 | -0.51 | -0.61 | -0.5975 | -0.7825 | -1.275 | 0.187019 | 0.081403 | 0.007166 | 0.999971 | 1 | 0.582714 |
| Tmem114 | Q9D563 | 2.19 | -2.17 | -1.99 | -0.44 | -1.67 | -1.23 | -0.98 | -1.72 | -3.11 | -0.99 | -1.34 | -1.08 | -1.79 | -1.54 | -1.53 | -0.85 | -0.7975 | -1.0275 | -0.825 | 0.394412 | 0.324557 | 0.381826 | 0.999971 | 1 | 0.871816 |
| Tmem115 | Q9WUH1 | -0.06 | -0.27 | 0.46 | 1.01 | -0.22 | -0.26 | -0.09 | 0.89 | -0.39 | -0.16 | 0.29 | 0.29 | -0.1 | -0.06 | 0.56 | 0.86 | -0.205 | -0.2775 | 0.03 | 0.593574 | 0.412339 | 0.934024 | 0.999971 | 1 | 0.995387 |
| Tmem117 | Q8BH18 | -6.39 | -9.36 | -10.54 | -7.94 | -9.65 | -7.58 | -7.8 | -7.41 | -9.38 | -6.96 | -12.72 | -8.49 | -9.76 | -9.65 | -10.18 | -8.8 | 0.4475 | -0.83 | -1.04 | 0.631258 | 0.543978 | 0.239647 | 0.999971 | 1 | 0.83365 |
| Tmem120a | Q8C1E7 | -0.53 | -0.03 | 0.02 | 0.45 | -0.03 | -0.09 | 0.2 | 0.53 | -0.21 | -1.68 | 0.13 | 0.16 | 0.58 | 0.13 | 1.12 | 0.85 | 0.175 | -0.3775 | 0.6925 | 0.515632 | 0.412719 | 0.043693 | 0.999971 | 1 | 0.702317 |
| Tmem121b | Q99MX7 | -0.94 | 1.23 | 0.19 | -0.2 | 0.44 | 0.71 | -0.2 | 0.03 | 0.4 | -0.18 | -0.17 | -1.98 | 0.98 | 0.8 | 0.28 | -0.11 | 0.175 | -0.5525 | 0.4175 | 0.706856 | 0.390572 | 0.396153 | 0.999971 | 1 | 0.871816 |
| Tmem126a | Q9D8Y1 | -0.9 | -0.63 | -0.63 | -0.69 | -0.74 | -0.4 | -0.57 | -0.72 | -0.91 | -1.21 | -0.71 | -0.49 | -0.81 | -0.81 | -0.84 | -0.9 | 0.105 | -0.1175 | -0.1275 | 0.564012 | 0.581824 | 0.456154 | 0.999971 | 1 | 0.887217 |
| Tmem126b | Q9D1R1 | -0.38 | 0.41 | 0.12 | 0.62 | 0.22 | 0.16 | 0.02 | 0.87 | 0.13 | -0.66 | 0.12 | 0.55 | 0.55 | 0.41 | 0.98 | 1.1 | 0.125 | -0.1575 | 0.5675 | 0.675094 | 0.636675 | 0.074362 | 0.999971 | 1 | 0.746069 |
| Tmem127 | G3UXI2 | 0.01 | -0.26 | -1.57 | -1.47 | -0.45 | 0.15 | -0.94 | -1.21 | -0.15 | 0.48 | -0.72 | -1.98 | 0.37 | -0.1 | -0.92 | -0.71 | 0.21 | 0.23 | 0.4825 | 0.657513 | 0.706281 | 0.319203 | 0.999971 | 1 | 0.853893 |
| Tmem130 | Q6NXM3 | 1.05 | 0.8 | 0.47 | 0.69 | 1.62 | 2.18 | 0.01 | 0.58 | 0.8 | -0.19 | 0.18 | 1.18 | 0.11 | 2.31 | 0.73 | 0.23 | 0.345 | -0.26 | 0.0925 | 0.471652 | 0.438102 | 0.848824 | 0.999971 | 1 | 0.981643 |
| Tmem131 | O70472 | 1.71 | 0.12 | -0.18 | -0.38 | 0.04 | 0.04 | 0.62 | -1.42 | -0.23 | 1.19 | -0.71 | 0.62 | 0.16 | 0.81 | -0.92 | 0.19 | -0.4975 | -0.1 | -0.2575 | 0.408038 | 0.864575 | 0.640811 | 0.999971 | 1 | 0.93301 |
| Tmem132a | Q922P8 | -0.35 | -0.36 | -0.7 | -0.54 | -0.34 | 0.22 | -0.85 | 0.1 | -0.12 | 0.08 | -0.5 | -0.49 | 0.1 | 0.41 | -0.32 | 0.17 | 0.27 | 0.23 | 0.5775 | 0.338566 | 0.293144 | 0.025395 | 0.999971 | 1 | 0.678474 |
| Tmem132b | F7BAB2 | 0.05 | -0.25 | -0.54 | -0.71 | -0.28 | -0.29 | -0.43 | -0.85 | -0.46 | -0.31 | -0.45 | -0.54 | 0.02 | -0.45 | -1.03 | -0.72 | -0.1 | -0.0775 | -0.1825 | 0.683805 | 0.721093 | 0.534779 | 0.999971 | 1 | 0.910077 |
| Tmem132c | Q8CEF9 | 0.65 | 1.38 | 0.62 | 0.11 | 1.55 | 1.23 | -0.64 | 0.88 | 1.37 | 0.27 | 0.34 | -1.07 | 1.91 | 0.86 | 0.29 | 0.84 | 0.065 | -0.4625 | 0.285 | 0.898409 | 0.388196 | 0.491295 | 0.999971 | 1 | 0.899128 |
| Tmem132d | A0A0R4J0I | -0.33 | -0.45 | -0.63 | -0.53 | -0.62 | -0.15 | -0.6 | -0.49 | -0.79 | -0.56 | -0.52 | -0.42 | -0.83 | -1.12 | -0.83 | -0.55 | 0.02 | -0.0875 | -0.3475 | 0.916866 | 0.622652 | 0.103258 | 0.999971 | 1 | 0.78268 |
| Tmem132e | Q6IEE6 | -0.14 | -1.27 | -0.65 | -0.79 | -0.88 | -1.21 | -0.6 | -0.85 | -1 | -0.39 | -0.93 | -0.7 | -1.13 | -0.9 | -0.67 | -1.14 | -0.1725 | -0.0425 | -0.2475 | 0.541688 | 0.880654 | 0.380695 | 0.999971 | 1 | 0.871816 |
| Tmem135 | Q9CYV5 | -1.72 | -0.53 | 0.06 | 0.19 | -0.44 | -0.13 | -0.17 | 0.55 | -0.5 | -1.64 | -0.03 | -0.02 | 0.48 | -0.7 | -0.23 | 0.18 | 0.4525 | -0.0475 | 0.4325 | 0.332886 | 0.929324 | 0.371958 | 0.999971 | 1 | 0.869526 |
| Tmem141 | A2AJB2 | 0.66 | 0.04 | 0.59 | -0.32 | 0.01 | 0.71 | 1 | 1 | -0.66 | 0.4 | 0.53 | 0.53 | 0.65 | -0.6 | 0.23 | 0.03 | 0.4375 | -0.0425 | -0.165 | 0.206774 | 0.906513 | 0.634882 | 0.999971 | 1 | 0.93301 |
| Tmem143 | A0A1B0GT | 0.62 | -0.83 | 0.13 | 0.2 | -0.92 | -0.86 | -0.24 | 0.37 | -1.34 | 0.08 | -0.1 | 0.02 | -2.28 | -1.15 | 0.02 | -0.01 | -0.4425 | -0.365 | -0.885 | 0.296821 | 0.408016 | 0.150173 | 0.999971 | 1 | 0.793248 |
| Tmem144 | Q8VEH0 | -0.26 | -0.22 | 0.39 | -0.04 | 0.06 | 0.53 | 0.35 | 1.25 | -0.08 | -0.4 | 0.15 | 0.02 | 0.74 | 0.86 | 0.51 | 1.1 | 0.58 | -0.045 | 0.835 | 0.081421 | 0.842516 | 0.005493 | 0.999971 | 1 | 0.582714 |
| Tmem145 | E9PUE6 | 0.08 | 0.96 | -0.02 | -0.06 | 0.85 | 1.02 | 0.55 | 0.09 | 0.49 | 0.46 | -0.27 | -0.51 | 0.26 | 0.83 | -0.32 | -0.55 | 0.3875 | -0.1975 | -0.185 | 0.244078 | 0.571793 | 0.628582 | 0.999971 | 1 | 0.932176 |
| Tmem147 | A0A140LIQ | 2.55 | 1.47 | 1.01 | 1.29 | -0.55 | 0.48 | 0.14 | 0.64 | -0.08 | 0.69 | 0.75 | -0.49 | 1.57 | 0.67 | 0.76 | 1.08 | -1.4025 | -1.3625 | -0.56 | 0.007174 | 0.011075 | 0.166499 | 0.999971 | 1 | 0.800215 |
| Tmem14c | Q9CQN6 | -0.25 | -1.33 | -1.46 | -1.67 | -1.41 | -0.56 | -1.91 | -1.62 | -1.95 | -1.27 | -1.9 | -1.87 | -1.14 | -1.16 | -2.13 | -1.68 | -0.1975 | -0.57 | -0.35 | 0.631528 | 0.130044 | 0.371706 | 0.999971 | 1 | 0.869526 |
| Tmem151a | Q6GQT5 | -1.29 | -0.78 | -1.31 | -1.27 | -1.64 | -0.87 | -1.42 | -1.01 | -1.18 | -1.83 | -1.57 | -1.02 | -1.09 | -0.8 | -1.28 | -1.08 | -0.0725 | -0.2375 | 0.1 | 0.770599 | 0.354747 | 0.639074 | 0.999971 | 1 | 0.93301 |
| Tmem151b | Q68FE7 | 0.48 | -1.04 | -1.31 | -1.39 | -1.24 | -0.48 | -1.13 | -1.14 | -1.01 | -0.27 | -1.66 | -1.87 | -0.65 | -1.1 | -1.44 | -0.87 | -0.1825 | -0.3875 | -0.2 | 0.681987 | 0.468745 | 0.653209 | 0.999971 | 1 | 0.934005 |
| Tmem160 | Q9D938 | 0.74 | -1.38 | -0.62 | -0.78 | -1.98 | -1.15 | 0.97 | -0.77 | -2.19 | -0.27 | -0.83 | -1.23 | -2 | -1.38 | -1.08 | -0.84 | -0.2225 | -0.62 | -0.815 | 0.748464 | 0.283207 | 0.11619 | 0.999971 | 1 | 0.78268 |
| Tmem161b | Q8C2L6 | 0.16 | 0.54 | 0.61 | 0.39 | -0.29 | 1.23 | 0.19 | 0.74 | -0.47 | -0.51 | 0.64 | 0.69 | -0.09 | 1.64 | 0.42 | 1.16 | 0.0425 | -0.3375 | 0.3575 | 0.900714 | 0.340056 | 0.363272 | 0.999971 | 1 | 0.869526 |
| Tmem163 | Q8C996 | 0.02 | -1.45 | -0.77 | -0.21 | -0.94 | -1.19 | 0.29 | -0.71 | -0.5 | -0.21 | -0.74 | -0.95 | -0.72 | -1.18 | -1.08 | -0.73 | -0.035 | 0.0025 | -0.325 | 0.935788 | 0.994402 | 0.358224 | 0.999971 | 1 | 0.869102 |
| Tmem164 | Q6PHN7 | 3.32 | -0.61 | -0.1 | 0.85 | -0.01 | 1.01 | 0.08 | 1 | 0.13 | 2.7 | 0.05 | 0.07 | 0.91 | 0.84 | 1.21 | 1.54 | -0.345 | -0.1275 | 0.26 | 0.676231 | 0.896533 | 0.745261 | 0.999971 | 1 | 0.961064 |
| Tmem165 | P52875 | 0.25 | 0.97 | -0.04 | 0.4 | 0.93 | 1.04 | -0.12 | -0.04 | 0.37 | 0.52 | -0.5 | 0 | 0.61 | 1.1 | 0.07 | 0.94 | 0.0575 | -0.2975 | 0.285 | 0.874664 | 0.356705 | 0.377891 | 0.999971 | 1 | 0.871417 |
| Tmem167 | E0CY07 | -0.89 | -0.5 | -1.36 | -1.63 | 0.29 | 0.07 | -1.1 | -0.89 | -0.29 | -1.43 | -1.08 | -0.06 | 0.12 | 0.38 | 0.3 | -0.71 | 0.6875 | 0.38 | 1.1175 | 0.117822 | 0.347357 | 0.010411 | 0.999971 | 1 | 0.613095 |
| Tmem168 | A0A0N4SW | -0.66 | -0.01 | 0.77 | 0.86 | 0.33 | 0.05 | 0.48 | 0.66 | 0.45 | -0.24 | 0.19 | 0.58 | 0.92 | -0.22 | 0.54 | 0.65 | 0.14 | 0.005 | 0.2325 | 0.705612 | 0.989676 | 0.576871 | 0.999971 | 1 | 0.918194 |
| Tmem169 | Q8BG50 | -1.74 | -1.71 | -0.61 | -0.85 | -2.54 | -2.21 | -1.3 | -0.77 | -1.78 | -1.59 | -0.63 | -0.73 | -2.88 | -1.7 | -0.42 | -0.33 | -0.4775 | 0.045 | -0.105 | 0.322134 | 0.909709 | 0.863677 | 0.999971 | 1 | 0.983079 |
| Tmem175 | Q9CXY1 | -0.11 | 1.17 | 0.57 | 0.07 | 0.6 | 0.93 | 3.3 | -0.94 | 0.55 | 0.63 | -0.77 | -0.44 | 0.55 | 1.01 | 0.23 | -1.76 | 0.5475 | -0.4325 | -0.4175 | 0.513443 | 0.329748 | 0.503342 | 0.999971 | 1 | 0.904824 |
| Tmem177 | Q8BPE4 | 0.56 | 0.15 | 0.43 | 0.73 | -0.07 | 0.56 | 0.51 | 1.51 | 0.01 | 0.06 | 0.41 | 0.7 | -0.42 | 0.6 | 1.09 | 0.75 | 0.16 | -0.1725 | 0.0375 | 0.64417 | 0.470448 | 0.913038 | 0.999971 | 1 | 0.992618 |
| Tmem178a | Q9CZ16 | 0.11 | -0.25 | -0.41 | -0.62 | -0.48 | -0.24 | 0.31 | -0.86 | -1.02 | -0.64 | -0.22 | -1.61 | -0.23 | -0.75 | -1.66 | -1.35 | -0.025 | -0.58 | -0.705 | 0.933175 | 0.108105 | 0.069422 | 0.999971 | 1 | 0.733937 |
| Tmem178b | M0QWJ9 | -0.15 | 1.04 | -1.47 | -1.47 | 2.96 | 1.13 | -1.55 | -1.41 | 1.59 | 1.04 | -1.49 | -1.24 | 1.78 | 0.85 | -1.3 | -1.09 | 0.795 | 0.4875 | 0.5725 | 0.478401 | 0.587859 | 0.513137 | 0.999971 | 1 | 0.905815 |
| Tmem181a | Q3U3W2 | 0.01 | 0.61 | 0.1 | 0.38 | 0.8 | 0.62 | 0.02 | 0.94 | 0.93 | 0.69 | 0 | 0.52 | 2.77 | 1.04 | 0.55 | 1.3 | 0.32 | 0.26 | 1.14 | 0.247723 | 0.333496 | 0.034707 | 0.999971 | 1 | 0.69205 |
| Tmem184b | A0A2R8VI0 | 0.02 | -0.09 | -0.82 | -0.41 | 0.16 | -0.99 | -1.21 | 0.34 | -0.63 | -0.37 | -0.72 | -0.49 | 0.1 | -0.1 | 0.21 | 0.18 | -0.1 | -0.2275 | 0.4225 | 0.809673 | 0.348052 | 0.101407 | 0.999971 | 1 | 0.78268 |
| Tmem185a | Q3TYW3 | 1.7 | 0.1 | -0.57 | 0.12 | -0.35 | 0.31 | -0.55 | 1.08 | 0.37 | 0.91 | -0.5 | -0.21 | 0.65 | 0.22 | 0.52 | 0.88 | -0.215 | -0.195 | 0.23 | 0.700207 | 0.715019 | 0.625965 | 0.999971 | 1 | 0.930479 |
| Tmem186 | Q9CR76 | 0.68 | 0.5 | 0.1 | 0.04 | -0.09 | -0.36 | 0.11 | 0.12 | 0.06 | 0.43 | -0.68 | -0.56 | 0.3 | 0.21 | -0.45 | -0.37 | -0.385 | -0.5175 | -0.4075 | 0.120822 | 0.12174 | 0.155099 | 0.999971 | 1 | 0.795578 |
| Tmem19 | A0A1W2P7 | -0.21 | -0.08 | -0.3 | -0.11 | -0.02 | 0.26 | -0.2 | 0.37 | 0.55 | 0.49 | 0.67 | 0.69 | 1.08 | 0.19 | 0.08 | 0.85 | 0.2775 | 0.775 | 0.725 | 0.186376 | 0.001166 | 0.023646 | 0.999971 | 0.590539 | 0.663563 |
| Tmem198 | Q8BG75 | -0.57 | -1.36 | -0.69 | -0.47 | -1.32 | -1.22 | -0.43 | -0.12 | -1.93 | -1.23 | -0.19 | -0.39 | -1.62 | -1.2 | -0.71 | -0.37 | 3.14E-16 | -0.1625 | -0.2025 | 1 | 0.703866 | 0.553123 | 1 | 1 | 0.913595 |
| Tmem198b | Q8CFU0 | 1.4 | 0.43 | 1.34 | -6.83 | 0.42 | -0.15 | 0.93 | -5.17 | 0.25 | 0.9 | 1.41 | 0.49 | -0.53 | 0.63 | -0.61 | -4.33 | -0.0775 | 1.6775 | -0.295 | 0.971224 | 0.359877 | 0.882674 | 0.999971 | 1 | 0.986576 |
| Tmem199 | Q5SYH2 | -0.82 | -2.08 | 1.66 | 1.48 | -0.73 | -2.33 | 0.85 | 1.73 | -1.62 | -1.33 | 1.62 | 0.73 | -1.08 | -1.47 | 1.96 | 1.2 | -0.18 | -0.21 | 0.0925 | 0.874012 | 0.845931 | 0.933269 | 0.999971 | 1 | 0.995138 |
| Tmem200a | Q8C817 | -2 | -2.18 | -1.35 | -3.46 | -1.88 | -2.25 | -0.31 | -2.59 | -1.86 | -2.61 | -0.83 | -1.29 | -1.9 | -1.99 | -1.77 | -3.44 | 0.49 | 0.6 | -0.0275 | 0.43091 | 0.285251 | 0.959607 | 0.999971 | 1 | 0.998721 |
| Tmem200c | J3QK46 | -1.23 | -0.63 | -0.93 | -1.68 | -2.29 | -1.56 | -2.07 | -1.98 | -2.26 | -1.28 | -1.87 | -1.71 | -0.27 | -1.63 | -1.88 | -1.6 | -0.8575 | -0.6625 | -0.2275 | 0.013927 | 0.056237 | 0.580394 | 0.999971 | 1 | 0.91995 |
| Tmem201 | A2A8U2-2 | -0.72 | -0.58 | -1.76 | -1.95 | -0.86 | -0.25 | -0.91 | -1.73 | -0.84 | -0.92 | -1.54 | -1.76 | -0.97 | -0.32 | -1.87 | -1.77 | 0.315 | -0.0125 | 0.02 | 0.478254 | 0.975107 | 0.966261 | 0.999971 | 1 | 0.998721 |
| Tmem205 | A0A1L1SSA | 0.93 | 0.88 | 0.3 | -0.37 | 1.06 | 0.84 | 0.24 | 0.36 | 1.04 | 1.05 | 0.66 | 0.73 | 1.49 | 1.2 | 0.18 | 0.58 | 0.19 | 0.435 | 0.4275 | 0.594081 | 0.200042 | 0.307841 | 0.999971 | 1 | 0.847402 |
| Tmem209 | Q8BRG8-2 | -1.16 | -1.05 | -0.41 | -0.53 | -1.05 | -0.94 | -0.65 | 0.17 | -0.74 | -1.24 | -0.45 | 0.62 | -1.01 | -0.69 | -0.3 | -0.09 | 0.17 | 0.335 | 0.265 | 0.610736 | 0.428196 | 0.371856 | 0.999971 | 1 | 0.869526 |
| Tmem214 | Q8BM55 | -0.38 | 0.08 | 0.08 | 0.14 | 0.05 | -0.1 | 0.08 | 0.05 | 0.12 | -0.66 | -0.11 | 0.02 | 0.77 | -0.07 | -0.16 | -0.23 | 0.04 | -0.1375 | 0.0975 | 0.835448 | 0.572525 | 0.727696 | 0.999971 | 1 | 0.956041 |
| Tmem222 | Q8BVA2 | -0.72 | -0.34 | -0.91 | -0.96 | -0.45 | -0.99 | -0.94 | -0.96 | -0.45 | -1.17 | -1 | -0.56 | -0.33 | -1.02 | -0.74 | -1.22 | -0.1025 | -0.0625 | -0.095 | 0.656422 | 0.801867 | 0.717943 | 0.999971 | 1 | 0.953282 |
| Tmem223 | Q9CQE2 | -1.83 | -0.39 | -1.19 | -1.14 | -0.26 | -0.75 | -0.85 | -0.5 | -0.56 | -0.97 | -1.01 | -0.65 | -0.29 | 0.38 | -0.27 | -0.73 | 0.5475 | 0.34 | 0.91 | 0.118195 | 0.301079 | 0.031997 | 0.999971 | 1 | 0.681356 |
| Tmem229a | B9EJI9 | 3.08 | -1.25 | -0.12 | -0.28 | -1.65 | -0.6 | -0.37 | -0.3 | -1.35 | 2.18 | 0.01 | -0.7 | -1.01 | -0.71 | -0.52 | -0.25 | -1.0875 | -0.3225 | -0.98 | 0.240894 | 0.767655 | 0.271133 | 0.999971 | 1 | 0.843794 |
| Tmem230 | Q8CIB6 | 0.77 | 0.46 | 1.37 | 1.63 | 0.35 | 0.39 | 1.73 | 1.81 | 0.28 | 0.21 | 1.3 | 1.71 | 1.96 | 0.86 | 1.35 | 1.02 | 0.0125 | -0.1825 | 0.24 | 0.978065 | 0.67662 | 0.505424 | 0.999971 | 1 | 0.905277 |
| Tmem231 | Q3U284 | -2.25 | -0.67 | -0.19 | -0.51 | -1.13 | -1.96 | -0.44 | 0.45 | -1.13 | -2.16 | -1.62 | -0.88 | -1.27 | -0.75 | -0.09 | -0.41 | 0.135 | -0.5425 | 0.275 | 0.829198 | 0.296969 | 0.576577 | 0.999971 | 1 | 0.918194 |
| Tmem237 | Q3V0J1-3 | 5.32 | 0.43 | -0.05 | 0.21 | 0.26 | 0.68 | -0.03 | 0.18 | 0.18 | 4.25 | -0.5 | -0.43 | 0.6 | 0.81 | -0.62 | -0.43 | -1.205 | -0.6025 | -1.3875 | 0.310332 | 0.695115 | 0.261734 | 0.999971 | 1 | 0.842262 |
| Tmem240 | B2RWJ3 | -2.46 | -2.47 | -1.91 | -1.73 | -2.34 | -2.86 | -1.74 | -1.83 | -2.07 | -2.17 | -0.83 | -0.9 | -3.98 | -2.55 | -1.2 | -1.21 | -0.05 | 0.65 | -0.0925 | 0.876703 | 0.126446 | 0.882849 | 0.999971 | 1 | 0.986576 |
| Tmem243 | A0A0G2JEI | -2.31 | 0.76 | -0.17 | -0.27 | -0.26 | -0.31 | -0.43 | -0.61 | -0.25 | -0.96 | -0.65 | -0.37 | 1.12 | -0.01 | -0.19 | -0.56 | 0.095 | -0.06 | 0.5875 | 0.873104 | 0.921701 | 0.393242 | 0.999971 | 1 | 0.871816 |
| Tmem245 | B1AZA5 | 0.91 | -0.02 | -0.37 | -0.28 | -0.05 | 0.75 | 0.3 | -0.64 | -0.17 | 1.16 | 0.68 | -0.76 | -0.1 | 0.4 | -0.68 | -0.4 | 0.03 | 0.1675 | -0.255 | 0.939664 | 0.730779 | 0.490014 | 0.999971 | 1 | 0.898612 |
| Tmem25 | A0A1N9MY | 1.15 | -0.25 | 0.06 | -0.84 | -0.26 | 0.24 | 1.37 | -1.51 | -1.17 | -0.63 | -1.22 | -1.68 | -1.74 | -1.27 | -1.83 | -3 | -0.07 | -1.205 | -1.99 | 0.915519 | 0.022748 | 0.003832 | 0.999971 | 1 | 0.582714 |
| Tmem254 | V9GX22 | 0.04 | -0.5 | -1.59 | -2.05 | -0.23 | -0.04 | -1.06 | -2.36 | -0.72 | 0.26 | -1.09 | -1.94 | 0.29 | 0.13 | -0.6 | -0.77 | 0.1025 | 0.1525 | 0.7875 | 0.874345 | 0.802143 | 0.147726 | 0.999971 | 1 | 0.793248 |
| Tmem256 | F6YWB5 | 0.5 | 0.74 | 1.54 | 1.11 | 0.76 | 0.57 | 0.08 | 1.61 | 0.48 | 1.04 | 1.13 | 1.23 | 0.73 | 0.75 | 1.59 | 2.27 | -0.2175 | -0.0025 | 0.3625 | 0.568746 | 0.99317 | 0.391909 | 0.999971 | 1 | 0.871816 |
| Tmem256_ | Q5F285 | -2.81 | -3.62 | -3.18 | -2.99 | -3.83 | -4.93 | -2.5 | -3.62 | -4.31 | -3.33 | -3.8 | -3.78 | -4.2 | -3.43 | -3.18 | -2.95 | -0.57 | -0.655 | -0.29 | 0.263357 | 0.041328 | 0.382404 | 0.999971 | 1 | 0.871816 |
| Tmem258 | P61166 | 3.07 | 0.55 | -0.28 | -0.42 | -0.45 | 0.36 | 2.06 | -0.28 | 0.58 | 2.83 | -0.47 | 0.27 | -0.53 | 0.32 | -0.37 | -0.95 | -0.3075 | 0.0725 | -1.1125 | 0.729611 | 0.940296 | 0.171985 | 0.999971 | 1 | 0.804075 |
| Tmem259 | Q8CIV2-2 | -0.79 | -0.18 | 0.26 | -0.44 | -0.16 | -0.36 | 2.16 | 0.02 | 0.55 | -0.43 | 1.05 | 1.06 | 0.13 | -0.98 | -0.06 | -0.87 | 0.7025 | 0.845 | -0.1575 | 0.241237 | 0.059093 | 0.655989 | 0.999971 | 1 | 0.934061 |
| Tmem260 | A0A2I3BQ | -0.02 | 0.26 | -1.02 | -0.73 | 0.12 | -0.23 | 1.12 | -0.84 | 0.04 | -0.27 | -1.65 | -0.8 | -0.81 | 0.42 | -1.44 | -1.15 | 0.42 | -0.2925 | -0.3675 | 0.385966 | 0.5195 | 0.445762 | 0.999971 | 1 | 0.884152 |
| Tmem263 | Q9DAM7 | 0.17 | -0.04 | 0.2 | 0.71 | 0.37 | -0.26 | 0.72 | 0.65 | 0.43 | -0.12 | 0.01 | 0.4 | 0.62 | -0.19 | 1.77 | 0.55 | 0.11 | -0.08 | 0.4275 | 0.702573 | 0.740354 | 0.317211 | 0.999971 | 1 | 0.853841 |
| Tmem266 | Q8BZB3 | -0.12 | -2.3 | -1.37 | -1.38 | -1.67 | -2.19 | -1.18 | -0.96 | -2.04 | -0.38 | -1.18 | -1.97 | -1.74 | -1.97 | -0.99 | -1.04 | -0.2075 | -0.1 | -0.1425 | 0.671878 | 0.855328 | 0.765739 | 0.999971 | 1 | 0.962465 |
| Tmem29 | E9PZH0 | 3.9 | 2.25 | 3.77 | 2.86 | 2.76 | 2.84 | 4 | 3.54 | 3.07 | 2.28 | 3.33 | 3.55 | 1.78 | 0.91 | 2.79 | 0.85 | 0.09 | -0.1375 | -1.6125 | 0.84454 | 0.761182 | 0.016233 | 0.999971 | 1 | 0.613095 |
| Tmem30a | Q8VEK0 | -1.16 | -1.38 | -0.89 | -0.82 | -0.97 | -1.27 | -0.87 | -0.76 | -0.67 | -1.08 | -0.96 | -0.8 | -0.78 | -1.16 | -0.56 | -0.95 | 0.095 | 0.185 | 0.2 | 0.661298 | 0.380889 | 0.382607 | 0.999971 | 1 | 0.871816 |
| Tmem33 | A0A0R4J1Z | 0.78 | 0.58 | 1.37 | 0.78 | 0.78 | 1.21 | 0.46 | 1.54 | 0.23 | 1.17 | 1.57 | 0.86 | 1.03 | 1.44 | 1.98 | 1.94 | 0.12 | 0.08 | 0.72 | 0.690902 | 0.808358 | 0.034666 | 0.999971 | 1 | 0.69205 |
| Tmem35a | Q9D328 | -0.76 | 0.05 | 0.03 | 0.47 | 0.34 | 0.68 | -0.01 | 0.69 | 0.88 | 0.3 | 0.51 | 0.35 | 0.31 | 0.43 | 0.97 | 0.31 | 0.4775 | 0.5625 | 0.5575 | 0.15037 | 0.085539 | 0.097608 | 0.999971 | 1 | 0.78268 |
| Tmem38a | Q3TMP8 | 3.87 | 1.29 | 0.96 | 0.18 | 0.64 | 3.56 | 1.48 | 1.29 | 1.48 | 3.57 | 1.59 | -0.36 | 0.18 | 1.74 | 0.56 | 0.66 | 0.1675 | -0.005 | -0.79 | 0.854195 | 0.996078 | 0.326318 | 0.999971 | 1 | 0.856472 |
| Tmem38b | Q9DAV9 | 2.65 | -0.07 | -0.29 | -0.25 | -0.05 | 0.99 | -0.31 | -0.42 | -1.05 | 1.29 | 0.49 | 0.18 | -1.25 | -0.97 | -0.11 | -0.24 | -0.4575 | -0.2825 | -1.1525 | 0.523264 | 0.71905 | 0.124293 | 0.999971 | 1 | 0.78268 |

| | | | | | | | | | | | | | | | | | | | | | | | | | | |
|---|---|---|---|---|---|---|---|---|---|---|---|---|---|---|---|---|---|---|---|---|---|---|---|---|---|---|
| Tmem41a | Q9D8U2 | -0.88 | -0.57 | -2.02 | -1.09 | -0.49 | -0.56 | -1.85 | -0.05 | -0.55 | -0.67 | -1.57 | -1.78 | -1.02 | 0.13 | -0.83 | -0.45 | 0.4025 | -0.0025 | 0.5975 | 0.397308 | 0.995242 | 0.149403 | 0.999971 | 1 | 0.793248 |
| Tmem43 | Q9DBS1 | -0.36 | -0.06 | -0.25 | -0.42 | 0.23 | -0.18 | -0.52 | 0.06 | 0.35 | 0.09 | -0.36 | 0.2 | 0.44 | -0.12 | 0.23 | -0.24 | 0.17 | 0.3425 | 0.35 | 0.453427 | 0.138561 | 0.138625 | 0.999971 | 1 | 0.784948 |
| Tmem47 | Q9JJG6 | -1.77 | -0.8 | -0.71 | -0.42 | -0.26 | -0.51 | 1.09 | -1.07 | -0.2 | -1.54 | -0.97 | -0.52 | -0.89 | -1.34 | -1.26 | -2.38 | 0.7375 | 0.1175 | -0.5425 | 0.168337 | 0.76737 | 0.211594 | 0.999971 | 1 | 0.823077 |
| Tmem50b | Q9D1X9 | 1.7 | 0.85 | 2.32 | 1.39 | 0.37 | 0.81 | 1.82 | 1.02 | 0.8 | 3.18 | 1.56 | 2.94 | 1.59 | 0.75 | 0.93 | 1.78 | -0.56 | 0.555 | -0.3025 | 0.196029 | 0.361202 | 0.437291 | 0.999971 | 1 | 0.882419 |
| Tmem51 | Q99LG1 | 0.01 | -1.44 | -1 | -0.48 | -0.64 | -0.69 | 0.65 | -1.32 | -0.8 | -0.63 | -0.64 | -1.2 | -0.79 | -1.85 | -1.35 | -1.59 | 0.2275 | -0.09 | -0.6675 | 0.639817 | 0.790587 | 0.103122 | 0.999971 | 1 | 0.78268 |
| Tmem59l | Q7TNI2 | -0.89 | 1.79 | -0.74 | -1.17 | 0.22 | 0.84 | -0.28 | -0.48 | 0.41 | -0.73 | -0.32 | -0.32 | 0.6 | 0.87 | -1.61 | -0.44 | 0.3275 | 0.0125 | 0.1075 | 0.630431 | 0.984941 | 0.893032 | 0.999971 | 1 | 0.98828 |
| Tmem62 | Q8BXJ9 | -0.93 | -0.83 | 0.54 | 0.37 | -1.32 | -1.12 | 0 | 0.83 | -0.61 | -0.85 | 1.43 | 0.84 | -1.83 | -0.34 | 0.93 | 1.06 | -0.19 | 0.415 | 0.1675 | 0.744105 | 0.508404 | 0.811943 | 0.999971 | 1 | 0.974279 |
| Tmem63b | Q3TWI9 | -1.23 | -0.58 | -0.48 | -0.59 | -0.34 | -0.22 | -0.26 | -0.15 | -0.39 | -0.59 | -0.05 | -0.06 | 0.25 | -0.23 | -0.18 | 0.02 | 0.4775 | 0.4475 | 0.685 | 0.054503 | 0.095034 | 0.017311 | 0.999971 | 1 | 0.617356 |
| Tmem63b_ | D3Z2D8 | -1.12 | -1.18 | -0.71 | -0.3 | 0.45 | -1.26 | 0.35 | -0.57 | 0.84 | 0.14 | -0.28 | 0.24 | -0.05 | -0.42 | 0.02 | -0.22 | 0.57 | 1.0625 | 0.66 | 0.206712 | 0.007732 | 0.027025 | 0.999971 | 1 | 0.678474 |
| Tmem63c | Q8CBX0 | 0.21 | 0.13 | 0 | 0 | 0.28 | 0.31 | -0.15 | 0.43 | 0.2 | 0.11 | 0.13 | 0.47 | 0.58 | 0.17 | 0.46 | 0.56 | 0.1325 | 0.1425 | 0.3575 | 0.507583 | 0.425481 | 0.078398 | 0.999971 | 1 | 0.754157 |
| Tmem65 | Q4VAE3 | -0.56 | -0.58 | -0.14 | 0.01 | -0.73 | -0.18 | -0.09 | -0.17 | -0.56 | -0.48 | -0.25 | -0.12 | -0.29 | -0.33 | -0.26 | 0.11 | 0.025 | -0.035 | 0.125 | 0.91749 | 0.873893 | 0.579195 | 0.999971 | 1 | 0.919302 |
| Tmem67 | A2AJP5 | -1.72 | -3.93 | 0.26 | -0.46 | -3.39 | -4.48 | 0.22 | -0.77 | -4.73 | -3.37 | -1.03 | -1.1 | -4.89 | -3.31 | -0.47 | -0.28 | -0.6425 | -1.095 | -0.775 | 0.615806 | 0.356557 | 0.55228 | 0.999971 | 1 | 0.913595 |
| Tmem68 | Q9D850 | -0.28 | -0.14 | -0.44 | -0.14 | 0.15 | -0.12 | -0.36 | -0.09 | 0.01 | -0.05 | -0.13 | 0.07 | 0.26 | 0.05 | -0.13 | -0.15 | 0.145 | 0.225 | 0.2575 | 0.45748 | 0.206723 | 0.196447 | 0.999971 | 1 | 0.812891 |
| Tmem69 | Q3KQJ0 | 0.81 | 1.42 | 1.99 | 1.91 | 0.9 | 2.16 | 1.19 | 1.65 | 1.44 | 0.29 | 0.89 | 1.48 | -0.89 | 1.06 | 0.74 | 1.93 | -0.0575 | -0.5075 | -0.8225 | 0.877962 | 0.202355 | 0.189937 | 0.999971 | 1 | 0.810547 |
| Tmem74b | A2AU34 | 0.86 | -1.3 | -1.51 | -1.88 | 0.66 | -0.55 | -1.56 | -1.63 | -1.11 | 0.32 | -0.96 | -1.78 | 0.1 | 0.82 | -0.24 | -1.32 | 0.1875 | 0.075 | 0.7975 | 0.79952 | 0.91337 | 0.266328 | 0.999971 | 1 | 0.842677 |
| Tmem87a | A2AQJ5 | 0.53 | 1.83 | 1.53 | 1.49 | 1.24 | 1.82 | 1.45 | 1.37 | 1.25 | 1.22 | 1.72 | 1.55 | 1.18 | 1.51 | 1.53 | 1.37 | 0.125 | 0.09 | 0.0525 | 0.69046 | 0.772728 | 0.861985 | 0.999971 | 1 | 0.982963 |
| Tmem87b | Z4YML9 | -0.92 | 2.27 | 0.91 | -0.35 | 2.13 | 1.9 | 0.69 | 0.79 | 2.42 | -0.67 | 1.37 | 1.74 | 1.73 | 1.94 | 1.42 | 0.32 | 0.9 | 0.7375 | 0.875 | 0.233676 | 0.41037 | 0.244034 | 0.999971 | 1 | 0.837316 |
| Tmem88b | Q3TYP4 | 0.39 | 1.11 | 1.49 | 0.8 | 0.23 | 0.35 | 0.34 | 1.03 | 0.99 | 0.8 | 0.32 | 0.96 | 1.24 | 0.75 | 1.16 | 1.04 | -0.46 | -0.18 | 0.1 | 0.15547 | 0.540194 | 0.717051 | 0.999971 | 1 | 0.952711 |
| Tmem9 | Q9CR23 | 0.67 | -0.05 | -0.12 | -0.89 | 0.04 | 0.25 | 0.07 | -0.15 | 0.02 | 0.56 | -0.2 | -0.65 | 1.12 | 0.68 | -0.44 | 0.08 | 0.15 | 0.03 | 0.4575 | 0.650042 | 0.938766 | 0.314575 | 0.999971 | 1 | 0.85151 |
| Tmem94 | A0A668KL5 | -0.78 | -0.76 | -1.47 | -0.87 | -0.8 | -0.58 | -1.06 | -0.98 | -0.81 | -1.01 | -1.03 | -1.25 | -0.31 | -0.76 | -1.24 | -0.85 | 0.115 | -0.055 | 0.18 | 0.62646 | 0.80903 | 0.515602 | 0.999971 | 1 | 0.905815 |
| Tmem97 | Q8VD00 | -0.58 | 1.46 | 0 | 0.91 | 0.52 | 2.2 | -0.08 | 0.36 | 0.92 | 1.85 | 0.27 | -0.27 | 0.44 | 2.12 | 0.59 | 0.5 | 0.3025 | 0.245 | 0.465 | 0.625634 | 0.680638 | 0.415598 | 0.999971 | 1 | 0.876885 |
| Tmem9b | Q9JJR8 | 0.48 | 1.25 | -0.06 | -0.1 | 1.24 | 1.19 | -0.13 | -0.41 | 1.63 | 0.56 | 0.22 | -0.06 | 2.5 | 0.71 | 0.25 | 0.42 | 0.08 | 0.195 | 0.5775 | 0.871931 | 0.671203 | 0.315861 | 0.999971 | 1 | 0.85297 |
| Tmf1 | B9EKI3 | 0.2 | 0.42 | 0.04 | 0.59 | 0.71 | 0.24 | -0.09 | 0.63 | 0.34 | 0 | 0.33 | 0.38 | 0.34 | 0.49 | 0.48 | 0.42 | 0.06 | -0.05 | 0.12 | 0.810036 | 0.804546 | 0.537224 | 0.999971 | 1 | 0.910077 |
| Tmlhe | Q91ZE0 | -0.58 | -0.37 | -1.23 | -0.49 | -1.12 | -0.9 | -0.67 | -0.35 | -1.02 | -0.59 | -1.17 | -0.91 | -1.47 | -1.19 | -1.2 | -0.53 | -0.0925 | -0.255 | -0.43 | 0.734299 | 0.327758 | 0.163601 | 0.999971 | 1 | 0.800215 |
| Tmod1 | P49813 | 0.33 | -0.35 | -1.23 | -0.56 | -0.7 | -0.74 | -1.72 | -1.2 | -1.03 | 0.21 | -1.24 | -1.39 | -0.57 | -0.06 | -1.47 | -0.93 | -0.6375 | -0.41 | -0.305 | 0.124956 | 0.381702 | 0.470924 | 0.999971 | 1 | 0.892857 |
| Tmod2 | Q9JKK7 | -1.38 | -0.93 | -1.42 | -1.32 | -0.94 | -1.37 | -0.89 | -1.43 | -0.95 | -1.44 | -1.04 | -0.84 | -0.69 | -1.12 | -1.22 | -1.1 | 0.105 | 0.195 | 0.23 | 0.639334 | 0.377493 | 0.295644 | 0.999971 | 1 | 0.847402 |
| Tmod3 | Q9JHJ0 | 0.41 | 0.39 | 0.18 | 0.35 | 1.12 | 0.52 | 0.48 | 0.12 | 0.41 | 0.6 | 0.31 | 0.72 | 0.22 | 0.86 | 0.39 | 0.13 | 0.2275 | 0.1775 | 0.0675 | 0.363945 | 0.335534 | 0.756721 | 0.999971 | 1 | 0.961116 |
| Tmpo | Q61033 | 2.03 | 1.1 | 0.56 | 0.61 | 0.59 | 1.04 | 0.42 | 0.61 | 1.02 | 1.55 | 0.42 | 0.6 | 0.22 | 1.24 | 0.19 | 0.63 | -0.41 | -0.1775 | -0.505 | 0.269544 | 0.662959 | 0.229937 | 0.999971 | 1 | 0.828687 |
| Tmpo_1 | Q61029 | 0.36 | -1.26 | -0.33 | -0.62 | -0.96 | -1.08 | -0.98 | -0.89 | -1.31 | -0.37 | -1.08 | -0.56 | -1.79 | -0.91 | -0.53 | -0.68 | -0.515 | -0.3675 | -0.515 | 0.15118 | 0.355269 | 0.237689 | 0.999971 | 1 | 0.833594 |
| Tmppe | D3Z286 | 0.61 | 0.35 | 0.38 | 0.42 | 0.19 | 0.27 | 0.05 | 0.51 | 0.19 | 0.26 | 0.7 | 0.15 | 0.25 | 0.46 | 0.33 | 0.02 | -0.185 | -0.115 | -0.175 | 0.33211 | 0.561829 | 0.355413 | 0.999971 | 1 | 0.868793 |
| Tmsb10 | Q6ZWY8 | 2.11 | 0.31 | -1.91 | -0.46 | 0.37 | 0.61 | -0.93 | 0.1 | 0.67 | -0.17 | -0.25 | -1.04 | 1.27 | 1.56 | -0.45 | 0.31 | 0.025 | -0.21 | 0.66 | 0.975376 | 0.798115 | 0.449322 | 0.999971 | 1 | 0.885522 |
| Tmsb15l | Q8C0W0 | 0.14 | -0.13 | 0.19 | 0.2 | 0.68 | 0.05 | 0.8 | 0.46 | 0.59 | -0.72 | 0.31 | 0.54 | 1.01 | -0.29 | 0.15 | 0.18 | 0.3975 | 0.08 | 0.1625 | 0.103111 | 0.801657 | 0.583281 | 0.999971 | 1 | 0.920364 |
| Tmsb4x | P20065-2 | 0.15 | -1.52 | -1.33 | -0.6 | -1.24 | -0.53 | 0.47 | -0.99 | -1.34 | -0.39 | -0.9 | -1.74 | -2.7 | -1.86 | -1.36 | -1.45 | 0.2525 | -0.2675 | -1.0175 | 0.614039 | 0.557327 | 0.050419 | 0.999971 | 1 | 0.722519 |
| Tmtc1 | Q3UV71 | 1.66 | -1.11 | 1.12 | 0.1 | -1.38 | -1.39 | 0.19 | -0.34 | -1.68 | 0.52 | 0.27 | -0.09 | -0.75 | -0.91 | 0.89 | 1.12 | -1.1725 | -0.6875 | -0.355 | 0.102374 | 0.349034 | 0.629865 | 0.999971 | 1 | 0.932649 |
| Tmtc3 | G5E8C4 | 0.46 | -0.51 | -0.2 | -0.25 | 0.06 | -0.52 | -0.17 | 0.07 | -0.28 | 0.59 | 0.11 | 0.34 | -0.14 | 0.03 | -0.27 | 0.26 | -0.015 | 0.315 | 0.095 | 0.955398 | 0.291675 | 0.716091 | 0.999971 | 1 | 0.952576 |
| Tmub1 | Q9JMG3-2 | 0.41 | -0.3 | -0.31 | 0.14 | -0.51 | -0.22 | -0.37 | 0.38 | -0.52 | 0.79 | 0.3 | 1.26 | 0.23 | 0.52 | 0.71 | 0.54 | -0.165 | 0.4725 | 0.515 | 0.558464 | 0.258777 | 0.054597 | 0.999971 | 1 | 0.725348 |
| Tmx1 | Q8VBT0 | 0.03 | -1.08 | -0.38 | -0.12 | -0.56 | -1.26 | 0.25 | -0.76 | -0.84 | -0.37 | -0.43 | -0.52 | -0.77 | -0.93 | -0.62 | -0.49 | -0.195 | -0.1525 | -0.315 | 0.614158 | 0.590457 | 0.278118 | 0.999971 | 1 | 0.843812 |
| Tmx2 | Q9D710 | -0.31 | 0.13 | -0.22 | -0.64 | -0.06 | 0.14 | -0.02 | -0.38 | 0.27 | -0.11 | -0.18 | -0.32 | -0.28 | 0.06 | -0.29 | -0.76 | 0.18 | 0.175 | -0.0575 | 0.442255 | 0.463778 | 0.822995 | 0.999971 | 1 | 0.975901 |
| Tmx3 | Q8BXZ1 | -0.09 | 0.06 | -0.06 | 0.17 | -0.03 | -0.09 | -0.29 | 0.2 | -0.07 | -0.16 | -0.17 | 0.07 | 0.39 | 0.13 | 0.07 | 0.2 | -0.0725 | -0.1025 | 0.1775 | 0.699878 | 0.547695 | 0.329617 | 0.999971 | 1 | 0.856787 |
| Tmx4 | Q8C0L0 | 0.19 | 0.93 | 0.9 | 1.04 | 0.41 | 0.38 | 1.35 | 0.41 | 0.27 | 0.07 | 0.26 | 0.71 | 0.83 | 0.6 | 0.59 | 0.96 | -0.1275 | -0.4375 | -0.02 | 0.683553 | 0.118322 | 0.934958 | 0.999971 | 1 | 0.995618 |
| Tnc | Q80YX1 | -0.02 | 0.34 | 0.65 | 0.14 | 0.52 | 0.63 | 1.25 | 0.43 | 0.74 | 0.43 | 1.22 | 0.99 | 0.63 | 0.15 | 0.74 | 0.91 | 0.43 | 0.5675 | 0.33 | 0.123672 | 0.045757 | 0.205393 | 0.999971 | 1 | 0.816688 |
| Tnfaip1 | O70479 | 0.28 | 2.15 | 0.28 | 0.14 | -0.13 | 1.9 | 0.24 | -0.42 | 0.3 | 0.85 | 0.16 | -0.17 | 0.31 | 0.88 | 0.15 | -0.95 | -0.315 | -0.4275 | -0.615 | 0.625933 | 0.39412 | 0.292364 | 0.999971 | 1 | 0.847402 |
| Tnfaip3 | Q60769 | 1.81 | 0.81 | 0.4 | 0.6 | 0.27 | 0.03 | 1.56 | 1.03 | 0.46 | 1.04 | 0.06 | -0.93 | 0.45 | 0.69 | -0.42 | -1.19 | -0.1825 | -0.7475 | -1.0225 | 0.681427 | 0.150209 | 0.064564 | 0.999971 | 1 | 0.726597 |
| Tnfaip8 | D3Z325 | 1.02 | 1.32 | -0.08 | -0.06 | 0.32 | 1.56 | -0.05 | 0.21 | 0.64 | 2 | -0.03 | 0.09 | 0.86 | 1.61 | -0.01 | 0.94 | -0.04 | 0.125 | 0.3 | 0.932927 | 0.818899 | 0.520854 | 0.999971 | 1 | 0.906972 |
| Tnfaip8l3 | Q3TBL6 | 0.86 | -0.32 | 0.92 | 0.53 | 0.27 | 1.28 | 0.42 | 1.15 | 0.53 | 0.84 | 0.31 | 0.24 | 0.29 | 0.61 | 0.67 | 0.79 | 0.2825 | -0.0175 | 0.0925 | 0.453948 | 0.956102 | 0.766394 | 0.999971 | 1 | 0.962472 |
| Tnfrsf21 | Q9EPU5 | 0.97 | 0.45 | 0.34 | 0.35 | 0.19 | 0.29 | 0.13 | 0.14 | 0.16 | 0.75 | -0.28 | -0.7 | 0.43 | -0.09 | 0.4 | 0.24 | -0.34 | -0.545 | -0.2825 | 0.127096 | 0.13677 | 0.240039 | 0.999971 | 1 | 0.834057 |
| Tnik | B2RQ80 | -0.25 | -0.85 | -0.38 | -0.56 | -1.11 | -0.82 | -0.17 | -0.7 | -0.61 | -0.25 | -0.51 | -0.46 | -0.89 | -0.95 | -0.93 | -0.6 | -0.19 | 0.0525 | -0.3325 | 0.470705 | 0.79549 | 0.135323 | 0.999971 | 1 | 0.784948 |
| Tnip1 | D3Z2W0 | -0.15 | 2.99 | 0.48 | 0.57 | 1.44 | 1.46 | 0.01 | 0.73 | 0.76 | 1.09 | 0.28 | 0.59 | 1.34 | 0.76 | 0.35 | 0.45 | -0.0625 | -0.2925 | -0.2475 | 0.928613 | 0.654546 | 0.708547 | 0.999971 | 1 | 0.95049 |
| Tnk2 | O54967-2 | 0.51 | -0.07 | 0.01 | 0.06 | -0.36 | -0.29 | 0.15 | -0.3 | 0 | 0.27 | -0.28 | -0.19 | -0.08 | -0.57 | -0.45 | 0.05 | -0.3275 | -0.1775 | -0.39 | 0.161334 | 0.427047 | 0.122245 | 0.999971 | 1 | 0.78268 |
| Tnks | Q6PFX9 | 1.7 | -0.66 | -0.52 | -0.28 | -0.21 | -0.03 | -1.05 | -0.11 | -0.39 | 1.8 | -0.79 | -0.59 | -0.11 | -0.33 | -0.5 | 0.14 | -0.41 | -0.0525 | -0.26 | 0.464505 | 0.943557 | 0.623556 | 0.999971 | 1 | 0.930282 |
| Tnks1bp1 | P58871 | 1.11 | 0.25 | 0.76 | 0.42 | 0.21 | 0.54 | 0.78 | 0.28 | 0.14 | 0.82 | 0.51 | 0.4 | 0.02 | 0.11 | 0.33 | 0.54 | -0.1825 | -0.1675 | -0.385 | 0.482371 | 0.522815 | 0.151725 | 0.999971 | 1 | 0.795578 |
| Tnpo1 | Q8BFY9-2 | -0.47 | 1.06 | 0.34 | -0.01 | 1.1 | 0.78 | 0.2 | 0.14 | 1.17 | -0.25 | 0.08 | 0.53 | 1.29 | 0.83 | 0.3 | 0.19 | 0.325 | 0.1525 | 0.4225 | 0.405832 | 0.719629 | 0.30051 | 0.999971 | 1 | 0.847402 |
| Tnpo2 | Q99LG2 | 0.02 | 0.9 | 0.76 | 0.29 | 1.09 | 0.91 | 0.16 | 0.27 | 1.13 | 0.26 | 0.57 | -0.07 | 1.25 | 1.11 | 0.41 | 0.56 | 0.115 | -0.02 | 0.34 | 0.713864 | 0.951177 | 0.273607 | 0.999971 | 1 | 0.843794 |
| Tnpo3 | Q6P2B1 | -0.59 | 0.52 | 0.92 | 0.62 | 0.77 | 0.45 | 0.94 | 0.9 | 0.76 | -0.05 | 0.85 | 0.6 | 1.04 | 0.51 | 0.87 | 0.76 | 0.3975 | 0.1725 | 0.4275 | 0.266139 | 0.648287 | 0.235483 | 0.999971 | 1 | 0.832918 |
| Tnr | Q8BYI9 | 0.83 | 1.13 | 0.56 | 0.46 | 1 | 0.96 | 0.28 | 0.32 | 0.86 | 0.44 | 0.02 | 0.44 | 0.82 | 0.45 | 0.34 | 0.36 | -0.105 | -0.305 | -0.2525 | 0.695998 | 0.247247 | 0.28459 | 0.999971 | 1 | 0.844871 |
| Tnrc18 | Q80WC3 | 0.69 | 1.38 | 1.54 | 1.7 | 1.58 | 1.72 | 2.49 | 0.75 | 1.19 | 1.32 | 0.5 | -0.16 | 2.2 | 2.19 | -0.39 | -0.5 | 0.3075 | -0.615 | -0.4525 | 0.451233 | 0.143884 | 0.533362 | 0.999971 | 1 | 0.910069 |
| Tnrc6b | Q8BKI2 | 1.56 | 0.39 | 1.31 | 1.41 | 0.56 | 0.74 | 2.28 | 1.08 | 0.89 | 1.42 | 1.35 | 1.08 | 0 | 0.26 | 0.65 | 0.68 | -0.0025 | 0.0175 | -0.77 | 0.995473 | 0.953309 | 0.034984 | 0.999971 | 1 | 0.69205 |
| Tnrc6c | B1ATC3 | 1.69 | 0.22 | 0.07 | 0.31 | 0.68 | 0.47 | 0.52 | 0.11 | 1.22 | 0.11 | -0.15 | -0.53 | -0.16 | -0.61 | -0.02 | -0.04 | -0.1275 | -0.41 | -0.78 | 0.73824 | 0.41789 | 0.069699 | 0.999971 | 1 | 0.733937 |
| Tns1 | A0A087WC | 0.52 | 0.37 | 0.69 | 0.5 | 2.1 | 0.56 | 0.44 | -0.26 | 0.53 | 1.37 | 1.05 | 1.55 | -0.4 | 0.66 | 0.25 | 0.01 | 0.19 | 0.605 | -0.39 | 0.686301 | 0.040112 | 0.154917 | 0.999971 | 1 | 0.795578 |
| Tns2 | A0A2R8VH | 0.29 | 0.7 | -0.99 | 0.34 | 1.33 | 0.06 | 0.17 | -0.89 | 0.15 | 0.71 | -0.83 | -1.03 | -0.82 | 0.2 | -1.03 | -0.78 | 0.0825 | -0.335 | -0.6925 | 0.87837 | 0.519622 | 0.138435 | 0.999971 | 1 | 0.784948 |
| Tns3 | A0A5F8MP | 0.59 | -0.06 | 0.35 | 0.28 | -0.21 | 0.25 | 0.11 | -0.05 | -0.04 | 0.59 | -0.39 | -0.05 | 0.14 | 0.02 | 0.22 | -0.04 | -0.265 | -0.2625 | -0.205 | 0.23712 | 0.334901 | 0.32766 | 0.999971 | 1 | 0.856472 |
| Toe1 | Q9D2E2 | -0.15 | 1.3 | 1.57 | 2.29 | 1.41 | 0.66 | 0.89 | 1.1 | 0.63 | 0.54 | 1.51 | 0.73 | 0.08 | 0.8 | 0.81 | 0.82 | -0.2375 | -0.4 | -0.625 | 0.635089 | 0.448501 | 0.236094 | 0.999971 | 1 | 0.832918 |
| Togaram1 | A0A1Y7VLS | -3.06 | -0.27 | 0.08 | -0.56 | 1.79 | -0.22 | 0.26 | -0.38 | 0.33 | -2.24 | 0.47 | 0.05 | 0.11 | -0.54 | -0.3 | -0.3 | 1.315 | 0.605 | 0.695 | 0.1197 | 0.490073 | 0.308492 | 0.999971 | 1 | 0.847402 |
| Tollip | Q9QZ06 | -0.79 | -0.95 | -0.64 | -0.55 | -0.92 | -0.93 | -0.47 | -0.61 | -1.09 | -0.96 | -0.32 | -0.45 | -1.22 | -0.82 | -0.5 | -0.38 | 1.57E-16 | 0.0275 | 0.0025 | 1 | 0.908253 | 0.991707 | 1 | 1 | 0.99925 |
| Tom1 | O88746 | -0.67 | -0.55 | -0.36 | -0.31 | -0.82 | -0.87 | -0.52 | -0.25 | -0.83 | -0.67 | -0.67 | -0.29 | -0.53 | -0.62 | -0.07 | 0.09 | -0.1425 | -0.1425 | 0.19 | 0.511477 | 0.477333 | 0.419651 | 0.999971 | 1 | 0.87788 |
| Tom1l1 | Q8BZR6 | 0.46 | 1.05 | 0.05 | -0.6 | 1.12 | 1.03 | -1.32 | 0.01 | 0.57 | 0.6 | -0.47 | -0.14 | 1.93 | 0.9 | 0.24 | -0.29 | -0.03 | -0.1 | 0.455 | 0.960563 | 0.810429 | 0.412933 | 0.999971 | 1 | 0.876042 |
| Tom1l2 | Q5SRX1 | -0.9 | -1.38 | -0.96 | -0.84 | -1.28 | -1.46 | -0.9 | -1.1 | -1.56 | -1.39 | -1.14 | -1.33 | -1.49 | -1.3 | -0.81 | -0.67 | -0.165 | -0.335 | -0.0475 | 0.455116 | 0.124599 | 0.853023 | 0.999971 | 1 | 0.98187 |
| Tom1l2_1 | Q5SRX1-2 | -3.14 | -5.47 | -4.48 | -5.88 | -2.94 | -6.83 | -4.3 | -5.56 | -4.31 | -6.35 | -4.39 | -4.4 | -4.75 | -4.44 | -5.02 | -4.7 | -0.165 | -0.12 | 0.015 | 0.858275 | 0.866635 | 0.978983 | 0.999971 | 1 | 0.99925 |
| Tomm20 | Q9DCC8 | 1.08 | 1.09 | 0.98 | 1.31 | 0.66 | 1.03 | 1.14 | 1.36 | 0.59 | 1.17 | 0.93 | 0.77 | 1.1 | 0.68 | 1.05 | 1.11 | -0.0675 | -0.25 | -0.13 | 0.750155 | 0.226236 | 0.502723 | 0.999971 | 1 | 0.904573 |
| Tomm22 | Q9CPQ3 | -0.89 | -1.1 | -0.82 | -1.06 | -1.05 | -1.29 | -0.95 | -0.64 | -1.22 | -0.84 | -0.62 | -0.77 | -1.02 | -0.77 | -0.75 | -0.48 | -0.015 | 0.105 | 0.2125 | 0.94157 | 0.600131 | 0.290228 | 0.999971 | 1 | 0.847402 |
| Tomm34 | Q9CYG7 | 1.18 | 1.4 | 1.12 | 1.15 | 1.21 | 2.15 | 1.3 | 1.5 | 1.65 | 1.09 | 1.26 | 1.11 | 0.71 | 1.58 | 1.17 | 1.04 | 0.3275 | 0.065 | -0.0875 | 0.211645 | 0.744497 | 0.704607 | 0.999971 | 1 | 0.950182 |
| Tomm34_1 | Q9CYG7-2 | -0.07 | 1.03 | 0.83 | 0.97 | 1.21 | 1.46 | 1.18 | 1.3 | 1.57 | -0.14 | 1.28 | 1.48 | 0.47 | 0.61 | 0.46 | 1.48 | 0.5975 | 0.3575 | 0.065 | 0.057626 | 0.434311 | 0.852354 | 0.999971 | 1 | 0.98187 |
| Tomm40 | Q9QYA2 | 0.79 | -0.32 | -0.34 | -0.36 | -0.3 | -0.57 | -0.24 | -0.37 | -0.18 | -0.31 | -0.4 | -0.2 | 0.02 | -0.27 | -0.51 | -0.36 | -0.3125 | -0.215 | -0.2225 | 0.312867 | 0.474588 | 0.479444 | 0.999971 | 1 | 0.894438 |
| Tomm40l | Q9CZR3 | 1.32 | 0.46 | 0.76 | 0.39 | 0.17 | 0.45 | 0.28 | 0.94 | 0.19 | 1.02 | 0.43 | 0.24 | 0.17 | 0.77 | 0.69 | 0.95 | -0.2725 | -0.2625 | -0.0875 | 0.351985 | 0.383463 | 0.758884 | 0.999971 | 1 | 0.961116 |
| Tomm5 | B1AXP6 | -1.9 | -3.51 | -1.19 | -1.6 | -2.42 | -3.38 | -1.52 | -2.44 | -3.44 | -1.58 | -1.73 | -2.21 | -3.09 | -3 | -2.44 | -2.04 | -0.39 | -0.19 | -0.5925 | 0.50689 | 0.754327 | 0.274664 | 0.999971 | 1 | 0.843794 |
| Tomm6 | Q9CQN3 | -0.55 | -0.79 | -1.18 | -0.77 | -0.44 | -1.39 | -1.27 | -0.18 | -1.64 | -1.27 | -0.92 | -1.1 | -0.26 | -1.36 | -0.4 | -0.63 | 0.0025 | -0.41 | 0.16 | 0.993903 | 0.108081 | 0.584575 | 0.999971 | 1 | 0.920364 |
| Tomm7 | Q9D173 | -1.81 | -1.36 | -1.1 | -0.82 | -1.22 | -2.51 | -1.28 | -0.85 | -1.3 | -1.03 | -0.92 | -1.47 | -0.44 | -1.1 | -1.9 | -0.53 | -0.1925 | 0.0925 | 0.28 | 0.632012 | 0.727489 | 0.471603 | 0.999971 | 1 | 0.892857 |
| Tomm70 | Q9CZW5 | -0.13 | -0.18 | -0.76 | -0.98 | -0.23 | -0.75 | -0.47 | -1 | -0.2 | -0.65 | -0.63 | -0.53 | -0.09 | -0.43 | -1.05 | -1.44 | -0.1 | 0.01 | -0.24 | 0.725238 | 0.969067 | 0.512253 | 0.999971 | 1 | 0.905815 |
| Top1 | Q04750 | 0.23 | -0.54 | -0.18 | -0.11 | -0.32 | -0.58 | -0.49 | -0.23 | -0.41 | -0.29 | -0.18 | -0.49 | -0.31 | -0.54 | -0.29 | -0.15 | -0.255 | -0.1925 | -0.1725 | 0.265688 | 0.38237 | 0.444107 | 0.999971 | 1 | 0.883802 |
| Top2b | Q64511 | -0.75 | -1.09 | -0.96 | -0.97 | -0.96 | -1.43 | -0.97 | -1.16 | -0.66 | -0.8 | -0.89 | -1.01 | -0.27 | -1.08 | -1.08 | -1.03 | -0.1875 | 0.1025 | 0.0775 | 0.347426 | 0.566798 | 0.750393 | 0.999971 | 1 | 0.961064 |
| Top3b | Q9Z321 | 0.41 | 0.73 | 1.23 | 0.9 | 0.46 | 0.79 | 0.5 | 1.32 | 0.25 | 0.73 | 0.86 | 1.21 | -0.1 | 0.88 | 1.11 | 1.25 | -0.05 | -0.055 | -0.0325 | 0.857271 | 0.842766 | 0.924961 | 0.999971 | 1 | 0.99401 |
| Tor1a | Q9ER39 | -0.44 | 0.15 | 0.34 | 0.2 | -0.03 | -0.03 | -0.17 | 0.17 | 0.06 | 0.05 | 0.07 | 0.72 | 0.15 | 0.09 | 0.18 | 0.53 | -0.0775 | 0.1625 | 0.175 | 0.732532 | 0.536794 | 0.463921 | 0.999971 | 1 | 0.890905 |
| Tor1aip1 | Q921T2-2 | -1.42 | -1.16 | 0.41 | 0.45 | -0.57 | -0.98 | 0.73 | -0.2 | -0.3 | -1.4 | 0.08 | 0.41 | -0.71 | -1.06 | 0.02 | 0.09 | 0.175 | 0.1275 | 0.015 | 0.757933 | 0.827484 | 0.977366 | 0.999971 | 1 | 0.99925 |
| Tor1aip2 | Q8BYU6 | 0.47 | -0.52 | -1.19 | -0.81 | 0 | 0.21 | -0.83 | -0.23 | -0.35 | 0.42 | -0.34 | -1.05 | 0.42 | 0.25 | -0.63 | -0.38 | 0.3 | 0.1825 | 0.4275 | 0.462016 | 0.679466 | 0.316755 | 0.999971 | 1 | 0.853713 |
| Tor1aip2_1 | Q9ER81 | -2.56 | 0.71 | 2.11 | 2.53 | -0.79 | -1.16 | 1.23 | 2.04 | -1.18 | -1.24 | 2.87 | 1.79 | -0.01 | -0.88 | 0.9 | 2.01 | -0.3675 | -0.1375 | -0.1925 | 0.766326 | 0.921248 | 0.868882 | 0.999971 | 1 | 0.983462 |
| Tor1b | Q9ER41 | -2.05 | -1.54 | -0.79 | -0.75 | -0.4 | -0.89 | -0.25 | -0.46 | -0.28 | -2.11 | -0.59 | -0.1 | -0.79 | -0.79 | -0.15 | -0.27 | 0.7825 | 0.5125 | 0.7825 | 0.043619 | 0.334665 | 0.049636 | 0.999971 | 1 | 0.715795 |
| Tor2a | Q8R1J9 | -0.96 | -1.22 | -0.8 | -0.47 | -1.42 | -1.6 | -0.53 | -0.13 | -2.27 | -1.09 | -0.33 | -0.87 | -1.3 | -1.09 | -0.3 | 0.13 | -0.0575 | -0.2775 | 0.2225 | 0.877562 | 0.511643 | 0.542828 | 0.999971 | 1 | 0.911251 |
| Tor3a | Q9ER38 | -1.28 | 0.32 | 0.83 | -0.53 | -0.5 | -0.06 | -0.91 | 0.53 | 0.14 | -0.71 | 0.72 | 1.08 | -1.43 | -0.09 | 1.51 | 1.1 | -0.07 | 0.4725 | 0.4375 | 0.892046 | 0.409331 | 0.55343 | 0.999971 | 1 | 0.913595 |
| Tox2 | E9Q0B9 | 3.08 | 0.75 | -1.56 | -2.77 | -0.19 | 0.71 | 0.4 | -2.75 | -0.06 | 1 | -2.19 | -1.26 | -0.64 | -0.33 | -2.62 | -2.87 | -0.3325 | -0.5025 | -1.49 | 0.804704 | 0.703315 | 0.266926 | 0.999971 | 1 | 0.842677 |
| Tox4 | Q8BU11 | 2.09 | 1.62 | 1.98 | 2.43 | 1.39 | 1.54 | 2.98 | 1.35 | 1.29 | 2.02 | 1.5 | 0.87 | 0.86 | 1.16 | 1.4 | 1.31 | -0.215 | -0.61 | -0.8475 | 0.598356 | 0.067924 | 0.006038 | 0.999971 | 1 | 0.582714 |
| Tp53bp1 | P70399-5 | 0.95 | 0.64 | 1.41 | 1.32 | 0.96 | 0.37 | 1.43 | 0.93 | 0.98 | 0.44 | 1.11 | 1.56 | 0.41 | 0.42 | 1.26 | 1.21 | -0.1575 | -0.0575 | -0.255 | 0.591867 | 0.84741 | 0.411102 | 0.999971 | 1 | 0.875847 |
| Tpbg | Q9Z0L0 | -0.19 | -0.72 | -1.29 | -1.35 | -0.3 | -0.98 | -1.24 | -1.52 | 0.07 | -0.66 | -1.26 | -1.06 | -0.37 | -1.37 | -1.21 | -1.03 | -0.1225 | 0.16 | -0.1075 | 0.739 | 0.680146 | 0.756336 | 0.999971 | 1 | 0.961116 |
| Tpbgl | Q8C013 | 0.91 | -1.15 | -2.41 | -1.29 | -1.31 | -1.52 | -1.25 | -1.2 | -0.68 | 1.23 | -1.48 | -0.23 | -0.99 | -0.88 | -1.61 | -0.11 | -0.335 | 0.695 | 0.0875 | 0.59895 | 0.401303 | 0.898492 | 0.999971 | 1 | 0.989489 |
| Tpcn1 | Q9EQJ0 | 2.21 | -0.66 | 0.68 | 0.64 | -0.91 | -0.1 | 2.74 | 0.69 | -1.44 | 1.03 | -0.03 | -0.52 | -1.72 | -1.87 | -0.48 | -1.08 | -0.1125 | -0.9575 | -2.005 | 0.897778 | 0.200737 | 0.009127 | 0.999971 | 1 | 0.613095 |
| Tpd52 | Q62393-3 | -1.71 | -2.12 | -1.66 | -1.37 | -1.86 | -1.58 | -1.31 | -1.28 | -1.71 | -1.76 | -0.95 | -1.16 | -2.31 | -1.93 | -1.35 | -1.37 | 0.2075 | 0.32 | -0.025 | 0.39531 | 0.256956 | 0.931595 | 0.999971 | 1 | 0.995069 |
| Tpd52l1 | B2RUC1 | 0.35 | 0.46 | 0.38 | 0.46 | 0.66 | 0.75 | 0.23 | 0.83 | 0.67 | 0.33 | 0.45 | 0.44 | 0.11 | 0.53 | 0.45 | 0.72 | 0.205 | 0.06 | 0.04 | 0.313971 | 0.720025 | 0.837326 | 0.999971 | 1 | 0.979416 |
| Tpd52l2 | A2AUD5 | -0.58 | -1.74 | -3.34 | -1.82 | -2.65 | -2.36 | -2.84 | -2.5 | -3.03 | -0.99 | -3.61 | -3.53 | -2.37 | -1.69 | -2.92 | -2.13 | -0.7175 | -0.92 | -0.4075 | 0.199076 | 0.24463 | 0.481359 | 0.999971 | 1 | 0.89485 |
| Tpd52l2_1 | Q9CYZ2 | 0.32 | 0.32 | -0.51 | -0.42 | 0.3 | 0.4 | -0.73 | -0.3 | 0.23 | 0.55 | -0.22 | -0.12 | 0.01 | 0.24 | -0.35 | -0.38 | -0.01 | 0.1825 | -0.0475 | 0.97683 | 0.542075 | 0.868084 | 0.999971 | 1 | 0.983462 |
| Tpgs1 | A0A6Q6PK | -1.32 | -0.26 | 0.43 | -0.67 | -0.62 | -0.31 | -0.23 | -0.09 | -0.89 | -0.97 | 0.46 | 0.14 | -0.51 | -0.11 | 0.55 | 0.19 | 0.1425 | 0.14 | 0.485 | 0.702573 | 0.771732 | 0.256718 | 0.999971 | 1 | 0.839012 |
| Tpgs2 | E9PUD7 | -1.63 | -3.55 | -1.74 | -1.53 | -2.89 | -3.1 | -1.77 | -0.7 | -3.12 | -2.16 | -1.99 | -2.18 | -3.24 | -2.33 | -1.94 | -1.18 | -0.0025 | -0.25 | -0.06 | 0.996999 | 0.624389 | 0.918894 | 1 | 1 | 0.992633 |
| Tph2 | Q8CGV2 | -1 | -0.96 | -0.85 | -0.52 | -0.96 | -1.26 | -0.95 | -0.94 | -0.84 | -0.6 | -1.23 | -1.05 | -1.07 | -0.44 | -0.32 | -1.05 | -0.195 | -0.0975 | 0.1125 | 0.333106 | 0.654369 | 0.65854 | 0.999971 | 1 | 0.934638 |
| Tpi1 | P17751-2 | 0.1 | 1.08 | 0.47 | -0.42 | 1.01 | 0.89 | 0.7 | -0.18 | 0.72 | -0.3 | 0.25 | 0.5 | 1.75 | 0.49 | 0.14 | -0.5 | 0.2975 | -0.015 | 0.1625 | 0.461056 | 0.967914 | 0.757749 | 0.999971 | 1 | 0.961116 |
| Tpk1 | Q9R0M5 | -0.94 | 0.22 | -0.36 | 0.01 | -0.02 | 0.43 | 0.01 | 0.4 | 0.2 | -1.1 | -0.67 | 0.1 | 0.07 | 0.6 | -0.02 | 0.1 | 0.4725 | -0.1 | 0.455 | 0.133076 | 0.796823 | 0.154564 | 0.999971 | 1 | 0.795578 |
| Tpm1 | F8WID5 | 1.03 | -0.49 | -0.43 | 1.64 | 9.04 | 2.22 | 0.34 | -2.11 | 2.6 | 7.46 | 4.48 | 8.56 | -3.85 | 1.09 | 0.77 | 0.04 | 1.935 | 5.3375 | -0.925 | 0.382843 | 0.002908 | 0.419695 | 0.999971 | 0.918989 | 0.87788 |
| Tpm1_1 | Q8BP43 | 1.95 | 1.19 | 0.63 | 1.76 | 1.7 | 1 | 0.91 | 0.21 | 1.06 | 2.04 | 0.46 | 1.19 | -0.58 | 1.16 | 1.35 | 1.04 | -0.4275 | -0.195 | -0.64 | 0.308508 | 0.644583 | 0.220672 | 0.999971 | 1 | 0.827096 |
| Tpm1_2 | G5E8R0 | 0.82 | -0.19 | -0.58 | 0.27 | -0.04 | 0.46 | -0.46 | -0.26 | -0.16 | 0.67 | -0.26 | -0.85 | -0.23 | -0.01 | -0.72 | -0.15 | -0.155 | -0.23 | -0.3575 | 0.662698 | 0.582266 | 0.304796 | 0.999971 | 1 | 0.847402 |
| Tpm1_3 | E9Q452 | 1.66 | -0.99 | -0.89 | -0.16 | 0.43 | -0.23 | 1.52 | -1.15 | -0.9 | 1.94 | -0.29 | 0.33 | -1.93 | -0.82 | -0.96 | -1.29 | 0.2375 | 0.365 | -1.155 | 0.751824 | 0.643481 | 0.083172 | 0.999971 | 1 | 0.755254 |
| Tpm1_4 | P58771 | -1.01 | -0.43 | -0.03 | 0.81 | -1.15 | -0.4 | 0.16 | 1.85 | -0.57 | -0.85 | 1.57 | 0.7 | -0.2 | -0.47 | -0.33 | 2.67 | 0.28 | 0.3775 | 0.5825 | 0.678798 | 0.549559 | 0.452093 | 0.999971 | 1 | 0.887217 |
| Tpm3 | P21107-2 | -0.13 | -0.4 | -1.23 | -1.22 | -0.12 | -0.37 | -0.58 | -1.75 | 0.04 | 0.1 | -1.18 | -1.18 | -0.34 | -0.32 | -1.43 | -1.5 | 0.04 | 0.19 | -0.1525 | 0.926266 | 0.663366 | 0.71259 | 0.999971 | 1 | 0.951793 |
| Tpm3_1 | D3Z6I8 | -0.11 | 0.42 | 0.88 | 0.74 | 0.3 | 0.65 | 0.89 | 0.25 | 0.73 | 0.33 | 0.77 | 0.06 | 1.08 | 0.06 | 0.49 | 0.51 | 0.04 | -0.01 | 0.0525 | 0.887022 | 0.972346 | 0.865327 | 0.999971 | 1 | 0.983318 |
| Tpm3_2 | E9Q7Q3 | -1.15 | -0.8 | -0.86 | -0.46 | -0.89 | -1.26 | -0.59 | -0.24 | -1.77 | -1.21 | -0.72 | -1.59 | -1.69 | -1.11 | -1.11 | -0.65 | 0.0725 | -0.505 | -0.3225 | 0.79308 | 0.10363 | 0.258623 | 0.999971 | 1 | 0.839012 |
| Tpm4 | Q6IRU2 | -0.47 | -1.29 | -0.73 | -0.2 | -0.59 | -1.07 | -0.1 | -1.12 | -0.95 | -0.56 | -0.48 | -0.9 | -2.19 | -0.97 | -0.46 | -1.02 | -0.0475 | -0.05 | -0.4875 | 0.886171 | 0.856459 | 0.257199 | 0.999971 | 1 | 0.839012 |
| Tpmt | A0A0R4J01 | -0.79 | 0.88 | 1.11 | 0.09 | 1.34 | 0.55 | 1.75 | 0.18 | 1.56 | -0.34 | 1.49 | 2.45 | 2.4 | 0.57 | 0.43 | -0.48 | 0.6325 | 0.9675 | 0.4075 | 0.241869 | 0.170037 | 0.548793 | 0.999971 | 1 | 0.911727 |
| Tpp1 | O89023 | -1.08 | 0.13 | 0.43 | -0.74 | 0.26 | 0.06 | 0.57 | -0.72 | -0.23 | -1.72 | 0.08 | -0.3 | 0 | -0.44 | -0.09 | -0.78 | 0.3575 | -0.2275 | -0.0125 | 0.410511 | 0.650708 | 0.974016 | 0.999971 | 1 | 0.99925 |
| Tpp2 | Q64514-2 | -0.29 | 0.42 | -0.1 | -0.57 | 0.22 | 0.27 | 0.18 | -0.12 | 0.55 | 0.09 | 0.35 | 0.18 | 0.94 | 0.36 | -0.02 | 0.07 | 0.2725 | 0.4275 | 0.4725 | 0.29761 | 0.121552 | 0.150489 | 0.999971 | 1 | 0.793248 |
| Tppp | Q7TQD2 | 0.32 | 0.15 | 0.38 | 0.44 | -0.19 | 0.55 | 0.57 | 0.15 | -0.06 | 0.09 | -0.18 | 0.04 | -0.13 | 0.2 | -0.09 | 0.2 | -0.0525 | -0.35 | -0.2775 | 0.819468 | 0.066293 | 0.158068 | 0.999971 | 1 | 0.797252 |
| Tppp3 | Q9CRB6 | -0.24 | -0.79 | 0.05 | 0.19 | -0.35 | 0.15 | 0.04 | 0.82 | 0.3 | 0.36 | 0.31 | 0.4 | 0.1 | 0.32 | 0.97 | 0.24 | 0.3625 | 0.54 | 0.605 | 0.283768 | 0.052318 | 0.0697 | 0.999971 | 1 | 0.733937 |
| Tpr | Q7M739 | 1.14 | 0.52 | 0.53 | 0.83 | 0.39 | 0.37 | 1.09 | 0.21 | 0.35 | 0.64 | 0.52 | 0.41 | -0.2 | 0.3 | 0.12 | 0.01 | -0.24 | -0.275 | -0.6975 | 0.378147 | 0.209125 | 0.011687 | 0.999971 | 1 | 0.613095 |
| Tprg1l | Q9DBS2 | -1.16 | -1.34 | -0.61 | -0.67 | -1.1 | -1.23 | -1.04 | -0.38 | -1.1 | -0.97 | -1.09 | -1.22 | -0.9 | -1.05 | -0.6 | -0.74 | 0.0075 | -0.15 | 0.1225 | 0.978464 | 0.510652 | 0.611089 | 0.999971 | 1 | 0.92673 |
| Tprkb | Q8QZZ7 | 0.43 | 0.87 | 2.71 | 2.39 | 0.51 | 0.9 | 4.22 | 1.13 | 0.5 | 0.29 | 1.57 | 1.11 | -0.4 | 0.88 | 1.75 | -0.22 | 0.09 | -0.7325 | -1.0975 | 0.921357 | 0.228639 | 0.134557 | 0.999971 | 1 | 0.784948 |
| Tprn | A2AI08 | -0.46 | -0.2 | -1.02 | -1.69 | -0.86 | -1 | -0.81 | -0.94 | -0.16 | -0.03 | -0.7 | -0.73 | 0.52 | -0.7 | -1.27 | -1.06 | -0.06 | 0.4375 | 0.215 | 0.856457 | 0.252853 | 0.658402 | 0.999971 | 1 | 0.934638 |
| Tpst1 | O70281 | -0.98 | -0.2 | -0.41 | -1.29 | 1.05 | -1.26 | -0.97 | -0.09 | 0.59 | -1.67 | 0.49 | 2.76 | 1.46 | 0.81 | 0.95 | -0.3 | 0.4025 | 1.2625 | 1.45 | 0.456501 | 0.162359 | 0.00753 | 0.999971 | 1 | 0.582714 |
| Tpt1 | P63028 | -0.19 | -0.48 | 1.9 | 1.48 | -0.13 | -0.87 | 1.41 | 1.96 | -0.17 | -0.78 | 1.9 | 1.29 | -0.56 | -0.65 | 1.77 | 1.38 | -0.085 | -0.1175 | -0.1925 | 0.91495 | 0.880154 | 0.805938 | 0.999971 | 1 | 0.971988 |
| Tra2a | A0A0N4SV | -0.82 | -0.46 | -0.69 | -0.85 | -0.01 | -1.15 | -0.78 | -0.9 | 0.36 | -0.81 | -0.87 | -0.68 | 0.5 | -0.89 | -1.29 | -0.64 | -0.005 | 0.205 | 0.125 | 0.985616 | 0.511071 | 0.743544 | 0.999971 | 1 | 0.961064 |
| Tra2b | P62996 | -0.04 | -1.16 | -0.58 | -0.81 | -0.85 | -0.82 | -0.31 | -1.11 | -1.15 | -0.42 | -1.02 | -1.12 | -1.28 | -0.75 | -0.87 | -1 | -0.125 | -0.28 | -0.3275 | 0.675933 | 0.361319 | 0.257989 | 0.999971 | 1 | 0.839012 |
| Trabd | E9Q7A5 | -0.31 | -1.38 | -0.42 | -0.78 | -1.5 | -1.38 | -0.9 | -0.63 | -1.01 | -0.23 | -0.82 | -0.88 | -0.83 | -0.63 | -1.21 | -0.91 | -0.38 | -0.0125 | -0.1725 | 0.252087 | 0.967093 | 0.548099 | 0.999971 | 1 | 0.911727 |
| Traf2 | P39429 | 0.9 | 0.41 | 0.17 | 0.43 | -0.53 | 0.11 | 0.19 | 0.39 | -0.72 | 0.08 | 0.32 | 0.39 | -0.73 | -0.29 | -0.16 | 0.1 | -0.4375 | -0.46 | -0.7475 | 0.131906 | 0.155643 | 0.016365 | 0.999971 | 1 | 0.613095 |
| Traf3 | Q3UHJ1 | 0.94 | 0.55 | 0.88 | 0.69 | 0.22 | 0.68 | 0.91 | 1.08 | 0.18 | 0.54 | 0.68 | 1.6 | 0.77 | 0.7 | 0.95 | 0.66 | -0.0425 | -0.015 | 0.005 | 0.859252 | 0.962297 | 0.978385 | 0.999971 | 1 | 0.99925 |
| Traf6 | P70196 | -0.44 | -0.3 | -0.16 | 0.5 | 0.15 | -0.21 | -0.38 | 0.42 | -0.33 | -0.03 | -0.01 | 0.9 | 0.13 | -0.14 | 0.52 | 0.81 | 0.095 | 0.2325 | 0.43 | 0.741708 | 0.494362 | 0.18077 | 0.999971 | 1 | 0.808532 |
| Traf7 | F8WJF7 | -0.7 | -1.5 | 0.33 | -0.4 | -0.64 | -0.42 | 0.08 | -0.41 | -0.32 | 0.12 | -0.15 | 0 | -0.8 | -0.24 | -0.56 | -0.38 | 0.22 | 0.48 | 0.0725 | 0.577495 | 0.225972 | 0.850198 | 0.999971 | 1 | 0.98187 |
| Trafd1 | Q3UDK1-2 | -0.33 | 0.4 | 0.32 | -0.4 | 1.03 | 0.04 | 1.97 | 0.25 | 1.5 | 0.13 | 1.39 | 1.04 | 0.72 | -0.36 | 0.02 | -0.21 | 0.825 | 1.0175 | 0.045 | 0.097392 | 0.020544 | 0.888501 | 0.999971 | 1 | 0.987485 |
| Trak1 | A0A5F8MP | -0.65 | -0.22 | 0.12 | -0.2 | -0.34 | -0.37 | 0.81 | -0.15 | -0.41 | -0.74 | -0.47 | -1.47 | 0.45 | -0.54 | -0.2 | -0.78 | 0.225 | -0.535 | -0.03 | 0.490592 | 0.100558 | 0.924053 | 0.999971 | 1 | 0.993608 |
| Trak2 | Q6P9N8 | 0.05 | -0.01 | -0.76 | -0.53 | -0.27 | -0.1 | -0.64 | -0.78 | -0.26 | 0.28 | -0.36 | -0.23 | 0.59 | -0.02 | -0.66 | -0.61 | -0.135 | 0.17 | 0.1375 | 0.621901 | 0.525301 | 0.69476 | 0.999971 | 1 | 0.947174 |
| Tram1 | Q91V04 | -2.32 | -0.35 | 1.18 | -0.74 | -0.87 | -0.12 | 0.93 | -0.89 | -1.3 | -2.83 | 0.88 | 0.02 | -1.46 | -0.3 | 0.78 | -0.61 | 0.32 | -0.25 | 0.16 | 0.672556 | 0.79765 | 0.835589 | 0.999971 | 1 | 0.979185 |
| Tram1l1 | Q8QZR0 | 0.32 | 0.1 | 0.85 | 2.43 | 0.42 | 0.45 | 2.31 | 1.5 | 0.34 | 1.12 | 0.61 | 0.98 | 1.05 | 0.4 | 1.37 | 0.29 | 0.245 | -0.1625 | -0.1475 | 0.699474 | 0.753112 | 0.78515 | 0.999971 | 1 | 0.966747 |
| Trap1 | Q9CQN1 | -1.22 | -0.82 | -1.31 | -1.33 | -0.75 | -0.99 | -1.1 | -1.09 | -0.68 | -1.23 | -0.82 | -0.86 | -0.19 | -0.73 | -1.12 | -0.95 | 0.1875 | 0.2725 | 0.4225 | 0.364017 | 0.221122 | 0.130123 | 0.999971 | 1 | 0.784948 |
| Trappc1 | Q5NCF2 | -1.71 | -0.68 | -0.71 | -0.88 | -0.82 | -0.54 | -0.77 | -0.75 | -0.75 | -0.4 | -0.89 | -0.68 | -0.78 | -0.42 | -0.69 | -0.51 | 0.275 | 0.315 | 0.395 | 0.322363 | 0.275451 | 0.174602 | 0.999971 | 1 | 0.808161 |
| Trappc10 | Q3TLI0 | -0.69 | -0.03 | -0.62 | -0.91 | -0.09 | -0.51 | -0.46 | -0.95 | 0.11 | -0.47 | -0.55 | -0.77 | 0.27 | -0.4 | -0.56 | -0.65 | 0.06 | 0.1425 | 0.2275 | 0.827336 | 0.613161 | 0.444545 | 0.999971 | 1 | 0.88409 |
| Trappc11 | B2RXC1 | -1.31 | 0.01 | 0.32 | -0.14 | -0.37 | -0.16 | 0.69 | -0.11 | -0.14 | -1.17 | 0.1 | -0.18 | -0.44 | -0.45 | 0.07 | -0.32 | 0.2925 | -0.0675 | -0.005 | 0.477745 | 0.87537 | 0.989123 | 0.999971 | 1 | 0.99925 |
| Trappc12 | Q8K2L8 | 0.14 | 0.29 | 0.6 | 0.59 | 0.21 | 0.4 | 0.28 | 0.34 | 0.34 | 0.38 | 0.64 | 0.61 | 0.35 | 0.37 | 0.77 | 1.1 | -0.0975 | 0.0875 | 0.2425 | 0.609011 | 0.65545 | 0.334357 | 0.999971 | 1 | 0.858827 |
| Trappc13 | Q3TIR1 | 0.36 | -0.16 | 0.77 | 0.66 | -0.45 | 0.21 | 1.09 | 0.4 | -0.47 | 0.2 | 0.61 | -0.15 | -0.9 | 0.08 | 0.17 | 0.05 | -0.095 | -0.36 | -0.5575 | 0.796546 | 0.271325 | 0.115763 | 0.999971 | 1 | 0.78268 |
| Trappc14 | Q3UTZ3 | 0.46 | -0.6 | -0.36 | -0.03 | -0.36 | -0.06 | -0.19 | -0.12 | -0.27 | 0.36 | 0 | -0.22 | -0.17 | -0.05 | -0.1 | 0.13 | -0.05 | 0.1 | 0.085 | 0.848302 | 0.725086 | 0.746056 | 0.999971 | 1 | 0.961064 |
| Trappc2 | Q9CQP2 | -2.89 | 0.12 | -0.52 | -1.23 | 0.29 | -0.56 | -0.76 | -0.64 | -0.03 | -2.73 | -0.5 | -0.21 | 0.38 | -0.98 | -0.2 | -0.89 | 0.7125 | 0.2625 | 0.7075 | 0.274771 | 0.748352 | 0.297696 | 0.999971 | 1 | 0.847402 |
| Trappc2l | Q9JME7 | -0.34 | -0.18 | 0.17 | 0.05 | -0.41 | 0.14 | -0.21 | -0.08 | -0.07 | -0.16 | 0.11 | -0.26 | -0.2 | 0.26 | 0.22 | 0.46 | -0.065 | -0.02 | 0.26 | 0.759382 | 0.918537 | 0.262226 | 0.999971 | 1 | 0.842262 |
| Trappc3 | O55013 | 0.11 | 0.6 | -0.62 | -0.02 | 0.29 | 0.7 | -0.28 | -0.35 | 0.66 | 0.23 | 0.21 | -0.75 | 0.33 | 0.12 | -0.27 | 0.04 | 0.0725 | 0.07 | 0.0375 | 0.834874 | 0.852829 | 0.897783 | 0.999971 | 1 | 0.989101 |
| Trappc4 | Q9ES56 | -0.7 | -0.19 | -0.61 | -1.08 | -0.22 | -0.22 | -0.73 | -0.74 | -0.42 | -0.84 | -0.66 | -0.5 | -0.2 | 0.12 | -0.5 | -0.68 | 0.1675 | 0.04 | 0.33 | 0.523293 | 0.865803 | 0.24586 | 0.999971 | 1 | 0.838611 |
| Trappc5 | Q9CQA1 | 1.46 | 0.19 | 0.3 | 0.63 | -0.23 | 0.57 | 0.2 | 0.41 | 0.1 | 0.86 | 0.43 | 0.42 | 0 | 0.01 | 0.47 | 0.98 | -0.4075 | -0.1925 | -0.28 | 0.242175 | 0.559848 | 0.446548 | 0.999971 | 1 | 0.884555 |
| Trappc6b | Q9D289 | -0.74 | 0.27 | 0.16 | -0.49 | 0.04 | 0.4 | 0.18 | 0 | 0.05 | -0.19 | 0.53 | 0.61 | 0.47 | 0.33 | 0.09 | 0.22 | 0.355 | 0.45 | 0.4775 | 0.22326 | 0.17803 | 0.111512 | 0.999971 | 1 | 0.78268 |
| Trappc8 | A0A286YC | 1.31 | 0.21 | 0.6 | 0.31 | 0.11 | 0.22 | 0.56 | 0.31 | 0.17 | 0.83 | 0.6 | 0.19 | -0.1 | 0.23 | 0.43 | 0.34 | -0.3075 | -0.16 | -0.3825 | 0.290864 | 0.600262 | 0.206088 | 0.999971 | 1 | 0.817241 |
| Trappc9 | Q3U0M1 | -0.21 | -0.08 | 0.18 | 0.14 | -0.03 | 0.26 | 0.11 | 0.28 | 0.17 | -0.14 | 0.14 | -0.12 | 0.46 | 0.04 | 0.3 | 0.44 | 0.1475 | 0.005 | 0.3025 | 0.439659 | 0.978732 | 0.149563 | 0.999971 | 1 | 0.793248 |
| Trex1 | Q91XB0 | 0.22 | -0.16 | -1.81 | -1.33 | 0.06 | 0.57 | -0.61 | -0.88 | -0.09 | -0.09 | -0.56 | -0.69 | 0.74 | 0.49 | -0.28 | -0.45 | 0.555 | 0.4125 | 0.895 | 0.314098 | 0.393024 | 0.111326 | 0.999971 | 1 | 0.78268 |
| Trh | Q62361 | 1.44 | -2.65 | 4.6 | 3.24 | 0.61 | 0.34 | 4.46 | 3.97 | 3.11 | -0.26 | 0.64 | 4.84 | -0.06 | 0.11 | 4.39 | -1.65 | 0.6875 | 0.425 | -0.96 | 0.685902 | 0.807916 | 0.59789 | 0.999971 | 1 | 0.92377 |
| Trhde | Q8K093 | -0.3 | -0.75 | -0.19 | -0.7 | -1.14 | -0.42 | -0.28 | -0.47 | -1.16 | -0.63 | -0.23 | -0.63 | -0.89 | -0.69 | -0.74 | -0.58 | -0.0925 | -0.1775 | -0.24 | 0.723753 | 0.499142 | 0.266943 | 0.999971 | 1 | 0.842677 |
| Trhr | P21761 | -0.92 | 0.57 | -1.14 | -2.25 | 1.22 | 0.78 | -1.38 | -1.4 | 1.1 | -0.1 | -1.48 | -1.13 | 1.94 | 0.73 | -1.35 | -1.75 | 0.74 | 0.5325 | 0.8275 | 0.374462 | 0.481116 | 0.388576 | 0.999971 | 1 | 0.871816 |
| Triap1 | Q9D8Z2 | 0.44 | 1.46 | 0.25 | 1.2 | 1.05 | 1.7 | 0.12 | 0.73 | 1.92 | -0.5 | 0.85 | 0.44 | 0.96 | 1.09 | 0.35 | 1.05 | 0.0625 | -0.16 | 0.025 | 0.880942 | 0.766177 | 0.940875 | 0.999971 | 1 | 0.996218 |
| Tril | Q9DBY4 | -0.05 | -0.58 | -1.9 | -1.44 | -1.22 | -0.83 | -1.43 | -0.82 | -1.76 | -1.02 | -0.75 | -1.28 | -1.38 | -1.31 | -1.18 | -1.97 | -0.0825 | -0.21 | -0.4675 | 0.844295 | 0.637476 | 0.291685 | 0.999971 | 1 | 0.847402 |
| Trim2 | Q9ESN6 | 0.07 | -0.57 | -0.55 | -0.49 | -0.45 | -0.29 | -0.28 | -0.56 | -0.48 | 0.03 | -0.8 | -0.67 | -0.75 | -0.23 | -0.42 | -0.79 | -0.01 | -0.095 | -0.1625 | 0.962764 | 0.715217 | 0.500732 | 0.999971 | 1 | 0.904223 |
| Trim23 | Q8BGX0-3 | -1.39 | -0.22 | 1.17 | 0.42 | -0.45 | 0.22 | 1.09 | 0.64 | -0.53 | -0.72 | 1.75 | 2.41 | 0.07 | -0.14 | 1.76 | 2.83 | 0.38 | 0.7325 | 1.135 | 0.51697 | 0.408294 | 0.18178 | 0.999971 | 1 | 0.808532 |

| | | | | | | | | | | | | | | | | | | | | | | | | | | |
|---|---|---|---|---|---|---|---|---|---|---|---|---|---|---|---|---|---|---|---|---|---|---|---|---|---|---|
| Trim24 | Q64127 | 2.93 | 1.57 | 0.38 | 1.18 | 1.44 | 1.1 | 1.17 | 0.13 | 1.51 | 2.23 | 0.18 | -0.31 | 0.03 | 0.46 | -0.11 | -0.36 | -0.555 | -0.6125 | -1.51 | 0.33153 | 0.405067 | 0.016403 | 0.999971 | 1 | 0.613095 |
| Trim25 | Q61510 | 0.9 | 0.59 | 0.83 | 0.83 | 1.23 | -0.03 | 0.15 | 0.81 | 0.54 | 0.38 | 0.49 | 1 | 0.91 | 1.15 | 0.53 | 0.97 | -0.2475 | -0.185 | 0.1025 | 0.42815 | 0.375124 | 0.617138 | 0.999971 | 1 | 0.928457 |
| Trim26 | J3QN94 | -0.15 | 0.76 | 0.8 | 0.26 | 0.47 | 0.28 | -0.5 | 1.63 | -0.23 | -0.36 | 0.36 | 1.19 | 0.95 | -0.07 | 0.76 | 0.77 | 0.0525 | -0.1775 | 0.185 | 0.909551 | 0.660227 | 0.5705 | 0.999971 | 1 | 0.91816 |
| Trim28 | Q62318 | 0.7 | 1.27 | 0.62 | 0.52 | 1.04 | 1.21 | 0.64 | 0.52 | 1.06 | 0.69 | 0.56 | 0.54 | 1.13 | 1.21 | 0.53 | 0.73 | 0.075 | -0.065 | 0.1225 | 0.771985 | 0.785131 | 0.637707 | 0.999971 | 1 | 0.93301 |
| Trim3 | Q9R1R2 | -0.23 | 0.11 | -0.03 | 0.08 | 1.03 | 0.31 | 0.22 | 0.37 | 1.33 | 0.16 | 0.16 | 0.29 | 2.48 | 0.09 | 0.21 | 0.09 | 0.5 | 0.5025 | 0.735 | 0.058733 | 0.122373 | 0.20157 | 0.999971 | 1 | 0.815758 |
| Trim32 | Q8CH72 | -0.19 | 0.95 | 0.5 | 0.12 | 0.56 | 0.9 | -0.04 | 0.44 | 0.94 | 0.16 | 0.37 | -0.26 | 1.56 | 0.9 | 0.4 | 0.58 | 0.12 | -0.0425 | 0.515 | 0.705812 | 0.902082 | 0.16586 | 0.999971 | 1 | 0.800215 |
| Trim33 | E9QME5 | 1.84 | 2.04 | 1.96 | 2.49 | 1.95 | 1.55 | 1.54 | 2.09 | 1.67 | 1.56 | 2.19 | 2.14 | 1.14 | 1.8 | 1.97 | 2.25 | -0.3 | -0.1925 | -0.2925 | 0.222787 | 0.437051 | 0.325021 | 0.999971 | 1 | 0.856472 |
| Trim35 | Q8C006 | 1.65 | 0.34 | 1.75 | 1.75 | 0.31 | 0.63 | 1.98 | 1.87 | 0.8 | 0.87 | 1.53 | 0.7 | 0.28 | 0.5 | 1.06 | 1.48 | -0.175 | -0.3975 | -0.5425 | 0.730729 | 0.311544 | 0.216168 | 0.999971 | 1 | 0.826893 |
| Trim36 | Q80WG7 | 0.71 | -1.24 | -0.47 | -0.47 | 0.1 | -1.37 | -0.01 | -0.21 | 0.28 | -0.27 | -0.49 | -0.5 | 0.78 | -0.81 | -0.14 | -1.41 | -0.005 | 0.1225 | -0.0275 | 0.991828 | 0.771231 | 0.961334 | 0.999971 | 1 | 0.998721 |
| Trim37 | F6XMH5 | -4.6 | -0.84 | -0.47 | -5.11 | 0.53 | -2.66 | -0.39 | -3.45 | 0.43 | -4.95 | 1.3 | 1.05 | 0.28 | -2.61 | -1.75 | -2.65 | 1.2625 | 2.2125 | 1.0725 | 0.366469 | 0.218308 | 0.399351 | 0.999971 | 1 | 0.871816 |
| Trim44 | F7B7X2 | 2.63 | 2.09 | 1.09 | 0.35 | 1.93 | 1.69 | 1.02 | 1.54 | 2.05 | 2.97 | 0.31 | 1.46 | 1.5 | 2.34 | 0.87 | -0.02 | 0.005 | 0.1575 | -0.3675 | 0.992076 | 0.818885 | 0.574146 | 0.999971 | 1 | 0.91816 |
| Trim46 | Q7TNM2 | 0.36 | 0.47 | 0.87 | 1.14 | 0.43 | 0.91 | 1.18 | 1.47 | 0.53 | 0.38 | 1.18 | 0.9 | 0.36 | 1.05 | 1.39 | 1.46 | 0.2875 | 0.0375 | 0.355 | 0.343648 | 0.890452 | 0.27645 | 0.999971 | 1 | 0.843794 |
| Trim47 | Q8C0E3 | 0.02 | -0.63 | 0.86 | 0.64 | 1.04 | -0.22 | 1.81 | 0.58 | -0.35 | -0.05 | 0.8 | 0.96 | 1.21 | -0.68 | 0.56 | -0.17 | 0.58 | 0.1175 | 0.0075 | 0.265992 | 0.788796 | 0.987908 | 0.999971 | 1 | 0.99925 |
| Trim56 | Q80VI1 | 1.26 | 0.96 | 1.08 | 1.01 | 0.14 | 1.55 | -0.66 | 0.35 | -0.43 | 1.11 | 0.29 | -0.12 | 0.09 | 1.97 | 0.55 | 0.37 | -0.7325 | -0.865 | -0.3325 | 0.119615 | 0.028941 | 0.421504 | 0.999971 | 1 | 0.87788 |
| Trim65 | Q8BFW4 | -0.95 | -0.18 | -0.35 | -0.12 | -1.12 | -0.9 | -0.04 | -0.17 | -0.72 | -0.33 | 0.07 | -0.22 | -0.53 | -0.62 | 0.33 | 0.15 | -0.1575 | 0.1 | 0.2325 | 0.632327 | 0.710685 | 0.461784 | 0.999971 | 1 | 0.890095 |
| Trim66 | E9PZP2 | -1.44 | -0.16 | -0.19 | -0.41 | 0.5 | -1.01 | -0.37 | -0.53 | 0.75 | -0.84 | 0.3 | -0.42 | 0.78 | -0.19 | -1.2 | -0.27 | 0.1975 | 0.4975 | 0.33 | 0.635782 | 0.277731 | 0.490627 | 0.999971 | 1 | 0.898612 |
| Trim72 | Q1XH17 | -0.78 | -0.04 | -0.46 | 0.78 | -1.05 | -1.45 | 1.21 | 0.11 | -0.66 | -0.49 | 0.5 | 0.92 | 0.23 | -0.89 | -0.1 | 2.29 | -0.17 | 0.1925 | 0.5075 | 0.786511 | 0.688107 | 0.466854 | 0.999971 | 1 | 0.891477 |
| Trim9 | Q8C7M3 | -2.81 | -1.31 | -1.27 | -1.3 | -0.32 | -1.32 | -1.01 | -1.12 | -0.93 | -3.41 | -0.17 | -0.37 | -1.1 | -1.46 | -0.75 | -1.41 | 0.73 | 0.4525 | 0.4925 | 0.106069 | 0.55567 | 0.234995 | 0.999971 | 1 | 0.832918 |
| Trim9_1 | A0A6I8MW | -0.15 | -0.49 | -0.37 | -0.04 | -0.9 | -0.42 | -0.2 | -0.76 | -1.21 | -0.73 | 0.06 | -1.04 | -1.76 | -0.53 | -0.72 | -0.32 | -0.3075 | -0.4675 | -0.57 | 0.200834 | 0.152108 | 0.115839 | 0.999971 | 1 | 0.78268 |
| Trim9_2 | E9QLH4 | 0.04 | 0.31 | -0.14 | -0.23 | -0.11 | 0.16 | -0.2 | 0.07 | -0.19 | 0 | 0 | -0.35 | -0.04 | -0.11 | -0.24 | 0.06 | -0.015 | -0.13 | -0.0775 | 0.940646 | 0.520334 | 0.695089 | 0.999971 | 1 | 0.947174 |
| Trio | Q0KL02-3 | -0.8 | -0.43 | -0.99 | -1.07 | -0.5 | -0.68 | -0.1 | -1.24 | -0.36 | -0.56 | -0.92 | -0.84 | -0.43 | -0.75 | -1.18 | -1.24 | 0.1925 | 0.1525 | -0.0775 | 0.510231 | 0.509597 | 0.767675 | 0.999971 | 1 | 0.962472 |
| Trio_1 | A0A2I3BQ1 | 1.75 | 0.08 | -1.62 | -2.42 | -0.65 | -0.28 | -0.24 | -1.82 | -0.57 | 1.31 | -1.09 | -3.41 | -3.29 | -0.61 | -0.06 | -2.16 | -0.195 | -0.3875 | -0.9775 | 0.827321 | 0.748003 | 0.367233 | 0.999971 | 1 | 0.869526 |
| Triobp | Q99KW3-4 | 3.23 | 0.35 | 0.33 | 1.13 | 0.82 | 0.6 | 1.27 | 0.03 | 0.83 | 2.66 | 0.09 | 0.35 | 0.04 | 0.53 | 0.39 | 0.07 | -0.58 | -0.2775 | -1.0025 | 0.391238 | 0.732636 | 0.13903 | 0.999971 | 1 | 0.784948 |
| Trip11 | E9Q512 | 1.48 | 1.62 | 1.85 | 1.85 | 1.5 | 1.28 | 1.35 | 2.01 | 1.51 | 1.44 | 1.7 | 1.97 | 1.05 | 1.5 | 1.71 | 2.13 | -0.165 | -0.045 | -0.1025 | 0.475464 | 0.823626 | 0.699764 | 0.999971 | 1 | 0.948208 |
| Trip12 | G5E870 | -0.02 | 0.2 | -0.39 | -0.37 | 0.17 | -0.07 | -0.32 | -0.01 | 0.49 | 0.11 | -0.25 | -0.18 | 0.14 | 0.01 | -0.15 | -0.5 | 0.0875 | 0.1875 | 0.02 | 0.69131 | 0.456544 | 0.932185 | 0.999971 | 1 | 0.995069 |
| Trip4 | Q9QXN3-2 | 0.81 | 0.44 | 1.02 | 0.53 | 0.79 | 0.85 | 1.96 | 0.88 | 0.76 | 1.01 | 1.98 | 2.32 | 0.72 | 0.66 | 1.51 | 0.88 | 0.42 | 0.8175 | 0.2425 | 0.204072 | 0.058938 | 0.363266 | 0.999971 | 1 | 0.869526 |
| Trir | Q9D735 | 4.18 | 2.19 | 4.39 | 5.1 | 2.08 | 2.68 | 4.82 | 3.31 | 2.31 | 3.36 | 3.03 | 2.1 | 0.93 | 1.76 | 3.12 | 2.64 | -0.7425 | -1.265 | -1.8525 | 0.348962 | 0.072192 | 0.027477 | 0.999971 | 1 | 0.678474 |
| Trit1 | Q80UN9 | -1.77 | -0.38 | -0.92 | 1.23 | -0.97 | -0.76 | -1.49 | -0.64 | -0.81 | -1.02 | -0.99 | -1.46 | -0.85 | -0.99 | -1.32 | 0.32 | -0.505 | -0.61 | -0.25 | 0.410635 | 0.319755 | 0.70557 | 0.999971 | 1 | 0.950182 |
| Trmt1 | H3BKS8 | 1.22 | 1.82 | 1.79 | 0.98 | 1.82 | 1.63 | 1.39 | 1.21 | 1.38 | 1.05 | 1.92 | 1.54 | 1.01 | 1.18 | 1.46 | 1.21 | 0.06 | 0.02 | -0.2375 | 0.823238 | 0.944594 | 0.36467 | 0.999971 | 1 | 0.869526 |
| Trmt10c | Q3UFY8 | -0.58 | -0.73 | 0.09 | 0.06 | -0.58 | -0.69 | 0.01 | 0.44 | -1.06 | -0.61 | 0.03 | -0.24 | -0.6 | -0.54 | 0.44 | 0.48 | 0.085 | -0.18 | 0.235 | 0.800929 | 0.577216 | 0.51752 | 0.999971 | 1 | 0.905815 |
| Trmt11 | E9QKG3 | -0.23 | 0.03 | 0.11 | 0.36 | -0.6 | -0.92 | -0.73 | 0.1 | -0.55 | -0.86 | -0.57 | -0.75 | 0.03 | -0.44 | -0.36 | 0.22 | -0.605 | -0.75 | -0.205 | 0.049795 | 0.004256 | 0.393762 | 0.999971 | 0.969702 | 0.871816 |
| Trmt112 | Q9DCG9 | 1.6 | 1.5 | 0.13 | 0.39 | 1.24 | 1.53 | -0.18 | 0.29 | 1.04 | 1.56 | -0.03 | -0.06 | 0.89 | 1.79 | 0.26 | 0.61 | -0.185 | -0.2775 | -0.0175 | 0.716852 | 0.591312 | 0.97009 | 0.999971 | 1 | 0.998787 |
| Trmt1l | A0A0R4J0L | -0.16 | 1.62 | -0.93 | -0.44 | 0.98 | 1.48 | -0.72 | -1.47 | 1.43 | 1.35 | -1.59 | -1.08 | 0.95 | 1.38 | -2.73 | -0.82 | 0.045 | 0.005 | -0.3275 | 0.955101 | 0.995432 | 0.737629 | 0.999971 | 1 | 0.959801 |
| Trmt44 | Q9D2Q2 | -2.05 | 0.83 | 0.42 | -0.64 | 0.82 | 0.07 | 0.44 | -0.54 | 1.48 | -1.58 | -0.21 | -0.13 | -0.74 | -0.94 | -0.42 | -1.37 | 0.5575 | 0.25 | -0.5075 | 0.394552 | 0.758486 | 0.417954 | 0.999971 | 1 | 0.877685 |
| Trmt6 | Q8CE96 | 1.92 | 2.36 | 2.65 | 2.36 | 2.07 | 1.59 | 2.56 | 1.85 | 1.89 | 2.15 | 2.51 | 2.03 | 1.32 | 1.86 | 1.76 | 1.2 | -0.305 | -0.1775 | -0.7875 | 0.281337 | 0.455598 | 0.01118 | 0.999971 | 1 | 0.613095 |
| Trmt61a | Q80XC2 | 3.23 | 2.83 | 2.19 | 1.69 | 1.49 | 2.21 | 4.04 | 1.05 | 1.86 | 2.61 | 1.87 | 1.85 | 2.17 | 2.19 | 1.78 | 1.37 | -0.2875 | -0.4375 | -0.6075 | 0.670228 | 0.264659 | 0.136088 | 0.999971 | 1 | 0.784948 |
| Trmu | Q9DAT5 | 0.81 | 0.23 | 0.82 | 0.82 | -0.07 | 0.51 | 0.74 | 0.86 | 0.15 | 0.46 | 0.32 | 0.46 | 0.13 | 0.26 | 0.45 | 0.67 | -0.16 | -0.3225 | -0.2925 | 0.560042 | 0.151562 | 0.221329 | 0.999971 | 1 | 0.8275 |
| Trnt1 | Q8K1J6 | -0.3 | 1.12 | 0.84 | 1.13 | 0.84 | 1.01 | 0.87 | 0.94 | 1.17 | 0.56 | 1.21 | 1.01 | 1.16 | 1.25 | 0.91 | 1.24 | 0.2175 | 0.29 | 0.4425 | 0.52516 | 0.43172 | 0.220741 | 0.999971 | 1 | 0.827096 |
| Tro | Q6DIC6 | 1.12 | 0.29 | -0.13 | 0.29 | 0.95 | 0.2 | 2.37 | -0.95 | 0.36 | -1.54 | 0.11 | -0.01 | -0.42 | 0.79 | -0.17 | -0.1 | 0.25 | -0.6625 | -0.3675 | 0.711337 | 0.18514 | 0.326354 | 0.999971 | 1 | 0.856472 |
| Trp53bp2 | E9QJU8 | 0.57 | -0.83 | -0.38 | -1.23 | -0.47 | -0.69 | -0.53 | -0.77 | -0.52 | -0.11 | -0.25 | -0.07 | -0.84 | -0.48 | -0.9 | -0.79 | -0.1475 | 0.23 | -0.285 | 0.69854 | 0.55538 | 0.465567 | 0.999971 | 1 | 0.891255 |
| Trp53i11 | A2AGS6 | -0.47 | -0.47 | -0.1 | -0.2 | 0.24 | -0.73 | 0.56 | 0.28 | 0.43 | -0.26 | 0.71 | 0.92 | 2.97 | -1.14 | 0.13 | 0.15 | 0.3975 | 0.76 | 0.8375 | 0.213312 | 0.024858 | 0.303119 | 0.999971 | 1 | 0.847402 |
| Trp53rkb | Q543M9 | 0.25 | 0.52 | 1.79 | 1.26 | 0.2 | 0.35 | 1.08 | 1.67 | 0.25 | 0.17 | 1.95 | 1.79 | 0.26 | 0.46 | 1.93 | 1.92 | -0.13 | 0.085 | 0.1875 | 0.776915 | 0.87709 | 0.724086 | 0.999971 | 1 | 0.955315 |
| Trpc1 | A0A087WS | -1.1 | -0.76 | -0.15 | -0.68 | -1.19 | -0.58 | -1.16 | -0.09 | -0.84 | -0.05 | 0.45 | -1.08 | -1.37 | -0.27 | -0.49 | -0.42 | -0.0825 | 0.2925 | 0.035 | 0.801438 | 0.4603 | 0.912807 | 0.999971 | 1 | 0.992618 |
| Trpc4 | Q9QUQ5 | -1.04 | -0.61 | 0.32 | 0.32 | -0.9 | -0.18 | 0.05 | 0.48 | -0.82 | -0.45 | 0.64 | 0.63 | -0.01 | -0.07 | 0.96 | 1.21 | 0.115 | 0.2525 | 0.775 | 0.786349 | 0.597967 | 0.110375 | 0.999971 | 1 | 0.78268 |
| Trpc4ap | Q9JLV2-2 | 0.38 | 0.32 | 0.91 | 1.11 | 0.34 | 0.73 | 0.61 | 1.71 | 0.38 | 0.59 | 0.95 | 1.42 | 0.85 | 0.9 | 1.65 | 1.8 | 0.1675 | 0.155 | 0.62 | 0.634847 | 0.614832 | 0.07858 | 0.999971 | 1 | 0.754157 |
| Trpc6 | Q3UZG1 | 0.29 | -1.06 | -0.92 | -0.02 | 0.47 | 0.47 | -0.8 | -0.03 | -1.5 | 0.14 | -0.49 | -0.83 | 1.3 | -0.2 | -0.74 | -0.03 | 0.455 | -0.2425 | 0.51 | 0.299188 | 0.592611 | 0.328275 | 0.999971 | 1 | 0.856472 |
| Trpm3 | Q5F4S7 | -1.13 | -1.55 | -0.09 | 0.02 | -1.16 | -0.94 | -0.45 | 0.33 | -0.99 | -1.28 | -0.6 | -0.33 | -0.7 | -0.54 | -0.22 | -0.37 | 0.1325 | -0.1125 | 0.23 | 0.780302 | 0.788596 | 0.555727 | 0.999971 | 1 | 0.914901 |
| Trpm7 | Q923J1 | -1.7 | -2.38 | -0.57 | -0.85 | -1.33 | -2.95 | -1.66 | -0.89 | -4 | -2.51 | -1.55 | -1.11 | -2.98 | -2.5 | -1.25 | -0.48 | -0.3325 | -0.9175 | -0.4275 | 0.553493 | 0.208433 | 0.510725 | 0.999971 | 1 | 0.905815 |
| Trpv2 | Q9WTR1 | -1.05 | -1.04 | -0.4 | -0.46 | -1.29 | -0.99 | -0.45 | -0.44 | -1.25 | -0.85 | -0.31 | -0.83 | -0.98 | -0.87 | -0.69 | -0.03 | -0.055 | -0.0725 | 0.095 | 0.848423 | 0.794046 | 0.743848 | 0.999971 | 1 | 0.961064 |
| Trrap | E9QLK7 | -0.12 | -1.78 | -0.93 | -0.94 | -1.4 | -2.8 | 0.66 | -1.63 | -1.96 | -2.31 | -1.97 | -0.2 | -3.59 | -2.64 | -0.73 | -3.75 | -0.35 | -0.6675 | -1.735 | 0.628023 | 0.238349 | 0.032969 | 0.999971 | 1 | 0.685881 |
| Trrap_1 | E9PWT1 | 0.13 | 0.77 | 1.01 | 1.06 | 0.79 | 0.69 | 0.68 | 0.6 | 0.7 | 0.37 | 0.9 | 0.9 | 0.54 | 1.02 | 0.92 | 1.07 | -0.0525 | -0.025 | 0.145 | 0.831565 | 0.925255 | 0.589738 | 0.999971 | 1 | 0.922012 |
| Trub1 | Q8C0D0 | 1.2 | 0.22 | 1.11 | 2.18 | 0.51 | -0.26 | 1.42 | 1.55 | 0.27 | 0.64 | 0.86 | 1.05 | -0.45 | 0.36 | 1.26 | 1.23 | -0.3725 | -0.4725 | -0.5775 | 0.493789 | 0.272369 | 0.290942 | 0.999971 | 1 | 0.847402 |
| Try10 | Q792Z1 | -0.88 | 2.79 | 3.4 | 1.04 | 0.6 | 4.21 | 1.94 | 2.1 | 1.85 | 0.86 | 2.96 | 1.29 | 0.6 | 2.1 | 1.55 | 2.99 | 0.625 | 0.1525 | 0.2225 | 0.568004 | 0.873493 | 0.818777 | 0.999971 | 1 | 0.975596 |
| Tsc1 | Q9EP53-3 | 0.32 | -0.26 | 0.03 | 0.06 | -0.36 | -0.11 | -0.07 | -0.05 | -0.23 | 0.26 | 0.11 | -0.28 | -0.44 | -0.53 | -0.19 | -0.2 | -0.185 | -0.0725 | -0.3775 | 0.363288 | 0.740421 | 0.090464 | 0.999971 | 1 | 0.771991 |
| Tsc2 | A0A2I3BPP | -0.3 | -0.31 | 0.14 | -0.28 | -0.37 | -0.42 | -0.02 | -0.16 | -0.27 | -0.21 | -0.03 | -0.36 | -0.33 | -0.35 | -0.1 | 0.04 | -0.055 | -0.03 | 0.0025 | 0.784715 | 0.875009 | 0.99012 | 0.999971 | 1 | 0.99925 |
| Tsc22d1 | P62500-2 | 1.69 | 1.5 | 2.31 | 3.25 | 1.63 | 1.93 | 1.21 | 3.53 | 2.38 | 1.84 | 3.06 | 2.28 | 2.75 | 1.94 | 2.68 | 2.82 | -0.1125 | 0.2025 | 0.36 | 0.847967 | 0.648862 | 0.403433 | 0.999971 | 1 | 0.873348 |
| Tsc22d1_1 | D3Z0V7 | 1.17 | 0.32 | 0.5 | 1.26 | 0.19 | 0.05 | 0.5 | 1.47 | 0.29 | -0.11 | 1.17 | 1.46 | 0.93 | 0.39 | 1.85 | 1.47 | -0.26 | -0.11 | 0.3475 | 0.50293 | 0.791881 | 0.375863 | 0.999971 | 1 | 0.870465 |
| Tsc22d2 | A0A0G2JFZ | 0.89 | 1.56 | 0.75 | 0.41 | 2.21 | 0.97 | 1.24 | 0.37 | 1.3 | 0.59 | 1.84 | 1.36 | 2.37 | 0.47 | 1.45 | 1.23 | 0.295 | 0.37 | 0.4775 | 0.496787 | 0.30329 | 0.288781 | 0.999971 | 1 | 0.847402 |
| Tsc22d3 | Q9Z2S7-3 | 0.41 | 1.23 | 0.17 | 0.42 | 0.41 | 0.56 | -0.41 | 0.46 | 0.29 | -0.69 | 0.53 | 0.42 | 1.32 | 1.14 | 0.47 | 0.89 | -0.3025 | -0.42 | 0.3975 | 0.361701 | 0.257571 | 0.212923 | 0.999971 | 1 | 0.823987 |
| Tsc22d4 | Q9EQN3 | -0.1 | 0.18 | 0.04 | 0.01 | -0.23 | -0.67 | -0.08 | 0.41 | 0.13 | 0.44 | -0.64 | -0.41 | 0.28 | 0.01 | -0.24 | -0.09 | -0.175 | -0.1525 | -0.0425 | 0.499158 | 0.576847 | 0.824297 | 0.999971 | 1 | 0.976309 |
| Tsen34 | H7BX14 | -0.1 | 1.63 | 1.15 | 1.17 | 1.21 | 0.31 | 0.42 | 0.6 | 0.47 | -0.13 | 0.81 | 1.15 | -0.26 | 0.88 | 1.03 | 0.83 | -0.3275 | -0.3875 | -0.3425 | 0.425115 | 0.385503 | 0.451574 | 0.999971 | 1 | 0.886993 |
| Tsfm | Q9CZR8 | -0.82 | -0.02 | 0.52 | 0.61 | -0.4 | -0.4 | -0.02 | 0.62 | -0.37 | -0.42 | 0.56 | -0.11 | -0.19 | 0.13 | 0.58 | 0.63 | -0.1225 | -0.1575 | 0.215 | 0.754363 | 0.683461 | 0.566224 | 0.999971 | 1 | 0.917959 |
| Tsg101 | Q61187 | -0.4 | -0.32 | -0.55 | -0.54 | -0.59 | -0.33 | -0.75 | -0.5 | -0.27 | -0.57 | -0.48 | -0.46 | 0.1 | -0.23 | -0.58 | -0.12 | -0.09 | 0.0075 | 0.245 | 0.621956 | 0.964724 | 0.255077 | 0.999971 | 1 | 0.839012 |
| Tsga10 | Q6NY15-2 | -0.03 | 0.68 | 0.33 | 1.09 | -0.59 | -0.12 | -0.08 | 0.38 | 0.73 | 0.98 | -0.59 | -1.04 | 0.89 | 0.53 | -1.19 | -1.23 | -0.62 | -0.4975 | -0.7675 | 0.075419 | 0.343179 | 0.193482 | 0.999971 | 1 | 0.810547 |
| Tshz3 | A0A0R4J01 | -3.7 | -0.77 | -0.4 | -1.3 | 1.31 | -0.7 | 1.27 | -1.48 | 0.65 | -2.65 | 0.98 | 1.66 | 1.17 | -2.59 | -1.48 | -0.72 | 1.6425 | 1.7025 | 0.6375 | 0.100324 | 0.144803 | 0.516488 | 0.999971 | 1 | 0.905815 |
| Tsn | Q62348 | 0.86 | 0.13 | 0.55 | 0.8 | 0.55 | -0.42 | 0.42 | 0.84 | -0.11 | -0.09 | 0.36 | 0.34 | -0.52 | -0.01 | 0.42 | 0.53 | -0.2375 | -0.46 | -0.48 | 0.46455 | 0.083188 | 0.136168 | 0.999971 | 1 | 0.784948 |
| Tsnax | Q9QZE7 | 0.46 | 0.83 | 0.35 | 0.44 | 0.75 | 0.52 | 0.33 | 0.48 | 0.69 | 0.23 | -0.18 | 0.03 | 0.37 | 0.49 | 0.1 | 0.4 | -1.23E-16 | -0.3275 | -0.18 | 1 | 0.200618 | 0.373168 | 1 | 1 | 0.869526 |
| Tspan13 | Q9D8C2 | -0.21 | 2.5 | 0.71 | 1.6 | 2.36 | 2.44 | 0.93 | 2.3 | 2.52 | -0.93 | 0.66 | 1.9 | 2.13 | 1.17 | 1.09 | 1.8 | 0.8575 | -0.1125 | 0.3975 | 0.192302 | 0.896476 | 0.499668 | 0.999971 | 1 | 0.903837 |
| Tspan14 | Q8QZY6 | -0.59 | 0.04 | -0.05 | -0.26 | -0.17 | 0.68 | -0.32 | 0.16 | 0.07 | 0.13 | -0.12 | 0 | 0.62 | 0.21 | -0.23 | 0 | 0.3025 | 0.235 | 0.365 | 0.29331 | 0.263437 | 0.176831 | 0.999971 | 1 | 0.808532 |
| Tspan15 | F7BWT7 | -1.86 | -0.1 | -0.96 | -1.56 | -0.57 | -0.51 | -2.32 | -0.49 | 0.01 | -0.64 | -0.58 | -0.92 | -0.15 | -0.36 | 0.02 | -1.1 | 0.1475 | 0.5875 | 0.7225 | 0.787231 | 0.180977 | 0.123769 | 0.999971 | 1 | 0.78268 |
| Tspan2 | Q9D1X8 | 0.33 | 1.94 | 2.66 | 0.95 | 1.47 | 1.84 | 2.03 | 1.73 | 2.21 | 0.6 | 1.87 | 2.32 | 3.56 | 0.55 | 2.17 | 0.8 | 0.2975 | 0.28 | 0.3 | 0.549668 | 0.64136 | 0.701659 | 0.999971 | 1 | 0.949019 |
| Tspan3 | Q9QY33 | -2.22 | 0.06 | -0.38 | -0.88 | 0.22 | 0.34 | 0.01 | -0.24 | 0.26 | -2.81 | -1.29 | -0.84 | 1.77 | -0.74 | -0.78 | -1.13 | 0.9375 | -0.315 | 0.635 | 0.075313 | 0.668274 | 0.407422 | 0.999971 | 1 | 0.874911 |
| Tspan31 | Q9CQ88 | 0.9 | 1.46 | 1.55 | -1 | 1.49 | 1.42 | 2.43 | 0.93 | 2.77 | 0.53 | 1.17 | 2.78 | 0.3 | 0.24 | -0.16 | -0.51 | 0.84 | 1.085 | -0.76 | 0.193043 | 0.1718 | 0.206328 | 0.999971 | 1 | 0.817242 |
| Tspan33 | Q8R3S2-2 | 0.02 | 1.11 | 0.55 | -1.82 | 0.23 | 1.96 | -0.5 | 1.18 | 0.61 | 0.16 | -0.56 | 0.44 | 1.13 | 1.26 | 0.56 | 1.16 | 0.7525 | 0.1975 | 1.0625 | 0.329996 | 0.753542 | 0.103434 | 0.999971 | 1 | 0.78268 |
| Tspan4 | A0A668KLD | 1.89 | 2.39 | 1.62 | 2.47 | 2.38 | 2.32 | 1.62 | 2.03 | 2.66 | 2.31 | 0.94 | 1.73 | 2.78 | 2.67 | 2.02 | 2.07 | -0.005 | -0.1825 | 0.2925 | 0.985836 | 0.656701 | 0.334818 | 0.999971 | 1 | 0.858827 |
| Tspan5 | D3Z641 | -0.36 | -0.52 | 0.27 | 0.62 | 0.34 | 0.17 | -0.32 | 0.19 | -0.16 | -0.13 | 0.37 | 0.06 | 0.58 | 0.06 | 0.85 | 0.77 | 0.0925 | 0.0325 | 0.5625 | 0.764987 | 0.91397 | 0.109053 | 0.999971 | 1 | 0.78268 |
| Tspan6 | Q99L96 | -0.13 | 0.34 | -0.12 | -0.12 | 0.91 | 0.9 | -0.07 | -0.2 | 1.1 | 0.65 | -0.64 | -0.62 | 1.52 | 1.05 | 0.03 | -0.86 | 0.3925 | 0.13 | 0.4425 | 0.245511 | 0.765304 | 0.391859 | 0.999971 | 1 | 0.871816 |
| Tspan7 | Q62283 | -0.78 | -1.36 | 0.12 | -0.42 | -1.37 | -0.95 | 0.14 | -0.39 | -1.2 | -0.88 | -0.4 | -1.09 | -1.1 | -1.41 | -0.53 | -0.34 | -0.0325 | -0.2825 | -0.235 | 0.939427 | 0.430495 | 0.544954 | 0.999971 | 1 | 0.911398 |
| Tspan8 | Q8R3G9 | -0.02 | 0.65 | -1.73 | 0.76 | 0.54 | 0.32 | 0.42 | -1.49 | 0.51 | 1.08 | 1.1 | 0.49 | -0.02 | 0.41 | -0.45 | 1.05 | 0.0325 | 0.88 | 0.3325 | 0.961764 | 0.138989 | 0.583497 | 0.999971 | 1 | 0.920364 |
| Tspan9 | Q8BJU2 | -0.84 | -0.58 | -0.54 | -0.57 | -0.16 | -0.38 | -0.4 | -0.76 | -0.24 | -0.83 | -1.22 | -0.66 | 0.01 | -0.65 | -1.16 | -1.38 | 0.2075 | -0.105 | -0.1625 | 0.315546 | 0.667693 | 0.613789 | 0.999971 | 1 | 0.927364 |
| Tspoap1 | Q5NCP6 | 0.71 | 0.69 | 1.09 | 1.02 | 0.12 | 0.81 | 0.46 | 1.28 | -0.33 | 0.37 | 1.1 | 1.67 | -0.17 | 0.63 | 1.55 | 1.77 | -0.21 | -0.175 | 0.0675 | 0.465345 | 0.68174 | 0.876158 | 0.999971 | 1 | 0.985129 |
| Tspyl1 | O88852 | 0.84 | 1.39 | 2.87 | 0.2 | 1.5 | 1.85 | 1.55 | 0.5 | 1.21 | 1.1 | -1.92 | -0.22 | 1.69 | 1.8 | 2.3 | 2.75 | 0.025 | -1.2825 | 0.81 | 0.965912 | 0.151825 | 0.178942 | 0.999971 | 1 | 0.808532 |
| Tspyl4 | Q8VD63 | 0.54 | 1.24 | 1.37 | 1.69 | 0.9 | 0.62 | 1.46 | 0.71 | 0.79 | 0.59 | 1.3 | 1.31 | -0.38 | 0.59 | 1.03 | 0.8 | -0.2875 | -0.2125 | -0.7 | 0.368351 | 0.496832 | 0.093414 | 0.999971 | 1 | 0.776565 |
| Tspyl5 | Q69ZB3 | 2.21 | 0.68 | 1.87 | 2.26 | 0.6 | 1.34 | 1.82 | 1.97 | 0.54 | 2.1 | 1.62 | 1.1 | 0.93 | 1.05 | 1.27 | 1.78 | -0.3225 | -0.415 | -0.4975 | 0.481095 | 0.386414 | 0.231182 | 0.999971 | 1 | 0.830179 |
| Tsr2 | Q8C8T8 | 0.71 | 0.7 | 0.52 | 0.58 | 0.78 | 1.27 | 0.22 | 0.99 | 0.4 | 0.39 | 0.19 | -0.08 | 1.28 | 1.52 | 0.54 | 1 | 0.1875 | -0.4025 | 0.4575 | 0.466494 | 0.055962 | 0.090309 | 0.999971 | 1 | 0.771991 |
| Tsr2_1 | H7BWY8 | 1.38 | 0.74 | 2.42 | 1.8 | 0.81 | 1.97 | 0.59 | 0.53 | 0.46 | 2.08 | 2.48 | 1.15 | 0.68 | 1.76 | 2.69 | 1.55 | -0.61 | -0.0425 | 0.085 | 0.205166 | 0.936543 | 0.866119 | 0.999971 | 1 | 0.983462 |
| Tssc4 | Q9JHE7 | 2.8 | 1.3 | 1.66 | 0.77 | 1.16 | 1.41 | 1.73 | 1.04 | 0.95 | 1.83 | 0.93 | 1.13 | -0.18 | 0.58 | 1.04 | 1.03 | -0.2975 | -0.4225 | -1.015 | 0.495886 | 0.362063 | 0.060704 | 0.999971 | 1 | 0.725348 |
| Tst | P52196 | -0.93 | -0.59 | 0.17 | -0.31 | -0.56 | -0.67 | 0.55 | -0.17 | -0.61 | -0.97 | 0.1 | -0.09 | -0.11 | -0.75 | -0.2 | -0.47 | 0.2025 | 0.0225 | 0.0325 | 0.570427 | 0.946397 | 0.909906 | 0.999971 | 1 | 0.991822 |
| Tstd3 | Q9D0B5 | -0.28 | -0.12 | -0.23 | -0.2 | -0.26 | -0.54 | -0.71 | -0.02 | -0.59 | -0.12 | -0.33 | -0.12 | -0.49 | -0.2 | -0.28 | 0.08 | -0.175 | -0.0825 | -0.015 | 0.409891 | 0.655656 | 0.937273 | 0.999971 | 1 | 0.995618 |
| Ttbk1 | A0A286YDI | -0.08 | 0.29 | 0 | 0.04 | 0.4 | 0.4 | 0.02 | 0.28 | 0.13 | 0.75 | -0.29 | -0.1 | 1 | 0.77 | -0.12 | 0.02 | 0.2125 | 0.06 | 0.355 | 0.277841 | 0.818307 | 0.251525 | 0.999971 | 1 | 0.839012 |
| Ttbk1_1 | Q6PCN3 | 0.95 | 0.18 | -0.21 | -0.53 | 0.02 | 0.14 | -0.23 | -0.7 | 0.2 | 0.61 | -0.75 | -0.56 | 0.12 | -0.1 | -0.62 | -0.61 | -0.29 | -0.2225 | -0.4 | 0.429746 | 0.606389 | 0.285406 | 0.999971 | 1 | 0.845133 |
| Ttbk2 | A2AW15 | 1.27 | 0.9 | 0.12 | 0.08 | 0.8 | 0.69 | -0.41 | -0.01 | 0.8 | 1.11 | 0.09 | 0.14 | 0.38 | 0.62 | 0.09 | 0.09 | -0.325 | -0.0575 | -0.2975 | 0.419577 | 0.878119 | 0.369119 | 0.999971 | 1 | 0.869526 |
| Ttc1 | Q91Z38 | 0.6 | -0.04 | 0.15 | 0.46 | 0.07 | 0.09 | 0.1 | 0.29 | -0.15 | 0.64 | 0.36 | 0.49 | -0.44 | 0.11 | 0.14 | 0.42 | -0.155 | 0.0425 | -0.235 | 0.461168 | 0.865203 | 0.372243 | 0.999971 | 1 | 0.869526 |
| Ttc13 | A0A338P6I | -1.57 | -0.42 | 1.29 | -0.6 | -0.5 | -0.37 | 0.47 | 0.71 | -0.25 | -1.03 | 0.85 | 2.05 | -0.25 | -0.44 | 0.38 | 0.37 | 0.4025 | 0.73 | 0.34 | 0.512958 | 0.379354 | 0.560562 | 0.999971 | 1 | 0.91591 |
| Ttc14 | A0A0G2JES | 0.02 | 0.02 | -0.11 | 0.42 | 0.67 | 0.18 | -0.33 | 0.19 | -0.52 | -0.35 | 0.26 | 0.46 | 1.97 | 0.38 | 0.18 | 1.35 | 0.09 | -0.125 | 0.8825 | 0.728417 | 0.653949 | 0.058546 | 0.999971 | 1 | 0.725348 |
| Ttc19 | Q8CC21 | 0.01 | 0.11 | 0.14 | -0.03 | 0.52 | 0.3 | 0.9 | 0.35 | 0.54 | 0.14 | 0.54 | -0.07 | 1.4 | 0.06 | -0.09 | -0.26 | 0.46 | 0.23 | 0.22 | 0.044014 | 0.28237 | 0.557494 | 0.999971 | 1 | 0.915051 |
| Ttc21b | Q0HA38 | 1.34 | 1.91 | 0.34 | -0.23 | 1.97 | 1.67 | 0.17 | 0.23 | 2.13 | 2.11 | 0.35 | 1.04 | 3.16 | 1.75 | 0.16 | 0.07 | 0.17 | 0.5675 | 0.445 | 0.782299 | 0.354313 | 0.577204 | 0.999971 | 1 | 0.918236 |
| Ttc26 | Q8BS45 | -0.84 | -0.28 | 0.09 | 0.06 | -0.24 | -0.11 | -0.7 | -0.22 | -0.37 | -0.6 | -0.76 | -0.29 | -0.19 | -0.16 | 0.06 | -0.05 | -0.075 | -0.2625 | 0.1575 | 0.782521 | 0.330949 | 0.535945 | 0.999971 | 1 | 0.910077 |
| Ttc27 | Q8CD92-2 | -0.66 | 0.77 | 0.8 | 0.67 | 1.03 | 0.71 | 0.4 | 0.91 | 0.83 | -0.54 | 0.72 | 0.57 | 0.97 | 0.51 | 1 | 1.06 | 0.3675 | -3.14E-16 | 0.49 | 0.331916 | 1 | 0.203052 | 0.999971 | 1 | 0.81617 |
| Ttc3 | G5E8T2 | 0.49 | 0.88 | 0.94 | 1.01 | 0.73 | 1.02 | 1.17 | 1.28 | 0.54 | 0.75 | 0.91 | 1 | 0.69 | 1.04 | 1.59 | 1.08 | 0.22 | -0.03 | 0.27 | 0.319418 | 0.883288 | 0.293742 | 0.999971 | 1 | 0.847402 |
| Ttc30b | Q9CY00 | 2.27 | 0.1 | 0.32 | 1.08 | 0.02 | 0.54 | 0.5 | 0.8 | 0 | 1.63 | 0.14 | 0.1 | -0.59 | 0.21 | 0.63 | 0.57 | -0.4775 | -0.475 | -0.7375 | 0.334315 | 0.418774 | 0.182128 | 0.999971 | 1 | 0.808532 |
| Ttc33 | D3Z767 | 1.79 | 2.9 | 2.32 | 2.87 | 2.41 | 3.11 | 2.3 | 2.47 | 2.05 | 1.52 | 3.16 | 2.94 | 2.16 | 2.42 | 2.64 | 2.18 | 0.1025 | -0.0525 | -0.12 | 0.750592 | 0.904909 | 0.687119 | 0.999971 | 1 | 0.945467 |
| Ttc37 | F8VPK0 | 0.12 | 0.3 | 0.17 | 0.16 | 0.21 | 0.35 | 0.01 | 0.52 | 0.16 | 0.15 | 0.34 | 0.32 | 0.17 | 0.43 | 0.52 | 0.59 | 0.085 | 0.055 | 0.24 | 0.650006 | 0.735996 | 0.205552 | 0.999971 | 1 | 0.816688 |
| Ttc38 | A3KMP2 | 0.67 | 0.73 | 0.29 | 0.07 | 0.68 | 0.95 | 0.33 | 1.05 | 0.77 | 0.99 | 0.01 | 0.2 | 0.63 | 0.56 | 0.25 | 0.47 | 0.3125 | 0.0525 | 0.0375 | 0.235476 | 0.856547 | 0.865609 | 0.999971 | 1 | 0.983318 |
| Ttc39b | Q8BYY4 | 0.37 | 1.35 | 1.12 | 1.11 | 0.27 | 0.96 | 0.26 | 0.43 | 0.88 | 1.1 | 1.1 | 1.1 | 0.34 | 1.29 | 0.55 | 0.5 | -0.5075 | 0.0575 | -0.3175 | 0.100718 | 0.816157 | 0.316478 | 0.999971 | 1 | 0.85361 |
| Ttc39c | A0A3Q4EG | -0.14 | -0.06 | -0.11 | -0.3 | 0.45 | 0.19 | 0.11 | 0.14 | 0.16 | -0.22 | -0.45 | -0.45 | 0.36 | 0.22 | 0.05 | -0.21 | 0.375 | -0.0875 | 0.2575 | 0.058914 | 0.669183 | 0.211431 | 0.999971 | 1 | 0.823077 |
| Ttc5 | Q99LG4 | -0.38 | 0.51 | 1.15 | 0.67 | 1.11 | 0.63 | 2.08 | 0.94 | 0.79 | 0 | 1.79 | 1.38 | 1.06 | 0.72 | 0.78 | 0.68 | 0.7025 | 0.5025 | 0.3225 | 0.124261 | 0.303072 | 0.343115 | 0.999971 | 1 | 0.861474 |
| Ttc7b | A0A1Y7VL4 | -1.11 | -1.03 | -1.08 | -1.1 | -0.98 | -0.94 | -1.17 | -0.96 | -1.06 | -1.04 | -1.13 | -0.94 | -0.34 | -0.75 | -0.98 | -0.78 | 0.0675 | 0.0375 | 0.3675 | 0.682341 | 0.811648 | 0.089712 | 0.999971 | 1 | 0.771991 |
| Ttc8 | Q8VD72 | -2.01 | -0.27 | 0.16 | 0.4 | 0.43 | 0.35 | 0.36 | -0.07 | 0.13 | -1.73 | 0.06 | 0.71 | -0.56 | -0.31 | 0.88 | 0.84 | 0.6975 | 0.2225 | 0.6425 | 0.198182 | 0.747922 | 0.304463 | 0.999971 | 1 | 0.847402 |
| Ttc9 | Q3V038 | 0.44 | -0.43 | -1.3 | -1.1 | -0.66 | -0.36 | -0.16 | -1.14 | -0.47 | -0.21 | -1.1 | -0.94 | -0.49 | -0.81 | -1.38 | -1.41 | 0.0175 | -0.0825 | -0.425 | 0.966976 | 0.844943 | 0.335497 | 0.999971 | 1 | 0.858827 |
| Ttc9b | Q9D6E4 | 0.78 | -0.25 | 0.45 | 0.46 | -0.56 | 0.05 | 0.27 | 0.53 | -0.45 | 0.97 | 0.61 | 0.43 | -0.36 | -0.19 | 0.93 | 1.17 | -0.2875 | 0.03 | 0.0275 | 0.380698 | 0.934063 | 0.947955 | 0.999971 | 1 | 0.997651 |
| Ttc9c | Q810A3 | -0.63 | -1.05 | -0.22 | -1.19 | -0.54 | -0.63 | -0.38 | -1.49 | -0.87 | -1.18 | -1.4 | 0.08 | -0.57 | -0.54 | -1.07 | -0.9 | 0.0125 | -0.07 | 0.0025 | 0.969815 | 0.853931 | 0.992695 | 0.999971 | 1 | 0.99925 |
| Tti2 | A0A1B0GS. | 0.52 | 0.13 | -0.36 | -0.2 | -0.34 | 0.21 | -1.53 | -0.9 | 0 | 0.65 | -1.26 | -0.24 | -0.1 | 0.49 | -0.09 | -0.67 | -0.6625 | -0.235 | -0.115 | 0.12638 | 0.579556 | 0.713343 | 0.999971 | 1 | 0.951793 |
| Ttl | P38585 | -0.07 | 0.67 | -0.77 | -0.49 | 0.65 | 0.4 | -0.22 | -0.59 | 0.31 | -0.08 | 0.13 | -0.15 | 0.2 | 0.65 | -0.27 | -0.01 | 0.225 | 0.2175 | 0.3075 | 0.580167 | 0.514598 | 0.404167 | 0.999971 | 1 | 0.87384 |
| Ttll1 | Q91V51 | -0.7 | -0.15 | 0.06 | -0.14 | -0.47 | -0.19 | -0.39 | 0.2 | -0.36 | -0.56 | -0.19 | -0.18 | -0.19 | -0.28 | 0.22 | 0.1 | 0.02 | -0.09 | 0.195 | 0.936059 | 0.68931 | 0.419496 | 0.999971 | 1 | 0.87788 |
| Ttll12 | Q3UDE2 | 0.08 | 0.12 | -0.67 | -0.63 | 0.15 | 0.35 | -0.31 | -0.53 | 0.44 | 0.31 | -0.69 | -0.46 | 0.3 | 0.77 | -0.28 | -0.24 | 0.19 | 0.175 | 0.4125 | 0.536437 | 0.617749 | 0.232448 | 0.999971 | 1 | 0.831121 |
| Ttll7 | E9Q6C6 | 0.18 | 0.63 | 0.94 | 1.02 | 0.82 | 0.78 | 0.87 | 0.9 | 1.07 | 0.72 | 1.07 | 0.99 | 0.97 | 0.77 | 0.87 | 0.56 | 0.15 | 0.27 | 0.1 | 0.519753 | 0.275769 | 0.681047 | 0.999971 | 1 | 0.943097 |
| Ttn | A0A5K1VV | 0.26 | 0.34 | -0.48 | -0.24 | 0.23 | 0.66 | -0.59 | -0.25 | 0.2 | 0.56 | -0.5 | -0.73 | -0.18 | 0.16 | -0.61 | -0.11 | 0.0425 | -0.0875 | -0.155 | 0.899047 | 0.804238 | 0.573517 | 0.999971 | 1 | 0.91816 |
| Ttpa | Q8BWP5 | 0.64 | -0.04 | -2.46 | -2.6 | 0.23 | -0.24 | -2.79 | -1.12 | -0.77 | -0.16 | -2.99 | -3.24 | -0.92 | -0.43 | -1.82 | -1.36 | 0.135 | -0.675 | -0.0175 | 0.886885 | 0.513954 | 0.982379 | 0.999971 | 1 | 0.99925 |
| Ttpal | Q9D3D0 | -0.68 | -0.48 | -0.64 | -0.44 | -0.04 | -0.18 | -0.77 | -0.64 | -0.69 | -0.43 | 0.68 | -0.58 | -0.41 | -0.43 | -0.73 | -0.25 | 0.1525 | 0.305 | 0.105 | 0.505467 | 0.357352 | 0.579075 | 0.999971 | 1 | 0.919302 |
| Ttr | P07309 | 0.46 | 3.53 | 1.67 | 2.9 | 3.97 | 1.68 | 2.17 | 1.39 | 2.59 | -0.07 | 0.48 | 1.55 | 2.52 | 1.4 | 1.91 | 1.57 | 0.1625 | -1.0025 | -0.29 | 0.839506 | 0.238927 | 0.660876 | 0.999971 | 1 | 0.93601 |
| Ttyh1 | Q9D3A9 | -0.72 | -1.15 | -2.6 | -1.77 | -0.84 | -0.93 | -2.64 | -2.55 | -0.83 | -0.87 | -2.4 | -2.21 | -0.41 | -0.81 | -2.49 | -1.7 | -0.18 | -0.0175 | 0.2075 | 0.759099 | 0.974261 | 0.7158 | 0.999971 | 1 | 0.952576 |
| Ttyh2 | Q3TH73 | 0.02 | -0.53 | -0.23 | -1.1 | -0.97 | -0.77 | -0.56 | -0.86 | -0.27 | 0.2 | -0.37 | -0.4 | -0.64 | -0.68 | -0.48 | -0.58 | -0.33 | 0.25 | -0.135 | 0.248134 | 0.3983 | 0.615148 | 0.999971 | 1 | 0.927914 |
| Ttyh3 | Q6P5F7 | -0.21 | -2.35 | -1.11 | -1.08 | -2.61 | -1.87 | -0.75 | -1.34 | -1.76 | -0.52 | -1.06 | -1.25 | -2.85 | -1.87 | -1.31 | -0.6 | -0.455 | 0.04 | -0.47 | 0.412502 | 0.933095 | 0.436087 | 0.999971 | 1 | 0.882419 |
| Tub | P50586 | 0.31 | -0.29 | -0.69 | -0.21 | -0.07 | -0.26 | -0.11 | -0.46 | -0.48 | -0.45 | -0.37 | -0.61 | -0.29 | -0.52 | -0.5 | -0.52 | -0.005 | -0.2575 | -0.2375 | 0.984098 | 0.301845 | 0.344718 | 0.999971 | 1 | 0.862646 |
| Tuba1a | P68369 | -0.27 | 0.14 | -0.09 | -0.27 | -0.09 | -0.03 | 0.18 | -0.24 | 0.36 | -0.05 | -0.05 | 0.07 | 0.14 | 0.15 | 0 | -0.06 | 0.0775 | 0.205 | 0.18 | 0.690795 | 0.308421 | 0.343617 | 0.999971 | 1 | 0.86226 |
| Tuba1b | P05213 | -0.03 | 1.8 | 0.47 | 0.15 | 1.19 | 1.79 | 0.17 | 0.29 | 1.3 | 0.57 | 0.25 | -0.01 | 1.32 | 2.24 | 0.21 | 0.22 | 0.2625 | -0.07 | 0.4 | 0.617162 | 0.881701 | 0.501249 | 0.999971 | 1 | 0.904273 |
| Tuba3b | P05214 | 2.65 | 2.13 | 3.39 | 1.86 | 1.67 | 2.7 | 2.11 | 1.6 | 1.05 | 2.3 | 3.14 | 1.8 | 1.01 | 1.83 | 2.62 | 2.94 | -0.4875 | -0.435 | -0.4075 | 0.245474 | 0.407534 | 0.431046 | 0.999971 | 1 | 0.881611 |
| Tuba4a | A0A0A0MC | 1.19 | 1.76 | 1.76 | 0.81 | 1.27 | 1.37 | 1.14 | 1.96 | 0.75 | 1.05 | 2.26 | 1.87 | 1.72 | 1.38 | 2.18 | 1.85 | 0.055 | 0.1025 | 0.4025 | 0.855737 | 0.799575 | 0.197248 | 0.999971 | 1 | 0.812891 |
| Tuba8 | Q9JJZ2 | 0.48 | 2.42 | 1.67 | 1.47 | 1.47 | 1.92 | 1.25 | 0.85 | 1.17 | 0.59 | 1.24 | 0.93 | 1.17 | 1.68 | 1.09 | 1.3 | -0.1375 | -0.5275 | -0.2 | 0.750827 | 0.217315 | 0.621569 | 0.999971 | 1 | 0.929641 |
| Tubal3 | Q3UX10 | -4.45 | 3.85 | 3.44 | -1.59 | 1.83 | 3.92 | 2.83 | 0.55 | 2.56 | -4.29 | 3.36 | 1.14 | 2.37 | 1.31 | 1.74 | -0.2 | 1.97 | 0.38 | 0.9925 | 0.312505 | 0.872035 | 0.594149 | 0.999971 | 1 | 0.923443 |
| Tubb1 | A2AQ07 | 0.43 | -1.17 | -2.37 | -1.84 | -1.78 | -0.7 | -1.93 | -2.96 | -2.22 | -0.09 | -3.34 | -2.91 | -3.57 | 0.43 | -2.86 | -2.18 | -0.605 | -0.9025 | -0.8075 | 0.391662 | 0.305404 | 0.406052 | 0.999971 | 1 | 0.874911 |
| Tubb2a | Q7TMM9 | -0.86 | -0.73 | -1.27 | -1.07 | -0.49 | -0.99 | -0.72 | -1.36 | -0.46 | -1.02 | -1.02 | -0.76 | -0.56 | -0.89 | -1.05 | -0.88 | 0.0925 | 0.1675 | 0.1375 | 0.711878 | 0.452898 | 0.516372 | 0.999971 | 1 | 0.905815 |
| Tubb2b | Q9CWF2 | -1.05 | -1.03 | -1.4 | -1.83 | -0.61 | -1.19 | -1.76 | -2.15 | -0.71 | -1.71 | -0.95 | 0.6 | -0.64 | -1.36 | -0.12 | 0.36 | -0.1 | 0.635 | 0.8875 | 0.788909 | 0.211018 | 0.049083 | 0.999971 | 1 | 0.714598 |
| Tubb3 | Q9ERD7 | -0.27 | 0.18 | 0.12 | -0.1 | -0.03 | 0 | 0.57 | 0.03 | 0.12 | -0.13 | 0.03 | -0.27 | 0.11 | 0.22 | 0.32 | -0.02 | 0.16 | -0.045 | 0.175 | 0.474429 | 0.816236 | 0.375586 | 0.999971 | 1 | 0.870465 |
| Tubb4a | Q9D6F9 | -0.66 | 0.63 | 0.39 | -0.11 | 0.22 | 0.64 | 0.13 | -0.36 | 0 | -0.53 | 0.36 | 0.37 | -0.08 | 0.35 | 0.12 | 0.03 | 0.095 | -0.0125 | 0.0425 | 0.784279 | 0.971467 | 0.889893 | 0.999971 | 1 | 0.98799 |
| Tubb4b | P68372 | -0.59 | 0.15 | 0.14 | 0.14 | 0.03 | -0.03 | 0.11 | 0.01 | -0.13 | -0.44 | -0.02 | -0.22 | -0.29 | 0.02 | 0.24 | 0.2 | 0.07 | -0.1625 | 0.0825 | 0.757259 | 0.497064 | 0.740934 | 0.999971 | 1 | 0.960899 |
| Tubb5 | P99024 | -0.33 | -0.74 | -1.95 | -1.22 | -0.81 | -0.55 | -1.19 | -2.03 | -0.83 | -0.3 | -2.12 | -1.68 | -1.18 | -0.53 | -1.93 | -1.4 | -0.085 | -0.1725 | -0.2 | 0.848916 | 0.731411 | 0.642988 | 0.999971 | 1 | 0.93301 |
| Tubb6 | Q922F4 | -0.55 | 0.34 | 0.12 | 0.82 | -0.12 | 0.52 | 0.88 | 0.73 | -0.05 | -0.02 | 0.11 | 0.03 | -0.61 | 0.1 | -0.42 | 0.99 | 0.32 | -0.165 | -0.1675 | 0.375935 | 0.580155 | 0.699693 | 0.999971 | 1 | 0.948208 |
| Tubg1 | P83887 | 0.02 | 0.43 | 0.39 | 0.24 | 0.48 | 0.52 | 0.44 | 0.39 | 0.72 | 0.22 | 0.51 | 0.38 | 0.65 | 0.24 | 0.36 | 0.53 | 0.1875 | 0.1875 | 0.175 | 0.308429 | 0.353551 | 0.379228 | 0.999971 | 1 | 0.871816 |
| Tubg2 | Q8VCK3 | 0.37 | 0.32 | 0.13 | 0.5 | 0.52 | -0.18 | 0.56 | 0.23 | 0.26 | 0.02 | 0.21 | 0.03 | -0.59 | -0.33 | -0.23 | 0.03 | -0.0475 | -0.2 | -0.61 | 0.834309 | 0.272744 | 0.014722 | 0.999971 | 1 | 0.613095 |
| Tubgcp2 | A0A1B0GR | -0.34 | 0.14 | -0.59 | -0.85 | -0.08 | -0.26 | -0.5 | -0.86 | -0.08 | -0.33 | -0.2 | -0.34 | -0.14 | -0.31 | -0.71 | -0.66 | -0.015 | 0.1725 | -0.045 | 0.95783 | 0.490741 | 0.868314 | 0.999971 | 1 | 0.983462 |
| Tubgcp3 | P58854 | -0.33 | 1.19 | 0.53 | 0.05 | 1.23 | 0.71 | 0.55 | 0.27 | 0.87 | -0.04 | 0.84 | 1.33 | 1.78 | 0.51 | 0.61 | 0.17 | 0.33 | 0.39 | 0.4075 | 0.387042 | 0.359992 | 0.378038 | 0.999971 | 1 | 0.871417 |
| Tubgcp4 | Q9D4F8 | -0.42 | -0.33 | -0.43 | -0.55 | -0.48 | -0.37 | -0.56 | -0.53 | -0.84 | -1.49 | -0.68 | 0.18 | -0.47 | 0.05 | -0.21 | 0.06 | -0.0525 | -0.275 | 0.29 | 0.751908 | 0.43099 | 0.16493 | 0.999971 | 1 | 0.800215 |
| Tubgcp5 | Q8BKN5-3 | 0.91 | 0.62 | 1.19 | 0.23 | 0.66 | 0.15 | 1.46 | 0.76 | 0.22 | 1.05 | 0.74 | 1.1 | 0.79 | -0.05 | 0.71 | 0.65 | 0.02 | 0.04 | -0.2125 | 0.952536 | 0.892731 | 0.476142 | 0.999971 | 1 | 0.894333 |
| Tubgcp6 | E9Q6P4 | -0.21 | 0.67 | 0.08 | 1.33 | 0.78 | 0.52 | 0.37 | 0.92 | 0.3 | -0.04 | 0.82 | 1.12 | 0.33 | 0.2 | 0.62 | 0.94 | 0.18 | 0.0825 | 0.055 | 0.614746 | 0.840305 | 0.881324 | 0.999971 | 1 | 0.986007 |
| Tufm | Q8BFR5 | -0.53 | -0.22 | -1.2 | -1.3 | -0.43 | -0.31 | -1.08 | -1.38 | -0.36 | -0.47 | -1.31 | -1.11 | -0.04 | -0.38 | -1.26 | -1.33 | 0.0125 | -7.85E-17 | 0.06 | 0.97212 | 1 | 0.879874 | 0.999971 | 1 | 0.985441 |
| Tulp3 | O88413 | -1.63 | -1.51 | -0.29 | -0.12 | -1.47 | -1.86 | -0.91 | -0.09 | -1.16 | -2.35 | -0.57 | -0.72 | -1.38 | -2.39 | -0.93 | 0.26 | -0.195 | -0.3125 | -0.2225 | 0.703441 | 0.555393 | 0.719603 | 0.999971 | 1 | 0.954401 |
| Tulp4 | Q9JIL5 | -0.74 | -0.5 | -0.5 | -0.63 | -0.2 | -0.25 | -0.86 | -0.25 | -0.67 | -0.07 | -0.95 | -0.21 | -0.5 | -0.12 | -0.58 | -0.38 | 0.2025 | 0.1175 | 0.1975 | 0.358243 | 0.628635 | 0.307896 | 0.999971 | 1 | 0.847402 |
| Tusc2 | Q9WVF8 | -1.01 | -0.07 | -0.98 | -0.84 | 0.55 | -0.72 | -0.5 | -0.67 | 0.17 | -1.24 | -0.7 | -0.7 | 0.18 | -0.64 | -0.28 | 0.03 | 0.39 | 0.1075 | 0.5475 | 0.298192 | 0.764632 | 0.092279 | 0.999971 | 1 | 0.775144 |
| Tusc3 | Q8BTV1 | -0.75 | 2.13 | 0.83 | 0.71 | 1.16 | 1.33 | 1.29 | 0.82 | -0.34 | -0.52 | 1.12 | 0.92 | 3.44 | 0.78 | 0.97 | 1.09 | 0.42 | -0.435 | 0.84 | 0.453693 | 0.516012 | 0.294991 | 0.999971 | 1 | 0.847402 |
| Tut1 | Q8R3F9 | 3.35 | 2.9 | 2.1 | 2.52 | 2.89 | 3.11 | 1.27 | 2.08 | 2 | 2.93 | 1.97 | 2.29 | 1.64 | 3.05 | 2.69 | 3.04 | -0.38 | -0.42 | -0.1125 | 0.422368 | 0.241584 | 0.782575 | 0.999971 | 1 | 0.965859 |
| Tut4 | B2RX14 | -1.04 | 0.79 | 0.33 | -0.23 | 0.83 | 0.24 | -0.17 | -0.84 | 0.75 | -0.93 | -0.35 | 0.17 | 0.75 | 0.2 | -0.32 | -0.04 | 0.0525 | -0.0525 | 0.185 | 0.914653 | 0.915801 | 0.668904 | 0.999971 | 1 | 0.938711 |
| Tut7 | E9PUA2 | 0.69 | 1.16 | 0.49 | 0.39 | 0.56 | 0.79 | 0.07 | 0.44 | 0.57 | 0.96 | 0.74 | 0.64 | 0.65 | 1.2 | 0.56 | 0.73 | -0.2175 | 0.045 | 0.1025 | 0.401641 | 0.84316 | 0.684323 | 0.999971 | 1 | 0.944176 |
| Twf1 | Q91YR1 | 0.08 | 1.47 | 0.32 | 0.17 | 1.07 | 1.65 | 0.25 | 0.36 | 1.25 | 0.7 | 0.13 | 0.25 | 1.44 | 1.26 | 0.13 | 0.31 | 0.3225 | 0.0725 | 0.275 | 0.465105 | 0.854536 | 0.533808 | 0.999971 | 1 | 0.910077 |
| Twf2 | Q9Z0P5 | -0.7 | -0.35 | -0.74 | -1.09 | -0.08 | -0.61 | -0.35 | -0.7 | -0.49 | -0.91 | -0.59 | -0.72 | -0.3 | -0.74 | -0.87 | -0.77 | 0.285 | 0.0425 | 0.05 | 0.252581 | 0.845485 | 0.830897 | 0.999971 | 1 | 0.978535 |
| Txlna | Q6PAM1 | 3.66 | 1.26 | 1.93 | 2.01 | 0.77 | 1.35 | 2.46 | 1.08 | 1.08 | 2.7 | 1.68 | 1.72 | -0.19 | 0.74 | 0.83 | 0.95 | -0.8 | -0.42 | -1.6325 | 0.188413 | 0.463107 | 0.012541 | 0.999971 | 1 | 0.613095 |
| Txlng | Q8BHN1-3 | 1.03 | 0.61 | 0.21 | -0.18 | 0.34 | 0.13 | 0.06 | 0.02 | 0.4 | 0.99 | 0.2 | 0.27 | -0.68 | 0.9 | 0.01 | 0.35 | -0.28 | 0.0475 | -0.2725 | 0.337551 | 0.881399 | 0.504183 | 0.999971 | 1 | 0.905265 |

| | | | | | | | | | | | | | | | | | | | | | | | | | | |
|---|---|---|---|---|---|---|---|---|---|---|---|---|---|---|---|---|---|---|---|---|---|---|---|---|---|---|
| Txn | P10639 | -0.58 | 0.44 | 0.68 | 0.59 | 0.63 | 0.58 | 0.51 | 0.92 | 0.98 | -0.14 | 0.81 | 0.41 | 0.36 | 0.79 | 1.06 | 0.97 | 0.3775 | 0.2325 | 0.5125 | 0.243855 | 0.537094 | 0.147318 | 0.999971 | 1 | 0.793248 |
| Txn2 | G3UZY2 | -1.98 | 1.29 | -1.97 | -2.16 | 0.56 | 0.81 | -1.07 | -2.45 | 0.91 | -1.43 | -1.35 | -2.35 | 1.8 | 0.73 | -2.25 | -2.74 | 0.6675 | 0.15 | 0.59 | 0.512621 | 0.877666 | 0.635259 | 0.999971 | 1 | 0.93301 |
| Txndc12 | Q9CQU0 | -0.89 | 0.08 | -0.29 | -0.09 | 0.25 | -0.07 | 0.64 | -0.6 | 0.1 | -0.62 | -0.48 | -0.15 | 0.76 | -0.3 | -0.32 | -0.51 | 0.3525 | 0.01 | 0.205 | 0.307994 | 0.971544 | 0.563721 | 0.999971 | 1 | 0.917087 |
| Txndc15 | Q6P6J9 | -0.16 | -0.55 | -1.49 | -0.85 | -0.36 | -0.89 | -1.29 | -0.78 | -0.52 | -0.32 | -1.46 | -1.28 | -0.03 | -0.48 | -0.61 | -0.58 | -0.0675 | -0.1325 | 0.3375 | 0.841225 | 0.729992 | 0.301047 | 0.999971 | 1 | 0.847402 |
| Txndc16 | Q7TN22 | -0.72 | 0.55 | -0.63 | -0.59 | -0.28 | 0.4 | -0.85 | -0.44 | -0.44 | -0.14 | -0.8 | -0.56 | -0.04 | -0.12 | -0.88 | -0.39 | 0.055 | -0.1375 | -0.01 | 0.885725 | 0.677859 | 0.977181 | 0.999971 | 1 | 0.99925 |
| Txndc17 | Q9CQM5 | 0.46 | 1.55 | 1.26 | 1.51 | 1.5 | 2.09 | 1.54 | 1.98 | 1.58 | 1.25 | 1.84 | 1.59 | 0.41 | 1.61 | 1.73 | 1.37 | 0.5825 | 0.37 | 0.085 | 0.080854 | 0.225263 | 0.822904 | 0.999971 | 1 | 0.975901 |
| Txndc5 | Q91W90 | -0.04 | 0.19 | 0.18 | 0.51 | 0.31 | 0.17 | 0.93 | 0.73 | 0.32 | 0.02 | 0.29 | 0.49 | 0.25 | 0.23 | 0.62 | 0.28 | 0.325 | 0.07 | 0.135 | 0.202327 | 0.729169 | 0.511602 | 0.999971 | 1 | 0.905815 |
| Txndc9 | Q9CQ79 | -0.24 | 0.79 | 0.13 | 0.28 | 0.7 | 1.04 | 0.04 | 0.51 | 0.65 | 0.54 | 0.63 | 0.57 | 0.95 | 1.29 | 0.74 | 0.72 | 0.3325 | 0.3575 | 0.685 | 0.292535 | 0.167423 | 0.030355 | 0.999971 | 1 | 0.678474 |
| Txnip | Q8BG60-2 | 0.29 | 0.58 | -0.31 | -0.89 | 0.06 | 0.39 | -0.58 | 1.23 | 0.63 | 0.23 | -0.05 | 0.55 | 0.44 | 1.21 | -0.66 | -0.46 | 0.3575 | 0.4225 | 0.215 | 0.45078 | 0.253693 | 0.670269 | 0.999971 | 1 | 0.938809 |
| Txnl1 | A0A498WC | -0.42 | 0.52 | -0.07 | -0.68 | 0.63 | 0.55 | 0.41 | 0.03 | 0.7 | -0.29 | 0.04 | 0.46 | 1.42 | 0.72 | 0.08 | -0.18 | 0.5675 | 0.39 | 0.6725 | 0.085634 | 0.266443 | 0.136019 | 0.999971 | 1 | 0.784948 |
| Txnl4a | P83877 | 0.28 | 1.43 | 0.98 | 0.4 | 2.01 | 0.51 | 0.63 | 0.85 | 2.2 | 0.31 | 0.93 | 1.11 | 3.07 | 1.21 | 0.73 | 0.82 | 0.2275 | 0.365 | 0.685 | 0.58569 | 0.425016 | 0.241465 | 0.999971 | 1 | 0.835635 |
| Txnl4b | Q8BUH1 | 0.03 | -0.07 | 0.63 | 0.78 | -0.2 | 1.24 | 0.54 | 1.2 | -0.74 | 0.07 | -0.66 | 0.29 | -0.09 | 0.55 | -0.14 | 0.08 | 0.3525 | -0.6025 | -0.2425 | 0.371515 | 0.098162 | 0.397633 | 0.999971 | 1 | 0.871816 |
| Txnrd1 | Q9JMH6 | 0.5 | 0.59 | 0.66 | 0.66 | 0.37 | 0.98 | 0.62 | 0.9 | 0.6 | 0.7 | 0.95 | 1.12 | 0.8 | 1.06 | 1.3 | 0.83 | 0.115 | 0.24 | 0.395 | 0.571551 | 0.223085 | 0.063902 | 0.999971 | 1 | 0.725348 |
| Txnrd2 | J3QMN4 | -0.1 | -0.35 | -0.84 | -0.76 | -0.53 | -0.14 | -0.59 | -0.49 | -0.59 | -0.07 | -0.93 | -1.07 | -0.38 | 0.06 | -0.58 | -0.35 | 0.075 | -0.1525 | 0.2 | 0.751499 | 0.605012 | 0.431123 | 0.999971 | 1 | 0.881611 |
| Txnrd3 | Q99MD6 | 0.92 | -0.58 | 0.1 | -0.11 | -0.11 | -0.41 | 0.42 | -0.09 | -0.76 | 0.99 | -0.41 | 0.45 | -0.41 | -0.11 | 0.18 | 0.47 | -0.13 | -0.015 | -0.05 | 0.711956 | 0.974793 | 0.88904 | 0.999971 | 1 | 0.987653 |
| Tyk2 | Q9R117 | 1.1 | -1.83 | -0.25 | -0.43 | -2.87 | -1.02 | -0.04 | -0.81 | -1 | 0.93 | 0.89 | 0.24 | -2.19 | -1.24 | -0.28 | -0.24 | -0.8325 | 0.6175 | -0.635 | 0.292077 | 0.377632 | 0.366945 | 0.999971 | 1 | 0.869526 |
| Tyro3 | P55144-3 | -0.26 | -0.21 | -0.57 | -0.26 | 0.66 | -0.35 | -0.7 | -0.41 | -0.48 | -0.63 | -0.94 | -1.01 | 0.81 | -0.62 | -0.55 | -0.54 | 0.125 | -0.44 | 0.1 | 0.690211 | 0.054325 | 0.775399 | 0.999971 | 1 | 0.96393 |
| Tyw1 | Q8BJM7 | -1.15 | -0.62 | 0.12 | 0.03 | -0.05 | -0.19 | -0.27 | -0.16 | -0.63 | -0.49 | 0.33 | 0.2 | -0.94 | -0.05 | 0.89 | 0.43 | 0.2375 | 0.2575 | 0.4875 | 0.447798 | 0.495465 | 0.306328 | 0.999971 | 1 | 0.847402 |
| Tyw3 | Q8BSA9 | 4.52 | 0.08 | 2.14 | 1.62 | 0.36 | 1.36 | 2.01 | 0.93 | -0.39 | 2.25 | 0.75 | -0.19 | -0.51 | 0.09 | 0.95 | -0.27 | -0.925 | -1.485 | -2.025 | 0.309401 | 0.159318 | 0.043119 | 0.999971 | 1 | 0.701725 |
| Tyw5 | A2RSX7 | 0.24 | 1.08 | 0.24 | -0.14 | 1.11 | 0.92 | 0.57 | 0.18 | 0.92 | 0.61 | 1.08 | 1.12 | 1.36 | 0.79 | 0.24 | -0.16 | 0.34 | 0.5775 | 0.2025 | 0.31629 | 0.075254 | 0.616559 | 0.999971 | 1 | 0.928229 |
| U2af1 | Q9D883 | 1.45 | 2.14 | 1.84 | 2.06 | 2.16 | 2.46 | 1.67 | 2.02 | 2.73 | 1.82 | 1.96 | 1.54 | 1.12 | 1.87 | 2.15 | 1.63 | 0.205 | 0.14 | -0.18 | 0.425178 | 0.647193 | 0.529253 | 0.999971 | 1 | 0.910069 |
| U2af1l4 | Q8BGJ9 | 2.45 | 1.73 | 1.4 | 1.79 | 1.61 | 1.25 | 1.88 | 1.53 | 2.31 | 2.55 | 2.69 | 1.53 | 0.47 | 1.89 | 1.68 | 2.03 | -0.275 | 0.4275 | -0.325 | 0.328562 | 0.226118 | 0.426803 | 0.999971 | 1 | 0.880793 |
| U2af2 | Q80XR5 | 1.04 | 1.35 | 0.93 | 1.23 | 1.28 | 1.22 | 0.97 | 1.15 | 1.33 | 1.21 | 0.65 | 0.56 | 0.71 | 1.63 | 0.95 | 1.08 | 0.0175 | -0.2 | -0.045 | 0.925244 | 0.422114 | 0.855329 | 0.999971 | 1 | 0.982398 |
| U2surp | Q6NV83-2 | 1.06 | 1.16 | 1.04 | 1.11 | 1.24 | 0.91 | 0.82 | 1.22 | 0.88 | 0.72 | 0.83 | 0.43 | 1.24 | 1.51 | 0.98 | 1 | -0.045 | -0.3775 | 0.09 | 0.807147 | 0.059494 | 0.642871 | 0.999971 | 1 | 0.93301 |
| Uaca | A0A1L1STY | -0.44 | -0.04 | 1.38 | 1.29 | -1.65 | 0.16 | 0.42 | 1.64 | 0.45 | 0.44 | 1.13 | 2 | -2.19 | 1.44 | 1.87 | 1.57 | -0.405 | 0.4575 | 0.125 | 0.587479 | 0.412506 | 0.89551 | 0.999971 | 1 | 0.988571 |
| Uap1 | A0A0R4J1F | 0.59 | 0.42 | 0.38 | 0.49 | 0.21 | 0.62 | 0.56 | 0.52 | 0.23 | 0.67 | 0.64 | -0.03 | -0.05 | 0.62 | 0.33 | 0.9 | 0.0075 | -0.0925 | -0.02 | 0.966847 | 0.671673 | 0.933929 | 0.999971 | 1 | 0.995387 |
| Uap1l1 | Q3TW96 | 0.32 | 0.94 | 0.77 | 0.71 | 0.87 | 0.83 | 1.25 | 0.37 | 0.49 | 0.39 | 1.49 | 0.55 | 0.7 | 0.91 | 0.84 | 1 | 0.145 | 0.045 | 0.1775 | 0.566364 | 0.879124 | 0.392104 | 0.999971 | 1 | 0.871816 |
| Uba1 | Q02053 | 0.47 | 0.72 | 0.76 | 0.74 | 0.55 | 0.35 | 0.55 | 0.59 | 0.45 | 0.39 | 0.75 | 0.89 | 0.41 | 0.54 | 0.63 | 0.81 | -0.1625 | -0.0525 | -0.075 | 0.36327 | 0.788053 | 0.684601 | 0.999971 | 1 | 0.944227 |
| Uba2 | Q9Z1F9 | 2.73 | 2.89 | 3.07 | 3.1 | 2.36 | 2.57 | 3.45 | 2.66 | 2.67 | 2.83 | 3.23 | 3.47 | 1.69 | 2.53 | 2.78 | 2.53 | -0.1875 | 0.1025 | -0.565 | 0.496533 | 0.664116 | 0.063773 | 0.999971 | 1 | 0.725348 |
| Uba3 | Q8C878 | 0.61 | 1.56 | 0.9 | 0.89 | 1.28 | 0.83 | 0.74 | 1.02 | 1.25 | 0.93 | 0.76 | 0.48 | 1.27 | 1.09 | 0.74 | 0.98 | -0.0225 | -0.135 | 0.03 | 0.930561 | 0.624807 | 0.906699 | 0.999971 | 1 | 0.991405 |
| Uba5 | Q8VE47 | -0.11 | 0.14 | 0.02 | 0.07 | 0.15 | -0.05 | 0.64 | -0.11 | 0.13 | -0.46 | -0.05 | -0.07 | 0.25 | 0.23 | -0.02 | 0.03 | 0.1275 | -0.1425 | 0.0925 | 0.568146 | 0.468853 | 0.599711 | 0.999971 | 1 | 0.92422 |
| Uba6 | Q8C7R4 | -0.5 | -0.11 | 0.11 | 0.13 | -0.11 | -0.14 | 0.11 | 0.3 | -0.16 | -0.37 | 0.09 | 0.21 | -0.09 | 0.2 | 0.26 | 0.36 | 0.1325 | 0.035 | 0.275 | 0.554252 | 0.879209 | 0.232547 | 0.999971 | 1 | 0.831128 |
| Ubac1 | Q8VDI7 | 0.08 | 0.54 | 0.43 | 1.15 | 0.78 | 0.2 | 0.29 | 1.04 | 0.77 | 0.54 | 0.77 | 0.93 | 1.56 | 0.12 | 0.7 | 1.28 | 0.0275 | 0.2025 | 0.365 | 0.928303 | 0.442292 | 0.347014 | 0.999971 | 1 | 0.864164 |
| Ubac2 | Q8R1K1 | -1.17 | 0.02 | -0.32 | -0.77 | -0.02 | 0.13 | -0.83 | -0.58 | -0.17 | -0.91 | 0.01 | 0.19 | 0.02 | -0.07 | -0.48 | -0.77 | 0.235 | 0.34 | 0.235 | 0.497048 | 0.343615 | 0.471175 | 0.999971 | 1 | 0.892857 |
| Ubald1 | Q6P3B2 | 1.79 | 1.62 | 1.19 | 1.97 | 1.94 | 0.97 | 2.21 | 0.76 | 2.32 | 0.06 | 1.14 | 0.19 | -3.86 | -0.35 | -0.94 | -1.89 | -0.1725 | -0.715 | -3.4025 | 0.651315 | 0.185726 | 0.001064 | 0.999971 | 1 | 0.387721 |
| Ubap1 | Q8BH48 | -0.39 | -0.37 | -0.38 | -0.32 | -0.08 | -0.63 | -0.71 | -0.16 | -0.34 | -1.06 | -0.77 | -0.24 | 0.08 | 0 | -0.29 | -0.22 | -0.03 | -0.2375 | 0.2575 | 0.886926 | 0.31273 | 0.168171 | 0.999971 | 1 | 0.800215 |
| Ubap2 | A2AMY5 | 2.09 | 2.12 | 1.7 | 1.75 | 2.48 | 2.33 | 1.88 | 1.76 | 2.64 | 1.82 | 1.97 | 2.08 | 2.59 | 2.23 | 1.66 | 1.54 | 0.1975 | 0.2125 | 0.09 | 0.417605 | 0.387566 | 0.7523 | 0.999971 | 1 | 0.961116 |
| Ubap2l | Q80X50 | 1.16 | 1.6 | 1.9 | 1.58 | 1.46 | 1.68 | 2.29 | 1.8 | 1.68 | 1.1 | 2.05 | 1.95 | 1.42 | 1.44 | 1.81 | 2.27 | 0.2475 | 0.135 | 0.175 | 0.348654 | 0.628326 | 0.523207 | 0.999971 | 1 | 0.907797 |
| Ubash3b | Q8BGG7 | 0.57 | -0.22 | -0.36 | -0.32 | 0.08 | -0.38 | -0.3 | -0.33 | -0.48 | -0.02 | 0.31 | 0.33 | 0.01 | -0.8 | -0.35 | -0.31 | -0.15 | 0.1175 | -0.28 | 0.574686 | 0.694652 | 0.345434 | 0.999971 | 1 | 0.863502 |
| Ube2a | A0A0J9YUN | 0.77 | 1.19 | 0.6 | 0.82 | 1.07 | 1.38 | 0.36 | 0.76 | 0.93 | 0.92 | 0.65 | 1.21 | 0.62 | 1.48 | 0.91 | 1.4 | 0.0475 | 0.0825 | 0.2575 | 0.860432 | 0.700059 | 0.340171 | 0.999971 | 1 | 0.860511 |
| Ube2b | P63147 | 0.4 | 0.97 | 1.1 | 0.98 | 0.23 | -0.14 | 0.37 | 1.11 | -0.21 | 0.33 | 1 | 1.41 | -0.37 | 0.91 | 0.99 | 1.42 | -0.47 | -0.23 | -0.125 | 0.155904 | 0.547974 | 0.754687 | 0.999971 | 1 | 0.961116 |
| Ube2d1 | P61080 | -2.16 | 1.19 | 0.68 | -0.03 | 1.36 | 1.63 | 0.55 | 1.04 | 2.2 | -0.56 | 1.27 | 0.84 | 2.69 | 2.09 | 0.9 | 0.79 | 1.225 | 1.0175 | 1.6975 | 0.106887 | 0.248088 | 0.055151 | 0.999971 | 1 | 0.725348 |
| Ube2d3 | A0A0G2JE3 | 2.66 | 0.63 | 1.49 | 1.43 | 0.81 | 0.54 | 0.81 | 1.78 | 0.42 | 1.53 | 1.63 | -0.53 | 0.63 | 1.66 | 1.41 | 1.72 | -0.5675 | -0.79 | -0.1975 | 0.24362 | 0.214479 | 0.667543 | 0.999971 | 1 | 0.938453 |
| Ube2e2 | D3YXD1 | 2.34 | 2.16 | 2.83 | 3.07 | 1.97 | 2.09 | 2.8 | 2.75 | 2.04 | 2.02 | 2.74 | 1.96 | 1.07 | 2.34 | 2.39 | 2.46 | -0.1975 | -0.41 | -0.535 | 0.526609 | 0.182833 | 0.184416 | 0.999971 | 1 | 0.809025 |
| Ube2e3 | P52483 | 1.48 | 4.27 | 3.94 | 4.34 | 3.23 | 4.38 | 3.65 | 4.08 | 3.96 | 3.46 | 5.09 | 3.88 | 2.36 | 3.85 | 3.87 | 4.71 | 0.3275 | 0.59 | 0.19 | 0.621327 | 0.407162 | 0.801893 | 0.999971 | 1 | 0.971467 |
| Ube2f | Q9CY34 | 1.02 | 2.24 | 1.51 | 0.76 | 2.29 | 1.37 | 1.52 | 0.91 | 2.18 | 0.8 | 1.49 | 2.2 | 2.3 | 0.97 | 1.23 | 0.7 | 0.14 | 0.285 | -0.0825 | 0.734924 | 0.522467 | 0.854512 | 0.999971 | 1 | 0.982125 |
| Ube2g1 | P62254 | -0.17 | 0.53 | 0.99 | 1.03 | 0.38 | 0.24 | 0.59 | 1.45 | 0.22 | 0.46 | 1.22 | 1.17 | 0.64 | 1.1 | 0.97 | 1.48 | 0.07 | 0.1725 | 0.4525 | 0.852442 | 0.639735 | 0.192763 | 0.999971 | 1 | 0.810547 |
| Ube2g2 | P60605 | -1.54 | -0.34 | -1.05 | -1.04 | -0.42 | -0.51 | -1.48 | -0.83 | -0.23 | -0.75 | -1.24 | -1.17 | 0.11 | -0.06 | -0.53 | -0.64 | 0.1825 | 0.145 | 0.7125 | 0.594999 | 0.668092 | 0.045132 | 0.999971 | 1 | 0.710102 |
| Ube2h | P62257 | -0.62 | 0.05 | 0.07 | 0.5 | 0.14 | 0.11 | 0.17 | 0.47 | 0.64 | 0.27 | 0.61 | 0.82 | 0.63 | 0.35 | 0.63 | 0.58 | 0.2225 | 0.585 | 0.5475 | 0.413393 | 0.058195 | 0.063374 | 0.999971 | 1 | 0.725348 |
| Ube2i | P63280 | -0.36 | 0.83 | 1.41 | 1.09 | 0.94 | 0.6 | 1.84 | 1.56 | 1.05 | 0.3 | 1.92 | 1.84 | 0.4 | 1.35 | 1.42 | 1.36 | 0.4925 | 0.535 | 0.39 | 0.290146 | 0.305254 | 0.378666 | 0.999971 | 1 | 0.871816 |
| Ube2j1 | Q9JJZ4 | 1.85 | -0.53 | -0.13 | 0.15 | -0.71 | 0.37 | 1.52 | 0.07 | -0.8 | 0.99 | -0.35 | -0.42 | -0.29 | 0.01 | -0.03 | -0.1 | -0.0225 | -0.48 | -0.4375 | 0.971706 | 0.432658 | 0.384538 | 0.999971 | 1 | 0.871816 |
| Ube2j2 | B1ASK8 | 1.85 | 0.24 | 1.23 | 1.77 | 0.42 | 0.56 | 1.31 | 0.27 | 0.55 | 1.71 | 0.48 | 0.03 | -0.21 | 0.25 | 0.28 | -0.05 | -0.6325 | -0.58 | -1.205 | 0.15328 | 0.249049 | 0.010544 | 0.999971 | 1 | 0.613095 |
| Ube2k | P61087 | 0.95 | 0.63 | 0.83 | 0.43 | 0.9 | 0.96 | 0.44 | 0.73 | 0.91 | 1.01 | 0.81 | 0.24 | 0.7 | 0.95 | 0.84 | 0.56 | 0.0475 | 0.0325 | 0.0525 | 0.823018 | 0.891347 | 0.794188 | 0.999971 | 1 | 0.970141 |
| Ube2l3 | P68037 | 0.25 | 0.54 | 0.78 | 0.52 | 0.5 | 0.76 | 0.85 | 1.62 | 0.85 | 0.61 | 1.05 | 1.04 | 0.5 | 0.98 | 1.31 | 1.06 | 0.41 | 0.365 | 0.44 | 0.167626 | 0.098191 | 0.090116 | 0.999971 | 1 | 0.771991 |
| Ube2m | P61082 | 1.29 | 0.67 | 0.54 | 0.45 | 0.78 | 1.14 | 1.37 | 0.28 | 0.68 | 1.08 | 0.41 | 0.5 | 1.29 | 0.71 | 0.53 | 0.65 | 0.155 | -0.07 | 0.0575 | 0.618718 | 0.789422 | 0.833322 | 0.999971 | 1 | 0.979036 |
| Ube2n | P61089 | 0.96 | 1.02 | 1.12 | 1.2 | 0.91 | 0.99 | 1.18 | 0.72 | 1.15 | 0.79 | 0.77 | 0.94 | 1.04 | 0.93 | 0.46 | 0.55 | -0.125 | -0.1625 | -0.33 | 0.501746 | 0.3717 | 0.135721 | 0.999971 | 1 | 0.784948 |
| Ube2o | Q6ZPJ3 | 0.02 | 0.47 | 0.44 | 0.38 | 0.36 | 0.46 | 0.4 | 0.38 | 0.7 | 0.45 | 0.59 | 0.52 | 0.79 | 0.57 | 0.54 | 0.67 | 0.0725 | 0.2375 | 0.315 | 0.692151 | 0.219369 | 0.122177 | 0.999971 | 1 | 0.78268 |
| Ube2q1 | Q7TSS2 | 0.64 | 0.47 | 0.09 | 0.35 | 0.34 | 1.16 | 0.3 | 0.46 | 0.73 | 0.82 | 0.18 | 0.36 | 0.49 | 0.52 | 0.09 | 0.6 | 0.1775 | 0.135 | 0.0375 | 0.495564 | 0.556324 | 0.859901 | 0.999971 | 1 | 0.982702 |
| Ube2q2 | D3Z647 | -0.82 | 1.45 | 1.33 | 0.82 | 1.51 | 0.5 | 0.88 | 0.67 | 1.32 | -0.8 | 1.3 | 1.36 | 0.58 | 1.4 | 1.23 | 0.91 | 0.195 | 0.1 | 0.335 | 0.710637 | 0.883021 | 0.518842 | 0.999971 | 1 | 0.905901 |
| Ube2ql1 | E9QMD2 | 2.46 | 1.15 | 1.41 | 1.5 | 0.85 | 1.85 | 2.46 | 1.33 | 0.75 | 2.13 | 1.41 | 0.3 | -0.86 | -0.04 | 0.85 | -0.85 | -0.0075 | -0.4825 | -1.855 | 0.985906 | 0.311709 | 0.00322 | 0.999971 | 1 | 0.577271 |
| Ube2r2 | Q6ZWZ2 | 2.39 | 0.99 | 1 | 1.37 | 1.33 | 1.41 | 0.94 | 1.34 | 1.23 | 1.71 | 1.17 | 0.8 | 0.69 | 0.9 | 1.19 | 1.44 | -0.1825 | -0.21 | -0.3825 | 0.596709 | 0.572544 | 0.304479 | 0.999971 | 1 | 0.847402 |
| Ube2v1 | Q9CZY3 | 5.52 | 1.12 | 1.67 | 2.1 | 2.14 | 1.62 | 2.44 | 0.34 | 2.78 | 5.28 | 1.12 | 0.17 | -2.3 | 1.89 | 1.65 | 0.68 | -0.9675 | -0.265 | -2.1225 | 0.336499 | 0.843086 | 0.112166 | 0.999971 | 1 | 0.78268 |
| Ube2v2 | Q9D2M8 | -0.16 | 0.02 | -0.02 | -0.72 | 0.22 | -0.28 | 0.2 | -0.52 | 0.08 | -0.28 | 0.83 | 0.63 | 0.05 | -0.29 | -0.3 | -1 | 0.125 | 0.535 | -0.165 | 0.645012 | 0.112652 | 0.574629 | 0.999971 | 1 | 0.91816 |
| Ube2w | Q8VDW4 | 1.48 | 1.35 | 0.24 | 0.37 | 1.38 | 0.88 | -0.81 | 1.74 | 0.79 | 1.14 | -0.48 | 1.81 | 0.21 | 1.26 | 0.6 | 1.86 | -0.0625 | -0.045 | 0.1225 | 0.916042 | 0.933044 | 0.788783 | 0.999971 | 1 | 0.96741 |
| Ube2z | Q3UE37 | -0.09 | 0.73 | 0.86 | 0.73 | 1.05 | 0.2 | 0.96 | 0.85 | 1.36 | -0.27 | 1.09 | 0.64 | 1.35 | -0.07 | 1.25 | 1.02 | 0.2075 | 0.1475 | 0.33 | 0.494478 | 0.714032 | 0.395591 | 0.999971 | 1 | 0.871816 |
| Ube3a | O08759 | 0.33 | 0.64 | 0.76 | 0.36 | 0.65 | 0.55 | 0.91 | 0.64 | 0.62 | 0.25 | 0.88 | 1.03 | 0.52 | 0.27 | 0.9 | 0.58 | 0.165 | 0.1725 | 0.045 | 0.40563 | 0.468012 | 0.834471 | 0.999971 | 1 | 0.979036 |
| Ube3b | Q9ES34 | -0.86 | -0.47 | -0.69 | -0.93 | -0.54 | -0.44 | -0.73 | -0.26 | -0.67 | -0.64 | -0.37 | -0.23 | -0.71 | -0.33 | -0.26 | -0.46 | 0.245 | 0.26 | 0.2975 | 0.240848 | 0.217387 | 0.164299 | 0.999971 | 1 | 0.800215 |
| Ube3c | Q80U95 | -0.45 | 0.28 | -0.16 | -0.1 | 0.21 | 0.12 | -0.12 | -0.22 | 0.08 | -0.11 | -0.29 | -0.2 | 0.34 | 0.3 | -0.05 | -0.05 | 0.105 | -0.0225 | 0.2425 | 0.639505 | 0.91614 | 0.299178 | 0.999971 | 1 | 0.847402 |
| Ube4a | G3X9Y5 | 0.42 | 0.65 | 0.71 | 0.41 | 0.64 | 0.78 | 1.11 | 0.75 | 0.91 | 0.45 | 0.94 | 0.86 | 0.68 | 0.41 | 0.72 | 0.17 | 0.2725 | 0.2425 | -0.0525 | 0.181044 | 0.235428 | 0.798344 | 0.999971 | 1 | 0.970479 |
| Ube4b | A0A571BD! | 0.5 | 0.66 | 1.06 | 1.04 | 0.74 | 0.56 | 1.43 | 0.92 | 0.67 | 0.65 | 1 | 1.1 | 0.77 | 0.67 | 0.82 | 0.74 | 0.0975 | 0.04 | -0.065 | 0.706553 | 0.856178 | 0.747527 | 0.999971 | 1 | 0.961064 |
| Ubfd1 | Q78JW9 | 0.21 | 0.14 | -0.12 | 0.13 | 0.22 | 0.15 | 0.07 | 0.29 | 0.3 | 0.02 | 0.27 | 0.4 | 0.71 | 0.48 | 0.34 | 0.61 | 0.0925 | 0.1575 | 0.445 | 0.597873 | 0.392552 | 0.036144 | 0.999971 | 1 | 0.700096 |
| Ubiad1 | Q9DC60 | 0.2 | -0.33 | 0.62 | 0.4 | -0.59 | -0.12 | -0.17 | -0.01 | -0.56 | 0.85 | 0.31 | 0.01 | -0.53 | 0.54 | 0.18 | 1.36 | -0.445 | -0.07 | 0.165 | 0.116673 | 0.842109 | 0.695831 | 0.999971 | 1 | 0.947174 |
| Ubl3 | Q9Z2M6 | -0.36 | -0.29 | -1.07 | -0.99 | -0.49 | -0.42 | -0.54 | -1.27 | 1.18 | -0.04 | -1.35 | -0.77 | 0.59 | -0.54 | -0.74 | -0.9 | -0.0025 | 0.4325 | 0.28 | 0.993231 | 0.431278 | 0.469943 | 0.999971 | 1 | 0.892481 |
| Ubl4a | P21126 | -1.92 | -3.04 | 0.63 | 0.07 | -1.51 | -3.55 | 0.85 | 0.47 | -2.4 | -2.96 | 0.55 | 1.12 | -1.91 | -2.48 | 0.17 | -0.01 | 0.13 | 0.1425 | 0.0075 | 0.912174 | 0.905285 | 0.993832 | 0.999971 | 1 | 0.999535 |
| Ubl5 | E0CY91 | 1.24 | 1.6 | 1.13 | 1.39 | 1.76 | 1.56 | 0.28 | 1.52 | 1.84 | 1.94 | 0.76 | 0.66 | 2.33 | 1.84 | 0.99 | 1.02 | -0.06 | -0.04 | 0.205 | 0.862696 | 0.909105 | 0.550986 | 0.999971 | 1 | 0.913169 |
| Ubl7 | Q91W67 | 1.61 | 0.64 | 0.33 | 0.55 | 0.54 | 0.8 | 0.62 | 0.49 | 0.44 | 0.64 | 0.47 | 0.3 | -0.12 | 0.47 | 0.26 | 0.54 | -0.17 | -0.32 | -0.495 | 0.574273 | 0.301759 | 0.150552 | 0.999971 | 1 | 0.793248 |
| Ublcp1 | Q8BGR9 | 0.28 | 2.01 | 2.38 | 2.47 | 1.6 | 1.07 | 2.63 | 1.52 | 1.47 | 0.57 | 1.73 | 1.4 | 1.44 | 1.39 | 1.42 | 1.25 | -0.08 | -0.4925 | -0.41 | 0.885989 | 0.363626 | 0.402273 | 0.999971 | 1 | 0.872309 |
| Ubn1 | Q4G0F8-2 | -0.29 | -0.49 | -0.6 | -0.49 | -0.3 | -0.66 | -0.68 | -0.81 | -0.23 | -0.43 | 0.03 | -0.58 | -1.27 | -0.64 | -0.57 | -0.33 | -0.145 | 0.165 | -0.235 | 0.458178 | 0.419432 | 0.346553 | 0.999971 | 1 | 0.864164 |
| Ubp1 | Q811S7 | -0.26 | 0.28 | 0.08 | -0.46 | 0.18 | -0.21 | 0.21 | -0.09 | 0.33 | -0.4 | 0.17 | 0.38 | 0.35 | -0.17 | -0.27 | -0.06 | 0.1125 | 0.21 | 0.0525 | 0.630272 | 0.43538 | 0.830907 | 0.999971 | 1 | 0.978535 |
| Ubqln1 | Q8R317 | 0.11 | 0.93 | -0.14 | 0.48 | 0.53 | 0.44 | -0.19 | 0.32 | 0.34 | -0.13 | 0.16 | 0.6 | 0.37 | 0.68 | 0.09 | 0.43 | -0.07 | -0.1025 | 0.0475 | 0.811704 | 0.724355 | 0.86488 | 0.999971 | 1 | 0.983225 |
| Ubqln1_1 | Q8R317-2 | -0.99 | 0.38 | -0.37 | -0.36 | -0.21 | -0.15 | -0.59 | 0.3 | 0.18 | -0.92 | -0.09 | 0.6 | -0.26 | 0.55 | -0.09 | -0.39 | 0.1725 | 0.2775 | 0.2875 | 0.606874 | 0.501409 | 0.41448 | 0.999971 | 1 | 0.876329 |
| Ubqln2 | Q9QZM0 | -0.38 | 0.7 | 0.13 | 0.22 | 0.61 | 0.25 | -0.05 | 0.42 | 0.51 | -0.32 | 0.43 | 0.53 | 1.04 | 0.54 | 0.41 | 0.49 | 0.14 | 0.12 | 0.4525 | 0.61703 | 0.696442 | 0.132392 | 0.999971 | 1 | 0.784948 |
| Ubqln4 | Q99NB8 | -1.31 | 1.87 | 0.7 | 0.93 | 1.81 | 1.01 | 0.66 | 1.18 | 1.63 | -1.25 | 1.29 | 1.55 | 2.23 | 0.7 | 1.33 | 1.47 | 0.6175 | 0.2575 | 0.885 | 0.351628 | 0.766758 | 0.209539 | 0.999971 | 1 | 0.822138 |
| Ubr1 | O70481 | 0.15 | 0.48 | 0.17 | 0.04 | 0.33 | 0.42 | -0.36 | 0.34 | 0.17 | 0.33 | -0.07 | 0.23 | 0.39 | 0.51 | 0.31 | 0.36 | -0.0275 | -0.045 | 0.1825 | 0.908154 | 0.812329 | 0.3298 | 0.999971 | 1 | 0.856787 |
| Ubr2 | Q6WKZ8-2 | -0.13 | 0.12 | 0.55 | 0.22 | 0.11 | 0.19 | 0.34 | 0.5 | 0.37 | 0.18 | 0.37 | 0.49 | 0.68 | 0.35 | 0.29 | 0.43 | 0.095 | 0.1625 | 0.2475 | 0.657731 | 0.436028 | 0.265684 | 0.999971 | 1 | 0.842677 |
| Ubr3 | Q5U430 | 0.13 | 0.46 | -0.11 | 0.15 | 0.52 | 0.67 | 0.2 | 0.14 | 0.51 | 0.39 | 0.09 | 0.09 | 0.59 | 0.49 | 0.25 | 0.21 | 0.225 | 0.1125 | 0.2275 | 0.316802 | 0.589787 | 0.283501 | 0.999971 | 1 | 0.844538 |
| Ubr4 | A2AN08-3 | 0.01 | 0.26 | 0.24 | 0.16 | 0.15 | 0.41 | 0.25 | 0.34 | 0.25 | 0.23 | 0.3 | 0.08 | 0.42 | 0.37 | 0.37 | 0.28 | 0.12 | 0.0475 | 0.1925 | 0.490701 | 0.775048 | 0.267763 | 0.999971 | 1 | 0.842677 |
| Ubr5 | A0A2I3BQS | 1.86 | 1.54 | 1.65 | 1.48 | 1.34 | 1.81 | 1.33 | 1.64 | 1.71 | 2.27 | 1.56 | 1.32 | 1.1 | 1.54 | 1.23 | 1.43 | -0.1025 | 0.0825 | -0.3075 | 0.613539 | 0.737876 | 0.139672 | 0.999971 | 1 | 0.784948 |
| Ubr7 | Q8BU04 | 2.15 | 2.44 | 2.05 | 2.12 | 1.92 | 2.2 | 2.08 | 1.68 | 1.97 | 1.59 | 2.26 | 2.26 | 0.75 | 1.39 | 1.68 | 1.57 | -0.22 | -0.17 | -0.8425 | 0.287952 | 0.449075 | 0.008376 | 0.999971 | 1 | 0.602121 |
| Ubtd2 | Q6PGH0 | -0.22 | -0.44 | -1.51 | -0.83 | -0.51 | -0.24 | -1.79 | -0.3 | -0.24 | 0.12 | -0.87 | -1.26 | 0.08 | -0.12 | -1.35 | -0.57 | 0.04 | 0.1875 | 0.26 | 0.926722 | 0.642865 | 0.527075 | 0.999971 | 1 | 0.909459 |
| Ubtf | A2AWT6 | -0.21 | -0.81 | -0.78 | -0.28 | -0.64 | -0.82 | -0.75 | -0.93 | -0.58 | -0.28 | -1.1 | -1.36 | -0.55 | -0.7 | -1.11 | -0.76 | -0.265 | -0.31 | -0.26 | 0.241577 | 0.316444 | 0.285791 | 0.999971 | 1 | 0.845133 |
| Ubxn1 | Q922Y1 | 0.99 | 0.51 | 0.94 | 0.89 | 0.5 | 0.93 | 0.54 | 0.58 | 0.64 | 1.29 | 0.38 | 0.59 | 0.34 | 1.14 | 0.62 | 0.72 | -0.195 | -0.1075 | -0.1275 | 0.350163 | 0.669348 | 0.590519 | 0.999971 | 1 | 0.922217 |
| Ubxn2a | Q99KJ0 | -1.45 | -0.11 | -0.55 | -0.5 | -0.52 | -0.52 | -0.37 | -0.54 | -0.65 | -1.51 | -0.38 | -0.18 | -0.74 | -0.33 | 0.24 | -0.18 | 0.165 | -0.0275 | 0.4 | 0.58046 | 0.944047 | 0.26424 | 0.999971 | 1 | 0.842533 |
| Ubxn2b | Q0KL01 | -0.34 | 0.02 | 0.09 | 0.05 | 0.21 | -0.21 | 0.22 | 0.26 | 0.16 | -0.47 | 0.35 | 0.35 | 0.2 | -0.54 | 0.33 | 0.22 | 0.165 | 0.1425 | 0.0975 | 0.427102 | 0.565952 | 0.699039 | 0.999971 | 1 | 0.947812 |
| Ubxn4 | A0A0R4J07 | 0.49 | -0.78 | -0.3 | -0.55 | -0.74 | -0.19 | 0.06 | -0.36 | -0.63 | 0.26 | -0.9 | -0.49 | -0.44 | -0.13 | -0.3 | -0.8 | -0.0225 | -0.155 | -0.1325 | 0.944633 | 0.670623 | 0.675698 | 0.999971 | 1 | 0.941576 |
| Ubxn6 | Q99PL6 | 0.89 | -0.71 | -0.73 | -0.5 | -0.78 | -0.27 | 0.01 | -0.82 | -1.02 | -0.05 | -0.65 | -0.64 | -1 | -0.78 | -0.71 | -0.54 | -0.2025 | -0.3275 | -0.495 | 0.627896 | 0.439595 | 0.220553 | 0.999971 | 1 | 0.827096 |
| Ubxn7 | G5E8R8 | 1.42 | 1.39 | 0.74 | 1.37 | 1.26 | 1.38 | 1.11 | 1.32 | 1.21 | 1.41 | 0.76 | 1.03 | 1.44 | 1.82 | 0.82 | 1.59 | 0.0375 | -0.1275 | 0.1875 | 0.863723 | 0.602551 | 0.514638 | 0.999971 | 1 | 0.905815 |
| Uchl1 | Q9R0P9 | 0 | -0.92 | -0.99 | -0.72 | -0.4 | -0.17 | -0.72 | -0.43 | -0.62 | -0.24 | -1.03 | -1.16 | -0.39 | 0.28 | -0.32 | -0.49 | 0.2275 | -0.105 | 0.4275 | 0.411958 | 0.737164 | 0.173349 | 0.999971 | 1 | 0.807307 |
| Uchl3 | Q9JKB1 | -0.18 | 0.85 | 1.06 | 0.38 | 0.87 | 0.64 | 0.94 | 0.33 | 0.81 | 0.61 | 0.24 | 0.01 | 0.87 | 1.15 | 0.29 | 0.42 | 0.1675 | -0.11 | 0.155 | 0.594613 | 0.738378 | 0.648262 | 0.999971 | 1 | 0.933842 |
| Uchl5 | Q9WUP7-2 | -0.51 | -0.79 | -0.42 | 0.21 | -0.74 | -1.44 | -0.9 | -0.03 | -0.33 | -1.68 | 0.3 | -0.16 | -0.82 | -0.4 | 0.16 | 0.2 | -0.4 | -0.09 | 0.1625 | 0.274187 | 0.840852 | 0.618663 | 0.999971 | 1 | 0.928581 |
| Uchl5_1 | Q9WUP7 | 0.01 | 0.39 | -1.47 | -2 | 0.45 | 0.37 | -1.12 | -1.27 | 0.38 | 0.2 | -1.45 | -1.59 | 0.91 | 0.59 | -1.48 | -1.35 | 0.375 | 0.1525 | 0.435 | 0.578561 | 0.829401 | 0.574512 | 0.999971 | 1 | 0.91816 |
| Uck1 | P52623 | 0.28 | 0.93 | 1.13 | 1.3 | 1.01 | 1.17 | 0.86 | 1.43 | 1.14 | 1.1 | 1 | 1.33 | 1.01 | 1.28 | 1.02 | 1.2 | 0.2075 | 0.2325 | 0.2175 | 0.454155 | 0.376559 | 0.409362 | 0.999971 | 1 | 0.875804 |
| Uckl1 | Q91YL3 | 0.76 | 0.58 | 0.22 | 0.1 | 0.81 | 0.08 | 0.9 | 0.23 | 0.92 | 0.25 | 0.79 | 1.14 | 0.71 | 0.22 | 0.1 | -0.01 | 0.09 | 0.36 | -0.16 | 0.743227 | 0.196057 | 0.526901 | 0.999971 | 1 | 0.909459 |
| Ucp2 | D3YXM5 | 0.8 | -0.32 | -0.36 | -0.85 | -0.64 | -0.48 | -0.81 | -0.52 | -1.75 | -0.43 | 0.23 | -0.73 | -0.63 | -1.35 | -1.77 | -0.51 | -0.43 | -0.4875 | -0.8825 | 0.23999 | 0.345905 | 0.068626 | 0.999971 | 1 | 0.733937 |
| Ucp3 | P56501 | -0.41 | 0.08 | -1.09 | -1.34 | -0.63 | -0.57 | -1.51 | -0.47 | -0.61 | -0.47 | -1.37 | -0.61 | 0.02 | -0.53 | -0.97 | -0.7 | -0.105 | -0.075 | 0.145 | 0.786592 | 0.840082 | 0.699075 | 0.999971 | 1 | 0.947812 |
| Uevld | Q3U1V6 | -0.19 | 0.84 | -0.06 | -0.26 | 0.74 | 1.01 | 0.09 | -0.13 | 0.55 | 0.19 | -0.16 | -0.22 | 1.24 | 0.87 | -0.1 | 0.02 | 0.345 | 0.0075 | 0.425 | 0.35258 | 0.980979 | 0.301183 | 0.999971 | 1 | 0.847402 |
| Ufc1 | Q9CR09 | -1.05 | 0.04 | -2.73 | -2.28 | 0.46 | -0.18 | -2.15 | -2.61 | 0.58 | -0.66 | -2.34 | -1.95 | 0.58 | 0.32 | -2 | -2.35 | 0.385 | 0.4125 | 0.6425 | 0.660186 | 0.619123 | 0.474604 | 0.999971 | 1 | 0.894044 |
| Ufd1 | P70362 | 0.88 | 0.2 | 0.27 | 0.54 | 0.04 | 0.24 | 0.93 | -0.12 | 0.04 | 0.74 | 0.24 | 0.27 | -0.03 | 0.39 | 0.03 | 0.09 | -0.2 | -0.15 | -0.3525 | 0.496134 | 0.541336 | 0.140845 | 0.999971 | 1 | 0.784967 |
| Ufl1 | Q8CCJ3 | -0.22 | -0.3 | 0.26 | -0.03 | -0.2 | -0.22 | 0.4 | 0.11 | -0.03 | -0.25 | 0.18 | 0.35 | -0.1 | -0.36 | 0.33 | -0.04 | 0.095 | 0.135 | 0.03 | 0.681295 | 0.54438 | 0.895635 | 0.999971 | 1 | 0.988571 |
| Ufm1 | P61961 | 0.16 | 0.12 | 0.9 | 0.89 | 0 | 0.23 | 0.19 | 0.52 | 0.05 | 0.36 | 0.33 | 0.57 | -0.04 | 0.23 | 0.59 | 1.18 | -0.2825 | -0.19 | -0.0275 | 0.301746 | 0.477149 | 0.935426 | 0.999971 | 1 | 0.995618 |
| Ufsp1 | Q9CZP0 | 0.39 | 1.07 | 0.45 | -0.7 | 0.44 | 0.73 | -0.08 | -0.11 | -1.54 | 0.09 | 0.51 | -0.04 | -0.97 | 0.4 | 0.34 | 0.46 | -0.0575 | -0.5475 | -0.245 | 0.886244 | 0.322278 | 0.605408 | 0.999971 | 1 | 0.92451 |
| Ufsp2 | Q99K23 | -0.79 | -0.15 | -1.09 | -0.8 | -0.22 | -0.09 | -1.24 | -0.81 | -0.01 | -0.6 | -1.12 | -1.16 | 0.21 | -0.34 | -0.89 | -0.74 | 0.1175 | -0.015 | 0.2675 | 0.723952 | 0.964042 | 0.411651 | 0.999971 | 1 | 0.875847 |
| Ugcg | O88693 | 0.24 | 0.8 | 1.07 | 1 | 0.46 | -0.05 | 0.38 | 0.88 | 0.26 | 0.21 | 0.87 | 0.36 | 1.11 | 1.15 | 0.65 | 1.51 | -0.36 | -0.3525 | 0.3275 | 0.22358 | 0.202029 | 0.25424 | 0.999971 | 1 | 0.839012 |
| Ugdh | O70475 | 0.27 | 0.84 | 0.07 | 0.21 | 0.68 | 0.69 | -0.11 | -0.07 | 0.92 | 0.74 | -0.27 | 0.23 | 0.72 | 0.9 | 0.39 | 0.52 | -0.05 | 0.0575 | 0.285 | 0.863987 | 0.857079 | 0.250507 | 0.999971 | 1 | 0.839012 |
| Uggt1 | Q6P5E4 | -0.93 | -0.81 | -0.33 | -0.33 | -0.6 | -0.8 | -0.12 | -0.32 | -0.36 | -0.7 | -0.16 | 0.01 | -0.48 | -0.45 | -0.25 | -0.05 | 0.14 | 0.2975 | 0.2925 | 0.574433 | 0.247727 | 0.219871 | 0.999971 | 1 | 0.827096 |
| Uggt2 | E9Q4X2 | 4.24 | 2.28 | -0.11 | 0.11 | 1.09 | 2.16 | -0.22 | 0.03 | 1.16 | 2.96 | 0.1 | 0.45 | 0.62 | 1.05 | -0.4 | 1.11 | -0.865 | -0.4625 | -1.035 | 0.412288 | 0.670565 | 0.300912 | 0.999971 | 1 | 0.847402 |
| Ugp2 | Q91ZJ5 | -0.78 | -0.15 | -0.53 | -1.09 | -0.3 | -0.24 | -0.51 | -0.77 | -0.06 | -0.81 | -0.32 | -0.25 | 0.13 | -0.61 | -0.69 | -0.9 | 0.1825 | 0.2775 | 0.12 | 0.483943 | 0.32256 | 0.696804 | 0.999971 | 1 | 0.947174 |
| Ugt1a7 | Q6ZQM8 | 0.59 | 0.41 | 0.73 | 0.43 | -0.33 | 0.67 | 0.13 | -0.21 | 0.1 | 1.45 | -0.54 | -1.75 | -0.1 | -0.11 | -0.21 | -0.01 | -0.475 | -0.725 | -0.6475 | 0.094659 | 0.257691 | 0.004711 | 0.999971 | 1 | 0.582714 |
| Ugt8 | Q64676 | -0.89 | -1.04 | -1.75 | -1.62 | -0.59 | -0.79 | -0.72 | -2.45 | -0.13 | -0.25 | -2.37 | -1.35 | 0.06 | 0.43 | -1 | -0.98 | 0.1875 | 0.3 | 0.9525 | 0.682865 | 0.571798 | 0.040035 | 0.999971 | 1 | 0.700838 |
| Uhmk1 | P97343 | -1.26 | 0.04 | 0.43 | -0.13 | -0.62 | 0.13 | 0.84 | 0.72 | -0.4 | -0.96 | 0.81 | 0.58 | -0.11 | -0.12 | 0.59 | 0.46 | 0.4975 | 0.2375 | 0.435 | 0.297336 | 0.645259 | 0.284827 | 0.999971 | 1 | 0.845133 |
| Uhrf1bp1 | B2KF50 | -0.37 | -0.58 | -0.3 | -0.31 | -0.74 | -0.56 | -0.61 | -0.31 | -0.46 | -0.46 | -0.21 | -0.48 | -0.67 | -0.73 | -0.46 | -0.36 | -0.165 | -0.0125 | -0.165 | 0.382434 | 0.942245 | 0.381499 | 0.999971 | 1 | 0.871816 |
| Uhrf1bp1l | A2RSJ4 | 0.09 | -0.17 | -0.38 | -0.34 | -0.59 | -0.43 | -0.25 | -0.58 | -0.08 | -0.06 | 0.02 | -0.41 | -0.88 | -0.56 | -0.74 | -0.73 | -0.2625 | 0.0675 | -0.5275 | 0.202165 | 0.733913 | 0.021318 | 0.999971 | 1 | 0.640714 |
| Ulk1 | A0A0R4J0E | -0.71 | -0.69 | 0.35 | 0.64 | -0.27 | -0.42 | 0.79 | 0.29 | -0.35 | -0.76 | 0.08 | -0.56 | 0.14 | -0.9 | -0.27 | -0.55 | 0.2 | -0.295 | -0.2925 | 0.638134 | 0.446487 | 0.468068 | 0.999971 | 1 | 0.89236 |
| Ulk2 | Q9QY01 | -1.3 | -0.67 | -0.98 | -0.87 | -0.63 | -1.21 | -0.79 | -0.98 | -1.01 | -1.11 | -0.72 | -0.99 | -0.7 | -0.78 | -0.88 | -0.76 | 0.0525 | -0.0025 | 0.175 | 0.814356 | 0.990284 | 0.38793 | 0.999971 | 1 | 0.871816 |
| Ulk3 | Q3U3Q1 | -0.53 | 0.42 | 0.78 | 0.75 | 0.2 | 0.52 | 0.41 | 0.81 | 0.1 | -0.41 | 0.73 | -0.96 | 0.49 | 0.06 | 0.81 | 0.74 | 0.13 | -0.49 | 0.17 | 0.695182 | 0.289526 | 0.625206 | 0.999971 | 1 | 0.930479 |
| Umad1 | E0CZ61 | 0.83 | 0.22 | 3.18 | 2.5 | 0.14 | -0.1 | 1.3 | 2.13 | -0.34 | -0.73 | 2.84 | 2.85 | 0.14 | -0.39 | 3.04 | 3.4 | -0.815 | -0.5275 | -0.135 | 0.311393 | 0.626606 | 0.89931 | 0.999971 | 1 | 0.989654 |
| Umps | P13439 | -0.3 | 1.16 | 0.67 | 0.44 | 1.43 | 0.64 | 0.7 | 0.32 | 1.02 | -0.14 | 0.74 | 1.14 | 0.76 | 0.56 | 0.36 | 0.29 | 0.28 | 0.1975 | -1.28E-16 | 0.459022 | 0.625426 | 1 | 0.999971 | 1 | 1 |
| Unc119 | Q9Z2R6 | 0.69 | 0.18 | 2.17 | 1.94 | 0.08 | 0.41 | 1.05 | 2.66 | 0.26 | 1.07 | 2.61 | 1.95 | 0.63 | 0.85 | 2.79 | 1.95 | -0.195 | 0.2275 | 0.31 | 0.773867 | 0.724224 | 0.627093 | 0.999971 | 1 | 0.930921 |
| Unc119b | Q8C4B4 | -0.78 | 0.15 | 0.15 | 0.23 | 0.09 | 0.14 | 0.16 | 0.48 | 0 | -0.28 | 0.39 | 0.59 | 0.34 | 0.2 | 0.58 | 0.65 | 0.28 | 0.2375 | 0.505 | 0.320983 | 0.455557 | 0.097188 | 0.999971 | 1 | 0.78268 |
| Unc13a | H3BJZ7 | -1.01 | -0.77 | -0.72 | -1.09 | -1.11 | -0.59 | -0.5 | -1.1 | -0.86 | -0.81 | -0.69 | -1.01 | -0.51 | -1.03 | -1.1 | -1.26 | 0.0725 | 0.055 | -0.0775 | 0.749137 | 0.762635 | 0.733104 | 0.999971 | 1 | 0.956962 |
| Unc13b | F6X605 | -5.97 | -2.78 | -2.77 | -1.44 | -2.63 | -4.67 | -2.54 | -2.49 | -3.68 | -3.54 | -2.53 | -1.74 | -3.87 | -3.92 | -1.74 | -2.03 | 0.1575 | 0.3675 | 0.35 | 0.872406 | 0.702577 | 0.728647 | 0.999971 | 1 | 0.956041 |
| Unc13b_1 | A0A140LH) | 1.54 | 0.01 | 0.56 | 0.22 | -0.31 | -0.33 | 0.38 | 0.04 | -0.34 | 0.99 | 0.05 | 1.2 | -0.41 | -0.56 | -0.71 | -0.17 | -0.6375 | -0.1075 | -1.045 | 0.110067 | 0.819178 | 0.015029 | 0.999971 | 1 | 0.613095 |
| Unc13c | Q8K0T7 | -1.2 | -2.03 | -1.95 | -1.77 | -1.17 | -1.76 | -1.63 | -1.82 | -0.91 | -1.44 | -1.71 | -1.79 | -0.9 | -1.41 | -0.94 | -2.36 | 0.1425 | 0.275 | 0.335 | 0.58834 | 0.348529 | 0.384059 | 0.999971 | 1 | 0.871816 |
| Unc45a | Q99KD5 | -0.25 | 0.31 | 0.3 | 0.2 | 1.17 | 0.99 | -0.13 | 1.25 | 0.35 | 0.43 | 0.38 | 0.83 | 1.23 | 1.91 | 0.99 | 1.54 | 0.68 | 0.3575 | 1.2775 | 0.074033 | 0.125408 | 0.000923 | 0.999971 | 1 | 0.365812 |
| Unc5a | Q8K1S4-2 | -1.69 | -1.28 | -0.21 | -0.33 | -1.06 | -1.22 | 0.42 | -0.62 | -1.01 | -1.24 | -0.65 | -0.77 | -0.32 | -1.39 | -0.93 | -0.78 | 0.2575 | -0.04 | 0.0225 | 0.595299 | 0.914589 | 0.955636 | 0.999971 | 1 | 0.998056 |
| Unc5b | Q8K1S3-2 | 0.1 | -0.09 | -0.2 | 0.09 | -0.55 | -0.23 | -0.83 | -0.89 | -0.53 | 0.21 | -0.77 | -0.85 | -0.17 | -0.22 | -0.1 | 0.07 | -0.6 | -0.46 | -0.08 | 0.019761 | 0.115866 | 0.656338 | 0.999971 | 1 | 0.934061 |
| Unc5c | O08747-2 | -0.31 | 0.16 | -0.74 | -1.34 | 0.09 | 0.45 | -0.96 | -1.09 | 0.35 | -0.18 | -0.76 | -1.49 | 0.63 | 0.13 | -0.77 | -1.5 | 0.18 | 0.0375 | 0.18 | 0.699828 | 0.937094 | 0.733512 | 0.999971 | 1 | 0.957221 |
| Unc5d | B7ZMT2 | -0.39 | -0.56 | -1.64 | -2.06 | -0.35 | -0.09 | -1.5 | -2.06 | -0.21 | -0.07 | -1.13 | -2.15 | 0.26 | -0.39 | -1.63 | -2.04 | 0.1625 | 0.2725 | 0.2125 | 0.775225 | 0.640701 | 0.729765 | 0.999971 | 1 | 0.956041 |
| Unc79 | E5CYJ9 | 0.18 | -0.48 | -0.31 | -0.71 | -0.06 | 0.28 | -0.97 | -0.74 | 0.07 | 0.38 | -0.63 | -1.02 | 0.57 | 0.44 | -1.05 | -0.37 | -0.0425 | 0.03 | 0.2275 | 0.90119 | 0.933986 | 0.576969 | 0.999971 | 1 | 0.918194 |
| Unc80 | A0A1D5RL\ | -0.89 | -0.72 | -0.85 | -0.3 | -0.56 | -0.72 | -0.58 | -0.3 | -0.59 | -0.68 | -0.53 | -0.63 | -0.29 | -0.83 | -0.7 | -0.36 | 0.15 | 0.0825 | 0.145 | 0.483304 | 0.674658 | 0.529601 | 0.999971 | 1 | 0.910069 |
| Unc93b1 | E9PYK0 | -1.18 | 0.52 | 0.12 | 0.05 | 1.47 | -0.05 | 0.9 | 0.39 | 2.08 | -0.86 | 0.49 | 0.71 | 1.42 | -0.99 | -0.09 | -0.86 | 0.8 | 0.7275 | -0.0075 | 0.109932 | 0.277045 | 0.990128 | 0.999971 | 1 | 0.99925 |
| Unk | Q8BL48-2 | 0.8 | 0.79 | 0.46 | 0.44 | 0.83 | 0.72 | 1.35 | 0.53 | 0.94 | 0.95 | 1.03 | 1.04 | 2.01 | 0.64 | 0.62 | -0.08 | 0.235 | 0.3675 | 0.175 | 0.33414 | 0.064519 | 0.682195 | 0.999971 | 1 | 0.943204 |
| Unkl | E0CYD7 | 0.24 | 0.74 | 1.21 | 1.32 | 0.67 | 0.93 | 1.31 | 1.92 | 0.68 | 0.47 | 1.09 | 1.31 | 0.3 | 0.83 | 1.12 | 0.8 | 0.33 | 0.01 | -0.115 | 0.369238 | 0.974664 | 0.708852 | 0.999971 | 1 | 0.950592 |
| Upf1 | Q9EPU0-2 | 0.27 | 0.82 | 1.1 | 0.96 | 0.85 | 0.75 | 0.88 | 1.19 | 0.74 | 0.37 | 1 | 1.17 | 0.64 | 0.78 | 1.15 | 1.2 | 0.13 | 0.0325 | 0.155 | 0.588864 | 0.903995 | 0.54705 | 0.999971 | 1 | 0.911483 |
| Upf2 | A2AT37 | 0.25 | -0.36 | 0.6 | 0.79 | 0.27 | -0.43 | 0.91 | 0.54 | 0.2 | -0.2 | 0.24 | 0.32 | 0.11 | -0.31 | 0.22 | 0.24 | 0.0025 | -0.18 | -0.255 | 0.994581 | 0.538203 | 0.39702 | 0.999971 | 1 | 0.871816 |
| Upf3b | Q3ULL6 | 0.23 | 0.24 | 0.02 | 0.31 | 0 | 0.29 | 0.05 | 0.43 | 0.26 | 0.27 | 0.3 | 0.27 | -0.22 | 0.19 | 0.22 | 0.12 | -0.0075 | 0.075 | -0.1225 | 0.968204 | 0.646209 | 0.522038 | 0.999971 | 1 | 0.907467 |
| Upp1 | P52624 | -0.08 | 0.48 | 0.24 | -0.03 | 0.51 | 0.32 | 0.11 | 1.07 | 0.04 | 0.48 | -0.03 | 0.58 | -0.36 | -0.22 | 0.38 | 0.62 | 0.35 | 0.115 | -0.0475 | 0.208786 | 0.625626 | 0.866924 | 0.999971 | 1 | 0.983462 |
| Uprt | B1AVZ0 | 0.69 | 1.29 | 0.87 | 0.74 | 0.88 | 0.83 | 0.9 | 0.92 | 0.52 | 0.45 | 0.65 | 0.77 | 0.22 | 0.91 | 0.59 | 0.25 | -0.015 | -0.3 | -0.405 | 0.939495 | 0.166957 | 0.123744 | 0.999971 | 1 | 0.78268 |
| Uqcc1 | Q9CWU6 | -0.96 | -1.17 | -0.85 | -0.71 | -1.06 | -0.99 | -1.38 | -0.73 | -1.19 | -0.87 | -0.88 | -0.88 | -1.23 | -0.72 | -0.54 | -0.51 | -0.1175 | -0.0325 | 0.1725 | 0.585178 | 0.862883 | 0.462917 | 0.999971 | 1 | 0.890653 |
| Uqcc2 | Q9CQY6 | -0.6 | -0.7 | -1.24 | -0.89 | -0.85 | -0.46 | -1.05 | -1.22 | -1.17 | -0.32 | -1.08 | -1.13 | -0.24 | -0.14 | -0.64 | -0.74 | -0.0375 | -0.0675 | 0.4175 | 0.878712 | 0.7999 | 0.107178 | 0.999971 | 1 | 0.78268 |
| Uqcc3 | Q8K2T4 | -1.55 | -0.62 | -0.93 | 0.16 | 0.12 | -0.18 | -0.94 | -0.55 | -0.18 | -0.35 | 0.09 | 0 | 0.12 | 0.1 | -0.7 | 0.14 | 0.3475 | 0.625 | 0.65 | 0.399707 | 0.110692 | 0.126415 | 0.999971 | 1 | 0.783193 |
| Uqcr10 | Q8R1I1 | -1.75 | -0.06 | -0.68 | -3.16 | -0.14 | 0.09 | 1.4 | -2.97 | -0.03 | -1.73 | 0.26 | 0.35 | -0.07 | -0.37 | -2.66 | -3.26 | 1.0075 | 1.125 | -0.1775 | 0.336874 | 0.163621 | 0.850432 | 0.999971 | 1 | 0.98187 |
| Uqcr11 | Q9CPX8 | -3.06 | -1.25 | -0.74 | -0.52 | -0.69 | -1.79 | -1.48 | -1.7 | -1.66 | -2.37 | -1.87 | -1.66 | -1.9 | -0.5 | -0.84 | -1.12 | -0.0225 | -0.4975 | 0.3025 | 0.968761 | 0.380943 | 0.612981 | 0.999971 | 1 | 0.927364 |
| Uqcrb | Q9CQB4 | -0.34 | -0.79 | -1.32 | -0.85 | -1.05 | -0.81 | -0.55 | -1.49 | -1.14 | -0.49 | -1.42 | -1.6 | -1.02 | -0.73 | -1.77 | -0.95 | -0.15 | -0.3375 | -0.2925 | 0.611907 | 0.304183 | 0.355981 | 0.999971 | 1 | 0.868793 |
| Uqcrc1 | Q9CZ13 | -1.6 | -1.24 | -0.65 | -0.47 | -1.08 | -1.35 | -0.76 | -0.43 | -1.14 | -1.68 | -0.72 | -0.73 | -1.09 | -1.21 | -0.63 | -0.22 | 0.085 | -0.0775 | 0.2025 | 0.795954 | 0.820788 | 0.558574 | 0.999971 | 1 | 0.915051 |
| Uqcrc2 | Q9DB77 | -0.75 | -1.07 | -1.1 | -0.99 | -1.26 | -1.22 | -1.18 | -1.08 | -1.6 | -1.28 | -1.07 | -0.81 | -1.62 | -1.08 | -0.95 | -0.69 | -0.2075 | -0.2125 | -0.1075 | 0.254267 | 0.354621 | 0.660369 | 0.999971 | 1 | 0.935582 |
| Uqcrfs1 | Q9CR68 | -1.14 | -1.6 | -1.33 | -2.05 | -1.49 | -1.32 | -1.54 | -2.12 | -1.72 | -0.86 | -1.86 | -2.16 | -1.8 | -1.37 | -1.97 | -1.9 | -0.0875 | -0.12 | -0.23 | 0.754258 | 0.723777 | 0.391263 | 0.999971 | 1 | 0.871816 |
| Uqcrh | P99028 | 0.81 | 0.77 | 1.29 | 1.07 | 0.44 | 0.76 | 0.92 | 0.85 | 0.45 | 1.24 | 0.45 | 1.12 | 0.59 | 0.62 | 0.86 | 1.53 | -0.2425 | -0.17 | -0.085 | 0.268812 | 0.525198 | 0.753577 | 0.999971 | 1 | 0.961116 |

| | | | | | | | | | | | | | | | | | | | | | | | | | | |
|---|---|---|---|---|---|---|---|---|---|---|---|---|---|---|---|---|---|---|---|---|---|---|---|---|---|---|
| Uqcrq | Q9CQ69 | -0.19 | -1.24 | -0.99 | -0.66 | -1.11 | -1.53 | -0.7 | -0.85 | -1.4 | -0.55 | -0.71 | -0.78 | -1.71 | -1.16 | -0.91 | -1.26 | -0.2775 | -0.09 | -0.49 | 0.366001 | 0.765303 | 0.121913 | 0.999971 | 1 | 0.78268 |
| Urb1 | E9PU96 | 0.02 | -1.03 | -0.97 | -1.01 | -1.25 | -1.41 | -0.55 | -1.28 | -1.3 | -0.49 | -0.45 | -0.24 | -0.96 | -1.12 | -1.17 | -0.73 | -0.375 | 0.1275 | -0.2475 | 0.263789 | 0.710489 | 0.400386 | 0.999971 | 1 | 0.871816 |
| Urb2 | E9Q7L1 | -1.15 | -1.09 | -0.39 | -0.05 | -1.67 | -2.47 | -0.4 | -0.41 | -1.74 | -1.46 | 0.48 | 0.46 | -2.32 | -0.67 | -0.41 | -1.37 | -0.5675 | 0.105 | -0.5225 | 0.300093 | 0.861921 | 0.285554 | 0.999971 | 1 | 0.845133 |
| Urgcp | Q5NCI0-2 | 0.3 | 1 | 0.05 | 0.07 | 1.5 | 0.78 | -0.25 | 0.21 | 1.21 | 0.67 | -0.08 | 0.48 | 1.89 | 1.16 | 0.38 | 0.59 | 0.205 | 0.215 | 0.65 | 0.624469 | 0.535056 | 0.122415 | 0.999971 | 1 | 0.78268 |
| Uri1 | A0A0U1RP | -0.02 | 0.26 | 0.42 | 1.05 | -0.44 | -0.77 | 0 | 0.73 | -0.49 | -0.57 | 0.69 | 0.6 | -1.3 | -0.16 | 0.57 | 0.55 | -0.5475 | -0.37 | -0.5125 | 0.175739 | 0.359226 | 0.285711 | 0.999971 | 1 | 0.845133 |
| Urm1 | Q9D2P4 | -1.17 | 0.08 | -0.67 | -1.02 | -0.37 | -0.32 | -0.76 | -1.05 | -0.07 | -1.02 | -0.53 | -0.29 | -0.64 | 0.09 | -0.6 | -0.6 | 0.07 | 0.2175 | 0.2575 | 0.830926 | 0.529006 | 0.443259 | 0.999971 | 1 | 0.883386 |
| Urod | P70697 | -0.03 | 0.47 | 0.03 | 0.13 | 0.83 | 0 | -0.34 | -0.01 | 0.54 | -0.15 | 0.1 | 0.24 | 1.11 | 0.3 | 0.09 | 0.41 | -0.03 | 0.0325 | 0.3275 | 0.916528 | 0.883415 | 0.242592 | 0.999971 | 1 | 0.837316 |
| Uros | P51163 | -0.01 | 0.13 | -0.38 | -0.15 | -0.47 | -0.28 | -0.59 | 0.77 | -0.05 | -0.17 | -0.28 | -0.56 | -0.05 | 0.06 | 0.47 | 0.12 | -0.04 | -0.1625 | 0.2525 | 0.903132 | 0.435102 | 0.24657 | 0.999971 | 1 | 0.839012 |
| Usb1 | A0A0R4J0E | 1.34 | 0.44 | 1.26 | 0.98 | 0.27 | 1.02 | 0.22 | 1.41 | 0.03 | 1.5 | 0.38 | 0.87 | 0.22 | 1.37 | 0.6 | 1.27 | -0.275 | -0.31 | -0.14 | 0.439379 | 0.409801 | 0.681305 | 0.999971 | 1 | 0.943107 |
| Use1 | Q9CQ56 | -0.64 | -0.15 | -0.22 | -0.51 | 0.17 | -0.29 | -0.46 | -0.21 | -0.2 | -0.29 | -0.18 | -0.32 | -0.21 | -0.42 | -0.36 | -0.45 | 0.1825 | 0.1325 | 0.02 | 0.417285 | 0.483854 | 0.917655 | 0.999971 | 1 | 0.992633 |
| Usf1 | Q61069 | 0.83 | 2.48 | 1.58 | 1.95 | 2.97 | 1.38 | 2.07 | 1.77 | 3.24 | -0.79 | 1.78 | 2.29 | 1.8 | -0.03 | 0.74 | 1.1 | 0.3375 | -0.08 | -0.8075 | 0.464618 | 0.923976 | 0.120179 | 0.999971 | 1 | 0.78268 |
| Usf2 | F6PWU6 | -0.31 | 2.28 | 1.21 | 0.42 | 2.69 | 2.62 | 2.81 | 1.53 | 2.59 | 0.33 | 0.92 | 1.26 | 1.28 | 0.54 | 0.67 | 0.45 | 1.5125 | 0.375 | -0.165 | 0.025973 | 0.578216 | 0.76035 | 0.999971 | 1 | 0.961116 |
| Uso1 | Q9Z1Z0 | 0.32 | 0.96 | 0.91 | 0.95 | 1.06 | 1.17 | 1.01 | 1.12 | 1.15 | 0.48 | 1.03 | 1.49 | 1.06 | 1.1 | 1.33 | 1.43 | 0.305 | 0.2525 | 0.445 | 0.170965 | 0.372934 | 0.071966 | 0.999971 | 1 | 0.739457 |
| Usp10 | P52479-2 | 0.95 | 1.16 | 1.08 | 1.3 | 1.51 | 0.74 | 1.27 | 0.96 | 1.22 | 0.97 | 1.14 | 1.4 | 1.11 | 0.99 | 1.04 | 0.88 | -0.0025 | 0.06 | -0.1175 | 0.99114 | 0.744329 | 0.508306 | 0.999971 | 1 | 0.905815 |
| Usp11 | Q99K46 | 0.09 | 0.79 | 1 | 0.98 | 0.83 | 0.53 | 1.01 | 1.13 | 0.65 | 0 | 1.04 | 1.23 | 0.91 | 0.81 | 1.3 | 1.18 | 0.16 | 0.015 | 0.335 | 0.55701 | 0.964947 | 0.226776 | 0.999971 | 1 | 0.828236 |
| Usp12 | Q9D9M2 | 0.48 | 2.72 | 1.39 | 0.07 | 3.09 | 2.72 | 1.05 | 0.69 | 3.17 | 0.64 | 1.01 | 1.3 | 4.2 | 1.49 | 0.86 | 0.67 | 0.7225 | 0.365 | 0.64 | 0.35092 | 0.623513 | 0.484444 | 0.999971 | 1 | 0.896402 |
| Usp13 | J3QSN2 | 0.26 | 0.35 | 0.16 | -0.74 | 0.12 | -0.05 | 0.14 | -0.3 | 0.42 | 0 | 0.05 | 0.05 | 0.93 | -0.05 | -0.36 | -0.55 | -0.03 | 0.1225 | -0.015 | 0.916232 | 0.667025 | 0.969879 | 0.999971 | 1 | 0.998787 |
| Usp14 | Q9JMA1 | 0.14 | 0.11 | 0.5 | 0.4 | -0.11 | 0.16 | 0.49 | 0.6 | 0.03 | -0.05 | 0.59 | 0.54 | -0.07 | 0.33 | 0.69 | 0.6 | -0.0025 | -0.01 | 0.1 | 0.991222 | 0.965081 | 0.669546 | 0.999971 | 1 | 0.938711 |
| Usp15 | Q8R5H1 | -0.35 | 0.45 | 0.51 | 0.27 | 0.77 | 0.43 | 0.91 | 0.31 | 1.02 | -0.18 | 0.82 | 1.02 | 1.18 | 0.44 | 0.43 | 0.39 | 0.385 | 0.45 | 0.39 | 0.169259 | 0.212948 | 0.197862 | 0.999971 | 1 | 0.813335 |
| Usp19 | E9Q9M5 | 0.42 | 1.08 | 0.15 | 0.09 | 0.67 | 1.08 | 0.31 | 0.28 | 0.74 | 0.38 | 0.42 | 0.53 | 1.08 | 1.03 | 0.09 | 0.32 | 0.15 | 0.0825 | 0.195 | 0.621559 | 0.752985 | 0.565363 | 0.999971 | 1 | 0.917745 |
| Usp20 | Q8C6M1 | -0.22 | -0.31 | 0.14 | -0.31 | -0.5 | -0.14 | -0.04 | 0.83 | 0.03 | 0.01 | 0.37 | -0.13 | -0.09 | -0.21 | 0.72 | 0.17 | 0.2125 | 0.245 | 0.3225 | 0.495647 | 0.24638 | 0.232124 | 0.999971 | 1 | 0.831121 |
| Usp22 | Q5DU02 | -2.5 | -4.93 | 1.92 | 1.76 | -4.81 | -3.45 | 1.85 | 1.71 | -4.86 | -3.67 | 1.74 | 1.54 | -8.12 | -5.08 | 1.35 | 0.77 | -0.2375 | -0.375 | -1.8325 | 0.911133 | 0.861093 | 0.473759 | 0.999971 | 1 | 0.894044 |
| Usp24 | E9PV45 | 0.16 | 0.24 | 0.42 | 0.39 | 0.15 | 0.27 | 0.39 | 0.46 | 0.15 | 0.21 | 0.16 | 0.2 | 0.4 | 0.48 | 0.68 | 0.64 | 0.015 | -0.1225 | 0.2475 | 0.932124 | 0.458543 | 0.184454 | 0.999971 | 1 | 0.809025 |
| Usp25 | A0A5F8MP | -0.48 | 0.06 | 0.2 | 0.12 | 0.24 | 0.33 | 0.09 | 0.14 | -0.03 | -0.18 | 0.16 | 0.23 | 0.32 | 0.15 | 0.26 | 0.44 | 0.225 | 0.07 | 0.3175 | 0.305028 | 0.751378 | 0.163936 | 0.999971 | 1 | 0.800215 |
| Usp30 | Q3UN04 | 0.04 | -0.17 | -0.33 | -0.13 | -0.2 | -0.37 | 0.5 | 0.37 | 0.36 | 0.63 | -0.51 | -1.32 | 0.27 | 0.63 | -0.07 | -1.14 | 0.2225 | -0.0625 | 0.07 | 0.388801 | 0.883479 | 0.85282 | 0.999971 | 1 | 0.98187 |
| Usp31 | E9Q6Y8 | 0.21 | -0.36 | -0.47 | -0.59 | -0.58 | -0.31 | -0.05 | -0.78 | -0.46 | -0.16 | -0.45 | -0.77 | -0.46 | -0.49 | -0.83 | -0.94 | -0.1275 | -0.1575 | -0.3775 | 0.62758 | 0.524758 | 0.149925 | 0.999971 | 1 | 0.793248 |
| Usp32 | F8VPZ3 | 0.09 | -0.58 | -0.41 | -0.39 | -0.64 | -0.42 | 0.16 | -0.38 | -0.39 | -0.13 | -0.24 | -0.37 | -0.22 | -0.37 | -0.34 | -0.23 | 0.0025 | 0.04 | 0.0325 | 0.992027 | 0.845858 | 0.873818 | 0.999971 | 1 | 0.984109 |
| Usp33 | Q8R5K2-2 | -0.78 | -0.12 | -0.7 | -1.02 | -0.32 | 0.24 | -1.09 | -0.64 | -0.33 | -0.37 | -0.62 | -0.78 | -0.1 | 0.22 | -0.46 | -0.67 | 0.2025 | 0.13 | 0.4025 | 0.550984 | 0.600105 | 0.185571 | 0.999971 | 1 | 0.809025 |
| Usp34 | F6WJB7 | 0.28 | 0.72 | 0.75 | 0.6 | 1.04 | 0.73 | 0.47 | 0.51 | 0.66 | 0.95 | 0.19 | -0.02 | 0.44 | 1.07 | 0.5 | 0.75 | 0.1 | -0.1425 | 0.1025 | 0.645019 | 0.593591 | 0.647069 | 0.999971 | 1 | 0.933613 |
| Usp35 | M0QWN7 | 0.56 | 0.56 | 0.05 | 0.44 | 0.38 | 0.34 | 0.69 | 0.28 | 0.44 | 0.18 | 0.12 | 0.14 | -0.17 | 0.53 | 0.04 | -0.44 | 0.02 | -0.1825 | -0.4125 | 0.922417 | 0.36746 | 0.141317 | 0.999971 | 1 | 0.784967 |
| Usp38 | Q8BW70 | -0.51 | 0.79 | 1.04 | 0 | 0.83 | 0.63 | 0.44 | 0.79 | 0.74 | -0.02 | 1.1 | 1.08 | 0.5 | 0.51 | 0.4 | 0.52 | 0.3425 | 0.395 | 0.1525 | 0.353323 | 0.360975 | 0.666659 | 0.999971 | 1 | 0.937557 |
| Usp39 | Q3TIX9 | 0.79 | 0.94 | 0.85 | 0.79 | 0.81 | 0.38 | 0.42 | 0.3 | 0.64 | 0.93 | 0.38 | 0.63 | 0.29 | 0.88 | 0.46 | 0.8 | -0.365 | -0.1975 | -0.235 | 0.079264 | 0.301195 | 0.263252 | 0.999971 | 1 | 0.842533 |
| Usp4 | P35123 | 0.52 | 0.95 | 0.66 | 0.3 | 0.39 | 0.56 | 0.83 | 0.77 | 0.51 | 0.59 | 1.05 | 0.89 | 0.56 | 0.73 | 0.86 | 0.36 | 0.03 | 0.1525 | 0.02 | 0.889451 | 0.502029 | 0.927303 | 0.999971 | 1 | 0.99417 |
| Usp43 | A6PWR8 | 1.24 | -0.97 | -0.51 | -0.47 | -1.47 | -0.52 | 0.71 | -0.7 | -0.91 | 0.98 | -0.26 | -0.98 | -1.85 | -1.37 | -1.33 | -0.26 | -0.3175 | -0.115 | -1.025 | 0.602323 | 0.850459 | 0.086911 | 0.999971 | 1 | 0.761242 |
| Usp45 | Q8K387 | 4.4 | 0.24 | 0.97 | 1.32 | 0.28 | 0.88 | 1.4 | 1.08 | -0.44 | 2.66 | 0.73 | 0.17 | -2.29 | -0.47 | -0.03 | 0.21 | -0.8225 | -0.9525 | -2.3775 | 0.345873 | 0.363401 | 0.03381 | 0.999971 | 1 | 0.689803 |
| Usp46 | P62069 | -0.92 | -0.34 | -0.23 | -0.37 | -0.37 | -0.79 | -0.5 | -0.39 | -0.28 | -0.92 | -0.33 | -0.29 | -0.19 | -0.75 | -0.55 | -0.43 | -0.0475 | 0.01 | -0.015 | 0.832108 | 0.967608 | 0.948392 | 0.999971 | 1 | 0.997766 |
| Usp47 | A0A1L1SV7 | 0.94 | 0.47 | 1.12 | 0.89 | 0.4 | 0.78 | 1.2 | 1.06 | 0.56 | 0.87 | 0.98 | 0.87 | 0.49 | 0.62 | 1.22 | 0.99 | 0.005 | -0.035 | -0.025 | 0.984098 | 0.868071 | 0.919361 | 0.999971 | 1 | 0.992633 |
| Usp48 | A2ALR9 | 2.71 | 1.31 | 0.65 | 0.42 | 0.67 | 1.16 | -0.17 | 0.94 | 0.73 | 1.61 | 0.76 | 1.36 | 0.71 | 1.16 | 0.81 | 0.63 | -0.6225 | -0.1575 | -0.445 | 0.270585 | 0.762326 | 0.376753 | 0.999971 | 1 | 0.870465 |
| Usp5 | Q3U4W8 | -0.21 | -0.38 | 0.68 | 0.42 | -0.14 | -0.3 | 0.37 | 0.22 | -0.16 | -0.32 | 0.45 | 0.63 | -0.42 | -0.34 | 0.61 | 0.4 | -0.09 | 0.0225 | -0.065 | 0.767089 | 0.946946 | 0.854818 | 0.999971 | 1 | 0.982125 |
| Usp53 | P15975 | 2.51 | 0.17 | 0.42 | 0.63 | -0.28 | -0.02 | 0.86 | -0.26 | -0.51 | 1.54 | -0.37 | -0.24 | -1.31 | -1.52 | -0.91 | -0.39 | -0.8575 | -0.8275 | -1.965 | 0.144712 | 0.229056 | 0.005503 | 0.999971 | 1 | 0.582714 |
| Usp54 | Q8BL06 | -0.37 | -0.69 | -0.94 | -1 | -0.54 | 0.21 | -1.1 | -0.09 | -0.3 | -0.46 | -0.99 | -0.64 | 0.06 | -0.47 | -0.69 | -0.18 | 0.37 | 0.1525 | 0.43 | 0.267508 | 0.525817 | 0.110008 | 0.999971 | 1 | 0.78268 |
| Usp6nl | Q80XC3 | -2.78 | -1.92 | -0.49 | -1.32 | -2.82 | -2.13 | -1.03 | -1.03 | -2.91 | -1.79 | -0.45 | -0.47 | -3.74 | -2.71 | -0.77 | -1.23 | -0.125 | 0.2225 | -0.485 | 0.834095 | 0.749534 | 0.52525 | 0.999971 | 1 | 0.909237 |
| Usp7 | F8VPX1 | 0.34 | 0.55 | 0.74 | 0.68 | 0.64 | 0.5 | 0.53 | 0.62 | 0.66 | 0.18 | 0.54 | 0.87 | 0.88 | 0.37 | 0.56 | 0.92 | -0.005 | -0.015 | 0.105 | 0.977494 | 0.944012 | 0.619927 | 0.999971 | 1 | 0.929296 |
| Usp8 | Q80U87 | 2.38 | 0.54 | -0.18 | -0.25 | 0.3 | 0.89 | 0.39 | -0.39 | 0.25 | 1.42 | -0.36 | -0.5 | 0.07 | 0.3 | -0.4 | -0.58 | -0.325 | -0.42 | -0.775 | 0.59566 | 0.544504 | 0.211874 | 0.999971 | 1 | 0.823112 |
| Usp9x | Q4FE56 | -0.1 | 0.13 | 0.16 | 0.05 | 0.22 | 0.36 | 0.19 | 0.27 | 0.43 | 0.12 | 0.35 | 0.13 | 0.66 | 0.25 | 0.33 | 0.3 | 0.2 | 0.1975 | 0.325 | 0.253093 | 0.277438 | 0.105692 | 0.999971 | 1 | 0.78268 |
| Usp9x_1 | G3UY52 | 0.77 | 1.95 | -0.58 | -0.97 | 1.38 | 2.33 | -0.58 | -0.75 | 1.67 | 2.63 | -1.2 | -2.41 | 2.63 | 2.41 | -1.2 | -1.12 | 0.3025 | -0.12 | 0.3875 | 0.737452 | 0.921418 | 0.729817 | 0.999971 | 1 | 0.956041 |
| Ust | Q8BUB6 | -0.59 | 0.92 | -3.95 | -0.32 | 1.71 | 0.08 | -8.7 | -1.59 | 0.61 | -0.05 | -6.29 | -9.22 | 0.61 | 1.69 | -2.69 | 0.25 | -1.14 | -2.7525 | 0.95 | 0.610947 | 0.25691 | 0.453643 | 0.999971 | 1 | 0.887217 |
| Utp11 | Q9CZJ1 | -1.96 | -1.34 | -1.55 | -2.04 | -1.26 | -2.14 | -1.6 | -1.9 | -1.04 | -1.84 | -1.4 | -0.96 | -0.7 | -2.35 | -1.5 | -1.75 | -0.0025 | 0.4125 | 0.1475 | 0.992649 | 0.161304 | 0.691338 | 0.999971 | 1 | 0.946835 |
| Utp14a | Q640M1 | 0.56 | -1.19 | -1.58 | -1.35 | -1.23 | -1.58 | -1.88 | -1.75 | -1.35 | 0.18 | -1.75 | -2 | -1.9 | -1.37 | -1.73 | -1.76 | -0.72 | -0.34 | -0.8 | 0.155536 | 0.594285 | 0.116358 | 0.999971 | 1 | 0.78268 |
| Utp15 | Q8C7V3 | -0.37 | -1.41 | -0.61 | -0.56 | -1.61 | -1.4 | -0.79 | -0.97 | -1.53 | -1.01 | -0.78 | -0.82 | -1.8 | -1.64 | -0.48 | -1.45 | -0.455 | -0.2975 | -0.605 | 0.160688 | 0.329763 | 0.125362 | 0.999971 | 1 | 0.78268 |
| Utp18 | Q5SSI6 | 2.47 | -0.47 | 0.41 | 1.08 | 0.35 | 0.21 | 0.22 | -0.11 | -0.29 | 1.5 | -0.45 | -0.18 | -0.41 | -0.64 | -0.96 | 0.97 | -0.705 | -0.7275 | -1.1325 | 0.239992 | 0.314127 | 0.123866 | 0.999971 | 1 | 0.78268 |
| Utp20 | E9QK83 | -0.1 | -0.62 | -0.63 | -0.93 | -0.7 | -1.37 | -1.12 | -1 | -0.83 | -1.21 | -0.98 | -0.56 | -1.25 | -1.27 | -1.18 | -1.08 | -0.4775 | -0.325 | -0.625 | 0.082628 | 0.210516 | 0.018922 | 0.999971 | 1 | 0.635388 |
| Utp3 | Q9JI13-2 | -0.69 | -0.69 | -0.09 | -0.48 | -1.09 | -0.71 | 0.05 | -0.76 | -1.36 | -0.64 | 0.04 | -0.04 | 0.08 | -0.69 | -0.25 | -0.33 | -0.14 | -0.0125 | 0.19 | 0.632532 | 0.971343 | 0.444102 | 0.999971 | 1 | 0.883802 |
| Utp4 | Q8R2N2 | -0.62 | 0.5 | 0.86 | -0.95 | 1.07 | -0.5 | 0.72 | 0 | 1.39 | -0.72 | 0.59 | 0.78 | 2.17 | -0.22 | -0.4 | -0.74 | 0.375 | 0.5625 | 0.255 | 0.475235 | 0.339604 | 0.723298 | 0.999971 | 1 | 0.955258 |
| Utp6 | Q8VCY6 | -1.25 | -0.63 | 0.33 | 0.04 | -0.12 | -0.7 | -0.32 | 0.96 | -0.52 | -1.03 | -0.6 | 0.26 | 0.25 | -0.03 | -0.2 | 0.09 | 0.3325 | -0.095 | 0.405 | 0.48429 | 0.822091 | 0.276672 | 0.999971 | 1 | 0.843794 |
| Utrn | E9Q6R7 | 0.74 | 0.15 | -0.06 | 0.36 | 0.61 | 0.21 | 0.18 | -0.51 | 0.17 | 0.88 | -0.17 | 0.33 | -0.46 | 0.36 | 0.14 | 0.12 | -0.175 | 0.005 | -0.2575 | 0.560342 | 0.98617 | 0.346858 | 0.999971 | 1 | 0.864164 |
| Uvrag | Q8K245 | 0.28 | -0.16 | 0.56 | 0.03 | -0.27 | 0.04 | 0.42 | 0.05 | -0.2 | 0.28 | 0.2 | 0.26 | -0.29 | -0.04 | 0.18 | 0.3 | -0.1175 | -0.0425 | -0.14 | 0.629677 | 0.8525 | 0.5601 | 0.999971 | 1 | 0.915694 |
| Uxs1 | Q91XL3 | -0.95 | -0.52 | -0.12 | 0.43 | -0.86 | -1.52 | -0.19 | 0.94 | -1.46 | -1.62 | 0.37 | 0.08 | -0.95 | -1.16 | 0.2 | -0.15 | -0.1175 | -0.3675 | -0.225 | 0.831573 | 0.507509 | 0.591189 | 0.999971 | 1 | 0.922461 |
| Vac14 | Q80WQ2 | -0.73 | 0.02 | 0.44 | 0.33 | 0.37 | 0.02 | -0.21 | 0.18 | 0.27 | -0.38 | 0.28 | 0.47 | 0.74 | 0.15 | 0.68 | 0.36 | 0.075 | 0.145 | 0.4675 | 0.802495 | 0.655705 | 0.151889 | 0.999971 | 1 | 0.795578 |
| Vamp1 | D3YTU0 | -2.37 | -3.08 | -0.55 | -0.29 | -3.62 | -3.17 | -1.14 | -0.49 | -3.81 | -3.53 | -0.45 | -0.24 | -4.49 | -3.24 | -0.82 | 0.02 | -0.5325 | -0.435 | -0.56 | 0.564378 | 0.68278 | 0.617693 | 0.999971 | 1 | 0.928581 |
| Vamp2 | B0QZN5 | -2.62 | -1.56 | -0.9 | -0.67 | -1.64 | -1.55 | -1.09 | -0.84 | -1.35 | -2.13 | -0.88 | -1.32 | -1.44 | -1.58 | -0.98 | -0.5 | 0.1575 | 0.0175 | 0.3125 | 0.725642 | 0.970655 | 0.510366 | 0.999971 | 1 | 0.905815 |
| Vamp3 | P63024 | 0.06 | -0.4 | -1.97 | -1.84 | -0.36 | -1.37 | -1.84 | 1.6 | -0.34 | -0.21 | -1.24 | -0.15 | -0.51 | -1.19 | -1.79 | -2.09 | 0.545 | 0.5525 | -0.3575 | 0.513422 | 0.311756 | 0.533531 | 0.999971 | 1 | 0.910069 |
| Vamp4 | O70480 | 0.11 | -0.78 | 0.69 | 0.1 | -0.52 | -0.23 | 1.63 | 0.67 | -0.46 | -0.06 | 1.83 | 2.32 | -2.01 | 0.05 | 0.86 | 0.55 | 0.3575 | 0.8775 | -0.1675 | 0.503788 | 0.221285 | 0.79655 | 0.999971 | 1 | 0.970479 |
| Vamp7 | P70280 | -0.57 | -0.63 | -0.3 | -0.56 | -0.36 | -0.14 | -0.14 | -0.02 | -0.31 | -0.65 | 0.11 | -0.68 | -0.05 | -0.22 | 0.43 | 0.16 | 0.35 | 0.1325 | 0.595 | 0.079107 | 0.572054 | 0.018505 | 0.999971 | 1 | 0.635388 |
| Vamp8 | O70404 | 1.02 | 0.07 | 1.69 | 2.02 | 0.08 | 0.94 | 1.01 | 1.47 | 0.19 | 1.38 | 1.8 | 1.24 | 0.05 | 0.32 | 1.77 | 0.08 | -0.325 | -0.0475 | -0.645 | 0.506421 | 0.925911 | 0.257509 | 0.999971 | 1 | 0.839012 |
| Vangl2 | D3YY75 | -1.03 | -0.79 | -1.22 | -1.57 | -0.73 | -0.73 | -1.06 | -1.59 | -0.24 | -0.86 | -1.27 | -1.06 | 0.27 | -0.68 | -1.38 | -1.54 | 0.125 | 0.295 | 0.32 | 0.654334 | 0.320682 | 0.455486 | 0.999971 | 1 | 0.887217 |
| Vapa | Q9WV55 | -0.17 | -0.76 | -0.68 | -0.48 | -0.61 | -0.69 | -0.03 | -0.47 | -0.06 | -0.48 | -0.2 | -0.08 | -0.2 | -0.85 | -0.39 | -0.76 | 0.0725 | 0.3175 | -0.0275 | 0.756648 | 0.156237 | 0.907642 | 0.999971 | 1 | 0.991405 |
| Vapb | Q8BH80 | 0.42 | -0.53 | -0.75 | -0.42 | -0.74 | -0.23 | -0.68 | -0.45 | -0.41 | 0.31 | -0.9 | -0.6 | -0.67 | -0.2 | -0.68 | -0.45 | -0.205 | -0.08 | -0.18 | 0.489293 | 0.822139 | 0.541941 | 0.999971 | 1 | 0.911119 |
| Vars1 | Q9Z1Q9 | 0.64 | 1.45 | 1.68 | 1.51 | 1.53 | 1.28 | 1.52 | 1.44 | 1.61 | 0.86 | 1.62 | 1.56 | 1.4 | 1.57 | 1.59 | 1.84 | 0.1225 | 0.0925 | 0.28 | 0.641657 | 0.760657 | 0.313907 | 0.999971 | 1 | 0.850589 |
| Vars2 | Q3U2A8 | -0.44 | -0.13 | -0.39 | -0.5 | -0.22 | -0.33 | -0.58 | -0.22 | -0.28 | -0.44 | -0.41 | -0.05 | 0.43 | -0.49 | -0.43 | -0.18 | 0.0275 | 0.07 | 0.1975 | 0.883441 | 0.707652 | 0.443066 | 0.999971 | 1 | 0.883386 |
| Vash1 | Q8C1W1 | 1.41 | 0.19 | -0.09 | -0.16 | 0.45 | 0.52 | 0.12 | 0.68 | 0.34 | 1.77 | 0.16 | 0.13 | -0.27 | 0.1 | 0.1 | 0.33 | 0.105 | 0.2625 | -0.2725 | 0.778083 | 0.602341 | 0.47371 | 0.999971 | 1 | 0.894044 |
| Vasn | Q9CZT5 | -2.69 | 0.59 | 0.8 | -0.87 | -0.23 | 0.5 | 0.14 | 0.73 | 0.77 | -2.1 | 0.56 | 1.53 | 0.96 | -0.29 | 0.33 | 0.28 | 0.8275 | 0.7325 | 0.8625 | 0.287346 | 0.477483 | 0.27613 | 0.999971 | 1 | 0.843794 |
| Vasp | P70460 | 0.61 | 0.4 | -0.64 | 0.88 | 1.07 | 0.13 | -0.16 | 0.12 | 0.6 | 0.68 | 0.39 | 0.44 | 0.77 | 0.85 | 0.28 | 0.25 | -0.0225 | 0.215 | 0.225 | 0.955884 | 0.527894 | 0.536855 | 0.999971 | 1 | 0.910077 |
| Vat1 | Q62465 | -0.46 | -1.09 | -1.14 | -1.46 | -0.39 | -0.72 | -1.12 | -0.68 | -0.24 | -0.32 | -1.27 | -0.71 | -0.03 | -0.48 | -0.2 | -0.96 | 0.31 | 0.4025 | 0.62 | 0.277052 | 0.22621 | 0.065045 | 0.999971 | 1 | 0.727266 |
| Vat1l | Q80TB8 | -0.06 | -1.36 | -1.12 | -0.8 | -0.55 | -0.81 | -0.83 | -0.5 | -0.07 | -0.83 | -1.57 | -0.79 | -0.46 | -0.19 | -0.01 | -1.24 | 0.1625 | 0.02 | 0.36 | 0.59454 | 0.960056 | 0.355234 | 0.999971 | 1 | 0.868793 |
| Vav1 | Q8VDU4 | 0.28 | 0 | 0.65 | 0.37 | 0.4 | 0.09 | -0.09 | 1.86 | -0.37 | -0.48 | 1.12 | 0.73 | 0.11 | 0.49 | 0.69 | 0.26 | 0.24 | -0.075 | 0.0625 | 0.585697 | 0.852235 | 0.782649 | 0.999971 | 1 | 0.965859 |
| Vav2 | Q60992 | -0.23 | -0.97 | -0.18 | 1.23 | -0.94 | -0.97 | -0.57 | 0.13 | -1.04 | -0.24 | 1.02 | 0.54 | -0.64 | -1.12 | 0.74 | -0.21 | -0.55 | 0.1075 | -0.27 | 0.278103 | 0.855966 | 0.630294 | 0.999971 | 1 | 0.9327 |
| Vav3 | Q9R0C8 | -1.45 | -0.54 | -0.56 | -0.47 | -0.26 | -0.8 | -0.14 | -0.48 | -0.61 | -1.26 | -0.78 | -0.72 | -0.45 | -0.54 | -0.86 | -0.71 | 0.335 | -0.0875 | 0.115 | 0.261597 | 0.759694 | 0.671538 | 0.999971 | 1 | 0.939561 |
| Vbp1 | P61759 | 0.53 | 0.96 | 0.64 | 0.68 | 0.93 | 1.02 | 1.72 | 0.81 | 0.95 | 0.51 | 0.9 | 1.05 | 1.6 | 0.72 | 0.91 | 0.11 | 0.4175 | 0.15 | 0.1325 | 0.123676 | 0.464622 | 0.682449 | 0.999971 | 1 | 0.943204 |
| Vcam1 | Q3UPN1 | -0.33 | -0.07 | -0.42 | -0.52 | -0.08 | -0.27 | -0.23 | -0.99 | 0.11 | -0.1 | -0.19 | -0.07 | 0.83 | 0.05 | -0.27 | -0.36 | -0.0575 | 0.2725 | 0.3975 | 0.81965 | 0.162793 | 0.20407 | 0.999971 | 1 | 0.81617 |
| Vcan | E9PYH0 | 0.16 | 0.67 | 0.36 | 0.37 | 0.53 | 0.4 | 0.74 | 0.36 | 0.74 | -0.07 | 0.15 | 0.33 | 0.47 | -0.13 | 0.31 | -0.12 | 0.1175 | -0.1025 | -0.2575 | 0.554275 | 0.663268 | 0.273018 | 0.999971 | 1 | 0.843794 |
| Vcl | Q64727 | 0.68 | 0.18 | 0.14 | 0.72 | 1.47 | 0.8 | 0.59 | 0.7 | 0.51 | 1.14 | 0.64 | 0.94 | 0.03 | 0.53 | 0.33 | 0.51 | 0.46 | 0.3775 | -0.08 | 0.116894 | 0.144253 | 0.731293 | 0.999971 | 1 | 0.956041 |
| Vcp | Q01853 | -1.14 | -0.04 | 0.12 | 0.02 | 0.1 | -0.24 | 0.14 | 0.06 | 0.25 | -0.58 | 0.29 | 0.4 | 0.1 | -0.14 | 0.22 | 0.28 | 0.275 | 0.35 | 0.375 | 0.389076 | 0.347966 | 0.252194 | 0.999971 | 1 | 0.839012 |
| Vcpip1 | A0A0R4J0N | 0.3 | 0.18 | 0.07 | 0.03 | 0.03 | 0.32 | 0.56 | -0.27 | 0.09 | 0.31 | -0.1 | -0.21 | -0.07 | 0.15 | -0.24 | -0.23 | 0.015 | -0.1225 | -0.2425 | 0.947618 | 0.525239 | 0.211007 | 0.999971 | 1 | 0.822884 |
| Vcpkmt | A0A571BD | -1.38 | -2.36 | -4.45 | -3.81 | -0.53 | -1 | -2.26 | -2.34 | -1.2 | -1.66 | -2.24 | 0.14 | -1.34 | -4.11 | -7.67 | 0.93 | 1.4675 | 1.76 | -0.0475 | 0.076391 | 0.047623 | 0.978381 | 0.999971 | 1 | 0.99925 |
| Vdac1 | Q60932-2 | -0.75 | -0.25 | -0.98 | -0.97 | -0.17 | -0.53 | -0.63 | -0.78 | -0.18 | -0.74 | -0.77 | -0.71 | 0.37 | -0.61 | -0.69 | -0.81 | 0.21 | 0.1375 | 0.3025 | 0.401372 | 0.582491 | 0.361334 | 0.999971 | 1 | 0.869526 |
| Vdac2 | Q60930 | -0.9 | -0.37 | 0.07 | -0.54 | -0.21 | -0.52 | 0.14 | -0.47 | -0.07 | -0.98 | -0.23 | -0.04 | -0.27 | -0.83 | -0.38 | -0.67 | 0.17 | 0.105 | -0.1025 | 0.534707 | 0.731243 | 0.69723 | 0.999971 | 1 | 0.94731 |
| Vdac2_1 | G3UX26 | 1.2 | 1.73 | 1.8 | 2.35 | 1.2 | 1.13 | 0.37 | 2.08 | 0.79 | 1.05 | 1.43 | 1.48 | 1.16 | 1.35 | 1.36 | 2.44 | -0.575 | -0.5825 | -0.1925 | 0.177525 | 0.075631 | 0.600925 | 0.999971 | 1 | 0.92426 |
| Vdac3 | J3QMG3 | -2.12 | 0.17 | -0.35 | -0.94 | 0.23 | -0.06 | 0.12 | -0.52 | 0.39 | -1.96 | -0.29 | -0.04 | 0.66 | -0.7 | -0.34 | -0.68 | 0.7525 | 0.335 | 0.545 | 0.14552 | 0.609161 | 0.327905 | 0.999971 | 1 | 0.856472 |
| Vezt | D3Z4E6 | 1.58 | -0.11 | -0.67 | -0.7 | 0.23 | -0.3 | -0.71 | -0.8 | -0.07 | 0.82 | -0.62 | -0.25 | 0.03 | -0.27 | -0.75 | -0.9 | -0.42 | -0.055 | -0.4975 | 0.443483 | 0.922862 | 0.362803 | 0.999971 | 1 | 0.869526 |
| Vgf | Q0VGU4 | 0.66 | -1.19 | -0.25 | -0.27 | -0.53 | -0.75 | -0.18 | -0.59 | -0.43 | -0.5 | -0.34 | -0.3 | -1.45 | -1.24 | -0.41 | -0.66 | -0.25 | -0.13 | -0.6775 | 0.517973 | 0.726011 | 0.138407 | 0.999971 | 1 | 0.784948 |
| Vhl | P40338 | 1.49 | 0.92 | 0.82 | 1.16 | -0.32 | 0.78 | 0.6 | 1.79 | 0.24 | 1.25 | 1.2 | 1.46 | -0.18 | 1.42 | 1.01 | 0.77 | -0.385 | -0.06 | -0.3425 | 0.383806 | 0.848251 | 0.357108 | 0.999971 | 1 | 0.868793 |
| Vill | G5E8H0 | 2.25 | 1.23 | -0.27 | 0.03 | 1.38 | 0.08 | 0.35 | 0.66 | 1.85 | 0.11 | -0.23 | -0.16 | 1.83 | 0.36 | 0.46 | 0.42 | -0.1925 | -0.4175 | -0.0425 | 0.7439 | 0.550273 | 0.945251 | 0.999971 | 1 | 0.997427 |
| Vim | P20152 | -0.59 | -0.42 | -0.05 | 0.13 | 1.91 | 0.16 | 0.32 | -0.72 | 0.44 | 0.59 | 0.39 | 1.97 | -0.97 | -0.07 | 0.46 | -0.43 | 0.65 | 1.08 | -0.02 | 0.238149 | 0.022177 | 0.953133 | 0.999971 | 1 | 0.998056 |
| Vip | A0A0R4J00 | -1.4 | -1.6 | -0.59 | -1.22 | -0.84 | -1.04 | -0.71 | -0.72 | -1.98 | -1.63 | -0.65 | -1.22 | -1.4 | -1.04 | -0.69 | -0.09 | 0.375 | -0.1675 | 0.3975 | 0.168612 | 0.637291 | 0.273126 | 0.999971 | 1 | 0.843794 |
| Vipas39 | Q8BGQ1 | -0.93 | -0.18 | 0.16 | -0.29 | 0.55 | -0.22 | 0.11 | 0 | 0.55 | -0.59 | 0.3 | 0.59 | 1.92 | 0.18 | 0.56 | 0.44 | 0.42 | 0.5225 | 1.085 | 0.173484 | 0.162353 | 0.030168 | 0.999971 | 1 | 0.678474 |
| Virma | E9PZY8 | 4.64 | 0.93 | 0.25 | 0.48 | 0.33 | 1.31 | 0.6 | 0.03 | 0.45 | 3.37 | 0.16 | 0.36 | -1.49 | -0.69 | -0.06 | -0.01 | -1.0075 | -0.49 | -2.1375 | 0.306257 | 0.671733 | 0.052344 | 0.999971 | 1 | 0.723916 |
| Vkorc1 | Q9CRC0 | 1.9 | -0.22 | 0.37 | 1.13 | -0.37 | -0.3 | 1.53 | 1.02 | -0.42 | 1.41 | 0.22 | -1.2 | -1.34 | -0.29 | -0.52 | -1.58 | -0.325 | -0.7925 | -1.7275 | 0.593604 | 0.247309 | 0.008239 | 0.999971 | 1 | 0.602121 |
| Vkorc1l1 | Q6TEK5 | -0.51 | 0.23 | -0.14 | 0.01 | 0.12 | 0.47 | 1.36 | 0.05 | 0.53 | 0.11 | 0.22 | -0.26 | 1.21 | 0.88 | -0.03 | -0.26 | 0.6025 | 0.2525 | 0.5525 | 0.100635 | 0.328539 | 0.165529 | 0.999971 | 1 | 0.800215 |
| Vldlr | Q91YY0 | -0.5 | -0.09 | 0.06 | -0.32 | 0 | 0.29 | 0.14 | 0.43 | 0 | -0.34 | 0.33 | 0.17 | 0.46 | -0.07 | 0.3 | -0.01 | 0.4275 | 0.2525 | 0.3825 | 0.06483 | 0.282252 | 0.11081 | 0.999971 | 1 | 0.78268 |
| Vma21 | Q78T54 | -1.63 | 0.03 | -0.14 | -0.99 | -0.37 | -0.16 | -0.45 | -1.01 | -0.23 | -1.34 | -0.17 | -0.13 | 0.18 | -1.7 | -0.67 | -0.81 | 0.185 | 0.215 | -0.0675 | 0.651222 | 0.63925 | 0.893854 | 0.999971 | 1 | 0.988303 |
| Vmp1 | Q99KU0 | -1.29 | -0.31 | 0.02 | -0.08 | 0.09 | 0.15 | -0.27 | 0.5 | 0.16 | -1.15 | 0.54 | 0.57 | 0.5 | -0.36 | 0.6 | 0.89 | 0.5325 | 0.445 | 0.8225 | 0.139698 | 0.358415 | 0.05934 | 0.999971 | 1 | 0.725348 |
| Vopp1 | A0A0N4SV | 1.44 | -1 | 1.39 | 1.23 | -1.19 | -0.31 | 2.15 | 0.55 | 0.02 | 1.34 | 1.44 | 1.11 | -1.53 | -1.02 | 0.28 | -0.86 | -0.465 | 0.2125 | -1.5475 | 0.578356 | 0.731647 | 0.036472 | 0.999971 | 1 | 0.700096 |
| Vps11 | Q91W86 | 0.16 | 0.29 | 0.73 | -0.07 | 0.24 | 0.47 | 0.87 | 0.22 | 0.3 | 0.47 | 0.84 | 0.78 | 0.67 | 0.31 | 0.39 | 0.37 | 0.1725 | 0.32 | 0.1575 | 0.500669 | 0.207197 | 0.493985 | 0.999971 | 1 | 0.901531 |
| Vps13a | A0A494B9( | -1.7 | 0.3 | -0.3 | 0.5 | 0.86 | -0.09 | -0.65 | -0.47 | 0.34 | -1.09 | 1.36 | 0.24 | -0.42 | -0.08 | -0.63 | 0.55 | 0.2125 | 0.5125 | 0.155 | 0.700807 | 0.436016 | 0.765159 | 0.999971 | 1 | 0.962405 |
| Vps13a_1 | Q5H8C4 | -0.59 | -0.17 | -0.16 | -0.11 | -0.57 | -0.05 | -0.34 | -0.25 | -0.53 | -0.43 | -0.31 | -0.53 | -0.62 | -0.18 | -0.3 | -0.03 | -0.045 | -0.1925 | -0.025 | 0.828438 | 0.319412 | 0.907667 | 0.999971 | 1 | 0.991405 |
| Vps13b | Q80TY5 | -0.3 | 0.44 | 1.55 | 1.3 | -0.12 | -0.66 | 1.2 | 1.76 | -1.21 | -0.39 | 2.02 | 0.82 | -0.17 | -2.43 | 0.42 | 0.89 | -0.2025 | -0.4375 | -1.07 | 0.751881 | 0.561817 | 0.185788 | 0.999971 | 1 | 0.809025 |
| Vps13c | Q8BX70 | -0.54 | -0.92 | -0.47 | -0.47 | -0.92 | -0.62 | -0.57 | -0.46 | -0.89 | -0.81 | -0.51 | -0.59 | -0.98 | -0.66 | -0.39 | -0.13 | -0.0425 | -0.1 | 0.06 | 0.833733 | 0.613456 | 0.805712 | 0.999971 | 1 | 0.971844 |
| Vps13d | B1ART2 | -0.85 | -0.6 | -0.38 | -0.47 | -0.39 | -0.22 | -0.13 | -0.23 | -0.73 | -0.67 | -0.35 | -0.21 | -0.16 | -0.33 | -0.36 | -0.16 | 0.3325 | 0.085 | 0.3225 | 0.102076 | 0.685364 | 0.112313 | 0.999971 | 1 | 0.78268 |
| Vps16 | G3X8X7 | -0.23 | 0.49 | 0.42 | 0.38 | 0.25 | 0.32 | 0.04 | 0.47 | 0.88 | 0.26 | 0.53 | 0.08 | 1.09 | 0.13 | 0.37 | 0.41 | 0.005 | 0.1725 | 0.235 | 0.982496 | 0.515332 | 0.412569 | 0.999971 | 1 | 0.875881 |
| Vps18 | Q8R307 | -0.19 | 0.3 | -0.19 | -0.25 | 0.3 | 0.24 | 0.15 | 0.04 | 0.75 | -0.03 | 0.24 | 0.37 | 0.92 | 0.02 | -0.04 | -0.16 | 0.265 | 0.415 | 0.2675 | 0.205923 | 0.108743 | 0.370336 | 0.999971 | 1 | 0.869526 |
| Vps25 | Q9CQ80 | -0.17 | -0.75 | -0.51 | -0.34 | -0.43 | -0.56 | -0.87 | -0.07 | -0.43 | -0.68 | -0.82 | -0.68 | -0.55 | -0.23 | -0.65 | -0.37 | -0.04 | -0.21 | -0.0075 | 0.867436 | 0.311498 | 0.971237 | 0.999971 | 1 | 0.99925 |
| Vps26a | P40336 | -0.49 | 0.29 | -0.32 | -0.82 | 0.99 | -0.21 | 0.24 | -0.54 | 1.24 | -0.68 | 0.51 | 0.57 | 2.1 | 0 | -0.29 | -0.27 | 0.455 | 0.745 | 0.72 | 0.260737 | 0.116245 | 0.227567 | 0.999971 | 1 | 0.828598 |
| Vps26b | Q8C0E2 | -0.37 | -0.71 | -0.63 | -0.43 | -0.46 | -0.39 | -0.69 | -0.41 | -0.46 | -0.52 | -0.38 | -0.82 | -0.49 | -0.4 | -0.55 | -0.24 | 0.0475 | -0.01 | 0.115 | 0.79462 | 0.957654 | 0.533312 | 0.999971 | 1 | 0.910069 |
| Vps26c | O35075 | 0.53 | 0.37 | 0.85 | 0.85 | 0.35 | 0.65 | 0.35 | 0.83 | 0.13 | 0.62 | 0.93 | 0.03 | -0.03 | 1.01 | 0.71 | 0.95 | -0.105 | -0.2225 | 0.01 | 0.628242 | 0.408103 | 0.971756 | 0.999971 | 1 | 0.99925 |
| Vps28 | Q9D1C8 | -0.06 | -0.49 | -0.22 | -0.15 | -0.58 | -0.78 | -0.19 | -0.09 | -0.81 | -0.4 | 0.11 | -0.19 | -0.36 | -0.54 | 0.02 | 0.31 | -0.18 | -0.0925 | 0.0875 | 0.435922 | 0.704619 | 0.720554 | 0.999971 | 1 | 0.954713 |
| Vps29 | D3Z645 | -0.3 | -0.67 | -1.19 | -1.65 | -0.71 | -0.61 | -0.17 | -1.32 | 0.36 | -0.91 | -1.06 | -0.83 | -0.56 | -0.73 | -1.1 | -1.59 | 0.25 | 0.3425 | -0.0425 | 0.502575 | 0.423466 | 0.907111 | 0.999971 | 1 | 0.991405 |
| Vps33a | Q9D2N9 | -0.65 | -0.2 | -0.26 | -0.01 | -0.04 | -0.32 | -0.42 | -0.08 | -0.1 | -0.63 | -0.18 | -0.25 | -0.22 | -0.61 | 0 | 0.05 | 0.065 | -0.01 | 0.085 | 0.75968 | 0.9636 | 0.720576 | 0.999971 | 1 | 0.954713 |
| Vps33b | P59016 | -0.66 | -0.68 | -0.55 | -0.61 | -0.8 | -0.61 | -0.58 | -0.61 | -0.73 | -0.66 | -0.65 | -0.51 | -0.59 | -0.58 | -0.28 | -0.52 | -0.025 | -0.0125 | 0.1325 | 0.879397 | 0.937663 | 0.448345 | 0.999971 | 1 | 0.885289 |
| Vps35 | Q9EQH3 | -0.35 | -0.05 | 0.67 | -0.03 | -0.1 | -0.08 | 1.05 | 0.02 | 0.04 | -0.08 | 0.69 | -0.38 | 0.17 | -0.16 | 0.39 | 0.22 | 0.1625 | 0.0075 | 0.095 | 0.640711 | 0.981009 | 0.72229 | 0.999971 | 1 | 0.955258 |
| Vps35l | Q8BWQ6 | 0 | 0.52 | 0.31 | 0.25 | 0.34 | 0.5 | 0.27 | 0.38 | 0.41 | 0.45 | 0.26 | 0.16 | 0.47 | 0.73 | 0.21 | 0.49 | 0.1025 | 0.05 | 0.205 | 0.587691 | 0.791632 | 0.33312 | 0.999971 | 1 | 0.857921 |
| Vps36 | Q91XD6 | -0.22 | 0.02 | -0.09 | -0.33 | -0.04 | 0.14 | 0.17 | -0.03 | 0.05 | 0.16 | -0.46 | -0.13 | -0.08 | 0.02 | -0.21 | -0.21 | 0.215 | 0.06 | 0.035 | 0.244627 | 0.770453 | 0.844144 | 0.999971 | 1 | 0.979814 |
| Vps37a | Q8CHS8 | -0.85 | -1.22 | -1.26 | -0.74 | -1.1 | -1.4 | -1.02 | -0.65 | -1.09 | -1.47 | -0.84 | -0.73 | -1.44 | -1.12 | -0.53 | -0.28 | -0.025 | -0.015 | 0.175 | 0.915814 | 0.950123 | 0.568653 | 0.999971 | 1 | 0.91816 |
| Vps37b | Q8R0J7 | -0.31 | -0.25 | -0.58 | -0.46 | -0.55 | -0.82 | -0.71 | -0.33 | -0.56 | -0.34 | -0.78 | -0.5 | -0.28 | -0.08 | -0.48 | -0.83 | -0.2025 | -0.145 | -0.0175 | 0.311288 | 0.440486 | 0.936807 | 0.999971 | 1 | 0.995618 |
| Vps37c | Q8R105 | -0.71 | -0.73 | -0.48 | -0.16 | -0.76 | -0.85 | -0.41 | -0.08 | -0.87 | -0.84 | -0.09 | -0.45 | -0.56 | -0.71 | -0.6 | -0.23 | -0.005 | -0.0425 | -0.005 | 0.983959 | 0.866058 | 0.981506 | 0.999971 | 1 | 0.99925 |
| Vps37d | Q810I0 | -0.55 | -5.4 | -0.53 | -1.24 | -3.58 | -4.65 | -0.43 | -0.86 | -3.4 | -2.31 | -0.41 | -0.49 | -4.53 | -4.19 | -0.84 | -0.97 | -0.45 | 0.2775 | -0.7025 | 0.745071 | 0.82209 | 0.609877 | 0.999971 | 1 | 0.92673 |
| Vps39 | Q8R5L3-2 | -0.26 | -0.06 | 0.05 | 0.02 | -0.1 | -0.09 | -0.17 | 0.13 | -0.16 | -0.06 | -0.05 | 0.04 | 0.29 | -0.02 | 0.14 | 0.22 | 0.005 | 0.005 | 0.22 | 0.977521 | 0.9763 | 0.238907 | 0.999971 | 1 | 0.833594 |
| Vps41 | Q5KU39 | -0.86 | 0.19 | -0.28 | -0.53 | 0.4 | -0.26 | -0.27 | -0.32 | 0.61 | -0.55 | -0.24 | 0.07 | 0.94 | -0.53 | -0.28 | -0.29 | 0.2575 | 0.3425 | 0.33 | 0.387454 | 0.315865 | 0.401404 | 0.999971 | 1 | 0.871816 |
| Vps45 | P97390 | -0.01 | 0.04 | -0.25 | -0.24 | 0 | 0.24 | -0.4 | -0.3 | -0.09 | -0.07 | -0.29 | -0.52 | 0.42 | 0.23 | -0.43 | 0.1 | -3.93E-17 | -0.1275 | 0.195 | 1 | 0.509214 | 0.414511 | 1 | 1 | 0.876329 |
| Vps4a | Q8VEJ9 | -0.14 | -0.22 | 0.16 | 0.09 | -0.06 | 0.05 | 0.44 | 0.09 | -0.13 | -0.02 | 0.02 | 0.29 | -0.11 | 0.13 | 0.24 | 0.1 | 0.1575 | 0.0675 | 0.1175 | 0.438551 | 0.722429 | 0.536089 | 0.999971 | 1 | 0.910077 |
| Vps4b | P46467 | 0.64 | 0.54 | 0.7 | 0.49 | 0.66 | 1.29 | 1.07 | 0.66 | 0.74 | 0.75 | 0.84 | 0.92 | 1.02 | 0.4 | 0.63 | 0.41 | 0.3275 | 0.22 | 0.0225 | 0.149275 | 0.199258 | 0.913305 | 0.999971 | 1 | 0.992618 |
| Vps50 | Q8CI71 | -1.41 | -0.63 | -0.2 | -0.23 | -0.62 | -0.91 | -0.12 | -0.42 | -0.68 | -1.28 | 0.19 | 0.16 | -1.29 | -0.84 | -0.25 | -0.24 | 0.1 | 0.215 | -0.0375 | 0.760377 | 0.619585 | 0.919036 | 0.999971 | 1 | 0.992633 |
| Vps51 | Q3UVL4 | 0.19 | 0.69 | -0.25 | -0.33 | 0.49 | 0.44 | -0.38 | -0.48 | 0.66 | 0.68 | -0.2 | -0.38 | 0.18 | 0.38 | -0.55 | -0.32 | -0.0575 | 0.115 | -0.1525 | 0.86757 | 0.74817 | 0.637125 | 0.999971 | 1 | 0.93301 |
| Vps52 | Q8C754 | 2.22 | 0.19 | -0.22 | -0.3 | 0.16 | 0 | -0.34 | -0.41 | 0.12 | 0.9 | -0.2 | -0.29 | 0.04 | 0.08 | -0.46 | -0.04 | -0.62 | -0.34 | -0.5675 | 0.283469 | 0.573191 | 0.322625 | 0.999971 | 1 | 0.85548 |
| Vps53 | Q8CCB4 | 0.06 | 0.16 | 0.25 | -0.18 | 0.13 | -0.14 | 0.3 | -0.37 | 0.13 | -0.17 | 0.43 | 0.18 | -0.47 | -0.4 | 0.05 | -0.27 | -0.0925 | 0.07 | -0.345 | 0.674125 | 0.732761 | 0.118084 | 0.999971 | 1 | 0.78268 |
| Vps54 | Q5SRW8 | 1.96 | 0.83 | -1.8 | -0.47 | 0.94 | 1.53 | 0.04 | 0.84 | 1.14 | 1.92 | 1.26 | 1.11 | 0.04 | 0.93 | -1.7 | 0.3 | 0.7075 | 1.2275 | -0.2375 | 0.376447 | 0.13216 | 0.789008 | 0.999971 | 1 | 0.96741 |
| Vps8 | D3YUP0 | 0.09 | 0.01 | -0.03 | -0.24 | -0.34 | 0.03 | -0.11 | -0.16 | -0.03 | -0.12 | -0.14 | -0.19 | -0.39 | -0.22 | 0.22 | -0.05 | -0.1025 | -0.0775 | -0.0675 | 0.575211 | 0.645252 | 0.741494 | 0.999971 | 1 | 0.96105 |
| Vps9d1 | Q8C190 | 0.91 | 0.3 | -0.43 | 0.28 | 0.07 | 0.33 | -0.54 | -0.3 | 0.12 | 0.63 | -0.21 | -0.25 | 0.57 | 0.73 | -0.43 | 0.02 | -0.375 | -0.1925 | -0.0425 | 0.278313 | 0.572815 | 0.908709 | 0.999971 | 1 | 0.991707 |
| Vrk1 | A0A1Y7VM | 2.42 | 0.79 | 0.77 | 1.05 | 0.74 | 0.26 | 0.06 | 0.85 | 0.17 | 0.6 | 0.2 | 0.56 | 0.96 | 0.41 | 1.29 | 0.98 | -0.78 | -0.875 | -0.3475 | 0.086858 | 0.050386 | 0.409799 | 0.999971 | 1 | 0.875847 |
| Vrk2 | Q8BN21-2 | 1.01 | 0.24 | -0.41 | 0 | -0.43 | 0.23 | -1.41 | 0.27 | -1.25 | 0.54 | -0.76 | -0.99 | -0.99 | 0.15 | -1.02 | -0.06 | -0.545 | -0.825 | -0.69 | 0.256762 | 0.105118 | 0.118408 | 0.999971 | 1 | 0.78268 |
| Vrk3 | Q8K3G5 | -0.4 | 0.81 | 1.44 | 0.74 | 0.89 | 0.77 | 0.04 | 0.99 | 0.01 | -0.4 | 0.19 | 1.32 | 0.76 | 0.41 | 0.79 | 1.32 | 0.025 | -0.3675 | 0.1725 | 0.952164 | 0.46551 | 0.673405 | 0.999971 | 1 | 0.94048 |
| Vsnl1 | P62761 | -0.56 | -0.57 | -0.79 | -1.01 | -0.71 | -0.48 | -0.91 | -0.85 | -0.77 | -0.81 | -0.77 | -0.73 | -0.5 | -0.59 | -0.97 | -0.71 | -0.005 | -0.0375 | 0.04 | 0.980115 | 0.834917 | 0.84454 | 0.999971 | 1 | 0.979814 |
| Vstm2a | A0A140LHI | -0.9 | 0.46 | -0.39 | -0.73 | 1.96 | 0 | 1.27 | 0.02 | 1.49 | -1.02 | 1.09 | 1.5 | 1.61 | 0.1 | -0.18 | -1.16 | 1.2025 | 1.155 | 0.4825 | 0.045364 | 0.089314 | 0.425002 | 0.999971 | 1 | 0.880275 |
| Vstm2b | Q9JME9 | 0.72 | 1.48 | -0.26 | 0.68 | 2.27 | 1.6 | 0.08 | 0.4 | 1.1 | 0.68 | 0.24 | 0.23 | 2 | 0.83 | -0.1 | 0.82 | 0.4325 | -0.0925 | 0.2325 | 0.456564 | 0.81545 | 0.654678 | 0.999971 | 1 | 0.934061 |
| Vstm2l | A2ACD2 | 2.08 | 0.7 | -0.05 | 0.88 | 0.79 | 1.43 | 0.3 | 0.91 | 1.13 | 2.05 | -0.21 | -0.15 | 1.93 | 1.95 | 0.95 | 0.38 | -0.045 | -0.1975 | 0.4 | 0.922976 | 0.758373 | 0.466353 | 0.999971 | 1 | 0.891431 |
| Vta1 | Q9CR26 | 0.48 | -0.64 | 0.37 | 0.08 | -0.62 | -0.64 | 1.04 | 0.13 | -0.34 | 0 | 0.54 | 0.2 | -0.08 | -1 | -0.01 | -0.28 | -0.095 | 0.0275 | -0.415 | 0.830031 | 0.930437 | 0.238686 | 0.999971 | 1 | 0.833594 |
| Vti1a | A0A286YDI | 1.5 | 0.02 | 1.51 | 0.27 | 1.02 | 0.17 | 1.15 | 0.11 | 0.86 | 0.47 | 0.66 | 0.38 | 0.83 | 0.06 | 0.26 | 0.01 | -0.2125 | -0.2325 | -0.535 | 0.640166 | 0.559221 | 0.221203 | 0.999971 | 1 | 0.8275 |
| Vti1a_1 | Q5FWJ7 | -0.51 | -0.21 | -0.84 | -0.51 | -0.3 | 0.18 | -0.91 | -0.31 | -0.14 | -0.46 | -0.63 | -0.76 | 0 | 0.44 | -0.51 | -0.3 | 0.1825 | 0.02 | 0.425 | 0.512233 | 0.928825 | 0.135546 | 0.999971 | 1 | 0.784948 |
| Vti1b | Q91XH6 | 0.69 | 0.72 | 0.23 | 0.62 | 0.69 | 0.91 | 0.23 | 0.94 | 0.6 | 1.1 | 0.47 | 0.68 | 1.15 | 1.22 | 0.69 | 1.11 | 0.1275 | 0.1475 | 0.4775 | 0.590277 | 0.505651 | 0.049269 | 0.999971 | 1 | 0.715024 |
| Vtn | P29788 | 2.16 | 1.69 | 0.32 | 1.7 | 2.23 | 1.43 | 1.86 | 0.93 | 1.42 | 2.28 | 1.32 | 1.82 | 2 | 1.82 | 1.3 | 0.61 | 0.145 | 0.2425 | -0.035 | 0.751007 | 0.576737 | 0.940909 | 0.999971 | 1 | 0.996218 |
| Vwa1 | Q8R2Z5 | 0.93 | 0.53 | -0.49 | 0.63 | 1.19 | 0.62 | -0.61 | -0.64 | 0.08 | 1 | -0.37 | -0.05 | 0.21 | 0.22 | 0.34 | 0.15 | -0.26 | -0.235 | -0.17 | 0.61268 | 0.567853 | 0.593053 | 0.999971 | 1 | 0.923349 |
| Vwa5a | Q99KC8 | -0.25 | 0.01 | 0.23 | -0.15 | 0.52 | 0.33 | 0.17 | -0.3 | 0.31 | 0.11 | 0.04 | 0.1 | 0.06 | 0.15 | 0.31 | -0.49 | 0.22 | 0.18 | 0.0475 | 0.36719 | 0.346044 | 0.841684 | 0.999971 | 1 | 0.979814 |
| Vwa5b2 | E0CZ01 | -1.78 | -3.69 | 0.06 | -0.02 | -2.28 | -2.52 | -0.5 | -0.27 | -3.08 | -1.59 | -0.23 | -0.68 | -4.03 | -2.49 | 0.41 | 0.19 | -0.035 | -0.0375 | -0.1225 | 0.970538 | 0.96936 | 0.92125 | 0.999971 | 1 | 0.992702 |
| Vwa8 | Q8CC88 | -0.29 | -0.73 | -0.69 | -0.56 | -0.61 | -0.62 | -0.68 | -0.37 | -0.88 | -0.56 | -0.79 | -0.69 | -0.5 | -0.6 | -0.66 | -0.6 | -0.0025 | -0.1625 | -0.0225 | 0.98944 | 0.393409 | 0.90176 | 0.999971 | 1 | 0.990759 |
| Vwc2l | Q505H4-3 | -0.23 | -1.21 | -0.8 | -0.84 | 0.2 | -0.84 | -1.06 | 0.05 | -0.94 | 0.32 | -1.5 | -0.36 | -2.07 | 0.68 | -0.53 | -1.02 | 0.3575 | 0.15 | 0.035 | 0.340238 | 0.720909 | 0.949821 | 0.999971 | 1 | 0.998038 |
| Vwf | E9QPU1 | 2.48 | 0.32 | 0.71 | 1.2 | 4.75 | 1.64 | 1.29 | 1.05 | 1.83 | 2.3 | 0.39 | 1.73 | -0.78 | 1.75 | 1.01 | 0.55 | 1.005 | 0.385 | -0.545 | 0.271992 | 0.508327 | 0.408437 | 0.999971 | 1 | 0.875014 |
| Vxn | Q8BG31 | 2.36 | 0.47 | 0.87 | 1.3 | 0.58 | 1.36 | 1.86 | 1.06 | 0.54 | 2.07 | 1.51 | 0.51 | 0.26 | 1.14 | 0.78 | 1.49 | -0.035 | -0.0925 | -0.3325 | 0.938859 | 0.858472 | 0.47192 | 0.999971 | 1 | 0.892857 |
| Wac | Q924H7-3 | 2.33 | 2.27 | 1.16 | 1.1 | 1.96 | 1.71 | 1.04 | 0.75 | 1.86 | 2.59 | -0.87 | 0.41 | 0.95 | 2.47 | 0.66 | 1.68 | -0.35 | -0.7175 | -0.275 | 0.412082 | 0.359375 | 0.578801 | 0.999971 | 1 | 0.919236 |

| | | | | | | | | | | | | | | | | | | | | | | | | | | |
|---|---|---|---|---|---|---|---|---|---|---|---|---|---|---|---|---|---|---|---|---|---|---|---|---|---|---|
| Wapl | B7ZP47 | 2.49 | 2.89 | 3.16 | 3.58 | 2.61 | 2.71 | 3.31 | 3.01 | 2.85 | 2.62 | 3.25 | 2.77 | 0.83 | 2.44 | 2.68 | 2.23 | -0.12 | -0.1575 | -0.985 | 0.68057 | 0.577179 | 0.051678 | 0.999971 | 1 | 0.723736 |
| Wars1 | P32921-2 | -0.32 | 0.58 | -0.03 | 0.18 | 0.74 | 0.71 | 0.47 | 0.28 | 0.8 | 0.28 | 0.4 | 0.49 | 0.6 | 0.66 | 0.17 | -0.11 | 0.4475 | 0.39 | 0.2275 | 0.097617 | 0.140596 | 0.424889 | 0.999971 | 1 | 0.880275 |
| Wars2 | Q9CYK1 | -0.45 | -0.41 | -0.26 | -0.29 | -0.39 | -0.54 | -0.26 | -0.34 | -0.53 | -0.76 | -0.46 | -0.15 | -0.53 | -0.76 | -0.39 | -0.23 | -0.03 | -0.1225 | -0.125 | 0.85979 | 0.53057 | 0.51617 | 0.999971 | 1 | 0.905815 |
| Wasf1 | Q8R5H6 | -1.19 | -2.56 | -1.23 | -1.67 | -2.49 | -2.5 | -1.1 | -1.63 | -2.36 | -2.18 | -1.02 | -1.19 | -3.24 | -2.68 | -1.61 | -1.78 | -0.2675 | -0.025 | -0.665 | 0.548297 | 0.954689 | 0.179105 | 0.999971 | 1 | 0.808532 |
| Wasf2 | Q8BH43 | 0.88 | 0.92 | 0.33 | 0.45 | 0.91 | 0.95 | 0.02 | 0.08 | 0.86 | 1.54 | 0.31 | 0.35 | 0.77 | 1.22 | 0.39 | 0.79 | -0.155 | 0.12 | 0.1475 | 0.611121 | 0.712261 | 0.564266 | 0.999971 | 1 | 0.917087 |
| Wasf3 | A0A571BF7 | -0.1 | -0.63 | -1.75 | -0.79 | -1.31 | -0.81 | -0.98 | -0.57 | -0.94 | -0.32 | -0.81 | -1.52 | -1.01 | -1.09 | -0.57 | -0.69 | -0.1 | -0.08 | -0.0225 | 0.785493 | 0.843496 | 0.949915 | 0.999971 | 1 | 0.998038 |
| Wasf3_1 | Q8VHI6 | 0.31 | -0.68 | -0.67 | -0.1 | -1.01 | -0.84 | -0.45 | -0.63 | -0.97 | -0.17 | -0.62 | -0.85 | -1.23 | -0.95 | -0.71 | -0.69 | -0.4475 | -0.3675 | -0.61 | 0.140897 | 0.247308 | 0.057682 | 0.999971 | 1 | 0.725348 |
| Washc1 | Q8VDD8 | -0.23 | -0.02 | 0.37 | 0.78 | 0.32 | -0.74 | 1.26 | 0.41 | 0.26 | -0.87 | 0.77 | 0.39 | 0.22 | -1.35 | 0.54 | 0.6 | 0.0875 | -0.0875 | -0.2225 | 0.842162 | 0.826999 | 0.642265 | 0.999971 | 1 | 0.93301 |
| Washc2 | Q6PGL7 | 0.79 | -0.19 | 0.22 | 0.35 | -0.23 | -0.26 | -0.12 | 0.39 | -0.19 | 0.26 | -0.05 | 0.33 | -0.31 | -0.27 | 0.2 | 0.76 | -0.3475 | -0.205 | -0.1975 | 0.221443 | 0.437033 | 0.545684 | 0.999971 | 1 | 0.911398 |
| Washc3 | S4R287 | 0.27 | 0.94 | 0.12 | 0.75 | 0.7 | 0.22 | 0.07 | 0.93 | 0.5 | 0.48 | 0.24 | 0.29 | 0.99 | 0.41 | 0.25 | 1.05 | -0.04 | -0.1425 | 0.155 | 0.890653 | 0.551324 | 0.598641 | 0.999971 | 1 | 0.92377 |
| Washc4 | Q3UMB9 | 0.34 | 0.25 | 0.5 | 0.33 | 0.38 | 0.33 | 0.73 | 0.4 | 0.41 | 0.31 | 0.7 | 0.78 | 0.46 | 0.15 | 0.43 | 0.38 | 0.105 | 0.195 | -7.85E-17 | 0.567269 | 0.314903 | 1 | 0.999971 | 1 | 1 |
| Washc5 | Q8C2E7 | 0.26 | 0.23 | 0.22 | 0.16 | 0.32 | 0.1 | 0 | 0.07 | 0.33 | 0.18 | 0.2 | 0.03 | 0.08 | 0.33 | 0.02 | 0.3 | -0.095 | -0.0325 | -0.035 | 0.577419 | 0.841984 | 0.839364 | 0.999971 | 1 | 0.979728 |
| Wasl | Q91YD9 | 0.38 | 0.69 | 0.15 | 0.42 | 0.46 | 0.54 | 0.32 | 0.53 | 0.23 | 0.42 | 0.08 | 0.24 | 0.1 | 0.38 | 0.34 | 0.34 | 0.0525 | -0.1675 | -0.12 | 0.782186 | 0.392084 | 0.539434 | 0.999971 | 1 | 0.910077 |
| Wbp11 | Q923D5 | 1.25 | 1.45 | 1.5 | 1.84 | 1.38 | 1.36 | 1.3 | 1.53 | 1.48 | 1.69 | 1.48 | 1.37 | 1.04 | 1.63 | 1.45 | 1.77 | -0.1175 | -0.005 | -0.0375 | 0.550503 | 0.979645 | 0.873744 | 0.999971 | 1 | 0.984109 |
| Wbp2 | P97765 | -0.46 | -0.2 | -0.17 | -0.19 | -0.02 | -0.26 | -0.03 | 0.18 | -0.47 | -0.61 | 0.38 | -0.16 | -0.28 | -0.03 | 0.31 | 0.17 | 0.2225 | 0.04 | 0.2975 | 0.249773 | 0.874849 | 0.16723 | 0.999971 | 1 | 0.800215 |
| Wbp4 | Q61048 | 4.74 | -0.46 | 1.52 | 2.02 | 1.43 | 0.1 | 1.42 | 1.98 | 0.92 | 1.35 | 1.59 | 1.88 | 0.19 | -0.29 | 1.9 | 2.14 | -0.7225 | -0.52 | -0.97 | 0.48504 | 0.599004 | 0.388246 | 0.999971 | 1 | 0.871816 |
| Wdcp | A0A1Y7VJP | -0.92 | -1.42 | -1.51 | -0.91 | -2.49 | 1.42 | -1.33 | -0.42 | -2.43 | -2.48 | -2.05 | -1.98 | -2.39 | 0.24 | -1.48 | -0.75 | 0.485 | -1.045 | 0.095 | 0.526556 | 0.001751 | 0.859034 | 0.999971 | 0.693667 | 0.982702 |
| Wdfy1 | E9Q4P1 | -1.04 | -5.34 | -4.49 | -3.64 | -5.32 | -4.64 | -2.74 | -3.59 | 5.62 | 5.15 | 4.76 | 4.64 | -7.16 | -4.71 | -4.94 | -4.24 | -0.445 | 8.67 | -1.635 | 0.649263 | 5.90E-06 | 0.133459 | 0.999971 | 0.026904 | 0.784948 |
| Wdfy2 | Q8BUB4 | 1.16 | 0.38 | 0.71 | 1.18 | -0.31 | 0.26 | 0.09 | -0.53 | -0.79 | 0.65 | 0.53 | -0.18 | -0.81 | 0.14 | 0.26 | 0.9 | -0.98 | -0.805 | -0.735 | 0.006454 | 0.057165 | 0.085064 | 0.999971 | 1 | 0.758588 |
| Wdfy3 | Q6VNB8 | -0.43 | -0.69 | -0.32 | -0.59 | -0.63 | -0.63 | -0.29 | -0.57 | -0.69 | -0.71 | -0.58 | -0.53 | -0.35 | -0.64 | -0.33 | -0.57 | -0.0225 | -0.12 | 0.035 | 0.904003 | 0.494639 | 0.850989 | 0.999971 | 1 | 0.98187 |
| Wdr1 | O88342 | -1.4 | -1.19 | -0.3 | -0.57 | -1.12 | -0.9 | 0.03 | -0.24 | -1.25 | -1.43 | -0.2 | -0.22 | -1.7 | -1.17 | -0.5 | -0.31 | 0.3075 | 0.09 | -0.055 | 0.408129 | 0.822205 | 0.888924 | 0.999971 | 1 | 0.987646 |
| Wdr11 | G5E8J3 | 0.48 | 0.49 | 0.24 | 0.16 | 0.6 | 0.62 | 0.19 | 0.37 | 0.44 | 0.83 | 0.03 | 0.13 | 0.55 | 0.66 | 0.35 | 0.01 | 0.1025 | 0.015 | 0.05 | 0.601813 | 0.948515 | 0.815671 | 0.999971 | 1 | 0.974918 |
| Wdr12 | Q9JJA4 | -0.57 | 0.22 | 1.76 | 1.03 | -0.04 | -0.15 | 3.45 | 0.36 | -0.2 | -0.01 | 0.65 | 0.12 | -1.02 | 0.04 | 0.47 | -0.38 | 0.295 | -0.47 | -0.8325 | 0.740457 | 0.359158 | 0.154733 | 0.999971 | 1 | 0.795578 |
| Wdr13 | Q91V09 | -0.6 | -0.04 | -0.25 | -0.73 | 0.07 | -0.31 | -0.09 | -0.35 | 0.18 | -0.6 | 0.05 | 0.09 | 0.03 | -0.45 | -0.38 | -0.65 | 0.235 | 0.335 | 0.0425 | 0.31456 | 0.219958 | 0.862087 | 0.999971 | 1 | 0.982963 |
| Wdr17 | H3BLI8 | 0.72 | 0.32 | 0.68 | 0.5 | 0.05 | 0.29 | -0.35 | 0.61 | -0.09 | 0.7 | -0.22 | 0.13 | 0.65 | 0.75 | -0.01 | -0.12 | -0.405 | -0.425 | -0.2375 | 0.131968 | 0.115512 | 0.379367 | 0.999971 | 1 | 0.871816 |
| Wdr18 | Q4VBE8 | 0.49 | 1.54 | 1.69 | 1.7 | 1.56 | 1.75 | 2.88 | 1.33 | 1.94 | 1.02 | 1.18 | 1.1 | 2.36 | 1.68 | 1.61 | 1.57 | 0.525 | -0.045 | 0.45 | 0.238451 | 0.8986 | 0.210791 | 0.999971 | 1 | 0.822884 |
| Wdr19 | Q3UGF1-3 | 0.38 | 0.17 | 0.5 | 1.26 | 0.12 | 0.23 | 0.87 | 0.53 | 0.1 | 0.15 | -0.11 | -0.06 | 0.11 | 0.21 | 0.28 | -0.02 | -0.14 | -0.5575 | -0.4325 | 0.642212 | 0.061057 | 0.132632 | 0.999971 | 1 | 0.784948 |
| Wdr20 | Q3UWE6 | 0.2 | 0.26 | -0.92 | -1 | -0.23 | 0.33 | -0.7 | -1.13 | 0.21 | -0.24 | -1.25 | -0.9 | 0.78 | -0.03 | -1.27 | -0.91 | -0.0675 | -0.18 | 0.0075 | 0.877657 | 0.689117 | 0.988708 | 0.999971 | 1 | 0.99925 |
| Wdr24 | Q8CFJ9 | 0.36 | 0.98 | 0.65 | 1.11 | 0.38 | 0.86 | 1.89 | 0.39 | 0.55 | 0.94 | 0.55 | 0.11 | 0.26 | 0.84 | 0.31 | 0.29 | 0.105 | -0.2375 | -0.35 | 0.782181 | 0.373978 | 0.182636 | 0.999971 | 1 | 0.808532 |
| Wdr25 | A0A1Y7VKV | 5.11 | 2.27 | 1.92 | 2.51 | 0.03 | 0.96 | 3.37 | 3.33 | 1.87 | 3.02 | 1.8 | 1.8 | -2.38 | 1.18 | 1.55 | 0.9 | -1.03 | -0.83 | -2.64 | 0.316392 | 0.265722 | 0.029981 | 0.999971 | 1 | 0.678474 |
| Wdr26 | A0A494BB1 | -0.72 | -0.01 | -0.84 | -0.66 | -0.01 | 0.02 | -0.27 | -0.61 | -0.29 | -0.36 | -0.36 | -0.47 | 0.01 | -0.1 | -0.97 | -0.7 | 0.34 | 0.1875 | 0.1175 | 0.213279 | 0.417778 | 0.703233 | 0.999971 | 1 | 0.949575 |
| Wdr3 | Q8BHB4 | 0.65 | 0.08 | -0.74 | -0.71 | -0.17 | 0.15 | -0.84 | -0.98 | -0.16 | 0.76 | -0.62 | -0.76 | 0.82 | 0.53 | -1.07 | -1.01 | -0.28 | -0.015 | -0.0025 | 0.500035 | 0.973513 | 0.996389 | 0.999971 | 1 | 1 |
| Wdr33 | Q8K4P0 | 2.33 | 1.53 | 1.3 | 2.11 | 1.46 | 2.3 | 1.77 | 1.36 | 1.56 | 2.47 | 1.38 | 1.16 | 0.31 | 1.62 | 1.49 | 1.19 | -0.095 | -0.175 | -0.665 | 0.768536 | 0.634648 | 0.100301 | 0.999971 | 1 | 0.78268 |
| Wdr35 | Q8BND3 | -0.1 | 0.68 | 0.2 | 0.39 | 0.35 | 0.44 | 0.07 | 0.78 | 0.29 | 0.27 | 0.32 | 0.52 | 0.84 | 0.59 | 0.62 | 1.04 | 0.1175 | 0.0575 | 0.48 | 0.638196 | 0.790679 | 0.064622 | 0.999971 | 1 | 0.726597 |
| Wdr36 | Q3TA68 | -0.83 | -1.16 | -1.18 | -0.09 | -1.37 | -1.57 | -1.14 | -0.96 | -1.06 | -0.78 | -0.43 | 0.24 | -1.08 | -1.08 | -0.05 | -0.45 | -0.445 | 0.3075 | 0.15 | 0.159264 | 0.413988 | 0.672345 | 0.999971 | 1 | 0.93984 |
| Wdr37 | Q8CBE3 | 0.42 | -0.12 | -0.02 | -0.6 | -0.13 | 0.14 | 0.28 | -0.49 | 0.23 | -0.26 | 0.25 | -0.08 | 0.32 | -0.42 | -0.28 | -0.96 | 0.03 | 0.115 | -0.255 | 0.915669 | 0.664122 | 0.454077 | 0.999971 | 1 | 0.887217 |
| Wdr41 | Q3UDP0 | -0.05 | 0.29 | 0.21 | 0.34 | 0.11 | 0.11 | -0.14 | 0.54 | 0.14 | 0.03 | 0.1 | 0.46 | 0.71 | 0.53 | 0.74 | 0.5 | -0.0425 | -0.015 | 0.4225 | 0.842425 | 0.937128 | 0.043416 | 0.999971 | 1 | 0.701725 |
| Wdr43 | Q6ZQL4 | -0.96 | 0.01 | -1.04 | -0.75 | -0.23 | -0.34 | -1.54 | -1.11 | -0.34 | 0.36 | -0.52 | -0.68 | 0.31 | -0.46 | -0.5 | -0.31 | -0.12 | 0.39 | 0.445 | 0.753068 | 0.25739 | 0.176276 | 0.999971 | 1 | 0.808532 |
| Wdr44 | Q6NVE8 | -0.1 | -0.05 | 0.03 | -0.3 | 0.14 | 0.04 | 0.07 | -0.15 | -0.08 | -0.19 | -0.07 | -0.01 | 0.58 | 0.12 | 0.09 | -0.12 | 0.13 | 0.0175 | 0.2725 | 0.469508 | 0.916987 | 0.222308 | 0.999971 | 1 | 0.827753 |
| Wdr45 | Q91VM3 | -1.02 | 0.23 | 0.75 | -0.54 | 0.59 | 0.69 | 0.26 | 0.28 | 0.58 | -0.7 | 0.89 | 0.77 | 1.14 | 0.19 | 0.74 | -0.03 | 0.6 | 0.53 | 0.655 | 0.151876 | 0.306905 | 0.168226 | 0.999971 | 1 | 0.800215 |
| Wdr45b | Q9CR39 | -2.98 | -0.48 | -0.62 | -1.9 | -0.43 | -1.17 | -0.61 | -1.2 | -0.98 | -1.86 | -0.55 | -0.42 | -0.66 | -1.39 | -0.77 | -1.81 | 0.6425 | 0.5425 | 0.3375 | 0.276291 | 0.390061 | 0.57283 | 0.999971 | 1 | 0.91816 |
| Wdr46 | Q9Z0H1 | 2.16 | -1.15 | -0.69 | -0.65 | -1.04 | -0.63 | -0.82 | -0.83 | -1.13 | 0.59 | -0.21 | 0.04 | -1.88 | -1.34 | -0.86 | 0.27 | -0.7475 | -0.095 | -0.87 | 0.29393 | 0.900321 | 0.291069 | 0.999971 | 1 | 0.847402 |
| Wdr47 | Q8CGF6 | -1.14 | -0.65 | -0.88 | -1.32 | -0.51 | -0.74 | -0.2 | -1.19 | -0.24 | -0.87 | -0.64 | -0.59 | -0.16 | -0.84 | -1.2 | -1.49 | 0.3375 | 0.4125 | 0.075 | 0.236907 | 0.101622 | 0.817098 | 0.999971 | 1 | 0.974929 |
| Wdr48 | Q8BH57 | -0.47 | -0.7 | -0.67 | -0.77 | -0.7 | -0.45 | -0.44 | -0.7 | -0.53 | -0.29 | -0.69 | -0.65 | -0.54 | -0.64 | -0.72 | -0.46 | 0.08 | 0.1125 | 0.0625 | 0.655971 | 0.539158 | 0.72118 | 0.999971 | 1 | 0.954937 |
| Wdr5 | P61965 | 1.57 | 1.27 | 1.68 | 1.58 | 1.16 | 1.32 | 1.11 | 1.5 | 0.93 | 1.25 | 1.17 | 1.19 | 0.66 | 0.9 | 1.53 | 1.76 | -0.2525 | -0.39 | -0.3125 | 0.208917 | 0.057602 | 0.293164 | 0.999971 | 1 | 0.847402 |
| Wdr54 | Q9R0D8 | -0.11 | 0.23 | -0.38 | -0.03 | -0.09 | -0.01 | -0.32 | 0.06 | 0.06 | 0.11 | -0.62 | -0.37 | -0.45 | 0.42 | -0.13 | 0.23 | -0.0175 | -0.1325 | 0.09 | 0.931844 | 0.590278 | 0.726292 | 0.999971 | 1 | 0.956041 |
| Wdr59 | A0A1D5RLJ | 0.8 | 1.38 | 1.43 | 1.09 | 0.74 | 1.08 | 1.05 | 1.08 | 0.96 | 1.15 | 1.14 | 0.67 | 0.9 | 1.36 | 1.03 | 1.32 | -0.1875 | -0.195 | -0.0225 | 0.393425 | 0.392301 | 0.920435 | 0.999971 | 1 | 0.992702 |
| Wdr6 | Q99ME2 | -0.51 | 0.22 | -0.21 | -0.07 | 0.42 | 0.41 | -0.12 | 0.76 | 0.68 | 0.25 | 0.92 | 0.08 | 0.8 | 1.04 | 1.05 | 0.82 | 0.51 | 0.625 | 1.07 | 0.07651 | 0.040309 | 0.000795 | 0.999971 | 1 | 0.365812 |
| Wdr61 | Q9ERF3 | -1.51 | -0.19 | 0.37 | 0.56 | 1.01 | -0.91 | 0.43 | 0.76 | 0.4 | -1.71 | 0.42 | 0.77 | 1.11 | -1.02 | 0.78 | 0.88 | 0.515 | 0.1625 | 0.63 | 0.385024 | 0.808421 | 0.322833 | 0.999971 | 1 | 0.85548 |
| Wdr7 | Q920I9 | -0.25 | -0.7 | -0.63 | -0.78 | -0.94 | -0.44 | -0.58 | -0.65 | -0.78 | -0.36 | -0.74 | -1.1 | -0.68 | -0.5 | -0.66 | -0.68 | -0.0625 | -0.155 | -0.04 | 0.765345 | 0.502207 | 0.835092 | 0.999971 | 1 | 0.979036 |
| Wdr73 | Q9CWR1 | -0.63 | 0.2 | 0.48 | 0.55 | -0.25 | 0.18 | 0.57 | 0.25 | 0.39 | -1.7 | 0.53 | 0.61 | 0.31 | -0.73 | 0.16 | 0.28 | 0.0375 | -0.1925 | -0.145 | 0.906921 | 0.735609 | 0.687553 | 0.999971 | 1 | 0.945573 |
| Wdr75 | Q3U821 | -0.83 | -0.95 | -1.34 | -1.54 | -1.36 | -1.63 | -1.38 | -1.65 | -1.69 | -1.25 | -0.93 | -1.13 | -0.49 | -1.31 | -0.86 | -0.84 | -0.34 | -0.085 | 0.29 | 0.155473 | 0.73925 | 0.282567 | 0.999971 | 1 | 0.844538 |
| Wdr77 | Q99J09 | -0.05 | 0.69 | 0.51 | 0.03 | 0.47 | 0.76 | 1.12 | 0.92 | 0.8 | 0.33 | 0.65 | 1.35 | 0.12 | 1.06 | 0.72 | 0.28 | 0.5225 | 0.4875 | 0.25 | 0.064774 | 0.120273 | 0.4016 | 0.999971 | 1 | 0.871816 |
| Wdr81 | K4DI77 | 0.54 | -0.03 | -0.03 | -0.28 | -0.03 | 0.15 | 0.08 | 0.03 | -0.19 | 0.29 | 0.11 | -0.02 | 0.28 | 0.05 | -0.07 | 0.11 | 0.0075 | -0.0025 | 0.0425 | 0.97291 | 0.991473 | 0.852362 | 0.999971 | 1 | 0.98187 |
| Wdr82 | Q8BFQ4 | 1.35 | 2.06 | 1.39 | 1.51 | 1.63 | 1.88 | 1.64 | 2.08 | 1.73 | 1.52 | 1.69 | 1.63 | 1.85 | 1.57 | 1.65 | 1.4 | 0.23 | 0.065 | 0.04 | 0.337204 | 0.762246 | 0.861231 | 0.999971 | 1 | 0.982963 |
| Wdr89 | Q9D0R9 | 1.4 | 0.51 | 1.22 | 0.72 | -0.22 | 0.06 | -0.14 | 0.78 | 0.21 | 1.62 | 0.21 | -0.02 | 1.23 | 0.57 | 0.62 | 1.61 | -0.8425 | -0.4575 | 0.045 | 0.023274 | 0.282494 | 0.8905 | 0.999971 | 1 | 0.988184 |
| Wdr91 | Q7TMQ7 | -1.36 | 0.04 | 0.4 | 0.07 | 0.49 | -0.15 | 0.25 | 0.37 | 0.86 | -0.97 | 0.73 | 0.53 | 1.72 | -0.78 | 0.49 | 0.44 | 0.4525 | 0.5 | 0.68 | 0.272593 | 0.362253 | 0.26655 | 0.999971 | 1 | 0.842677 |
| Wdsub1 | Q9D0I6 | 0.71 | 1.14 | -0.95 | -0.51 | 0.9 | 1.06 | -0.27 | -0.67 | 0.8 | 0.83 | -1.21 | -1.12 | 0.52 | 0.8 | -0.89 | -0.35 | 0.1575 | -0.2725 | -0.0775 | 0.791928 | 0.693342 | 0.893085 | 0.999971 | 1 | 0.98828 |
| Wdtc1 | Q80ZK9 | 1.5 | 0.9 | 0.79 | 0.46 | 0.46 | 1.15 | 0.72 | 0.75 | 0.21 | 1.54 | 0.38 | 0.37 | 0.37 | 1 | 0.68 | 0.44 | -0.1425 | -0.2875 | -0.29 | 0.608761 | 0.441357 | 0.310547 | 0.999971 | 1 | 0.847402 |
| Wfs1 | P56695 | -0.08 | -1.57 | -0.69 | -0.33 | -0.98 | -1.29 | 0.25 | -0.36 | -1.1 | -0.52 | -0.67 | -0.48 | -1.57 | -0.78 | -0.4 | -0.77 | 0.0725 | -0.025 | -0.2125 | 0.870256 | 0.942932 | 0.591514 | 0.999971 | 1 | 0.922652 |
| Whrn | Q80VW5-4 | -0.84 | -0.21 | -0.63 | -0.56 | -0.68 | 0.1 | -1.03 | -0.42 | -0.44 | -0.09 | -0.16 | -0.1 | 0.57 | 0.36 | -0.48 | 0.16 | 0.0525 | 0.3625 | 0.7125 | 0.853935 | 0.104291 | 0.02902 | 0.999971 | 1 | 0.678474 |
| Wipf2 | Q6PEV3 | 1.15 | 0.34 | 0.12 | 0.15 | 0.45 | 0.48 | 0.05 | -0.11 | 0.01 | 0.8 | -0.17 | -0.45 | 0.45 | 0.22 | -0.37 | 0.02 | -0.2225 | -0.3925 | -0.36 | 0.455701 | 0.285031 | 0.25731 | 0.999971 | 1 | 0.839012 |
| Wipf3 | E9Q2D0 | 0.53 | 0.62 | 0.58 | 0.47 | 0.23 | 0.52 | 0.29 | 0.39 | -0.11 | 0.3 | 0.12 | 0.05 | -0.2 | 0.04 | 0.3 | 0.39 | -0.1925 | -0.46 | -0.4175 | 0.271789 | 0.024379 | 0.06051 | 0.999971 | 1 | 0.725348 |
| Wipi1 | Q8R3E3-2 | -0.26 | 0.34 | 0.22 | -0.53 | 0.35 | 0.58 | -0.38 | 0.61 | 0.43 | -0.47 | 0.24 | 0.23 | 0.66 | 0.31 | 0.01 | -0.01 | 0.3475 | 0.165 | 0.3 | 0.283254 | 0.577775 | 0.292936 | 0.999971 | 1 | 0.847402 |
| Wipi2 | Q80W47 | 0.38 | 0.39 | 0.65 | 0.19 | -0.19 | 1.02 | 0.79 | 0.61 | 0.13 | 0.07 | 0.5 | 0.16 | 0.18 | 0.53 | 0.76 | 0.45 | 0.155 | -0.1875 | 0.0775 | 0.597013 | 0.346596 | 0.708644 | 0.999971 | 1 | 0.95049 |
| Wiz | G5E8J8 | 0.5 | 0.62 | 0.54 | 1.71 | 0.72 | -0.06 | -0.08 | 0.77 | 0.27 | 0.94 | 0.07 | 0.28 | 1.01 | 0.72 | 0.86 | 1.33 | -0.505 | -0.4525 | 0.1375 | 0.189038 | 0.209262 | 0.669951 | 0.999971 | 1 | 0.938809 |
| Wls | Q6DID7 | -1.12 | 0.2 | 0.28 | 0.21 | 0.16 | 0.75 | -0.17 | 0.25 | -0.02 | -1.07 | 0.45 | 0.64 | 0.95 | 0.52 | 0.73 | 0.77 | 0.355 | 0.1075 | 0.85 | 0.356699 | 0.822162 | 0.03399 | 0.999971 | 1 | 0.689944 |
| Wnk1 | P83741-4 | 0.37 | -0.39 | 0.4 | 0.25 | -0.3 | -0.39 | 0.31 | -0.03 | -0.3 | 0.11 | 0.39 | 0.08 | -0.45 | -0.47 | 0.37 | 0.19 | -0.26 | -0.0875 | -0.2475 | 0.339478 | 0.733599 | 0.412501 | 0.999971 | 1 | 0.875881 |
| Wnk1_1 | F6V9V5 | -0.58 | -0.44 | -0.35 | -0.17 | -0.33 | -0.68 | -0.26 | 1.07 | -0.78 | -1.56 | -0.39 | -0.65 | 1.21 | 0.12 | 1.39 | 0.36 | 0.335 | -0.46 | 1.155 | 0.388937 | 0.127725 | 0.006055 | 0.999971 | 1 | 0.582714 |
| Wnk2 | E9QJQ9 | 0.1 | -0.51 | -0.36 | -0.41 | -0.72 | -0.57 | -0.41 | -0.27 | -0.58 | -0.01 | -0.33 | -0.33 | -0.63 | -0.6 | -0.37 | -0.29 | -0.1975 | -0.0175 | -0.1775 | 0.368776 | 0.936414 | 0.409178 | 0.999971 | 1 | 0.875789 |
| Wnk3 | V9GWU7 | -1.19 | -2.27 | -1.57 | -1.71 | -1.72 | -2.8 | -1.49 | -1.57 | -2.28 | -1.96 | -1.07 | -1.16 | -2.24 | -2.14 | -1.51 | -1.85 | -0.21 | 0.0675 | -0.25 | 0.571577 | 0.852748 | 0.399164 | 0.999971 | 1 | 0.871816 |
| Wrnip1 | Q91XU0 | 1.19 | 1.29 | 1.14 | 1.38 | 1.64 | 0.88 | 1.12 | 0.95 | 1.33 | 1.29 | 1.33 | 1.31 | 1.04 | 0.99 | 0.97 | 1.38 | -0.1025 | 0.065 | -0.155 | 0.646663 | 0.685652 | 0.411076 | 0.999971 | 1 | 0.875847 |
| Wsb2 | O54929 | -0.22 | 0.07 | -0.52 | -0.76 | -0.03 | 0.59 | -0.77 | -0.51 | 0.03 | 0.07 | -0.4 | -1.36 | -0.17 | 0.56 | -0.6 | 0.14 | 0.1775 | -0.0575 | 0.34 | 0.608847 | 0.875859 | 0.290025 | 0.999971 | 1 | 0.847402 |
| Wtap | E0CYH0 | 1.4 | 0.97 | 0.25 | 0.04 | 0.84 | 0.89 | 2.43 | -1.03 | 1.16 | 1.13 | -0.43 | -0.82 | 0.91 | 1.03 | -0.58 | -1.48 | 0.1175 | -0.405 | -0.695 | 0.866765 | 0.475723 | 0.282296 | 0.999971 | 1 | 0.844538 |
| Wwc1 | Q5SXA9 | 0.07 | -0.53 | -1.09 | -1.39 | -0.61 | -0.43 | -0.94 | -0.99 | -0.41 | 0.26 | -0.65 | -1.21 | -0.53 | -0.55 | -1.38 | -1.11 | -0.0075 | 0.2325 | -0.1575 | 0.982596 | 0.584575 | 0.674887 | 0.999971 | 1 | 0.941083 |
| Wwc2 | Q6NXJ0 | 0.39 | 0.57 | -0.01 | 0.05 | 0.5 | 0.53 | 0.29 | 0.54 | 0.27 | 0.44 | 0.42 | 0.33 | 0.51 | 0.48 | -0.02 | 0.47 | 0.215 | 0.115 | 0.11 | 0.309329 | 0.566219 | 0.63163 | 0.999971 | 1 | 0.932858 |
| Wwox | Q91WL8 | -0.57 | -0.93 | -0.54 | -0.33 | -0.78 | -0.81 | -0.81 | -0.08 | -0.56 | -0.22 | -1.07 | -0.18 | -0.48 | -0.19 | -0.25 | -0.26 | -0.0275 | 0.085 | 0.2975 | 0.911614 | 0.745869 | 0.161586 | 0.999971 | 1 | 0.800215 |
| Wwp1 | Q8BZZ3 | 1.1 | -0.01 | -0.54 | -0.11 | -0.27 | -0.02 | 0.36 | -0.53 | 0.12 | 0.72 | -0.38 | -0.76 | 0.18 | -0.33 | -0.92 | -0.82 | -0.225 | -0.185 | -0.5825 | 0.560684 | 0.680573 | 0.181126 | 0.999971 | 1 | 0.808532 |
| Wwp2 | Q9DBH0 | -0.4 | -1.08 | -0.53 | -0.67 | -0.6 | -0.89 | -0.04 | -0.59 | -0.58 | -0.86 | -0.57 | -0.97 | -0.9 | -1.62 | -0.68 | -1.32 | 0.14 | -0.075 | -0.46 | 0.587518 | 0.733142 | 0.121601 | 0.999971 | 1 | 0.78268 |
| Xab2 | Q9DCD2 | 1.31 | 1.44 | 1.18 | 1.75 | 0.98 | 0.84 | 0.59 | 1.31 | 1.23 | 0.85 | 1.14 | 1.29 | 1.06 | 1.31 | 1.17 | 1.71 | -0.49 | -0.2925 | -0.1075 | 0.058942 | 0.179641 | 0.638897 | 0.999971 | 1 | 0.93301 |
| Xdh | Q00519 | 0.31 | 1.49 | 0.13 | 0.65 | 2.88 | 1.2 | 0.32 | 0.15 | 0.44 | 0.02 | -0.43 | 0.8 | 0.66 | 1.14 | -0.32 | -0.19 | 0.4925 | -0.4375 | -0.3225 | 0.443044 | 0.27695 | 0.466268 | 0.999971 | 1 | 0.891431 |
| Xiap | Q60989 | -0.44 | -0.23 | -0.17 | 0.18 | -0.26 | -0.5 | -0.26 | 0.2 | -0.26 | -0.01 | -0.19 | -0.18 | -0.71 | 0.08 | 0.03 | 0.04 | -0.04 | 0.005 | 0.025 | 0.863061 | 0.979624 | 0.922135 | 0.999971 | 1 | 0.993069 |
| Xk | Q9QXY7 | 0.54 | -0.24 | 0.02 | 0.29 | -0.65 | -0.46 | 0.98 | -1.1 | -1.67 | 0.01 | 0.39 | -0.71 | -0.63 | -1.19 | -0.38 | -0.44 | -0.46 | -0.6475 | -0.8125 | 0.322166 | 0.179776 | 0.01399 | 0.999971 | 1 | 0.613095 |
| Xkr4 | Q5GH67 | 0.13 | -2.28 | -0.67 | -1.42 | -1.95 | -1.77 | 1.05 | -1.74 | -1.94 | -0.56 | -0.89 | -1.18 | -1.89 | -1.7 | -0.91 | -0.97 | -0.0425 | -0.0825 | -0.3075 | 0.957314 | 0.880396 | 0.564843 | 0.999971 | 1 | 0.917442 |
| Xkr6 | E9Q6C8 | 5.8 | 0.27 | 0.49 | 0.76 | -0.03 | 0.03 | 0.89 | 0.69 | 0.03 | 3.13 | 0.35 | 0.24 | 0.95 | 0.06 | 0.25 | 0.38 | -1.435 | -0.8925 | -1.42 | 0.250355 | 0.515874 | 0.25379 | 0.999971 | 1 | 0.839012 |
| Xkr7 | Q5GH64 | 0.52 | -0.56 | -0.62 | 0.26 | -0.03 | 0.01 | -0.75 | -0.5 | -0.59 | 0.31 | -1.08 | -1.44 | -0.63 | -0.19 | -0.89 | -0.58 | -0.2175 | -0.6 | -0.4725 | 0.52608 | 0.204657 | 0.169876 | 0.999971 | 1 | 0.802438 |
| Xpc | A0A0U1RP | -0.49 | 0.37 | 1.88 | 2.19 | 0.38 | -1.2 | 2.35 | 1.93 | -1.43 | -0.29 | 2.03 | 1.26 | -0.57 | -1.11 | 1.6 | 2.13 | -0.1225 | -0.595 | -0.475 | 0.893775 | 0.515095 | 0.604903 | 0.999971 | 1 | 0.924454 |
| Xpnpep1 | Q3UE92 | 0.01 | 0.39 | 0.68 | 0.44 | 0.3 | 0.46 | 0.75 | 0.52 | 0.45 | 0.19 | 0.56 | 0.39 | 0.21 | 0.05 | 0.34 | 0.28 | 0.1275 | 0.0175 | -0.16 | 0.556259 | 0.932663 | 0.446407 | 0.999971 | 1 | 0.884467 |
| Xpnpep3 | B7ZMP1 | -1.07 | -0.97 | -0.99 | -1.61 | -0.51 | -1.21 | -0.77 | -1.13 | -1.06 | -1.2 | -1 | -0.78 | -1.13 | -0.91 | -1.26 | -1.44 | 0.255 | 0.15 | -0.025 | 0.322779 | 0.495703 | 0.912734 | 0.999971 | 1 | 0.992618 |
| Xpo1 | Q6P5F9 | -0.54 | 0.41 | 0.35 | -0.03 | 0.58 | 0.31 | 0.2 | 0.16 | 0.5 | -0.31 | 0.25 | 0.68 | 0.58 | 0.24 | 0.6 | 0.27 | 0.265 | 0.2325 | 0.375 | 0.324921 | 0.462553 | 0.177631 | 0.999971 | 1 | 0.808532 |
| Xpo4 | A0A0R4J25 | -0.61 | 0.01 | 0 | -0.06 | -0.03 | -0.25 | 0.05 | 0.16 | 0.01 | -0.1 | 0.53 | 0.93 | -0.06 | -0.02 | 0.15 | 0.11 | 0.1475 | 0.5075 | 0.21 | 0.503987 | 0.109859 | 0.330459 | 0.999971 | 1 | 0.857107 |
| Xpo5 | Q924C1 | 0.44 | 1.34 | 1.88 | 1.52 | 0.89 | 1.25 | 1.45 | 1.61 | 1.15 | 0.7 | 1.76 | 1.23 | 0.88 | 1.39 | 1.71 | 1.59 | 0.005 | -0.085 | 0.0975 | 0.98825 | 0.81648 | 0.781658 | 0.999971 | 1 | 0.965859 |
| Xpo6 | Q924Z6 | -0.32 | 1.43 | 0.74 | 1.2 | 1.04 | 0.73 | 0.73 | 1.28 | 0.98 | 0.52 | 0.8 | 0.6 | 1.18 | 0.64 | 1 | 1.12 | 0.1825 | -0.0375 | 0.2225 | 0.644718 | 0.922833 | 0.573101 | 0.999971 | 1 | 0.91816 |
| Xpo7 | Q9EPK7 | -0.01 | 0.29 | -0.04 | -0.12 | 0.02 | 0.22 | -0.18 | -0.27 | -0.04 | -0.16 | 0.12 | -0.06 | -0.1 | 0.02 | -0.03 | -0.35 | -0.0825 | -0.065 | -0.145 | 0.680985 | 0.717931 | 0.453692 | 0.999971 | 1 | 0.887217 |
| Xpot | Q9CRT8 | 0.27 | 0.74 | 1.03 | 0.77 | 0.77 | 0.58 | 0.88 | 0.92 | 0.81 | 0.63 | 1.1 | 0.07 | 0.59 | 0.62 | 1.12 | 0.94 | 0.085 | -0.05 | 0.115 | 0.699873 | 0.859255 | 0.631505 | 0.999971 | 1 | 0.932858 |
| Xpr1 | Q9Z0U0 | -1.39 | -0.21 | 0.06 | -0.44 | -0.25 | -0.05 | -0.61 | -0.02 | 0.06 | -0.99 | -0.08 | -0.06 | 0.56 | -0.35 | -0.02 | -0.35 | 0.2625 | 0.2275 | 0.455 | 0.447778 | 0.557852 | 0.240315 | 0.999971 | 1 | 0.834057 |
| Xrcc1 | Q60596 | 0.56 | 1.56 | 0.91 | 1.03 | 1.29 | 0.52 | 0.62 | 1.16 | 1.09 | 1.49 | 1.18 | 0.81 | 1.24 | 1.44 | 1.03 | 1.75 | -0.1175 | 0.1275 | 0.35 | 0.689803 | 0.636931 | 0.225503 | 0.999971 | 1 | 0.828225 |
| Xrcc4 | A0A0R4J1Y | 0.26 | 1.21 | 0.71 | 0.41 | 1.47 | 0.84 | 0.87 | 0.68 | 1.34 | 0.22 | 1.03 | 1.41 | 1.48 | 0.82 | 0.73 | 0.19 | 0.3175 | 0.3525 | 0.1575 | 0.283232 | 0.315803 | 0.64118 | 0.999971 | 1 | 0.93301 |
| Xrcc5 | P27641 | 1.22 | 2.13 | 0.03 | 2.39 | 1.46 | 1.19 | -0.55 | 5.14 | 1.34 | 1.31 | -1.56 | -0.59 | 1.38 | 2 | 0.17 | 1.38 | 0.3675 | -1.3175 | -0.21 | 0.752817 | 0.131577 | 0.727239 | 0.999971 | 1 | 0.956041 |
| Xrcc6 | A0A0R4J18 | 0.11 | -0.66 | 1.4 | 0.98 | -0.59 | -1.02 | 0.6 | 1.62 | -0.95 | -0.01 | 0.68 | 1.09 | -1.39 | -0.48 | 1.29 | 1.06 | -0.305 | -0.255 | -0.3375 | 0.656312 | 0.66642 | 0.637352 | 0.999971 | 1 | 0.93301 |
| Xrn1 | A0A087WC | 1.58 | 0.48 | 0.65 | 0.87 | 0.68 | 0.33 | 0.92 | 0.52 | 0.58 | 0.85 | 0.53 | 0.62 | 0.37 | -0.13 | 0.4 | 0.55 | -0.2825 | -0.25 | -0.5975 | 0.336464 | 0.366414 | 0.068664 | 0.999971 | 1 | 0.733937 |
| Xrn2 | Q9DBR1-2 | 1.45 | 2.15 | 1.62 | 1.75 | 1.83 | 1.95 | 1.41 | 1.79 | 2.07 | 1.9 | 1.29 | 1.46 | 1.46 | 1.99 | 1.52 | 1.86 | 0.0025 | -0.0625 | -0.035 | 0.991255 | 0.808908 | 0.881066 | 0.999971 | 1 | 0.986006 |
| Xxylt1 | Q3U4G3 | 0.01 | 0.13 | -0.3 | 0.3 | 0.11 | -0.12 | -0.72 | 0.53 | 0.31 | -0.18 | -0.01 | 0.11 | -0.7 | -0.01 | -0.05 | 0.01 | -0.085 | 0.0225 | -0.2225 | 0.776286 | 0.914571 | 0.37437 | 0.999971 | 1 | 0.87031 |
| Xylb | Q3TNA1 | 3.62 | 1.66 | -0.04 | -0.41 | 0.88 | 1.91 | -0.19 | 0.26 | 0.5 | 2.18 | 0.35 | -0.2 | -2.03 | 0.21 | -0.21 | 0.31 | -0.4925 | -0.5 | -1.6375 | 0.594982 | 0.60146 | 0.114915 | 0.999971 | 1 | 0.78268 |
| Yae1d1 | Q9DAY6 | -0.68 | 0.92 | 1.15 | 1.16 | 1.41 | 0.77 | 2.56 | 0.23 | 0.09 | -0.76 | 0.32 | -0.12 | 1.48 | -0.18 | 0.35 | -0.54 | 0.605 | -0.755 | -0.36 | 0.334676 | 0.134658 | 0.534392 | 0.999971 | 1 | 0.910077 |
| Yaf2 | Q99LW6 | 2.98 | 0.55 | 0.45 | 2.38 | 0.64 | 1.34 | 3.65 | 1.56 | 1.21 | 2.29 | -1.47 | -1.63 | -2.22 | 0.36 | -0.94 | -1.81 | 0.2075 | -1.49 | -2.7425 | 0.800013 | 0.180897 | 0.006154 | 0.999971 | 1 | 0.582714 |
| Yap1 | P46938 | -0.58 | 0.64 | 0.12 | 0.35 | 2.02 | 0.61 | 0.28 | 0.1 | 1.52 | -0.55 | -0.26 | 0.52 | 1.55 | -0.49 | 0.23 | 0.1 | 0.62 | 0.175 | 0.215 | 0.212358 | 0.724689 | 0.649489 | 0.999971 | 1 | 0.933842 |
| Yars | A2A7S7 | 0.5 | 1.36 | 1.44 | 1.14 | 1.26 | 1.21 | 1.47 | 1.32 | 1.47 | 0.93 | 1.65 | 1.95 | 1.29 | 1.09 | 1.47 | 1.56 | 0.205 | 0.39 | 0.2425 | 0.4192 | 0.226176 | 0.364948 | 0.999971 | 1 | 0.869526 |
| Yars1 | Q91WQ3 | 2.64 | 0.67 | 2.37 | 2.23 | 1.32 | 0.87 | 2.68 | 1.53 | 0.97 | 2.36 | 0.73 | 0.8 | -0.56 | 0.13 | 0.47 | 1.66 | -0.3775 | -0.7625 | -1.5525 | 0.491349 | 0.185139 | 0.025481 | 0.999971 | 1 | 0.678474 |
| Yars2 | Q8BYL4 | -0.51 | -0.58 | -0.15 | -0.43 | -0.69 | -0.82 | 0.07 | -0.35 | -0.55 | -0.81 | 0.21 | 0.07 | -0.41 | -0.63 | -0.35 | -0.38 | -0.03 | 0.1475 | -0.025 | 0.903926 | 0.597363 | 0.893317 | 0.999971 | 1 | 0.98828 |
| Ybx1 | P62960 | 0.01 | -0.46 | 0.14 | 0.5 | -0.3 | -0.69 | 1.84 | 0.03 | 0.08 | 0.58 | 0.15 | 0.31 | -0.83 | -0.5 | 0.53 | 0.3 | 0.1725 | 0.2325 | -0.1725 | 0.752458 | 0.368569 | 0.641754 | 0.999971 | 1 | 0.93301 |
| Ybx3 | Q9JKB3 | 1.75 | 0.65 | 0.17 | 0.65 | 0.83 | 0.71 | 0.08 | 0.45 | 1.08 | 1.99 | -0.34 | 0.07 | 0.04 | 1.17 | 0.56 | 0.63 | -0.2875 | -0.105 | -0.205 | 0.437764 | 0.854419 | 0.603131 | 0.999971 | 1 | 0.924454 |
| Ydjc | Q14BV6 | 0.57 | 1.15 | 0.19 | 0.58 | 0.88 | 0.48 | 0 | 0.75 | 0.44 | 0.76 | 0.45 | 0.38 | 1.86 | 0.35 | 0.71 | 0.75 | -0.095 | -0.115 | 0.295 | 0.743388 | 0.639434 | 0.435549 | 0.999971 | 1 | 0.882419 |
| Yeats4 | Q9CR11 | -0.64 | -0.78 | -0.2 | 0.16 | 0.08 | -1.6 | -0.08 | -0.63 | -0.38 | -1.21 | -0.5 | -0.32 | 0.3 | -0.6 | -0.57 | -0.84 | -0.1925 | -0.2375 | -0.0625 | 0.643803 | 0.442432 | 0.849561 | 0.999971 | 1 | 0.98187 |
| Yes1 | Q04736 | -0.29 | -0.33 | -0.38 | -0.02 | -0.71 | -0.26 | -0.72 | -0.01 | -0.3 | -0.62 | -0.62 | -0.43 | 0.18 | -0.11 | 0.23 | -0.02 | -0.17 | -0.2375 | 0.325 | 0.467767 | 0.214028 | 0.108819 | 0.999971 | 1 | 0.78268 |
| Yif1b | Q9CX30-2 | 0.02 | -0.43 | 0.33 | 0 | -0.38 | -0.47 | 0.39 | 0.08 | -0.65 | -0.26 | -0.2 | -0.21 | -1.04 | -0.35 | -0.31 | 0.57 | -0.075 | -0.31 | -0.2625 | 0.783883 | 0.194689 | 0.470134 | 0.999971 | 1 | 0.892481 |
| Yipf1 | A2A8S2 | 4.56 | -0.04 | -0.4 | -0.17 | -0.01 | 0.64 | -0.19 | 1.32 | 1.34 | 3.28 | 0.23 | 0.52 | 0.73 | 0.29 | 0.58 | 0.41 | -0.5475 | 0.355 | -0.485 | 0.622622 | 0.773787 | 0.651606 | 0.999971 | 1 | 0.933842 |
| Yipf3 | Q3UDR8 | 0.89 | 0.26 | 1.08 | 1.68 | 0.77 | 0.94 | 0.88 | 1.76 | 0.69 | 1.33 | 1.01 | 1.62 | 0.19 | 0.73 | 1.91 | 2.48 | 0.11 | 0.185 | 0.35 | 0.761067 | 0.599017 | 0.531933 | 0.999971 | 1 | 0.910069 |
| Yipf4 | Q8C407 | -1.17 | 1.13 | 1.95 | 1.91 | 1.04 | 1.07 | 1.71 | 2.57 | 1.02 | -1.02 | 1.46 | 2.57 | 2.34 | 1.33 | 2.15 | 1.98 | 0.6425 | 0.0525 | 0.995 | 0.393229 | 0.955628 | 0.175921 | 0.999971 | 1 | 0.808532 |
| Yipf5 | Q9EQQ2 | -7.85 | -0.42 | -0.03 | -2.14 | 0.5 | -0.84 | 0.67 | -1.4 | 1.71 | -6.13 | -0.05 | 0.9 | -0.01 | -2.72 | -0.39 | -2.9 | 2.3425 | 1.7175 | 1.105 | 0.182457 | 0.455131 | 0.529308 | 0.999971 | 1 | 0.910069 |
| Yipf6 | Q8BR70 | 1.55 | 0.18 | -0.16 | 0.22 | -0.11 | -0.03 | 0.45 | 0.4 | -0.26 | 1.75 | 0 | -0.03 | -1.53 | 0.08 | 0.06 | 0.67 | -0.27 | -0.0825 | -0.6275 | 0.492675 | 0.881411 | 0.276496 | 0.999971 | 1 | 0.843794 |
| Yjefn3 | F6W8I0 | -0.36 | -0.35 | -0.65 | -0.68 | -0.64 | 0.45 | -0.13 | -0.24 | -0.27 | 0.41 | -0.19 | -0.99 | 0.37 | 0.3 | -0.42 | 0.32 | 0.37 | 0.25 | 0.6525 | 0.185074 | 0.423454 | 0.022851 | 0.999971 | 1 | 0.652043 |
| Yju2 | Q9D6J3 | 1.67 | 1.41 | -0.69 | 0.37 | 0.71 | 0.74 | -0.74 | -0.29 | 0.66 | 1.56 | 1.64 | 2.12 | 0 | 1.16 | -0.2 | 0.21 | -0.585 | 0.805 | -0.3975 | 0.340311 | 0.182766 | 0.489337 | 0.999971 | 1 | 0.898346 |
| Ykt6 | Q9CQW1 | -0.25 | -0.22 | -1.22 | -0.79 | -0.25 | -0.3 | -0.78 | -1.28 | -0.08 | -0.17 | -0.93 | -1.28 | -0.05 | -0.43 | -1.12 | -0.9 | -0.0325 | 0.005 | -0.005 | 0.92305 | 0.98914 | 0.988142 | 0.999971 | 1 | 0.99925 |
| Ylpm1 | D3YWX2 | 0.43 | -0.37 | 0.44 | 0.33 | -0.41 | -0.34 | 0.62 | 0.28 | -0.41 | 0.16 | 0.47 | 0.38 | -0.57 | -0.31 | -0.04 | 0.16 | -0.17 | -0.0575 | -0.3975 | 0.59588 | 0.841989 | 0.167258 | 0.999971 | 1 | 0.800215 |
| Yme1l1 | O88967 | 0.06 | 0.02 | 0.01 | 0.01 | -0.37 | -0.29 | -0.13 | -0.08 | -0.26 | 0.12 | -0.16 | 0 | -0.13 | 0.02 | 0.01 | 0.04 | -0.2425 | -0.1 | -0.04 | 0.173265 | 0.560391 | 0.80452 | 0.999971 | 1 | 0.971467 |
| Yod1 | Q8CB27 | 1.12 | 0.38 | 0.5 | 0.85 | 0.39 | 0.67 | 0.22 | 1.04 | 0.58 | 1.16 | 0.33 | 0 | 0.17 | 0.38 | 0.21 | 0.96 | -0.1325 | -0.195 | -0.2825 | 0.621605 | 0.525862 | 0.309164 | 0.999971 | 1 | 0.847402 |
| Ypel1 | A0A338P71 | -2.29 | 0.74 | 0.03 | -4.56 | 0.43 | -0.36 | 3.09 | -3.95 | 1.9 | -4.62 | 1.14 | 0.3 | -0.79 | -4.5 | -2.21 | -5.6 | 1.3225 | 1.2 | -1.755 | 0.436524 | 0.485395 | 0.243207 | 0.999971 | 1 | 0.837316 |
| Ypel5 | P62700 | -0.19 | 1.49 | 0.7 | 0.4 | 1.33 | 1 | 0.35 | 0.43 | 1.08 | 0.56 | 0.95 | 0.94 | 1.81 | 1.08 | 0.37 | 0.55 | 0.1775 | 0.2825 | 0.3525 | 0.66028 | 0.43988 | 0.439825 | 0.999971 | 1 | 0.883105 |
| Yrdc | Q3U5F4 | 1.32 | 0.65 | -1.11 | 0.24 | -0.39 | 0.58 | -1.12 | -0.38 | -0.53 | 0.1 | -0.41 | -0.91 | -0.22 | 0.22 | 0.13 | 0.23 | -0.6025 | -0.7125 | -0.185 | 0.304565 | 0.190017 | 0.705306 | 0.999971 | 1 | 0.950182 |
| Ythdc1 | E9Q5K9-3 | 0.81 | -0.16 | 0.5 | 0.44 | -0.46 | -0.03 | 1.18 | 0.44 | -0.14 | 0.18 | 2.56 | 0.91 | -0.6 | -0.17 | 0.41 | -0.02 | -0.115 | 0.48 | -0.4925 | 0.768092 | 0.420918 | 0.127634 | 0.999971 | 1 | 0.783863 |
| Ythdc2 | B2RR83 | 0.63 | 1.4 | 0.72 | 0.68 | 1.33 | 0.76 | 0.75 | 0.54 | 1.51 | 1.06 | 1.25 | 1.17 | 1.42 | 0.94 | 0.62 | 1.02 | -0.0125 | 0.39 | 0.1425 | 0.962822 | 0.127125 | 0.596107 | 0.999971 | 1 | 0.92377 |
| Ythdf1 | A2AWN8 | 2.58 | 2.01 | 0.69 | 0.33 | 1.36 | 1.95 | 1.17 | 0.56 | 1.63 | 1.88 | 0.75 | 1.06 | 0.32 | 1.15 | 0.7 | 0.31 | -0.1425 | -0.0725 | -0.7825 | 0.797766 | 0.894655 | 0.162355 | 0.999971 | 1 | 0.800215 |
| Ythdf2 | Q91YT7 | 0.34 | 1.48 | 2.52 | 2.82 | 2.18 | 0.53 | 2.51 | 3 | 1.44 | -0.45 | 3.24 | 2.61 | 0.35 | -1.11 | 2.45 | 3.38 | 0.265 | -0.08 | -0.5225 | 0.706603 | 0.928301 | 0.616463 | 0.999971 | 1 | 0.928229 |
| Ythdf3 | Q8BYK6-3 | 0.69 | 1.87 | 1.18 | 1.15 | 1.68 | 1.54 | 1.4 | 1.09 | 1.83 | 1.01 | 1.35 | 1.41 | 1.45 | 1.6 | 0.97 | 1.23 | 0.205 | 0.1775 | 0.09 | 0.481422 | 0.561325 | 0.757916 | 0.999971 | 1 | 0.961116 |
| Ywhab | Q9CQV8 | -0.18 | -0.58 | -0.37 | 0.09 | -1.05 | -0.8 | -0.54 | -0.13 | -1.14 | -0.46 | -0.58 | -0.3 | -1.35 | -0.5 | -0.25 | 0.24 | -0.37 | -0.36 | -0.205 | 0.185436 | 0.182608 | 0.567591 | 0.999971 | 1 | 0.91816 |
| Ywhae | P62259 | -0.66 | 0 | 0.81 | 0.36 | -0.2 | 0.14 | 0.12 | 1.09 | -0.15 | -0.46 | 0.8 | -1.19 | -0.02 | 0.05 | 1.25 | 1.05 | 0.16 | -0.3775 | 0.455 | 0.689475 | 0.442512 | 0.30529 | 0.999971 | 1 | 0.847402 |
| Ywhag | P61982 | 0.16 | 0.29 | 0.32 | 0.27 | -0.22 | 0.2 | 1.13 | 0.11 | -0.04 | -0.01 | 0.45 | 0.49 | 0.38 | 0.05 | 0.46 | 0.6 | 0.045 | -0.0375 | 0.1125 | 0.88071 | 0.851711 | 0.558463 | 0.999971 | 1 | 0.915051 |
| Ywhah | P68510 | 0.1 | -0.05 | 0.53 | 0.66 | -0.36 | 0.51 | 0.48 | 0.53 | -0.17 | 0.05 | 0.78 | 0.7 | 0.25 | -0.05 | 0.74 | 1.23 | -0.02 | 0.03 | 0.2325 | 0.944557 | 0.919883 | 0.485799 | 0.999971 | 1 | 0.897483 |
| Ywhaq | P68254 | 0.36 | 0.6 | -0.38 | 0.17 | 0.44 | 0.33 | -0.5 | -0.08 | 0.58 | 0.76 | -0.15 | -0.81 | 0.86 | 0.61 | 0.1 | 0.18 | -0.14 | -0.0925 | 0.25 | 0.648603 | 0.816903 | 0.396864 | 0.999971 | 1 | 0.871816 |
| Ywhaz | P63101 | -0.11 | 0.93 | 0.03 | 0.66 | 0.25 | 0.09 | 0.25 | 0.2 | 0.07 | 0.39 | 0.48 | -0.47 | 0.46 | 1.15 | 0.48 | 0.79 | -0.18 | -0.26 | 0.3425 | 0.511556 | 0.436949 | 0.278656 | 0.999971 | 1 | 0.843812 |
| Yy1 | Q00899 | 3.25 | 3.53 | 3.09 | 3.59 | 3.19 | 3 | 2.93 | 3.07 | 2.8 | 3.31 | 2.99 | 2.84 | 2.82 | 3.18 | 3.78 | 2.62 | -0.3175 | -0.38 | -0.265 | 0.128351 | 0.099454 | 0.376278 | 0.999971 | 1 | 0.870465 |
| Zbed3 | Q9D0L1 | 2.48 | -0.29 | 1.25 | 1.46 | -0.26 | 0.46 | 2.6 | 1.4 | 0.27 | 1.97 | 0.57 | -0.02 | -3 | -1.81 | -0.3 | -1.1 | -0.175 | -0.5275 | -2.7775 | 0.817618 | 0.432458 | 0.004216 | 0.999971 | 1 | 0.582714 |
| Zbtb11 | G5E8B9 | 1.92 | 0.85 | 0.47 | 0.2 | 0.23 | 1.04 | 0.29 | 0.26 | 0.06 | 1.87 | 0.7 | 0.8 | 0.61 | 0.95 | 0.11 | 0.65 | -0.405 | -0.0025 | -0.28 | 0.331972 | 0.995978 | 0.488758 | 0.999971 | 1 | 0.898346 |
| Zbtb16 | Q3UQ17 | 0.75 | 0.16 | 1.71 | 0.02 | 0.99 | -0.27 | 1.3 | 0.2 | 1.15 | 0.37 | 1.11 | 0.34 | 1.26 | 0.07 | 0.06 | 0.47 | -0.105 | 0.0825 | -0.195 | 0.829993 | 0.845298 | 0.665159 | 0.999971 | 1 | 0.93697 |
| Zbtb18 | A0A0A6YY3 | 2.98 | -0.84 | 1.97 | 1.72 | -2.06 | 1.36 | 1.27 | 1.98 | -1.52 | 2.12 | 1.52 | 1.29 | -3.44 | -1.58 | 1.9 | 2.75 | -0.82 | -0.605 | -1.55 | 0.458646 | 0.56122 | 0.310885 | 0.999971 | 1 | 0.84774 |
| Zbtb20 | Q8K0L9 | 0.87 | 0.23 | 2.45 | 1.41 | -0.35 | 0.74 | 2.08 | 1.26 | -0.7 | 0.76 | 0.66 | -2.22 | -1.53 | 0.09 | 1.4 | 1.23 | -0.3075 | -1.615 | -0.9425 | 0.627558 | 0.060211 | 0.225796 | 0.999971 | 1 | 0.828225 |
| Zbtb38 | F6Z595 | 1.57 | -0.13 | 0.43 | 0.73 | 1.1 | 0.6 | 0.12 | 0.04 | 0.48 | 2.5 | -0.24 | -0.47 | -0.77 | 0.38 | -0.02 | -0.04 | -0.185 | -0.0825 | -0.7625 | 0.653789 | 0.905364 | 0.089425 | 0.999971 | 1 | 0.771991 |
| Zbtb45 | Q52KG4 | 1.02 | 1.87 | 0.49 | 0.09 | 0.48 | -0.4 | 0.2 | 0.74 | 0.95 | 0.77 | 0.67 | 0.28 | 1.64 | 1.47 | 0.81 | 0.22 | -0.6125 | -0.2 | 0.1675 | 0.179281 | 0.614297 | 0.722996 | 0.999971 | 1 | 0.955258 |
| Zbtb7a | O88939-2 | 0.42 | 0.92 | 1.96 | 1.32 | 0.87 | -0.15 | 3.29 | 1.57 | 0.56 | 0.97 | 2.14 | 0.48 | 0.25 | 0.53 | 1.11 | -0.15 | 0.24 | -0.1175 | -0.72 | 0.738191 | 0.803251 | 0.100606 | 0.999971 | 1 | 0.78268 |
| Zbtb8os | B2KGA7 | -0.95 | 0.19 | 0 | -0.68 | 1.6 | 1.89 | 1.25 | 1.57 | 0.39 | -1.08 | -0.09 | -0.28 | 2.51 | 1.1 | 1.8 | 1.16 | 1.9375 | 0.095 | 2.0025 | 0.000161 | 0.809267 | 0.000897 | 0.244833 | 1 | 0.365812 |
| Zc2hc1a | Q8BJH1 | 1.58 | -0.01 | 0.28 | 0.39 | 0.86 | 1.22 | 0.5 | 0.56 | 0.99 | 1.58 | 0.15 | 0.25 | 1.15 | 1.32 | 0.47 | 0.64 | 0.225 | 0.1825 | 0.335 | 0.552249 | 0.690795 | 0.400086 | 0.999971 | 1 | 0.871816 |
| Zc3h11a | Q6NZF1 | 0.23 | 0.42 | 0.75 | 1.26 | 0.41 | -0.06 | 0.57 | 0.64 | 0.11 | 0.18 | 0.65 | 0.84 | -0.03 | 0.32 | 1.04 | 1.13 | -0.275 | -0.22 | -0.05 | 0.352075 | 0.464881 | 0.887687 | 0.999971 | 1 | 0.987485 |
| Zc3h12b | G3X9I7 | -0.96 | -0.11 | -2.41 | -1.15 | -0.56 | -0.2 | -1.58 | -1.76 | -0.67 | -1.52 | -0.85 | -1.99 | -0.3 | -0.07 | -1.48 | -2.62 | 0.1325 | -0.1 | 0.04 | 0.812635 | 0.848556 | 0.953416 | 0.999971 | 1 | 0.998056 |
| Zc3h13 | E9Q784 | -4.67 | -1.1 | 0.7 | 0.85 | -0.29 | -2.3 | 0.94 | -0.59 | -1.05 | -5.72 | 0.73 | -0.24 | -0.19 | -4.62 | 0.08 | 0.96 | 0.495 | -0.515 | 0.1125 | 0.701058 | 0.764387 | 0.943534 | 0.999971 | 1 | 0.99692 |
| Zc3h14 | Q8BJ05-2 | 0.75 | 0.76 | 1.28 | 0.62 | 0.54 | 0.5 | 1.04 | 0.78 | 0.44 | 0.49 | 0.86 | 1.42 | 0.38 | 0.64 | 0.94 | 0.55 | -0.1375 | -0.05 | -0.225 | 0.554089 | 0.859505 | 0.336891 | 0.999971 | 1 | 0.858827 |
| Zc3h15 | Q3TIV5 | 0.75 | 1.05 | 1.58 | 1.96 | 1.8 | 0.92 | 1.57 | 1.64 | 1.17 | 1.05 | 1.66 | 1.57 | 0.54 | 1.22 | 1.59 | 1.65 | 0.1475 | 0.0275 | -0.085 | 0.65763 | 0.929763 | 0.814902 | 0.999971 | 1 | 0.974918 |
| Zc3h18 | H3BIW0 | 1.4 | 0.78 | 0.95 | 1.62 | 1.06 | 0.76 | 0.7 | 1.21 | 0.91 | 1.05 | 1.12 | 1.2 | 0.54 | 0.85 | 1.33 | 1.46 | -0.255 | -0.1175 | -0.1425 | 0.331531 | 0.621231 | 0.635273 | 0.999971 | 1 | 0.93301 |

| | | | | | | | | | | | | | | | | | | | | | | | | | | |
|---|---|---|---|---|---|---|---|---|---|---|---|---|---|---|---|---|---|---|---|---|---|---|---|---|---|---|
| Zc3h3 | Q8CHP0 | 2.37 | 1.49 | 0.8 | 1.78 | 1.03 | 0.97 | 0.36 | 0.97 | 1.48 | 1.92 | 0.82 | 0.19 | 1.24 | 1.19 | 0.12 | 1.96 | -0.7775 | -0.5075 | -0.4825 | 0.052758 | 0.29611 | 0.317806 | 0.999971 | 1 | 0.853893 |
| Zc3h4 | E9Q8K8 | 2.82 | 1.56 | 1.28 | 1.64 | 1.87 | 1.49 | 1.5 | 1.52 | 1.47 | 1.92 | 1.29 | 1.45 | 1.05 | 1.25 | 1.43 | 1.33 | -0.23 | -0.2925 | -0.56 | 0.513004 | 0.423795 | 0.132234 | 0.999971 | 1 | 0.784948 |
| Zc3h6 | A2AP88 | 0.65 | -0.53 | -0.78 | -0.48 | -0.44 | -0.98 | 0.1 | -0.01 | 0.8 | 0.73 | 1.02 | 1.1 | 0.42 | -0.4 | 0.29 | -0.31 | -0.0475 | 1.1975 | 0.285 | 0.902145 | 0.005468 | 0.448893 | 0.999971 | 1 | 0.885351 |
| Zc3h7b | F8VPP8 | 0.02 | 1.13 | 0.79 | 0.36 | 1.09 | 1.03 | 1.34 | 0.51 | 1.26 | 0.28 | 1.32 | 1.22 | 0.92 | 0.92 | 0.82 | 0.38 | 0.4175 | 0.445 | 0.185 | 0.19517 | 0.216355 | 0.526011 | 0.999971 | 1 | 0.909459 |
| Zc3hc1 | H3BKM2 | 0.52 | 1.38 | 0.59 | 0.66 | 1.52 | 0.91 | -0.06 | 0.11 | 1.4 | 0.39 | -0.08 | 0.29 | 0.1 | 0.83 | 0.69 | 0.39 | -0.1675 | -0.2875 | -0.285 | 0.67629 | 0.439289 | 0.315957 | 0.999971 | 1 | 0.85297 |
| Zcchc10 | F6QZB1 | 0.07 | 0.45 | -0.74 | -0.12 | 0.34 | -0.95 | -1.16 | -0.39 | 0.16 | -0.35 | -0.33 | -1.11 | 0.81 | 0.07 | -1.06 | -0.87 | -0.455 | -0.3225 | -0.1775 | 0.272214 | 0.37442 | 0.705746 | 0.999971 | 1 | 0.950182 |
| Zcchc17 | Q9ESX4-2 | 1.68 | 1.63 | 0.77 | 1.25 | 0.77 | 0.39 | -0.7 | 0 | 1.41 | 1.46 | 0.41 | 1.52 | 1.2 | 2.01 | 0.64 | 0.7 | -1.2175 | -0.1325 | -0.195 | 0.008696 | 0.694147 | 0.601078 | 0.999971 | 1 | 0.92426 |
| Zcchc18 | Q8VD24 | -1.47 | 0.88 | 0.69 | 0.83 | 0.24 | 0.79 | 1.28 | 1.48 | 0.63 | -1.08 | 0.93 | 0.6 | 1.02 | 0.49 | 1.16 | 0.83 | 0.715 | 0.0375 | 0.6425 | 0.237174 | 0.955008 | 0.254531 | 0.999971 | 1 | 0.839012 |
| Zcchc8 | Q9CYA6 | 2.94 | 2.01 | 1.17 | 1.13 | 3.89 | 2.6 | 0.17 | 0.92 | 0.87 | 1.7 | 0.87 | 0.48 | 3.94 | 2.97 | 0.75 | 1.33 | 0.0825 | -0.8325 | 0.435 | 0.921921 | 0.103048 | 0.5736 | 0.999971 | 1 | 0.91816 |
| Zcrb1 | Q9CZ96-2 | 1.95 | 1.54 | 1.95 | 2.54 | 1.13 | 2.1 | 1.85 | 2.42 | 1.31 | 1.86 | 1.93 | 1.59 | 1.25 | 1.74 | 2.28 | 2.27 | -0.12 | -0.3225 | -0.11 | 0.724528 | 0.248861 | 0.734204 | 0.999971 | 1 | 0.957712 |
| Zdhhc13 | Q9CWU2 | 1.32 | 1.6 | 1.31 | 0.46 | 0.74 | 1.19 | 0.33 | 0.97 | 0.43 | 1.31 | 0.89 | 0.83 | 1.39 | 1.46 | 1.05 | 1.52 | -0.365 | -0.3075 | 0.1825 | 0.261134 | 0.335518 | 0.524477 | 0.999971 | 1 | 0.908589 |
| Zdhhc14 | Q8BQQ1 | 4.65 | -0.06 | -0.95 | -0.95 | -0.95 | 0.13 | -1.52 | -0.44 | -0.12 | 4.01 | -1.46 | -1.33 | -1.6 | 0.5 | -1.35 | -0.81 | -1.3675 | -0.3975 | -1.4875 | 0.284597 | 0.810467 | 0.258667 | 0.999971 | 1 | 0.839012 |
| Zdhhc17 | Q80TN5 | 1.38 | 1.07 | 1.24 | 1.28 | 1.25 | 1.36 | 1.59 | 0.89 | 1.32 | 1.5 | 1.2 | 1.4 | 1.92 | 1.45 | 1.43 | 1.54 | 0.03 | 0.1125 | 0.3425 | 0.886402 | 0.518879 | 0.106497 | 0.999971 | 1 | 0.78268 |
| Zdhhc18 | Q5Y5T2 | 1.81 | -1.19 | 0.27 | 1.27 | -2.1 | 0.05 | 0.27 | 0.71 | -2.05 | 1.39 | -0.54 | 0.08 | -2.16 | 0.3 | 0.57 | 0.51 | -0.8075 | -0.82 | -0.735 | 0.336234 | 0.362184 | 0.39041 | 0.999971 | 1 | 0.871816 |
| Zdhhc2 | P59267 | -0.3 | -1 | -1.68 | -1.08 | -1.18 | -0.66 | -1.26 | -0.96 | -0.66 | 0.12 | -0.82 | -0.92 | -1.47 | -1.07 | -1.14 | -1.11 | 7.85E-17 | 0.445 | -0.1825 | 1 | 0.237286 | 0.553425 | 1 | 1 | 0.913595 |
| Zdhhc20 | Q5Y5T1-2 | -0.38 | 0.34 | -0.9 | -0.82 | -0.65 | -0.21 | -1.35 | 1.03 | 0.07 | -0.52 | -0.97 | -0.66 | -0.64 | 0.66 | -1.14 | -0.76 | 0.145 | -0.08 | -0.03 | 0.784699 | 0.820157 | 0.947313 | 0.999971 | 1 | 0.997651 |
| Zdhhc22 | A0PK84 | -0.67 | -0.64 | -0.35 | -0.85 | -0.19 | -0.68 | 0.47 | 1.17 | -0.75 | 0.08 | 0.33 | 0.86 | -0.81 | -0.09 | -0.14 | -0.45 | 0.82 | 0.7575 | 0.255 | 0.06484 | 0.053228 | 0.290761 | 0.999971 | 1 | 0.847402 |
| Zdhhc3 | A0A571BEV | -0.08 | -0.84 | 2.96 | 3.02 | 0.66 | -1.68 | 0.45 | 2.61 | -1.36 | -0.47 | 3.43 | 2.15 | 0.22 | -0.44 | 2.84 | 3.65 | -0.755 | -0.3275 | 0.3025 | 0.529783 | 0.807793 | 0.810138 | 0.999971 | 1 | 0.973855 |
| Zdhhc5 | Q8VDZ4 | -0.12 | -2.11 | -1.62 | -1.74 | -2.04 | -1.83 | -1.06 | -2.51 | -1.78 | -0.66 | -1.4 | -1.9 | -1.79 | -1.94 | -2.55 | -2.34 | -0.4625 | -0.0375 | -0.7575 | 0.361704 | 0.938429 | 0.116109 | 0.999971 | 1 | 0.78268 |
| Zdhhc6 | Q9CPV7 | -0.85 | -0.43 | 0.81 | 0.97 | -0.24 | -0.59 | -0.05 | 1.86 | -0.47 | -1.54 | 2.47 | 2.42 | 0.13 | -1.24 | 1.24 | 2.56 | 0.12 | 0.595 | 0.5475 | 0.852557 | 0.556824 | 0.515676 | 0.999971 | 1 | 0.905815 |
| Zdhhc8 | Q5Y5T5 | -1.28 | -1.93 | -2.17 | -1.66 | -1.37 | -1.92 | -1.81 | -1.77 | -1.69 | -0.85 | -1.15 | -1.74 | -1.59 | -1.88 | -1.88 | -1.56 | 0.0425 | 0.4025 | 0.0325 | 0.866257 | 0.198178 | 0.893437 | 0.999971 | 1 | 0.98828 |
| Zdhhc9 | P59268 | -0.76 | -0.31 | 0.61 | 0.34 | -0.76 | -0.13 | 0.41 | 0.23 | -1.06 | -0.09 | 0.46 | 0.21 | -0.48 | 0.03 | 0.61 | 1.49 | -0.0325 | -0.09 | 0.4425 | 0.933206 | 0.834891 | 0.376542 | 0.999971 | 1 | 0.870465 |
| Zeb1 | Q64318 | -0.16 | 0.69 | 0.74 | 1.01 | 1.02 | 0.97 | 0.58 | 0.67 | 0.86 | 0.56 | 0.58 | 1.3 | 0.69 | 0.81 | 0.7 | 0.69 | 0.24 | 0.255 | 0.1525 | 0.414382 | 0.421295 | 0.58074 | 0.999971 | 1 | 0.920178 |
| Zeb2 | A0A0J9YVF | 0.73 | -0.68 | 2.47 | 2.1 | -0.48 | -0.42 | 2.13 | 2.45 | -0.93 | 0.73 | 2 | 1.56 | -3.12 | -0.37 | 1.97 | 2.32 | -0.235 | -0.315 | -0.955 | 0.805859 | 0.718456 | 0.465442 | 0.999971 | 1 | 0.891255 |
| Zer1 | Q80ZJ6 | -0.37 | 0.17 | 1.12 | 0.58 | 0.13 | 0.23 | 0.88 | 0.35 | 0.47 | -0.01 | 1.09 | -0.6 | 0.06 | -0.15 | 0.59 | 0.66 | 0.0225 | -0.1375 | -0.085 | 0.948769 | 0.760555 | 0.815472 | 0.999971 | 1 | 0.974918 |
| Zfa | P23607 | 0.26 | -0.42 | -0.27 | -0.62 | -0.89 | -0.36 | -0.14 | 0.2 | -1.24 | -0.61 | -0.65 | -1.34 | -2.55 | -0.49 | 0.13 | -0.88 | -0.035 | -0.6975 | -0.685 | 0.908129 | 0.03354 | 0.23763 | 0.999971 | 1 | 0.833594 |
| Zfand1 | Q8BFR6 | 1.02 | -0.23 | 0.83 | -0.34 | -0.1 | -0.76 | 0.83 | 0.54 | -0.01 | 0.45 | 0.75 | -0.03 | -1.11 | -0.84 | 0.19 | -0.73 | -0.1925 | -0.03 | -0.9425 | 0.681164 | 0.937956 | 0.051809 | 0.999971 | 1 | 0.723736 |
| Zfand2b | Q91X58 | 0.43 | -0.32 | -0.09 | -0.21 | -0.94 | -0.19 | -0.3 | -0.01 | -0.77 | 0.47 | 0.05 | -0.05 | -0.12 | 0.16 | -0.22 | -0.07 | -0.3125 | -0.0275 | -0.015 | 0.278403 | 0.929442 | 0.947011 | 0.999971 | 1 | 0.997651 |
| Zfand5 | O88878 | -0.08 | 0.4 | 0.66 | 0.58 | 0.1 | 0.17 | -0.07 | 0.53 | 0.38 | 0.88 | 0.69 | -0.05 | 0.21 | 1.05 | 0.65 | 1.07 | -0.2075 | 0.085 | 0.355 | 0.399109 | 0.759665 | 0.224263 | 0.999971 | 1 | 0.828225 |
| Zfand6 | Q9DCH6 | 2.51 | 1.85 | 0.7 | 0.92 | 1.66 | 1.67 | 0.59 | 1.35 | 1.27 | 2.27 | 0.72 | -0.08 | 1.78 | 1.97 | 0.22 | 0.97 | -0.1775 | -0.45 | -0.26 | 0.700307 | 0.457417 | 0.630042 | 0.999971 | 1 | 0.932649 |
| Zfc3h1 | B2RT41 | 2.43 | 0.42 | -0.23 | -0.01 | 0.74 | 0.8 | -0.43 | 0.31 | 0.76 | 2.3 | -0.61 | 0.8 | 2.4 | 1.42 | 0.08 | -0.22 | -0.2975 | 0.16 | 0.2675 | 0.628497 | 0.835515 | 0.73072 | 0.999971 | 1 | 0.956041 |
| Zfp180 | Q6NZI9 | 2.15 | 1.54 | 1.52 | 2.17 | 0.99 | 0.87 | 1.69 | 2.13 | 0.67 | 1.32 | 1.83 | 0.65 | -0.31 | 0.99 | 2.12 | 1.91 | -0.425 | -0.7275 | -0.6675 | 0.237483 | 0.054801 | 0.234778 | 0.999971 | 1 | 0.832918 |
| Zfp219 | Q4VA98 | 1.24 | -0.09 | 1.92 | 2.27 | 0.17 | -1.11 | 2.39 | 2.18 | 0.17 | -0.65 | 1.73 | 0.62 | -0.89 | -0.83 | 2.27 | 0.89 | -0.4275 | -0.8675 | -0.975 | 0.630619 | 0.209445 | 0.257745 | 0.999971 | 1 | 0.839012 |
| Zfp280d | V9GWW1 | 2.54 | 1.18 | 0.06 | 0.73 | 1.45 | 0.39 | -0.24 | 1 | 1.9 | -0.17 | 0.25 | 1.23 | -2.73 | 0.35 | -0.07 | 1.4 | -0.4775 | -0.325 | -1.39 | 0.423493 | 0.615678 | 0.155292 | 0.999971 | 1 | 0.795578 |
| Zfp324 | Q78F42 | 6.18 | 1.6 | 0.75 | -0.6 | -0.42 | 4.19 | 1.5 | 0.5 | 0.17 | 7.39 | 0.82 | -0.82 | -1.14 | 4.98 | 0.67 | 0.79 | -0.54 | -0.0925 | -0.6575 | 0.732083 | 0.965095 | 0.706112 | 0.999971 | 1 | 0.950182 |
| Zfp326 | A0A0R4J09 | -0.75 | 0.27 | 0.41 | 0.77 | 0.07 | 0.21 | 0.45 | 0.79 | 0.06 | -0.12 | 0.54 | 0.39 | -0.14 | 0.15 | 0.6 | 0.63 | 0.205 | 0.0425 | 0.135 | 0.566709 | 0.903964 | 0.713021 | 0.999971 | 1 | 0.951793 |
| Zfp346 | E9PYN2 | 0.98 | 0.09 | -0.23 | 0.04 | -0.31 | -0.05 | 0.77 | -0.32 | 0.01 | 1.15 | -0.6 | -0.03 | 0.1 | 0.05 | -0.25 | -0.32 | -0.1975 | -0.0875 | -0.325 | 0.5854 | 0.838132 | 0.286692 | 0.999971 | 1 | 0.846151 |
| Zfp362 | E9QA68 | 1.9 | -1.17 | 1.99 | 0.59 | -0.89 | -1.03 | 2.44 | 1 | -1.69 | 0.59 | 2.47 | 3.17 | -1.83 | -1.21 | 1.41 | 0.25 | -0.4475 | 0.3075 | -1.1725 | 0.653061 | 0.794369 | 0.228498 | 0.999971 | 1 | 0.828687 |
| Zfp36l2 | P23949 | 1.55 | 0.96 | 0 | 0.4 | 0.95 | 0.33 | 0.44 | 0.86 | 0.23 | 1.81 | 0.03 | -1.15 | -1.52 | 0.86 | -0.38 | 0.33 | -0.0825 | -0.4975 | -0.905 | 0.819513 | 0.44266 | 0.13609 | 0.999971 | 1 | 0.784948 |
| Zfp37 | A0A1Y7VJF | -0.13 | -0.19 | 1.12 | 1.53 | -0.67 | -0.74 | 1.21 | 1.59 | -0.36 | -0.35 | 1.62 | 0.18 | -1.34 | -0.2 | 0.76 | 0.46 | -0.235 | -0.31 | -0.6625 | 0.730779 | 0.600708 | 0.275127 | 0.999971 | 1 | 0.843794 |
| Zfp385a | A0A2R8VH | -0.18 | 0.72 | 0.96 | 0.4 | 0.74 | 0.77 | 1.07 | 1.33 | 1.02 | 0.23 | 0.84 | 1.42 | 1.78 | 0.49 | 1.22 | 0.7 | 0.5025 | 0.4025 | 0.5725 | 0.114276 | 0.26205 | 0.147411 | 0.999971 | 1 | 0.793248 |
| Zfp385b | G5E8Y8 | -0.15 | -0.27 | -0.18 | 0.58 | -0.74 | -0.85 | 0.4 | 1.44 | 0 | -0.03 | 0.34 | 0.65 | -0.24 | -0.59 | -0.1 | 1.29 | 0.0675 | 0.245 | 0.095 | 0.898404 | 0.377717 | 0.826336 | 0.999971 | 1 | 0.976804 |
| Zfp428 | H3BL05 | 1.38 | 2.57 | 1.44 | 1.77 | 2.72 | 1.98 | 2.02 | 1.27 | 2.57 | 1.58 | 0.8 | -0.8 | 3.56 | 2.45 | 0.28 | 0.43 | 0.2075 | -0.7525 | -0.11 | 0.595331 | 0.29515 | 0.885062 | 0.999971 | 1 | 0.98711 |
| Zfp512b | Q6PHP4 | 1.29 | -0.83 | -1.93 | -0.23 | -0.01 | -0.03 | -1.26 | -2.39 | -0.5 | 0.23 | -2.08 | -1.3 | -1.28 | -0.18 | -2.68 | -1.24 | -0.4975 | -0.4875 | -0.92 | 0.534213 | 0.525463 | 0.247118 | 0.999971 | 1 | 0.839012 |
| Zfp532 | S4R2I9 | -0.39 | -0.63 | -0.46 | -0.17 | -0.17 | -0.56 | -0.25 | -0.26 | -0.38 | -0.18 | -0.22 | 0.04 | 0.13 | -0.54 | -0.46 | -0.45 | 0.1025 | 0.2275 | 0.0825 | 0.597931 | 0.251437 | 0.713715 | 0.999971 | 1 | 0.952107 |
| Zfp638 | A0A0N4SV: | 0.97 | 0.51 | 0.36 | 0.69 | 0.56 | 0.73 | 0.46 | 0.36 | 0.34 | 0.64 | 0.32 | 0.2 | 0.25 | 0.66 | 0.46 | 0.87 | -0.105 | -0.2575 | -0.0725 | 0.613382 | 0.237736 | 0.750379 | 0.999971 | 1 | 0.961064 |
| Zfp651 | A0A1L1STN | -1.32 | 1.22 | 2.68 | 2.71 | 0.99 | -0.55 | 2.92 | 2.83 | -0.2 | -0.52 | 2.57 | 2.39 | 0.56 | -0.21 | 1.67 | 1.74 | 0.225 | -0.2625 | -0.3825 | 0.840776 | 0.815697 | 0.68716 | 0.999971 | 1 | 0.945467 |
| Zfp708 | A0A087WF | 0.32 | 1.32 | 5.84 | 1.67 | 0.08 | 0.17 | 7.31 | 4.17 | 1.2 | -3.32 | 8.48 | 9.76 | 2.24 | -1.08 | 5.15 | 2.64 | 0.645 | 1.7425 | -0.05 | 0.732302 | 0.559653 | 0.97453 | 0.999971 | 1 | 0.99925 |
| Zfp947 | Q8BIQ6 | 2.26 | 2.19 | 1.78 | 3.15 | 1.25 | 3.66 | 1.98 | 0.86 | 2.42 | 3.99 | 1.62 | 0.71 | 1.67 | 3.11 | 1.68 | 2.61 | -0.4075 | -0.16 | -0.0775 | 0.517377 | 0.815723 | 0.85827 | 0.999971 | 1 | 0.982702 |
| Zfpl1 | A0A494B9Y | -1.33 | -0.66 | 2 | 1.93 | -1.32 | -1.05 | 1 | 2.18 | -1.07 | -1.75 | 1.25 | 0.99 | -1.38 | -1.91 | 1.52 | 1.65 | -0.2825 | -0.63 | -0.515 | 0.792589 | 0.543734 | 0.652694 | 0.999971 | 1 | 0.934005 |
| Zfpm1 | O35615 | 2.32 | 0.9 | 1.16 | 2.53 | 1.34 | 1.43 | 1.49 | 2.5 | -0.08 | 2.04 | 1.58 | 0.6 | -1.06 | 1.19 | 1.52 | 2.06 | -0.0375 | -0.6925 | -0.8 | 0.934869 | 0.251215 | 0.285181 | 0.999971 | 1 | 0.845133 |
| Zfr | O88532 | 0.77 | 1 | 0.96 | 0.61 | 0.81 | 0.93 | 1.3 | 0.54 | 0.84 | 0.84 | 0.85 | 0.67 | 0.75 | 1.01 | 0.58 | 0.6 | 0.06 | -0.035 | -0.1 | 0.788619 | 0.842707 | 0.613594 | 0.999971 | 1 | 0.927364 |
| Zfr2 | E9Q5M4 | 2.95 | 2.67 | 3.37 | 2.51 | 2.47 | 2.93 | 3.45 | 3.13 | -2.39 | -1.36 | -2.02 | -2.58 | -4.25 | -3.07 | -2.13 | -2.51 | 0.12 | -4.9625 | -5.865 | 0.680615 | 2.39E-07 | 9.59E-07 | 0.999971 | 0.00218 | 0.004371 |
| Zfyve1 | Q810J8 | -0.23 | -0.44 | -0.16 | -0.69 | -0.18 | -0.02 | 0.04 | -0.17 | -0.02 | -0.22 | 0.23 | 0.35 | 0.3 | -0.31 | 0.02 | -0.46 | 0.2975 | 0.465 | 0.2675 | 0.151781 | 0.057132 | 0.284247 | 0.999971 | 1 | 0.844538 |
| Zfyve16 | Q80U44 | 1.28 | 1.42 | 1.01 | 0.39 | 1.13 | 1.45 | 0.58 | 0.67 | 1.21 | 1.36 | 1.17 | 0.47 | 0.96 | 1.87 | 1.25 | 0.24 | -0.0675 | 0.0275 | 0.055 | 0.828101 | 0.928721 | 0.88846 | 0.999971 | 1 | 0.987485 |
| Zfyve19 | Q9DAZ9 | 0.92 | -0.71 | -0.12 | 0.44 | -0.53 | -0.25 | 0.33 | 0.41 | -0.54 | 0.47 | -0.11 | -0.49 | -0.91 | -0.44 | -0.14 | -0.19 | -0.1425 | -0.3 | -0.5525 | 0.722759 | 0.46542 | 0.171763 | 0.999971 | 1 | 0.804075 |
| Zfyve21 | A0A1Y7VN | -0.72 | -0.48 | -0.89 | 2.34 | 0.42 | -0.34 | -1.2 | 1.09 | 0.12 | 0.09 | -1.82 | -1.48 | -0.4 | 0.36 | -1.37 | 1.56 | -0.07 | -0.835 | -0.025 | 0.931594 | 0.329417 | 0.977356 | 0.999971 | 1 | 0.99925 |
| Zfyve26 | Q5DU37 | 0.44 | 0.63 | 2.02 | 0.87 | 1.01 | 0.41 | 2.26 | 1.22 | 0.94 | 0.39 | 2.69 | 2.68 | 1.43 | 0.44 | 1.2 | 0.79 | 0.235 | 0.685 | -0.025 | 0.631369 | 0.295812 | 0.950056 | 0.999971 | 1 | 0.998038 |
| Zfyve27 | Q3TXX3 | -0.86 | -0.79 | 0.09 | -0.05 | 0.18 | -0.87 | 0.06 | 0.05 | 0.55 | -0.43 | 0.37 | 0.5 | 0.11 | -1.67 | 0.14 | -0.14 | 0.2575 | 0.65 | 0.0125 | 0.458944 | 0.078965 | 0.978536 | 0.999971 | 1 | 0.99925 |
| Zfyve28 | Q6ZPK7 | 0 | 0.29 | 0.06 | 0.78 | -0.32 | -0.31 | 0.44 | 0.3 | -0.63 | -0.15 | 0.06 | 0.42 | -0.04 | -0.31 | 0.64 | 0.69 | -0.255 | -0.3575 | -0.0375 | 0.377019 | 0.241316 | 0.904038 | 0.999971 | 1 | 0.991091 |
| Zfyve9 | A2A8R0 | -0.38 | 1.49 | 1.57 | 1.56 | 1.22 | 0.91 | 1.38 | 1.7 | 1.13 | -0.34 | 1.73 | 1.72 | 0.99 | 0.64 | 1.32 | 1.45 | 0.2425 | -1.57E-16 | 0.04 | 0.611146 | 1 | 0.933421 | 0.999971 | 1 | 0.995138 |
| Zgpat | Q8VDM1 | 2.51 | 2.88 | 0.93 | 1.25 | 3.36 | 2.52 | 1.02 | 1.42 | 3.01 | 2.15 | 1.51 | 1.66 | 3.99 | 2.39 | 1.53 | 1.48 | 0.1875 | 0.19 | 0.455 | 0.772156 | 0.725136 | 0.510951 | 0.999971 | 1 | 0.905815 |
| Zhx2 | Q8C0C0 | 2.12 | 1.62 | 1.68 | 1.54 | 1.17 | 1.79 | 1.57 | 1.49 | 0.64 | 1.89 | 0.91 | 1.32 | 0.6 | 1.62 | 1.46 | 0.41 | -0.235 | -0.55 | -0.7175 | 0.3098 | 0.100874 | 0.054204 | 0.999971 | 1 | 0.725348 |
| Zhx3 | Q8C0Q2 | 1.26 | 1.69 | 1.31 | 1.67 | 1.82 | 1.59 | 0.72 | 1.14 | 1.62 | 1.54 | 1.05 | 1.48 | 1.76 | 2.49 | 1.4 | 2.17 | -0.165 | -0.06 | 0.4725 | 0.564618 | 0.780403 | 0.118886 | 0.999971 | 1 | 0.78268 |
| Zkscan1 | Q8BGS3-2 | 4.24 | 0.79 | 1.17 | 1.9 | 1.07 | 2.15 | 2.23 | 1.41 | 1.53 | 3.16 | 0.83 | 0.95 | -0.48 | 0.23 | 0.83 | -0.07 | -0.31 | -0.4075 | -1.8975 | 0.677569 | 0.633794 | 0.029047 | 0.999971 | 1 | 0.678474 |
| Zkscan16 | A2ALW2 | 0.24 | -1.25 | -0.92 | -0.66 | -1.03 | -0.88 | -0.8 | -0.56 | -0.85 | -0.53 | -1.34 | -1.13 | -0.41 | -0.54 | -0.6 | -0.78 | -0.17 | -0.315 | 0.065 | 0.611967 | 0.388678 | 0.843438 | 0.999971 | 1 | 0.979814 |
| Zmat2 | Q9CPW7 | 1.95 | 1.98 | 2.53 | 3.32 | 2.01 | 2.28 | 2.36 | 3.35 | 2.03 | 1.71 | 2.88 | 2.68 | 2.09 | 2.15 | 3.28 | 3.36 | 0.055 | -0.12 | 0.275 | 0.893991 | 0.766697 | 0.541206 | 0.999971 | 1 | 0.910733 |
| Zmat3 | B9EI21 | 2.76 | 1.07 | 2.01 | 0.37 | 0.9 | 1.16 | 1.66 | 0.55 | 0.81 | 1.75 | 0.58 | 1.66 | -0.25 | 0.61 | 1.72 | 0.39 | -0.485 | -0.3525 | -0.935 | 0.371138 | 0.530762 | 0.150248 | 0.999971 | 1 | 0.793248 |
| Zmiz1 | Q6P1E1-2 | 0.77 | 3.53 | 1.96 | 1.97 | 3.4 | 3.29 | 1.53 | 1.86 | 2.75 | 1.6 | 1.92 | 1.95 | 3 | 2.54 | 1.88 | 2.12 | 0.4625 | -0.0025 | 0.3275 | 0.497815 | 0.996486 | 0.566962 | 0.999971 | 1 | 0.917976 |
| Zmpste24 | Q80W54 | -0.92 | -0.03 | 0.05 | 0.13 | 0.15 | 0.1 | 0.22 | 0.56 | -0.18 | -0.42 | -0.12 | 0.15 | 0.09 | 0.19 | 0.52 | 0.48 | 0.45 | 0.05 | 0.5125 | 0.135702 | 0.860257 | 0.096736 | 0.999971 | 1 | 0.78268 |
| Zmym2 | Q9CU65 | 1.55 | 1.72 | 2.5 | 2.32 | 1.94 | 1.62 | 2.55 | 1.96 | 1.73 | 0.56 | 2.39 | 1.97 | 0.43 | 0.53 | 1.64 | 0.99 | -0.005 | -0.36 | -1.125 | 0.986974 | 0.41301 | 0.010633 | 0.999971 | 1 | 0.613095 |
| Zmym3 | B1AXS8 | 0.79 | 0.48 | 1.12 | 1.17 | 0.15 | 0.34 | 1.47 | 0.95 | 0.35 | 0.57 | 0.56 | 0.71 | -0.05 | 0.57 | 0.87 | 0.62 | -0.1625 | -0.3425 | -0.3875 | 0.632658 | 0.143308 | 0.179048 | 0.999971 | 1 | 0.808532 |
| Zmym4 | A2A791-2 | 1.16 | 0.76 | 1.01 | 1.08 | 0.82 | -0.02 | 1.2 | 0.59 | 0.54 | 1.3 | 0.35 | 0.45 | 0.03 | 0.23 | 0.53 | 0.26 | -0.355 | -0.3425 | -0.74 | 0.231798 | 0.204645 | 0.004373 | 0.999971 | 1 | 0.582714 |
| Zmynd11 | Q8R5C8 | 0.37 | -0.98 | -0.19 | -0.14 | -1.06 | -0.58 | 0.21 | -0.51 | -1.04 | -0.6 | -0.26 | -0.14 | -1.65 | -1.59 | -0.36 | -1.12 | -0.25 | -0.275 | -0.945 | 0.505858 | 0.428218 | 0.037077 | 0.999971 | 1 | 0.700096 |
| Zmynd8 | A2A482 | -0.47 | -0.74 | -0.35 | -0.5 | -0.43 | -0.36 | -0.68 | -0.32 | -0.48 | -0.54 | -0.27 | -0.31 | -0.29 | -0.57 | -0.09 | -0.09 | 0.0675 | 0.115 | 0.255 | 0.717957 | 0.524009 | 0.223098 | 0.999971 | 1 | 0.827753 |
| Znf148 | Q61624 | 0.31 | 1.4 | 1.34 | 1.47 | 1.18 | 1.36 | 0.56 | 0.75 | 1.43 | 0.52 | 0.98 | 1.15 | -0.22 | 0.83 | 0.71 | 1.62 | -0.1675 | -0.11 | -0.395 | 0.614754 | 0.741591 | 0.380795 | 0.999971 | 1 | 0.871816 |
| Znf207 | Q9JMD0-2 | 2.08 | 2.63 | 0.22 | 0.55 | 2.56 | 2.1 | 3.24 | 0.31 | 2.45 | 1.98 | -0.35 | 0.15 | 1.52 | 2.42 | 0.26 | 0.57 | 0.6825 | -0.3125 | -0.1775 | 0.386491 | 0.701538 | 0.797204 | 0.999971 | 1 | 0.970479 |
| Znf22 | Q9ERU3 | -2.56 | 2.62 | -2.67 | -3.51 | 3.48 | 0.89 | -1.09 | -3.05 | 3.76 | 0.36 | -1.18 | -2.03 | 4.86 | 0.17 | -2.82 | -3.57 | 1.5875 | 1.7575 | 1.19 | 0.375599 | 0.31417 | 0.574068 | 0.999971 | 1 | 0.91816 |
| Znf280c | Q6P3Y5-2 | -0.31 | -0.84 | -1 | -0.8 | -1.85 | -0.98 | -2.37 | -1.06 | -1.54 | -0.59 | -1.43 | -1.11 | -1.43 | -2.12 | -1.3 | -0.7 | -0.8275 | -0.43 | -0.65 | 0.042603 | 0.145186 | 0.074115 | 0.999971 | 1 | 0.746069 |
| Znf318 | Q99PP2 | 2.15 | 0.71 | 2.58 | 1.91 | 0.95 | 0.6 | 0.75 | 1.82 | 0.47 | 0.99 | 1.36 | 2.2 | 0.76 | 0.59 | 2.08 | 1.99 | -0.8075 | -0.5825 | -0.4825 | 0.103017 | 0.266605 | 0.365825 | 0.999971 | 1 | 0.869526 |
| Znf32 | Q80V23 | 1.61 | -0.39 | 2.1 | 2.22 | -0.94 | -0.47 | 2.18 | 2.89 | -1.26 | 0.48 | 2.13 | 1.37 | -2.14 | -0.75 | 1.82 | 1.44 | -0.47 | -0.705 | -1.2925 | 0.64235 | 0.419722 | 0.216507 | 0.999971 | 1 | 0.826893 |
| Znf330 | Q922H9 | 1.41 | 0.74 | 1.91 | 1.21 | 0.17 | 0.72 | 2.11 | 1.88 | 0.26 | 0.98 | 2.01 | 1.31 | 0.11 | 0.56 | 1.88 | 0.97 | -0.0975 | -0.1775 | -0.4375 | 0.841169 | 0.671348 | 0.318323 | 0.999971 | 1 | 0.853893 |
| Znf365 | Q8BG89 | -0.82 | -1.4 | -1.59 | -1.13 | -1.44 | -1.6 | -1.54 | -1.08 | -1.78 | -1.34 | -1.23 | -1.18 | -1.03 | -1.2 | -1.41 | -0.84 | -0.18 | -0.1475 | 0.115 | 0.456649 | 0.550299 | 0.634503 | 0.999971 | 1 | 0.93301 |
| Znf414 | Q9DCK4 | 5.06 | 2.11 | 3.65 | 4.21 | 2.57 | 3.25 | 5.2 | 1.48 | 2.92 | 4.72 | 1.57 | 1.28 | -0.48 | 0.91 | 0.63 | 0.79 | -0.6325 | -1.135 | -3.295 | 0.485363 | 0.229882 | 0.000634 | 0.999971 | 1 | 0.360999 |
| Znf48 | Q3US17 | 3.03 | 0.31 | 2.24 | 2.74 | 0.52 | 1.1 | 2.72 | 1.8 | 1.48 | 2.44 | 2.15 | 2.05 | -1.47 | -0.27 | 1.14 | 0.79 | -0.545 | -0.05 | -2.0325 | 0.444067 | 0.932644 | 0.024853 | 0.999971 | 1 | 0.67615 |
| Znf512 | Q69Z99 | 0.36 | -1.54 | -1.24 | -0.67 | -1.5 | -1.31 | -0.96 | -1.05 | -1.36 | -0.08 | -0.93 | -1.27 | -2 | -1.32 | -1 | -1.31 | -0.4325 | -0.1375 | -0.635 | 0.312012 | 0.773777 | 0.175529 | 0.999971 | 1 | 0.808532 |
| Znf598 | Q80YR4-2 | 0.44 | 0.7 | 0.18 | 0.16 | 0.49 | 0.48 | 0.82 | 0.33 | 0.67 | 0.27 | 0.76 | 0.93 | 0.51 | 0.48 | 0.15 | 0.4 | 0.16 | 0.2875 | 0.015 | 0.459169 | 0.225217 | 0.941889 | 0.999971 | 1 | 0.996889 |
| Znf622 | Q91VY9 | 1.12 | 1.07 | 0.76 | 0.26 | 1.08 | 1.17 | 0.93 | 0.83 | 0.49 | 1.27 | 1.14 | 1.49 | 0.59 | 1.02 | 1.01 | 0.5 | 0.2 | 0.295 | -0.0225 | 0.419581 | 0.338837 | 0.931866 | 0.999971 | 1 | 0.995069 |
| Znf668 | Q8K2R5 | 0.36 | 0.72 | 0.15 | 0.19 | -0.09 | 0.62 | 0.47 | 0.74 | -0.1 | 0.66 | 1.08 | 0.71 | 0.26 | 0.24 | 1.16 | 0.29 | 0.08 | 0.2325 | 0.1325 | 0.75131 | 0.431913 | 0.63431 | 0.999971 | 1 | 0.93301 |
| Znf687 | Q9D2D7 | -1.63 | 0.92 | -0.74 | 0.02 | -0.75 | 0.24 | -0.63 | -0.55 | -0.58 | -1.42 | -1.14 | -1.73 | 0.9 | 0.56 | -1.13 | -1.37 | -0.065 | -0.86 | 0.0975 | 0.904249 | 0.144844 | 0.891818 | 0.999971 | 1 | 0.988225 |
| Znf704 | Q9ERQ3 | -1.26 | -0.93 | 0.18 | -0.64 | -1.57 | -1.56 | -0.31 | 0.91 | -1.07 | -1.6 | -0.83 | -1.25 | -1.02 | -1.49 | -0.18 | 0 | 0.03 | -0.525 | -0.01 | 0.960679 | 0.15293 | 0.981892 | 0.999971 | 1 | 0.99925 |
| Znf706 | Q9D115 | 3.88 | 3.7 | 2 | 1.74 | 3.11 | 3.65 | 2.24 | 3.63 | 2.83 | 3.05 | 2.19 | -2.03 | 3.31 | 3.59 | 2.83 | 1.16 | 0.3275 | -1.32 | -0.1075 | 0.583436 | 0.281509 | 0.878829 | 0.999971 | 1 | 0.985441 |
| Znf771 | Q8BJ90 | 0.67 | 1.74 | -0.46 | 0.94 | 1.66 | 1.28 | 0.36 | 0.4 | 1.48 | 2.37 | -0.43 | 0.18 | 1.74 | 1.48 | 0.4 | 0.25 | 0.2025 | 0.1775 | 0.245 | 0.695903 | 0.802196 | 0.654055 | 0.999971 | 1 | 0.934061 |
| Znf787 | Q8BIF9 | 7.12 | 0.63 | 4.19 | 5.59 | 1.44 | 3.52 | 5.22 | 3.18 | -0.79 | 4.96 | 2.87 | 2.08 | -5.81 | -1.14 | -1.68 | -0.27 | -1.0425 | -2.1025 | -6.6075 | 0.466297 | 0.219542 | 0.003049 | 0.999971 | 1 | 0.577271 |
| Znf800 | Q0VEE6 | -1.24 | -1.87 | -1.34 | -0.97 | -1.55 | -1.68 | -1.86 | -1.23 | -2.15 | -1.21 | -1.31 | -1.7 | -1.7 | -1.51 | -0.89 | -1 | -0.225 | -0.2375 | 0.08 | 0.390061 | 0.42881 | 0.779969 | 0.999971 | 1 | 0.965847 |
| Znf830 | Q8R1N0 | -1.08 | 1.41 | 2.08 | 0.96 | 1.99 | -0.35 | 1.56 | 1.32 | 1 | -1.46 | 1.71 | 1.41 | -1.34 | -0.46 | 1.73 | 1.12 | 0.2875 | -0.1775 | -0.58 | 0.708663 | 0.842917 | 0.515018 | 0.999971 | 1 | 0.905815 |
| Znfx1 | Q8R151 | 0.48 | 0.93 | 0.93 | 0.68 | 0.61 | 0.61 | 0.81 | 0.88 | 0.51 | 0.53 | 0.98 | 0.95 | 0.61 | 0.64 | 0.92 | 1.09 | -0.0275 | -0.0125 | 0.06 | 0.886863 | 0.953204 | 0.775671 | 0.999971 | 1 | 0.96393 |
| Znhit2 | Q9QY66 | 2.91 | 2.14 | 1.1 | 1.02 | 3.69 | 1.99 | 0.08 | 1.27 | 3.03 | 2.75 | 1.37 | 1.19 | 3.26 | -0.62 | 0.66 | 1.09 | -0.035 | 0.2925 | -0.695 | 0.964653 | 0.627172 | 0.412726 | 0.999971 | 1 | 0.87601 |
| Znrd2 | P56873 | -0.83 | 0.27 | 1.69 | 0.18 | -0.07 | 0.2 | 0.69 | 1 | -0.36 | 0.8 | 0.95 | 0.73 | -0.58 | 0.37 | 1.41 | 0.96 | 0.1275 | 0.2025 | 0.2125 | 0.80885 | 0.715126 | 0.730129 | 0.999971 | 1 | 0.956041 |
| Znrf1 | E9PZB7 | 0.07 | -0.5 | -1.24 | -0.54 | -0.53 | -0.55 | -1.47 | -0.67 | -0.4 | 0.41 | -0.5 | -0.26 | -0.46 | -0.19 | -1.02 | -0.44 | -0.2525 | 0.365 | 0.025 | 0.470553 | 0.29334 | 0.938182 | 0.999971 | 1 | 0.99575 |
| Znrf2 | Q71FD5 | 2.56 | -0.6 | -0.11 | 0.19 | -0.65 | -0.24 | 0.65 | -0.29 | -0.33 | 1.09 | 0.05 | -0.23 | -0.24 | -0.09 | -0.56 | -1.74 | -0.6425 | -0.365 | -1.1675 | 0.359413 | 0.606967 | 0.131679 | 0.999971 | 1 | 0.784948 |
| Zpr1 | Q62384 | 0.47 | 0.92 | 0.69 | 0.59 | 0.92 | 0.74 | 0.74 | 0.87 | 0.62 | 0.82 | 0.96 | 1.09 | 0.63 | 0.94 | 0.98 | 0.77 | 0.15 | 0.205 | 0.1625 | 0.419453 | 0.310353 | 0.407099 | 0.999971 | 1 | 0.874911 |
| Zranb2 | D3Z4U0 | 1.43 | 1.64 | 1.45 | 1.18 | 1.63 | 1.27 | 2 | 1.3 | 2.1 | 1.49 | 1.29 | 0.87 | 1.24 | 1.28 | 0.83 | 0.91 | 0.125 | 0.0125 | -0.36 | 0.593058 | 0.964995 | 0.105206 | 0.999971 | 1 | 0.78268 |
| Zscan18 | E9PUD6 | 2.75 | 0.59 | 0.89 | 1.24 | 0.78 | 0.07 | 0.18 | 1.31 | 1.35 | 2.76 | 1.28 | 0.54 | 1.19 | 1.2 | 0.36 | 1.49 | -0.7825 | 0.115 | -0.3075 | 0.155667 | 0.850865 | 0.54268 | 0.999971 | 1 | 0.911199 |
| Zscan26 | F8WJ31 | 1.4 | 2.13 | 0.74 | 1.61 | 1.31 | 1.01 | 1.73 | 0.6 | 1.36 | 1.76 | 0.53 | 0.87 | 0.77 | 1.09 | 0.43 | 0.5 | -0.3075 | -0.34 | -0.7725 | 0.407986 | 0.384341 | 0.039338 | 0.999971 | 1 | 0.700838 |
| Zswim8 | Q3UHH1-3 | -0.39 | 0.11 | 0.01 | -0.34 | 0.07 | -0.22 | -0.1 | -0.23 | 0.22 | -0.04 | -0.04 | 0.3 | -0.12 | 0.08 | -0.2 | -0.03 | 0.0325 | 0.2625 | 0.085 | 0.871354 | 0.219344 | 0.670753 | 0.999971 | 1 | 0.93891 |
| Zw10 | O54692 | -0.3 | 0.18 | -0.04 | 0.07 | 0.03 | 0.04 | -0.47 | -0.03 | -0.04 | -0.08 | 0.16 | 0.14 | 0.3 | 0.34 | 0.15 | 0.22 | -0.085 | 0.0675 | 0.275 | 0.686902 | 0.716991 | 0.16301 | 0.999971 | 1 | 0.800215 |
| Zwint | Q9CQU5 | 2.35 | 0.34 | 0.44 | 0.71 | 0.23 | 1.06 | 1.72 | 0.75 | 0.04 | 1.19 | 0.43 | 0.54 | 0.2 | 0.92 | 0.61 | 0.32 | -0.02 | -0.41 | -0.4475 | 0.969308 | 0.414324 | 0.349247 | 0.999971 | 1 | 0.865192 |
| Zyg11b | Q3UFS0 | -0.61 | 0.61 | -0.03 | -0.25 | 0.55 | 0.38 | -0.24 | -0.17 | 0.53 | -0.09 | -0.02 | 0.19 | 0.71 | 0.44 | -0.3 | 0.04 | 0.2 | 0.2225 | 0.2925 | 0.541671 | 0.464694 | 0.396164 | 0.999971 | 1 | 0.871816 |
| Zyx | Q62523 | 0.77 | 0.56 | 0.77 | 0.79 | 1.42 | 1.14 | 0.63 | 1.07 | 1.47 | 1.17 | 0.96 | 0.76 | 1.94 | 0.72 | 1.12 | 0.88 | 0.3425 | 0.3675 | 0.4425 | 0.141945 | 0.105859 | 0.152155 | 0.999971 | 1 | 0.795578 |
| Zzef1 | E9Q5W5 | 0.58 | 0.71 | 0.21 | -0.06 | 0.39 | 0.63 | -0.25 | 0.03 | 0.69 | 0.87 | 0.22 | 0.34 | 0.82 | 1.05 | 0.15 | 0.34 | -0.16 | 0.17 | 0.23 | 0.568498 | 0.51067 | 0.430626 | 0.999971 | 1 | 0.881611 |

**GSEA cortex**

** GSEA mouse MSigDB canonical pathways: m2.cp.v2023.1.Mm.symbols*

**significant pathways (FDR<0.25) are marked in yellow*

| CX MNK1KO / WT | | | | CX MNK2KO / WT | | | | CX MNKDKO / WT | | | |
|---|---|---|---|---|---|---|---|---|---|---|---|
| canonical pathway | NES | NOM p-val | FDR q-val | canonical pathway | NES | NOM p-val | FDR q-val | canonical pathway | NES | NOM p-val | FDR q-val |
| REACTOME_FORMATION | 2.03725 | 0 | 0.01586 | REACTOME_FORMATION_ | 1.565585 | 0.015556 | 0.36954 | REACTOME_FORMATION_ | 2.030022 | 0 | 0.079391 |
| REACTOME_ACTIVATION | 1.880446 | 0.001102 | 0.052729 | REACTOME_ACTIVATION_ | 1.362647 | 0.097606 | 0.686704 | REACTOME_ACTIVATION_ | 2.046375 | 4.06E-04 | 0.129565 |
| REACTOME_TRANSLATI | 1.690334 | 3.36E-04 | 0.186555 | REACTOME_TRANSLATION | 1.399665 | 0.023346 | 0.616157 | REACTOME_TRANSLATION | 1.846749 | 0 | 0.172686 |
| REACTOME_NONSENSE_ | 2.000028 | 0 | 0.024238 | REACTOME_NONSENSE_M | 1.523747 | 0.0272 | 0.445504 | REACTOME_NONSENSE_ | 1.864469 | 4.71E-04 | 0.177585 |
| REACTOME_RHO_GTPA | -1.94442 | 1.04E-03 | 0.370116 | REACTOME_RHO_GTPASE | -1.87132 | 7.71E-04 | 0.138072 | REACTOME_RHO_GTPASE | -1.77925 | 0.002181 | 0.178952 |
| WP_CYTOPLASMIC_RIBO | 1.987372 | 1.81E-04 | 0.024618 | WP_CYTOPLASMIC_RIBOS | 1.478443 | 0.033279 | 0.505782 | WP_CYTOPLASMIC_RIBOS | 1.885397 | 4.49E-04 | 0.183386 |
| REACTOME_EUKARYOTI | 1.961733 | 0 | 0.027057 | REACTOME_EUKARYOTIC_ | 1.586985 | 0.008583 | 0.363702 | REACTOME_EUKARYOTIC_ | 1.901042 | 0 | 0.211881 |
| REACTOME_EPIGENETIC | -1.57303 | 0.027465 | 0.437501 | REACTOME_EPIGENETIC_R | -1.10368 | 0.326811 | 0.868273 | REACTOME_EPIGENETIC_ | -1.77959 | 0.003359 | 0.237636 |
| REACTOME_PLATELET_A | 1.275385 | 0.15758 | 0.793899 | REACTOME_PLATELET_AG | 0.937522 | 0.529756 | 0.967624 | REACTOME_PLATELET_AG | -1.73431 | 0.004446 | 0.258368 |
| WP_TGFBETA_SIGNALIN | -1.63026 | 2.95E-02 | 0.426965 | WP_TGFBETA_SIGNALING_ | -1.08004 | 0.363324 | 0.883869 | WP_TGFBETA_SIGNALING | -1.80179 | 0.001233 | 0.259602 |
| REACTOME_RUNX1_REG | -1.84279 | 0.004265 | 0.503295 | REACTOME_RUNX1_REGU | -1.53353 | 0.043108 | 0.583698 | REACTOME_RUNX1_REGU | -1.70599 | 0.008219 | 0.260255 |
| REACTOME_NONHOMO | -1.79293 | 7.13E-03 | 0.515077 | REACTOME_NONHOMOLO | -1.92679 | 5.79E-04 | 0.123981 | REACTOME_NONHOMOL | -1.71584 | 0.00605 | 0.270545 |
| REACTOME_CITRIC_ACID | 1.621875 | 0.031439 | 0.254331 | REACTOME_CITRIC_ACID_ | 1.50609 | 0.069173 | 0.467476 | REACTOME_CITRIC_ACID_ | 1.764501 | 0.009613 | 0.296197 |
| BIOCARTA_VDR_PATHW | -1.70266 | 2.00E-02 | 0.515924 | BIOCARTA_VDR_PATHWAY | -1.37036 | 0.115415 | 0.680381 | BIOCARTA_VDR_PATHWA | -1.8209 | 0.002799 | 0.390858 |
| REACTOME_FOXO_MED | -0.91109 | 0.569936 | 0.952196 | REACTOME_FOXO_MEDIA | -1.00846 | 0.45377 | 0.991491 | REACTOME_FOXO_MEDIA | -1.63594 | 0.018824 | 0.396437 |
| REACTOME_NONSENSE_ | 1.97281 | 0 | 0.026241 | REACTOME_NONSENSE_M | 1.46745 | 0.028805 | 0.502116 | REACTOME_NONSENSE_ | 1.689707 | 0.001654 | 0.407544 |
| REACTOME_COPI_INDEP | 1.311902 | 0.11237 | 0.713792 | REACTOME_COPI_INDEPE | 1.303251 | 0.136225 | 0.727944 | REACTOME_COPI_INDEPE | 1.706066 | 0.007602 | 0.407662 |
| REACTOME_CHOLESTER | 1.50879 | 0.048542 | 0.407597 | REACTOME_CHOLESTEROL | 1.251743 | 0.196001 | 0.773661 | REACTOME_CHOLESTERO | 1.673218 | 0.020371 | 0.411628 |
| REACTOME_BUTYRATE_ | 0.872135 | 0.621181 | 0.980687 | REACTOME_BUTYRATE_RE | 0.88707 | 0.593875 | 1 | REACTOME_BUTYRATE_R | -1.62258 | 0.016069 | 0.412676 |
| REACTOME_DNA_REPAI | -1.47702 | 0.01572 | 0.508199 | REACTOME_DNA_REPAIR | -1.60532 | 0.002017 | 0.705637 | REACTOME_DNA_REPAIR | -1.64127 | 8.30E-04 | 0.416997 |
| REACTOME_DNA_DOUB | -1.63895 | 0.011537 | 0.536297 | REACTOME_DNA_DOUBLE | -1.76441 | 0.00236 | 0.26655 | REACTOME_DNA_DOUBL | -1.64146 | 0.006481 | 0.468216 |
| REACTOME_POSITIVE_E | -1.02317 | 0.407847 | 0.869478 | REACTOME_POSITIVE_EPI | -0.57526 | 0.955637 | 1 | REACTOME_POSITIVE_EPI | -1.58139 | 0.027231 | 0.482048 |
| REACTOME_MET_PROM | 1.459251 | 0.071429 | 0.474245 | REACTOME_MET_PROMO | 2.07757 | 4.15E-04 | 0.006248 | REACTOME_MET_PROMC | -1.54082 | 0.035364 | 0.485495 |
| WP_CHOLESTEROL_MET | 1.731039 | 0.005724 | 0.144572 | WP_CHOLESTEROL_META | 1.409934 | 0.082571 | 0.59684 | WP_CHOLESTEROL_META | 1.622111 | 0.015816 | 0.487879 |
| REACTOME_SIGNALLING | 0.868639 | 0.637072 | 0.982782 | REACTOME_SIGNALLING_T | 0.998429 | 0.437461 | 0.920315 | REACTOME_SIGNALLING_ | 1.633789 | 0.021477 | 0.492286 |
| REACTOME_KSRP_KHSR | -0.88869 | 0.60818 | 0.929108 | REACTOME_KSRP_KHSRP_ | 0.883498 | 0.601854 | 1 | REACTOME_KSRP_KHSRP_ | -1.57158 | 0.027484 | 0.493332 |
| REACTOME_COLLAGEN_ | 1.709907 | 0.01152 | 0.165944 | REACTOME_COLLAGEN_BI | 2.154113 | 2.10E-04 | 0.002751 | REACTOME_COLLAGEN_B | -1.55005 | 0.032072 | 0.497267 |
| REACTOME_NUCLEOTID | -1.15091 | 0.22147 | 0.767604 | REACTOME_NUCLEOTIDE_ | -1.34508 | 0.078034 | 0.713106 | REACTOME_NUCLEOTIDE_ | -1.54232 | 0.01448 | 0.504351 |
| WP_MECHANISMS_ASS | -1.42625 | 0.012566 | 0.585766 | WP_MECHANISMS_ASSOC | -1.28279 | 0.057563 | 0.7243 | WP_MECHANISMS_ASSO | -1.55438 | 0.003223 | 0.50702 |
| WP_GPCRS_OTHER | -1.35764 | 0.126527 | 0.649127 | WP_GPCRS_OTHER | -1.41954 | 0.08862 | 0.563831 | WP_GPCRS_OTHER | -1.58178 | 0.027839 | 0.517283 |
| REACTOME_TRISTETRAP | 0.827057 | 0.693612 | 1 | REACTOME_TRISTETRAPR | 0.908755 | 0.565024 | 0.98057 | REACTOME_TRISTETRAPR | -1.55762 | 0.03256 | 0.523402 |
| WP_GPCRS_NONODORA | -1.32923 | 0.086947 | 0.680599 | WP_GPCRS_NONODORAN | -1.29277 | 0.106651 | 0.71209 | WP_GPCRS_NONODORAN | -1.49242 | 0.027156 | 0.53502 |
| REACTOME_ESTROGEN_ | 0.991009 | 0.46256 | 0.95541 | REACTOME_ESTROGEN_D | -0.91632 | 0.587396 | 1 | REACTOME_ESTROGEN_D | -1.51613 | 0.033867 | 0.542885 |
| REACTOME_MRNA_SPLI | -1.50808 | 0.003863 | 0.501424 | REACTOME_MRNA_SPLICI | -1.20087 | 0.108152 | 0.776795 | REACTOME_MRNA_SPLIC | -1.49472 | 0.004442 | 0.545504 |
| WP_DELTANOTCH_SIGN | 0.881389 | 0.641398 | 0.974014 | WP_DELTANOTCH_SIGNAL | -0.92165 | 0.584454 | 1 | WP_DELTANOTCH_SIGNA | -1.50404 | 0.027744 | 0.548667 |
| REACTOME_ERK_MAPK_ | 1.473833 | 0.067224 | 0.442872 | REACTOME_ERK_MAPK_T | 0.756483 | 0.765931 | 1 | REACTOME_ERK_MAPK_T | 1.591535 | 0.034347 | 0.549716 |
| REACTOME_FORMATION | -0.81597 | 0.748063 | 0.970468 | REACTOME_FORMATION_ | -1.26436 | 0.148386 | 0.73318 | REACTOME_FORMATION_ | -1.48417 | 0.041092 | 0.55026 |
| REACTOME_PROCESSIN | -1.55307 | 0 | 0.467231 | REACTOME_PROCESSING_ | -1.34767 | 0.01823 | 0.71995 | REACTOME_PROCESSING_ | -1.50788 | 0.001942 | 0.5555 |
| REACTOME_RUNX1_INT | -1.16177 | 0.25798 | 0.741172 | REACTOME_RUNX1_INTER | -1.51454 | 0.04136 | 0.592172 | REACTOME_RUNX1_INTE | -1.51889 | 0.043175 | 0.555726 |
| WP_DYSREGULATED_MI | 1.360839 | 0.117366 | 0.670638 | WP_DYSREGULATED_MIR | 1.590958 | 0.04497 | 0.365659 | WP_DYSREGULATED_MIR | -1.58242 | 0.028751 | 0.557149 |
| REACTOME_DUAL_INCIS | -1.19574 | 0.20943 | 0.742141 | REACTOME_DUAL_INCISIO | -1.46153 | 0.046693 | 0.521242 | REACTOME_DUAL_INCISI | -1.49639 | 0.040799 | 0.560205 |
| REACTOME_HSF1_DEPE | 1.022819 | 0.419508 | 0.959538 | REACTOME_HSF1_DEPEND | -1.3424 | 0.12234 | 0.707231 | REACTOME_HSF1_DEPEN | -1.47508 | 0.053519 | 0.569726 |
| REACTOME_COLLAGEN_ | 1.685491 | 0.01648 | 0.185732 | REACTOME_COLLAGEN_FC | 2.147013 | 4.13E-04 | 0.002843 | REACTOME_COLLAGEN_F | -1.46736 | 0.05544 | 0.583978 |
| REACTOME_COLLAGEN_ | 1.757987 | 0.008895 | 0.118669 | REACTOME_COLLAGEN_DE | 2.10213 | 4.22E-04 | 0.004724 | REACTOME_COLLAGEN_D | -1.45781 | 0.065939 | 0.606696 |
| REACTOME_CHROMATI | -1.14453 | 0.214732 | 0.770889 | REACTOME_CHROMATIN_ | -1.1612 | 0.20525 | 0.819015 | REACTOME_CHROMATIN_ | -1.40092 | 0.041568 | 0.623161 |
| REACTOME_HDACS_DEA | -1.3618 | 0.120905 | 0.651951 | REACTOME_HDACS_DEAC | -0.77756 | 0.770366 | 1 | REACTOME_HDACS_DEAC | -1.40301 | 0.093451 | 0.628602 |
| REACTOME_MRNA_DEC | 0.84861 | 0.66673 | 1 | REACTOME_MRNA_DECAY | -0.93356 | 0.554119 | 1 | REACTOME_MRNA_DECA | -1.40599 | 0.092703 | 0.630392 |
| REACTOME_ASSEMBLY_ | 1.91743 | 0.00189 | 0.037501 | REACTOME_ASSEMBLY_OF | 2.23117 | 0 | 0.001058 | REACTOME_ASSEMBLY_O | -1.37079 | 0.112001 | 0.631327 |
| REACTOME_SUMOYLATI | -1.06995 | 0.352866 | 0.809408 | REACTOME_SUMOYLATIO | -1.14634 | 0.274707 | 0.812621 | REACTOME_SUMOYLATIO | -1.41625 | 0.08193 | 0.63219 |
| REACTOME_TRANSPORT | -0.91188 | 0.620903 | 0.955492 | REACTOME_TRANSPORT_ | -0.93029 | 0.579411 | 1 | REACTOME_TRANSPORT_ | -1.38488 | 0.050963 | 0.63577 |
| REACTOME_REGULATIO | -0.9078 | 0.583388 | 0.945102 | REACTOME_REGULATION_ | 0.724047 | 0.834716 | 1 | REACTOME_REGULATION | -1.40798 | 0.081247 | 0.636926 |
| REACTOME_G2_M_DNA | -1.70844 | 0.014166 | 0.594058 | REACTOME_G2_M_DNA_D | -1.50397 | 0.047537 | 0.517898 | REACTOME_G2_M_DNA_ | -1.37187 | 0.10079 | 0.638862 |
| REACTOME_GPVI_MEDI | 0.805593 | 0.735919 | 0.997564 | REACTOME_GPVI_MEDIAT | 0.921974 | 0.547474 | 0.981586 | REACTOME_GPVI_MEDIA | -1.41058 | 0.077687 | 0.641129 |
| REACTOME_SIGNALING_ | -0.6624 | 0.896649 | 1 | REACTOME_SIGNALING_B | -1.02468 | 0.428625 | 0.9559 | REACTOME_SIGNALING_B | -1.38027 | 0.106582 | 0.641788 |
| REACTOME_CELLULAR_ | -0.73357 | 0.9317 | 1 | REACTOME_CELLULAR_RE | -0.90958 | 0.627531 | 1 | REACTOME_CELLULAR_RE | -1.39276 | 0.046365 | 0.643235 |
| REACTOME_PROCESSIN | -1.53492 | 0.044563 | 0.46941 | REACTOME_PROCESSING_ | -1.44126 | 0.066101 | 0.537634 | REACTOME_PROCESSING_ | -1.37674 | 0.097598 | 0.643604 |
| WP_NEURAL_CREST_DIF | -1.00441 | 0.43353 | 0.889533 | WP_NEURAL_CREST_DIFFE | -1.19882 | 0.212139 | 0.765273 | WP_NEURAL_CREST_DIFF | -1.38902 | 0.083849 | 0.645231 |
| BIOCARTA_HER2_PATHV | -0.83769 | 0.671119 | 0.965593 | BIOCARTA_HER2_PATHWA | 0.397209 | 0.997703 | 1 | BIOCARTA_HER2_PATHW | -1.38547 | 0.103781 | 0.646556 |
| REACTOME_DEGRADATI | 1.789369 | 0.005235 | 0.09825 | REACTOME_DEGRADATION | 2.276743 | 0 | 9.25E-04 | REACTOME_DEGRADATIO | -1.4164 | 0.066038 | 0.647726 |
| REACTOME_INTEGRIN_S | 1.063829 | 0.37085 | 1 | REACTOME_INTEGRIN_SIG | -0.8354 | 0.703838 | 1 | REACTOME_INTEGRIN_SI | -1.37251 | 0.101376 | 0.648496 |
| REACTOME_RAC3_GTPA | -0.65324 | 0.981053 | 1 | REACTOME_RAC3_GTPASE | 1.355795 | 0.081419 | 0.697161 | REACTOME_RAC3_GTPAS | -1.43423 | 0.036544 | 0.656252 |
| REACTOME_INTEGRIN_ | 2.097309 | 0 | 0.027884 | REACTOME_INTEGRIN_CEL | 2.202235 | 0 | 0.001243 | REACTOME_INTEGRIN_CE | -1.42902 | 0.061339 | 0.661248 |
| REACTOME_TRANSCRIPT | -1.01797 | 0.410766 | 0.872711 | REACTOME_TRANSCRIPTIC | -1.2564 | 0.141655 | 0.73976 | REACTOME_TRANSCRIPTI | -1.4166 | 0.059233 | 0.663905 |
| REACTOME_IMMUNORE | 0.914162 | 0.577978 | 0.955271 | REACTOME_IMMUNOREG | 1.090197 | 0.337202 | 0.882776 | REACTOME_IMMUNOREG | -1.43669 | 0.067832 | 0.664515 |
| WP_FIBRIN_COMPLEME | 0.656221 | 0.91597 | 1 | WP_FIBRIN_COMPLEMENT | -1.16453 | 0.260836 | 0.816041 | WP_FIBRIN_COMPLEMEN | -1.35891 | 0.105576 | 0.668288 |
| REACTOME_CYTOPROTE | 0.789522 | 0.797688 | 1 | REACTOME_CYTOPROTECT | -1.64372 | 0.009971 | 0.707988 | REACTOME_CYTOPROTEC | -1.42334 | 0.058189 | 0.668737 |
| REACTOME_DNA_DOUB | -1.13748 | 0.278019 | 0.776857 | REACTOME_DNA_DOUBLE | -1.39246 | 0.088717 | 0.629122 | REACTOME_DNA_DOUBL | -1.41882 | 0.076126 | 0.671393 |
| REACTOME_NON_INTEG | 1.521135 | 0.049736 | 0.406799 | REACTOME_NON_INTEGRI | 2.091786 | 2.09E-04 | 0.005304 | REACTOME_NON_INTEGR | -1.43936 | 0.076436 | 0.67216 |
| REACTOME_SUMOYLATI | -1.14318 | 0.248342 | 0.766645 | REACTOME_SUMOYLATIO | -1.29671 | 0.126174 | 0.709965 | REACTOME_SUMOYLATIO | -1.35369 | 0.094802 | 0.677885 |
| WP_GLUTATHIONE_AND | -1.39022 | 0.094759 | 0.627407 | WP_GLUTATHIONE_AND_ | -0.92803 | 0.567537 | 1 | WP_GLUTATHIONE_AND_ | -1.33525 | 0.123926 | 0.70788 |
| REACTOME_ACTIVATION | -1.63329 | 0.02384 | 0.456218 | REACTOME_ACTIVATION_ | -1.57171 | 0.027692 | 0.724862 | REACTOME_ACTIVATION_ | -1.32953 | 0.128767 | 0.708635 |
| WP_NOTCH_SIGNALING | 0.758008 | 0.792992 | 1 | WP_NOTCH_SIGNALING_P | -1.28976 | 0.162662 | 0.710657 | WP_NOTCH_SIGNALING_ | -1.33717 | 0.132438 | 0.711553 |
| REACTOME_REGULATIO | -1.25538 | 0.152962 | 0.720221 | REACTOME_REGULATION_ | -1.35452 | 0.093629 | 0.709418 | REACTOME_REGULATION | -1.33045 | 0.111266 | 0.716352 |
| REACTOME_CYTOSOLIC_ | -1.61402 | 0.030647 | 0.439745 | REACTOME_CYTOSOLIC_SE | -1.09834 | 0.342569 | 0.864042 | REACTOME_CYTOSOLIC_S | -1.32483 | 0.142045 | 0.71712 |
| REACTOME_HOMOLOGY | -1.59146 | 0.027051 | 0.441366 | REACTOME_HOMOLOGY_ | -1.33087 | 0.114916 | 0.705276 | REACTOME_HOMOLOGY_ | -1.33774 | 0.117698 | 0.721447 |
| REACTOME_CARGO_CO | 1.147502 | 0.278679 | 0.97 | REACTOME_CARGO_CONC | -1.22357 | 0.205633 | 0.752917 | REACTOME_CARGO_CON | 1.51241 | 0.040854 | 0.727361 |
| REACTOME_RAC1_GTPA | -0.89754 | 0.701461 | 0.926149 | REACTOME_RAC1_GTPASE | 0.87673 | 0.70664 | 1 | REACTOME_RAC1_GTPAS | -1.33875 | 0.05509 | 0.729685 |
| REACTOME_SYNTHESIS_ | 0.974385 | 0.489646 | 0.951464 | REACTOME_SYNTHESIS_OF | 0.974145 | 0.47902 | 0.936369 | REACTOME_SYNTHESIS_O | 1.403228 | 0.085017 | 0.741913 |
| REACTOME_RNA_POLYN | -0.89045 | 0.6404 | 0.939647 | REACTOME_RNA_POLYME | -1.15277 | 0.235709 | 0.814635 | REACTOME_RNA_POLYME | -1.31563 | 0.099947 | 0.74465 |

| | | | | | | | | | | | |
|---|---|---|---|---|---|---|---|---|---|---|---|
| BIOCARTA_ERK_PATHW/ | 1.271617 | 0.175845 | 0.778851 | BIOCARTA_ERK_PATHWAY | -0.6939 | 0.859133 | 1 | BIOCARTA_ERK_PATHWAY | 1.415012 | 0.09235 | 0.747636 |
| REACTOME_ASPARAGIN | 1.517258 | 0.003172 | 0.406266 | REACTOME_ASPARAGINE_ | 1.01277 | 0.412723 | 0.906119 | REACTOME_ASPARAGINE_ | 1.407596 | 0.008611 | 0.75062 |
| REACTOME_INSULIN_RE | 1.083618 | 0.352425 | 0.996916 | REACTOME_INSULIN_RECE | 0.594172 | 0.943273 | 1 | REACTOME_INSULIN_REC | 1.516076 | 0.054685 | 0.759404 |
| REACTOME_NUCLEAR_E | 1.115569 | 0.317382 | 0.986856 | REACTOME_NUCLEAR_EVE | 0.668393 | 0.882097 | 1 | REACTOME_NUCLEAR_EV | 1.416776 | 0.082795 | 0.768872 |
| REACTOME_PEPTIDE_LI( | 1.391596 | 0.074054 | 0.641587 | REACTOME_PEPTIDE_LIGA | 1.290004 | 0.140494 | 0.731727 | REACTOME_PEPTIDE_LIG/ | 1.389854 | 0.064018 | 0.769489 |
| REACTOME_METABOLIS | -1.22635 | 0.190538 | 0.747138 | REACTOME_METABOLISM_ | -1.36744 | 0.10506 | 0.67435 | REACTOME_METABOLISM | -1.30721 | 0.150534 | 0.769869 |
| REACTOME_NOREPINEP | 0.647122 | 0.903318 | 1 | REACTOME_NOREPINEPHF | 0.941905 | 0.512858 | 0.963619 | REACTOME_NOREPINEPH | -1.30416 | 0.163421 | 0.771476 |
| WP_FATTY_ACID_BIOSY | -1.07845 | 0.346969 | 0.796465 | WP_FATTY_ACID_BIOSYNT | -0.92392 | 0.568898 | 1 | WP_FATTY_ACID_BIOSYN | -1.30103 | 0.16016 | 0.773203 |
| REACTOME_DNA_REPLI( | 1.009345 | 0.433448 | 0.965452 | REACTOME_DNA_REPLICA | 1.20259 | 0.191596 | 0.832159 | REACTOME_DNA_REPLIC/ | 1.421159 | 0.038623 | 0.781075 |
| REACTOME_DEADENYLA | -1.11759 | 0.272748 | 0.808533 | REACTOME_DEADENYLATI | -0.88478 | 0.659671 | 1 | REACTOME_DEADENYLAT | -1.28211 | 0.130272 | 0.787927 |
| REACTOME_MITOCHON | 0.834356 | 0.778549 | 0.998819 | REACTOME_MITOCHONDF | 0.807184 | 0.803061 | 1 | REACTOME_MITOCHOND | 1.425913 | 0.02531 | 0.793241 |
| REACTOME_ASSEMBLY_ | 0.992597 | 0.462943 | 0.954775 | REACTOME_ASSEMBLY_OF | 1.161884 | 0.23762 | 0.873511 | REACTOME_ASSEMBLY_O | 1.377511 | 0.04957 | 0.794255 |
| REACTOME_PHASE_II_C | 1.637782 | 0.011113 | 0.24161 | REACTOME_PHASE_II_CON | 0.976545 | 0.465988 | 0.943385 | REACTOME_PHASE_II_CO | 1.520062 | 0.018541 | 0.795124 |
| REACTOME_DDX58_IFIH | -0.9463 | 0.522426 | 0.910915 | REACTOME_DDX58_IFIH1_ | -0.88627 | 0.613444 | 1 | REACTOME_DDX58_IFIH1 | -1.29056 | 0.160396 | 0.79559 |
| REACTOME_ER_TO_GOL | 1.405176 | 0.024483 | 0.604608 | REACTOME_ER_TO_GOLGI | 1.078497 | 0.305479 | 0.878339 | REACTOME_ER_TO_GOLG | 1.432887 | 0.006735 | 0.797418 |
| REACTOME_INWARDLY_ | -1.68494 | 0.017712 | 0.502986 | REACTOME_INWARDLY_RE | -1.53444 | 0.034695 | 0.621212 | REACTOME_INWARDLY_R | -1.28231 | 0.167913 | 0.797962 |
| REACTOME_CD209_DC_ | -0.78318 | 0.748886 | 0.987325 | REACTOME_CD209_DC_SI( | 0.54207 | 0.963991 | 1 | REACTOME_CD209_DC_SI | -1.28414 | 0.172304 | 0.801016 |
| BIOCARTA_TNFR1_PATH | -1.24197 | 0.193281 | 0.741264 | BIOCARTA_TNFR1_PATHW | 0.70218 | 0.837832 | 1 | BIOCARTA_TNFR1_PATHW | -1.25428 | 0.198669 | 0.803626 |
| REACTOME_REGULATIO | -1.06988 | 0.319917 | 0.802622 | REACTOME_REGULATION_ | -1.1209 | 0.257453 | 0.838305 | REACTOME_REGULATION_ | -1.29138 | 0.103266 | 0.803738 |
| REACTOME_GOLGI_ASS( | 0.810823 | 0.778224 | 1 | REACTOME_GOLGI_ASSOC | 0.682339 | 0.923544 | 1 | REACTOME_GOLGI_ASSO( | -1.24489 | 0.171251 | 0.804692 |
| REACTOME_TRANSPORT | -1.59487 | 0.025751 | 0.462409 | REACTOME_TRANSPORT_( | -1.48558 | 0.039167 | 0.538562 | REACTOME_TRANSPORT_ | -1.28539 | 0.147974 | 0.806838 |
| REACTOME_DEACTIVATI | -0.86323 | 0.644383 | 0.95896 | REACTOME_DEACTIVATIOI | -1.38739 | 0.099104 | 0.631184 | REACTOME_DEACTIVATIO | -1.25576 | 0.195372 | 0.807293 |
| REACTOME_TRANSCRIPT | -1.149 | 0.208411 | 0.765125 | REACTOME_TRANSCRIPTIC | -1.22678 | 0.133936 | 0.761698 | REACTOME_TRANSCRIPTI( | -1.24578 | 0.136464 | 0.810479 |
| REACTOME_INTRINSIC_I | 0.828887 | 0.721933 | 1 | REACTOME_INTRINSIC_PA | 0.745588 | 0.82678 | 1 | REACTOME_INTRINSIC_P/ | 1.463957 | 0.057479 | 0.810967 |
| REACTOME_DRUG_ADM | 0.919905 | 0.578529 | 0.949912 | REACTOME_DRUG_ADME | -0.95485 | 0.532422 | 1 | REACTOME_DRUG_ADME | -1.23658 | 0.181793 | 0.811462 |
| BIOCARTA_P38MAPK_P/ | -0.95224 | 0.510117 | 0.910216 | BIOCARTA_P38MAPK_PAT | -1.07996 | 0.356114 | 0.876832 | BIOCARTA_P38MAPK_PAT | -1.25933 | 0.190674 | 0.812173 |
| WP_MICROGLIA_PATHO | -1.52935 | 0.042051 | 0.46239 | WP_MICROGLIA_PATHOGI | -0.85156 | 0.676973 | 1 | WP_MICROGLIA_PATHOG | -1.24958 | 0.187708 | 0.813786 |
| REACTOME_FLT3_SIGNA | 0.726555 | 0.821632 | 1 | REACTOME_FLT3_SIGNALI | -1.13441 | 0.307105 | 0.814526 | REACTOME_FLT3_SIGNAL | -1.26123 | 0.19863 | 0.81428 |
| REACTOME_ECM_PROTI | 1.936779 | 0 | 0.032647 | REACTOME_ECM_PROTEC | 2.164128 | 2.06E-04 | 0.002557 | REACTOME_ECM_PROTEC | -1.25612 | 0.181316 | 0.815648 |
| REACTOME_NCAM_SIGN | 1.55138 | 0.033854 | 0.360326 | REACTOME_NCAM_SIGNA | 2.110592 | 0 | 0.004494 | REACTOME_NCAM_SIGN/ | -1.23107 | 0.213746 | 0.816219 |
| REACTOME_SUMOYLATI | -1.2825 | 0.134531 | 0.720701 | REACTOME_SUMOYLATIOI | -1.31441 | 0.117693 | 0.669963 | REACTOME_SUMOYLATIO | -1.23974 | 0.185299 | 0.816787 |
| WP_OXIDATIVE_STRESS_ | -1.26792 | 0.129069 | 0.716126 | WP_OXIDATIVE_STRESS_A | -1.5583 | 0.011776 | 0.724388 | WP_OXIDATIVE_STRESS_/ | -1.24636 | 0.153786 | 0.817591 |
| REACTOME_GABA_SYNT | 0.710563 | 0.850347 | 1 | REACTOME_GABA_SYNTHI | 0.975925 | 0.479038 | 0.935768 | REACTOME_GABA_SYNTH | -1.22822 | 0.218415 | 0.818669 |
| REACTOME_CHROMOSC | -1.36614 | 0.096019 | 0.672709 | REACTOME_CHROMOSOM | -0.85865 | 0.681493 | 1 | REACTOME_CHROMOSON | -1.23693 | 0.187799 | 0.819091 |
| REACTOME_SENSORY_P | -1.21915 | 0.188271 | 0.73758 | REACTOME_SENSORY_PER | -1.05021 | 0.379777 | 0.929765 | REACTOME_SENSORY_PEI | -1.23212 | 0.192865 | 0.820781 |
| WP_KIT_RECEPTOR_SIGI | -1.04389 | 0.382108 | 0.843926 | WP_KIT_RECEPTOR_SIGN/ | -0.81287 | 0.772445 | 1 | WP_KIT_RECEPTOR_SIGN/ | -1.22552 | 0.189312 | 0.820981 |
| REACTOME_DOWNREGU | -0.89868 | 0.592003 | 0.933305 | REACTOME_DOWNREGUL/ | -1.09136 | 0.344886 | 0.86608 | REACTOME_DOWNREGUL | -1.22286 | 0.225755 | 0.823146 |
| REACTOME_RAC2_GTPA | -0.85235 | 0.73049 | 0.955781 | REACTOME_RAC2_GTPASE | 1.053212 | 0.361491 | 0.8744 | REACTOME_RAC2_GTPAS | -1.26155 | 0.13394 | 0.823283 |
| REACTOME_BIOLOGICAL | 1.878272 | 3.53E-04 | 0.049824 | REACTOME_BIOLOGICAL_( | 1.425181 | 0.046591 | 0.574811 | REACTOME_BIOLOGICAL_ | 1.480575 | 0.01248 | 0.824744 |
| REACTOME_SUMOYLATI | -1.29199 | 0.140827 | 0.716772 | REACTOME_SUMOYLATIOI | -1.0548 | 0.37433 | 0.943179 | REACTOME_SUMOYLATIO | -1.27019 | 0.160722 | 0.828817 |
| WP_P38_MAPK_SIGNAL | -1.11005 | 0.308212 | 0.808571 | WP_P38_MAPK_SIGNALIN( | -1.25452 | 0.180312 | 0.713652 | WP_P38_MAPK_SIGNALIN | -1.2618 | 0.180668 | 0.832842 |
| REACTOME_REGULATIO | -0.93827 | 0.548181 | 0.917813 | REACTOME_REGULATION_ | -0.93062 | 0.57154 | 1 | REACTOME_REGULATION_ | -1.26673 | 0.142538 | 0.832943 |
| REACTOME_CDK_MEDIA | 1.078915 | 0.324505 | 1 | REACTOME_CDK_MEDIATE | 1.321109 | 0.111601 | 0.701251 | REACTOME_CDK_MEDIAT | 1.433005 | 0.038099 | 0.83313 |
| REACTOME_SEROTONIN | -0.52409 | 0.981068 | 1 | REACTOME_SEROTONIN_N | 0.754813 | 0.775187 | 1 | REACTOME_SEROTONIN_ | -1.26241 | 0.192367 | 0.841061 |
| REACTOME_PYRUVATE_ | 1.275217 | 0.145354 | 0.784406 | REACTOME_PYRUVATE_M | 1.247277 | 0.178212 | 0.779364 | REACTOME_PYRUVATE_N | 1.466622 | 0.038043 | 0.843386 |
| BIOCARTA_EIF4_PATHW | 1.481367 | 0.059466 | 0.431458 | BIOCARTA_EIF4_PATHWAY | 0.919152 | 0.541771 | 0.976762 | BIOCARTA_EIF4_PATHWA | 1.359173 | 0.113256 | 0.845941 |
| REACTOME_HSP90_CHA | 1.381645 | 0.075451 | 0.653672 | REACTOME_HSP90_CHAPE | 1.124515 | 0.284556 | 0.902886 | REACTOME_HSP90_CHAP | 1.43818 | 0.045219 | 0.848575 |
| REACTOME_PEROXISOM | 1.210986 | 0.185374 | 0.83392 | REACTOME_PEROXISOMA | 0.851977 | 0.686757 | 1 | REACTOME_PEROXISOMA | 1.352408 | 0.072521 | 0.849792 |
| REACTOME_JNK_C_JUN_ | -1.40361 | 0.104513 | 0.601736 | REACTOME_JNK_C_JUN_K | 0.586043 | 0.932065 | 1 | REACTOME_JNK_C_JUN_I | -1.21415 | 0.246456 | 0.850044 |
| REACTOME_APOPTOTIC_ | -0.79111 | 0.781142 | 0.982187 | REACTOME_APOPTOTIC_C | -0.5599 | 0.983229 | 1 | REACTOME_APOPTOTIC_( | -1.20561 | 0.224182 | 0.850493 |
| WP_TYROBP_CAUSAL_N | -0.85781 | 0.661874 | 0.952443 | WP_TYROBP_CAUSAL_NET | -1.05506 | 0.387934 | 0.949904 | WP_TYROBP_CAUSAL_NE | -1.2031 | 0.237952 | 0.852274 |
| REACTOME_TOLL_LIKE_I | -1.40406 | 0.034799 | 0.619117 | REACTOME_TOLL_LIKE_RE | -1.26069 | 0.097085 | 0.735169 | REACTOME_TOLL_LIKE_RI | -1.20723 | 0.168359 | 0.852314 |
| REACTOME_SIGNALING_ | -0.99175 | 0.469556 | 0.897526 | REACTOME_SIGNALING_B | -0.95766 | 0.515307 | 1 | REACTOME_SIGNALING_B | -1.21059 | 0.244186 | 0.856018 |
| REACTOME_GENERIC_TI | -0.99921 | 0.447792 | 0.899729 | REACTOME_GENERIC_TRA | -1.25633 | 0.032339 | 0.728825 | REACTOME_GENERIC_TR/ | -1.20772 | 0.085422 | 0.859025 |
| REACTOME_SIGNALING_ | 0.646021 | 0.95537 | 1 | REACTOME_SIGNALING_B | -0.57856 | 0.986021 | 1 | REACTOME_SIGNALING_B | 1.343661 | 0.081955 | 0.862579 |
| BIOCARTA_FAS_PATHW/ | -1.28618 | 0.162271 | 0.72225 | BIOCARTA_FAS_PATHWAY | -0.73003 | 0.823644 | 1 | BIOCARTA_FAS_PATHWAY | -1.19703 | 0.249661 | 0.86871 |
| REACTOME_GENE_EXPR | -1.05731 | 0.249787 | 0.808311 | REACTOME_GENE_EXPRES | -1.27122 | 0.013769 | 0.731218 | REACTOME_GENE_EXPRE! | -1.18684 | 0.076735 | 0.884919 |
| REACTOME_ACTIVATION | 1.262149 | 0.191372 | 0.795911 | REACTOME_ACTIVATION_( | 0.713504 | 0.811388 | 1 | REACTOME_ACTIVATION_ | 1.332804 | 0.131042 | 0.885383 |
| REACTOME_CDC42_GTP | -0.68922 | 0.963357 | 1 | REACTOME_CDC42_GTPAS | 0.637723 | 0.976687 | 1 | REACTOME_CDC42_GTPA | -1.18879 | 0.207422 | 0.885452 |
| REACTOME_EXTRACELLU | 2.059572 | 0 | 0.018745 | REACTOME_EXTRACELLUL/ | 2.519486 | 0 | 0 | REACTOME_EXTRACELLUL | -1.18448 | 0.199318 | 0.88623 |
| REACTOME_GABA_B_RE | -1.43466 | 0.067491 | 0.599043 | REACTOME_GABA_B_RECE | -1.245 | 0.167669 | 0.716698 | REACTOME_GABA_B_REC | -1.19038 | 0.238523 | 0.887421 |
| REACTOME_TRANSPORT | 1.506436 | 0.008696 | 0.40382 | REACTOME_TRANSPORT_T | 1.10052 | 0.263379 | 0.887048 | REACTOME_TRANSPORT_ | 1.438551 | 0.00593 | 0.889169 |
| REACTOME_PKMTS_ME | -1.34601 | 0.115622 | 0.655818 | REACTOME_PKMTS_METH | -1.51354 | 0.042132 | 0.507083 | REACTOME_PKMTS_METI | -1.17728 | 0.262019 | 0.890702 |
| WP_WNT_SIGNALING_P | 0.728598 | 0.914402 | 1 | WP_WNT_SIGNALING_PAT | -1.03921 | 0.382364 | 0.940075 | WP_WNT_SIGNALING_PA | -1.17972 | 0.219635 | 0.8974 |
| REACTOME_GENE_SILEN | -1.24344 | 0.168265 | 0.748693 | REACTOME_GENE_SILENCI | -1.13352 | 0.276604 | 0.809825 | REACTOME_GENE_SILENC | -1.17762 | 0.246688 | 0.897597 |
| REACTOME_SUMOYLATI | -0.94564 | 0.532431 | 0.907131 | REACTOME_SUMOYLATIOI | -0.93817 | 0.551204 | 1 | REACTOME_SUMOYLATIO | -1.15974 | 0.265641 | 0.905516 |
| WP_MRNA_PROCESSING | -0.89881 | 0.808896 | 0.938129 | WP_MRNA_PROCESSING | -0.9544 | 0.609602 | 1 | WP_MRNA_PROCESSING | -1.16241 | 0.140777 | 0.910591 |
| REACTOME_INOSITOL_P | -0.66741 | 0.936518 | 1 | REACTOME_INOSITOL_PH( | 0.76185 | 0.808162 | 1 | REACTOME_INOSITOL_PH | -1.16006 | 0.265938 | 0.912141 |
| REACTOME_ESR_MEDIA | 1.135938 | 0.244628 | 0.984113 | REACTOME_ESR_MEDIATE | -0.91478 | 0.617541 | 1 | REACTOME_ESR_MEDIATI | -1.16688 | 0.225293 | 0.917004 |
| WP_ESTROGEN_SIGNALI | -1.23947 | 0.152405 | 0.725682 | WP_ESTROGEN_SIGNALIN( | -1.15636 | 0.22592 | 0.810172 | WP_ESTROGEN_SIGNALIN | -1.16249 | 0.248636 | 0.91834 |
| REACTOME_RHOB_GTP/ | 0.543792 | 0.996969 | 1 | REACTOME_RHOB_GTPASI | 0.939264 | 0.538925 | 0.966957 | REACTOME_RHOB_GTPAS | -1.16689 | 0.244507 | 0.925294 |
| REACTOME_GPCR_LIGAI | -1.20234 | 0.143091 | 0.740656 | REACTOME_GPCR_LIGANE | -1.19765 | 0.15293 | 0.760276 | REACTOME_GPCR_LIGANI | -1.16253 | 0.222276 | 0.92639 |
| REACTOME_SUMOYLATI | -1.25586 | 0.11349 | 0.731248 | REACTOME_SUMOYLATIOI | -1.32168 | 0.064548 | 0.682553 | REACTOME_SUMOYLATIO | -1.1358 | 0.263623 | 0.931979 |
| REACTOME_CLASS_A_1_ | -1.17556 | 0.190348 | 0.760082 | REACTOME_CLASS_A_1_RI | -0.8828 | 0.687488 | 1 | REACTOME_CLASS_A_1_R | -1.1454 | 0.259571 | 0.931981 |
| REACTOME_NUCLEAR_P | -1.34769 | 0.115318 | 0.666374 | REACTOME_NUCLEAR_POI | -1.01596 | 0.432956 | 0.972193 | REACTOME_NUCLEAR_PO | -1.14254 | 0.289685 | 0.935765 |
| REACTOME_CELL_CELL_ | -1.12192 | 0.280269 | 0.811102 | REACTOME_CELL_CELL_JU | 1.050636 | 0.375103 | 0.877142 | REACTOME_CELL_CELL_JU | -1.13862 | 0.290096 | 0.936088 |
| WP_FAS_PATHWAY_ANI | -0.97623 | 0.4774 | 0.908762 | WP_FAS_PATHWAY_AND_ | -0.74552 | 0.822409 | 1 | WP_FAS_PATHWAY_AND_ | -1.13622 | 0.299133 | 0.937823 |
| REACTOME_ADHERENS_ | -0.83165 | 0.710064 | 0.969613 | REACTOME_ADHERENS_JU | 0.952686 | 0.502808 | 0.949774 | REACTOME_ADHERENS_JI | -1.14569 | 0.29493 | 0.938605 |
| REACTOME_SIGNALING_ | 1.424691 | 0.068708 | 0.56108 | REACTOME_SIGNALING_B' | 1.878707 | 0.001685 | 0.078986 | REACTOME_SIGNALING_B | -1.13966 | 0.288073 | 0.939603 |
| REACTOME_VISUAL_PH( | -1.32808 | 0.115468 | 0.639914 | REACTOME_VISUAL_PHOT | -1.11397 | 0.296631 | 0.847355 | REACTOME_VISUAL_PHOT | -1.14809 | 0.283741 | 0.944801 |
| REACTOME_FACTORS_IN | -0.8783 | 0.691684 | 0.940977 | REACTOME_FACTORS_INV | -0.86044 | 0.723311 | 1 | REACTOME_FACTORS_IN\ | -1.14601 | 0.257799 | 0.945278 |
| REACTOME_GLOBAL_GE | -0.9094 | 0.607421 | 0.951511 | REACTOME_GLOBAL_GEN( | -1.14466 | 0.245121 | 0.810416 | REACTOME_GLOBAL_GEN | -1.12738 | 0.293283 | 0.957817 |
| REACTOME_SIGNALING_ | 0.957303 | 0.516032 | 0.950623 | REACTOME_SIGNALING_B' | 0.641005 | 0.916192 | 1 | REACTOME_SIGNALING_B | 1.065698 | 0.35416 | 0.969691 |
| REACTOME_S_PHASE | 0.629879 | 0.988411 | 1 | REACTOME_S_PHASE | 0.662886 | 0.978839 | 1 | REACTOME_S_PHASE | 0.642285 | 0.996263 | 0.972452 |

| | | | | | | | | | | | |
|---|---|---|---|---|---|---|---|---|---|---|---|
| REACTOME_OPIOID_SIG | -0.82018 | 0.778783 | 0.966272 | REACTOME_OPIOID_SIGNA | -0.52797 | 0.998608 | 1 | REACTOME_OPIOID_SIGN | 0.651438 | 0.977634 | 0.973526 |
| WP_ALZHEIMERS_DISEA | 0.836316 | 0.737166 | 1 | WP_ALZHEIMERS_DISEASE | -0.60667 | 0.9835 | 1 | WP_ALZHEIMERS_DISEASI | 0.660842 | 0.970329 | 0.973982 |
| REACTOME_PLASMA_LII | 0.984177 | 0.475563 | 0.9551 | REACTOME_PLASMA_LIPO | -1.14457 | 0.26184 | 0.802726 | REACTOME_PLASMA_LIPO | -1.12143 | 0.308902 | 0.974032 |
| REACTOME_ABC_FAMILY | -1.27575 | 0.102129 | 0.702696 | REACTOME_ABC_FAMILY_ | 0.927492 | 0.577496 | 0.983438 | REACTOME_ABC_FAMILY_ | 0.64366 | 0.995054 | 0.974983 |
| WP_PRIMARY_FOCAL_SI | 1.314465 | 0.103486 | 0.724422 | WP_PRIMARY_FOCAL_SEG | 1.727188 | 0.009388 | 0.220767 | WP_PRIMARY_FOCAL_SEC | 0.654138 | 0.974186 | 0.975169 |
| REACTOME_BRANCHED_ | -0.61335 | 0.947888 | 1 | REACTOME_BRANCHED_CI | -0.80414 | 0.727238 | 1 | REACTOME_BRANCHED_C | 0.631507 | 0.943468 | 0.975223 |
| REACTOME_NEUREXINS_ | -0.68553 | 0.911266 | 1 | REACTOME_NEUREXINS_A | -0.4997 | 0.994599 | 1 | REACTOME_NEUREXINS_/ | 0.663238 | 0.938247 | 0.975672 |
| REACTOME_DEVELOPMI | -1.10727 | 0.199798 | 0.801286 | REACTOME_DEVELOPMEN | 1.484779 | 0.005077 | 0.511865 | REACTOME_DEVELOPMEN | 1.066036 | 0.23125 | 0.97762 |
| REACTOME_HEDGEHOG | 0.648813 | 0.985984 | 1 | REACTOME_HEDGEHOG_C | 0.872846 | 0.685622 | 1 | REACTOME_HEDGEHOG_( | 0.66386 | 0.990674 | 0.978672 |
| REACTOME_APOPTOSIS | 0.588923 | 0.993825 | 1 | REACTOME_APOPTOSIS | 0.63659 | 0.979608 | 1 | REACTOME_APOPTOSIS | 1.067601 | 0.308782 | 0.980868 |
| BIOCARTA_AT1R_PATHW | 0.796588 | 0.74243 | 1 | BIOCARTA_AT1R_PATHWA | -0.75896 | 0.794572 | 1 | BIOCARTA_AT1R_PATHW/ | 0.665259 | 0.904079 | 0.981126 |
| REACTOME_SELECTIVE_/ | 1.113136 | 0.286915 | 0.988913 | REACTOME_SELECTIVE_AU | 0.91369 | 0.578357 | 0.983789 | REACTOME_SELECTIVE_AI | 0.681992 | 0.965842 | 0.981906 |
| REACTOME_DOWNSTRE | 0.838992 | 0.689383 | 1 | REACTOME_DOWNSTREAN | 1.210432 | 0.219614 | 0.840492 | REACTOME_DOWNSTREA | 1.060151 | 0.367065 | 0.981923 |
| REACTOME_SIGNALING_ | 0.784188 | 0.810536 | 1 | REACTOME_SIGNALING_BY | 0.443171 | 0.998991 | 1 | REACTOME_SIGNALING_B | 0.677682 | 0.955759 | 0.981967 |
| REACTOME_PLATELET_S | -1.47434 | 0.070135 | 0.496659 | REACTOME_PLATELET_SEN | 0.65007 | 0.881943 | 1 | REACTOME_PLATELET_SE | 0.672785 | 0.886466 | 0.982401 |
| BIOCARTA_IGF1_PATHW | 1.257466 | 0.196456 | 0.78588 | BIOCARTA_IGF1_PATHWAY | 1.23512 | 0.221988 | 0.801407 | BIOCARTA_IGF1_PATHWA | 1.06937 | 0.366513 | 0.983361 |
| REACTOME_HEDGEHOG | 0.986929 | 0.468593 | 0.951651 | REACTOME_HEDGEHOG_C | 0.887246 | 0.615874 | 1 | REACTOME_HEDGEHOG_( | 0.666344 | 0.96549 | 0.983832 |
| REACTOME_RHOBTB_G1 | 1.182318 | 0.233609 | 0.904636 | REACTOME_RHOBTB_GTP/ | 1.034811 | 0.394471 | 0.878861 | REACTOME_RHOBTB_GTP | 0.682396 | 0.933645 | 0.985111 |
| REACTOME_INSULIN_RE | -0.65166 | 0.924724 | 1 | REACTOME_INSULIN_RECE | -0.59405 | 0.960161 | 1 | REACTOME_INSULIN_REC | 0.602508 | 0.958296 | 0.985497 |
| REACTOME_APC_C_MEI | 1.068829 | 0.343694 | 1 | REACTOME_APC_C_MEDI/ | 1.227723 | 0.175026 | 0.7995 | REACTOME_APC_C_MEDI | 1.070954 | 0.314815 | 0.986715 |
| REACTOME_RAS_PROCE | 1.083732 | 0.35678 | 1 | REACTOME_RAS_PROCESS | -0.61593 | 0.930778 | 1 | REACTOME_RAS_PROCES! | 1.056465 | 0.377348 | 0.987232 |
| REACTOME_TRANSCRIPT | 0.996671 | 0.453282 | 0.950595 | REACTOME_TRANSCRIPTIC | 0.670055 | 0.943377 | 1 | REACTOME_TRANSCRIPTI | 0.682826 | 0.957668 | 0.988286 |
| REACTOME_TRANSFERR | 0.950868 | 0.525205 | 0.958247 | REACTOME_TRANSFERRIN | 0.645256 | 0.897165 | 1 | REACTOME_TRANSFERRIN | 1.001384 | 0.440793 | 0.989188 |
| REACTOME_METABOLIS | 1.677806 | 0.004893 | 0.189697 | REACTOME_METABOLISM_ | 0.991934 | 0.451482 | 0.915981 | REACTOME_METABOLISM | 1.072741 | 0.313348 | 0.989297 |
| REACTOME_SIGNALING_ | 0.710899 | 0.873712 | 1 | REACTOME_SIGNALING_BY | -0.55326 | 0.982762 | 1 | REACTOME_SIGNALING_B | 0.58292 | 0.976467 | 0.989353 |
| WP_SPLICING_FACTOR_ | -0.96717 | 0.488805 | 0.912202 | WP_SPLICING_FACTOR_NC | -0.82455 | 0.746792 | 1 | WP_SPLICING_FACTOR_N | 0.980787 | 0.470819 | 0.990503 |
| BIOCARTA_MCALPAIN_F | 1.170356 | 0.265935 | 0.92728 | BIOCARTA_MCALPAIN_PAT | 1.669477 | 0.031904 | 0.271098 | BIOCARTA_MCALPAIN_PA | 0.683558 | 0.877852 | 0.991237 |
| REACTOME_EPH_EPHRII | 0.602479 | 0.953188 | 1 | REACTOME_EPH_EPHRIN_ | -0.44498 | 0.997351 | 0.997782 | REACTOME_EPH_EPHRIN_ | 0.564119 | 0.976612 | 0.991622 |
| REACTOME_MITOCHON | 0.478595 | 0.986528 | 1 | REACTOME_MITOCHONDR | 0.839464 | 0.650126 | 1 | REACTOME_MITOCHOND | 0.684373 | 0.879234 | 0.994148 |
| REACTOME_SIGNALING_ | 1.043162 | 0.393805 | 0.996893 | REACTOME_SIGNALING_BY | 0.74857 | 0.817224 | 1 | REACTOME_SIGNALING_B | 0.997881 | 0.445452 | 0.99427 |
| REACTOME_REGULATIO | 1.286427 | 0.119254 | 0.780719 | REACTOME_REGULATION_ | 1.316162 | 0.118154 | 0.706979 | REACTOME_REGULATION | 1.127653 | 0.235342 | 0.994538 |
| REACTOME_FC_EPSILON | 0.746294 | 0.915883 | 1 | REACTOME_FC_EPSILON_F | 1.039264 | 0.37462 | 0.885727 | REACTOME_FC_EPSILON_ | 0.706605 | 0.981193 | 0.994854 |
| REACTOME_RET_SIGNAI | 1.283321 | 0.154539 | 0.77251 | REACTOME_RET_SIGNALIN | 1.598566 | 0.037037 | 0.359904 | REACTOME_RET_SIGNALII | 0.694565 | 0.897333 | 0.995796 |
| BIOCARTA_STRESS_PATI | -1.03173 | 0.407875 | 0.869435 | BIOCARTA_STRESS_PATHW | 0.762229 | 0.75671 | 1 | BIOCARTA_STRESS_PATHV | 0.479676 | 0.989978 | 0.995819 |
| REACTOME_CA2_PATHW | 0.924852 | 0.558044 | 0.954099 | REACTOME_CA2_PATHWA | -1.03585 | 0.404823 | 0.944814 | REACTOME_CA2_PATHW/ | 0.981185 | 0.47644 | 0.996103 |
| REACTOME_REGULATIO | 0.605498 | 0.942911 | 1 | REACTOME_REGULATION_ | -0.95751 | 0.506807 | 1 | REACTOME_REGULATION | 1.001505 | 0.437236 | 0.996255 |
| BIOCARTA_PYK2_PATHW | -0.58812 | 0.961596 | 1 | BIOCARTA_PYK2_PATHWA | -0.70745 | 0.865305 | 1 | BIOCARTA_PYK2_PATHW/ | 0.505109 | 0.990407 | 0.996383 |
| REACTOME_L1CAM_INT | 1.023263 | 0.411969 | 0.963151 | REACTOME_L1CAM_INTER | 0.796559 | 0.793019 | 1 | REACTOME_L1CAM_INTEI | 0.685715 | 0.968504 | 0.996595 |
| REACTOME_AURKA_ACT | -0.43622 | 0.999339 | 0.998502 | REACTOME_AURKA_ACTIV | 1.04263 | 0.383719 | 0.885791 | REACTOME_AURKA_ACTIV | 1.044675 | 0.374067 | 0.996998 |
| REACTOME_SYNTHESIS_ | 0.81546 | 0.711494 | 1 | REACTOME_SYNTHESIS_OI | 0.909456 | 0.557842 | 0.982812 | REACTOME_SYNTHESIS_O | 0.707529 | 0.863063 | 0.997605 |
| REACTOME_GOLGI_TO_ | 1.127399 | 0.244588 | 0.978263 | REACTOME_GOLGI_TO_ER | 1.177458 | 0.194227 | 0.843635 | REACTOME_GOLGI_TO_EI | 1.073014 | 0.287671 | 0.998117 |
| BIOCARTA_GLEEVEC_PA | 0.812954 | 0.709808 | 1 | BIOCARTA_GLEEVEC_PATH | 0.433068 | 0.992166 | 1 | BIOCARTA_GLEEVEC_PATI | 0.710489 | 0.853816 | 0.998147 |
| BIOCARTA_PDGF_PATHV | 0.731061 | 0.825779 | 1 | BIOCARTA_PDGF_PATHWA | -0.83753 | 0.694455 | 1 | BIOCARTA_PDGF_PATHW | 0.695712 | 0.883834 | 0.99839 |
| REACTOME_SIGNALING_ | -0.69599 | 0.907521 | 1 | REACTOME_SIGNALING_BY | -0.6299 | 0.956163 | 1 | REACTOME_SIGNALING_B | 0.514676 | 0.996925 | 0.998455 |
| REACTOME_MAPK_FAM | 1.023786 | 0.401715 | 0.976974 | REACTOME_MAPK_FAMILY | 0.779922 | 0.943042 | 1 | REACTOME_MAPK_FAMIL | 0.713584 | 0.998912 | 0.99853 |
| WP_PROTEASOME_DEG | -0.90535 | 0.598805 | 0.935932 | WP_PROTEASOME_DEGRA | -0.71428 | 0.904412 | 1 | WP_PROTEASOME_DEGR/ | 0.699004 | 0.927246 | 0.998915 |
| BIOCARTA_GPCR_PATHV | -0.54297 | 0.980306 | 1 | BIOCARTA_GPCR_PATHWA | -0.82438 | 0.694026 | 1 | BIOCARTA_GPCR_PATHW | -0.33886 | 0.999692 | 0.999895 |
| REACTOME_SMOOTH_N | 2.095553 | 1.89E-04 | 0.014507 | REACTOME_SMOOTH_MU | 2.23465 | 0 | 0.001176 | REACTOME_SMOOTH_MU | -0.88747 | 0.620323 | 1 |
| REACTOME_RHO_GTPAS | 2.047715 | 1.86E-04 | 0.017034 | REACTOME_RHO_GTPASES | 2.209499 | 0 | 0.001241 | REACTOME_RHO_GTPASE | -0.45823 | 0.990926 | 1 |
| REACTOME_COMPLEME | 1.871256 | 0.002273 | 0.05027 | REACTOME_COMPLEMENT | 1.450704 | 0.094501 | 0.525419 | REACTOME_COMPLEMEN | -0.95891 | 0.519225 | 1 |
| REACTOME_PHASE_I_FU | 1.840742 | 0.002778 | 0.066004 | REACTOME_PHASE_I_FUN | 1.630544 | 0.021846 | 0.325468 | REACTOME_PHASE_I_FUN | 1.205816 | 0.187153 | 1 |
| REACTOME_RESPONSE_ | 1.798976 | 5.37E-04 | 0.095028 | REACTOME_RESPONSE_TC | 1.275612 | 0.123618 | 0.740893 | REACTOME_RESPONSE_T( | -0.98058 | 0.503002 | 1 |
| WP_OXIDATIVE_STRESS_ | 1.758821 | 0.007422 | 0.124298 | WP_OXIDATIVE_STRESS_R | -0.67299 | 0.899329 | 1 | WP_OXIDATIVE_STRESS_F | 0.901859 | 0.597218 | 1 |
| REACTOME_REGULATIO | 1.656866 | 0.007111 | 0.216007 | REACTOME_REGULATION_ | 1.612106 | 0.011934 | 0.353559 | REACTOME_REGULATION | -0.82697 | 0.747113 | 1 |
| WP_METAPATHWAY_BI | 1.631887 | 0.01252 | 0.244089 | WP_METAPATHWAY_BIOT | 1.183106 | 0.220864 | 0.832484 | WP_METAPATHWAY_BIOT | 1.171471 | 0.203804 | 1 |
| BIOCARTA_MTOR_PATH | 1.550841 | 0.037663 | 0.350919 | BIOCARTA_MTOR_PATHW | 1.145075 | 0.297782 | 0.884687 | BIOCARTA_MTOR_PATHW | 1.135572 | 0.295699 | 1 |
| REACTOME_DISASSEMB | 1.572992 | 0.036513 | 0.351786 | REACTOME_DISASSEMBLY_ | 1.185451 | 0.240902 | 0.832538 | REACTOME_DISASSEMBLY | 0.863328 | 0.653726 | 1 |
| BIOCARTA_IGF1MTOR_F | 1.565579 | 0.040882 | 0.359024 | BIOCARTA_IGF1MTOR_PAT | 1.339213 | 0.149928 | 0.684472 | BIOCARTA_IGF1MTOR_PA | 1.19328 | 0.236763 | 1 |
| WP_FOCAL_ADHESION | 1.558832 | 0.006217 | 0.364024 | WP_FOCAL_ADHESION | 2.258373 | 0 | 8.77E-04 | WP_FOCAL_ADHESION | -0.90816 | 0.643803 | 1 |
| REACTOME_TGF_BETA_I | 1.54002 | 0.042164 | 0.367519 | REACTOME_TGF_BETA_RE | 1.995003 | 0.00146 | 0.020409 | REACTOME_TGF_BETA_RI | -0.81692 | 0.708075 | 1 |
| WP_TRYPTOPHAN_MET/ | 1.552856 | 0.033421 | 0.367809 | WP_TRYPTOPHAN_METAB | 1.241386 | 0.196483 | 0.789737 | WP_TRYPTOPHAN_METAI | 1.096716 | 0.30925 | 1 |
| REACTOME_GLUTATHIO | 1.502735 | 0.044757 | 0.403892 | REACTOME_GLUTATHIONE | -0.82569 | 0.707884 | 1 | REACTOME_GLUTATHION | 1.001763 | 0.447789 | 1 |
| BIOCARTA_TFF_PATHW/ | 1.498037 | 0.064314 | 0.406534 | BIOCARTA_TFF_PATHWAY | 1.089106 | 0.344872 | 0.880138 | BIOCARTA_TFF_PATHWAY | 1.130845 | 0.296668 | 1 |
| REACTOME_ROLE_OF_P | 1.510145 | 0.055081 | 0.414669 | REACTOME_ROLE_OF_PHC | -1.00504 | 0.462185 | 0.989449 | REACTOME_ROLE_OF_PH | -0.90923 | 0.583097 | 1 |
| REACTOME_COPI_MEDI/ | 1.487672 | 0.01938 | 0.424171 | REACTOME_COPI_MEDIAT | 1.068519 | 0.325105 | 0.875764 | REACTOME_COPI_MEDIAT | 1.27378 | 0.087838 | 1 |
| REACTOME_ADRENALIN | -1.55299 | 0.040319 | 0.442891 | REACTOME_ADRENALINE_ | -1.33461 | 0.130852 | 0.70594 | REACTOME_ADRENALINE_ | -0.83963 | 0.684973 | 1 |
| REACTOME_PRESYNAPT | -1.57724 | 3.81E-02 | 0.452492 | REACTOME_PRESYNAPTIC_ | -1.48479 | 0.066895 | 0.519091 | REACTOME_PRESYNAPTIC | -0.81094 | 0.714218 | 1 |
| REACTOME_POTASSIUM | -1.68077 | 0.008722 | 0.453404 | REACTOME_POTASSIUM_( | -1.84142 | 9.72E-04 | 0.139027 | REACTOME_POTASSIUM_ | -1.07326 | 0.354291 | 1 |
| REACTOME_G_PROTEIN | -1.50411 | 0.049633 | 0.470454 | REACTOME_G_PROTEIN_B | -1.32346 | 0.135078 | 0.68963 | REACTOME_G_PROTEIN_I | -0.81643 | 0.711515 | 1 |
| REACTOME_G_ALPHA_Z | -1.49667 | 0.053191 | 0.471826 | REACTOME_G_ALPHA_Z_S | -1.32361 | 0.123367 | 0.703379 | REACTOME_G_ALPHA_Z_! | -0.98589 | 0.488935 | 1 |
| REACTOME_G_BETA_GA | -1.63853 | 0.027694 | 0.484038 | REACTOME_G_BETA_GAM | -1.44739 | 0.078537 | 0.534043 | REACTOME_G_BETA_GAN | -0.84634 | 0.66886 | 1 |
| REACTOME_REGULATIO | -1.50515 | 4.69E-02 | 0.487974 | REACTOME_REGULATION_ | -0.99682 | 0.458509 | 1 | REACTOME_REGULATION | -0.99032 | 0.483852 | 1 |
| REACTOME_MUSCLE_CC | 1.431518 | 0.027851 | 0.549783 | REACTOME_MUSCLE_CON | 1.855598 | 6.06E-04 | 0.092988 | REACTOME_MUSCLE_CON | -0.76617 | 0.865063 | 1 |
| WP_FOCAL_ADHESION_ | 1.419392 | 0.019932 | 0.567696 | WP_FOCAL_ADHESION_PI3 | 1.678641 | 0.001572 | 0.274626 | WP_FOCAL_ADHESION_PI | -0.88798 | 0.69818 | 1 |
| REACTOME_DNA_DAMA | -1.42656 | 0.078599 | 0.604991 | REACTOME_DNA_DAMAGI | -1.0927 | 0.33875 | 0.876412 | REACTOME_DNA_DAMAG | -0.99773 | 0.470141 | 1 |
| REACTOME_RECOGNITI( | -1.4134 | 0.096913 | 0.607543 | REACTOME_RECOGNITION | -1.14434 | 0.298029 | 0.795635 | REACTOME_RECOGNITION | -1.04498 | 0.417378 | 1 |
| REACTOME_G_ALPHA_S | -1.38823 | 0.073708 | 0.615996 | REACTOME_G_ALPHA_S_S | -1.27711 | 0.128586 | 0.73331 | REACTOME_G_ALPHA_S_! | -1.09922 | 0.333198 | 1 |
| REACTOME_G_BETA_GA | -1.32852 | 0.134291 | 0.652716 | REACTOME_G_BETA_GAM | -1.33968 | 0.127647 | 0.701931 | REACTOME_G_BETA_GAN | -0.69969 | 0.847664 | 1 |
| REACTOME_RECEPTOR_ | 1.384766 | 0.10579 | 0.654493 | REACTOME_RECEPTOR_TY | 0.557713 | 0.957545 | 1 | REACTOME_RECEPTOR_T | 0.992129 | 0.456877 | 1 |
| REACTOME_TRANSLESI( | -1.34113 | 0.129457 | 0.656178 | REACTOME_TRANSLESION | -1.1407 | 0.293814 | 0.800552 | REACTOME_TRANSLESION | -0.91046 | 0.583033 | 1 |
| REACTOME_OXIDATIVE_ | -1.3646 | 0.110028 | 0.659745 | REACTOME_OXIDATIVE_ST | 1.149954 | 0.275756 | 0.883304 | REACTOME_OXIDATIVE_S | -0.92718 | 0.570148 | 1 |
| REACTOME_THROMBOX | -1.3172 | 0.141044 | 0.661092 | REACTOME_THROMBOXAI | -0.92244 | 0.558929 | 1 | REACTOME_THROMBOXA | -0.77801 | 0.752917 | 1 |
| REACTOME_FCGAMMA_ | 1.372073 | 0.062075 | 0.664576 | REACTOME_FCGAMMA_RI | 0.885681 | 0.636843 | 1 | REACTOME_FCGAMMA_R | -0.64871 | 0.957159 | 1 |
| REACTOME_AGGREPHA( | 1.356127 | 0.106917 | 0.665621 | REACTOME_AGGREPHAGY | 0.714442 | 0.845535 | 1 | REACTOME_AGGREPHAGY | 0.768583 | 0.82577 | 1 |

| | | | | | | | | | | | |
|---|---|---|---|---|---|---|---|---|---|---|---|
| WP_GPCRS_CLASS_A_RH | -1.329 | 0.118739 | 0.665931 | WP_GPCRS_CLASS_A_RHO | -1.19231 | 0.224802 | 0.760924 | WP_GPCRS_CLASS_A_RH( | -1.08858 | 0.354356 | 1 |
| REACTOME_VEGFR2_MI | 1.373718 | 0.097124 | 0.670626 | REACTOME_VEGFR2_MED | 1.470035 | 0.075565 | 0.506028 | REACTOME_VEGFR2_MED | 0.884283 | 0.626313 | 1 |
| REACTOME_CELL_EXTRA | 1.357794 | 0.126918 | 0.670663 | REACTOME_CELL_EXTRACI | 1.733791 | 0.017402 | 0.219839 | REACTOME_CELL_EXTRAC | -0.67271 | 0.87217 | 1 |
| REACTOME_SIGNALING_ | 1.36625 | 0.073047 | 0.674196 | REACTOME_SIGNALING_B' | 1.135191 | 0.262416 | 0.89588 | REACTOME_SIGNALING_B | 1.176832 | 0.177988 | 1 |
| BIOCARTA_NFAT_PATHV | 1.361687 | 0.100884 | 0.679085 | BIOCARTA_NFAT_PATHWA | 1.158855 | 0.263003 | 0.868387 | BIOCARTA_NFAT_PATHW. | 0.933962 | 0.540462 | 1 |
| REACTOME_TELOMERE_ | -1.28052 | 0.157676 | 0.699982 | REACTOME_TELOMERE_M | -0.80893 | 0.74491 | 1 | REACTOME_TELOMERE_N | -1.04623 | 0.411712 | 1 |
| REACTOME_SIGNALING_ | 1.335048 | 0.092478 | 0.700333 | REACTOME_SIGNALING_B' | 1.171513 | 0.224419 | 0.848335 | REACTOME_SIGNALING_B | 1.194714 | 0.186298 | 1 |
| REACTOME_AMINE_LIG/ | -1.71628 | 0.016856 | 0.701106 | REACTOME_AMINE_LIGAN | -1.48621 | 0.059207 | 0.56052 | REACTOME_AMINE_LIGAI | -0.95905 | 0.52393 | 1 |
| REACTOME_RHOH_GTP/ | 1.337284 | 0.116345 | 0.702549 | REACTOME_RHOH_GTPAS | 1.474681 | 0.067132 | 0.504604 | REACTOME_RHOH_GTPAS | -1.02552 | 0.431049 | 1 |
| REACTOME_ACTIVATION | -1.28183 | 0.152859 | 0.709089 | REACTOME_ACTIVATION_( | -1.48437 | 0.054514 | 0.499846 | REACTOME_ACTIVATION_ | 0.921798 | 0.560574 | 1 |
| REACTOME_ADP_SIGNA | -1.29836 | 0.156363 | 0.709312 | REACTOME_ADP_SIGNALLI | -0.91242 | 0.578844 | 1 | REACTOME_ADP_SIGNALL | -0.69951 | 0.841673 | 1 |
| WP_TRANSLATION_FAC' | 1.338444 | 0.104578 | 0.709448 | WP_TRANSLATION_FACTO | 1.199925 | 0.211626 | 0.816169 | WP_TRANSLATION_FACT( | 1.004119 | 0.442544 | 1 |
| REACTOME_PLATELET_A | 1.340225 | 0.036528 | 0.713951 | REACTOME_PLATELET_ACT | 1.016457 | 0.406108 | 0.899982 | REACTOME_PLATELET_AC | -1.00256 | 0.471843 | 1 |
| REACTOME_FCERI_MED | -1.26344 | 0.16816 | 0.718355 | REACTOME_FCERI_MEDIA' | -0.76459 | 0.791276 | 1 | REACTOME_FCERI_MEDIA | -0.88278 | 0.622608 | 1 |
| REACTOME_LDL_CLEAR/ | 1.312911 | 0.151354 | 0.72004 | REACTOME_LDL_CLEARAN | 1.206641 | 0.244355 | 0.835683 | REACTOME_LDL_CLEARAI | 0.864418 | 0.649756 | 1 |
| BIOCARTA_INTEGRIN_P/ | 1.317096 | 0.126351 | 0.725243 | BIOCARTA_INTEGRIN_PATI | 1.611432 | 0.034288 | 0.343608 | BIOCARTA_INTEGRIN_PAT | 0.923021 | 0.572866 | 1 |
| REACTOME_POST_TRAN | 1.319429 | 0.124207 | 0.727204 | REACTOME_POST_TRANSL | 1.140265 | 0.286604 | 0.893466 | REACTOME_POST_TRANS | 1.267513 | 0.145875 | 1 |
| REACTOME_BASE_EXCIS | -1.21725 | 0.211616 | 0.733037 | REACTOME_BASE_EXCISIO | -0.7337 | 0.823652 | 1 | REACTOME_BASE_EXCISI( | -0.8185 | 0.699875 | 1 |
| REACTOME_SUMOYLATI | -1.23988 | 0.153761 | 0.736181 | REACTOME_SUMOYLATIOI | -1.46462 | 0.032277 | 0.52941 | REACTOME_SUMOYLATIO | -0.92801 | 0.584489 | 1 |
| REACTOME_G_PROTEIN_ | -1.22277 | 0.206976 | 0.736512 | REACTOME_G_PROTEIN_A | -0.97592 | 0.491141 | 1 | REACTOME_G_PROTEIN_/ | -0.875 | 0.631464 | 1 |
| WP_IL5_SIGNALING_PAT | -1.19427 | 0.185419 | 0.736978 | WP_IL5_SIGNALING_PATH | -0.587 | 0.98749 | 1 | WP_IL5_SIGNALING_PATH | -0.74277 | 0.848656 | 1 |
| REACTOME_PLATELET_C | 1.319464 | 0.152365 | 0.738083 | REACTOME_PLATELET_CAL | -0.80368 | 0.728852 | 1 | REACTOME_PLATELET_CA | -0.8198 | 0.704897 | 1 |
| REACTOME_BILE_ACID_ | 1.302678 | 0.148148 | 0.738606 | REACTOME_BILE_ACID_AN | 0.83776 | 0.663609 | 1 | REACTOME_BILE_ACID_AI | -0.97236 | 0.504807 | 1 |
| REACTOME_EGFR_DOW | -1.22488 | 0.208209 | 0.740682 | REACTOME_EGFR_DOWNF | -0.95514 | 0.525375 | 1 | REACTOME_EGFR_DOWN | -0.59933 | 0.943078 | 1 |
| REACTOME_NEGATIVE_ | -1.2116 | 0.22103 | 0.741224 | REACTOME_NEGATIVE_RE | -0.98104 | 0.47923 | 1 | REACTOME_NEGATIVE_RI | -0.71924 | 0.826204 | 1 |
| WP_ENDOCHONDRAL_C | -1.20781 | 0.21317 | 0.743246 | WP_ENDOCHONDRAL_OSS | -1.15988 | 0.269135 | 0.81496 | WP_ENDOCHONDRAL_OS | -0.90695 | 0.596935 | 1 |
| WP_EXERCISEINDUCED_ | -1.16351 | 0.244783 | 0.744357 | WP_EXERCISEINDUCED_CI | 0.717694 | 0.850621 | 1 | WP_EXERCISEINDUCED_C | -1.0655 | 0.385267 | 1 |
| REACTOME_AMINO_ACI | -1.19775 | 0.228832 | 0.745686 | REACTOME_AMINO_ACID_ | 0.842614 | 0.667084 | 1 | REACTOME_AMINO_ACID | -0.88632 | 0.61922 | 1 |
| BIOCARTA_ECM_PATHW | 1.320114 | 0.143582 | 0.746869 | BIOCARTA_ECM_PATHWA' | 1.75156 | 0.015962 | 0.211785 | BIOCARTA_ECM_PATHWA | 0.760377 | 0.788698 | 1 |
| REACTOME_ADP_SIGNA | -1.20328 | 0.228754 | 0.7479 | REACTOME_ADP_SIGNALLI | -1.27077 | 0.182342 | 0.721035 | REACTOME_ADP_SIGNALL | -0.62687 | 0.913453 | 1 |
| REACTOME_SLC_MEDIA' | -1.164 | 0.168183 | 0.751608 | REACTOME_SLC_MEDIATE | -1.27532 | 0.070133 | 0.727862 | REACTOME_SLC_MEDIATI | -1.05159 | 0.379613 | 1 |
| WP_IL9_SIGNALING_PAT | -1.1667 | 0.257675 | 0.751824 | WP_IL9_SIGNALING_PATH | 1.254199 | 0.199316 | 0.793302 | WP_IL9_SIGNALING_PATH | -0.78427 | 0.745307 | 1 |
| WP_IL2_SIGNALING_PAT | -1.16879 | 0.2125 | 0.754166 | WP_IL2_SIGNALING_PATH | 1.050159 | 0.375729 | 0.873366 | WP_IL2_SIGNALING_PATH | 0.914932 | 0.624294 | 1 |
| REACTOME_CHAPERONI | -1.22644 | 0.184801 | 0.758694 | REACTOME_CHAPERONIN_ | -0.7721 | 0.805208 | 1 | REACTOME_CHAPERONIN | -0.59486 | 0.963243 | 1 |
| REACTOME_TNFR1_INDI | -1.16962 | 0.264107 | 0.760717 | REACTOME_TNFR1_INDUC | 0.624474 | 0.905411 | 1 | REACTOME_TNFR1_INDU( | -0.92815 | 0.564799 | 1 |
| BIOCARTA_MAPK_PATH | -1.18072 | 0.205285 | 0.762373 | BIOCARTA_MAPK_PATHW. | 0.843178 | 0.705484 | 1 | BIOCARTA_MAPK_PATHW. | -0.64491 | 0.945559 | 1 |
| BIOCARTA_TOLL_PATHV | -1.1776 | 0.258346 | 0.762996 | BIOCARTA_TOLL_PATHWA | 0.760412 | 0.749894 | 1 | BIOCARTA_TOLL_PATHW/ | -0.74632 | 0.78568 | 1 |
| REACTOME_CLASS_B_2_ | -1.18316 | 0.235064 | 0.764285 | REACTOME_CLASS_B_2_SI | -1.54688 | 0.031974 | 0.608624 | REACTOME_CLASS_B_2_S | -0.90837 | 0.593037 | 1 |
| REACTOME_MAPK_TAR( | -1.1709 | 0.253687 | 0.765895 | REACTOME_MAPK_TARGE | 0.647289 | 0.909691 | 1 | REACTOME_MAPK_TARGI | 1.230528 | 0.188998 | 1 |
| REACTOME_METABOLIS | 1.292727 | 0.021572 | 0.766503 | REACTOME_METABOLISM_ | 1.089052 | 0.233073 | 0.874235 | REACTOME_METABOLISM | 1.097387 | 0.133739 | 1 |
| BIOCARTA_INSULIN_PAT | 1.286236 | 0.179649 | 0.77111 | BIOCARTA_INSULIN_PATH' | 1.278426 | 0.191809 | 0.741208 | BIOCARTA_INSULIN_PATH | 1.149609 | 0.280759 | 1 |
| REACTOME_RNA_POLYN | 1.256405 | 0.197394 | 0.771758 | REACTOME_RNA_POLYME | 1.103905 | 0.337553 | 0.88246 | REACTOME_RNA_POLYMI | 1.07441 | 0.360334 | 1 |
| WP_SPHINGOLIPID_MET | 1.274238 | 0.173594 | 0.778291 | WP_SPHINGOLIPID_METAI | 1.349307 | 0.14348 | 0.695409 | WP_SPHINGOLIPID_META | 0.82705 | 0.694975 | 1 |
| REACTOME_GAMMA_C/ | 1.256574 | 0.190531 | 0.780114 | REACTOME_GAMMA_CAR | 1.011111 | 0.428183 | 0.906192 | REACTOME_GAMMA_CAF | 0.834218 | 0.690357 | 1 |
| WP_SPHINGOLIPID_MET | 1.268731 | 0.182319 | 0.780141 | WP_SPHINGOLIPID_METAI | 1.347733 | 0.139217 | 0.689432 | WP_SPHINGOLIPID_META | 0.825807 | 0.707257 | 1 |
| REACTOME_MYD88_MA | -1.13368 | 0.242175 | 0.780759 | REACTOME_MYD88_MAL_ | 0.722711 | 0.895982 | 1 | REACTOME_MYD88_MAL_ | 0.776157 | 0.882254 | 1 |
| REACTOME_SIGNALING_ | 1.24436 | 0.056774 | 0.782274 | REACTOME_SIGNALING_B' | 1.384746 | 0.017362 | 0.639522 | REACTOME_SIGNALING_B | -0.89468 | 0.724495 | 1 |
| REACTOME_HEMOSTASI | 1.245199 | 0.051649 | 0.787661 | REACTOME_HEMOSTASIS | 0.958481 | 0.564949 | 0.95088 | REACTOME_HEMOSTASIS | -0.94554 | 0.624174 | 1 |
| REACTOME_NERVOUS_S | 1.249499 | 0.072069 | 0.789004 | REACTOME_NERVOUS_SYS | 1.434095 | 0.01882 | 0.560475 | REACTOME_NERVOUS_SY | 0.907966 | 0.75615 | 1 |
| REACTOME_BINDING_AI | -1.0983 | 0.338776 | 0.791656 | REACTOME_BINDING_ANC | -1.17813 | 0.265903 | 0.785038 | REACTOME_BINDING_ANI | -1.00475 | 0.471024 | 1 |
| WP_CALCIUM_REGULAT | -1.08676 | 0.28184 | 0.791694 | WP_CALCIUM_REGULATIO | -1.32732 | 0.053923 | 0.703952 | WP_CALCIUM_REGULATI( | -0.68062 | 0.960855 | 1 |
| REACTOME_SIGNALING_ | 1.2606 | 0.153957 | 0.792392 | REACTOME_SIGNALING_B' | 1.7706 | 0.009083 | 0.191732 | REACTOME_SIGNALING_B | -0.77181 | 0.808443 | 1 |
| REACTOME_AMINO_ACI | -1.06857 | 0.344088 | 0.793174 | REACTOME_AMINO_ACIDS | -0.70998 | 0.899234 | 1 | REACTOME_AMINO_ACID | -0.82975 | 0.71931 | 1 |
| REACTOME_RNA_POLYN | 1.257682 | 0.187416 | 0.79439 | REACTOME_RNA_POLYME | 1.107407 | 0.33175 | 0.897258 | REACTOME_RNA_POLYMI | 1.07888 | 0.356589 | 1 |
| REACTOME_REGULATIO | 1.245484 | 0.203041 | 0.795429 | REACTOME_REGULATION_ | 1.088606 | 0.343315 | 0.863753 | REACTOME_REGULATION_ | 1.112129 | 0.312916 | 1 |
| REACTOME_THROMBIN_ | -1.09455 | 0.330087 | 0.796131 | REACTOME_THROMBIN_SI | -0.86956 | 0.640192 | 1 | REACTOME_THROMBIN_S | -0.63871 | 0.915117 | 1 |
| REACTOME_TAK1_DEPE | -1.08977 | 0.328643 | 0.796635 | REACTOME_TAK1_DEPEND | -0.65591 | 0.922432 | 1 | REACTOME_TAK1_DEPEN | -0.73799 | 0.820543 | 1 |
| REACTOME_RND1_GTP/ | 1.236061 | 0.184529 | 0.796733 | REACTOME_RND1_GTPASI | 1.522596 | 0.05583 | 0.437405 | REACTOME_RND1_GTPAS | 1.129143 | 0.271275 | 1 |
| REACTOME_RAB_GERAN | -1.06936 | 0.330651 | 0.797442 | REACTOME_RAB_GERANYI | -0.77015 | 0.852578 | 1 | REACTOME_RAB_GERANY | -0.77587 | 0.805597 | 1 |
| REACTOME_SYNTHESIS_ | -1.09847 | 0.320843 | 0.798724 | REACTOME_SYNTHESIS_OI | -0.92978 | 0.558852 | 1 | REACTOME_SYNTHESIS_O | -1.07655 | 0.378993 | 1 |
| REACTOME_SNRNP_ASS | -1.08678 | 0.314362 | 0.798904 | REACTOME_SNRNP_ASSEI | -1.04709 | 0.38026 | 0.933623 | REACTOME_SNRNP_ASSEI | -0.76812 | 0.812713 | 1 |
| REACTOME_RNA_POLYN | -1.10546 | 0.288759 | 0.799262 | REACTOME_RNA_POLYME | -1.22359 | 0.169797 | 0.763001 | REACTOME_RNA_POLYMI | -0.90797 | 0.608807 | 1 |
| REACTOME_INTERLEUKI | -1.0794 | 0.326594 | 0.800459 | REACTOME_INTERLEUKIN_ | 0.87925 | 0.634101 | 1 | REACTOME_INTERLEUKIN | -0.62655 | 0.955956 | 1 |
| REACTOME_ACTIVATED_ | -1.09072 | 0.353401 | 0.801078 | REACTOME_ACTIVATED_T/ | 1.077878 | 0.360852 | 0.874588 | REACTOME_ACTIVATED_T | -0.82368 | 0.691545 | 1 |
| REACTOME_MHC_CLASS | 1.232315 | 0.137919 | 0.80231 | REACTOME_MHC_CLASS_I | 0.901928 | 0.62927 | 0.987026 | REACTOME_MHC_CLASS_ | 1.140667 | 0.192608 | 1 |
| REACTOME_GLUCAGON_ | -1.06112 | 0.369135 | 0.803043 | REACTOME_GLUCAGON_T | -1.51438 | 0.046368 | 0.559899 | REACTOME_GLUCAGON_' | 1.075015 | 0.359748 | 1 |
| REACTOME_MYD88_IND | -1.09931 | 0.280655 | 0.80374 | REACTOME_MYD88_INDEF | 0.847211 | 0.709148 | 1 | REACTOME_MYD88_INDE | 0.926727 | 0.609895 | 1 |
| REACTOME_VASOPRESS | -1.10894 | 0.311454 | 0.803933 | REACTOME_VASOPRESSIN_ | -1.52299 | 0.041825 | 0.591767 | REACTOME_VASOPRESSIN | 0.981203 | 0.478339 | 1 |
| REACTOME_CELLULAR_S | -1.06285 | 0.344141 | 0.804362 | REACTOME_CELLULAR_SEI | 0.835746 | 0.707849 | 1 | REACTOME_CELLULAR_SE | -0.97074 | 0.512055 | 1 |
| REACTOME_ANTIGEN_P | -1.11378 | 0.271775 | 0.804395 | REACTOME_ANTIGEN_PRC | 0.749621 | 0.861907 | 1 | REACTOME_ANTIGEN_PR( | -0.60895 | 0.976033 | 1 |
| REACTOME_GLUCONEO( | 1.236172 | 0.194237 | 0.804971 | REACTOME_GLUCONEOGE | 1.050157 | 0.379137 | 0.868169 | REACTOME_GLUCONEOGI | -0.7566 | 0.800808 | 1 |
| BIOCARTA_HIVNEF_PATI | -1.08018 | 0.336807 | 0.805225 | BIOCARTA_HIVNEF_PATHV | 0.649479 | 0.930715 | 1 | BIOCARTA_HIVNEF_PATH' | -0.89031 | 0.623926 | 1 |
| WP_ONECARBON_META | -1.11597 | 0.299957 | 0.805515 | WP_ONECARBON_METAB( | -1.09955 | 0.332295 | 0.867447 | WP_ONECARBON_METAB | -0.96813 | 0.512648 | 1 |
| REACTOME_AQUAPORIN | -1.12088 | 0.285927 | 0.805999 | REACTOME_AQUAPORIN_I | -1.59471 | 0.021146 | 0.675754 | REACTOME_AQUAPORIN_ | 0.9662 | 0.497605 | 1 |
| REACTOME_TOLL_LIKE_I | -1.09962 | 0.297715 | 0.810576 | REACTOME_TOLL_LIKE_RE | 0.686758 | 0.939555 | 1 | REACTOME_TOLL_LIKE_RI | 0.764283 | 0.895037 | 1 |
| REACTOME_SIGNALING_ | 1.227781 | 0.174489 | 0.811205 | REACTOME_SIGNALING_B' | 1.713913 | 0.009683 | 0.23268 | REACTOME_SIGNALING_B | 0.958745 | 0.51098 | 1 |
| REACTOME_MAP2K_AN | 1.225052 | 0.188927 | 0.813153 | REACTOME_MAP2K_AND_ | 0.92245 | 0.553712 | 0.984663 | REACTOME_MAP2K_AND_ | -0.97355 | 0.5058 | 1 |
| REACTOME_TOLL_LIKE_I | -1.07062 | 0.326225 | 0.814401 | REACTOME_TOLL_LIKE_RE | 0.763107 | 0.871027 | 1 | REACTOME_TOLL_LIKE_RI | 0.868055 | 0.75073 | 1 |
| REACTOME_CELL_SURF/ | 1.217299 | 0.181059 | 0.818016 | REACTOME_CELL_SURFACI | 1.584074 | 0.026716 | 0.359633 | REACTOME_CELL_SURFAC | -0.99619 | 0.473026 | 1 |
| REACTOME_RND3_GTP/ | 1.218988 | 0.194254 | 0.819942 | REACTOME_RND3_GTPASI | 1.322058 | 0.135428 | 0.708239 | REACTOME_RND3_GTPAS | 1.12263 | 0.273362 | 1 |
| REACTOME_NEUTROPHI | 1.220147 | 0.073319 | 0.823814 | REACTOME_NEUTROPHIL_ | -1.12452 | 0.160604 | 0.833392 | REACTOME_NEUTROPHIL_ | -0.8028 | 0.896083 | 1 |
| WP_EUKARYOTIC_TRAN | -1.03625 | 0.395578 | 0.861785 | WP_EUKARYOTIC_TRANSC | -1.4142 | 0.067803 | 0.566321 | WP_EUKARYOTIC_TRANS( | -0.89814 | 0.610256 | 1 |
| REACTOME_REGULATIO | -1.0214 | 0.414488 | 0.868472 | REACTOME_REGULATION_ | -1.11962 | 0.276187 | 0.834975 | REACTOME_REGULATION_ | -0.7155 | 0.884501 | 1 |
| REACTOME_TRANSPORT | -1.02411 | 0.364138 | 0.873269 | REACTOME_TRANSPORT_( | -1.25243 | 0.027409 | 0.700609 | REACTOME_TRANSPORT_ | -0.83218 | 0.875748 | 1 |

| | | | | | | | | | | | |
|---|---|---|---|---|---|---|---|---|---|---|---|
| REACTOME_TRANSCRIPT | -1.01472 | 0.424066 | 0.876481 | REACTOME_TRANSCRIPTIC | -0.83554 | 0.71072 | 1 | REACTOME_TRANSCRIPTI | -1.02987 | 0.428162 | 1 |
| REACTOME_DUAL_INCIS | -1.02445 | 0.414608 | 0.879088 | REACTOME_DUAL_INCISIO | -1.00053 | 0.45767 | 0.99813 | REACTOME_DUAL_INCISI | -0.85877 | 0.648196 | 1 |
| REACTOME_RMTS_MET | 1.196945 | 0.222449 | 0.880812 | REACTOME_RMTS_METHY | 1.071142 | 0.355268 | 0.873187 | REACTOME_RMTS_METHY | 0.837394 | 0.713443 | 1 |
| REACTOME_DNA_DAMA | -1.02518 | 0.404649 | 0.883733 | REACTOME_DNA_DAMAGI | -1.10454 | 0.321234 | 0.872996 | REACTOME_DNA_DAMAG | -1.10954 | 0.338106 | 1 |
| REACTOME_G2_M_CHE | -0.997 | 0.450825 | 0.887195 | REACTOME_G2_M_CHECK | -0.92653 | 0.588235 | 1 | REACTOME_G2_M_CHECK | -0.89904 | 0.635339 | 1 |
| WP_IL3_SIGNALING_PAT | -0.96204 | 0.516092 | 0.892352 | WP_IL3_SIGNALING_PATH | 0.546276 | 0.997168 | 1 | WP_IL3_SIGNALING_PATH | -0.81699 | 0.770544 | 1 |
| REACTOME_RESPIRATOI | -0.9973 | 0.455039 | 0.892682 | REACTOME_RESPIRATORY_ | -1.31019 | 0.079105 | 0.672902 | REACTOME_RESPIRATORY | -0.95808 | 0.540604 | 1 |
| REACTOME_SIGNAL_AM | -0.99873 | 0.440537 | 0.894652 | REACTOME_SIGNAL_AMPL | -0.81031 | 0.736721 | 1 | REACTOME_SIGNAL_AMP | -0.53147 | 0.976506 | 1 |
| BIOCARTA_MET_PATHW | 1.186874 | 0.240338 | 0.894909 | BIOCARTA_MET_PATHWAY | 1.355104 | 0.128435 | 0.688007 | BIOCARTA_MET_PATHWA | 1.097023 | 0.310599 | 1 |
| REACTOME_G_ALPHA_I_ | -1.00444 | 0.431221 | 0.896112 | REACTOME_G_ALPHA_I_SI | -0.85326 | 0.780418 | 1 | REACTOME_G_ALPHA_I_S | -0.86868 | 0.710273 | 1 |
| REACTOME_COMPLEX_I_ | -0.96266 | 0.507675 | 0.896277 | REACTOME_COMPLEX_I_B | -1.21278 | 0.176355 | 0.762533 | REACTOME_COMPLEX_I_I | -0.77092 | 0.814516 | 1 |
| WP_FOLIC_ACID_NETW | 1.190749 | 0.252475 | 0.896718 | WP_FOLIC_ACID_NETWOR | 0.921782 | 0.539141 | 0.977801 | WP_FOLIC_ACID_NETWOI | -0.85042 | 0.665173 | 1 |
| REACTOME_RHOU_GTP/ | 1.18826 | 0.225854 | 0.898047 | REACTOME_RHOU_GTPAS | 1.286655 | 0.152428 | 0.733426 | REACTOME_RHOU_GTPAS | 1.17144 | 0.211991 | 1 |
| REACTOME_INTRA_GOL | -0.96348 | 0.494063 | 0.899606 | REACTOME_INTRA_GOLGI_ | -0.5065 | 0.992982 | 1 | REACTOME_INTRA_GOLG | 0.78065 | 0.804194 | 1 |
| REACTOME_CLATHRIN_I | -0.96664 | 0.522179 | 0.901744 | REACTOME_CLATHRIN_ME | -0.75571 | 0.936069 | 1 | REACTOME_CLATHRIN_M | -1.03192 | 0.418273 | 1 |
| REACTOME_RESPIRATOI | -0.98036 | 0.479243 | 0.901953 | REACTOME_RESPIRATORY_ | -1.31607 | 0.067856 | 0.676947 | REACTOME_RESPIRATORY | -0.96787 | 0.520741 | 1 |
| BIOCARTA_DEATH_PATH | -0.96458 | 0.493272 | 0.902109 | BIOCARTA_DEATH_PATHW | 0.848799 | 0.645181 | 1 | BIOCARTA_DEATH_PATHV | 0.903109 | 0.590094 | 1 |
| REACTOME_BIOSYNTHE | 1.180731 | 0.216415 | 0.902456 | REACTOME_BIOSYNTHESIS | -1.0431 | 0.388802 | 0.940522 | REACTOME_BIOSYNTHESI | -1.00734 | 0.464586 | 1 |
| REACTOME_SIGNALING_ | -1.0045 | 0.42975 | 0.902712 | REACTOME_SIGNALING_BY | 1.063904 | 0.325065 | 0.879111 | REACTOME_SIGNALING_B | 0.760297 | 0.975793 | 1 |
| WP_WNT_SIGNALING_P | -0.98159 | 0.472955 | 0.904363 | WP_WNT_SIGNALING_PAT | -0.69972 | 0.922187 | 1 | WP_WNT_SIGNALING_PA | -1.0214 | 0.436193 | 1 |
| REACTOME_ROS_AND_F | -0.98546 | 0.470512 | 0.904834 | REACTOME_ROS_AND_RN | -0.85677 | 0.657241 | 1 | REACTOME_ROS_AND_RN | -0.69588 | 0.85396 | 1 |
| REACTOME_SUMOYLATI | -0.94957 | 0.515 | 0.906841 | REACTOME_SUMOYLATIOI | -0.98717 | 0.474223 | 1 | REACTOME_SUMOYLATIO | -1.00532 | 0.455932 | 1 |
| REACTOME_TRANSPORT | -0.96686 | 0.480026 | 0.907067 | REACTOME_TRANSPORT_( | -0.81943 | 0.758375 | 1 | REACTOME_TRANSPORT_ | -1.00664 | 0.458101 | 1 |
| REACTOME_NUCLEAR_E | -0.98236 | 0.479867 | 0.9083 | REACTOME_NUCLEAR_EN\ | -0.91548 | 0.595528 | 1 | REACTOME_NUCLEAR_EN | -0.77434 | 0.798811 | 1 |
| REACTOME_ANTIGEN_P | -0.95031 | 0.515754 | 0.91035 | REACTOME_ANTIGEN_PRE | -0.88955 | 0.612029 | 1 | REACTOME_ANTIGEN_PRI | -0.59414 | 0.936456 | 1 |
| REACTOME_SIALIC_ACID | -0.9857 | 0.466807 | 0.910494 | REACTOME_SIALIC_ACID_I | -1.21586 | 0.230948 | 0.76122 | REACTOME_SIALIC_ACID_ | -1.09008 | 0.368413 | 1 |
| REACTOME_RNA_POLYM | -0.96949 | 0.483071 | 0.911129 | REACTOME_RNA_POLYME | -1.42056 | 0.070237 | 0.577152 | REACTOME_RNA_POLYMI | -1.02713 | 0.432292 | 1 |
| REACTOME_MITOTIC_PI | -0.97093 | 0.496082 | 0.912818 | REACTOME_MITOTIC_PRO | -0.95312 | 0.531219 | 1 | REACTOME_MITOTIC_PRO | -0.75754 | 0.839111 | 1 |
| WP_DRAVET_SYNDROM | -0.94181 | 0.556138 | 0.912858 | WP_DRAVET_SYNDROME_ | -1.5521 | 0.013455 | 0.633422 | WP_DRAVET_SYNDROME_ | -1.05008 | 0.391047 | 1 |
| BIOCARTA_FMLP_PATH\ | -0.97293 | 0.490489 | 0.912941 | BIOCARTA_FMLP_PATHW | 0.612017 | 0.927079 | 1 | BIOCARTA_FMLP_PATHW | -0.88761 | 0.616432 | 1 |
| REACTOME_TP53_REGU | -0.95277 | 0.528539 | 0.914472 | REACTOME_TP53_REGULA | -0.82412 | 0.774942 | 1 | REACTOME_TP53_REGUL/ | -0.67272 | 0.932674 | 1 |
| REACTOME_RUNX1_REG | 1.175042 | 0.214208 | 0.916774 | REACTOME_RUNX1_REGU | 0.924111 | 0.560772 | 0.984318 | REACTOME_RUNX1_REGU | 1.143391 | 0.220972 | 1 |
| REACTOME_GLYOXYLAT | -0.93517 | 0.530956 | 0.921447 | REACTOME_GLYOXYLATE_ | -0.90387 | 0.591611 | 1 | REACTOME_GLYOXYLATE_ | -0.78937 | 0.753068 | 1 |
| REACTOME_PURINE_CA | 1.169238 | 0.275384 | 0.923323 | REACTOME_PURINE_CATA | 0.732195 | 0.799163 | 1 | REACTOME_PURINE_CAT/ | 1.007488 | 0.436578 | 1 |
| REACTOME_PCP_CE_PA | 1.166725 | 0.217774 | 0.924975 | REACTOME_PCP_CE_PATH | 0.993439 | 0.454806 | 0.921019 | REACTOME_PCP_CE_PATH | 0.748586 | 0.926144 | 1 |
| REACTOME_SIGNALING_ | -0.88796 | 0.798184 | 0.926081 | REACTOME_SIGNALING_BY | -0.82732 | 0.926713 | 1 | REACTOME_SIGNALING_B | -1.00609 | 0.466495 | 1 |
| WP_APOPTOSIS | -0.93123 | 0.546743 | 0.927358 | WP_APOPTOSIS | -0.86861 | 0.654636 | 1 | WP_APOPTOSIS | -0.83555 | 0.710393 | 1 |
| REACTOME_RAB_GEFS_ | -0.92684 | 0.584165 | 0.92903 | REACTOME_RAB_GEFS_EX | -1.15252 | 0.22337 | 0.807147 | REACTOME_RAB_GEFS_EX | -0.87069 | 0.680623 | 1 |
| REACTOME_TNFR1_IND | -0.89777 | 0.586073 | 0.930616 | REACTOME_TNFR1_INDUC | -0.61535 | 0.948454 | 1 | REACTOME_TNFR1_INDU | -0.78943 | 0.746494 | 1 |
| REACTOME_CARGO_REC | -0.8899 | 0.660581 | 0.930979 | REACTOME_CARGO_RECO | 0.666938 | 0.963496 | 1 | REACTOME_CARGO_RECO | -0.96135 | 0.52643 | 1 |
| REACTOME_TRANSCRIPT | -0.92798 | 0.646229 | 0.931319 | REACTOME_TRANSCRIPTIC | -1.18619 | 0.117536 | 0.773811 | REACTOME_TRANSCRIPTI | -1.09152 | 0.297105 | 1 |
| REACTOME_TRANSPORT | -0.88321 | 0.676145 | 0.933374 | REACTOME_TRANSPORT_( | -1.00835 | 0.449298 | 0.98485 | REACTOME_TRANSPORT_ | 0.89115 | 0.697865 | 1 |
| REACTOME_G_ALPHA_1 | -0.89036 | 0.650619 | 0.934803 | REACTOME_G_ALPHA_12_ | -0.92887 | 0.579162 | 1 | REACTOME_G_ALPHA_12_ | 0.89259 | 0.671932 | 1 |
| REACTOME_RAP1_SIGN/ | -0.9075 | 0.577304 | 0.935203 | REACTOME_RAP1_SIGNAL | -0.76586 | 0.77027 | 1 | REACTOME_RAP1_SIGNAI | -0.66167 | 0.873934 | 1 |
| REACTOME_CLASS_I_MI | -0.90295 | 0.744046 | 0.937261 | REACTOME_CLASS_I_MHC | -0.73646 | 0.991785 | 1 | REACTOME_CLASS_I_MH | -0.82559 | 0.844339 | 1 |
| REACTOME_SIGNALING_ | 0.966723 | 0.522594 | 0.937948 | REACTOME_SIGNALING_BY | 1.346207 | 0.07867 | 0.683543 | REACTOME_SIGNALING_B | -0.64135 | 0.972402 | 1 |
| REACTOME_DETOXIFICA | 0.967738 | 0.492031 | 0.938865 | REACTOME_DETOXIFICATI | 0.656213 | 0.907824 | 1 | REACTOME_DETOXIFICAT | -0.74375 | 0.818168 | 1 |
| REACTOME_VOLTAGE_G | -0.92149 | 0.570007 | 0.938966 | REACTOME_VOLTAGE_GAT | -1.61293 | 0.01773 | 0.77277 | REACTOME_VOLTAGE_GA | -0.56163 | 0.967765 | 1 |
| REACTOME_REGULATIO | -0.90029 | 0.607081 | 0.939288 | REACTOME_REGULATION_ | -1.18375 | 0.224808 | 0.773731 | REACTOME_REGULATION_ | -0.83151 | 0.705166 | 1 |
| REACTOME_NEURONAL_ | -0.9194 | 0.710782 | 0.939371 | REACTOME_NEURONAL_S\ | -1.21203 | 0.073782 | 0.755583 | REACTOME_NEURONAL_S | -0.78126 | 0.916542 | 1 |
| REACTOME_GLI3_IS_PR | 1.161135 | 0.23123 | 0.939378 | REACTOME_GLI3_IS_PROC | 1.202657 | 0.198427 | 0.840508 | REACTOME_GLI3_IS_PRO | 0.882445 | 0.668885 | 1 |
| REACTOME_E3_UBIQUIT | 0.968796 | 0.495546 | 0.939678 | REACTOME_E3_UBIQUITIN | -1.20025 | 0.216031 | 0.769515 | REACTOME_E3_UBIQUITII | 0.75258 | 0.856762 | 1 |
| WP_CHEMOKINE_SIGNA | -0.90769 | 0.649575 | 0.940001 | WP_CHEMOKINE_SIGNALII | -0.56469 | 0.9996 | 1 | WP_CHEMOKINE_SIGNALI | -0.56828 | 0.997705 | 1 |
| WP_G13_SIGNALING_P/ | 0.963289 | 0.508336 | 0.940405 | WP_G13_SIGNALING_PATI | 1.18902 | 0.228419 | 0.828479 | WP_G13_SIGNALING_PAT | 0.888813 | 0.631789 | 1 |
| REACTOME_PYRUVATE_ | 0.968924 | 0.495837 | 0.943391 | REACTOME_PYRUVATE_M | -0.69866 | 0.863418 | 1 | REACTOME_PYRUVATE_M | -0.52538 | 0.975528 | 1 |
| REACTOME_GAP_JUNCT | -0.87512 | 0.624249 | 0.944153 | REACTOME_GAP_JUNCTIO | -1.45743 | 0.069989 | 0.517301 | REACTOME_GAP_JUNCTIC | 0.717878 | 0.840542 | 1 |
| REACTOME_METABOLIS | 0.963299 | 0.525477 | 0.944394 | REACTOME_METABOLISM_ | 0.846153 | 0.756688 | 1 | REACTOME_METABOLISM | -0.77657 | 0.858514 | 1 |
| REACTOME_SIGNALING_ | 1.157554 | 0.202102 | 0.945602 | REACTOME_SIGNALING_BY | -0.82039 | 0.844565 | 1 | REACTOME_SIGNALING_B | -0.998 | 0.473621 | 1 |
| REACTOME_ARACHIDON | 0.969298 | 0.499443 | 0.94637 | REACTOME_ARACHIDONIC | 1.189848 | 0.242779 | 0.83367 | REACTOME_ARACHIDONI | 1.047133 | 0.382465 | 1 |
| REACTOME_CELL_DEATI | 0.956952 | 0.518632 | 0.947701 | REACTOME_CELL_DEATH_ | 0.592337 | 0.981672 | 1 | REACTOME_CELL_DEATH_ | 0.855604 | 0.728357 | 1 |
| REACTOME_EPHB_MEDI | 0.919392 | 0.572613 | 0.947742 | REACTOME_EPHB_MEDIAT | 0.677156 | 0.904067 | 1 | REACTOME_EPHB_MEDIA | 1.095049 | 0.301779 | 1 |
| WP_SPINAL_CORD_INJU | 0.974037 | 0.492651 | 0.948314 | WP_SPINAL_CORD_INJURY | 1.000947 | 0.433861 | 0.917573 | WP_SPINAL_CORD_INJUR | 0.912937 | 0.601193 | 1 |
| REACTOME_SIGNALING_ | 0.969966 | 0.492821 | 0.948437 | REACTOME_SIGNALING_BY | 0.719784 | 0.822673 | 1 | REACTOME_SIGNALING_B | 0.7284 | 0.837001 | 1 |
| REACTOME_FATTY_ACYI | 0.924266 | 0.5582 | 0.948505 | REACTOME_FATTY_ACYL_( | -0.83812 | 0.698024 | 1 | REACTOME_FATTY_ACYL_ | -0.99055 | 0.477812 | 1 |
| REACTOME_TCF_DEPEN | 0.971205 | 0.504536 | 0.948778 | REACTOME_TCF_DEPENDE | 0.868768 | 0.717439 | 1 | REACTOME_TCF_DEPEND | -0.87339 | 0.693927 | 1 |
| REACTOME_STIMULI_SE | -0.90816 | 0.596314 | 0.949524 | REACTOME_STIMULI_SENS | -0.73228 | 0.888975 | 1 | REACTOME_STIMULI_SEN | -0.82332 | 0.738196 | 1 |
| WP_MECP2_AND_ASSO | 0.990008 | 0.470272 | 0.949621 | WP_MECP2_AND_ASSOCI/ | 0.996053 | 0.449803 | 0.922715 | WP_MECP2_AND_ASSOCI | -1.09798 | 0.349692 | 1 |
| REACTOME_CARGO_TR/ | 0.995352 | 0.456718 | 0.950318 | REACTOME_CARGO_TRAFI | 0.729834 | 0.841139 | 1 | REACTOME_CARGO_TRAF | -0.68009 | 0.891904 | 1 |
| REACTOME_HEDGEHOG | 0.998165 | 0.453306 | 0.950364 | REACTOME_HEDGEHOG_L | 1.008825 | 0.427359 | 0.908312 | REACTOME_HEDGEHOG_I | 1.022935 | 0.396565 | 1 |
| REACTOME_NUCLEAR_E | 1.00228 | 0.452575 | 0.95083 | REACTOME_NUCLEAR_EVE | 1.137986 | 0.27118 | 0.893734 | REACTOME_NUCLEAR_EV | -0.83418 | 0.721936 | 1 |
| REACTOME_G1_S_TRAN | 0.971873 | 0.5 | 0.950874 | REACTOME_G1_S_TRANSI | 0.65096 | 0.965269 | 1 | REACTOME_G1_S_TRANSI | 0.733728 | 0.936903 | 1 |
| REACTOME_ANCHORIN( | -0.85203 | 0.707257 | 0.951757 | REACTOME_ANCHORING_( | -0.90094 | 0.624364 | 1 | REACTOME_ANCHORING_ | -0.79799 | 0.771174 | 1 |
| WP_OXIDATIVE_DAMAG | 0.924386 | 0.558227 | 0.951772 | WP_OXIDATIVE_DAMAGE_ | 1.054906 | 0.374342 | 0.874446 | WP_OXIDATIVE_DAMAGE | 0.897218 | 0.608056 | 1 |
| WP_MITOCHONDRIAL_L | 0.987897 | 0.476109 | 0.95198 | WP_MITOCHONDRIAL_LOI | 0.499353 | 0.98147 | 1 | WP_MITOCHONDRIAL_LO | -0.53066 | 0.968192 | 1 |
| REACTOME_CELLULAR_I | 0.921731 | 0.694508 | 0.952017 | REACTOME_CELLULAR_RE | -0.90295 | 0.786252 | 1 | REACTOME_CELLULAR_RE | -1.0387 | 0.391816 | 1 |
| REACTOME_GPER1_SIGI | -0.85593 | 0.658116 | 0.952128 | REACTOME_GPER1_SIGNA | -1.03277 | 0.410755 | 0.941561 | REACTOME_GPER1_SIGN/ | -0.67894 | 0.87988 | 1 |
| REACTOME_REGULATIO | 0.990516 | 0.461496 | 0.952472 | REACTOME_REGULATION_ | 1.06138 | 0.361785 | 0.875901 | REACTOME_REGULATION_ | -0.85422 | 0.687754 | 1 |
| REACTOME_METABOLIS | 0.94231 | 0.530682 | 0.952477 | REACTOME_METABOLISM_ | -1.05091 | 0.402034 | 0.949573 | REACTOME_METABOLISM | -0.62801 | 0.909978 | 1 |
| REACTOME_SCF_SKP2_I | 0.957909 | 0.513139 | 0.95279 | REACTOME_SCF_SKP2_ME | 0.904574 | 0.587889 | 0.983867 | REACTOME_SCF_SKP2_MI | -0.51411 | 0.995615 | 1 |
| REACTOME_PEPTIDE_H( | 0.976676 | 0.483729 | 0.952805 | REACTOME_PEPTIDE_HOR | 1.294068 | 0.156891 | 0.737803 | REACTOME_PEPTIDE_HOF | 1.279599 | 0.137455 | 1 |
| REACTOME_ADAPTIVE_I | 0.930372 | 0.685092 | 0.953059 | REACTOME_ADAPTIVE_IM | 0.905975 | 0.760181 | 0.984085 | REACTOME_ADAPTIVE_IM | -0.77432 | 0.953448 | 1 |
| WP_FACTORS_AND_PAT | 0.920033 | 0.562217 | 0.953145 | WP_FACTORS_AND_PATH\ | 1.063472 | 0.369942 | 0.874909 | WP_FACTORS_AND_PATH | 0.715977 | 0.863326 | 1 |
| WP_PURINE_METABOLI | 0.953801 | 0.542903 | 0.953268 | WP_PURINE_METABOLISM | -0.67691 | 0.988119 | 1 | WP_PURINE_METABOLISN | 1.024195 | 0.392275 | 1 |
| REACTOME_RHOC_GTP/ | 0.916051 | 0.589571 | 0.953556 | REACTOME_RHOC_GTPASI | 1.065198 | 0.344968 | 0.880648 | REACTOME_RHOC_GTPAS | -0.83495 | 0.72788 | 1 |

| | | | | | | | | | | | |
|---|---|---|---|---|---|---|---|---|---|---|---|
| BIOCARTA_BAD_PATHW | 0.974961 | 0.484256 | 0.953906 | BIOCARTA_BAD_PATHWAY | 0.656678 | 0.883932 | 1 | BIOCARTA_BAD_PATHWA | 1.101857 | 0.324693 | 1 |
| REACTOME_TRANSCRIPT | 0.909448 | 0.605093 | 0.954202 | REACTOME_TRANSCRIPTIO | 0.964069 | 0.496702 | 0.938761 | REACTOME_TRANSCRIPTI | -0.79286 | 0.782406 | 1 |
| REACTOME_MITOTIC_G | 0.980335 | 0.486364 | 0.954272 | REACTOME_MITOTIC_G1_ | -0.68585 | 0.961801 | 1 | REACTOME_MITOTIC_G1_ | 0.732848 | 0.94536 | 1 |
| REACTOME_DAP12_INTI | 1.002624 | 0.453218 | 0.954333 | REACTOME_DAP12_INTER | 0.816655 | 0.678774 | 1 | REACTOME_DAP12_INTER | -0.63808 | 0.900189 | 1 |
| REACTOME_SYNTHESIS_ | 0.935127 | 0.542216 | 0.95438 | REACTOME_SYNTHESIS_OI | 0.538252 | 0.976341 | 1 | REACTOME_SYNTHESIS_O | 0.862086 | 0.650807 | 1 |
| REACTOME_ORC1_REM | 0.998177 | 0.454997 | 0.954937 | REACTOME_ORC1_REMOV | 1.030129 | 0.395775 | 0.88803 | REACTOME_ORC1_REMO | 0.834553 | 0.758919 | 1 |
| REACTOME_COPII_MEDI | 0.98146 | 0.487027 | 0.95508 | REACTOME_COPII_MEDIAT | -0.89466 | 0.64337 | 1 | REACTOME_COPII_MEDIA | 1.297888 | 0.088567 | 1 |
| REACTOME_INTRA_GOL | 0.910364 | 0.660021 | 0.955131 | REACTOME_INTRA_GOLGI_ | 1.133593 | 0.223578 | 0.879283 | REACTOME_INTRA_GOLG | 0.905007 | 0.732919 | 1 |
| REACTOME_TRANSPORT | 0.942701 | 0.530507 | 0.955173 | REACTOME_TRANSPORT_ | -0.74051 | 0.8117 | 1 | REACTOME_TRANSPORT_ | -0.99563 | 0.47696 | 1 |
| REACTOME_UNBLOCKIN | -0.85868 | 0.646707 | 0.955178 | REACTOME_UNBLOCKING_ | -1.14833 | 0.284366 | 0.813828 | REACTOME_UNBLOCKING | -0.61772 | 0.922058 | 1 |
| REACTOME_CELLULAR_ | 0.912914 | 0.595544 | 0.955182 | REACTOME_CELLULAR_RE | 1.058893 | 0.354193 | 0.872816 | REACTOME_CELLULAR_RE | -0.73 | 0.867554 | 1 |
| REACTOME_ION_TRANS | 0.925687 | 0.559172 | 0.955407 | REACTOME_ION_TRANSPO | -0.95363 | 0.529849 | 1 | REACTOME_ION_TRANSP | 0.71538 | 0.898435 | 1 |
| REACTOME_P75_NTR_R | 0.977147 | 0.487161 | 0.955636 | REACTOME_P75_NTR_REC | 0.715125 | 0.903725 | 1 | REACTOME_P75_NTR_RE | -0.64767 | 0.953951 | 1 |
| REACTOME_STABILIZATI | 0.932042 | 0.5544 | 0.955784 | REACTOME_STABILIZATION | 0.99215 | 0.446929 | 0.920075 | REACTOME_STABILIZATIO | 1.078672 | 0.327147 | 1 |
| WP_FATTY_ACID_BETAC | 0.930546 | 0.555227 | 0.956294 | WP_FATTY_ACID_BETAOXI | 0.647958 | 0.925972 | 1 | WP_FATTY_ACID_BETAOX | -0.77259 | 0.783936 | 1 |
| REACTOME_CHONDROIT | 0.978319 | 0.487112 | 0.956324 | REACTOME_CHONDROITIN | 0.744257 | 0.811852 | 1 | REACTOME_CHONDROITI | -0.92649 | 0.569859 | 1 |
| REACTOME_CLASS_I_PE | 0.91113 | 0.576329 | 0.956553 | REACTOME_CLASS_I_PERC | 1.116369 | 0.314095 | 0.908934 | REACTOME_CLASS_I_PER | 0.968376 | 0.49322 | 1 |
| REACTOME_BETA_CATE | 0.944868 | 0.548263 | 0.956585 | REACTOME_BETA_CATENII | 0.730049 | 0.926902 | 1 | REACTOME_BETA_CATEN | -0.50274 | 0.999507 | 1 |
| REACTOME_TRANS_GOL | 0.905913 | 0.620665 | 0.956956 | REACTOME_TRANS_GOLGI | 0.781536 | 0.824497 | 1 | REACTOME_TRANS_GOLG | -1.0279 | 0.420018 | 1 |
| REACTOME_DOWNSTRE | 0.927728 | 0.577614 | 0.957014 | REACTOME_DOWNSTREAM | 1.225042 | 0.169068 | 0.799762 | REACTOME_DOWNSTREA | 0.857249 | 0.752915 | 1 |
| WP_ALPHA_6_BETA_4_I | 0.935484 | 0.548778 | 0.957166 | WP_ALPHA_6_BETA_4_INT | 1.579593 | 0.028274 | 0.35946 | WP_ALPHA_6_BETA_4_IN | 1.218336 | 0.161168 | 1 |
| REACTOME_INTERLEUKI | -0.85979 | 0.648997 | 0.957439 | REACTOME_INTERLEUKIN_ | 0.841243 | 0.653137 | 1 | REACTOME_INTERLEUKIN | -0.86611 | 0.645222 | 1 |
| REACTOME_METABOLIS | 0.949815 | 0.543177 | 0.95747 | REACTOME_METABOLISM_ | -0.71299 | 0.939654 | 1 | REACTOME_METABOLISM | 1.026769 | 0.394652 | 1 |
| REACTOME_A_TETRASA | 1.003099 | 0.446686 | 0.957518 | REACTOME_A_TETRASACC | 1.09907 | 0.334244 | 0.885347 | REACTOME_A_TETRASAC | -0.87824 | 0.62171 | 1 |
| BIOCARTA_CREB_PATHW | 0.998776 | 0.455023 | 0.957648 | BIOCARTA_CREB_PATHWA | -0.78158 | 0.767304 | 1 | BIOCARTA_CREB_PATHW | 1.026522 | 0.412948 | 1 |
| WP_MICRORNAS_IN_CA | 1.152518 | 0.234116 | 0.957839 | WP_MICRORNAS_IN_CARD | 1.576254 | 0.024006 | 0.356673 | WP_MICRORNAS_IN_CAR | 0.835664 | 0.777687 | 1 |
| REACTOME_DECTIN_1_ | 0.906844 | 0.606974 | 0.957926 | REACTOME_DECTIN_1_ME | 1.10682 | 0.297585 | 0.879279 | REACTOME_DECTIN_1_M | 0.975673 | 0.488012 | 1 |
| REACTOME_ACTIVATION | 1.021804 | 0.406125 | 0.957979 | REACTOME_ACTIVATION_ | 1.228094 | 0.170107 | 0.807006 | REACTOME_ACTIVATION_ | 1.221364 | 0.138215 | 1 |
| REACTOME_G1_S_DNA_ | 0.948285 | 0.534446 | 0.958119 | REACTOME_G1_S_DNA_D | 0.887864 | 0.629078 | 1 | REACTOME_G1_S_DNA_D | 0.855502 | 0.72693 | 1 |
| REACTOME_MITOTIC_G | 0.925918 | 0.615531 | 0.95842 | REACTOME_MITOTIC_G2_ | 0.88028 | 0.701248 | 1 | REACTOME_MITOTIC_G2_ | 1.113288 | 0.205128 | 1 |
| REACTOME_SPHINGOLIF | 0.945511 | 0.537792 | 0.958594 | REACTOME_SPHINGOLIPID | 1.208133 | 0.189195 | 0.83942 | REACTOME_SPHINGOLIPI | 0.954331 | 0.529935 | 1 |
| REACTOME_KEAP1_NFE | 0.981724 | 0.483246 | 0.958653 | REACTOME_KEAP1_NFE2L | 1.095906 | 0.308283 | 0.882945 | REACTOME_KEAP1_NFE2 | -0.7167 | 0.897755 | 1 |
| REACTOME_SEPARATIO | 0.942791 | 0.573853 | 0.95879 | REACTOME_SEPARATION_ | 1.0281 | 0.382899 | 0.884196 | REACTOME_SEPARATION_ | 1.075601 | 0.274118 | 1 |
| REACTOME_FATTY_ACID | 0.938751 | 0.572259 | 0.959115 | REACTOME_FATTY_ACID_ | 0.851944 | 0.735926 | 1 | REACTOME_FATTY_ACID_ | -0.56281 | 0.996136 | 1 |
| WP_GLYCEROLIPIDS_AN | 0.93211 | 0.542766 | 0.959348 | WP_GLYCEROLIPIDS_AND_ | 0.701049 | 0.852632 | 1 | WP_GLYCEROLIPIDS_AND | -0.99848 | 0.472684 | 1 |
| WP_ADIPOGENESIS_GEN | 1.145939 | 0.251365 | 0.959657 | WP_ADIPOGENESIS_GENE | 1.106852 | 0.301623 | 0.885738 | WP_ADIPOGENESIS_GENE | -0.88993 | 0.634306 | 1 |
| WP_MYOMETRIAL_RELA | 1.00812 | 0.428319 | 0.959822 | WP_MYOMETRIAL_RELAX | 1.061257 | 0.336712 | 0.870849 | WP_MYOMETRIAL_RELAX | -0.80289 | 0.815681 | 1 |
| REACTOME_METABOLIS | 0.935737 | 0.603898 | 0.960247 | REACTOME_METABOLISM_ | -0.84556 | 0.827276 | 1 | REACTOME_METABOLISM | -1.03855 | 0.399683 | 1 |
| REACTOME_OVARIAN_T | 0.93692 | 0.543391 | 0.960646 | REACTOME_OVARIAN_TU | 0.694877 | 0.843262 | 1 | REACTOME_OVARIAN_TU | 0.762242 | 0.794225 | 1 |
| REACTOME_CYCLIN_A_ | 1.003467 | 0.440145 | 0.96104 | REACTOME_CYCLIN_A_CD | 0.729785 | 0.877979 | 1 | REACTOME_CYCLIN_A_CD | -0.49991 | 0.996023 | 1 |
| REACTOME_GLUCOSE_ | 1.006323 | 0.435205 | 0.961074 | REACTOME_GLUCOSE_ME | 0.915143 | 0.578492 | 0.983941 | REACTOME_GLUCOSE_ME | -0.81935 | 0.760383 | 1 |
| REACTOME_AUF1_HNRN | 1.009077 | 0.435775 | 0.961467 | REACTOME_AUF1_HNRNP | 1.133669 | 0.259896 | 0.88631 | REACTOME_AUF1_HNRNP | 0.884124 | 0.661094 | 1 |
| REACTOME_CYTOKINE_ | -0.85981 | 0.841592 | 0.962296 | REACTOME_CYTOKINE_SIG | 0.912458 | 0.667577 | 0.982967 | REACTOME_CYTOKINE_SI | -0.73222 | 0.949573 | 1 |
| REACTOME_NEUROTRA | -0.82858 | 0.828143 | 0.962304 | REACTOME_NEUROTRANS | -0.96645 | 0.522041 | 1 | REACTOME_NEUROTRAN | -0.73513 | 0.910865 | 1 |
| REACTOME_DEUBIQUITI | 0.945554 | 0.580417 | 0.96241 | REACTOME_DEUBIQUITIN | 0.83603 | 0.813903 | 1 | REACTOME_DEUBIQUITIN | -0.6875 | 0.970295 | 1 |
| REACTOME_EFFECTS_O | 1.147194 | 0.285796 | 0.962851 | REACTOME_EFFECTS_OF_ | -0.75669 | 0.800345 | 1 | REACTOME_EFFECTS_OF_ | -0.91734 | 0.572516 | 1 |
| REACTOME_CELLULAR_ | 1.004306 | 0.439328 | 0.963044 | REACTOME_CELLULAR_RE | -0.86653 | 0.76861 | 1 | REACTOME_CELLULAR_RE | -1.09389 | 0.313869 | 1 |
| WP_WNT_SIGNALING_P | -0.86355 | 0.660261 | 0.963122 | WP_WNT_SIGNALING_PAT | -0.90158 | 0.605691 | 1 | WP_WNT_SIGNALING_PA | -0.79413 | 0.75058 | 1 |
| WP_OXIDATIVE_PHOSP | -0.8297 | 0.722562 | 0.964467 | WP_OXIDATIVE_PHOSPHO | -1.25623 | 0.156701 | 0.718248 | WP_OXIDATIVE_PHOSPH | -0.81367 | 0.740088 | 1 |
| REACTOME_TCR_SIGNA | 0.901871 | 0.63331 | 0.964523 | REACTOME_TCR_SIGNALIN | 1.233041 | 0.153502 | 0.799403 | REACTOME_TCR_SIGNALI | 0.934673 | 0.587783 | 1 |
| REACTOME_SIGNALING_ | -0.8444 | 0.692291 | 0.964662 | REACTOME_SIGNALING_B | -0.93306 | 0.553861 | 1 | REACTOME_SIGNALING_B | -0.72661 | 0.836841 | 1 |
| BIOCARTA_BARRESTIN_ | 0.899105 | 0.590658 | 0.965008 | BIOCARTA_BARRESTIN_SR | -0.88724 | 0.610347 | 1 | BIOCARTA_BARRESTIN_SR | 1.009849 | 0.430888 | 1 |
| REACTOME_TRANSPORT | -0.83111 | 0.719518 | 0.96609 | REACTOME_TRANSPORT_ | -0.80447 | 0.762261 | 1 | REACTOME_TRANSPORT_ | -1.0067 | 0.458694 | 1 |
| REACTOME_ENERGY_DE | -0.82231 | 0.722564 | 0.966411 | REACTOME_ENERGY_DEPE | 0.748437 | 0.800625 | 1 | REACTOME_ENERGY_DEP | 0.79881 | 0.787255 | 1 |
| REACTOME_CLEC7A_DE | 1.013408 | 0.425536 | 0.966641 | REACTOME_CLEC7A_DECT | 1.119387 | 0.271719 | 0.905925 | REACTOME_CLEC7A_DECT | 0.869408 | 0.740655 | 1 |
| REACTOME_RAB_REGUL | -0.8638 | 0.735638 | 0.967498 | REACTOME_RAB_REGULAT | -1.05026 | 0.362686 | 0.936924 | REACTOME_RAB_REGULA | -0.69554 | 0.940276 | 1 |
| REACTOME_NRAGE_SIG | 0.899419 | 0.609064 | 0.967636 | REACTOME_NRAGE_SIGNA | 0.501236 | 0.994066 | 1 | REACTOME_NRAGE_SIGN | -0.81729 | 0.728633 | 1 |
| REACTOME_SULFUR_AM | 1.023341 | 0.428816 | 0.968049 | REACTOME_SULFUR_AMIN | 1.115881 | 0.314031 | 0.903409 | REACTOME_SULFUR_AMI | 1.162183 | 0.254693 | 1 |
| REACTOME_EPH_EPHRI | 1.011452 | 0.433064 | 0.968292 | REACTOME_EPH_EPHRIN_ | 0.565282 | 0.991154 | 1 | REACTOME_EPH_EPHRIN_ | 0.815535 | 0.804564 | 1 |
| REACTOME_TRANSMISS | -0.81475 | 0.903632 | 0.968438 | REACTOME_TRANSMISSIO | -1.08073 | 0.28161 | 0.888882 | REACTOME_TRANSMISSIO | -0.86822 | 0.741024 | 1 |
| REACTOME_TNF_SIGNA | -0.82316 | 0.748277 | 0.96919 | REACTOME_TNF_SIGNALI | -1.1016 | 0.308584 | 0.867798 | REACTOME_TNF_SIGNALI | -0.92703 | 0.575812 | 1 |
| REACTOME_PROCESSIN | -0.838 | 0.692025 | 0.969612 | REACTOME_PROCESSING_ | -1.1969 | 0.23071 | 0.753998 | REACTOME_PROCESSING_ | -0.74088 | 0.814705 | 1 |
| REACTOME_GABA_RECE | -0.84002 | 0.722838 | 0.969795 | REACTOME_GABA_RECEPT | -0.8979 | 0.619121 | 1 | REACTOME_GABA_RECEP | -0.83572 | 0.709431 | 1 |
| BIOCARTA_CHEMICAL_P | -0.81163 | 0.715032 | 0.970144 | BIOCARTA_CHEMICAL_PAT | -0.74918 | 0.802652 | 1 | BIOCARTA_CHEMICAL_PA | 0.843375 | 0.67456 | 1 |
| REACTOME_REGULATED | 1.00939 | 0.442441 | 0.970253 | REACTOME_REGULATED_ | 1.029632 | 0.404457 | 0.884529 | REACTOME_REGULATED_ | 0.988513 | 0.462236 | 1 |
| REACTOME_UBIQUITIN_ | 1.013678 | 0.4275 | 0.970699 | REACTOME_UBIQUITIN_M | 1.150475 | 0.259221 | 0.889338 | REACTOME_UBIQUITIN_M | 1.141135 | 0.23399 | 1 |
| REACTOME_SIGNALING_ | 0.887885 | 0.668544 | 0.970735 | REACTOME_SIGNALING_B | 0.86585 | 0.692958 | 1 | REACTOME_SIGNALING_B | 0.782065 | 0.902332 | 1 |
| REACTOME_TRAFFICKIN | 1.016417 | 0.434019 | 0.971437 | REACTOME_TRAFFICKING_ | -0.81206 | 0.741412 | 1 | REACTOME_TRAFFICKING | -0.57277 | 0.964987 | 1 |
| WP_GLYCOLYSIS_AND_ | 1.014856 | 0.431114 | 0.97168 | WP_GLYCOLYSIS_AND_GL | -0.82203 | 0.735294 | 1 | WP_GLYCOLYSIS_AND_GL | -0.80445 | 0.746252 | 1 |
| REACTOME_METABOLIS | 1.134753 | 0.271832 | 0.972562 | REACTOME_METABOLISM_ | 1.202505 | 0.214801 | 0.824019 | REACTOME_METABOLISM | 1.248348 | 0.136936 | 1 |
| REACTOME_APC_C_CDH | 1.023492 | 0.411359 | 0.972711 | REACTOME_APC_C_CDH1_ | 1.251862 | 0.159533 | 0.782371 | REACTOME_APC_C_CDH1 | 1.269724 | 0.109701 | 1 |
| REACTOME_PROTEIN_L | 0.884258 | 0.67532 | 0.973356 | REACTOME_PROTEIN_LOC | 0.995169 | 0.44258 | 0.920626 | REACTOME_PROTEIN_LO | 1.00597 | 0.426972 | 1 |
| REACTOME_GLUCAGON_ | -0.83204 | 0.703118 | 0.973423 | REACTOME_GLUCAGON_L | -1.51407 | 0.040488 | 0.531737 | REACTOME_GLUCAGON_ | 0.821382 | 0.725191 | 1 |
| WP_ERBB_SIGNALING_F | 0.888145 | 0.608973 | 0.973485 | WP_ERBB_SIGNALING_PAT | -0.87374 | 0.648208 | 1 | WP_ERBB_SIGNALING_PA | -1.01534 | 0.449743 | 1 |
| REACTOME_SIGNALING_ | 1.126576 | 0.276052 | 0.973692 | REACTOME_SIGNALING_B | 1.695924 | 0.0102 | 0.253623 | REACTOME_SIGNALING_B | -1.00788 | 0.458084 | 1 |
| REACTOME_RHO_GTPAS | 0.889297 | 0.610643 | 0.973915 | REACTOME_RHO_GTPASES | -0.66568 | 0.911279 | 1 | REACTOME_RHO_GTPASE | 0.754147 | 0.830769 | 1 |
| BIOCARTA_HDAC_PATH | 1.124533 | 0.312819 | 0.974008 | BIOCARTA_HDAC_PATHW | 0.601937 | 0.929926 | 1 | BIOCARTA_HDAC_PATHW | -0.64658 | 0.901296 | 1 |
| BIOCARTA_RHO_PATHW | 1.132182 | 0.305392 | 0.974913 | BIOCARTA_RHO_PATHWAY | 1.384454 | 0.127646 | 0.629449 | BIOCARTA_RHO_PATHWA | -0.59319 | 0.93664 | 1 |
| REACTOME_THE_CITRIC | 0.884587 | 0.71463 | 0.975895 | REACTOME_THE_CITRIC_A | -0.92339 | 0.641252 | 1 | REACTOME_THE_CITRIC_ | -0.64769 | 0.987544 | 1 |
| REACTOME_IRON_UPTA | 0.889809 | 0.616645 | 0.976014 | REACTOME_IRON_UPTAKE | 0.569478 | 0.973777 | 1 | REACTOME_IRON_UPTAK | -0.57692 | 0.973112 | 1 |
| REACTOME_GLYCOSAMI | 0.892126 | 0.639547 | 0.976796 | REACTOME_GLYCOSAMINO | 0.7916 | 0.787873 | 1 | REACTOME_GLYCOSAMIN | -0.75903 | 0.828314 | 1 |
| BIOCARTA_RACCYCD_PA | 1.028322 | 0.416442 | 0.977055 | BIOCARTA_RACCYCD_PATH | -0.6252 | 0.932657 | 1 | BIOCARTA_RACCYCD_PAT | 0.735263 | 0.825235 | 1 |
| REACTOME_SEALING_O | 0.881498 | 0.622084 | 0.97711 | REACTOME_SEALING_OF_ | -0.8269 | 0.712279 | 1 | REACTOME_SEALING_OF_ | 0.834956 | 0.711091 | 1 |
| REACTOME_MITOTIC_SF | 0.875845 | 0.671728 | 0.978088 | REACTOME_MITOTIC_SPIN | 1.16152 | 0.233808 | 0.866882 | REACTOME_MITOTIC_SPI | 1.098349 | 0.270282 | 1 |

| | | | | | | | | | | | |
|---|---|---|---|---|---|---|---|---|---|---|---|
| REACTOME_PROGRAMN | 0.890266 | 0.686386 | 0.978237 | REACTOME_PROGRAMME | 0.761102 | 0.883418 | 1 | REACTOME_PROGRAMMI | 1.091823 | 0.258804 | 1 |
| REACTOME_ANTIGEN_A | 0.878387 | 0.627083 | 0.978356 | REACTOME_ANTIGEN_ACT | -0.94513 | 0.529434 | 1 | REACTOME_ANTIGEN_AC | -1.08641 | 0.361759 | 1 |
| REACTOME_RHOA_GTP/ | 0.876947 | 0.705965 | 0.978655 | REACTOME_RHOA_GTPASI | 1.175255 | 0.189124 | 0.843339 | REACTOME_RHOA_GTPAS | -0.90543 | 0.64598 | 1 |
| REACTOME_NUCLEOTID | 0.892608 | 0.605993 | 0.978979 | REACTOME_NUCLEOTIDE_ | 0.76304 | 0.774012 | 1 | REACTOME_NUCLEOTIDE_ | 1.154805 | 0.249534 | 1 |
| WP_INTEGRINMEDIATED | 1.119362 | 0.261519 | 0.979356 | WP_INTEGRINMEDIATED_ | 1.345623 | 0.085958 | 0.674856 | WP_INTEGRINMEDIATED_ | 0.775426 | 0.912197 | 1 |
| REACTOME_METABOLIS | 1.13504 | 0.242506 | 0.97953 | REACTOME_METABOLISM_ | 0.934197 | 0.550669 | 0.968388 | REACTOME_METABOLISM | -0.57462 | 0.992205 | 1 |
| REACTOME_AZATHIOPR | -0.80441 | 0.723664 | 0.979801 | REACTOME_AZATHIOPRIN | 0.686468 | 0.860657 | 1 | REACTOME_AZATHIOPRIN | -0.98449 | 0.497127 | 1 |
| REACTOME_COPI_DEPEI | 0.871031 | 0.690425 | 0.980125 | REACTOME_COPI_DEPEND | 0.756663 | 0.870307 | 1 | REACTOME_COPI_DEPENI | -0.86714 | 0.681672 | 1 |
| WP_SIDS_SUSCEPTIBILIT | 1.028966 | 0.41308 | 0.980169 | WP_SIDS_SUSCEPTIBILITY_ | 0.976474 | 0.481118 | 0.938865 | WP_SIDS_SUSCEPTIBILITY_ | -1.04489 | 0.415964 | 1 |
| BIOCARTA_EDG1_PATHV | 1.030304 | 0.415657 | 0.980766 | BIOCARTA_EDG1_PATHWA | -0.59042 | 0.953205 | 1 | BIOCARTA_EDG1_PATHW. | 0.760899 | 0.788462 | 1 |
| WP_MAPK_CASCADE | 0.873179 | 0.63916 | 0.981363 | WP_MAPK_CASCADE | 1.451785 | 0.086665 | 0.533331 | WP_MAPK_CASCADE | -0.91139 | 0.582806 | 1 |
| WP_INSULIN_SIGNALING | -0.79598 | 0.904797 | 0.982237 | WP_INSULIN_SIGNALING | 1.293065 | 0.082104 | 0.731247 | WP_INSULIN_SIGNALING | -0.62837 | 0.989673 | 1 |
| WP_GLYCOGEN_METAB | 1.023797 | 0.418583 | 0.982271 | WP_GLYCOGEN_METABOL | 0.686829 | 0.873561 | 1 | WP_GLYCOGEN_METABO | 1.253929 | 0.158921 | 1 |
| REACTOME_SIGNALING_ | 1.112862 | 0.316633 | 0.982404 | REACTOME_SIGNALING_B\ | 0.419237 | 0.996022 | 1 | REACTOME_SIGNALING_E | 0.927667 | 0.551044 | 1 |
| REACTOME_REGULATIO | 1.128317 | 0.273402 | 0.982601 | REACTOME_REGULATION_ | 1.098385 | 0.303875 | 0.881184 | REACTOME_REGULATION_ | 1.085638 | 0.288813 | 1 |
| WP_MAPK_SIGNALING_ | -0.80049 | 0.875524 | 0.9828 | WP_MAPK_SIGNALING_PA | 1.00745 | 0.423853 | 0.907575 | WP_MAPK_SIGNALING_P/ | -1.09445 | 0.313527 | 1 |
| REACTOME_DEGRADATI | 1.138144 | 0.254783 | 0.983375 | REACTOME_DEGRADATION | 1.072752 | 0.338178 | 0.873636 | REACTOME_DEGRADATIO | 0.983695 | 0.469244 | 1 |
| BIOCARTA_NOS1_PATHV | -0.7698 | 0.771015 | 0.983664 | BIOCARTA_NOS1_PATHW/ | -1.47918 | 0.064316 | 0.498456 | BIOCARTA_NOS1_PATHW | -0.66582 | 0.881934 | 1 |
| REACTOME_DEATH_REC | 1.035664 | 0.38819 | 0.98372 | REACTOME_DEATH_RECEF | -0.78283 | 0.888756 | 1 | REACTOME_DEATH_RECE | -0.8124 | 0.795066 | 1 |
| WP_SIGNAL_TRANSDUC | -0.79758 | 0.73085 | 0.983759 | WP_SIGNAL_TRANSDUCTI( | -0.77305 | 0.760935 | 1 | WP_SIGNAL_TRANSDUCTI | -1.07765 | 0.379599 | 1 |
| BIOCARTA_IGF1R_PATH\ | 1.108627 | 0.31946 | 0.984306 | BIOCARTA_IGF1R_PATHW/ | 0.687317 | 0.859798 | 1 | BIOCARTA_IGF1R_PATHW | 0.901165 | 0.589 | 1 |
| REACTOME_RHOBTB1_( | 1.041673 | 0.402083 | 0.984749 | REACTOME_RHOBTB1_GTI | 0.955824 | 0.496676 | 0.949589 | REACTOME_RHOBTB1_GT | -0.78901 | 0.744991 | 1 |
| REACTOME_CARBOXYTE | 1.032296 | 0.421142 | 0.98478 | REACTOME_CARBOXYTERN | -0.81855 | 0.707854 | 1 | REACTOME_CARBOXYTER | 1.144907 | 0.285903 | 1 |
| REACTOME_CTLA4_INHI | 1.030525 | 0.41314 | 0.985501 | REACTOME_CTLA4_INHIBI | 0.770441 | 0.75557 | 1 | REACTOME_CTLA4_INHIB | 1.139931 | 0.27622 | 1 |
| REACTOME_SIGNALING_ | 1.033625 | 0.412307 | 0.985601 | REACTOME_SIGNALING_B\ | 0.688258 | 0.862683 | 1 | REACTOME_SIGNALING_E | 1.088966 | 0.332163 | 1 |
| REACTOME_RHO_GTPAS | 1.119678 | 0.257479 | 0.985891 | REACTOME_RHO_GTPASES | 0.926275 | 0.57302 | 0.982482 | REACTOME_RHO_GTPASE | 1.026938 | 0.395243 | 1 |
| REACTOME_PTEN_REGU | -0.79146 | 0.865252 | 0.986006 | REACTOME_PTEN_REGULA | 0.80491 | 0.81339 | 1 | REACTOME_PTEN_REGUL | -0.53136 | 0.998646 | 1 |
| REACTOME_MTORC1_M | -0.7707 | 0.785111 | 0.986531 | REACTOME_MTORC1_MEI | 0.964289 | 0.487672 | 0.94266 | REACTOME_MTORC1_ME | 1.159547 | 0.256993 | 1 |
| REACTOME_HEPARAN_S | 1.109907 | 0.319165 | 0.986745 | REACTOME_HEPARAN_SUI | 1.038139 | 0.390364 | 0.878943 | REACTOME_HEPARAN_SU | 0.717842 | 0.884116 | 1 |
| REACTOME_PHOSPHOLI | 1.023862 | 0.402418 | 0.987417 | REACTOME_PHOSPHOLIPI | 0.847107 | 0.781244 | 1 | REACTOME_PHOSPHOLIPI | 1.031233 | 0.370203 | 1 |
| REACTOME_POSTMITOT | -0.78557 | 0.760163 | 0.987673 | REACTOME_POSTMITOTIC_ | -1.01896 | 0.437757 | 0.968853 | REACTOME_POSTMITOTI( | -1.1065 | 0.337183 | 1 |
| BIOCARTA_PTEN_PATHV | 1.036061 | 0.415644 | 0.987915 | BIOCARTA_PTEN_PATHWA | 0.738659 | 0.7883 | 1 | BIOCARTA_PTEN_PATHW/ | -0.98298 | 0.495346 | 1 |
| REACTOME_MRNA_CAP | 0.86508 | 0.642209 | 0.98823 | REACTOME_MRNA_CAPPII | -0.70177 | 0.867038 | 1 | REACTOME_MRNA_CAPPI | 0.721747 | 0.858914 | 1 |
| WP_ONECARBON_MET/ | -0.77719 | 0.808649 | 0.988794 | WP_ONECARBON_METAB( | 0.969569 | 0.473214 | 0.936117 | WP_ONECARBON_METAB | -0.74908 | 0.819913 | 1 |
| WP_PPAR_SIGNALING_F | 1.038916 | 0.396 | 0.988954 | WP_PPAR_SIGNALING_PAT | 1.149039 | 0.266749 | 0.878768 | WP_PPAR_SIGNALING_PA | 0.889813 | 0.643948 | 1 |
| REACTOME_EXTRA_NUC | 1.04186 | 0.386519 | 0.989873 | REACTOME_EXTRA_NUCL | 0.714513 | 0.882026 | 1 | REACTOME_EXTRA_NUCL | 0.789883 | 0.833273 | 1 |
| REACTOME_HATS_ACET | -0.77091 | 0.771875 | 0.990534 | REACTOME_HATS_ACETYL | 0.741888 | 0.783043 | 1 | REACTOME_HATS_ACETYI | -1.10836 | 0.344979 | 1 |
| BIOCARTA_BIOPEPTIDES | 1.036837 | 0.401046 | 0.990782 | BIOCARTA_BIOPEPTIDES_P | 1.424508 | 0.092414 | 0.565906 | BIOCARTA_BIOPEPTIDES_I | 1.087594 | 0.327879 | 1 |
| REACTOME_CARDIAC_C( | -0.77857 | 0.847124 | 0.990816 | REACTOME_CARDIAC_CON | -1.16669 | 0.210856 | 0.817092 | REACTOME_CARDIAC_COI | -0.82123 | 0.755962 | 1 |
| REACTOME_DEADENYLA | -0.76225 | 0.79373 | 0.99119 | REACTOME_DEADENYLATI | 0.539771 | 0.976093 | 1 | REACTOME_DEADENYLAT | -0.62569 | 0.932014 | 1 |
| REACTOME_IRS_MEDIAT | -0.77313 | 0.773124 | 0.991219 | REACTOME_IRS_MEDIATEI | -0.65965 | 0.900747 | 1 | REACTOME_IRS_MEDIATE | -0.64341 | 0.905935 | 1 |
| REACTOME_GLYCEROPH | 1.095725 | 0.302707 | 0.991639 | REACTOME_GLYCEROPHOS | 0.850576 | 0.717448 | 1 | REACTOME_GLYCEROPHC | 0.915352 | 0.641001 | 1 |
| REACTOME_PINK1_PRKI | 0.86222 | 0.636449 | 0.99191 | REACTOME_PINK1_PRKN_ | 0.653652 | 0.88151 | 1 | REACTOME_PINK1_PRKN_ | -0.55802 | 0.960671 | 1 |
| REACTOME_SENESCENC | 1.042746 | 0.403634 | 0.992484 | REACTOME_SENESCENCE_ | -0.76207 | 0.801025 | 1 | REACTOME_SENESCENCE_ | 1.151517 | 0.254879 | 1 |
| WP_AMINO_ACID_MET/ | 0.85595 | 0.713445 | 0.993855 | WP_AMINO_ACID_METAB | 0.596457 | 0.990746 | 1 | WP_AMINO_ACID_METAE | -0.65376 | 0.960954 | 1 |
| REACTOME_RHOQ_GTP/ | 1.092869 | 0.319631 | 0.995628 | REACTOME_RHOQ_GTPAS | 1.3282 | 0.116966 | 0.698982 | REACTOME_RHOQ_GTPAS | -0.72487 | 0.867914 | 1 |
| REACTOME_CD28_CO_S | 1.090975 | 0.340122 | 0.995929 | REACTOME_CD28_CO_STII | 0.701385 | 0.853886 | 1 | REACTOME_CD28_CO_STI | 1.155326 | 0.253198 | 1 |
| REACTOME_MAJOR_PAT | 1.096291 | 0.268584 | 0.996661 | REACTOME_MAJOR_PATH | 1.095366 | 0.273321 | 0.878389 | REACTOME_MAJOR_PATH | -1.06853 | 0.34865 | 1 |
| WP_TCA_CYCLE | 0.462095 | 0.997423 | 0.996845 | WP_TCA_CYCLE | -1.02639 | 0.418825 | 0.956913 | WP_TCA_CYCLE | -0.59122 | 0.958968 | 1 |
| REACTOME_INTERCONV | 1.06122 | 0.385376 | 0.996947 | REACTOME_INTERCONVER | -0.9436 | 0.530883 | 1 | REACTOME_INTERCONVEI | -0.8513 | 0.658487 | 1 |
| REACTOME_MITOPHAGY | 0.856022 | 0.658632 | 0.996975 | REACTOME_MITOPHAGY | 0.592439 | 0.952689 | 1 | REACTOME_MITOPHAGY | -0.51754 | 0.981802 | 1 |
| REACTOME_RHO_GTPAS | 1.101647 | 0.313976 | 0.997363 | REACTOME_RHO_GTPASES | 0.983328 | 0.47284 | 0.932368 | REACTOME_RHO_GTPASE | -0.72264 | 0.850463 | 1 |
| REACTOME_SWITCHING | 1.062797 | 0.350476 | 0.997427 | REACTOME_SWITCHING_C | 1.197755 | 0.204217 | 0.815404 | REACTOME_SWITCHING_( | 1.195855 | 0.162117 | 1 |
| REACTOME_DEGRADATI | 1.103426 | 0.299144 | 0.997693 | REACTOME_DEGRADATION | 1.283445 | 0.145931 | 0.734327 | REACTOME_DEGRADATIO | 1.143913 | 0.229632 | 1 |
| REACTOME_REGULATIO | 0.857013 | 0.726292 | 0.997916 | REACTOME_REGULATION_ | 0.81244 | 0.782069 | 1 | REACTOME_REGULATION_ | -0.7898 | 0.816732 | 1 |
| REACTOME_RHO_GTPAS | 1.081568 | 0.285617 | 0.998052 | REACTOME_RHO_GTPASE_ | 1.298735 | 0.064822 | 0.732516 | REACTOME_RHO_GTPASE | -0.9963 | 0.488929 | 1 |
| REACTOME_NUCLEOTID | 0.806835 | 0.741671 | 0.998132 | REACTOME_NUCLEOTIDE_ | 0.884882 | 0.597971 | 1 | REACTOME_NUCLEOTIDE_ | 0.856547 | 0.683458 | 1 |
| REACTOME_ACTIVATION | 0.725793 | 0.86554 | 0.998624 | REACTOME_ACTIVATION_( | -0.66193 | 0.927997 | 1 | REACTOME_ACTIVATION_ | -0.40872 | 0.998995 | 1 |
| WP_ESC_PLURIPOTENCY | 1.0976 | 0.304126 | 0.998733 | WP_ESC_PLURIPOTENCY_F | 0.880303 | 0.628158 | 1 | WP_ESC_PLURIPOTENCY_ | 0.806882 | 0.828229 | 1 |
| REACTOME_FORMATION | 0.857945 | 0.668832 | 0.999031 | REACTOME_FORMATION_( | -1.03402 | 0.410128 | 0.944175 | REACTOME_FORMATION_ | 0.72303 | 0.864865 | 1 |
| REACTOME_RHOBTB2_( | 1.077831 | 0.356895 | 0.999042 | REACTOME_RHOBTB2_GTI | 0.946256 | 0.512448 | 0.959694 | REACTOME_RHOBTB2_GT | 1.149355 | 0.262583 | 1 |
| REACTOME_MITOCHON | 0.46346 | 0.997427 | 0.999061 | REACTOME_MITOCHONDR | -0.9365 | 0.552156 | 1 | REACTOME_MITOCHOND | -1.0141 | 0.44401 | 1 |
| REACTOME_MAPK6_MA | 0.835482 | 0.74504 | 0.999452 | REACTOME_MAPK6_MAPK | 1.134123 | 0.258723 | 0.892148 | REACTOME_MAPK6_MAP | 0.885533 | 0.675395 | 1 |
| REACTOME_INTEGRATIC | 0.832465 | 0.75522 | 0.999816 | REACTOME_INTEGRATION_ | -0.76065 | 0.878664 | 1 | REACTOME_INTEGRATION | -0.56035 | 0.989122 | 1 |
| REACTOME_SPHINGOLIF | 1.049671 | 0.395141 | 1 | REACTOME_SPHINGOLIPID | 1.737207 | 0.016383 | 0.224926 | REACTOME_SPHINGOLIPII | 1.023133 | 0.413555 | 1 |
| REACTOME_CELL_JUNCT | 0.6835 | 0.941091 | 1 | REACTOME_CELL_JUNCTIC | 1.673977 | 0.010468 | 0.272772 | REACTOME_CELL_JUNCTI( | -0.94073 | 0.555156 | 1 |
| REACTOME_CELL_CELL_ | 0.726179 | 0.909838 | 1 | REACTOME_CELL_CELL_C( | 1.656672 | 0.010814 | 0.284438 | REACTOME_CELL_CELL_C( | -0.70251 | 0.910752 | 1 |
| WP_COMPREHENSIVE_II | 1.084816 | 0.32201 | 1 | WP_COMPREHENSIVE_IL1 | 1.539126 | 0.028015 | 0.41883 | WP_COMPREHENSIVE_IL1 | -0.7333 | 0.871882 | 1 |
| BIOCARTA_RAC1_PATHV | 0.841363 | 0.674929 | 1 | BIOCARTA_RAC1_PATHWA | 1.483259 | 0.07442 | 0.504325 | BIOCARTA_RAC1_PATHW/ | -0.82665 | 0.695356 | 1 |
| REACTOME_CYCLIN_D_/ | 0.770026 | 0.769726 | 1 | REACTOME_CYCLIN_D_AS! | -1.42411 | 0.081936 | 0.581573 | REACTOME_CYCLIN_D_AS | -1.07777 | 0.371842 | 1 |
| REACTOME_RND2_GTP/ | 1.069814 | 0.353335 | 1 | REACTOME_RND2_GTPASI | 1.389783 | 0.099461 | 0.635081 | REACTOME_RND2_GTPAS | 0.877662 | 0.664601 | 1 |
| REACTOME_RNA_POLYN | 0.762363 | 0.852157 | 1 | REACTOME_RNA_POLYME | -1.55694 | 0.017305 | 0.665458 | REACTOME_RNA_POLYMI | -0.95657 | 0.534145 | 1 |
| BIOCARTA_AGR_PATHW | -0.71464 | 0.865649 | 1 | BIOCARTA_AGR_PATHWAY | 1.335569 | 0.138231 | 0.685738 | BIOCARTA_AGR_PATHWA | -0.54228 | 0.976879 | 1 |
| REACTOME_GAP_JUNCT | 0.749883 | 0.826233 | 1 | REACTOME_GAP_JUNCTIO | -1.31704 | 0.123443 | 0.686523 | REACTOME_GAP_JUNCTI( | 0.739261 | 0.876433 | 1 |
| BIOCARTA_BCR_PATHW | -0.50643 | 0.988891 | 1 | BIOCARTA_BCR_PATHWAY | -1.25445 | 0.193356 | 0.703589 | BIOCARTA_BCR_PATHWA | -0.78761 | 0.752469 | 1 |
| WP_REGULATION_OF_A | 1.054796 | 0.348748 | 1 | WP_REGULATION_OF_ACT | 1.303654 | 0.089998 | 0.736729 | WP_REGULATION_OF_AC | -0.64932 | 0.979334 | 1 |
| REACTOME_KINESINS | 0.757888 | 0.824135 | 1 | REACTOME_KINESINS | -1.23401 | 0.187682 | 0.745745 | REACTOME_KINESINS | -0.85677 | 0.665762 | 1 |
| REACTOME_PROCESSING | -0.73225 | 0.830802 | 1 | REACTOME_PROCESSING_ | -1.22017 | 0.219327 | 0.755344 | REACTOME_PROCESSING_ | -0.90836 | 0.586016 | 1 |
| REACTOME_TP53_REGU | 0.603669 | 0.968635 | 1 | REACTOME_TP53_REGULA | -1.20751 | 0.20865 | 0.762363 | REACTOME_TP53_REGUL/ | -0.61728 | 0.94634 | 1 |
| REACTOME_SIGNAL_TR/ | 0.501771 | 0.98448 | 1 | REACTOME_SIGNAL_TRAN | 1.259447 | 0.197942 | 0.784926 | REACTOME_SIGNAL_TRAN | 1.234538 | 0.19628 | 1 |
| REACTOME_SIGNALING_ | 0.848494 | 0.926562 | 1 | REACTOME_SIGNALING_B\ | 1.253271 | 0.031451 | 0.786947 | REACTOME_SIGNALING_E | -0.85807 | 0.853188 | 1 |
| REACTOME_FORMATION | 0.799737 | 0.785293 | 1 | REACTOME_FORMATION_( | -1.15753 | 0.240734 | 0.814546 | REACTOME_FORMATION_ | -0.53427 | 0.989359 | 1 |
| REACTOME_FCERI_MED | 0.837745 | 0.756972 | 1 | REACTOME_FCERI_MEDIA | 1.200951 | 0.189756 | 0.820939 | REACTOME_FCERI_MEDIA | 0.797761 | 0.865763 | 1 |
| REACTOME_ACTIVATION | 0.768155 | 0.841153 | 1 | REACTOME_ACTIVATION_( | 1.08568 | 0.319903 | 0.867094 | REACTOME_ACTIVATION_ | 1.018969 | 0.408619 | 1 |

| | | | | | | | | | | | |
|---|---|---|---|---|---|---|---|---|---|---|---|
| REACTOME_SYNTHESIS_ | 0.839459 | 0.670474 | 1 | REACTOME_SYNTHESIS_OI | 1.083608 | 0.349551 | 0.867771 | REACTOME_SYNTHESIS_O | -0.81829 | 0.713771 | 1 |
| REACTOME_UCH_PROTE | 1.046732 | 0.374507 | 1 | REACTOME_UCH_PROTEIN | 1.08895 | 0.313339 | 0.86857 | REACTOME_UCH_PROTEII | 0.901372 | 0.665274 | 1 |
| REACTOME_O_LINKED_( | 1.06488 | 0.373686 | 1 | REACTOME_O_LINKED_GL | 1.077203 | 0.355178 | 0.871001 | REACTOME_O_LINKED_GI | 1.100638 | 0.312168 | 1 |
| REACTOME_NUCLEAR_E | -0.71126 | 0.938744 | 1 | REACTOME_NUCLEAR_ENV | -1.09192 | 0.306037 | 0.87162 | REACTOME_NUCLEAR_EN | -0.7653 | 0.827141 | 1 |
| REACTOME_RHOV_GTPA | 1.045884 | 0.390289 | 1 | REACTOME_RHOV_GTPASI | 1.074384 | 0.342138 | 0.874134 | REACTOME_RHOV_GTPAS | 0.856068 | 0.690117 | 1 |
| REACTOME_SIGNALING_ | 0.706266 | 0.845847 | 1 | REACTOME_SIGNALING_BY | 1.056556 | 0.389359 | 0.8747 | REACTOME_SIGNALING_B | 0.880211 | 0.623077 | 1 |
| REACTOME_AMPLIFICAT | 0.789798 | 0.80373 | 1 | REACTOME_AMPLIFICATIO | 1.037072 | 0.384913 | 0.877083 | REACTOME_AMPLIFICATI | -0.64513 | 0.94646 | 1 |
| REACTOME_G_PROTEIN_ | 0.514286 | 0.993934 | 1 | REACTOME_G_PROTEIN_M | 1.093112 | 0.323938 | 0.879563 | REACTOME_G_PROTEIN_I | -0.49567 | 0.992671 | 1 |
| WP_IL6_SIGNALING_PAT | 0.808807 | 0.814682 | 1 | WP_IL6_SIGNALING_PATH | 1.042044 | 0.376719 | 0.882339 | WP_IL6_SIGNALING_PATH | -0.73808 | 0.882531 | 1 |
| REACTOME_CROSS_PRE | 0.777836 | 0.81951 | 1 | REACTOME_CROSS_PRESE | 1.038377 | 0.383556 | 0.883365 | REACTOME_CROSS_PRESI | 1.245438 | 0.140384 | 1 |
| REACTOME_INTERLEUKI | 0.727957 | 0.937062 | 1 | REACTOME_INTERLEUKIN_ | 1.026503 | 0.397533 | 0.88412 | REACTOME_INTERLEUKIN | 0.715593 | 0.966995 | 1 |
| REACTOME_RHOF_GTPA | 0.675758 | 0.915821 | 1 | REACTOME_RHOF_GTPASE | 1.112342 | 0.307202 | 0.887522 | REACTOME_RHOF_GTPAS | -0.77244 | 0.786802 | 1 |
| REACTOME_ION_HOME | 0.783829 | 0.791139 | 1 | REACTOME_ION_HOMEOS | -1.08289 | 0.331435 | 0.888653 | REACTOME_ION_HOMEO | -0.9329 | 0.559743 | 1 |
| REACTOME_PLASMA_LII | 1.098258 | 0.32859 | 1 | REACTOME_PLASMA_LIPO | 1.106852 | 0.324218 | 0.892397 | REACTOME_PLASMA_LIP | 0.788305 | 0.771736 | 1 |
| REACTOME_C_TYPE_LEC | 0.843234 | 0.761643 | 1 | REACTOME_C_TYPE_LECTI | 1.020334 | 0.411323 | 0.893067 | REACTOME_C_TYPE_LECT | -0.7128 | 0.921511 | 1 |
| REACTOME_RHOD_GTP | 0.470948 | 0.999095 | 1 | REACTOME_RHOD_GTPASI | 1.112356 | 0.292553 | 0.894299 | REACTOME_RHOD_GTPAS | -0.76484 | 0.813226 | 1 |
| REACTOME_RETROGRAI | 0.598286 | 0.980099 | 1 | REACTOME_RETROGRADE_ | 1.021487 | 0.406652 | 0.894447 | REACTOME_RETROGRADE | -0.42685 | 0.999041 | 1 |
| REACTOME_MACROAUT | 1.086542 | 0.292147 | 1 | REACTOME_MACROAUTOI | 1.124016 | 0.251056 | 0.897277 | REACTOME_MACROAUTO | -0.58073 | 0.993945 | 1 |
| REACTOME_RHO_GTPAS | 0.708578 | 0.999036 | 1 | REACTOME_RHO_GTPASE_ | 1.113391 | 0.191757 | 0.897742 | REACTOME_RHO_GTPASE | -0.81035 | 0.894496 | 1 |
| WP_TOLLLIKE_RECEPTOI | 0.797561 | 0.731841 | 1 | WP_TOLLLIKE_RECEPTOR_ | 1.113717 | 0.317011 | 0.903667 | WP_TOLLLIKE_RECEPTOR_ | -0.9811 | 0.491944 | 1 |
| REACTOME_TRANSCRIPT | 0.819076 | 0.706993 | 1 | REACTOME_TRANSCRIPTIC | -1.06626 | 0.377731 | 0.917242 | REACTOME_TRANSCRIPTI | 0.881072 | 0.628008 | 1 |
| REACTOME_ASYMMETR | 1.047637 | 0.377001 | 1 | REACTOME_ASYMMETRIC_ | 1.000984 | 0.431923 | 0.922371 | REACTOME_ASYMMETRIC | 0.94616 | 0.540084 | 1 |
| REACTOME_THE_ROLE_ | 1.051182 | 0.368142 | 1 | REACTOME_THE_ROLE_OF | 0.983485 | 0.46413 | 0.936673 | REACTOME_THE_ROLE_O | 1.188355 | 0.175521 | 1 |
| REACTOME_TNFR2_NON | 0.757597 | 0.86266 | 1 | REACTOME_TNFR2_NON_( | 0.980144 | 0.467463 | 0.937312 | REACTOME_TNFR2_NON_ | 0.807166 | 0.828051 | 1 |
| REACTOME_UB_SPECIFI | 1.063338 | 0.330005 | 1 | REACTOME_UB_SPECIFIC_ | 0.967533 | 0.511891 | 0.937629 | REACTOME_UB_SPECIFIC_ | 0.933109 | 0.63458 | 1 |
| REACTOME_NEUROTRAI | -0.63014 | 0.977833 | 1 | REACTOME_NEUROTRANS | -1.04188 | 0.391628 | 0.937693 | REACTOME_NEUROTRAN | -0.99694 | 0.470929 | 1 |
| REACTOME_RIPK1_MED | 1.046682 | 0.392628 | 1 | REACTOME_RIPK1_MEDIAT | 0.969855 | 0.477496 | 0.939901 | REACTOME_RIPK1_MEDIA | 0.755841 | 0.803515 | 1 |
| REACTOME_PLATELET_H | 0.748023 | 0.872129 | 1 | REACTOME_PLATELET_HO | -1.05089 | 0.375196 | 0.942104 | REACTOME_PLATELET_HC | 0.71588 | 0.936709 | 1 |
| REACTOME_DOWNSTRE | 0.817051 | 0.786191 | 1 | REACTOME_DOWNSTREAN | 0.970231 | 0.479844 | 0.943448 | REACTOME_DOWNSTREA | 0.92898 | 0.602174 | 1 |
| BIOCARTA_IL2RB_PATHV | 0.850943 | 0.663507 | 1 | BIOCARTA_IL2RB_PATHWA | 0.9553 | 0.501161 | 0.946618 | BIOCARTA_IL2RB_PATHW | 0.874866 | 0.636683 | 1 |
| REACTOME_SEMAPHOR | 0.671285 | 0.938658 | 1 | REACTOME_SEMAPHORIN_ | 0.957624 | 0.494303 | 0.948884 | REACTOME_SEMAPHORIN | 1.161969 | 0.216236 | 1 |
| REACTOME_DEGRADATI | 1.043907 | 0.381828 | 1 | REACTOME_DEGRADATION | 0.945706 | 0.520618 | 0.956808 | REACTOME_DEGRADATIO | 0.892425 | 0.643218 | 1 |
| REACTOME_MYOGENES | 1.072242 | 0.371736 | 1 | REACTOME_MYOGENESIS | 0.947351 | 0.504205 | 0.960938 | REACTOME_MYOGENESIS | 0.860703 | 0.648878 | 1 |
| REACTOME_SIGNALING_ | 1.068718 | 0.321 | 1 | REACTOME_SIGNALING_BY | 0.934705 | 0.58565 | 0.971267 | REACTOME_SIGNALING_B | -0.82939 | 0.807274 | 1 |
| REACTOME_DNA_REPLI | 0.777352 | 0.860589 | 1 | REACTOME_DNA_REPLICAT | 0.920186 | 0.582225 | 0.978044 | REACTOME_DNA_REPLICA | 0.989963 | 0.468764 | 1 |
| REACTOME_REGULATIO | 0.498867 | 0.996896 | 1 | REACTOME_REGULATION_ | 0.9103 | 0.57572 | 0.9847 | REACTOME_REGULATION | 0.85471 | 0.720583 | 1 |
| REACTOME_EPHRIN_SIG | 0.595787 | 0.944754 | 1 | REACTOME_EPHRIN_SIGNA | 0.383511 | 0.998948 | 0.999479 | REACTOME_EPHRIN_SIGN | 0.89779 | 0.589658 | 1 |
| REACTOME_DARPP_32_ | -0.56796 | 0.961929 | 1 | REACTOME_DARPP_32_EV | -0.46304 | 0.991171 | 0.999865 | REACTOME_DARPP_32_E | 1.136998 | 0.291892 | 1 |
| BIOCARTA_AKT_PATHW | 0.660415 | 0.887364 | 1 | BIOCARTA_AKT_PATHWAY | 0.685065 | 0.846399 | 1 | BIOCARTA_AKT_PATHWA | -0.6815 | 0.854018 | 1 |
| BIOCARTA_CASPASE_PA | 0.602699 | 0.929691 | 1 | BIOCARTA_CASPASE_PATH | 0.50768 | 0.974559 | 1 | BIOCARTA_CASPASE_PATI | -0.73116 | 0.810664 | 1 |
| BIOCARTA_CERAMIDE_F | -0.59644 | 0.955365 | 1 | BIOCARTA_CERAMIDE_PAT | 0.608019 | 0.929774 | 1 | BIOCARTA_CERAMIDE_PA | 1.002839 | 0.446014 | 1 |
| BIOCARTA_CXCR4_PATH | 0.818967 | 0.700076 | 1 | BIOCARTA_CXCR4_PATHW | -0.65754 | 0.897335 | 1 | BIOCARTA_CXCR4_PATHW | 0.900396 | 0.580491 | 1 |
| BIOCARTA_EGF_PATHW | 0.770279 | 0.777348 | 1 | BIOCARTA_EGF_PATHWAY | -0.83344 | 0.692765 | 1 | BIOCARTA_EGF_PATHWA | 0.779765 | 0.777035 | 1 |
| BIOCARTA_FCER1_PATH | 0.630272 | 0.936636 | 1 | BIOCARTA_FCER1_PATHW | -0.51078 | 0.987978 | 1 | BIOCARTA_FCER1_PATHW | -0.40573 | 0.997901 | 1 |
| BIOCARTA_GH_PATHWA | -0.65402 | 0.908153 | 1 | BIOCARTA_GH_PATHWAY | -0.72469 | 0.824155 | 1 | BIOCARTA_GH_PATHWAY | 0.775043 | 0.781548 | 1 |
| BIOCARTA_KERATINOCY | -0.73473 | 0.858307 | 1 | BIOCARTA_KERATINOCYTE | -0.53628 | 0.986146 | 1 | BIOCARTA_KERATINOCYTI | 1.002535 | 0.429972 | 1 |
| BIOCARTA_MITOCHOND | 0.770647 | 0.764616 | 1 | BIOCARTA_MITOCHONDRI | 0.817224 | 0.688586 | 1 | BIOCARTA_MITOCHONDR | 1.189093 | 0.243836 | 1 |
| BIOCARTA_PAR1_PATHV | 0.808389 | 0.708047 | 1 | BIOCARTA_PAR1_PATHWA | -0.78914 | 0.744531 | 1 | BIOCARTA_PAR1_PATHW | -0.5763 | 0.946044 | 1 |
| BIOCARTA_RAS_PATHW | 0.72782 | 0.823217 | 1 | BIOCARTA_RAS_PATHWAY | -0.64508 | 0.911499 | 1 | BIOCARTA_RAS_PATHWA | 0.822822 | 0.712321 | 1 |
| BIOCARTA_TCR_PATHW | 0.632571 | 0.92906 | 1 | BIOCARTA_TCR_PATHWAY | -0.79656 | 0.748348 | 1 | BIOCARTA_TCR_PATHWA | -0.3788 | 0.998797 | 1 |
| BIOCARTA_TPO_PATHW | 0.696284 | 0.858586 | 1 | BIOCARTA_TPO_PATHWAY | -0.82304 | 0.703269 | 1 | BIOCARTA_TPO_PATHWA | 0.843406 | 0.677321 | 1 |
| BIOCARTA_VIP_PATHW | -0.66842 | 0.891327 | 1 | BIOCARTA_VIP_PATHWAY | -0.75632 | 0.789789 | 1 | BIOCARTA_VIP_PATHWAY | -0.37829 | 0.99906 | 1 |
| REACTOME_ACETYLCHO | 0.743975 | 0.804624 | 1 | REACTOME_ACETYLCHOLII | 0.880861 | 0.602612 | 1 | REACTOME_ACETYLCHOLI | -1.06405 | 0.391385 | 1 |
| REACTOME_ANTIGEN_P | 0.731865 | 0.986525 | 1 | REACTOME_ANTIGEN_PRC | 0.758243 | 0.964518 | 1 | REACTOME_ANTIGEN_PR | -0.76107 | 0.921287 | 1 |
| REACTOME_ANTIVIRAL_ | 0.712357 | 0.854707 | 1 | REACTOME_ANTIVIRAL_M | -0.68056 | 0.898276 | 1 | REACTOME_ANTIVIRAL_M | -0.85092 | 0.670834 | 1 |
| REACTOME_APOPTOTIC_ | -0.63591 | 0.962575 | 1 | REACTOME_APOPTOTIC_E | 0.493367 | 0.994906 | 1 | REACTOME_APOPTOTIC_I | 0.755542 | 0.854017 | 1 |
| REACTOME_CA_DEPEND | -0.49218 | 0.991008 | 1 | REACTOME_CA_DEPENDEI | 0.573609 | 0.960825 | 1 | REACTOME_CA_DEPENDE | -0.48473 | 0.989734 | 1 |
| REACTOME_CELL_CYCLE | 0.631993 | 1 | 1 | REACTOME_CELL_CYCLE | -0.76131 | 0.99177 | 1 | REACTOME_CELL_CYCLE | -0.75084 | 0.95944 | 1 |
| REACTOME_CELL_CYCLE | -0.63264 | 0.997389 | 1 | REACTOME_CELL_CYCLE_C | 0.716913 | 0.970322 | 1 | REACTOME_CELL_CYCLE_ | -0.76362 | 0.892272 | 1 |
| REACTOME_COSTIMULA | 0.730087 | 0.866177 | 1 | REACTOME_COSTIMULATI | 0.742093 | 0.832632 | 1 | REACTOME_COSTIMULAT | 1.155432 | 0.232597 | 1 |
| REACTOME_DAP12_SIGI | 0.836652 | 0.679163 | 1 | REACTOME_DAP12_SIGNA | 0.835987 | 0.656759 | 1 | REACTOME_DAP12_SIGNA | -0.6425 | 0.898292 | 1 |
| REACTOME_DOPAMINE_ | 0.5685 | 0.971348 | 1 | REACTOME_DOPAMINE_N | -0.91555 | 0.575187 | 1 | REACTOME_DOPAMINE_N | -0.97609 | 0.503969 | 1 |
| REACTOME_DOWNREGU | 0.737226 | 0.802225 | 1 | REACTOME_DOWNREGULA | -0.67383 | 0.871393 | 1 | REACTOME_DOWNREGUL | 0.897472 | 0.587854 | 1 |
| REACTOME_DOWNSTRE | -0.68186 | 0.872303 | 1 | REACTOME_DOWNSTREAN | -0.80657 | 0.721888 | 1 | REACTOME_DOWNSTREA | -0.57061 | 0.95313 | 1 |
| REACTOME_ENDOSOMA | -0.75294 | 0.819806 | 1 | REACTOME_ENDOSOMAL_ | -0.78041 | 0.775462 | 1 | REACTOME_ENDOSOMAL | -0.56299 | 0.968778 | 1 |
| REACTOME_FORMATION | 0.497099 | 0.991852 | 1 | REACTOME_FORMATION_ | -0.8975 | 0.606236 | 1 | REACTOME_FORMATION_ | -1.02456 | 0.437023 | 1 |
| REACTOME_G_ALPHA_C | 0.806965 | 0.826396 | 1 | REACTOME_G_ALPHA_Q_S | -0.804 | 0.849473 | 1 | REACTOME_G_ALPHA_Q_ | 1.001665 | 0.452445 | 1 |
| REACTOME_GLUTAMAT | -0.46359 | 0.995511 | 1 | REACTOME_GLUTAMATE_ | 0.805416 | 0.709631 | 1 | REACTOME_GLUTAMATE_ | -0.87639 | 0.635511 | 1 |
| REACTOME_GLYCOLYSIS | 0.82419 | 0.75762 | 1 | REACTOME_GLYCOLYSIS | 0.657266 | 0.949805 | 1 | REACTOME_GLYCOLYSIS | -1.08277 | 0.34861 | 1 |
| REACTOME_GLYCOSPHII | 0.712954 | 0.878617 | 1 | REACTOME_GLYCOSPHING | -0.88248 | 0.634066 | 1 | REACTOME_GLYCOSPHIN | -0.92473 | 0.572258 | 1 |
| REACTOME_INTERFERO | 0.613782 | 0.967192 | 1 | REACTOME_INTERFERON_ | -0.58955 | 0.978409 | 1 | REACTOME_INTERFERON_ | -0.90739 | 0.599533 | 1 |
| REACTOME_INTERLEUKI | -0.62254 | 0.945911 | 1 | REACTOME_INTERLEUKIN_ | -0.63264 | 0.937476 | 1 | REACTOME_INTERLEUKIN | -0.83269 | 0.695173 | 1 |
| REACTOME_INTRACELLU | -0.67412 | 0.997565 | 1 | REACTOME_INTRACELLUL | 0.728739 | 0.971845 | 1 | REACTOME_INTRACELLUL | 0.685976 | 0.994203 | 1 |
| REACTOME_INTRAFLAGI | -0.71035 | 0.910706 | 1 | REACTOME_INTRAFLAGELI | -0.67382 | 0.935209 | 1 | REACTOME_INTRAFLAGEL | 1.049958 | 0.364961 | 1 |
| REACTOME_ION_CHANN | 0.685386 | 0.97282 | 1 | REACTOME_ION_CHANNEI | -0.89063 | 0.681604 | 1 | REACTOME_ION_CHANNE | 0.863895 | 0.773246 | 1 |
| REACTOME_LYSOSOME_ | 0.840045 | 0.698538 | 1 | REACTOME_LYSOSOME_VI | 0.829405 | 0.689005 | 1 | REACTOME_LYSOSOME_V | -0.72197 | 0.84338 | 1 |
| REACTOME_M_PHASE | 0.759436 | 0.977707 | 1 | REACTOME_M_PHASE | 0.839268 | 0.860483 | 1 | REACTOME_M_PHASE | 0.982146 | 0.517016 | 1 |
| REACTOME_MEMBRANE | 0.727639 | 0.99937 | 1 | REACTOME_MEMBRANE_T | -0.72409 | 0.999578 | 1 | REACTOME_MEMBRANE_ | -0.65111 | 0.998772 | 1 |
| REACTOME_METABOLIS | 0.748868 | 0.964762 | 1 | REACTOME_METABOLISM_ | -0.77362 | 0.950694 | 1 | REACTOME_METABOLISM | 0.847487 | 0.900981 | 1 |
| REACTOME_MITOTIC_M | 0.84741 | 0.812074 | 1 | REACTOME_MITOTIC_MET | 0.839198 | 0.819759 | 1 | REACTOME_MITOTIC_ME | 0.875393 | 0.834691 | 1 |
| REACTOME_MITOTIC_PI | 0.712857 | 0.962155 | 1 | REACTOME_MITOTIC_PRO | 0.791969 | 0.859956 | 1 | REACTOME_MITOTIC_PRO | 0.947699 | 0.588235 | 1 |
| REACTOME_MRNA_SPLI | -0.55214 | 0.990714 | 1 | REACTOME_MRNA_SPLICII | 0.574753 | 0.97854 | 1 | REACTOME_MRNA_SPLICI | 0.717269 | 0.908741 | 1 |
| REACTOME_MTOR_SIGN | -0.69605 | 0.910691 | 1 | REACTOME_MTOR_SIGNAI | 0.796237 | 0.758138 | 1 | REACTOME_MTOR_SIGNA | 0.965497 | 0.503009 | 1 |
| REACTOME_NEDDYLATI | 0.751141 | 0.960075 | 1 | REACTOME_NEDDYLATION | 0.709805 | 0.978732 | 1 | REACTOME_NEDDYLATIOI | -0.69566 | 0.965848 | 1 |

| | | | | | | | | | | | |
|---|---|---|---|---|---|---|---|---|---|---|---|
| REACTOME_NEGATIVE_I | 0.788226 | 0.735025 | 1 | REACTOME_NEGATIVE_RE | 0.484684 | 0.983757 | 1 | REACTOME_NEGATIVE_RE | 0.905189 | 0.575602 | 1 |
| REACTOME_NEGATIVE_I | 0.705336 | 0.881086 | 1 | REACTOME_NEGATIVE_RE | -0.99413 | 0.465553 | 1 | REACTOME_NEGATIVE_RE | 0.756776 | 0.834092 | 1 |
| REACTOME_NEGATIVE_I | 0.571306 | 0.991797 | 1 | REACTOME_NEGATIVE_RE | 0.586951 | 0.988571 | 1 | REACTOME_NEGATIVE_RE | 1.045184 | 0.360905 | 1 |
| REACTOME_NETRIN_1_S | -0.52605 | 0.982093 | 1 | REACTOME_NETRIN_1_SIG | 0.667031 | 0.874576 | 1 | REACTOME_NETRIN_1_SI | 1.138849 | 0.28736 | 1 |
| REACTOME_NOD1_2_SI | 0.655279 | 0.900096 | 1 | REACTOME_NOD1_2_SIGN | 0.613909 | 0.924602 | 1 | REACTOME_NOD1_2_SIGI | -0.86718 | 0.645875 | 1 |
| REACTOME_NUCLEOTID | 0.777648 | 0.751586 | 1 | REACTOME_NUCLEOTIDE_ | -0.72338 | 0.823004 | 1 | REACTOME_NUCLEOTIDE_ | 0.810137 | 0.716592 | 1 |
| REACTOME_ORGANELLE | 0.694283 | 0.984375 | 1 | REACTOME_ORGANELLE_E | -0.89692 | 0.693882 | 1 | REACTOME_ORGANELLE_ | 0.859987 | 0.82044 | 1 |
| REACTOME_PEROXISOM | 0.779972 | 0.773487 | 1 | REACTOME_PEROXISOMAI | 0.743485 | 0.7945 | 1 | REACTOME_PEROXISOMA | 0.873472 | 0.644451 | 1 |
| REACTOME_PI_METABO | 0.734331 | 0.896764 | 1 | REACTOME_PI_METABOLI: | 0.843598 | 0.712861 | 1 | REACTOME_PI_METABOLI | 1.009563 | 0.425043 | 1 |
| REACTOME_PROTEIN_PI | 0.767188 | 0.865323 | 1 | REACTOME_PROTEIN_PRO | -0.49328 | 0.99961 | 1 | REACTOME_PROTEIN_PR( | 0.781741 | 0.886186 | 1 |
| REACTOME_PROTEIN_U | 0.807989 | 0.777309 | 1 | REACTOME_PROTEIN_UBI( | -0.99283 | 0.46502 | 1 | REACTOME_PROTEIN_UBI | 0.84544 | 0.752076 | 1 |
| REACTOME_RAF_ACTIV/ | 1.044984 | 0.387585 | 1 | REACTOME_RAF_ACTIVATI | 0.651302 | 0.925673 | 1 | REACTOME_RAF_ACTIVAT | 1.130299 | 0.263875 | 1 |
| REACTOME_RECRUITME | 0.757608 | 0.860188 | 1 | REACTOME_RECRUITMENT | 0.789956 | 0.794383 | 1 | REACTOME_RECRUITMEN | 1.01073 | 0.42609 | 1 |
| REACTOME_RECYCLING_ | 1.075442 | 0.349221 | 1 | REACTOME_RECYCLING_P/ | -0.67135 | 0.923476 | 1 | REACTOME_RECYCLING_F | 0.743797 | 0.867308 | 1 |
| REACTOME_REGULATIO | 0.486252 | 0.987421 | 1 | REACTOME_REGULATION_ | -0.68083 | 0.874019 | 1 | REACTOME_REGULATION_ | -0.55111 | 0.955882 | 1 |
| REACTOME_RESOLUTIOI | 0.826975 | 0.768642 | 1 | REACTOME_RESOLUTION_ | 0.783044 | 0.835739 | 1 | REACTOME_RESOLUTION_ | 0.838058 | 0.799244 | 1 |
| REACTOME_RHO_GTPAS | -0.44627 | 0.996293 | 1 | REACTOME_RHO_GTPASES | -0.82259 | 0.70441 | 1 | REACTOME_RHO_GTPASE | 0.734202 | 0.832107 | 1 |
| REACTOME_RHOG_GTP/ | -0.59088 | 0.99362 | 1 | REACTOME_RHOG_GTPAS | 0.792212 | 0.799104 | 1 | REACTOME_RHOG_GTPAS | -0.86669 | 0.681979 | 1 |
| REACTOME_RNA_POLYN | -0.73157 | 0.848632 | 1 | REACTOME_RNA_POLYME | -0.7521 | 0.819388 | 1 | REACTOME_RNA_POLYMI | -0.82608 | 0.705123 | 1 |
| REACTOME_SIGNALING_ | 0.820223 | 0.725828 | 1 | REACTOME_SIGNALING_BY | -0.75618 | 0.82938 | 1 | REACTOME_SIGNALING_B | -0.83269 | 0.701934 | 1 |
| REACTOME_SIGNALING_ | 0.74971 | 0.919585 | 1 | REACTOME_SIGNALING_BY | 0.769913 | 0.891821 | 1 | REACTOME_SIGNALING_B | -0.48535 | 1 | 1 |
| REACTOME_SIGNALING_ | -0.67452 | 0.911803 | 1 | REACTOME_SIGNALING_BY | 0.58335 | 0.956989 | 1 | REACTOME_SIGNALING_B | 0.839924 | 0.701153 | 1 |
| REACTOME_SIGNALING_ | -0.73888 | 0.824705 | 1 | REACTOME_SIGNALING_BY | -0.63033 | 0.927461 | 1 | REACTOME_SIGNALING_B | -0.61609 | 0.934773 | 1 |
| REACTOME_SYNAPTIC_/ | 0.611384 | 0.9383 | 1 | REACTOME_SYNAPTIC_AD | -0.85027 | 0.656662 | 1 | REACTOME_SYNAPTIC_AD | 0.84066 | 0.69213 | 1 |
| REACTOME_SYNTHESIS_ | 0.519683 | 0.992852 | 1 | REACTOME_SYNTHESIS_OI | 0.837491 | 0.690764 | 1 | REACTOME_SYNTHESIS_O | 0.971367 | 0.484518 | 1 |
| REACTOME_TBC_RABGA | 0.769042 | 0.790023 | 1 | REACTOME_TBC_RABGAPS | 0.62788 | 0.933778 | 1 | REACTOME_TBC_RABGAP | 1.150942 | 0.248417 | 1 |
| REACTOME_TP53_REGU | 0.766617 | 0.765075 | 1 | REACTOME_TP53_REGULA | -0.69492 | 0.847908 | 1 | REACTOME_TP53_REGUL/ | -1.03563 | 0.428996 | 1 |
| REACTOME_TRAFFICKIN | 0.638594 | 0.911003 | 1 | REACTOME_TRAFFICKING_ | -0.94093 | 0.53545 | 1 | REACTOME_TRAFFICKING_ | -0.70595 | 0.834825 | 1 |
| REACTOME_VESICLE_MI | 0.755426 | 0.998446 | 1 | REACTOME_VESICLE_MED | -0.83837 | 0.96737 | 1 | REACTOME_VESICLE_MED | -0.69312 | 0.9956 | 1 |
| WP_EGFR1_SIGNALING_ | 0.567677 | 1 | 1 | WP_EGFR1_SIGNALING_P/ | 0.814753 | 0.843177 | 1 | WP_EGFR1_SIGNALING_P | -0.46567 | 1 | 1 |
| WP_ELECTRON_TRANSP | 0.817384 | 0.788605 | 1 | WP_ELECTRON_TRANSPOF | -0.94386 | 0.556256 | 1 | WP_ELECTRON_TRANSPO | -0.85886 | 0.704022 | 1 |
| WP_EPO_RECEPTOR_SIG | -0.64926 | 0.917906 | 1 | WP_EPO_RECEPTOR_SIGN | 0.698123 | 0.841033 | 1 | WP_EPO_RECEPTOR_SIGN | 1.039833 | 0.392999 | 1 |
| WP_G_PROTEIN_SIGNAI | 0.662476 | 0.97652 | 1 | WP_G_PROTEIN_SIGNALIN | -0.76371 | 0.888621 | 1 | WP_G_PROTEIN_SIGNALII | -0.72505 | 0.90562 | 1 |
| WP_GLUTATHIONE_MET | 0.590356 | 0.950851 | 1 | WP_GLUTATHIONE_METAI | 0.727016 | 0.81115 | 1 | WP_GLUTATHIONE_META | 0.782455 | 0.769401 | 1 |
| WP_IL1_SIGNALING_PA1 | 0.685312 | 0.891847 | 1 | WP_IL1_SIGNALING_PATH | 0.87292 | 0.620675 | 1 | WP_IL1_SIGNALING_PATH | 0.895187 | 0.599087 | 1 |
| WP_IL7_SIGNALING_PAT | -0.58643 | 0.973781 | 1 | WP_IL7_SIGNALING_PATH | 0.867823 | 0.632099 | 1 | WP_IL7_SIGNALING_PATH | 0.906612 | 0.598162 | 1 |
| WP_P53_SIGNALING | 1.049565 | 0.388647 | 1 | WP_P53_SIGNALING | 0.632601 | 0.915312 | 1 | WP_P53_SIGNALING | 0.987979 | 0.452677 | 1 |
| WP_PARKINSONS_DISEA | 0.807719 | 0.725869 | 1 | WP_PARKINSONS_DISEASE | -0.71307 | 0.87088 | 1 | WP_PARKINSONS_DISEAS | 1.189297 | 0.21242 | 1 |
| WP_SELENIUM_METABO | 0.54309 | 0.983619 | 1 | WP_SELENIUM_METABOLI | 0.704922 | 0.858921 | 1 | WP_SELENIUM_METABOL | -0.59332 | 0.955527 | 1 |

**GSEA cluster cortex**

** NbClust results. Method=ward.D2, distance=euclidean*

| | | NES | | |
|---|---|---|---|---|
| **cluster** | **ID** | **MNK1KO** | **MNK2KO** | **MNKDKO** |
| 1 | REACTOME_FORMATION_OF_A_POO | 2.03725 | 1.565585 | 2.030022 |
| 1 | REACTOME_ACTIVATION_OF_THE_MI | 1.880446 | 1.362647 | 2.046375 |
| 1 | REACTOME_TRANSLATION | 1.690334 | 1.399665 | 1.846749 |
| 1 | REACTOME_NONSENSE_MEDIATED_D | 2.000028 | 1.523747 | 1.864469 |
| 1 | WP_CYTOPLASMIC_RIBOSOMAL_PRO | 1.987372 | 1.478443 | 1.885397 |
| 1 | REACTOME_EUKARYOTIC_TRANSLATI | 1.961733 | 1.586985 | 1.901042 |
| 1 | REACTOME_NONSENSE_MEDIATED_D | 1.97281 | 1.46745 | 1.689707 |
| 1 | WP_CHOLESTEROL_METABOLISM_WI | 1.731039 | 1.409934 | 1.622111 |
| 1 | REACTOME_PHASE_II_CONJUGATION_ | 1.637782 | 0.976545 | 1.520062 |
| 1 | REACTOME_BIOLOGICAL_OXIDATIONS | 1.878272 | 1.425181 | 1.480575 |
| 1 | WP_PRIMARY_FOCAL_SEGMENTAL_G | 1.314465 | 1.727188 | 0.654138 |
| 1 | REACTOME_METABOLISM_OF_STERO | 1.677806 | 0.991934 | 1.072741 |
| 1 | BIOCARTA_ECM_PATHWAY | 1.320114 | 1.75156 | 0.760377 |
| 1 | REACTOME_SPHINGOLIPID_DE_NOVO | 1.049671 | 1.737207 | 1.023133 |
| 1 | REACTOME_SIGNALING_BY_TGFB_FA | 1.227781 | 1.713913 | 0.958745 |
| 1 | REACTOME_PHASE_I_FUNCTIONALIZA | 1.840742 | 1.630544 | 1.205816 |
| 1 | WP_METAPATHWAY_BIOTRANSFORM | 1.631887 | 1.183106 | 1.171471 |
| 1 | WP_OXIDATIVE_STRESS_RESPONSE | 1.758821 | -0.67299 | 0.901859 |
| 2 | REACTOME_RHO_GTPASES_ACTIVATE | -1.94442 | -1.87132 | -1.77925 |
| 2 | REACTOME_NONHOMOLOGOUS_END | -1.79293 | -1.92679 | -1.71584 |
| 2 | REACTOME_POTASSIUM_CHANNELS | -1.68077 | -1.84142 | -1.07326 |
| 3 | REACTOME_MET_PROMOTES_CELL_M | 1.459251 | 2.07757 | -1.54082 |
| 3 | REACTOME_COLLAGEN_BIOSYNTHESI | 1.709907 | 2.154113 | -1.55005 |
| 3 | REACTOME_COLLAGEN_FORMATION | 1.685491 | 2.147013 | -1.46736 |
| 3 | REACTOME_COLLAGEN_DEGRADATIO | 1.757987 | 2.10213 | -1.45781 |
| 3 | REACTOME_ASSEMBLY_OF_COLLAGEI | 1.91743 | 2.23117 | -1.37079 |
| 3 | REACTOME_DEGRADATION_OF_THE_ | 1.789369 | 2.276743 | -1.4164 |
| 3 | REACTOME_INTEGRIN_CELL_SURFACE | 2.097309 | 2.202235 | -1.42902 |
| 3 | REACTOME_NON_INTEGRIN_MEMBRA | 1.521135 | 2.091786 | -1.43936 |
| 3 | REACTOME_ECM_PROTEOGLYCANS | 1.936779 | 2.164128 | -1.25612 |
| 3 | REACTOME_NCAM_SIGNALING_FOR_ | 1.55138 | 2.110592 | -1.23107 |
| 3 | REACTOME_EXTRACELLULAR_MATRIX | 2.059572 | 2.519486 | -1.18448 |
| 3 | REACTOME_SIGNALING_BY_PDGF | 1.424691 | 1.878707 | -1.13966 |
| 3 | WP_FOCAL_ADHESION | 1.558832 | 2.258373 | -0.90816 |
| 3 | REACTOME_SMOOTH_MUSCLE_CONT | 2.095553 | 2.23465 | -0.88747 |
| 3 | REACTOME_RHO_GTPASES_ACTIVATE | 2.047715 | 2.209499 | -0.45823 |
| 3 | REACTOME_TGF_BETA_RECEPTOR_SI | 1.54002 | 1.995003 | -0.81692 |
| 3 | REACTOME_MUSCLE_CONTRACTION | 1.431518 | 1.855598 | -0.76617 |
| 3 | REACTOME_SIGNALING_BY_TGF_BETA | 1.2606 | 1.7706 | -0.77181 |
| 3 | REACTOME_CELL_EXTRACELLULAR_M | 1.357794 | 1.733791 | -0.67271 |
| 3 | REACTOME_REGULATION_OF_INSULI | 1.656866 | 1.612106 | -0.82697 |
| 3 | REACTOME_COMPLEMENT_CASCADE | 1.871256 | 1.450704 | -0.95891 |
| 3 | REACTOME_RESPONSE_TO_ELEVATED | 1.798976 | 1.275612 | -0.98058 |
