## Supplementary Table 3 for "Distinct roles for MNK1 and MNK2 in social and cognitive behavior through kinase-specific regulation of the synaptic proteome and phosphoproteome"

**TMT synaptoneurosomes all proteins**

| protein names | | normalized protein expression | | | | | | | | | | | | | | | | LogFC over WT | | | p-value / WT | | | adjusted p-value / WT | | |
|---|---|---|---|---|---|---|---|---|---|---|---|---|---|---|---|---|---|---|---|---|---|---|---|---|---|---|
| id | GeneName | SYP_WT | SYP_WT | SYP_WT | SYP_WT | SYP_MNK1 | SYP_MNK1 | SYP_MNK1 | SYP_MNK1 | SYP_MNK2 | SYP_MNK2 | SYP_MNK2 | SYP_MNK2 | SYP_MNKD | SYP_MNKD | SYP_MNKD | SYP_MNKD | logFC.SYP_ | logFC.SYP_ | logFC.SYP_ | P.Value.SY | P.Value.SY | P.Value.SY | adj.P.Val.S | adj.P.Val.S | adj.P.Val.SYP_MNKDKO.over.SYP_WT |
| 0610009B2 | Q8R3W2 | -1.7 | -0.8 | -0.54 | 0.12 | -0.68 | -1.78 | 0.14 | -0.26 | 1 | -0.33 | -0.09 | -1.53 | 0.6 | 0.42 | -0.44 | -0.46 | 0.085 | 0.4925 | 0.76 | 0.868912 | 0.412619 | 0.114639 | 1 | 0.996511 | 0.717287 |
| 0610010K1 | D3Z687 | -0.08 | 1.15 | -0.72 | -0.68 | 1.35 | 0.08 | -0.1 | -0.82 | -0.52 | -0.28 | -0.78 | 0.63 | -0.08 | -0.53 | -0.15 | -0.16 | 0.21 | -0.155 | -0.1475 | 0.715284 | 0.755239 | 0.730327 | 1 | 0.996511 | 0.958635 |
| 0610012G | A0A0J9YTR | 2.02 | 1.73 | 1.04 | 1.16 | 0.83 | 1.64 | 0.97 | 1.11 | 1.26 | -0.46 | 1.27 | 2.49 | 2.92 | -1.21 | 1.15 | 1.69 | -0.35 | -0.3475 | -0.35 | 0.250757 | 0.563536 | 0.666256 | 1 | 0.996511 | 0.950424 |
| 1110059G1 | Q8K039 | -3.59 | -4.54 | -6.47 | -10.05 | -5.5 | -5.18 | -13.84 | -10.44 | -3.43 | -3.02 | -6.47 | -3.71 | -2.02 | -5.38 | -6.03 | -8.29 | -2.5775 | 2.005 | 0.7325 | 0.270406 | 0.192102 | 0.669877 | 1 | 0.996511 | 0.952558 |
| 1110065P2 | V9GXF1 | -2.02 | -2.04 | -0.23 | -0.42 | -2.95 | -0.9 | -0.67 | -0.06 | 1.56 | 0.24 | 0.01 | -1.75 | 2.45 | -1.55 | -0.71 | -0.36 | 0.0325 | 1.1925 | 1.135 | 0.964067 | 0.144061 | 0.226475 | 1 | 0.996511 | 0.787447 |
| 1500009L1 | A0A1W2P6 | 1.91 | 2.18 | 2.32 | 2.16 | 1.36 | 2.03 | 1.93 | 2.52 | 1.23 | 1.84 | 2.38 | 2.22 | 2.11 | 0.24 | 2.73 | 2.76 | -0.1825 | -0.225 | -0.1825 | 0.491004 | 0.432424 | 0.741729 | 1 | 0.996511 | 0.958881 |
| 1600014C1 | Q8WUR0 | -2.03 | -2.22 | 1.06 | -0.42 | -2.02 | -1.57 | 0.1 | 0.51 | -2.23 | -0.55 | -0.75 | -1.84 | -2.01 | -1.78 | -0.72 | -0.06 | 0.1575 | -0.44 | -0.24 | 0.85914 | 0.579953 | 0.766232 | 1 | 0.996511 | 0.964134 |
| 1700025G | Q8K207 | 1.31 | 1.35 | 1.63 | 1.6 | 1.16 | 1.42 | 1.06 | 1.46 | 0.2 | -0.37 | 1.63 | 1.58 | 0.67 | -2.48 | 1.39 | 1.39 | -0.1975 | -0.7125 | -1.23 | 0.279916 | 0.159826 | 0.165576 | 1 | 0.996511 | 0.758837 |
| 1700037H | H7BXB9 | -0.71 | -0.82 | -0.25 | -0.26 | -0.38 | -0.93 | 0.11 | 0.08 | -0.98 | 0.04 | -0.14 | -0.63 | -0.63 | -0.57 | -0.17 | -0.16 | 0.23 | 0.0825 | 0.1275 | 0.43159 | 0.77343 | 0.592333 | 1 | 0.996511 | 0.929843 |
| 2210016F1 | G3X8U3 | -0.69 | -0.8 | -0.42 | -0.36 | -0.76 | -1.19 | -0.46 | -0.32 | -1.12 | 0.21 | -0.54 | -1.07 | -1.41 | -0.66 | -0.6 | -0.44 | -0.115 | -0.0625 | -0.21 | 0.628967 | 0.847821 | 0.43894 | 1 | 0.996511 | 0.883205 |
| 2310061I | B8JJ66 | -0.06 | -0.19 | -0.69 | -0.89 | -0.02 | -0.22 | -1.37 | -0.9 | 0.35 | -1.13 | -0.85 | -0.04 | 0.02 | -0.66 | -1.37 | -0.6 | -0.17 | 0.04 | -0.195 | 0.633095 | 0.916728 | 0.575755 | 1 | 0.996755 | 0.926728 |
| 2410002F2 | Q3TE80 | -1.46 | -1.31 | -2.08 | -2.45 | -0.94 | -1.97 | -1.12 | -2.3 | -1.77 | -2.6 | -2.28 | -1.57 | -1.74 | -1.81 | -1.86 | -1.85 | 0.2425 | -0.23 | 0.01 | 0.547536 | 0.514996 | 0.972105 | 1 | 0.996511 | 0.992865 |
| 2410004B1 | Q9CWU4 | -2.24 | -2.69 | -2.76 | -2.62 | -3.2 | -2.34 | -4.1 | -3.86 | -0.75 | -1.45 | -2.97 | -2 | -0.01 | -2.88 | -2.94 | -3.49 | -0.7975 | 0.785 | 0.2475 | 0.066651 | 0.111303 | 0.731411 | 1 | 0.996511 | 0.958635 |
| 2900026A | A0A1D5RLI | -0.17 | 0.64 | -0.82 | -1.11 | 0.54 | 0.07 | -0.02 | -0.02 | 0.84 | -0.15 | 0.18 | 0.09 | 1.16 | -0.3 | 2.22 | -0.83 | 0.5075 | 0.605 | 0.9275 | 0.214396 | 0.174701 | 0.219859 | 1 | 0.996511 | 0.787447 |
| 2900026A | A0A1B0GR | 0.57 | 0.32 | 0.99 | 1.24 | 0.2 | 0.64 | 0.69 | 1.35 | 1.05 | 0.64 | 0.79 | 0.56 | 1.39 | 0.27 | 0.77 | 0.59 | -0.06 | -0.02 | -0.025 | 0.846374 | 0.93728 | 0.937626 | 1 | 0.996755 | 0.989767 |
| 3110082I1 | Q9CXL3 | -2.06 | -3.28 | -1.08 | -0.49 | -3.51 | -3.16 | -1.17 | -1.52 | -4.99 | 0.87 | -0.93 | -3.24 | -2.41 | -3.66 | -0.49 | -0.65 | -0.6125 | -0.345 | -0.075 | 0.43068 | 0.786702 | 0.931494 | 1 | 0.996511 | 0.989713 |
| 4931406C | Q91V76 | 1.23 | 0.12 | 1.26 | 0.82 | 0.8 | 1.05 | 1.63 | 1.12 | -0.22 | 0.49 | 1.4 | 1.04 | -0.55 | 0.5 | 0.88 | 1.05 | 0.2925 | -0.18 | -0.3875 | 0.361386 | 0.669161 | 0.374327 | 1 | 0.996511 | 0.859424 |
| 4933402D | Q9D4D3 | 0.36 | 1.34 | 1.06 | 0.77 | 0.64 | 0.12 | 0.72 | 0.53 | 0.4 | 0.13 | 0.82 | 0.81 | 0.05 | -0.23 | 0.43 | 0.44 | -0.38 | -0.3425 | -0.71 | 0.166026 | 0.242512 | 0.031741 | 1 | 0.996511 | 0.641974 |
| 4933427D1 | A0A0N4SU | 1.74 | 0.11 | 1.34 | 1.16 | 0.3 | 1.65 | 0.73 | 0.84 | 0.72 | 0.23 | 0.93 | 1.62 | 2.72 | 0.43 | 1.66 | 2.13 | -0.2075 | -0.2125 | 0.6475 | 0.62334 | 0.621625 | 0.259406 | 1 | 0.996511 | 0.812337 |
| 4933434E2 | H7BX33 | 0.16 | 0.11 | 0.71 | 0.6 | 0.42 | 0.24 | 0.99 | 0.58 | 0.45 | 0.96 | 0.73 | 0.27 | -0.35 | -0.66 | -0.61 | 0.91 | 0.1625 | 0.2075 | -0.5725 | 0.499525 | 0.401245 | 0.16456 | 1 | 0.996511 | 0.758837 |
| 5031439G | B1APX2 | 0.27 | 0.38 | 0.11 | -0.02 | 0.56 | 0.22 | 0.17 | -0.1 | 0.52 | 0.17 | 0.24 | 0.56 | 0.4 | 0.32 | 0.06 | 0.1 | 0.0275 | 0.1875 | 0.035 | 0.888636 | 0.33621 | 0.856773 | 1 | 0.996511 | 0.976225 |
| 5730455P1 | Q9CYI0 | -0.32 | 0.25 | -0.64 | -1.78 | 0.67 | -0.33 | -1.69 | -2.05 | 0.2 | -1.26 | -1.15 | -0.12 | -0.84 | 0.41 | -0.57 | -0.92 | -0.2275 | 0.04 | 0.1425 | 0.741579 | 0.938788 | 0.772066 | 1 | 0.996755 | 0.965933 |
| 6030458C1 | Q8BGC1 | -1.14 | -0.35 | -1.34 | -1.28 | 0.45 | -1.05 | -0.76 | -1.37 | 0.28 | -0.86 | -1.21 | -0.97 | -0.63 | 0.24 | -1.4 | -1.51 | 0.345 | 0.3375 | 0.2025 | 0.430618 | 0.394534 | 0.648411 | 1 | 0.996511 | 0.946083 |
| 6820408C1 | Q8BJX2 | 0.14 | 0.42 | 0.21 | -0.21 | 0.87 | -0.12 | -0.26 | -0.23 | -0.19 | 0.17 | 0.03 | -0.59 | -0.49 | 0.82 | 0.03 | 0.07 | -0.075 | -0.285 | -0.0325 | 0.801626 | 0.253009 | 0.916317 | 1 | 0.996511 | 0.986556 |
| 8030462N | Q0VAW6 | 0.16 | 0.35 | -0.25 | 0.15 | -0.09 | 0.28 | -1.39 | -0.78 | 0.24 | -0.09 | -0.71 | 0.54 | -0.46 | -0.09 | -1.23 | -0.77 | -0.5975 | -0.1075 | -0.74 | 0.135983 | 0.722296 | 0.029004 | 1 | 0.996511 | 0.637553 |
| A830018L1 | B7ZN57 | -0.57 | 0.2 | 0.22 | 0.44 | 0.75 | -0.74 | 0.19 | 0.31 | 0.62 | 0.67 | 0.1 | 1.02 | -0.69 | 0.99 | -0.85 | 0.07 | 0.055 | 0.53 | -0.1925 | 0.880484 | 0.102856 | 0.669881 | 1 | 0.996511 | 0.952558 |
| Aaas | P58742 | 0.3 | 0.34 | 0.03 | 0.42 | 0.86 | 0.22 | 1.08 | 0.64 | -0.1 | -0.09 | 0.21 | 0.17 | 0.47 | 1.09 | 0.62 | 0.49 | 0.4275 | -0.225 | 0.395 | 0.083581 | 0.237794 | 0.098884 | 1 | 0.996511 | 0.698156 |
| Aacs | Q9D2R0 | -0.34 | -0.43 | -0.22 | -0.42 | -0.04 | -0.39 | -0.51 | 0.01 | -0.25 | 0.25 | 0.25 | 0.17 | -0.06 | 0.09 | 0.12 | 0.22 | 0.12 | 0.4575 | 0.445 | 0.513658 | 0.035742 | 0.029883 | 1 | 0.996511 | 0.640893 |
| Aagab | Q8R2R3 | -1.58 | -0.66 | -1.18 | -0.58 | -1.4 | -1.16 | -0.82 | -0.86 | -0.82 | -0.11 | -0.55 | -1.08 | -0.73 | -0.66 | -0.91 | -0.93 | -0.06 | 3.60E-01 | 0.1925 | 0.827743 | 0.271113 | 0.48072 | 1 | 0.996511 | 0.893906 |
| Aak1 | Q3UHJ0 | 0.09 | 0.3 | 0.36 | 0.3 | 0.11 | 0.11 | 0.21 | 0.02 | -0.12 | -0.13 | 0 | 0.15 | -0.21 | 0.2 | -0.03 | -0.16 | -0.15 | -0.2875 | -0.3125 | 0.327326 | 0.115407 | 0.124248 | 1 | 0.996511 | 0.721448 |
| Aak1 | A0A571BEI | -0.69 | -0.33 | -0.08 | 0.16 | -0.75 | -0.84 | 0.49 | 0.16 | 0.53 | 0.44 | 0.22 | -0.46 | 0.87 | -0.95 | -0.2 | 0.21 | 1.96E-16 | 0.4175 | 0.2175 | 1 | 0.18228 | 0.59498 | 1 | 0.996511 | 0.929843 |
| Aak1 | A0A571BD | 1.3 | 0.69 | -1.14 | 0.39 | 0.68 | 0.73 | 0.27 | 0.64 | 0.77 | 0.89 | -0.2 | 1.84 | 0.08 | 0.39 | 0.85 | 0.64 | 0.27 | 0.515 | 0.18 | 0.584401 | 0.4084 | 0.723468 | 1 | 0.996511 | 0.958398 |
| Aamdc | D3Z442 | -0.99 | -0.38 | -0.18 | -0.3 | -0.07 | -1.14 | -0.17 | 0.19 | -0.8 | -0.58 | -0.01 | -0.84 | -0.5 | 0.05 | -0.3 | -0.64 | 0.165 | -0.095 | 0.115 | 0.621901 | 0.731771 | 0.662845 | 1 | 0.996511 | 0.94972 |
| Aamp | J3QN89 | -0.31 | -0.36 | -1.25 | -1.16 | -0.43 | -0.43 | -1.79 | -1.31 | -0.33 | -0.94 | -1.25 | -0.2 | -0.42 | -0.34 | -1.01 | -1.11 | -0.22 | 0.09 | 0.05 | 0.583464 | 0.796022 | 0.877809 | 1 | 0.996511 | 0.979626 |
| Aar2 | Q9D2V5 | 0.29 | 0.71 | 0.44 | 0.64 | 0.96 | 0.14 | 0.45 | 0.25 | 0.5 | 0.8 | 0.37 | 0.64 | 0.04 | 0.78 | -0.21 | 0.77 | -0.07 | 0.0575 | -0.175 | 0.757505 | 0.762691 | 0.547981 | 1 | 0.996511 | 0.912571 |
| Aars1 | Q8BGQ7 | -0.21 | -0.27 | -0.19 | -0.34 | -0.71 | -0.38 | -0.34 | -0.31 | -0.3 | -0.11 | -0.27 | -0.21 | -0.43 | -0.22 | -0.66 | -0.73 | -0.1825 | 0.03 | -0.2575 | 0.273921 | 0.848197 | 0.206709 | 1 | 0.996511 | 0.785338 |
| Aars2 | Q14CH7 | 0.11 | 0.22 | 0.28 | 0.33 | 0.01 | 0.25 | 0.33 | 0.45 | 0.98 | 0.67 | 0.65 | 0.09 | 0.88 | 0.74 | 0.29 | 0.26 | 0.025 | 0.3625 | 0.3075 | 0.8783 | 0.13502 | 0.180994 | 1 | 0.996511 | 0.775542 |
| Aarsd1 | Q3THG9 | -1.38 | -0.89 | -0.85 | -1.07 | -0.68 | -1.41 | -0.69 | -0.86 | -1.3 | -0.34 | -0.68 | -1.37 | -0.97 | -0.39 | -0.91 | -0.75 | 0.1375 | 0.125 | 0.2925 | 0.553205 | 0.664035 | 0.215867 | 1 | 0.996511 | 0.787447 |
| Aasdhppt | Q9CQF6 | -1.92 | -2.01 | 0.37 | -0.72 | -2.05 | -2.53 | -0.74 | -0.28 | -2.5 | -0.1 | -0.37 | -1.96 | -2.16 | -2.25 | -0.7 | -0.49 | -0.33 | -0.1625 | -0.33 | 0.639897 | 0.82599 | 0.622305 | 1 | 0.996511 | 0.938628 |
| Aass | Q99K67 | 1.93 | 1.13 | 0.91 | 1.16 | 1 | 1.66 | -0.28 | 0.87 | 1.14 | 0.56 | 1.04 | 1.26 | 1.05 | 0.6 | 0.12 | 0.21 | -0.47 | -0.2825 | -0.7875 | 0.292226 | 0.330971 | 0.03214 | 1 | 0.996511 | 0.641974 |
| Aatf | Q9JKX4 | 1.01 | 1.15 | 1.69 | 1.65 | 0.65 | 0.35 | 1.45 | 1.39 | 0.4 | 0.6 | 1.98 | 0.77 | 0.18 | 1.04 | 2.11 | 2.35 | -0.415 | -0.4375 | 0.045 | 0.212524 | 0.269531 | 0.927571 | 1 | 0.996511 | 0.989337 |
| Aatk | B1AZF9 | 0.18 | 0.45 | 0.63 | 0.22 | 0.61 | 0.21 | 0.91 | 0.44 | -0.16 | 0.16 | 0.59 | 0.4 | -0.06 | 0.5 | 0.51 | 0.73 | 0.1725 | -0.1225 | 0.05 | 0.419352 | 0.592089 | 0.833942 | 1 | 0.996511 | 0.975371 |
| Abat | P61922 | 0.19 | -0.07 | 0.23 | 1.17 | -0.16 | -0.12 | 1.18 | 0.66 | -0.15 | 0.67 | 0.39 | -0.2 | -0.43 | 0.01 | 0.39 | 0.34 | 0.01 | -0.2025 | -0.3025 | 0.979974 | 0.555123 | 0.375626 | 1 | 0.996511 | 0.859611 |
| Abca1 | P41233 | 0.94 | 0.83 | 0.3 | 0.29 | 0.94 | 0.86 | 0.31 | 0.04 | 0.52 | -0.08 | 0.12 | 0.74 | 0.32 | 0.19 | 0.17 | 0.57 | -0.0525 | -0.265 | -0.2775 | 0.851186 | 0.338903 | 0.258631 | 1 | 0.996511 | 0.812074 |
| Abca2 | A2AJ26 | -0.64 | -0.69 | -0.13 | -0.48 | 0.02 | -0.66 | -0.22 | -0.24 | -0.76 | 0.02 | 0.02 | -0.26 | -0.5 | -0.67 | 0.04 | -0.01 | 0.21 | 0.24 | 0.2 | 0.342893 | 0.345681 | 0.433803 | 1 | 0.996511 | 0.879859 |
| Abca3 | Q8R420 | 0.14 | 0.08 | 0.65 | 0.82 | 0.87 | 0.68 | 0.92 | 0.7 | 0.23 | 0.52 | 0.77 | 0.61 | 0.02 | 0.51 | 0.79 | 0.93 | 0.37 | 0.11 | 0.14 | 0.1173 | 0.652035 | 0.630686 | 1 | 0.996511 | 0.941132 |
| Abca5 | Q8K448 | -0.15 | 0.32 | 0.44 | 0.14 | 0.63 | -0.22 | 0.21 | 0.08 | -0.39 | -0.02 | 0.18 | -0.3 | -0.65 | 0.49 | -0.04 | 0.13 | -0.0125 | -0.32 | -0.205 | 0.957614 | 0.171323 | 0.482848 | 1 | 0.996511 | 0.893925 |
| Abca6 | Q8K441 | -1.67 | -0.72 | -1.86 | -2.84 | -0.45 | -1.5 | -3.14 | -3.45 | -2.55 | -2.25 | -2.53 | -2.57 | -2.57 | -0.75 | -2.9 | -2.78 | -0.3625 | -0.7025 | -0.4775 | 0.630643 | 0.12103 | 0.441599 | 1 | 0.996511 | 0.883549 |
| Abca7 | E9Q6G4 | 1.52 | 2.36 | 1.9 | 2.17 | -1.69 | -1.92 | -4 | -4.2 | 1.38 | 2.95 | 1.61 | 1.4 | 1.01 | 2.03 | 1.39 | 1.71 | -4.94 | -0.1525 | -0.4525 | 3.66E-05 | 0.70331 | 0.154826 | 0.0835 | 0.996511 | 0.750763 |
| Abca8b | Q8K440-2 | 1.08 | 1.11 | 0.82 | 0.81 | 0.87 | 0.8 | 1.65 | 0.79 | 1 | 1.11 | 1.03 | 0.93 | 0.26 | 0.78 | 0.84 | 1.79 | 0.0725 | 0.0625 | -0.0375 | 0.762402 | 0.712237 | 0.90992 | 1 | 0.996511 | 0.986458 |
| Abca9 | Q8K449 | -1.25 | -1.14 | -2.01 | -0.85 | -1.4 | -0.94 | -2.13 | -1.58 | -0.96 | -1.36 | -1.36 | -1.19 | -1.37 | -0.89 | -2.17 | -2.18 | -0.2 | 0.095 | -0.34 | 0.557543 | 0.733031 | 0.393675 | 1 | 0.996511 | 0.865775 |
| Abcb10 | Q9JI39 | -0.3 | -0.08 | 0.06 | 0.27 | 0.26 | -0.23 | 0.29 | 0.22 | -0.19 | 0.39 | 0.46 | -0.51 | -0.32 | 0.05 | -0.27 | -0.1 | 0.1475 | 0.05 | -0.1475 | 0.474105 | 0.856228 | 0.482929 | 1 | 0.996511 | 0.893925 |
| Abcb11 | J3QNY6 | -2.29 | -2.83 | -1.49 | -0.51 | -2.93 | -1.23 | -1.59 | -0.18 | -1.9 | -2.98 | -1.44 | -3.5 | -0.85 | -2.01 | -2.38 | -2.06 | 0.2975 | -0.675 | -0.045 | 0.667241 | 0.302829 | 0.935744 | 1 | 0.996511 | 0.989767 |
| Abcb1a | P21447 | 0.04 | 0.09 | 0.4 | 0.46 | 0.14 | -0.07 | 0.15 | -0.11 | 0.05 | 0.29 | 0.59 | 0.15 | 0.3 | -0.64 | 0.23 | 0.33 | -0.22 | 0.0225 | -0.1925 | 0.230529 | 0.912485 | 0.493764 | 1 | 0.996744 | 0.89758 |
| Abcb6 | Q9DC29 | -0.85 | -0.51 | -0.39 | -0.85 | -0.39 | -0.39 | -0.47 | -1.01 | -0.22 | -1.08 | -0.65 | -0.45 | 0.34 | -0.77 | -0.8 | -0.49 | 0.085 | 0.05 | 0.22 | 0.694861 | 0.837296 | 0.47462 | 1 | 0.996511 | 0.891711 |
| Abcb7 | Q61102 | 0.12 | 0.18 | 0.32 | 0.5 | 0.15 | 0.32 | 0.24 | 0.17 | 0.36 | -0.03 | 0.35 | -0.09 | 0.32 | -0.07 | -0.05 | 0.28 | -0.06 | -0.1325 | -0.16 | 0.70479 | 0.508988 | 0.433692 | 1 | 0.996511 | 0.879859 |
| Abcb8 | Q9CXJ4 | -0.03 | 0.12 | -0.21 | 0.05 | -0.25 | -0.1 | 0 | -0.17 | 0.29 | -0.38 | -0.04 | -0.11 | 0.34 | 0.38 | -0.16 | 0.06 | -0.1125 | -0.0425 | 0.1725 | 0.477423 | 0.833998 | 0.413217 | 1 | 0.996511 | 0.874811 |
| Abcb9 | Q9JJ59 | -0.13 | 0.28 | 0.23 | 0.28 | 0.26 | 0.1 | 0.45 | 0.2 | -0.36 | -0.1 | 0.31 | -0.03 | -0.05 | 0.04 | 0.16 | 0.57 | 0.0875 | -0.21 | 0.015 | 0.616427 | 0.335487 | 0.945592 | 1 | 0.996511 | 0.992376 |
| Abcc1 | O35379 | 0.41 | 1.08 | 0.31 | 0.57 | 1.14 | 0.99 | 0.5 | 0.7 | 0.68 | 1.08 | 0.49 | 0.51 | 0.76 | 1.14 | 0.29 | 0.37 | 0.24 | 0.0975 | 0.0475 | 0.330246 | 0.691355 | 0.865441 | 1 | 0.996511 | 0.977956 |
| Abcc4 | E9Q236 | 0.26 | 0.05 | 0.76 | 0.21 | 0.24 | 0.41 | 1.33 | 0.37 | -0.2 | -0.27 | 0.94 | 0.45 | -0.41 | -0.08 | 0.98 | 0.62 | 0.2675 | -9.00E-02 | -0.0425 | 0.373068 | 0.781788 | 0.903375 | 1 | 0.996511 | 0.985087 |
| Abcc5 | Q9R1X5 | -0.77 | -0.17 | 0.98 | 0.54 | -0.8 | -0.22 | -0.28 | 0.27 | 2.5 | 0.4 | 0.5 | -0.6 | 3.85 | -0.67 | 1.47 | 1.42 | -0.4025 | 0.555 | 1.3725 | 0.347358 | 0.42841 | 0.152588 | 1 | 0.996511 | 0.750422 |
| Abcc8 | B2RUS7 | 0.58 | 0.92 | 0.84 | 0.69 | 1.11 | 0.59 | 0.19 | 0.38 | 0.57 | 0.84 | 0.66 | 0.56 | 0.72 | 0.65 | 0.33 | 0.57 | -0.19 | -0.1 | -0.19 | 0.420819 | 0.568005 | 0.333016 | 1 | 0.996511 | 0.845601 |
| Abcd1 | P48410 | 0.76 | 0.56 | 0.69 | 0.38 | 1.26 | 0.57 | 0.74 | 0.51 | 0.82 | 1.4 | 0.69 | 1.23 | 0.47 | 1.06 | 0.02 | 0.8 | 0.1725 | 0.4375 | -0.01 | 0.4319 | 0.076612 | 0.969961 | 1 | 0.996511 | 0.992767 |
| Abcd2 | Q61285 | -0.17 | 0.59 | 0.97 | 0.8 | 1.04 | 0.53 | 1.33 | 0.89 | 0.52 | 0.88 | 1.16 | 0.51 | 0.71 | 0.81 | 1.01 | 1.12 | 0.4 | 0.22 | 0.365 | 0.204725 | 0.472781 | 0.223677 | 1 | 0.996511 | 0.787447 |
| Abcd3 | P55096 | 0 | 0.33 | 0.65 | 0.51 | 0.61 | 0.33 | 0.96 | 0.47 | 1.09 | 0.93 | 0.37 | 0.36 | 1.33 | 1.05 | 0.84 | 0.6 | 0.22 | 0.315 | 0.5825 | 0.329325 | 0.239478 | 0.039049 | 1 | 0.996511 | 0.658454 |
| Abcd4 | O89016 | -0.53 | -0.42 | 0.65 | 0.19 | 0.28 | -0.04 | 0.26 | 0.39 | -0.1 | -0.2 | 0.38 | 0.03 | -0.36 | 0.28 | 1.05 | 1.74 | 0.25 | 0.055 | 0.705 | 0.400432 | 0.857935 | 0.180965 | 1 | 0.996511 | 0.775542 |
| Abce1 | P61222 | -0.93 | -0.6 | -1.42 | -1.51 | -0.81 | -0.94 | -1.5 | -1.5 | -0.77 | -0.78 | -1.38 | -0.69 | -1.04 | -0.1 | -1.88 | -1.71 | -0.0725 | 0.21 | -0.0675 | 0.798092 | 0.459519 | 0.87643 | 1 | 0.996511 | 0.979626 |
| Abcf1 | Q6P542 | -1.25 | -1.46 | -0.63 | -0.77 | -1.16 | -1.02 | -0.86 | -0.86 | -1.59 | -0.42 | -0.56 | -1.17 | -1.31 | -1.22 | -0.61 | -0.78 | 0.0525 | 0.0925 | 0.0475 | 0.818611 | 0.781256 | 0.865068 | 1 | 0.996511 | 0.977944 |
| Abcf2 | Q99LE6 | 0.27 | -0.27 | -0.52 | -0.61 | -0.17 | -0.12 | -0.67 | -0.48 | -0.23 | -0.02 | -0.73 | 0 | -0.44 | 0.07 | -0.86 | -0.91 | -0.0775 | 0.0375 | -0.2525 | 0.756309 | 0.891337 | 0.425554 | 1 | 0.996511 | 0.877887 |
| Abcf3 | Q8K268 | 0.14 | 0.27 | 0.56 | 0.39 | -0.05 | 0.11 | 0.22 | 0.31 | -0.23 | 0.4 | 0.32 | 0.34 | 0.18 | 0.05 | 0.01 | 0.13 | -0.1925 | -0.1325 | -0.2475 | 0.277157 | 0.538568 | 0.200084 | 1 | 0.996511 | 0.78004 |
| Abcg1 | Q64343 | 0.46 | 0.51 | 0.7 | 0.46 | 0.29 | 0.81 | -0.07 | 0.43 | 1.36 | -0.97 | 0.03 | 0.2 | 1.9 | -0.49 | 0.7 | 0.52 | -0.1675 | -0.3775 | 0.125 | 0.446062 | 0.413367 | 0.787982 | 1 | 0.996511 | 0.968587 |
| Abcg2 | S4R2E1 | 0.24 | 0.73 | 0.62 | 0.44 | 0.71 | 0.37 | 0.5 | 0.3 | 0.39 | 0.32 | 0.77 | 0.58 | 0.21 | 0.71 | 0.39 | 0.58 | -0.0375 | 0.0075 | -0.035 | 0.837804 | 0.969796 | 0.868176 | 1 | 0.996811 | 0.978872 |
| Abcg4 | Q91WA9 | 1.01 | 0.69 | 0.39 | 1.22 | 0.7 | 0.76 | 0.35 | 1.23 | 2.37 | 2.58 | 1 | 1.48 | 1.95 | 2.87 | 1.59 | 1.45 | -0.0675 | 1.03 | 1.1375 | 0.798525 | 0.027579 | 0.011431 | 1 | 0.996511 | 0.566767 |
| Abhd10 | Q6PE15-2 | 0.49 | 0.31 | -0.01 | 0.16 | -0.05 | 0.24 | 0.08 | 0.07 | 0.25 | 0.28 | -0.15 | 0.21 | 0.19 | 0.73 | -0.54 | -0.51 | -0.1525 | -0.09 | -0.27 | 0.38821 | 0.650511 | 0.418699 | 1 | 0.996511 | 0.87558 |
| Abhd11 | Q8K4F5 | 0.4 | 0.65 | 1.37 | 1.85 | 0.24 | 0.33 | 0.6 | 1.26 | 0.45 | 0.85 | 1.16 | 0.27 | 0.18 | 0.02 | 0.89 | 0.89 | -0.46 | -0.385 | -0.5725 | 0.250164 | 0.31949 | 0.168673 | 1 | 0.996511 | 0.759484 |
| Abhd12 | Q99LR1 | 0.25 | -0.17 | -0.85 | -0.78 | 0.18 | 0.29 | -0.64 | -0.64 | -0.16 | -0.5 | -0.86 | 0.4 | 0.05 | -0.6 | 0.09 | -0.79 | 0.185 | 0.1075 | 0.075 | 0.6003 | 0.76815 | 0.827472 | 1 | 0.996511 | 0.973757 |
| Abhd14b | E9QN99 | 0.19 | -0.55 | -1.45 | -1.2 | -0.69 | 0.15 | -1.07 | -1.49 | -1 | -0.98 | -1.76 | -0.52 | -0.84 | -0.32 | -1.6 | -1.47 | -0.0225 | -0.3125 | -0.305 | 0.961743 | 0.468764 | 0.500371 | 1 | 0.996511 | 0.900792 |
| Abhd16a | Q9Z1Q2 | -0.16 | -0.13 | 0.28 | 0.2 | 0.26 | -0.18 | 0.49 | 0.33 | 0.14 | 0.06 | 0.26 | -0.31 | 0.75 | -0.14 | 0.42 | 0.29 | 0.1775 | -0.01 | 0.2825 | 0.408261 | 0.961942 | 0.275658 | 1 | 0.996755 | 0.822084 |
| Abhd17a | Q99JW1 | -0.16 | -0.27 | 0.64 | 0.48 | -0.26 | 0.33 | 0.05 | 0.39 | 2.17 | 0.07 | 0.52 | -0.04 | 3.15 | -1.72 | 0.64 | 0.26 | -0.045 | 0.5075 | 0.41 | 0.870363 | 0.341525 | 0.65736 | 1 | 0.996511 | 0.948659 |
| Abhd17b | Q7M759 | -0.02 | 0.42 | -0.1 | -0.13 | 0.63 | 0.4 | 0.57 | 0.18 | 0.26 | -1.21 | -0.08 | 0.52 | 0.17 | 0.71 | 0.02 | 0.29 | 0.4025 | -0.17 | 0.255 | 0.067753 | 0.661708 | 0.297392 | 1 | 0.996511 | 0.826603 |
| Abhd17c | Q8VCV1 | 0.47 | 0.85 | 0.65 | 1.18 | 0.77 | 0.6 | 0.81 | 1.03 | 1.19 | 2.65 | 0.76 | 0.62 | 0.39 | 1.56 | 0.47 | 0.84 | 0.015 | 0.5175 | 0.0275 | 0.941885 | 0.277226 | 0.930426 | 1 | 0.996511 | 0.989713 |
| Abhd3 | Q91ZH7 | -0.45 | -1.13 | -1.37 | -1.13 | -0.85 | 0.14 | -0.43 | -0.99 | -0.17 | -0.92 | -1.17 | 0.14 | 0.08 | -0.71 | -0.82 | -0.66 | 0.4875 | 0.49 | 0.4925 | 0.149241 | 0.194349 | 0.126311 | 1 | 0.996511 | 0.721505 |
| Abhd4 | Q3U7M5 | 0.34 | 0.31 | 0.27 | 0.77 | 0.46 | 0.45 | 1.03 | 0.79 | 0.55 | 0.47 | 0.77 | 0.38 | 0.06 | 0.64 | 0.15 | 0.49 | 0.26 | 0.12 | -0.0875 | 0.237077 | 0.544585 | 0.700784 | 1 | 0.996511 | 0.956416 |
| Abhd5 | Q9DBL9 | 0 | 0.19 | -0.18 | 0.44 | -0.33 | 0.13 | 0.4 | 0.16 | 0.49 | 0.43 | -0.04 | 0.08 | 0.01 | -0.24 | -0.45 | 0.46 | -0.0225 | 0.1275 | -0.1675 | 0.919933 | 0.569374 | 0.530186 | 1 | 0.996511 | 0.908663 |
| Abhd6 | Q8R2Y0-2 | 1.08 | 0.2 | 0.6 | 1.3 | 0.48 | 0.95 | 0.92 | 1.47 | 1.08 | 0.81 | 1.02 | 1.41 | 0.52 | 1.09 | 0.96 | 0.77 | 0.16 | 0.285 | 0.04 | 0.612144 | 0.333421 | 0.890908 | 1 | 0.996511 | 0.98183 |
| Abi1 | B7ZCU3 | 1.38 | 1.55 | 1.29 | 1.26 | 1.56 | 1.13 | 1.07 | 1.29 | -0.92 | -1.4 | 1.71 | 1.5 | -0.34 | -1.63 | 0.95 | 1.86 | -0.1075 | -1.1475 | -1.16 | 0.545031 | 0.144223 | 0.121712 | 1 | 0.996511 | 0.721448 |
| Abi1 | B7ZCU4 | 1.33 | 1.04 | 1.32 | 1.13 | 1 | 1.35 | 0.68 | 1.15 | 0.99 | 0.71 | 1.18 | 1.26 | 0.69 | 1.66 | 0.98 | 0.65 | -0.16 | -0.17 | -0.21 | 0.419922 | 0.39384 | 0.443498 | 1 | 0.996511 | 0.883549 |
| Abi2 | A0A087WF | 0.98 | 0.44 | 0.66 | 0.71 | 0.21 | 1.14 | 0.34 | 0.33 | 1.7 | -0.38 | 0.56 | 1.04 | 1.86 | -0.12 | 0.44 | 0.11 | -0.1925 | 0.0325 | -0.125 | 0.453031 | 0.939334 | 0.774206 | 1 | 0.996755 | 0.966062 |
| Abi2 | A0A087WF | -4.4 | -3.92 | -4.41 | -1.45 | -5.51 | -3.5 | -4.2 | -2.3 | -0.54 | -3.09 | -2.61 | -3.7 | 1.13 | -7.74 | -2.33 | -3.58 | -0.3325 | 1.06 | 0.415 | 0.705643 | 0.252325 | 0.81169 | 1 | 0.996511 | 0.972642 |
| Abi3 | Q8BYZ1 | -0.94 | -1.6 | -0.79 | -0.66 | -1.08 | -1.22 | -0.93 | -0.78 | -1.65 | -0.51 | -0.59 | -1.34 | -1.7 | -1.56 | -1.53 | -1.56 | -0.005 | -0.025 | -0.59 | 0.983612 | 0.942009 | 0.038354 | 1 | 0.996755 | 0.658454 |
| Abl1 | P00520-2 | -0.21 | 0.16 | 0.51 | 0.59 | -0.7 | 0.13 | 0.15 | 0.47 | 1.07 | 0.32 | 0.58 | 0.12 | 1.9 | -0.05 | 0.94 | 0.54 | -0.25 | 0.26 | 0.57 | 0.42319 | 0.374248 | 0.205353 | 1 | 0.996511 | 0.783452 |
| Abl2 | F8VQH0 | -0.02 | 0.17 | 0.14 | -0.03 | -0.09 | 0.17 | -0.16 | 0.05 | 0.69 | 0.04 | 0.19 | 0.43 | 0.77 | 0.03 | 0.29 | -0.07 | -0.0725 | 0.2725 | 0.19 | 0.645381 | 0.197936 | 0.429471 | 1 | 0.996511 | 0.879165 |
| Ablim1 | E9Q9C0 | 0.3 | 0.34 | 0.87 | 0.96 | 0.12 | 0.38 | 0.73 | 0.89 | 0.64 | 1.23 | 0.89 | 0.46 | 0.92 | 0.46 | 0.68 | 0.89 | -0.0875 | 0.1875 | 0.12 | 0.732362 | 0.477115 | 0.621452 | 1 | 0.996511 | 0.938628 |
| Ablim2 | Q8BL65 | 0.78 | 0.63 | 0.51 | 0.78 | 0.4 | 0.75 | 0.25 | 0.55 | 0.48 | 0.7 | 0.5 | 0.6 | 0.58 | -0.06 | 0.33 | 0.02 | -0.1875 | -0.105 | -0.4575 | 0.298951 | 0.532432 | 0.058729 | 1 | 0.996511 | 0.661738 |
| Ablim2 | E9Q0W6 | 1.58 | 1.23 | 1.48 | 1.68 | 1.03 | 0.47 | 1.17 | 0.97 | 0.67 | 0.99 | 1.03 | 1.19 | 0.51 | 1.07 | 1.59 | 1.34 | -0.5825 | -0.5225 | -0.365 | 0.020212 | 0.025132 | 0.205306 | 1 | 0.996511 | 0.783452 |
| Ablim2 | Q8BL65-4 | 1.98 | 2.76 | 1.1 | 1.64 | 1.28 | 0.25 | 0.7 | 0.49 | 1.72 | 0.08 | 0.95 | 1.99 | 1.38 | 1.38 | 1.38 | 1.57 | -1.19 | -0.685 | -0.4425 | 0.013294 | 0.204486 | 0.225849 | 1 | 0.996511 | 0.787447 |
| Ablim3 | Q69ZX8 | 0.7 | 0.37 | 0.36 | 0.28 | 0.03 | 0.72 | 0.21 | -0.14 | 0.76 | 0.23 | 0.53 | 1.05 | 1.32 | -0.21 | 0.24 | 0.55 | -0.2225 | 0.215 | 0.0475 | 0.342948 | 0.361483 | 0.887717 | 1 | 0.996511 | 0.980447 |
| Abr | A0A571BG | -0.36 | -0.47 | 0.42 | 0.88 | -1.52 | -0.55 | -0.21 | 0.89 | 1.42 | 2.7 | 0.07 | 0.45 | 0.84 | 0.08 | 1.23 | 0.94 | -0.465 | 1.0425 | 0.655 | 0.402453 | 0.116671 | 0.121966 | 1 | 0.996511 | 0.721448 |
| Abr | Q5SSL4 | 0.37 | 0.31 | 0.03 | -0.02 | 0.39 | 0.62 | 0.2 | 0.09 | 0 | 0.04 | 0.22 | 0.59 | -0.15 | 0.04 | -0.07 | -0.08 | 0.1525 | 0.04 | -0.2375 | 0.431996 | 0.84875 | 0.225111 | 1 | 0.996511 | 0.787447 |
| Abr | Q5SSL4-4 | 0.35 | -0.28 | 0.07 | -0.16 | 0.51 | 0.52 | 0.22 | 1.21 | 0.16 | 0.43 | -0.53 | -0.88 | 0.15 | 1.02 | -0.17 | -0.03 | 0.62 | -0.2 | 0.2475 | 0.039091 | 0.549283 | 0.434186 | 1 | 0.996511 | 0.880349 |
| Abracl | A0A1W2P7 | -3.53 | -4.09 | -2.88 | -2.86 | -3.6 | -3.65 | -2.19 | -2.64 | -5.58 | -3.02 | -3.14 | -3.15 | -4.85 | -5.63 | -3.1 | -2.43 | 0.32 | -0.3825 | -0.6625 | 0.469441 | 0.546419 | 0.372605 | 1 | 0.996511 | 0.859424 |
| Abraxas2 | Z4YJY0 | 0.27 | 0.03 | -0.31 | -0.06 | -0.39 | 0.21 | -1.03 | -0.22 | 0.14 | 0.25 | 0.06 | 0.09 | 0.36 | -0.03 | -0.36 | -0.56 | -0.34 | 0.1525 | -0.13 | 0.253126 | 0.423368 | 0.622468 | 1 | 0.996511 | 0.938628 |
| Abtb2 | Q7TQI7 | -0.15 | 0.47 | 0.29 | 0.81 | 0.21 | 0.36 | 0.47 | 0.3 | 0.45 | 0.36 | 0.17 | 0.16 | 0.14 | 0.42 | 0.13 | 0.93 | -0.02 | -0.07 | 0.05 | 0.929915 | 0.770851 | 0.863346 | 1 | 0.996511 | 0.977235 |
| Acaa1a | Q921H8 | 0.43 | 0.44 | 0.61 | 0.72 | 0.54 | 0.2 | 1.04 | 0.78 | 0.71 | 1.25 | 0.71 | 0.66 | 0.64 | 1.23 | 0.91 | 0.98 | 0.09 | 0.2825 | 0.39 | 0.679475 | 0.188971 | 0.082536 | 1 | 0.996511 | 0.68865 |
| Acaa2 | Q8BWT1 | 1.07 | 0.89 | 0.39 | 0.8 | 0.72 | 0.91 | 0.88 | 0.5 | 0.41 | 0.24 | 0.5 | 0.74 | 0.28 | 0.28 | 0.23 | 0.45 | -0.035 | -0.315 | -0.4775 | 0.86323 | 0.171237 | 0.046059 | 1 | 0.996511 | 0.658454 |
| Acaca | Q5SWU9 | 0.27 | 0.25 | 0.5 | 0.49 | 0.58 | 0.43 | 0.35 | 0.48 | 0.4 | 0.51 | 0.64 | 0.35 | 0.13 | 0.79 | 0.34 | 0.32 | 0.0825 | 0.0975 | 0.0175 | 0.594155 | 0.571513 | 0.934217 | 1 | 0.996511 | 0.989767 |
| Acad11 | A0A0R4J0I | -0.32 | 0.24 | 0.14 | 0.3 | 0.55 | 0.3 | 0.83 | 0.23 | 1.09 | 0.55 | 0.27 | 0.55 | 1.46 | 0.41 | 0.43 | 0.32 | 0.3875 | 0.525 | 0.565 | 0.105112 | 0.058482 | 0.098148 | 1 | 0.996511 | 0.698156 |
| Acad12 | D3Z7X0 | 0.22 | 0.5 | 0.59 | 1.2 | 0.4 | -0.01 | 1.87 | 0.68 | -0.07 | 0.15 | 1.33 | -0.26 | 0.2 | 0.2 | 0.01 | 0.87 | 0.1075 | -0.34 | -0.3075 | 0.800425 | 0.400215 | 0.312098 | 1 | 0.996511 | 0.834523 |
| Acad8 | D3YTT4 | -0.16 | -0.02 | 0.02 | 0.34 | -0.12 | -0.07 | 0.22 | 0.14 | -0.29 | 0.01 | 0.33 | -0.45 | -0.44 | 0.32 | -0.1 | -0.29 | -0.0025 | -0.145 | -0.1725 | 0.988859 | 0.537953 | 0.472496 | 1 | 0.996511 | 0.891144 |
| Acad9 | Q8JZN5 | 0.1 | 0.36 | 0.3 | 0.37 | -0.01 | 0.35 | 0.28 | 0.29 | 0.32 | -0.03 | 0.49 | 0.11 | 0.45 | 0.16 | 0.03 | 0.07 | -0.055 | -0.06 | -0.105 | 0.735343 | 0.751136 | 0.584796 | 1 | 0.996511 | 0.927734 |
| Acadl | A0A0R4J08 | 0.23 | -0.07 | 0.5 | 1.03 | 0.27 | 0.26 | 1.06 | 0.78 | 0.03 | 0.34 | 0.83 | 0.07 | -0.02 | -0.18 | 0.44 | 0.8 | 0.17 | -0.105 | -0.1625 | 0.578016 | 0.729321 | 0.621668 | 1 | 0.996511 | 0.938628 |
| Acadm | P45952 | 0.9 | 0.69 | 0.39 | 0.68 | 0.72 | 0.68 | 0.88 | 0.54 | 0.87 | 0.02 | 0.67 | 0.71 | 0.48 | 0.89 | 0.32 | 0.57 | 0.04 | -0.0975 | -0.1 | 0.819192 | 0.687806 | 0.642766 | 1 | 0.996511 | 0.94365 |
| Acads | Q07417 | 1.06 | 0.82 | 0.97 | 1.44 | 0.54 | 0.94 | 1.62 | 1.62 | 0.61 | 1.1 | 1.62 | 1.02 | 0.4 | -0.25 | 0.93 | 1.04 | 0.1075 | 0.015 | -0.5425 | 0.717664 | 0.954612 | 0.124694 | 1 | 0.996755 | 0.721448 |
| Acadsb | Q9DBL1 | 0.1 | -0.1 | 0.1 | 0.46 | -0.29 | -0.16 | 0.59 | 0.08 | -0.34 | 0.3 | 0.37 | -0.07 | -0.14 | -0.06 | 0.06 | 0.41 | -0.085 | -0.075 | -0.0725 | 0.725649 | 0.748493 | 0.741873 | 1 | 0.996511 | 0.958932 |
| Acadvl | P50544 | 0.78 | 0.81 | 0.2 | 0.26 | 0.71 | 0.68 | 0.54 | 0.17 | 0.39 | -0.05 | 0.48 | 0.6 | 0.25 | 0.31 | 0.09 | 0.22 | 0.0125 | -0.1575 | -0.295 | 0.955899 | 0.52163 | 0.204035 | 1 | 0.996511 | 0.782817 |
| Acan | Q61282 | -0.03 | -0.02 | -1.03 | -0.83 | -0.56 | -0.2 | -1.04 | -0.91 | -0.18 | -0.65 | -1.04 | 0 | -0.22 | -0.23 | -0.98 | -1.07 | -0.2 | 0.01 | -0.1475 | 0.534219 | 0.976998 | 0.67427 | 1 | 0.997808 | 0.952944 |
| Acap2 | D4AFX6 | -1.12 | -0.29 | -0.12 | -0.27 | -1.75 | -0.09 | -0.84 | -0.44 | -1.13 | -1.55 | -0.3 | -0.68 | 0.62 | -0.4 | 0.94 | -0.12 | -0.33 | -0.465 | 0.71 | 0.418185 | 0.203393 | 0.086688 | 1 | 0.996511 | 0.691619 |
| Acap2 | A0A338P6I | 1.04 | 0.96 | 0.53 | 0.33 | 1.2 | 0.79 | 1.1 | 0.52 | 0.28 | 0.07 | 0.6 | 1.15 | 0.14 | 0.66 | 0.4 | 0.8 | 0.1875 | -0.19 | -0.215 | 0.45034 | 0.527398 | 0.409539 | 1 | 0.996511 | 0.87457 |
| Acap3 | A2ADA6 | -0.45 | -0.33 | 0.01 | 0.13 | -0.35 | -0.35 | -0.21 | -0.17 | -0.16 | 0.43 | -0.12 | -0.17 | 0.27 | -0.35 | -0.01 | 0.18 | -0.11 | 0.155 | 0.1825 | 0.558414 | 0.508571 | 0.446278 | 1 | 0.996511 | 0.884817 |
| Acat1 | Q8QZT1 | 1.11 | 0.59 | 0.93 | 0.75 | 0.27 | 1.02 | 1.07 | 1.14 | 0.69 | 0.25 | 1.27 | 1.2 | 0.74 | 0.43 | 0.94 | 0.97 | 0.03 | 0.0075 | -0.075 | 0.902775 | 0.978383 | 0.73286 | 1 | 0.998749 | 0.958635 |
| Acat2 | Q8CAY6 | 0.09 | 0.12 | 0.33 | 0.45 | 0.16 | 0 | 0.79 | 0.67 | -0.03 | 0.33 | 0.58 | 0.34 | -0.19 | 0.19 | 0.53 | 0.53 | 0.1575 | 0.0575 | 0.0175 | 0.499187 | 0.775297 | 0.940357 | 1 | 0.996511 | 0.990894 |
| Acbd3 | A0A0R4J07 | -0.81 | -0.71 | -1.12 | -0.96 | -0.36 | -1.02 | -0.63 | -0.94 | -0.75 | -0.6 | -0.93 | -0.52 | -0.93 | -0.63 | -0.73 | -1.09 | 0.1625 | 0.2 | 0.055 | 0.438376 | 0.300899 | 0.785355 | 1 | 0.996511 | 0.967642 |
| Acbd5 | A0A2I3BRB | -0.21 | 0.4 | 0.4 | 0.29 | 0.1 | 0.25 | 1.01 | 0.27 | 1.02 | 0.14 | 0.69 | 0.32 | 1.47 | 0.59 | 1.11 | 0.79 | 0.1875 | 0.3225 | 0.77 | 0.477346 | 0.239877 | 0.017171 | 1 | 0.996511 | 0.602211 |
| Acbd6 | Q9D061 | -0.71 | -0.33 | 0.1 | -0.1 | -0.49 | -0.87 | 0.04 | -0.06 | -1.02 | -0.2 | -0.04 | -0.66 | -0.85 | -0.31 | -0.19 | -0.18 | -0.085 | -0.22 | -0.1225 | 0.758717 | 0.457562 | 0.643245 | 1 | 0.996511 | 0.943874 |
| Ace | P09470 | 0.12 | 0.43 | 0.48 | 0.12 | 0.11 | -0.31 | 0.59 | -0.15 | -1.01 | -0.68 | -0.08 | 0.12 | -0.53 | -0.69 | 0.65 | 0.47 | -0.2275 | -0.7 | -0.3125 | 0.349125 | 0.03627 | 0.38561 | 1 | 0.996511 | 0.861533 |
| Ace2 | Q8R0I0 | -0.4 | 0.17 | 0.06 | -0.23 | 0.04 | -0.22 | 0.4 | -0.39 | 0.66 | -0.22 | 0.31 | -0.01 | 1.34 | 0.23 | 0.05 | 0.38 | 0.0575 | 0.285 | 0.6 | 0.806029 | 0.279772 | 0.089788 | 1 | 0.996511 | 0.69217 |
| Ache | P21836 | 1.34 | 0.9 | 0.47 | 1.25 | 0.73 | 0.92 | 1.06 | 0.94 | 0.91 | 1.28 | 0.36 | 1.26 | 0.54 | -0.32 | 1.17 | 0.85 | -0.0775 | -0.0375 | -0.43 | 0.735606 | 0.899935 | 0.262646 | 1 | 0.996511 | 0.81524 |
| Acin1 | Q9JIX8-4 | 0.78 | -0.23 | -0.58 | -0.59 | -1.09 | 0.51 | -0.91 | 0.05 | 0.46 | -0.34 | -0.57 | -0.86 | 1.37 | -0.52 | -0.33 | -1.15 | -0.205 | -0.1725 | -0.0025 | 0.660844 | 0.674672 | 0.99654 | 1 | 0.996511 | 0.999143 |
| Acly | Q91V92 | 0.16 | -0.27 | 0.11 | 0.9 | -0.21 | -0.61 | -0.87 | -0.14 | 0.62 | 1.81 | -0.62 | -0.12 | 1.75 | 0.3 | 0.46 | 0.06 | -0.6825 | 0.1975 | 0.4175 | 0.044984 | 0.714288 | 0.343827 | 1 | 0.996511 | 0.848473 |
| Acly | Q3V117 | 0.64 | 0.23 | 0.39 | 0.34 | 0.11 | 0.52 | 0.17 | 0.49 | 0.13 | 0.16 | 0.4 | 0.63 | 0.08 | 0.07 | 0.36 | 0.39 | -0.0775 | -0.07 | -0.175 | 0.670605 | 0.723265 | 0.380336 | 1 | 0.996511 | 0.860984 |
| Aco1 | P28271 | 0.67 | 0.5 | 0.1 | 0.15 | 0.57 | 0.65 | 0.26 | 0.27 | 0.73 | 0.07 | 0.15 | 0.69 | 0.4 | 0.52 | -0.06 | -0.05 | 0.0825 | 0.055 | -0.1525 | 0.684876 | 0.823633 | 0.533125 | 1 | 0.996511 | 0.911443 |
| Aco2 | Q99KI0 | 0.8 | 0.75 | 0.67 | 0.91 | 0.44 | 0.61 | 0.8 | 0.95 | 0.37 | 0.68 | 0.84 | 0.6 | 0.11 | 0.65 | 0.41 | 0.5 | -0.0825 | -0.16 | -0.365 | 0.634854 | 0.380631 | 0.089319 | 1 | 0.996511 | 0.69217 |
| Acot1 | O55137 | 0.24 | -0.4 | 2.28 | 3.16 | -0.36 | 0.25 | 2.68 | 3.63 | -0.53 | 2.85 | 2.09 | -0.21 | -0.87 | 0.04 | 1.68 | 2.57 | 0.23 | -0.27 | -0.465 | 0.839816 | 0.799668 | 0.651006 | 1 | 0.996511 | 0.947407 |
| Acot11 | Q8VHQ9 | -0.46 | -0.36 | -0.44 | -0.37 | -0.33 | 0.33 | -0.49 | -0.46 | 0.41 | -0.5 | -0.38 | -0.32 | 1.32 | -0.55 | -0.26 | -0.21 | 0.17 | 0.21 | 0.4825 | 0.444175 | 0.385747 | 0.254201 | 1 | 0.996511 | 0.809524 |
| Acot13 | Q9CQR4 | 0.68 | 1.07 | 0.53 | 1.4 | 0.38 | 0.44 | 0.5 | 1.79 | 1.07 | 0.84 | 1.17 | 0.24 | 0.87 | 1.03 | 0.13 | -0.54 | -0.1425 | -0.09 | -0.5475 | 0.703077 | 0.760002 | 0.191996 | 1 | 0.996511 | 0.777117 |
| Acot2 | Q9QYR9 | 0.09 | 0.55 | 0.37 | 0.47 | 0.56 | 0.27 | 0.85 | 0.56 | 0.15 | 0.04 | 0.48 | -0.08 | -0.29 | 0.39 | 0.1 | 0.25 | 0.19 | -0.2225 | -0.2575 | 0.33738 | 0.29164 | 0.271058 | 1 | 0.996511 | 0.822084 |
| Acot7 | Q91V12-2 | -0.24 | -0.23 | -0.38 | -0.79 | -0.13 | -0.04 | -0.18 | -1.02 | 0.35 | -0.8 | -0.37 | 0.15 | 0.24 | 0.03 | -0.68 | -0.85 | 0.0675 | 0.2425 | 0.095 | 0.801986 | 0.424351 | 0.757396 | 1 | 0.996511 | 0.962481 |
| Acot8 | P58137 | -0.54 | -0.22 | -0.19 | -0.11 | 0.25 | -0.28 | 0.33 | -0.25 | -0.6 | 0 | 0.01 | -0.39 | -0.28 | 0.37 | 0.1 | 0.05 | 0.2775 | 0.02 | 0.325 | 0.215562 | 0.926456 | 0.159283 | 1 | 0.996755 | 0.754511 |
| Acot9 | Q9R0X4 | 0.72 | 0.39 | 1.06 | 1.12 | 0.48 | 1.17 | 0.8 | 1.38 | -0.22 | 0.38 | 1.16 | 0.62 | -0.5 | 0.06 | 0.56 | 0.63 | 0.135 | -0.3375 | -0.635 | 0.61657 | 0.32151 | 0.072749 | 1 | 0.996511 | 0.668066 |
| Acox1 | Q9R0H0-2 | -0.35 | 0.19 | -0.03 | -0.23 | 0.83 | 0.08 | 1.28 | -0.05 | 0.43 | 0.35 | 0.46 | 0.46 | 0.26 | 0.92 | 0.52 | 0.36 | 0.64 | 0.53 | 0.62 | 0.078029 | 0.017299 | 0.023717 | 1 | 0.996511 | 0.624734 |
| Acox3 | Q9EPL9 | 0.25 | 0.76 | 0.47 | 0.17 | 0.94 | 0.73 | 1.13 | 2.11 | 0.24 | 0.44 | 1.9 | 0.47 | 0.48 | 0.75 | 0.6 | 2.93 | 0.815 | 0.35 | 0.7775 | 0.031622 | 0.379471 | 0.182777 | 1 | 0.996511 | 0.776002 |
| Acp1 | Q561M1 | 2.01 | 0.09 | 1.11 | -0.39 | 0.87 | 2.23 | 2.59 | 1.82 | -2.93 | -1.97 | 1.2 | 1.56 | -2.81 | -2.37 | 2.08 | 1.69 | 1.1725 | -1.24 | -1.0575 | 0.075721 | 0.284584 | 0.407303 | 1 | 0.996511 | 0.874084 |
| Acp1 | Q9D358 | 0.44 | 0.03 | 0.65 | 1.54 | -0.14 | 0.66 | 1.1 | 1.48 | 0.64 | 0.86 | 0.68 | 0.53 | 0.88 | 0.09 | 0.48 | 0.59 | 0.11 | 0.0125 | -0.155 | 0.802887 | 0.969301 | 0.663616 | 1 | 0.996755 | 0.949825 |
| Acp2 | A0A5F8MP | -1.03 | -0.55 | -0.72 | -0.84 | -0.32 | -0.61 | -0.42 | -1.13 | -0.94 | -0.54 | -0.8 | -0.62 | -0.98 | 0.02 | -1.33 | -1.13 | 0.165 | 0.06 | -0.07 | 0.474615 | 0.754321 | 0.82826 | 1 | 0.996511 | 0.973757 |
| Acp6 | Q8BP40 | 0.03 | 0.13 | -0.08 | 0.4 | 0.05 | -0.19 | 0.16 | 0.09 | -0.11 | 0.5 | 0.02 | -0.27 | -0.44 | 0.51 | -0.44 | -0.77 | -0.0925 | -0.085 | -0.405 | 0.601802 | 0.710498 | 0.207808 | 1 | 0.996511 | 0.785338 |
| Acsbg1 | Q99PU5 | 0.36 | 0.29 | 1.1 | 1.18 | 0.54 | 0.23 | 1.96 | 1.28 | 0.5 | 0.44 | 1.45 | 0.68 | -0.31 | 0.43 | 0.75 | 1.15 | 0.27 | 0.035 | -0.2275 | 0.530252 | 0.915417 | 0.552465 | 1 | 0.996755 | 0.91565 |
| Acsf2 | Q8VCW8 | 1.75 | 0.77 | 1.25 | 1.14 | 0.79 | 1.67 | 1.14 | 0.98 | -0.09 | 0.51 | 1.29 | 1.4 | 0.11 | -0.55 | 1.05 | 1.2 | -0.0825 | -0.45 | -0.775 | 0.768506 | 0.265754 | 0.103001 | 1 | 0.996511 | 0.700625 |
| Acsf3 | Q3URE1 | 0.55 | 0.76 | 1.16 | 1.23 | 0.57 | 0.54 | 0.82 | 1.23 | 0.52 | 0.65 | 1.2 | 0.36 | 0.66 | 0.76 | 1.15 | 1.01 | -0.135 | -0.2425 | -0.03 | 0.580838 | 0.367749 | 0.89948 | 1 | 0.996511 | 0.983509 |
| Acsl1 | D3Z041 | 0.1 | 0.24 | 0.58 | 0.45 | 0.4 | -0.01 | 0.32 | 0.09 | -0.38 | 0.07 | 0.43 | -0.07 | -0.39 | -0.1 | 0.28 | 0.06 | -0.1425 | -0.33 | -0.38 | 0.449822 | 0.177609 | 0.118125 | 1 | 0.996511 | 0.720612 |
| Acsl3 | Q9CZW4 | 0.55 | 0.71 | 0.59 | 0.54 | 0.88 | 0.5 | 1.62 | 0.81 | 1.08 | 0.81 | 0.73 | 0.75 | 0.69 | 0.87 | 0.85 | 0.96 | 0.355 | 0.245 | 0.245 | 0.183283 | 0.172517 | 0.180938 | 1 | 0.996511 | 0.775542 |
| Acsl4 | Q9QUJ7-2 | -0.34 | -0.72 | -0.31 | -0.43 | -0.1 | -0.18 | 0.13 | -0.18 | -1.76 | -0.99 | -0.32 | -0.36 | -1.28 | -1.47 | 0.08 | 0.25 | 0.3675 | -0.4075 | -0.155 | 0.056633 | 0.256426 | 0.722091 | 1 | 0.996511 | 0.958398 |
| Acsl5 | Q8JZR0 | -0.11 | 0.13 | 0.65 | 0.22 | 0.47 | -0.1 | -0.06 | 0.15 | -0.68 | -0.64 | 0.24 | -0.09 | -0.89 | -0.25 | 0.25 | 0.53 | -0.1075 | -0.515 | -0.3125 | 0.63677 | 0.097715 | 0.380349 | 1 | 0.996511 | 0.860984 |
| Acsl6 | Q5ICG5 | -0.8 | -0.6 | -0.41 | -0.34 | -1.17 | 0.19 | -0.73 | -1.37 | 0.48 | 0.49 | -0.33 | 0.18 | 1.13 | -0.8 | 0.23 | -1.02 | -0.2325 | 0.7425 | 0.4225 | 0.509907 | 0.013442 | 0.385458 | 1 | 0.996511 | 0.861533 |
| Acsl6 | Q91WC3 | -0.01 | -0.06 | 0.24 | 0.35 | 0.14 | 0.19 | 0.75 | 0.18 | -0.01 | 0.14 | 0.29 | 0 | -0.29 | 0.05 | 0.03 | 0.36 | 0.185 | -0.025 | -0.0925 | 0.379875 | 0.891717 | 0.672473 | 1 | 0.996511 | 0.952944 |
| Acss1 | Q99NB1 | 0.67 | 0.8 | 0.39 | 0.74 | 0.68 | 0.89 | 1.18 | 0.54 | 0.45 | -0.39 | 1 | 0.63 | 0.16 | 0.02 | 0.14 | 0.31 | 0.1725 | -0.2275 | -0.4925 | 0.397541 | 0.470873 | 0.02629 | 1 | 0.996511 | 0.633075 |

| | | | | | | | | | | | | | | | | | | | | | | | | | | |
|---|---|---|---|---|---|---|---|---|---|---|---|---|---|---|---|---|---|---|---|---|---|---|---|---|---|---|
| Acss2 | A2AQN4 | -0.62 | -0.49 | -0.26 | -0.37 | -0.32 | -0.75 | -0.23 | -0.39 | -1.06 | -0.69 | -0.13 | -0.46 | -0.71 | -0.81 | -0.62 | -0.54 | 0.0125 | -0.15 | -0.235 | 0.945122 | 0.534245 | 0.221207 | 1 | 0.996511 | 0.787447 |
| Actb | P60710 | 0.97 | 0.51 | 0.54 | 0.69 | 0.72 | 0.85 | 0.72 | 1.09 | 0.42 | 0.28 | 0.73 | 0.92 | -0.06 | 0.36 | 0.3 | 0.64 | 0.1675 | -0.09 | -0.3675 | 0.367048 | 0.680847 | 0.130087 | 1 | 0.996511 | 0.723659 |
| Actbl2 | Q8BFZ3 | 0.83 | 0.73 | 1.62 | 0.03 | 2.55 | 1.63 | 2.11 | 1.02 | -1.77 | -0.95 | 1.37 | 1.06 | -1.36 | -0.66 | 2.02 | 2.37 | 1.025 | -0.875 | -0.21 | 0.039658 | 0.264172 | 0.813977 | 1 | 0.996511 | 0.972768 |
| Actc1 | P68033 | -0.01 | 0.19 | -0.11 | 0.31 | 0.26 | -0.03 | -0.01 | 0.24 | 0.57 | 1.21 | 0.19 | -0.02 | 0.59 | 1 | -0.83 | -0.96 | 0.02 | 0.3925 | -0.145 | 0.908028 | 0.204826 | 0.759964 | 1 | 0.996511 | 0.963123 |
| Actg1 | P63260 | -0.11 | 0.39 | 0.02 | 0.21 | 0.64 | -0.45 | 0.93 | 0.21 | 0.7 | 0.66 | -0.12 | -0.36 | -0.47 | 1.07 | 0.25 | 0.55 | 0.205 | 0.0925 | 0.2225 | 0.518438 | 0.756733 | 0.518103 | 1 | 0.996511 | 0.90668 |
| Actl6a | Q9Z2N8 | -0.72 | -1.74 | -1.25 | -0.93 | -1.09 | -1.17 | -1.09 | -0.89 | -0.79 | -0.91 | -1.29 | -1.1 | -1.33 | -0.16 | -1.61 | -1.19 | 0.1 | 0.1375 | 0.0875 | 0.683224 | 0.606514 | 0.81661 | 1 | 0.996511 | 0.972768 |
| Actl6b | Q99MR0 | 0.03 | -1.15 | -1.46 | -0.93 | -1.13 | -0.72 | -1.25 | -1.66 | -0.66 | -0.24 | -1.57 | -0.61 | -0.46 | -0.7 | -1.26 | -1.55 | -0.3125 | 0.1075 | -0.115 | 0.395 | 0.792387 | 0.770698 | 1 | 0.996511 | 0.965933 |
| Actn1 | A1BN54 | 1.93 | 0.83 | 1.14 | 1.03 | 1.01 | 1.64 | 1.01 | 1.12 | 0.38 | 0.36 | 1.11 | 1.19 | 0.12 | 0.74 | 0.62 | 0.79 | -0.0375 | -0.4725 | -0.665 | 0.895527 | 0.175463 | 0.050038 | 1 | 0.996511 | 0.658454 |
| Actn1 | Q7TPR4 | 9.06 | 1.26 | 5.22 | 0.72 | 2.45 | 8.65 | 8.61 | 4.93 | 0.46 | 0.84 | 5.66 | 6.65 | 0.55 | -0.22 | 5.44 | 7.38 | 2.095 | -0.6625 | -0.7775 | 0.352868 | 0.768224 | 0.744064 | 1 | 0.996511 | 0.959371 |
| Actn2 | Q9JI91 | 1.6 | 0.98 | 1.74 | 1.8 | 0.93 | 1.03 | 2.1 | 1.9 | 0.96 | 0.69 | 1.48 | 1.29 | 0.9 | 0.7 | 1.69 | 1.47 | -0.04 | -0.425 | -0.34 | 0.906406 | 0.144634 | 0.288793 | 1 | 0.996511 | 0.824705 |
| Actn4 | A0A1L1SV2 | 1.06 | 0.77 | 0.86 | 0.91 | 1.25 | 1.07 | 0.73 | 0.93 | 0.47 | 0.83 | 1.04 | 0.98 | 0.2 | 1.02 | 0.48 | 0.55 | 0.095 | -0.07 | -0.3375 | 0.589648 | 0.720212 | 0.162564 | 1 | 0.996511 | 0.757516 |
| Actn4 | A0A1L1SVJ | 2.49 | 2.49 | 0.48 | 1.63 | 2.35 | 1.87 | 0.1 | 1.26 | 1.81 | -0.36 | 1 | 2.55 | 2.27 | -3.11 | 1.07 | 0.29 | -0.3775 | -0.5225 | -1.6425 | 0.546541 | 0.470284 | 0.165666 | 1 | 0.996511 | 0.758837 |
| Actn4 | E9Q2W9 | 0.4 | 0.62 | 0.78 | 0.81 | 1.22 | 0.47 | 0.87 | 1.1 | 0.75 | 0.63 | 1.19 | 0.34 | 0.58 | 0.91 | 0.29 | 0.72 | 0.2625 | 0.075 | -0.0275 | 0.243452 | 0.746483 | 0.898268 | 1 | 0.996511 | 0.983509 |
| Actr10 | Q9QZB7 | 0.02 | 0.1 | 0.66 | 0.46 | 0.23 | -0.03 | 0.54 | 0.58 | -0.08 | 0.16 | 0.61 | 0.14 | -0.65 | 0.17 | 0.46 | 0.62 | 0.02 | -0.1025 | -0.16 | 0.930099 | 0.667806 | 0.624884 | 1 | 0.996511 | 0.939167 |
| Actr1a | P61164 | 0.15 | 0.11 | 0.38 | 0.45 | 0.01 | 0.27 | 0.1 | 0.53 | 0.78 | 1.26 | 0.5 | 0.41 | 0.48 | 0.85 | 0.3 | 0.18 | -0.045 | 0.465 | 0.18 | 0.807108 | 0.077168 | 0.421786 | 1 | 0.996511 | 0.875736 |
| Actr1b | Q8R5C5 | 0.7 | 0.01 | 0.04 | 0.49 | 0.23 | 0.53 | 0.35 | 0.48 | 0.33 | 0.44 | 0.16 | 0.77 | -0.09 | 0.02 | -0.13 | 0.17 | 0.0875 | 0.115 | -0.3175 | 0.679327 | 0.634737 | 0.188362 | 1 | 0.996511 | 0.776843 |
| Actr2 | P61161 | 1.01 | 0.46 | 0.35 | 0.48 | 0.74 | 1.03 | 0.64 | 0.7 | 0.11 | 0.26 | 0.56 | 0.75 | -0.43 | 0.4 | 0 | 0.23 | 0.2025 | -0.155 | -0.525 | 0.332419 | 0.516551 | 0.070697 | 1 | 0.996511 | 0.668066 |
| Actr3 | Q99JY9 | 1.32 | 0.48 | 0.37 | 1.05 | 0.64 | 1.22 | 0.6 | 1.3 | 0.22 | -0.49 | 0.71 | 1.09 | -0.42 | 0.01 | 0.26 | 0.25 | 0.135 | -0.4225 | -0.78 | 0.646863 | 0.298812 | 0.024485 | 1 | 0.996511 | 0.628313 |
| Actr3b | Q641P0 | 0.32 | -0.39 | 0.16 | 0.41 | -0.01 | 0.27 | 0.18 | 0.5 | -0.09 | -0.14 | 0.28 | -0.33 | -0.72 | -0.36 | -0.53 | -0.31 | 0.11 | -0.195 | -0.605 | 0.632095 | 0.435224 | 0.031027 | 1 | 0.996511 | 0.641481 |
| Actr6 | A0A0R4J00 | -2.01 | -1.75 | -2.37 | -1.53 | -2.75 | -3.05 | -3.22 | -0.81 | -3.13 | -1.37 | -2.89 | -2.29 | -2.71 | -4.33 | -1.89 | -1.46 | -0.5425 | -0.505 | -0.6825 | 0.327951 | 0.239805 | 0.275602 | 1 | 0.996511 | 0.822084 |
| Acvr1 | P37172 | -0.28 | -0.12 | -1.84 | -0.94 | -0.15 | 0.51 | -1.33 | -1.2 | -0.43 | -0.29 | -0.93 | 0.15 | -0.59 | -0.06 | -0.89 | -0.5 | 0.2525 | 0.42 | 0.285 | 0.641748 | 0.338224 | 0.492821 | 1 | 0.996511 | 0.897531 |
| Acvr1b | Q61271 | 0.62 | 0.36 | 0.46 | 0.85 | 0.96 | 0.94 | 1.12 | 1.11 | -0.28 | 1.8 | 0.55 | 0.4 | -0.09 | 0.52 | 0.61 | 1.16 | 0.46 | 0.045 | -0.0225 | 0.023008 | 0.915281 | 0.93873 | 1 | 0.996755 | 0.989956 |
| Acvr1c | Q8K348 | 0.59 | 0.96 | -0.59 | 1.01 | 0.91 | 0 | -0.15 | 0.61 | 0.46 | 0.19 | -0.07 | -0.32 | -0.41 | 0.91 | 0.04 | 0.79 | -0.15 | -0.4275 | -0.16 | 0.722436 | 0.292573 | 0.728184 | 1 | 0.996511 | 0.958635 |
| Acvr2a | P27038 | 0.07 | 0.18 | 0.84 | 1.12 | 0.08 | 0.42 | 1.21 | 0.94 | -0.55 | 2.12 | 0.79 | 0.35 | -0.5 | -0.84 | 0.61 | 0.99 | 0.11 | 0.125 | -0.4875 | 0.751293 | 0.824333 | 0.319142 | 1 | 0.996511 | 0.838969 |
| Acy1 | A0A0R4J05 | -0.03 | -0.12 | 0.07 | -0.11 | -0.21 | -0.5 | 0.01 | -0.23 | -0.29 | 0.2 | -0.45 | -0.29 | -0.11 | -0.3 | -0.28 | -0.5 | -0.185 | -0.16 | -0.25 | 0.287938 | 0.428822 | 0.190675 | 1 | 0.996511 | 0.776865 |
| Acy3 | Q91XE4 | 0.67 | 0.34 | -1.08 | -1.38 | 0.55 | 0.18 | -0.99 | -1.3 | -0.13 | -0.21 | -1.14 | 0.13 | -0.42 | 0.36 | -1.67 | -0.97 | -0.0275 | 0.025 | -0.3125 | 0.964406 | 0.962744 | 0.61077 | 1 | 0.996755 | 0.935047 |
| Acyp1 | P56376 | -0.76 | -0.42 | -0.14 | -0.34 | -0.53 | -0.61 | -0.58 | -0.96 | -0.8 | -2.38 | -0.46 | -0.71 | -0.76 | -0.7 | -0.88 | -0.58 | -0.255 | -0.6725 | -0.315 | 0.217382 | 0.145244 | 0.151275 | 1 | 0.996511 | 0.749304 |
| Acyp2 | P56375 | 0.21 | -0.16 | -0.18 | -0.62 | -0.73 | 0.19 | -0.65 | -0.09 | 0.84 | -1.03 | 0.11 | -0.03 | 0.72 | -0.87 | -0.69 | -0.86 | -0.1325 | 0.16 | -0.2375 | 0.640176 | 0.691624 | 0.560414 | 1 | 0.996511 | 0.919045 |
| Ada | P03958 | 0.59 | 0.56 | -1.74 | -1.79 | 0.47 | 0.35 | -1.1 | -1.69 | -0.4 | -0.57 | -1.27 | 0.45 | 0.07 | 0.13 | -1.22 | -1.54 | 0.1025 | 0.1475 | -0.045 | 0.895123 | 0.83161 | 0.950538 | 1 | 0.996511 | 0.992476 |
| Adal | Q80SY6 | -0.28 | 0.26 | -0.56 | -0.61 | 0.33 | -0.21 | -1.2 | -0.32 | 0.1 | -0.37 | -0.55 | 0.02 | -0.23 | 0.08 | -0.33 | -1.16 | -0.0525 | 0.0975 | -0.1125 | 0.883574 | 0.717763 | 0.736723 | 1 | 0.996511 | 0.958635 |
| Adam10 | O35598 | -0.16 | 0.09 | -0.14 | -0.22 | 0.14 | 0.16 | 0.15 | -0.54 | 0.27 | -0.02 | -0.19 | 0.24 | 0.4 | 0.01 | 0.19 | 0.09 | 0.085 | 0.1825 | 0.28 | 0.690181 | 0.345023 | 0.161958 | 1 | 0.996511 | 0.757516 |
| Adam11 | Q7TQG7 | 0.34 | -0.32 | -0.8 | -0.54 | 0.05 | 0.85 | -0.48 | -0.85 | 0.46 | -0.78 | -0.79 | 0.4 | 0.18 | -0.06 | -1.1 | -0.77 | 0.2225 | 0.1525 | -0.1075 | 0.594926 | 0.709135 | 0.775632 | 1 | 0.996511 | 0.966799 |
| Adam15 | O88839-2 | 0.31 | 0.3 | 0.19 | 0.91 | 0.24 | 0.67 | 1.15 | 0.66 | 0.62 | 1.34 | 0.87 | 0.6 | 1.74 | 1.09 | 1.17 | 0.67 | 0.2525 | 0.43 | 0.74 | 0.339918 | 0.122253 | 0.02945 | 1 | 0.996511 | 0.637553 |
| Adam17 | Q9Z0F8 | -0.16 | 0.36 | -0.02 | -0.12 | 0.14 | 0.29 | -1.31 | -0.45 | -0.19 | 0.19 | -0.4 | 0.17 | 0.65 | 0.04 | -0.34 | -0.15 | -0.3475 | -0.0725 | 0.035 | 0.353709 | 0.745031 | 0.896743 | 1 | 0.996511 | 0.983509 |
| Adam22 | Q9R1V6-14 | 0.26 | 0.34 | 0.42 | 0.16 | 0.63 | 0.49 | 0.18 | 0.18 | -0.14 | 0.12 | 0.13 | 0.28 | 0.09 | 0.26 | 0.47 | 0.32 | 0.075 | -0.1975 | -0.01 | 0.66982 | 0.277231 | 0.95637 | 1 | 0.996511 | 0.992636 |
| Adam23 | Q9R1V7-2 | 0.86 | 0.69 | 0.76 | 0.73 | 0.45 | 0.82 | 0.68 | 0.36 | 0.95 | 0.21 | 0.28 | 0.7 | 0.98 | -0.12 | 0.21 | 0.17 | -0.1825 | -0.225 | -0.45 | 0.291184 | 0.316857 | 0.116412 | 1 | 0.996511 | 0.718608 |
| Adam9 | Q61072 | -0.02 | 0.23 | -0.28 | 0.01 | 0.4 | 0.25 | -0.02 | 0.02 | 0.46 | 0.68 | -0.01 | 0.45 | 0.21 | 0.11 | -0.08 | -0.22 | 0.1775 | 0.41 | 0.02 | 0.351537 | 0.086766 | 0.922103 | 1 | 0.996511 | 0.987447 |
| Adap1 | E9PY16 | 0.48 | -0.04 | 0.79 | 1.09 | 0.37 | 0.47 | 1.19 | 1.29 | -0.49 | 0.02 | 1.11 | 0.61 | -1.04 | -0.25 | 0.62 | 0.65 | 0.25 | -0.2675 | -0.585 | 0.456063 | 0.513554 | 0.21094 | 1 | 0.996511 | 0.786269 |
| Adar | Q99MU3-5 | 0.2 | -1.34 | -0.89 | -1.11 | -0.52 | -0.13 | -0.41 | -0.53 | -1.25 | -0.92 | -0.44 | -0.58 | -0.94 | -0.73 | 0.2 | -0.25 | 0.3875 | -0.0125 | 0.355 | 0.274535 | 0.973276 | 0.395614 | 1 | 0.997515 | 0.866946 |
| Adarb1 | Q91ZS8-4 | 0.23 | -0.77 | -0.99 | -0.59 | -0.56 | -0.2 | -0.23 | -0.46 | -0.97 | -0.68 | -0.84 | -1.16 | -0.8 | -0.69 | -0.63 | -0.49 | 1.68E-01 | -0.3825 | -0.1225 | 0.557334 | 0.21407 | 0.674606 | 1 | 0.996511 | 0.952944 |
| Adat2 | Q6P6J0 | -0.44 | -0.47 | 0.33 | 0.41 | -0.23 | -0.05 | -0.16 | 0.43 | 0 | 0.36 | 0.07 | -0.1 | -0.57 | 0.34 | -0.67 | -0.49 | 0.04 | 0.125 | -0.305 | 0.887712 | 0.648891 | 0.374106 | 1 | 0.996511 | 0.859424 |
| Adat3 | Q6PAT0 | 6.99 | 3.15 | 5.06 | 3.45 | 4.27 | 6.17 | 5.95 | 8.93 | 1.83 | 2.65 | 6.22 | 6.41 | 3.49 | -0.04 | 7.06 | 7.71 | 1.6675 | -0.385 | -0.1075 | 0.178966 | 0.77151 | 0.951509 | 1 | 0.996511 | 0.992476 |
| Adck1 | Q9D0L4 | 0.01 | -0.43 | 0.26 | -0.53 | -0.08 | 0.33 | 0.74 | 0.05 | -0.54 | -0.79 | 0.74 | 0.03 | -0.7 | -0.54 | 0.34 | 0.61 | 0.4325 | 0.0325 | 0.1 | 0.131926 | 0.930777 | 0.784796 | 1 | 0.996755 | 0.967642 |
| Adck2 | E9QL72 | -1.14 | -1.29 | -1.08 | -1.35 | -1.01 | -1.34 | -2.29 | -2.59 | -1.05 | 0.36 | -1.85 | -1.21 | -1.16 | -0.05 | -1.59 | -2.57 | -0.5925 | 0.2775 | -0.1275 | 0.131409 | 0.535645 | 0.795476 | 1 | 0.996511 | 0.969727 |
| Adck5 | E9PUK2 | -0.02 | 1.61 | -0.23 | 0.2 | 0.21 | -0.76 | -0.39 | -0.02 | 1 | 0.19 | 0.03 | -0.02 | -0.68 | 0.4 | 0.15 | -0.22 | -0.63 | -0.09 | -0.4775 | 0.173996 | 0.841704 | 0.306025 | 1 | 0.996511 | 0.832533 |
| Adcy1 | O88444 | -0.16 | 0.02 | 0.29 | 0.13 | 0.53 | 0.18 | 0.46 | -0.14 | 0.07 | -0.2 | 0.23 | 0.01 | 0.11 | 0.34 | -0.1 | 0.11 | 0.1875 | -0.0425 | 0.045 | 0.379584 | 0.82158 | 0.820926 | 1 | 0.996511 | 0.972768 |
| Adcy2 | Q3V1Q3 | -0.11 | 0.18 | 0.61 | 0.18 | 0.54 | 0.21 | 0.58 | 0.46 | 0.14 | 0.27 | 0.68 | 0.18 | -0.01 | 0.27 | 0.16 | 0.43 | 0.2325 | 0.1025 | -0.0025 | 0.268229 | 0.653288 | 0.991067 | 1 | 0.996511 | 0.996154 |
| Adcy3 | B8JK56 | 0.36 | 0.7 | 0.62 | 0.37 | 0.63 | 0.86 | 0.97 | 0.45 | 0.73 | 0.37 | 0.76 | 0.37 | 0.37 | 0.28 | 0.51 | 0.56 | 0.215 | 0.045 | -0.0825 | 0.266932 | 0.816056 | 0.662232 | 1 | 0.996511 | 0.949645 |
| Adcy5 | P84309 | 2.38 | 0.58 | 1.75 | 2.26 | 0.28 | 1 | 2.32 | 1.93 | 1.03 | 1.44 | 1.1 | 2.05 | 0.65 | 0.38 | 2.69 | 2.06 | -0.36 | -0.3375 | -0.2975 | 0.528513 | 0.455748 | 0.63897 | 1 | 0.996511 | 0.942804 |
| Adcy6 | A0A2I3BRC | 0.69 | 1.03 | 1.16 | 1.54 | 1.12 | 0.55 | 1.69 | 1.4 | 0.53 | 0.39 | 1.61 | 0.93 | 0.34 | 1.75 | 0.8 | 1.45 | 0.085 | -0.24 | -0.02 | 0.775808 | 0.466413 | 0.955457 | 1 | 0.996511 | 0.992482 |
| Adcy8 | A0A2I3BQ4 | 1.28 | 1.04 | 1.14 | 0.75 | 1.16 | 1.43 | 1.49 | 1.22 | 0.82 | 0.63 | 1.36 | 1.31 | 0.51 | 0.89 | 0.97 | 1.88 | 0.2725 | -0.0225 | 0.01 | 0.159486 | 0.925282 | 0.974911 | 1 | 0.996755 | 0.992977 |
| Adcy9 | P51830 | 1.22 | 1.19 | 1.68 | 1.11 | 1.02 | 1.04 | 1.42 | 1.2 | 0.87 | 0.79 | 1.22 | 0.93 | 0.77 | 0.43 | 1.12 | 1.18 | -0.13 | -0.3475 | -0.425 | 0.509579 | 0.117905 | 0.116306 | 1 | 0.996511 | 0.718608 |
| Adcyap1 | O70176 | 1.4 | 2.08 | -0.1 | 0.7 | 2.1 | 2.32 | 1.39 | 0.44 | 1.19 | 2.48 | 1.4 | 2.58 | 0.87 | 2.81 | 1.04 | 1.5 | 0.5425 | 0.8925 | 0.535 | 0.359525 | 0.128673 | 0.375072 | 1 | 0.996511 | 0.859611 |
| Adcyap1r1 | E9Q968 | 0.13 | 0.45 | 0.38 | 0.11 | 0.11 | 0.21 | 0.46 | 0.53 | 0.34 | 0.32 | 0.05 | 0.29 | 0.33 | 0.05 | 0.06 | 0.06 | 0.06 | -0.0175 | -0.1425 | 0.737256 | 0.921638 | 0.458229 | 1 | 0.996755 | 0.887902 |
| Adcyap1r1 | E9Q3E8 | -0.12 | 2.3 | -0.41 | 0.02 | -0.16 | 0.28 | 0.12 | 0.08 | -0.3 | -0.25 | -0.58 | -0.13 | 0.23 | 0.26 | 0.39 | 0.31 | -0.3675 | -0.7625 | -0.15 | 0.528567 | 0.211273 | 0.794289 | 1 | 0.996511 | 0.969367 |
| Add1 | E9Q1K3 | 1 | 0.4 | 0.83 | 0.6 | 1.7 | 0.02 | 0.75 | 0.69 | 0.11 | 1.77 | -0.04 | 0.08 | 0.74 | 1.22 | 1.58 | 1.1 | 0.0825 | -0.2275 | 0.4525 | 0.81615 | 0.596981 | 0.098416 | 1 | 0.996511 | 0.698156 |
| Add1 | F8WGR0 | 0.78 | 0.12 | 0.68 | 1.02 | -0.17 | 0.56 | 0.48 | 1.03 | 0.56 | 1.16 | 0.74 | 0.34 | 0.48 | -0.07 | 0.27 | 0.34 | -0.175 | 0.05 | -0.395 | 0.572983 | 0.85446 | 0.147274 | 1 | 0.996511 | 0.746406 |
| Add1 | Q9QYC0 | -0.86 | -0.26 | -1.29 | -2.21 | -1.2 | -0.86 | -1.95 | -2.33 | -1.44 | -1.8 | -0.77 | -0.77 | -1.01 | -0.96 | -1.74 | -1.18 | -0.43 | -0.04 | -0.0675 | 0.391516 | 0.929787 | 0.874214 | 1 | 0.996755 | 0.979626 |
| Add2 | Q9QYB8 | 1.07 | 0.69 | 1.07 | 0.74 | 0.68 | 0.94 | 0.97 | 0.98 | 0.6 | 0.34 | 1.14 | 0.5 | 0.39 | 0.93 | 0.72 | 0.83 | -5.9E-17 | -0.2475 | -0.175 | 1 | 0.303511 | 0.419448 | 1 | 0.996511 | 0.87558 |
| Add3 | Q9QYB5-2 | 0.27 | 0.24 | 0.56 | 0.52 | 0.23 | 0.33 | 0.55 | 0.56 | 0.28 | 0.2 | 0.66 | 0.15 | 0.43 | -0.31 | 0.33 | 0.25 | 0.02 | -0.075 | -0.2225 | 0.906135 | 0.701944 | 0.346546 | 1 | 0.996511 | 0.851096 |
| Add3 | Q9QYB5 | -1.22 | -1.18 | 0.33 | -0.06 | -0.83 | -1.23 | -1.23 | 0.49 | -0.68 | 0.57 | 0.71 | -1.53 | -0.98 | -0.86 | -1.03 | -0.79 | -0.1675 | 0.3 | -0.3825 | 0.748478 | 0.623684 | 0.335532 | 1 | 0.996511 | 0.847442 |
| Adgb | D3YZ77 | -1.12 | -1.03 | -0.8 | -0.4 | -1.41 | -0.76 | -0.48 | 0.02 | 0.61 | 0.02 | 0.04 | -1.69 | -1.72 | -0.9 | -0.58 | -0.01 | 0.18 | 0.5825 | 0.035 | 0.587858 | 0.251875 | 0.926746 | 1 | 0.996511 | 0.989128 |
| Adgra1 | Q8C4G9 | -0.12 | 0.65 | 0.31 | 0.79 | 1.16 | 0.38 | 0.25 | 0.76 | 0.95 | 1.44 | 0.74 | 0.35 | 0.76 | 0.95 | 0.53 | 0.77 | 0.23 | 0.4625 | 0.345 | 0.434582 | 0.159056 | 0.19534 | 1 | 0.996511 | 0.78004 |
| Adgra2 | J3QMG7 | -2.28 | -0.3 | -1.02 | -1.03 | 0.25 | -2.61 | -3.17 | 0.87 | 3.67 | 5.63 | -1.49 | -2.22 | 4.57 | 2.79 | -1.61 | -3.05 | -0.0075 | 2.555 | 1.8325 | 0.993882 | 0.171009 | 0.280049 | 1 | 0.996511 | 0.822084 |
| Adgrb1 | Q3UHD1 | 1.15 | 1.02 | 1.4 | 1.15 | 1.13 | 1.17 | 1.04 | 1.17 | 1.37 | 0.94 | 1.36 | 1.27 | 1.58 | 1.27 | 0.73 | 1.06 | -0.0525 | 0.055 | -0.02 | 0.734498 | 0.769757 | 0.932538 | 1 | 0.996511 | 0.989713 |
| Adgrb2 | H7BX74 | 0.63 | 0.84 | 0.56 | 0.56 | 2.41 | 0.82 | 0.47 | 0.54 | 1.34 | 0.65 | 0.85 | 0.76 | 1.6 | 1.16 | 0.62 | 0.65 | 0.4125 | 0.2525 | 0.36 | 0.351812 | 0.247869 | 0.201858 | 1 | 0.996511 | 0.781867 |
| Adgrb3 | Q80ZF8 | 1.23 | 1.08 | 1 | 0.97 | 0.94 | 1.23 | 0.69 | 0.85 | 0.54 | 0.78 | 1.11 | 1.27 | 0.66 | 0.38 | 0.93 | 0.81 | -0.1425 | -0.145 | -0.375 | 0.426319 | 0.50451 | 0.087345 | 1 | 0.996511 | 0.691619 |
| Adgre1 | Q61549 | 0.33 | -0.54 | -3.92 | -4 | -0.07 | 0.03 | -5.76 | -5.34 | 0.07 | -3 | -3.42 | 0.43 | 0.88 | -1.95 | -4.33 | -5.22 | -0.7525 | 0.5525 | -0.6225 | 0.667734 | 0.683644 | 0.693136 | 1 | 0.996511 | 0.955729 |
| Adgrf5 | G5E8Q8 | -0.1 | 0.38 | -1.38 | -2.11 | 0.81 | -0.28 | -1.11 | -2.12 | -0.44 | -1.19 | -0.89 | -0.05 | -0.47 | -0.57 | -1.61 | -1.6 | 0.1275 | 0.16 | -0.26 | 0.867314 | 0.780986 | 0.665524 | 1 | 0.996511 | 0.949825 |
| Adgrg1 | Q8K209 | 1.14 | 1.57 | 0.8 | 0.76 | 1.61 | 1.18 | 0.92 | 0.64 | 1.28 | 0.47 | 0.2 | 0.98 | 0.97 | 1.45 | 0.6 | 1.07 | 0.02 | -0.335 | -0.045 | 0.943307 | 0.29752 | 0.871351 | 1 | 0.996511 | 0.979293 |
| Adgrl1 | H7BX15 | 2.27 | 1.72 | 2.07 | 1.95 | -0.36 | 2.67 | -0.58 | 0.83 | 2.24 | 0.45 | 1.54 | 1.94 | 3.04 | -4.65 | 1.34 | 1.13 | -1.3625 | -0.46 | -1.7875 | 0.072908 | 0.259124 | 0.250768 | 1 | 0.996511 | 0.807201 |
| Adgrl1 | E9Q3V9 | 0.27 | 0.53 | 0.31 | 0.57 | 0.54 | 0.36 | 0.42 | 0.52 | 0.5 | 0.36 | 0.51 | 0.66 | 0.2 | 0.6 | 0.25 | 0.07 | 0.04 | 0.0875 | -0.14 | 0.796796 | 0.614414 | 0.49366 | 1 | 0.996511 | 0.89758 |
| Adgrl2 | A0A0G2JFV | 1.3 | 1.13 | 0.97 | 0.97 | 1.45 | 1.43 | 1.26 | 1.1 | 1.08 | 0.68 | 1.17 | 1.49 | 0.91 | 1.15 | 0.82 | 0.91 | 0.2175 | 0.0125 | -0.145 | 0.217931 | 0.954877 | 0.446174 | 1 | 0.996755 | 0.884817 |
| Adgrl3 | D3YTW7 | 0.99 | 0.83 | 0.8 | 1 | 0.68 | 1.08 | 0.87 | 1.11 | 1.2 | 0.75 | 1.11 | 1.29 | 1.1 | 0.78 | 0.61 | 0.68 | 0.03 | 0.1825 | -0.1125 | 0.858194 | 0.345443 | 0.564973 | 1 | 0.996511 | 0.920315 |
| Adgrl3 | D3Z6H9 | 0.07 | 0.33 | -1.32 | -1.8 | 0.09 | 0.74 | -0.94 | -0.55 | 0.67 | -2.34 | -0.15 | 0.06 | 1.87 | -0.25 | -1.64 | -1.56 | 0.515 | 0.24 | 0.285 | 0.387694 | 0.75234 | 0.744318 | 1 | 0.996511 | 0.959371 |
| Adh5 | P28474 | 0.68 | -0.82 | -0.8 | -0.81 | -0.57 | 0.54 | -1.04 | -1 | 0.51 | -0.39 | -1.02 | 0.44 | 0.02 | 0.38 | -1.03 | -0.85 | -0.08 | 0.3225 | 0.0675 | 0.868798 | 0.5124 | 0.886456 | 1 | 0.996511 | 0.979767 |
| Adhfe1 | A0A087WF | 0.08 | 0.4 | 1.29 | 0.98 | 0.3 | 0.23 | 1.01 | 0.8 | -0.22 | 0.69 | 1.16 | 0.52 | 0.27 | -0.51 | 0.61 | 1 | -0.1025 | -0.15 | -0.345 | 0.753012 | 0.695566 | 0.405119 | 1 | 0.996511 | 0.87246 |
| Adi1 | Q99JT9 | 0.83 | 0.73 | 0.51 | 0.27 | 0.44 | -0.07 | -0.19 | 0 | 0.25 | 0.27 | -0.31 | 0.4 | -0.2 | 0.95 | -0.23 | -0.47 | -0.54 | -0.4325 | -0.5725 | 0.029915 | 0.089884 | 0.121025 | 1 | 0.996511 | 0.721448 |
| Adipoq | Q60994 | -2.26 | -2.65 | -3.05 | -3.05 | -0.32 | -2.48 | -0.98 | -2.28 | -2.37 | -1.03 | -3.31 | -1.83 | -2.06 | -2.5 | -1.99 | -3.09 | 1.2375 | 0.6175 | 0.3425 | 0.036019 | 0.220886 | 0.303432 | 1 | 0.996511 | 0.831662 |
| Adipor1 | Q91VH1 | -0.11 | 0.56 | 0.66 | 0.77 | 0.91 | 0.18 | 1.45 | 0.65 | 0.18 | 1.12 | 0.91 | 0.32 | 0.02 | -0.03 | 0.44 | 0.89 | 0.3275 | 0.1625 | -0.14 | 0.325138 | 0.597589 | 0.646615 | 1 | 0.996511 | 0.945037 |
| Adk | P55264-2 | -1.61 | -2.53 | -1.22 | -1.43 | -2.65 | -1.1 | -0.95 | -1.34 | -2.38 | -1.3 | -1.26 | -1.99 | -1.56 | -2.73 | -1.64 | -1.42 | 0.1875 | -0.035 | -0.14 | 0.678598 | 0.927087 | 0.728204 | 1 | 0.996755 | 0.958635 |
| Adnp | Q9Z103 | 0.67 | -0.66 | 0.88 | 0.26 | -0.69 | 0.22 | 1.01 | 0.43 | -0.04 | 0.13 | 0.88 | -0.04 | -0.04 | 0.11 | 1.81 | 1.78 | -0.045 | -0.055 | 0.6275 | 0.92137 | 0.887637 | 0.283522 | 1 | 0.996511 | 0.822669 |
| Ado | Q6PDY2 | -0.22 | 0.26 | 0.42 | 0.08 | 0.15 | -0.19 | -0.17 | 0.09 | 0.21 | 0.82 | 0.18 | 0.05 | 0.08 | 0.44 | 0.17 | -0.03 | -0.165 | 0.18 | 0.03 | 0.412601 | 0.469943 | 0.891777 | 1 | 0.996511 | 0.982213 |
| Adora1 | Q60612 | 0.85 | 1.05 | 0.46 | 0.36 | -0.6 | 1.97 | -0.97 | -1 | 3.16 | -2.56 | 0.02 | 0.24 | 3.63 | -3.73 | -0.4 | -0.28 | -0.83 | -0.465 | -0.875 | 0.232074 | 0.662364 | 0.520712 | 1 | 0.996511 | 0.907667 |
| Adora2a | Q60613 | 1.19 | -0.1 | 2.02 | 1.31 | -1.18 | -0.26 | 0.48 | 1.43 | 1.93 | 1.5 | 0.27 | 1.03 | 2.57 | 0.14 | 1.88 | 1.4 | -0.9875 | 0.0775 | 0.3925 | 0.151439 | 0.88254 | 0.530302 | 1 | 0.996511 | 0.908663 |
| Adpgk | Q3UDS7 | -0.17 | 0.05 | -0.88 | -0.4 | 0.53 | -0.07 | -0.17 | -0.73 | -0.14 | 0.15 | -0.46 | -0.12 | -0.04 | 0.3 | 0.01 | 0.09 | 0.24 | 0.2075 | 0.44 | 0.459373 | 0.426458 | 0.102978 | 1 | 0.996511 | 0.700625 |
| Adprh | P54923 | -0.64 | -0.64 | -1.24 | -1.13 | -0.54 | -0.61 | -0.86 | -1.06 | -0.97 | -0.53 | -1.18 | -0.51 | -1 | -0.53 | -1 | -1.16 | 0.145 | 0.115 | -0.01 | 0.518046 | 0.650142 | 0.967454 | 1 | 0.996511 | 0.9927 |
| Adprm | Q99KS6-2 | -2.07 | -1.69 | -2.57 | -1.47 | -1.36 | -1.88 | -2.35 | -2.07 | -1.51 | -2.05 | -1.13 | -1.65 | -1.55 | -1.8 | -2.5 | -1.53 | 0.035 | 0.365 | 0.105 | 0.910951 | 0.258186 | 0.753232 | 1 | 0.996511 | 0.961479 |
| Adprs | Q8CG72 | -1.14 | -0.93 | -0.93 | -0.53 | -1.35 | -0.99 | -0.96 | -0.97 | -0.41 | -1.1 | -1.02 | -1.13 | -0.34 | -0.87 | -1.34 | -1.17 | -0.185 | -0.0325 | -0.0475 | 0.354679 | 0.892224 | 0.863284 | 1 | 0.996511 | 0.977235 |
| Adra1a | P97718 | 0.88 | 0.88 | 0.89 | 0.91 | 0.59 | 0.3 | -0.31 | 0.44 | 1.89 | 0.93 | 0.57 | 0.94 | 1.46 | 1.27 | 0.75 | 0.13 | -0.635 | 0.1925 | 0.0125 | 0.017515 | 0.514004 | 0.96763 | 1 | 0.996511 | 0.9927 |
| Adra1b | P97717 | -0.64 | 0.18 | -0.26 | -0.14 | 1.08 | -0.24 | 0.11 | -0.37 | 0.13 | -0.37 | 0.28 | 0.37 | 0.78 | 1.29 | 0.13 | -0.53 | 0.36 | 0.3175 | 0.6325 | 0.32415 | 0.237361 | 0.150975 | 1 | 0.996511 | 0.749304 |
| Adra2a | Q01338 | -0.23 | 0.73 | 1.14 | 1.49 | -0.54 | 0.49 | 0.51 | 1.14 | 1.29 | 3.03 | 1.1 | 0.15 | 1.88 | 0.66 | 1.55 | 0.75 | -0.3825 | 0.61 | 0.4275 | 0.426512 | 0.35662 | 0.356005 | 1 | 0.996511 | 0.8553 |
| Adra2c | Q01337 | -0.18 | -0.12 | 1.83 | 1.24 | -2 | -0.85 | 0.93 | 1.2 | 0.99 | 2.66 | 0.4 | 0.68 | 2.79 | 1.22 | 0.33 | 2.3 | -0.8725 | 0.49 | 0.9675 | 0.303071 | 0.458868 | 0.177726 | 1 | 0.996511 | 0.772068 |
| Adrbk2 | Q3UYH7 | 0.17 | -0.25 | 0.25 | 0.17 | -0.22 | 0.24 | 0.7 | 0.34 | -0.06 | -0.3 | 0.38 | -0.24 | 0.29 | -0.39 | 0.11 | -0.12 | 0.18 | -0.14 | -0.1125 | 0.455859 | 0.539038 | 0.627212 | 1 | 0.996511 | 0.939737 |
| Adrm1 | Q9JKV1 | -1.62 | -1.31 | 0.76 | 0.56 | -1.36 | -1.13 | 0.28 | 0.78 | -0.1 | 0.21 | 0.92 | -1.54 | 0.01 | -0.75 | 0.9 | 0.55 | 0.045 | 0.275 | 0.58 | 0.951001 | 0.707749 | 0.384296 | 1 | 0.996511 | 0.860984 |
| Adsl | P54822 | 0.02 | 0.04 | 0.48 | 0.6 | -0.22 | -0.28 | 0.66 | 0.68 | -0.08 | 0.15 | 0.21 | -0.01 | -0.53 | 0.26 | 0.04 | 0.14 | -0.075 | -0.2175 | -0.3075 | 0.804101 | 0.311522 | 0.254964 | 1 | 0.996511 | 0.809524 |
| Adss1 | P28650 | -1.93 | -2.26 | -2.69 | -3.21 | -2.27 | -2.66 | -2.72 | -2.82 | -2.01 | -1.47 | -2.46 | -1.97 | -1.21 | -2.89 | -1.09 | -3.38 | -0.095 | 0.545 | 0.38 | 0.751837 | 0.134723 | 0.525492 | 1 | 0.996511 | 0.908107 |
| Adss2 | P46664 | -0.11 | -0.35 | -0.02 | 0.26 | 0.17 | 0.12 | 0.29 | 0.03 | -0.68 | -0.15 | 0.01 | -0.35 | -0.86 | 0.27 | -0.22 | 0.12 | 0.2075 | -0.2375 | -0.1175 | 0.27094 | 0.313119 | 0.692102 | 1 | 0.996511 | 0.955729 |
| Afap1 | Q80YS6 | -0.38 | -0.07 | 0.66 | 0.53 | -0.56 | -0.07 | 0.01 | 0.12 | 0.73 | 1.48 | 0.94 | 0.18 | 0.67 | -0.61 | 0.23 | -0.29 | -0.31 | 0.6475 | -0.185 | 0.300072 | 0.096831 | 0.617467 | 1 | 0.996511 | 0.937152 |
| Afap1l1 | Q8BZI0 | 0.31 | 0.19 | -0.11 | -0.39 | -0.05 | -0.17 | -0.61 | -0.55 | 1.05 | -0.35 | -0.3 | 0.34 | 1.26 | -0.28 | -0.87 | 0.88 | -0.345 | 0.185 | 0.2475 | 0.158109 | 0.605543 | 0.613578 | 1 | 0.996511 | 0.935813 |
| Afap1l2 | Q5DTU0 | -0.06 | 1.17 | 0.37 | 0.43 | 1.43 | 0.61 | 0.16 | 0.09 | 0.13 | 0.45 | 0.37 | -0.22 | -0.02 | 1.51 | -0.05 | 0.3 | 0.095 | -0.295 | -0.0425 | 0.802747 | 0.341934 | 0.920498 | 1 | 0.996511 | 0.98706 |
| Afdn | Q9QZQ1 | 0.71 | 0.7 | 0.3 | 0.17 | 0.76 | 0.72 | 0.26 | 0.23 | 0.86 | 0.12 | 0.6 | 1.1 | 0.96 | 0.56 | 0.33 | 0.35 | 0.0225 | 0.2 | 0.08 | 0.919154 | 0.462418 | 0.739588 | 1 | 0.996511 | 0.958796 |
| Afdn | E9PYX7 | -0.26 | 0.43 | -0.21 | -0.53 | 0.26 | 0.11 | -0.44 | -0.24 | -0.09 | -0.24 | 0.5 | 0.12 | -0.54 | 0.49 | -0.22 | -0.25 | 0.065 | 0.215 | 0.0125 | 0.806948 | 0.44011 | 0.967682 | 1 | 0.996511 | 0.9927 |
| Aff4 | Q9ESC8 | -1.36 | -2.36 | -1.15 | -0.92 | -1.95 | -1.39 | -1.45 | -1.5 | -0.65 | 0.05 | -1.28 | -2.45 | -0.25 | -0.01 | 0.8 | -0.61 | -0.125 | 0.365 | 1.43 | 0.707468 | 0.526036 | 0.007342 | 1 | 0.996511 | 0.552614 |
| Afg1l | Q3V384 | 0.44 | 0.4 | 0.69 | 0.33 | 0.1 | 0.49 | 1.32 | 0.87 | -0.32 | -0.42 | 0.87 | 0.53 | -0.65 | -0.15 | 0.34 | 1.24 | 0.23 | -0.3 | -0.27 | 0.415086 | 0.369102 | 0.501416 | 1 | 0.996511 | 0.901002 |
| Afg3l1 | Q920A7 | 0.15 | 0.14 | 0.27 | 0.14 | 0.53 | 0.11 | 0.51 | 0.25 | 1.83 | 0.63 | 1.11 | 0.89 | 2.32 | 1.05 | 0.92 | 0.3 | 0.175 | 0.94 | 0.9725 | 0.304229 | 0.007363 | 0.038664 | 1 | 0.996511 | 0.658454 |
| Afg3l2 | Q8JZQ2 | 0.28 | 0.47 | 0.76 | 0.95 | -0.05 | 0.58 | 0.48 | 0.88 | 0.27 | 0.87 | 0.93 | 0.26 | 0.29 | 0.2 | 0.42 | 0.43 | -0.1425 | -0.0325 | -0.28 | 0.579545 | 0.899326 | 0.213075 | 1 | 0.996511 | 0.786931 |
| Afm | O89020 | -3.04 | -3.24 | -2.53 | -2.99 | -2.53 | -3.15 | -3.07 | -2.76 | -3.71 | -1.49 | -2.87 | -2.83 | -3.38 | -2.31 | -3.73 | -3.54 | 0.0725 | 0.225 | -0.29 | 0.750718 | 0.622326 | 0.416106 | 1 | 0.996511 | 0.874945 |
| Aftph | Q80WT5-2 | -0.58 | -0.56 | -0.17 | -0.28 | -0.68 | -0.59 | -0.4 | -0.32 | -0.59 | -0.47 | -0.27 | -0.38 | -0.5 | -0.09 | -0.23 | -0.34 | -0.1 | -0.03 | 0.1075 | 0.578018 | 0.870842 | 0.594784 | 1 | 0.996511 | 0.929843 |
| Aga | Q64191 | -0.29 | -0.7 | -0.47 | -0.5 | -0.72 | -0.92 | 0.14 | -0.78 | -0.63 | -0.45 | -0.48 | 0.25 | -0.21 | -1.08 | -0.75 | -0.22 | -0.08 | 0.1625 | -0.075 | 0.761089 | 0.505515 | 0.772143 | 1 | 0.996511 | 0.965933 |
| Agap1 | A0A087WF | 1.02 | 0.86 | 0.11 | 0.25 | 0.74 | 0.93 | 0.35 | 0.42 | 1.61 | 1.04 | 0.23 | 1.23 | 1.06 | 1.67 | 0.95 | 0.12 | 0.05 | 0.4675 | 0.39 | 0.852135 | 0.214045 | 0.318955 | 1 | 0.996511 | 0.838969 |
| Agap1 | A0A1D5RLI | 1.37 | 0.48 | 0.56 | 0.65 | -0.04 | 1.06 | 0.12 | 0.6 | 1.64 | 1.45 | 0.84 | 0.88 | 2.52 | 1.16 | 1.76 | -0.14 | -0.33 | 0.4375 | 0.56 | 0.311879 | 0.163432 | 0.324973 | 1 | 0.996511 | 0.842881 |
| Agap1 | E9Q3K3 | 0.43 | 0.35 | 0.35 | 0.73 | 0.16 | 0.29 | 0.21 | 0.39 | 0.88 | 0.62 | 0.47 | 0.31 | 0.57 | 0.55 | 0.21 | 0.25 | -0.2025 | 1.05E-01 | -0.07 | 0.234596 | 0.603084 | 0.72611 | 1 | 0.996511 | 0.958398 |
| Agap2 | Q3UHD9 | 0.4 | -0.1 | 0.28 | 0.53 | 0.03 | 0.56 | 0.29 | 0.64 | 0.95 | 0.28 | 0.58 | 0.24 | 1.26 | 0.12 | 0.68 | -0.07 | 0.1025 | 0.235 | 0.22 | 0.640697 | 0.342192 | 0.512878 | 1 | 0.996511 | 0.904133 |
| Agap3 | Q8VHH5 | 0.91 | 0.64 | 0.97 | 1.23 | 0.53 | 1.06 | 0.82 | 1.49 | 1.01 | 1.22 | 1.14 | 1.2 | 0.71 | 0.93 | 0.61 | 0.5 | 0.0375 | 0.205 | -0.25 | 0.880174 | 0.293417 | 0.253316 | 1 | 0.996511 | 0.809524 |
| Agap3 | A0A1D5RM | 0.05 | -0.38 | -1.1 | -0.64 | -1.01 | 0.44 | -0.17 | -0.77 | 0.11 | -1.04 | -1 | -0.05 | 0.31 | 0 | -1.32 | -0.96 | 0.14 | 0.0225 | 0.025 | 0.715808 | 0.952163 | 0.953883 | 1 | 0.996755 | 0.992476 |
| Agbl4 | E4W7Y4 | -0.19 | 0.21 | 0.33 | -0.1 | 0.51 | 0.24 | 0.6 | 0.88 | -0.74 | -0.83 | 0.46 | 0.05 | -0.31 | 0.03 | 0.22 | 0.38 | 0.495 | -0.3275 | 0.0175 | 0.039879 | 0.339119 | 0.940556 | 1 | 0.996511 | 0.990894 |
| Agfg1 | A0A087WN | 0.32 | -0.28 | 0.06 | 0.26 | -0.03 | 0.22 | -0.16 | 0.16 | 0.78 | -0.12 | 0.04 | -0.14 | 0.72 | 0.62 | -0.21 | -0.47 | -0.0425 | 0.05 | 0.075 | 0.828517 | 0.853646 | 0.82022 | 1 | 0.996511 | 0.972768 |
| Agfg2 | Q3U2K8 | 0.36 | -0.18 | -0.23 | -0.11 | -0.44 | -0.07 | -0.28 | -0.35 | -0.07 | -0.39 | -0.16 | -0.03 | -0.25 | 0.14 | -0.47 | -0.62 | -0.245 | -0.1225 | -0.26 | 0.226781 | 0.552911 | 0.310465 | 1 | 0.996511 | 0.834523 |
| Agk | Q9ESW4 | 0.57 | 0.38 | -0.3 | -0.12 | 0.22 | 0.53 | -0.13 | -0.34 | 0.63 | -0.64 | -0.1 | 0.59 | 0.4 | 1.04 | -0.93 | -0.51 | -0.0625 | -0.0125 | -0.1325 | 0.82523 | 0.97207 | 0.77399 | 1 | 0.997515 | 0.966062 |
| Agl | F8VPN4 | 0.28 | -0.06 | 0.1 | 0.16 | 0.73 | 0.56 | 0.5 | 0.3 | -0.16 | -0.44 | 0.15 | 0.05 | -0.52 | 0.07 | -0.13 | 0.27 | 0.4025 | -0.22 | -0.1975 | 0.038028 | 0.285703 | 0.397548 | 1 | 0.996511 | 0.867334 |
| Ago1 | A0A6I8MX: | -0.37 | -0.72 | 0.04 | 0.48 | -0.53 | -0.71 | -0.29 | -0.05 | -0.48 | -0.07 | -0.01 | -0.23 | -1.01 | -0.78 | -0.1 | -0.19 | -0.2525 | -0.055 | -0.3775 | 0.402692 | 0.849035 | 0.286053 | 1 | 0.996511 | 0.823465 |
| Ago2 | Q8CJG0 | -0.56 | -0.88 | -1.18 | -0.94 | -0.77 | -0.49 | -0.63 | -1.05 | -0.27 | -0.8 | -1.17 | -0.5 | 0.1 | -0.36 | -0.91 | -0.78 | 0.155 | 0.205 | 0.4025 | 0.457705 | 0.429132 | 0.177236 | 1 | 0.996511 | 0.772068 |
| Ago3 | Q8CJF9 | -1.69 | -1.51 | -0.58 | -0.55 | -1.36 | -1.25 | -0.2 | -0.24 | -1.03 | 0.31 | -0.24 | -1.38 | -0.33 | -0.52 | -0.16 | 0.55 | 0.32 | 0.4975 | 0.9675 | 0.442518 | 0.292886 | 0.027338 | 1 | 0.996511 | 0.635993 |
| Agpat1 | A0A0R4J26 | 2.02 | 1.11 | 1.36 | 0.27 | 1.72 | 1.63 | 2.16 | 1.58 | 0.17 | 0.91 | 1.35 | 1.8 | 0.03 | 0.51 | 1.75 | 2.55 | 0.5825 | -0.1325 | 0.02 | 0.139988 | 0.778021 | 0.974281 | 1 | 0.996511 | 0.992977 |
| Agpat3 | Q9D517 | 0.39 | -0.04 | -0.19 | -0.23 | 0.29 | 0.18 | 0.48 | -0.42 | 0.38 | -0.12 | -0.3 | 0.48 | 0.48 | 0.4 | -0.06 | 0.06 | 0.15 | 0.1275 | 0.2375 | 0.555722 | 0.622342 | 0.325823 | 1 | 0.996511 | 0.843144 |
| Agpat4 | Q8K4X7 | 0.12 | 0.09 | 0.08 | 0.2 | 0.46 | 0.19 | 0.42 | 0.14 | 0.13 | 0.23 | 0.19 | 0.35 | 0.51 | 0.39 | 0.79 | 0.86 | 0.18 | 0.1025 | 0.515 | 0.262282 | 0.51828 | 0.023259 | 1 | 0.996511 | 0.623925 |
| Agpat5 | Q9D1E8 | 1.74 | 1.08 | 1.42 | 0.51 | 1.28 | 1.87 | 1.62 | 0.78 | 1.77 | 0.77 | 1.34 | 1.79 | 1.6 | 1.36 | 1.54 | 1.7 | 0.2 | 0.23 | 0.3625 | 0.561608 | 0.51499 | 0.231237 | 1 | 0.996511 | 0.788863 |
| Agps | A2AL50 | -0.14 | 0.29 | 0.11 | 0.81 | 0.73 | 0.45 | 0.64 | 0.28 | 0.3 | 0.76 | 0.16 | 0.28 | 0.25 | 0.75 | 0.32 | 0.65 | 0.2575 | 0.1075 | 0.225 | 0.300333 | 0.680549 | 0.402488 | 1 | 0.996511 | 0.870698 |
| Agrn | M0QWP1 | -0.24 | -0.47 | -0.66 | -0.67 | -0.08 | -0.29 | -0.55 | -0.9 | -0.21 | -0.48 | -0.65 | -0.03 | 0.1 | -0.46 | 0.49 | -0.25 | 0.055 | 0.1675 | 0.48 | 0.807073 | 0.439132 | 0.092496 | 1 | 0.996511 | 0.695968 |
| Agt | Q3UTR7 | -0.79 | -0.56 | -2.29 | -2.14 | -0.45 | -1.83 | -1.42 | -1.89 | -0.88 | -0.7 | -1.97 | -0.36 | -1.92 | 0.16 | -1.5 | -1.58 | 0.0475 | 0.4675 | 0.235 | 0.926279 | 0.384705 | 0.692895 | 1 | 0.996511 | 0.955729 |
| Agtpbp1 | Q641K1-2 | 1.26 | 0.96 | 0.3 | -0.02 | 0.65 | 1.24 | 0.01 | 0.2 | 1.34 | 0.23 | 0.36 | 1.13 | 0.79 | 0.81 | -0.21 | -0.21 | -0.1 | 0.14 | -0.33 | 0.793229 | 0.718716 | 0.417962 | 1 | 0.996511 | 0.87558 |
| Ahctf1 | Q8CJF7 | 0.33 | 0.51 | 0.63 | 0.95 | 0.35 | 0.14 | 0.79 | 0.85 | 0.42 | 0.96 | 0.52 | -0.04 | 0.5 | 1.21 | 1.17 | 1.31 | -0.0725 | -0.14 | 0.4425 | 0.756917 | 0.59574 | 0.11209 | 1 | 0.996511 | 0.713445 |
| Ahcy | P50247 | -0.62 | -0.75 | -0.46 | -0.35 | -0.54 | -0.75 | -0.41 | -0.4 | -1.14 | -0.21 | -0.58 | -0.73 | -1.08 | -1 | -0.82 | -0.93 | 0.02 | -0.12 | -0.4125 | 0.907119 | 0.618575 | 0.050567 | 1 | 0.996511 | 0.658454 |
| Ahcyl1 | Q80SW1 | 0.49 | 0.58 | -0.37 | -0.1 | 0.72 | 0.51 | 0.28 | -0.21 | 0.26 | -0.14 | -0.37 | 0.62 | 0.25 | 0.31 | -0.1 | -0.11 | 0.175 | -0.0575 | -0.0625 | 0.5656 | 0.856537 | 0.821689 | 1 | 0.996511 | 0.972768 |
| Ahcyl2 | Q68FL4-2 | 0.34 | -1.42 | -1.19 | -1.61 | 0.66 | -1.67 | -1.47 | -2.38 | -0.83 | -1.24 | -2.72 | -1.47 | 1.01 | 0.19 | -1.12 | -1.61 | -0.245 | -0.595 | 0.5875 | 0.733071 | 0.300885 | 0.397128 | 1 | 0.996511 | 0.867334 |
| Ahcyl2 | F8WGT1 | 0.13 | -0.11 | -0.08 | 0.11 | 0.54 | 0.28 | 0.5 | 0.13 | 0.07 | 0.27 | -0.05 | 0.27 | -0.16 | -0.06 | 0.43 | 0.58 | 0.35 | 0.1275 | 0.185 | 0.064054 | 0.470346 | 0.437587 | 1 | 0.996511 | 0.883161 |
| Ahdc1 | Q6PAL7 | -0.35 | -0.82 | -0.96 | -0.7 | -0.85 | -0.47 | -0.61 | -0.52 | -0.54 | -0.71 | -0.8 | -0.47 | -0.11 | -0.57 | -0.56 | -0.44 | 0.095 | 0.0775 | 0.2875 | 0.623994 | 0.699824 | 0.213428 | 1 | 0.996511 | 0.786931 |
| Ahi1 | E9QP54 | -0.35 | -0.02 | -1.48 | -0.86 | -0.8 | -0.45 | -0.27 | -0.94 | -0.39 | -0.05 | -1.45 | -0.87 | 0.55 | -0.78 | 0.35 | 0.04 | 0.0625 | -0.0125 | 0.7175 | 0.854871 | 0.976126 | 0.110822 | 1 | 0.997588 | 0.712422 |
| Ahnak | E9Q616 | -0.16 | -0.51 | -0.32 | 0.22 | -0.44 | -0.5 | -0.06 | -0.66 | -0.23 | 0.13 | -0.14 | -0.19 | -0.42 | 0 | -0.47 | -0.26 | -0.2225 | 0.085 | -0.095 | 0.332235 | 0.693806 | 0.682698 | 1 | 0.996511 | 0.955729 |
| Ahnak2 | **A0A7N9VR** | **0.66** | 0.68 | 0.88 | 0.78 | **1.1** | 0.08 | 1.3 | 0.3 | 1.14 | 1.27 | 1.19 | 1.24 | **1.19** | 1.28 | 1.29 | 1.24 | -0.055 | 0.46 | 0.5 | 0.854448 | 0.016502 | 0.015349 | 1 | 0.996511 | 0.589387 |
| Ahsa1 | Q8BK64 | 0.17 | 0.19 | -0.07 | -0.28 | 0.1 | -0.22 | -0.15 | -0.33 | -0.75 | 0.2 | -0.32 | 0.08 | -0.8 | -0.44 | -0.26 | -0.33 | -0.1525 | -0.2 | -0.46 | 0.419955 | 0.451442 | 0.058993 | 1 | 0.996511 | 0.661738 |
| Ahsa2 | F6S1R2 | 0.09 | -0.2 | -0.52 | -0.07 | 0.24 | 0.03 | 0.02 | -0.3 | -0.06 | -0.66 | -0.1 | -0.11 | -0.41 | 0.17 | -0.12 | -0.14 | 0.1725 | -0.0575 | 0.05 | 0.404349 | 0.79991 | 0.823367 | 1 | 0.996511 | 0.972768 |
| Ahsg | P29699 | -4.03 | -4.47 | -4.1 | -4.77 | -4 | -4.6 | -4.78 | -4.18 | -4.51 | -2.46 | -4.6 | -4.15 | -3.34 | -2.33 | -4.78 | -4.75 | -0.0475 | 0.4125 | 0.5425 | 0.854399 | 0.411747 | 0.353456 | 1 | 0.996511 | 0.855029 |
| Aida | Q8C4Q6 | 0.72 | 0.25 | 0.59 | 1.35 | 0.15 | 0.5 | 0.99 | 1.31 | 0.34 | 1.32 | 0.73 | 0.41 | -0.12 | 0.53 | 0.75 | 0.65 | 0.01 | -0.0275 | -0.275 | 0.976115 | 0.931617 | 0.388031 | 1 | 0.996755 | 0.864251 |
| Aif1 | O70200 | 1.02 | 0.48 | -0.4 | 0.05 | -0.26 | 0.77 | -0.56 | -0.05 | 0.77 | -0.69 | 0.01 | 0.54 | 0.87 | 0.08 | -0.82 | -0.47 | -0.3125 | -0.13 | -0.3725 | 0.435366 | 0.758001 | 0.417951 | 1 | 0.996511 | 0.87558 |
| Aifm1 | B1AU25 | 0.46 | 0.57 | 0.31 | 0.45 | 0.48 | 0.64 | 0.48 | 0.34 | 0.3 | -0.2 | 0.69 | 0.44 | 0.26 | 0.2 | 0.02 | 0.27 | 0.0375 | -0.14 | -0.26 | 0.806187 | 0.543583 | 0.165551 | 1 | 0.996511 | 0.758837 |
| Aifm2 | Q8BUE4 | -0.46 | -0.28 | 0.8 | 0.68 | 0.64 | -0.3 | 0.82 | 0.81 | 0.31 | 0.3 | 0.79 | -0.43 | -0.17 | 0.31 | 0.65 | 0.87 | 0.3075 | 0.0575 | 0.23 | 0.445864 | 0.883601 | 0.553033 | 1 | 0.996511 | 0.916262 |
| Aifm3 | Q3TY86 | 0.58 | 1.09 | 0.81 | 0.57 | 1.06 | 1.01 | -0.38 | 0.47 | 0.83 | 0.27 | 0.48 | 0.76 | 0.88 | 0.7 | 0.51 | 0.65 | -0.2225 | -0.1775 | -0.0775 | 0.522371 | 0.422677 | 0.707669 | 1 | 0.996511 | 0.957213 |
| Aig1 | H7BX91 | -4.08 | -6.79 | -3.4 | -11.8 | -8.27 | -1.36 | -5.25 | -4.1 | 5.02 | -2.27 | -2.4 | -5.82 | 8.79 | -8.14 | 4.73 | -6.34 | 1.7725 | 5.15 | 6.2775 | 0.413701 | 0.079003 | 0.14557 | 1 | 0.996511 | 0.745321 |
| Aimp1 | P31230 | -1.63 | -1.78 | -2.16 | -1.96 | -1.66 | -1.76 | -2.11 | -2.03 | -1.71 | -1.21 | -2.12 | -1.61 | -1.53 | -1.14 | -1.76 | -2.16 | -0.0075 | 0.22 | 0.235 | 0.969041 | 0.379975 | 0.390837 | 1 | 0.996511 | 0.865375 |
| Aimp2 | Q8R010 | -2.31 | -2.76 | -2.89 | -2.8 | -2.04 | -2.58 | -3.05 | -2.8 | -2.02 | -1.37 | -2.5 | -2.24 | -1.53 | -1.39 | -2.22 | -2.9 | 0.0725 | 0.6575 | 0.68 | 0.780631 | 0.043793 | 0.089672 | 1 | 0.996511 | 0.69217 |
| Aip | O08915 | 1.03 | -0.19 | -0.26 | -0.09 | 0.17 | 0.82 | -0.49 | -0.12 | -0.34 | 0.55 | -0.34 | 0.74 | -0.11 | -0.07 | 0 | -0.33 | -0.0275 | 0.03 | -0.25 | 0.943649 | 0.94016 | 0.441956 | 1 | 0.996755 | 0.883549 |
| Ajap1 | A2ALI5 | 5.02 | 4.08 | 0.09 | 0.07 | 4.87 | 4.59 | -0.45 | -0.82 | 0.48 | -1.03 | -0.04 | 5.31 | -0.38 | 1.06 | 0.3 | 0.11 | -0.2675 | -1.135 | -2.0425 | 0.882299 | 0.513397 | 0.114182 | 1 | 0.996511 | 0.717287 |
| Ajm1 | A2AJA9 | 1.3 | 1.18 | 1.29 | 1.35 | 0.94 | 1.35 | 0.73 | 1.04 | 1.06 | 0.27 | 1.37 | 1.18 | 0.85 | 1.02 | 0.64 | 0.95 | -0.265 | -0.31 | -0.415 | 0.165211 | 0.257419 | 0.044187 | 1 | 0.996511 | 0.658454 |
| Ak1 | Q9R0Y5 | -0.87 | -0.73 | -0.15 | 0.31 | -0.51 | -1.12 | -0.28 | 0.13 | -0.68 | 0.43 | -0.25 | -0.63 | -1.29 | -0.13 | -0.58 | -0.75 | -0.085 | 0.0775 | -0.3275 | 0.813935 | 0.831233 | 0.372325 | 1 | 0.996511 | 0.859424 |
| Ak2 | Q9WTP6 | 0.97 | 0.37 | 0.06 | 0.19 | 0.13 | 0.65 | 0.14 | -0.22 | 0.62 | 0.76 | 0.42 | 0.83 | 0.14 | 0.83 | -0.29 | -0.07 | -0.2225 | 0.26 | -0.245 | 0.425836 | 0.305765 | 0.453065 | 1 | 0.996511 | 0.885993 |
| Ak3 | Q9WTP7 | 0.87 | 0.74 | 1.2 | 1.46 | 0.65 | 0.86 | 1.43 | 1.52 | 0.53 | 0.75 | 1.54 | 0.6 | 0.29 | 0.72 | 1.06 | 1.17 | 0.0475 | -0.2125 | -0.2575 | 0.861861 | 0.474207 | 0.36596 | 1 | 0.996511 | 0.858761 |

| | | | | | | | | | | | | | | | | | | | | | | | | | | |
|---|---|---|---|---|---|---|---|---|---|---|---|---|---|---|---|---|---|---|---|---|---|---|---|---|---|---|
| Ak4 | Q9WUR9 | 0.35 | 0.58 | 0.52 | 0.43 | 0.37 | 0.4 | 0.6 | 0.44 | 0.07 | 0.47 | 0.73 | 0.17 | 0.07 | 0.41 | 0.09 | 0.34 | -0.0175 | -0.11 | -0.2425 | 0.906452 | 0.59371 | 0.21111 | 1 | 0.996511 | 0.786269 |
| Ak5 | Q920P5 | -0.66 | -0.56 | -0.65 | -0.63 | -0.36 | -0.67 | -0.78 | -0.71 | -0.29 | -0.23 | -0.67 | -0.49 | -0.03 | 0.01 | -0.74 | -0.78 | -0.005 | 0.205 | 0.24 | 0.974897 | 0.259842 | 0.352139 | 1 | 0.996511 | 0.854088 |
| Akap1 | O08715 | 0.28 | 0.24 | 0.03 | 0.18 | 0.1 | 0.49 | -0.06 | 0.22 | 0.67 | 0.38 | 0.29 | 0.4 | 0.78 | 0.26 | 0.28 | 0.3 | 0.005 | 0.2525 | 0.2225 | 0.97734 | 0.169367 | 0.287494 | 1 | 0.996511 | 0.824705 |
| Akap10 | O88845 | 0.71 | 0.63 | 0.82 | 0.24 | 0.78 | 0.72 | 0.69 | 0.7 | -0.16 | -0.14 | 0.7 | 0.7 | 0.01 | -0.16 | 0.54 | 0.57 | 0.1225 | -0.325 | -0.36 | 0.492339 | 0.274144 | 0.182619 | 1 | 0.996511 | 0.776002 |
| Akap11 | E9Q777 | -0.31 | -0.01 | 0.18 | -0.02 | 1.49 | -0.33 | -0.33 | -0.07 | 0.11 | 0.03 | -0.17 | 0.04 | -1.09 | 1.17 | -0.56 | 0.03 | 0.23 | 0.0425 | -0.0725 | 0.588028 | 0.815465 | 0.876285 | 1 | 0.996511 | 0.979626 |
| Akap12 | Q9WTQ5 | 0.37 | 0.44 | 0.54 | 0.63 | 0.16 | 0.13 | 0.51 | 0.76 | 1.05 | 1.18 | 0.73 | 0.84 | 0.89 | 0.92 | 1.08 | 0.79 | -0.105 | 0.455 | 0.425 | 0.595244 | 0.031553 | 0.038052 | 1 | 0.996511 | 0.658454 |
| Akap13 | E9Q394-2 | -2.73 | -1.28 | -1.57 | -1.01 | -1.4 | -2.9 | -2.13 | -2.08 | -1.52 | -0.57 | -0.93 | -2.3 | -0.89 | -2.5 | -0.42 | -2.1 | -0.48 | 0.3175 | 0.17 | 0.305628 | 0.528428 | 0.766165 | 1 | 0.996511 | 0.964134 |
| Akap5 | H3BIV5 | 2 | 1.25 | 0.96 | 1.68 | 0.86 | 1.07 | 0.93 | 1.2 | 1.65 | 1 | 0.86 | 1.38 | 1.42 | 1.66 | 0.99 | 0.59 | -0.4575 | -0.25 | -0.3075 | 0.098262 | 0.411691 | 0.366963 | 1 | 0.996511 | 0.858904 |
| Akap6 | E9Q9K8 | 0.72 | 0.94 | 0.62 | 0.07 | 0.73 | 0.71 | 0.82 | 0.3 | 0.25 | 0.81 | 0.35 | 1.19 | 0.93 | 0.66 | 0.83 | 0.67 | 0.0525 | 0.0625 | 0.185 | 0.823716 | 0.831357 | 0.442863 | 1 | 0.996511 | 0.883549 |
| Akap7 | Q7TN79 | 1.39 | 1.59 | 1.26 | 1.1 | 1.39 | 1.34 | 1.02 | 0.89 | 1.87 | 1.9 | 1.13 | 1.45 | 1.7 | 2.1 | 1.55 | 0.76 | -0.175 | 0.2525 | 0.1925 | 0.3818 | 0.30671 | 0.538241 | 1 | 0.996511 | 0.912212 |
| Akap8 | Q9DBR0 | 0.63 | -1.92 | -2.76 | -2.72 | -1.82 | -0.07 | -3.72 | -2.64 | -1.53 | -1.89 | -2.51 | -0.59 | -1.1 | -1.64 | -2.06 | -3.34 | -0.37 | 0.0625 | -0.3425 | 0.710875 | 0.938237 | 0.683475 | 1 | 0.996755 | 0.955729 |
| Akap8l | A0A494B9I | 0.14 | -1.13 | -0.7 | -0.73 | -1.68 | -0.47 | -1.09 | -0.19 | -0.48 | -0.9 | -0.63 | -1.48 | 0.76 | -0.72 | -0.3 | -1.25 | -0.2525 | -0.2675 | 0.2275 | 0.534484 | 0.441337 | 0.632026 | 1 | 0.996511 | 0.941132 |
| Akap9 | E9QQ10 | -0.72 | -0.96 | -1.62 | -1.45 | -0.85 | -1.09 | -1.26 | -1.56 | -0.75 | -0.56 | -1.27 | -1 | -0.64 | -0.63 | -1.07 | -1.29 | -0.0025 | 0.2925 | 0.28 | 0.992453 | 0.302227 | 0.338757 | 1 | 0.996511 | 0.847578 |
| Akr1a1 | Q9JII6 | 0.47 | 0.06 | -0.08 | -0.65 | -0.34 | 0.45 | -0.53 | -0.29 | 0.41 | -1.02 | -0.3 | 0.23 | 0.82 | -0.7 | -0.46 | -0.82 | -0.1275 | -0.12 | -0.24 | 0.683301 | 0.7535 | 0.573691 | 1 | 0.996511 | 0.925747 |
| Akr1b1 | P45376 | 0 | -0.91 | -0.64 | -0.76 | -0.57 | 0.01 | -0.51 | -0.75 | -1.46 | -0.55 | -0.68 | -0.04 | -1.44 | -1.09 | -0.82 | -0.95 | 0.1225 | -0.105 | -0.4975 | 0.646383 | 0.76368 | 0.089356 | 1 | 0.996511 | 0.69217 |
| Akr1b10 | G5E895 | 1.22 | 0.59 | 0.89 | 0 | 1.14 | 1.43 | 0.76 | 0.74 | -1.05 | -0.98 | 0.96 | 0.7 | -1.27 | -0.85 | 0.61 | 1 | 0.3425 | -0.7675 | -0.8025 | 0.278095 | 0.186328 | 0.177241 | 1 | 0.996511 | 0.772068 |
| Akr1b7 | P21300 | -1.02 | -0.74 | -1.49 | -0.81 | -0.37 | -1.87 | -0.99 | -0.48 | -2.07 | -1.78 | -0.53 | -2.55 | -2.69 | 0.31 | -1.29 | -2.25 | 0.0875 | -0.7175 | -0.465 | 0.8104 | 0.12817 | 0.46499 | 1 | 0.996511 | 0.888267 |
| Akr1e2 | Q9DCT1 | -0.66 | -0.7 | -1.19 | -1.2 | -0.29 | -0.8 | -0.92 | -1.08 | -1.05 | -1.24 | -1.1 | -0.9 | -1.29 | -0.49 | -1.5 | -1.43 | 0.165 | -0.135 | -0.24 | 0.50051 | 0.522455 | 0.421122 | 1 | 0.996511 | 0.875639 |
| Akr7a2 | Q8CG76 | -0.02 | 0.13 | 0.17 | 0.29 | -0.07 | -0.14 | -0.02 | 0.06 | -0.14 | -0.06 | -0.02 | -0.39 | 0.17 | -0.49 | -0.07 | -0.27 | -0.185 | -0.295 | -0.3075 | 0.240429 | 0.120607 | 0.171151 | 1 | 0.996511 | 0.762774 |
| Akt1 | P31750 | -0.61 | -0.75 | -0.99 | -1.37 | -0.67 | -0.83 | -1.06 | -1.29 | -0.82 | -1.52 | -1.21 | -0.94 | -0.46 | -1.09 | -1.47 | -1.49 | -0.0325 | -0.1925 | -0.1975 | 0.888917 | 0.450521 | 0.521303 | 1 | 0.996511 | 0.907667 |
| Akt1s1 | Q9D1F4 | 2.17 | 0.9 | 0.45 | 0.28 | 0.81 | 1.78 | 1.09 | 0.24 | 0.94 | 0.39 | 0.61 | 1.74 | 0.37 | 1.14 | -0.13 | 0.22 | 0.03 | -0.03 | -0.55 | 0.951421 | 0.950598 | 0.263958 | 1 | 0.996755 | 0.816045 |
| Akt2 | Q60823 | -0.43 | -0.38 | 0.21 | 0.21 | 0.17 | 0.09 | 0.73 | 0.08 | -0.06 | 0.05 | -0.01 | -0.15 | -0.22 | 0.15 | -0.24 | -0.09 | 0.365 | 0.055 | -0.0025 | 0.167658 | 0.802822 | 0.991636 | 1 | 0.996511 | 0.996154 |
| Akt3 | Q9WUA6 | -0.46 | -0.37 | -0.22 | 0.44 | -0.1 | -0.51 | -0.23 | 0.38 | -0.46 | 0.28 | 0.02 | -0.2 | -0.57 | 0.03 | -0.12 | -0.23 | 0.0375 | 0.0625 | -0.07 | 0.893005 | 0.818555 | 0.792947 | 1 | 0.996511 | 0.96933 |
| Aktip | D3Z632 | -2.33 | -2.37 | -1.07 | -1.44 | -1.82 | -2.18 | -0.98 | -1.37 | -2.78 | -0.34 | -1.14 | -2.54 | -2.94 | -1.16 | -1.41 | -1.58 | 0.215 | 0.1025 | 0.03 | 0.588666 | 0.86629 | 0.950293 | 1 | 0.996511 | 0.992476 |
| Alad | P10518 | -0.59 | -0.76 | -0.85 | -0.73 | -0.7 | -0.75 | -1.2 | -0.97 | -0.82 | -0.78 | -1.2 | -0.72 | -0.99 | -0.07 | -1.46 | -1.19 | -0.1725 | -0.1475 | -0.195 | 0.338742 | 0.431131 | 0.539795 | 1 | 0.996511 | 0.912212 |
| Alb | P07724 | -5.88 | -7.07 | -4.97 | -4.85 | -5.67 | -5.72 | -4.64 | -5.03 | -6.39 | -3.03 | -4.78 | -5.44 | -5.51 | -4.33 | -5.05 | -5.51 | 0.4275 | 0.7825 | 0.5925 | 0.428209 | 0.33686 | 0.289725 | 1 | 0.996511 | 0.825111 |
| Alcam | E9Q3Q6 | 1.15 | 0.56 | 0.31 | 0.95 | 0.91 | 1.5 | 0.61 | 0.99 | 1.47 | 1.02 | 0.53 | 1.66 | 1.11 | 1.52 | 0.62 | 0.57 | 0.26 | 0.4275 | 0.2125 | 0.349349 | 0.200022 | 0.491847 | 1 | 0.996511 | 0.897531 |
| Aldh16a1 | D3Z0B9 | 1.59 | 0.41 | 1.41 | 0.48 | 0.21 | 0.57 | -0.9 | -0.06 | 0.52 | 0.42 | -0.18 | 0.16 | 2.14 | 0.78 | 1.3 | -0.63 | -1.0175 | -0.7425 | -0.075 | 0.033864 | 0.053715 | 0.900816 | 1 | 0.996511 | 0.983757 |
| Aldh18a1 | Q9Z110 | 0.49 | 0.55 | 0.71 | 0.58 | 0 | 0.53 | 0.26 | 0.58 | 0.48 | 0.78 | 0.64 | 0.45 | 0.22 | 0.07 | 0.14 | 0.36 | -0.24 | 0.005 | -0.385 | 0.214844 | 0.976372 | 0.052966 | 1 | 0.997588 | 0.661271 |
| Aldh18a1 | Q9Z110-2 | -0.64 | -0.96 | -0.3 | 0 | -1.15 | -1.17 | 0.34 | -0.48 | -0.68 | -0.8 | 0.51 | -0.57 | -1.26 | 0.47 | -1.14 | -1.08 | -0.14 | 0.09 | -0.2775 | 0.720663 | 0.801498 | 0.529224 | 1 | 0.996511 | 0.908663 |
| Aldh1a1 | P24549 | 1.46 | 0.51 | 0.55 | 0.85 | 1.04 | 1.04 | 0.57 | 1.28 | 1.17 | -0.22 | 0.46 | 1.41 | 0.23 | 1.18 | 0.2 | 0.12 | 0.14 | -0.1375 | -0.41 | 0.607637 | 0.737121 | 0.240874 | 1 | 0.996511 | 0.795697 |
| Aldh1a2 | Q62148 | -2.5 | -2.97 | -1.74 | -1.63 | -2.79 | -2.09 | -1.24 | -1.88 | -2.5 | -0.47 | -1.53 | -2.53 | -2.03 | -1.59 | -2.26 | -1.48 | 0.21 | 0.4525 | 0.37 | 0.621752 | 0.410629 | 0.321815 | 1 | 0.996511 | 0.840788 |
| Aldh1a7 | O35945 | -0.2 | 0.15 | -0.09 | -0.88 | -0.72 | 0.06 | -0.06 | -0.69 | 1.79 | 0.23 | 0.03 | 0.43 | 0.32 | 0.94 | -1.25 | -0.48 | -0.0975 | 0.875 | 0.1375 | 0.745108 | 0.068859 | 0.779592 | 1 | 0.996511 | 0.967289 |
| Aldh1b1 | Q9CZS1 | 0.65 | 0.37 | 0.39 | 0.91 | -0.11 | 0.58 | 0.58 | 0.7 | 0.72 | 0.43 | 0.55 | 0.51 | 0.42 | 0.79 | 0.18 | 0.04 | -0.1425 | -0.0275 | -0.2225 | 0.556498 | 0.887563 | 0.373953 | 1 | 0.996511 | 0.859424 |
| Aldh1l1 | Q8R0Y6 | 0.5 | -0.2 | -0.24 | 0.25 | 0.43 | 0.79 | 0.27 | 0.02 | 0.03 | -0.18 | 0 | 0.03 | -0.13 | 0.17 | -0.51 | -0.15 | 0.3 | -0.1075 | -0.2325 | 0.254333 | 0.630983 | 0.377536 | 1 | 0.996511 | 0.860984 |
| Aldh1l2 | Q8K009 | 0.38 | 0.45 | 0.49 | 0.72 | 0.24 | 0.67 | 0.45 | 0.66 | -0.25 | -0.64 | 0.33 | 0.14 | -0.48 | -0.41 | -0.55 | -0.57 | -0.005 | -0.615 | -1.0125 | 0.977145 | 0.034527 | 0.000326 | 1 | 0.996511 | 0.270112 |
| Aldh2 | P47738 | 0.55 | 0.36 | 0.49 | 0.73 | 0.18 | 0.38 | 0.89 | 0.68 | 0.64 | 0.27 | 0.7 | 0.12 | 0.76 | 0.17 | 1.04 | 0.36 | 4.91E-17 | -0.1 | 0.05 | 1 | 0.629706 | 0.839041 | 1 | 0.996511 | 0.976218 |
| Aldh3a2 | B1AV77 | 1.13 | 1.38 | 0.88 | 1.08 | 1.36 | 1.03 | 1.23 | 0.95 | 1.19 | 0.71 | 0.67 | 1.03 | 0.61 | 1.14 | 1 | 1.12 | 0.025 | -0.2175 | -0.15 | 0.890934 | 0.309638 | 0.491442 | 1 | 0.996511 | 0.897531 |
| Aldh3b1 | Q80VQ0 | 1.66 | 1.66 | 0.83 | 0.83 | 1.79 | 1.11 | 1.22 | 0.44 | 0.76 | 0.77 | 0.08 | 1.14 | 0.13 | 0.57 | 1.09 | 0.95 | -0.105 | -0.5575 | -0.56 | 0.765725 | 0.11404 | 0.11361 | 1 | 0.996511 | 0.716416 |
| Aldh4a1 | Q8CHT0 | 0.94 | 1.01 | 0.83 | 1.13 | 0.6 | 0.76 | 0.94 | 0.75 | 1.01 | 0.56 | 0.97 | 0.66 | 0.92 | 0.44 | 0.8 | 0.91 | -0.215 | -0.1775 | -0.21 | 0.197721 | 0.356714 | 0.304461 | 1 | 0.996511 | 0.831662 |
| Aldh5a1 | Q8BWF0 | 1.51 | 1.4 | 0.8 | 0.68 | 1.01 | 1.7 | 1.03 | 0.85 | 0.61 | -0.24 | 0.89 | 1.19 | 0.78 | -0.4 | 0.52 | 0.88 | 0.05 | -0.485 | -0.6525 | 0.859997 | 0.203628 | 0.09353 | 1 | 0.996511 | 0.698156 |
| Aldh6a1 | Q9EQ20 | 1.87 | 0.8 | 0.67 | 1.12 | 0.68 | 1.76 | 1.24 | 0.93 | 0.72 | 0.74 | 0.81 | 1.13 | 0.55 | 0.36 | 0.48 | 0.82 | 0.0375 | -0.265 | -0.5625 | 0.912889 | 0.377681 | 0.087336 | 1 | 0.996511 | 0.691619 |
| Aldh7a1 | Q9DBF1-2 | 0.59 | 0.51 | 0.11 | 0.27 | 0.73 | 0.51 | 0.7 | 0.17 | -0.1 | 0.13 | 0.09 | 0.21 | -0.29 | 0.17 | -0.21 | 0.14 | 0.1575 | -0.2875 | -0.4175 | 0.44292 | 0.151431 | 0.07972 | 1 | 0.996511 | 0.681818 |
| Aldh9a1 | Q9JLJ2 | -0.23 | -0.47 | -0.31 | -0.25 | 0.02 | -0.03 | 0.56 | -0.02 | -0.37 | 0.07 | -0.43 | -0.41 | -0.32 | -0.19 | -0.39 | -0.13 | 0.4475 | 0.03 | 0.0575 | 0.041381 | 0.873778 | 0.745815 | 1 | 0.996511 | 0.960079 |
| Aldoa | A6ZI44 | -0.11 | -0.8 | -0.16 | 0.88 | -0.54 | -0.96 | 0.68 | 0.61 | 0.61 | 1.36 | 0.97 | 0.38 | 0.46 | 0.68 | 0.53 | 1.38 | -0.005 | 0.8775 | 0.81 | 0.991964 | 0.048837 | 0.06511 | 1 | 0.996511 | 0.66433 |
| Aldoa | P05064 | -0.49 | -0.73 | -0.8 | -0.68 | -0.75 | -0.42 | -0.74 | -0.64 | -0.45 | -0.5 | -0.71 | -0.26 | -0.8 | -0.55 | -1.01 | -1.09 | 0.0375 | 0.195 | -0.1875 | 0.816932 | 0.295508 | 0.36653 | 1 | 0.996511 | 0.858904 |
| Aldoc | P05063 | 0.87 | 0.13 | 0.16 | 0.61 | 0.56 | 0.35 | 0.39 | 0.56 | 0.5 | 0.35 | 0.29 | 0.26 | -0.29 | 0.56 | -0.27 | 0.08 | 0.0225 | -0.0925 | -0.4225 | 0.916314 | 0.679934 | 0.165252 | 1 | 0.996511 | 0.758837 |
| Alg1 | Q921Q3 | 2.03 | 1.4 | 0.66 | 0.57 | 1.51 | 1.81 | 0.78 | 0.44 | 1.92 | 0.89 | 0.17 | 1.63 | 1.68 | 1.79 | 1.05 | 0.86 | -0.03 | -0.0125 | 0.18 | 0.945109 | 0.979472 | 0.653094 | 1 | 0.998773 | 0.947633 |
| Alg10b | Q3UGP8 | -0.61 | 0.21 | -0.59 | -0.55 | 0.73 | -0.24 | 0.47 | -0.11 | -0.29 | -0.17 | -0.3 | -0.49 | -0.86 | 0.51 | -0.01 | 0.16 | 0.5975 | 0.0725 | 0.335 | 0.075124 | 0.761011 | 0.351348 | 1 | 0.996511 | 0.853984 |
| Alg11 | Q3TZM9 | -0.15 | 0.19 | -1.32 | -1.16 | 0.65 | 0 | 0.22 | -0.93 | 0.11 | -2.33 | -1.3 | -0.19 | 0 | 0.06 | -0.8 | -0.49 | 0.595 | -0.3175 | 0.3025 | 0.221372 | 0.607333 | 0.465502 | 1 | 0.996511 | 0.888369 |
| Alg13 | Q9D8C3 | -1.28 | 0.53 | 1.01 | 1.33 | 2.7 | 5.22 | 1.33 | 0.45 | 0 | -0.15 | 0.61 | -0.94 | 0.36 | 2.68 | 0.44 | 1.44 | 2.0275 | -0.5175 | 0.8325 | 0.085418 | 0.405247 | 0.266702 | 1 | 0.996511 | 0.818426 |
| Alg2 | Q9DBE8 | -0.36 | -0.66 | -0.32 | -0.56 | -0.21 | -0.43 | 0.44 | -0.28 | -0.85 | -0.39 | -0.3 | -0.5 | -0.55 | -0.04 | 0.21 | 0.2 | 0.355 | -0.035 | 0.43 | 0.146864 | 0.85892 | 0.096256 | 1 | 0.996511 | 0.698156 |
| Alg3 | A0A338P7I | 0.1 | 1 | 0.31 | -0.34 | 0.02 | 1.28 | 1.66 | 0.72 | 0.75 | 1.56 | 1.12 | 1.15 | -0.04 | 1.19 | 1.38 | 1.3 | 0.6525 | 0.8775 | 0.69 | 0.150563 | 0.023026 | 0.125218 | 1 | 0.996511 | 0.721448 |
| Alg5 | Q9DB25 | 0 | 0.23 | -0.22 | -0.11 | 0.45 | -0.34 | 0.03 | -0.32 | -0.1 | 0.1 | -0.38 | -0.13 | 0.12 | 0.64 | 0.17 | 0.36 | -0.02 | -0.1025 | 0.3475 | 0.930308 | 0.597529 | 0.125307 | 1 | 0.996511 | 0.721448 |
| Alg9 | Q8VDI9 | 0.17 | 0.1 | -0.74 | -0.23 | -0.39 | 0.86 | -1.57 | -0.81 | 1.83 | -0.46 | -0.94 | -0.22 | 2.94 | -0.72 | 0.14 | -0.26 | -0.3025 | 0.2275 | 0.7 | 0.554102 | 0.701883 | 0.373274 | 1 | 0.996511 | 0.859424 |
| Alk | P97793 | -0.06 | 0.01 | -0.66 | 0.06 | -0.32 | -0.03 | -0.23 | 0.14 | 1.12 | 0.56 | -0.48 | -0.18 | 0.91 | 0.33 | -0.05 | -0.51 | 0.0525 | 0.4175 | 0.3325 | 0.811709 | 0.293602 | 0.346397 | 1 | 0.996511 | 0.851096 |
| Alkbh3 | Q8K1E6 | -0.38 | -0.5 | -1.09 | -0.81 | -2.66 | -0.03 | -4.46 | -1.55 | -1.81 | -4.94 | -2.49 | -0.44 | -0.02 | -8.36 | -2.8 | -2.54 | -1.48 | -1.725 | -2.735 | 0.110332 | 0.072177 | 0.108515 | 1 | 0.996511 | 0.707441 |
| Alkbh5 | Q3TSG4 | -0.9 | -0.86 | -2.72 | -2.21 | -0.97 | -1.35 | -2.39 | -2.03 | -0.34 | -0.73 | -2.51 | -0.73 | -0.93 | -0.82 | -2.2 | -2.21 | -0.0125 | 0.595 | 0.1325 | 0.980912 | 0.349272 | 0.812852 | 1 | 0.996511 | 0.972768 |
| Alkbh6 | Q8K2U2 | -1.79 | -1.61 | 0.48 | -0.74 | -1.6 | -2.03 | 0.17 | 0.53 | -1.55 | -0.2 | 0.16 | -1.62 | -1 | -1.31 | 0.21 | 0.52 | 0.1825 | 0.1125 | 0.52 | 0.805686 | 0.858734 | 0.415504 | 1 | 0.996511 | 0.874945 |
| Alkbh7 | Q9D6Z0 | -0.06 | -0.72 | -0.92 | -0.59 | -0.68 | 0.35 | -0.22 | 0.09 | 0.38 | -1.36 | -0.45 | -0.99 | -0.88 | -2.11 | -0.62 | -1.66 | 0.4575 | -0.0325 | -0.745 | 0.139735 | 0.935412 | 0.076859 | 1 | 0.996755 | 0.67615 |
| Alkbh8 | Q80Y20-2 | 0.81 | 0.3 | 1 | 0.66 | -0.6 | 0.71 | 1.03 | 0.9 | 0.26 | -0.75 | 0.92 | 0.78 | -0.89 | -0.07 | 1.1 | 0.95 | -0.1825 | -0.39 | -0.42 | 0.635888 | 0.331871 | 0.374084 | 1 | 0.996511 | 0.859424 |
| Alox5ap | A0A0J9YUC | 0.4 | 0.91 | 0.73 | 0.83 | 0.54 | 0.66 | 1.33 | -0.91 | 0.21 | 3.85 | 0.75 | 0.68 | -0.6 | 0.32 | 0.36 | 1.71 | -0.3125 | 0.655 | -0.27 | 0.494559 | 0.400777 | 0.561403 | 1 | 0.996511 | 0.91997 |
| Alpl | P09242 | 0.1 | 0.18 | -0.25 | 0.02 | 0.13 | -0.27 | -1.19 | -0.54 | -0.21 | -1.07 | -0.45 | -0.2 | 0.01 | -0.19 | 0.02 | -0.64 | -0.48 | -0.495 | -0.2125 | 0.128075 | 0.073043 | 0.361456 | 1 | 0.996511 | 0.858475 |
| Als2 | Q920R0 | -0.25 | 0.88 | 0.47 | 0.16 | 1.01 | -0.1 | 0.05 | 0.01 | 0.86 | 0.23 | 0.34 | 0.36 | 0.96 | 1.29 | 0.5 | 0.43 | -0.0725 | 0.1325 | 0.48 | 0.831144 | 0.646429 | 0.1591 | 1 | 0.996511 | 0.754492 |
| Alyref | O08583 | 0.99 | -0.96 | -0.63 | -0.63 | -0.97 | 0.11 | -0.3 | -0.09 | -1.3 | -2.66 | -0.6 | -1.1 | -1.46 | -1.23 | -0.03 | -0.44 | -0.005 | -1.1075 | -0.4825 | 0.991365 | 0.080271 | 0.359802 | 1 | 0.996511 | 0.856645 |
| Amacr | O09174 | 1.4 | 1.07 | 0.65 | 0.78 | 1.01 | 1.09 | 0.98 | 0.9 | 0.94 | 0.54 | 0.76 | 1.4 | 0.51 | 0.87 | 0.64 | 0.77 | 0.02 | -0.065 | -0.2775 | 0.921377 | 0.80624 | 0.245849 | 1 | 0.996511 | 0.802442 |
| Ambra1 | A2AH22-7 | -2.18 | -2.23 | -1.89 | -2.42 | -2.5 | -1.51 | -2.13 | -2.18 | -0.22 | -1.43 | -2.18 | -2.44 | 1.83 | -2.66 | -1.42 | -1.98 | 0.1 | 0.6125 | 1.1225 | 0.687077 | 0.220395 | 0.23248 | 1 | 0.996511 | 0.788994 |
| Amdhd2 | F6UP77 | 4.49 | 2.36 | 2.31 | 0.14 | 4.05 | 4.62 | 2.34 | 1.57 | -0.75 | -0.44 | 2.2 | 4.52 | -0.78 | 0.13 | 2.12 | 2.52 | 0.82 | -0.9425 | -1.3275 | 0.431257 | 0.494915 | 0.232062 | 1 | 0.996511 | 0.788863 |
| Amer2 | A0A0R4J03 | -0.46 | -0.46 | 0.14 | -0.01 | -0.31 | -0.37 | -0.18 | -0.38 | -0.52 | 0.29 | -0.08 | -0.48 | -0.47 | -0.1 | -0.25 | 0.02 | -0.1125 | 1.18E-16 | -0.0025 | 0.570238 | 1 | 0.991361 | 1 | 1 | 0.996154 |
| Amer3 | Q6NS69 | -0.83 | 0.98 | -0.62 | -0.63 | 0.15 | -1.29 | -1.57 | -0.77 | 1.4 | 0.51 | -0.01 | 0.65 | 3.21 | 0.31 | -0.01 | -0.11 | -0.595 | 0.9125 | 1.125 | 0.271575 | 0.08392 | 0.187716 | 1 | 0.996511 | 0.776843 |
| Amfr | Q9R049-2 | 0.01 | -0.21 | -0.36 | -0.51 | 0.41 | -0.18 | 0.17 | -0.25 | 0.88 | -0.28 | -0.52 | 0.16 | 1.17 | 1.3 | 0.53 | 0.13 | 0.305 | 0.3275 | 1.05 | 0.18169 | 0.329266 | 0.007395 | 1 | 0.996511 | 0.552614 |
| Amigo1 | Q80ZD8 | 0.69 | 0.91 | 0.67 | 1.08 | 1.26 | 0.75 | 0.82 | 0.67 | 0.91 | 0.84 | 0.49 | 0.85 | 0.85 | 1.72 | 0.2 | 0.7 | 0.0375 | -0.065 | 0.03 | 0.849843 | 0.735526 | 0.92823 | 1 | 0.996511 | 0.98946 |
| Amigo2 | Q80ZD9 | 0.84 | 2.31 | -0.07 | 0.67 | 2.05 | 0.64 | 0.77 | 0.34 | 1.01 | 1.25 | 0.66 | 1.34 | 0.24 | 3.15 | 0.68 | 1.12 | 0.0125 | 0.1275 | 0.36 | 0.982535 | 0.793225 | 0.626997 | 1 | 0.996511 | 0.939737 |
| Amot | Q8VHG2-2 | 0.99 | 0.5 | 1.16 | 1.13 | 0.66 | 1.52 | 1.16 | 0.99 | 1.57 | 1.41 | 1.23 | 1.12 | 2.29 | 0.27 | 1.19 | 1.29 | 0.1375 | 0.3875 | 0.315 | 0.582773 | 0.105421 | 0.461385 | 1 | 0.996511 | 0.887902 |
| Amotl1 | A0A1Y7VM | 1.31 | 1.4 | 0.42 | 0.24 | 0.99 | 1.3 | 0.48 | 0.37 | 2.06 | 0.13 | 0.51 | 2.07 | 2.25 | 1.15 | 0.13 | 0.26 | -0.0575 | 0.35 | 0.105 | 0.871292 | 0.525955 | 0.843377 | 1 | 0.996511 | 0.976225 |
| Amotl2 | B8JK87 | -2.18 | -1.05 | -1.24 | 1.45 | -2.24 | -1.68 | -0.58 | 0.04 | 1.69 | -3.95 | 1.32 | -1.83 | 1.74 | -1.42 | 1.74 | 0.84 | -0.36 | 0.0625 | 1.48 | 0.669451 | 0.963985 | 0.150583 | 1 | 0.996755 | 0.749304 |
| Ampd2 | A2AE27 | 0.73 | 0.71 | -0.12 | -0.14 | 0.67 | 0.4 | -0.08 | -0.06 | 0.07 | -0.84 | -0.27 | 0.58 | -0.16 | 0.52 | 0.03 | -0.38 | -0.0625 | -0.41 | -0.2925 | 0.836704 | 0.289563 | 0.370185 | 1 | 0.996511 | 0.859245 |
| Ampd3 | A0A1L1SRX | 1.37 | 0.08 | 0.18 | 1.19 | 0.86 | 0.61 | 0.32 | 0.68 | 0.1 | -0.04 | 0.27 | 0.96 | 0.01 | 0.78 | 0.05 | 0.27 | -0.0875 | -0.3825 | -0.4275 | 0.797658 | 0.33588 | 0.267314 | 1 | 0.996511 | 0.818895 |
| Amph | Q7TQF7 | 0.58 | 0.65 | -0.12 | 0.11 | 0.71 | 0.4 | -0.67 | -0.75 | 0.68 | 0.1 | -0.48 | 0.54 | 0.85 | 0.54 | -0.48 | -0.67 | -0.3825 | -0.095 | -0.245 | 0.343085 | 0.767329 | 0.546646 | 1 | 0.996511 | 0.912406 |
| Amt | Q8CFA2 | 0.11 | 0.11 | 0.33 | 0.54 | 0 | 0.17 | 1.44 | 0.45 | -0.17 | 0.18 | 1.23 | 0 | 0.63 | 0.08 | 0.71 | 0.52 | 0.2425 | 0.0375 | 0.2125 | 0.468021 | 0.909278 | 0.353828 | 1 | 0.996744 | 0.855029 |
| Amz2 | B1AT59 | -0.08 | -0.57 | -0.63 | -0.63 | -0.78 | -0.16 | -0.71 | -0.62 | 0.1 | -1.58 | -0.41 | -0.08 | 0.01 | -1.98 | -0.76 | -0.25 | -0.09 | -0.015 | -0.2675 | 0.68054 | 0.968901 | 0.545513 | 1 | 0.996755 | 0.912258 |
| Anapc1 | P53995 | -0.75 | -0.68 | -0.67 | -0.67 | -0.42 | -1.06 | -0.65 | -0.88 | -1.05 | -0.55 | -0.73 | -0.71 | -1.06 | -0.73 | -0.89 | -0.63 | -0.06 | -0.0675 | -0.135 | 0.743951 | 0.703201 | 0.467564 | 1 | 0.996511 | 0.889213 |
| Anapc13 | Q8R034 | -0.98 | -0.13 | -0.93 | -1.31 | -0.25 | -1.83 | -1.55 | -0.91 | -0.97 | -1.81 | -0.9 | -1.65 | -0.9 | 0.5 | -0.64 | -0.44 | -0.2975 | -0.495 | 0.4675 | 0.471575 | 0.167931 | 0.243039 | 1 | 0.996511 | 0.798789 |
| Anapc16 | Q9CPV2 | -1.05 | -0.81 | -0.7 | -0.54 | -1.3 | -1.66 | -0.66 | -1.37 | -1.58 | -0.52 | -1.16 | -0.43 | -0.88 | -0.67 | -1.62 | -1.46 | -0.4725 | -0.1475 | -0.3825 | 0.085085 | 0.624396 | 0.186828 | 1 | 0.996511 | 0.776843 |
| Anapc2 | Q8BZQ7 | -0.76 | -0.82 | 0.03 | -0.22 | -0.58 | -1.1 | -0.04 | -0.16 | -0.87 | 0.23 | -0.06 | -0.89 | -1.04 | -0.5 | -0.18 | 0.09 | -0.0275 | 0.045 | 0.035 | 0.929654 | 0.896488 | 0.91403 | 1 | 0.996511 | 0.986458 |
| Anapc4 | Q91W96 | -0.54 | -0.25 | -0.44 | -0.5 | -0.33 | -0.77 | 0.07 | -0.54 | -0.37 | -1.13 | -0.68 | -0.69 | -0.92 | 0 | -1.05 | -0.26 | 0.04 | -0.285 | -0.125 | 0.852583 | 0.20049 | 0.658879 | 1 | 0.996511 | 0.949645 |
| Anapc5 | A0A0G2JEC | -0.9 | -0.46 | -0.35 | -0.79 | -0.71 | -1.05 | -0.32 | -0.95 | -0.64 | -0.28 | -0.5 | -0.74 | -0.59 | -0.04 | -0.54 | -0.66 | -0.1325 | 0.085 | 0.1675 | 0.565834 | 0.684771 | 0.481674 | 1 | 0.996511 | 0.893906 |
| Anapc7 | Q9WVM3 | 0.72 | 0.37 | -0.07 | 0.21 | 0.91 | 0.34 | -0.23 | -0.22 | 0.88 | 0.53 | -0.01 | 0.78 | 0.79 | -0.1 | 0.03 | -0.15 | -0.1075 | 0.2375 | -0.165 | 0.731389 | 0.395001 | 0.573735 | 1 | 0.996511 | 0.925747 |
| Angel2 | Q8K1C0-3 | -0.3 | -0.1 | -0.71 | 0.12 | -0.93 | -0.01 | -0.31 | -0.21 | 0.17 | -1.13 | -0.56 | -0.44 | 0.38 | -0.98 | -1.13 | -0.76 | -0.1175 | -0.2425 | -0.375 | 0.666156 | 0.456218 | 0.332993 | 1 | 0.996511 | 0.845601 |
| Ank1 | A0A0R4J1N | 0.59 | 0.19 | 0.76 | 1.5 | 0.42 | 0.73 | 0.78 | 1.25 | 0.12 | 0.09 | 0.52 | -0.17 | 0.1 | 0.41 | 0.37 | 0.57 | 0.035 | -0.62 | -0.3975 | 0.911952 | 0.073418 | 0.211745 | 1 | 0.996511 | 0.786269 |
| Ank2 | S4R2D6 | 1.5 | 0.88 | 1.28 | 1.11 | 0.3 | 1.51 | 1.06 | 2.01 | 1.2 | 0.71 | 1.72 | 1.82 | 0.82 | 0.92 | 1.91 | 1.91 | 0.0275 | 0.17 | 0.1975 | 0.940386 | 0.568605 | 0.555049 | 1 | 0.996511 | 0.916796 |
| Ank2 | Q8C8R3-7 | 0.57 | 0.72 | 1.64 | 1.86 | -1.02 | 1.08 | 0.12 | 1.75 | 2.17 | 1.4 | 1.41 | 0.3 | 3.04 | -0.14 | 1.73 | 1.14 | -0.715 | 0.1225 | 0.245 | 0.269221 | 0.794814 | 0.714949 | 1 | 0.996511 | 0.958106 |
| Ank2 | Q8C8R3-2 | 1.52 | 1.26 | 1.61 | 1.32 | 1.43 | 1.1 | 1.39 | 1.74 | 1.18 | 0.86 | 1.76 | 1.4 | 0.95 | 1.67 | 1.4 | 1.48 | -0.0125 | -0.1275 | -0.0525 | 0.948071 | 0.591768 | 0.813708 | 1 | 0.996511 | 0.972768 |
| Ank2 | Q8C8R3-4 | 0.75 | 0.79 | 0.65 | 0.8 | 0.69 | 0.69 | 0.7 | 0.8 | 1.01 | 0.64 | 0.79 | 0.89 | 1.19 | 0.58 | 0.86 | 0.49 | -0.0275 | 0.085 | 0.0325 | 0.844423 | 0.61408 | 0.880116 | 1 | 0.996511 | 0.979626 |
| Ank3 | S4R2K9 | 1.05 | 0.69 | 0.88 | 0.94 | 0.6 | 0.9 | 0.71 | 0.75 | 0.8 | 0.69 | 0.84 | 0.77 | 0.74 | 0.71 | 0.68 | 0.68 | -0.15 | -0.115 | -0.1875 | 0.361222 | 0.494297 | 0.301876 | 1 | 0.996511 | 0.830681 |
| Ank3 | A0A087WN | 1.39 | 0.96 | 0.62 | 0.88 | 0.81 | 0.79 | 0.23 | 0.23 | 0.96 | 0.96 | 0.67 | 1.1 | 0.9 | 1.41 | 0.38 | 0.18 | -0.4475 | -0.04 | -0.245 | 0.094016 | 0.856145 | 0.457469 | 1 | 0.996511 | 0.887902 |
| Ank3 | A0A087WR | 0.65 | 0.94 | -0.07 | -0.28 | 1.58 | 0.69 | -0.9 | 0.22 | 1.67 | 1.77 | -0.86 | 1.4 | 1.52 | 0.27 | -1.47 | -0.48 | 0.0875 | 0.685 | -0.35 | 0.871922 | 0.292505 | 0.583602 | 1 | 0.996511 | 0.927734 |
| Ank3 | G5E8K2 | -0.34 | -0.33 | -0.78 | -0.37 | -0.16 | -0.75 | -0.03 | -0.16 | -0.32 | -1.38 | -0.29 | -0.53 | -0.51 | -0.25 | -0.81 | -0.53 | 0.18 | -0.175 | -0.07 | 0.419623 | 0.547021 | 0.743561 | 1 | 0.996511 | 0.959321 |
| Ankfn1 | F6X7B3 | 0.79 | 0.33 | 1.88 | 2.33 | 0.45 | 0.55 | 1.76 | 3.26 | -0.1 | 1.94 | 1.42 | -0.25 | 0.29 | -0.39 | 0.45 | 3.82 | 0.1725 | -0.58 | -0.29 | 0.81271 | 0.386752 | 0.758758 | 1 | 0.996511 | 0.963116 |
| Ankfy1 | Q810B6 | -0.2 | -0.35 | -0.22 | -0.24 | -0.21 | -0.36 | -0.22 | -0.24 | -0.58 | -0.18 | -0.22 | -0.32 | -0.64 | -0.24 | -0.54 | -0.46 | -0.005 | -0.0725 | -0.2175 | 0.97175 | 0.674361 | 0.249187 | 1 | 0.996511 | 0.804842 |
| Ankh | Q9JHZ2 | -2.06 | -1.75 | -1.27 | -1.66 | -1.26 | -1.93 | -1.28 | -1.33 | -1.41 | -1.49 | -1.21 | -2 | -1.34 | -0.75 | -1.6 | -1.26 | 0.235 | 0.1575 | 0.4475 | 0.347188 | 0.541538 | 0.120862 | 1 | 0.996511 | 0.721448 |
| Ankhd1 | E9Q1M6 | -2.08 | -1.53 | -1.35 | -1.38 | -2.22 | -1.7 | -1.05 | -0.87 | -1.26 | -0.76 | -1.22 | -1.89 | -0.51 | -2.22 | -1.51 | -1.46 | 0.125 | 0.3025 | 0.16 | 0.714909 | 0.320115 | 0.674844 | 1 | 0.996511 | 0.952944 |
| Ankib1 | Q6ZPS6 | 0.56 | 0.64 | 0.6 | 0.42 | 0.65 | 0.6 | 0.46 | 0.21 | 1 | 0.86 | 0.56 | 0.93 | 1.21 | 0.67 | 0.68 | 0.57 | -0.075 | 0.2825 | 0.2275 | 0.653204 | 0.137799 | 0.295996 | 1 | 0.996511 | 0.825612 |
| Ankle2 | Q6P1H6 | 0.41 | 0.43 | -0.47 | -0.04 | 0.55 | 0.4 | -0.55 | -0.43 | 0.41 | 0.29 | -0.68 | 0.46 | 0.22 | 1.12 | -0.14 | -0.51 | -0.09 | 0.0375 | 0.09 | 0.792203 | 0.911883 | 0.820132 | 1 | 0.996744 | 0.972768 |
| Ankmy1 | E0CYY3 | -2.24 | -2.33 | 0.07 | -0.65 | -3.88 | -1.36 | 0.99 | -0.56 | 0.14 | -0.59 | 1.28 | -0.93 | 1.39 | -4.02 | -0.12 | -0.38 | 0.085 | 1.2625 | 0.505 | 0.935791 | 0.099088 | 0.662787 | 1 | 0.996511 | 0.94972 |
| Ankmy2 | Q3TPE9 | 0.62 | -0.47 | 0 | 0.28 | 0.07 | -0.3 | 0.98 | 0.44 | -0.03 | 0.02 | 0.11 | 0.07 | 0.48 | 0.04 | 0.69 | 0.77 | 0.19 | -0.065 | 0.3875 | 0.584565 | 0.798304 | 0.21378 | 1 | 0.996511 | 0.786931 |
| Ankrd13a | Q80UP5 | -0.7 | -0.86 | 0.43 | -0.15 | -0.8 | -0.5 | 0.14 | -0.17 | -0.22 | 0.72 | 0.07 | -0.6 | -0.09 | -0.6 | 0.22 | 0.06 | -0.0125 | 0.3125 | 0.2175 | 0.970957 | 0.429678 | 0.529657 | 1 | 0.996511 | 0.908663 |
| Ankrd13c | Q3UX43 | -0.14 | -0.17 | -0.08 | -0.26 | 0.57 | -0.23 | 0.15 | -0.14 | -0.81 | -0.76 | 0.01 | -0.22 | -0.84 | -0.45 | -0.24 | -0.36 | 0.25 | -0.2825 | -0.31 | 0.256259 | 0.249595 | 0.149195 | 1 | 0.996511 | 0.748169 |
| Ankrd13d | Q6PD24 | 0.6 | 0.1 | 0.27 | 0.13 | -0.01 | 0.44 | 0.41 | 0.42 | 0.11 | 0.04 | 0.39 | 0.56 | 0.13 | 0.49 | 0.65 | 0.24 | 0.04 | 3.93E-17 | 0.1025 | 0.836724 | 1 | 0.639143 | 1 | 1 | 0.942804 |
| Ankrd17 | A0A0G2JD: | -2.51 | -2.65 | -2.42 | -2.4 | -2.28 | -2.55 | -2.11 | -2.46 | -2.28 | -1.27 | -2.1 | -2.24 | -2.26 | -1.88 | -2.62 | -2.7 | 0.145 | 0.5225 | 0.13 | 0.39813 | 0.073235 | 0.586316 | 1 | 0.996511 | 0.928178 |
| Ankrd27 | Q3UMR0-3 | -0.2 | 0.17 | 0.22 | 0.24 | 0.96 | 0.05 | -0.12 | 0.08 | 0.24 | 1.45 | 0.45 | -0.08 | 0.1 | 0.69 | 0.31 | 0.13 | 0.135 | 0.4075 | 0.2 | 0.619629 | 0.251935 | 0.376956 | 1 | 0.996511 | 0.860984 |
| Ankrd28 | Q505D1 | -0.15 | -0.1 | -0.28 | -0.48 | -0.07 | -0.23 | -0.37 | -0.69 | -0.19 | -0.23 | -0.32 | -0.06 | -0.5 | -0.18 | -0.5 | -0.5 | -0.0875 | 0.0525 | -0.1675 | 0.652694 | 0.762989 | 0.39329 | 1 | 0.996511 | 0.865775 |
| Ankrd29 | D3YTU5 | 0.86 | 0.78 | 1.02 | 0.97 | 0.58 | 0.43 | 0.77 | 0.73 | 0.1 | 0.81 | 0.64 | 0.22 | -0.01 | 0.47 | 0.8 | 0.54 | -0.28 | -0.465 | -0.4575 | 0.105139 | 0.05663 | 0.068163 | 1 | 0.996511 | 0.664936 |
| Ankrd33b | Q3U0L2 | 0.96 | 0.44 | 1.44 | 1.21 | 0.8 | 0.68 | 0.05 | 1.26 | 1.09 | 0.62 | 0.85 | 0.32 | 1.53 | 1.12 | 0.18 | 0.42 | -0.315 | -0.2925 | -0.2 | 0.340924 | 0.314672 | 0.592235 | 1 | 0.996511 | 0.929843 |
| Ankrd34a | B2RW11 | 0.15 | -0.16 | 1.22 | 0.41 | -0.68 | 0.33 | 0.48 | 1.02 | 1.49 | 0.21 | 1.18 | 0.31 | 2.43 | -0.25 | 1.02 | 0.74 | -0.1175 | 0.3925 | 0.58 | 0.78578 | 0.354528 | 0.328952 | 1 | 0.996511 | 0.844943 |
| Ankrd34b | Q3UUF8 | 0.24 | 0.33 | 1.33 | 1.28 | 0.31 | 0.06 | 1.4 | 1.16 | 0.67 | 1.05 | 1.29 | 0.5 | 1.05 | 0.71 | 1.17 | 1.17 | -0.0625 | 0.0825 | 0.23 | 0.879329 | 0.808698 | 0.479968 | 1 | 0.996511 | 0.893906 |
| Ankrd40 | Q5SUE8-2 | 0.11 | 0.14 | -0.57 | -0.19 | -0.13 | 0.46 | -0.11 | -0.31 | 0.8 | -0.02 | -0.4 | 0.51 | 0.42 | 0.76 | -0.32 | -0.15 | 0.105 | 0.35 | 0.305 | 0.673191 | 0.285783 | 0.338599 | 1 | 0.996511 | 0.847578 |
| Ankrd44 | E9QNQ2 | -0.41 | 0.56 | 0.14 | 0.6 | 0.32 | 0.09 | 0.46 | 0.51 | 0.42 | 1.16 | 0.7 | 0.16 | 0.42 | 0.58 | 0.37 | 0.64 | 0.1225 | 0.3875 | 0.28 | 0.641516 | 0.244173 | 0.31287 | 1 | 0.996511 | 0.835247 |
| Ankrd45 | Q810N6 | 1.44 | -0.06 | 0.47 | 0.18 | 0.92 | 0.25 | -0.86 | -0.05 | -0.02 | 0.27 | -1.03 | -0.27 | 0.93 | 0.8 | -0.41 | -0.79 | -0.4425 | -0.77 | -0.375 | 0.348841 | 0.088452 | 0.464873 | 1 | 0.996511 | 0.888267 |
| Ankrd46 | Q8BTZ5 | -0.06 | -0.02 | -2.86 | -2.1 | 0.39 | -0.04 | -1.63 | -3.11 | 0.9 | -0.33 | -2.77 | 0.21 | 0.84 | 1.75 | -2.04 | -2.04 | 1.1625 | 0.7625 | 0.8875 | 0.866453 | 0.439019 | 0.423162 | 1 | 0.996511 | 0.875782 |
| Ankrd49 | Q8VE42 | 0.55 | 0.51 | -0.05 | -0.19 | 0.69 | 0.65 | -0.37 | -0.1 | -0.01 | -0.39 | -0.18 | 0.35 | 0.65 | -0.18 | -0.07 | -0.48 | 0.0125 | -0.2625 | -0.2225 | 0.968852 | 0.334123 | 0.480316 | 1 | 0.996511 | 0.893906 |
| Ankrd50 | F7BE84 | 0.85 | 0.29 | -0.03 | 0.23 | 0.48 | 0.01 | 0.33 | 0.3 | 0.72 | 0.22 | 0.31 | 0.16 | 1.14 | 0.49 | -0.05 | 0.06 | -0.055 | 0.0175 | 0.075 | 0.810594 | 0.943735 | 0.820554 | 1 | 0.996755 | 0.972768 |
| Ankrd52 | Q8BTI7 | -0.81 | -0.58 | -0.74 | -0.14 | 0.07 | -0.35 | -0.2 | -0.53 | -0.3 | -0.34 | -0.51 | -0.69 | -1.17 | -0.11 | -0.09 | 0.46 | 0.315 | 0.1075 | 0.34 | 0.177386 | 0.619419 | 0.365028 | 1 | 0.996511 | 0.858475 |
| Ankrd54 | Q91WK7 | -1.8 | -1.06 | -0.96 | -1.38 | -0.71 | -1.93 | -1.45 | -1.62 | -0.75 | -0.27 | -1.5 | -1.7 | -0.27 | -0.72 | -1.37 | -1.06 | -0.1275 | 0.245 | 0.445 | 0.686674 | 0.513594 | 0.177656 | 1 | 0.996511 | 0.772068 |
| Ankrd63 | A2ARS0 | 1.32 | 0.17 | 0.38 | 0.97 | -0.47 | -0.06 | 0.82 | 0.58 | 0.67 | 0.67 | -0.36 | 0.66 | 0.42 | 0.34 | 0.77 | 0.19 | -0.4925 | -0.3 | -0.28 | 0.213605 | 0.413421 | 0.368354 | 1 | 0.996511 | 0.859245 |
| Anks1 | A0A3B2W4 | 0.9 | 0.74 | 1.03 | 0.72 | 1.36 | 0.47 | 0.62 | 0.39 | 0.35 | 0.86 | 0.86 | 0.84 | 0.18 | 0.73 | 0.49 | 0.97 | -0.1375 | -0.12 | -0.255 | 0.580699 | 0.546969 | 0.282953 | 1 | 0.996511 | 0.822595 |
| Anks1b | A0A0R4J2A | 0.2 | 0.47 | 0.9 | -0.16 | 0.98 | 0.61 | -0.5 | -0.45 | 0.77 | 0.65 | 0.59 | 0.24 | 1.09 | 1.78 | 0.58 | 1.47 | -0.1925 | 0.21 | 0.8775 | 0.640709 | 0.440565 | 0.028548 | 1 | 0.996511 | 0.637553 |
| Anks1b | A0A0R4J2A | 2.08 | 1.56 | 2.49 | 1.13 | 2.07 | 2.27 | 2.92 | 2.75 | -3.66 | -2.96 | 3.18 | 2 | -2.31 | -4.08 | 2.71 | 2.19 | 0.6875 | -2.125 | -2.1875 | 0.073238 | 0.202923 | 0.17192 | 1 | 0.996511 | 0.76392 |
| Anks1b | S4R286 | 0.78 | 0.72 | 0.81 | 1.01 | 0.97 | 0.99 | 0.98 | 0.87 | 0.8 | 0.25 | 1.07 | 0.78 | 0.36 | 0.39 | 0.75 | 0.75 | 0.1225 | -0.105 | -0.2675 | 0.416777 | 0.6353 | 0.19583 | 1 | 0.996511 | 0.78004 |
| Anks1b | S4R1Q0 | 1.02 | 0.79 | 2.17 | 1.89 | 1.1 | 0.93 | 2.03 | 2.02 | 0.9 | 1.92 | 2.19 | 1.01 | 0.57 | 1.25 | 1.35 | 1.24 | 0.0525 | 0.0375 | -0.365 | 0.899676 | 0.9317 | 0.338315 | 1 | 0.996755 | 0.847578 |
| Anks1b | S4R2Q2 | 2.96 | 1.81 | 3.12 | 3.87 | 1.46 | 2.94 | 1.91 | 4.32 | 3.02 | 4.61 | 3.45 | 2.93 | 3.08 | 2.61 | 3.08 | 3.35 | -0.2825 | 0.5625 | 0.09 | 0.684632 | 0.306098 | 0.83453 | 1 | 0.996511 | 0.975487 |
| Anks3 | A0A2R8VH | -0.32 | -0.11 | 0.17 | -0.61 | -0.97 | -1.49 | -1.43 | -0.69 | 0.53 | 0.81 | -0.25 | -0.86 | 2.92 | -0.45 | 0.82 | -0.96 | -0.9275 | 0.275 | 0.8 | 0.006143 | 0.493528 | 0.32772 | 1 | 0.996511 | 0.844684 |
| Ankzf1 | J3QM81 | -0.58 | -0.61 | -4.11 | -4.99 | -0.52 | -1.3 | -4.07 | -3.17 | 0.04 | -1.9 | -5.2 | -0.94 | -0.18 | 0.62 | -6.79 | -4.78 | 0.3075 | 0.5725 | -0.21 | 0.808345 | 0.693858 | 0.911329 | 1 | 0.996511 | 0.986458 |
| Anln | Q8K298 | 1.39 | 1.27 | 1.23 | 0.75 | 1.37 | 0.85 | -0.3 | 0.71 | 0.96 | 0.47 | 0.72 | 0.82 | 0.28 | 0.56 | 0.92 | 0.99 | -0.5025 | -0.4175 | -0.4725 | 0.187567 | 0.079057 | 0.086473 | 1 | 0.996511 | 0.691619 |
| Ano10 | Q8BH79-2 | 0.37 | 0.24 | 0.57 | 0.61 | 0.91 | 0.27 | 0.63 | 0.63 | -0.02 | 0.76 | 0.42 | 0.41 | -0.16 | 0.47 | 0.98 | 0.81 | 0.1625 | -0.055 | 0.0775 | 0.411394 | 0.802276 | 0.785877 | 1 | 0.996511 | 0.967642 |
| Ano3 | A2AHL1 | 0.38 | 0.98 | 0.91 | 0.52 | 0.67 | 0.08 | 1.28 | 0.59 | 0.48 | -0.36 | 0.82 | 0.12 | 0.1 | 0.96 | 0.79 | 1.07 | -0.0425 | -0.4325 | 0.0325 | 0.882313 | 0.172351 | 0.908222 | 1 | 0.996511 | 0.986458 |
| Ano6 | A0A2I3BPX | 1.54 | 1.12 | 0.79 | 1.32 | 1.01 | 0.92 | 1.14 | 1.18 | 0.81 | 0.79 | 0.94 | 1.31 | -0.19 | 0.73 | 0.71 | 0.62 | -0.13 | -0.23 | -0.725 | 0.524936 | 0.33411 | 0.031387 | 1 | 0.996511 | 0.641974 |
| Ano8 | A0A1L1SR9 | -0.03 | 0.19 | 0.41 | 0.22 | 0.08 | 0.34 | 0.3 | 0.38 | 0.16 | 0.4 | 0.23 | -0.28 | 0.42 | 0.19 | 0.5 | 0.73 | 0.0775 | -0.07 | 0.2625 | 0.645377 | 0.742189 | 0.22275 | 1 | 0.996511 | 0.787447 |
| Anp32a | Q3Z7M9 | -1.78 | -2.53 | -2.59 | -2.47 | -1.98 | -2.89 | -2.47 | -2.65 | -2.81 | -1.02 | -2.47 | -1.3 | -2.69 | -1.71 | -2.72 | -3 | -0.155 | 0.4425 | -0.1875 | 0.575826 | 0.337121 | 0.584547 | 1 | 0.996511 | 0.927734 |
| Anp32b | Q9EST5 | -1.84 | -2.82 | -4.65 | -5.75 | -3.13 | -2.48 | -5.35 | -5.49 | -2.19 | -2.54 | -4.77 | -1.71 | -2.55 | -2.82 | -5.91 | -5.85 | -0.3475 | 0.9625 | -0.5175 | 0.740679 | 0.433569 | 0.651296 | 1 | 0.996511 | 0.947503 |
| Anp32e | P97822-2 | -4.26 | -4.15 | -5 | -5.92 | -3.99 | -4.86 | -5.86 | -6.14 | -4.71 | -3.41 | -5.5 | -3.62 | -4.64 | -3.24 | -6.35 | -6.11 | -0.38 | 0.5225 | -0.2525 | 0.520255 | 0.382957 | 0.736926 | 1 | 0.996511 | 0.958635 |
| Anpep | P97449 | -1.16 | -0.85 | -0.93 | -0.7 | -0.76 | -1.18 | -0.79 | -1.42 | -0.84 | -0.73 | -0.98 | -0.84 | -0.81 | -0.87 | -0.64 | -0.64 | -0.1275 | 0.0625 | 0.12 | 0.553166 | 0.725711 | 0.528917 | 1 | 0.996511 | 0.908663 |
| Antkmt | Q501J2 | -0.04 | 0.76 | 0.11 | 0.13 | 0.48 | -0.23 | 0.81 | 0.38 | 0.67 | 0.41 | 0.08 | 0.28 | 0.56 | 1.02 | -0.14 | 0.31 | 0.12 | 0.3 | 0.1975 | 0.67257 | 0.233553 | 0.528795 | 1 | 0.996511 | 0.908663 |
| Antxr1 | Q9CZ52 | 0.39 | 0.45 | 0.92 | 1.47 | 0.16 | 0.83 | 0.8 | 0.69 | 0.21 | -1.1 | 0.6 | 0.28 | 0.36 | -0.68 | 0.69 | 1.28 | -0.1875 | -0.81 | -0.395 | 0.529156 | 0.085701 | 0.395115 | 1 | 0.996511 | 0.866783 |
| Anxa1 | P10107 | -0.18 | 1.34 | -0.7 | 0.23 | -1.22 | -0.47 | -0.51 | -1.07 | -0.23 | -0.39 | -1.17 | 2.94 | -0.21 | 3.87 | -0.98 | -0.73 | -0.99 | 0.115 | 0.315 | 0.049064 | 0.898916 | 0.772458 | 1 | 0.996511 | 0.965933 |
| Anxa11 | P97384 | 0.3 | 0.81 | 0.54 | 0.11 | 1.33 | 0.18 | 0.34 | 0.1 | -0.5 | -0.35 | 0.26 | 0.14 | -0.59 | 0.51 | 0.4 | 0.66 | 0.0475 | -0.5525 | -0.195 | 0.880679 | 0.05789 | 0.553699 | 1 | 0.996511 | 0.916696 |
| Anxa2 | P07356 | -0.06 | -0.45 | -0.94 | 0.27 | -0.41 | -0.05 | -0.33 | -0.81 | -0.06 | -0.62 | -0.92 | -0.47 | 0.03 | -0.18 | -1.02 | -0.46 | -0.105 | -0.2225 | -0.1125 | 0.728351 | 0.489213 | 0.744851 | 1 | 0.996511 | 0.959729 |
| Anxa3 | O35639 | 0.99 | 0.62 | 0.02 | 0.19 | 0.52 | 0.55 | -0.2 | -0.14 | 0.56 | 0.39 | 0.21 | 0.6 | 0.29 | 0.49 | -0.13 | -0.04 | -0.2725 | -0.015 | -0.3025 | 0.371374 | 0.953343 | 0.299078 | 1 | 0.996755 | 0.828257 |
| Anxa4 | P97429 | 0.94 | 1.13 | 0.14 | 0.4 | 0.83 | 0.85 | 0.3 | -0.13 | 0.46 | -0.3 | -0.4 | 0.67 | 0.25 | 0.44 | -0.03 | 0.24 | -0.18 | -0.545 | -0.4275 | 0.558531 | 0.143308 | 0.143845 | 1 | 0.996511 | 0.742777 |
| Anxa5 | P48036 | 0.27 | 0.33 | 0.5 | 0.42 | 0.77 | 0.06 | 0.42 | 0.29 | -0.09 | -0.12 | 0.31 | 0.22 | -0.74 | -0.17 | -0.17 | 0.72 | 0.005 | -0.3 | -0.47 | 0.97935 | 0.128393 | 0.160087 | 1 | 0.996511 | 0.755194 |
| Anxa6 | F8WIT2 | 1.25 | 1.09 | 1.48 | 2.74 | 0.75 | 0.47 | 1.26 | 2.87 | 1.46 | 2.75 | 1.52 | 0.64 | 0.72 | 2.11 | 0.95 | 1.31 | -0.3025 | -0.0475 | -0.3675 | 0.614976 | 0.928709 | 0.429022 | 1 | 0.996755 | 0.879165 |
| Anxa6 | P14824 | 0.73 | 0.6 | 1.14 | 1.21 | 0.92 | 0.81 | 1.16 | 1.35 | 0.79 | 1.06 | 1.28 | 0.99 | 0.36 | 0.54 | 1.11 | 1.13 | 0.14 | 0.11 | -0.135 | 0.524987 | 0.61787 | 0.621526 | 1 | 0.996511 | 0.938628 |
| Anxa7 | Q07076 | 0.69 | 0.48 | 0.69 | 0.62 | 0.32 | 0.73 | 0.6 | 0.52 | 0.23 | -0.18 | 0.52 | 0.38 | 0.69 | -0.38 | 0.5 | 0.58 | -0.0775 | -0.3825 | -0.2725 | 0.631939 | 0.089785 | 0.333343 | 1 | 0.996511 | 0.845601 |
| Aoc3 | O70423-2 | -2.28 | -1.35 | -3.88 | -3.27 | -1.29 | -2.3 | -2.73 | -3.6 | -2.13 | -3.86 | -2.83 | -2.14 | -2.75 | -1.8 | -3.51 | -3.09 | 0.215 | -0.045 | -0.0925 | 0.747721 | 0.943049 | 0.879166 | 1 | 0.996755 | 0.979626 |
| Aox1 | Q3X8P9 | -0.44 | -0.75 | -0.62 | -0.38 | -0.09 | -0.65 | -1.12 | -0.58 | -0.62 | 0.38 | -0.9 | -0.87 | -1.15 | -0.33 | -0.2 | -0.26 | -0.0625 | 0.045 | 0.0625 | 0.796418 | 0.8861 | 0.813912 | 1 | 0.996511 | 0.972768 |
| Ap1b1 | Q5SVG5 | -0.22 | -0.22 | 0.32 | 0.35 | -0.11 | -0.33 | 0.52 | 0.49 | -0.47 | 0.11 | 0.36 | -0.39 | -0.99 | -0.03 | 0.04 | -0.05 | 0.085 | -0.155 | -0.315 | 0.755645 | 0.570998 | 0.315776 | 1 | 0.996511 | 0.838085 |
| Ap1g1 | Q8CBB7 | -0.19 | -0.5 | 0.34 | 0.39 | -0.19 | -0.15 | 0.51 | 0.49 | -0.62 | 0.31 | 0.46 | -0.3 | -1.01 | -0.38 | -0.07 | 0.15 | 0.155 | -0.0475 | -0.3375 | 0.595405 | 0.885689 | 0.326068 | 1 | 0.996511 | 0.843537 |
| Ap1m1 | P35585 | -0.02 | 0.24 | -0.24 | -0.18 | 0.54 | -0.16 | 0.12 | 0.07 | -0.25 | -0.48 | -0.03 | -0.01 | -0.51 | 0.32 | -0.44 | -0.28 | 0.1925 | -0.1425 | -0.1775 | 0.369875 | 0.486787 | 0.485002 | 1 | 0.996511 | 0.894618 |

| | | | | | | | | | | | | | | | | | | | | | | | | | | |
|---|---|---|---|---|---|---|---|---|---|---|---|---|---|---|---|---|---|---|---|---|---|---|---|---|---|---|
| Ap1s1 | P61967 | 0.62 | -0.31 | -0.76 | -0.97 | -0.12 | 0.53 | -0.62 | -0.97 | 0.33 | -0.6 | -0.68 | 0.28 | -0.51 | 0.55 | -1.4 | -1.02 | 0.06 | 0.1875 | -0.24 | 0.893238 | 0.659467 | 0.641355 | 1 | 0.996511 | 0.943421 |
| Ap1s2 | Q9DB50 | -0.31 | -0.48 | 0.12 | 0.43 | 0.2 | -0.54 | 0.53 | 0.24 | -0.25 | 0.83 | 0.35 | -0.65 | -0.67 | 0.66 | -0.18 | 0.58 | 0.1675 | 0.13 | 0.1575 | 0.585136 | 0.728458 | 0.672899 | 1 | 0.996511 | 0.952944 |
| Ap2a1 | P17426-2 | 1.29 | 0.26 | 0.83 | 0.88 | 0.57 | 1.33 | 1.47 | 1.01 | -0.06 | 0.68 | 0.91 | 0.65 | -0.28 | 0.45 | 0.71 | 0.92 | 0.28 | -0.27 | -0.365 | 0.349427 | 0.384321 | 0.296194 | 1 | 0.996511 | 0.825612 |
| Ap2a2 | P17427 | 1.66 | 0.15 | 0.44 | 0.32 | 0.42 | 1.93 | 0.66 | 0.44 | -0.16 | -0.19 | 0.34 | 1.42 | -0.76 | -0.11 | -0.12 | 0.01 | 0.22 | -0.29 | -0.8875 | 0.636418 | 0.547075 | 0.040315 | 1 | 0.996511 | 0.658454 |
| Ap2b1 | Q9DBG3 | 1.08 | 0.37 | 0.37 | 0.46 | 0.51 | 1.42 | 1.25 | 0.69 | 0.51 | 0.06 | 0.48 | 0.8 | -0.09 | 0.45 | 0.18 | 0.33 | 3.98E-01 | -0.1075 | -0.3525 | 0.180817 | 0.670901 | 0.17315 | 1 | 0.996511 | 0.764376 |
| Ap2m1 | A0A338P6T | -2.52 | -0.58 | -3.54 | -4.76 | 0.86 | -1.12 | -2.64 | -4.13 | -1.31 | -4 | -3.32 | -2.08 | -1.15 | -0.89 | -4.33 | -2.77 | 1.0925 | 0.1725 | 0.565 | 0.389157 | 0.857889 | 0.59833 | 1 | 0.996511 | 0.930093 |
| Ap2m1 | P84091 | -0.04 | 0 | -0.05 | -0.07 | 0.21 | 0.38 | 0.15 | 0.14 | 0.12 | -0.66 | 0.11 | -0.23 | -0.14 | 0.18 | 0.36 | -0.24 | 0.26 | -0.125 | 0.08 | 0.09867 | 0.575483 | 0.695584 | 1 | 0.996511 | 0.955729 |
| Ap2s1 | P62743 | 1.02 | 1.02 | 1.57 | 1.16 | 1.47 | 1.05 | 1.92 | 1.71 | -1.48 | -2.06 | 1.67 | 0.79 | -1.96 | -1.72 | 1.05 | 1.59 | 0.345 | -1.4625 | -1.4525 | 0.179202 | 0.10054 | 0.109816 | 1 | 0.996511 | 0.709835 |
| Ap3b1 | Q9Z1T1 | -0.62 | -0.53 | 0.18 | 0.14 | -0.91 | -0.63 | 0.07 | -0.05 | -0.31 | 0.13 | 0.2 | -0.56 | -0.43 | -0.14 | -0.44 | -0.45 | -0.1725 | 0.0725 | -0.1575 | 0.582579 | 0.802078 | 0.543596 | 1 | 0.996511 | 0.912212 |
| Ap3b2 | Q9JME5 | 0.79 | 0.68 | 0.96 | 1.02 | 0.7 | 0.85 | 0.64 | 0.88 | 0.9 | 1.28 | 0.96 | 0.84 | 0.4 | 1.12 | 0.66 | 0.63 | -0.095 | 0.1325 | -0.16 | 0.556222 | 0.48261 | 0.474743 | 1 | 0.996511 | 0.891741 |
| Ap3d1 | O54774 | 0.02 | 0.3 | 0.69 | 0.26 | 0.54 | 0.19 | 0.58 | 0.42 | -0.67 | -0.53 | 0.62 | 0.08 | -0.57 | -0.85 | 0.46 | 0.52 | 0.115 | -0.4425 | -0.4275 | 0.565366 | 0.198782 | 0.266509 | 1 | 0.996511 | 0.818107 |
| Ap3m1 | Q9JKC8 | 0.45 | 0.29 | 0.01 | -0.03 | 0.57 | 0.09 | 0.15 | -0.04 | 0.13 | -0.12 | 0.1 | 0.74 | -0.72 | 0.19 | -0.38 | -0.26 | 0.0125 | 0.0325 | -0.4725 | 0.951381 | 0.89369 | 0.088683 | 1 | 0.996511 | 0.691619 |
| Ap3m2 | Q8R2R9 | -0.3 | -0.06 | -0.06 | -0.18 | 0.32 | -0.02 | -0.05 | -0.04 | -0.34 | -0.36 | 0 | -0.29 | -0.51 | 0.21 | -0.27 | -0.14 | 0.2025 | -0.0975 | -0.0275 | 0.238416 | 0.579183 | 0.898261 | 1 | 0.996511 | 0.983509 |
| Ap3s1 | Q9DCR2 | 0.3 | 0.07 | 0.27 | 0.37 | 0.23 | 0.14 | 0.42 | 0.51 | -0.52 | 0.27 | 0.55 | 0.44 | -0.6 | -0.66 | -0.05 | 0.14 | 0.0725 | -0.0675 | -0.545 | 0.660945 | 0.800634 | 0.050091 | 1 | 0.996511 | 0.658454 |
| Ap3s2 | Q8BSZ2 | -0.16 | -0.09 | -0.29 | -0.05 | 0.15 | -0.09 | -0.41 | 0.09 | -0.24 | -0.84 | -0.14 | -0.05 | -0.7 | -0.3 | -0.42 | -0.21 | 0.0825 | -0.17 | -0.26 | 0.650173 | 0.452013 | 0.200814 | 1 | 0.996511 | 0.780477 |
| Ap4b1 | A0A0G2JFY | -0.88 | -0.66 | -0.08 | 0.13 | 0 | -1.66 | -0.6 | -0.51 | -0.24 | -0.27 | -0.68 | -1.04 | -1.54 | 0.28 | -0.28 | -0.19 | -0.32 | -0.185 | -0.06 | 0.430369 | 0.551888 | 0.889846 | 1 | 0.996511 | 0.98149 |
| Ap4e1 | Q80V94 | -0.25 | 0.11 | -0.5 | -0.4 | 0.45 | -0.24 | -0.33 | -0.44 | -0.11 | -0.48 | -0.53 | -0.28 | -0.17 | 0.37 | -0.95 | -0.62 | 0.12 | -0.09 | -0.0825 | 0.636688 | 0.667764 | 0.79787 | 1 | 0.996511 | 0.969727 |
| Ap4m1 | Q9JKC7 | -0.45 | -0.33 | 0.02 | 0.85 | -0.22 | -0.13 | 0.46 | 0.26 | 0.98 | 0.38 | 0.19 | -0.27 | 0.85 | -0.34 | 0.19 | 0.03 | 0.07 | 0.2975 | 0.16 | 0.829985 | 0.439272 | 0.671461 | 1 | 0.996511 | 0.952631 |
| Ap4s1 | Q9WVL1 | 1.63 | 1.15 | 1.02 | 1.36 | 0.95 | 1.04 | 0.21 | 0.74 | 1.4 | 0.75 | 1.33 | 1.71 | 1.78 | 0.47 | 1.02 | 0.94 | -0.555 | 0.0075 | -0.2375 | 0.045711 | 0.976956 | 0.453248 | 1 | 0.997808 | 0.885993 |
| Ap5s1 | A2ANC4 | -2.04 | -2.91 | -2.34 | -2.8 | -2.1 | -2.1 | -1.23 | -2.48 | -3.99 | -2.49 | -3.72 | -6.21 | -4.1 | -2.69 | -3.01 | -1.64 | 0.545 | -1.58 | -0.3375 | 0.121285 | 0.054617 | 0.511607 | 1 | 0.996511 | 0.903991 |
| Ap5z1 | Q3U829 | -1.37 | -0.57 | -0.48 | -0.61 | -1.63 | -0.4 | -0.12 | -0.54 | 0.5 | -0.23 | -0.43 | -0.82 | -1.2 | -1.26 | -0.8 | -0.67 | 0.085 | 0.5125 | -0.225 | 0.819023 | 0.158128 | 0.422248 | 1 | 0.996511 | 0.875782 |
| Apaf1 | G3XA09 | -0.69 | -0.43 | -1.09 | -0.7 | -0.77 | -0.4 | -2.28 | -0.79 | -1.1 | 0.78 | -1.11 | 0.47 | -0.36 | 0.79 | 1.19 | -0.32 | -0.3325 | 0.4875 | 1.0525 | 0.428291 | 0.330587 | 0.025983 | 1 | 0.996511 | 0.633075 |
| Apba1 | B2RUJ5 | 0.13 | 0.59 | 0.43 | 0.38 | 0.59 | 0.12 | 0.14 | 0.27 | 0.9 | 0.98 | 0.6 | 0.71 | 0.67 | 1.21 | 0.29 | 0.27 | -0.1025 | 0.415 | 0.2275 | 0.584339 | 0.051427 | 0.402763 | 1 | 0.996511 | 0.870698 |
| Apba2 | P98084 | -0.21 | -0.02 | 0.22 | 0.44 | -0.77 | 0.11 | 0.54 | 0.25 | 0.58 | 0.44 | 0.35 | 0.6 | 1.02 | 0.12 | 0.35 | 0.38 | -0.075 | 0.385 | 0.36 | 0.809351 | 0.083667 | 0.198695 | 1 | 0.996511 | 0.78004 |
| Apba3 | O88888 | 0.32 | 0.54 | -0.04 | 0.42 | 0.75 | -0.1 | -0.08 | -0.71 | 0.39 | 1.34 | 0.21 | 0.2 | 0.14 | 1.02 | 0.03 | -0.05 | -0.345 | 0.225 | -0.025 | 0.294838 | 0.46643 | 0.932013 | 1 | 0.996511 | 0.989713 |
| Apbb1 | Q9QXJ1 | 0.1 | 0.62 | 0.25 | 0.55 | 0.7 | 0.48 | 0.76 | 0.58 | 0.57 | -0.07 | 0.61 | 0.42 | 0.38 | 1.45 | 0.37 | 0.34 | 0.25 | 0.0025 | 0.255 | 0.193028 | 0.991346 | 0.417853 | 1 | 0.999285 | 0.87558 |
| Apbb1ip | Q8R5A3 | -0.74 | -1.49 | -0.53 | -0.66 | -0.34 | -0.98 | -0.57 | -0.41 | 0.05 | 1.47 | -0.28 | -1.02 | 1.63 | -0.7 | -0.97 | -0.63 | 0.28 | 0.91 | 0.6875 | 0.308743 | 0.109853 | 0.261519 | 1 | 0.996511 | 0.81412 |
| Apbb2 | E9PWH3 | 0.58 | 0.69 | 1.28 | 1.24 | 0.41 | 0.89 | 0.79 | 1.87 | 0.69 | 1.17 | 1.18 | 0.64 | 0.79 | 0.63 | 1.78 | 1.99 | 0.0425 | -0.0275 | 0.35 | 0.90246 | 0.914201 | 0.367563 | 1 | 0.996755 | 0.859245 |
| Apc | B2RUG9 | 0.39 | 0.62 | 0.69 | 0.53 | 0.27 | 0.47 | 0.27 | 0.73 | 0.9 | 0.76 | 0.85 | 0.84 | 1.06 | 0.25 | 0.88 | 0.44 | -0.1225 | 0.28 | 0.1 | 0.491049 | 0.111393 | 0.676536 | 1 | 0.996511 | 0.953006 |
| Apc2 | G5E832 | 0.21 | 0.32 | 0.89 | 0.74 | 0.34 | 0.29 | 0.41 | 0.55 | 1.47 | 1.09 | 0.71 | 0.46 | 1.69 | 1.05 | 1.13 | 0.93 | -0.1425 | 0.3925 | 0.66 | 0.490646 | 0.192663 | 0.031062 | 1 | 0.996511 | 0.641481 |
| Apeh | A0A0R4J10 | -0.41 | -0.91 | -0.32 | 0.26 | -0.76 | -0.18 | -0.24 | -0.07 | 0.03 | 0.03 | -0.11 | -0.37 | 0.04 | 0.09 | -0.24 | -0.22 | 0.0325 | 0.24 | 0.2625 | 0.909445 | 0.389283 | 0.355305 | 1 | 0.996511 | 0.855029 |
| Apex1 | P28352 | -5.42 | -5.88 | -5.06 | -4.64 | -5.28 | -5.66 | -4.49 | -5.03 | -5.3 | -3.06 | -4.59 | -5.08 | -4.56 | -4.06 | -5.2 | -5.35 | 0.135 | 0.7425 | 0.4575 | 0.698159 | 0.183503 | 0.255715 | 1 | 0.996511 | 0.809552 |
| Aph1b | F8WHK7 | 0.45 | 0.04 | 1.05 | 1.73 | 0.62 | 0.12 | 0.43 | 1.22 | 0.95 | 1.9 | 0.83 | 0.16 | 0.31 | 1.55 | 0.55 | 0.24 | -0.22 | 0.1425 | -0.155 | 0.593725 | 0.766833 | 0.731591 | 1 | 0.996511 | 0.958635 |
| Api5 | O35841 | -1.44 | -2.65 | -2.43 | -2.33 | -2.29 | -2.19 | -2.7 | -2.41 | -2.12 | -1.2 | -2.33 | -2.15 | -2.36 | -1.9 | -2.64 | -2.53 | -0.185 | 0.2625 | -0.145 | 0.526545 | 0.471373 | 0.651044 | 1 | 0.996511 | 0.947407 |
| Apip | Q9WVQ5 | -1.43 | -1.59 | -1.31 | -1.42 | -1.53 | -1.38 | -0.84 | -1.34 | -1.98 | -1.27 | -1.2 | -1.06 | -1.72 | -2.01 | -1.75 | -1.14 | 0.165 | 0.06 | -0.2175 | 0.409363 | 0.803318 | 0.365626 | 1 | 0.996511 | 0.858622 |
| Aplf | Q9D842-2 | -2.89 | -3.61 | -1.94 | -2.23 | -5.36 | -2.95 | -3.76 | -2.46 | -2.74 | 2.39 | -4.05 | -1.61 | -1.46 | -5.08 | -1.65 | -2.33 | -0.965 | 1.165 | 0.0375 | 0.17366 | 0.377058 | 0.963705 | 1 | 0.996511 | 0.9927 |
| Aplp1 | Q03157 | -0.36 | -0.44 | -1.11 | -1.63 | -0.37 | -0.57 | -1.05 | -1.23 | 0.17 | -0.75 | -1.09 | -0.46 | 0.56 | -0.26 | -0.15 | -1.2 | 0.08 | 0.3525 | 0.6225 | 0.818076 | 0.3733 | 0.185691 | 1 | 0.996511 | 0.776843 |
| Aplp2 | Q60709 | -0.11 | 0.11 | -0.37 | -0.35 | 0.23 | -0.01 | -0.09 | -0.43 | -0.7 | -0.59 | -0.12 | 0.15 | -0.68 | -0.38 | -0.4 | -0.47 | 0.105 | -0.135 | -0.3025 | 0.614321 | 0.595095 | 0.155522 | 1 | 0.996511 | 0.750763 |
| Aplp2 | Q06335 | 0.86 | 1.78 | -0.76 | 0.35 | 2.42 | 0.73 | 0.05 | 0.73 | 0 | 0.63 | 0.81 | 0.51 | 0.52 | 0.95 | 0.66 | 1.1 | 0.425 | -0.07 | 0.25 | 0.52727 | 0.892191 | 0.624899 | 1 | 0.996511 | 0.939167 |
| Apmap | Q9D7N9 | 0.23 | 0.38 | 0.42 | 0.4 | 0.78 | 0.32 | 0.78 | 0.43 | 0.37 | 0.08 | 0.2 | 0.18 | 0.44 | 0.56 | 0.66 | 0.71 | 0.22 | -0.15 | 0.235 | 0.230619 | 0.369651 | 0.201385 | 1 | 0.996511 | 0.781032 |
| Apoa1 | Q00623 | -4.79 | -5.82 | -3.91 | -4.62 | -4.94 | -5.02 | -4.22 | -4.35 | -5.46 | -2 | -3.86 | -4.56 | -4.32 | -3.14 | -4.42 | -4.67 | 0.1525 | 0.815 | 0.6475 | 0.714889 | 0.296884 | 0.206497 | 1 | 0.996511 | 0.785338 |
| Apoa2 | P09813 | -4.1 | -4.13 | -1.2 | -3.57 | -5.21 | -3.09 | -4.68 | -3.74 | -3.87 | -0.64 | -1.62 | -3.68 | -0.93 | -4.05 | -2.67 | -1.27 | -0.93 | 0.7975 | 1.02 | 0.241576 | 0.409604 | 0.272266 | 1 | 0.996511 | 0.822084 |
| Apoa4 | P06728 | -5.57 | -6.36 | -4.55 | -4.7 | -5.57 | -6.04 | -4.77 | -5.33 | -5.86 | -2.36 | -4.51 | -5.52 | -4.59 | -3.96 | -5.29 | -5.48 | -0.1325 | 0.7325 | 0.465 | 0.774135 | 0.375955 | 0.371376 | 1 | 0.996511 | 0.859424 |
| Apob | E9Q1Y3 | -2.78 | -4.55 | -1.51 | -2.03 | -4.94 | -4.35 | -1.02 | -2.26 | -3.16 | -1.49 | -1.44 | -3.09 | -2.97 | -2.52 | -1.74 | -2.23 | -0.425 | 0.4225 | 0.3525 | 0.675832 | 0.572999 | 0.590222 | 1 | 0.996511 | 0.929142 |
| Apoc1 | P34928 | -4.22 | -4.01 | -2.41 | -2.69 | -4.65 | -4.37 | -4.55 | -3.11 | -5.49 | -2.53 | -2.57 | -3.24 | -3.41 | -4.41 | -3.12 | -4.15 | -0.8375 | -0.125 | -0.44 | 0.143243 | 0.868508 | 0.399186 | 1 | 0.996511 | 0.868093 |
| Apoc3 | E9QP56 | -1.28 | -1.26 | -2.82 | -2.79 | -0.99 | -1.57 | -3.65 | -2.94 | -0.58 | -1.94 | -2.57 | -1.67 | -0.37 | -1.41 | -3.56 | -3.92 | -0.25 | 0.3475 | -0.2775 | 0.715727 | 0.539082 | 0.748495 | 1 | 0.996511 | 0.960744 |
| Apod | P51910 | -0.3 | -0.67 | -0.79 | -1.17 | 0.06 | -1.35 | 0 | -0.94 | -0.55 | -0.3 | -0.78 | -0.63 | -0.46 | 0.44 | -0.91 | -1.03 | 0.175 | 0.1675 | 0.2425 | 0.641709 | 0.48286 | 0.518202 | 1 | 0.996511 | 0.90668 |
| Apoe | P08226 | -1.07 | -1.45 | -1.21 | -1.42 | -1.1 | -1.25 | -1.11 | -1.37 | -1.09 | -1.26 | -1.22 | -0.55 | -0.94 | -1.01 | -1.19 | -1.47 | 0.08 | 0.2575 | 0.135 | 0.63302 | 0.266624 | 0.521945 | 1 | 0.996511 | 0.907667 |
| Apoh | Q01339 | -2.78 | -3.46 | -2.27 | -2.11 | -3.02 | -2.69 | -2.68 | -1.93 | -2.76 | -0.75 | -2.53 | -2.83 | -2.2 | -1.99 | -2.08 | -2.43 | 0.075 | 0.4375 | 0.48 | 0.837015 | 0.423292 | 0.163588 | 1 | 0.996511 | 0.758352 |
| Apoo | Q9DCZ4-2 | 2.1 | 2.34 | 0.62 | 0.9 | 2.04 | 2.13 | 0.92 | 0.76 | 1.62 | 0.5 | 0.73 | 1.87 | 1.96 | 1.83 | 0.39 | 0.69 | -0.0275 | -0.31 | -0.2725 | 0.957426 | 0.54317 | 0.616177 | 1 | 0.996511 | 0.93691 |
| Apool | Q78IK4 | -0.16 | -0.17 | -0.51 | -0.61 | -0.38 | -0.07 | -0.32 | -0.63 | -0.49 | -0.89 | -0.2 | -0.12 | -0.08 | -0.37 | -0.49 | -0.17 | 0.0125 | -0.0625 | 0.085 | 0.9495 | 0.792873 | 0.684568 | 1 | 0.996511 | 0.955729 |
| App | P12023-2 | -2.3 | -3.18 | 0.62 | 0.53 | -1 | -3.2 | 0.87 | 0.98 | -2.42 | 1.99 | 0.37 | -3.44 | -0.7 | 0.16 | 2.72 | 0.44 | 0.495 | 0.2075 | 1.7375 | 0.690251 | 0.883381 | 0.136503 | 1 | 0.996511 | 0.732247 |
| App | A0A2I3BQ2 | -1.02 | -0.91 | -0.74 | -0.71 | -0.74 | -1.2 | -0.62 | -0.86 | -1.01 | -0.68 | -0.85 | -0.77 | -0.89 | -0.7 | -0.46 | -0.77 | -0.01 | 0.0175 | 0.14 | 0.957072 | 0.91987 | 0.472322 | 1 | 0.996755 | 0.891144 |
| Appbp2 | Q9DAX9 | -0.21 | -0.37 | -1.23 | -1.83 | 0.61 | -0.43 | -1.31 | -0.96 | 0.49 | -0.39 | -1.09 | -0.14 | 0.05 | -0.35 | -0.72 | -1.11 | 0.3875 | 0.6275 | 0.3775 | 0.463457 | 0.204028 | 0.392566 | 1 | 0.996511 | 0.865775 |
| Appl1 | Q8K3H0 | 0.03 | -0.21 | 0.06 | 0.07 | -0.23 | -0.01 | -0.09 | 0.34 | -0.03 | 0.27 | 0.24 | -0.17 | -0.15 | -0.01 | -0.16 | 0.02 | 0.015 | 0.09 | -0.0625 | 0.934275 | 0.630786 | 0.725058 | 1 | 0.996511 | 0.958398 |
| Appl2 | Q8K3G9 | 0.94 | 0.89 | 0.95 | 1 | 0.95 | 0.71 | 0.75 | 0.94 | 0.14 | 0.33 | 0.69 | 0.71 | -0.12 | 0.26 | 0.46 | 0.98 | -0.1075 | -0.4775 | -0.55 | 0.471733 | 0.034734 | 0.059152 | 1 | 0.996511 | 0.661738 |
| Aprt | P08030 | -1.12 | -0.59 | -0.59 | -0.57 | -0.62 | -0.89 | -0.89 | -0.8 | -0.23 | -0.05 | -0.65 | -0.84 | -0.12 | 0.08 | -0.78 | -0.92 | -0.0825 | 0.275 | 0.2825 | 0.662855 | 0.288378 | 0.350052 | 1 | 0.996511 | 0.85327 |
| Aptx | Q7TQC5-4 | -2.52 | -2.1 | -2.87 | -3 | -1.53 | -2.14 | -2.79 | -2.4 | -2.14 | -1.29 | -1.99 | -1.21 | -1.47 | -0.81 | -2.58 | -2.67 | 0.4075 | 0.965 | 0.74 | 0.230291 | 0.012828 | 0.137348 | 1 | 0.996511 | 0.732462 |
| Aqp1 | Q02013 | -1.47 | -1.15 | -0.31 | -0.07 | 0.1 | -1.06 | -0.67 | -0.14 | -0.41 | -0.1 | -0.76 | -1.4 | 0.1 | -0.28 | -0.16 | -0.1 | 0.3075 | 0.0825 | 0.64 | 0.449497 | 0.841891 | 0.088938 | 1 | 0.996511 | 0.691619 |
| Aqp11 | A0A0X1KG | 0.59 | 0.38 | 0.64 | -0.9 | 0.32 | 0.65 | -0.45 | -1.28 | 0.88 | 1.27 | -0.15 | -0.59 | 2.22 | -0.47 | -0.52 | 0.12 | -0.3675 | 0.175 | 0.16 | 0.484777 | 0.739177 | 0.812065 | 1 | 0.996511 | 0.972642 |
| Aqp4 | A0A5F8MC | -0.66 | 1.89 | 1.44 | 1.64 | 0.96 | 1.12 | 0.87 | 1.46 | 0.75 | 1.88 | 0.98 | 0.83 | 1.33 | 2.14 | 1.97 | 2.72 | 0.025 | 0.0325 | 0.9625 | 0.963737 | 0.955965 | 0.134366 | 1 | 0.996755 | 0.731111 |
| Aqp4 | A0A0R4J0Z | 0.13 | 0.6 | -0.03 | 0.24 | 0.5 | 0.04 | -0.14 | -0.02 | 0.47 | 0.45 | 0.46 | 0.57 | -0.68 | 0.74 | -0.14 | -0.4 | -0.14 | 0.2525 | -0.355 | 0.525765 | 0.21106 | 0.307061 | 1 | 0.996511 | 0.832533 |
| Aqr | Q8CFQ3 | -0.2 | -0.93 | -0.98 | -0.92 | -1.37 | -0.81 | -0.92 | -0.79 | -1.13 | -0.41 | -1.03 | -1.18 | -0.45 | -0.79 | -0.43 | -0.82 | -0.215 | -0.18 | 0.135 | 0.389971 | 0.514586 | 0.59012 | 1 | 0.996511 | 0.929142 |
| Araf | P04627 | -0.73 | -1.02 | -1.38 | -0.81 | -0.36 | -0.48 | -0.94 | -1.1 | -0.68 | 0.96 | -1.1 | -0.48 | 0.16 | -0.11 | -0.84 | -1.57 | 0.265 | 0.66 | 0.395 | 0.294 | 0.162306 | 0.337556 | 1 | 0.996511 | 0.847578 |
| Araf | Q8CAD1 | -0.52 | -0.42 | -0.75 | -0.41 | 0.41 | 0.29 | 0.33 | 0.67 | 1.46 | -0.29 | -0.12 | -0.22 | 0.05 | -0.32 | -0.61 | -0.9 | 0.95 | 0.7325 | 0.08 | 0.000384 | 0.099456 | 0.750537 | 0.544245 | 0.996511 | 0.961479 |
| Arap1 | Q4LDD4-2 | 0.39 | 0.25 | 0.19 | 0.19 | 0.21 | 0.13 | -0.23 | 0.16 | 0.35 | 0.04 | 0.15 | 0.2 | 0.27 | -0.19 | 0.27 | 0.36 | -0.1875 | -0.07 | -0.0775 | 0.278711 | 0.673094 | 0.699391 | 1 | 0.996511 | 0.955729 |
| Arap2 | E9QP44 | 0.34 | 0.44 | 0.64 | 0.45 | 0.65 | 0.63 | 0.71 | 0.99 | 0.44 | 0.47 | 1.17 | 0.46 | 0.33 | 0.79 | 0.38 | 0.89 | 0.2775 | 0.1675 | 0.13 | 0.11686 | 0.462376 | 0.543911 | 1 | 0.996511 | 0.912212 |
| Arc | Q9WV31 | 0.42 | 0.97 | 1.23 | 1.15 | 1.14 | 0.44 | 0.75 | 0.93 | 0.16 | 0.73 | 0.93 | 0.26 | -0.2 | 0.76 | 0.83 | 0.62 | -0.1275 | -0.4225 | -0.44 | 0.609663 | 0.149265 | 0.179841 | 1 | 0.996511 | 0.775542 |
| Arcn1 | Q5XJY5 | -0.98 | -1.19 | -1.51 | -1.6 | -0.78 | -1.31 | -1.55 | -1.87 | -1.15 | -1.12 | -1.45 | -0.83 | -0.89 | -0.77 | -1.34 | -1.43 | -0.0575 | 0.1825 | 0.2125 | 0.834574 | 0.427741 | 0.406417 | 1 | 0.996511 | 0.87356 |
| Arel1 | Q8CHG5 | -0.13 | -0.02 | 0.41 | -0.05 | 0.01 | -0.37 | 0.46 | 0.19 | -0.12 | -0.22 | 0.08 | -0.37 | -0.51 | -0.5 | 0.26 | 0.76 | 0.02 | -0.21 | -0.05 | 0.931005 | 0.313855 | 0.881325 | 1 | 0.996511 | 0.979767 |
| Arf1 | P84078 | -0.6 | -0.39 | -0.57 | -0.67 | -0.23 | -0.62 | -0.23 | -0.79 | -0.58 | -1.24 | -0.51 | -0.61 | -1.21 | 0.17 | -1.34 | -1.05 | 0.09 | -0.1775 | -0.3 | 0.640118 | 0.424882 | 0.401817 | 1 | 0.996511 | 0.870698 |
| Arf2 | Q8BSL7 | -1.55 | -0.36 | -0.47 | -0.98 | 0.07 | -1.33 | -0.71 | -1.49 | -0.35 | -1.47 | -1.04 | -1.69 | -2.12 | -0.76 | -2.64 | -1.31 | -0.025 | -0.2975 | -0.8675 | 0.952382 | 0.449166 | 0.091703 | 1 | 0.996511 | 0.693639 |
| Arf4 | P61750 | 0.03 | 0.38 | 1.22 | 1.11 | 0.43 | 0.15 | 1.19 | 0.94 | 0.13 | 1.43 | 1.33 | 0.41 | -0.38 | 0.65 | 1.04 | 1.25 | -0.0075 | 0.14 | -0.045 | 0.983198 | 0.735468 | 0.918184 | 1 | 0.996511 | 0.986868 |
| Arf5 | P84084 | -0.43 | -0.75 | 0.19 | 0.42 | -0.63 | -0.16 | 0 | 0.51 | 0.06 | 0.4 | 0.43 | 0.07 | -1.16 | 0.02 | -0.31 | -0.21 | 0.0725 | 0.3825 | -0.2725 | 0.833808 | 0.218397 | 0.464867 | 1 | 0.996511 | 0.888267 |
| Arf6 | P62331 | 0.23 | 0.35 | 0.1 | 0.46 | 0.11 | 0.3 | 0.34 | 0.41 | 0.41 | -0.08 | 0.39 | -0.2 | -0.09 | 0.77 | -0.3 | 0.03 | 0.005 | -0.155 | -0.1825 | 0.975277 | 0.478861 | 0.504589 | 1 | 0.996511 | 0.901002 |
| Arfgap1 | V9GXM1 | 0.33 | 0.8 | 0.86 | 1.06 | 0.87 | 0.3 | 0.94 | 0.92 | -0.01 | 0.37 | 1 | 0.31 | 0.01 | 0.44 | 0.79 | 0.59 | -0.005 | -0.345 | -0.305 | 0.983011 | 0.23155 | 0.254295 | 1 | 0.996511 | 0.809524 |
| Arfgap2 | Q99K28 | -1.03 | -0.89 | -0.43 | -0.71 | -1 | -1.15 | -0.63 | -0.68 | -1.61 | -1.04 | -0.44 | -0.76 | -1.02 | -1.2 | -0.37 | -0.5 | -0.1 | -0.1975 | -0.0075 | 0.634194 | 0.500442 | 0.9774 | 1 | 0.996511 | 0.993727 |
| Arfgap3 | A0A0R4J0T | 0.03 | 0.11 | 0.35 | 0.36 | -0.08 | 0.05 | -0.65 | 0.01 | -0.11 | 0.14 | 0.27 | 0.24 | 0.71 | -0.41 | -0.12 | -0.53 | -0.38 | -0.0775 | -0.3 | 0.09938 | 0.673957 | 0.336867 | 1 | 0.996511 | 0.847578 |
| Arfgef1 | G3X9K3 | -0.34 | -0.37 | -0.48 | -0.37 | -0.35 | -0.31 | -0.48 | -0.55 | -0.18 | -0.04 | -0.35 | -0.06 | -0.41 | 0 | -0.73 | -0.61 | -0.0325 | 0.2325 | -0.0475 | 0.824041 | 0.181566 | 0.826406 | 1 | 0.996511 | 0.973235 |
| Arfgef2 | A2A5R2 | 0.88 | 0.37 | 0.63 | 0.51 | 0.19 | 0.99 | 0.3 | 0.4 | 0.55 | 0.23 | 0.71 | 0.74 | 0.73 | -0.12 | 0.48 | 0.52 | -0.1275 | -0.04 | -0.195 | 0.5805 | 0.845488 | 0.438682 | 1 | 0.996511 | 0.883205 |
| Arfgef3 | Q3UGY8 | -0.31 | -0.31 | 0.01 | -0.06 | -0.23 | -0.41 | -0.09 | -0.2 | -0.57 | -0.1 | 0.01 | -0.44 | -0.62 | -0.32 | -0.35 | -0.29 | -0.065 | -0.1075 | -0.2275 | 0.693924 | 0.604725 | 0.251094 | 1 | 0.996511 | 0.807507 |
| Arfip1 | G5E8V9-2 | -0.12 | 0.11 | -0.15 | 0 | 0.31 | -0.21 | -0.09 | -0.03 | -0.01 | 0.36 | -0.15 | -0.05 | -0.14 | 0.84 | -0.28 | -0.32 | 0.035 | 0.0775 | 0.065 | 0.841626 | 0.679104 | 0.825965 | 1 | 0.996511 | 0.973155 |
| Arfip2 | A0A1B0GS | -0.51 | -0.11 | -0.25 | -0.11 | -0.59 | -0.68 | -0.15 | -0.18 | 0.2 | 0.36 | -0.35 | -0.45 | 0.37 | 0.04 | -0.38 | -0.83 | -0.155 | 0.185 | 0.045 | 0.445535 | 0.459636 | 0.877672 | 1 | 0.996511 | 0.979626 |
| Arfrp1 | Q8BXL7 | -0.98 | -1.05 | -1.09 | -1.18 | -0.09 | -0.87 | -0.71 | -1.27 | -0.93 | -1.36 | -1.63 | -0.82 | -1.84 | -0.34 | -1.55 | -0.87 | 0.34 | -0.11 | -0.075 | 0.210392 | 0.630011 | 0.825865 | 1 | 0.996511 | 0.973155 |
| Arg2 | O08691 | 0.3 | 0.7 | -0.1 | 0.47 | 0.31 | 0.49 | 0.75 | 0.53 | 0.92 | 0.7 | 0.1 | 0.59 | 0.39 | 0.76 | -0.26 | 0.15 | 0.1775 | 0.235 | -0.0825 | 0.421513 | 0.379163 | 0.775927 | 1 | 0.996511 | 0.966799 |
| Arglu1 | Q3UL36 | 0.51 | -1.08 | -0.33 | -0.79 | -0.18 | -0.24 | -1.73 | 0.08 | 0.26 | -0.21 | -0.91 | -0.95 | 1.4 | -0.13 | 0.7 | -0.83 | -0.095 | -0.03 | 0.7075 | 0.8481 | 0.944222 | 0.22085 | 1 | 0.996755 | 0.787447 |
| Arhgap1 | A2AH25 | 1.69 | 0.68 | 0.78 | 0.42 | 0.61 | 1.75 | 0.57 | 0.75 | -0.31 | 0.34 | 0.87 | 1.83 | -0.14 | -0.36 | 0.58 | 0.55 | 0.0275 | -0.21 | -0.735 | 0.941327 | 0.671949 | 0.06767 | 1 | 0.996511 | 0.664936 |
| Arhgap1 | Q5FWK3 | 0.47 | 0.83 | 0.82 | 0.8 | 0.68 | -0.03 | 0.94 | 1.16 | 1.43 | 1.19 | 1.09 | 0.82 | 0.67 | 1.43 | 0.25 | 0.25 | -0.0425 | 0.4025 | -0.08 | 0.877859 | 0.072624 | 0.79224 | 1 | 0.996511 | 0.96933 |
| Arhgap10 | Q6Y5D8-3 | 0.44 | 0.46 | -0.2 | -0.61 | 0.01 | -0.2 | -1.1 | -1.28 | 0.33 | -0.22 | -1.52 | 0.24 | 0.93 | -1.82 | -0.37 | -0.08 | -0.665 | -0.315 | -0.3575 | 0.116849 | 0.506595 | 0.538401 | 1 | 0.996511 | 0.912212 |
| Arhgap12 | B2RUJ8 | -0.1 | 0.03 | 0.39 | 0.07 | 0.56 | 0.07 | 0.52 | 0.33 | -0.28 | -0.52 | 0.31 | -0.23 | -0.18 | -1.1 | 0.44 | 0.33 | 0.2725 | -0.2775 | -0.225 | 0.178154 | 0.255091 | 0.537588 | 1 | 0.996511 | 0.912212 |
| Arhgap17 | Q3UIA2-2 | 0.05 | -0.61 | -0.21 | 0.23 | -0.76 | -0.13 | -0.75 | -0.28 | 1.19 | 0.12 | -0.29 | 0.22 | 1.46 | -0.78 | -0.54 | -0.3 | -0.345 | 0.445 | 0.095 | 0.198405 | 0.230486 | 0.850511 | 1 | 0.996511 | 0.976225 |
| Arhgap18 | Q8K0Q5 | -0.9 | -0.58 | 0.16 | 0.08 | 0.09 | 0.08 | -0.38 | 0.37 | -0.38 | 0.59 | 0.21 | -0.16 | -0.2 | 0.27 | 0.08 | 1.4 | 0.35 | 0.375 | 0.6975 | 0.260285 | 0.276069 | 0.121259 | 1 | 0.996511 | 0.721448 |
| Arhgap20 | Q6IFT4 | 0.27 | -0.2 | 0.04 | 0.18 | -0.44 | 0.78 | -0.09 | -0.24 | 1.38 | -0.17 | -0.14 | 0.17 | 2.78 | 0.65 | -0.11 | -0.21 | -0.07 | 0.2375 | 0.705 | 0.808386 | 0.523337 | 0.287965 | 1 | 0.996511 | 0.824705 |
| Arhgap21 | B7ZCJ1 | 1.1 | 1 | 1.25 | 1.04 | 0.95 | 1.19 | 0.93 | 1 | 0.75 | 1.31 | 1.24 | 1.13 | 0.64 | 0.93 | 1.25 | 1.17 | -0.08 | 0.01 | -0.1 | 0.602746 | 0.958399 | 0.632533 | 1 | 0.996755 | 0.941132 |
| Arhgap23 | B1AQY2 | 0.1 | 0.42 | 0.34 | 0.18 | 0.27 | 0.09 | 0.16 | 0.26 | 0.31 | 0.17 | 0.47 | 0.37 | 0.35 | 0.46 | 0.43 | 0.29 | -0.065 | 0.07 | 0.1225 | 0.674755 | 0.685066 | 0.496572 | 1 | 0.996511 | 0.897918 |
| Arhgap25 | Q8BYW1 | -0.28 | 0.3 | 0.9 | 0.59 | -0.08 | -0.11 | 0.03 | 0.64 | -0.11 | 0.28 | 0.54 | 0.04 | -0.12 | 0.11 | 0.26 | 0.06 | -0.2575 | -0.19 | -0.3 | 0.406159 | 0.525619 | 0.304216 | 1 | 0.996511 | 0.831662 |
| Arhgap26 | F6T836 | 0.96 | 0.9 | 0.57 | 0.33 | 0.96 | 0.92 | 0.54 | 0.35 | 0.29 | 0.46 | 0.4 | 1.16 | -0.01 | 0.55 | 0.2 | 0.33 | 0.0025 | -0.1125 | -0.4225 | 0.991279 | 0.671599 | 0.095828 | 1 | 0.996511 | 0.698156 |
| Arhgap27 | A2AB59 | -0.22 | -0.39 | -0.66 | -0.28 | -0.69 | -0.41 | -0.77 | -0.48 | 0.64 | -0.05 | -0.1 | -0.89 | 0.69 | -0.07 | -0.18 | -0.59 | -0.2 | 0.2875 | 0.35 | 0.275978 | 0.389342 | 0.259197 | 1 | 0.996511 | 0.812074 |
| Arhgap29 | A0A0G2JEY | 0.87 | 1.37 | 0.87 | 1.2 | 1.69 | 0.53 | 1.34 | 0.85 | 1.11 | 1.32 | 0.57 | 0.79 | 0.2 | 2 | 0.19 | 0.31 | 0.025 | -0.13 | -0.4025 | 0.930486 | 0.586391 | 0.36854 | 1 | 0.996511 | 0.859245 |
| Arhgap31 | A6X8Z5 | -1.14 | -0.36 | -0.38 | -0.07 | -0.24 | -0.95 | 0.15 | 0.23 | -1.19 | -0.56 | 0.4 | -1.09 | -0.65 | -0.1 | -0.2 | -0.05 | 0.285 | -0.1225 | 0.2375 | 0.413517 | 0.765217 | 0.414358 | 1 | 0.996511 | 0.874945 |
| Arhgap32 | S4R2G6 | 0.13 | 0.13 | 0.45 | 1.14 | 0.36 | 0.66 | 0.32 | 0.64 | 0.4 | 2.58 | 0.31 | 0.46 | 1.74 | 0.54 | 1.16 | 0.94 | 0.0325 | 0.475 | 0.6325 | 0.90143 | 0.399396 | 0.09382 | 1 | 0.996511 | 0.698156 |
| Arhgap32 | Q811P8 | 0.46 | 0.57 | 0.9 | 0.77 | 0.15 | 0.47 | 0.47 | 0.78 | 0.7 | 0.51 | 1.03 | 0.68 | 0.57 | 0.66 | 0.39 | 0.43 | -0.2075 | 0.055 | -0.1625 | 0.307671 | 0.781349 | 0.407221 | 1 | 0.996511 | 0.874084 |
| Arhgap33 | A0A140LH | 0.27 | 0.49 | 0.03 | 0.52 | 0.06 | 0.65 | 0.57 | 0.57 | 1.62 | 0.82 | 0.66 | 0.64 | 1.85 | 0.8 | 0.52 | 0.36 | 0.135 | 0.6075 | 0.555 | 0.518641 | 0.050025 | 0.140486 | 1 | 0.996511 | 0.737129 |
| Arhgap35 | Q91YM2 | 0.67 | 0.24 | 0.6 | 0.3 | 0.29 | 0.72 | 0.71 | 0.55 | 0.32 | 0.18 | 0.71 | 0.8 | -0.08 | 0.24 | 0.36 | 0.48 | 0.115 | 0.05 | -0.2025 | 0.543279 | 0.821001 | 0.35957 | 1 | 0.996511 | 0.856645 |
| Arhgap39 | P59281-2 | -2.86 | 0.87 | -2.47 | -4.22 | -4.62 | -1.38 | -6.64 | -2.89 | 2.7 | 2.65 | -5.42 | -1.06 | 4.52 | -2.75 | -0.49 | -1.06 | -1.7125 | 1.8875 | 2.225 | 0.241282 | 0.350648 | 0.209389 | 1 | 0.996511 | 0.786218 |
| Arhgap39 | H7BX46 | 0.34 | 0.69 | 0.45 | 0.26 | 0.58 | 0.36 | 0.37 | 0.37 | 0.9 | 0.52 | 0.72 | 0.53 | 1.24 | 0.58 | 0.63 | 0.37 | -0.015 | 0.2325 | 0.27 | 0.927519 | 0.23774 | 0.287832 | 1 | 0.996511 | 0.824705 |
| Arhgap42 | B2RQE8 | -1.07 | -1.48 | 0.26 | -0.88 | -2.58 | -0.93 | -1.56 | -1.43 | -0.87 | -0.65 | -0.98 | -1.21 | -0.17 | -2.26 | -0.85 | -0.33 | -0.8325 | -0.135 | -0.11 | 0.105256 | 0.721006 | 0.843405 | 1 | 0.996511 | 0.976225 |
| Arhgap44 | Q5SSM3 | -0.38 | -0.35 | 0.59 | 0.6 | -0.42 | -0.15 | 0.51 | 0.39 | 0.03 | 0.75 | 0.55 | -0.38 | 0.18 | 0.21 | 0.32 | 0.36 | -0.0325 | 0.1225 | 0.1525 | 0.92423 | 0.73859 | 0.608176 | 1 | 0.996511 | 0.93378 |
| Arhgap5 | E9PYT0 | 1.02 | 0.63 | 0.4 | 0.17 | 0.83 | 0.92 | 0.69 | 0.39 | -0.25 | 0.14 | 0.5 | 1.15 | -0.39 | -0.36 | 0.13 | 0.22 | 0.1525 | -0.17 | -0.655 | 0.5185 | 0.622383 | 0.034559 | 1 | 0.996511 | 0.651434 |
| Arhgap6 | G3UZI7 | -0.17 | -0.57 | -1.79 | -1.77 | -1.23 | -1.05 | -1.3 | -1.03 | -0.07 | -1.75 | -2.03 | -0.34 | 0.68 | -0.94 | -1.63 | -0.46 | -0.0775 | 0.0275 | 0.4875 | 0.845205 | 0.962869 | 0.415526 | 1 | 0.996755 | 0.874945 |
| Arhgdia | Q99PT1 | 1.17 | 0.94 | 0.84 | 0.87 | 0.85 | 0.89 | 0.61 | 0.76 | 1.07 | 0.96 | 0.6 | 0.91 | 0.6 | 1.93 | 0.57 | 0.2 | -0.1775 | -0.07 | -0.13 | 0.284159 | 0.707585 | 0.731058 | 1 | 0.996511 | 0.958635 |
| Arhgdib | Q61599 | -0.84 | -1.29 | -0.47 | 0.19 | -1.89 | -1.59 | -0.96 | 0.06 | -1.33 | 1.13 | -0.16 | -1.31 | -1.91 | -0.77 | -0.44 | -0.36 | -0.4925 | 0.185 | -0.2675 | 0.331236 | 0.760899 | 0.555652 | 1 | 0.996511 | 0.917092 |
| Arhgef1 | E9PUF7 | 0.6 | 0.57 | -0.09 | 0.05 | 0.43 | 0.47 | -0.04 | 0.18 | 0.55 | 0.34 | 0.01 | 0.27 | 0.23 | 0.42 | -0.31 | 0.14 | -0.0225 | 0.01 | -0.1625 | 0.922653 | 0.966415 | 0.540865 | 1 | 0.996755 | 0.912212 |
| Arhgef10 | Q8C033 | -0.17 | 0.09 | 0.33 | 0.09 | -0.54 | -0.3 | 0.11 | -0.36 | 0.77 | -0.22 | 0.09 | -0.28 | 1.12 | -1.11 | 0.2 | 0.21 | -0.3575 | 0.005 | 0.02 | 0.105503 | 0.985492 | 0.964118 | 1 | 0.998892 | 0.9927 |
| Arhgef10l | A2AWP8-3 | -0.04 | 0.05 | 0.02 | 0.04 | -0.24 | -0.06 | -0.26 | -0.22 | 0.13 | -0.34 | -0.1 | 0.04 | 0.14 | -0.02 | 0.02 | -0.22 | -0.2125 | -0.085 | -0.0375 | 0.159788 | 0.630422 | 0.831275 | 1 | 0.996511 | 0.973982 |
| Arhgef11 | Q68FM7 | 0.53 | 0.43 | 1.35 | 1.05 | 0.59 | 0.48 | 0.99 | 1.35 | 0.71 | 0.57 | 1.37 | 0.61 | 0.32 | 0.55 | 0.77 | 1.18 | 0.0125 | -0.025 | -0.135 | 0.966049 | 0.932329 | 0.65243 | 1 | 0.996755 | 0.947633 |
| Arhgef12 | Q8R4H2 | 1.42 | 1.25 | 0.7 | 0.84 | 1.53 | 1.41 | -0.2 | 0.66 | 2.3 | -1.12 | 0.72 | 1.66 | 2.05 | 1.18 | -0.56 | -0.26 | -0.2025 | -0.1625 | -0.45 | 0.622102 | 0.815049 | 0.449485 | 1 | 0.996511 | 0.884817 |
| Arhgef12 | F8VQN6 | 0.45 | 0.52 | 0.9 | 0.45 | 0.56 | 0.38 | 0.76 | 0.74 | 0.45 | 0.58 | 0.99 | 0.52 | 0.23 | 0.58 | 0.84 | 0.65 | 0.03 | 0.055 | -0.005 | 0.869721 | 0.790857 | 0.981752 | 1 | 0.996511 | 0.994858 |
| Arhgef17 | Q80U35 | 0.17 | 0.4 | 0.74 | 0.46 | 0.37 | 0.24 | 0.18 | 0.53 | -0.14 | -0.48 | 0.68 | 0.26 | 0.14 | -0.23 | 0.56 | 0.39 | -0.1125 | -0.3625 | -0.2275 | 0.543964 | 0.227588 | 0.364062 | 1 | 0.996511 | 0.858475 |
| Arhgef18 | A0A5F8MP | 0.14 | 0.36 | 0.38 | 0.48 | 0.18 | 0.05 | 0.29 | 0.59 | 0.32 | 0.61 | 0.28 | 0.41 | 0.15 | 0.59 | -0.21 | 0.26 | -0.0625 | 0.065 | -0.1425 | 0.729793 | 0.71085 | 0.533942 | 1 | 0.996511 | 0.911825 |
| Arhgef19 | A2A832 | -1.41 | -1.89 | -2.41 | 2.33 | -3.71 | -2.46 | -1.9 | -0.56 | 0.23 | 0.55 | 0 | -1.42 | -0.31 | -1.75 | -0.62 | -2.18 | -1.3125 | 0.685 | -0.37 | 0.263836 | 0.516754 | 0.723399 | 1 | 0.996511 | 0.958398 |
| Arhgef2 | H3BJU7 | -0.69 | -0.44 | -0.3 | -0.59 | -0.38 | -0.9 | -0.5 | -0.48 | -0.58 | -0.24 | -0.21 | -0.21 | -0.3 | -0.4 | -0.63 | -0.46 | -0.06 | 0.195 | 0.0575 | 0.746129 | 0.308735 | 0.761207 | 1 | 0.996511 | 0.963123 |
| Arhgef25 | A0A1W2P6 | 0.31 | -0.13 | 0.48 | 0.6 | -0.39 | 0.35 | 0.4 | 0.76 | 0.54 | 0.91 | 0.81 | 0.17 | -0.54 | 0.81 | 0.05 | 0.08 | -0.035 | 0.2925 | -0.215 | 0.903793 | 0.265136 | 0.512769 | 1 | 0.996511 | 0.904116 |
| Arhgef26 | D3YYY8 | -0.34 | -0.03 | 0.12 | 0.23 | -0.29 | 0.04 | -0.11 | -0.06 | -0.15 | 0.41 | -0.05 | 0.05 | 0.15 | 0.12 | 0.21 | 0.52 | -0.1 | 0.07 | 0.255 | 0.590078 | 0.744488 | 0.247451 | 1 | 0.996511 | 0.803677 |
| Arhgef28 | A0A286YD | 0.05 | 0.67 | -0.13 | -0.14 | 0.65 | 0.24 | 0.14 | -0.23 | 0.33 | 0.7 | -0.07 | 0.12 | 0.3 | 0.28 | 0.04 | -0.08 | 0.0875 | 0.1575 | 0.0225 | 0.745592 | 0.561288 | 0.927457 | 1 | 0.996511 | 0.989331 |
| Arhgef4 | E9PYK8 | -0.91 | -0.71 | -0.83 | -1.12 | -0.23 | -0.83 | -0.47 | -0.93 | -0.74 | -0.39 | -0.93 | -0.38 | -0.55 | 0.76 | -1.54 | -1.19 | 0.2775 | 0.2825 | 0.2625 | 0.211048 | 0.194178 | 0.588621 | 1 | 0.996511 | 0.929131 |
| Arhgef40 | Q3UPH7-3 | -0.41 | -0.04 | 0.32 | 0.3 | 0.32 | -0.11 | -0.2 | 0.42 | 0.01 | 0.03 | 0.1 | -0.03 | 0.18 | -0.48 | 0.11 | 0.38 | 0.065 | -0.015 | 0.005 | 0.790828 | 0.944193 | 0.98548 | 1 | 0.996755 | 0.996154 |
| Arhgef6 | F6WMJ3 | 0.86 | 0.74 | 0.11 | -0.16 | 0.62 | 0.64 | 0.07 | -0.02 | 1.19 | -0.66 | -0.27 | 0.65 | 1.08 | 0.74 | -0.08 | -0.06 | -0.06 | -0.16 | 0.0325 | 0.841386 | 0.727867 | 0.930461 | 1 | 0.996511 | 0.989713 |
| Arhgef7 | Q9ES28 | 0.17 | -0.03 | 0.94 | 0.85 | -0.11 | 0.35 | 0.58 | 0.94 | -0.04 | 0.63 | 0.98 | 0.24 | -0.23 | -0.2 | 0.31 | 0.31 | -0.0425 | -0.03 | -0.435 | 0.894284 | 0.927016 | 0.170883 | 1 | 0.996755 | 0.762322 |
| Arhgef7 | Q9ES28-5 | -3.69 | -5.72 | -3.2 | -2.71 | -5.37 | -3.93 | -5.54 | -1.36 | -2.96 | 1.07 | -2.65 | -3.67 | -1.81 | -3.96 | -0.76 | -3.91 | -0.22 | 1.7775 | 1.22 | 0.834055 | 0.140008 | 0.210355 | 1 | 0.996511 | 0.786218 |
| Arhgef9 | B1AXI4 | -0.03 | -0.23 | -0.09 | 0.08 | 0.55 | 0.91 | 0.48 | 0.5 | 1.09 | 0.62 | 0.69 | 0.49 | 0.79 | 0.76 | 0.34 | 0.58 | 0.6775 | 0.79 | 0.685 | 0.003409 | 0.003073 | 0.006299 | 1 | 0.996511 | 0.552614 |
| Arid1a | E9QAQ7 | -2.43 | -3.14 | -2.8 | -3.15 | -3.57 | -2.91 | -3.31 | -2.78 | -0.39 | -0.92 | -2.48 | -2.81 | 0.54 | -2.45 | -0.02 | -3.3 | -0.2625 | 1.23 | 1.5725 | 0.324805 | 0.054061 | 0.091377 | 1 | 0.996511 | 0.693316 |
| Arid1b | E9Q6R4 | -2.65 | -3.08 | -1.79 | -1.79 | -2.82 | -3.07 | -2.15 | -2.16 | -2.53 | -0.85 | -1.87 | -2.73 | -2.29 | -1.61 | -2.36 | -2.05 | -0.2225 | 0.3325 | 0.25 | 0.560403 | 0.507852 | 0.492751 | 1 | 0.996511 | 0.897531 |
| Arid2 | E9Q7E2 | 0.85 | 0.54 | 0.69 | 0.66 | 0.36 | 0.66 | -0.02 | 0.02 | 1.47 | 0.24 | 0.33 | 0.61 | 1.87 | 1.28 | 0.91 | 0.7 | -0.43 | -0.0225 | 0.505 | 0.059902 | 0.939155 | 0.10215 | 1 | 0.996755 | 0.700524 |
| Arid4a | F8VPQ2 | -1.79 | -2.5 | -1.23 | -1.73 | -2.86 | -2.38 | -1.19 | -1.11 | -2.77 | -0.53 | -2.82 | -2.39 | -1.52 | -1.32 | -0.76 | -0.62 | -0.0725 | -0.315 | 0.7575 | 0.877641 | 0.573144 | 0.049579 | 1 | 0.996511 | 0.658454 |
| Arid4b | Z4YL44 | -0.48 | -7.99 | -9.97 | -6.55 | -8.72 | -2.15 | -8.15 | -9.4 | -0.78 | -3.58 | -8.43 | -3.59 | 0.67 | -3.44 | -3.23 | -5.75 | -0.8575 | 2.1525 | 3.31 | 0.716416 | 0.363284 | 0.151698 | 1 | 0.996511 | 0.750178 |
| Arih1 | Q9Z1K5 | -0.65 | -0.83 | -0.28 | -0.89 | -0.87 | -0.69 | -0.94 | -0.63 | -1.29 | -1.13 | -0.4 | -0.42 | -1.35 | -1.38 | -0.34 | -0.46 | -0.12 | -0.1475 | -0.22 | 0.537981 | 0.603901 | 0.495731 | 1 | 0.996511 | 0.89758 |
| Arih2 | Q9Z1K6 | -1.12 | -1.25 | -1.01 | -0.64 | -0.5 | -0.95 | -1.38 | -1.17 | -0.71 | -0.03 | -0.81 | -0.76 | -0.94 | -0.62 | -0.6 | -0.86 | 0.005 | 0.4275 | 0.25 | 0.983621 | 0.113876 | 0.2589 | 1 | 0.996511 | 0.812074 |
| Arl1 | F8WIB1 | -1.91 | -1.41 | -2.68 | -2.57 | -1.43 | -1.76 | -2.27 | -2.94 | -1.33 | -2.37 | -2.35 | -1.66 | -1.96 | -0.99 | -3.72 | -2.96 | 0.0425 | 0.215 | -0.265 | 0.918626 | 0.575154 | 0.664665 | 1 | 0.996511 | 0.949825 |
| Arl10 | Q9QXJ4 | 0.75 | 1.04 | 0.7 | 0.99 | 1.01 | 1.03 | 1.25 | 0.97 | 0.38 | -0.35 | 1.35 | 0.61 | 0.43 | 0.73 | 1.5 | 1.58 | 0.195 | -0.3725 | 0.19 | 0.25406 | 0.306715 | 0.540786 | 1 | 0.996511 | 0.912212 |
| Arl13b | Q640N2 | -2.02 | -1.48 | -0.64 | -0.68 | -0.83 | -1.21 | -0.84 | -0.05 | -0.49 | 0.96 | -1.05 | -2.03 | -2.14 | -1.16 | -0.16 | 0.31 | 0.4725 | 0.5525 | 0.4175 | 0.246923 | 0.402885 | 0.483143 | 1 | 0.996511 | 0.893925 |
| Arl15 | Q8BGR6 | 1.09 | 0.89 | 0.49 | 0.6 | 1.32 | 1.25 | 0.39 | 0.19 | 0.21 | 0.33 | 0.02 | 0.85 | -0.37 | 0.61 | 0.18 | 0.18 | 0.02 | -0.415 | -0.6175 | 0.949322 | 0.121864 | 0.043943 | 1 | 0.996511 | 0.658454 |
| Arl2 | Q9D0J4 | -0.19 | 0.27 | -0.6 | -0.34 | 0.53 | -0.4 | 0.41 | -0.38 | -0.37 | -0.95 | -0.38 | 0 | -0.57 | 0.29 | -0.64 | -0.07 | 0.255 | -0.21 | -0.0325 | 0.41353 | 0.460751 | 0.913371 | 1 | 0.996511 | 0.986458 |
| Arl2bp | Q9D385 | -1.66 | -1.09 | -1.1 | -1.01 | -1.5 | -1.55 | -0.49 | -1.02 | -1.82 | -0.7 | -1.31 | -1.31 | -1.86 | -1.71 | -1.41 | -1.28 | 0.075 | -0.07 | -0.35 | 0.795841 | 0.805733 | 0.168996 | 1 | 0.996511 | 0.759484 |
| Arl3 | Q9WUL7 | -1.9 | -1.77 | -1.97 | -2.03 | -1.55 | -1.75 | -1.39 | -2 | -1.36 | -2.26 | -2.1 | -1.78 | -1.37 | -1.17 | -2.18 | -2.45 | 0.245 | 0.0425 | 0.125 | 0.207901 | 0.856925 | 0.697717 | 1 | 0.996511 | 0.955729 |
| Arl4c | P61208 | 1.33 | 0.71 | 0.6 | 1.87 | 0.16 | 1.32 | -0.24 | 1.35 | 2.07 | 3.15 | 0.55 | 2.18 | 0.9 | 1.29 | -1.11 | -0.91 | -0.48 | 0.86 | -1.085 | 0.317548 | 0.153732 | 0.108937 | 1 | 0.996511 | 0.707444 |
| Arl5a | Q80ZU0 | -1.09 | -0.61 | -0.21 | -0.09 | -0.22 | -0.87 | -0.2 | -0.42 | -0.99 | -0.21 | -0.2 | -1.06 | -1.41 | -0.38 | -0.72 | -0.43 | 0.0725 | -0.115 | -0.235 | 0.7944 | 0.725256 | 0.483736 | 1 | 0.996511 | 0.894091 |
| Arl6 | O88848 | -0.36 | 0.01 | 0.23 | -0.59 | 0.27 | -0.28 | 0.81 | -0.04 | -0.17 | -1.29 | 0.08 | -0.2 | -0.53 | 0.52 | 0.12 | 0.29 | 0.3675 | -0.2175 | 0.2775 | 0.236814 | 0.537705 | 0.371296 | 1 | 0.996511 | 0.859424 |
| Arl6ip1 | Q9JKW0 | 1.23 | 0.67 | 1.57 | 0.93 | 0.23 | 2.29 | -0.12 | 0.66 | 0.55 | 2.09 | 0.94 | 0.94 | 1.77 | 0.39 | 1.36 | 0.92 | -0.335 | 0.03 | 0.01 | 0.525598 | 0.93581 | 0.977266 | 1 | 0.996755 | 0.993727 |
| Arl6ip4 | Q9JM93 | 0.44 | -0.18 | 0.17 | 0.34 | -2.54 | 0.05 | -1.37 | 0.15 | 1.24 | 0.61 | 0.12 | -0.26 | 2.27 | 1.12 | -0.81 | -1.82 | -1.12 | 0.235 | -0.0025 | 0.088236 | 0.502167 | 0.997621 | 1 | 0.996511 | 0.999705 |
| Arl6ip5 | Q8R5J9 | 1.03 | 1.27 | 1.42 | 1.18 | 1.22 | 0.88 | 1.73 | 1.76 | -0.67 | -0.95 | 1.37 | 0.93 | -0.84 | -0.28 | 1.28 | 1.45 | 0.1725 | -1.055 | -0.8225 | 0.483107 | 0.076794 | 0.148223 | 1 | 0.996511 | 0.746406 |
| Arl6ip6 | Q8BH07 | 3.37 | 1.81 | 1.94 | 2.15 | 1.26 | 2.47 | 2.32 | 3.04 | 1.52 | 0.82 | 1.85 | 1.93 | 2.71 | 0.9 | 2.92 | 3.66 | -0.045 | -0.7875 | 0.23 | 0.924857 | 0.08612 | 0.714575 | 1 | 0.996511 | 0.958106 |
| Arl8a | Q8VEH3 | -0.15 | 0.56 | 0.39 | 0.3 | 0.97 | 0.18 | 1.4 | 0.49 | -0.05 | -1.13 | 0.68 | 0.2 | -0.33 | 0.32 | 0.58 | 1.03 | 0.485 | -0.35 | 0.125 | 0.138368 | 0.386201 | 0.702806 | 1 | 0.996511 | 0.956859 |
| Arl8b | Q9CQW2 | 0.97 | 0.49 | 0.24 | 0.37 | 1.19 | 0.82 | 0.85 | 0.38 | 1 | -0.13 | 0.4 | 1.49 | -0.09 | 0.6 | -0.15 | 0.26 | 0.2925 | 0.1725 | -0.3625 | 0.250769 | 0.646667 | 0.191948 | 1 | 0.996511 | 0.777117 |
| Armc1 | Q9D7A8 | -0.08 | -0.31 | 0.09 | 0.18 | -0.57 | -0.03 | 0.08 | 0.27 | 0.76 | -0.27 | 0.14 | -0.14 | 0.49 | -0.15 | -0.47 | -0.5 | -0.0325 | 0.1525 | -0.1275 | 0.887357 | 0.574409 | 0.646012 | 1 | 0.996511 | 0.944749 |
| Armc10 | Q9D0L7 | 0.97 | 0.73 | -0.05 | 0.26 | 0.64 | 1.23 | -0.19 | 0.26 | 0.15 | 0.23 | 0.27 | 1.16 | 0.25 | -0.54 | -0.12 | -0.27 | 0.0075 | -0.025 | -0.6475 | 0.983442 | 0.939234 | 0.053453 | 1 | 0.996755 | 0.661738 |
| Armc6 | Q8BNU0 | -0.11 | -0.22 | 0.24 | 0.01 | -0.44 | -0.04 | 0.5 | -0.04 | 0.28 | 0.59 | 0.3 | 0.33 | -0.16 | 0.3 | -0.07 | 0.56 | 0.015 | 0.395 | 0.1775 | 0.948924 | 0.057285 | 0.458295 | 1 | 0.996511 | 0.887902 |
| Armc8 | Q9DBR3-2 | 1.39 | 0.95 | 0.14 | 0.45 | 0.89 | 1.64 | -0.35 | -0.15 | 0.26 | 1 | -0.45 | 1.54 | 0.49 | 0.05 | -0.22 | -0.37 | -0.225 | -0.145 | -0.745 | 0.6537 | 0.76304 | 0.050303 | 1 | 0.996511 | 0.658454 |

| | | | | | | | | | | | | | | | | | | | | | | | | | | |
|---|---|---|---|---|---|---|---|---|---|---|---|---|---|---|---|---|---|---|---|---|---|---|---|---|---|---|
| Armc8 | G3X920 | -0.43 | -0.34 | -0.58 | -0.82 | 0.18 | -0.36 | -0.33 | -0.62 | -0.59 | -0.18 | -0.59 | -0.24 | -0.58 | 0.06 | -0.9 | -0.57 | 0.26 | 0.1425 | 0.045 | 0.258082 | 0.484954 | 0.861027 | 1 | 0.996511 | 0.977204 |
| Armc9 | Q9D2I5-2 | -0.41 | -0.2 | -0.27 | -0.09 | 0.17 | -0.66 | 0.13 | -0.23 | -0.64 | 0.17 | 0.14 | -0.02 | -0.47 | -0.42 | 0.05 | -0.09 | 0.095 | 0.155 | 0.01 | 0.67504 | 0.509084 | 0.961108 | 1 | 0.996511 | 0.9927 |
| Armcx2 | Q6A058 | -0.39 | -0.17 | -0.17 | 0.25 | -0.35 | -0.29 | -0.42 | -0.18 | 1.45 | 0.9 | -0.2 | 0.04 | 2.38 | -0.17 | -0.11 | -0.18 | -0.19 | 0.6675 | 0.6 | 0.319347 | 0.114015 | 0.326813 | 1 | 0.996511 | 0.843799 |
| Armcx3 | Q8BHS6 | -0.6 | -0.65 | -0.44 | -0.84 | -0.12 | -0.62 | -0.25 | -0.51 | -1.26 | -0.97 | -0.43 | -0.71 | -0.9 | -0.81 | -0.56 | -0.75 | 0.2575 | -0.21 | -0.1225 | 0.185855 | 0.370269 | 0.521018 | 1 | 0.996511 | 0.907667 |
| Armcx4 | E9PWM3 | 1.29 | 1.36 | -0.69 | -0.2 | 1.26 | 0.95 | -0.29 | 0.01 | 1.12 | 1.68 | -0.38 | 0.52 | 0.91 | 1.21 | -0.5 | 0.01 | 0.0425 | 0.295 | -0.0325 | 0.941884 | 0.638416 | 0.956714 | 1 | 0.996511 | 0.992636 |
| Armh3 | Q6PD19 | 0.18 | 0.12 | 0.12 | -0.01 | 0.21 | 0.24 | 0.67 | 0.16 | -0.7 | -0.43 | 0.59 | 0.11 | -0.42 | -0.16 | 0.35 | 0.67 | 0.2175 | -0.21 | 0.0075 | 0.232445 | 0.485571 | 0.978025 | 1 | 0.996511 | 0.993727 |
| Armt1 | A6H630 | 0.98 | 0.51 | 0.35 | -0.08 | 0.84 | 0.9 | 0.62 | 0.23 | -0.25 | 1.28 | 0.11 | 0.82 | -0.33 | 0.04 | -0.01 | 0.19 | 0.2075 | 0.05 | -0.4675 | 0.452757 | 0.898347 | 0.110139 | 1 | 0.996511 | 0.710755 |
| Arnt2 | Q61324-2 | -1.09 | -0.93 | -1.22 | -1.29 | -0.75 | -1.25 | -1.3 | -1.41 | -0.98 | -1.17 | -0.93 | -0.94 | -1.16 | -0.62 | -1.35 | -1.52 | -0.045 | 0.1275 | -0.03 | 0.821758 | 0.466273 | 0.902981 | 1 | 0.996511 | 0.985011 |
| Arntl | A0A1B0GS | -4.58 | -4.58 | -2.26 | -2.24 | -5.03 | -5.07 | -3.18 | -3.95 | -4.75 | -0.99 | -2.25 | -5.06 | -1.17 | -2.76 | -1.67 | -1.73 | -0.8925 | 0.1525 | 1.5825 | 0.245018 | 0.886596 | 0.042832 | 1 | 0.996511 | 0.658454 |
| Arpc1a | Q9R0Q6 | -0.51 | -0.62 | 0.51 | 0.93 | 0.05 | -0.26 | 0.5 | 0.82 | -1.01 | -0.66 | 0.67 | -0.98 | -1.74 | -0.9 | 0.32 | 0.1 | 0.2 | -0.5725 | -0.6325 | 0.637388 | 0.279977 | 0.278062 | 1 | 0.996511 | 0.822084 |
| Arpc1b | Q91Z25 | -0.07 | -0.13 | -0.28 | -0.24 | -0.09 | -0.34 | -0.15 | -0.17 | -0.33 | -0.36 | -0.22 | 0.02 | -0.6 | -0.03 | -0.51 | -0.6 | -0.0075 | -0.0425 | -0.255 | 0.959853 | 0.806171 | 0.236974 | 1 | 0.996511 | 0.792722 |
| Arpc2 | Q9CVB6 | 1.65 | 0.74 | 1.22 | 1.88 | 1.05 | 1.61 | 1.48 | 2.2 | 1 | 0.81 | 1.47 | 1.48 | -0.01 | -0.19 | 0.71 | 1.06 | 0.2125 | -0.1825 | -0.98 | 0.530693 | 0.555993 | 0.027534 | 1 | 0.996511 | 0.635993 |
| Arpc3 | Q9JM76 | 0.44 | -0.05 | 0.57 | 0.49 | -0.47 | 0.49 | 0.34 | 0.63 | 0.26 | -0.06 | 0.79 | 0 | -0.67 | 0.7 | 0.12 | -0.3 | -0.115 | -0.115 | -0.4 | 0.687817 | 0.65849 | 0.244477 | 1 | 0.996511 | 0.799771 |
| Arpc4 | P59999 | 0.53 | 0.29 | -0.2 | -0.77 | 1.37 | 0.28 | -0.24 | -0.14 | -1.09 | -1.64 | -0.3 | 0.01 | -2.62 | -0.79 | -1.47 | -0.69 | 0.355 | -0.7175 | -1.355 | 0.425132 | 0.134162 | 0.021612 | 1 | 0.996511 | 0.621926 |
| Arpc5 | Q9CPW4 | -0.24 | -0.32 | 0.11 | 0.16 | 0.68 | -0.48 | -0.05 | 0.29 | 0 | -0.1 | 0.11 | -0.62 | -0.45 | 0.15 | -0.12 | -0.26 | 0.1825 | -0.08 | -0.0975 | 0.517466 | 0.731922 | 0.664488 | 1 | 0.996511 | 0.949825 |
| Arpc5l | Q9D898 | 0.58 | 0.16 | -0.05 | -0.66 | 0.68 | 0.27 | -0.31 | -0.5 | -0.3 | -0.54 | -0.43 | -0.16 | -0.98 | 0.31 | -0.85 | -0.56 | 0.0275 | -0.365 | -0.5275 | 0.938721 | 0.218143 | 0.187506 | 1 | 0.996511 | 0.776843 |
| Arpin | Q9D0A3 | -0.12 | -0.8 | -0.27 | -0.29 | -0.22 | -0.84 | 0.07 | -0.47 | -0.65 | -0.43 | -0.37 | -0.52 | -0.75 | -0.28 | -0.53 | -0.29 | 0.005 | -0.1225 | -0.0925 | 0.984283 | 0.555852 | 0.689037 | 1 | 0.996511 | 0.955729 |
| Arpp19 | P56212 | -0.58 | -0.8 | -1.51 | -1.69 | -1.16 | -0.96 | -1.65 | -1.56 | -0.45 | -0.63 | -1.18 | -0.48 | -0.6 | -0.23 | -1.75 | -1.88 | -0.1875 | 0.46 | 0.03 | 0.549559 | 0.174028 | 0.948308 | 1 | 0.996511 | 0.992476 |
| Arpp21 | Q9DCB4 | 0.46 | 0.8 | -0.08 | 0.17 | -0.22 | -0.07 | 0.02 | -0.16 | 1.37 | -0.07 | -0.05 | 1.17 | 1.67 | 1.33 | 0.42 | 0.04 | -0.445 | 0.2675 | 0.5275 | 0.06977 | 0.519044 | 0.218711 | 1 | 0.996511 | 0.787447 |
| Arpp21 | Q9DCB4-2 | -0.64 | -2.36 | -1.6 | -2.57 | -3.79 | -0.95 | -4.34 | -1.92 | 1.29 | 0.71 | -1.99 | -0.93 | 1.83 | -1.65 | -2.68 | -1.96 | -0.9575 | 1.5625 | 0.6775 | 0.259777 | 0.074003 | 0.495722 | 1 | 0.996511 | 0.89758 |
| Arrb1 | Q8BWG8 | -0.59 | -0.18 | -0.11 | -0.02 | -0.39 | -0.27 | -0.06 | -0.06 | -0.63 | -0.26 | -0.17 | -0.6 | -1.13 | -0.13 | -0.49 | -0.43 | 0.03 | -0.19 | -0.32 | 0.874024 | 0.383922 | 0.254865 | 1 | 0.996511 | 0.809524 |
| Arrb2 | Q91YI4 | -0.41 | -1.02 | -0.54 | -0.51 | -1.08 | -0.73 | 0.05 | 0.32 | -1.18 | 0.54 | -0.07 | -1.18 | -1.25 | -0.8 | -0.92 | -0.35 | 0.26 | 0.1475 | -0.21 | 0.455178 | 0.729111 | 0.427659 | 1 | 0.996511 | 0.879165 |
| Arrdc1 | A2AIS9 | 0.03 | -0.64 | 0.94 | 0.78 | -0.53 | -0.24 | 0.59 | 0.89 | -0.25 | 0.01 | 1.05 | -0.06 | 0.06 | 0.55 | 0.2 | 0.57 | -0.1 | -0.09 | 0.0675 | 0.828104 | 0.838266 | 0.857813 | 1 | 0.996511 | 0.976225 |
| Arsa | P50428 | -0.45 | -0.39 | -0.15 | -0.01 | -0.69 | -0.16 | -0.21 | -0.68 | -0.94 | 0.35 | -0.22 | -0.61 | -0.43 | -1.22 | -0.88 | -0.48 | -0.185 | -0.105 | -0.5025 | 0.382864 | 0.728198 | 0.068215 | 1 | 0.996511 | 0.664936 |
| Arsb | A0A0R4J13 | -0.52 | -0.55 | -0.16 | 0.31 | -0.59 | -0.46 | 0.84 | 0.34 | -0.58 | -0.22 | 0.58 | -0.55 | -0.5 | -0.3 | -0.28 | -0.04 | 0.2625 | 0.0375 | -0.05 | 0.489686 | 0.91038 | 0.843722 | 1 | 0.996744 | 0.976225 |
| Arsg | Q3TYD4 | 0.59 | 0.78 | -0.4 | -0.56 | 2.52 | 0.16 | 0.74 | 0.09 | 0.17 | -1.65 | 0.72 | -0.35 | 0.74 | 1.26 | -1.07 | -0.4 | 0.775 | -0.38 | 0.03 | 0.220394 | 0.504811 | 0.958612 | 1 | 0.996511 | 0.992636 |
| Arsj | Q8BM89 | -5.14 | -2.98 | -3.13 | -3.93 | -2.63 | -4.69 | -3.96 | -4.05 | -1.72 | 0.03 | -3.7 | -4.51 | 1.16 | 0.04 | -3.14 | -3.89 | -0.0375 | 1.32 | 2.3375 | 0.950043 | 0.21706 | 0.07284 | 1 | 0.996511 | 0.668066 |
| Arvcf | P98203 | 0.75 | 0.76 | 0.91 | 0.84 | 0.93 | 0.79 | 0.74 | 0.64 | 0.99 | 0.25 | 0.91 | 0.99 | 0.85 | 0.75 | 0.87 | 0.91 | -0.04 | -0.03 | 3.00E-02 | 0.788173 | 0.891646 | 0.858435 | 1 | 0.996511 | 0.976225 |
| Arxes2 | C0HK80 | 0.3 | 0.91 | 0.78 | -0.35 | 1.77 | 0.65 | 1.41 | 0.64 | -1.69 | -1.28 | 0.57 | 0.51 | -1.4 | -1.46 | 1.49 | 1.74 | 0.7075 | -0.8825 | -0.3175 | 0.091139 | 0.167693 | 0.704035 | 1 | 0.996511 | 0.957084 |
| As3mt | Q91WU5 | -0.26 | -0.57 | 0.12 | 0.77 | -0.24 | -0.37 | -0.13 | 0.51 | -0.5 | 0.95 | 0.2 | -0.2 | -0.51 | -0.21 | 0.62 | 0.56 | -0.0725 | 0.0975 | 0.1 | 0.829862 | 0.810594 | 0.798009 | 1 | 0.996511 | 0.969727 |
| Asah1 | Q9WV54 | -0.02 | -0.39 | -1 | -0.33 | -0.26 | -0.11 | -1.17 | -1.15 | -0.35 | -0.56 | -0.38 | -0.33 | -0.93 | -0.54 | -1.57 | -0.82 | -0.2375 | 0.03 | -5.30E-01 | 0.488849 | 0.900249 | 0.113664 | 1 | 0.996511 | 0.716416 |
| Asah2 | B7ZN55 | 0.4 | 0.69 | 0.4 | 0.37 | 0.43 | 0.07 | 1.54 | 0.87 | 0.28 | 0.29 | 0.6 | -0.1 | 0.23 | 0.27 | 0.92 | 1.19 | 0.2625 | -0.1975 | 0.1875 | 0.419139 | 0.352666 | 0.499051 | 1 | 0.996511 | 0.899595 |
| Asap1 | H3BJY2 | -0.22 | -0.38 | 1.8 | 0.94 | -0.91 | -0.79 | -0.27 | 1.25 | -1.13 | 0.93 | 1.31 | -1.09 | -0.51 | -3.82 | 0.23 | -0.71 | -0.715 | -0.53 | -1.7375 | 0.288133 | 0.486207 | 0.087877 | 1 | 0.996511 | 0.691619 |
| Asap1 | E9QMJ1 | 0.39 | 0.04 | 0.49 | 0.68 | 0.18 | 0.45 | 0.05 | 0.61 | 0.14 | 0.33 | 0.51 | 0.07 | -0.07 | 0.32 | -0.2 | -0.05 | -0.0775 | -0.1375 | -0.4 | 0.715443 | 0.517698 | 0.100723 | 1 | 0.996511 | 0.698156 |
| Asap2 | D3YX85 | 1.01 | 0.52 | -0.02 | -3.32 | -1.81 | 0.66 | -4.58 | -3.46 | 3.13 | -3.17 | -1.08 | 0.22 | 3.96 | -0.32 | 2.53 | -2.28 | -1.845 | 0.2275 | 1.425 | 0.192962 | 0.876678 | 0.360443 | 1 | 0.996511 | 0.85741 |
| Asap2 | E9PX52 | -0.06 | 0.34 | 0.81 | 0.75 | 0.39 | -0.04 | 0 | 0.31 | 0.24 | 0.89 | 0.72 | 0.15 | 0.06 | 0.62 | 0.26 | 0.27 | -0.295 | 0.04 | -0.1575 | 0.246359 | 0.887339 | 0.551322 | 1 | 0.996511 | 0.914255 |
| Asb13 | Q8VBX0 | -0.28 | 0.35 | 1.98 | -0.13 | 0.7 | -0.69 | -0.03 | -0.34 | -0.37 | 0.31 | -0.18 | -0.18 | -0.13 | -0.06 | 1.63 | 0.64 | -0.57 | -0.585 | 0.04 | 0.312771 | 0.261262 | 0.947162 | 1 | 0.996511 | 0.992376 |
| Asb3 | A0A0N4SV | 0.33 | -0.14 | -1.2 | -1.94 | -1.29 | 1.27 | -0.88 | -2 | 0.14 | -2.76 | -1.2 | 0.84 | 2.45 | -1.83 | -1.78 | -1.25 | 0.0125 | -0.0075 | 0.135 | 0.987296 | 0.992974 | 0.895187 | 1 | 0.999285 | 0.983509 |
| Asb6 | Q91ZU1 | -0.59 | -0.37 | 0.1 | -2.04 | 0.02 | -0.74 | -1.84 | -1.17 | 0 | 2.42 | -0.87 | -0.63 | -0.65 | 0.39 | -1.4 | 0.17 | -0.2075 | 0.955 | 0.3525 | 0.708638 | 0.2514 | 0.539075 | 1 | 0.996511 | 0.912212 |
| Ascc1 | Q9D8Z1 | -0.07 | -0.03 | 0.1 | -0.4 | -0.19 | -0.81 | -0.13 | -0.14 | -0.19 | 0.12 | -0.22 | -0.11 | -0.07 | -0.09 | -0.08 | 0.38 | -0.2175 | 3.8E-17 | 0.135 | 0.336238 | 1 | 0.530666 | 1 | 1 | 0.908944 |
| Ascc2 | Q91WR3-2 | -1.7 | -0.5 | -0.99 | -1.12 | -1.85 | -0.86 | -0.93 | -0.85 | -0.96 | 0.45 | 0.1 | -1.35 | -0.94 | -0.92 | -0.82 | -0.04 | -0.045 | 0.6375 | 0.3975 | 0.893271 | 0.191622 | 0.248712 | 1 | 0.996511 | 0.804385 |
| Ascc3 | E9PZJ8 | -0.63 | -0.6 | 0.16 | -0.21 | -0.36 | -0.72 | 0.24 | 0.02 | -0.37 | 0.47 | 0.19 | -0.16 | -0.3 | -0.29 | 0.14 | 0.23 | 0.115 | 0.3525 | 0.265 | 0.686738 | 0.224238 | 0.325166 | 1 | 0.996511 | 0.842881 |
| Asf1a | Q9CQE6 | -0.18 | -0.1 | -0.88 | -0.22 | 0.39 | -0.15 | -1.05 | -0.09 | -0.67 | 0.32 | -0.24 | 0.05 | -0.31 | 0.71 | -0.65 | -1.01 | 0.12 | 0.21 | 0.03 | 0.723894 | 0.472397 | 0.939785 | 1 | 0.996511 | 0.990645 |
| Ash2l | Q91X20 | -2.4 | -2.12 | -2.47 | -2.97 | -1.61 | -2.92 | -3.01 | -3.17 | -1.92 | -0.83 | -2.81 | -2.77 | -1.35 | -0.99 | -2.55 | -2.9 | -0.1875 | 0.4075 | 0.5425 | 0.624433 | 0.392919 | 0.261429 | 1 | 0.996511 | 0.81412 |
| Asic1 | Q6NXK8 | 0.51 | 0.31 | 0.31 | 0.63 | 0.44 | 0.44 | 0.66 | 0.62 | 0.6 | 1.34 | 0.75 | 0.53 | 0.52 | 0.95 | 0.67 | 0.6 | 0.1 | 0.365 | 0.245 | 0.537354 | 0.141855 | 0.22918 | 1 | 0.996511 | 0.787447 |
| Asic2 | Q925H0-2 | 0.06 | 0.1 | 0.16 | -0.14 | 0.47 | 0.68 | 0.24 | 0.05 | -1.33 | -0.28 | 0.15 | 0.65 | -1.59 | -2.84 | 0.01 | 0.17 | 0.315 | -0.2475 | -1.1075 | 0.124927 | 0.546809 | 0.117029 | 1 | 0.996511 | 0.718608 |
| Asic2 | Q925H0 | -0.62 | -0.37 | -0.87 | -0.57 | -0.61 | -0.82 | -0.25 | -1.29 | -0.19 | -0.01 | -1.01 | -0.65 | -0.6 | 0.45 | -0.21 | -0.68 | -0.135 | 0.1425 | 0.3475 | 0.594802 | 0.594642 | 0.25517 | 1 | 0.996511 | 0.809524 |
| Asic4 | D3YYS9 | -0.09 | -0.17 | 0.86 | 1.32 | -0.14 | -0.27 | 1.01 | 1.27 | 0.66 | 1.49 | 0.85 | 0.11 | 0.55 | -0.33 | 0.88 | 1.64 | -0.0125 | 0.2975 | 0.205 | 0.979829 | 0.501721 | 0.688509 | 1 | 0.996511 | 0.955729 |
| Asl | Q91YI0 | -0.11 | -0.25 | -0.07 | 0.08 | -0.35 | -0.45 | -0.12 | 0.04 | -0.73 | 0.46 | -0.24 | -0.33 | -0.96 | -0.69 | -0.28 | -0.21 | -0.1325 | -0.1225 | -0.4475 | 0.461917 | 0.653731 | 0.081212 | 1 | 0.996511 | 0.687007 |
| Asns | Q61024 | -1.26 | -1.51 | -0.98 | -1.02 | -1.85 | -1.26 | -0.75 | -0.69 | -1.03 | -0.43 | -1.06 | -1.24 | -1.16 | -1.17 | -1.48 | -1.72 | 0.055 | 0.2525 | -0.19 | 0.8522 | 0.31139 | 0.414188 | 1 | 0.996511 | 0.874945 |
| Aspa | Q8R3P0 | 4.25 | 2.28 | 0.41 | -0.63 | 2.5 | 3.75 | 0 | -0.75 | -0.08 | -1.62 | -0.18 | 3.3 | -0.51 | -0.29 | -0.11 | 0.44 | -0.2025 | -1.2225 | -1.695 | 0.880276 | 0.373193 | 0.110954 | 1 | 0.996511 | 0.712422 |
| Asph | A2AL85 | 0.08 | 0.45 | 0.39 | 0.31 | 1.06 | 0.36 | 1.43 | 0.72 | 0.26 | 0.05 | 0.25 | 0.19 | 0.22 | 1.03 | 0.98 | 1.3 | 0.585 | -0.12 | 0.575 | 0.044163 | 0.489271 | 0.058283 | 1 | 0.996511 | 0.661738 |
| Asph | Q8BSY0-3 | -0.28 | -0.03 | 0.87 | 0.74 | 0.82 | -0.4 | 1.24 | 1.23 | -0.63 | 0.51 | 1.63 | -0.33 | -0.34 | 0.39 | 1.34 | 1.8 | 0.3975 | -0.03 | 0.4725 | 0.384579 | 0.955324 | 0.372711 | 1 | 0.996755 | 0.859424 |
| Asphd1 | Q2TA57 | 0.3 | 0.36 | -0.03 | -0.6 | -0.34 | -0.2 | -0.8 | -0.43 | -0.34 | -0.78 | -0.63 | -0.44 | -0.43 | -0.14 | -0.74 | -0.73 | -0.45 | -0.555 | -0.5175 | 0.114279 | 0.05808 | 0.09373 | 1 | 0.996511 | 0.698156 |
| Asphd2 | Q80VP9 | 1.41 | 0.32 | 0.29 | 0.08 | 0.36 | 1.31 | 1.63 | 0.2 | -0.15 | -1.52 | 0.26 | 0.94 | -0.78 | -0.3 | 0.07 | 0.3 | 0.35 | -0.6425 | -0.7025 | 0.426543 | 0.2638 | 0.086931 | 1 | 0.996511 | 0.691619 |
| Aspscr1 | Q8VBT9 | 0.23 | 0.33 | -0.28 | -0.5 | 0.09 | 0.73 | -0.67 | -0.48 | 0.01 | -0.12 | -0.15 | 0.33 | -0.19 | -0.01 | -0.46 | -0.46 | -0.0275 | 0.0725 | -0.225 | 0.938639 | 0.772854 | 0.393607 | 1 | 0.996511 | 0.865775 |
| Asrgl1 | Q8C0M9 | 0.01 | -0.33 | -0.48 | -0.23 | -0.35 | 0.1 | -0.15 | -0.28 | -0.08 | -0.51 | -0.46 | -0.24 | -0.27 | -0.43 | -0.6 | -0.48 | 0.0875 | -0.065 | -0.1875 | 0.63787 | 0.74065 | 0.349885 | 1 | 0.996511 | 0.85327 |
| Ass1 | P16460 | 0.11 | -0.19 | -0.14 | -0.37 | -0.37 | -0.27 | 0.11 | -0.22 | -0.21 | 0.14 | -0.56 | -0.28 | -0.3 | 0.1 | -0.18 | -0.55 | -0.04 | -0.08 | -0.085 | 0.829084 | 0.710594 | 0.699074 | 1 | 0.996511 | 0.955729 |
| Astn1 | Q61137-2 | 0.53 | 0.57 | 0.99 | 1.08 | 0.52 | 0.74 | 0.91 | 1.01 | 0.84 | 0.76 | 1.17 | 0.97 | 0.33 | 0.42 | 0.69 | 0.83 | 0.0025 | 0.1425 | -0.225 | 0.990353 | 0.50365 | 0.338591 | 1 | 0.996511 | 0.847578 |
| Astn2 | Q80Z10 | 0.75 | 0.71 | 0.26 | 0.19 | 0.55 | 1.13 | 0.07 | 0.13 | 0.16 | -0.42 | 0.33 | 1.02 | 0.07 | 0.07 | -0.16 | 0.28 | -0.0075 | -0.205 | -0.4125 | 0.979084 | 0.537605 | 0.090173 | 1 | 0.996511 | 0.69238 |
| Asxl2 | D3Z112 | -2.38 | -1.06 | 0.96 | 1.44 | -0.4 | -3.31 | 0.52 | 3.97 | -1.65 | 1.28 | 0.61 | -0.96 | -1.49 | -0.38 | -0.8 | -0.06 | 0.455 | 0.08 | -0.4225 | 0.770435 | 0.936536 | 0.62044 | 1 | 0.996755 | 0.93807 |
| Atad1 | Q9D5T0 | 0.52 | -0.06 | 0.01 | -0.45 | 0.46 | 0.89 | 0.16 | -0.02 | -0.42 | 0.15 | 0.4 | 0.59 | -0.18 | 0.09 | 0.16 | 0.1 | 0.3675 | 0.175 | 0.0375 | 0.217204 | 0.565687 | 0.879736 | 1 | 0.996511 | 0.979626 |
| Atad2b | E9Q166 | 1.07 | 0.5 | 0.73 | 0.56 | 0.25 | 0.02 | 0.02 | 0.29 | 0.65 | 0.8 | 1.13 | 0.77 | 1.1 | 0.33 | 0.8 | 1.3 | -0.57 | 0.1225 | 0.1675 | 0.013482 | 0.560214 | 0.540785 | 1 | 0.996511 | 0.912212 |
| Atad3 | Q925I1 | 1.13 | 1.06 | 0.96 | 1.08 | 0.65 | 1.09 | 0.53 | 0.94 | 1.04 | 0.82 | 0.92 | 0.94 | 1.03 | 1.02 | 0.58 | 0.55 | -0.255 | -0.1275 | -0.2625 | 0.179595 | 0.427827 | 0.216536 | 1 | 0.996511 | 0.787447 |
| Atat1 | Q8K341-4 | 0.7 | 1.1 | 0.86 | 0.47 | 1.04 | 0.95 | 0.64 | 0.56 | 1.34 | 1.1 | 0.99 | 1.06 | 1.38 | 1.7 | 0.67 | 0.66 | 0.015 | 0.34 | 0.32 | 0.942019 | 0.11889 | 0.308599 | 1 | 0.996511 | 0.834095 |
| Atcay | Q8BHE3 | 1.36 | -0.01 | 0.37 | 1.31 | -0.67 | 1.21 | 0.83 | 1.52 | 1.46 | 0.44 | 1.08 | 1.83 | 0.53 | 0.17 | 0.33 | 0.2 | -0.035 | 0.445 | -0.45 | 0.948705 | 0.314895 | 0.21998 | 1 | 0.996511 | 0.787447 |
| Ate1 | A0A1L1SQ | -0.45 | -0.85 | -0.58 | -0.75 | -1.18 | -0.58 | -0.66 | -0.52 | -0.09 | -0.34 | -0.57 | -0.26 | 0.18 | -0.9 | -0.5 | -0.68 | -0.0775 | 3.43E-01 | 0.1825 | 0.707018 | 0.09963 | 0.50911 | 1 | 0.996511 | 0.902804 |
| Atf2 | P16951 | -3.78 | -2.73 | -1.95 | -3.01 | -1.83 | -3.22 | -2.13 | -3.54 | -1.37 | -2.23 | -2.02 | -2.07 | -3.94 | -0.62 | -1.03 | -2.63 | 0.1875 | 0.945 | 0.8125 | 0.717121 | 0.041663 | 0.30655 | 1 | 0.996511 | 0.832533 |
| Atf7ip | Q7TT18 | -1.05 | -2.52 | -1.75 | -0.26 | -1.98 | -1.82 | -0.97 | 0.02 | -2.24 | 1.66 | -0.25 | -2.05 | -1.75 | -1.15 | 0.31 | -0.71 | 0.2075 | 0.675 | 0.57 | 0.733373 | 0.474046 | 0.351467 | 1 | 0.996511 | 0.853984 |
| Atg101 | Q9D8Z6 | 0.11 | 0.64 | 0.54 | 0.64 | 0.26 | 0.02 | 0.1 | 0.42 | 0.47 | 0.37 | 0.57 | 0.21 | 0.71 | 1 | 0.66 | 0.57 | -0.2825 | -0.0775 | 0.2525 | 0.166447 | 0.697286 | 0.254962 | 1 | 0.996511 | 0.809524 |
| Atg12 | Q9CQY1 | -2.01 | -1.98 | -1.14 | -1.08 | -2.05 | -1.3 | -1.12 | -1.17 | -1.52 | -0.53 | -0.76 | -2.15 | -2 | -0.92 | -1.42 | -2 | 0.1425 | 0.3125 | -0.0325 | 0.664119 | 0.470241 | 0.928011 | 1 | 0.996511 | 0.98946 |
| Atg13 | Q91YI1-2 | 0.13 | 1.02 | 1.76 | 1.51 | 0.49 | 0.01 | 1.31 | 1.55 | 0.18 | 2.96 | 1.42 | 0.59 | 0.35 | 0.07 | 1.71 | 2.5 | -0.265 | 0.1825 | 0.0525 | 0.576558 | 0.77901 | 0.932303 | 1 | 0.996511 | 0.989713 |
| Atg14 | Q8CDJ3 | -1.33 | -1.74 | -0.59 | -0.86 | -1.52 | -0.99 | 0.52 | -0.42 | -0.55 | 0.37 | -0.53 | -1.06 | 0.39 | -1.04 | -0.65 | 0.42 | 0.5275 | 0.6875 | 0.91 | 0.278178 | 0.096567 | 0.058471 | 1 | 0.996511 | 0.661738 |
| Atg16l1 | Q8C0J2 | 0.3 | 0.17 | 0.13 | 0.31 | -0.42 | -0.02 | -0.55 | 0 | 0.52 | 0.58 | 0.15 | 0.02 | 0.2 | -0.3 | -0.59 | -0.75 | -0.475 | 0.09 | -0.5875 | 0.030255 | 0.64905 | 0.039496 | 1 | 0.996511 | 0.658454 |
| Atg2a | Q6P4T0 | -0.1 | 0.29 | 0.31 | 0.03 | 0.49 | -0.17 | -0.23 | 0.01 | 0.43 | 0.68 | 0.18 | -0.09 | 0.32 | 0.72 | 0.17 | 0.37 | -0.1075 | 0.1675 | 0.2625 | 0.622019 | 0.467762 | 0.234404 | 1 | 0.996511 | 0.790657 |
| Atg2b | Q80XK6 | 0.4 | 0.36 | 0.17 | 0.03 | 0.27 | 0.16 | 0.05 | -0.02 | 0.23 | 0.84 | 0.07 | 0.33 | 0.19 | 0.6 | 0.01 | 0.14 | -0.125 | 0.1275 | -5.00E-03 | 0.455586 | 0.571114 | 0.981094 | 1 | 0.996511 | 0.994858 |
| Atg3 | Q9CPX6 | -0.51 | -0.15 | 0.11 | -0.37 | 0.17 | -0.59 | 0.32 | -0.03 | -0.85 | -0.32 | 0.02 | -0.28 | -1.52 | -0.64 | -0.29 | 0.24 | 0.1975 | -0.1275 | -0.3225 | 0.441875 | 0.611815 | 0.409438 | 1 | 0.996511 | 0.87457 |
| Atg4b | A0A0R4J06 | 0.1 | 0.57 | -1.7 | -1.4 | 0.48 | 0.01 | -0.92 | -1.8 | 0.36 | -1.34 | -1.72 | 0.06 | -0.35 | 1.23 | -2.05 | -0.99 | 0.05 | -0.0525 | 0.0675 | 0.94147 | 0.939079 | 0.932372 | 1 | 0.996755 | 0.989713 |
| Atg4c | Q811C2 | -0.51 | 0.47 | 0.49 | 0.37 | 0.12 | 0.12 | -1.57 | -1.79 | 0.5 | 0.51 | 0.52 | -0.11 | 0.7 | 0.28 | 0.07 | 0.7 | -0.985 | 0.15 | 0.2325 | 0.089616 | 0.611802 | 0.44628 | 1 | 0.996511 | 0.884817 |
| Atg5 | Q99J83 | -0.01 | -0.07 | 1.1 | 0.93 | 0.52 | 0.06 | 0.4 | 1.27 | -0.14 | -0.35 | 0.77 | -0.16 | -0.79 | -0.26 | -0.05 | -0.19 | 0.075 | -0.4575 | -0.81 | 0.843147 | 0.250873 | 0.041142 | 1 | 0.996511 | 0.658454 |
| Atg7 | A0A0A0MC | -0.01 | 0.13 | 0.33 | 0.22 | 0.28 | -0.16 | 0.26 | 0.22 | -0.26 | 0.3 | 0.1 | -0.12 | -0.69 | 0.18 | 0.04 | 0.12 | -0.0175 | -0.1625 | -0.255 | 0.920373 | 0.415262 | 0.321166 | 1 | 0.996511 | 0.839916 |
| Atg9a | Q68FE2 | 0.43 | 0.17 | 0.2 | 0.02 | 0.39 | 0.64 | 0.13 | 0.31 | 0.5 | -0.36 | 0.54 | 0.41 | 0.77 | -0.39 | 0 | 0.14 | 0.1625 | 0.0675 | -0.075 | 0.379273 | 0.788653 | 0.786931 | 1 | 0.996511 | 0.968024 |
| Atic | Q9CWJ9 | -0.26 | -0.51 | -0.11 | -0.03 | -0.54 | -0.22 | -0.09 | -0.03 | -0.21 | 0.12 | -0.22 | -0.24 | -0.52 | -0.09 | -0.46 | -0.23 | 0.0075 | 0.09 | -0.0975 | 0.968878 | 0.640937 | 0.63902 | 1 | 0.996511 | 0.942804 |
| Atl1 | Q8BH66 | 0.67 | 0.49 | 0.81 | 0.73 | 0.99 | 0.81 | 1 | 0.96 | 0.52 | 0.19 | 0.8 | 0.71 | 0.64 | 0.86 | 0.99 | 1.1 | 0.265 | -0.12 | 0.2225 | 0.110217 | 0.553353 | 0.268502 | 1 | 0.996511 | 0.820082 |
| Atl2 | Q6PA06 | -0.16 | -0.13 | -0.42 | -0.32 | 0.03 | -0.11 | -0.16 | -0.36 | -0.2 | -0.2 | -0.48 | -0.18 | -0.15 | 0.26 | -0.12 | -0.43 | 0.1075 | -0.0075 | 0.1475 | 0.517579 | 0.965416 | 0.495049 | 1 | 0.996755 | 0.89758 |
| Atl3 | Q91YH5 | 0.19 | 0.29 | -0.15 | -0.5 | 0.48 | -0.01 | 0.01 | -0.51 | 0.25 | -0.25 | -0.3 | 0.36 | 0.25 | 0.48 | 0.09 | 0.01 | 0.035 | 0.0575 | 0.25 | 0.898538 | 0.8278 | 0.320141 | 1 | 0.996511 | 0.839402 |
| Atox1 | O08997 | 0.16 | 0.06 | -1.97 | -1.83 | -0.14 | -0.34 | -1.45 | -2.06 | -0.09 | -1.2 | -1.92 | 0.3 | -0.3 | 0.77 | -2.38 | -2.63 | -0.1025 | 0.1675 | -0.24 | 0.878477 | 0.811597 | 0.790723 | 1 | 0.996511 | 0.969105 |
| Atp11a | E9Q3G7 | 1.55 | 0.68 | 0.91 | 0.56 | 1.05 | 1.57 | 0.97 | 0.54 | 0.54 | 0.48 | 0.48 | 1.4 | 0.75 | 0.71 | 1.04 | 0.85 | 0.1075 | -0.2 | -0.0875 | 0.72312 | 0.532378 | 0.738143 | 1 | 0.996511 | 0.958796 |
| Atp11b | Q6DFW5 | -0.09 | -0.11 | -1.25 | -1.46 | 0.37 | -0.02 | -0.91 | -1.41 | -0.79 | -1.24 | -0.92 | 0.01 | -0.76 | -0.44 | -1.17 | -0.86 | 0.235 | -0.0075 | -0.08 | 0.642751 | 0.985982 | 0.834841 | 1 | 0.998892 | 0.975487 |
| Atp11c | E9QKK8 | -0.12 | 0.41 | 0.29 | 0.25 | 0.4 | 0.16 | 0.77 | 0.42 | -0.4 | 0.28 | 0.15 | 0.1 | -0.23 | -0.09 | 0.03 | 0.8 | 0.23 | -0.175 | -0.08 | 0.272802 | 0.441676 | 0.774149 | 1 | 0.996511 | 0.966062 |
| Atp13a1 | Q9EPE9 | -0.34 | -0.24 | -0.24 | -0.26 | 0.29 | -0.17 | 0.27 | -0.32 | -0.6 | -0.42 | -0.39 | -0.1 | -0.45 | -0.01 | 0.32 | 0.34 | 0.2875 | -0.1075 | 0.32 | 0.162458 | 0.546499 | 0.189115 | 1 | 0.996511 | 0.776843 |
| Atp13a2 | E9PYX5 | 0.09 | 0.39 | 1.23 | 1.59 | 0.47 | 0 | 1.54 | 1.68 | 0.26 | 0.8 | 1.98 | 0.34 | -0.13 | 0.68 | 1.48 | 1.77 | 0.0975 | 0.02 | 0.125 | 0.844695 | 0.96762 | 0.808039 | 1 | 0.996755 | 0.971953 |
| Atp13a3 | Q5XF89 | -0.71 | 0.07 | 0.56 | -0.42 | -0.27 | -1.14 | 0.38 | -1.49 | -0.52 | -0.72 | -0.69 | -0.83 | -0.96 | 0.03 | -1.26 | -0.53 | -0.505 | -0.565 | -0.555 | 0.29934 | 0.082191 | 0.173585 | 1 | 0.996511 | 0.764376 |
| Atp13a4 | A0A0R4J0 | -0.09 | -0.31 | -0.21 | 0.83 | -0.4 | -0.76 | 0.81 | 1.11 | 0.15 | 0.32 | 0.3 | -0.02 | -1.29 | -0.35 | 0.07 | 1.02 | 0.135 | 0.1325 | -0.1925 | 0.781212 | 0.643375 | 0.705918 | 1 | 0.996511 | 0.957084 |
| Atp1a1 | Q8VDN2 | 0.86 | 1.02 | 1.32 | 1.29 | 1.31 | 0.86 | 1.02 | 1.25 | 0.51 | 0.84 | 1.36 | 0.8 | 0.4 | 0.86 | 0.66 | 0.83 | -0.0125 | -0.245 | -0.435 | 0.947582 | 0.31704 | 0.064631 | 1 | 0.996511 | 0.66433 |
| Atp1a2 | Q6PIE5 | 0.93 | 0.73 | 0.53 | 1 | 0.85 | 1 | 0.66 | 0.73 | 0.75 | 0.64 | 0.46 | 1.02 | 0.22 | 0.79 | -0.04 | 0.54 | 0.0125 | -0.08 | -0.42 | 0.943617 | 0.696744 | 0.114967 | 1 | 0.996511 | 0.717287 |
| Atp1a3 | A0A0G2JG | 0.38 | 0.58 | 0.67 | 0.94 | 0.65 | 0.4 | 0.87 | 0.9 | 0.46 | 0.6 | 0.7 | 0.53 | 0.05 | 0.81 | 0.42 | 0.62 | 0.0625 | -0.07 | -0.1675 | 0.752966 | 0.708866 | 0.489841 | 1 | 0.996511 | 0.897531 |
| Atp1b1 | P14094 | 1.78 | 1.35 | 1.76 | 2.14 | 1.22 | 1.79 | 1.86 | 2.11 | 1.27 | 1.36 | 1.75 | 1.87 | 0.96 | 0.79 | 1.61 | 1.82 | -0.0125 | -0.195 | -0.4625 | 0.961237 | 0.432344 | 0.157721 | 1 | 0.996511 | 0.751678 |
| Atp1b2 | P14231 | 2.2 | 1.77 | 1.38 | 1.78 | 1.77 | 1.89 | 1.88 | 1.93 | 1.41 | 1.71 | 1.55 | 2.17 | 0.95 | 1.43 | 1.3 | 1.56 | 0.085 | -0.0725 | -0.4725 | 0.676196 | 0.777373 | 0.083397 | 1 | 0.996511 | 0.689531 |
| Atp1b3 | P97370 | 0.71 | 0.72 | -0.44 | -0.47 | 0.55 | 0.72 | -0.36 | -0.67 | 0.37 | -0.49 | -0.48 | 0.79 | 0.38 | 0 | -0.72 | -0.52 | -0.07 | -0.0825 | -0.345 | 0.87534 | 0.850782 | 0.403476 | 1 | 0.996511 | 0.870698 |
| Atp23 | G3UW46 | 0.24 | -0.02 | -0.4 | -0.33 | 0.98 | 1.84 | -0.14 | -0.31 | -0.06 | -1.55 | -0.21 | 0.21 | 0.8 | 0.14 | -1.15 | -0.44 | 0.72 | -0.275 | -0.035 | 0.164438 | 0.498384 | 0.933696 | 1 | 0.996511 | 0.989767 |
| Atp2a2 | O55143 | 0.26 | 0.18 | 0.57 | 0.67 | 0.49 | 0.29 | 0.62 | 0.62 | 0.13 | 0.34 | 0.23 | 0.08 | 0.31 | 0.58 | 0.39 | 0.72 | 0.085 | -0.225 | 0.08 | 0.646198 | 0.254146 | 0.703253 | 1 | 0.996511 | 0.956919 |
| Atp2a3 | B1ATS5 | -0.73 | -1.4 | -0.47 | -0.45 | -0.8 | -1.5 | -0.4 | -0.55 | -0.27 | 0.19 | -0.21 | -1.19 | 0.61 | -0.32 | -0.01 | -0.13 | -0.05 | 0.3925 | 0.8 | 0.876489 | 0.28952 | 0.027704 | 1 | 0.996511 | 0.635993 |
| Atp2b1 | G5E829 | 0.62 | 0.64 | 0.36 | 0.44 | 0.69 | 0.66 | 0.31 | 0.23 | 0.31 | -0.29 | 0.34 | 0.78 | 0.38 | -0.32 | 0.05 | 0.1 | -0.0425 | -0.23 | -0.4625 | 0.81453 | 0.373987 | 0.055615 | 1 | 0.996511 | 0.661738 |
| Atp2b1 | A0A1W2P7 | 0.24 | 0.44 | 1.02 | 1.2 | 0.11 | 0.1 | 0.98 | 0.95 | 1.07 | 0.76 | 1.01 | 0.56 | 0.56 | 1.06 | 0.16 | 0.54 | -0.19 | 0.125 | -0.145 | 0.566809 | 0.648003 | 0.636905 | 1 | 0.996511 | 0.942665 |
| Atp2b2 | F8WHB1 | 1.04 | 1.14 | 1.49 | 1.37 | 1.13 | 1.06 | 1.14 | 1.27 | 0.74 | 0.59 | 1.4 | 1.06 | 0.57 | 0.56 | 0.68 | 0.98 | -0.11 | -0.3125 | -0.5625 | 0.516218 | 0.209019 | 0.021375 | 1 | 0.996511 | 0.621926 |
| Atp2b2 | S4R1C4 | -0.31 | 0.26 | 0.7 | 1.15 | 0.57 | -0.37 | 0.6 | 0.7 | 0.1 | 0.16 | 0.7 | 0.15 | -0.08 | 0.67 | 0.84 | 1.21 | -0.075 | -0.1725 | 0.21 | 0.843322 | 0.612463 | 0.601134 | 1 | 0.996511 | 0.930802 |
| Atp2b3 | Q0VF55 | 0.54 | 0.56 | 0.77 | 0.75 | 0.57 | 0.67 | 0.5 | 0.62 | 0.45 | 0.37 | 0.58 | 0.46 | 0.47 | 0.18 | 0.39 | 0.7 | -0.065 | -0.19 | -0.22 | 0.66345 | 0.263398 | 0.277408 | 1 | 0.996511 | 0.822084 |
| Atp2b3 | A2ALL9 | 0.46 | 1.12 | 0.69 | 0.54 | 1.91 | 0.47 | 0.77 | 0.39 | 0.39 | 0.14 | 0.62 | 0.54 | 0.36 | 1.05 | 0.36 | 0.79 | 0.1825 | -0.28 | -0.0625 | 0.618482 | 0.222619 | 0.807151 | 1 | 0.996511 | 0.97178 |
| Atp2b4 | E9Q828 | 1.72 | 1.59 | 1.34 | 1.61 | 1.69 | 1.97 | 1.51 | 1.67 | 1.22 | 1.11 | 1.55 | 1.9 | 0.85 | 0.74 | 1.42 | 1.37 | 0.145 | -0.12 | -0.47 | 0.41487 | 0.602085 | 0.071342 | 1 | 0.996511 | 0.668066 |
| Atp2c1 | H3BL44 | -0.3 | -0.37 | -0.61 | -0.56 | -0.07 | -0.25 | -0.43 | -0.54 | -0.38 | 0.28 | -0.32 | -0.34 | -0.36 | -0.49 | -0.17 | -0.2 | 0.1375 | 0.27 | 0.155 | 0.441823 | 0.228967 | 0.415876 | 1 | 0.996511 | 0.874945 |
| Atp5c1 | A2AKU9 | 1.01 | 0.63 | 0.98 | 1.54 | 0.58 | 1.39 | 0.67 | 1.66 | 0.64 | 1.15 | 1.15 | 0.73 | 1.34 | -0.42 | 0.34 | 0.51 | 0.035 | -0.1225 | -0.5975 | 0.912511 | 0.630933 | 0.15489 | 1 | 0.996511 | 0.750763 |
| Atp5c1 | Q8C2Q8 | -0.33 | 0.75 | 0.08 | -0.93 | -0.43 | 0.27 | -0.79 | -0.51 | 1.11 | 0.06 | -0.87 | -0.15 | -0.15 | 0.18 | -1.19 | 1.03 | -0.2575 | 0.145 | 0.075 | 0.520255 | 0.77325 | 0.888649 | 1 | 0.996511 | 0.98074 |
| Atp5f1a | Q03265 | 0.88 | 0.82 | 0.65 | 1.09 | 0.71 | 0.64 | 0.89 | 1.15 | 0.87 | 0.51 | 0.98 | 0.7 | 0.37 | 0.91 | 0.24 | 0.63 | -0.0125 | -0.095 | -0.3225 | 0.946492 | 0.623674 | 0.172753 | 1 | 0.996511 | 0.76428 |
| Atp5f1b | P56480 | -0.28 | 0.08 | 0.29 | 0.7 | -0.02 | -0.36 | 0.47 | 0.6 | 0.07 | 0.73 | 0.42 | -0.44 | -0.38 | 0.8 | -0.07 | 0.24 | -0.025 | -0.0025 | -0.05 | 0.93348 | 0.993824 | 0.878902 | 1 | 0.999285 | 0.979626 |
| Atp5f1d | Q9D3D9 | -1.51 | -0.37 | -1.3 | -1.34 | -0.32 | -1.61 | -1.42 | -1.74 | -1.52 | -2.39 | -1.12 | -1.51 | -1.63 | -2.3 | -1.82 | -1.71 | -0.1425 | -0.505 | -0.735 | 0.7167 | 0.18714 | 0.038438 | 1 | 0.996511 | 0.658454 |
| Atp5f1e | P56382 | 0.12 | -0.29 | -0.2 | -0.5 | -0.78 | 0.53 | -0.25 | -0.12 | 1.52 | -1.05 | 0.37 | -0.38 | 2.69 | -0.6 | 0.83 | -0.68 | 0.0625 | 0.3325 | 0.7775 | 0.83337 | 0.530487 | 0.299737 | 1 | 0.996511 | 0.829161 |
| Atp5j2 | F8WHP8 | 0.29 | 0.59 | 0.38 | 0.64 | 0.73 | 0.46 | 0.81 | 0.63 | 0.3 | -0.8 | 0.77 | 0.28 | -0.14 | 1.19 | -0.19 | 0.22 | 0.1825 | -0.3375 | -0.205 | 0.292917 | 0.332899 | 0.542668 | 1 | 0.996511 | 0.912212 |
| Atp5mc1 | Q9CR84 | 1.5 | 0.78 | 1.61 | 1.22 | -0.44 | 3 | 0.05 | 0.93 | 4.27 | -1.55 | 1.46 | 0.95 | 5.09 | -1.31 | 1.51 | 1.43 | -0.3925 | 0.005 | 0.4025 | 0.582636 | 0.996313 | 0.733876 | 1 | 0.999713 | 0.958635 |
| Atp5me | Q06185 | 0.53 | 1.73 | -0.61 | -0.72 | 1.26 | 0.98 | -0.1 | -0.34 | 1.29 | -2.39 | 0.09 | 0.32 | 0.39 | 1.18 | -1.56 | -0.51 | 0.2175 | -0.405 | -0.3575 | 0.731837 | 0.645538 | 0.632967 | 1 | 0.996511 | 0.941132 |
| Atp5mg | Q9CPQ8 | 2 | 0.6 | 0.29 | 1.26 | -0.04 | 1.34 | 0.97 | 1.63 | 1.55 | -0.39 | 1.19 | 1.82 | 0.42 | 0.85 | -0.29 | 0.16 | -0.0625 | 0.005 | -0.7525 | 0.897658 | 0.993032 | 0.105572 | 1 | 0.999285 | 0.703417 |
| Atp5mj | P56379 | 0.48 | 0.79 | 0.04 | 0 | 0.32 | 0.71 | 0.67 | 0.24 | -1.05 | -2.26 | 0.52 | 0.58 | -0.75 | -2.78 | 0.85 | 0.84 | 0.1575 | -0.88 | -0.7875 | 0.515788 | 0.197169 | 0.335074 | 1 | 0.996511 | 0.847354 |
| Atp5mk | Q78IK2 | 0.32 | 0.72 | 0.56 | 0.64 | 0.78 | 0.33 | 0.74 | 0.6 | -0.25 | 0.7 | 0.64 | 0.05 | -0.08 | 0.64 | 0.12 | 0.57 | 0.0525 | -0.275 | -0.2475 | 0.769939 | 0.313465 | 0.309413 | 1 | 0.996511 | 0.834523 |
| Atp5pb | Q9CQQ7 | 0.5 | 0.73 | 0.5 | 0.36 | 0.53 | 0.56 | 0.29 | 0.39 | 0.2 | 0.14 | 0.6 | 0.18 | 0.14 | 0.69 | -0.03 | 0.07 | -0.08 | -0.2425 | -0.305 | 0.621188 | 0.222353 | 0.199784 | 1 | 0.996511 | 0.78004 |
| Atp5pd | Q9DCX2 | 1.62 | 1.38 | 1.07 | 1.61 | 1.07 | 1.12 | 1.28 | 1.55 | 1.3 | 1.54 | 1.31 | 1.46 | 0.92 | 1.59 | 0.95 | 0.89 | -0.165 | -0.0175 | -0.3325 | 0.420843 | 0.928439 | 0.20127 | 1 | 0.996755 | 0.780919 |
| Atp5pf | P97450 | 1.41 | 1.07 | 0.96 | 1.52 | 0.61 | 0.96 | 0.82 | 1.57 | 1.38 | 0.99 | 1.46 | 1.56 | 1.09 | 1.34 | 0.57 | 0.59 | -0.25 | 0.1075 | -0.3425 | 0.342393 | 0.62752 | 0.211792 | 1 | 0.996511 | 0.786269 |
| Atp5po | Q9DB20 | 0.39 | 0.26 | 0.35 | 0.67 | 0.17 | 0.34 | 0.19 | 0.64 | 0.33 | 0.36 | 0.29 | 0.22 | 0.2 | 0.15 | 0.25 | 0.18 | -0.0825 | -0.1175 | -0.2225 | 0.653968 | 0.497131 | 0.238874 | 1 | 0.996511 | 0.793481 |
| Atp6ap1 | Q9R1Q9 | 0.41 | 0.69 | 0.86 | 0.45 | 1.08 | 0.45 | 0.86 | 0.92 | -0.05 | 0.43 | 0.68 | 0.45 | 0.11 | 0.41 | 0.72 | 0.9 | 0.225 | -0.225 | -0.0675 | 0.284029 | 0.326231 | 0.780328 | 1 | 0.996511 | 0.9676 |
| Atp6ap2 | Q9CYN9 | 0.2 | 0.9 | 1.14 | 0.64 | 0.29 | 0.53 | 0.74 | 0.88 | -0.61 | -1.07 | 0.97 | 0.31 | -0.29 | -0.5 | 0.84 | 1.05 | -0.11 | -0.82 | -0.445 | 0.662385 | 0.107689 | 0.305874 | 1 | 0.996511 | 0.832533 |
| Atp6v0a1 | K3W4T3 | 0.8 | 0.71 | 1.28 | 1.42 | 0.58 | 0.93 | 0.92 | 1.52 | 0.61 | 1.01 | 1.35 | 0.94 | 0.4 | 0.29 | 0.82 | 1.13 | -0.065 | -0.075 | -0.3925 | 0.808866 | 0.76776 | 0.186723 | 1 | 0.996511 | 0.776843 |
| Atp6v0a1 | Q9Z1G4 | 0.09 | 1.58 | 0.98 | 0.26 | 2.01 | 1.15 | 1.19 | 0.86 | 0.86 | 1.19 | 1.49 | 0.44 | 0.41 | 0.28 | 1.17 | 1.36 | 0.575 | 0.2675 | 0.0775 | 0.174753 | 0.502652 | 0.852803 | 1 | 0.996511 | 0.976225 |
| Atp6v0a2 | P15920 | -0.69 | -0.77 | -0.7 | -0.73 | -0.52 | -0.58 | -0.79 | -0.57 | -0.76 | -0.34 | -0.75 | -0.53 | 0.07 | -0.57 | -0.64 | -0.75 | 0.1075 | 0.1275 | 0.25 | 0.46754 | 0.471592 | 0.293716 | 1 | 0.996511 | 0.825612 |
| Atp6v0c | E9Q9C5 | -0.72 | 0.43 | 0.58 | 0.05 | 0.68 | 0.23 | 0.34 | -0.03 | 0.32 | 0.02 | 0.46 | -0.25 | 0.58 | -0.3 | 0.13 | 0.28 | 0.22 | 0.0525 | 0.0875 | 0.49616 | 0.873192 | 0.79887 | 1 | 0.996511 | 0.969753 |
| Atp6v0d1 | P51863 | 0.91 | 0.42 | 0.3 | 0.35 | 0.5 | 0.82 | 0.07 | 0.09 | 1.49 | 0.57 | 0.09 | 1.03 | -0.03 | 0.69 | -0.17 | 0.34 | -0.125 | 0.3 | -0.2875 | 0.609298 | 0.374894 | 0.295521 | 1 | 0.996511 | 0.825612 |
| Atp6v1a | P50516 | 1.72 | 1.38 | 1.34 | 0.93 | 1.23 | 1.65 | 1.41 | 1.32 | 1.33 | 0.76 | 1.22 | 1.84 | 0.64 | 0.93 | 1.01 | 1.22 | 0.06 | -0.055 | -0.3925 | 0.777747 | 0.846988 | 0.129159 | 1 | 0.996511 | 0.723655 |
| Atp6v1b2 | P62814 | 0.1 | 0.29 | 0.63 | 0.63 | 0.21 | 0.11 | 0.4 | 0.69 | 0.27 | 0.57 | 0.6 | 0.27 | 0.06 | 0.4 | 0.31 | 0.39 | -0.06 | 0.015 | -0.1225 | 0.776265 | 0.941807 | 0.56528 | 1 | 0.996755 | 0.920487 |
| Atp6v1c1 | Q9Z1G3 | 1.37 | 1.08 | 0.91 | 0.82 | 0.99 | 1.18 | 1.66 | 0.86 | 0.89 | 0.65 | 0.84 | 1.24 | 0.35 | 1.17 | 0.75 | 0.84 | 0.1275 | -0.14 | -0.2675 | 0.585395 | 0.517266 | 0.290216 | 1 | 0.996511 | 0.825134 |
| Atp6v1d | P57746 | 0.75 | 1.04 | 0.95 | 0.8 | 0.85 | 0.66 | 0.9 | 0.9 | 0.73 | 0.51 | 0.9 | 0.75 | 0.11 | 0.53 | 0.56 | 0.77 | -0.0575 | -0.1625 | -0.3925 | 0.712623 | 0.366976 | 0.090175 | 1 | 0.996511 | 0.69238 |
| Atp6v1e1 | P50518 | 3.21 | 2.46 | 2.19 | 1.43 | 2.93 | 3.03 | 2.29 | 2.33 | 0.52 | 0.52 | 2.22 | 3.26 | 0.46 | 0.3 | 1.89 | 2.29 | 0.3225 | -0.6925 | -1.0875 | 0.421265 | 0.335827 | 0.084449 | 1 | 0.996511 | 0.690427 |
| Atp6v1f | Q9D1K2 | 1.49 | 1.93 | 2.31 | 1.67 | 1.73 | 1.49 | 3.05 | 2.48 | 0.98 | 1.26 | 2.36 | 1.58 | 0.81 | 1.72 | 2.11 | 2.77 | 0.3375 | -0.305 | 0.0025 | 0.383903 | 0.38368 | 0.995309 | 1 | 0.996511 | 0.998317 |
| Atp6v1g1 | Q9CR51 | -0.5 | -0.23 | 0.08 | 0.07 | 0.15 | -0.51 | -0.22 | 0.06 | -0.1 | 0.66 | 0.39 | -0.14 | 0.62 | 0.75 | 0.14 | 0.27 | 0.015 | 0.3475 | 0.59 | 0.946791 | 0.202015 | 0.033453 | 1 | 0.996511 | 0.644622 |
| Atp6v1g2 | Q9WTT4 | 0.92 | 0.29 | 0.79 | 1.6 | -0.24 | 0.34 | 0.18 | 1.55 | 1.59 | 1.36 | 0.79 | 1.01 | 1.04 | -0.07 | 0.55 | 0.3 | -0.4425 | 0.2875 | -0.445 | 0.327325 | 0.384771 | 0.228875 | 1 | 0.996511 | 0.787447 |
| Atp6v1h | A0A0A6YW | -4.2 | -5.13 | -5.6 | -5.83 | -3.93 | -3.28 | -5.09 | -6.21 | -5.08 | -2.57 | -4.4 | -4.14 | -4.41 | -3.59 | -4.99 | -5.24 | 0.5625 | 1.1425 | 0.6325 | 0.412273 | 0.079798 | 0.211382 | 1 | 0.996511 | 0.786269 |
| Atp6v1h | Q8BVE3 | -0.33 | -0.06 | 0.39 | 0.56 | 0.03 | -0.12 | 0.49 | 0.64 | -0.09 | 0.22 | 0.53 | -0.25 | -0.37 | 0.16 | 0.1 | 0.2 | 0.12 | -0.0375 | -0.1175 | 0.666262 | 0.893268 | 0.66326 | 1 | 0.996511 | 0.949825 |
| Atp7a | A2AG68 | 0.38 | 0.47 | 0.78 | 0.25 | 0.1 | -0.59 | 1.37 | 0.27 | -1.08 | 1.01 | 0.62 | -0.15 | -0.79 | -0.49 | 0.63 | 0.64 | -0.1825 | -0.37 | -0.4725 | 0.648085 | 0.41736 | 0.234111 | 1 | 0.996511 | 0.790641 |
| Atp8a1 | P70704 | 1 | 0.81 | 0.55 | 0.55 | 0.82 | 1.07 | 0.64 | 0.65 | 0.46 | 0.25 | 0.52 | 0.96 | 0.24 | 0.31 | 0.48 | 0.51 | 0.0675 | -0.18 | -0.3425 | 0.720895 | 0.425612 | 0.1098 | 1 | 0.996511 | 0.709835 |
| Atp8a2 | P98200 | 0.64 | 0.84 | 0.86 | 0.6 | 1.21 | 0.99 | 1.03 | 0.65 | 0.05 | 0.31 | 0.69 | 0.83 | 0.19 | 0.12 | 0.84 | 1.13 | 0.235 | -0.265 | -0.165 | 0.213911 | 0.257877 | 0.556231 | 1 | 0.996511 | 0.917092 |
| Atp9a | A2AQC3 | -0.17 | 0.03 | -0.14 | -0.24 | 0.4 | -0.11 | -0.18 | -0.49 | 0.12 | -0.4 | -0.14 | 0 | 0.66 | 0.41 | 0.25 | 0.35 | 0.035 | 0.025 | 0.5475 | 0.872251 | 0.893081 | 0.01571 | 1 | 0.996511 | 0.589387 |
| Atp9b | A0A286YC | 0.49 | 0.77 | -0.09 | 0.08 | 0.55 | 0.41 | -0.36 | -0.08 | 0.86 | 0.5 | 0 | 0.61 | 0.59 | 0.9 | -0.18 | 0 | -0.1825 | 0.18 | 0.015 | 0.532058 | 0.523193 | 0.963023 | 1 | 0.996511 | 0.9927 |
| Atpaf1 | H3BLL2 | 0.01 | 0.14 | 0.37 | 0.49 | -0.13 | 0.17 | 0.02 | 0.45 | 0.78 | -0.7 | 0.22 | 0.11 | 0.69 | -0.03 | -0.74 | -0.33 | -0.125 | -0.15 | -0.355 | 0.533558 | 0.644649 | 0.291946 | 1 | 0.996511 | 0.825497 |
| Atpaf2 | Q91YY4 | -0.26 | 0.22 | 0 | -0.17 | 0.69 | 0.03 | 0.68 | 0.22 | -0.14 | -0.33 | 0.58 | 0 | -0.42 | 0.6 | -0.15 | 0.38 | 0.4575 | 0.08 | 0.155 | 0.063823 | 0.746815 | 0.581141 | 1 | 0.996511 | 0.927734 |
| Atpif1 | E9PV44 | 2.07 | 0.42 | 0.97 | 1.95 | -0.81 | 1.39 | -0.08 | 1.71 | 2.98 | 1.22 | 1.19 | 2.44 | 2.34 | -0.08 | 0.93 | 0.48 | -0.8 | 0.605 | -0.435 | 0.240569 | 0.290169 | 0.473414 | 1 | 0.996511 | 0.891467 |
| Atpsckmt | E0CXY7 | -0.21 | -0.14 | 0.18 | 0.47 | 0.46 | -0.15 | 0.46 | 0.31 | 0.15 | 0.78 | 0.71 | -0.11 | 0.29 | 1.69 | 0.38 | 0.36 | 0.195 | 0.3075 | 0.605 | 0.410011 | 0.288395 | 0.124921 | 1 | 0.996511 | 0.721448 |
| Atraid | G3UXG8 | -3.3 | -3.7 | -2.17 | -4.5 | -6.26 | -1.5 | -2.75 | -3.21 | -4.48 | -5.4 | -3.12 | -2.79 | -1.35 | -4.65 | -1.76 | -2.67 | -0.0125 | -0.53 | 0.81 | 0.990067 | 0.460213 | 0.322185 | 1 | 0.996511 | 0.840788 |
| Atrn | Q9WU60 | -0.62 | -0.42 | -0.48 | -0.2 | -0.62 | -0.26 | -0.11 | -0.57 | 0.24 | -0.33 | -0.31 | -0.32 | 0.52 | -0.09 | 0 | -0.12 | 0.04 | 0.25 | 0.5075 | 0.832919 | 0.250943 | 0.045564 | 1 | 0.996511 | 0.658454 |
| Atrnl1 | Q6A051 | -0.14 | -0.54 | -0.49 | -0.29 | -0.53 | 0.16 | -0.51 | -0.75 | 0.65 | -0.3 | -0.54 | -0.09 | 1.63 | -1.12 | -0.28 | -0.41 | -0.0425 | 0.295 | 0.32 | 0.856452 | 0.314339 | 0.564127 | 1 | 0.996511 | 0.920254 |
| Atrx | Q61687 | 0.97 | 0.71 | 0.85 | 0.71 | 0.44 | 0.58 | 0.37 | 0.33 | 0.77 | 0.59 | 0.72 | 0.87 | 0.74 | 0.68 | 0.96 | 1.06 | -0.38 | -0.0725 | 0.05 | 0.03348 | 0.666473 | 0.79122 | 1 | 0.996511 | 0.969105 |
| Atxn1 | J3QPR1 | -3.48 | -3.62 | -2.2 | -2.18 | -4.05 | -3.71 | -2.71 | -2.86 | -2.93 | -0.95 | -2.81 | -2.37 | -1.23 | -2.34 | -3.06 | -3.1 | -0.4625 | 0.605 | 0.4375 | 0.342048 | 0.292109 | 0.429804 | 1 | 0.996511 | 0.879165 |

| | | | | | | | | | | | | | | | | | | | | | | | | | | |
|---|---|---|---|---|---|---|---|---|---|---|---|---|---|---|---|---|---|---|---|---|---|---|---|---|---|---|
| Atxn10 | P28658 | -0.69 | -0.82 | -0.96 | -0.91 | -0.33 | -0.82 | -0.68 | -0.8 | -0.89 | -0.48 | -0.97 | -0.53 | -1.01 | -0.7 | -1.08 | -0.71 | 0.1875 | 0.1275 | -0.03 | 0.302983 | 0.513385 | 0.875416 | 1 | 0.996511 | 0.979626 |
| Atxn1l | P0C7T6 | -0.1 | -0.69 | -0.37 | -0.13 | -0.55 | -1.15 | -0.76 | -0.23 | -0.03 | 0.74 | -0.65 | -0.49 | 1.1 | 1.51 | -0.19 | -0.54 | -0.35 | 0.215 | 0.7925 | 0.183962 | 0.52763 | 0.126276 | 1 | 0.996511 | 0.721505 |
| Atxn2 | E9QM77 | -2.41 | -2.73 | -1.7 | -1.75 | -2.59 | -2.96 | -1.88 | -2.03 | -1.77 | -0.49 | -1.57 | -2.32 | -1.27 | -1.56 | -1.39 | -1.72 | -0.2175 | 0.61 | 0.6625 | 0.531086 | 0.184315 | 0.044338 | 1 | 0.996511 | 0.658454 |
| Atxn2 | F6U2C2 | -0.6 | -0.92 | -4.02 | -4.31 | -0.33 | -1.12 | -4.45 | -4.14 | -0.48 | -1.79 | -3.02 | -0.69 | -1.02 | -0.76 | -3.04 | -3.47 | -0.0475 | 0.9675 | 0.39 | 0.970482 | 0.360035 | 0.71769 | 1 | 0.996511 | 0.958398 |
| Atxn2l | Q7TQH0-2 | -1.9 | -2.02 | -2.01 | -2.3 | -2.09 | -2.06 | -2.07 | -2.39 | -1.61 | -1.23 | -2 | -1.77 | -1.05 | -1.56 | -2.06 | -2.42 | -0.095 | 0.405 | 0.285 | 0.578947 | 0.093498 | 0.380844 | 1 | 0.996511 | 0.860984 |
| Atxn3 | Q9CVD2 | -0.33 | -0.77 | -0.11 | 0.33 | -0.63 | -0.41 | -0.15 | 0.15 | 1.33 | 0.18 | -0.17 | -0.4 | 1.95 | -1.21 | 0.11 | -0.51 | -0.04 | 0.455 | 0.305 | 0.888554 | 0.299481 | 0.641679 | 1 | 0.996511 | 0.943421 |
| Atxn7l3b | Q3UD01 | -0.74 | -0.21 | -0.87 | -0.59 | 0.16 | -0.42 | -0.49 | -1.32 | 0.36 | -0.43 | -0.5 | -0.12 | -0.45 | 0.45 | -1.67 | -1.67 | 0.085 | 0.43 | -0.2325 | 0.795391 | 0.126916 | 0.643304 | 1 | 0.996511 | 0.943874 |
| Auh | A0A0R4J02 | 0.93 | 0.48 | 0.55 | 0.52 | 0.21 | 0.85 | 0.33 | 0.44 | 0.27 | 0.27 | 0.3 | 0.32 | 0.05 | 0.35 | -0.12 | 0.15 | -0.1625 | -0.33 | -0.5125 | 0.435434 | 0.088013 | 0.032071 | 1 | 0.996511 | 0.641974 |
| Aup1 | P70295 | 0.7 | 0.77 | -0.22 | 0.35 | 0.69 | 0.17 | 0.15 | 0.45 | 1.05 | 0.8 | 0.24 | 0.95 | 0.95 | 1.27 | 0.34 | 0.18 | -0.035 | 0.36 | 0.285 | 0.895546 | 0.244905 | 0.413626 | 1 | 0.996511 | 0.874945 |
| Aurkaip1 | Q9DCJ7 | -0.44 | -0.46 | -0.96 | -0.26 | -0.96 | -0.15 | 0.76 | -0.41 | 0.12 | 1.19 | 0.77 | -1.53 | 0.03 | 1.82 | 0.03 | -1.03 | 0.34 | 0.6675 | 0.7425 | 0.371068 | 0.25865 | 0.208998 | 1 | 0.996511 | 0.786218 |
| Aven | A2AGL5 | -1.6 | -0.24 | 1.47 | 0.9 | -0.09 | -0.9 | 0.52 | 1.11 | 0.15 | 1.61 | 1.47 | -1.58 | -0.61 | -2.32 | 1.13 | 1.48 | 0.0275 | 0.28 | -0.2125 | 0.969764 | 0.75743 | 0.830228 | 1 | 0.996511 | 0.973939 |
| Avl9 | Q80U56 | 1.09 | 0.87 | 0.42 | 0.17 | 0.6 | 1.16 | 0.69 | 0.31 | 0.09 | -0.55 | 0.62 | 1 | 0.13 | 0.12 | 0.14 | 0.46 | 0.0525 | -0.3475 | -0.425 | 0.849848 | 0.374426 | 0.123613 | 1 | 0.996511 | 0.721448 |
| AW209491 | Q91WD4 | -0.54 | -1.29 | -1.1 | -1.59 | -1.4 | -1.02 | -2.28 | -1.44 | 0.78 | -4.35 | -0.99 | -1 | 0.8 | -1.45 | -0.21 | -1.48 | -0.405 | -0.26 | 0.545 | 0.246835 | 0.791735 | 0.332484 | 1 | 0.996511 | 0.845601 |
| AW551984 | Q8BGF0 | 0.28 | 0.74 | -0.43 | 0.2 | 0.48 | 0.17 | 0.85 | 0.18 | 0.25 | 0.71 | 0.15 | 0.76 | 0.31 | 0.26 | 1.19 | 0.65 | 0.2225 | 0.27 | 0.405 | 0.450217 | 0.371095 | 0.235489 | 1 | 0.996511 | 0.791493 |
| Axl | Q6PE80 | 1.64 | 0.42 | 0.68 | 0.33 | 1.28 | 0.7 | 0.41 | 0.85 | -0.17 | -0.25 | 0.56 | 0.9 | 0.23 | 0.23 | -0.35 | 0.51 | 0.0425 | -0.5075 | -0.6125 | 0.900193 | 0.21956 | 0.106206 | 1 | 0.996511 | 0.703417 |
| Azi2 | Q9QYP6 | -0.22 | -0.44 | 0.57 | 0.26 | -1.51 | 0.08 | -0.7 | -0.23 | 1.96 | 1.02 | 0.13 | 0.02 | 2.74 | -0.7 | 0.6 | 0.48 | -0.6325 | 0.74 | 0.7375 | 0.133977 | 0.144882 | 0.296629 | 1 | 0.996511 | 0.825612 |
| B2m | P01887 | 2.36 | 0.84 | 0.88 | 0.06 | 0.56 | 1.51 | 1.36 | 0.77 | 1.04 | 0.52 | 0.84 | 1.5 | 1.49 | 0.45 | 1.36 | 2.01 | 0.015 | -0.06 | 0.2925 | 0.975618 | 0.901883 | 0.588626 | 1 | 0.996609 | 0.929131 |
| B3galnt1 | Q920V1 | -1.83 | -2.25 | -1.34 | -0.08 | -1.73 | -1.93 | -0.79 | -1.11 | -1.54 | -0.2 | -0.55 | -1.39 | -1.54 | -1.54 | -0.81 | 0.12 | -0.015 | 0.455 | 0.4325 | 0.975951 | 0.399585 | 0.451514 | 1 | 0.996511 | 0.885538 |
| B3galt5 | Q9JI67 | -0.55 | -0.31 | -1.66 | -1.18 | -2.52 | -0.08 | -0.72 | -1.21 | 1.16 | 0.6 | -1.51 | -1.16 | 1.93 | -0.95 | 0.86 | -0.98 | -0.2075 | 0.6975 | 1.14 | 0.707289 | 0.305236 | 0.133055 | 1 | 0.996511 | 0.728417 |
| B3galt6 | Q91Z92 | -1.67 | -2.04 | -2.36 | -1.45 | -0.97 | -1.14 | -1.71 | -1.91 | -1.17 | -2.23 | -2.33 | -1.94 | -2.16 | -1.33 | -1.8 | -1.49 | 0.4475 | -0.0375 | 0.185 | 0.160709 | 0.909129 | 0.526961 | 1 | 0.996744 | 0.90845 |
| B3gat3 | P58158 | -1.3 | -3.12 | -1.33 | -0.1 | -2.31 | -0.49 | -1.34 | -0.94 | -1.37 | -1.33 | -1.13 | -2.62 | -0.3 | -2.81 | -0.36 | -0.33 | 0.1925 | -0.15 | 0.5125 | 0.772722 | 0.816822 | 0.522652 | 1 | 0.996511 | 0.907667 |
| B3glct | Q8BHT6 | -0.41 | 0.15 | -0.71 | -0.72 | 0.56 | -0.27 | 0.2 | -0.2 | -0.24 | 0.19 | -0.56 | -0.21 | -0.78 | 0.63 | 0.32 | 0.18 | 0.495 | 0.2175 | 0.51 | 0.107885 | 0.430531 | 0.181838 | 1 | 0.996511 | 0.776002 |
| B4galnt1 | Q09200 | -0.86 | -0.76 | -0.7 | -0.45 | -0.49 | -0.73 | 0.11 | -0.16 | -0.71 | 0.25 | -0.59 | -1.13 | -0.65 | -0.23 | 0.26 | 0.21 | 0.375 | 0.1475 | 0.59 | 0.123318 | 0.632159 | 0.046048 | 1 | 0.996511 | 0.658454 |
| B4galnt4 | Q766D5 | -3.81 | -2.29 | -3.04 | -2.68 | -4.61 | -3.82 | -6.99 | -3.44 | -6.84 | -1.79 | -4.3 | -2.08 | 1.55 | -9.06 | -2.41 | -0.87 | -1.76 | -0.7975 | 0.2575 | 0.046949 | 0.471569 | 0.899122 | 1 | 0.996511 | 0.983509 |
| B4galt3 | Q91YY2 | -0.26 | 0.04 | -0.76 | -0.74 | -0.75 | 0.07 | -0.39 | -0.65 | -0.21 | -1.62 | -1.45 | -0.12 | 0.15 | -0.24 | -0.8 | -0.28 | 7.85E-17 | -0.42 | 0.1375 | 1 | 0.331728 | 0.639287 | 1 | 0.996511 | 0.942804 |
| B4galt7 | D3Z065 | -0.75 | -1.09 | -0.1 | -0.37 | -2.54 | 0.24 | -1.61 | 0.32 | 1.31 | 0.09 | 0.81 | -0.66 | 2.36 | -1.58 | -0.86 | -1.61 | -0.32 | 0.965 | 0.155 | 0.635074 | 0.058358 | 0.858533 | 1 | 0.996511 | 0.976225 |
| B4gat1 | Q8BWP8 | -2.3 | -2.34 | -1.39 | -1.57 | -1.13 | -1.63 | -1.33 | -1.36 | -2.01 | -0.58 | -1.28 | -2.39 | -1.6 | -1.57 | -1.08 | -1.27 | 0.5375 | 0.335 | 0.52 | 0.074702 | 0.457247 | 0.10173 | 1 | 0.996511 | 0.698697 |
| B630019K0 | Q7TNS5 | -0.05 | 0.86 | -0.96 | -0.56 | 0.64 | -0.14 | -0.91 | -0.87 | 0.35 | -1.05 | -0.35 | 0 | -0.13 | 0.29 | -0.36 | -0.82 | -0.1425 | -0.085 | -0.0775 | 0.773973 | 0.854249 | 0.858069 | 1 | 0.996511 | 0.976225 |
| B9d1 | Q5NCM9 | 0.4 | -0.28 | 1.21 | 1.22 | -0.23 | 0.27 | 1.28 | 1.48 | 0.64 | 1.99 | 1.09 | 0.9 | 0.86 | -2.05 | 0.8 | 1.52 | 0.0625 | 0.5175 | -0.355 | 0.900915 | 0.25778 | 0.653676 | 1 | 0.996511 | 0.947633 |
| Baalc | Q8VHV1 | 1 | 1.16 | 1.65 | 1.24 | 1.65 | 0.78 | 1.02 | 1.53 | -0.8 | 0.65 | 1.39 | 0.98 | -0.78 | -1.19 | 1.63 | 1.86 | -0.0175 | -0.7075 | -0.8825 | 0.945896 | 0.154276 | 0.246269 | 1 | 0.996511 | 0.802442 |
| Babam1 | Q3UI43 | -0.19 | 0.11 | -0.33 | -0.62 | 0.43 | -0.27 | -0.91 | -0.73 | -0.02 | 0.43 | -0.55 | 0.01 | 0.3 | 0.69 | -0.42 | -0.98 | -1.13E-01 | 0.225 | 0.155 | 0.731032 | 0.410927 | 0.691767 | 1 | 0.996511 | 0.955729 |
| Babam2 | Q8K3W0 | -0.39 | -0.37 | 0.11 | -0.08 | -0.4 | -0.06 | -0.25 | 0.06 | -0.73 | 0.63 | 0.03 | -0.3 | -0.44 | -0.48 | 0.1 | -0.2 | 0.02 | 0.09 | -0.0725 | 0.917827 | 0.774159 | 0.749504 | 1 | 0.996511 | 0.961298 |
| Bace1 | P56818 | 0.18 | 0.52 | 0.6 | 0.16 | -0.32 | 0.44 | -0.46 | -0.03 | 1.6 | 0.79 | 0.23 | 0.59 | 1.58 | 0.1 | 0.8 | 0.12 | -0.4575 | 0.4375 | 0.285 | 0.087714 | 0.187177 | 0.439978 | 1 | 0.996511 | 0.883445 |
| Bad | Q61337 | -0.7 | -0.53 | -1.13 | -1.13 | -0.68 | -1.17 | -2.32 | -1.72 | -0.13 | -1.13 | -1.02 | -0.73 | 0.35 | -0.65 | -0.99 | -1.09 | -0.6 | 0.12 | 0.2775 | 0.129791 | 0.672382 | 0.446234 | 1 | 0.996511 | 0.884817 |
| Bad | Q3U9H3 | 0.34 | 0.73 | 0.41 | -0.05 | 1.37 | 0.1 | -0.7 | 0.17 | 0.86 | -0.05 | 0.34 | 0.98 | 0.63 | -0.05 | -0.52 | 0.86 | -0.1225 | 0.175 | -0.1275 | 0.774336 | 0.556917 | 0.717586 | 1 | 0.996511 | 0.958398 |
| Bag1 | F6TCF9 | -0.62 | -0.25 | 0.7 | 0.62 | -0.28 | -0.98 | 0.44 | 0.7 | -0.98 | 0.64 | 0.36 | -0.41 | -0.66 | -0.1 | 0.87 | 0.73 | -0.1425 | -0.21 | 0.0975 | 0.758897 | 0.650613 | 0.831422 | 1 | 0.996511 | 0.973982 |
| Bag2 | Q91YN9 | -0.06 | 0.04 | 0.14 | -0.16 | 0.11 | 0.3 | -1 | -0.53 | 0.7 | -1.06 | -0.12 | -0.06 | 1.03 | -0.66 | 0.12 | -0.58 | -0.27 | -0.125 | -0.0125 | 0.383404 | 0.727151 | 0.974131 | 1 | 0.996511 | 0.992977 |
| Bag3 | Q9JLV1 | -0.42 | -0.52 | -0.43 | -0.27 | -1.33 | -0.26 | -0.8 | -0.34 | 0.94 | -0.12 | -0.3 | -0.24 | 1.5 | -0.54 | -0.25 | -0.34 | -0.2725 | 0.48 | 0.5025 | 0.311233 | 0.13911 | 0.281473 | 1 | 0.996511 | 0.822084 |
| Bag4 | Q8CI61 | 1 | 1.12 | 2.83 | -2.12 | 1.37 | 1.05 | 2.41 | 0.95 | -1.14 | -1.97 | 1.66 | 1.57 | -0.68 | -1.08 | 2.74 | 2.49 | 0.7375 | -0.6775 | 0.16 | 0.456088 | 0.588886 | 0.900958 | 1 | 0.996511 | 0.983757 |
| Bag5 | Q8CI32 | -0.28 | -0.25 | 0.45 | 0.64 | -0.33 | -0.33 | 0.93 | 0.82 | -0.08 | 1.03 | 0.56 | -0.17 | 0.16 | -0.46 | 0.94 | 0.85 | 0.1325 | 0.195 | 0.2325 | 0.739706 | 0.59082 | 0.556634 | 1 | 0.996511 | 0.917092 |
| Bag6 | Q3UF95 | 0.01 | 0.16 | -0.01 | -0.08 | 0.25 | -0.27 | -0.31 | -0.28 | -0.55 | -0.17 | -0.34 | 0.12 | -0.5 | -0.54 | -0.16 | -0.26 | -0.1725 | -0.255 | -0.385 | 0.364075 | 0.223722 | 0.064956 | 1 | 0.996511 | 0.66433 |
| Baiap2 | Q8BKX1-2 | 0.84 | 1 | 1.38 | 1.04 | 1.13 | 0.7 | 1.55 | 1.36 | 0.47 | 0.49 | 1.44 | 1.2 | 0.59 | 0.58 | 1.45 | 1.29 | 0.12 | -0.165 | -0.0875 | 0.609903 | 0.564085 | 0.752765 | 1 | 0.996511 | 0.961479 |
| Baiap3 | S4R1K2 | 0.12 | 0.71 | -1.73 | 0.04 | -0.41 | 0.29 | 0.43 | 0.24 | 1.18 | 1.96 | -1.1 | 0.13 | 0.58 | -0.36 | 2.3 | 0.75 | 0.3525 | 0.7575 | 1.0325 | 0.499148 | 0.33745 | 0.161214 | 1 | 0.996511 | 0.757375 |
| Bak1 | O08734 | -0.8 | -0.64 | -0.97 | -1.24 | 0.06 | -0.42 | -0.52 | -0.53 | -0.31 | -1.13 | -0.45 | -0.93 | 0.5 | -0.34 | -0.79 | -0.64 | 0.56 | 0.2075 | 0.595 | 0.027239 | 0.423585 | 0.091438 | 1 | 0.996511 | 0.693316 |
| Banf1 | O54962 | 0.98 | 1.73 | 1.72 | 1.41 | 1.21 | 0.67 | 2.25 | 1.41 | 0.7 | 1.26 | 1.44 | 0.9 | 1.27 | 1.53 | 1.78 | 1.76 | -0.075 | -0.385 | 0.125 | 0.833613 | 0.167224 | 0.616392 | 1 | 0.996511 | 0.936924 |
| Basp1 | Q91XV3 | 1.2 | 1.09 | 1.55 | 1.79 | 0.41 | 1.5 | 0.23 | 1.44 | 2.34 | 2.09 | 1.4 | 1.31 | 2.81 | 1.31 | 1.72 | 0.93 | -0.5125 | 0.3775 | 0.285 | 0.174762 | 0.236869 | 0.500787 | 1 | 0.996511 | 0.900975 |
| Bax | A0A1B0GT | 0.45 | 0.02 | 0 | 0.25 | 0 | 0.12 | 0.39 | -0.32 | -0.43 | 0.04 | 0.09 | 0.46 | -0.17 | 0.18 | -0.54 | 0.34 | -0.1325 | -0.14 | -0.2275 | 0.532304 | 0.562146 | 0.382237 | 1 | 0.996511 | 0.860984 |
| Baz1b | Q9Z277 | 1.26 | 0.91 | 1.06 | 0.96 | 0.9 | 0.97 | 0.98 | 0.92 | 0.66 | 0.02 | 1.23 | 0.98 | 0.72 | 0.77 | 1.35 | 1.49 | -0.105 | -0.325 | 0.035 | 0.496027 | 0.270469 | 0.887287 | 1 | 0.996511 | 0.980329 |
| Baz2a | F8VPM0 | 1.13 | 1.03 | 1.67 | 2.37 | -0.33 | 1.39 | -0.7 | 1.06 | 2.18 | 1.65 | 1.42 | 0.57 | 2.44 | 0.38 | 1.58 | 0.95 | -1.195 | -0.095 | -0.2125 | 0.053998 | 0.825302 | 0.673203 | 1 | 0.996511 | 0.952944 |
| Bbln | P58686 | -0.96 | 0.23 | 0.29 | 0.66 | 0.07 | -1.09 | 0.53 | 0.24 | -0.51 | 0.31 | 0.27 | -0.15 | -0.67 | 0.54 | -0.01 | 0.27 | -0.1175 | -0.075 | -0.0225 | 0.800267 | 0.845907 | 0.957021 | 1 | 0.996511 | 0.992636 |
| Bbs1 | Q3V3N7 | -0.05 | -0.37 | -0.34 | 0.12 | -0.11 | -0.58 | 0.04 | -0.28 | -0.54 | -0.33 | -0.27 | -0.19 | -1.19 | -0.32 | -0.31 | -0.32 | -0.0725 | -0.1725 | -0.375 | 0.727119 | 0.384613 | 0.191488 | 1 | 0.996511 | 0.777117 |
| Bbs2 | Q9CWF6 | 0.09 | -0.44 | -0.85 | -0.77 | -0.56 | -0.77 | -0.2 | -0.64 | -0.2 | -0.5 | -0.87 | -0.28 | -0.45 | -0.22 | -0.94 | -0.33 | -0.03 | 0.09 | -0.0075 | 0.845186 | 0.913005 | 0.978963 | 1 | 0.996755 | 0.994274 |
| Bbs4 | Q8C1Z7 | 0.53 | 0.4 | -0.18 | -0.75 | 0.34 | 0.24 | 0.91 | -0.01 | 0.03 | -0.76 | -0.38 | 0.72 | 0.44 | 0.75 | 0.53 | 0.32 | 0.37 | -0.0975 | 0.51 | 0.293994 | 0.812707 | 0.1327 | 1 | 0.996511 | 0.728417 |
| Bbs5 | A2AUC6 | -1.16 | -0.19 | -0.01 | 0.06 | -0.47 | -0.56 | 0.25 | -0.12 | -2.37 | 1.33 | 0.65 | -2.34 | -2.2 | -2.34 | -0.29 | -0.1 | 0.1 | -0.3575 | -0.9075 | 0.761477 | 0.697011 | 0.161206 | 1 | 0.996511 | 0.757375 |
| Bbs7 | E9Q0R0 | -0.54 | -0.72 | 0.03 | -0.7 | -0.67 | -1.02 | 0.09 | -0.42 | -0.58 | 0.04 | -0.24 | -0.71 | -0.51 | -0.26 | -0.03 | 0.2 | -0.0225 | 0.11 | 0.3325 | 0.938489 | 0.676446 | 0.223993 | 1 | 0.996511 | 0.787447 |
| Bbs9 | Q811G0-2 | 0.58 | 1 | 0.16 | -0.07 | 1.3 | 0.32 | 0.45 | 0.55 | 0.5 | -1.2 | 0.13 | 0.6 | -0.41 | -0.13 | -0.12 | 0.56 | 0.2375 | -0.41 | -0.4425 | 0.460513 | 0.373324 | 0.190464 | 1 | 0.996511 | 0.776843 |
| BC004004 | A0A3Q4L2 | -0.79 | 0.64 | 0.42 | -0.87 | 0.87 | 0.18 | 0.62 | 0.6 | 1.47 | 0.66 | -0.21 | 0.54 | 1.52 | 1.86 | -0.14 | 0.99 | 0.7175 | 0.765 | 1.2075 | 0.098591 | 0.144148 | 0.050737 | 1 | 0.996511 | 0.658454 |
| BC024978 | D3Z7G4 | -0.01 | -0.05 | 0.43 | 0.72 | -0.35 | -0.7 | 0.71 | 1.06 | -2.09 | 0.49 | 0.72 | -0.12 | -0.85 | -2.1 | 0.68 | 0.53 | -0.0925 | -0.5225 | -0.7075 | 0.829217 | 0.401043 | 0.272248 | 1 | 0.996511 | 0.822084 |
| Bcam | Q9R069 | -0.76 | -1.1 | -0.73 | -0.66 | -0.95 | -0.88 | -0.37 | -0.43 | -1.12 | -0.42 | -0.69 | -0.87 | -0.63 | -0.75 | -0.27 | -0.46 | 0.155 | 0.0375 | 0.285 | 0.463615 | 0.862441 | 0.189003 | 1 | 0.996511 | 0.776843 |
| Bcan | Q61361 | -0.72 | -1.35 | -1.09 | -1.06 | -1.13 | -0.99 | -1.02 | -1.09 | -1.64 | -0.73 | -1.11 | -1.1 | -1.24 | -1.18 | -1.03 | -1.49 | -0.0025 | -0.09 | -0.18 | 0.988846 | 0.720248 | 0.412604 | 1 | 0.996511 | 0.874811 |
| Bcap29 | Q61334 | 1.33 | 1.04 | 0.66 | 0.54 | 1.69 | 1.02 | 0.51 | 0.71 | 0.58 | 1.79 | 0.36 | 1.08 | 1.01 | -1.48 | 1.18 | 0.73 | 0.09 | 0.06 | -0.5325 | 0.772054 | 0.866258 | 0.380223 | 1 | 0.996511 | 0.860984 |
| Bcap31 | Q61335 | 0.33 | 0.57 | 0.52 | 0.64 | 0.49 | 0.17 | 0.43 | 0.63 | 0.34 | 0.78 | 0.3 | 0.25 | 1.06 | 0.39 | 1.37 | 0.89 | -0.135 | -0.1375 | 0.3625 | 0.4508 | 0.465304 | 0.177168 | 1 | 0.996511 | 0.772068 |
| Bcar1 | Q61140 | 0.4 | 0.48 | 0.4 | 0.68 | 0.5 | 0.09 | 0.59 | 0.46 | 0.67 | 0.76 | 0.42 | 0.23 | 1.01 | 0.65 | 0.33 | 0.37 | -0.08 | 0.03 | 0.1 | 0.651713 | 0.87613 | 0.652435 | 1 | 0.996511 | 0.947633 |
| Bcar3 | Q9QZK2 | 0.61 | 0.32 | 0.32 | 0.21 | 0.25 | -0.06 | 0.1 | 0.23 | 0.7 | 0.24 | 0.41 | 0.09 | 0.55 | 0.92 | 0.21 | 0.5 | -0.235 | -0.005 | 0.18 | 0.183909 | 0.98038 | 0.422454 | 1 | 0.998773 | 0.875782 |
| Bcas1 | Q80YN3 | 3.42 | 2.38 | 1.23 | 0.94 | 2.31 | 2.89 | 0.39 | 0.82 | 1.39 | 0.71 | 1.07 | 3.32 | 1.78 | 0.18 | 1.89 | 1.29 | -0.39 | -0.37 | -0.7075 | 0.602648 | 0.618659 | 0.278963 | 1 | 0.996511 | 0.822084 |
| Bcas2 | Q9D287 | 0.72 | -0.24 | -0.44 | -0.87 | -0.16 | -0.07 | -0.73 | -0.3 | -0.48 | 0.08 | -0.8 | -0.04 | -0.55 | 0.23 | -0.36 | -0.62 | -0.1075 | -0.1025 | -0.1175 | 0.760476 | 0.786888 | 0.756262 | 1 | 0.996511 | 0.962333 |
| Bcas3 | B1AR74 | 0.3 | 0.32 | 0.38 | 0.33 | 0.25 | 0.27 | 0.2 | 0.33 | -0.01 | -0.02 | 0.35 | 0.45 | -0.12 | -0.11 | 0.1 | -0.05 | -0.07 | -0.14 | -0.3775 | 0.613592 | 0.455842 | 0.049012 | 1 | 0.996511 | 0.658454 |
| Bcas3 | Q8CCN5 | 0.29 | 0.74 | 0.22 | 0.49 | 0.18 | -0.2 | -0.21 | -0.31 | 1.64 | 0.58 | -0.08 | 0.43 | 1.23 | 1.55 | 0.04 | -0.63 | -0.57 | 0.2075 | 0.1125 | 0.016178 | 0.576498 | 0.818014 | 1 | 0.996511 | 0.972768 |
| Bcat1 | Q8CBC8 | -0.86 | -0.57 | 0.3 | 0.45 | -0.87 | -0.7 | 0.36 | 0.3 | -0.75 | 0.13 | 0.15 | -0.62 | -0.74 | -0.07 | 0.24 | 0.38 | -0.0575 | -0.1025 | 0.1225 | 0.89288 | 0.790698 | 0.75607 | 1 | 0.996511 | 0.962333 |
| Bcat2 | O35855 | 0.24 | 0.26 | -0.35 | -0.25 | -0.01 | -0.05 | 0.3 | -0.17 | -0.4 | -0.61 | 0.15 | 0.04 | -0.89 | 0.05 | -0.39 | -0.54 | 0.0425 | -0.18 | -0.4175 | 0.843117 | 0.494946 | 0.154386 | 1 | 0.996511 | 0.750763 |
| Bccip | Q9CWI3 | -4.39 | -5.22 | -2.1 | -3.06 | -5.51 | -3.99 | -3.66 | -2.82 | -3.34 | -1.32 | -2.36 | -4.6 | -2.17 | -4.5 | -2.25 | -3.37 | -0.3025 | 0.7875 | 0.62 | 0.707739 | 0.38586 | 0.444519 | 1 | 0.996511 | 0.884411 |
| Bckdha | Q3U3J1 | 0.51 | 0.32 | 0.76 | 0.81 | -0.01 | 0.38 | 1.06 | 0.62 | 0.82 | 0.6 | 1.02 | 0.27 | 0.94 | 0.71 | 0.68 | 0.94 | -0.0875 | 0.0775 | 0.2175 | 0.737242 | 0.736225 | 0.294963 | 1 | 0.996511 | 0.825612 |
| Bckdhb | Q6P3A8 | -0.52 | -0.62 | -1.66 | -1.36 | -0.67 | -0.42 | -1.45 | -2.01 | -0.18 | -0.97 | -1.77 | -0.73 | -0.1 | -0.34 | -2.05 | -1.33 | -0.0975 | 0.1275 | 0.085 | 0.820371 | 0.757124 | 0.863778 | 1 | 0.996511 | 0.977235 |
| Bckdk | A0A0U1RN | 0.19 | 0.55 | -0.15 | -0.16 | 0.26 | 0.15 | -0.49 | -0.17 | 0.89 | 0.23 | 0.46 | 0.11 | 0.98 | 1.17 | -0.15 | -0.08 | -0.17 | 0.315 | 0.3725 | 0.503337 | 0.245775 | 0.335448 | 1 | 0.996511 | 0.847442 |
| Bcl10 | Q9Z0H7 | -0.4 | 0.2 | -0.84 | -1.47 | 0.5 | -0.61 | -0.96 | -1.03 | -0.46 | 0.08 | -0.61 | -0.69 | -0.85 | -0.24 | -1.63 | -1.03 | 0.1025 | 0.2075 | -0.31 | 0.825123 | 0.587207 | 0.479489 | 1 | 0.996511 | 0.893468 |
| Bcl11a | Q5STS9 | -0.63 | -0.7 | -0.57 | -0.75 | -0.38 | 0.01 | -0.33 | -0.57 | -0.65 | 0.16 | -0.53 | -0.43 | 0.24 | -0.21 | -0.09 | -0.14 | 0.345 | 0.3 | 0.6125 | 0.076305 | 0.197924 | 0.009377 | 1 | 0.996511 | 0.563386 |
| Bcl11b | A0A0R4J1E | -0.15 | -0.89 | -0.33 | -0.42 | -1.24 | 0.17 | -0.6 | -0.85 | 1.81 | -1.47 | -0.95 | -0.55 | 2.48 | -1.15 | 0.61 | -0.28 | -0.1825 | 0.1575 | 0.8625 | 0.580898 | 0.815765 | 0.24922 | 1 | 0.996511 | 0.804842 |
| Bcl2 | P10417-2 | -1.43 | -0.5 | -1 | -0.84 | -0.21 | -0.91 | -0.44 | 0.04 | -1.07 | -1.99 | -0.45 | -0.97 | -1.37 | -0.6 | -2.1 | -0.3 | 0.5625 | -0.1775 | -0.15 | 0.072926 | 0.627652 | 0.726143 | 1 | 0.996511 | 0.958398 |
| Bcl2l1 | A2AHX9 | -0.28 | 0.07 | 0.01 | -0.07 | 0.5 | -0.22 | 0.13 | -0.09 | -0.05 | 0.44 | 0.13 | 0.05 | -0.27 | 0.6 | -0.34 | -0.06 | 0.1475 | 0.21 | 0.05 | 0.479288 | 0.28326 | 0.8461 | 1 | 0.996511 | 0.976225 |
| Bcl2l13 | P59017 | 1.36 | 1.01 | -0.08 | -0.25 | 0.68 | 1.43 | -0.23 | -0.24 | 1.05 | -1.5 | -0.03 | 1.27 | 1.2 | 0.56 | -0.95 | -0.4 | -0.1 | -0.3125 | -0.4075 | 0.847678 | 0.647982 | 0.484068 | 1 | 0.996511 | 0.894252 |
| Bcl7a | Q9CXE2 | -0.66 | -1.39 | -1.03 | -1.45 | -1.59 | -1.27 | -1.18 | -1.57 | -1.47 | -0.56 | -0.93 | -1.14 | -1 | -0.83 | -2.01 | -1.45 | -0.27 | 0.1075 | -0.19 | 0.259308 | 0.699287 | 0.563374 | 1 | 0.996511 | 0.920024 |
| Bcl7b | Q921K9 | -0.92 | -1.27 | -1.43 | -1.36 | -1.06 | -1.16 | -1.11 | -1.68 | -1 | -1.11 | -1.31 | -1.03 | -1.22 | -0.77 | -1.49 | -1.65 | -0.0075 | 0.1375 | 0.305 | 0.971586 | 0.491877 | 0.882918 | 1 | 0.996511 | 0.979767 |
| Bclaf1 | Q8K019-2 | 1.27 | -0.09 | -0.12 | -0.24 | -1.05 | 0.75 | -0.62 | 0.58 | 0.52 | 0.11 | -0.22 | -0.55 | 2.05 | 0.01 | 1 | -0.47 | -0.29 | -0.24 | 0.4425 | 0.582532 | 0.55678 | 0.472754 | 1 | 0.996511 | 0.891144 |
| Bcr | Q6PAJ1 | 0.1 | -0.11 | 0.13 | 0.32 | 0.08 | -0.22 | 0.36 | 0.13 | 0.22 | -0.05 | 0.1 | 0.02 | -0.05 | 0.17 | -0.07 | 0.04 | -0.0225 | -0.0375 | -0.0875 | 0.904712 | 0.831347 | 0.64012 | 1 | 0.996511 | 0.942804 |
| Bcs1l | Q9CZP5 | -0.25 | 0.22 | 0.24 | 0.26 | -0.23 | -0.19 | 0.43 | 0.25 | -0.61 | 0.2 | 0.29 | -0.39 | -0.86 | -0.7 | -0.27 | 0.14 | -0.0525 | -0.245 | -0.54 | 0.815984 | 0.373331 | 0.078719 | 1 | 0.996511 | 0.680689 |
| Bdh1 | Q80XN0 | 0.24 | -0.09 | -0.14 | 0.08 | 0 | 0.44 | 0.28 | 0.15 | -0.12 | -0.58 | 0.25 | 0.36 | -0.19 | -0.21 | -0.54 | -0.22 | 0.195 | -0.045 | -0.3125 | 0.284563 | 0.858145 | 0.133263 | 1 | 0.996511 | 0.728417 |
| Bdnf | H9H9S8 | 0.75 | 1.14 | 1.01 | 0.96 | -0.83 | 1.24 | -1 | -0.01 | 1.13 | -0.51 | 0.2 | 0.58 | 1.78 | -2.48 | 0.51 | 0.81 | -1.115 | -0.615 | -0.81 | 0.042684 | 0.104464 | 0.342849 | 1 | 0.996511 | 0.848473 |
| Becn1 | O88597 | 0.11 | 0.32 | 0.55 | 0.06 | 0.14 | 0.29 | 0.68 | 0.27 | 0.33 | 0.61 | 0.35 | -0.07 | 0.45 | -0.15 | -0.05 | 0.63 | 0.085 | 0.045 | -0.04 | 0.666806 | 0.836443 | 0.874105 | 1 | 0.996511 | 0.979626 |
| Begain | F8WIG2 | 1.45 | 1.31 | 1.68 | 1.47 | 0.65 | 1.44 | 0.61 | 1.06 | 1.36 | 1.42 | 1.54 | 1.46 | 1.36 | 0.45 | 1.22 | 0.76 | -0.5375 | -0.0325 | -0.53 | 0.041327 | 0.845916 | 0.062902 | 1 | 0.996511 | 0.66433 |
| Bet1 | O35623 | -0.58 | -0.11 | -1.02 | -0.94 | 0.27 | 0.01 | -0.23 | -0.59 | 0.49 | -1.6 | -0.29 | 0.23 | 0.68 | 1.12 | -0.92 | -0.48 | 0.5275 | 0.37 | 0.7625 | 0.086906 | 0.445113 | 0.144675 | 1 | 0.996511 | 0.743156 |
| Bex2 | Q9WTZ8 | -0.73 | 0.31 | -2.78 | -3.96 | -0.81 | -0.37 | -3.99 | -2.91 | -0.36 | -1.57 | -2.91 | -0.2 | -1.3 | -0.27 | -2.79 | -4.05 | -0.23 | 0.53 | -0.3125 | 0.842586 | 0.611488 | 0.783626 | 1 | 0.996511 | 0.967642 |
| Bfar | A0A0R4J04 | -0.38 | 0.13 | 0.1 | -0.25 | -0.4 | 0.25 | -0.21 | -0.98 | 0.02 | -0.12 | 0.02 | -0.16 | -0.6 | 0.42 | 0.05 | 0.94 | -0.4 | -0.2125 | 0.275 | 0.166722 | 0.369256 | 0.447547 | 1 | 0.996511 | 0.884817 |
| Bgn | P28653 | -1.48 | -1.47 | 0.74 | 0.37 | -0.56 | -1.72 | 0.05 | 0.1 | -1.67 | -1.08 | 0.49 | -0.94 | -1.62 | -1.07 | 0.06 | 1.1 | -0.0725 | -0.34 | 0.0775 | 0.912452 | 0.619797 | 0.91913 | 1 | 0.996511 | 0.986868 |
| Bhlhe23 | Q8BGW3 | 1.32 | 2.16 | 1.28 | 0.96 | 1.74 | 1.47 | 0.36 | 1.2 | -0.81 | -0.12 | 1.38 | 1.73 | 0.68 | 1.21 | 1.9 | 0.97 | -0.2375 | -0.885 | -0.24 | 0.530691 | 0.166801 | 0.510659 | 1 | 0.996511 | 0.90359 |
| Bicd1 | G3UXK5 | 0.51 | 0.53 | 0.88 | 1.31 | 0.79 | 0.31 | 1.09 | 0.56 | 0.53 | 1.19 | 0.49 | 0.43 | 0.36 | 1.31 | 1.26 | 0.55 | -0.12 | -0.1475 | 0.0625 | 0.645917 | 0.59226 | 0.842791 | 1 | 0.996511 | 0.976225 |
| Bicd2 | Q921C5 | 0.61 | 0.83 | 0.63 | 0.52 | 1.66 | 0.44 | 0.75 | 0.64 | 1.27 | 0.77 | 0.87 | 0.81 | 0.95 | 1.09 | 0.58 | 0.83 | 0.225 | 0.2825 | 0.215 | 0.431761 | 0.162194 | 0.290926 | 1 | 0.996511 | 0.825497 |
| Bicra | F8VPZ9 | -0.05 | 0.25 | 0.38 | -0.15 | -1.55 | -0.79 | -1.49 | 0.11 | 1.06 | 1.89 | 0.67 | -0.48 | 1.85 | 1.7 | 0.77 | -2.19 | -1.0375 | 0.6775 | 0.425 | 0.023688 | 0.178502 | 0.619886 | 1 | 0.996511 | 0.93807 |
| Bicral | Q8CHH5 | 0.1 | 0.05 | 0.26 | -0.5 | -0.76 | -0.18 | 0.51 | 0.59 | -0.48 | 0.29 | -0.77 | -0.1 | -0.58 | -0.04 | -0.21 | 0.63 | 0.0625 | -0.2425 | -0.0275 | 0.85634 | 0.41565 | 0.9296 | 1 | 0.996511 | 0.989687 |
| Bid | P70444 | -0.06 | 0.15 | -0.74 | -0.39 | 0.69 | -0.19 | -0.26 | -0.56 | -0.64 | -0.12 | -0.3 | 0.15 | -0.2 | 1.4 | -0.26 | -0.43 | 0.18 | 0.0325 | 0.3875 | 0.581745 | 0.904485 | 0.393845 | 1 | 0.996744 | 0.865775 |
| Bin1 | O08539 | 1.31 | 1.11 | 0.8 | 1 | 0.89 | 1.29 | 1 | 0.68 | 0.74 | 0.09 | 0.78 | 1.3 | 0.45 | 1.09 | 0.58 | 0.61 | -0.09 | -0.3275 | -3.73E-01 | 0.653205 | 0.263462 | 0.123737 | 1 | 0.996511 | 0.721448 |
| Bin2 | S4R270 | -0.7 | -2.15 | -1.48 | -1.02 | -2.61 | -0.67 | -1.88 | -1.07 | -0.99 | -1.03 | -0.99 | -1.63 | -0.19 | -2.53 | -1.62 | -1.78 | -0.22 | 0.1775 | -0.1925 | 0.657348 | 0.61005 | 0.721145 | 1 | 0.996511 | 0.958398 |
| Bin3 | Q9JI08 | -0.19 | 0.35 | -0.42 | -0.22 | 0.67 | 0.13 | -1.08 | -0.65 | 0.82 | 0.49 | -0.34 | 0.33 | -0.15 | 0.78 | -0.61 | -0.85 | -0.1125 | 0.445 | -0.0875 | 0.779059 | 0.163968 | 0.820068 | 1 | 0.996511 | 0.972768 |
| Birc2 | Q62210 | -0.16 | -0.41 | -0.52 | -0.48 | -0.26 | -0.4 | -0.15 | -0.43 | 0.75 | 0.39 | -0.4 | 0.2 | 0.38 | 0.42 | -0.28 | -0.12 | 0.0825 | 0.6275 | 0.4925 | 0.615257 | 0.042349 | 0.061934 | 1 | 0.996511 | 0.662517 |
| Birc6 | O88738 | -0.56 | -0.58 | -0.54 | -0.49 | -0.63 | -0.51 | -0.6 | -0.38 | 0.08 | -0.06 | -0.25 | -0.46 | 0.42 | -0.39 | -0.57 | -0.56 | 0.0125 | 0.37 | 0.2675 | 0.931058 | 0.069153 | 0.324615 | 1 | 0.996511 | 0.842881 |
| Bivm | Q8CBX9 | 2.83 | 1.92 | -0.52 | -0.94 | 0.84 | 4.6 | -0.31 | -0.52 | -0.72 | -0.78 | -0.22 | 3.03 | 1.84 | -2.48 | -0.14 | -0.23 | 0.33 | -0.495 | -1.075 | 0.805915 | 0.670696 | 0.356988 | 1 | 0.996511 | 0.855416 |
| Bles03 | Q8VD62 | 0 | -1.04 | -0.82 | -0.5 | -1.29 | 0.18 | 0.29 | -0.85 | -0.96 | -0.37 | -0.74 | -0.5 | -1.06 | -0.65 | -0.77 | -1.12 | 0.1725 | -0.0525 | -0.31 | 0.683108 | 0.848494 | 0.277322 | 1 | 0.996511 | 0.822084 |
| Blmh | Q8R016 | -0.21 | -0.26 | -0.06 | -0.13 | -0.23 | -0.37 | 0.08 | 0.05 | -0.29 | -0.04 | -0.21 | -0.11 | -0.41 | -0.13 | -0.15 | -0.12 | 0.0475 | 0.0025 | -0.0375 | 0.780756 | 0.987658 | 0.832824 | 1 | 0.999285 | 0.975129 |
| Bloc1s1 | O55102 | -0.72 | 0.15 | 0.2 | 0.5 | 0.19 | -0.3 | 0.77 | 0.35 | -0.22 | 0.26 | 0.5 | -0.47 | -1.25 | 0.78 | 0.32 | 0.64 | 0.22 | -0.015 | 0.09 | 0.514153 | 0.964635 | 0.857111 | 1 | 0.996755 | 0.976225 |
| Bloc1s2 | Q9CWG9 | -0.75 | 0.08 | -0.98 | -0.87 | -0.13 | -0.23 | -0.65 | -1.16 | 0.3 | -0.5 | -1.04 | -0.2 | 0.6 | 0.05 | -0.38 | -1.18 | 0.0875 | 0.27 | 0.4025 | 0.789623 | 0.461072 | 0.356961 | 1 | 0.996511 | 0.855416 |
| Bloc1s4 | Q8VED2 | -0.92 | -0.22 | -0.61 | 0.53 | 0.17 | -0.21 | -0.04 | 0.02 | 1.39 | 0.71 | 0.34 | -0.5 | 1.03 | 0.2 | 0.39 | 0.33 | 0.29 | 0.79 | 0.7925 | 0.371345 | 0.120923 | 0.051333 | 1 | 0.996511 | 0.658454 |
| Blvra | Q9CY64 | 0.11 | -0.14 | 0.14 | -0.07 | -0.29 | -0.06 | -0.02 | 0.3 | -0.74 | 0 | -0.12 | 0.03 | -1.06 | -0.44 | -0.42 | -0.21 | -0.0275 | -0.2175 | -0.5425 | 0.880253 | 0.350489 | 0.044519 | 1 | 0.996511 | 0.658454 |
| Blvrb | Q923D2 | -0.17 | -1.61 | 0.22 | 1.07 | 1.46 | 0.44 | -0.32 | 1.02 | 0.67 | -0.02 | 0.15 | -1.45 | -1.71 | -0.7 | 0.09 | -0.26 | 0.0425 | -0.04 | 0.3325 | 0.951633 | 0.951382 | 0.635086 | 1 | 0.996755 | 0.941407 |
| Bmerb1 | Q8R1Y2 | 0.71 | 1.44 | 0.91 | 0.85 | 1.68 | 0.72 | 1.63 | 0.52 | 0.86 | 1.48 | 1 | 1.05 | 1.1 | 2.17 | 1.64 | 1.69 | 0.16 | 0.12 | 0.6725 | 0.631536 | 0.615342 | 0.041794 | 1 | 0.996511 | 0.658454 |
| Bmp2k | Q91Z96 | 0.71 | 0.96 | 0.35 | 0.38 | 0.58 | 0.92 | 0.18 | 0.61 | 2.66 | 0.49 | 0.58 | 1.02 | 2.79 | 1.77 | 0.78 | 0.49 | -0.0275 | 0.5875 | 0.8575 | 0.904579 | 0.249637 | 0.116577 | 1 | 0.996511 | 0.718608 |
| Bmpr1a | Q60607 | 1.75 | 1.32 | 1.09 | 1.08 | 1.84 | 1.42 | 0.83 | 1.3 | 1.12 | 1.77 | 0.83 | 2.19 | 0.59 | 0.97 | 1.48 | 1.53 | 0.0375 | 0.1675 | -0.1675 | 0.888243 | 0.625469 | 0.566859 | 1 | 0.996511 | 0.921806 |
| Bmpr1b | P36898 | -0.01 | 0.25 | -0.1 | 0.41 | -0.11 | 0.52 | 0.12 | 0.24 | -0.18 | 1.32 | 0.21 | 0.96 | 0.7 | 0.62 | 0.45 | 0.95 | 0.055 | 0.44 | 0.5425 | 0.789642 | 0.234661 | 0.029415 | 1 | 0.996511 | 0.637553 |
| Bmpr2 | O35607 | 0.3 | 0.54 | 0.63 | 0.65 | 0.68 | 0.53 | 0.37 | 0.61 | 0.64 | 0.78 | 0.77 | 0.6 | 0.33 | 0.67 | 0.61 | 0.56 | 0.0175 | 0.1675 | 0.0125 | 0.914626 | 0.338814 | 0.947107 | 1 | 0.996511 | 0.992376 |
| Bms1 | Q6PGF5 | 1.21 | 1.15 | 1.89 | 1.55 | 0.97 | 0.46 | 1.21 | 1.34 | 0.94 | 1.25 | 1.51 | 1.25 | 0.98 | 1.52 | 2.12 | 1.98 | -0.455 | -0.2125 | 0.2 | 0.114641 | 0.379835 | 0.532334 | 1 | 0.996511 | 0.910774 |
| Bmt2 | A0A0N4SU | -0.12 | -0.14 | -0.34 | 0.04 | -0.91 | 0.33 | -1.1 | -1.08 | 0.32 | -1.05 | -0.75 | -0.14 | 0.89 | -0.29 | 0.05 | -0.88 | -0.55 | -0.265 | 0.0825 | 0.13269 | 0.41384 | 0.823702 | 1 | 0.996511 | 0.972768 |
| Bnip1 | Q6QD59 | -0.13 | -0.07 | -0.33 | -0.09 | 0.29 | -0.05 | 0.38 | 0.29 | -0.1 | 0.8 | -0.24 | -0.18 | 0.02 | 0.83 | 0.19 | 0.3 | 0.3825 | 0.225 | 0.49 | 0.045408 | 0.411568 | 0.057744 | 1 | 0.996511 | 0.661738 |
| Bnip2 | Q91VL0 | 0.21 | 0.12 | 1.04 | 1.28 | 0.14 | 0.61 | 1.64 | 1.55 | 0.89 | 0.27 | 1.21 | 0.59 | 0.27 | -0.65 | 0.62 | 0.71 | 0.3225 | 0.0775 | -0.425 | 0.467479 | 0.82385 | 0.314391 | 1 | 0.996511 | 0.836924 |
| Bnip3 | O55003 | 0.18 | 0.14 | 0.78 | 0.56 | -1.04 | 0.34 | -0.67 | 0.22 | 2.42 | 0.77 | 0.4 | 0.41 | 3.72 | -0.5 | 0.44 | -0.22 | -0.7025 | 0.585 | 0.445 | 0.075266 | 0.236603 | 0.617467 | 1 | 0.996511 | 0.937152 |
| Bnip3l | Q9Z2F7 | -0.68 | 0.78 | -0.95 | -0.97 | 0.89 | 0.25 | -0.66 | -0.45 | -0.77 | -1.63 | -0.44 | -0.26 | 0.3 | 0.72 | -0.85 | -1.19 | 0.4625 | -0.32 | 0.2 | 0.370416 | 0.511904 | 0.725599 | 1 | 0.996511 | 0.958398 |
| Boc | D3Z763 | -2.85 | -3.45 | -1.51 | -1.95 | -3.38 | -1.74 | -2.84 | -2.97 | -2.44 | 0.63 | -1.62 | -2.63 | -1.9 | -2.02 | -1.09 | -2.25 | -0.2925 | 0.925 | 0.625 | 0.574041 | 0.257338 | 0.209559 | 1 | 0.996511 | 0.786218 |
| Bod1 | Q5SQY2 | -1.32 | -2.17 | 0.19 | -1.38 | -1.5 | -1.25 | 0.12 | -0.7 | -2.8 | -2.36 | -0.47 | -1.56 | -2.37 | -1.99 | -0.74 | -0.12 | 0.3375 | -0.6275 | -0.135 | 0.549709 | 0.346561 | 0.837178 | 1 | 0.996511 | 0.975831 |
| Bod1l | E9Q6J5-2 | -2.03 | -2.34 | -2.8 | -3.18 | -2.81 | -1.66 | -3.35 | -3.35 | 0.01 | -1.59 | -2.97 | -2.24 | 0.15 | -2.58 | -2.72 | -3.43 | -0.2 | 0.89 | 0.4425 | 0.65024 | 0.179374 | 0.557829 | 1 | 0.996511 | 0.917135 |
| Bok | O35425 | 0.47 | 0.3 | -1.14 | -0.87 | -0.48 | 0.3 | -1.33 | -0.45 | -0.17 | -0.09 | -1.21 | 0.17 | -0.2 | 0.47 | -1 | -1.44 | -0.18 | -0.015 | -0.2325 | 0.711979 | 0.974752 | 0.669941 | 1 | 0.997515 | 0.952558 |
| Bola1 | Q9D8S9 | 1.88 | 0.68 | 1.14 | 1.98 | 0.4 | 1.26 | 1.06 | 1.61 | 1.44 | 1.63 | 1.16 | 2.2 | 0.95 | 0.85 | 0.75 | 0.51 | -0.3375 | 0.1875 | -0.655 | 0.386182 | 0.613278 | 0.072149 | 1 | 0.996511 | 0.668066 |
| Bola2 | Q8BGS2 | -1.49 | -1.36 | -0.48 | -0.48 | -0.76 | -1.44 | 0.11 | -0.21 | -1.01 | 0.27 | -0.38 | -1.03 | -0.76 | -0.41 | -0.04 | -0.31 | 0.3775 | 0.45 | 0.5725 | 0.369719 | 0.309641 | 0.099891 | 1 | 0.996511 | 0.698156 |
| Bola3 | E9Q705 | 1.35 | 0.63 | -0.79 | -1.19 | -0.41 | 1.14 | -1.31 | -1.31 | 1.94 | -1.42 | -0.75 | 1.38 | 1.96 | 1.09 | -0.84 | -0.92 | -0.4725 | 0.2875 | 0.3225 | 0.533406 | 0.75182 | 0.701526 | 1 | 0.996511 | 0.956791 |
| Bop1 | P97452 | 0.2 | 0.56 | 1.02 | 0.33 | 0.4 | -0.21 | 0.75 | 0.01 | 0.27 | 0.68 | 0.96 | 0.36 | 0.92 | 0.1 | 0.47 | 1.19 | -0.29 | 0.04 | 0.1425 | 0.31768 | 0.877286 | 0.648166 | 1 | 0.996511 | 0.945938 |
| Borcs5 | E9Q2W8 | 0.09 | 0.33 | 0.76 | 0.68 | 0.39 | 0.09 | 0.77 | 0.66 | 0.07 | 1.3 | 0.61 | 0.14 | 0.56 | 0.28 | 0.71 | 0.87 | 0.0125 | 0.065 | 0.14 | 0.957564 | 0.840542 | 0.562487 | 1 | 0.996511 | 0.920024 |
| Borcs6 | Q9D6W8 | -0.37 | 0.04 | -0.28 | -0.14 | -0.03 | -0.5 | -0.33 | 0.12 | -0.05 | 1.34 | -0.44 | -0.36 | -0.2 | 0.07 | -0.28 | -0.25 | 0.0025 | 0.31 | 0.0225 | 0.990005 | 0.451474 | 0.907377 | 1 | 0.996511 | 0.986458 |
| Borcs7 | Q9CRC6 | 0.06 | 0.04 | -0.08 | 0.87 | 0.81 | -0.32 | 0.37 | 0.66 | -0.26 | 1.46 | 0.64 | 0.12 | -0.05 | 1.49 | 0.46 | 0.74 | 0.1575 | 0.2675 | 0.4375 | 0.629105 | 0.52007 | 0.266312 | 1 | 0.996511 | 0.818107 |
| Borcs8 | Q9D6Y4 | -0.05 | 0.46 | 0.69 | 0.72 | 0.56 | -0.37 | 1.14 | 0.89 | -1.09 | 1.04 | 0.71 | 0.05 | -0.03 | 0.2 | 0.75 | 0.73 | 0.1 | -0.2775 | -0.0425 | 0.781094 | 0.559112 | 0.88068 | 1 | 0.996511 | 0.979767 |
| Bpgm | P15327 | 2.48 | 0.07 | 0.63 | 1.15 | 0.14 | 3.34 | 0.39 | 1.08 | 1.52 | 0.11 | 0.01 | 1.28 | 1.34 | 0.79 | 0.09 | 0.25 | 0.155 | -0.3525 | -0.465 | 0.847249 | 0.556379 | 0.402519 | 1 | 0.996511 | 0.870698 |
| Bphl | Q8R164 | 1.18 | 1.01 | 1.95 | 1.38 | 0.79 | 1.06 | 2.19 | 1.96 | 0.02 | 0.65 | 1.82 | 0.87 | -0.42 | 0.11 | 1.47 | 2.09 | 0.12 | -0.54 | -0.5675 | 0.750741 | 0.206288 | 0.332364 | 1 | 0.996511 | 0.845601 |
| Bpnt1 | Q9Z0S1 | 0.01 | 0.6 | 0.35 | 0.35 | 1.01 | 0.87 | 0.5 | 0.43 | 1.16 | 0.06 | 0.34 | 1.62 | 1.03 | -2.86 | 0.02 | -0.71 | 0.375 | 0.4675 | -0.9575 | 0.105386 | 0.226629 | 0.224997 | 1 | 0.996511 | 0.787447 |
| Bpnt1 | A0A1B0GR | -0.65 | -0.79 | -0.48 | -0.58 | -0.99 | -0.47 | -0.38 | -0.3 | -0.68 | -0.57 | -0.37 | -0.72 | 0.04 | -1.45 | -0.28 | -0.83 | 0.09 | 0.04 | -0.005 | 0.656218 | 0.818696 | 0.988089 | 1 | 0.996511 | 0.996154 |
| Bpnt2 | Q80V26 | -1.5 | -3.21 | -2.29 | -2.07 | -1.59 | -1.04 | -0.91 | -2.24 | -3.61 | -2.81 | -2.18 | -1.67 | -2.81 | -2.55 | -1.01 | -1.79 | 0.8225 | -0.3 | 0.2275 | 0.087098 | 0.560096 | 0.652638 | 1 | 0.996511 | 0.947633 |
| Bptf | A2A654 | -0.62 | -1.65 | -0.04 | 0.03 | -1.83 | -0.58 | -0.19 | 0.25 | -0.27 | 0.79 | 0.19 | -1.29 | 0.88 | -2.33 | 0.55 | 0.16 | -0.0175 | 0.425 | 0.385 | 0.974283 | 0.440801 | 0.609123 | 1 | 0.996511 | 0.934202 |
| Braf | P28028-1 | -0.25 | -0.15 | -0.34 | -0.42 | -0.4 | -0.22 | -0.17 | -0.56 | -0.59 | -0.64 | -0.55 | -0.24 | -0.71 | -1.14 | -0.77 | -0.96 | -0.0475 | -0.215 | -0.605 | 0.772976 | 0.244484 | 0.01076 | 1 | 0.996511 | 0.563386 |
| Braf | P28028 | -0.97 | 0.49 | 0.13 | 0.42 | -0.14 | -0.66 | 0.49 | 0.21 | 0.57 | 0.65 | 0.21 | -0.17 | -0.09 | 0.86 | -0.05 | 0.05 | -0.0425 | 0.2975 | 0.175 | 0.914455 | 0.435175 | 0.657604 | 1 | 0.996511 | 0.948659 |
| Brap | Q99MP8 | -0.95 | -0.7 | -0.81 | -1.21 | -0.77 | -0.82 | -1.03 | -1.37 | -0.6 | -0.31 | -1.21 | -0.86 | -0.31 | -0.77 | -1.17 | -1.25 | -0.08 | 0.1725 | 0.2425 | 0.698104 | 0.48915 | 0.873819 | 1 | 0.996511 | 0.979626 |
| Brat1 | E9QLK3 | -0.97 | -0.95 | -0.71 | -0.46 | -0.81 | -1.54 | -0.76 | -1.26 | -0.65 | -0.34 | -0.88 | -1.16 | -0.78 | -1.13 | -1.34 | -1.04 | -0.32 | 0.015 | -0.3 | 0.202242 | 0.949927 | 0.193495 | 1 | 0.996755 | 0.777907 |
| Brawnin | Q8BTC1 | 1.73 | 1.59 | 2.41 | 2.44 | 1.28 | 1.58 | 1.98 | 2.38 | 0.96 | 1.74 | 1.7 | 1.1 | 0.65 | 1.36 | 1.2 | 1.47 | -0.2375 | -0.6675 | -0.8725 | 0.465014 | 0.053168 | 0.016597 | 1 | 0.996511 | 0.591538 |
| Brcc3 | P46737 | -1.23 | -1.65 | -1.45 | -1.07 | -1.17 | -1.12 | -1 | -1.22 | -1.93 | -1 | -1.11 | -1.11 | -1.88 | -1.12 | -1.61 | -1.13 | 0.2225 | 0.0625 | -0.085 | 0.237685 | 0.814997 | 0.741113 | 1 | 0.996511 | 0.958796 |
| Brd1 | G5E8P1 | 1.34 | 0.99 | 1.79 | 1.53 | 0.52 | 1.52 | 1.3 | 1.35 | 0.7 | -2.34 | 1.35 | 0.74 | 1.91 | -2.35 | 1.82 | 2.07 | -0.24 | -1.3 | -0.55 | 0.40423 | 0.118073 | 0.574747 | 1 | 0.996511 | 0.926139 |
| Brd2 | Q7JJ13 | 1.6 | 1.61 | 1.09 | 0.81 | -2.17 | 1.18 | 0.65 | 0.92 | 3.03 | 0.37 | 0.67 | 0.12 | 2.8 | 0.49 | 1.46 | 1 | -1.1325 | -0.23 | 0.16 | 0.146006 | 0.719415 | 0.748331 | 1 | 0.996511 | 0.960744 |
| Brd3 | Q8K2F0 | 1.28 | -0.19 | 0.71 | 1.12 | -1.53 | 0.78 | 0.8 | 0.54 | 1.04 | 1.14 | 0.66 | 0.62 | 1.15 | -1.26 | 1.4 | 1.11 | -0.5825 | 0.135 | -0.13 | 0.341041 | 0.699136 | 0.839758 | 1 | 0.996511 | 0.976219 |
| Brd4 | Q3UH70 | 1.01 | 0.04 | -0.48 | -0.29 | -0.04 | 0.2 | -0.46 | -0.34 | -0.39 | -0.01 | -0.65 | 0.05 | 0.2 | 0.16 | 0.38 | -0.51 | -0.23 | -0.02 | -0.0125 | 0.515641 | 0.385853 | 0.97341 | 1 | 0.996511 | 0.992946 |
| Brd7 | O88665 | 0.38 | 0.4 | 0.78 | 1.94 | 0.57 | 0.4 | 0.56 | 1.56 | -0.07 | 1.69 | 0.5 | 0.54 | 0.24 | -0.19 | 1.41 | 0.74 | -0.1025 | -0.21 | -0.325 | 0.809115 | 0.666723 | 0.497102 | 1 | 0.996511 | 0.897918 |
| Brd8 | Q8R3B7-2 | -2.27 | -2.04 | -1.6 | -2.34 | -3.48 | -1.62 | -2.51 | -1.98 | -0.51 | -0.71 | -2.04 | -2.62 | 0.21 | -3.78 | -1.57 | -1.8 | -0.335 | 0.5925 | 0.3275 | 0.424505 | 0.256291 | 0.665143 | 1 | 0.996511 | 0.949825 |
| Brd9 | A0A1Y7VJ6 | -0.34 | -2.11 | -0.94 | -2.14 | -2.05 | -1.14 | -0.97 | -0.59 | -0.93 | -0.17 | -1.47 | -2.62 | 0.55 | -0.14 | -0.82 | -0.51 | 0.195 | 0.085 | 1.1525 | 0.698047 | 0.891449 | 0.043415 | 1 | 0.996511 | 0.658454 |
| Bri3bp | Q8BXV2 | 1.2 | -0.64 | -0.08 | -2.02 | 0.11 | 1.71 | 1.35 | -0.06 | -2.02 | -0.66 | -0.28 | 1.02 | -1.08 | -1.26 | 1.2 | 1.46 | 1.1625 | -0.1 | 0.465 | 0.135567 | 0.903576 | 0.601809 | 1 | 0.996609 | 0.930802 |
| Brinp1 | Q920P3 | 0.76 | 0.53 | 1.32 | 0.73 | 0.81 | 1.18 | 1.41 | 1.18 | 0.13 | 0.55 | 1.5 | 1.05 | -0.13 | 0.1 | 0.81 | 1.23 | 0.31 | -0.0275 | -0.3325 | 0.200362 | 0.935154 | 0.359734 | 1 | 0.996755 | 0.856645 |
| Brinp2 | Q6DFY8 | 0.45 | 0.81 | 1.31 | 0.6 | 0.78 | 0.6 | 0.86 | 0.9 | 0.41 | -0.19 | 1.21 | 0.87 | 0.12 | 0.45 | 0.4 | 0.6 | -0.0075 | -0.2175 | -0.4 | 0.973001 | 0.538261 | 0.132991 | 1 | 0.996511 | 0.728417 |
| Brinp3 | Q499E0 | 1.04 | 1.2 | 0.92 | 0.79 | 1.27 | 1.11 | 0.91 | 1.15 | 0.52 | -0.94 | 0.84 | 1.26 | 1.17 | 0.25 | 0.45 | 1.07 | 0.1225 | -0.5675 | -0.2525 | 0.475017 | 0.23462 | 0.359314 | 1 | 0.996511 | 0.856645 |

| | | | | | | | | | | | | | | | | | | | | | | | | | | |
|---|---|---|---|---|---|---|---|---|---|---|---|---|---|---|---|---|---|---|---|---|---|---|---|---|---|---|
| Brix1 | Q9DCA5 | 0.7 | -0.06 | 0.38 | -0.73 | -0.29 | 0.9 | 0.35 | -0.53 | 2.94 | -1.94 | -0.29 | -0.33 | 4.84 | -0.01 | -0.74 | 0.3 | 0.035 | 0.0225 | 1.025 | 0.933443 | 0.981307 | 0.388662 | 1 | 0.998773 | 0.86481 |
| Brk1 | Q91VR8 | 2.12 | 0.47 | 1.37 | 0.32 | 0.7 | 2.31 | 1.38 | 1.51 | 0.85 | 0.07 | 1.73 | 2.11 | 0.01 | -0.62 | 2.66 | 1.03 | 0.405 | 0.12 | -0.3 | 0.420948 | 0.833578 | 0.69036 | 1 | 0.996511 | 0.955729 |
| Brms1l | Q3U1T3 | -1.52 | -1.24 | 1.04 | 1.14 | -1.28 | -0.67 | 0.89 | 1.42 | -0.5 | -0.7 | 0.99 | -0.69 | -1.68 | -0.85 | 0.46 | 0.47 | 0.235 | -0.08 | -0.255 | 0.785421 | 0.914598 | 0.750756 | 1 | 0.996755 | 0.961479 |
| Brox | Q8K2Q7 | -0.95 | -0.8 | -0.33 | -0.76 | -0.99 | -0.86 | -1.23 | -0.65 | -0.13 | -0.63 | -0.73 | -0.61 | 0.13 | -0.53 | -0.76 | -1.19 | -0.2225 | 0.185 | 0.1225 | 0.302717 | 0.418597 | 0.697758 | 1 | 0.996511 | 0.955729 |
| Brpf1 | A0A0N4SU | 6.04 | 5.17 | 5.28 | 3.17 | 5.33 | 5.49 | 3.85 | 3.75 | 1.22 | 1.6 | 4.63 | 5.26 | 1.36 | 1.3 | 4.97 | 5.78 | -0.31 | -1.7375 | -1.5625 | 0.658498 | 0.133699 | 0.209599 | 1 | 0.996511 | 0.786218 |
| Brsk1 | Q5RJI5 | 1.06 | 0.77 | 0.87 | 0.55 | 0.69 | 1.05 | 0.93 | 0.84 | 0.6 | 0.7 | 0.92 | 1.23 | 0.77 | 0.6 | 0.76 | 0.69 | 0.065 | 0.05 | -0.1075 | 0.715051 | 0.81645 | 0.575602 | 1 | 0.996511 | 0.926728 |
| Brsk2 | G3UWV4 | -0.11 | 0.28 | 0.25 | 0.83 | -0.56 | -0.48 | -0.42 | 0.48 | 0.63 | 1.05 | 0.21 | -0.3 | 1.6 | 0.46 | 0.44 | -0.4 | -0.5575 | 0.085 | 0.2125 | 0.097337 | 0.804207 | 0.624978 | 1 | 0.996511 | 0.939167 |
| Brsk2 | Q69Z98 | 0.03 | 0.59 | 0.92 | 0.79 | 0.7 | 0.46 | 0.5 | 0.79 | 0.4 | 1.08 | 0.79 | 0.02 | 0.53 | 0.7 | 0.49 | 0.6 | 0.03 | -0.01 | -0.0025 | 0.896499 | 0.973983 | 0.991691 | 1 | 0.997515 | 0.996154 |
| Brwd1 | Q921C3 | 1.09 | 1.82 | 1.52 | 1.25 | 1.51 | 1.03 | -0.85 | 0.62 | 2.63 | -1.66 | 1.65 | 1.75 | 2.65 | 2.11 | 0.88 | 2.18 | -0.8425 | -0.3275 | 0.535 | 0.115346 | 0.705363 | 0.200234 | 1 | 0.996511 | 0.78004 |
| Bscl2 | A0A0R4J22 | 0.51 | 0.86 | -0.22 | -0.74 | 1.34 | 1.32 | 0.37 | -0.6 | 0.04 | -1.03 | -0.38 | 1.04 | 1.27 | 0.09 | 0.17 | 0.72 | 0.505 | -0.185 | 0.46 | 0.359974 | 0.723965 | 0.30151 | 1 | 0.996511 | 0.830308 |
| Bsdc1 | Q80Y55 | 0.13 | 0.41 | 0.17 | -0.15 | 0.36 | 0.4 | -0.11 | -0.16 | 0.98 | -0.22 | 0.28 | 0.41 | 1.38 | 0.31 | 0.25 | 0.02 | -0.0175 | 0.2225 | 0.35 | 0.934859 | 0.440314 | 0.302147 | 1 | 0.996511 | 0.830699 |
| Bsg | K3W4Q8 | 0.32 | 0.27 | 0.47 | 0.35 | -0.09 | 0.73 | 0.56 | 0.23 | 1.47 | 1.13 | 0.62 | 0.42 | 2.2 | 0.55 | 0.92 | 0.26 | 0.005 | 0.5575 | 0.63 | 0.981268 | 0.058349 | 0.152372 | 1 | 0.996511 | 0.750422 |
| Bsn | O88737 | 1.38 | 1.1 | 1.24 | 1.24 | 0.95 | 1.27 | 0.75 | 1 | 1.27 | 0.88 | 1.2 | 1.3 | 1.41 | 0.95 | 1.03 | 0.84 | -0.2475 | -0.0775 | -0.1825 | 0.175831 | 0.666996 | 0.377063 | 1 | 0.996511 | 0.860984 |
| Btaf1 | E9QAE3 | -0.63 | -0.37 | -0.43 | -0.67 | 0.25 | -1.18 | -0.72 | -1.15 | -0.55 | 0.88 | -0.56 | -0.56 | -1.39 | -0.66 | -0.99 | -0.48 | -0.175 | 0.3275 | -0.355 | 0.600672 | 0.371801 | 0.17675 | 1 | 0.996511 | 0.771503 |
| Btbd1 | P58544 | -1.71 | -1 | 0.64 | -0.89 | -0.73 | -0.78 | 0.47 | -0.51 | 2.3 | -0.67 | 0.07 | -1.07 | 2.58 | 0.11 | -0.24 | 0.17 | 0.3525 | 0.8975 | 1.395 | 0.509696 | 0.287227 | 0.085798 | 1 | 0.996511 | 0.691619 |
| Btbd10 | E9Q6L8 | -0.18 | 0.62 | 1.05 | 0.58 | 0.15 | -0.12 | 0.25 | 0.81 | 0.96 | 1.06 | 0.64 | 0.46 | 1.29 | 1.15 | 0.73 | 0.22 | -0.245 | 0.2625 | 0.33 | 0.445817 | 0.38895 | 0.357002 | 1 | 0.996511 | 0.855416 |
| Btbd11 | Q6GQW0 | 0.76 | 0.94 | 1.03 | 0.97 | 1.21 | 0.79 | 0.85 | 1 | 0.83 | 1.25 | 0.95 | 0.68 | 0.17 | 1 | 0.21 | 0.9 | 0.0375 | 0.0025 | -0.355 | 0.821696 | 0.989528 | 0.192243 | 1 | 0.999285 | 0.77733 |
| Btbd17 | Q9DB72 | -0.54 | -0.19 | -0.12 | -0.13 | 0.22 | -0.52 | 0.31 | -0.17 | -0.4 | -0.71 | -0.29 | -0.31 | -0.86 | 0.07 | -0.55 | -0.6 | 0.205 | -0.1825 | -0.24 | 0.390645 | 0.358173 | 0.355562 | 1 | 0.996511 | 0.855029 |
| Btbd8 | Q3TEG0 | 0.14 | 0.03 | 0.37 | 0.58 | 0 | -0.15 | -0.33 | -0.08 | -1.4 | -1.41 | 0.03 | 0.76 | -0.46 | -0.87 | 0.06 | 0.42 | -0.42 | -0.785 | -0.4925 | 0.045513 | 0.153151 | 0.146238 | 1 | 0.996511 | 0.746406 |
| Btbd9 | Q8C726 | -1.26 | -0.74 | -1.78 | -2.21 | -0.69 | -1.47 | -1.77 | -2.75 | -0.14 | 0.45 | -2.15 | -1.01 | -0.04 | 1.45 | -2.3 | -2.47 | -0.1725 | 0.785 | 0.6575 | 0.726275 | 0.209929 | 0.4688 | 1 | 0.996511 | 0.8901 |
| Btd | A0A0R4J13 | 0.43 | 0.74 | -4.11 | -3.94 | 0.54 | 0.58 | -4.06 | -4.87 | 0.13 | -0.65 | -4.3 | 1 | 0.37 | 0.02 | -2.37 | -3.58 | -0.2325 | 0.765 | 0.33 | 0.895014 | 0.630412 | 0.820564 | 1 | 0.996511 | 0.972768 |
| Btf3 | Q64152 | -4.33 | -4.66 | -3.15 | -3.32 | -4.17 | -4.82 | -3.2 | -3.22 | -3.54 | -1.9 | -3.31 | -4.18 | -3.07 | -3.77 | -3.12 | -3.7 | 0.0125 | 0.6325 | 0.45 | 0.980018 | 0.27655 | 0.278694 | 1 | 0.996511 | 0.822084 |
| Btf3l4 | Q9CQH7 | -4 | -4.24 | -3.22 | -3.53 | -3.57 | -4.48 | -3.57 | -3.83 | -4.11 | -1.52 | -3.57 | -3.51 | -4.14 | -3.09 | -3.37 | -3.83 | -0.115 | 0.57 | 0.14 | 0.711212 | 0.32737 | 0.673617 | 1 | 0.996511 | 0.952944 |
| Bub3 | A0A140LH | -5.47 | -6.42 | -4.81 | -6.04 | -6.14 | -6.25 | -5.91 | -5.46 | -5.9 | -3.67 | -5.01 | -5.2 | -4.12 | -5.35 | -4.46 | -5.59 | -0.255 | 0.74 | 0.805 | 0.50042 | 0.193691 | 0.112085 | 1 | 0.996511 | 0.713445 |
| Bud13 | Q8R149 | -1.36 | -2.2 | -0.67 | -0.45 | -1.27 | -1.6 | 0.04 | 0.43 | -1.16 | -0.87 | -0.36 | -1 | -0.14 | 0.77 | -0.1 | 0.65 | 0.57 | 0.3225 | 1.465 | 0.337344 | 0.440003 | 0.008398 | 1 | 0.996511 | 0.563386 |
| Bud31 | Q6PGH1 | -1.4 | -1.55 | -1.82 | -1.83 | -1.82 | -1.39 | -1.85 | -1.3 | -0.06 | -3.09 | -1.89 | -1.74 | 0.12 | -1 | -2.17 | -2.75 | 0.06 | -0.045 | 0.2 | 0.772319 | 0.93817 | 0.7364 | 1 | 0.996755 | 0.958635 |
| Bysl | O54825 | -1.11 | -1.56 | -0.15 | -0.85 | -1.52 | -1.73 | -1.34 | -0.8 | -2.95 | -2.98 | -0.53 | -1.25 | -2.93 | -4.09 | -1.58 | -0.88 | -0.43 | -1.01 | -1.4525 | 0.231633 | 0.133419 | 0.063997 | 1 | 0.996511 | 0.66433 |
| Bzw1 | A0A087WC | -1.63 | -1.94 | -2.32 | -2.34 | -2.01 | -1.9 | -2.63 | -2.68 | -2.11 | -1.4 | -2.48 | -1.69 | -1.81 | -1.63 | -2.89 | -3.28 | -0.2475 | 0.1375 | -0.345 | 0.372354 | 0.646232 | 0.419212 | 1 | 0.996511 | 0.87558 |
| Bzw2 | Q91VK1 | -1.79 | -1.99 | -1.38 | -1.39 | -1.58 | -1.75 | -1.18 | -1.12 | -2.13 | -0.82 | -1.26 | -1.85 | -1.5 | -1.15 | -1.52 | -1.63 | 0.23 | 0.1225 | 0.1875 | 0.338395 | 0.710512 | 0.421903 | 1 | 0.996511 | 0.875736 |
| C030006K1 | Q8VE95 | 2.93 | 1.21 | 0.71 | 1.21 | 0.52 | 2.48 | 0.31 | 1.17 | 0.79 | 1.82 | 1.02 | 2.05 | 0.69 | 1.05 | 0.21 | -0.14 | -0.395 | -0.095 | -1.0625 | 0.533688 | 0.857932 | 0.064355 | 1 | 0.996511 | 0.66433 |
| C1qa | P98086 | -3.88 | -5.07 | -3.67 | -3.52 | -4.13 | -4.08 | -2.58 | -3.47 | -3.8 | -2.14 | -3.92 | -3.61 | -3.1 | -3.51 | -2.74 | -3.35 | 0.47 | 0.6675 | 0.86 | 0.329966 | 0.209427 | 0.047129 | 1 | 0.996511 | 0.658454 |
| C1qb | P14106 | -3.11 | -3.38 | -2.18 | -1.34 | -2.7 | -3.62 | -1.14 | -1.87 | -4.47 | -0.77 | -1.45 | -2.75 | -2.99 | -3.43 | -1.19 | -0.93 | 0.17 | 0.1425 | 0.3675 | 0.791999 | 0.866266 | 0.607029 | 1 | 0.996511 | 0.932961 |
| C1qbp | Q8R5L1 | 0.42 | 0.34 | 0.77 | 1.76 | -0.13 | 0.24 | 0.88 | 1.55 | 0.13 | 0.57 | 0.94 | 0.4 | -1.23 | 0.07 | 0.15 | 0.34 | -0.1875 | -0.3125 | -0.99 | 0.683571 | 0.394003 | 0.055086 | 1 | 0.996511 | 0.661738 |
| C1qc | Q02105 | -2.38 | -3.47 | -4.21 | -3.53 | -3.06 | -2.55 | -3.31 | -3.88 | -2.58 | -2.45 | -3.82 | -2.46 | -2.31 | -2.85 | -2.99 | -3.66 | 0.1975 | 0.57 | 0.445 | 0.653402 | 0.246506 | 0.330978 | 1 | 0.996511 | 0.845601 |
| C1ql3 | Q9ESN4 | 0.15 | 0.6 | 0.47 | 0.35 | 0.66 | 0.45 | 0.94 | 0.57 | 0.15 | -0.18 | 0.25 | 0.59 | 0.21 | 0.81 | 0.38 | 0.44 | 0.2625 | -0.19 | 0.0675 | 0.178268 | 0.40142 | 0.752806 | 1 | 0.996511 | 0.961479 |
| C1qtnf4 | Q8R066 | 0.43 | 0.45 | 0.18 | 0.32 | 0.71 | 0.52 | 0.59 | 0.52 | -0.12 | 1.16 | 0.1 | 0.67 | 0.11 | -0.07 | 0.85 | 0.77 | 0.24 | 0.1075 | 0.07 | 0.138302 | 0.721685 | 0.792676 | 1 | 0.996511 | 0.96933 |
| C2cd2 | E9Q3C1 | 0.27 | 0.39 | 0.46 | 0.26 | 0.64 | 0.56 | 0.14 | 0.17 | 0.57 | -0.45 | 0.12 | 0.39 | 1.56 | 0.79 | 0.75 | 0.58 | 0.0325 | -0.1875 | 0.575 | 0.858334 | 0.462028 | 0.047342 | 1 | 0.996511 | 0.658454 |
| C2cd2l | Q80X80 | 0.35 | 0.01 | 0.08 | 0.32 | -0.1 | 0.16 | 0.6 | 0.38 | 0.13 | 0.16 | 0.04 | 0.36 | 0.66 | 0.45 | 0.54 | 0.34 | 0.07 | -0.0175 | 0.3075 | 0.731643 | 0.921428 | 0.12943 | 1 | 0.996755 | 0.723655 |
| C2cd4c | Q5HZI2 | -0.14 | 0.15 | 0.5 | 0.4 | -0.02 | -0.22 | 0.44 | 0.24 | 0.77 | 0.43 | 0.64 | 0.32 | 1.06 | 0.69 | 0.61 | 0.61 | -0.1175 | 0.3125 | 0.515 | 0.604133 | 0.171537 | 0.045606 | 1 | 0.996511 | 0.658454 |
| C2cd5 | Q7TPS5 | 1.63 | 2.17 | 1.23 | 1.9 | 5.79 | 0.2 | 0.09 | 0.73 | 5.68 | 2 | 1.41 | 2.98 | 3 | 3.15 | 3.12 | 1.19 | -0.03 | 1.285 | 0.8825 | 0.980647 | 0.165667 | 0.095507 | 1 | 0.996511 | 0.698156 |
| C2cd5 | A0A0N4SW | 0.65 | 0.73 | 0.87 | 0.71 | 0.86 | 0.57 | 0.91 | 0.81 | 0.37 | 0.6 | 0.66 | 0.64 | 0.47 | 0.47 | 0.69 | 0.99 | 0.0475 | -0.1725 | -0.085 | 0.760999 | 0.313667 | 0.671161 | 1 | 0.996511 | 0.952631 |
| C3 | P01027 | -3.81 | -4.8 | -3.86 | -4.47 | -3.4 | -4.48 | -3.28 | -4.09 | -4.64 | -2.17 | -3.9 | -3.62 | -4.07 | -3.53 | -4.26 | -4.23 | 0.4225 | 0.6525 | 0.2125 | 0.256939 | 0.237651 | 0.492837 | 1 | 0.996511 | 0.897531 |
| C4b | P01029 | -1.07 | -0.8 | -0.57 | -0.2 | -0.48 | -1.03 | 0.28 | -0.08 | -0.63 | 1.16 | -0.22 | 0.07 | -1.16 | 0.36 | -1.07 | 0.35 | 0.3325 | 0.755 | 0.28 | 0.324265 | 0.090294 | 0.528634 | 1 | 0.996511 | 0.908663 |
| C9orf72 | Q6DFW0 | -0.95 | -0.55 | 0.67 | 0.7 | -0.43 | -0.7 | 0.94 | 0.13 | -0.47 | 0.96 | 0.67 | -0.99 | -0.49 | -1.54 | 0.29 | 0.23 | 0.0175 | 0.075 | -0.345 | 0.972721 | 0.896269 | 0.537592 | 1 | 0.996511 | 0.912212 |
| Ca1 | P13634 | 0.72 | -0.94 | 0.8 | 2 | -0.61 | 1.7 | 0.43 | 2.37 | -1.16 | -1.14 | -0.45 | -1.64 | 0.03 | -0.81 | -0.71 | -0.79 | 0.3275 | -1.7425 | -1.215 | 0.686776 | 0.016573 | 0.063452 | 1 | 0.996511 | 0.66433 |
| Ca10 | P61215 | 0.7 | 1.14 | 0.81 | 1.18 | 0.7 | 1.06 | 0.38 | 1.27 | 1.27 | 2.05 | 1.28 | 1.12 | 1.24 | 0.25 | 0.51 | 0.46 | -0.105 | 0.4725 | -0.3425 | 0.668639 | 0.097379 | 0.227181 | 1 | 0.996511 | 0.787447 |
| Ca11 | O70354 | 2.67 | 1.05 | 0.6 | 0.39 | 0.7 | 2.8 | 0.7 | 0.08 | 1.24 | 0 | 0.5 | 2.19 | 1.34 | 0.83 | -0.08 | 0.02 | -0.1075 | -0.195 | -0.65 | 0.880096 | 0.761522 | 0.271002 | 1 | 0.996511 | 0.822084 |
| Ca14 | Q9WVT6 | 1.87 | 1.31 | 1.54 | 0.99 | 0.98 | 1.21 | 1.18 | 0.63 | 0.94 | 0.11 | 0.63 | 1.26 | -0.06 | 0.21 | 0.84 | 0.89 | -0.4275 | -0.6925 | -0.9575 | 0.10641 | 0.04964 | 0.012226 | 1 | 0.996511 | 0.571403 |
| Ca2 | P00920 | 1.2 | 0.17 | 0.76 | 1.13 | 0.54 | 1.44 | 1.32 | 1.55 | -0.5 | -1.1 | 0.86 | 0.31 | -0.71 | -0.23 | 0.48 | 0.59 | 0.3975 | -0.9225 | -0.7825 | 0.236712 | 0.07287 | 0.063994 | 1 | 0.996511 | 0.66433 |
| Ca4 | Q64444 | 0.88 | 1.14 | 1.88 | 1.57 | 0.91 | 0.98 | 1.5 | 1.63 | 0.62 | 1.54 | 1.91 | 1.06 | 0.04 | 0.4 | 1.03 | 1.85 | -0.1125 | -0.085 | -0.5375 | 0.696999 | 0.808797 | 0.234733 | 1 | 0.996511 | 0.791172 |
| Ca5b | Q9QZA0 | 0.55 | 0.56 | -0.1 | 0.12 | 0.14 | 0.88 | 0.12 | -0.19 | 1 | -0.91 | 0.16 | 1.11 | 0.73 | 0.76 | -0.5 | -0.73 | -0.045 | 0.0575 | -0.2175 | 0.873294 | 0.901369 | 0.59915 | 1 | 0.996609 | 0.930565 |
| Ca7 | Q9ERQ8 | 1.19 | 0.8 | 0.96 | 1.11 | 0.94 | 0.44 | -0.05 | 1.18 | 1.2 | 0.75 | 0.45 | 0.7 | 0.06 | 1.69 | 0.05 | 0.14 | -0.3875 | -0.24 | -0.53 | 0.201171 | 0.286315 | 0.204793 | 1 | 0.996511 | 0.783452 |
| Ca8 | P28651 | -0.67 | -0.45 | 0.15 | -0.12 | 0.35 | -0.53 | 0.09 | 0.25 | 0.08 | 0.35 | -0.17 | -0.53 | -0.02 | 0.7 | -0.03 | 0.32 | 0.3125 | 0.205 | 0.515 | 0.270143 | 0.462111 | 0.086564 | 1 | 0.996511 | 0.691619 |
| Caap1 | Q8VDY9 | -2.31 | -2.3 | 0.84 | 2.96 | -3.25 | -2.75 | -3.43 | 1.6 | -0.85 | 0.54 | 0.46 | -1.61 | -2.48 | -1.22 | -0.74 | 0.24 | -1.755 | -0.1625 | -0.8475 | 0.280879 | 0.896072 | 0.504368 | 1 | 0.996511 | 0.901002 |
| Cab39 | Q06138 | 0.43 | -0.18 | 0.36 | 0.5 | 0.06 | 0.51 | 0.06 | 0.92 | -0.24 | -0.22 | 0.33 | 0.32 | -0.47 | -0.26 | 0.25 | -0.04 | 0.11 | -0.23 | -0.4075 | 0.680124 | 0.364864 | 0.132089 | 1 | 0.996511 | 0.726871 |
| Cab39l | Q9DB16 | -0.29 | -0.14 | -1.05 | -1.11 | -0.59 | -0.73 | -1.43 | -1.21 | -1.24 | -0.71 | -1.39 | -0.5 | -0.9 | -0.32 | -1.4 | -0.97 | -0.3425 | -0.3125 | -0.25 | 0.292947 | 0.352982 | 0.465372 | 1 | 0.996511 | 0.888369 |
| Cabin1 | G3X8Q1 | -0.05 | 0.16 | -0.54 | -0.17 | -0.78 | 0.19 | -1.41 | -0.67 | 0.86 | -0.19 | -0.27 | 0.2 | 1.81 | -0.96 | -0.12 | -0.96 | -0.5175 | 0.3 | 0.0925 | 0.161733 | 0.334842 | 0.879924 | 1 | 0.996511 | 0.979626 |
| Cables1 | Q3U148 | 1.3 | 0.93 | 0.61 | 0.74 | 1.28 | 0.82 | 0.31 | 0.5 | 0.93 | 0.77 | 0.45 | 1.29 | 0.43 | 0.5 | 0.19 | 0.59 | -0.1675 | -0.035 | -0.4675 | 0.534613 | 0.889726 | 0.060817 | 1 | 0.996511 | 0.661738 |
| Cables2 | Q8K3M5 | -0.23 | -0.27 | -0.06 | 0.18 | -0.05 | -0.26 | 1.3 | 0.11 | -0.66 | 0.11 | 0.36 | 0 | -0.41 | -0.64 | 1.13 | 0.7 | 0.37 | 0.0475 | 0.29 | 0.307171 | 0.854962 | 0.495883 | 1 | 0.996511 | 0.89758 |
| Cabp1 | D3Z1M4 | -1.82 | -1.08 | -0.97 | -2.1 | -0.59 | -0.99 | -1.05 | -1.44 | 0.72 | -1.29 | -0.8 | -1.34 | 1.81 | -0.88 | -0.24 | -1.35 | 0.475 | 0.815 | 1.3275 | 0.163229 | 0.140544 | 0.078176 | 1 | 0.996511 | 0.677244 |
| Cacfd1 | A6PWX1 | -0.04 | 0.33 | 0.37 | 0.21 | 1.31 | 1.61 | 0.79 | 0.87 | 0.53 | 0.28 | 0.13 | 0.87 | 1.36 | -0.17 | 0.4 | 1.1 | 0.9275 | 0.235 | 0.455 | 0.003191 | 0.306432 | 0.221263 | 1 | 0.996511 | 0.787447 |
| Cachd1 | Z4YK18 | -0.69 | 0.46 | 0.09 | 0.04 | 0.82 | -0.3 | 0.79 | 0.27 | -0.23 | 0.48 | 0.19 | -0.8 | -0.06 | 1 | 0.36 | 0.89 | 0.42 | -0.065 | 0.5725 | 0.243023 | 0.856096 | 0.122761 | 1 | 0.996511 | 0.721448 |
| Cacna1a | A0A571BE | 0.8 | 0.9 | 1.02 | 0.84 | 0.83 | 1.02 | 0.58 | 0.73 | 0.79 | 0.32 | 0.93 | 0.66 | 1.08 | 0.26 | 0.71 | 0.77 | -0.1 | -0.215 | -0.185 | 0.544357 | 0.283815 | 0.418254 | 1 | 0.996511 | 0.87558 |
| Cacna1a | A0A571BE | -0.49 | -0.68 | 1.08 | 1.49 | 0.13 | -1.23 | 1.1 | 0.89 | -1.11 | 0.48 | 1.6 | -0.77 | -2.22 | 0 | 1.15 | 1.96 | -0.1275 | -0.3 | -0.1275 | 0.853162 | 0.690123 | 0.893165 | 1 | 0.996511 | 0.982875 |
| Cacna1b | O55017-2 | 1.23 | 1.2 | 1.52 | 1.39 | 1.1 | 1.37 | 1.3 | 1.54 | 1.24 | 1.23 | 1.65 | 1.31 | 1.47 | 0.57 | 1.92 | 1.51 | -0.0075 | 0.0225 | 0.0325 | 0.964829 | 0.903261 | 0.915192 | 1 | 0.996609 | 0.986556 |
| Cacna1c | A0A087WF | 1.67 | 1.69 | 0.97 | 0.89 | 1.63 | 1.43 | 0.78 | 0.72 | 1.41 | 0.36 | 0.99 | 1.44 | 1.68 | 1.68 | 0.48 | 0.64 | -0.165 | -0.255 | -0.185 | 0.598316 | 0.448154 | 0.629048 | 1 | 0.996511 | 0.940595 |
| Cacna1d | A0A589Q4 | 0.51 | 0.2 | 0.58 | 1.18 | 0.34 | 0.34 | 0.1 | 0.23 | 0.49 | 0.77 | 0.68 | 0.73 | 0.03 | 1.07 | 0.43 | 0.5 | -0.365 | 0.05 | -0.11 | 0.141588 | 0.835757 | 0.720957 | 1 | 0.996511 | 0.958398 |
| Cacna1e | A0A087WS | 0.99 | 1.21 | 1.01 | 0.83 | 1.05 | 0.86 | 0.85 | 0.75 | 0.68 | 0.69 | 1 | 0.92 | 0.77 | 0.84 | 0.86 | 0.55 | -0.1325 | -0.1875 | -0.255 | 0.420361 | 0.311492 | 0.195558 | 1 | 0.996511 | 0.78004 |
| Cacna1e | A0A571BF | 0.72 | 0.44 | 1.79 | 1.54 | -0.35 | 0.23 | 0.48 | 0.97 | 0.88 | 1.56 | 0.88 | 0.34 | 1.7 | -1.85 | 2.01 | 1.39 | -0.79 | -0.2075 | -0.31 | 0.075048 | 0.598853 | 0.718168 | 1 | 0.996511 | 0.958398 |
| Cacna1g | Q5SUG2 | 0.25 | 0.31 | 0.52 | 1.08 | 0.08 | 1.66 | 0.24 | 0.77 | 1.83 | 0.22 | 0.22 | 0.1 | 3.2 | 0.34 | 1.02 | 0.8 | 0.1475 | 0.0525 | 0.8 | 0.700805 | 0.902786 | 0.210271 | 1 | 0.996609 | 0.786218 |
| Cacna1h | E9Q6P9 | -0.24 | -0.36 | 0.06 | 0.82 | -0.07 | -0.31 | -0.33 | 0.63 | -0.27 | 2.62 | 0.32 | -0.09 | 1.29 | -0.26 | 1.3 | 0.12 | -0.09 | 0.575 | 0.5425 | 0.789334 | 0.392249 | 0.251921 | 1 | 0.996511 | 0.80892 |
| Cacna1i | E9Q7P2 | 0.41 | 0.97 | 0.95 | 0.87 | 0.58 | 0.72 | 0.07 | 0.4 | 1.48 | 0.71 | 0.88 | 0.73 | 1.57 | 0.75 | 0.62 | 0.35 | -0.3575 | 0.15 | 0.0225 | 0.127136 | 0.549206 | 0.941179 | 1 | 0.996511 | 0.991085 |
| Cacna2d1 | O08532-5 | 1.12 | 1.44 | 1.24 | 1.28 | 1.31 | 0.96 | 1.3 | 1.14 | 0.75 | 0.8 | 1.22 | 1.25 | 0.24 | 1.22 | 0.81 | 0.81 | -0.0925 | -0.265 | -0.5 | 0.57608 | 0.205882 | 0.069191 | 1 | 0.996511 | 0.667373 |
| Cacna2d2 | E9Q683 | 1.29 | 0.84 | 0.67 | 1.11 | 0.8 | 0.85 | 1.27 | 0.8 | 0.17 | 0.44 | 0.49 | 1.29 | -0.37 | -0.21 | 1.13 | 0.96 | -0.0475 | -0.38 | -0.6 | 0.819457 | 0.209212 | 0.157233 | 1 | 0.996511 | 0.751678 |
| Cacna2d3 | Q9Z1L5 | 0.67 | 0.6 | 1.19 | 0.92 | 0.49 | 0.71 | 1.75 | 1.14 | 0.25 | 0.14 | 1.17 | 0.79 | -0.02 | 0.1 | 1.06 | 1.24 | 0.1775 | -0.2575 | -0.25 | 0.563975 | 0.378893 | 0.479372 | 1 | 0.996511 | 0.893468 |
| Cacnb1 | Q8R3Z5-3 | -0.1 | -0.76 | -0.02 | 0.76 | 0.32 | 0.17 | -0.63 | 0.5 | 1.19 | 1.99 | 0.44 | 0.14 | 1.66 | 0.93 | 0.65 | 0.67 | 0.12 | 0.97 | 1.0075 | 0.751877 | 0.071502 | 0.02511 | 1 | 0.996511 | 0.633075 |
| Cacnb1 | A2A543 | 1.34 | 1.21 | 1.34 | 1.62 | 1.45 | 1.27 | 1.87 | 1.84 | 1.01 | 1.3 | 1.89 | 1.57 | 1.05 | 0.59 | 1.79 | 1.71 | 0.23 | 0.065 | -0.0925 | 0.274646 | 0.783412 | 0.764366 | 1 | 0.996511 | 0.964134 |
| Cacnb2 | Q8CC27 | 0.83 | 0.63 | 0.75 | 1.6 | 0.67 | 0.68 | 0.58 | 1.16 | 0.64 | 0.9 | 0.64 | 0.61 | 0.59 | 0.06 | 1.21 | 1.25 | -0.18 | -0.255 | -0.175 | 0.500552 | 0.326753 | 0.623898 | 1 | 0.996511 | 0.938628 |
| Cacnb3 | P54285 | 1.37 | 0.9 | 0.62 | 0.7 | 1.08 | 1.1 | 0.41 | 0.52 | 0.76 | 0.37 | 0.7 | 1.33 | 0.71 | 0.95 | 0.64 | 0.41 | -0.12 | -0.1075 | -0.22 | 0.642967 | 0.696717 | 0.371667 | 1 | 0.996511 | 0.859424 |
| Cacnb4 | Q8R0S4 | 0.87 | 0.56 | 1.36 | 1.4 | 0.52 | 0.81 | 1.38 | 1.38 | 0.89 | 0.9 | 1.5 | 0.68 | 0.91 | 0.6 | 1.28 | 1.29 | -0.025 | -0.055 | -0.0275 | 0.932219 | 0.84459 | 0.922088 | 1 | 0.996511 | 0.987447 |
| Cacng2 | O88602 | 1.42 | 1.08 | 1.36 | 1.63 | 1.36 | 1.33 | 1.2 | 1.75 | 0.39 | 0.93 | 1.91 | 1.29 | 0.43 | 0.31 | 0.91 | 0.87 | 0.0375 | -0.2425 | -0.7425 | 0.849855 | 0.476121 | 0.010544 | 1 | 0.996511 | 0.563386 |
| Cacng3 | Q9JJV5 | 2.66 | 0.17 | 4.07 | 1.15 | 0.72 | 2.57 | 4.84 | 3.15 | -0.46 | 0.62 | 4.12 | 1.76 | -1.02 | -1.36 | 4.47 | 4.94 | 0.8075 | -0.5025 | -0.255 | 0.462339 | 0.668084 | 0.880202 | 1 | 0.996511 | 0.979626 |
| Cacng4 | Q9JJV4 | 0.79 | 0.35 | 0.7 | 1.1 | -0.25 | -0.75 | 1.14 | 0.84 | 0.09 | 0.48 | 1.03 | 0.46 | -0.24 | -0.38 | 0.79 | 1.15 | -0.49 | -0.22 | -0.405 | 0.283603 | 0.416022 | 0.319621 | 1 | 0.996511 | 0.838969 |
| Cacng7 | P62956 | -0.35 | 0.01 | 0.84 | 0.3 | 0.94 | 0.98 | 1.21 | 0.31 | -0.58 | -0.57 | 1.16 | 0.15 | -0.32 | -0.18 | 0.81 | 0.87 | 0.66 | -0.16 | 0.095 | 0.059823 | 0.724429 | 0.808254 | 1 | 0.996511 | 0.971953 |
| Cacng8 | S4R2F1 | 0.89 | 0.26 | 0.66 | 1.3 | 0.16 | 0.99 | 0.79 | 1.09 | 1.01 | 0.49 | 0.76 | 1 | 1.64 | -0.21 | 1.27 | 0.95 | -0.02 | 0.0375 | 0.135 | 0.946663 | 0.888068 | 0.755557 | 1 | 0.996511 | 0.962333 |
| Cactin | Q9CS00 | -0.7 | -0.32 | 1.05 | 1.61 | -0.28 | -0.03 | 0.4 | 2 | -0.49 | 0.2 | 1.41 | -0.73 | -1.2 | -0.22 | 0.57 | 1.7 | 0.1125 | -0.3125 | -0.1975 | 0.86869 | 0.640117 | 0.791043 | 1 | 0.996511 | 0.969105 |
| Cacul1 | E9PV69 | -0.15 | -0.55 | -2.64 | -1.76 | -0.98 | -0.51 | -3.32 | -1.88 | -0.73 | 0.15 | -0.92 | -0.11 | -0.29 | -0.96 | -1.29 | -2.73 | -0.3975 | 0.8725 | -0.0425 | 0.603353 | 0.154001 | 0.951289 | 1 | 0.996511 | 0.992476 |
| Cacybp | Q9CXW3 | -0.11 | -0.4 | -1.98 | -2.09 | -0.06 | -0.44 | -1.59 | -1.83 | -1.59 | -2.04 | -1.76 | -0.1 | -1.27 | -1.18 | -2.02 | -1.99 | 0.165 | -0.2275 | -0.47 | 0.788332 | 0.713802 | 0.381041 | 1 | 0.996511 | 0.860984 |
| Cad | E9QAI5 | 0.34 | 0.01 | -0.33 | -0.24 | 0.32 | 0.36 | 0.05 | -0.18 | 0.31 | -0.03 | -0.23 | 0.38 | -0.06 | 0.68 | -0.25 | -0.13 | 0.1925 | 0.1625 | 0.115 | 0.391782 | 0.498464 | 0.681722 | 1 | 0.996511 | 0.955729 |
| Cadm1 | E0CY16 | -0.06 | 0.49 | 0.59 | 0.65 | 0.17 | 0.8 | 1.29 | 0.8 | 0.63 | 0.2 | 0.25 | -0.39 | -0.33 | 0.65 | -0.12 | -0.55 | 0.3475 | -0.245 | -0.505 | 0.240595 | 0.390223 | 0.136033 | 1 | 0.996511 | 0.731656 |
| Cadm1 | Q8R5M8-4 | 0.72 | 0.53 | -0.38 | -0.13 | 0.06 | 0.68 | -0.2 | -0.12 | 0.53 | 0.28 | -0.48 | 0.84 | -0.02 | 0.07 | -0.37 | -0.61 | -0.08 | 0.1075 | -0.4175 | 0.803691 | 0.77318 | 0.206378 | 1 | 0.996511 | 0.785338 |
| Cadm2 | G3UZM4 | 0.91 | 0.62 | 1.35 | 1.76 | 0.25 | 1.05 | 0.3 | 1.26 | 1.22 | 1.77 | 0.93 | 1.03 | 0.73 | 0.43 | 0.25 | 0.63 | -0.445 | 0.0775 | -0.65 | 0.220487 | 0.805403 | 0.047887 | 1 | 0.996511 | 0.658454 |
| Cadm3 | K4DI58 | 0.68 | 0.54 | 0.82 | 1.05 | 0.02 | 1.22 | 0.4 | 0.59 | 0.95 | 0.98 | 0.79 | 0.83 | 1.5 | 0.16 | 1.06 | 0.83 | -0.215 | 0.115 | 0.115 | 0.445819 | 0.532598 | 0.71129 | 1 | 0.996511 | 0.957874 |
| Cadm4 | Q8R464 | 0.86 | 0.95 | 0.49 | 0.79 | 1.11 | 1.11 | 0.42 | 0.48 | 0.74 | 1.04 | 0.53 | 1.21 | 0.77 | 0.87 | 1.05 | 0.8 | 0.0075 | 0.1075 | 0.1 | 0.974345 | 0.626283 | 0.606048 | 1 | 0.996511 | 0.932425 |
| Cadps | Q80TJ1 | 0.63 | 0.35 | 0.56 | 0.73 | 0.13 | 0.82 | 0.72 | 0.66 | 0.06 | 0.61 | 0.54 | 0.37 | -0.19 | 0.1 | 0.16 | 0.31 | 0.015 | -0.1725 | -0.4725 | 0.941534 | 0.393652 | 0.039448 | 1 | 0.996511 | 0.658454 |
| Cadps | Q80TJ1-4 | -0.57 | 0.55 | 0.4 | -0.12 | 0.87 | -0.54 | -0.79 | -0.09 | 0.1 | -1.01 | -0.57 | -0.65 | -1.43 | -0.14 | -1.62 | -0.54 | -0.2025 | -0.5975 | -0.9975 | 0.630743 | 0.106898 | 0.038311 | 1 | 0.996511 | 0.658454 |
| Cadps | Q80TJ1-2 | 0.84 | 1.05 | 1.02 | 1.06 | 1.11 | 0.95 | 1.34 | 1.19 | 1.01 | 1.36 | 1.09 | 0.47 | 0.23 | 1.25 | 0.55 | 0.86 | 0.155 | -0.01 | -0.27 | 0.344388 | 0.964845 | 0.305667 | 1 | 0.996755 | 0.832533 |
| Cadps | A0A286YE | 0.25 | -0.3 | 0.7 | 1.43 | -0.23 | -0.14 | 1.49 | 0.68 | 0.67 | -0.34 | 0.79 | 0.46 | 0.15 | 0.53 | 0.74 | -0.07 | -0.07 | -0.125 | -0.1825 | 0.888981 | 0.76772 | 0.646052 | 1 | 0.996511 | 0.944749 |
| Cadps2 | E9Q835 | 1.19 | 1.04 | 1.41 | 1.37 | 0.66 | 1.18 | 0.99 | 1.08 | 0.69 | 0.62 | 1.09 | 0.73 | 0.5 | 0.76 | 0.65 | 0.91 | -0.275 | -0.47 | -0.5475 | 0.160649 | 0.034432 | 0.019041 | 1 | 0.996511 | 0.604671 |
| Cadps2 | Q8BYR5-4 | -0.89 | -0.12 | -0.92 | -1.75 | -1.52 | -0.38 | 0.78 | 0.13 | -0.4 | -0.83 | -0.76 | -0.45 | -1.26 | -0.34 | -0.37 | 0.24 | 0.6725 | 0.31 | 0.4875 | 0.234509 | 0.379414 | 0.277232 | 1 | 0.996511 | 0.822084 |
| Calb1 | P12658 | -1.21 | -0.68 | -0.63 | -1.17 | -0.28 | -0.85 | -0.17 | -0.74 | -1.93 | -1.31 | -0.58 | -0.79 | -1.41 | -1.03 | -0.59 | -0.39 | 0.4125 | -0.23 | 0.0675 | 0.116954 | 0.498561 | 0.817229 | 1 | 0.996511 | 0.972768 |
| Calb2 | Q08331 | 1.06 | 0.83 | 0.01 | -0.37 | 0.67 | 0.78 | 0.91 | 0.06 | -0.25 | -0.86 | -0.05 | 1.59 | -0.12 | -0.54 | 1.06 | 1.11 | 0.2225 | -0.275 | -0.005 | 0.549953 | 0.633144 | 0.992017 | 1 | 0.996511 | 0.996154 |
| Calcoco1 | Q8CGU1 | 0.91 | 0.55 | 1.18 | 1.41 | 0.06 | 0.63 | 0.64 | 0.8 | 1.03 | 1.24 | 0.99 | 0.4 | 1.07 | 0.26 | 0.67 | 0.68 | -0.48 | -0.0975 | -0.3425 | 0.088482 | 0.721635 | 0.228835 | 1 | 0.996511 | 0.787447 |
| Cald1 | Q8VCQ8 | -2.91 | -4.8 | -2.78 | -3.17 | -4.6 | -3.87 | -4.7 | -4.26 | -3.4 | -2.05 | -2.41 | -3.87 | -2.86 | -4.35 | -2.98 | -4.29 | -0.9425 | 0.4825 | -0.205 | 0.070983 | 0.415293 | 0.719948 | 1 | 0.996511 | 0.958398 |
| Cald1 | E9QA15 | -2.18 | -3.07 | -3.12 | -2.98 | -3.17 | -2.72 | -4.26 | -3.86 | -2.43 | -1.84 | -2.45 | -2.48 | -1.85 | -3.14 | -3.33 | -3.43 | -0.665 | 0.5375 | -0.1 | 0.114037 | 0.083178 | 0.808051 | 1 | 0.996511 | 0.971953 |
| Calhm2 | D3Z2E8 | -0.19 | 0.15 | 0.64 | 0.82 | 0.76 | 0.32 | 0.04 | 0.51 | -0.1 | 0.48 | 0.54 | 0.12 | -0.83 | 0.44 | -0.59 | 0.14 | 0.0525 | -0.095 | -0.565 | 0.850845 | 0.740358 | 0.152659 | 1 | 0.996511 | 0.750422 |
| Calhm5 | Q8R100 | -0.48 | 0.39 | 0.07 | -0.6 | 1.72 | 0.43 | 0.87 | 0.54 | -0.35 | 0.18 | -0.46 | -0.71 | -0.62 | 0.87 | -0.54 | 0.6 | 1.045 | -0.18 | 0.2325 | 0.016482 | 0.557534 | 0.589889 | 1 | 0.996511 | 0.929142 |
| Calm3 | P0DP28 | 2.67 | 3.48 | 3.05 | 2.12 | 4.03 | 3.89 | 2.9 | 2.53 | 0.75 | 0.22 | 3.16 | 2.85 | 1.44 | 1.53 | 2.25 | 3.64 | 0.5075 | -1.085 | -0.615 | 0.264443 | 0.157724 | 0.273371 | 1 | 0.996511 | 0.822084 |
| Calr | P14211 | 1.46 | 0.19 | 0.67 | -0.58 | 1.12 | 2 | 2.37 | 0.98 | -0.18 | -1.14 | 0.85 | 1.61 | 0.92 | -0.64 | 2.71 | 2.65 | 1.1825 | -0.15 | 0.975 | 0.040895 | 0.823277 | 0.251976 | 1 | 0.996511 | 0.80892 |
| Calu | Q6XLQ8 | 0.43 | 1.09 | 0.61 | 0.3 | 1.66 | 0.38 | 1.52 | 0.64 | -0.1 | 0.03 | 0.35 | 0.85 | 0.03 | 1.43 | 1.36 | 1.27 | 0.4425 | -0.325 | 0.415 | 0.225273 | 0.2709 | 0.276861 | 1 | 0.996511 | 0.822084 |
| Calu | O35887 | 2.27 | 2.29 | 1.12 | 0.25 | 3.1 | 2.29 | 2.34 | 0.96 | -0.01 | 0.68 | 0.71 | 2.64 | 0.14 | 1.29 | 1.87 | 1.84 | 0.69 | -0.4775 | -0.1975 | 0.272831 | 0.492515 | 0.736608 | 1 | 0.996511 | 0.958635 |
| Caly | Q9DCA7 | -2.76 | -4.16 | -2.05 | -2.27 | -3.78 | -1.5 | -2.42 | -2.72 | -0.44 | -2.23 | -2 | -2.99 | 1.03 | -4.46 | -2.06 | -1.72 | 0.205 | 0.895 | 1.0075 | 0.736688 | 0.193796 | 0.366879 | 1 | 0.996511 | 0.858904 |
| Camk1 | Q91YS8 | 0.63 | 0.84 | 1.38 | 0.83 | 0.71 | 0.86 | 1.54 | 1.24 | 1.05 | 0.55 | 1.62 | 0.88 | 0.04 | 1.21 | 1.57 | 1.5 | 0.1675 | 0.105 | 0.16 | 0.519562 | 0.714276 | 0.674994 | 1 | 0.996511 | 0.952944 |
| Camk1d | Q8BW96 | 0.98 | 0.99 | 0.27 | 0.29 | 0.87 | 0.8 | 0.62 | 0.12 | 0.54 | 0.19 | 0.16 | 0.81 | 0.34 | 0.58 | 0.09 | 0.1 | -0.03 | -0.2075 | -0.355 | 0.911602 | 0.451582 | 0.199042 | 1 | 0.996511 | 0.78004 |
| Camk1d | Q8BW96-2 | -1.3 | -0.86 | -1.59 | -1.74 | 0.88 | -0.63 | -0.1 | -1.4 | -0.82 | -3.23 | -1.89 | -1.66 | -2.38 | -1.66 | -2.54 | -1.12 | 1.06 | -0.5275 | -0.5525 | 0.051018 | 0.305899 | 0.164634 | 1 | 0.996511 | 0.758837 |
| Camk1g | F6YRI7 | 0.28 | 0.51 | 0.64 | 1.19 | 0.49 | 0.35 | 1.07 | 0.99 | 0.73 | 0.68 | 1.26 | 0.49 | 0.12 | 1.48 | 0.32 | 0.6 | 0.07 | 0.135 | -0.025 | 0.795258 | 0.619209 | 0.943562 | 1 | 0.996511 | 0.991609 |
| **Camk2a** | **P11798** | **0.73** | **1.15** | **0.49** | **1.72** | **0.22** | **0.35** | **-0.26** | **2.14** | **0.72** | **2.22** | **0.85** | **1.67** | **1.58** | **-0.09** | **0.26** | **0.37** | -0.41 | 0.3425 | -0.4925 | 0.457048 | 0.426233 | 0.270984 | 1 | 0.996511 | 0.822084 |
| Camk2a | F8WIS9 | 0.19 | 0.2 | -0.39 | -0.31 | 0.23 | 0.12 | -0.47 | -0.49 | 0.39 | -0.26 | -0.72 | 0.46 | 0.64 | 0.32 | -0.59 | -0.35 | -0.075 | 0.045 | 0.0825 | 0.771026 | 0.888815 | 0.802907 | 1 | 0.996511 | 0.971091 |
| Camk2b | P28652 | 0.05 | 0.65 | 0.62 | 0.35 | 0.41 | 0.12 | 0.49 | 0.27 | 0.46 | 0.43 | 0.52 | 0.58 | 1.08 | 0.82 | 0.39 | 0.79 | -0.095 | 0.08 | 0.3525 | 0.631185 | 0.684571 | 0.165438 | 1 | 0.996511 | 0.758837 |
| Camk2b | Q5SVI1 | 0.91 | 1.53 | 1.07 | 1.21 | 1.2 | 0.12 | 1.48 | 0.38 | 0.76 | 1.39 | 1.1 | 0.95 | 0.97 | 1.22 | 1.33 | 0.62 | -0.385 | -0.13 | -0.145 | 0.273651 | 0.563628 | 0.554337 | 1 | 0.996511 | 0.916796 |
| Camk2d | E9Q1T1 | 0.32 | 0.74 | -0.22 | 0 | 0.69 | 0.32 | 0.21 | 0.24 | 0.5 | 0.36 | 0.03 | 0.78 | 0.96 | 0.55 | 0.52 | 0.39 | 0.155 | 0.2075 | 0.395 | 0.537716 | 0.45722 | 0.165418 | 1 | 0.996511 | 0.758837 |
| Camk2d | F6RWZ9 | -0.95 | 1.1 | -0.71 | 0.63 | 0.86 | -0.45 | 0.49 | 0.42 | 1.17 | 0.62 | -0.5 | 0.91 | 1.21 | 0.64 | -0.36 | 0.94 | 0.3125 | 0.5325 | 0.59 | 0.556716 | 0.363943 | 0.307473 | 1 | 0.996511 | 0.832533 |
| Camk2d | A0A0G2JG | -6.22 | -5.38 | -7.58 | -5.93 | -5.78 | -5.34 | -5.87 | -6.56 | -3.76 | -7.3 | -6.17 | -5.83 | -5.4 | -6.13 | -5.47 | -5.51 | 0.39 | 0.5125 | 0.65 | 0.435685 | 0.522001 | 0.188444 | 1 | 0.996511 | 0.776843 |
| Camk2g | A0A286YC | -2.83 | -1.57 | -1.57 | -1.33 | -1.14 | -3.2 | -1.99 | -2.44 | -1.31 | -1.34 | -1.88 | -2.45 | -0.88 | 2.42 | -2.62 | -1.69 | -0.3675 | 0.08 | 1.1325 | 0.474226 | 0.846193 | 0.286945 | 1 | 0.996511 | 0.824333 |
| Camk2g | A0A286YC | 0.43 | 0.67 | 1.72 | 0.54 | 0.69 | -0.05 | 1.05 | 0.66 | -0.47 | -2.23 | 0.91 | 0.57 | 0.2 | -0.24 | 0.95 | 1.17 | -0.2525 | -1.145 | -0.32 | 0.489355 | 0.127506 | 0.456258 | 1 | 0.996511 | 0.887169 |
| Camk2g | Q923T9-2 | -1.09 | 0.19 | -0.24 | -0.13 | -3.22 | 0.94 | -2 | -1.49 | 4.39 | -4.18 | -0.37 | -0.23 | 6.07 | -2.28 | 0.72 | -0.85 | -1.125 | 0.22 | 1.2325 | 0.196094 | 0.889474 | 0.458821 | 1 | 0.996511 | 0.887902 |
| Camk2g | A0A2I3BQ | -0.26 | 0.51 | 0.46 | 0.13 | 0.27 | -0.09 | 0.31 | -0.16 | 0.22 | -0.38 | 0.34 | 0.08 | 0.63 | -0.05 | -0.12 | -0.03 | -0.1275 | -0.145 | -0.1025 | 0.587813 | 0.577244 | 0.708308 | 1 | 0.996511 | 0.957562 |
| Camk2g | Q923T9-3 | -0.71 | 0.35 | 1.11 | 1.55 | 0.25 | -0.79 | 0.55 | 1.32 | -0.2 | 1.56 | 0.89 | 0.26 | -0.03 | 0.56 | 1.28 | 1.25 | -0.2425 | 0.0525 | 0.19 | 0.688219 | 0.927212 | 0.72632 | 1 | 0.996755 | 0.958398 |
| Camk2n1 | Q6QWF9 | 1.28 | 1.61 | 0.56 | 0.22 | 0.99 | 0.87 | 0.89 | 0.32 | 0.78 | -0.38 | 0.76 | 1.73 | 1.11 | 1.87 | 0.15 | 0.39 | -0.15 | -0.195 | -0.0375 | 0.662092 | 0.697329 | 0.936609 | 1 | 0.996511 | 0.989767 |
| Camk4 | Q8BGR3 | -0.42 | -1.28 | -0.37 | -0.69 | -1.23 | -0.74 | -0.1 | -0.91 | -1.71 | -0.82 | -0.62 | -0.41 | -1.25 | -1.95 | -0.64 | -0.79 | -0.055 | -0.2 | -0.4675 | 0.859763 | 0.568554 | 0.212251 | 1 | 0.996511 | 0.786366 |
| Camkk1 | Q8VBY2 | 0.38 | 0.41 | 0.59 | 0.72 | 0.18 | 0.06 | 0.07 | 0.53 | -0.13 | 0.66 | 0.14 | 0.22 | -0.23 | -0.3 | -0.07 | 0.19 | -0.315 | -0.3025 | -0.6275 | 0.108904 | 0.192439 | 0.01171 | 1 | 0.996511 | 0.566767 |
| Camkk2 | Q8C078 | -0.23 | -0.19 | 0.21 | 0.43 | -0.16 | -0.26 | 0.25 | 0.7 | -0.27 | 0.5 | 0.25 | 0.12 | -0.51 | 0.22 | -0.47 | 0.03 | 0.0775 | 0.095 | -0.2375 | 0.778639 | 0.704777 | 0.385848 | 1 | 0.996511 | 0.861533 |
| Camkmt | Q3U2J5 | -1.02 | -0.58 | -0.4 | -0.61 | -0.39 | -1.04 | -0.63 | -0.75 | -0.69 | -0.46 | -0.66 | -1.26 | -0.75 | -0.02 | -1.55 | -0.81 | -0.05 | -0.115 | -0.13 | 0.815592 | 0.63746 | 0.702769 | 1 | 0.996511 | 0.956859 |
| Camkv | Q3UHL1 | 0.37 | -0.42 | 0.28 | 0.58 | -0.38 | 0.59 | 0.82 | 0.73 | -0.41 | 0.19 | 0.62 | 0.29 | -0.99 | -1.14 | 0.41 | 0.21 | 0.2375 | -0.03 | -5.80E-01 | 0.491367 | 0.9225 | 0.203135 | 1 | 0.996755 | 0.782817 |
| Camlg | P49070 | 0.01 | 0.03 | 0.2 | 0.46 | 0.28 | 0.32 | 0.62 | 1.57 | 0.02 | 0.64 | 0.46 | 0.11 | 0.94 | 0.36 | 1.66 | 0.95 | 0.5225 | 0.1325 | 0.8025 | 0.122295 | 0.547174 | 0.023645 | 1 | 0.996511 | 0.624626 |
| Camsap1 | A0A0A0M | 0.44 | 0.59 | 0.45 | 0.52 | 0.33 | 0.51 | 0.28 | 0.5 | 1.07 | 0.69 | 0.5 | 0.94 | 1.03 | 0.72 | 0.23 | 0.26 | -0.095 | 0.3 | 0.06 | 0.524953 | 0.14021 | 0.799976 | 1 | 0.996511 | 0.970393 |
| Camsap2 | A0A0A6YY | -0.3 | -0.52 | -0.18 | -0.48 | -0.68 | -0.76 | -0.71 | -0.69 | -0.51 | -0.02 | -0.31 | -0.24 | -0.21 | -0.37 | -0.54 | -0.82 | -0.34 | 0.1 | -0.115 | 0.050081 | 0.597014 | 0.588077 | 1 | 0.996511 | 0.929131 |
| Camsap3 | Q80VC9-14 | 0.01 | 0.15 | 0.43 | 0.53 | 0.2 | -0.03 | 0.49 | 0.42 | 0.5 | 0.56 | 0.62 | 0.33 | 0.36 | 0.35 | 0.54 | 0.36 | -0.01 | 0.2225 | 0.1225 | 0.960277 | 0.264492 | 0.54056 | 1 | 0.996511 | 0.912212 |
| Camsap3 | Q80VC9-5 | 1.6 | 2.42 | 4.34 | 2.67 | 1.79 | 1.48 | 2.8 | 2.8 | 3.32 | 4.34 | 3.18 | 1.43 | 2.4 | 1.26 | 1.37 | 4.13 | -0.54 | 0.31 | -0.4675 | 0.386871 | 0.682853 | 0.559119 | 1 | 0.996511 | 0.91833 |
| Camta1 | A2A891-5 | -2.88 | -1.41 | -0.8 | -1.79 | -2.05 | -3.13 | -2.4 | -1.94 | -2.17 | 0.57 | -1.4 | -2 | 2.24 | -0.78 | -0.68 | -1.24 | -0.66 | 0.47 | 1.605 | 0.189108 | 0.505372 | 0.076154 | 1 | 0.996511 | 0.674504 |
| Camta2 | B0QZH6 | -1.03 | -0.17 | -0.34 | -0.35 | -0.44 | -0.86 | 0.15 | -0.33 | 0.11 | -0.32 | -0.29 | -0.13 | 0.42 | 0.33 | 0.08 | 0 | 0.1025 | 0.315 | 0.68 | 0.718638 | 0.214011 | 0.021918 | 1 | 0.996511 | 0.621926 |
| Cand1 | Q6ZQ38 | -0.5 | -1 | -0.24 | -0.2 | -0.77 | -0.63 | -0.21 | -0.2 | -1.03 | -0.03 | -0.39 | -0.61 | -1.29 | -0.81 | -0.61 | -0.32 | 0.0325 | -0.03 | -0.2725 | 0.895551 | 0.917026 | 0.36154 | 1 | 0.996755 | 0.858475 |
| Cand2 | Q6ZQ73 | 0.32 | -0.81 | -0.1 | -0.01 | 0.08 | 0.44 | 0.47 | 0.12 | -0.7 | 0.31 | 0.35 | 0.06 | -0.71 | -0.4 | 0.17 | 0.36 | 0.4275 | 0.155 | 0.005 | 0.135284 | 0.647006 | 0.988362 | 1 | 0.996511 | 0.996154 |
| Cant1 | Q8VCF1-3 | -2.01 | -2.43 | -1.2 | -1.04 | -1.79 | -2.06 | 0.07 | -1.06 | -1.33 | -0.67 | -0.73 | -1.17 | -1.5 | -0.79 | -0.79 | 0.49 | 0.46 | 0.695 | 1.0225 | 0.398022 | 0.080253 | 0.064713 | 1 | 0.996511 | 0.66433 |
| Canx | P35564 | 0.34 | 0.18 | 0.06 | 0.24 | 0.48 | 0.34 | 0.45 | 0.19 | 0.29 | 0.37 | 0.08 | 0.74 | 0.35 | 1.01 | 0.94 | 0.31 | 0.16 | 0.165 | 0.4475 | 0.31894 | 0.417601 | 0.085695 | 1 | 0.996511 | 0.691619 |
| Cap1 | P40124 | 0.95 | 0.12 | 0.45 | 0.09 | 0.19 | 0.83 | 0.49 | 0.6 | -0.31 | -0.25 | 0.37 | 0.69 | -0.44 | -0.25 | 0.31 | 0.2 | 0.125 | -0.2775 | -0.4475 | 0.62171 | 0.390846 | 0.144342 | 1 | 0.996511 | 0.743156 |
| Cap2 | Q9CYT6 | 0.95 | 0.49 | 1.06 | 0.56 | 0.87 | 0.81 | 0.83 | 0.96 | -0.18 | 0.38 | 0.92 | 0.64 | -0.33 | 0.26 | 0.25 | 0.49 | 0.1025 | -0.325 | -0.5975 | 0.584931 | 0.271215 | 0.041509 | 1 | 0.996511 | 0.658454 |
| Capg | Q99LB4 | -1.28 | -2.07 | -0.69 | -0.66 | -2.24 | -2.77 | -0.65 | -0.71 | -3.06 | -0.5 | -0.5 | -1.82 | -2.69 | -3.17 | -0.78 | -0.98 | -0.4175 | -0.295 | -0.73 | 0.476254 | 0.645444 | 0.262916 | 1 | 0.996511 | 0.81524 |
| Capn1 | O35350 | -0.21 | -0.18 | -0.16 | 0.01 | -0.03 | -0.01 | -0.24 | 0.09 | 0.28 | -0.12 | -0.29 | -0.19 | 0.8 | 0.27 | -0.61 | -0.31 | 0.0875 | 0.055 | 0.1725 | 0.574342 | 0.773536 | 0.595677 | 1 | 0.996511 | 0.929911 |
| Capn10 | Q9ESK3 | -0.37 | -0.91 | -0.49 | -0.35 | -1.53 | -1.76 | -0.21 | -0.05 | 2.12 | -0.18 | -0.48 | -0.76 | 1.38 | 0.26 | -0.49 | -0.32 | -0.3575 | 0.705 | 0.7375 | 0.415508 | 0.270147 | 0.108166 | 1 | 0.996511 | 0.706427 |
| Capn2 | O08529 | -0.14 | 0.1 | 0.04 | 0.24 | 0.01 | -0.06 | 0.1 | -0.1 | -0.16 | 0.38 | -0.07 | -0.09 | -0.35 | 0.09 | -0.29 | -0.13 | -0.0725 | -0.045 | -0.23 | 0.64525 | 0.819881 | 0.260459 | 1 | 0.996511 | 0.814069 |
| Capn5 | O08688 | 0.41 | 0.33 | 0.88 | 0.86 | 0.84 | 0.61 | 1.64 | 0.81 | -0.02 | 0.49 | 0.98 | 0.56 | 0.04 | 0.24 | 0.79 | 0.79 | 0.355 | -0.1175 | -0.155 | 0.218045 | 0.662368 | 0.564958 | 1 | 0.996511 | 0.920315 |
| Capn7 | Q9R1S8 | 0.11 | 0.18 | 0.11 | 0 | -0.38 | 0.24 | -0.48 | -0.18 | 0.24 | 0.02 | -0.15 | 0.29 | 0.09 | -0.25 | -0.61 | -0.3 | -0.3 | -7.9E-17 | -0.3675 | 0.155451 | 1 | 0.10463 | 1 | 1 | 0.702979 |
| Capns1 | O88456 | 0.24 | 0.23 | 0.61 | 0.59 | 0.05 | 0.29 | 0.72 | 0.51 | 0.48 | 0.1 | 0.56 | 0.41 | 0.36 | 0.64 | 0.03 | 0.57 | -0.025 | -0.03 | -0.0175 | 0.904113 | 0.87908 | 0.9372 | 1 | 0.996511 | 0.989767 |
| Caprin1 | Q60865 | -2.32 | -2.32 | -2.15 | -2.35 | -1.77 | -2.48 | -1.9 | -2.26 | -2.79 | -1.41 | -2.05 | -1.92 | -1.97 | -1.89 | -1.99 | -1.96 | 0.1825 | 0.2425 | 0.3325 | 0.37671 | 0.421452 | 0.074569 | 1 | 0.996511 | 0.671911 |
| Capza1 | Q5RKN9 | 0.34 | 0.49 | 0.22 | 0.56 | 0.59 | 0.58 | 0.59 | 0.55 | 0.35 | 0.79 | 0.57 | 0.65 | -0.03 | 0.63 | 0.2 | 0.51 | 0.175 | 0.1875 | -0.075 | 0.264934 | 0.320329 | 0.733142 | 1 | 0.996511 | 0.958635 |
| Capza2 | P47754 | 0.27 | -0.04 | 0.45 | 0.41 | 0.36 | 0.61 | 0.89 | 0.84 | -0.62 | -0.12 | 0.47 | 0.18 | -0.97 | -0.34 | 0.23 | 0.63 | 0.4025 | -0.295 | -0.385 | 0.068952 | 0.29582 | 0.301549 | 1 | 0.996511 | 0.830308 |
| Capzb | P47757-2 | 1.43 | 0.98 | 1.04 | 1.44 | 0.96 | 1.12 | 1.07 | 1.52 | 1.56 | 1.38 | 1.18 | 1.4 | 1.01 | 1.55 | 0.73 | 0.68 | -0.055 | 0.1575 | -0.23 | 0.788082 | 0.433409 | 0.391476 | 1 | 0.996511 | 0.865375 |

| | | | | | | | | | | | | | | | | | | | | | | | | | | | |
|---|---|---|---|---|---|---|---|---|---|---|---|---|---|---|---|---|---|---|---|---|---|---|---|---|---|---|---|
| Capzb | P47757 | 0.4 | 0.64 | 0.6 | 1.84 | 0.03 | 0.13 | 0.55 | 0.78 | 0.95 | 2.72 | 1.05 | 0.19 | 0.74 | 1.22 | 1.01 | 1.06 | -0.4975 | 0.3575 | 0.1375 | 0.187366 | 0.538928 | 0.689026 | | 1 | 0.996511 | 0.955729 |
| Card19 | Q9D1I2 | 0.07 | 0.18 | 0.75 | 1.23 | -0.59 | 0.11 | 0.98 | 0.72 | 0.45 | 0.72 | 0.67 | -0.03 | 0.93 | -0.37 | 0.58 | 1.56 | -0.2525 | -0.105 | 0.1175 | 0.546586 | 0.742999 | 0.797326 | | 1 | 0.996511 | 0.969727 |
| Carhsp1 | Q9CR86 | 0.12 | -0.24 | -0.68 | -0.27 | -0.5 | -0.73 | -0.51 | -0.72 | -0.08 | -1.82 | -0.77 | -0.12 | 0.19 | -0.21 | -0.98 | -1.13 | -0.3475 | -0.43 | -0.265 | 0.118549 | 0.316836 | 0.458134 | | 1 | 0.996511 | 0.887902 |
| Carm1 | D3YUP1 | -0.53 | -0.24 | -1.02 | -1.21 | -0.17 | -0.4 | -1.15 | -1.32 | -0.49 | -1.77 | -1.04 | -0.68 | -0.7 | -0.32 | -1.48 | -1.64 | -0.01 | -0.245 | -0.285 | 0.976846 | 0.492335 | 0.456076 | | 1 | 0.996511 | 0.887169 |
| Carmil1 | D3Z030 | 0.16 | 0.2 | 0.83 | 0.65 | 0.45 | 0.19 | 1.04 | 0.82 | 0.11 | 0.96 | 1.01 | 0.35 | 0.17 | 0.29 | 0.73 | 0.81 | 0.165 | 0.1475 | 0.04 | 0.531215 | 0.611204 | 0.877236 | | 1 | 0.996511 | 0.979626 |
| Carmil2 | S0DHL8 | 0.86 | 0.95 | 0.47 | 0.65 | 1.1 | 0.4 | 0.85 | 0.69 | 0.92 | -0.33 | 0.46 | 0.49 | 1.55 | 0.54 | 0.88 | 0.23 | 0.0275 | -0.3475 | 0.0675 | 0.895767 | 0.251534 | 0.828735 | | 1 | 0.996511 | 0.973757 |
| Carmil3 | Q3UFQ8 | 0.31 | 0.21 | -0.09 | 0.33 | -0.12 | 0.59 | 0.67 | 0.44 | 0.55 | 0.21 | 0.17 | 0.38 | 0.59 | 0.59 | 0.32 | 0.38 | 0.205 | 0.1375 | 0.28 | 0.372966 | 0.472882 | 0.172462 | | 1 | 0.996511 | 0.76392 |
| Carnmt1 | Q80UY1 | -1.79 | -2.6 | -0.3 | -0.42 | -3.39 | -1.01 | -1.39 | -0.96 | -0.06 | -0.74 | -0.19 | -0.84 | 1.73 | -1.13 | -0.78 | -0.55 | -0.41 | 0.82 | 1.095 | 0.574506 | 0.157462 | 0.1799 | | 1 | 0.996511 | 0.775542 |
| Cars1 | Q9ER72 | -0.55 | -0.97 | -0.82 | -0.8 | -1.07 | -0.75 | -0.98 | -0.57 | -1.12 | -0.83 | -0.66 | -0.45 | -1.08 | -0.71 | -1.04 | -0.98 | -0.0575 | 0.02 | -0.1675 | 0.756025 | 0.923889 | 0.397714 | | 1 | 0.996755 | 0.867334 |
| Cars2 | Q8BYM8 | 1.5 | 1.83 | 0.36 | 0.53 | 0.69 | 1.48 | 0.27 | 0.45 | 0.62 | 0.02 | 0.37 | 1.58 | 0.3 | 1.16 | -0.17 | 0.11 | -0.3325 | -0.4075 | -0.705 | 0.437392 | 0.387765 | 0.133657 | | 1 | 0.996511 | 0.729871 |
| Cartpt | P56388-2 | -1.22 | -0.78 | -1.36 | -0.5 | -0.35 | -2.13 | 0.06 | -0.71 | -0.78 | 0.34 | -1.23 | -0.63 | -0.51 | 0.17 | 1.61 | 0.86 | 0.1825 | 0.39 | 1.4975 | 0.70326 | 0.311379 | 0.01033 | | 1 | 0.996511 | 0.563386 |
| Casc3 | Q8K3W3 | -1.71 | -1.76 | -1.29 | -1.76 | -2.27 | -2.21 | -2.07 | -1.48 | -1.94 | 0.18 | -1.58 | -1.92 | -0.29 | -1.52 | -0.76 | -1.67 | -0.3775 | 0.315 | 0.57 | 0.13117 | 0.519671 | 0.124287 | | 1 | 0.996511 | 0.721448 |
| Casd1 | Q7TN73 | -0.29 | -0.59 | -0.32 | 0.12 | 0.24 | -2.74 | 0.44 | -0.82 | -0.26 | -0.73 | -0.93 | -0.93 | -0.47 | 1.27 | -0.49 | 0.41 | -0.45 | -0.4425 | 0.45 | 0.509596 | 0.09516 | 0.304486 | | 1 | 0.996511 | 0.831662 |
| Cask | B1AUH5 | 0.71 | 1.02 | 1.24 | 2.51 | 1.77 | 0.65 | 1.34 | 2.77 | 0.59 | 3.2 | 2.14 | 0.77 | 0.15 | -1.07 | 1.97 | 1.31 | 0.2625 | 0.305 | -0.78 | 0.632205 | 0.648897 | 0.285545 | | 1 | 0.996511 | 0.822669 |
| Cask | O70589 | 0.75 | 0.48 | 0.93 | 0.87 | 0.75 | 0.79 | 1.17 | 1.03 | 0.29 | 0.83 | 0.95 | 0.8 | -0.06 | 0.29 | 0.89 | 0.82 | 0.1775 | -0.04 | -0.2725 | 0.347345 | 0.853287 | 0.328886 | | 1 | 0.996511 | 0.844943 |
| Caskin1 | Q6P9K8 | 1.29 | 0.92 | 0.93 | 1.05 | 0.48 | 1.08 | 0.36 | 0.74 | 1.15 | 1.15 | 0.81 | 1.11 | 1.01 | 1.14 | 0.74 | 0.37 | -0.3825 | 0.0075 | -0.2325 | 0.095494 | 0.967218 | 0.33302 | | 1 | 0.996755 | 0.845601 |
| Caskin2 | Q8VHK1 | -0.03 | 0.46 | 0.4 | 0.39 | 0.13 | -0.06 | 0.53 | 0.35 | -0.3 | 0.99 | 0.45 | -0.06 | -0.26 | -1.02 | 0.85 | 0.52 | -0.0675 | -0.035 | -0.2825 | 0.739664 | 0.910238 | 0.501615 | | 1 | 0.996744 | 0.901002 |
| Casp1 | P29452 | -1.42 | -2.28 | -1.31 | -0.59 | -0.68 | -0.84 | -0.32 | -0.07 | -0.86 | 3.21 | -1.24 | -1.82 | -0.76 | -1.15 | 0.05 | -0.92 | 0.9225 | 1.2225 | 0.705 | 0.032423 | 0.27342 | 0.116867 | | 1 | 0.996511 | 0.718608 |
| Casp3 | P70677 | -0.7 | -1.19 | -1.67 | -1.44 | -0.7 | -1.01 | -1.41 | -1.23 | -0.45 | -0.79 | -1.52 | -0.07 | -1.01 | -0.72 | -1.61 | -1.57 | 0.1625 | 0.5425 | 0.0225 | 0.544526 | 0.159708 | 0.942178 | | 1 | 0.996511 | 0.991114 |
| Casp6 | O08738 | -2.44 | -2.35 | -1.7 | -2.02 | -0.67 | -0.88 | -1.23 | -1.9 | -1.1 | -0.62 | -1.37 | -0.79 | -3.22 | -0.42 | -1.92 | -1.72 | 0.9575 | 1.1575 | 0.3075 | 0.01312 | 0.001567 | 0.580307 | | 1 | 0.996511 | 0.927734 |
| Casp7 | P97864 | -0.85 | -1.31 | -0.64 | -0.29 | -2.35 | -0.9 | -1.26 | -0.9 | 0.6 | 0.29 | -0.83 | -0.88 | 1.63 | -2.07 | -0.75 | -1.39 | -0.58 | 0.5675 | 0.1275 | 0.156249 | 0.195096 | 0.865407 | | 1 | 0.996511 | 0.977956 |
| Casp8 | A0A087WC | -1.92 | -2.31 | -0.39 | 0.32 | -1.55 | -2.47 | -1.1 | 0.16 | -3.24 | 0.38 | -0.23 | -1.69 | -3.09 | -3.49 | -0.73 | -0.1 | -0.165 | -0.12 | -0.7775 | 0.825702 | 0.89601 | 0.417428 | | 1 | 0.996511 | 0.87558 |
| Casq2 | E9PZ67 | -1.03 | -0.83 | -0.04 | -0.73 | -0.14 | -1.13 | -0.6 | -0.39 | -0.3 | 0.91 | -0.04 | -1 | 1.11 | -0.43 | 0.09 | -0.3 | 0.0925 | 0.55 | 0.775 | 0.757438 | 0.218931 | 0.077145 | | 1 | 0.996511 | 0.676667 |
| Cast | Q8C281 | -0.85 | -0.85 | -0.43 | -0.39 | -1.37 | -0.87 | -1.27 | -0.77 | -0.89 | -0.33 | -0.87 | -0.64 | -0.66 | -0.73 | -1.06 | -1.1 | -0.44 | -0.0525 | -0.2575 | 0.070813 | 0.811296 | 0.26158 | | 1 | 0.996511 | 0.81412 |
| Castor2 | Q8CAB8 | -1.16 | -0.6 | -0.85 | -1.39 | -1.08 | -1.09 | -0.93 | -1.1 | 0.23 | -0.91 | -1.09 | -0.82 | 0.43 | 0.03 | -1.13 | -0.83 | -0.05 | 0.3525 | 0.625 | 0.809479 | 0.315124 | 0.136217 | | 1 | 0.996511 | 0.731656 |
| Cat | P24270 | 0.07 | -0.02 | 0.61 | 0.8 | 0.6 | 0.69 | 1.18 | 0.83 | 0.72 | 0.41 | 0.6 | 0.28 | 0.91 | 0.75 | 0.81 | 0.84 | 0.46 | 0.1375 | 0.4625 | 0.093463 | 0.582131 | 0.082627 | | 1 | 0.996511 | 0.68865 |
| Cav1 | P49817 | -0.42 | -0.44 | 0.08 | -0.37 | -0.3 | -0.5 | 0.08 | -0.33 | -1.14 | -0.5 | 0.51 | -0.15 | -0.18 | -1.69 | 0.21 | -0.03 | 0.025 | -0.0325 | -0.135 | 0.902636 | 0.927386 | 0.752074 | | 1 | 0.996755 | 0.961479 |
| Cavin1 | O54724 | -0.57 | -0.54 | -0.3 | -0.29 | -0.34 | -0.52 | -0.49 | -0.43 | -0.54 | -0.12 | -0.13 | -0.33 | -0.47 | -0.41 | -0.64 | -0.46 | -0.02 | 0.145 | -0.07 | 0.896865 | 0.442812 | 0.699462 | | 1 | 0.996511 | 0.955729 |
| Cavin2 | Q63918 | -0.45 | -1.26 | -0.37 | -0.11 | -0.8 | -0.73 | -0.13 | -0.7 | -0.39 | -0.36 | 0.1 | -0.76 | -0.14 | -0.89 | -0.47 | 0.09 | -0.0425 | 0.195 | 0.195 | 0.884346 | 0.531499 | 0.558879 | | 1 | 0.996511 | 0.918101 |
| Cavin3 | Q91VJ2 | -2.61 | -3.36 | -3.43 | -3.14 | -2.29 | -2.61 | -3.57 | -3.39 | -2.56 | -1.62 | -3.18 | -2.61 | -1.44 | -2.99 | -3.12 | -3.26 | 0.17 | 0.6425 | 0.4325 | 0.624924 | 0.104148 | 0.338764 | | 1 | 0.996511 | 0.847578 |
| Cbarp | Q66L44 | 0.36 | 0.59 | 0.17 | 0.03 | 0.85 | 0.76 | -0.11 | -0.09 | 2.42 | -0.06 | 0.31 | 0.79 | 3.07 | 0.93 | 1.09 | 0.28 | 0.065 | 0.5775 | 1.055 | 0.82225 | 0.283513 | 0.08943 | | 1 | 0.996511 | 0.69217 |
| Cbarp | E9Q6U0 | -1.05 | -2.27 | -1.31 | -0.73 | -1.87 | -0.01 | -0.56 | -0.26 | -1.62 | 0.21 | -0.22 | -1.09 | -1.97 | 0.17 | -0.75 | -0.3 | 0.665 | 0.66 | 0.6275 | 0.198497 | 0.204646 | 0.251977 | | 1 | 0.996511 | 0.80892 |
| Cbfa2t2 | A2AKD9 | -0.73 | -2.47 | -1.38 | -3.96 | -2.53 | -2.16 | -1.91 | -2.87 | -1.52 | -1.56 | -2.56 | -2.53 | -1.1 | -3.28 | -1.01 | -2.33 | -0.2325 | 0.0925 | 0.205 | 0.72882 | 0.893979 | 0.798441 | | 1 | 0.996511 | 0.969727 |
| Cbfb | Q08024-2 | -0.29 | -0.42 | -0.98 | -0.8 | 0.14 | -0.46 | -0.8 | -0.57 | -0.42 | -0.46 | -0.58 | -0.5 | -0.93 | 0.78 | -0.85 | -1.2 | 0.2 | 0.1325 | 0.0725 | 0.457207 | 0.529908 | 0.87231 | | 1 | 0.996511 | 0.979626 |
| Cbl | P22682 | 0.07 | -0.12 | -0.13 | 0.31 | -0.38 | -0.4 | -0.16 | -0.05 | -0.25 | 0.01 | -0.21 | -0.1 | -0.15 | -0.21 | -0.09 | -0.19 | -0.28 | -0.17 | -0.1925 | 0.145363 | 0.364009 | 0.319146 | | 1 | 0.996511 | 0.838969 |
| Cblb | B9EKI5 | 0.25 | 0.39 | -0.92 | -0.26 | 0.68 | 0.19 | -0.09 | -0.29 | 0.86 | 0.26 | -0.29 | -0.27 | 0.53 | 0.83 | -0.77 | -0.5 | 0.2575 | 0.275 | 0.1575 | 0.468637 | 0.483506 | 0.732982 | | 1 | 0.996511 | 0.958635 |
| Cbll1 | E9QM14 | -0.26 | -0.95 | -0.41 | -0.24 | -0.38 | -1.06 | -0.64 | -0.09 | -0.71 | 1.47 | -0.57 | -0.79 | 0.23 | -0.13 | 1.32 | -0.16 | -0.0775 | 0.315 | 0.78 | 0.774608 | 0.552499 | 0.063405 | | 1 | 0.996511 | 0.66433 |
| Cbln1 | Q9R171 | 0.26 | 0.63 | -0.93 | -1.03 | 1.37 | 1.01 | -0.12 | -0.73 | 0.71 | -1.17 | -0.08 | 0.11 | 0.63 | 1.38 | -0.44 | -1.09 | 0.65 | 0.16 | 0.3875 | 0.285755 | 0.76346 | 0.543407 | | 1 | 0.996511 | 0.912212 |
| Cbln4 | Q8BME9 | -0.61 | 0.83 | -0.15 | 0.26 | -0.05 | -1.24 | 0.41 | 0 | -1.06 | -0.27 | 0.12 | -0.63 | -1.15 | 1.17 | -0.05 | 0.28 | -0.3025 | -0.5425 | -0.02 | 0.495582 | 0.180168 | 0.969703 | | 1 | 0.996511 | 0.992767 |
| Cbr1 | P48758 | 0.33 | -0.35 | -0.15 | -0.53 | -0.18 | 0.1 | 0.39 | -0.04 | 0.04 | -1.65 | 0.03 | 0.29 | 0.34 | -0.55 | 0.21 | 0 | 0.2425 | -0.1475 | 0.175 | 0.323295 | 0.745974 | 0.547037 | | 1 | 0.996511 | 0.912571 |
| Cbr2 | P08074 | 0 | -0.15 | -0.38 | 0.86 | -0.83 | -0.86 | -0.58 | -0.1 | -0.43 | -0.81 | -0.09 | -0.01 | -1.12 | -0.97 | -0.39 | -0.69 | -0.675 | -0.4175 | -0.875 | 0.058377 | 0.221559 | 0.022131 | | 1 | 0.996511 | 0.621926 |
| Cbr3 | Q8K354 | 0.15 | -0.02 | -0.7 | -0.31 | 0.01 | -0.23 | -0.61 | -0.14 | 0.08 | -0.43 | -0.62 | -0.22 | -0.13 | 0.31 | -0.99 | -0.83 | -0.0225 | -0.0775 | -0.19 | 0.926111 | 0.764994 | 0.593607 | | 1 | 0.996511 | 0.929843 |
| Cbr4 | Q91VT4 | 1.6 | 0.77 | 0.71 | 0.54 | 0.34 | 1.22 | 0.82 | 0.78 | -0.06 | -0.06 | 1.14 | 1.14 | -0.06 | 0.15 | 0.31 | 0.78 | -0.115 | -0.365 | -0.61 | 0.698755 | 0.374484 | 0.073542 | | 1 | 0.996511 | 0.669435 |
| Cbs | Q91WT9-2 | 0.28 | -0.72 | -0.26 | -0.23 | -0.14 | 0.23 | -0.46 | -0.06 | -0.15 | 0.21 | 0.11 | -0.07 | -0.49 | -0.04 | -0.15 | -0.44 | 0.125 | 0.2575 | -0.0475 | 0.630456 | 0.309803 | 0.855366 | | 1 | 0.996511 | 0.976225 |
| Cbwd1 | Q8VEH6 | -0.64 | -0.44 | 0.5 | 0.92 | -0.66 | -0.72 | 0.75 | 0.77 | -0.44 | 0.69 | 0.61 | -0.05 | -0.48 | -1.39 | -0.45 | 0.28 | -0.05 | 0.1175 | -0.595 | 0.922777 | 0.787547 | 0.231195 | | 1 | 0.996511 | 0.788863 |
| Cbx1 | P83917 | 0.27 | -0.13 | -0.17 | 0.18 | 0.68 | 0.37 | 0.27 | -0.13 | 0.15 | -1.16 | -0.34 | 0.77 | -0.41 | -0.3 | -0.24 | 0.29 | 0.26 | -0.1825 | -0.2025 | 0.261705 | 0.652473 | 0.395902 | | 1 | 0.996511 | 0.867334 |
| Cbx3 | Q9DCC5 | 0.49 | -0.34 | -1.4 | -0.66 | -0.73 | -0.31 | -1.27 | -0.83 | -0.05 | -1.74 | 0.5 | -0.05 | -0.52 | 0.04 | -1.56 | -1.07 | -0.3075 | 0.1425 | -0.3 | 0.462711 | 0.80404 | 0.543247 | | 1 | 0.996511 | 0.912212 |
| Cbx5 | Q61686 | -0.05 | -0.37 | 0.09 | 0.34 | -0.35 | 0.11 | 0.31 | 0.3 | 0.05 | 0.08 | -0.14 | -0.12 | 0.1 | -0.44 | 0.08 | 0.28 | 0.09 | -0.035 | 0.0025 | 0.699703 | 0.864314 | 0.991971 | | 1 | 0.996511 | 0.996154 |
| Cbx8 | Q9QXV1 | 0.41 | -0.64 | 0.02 | 0.7 | -0.99 | -0.49 | 0.03 | -0.22 | 0.2 | 0.51 | -0.18 | -0.2 | 0.67 | 0.21 | 0.57 | 0.68 | -0.54 | -0.04 | 0.41 | 0.148501 | 0.904294 | 0.218258 | | 1 | 0.996744 | 0.787447 |
| Cby1 | Q9D1C2 | 0.56 | 0.23 | -0.33 | -0.6 | 0.06 | -0.18 | 0.1 | -0.78 | -0.14 | 0.52 | -1.29 | 0.59 | 1.78 | 0.26 | 0.02 | 0.81 | -0.165 | -0.045 | 0.7525 | 0.613238 | 0.92456 | 0.118849 | | 1 | 0.996755 | 0.721448 |
| Cc2d1a | Q8K1A6 | 0.19 | 0.45 | 0.28 | 0.56 | 0.62 | 0.58 | 0.36 | 0.76 | 0.39 | 0.84 | 0.7 | 0.42 | 0.32 | 0.73 | 0.4 | 0.56 | 0.21 | 0.2175 | 0.1325 | 0.237804 | 0.276454 | 0.502576 | | 1 | 0.996511 | 0.901002 |
| Cc2d1a | E9PX94 | 0.73 | 1.85 | -0.33 | 0.49 | 0.97 | 1.27 | -0.21 | -0.08 | 1.44 | 2.22 | 0 | 2.11 | 0.75 | 1.21 | 0.07 | -0.04 | -0.1975 | 0.7575 | -0.1875 | 0.713113 | 0.244032 | 0.709289 | | 1 | 0.996511 | 0.957656 |
| Cc2d1b | Q8BRN9 | 0.4 | 0.41 | 0.21 | 0.53 | 0.2 | 0.52 | 0.03 | 0.23 | 1.41 | 1.06 | 0.2 | 0.48 | 1.81 | -0.35 | 0.48 | 0.31 | -0.1425 | 0.4 | 0.175 | 0.417623 | 0.192621 | 0.688479 | | 1 | 0.996511 | 0.955729 |
| Cc2d2a | Q8CFW7-2 | -1 | 0.5 | 1.31 | 0.93 | -3.29 | -0.89 | -1.68 | -1.31 | 1.2 | -0.39 | -1.76 | -1.34 | 2.37 | -0.97 | 5.33 | -0.87 | -2.2275 | -1.0075 | 1.03 | 0.008233 | 0.203185 | 0.472016 | | 1 | 0.996511 | 0.891144 |
| Ccar1 | Q8CH18-3 | -1.95 | -2.39 | -2.18 | -3.06 | -1.65 | -2.42 | -2.3 | -3.26 | -2.26 | -1.03 | -2.89 | -2.1 | -1.91 | -1.68 | -0.39 | -2.72 | -0.0125 | 3.25E-01 | 0.72 | 0.974176 | 0.457266 | 0.176331 | | 1 | 0.996511 | 0.771503 |
| Ccar2 | Q8VDP4 | -1.92 | -3.29 | -2.73 | -2.68 | -3.4 | -2.99 | -3.19 | -2.75 | -2.85 | -1.35 | -2.29 | -3.31 | -2.39 | -2.71 | -2.89 | -3.26 | -0.4275 | 0.205 | -0.1575 | 0.190803 | 0.66698 | 0.641481 | | 1 | 0.996511 | 0.943421 |
| Ccdc106 | E9QPN6 | -0.5 | -0.52 | -0.49 | 0.63 | -1.7 | -0.25 | 0.57 | 0.68 | 1.3 | 1.01 | 0.1 | -1.18 | 0.15 | -0.33 | 0.33 | 1.31 | 0.045 | 0.5275 | 0.585 | 0.936497 | 0.371163 | 0.192636 | | 1 | 0.996511 | 0.777539 |
| Ccdc107 | Q9DCC3 | -0.49 | -1.02 | 0.09 | 0.58 | -0.84 | -0.83 | -0.71 | -0.51 | 2.75 | 2.66 | -1.2 | -0.31 | 2.83 | 3.39 | 1.32 | 1.45 | -0.5125 | 1.185 | 2.4575 | 0.161375 | 0.240094 | 0.001997 | | 1 | 0.996511 | 0.419079 |
| Ccdc115 | Q8VE99 | 0.61 | -0.6 | -0.04 | -0.04 | 0.16 | -0.99 | -0.17 | 0.35 | -0.08 | 0.02 | -0.49 | -0.38 | 0.2 | 0.34 | 1.48 | 0.92 | -0.145 | -0.215 | 0.7525 | 0.694659 | 0.459089 | 0.071072 | | 1 | 0.996511 | 0.668066 |
| Ccdc12 | Q8R344 | -1.72 | -2.49 | -2.05 | -1.95 | -3.76 | -1.89 | -2.65 | -1.97 | -1.82 | -1.99 | -1.91 | -2.3 | -0.03 | -2.7 | -1.3 | -2.49 | -0.515 | 0.0475 | 0.4225 | 0.252525 | 0.833731 | 0.476664 | | 1 | 0.996511 | 0.892174 |
| Ccdc120 | A2AEV7 | 0.27 | 0.42 | 0.02 | 0.51 | 1.32 | 0.86 | -0.08 | 0.58 | 0.25 | 1.02 | 0.43 | 0.7 | 0.85 | -0.69 | -0.29 | 0.37 | 0.365 | 0.295 | -0.245 | 0.255018 | 0.222942 | 0.495894 | | 1 | 0.996511 | 0.89758 |
| Ccdc124 | Q9D8X2 | -1.9 | -2.07 | -1.02 | -1.16 | -1.7 | -2.02 | -1.59 | -1.34 | -0.92 | -1.14 | -1.21 | -1.76 | -0.2 | -0.63 | -0.92 | -1.39 | -0.125 | 0.28 | 0.7525 | 0.674602 | 0.389666 | 0.060939 | | 1 | 0.996511 | 0.661738 |
| Ccdc127 | E0CYX9 | -0.16 | 0.17 | 0.4 | 0.2 | -0.02 | 0.16 | -0.11 | 0.03 | 0.1 | -0.04 | 0.21 | -0.22 | 0.34 | 0.49 | -0.09 | 0.27 | -0.1375 | -0.14 | 0.1 | 0.444367 | 0.487845 | 0.651317 | | 1 | 0.996511 | 0.947503 |
| Ccdc134 | Q8C7V8 | -0.26 | -0.16 | -0.17 | -0.06 | 0.32 | -0.46 | 0.81 | 0.32 | -0.44 | 0.51 | -0.34 | -0.07 | -0.31 | 0.69 | 0.69 | 1.12 | 0.41 | 0.0775 | 0.71 | 0.156741 | 0.750502 | 0.047187 | | 1 | 0.996511 | 0.658454 |
| Ccdc136 | D3Z165 | -0.01 | 0.58 | 0.32 | 0.28 | 0.77 | -0.1 | 0.72 | 0.27 | 0.64 | -0.14 | 0.36 | 0.12 | 0.25 | 1.48 | 0.21 | 0.65 | 0.1225 | -0.0475 | 0.355 | 0.626624 | 0.840127 | 0.289794 | | 1 | 0.996511 | 0.825111 |
| Ccdc137 | Q8R0K4 | 0.35 | -0.48 | -0.24 | 0.17 | -0.92 | 0.02 | -0.24 | 0.31 | 0.16 | 0.58 | -0.15 | -1.08 | 1.06 | -0.38 | -0.1 | -0.2 | -0.1575 | -0.0725 | 0.145 | 0.621881 | 0.850585 | 0.696239 | | 1 | 0.996511 | 0.955729 |
| Ccdc141 | E9Q8Q6 | 0.01 | 0.3 | -0.39 | -0.01 | 0.36 | -0.2 | -0.07 | -0.81 | 0.56 | 0.49 | -0.43 | 0.17 | 0.48 | 0.61 | 0.02 | -0.12 | -0.1575 | 0.22 | 0.27 | 0.580375 | 0.438737 | 0.308166 | | 1 | 0.996511 | 0.833667 |
| Ccdc149 | A0A1D5RN | -0.57 | -0.77 | -2.18 | -1.53 | -0.81 | -0.41 | -2.25 | -2.05 | 0.18 | -1.91 | -1.69 | -0.09 | 0.14 | -0.57 | -0.98 | -1.5 | -0.1175 | 0.385 | 0.535 | 0.826869 | 0.524587 | 0.277624 | | 1 | 0.996511 | 0.822084 |
| Ccdc167 | Q9D162 | 2.8 | 2.42 | 1.03 | 0.49 | 2.53 | 2.44 | 0.77 | 0.69 | 3.65 | 0.91 | 0.66 | 2.75 | 4.18 | 3.86 | 1.74 | 1.71 | -0.0775 | 0.3075 | 1.1875 | 0.909064 | 0.708901 | 0.153669 | | 1 | 0.996511 | 0.750763 |
| Ccdc177 | Q3UHB8 | 0.68 | 0.83 | 0.94 | 0.56 | 0.27 | 0.59 | 0.51 | 0.42 | 0.59 | 0.66 | 0.68 | 0.33 | 0.86 | 0.16 | 0.41 | 0.47 | -0.305 | -0.1875 | -0.2775 | 0.092447 | 0.315996 | 0.226122 | | 1 | 0.996511 | 0.787447 |
| Ccdc184 | Q8BMK5 | -0.67 | -0.21 | -0.72 | -0.41 | 0.27 | -0.31 | -1.28 | -0.87 | 0.36 | -0.64 | -0.32 | -1.45 | 0.29 | 0.27 | -0.98 | -0.11 | -0.045 | -0.01 | 0.37 | 0.896094 | 0.978979 | 0.272827 | | 1 | 0.998749 | 0.822084 |
| Ccdc186 | Q8C9S4 | -1.25 | -1.15 | -0.27 | -0.66 | -1.94 | -1.15 | -1.3 | -1 | -0.74 | -0.19 | -0.65 | -0.98 | -0.08 | -0.77 | -0.68 | -0.82 | -0.515 | 0.1925 | 0.245 | 0.118094 | 0.512504 | 0.421079 | | 1 | 0.996511 | 0.875639 |
| Ccdc22 | Q9JIG7 | -0.01 | 0.12 | 0.14 | -0.04 | 0.56 | -0.3 | 0.09 | 0.2 | 0.17 | 0 | 0.23 | 0.04 | 0.07 | 0.73 | 0.01 | 0.15 | 0.085 | 0.0575 | 0.1875 | 0.687623 | 0.72311 | 0.407644 | | 1 | 0.996511 | 0.874084 |
| Ccdc25 | Q78PG9 | -0.24 | -1.33 | -0.77 | -0.72 | -1.07 | -0.1 | -0.56 | -0.65 | -1.19 | 0.99 | -0.58 | 0.15 | -0.63 | -1.2 | -0.66 | -0.54 | 0.17 | 0.6075 | 0.0075 | 0.570999 | 0.232132 | 0.979099 | | 1 | 0.996511 | 0.994274 |
| Ccdc28a | Q3UFK1 | -0.09 | 0.38 | 0.23 | 0.48 | 0.97 | -0.43 | 0.19 | 0.42 | -0.22 | -0.53 | 0.25 | -0.12 | -2.51 | -1.49 | -1.89 | -0.02 | 0.0375 | -0.405 | -1.7275 | 0.903792 | 0.110691 | 0.007519 | | 1 | 0.996511 | 0.552614 |
| Ccdc28b | Q8CEG5 | -6.5 | -7.69 | -7.85 | -8.77 | -7.58 | -4.38 | -5.05 | -7.09 | -8.52 | -14.08 | -4.92 | -5.9 | -9.04 | -8.66 | -8.11 | -6.34 | 1.6775 | -0.6525 | -0.335 | 0.065973 | 0.728912 | 0.62845 | | 1 | 0.996511 | 0.940342 |
| Ccdc32 | Q3UHY7 | 0.63 | 0.7 | -0.79 | -0.9 | 0.42 | 0.37 | -1.13 | -1.19 | 0.55 | -0.39 | -0.61 | 1.15 | 0.9 | 2.03 | -1.49 | -1.62 | -0.2925 | 0.265 | 0.045 | 0.612164 | 0.633875 | 0.960054 | | 1 | 0.996511 | 0.992636 |
| Ccdc33 | A0A1L1SR7 | 0.14 | 0.71 | 0.33 | 0.83 | 0.19 | -0.71 | 0.88 | 0.24 | -0.69 | -0.91 | 0.04 | -0.2 | -0.59 | 0.4 | 0.42 | 0.11 | -0.3525 | -0.9425 | -0.4175 | 0.32895 | 0.008549 | 0.186594 | | 1 | 0.996511 | 0.776843 |
| Ccdc43 | Q9CR29 | -2.71 | -2.65 | -3.04 | -3.92 | -3.12 | -2.88 | -3.94 | -4 | -2.38 | -1.34 | -3.66 | -2.42 | -1.78 | -2.3 | -3.4 | -3.5 | -0.405 | 0.63 | 0.335 | 0.310755 | 0.243407 | 0.491842 | | 1 | 0.996511 | 0.897531 |
| Ccdc47 | Q9D024 | -0.19 | -0.21 | -0.07 | -0.16 | 0.38 | -0.24 | 0.79 | 0.31 | -0.33 | -0.27 | -0.23 | -0.14 | 0.27 | -0.18 | 1.06 | 1 | 0.4675 | -0.085 | 0.695 | 0.070879 | 0.588565 | 0.048645 | | 1 | 0.996511 | 0.658454 |
| Ccdc50 | A0A668KL3 | 0.13 | -0.25 | -0.29 | -0.32 | -0.14 | -0.53 | -1.02 | -0.88 | -0.06 | 0.55 | -0.49 | 0.12 | -0.19 | 0.36 | -0.4 | -0.64 | -0.46 | 0.2125 | -0.035 | 0.081517 | 0.421764 | 0.894755 | | 1 | 0.996511 | 0.983453 |
| Ccdc51 | Q3URS9-2 | 0.13 | 0.3 | 0.49 | 0.72 | -0.05 | 0.25 | -0.31 | 0.2 | 0.28 | -0.57 | 0.42 | -0.07 | 0.78 | 0.09 | -0.29 | -0.05 | -0.3875 | -0.395 | -0.2775 | 0.091531 | 0.168784 | 0.338877 | | 1 | 0.996511 | 0.847578 |
| Ccdc59 | Q32M26 | 0.02 | -0.26 | 0.94 | 1.06 | 1.42 | -0.19 | -1.24 | 0.38 | 1.09 | 0.94 | 0.47 | -0.42 | 2.09 | 0 | 1.41 | 0.17 | -0.3475 | 0.08 | 0.4775 | 0.559337 | 0.857542 | 0.398646 | | 1 | 0.996511 | 0.867334 |
| Ccdc6 | D3YZP9 | 0.38 | 0.31 | 0.11 | 0.18 | 0.09 | 0.08 | -0.05 | 0.01 | 1.15 | 0.36 | -0.24 | 0.39 | 1.25 | 0.73 | 0.01 | -0.26 | -0.2125 | 0.17 | 0.1875 | 0.176083 | 0.571678 | 0.592153 | | 1 | 0.996511 | 0.929843 |
| Ccdc85a | F6TRJ8 | -0.36 | -0.07 | 0.67 | 0.47 | -0.28 | -0.21 | -0.25 | 1.01 | -0.95 | 1.03 | 1.1 | -0.16 | 0.35 | -0.02 | 1.31 | 0.27 | -0.11 | 0.0775 | 0.3 | 0.769648 | 0.8791 | 0.42291 | | 1 | 0.996511 | 0.875782 |
| Ccdc85a | E0CX32 | -0.37 | 0.36 | 0.21 | 0.43 | -0.05 | -0.08 | -0.5 | 0.1 | 0.58 | 0.19 | 0.27 | -0.04 | 1.05 | -0.39 | 0.44 | -0.08 | -0.29 | 0.0925 | 0.0975 | 0.244688 | 0.708883 | 0.786342 | | 1 | 0.996511 | 0.967714 |
| Ccdc85b | Q6PDY0 | -1.23 | -1.55 | -1.75 | -1.64 | -1.35 | -0.92 | -1.44 | -1.72 | -1.79 | -1.2 | -1.8 | -1.38 | -1.62 | -0.96 | -2.46 | -2.02 | 0.185 | 1.96E-16 | -0.2225 | 0.415228 | 1 | 0.516137 | | 1 | 1 | 0.905674 |
| Ccdc85c | A0A217FL8 | -0.3 | -0.71 | 0.3 | 0.45 | -1.01 | -0.64 | -0.12 | 0.13 | 0.82 | 0.1 | 0.29 | -0.45 | 2.09 | -0.46 | 0.09 | 0.15 | -0.345 | 0.255 | 0.5325 | 0.346506 | 0.491068 | 0.362612 | | 1 | 0.996511 | 0.858475 |
| Ccdc88a | Q5SNZ0-2 | 0.31 | 0.49 | 0.53 | 0.5 | 0.46 | 0.43 | 0.25 | 0.38 | 0.61 | 0.84 | 0.59 | 0.7 | 0.89 | 0.81 | 0.44 | 0.09 | -0.0775 | 0.2275 | 0.1 | 0.602343 | 0.188447 | 0.669732 | | 1 | 0.996511 | 0.952558 |
| Ccdc88c | Q6VGS5 | 0.85 | -0.18 | -1.45 | -1.43 | 0.64 | 0.68 | -2.08 | -1.69 | 0.94 | -0.53 | -1.71 | 0.99 | 1.26 | 0.97 | -1.05 | -2.02 | -0.06 | 0.475 | 0.3425 | 0.942452 | 0.542793 | 0.694444 | | 1 | 0.996511 | 0.955729 |
| Ccdc9 | D3YW42 | -0.62 | -0.79 | -0.28 | -0.27 | -0.72 | -1.22 | -1.15 | -0.61 | -0.72 | 0.47 | -0.53 | -0.78 | 0.07 | -0.35 | -0.27 | -0.71 | -0.435 | 0.1 | 0.175 | 0.077329 | 0.754641 | 0.477845 | | 1 | 0.996511 | 0.892279 |
| Ccdc90b | Q8C3X2 | 0.31 | 0.66 | 0.64 | 0.67 | 0.16 | 0.44 | 0.81 | 0.44 | 0.3 | 0.47 | 0.94 | 0.12 | 0.75 | 0.21 | 0.45 | 1.1 | -0.1075 | -0.1125 | 0.0575 | 0.584526 | 0.625474 | 0.8157 | | 1 | 0.996511 | 0.972768 |
| Ccdc91 | Q9D8L5 | 0.33 | 0.45 | -0.64 | -0.81 | 0.2 | 0.2 | -0.64 | -0.68 | 0.18 | -0.75 | -0.7 | 0.19 | -0.19 | 0.2 | -0.91 | -1.12 | -0.0625 | -0.1025 | -0.3375 | 0.871707 | 0.797692 | 0.437245 | | 1 | 0.996511 | 0.883161 |
| Ccdc92 | Q8VDN4 | 0.19 | 0.22 | 0.61 | 0.3 | 0.18 | 0.03 | 0.24 | 0.48 | 0.51 | 0.78 | 0.55 | 0.06 | 0.26 | 0.72 | 0.43 | 0.27 | -0.0975 | 0.145 | 0.09 | 0.590336 | 0.510283 | 0.663526 | | 1 | 0.996511 | 0.949825 |
| Ccdc92b | Q5SUE3 | 1.11 | 2.22 | 0.84 | 1.74 | 1.97 | 1.16 | 0.2 | 0.87 | 1.06 | 1.23 | 1.02 | 1.77 | 0.95 | 1.15 | 0.87 | 0.72 | -0.4275 | -0.2075 | -0.555 | 0.352933 | 0.556123 | 0.117689 | | 1 | 0.996511 | 0.718913 |
| Ccdc93 | E9QAD4 | 1.31 | -1.01 | -1.12 | 0.35 | -0.42 | 0.43 | -1.25 | -0.92 | 0.02 | -0.57 | -0.22 | 0.61 | -0.23 | 0.39 | -1.05 | -0.8 | -0.4225 | 0.0775 | -0.305 | 0.504861 | 0.893808 | 0.618266 | | 1 | 0.996511 | 0.937583 |
| Ccdc93 | Q7TQK5 | -0.06 | 0.06 | -0.01 | 0.09 | -0.05 | -0.13 | -0.58 | -0.24 | 0.23 | 0.07 | -0.17 | -0.21 | 0.44 | -0.02 | -0.17 | -0.33 | -0.27 | -0.04 | -0.04 | 0.143929 | 0.822099 | 0.855849 | | 1 | 0.996511 | 0.976225 |
| Ccdc97 | Q9DBT3 | -1.72 | -0.94 | -1.59 | -1.73 | -1.92 | -1.87 | -1.68 | -1.81 | -1.27 | -1.49 | -1.5 | -1.45 | 0.03 | -1.11 | -2.22 | -2.09 | -0.325 | 0.0675 | 0.1475 | 0.16305 | 0.767843 | 0.774997 | | 1 | 0.996511 | 0.966519 |
| Cck | P09240 | 0.73 | 1.04 | 0.36 | 0.75 | 0.77 | 1.69 | 0.61 | 0.39 | 0.72 | 0.77 | 0.88 | 1.18 | 1.07 | 1.66 | 0.86 | 1.18 | 0.145 | 0.1675 | 0.4725 | 0.64462 | 0.442032 | 0.08801 | | 1 | 0.996511 | 0.691619 |
| Cckbr | P56481 | 1.75 | 1.57 | 2.11 | 0.36 | 2.65 | 2.22 | 0.65 | 1.58 | -0.75 | 1.28 | 1.97 | 2.71 | -0.45 | -1.15 | 1.85 | 1.6 | 0.3275 | -0.145 | -0.985 | 0.541185 | 0.848024 | 0.214238 | | 1 | 0.996511 | 0.786931 |
| Ccm2 | Q8K2Y9-2 | 0.55 | -0.36 | -0.04 | -0.01 | -0.49 | -0.08 | -0.22 | -0.21 | 0.28 | -0.17 | -0.3 | -0.61 | 0.4 | -0.27 | -0.22 | -0.43 | -0.285 | -0.235 | -0.165 | 0.232302 | 0.406855 | 0.562588 | | 1 | 0.996511 | 0.920024 |
| Ccn2 | P29268 | -0.25 | -0.44 | -1.12 | -0.29 | -0.57 | 0.3 | -0.99 | -0.83 | -0.72 | -1.17 | -0.89 | -0.05 | 0.09 | -0.78 | -0.29 | -0.69 | 0.0025 | -0.1825 | 0.1075 | 0.994125 | 0.565255 | 0.719334 | | 1 | 0.996511 | 0.958398 |
| Ccn3 | Q64299 | -3.03 | -3.98 | -3.67 | -2.64 | -3.35 | -2.25 | -2.35 | -2.49 | -2.17 | -0.63 | -3.25 | -2.14 | -2.07 | -2.61 | -2.09 | -2.33 | 0.72 | 1.2825 | 1.055 | 0.082257 | 0.047893 | 1.07E-02 | | 1 | 0.996511 | 0.563386 |
| Ccnk | Q3U3M5 | -1.45 | -2.59 | -3.04 | -2.47 | -2.59 | -2.16 | -3.51 | -2.98 | -2.05 | -2.57 | -2.55 | -2.56 | -0.96 | -1.93 | -3.22 | -3.6 | -0.4225 | -0.045 | -0.04 | 0.323384 | 0.898093 | 0.949504 | | 1 | 0.996511 | 0.992476 |
| Ccnl2 | Q9JJA7 | -0.65 | -0.13 | -1.82 | -2.72 | -0.31 | -0.07 | -1.55 | -1.68 | 0.26 | -0.97 | -1.35 | -0.15 | 0.08 | 0.19 | -1.24 | -0.66 | 0.4275 | 0.7775 | 0.9225 | 0.516049 | 0.238175 | 0.15931 | | 1 | 0.996511 | 0.754511 |
| Ccnt1 | Q9QWV9 | -1.05 | -1.03 | -6.01 | -8.37 | -0.84 | -1.65 | -6.87 | -6.88 | -0.76 | -5.13 | -6.73 | -1.26 | -1.92 | -0.14 | -6.15 | -6.11 | 0.055 | 0.645 | 0.535 | 0.979957 | 0.758373 | 0.800103 | | 1 | 0.996511 | 0.970393 |
| Ccnt2 | Q7TQK0-2 | -3.79 | -5.6 | -4.21 | -4.55 | -6.57 | -4.54 | -4.54 | -4.37 | -4.13 | -0.16 | -3.93 | -3.14 | -2.54 | -4.85 | -2.88 | -4.12 | -0.4675 | 1.6975 | 0.94 | 0.439477 | 0.086288 | 0.147594 | | 1 | 0.996511 | 0.746406 |
| Ccny | Q8BGU5 | 0.48 | 0.74 | 0.7 | 0.84 | 0.66 | 0.27 | 0.61 | 0.63 | 0.48 | 0.84 | 0.71 | 0.39 | 0.37 | 1.07 | 0.52 | 0.57 | -0.1475 | -0.085 | -0.0575 | 0.398619 | 0.652076 | 0.794487 | | 1 | 0.996511 | 0.969367 |
| Ccnyl1 | D3YUJ3 | 1.44 | 1.47 | 0.91 | 0.71 | 1.69 | 0.94 | 0.9 | 0.69 | 1.83 | 0.97 | 0.91 | 1.71 | 1.46 | 2.04 | 0.99 | 0.95 | -0.0775 | 0.2225 | 0.2275 | 0.789768 | 0.480117 | 0.488067 | | 1 | 0.996511 | 0.897531 |
| Ccp110 | Q7TSH4 | -0.46 | 0.12 | 1.08 | 0.49 | -0.77 | 0.09 | 0.32 | 0.41 | 0.79 | 1.2 | 0.67 | 0.07 | 2.01 | -0.74 | 0.77 | 0.85 | -0.295 | 0.375 | 0.415 | 0.464979 | 0.342388 | 0.492278 | | 1 | 0.996511 | 0.897531 |
| Ccs | A0A494BB! | 0.2 | -0.09 | -0.52 | -0.27 | -0.55 | 0.36 | -0.59 | 0.11 | 0.01 | 0.1 | -0.32 | 0 | 0.65 | -1.06 | -0.33 | -0.4 | 0.0025 | 0.1175 | -0.115 | 0.992971 | 0.590447 | 0.759393 | | 1 | 0.996511 | 0.963123 |
| Ccsap | Q8QZT2 | -1.15 | -1.57 | -1.34 | -1.44 | -2.65 | -1.7 | -3.08 | -2.05 | -2.56 | -0.92 | -2.2 | -0.99 | -1.09 | -1.68 | -1.87 | -2.42 | -0.995 | -0.2925 | -0.39 | 0.011171 | 0.479512 | 0.216918 | | 1 | 0.996511 | 0.787447 |
| Ccser1 | Q8C0C4-3 | 0.46 | 0.9 | 1.01 | 0.39 | 0.64 | -0.27 | 0.57 | 0.81 | -0.05 | -0.11 | 0.86 | 0.67 | -0.42 | -0.49 | 0.22 | -0.15 | -0.2525 | -0.3475 | -0.9 | 0.391041 | 0.263144 | 0.00623 | | 1 | 0.996511 | 0.552614 |
| Ccser2 | Q3UHI0 | -0.22 | 0.69 | 0.95 | 0.56 | 0.8 | -0.23 | 0.25 | 0.75 | -0.62 | -1.08 | 0.79 | 0.36 | 0.12 | 0.71 | 0.89 | 1.06 | -0.1025 | -0.6325 | 0.2 | 0.761962 | 0.199478 | 0.546596 | | 1 | 0.996511 | 0.912406 |
| Ccsmst1 | Q6RUT7 | -1.48 | -0.92 | -0.9 | -1.32 | -1.13 | -1.06 | -0.75 | -1.08 | -1.49 | -0.69 | -0.93 | -1.61 | -1.61 | -0.47 | -1.37 | -0.86 | 0.15 | -0.025 | 0.0775 | 0.46345 | 0.928073 | 0.799871 | | 1 | 0.996755 | 0.970393 |
| Cct2 | P80314 | -0.66 | -0.63 | -0.56 | -0.5 | -0.66 | -0.81 | -0.57 | -0.67 | -0.59 | -0.51 | -0.69 | -0.61 | -1.1 | -0.46 | -0.8 | -0.63 | -0.09 | -0.0125 | -0.16 | 0.539901 | 0.936124 | 0.441587 | | 1 | 0.996755 | 0.883549 |
| Cct3 | P80318 | -0.11 | -0.22 | -0.14 | 0.05 | -0.23 | -0.18 | -0.28 | 0.14 | -0.13 | 0.49 | 0.01 | -0.01 | -0.37 | -0.06 | -0.12 | -0.01 | -0.0325 | 0.195 | -0.035 | 0.845192 | 0.340011 | 0.848867 | | 1 | 0.996511 | 0.976225 |
| Cct4 | P80315 | -0.12 | -0.91 | -0.71 | -0.37 | -0.96 | -0.17 | -0.59 | -0.37 | -0.11 | 0.1 | -0.58 | -0.17 | -0.5 | -0.23 | -0.74 | -0.96 | 0.005 | 0.3375 | -0.08 | 0.984309 | 0.199536 | 0.761588 | | 1 | 0.996511 | 0.963123 |
| Cct5 | P80316 | -0.27 | -0.49 | -0.53 | -0.54 | -0.28 | -0.38 | -0.32 | -0.47 | -0.56 | -0.24 | -0.62 | -0.21 | -0.83 | -0.41 | -0.54 | -0.54 | 0.095 | 0.05 | -0.1225 | 0.532541 | 0.78735 | 0.520761 | | 1 | 0.996511 | 0.907667 |
| Cct6a | Q52KG9 | 0.37 | -0.03 | -0.25 | -0.23 | 0.34 | 0.15 | -0.34 | -0.11 | -0.77 | 0.3 | -0.3 | 0.36 | -0.87 | -0.35 | -0.31 | -0.38 | 0.045 | -0.0675 | -0.4425 | 0.84314 | 0.826419 | 0.088016 | | 1 | 0.996511 | 0.691619 |
| Cct7 | P80313 | -0.33 | -0.64 | -0.46 | -0.44 | -0.58 | -0.34 | -0.23 | -0.24 | -0.96 | -0.61 | -0.32 | -0.38 | -1.11 | -1.09 | -0.56 | -0.28 | 0.12 | -0.1 | -0.2925 | 0.468743 | 0.62767 | 0.258512 | | 1 | 0.996511 | 0.812074 |
| Cct8 | P42932 | -0.25 | -0.48 | -0.51 | -0.37 | -0.43 | -0.24 | -0.82 | -0.59 | -0.85 | -0.07 | -0.57 | -0.3 | -0.77 | -0.72 | -0.61 | -0.82 | -0.1175 | -0.045 | -0.3275 | 0.521158 | 0.836072 | 0.088215 | | 1 | 0.996511 | 0.691619 |
| Ccz1 | Q8C1Y8 | 0.17 | 0.19 | -1.01 | -1.11 | 0.68 | 0.09 | -0.45 | -1.08 | -0.12 | -1.07 | -0.78 | 0.17 | 0.21 | 0.37 | -1.21 | -0.67 | 0.25 | -0.01 | 0.115 | 0.60575 | 0.981589 | 0.812456 | | 1 | 0.998773 | 0.972642 |
| Cd109 | Q8R422 | 0.46 | 0.99 | 0.33 | 0.32 | 1.31 | 0.55 | 0.92 | 0.5 | 0.63 | 0.06 | 0.56 | 0.74 | 0.33 | 1.42 | 0.06 | 0.56 | 0.295 | -0.0275 | 0.0675 | 0.267883 | 0.910244 | 0.840667 | | 1 | 0.996744 | 0.976225 |
| Cd151 | O35566 | 3.98 | 1.41 | 0.69 | 0.5 | 1.81 | 3.9 | 1.75 | 1.26 | 1.67 | 1.71 | 1.41 | 2.75 | 2.01 | 1.69 | 2 | 2.43 | 0.535 | 0.24 | 0.3875 | 0.553603 | 0.757073 | 0.602728 | | 1 | 0.996511 | 0.931061 |
| Cd163 | Q2VLH6 | -0.54 | 1.42 | -1.68 | -0.73 | 1.77 | 0.9 | -1.29 | -1.63 | 1.31 | -1.75 | -0.26 | 0.6 | -0.35 | 1.03 | -0.96 | -0.14 | 0.32 | 0.3575 | 0.2775 | 0.735947 | 0.670281 | 0.69316 | | 1 | 0.996511 | 0.955729 |
| Cd180 | Q62192 | -0.72 | -0.48 | 0.35 | 0.09 | 0.26 | -0.59 | -0.34 | 0.1 | -0.67 | 0.09 | 0.36 | -0.33 | -1.14 | -0.54 | -1.18 | -0.59 | 0.0475 | 0.0525 | -0.6725 | 0.878416 | 0.874702 | 0.055161 | | 1 | 0.996511 | 0.661738 |
| Cd200 | Q80VX2 | 0.46 | 0.73 | 1.14 | 0.79 | 0.74 | 0.7 | 1.29 | 1 | -0.47 | 0.6 | 1.25 | 1.1 | -0.43 | -0.91 | 1.21 | 0.9 | 0.1525 | -0.16 | -0.5875 | 0.492827 | 0.687288 | 0.254573 | | 1 | 0.996511 | 0.809524 |
| Cd276 | Q8VE98 | 0.63 | 1.94 | 0.67 | 0.71 | 1.02 | 1.01 | 0.83 | 1.04 | 1.63 | 0.26 | 0.78 | 1.5 | 0.52 | 1.75 | 0.22 | 1.01 | -0.0125 | 0.055 | -0.1125 | 0.968361 | 0.897606 | 0.797357 | | 1 | 0.996511 | 0.969727 |
| Cd2ap | Q9JLQ0 | 0.34 | -0.24 | -0.17 | 0.98 | -0.4 | -0.46 | -0.29 | 0.55 | 0.34 | 1.28 | -0.21 | 0.29 | -0.26 | 0.23 | -0.13 | -0.26 | -0.3775 | 0.1975 | -0.3325 | 0.301503 | 0.624822 | 0.304789 | | 1 | 0.996511 | 0.831662 |
| Cd2bp2 | Q9CWK3 | -2.37 | -3.41 | -3.31 | -3.96 | -2.76 | -3.65 | -2.36 | -3.17 | -3.37 | -1.33 | -2.71 | -2.63 | -2.08 | -1.84 | -0.91 | -2.17 | 0.2775 | 0.7525 | 1.5125 | 0.499556 | 0.159582 | 0.005582 | | 1 | 0.996511 | 0.552614 |
| Cd34 | Q64314 | 1.22 | 0.81 | -1.61 | -1.56 | 0.44 | 1.3 | -1.68 | -1.87 | 0.65 | -0.27 | -1.85 | 0.83 | 0.7 | 0.17 | -2.44 | -2.5 | -0.1675 | 0.125 | -0.7325 | 0.863908 | 0.886571 | 0.476701 | | 1 | 0.996511 | 0.892174 |
| Cd36 | Q08857 | -0.96 | -0.12 | -0.42 | 1.4 | 0.1 | -0.58 | 1.06 | 1.39 | -0.25 | -0.83 | -0.09 | 0.39 | -0.9 | -0.1 | 0.17 | 0.83 | 0.5175 | -0.17 | 0.025 | 0.411199 | 0.745807 | 0.964991 | | 1 | 0.996511 | 0.9927 |
| Cd38 | P56528 | 1.07 | 2.7 | 0.2 | -0.21 | 2.47 | 0.76 | 2.83 | 0.39 | 0.4 | -0.78 | 0.76 | 0.73 | -0.4 | 0.08 | 0.92 | 0.86 | 0.6725 | -0.6625 | -0.575 | 0.409976 | 0.338977 | 0.390674 | | 1 | 0.996511 | 0.865375 |
| Cd44 | A2APM2 | 1.13 | 1.63 | 0.6 | 0.32 | 1.64 | 0.86 | 0.91 | 0.08 | 0.83 | 0.36 | 0.81 | 1.67 | 0.78 | 1.45 | 1.01 | 1.36 | -0.0475 | -0.0025 | 0.23 | 0.906742 | 0.994796 | 0.494525 | | 1 | 0.999285 | 0.89758 |
| Cd47 | A0A2R8VK | -1.13 | 0.04 | 0.36 | -0.08 | 1.13 | 0.07 | 0.9 | 0.24 | -0.75 | 0.15 | -0.08 | -0.93 | -0.92 | 0.61 | -0.39 | -1.11 | 0.7875 | -0.2 | -0.25 | 0.069801 | 0.616483 | 0.599293 | | 1 | 0.996511 | 0.930565 |
| Cd47 | A0A2R8W6 | 0.45 | 0.83 | 0.95 | 0.9 | 1.34 | 0.51 | 1.22 | 0.82 | 0.46 | 0.7 | 0.79 | 0.52 | 0.1 | 1.22 | 0.65 | 0.72 | 0.19 | -0.165 | -0.11 | 0.433043 | 0.401245 | 0.692782 | | 1 | 0.996511 | 0.955729 |
| Cd55 | Q61475 | -1.34 | -0.86 | 0.3 | -0.4 | -2.05 | -0.18 | 0.81 | 0.09 | -0.45 | 0.3 | 0.21 | -0.81 | -1.16 | -1.79 | 0.1 | 0.17 | 0.2425 | 0.3875 | -0.095 | 0.705127 | 0.365373 | 0.862789 | | 1 | 0.996511 | 0.977235 |
| Cd63 | A0A1W2P8 | -0.97 | 0.2 | -0.53 | -1.47 | 0.53 | -0.38 | 0.41 | -1.18 | 0.79 | -3.29 | -0.07 | -0.5 | 0.29 | 0.82 | 0.28 | -0.03 | 0.5375 | -0.075 | 1.0325 | 0.291049 | 0.930267 | 0.023629 | | 1 | 0.996755 | 0.624626 |
| Cd81 | P35762 | 0.3 | 0.63 | 0.11 | 0.25 | 0.19 | 0.36 | 0.75 | 0.19 | 0.27 | -0.5 | 0.07 | 0.4 | -0.05 | -0.42 | -0.07 | 0.13 | 0.05 | -0.2625 | -0.425 | 0.805861 | 0.309912 | 0.073839 | | 1 | 0.996511 | 0.670574 |
| Cd82 | P40237 | 1.26 | 1.17 | 1.64 | 0.94 | 0.83 | 1.18 | 1.43 | 0.71 | -0.04 | 0.11 | 0.6 | 0.55 | 0.19 | -0.91 | 1.31 | 1.68 | -0.215 | -0.9475 | -0.685 | 0.375238 | 0.003597 | 0.238776 | | 1 | 0.996511 | 0.793481 |
| Cd9 | P40240 | 0.86 | -0.13 | 1.4 | 1.71 | -0.21 | 0.48 | 1.27 | 2.13 | 1 | 2.27 | 0.92 | 0.56 | 0.04 | 0.16 | 1.21 | 0.99 | -0.0425 | 0.2275 | -0.36 | 0.942507 | 0.657419 | 0.450364 | | 1 | 0.996511 | 0.884817 |
| Cd93 | O89103 | -1.05 | -1.56 | 0 | -0.18 | -0.8 | -1.26 | 0.04 | -1.23 | -1.96 | -1.08 | -0.42 | -1.03 | -1.55 | -1.64 | 0.39 | -0.67 | -0.115 | -0.425 | -0.17 | 0.795882 | 0.363665 | 0.758848 | | 1 | 0.996511 | 0.963116 |
| Cd99 | Q8VCN6 | -2.41 | -1.98 | -1.21 | -1.21 | -1.92 | -2.18 | -1.21 | -1.59 | -2.84 | -1.14 | -0.88 | -2.2 | -2.28 | -1.16 | -1.67 | -1.57 | -0.0225 | -0.0625 | 0.0325 | 0.948709 | 0.9019 | 0.929841 | | 1 | 0.996609 | 0.989687 |
| Cd99l2 | A2AP74 | -0.29 | 0.81 | 0.04 | 0.34 | 0.18 | -0.01 | 0.01 | -0.37 | 1.89 | 0.98 | 0.8 | 1.29 | 0.89 | 1.06 | -0.42 | -0.63 | -0.2725 | 1.015 | -2.4E-16 | 0.322901 | 0.013197 | 1 | | 1 | 0.996511 | 1 |
| Cda | P56389 | 0.56 | 1.33 | 0.03 | 0.48 | 1.01 | 0.68 | 0.39 | -0.2 | 1.32 | -0.73 | -1.1 | 0.9 | 1.13 | 0.58 | -0.29 | -2.15 | -0.13 | -0.5025 | -0.7825 | 0.716592 | 0.411732 | 0.278601 | | 1 | 0.996511 | 0.822084 |
| Cdc123 | Q8CII2 | -3.86 | -2.27 | -2.41 | -3.41 | -1.99 | -4.27 | -3.18 | -3.45 | -2.16 | -4.46 | -3.54 | -3.21 | -2.14 | -3.68 | -3.77 | -2.6 | -0.235 | -0.355 | -0.06 | 0.675114 | 0.533309 | 0.907731 | | 1 | 0.996511 | 0.986458 |
| Cdc16 | Q8R349 | -0.78 | -1.12 | -1 | -1.12 | -0.35 | -0.97 | -0.83 | -1.29 | -1.19 | -0.77 | -0.77 | -1.11 | -1.39 | -0.76 | -0.98 | -0.89 | 0.145 | 0.045 | 2.36E-16 | 0.533865 | 0.815003 | 1 | | 1 | 0.996511 | 1 |
| Cdc23 | G3X8W7 | 0.42 | -0.98 | -0.03 | -1.11 | -0.58 | 0.31 | -0.04 | -0.49 | -1.34 | -1.05 | 0.16 | 0.35 | -1.13 | -1.23 | -0.14 | 0.31 | 0.225 | -0.045 | -0.1225 | 0.577405 | 0.931239 | 0.804174 | | 1 | 0.996755 | 0.971404 |
| Cdc26 | Q99JP4 | 0.82 | 0.93 | 1.13 | 1.65 | 0.3 | 0.36 | 1.36 | 1.54 | 0.64 | 1.59 | 1.26 | 1.01 | 0.46 | 0.13 | 1.08 | 0.88 | -0.2425 | -0.0075 | -0.495 | 0.50425 | 0.978858 | 0.121474 | | 1 | 0.998749 | 0.721448 |
| Cdc27 | A2A6Q5 | -0.23 | 0.24 | -0.31 | -0.27 | 0.08 | -0.39 | -0.57 | -0.53 | 0.17 | 0.89 | -0.16 | -0.09 | -0.03 | 0.13 | -0.67 | -0.54 | -0.21 | 0.345 | -0.135 | 0.353543 | 0.243549 | 0.607634 | | 1 | 0.996511 | 0.933262 |
| Cdc34 | Q8CFI2 | -1.19 | -0.77 | -0.31 | -0.96 | -1.05 | -0.99 | -0.81 | -0.89 | -1.03 | -0.09 | -0.49 | -0.82 | -0.98 | 0.01 | -0.87 | -0.42 | -0.1275 | 0.2 | 0.2425 | 0.563147 | 0.492448 | 0.435169 | | 1 | 0.996511 | 0.881557 |
| Cdc37 | Q61081 | 0.29 | 0.14 | 0.38 | 0.26 | 0.49 | 0.55 | 0.29 | 0.32 | 0.06 | 0.11 | 0.25 | 0.24 | -0.12 | 0.71 | -0.02 | 0.06 | 0.145 | -0.1025 | -0.11 | 0.354458 | 0.529551 | 0.642481 | | 1 | 0.996511 | 0.94365 |
| Cdc37l1 | Q9CZP7-2 | -0.93 | -0.3 | -0.34 | -0.62 | -0.69 | -0.63 | -0.78 | -0.62 | -0.22 | 0.08 | -0.63 | -0.76 | 0.02 | 0.06 | -0.62 | -0.86 | -0.1325 | 0.165 | 0.1975 | 0.490697 | 0.531407 | 0.501192 | | 1 | 0.996511 | 0.901002 |
| Cdc40 | Q9DC48 | 0.67 | 0.25 | 0.33 | 0.28 | -0.14 | 0.46 | 0.08 | 0.61 | 0.14 | 0.72 | 0.12 | 0.01 | 0.23 | -0.01 | 0.5 | 0.16 | -0.13 | -0.135 | -0.1625 | 0.559802 | 0.549146 | 0.437977 | | 1 | 0.996511 | 0.883161 |
| Cdc42 | P60766 | -0.1 | -0.62 | -0.14 | 0.21 | -0.64 | -0.04 | -0.28 | -0.1 | 0.25 | 0.63 | -0.02 | 0.12 | -0.78 | -0.39 | -0.35 | -0.46 | -0.1025 | 0.4075 | -0.3325 | 0.664927 | 0.124935 | 0.185492 | | 1 | 0.996511 | 0.776843 |
| Cdc42 | P60766-1 | 0.11 | 0.24 | 0.73 | 0.54 | 0.55 | 0.51 | 0.33 | 0.36 | 0.63 | 0.86 | 0.22 | 0.32 | 0.11 | 0.55 | -0.12 | 0.73 | 0.0325 | 0.1025 | -0.0875 | 0.864008 | 0.662426 | 0.743605 | | 1 | 0.996511 | 0.959321 |
| Cdc42bpa | D3YYN8 | 0.61 | 0.21 | 0.41 | 0.22 | 0.29 | 0.7 | 0.49 | 0.29 | 0.18 | 0.03 | 0.58 | 0.51 | -0.03 | 0.21 | 0.37 | 0.57 | 0.08 | -0.0375 | -0.0825 | 0.65965 | 0.856132 | 0.700493 | | 1 | 0.996511 | 0.956416 |

| | | | | | | | | | | | | | | | | | | | | | | | | | | |
|---|---|---|---|---|---|---|---|---|---|---|---|---|---|---|---|---|---|---|---|---|---|---|---|---|---|---|
| Cdc42bpb | Q7TT50 | 0.55 | -0.05 | 0.41 | 0.34 | 0.02 | 0.42 | 0.26 | 0.45 | 0.06 | 0.43 | 0.52 | 0.28 | -0.27 | 0.08 | 0.12 | -0.05 | -0.025 | 0.01 | -0.3425 | 0.898792 | 0.961547 | 0.133311 | 1 | 0.996755 | 0.728417 |
| Cdc42ep1 | A0A0R4J0S | 0.6 | 1.08 | 1.06 | 0.81 | 0.86 | 0.24 | 0.59 | 0.06 | 0.32 | 0.12 | 0.48 | 0.11 | -0.01 | 0.32 | 0.59 | 0.91 | -0.45 | -0.63 | -0.435 | 0.078444 | 0.010364 | 0.117526 | 1 | 0.996511 | 0.718609 |
| Cdc42ep2 | Q8JZX9 | 1.55 | 2.67 | 0.82 | -0.59 | 2.7 | 2.49 | -0.27 | 0.37 | 1.18 | 0.83 | -0.24 | 1.72 | 0.59 | 1.81 | -0.31 | 0.32 | 0.21 | -0.24 | -0.51 | 0.817649 | 0.741169 | 0.49513 | 1 | 0.996511 | 0.89758 |
| Cdc42ep4 | Q9JM96 | 0.08 | 0.35 | 0.41 | 0.19 | -0.2 | 0.34 | 0.3 | 0.13 | 0.35 | 0.2 | 0.76 | 0.59 | 0.85 | 0.51 | 0.48 | -0.24 | -0.115 | 0.2175 | 0.1425 | 0.539864 | 0.286863 | 0.596576 | 1 | 0.996511 | 0.929911 |
| Cdc42se2 | Q8BGH7 | 0.34 | 1.02 | 0.3 | 0.43 | 0.73 | 0.81 | 0.69 | 0.58 | 1.69 | 0.59 | 0.44 | 0.71 | 1.46 | 2.09 | 0.18 | 0.47 | 0.18 | 0.335 | 0.5275 | 0.391037 | 0.321454 | 0.257393 | 1 | 0.996511 | 0.812003 |
| Cdc5l | Q6A068 | 1.18 | -0.44 | -0.09 | 0.28 | -0.48 | 0.47 | 0 | 1.11 | -0.02 | 0.14 | 0.27 | -0.49 | 0.47 | 0.27 | 0.9 | -0.02 | 0.0425 | -0.2575 | 0.1725 | 0.925095 | 0.496338 | 0.657183 | 1 | 0.996511 | 0.948659 |
| Cdc73 | Q8JZM7 | 0.05 | -2.09 | -1.11 | -0.47 | -2.08 | -1.03 | -1.04 | -0.35 | -1.06 | -0.33 | -0.61 | -1.86 | 0 | -0.77 | 0.28 | -1 | -0.22 | -0.06 | 0.5325 | 0.682277 | 0.909259 | 0.315118 | 1 | 0.996744 | 0.837638 |
| Cdh10 | P70408 | 0.68 | 0.99 | 1.09 | 0.98 | 1.08 | 0.76 | 1.04 | 0.89 | 0.87 | 1.4 | 1.21 | 1 | 0.34 | 1.3 | 0.63 | 0.75 | 0.0075 | 0.185 | -0.18 | 0.964533 | 0.361805 | 0.481997 | 1 | 0.996511 | 0.893925 |
| Cdh11 | P55288 | 0.61 | 1.13 | 1.01 | 1.03 | 1.17 | 0.91 | 1.02 | 0.88 | 0.8 | 0.8 | 1.31 | 1.03 | 0.01 | 0.34 | 0.42 | 0.87 | 0.05 | 0.04 | -0.535 | 0.779622 | 0.848887 | 0.05449 | 1 | 0.996511 | 0.661738 |
| Cdh12 | Q5RJH3 | 0.36 | 0.77 | 1.1 | 0.81 | 1.08 | 0.26 | 0.7 | 0.67 | 0.25 | 0.83 | 1.04 | 0.71 | -0.38 | 0.34 | 0.07 | 1.03 | -0.0825 | -0.0525 | -0.495 | 0.733315 | 0.833401 | 0.163845 | 1 | 0.996511 | 0.758352 |
| Cdh13 | Q9WTR5 | 1.02 | 0.32 | 0.69 | 0.69 | 0.38 | 1.25 | 2.83 | 1 | 0.65 | 0.03 | 1.08 | 1.52 | 0.2 | 0.5 | 0.68 | 1.32 | 0.685 | 0.14 | -0.005 | 0.193657 | 0.684325 | 0.986341 | 1 | 0.996511 | 0.996154 |
| Cdh18 | E9Q9Q6 | 1.86 | 1.66 | 1.55 | 1.19 | 1.77 | 1.93 | 1.58 | 1.55 | 1.26 | 0.63 | 1.75 | 2.07 | 1.08 | -0.08 | 0.79 | 1.99 | 0.1425 | -0.1375 | -0.62 | 0.481582 | 0.686448 | 0.172322 | 1 | 0.996511 | 0.76392 |
| Cdh19 | E9Q3A7 | 0.87 | 1.11 | 0.67 | 0.74 | 1.65 | 0.51 | 0.05 | 1.24 | 1.12 | 1.2 | 0.71 | 0.87 | 0.99 | 1.93 | 0.62 | 0.94 | 0.015 | 0.1275 | 0.2725 | 0.966402 | 0.526586 | 0.387926 | 1 | 0.996511 | 0.864251 |
| Cdh2 | D3YYT0 | 0.62 | 1.08 | 0.96 | 1.04 | 1.13 | 0.63 | 1.06 | 0.88 | 0.66 | 0.87 | 1.09 | 0.98 | 0.2 | 0.67 | 0.44 | 0.86 | -2.4E-16 | -0.025 | -0.3825 | 1 | 0.897152 | 0.116187 | 1 | 0.996511 | 0.718608 |
| Cdh20 | Q9Z0M3 | 0.15 | 1.19 | 0.98 | 0.93 | 1.06 | 0.48 | 1.35 | 1.32 | 0.78 | 1.14 | 1.14 | 0.76 | -0.32 | 0.75 | 1.35 | 1.21 | 0.24 | 0.1425 | -0.065 | 0.43483 | 0.597881 | 0.87745 | 1 | 0.996511 | 0.979626 |
| Cdh22 | Q9WTP5 | 1.81 | 1.57 | 2.33 | 1.62 | 2.06 | 1.55 | 0.6 | 1.54 | -0.65 | 0.7 | 1.85 | 1.33 | -1.36 | -1.37 | 1.4 | 1.75 | -0.395 | -1.025 | -1.7275 | 0.262897 | 0.078143 | 0.051801 | 1 | 0.996511 | 0.658454 |
| Cdh4 | P39038 | 1.61 | 1.68 | 0.89 | 1.19 | 1.79 | 1.52 | 1.37 | 0.95 | 1.18 | 0.73 | 1.05 | 1.68 | 0.56 | 1.04 | 0.92 | 1.03 | 0.065 | -0.1825 | -0.455 | 0.805167 | 0.523152 | 0.096665 | 1 | 0.996511 | 0.698156 |
| Cdh5 | P55284 | -1.4 | -0.92 | -0.49 | 0.13 | -0.27 | -1.3 | -0.49 | -0.87 | -0.78 | 0.54 | 0.25 | -0.96 | -0.36 | -0.55 | 0.42 | 0.22 | -0.0625 | 0.4325 | 0.6025 | 0.868106 | 0.362693 | 0.145712 | 1 | 0.996511 | 0.745321 |
| Cdh6 | P97326 | 1.33 | 1.58 | 2.43 | 2.07 | 1.29 | 0.96 | 1.81 | 1.66 | 1.39 | 1.95 | 2.26 | 1.35 | 0.52 | 1.6 | 1.05 | 1.77 | -0.4225 | -0.115 | -0.6175 | 0.19338 | 0.727651 | 0.120952 | 1 | 0.996511 | 0.721448 |
| Cdh7 | Q8BM92 | 1.85 | 2.01 | 1.17 | 0.74 | 2.15 | 1.75 | 1.24 | 0.99 | 1.49 | -0.66 | 1.47 | 1.9 | 0.83 | 1.91 | 0.48 | 1.44 | 0.09 | -0.3925 | -0.2775 | 0.810441 | 0.515711 | 0.510292 | 1 | 0.996511 | 0.903419 |
| Cdh8 | P97291 | 1.3 | 1.56 | 1.21 | 0.94 | 1.83 | 1.42 | 1.22 | 1.16 | 0.39 | 0.83 | 1.46 | 1.75 | -0.59 | 0.12 | 0.16 | 1.22 | 0.155 | -0.145 | -1.025 | 0.489749 | 0.662079 | 0.024101 | 1 | 0.996511 | 0.626254 |
| Cdh9 | F8WHU6 | 1.32 | 1.14 | 2.41 | 0.85 | 1.28 | 1.62 | 3.28 | 1.87 | 0.42 | 0.21 | 2.46 | 1.54 | 0 | 1 | 2.66 | 3.19 | 0.5825 | -0.2725 | 0.2825 | 0.273626 | 0.637081 | 0.701646 | 1 | 0.996511 | 0.956791 |
| Cdip1 | Q9DB75 | -0.27 | 0.05 | -0.18 | 0.48 | 0.2 | -0.51 | -0.94 | 0.01 | -0.13 | 1.35 | -0.45 | -0.13 | 0.89 | 1.18 | -0.17 | -0.62 | -0.33 | 0.14 | 0.3 | 0.294977 | 0.736812 | 0.497035 | 1 | 0.996511 | 0.897918 |
| Cdipt | Q8VDP6 | -2.1 | -1.28 | -1.16 | -1.59 | -0.35 | -1.66 | -0.83 | -1.67 | -2.04 | -1.41 | -1.55 | -1.91 | -1.95 | -1.5 | -1.62 | -1.05 | 0.405 | -0.195 | 0.0025 | 0.28976 | 0.479695 | 0.993236 | 1 | 0.996511 | 0.996845 |
| Cdk11b | A2A9P6 | -1.05 | -2.14 | -1.19 | -1.77 | -1.56 | -2.12 | -0.94 | -1.07 | -2.11 | -0.45 | -1.86 | -2.03 | -1.18 | -0.5 | -0.1 | -0.86 | 0.115 | -0.075 | 0.8775 | 0.746189 | 0.864596 | 0.029322 | 1 | 0.996511 | 0.637553 |
| Cdk13 | Q69ZA1 | 1.42 | -0.25 | -1.68 | -1.23 | -0.59 | 0.27 | -1.39 | -1.35 | 0.12 | -3.86 | -1.48 | 0.4 | 1.09 | 0.69 | -2.13 | -2.22 | -0.33 | -0.77 | -0.2075 | 0.64636 | 0.479057 | 0.836378 | 1 | 0.996511 | 0.975831 |
| Cdk14 | O35495 | 0.83 | -0.25 | 0.69 | 0.67 | 0.35 | 1.41 | 0.25 | 0.91 | -1.27 | -0.04 | 0.6 | 0.77 | -0.88 | -1.39 | 0.55 | 0.78 | 0.245 | -0.47 | -0.72 | 0.49192 | 0.349689 | 0.207053 | 1 | 0.996511 | 0.785338 |
| Cdk16 | Q04735 | 0.51 | 0.79 | 1.16 | 0.96 | 0.69 | 0.66 | 0.74 | 0.73 | 1.36 | 3.2 | 1.1 | 1.03 | 1.34 | 1.39 | 0.67 | 0.89 | -0.15 | 0.8175 | 0.2175 | 0.420143 | 0.126374 | 0.402486 | 1 | 0.996511 | 0.870698 |
| Cdk17 | Q8K0D0 | 0.02 | 0.8 | 0.89 | 0.31 | 0.63 | -0.05 | 0.44 | 0.12 | -0.01 | 0.22 | 0.65 | 1.04 | 0.6 | 0.36 | 0.73 | 0.6 | -0.22 | -0.03 | 0.0675 | 0.414765 | 0.923566 | 0.789375 | 1 | 0.996755 | 0.968677 |
| Cdk18 | Q04899 | 0.02 | -0.04 | 0.21 | 0.36 | -0.02 | -0.72 | 0.42 | 0.05 | -0.09 | 0.45 | -0.14 | 0.12 | 0.11 | 0.07 | 0.2 | -0.18 | -0.205 | -0.0525 | -0.0875 | 0.443655 | 0.80004 | 0.654724 | 1 | 0.996511 | 0.948172 |
| Cdk19 | Q8BWD8 | -1.27 | -2.04 | 1.13 | 0.59 | -1.17 | -1.44 | 1.38 | 1.21 | -1.81 | 0.88 | 1.09 | -1.38 | -1.6 | -1.58 | 0.86 | 0.5 | 0.3925 | 0.0925 | -0.0575 | 0.682444 | 0.922802 | 0.948829 | 1 | 0.996755 | 0.992476 |
| Cdk20 | Q9JHU3 | -0.31 | -0.76 | -0.56 | 0.46 | -1.24 | -0.76 | -0.74 | -0.85 | 0.12 | 0.26 | -0.17 | -0.56 | -0.18 | 0 | -0.35 | -0.13 | -0.605 | 0.205 | 0.1275 | 0.064079 | 0.529882 | 0.66454 | 1 | 0.996511 | 0.949825 |
| Cdk2ap1 | F2Z4B3 | -2.24 | -2.37 | -2.81 | -3.38 | -0.99 | -1.94 | -3.19 | -3.64 | -2.15 | -0.21 | -3.55 | -2.79 | -1.33 | -1.18 | -1.75 | -2.55 | 0.26 | 0.525 | 0.9975 | 0.665057 | 0.455138 | 0.028669 | 1 | 0.996511 | 0.637553 |
| Cdk4 | E9Q9E4 | -0.07 | 0.16 | -0.57 | 0.43 | -0.42 | -0.51 | -0.62 | -0.65 | -0.23 | 0.36 | -0.04 | -0.26 | -0.3 | 0.05 | -1.31 | -0.73 | -0.5375 | -0.03 | -0.56 | 0.045851 | 0.911767 | 0.143517 | 1 | 0.996744 | 0.742733 |
| Cdk5 | P49615 | 1.09 | 0.1 | -0.28 | -0.24 | 0.37 | 1.08 | 0.73 | -0.3 | -0.05 | -0.45 | 0.04 | 1.03 | -0.58 | 0.84 | -0.09 | -0.14 | 0.3025 | -0.025 | -0.16 | 0.466346 | 0.952993 | 0.702847 | 1 | 0.996755 | 0.956859 |
| Cdk5r1 | P61809 | -0.72 | -0.68 | -0.3 | -0.16 | -0.23 | -0.87 | -0.41 | -0.64 | -0.68 | 0.32 | -0.4 | -0.68 | -0.26 | 0.27 | -0.53 | -0.65 | -0.0725 | 0.105 | 0.1725 | 0.74237 | 0.713054 | 0.529591 | 1 | 0.996511 | 0.908663 |
| Cdk5r2 | O35926 | 0.35 | -0.47 | -0.55 | -0.74 | -0.41 | 0.11 | -0.68 | -0.66 | 0.49 | -0.26 | -0.19 | 0.14 | 0.7 | 0.8 | -1.05 | -1.27 | -0.0575 | 3.98E-01 | 0.1475 | 0.848319 | 0.210057 | 0.791278 | 1 | 0.996511 | 0.969105 |
| Cdk5rap3 | Q99LM2 | -0.58 | -0.42 | -0.44 | -0.48 | -0.49 | -0.44 | -0.06 | -0.49 | 0.18 | -0.22 | -0.7 | -0.49 | 0.48 | -0.14 | 0.02 | -0.06 | 0.11 | 0.1725 | 0.555 | 0.513432 | 0.454812 | 0.022954 | 1 | 0.996511 | 0.623147 |
| Cdk7 | A0A286YD( | -4.72 | -4.54 | -2.43 | -1.97 | -4.02 | -5 | -2.2 | -2.46 | -4.09 | -1.36 | -1.57 | -4.21 | -2.29 | -2.27 | -2.49 | -1.6 | -0.005 | 0.6075 | 1.2525 | 0.995424 | 0.525692 | 0.088256 | 1 | 0.996511 | 0.691619 |
| Cdk9 | Q99J95 | -2.28 | -3.02 | -3.07 | -3.17 | -2.37 | -2.95 | -2.65 | -3.02 | -2.89 | -2.63 | -3.09 | -2.12 | -2.43 | -1.68 | -2.96 | -3.21 | 0.1375 | 0.2025 | 0.315 | 0.601503 | 0.503864 | 0.421186 | 1 | 0.996511 | 0.875639 |
| Cdkal1 | Q91WE6-2 | -0.02 | 0.37 | -0.26 | 0.04 | 1.24 | -0.66 | -0.23 | -0.41 | 0.47 | 0.35 | -0.48 | 0.19 | 1.31 | 1.16 | 0.27 | 0.33 | -0.0475 | 0.1 | 0.735 | 0.909684 | 0.707704 | 0.039664 | 1 | 0.996511 | 0.658454 |
| Cdkl5 | A0A0G2JG\ | 1.39 | 0.56 | 1.13 | 0.81 | 0.39 | 1.47 | 0.53 | 0.99 | 0.68 | 0.54 | 1.07 | 1.07 | 0.31 | 0.5 | 0.45 | 0.64 | -0.1275 | -0.1325 | -0.4975 | 0.674161 | 0.599029 | 0.05977 | 1 | 0.996511 | 0.661738 |
| Cdkn1b | P46414 | -1.98 | -1.59 | -1.8 | -2.29 | -1.58 | -2.05 | -2.14 | -1.74 | -1.13 | -1.51 | -1.04 | -1.57 | 0.07 | -3.89 | -0.66 | -1.79 | 0.0375 | 0.6025 | 0.3475 | 0.865223 | 0.026894 | 0.661651 | 1 | 0.996511 | 0.949645 |
| Cdkn2aip | Q8BI72 | -2.06 | -3.15 | -1.87 | -1.8 | -3.21 | -2.68 | -2.13 | -2.25 | -2.74 | -1.12 | -1.73 | -2.8 | -2.09 | -2.68 | -2.18 | -2.06 | -0.3475 | 0.1225 | -0.0325 | 0.370844 | 0.799193 | 0.924973 | 1 | 0.996511 | 0.988301 |
| Cdkn2c | Q60772 | 0.01 | -0.24 | -2.05 | -2.09 | 0.26 | -0.42 | -2.41 | -2.26 | -1.57 | -1.41 | -1.86 | -0.17 | -0.92 | -0.46 | -2.24 | -2.37 | -0.115 | -0.16 | -0.405 | 0.884084 | 0.796727 | 0.551459 | 1 | 0.996511 | 0.914317 |
| Cdr2l | A2A6T1 | -0.95 | -0.76 | -1.23 | -0.99 | -0.39 | -0.84 | -0.59 | -1.53 | -0.36 | -0.58 | -1.11 | -0.65 | -0.3 | -0.41 | -0.91 | -0.73 | 0.145 | 0.3075 | 0.395 | 0.596749 | 0.188846 | 0.101407 | 1 | 0.996511 | 0.698697 |
| Cds1 | P98191 | -0.85 | -0.66 | -0.03 | -0.14 | 0.27 | -0.81 | -0.16 | 0.47 | -0.59 | 1.34 | -0.12 | -0.95 | 0.1 | 0.49 | 0.77 | 0.81 | 0.3625 | 0.34 | 0.9625 | 0.29615 | 0.504466 | 0.00697 | 1 | 0.996511 | 0.552614 |
| Cds2 | Q99L43 | -0.46 | -0.55 | -1.41 | -0.71 | -1.12 | 0.24 | -0.82 | -0.62 | 0.53 | -1.09 | -1.16 | -0.65 | 1.14 | -1.24 | -0.22 | -1.03 | 0.2025 | 0.19 | 0.445 | 0.565621 | 0.655131 | 0.417019 | 1 | 0.996511 | 0.875537 |
| Cdv3 | Q4VAA2 | -0.14 | -0.26 | 0.48 | 0.51 | -0.91 | -0.25 | 0.22 | 0.33 | -0.32 | 0.44 | 0.56 | 0.03 | -0.05 | -0.4 | 0.37 | -0.01 | -3.00E-01 | 0.03 | -0.17 | 0.383461 | 0.918405 | 0.544736 | 1 | 0.996755 | 0.912212 |
| Cebpz | A0A0R4J04 | 0.55 | 2.78 | 7.67 | 8.1 | 1.29 | 1.15 | 8.94 | 8.88 | -0.39 | 12.49 | 8.5 | 1.54 | 0.7 | 1.25 | 8.7 | 12.02 | 0.29 | 0.76 | 0.8925 | 0.910291 | 0.808893 | 0.763861 | 1 | 0.996511 | 0.964134 |
| Cebpzos | Q8BTE5 | 0.53 | 1.57 | 0.06 | 0.42 | 0.79 | 0.5 | 0.38 | -0.34 | -0.29 | -0.58 | 0.37 | 0.46 | 0.21 | 0.99 | -0.42 | 0.27 | -0.3125 | -0.655 | -0.3825 | 0.423413 | 0.124101 | 0.369187 | 1 | 0.996511 | 0.859245 |
| Celf1 | A0A0R4J0T | -0.16 | -1.81 | -1.84 | -2.07 | -1.36 | -0.7 | -1.57 | -1.81 | -2.89 | -1.39 | -1.54 | -0.86 | -1.8 | -1.85 | -1.43 | -2.04 | 0.11 | -0.2 | -0.31 | 0.813083 | 0.725415 | 0.483174 | 1 | 0.996511 | 0.893925 |
| Celf2 | V9GX43 | -1.38 | -2.81 | -1.89 | -2.01 | -2.64 | -1.75 | -1.94 | -2.11 | -1.28 | -1.36 | -1.96 | -2.25 | -0.32 | -2.78 | -1.53 | -2.53 | -0.0875 | 0.31 | 0.2325 | 0.797383 | 0.407834 | 0.690414 | 1 | 0.996511 | 0.955729 |
| Celf3 | A0A0G2JGI | -1.09 | -1.85 | -1.68 | -2.2 | -1.45 | -1.93 | -2.15 | -2.27 | -1.75 | -1.38 | -1.75 | -1.07 | -1.08 | -1.3 | -1.61 | -1.78 | -0.245 | 0.2175 | 0.2625 | 0.413509 | 0.464181 | 0.384072 | 1 | 0.996511 | 0.860984 |
| Celf4 | Q7TSY6-4 | -2.15 | -2.4 | -1.81 | -1.69 | -2.32 | -2.52 | -1.46 | -1.81 | -1.76 | -1.17 | -1.52 | -2.22 | -0.48 | -1.79 | -0.74 | -1.23 | -0.015 | 0.345 | 0.9525 | 0.958777 | 0.244991 | 0.017893 | 1 | 0.996511 | 0.602211 |
| Celf5 | D3Z580 | -2.2 | -3.36 | -2.21 | -3.08 | -3.79 | -2.04 | -2.87 | -2.57 | -0.31 | -1.22 | -2.6 | -2.18 | 1.61 | -2.21 | -2.75 | -2.34 | -0.105 | 1.135 | 1.29 | 0.811983 | 0.064043 | 0.197687 | 1 | 0.996511 | 0.78004 |
| Celf6 | D6RHP1 | -1.32 | -1 | 0.81 | -1.1 | -1.17 | -0.85 | -0.31 | 0.18 | -0.95 | 0.37 | -0.66 | -0.87 | -0.34 | -0.38 | -0.19 | 0.01 | 0.115 | 0.125 | 0.4275 | 0.827812 | 0.81549 | 0.374437 | 1 | 0.996511 | 0.859424 |
| Celsr2 | Q9R0M0 | -0.05 | 0.14 | 0.11 | 0.09 | 0.38 | 0.22 | 0.1 | 0.11 | 0.44 | -0.15 | 0.24 | 0.31 | 0.67 | 0.28 | 0.5 | 0.31 | 0.13 | 0.1375 | 0.3675 | 0.400261 | 0.477147 | 0.072901 | 1 | 0.996511 | 0.668066 |
| Cemip2 | Q5FWI3 | -0.14 | 0.45 | -0.33 | -0.19 | 0.96 | -0.13 | 0.3 | -0.12 | 0.34 | -0.62 | 0.08 | 0.5 | 0.45 | 0.54 | -0.14 | 0.14 | 0.305 | 0.1275 | 0.3 | 0.331948 | 0.677943 | 0.267772 | 1 | 0.996511 | 0.819502 |
| Cend1 | Q9JKC6 | 1.55 | -0.03 | 0.86 | 1.52 | -1.28 | 1.21 | -0.49 | 1.12 | 3.05 | 1.08 | 0.87 | 1.64 | 3.59 | -0.43 | 1.23 | 0.29 | -0.835 | 0.685 | 0.195 | 0.221824 | 0.246662 | 0.819597 | 1 | 0.996511 | 0.972768 |
| Cenpa | A0A0G2JGI | 1.43 | 1.29 | 2.09 | 2.1 | 0.69 | 1.09 | 1.02 | 1.9 | -1.2 | -3.57 | 2.2 | 1.17 | -0.17 | -1.89 | 2.32 | 2.54 | -0.5525 | -2.0775 | -1.0275 | 0.117702 | 0.10354 | 0.304386 | 1 | 0.996511 | 0.831662 |
| Cenpv | Q9CXS4 | 1.72 | 1.08 | 1.42 | 1.36 | 1.03 | 1.24 | 0.99 | 1.17 | 0.86 | 0.68 | 1.15 | 1.03 | 1.47 | 0.75 | 1.56 | 1.3 | -0.2875 | -0.465 | -0.125 | 0.147266 | 0.050695 | 0.626509 | 1 | 0.996511 | 0.939737 |
| Cep112 | Z4YKK6 | -0.32 | 0.1 | 0.78 | 0.28 | 0.12 | -0.37 | -0.42 | 0.43 | -0.42 | -0.34 | 1.05 | -0.29 | -0.5 | 0.14 | -0.01 | 1.47 | -0.27 | -0.21 | 0.065 | 0.383665 | 0.602435 | 0.885661 | 1 | 0.996511 | 0.979767 |
| Cep128 | Q8BI22 | -1.05 | -1.46 | -0.02 | -1.19 | -1.47 | -0.84 | -1.9 | -1.56 | 1.72 | -4.27 | -0.72 | -0.73 | 2.34 | -1.85 | 1.45 | -1.84 | -0.5125 | -0.07 | 0.955 | 0.187216 | 0.951007 | 0.360895 | 1 | 0.996755 | 0.857677 |
| Cep131 | B1AXI9 | -0.59 | -0.28 | 1.32 | 0.62 | -0.8 | -0.49 | 0.53 | 0.54 | 0.71 | 2.32 | 0.98 | -0.27 | 1.83 | -0.26 | 0.89 | 0.84 | -0.3225 | 0.6675 | 0.5575 | 0.533423 | 0.305062 | 0.334908 | 1 | 0.996511 | 0.847169 |
| Cep135 | Q6P5D4 | 0.71 | 1.15 | 0.86 | 0.4 | 0.27 | 0.89 | 1.1 | 0.22 | 1.05 | -0.09 | 0.65 | 0.38 | 1.1 | 1.68 | 0.44 | 1.06 | -0.16 | -0.2825 | 0.29 | 0.564408 | 0.349239 | 0.358825 | 1 | 0.996511 | 0.856645 |
| Cep170 | Q6A065 | 0.75 | 0.82 | 0.83 | 0.65 | 0.41 | 0.44 | 0.42 | 0.5 | 0.87 | 0.6 | 0.69 | 1 | 0.96 | 0.6 | 0.76 | 0.31 | -0.32 | 0.0275 | -0.105 | 0.046374 | 0.873683 | 0.611874 | 1 | 0.996511 | 0.935047 |
| Cep170 | A0A0A6YX! | 1.52 | 0.39 | 0.35 | 0.49 | 2.11 | 1.32 | 0.82 | -0.59 | 0.74 | 1.75 | -0.04 | 1.58 | -0.41 | 2.85 | 0.72 | 0.68 | 0.2275 | 0.32 | 0.2725 | 0.69499 | 0.499062 | 0.685582 | 1 | 0.996511 | 0.955729 |
| Cep170b | Q80U49 | 0.05 | 0.26 | 0.04 | -0.25 | -0.1 | -0.07 | -0.42 | -0.38 | 0.34 | 0.13 | 0.09 | 0.29 | 0.88 | 0.35 | 0.07 | -0.61 | -0.2675 | 0.1875 | 0.1475 | 0.16808 | 0.322313 | 0.657178 | 1 | 0.996511 | 0.948659 |
| Cep41 | Q99NF3 | -0.81 | -1.04 | -1.18 | -0.65 | -0.82 | -0.65 | -0.84 | -0.97 | -0.91 | -0.16 | -0.96 | -0.74 | -0.98 | -0.6 | -1.14 | -1.04 | 0.1 | 0.2275 | -0.02 | 0.582043 | 0.36358 | 0.927095 | 1 | 0.996511 | 0.989293 |
| Cep76 | Q0VEJ0 | -0.41 | 0.13 | -4.79 | -2.47 | 0.14 | -0.44 | -3.42 | -4.53 | -0.53 | -2.1 | -3.31 | -0.51 | -0.34 | 0.55 | -3.35 | -4.75 | -0.1775 | 0.2725 | -0.0875 | 0.900622 | 0.816173 | 0.953038 | 1 | 0.996511 | 0.992476 |
| Cep97 | Q9CZ62 | -0.06 | 0.06 | -0.74 | -0.55 | 0.4 | -0.4 | -0.78 | -0.99 | -0.1 | -0.81 | -0.72 | 0.11 | -0.39 | 1.09 | -0.6 | -0.7 | -0.12 | -0.0575 | 0.1725 | 0.730776 | 0.849225 | 0.694113 | 1 | 0.996511 | 0.955729 |
| Cept1 | A0A0A6YXI | -0.8 | -0.56 | 0.38 | 0.33 | 0.12 | -0.81 | 0.84 | 1 | -0.93 | -0.48 | 0.2 | -0.62 | -0.65 | -0.17 | 0.61 | 0.83 | 0.45 | -0.295 | 0.3175 | 0.355803 | 0.437564 | 0.472514 | 1 | 0.996511 | 0.891144 |
| Cerk | Q8K4Q7 | -0.76 | -1.19 | -1.48 | 0.46 | -1.39 | -1.54 | -0.73 | 0.13 | -0.09 | -0.56 | -0.68 | -1.13 | 1.38 | -0.8 | 0.66 | -0.43 | -0.14 | 0.1275 | 0.945 | 0.790331 | 0.776958 | 0.143262 | 1 | 0.996511 | 0.742711 |
| Cers1 | P27545 | 1.25 | 0.73 | 1.16 | 1.19 | 1.69 | 1.64 | 2.04 | 1.96 | 1.52 | 1.06 | 1.29 | 1.79 | 1.23 | 1.58 | 2.56 | 2.71 | 0.75 | 0.3325 | 0.9375 | 0.003629 | 0.171738 | 0.031534 | 1 | 0.996511 | 0.641974 |
| Cers2 | Q924Z4 | -0.21 | -0.31 | 1.48 | 0.54 | 0.43 | 0.12 | 1.26 | 1.15 | 0.69 | 0.4 | 1.68 | 0.65 | 0.85 | 0.91 | 2.45 | 2.26 | 0.365 | 0.48 | 1.2425 | 0.438612 | 0.321483 | 0.047222 | 1 | 0.996511 | 0.658454 |
| Cers4 | Q9D6J1 | -0.52 | 0.25 | -0.34 | -0.41 | 0.28 | -0.1 | -0.63 | -0.07 | 0.89 | 0.39 | -0.65 | -0.26 | 2.26 | 0.35 | 0.37 | 0.05 | 0.125 | 0.3475 | 1.0125 | 0.636378 | 0.361029 | 0.06859 | 1 | 0.996511 | 0.665034 |
| Cers6 | H3BL08 | 0.08 | 0.47 | 0.77 | 0.46 | 0.95 | 0.34 | 1.54 | 2.09 | -1.92 | -5.54 | 0.84 | -0.19 | -1.53 | -1.21 | 1.91 | 1.52 | 0.785 | -2.1475 | -0.2725 | 0.066296 | 0.115857 | 0.739406 | 1 | 0.996511 | 0.958796 |
| Cert1 | Q9EQG9 | -0.86 | -1.12 | -0.82 | -0.94 | -1.01 | -1.15 | -1.13 | -1.08 | -1.45 | -0.69 | -0.85 | -0.92 | -0.73 | -1.26 | -0.91 | -0.87 | -0.1575 | -0.0425 | -0.0075 | 0.308257 | 0.844554 | 0.969885 | 1 | 0.996511 | 0.992767 |
| Ces1c | P23953 | -5.41 | -6.28 | -3.04 | -3.14 | -5 | -5.11 | -2.82 | -2.93 | -5.44 | -1.49 | -2.8 | -5.06 | -4.62 | -4.38 | -3 | -2.82 | 0.5025 | 0.77 | 0.7625 | 0.590376 | 0.495694 | 0.377308 | 1 | 0.996511 | 0.860984 |
| Cetn2 | Q9R1K9 | 0.91 | 1.14 | 0.42 | 0.65 | 1 | 0.68 | 0.48 | 0.37 | 0.87 | -0.06 | 0.68 | 1.22 | 0.68 | 1.24 | 0.71 | 0.81 | -0.1475 | -0.1025 | 0.08 | 0.523855 | 0.744636 | 0.740806 | 1 | 0.996511 | 0.958796 |
| Cetn3 | A0A286YDI | 2.09 | 2.96 | 1.36 | 0.25 | 3.09 | 1.83 | 1.5 | 0.59 | 1.41 | -0.22 | 0.94 | 2.23 | 0.51 | 2.98 | 0.81 | 1.21 | 0.0875 | -0.575 | -0.2875 | 0.900434 | 0.420092 | 0.691558 | 1 | 0.996511 | 0.955729 |
| Cetn4 | Q8K4K1-3 | -0.3 | 1.64 | -1.55 | -0.53 | -0.15 | -1.35 | -0.06 | -1 | 0.02 | -0.85 | -0.85 | 0.74 | 0.93 | 1.4 | -1.15 | -0.92 | -0.455 | -0.05 | 0.25 | 0.502653 | 0.942935 | 0.764677 | 1 | 0.996755 | 0.964134 |
| Cfap20 | Q8BTU1 | -0.73 | -0.56 | -1.6 | -1.49 | -0.85 | -1.32 | -0.48 | -1.36 | -1.13 | -1.86 | -1.42 | -0.85 | -0.39 | -0.94 | -0.86 | -1.95 | 0.0925 | -0.22 | 0.06 | 0.777708 | 0.518354 | 0.882082 | 1 | 0.996511 | 0.979767 |
| Cfap298 | Q8BL95 | 0.55 | 0.48 | 0.85 | -0.4 | 0.87 | 0.53 | 0.47 | 0.57 | 2.19 | 0.92 | 0.3 | 0.05 | 2.48 | 1.94 | 0.03 | 0.3 | 0.24 | 0.495 | 0.8175 | 0.409505 | 0.344611 | 0.199586 | 1 | 0.996511 | 0.78004 |
| Cfap300 | Q8CC70 | -0.27 | -0.8 | -0.44 | -0.28 | -0.64 | -0.32 | 0.15 | -0.9 | 0.02 | -0.16 | -0.91 | 0.01 | 0.08 | -2.78 | -0.28 | -1.85 | 0.02 | 0.1875 | -0.76 | 0.939796 | 0.491847 | 0.24304 | 1 | 0.996511 | 0.798789 |
| Cfap36 | Q8C6E0 | -1.03 | -0.8 | 0.42 | 0.13 | -1.2 | -0.87 | 0.1 | 0.11 | -0.41 | 0.29 | 0.04 | -0.91 | -0.26 | -0.98 | 0.19 | -0.3 | -0.145 | 0.0725 | -0.0175 | 0.749183 | 0.862203 | 0.965871 | 1 | 0.996511 | 0.9927 |
| Cfap410 | Q8C6G1 | -1.53 | -0.95 | -0.23 | -1.01 | -1.62 | -1.2 | -0.33 | -1.22 | -1.09 | -1.09 | -0.35 | -0.58 | -0.89 | -1.06 | -1.25 | -0.33 | -0.1625 | 0.1525 | 0.0475 | 0.657201 | 0.64081 | 0.887098 | 1 | 0.996511 | 0.980239 |
| Cfap418 | B1AV75 | -0.76 | -1.19 | -0.63 | -0.74 | -1.55 | -0.55 | -1.26 | -1.42 | -0.59 | -0.68 | -0.89 | -0.66 | 0.32 | -1.63 | -0.21 | -1.14 | -0.365 | 0.125 | 0.165 | 0.189035 | 0.523417 | 0.70549 | 1 | 0.996511 | 0.957084 |
| Cfap74 | G3UWF6 | -4.1 | -3.76 | -2.27 | -2.87 | -3 | -3.8 | -2.89 | -3.36 | -2.37 | -1.34 | -2.18 | -3.94 | -1.35 | -2.89 | -2.75 | -2.56 | -0.0125 | 0.7925 | 0.8625 | 0.977014 | 0.226882 | 0.117003 | 1 | 0.996511 | 0.718608 |
| Cfdp1 | O88271 | -2.32 | -2.61 | -3.36 | -3.3 | -2.25 | -2.92 | -3.38 | -3.56 | -4.6 | -1.6 | -3.26 | -2 | -3.26 | -2.82 | -3.4 | -3.75 | -0.13 | 0.0325 | -0.41 | 0.726548 | 0.960738 | 0.229108 | 1 | 0.996755 | 0.787447 |
| Cfh | E9Q8I0 | -2.69 | -2.98 | -1.85 | -1.75 | -2.81 | -2.73 | -1.87 | -1.46 | -2 | -1.03 | -1.5 | -2.61 | -2.37 | -0.85 | -2.02 | -1.51 | 0.1 | 0.5325 | 0.63 | 0.812575 | 0.237788 | 0.166645 | 1 | 0.996511 | 0.759484 |
| Cfl1 | P18760 | 0.4 | -0.64 | 0.13 | 0.42 | -0.46 | 0.95 | 0.03 | 0.44 | -0.08 | -0.25 | 0.14 | 0.24 | -0.22 | -0.18 | -0.36 | -0.17 | 0.1625 | -0.065 | -0.31 | 0.662788 | 0.818284 | 0.276742 | 1 | 0.996511 | 0.822084 |
| Cfl2 | P45591 | 7.57 | 3.34 | -0.03 | 0.1 | 4.74 | 7.58 | 0.3 | 0.15 | 0.29 | -0.87 | -0.04 | 6.74 | -0.8 | 0.17 | -0.44 | -0.01 | 0.4475 | -1.215 | -3.015 | 0.843412 | 0.588989 | 0.086441 | 1 | 0.996511 | 0.691619 |
| Cgn | D3YUW7 | 0.17 | 0.57 | -0.4 | -0.62 | -0.58 | 0.13 | -1.4 | -0.65 | 1.56 | 1.83 | -1.47 | 0.15 | 1.74 | 0.51 | 0.49 | -1.32 | -0.555 | 0.5875 | 0.425 | 0.180612 | 0.430002 | 0.503224 | 1 | 0.996511 | 0.901002 |
| Cgnl1 | B2RSU6 | -0.41 | -0.34 | 1.09 | 0.77 | 0.11 | -0.53 | 1.27 | -0.02 | 0.17 | 1.73 | 0.46 | -0.2 | 0.6 | -0.18 | 1.12 | 0.93 | -0.07 | 0.2625 | 0.34 | 0.888105 | 0.619737 | 0.459061 | 1 | 0.996511 | 0.887902 |
| Cgref1 | E9QPK1 | -0.02 | -0.74 | -1.05 | -0.67 | 0.85 | -0.04 | -0.92 | -1.18 | 0.13 | 0.78 | -0.97 | 0.27 | 0.85 | 1.44 | -0.16 | -0.34 | 0.2975 | 0.6725 | 1.0675 | 0.533054 | 0.126163 | 0.038816 | 1 | 0.996511 | 0.658454 |
| Chac2 | Q9CQG1 | -1.06 | -0.77 | 0.11 | 0.01 | -1.21 | -0.85 | -0.03 | 0.37 | -0.77 | -0.49 | -0.28 | -0.76 | -0.9 | -0.3 | -0.67 | -0.29 | -0.0025 | -0.1475 | -0.1125 | 0.995393 | 0.640036 | 0.732625 | 1 | 0.996511 | 0.958635 |
| Chadl | E9Q7T7 | -1.33 | -1.49 | -1.4 | -1.79 | -0.87 | -1.94 | -0.25 | -1.45 | -0.25 | -0.33 | -1.55 | -1.32 | -0.15 | 0.46 | -0.06 | -0.53 | 0.375 | 0.64 | 1.4325 | 0.316147 | 0.090265 | 0.000389 | 1 | 0.996511 | 0.270112 |
| Champ1 | Q8K327 | -0.02 | -0.94 | -0.79 | -0.28 | -0.83 | -0.78 | -0.53 | -0.66 | -0.61 | -0.15 | -0.79 | -0.77 | -0.74 | -0.6 | -0.17 | -0.6 | -0.1925 | -0.0725 | -0.02 | 0.43279 | 0.792642 | 0.941339 | 1 | 0.996511 | 0.991085 |
| Chat | Q8BQV2 | -0.24 | -0.24 | -0.07 | 0.35 | 0.17 | -0.1 | 0.27 | 0.31 | 0.19 | -0.82 | -0.29 | 0 | -0.15 | 0.43 | -0.06 | 0.2 | 0.2125 | -0.18 | 0.155 | 0.304213 | 0.517161 | 0.512627 | 1 | 0.996511 | 0.904116 |
| Chchd1 | Q9CQA6 | -1.39 | -2.48 | -3.88 | -3.05 | -1.38 | -0.99 | -2.57 | -3.2 | -2.53 | -1.63 | -2.8 | -1.54 | -2.3 | -1.99 | -3.03 | -4.19 | 0.665 | 0.575 | -0.1775 | 0.331786 | 0.322127 | 0.78569 | 1 | 0.996511 | 0.967642 |
| Chchd10 | Q7TNL9 | 0.55 | -2.08 | -2.48 | -3.06 | -2.9 | 0.51 | -2.49 | -2.62 | -0.61 | -2.45 | -2.79 | 1.06 | -0.77 | -2.74 | -3.09 | -3.5 | -0.1075 | 0.57 | -0.7575 | 0.915461 | 0.598454 | 0.40921 | 1 | 0.996511 | 0.87457 |
| Chchd2 | Q9D1L0 | -1.92 | -2.53 | -0.58 | -0.41 | -3.13 | -2.41 | -0.41 | -0.37 | -2.33 | -0.07 | -0.52 | -1.89 | -2.3 | -1.65 | -0.46 | -0.47 | -0.22 | 0.1575 | 0.14 | 0.779984 | 0.816811 | 0.823824 | 1 | 0.996511 | 0.972768 |
| Chchd3 | Q9CRB9 | 1.72 | 1.16 | 0.65 | 1.02 | 0.27 | 1.31 | -0.01 | 0.5 | 1.61 | 0.55 | 0.66 | 1.42 | 1.64 | 0.17 | 0.78 | 0.13 | -0.62 | -0.0775 | -0.4575 | 0.101963 | 0.82066 | 0.272742 | 1 | 0.996511 | 0.822084 |
| Chchd3 | D3Z0L4 | 15.87 | 10.82 | 8.29 | 4.03 | 13.13 | 16.44 | 9.31 | 8.32 | -1.67 | -0.01 | 8.45 | 15.19 | -2.57 | 2.91 | 9.73 | 11.36 | 2.0475 | -4.2625 | -4.395 | 0.464949 | 0.31643 | 0.24052 | 1 | 0.996511 | 0.795658 |
| Chchd4 | Q8VEA4 | -4.35 | -3.97 | -3.06 | -2.96 | -3.27 | -3.37 | -3.07 | -3.02 | -5.15 | -1.98 | -2.47 | -3.61 | -2.7 | -4.29 | -3.17 | -3.15 | 0.4025 | 0.2825 | 0.2575 | 0.255572 | 0.69225 | 0.574101 | 1 | 0.996511 | 0.925754 |
| Chchd5 | Q9CQP3 | -0.32 | -0.33 | 0.39 | 1.34 | -0.41 | 0.12 | 1.17 | 1.02 | 0.11 | 0.92 | 0.9 | -0.2 | 0.53 | -0.48 | 0.42 | 0.7 | 0.205 | 0.1625 | 0.0225 | 0.684331 | 0.722299 | 0.959972 | 1 | 0.996511 | 0.992636 |
| Chchd6 | Q91VN4 | 0.56 | 0.32 | 0.57 | 0.65 | -0.01 | 0.65 | 0.07 | 0.33 | 1.09 | 0.55 | 0.58 | 0.66 | 1.22 | 0.17 | 0.41 | 0.34 | -0.265 | 0.195 | 0.01 | 0.205337 | 0.334911 | 0.970351 | 1 | 0.996511 | 0.992767 |
| Chchd7 | Q8K2Q5 | -0.44 | 0.78 | -0.78 | -1.88 | -0.83 | -0.14 | -1.29 | -1.13 | 0.08 | -1.24 | -0.67 | 1.3 | -0.32 | 1.38 | 0.01 | -0.51 | -0.2675 | 0.4475 | 0.72 | 0.63091 | 0.530142 | 0.274009 | 1 | 0.996511 | 0.822084 |
| Chd2 | E9PZM4 | 0.35 | 0.25 | 0.72 | 0.16 | 0.36 | 0.27 | 0 | 0.06 | 0.17 | 0.9 | 0.56 | -0.24 | 0.6 | 0.98 | 1.2 | 0.48 | -0.1975 | -0.0225 | 0.445 | 0.311395 | 0.937132 | 0.095496 | 1 | 0.996755 | 0.698156 |
| Chd3 | B1AR17 | -1.04 | -1.14 | -0.91 | -1.16 | -1 | -1.37 | -0.89 | -1.31 | -0.63 | -0.57 | -1.1 | -1.05 | -0.53 | -0.48 | -0.89 | -0.58 | -0.08 | 0.225 | 0.4425 | 0.653504 | 0.277037 | 0.040579 | 1 | 0.996511 | 0.658454 |
| Chd4 | E9QAS4 | -1.27 | -1.84 | -1.4 | -1.55 | -1.26 | -1.76 | -1.3 | -1.77 | -1.77 | -1.11 | -1.47 | -1.79 | -1.31 | -1.56 | -1.07 | -1.13 | -0.0075 | -0.02 | 0.2475 | 0.97187 | 0.931305 | 0.273015 | 1 | 0.996755 | 0.822084 |
| Chd5 | E9PYU4 | -0.93 | -2.37 | -0.49 | -1.33 | -1.14 | -1.16 | 0.33 | -1.83 | -1.42 | -0.37 | -0.6 | -0.61 | -0.68 | -1.25 | 0.24 | 0.17 | 0.33 | 0.53 | 0.9 | 0.557161 | 0.245554 | 0.100125 | 1 | 0.996511 | 0.698156 |
| Chd6 | A3KFM7-2 | 0.49 | 0.54 | -0.78 | -0.19 | 0.06 | 0.12 | 0.16 | -0.22 | 1.1 | -1.83 | -0.28 | 0.67 | 1.76 | 0.84 | -0.45 | -0.2 | 0.015 | -0.1 | 0.4725 | 0.962316 | 0.879013 | 0.40107 | 1 | 0.996511 | 0.870023 |
| Chd8 | Q09XV5 | -0.04 | -0.4 | -1.67 | -2.03 | -0.39 | -0.53 | -1.91 | -2.05 | -0.16 | -1.5 | -2.19 | -0.1 | 0.41 | 0.11 | -1.48 | -1.59 | -0.185 | 0.0475 | 0.3975 | 0.756611 | 0.941127 | 0.543705 | 1 | 0.996755 | 0.912212 |
| Chd9 | Q8BYH8-2 | -0.43 | 0.47 | 1.19 | -0.41 | -0.43 | -0.48 | -1.69 | -1.86 | 0.23 | -4.08 | 0.59 | 0.03 | 0.33 | 0.42 | 0.59 | -0.77 | -1.32 | -1.0125 | -0.0625 | 0.026549 | 0.346069 | 0.893589 | 1 | 0.996511 | 0.983001 |
| Chdh | Q8BJ64 | 0.65 | 1.53 | 0.73 | 1.04 | 1.09 | 1.11 | 0.78 | 0.83 | 1.24 | -0.04 | 0.39 | 0.77 | 1.47 | 0.87 | 0.7 | 1.12 | -0.035 | -0.3975 | 0.0525 | 0.881424 | 0.254256 | 0.851583 | 1 | 0.996511 | 0.976225 |
| Cherp | G5E8I8 | 0.19 | -0.8 | -0.96 | -0.99 | -0.63 | -0.77 | -0.65 | -0.5 | -0.88 | -1.03 | -0.92 | -1.13 | -0.56 | 0.06 | 0.12 | -0.82 | 0.0025 | -0.35 | 0.34 | 0.993028 | 0.25249 | 0.355746 | 1 | 0.996511 | 0.855029 |
| Chga | P26339 | -1.15 | -1.72 | -2 | -2.51 | -1.48 | -1.25 | -2.19 | -2.31 | -0.59 | -1.08 | -1.66 | -0.59 | -0.69 | -0.2 | -1.35 | -1.94 | 0.0375 | 0.865 | 0.8 | 0.918686 | 0.041591 | 0.101999 | 1 | 0.996511 | 0.700016 |
| Chgb | P16014 | 0.35 | 0.28 | -0.14 | -0.24 | 0.34 | 0.41 | -0.01 | -0.05 | 0.43 | 0.22 | -0.04 | 0.75 | 0.57 | 0.88 | 0.59 | 0.33 | 0.11 | 0.2775 | 0.53 | 0.611466 | 0.279827 | 0.043994 | 1 | 0.996511 | 0.658454 |
| Chic1 | Q8CBW7 | -0.39 | 0.1 | 0.03 | 0.54 | 0.53 | -0.23 | 0.25 | -0.27 | 0.09 | -0.02 | -0.04 | -0.17 | 0.53 | 1.14 | 0.34 | 0.48 | -7.9E-17 | -0.105 | 0.5525 | 1 | 0.649745 | 0.075411 | 1 | 0.996511 | 0.673156 |
| Chid1 | A0A0R4J24 | -1.52 | -1.5 | -1.97 | -2.08 | -0.84 | -1.23 | -0.63 | -1.89 | -1.4 | -2.38 | -2.21 | -1.77 | -0.32 | -0.96 | -0.75 | -1.26 | 0.62 | -0.1725 | 0.945 | 0.072629 | 0.541901 | 0.00678 | 1 | 0.996511 | 0.552614 |
| Chka | O54804-2 | -0.3 | -0.21 | -0.37 | -0.3 | -0.26 | -0.05 | -0.13 | -0.6 | -1.12 | 0.5 | -0.67 | -0.6 | -0.65 | -1.13 | -1.1 | -1.12 | 0.035 | -0.1775 | -0.705 | 0.84126 | 0.605106 | 0.005343 | 1 | 0.996511 | 0.552614 |
| Chkb | O55229 | 0.89 | 1.04 | -0.14 | -0.19 | 0.96 | 0.5 | -0.23 | -0.2 | 0.7 | -0.5 | 0.02 | 0.97 | 0.59 | 1.12 | -0.29 | -0.17 | -0.1425 | -0.1025 | -0.0875 | 0.729234 | 0.816073 | 0.843204 | 1 | 0.996511 | 0.976225 |
| Chl1 | P70232-2 | 2.02 | 2.58 | 1.49 | 2.12 | 1.74 | 3.72 | 2.31 | 3.1 | 2.56 | 3.15 | 1.87 | 3.19 | 2.63 | 1.18 | 3.56 | 3.51 | 0.665 | 0.64 | 0.6675 | 0.16935 | 0.111595 | 0.247251 | 1 | 0.996511 | 0.803366 |
| Chl1 | P70232 | -0.35 | -0.05 | -0.43 | -0.31 | -0.09 | -0.33 | -0.08 | -0.07 | -0.24 | -0.32 | -0.1 | -0.08 | -0.19 | 0.44 | -0.24 | -0.52 | 0.1425 | 0.1 | 0.1575 | 0.392329 | 0.568354 | 0.535885 | 1 | 0.996511 | 0.912212 |
| Chm | A2AD03 | 0.17 | -0.49 | -0.13 | -0.33 | -0.29 | 0.27 | -0.68 | -0.43 | -0.89 | -0.44 | -0.3 | 0.25 | -0.59 | -1.12 | -0.38 | -0.68 | -0.0875 | -0.15 | -0.4975 | 0.733474 | 0.60161 | 0.069703 | 1 | 0.996511 | 0.668066 |
| Chml | Q9QZD5 | -0.64 | -0.22 | -0.02 | 0.38 | -0.78 | -1.3 | -0.79 | -0.23 | -0.62 | 0.38 | 0.29 | -0.31 | -1.22 | 0.4 | -0.1 | -0.62 | -0.65 | 0.06 | -0.26 | 0.056501 | 0.851567 | 0.514445 | 1 | 0.996511 | 0.90567 |
| Chmp1a | Q921W0 | -0.38 | -0.19 | 0.41 | 0.39 | -0.08 | -0.83 | -0.15 | 0.5 | 0.25 | 0.73 | 0.52 | -0.34 | 1.6 | -0.44 | 0.48 | 0.19 | -0.1975 | 0.2325 | 0.4 | 0.552842 | 0.460974 | 0.381511 | 1 | 0.996511 | 0.860984 |
| Chmp1b1 | Q99LU0 | -0.74 | -0.37 | 0.2 | -0.76 | -0.3 | -0.7 | -0.72 | -0.65 | -0.41 | -1.85 | -0.09 | -0.4 | 0.6 | -0.49 | 0.11 | -0.13 | -0.175 | -0.27 | 0.44 | 0.499284 | 0.534311 | 0.199199 | 1 | 0.996511 | 0.78004 |
| Chmp1b2 | Q9CQD4 | -1.03 | -0.85 | -1.37 | -2.67 | -0.86 | -1.19 | -2.91 | -1.9 | -0.31 | -3.52 | -1.45 | -0.83 | 1.23 | -1.76 | -0.14 | -3.16 | -0.235 | -0.0475 | 0.5225 | 0.676342 | 0.948712 | 0.578421 | 1 | 0.996755 | 0.927198 |
| Chmp2a | Q9DB34 | -0.55 | -0.45 | 0.16 | 0.08 | -0.62 | -0.65 | -0.36 | 0.05 | -0.32 | 0.08 | 0.49 | -0.45 | 0.97 | -0.59 | 0.56 | 0.3 | -0.205 | 0.14 | 0.5 | 0.428 | 0.629607 | 0.198717 | 1 | 0.996511 | 0.78004 |
| Chmp2b | Q8BJF9 | -0.59 | -0.39 | -0.28 | -0.53 | -1.19 | -0.77 | -1.44 | -0.83 | -0.25 | -0.73 | -0.76 | -0.38 | 0.86 | -0.61 | -0.57 | -1.1 | -0.61 | -0.0825 | 0.0925 | 0.014484 | 0.676475 | 0.821956 | 1 | 0.996511 | 0.972768 |
| Chmp3 | Q9CQ10 | 0.16 | 0.25 | 0.21 | 0.68 | -0.23 | 0.11 | -0.06 | 0.54 | 0.26 | -0.63 | 0.31 | 0.51 | 0.58 | 0.06 | 0.29 | 0.01 | -0.235 | -0.2125 | -0.09 | 0.312629 | 0.469777 | 0.69 | 1 | 0.996511 | 0.955729 |
| Chmp4b | Q9D8B3 | -0.16 | 0.65 | -0.02 | -0.58 | 0.3 | 0.04 | -0.63 | -0.64 | 0.21 | 0.18 | -0.45 | 0.16 | 1.04 | 0.5 | -0.23 | -0.45 | -0.205 | 0.0525 | 0.2425 | 0.548342 | 0.863319 | 0.557657 | 1 | 0.996511 | 0.917092 |
| Chmp5 | Q9D7S9 | 0.13 | 0.37 | 0.51 | 0.36 | 0.04 | -0.19 | 0.02 | 0.22 | 0.09 | 0.78 | 0.23 | 0.1 | 0.21 | 0.43 | 0.06 | 0.39 | -0.32 | -0.0425 | -0.07 | 0.085511 | 0.846134 | 0.715851 | 1 | 0.996511 | 0.958106 |
| Chmp6 | P0C0A3 | 1.3 | 0.93 | 1.25 | 1.25 | 0.49 | 1.63 | 1.66 | 1.64 | 0.57 | 0.56 | 1.54 | 1.39 | 0.48 | -0.02 | 1.16 | 1.54 | 0.1725 | -0.1675 | -0.3925 | 0.567298 | 0.561223 | 0.285315 | 1 | 0.996511 | 0.822669 |
| Chmp7 | Q8R1T1 | -0.28 | -0.15 | -0.35 | -0.42 | 0.17 | -0.13 | -0.18 | -0.28 | 0.4 | -0.11 | -0.03 | -0.12 | 0.31 | 0.62 | -0.67 | -0.5 | 0.195 | 0.335 | 0.24 | 0.263551 | 0.108302 | 0.464024 | 1 | 0.996511 | 0.888267 |
| Chn1 | Q91V57 | -0.82 | -0.57 | -0.32 | -0.23 | -0.2 | -0.49 | -0.58 | -0.4 | -0.65 | -0.26 | -0.2 | -0.56 | -0.73 | -0.49 | -0.41 | -0.43 | 0.0675 | 0.0675 | -0.03 | 0.726942 | 0.75254 | 0.886423 | 1 | 0.996511 | 0.979767 |
| Chn2 | Q3V2R3 | 0.02 | -0.08 | -0.63 | -0.23 | -0.36 | -0.04 | -0.24 | -0.07 | 0.13 | -0.48 | -0.42 | -0.24 | 0.23 | 0.33 | -1.17 | -0.73 | 0.0525 | -0.0225 | -0.105 | 0.789471 | 0.922092 | 0.784144 | 1 | 0.996755 | 0.967642 |
| Chordc1 | Q9D1P4 | -0.99 | -1.21 | -0.83 | -0.59 | -1.07 | -0.81 | -0.83 | -0.48 | -1.33 | -0.25 | -0.62 | -0.93 | -1.32 | -1.04 | -0.83 | -0.99 | 0.1075 | 0.1225 | -0.14 | 0.607098 | 0.660151 | 0.525896 | 1 | 0.996511 | 0.908107 |
| Chp1 | P61022 | 1.37 | 1.08 | 1.18 | 1.42 | 0.68 | 1.23 | 1.72 | 1.75 | 0.79 | 0.6 | 1.53 | 1.69 | 0.9 | 0.42 | 1.28 | 1.28 | 0.0825 | -0.11 | -0.2925 | 0.760288 | 0.705084 | 0.264136 | 1 | 0.996511 | 0.816045 |
| Chpf | Q6IQX7 | -0.74 | -0.45 | 0.72 | 1.32 | -0.67 | -0.62 | -0.1 | 0.79 | -0.11 | 0.08 | 0.32 | -0.39 | -0.09 | -0.61 | 1.98 | 1.5 | -0.3625 | -0.2375 | 0.4825 | 0.510037 | 0.619635 | 0.504773 | 1 | 0.996511 | 0.901002 |
| Chpf2 | Q3UU43 | -0.42 | -1.94 | -2.52 | -2.82 | -0.94 | -0.07 | -2.66 | -4.44 | -0.8 | 1.58 | -3.09 | -1.59 | 0.78 | -0.05 | -0.05 | -0.68 | -0.1025 | 0.95 | 1.925 | 0.917611 | 0.354542 | 0.007503 | 1 | 0.996511 | 0.552614 |
| Chpt1 | Q8C025 | -0.27 | 0.29 | -0.02 | -0.18 | 0.78 | -0.65 | 1.2 | 0.13 | 2.98 | 0.59 | 0.04 | -0.55 | 2.9 | 5.73 | 0.73 | 0.81 | 0.41 | 0.81 | 2.5875 | 0.318394 | 0.273565 | 0.034598 | 1 | 0.996511 | 0.651434 |
| Chrm1 | P12657 | 0.54 | 0.95 | 0.56 | 0.67 | 0.87 | 0.82 | 0.14 | 0.19 | 1.67 | 0.7 | 0.39 | 1 | 1.96 | 1.48 | 1.55 | 0.34 | -0.175 | 0.26 | 0.6525 | 0.464849 | 0.390981 | 0.094208 | 1 | 0.996511 | 0.698156 |
| Chrm2 | Q9ERZ4 | 0.74 | 0.64 | 1.58 | 2.08 | -0.49 | 0.59 | 1.32 | 1.87 | 1.64 | 2.85 | 1.64 | 1.47 | 2.06 | 0.16 | 2.21 | 1.77 | -0.4375 | 0.64 | 0.29 | 0.445743 | 0.173739 | 0.594773 | 1 | 0.996511 | 0.929843 |
| Chrm3 | Q9ERZ3 | 0.3 | 0.58 | 1.39 | 1.05 | 1.08 | 0.3 | 0.95 | 1.1 | -0.36 | 0.97 | 2.07 | 0.04 | -0.37 | 0.36 | 0.93 | 1.42 | 0.0275 | -0.15 | -0.245 | 0.927852 | 0.784109 | 0.574675 | 1 | 0.996511 | 0.926139 |
| Chrm4 | P32211 | 0.23 | 0.15 | 0.72 | 0.43 | -0.15 | 0.11 | 0.37 | 0.39 | 1.04 | 0.75 | 0.41 | 0.57 | 1.28 | 0.11 | 0.22 | 0.56 | -0.2025 | 0.31 | 0.16 | 0.344836 | 0.186268 | 0.601883 | 1 | 0.996511 | 0.930802 |
| Chrna4 | B7ZBU7 | 0.6 | 1.37 | 0.27 | 0.64 | 1.45 | 0.64 | 0.61 | 0.3 | 0.84 | 0.57 | 0.33 | 0.91 | 0.48 | 1.27 | 0.7 | 0.33 | 0.03 | -0.0575 | -0.025 | 0.927205 | 0.837129 | 0.9372 | 1 | 0.996511 | 0.989767 |
| Chrnb2 | A0A0G2JG( | -0.29 | -0.5 | 0.16 | 0.47 | 0.11 | -0.75 | 1.03 | 0.54 | -0.16 | -0.8 | 0.14 | -0.42 | -0.92 | -0.49 | 0.14 | 0.34 | 0.2725 | -0.27 | -0.1925 | 0.512012 | 0.381836 | 0.593984 | 1 | 0.996511 | 0.929843 |
| Chst2 | Q80WV3 | 0.23 | 0.14 | -0.2 | -0.18 | 0.28 | -0.11 | 0.7 | 0 | 0.59 | -0.27 | -0.12 | 0.37 | 1.27 | 0.94 | 0.02 | -0.18 | 0.22 | 0.145 | 0.515 | 0.353013 | 0.56939 | 0.179495 | 1 | 0.996511 | 0.775542 |
| Chsy3 | Q5DTK1 | -1.64 | -3.6 | 0.16 | -0.47 | -3.64 | -1.47 | -0.86 | 0.16 | -1.69 | 0.97 | -0.29 | -3.57 | -0.12 | -2.45 | -0.1 | 0.34 | -0.065 | 0.2425 | 0.805 | 0.949788 | 0.832263 | 0.398203 | 1 | 0.996511 | 0.867334 |
| Chtop | E9PW20 | 1.12 | 0.48 | 0.41 | 0.58 | 0.24 | 0.56 | 0.19 | 1 | -0.03 | 0.98 | 0.47 | -0.77 | 0.83 | 0.17 | 0.79 | 0.5 | -0.15 | -0.485 | -0.075 | 0.562541 | 0.234774 | 0.768225 | 1 | 0.996511 | 0.965206 |
| Chuk | E9Q605 | -0.49 | -0.67 | 0.06 | -0.24 | -0.2 | -0.52 | -0.01 | -0.41 | 0.04 | -0.26 | -0.17 | -0.32 | -0.14 | 0.03 | -0.67 | -0.25 | 0.05 | 0.1575 | 0.0775 | 0.819332 | 0.473424 | 0.757822 | 1 | 0.996511 | 0.962717 |
| Ciao1 | Q99KN2 | -1.27 | -1.54 | -1.57 | -0.9 | -1.95 | -0.92 | -1.17 | -1.45 | -0.48 | -0.68 | -1.07 | -0.82 | 0.34 | -1.29 | 1.6 | -1.4 | -0.0525 | 0.5575 | 1.1325 | 0.848362 | 0.036672 | 0.116841 | 1 | 0.996511 | 0.718608 |
| Ciao2a | Q9DCL2 | 0.56 | 0.48 | -0.48 | -0.64 | 0.8 | -0.17 | -0.6 | -0.29 | 2.07 | 0.62 | -0.59 | 1.51 | 1.37 | 1.75 | -0.49 | -0.72 | -0.045 | 0.9225 | 0.4975 | 0.91248 | 0.1533 | 0.447109 | 1 | 0.996511 | 0.884817 |

| | | | | | | | | | | | | | | | | | | | | | | | | | | | |
|---|---|---|---|---|---|---|---|---|---|---|---|---|---|---|---|---|---|---|---|---|---|---|---|---|---|---|---|
| Ciao2b | Q9D187 | 0.34 | 0.47 | 0.03 | -0.05 | 0.83 | -0.33 | -0.2 | -0.31 | 0.15 | -0.01 | -0.27 | 0.22 | -0.78 | 0.37 | -0.77 | -0.64 | -0.2 | -0.175 | -0.6525 | 0.513649 | 0.411329 | 0.062133 | | 1 | 0.996511 | 0.663872 |
| Ciao3 | Q7TMW6 | -0.21 | -0.19 | -0.6 | -0.56 | -0.17 | -0.31 | -0.09 | -0.25 | 0.14 | -0.62 | -0.78 | -0.22 | 0.47 | -0.57 | -0.52 | -0.1 | 0.185 | 0.02 | 0.21 | 0.297491 | 0.937524 | 0.466579 | | 1 | 0.996755 | 0.888757 |
| Ciapin1 | Q8WTY4 | -1.28 | -1.16 | -1.28 | -1.74 | -0.91 | -0.98 | -1.3 | -1.59 | -1.71 | -0.87 | -1.43 | -0.81 | -0.9 | -1 | -1.43 | -1.72 | 0.17 | 0.16 | 0.1025 | 0.455909 | 0.556468 | 0.693747 | | 1 | 0.996511 | 0.955729 |
| Cib1 | A0A0U1RP | 1.19 | 1.06 | 1.07 | 0.81 | 0.92 | 0.69 | 0.46 | 0.53 | 0.92 | 0.61 | 0.42 | 0.51 | 0.86 | 0.95 | 0.71 | -0.19 | -0.3825 | -0.4175 | -0.45 | 0.05556 | 0.053536 | 0.147574 | | 1 | 0.996511 | 0.746406 |
| Cibar1 | Q8BP22 | -0.34 | -0.05 | 0.08 | -0.37 | -0.25 | -0.36 | -0.99 | -0.13 | 0.58 | 0.32 | 0.06 | -0.24 | 1.13 | -0.66 | 0.44 | -0.01 | -0.2625 | 0.35 | 0.395 | 0.28656 | 0.164415 | 0.316212 | | 1 | 0.996511 | 0.838265 |
| Cic | Q924A2 | -3.24 | -3.39 | -2.99 | -3.03 | -4.64 | -3.59 | -3.36 | -3.49 | -3.44 | -0.95 | -2.6 | -2.95 | -1.78 | -2.61 | -2.92 | -3.84 | -0.6075 | 0.6775 | 0.375 | 0.073233 | 0.206009 | 0.379664 | | 1 | 0.996511 | 0.860984 |
| Cirbp | P60824 | -0.76 | -2.6 | -2.2 | -2.58 | -3.18 | -2.16 | -1.8 | -2.41 | -1.29 | -2.99 | -2.54 | -2.71 | 0.14 | -3.69 | -1.3 | -2.49 | -0.3525 | -0.3475 | 0.2 | 0.473903 | 0.518542 | 0.810743 | | 1 | 0.996511 | 0.972252 |
| Cisd1 | Q91WS0 | 3.21 | 1.08 | 3.9 | -0.23 | 2.22 | 3.37 | 4.98 | 3.1 | -0.23 | -0.19 | 3.78 | 2.61 | -0.28 | 0.44 | 4.07 | 5.27 | 1.4275 | -0.4975 | 0.385 | 0.177882 | 0.690315 | 0.79382 | | 1 | 0.996511 | 0.969367 |
| Cisd2 | Q9CQB5 | -0.23 | -0.49 | 0.37 | 0.17 | 0.16 | -0.2 | 0.75 | 0.33 | -0.57 | 0.19 | 0.02 | -0.32 | -0.26 | 0.32 | 0.78 | 0.82 | 0.305 | -0.125 | 0.46 | 0.292307 | 0.648516 | 0.179189 | | 1 | 0.996511 | 0.775542 |
| Cisd3 | Z4YKM2 | -0.83 | -1.23 | -0.65 | -1.03 | -1.09 | -0.53 | -1.39 | -1.52 | -0.73 | 0.83 | -1.36 | -0.97 | 0.02 | -0.72 | 0.56 | -0.55 | -0.1975 | 0.3775 | 0.7625 | 0.457836 | 0.426131 | 0.039734 | | 1 | 0.996511 | 0.658454 |
| Cit | E9QL53 | 1.08 | 0.92 | 0.91 | 1.06 | 0.9 | 1.06 | 0.74 | 0.87 | 1.13 | 0.94 | 0.93 | 0.97 | 1.02 | 1.03 | 0.84 | 0.79 | -0.1 | -2.7E-16 | -0.0725 | 0.516079 | 1 | 0.680827 | | 1 | 1 | 0.955729 |
| Ciz1 | Q3UUV0 | 1.42 | -0.55 | 0.22 | 1.39 | 0.69 | -0.42 | 0.38 | -1.05 | 1.27 | 2.71 | 0.31 | -0.01 | 0.44 | 2 | -0.22 | 1.15 | -0.72 | 0.45 | 0.2225 | 0.226314 | 0.52853 | 0.719743 | | 1 | 0.996511 | 0.958398 |
| Ckap4 | Q8BMK4 | -0.42 | -0.73 | -0.98 | -0.77 | 0.47 | -0.27 | 0.46 | -0.36 | -0.16 | -0.43 | -0.97 | 0.1 | 0.2 | -0.01 | 1.23 | 0.83 | 0.8 | 0.36 | 1.2875 | 0.013282 | 0.207551 | 0.002769 | | 1 | 0.996511 | 0.453261 |
| Ckap5 | A0A0R4J0K | 0.44 | 0.3 | 0.48 | 0.47 | 0.16 | 0.51 | 0.7 | 0.65 | 0.56 | 0.2 | 0.57 | 0.53 | 0.48 | 0.15 | 0.58 | 0.42 | 0.0825 | 0.0425 | -0.015 | 0.642124 | 0.805737 | 0.935293 | | 1 | 0.996511 | 0.989767 |
| Ckb | Q04447 | -0.42 | -0.49 | -1.16 | -0.76 | -0.65 | -0.56 | -0.75 | -1.08 | -0.52 | -1.21 | -1.23 | -0.68 | -0.88 | -0.35 | -1.46 | -1.29 | -0.0525 | -0.2025 | -0.2875 | 0.815016 | 0.452449 | 0.36391 | | 1 | 0.996511 | 0.858475 |
| Ckm | A0A0J9YKD | -0.78 | -0.27 | -3.62 | -3.09 | -0.44 | -0.29 | -2.91 | -3.09 | -1.19 | -1.92 | -3.14 | -0.76 | -1.18 | -1.4 | -3.8 | -3.52 | 0.2575 | 0.1875 | -0.535 | 0.799193 | 0.832007 | 0.582816 | | 1 | 0.996511 | 0.927734 |
| Ckmt1 | P30275 | 0.95 | 0.49 | -0.18 | 0.26 | 0.15 | 0.72 | -0.11 | 0.1 | 0.86 | 0.33 | 0.06 | 0.94 | 0.46 | 0.8 | -0.18 | -0.28 | -0.165 | 0.1675 | -0.18 | 0.578135 | 0.600593 | 0.607516 | | 1 | 0.996511 | 0.933262 |
| Ckmt2 | Q6P8J7 | -8.18 | -9.1 | -7 | -6.95 | -6.42 | -10.33 | -7.89 | -9 | -11.75 | -5.3 | -7.76 | -7.57 | -8.03 | -8.5 | -8.72 | -8.41 | -0.6025 | -0.2875 | -0.6075 | 0.4991 | 0.823027 | 0.2441 | | 1 | 0.996511 | 0.799567 |
| Clasp1 | A0A087WF | 0.4 | 0.38 | -0.37 | 0.98 | 0.61 | 0.75 | 0.78 | 0.17 | 1.06 | 0.53 | 1.31 | 0.77 | 1.84 | 1.5 | 1.44 | 0.57 | 0.23 | 0.57 | 0.99 | 0.460523 | 0.106382 | 0.026167 | | 1 | 0.996511 | 0.633075 |
| Clasp1 | E9QKH0 | -0.7 | -0.57 | -0.39 | -0.53 | -0.79 | -0.76 | -0.49 | -1.1 | -0.23 | 1.1 | -1.54 | -1.18 | -0.28 | -0.79 | -1.18 | -0.53 | -0.2375 | 0.085 | -0.1475 | 0.216878 | 0.876522 | 0.544758 | | 1 | 0.996511 | 0.912212 |
| Clasp1 | A0A087WC | 0.13 | 0.25 | 0.53 | 0.02 | 0.22 | 0.12 | -0.15 | 0.11 | 0.14 | -0.59 | 0.4 | 0.35 | 0.57 | -0.25 | 0.49 | 0.22 | -0.1575 | -0.1575 | 0.025 | 0.392876 | 0.562721 | 0.920029 | | 1 | 0.996511 | 0.98688 |
| Clasp2 | Q08EB5 | -8.49 | -7.6 | -2.65 | -2.53 | -5.58 | -8.22 | -4.23 | -4.85 | -8.12 | -2.48 | -3.14 | -8.15 | -5.18 | -5.8 | -1.19 | -4.05 | -0.4025 | -0.155 | 1.2625 | 0.803581 | 0.937276 | 0.457901 | | 1 | 0.996755 | 0.887902 |
| Clasp2 | Q8BRT1-5 | -1.18 | -1.08 | 0.39 | 0.28 | -0.97 | -1.68 | -0.21 | 0.12 | -1.89 | 0.5 | 0.24 | -1.46 | -1.85 | -1.78 | 0.16 | 0.52 | -0.2875 | -0.255 | -0.34 | 0.595155 | 0.703981 | 0.622982 | | 1 | 0.996511 | 0.938628 |
| Clasp2 | Q08EB6 | 0.74 | 0.64 | 0.62 | 0.49 | 0.56 | 0.8 | 0.53 | 0.47 | 0.59 | 0.77 | 0.79 | 1.03 | 0.64 | 0.52 | 0.6 | 0.55 | -0.0325 | 0.1725 | -0.045 | 0.835012 | 0.338998 | 0.791453 | | 1 | 0.996511 | 0.969105 |
| Clasp2 | F7DCH5 | 1.34 | 1.2 | 0.91 | 1.11 | 0.56 | 0.9 | 0.31 | 0.71 | 1.57 | 2.06 | 1.14 | 1.69 | 1.41 | 2.2 | 0.54 | 0.13 | -0.52 | 0.475 | -0.07 | 0.022663 | 0.072871 | 0.875035 | | 1 | 0.996511 | 0.979626 |
| Clasrp | Q8CFC7 | 2.16 | 0.79 | 1.05 | 1.2 | -0.6 | 0.91 | 0.07 | 1.77 | 1.68 | 2.21 | 0.63 | 0.41 | 2.31 | 2.34 | 2 | 0.04 | -0.7625 | -0.0675 | 0.3725 | 0.186358 | 0.889617 | 0.522296 | | 1 | 0.996511 | 0.907667 |
| Clcc1 | Q99LI2 | 0.15 | 0.53 | 1.01 | 0.95 | 0.87 | 0.5 | 1.41 | 1.04 | -0.05 | 0.79 | 0.83 | 0.34 | 0.36 | -0.3 | 1.59 | 1.5 | 0.295 | -0.1825 | 0.1275 | 0.306351 | 0.542151 | 0.78636 | | 1 | 0.996511 | 0.967714 |
| Clcn2 | A0A338P67 | -0.16 | 0.58 | 0.51 | 0.21 | 0.61 | 0.14 | 0.92 | 0.12 | -0.68 | 0.41 | -0.12 | -0.08 | -0.27 | 0.06 | 0.05 | 0.73 | 0.1625 | -0.4025 | -0.1425 | 0.54336 | 0.187569 | 0.622197 | | 1 | 0.996511 | 0.938628 |
| Clcn3 | Q8K4W8 | 0.07 | 0.19 | 0.39 | 0.68 | 0.13 | 0.95 | -0.56 | 0.56 | 0.41 | 0 | -0.46 | 0.47 | 0.39 | 1.6 | -0.19 | -0.08 | -0.0625 | -0.2275 | 0.0975 | 0.853449 | 0.407966 | 0.813495 | | 1 | 0.996511 | 0.972768 |
| Clcn3 | A0A6Q6Q7 | -0.28 | 0.4 | 0.56 | 0.15 | 0.6 | 0.02 | 0.11 | 0.07 | -0.2 | 0.12 | 0.39 | -0.15 | 0.01 | 0.65 | 0.11 | 0.18 | -0.0075 | -0.1675 | 0.03 | 0.975262 | 0.510131 | 0.908224 | | 1 | 0.996511 | 0.986458 |
| Clcn4 | Q3LRV6 | -0.04 | 0.28 | 0.42 | 0.37 | 0.63 | 0.04 | 0.4 | 0.45 | 0.11 | 0.1 | 0.41 | 0.09 | 0 | 0.51 | 0.19 | 0.11 | 0.1225 | -0.08 | -0.055 | 0.53697 | 0.671938 | 0.793084 | | 1 | 0.996511 | 0.96933 |
| Clcn5 | Q9WVD4 | -1.3 | -0.58 | -0.89 | -1.22 | -0.19 | -0.93 | -0.65 | -1.55 | -1.86 | -2.98 | -0.87 | -1.14 | -1.79 | -1.2 | -1.53 | -1.22 | 0.1675 | -0.715 | -0.4375 | 0.604868 | 0.152641 | 0.107773 | | 1 | 0.996511 | 0.706141 |
| Clcn6 | A2A7F6 | -0.02 | 0.21 | -0.18 | -0.17 | 0.25 | 0.15 | 0.16 | 0 | 0.26 | -0.01 | -0.25 | 0.17 | 0.23 | 0.14 | -0.05 | -0.13 | 0.18 | 0.0825 | 0.0875 | 0.288207 | 0.67622 | 0.655482 | | 1 | 0.996511 | 0.94818 |
| Clcn7 | F6SUM2 | 0.05 | -0.15 | -0.7 | -0.44 | -0.56 | 0.72 | 0 | -0.63 | 0.6 | -0.46 | -0.2 | 0.31 | 1.48 | -1.05 | -0.38 | -0.55 | 0.1925 | 0.3725 | 0.185 | 0.576127 | 0.231877 | 0.729981 | | 1 | 0.996511 | 0.958635 |
| Cldn1 | O88551 | 0.76 | 0.42 | 1.27 | 1.08 | 1.08 | 0.8 | 1.36 | -0.08 | 0.96 | 0.8 | 1.07 | 0.22 | 0.43 | 1.21 | 0.49 | 0.57 | -0.0925 | -0.12 | -0.2075 | 0.791473 | 0.667995 | 0.466077 | | 1 | 0.996511 | 0.888548 |
| Cldn10 | Q9Z0S6-6 | 0.89 | 1.35 | 1.09 | 0.67 | 0.92 | 1.17 | 0.97 | 0.89 | 0.81 | 1.22 | 1.08 | 1.09 | 1.13 | 0.37 | 0.65 | 0.2 | -0.0125 | 0.05 | -4.13E-01 | 0.948617 | 0.81276 | 0.156348 | | 1 | 0.996511 | 0.751493 |
| Cldn11 | Q60771 | 0.18 | -1.81 | 0.02 | -1.42 | -1.28 | -0.45 | -0.11 | -1.47 | -2.81 | -1.2 | -1.51 | -1.34 | -1.57 | -3.74 | -0.43 | 0.08 | -0.07 | -0.9575 | -0.6575 | 0.89848 | 0.122893 | 0.464815 | | 1 | 0.996511 | 0.888267 |
| Cldn12 | Q9ET43 | -1.91 | -1.21 | -3.34 | -3.04 | -1.22 | -1.45 | -2.87 | -2.92 | -1.84 | -1.43 | -2.77 | -1.86 | -2.51 | -0.73 | -3.57 | -3.54 | 0.26 | 0.4 | -0.2125 | 0.672223 | 0.456463 | 0.777402 | | 1 | 0.996511 | 0.967289 |
| Cldn3 | Q9Z0G9-2 | 4.55 | -0.89 | -1.61 | -2.2 | 1.47 | -1.38 | -2.46 | -3.25 | 3.72 | 4.37 | -2.87 | 8.52 | -3.5 | -4.66 | -0.96 | -3.99 | -1.3675 | 3.4725 | -3.24 | 0.420299 | 0.190765 | 0.062722 | | 1 | 0.996511 | 0.66433 |
| Cldn5 | O54942 | -2.39 | -2.13 | -0.32 | -1.86 | -3.44 | -0.88 | -4.1 | -2.69 | 0.97 | -0.64 | -1.05 | -2.13 | 2.08 | -3.46 | -1.55 | -1.8 | -1.1025 | 0.9625 | 0.4925 | 0.169715 | 0.204449 | 0.661899 | | 1 | 0.996511 | 0.949645 |
| Cldnd1 | Q9CQX5 | 0.77 | 0.81 | 1.43 | 0.88 | 1.03 | 1.1 | 2.33 | 1.1 | 0.14 | 0.21 | 1.69 | 1.07 | -1.24 | 0.81 | 1.6 | 1.75 | 0.4175 | -0.195 | -0.2425 | 0.237856 | 0.615808 | 0.707075 | | 1 | 0.996511 | 0.957157 |
| Clec16a | Z4YLG3 | 0.53 | -0.16 | 0.3 | 0.12 | 0.21 | 0.65 | 0.36 | 0.03 | -0.17 | -0.48 | 0.42 | 0.49 | -0.48 | -0.44 | 0.23 | 0.41 | 0.115 | -0.1325 | -0.2675 | 0.603481 | 0.645212 | 0.365442 | | 1 | 0.996511 | 0.858475 |
| Clec2l | P0C7M9 | -0.38 | -0.3 | -0.17 | 0.07 | -0.24 | -0.11 | -0.16 | -0.3 | -0.04 | -0.02 | -0.05 | -0.22 | -0.15 | 0.07 | -0.36 | -0.85 | -0.0075 | 0.1125 | -0.1275 | 0.963639 | 0.531165 | 0.615831 | | 1 | 0.996511 | 0.93691 |
| Clgn | P52194 | -0.16 | 0.15 | -0.58 | -1.14 | -0.39 | -0.23 | -0.41 | -2.01 | 0.49 | -0.32 | -0.6 | 1.12 | -0.01 | 0.56 | -0.2 | -0.27 | -0.3275 | 0.605 | 0.4525 | 0.489315 | 0.203898 | 0.203253 | | 1 | 0.996511 | 0.782817 |
| Clic1 | Q9Z1Q5 | -0.28 | -1.38 | -1.34 | -1.04 | -1.05 | -0.77 | -1.78 | -1.52 | -0.46 | -0.62 | -1.46 | -0.68 | -1.02 | -0.68 | -2.01 | -1.85 | -0.27 | 0.205 | -0.38 | 0.424763 | 0.545256 | 0.350338 | | 1 | 0.996511 | 0.853281 |
| Clic4 | Q9QYB1 | 0.62 | 0.42 | 0.73 | 0.02 | 0.69 | 0.05 | 1.16 | 0.13 | 0.16 | 0.08 | 0.37 | 0.32 | -0.22 | 0.57 | 0.66 | 0.91 | 0.06 | -0.215 | 0.0325 | 0.841921 | 0.325418 | 0.914299 | | 1 | 0.996511 | 0.986458 |
| Clic5 | A0A668KLE | -0.42 | 0.06 | -0.56 | -0.02 | 0.44 | -0.24 | 0.14 | -0.2 | -1.07 | -2.58 | -0.01 | -0.18 | -1.21 | -1.46 | -0.67 | 0.29 | 0.27 | -0.725 | -0.5275 | 0.271999 | 0.217038 | 0.210629 | | 1 | 0.996511 | 0.786218 |
| Clic6 | Q8BHB9 | -2.87 | -2.62 | -2.36 | -2.2 | -2.99 | -3.58 | -2.94 | -3.75 | -3.36 | -0.67 | -5.28 | -2.92 | -1.85 | -2.46 | -2.1 | -2.04 | -0.8025 | -0.545 | 0.4 | 0.012886 | 0.532873 | 0.117202 | | 1 | 0.996511 | 0.718609 |
| Clint1 | Q5SUH7 | 1.2 | -0.61 | 0.21 | -0.81 | -0.5 | -0.04 | -1.06 | -1.42 | -2.2 | -1.97 | -0.27 | 0.79 | 0.16 | -4.81 | 0.08 | -0.24 | -0.7525 | -0.91 | -1.2 | 0.163157 | 0.254797 | 0.312079 | | 1 | 0.996511 | 0.834523 |
| Clint1 | Q5SUH6 | 0.72 | 0.77 | -0.04 | 0.01 | 0.85 | 0.55 | 0.07 | 0.03 | 0.92 | 0.09 | 0.16 | 0.95 | 0.58 | 1.43 | 0.05 | -0.14 | 0.01 | 0.165 | 0.115 | 0.972897 | 0.610248 | 0.773903 | | 1 | 0.996511 | 0.966062 |
| Clip1 | D3Z3M7 | 0.02 | 0.59 | -0.24 | 0.22 | -0.89 | 0.79 | -1.09 | -0.39 | 1.31 | 2.55 | 1.02 | 0.04 | 0.57 | -1.8 | -0.05 | 0.65 | -0.5425 | 1.0825 | -0.305 | 0.22575 | 0.058215 | 0.582406 | | 1 | 0.996511 | 0.927734 |
| Clip1 | Q922J3 | 0.03 | 0.57 | 0.58 | 0.63 | -0.02 | -0.75 | -0.1 | -0.14 | -0.01 | 0.67 | 0.67 | -0.02 | 0.64 | 1.35 | 0.64 | -0.88 | -0.705 | -0.125 | -0.015 | 0.014798 | 0.635095 | 0.97408 | | 1 | 0.996511 | 0.992977 |
| Clip1 | D3Z2Z1 | 0.13 | -0.02 | 0.34 | -0.09 | 0.02 | -0.04 | 0.46 | -0.13 | -0.23 | 0.35 | 0.27 | 0.14 | -0.14 | 0.2 | 0.17 | 0.15 | -0.0125 | 0.0425 | 5.00E-03 | 0.949305 | 0.836254 | 0.979552 | | 1 | 0.996511 | 0.994502 |
| Clip2 | Q9Z0H8 | 0.83 | 0.5 | 0.47 | 0.39 | 0.52 | 0.61 | 0.5 | 0.06 | 0.54 | 0.99 | 0.39 | 0.99 | 0.38 | 0.79 | 0.48 | 0.27 | -0.125 | 0.18 | -0.0675 | 0.523165 | 0.422219 | 0.746113 | | 1 | 0.996511 | 0.960124 |
| Clip3 | B9EHT4 | 0.91 | 0.73 | 0.57 | 0.55 | 1.18 | 1.16 | 0.76 | 0.95 | 0.47 | 0.73 | 1 | 1.39 | 0.54 | 0.56 | 1.05 | 0.79 | 0.3225 | 0.2075 | 0.045 | 0.096008 | 0.399305 | 0.827764 | | 1 | 0.996511 | 0.973757 |
| Clip4 | A0A2R8VH | -1.06 | -0.29 | -0.84 | -1.06 | -0.24 | -0.93 | -1.06 | -1.51 | 0.38 | -0.89 | -1.3 | -0.7 | 0.42 | -0.07 | -0.98 | -1.27 | -0.1225 | 0.185 | 0.3375 | 0.697385 | 0.63418 | 0.424638 | | 1 | 0.996511 | 0.876965 |
| Clmn | Q8C5W0-4 | 0.02 | 0.31 | 0.48 | 0.36 | -0.12 | 0.12 | 0.44 | 0.29 | 0.85 | 0.64 | 0.5 | 0.28 | 1.19 | 0.25 | 0.76 | 0.6 | -0.11 | 0.275 | 0.4075 | 0.570644 | 0.197674 | 0.131507 | | 1 | 0.996511 | 0.726398 |
| Clmp | Q8R373 | 0.76 | 1.03 | 0.74 | 0.75 | 0.57 | 0.43 | -0.32 | 0.59 | 1.3 | 0.73 | 0.12 | 1.06 | 1.48 | 1.03 | 0.66 | 0 | -0.5025 | -0.0175 | -0.0275 | 0.063469 | 0.949762 | 0.932593 | | 1 | 0.996755 | 0.989713 |
| Cln6 | D6RE34 | 0.04 | 0.37 | 0.47 | -0.11 | 1.82 | -0.77 | 0.53 | 0.28 | 0.43 | -0.8 | 0.59 | 0.6 | -0.11 | 1.36 | 0.78 | 1.06 | 0.2725 | 0.0125 | 0.58 | 0.593537 | 0.971908 | 0.11936 | | 1 | 0.997515 | 0.721448 |
| Cln8 | Q9QUK3 | -0.29 | -0.18 | -0.56 | -0.11 | 0.26 | 0.5 | 0.94 | 0.16 | 0.84 | -0.76 | -0.39 | 0.43 | 2.08 | 0.94 | 0 | -0.78 | 0.75 | 0.315 | 0.845 | 0.008056 | 0.403026 | 0.166507 | | 1 | 0.996511 | 0.759484 |
| Clns1a | Q923F1 | -0.42 | -0.27 | -0.05 | -0.05 | -0.1 | -0.7 | 0.03 | 0 | -0.29 | -0.04 | 0.1 | -0.15 | -0.65 | 0.29 | -0.91 | -0.15 | 0.005 | 0.1025 | -0.1575 | 0.981697 | 0.581865 | 0.597599 | | 1 | 0.996511 | 0.930093 |
| Clock | A0A0J9YU6 | -4.12 | -2.33 | -4.21 | -2.65 | -3 | -4.13 | -3.58 | -3.43 | -1.4 | -0.78 | -3.28 | -2.65 | -0.62 | -0.12 | -2.37 | -4.1 | -0.2075 | 1.3 | 1.525 | 0.679061 | 0.084705 | 0.125296 | | 1 | 0.996511 | 0.721448 |
| Clpb | Q60649 | 0.17 | 0.18 | 0.03 | 0.43 | -0.19 | 0.21 | 0.09 | 0.18 | 0.44 | 0.38 | 0.44 | 0.28 | 0.72 | 0.11 | 0.14 | 0.07 | -0.13 | 0.1825 | 0.0575 | 0.460998 | 0.2988 | 0.797293 | | 1 | 0.996511 | 0.969727 |
| Clpp | O88696 | 1.09 | 1.1 | 0.5 | 0.76 | 0.6 | 1 | 0.64 | 0.45 | 0.74 | 0.64 | 0.36 | 0.84 | 0.38 | 1.11 | 0.39 | 0.78 | -0.19 | -0.2175 | -0.1975 | 0.382434 | 0.32991 | 0.450398 | | 1 | 0.996511 | 0.884817 |
| Clptm1 | Q8VBZ3 | -0.23 | -0.17 | 0.18 | 0.19 | 0.18 | -0.11 | 0.75 | 0.36 | 0.03 | 0.51 | 0.45 | 0.03 | 0.4 | 0.7 | 1.01 | 0.87 | 0.3025 | 0.2625 | 0.7525 | 0.212625 | 0.238499 | 0.007736 | | 1 | 0.996511 | 0.555152 |
| Clptm1l | Q8BXA5 | -0.03 | 0.05 | -0.25 | 0.21 | 0.03 | -0.2 | 0.37 | 0.18 | -0.67 | 0.1 | -0.02 | -0.23 | -0.35 | -0.38 | 0.43 | 0.4 | 0.1 | -0.2 | 0.03 | 0.604411 | 0.390762 | 0.911334 | | 1 | 0.996511 | 0.986458 |
| Clpx | Q9JHS4 | 0.04 | 0.08 | 0.57 | 0.76 | -0.4 | -0.01 | 0.53 | 0.56 | 0.56 | 0.38 | 0.69 | 0.04 | 0.51 | 0.3 | 0.43 | 0.43 | -0.1925 | 0.055 | 0.055 | 0.51495 | 0.82653 | 0.810588 | | 1 | 0.996511 | 0.972222 |
| Clstn1 | Q9EPL2-2 | -3.21 | -3.54 | -1.93 | -2.59 | -2.73 | -2.68 | -1.81 | -2.11 | -4.04 | -1.01 | -2.44 | -2.76 | -3.31 | -2.24 | -2.7 | -2.37 | 0.485 | 0.255 | 0.1625 | 0.241805 | 0.697553 | 0.693528 | | 1 | 0.996511 | 0.955729 |
| Clstn1 | Q9EPL2 | -0.59 | -0.83 | -1.08 | -1.22 | -0.3 | -0.62 | -0.65 | -1.03 | -0.87 | -1.35 | -0.57 | -0.34 | -0.61 | -1.18 | -1.03 | -0.9 | 0.28 | 0.1475 | 7.85E-17 | 0.235005 | 0.59226 | 1 | | 1 | 0.996511 | 1 |
| Clstn2 | Q9ER65-2 | -1.39 | -0.08 | -0.64 | -0.88 | -0.87 | -2.13 | -0.3 | -0.75 | -2.22 | -0.95 | -0.69 | -1.21 | -1.26 | -0.86 | -0.43 | -0.66 | -0.265 | -0.52 | -0.055 | 0.555364 | 0.227015 | 0.866778 | | 1 | 0.996511 | 0.978305 |
| Clstn3 | D3Z601 | -0.3 | -0.71 | -1.03 | -1.11 | -0.58 | -0.48 | -0.45 | -0.56 | -0.45 | -0.66 | -0.95 | -0.25 | 0.2 | -0.17 | -0.11 | -0.41 | 0.27 | 0.21 | 0.665 | 0.226822 | 0.423964 | 0.026428 | | 1 | 0.996511 | 0.633866 |
| Clta | B1AWD8 | 2.08 | 1.81 | 1.1 | 0.4 | 2.28 | 1.73 | 0.98 | 0.82 | -0.28 | -2.63 | 0.93 | 1.81 | 0.03 | -0.94 | 1.07 | 1.14 | 0.105 | -1.39 | -1.0225 | 0.823597 | 0.161286 | 0.100703 | | 1 | 0.996511 | 0.698156 |
| Clta | B1AWE0 | 1.72 | 1.31 | 1.84 | 2.27 | 1.27 | 1.37 | 1.31 | 2.17 | 0.45 | 0.39 | 1.55 | 1.34 | 0.6 | -0.34 | 2 | 1.79 | -0.255 | -0.8525 | -0.7725 | 0.392337 | 0.036208 | 0.175264 | | 1 | 0.996511 | 0.769808 |
| Cltb | Q6IRU5-2 | 0.37 | 1.4 | 1.2 | 1.56 | 0.66 | 0.38 | -0.05 | 1.08 | 1.98 | 2.91 | 1.44 | 1.68 | 1.59 | 0.91 | 1.77 | 1.04 | -0.615 | 0.87 | 0.195 | 0.10101 | 0.053921 | 0.567543 | | 1 | 0.996511 | 0.921806 |
| Cltb | Q6IRU5 | 1.31 | 1.14 | 1.83 | 2.63 | 1.4 | 0.85 | 1.42 | 2.22 | 1.21 | 2.49 | 1.91 | 1.38 | 1.14 | 1.6 | 2.13 | 1.76 | -0.255 | 0.02 | -0.07 | 0.539844 | 0.961953 | 0.85457 | | 1 | 0.996755 | 0.976225 |
| Cltc | Q68FD5 | 0.7 | 0.33 | 0.79 | 0.75 | 0.67 | 0.88 | 1.16 | 1.12 | 0.03 | -0.08 | 0.92 | 0.6 | -0.51 | -0.12 | 0.61 | 0.97 | 0.315 | -0.275 | -0.405 | 0.129216 | 0.329545 | 0.266432 | | 1 | 0.996511 | 0.818107 |
| Clu | Q06890 | -0.1 | 0.1 | 0.11 | -0.27 | 0.7 | 0.43 | -0.23 | 0.55 | -0.71 | -0.35 | 0.48 | 0.89 | -0.59 | -0.16 | 0.15 | 0.23 | 0.4025 | 0.1175 | -0.0525 | 0.120821 | 0.749817 | 0.829534 | | 1 | 0.996511 | 0.973939 |
| Cluh | A0A0R4J14 | -1.35 | -1.53 | -1.21 | -1.29 | -1.48 | -0.77 | -1.01 | -1.27 | -1.23 | -0.58 | -1.15 | -1.09 | -0.91 | -0.83 | -1.45 | -1.32 | 0.2125 | 0.3325 | 0.2175 | 0.306673 | 0.13499 | 0.333517 | | 1 | 0.996511 | 0.845601 |
| Clvs1 | Q9D4C9 | 0.51 | 0.1 | 0.36 | -0.02 | 0.28 | 0.43 | 0.33 | -0.03 | -0.19 | 0.13 | 0.34 | 0.62 | -0.23 | 0.49 | -0.2 | 0.37 | 0.015 | -0.0125 | -0.13 | 0.938001 | 0.958023 | 0.612309 | | 1 | 0.996755 | 0.935166 |
| Clvs2 | Q8BG92 | -0.37 | 0.11 | -0.09 | -0.04 | 0.14 | -0.21 | -0.34 | -0.26 | -0.24 | 0.01 | 0.06 | -0.16 | -0.38 | 0.44 | -0.51 | -0.36 | -0.07 | 0.015 | -0.105 | 0.708542 | 0.935154 | 0.693495 | | 1 | 0.996755 | 0.955729 |
| Clybl | Q8R4N0 | 0.91 | 1.01 | 0.88 | 1.23 | 0.73 | 0.7 | 1.09 | 1 | 0.65 | 1.45 | 0.63 | 0.62 | 0.6 | 0.94 | 0.63 | 0.68 | -0.1275 | -0.17 | -0.295 | 0.472116 | 0.493186 | 0.145284 | | 1 | 0.996511 | 0.745321 |
| Cmas | A0A0R4J0E | -4.48 | -3.42 | -3.1 | -3.9 | -3.6 | -5.1 | -3.44 | -4.41 | -3.61 | -1.53 | -3.3 | -2.09 | -4.4 | -2.34 | -3.38 | -3.8 | -0.4125 | 1.0925 | 0.245 | 0.375462 | 0.067026 | 0.621498 | | 1 | 0.996511 | 0.938628 |
| Cmbl | Q8R1G2 | -0.19 | -0.52 | -0.52 | -0.35 | -0.83 | -0.41 | 0.04 | -0.48 | -1.21 | -0.54 | -0.53 | -0.63 | -1.23 | -1.15 | 0.05 | -0.51 | -0.025 | -0.3325 | -0.315 | 0.909191 | 0.154461 | 0.335347 | | 1 | 0.996511 | 0.847442 |
| Cmc1 | Q9CPZ8 | 0.04 | 0.27 | -0.86 | -1.03 | 1 | 0.25 | -0.67 | -1.05 | 0.38 | -2.25 | -1.52 | -0.08 | -0.46 | 0.16 | -1.51 | -1.29 | 0.2775 | -0.4725 | -0.38 | 0.59543 | 0.46301 | 0.429419 | | 1 | 0.996511 | 0.879165 |
| Cmc2 | Q8K199 | 0.03 | -0.31 | -0.73 | -1.17 | -0.47 | -0.05 | -1.07 | -1.47 | -0.27 | -0.38 | -1.57 | -0.31 | 0.3 | 0.1 | -1.49 | -2.15 | -0.22 | -0.0875 | -0.265 | 0.572865 | 0.822868 | 0.659902 | | 1 | 0.996511 | 0.949645 |
| Cmc4 | Q61908 | -3.45 | -2.72 | -2.05 | -1.69 | -2.39 | -2.85 | -1.84 | -2.49 | -2.58 | -1.88 | -1.85 | -2.88 | -0.43 | -2.75 | -1.52 | -1.32 | 0.085 | 0.18 | 0.9725 | 0.837352 | 0.683237 | 0.11362 | | 1 | 0.996511 | 0.716416 |
| Cmip | Q9D486 | 1.12 | 0.57 | 0.81 | -0.04 | 0.27 | 1.33 | 0.61 | 0.12 | 0.26 | -0.61 | 0.55 | 1.13 | 0.8 | 0.19 | 0.59 | 0.78 | -0.0325 | -0.2825 | -0.025 | 0.926059 | 0.503018 | 0.932963 | | 1 | 0.996511 | 0.989767 |
| Cmpk1 | Q9DBP5 | 0.15 | 0.17 | 0.25 | 0.23 | -0.33 | -0.18 | 0.2 | 0.49 | -0.27 | -0.32 | 0.4 | 0.31 | -0.94 | -0.42 | -0.34 | -0.45 | -0.155 | -0.17 | -0.7375 | 0.474717 | 0.457475 | 0.005284 | | 1 | 0.996511 | 0.552614 |
| Cmpk2 | Q3U5Q7 | 0.17 | 0.03 | 0.86 | 1.55 | -0.26 | 0.08 | 0.68 | 1.28 | 0.17 | 1.03 | 1.12 | -0.26 | -0.02 | 0.19 | 0.87 | 0.52 | -0.2075 | -0.1375 | -0.2625 | 0.649093 | 0.763082 | 0.50471 | | 1 | 0.996511 | 0.901002 |
| Cmss1 | Q9CZT6 | -0.51 | -1.28 | -2.17 | -1.07 | -0.67 | -0.61 | -0.6 | -1.88 | -0.16 | -1.88 | -0.93 | -0.08 | -0.45 | 0.18 | -1.31 | -1.65 | 0.3175 | 0.495 | 0.45 | 0.47287 | 0.340108 | 0.381921 | | 1 | 0.996511 | 0.860984 |
| Cmtm4 | Q8CJ61 | 0.21 | 1.15 | 0.91 | 1.21 | 0.81 | 0.49 | 0.63 | 0.9 | 1 | 1.26 | 0.83 | 1.05 | 0.93 | 1.39 | 0.94 | 0.9 | -0.1625 | 0.165 | 0.17 | 0.531708 | 0.536192 | 0.545923 | | 1 | 0.996511 | 0.912321 |
| Cmtm5 | Q45TP8 | 0.2 | 0.04 | 0.62 | 0.28 | 0.69 | 0.37 | 0.95 | 0.3 | 0.47 | 0.58 | 0.64 | 0.19 | 0.38 | 1.05 | 0.64 | 0.36 | 0.2925 | 0.185 | 0.3225 | 0.203115 | 0.376988 | 0.202817 | | 1 | 0.996511 | 0.782817 |
| Cmtr1 | Q9DBC3 | -3.73 | -3.62 | -3.51 | -3.88 | -3.28 | -4.5 | -3.49 | -4 | -3.65 | -1.96 | -3.55 | -3.36 | -2.82 | -3.22 | -4.06 | -4.18 | -0.1325 | 0.555 | 0.115 | 0.644774 | 0.176373 | 0.735008 | | 1 | 0.996511 | 0.958635 |
| Cnbp | P53996-3 | -2.55 | -4.84 | -4.4 | -5.55 | -4.64 | -2.47 | -4.3 | -4.15 | -4.17 | -3.5 | -4.58 | -1.9 | -2.61 | -5.65 | -2.17 | -4.3 | 0.445 | 0.7975 | 0.6525 | 0.544457 | 0.325919 | 0.484238 | | 1 | 0.996511 | 0.894266 |
| Cndp2 | Q9D1A2 | -0.54 | -0.47 | -0.39 | -0.07 | -0.29 | -0.7 | 0.32 | -0.12 | -0.52 | -0.27 | -0.16 | -0.43 | -0.72 | 0.04 | -0.32 | -0.22 | 0.17 | 0.0225 | 0.0625 | 0.499006 | 0.905261 | 0.788026 | | 1 | 0.996744 | 0.968587 |
| Cnep1r1 | D3Z6J3 | 0.7 | 0.67 | 0.61 | 0.92 | 0.88 | 0.99 | 0.99 | 0.87 | 0.35 | 1 | 0.46 | 0.44 | 0.43 | 1.35 | 0.58 | 0.99 | 0.2075 | -0.1625 | 0.1125 | 0.191396 | 0.441303 | 0.656361 | | 1 | 0.996511 | 0.948332 |
| Cnih1 | D6RGU4 | -3.98 | -3.64 | -4.7 | -5.26 | -3.52 | -3.73 | -4.33 | -5.48 | -3.39 | -4.95 | -4.68 | -4.35 | -2.93 | -2.73 | -5.72 | -4.66 | 0.13 | 0.0525 | 0.385 | 0.804398 | 0.910295 | 0.599149 | | 1 | 0.996744 | 0.930565 |
| Cnih2 | A0A494B9S | 3.9 | 2.87 | 3.37 | 3.84 | 0.93 | 3.74 | 3.78 | 4.88 | 4.69 | 4.95 | 4.01 | 3.35 | 2.61 | 3.45 | 3.47 | 3.1 | -0.1625 | 0.755 | -0.3375 | 0.837236 | 0.09424 | 0.306691 | | 1 | 0.996511 | 0.832533 |
| Cnih3 | G3XA11 | -1.83 | -2.59 | -2.35 | -0.63 | -2.66 | -0.42 | -2.36 | -1.3 | -0.63 | 0.27 | -1.39 | -0.34 | 0.44 | -0.35 | -2.6 | -2.09 | 0.165 | 1.3275 | 0.7 | 0.788188 | 0.028618 | 0.367609 | | 1 | 0.996511 | 0.859245 |
| Cnih4 | A0A0A6YW | 0.2 | 1.17 | 0.33 | 0.2 | 1.17 | 0.03 | 0.65 | 0.31 | 0.2 | 0.08 | 0.18 | 0.59 | -0.34 | 0.71 | -0.19 | 0.41 | 0.065 | -0.2125 | -0.3275 | 0.843582 | 0.445482 | 0.349642 | | 1 | 0.996511 | 0.85327 |
| Cnksr2 | Q80YA9 | 0.14 | 0.27 | 0.1 | 0.12 | 0.23 | 0.24 | 0.29 | 0.26 | 0.27 | 0.32 | 0.63 | 0.64 | 0.55 | 0.58 | 0.01 | 0.02 | 0.0975 | 0.3075 | 0.1325 | 0.490624 | 0.108655 | 0.545286 | | 1 | 0.996511 | 0.912212 |
| Cnn1 | Q08091 | -3.39 | -2.94 | -4.85 | -3.76 | -3.18 | -3.4 | -3.63 | -4.05 | -2.61 | -4.55 | -4.05 | -3 | -2.3 | -3.55 | -4.14 | -4.32 | 0.17 | 0.1825 | 0.1575 | 0.686991 | 0.744972 | 0.780252 | | 1 | 0.996511 | 0.9676 |
| Cnn2 | Q08093 | -2.48 | -3 | -2.04 | -2.49 | -1.81 | -2.66 | -2.23 | -2.76 | -1.8 | -1.38 | -1.7 | -2.05 | -2.15 | -1.81 | -2.53 | -2.51 | 0.1375 | 0.77 | 0.2525 | 0.640571 | 0.015194 | 0.379241 | | 1 | 0.996511 | 0.860984 |
| Cnn3 | Q9DAW9 | -0.55 | -0.35 | -0.41 | -0.09 | -0.87 | -0.36 | -1.35 | -0.51 | 0.57 | -0.06 | -0.21 | -0.12 | 0.78 | -0.26 | -0.59 | -0.65 | -0.4225 | 0.395 | 0.17 | 0.120917 | 0.117352 | 0.623685 | | 1 | 0.996511 | 0.938628 |
| Cnnm1 | A0A286YDI | 1.04 | 1.28 | 1.13 | 0.95 | 1.01 | 0.88 | 1.11 | 1.01 | 0.89 | 1.29 | 1.22 | 1.08 | 0.48 | 1.18 | 0.94 | 1.09 | -0.0975 | 0.02 | -0.1775 | 0.531903 | 0.911267 | 0.430694 | | 1 | 0.996744 | 0.879165 |
| Cnnm2 | Q3TWN3 | 0.93 | 1.14 | 0.51 | 0.51 | 1 | 0.78 | 0.86 | 0.32 | 1.21 | -0.14 | 0.66 | 0.9 | 0.59 | 1.39 | 0.11 | 0.46 | -0.0325 | -0.115 | -0.135 | 0.8893 | 0.72611 | 0.674155 | | 1 | 0.996511 | 0.952944 |
| Cnnm3 | Q32NY4 | 1.5 | 1.15 | 0.8 | 0.56 | 1.49 | 1.51 | 1.25 | 0.72 | 0.92 | 0.75 | 0.71 | 1.67 | 0.53 | 0.8 | 0.93 | 0.64 | 0.24 | 0.01 | -0.2775 | 0.400009 | 0.97403 | 0.292763 | | 1 | 0.997515 | 0.825612 |
| Cnnm4 | Q69ZF7 | 0.22 | 0.61 | 1.2 | 1.06 | 1.3 | 0.02 | 1.09 | 1.36 | 0.45 | 1.07 | 1.57 | 0.87 | -0.09 | 0.9 | 1.26 | 1.19 | 0.17 | 0.2175 | 0.0425 | 0.64489 | 0.505152 | 0.909566 | | 1 | 0.996511 | 0.986458 |
| Cnot1 | Q6ZQ08 | -0.74 | -0.81 | -0.53 | -0.69 | -0.35 | -0.89 | -0.67 | -0.85 | -0.82 | 0.28 | -0.67 | -0.75 | -0.55 | -0.42 | -0.67 | -0.56 | 0.0025 | 0.2025 | 0.1425 | 0.988972 | 0.473385 | 0.427682 | | 1 | 0.996511 | 0.879165 |
| Cnot10 | Q8BH15-2 | -1.07 | -1.1 | -1.19 | -0.51 | -1.24 | -1.08 | -0.89 | -1.32 | -1.73 | 1.43 | -1.22 | -0.07 | -0.5 | -2.04 | 0.29 | -0.22 | -0.165 | 0.57 | 0.35 | 0.439673 | 0.394763 | 0.48198 | | 1 | 0.996511 | 0.893925 |
| Cnot10 | Q8BH15 | -0.86 | -1 | -0.83 | -0.86 | -0.8 | -0.69 | -1.03 | -1.16 | -1.17 | 0.03 | -0.76 | -0.57 | -0.7 | -1.41 | -0.79 | -0.66 | -0.0325 | 0.27 | -0.0025 | 0.846663 | 0.329166 | 0.99118 | | 1 | 0.996511 | 0.996154 |
| Cnot11 | Q9CWN7 | -1.3 | -1.26 | 0.03 | -0.03 | -1.36 | -1.62 | 0.14 | -0.28 | -0.66 | 0.01 | -0.04 | -1.03 | -0.73 | -0.62 | -0.6 | -0.04 | -0.14 | 0.21 | 0.1425 | 0.786609 | 0.622647 | 0.714849 | | 1 | 0.996511 | 0.958106 |
| Cnot2 | Q8C5L3-2 | -1.4 | -1.4 | -1.01 | -1.75 | -1.35 | -1.45 | -2.5 | -2.43 | -0.15 | -2.15 | -1.84 | -1.43 | 0.92 | -1.62 | -0.69 | -2.09 | -0.5425 | -0.0025 | 0.52 | 0.130864 | 0.995448 | 0.412884 | | 1 | 0.999285 | 0.874811 |
| Cnot3 | Q8K0V4 | -0.94 | -0.66 | -0.75 | -0.83 | -0.66 | -1.4 | -0.93 | -0.97 | -1.16 | -0.47 | -0.8 | -0.8 | -1.33 | -0.59 | -0.66 | -0.81 | -0.195 | -0.0125 | -0.0525 | 0.339003 | 0.950483 | 0.815737 | | 1 | 0.996755 | 0.972768 |
| Cnot4 | Q05BG1 | -0.81 | -0.56 | -0.4 | -0.63 | -0.52 | -0.8 | -0.4 | -0.73 | -0.69 | -0.4 | -0.82 | -0.66 | -0.65 | -0.33 | -0.7 | -1.08 | -0.0125 | -0.0425 | -0.09 | 0.942899 | 0.817801 | 0.688992 | | 1 | 0.996511 | 0.955729 |
| Cnot6l | Q8VEG6-2 | 1.93 | 1.21 | 0.17 | -0.25 | 2.55 | 1.7 | 0.08 | -0.24 | 0.45 | -1.01 | 0.04 | 2.5 | -1.11 | 0.74 | -0.7 | -0.01 | 0.2575 | -0.27 | -1.035 | 0.731235 | 0.736887 | 0.10569 | | 1 | 0.996511 | 0.703417 |
| Cnot7 | Q60809 | -1.54 | -1.16 | -1.74 | -2.27 | -0.68 | -1.44 | -1.61 | -2.27 | -1.03 | -2.26 | -1.69 | -1.5 | -1.18 | -0.68 | -2.57 | -1.92 | 0.1775 | 0.0575 | 0.09 | 0.64242 | 0.865398 | 0.841187 | | 1 | 0.996511 | 0.976225 |
| Cnot9 | Q9JKY0 | -1.41 | -1.01 | -1.43 | -1.27 | -0.78 | -1.4 | -1.12 | -1.68 | -1.32 | -1.33 | -1.44 | -1.39 | -1.59 | -0.58 | -1.94 | -1.45 | 0.035 | -0.09 | -0.11 | 0.880845 | 0.607401 | 0.726298 | | 1 | 0.996511 | 0.958398 |
| Cnp | P16330 | 0.86 | 1.07 | 1.49 | 1.08 | 1.09 | 0.19 | 1.47 | 0.87 | 0.21 | 0.49 | 1.04 | 0.43 | -0.23 | 0.43 | 1.16 | 1.31 | -0.22 | -0.5825 | -0.4575 | 0.466904 | 0.039348 | 0.238331 | | 1 | 0.996511 | 0.793335 |
| Cnpy2 | Q9QXT0 | -0.33 | -0.59 | -0.85 | -0.73 | -0.35 | -0.58 | -0.54 | -1.06 | -0.2 | -0.18 | -1.14 | -0.19 | 0.77 | 0.34 | 0.31 | -0.32 | -0.0075 | 0.1975 | 0.9 | 0.972022 | 0.480908 | 0.009043 | | 1 | 0.996511 | 0.563386 |
| Cnpy3 | Q9DAU1 | -0.35 | -0.57 | -0.75 | -1.15 | 0.31 | -0.77 | -0.56 | -1.3 | -1.28 | -0.43 | -1.2 | -0.76 | -0.71 | 0.22 | 0.06 | -0.09 | 0.125 | -0.2125 | 0.575 | 0.728728 | 0.447649 | 0.068873 | | 1 | 0.996511 | 0.666991 |
| Cnpy4 | Q8BQ47 | 0.75 | 1.2 | 1.05 | 0.88 | 1.22 | 0.92 | 1.82 | 1.38 | 0.72 | 0.69 | 0.68 | 1.43 | 0.52 | 1.3 | 1.35 | 1.24 | 0.365 | -0.09 | 0.1325 | 0.140687 | 0.705174 | 0.601326 | | 1 | 0.996511 | 0.930802 |
| Cnr1 | P47746 | -0.28 | -0.04 | 0.15 | 0.14 | -0.27 | 0.06 | 0.06 | 0.03 | -0.05 | -0.08 | -0.11 | -0.03 | 0.03 | 0.28 | -0.38 | -0.43 | -0.0225 | -0.06 | -0.1175 | 0.898525 | 0.731775 | 0.623319 | | 1 | 0.996511 | 0.938628 |
| Cnrip1 | Q5M8N0 | 0.43 | 0.01 | 0.31 | 0.25 | 0 | 0.49 | 0.42 | 0.33 | 0.11 | -0.25 | 0.04 | 0.01 | -0.22 | 0.35 | -0.48 | -0.4 | 0.06 | -0.2725 | -0.4375 | 0.743532 | 0.160682 | 0.100154 | | 1 | 0.996511 | 0.698156 |
| Cnst | Q8CBC4 | -0.12 | 0.24 | 0.26 | -0.01 | 0.37 | -0.55 | 0.01 | -0.07 | -0.54 | 0.38 | -0.43 | -0.22 | 0.29 | 0.09 | 0.45 | 0.02 | -0.1525 | -0.295 | 0.12 | 0.513563 | 0.255869 | 0.554919 | | 1 | 0.996511 | 0.916796 |
| Cntfr | O88507 | 0.35 | -0.07 | 0.32 | 0.41 | -0.04 | 0.57 | 0.41 | 0.23 | 0.07 | -0.13 | 0.48 | 0.51 | 0.09 | 0.32 | -0.08 | -0.81 | 0.04 | -0.02 | -0.3725 | 0.843214 | 0.929306 | 0.214378 | | 1 | 0.996755 | 0.786931 |
| Cntn1 | P12960 | 0.69 | 0.75 | 0.68 | 0.47 | 0.93 | 0.66 | 0.97 | 0.65 | 0.37 | 0.46 | 0.75 | 0.88 | 0.17 | 0.85 | 0.55 | 0.48 | 0.155 | -0.0325 | -0.135 | 0.356697 | 0.864831 | 0.525991 | | 1 | 0.996511 | 0.908107 |
| Cntn2 | Q61330 | 0.95 | 1.01 | 1.08 | 1.16 | 1.23 | 0.83 | 0.91 | 0.77 | 0.39 | 0.71 | 0.68 | 0.67 | 0.17 | 0.52 | 0.86 | 1.05 | -0.115 | -0.4375 | -0.4 | 0.497717 | 0.027138 | 0.120955 | | 1 | 0.996511 | 0.721448 |
| Cntn3 | Q07409 | 0.35 | 0.8 | 0.89 | 0.86 | 0.87 | 0.5 | 0.93 | 0.95 | 0.43 | 0.7 | 1.1 | 0.9 | 0.42 | 0.52 | 1.11 | 0.96 | 0.0875 | 0.0575 | 0.0275 | 0.660975 | 0.798837 | 0.91077 | | 1 | 0.996511 | 0.986458 |
| Cntn4 | Q69Z26 | 0.3 | 0.26 | 0.43 | 0.65 | 0.53 | 0.68 | 0.56 | 0.56 | 0.6 | 0.44 | 0.57 | 0.73 | 0.45 | 0.81 | 0.33 | 0.32 | 0.1725 | 0.175 | 0.0675 | 0.292755 | 0.335096 | 0.743796 | | 1 | 0.996511 | 0.959321 |
| Cntn5 | P68500 | 0.68 | 0.59 | 0.79 | 1.07 | 0.15 | 0.84 | 0.97 | 1.41 | 1.46 | 0.72 | 1.26 | 0.94 | 1.79 | 0.5 | 1.24 | 0.99 | 0.06 | 0.3125 | 0.3475 | 0.832261 | 0.194103 | 0.266285 | | 1 | 0.996511 | 0.818107 |
| Cntn6 | Q9JMB8 | 0.74 | 0.82 | 0.56 | 0.7 | 0.51 | 0.73 | 0.66 | 0.44 | 0.96 | 0.97 | 0.73 | 1.16 | 0.18 | 1.06 | 0.1 | 0.47 | -0.12 | 0.25 | -0.2525 | 0.446192 | 0.178051 | 0.33659 | | 1 | 0.996511 | 0.847578 |
| Cntnap1 | O54991 | 0.41 | 0.44 | 0.22 | 0.01 | 0.29 | 0.33 | 0.81 | -0.25 | 0.02 | -0.5 | 0 | 0.36 | -0.02 | 0.13 | 0.2 | 0.33 | 0.025 | -0.3 | -0.11 | 0.920405 | 0.220831 | 0.576298 | | 1 | 0.996511 | 0.927 |
| Cntnap2 | E9QNF7 | 1.25 | 0.88 | 0.63 | 0.71 | 1.2 | 1.36 | 0.69 | 0.61 | 0.79 | 0.54 | 0.56 | 1.01 | 0.82 | 0.91 | 0.93 | 0.6 | 0.0975 | -0.1425 | -0.0525 | 0.691724 | 0.514991 | 0.805545 | | 1 | 0.996511 | 0.971715 |
| Cntnap3 | E9PY62 | -0.12 | -0.19 | 2.51 | 1.26 | -0.15 | 0.11 | 3.35 | 2.79 | 1.69 | 0.59 | 2.85 | 0.31 | 1.9 | 0.43 | 2.76 | 3.23 | 0.66 | 0.495 | 1.215 | 0.511592 | 0.53169 | 0.155284 | | 1 | 0.996511 | 0.750763 |
| Cntnap4 | D3YWB9 | 0.36 | 0.68 | 0.32 | 0.38 | 0.58 | 0.5 | 0.54 | 0.35 | 0.6 | 0.31 | 0.53 | 0.48 | 0.37 | 0.85 | 0.11 | 0.31 | 0.0575 | 0.045 | -2.50E-02 | 0.719491 | 0.797247 | 0.911464 | | 1 | 0.996511 | 0.986458 |
| Cntnap5a | Q0V8T9 | 0.79 | 0.73 | -0.09 | 0.08 | 0.77 | 0.82 | -0.07 | 0.08 | 0.4 | -0.34 | 0.23 | 1.14 | 0.28 | 0.67 | 0.01 | 0.03 | 0.0225 | -0.02 | -0.13 | 0.943028 | 0.956519 | 0.654231 | | 1 | 0.996755 | 0.947657 |
| Cntnap5b | Q0V8T8 | 1.36 | 1.4 | 0.79 | 0.67 | 1.87 | 1.31 | 1.2 | 0.79 | 1.45 | 1.51 | 1.02 | 1.57 | 0.82 | 1.86 | 0.96 | 1.03 | 0.2375 | 0.3325 | 0.1125 | 0.425258 | 0.206538 | 0.718448 | | 1 | 0.996511 | 0.958398 |
| Cntnap5c | Q0V8T7 | 0.72 | 0.69 | 2.34 | 1.31 | 0.62 | 0.91 | 0.41 | 1.39 | -0.07 | 0.76 | 0.67 | 0.87 | 0.48 | -0.8 | 2.73 | 1.03 | -0.4325 | -0.7075 | -0.405 | 0.31133 | 0.118573 | 0.591125 | | 1 | 0.996511 | 0.929523 |
| Coa3 | Q9D2R6 | 0.59 | 0.74 | 0.76 | 1.54 | 0.53 | 0.89 | 0.8 | 1.52 | 0.76 | 0.7 | 0.97 | 0.59 | 0.43 | 1.43 | 0.37 | 0.45 | 0.0275 | -0.1525 | -0.2375 | 0.926375 | 0.548262 | 0.483833 | | 1 | 0.996511 | 0.894091 |
| COA4 | Q8BT51 | -0.95 | -2.67 | 0.26 | 1.19 | -2 | -0.97 | 0.85 | 0.82 | -2.37 | 1.14 | 0.64 | -1.03 | -2.34 | -2.79 | 0.31 | 0.21 | 0.2175 | 0.1375 | -0.61 | 0.824117 | 0.894641 | 0.563283 | | 1 | 0.996511 | 0.920024 |
| Coa6 | Q8BGD8 | -1.4 | -2.47 | -1.67 | -1.18 | -2.26 | -0.94 | -1.75 | -1.65 | -4.46 | -0.81 | -1.52 | -1.64 | -2.11 | -4.57 | -1.77 | -1.48 | 0.03 | -0.4275 | -0.8025 | 0.935784 | 0.58319 | 0.264435 | | 1 | 0.996511 | 0.816417 |
| Coa7 | Q921H9 | -0.04 | -0.17 | 0.28 | -0.02 | -0.47 | 0.42 | -0.46 | -0.03 | -0.44 | 0.5 | 0.26 | -0.08 | -0.24 | -0.76 | 0.09 | 0.27 | -0.1475 | 0.0475 | -0.1725 | 0.552212 | 0.848935 | 0.527698 | | 1 | 0.996511 | 0.908538 |
| Coa8 | E9Q6C3 | 0.92 | 1.61 | 0.7 | 0.57 | 2.03 | 1.19 | 1.82 | 1 | 1.69 | 1.46 | 0.58 | 1.93 | 0.24 | 1.92 | 0.72 | 0.77 | 0.56 | 0.465 | -0.0375 | 0.116388 | 0.223342 | 0.926749 | | 1 | 0.996511 | 0.989128 |
| Coasy | Q9DBL7 | -0.06 | 0.03 | 1.55 | 1.59 | 0.02 | -0.08 | 1.03 | 1.9 | -1.11 | 0.54 | 1.69 | -0.01 | -1.02 | -2.2 | 1.36 | 1.54 | -0.06 | -0.5 | -0.8575 | 0.91985 | 0.465975 | 0.362889 | | 1 | 0.996511 | 0.858475 |
| Cobl | G5E8P4 | 0.19 | -0.11 | 0.11 | 0.02 | 0.38 | 0.02 | -0.47 | -0.18 | 0.29 | 0.82 | 0.26 | 0.01 | 0.67 | 0.8 | -0.1 | -0.42 | -0.115 | 0.2925 | 0.185 | 0.596261 | 0.205717 | 0.556856 | | 1 | 0.996511 | 0.917092 |
| Cobll1 | B1AZ15 | -0.88 | -1.81 | -1.99 | -2.41 | -1.95 | -1.37 | -2.44 | -2.56 | -0.22 | -0.76 | -1.61 | -0.89 | -0.36 | -0.84 | -1.33 | -2.34 | -0.3075 | 0.9025 | 0.555 | 0.44763 | 0.052665 | 0.281441 | | 1 | 0.996511 | 0.822084 |
| Coch | Q62507 | 0.95 | -0.46 | 0.26 | -0.01 | 0.81 | 0.73 | 0.63 | -0.21 | -0.99 | 0.4 | -0.08 | 0.91 | -0.63 | -0.35 | 0.21 | 0.45 | 0.305 | -0.125 | -0.265 | 0.40885 | 0.789866 | 0.487734 | | 1 | 0.996511 | 0.897297 |
| Cog1 | Q9Z160 | -1.89 | -1.86 | -0.95 | -1.36 | -1.53 | -2.13 | -1.36 | -1.54 | -1.21 | -1.07 | -1.54 | -1.71 | -1.23 | -1.41 | -1.41 | -1.12 | -0.125 | 0.1325 | 0.2225 | 0.659963 | 0.638324 | 0.407266 | | 1 | 0.996511 | 0.874084 |
| Cog2 | Q921L5 | -2.77 | -2.24 | -0.96 | -2.18 | -1.15 | -2.73 | -1.86 | -2.42 | -2.9 | -1.59 | -2.16 | -2.37 | -2.54 | -2.01 | -1.08 | -1.83 | -0.0025 | -0.2175 | 0.1725 | 0.995826 | 0.625038 | 0.708301 | | 1 | 0.996511 | 0.957562 |
| Cog3 | E9QL65 | -1.76 | -1.86 | -1.3 | -1.03 | -1.44 | -2.01 | -1.21 | -1.37 | -1.77 | -0.61 | -1.34 | -1.77 | -2.01 | -1.12 | -1.7 | -1.17 | -0.02 | 0.115 | -0.0125 | 0.940476 | 0.730655 | 0.967001 | | 1 | 0.996511 | 0.9927 |
| Cog4 | G3UWH9 | -1.52 | -1.46 | -2.14 | -1.67 | -1.15 | -1.72 | -1.71 | -2.4 | -1.49 | -1.66 | -1.89 | -1.02 | -1.55 | -1.3 | -2.36 | -1.76 | -0.0475 | 0.1825 | -0.045 | 0.872891 | 0.488333 | 0.876769 | | 1 | 0.996511 | 0.979626 |
| Cog5 | Q8C0L8 | -1.17 | -1.45 | -1.36 | -1.42 | -1.18 | -1.23 | -1.24 | -1.74 | -1.36 | -1.13 | -1.71 | -1.38 | -0.66 | -0.67 | -1.73 | -1.15 | 0.0025 | -0.045 | 0.2975 | 0.989295 | 0.813972 | 0.305453 | | 1 | 0.996511 | 0.832258 |
| Cog6 | A0A0R4J0L | -1.6 | -1.54 | -1.22 | -1.55 | -1.19 | -1.43 | -0.38 | -1.48 | -0.99 | -0.71 | -1.17 | -1.25 | -0.07 | -0.65 | -1.39 | -0.9 | 0.3575 | 0.4475 | 0.725 | 0.214088 | 0.047754 | 0.036827 | | 1 | 0.996511 | 0.658454 |
| Cog7 | A0A0R4J0C | -0.37 | -0.71 | -1.4 | -1.53 | -0.47 | -0.45 | -0.98 | -1.84 | 0 | -1.25 | -1.43 | -0.1 | -0.27 | -0.22 | -1.77 | -1.18 | 0.0675 | 0.3075 | 0.1425 | 0.86692 | 0.490585 | 0.747691 | | 1 | 0.996511 | 0.960744 |
| Cog8 | Q9JJA2 | -2.76 | -2.85 | -1.72 | -1.89 | -1.31 | -2.53 | -1.16 | -1.66 | -0.75 | -1.53 | -1.58 | -2.27 | 0.39 | -0.99 | -1.88 | -1.47 | 0.64 | 0.7725 | 1.3175 | 0.135941 | 0.084773 | 0.032992 | | 1 | 0.996511 | 0.644622 |
| Coil | Q5SU73 | 1.63 | 1.52 | 1.69 | 1.84 | 0.66 | 1.02 | 1.04 | 1.88 | 1.29 | 1.18 | 1.48 | 1.18 | 1.38 | 1.2 | 1.61 | 1.57 | -0.52 | -0.3875 | -0.23 | 0.083556 | 0.048184 | 0.249196 | | 1 | 0.996511 | 0.804842 |
| Col12a1 | E9PX70 | -0.83 | -1.22 | -1.59 | -1.5 | -0.82 | -1.55 | -2.33 | -2.26 | -0.89 | -1.03 | -1.45 | -0.65 | -1.4 | -0.93 | -2.17 | -1.7 | -0.455 | 0.28 | -0.265 | 0.243932 | 0.296529 | 0.414798 | | 1 | 0.996511 | 0.874945 |

| | | | | | | | | | | | | | | | | | | | | | | | | | | |
|---|---|---|---|---|---|---|---|---|---|---|---|---|---|---|---|---|---|---|---|---|---|---|---|---|---|---|
| Col18a1 | E9QPX1 | -2.96 | -2.98 | -3.56 | -3.51 | -2.66 | -3.07 | -3.69 | -4.09 | -2.74 | -2.77 | -3.15 | -2.36 | -2.52 | -3.31 | -3.42 | -3.69 | -0.125 | 0.4975 | 0.0175 | 0.717821 | 0.075334 | 0.954884 | 1 | 0.996511 | 0.992482 |
| Col19a1 | F8WHV4 | -1.28 | -0.74 | -0.58 | -1.01 | -1.48 | -1.01 | -0.77 | -1.56 | -1.08 | -0.71 | -0.48 | -1.18 | -2.38 | 0.31 | -1.79 | -1.3 | -0.3025 | 0.04 | -0.3875 | 0.254068 | 0.871874 | 0.488442 | 1 | 0.996511 | 0.897531 |
| Col1a1 | P11087 | -6.49 | -7.24 | -7.02 | -7.74 | -6.5 | -6.67 | -7.49 | -7.38 | -6.21 | -7.08 | -6.24 | -6.15 | -4.45 | -6.53 | -7.84 | -7.4 | 0.1125 | 0.7025 | 0.5675 | 0.745704 | 0.062868 | 0.439471 | 1 | 0.996511 | 0.883205 |
| Col1a2 | Q01149 | -1.07 | -1.66 | -1.09 | -1.69 | -2.11 | -0.55 | -2.75 | -1.69 | -2.62 | -4.16 | -1.01 | -0.99 | -1.74 | -4.49 | -1.13 | -1.15 | -0.3975 | -0.8175 | -0.75 | 0.396873 | 0.265986 | 0.325523 | 1 | 0.996511 | 0.843085 |
| Col25a1 | V9GWX5 | -3.12 | -2.99 | -1.83 | -1.83 | -3.81 | -2.36 | -2.27 | -1.66 | 1.3 | -1.19 | -1.68 | -3.03 | 2.09 | -2.33 | -0.22 | -2.1 | -0.0825 | 1.2925 | 1.8025 | 0.876358 | 0.16569 | 0.090892 | 1 | 0.996511 | 0.693316 |
| Col3a1 | P08121 | -3.09 | -4.62 | -9.1 | -18.01 | -4.86 | -2.44 | -14.92 | -11.91 | 0.23 | -7.78 | -10.95 | -3.49 | 1.21 | -5.32 | -10.95 | -12.35 | 0.1725 | 3.2075 | 1.8525 | 0.965221 | 0.395537 | 0.647377 | 1 | 0.996511 | 0.945544 |
| Col4a1 | P02463 | -0.55 | -0.95 | -0.47 | -0.75 | -0.8 | -0.67 | -0.62 | -0.45 | -0.89 | 0.28 | -0.4 | -0.77 | -0.22 | -0.96 | -0.57 | -0.62 | 0.045 | 0.235 | 0.0875 | 0.799488 | 0.430576 | 0.704852 | 1 | 0.996511 | 0.957084 |
| Col4a2 | P08122 | -1.29 | -1.39 | -1.67 | -2.05 | -1.19 | -0.83 | -1.47 | -1.94 | -1 | -1.37 | -1.52 | -1.11 | -0.66 | -1.45 | -1.1 | -1.2 | 0.2425 | 0.35 | 0.4975 | 0.412558 | 0.16459 | 0.086718 | 1 | 0.996511 | 0.691619 |
| Col4a5 | Q63ZW6 | 2.17 | 2.13 | 0.02 | -1.93 | 1.52 | 2.21 | 1.34 | 1.72 | 1.15 | -0.85 | 2.62 | 2.29 | 0.88 | 1.27 | 1.99 | 2.98 | 1.1 | 0.705 | 1.1825 | 0.241126 | 0.535625 | 0.243125 | 1 | 0.996511 | 0.798789 |
| Col6a1 | Q04857 | -3.3 | -3.03 | -3.21 | -3.67 | -2.83 | -3.41 | -3.5 | -3.97 | -3.44 | -2.26 | -3.28 | -3.25 | -3.24 | -3.28 | -3.36 | -3.45 | -0.125 | 0.245 | -0.03 | 0.650667 | 0.431102 | 0.883603 | 1 | 0.996511 | 0.979767 |
| Col6a2 | Q02788 | -2.7 | -2.99 | -3.73 | -3.53 | -2.47 | -2.78 | -3.25 | -4.16 | -2.51 | -3.05 | -2.97 | -2.5 | -2.31 | -2.71 | -3.49 | -3.75 | 0.0725 | 0.48 | 0.1725 | 0.860296 | 0.123807 | 0.665165 | 1 | 0.996511 | 0.949825 |
| Col6a3 | J3QQ16 | -2.95 | -3.48 | -3.4 | -3.19 | -2.97 | -3.33 | -3.07 | -3.36 | -3.37 | -3.13 | -2.78 | -2.7 | -3.14 | -2.66 | -3.34 | -2.99 | 0.0725 | 0.26 | 0.2225 | 0.705073 | 0.27388 | 0.347433 | 1 | 0.996511 | 0.851503 |
| Colec12 | A0A452J8C | -0.21 | -0.25 | 0.28 | -0.22 | -0.3 | 0.17 | 0.28 | -0.49 | -0.39 | -0.1 | -0.78 | -0.04 | -0.29 | -0.18 | -0.41 | 0.34 | 0.015 | -0.2275 | -0.035 | 0.950055 | 0.354633 | 0.886218 | 1 | 0.996511 | 0.979767 |
| Colgalt1 | Q8K297 | -0.75 | -0.67 | -0.02 | -0.32 | 0.01 | -1.05 | 0.71 | -0.21 | -0.32 | -0.24 | -0.03 | -0.75 | 0.95 | 0.12 | 0.45 | 0.86 | 0.305 | 0.105 | 1.035 | 0.430584 | 0.675479 | 0.004568 | 1 | 0.996511 | 0.550241 |
| Commd1 | Q8VI86 | -0.89 | 0.18 | 1.42 | 1.88 | -0.49 | -1.21 | 1.04 | 1.89 | -1.05 | 1.84 | 1.57 | -0.7 | -0.17 | -0.35 | 1.66 | 1.33 | -0.34 | -0.2325 | -0.03 | 0.690108 | 0.791982 | 0.967269 | 1 | 0.996511 | 0.9927 |
| Commd10 | Q8JZY2 | -0.8 | -0.11 | -0.8 | -0.95 | -0.11 | -0.8 | -0.8 | -1.07 | -0.67 | -0.3 | -0.65 | -0.4 | -1.17 | 0.17 | -1.11 | -0.97 | -0.03 | 0.16 | -0.105 | 0.914957 | 0.507017 | 0.77232 | 1 | 0.996511 | 0.965933 |
| Commd2 | Q8BXC6 | -1.34 | -1.22 | -1.6 | -1.65 | -0.5 | -1.77 | -0.98 | -1.57 | -1.27 | -1.83 | -1.7 | -1.34 | -1.51 | -0.09 | -1.81 | -1.49 | 0.2475 | -0.0825 | 0.2275 | 0.423813 | 0.698753 | 0.560561 | 1 | 0.996511 | 0.919045 |
| Commd3 | Q63829 | -0.48 | -0.3 | -0.01 | -0.14 | -0.2 | -0.55 | -0.14 | -0.02 | -0.36 | 0.39 | -0.08 | -0.18 | -0.51 | 0.26 | -0.44 | 0.03 | 0.005 | 0.175 | 0.0675 | 0.979068 | 0.443846 | 0.784795 | 1 | 0.996511 | 0.967642 |
| Commd4 | Q9CQ02 | -2.63 | -2.27 | -1.73 | -3.52 | -1.94 | -1.83 | -2.22 | -3.26 | -2.5 | -3.23 | -2.41 | -2.75 | -2.61 | -1.93 | -3.42 | -2.71 | 0.225 | -0.185 | -0.13 | 0.62876 | 0.645772 | 0.776211 | 1 | 0.996511 | 0.966842 |
| Commd5 | A0A0R4J0U | -0.61 | -0.46 | -0.64 | -0.81 | -0.13 | -0.61 | -0.52 | -0.64 | -0.81 | -1.06 | -0.79 | -0.57 | -0.72 | -0.5 | -0.91 | -0.92 | 0.155 | -0.1775 | -0.1325 | 0.404094 | 0.350117 | 0.501587 | 1 | 0.996511 | 0.901002 |
| Commd6 | Q3V4B5 | -0.45 | -0.23 | -0.26 | 0.01 | -0.1 | -0.46 | -0.21 | -0.08 | -0.15 | -0.09 | -0.29 | -0.27 | -0.6 | 0.45 | -0.66 | 0.01 | 0.02 | 0.0325 | 0.0325 | 0.909651 | 0.853689 | 0.912323 | 1 | 0.996511 | 0.986458 |
| Commd7 | Q8BG94 | -0.3 | -0.27 | -0.28 | -0.04 | 0.27 | -0.35 | -0.73 | -0.3 | -0.07 | 0.57 | -0.53 | -0.62 | -0.09 | 1 | -0.55 | -0.75 | -0.055 | 0.06 | 0.125 | 0.813717 | 0.835281 | 0.745695 | 1 | 0.996511 | 0.96006 |
| Commd8 | Q9CZG3 | -1.13 | -0.58 | -0.46 | -0.12 | -0.14 | -1.19 | -0.23 | -0.21 | -0.89 | -0.23 | -0.05 | -0.97 | -0.82 | -0.1 | -0.49 | -0.41 | 0.13 | 0.0375 | 0.1175 | 0.685135 | 0.904972 | 0.673966 | 1 | 0.996744 | 0.952944 |
| Commd9 | Q8K2Q0 | 0.46 | 0.05 | -0.53 | -0.52 | 0.09 | -0.17 | -0.41 | -0.33 | -2.13 | -2.25 | -0.37 | 0.75 | -1.98 | -2.56 | -0.57 | -0.48 | -0.07 | -0.865 | -1.2625 | 0.795727 | 0.233797 | 0.038424 | 1 | 0.996511 | 0.658454 |
| Comt | O88587 | -1.42 | -1.29 | -0.36 | -0.33 | -0.66 | -1.47 | 0.71 | 0.14 | -2.04 | -2.32 | -0.08 | -1.46 | -1.73 | -1.03 | 0.05 | -0.07 | 0.53 | -0.625 | 0.155 | 0.317752 | 0.261294 | 0.747595 | 1 | 0.996511 | 0.960744 |
| Comtd1 | Q8BIG7 | -0.48 | -0.59 | -0.21 | -0.04 | -0.6 | -0.4 | -0.73 | -0.16 | 0.64 | 0.09 | -0.05 | -0.4 | 0.6 | -0.18 | -0.07 | -0.38 | -0.1425 | 0.4 | 0.3225 | 0.495857 | 0.159437 | 0.253172 | 1 | 0.996511 | 0.809524 |
| Copa | F8WHL2 | -1.66 | -2.16 | -2.11 | -2.4 | -1.7 | -1.8 | -1.86 | -2.55 | -1.68 | -0.96 | -2.18 | -1.28 | -1.36 | -1.56 | -2.05 | -2.02 | 0.105 | 0.5575 | 0.335 | 0.684547 | 0.098676 | 0.218616 | 1 | 0.996511 | 0.787447 |
| Copb1 | Q9JIF7 | -0.98 | -0.81 | -1.25 | -1.32 | -0.76 | -0.81 | -1.23 | -1.38 | -0.56 | -1 | -1.1 | -0.5 | -0.32 | -0.25 | -1.21 | -1.1 | 0.045 | 0.3 | 0.37 | 0.836741 | 0.20834 | 0.229501 | 1 | 0.996511 | 0.787447 |
| Copb2 | O55029 | -2.38 | -2.42 | -1.48 | -2.26 | -1.89 | -2.37 | -1.14 | -2.05 | -2.61 | -1.81 | -1.6 | -1.95 | -1.96 | -2.18 | -1.44 | -1.14 | 0.2725 | 0.1425 | 0.455 | 0.420819 | 0.650424 | 0.189574 | 1 | 0.996511 | 0.776843 |
| Cope | O89079 | -2.71 | -3.02 | -1.82 | -2.27 | -2.04 | -2.95 | -0.98 | -1.92 | -3.13 | -1.41 | -1.89 | -2.32 | -3.14 | -2.05 | -2.14 | -1.49 | 0.4825 | 0.2675 | 0.25 | 0.297565 | 0.53426 | 0.54967 | 1 | 0.996511 | 0.913178 |
| Copg1 | Q9QZE5 | -1.8 | -2.04 | -2.56 | -2.71 | -1.7 | -1.89 | -2.62 | -3.06 | -1.9 | -2.04 | -2.65 | -1.56 | -1.8 | -1.37 | -2.72 | -2.87 | -0.04 | 0.24 | 0.0875 | 0.912788 | 0.453177 | 0.828731 | 1 | 0.996511 | 0.973757 |
| Copg2 | Q9QXK3-4 | -0.05 | -0.85 | -0.76 | -1.16 | -0.3 | -0.27 | -0.64 | -0.76 | -1.25 | -0.79 | -0.79 | -0.05 | -0.97 | -1.22 | -0.87 | -0.6 | 0.2125 | -0.015 | -0.21 | 0.440182 | 0.964506 | 0.469215 | 1 | 0.996755 | 0.890107 |
| Coprs | Q9CQ13 | -1.06 | -1 | -0.14 | 0.86 | -0.82 | -0.65 | 0.26 | 0.82 | -0.72 | 1.44 | 0.68 | -0.96 | -1 | -1.22 | 0.52 | 1.28 | 0.2375 | 0.445 | 0.23 | 0.664303 | 0.508093 | 0.736988 | 1 | 0.996511 | 0.958635 |
| Cops2 | P61202-2 | 0.99 | 0.82 | 1.19 | 0.9 | 0.93 | 1.09 | 1.41 | 1.53 | -0.48 | 0.39 | 1.17 | 0.98 | -0.87 | -0.17 | 0.7 | 0.97 | 0.265 | -0.46 | -0.8175 | 0.198773 | 0.232451 | 0.072914 | 1 | 0.996511 | 0.668066 |
| Cops3 | O88543 | -0.54 | -0.71 | -0.31 | -0.43 | -0.78 | -0.44 | -0.53 | -0.22 | -0.72 | -0.18 | -0.46 | -0.55 | -0.74 | -0.95 | -0.68 | -0.31 | 0.005 | 0.02 | -0.1725 | 0.978448 | 0.918122 | 0.428168 | 1 | 0.996755 | 0.879165 |
| Cops4 | O88544 | -0.04 | -0.06 | -0.15 | -0.01 | -0.1 | -0.17 | -0.25 | 0.03 | -0.48 | -0.02 | -0.09 | -0.05 | -0.78 | -0.29 | -0.31 | -0.36 | -0.0575 | -0.095 | -0.37 | 0.696879 | 0.598618 | 0.082738 | 1 | 0.996511 | 0.68865 |
| Cops5 | O35864 | -0.52 | -0.3 | -0.94 | -0.6 | -0.34 | -0.4 | -0.52 | -0.62 | -0.37 | -1.53 | -0.63 | -0.52 | -0.65 | -0.76 | -1.24 | -1.05 | 0.12 | -0.1725 | -0.335 | 0.52634 | 0.568004 | 0.174018 | 1 | 0.996511 | 0.765074 |
| Cops6 | O88545 | -0.77 | -0.7 | -0.47 | -0.03 | -0.85 | -0.6 | -0.11 | -0.03 | -1.07 | -0.74 | -0.45 | -0.73 | -1.38 | -0.87 | -0.88 | -0.59 | 0.095 | -0.255 | -0.4375 | 0.720919 | 0.300385 | 0.12234 | 1 | 0.996511 | 0.721448 |
| Cops7a | Q9CZ04-2 | 0.47 | 0.6 | 0.32 | 0.29 | 0.26 | -0.01 | -0.31 | 0.22 | 0.21 | 0.91 | 0.04 | 0.2 | -0.12 | 0.23 | -0.03 | -0.43 | -0.38 | -0.08 | -0.5075 | 0.070668 | 0.735396 | 0.037327 | 1 | 0.996511 | 0.658454 |
| Cops7b | Q8BV13 | 0.05 | -0.27 | -0.05 | -0.21 | -0.59 | 0 | -0.44 | 0 | 0.3 | 0.29 | -0.07 | 0.31 | 0.12 | -0.59 | -0.37 | -0.58 | -0.1375 | 0.3275 | -0.235 | 0.499858 | 0.099892 | 0.317083 | 1 | 0.996511 | 0.838864 |
| Cops8 | Q8VBV7 | -0.48 | -0.07 | -0.7 | -0.2 | -0.33 | -0.28 | -0.27 | -0.48 | -0.15 | -0.25 | -0.31 | -0.32 | -0.65 | 0.54 | -1.04 | -1.2 | 0.0225 | 0.105 | -0.225 | 0.905077 | 0.598332 | 0.579362 | 1 | 0.996511 | 0.927734 |
| Cops9 | Q3U898 | -2.02 | -0.54 | -0.08 | -2.95 | -1.21 | -0.89 | -0.61 | -1.19 | -3.44 | 0.39 | -3.19 | -1.2 | -4.89 | -2.3 | -1.07 | -1.74 | 0.4225 | -0.4625 | -1.1025 | 0.499511 | 0.647397 | 0.266823 | 1 | 0.996511 | 0.818521 |
| Copz1 | P61924 | -0.72 | -1 | -0.21 | -0.84 | -0.28 | -0.91 | -0.27 | -0.48 | -3.4 | -1.85 | -0.19 | -0.17 | -3.33 | -3.38 | -0.39 | -0.31 | 0.2075 | -0.71 | -1.16 | 0.401466 | 0.337563 | 0.170618 | 1 | 0.996511 | 0.762322 |
| Copz2 | Q9JHH9 | -1.84 | -1.33 | -1.87 | -2.23 | -1.74 | -2.79 | -2.17 | -2.66 | -1.69 | -1.11 | -2.36 | -1.98 | -1.76 | -1.17 | -2.77 | -2.16 | -0.5225 | 0.0325 | -0.1475 | 0.110343 | 0.919515 | 0.695348 | 1 | 0.996755 | 0.955729 |
| Coq10a | A0A1W2P7 | -0.14 | 0.2 | -0.04 | 0.05 | 0.26 | -0.19 | 0.32 | 0.01 | 0.23 | 0.53 | 0.14 | -0.32 | 0.59 | 0.55 | 0.03 | -0.05 | 0.0825 | 0.1275 | 0.2625 | 0.651928 | 0.574315 | 0.26919 | 1 | 0.996511 | 0.820534 |
| Coq2 | F7CDY3 | 0.44 | 2.05 | -0.59 | -0.18 | 1.52 | 1.04 | 0.99 | -0.34 | 1.17 | 0.22 | 0.15 | 1.73 | 1.03 | 1.87 | -0.04 | 0.79 | 0.3725 | 0.3875 | 0.4825 | 0.564389 | 0.546022 | 0.457758 | 1 | 0.996511 | 0.887902 |
| Coq3 | Q8BMS4 | 1.51 | 1.36 | -0.04 | 0.47 | 0.5 | 1.11 | 0.63 | 0.07 | 0.69 | -0.91 | -0.12 | 0.4 | 0.82 | 0.55 | -0.11 | -0.61 | -0.2475 | -0.81 | -0.6625 | 0.54225 | 0.116311 | 0.175954 | 1 | 0.996511 | 0.771503 |
| Coq4 | Q8BGB8 | -0.86 | -0.14 | -0.86 | -0.19 | -0.67 | -0.47 | -0.17 | -0.72 | -0.2 | -0.2 | -0.59 | -0.48 | -0.14 | 0.54 | -0.76 | -0.55 | 0.005 | 0.145 | 0.285 | 0.983959 | 0.562784 | 0.421939 | 1 | 0.996511 | 0.875736 |
| Coq5 | Q9CXI0 | 0.32 | 0.76 | -0.1 | 0.06 | 0.42 | 0.01 | 0.14 | -0.07 | 0.27 | -0.47 | -0.01 | 0.15 | 0.22 | 0.56 | -0.49 | 0.05 | -1.35E-01 | -0.275 | -0.175 | 0.567938 | 0.315446 | 0.564925 | 1 | 0.996511 | 0.920315 |
| Coq6 | Q8R1S0 | 0.12 | 0.24 | 0.93 | 0.98 | -0.76 | -0.21 | -0.03 | 0.28 | -0.84 | -0.14 | 0.15 | -0.48 | -0.78 | -1.37 | 0.18 | -0.02 | -0.7475 | -0.895 | -1.065 | 0.036799 | 0.018036 | 0.026122 | 1 | 0.996511 | 0.633075 |
| Coq7 | P97478 | 0.75 | -1.13 | -0.42 | -0.2 | -1.23 | 0.21 | -0.79 | -0.97 | -1.18 | -0.15 | -0.73 | -0.89 | -2.46 | -1.2 | -1.79 | -0.57 | -0.445 | -0.4875 | -1.255 | 0.350593 | 0.266312 | 0.036812 | 1 | 0.996511 | 0.658454 |
| Coq8a | Q60936 | -0.02 | 0.4 | 0.49 | 0.69 | 0.28 | -0.34 | 1.11 | 0.88 | -0.3 | 0.07 | 0.72 | -0.12 | -0.41 | -0.25 | 0.47 | 0.78 | 0.0925 | -0.2975 | -0.2425 | 0.788875 | 0.304287 | 0.464362 | 1 | 0.996511 | 0.888267 |
| Coq8b | Q566J8 | -0.04 | 0.17 | -0.6 | -0.05 | -0.34 | 0.54 | 0.33 | -0.09 | 0.35 | 0.91 | 0.22 | 0.26 | 0.63 | 0.15 | 0.38 | 0.56 | 0.24 | 0.565 | 0.56 | 0.377703 | 0.048984 | 0.039799 | 1 | 0.996511 | 0.658454 |
| Coq9 | Q8K1Z0 | 0.69 | 0.83 | 0.7 | 0.81 | 0.42 | 0.64 | 0.48 | 0.58 | 0.31 | 0.03 | 0.71 | 0.24 | 0.31 | 0.56 | 0.64 | 0.64 | -0.2275 | -0.435 | -0.22 | 0.143865 | 0.052289 | 0.239442 | 1 | 0.996511 | 0.794419 |
| Coro1a | O89053 | 0.53 | -0.29 | 0.22 | 0.41 | -0.36 | 0.47 | -0.11 | 0.6 | -0.3 | 0.09 | 0.21 | -0.06 | -0.54 | -0.12 | -0.17 | -0.32 | -0.0675 | -0.2325 | -0.505 | 0.817301 | 0.344906 | 0.062269 | 1 | 0.996511 | 0.66433 |
| Coro1a | A0A0U1RP | -2.8 | -3.6 | -2.21 | -1.74 | -1.6 | -3.33 | -2.48 | -2.95 | -1.7 | -2.49 | -1.85 | -2.16 | -4.14 | -1.47 | -4.29 | -2.41 | -0.0025 | 0.5375 | -0.49 | 0.996045 | 0.218511 | 0.501286 | 1 | 0.996511 | 0.901002 |
| Coro1b | Q9WUM3 | 0.25 | -0.58 | 0.39 | 0.84 | -0.03 | 0.09 | 0.25 | 0.81 | 0.07 | 1.38 | 0.34 | 0.02 | -0.26 | 0.46 | 0.16 | 0.05 | 0.055 | 0.2275 | -0.1225 | 0.870831 | 0.584899 | 0.71451 | 1 | 0.996511 | 0.958106 |
| Coro1c | Q9WUM4 | 0.74 | 0 | 0.36 | 0.47 | 0.33 | 0.45 | 0.53 | 0.8 | 0.08 | 0.28 | 0.26 | 0.31 | -0.65 | 0.13 | -0.18 | -0.25 | 0.135 | -0.16 | -0.63 | 0.526424 | 0.447164 | 0.032723 | 1 | 0.996511 | 0.644622 |
| Coro2a | B1AVH5 | -0.17 | -0.22 | 0.55 | 0.38 | 0.36 | -0.05 | 0.15 | 0.08 | -0.12 | 0.09 | 0.5 | -0.28 | -0.01 | 0.72 | 0.18 | -0.24 | -7.9E-17 | -0.0875 | 0.0275 | 1 | 0.748235 | 0.926009 | 1 | 0.996511 | 0.988944 |
| Coro2b | Q8BH44 | 0.95 | 0.51 | 1.26 | 1.13 | 0.23 | 1.01 | 0.99 | 1.06 | 0.69 | 0.84 | 0.95 | 0.37 | 0.4 | 0.6 | 0.11 | 0.73 | -0.14 | -0.25 | -0.5025 | 0.599346 | 0.305296 | 0.068192 | 1 | 0.996511 | 0.664936 |
| Coro6 | Q920M5 | 0.38 | 0.47 | 1.85 | 1.08 | 0.62 | 0.46 | 0.75 | 1.44 | 0.88 | -0.29 | 1.39 | 0.09 | 1.14 | 0.72 | 1.58 | 2.22 | -0.1275 | -0.4275 | 0.47 | 0.738927 | 0.380413 | 0.30409 | 1 | 0.996511 | 0.831662 |
| Coro7 | Q9D2V7 | 0.14 | -0.77 | -0.68 | -0.09 | -0.47 | -0.38 | -0.33 | -0.17 | -0.53 | -0.47 | -0.53 | -0.62 | -0.81 | -0.11 | -0.95 | -1.06 | 0.0125 | -0.1875 | -0.3825 | 0.959252 | 0.457879 | 0.246077 | 1 | 0.996511 | 0.802442 |
| Cotl1 | Q9CQI6 | 0.54 | -0.66 | 0.28 | 0.36 | -0.6 | 0.24 | -0.37 | 0.48 | 0.27 | 1.06 | -0.1 | 0.13 | -0.13 | -0.28 | -0.21 | -0.74 | -0.1925 | 0.21 | -0.47 | 0.590111 | 0.56249 | 0.156436 | 1 | 0.996511 | 0.751493 |
| Cox10 | Q8CFY5 | 0.04 | 0.63 | -0.97 | -0.49 | 0.74 | -0.43 | -0.8 | 0.49 | 1.06 | -1.99 | -0.19 | 0.6 | 1.19 | 1.18 | -1.13 | -0.65 | 0.1975 | 0.0675 | 0.345 | 0.67358 | 0.921635 | 0.590982 | 1 | 0.996755 | 0.929458 |
| Cox11 | Q6P8I6 | 0.21 | 0.3 | -0.47 | -1.12 | 0.23 | 0.2 | -0.5 | -0.34 | 0.1 | 0.81 | 0.43 | -0.11 | 0.29 | 0.5 | -0.12 | 0.39 | 0.1675 | 0.5775 | 0.535 | 0.64668 | 0.149742 | 0.156833 | 1 | 0.996511 | 0.751493 |
| Cox14 | Q8BH51 | -0.7 | 0.23 | 1.15 | 1.43 | -0.5 | 0.5 | 1.26 | 1.25 | 2.19 | 1.83 | 1.47 | -0.83 | 4.47 | -1.32 | 1.43 | 1.19 | 0.1 | 0.6375 | 0.915 | 0.863567 | 0.409253 | 0.431451 | 1 | 0.996511 | 0.879165 |
| Cox15 | Q8BJ03-2 | -0.68 | -0.35 | -0.65 | -0.37 | 0.39 | -0.48 | 0.23 | -0.51 | -0.25 | -1.09 | -0.08 | -0.39 | 0 | 0.83 | -0.55 | -0.15 | 0.42 | 0.06 | 0.545 | 0.133035 | 0.816624 | 0.108216 | 1 | 0.996511 | 0.706427 |
| Cox17 | P56394 | -0.3 | -0.96 | -0.3 | -0.26 | -0.8 | -0.81 | -0.76 | -0.57 | -1.2 | 0.96 | -0.56 | -1.27 | 0.16 | -0.25 | -0.26 | -0.78 | -0.28 | -0.0625 | 0.1725 | 0.199676 | 0.901608 | 0.538614 | 1 | 0.996609 | 0.912212 |
| Cox18 | Q8VC74 | 0.03 | -0.93 | -0.8 | 0.07 | -2.88 | 0.56 | -1.08 | -1.09 | 1.85 | -0.79 | -0.48 | -0.21 | 3.37 | -1.76 | -1.18 | -0.07 | -0.715 | 0.5 | 0.4975 | 0.309601 | 0.414057 | 0.638533 | 1 | 0.996511 | 0.942804 |
| Cox19 | Q8K0C8 | -1.03 | -0.65 | -2.26 | -1.26 | -0.21 | -0.98 | -1.69 | -1.51 | -0.78 | -0.22 | -0.99 | -1.17 | -0.49 | 0.34 | -1.39 | -1.61 | 0.2025 | 0.51 | 0.5125 | 0.651155 | 0.20873 | 0.341015 | 1 | 0.996511 | 0.848473 |
| Cox20 | Q9D7J4 | -1.47 | -1.67 | -1.89 | -1.4 | -1.61 | -1.23 | -1.64 | -1.38 | -1.01 | -0.56 | -1.8 | -1.54 | 0.34 | -0.69 | -1.5 | -1.43 | 0.1425 | 0.38 | 0.7875 | 0.454702 | 0.23078 | 0.089341 | 1 | 0.996511 | 0.69217 |
| Cox4i1 | P19783 | 0.31 | 0.19 | 0.33 | 0.96 | 0.08 | 0.32 | 0.78 | 0.96 | 0.26 | 0.05 | 0.79 | 0.22 | 0.05 | 0.38 | 0.1 | 0.14 | 0.0875 | -0.1175 | -0.28 | 0.748403 | 0.648943 | 0.247005 | 1 | 0.996511 | 0.803237 |
| Cox4i1 | M0QWX7 | 2.05 | 3.64 | -0.79 | -1.21 | 2.91 | 2.9 | -2.85 | -3.11 | 2.63 | -2.26 | -1.62 | 1.92 | 3.77 | 1.03 | -1.97 | -2.1 | -0.96 | -0.755 | -0.74 | 0.603538 | 0.6197 | 0.647859 | 1 | 0.996511 | 0.945793 |
| Cox4i2 | Q91W29 | -0.95 | -0.78 | 2.12 | 2.88 | -0.7 | -0.79 | 1.53 | 2.03 | 0.28 | 2.87 | 2.38 | -1.45 | 2.84 | 0.31 | 2.62 | 2.3 | -0.3 | 0.2025 | 1.2 | 0.785541 | 0.871627 | 0.259061 | 1 | 0.996511 | 0.812074 |
| Cox5a | P12787 | 0.98 | 1.16 | 0.98 | 0.8 | 1.06 | 0.84 | 1.13 | 1.14 | 0.15 | 0 | 0.97 | 0.8 | -0.15 | 0.12 | 0.54 | 0.95 | 0.0625 | -0.5 | -0.615 | 0.700464 | 0.087974 | 0.048768 | 1 | 0.996511 | 0.658454 |
| Cox5b-ps | Q9D881 | 1.46 | 1.03 | 1.24 | 1.88 | 0.57 | 1.25 | 1.02 | 1.89 | 1.51 | 1.94 | 1.29 | 1.42 | 1.32 | 1.26 | 1.09 | 1.02 | -0.22 | 0.1375 | -0.23 | 0.500198 | 0.588828 | 0.343228 | 1 | 0.996511 | 0.848473 |
| Cox6a1 | Q9DCW5 | -1.12 | -1.24 | -0.4 | 0.23 | -1.37 | -1.27 | 0.91 | 0.15 | -1.39 | -1.37 | 0.15 | -1.36 | -3.89 | -0.39 | -0.39 | -0.31 | 0.2375 | -0.36 | -0.6125 | 0.691422 | 0.459188 | 0.478055 | 1 | 0.996511 | 0.892279 |
| Cox6a2 | P43023 | -1.06 | -1.14 | -0.11 | 0.44 | -1.56 | -1.24 | -0.08 | 0.14 | -0.57 | -0.62 | 0.38 | -1.09 | -0.35 | -2.08 | 0.11 | 0.67 | -0.2175 | -0.0075 | 0.055 | 0.678604 | 0.986995 | 0.931862 | 1 | 0.999285 | 0.989713 |
| Cox6b1 | P56391 | 1.4 | 0.53 | 0.49 | 0.59 | 0.76 | 1.37 | 0.14 | 0.33 | -0.25 | 0.27 | 0.35 | 1.03 | 0.3 | -0.12 | 0.16 | 0.16 | -0.1025 | -0.4025 | -0.6275 | 0.761623 | 0.251834 | 0.037798 | 1 | 0.996511 | 0.658454 |
| Cox6c | Q9CPQ1 | 2.74 | 1.52 | 2.12 | 2.16 | 1.22 | 1.88 | 1.76 | 2.8 | 1.43 | -0.26 | 2.23 | 2.41 | 0.7 | 0.6 | 1.53 | 1.99 | -0.22 | -0.6825 | -0.93 | 0.575947 | 0.276016 | 0.043224 | 1 | 0.996511 | 0.658454 |
| Cox7a1 | A0A140LIU | 0.56 | 1.1 | 0.8 | 1.51 | 1.71 | 1.12 | 1.57 | 1.18 | 0.14 | 0.33 | 1.46 | 0.85 | 0.77 | -0.14 | 1.23 | 1.48 | 0.4025 | -0.2975 | -0.1575 | 0.147789 | 0.40822 | 0.692805 | 1 | 0.996511 | 0.955729 |
| Cox7a2 | P48771 | 2.03 | 2.17 | 1.71 | 1.75 | 1.52 | 2.25 | 2.06 | 1.63 | 1.1 | 0.32 | 1.69 | 1.47 | 1.74 | 0.13 | 1.37 | 1.84 | -5.00E-02 | -0.77 | -0.645 | 0.825409 | 0.03779 | 0.129417 | 1 | 0.996511 | 0.723655 |
| Cox7a2l | E9PZS8 | 0.62 | 1.09 | 0.96 | 1.02 | 0.71 | 1.12 | 0.67 | 0.61 | 0.21 | 0.32 | 0.92 | 0.72 | 0.54 | -0.7 | 0.63 | 0.72 | -0.145 | -0.38 | -0.625 | 0.459768 | 0.124749 | 0.100296 | 1 | 0.996511 | 0.698156 |
| Cox7c | P17665 | 5.64 | 2.64 | 2.63 | 1.28 | 1.82 | 7.11 | 2.24 | 2.38 | 2.36 | 0.97 | 2.57 | 4.85 | 0.98 | 1.02 | 3.01 | 4.08 | 0.34 | -0.36 | -0.775 | 0.806199 | 0.743207 | 0.47573 | 1 | 0.996511 | 0.892174 |
| Cp | G3X9T8 | -1.1 | -1.35 | -0.94 | -0.68 | -1.05 | -1.55 | -0.74 | -1.08 | -1.65 | -0.9 | -0.84 | -0.94 | -1.38 | -1.67 | -1.27 | -1.17 | -0.0875 | -0.065 | -0.355 | 0.71129 | 0.800566 | 0.141062 | 1 | 0.996511 | 0.737973 |
| Cpd | O89001 | -0.94 | -1.64 | -1.34 | -1.31 | -1.17 | -1.14 | -1.68 | -1.39 | -1.65 | -1.24 | -1.49 | -1.6 | -0.96 | -1.2 | -1.41 | -1.7 | -0.0375 | -0.1875 | -0.01 | 0.861877 | 0.387232 | 0.967757 | 1 | 0.996511 | 0.9927 |
| Cpe | Q00493 | 1.13 | 1.25 | -0.17 | -0.23 | 1.47 | 0.96 | -0.1 | 0 | 0.38 | 0.59 | 0.04 | 1.45 | 0.07 | 0.52 | 0.42 | 0.13 | 0.0875 | 0.12 | -0.21 | 0.863722 | 0.798629 | 0.603739 | 1 | 0.996511 | 0.931517 |
| Cpeb2 | E9Q969 | 0.27 | -1.2 | -0.73 | -0.49 | -0.75 | 0.12 | -0.54 | -0.57 | -0.47 | -0.72 | -0.61 | 0.37 | 0.09 | 0.05 | -0.84 | -0.96 | 0.1025 | 0.18 | 0.1225 | 0.768605 | 0.638051 | 0.760489 | 1 | 0.996511 | 0.963123 |
| Cpeb3 | A0A0R4J10 | -0.94 | -0.63 | -1.3 | -1.17 | -0.66 | -0.85 | -0.9 | -1.21 | -1.03 | -1.2 | -1.18 | -0.57 | -0.73 | -0.17 | -1.07 | -1.25 | 0.105 | 0.015 | 0.205 | 0.623942 | 0.94941 | 0.492283 | 1 | 0.996755 | 0.897531 |
| Cpeb4 | Q5SU48 | -0.8 | -0.94 | -0.78 | -0.65 | -1.13 | -1.18 | -0.28 | -0.63 | -0.89 | -1.22 | -0.72 | -0.84 | -0.27 | 0.88 | -0.07 | -0.99 | -0.0125 | -0.125 | 0.68 | 0.9582 | 0.503536 | 0.100846 | 1 | 0.996511 | 0.698156 |
| Cplx1 | P63040 | 0.14 | 0.44 | 0.9 | 0.62 | 0.6 | -0.12 | 0.24 | 0.52 | 0.09 | 0.68 | 0.59 | 0.09 | 0.41 | 0.98 | 0.52 | 0.63 | -0.215 | -0.1625 | 0.11 | 0.386523 | 0.518539 | 0.649015 | 1 | 0.996511 | 0.946117 |
| Cplx2 | P84086 | 1.23 | 1.1 | 0.28 | 0.69 | 0.86 | 0.88 | 0.08 | 0.33 | 0.89 | 0.9 | 0.02 | 1.12 | 0.69 | 1.14 | 0.3 | 0.09 | -0.2875 | -0.0925 | -0.27 | 0.339462 | 0.775642 | 0.41099 | 1 | 0.996511 | 0.874811 |
| Cplx3 | Q8R1B5 | 0.03 | 0.81 | 1.01 | 0.72 | 0.87 | 0.6 | -0.18 | 0.61 | 0.65 | 1.49 | 1.11 | 0.35 | 0.78 | 0.72 | -0.05 | 0.21 | -0.1675 | 0.2575 | -0.2275 | 0.588744 | 0.440431 | 0.462094 | 1 | 0.996511 | 0.888262 |
| Cpm | Q80V42 | 1.08 | 1.03 | 1.39 | 0.59 | 1.28 | 0.72 | 0.84 | 0.98 | 0.48 | 0.73 | 1.17 | 0.88 | 0.24 | 0.89 | 1.84 | 1.67 | -0.0675 | -0.2075 | 0.1375 | 0.765015 | 0.406102 | 0.726463 | 1 | 0.996511 | 0.958398 |
| Cpne1 | Q8C166 | 0.09 | 0.02 | -0.3 | 0.06 | 0.17 | 0.34 | -0.1 | -0.14 | 1.05 | -1.22 | -0.21 | 0.12 | 0.97 | 0 | -0.44 | -0.28 | 0.1 | -0.0325 | 0.095 | 0.594489 | 0.942218 | 0.774744 | 1 | 0.996755 | 0.966468 |
| Cpne2 | P59108 | 0.74 | 1.18 | 0 | 0.07 | 1.04 | 0.98 | 0.81 | -0.16 | 0.97 | -1.04 | -0.05 | 1.08 | 0.77 | 1.28 | -0.39 | 0.55 | 0.17 | -0.2575 | 0.055 | 0.654431 | 0.628414 | 0.89764 | 1 | 0.996511 | 0.983509 |
| Cpne3 | Q8BT60 | 0.77 | 0.7 | 0.63 | 0.56 | 0.84 | 0.98 | 0.82 | 0.67 | -0.18 | 0.01 | 0.71 | 0.85 | -0.39 | -0.69 | 0.93 | 0.73 | 0.1625 | -0.3175 | -0.52 | 0.299344 | 0.2635 | 0.207178 | 1 | 0.996511 | 0.785338 |
| Cpne4 | Q8BLR2 | 0.58 | 0.29 | 0.58 | 0.61 | 0.64 | 0.81 | 1.4 | 0.88 | 0.3 | 0.22 | 0.72 | 0.6 | 0.07 | 0.28 | 0.76 | 0.89 | 0.4175 | -0.055 | -0.015 | 0.072308 | 0.777694 | 0.951085 | 1 | 0.996511 | 0.992476 |
| Cpne5 | E9Q1V2 | 0.23 | -0.02 | -0.41 | -0.46 | 1.61 | -0.62 | -0.79 | -0.82 | -0.4 | 0.61 | -0.23 | 0.52 | 1.57 | 2.03 | 0.23 | -0.39 | 0.01 | 0.29 | 1.025 | 0.985753 | 0.359894 | 0.08679 | 1 | 0.996511 | 0.691619 |
| Cpne5 | Q8JZW4 | -0.06 | 0.13 | 0.69 | 0.81 | 0.85 | -0.34 | 1.38 | 0.65 | -0.35 | 0.09 | 0.77 | 0.06 | -0.29 | 0.78 | 0.78 | 1.44 | 0.2425 | -0.25 | 0.285 | 0.542831 | 0.436585 | 0.483449 | 1 | 0.996511 | 0.893925 |
| Cpne6 | Q9Z140 | 0.18 | 0.17 | 0.54 | 0.52 | 0.51 | 0.36 | 0.54 | 0.65 | 0.41 | 1.64 | 0.58 | 0.29 | 0.25 | 0.42 | 0.88 | 0.85 | 0.1625 | 0.3775 | 0.2475 | 0.354527 | 0.264087 | 0.300799 | 1 | 0.996511 | 0.830276 |
| Cpne7 | Q0VE82 | 0.62 | 0.67 | 0.04 | -0.36 | 1.4 | 0.6 | 0.62 | 0.02 | 0.53 | 0.04 | -0.17 | 0.7 | 0.32 | 0.87 | 0.06 | 0.43 | 4.18E-01 | 0.0325 | 0.1775 | 0.264392 | 0.919092 | 0.569101 | 1 | 0.996755 | 0.922588 |
| Cpne8 | Q9DC53 | 0.36 | 0.46 | 0.24 | -0.17 | 1.16 | 0.16 | 0.77 | -0.5 | 0.61 | -0.06 | 0.39 | 0.67 | 0.51 | 1.58 | 0.85 | 0.4 | 0.175 | 0.18 | 0.6125 | 0.638539 | 0.464735 | 0.074567 | 1 | 0.996511 | 0.671911 |
| Cpne9 | A0A0R4J0J | 0.23 | 0.89 | 1.03 | 0.47 | 0.76 | 0.11 | 0.73 | 0.46 | 0.14 | 0.18 | 0.88 | 0.23 | 0.05 | 0.76 | 0.42 | 0.58 | -0.14 | -0.2975 | -0.2025 | 0.579803 | 0.288166 | 0.453169 | 1 | 0.996511 | 0.885993 |
| Cpox | P36552 | -0.17 | -0.03 | 0.03 | 0.09 | -0.38 | 0.09 | -0.29 | 0.2 | 0.2 | 0.23 | 0.23 | -0.09 | -0.17 | 0.28 | -0.57 | -0.33 | -0.075 | 0.1625 | -0.1775 | 0.694911 | 0.356491 | 0.45077 | 1 | 0.996511 | 0.885222 |
| Cpped1 | Q8BFS6 | -0.62 | -0.31 | -1.01 | -0.32 | -0.09 | -0.75 | -0.53 | -0.48 | -0.16 | 0.51 | -0.35 | -0.44 | -0.08 | 0.09 | -0.39 | -0.66 | 0.1025 | 0.455 | 0.305 | 0.661432 | 0.133926 | 0.263292 | 1 | 0.996511 | 0.81524 |
| Cpq | Q9WVJ3 | -0.55 | -0.19 | -0.67 | -0.54 | -0.6 | -0.23 | -0.48 | -0.88 | 0 | 0.01 | -0.33 | -0.46 | -0.06 | -0.33 | -1.36 | -0.83 | -0.06 | 0.2925 | -0.1575 | 0.766889 | 0.176559 | 0.617736 | 1 | 0.996511 | 0.937259 |
| Cpsf1 | Q9EPU4 | -0.75 | -2.09 | -1.89 | -1.53 | -1.16 | -1.42 | -0.91 | -1.55 | -1.57 | -1.14 | -1.54 | -1.79 | -0.75 | -0.83 | 0.07 | -0.66 | 0.305 | 0.055 | 1.0225 | 0.354183 | 0.865245 | 0.017629 | 1 | 0.996511 | 0.602211 |
| Cpsf2 | O35218 | -1.08 | -2.04 | -1.4 | -1.56 | -1.2 | -1.71 | -0.66 | -1.13 | -2 | -0.99 | -1.24 | -1.96 | -1.31 | -1.22 | 0.23 | -0.8 | 0.345 | -0.0275 | 0.745 | 0.258451 | 0.932201 | 0.084973 | 1 | 0.996755 | 0.691619 |
| Cpsf3 | Q9QXK7 | -2.75 | -4.02 | -1.39 | -1.67 | -2.76 | -3.7 | -1.19 | -1.62 | -4.24 | -0.64 | -1.47 | -3.83 | -2.62 | -2.7 | -1.23 | -1.28 | 0.14 | -0.0875 | 0.5 | 0.85092 | 0.92711 | 0.454871 | 1 | 0.996755 | 0.887169 |
| Cpsf4 | E0CXT7 | -0.4 | -0.85 | -1.14 | -1.04 | -0.65 | -0.72 | -0.49 | -1.01 | -1.47 | -1.5 | -1.04 | -0.91 | -1.26 | -0.91 | 0.21 | -0.15 | 0.14 | -0.3725 | 0.33 | 0.529692 | 0.156924 | 0.381503 | 1 | 0.996511 | 0.860984 |
| Cpsf6 | H3BJ30 | -3.62 | -5.15 | -3.8 | -3.39 | -4.32 | -4.61 | -3.77 | -3.78 | -2.14 | -1.83 | -3.68 | -4.09 | -2.05 | -0.56 | -2.37 | -3.58 | -0.13 | 1.055 | 1.85 | 0.756901 | 0.11949 | 0.020934 | 1 | 0.996511 | 0.621926 |
| Cpsf7 | Q8BTV2-2 | -3.07 | -4.5 | -4.08 | -4.53 | -3.54 | -3.74 | -3.79 | -4.06 | -3.48 | -2.49 | -3.98 | -3.51 | -2.95 | -3.28 | -2.27 | -4.1 | 0.2625 | 0.68 | 0.895 | 0.453856 | 0.146411 | 0.08881 | 1 | 0.996511 | 0.691619 |
| Cpt1a | P97742 | 0.21 | 0.47 | 0.88 | 1.08 | 0.84 | 0.76 | 1.35 | 1.09 | 0.48 | 0.9 | 1.22 | 0.47 | 0.4 | 0.31 | 0.8 | 1.05 | 0.35 | 0.1075 | -0.02 | 0.185297 | 0.701607 | 0.943367 | 1 | 0.996511 | 0.991596 |
| Cpt1c | Q8BGD5 | -0.67 | -0.46 | -0.44 | -0.29 | -0.18 | -0.5 | -0.06 | -0.12 | 0.03 | 0.31 | -0.2 | -0.64 | 0.4 | -0.1 | 0.3 | 0.02 | 0.25 | 0.34 | 0.62 | 0.176945 | 0.184521 | 0.013365 | 1 | 0.996511 | 0.585601 |
| Cpt2 | P52825 | 2.12 | 1.47 | 1.43 | 0.94 | 1.24 | 1.67 | 1.55 | 1.07 | 0.47 | 0.83 | 1.4 | 1.69 | 0.4 | 0.26 | 1 | 1.36 | -0.1075 | -0.3925 | -0.735 | 0.703378 | 0.289404 | 0.061745 | 1 | 0.996511 | 0.662517 |
| Cptp | Z4YLU2 | -1.81 | -1.17 | -2.17 | -2.46 | -0.63 | -1.4 | -1.34 | -2.27 | -2.03 | -2.91 | -1.62 | -1.98 | -1.75 | -1.19 | -1.95 | -1.01 | 0.4925 | -0.2325 | 0.4275 | 0.249531 | 0.54181 | 0.246188 | 1 | 0.996511 | 0.802442 |
| Cr1l | Q64735-2 | 0.45 | 0.28 | -0.52 | -0.63 | 0.65 | 0.68 | 0.12 | -0.28 | 0.06 | -0.7 | -0.64 | 0.62 | -0.1 | 0.18 | 0.2 | -0.06 | 0.3975 | -0.06 | 0.16 | 0.268244 | 0.880278 | 0.595417 | 1 | 0.996511 | 0.929911 |
| Crabp1 | P62965 | 0.22 | -0.69 | -1.54 | -0.71 | -0.51 | -0.56 | -0.76 | 0.87 | -0.31 | 0.57 | 0.2 | 0.48 | -0.56 | -0.34 | -0.69 | -1.62 | 0.44 | 0.915 | -0.1225 | 0.371399 | 0.042722 | 0.778135 | 1 | 0.996511 | 0.967289 |
| Cracd | E9Q5L4 | 1.04 | 0.97 | 0.69 | 0.79 | 0.58 | 1.16 | 0.6 | 1.06 | 1.65 | 1.21 | 1.04 | 1.2 | 1.71 | 1.61 | 0.83 | 0.75 | -0.0225 | 0.4025 | 0.3525 | 0.911698 | 0.072905 | 0.235097 | 1 | 0.996511 | 0.791437 |
| Cracdl | F7B0R9 | 0.03 | -0.23 | 1.86 | 1.83 | -0.48 | -0.85 | 3.4 | 3.22 | 0.05 | 0.86 | 3.21 | 0.02 | -0.06 | 0.96 | 1.82 | 3.12 | 0.45 | 0.1625 | 0.5875 | 0.695967 | 0.847919 | 0.465478 | 1 | 0.996511 | 0.888369 |
| Cracdl | E9Q3M9 | 0.54 | 0.33 | 0.4 | 0.65 | 0.37 | 0.07 | 0.1 | 0.43 | 0.79 | 1.09 | 0.37 | 0.49 | 1 | 0.98 | 0.4 | -0.14 | -0.2375 | 0.205 | 0.08 | 0.185805 | 0.355397 | 0.786915 | 1 | 0.996511 | 0.968024 |
| Cradd | O88843 | -1.49 | -0.12 | -1.48 | -0.65 | -0.51 | -1.82 | -0.49 | -0.85 | -1.03 | -1.71 | -1.61 | -1.09 | -2.13 | -1.07 | -2.65 | -1.61 | 0.0175 | -0.425 | -0.93 | 0.967394 | 0.265809 | 0.065165 | 1 | 0.996511 | 0.66433 |
| Crat | H7BX88 | 1.08 | 0.06 | 1.31 | 0.97 | -0.16 | 1.07 | 2.25 | 1.05 | 0 | 0.71 | 1.62 | 0.59 | -0.09 | 0.14 | 1.66 | 1.59 | 0.1975 | -0.125 | -0.03 | 0.704158 | 0.761972 | 0.952358 | 1 | 0.996511 | 0.992476 |
| Crbn | Q8C7D2-2 | -0.38 | -0.59 | -0.19 | -0.46 | -0.59 | -0.13 | 0.07 | -0.26 | -1.82 | -1.46 | -0.03 | -0.23 | -1.67 | -1.87 | -0.47 | 0 | 0.1775 | -0.48 | -0.5975 | 0.378401 | 0.279592 | 0.196956 | 1 | 0.996511 | 0.78004 |
| Creb1 | Q01147-1 | -1.75 | -2.42 | -1.74 | -2.36 | -1.5 | -2.4 | -2.32 | -2.33 | -2.45 | -0.87 | -1.92 | -2.04 | -1.88 | -1.46 | -1.22 | -1.87 | -0.07 | 0.2475 | 0.46 | 0.806121 | 0.51166 | 0.117087 | 1 | 0.996511 | 0.718608 |
| Crebbp | A0A0U1RQ | -2.52 | -2.71 | -1.98 | -2.01 | -2.61 | -2.25 | -1.97 | -1.81 | -1.13 | -1.17 | -1.76 | -2.15 | -0.9 | -1.77 | -2.12 | -1.88 | 0.145 | 0.7525 | 0.6375 | 0.583098 | 0.036022 | 0.078246 | 1 | 0.996511 | 0.677244 |
| Creg1 | K4DI63 | -1.76 | -2.02 | -2.05 | -2.48 | -5.18 | -1.29 | -2.21 | -2.67 | 0.24 | -1.67 | -1.79 | -2.52 | 2.24 | -4.85 | -2.61 | -2.15 | -0.76 | 0.6425 | 0.235 | 0.333642 | 0.269611 | 0.85929 | 1 | 0.996511 | 0.976627 |
| Creg2 | Q8BGC9 | -2.81 | -2.6 | 0.64 | 1.25 | -3.41 | -1.41 | -1.58 | 0.18 | 1.63 | 1.85 | 1.05 | -2.95 | 2.58 | -5.31 | 1.02 | 0.56 | -0.675 | 1.275 | 0.5925 | 0.563033 | 0.369165 | 0.742417 | 1 | 0.996511 | 0.958976 |
| Creld1 | Q91XD7 | -1.06 | -1.32 | -3.28 | -2.45 | -1.88 | -0.7 | -2.88 | -3.06 | -0.65 | -1.69 | -2.9 | -1.02 | 0.78 | -3.09 | -1.57 | -2.31 | -0.1025 | 0.4625 | 0.48 | 0.879886 | 0.484159 | 0.589116 | 1 | 0.996511 | 0.929131 |
| Creld2 | Q9CYA0 | -1.5 | -1.38 | -0.95 | -1.27 | -0.56 | -1.82 | -0.21 | -0.74 | -2.8 | -2.85 | -1.4 | -1.04 | -1.44 | -2.3 | 0.51 | -0.05 | 0.4425 | -0.7475 | 0.455 | 0.230273 | 0.127523 | 0.454243 | 1 | 0.996511 | 0.887073 |
| Crh | Q8CIT0 | 1.18 | 0.96 | 0.02 | 0.86 | 0.52 | 1.32 | 0.25 | 0.68 | 1.63 | 0.15 | 0.58 | 1.26 | 1.92 | 0.92 | 0.79 | 0.58 | -0.0625 | 0.15 | 0.2975 | 0.850229 | 0.708635 | 0.442824 | 1 | 0.996511 | 0.883549 |
| Crhbp | A0A217FL6 | -0.15 | 0.23 | 0.4 | -0.12 | -0.04 | 0.14 | 0.19 | 0.43 | -0.83 | -0.57 | 0.32 | -0.05 | -1.07 | -0.98 | 0.06 | 0.36 | 0.09 | -0.3725 | -0.4975 | 0.653257 | 0.231971 | 0.20942 | 1 | 0.996511 | 0.786218 |
| Crhr1 | P35347 | -0.22 | 0.26 | 1.21 | 0.66 | -0.32 | -0.55 | -0.52 | 0.48 | -0.58 | 0.53 | 1.2 | -0.93 | -0.53 | -0.38 | 0.02 | 0.36 | -0.705 | -0.4225 | -0.61 | 0.083924 | 0.437937 | 0.116665 | 1 | 0.996511 | 0.718608 |
| Crip1 | A0A0G2JEK | -7.97 | -10.56 | -5.28 | -4.81 | -9.98 | -9.04 | -5.08 | -4.72 | -7.82 | -4.87 | -4.79 | -8.43 | -5.38 | -10.35 | -4.63 | -5.86 | -0.05 | 0.6775 | 0.6 | 0.976359 | 0.645831 | 0.71609 | 1 | 0.996511 | 0.958106 |
| Crip2 | Q9DCT8 | -3.05 | -3.06 | -1.86 | -1.63 | -2.57 | -2.7 | -1.56 | -1.7 | -2.79 | -1.24 | -1.54 | -2.47 | -2.71 | -2.09 | -1.83 | -2.14 | 0.2675 | 0.39 | 0.2075 | 0.554146 | 0.438252 | 0.612883 | 1 | 0.996511 | 0.935333 |
| Cript | O70333 | -8.94 | -12.24 | -11.32 | -10.41 | -10.24 | -8.44 | -9.87 | -12.41 | -5.17 | -6.48 | -7.42 | -8.32 | -6.25 | -7.39 | -6.11 | -11.37 | 0.4875 | 3.88 | 2.9475 | 0.617848 | 0.001784 | 0.041707 | 1 | 0.996511 | 0.658454 |
| Crk | Q64010 | 1.22 | 0.96 | 0.42 | 0.2 | 0.79 | 0.85 | -0.11 | 0.18 | 0.52 | -0.37 | 0.33 | 1.24 | 0.15 | 0.76 | 0.17 | -0.08 | -0.2725 | -0.27 | -0.45 | 0.410572 | 0.494829 | 0.167513 | 1 | 0.996511 | 0.759484 |
| Crk | Q8JZR2 | 0.03 | -0.76 | -0.6 | -0.26 | -0.5 | -1.04 | -0.06 | -0.47 | 0.05 | -0.38 | 0.22 | -0.47 | 0.03 | 0.07 | -0.39 | -0.74 | -0.12 | 0.2525 | 0.14 | 0.66142 | 0.347353 | 0.620365 | 1 | 0.996511 | 0.93807 |
| Crkl | P47941 | 0.91 | 0.8 | 0.31 | 0.61 | 0.92 | 0.98 | 0.65 | 0.76 | 0.55 | 0.43 | 0.43 | 0.92 | 0.3 | 0.97 | 0.12 | 0.42 | 0.17 | -0.075 | -0.205 | 0.382285 | 0.728202 | 0.432189 | 1 | 0.996511 | 0.879165 |
| Crlf3 | Q9Z2L7 | -1.08 | -0.89 | -0.16 | 0.14 | -0.76 | -0.82 | -1.17 | -0.06 | -0.47 | 1.26 | -0.12 | -1.04 | 0.48 | -0.04 | 0.06 | -0.41 | -0.205 | 0.405 | 0.52 | 0.569182 | 0.449842 | 0.155296 | 1 | 0.996511 | 0.750763 |
| Crls1 | E9PUW6 | 0.51 | 1.09 | -2.17 | -1.07 | -1.48 | -0.72 | -1.59 | -1.67 | 1.89 | 1.2 | -2.04 | 2.47 | 2.12 | 2.67 | -1.38 | -2.17 | -0.955 | 1.29 | 0.72 | 0.197165 | 0.268944 | 0.574589 | 1 | 0.996511 | 0.926139 |
| Crmp1 | P97427 | -3.68 | -3.82 | -3.47 | -4.44 | -4.41 | -4.06 | -4.24 | -3.69 | -2.25 | -3.39 | -3.43 | -3.49 | -1.31 | -3.12 | -3.03 | -4.68 | -0.2475 | 0.7125 | 0.8175 | 0.365094 | 0.071503 | 0.233 | 1 | 0.996511 | 0.788994 |
| Crmp1 | Q6P1J1 | -0.25 | -0.26 | 0.29 | 0.33 | -0.39 | -0.33 | 0.47 | 0.52 | -0.21 | 0.12 | 0.66 | 0.13 | -0.62 | -0.11 | 0.15 | 0.1 | 0.04 | 0.1475 | -0.1475 | 0.892192 | 0.576418 | 0.582823 | 1 | 0.996511 | 0.927734 |
| Crnkl1 | P63154 | -0.69 | -0.89 | -1.29 | -1.44 | -1.44 | -1.11 | -1.46 | -2.07 | -1.19 | -1.98 | -1.81 | -1.26 | -0.99 | -0.88 | -1.37 | -2.02 | -0.4425 | -0.4825 | -0.2375 | 0.129353 | 0.108222 | 0.461997 | 1 | 0.996511 | 0.888262 |
| Crocc | Q8CJ40 | 0.35 | -0.35 | 0.1 | 0.48 | -0.39 | -0.1 | 0.32 | 0.43 | 0.23 | 0.94 | 0.46 | 0.06 | 0.89 | -0.11 | 0.99 | 0.66 | -0.08 | 0.2775 | 0.4625 | 0.767015 | 0.33154 | 0.168901 | 1 | 0.996511 | 0.759484 |
| Crocc | Q8CJ40-2 | -1.28 | 0.09 | -0.75 | -0.9 | 0.52 | -0.28 | -3.06 | -1.62 | 0.87 | -0.28 | -0.67 | -0.2 | 0.39 | 0.39 | -2.01 | -1.57 | -0.4 | 0.64 | 0.01 | 0.599741 | 0.150952 | 0.987463 | 1 | 0.996511 | 0.996154 |
| Crocc2 | F6XLV1 | 0.61 | 2.06 | 2.14 | 0.95 | 1.38 | 0.58 | 1.11 | 0.62 | 0.9 | 1.13 | 0.81 | 0.59 | 1.93 | 2.28 | 2.07 | 1.64 | -0.5175 | -0.5825 | 0.54 | 0.226272 | 0.158976 | 0.196347 | 1 | 0.996511 | 0.78004 |
| Crot | Q9DC50 | -0.18 | -0.14 | 0.22 | 0.27 | 0.36 | 0.22 | 0.49 | 0.31 | 0.68 | 0.9 | 0.5 | 0.6 | 0.59 | 0.48 | 0.26 | 0.22 | 0.3025 | 0.6275 | 0.345 | 0.115786 | 0.01072 | 0.123467 | 1 | 0.996511 | 0.721448 |
| Crppa | A0A1Y7VM | -0.68 | -0.44 | -0.39 | -0.19 | -0.57 | -0.84 | -0.59 | -0.51 | -0.83 | 0.53 | -0.42 | -0.91 | -1.41 | -0.52 | -0.58 | -0.66 | -0.2025 | 0.0175 | -0.3675 | 0.263239 | 0.958974 | 0.181617 | 1 | 0.996755 | 0.776002 |
| Crtac1 | Q8R555 | 0.7 | 0.78 | 0.49 | 0.72 | 0.7 | 0.68 | 0.91 | 0.72 | -0.16 | 0.43 | 0.49 | 0.94 | -0.3 | -0.43 | 0.66 | 0.64 | 0.08 | -0.2475 | -0.53 | 0.604435 | 0.346116 | 0.115058 | 1 | 0.996511 | 0.717287 |
| Crtc1 | Q68ED7 | -0.64 | -0.23 | -0.75 | -0.83 | -0.42 | -0.88 | -0.72 | -0.96 | -0.39 | -0.9 | -0.84 | -0.61 | -0.52 | -0.08 | -1.07 | -0.98 | -0.1325 | -0.0725 | -0.05 | 0.528792 | 0.738196 | 0.860061 | 1 | 0.996511 | 0.977204 |
| Crtc2 | Q3U182 | 1.23 | 0.59 | 0.15 | 0.58 | -0.2 | 0.55 | -1.38 | 0.33 | 1.24 | 0.54 | -0.62 | 0.9 | 2.23 | 0.09 | 2.85 | -0.82 | -0.8125 | -0.1225 | 0.45 | 0.09997 | 0.779004 | 0.580608 | 1 | 0.996511 | 0.927734 |
| Crtc3 | Q91X84 | -0.09 | 1.14 | 0.96 | 0.03 | 0.02 | -0.05 | 0.38 | 0.59 | -0.63 | 0.07 | 0.6 | 0.41 | -0.15 | -0.72 | 0.77 | 0.28 | -0.275 | -0.3975 | -0.465 | 0.424981 | 0.331578 | 0.29165 | 1 | 0.996511 | 0.825497 |
| Cry2 | Q9R194 | -1.6 | -1.12 | -1.05 | -0.94 | -1.53 | -1.05 | -0.98 | -0.76 | -1.13 | -0.15 | -0.85 | -1 | -0.75 | 0.03 | -1.06 | -1.29 | 0.0975 | 0.395 | 0.41 | 0.680014 | 0.177233 | 0.230788 | 1 | 0.996511 | 0.788863 |
| Cryab | P23927 | 0.36 | 0.33 | 0.82 | 0.65 | -0.21 | -0.23 | 0.29 | -0.03 | -1.36 | -0.23 | -0.59 | -1.43 | -0.83 | -0.57 | 0.15 | 0.46 | -0.585 | -1.4425 | -0.7375 | 0.016605 | 0.001508 | 0.047964 | 1 | 0.996511 | 0.658454 |
| Crybb1 | E9PYP8 | 0.05 | -0.41 | -0.47 | -0.1 | -0.02 | 0.03 | 0.07 | 0.32 | -0.11 | 0.34 | 0.23 | -0.13 | -0.56 | -0.31 | -1.14 | -0.73 | 0.3325 | 0.315 | -0.4525 | 0.102083 | 0.165062 | 0.096065 | 1 | 0.996511 | 0.698156 |
| Crybg3 | A0A338P6I | 0.55 | 0.17 | -1.13 | -0.69 | 0.21 | -0.13 | -0.77 | -0.58 | 2.31 | 0.13 | -1.35 | 0.72 | 1.62 | 3.75 | -0.46 | -0.99 | -0.0425 | 0.7275 | 1.255 | 0.918744 | 0.356999 | 0.241661 | 1 | 0.996511 | 0.796892 |

| | | | | | | | | | | | | | | | | | | | | | | | | | | | |
|---|---|---|---|---|---|---|---|---|---|---|---|---|---|---|---|---|---|---|---|---|---|---|---|---|---|---|---|
| Cryl1 | Q99KP3 | -1.77 | -2.04 | -0.73 | -1 | -1.96 | -1.88 | -0.72 | -0.85 | -2.59 | -1.01 | -0.84 | -1.9 | -2.43 | -1.94 | -1.13 | -1.08 | 0.0325 | -0.2 | -0.26 | 0.93879 | 0.677369 | 0.549116 | | 1 | 0.996511 | 0.912757 |
| Crym | O54983 | -0.75 | -0.85 | -0.95 | -0.78 | -0.62 | -0.99 | -0.65 | -0.65 | -0.54 | -0.53 | -1.01 | -0.68 | -0.95 | -0.02 | -0.43 | -1.17 | 0.105 | 0.1425 | 0.19 | 0.518006 | 0.445392 | 0.506672 | | 1 | 0.996511 | 0.902614 |
| Cryz | P47199 | -0.27 | -0.33 | -0.61 | -0.48 | -0.17 | -0.58 | -0.09 | -0.56 | -0.55 | -1.04 | -0.95 | -0.54 | -0.71 | -0.17 | -0.53 | -0.8 | 0.0725 | -0.3475 | -0.13 | 0.701851 | 0.111185 | 0.54798 | | 1 | 0.996511 | 0.912571 |
| Cryzl1 | Q921W4 | 0.32 | 0.26 | 0.21 | 0.34 | -0.08 | 0.04 | 0.12 | 0.28 | -0.22 | -0.35 | 0.15 | 0.14 | -0.48 | -0.52 | 0.06 | 0.05 | -0.1925 | -0.3525 | -0.505 | 0.228601 | 0.089134 | 0.041855 | | 1 | 0.996511 | 0.658454 |
| Cryzl2 | Q3UNZ8 | 0.49 | 0.16 | 0.32 | 0.44 | 0.23 | 0.39 | 1.36 | 0.41 | 0.05 | -0.03 | 0.6 | 0 | 0.4 | -0.01 | 0.63 | 1.24 | 0.245 | -0.1975 | 0.2125 | 0.38036 | 0.358693 | 0.468328 | | 1 | 0.996511 | 0.889423 |
| Cs | Q9CZU6 | 1.1 | 0.54 | 0.15 | 0.57 | 0.29 | 0.91 | 0.26 | 0.47 | 0.81 | 0.07 | 0.44 | 0.91 | 0.54 | 0.49 | -0.37 | -0.1 | -0.1075 | -0.0325 | -0.45 | 0.676022 | 0.908861 | 0.168159 | | 1 | 0.996744 | 0.759484 |
| Csad | Q9DBE0 | 0.13 | -0.18 | 0.1 | 0.1 | -0.13 | 0.13 | -0.77 | 0.37 | -0.2 | 0.02 | 0.07 | -0.17 | -0.32 | -0.32 | -0.14 | 0.23 | -0.1375 | -0.1075 | -0.175 | 0.604956 | 0.539354 | 0.410655 | | 1 | 0.996511 | 0.874811 |
| Csdc2 | Q91YQ3 | -0.37 | -0.34 | 1.16 | 1.8 | -0.54 | -0.37 | 0.73 | 1.42 | -0.3 | 2.05 | 1.27 | -0.17 | -0.33 | -0.09 | 1.41 | 1.1 | -0.2525 | 0.15 | -0.04 | 0.699258 | 0.834279 | 0.949664 | | 1 | 0.996511 | 0.992476 |
| Csde1 | Q91W50 | -0.93 | -1.59 | -1.34 | -1.61 | -1.44 | -1.13 | -1.43 | -1.47 | -1.19 | -0.5 | -1.4 | -0.93 | -0.96 | -0.8 | -1.39 | -1.67 | 3.14E-16 | 0.3625 | 0.1625 | 1 | 0.197897 | 0.560115 | | 1 | 0.996511 | 0.918806 |
| Cse1l | Q9ERK4 | 0.01 | -0.45 | -0.29 | -0.4 | -0.38 | -0.15 | -0.45 | -0.57 | -0.55 | -0.25 | -0.56 | 0.15 | -0.92 | -0.65 | -0.37 | -0.58 | -0.105 | -0.02 | -0.3475 | 0.564601 | 0.930348 | 0.126426 | | 1 | 0.996755 | 0.721505 |
| Csf1r | P09581 | -2.11 | -2.79 | -1.83 | -1.99 | -2.31 | -2.08 | -1.79 | -1.67 | -1.77 | -1.6 | -1.44 | -2.1 | -2.31 | -1.91 | -2.16 | -2.04 | 0.2175 | 0.4525 | 0.075 | 0.418706 | 0.120972 | 0.771482 | | 1 | 0.996511 | 0.965933 |
| Csk | P41241 | 0.06 | -0.41 | -0.74 | -1.14 | -0.08 | -0.07 | -0.25 | -0.65 | -0.63 | -1.82 | -0.75 | 0 | -0.67 | -0.43 | -0.76 | -0.83 | 0.295 | -0.2425 | -0.115 | 0.321972 | 0.576799 | 0.689531 | | 1 | 0.996511 | 0.955729 |
| Csl | Q80X68 | 1.38 | 1.72 | 3.05 | 2.75 | 0.74 | 1.03 | 0.93 | 2.71 | 1.14 | 1.23 | 1.83 | 1.27 | 1.16 | -0.26 | 2.23 | 2.91 | -0.8725 | -0.8575 | -0.715 | 0.143557 | 0.062261 | 0.336857 | | 1 | 0.996511 | 0.847578 |
| Csmd1 | Q923L3 | 0.14 | 0.11 | 0.65 | 0.61 | 0.14 | 0.55 | 0.72 | 0.58 | 0.25 | 0.63 | 0.96 | 0.34 | 0.41 | -0.29 | 0.16 | 0.56 | 0.12 | 0.1675 | -0.1675 | 0.583532 | 0.496825 | 0.530205 | | 1 | 0.996511 | 0.908663 |
| Csmd2 | V9GX34 | 0.82 | 0.47 | 0.64 | 0.47 | 0.92 | 1.15 | 0.61 | 0.52 | 0.5 | 0.09 | 0.56 | 0.72 | 0.72 | 0.6 | 0.41 | 0.6 | 0.2 | -0.1325 | -0.0175 | 0.331858 | 0.522141 | 0.925293 | | 1 | 0.996511 | 0.988527 |
| Csmd3 | E9PWJ0 | 0.38 | 0.38 | 0.07 | 0.03 | 0.44 | 0.72 | -0.2 | 0.7 | 0.58 | 0.01 | 0.38 | 0.49 | 0.8 | 0.59 | 0.02 | -0.09 | 0.2 | 0.15 | 0.115 | 0.42863 | 0.469689 | 0.664349 | | 1 | 0.996511 | 0.949825 |
| Csnk1a1 | E9Q4G7 | 2.18 | 0.4 | -0.17 | 0.75 | 0.68 | 1.83 | -1.6 | -1.4 | 2.06 | -2.13 | -0.42 | 1.94 | 1.82 | -1.75 | -0.08 | -0.79 | -0.9125 | -0.4275 | -0.99 | 0.311264 | 0.673837 | 0.245049 | | 1 | 0.996511 | 0.801068 |
| Csnk1a1 | Q8BK63-2 | -0.14 | -0.2 | -0.18 | -0.17 | 0.23 | -0.04 | 0.2 | 0.02 | -0.32 | -0.38 | -0.08 | -0.1 | -0.72 | 0.03 | -0.47 | -0.1 | 0.275 | -0.0475 | -0.1425 | 0.089616 | 0.772679 | 0.525522 | | 1 | 0.996511 | 0.908107 |
| Csnk1d | Q9DC28-2 | -1.1 | -0.6 | 0.76 | -0.36 | -1.22 | -1.01 | -0.31 | -0.01 | 0.56 | 0.72 | 0.02 | 0.06 | 1.11 | 0.57 | 0.39 | -0.18 | -0.3125 | 0.665 | 0.7975 | 0.495573 | 0.132382 | 0.103107 | | 1 | 0.996511 | 0.700625 |
| Csnk1d | Q9DC28 | 0.67 | 0.24 | 0.31 | 0.26 | 0.14 | 0.76 | 0.85 | 0.27 | 0.44 | 0.06 | 0.43 | 0.72 | 0.3 | -0.11 | 0.18 | 0.47 | 0.135 | 0.0425 | -0.16 | 0.552075 | 0.840571 | 0.461757 | | 1 | 0.996511 | 0.888262 |
| Csnk1e | Q9JMK2 | -0.31 | -0.11 | -0.55 | -0.55 | -0.63 | -0.36 | -0.64 | -0.62 | 0.52 | -0.84 | -0.34 | -0.08 | 0.73 | -0.27 | -0.98 | -1.26 | -0.1825 | 0.195 | -0.065 | 0.312291 | 0.529321 | 0.880801 | | 1 | 0.996511 | 0.979767 |
| Csnk1g1 | A0A0U1RP | 1.57 | 1.45 | 0.88 | 0.34 | 1.41 | 1.57 | 0.13 | 0.61 | 0.73 | 0.76 | 0.66 | 2.07 | 0.5 | 0.1 | 1.14 | 0.12 | -0.13 | -0.005 | -0.595 | 0.754614 | 0.990477 | 0.131466 | | 1 | 0.999285 | 0.726398 |
| Csnk1g2 | Q99K78 | 0.43 | 1.08 | 0.68 | 0.58 | 1.19 | 1.1 | 0.5 | -0.09 | 1.08 | 0.55 | 0.33 | 0.54 | 1.77 | 1.39 | 0.52 | 0.25 | -0.0175 | -0.0675 | 0.29 | 0.956464 | 0.778482 | 0.447288 | | 1 | 0.996511 | 0.884817 |
| Csnk1g3 | Q8C4X2 | 0.12 | 0.62 | 1.15 | 1.82 | 0.61 | 0.16 | 1.23 | 1.95 | -0.07 | 1.69 | 1.37 | 0.16 | -0.64 | 0.14 | 0.77 | 0.64 | 0.06 | -0.14 | -0.7 | 0.902884 | 0.790397 | 0.151648 | | 1 | 0.996511 | 0.750178 |
| Csnk2a1 | Q60737 | 0.98 | 1.08 | 0.92 | 1.04 | 1.4 | 0.87 | 1.11 | 1.12 | 0.59 | 0.91 | 1.04 | 1.07 | 0.39 | 0.99 | 0.85 | 1.02 | 0.12 | -0.1025 | -0.1925 | 0.482293 | 0.574218 | 0.368836 | | 1 | 0.996511 | 0.859245 |
| Csnk2a2 | O54833 | 1.48 | 1.75 | 1.89 | 1.64 | 1.9 | 1.56 | 1.67 | 1.9 | 0.61 | 0.88 | 1.93 | 1.42 | 0.22 | 0.53 | 1.6 | 1.67 | 0.0675 | -0.48 | -0.685 | 0.696601 | 0.146911 | 0.092607 | | 1 | 0.996511 | 0.695968 |
| Csnk2b | P67871 | 0.48 | 0.13 | 0.29 | 0.48 | 0.19 | 0.98 | 0.63 | 0.36 | -0.36 | 0.31 | 0.5 | 0.65 | -0.35 | -0.46 | 0.41 | 0.69 | 0.195 | -0.07 | -0.2725 | 0.380023 | 0.786639 | 0.384068 | | 1 | 0.996511 | 0.860984 |
| Cspg4 | Q8VHY0 | 0.74 | 1.05 | 0.25 | 0.6 | 1.08 | 0.83 | 0.54 | 0.5 | 0.94 | 0.34 | 0.61 | 1.19 | 0.43 | 1.19 | 0.31 | 0.49 | 0.0775 | 0.11 | -0.055 | 0.739756 | 0.68086 | 0.844004 | | 1 | 0.996511 | 0.976225 |
| Cspg5 | Q71M36-2 | -2.18 | -1.22 | -2.65 | -0.74 | -1.47 | -1.66 | -1.26 | -0.78 | -0.93 | -0.7 | -1.17 | -1.01 | -0.19 | -0.63 | -2.14 | -1.63 | 0.405 | 0.745 | 0.55 | 0.373007 | 0.106681 | 0.351563 | | 1 | 0.996511 | 0.853984 |
| Cspg5 | Q71M36 | -0.7 | 0.01 | 0.36 | 0.63 | -0.22 | -0.75 | 0.75 | 0.43 | -0.39 | 0.52 | 0.77 | 0.36 | -0.43 | 0.12 | 0.41 | 0.26 | -0.0225 | 0.24 | 0.015 | 0.956691 | 0.520952 | 0.964902 | | 1 | 0.996511 | 0.9927 |
| Cspp1 | D3Z378 | -1.99 | -2.15 | 0.53 | -0.12 | -1.28 | -1.32 | 0.6 | -0.67 | -2.13 | 0.32 | -0.24 | -2.17 | -2.46 | -1.7 | 1.09 | 0.51 | 0.265 | -0.1225 | 0.2925 | 0.717258 | 0.883795 | 0.763902 | | 1 | 0.996511 | 0.964134 |
| Csrp1 | P97315 | -0.68 | -1.03 | -0.62 | -0.62 | -0.99 | -0.57 | -0.95 | -0.95 | -1.83 | -0.49 | -0.7 | -1.06 | -1.3 | -1.7 | -0.72 | -0.61 | -0.1275 | -0.2825 | -0.345 | 0.491614 | 0.378079 | 0.254066 | | 1 | 0.996511 | 0.809524 |
| Csrp2 | P97314 | -2.28 | -2.55 | -3.14 | -2.43 | -2.45 | -3.27 | -3.42 | -3.59 | -2.87 | -2.29 | -3.11 | -2.6 | -2.11 | -2.93 | -3.3 | -2.85 | -0.5825 | -0.1175 | -0.1975 | 0.088195 | 0.668114 | 0.540408 | | 1 | 0.996511 | 0.912212 |
| Cst3 | P21460 | -0.05 | -1.23 | -1.34 | -1.46 | -1.04 | -0.42 | -0.56 | -1.14 | -0.11 | -2.03 | -1.19 | 0.38 | -0.69 | 0 | -1.33 | -1.35 | 0.23 | 0.2825 | 0.1775 | 0.523478 | 0.629022 | 0.684175 | | 1 | 0.996511 | 0.955729 |
| Cstb | Q62426 | -0.76 | -0.73 | -0.42 | -1.01 | -0.71 | -1.27 | 1.35 | -1 | -0.77 | -0.54 | -0.34 | -1.03 | -0.94 | -0.48 | -0.57 | 0.35 | 0.3225 | 0.06 | 0.32 | 0.56679 | 0.790775 | 0.314261 | | 1 | 0.996511 | 0.836924 |
| Cstf1 | Q99LC2 | -1.28 | -2.47 | -1.82 | -2.27 | -1.79 | -2.27 | -1.5 | -2.05 | -3.01 | -1.86 | -1.79 | -2.3 | -1.95 | -1.73 | -1.79 | -2.31 | 0.0575 | -0.28 | 0.015 | 0.851864 | 0.46015 | 0.960831 | | 1 | 0.996511 | 0.9927 |
| Cstf2 | Q8BIQ5-2 | -0.77 | -0.89 | -1.59 | -1.62 | -0.62 | -0.86 | -1.31 | -1.58 | -0.53 | -0.19 | -1.63 | -0.99 | -0.55 | -0.26 | -0.8 | -1.46 | 0.125 | 0.3825 | 0.45 | 0.68611 | 0.319173 | 0.209326 | | 1 | 0.996511 | 0.786218 |
| Cstf2 | F6ZKC7 | 0.55 | 1.44 | 0.07 | -0.8 | -0.01 | -0.61 | -1.41 | 0.84 | 1.49 | 2.61 | 0.01 | -1.29 | 0.01 | 1.05 | 0.76 | -1.04 | -0.6125 | 0.39 | -0.12 | 0.32817 | 0.657881 | 0.842745 | | 1 | 0.996511 | 0.976225 |
| Cstf2t | Q8C7E9 | -1.34 | -1.32 | -1.69 | -1.46 | -1.59 | -1.53 | -1.48 | -1.83 | -0.59 | -0.73 | -1.91 | -1.75 | 0.09 | -1.19 | -0.63 | -1.64 | -0.155 | 0.2075 | 0.61 | 0.370666 | 0.551707 | 0.129946 | | 1 | 0.996511 | 0.723659 |
| Cstf3 | Q99LI7 | -1.65 | -2.67 | -2.2 | -2.13 | -1.81 | -2.31 | -1.92 | -2.1 | -2.8 | -1.36 | -1.94 | -2.65 | -2.13 | -2.32 | -1.63 | -2.03 | 0.1275 | -0.025 | 0.135 | 0.61026 | 0.947422 | 0.627453 | | 1 | 0.996755 | 0.939823 |
| Cstpp1 | Q8BHR8 | 0.02 | -0.39 | 0.25 | 0.14 | 0.35 | -0.31 | 0.67 | -0.03 | -0.54 | 0.57 | 0.41 | 0.2 | -0.52 | 0.44 | 0.15 | 0.58 | 0.165 | 0.155 | 0.1575 | 0.536301 | 0.597193 | 0.597652 | | 1 | 0.996511 | 0.930093 |
| Ctbp1 | O88712 | 0.33 | 0.24 | 0.29 | 0.5 | -0.14 | 0.4 | 0.23 | 0.3 | 0.36 | 0.02 | 0.44 | 0.28 | -0.18 | 0.33 | -0.29 | -0.11 | -0.1425 | -0.065 | -0.4025 | 0.430962 | 0.714068 | 0.079291 | | 1 | 0.996511 | 0.681114 |
| Ctbp2 | Q91YZ2 | 0.02 | -0.2 | -0.03 | 0.13 | -0.51 | -0.28 | -0.18 | -0.31 | -0.34 | -0.29 | 0.02 | -0.53 | 0.13 | -0.57 | 0.26 | -0.04 | -0.3 | -0.265 | -0.035 | 0.088266 | 0.187502 | 0.882036 | | 1 | 0.996511 | 0.979767 |
| Ctbs | Q8R242 | 0.51 | 0.71 | -0.2 | -0.65 | 0.07 | 0.68 | 0.27 | -0.22 | -1.53 | 0.94 | -0.12 | 0.28 | -0.95 | -0.99 | -0.5 | -0.17 | 0.1075 | -0.2 | -0.745 | 0.761106 | 0.722575 | 0.067178 | | 1 | 0.996511 | 0.664936 |
| Ctc1 | Q5SUQ9-2 | -4.09 | -4.32 | -1.67 | -2.14 | -3.09 | -4.35 | -2.3 | -2.17 | -2.94 | -1.62 | -1.96 | -3.57 | -2.43 | -3.61 | -1.57 | -1.87 | 0.0775 | 0.5325 | 0.685 | 0.918424 | 0.474333 | 0.361952 | | 1 | 0.996511 | 0.858475 |
| Ctcf | Q61164 | 2.7 | 1.48 | 1.58 | 0.32 | 1.84 | 2.62 | 0.96 | 0.88 | -0.89 | 0.45 | 1.25 | 2.13 | -0.06 | -0.01 | 1.85 | 1.5 | 0.055 | -0.785 | -0.7 | 0.924572 | 0.298865 | 0.28791 | | 1 | 0.996511 | 0.824705 |
| Ctdnep1 | Q3TP92 | 0.71 | 1.16 | 1.62 | -0.09 | 1.18 | 0.79 | 0.38 | 1.24 | 0.33 | -0.31 | 0.54 | 0.67 | 0.7 | 0.18 | 1.02 | 0.74 | 0.0475 | -0.5425 | -0.19 | 0.90361 | 0.203407 | 0.62951 | | 1 | 0.996511 | 0.940703 |
| Ctdp1 | Q7TSG2 | -1.02 | -1.36 | -2 | -2.29 | -1.11 | -1.57 | -2.06 | -1.86 | -1.36 | -1.64 | -1.66 | -0.94 | -0.86 | -1.11 | -2.32 | -2.44 | 0.0175 | 0.2675 | -0.015 | 0.959332 | 0.430114 | 0.974487 | | 1 | 0.996511 | 0.992977 |
| Ctdsp1 | P58466 | 0.5 | 0.79 | -0.34 | 0.44 | 1.51 | 0.43 | 0.23 | 0.32 | 1.3 | 0.1 | 0.12 | 0.93 | 0.51 | 1.93 | 1 | 0.8 | 0.275 | 0.265 | 0.7125 | 0.460685 | 0.483475 | 0.088337 | | 1 | 0.996511 | 0.691619 |
| Ctdspl | G3UXN0 | -0.14 | 0.64 | -0.11 | -1.98 | 0.66 | -0.37 | -0.5 | -1.49 | 1.15 | 0.03 | -0.5 | 0.31 | 0.83 | 0.83 | -1.99 | -0.9 | -0.0275 | 0.645 | 0.09 | 0.96597 | 0.2995 | 0.910561 | | 1 | 0.996511 | 0.986458 |
| Ctdspl2 | Q8BG15-6 | 1.23 | 1.33 | 0.5 | 0.01 | 0.9 | 0.4 | 0.2 | 0.29 | 0.65 | -0.59 | 0.29 | 0.68 | 1.23 | 0.65 | 1.35 | 0.91 | -0.32 | -0.51 | 0.2675 | 0.357193 | 0.234879 | 0.450872 | | 1 | 0.996511 | 0.885233 |
| Cth | Q8VCN5 | -0.17 | -0.26 | -0.17 | -0.05 | 0.06 | -0.05 | 0.44 | -0.19 | 0.07 | 0.06 | 0.02 | -0.35 | -0.3 | 0.22 | -0.05 | 0.14 | 0.2275 | 0.1125 | 0.165 | 0.237311 | 0.531949 | 0.407695 | | 1 | 0.996511 | 0.874084 |
| Ctif | E9Q1U6 | -1.73 | -1.67 | -0.92 | -0.88 | -1.28 | -1.86 | -0.87 | -1.11 | -1.84 | 0.64 | -0.93 | -1.67 | -1.12 | -1.15 | -0.76 | -0.93 | 0.02 | 0.35 | 0.31 | 0.948273 | 0.538532 | 0.272834 | | 1 | 0.996511 | 0.822084 |
| Ctif | Q6PEE2 | 0.61 | -0.48 | 0.15 | -1.02 | 0.24 | 0.77 | -0.09 | -0.07 | 0.07 | -1.12 | -0.97 | -0.2 | 0.81 | -0.83 | -0.46 | 0.35 | 0.3975 | -0.37 | 0.1525 | 0.320317 | 0.404505 | 0.753011 | | 1 | 0.996511 | 0.961479 |
| Ctnna1 | P26231 | 0.41 | 0.59 | 0.78 | 0.9 | 0.36 | 0.49 | 1.51 | 0.88 | 0.79 | 0.74 | 0.84 | 0.66 | 0.44 | 0.63 | 0.49 | 0.53 | 0.14 | 0.0875 | -0.1475 | 0.622068 | 0.629649 | 0.448137 | | 1 | 0.996511 | 0.884817 |
| Ctnna2 | Q61301 | 1.19 | 1.14 | 1.33 | 1.33 | 1.43 | 1.23 | 1.35 | 1.36 | 1.04 | 1.09 | 1.54 | 1.38 | 0.19 | 0.82 | 0.67 | 1.27 | 0.095 | 0.015 | -0.51 | 0.520818 | 0.936252 | 0.074493 | | 1 | 0.996755 | 0.671911 |
| Ctnna2 | Q61301-2 | -0.88 | 1.72 | 0.58 | 1.02 | 1.91 | 1.96 | 0.66 | 0.39 | 1.71 | 0.23 | 0.57 | 0.54 | 0.14 | 1 | -0.37 | 0.66 | 0.62 | 0.1525 | -0.2525 | 0.334919 | 0.794919 | 0.662453 | | 1 | 0.996511 | 0.949645 |
| Ctnna3 | Q65CL1 | 1.16 | 1.74 | 3.18 | 0.93 | 0.21 | 2.19 | 0.36 | 0.4 | 1.91 | -0.59 | 1.74 | 0.66 | 2.88 | -2.59 | 2.02 | 2.65 | -0.9625 | -0.8225 | -0.5125 | 0.151278 | 0.257793 | 0.679071 | | 1 | 0.996511 | 0.955269 |
| Ctnnb1 | Q02248 | 0.91 | 0.81 | 0.97 | 0.59 | 1.05 | 1.29 | 0.99 | 0.74 | 0.44 | -0.02 | 1.26 | 1.09 | 0.07 | 0.24 | 0.37 | 0.96 | 0.1975 | -0.1275 | -0.41 | 0.297838 | 0.682597 | 0.122926 | | 1 | 0.996511 | 0.721448 |
| Ctnnbip1 | Q9JJN6 | 0.21 | 0.14 | 0.22 | 0.44 | 0.06 | 0.02 | 0.14 | 0.34 | -0.32 | 0.47 | 0.27 | 0.26 | -0.1 | 0.1 | 0.67 | 0.48 | -0.1125 | -0.0825 | 0.035 | 0.486895 | 0.708306 | 0.87951 | | 1 | 0.996511 | 0.979626 |
| Ctnnbl1 | Q9CWL8 | -1.42 | -4.33 | -3.7 | -3.6 | -3.24 | -2.72 | -3.41 | -3.14 | -3.36 | -2.14 | -2.8 | -3.55 | -2.71 | -2.55 | -1.86 | -2.84 | 0.135 | 0.3 | 0.7725 | 0.820518 | 0.645182 | 0.229632 | | 1 | 0.996511 | 0.787447 |
| Ctnnd1 | P30999-2 | 0.67 | 0.74 | 1.08 | 1.01 | 0.35 | 0.88 | 0.39 | 1.91 | 0.43 | 0.97 | 1.83 | 1.47 | 3.24 | 0.55 | 1.05 | 1.62 | 0.0075 | 0.3 | 0.74 | 0.98338 | 0.362178 | 0.199558 | | 1 | 0.996511 | 0.78004 |
| Ctnnd1 | G3X9V2 | 1.32 | 0.87 | 0.94 | 0.78 | 1.1 | 1.51 | 1.09 | 1.06 | 1.04 | 0.74 | 1.37 | 1.53 | 0.92 | 1 | 0.96 | 0.97 | 0.2125 | 0.1925 | -0.015 | 0.293587 | 0.431028 | 0.938147 | | 1 | 0.996511 | 0.989844 |
| Ctnnd2 | E9QKH8 | 0.96 | 0.89 | 1.38 | 1.1 | 0.83 | 1.17 | 1.27 | 1.28 | 0.51 | 0.82 | 1.67 | 1.24 | -0.49 | -0.8 | -0.01 | 1.34 | 0.055 | -0.0225 | -1.0725 | 0.772475 | 0.937057 | 0.040974 | | 1 | 0.996755 | 0.658454 |
| Ctnnd2 | A0A2I3BQ\ | -1.08 | -0.12 | -0.4 | -0.07 | -0.59 | -1.07 | -0.18 | -0.3 | -1.12 | 0.43 | -0.27 | -0.89 | -0.62 | 0.3 | -0.02 | 0.44 | -0.1175 | -0.045 | 0.4425 | 0.698716 | 0.910296 | 0.206921 | | 1 | 0.996744 | 0.785338 |
| Ctps1 | P70698 | -0.15 | 0.09 | 0.27 | -0.27 | -0.01 | 0.14 | 0.56 | -0.09 | -0.24 | -0.98 | 0.33 | 0.11 | -0.16 | -0.3 | 0.15 | 0.15 | 0.165 | -0.18 | -0.025 | 0.449782 | 0.569303 | 0.908678 | | 1 | 0.996511 | 0.986458 |
| Ctps2 | P70303 | -0.22 | -0.04 | -0.06 | -0.09 | 0.12 | -0.16 | 0.31 | -0.09 | -0.68 | -0.37 | -0.53 | -0.19 | -0.34 | -0.09 | -0.27 | 0.31 | 0.1475 | -0.34 | 0.005 | 0.391602 | 0.085686 | 0.980962 | | 1 | 0.996511 | 0.994858 |
| Ctr9 | Q62018 | -0.68 | -1.99 | -1.64 | -1.79 | -1.61 | -1.37 | -1.91 | -1.01 | -1.63 | -0.87 | -1.79 | -1.92 | -0.74 | -0.88 | -0.48 | -1.2 | 0.05 | -0.0275 | 0.7 | 0.881801 | 0.939633 | 0.059287 | | 1 | 0.996755 | 0.661738 |
| Ctsa | G3X8T3 | -0.45 | -0.67 | -1.03 | -0.49 | -0.87 | -0.22 | -0.13 | -0.37 | -0.52 | -0.17 | -0.08 | -0.28 | -0.4 | -1.14 | -0.53 | -0.72 | 0.2625 | 0.3975 | -0.0375 | 0.273372 | 0.082647 | 0.878305 | | 1 | 0.996511 | 0.979626 |
| Ctsb | P10605 | 0.33 | -0.63 | -1.41 | -1.47 | -0.49 | 0.28 | -0.79 | -1.6 | -1.13 | -1.03 | -1.11 | -0.06 | -0.83 | -0.94 | -1.82 | -1.58 | 0.145 | -0.0375 | -0.4975 | 0.783326 | 0.935398 | 0.293612 | | 1 | 0.996755 | 0.825612 |
| Ctsc | P97821 | -0.95 | -1.78 | -0.04 | 0.2 | -1.44 | -0.87 | 0.28 | 0.65 | -1.06 | 0.87 | 0.11 | -0.68 | -1.24 | -1.15 | -0.17 | 0.41 | 0.2975 | 0.4525 | 0.105 | 0.626153 | 0.438786 | 0.850232 | | 1 | 0.996511 | 0.976225 |
| Ctsd | P18242 | -0.32 | -0.76 | -0.65 | 0.63 | -0.84 | -0.3 | 0.14 | 0.65 | -0.4 | 0.44 | 0.42 | -0.25 | -0.58 | -0.55 | -1 | -0.39 | 0.1875 | 0.3275 | -0.355 | 0.656757 | 0.390286 | 0.313953 | | 1 | 0.996511 | 0.836827 |
| Ctsf | Q9R013 | -0.34 | 0.18 | -2.1 | -1.33 | -0.49 | 0.08 | -0.91 | -1.26 | 0.45 | -1.4 | -0.92 | -0.19 | 0.61 | -0.32 | -1.94 | -1.12 | 2.53E-01 | 0.3825 | 0.205 | 0.640056 | 0.526799 | 0.762788 | | 1 | 0.996511 | 0.963993 |
| Ctsh | P49935 | -1.95 | -1.23 | -1.69 | -0.47 | -0.99 | -1.93 | -0.73 | -1.52 | -0.79 | -0.72 | -0.68 | -1.88 | -0.62 | 0.04 | -0.79 | -1.46 | 0.0425 | 0.3175 | 0.6275 | 0.914804 | 0.450013 | 0.166497 | | 1 | 0.996511 | 0.759484 |
| Ctsl | P06797 | -1.02 | -0.26 | -0.85 | -0.17 | -0.56 | -0.5 | 0.98 | -0.54 | -0.46 | -0.87 | 0.42 | -0.18 | -0.74 | 0.12 | -1.07 | -0.68 | 0.42 | 0.3025 | -0.0175 | 0.318485 | 0.383716 | 0.957936 | | 1 | 0.996511 | 0.992636 |
| Ctso | Q8BM88 | 6.68 | 7.63 | 3.45 | 3.31 | 8.32 | 7.17 | 4.57 | 2.91 | 4.98 | 2.36 | 4.64 | 7.62 | 4.08 | 2.75 | 5.85 | 6.3 | 0.475 | -0.3675 | -0.5225 | 0.748123 | 0.790337 | 0.671614 | | 1 | 0.996511 | 0.952698 |
| Ctss | F6WR04 | -1.32 | -1.72 | -1.05 | -0.49 | -1.78 | -1.59 | -0.61 | -1.22 | -1.99 | -1.13 | -0.81 | -1.95 | -1.81 | -1.97 | -1.89 | -1.42 | -0.155 | -0.325 | -0.6275 | 0.660069 | 0.401407 | 0.06118 | | 1 | 0.996511 | 0.661738 |
| Ctsz | Q9WUU7 | 1.71 | 0.22 | -2.35 | -1.7 | 0.69 | 1.63 | -1.11 | -2.47 | -1.08 | -0.53 | -1.89 | 1.67 | -1.23 | -1.74 | -1.92 | -2.19 | 0.215 | 0.0725 | -1.24 | 0.853586 | 0.94618 | 0.170153 | | 1 | 0.996755 | 0.762229 |
| Cttn | Q60598 | 1.77 | 0.39 | 0.53 | 0.57 | 0.02 | 1.48 | 0.44 | 0.64 | 1.13 | 0.26 | 0.7 | 1.53 | 1.03 | -0.09 | 0.48 | 0.22 | -0.17 | 0.09 | -0.405 | 0.684442 | 0.823221 | 0.310955 | | 1 | 0.996511 | 0.834523 |
| Cttnbp2 | B9EJA2-5 | 0.96 | 0.81 | 0.86 | 0.88 | 0.79 | 0.89 | 0.56 | 0.77 | 1.05 | 0.84 | 0.83 | 0.97 | 0.43 | 1.27 | 0.34 | 0.29 | -0.125 | 0.045 | -0.295 | 0.416057 | 0.777402 | 0.276843 | | 1 | 0.996511 | 0.822084 |
| Cttnbp2nl | Q99LJ0 | 0.53 | 0.14 | -0.05 | 0.22 | -0.38 | 0.06 | -0.2 | -0.09 | 0.85 | 1.27 | 0.02 | 0.39 | 0.72 | 0.21 | 0.12 | -0.17 | -0.3625 | 0.4225 | 0.01 | 0.084495 | 0.187007 | 0.968478 | | 1 | 0.996511 | 0.992733 |
| Ctu1 | Q99J10 | -1.14 | -0.82 | -1.24 | -0.57 | -1.77 | -0.27 | -0.87 | -0.43 | 0.64 | -0.38 | -0.4 | -1.05 | 1.31 | -1.96 | -0.69 | -0.74 | 0.1075 | 0.645 | 0.4225 | 0.762249 | 0.108698 | 0.510272 | | 1 | 0.996511 | 0.903419 |
| Ctu2 | Q3U308 | -0.35 | 0.03 | -0.15 | 0.32 | -0.24 | -0.22 | -0.16 | 0.3 | 0.08 | -0.3 | 0.09 | -0.13 | -0.28 | 0.12 | -0.74 | -0.23 | -0.0425 | -0.0275 | -0.245 | 0.844275 | 0.89671 | 0.353336 | | 1 | 0.996511 | 0.855029 |
| Cuedc1 | Q8R3V6 | -0.55 | 0.18 | 0.26 | 0.58 | 0.22 | -0.32 | 0 | 0.53 | 0.73 | 0.38 | -0.12 | 0.03 | 0.42 | 0.24 | -0.18 | 0.31 | -0.01 | 0.1375 | 0.08 | 0.973107 | 0.657452 | 0.782436 | | 1 | 0.996511 | 0.967642 |
| Cuedc2 | A0A494BB! | -1.6 | -1.9 | -1.6 | -1.68 | -1.44 | -2.13 | -1.93 | -1.86 | -2.1 | -1.05 | -1.71 | -1.69 | -1.76 | -1.35 | -1.67 | -2.07 | -0.145 | 0.0575 | -0.0175 | 0.467564 | 0.819173 | 0.935723 | | 1 | 0.996511 | 0.989767 |
| Cul1 | Q9WTX6 | 0.03 | -0.15 | 0.01 | -0.08 | -0.02 | 0.06 | -0.3 | -0.01 | -0.15 | -0.12 | 0.04 | -0.04 | -0.16 | -0.2 | -0.02 | -0.1 | -0.02 | -0.02 | -0.0725 | 0.898207 | 0.899444 | 0.670888 | | 1 | 0.996511 | 0.952558 |
| Cul2 | Q9D4H8 | -0.24 | -0.19 | -0.23 | -0.41 | -0.21 | -0.32 | -0.28 | -0.15 | -0.09 | -0.68 | -0.09 | -0.1 | 0.01 | -0.15 | -0.32 | -0.1 | 0.0275 | 0.0275 | 0.1275 | 0.849926 | 0.891889 | 0.481078 | | 1 | 0.996511 | 0.893906 |
| Cul3 | Q9JLV5 | 0.39 | -0.22 | 0.61 | 0.79 | 0.11 | 0.36 | 0.82 | 0.92 | -0.29 | 0.49 | 0.63 | 0.4 | -0.72 | 0 | 0.59 | 0.38 | 0.16 | -0.085 | -0.33 | 0.586228 | 0.780744 | 0.368461 | | 1 | 0.996511 | 0.859245 |
| Cul4a | Q3TCH7 | -0.69 | -0.55 | -0.77 | -0.84 | -0.31 | -0.7 | -1.11 | -0.9 | -0.9 | -0.7 | -0.89 | -0.64 | -0.95 | -0.81 | -1.26 | -1.45 | -0.0425 | -0.07 | -0.405 | 0.83933 | 0.681003 | 0.085087 | | 1 | 0.996511 | 0.691619 |
| Cul4b | A2A432 | -1.08 | -0.86 | -0.43 | -0.55 | -1.18 | -1.11 | -0.93 | -0.64 | -1.02 | -0.41 | -0.65 | -1.26 | -1.19 | -0.64 | -1.16 | -1.09 | -0.235 | -0.105 | -0.29 | 0.293231 | 0.687044 | 0.240376 | | 1 | 0.996511 | 0.795658 |
| Cul5 | Q9D5V5 | -0.28 | -0.2 | -0.34 | -0.3 | -0.23 | -0.35 | -0.4 | -0.48 | -0.77 | -0.1 | -0.37 | -0.27 | -0.95 | -0.56 | -0.83 | -0.64 | -0.085 | -0.0975 | -0.465 | 0.561166 | 0.622904 | 0.029003 | | 1 | 0.996511 | 0.637553 |
| Cul7 | Q8VE73-3 | 1.05 | -0.16 | -0.9 | -0.35 | -1.71 | 0.1 | -0.52 | -0.07 | 4.78 | 1.12 | -0.91 | -0.97 | 6.51 | 1.36 | 1.94 | -0.38 | -0.46 | 1.095 | 2.4475 | 0.397854 | 0.396086 | 0.098725 | | 1 | 0.996511 | 0.698156 |
| Cul9 | E9QP09 | -0.35 | 0.07 | 0.2 | 0.11 | 0.07 | 0.02 | 0.31 | 0.34 | -0.19 | -0.39 | 0.27 | -0.16 | -0.05 | -0.01 | -0.04 | -0.04 | 0.1775 | -0.125 | -0.0425 | 0.356163 | 0.575994 | 0.827317 | | 1 | 0.996511 | 0.973757 |
| Custos | Q3UY34 | -2.16 | -3.61 | -1.84 | -1.72 | -4.19 | -2.9 | -2.68 | -2.62 | -2.71 | -1.03 | -2.01 | -2.85 | -2.58 | -2.81 | -1.63 | -2.66 | -0.765 | 0.1825 | -0.0875 | 0.170489 | 0.743054 | 0.855133 | | 1 | 0.996511 | 0.976225 |
| Cuta | D5MCW4 | -0.44 | -0.03 | -0.58 | -0.85 | -0.17 | -0.34 | -0.54 | -0.81 | -0.27 | -2.47 | -0.92 | 0.09 | -0.91 | 0.18 | -1.2 | -0.66 | 0.01 | -0.4175 | -0.1725 | 0.966232 | 0.452017 | 0.616939 | | 1 | 0.996511 | 0.937141 |
| Cutc | Q9D8X1 | -3.95 | -1.99 | -2.83 | -3.3 | -2.27 | -4.04 | -2.73 | -3.42 | -2.64 | -1.46 | -2.81 | -2.77 | -1.38 | -1.43 | -2.48 | -2.84 | -0.0975 | 0.5975 | 0.985 | 0.851567 | 0.241158 | 0.08283 | | 1 | 0.996511 | 0.688789 |
| Cux1 | H3BKT0 | -0.29 | -1.32 | -0.36 | -0.48 | -1.05 | -0.77 | 0.45 | -0.01 | -0.2 | -1.68 | -0.58 | -0.38 | 0.23 | -0.51 | 0.31 | 0.52 | 0.2675 | -0.0975 | 0.75 | 0.50482 | 0.804077 | 0.046643 | | 1 | 0.996511 | 0.658454 |
| Cux1 | H3BK24 | -1.74 | -2.4 | -0.77 | -0.49 | -2.05 | -1.5 | -0.98 | -0.88 | -1.48 | 0.28 | -1.01 | -2.08 | -1.02 | -2.15 | -0.57 | -0.47 | -0.0025 | 0.2775 | 0.2975 | 0.995824 | 0.651569 | 0.585556 | | 1 | 0.996511 | 0.928178 |
| Cwc15 | Q9JHS9 | -1.04 | -3.32 | -1.71 | -1.68 | -3.81 | -1.6 | -1.28 | -1.35 | -1.73 | -2.04 | -1 | -3.1 | -0.42 | -3.76 | -0.75 | -1.65 | -0.0725 | -0.03 | 0.2925 | 0.917689 | 0.95999 | 0.71759 | | 1 | 0.996755 | 0.958398 |
| Cwc22 | B1AYU7 | 0.82 | 0.14 | 0.09 | -0.19 | -1.29 | 0.17 | -1.02 | 0.03 | -0.06 | -0.46 | -0.35 | 0.14 | 1.08 | 0.02 | 0.12 | -1.22 | -0.7425 | -0.3975 | -0.215 | 0.091956 | 0.166184 | 0.659068 | | 1 | 0.996511 | 0.949645 |
| Cwc25 | Q9DBF7 | 0 | -0.72 | -0.49 | -1.21 | -1.1 | -1.41 | -0.98 | -1.37 | -1.28 | -0.91 | -0.41 | -0.79 | -0.82 | -1.29 | -1.03 | -1.83 | -0.61 | -0.2425 | -0.6375 | 0.051682 | 0.444365 | 0.08399 | | 1 | 0.996511 | 0.690249 |
| Cwc27 | Q3TKY6 | -1.74 | -1.28 | -2.03 | -1.79 | -2.33 | -2.02 | -2.15 | -1.75 | -1.73 | -1.74 | -2.03 | -1.58 | -1.24 | -0.59 | -1.24 | -1.99 | -0.3525 | -0.06 | 0.445 | 0.138347 | 0.785375 | 0.200142 | | 1 | 0.996511 | 0.78004 |
| Cx3cl1 | O35188 | -0.56 | -0.58 | -0.28 | -0.75 | -0.18 | -0.67 | 0.13 | -0.63 | -0.87 | -0.97 | -0.16 | -0.56 | -0.81 | 0.07 | -0.42 | -0.19 | 0.205 | -0.0975 | 0.205 | 0.38979 | 0.680797 | 0.414392 | | 1 | 0.996511 | 0.874945 |
| Cx3cr1 | Q9Z0D9 | -4 | -4.43 | -3.65 | -2.68 | -3.66 | -3.9 | -4.46 | -3.94 | -4.84 | -2.9 | -3.75 | -3.81 | -4.44 | -3.26 | -4.73 | -4.26 | -0.3 | -0.135 | -0.4825 | 0.446561 | 0.789786 | 0.311775 | | 1 | 0.996511 | 0.834523 |
| Cxadr | P97792 | 0.69 | 0.52 | 0.02 | 0.35 | 0.7 | 1.33 | 1.24 | 0.61 | 1.59 | -0.59 | 1.1 | 1.71 | 1.14 | 0.8 | 0.48 | 0.5 | 0.575 | 0.5575 | 0.335 | 0.041923 | 0.291652 | 0.197166 | | 1 | 0.996511 | 0.78004 |
| Cxcl12 | H7BX38 | 0.44 | -0.42 | -1.34 | -2.11 | 0.4 | 0.31 | -0.37 | 0.1 | -1.66 | -3.34 | 0.61 | 0.55 | -1.23 | -0.67 | -1.06 | -1.05 | 0.9675 | -0.1025 | -0.145 | 0.09701 | 0.917509 | 0.782648 | | 1 | 0.996755 | 0.967642 |
| Cxcl14 | Q9WUQ5 | -1.21 | -1.73 | -1.01 | -1.44 | -2.92 | 0.51 | -2.26 | -2.04 | 1.12 | -1.71 | -1.24 | -1.09 | 2.68 | -2.05 | -0.85 | -1.22 | -0.33 | 0.6175 | 0.9875 | 0.63757 | 0.315307 | 0.311763 | | 1 | 0.996511 | 0.834523 |
| Cxxc1 | Q9CWW7 | -4.64 | -8.3 | -3.29 | -2.52 | -7.69 | -9.31 | -4.05 | -3.54 | -5.68 | -0.29 | -3.66 | -5.06 | -2.97 | -13.83 | -3.45 | -3.65 | -1.46 | 1.015 | -1.2875 | 0.39924 | 0.52289 | 0.620355 | | 1 | 0.996511 | 0.93807 |
| Cyb561 | Q60720 | -0.46 | 0.38 | 0.01 | -1.84 | 1.21 | 0.6 | -0.37 | -0.57 | 0.65 | 4.63 | -1.5 | -0.58 | 0.87 | 0.95 | 1.19 | 0.75 | 0.695 | 1.2775 | 1.4175 | 0.254546 | 0.333953 | 0.013282 | | 1 | 0.996511 | 0.584789 |
| Cyb561d2 | A0A0A6YY: | 0.14 | -0.28 | -0.61 | 0.16 | -1.57 | -0.11 | 0.29 | 0.36 | 0.71 | 0.39 | 0.52 | 0.63 | -0.99 | -0.28 | 1.02 | 0.02 | -0.11 | 0.71 | 0.09 | 0.807957 | 0.012418 | 0.835555 | | 1 | 0.996511 | 0.975524 |
| Cyb5a | G5E850 | 0.3 | 0.51 | 0.8 | 0.41 | 0.73 | 0.7 | 1.59 | 0.65 | 0.47 | -0.36 | 0.74 | 0.58 | 0.37 | 0.56 | 0.92 | 1.44 | 0.4125 | -0.1475 | 0.3175 | 0.137391 | 0.60126 | 0.273695 | | 1 | 0.996511 | 0.822084 |
| Cyb5b | Q9CQX2 | 0.54 | 0.65 | -0.02 | 0.25 | 0.59 | 0.44 | 0.33 | 0.21 | 0.46 | -0.01 | 0.23 | 0.35 | 0.35 | 0.85 | -0.02 | -0.12 | 0.0375 | -0.0975 | -0.09 | 0.853247 | 0.658065 | 0.752666 | | 1 | 0.996511 | 0.961479 |
| Cyb5d2 | Q5SSH8 | 1.05 | 1.26 | 1.41 | 1.11 | 0.54 | 0.69 | 1.58 | 1.08 | -1.44 | 2.19 | 1.37 | 0.91 | -1.38 | -1.55 | 1.42 | 1.82 | -0.235 | -0.45 | -1.13 | 0.370646 | 0.531741 | 0.186733 | | 1 | 0.996511 | 0.776843 |
| Cyb5r1 | Q9DB73 | 0.15 | 0.28 | 1.08 | 1.73 | 0.25 | 0.08 | 1.06 | 1.76 | 0.18 | 1.18 | 1.62 | 0.06 | -0.12 | 0.27 | 1 | 0.99 | -0.0225 | -0.05 | -0.275 | 0.963686 | 0.91932 | 0.534893 | | 1 | 0.996755 | 0.912212 |
| Cyb5r3 | Q9DCN2 | 0.43 | -0.01 | 0.54 | 0.85 | -0.24 | 0.35 | 0.81 | 0.92 | 0.18 | 0.63 | 0.61 | 0.09 | -0.31 | 0.27 | 0.3 | 0.17 | 0.0075 | -0.075 | -0.345 | 0.980853 | 0.764911 | 0.203372 | | 1 | 0.996511 | 0.782817 |
| Cyb5r4 | Q3TDX8 | -0.5 | 0.14 | -1.16 | -1.19 | 0.3 | -0.46 | -1.07 | -1.64 | -0.27 | -0.75 | -0.95 | -0.25 | -0.71 | 0.66 | -1.15 | -1.06 | -0.04 | 0.1225 | 0.1125 | 0.933946 | 0.729334 | 0.818641 | | 1 | 0.996511 | 0.972768 |
| Cybc1 | Q3TYS2 | 0.66 | 0.41 | 1.47 | 0.76 | 0.43 | 0.2 | 0.75 | 1.2 | 0.15 | 0.29 | 1.28 | 0.65 | 0.66 | 0.67 | 2.1 | 1.68 | -0.18 | -0.2325 | 0.4525 | 0.56443 | 0.494805 | 0.288461 | | 1 | 0.996511 | 0.824705 |
| Cyc1 | Q9D0M3 | 1.31 | 1.42 | 1.12 | 0.61 | 1.26 | 1.56 | 1.07 | 0.99 | 0.93 | 0.03 | 1.11 | 1.2 | 0.73 | 1.04 | 0.67 | 1.12 | 0.105 | -0.2975 | -0.225 | 0.658761 | 0.368515 | 0.373307 | | 1 | 0.996511 | 0.859424 |
| Cycs | P62897 | 1.09 | 0.79 | 0.53 | 1.11 | 0.48 | 1.06 | 0.18 | 0.96 | 0.72 | 0.12 | 0.43 | 0.57 | 0.24 | 0.81 | -0.34 | -0.2 | -0.21 | -0.42 | -0.7525 | 0.424532 | 0.087557 | 0.033456 | | 1 | 0.996511 | 0.644622 |
| Cyfip1 | Q7TMB8 | 0.4 | 0.19 | 0.29 | 0.57 | 0.39 | 0.5 | 0.77 | 0.71 | -0.29 | 0.3 | 0.31 | 0.22 | -0.58 | -0.28 | -0.04 | 0.24 | 0.23 | -0.2275 | -0.5275 | 0.203837 | 0.292167 | 0.047855 | | 1 | 0.996511 | 0.658454 |
| Cyfip1 | A0A0R4J11 | 1.3 | 1.29 | 0.2 | 0.23 | 1.73 | 0.36 | 0.22 | 0.57 | 1.41 | 0.44 | 0.27 | 0.89 | 0.45 | 2.14 | 0.08 | 0.28 | -0.035 | -0.0025 | -0.0175 | 0.935662 | 0.994844 | 0.973419 | | 1 | 0.999285 | 0.992946 |
| Cyfip2 | Q5SQX6 | 0.69 | 0.37 | 0.67 | 0.7 | 0.53 | 0.68 | 0.43 | 0.57 | 0.73 | 0.81 | 0.65 | 0.61 | -0.18 | 0.7 | 0.18 | 0.24 | -0.055 | 0.0925 | -0.3725 | 0.729647 | 0.588599 | 0.143165 | | 1 | 0.996511 | 0.742711 |
| Cygb | Q9CX80 | -0.47 | -1.17 | -0.7 | -1.17 | -1.18 | -0.61 | -0.01 | -0.72 | -1.97 | -0.65 | -0.93 | -0.86 | -1.54 | -1.63 | -0.49 | -0.41 | 0.2475 | -0.225 | -0.14 | 0.413436 | 0.512182 | 0.702976 | | 1 | 0.996511 | 0.956859 |
| Cyhr1 | Q9QXA1 | -5.81 | -5.76 | -2.8 | -2.94 | -5.56 | -7.26 | -2.2 | -3.17 | -6.77 | -2.54 | -2.75 | -5.58 | -6.21 | -6.26 | -2.9 | -3.12 | -0.22 | -0.0825 | -0.295 | 0.862701 | 0.945318 | 0.792573 | | 1 | 0.996755 | 0.96933 |
| Cyld | Z4YJJ5 | -0.83 | -0.21 | 0.26 | 0.14 | 0.03 | -1.16 | 0.83 | 0.23 | -0.7 | 0.02 | 0.21 | -0.59 | -0.81 | 0.18 | 0.34 | 0.06 | 0.1425 | -0.105 | 0.1025 | 0.752038 | 0.750522 | 0.771283 | | 1 | 0.996511 | 0.965933 |
| Cyp20a1 | Q8BKE6 | 1.54 | 1.41 | 2.71 | 0.97 | 1.48 | 2.16 | 2.04 | 2.23 | -0.83 | 0.07 | 2.13 | 1.87 | 0.02 | -1.34 | 3.21 | 3.24 | 0.32 | -0.8475 | -0.375 | 0.417838 | 0.264331 | 0.729951 | | 1 | 0.996511 | 0.958635 |
| Cyp2d22 | Q9JKY7 | 0.39 | 0.41 | 0.92 | 0.95 | 0.63 | 0.74 | 1.56 | 1.49 | 0.61 | 1.08 | 1.01 | 0.38 | 0.66 | 0.78 | 1.07 | 1.11 | 0.4375 | 0.1025 | 0.2375 | 0.15661 | 0.683316 | 0.322141 | | 1 | 0.996511 | 0.840788 |
| Cyp2j6 | O54750 | -0.4 | 0.26 | 0.74 | -1 | 0.84 | -0.59 | 1.41 | 0.24 | -0.32 | 0.15 | -0.09 | -0.1 | -0.92 | 0.6 | 0.98 | 1.54 | 0.575 | 0.01 | 0.65 | 0.291527 | 0.978896 | 0.291916 | | 1 | 0.998749 | 0.825497 |
| Cyp2j9 | Q924D1 | -0.12 | 0.29 | -0.65 | -0.17 | 1.11 | 0.11 | 0.5 | -0.5 | -0.09 | 0.06 | -0.11 | 0.06 | 0.28 | 0.96 | 0 | 0.25 | 0.4675 | 0.1425 | 0.535 | 0.228745 | 0.54016 | 0.098131 | | 1 | 0.996511 | 0.698156 |
| Cyp2s1 | D3Z7F1 | 0.22 | 0.69 | 0.55 | 0.6 | -0.62 | -0.39 | 0.14 | 0.42 | -0.33 | -0.21 | -0.11 | 0.47 | -0.25 | -0.84 | 0.6 | 0.08 | -0.6275 | -0.56 | -0.6175 | 0.040668 | 0.038641 | 0.083676 | | 1 | 0.996511 | 0.689531 |
| Cyp46a1 | Q9WVK8 | -0.27 | -0.25 | -0.01 | -0.12 | 0.21 | -0.31 | 0.56 | 0.09 | -0.33 | 0.13 | -0.02 | -0.15 | -0.1 | 0.67 | 0.38 | 0.43 | 0.3 | 0.07 | 0.5075 | 0.187241 | 0.699484 | 0.045112 | | 1 | 0.996511 | 0.658454 |
| Cyp4f14 | Q9EP75 | 0.5 | 0.27 | 1.25 | 0.84 | 1.01 | 0.88 | 1.32 | 1.06 | 0.81 | 1.08 | 1.11 | 0.51 | 0.69 | 0.49 | 1.07 | 1.51 | 0.3525 | 0.1625 | 0.225 | 0.176619 | 0.552158 | 0.484616 | | 1 | 0.996511 | 0.894377 |
| Cyp4f16 | Q99N17 | -1.75 | 0.32 | -1.06 | -0.37 | 0.44 | -0.48 | -0.49 | -0.8 | -2.02 | -2.62 | -1.25 | -1.51 | -1.35 | -1.23 | -1.95 | -0.43 | 0.3825 | -1.135 | -0.525 | 0.435171 | 0.046987 | 0.315274 | | 1 | 0.996511 | 0.837729 |
| Cyp4x1 | A2A9Y7 | 0.95 | 1.03 | 1.18 | -0.19 | 1.25 | 1.16 | 1.1 | 0.77 | -0.15 | -0.54 | 1.01 | 0.97 | -0.48 | 0.45 | 1.26 | 1.57 | 0.3275 | -0.42 | -0.0425 | 0.325999 | 0.38419 | 0.934383 | | 1 | 0.996511 | 0.989767 |
| Cyp51a1 | Q8K0C4 | -0.24 | 0.11 | 0.09 | -0.19 | 0.29 | -0.26 | 0.49 | 0.08 | -0.22 | -0.79 | 0.16 | 0.36 | 0.1 | 0.41 | 0.94 | 0.65 | 0.2075 | -0.065 | 0.5825 | 0.340934 | 0.817108 | 0.035602 | | 1 | 0.996511 | 0.654189 |
| Cyp7b1 | Q60991 | -1.3 | -0.55 | 0.58 | 0.17 | -0.54 | -0.26 | 0.96 | 0.28 | -0.22 | -0.29 | 0.17 | -0.82 | 0.57 | -0.89 | 0.52 | 0.93 | 0.385 | -0.015 | 0.5575 | 0.439931 | 0.9725 | 0.311743 | | 1 | 0.997515 | 0.834523 |
| Cyria | Q8BHZ0 | -1.09 | -1.32 | -1.3 | -1.3 | -1.3 | -1.18 | -0.84 | -1.04 | -1.42 | -1.19 | -1.19 | -1.2 | -1.38 | -1.01 | -1.73 | -1.47 | 0.1625 | 0.0025 | -0.145 | 0.34596 | 0.987875 | 0.503551 | | 1 | 0.999285 | 0.901002 |
| Cyrib | A0A2I3BQ! | 0.63 | 2.14 | 1.08 | 0.61 | 2.44 | 0.51 | 1.48 | 0.66 | -0.58 | -0.46 | 1.14 | 0.87 | -3.18 | 0.28 | 0.99 | 0.55 | 0.1575 | -0.8725 | -1.455 | 0.76434 | 0.126139 | 0.139298 | | 1 | 0.996511 | 0.73493 |
| Cyrib | Q921M7 | 0.1 | -0.25 | -0.33 | -0.12 | -0.4 | 0.19 | -0.34 | -0.17 | 0.23 | -0.33 | -0.39 | -0.09 | 0.09 | -0.13 | -0.81 | -0.74 | -0.03 | 0.005 | -0.2475 | 0.878969 | 0.981112 | 0.367013 | | 1 | 0.998773 | 0.858904 |
| Cys1 | A0A0R4J21 | 1.12 | 1.26 | 1.8 | 1.33 | -0.92 | 1.97 | -1.92 | 0.47 | 3.02 | 0.02 | 1.47 | 1.14 | 3.67 | -1.17 | 0.92 | 0.39 | -1.4775 | 0.035 | -0.425 | 0.084292 | 0.952205 | 0.643903 | | 1 | 0.996755 | 0.944252 |
| Cystm1 | D3YVS7 | 2.08 | 0.93 | 1.46 | 1.42 | -0.01 | 0.5 | 0.66 | 0.36 | 2.05 | 0.05 | 1.04 | 1.46 | 2.03 | 0.21 | 0.67 | 0.88 | -1.095 | -0.3225 | -0.525 | 0.003575 | 0.483524 | 0.243441 | | 1 | 0.996511 | 0.79948 |
| Cyth1 | Q9QX11-2 | 0.27 | 0.34 | 0.29 | -0.01 | 0.63 | 0.19 | -0.24 | -0.33 | 0.14 | 0.32 | 0.19 | 0.32 | 0.12 | 0.84 | -0.14 | -0.15 | -0.16 | 0.02 | -0.055 | 0.522695 | 0.906138 | 0.838758 | | 1 | 0.996744 | 0.976176 |
| Cyth2 | A0A1B0GQ | -0.7 | 0.82 | -0.33 | 0.54 | 0.36 | -0.99 | -0.86 | 1.26 | 0.09 | 3.25 | 0.07 | -0.3 | 0.94 | 2.75 | -0.02 | -0.31 | -0.14 | 0.695 | 0.7575 | 0.811612 | 0.404444 | 0.298985 | | 1 | 0.996511 | 0.828257 |
| Cyth2 | Q99KH2 | 1.09 | 1.12 | 0.69 | 0.84 | 1.02 | 0.99 | 0.89 | 1.01 | 1.19 | 0.9 | 0.94 | 1.31 | 0.76 | 1.84 | 0.36 | 0.61 | 0.0425 | 0.15 | -0.0425 | 0.796711 | 0.450738 | 0.901076 | | 1 | 0.996511 | 0.983757 |
| Cyth3 | O08967 | -0.7 | -1.04 | -0.91 | -0.85 | -0.77 | -0.51 | -0.44 | -0.98 | -0.92 | 0.07 | -1.27 | -0.7 | -0.66 | -0.79 | -1.4 | -0.4 | 0.2 | 0.17 | 0.0625 | 0.29544 | 0.572786 | 0.806376 | | 1 | 0.996511 | 0.971715 |
| Czib | Q8BHG2 | -0.14 | -0.47 | -0.13 | -0.56 | -1.05 | -0.52 | -0.27 | -0.44 | -0.87 | -0.95 | -0.31 | -0.37 | -0.48 | -1.6 | -0.46 | -0.56 | -0.245 | -0.3 | -0.45 | 0.290419 | 0.216638 | 0.168779 | | 1 | 0.996511 | 0.759484 |
| D030056L2 | Q3U2A5 | -3.56 | -3.26 | -0.76 | -1.75 | -5.22 | -2.64 | -1.15 | -2.32 | 0.21 | 2.36 | -2.28 | -2.12 | 0.46 | -1.98 | -1.6 | -3.29 | -0.5 | 1.875 | 0.73 | 0.608881 | 0.12937 | 0.432403 | | 1 | 0.996511 | 0.879165 |
| D17h6s53e | Q9Z1R4 | -0.35 | -0.3 | 0.87 | 0.81 | -0.35 | -0.36 | 0.68 | 0.8 | 0.25 | 0.65 | 0.29 | -0.86 | 0.99 | -0.02 | 1.14 | 1.02 | -0.065 | -0.175 | 0.525 | 0.880425 | 0.693478 | 0.225352 | | 1 | 0.996511 | 0.787447 |
| D2hgdh | E9QN44 | -1.41 | -0.76 | -1.06 | -0.41 | -0.49 | -0.73 | 0.24 | -0.88 | -0.81 | -1.46 | -2.16 | -0.98 | 0.13 | 0.49 | -3.03 | -1.57 | 0.445 | -0.4425 | -0.085 | 0.188612 | 0.239996 | 0.910849 | | 1 | 0.996511 | 0.986458 |
| D3Ertd751 | Q8BGN2 | 0.16 | 0.18 | 0.78 | 0.65 | 0.07 | -0.53 | 1.11 | 0.26 | 0.13 | 0.83 | 0.18 | -0.06 | -0.13 | 0.12 | 0.27 | 0.46 | -0.215 | -0.1725 | -0.2625 | 0.553105 | 0.524023 | 0.292468 | | 1 | 0.996511 | 0.825497 |
| D430041D | A0A2R8VK | 0.58 | 0.73 | 0.9 | 0.9 | 0.61 | 0.59 | 0.59 | 0.8 | 0.76 | 0.7 | 0.83 | 0.68 | 0.75 | 0.47 | 0.55 | 0.61 | -0.13 | -0.035 | -0.1825 | 0.418927 | 0.833875 | 0.333477 | | 1 | 0.996511 | 0.845601 |
| D630045J1 | D3YTS3 | 0.42 | 0.63 | 0.71 | 0.51 | 0.97 | 0.48 | 0.74 | 0.55 | 0.6 | 0.32 | 0.67 | 0.45 | 0.38 | 0.94 | 0.45 | 0.64 | 0.1175 | -0.0575 | 0.035 | 0.508753 | 0.741708 | 0.863748 | | 1 | 0.996511 | 0.977235 |
| D6Wsu163 | Q91YN0 | -0.19 | -0.39 | -0.72 | -1.08 | -0.43 | -0.58 | -0.92 | -0.84 | -0.15 | -2.07 | -0.38 | -0.45 | -0.17 | 0.34 | -0.16 | -1.21 | -0.0975 | -0.1675 | 0.295 | 0.687119 | 0.712371 | 0.435941 | | 1 | 0.996511 | 0.88254 |
| D8Ertd738 | Q8R1F0 | 0.73 | -0.22 | -2.45 | -3.18 | -1.07 | 0.19 | -3.24 | -3.61 | 0.34 | -5.7 | -2.77 | -0.03 | 1.6 | -0.9 | -2.01 | -2.61 | -0.6525 | -0.76 | 0.3 | 0.575032 | 0.614474 | 0.797103 | | 1 | 0.996511 | 0.969727 |
| Daam1 | Q8BPM0 | 0.84 | 0.67 | 1.35 | 0.94 | 0.7 | 1.01 | 1.04 | 0.92 | 0.59 | 0.63 | 1.23 | 0.97 | 0.27 | 0.36 | 1.15 | 1.19 | -0.0325 | -0.095 | -0.2075 | 0.869494 | 0.690669 | 0.494917 | | 1 | 0.996511 | 0.89758 |
| Daam2 | Q80U19 | 1.41 | 1.1 | 1.3 | 0.88 | 1.86 | 1.67 | 0.88 | 0.95 | 0.43 | 0.5 | 1.33 | 1.7 | 0.21 | 0.6 | 0.63 | 0.89 | 0.1675 | -0.1825 | -0.59 | 0.551436 | 0.584915 | 0.027672 | | 1 | 0.996511 | 0.635993 |
| Dab1 | F2Z465 | 0.87 | 0.87 | 0.12 | 0.86 | 1.35 | 0.4 | -0.18 | 0.25 | 0.76 | 1 | 0.4 | 0.79 | -0.24 | 1.6 | 0.46 | 0.34 | -0.225 | 0.0575 | -0.14 | 0.533527 | 0.817025 | 0.733735 | | 1 | 0.996511 | 0.958635 |
| Dab2 | E9PX84 | -1.01 | -0.91 | -0.91 | -0.72 | -0.6 | -0.9 | -0.72 | -0.95 | -0.54 | -0.98 | -0.62 | -0.81 | -1.1 | -0.45 | -1.28 | -0.53 | 0.095 | 0.15 | 0.0475 | 0.560887 | 0.416893 | 0.848713 | | 1 | 0.996511 | 0.976225 |

| | | | | | | | | | | | | | | | | | | | | | | | | | | |
|---|---|---|---|---|---|---|---|---|---|---|---|---|---|---|---|---|---|---|---|---|---|---|---|---|---|---|
| Dab2ip | Q3UHC7 | 0.89 | 0.71 | 0.46 | 0.59 | 0.92 | 0.6 | 0.57 | 0.21 | 1.04 | 0.36 | 0.52 | 0.87 | 0.98 | 1.39 | 0.3 | 0.2 | -0.0875 | 0.035 | 0.055 | 0.667984 | 0.872903 | 0.858281 | 1 | 0.996511 | 0.976225 |
| Dact2 | Q7TN08 | -1.82 | -0.71 | 0.14 | 0.54 | -0.15 | 0.2 | -0.54 | -0.63 | 1.01 | 1.42 | 0.29 | -0.89 | 2.81 | 0.16 | 2.07 | -1.38 | 0.1825 | 0.92 | 1.3775 | 0.722311 | 0.189358 | 0.179299 | 1 | 0.996511 | 0.775542 |
| Dact3 | Q0PHV7 | 0.09 | -0.37 | 0.98 | 0.94 | -0.4 | 0.47 | 0.33 | 0.97 | 1.41 | 1.86 | 1.25 | 0.35 | 2.42 | 0.24 | 1.18 | 0.68 | -0.0675 | 0.8075 | 0.72 | 0.869101 | 0.08999 | 0.199983 | 1 | 0.996511 | 0.78004 |
| Dad1 | P61804 | -1.58 | -1.32 | -1.56 | -1.34 | -0.48 | -1.12 | -0.46 | -1.59 | -0.82 | -1.17 | -1.37 | -1.18 | -0.45 | -0.07 | -0.81 | -0.74 | 5.38E-01 | 0.315 | 0.9325 | 0.085674 | 0.125767 | 0.002714 | 1 | 0.996511 | 0.453261 |
| Dag1 | Q62165 | 0.93 | 1.06 | 0.84 | 1.12 | 1.12 | 0.99 | 0.84 | 0.91 | 0.84 | 1.44 | 1.11 | 1.12 | 0.67 | 1.42 | 0.8 | 0.67 | -0.0225 | 0.14 | -0.0975 | 0.884727 | 0.474709 | 0.676918 | 1 | 0.996511 | 0.953397 |
| Dagla | Q6WQJ1 | 1.45 | 1.56 | 1.8 | 1.52 | 1.51 | 1.11 | 1.23 | 1.13 | 0.44 | 0.82 | 1.32 | 1.23 | 0.33 | 0.78 | 0.7 | 0.98 | -0.3375 | -0.63 | -0.885 | 0.075692 | 0.027437 | 0.002427 | 1 | 0.996511 | 0.427013 |
| Daglb | Q91WC9 | 0.18 | 0.51 | 0.28 | 0.02 | 1.14 | 0.19 | -0.35 | -0.35 | -0.5 | -0.05 | -0.21 | -0.19 | -1.03 | 0.82 | -0.26 | 0 | -0.09 | -0.485 | -0.365 | 0.798211 | 0.031818 | 0.354484 | 1 | 0.996511 | 0.855029 |
| Dalrd3 | Q6PJN8 | -0.88 | -0.6 | 0.18 | -0.66 | -0.4 | -0.75 | -0.24 | -0.78 | -0.67 | -0.11 | -0.63 | -0.94 | -1 | 0.24 | -0.77 | -0.55 | -0.0525 | -0.0975 | -0.03 | 0.846916 | 0.742584 | 0.932049 | 1 | 0.996511 | 0.989713 |
| Dap | A0A087WR | 10.85 | 1.07 | 6.18 | 0.96 | 2.09 | 10.84 | 8.75 | 0.77 | 3.65 | 1.4 | 7.15 | 10.93 | 3.96 | 0.98 | 8.95 | 9.65 | 0.8475 | 1.0175 | 1.12 | 0.780863 | 0.717692 | 0.688121 | 1 | 0.996511 | 0.955729 |
| Dap3 | G3X9M0 | -0.4 | -0.32 | 0.15 | 0.27 | -0.56 | -0.39 | 0.01 | 0.26 | -0.27 | 0.06 | -0.03 | -0.74 | -0.43 | -0.01 | 0.16 | -0.12 | -0.095 | -0.17 | -0.025 | 0.715183 | 0.523258 | 0.918548 | 1 | 0.996511 | 0.986868 |
| Dapk1 | Q80YE7-2 | 0.5 | 0.22 | 0.57 | 0.64 | 0.56 | 0.44 | 0.37 | 0.79 | 0.33 | 0.39 | 0.46 | 0.43 | 0.3 | 0.26 | 0.44 | 0.46 | 0.0575 | -0.08 | -0.1175 | 0.74628 | 0.643523 | 0.533372 | 1 | 0.996511 | 0.911674 |
| Dapk3 | O54784 | -0.82 | -0.88 | -1.07 | -0.97 | -0.45 | -1.2 | -1.53 | -1.51 | -0.89 | -1.82 | -1.28 | -0.84 | -0.79 | -0.65 | -0.93 | -1.51 | -0.2375 | -0.2725 | -0.035 | 0.382234 | 0.30066 | 0.882588 | 1 | 0.996511 | 0.979767 |
| Dars1 | Q922B2 | -0.33 | -1.45 | -1.38 | -1.35 | -0.92 | -0.27 | -0.64 | -1.15 | -1.38 | -0.97 | -1.19 | -0.34 | -1.41 | -0.98 | -1.15 | -1.21 | 0.3825 | 0.1575 | -0.06 | 0.251764 | 0.648818 | 0.839222 | 1 | 0.996511 | 0.976218 |
| Dars2 | Q8BIP0 | 0.88 | 1.26 | 0.71 | 0.61 | 0.31 | 0.81 | 0.69 | 0.68 | -0.01 | -0.95 | 0.96 | 0.4 | -0.29 | -0.66 | -0.15 | 0.03 | -0.2425 | -0.765 | -1.1325 | 0.263068 | 0.087975 | 0.001204 | 1 | 0.996511 | 0.412464 |
| Daxx | O35613 | 1.18 | 1.08 | 1.08 | 1.07 | 0.32 | 0.29 | 1 | 1.06 | 0.42 | 1.51 | 1.33 | 1.02 | 0.76 | -0.12 | 1.53 | 1.02 | -0.435 | -0.0325 | -0.305 | 0.086628 | 0.900428 | 0.387659 | 1 | 0.996511 | 0.864251 |
| Dazap1 | Q3UGB5 | -3.82 | -4.06 | -3.39 | -3.11 | -3.38 | -3.81 | -3.11 | -3.34 | -3.89 | -1.4 | -3.36 | -3.08 | -3.96 | -3.48 | -2.38 | -3.53 | 0.185 | 0.6625 | 0.2575 | 0.491921 | 0.236253 | 0.510995 | 1 | 0.996511 | 0.90359 |
| Dbh | Q64237 | 0.51 | 1.06 | 0.73 | 0.76 | 0.67 | 0.17 | 0.74 | 0.6 | 0.74 | 0.68 | 0.7 | 0.81 | 0.5 | 1.18 | 0.8 | 0.57 | -0.22 | -0.0325 | -0.0025 | 0.294279 | 0.857852 | 0.991435 | 1 | 0.996511 | 0.996154 |
| Dbi | P31786 | -0.49 | -0.3 | -0.48 | 0 | -0.4 | -0.62 | -0.1 | 0.04 | -0.52 | -0.55 | -0.38 | -0.37 | -1.08 | -0.01 | -0.51 | -0.17 | 0.0475 | -0.1375 | -0.125 | 0.824227 | 0.464732 | 0.659748 | 1 | 0.996511 | 0.949645 |
| Dbn1 | Q9QXS6 | 1.56 | 1.09 | 1.3 | 1.55 | 0.99 | 1.29 | 0.86 | 1.43 | 1.1 | 1.16 | 1.43 | 1.27 | 0.74 | 0.89 | 0.92 | 0.91 | -0.2325 | -0.135 | -0.51 | 0.272587 | 0.484953 | 0.025555 | 1 | 0.996511 | 0.633075 |
| Dbn1 | A0A0R4J1E | 1.5 | 0.25 | 0.3 | 1.01 | 0.38 | 1.16 | 0.99 | 0.15 | 1.57 | 0.71 | 0.02 | 1.45 | 1 | 1.56 | -0.16 | -0.3 | -0.095 | 0.1725 | -0.24 | 0.79614 | 0.697417 | 0.636649 | 1 | 0.996511 | 0.942665 |
| Dbndd1 | H3BK17 | 0.8 | 0.59 | 0.09 | -0.02 | 3.48 | -0.52 | 0.63 | -0.17 | 2.71 | -0.81 | -0.58 | 0.85 | 1.75 | 1.45 | 0.13 | 0.8 | 0.49 | 0.1775 | 0.6675 | 0.561818 | 0.814399 | 0.119519 | 1 | 0.996511 | 0.721448 |
| Dbndd2 | Q9CRD4 | -0.06 | 0.05 | -1.28 | -0.13 | 0.24 | -1.2 | -2.02 | -1.98 | -1.74 | 0.38 | -2.23 | -0.64 | 0.06 | -1.38 | -1.49 | -1.59 | -0.885 | -0.7025 | -0.745 | 0.141528 | 0.264773 | 0.138074 | 1 | 0.996511 | 0.734193 |
| Dbnl | Q62418 | 0.35 | 0.04 | 0 | 0.56 | 0.19 | 0.26 | 0.22 | 0.63 | 0.34 | 0.46 | 0.31 | 0.34 | -0.4 | 0.1 | -0.1 | -0.13 | 0.0875 | 0.125 | -0.37 | 0.664358 | 0.520808 | 0.118914 | 1 | 0.996511 | 0.721448 |
| Dbr1 | Q923B1 | -2.98 | -2.57 | -2.93 | -2.99 | -3.45 | -2.94 | -3.12 | -3.1 | -3.45 | -2.05 | -3.52 | -2.54 | -2.14 | -2.72 | -2.9 | -3.01 | -0.285 | -0.0225 | 0.175 | 0.153626 | 0.950364 | 0.491952 | 1 | 0.996755 | 0.897531 |
| Dbt | P53395 | 0.68 | 0.79 | 0.59 | 0.69 | 0.55 | 0.82 | 0.51 | 0.36 | 0.21 | 0.21 | 0.66 | 0.33 | -0.11 | -0.1 | -0.05 | 0.07 | -0.1275 | -0.335 | -0.735 | 0.443097 | 0.090415 | 0.001955 | 1 | 0.996511 | 0.419079 |
| Dcaf1 | Q80TR8-4 | -2.26 | -3.13 | -1.74 | -1.78 | -3.2 | -2.91 | -2.45 | -2.29 | -0.96 | -2.68 | -2.3 | -2.22 | -0.56 | -2.29 | -3.04 | -2.21 | -0.485 | 0.1875 | 0.2025 | 0.209442 | 0.68685 | 0.721305 | 1 | 0.996511 | 0.958398 |
| Dcaf10 | A2AKB9-2 | -0.55 | 0.07 | -1.78 | -2.13 | 0.07 | -0.39 | -2.29 | -2.71 | -0.36 | 0.47 | -1.38 | -0.09 | -0.71 | -0.13 | -1.58 | -2.37 | -0.2325 | 0.7575 | -0.1 | 0.764811 | 0.222588 | 0.877556 | 1 | 0.996511 | 0.979626 |
| Dcaf11 | Q91VU6 | -0.97 | -1.07 | 0.97 | 1.09 | -0.2 | -1.18 | 1.23 | 0.74 | -1.5 | 0.61 | 1.3 | -1.22 | -1.16 | -1.15 | 1.04 | 0.8 | 0.1425 | -0.2075 | -0.1225 | 0.843351 | 0.80007 | 0.872297 | 1 | 0.996511 | 0.979626 |
| Dcaf13 | Q6PAC3 | 1.19 | 0.37 | 1.15 | 0.9 | -0.01 | 0.48 | 0.12 | 0.83 | 0.7 | 0.95 | 0.77 | 0.49 | 0.38 | 0.02 | 0.5 | 0.37 | -0.5475 | -0.175 | -0.585 | 0.071203 | 0.471435 | 0.04087 | 1 | 0.996511 | 0.658454 |
| Dcaf5 | Q80T85 | -4.69 | -6.37 | -1.79 | -0.3 | -7.24 | -5.47 | -1.66 | -0.98 | -2.82 | 0.84 | -1.63 | -4.16 | -1.67 | -5.76 | -1.16 | -0.98 | -0.55 | 1.345 | 0.895 | 0.76243 | 0.396067 | 0.574427 | 1 | 0.996511 | 0.926116 |
| Dcaf6 | Q9DC22 | -0.78 | -1 | -1.06 | -0.16 | -0.86 | -0.83 | -1 | -0.88 | -0.77 | 0.26 | -0.59 | -0.95 | -0.63 | -0.59 | -0.91 | -1.32 | -0.1425 | 0.2375 | -0.1125 | 0.538118 | 0.484006 | 0.693667 | 1 | 0.996511 | 0.955729 |
| Dcaf7 | P61963 | -0.16 | -0.23 | -0.23 | -0.07 | 0.06 | -0.32 | 0.1 | -0.12 | -0.2 | 0.18 | -0.26 | -0.15 | -0.09 | -0.21 | 0.05 | -0.16 | 0.1025 | 0.065 | 0.07 | 0.533736 | 0.712888 | 0.686771 | 1 | 0.996511 | 0.955729 |
| Dcaf8 | Q8N7N5 | -0.19 | -0.2 | 0.06 | -0.03 | -0.44 | -0.44 | 0.18 | -0.19 | -0.63 | 0.22 | 0.53 | -0.28 | -0.51 | -0.37 | 0.04 | 0.04 | -0.1325 | 5.00E-02 | -0.11 | 0.503154 | 0.85742 | 0.606352 | 1 | 0.996511 | 0.932425 |
| Dcakd | Q8BHC4 | 0.75 | 0.66 | 0.22 | -0.22 | 0.75 | 1.09 | 0.55 | 0.31 | 1.15 | -0.25 | 0.16 | 1.22 | 0.95 | 0.84 | 0.38 | 0.25 | 0.3225 | 0.2175 | 0.2525 | 0.26877 | 0.596785 | 0.403322 | 1 | 0.996511 | 0.870698 |
| Dcc | P70211-3 | 0.09 | 0.37 | 0.48 | 0.58 | 0.17 | 0.02 | 0.79 | 0.51 | 0.21 | 0.6 | 0.82 | 0.41 | -0.15 | 0.55 | 0.26 | 0.43 | -0.0075 | 0.13 | -0.1075 | 0.973302 | 0.541484 | 0.642602 | 1 | 0.996511 | 0.94365 |
| Dchs1 | E9PVD3 | 0.92 | 1.24 | 0.19 | 0.59 | 0.59 | 0.91 | 0.1 | 0.27 | 1.4 | 1.11 | 0.12 | 1.23 | 1.2 | 0.4 | 0.42 | 0.28 | -0.2675 | 0.23 | -0.16 | 0.365759 | 0.523986 | 0.615016 | 1 | 0.996511 | 0.936534 |
| Dclk1 | Q80VB6 | 1.17 | 1.94 | 0.84 | 0.82 | 2.28 | 1.44 | 1.27 | 1.47 | 2.47 | 2.16 | 1.18 | 2.31 | 1.52 | 3.64 | 1.45 | 1.35 | 0.4225 | 0.8375 | 0.7975 | 0.228867 | 0.050786 | 0.180396 | 1 | 0.996511 | 0.775542 |
| Dclk1 | A0A0G2JG( | 1.46 | 1.63 | 0.97 | 0.81 | 2.3 | 0.16 | 1.39 | 0.83 | 0.67 | -0.17 | 0.8 | 2.57 | 0.4 | 0.54 | 1.13 | 1.26 | -0.0475 | -0.25 | -0.385 | 0.917373 | 0.656983 | 0.222804 | 1 | 0.996511 | 0.787447 |
| Dclk1 | Q8CCN4 | 0.1 | 0.42 | -0.06 | 0.06 | 0.43 | 0.69 | -0.51 | 0.1 | 0.3 | -1.53 | -0.69 | -0.73 | 0.14 | 1.29 | -1.76 | -0.67 | 0.0475 | -0.7925 | -0.38 | 0.865833 | 0.060467 | 0.529855 | 1 | 0.996511 | 0.908663 |
| Dclk1 | H7BX36 | 0.32 | -1.16 | 0.23 | -0.78 | -2.49 | 0.26 | -1.07 | 0.4 | -0.62 | 0.31 | 0.04 | -0.26 | 1.94 | -2.95 | 0.35 | -0.1 | -0.3775 | 0.215 | 0.1575 | 0.59099 | 0.594301 | 0.870944 | 1 | 0.996511 | 0.979293 |
| Dclk1 | Q9JLM8 | 1.22 | 0.47 | -0.05 | 0.02 | 0.58 | 1.07 | 0.69 | 0.11 | 1.14 | 0.29 | 0.41 | 1.67 | 0.53 | 1.25 | -0.05 | -0.01 | 0.1975 | 0.4625 | 0.015 | 0.565848 | 0.282511 | 0.970436 | 1 | 0.996511 | 0.992767 |
| Dclk2 | Q6PGN3-3 | 0.53 | -0.63 | -0.19 | -0.36 | -0.57 | 0.45 | -0.28 | -0.3 | -0.2 | -0.22 | -0.32 | 0.42 | -0.31 | -0.64 | -0.21 | 0.04 | -0.0125 | 0.0825 | -0.1175 | 0.969067 | 0.786722 | 0.695141 | 1 | 0.996511 | 0.955729 |
| Dcn | P28654 | -0.79 | -0.16 | -1.14 | -0.68 | -0.45 | -0.72 | -1.08 | -0.87 | 0.14 | -1.75 | -1.36 | -0.34 | -0.11 | 1.26 | -2.09 | -0.56 | -0.0875 | -0.135 | 0.3175 | 0.730413 | 0.766594 | 0.630165 | 1 | 0.996511 | 0.941132 |
| Dcp1a | Q91YD3 | -0.94 | -0.67 | -0.04 | 0.17 | -0.8 | -1.22 | 0.13 | 0 | -1.11 | 0.07 | -0.13 | -1.01 | -1.15 | -0.6 | -0.42 | -0.45 | -0.1025 | -0.175 | -0.285 | 0.794148 | 0.649764 | 0.381481 | 1 | 0.996511 | 0.860984 |
| Dcp1b | B9EIX0 | -1.08 | -1.29 | -0.11 | 0 | -1.88 | -1.55 | -0.71 | 0.12 | -1.36 | 0.64 | -0.73 | -1.07 | -0.77 | -1.13 | -0.85 | -0.31 | -0.385 | -0.01 | -0.145 | 0.460228 | 0.984403 | 0.692179 | 1 | 0.998892 | 0.955729 |
| Dcp2 | Q9CYC6 | 0.03 | -0.05 | -1.41 | -2.03 | -0.49 | 0.28 | -1.52 | -1.52 | 0.46 | -1.56 | -1.7 | -0.16 | 1.46 | -1.5 | -1.26 | -1.66 | 0.0525 | 0.125 | 0.125 | 0.931596 | 0.851988 | 0.877014 | 1 | 0.996511 | 0.979626 |
| Dcps | Q9DAR7 | -4.01 | -3.63 | -3.96 | -4.39 | -3.23 | -4.05 | -4.25 | -4.36 | -3.27 | -0.28 | -3.58 | -3.46 | -2.66 | -2.21 | -4.36 | -4.51 | 0.025 | 1.35 | 0.5625 | 0.932998 | 0.092948 | 0.328219 | 1 | 0.996511 | 0.844932 |
| Dctn1 | E9Q586 | -1.64 | -0.47 | 0.94 | 0.16 | -0.97 | -0.52 | 1 | 0.77 | -1.19 | 0.2 | 0.83 | -0.74 | -0.72 | 0.62 | 0.29 | 1.35 | 0.3225 | 0.0275 | 0.6375 | 0.627555 | 0.966125 | 0.328676 | 1 | 0.996755 | 0.844943 |
| Dctn1 | E9Q3M3 | 0.23 | 0.14 | 0.21 | 0.34 | 0.4 | 0.08 | 0.26 | 0.23 | 0.61 | 0.7 | 0.28 | 0.22 | 0.32 | 0.8 | 0.21 | 0.09 | 0.0125 | 0.2225 | 0.125 | 0.934069 | 0.251994 | 0.565266 | 1 | 0.996511 | 0.920487 |
| Dctn2 | Q99KJ8 | 1.02 | 0.92 | 1.17 | 0.56 | 1.01 | 0.76 | 1.5 | 1 | 0.03 | -0.25 | 1.17 | 1.09 | -0.14 | 0.41 | 1.16 | 1.39 | 0.15 | -0.4075 | -0.2125 | 0.50895 | 0.29149 | 0.566469 | 1 | 0.996511 | 0.9216 |
| Dctn3 | Q9Z0Y1 | 0.28 | 0.4 | 0.78 | 0.69 | 0.76 | 0.02 | 0.11 | 0.38 | 0.47 | 1.05 | 0.57 | 0.26 | 0.06 | 0.74 | 0.34 | 0.17 | -0.22 | 0.05 | -0.21 | 0.342238 | 0.83131 | 0.378784 | 1 | 0.996511 | 0.860984 |
| Dctn4 | Q8CBY8-2 | -0.03 | -0.13 | -0.41 | -0.39 | 0.2 | -0.13 | -0.23 | -0.43 | -0.06 | -0.1 | -0.43 | 0.06 | -0.32 | 0.51 | -0.72 | -0.79 | 0.0925 | 0.1075 | -0.09 | 0.64038 | 0.584069 | 0.778644 | 1 | 0.996511 | 0.967289 |
| Dctn5 | Q9QZB9 | -0.39 | -0.16 | -1.09 | -0.85 | 0.53 | -0.35 | -0.85 | -0.99 | -0.16 | -0.21 | -1.15 | -0.1 | -0.05 | 0.91 | -1.23 | -0.95 | 0.2075 | 0.2175 | 0.2925 | 0.590922 | 0.51016 | 0.557158 | 1 | 0.996511 | 0.917092 |
| Dctn6 | D3Z6H3 | 0.39 | 0.27 | 0.48 | 0.58 | 0.12 | 0.14 | 0.22 | 0.58 | 0.85 | 0.84 | 0.69 | 0.36 | 1.01 | 0.38 | 0.64 | 0.18 | -0.165 | 0.255 | 0.1225 | 0.35799 | 0.201628 | 0.602612 | 1 | 0.996511 | 0.931041 |
| Dctpp1 | Q9QY93 | 0.31 | -0.5 | -0.21 | -0.69 | -0.55 | 0.69 | 0.06 | -0.15 | -0.17 | 0.04 | -0.24 | 0.13 | 0.5 | -0.73 | -0.08 | 0.1 | 0.285 | 0.2125 | 0.22 | 0.39692 | 0.415224 | 0.519606 | 1 | 0.996511 | 0.907667 |
| Dcun1d1 | Q9QZ73 | -0.63 | -0.57 | -0.49 | -0.64 | -0.8 | -0.42 | -0.48 | -0.54 | -1.21 | -0.94 | -0.44 | -0.79 | -1.15 | -1.35 | -0.96 | -1.23 | 0.0225 | -0.2625 | -0.59 | 0.885811 | 0.228832 | 0.009192 | 1 | 0.996511 | 0.563386 |
| Dcun1d2 | Q8BZJ7 | 0.06 | 0.47 | -0.42 | -0.68 | 0.37 | 0.15 | -0.45 | -0.46 | 0.31 | -1.6 | -0.71 | -0.27 | -0.2 | 0.53 | -1.4 | -0.81 | 0.045 | -0.425 | -0.3275 | 0.889262 | 0.356283 | 0.481363 | 1 | 0.996511 | 0.893906 |
| Dcun1d3 | Q8K0V2 | 0.55 | 0.63 | 0.37 | 0.76 | 0.21 | 0.66 | -0.29 | 0.81 | 1.08 | 1.26 | 0.63 | 0.69 | 0.7 | 0.96 | 0.21 | 0.29 | -0.23 | 0.3375 | -0.0375 | 0.400204 | 0.140796 | 0.87353 | 1 | 0.996511 | 0.979626 |
| Dcun1d4 | Q8C5X2 | -3.87 | -4.37 | -3.06 | -3.03 | -4.56 | -4.11 | -3.37 | -3.37 | -4.04 | -1.9 | -3.09 | -3.96 | -2.14 | -3.76 | -3.21 | -3.03 | -0.27 | 0.335 | 0.5475 | 0.518047 | 0.546392 | 0.238132 | 1 | 0.996511 | 0.79296 |
| Dcun1d5 | Q9CXV9 | -3.41 | -3.42 | -0.76 | -2 | -3.49 | -3.91 | -1.5 | -1.98 | -4.08 | -0.72 | -2.18 | -3 | -3.14 | -3.69 | -0.43 | -2.35 | -0.3225 | -0.0975 | -0.005 | 0.680449 | 0.909746 | 0.995359 | 1 | 0.996744 | 0.998317 |
| Dcx | O88809 | 0.27 | -0.58 | -0.83 | -1.19 | -0.24 | -0.06 | 0.04 | -0.49 | 1.51 | -0.12 | -0.18 | 1.24 | 1.28 | -0.67 | -0.5 | -0.36 | 0.395 | 1.195 | 0.52 | 0.242615 | 0.039544 | 0.32279 | 1 | 0.996511 | 0.840788 |
| Dcxr | Q91X52 | 0.71 | -0.04 | -0.93 | -0.44 | 0.02 | 0.15 | -1.47 | -1.03 | 0.03 | -0.16 | -0.86 | 0.02 | -0.08 | 0.09 | -1.3 | -1.25 | -0.4075 | -0.0675 | -0.46 | 0.412447 | 0.862342 | 0.347459 | 1 | 0.996511 | 0.851503 |
| Dda1 | Q9D9Z5 | -4.08 | -5.53 | -1.82 | -1.19 | -3.73 | -5.2 | -1.8 | -1.86 | -3.78 | -0.54 | -1.31 | -3.13 | -2.95 | -2.76 | -1.32 | -1.35 | 0.0075 | 0.965 | 1.06 | 0.994837 | 0.403063 | 0.297161 | 1 | 0.996511 | 0.826214 |
| Ddah1 | Q9CWS0 | 2.43 | 0.88 | 1.04 | 1.15 | 0.99 | 2.33 | 0.88 | 1.32 | 0.57 | 0.77 | 0.89 | 2.06 | 0.25 | -0.13 | 0.96 | 0.79 | 0.005 | -3.03E-01 | -0.9075 | 0.991161 | 0.515372 | 0.053855 | 1 | 0.996511 | 0.661738 |
| Ddah2 | Q99LD8 | 0.53 | -0.19 | 0.1 | 0.16 | 0 | 0.43 | -0.02 | 0.14 | -0.46 | -0.42 | -0.03 | 0.46 | -0.58 | -0.72 | 0.23 | 0.36 | -0.0125 | -0.2625 | -0.3275 | 0.952295 | 0.351676 | 0.320097 | 1 | 0.996511 | 0.839402 |
| Ddb1 | Q3U1J4 | -1.08 | -0.96 | -0.94 | -1.25 | -0.92 | -1.01 | -1.05 | -1.32 | -1.29 | -1.44 | -1.02 | -1.03 | -1.71 | -1.24 | -1.47 | -1.43 | -0.0175 | -0.1375 | -0.405 | 0.916545 | 0.466531 | 0.061802 | 1 | 0.996511 | 0.662517 |
| Ddc | O88533 | 0.95 | 0.38 | 0.66 | 0.81 | 0.24 | 0.04 | 1.24 | 0.61 | 0.74 | 0.53 | 0.4 | 0.87 | 0.1 | 0.66 | 0.62 | 0.68 | -0.1675 | -0.065 | -0.185 | 0.568165 | 0.753238 | 0.429296 | 1 | 0.996511 | 0.879165 |
| Ddhd1 | F8WIJ5 | 0.35 | 0.42 | 0.64 | 0.45 | 0.51 | 0.4 | 0.65 | 0.55 | 0.5 | 0.52 | 0.69 | 0.36 | 0.37 | 0.85 | 0.23 | 0.21 | 0.0625 | 0.0525 | -0.05 | 0.683213 | 0.757816 | 0.816804 | 1 | 0.996511 | 0.972768 |
| Ddhd2 | Q80Y98 | -0.2 | 0.1 | 0.03 | 0.07 | -0.32 | -0.2 | -0.86 | -0.44 | -0.25 | -0.37 | -0.33 | -0.1 | -0.2 | -0.2 | -0.55 | -0.6 | -0.455 | -0.2625 | -0.3875 | 0.041906 | 0.147715 | 0.076576 | 1 | 0.996511 | 0.675786 |
| Ddi2 | A2ADY9 | -0.13 | 0.03 | -0.33 | 0.04 | -0.17 | -0.09 | -0.19 | -0.19 | -0.25 | -0.38 | -0.42 | 0.04 | -0.3 | 0.29 | -0.82 | -0.59 | -0.0625 | -0.155 | -0.2575 | 0.690797 | 0.42707 | 0.36449 | 1 | 0.996511 | 0.858475 |
| Ddn | Q80TS7 | 2.25 | 1.02 | 0.98 | 0.93 | 0.86 | 2.15 | 0.87 | 1.16 | 0.89 | 1 | 1.23 | 1.83 | 0.84 | 0.6 | -0.04 | 0.29 | -0.035 | -0.0575 | -0.8725 | 0.932596 | 0.876419 | 0.038267 | 1 | 0.996511 | 0.658454 |
| Ddost | O54734 | -0.16 | -0.83 | -0.64 | -0.78 | 0.07 | 0.27 | 1 | -0.4 | 0.24 | -1.1 | -0.63 | 0.1 | 0.56 | -0.35 | 0.34 | 0.67 | 0.8375 | 0.255 | 0.9075 | 0.027167 | 0.466311 | 0.011862 | 1 | 0.996511 | 0.566767 |
| Ddr1 | Q03146-2 | 1.03 | 1.02 | 0.87 | 0.8 | 1.25 | 1.17 | 1.06 | 0.62 | -0.52 | 0.17 | 0.74 | 1.29 | -0.82 | -0.66 | 0.71 | 1.03 | 0.095 | -0.51 | -0.865 | 0.619694 | 0.200985 | 0.080668 | 1 | 0.996511 | 0.684354 |
| Ddrgk1 | Q80WW9 | -0.89 | -1.4 | 1.13 | 0.57 | -1.36 | -0.17 | 1.26 | 0.56 | 1.47 | 0.8 | 0.79 | -0.71 | 3.27 | -0.8 | 2.08 | 2.04 | 0.22 | 0.735 | 1.795 | 0.766418 | 0.300252 | 0.083603 | 1 | 0.996511 | 0.689531 |
| Ddt | O35215 | -0.77 | -1.12 | -0.22 | -0.28 | -1.31 | -0.57 | 0.18 | -0.48 | -1.64 | -1.21 | -0.41 | -0.97 | -1.9 | -1.44 | -1.08 | -0.88 | 0.0525 | -0.46 | -0.7275 | 0.882939 | 0.188959 | 0.044386 | 1 | 0.996511 | 0.658454 |
| Ddx1 | Q91VR5 | -2.22 | -2.57 | -2.21 | -2.33 | -1.96 | -2.5 | -1.82 | -2.41 | -2.34 | -1.69 | -2.28 | -1.87 | -2.67 | -1.59 | -2.07 | -2.2 | 0.16 | 0.2875 | 0.2 | 0.46014 | 0.208649 | 0.456811 | 1 | 0.996511 | 0.887902 |
| Ddx10 | Q80Y44 | 0.87 | -0.51 | 1.5 | 1.01 | -3.26 | 2.4 | -1.06 | -0.4 | 1.49 | 0.47 | 1.08 | -0.29 | 3.06 | -1.71 | 0.9 | 0.22 | -1.2975 | -0.03 | -0.1 | 0.263037 | 0.955287 | 0.917141 | 1 | 0.996755 | 0.986641 |
| Ddx17 | Q3U741 | -1.48 | -3.24 | -2.35 | -2 | -2.5 | -2.18 | -1.76 | -1.86 | -2.99 | -1.86 | -2.17 | -2.19 | -2.2 | -2.6 | -1.33 | -1.9 | 0.1925 | -0.035 | 0.26 | 0.619581 | 0.933439 | 0.553235 | 1 | 0.996755 | 0.916262 |
| Ddx18 | Q8K363 | 0.85 | 0.21 | 0.35 | 0.23 | 0.08 | 0.84 | 0.57 | 0.45 | 1.34 | -0.96 | 0.5 | -0.1 | 1.98 | 0.04 | -0.19 | 0.42 | 0.075 | -0.215 | 0.1525 | 0.750198 | 0.652436 | 0.750844 | 1 | 0.996511 | 0.961479 |
| Ddx19a | Q61655 | -0.09 | -0.45 | 0.81 | 0.83 | -0.52 | -0.47 | 0.22 | 0.82 | -0.39 | 0.58 | 0.55 | -0.14 | -0.57 | -0.24 | 0.46 | 0.54 | -0.2625 | -0.125 | -0.2275 | 0.540753 | 0.748746 | 0.577845 | 1 | 0.996511 | 0.927198 |
| Ddx19b | Q8R3C7 | 1.48 | 0.08 | -1.07 | -0.93 | -0.69 | 1.31 | -0.41 | -0.82 | 2.85 | -1.01 | -0.78 | 1.37 | 3.22 | -1.27 | -1.51 | -1.41 | -0.0425 | 0.7175 | -0.1325 | 0.951311 | 0.4718 | 0.908675 | 1 | 0.996511 | 0.986458 |
| Ddx20 | Q9JJY4 | -1.59 | -1.36 | -2.74 | -3.15 | 0.22 | -2.22 | -3.04 | -3.63 | -1.28 | -2.1 | -3.34 | -2.23 | -1.61 | -1.71 | -3.08 | -3.03 | 0.0425 | -0.0275 | -0.1475 | 0.960402 | 0.960742 | 0.78801 | 1 | 0.996755 | 0.968587 |
| Ddx21 | Q9JIK5 | 1.16 | 0.12 | 0.94 | 1.16 | -0.04 | 0.64 | 0.69 | 1.16 | 0.58 | 0.72 | 1.13 | 0.11 | -0.05 | 0.35 | 0.84 | 1.04 | -0.2325 | -0.21 | -0.3 | 0.497627 | 0.521883 | 0.397573 | 1 | 0.996511 | 0.867334 |
| Ddx23 | D3Z0M9 | 0.08 | -0.99 | -1.1 | -1.26 | -1.09 | -0.51 | -0.71 | -0.45 | -0.41 | -1.05 | -1.28 | -1.32 | 0.75 | -0.53 | 0.06 | -0.59 | 0.1275 | -0.1975 | 0.74 | 0.698364 | 0.586706 | 0.103643 | 1 | 0.996511 | 0.70231 |
| Ddx24 | F8WJA0 | 0.68 | 0.5 | 0.62 | 0.77 | -0.04 | 0.45 | 0.37 | 0.46 | 0.47 | 0.64 | 0.74 | 0.35 | 0.47 | 0.24 | 0.84 | 0.86 | -0.3325 | -0.0925 | -0.04 | 0.089135 | 0.600425 | 0.852536 | 1 | 0.996511 | 0.976225 |
| Ddx25 | Q9QY15 | -1.12 | -0.92 | -1.05 | -1.14 | -1.45 | -0.85 | -0.86 | -0.85 | -0.42 | 0.26 | -1.07 | -0.77 | -0.2 | -1.22 | -0.81 | -1.22 | 0.055 | 0.5575 | 0.195 | 0.776811 | 0.088156 | 0.477654 | 1 | 0.996511 | 0.89226 |
| Ddx27 | Q921N6 | 0.84 | 0.71 | 1.18 | 0.58 | 0.44 | 0.44 | 0.46 | 0.86 | 0.65 | 1.12 | 1.13 | 0.48 | 0.76 | 0.76 | 1.13 | 1.05 | -0.2775 | 0.0175 | 0.0975 | 0.187115 | 0.941351 | 0.652551 | 1 | 0.996755 | 0.947633 |
| Ddx28 | Q9CWT6 | -0.58 | -0.04 | -0.4 | -0.17 | 2.31 | -0.58 | -0.5 | 0.12 | 0.51 | -0.15 | -0.18 | -0.6 | 0.61 | 0.88 | -0.34 | -0.08 | 0.635 | 0.1925 | 0.565 | 0.324319 | 0.487515 | 0.102576 | 1 | 0.996511 | 0.700625 |
| Ddx39b | Q9Z1N5 | -0.89 | -3.4 | -3.29 | -2.53 | -2.95 | -2.02 | -2.47 | -2.31 | -3.7 | -4.06 | -2.8 | -3.21 | -3.79 | -3.3 | -2.82 | -3.49 | 0.09 | -0.915 | -0.8225 | 0.872125 | 0.146857 | 0.170635 | 1 | 0.996511 | 0.762322 |
| Ddx3x | Q62167 | -0.72 | -1.05 | -0.41 | -0.6 | -0.92 | -1.26 | -0.51 | -0.45 | -1.12 | 0.1 | -0.61 | -0.68 | -0.92 | -0.97 | -0.21 | -0.31 | -0.09 | 0.1175 | 0.0925 | 0.715322 | 0.690401 | 0.729102 | 1 | 0.996511 | 0.958635 |
| Ddx3y | Q62095 | -3.32 | 1.51 | 2.33 | -3.36 | -2.94 | 0.45 | -1.5 | 1.34 | -2.94 | -5.48 | 2.12 | 1.21 | 0.42 | -1.74 | 2.37 | -2.67 | 0.0475 | -0.5625 | 0.305 | 0.976459 | 0.788248 | 0.856324 | 1 | 0.996511 | 0.976225 |
| Ddx41 | A0A1S6GW | -0.7 | -0.67 | -0.1 | -0.19 | -0.9 | -0.94 | -0.73 | -0.41 | -0.13 | 0.46 | -0.46 | -0.72 | -0.68 | 1 | -0.18 | -0.51 | -0.33 | 0.2025 | 0.3225 | 0.161266 | 0.510736 | 0.424489 | 1 | 0.996511 | 0.876965 |
| Ddx42 | Q810A7 | -3.38 | -4.18 | -2.83 | -3.25 | -3.87 | -3.78 | -3.1 | -3.44 | -3.75 | -1.59 | -3.03 | -3.38 | -3.13 | -3.45 | -2.86 | -3.25 | -0.1375 | 0.4725 | 0.2375 | 0.672897 | 0.367298 | 0.459411 | 1 | 0.996511 | 0.887902 |
| Ddx46 | Q569Z5 | -1.38 | -2.94 | -1.43 | -1.4 | -2.81 | -2.32 | -1.56 | -1.28 | -2.13 | -1.47 | -1.25 | -2.5 | -1.79 | -1.82 | -1.05 | -1.56 | -0.205 | -0.05 | 0.2325 | 0.671454 | 0.911895 | 0.571392 | 1 | 0.996744 | 0.924332 |
| Ddx47 | F6QKD2 | -0.51 | -0.69 | -0.64 | -0.18 | -0.28 | -0.37 | -0.06 | -0.27 | -0.36 | -0.54 | -0.21 | -0.57 | -0.48 | -0.26 | 0.3 | 0.28 | 0.26 | 0.085 | 0.465 | 0.170355 | 0.664841 | 0.097978 | 1 | 0.996511 | 0.698156 |
| Ddx5 | Q8BTS0 | -0.22 | -1.95 | -1.39 | -1.04 | -2 | -1.03 | -1.35 | -0.76 | -1.71 | -1 | -1.21 | -1.47 | -1.02 | -1.72 | -0.71 | -1.36 | -0.135 | -0.1975 | -0.0525 | 0.749446 | 0.605724 | 0.897114 | 1 | 0.996511 | 0.983509 |
| Ddx50 | Q99MJ9 | 0.78 | 0.33 | 0.45 | 0.52 | 0.49 | 0.85 | 0.21 | 0.65 | 0.69 | -0.21 | 0.39 | -0.33 | 0.86 | 1.67 | 0.76 | 0.41 | 0.03 | -0.385 | 0.405 | 0.880063 | 0.18551 | 0.195984 | 1 | 0.996511 | 0.78004 |
| Ddx51 | Q6P9R1 | 1.12 | 0.79 | 0.22 | 0.32 | 0.57 | 0.49 | 0.77 | 0.36 | 0.87 | 0 | 0.75 | 0.37 | 1.19 | 0.7 | 0.6 | 0.11 | -0.065 | -0.115 | 0.0375 | 0.788303 | 0.698018 | 0.904752 | 1 | 0.996511 | 0.985525 |
| Ddx52 | Q8K301 | 1.79 | 1.53 | 2.21 | 0.74 | 1.26 | 1.16 | 1.84 | 0.99 | -3.44 | -6.2 | 2.51 | 1.63 | -3.53 | -3.57 | 1.61 | 2.17 | -0.255 | -2.9425 | -2.3975 | 0.469372 | 0.142471 | 0.119968 | 1 | 0.996511 | 0.721448 |
| Ddx54 | Q8K4L0 | 0.67 | 1.11 | 1.01 | 0.85 | 1.01 | 0.22 | 0.19 | 1.08 | 1.45 | 0.48 | 0.4 | 0.49 | 1.14 | 0.95 | 1.4 | 1.13 | -0.285 | -0.205 | 0.245 | 0.303469 | 0.470259 | 0.241371 | 1 | 0.996511 | 0.796473 |
| Ddx55 | E9Q9T6 | -3.17 | -0.76 | 0.08 | 0.52 | -3.21 | -2.38 | 0.57 | -0.37 | -2.23 | 1.37 | -0.15 | -3.05 | -1.9 | 0.22 | -0.75 | 0.23 | -0.515 | -0.1825 | 0.2825 | 0.634294 | 0.87515 | 0.745316 | 1 | 0.996511 | 0.959729 |
| Ddx56 | Q9D0R4 | 0.67 | 1.02 | 0.43 | 0.86 | 0.98 | 0.35 | 0.07 | -0.17 | -0.06 | -0.12 | 0.06 | 0.44 | -0.18 | 0.36 | 0.75 | 0.12 | -0.4375 | -0.665 | -0.4825 | 0.146581 | 0.013106 | 0.092857 | 1 | 0.996511 | 0.695968 |
| Ddx58 | A2AP29 | 0.36 | 2.89 | -0.54 | -0.78 | -0.18 | 1.9 | -0.91 | -0.75 | 2.33 | -1.23 | -1.42 | 2.6 | 1.37 | -0.81 | -1.68 | -0.88 | -0.4675 | 0.0875 | -0.9825 | 0.625638 | 0.943489 | 0.317589 | 1 | 0.996755 | 0.838864 |
| Ddx6 | P54823 | -1.39 | -2.17 | -1.97 | -2.1 | -1.69 | -1.44 | -1.76 | -2.08 | -1.76 | -1.44 | -1.86 | -1.47 | -1.83 | -1.23 | -2.03 | -1.88 | 0.165 | 0.275 | 0.165 | 0.493428 | 0.261941 | 0.550003 | 1 | 0.996511 | 0.913564 |
| Decr1 | Q9CQ62 | -0.04 | -0.25 | 0.23 | 0.5 | -0.41 | -0.12 | 0.21 | -0.07 | 0.47 | 0.02 | 0.23 | -0.43 | 0.67 | 0.29 | -0.24 | -0.02 | -0.2075 | -0.0375 | 0.065 | 0.372805 | 0.888499 | 0.814983 | 1 | 0.996511 | 0.972768 |
| Decr2 | Q9WV68 | -0.83 | -0.31 | 0.02 | -0.42 | 0.24 | -0.27 | 0.03 | -0.08 | -0.11 | -1.2 | -0.46 | -0.53 | 0.6 | -0.15 | 0.57 | 0.09 | 0.365 | -0.19 | 0.6625 | 0.133547 | 0.524316 | 0.037616 | 1 | 0.996511 | 0.658454 |
| Def6 | A0A0R4IZX | 1.95 | 1.3 | 0.79 | 0.85 | 1.07 | 0.11 | -0.56 | 0.08 | 1.76 | 2.15 | 0.63 | 0.84 | 2.2 | 2.32 | 0.7 | 1.13 | -1.0475 | 0.1225 | 0.365 | 0.027962 | 0.774832 | 0.429784 | 1 | 0.996511 | 0.879165 |
| Def8 | A0A0R4IZV | 0.79 | 1.08 | 1 | 0.99 | 0 | 1.11 | -1.52 | -0.19 | 2.38 | -0.08 | 0.46 | 1.35 | 2.9 | 1.26 | 0.76 | 0.1 | -1.115 | 0.0625 | 0.29 | 0.050158 | 0.901223 | 0.603521 | 1 | 0.996609 | 0.931339 |
| Degs1 | O09005 | 0.23 | -0.67 | 0.98 | 0.64 | 0.92 | -0.32 | 0.14 | 0.39 | 0.53 | 0.75 | 0.57 | 0.34 | -0.23 | 1.04 | 0.35 | 1.13 | -0.0125 | 0.2525 | 0.2775 | 0.975854 | 0.486105 | 0.543536 | 1 | 0.996511 | 0.912212 |
| Dek | Q7TNV0 | -2.92 | -3.78 | -1.88 | -1.77 | -3.74 | -3.85 | -2.47 | -2.62 | -3.57 | -0.66 | -1.95 | -3.06 | -3.57 | -3.38 | -2.95 | -2.3 | -0.5825 | 0.2775 | -0.4625 | 0.303675 | 0.703736 | 0.378685 | 1 | 0.996511 | 0.860984 |
| Dennd10 | Q9D8N2 | -0.76 | -0.5 | -1.63 | -1.32 | -0.14 | -0.98 | -2.03 | -1.89 | -0.67 | -0.74 | -1.73 | -0.86 | -0.6 | -0.12 | -1.23 | -1.91 | -0.2075 | 0.0525 | 0.0875 | 0.66214 | 0.880525 | 0.843001 | 1 | 0.996511 | 0.976225 |
| Dennd11 | Q3UHG7 | 0.58 | 0.7 | 0.26 | 0.34 | 0.46 | 0.32 | 0.32 | -0.08 | 1.07 | -1.14 | 0.17 | 0.72 | 0.87 | 1.15 | 0.08 | 0.31 | -0.215 | -0.265 | 0.1325 | 0.281338 | 0.572003 | 0.644434 | 1 | 0.996511 | 0.944749 |
| Dennd1a | Q8K382 | 0.52 | 0.36 | 0.79 | 0.64 | 0.44 | 0.48 | 0.71 | 0.67 | 0.81 | 0.79 | 0.7 | 0.23 | 0.55 | 0.53 | 0.47 | 0.43 | -0.0025 | 0.055 | -0.0825 | 0.988113 | 0.791816 | 0.65304 | 1 | 0.996511 | 0.947633 |
| Dennd1b | Q3U1T9-2 | 0.25 | -0.27 | -0.72 | -0.74 | -0.24 | -0.01 | -0.86 | -0.95 | 0.36 | 0.04 | -1.13 | 0.09 | 0.91 | 0.48 | -0.29 | -0.89 | -0.145 | 0.21 | 0.4225 | 0.65305 | 0.592249 | 0.349619 | 1 | 0.996511 | 0.85327 |
| Dennd2a | Q8C4S8 | -0.68 | -0.49 | -1.54 | -0.25 | -1.31 | -0.69 | -0.62 | -1.61 | -1.63 | -1.69 | -1.35 | -0.41 | -1.18 | -0.22 | -1.5 | -1.4 | -0.3175 | -0.53 | -0.335 | 0.382306 | 0.199277 | 0.403525 | 1 | 0.996511 | 0.870698 |
| Dennd4a | E9Q8V6 | 0.13 | 0.25 | 0.43 | 0.12 | 0.08 | -0.05 | 0.02 | -0.01 | 0 | 0.44 | 0.23 | 0.29 | 0.19 | 0.23 | 0.13 | -0.04 | -0.2225 | 0.0075 | -0.105 | 0.1662 | 0.966982 | 0.567335 | 1 | 0.996755 | 0.921806 |
| Dennd4b | A0A0R4J17 | 0.47 | 0.77 | 1.02 | 0.82 | 0.57 | 0.38 | 0.86 | 0.81 | 0.39 | 0.58 | 0.88 | 0.34 | -0.15 | 0.36 | 0.56 | 0.62 | -0.115 | -0.2225 | -0.4225 | 0.55914 | 0.305316 | 0.111765 | 1 | 0.996511 | 0.713445 |
| Dennd4c | A6H8H2-2 | 1.74 | 0.24 | 0.15 | 0.21 | -0.39 | -0.17 | -0.68 | -0.38 | 0.61 | 0.29 | 0.24 | 0.83 | 0.66 | 0.67 | -0.15 | 0.09 | -0.99 | -0.0925 | -0.2675 | 0.027031 | 0.814004 | 0.527268 | 1 | 0.996511 | 0.90845 |
| Dennd5a | A0A0R4J1E | -0.07 | 0 | 0.44 | 0.35 | 0.08 | -0.1 | 0.3 | 0.23 | -0.02 | 0.52 | 0.22 | 0.04 | 0.5 | -0.18 | 0.3 | 0.43 | -0.0525 | 0.01 | 0.0825 | 0.784632 | 0.962882 | 0.729947 | 1 | 0.996755 | 0.958635 |
| Dennd5b | A2RSQ0 | -0.29 | -0.77 | -0.12 | -0.45 | 0.31 | -0.51 | -0.17 | -0.07 | -0.55 | -0.16 | -0.12 | -0.57 | -0.66 | -0.19 | -0.52 | -0.37 | 0.2975 | 0.0575 | -0.0275 | 0.22785 | 0.794915 | 0.901061 | 1 | 0.996511 | 0.983757 |
| Dennd6a | Q8BH65 | -0.33 | -0.23 | 0.34 | 0.34 | -0.18 | -0.53 | -0.18 | 0.12 | -0.23 | 0.5 | 0.01 | -0.41 | -0.51 | 0.39 | -0.59 | 0.19 | -0.2225 | -0.0625 | -0.16 | 0.36539 | 0.823133 | 0.612581 | 1 | 0.996511 | 0.935166 |
| Dennd6b | Q9D9V7 | -0.22 | 0.24 | -0.31 | -0.23 | 0.22 | -0.13 | -0.47 | -0.49 | -0.42 | -0.9 | -0.36 | -0.07 | -0.64 | 0.55 | -0.93 | -0.64 | -0.0875 | -0.3075 | -0.285 | 0.703302 | 0.222094 | 0.423899 | 1 | 0.996511 | 0.876297 |
| Denr | Q9CQJ6 | -1.11 | -1.17 | -0.61 | -0.07 | -1 | -1.11 | -0.48 | -0.3 | -1.15 | 0.08 | -0.35 | -0.94 | -1.45 | -0.1 | -0.63 | -0.8 | 0.0175 | 0.15 | -0.005 | 0.955875 | 0.685032 | 0.989221 | 1 | 0.996511 | 0.996154 |
| Depdc5 | P61460-2 | 0.1 | 0.05 | 0.04 | -0.19 | 0.01 | 0.1 | 0.24 | -0.3 | 0.76 | -0.41 | -0.25 | 0.05 | 1.01 | 0.23 | -0.52 | -0.03 | 0.0125 | 0.0375 | 0.1725 | 0.943966 | 0.893161 | 0.60339 | 1 | 0.996511 | 0.931296 |
| Deptor | Q570Y9 | 1.11 | -0.35 | -0.17 | -0.16 | 0.37 | 0.8 | 1.83 | -0.22 | -0.56 | 0.06 | 0.23 | 0.52 | -0.83 | 0.41 | -0.37 | 0.2 | 0.5875 | -0.045 | -0.255 | 0.263898 | 0.908215 | 0.547372 | 1 | 0.996744 | 0.912571 |
| Dera | Q91YP3 | 0.72 | 0.82 | 0.98 | 0.33 | -1.36 | 0.51 | -0.67 | 0.25 | 0.91 | 0.7 | 0.32 | 0.67 | 2.25 | -0.42 | 0.18 | 0.8 | -1.03 | -0.0625 | -0.01 | 0.035863 | 0.777875 | 0.985331 | 1 | 0.996511 | 0.996154 |
| Derl1 | Q99J56 | 0.1 | 0.68 | 0.17 | -0.76 | 0.83 | -0.26 | 1.46 | 0.84 | -0.23 | -0.19 | 0.45 | 0.01 | -0.28 | 0.14 | 3.77 | 1.08 | 0.67 | -0.0375 | 1.13 | 0.150139 | 0.910448 | 0.211466 | 1 | 0.996744 | 0.786269 |
| Derl2 | Q3U957 | -0.34 | -0.07 | -0.36 | 1.26 | 1.2 | 0.34 | 1.83 | 1.79 | 0.87 | 1.16 | 1.04 | 0.37 | 0.47 | 1.51 | 1.72 | 1.98 | 1.1675 | 0.7375 | 1.2975 | 0.036042 | 0.09523 | 0.022204 | 1 | 0.996511 | 0.621926 |
| Derpc | P0CG14 | -2.52 | -3.64 | -1.53 | -1.79 | -3.2 | -3.78 | -2.37 | -2.24 | -3.63 | -1.1 | -2.16 | -3.28 | -3.12 | -3.31 | -2.39 | -2.61 | -0.5275 | -0.1725 | -0.4875 | 0.347583 | 0.799024 | 0.329642 | 1 | 0.996511 | 0.845345 |
| Des | P31001 | -0.47 | 0.15 | -1.01 | -0.83 | -0.81 | -0.86 | -0.7 | -1.43 | -0.75 | -0.54 | -0.65 | -0.59 | -0.67 | -0.63 | -0.93 | -0.86 | -0.41 | -0.0925 | -0.2325 | 0.196968 | 0.736391 | 0.423218 | 1 | 0.996511 | 0.875782 |
| Desi1 | Q9CQT7 | 0.05 | 0.66 | 0.04 | 0.02 | 0.54 | -0.18 | -0.16 | -0.45 | 0.37 | -0.05 | -0.29 | -0.09 | -0.16 | 0.75 | -0.64 | 0.05 | -0.255 | -0.2075 | -0.1925 | 0.355824 | 0.392798 | 0.564762 | 1 | 0.996511 | 0.920315 |
| Dffa | O54786-2 | -0.95 | -0.84 | -1.81 | -1.81 | -1.18 | -0.64 | -1.51 | -1.37 | -1.09 | -1.53 | -1.77 | -0.28 | -0.7 | -0.46 | -1.8 | -1.23 | 0.1775 | 0.185 | 0.305 | 0.581805 | 0.647656 | 0.439024 | 1 | 0.996511 | 0.883205 |
| Dgat1 | Q9Z2A7 | -0.06 | 0.15 | -0.62 | -0.61 | 0.24 | -0.2 | -0.99 | -0.8 | 0.07 | -0.47 | -0.91 | -0.08 | 0.24 | -0.27 | 0.18 | -0.3 | -0.1525 | -0.0625 | 0.2475 | 0.648453 | 0.834847 | 0.367524 | 1 | 0.996511 | 0.859245 |
| Dgcr2 | A0A0R4J1N | 0.12 | 0.25 | -1.36 | -0.74 | -0.36 | -0.06 | -2.13 | -1.54 | 0.56 | -1.98 | -1.53 | 0.31 | 0.44 | 0.51 | -1.95 | -1.76 | -0.59 | -0.2275 | -0.2575 | 0.31253 | 0.737841 | 0.714376 | 1 | 0.996511 | 0.958106 |
| Dgcr8 | Q9EQM6 | -0.43 | -0.96 | -0.4 | -0.74 | -0.42 | -0.9 | -1.36 | -0.69 | -1.25 | -0.13 | -1.42 | -0.88 | -0.93 | -0.73 | -0.87 | -0.81 | -0.21 | -0.2875 | -0.2025 | 0.4125 | 0.376298 | 0.334226 | 1 | 0.996511 | 0.846677 |
| Dgkb | Q6NS52-2 | -0.36 | -0.97 | 0.09 | 0.29 | 0.09 | -0.46 | 0.44 | 0.41 | -0.87 | 0.34 | -0.08 | -1.27 | -0.45 | 0.62 | -0.15 | 0.34 | 0.3575 | -0.2325 | 0.3275 | 0.305781 | 0.595991 | 0.378063 | 1 | 0.996511 | 0.860984 |
| Dgkb | Q6NS52 | 0.32 | -1.08 | -0.49 | -0.11 | -0.55 | 0.16 | 0.6 | 0.36 | -0.83 | -0.34 | -0.48 | -0.26 | -0.6 | -0.82 | 0.4 | 0.13 | 0.4825 | -0.1375 | 0.1175 | 0.213341 | 0.671201 | 0.769512 | 1 | 0.996511 | 0.965933 |
| Dgkd | E9PUQ8 | 1.01 | 0.28 | -0.61 | -0.13 | 0.02 | 1.08 | -0.37 | 0.02 | 0.19 | -0.87 | -0.4 | 0.08 | 0.44 | 0.46 | -1.19 | -1.16 | 0.05 | -0.3875 | -0.5 | 0.907992 | 0.348082 | 0.364641 | 1 | 0.996511 | 0.858475 |
| Dgke | Q9R1C6 | 0.02 | -0.01 | 0.09 | 0.05 | 0.57 | 0.1 | 0.1 | 0.21 | -0.46 | 0.09 | -0.1 | -0.06 | -0.05 | -0.44 | 0.88 | 0.6 | 0.2075 | -0.17 | 0.21 | 0.23754 | 0.363741 | 0.504993 | 1 | 0.996511 | 0.901074 |
| Dgkg | Q91WG7 | -0.46 | -0.58 | -0.83 | -0.67 | -0.57 | -0.26 | -0.11 | -0.45 | -0.56 | -1.51 | -0.82 | -0.6 | -0.21 | -0.73 | -0.37 | -0.28 | 0.2875 | -0.2375 | 0.2375 | 0.129748 | 0.363628 | 0.261531 | 1 | 0.996511 | 0.81412 |
| Dgkh | A0A2I3BQ4 | -0.38 | -1 | -0.31 | -0.03 | -0.99 | -0.26 | -0.01 | 0.15 | -1.05 | -0.59 | -0.13 | -0.93 | -0.79 | -0.8 | -0.83 | -0.53 | 0.1525 | -0.245 | -0.3075 | 0.633519 | 0.419028 | 0.238736 | 1 | 0.996511 | 0.793481 |
| Dgki | D3YWQ0-2 | -0.16 | -0.53 | -0.58 | -0.2 | -0.72 | -0.26 | -0.58 | -0.51 | -0.65 | -0.26 | -0.82 | -0.43 | -0.98 | -0.62 | -1.16 | -0.86 | -0.15 | -0.1725 | -0.5375 | 0.429773 | 0.417069 | 0.030713 | 1 | 0.996511 | 0.641481 |
| Dgkq | Q6P5E8 | -0.14 | -0.13 | 0.32 | 0.58 | 0.01 | 0.11 | 0.61 | 0.43 | 0.44 | -0.15 | 0.52 | 0.09 | 0.28 | 0.29 | 0.23 | 0.28 | 0.1325 | 0.0675 | 0.1125 | 0.585134 | 0.792744 | 0.619703 | 1 | 0.996511 | 0.93807 |
| Dgkz | Q80UP3 | 4.54 | 4.27 | 4.12 | 4.06 | 2.67 | 4.72 | 2.1 | 4.38 | 4.35 | 5.76 | 3.93 | 3.42 | 4.77 | 3.18 | 2.95 | 2.94 | -0.78 | 0.1175 | -0.7875 | 0.210232 | 0.806647 | 0.095182 | 1 | 0.996511 | 0.698156 |
| Dgkz | A2AHJ7 | -0.81 | -1.12 | -1.1 | -0.88 | -1.07 | -0.79 | -1.07 | -0.74 | -0.81 | -1.3 | -0.89 | -1.24 | -0.45 | -0.84 | -1.63 | -1.7 | 0.06 | -0.0825 | -0.1775 | 0.725653 | 0.676981 | 0.583761 | 1 | 0.996511 | 0.927734 |
| Dglucy | E9QMK9 | 1.02 | 0.29 | 0.36 | 0.59 | 0.58 | 0.84 | 0.71 | 0.83 | 0.17 | 0.29 | 0.46 | 0.48 | 0.23 | 0.29 | 0.32 | 0.43 | 0.175 | -0.215 | -0.2475 | 0.404973 | 0.338915 | 0.279307 | 1 | 0.996511 | 0.822084 |
| Dguok | Q9QX60-2 | 0.16 | 0.71 | -2.19 | -1.06 | -0.01 | 0.52 | -1.32 | -1.64 | 0.94 | -3.43 | -1.42 | 0.27 | 0.79 | 1.06 | -2.97 | -2.47 | -0.0175 | -0.315 | -0.3025 | 0.981328 | 0.764825 | 0.784983 | 1 | 0.996511 | 0.967642 |
| Dhcr24 | Q8VCH6 | -1.58 | -1.32 | -0.09 | -0.72 | -0.28 | -1.01 | 1.58 | -0.03 | -1.46 | -1.82 | 0.84 | -1.09 | -1.18 | -1.15 | 1.02 | 1.39 | 0.9925 | 0.045 | 0.9475 | 0.11687 | 0.942261 | 0.195408 | 1 | 0.996755 | 0.78004 |
| Dhcr7 | A0A140LIT | -1.09 | -0.59 | -1.48 | -1.27 | 0 | -0.48 | -1.16 | -1.36 | -0.55 | -0.64 | -1.46 | -0.48 | -0.22 | -0.34 | -1.08 | -1.24 | 0.3575 | 0.325 | 0.3875 | 0.324393 | 0.298683 | 0.252306 | 1 | 0.996511 | 0.80892 |
| Dhdds | A3KGL2 | -0.32 | -1.07 | -0.77 | -0.55 | -1.8 | -0.19 | -2.16 | -1.2 | 3.81 | -0.18 | -1.82 | -0.94 | 6.21 | -2.83 | 1.47 | -0.7 | -0.66 | 0.895 | 1.715 | 0.151006 | 0.434488 | 0.333236 | 1 | 0.996511 | 0.845601 |
| Dhdh | Q9DBB8 | -0.37 | -0.87 | -0.32 | -0.23 | -0.58 | -0.62 | -0.34 | 0.05 | -0.35 | -0.17 | -0.52 | -0.63 | 0.64 | -0.07 | 0 | -0.09 | 0.075 | 0.03 | 0.5675 | 0.744909 | 0.889163 | 0.050925 | 1 | 0.996511 | 0.658454 |
| Dhfr | P00375 | -0.25 | 0.16 | 0.91 | 1.18 | 0.21 | -0.32 | 1.5 | 1.08 | -0.15 | 0.98 | 0.78 | -0.07 | -0.04 | 0.54 | 0.57 | 0.74 | 0.1175 | -0.115 | -0.0475 | 0.8099 | 0.783003 | 0.896419 | 1 | 0.996511 | 0.983509 |
| Dhodh | O35435 | 0.57 | 0.75 | 1.04 | 1.18 | 0.32 | 0.37 | 1.25 | 1.03 | 0.31 | 0.6 | 0.67 | 0.29 | -0.2 | 0.45 | 0.44 | 0.99 | -0.1425 | -0.4175 | -0.465 | 0.608135 | 0.075091 | 0.141017 | 1 | 0.996511 | 0.737973 |
| Dhps | Q3TXU5 | -1.28 | -1.23 | -2.41 | -2.35 | -1.44 | -1.78 | -2.13 | -2.18 | -1.48 | -1.7 | -2.18 | -1.72 | -1.09 | -0.51 | -2.65 | -2.46 | -0.065 | 0.0475 | 0.14 | 0.854011 | 0.891914 | 0.804809 | 1 | 0.996511 | 0.971715 |
| Dhrs1 | Q99L04 | -0.47 | -0.68 | 0.58 | 0.68 | -0.74 | -0.14 | 0.35 | 0.88 | -0.34 | 1.47 | 0.72 | -0.38 | 0.15 | -0.01 | 0.64 | 0.62 | 0.06 | 0.34 | 0.3225 | 0.895685 | 0.523311 | 0.403867 | 1 | 0.996511 | 0.870999 |
| Dhrs11 | Q3U0B3 | 0.74 | 0.02 | 0.15 | 1.1 | 0.62 | 1.03 | 0.13 | 0.8 | 0.26 | -0.6 | 0.01 | 0.07 | -0.13 | 0.72 | -0.92 | 0.17 | 0.1425 | -0.5675 | -0.5425 | 0.649745 | 0.100141 | 0.207118 | 1 | 0.996511 | 0.785338 |
| Dhrs13 | Q5SS80 | -1.39 | -1.71 | -0.63 | -0.56 | -1.5 | -1.11 | -0.02 | -0.75 | -0.65 | -0.45 | -0.76 | -1.13 | -0.09 | -0.8 | 0.24 | -0.1 | 0.2275 | 0.325 | 0.885 | 0.572732 | 0.322256 | 0.03214 | 1 | 0.996511 | 0.641974 |
| Dhrs4 | Q99LB2 | 0.57 | 0.72 | 0.94 | 0.91 | 0.62 | 0.55 | 0.93 | 0.69 | 0.56 | 0.7 | 0.69 | 0.5 | 0.53 | 0.92 | 0.45 | 0.89 | -0.0875 | -0.1725 | -0.0875 | 0.612295 | 0.333918 | 0.676046 | 1 | 0.996511 | 0.952944 |

| | | | | | | | | | | | | | | | | | | | | | | | | | | | |
|---|---|---|---|---|---|---|---|---|---|---|---|---|---|---|---|---|---|---|---|---|---|---|---|---|---|---|---|
| Dhrs7 | A0A1Y7VN | 0.58 | 0.63 | 0.14 | 0.1 | 0.53 | 0.54 | 0.77 | -0.09 | -0.06 | 0.13 | -0.22 | 0.3 | 0.59 | 1.09 | -0.14 | 0.06 | 0.075 | -0.325 | 0.0375 | 0.760447 | 0.162535 | 0.905968 | | 1 | 0.996511 | 0.986321 |
| Dhrs7b | Z4YKT6 | 0.18 | 0.47 | 0.29 | 0.44 | 0.58 | 0.05 | 0.38 | 0.27 | 0.58 | 0.53 | 0.39 | 0.3 | 0.16 | 0.91 | 0.09 | 0.25 | -0.025 | 0.105 | 0.0075 | 0.887595 | 0.54256 | 0.975 | | 1 | 0.996511 | 0.992977 |
| Dhtkd1 | A2ATU0 | -2.61 | -2.63 | -1.27 | 0.07 | 0.48 | 0.69 | -1.4 | -0.6 | 0.77 | -1.5 | -0.38 | 0.44 | 0.68 | 0.42 | -0.22 | -2.58 | 1.4025 | 1.4425 | 1.185 | 0.082414 | 0.079361 | 0.202451 | | 1 | 0.996511 | 0.782817 |
| Dhx15 | O35286 | -1.51 | -1.61 | -1.1 | -1.31 | -1.55 | -1.7 | -1.3 | -1.01 | -2.12 | -0.86 | -1.38 | -1.76 | -1.71 | -1.69 | -1.25 | -1.33 | -0.0075 | -0.1475 | -0.1125 | 0.972092 | 0.623708 | 0.607359 | | 1 | 0.996511 | 0.933262 |
| Dhx16 | G3X8X0 | 0.45 | -1.44 | -0.2 | 0.44 | 0.4 | -0.59 | -0.55 | -0.06 | 1.68 | -0.36 | 0.07 | -0.62 | 0.97 | -0.79 | -0.44 | 0.14 | -0.0125 | 0.38 | 0.1575 | 0.978566 | 0.54604 | 0.772171 | | 1 | 0.996511 | 0.965933 |
| Dhx29 | Q6PGC1 | -1.56 | -1.7 | -1.67 | -1.7 | -1.38 | -1.28 | -1.63 | -1.64 | -1.12 | -0.98 | -1.51 | -1.4 | -1 | -1.15 | -1.38 | -1.41 | 0.175 | 0.405 | 0.4225 | 0.289471 | 0.054913 | 0.046004 | | 1 | 0.996511 | 0.658454 |
| Dhx30 | A0A0G2JGI | -0.25 | -0.22 | -0.54 | -0.35 | 0.09 | -0.23 | -0.29 | -0.11 | -0.42 | -0.51 | -0.43 | -0.26 | -0.43 | 0.12 | -0.31 | -0.1 | 0.205 | -0.065 | 0.16 | 0.239001 | 0.701346 | 0.4414 | | 1 | 0.996511 | 0.883549 |
| Dhx32 | D3Z4E3 | -1.94 | -2.22 | -1.53 | -1.74 | -1.77 | -2.96 | -1.5 | -2.27 | -2.17 | -0.66 | -2.08 | -2.14 | -1.18 | -1.71 | -1.74 | -2.19 | -0.2675 | 0.095 | 0.1525 | 0.440983 | 0.803651 | 0.583465 | | 1 | 0.996511 | 0.927734 |
| Dhx35 | Q8K1G9 | -0.34 | 0.03 | -0.27 | -0.63 | -1.12 | -1.26 | -0.38 | -0.29 | -0.69 | -0.07 | -0.27 | -0.42 | 0.34 | -0.86 | -0.21 | 0.49 | -0.46 | -0.06 | 0.2425 | 0.133929 | 0.788685 | 0.477542 | | 1 | 0.996511 | 0.892234 |
| Dhx36 | Q8VHK9 | -1.08 | -1.27 | -0.74 | -0.72 | -0.92 | -1.25 | -0.33 | -0.28 | -0.85 | -0.72 | -0.68 | -0.89 | -0.35 | -0.62 | -0.81 | -0.69 | 0.2575 | 0.1675 | 0.335 | 0.36197 | 0.402739 | 0.1506 | | 1 | 0.996511 | 0.749304 |
| Dhx38 | Q80X98 | -0.87 | -0.11 | -2.86 | -2.66 | -0.55 | -0.4 | -3.59 | -2.85 | -0.39 | -2.12 | -2.55 | -0.38 | -0.13 | -0.14 | -2.74 | -3.31 | -0.2225 | 0.265 | 0.045 | 0.813969 | 0.740571 | 0.962815 | | 1 | 0.996511 | 0.9927 |
| Dhx57 | Q6P5D3 | -1.06 | -1.35 | -0.77 | -0.92 | -0.95 | -1.52 | -0.7 | -1.12 | -1.66 | -0.2 | -0.93 | -1.15 | -1.13 | -0.86 | -0.38 | -0.75 | -0.0475 | 0.04 | 0.245 | 0.837127 | 0.902296 | 0.318821 | | 1 | 0.996609 | 0.838969 |
| Dhx8 | A2A4P0 | -1.7 | -1.24 | -1.31 | -0.85 | -1.69 | -1.17 | -1.37 | -1.31 | -0.22 | -0.52 | -1.5 | -2.38 | -1.03 | -1.37 | -1.43 | -1.64 | -0.11 | 0.12 | -0.0925 | 0.629675 | 0.805193 | 0.711493 | | 1 | 0.996511 | 0.957969 |
| Dhx9 | E9QNN1 | -0.16 | -0.92 | 0.45 | 0.66 | 0.28 | -1.02 | 0.78 | 0.76 | 0.37 | 1.03 | 0.67 | -0.44 | 0.08 | 0.56 | 1.47 | 1.18 | 0.1925 | 0.4 | 0.815 | 0.705983 | 0.380769 | 0.095701 | | 1 | 0.996511 | 0.698156 |
| Dhx9 | A0A087WF | 0.82 | 0.03 | -0.05 | 0.19 | -0.26 | 0.3 | 0.03 | 0.54 | -0.23 | -0.42 | 0.01 | -0.5 | 0.28 | 0.11 | 0.5 | -0.36 | -0.095 | -0.5325 | -0.115 | 0.724091 | 0.05893 | 0.689508 | | 1 | 0.996511 | 0.955729 |
| Diablo | Q9JIQ3 | 2.38 | 0.34 | -0.17 | -0.26 | -0.11 | 2.66 | -0.11 | -0.02 | 0.04 | -0.58 | -0.29 | 1.34 | 0.57 | 0.01 | -0.5 | 0.14 | 0.0325 | -0.445 | -0.5175 | 0.968755 | 0.51873 | 0.398891 | | 1 | 0.996511 | 0.867659 |
| Diaph1 | E9PV41 | 1.53 | 1.25 | 0.74 | 0.92 | 1.21 | 1.33 | 0.46 | 0.78 | 1.27 | 0.42 | 0.53 | 1.23 | 0.91 | 0.74 | 0.66 | 0.62 | -0.165 | -0.2475 | -0.3775 | 0.547659 | 0.408553 | 0.12886 | | 1 | 0.996511 | 0.723655 |
| Diaph2 | Q6W4W7 | 0.23 | -0.05 | 0.35 | 0.34 | 0.12 | -0.05 | 0.46 | 0.55 | -0.16 | -0.12 | 0.14 | -0.08 | -0.39 | 0.03 | 0.16 | 0.26 | 0.0525 | -0.2725 | -0.2025 | 0.794644 | 0.157149 | 0.371317 | | 1 | 0.996511 | 0.859424 |
| Dicer1 | F8VQ54 | -0.51 | -0.99 | -0.66 | -1.02 | 0.14 | -0.56 | -0.85 | -0.4 | -0.11 | -0.84 | -0.64 | 0.29 | 0 | -1.25 | -0.58 | -0.34 | 0.3775 | 0.47 | 0.2525 | 0.162271 | 0.136028 | 0.415654 | | 1 | 0.996511 | 0.874945 |
| Dido1 | Q8C9B9 | -1.43 | -2.58 | 3.47 | 1.7 | -3.07 | -2.31 | -1.55 | -2.12 | -2.21 | 1.46 | -1.86 | -2.27 | -1.53 | -2.65 | 3.84 | -2.04 | -2.5525 | -1.51 | -0.885 | 0.072544 | 0.322686 | 0.627477 | | 1 | 0.996511 | 0.939823 |
| Dimt1 | Q9D0D4 | -0.59 | -0.2 | -0.21 | -0.31 | -0.32 | -0.82 | -0.02 | -0.08 | -0.13 | -0.35 | -0.37 | -1.21 | -0.06 | 1.27 | -0.01 | -0.07 | 0.0175 | -0.1875 | 0.61 | 0.937918 | 0.493939 | 0.101659 | | 1 | 0.996511 | 0.698697 |
| Dip2a | F8WI56 | 0.17 | -0.05 | 0.08 | 0.4 | 0.28 | -0.06 | 0.28 | 0.17 | -0.14 | 0.49 | 0.28 | -0.08 | -0.46 | 0.08 | -0.1 | 0.19 | 0.0175 | -0.0125 | -0.2225 | 0.919759 | 0.9539 | 0.328784 | | 1 | 0.996755 | 0.844943 |
| Dip2b | B2RQC7 | 1.69 | 0.68 | 0.3 | 0.49 | 0.57 | 0.89 | 0.48 | 0.69 | 1.21 | 0.04 | 0.04 | 1.73 | -0.5 | 1.8 | -0.54 | -0.08 | -0.1325 | -0.035 | -0.62 | 0.676406 | 0.943113 | 0.303136 | | 1 | 0.996755 | 0.831227 |
| Dip2b | Q3UH60 | -0.26 | -0.23 | 0.04 | 0.06 | -0.01 | 0.02 | 0.14 | 0.06 | -0.25 | 0.02 | 0.08 | -0.12 | -0.46 | 0.21 | -0.46 | -0.42 | 0.15 | 0.03 | -0.185 | 0.352917 | 0.867355 | 0.429851 | | 1 | 0.996511 | 0.879165 |
| Dip2c | B2RQ71 | 0.47 | 0.29 | 0.31 | 0.41 | 0.42 | 0.39 | 0.03 | 0.19 | 0.69 | 0.07 | 0.37 | 0.02 | 0.19 | 0.78 | -0.13 | -0.21 | -0.1125 | -0.0825 | -0.2125 | 0.492444 | 0.690045 | 0.420142 | | 1 | 0.996511 | 0.87558 |
| Dip2c | B7ZMZ7 | 0.22 | -0.25 | 1.1 | 1.22 | 0.13 | -0.04 | 1.34 | 1.36 | 0.53 | 1.78 | 1.08 | -0.07 | 0.4 | 0.3 | 0.94 | 1.19 | 0.125 | 0.2575 | 0.135 | 0.794428 | 0.604238 | 0.735361 | | 1 | 0.996511 | 0.958635 |
| Dipk1b | Q99ML4 | -0.97 | -1.19 | -1.53 | -1.08 | -0.11 | -0.81 | -0.24 | -1.23 | -1.21 | -1.18 | -1.25 | -1.07 | -1.2 | -0.2 | -0.31 | -0.29 | 0.595 | 0.015 | 0.6925 | 0.06481 | 0.93598 | 0.034812 | | 1 | 0.996755 | 0.651434 |
| Dipk2a | Q3USZ8 | 0.42 | 0.61 | 2.37 | 3.29 | 1.3 | 0.89 | 3.65 | 3.72 | -0.11 | 1.91 | 3.43 | 0.97 | -1.28 | -0.46 | 2.44 | 4.32 | 0.7175 | -0.1225 | -0.4175 | 0.444039 | 0.894117 | 0.750208 | | 1 | 0.996511 | 0.961479 |
| Diras1 | Q91Z61 | 0.33 | 0.11 | -0.45 | -0.35 | 1.14 | -0.54 | -0.01 | -0.42 | -0.23 | -0.1 | -0.26 | -0.09 | 0.49 | 0.94 | -0.14 | -0.39 | 0.1325 | -0.08 | 0.315 | 0.741826 | 0.723291 | 0.38142 | | 1 | 0.996511 | 0.860984 |
| Diras2 | Q5PR73 | -0.3 | -0.26 | -0.09 | 0.06 | -0.11 | -0.43 | 0.01 | -0.22 | -0.27 | -0.11 | -0.15 | -0.3 | -0.58 | 0.28 | -0.8 | -0.84 | -0.04 | -0.06 | -0.3375 | 0.818575 | 0.726708 | 0.264071 | | 1 | 0.996511 | 0.816045 |
| Dis3 | Q9CSH3 | -3.5 | -3.78 | -2.56 | -2.71 | -4.2 | -3.78 | -2.64 | -3.26 | -2.38 | -2.08 | -2.47 | -3.73 | -1.02 | -3.61 | -1.65 | -1.87 | -0.3325 | 0.4725 | 1.1 | 0.438319 | 0.302807 | 0.084267 | | 1 | 0.996511 | 0.690427 |
| Dis3l2 | Q8CI75-2 | -0.59 | -0.51 | 0.06 | 0 | -0.37 | -0.72 | -0.01 | 0.09 | -0.81 | 0.46 | -0.09 | -0.51 | -0.77 | -0.08 | -0.25 | -0.32 | 0.0075 | 0.0225 | -0.095 | 0.976883 | 0.944351 | 0.710538 | | 1 | 0.996755 | 0.957819 |
| Disp2 | Q8CIP5 | 1.4 | -4.15 | -0.24 | 0.08 | -5.06 | 1.83 | -0.32 | -0.21 | 1.28 | 0.21 | -0.4 | -1.85 | 1.26 | 0.54 | 0.15 | -0.35 | -0.2125 | 0.5375 | 1.1275 | 0.899435 | 0.660891 | 0.323717 | | 1 | 0.996511 | 0.841517 |
| Dixdc1 | Q80Y83-7 | 0.87 | 1.11 | 0.67 | 0.8 | 1.15 | 1.01 | 1.09 | 1.03 | 0.98 | 0.19 | 1.05 | 1.19 | 1.42 | 0.5 | 1.28 | 0.96 | 0.2075 | -0.01 | 0.1775 | 0.216923 | 0.96951 | 0.492824 | | 1 | 0.996755 | 0.897531 |
| Dkc1 | Q9ESX5 | 1.89 | 1.42 | 1.2 | 1.03 | 1.54 | 1.69 | 1.32 | 1.09 | 0.73 | 0.17 | 1.29 | 1.32 | 0.71 | 1.28 | 1.34 | 1.51 | 0.025 | -0.5075 | -0.175 | 0.917817 | 0.148431 | 0.531596 | | 1 | 0.996511 | 0.910024 |
| Dkk3 | Q9QUN9 | -2.53 | -2.24 | -1.99 | -2.16 | -2.12 | -2.21 | -2.05 | -2.37 | -2.23 | -0.67 | -2.2 | -1.81 | -1.29 | -2.27 | -2.14 | -2.21 | 0.0425 | 0.5025 | 0.2525 | 0.811928 | 0.198375 | 0.37541 | | 1 | 0.996511 | 0.859611 |
| Dlat | Q8BMF4 | 0.62 | 0.76 | 0.05 | 0.42 | 0.3 | 0.62 | 0.08 | 0.17 | 0.62 | 0.3 | 0.05 | 0.37 | 0.37 | 0.78 | -0.25 | -0.14 | -0.17 | -0.1275 | -0.2725 | 0.444596 | 0.57813 | 0.372051 | | 1 | 0.996511 | 0.859424 |
| Dlc1 | A0A0R4J17 | -0.54 | 0.21 | 0.73 | 0.99 | -0.07 | -0.92 | 0.72 | 1 | -0.23 | -0.13 | 1 | -0.47 | -0.09 | -0.32 | 0.38 | 0.88 | -0.165 | -0.305 | -0.135 | 0.744476 | 0.49698 | 0.743676 | | 1 | 0.996511 | 0.959321 |
| Dld | O08749 | 1.59 | 1.43 | 1.43 | 1.49 | 0.91 | 1.3 | 1.12 | 1.49 | 0.72 | 1.34 | 1.39 | 1.51 | 0.89 | 0.37 | 0.87 | 1.39 | -0.28 | -0.245 | -0.605 | 0.140357 | 0.280616 | 0.034505 | | 1 | 0.996511 | 0.651434 |
| Dlg1 | E9Q9H0 | 0.58 | 0.57 | 1.16 | 1.35 | 0.58 | 0.61 | 0.91 | 1.28 | 0.62 | 1.19 | 1.23 | 0.57 | 0.34 | 0.1 | 0.87 | 0.73 | -0.07 | -0.0125 | -0.405 | 0.792134 | 0.964355 | 0.180193 | | 1 | 0.996755 | 0.775542 |
| Dlg1 | H7BWY4 | 2.44 | 1.91 | 1.03 | 1.47 | 1.06 | 1.59 | 1.65 | 1.23 | 1.27 | 1.28 | 1.06 | 1.71 | -0.05 | 1.64 | 1.58 | 1.42 | -0.33 | -0.3825 | -0.565 | 0.324917 | 0.264218 | 0.250822 | | 1 | 0.996511 | 0.807201 |
| Dlg1 | Q811D0-2 | -3.98 | -2.71 | -2.43 | -3.67 | -2.26 | -4.05 | -3.46 | -3.36 | -4.74 | -3.38 | -4.89 | -6.31 | -0.08 | -10.92 | -3.46 | -4.65 | -0.085 | -1.6325 | -1.58 | 0.861503 | 0.03042 | 0.44505 | | 1 | 0.996511 | 0.884817 |
| Dlg1 | D3Z3B8 | 2.01 | 0.66 | 1.59 | 2.23 | 0.45 | 2.18 | 1.12 | 2.3 | 2.54 | 1.67 | 1.63 | 2.2 | 2.4 | 0.94 | 1.42 | 1.41 | -0.11 | 0.3875 | -0.08 | 0.83173 | 0.33883 | 0.85559 | | 1 | 0.996511 | 0.976225 |
| Dlg2 | A0A338P6J | 1.53 | 1.21 | 1.12 | 1.37 | 1.24 | 1.51 | 1.23 | 1.4 | 0.89 | 0.84 | 1.34 | 1.41 | 0.63 | 1.08 | 0.89 | 1.04 | 0.0375 | -0.1875 | -0.3975 | 0.823123 | 0.392135 | 0.075306 | | 1 | 0.996511 | 0.673156 |
| Dlg2 | Q91XM9-6 | -0.91 | -1.34 | -0.48 | -0.12 | 0.91 | -0.9 | -0.04 | 0.28 | 0.03 | -3.14 | -0.08 | -1.51 | -4.09 | -0.34 | -1.55 | -0.46 | 0.775 | -0.4625 | -0.8975 | 0.099566 | 0.523442 | 0.290025 | | 1 | 0.996511 | 0.825134 |
| Dlg2 | A0A571BEI | 4.38 | 2.93 | 2.21 | -1.07 | 0.73 | 6.02 | -0.74 | 1.34 | 3.91 | -1.17 | 0.98 | 6.94 | 5.23 | -1.33 | 2.25 | 1.39 | -0.275 | 0.5525 | -0.2275 | 0.868403 | 0.769098 | 0.885184 | | 1 | 0.996511 | 0.979767 |
| Dlg3 | P70175 | 1.18 | 0.85 | 1 | 1.33 | 0.93 | 1.13 | 1.24 | 1.47 | 0.82 | 1.02 | 1.24 | 1.06 | 0.32 | 0.75 | 0.75 | 0.92 | 0.1025 | -0.055 | -0.405 | 0.595396 | 0.774278 | 0.090287 | | 1 | 0.996511 | 0.692659 |
| Dlg4 | A0A338P6I | 1.83 | 1.5 | 1.42 | 1.53 | 1.78 | 1.8 | 2.07 | 1.58 | 0.99 | 1.05 | 1.47 | 1.45 | 0.9 | 1.08 | 1.07 | 1.27 | 0.2375 | -0.33 | -0.49 | 0.209604 | 0.13164 | 0.029392 | | 1 | 0.996511 | 0.637553 |
| Dlg4 | Q62108-3 | -1.04 | -0.83 | 0.49 | -1.79 | -0.38 | -0.84 | -0.38 | -1.91 | -0.76 | 3.17 | -1.87 | -0.92 | -0.36 | 0.51 | 1.15 | -2.64 | -0.085 | 0.6975 | 0.4575 | 0.876378 | 0.526185 | 0.596771 | | 1 | 0.996511 | 0.929911 |
| Dlg5 | E9Q9I2 | 0.25 | 0.83 | 0.43 | 0.24 | 0.91 | 0.65 | 0.64 | 0.41 | 1.17 | 0.59 | 0.51 | 0.79 | 1.6 | 1.38 | 0.72 | 0.68 | 0.215 | 0.3275 | 0.6575 | 0.305712 | 0.1836 | 0.045268 | | 1 | 0.996511 | 0.658454 |
| Dlgap1 | Q9D415 | 1.08 | 0.96 | 1.34 | 1.3 | 0.88 | 1.26 | 0.82 | 1.45 | 0.98 | 1.23 | 1.3 | 0.75 | 1 | 0.88 | 0.95 | 1.03 | -0.0675 | -0.105 | -0.205 | 0.743935 | 0.606734 | 0.279893 | | 1 | 0.996511 | 0.822084 |
| Dlgap1 | D3Z508 | 0.09 | 1.24 | 0.44 | 1.66 | 0.9 | 1.28 | 1 | 1.39 | -0.11 | 0.98 | 1.56 | 0.72 | -0.34 | 1.96 | 0.91 | -0.32 | 0.285 | -0.07 | -0.305 | 0.43883 | 0.881051 | 0.61609 | | 1 | 0.996511 | 0.93691 |
| Dlgap1 | Q9D415-3 | 0.21 | 1.42 | 1 | 1.87 | -0.36 | 0.33 | 1.51 | 1.59 | 0.28 | -0.18 | 1.05 | 0.66 | -0.38 | 0.17 | 2.29 | 1.7 | -0.3575 | -0.6725 | -0.18 | 0.51471 | 0.134838 | 0.784097 | | 1 | 0.996511 | 0.967642 |
| Dlgap1 | Q3UVU8 | 0.08 | 0.49 | 1.55 | 0.93 | 1.04 | 0.24 | 1 | 1.04 | 0.01 | 0.7 | 1.63 | 0.55 | 0.02 | 1.34 | 0.18 | 1.16 | 0.0675 | -0.04 | -0.0875 | 0.849595 | 0.926734 | 0.841259 | | 1 | 0.996755 | 0.976225 |
| Dlgap1 | Q9D415-2 | -1.71 | -1.84 | -1.99 | -1.66 | -0.27 | -1.02 | -1.67 | -1.59 | -1.02 | -0.5 | -1.67 | -1.27 | -2.08 | -0.88 | -2.25 | -1.91 | 0.6625 | 0.685 | 0.02 | 0.067103 | 0.030668 | 0.950482 | | 1 | 0.996511 | 0.992476 |
| Dlgap2 | Q8BJ42 | 0.39 | 1.29 | 0.92 | 0 | 0.92 | -0.34 | 0.75 | 0.84 | 0.16 | 0.86 | 1.36 | -1.02 | 0.7 | 0.21 | 0.15 | 0 | -0.1075 | -0.31 | -0.385 | 0.782481 | 0.572298 | 0.257891 | | 1 | 0.996511 | 0.812074 |
| Dlgap2 | Q0VF59 | 1.69 | 1.24 | 1.65 | 1.37 | 1.19 | 1.77 | 1.2 | 1.29 | 1.3 | 1.48 | 1.73 | 1.56 | 1.25 | 1.46 | 1.45 | 1.26 | -0.125 | 0.03 | -0.1325 | 0.547652 | 0.877358 | 0.502822 | | 1 | 0.996511 | 0.901002 |
| Dlgap3 | B1AS06 | 1.89 | 1.21 | 1.65 | 1.43 | 1.41 | 1.74 | 1.77 | 1.37 | 0.94 | 1.11 | 1.56 | 1.47 | 0.83 | 1.16 | 1.39 | 1.55 | 0.0275 | -0.275 | -3.13E-01 | 0.895188 | 0.263218 | 0.229231 | | 1 | 0.996511 | 0.787447 |
| Dlgap4 | B7ZNS2 | 0.86 | 0.7 | 0.9 | 0.97 | 1.02 | 0.97 | 0.88 | 1 | 0.84 | 0.83 | 1.02 | 0.85 | 0.91 | 0.77 | 0.56 | 0.79 | 0.11 | 0.0275 | -0.1 | 0.46024 | 0.865702 | 0.58478 | | 1 | 0.996511 | 0.927734 |
| Dlgap4 | B1AZP2-2 | 0.09 | 0.5 | -0.21 | 0.15 | -0.1 | 0.22 | -0.82 | 0.67 | -0.26 | -0.25 | 0.66 | -0.35 | 0.73 | 0.37 | -0.31 | -0.14 | -0.14 | -0.1825 | 0.03 | 0.677385 | 0.531463 | 0.918717 | | 1 | 0.996511 | 0.986868 |
| Dlst | Q9D2G2 | -0.28 | -0.1 | 0.36 | 0.54 | -0.42 | -0.13 | 0.28 | 0.43 | -0.06 | -0.03 | 0.57 | -0.24 | -0.41 | -0.46 | -0.29 | 0.06 | -0.09 | -0.07 | -0.405 | 0.745467 | 0.798959 | 0.139827 | | 1 | 0.996511 | 0.735494 |
| Dmac1 | Q9CQ00 | 2.49 | 0.97 | 0.92 | 1.36 | 0.93 | 1.79 | 0.26 | 1.67 | 2.41 | 1.25 | 1.17 | 2.35 | 2.27 | 2.15 | 1.67 | 1.38 | -0.2725 | 0.36 | 0.4325 | 0.567878 | 0.448169 | 0.300908 | | 1 | 0.996511 | 0.830276 |
| Dmac2 | Q8CA44 | 2.76 | 1.25 | -0.14 | -0.5 | 0.66 | 3.14 | -0.75 | -0.77 | -0.37 | -0.08 | -0.53 | 2.62 | -0.44 | -1.09 | -0.62 | 0.4 | -0.2725 | -0.4325 | -1.28 | 0.797197 | 0.64887 | 0.106398 | | 1 | 0.996511 | 0.703417 |
| Dmac2l | Q9CRA7 | 2.99 | 1.27 | 1.04 | 0.65 | 1.46 | 2.81 | 1.28 | 0.69 | 0.55 | 0.98 | 0.72 | 2.39 | 0.36 | 1.32 | 0.36 | 0.68 | 0.0725 | -0.3275 | -0.8075 | 0.907239 | 0.594138 | 0.149125 | | 1 | 0.996511 | 0.748169 |
| Dmap1 | Q9JI44 | 0.08 | -0.71 | -1.17 | -0.96 | -0.22 | -0.37 | -1.74 | -0.57 | -0.42 | -0.92 | -1.23 | -0.29 | -0.64 | -0.56 | -0.81 | -0.69 | -0.035 | -0.025 | 0.015 | 0.932565 | 0.942072 | 0.959222 | | 1 | 0.996755 | 0.992636 |
| Dmd | P11531 | 0.91 | 0.46 | 0.57 | 0.61 | 0.83 | 0.68 | 0.56 | 0.69 | 1 | 0.53 | 0.72 | 0.67 | 0.98 | 0.74 | 0.58 | 0.63 | 0.0525 | 0.0925 | 0.095 | 0.753337 | 0.633342 | 0.635145 | | 1 | 0.996511 | 0.941407 |
| Dmtn | Q9WV69-2 | 0.44 | 0.87 | 0.49 | 0.73 | 0.29 | 0.57 | 0.71 | 1.13 | 0.17 | 0.52 | 0.64 | 0.24 | 0.12 | -0.1 | -0.2 | 0.16 | 0.0425 | -0.24 | -0.6375 | 0.849567 | 0.250589 | 0.010606 | | 1 | 0.996511 | 0.563386 |
| Dmtn | Q9WV69 | 1.92 | 0.99 | 0.49 | 0.87 | 1.05 | 1.66 | 0.07 | 0.67 | 0.82 | 0.12 | 0.61 | 1.69 | 0.5 | 1 | 0 | 0.04 | -0.205 | -0.2575 | -0.6825 | 0.629845 | 0.547529 | 0.09592 | | 1 | 0.996511 | 0.698156 |
| Dmwd | Q08274 | 0.04 | 0.08 | -0.15 | 0.33 | 0.06 | 0.07 | 0.26 | 0.16 | 0.48 | 0.66 | -0.09 | 0.32 | 0.41 | 0.87 | 0.21 | 0.19 | 0.0625 | 0.2675 | 0.345 | 0.707165 | 0.251011 | 0.160036 | | 1 | 0.996511 | 0.755194 |
| Dmwd | E9Q6J8 | 1.55 | 0.67 | 1.28 | 1.84 | 0.19 | 1.27 | 1.46 | 2.13 | 0.56 | 2.22 | 1.9 | 1 | 0.55 | 1.12 | 1.89 | 1.32 | -0.0725 | 0.085 | -0.115 | 0.869517 | 0.844776 | 0.753766 | | 1 | 0.996511 | 0.962025 |
| Dmxl1 | Q6PNC0 | 0.76 | 1.05 | 1.03 | 0.95 | 1.1 | 0.75 | 0.66 | 0.96 | 0.71 | 0.97 | 0.94 | 0.63 | 0.53 | 0.88 | 0.9 | 0.92 | -0.08 | -0.135 | -0.14 | 0.64265 | 0.452808 | 0.469806 | | 1 | 0.996511 | 0.890289 |
| Dmxl2 | Q8BPN8-2 | 0.56 | 0.52 | 0.73 | 0.69 | 0.63 | 0.6 | 0.6 | 0.71 | 0.38 | 0.29 | 0.72 | 0.48 | 0.18 | 0.47 | 0.49 | 0.41 | 0.01 | -0.1575 | -0.2375 | 0.944498 | 0.38232 | 0.208097 | | 1 | 0.996511 | 0.785665 |
| Dnaaf10 | Q8BGF3 | -0.15 | -0.22 | 0.02 | -0.48 | 0.5 | -1.01 | -0.66 | -0.61 | -0.14 | 0.69 | -0.5 | -0.67 | -0.53 | 0.42 | -0.48 | -0.1 | -0.2375 | 0.0525 | 0.035 | 0.482495 | 0.869724 | 0.896141 | | 1 | 0.996511 | 0.983509 |
| Dnaaf5 | B9EJR8 | 1.09 | 0.68 | 0.22 | 0.19 | 1.38 | 1.2 | 0.61 | -0.03 | 1.05 | 0.02 | 0.54 | 0.86 | -0.24 | 1.01 | 0.43 | 0.25 | 0.245 | 0.0725 | -0.1825 | 0.509242 | 0.816735 | 0.590532 | | 1 | 0.996511 | 0.929142 |
| Dnaaf9 | Q3V0A7 | 0.16 | -0.37 | -0.08 | 0.14 | -0.05 | -0.27 | 0.01 | -0.3 | -0.21 | -0.22 | 0.01 | -0.41 | -0.11 | 0.09 | -0.79 | -0.26 | -0.115 | -0.17 | -0.23 | 0.542174 | 0.40527 | 0.38026 | | 1 | 0.996511 | 0.860984 |
| Dnah2 | D3Z667 | -4.23 | -4.23 | -2.52 | -4.14 | -4.34 | -3.54 | -4.8 | -5.7 | -3.57 | -1.13 | -2.91 | -3.64 | -1.15 | -4.37 | -2 | -3.2 | -0.815 | 0.9675 | 1.1 | 0.173838 | 0.166229 | 0.163055 | | 1 | 0.996511 | 0.758224 |
| Dnaja1 | P63037 | -0.22 | 0.29 | 0.24 | 0.27 | -0.3 | 0.22 | 0.06 | 0.03 | 0.62 | 0.71 | 0.31 | 0.4 | 1.1 | 0.01 | 0.37 | 0.41 | -0.1425 | 0.365 | 0.3275 | 0.478205 | 0.098197 | 0.259679 | | 1 | 0.996511 | 0.812629 |
| Dnaja2 | Q9QYJ0 | 2.04 | 1.25 | 1.42 | 0.44 | 1.23 | 2.18 | 1.21 | 0.71 | 0.31 | -0.19 | 1.15 | 2.52 | 0.44 | 0.36 | 1.39 | 1.79 | 0.045 | -0.34 | -0.2925 | 0.915034 | 0.586828 | 0.526088 | | 1 | 0.996511 | 0.908107 |
| Dnaja3 | Q99M87-2 | -0.27 | 0.41 | -0.24 | 0.74 | 0.27 | -0.17 | 0.46 | 0.76 | -1.42 | -1.46 | 0.27 | 0.04 | -1.57 | -1.42 | 0.07 | 0.11 | 0.17 | -0.8025 | -0.8625 | 0.587867 | 0.128731 | 0.104668 | | 1 | 0.996511 | 0.702979 |
| Dnaja3 | Q99M87 | 0.67 | 0.85 | 0.66 | 1.11 | -0.03 | 0.96 | 0.73 | 0.81 | 0.75 | 0.53 | 0.78 | 0.7 | 1.02 | 0.27 | 0.09 | 0.76 | -0.205 | -0.1325 | -0.2875 | 0.430662 | 0.475301 | 0.295727 | | 1 | 0.996511 | 0.825612 |
| Dnaja4 | Q9JMC3 | 0.46 | 1.08 | -0.09 | -0.74 | 0.42 | 0.71 | -0.62 | -0.97 | 0.47 | -0.13 | -0.5 | 0.85 | 1.16 | 0.35 | -0.46 | -0.81 | -0.2925 | -0.005 | -0.1175 | 0.573813 | 0.991337 | 0.828514 | | 1 | 0.999285 | 0.973757 |
| Dnajb1 | Q9QYJ3 | 0.83 | 0.21 | 0.1 | 0 | -0.03 | 0.55 | -0.52 | -0.03 | 0.4 | 0.3 | -0.11 | 0.7 | 0.43 | 0.03 | -0.02 | -0.09 | -0.2925 | 0.0375 | -0.1975 | 0.325155 | 0.888373 | 0.442484 | | 1 | 0.996511 | 0.883549 |
| Dnajb11 | Q99KV1 | 0.5 | 0.45 | 0.79 | 0.43 | 1.02 | 0.65 | 1.54 | 1.32 | -0.58 | 0.07 | 0.75 | 0.79 | 0.25 | -0.26 | 1.96 | 1.8 | 0.59 | -0.285 | 0.395 | 0.028786 | 0.401365 | 0.455904 | | 1 | 0.996511 | 0.887169 |
| Dnajb12 | Q9QYI4 | 0.15 | -0.46 | -0.28 | -0.78 | -0.41 | 0.35 | -0.24 | -0.76 | 0.6 | -1.09 | -0.25 | -0.05 | 1.28 | -1.06 | 1.39 | 0.7 | 0.0775 | 0.145 | 0.92 | 0.796349 | 0.706612 | 0.123104 | | 1 | 0.996511 | 0.721448 |
| Dnajb14 | Q149L6 | 1.38 | 1.29 | -0.11 | 0.04 | 1.32 | 1.98 | 0.22 | 0.04 | 0.85 | 0.1 | -0.16 | 1.45 | 0.94 | 0.03 | 0.6 | 0.58 | 0.24 | -0.09 | -0.1125 | 0.667988 | 0.857694 | 0.78903 | | 1 | 0.996511 | 0.968677 |
| Dnajb2 | Q9QYI5 | 1.03 | 1 | 1.48 | 1.59 | 0.16 | 0.91 | 0.49 | 1.17 | 1.07 | 1.31 | 0.9 | 0.79 | 1.4 | 0.31 | 1.02 | 0.99 | -0.5925 | -0.2575 | -0.345 | 0.056169 | 0.269652 | 0.253086 | | 1 | 0.996511 | 0.809524 |
| Dnajb4 | Q9D832 | -0.24 | -0.21 | -0.09 | -0.16 | 0.03 | -0.21 | -0.25 | -0.01 | 0.34 | -0.36 | -0.07 | -0.12 | 0.34 | 0.38 | 0.16 | -0.21 | 0.065 | 0.1225 | 0.3425 | 0.668765 | 0.542856 | 0.118542 | | 1 | 0.996511 | 0.721448 |
| Dnajb5 | A2AG30 | 0.12 | 0.19 | -0.23 | -0.44 | -0.39 | 0.02 | -0.6 | -0.43 | 0.56 | -0.01 | -0.48 | 0.24 | 1.21 | 0.92 | -0.84 | -0.81 | -0.26 | 0.1675 | 0.21 | 0.259212 | 0.550759 | 0.691467 | | 1 | 0.996511 | 0.955729 |
| Dnajb6 | O54946 | 1.21 | 1.31 | 1.14 | 0.56 | 0.95 | 1.26 | 0.43 | 0.64 | 0.73 | 0.1 | 0.88 | 1.33 | 1.08 | 0.89 | 0.78 | 0.95 | -0.235 | -0.295 | -0.13 | 0.373075 | 0.352124 | 0.569707 | | 1 | 0.996511 | 0.923081 |
| Dnajb8 | A0A0N4SV | 1.94 | 2.87 | 1.6 | 1.81 | 2.23 | 1.72 | 1.02 | 1.19 | 2.71 | 2.23 | 1.33 | 1.84 | 2.51 | 2.75 | 2.05 | 1.76 | -0.515 | -0.0275 | 0.2125 | 0.191769 | 0.943637 | 0.55314 | | 1 | 0.996755 | 0.916262 |
| Dnajc1 | Q61712 | -0.43 | 0.18 | -1.41 | -0.44 | 0.02 | -0.48 | -0.48 | -0.76 | 0.39 | -0.92 | -1.1 | -0.67 | 0.46 | 0.6 | -0.84 | -0.64 | 0.1 | -0.05 | 0.42 | 0.776755 | 0.909767 | 0.37843 | | 1 | 0.996744 | 0.860984 |
| Dnajc10 | Q9DC23 | -0.77 | -0.55 | -1.4 | -1.26 | -0.04 | -0.75 | -0.64 | -1.31 | -0.79 | 0.3 | -1.45 | -0.45 | -0.09 | 0.01 | -0.18 | -0.7 | 0.31 | 0.3975 | 0.755 | 0.347783 | 0.332042 | 0.022112 | | 1 | 0.996511 | 0.621926 |
| Dnajc11 | Q5U458 | -0.24 | -0.25 | -0.14 | -0.09 | -0.38 | 0.02 | 0.04 | -0.05 | -0.48 | -0.34 | 0.1 | -0.37 | -0.49 | -0.34 | -0.56 | -0.2 | 0.0875 | -0.0925 | -0.2175 | 0.596142 | 0.629279 | 0.247123 | | 1 | 0.996511 | 0.803237 |
| Dnajc12 | D3Z0D9 | 0.46 | -0.29 | -0.35 | 0.39 | -0.47 | 0.32 | 1.16 | 0.18 | 0.53 | 0.49 | -0.87 | 0.52 | -0.16 | 1.23 | -0.04 | -0.75 | 0.245 | 0.115 | 0.0175 | 0.522687 | 0.769176 | 0.968552 | | 1 | 0.996511 | 0.992733 |
| Dnajc13 | G3X922 | -0.08 | 0.2 | -0.06 | -0.21 | 0.46 | -0.03 | 0.22 | -0.13 | 0.12 | -0.06 | 0.13 | 0.14 | -0.3 | 0.48 | -0.13 | -0.16 | 0.1675 | 0.12 | 0.01 | 0.398478 | 0.493351 | 0.96602 | | 1 | 0.996511 | 0.9927 |
| Dnajc14 | A0A1W2P6 | -0.28 | -1.25 | -1.5 | -0.29 | -3.36 | 0.6 | -2.59 | -1.1 | 3.35 | -3.25 | -0.63 | -0.4 | 5 | -1.32 | 0.39 | -1.6 | -0.7825 | 0.5975 | 1.4475 | 0.362891 | 0.634277 | 0.311572 | | 1 | 0.996511 | 0.834523 |
| Dnajc15 | Q78YY6 | -0.37 | -0.14 | -0.36 | -0.15 | -0.72 | 0.12 | -0.75 | -0.13 | 0.31 | 1.31 | -0.04 | 0.07 | 0.98 | 0.06 | 0.64 | 0.36 | -0.115 | 0.6675 | 0.765 | 0.63564 | 0.060209 | 0.011352 | | 1 | 0.996511 | 0.566767 |
| Dnajc16 | Q80TN4 | -0.09 | 0.2 | -0.3 | -0.33 | 0.7 | 0.26 | -0.02 | -0.27 | -0.05 | -0.97 | -0.37 | -0.17 | 0.7 | 0.25 | 0.59 | 0.68 | 0.2975 | -0.26 | 0.685 | 0.257937 | 0.328312 | 0.010647 | | 1 | 0.996511 | 0.563386 |
| Dnajc18 | Q9CZJ9 | -1.36 | -0.91 | 0.13 | 0.56 | -0.48 | -0.43 | 1.08 | 0.51 | -1.05 | 0.25 | 0.08 | -0.88 | 0.01 | -0.78 | 1.71 | 1.88 | 0.565 | -0.005 | 1.1 | 0.30881 | 0.992235 | 0.149862 | | 1 | 0.999285 | 0.748815 |
| Dnajc19 | Q9CQV7 | 1.17 | 1.11 | 0.61 | 0.36 | 1.12 | 0.94 | 0.26 | 0.37 | 0.97 | 0.6 | 0.6 | 1.13 | 1.03 | 1.42 | -0.11 | -0.09 | -0.14 | 0.0125 | -0.25 | 0.629878 | 0.961285 | 0.553946 | | 1 | 0.996755 | 0.916796 |
| Dnajc2 | E9Q9H2 | -0.29 | -0.52 | -2.2 | -2.5 | -0.59 | -0.67 | -2.46 | -2.44 | 0.17 | -1.22 | -2.19 | -0.79 | 0.72 | 0.05 | -2.08 | -2.57 | -0.1625 | 0.37 | 0.4075 | 0.816575 | 0.590401 | 0.64545 | | 1 | 0.996511 | 0.944749 |
| Dnajc24 | Q91ZF0 | -1.12 | -1.09 | -0.73 | -1.24 | -0.81 | -1.49 | -0.41 | -1.27 | -1.64 | -1.54 | -1.06 | -1.15 | -2.29 | -1.31 | -1.34 | -1.34 | 0.05 | -0.3025 | -0.525 | 0.853377 | 0.189787 | 0.091098 | | 1 | 0.996511 | 0.693316 |
| Dnajc25 | A2ALW5 | -1.21 | -0.76 | -2.15 | -1.44 | -0.27 | -0.95 | -0.43 | -0.55 | -0.64 | 0.19 | -2.39 | -0.43 | 0.43 | 0.01 | 1.01 | -0.57 | 0.84 | 0.5725 | 1.61 | 0.025573 | 0.333023 | 0.004115 | | 1 | 0.996511 | 0.548272 |
| Dnajc27 | Q8CFP6 | -1.83 | -1.18 | -1.32 | -0.95 | -1.74 | -1.48 | -1.01 | -1.18 | -0.85 | -0.78 | -0.93 | -1.1 | -0.33 | -1.24 | -1.24 | -1.17 | -0.0325 | 0.405 | 0.325 | 0.899147 | 0.106613 | 0.29862 | | 1 | 0.996511 | 0.828257 |
| Dnajc28 | Q8VCE1 | 0.7 | 0.82 | 1.25 | 0.44 | 1.08 | 0.48 | 0.93 | 1.18 | 0.35 | -1.48 | 0.6 | 0.49 | 0.29 | 0.63 | 0.64 | 2.13 | 0.115 | -0.8125 | 0.12 | 0.638639 | 0.121995 | 0.777317 | | 1 | 0.996511 | 0.967289 |
| Dnajc3 | Q91YW3 | -0.02 | 0.24 | 0.02 | 0.1 | 0.61 | -0.26 | 0.25 | 0.31 | -0.19 | 0.78 | -0.32 | -0.02 | 0.31 | 0.75 | 1.69 | 1.07 | 0.1425 | -0.0225 | 0.87 | 0.513124 | 0.93337 | 0.018478 | | 1 | 0.996755 | 0.602211 |
| Dnajc30 | P59041 | -0.04 | -0.04 | -0.21 | 0.18 | -0.21 | 0.35 | -0.2 | -0.07 | -0.23 | 1.05 | 0.43 | 0.19 | 0.13 | 0.13 | -0.18 | -0.17 | -0.005 | 0.3875 | 0.005 | 0.979154 | 0.202388 | 0.979311 | | 1 | 0.996511 | 0.994367 |
| Dnajc5 | P60904 | 1.89 | 1.47 | 2.24 | 2.01 | 0.64 | 1.86 | 1.2 | 1.76 | 2.43 | 0.95 | 2.08 | 1.72 | 2.55 | 0.39 | 1.88 | 1.67 | -0.5375 | -0.1075 | -0.28 | 0.118806 | 0.758299 | 0.540873 | | 1 | 0.996511 | 0.912212 |
| Dnajc6 | Q80TZ3-3 | -2.23 | -1.22 | -4.06 | -6.11 | -1.78 | -1.45 | -1.52 | -0.82 | 5.47 | -4.35 | -1.53 | -2.63 | 8.42 | -4.48 | 4.68 | -5.85 | 2.0125 | 2.645 | 4.0975 | 0.067118 | 0.238627 | 0.224436 | | 1 | 0.996511 | 0.787447 |
| Dnajc6 | Q80TZ3-2 | 0.38 | 0.89 | 0.43 | 0.5 | 1.06 | 0.44 | 0.67 | 0.49 | 0.5 | 0.52 | 0.37 | 0.5 | -0.32 | 1.46 | 0.03 | 0.41 | 0.115 | -0.0775 | -0.155 | 0.587486 | 0.674515 | 0.691967 | | 1 | 0.996511 | 0.955729 |
| Dnajc6 | Q80TZ3 | -0.84 | -0.76 | -0.72 | -0.67 | -0.57 | -0.68 | -0.37 | -0.71 | -1.07 | -0.73 | -0.49 | -0.58 | -0.99 | -0.68 | -0.56 | -1.07 | 0.165 | 0.03 | -0.0775 | 0.301148 | 0.874362 | 0.695147 | | 1 | 0.996511 | 0.955729 |
| Dnajc6 | B1B0B7 | -2.06 | -2.24 | -6.23 | -7.08 | -3.6 | -1.84 | -8.69 | -7.56 | -0.27 | -4.41 | -6.9 | -2.52 | 0.97 | -0.38 | -7.7 | -8.23 | -1.02 | 0.8775 | 0.5675 | 0.586055 | 0.611822 | 0.814896 | | 1 | 0.996511 | 0.972768 |
| Dnajc7 | Q9QYI3 | -0.77 | -1.41 | -0.94 | -1.19 | -1.21 | -0.64 | -1.36 | -1.3 | -1.17 | -0.5 | -1.21 | -0.84 | -0.63 | -0.86 | -1.04 | -1.42 | -0.05 | 0.1475 | 0.09 | 0.831301 | 0.54848 | 0.720918 | | 1 | 0.996511 | 0.958398 |
| Dnajc8 | Q6NZB0 | -2.68 | -3.48 | -3.16 | -3.47 | -3.72 | -3.54 | -3.65 | -3.6 | -3.79 | -2.14 | -3.24 | -3.13 | -3.09 | -3.29 | -3.12 | -4.04 | -0.43 | 0.1225 | -0.1875 | 0.073986 | 0.746343 | 0.540841 | | 1 | 0.996511 | 0.912212 |
| Dnajc9 | Q91WN1 | -4.75 | -5.17 | -4.2 | -5.2 | -3.99 | -4.54 | -3.92 | -4.84 | -3.78 | -1.79 | -4.8 | -3.87 | -2.61 | -3.2 | -4.05 | -4.04 | 0.5075 | 1.27 | 1.355 | 0.135142 | 0.066188 | 0.008355 | | 1 | 0.996511 | 0.563386 |
| Dnal1 | Q05A62-2 | -0.83 | -0.76 | -0.63 | -1.71 | -0.97 | -1.14 | 0.11 | -0.87 | -1.29 | -0.7 | -0.27 | -0.51 | -1.58 | -0.84 | -0.6 | -0.41 | 0.265 | 0.29 | 0.125 | 0.466493 | 0.386439 | 0.723504 | | 1 | 0.996511 | 0.958398 |
| Dnal4 | Q8CDT8 | 0.67 | 0.02 | 1.16 | -1.53 | -0.15 | 0.85 | -0.93 | 0.07 | -0.48 | -4.84 | 0.53 | 0.52 | 0.38 | -1.03 | 1 | 0.66 | -0.12 | -1.1475 | 0.1725 | 0.8485 | 0.373944 | 0.796772 | | 1 | 0.996511 | 0.969727 |
| Dner | Q8JZM4 | -0.32 | 0.03 | -0.71 | -0.65 | 0.05 | -0.13 | -0.81 | -0.41 | -0.02 | 0.5 | -0.79 | 0.05 | 0.37 | 0.66 | -0.81 | -0.72 | 0.0875 | 0.3475 | 0.2875 | 0.738728 | 0.291863 | 0.476161 | | 1 | 0.996511 | 0.892174 |
| Dnlz | Q9D113 | 0.5 | 0.37 | 0.01 | 0.39 | 0.65 | 0.41 | 0.34 | -0.11 | 0.67 | -0.95 | 0.67 | 0.44 | 0.63 | 1.37 | -0.87 | 0.14 | 0.005 | -0.11 | 0 | 0.981534 | 0.777132 | 1 | | 1 | 0.996511 | 1 |
| Dnm1 | P39053-4 | 0.44 | 0.29 | 0.53 | 0.98 | 0.33 | 0.3 | 0.44 | 1.01 | 0.48 | 0.61 | 0.59 | 0.42 | 0.15 | 0.57 | 0.3 | 0.22 | -0.04 | -0.035 | -0.25 | 0.866974 | 0.862977 | 0.280554 | | 1 | 0.996511 | 0.822084 |
| Dnm1 | A0A0J9YUN | 1.52 | 0.19 | -0.97 | -0.25 | 0.21 | 1.31 | -0.38 | -0.11 | 1.5 | -1.67 | -0.8 | 1.59 | -0.18 | 0.32 | -1.24 | -0.5 | 0.135 | 0.0325 | -0.5225 | 0.81789 | 0.970461 | 0.369711 | | 1 | 0.99727 | 0.859245 |
| Dnm1 | P39053-6 | 1.28 | 1.75 | 2.11 | 1.67 | 1.22 | 1.24 | 1.14 | 1.46 | 2.43 | 1.91 | 1.58 | 1.67 | 1.76 | 2.89 | 1.39 | 1.16 | -0.4375 | 0.195 | 0.0975 | 0.064168 | 0.478375 | 0.809442 | | 1 | 0.996511 | 0.972098 |
| Dnm1 | A0A0J9YUS | 1.17 | 1.58 | -2.62 | -1.88 | 1.32 | 1.44 | -2.46 | -3.15 | 1.84 | -1.85 | -2.38 | 1.59 | 2.04 | 1.87 | -2.24 | -2.8 | -0.275 | 0.2375 | 0.155 | 0.848691 | 0.8627 | 0.916979 | | 1 | 0.996511 | 0.986641 |
| Dnm1l | Q8K1M6-3 | -1.42 | -0.91 | -0.88 | 0.11 | -0.11 | -1.48 | -0.38 | -0.45 | -0.7 | -0.16 | 0.33 | -0.85 | -1.35 | 1.46 | 0.1 | 0.02 | 0.17 | 0.43 | 0.8325 | 0.6816 | 0.298792 | 0.190407 | | 1 | 0.996511 | 0.776843 |
| Dnm1l | E9PUD2 | 0.13 | -0.23 | -0.81 | -0.23 | 0.36 | -0.3 | 0.31 | -0.39 | -0.03 | -0.22 | 0.07 | -0.23 | -0.64 | 1.62 | 0.12 | -0.49 | 0.28 | 0.1825 | 0.4375 | 0.331856 | 0.449029 | 0.403584 | | 1 | 0.996511 | 0.870698 |
| Dnm1l | Q8K1M6 | -0.56 | -0.35 | -0.16 | -0.28 | -0.35 | -0.56 | -0.36 | -0.34 | -0.22 | 0.07 | -0.28 | -0.42 | -0.38 | -0.03 | -0.54 | -0.24 | -0.065 | 0.125 | 0.04 | 0.687183 | 0.515941 | 0.843068 | | 1 | 0.996511 | 0.976225 |
| Dnm2 | P39054-2 | 0.02 | -0.16 | 0.12 | 0.17 | -0.31 | -0.01 | 0.49 | -0.03 | 0.02 | 0.07 | -0.02 | -0.04 | -0.12 | 0.11 | -0.38 | -0.05 | -0.0025 | -0.03 | -0.1475 | 0.990482 | 0.854876 | 0.459453 | | 1 | 0.996511 | 0.887902 |
| Dnm3 | E9QLL2 | 0.44 | 0.3 | 0.33 | 0.32 | 0.43 | 0.26 | 0.54 | 0.27 | 0.06 | -0.19 | 0.11 | 0.13 | -0.11 | 0.41 | -0.08 | 0.09 | 0.0275 | -0.32 | -0.27 | 0.85449 | 0.080121 | 0.190106 | | 1 | 0.996511 | 0.776843 |
| Dnmbp | A0A1D5RM | 0.03 | 0.54 | 1.27 | 0.39 | -0.04 | 0.02 | -0.38 | 0.1 | 0.29 | 0.83 | 0.05 | 0.21 | 1.07 | 0.57 | 0.05 | -0.25 | -0.6325 | -0.2125 | -0.1975 | 0.049631 | 0.502777 | 0.606238 | | 1 | 0.996511 | 0.932425 |
| Dnmt1 | P13864 | -2.78 | -2.36 | -1.97 | -2.07 | -2.04 | -2.96 | -1.84 | -2.11 | -1.08 | -1.53 | -1.62 | -2.61 | -1.76 | -2.1 | -1.78 | -2.22 | 0.0575 | 0.585 | 0.33 | 0.849859 | 0.131827 | 0.207902 | | 1 | 0.996511 | 0.785338 |
| Dnmt3a | O88508 | 2.25 | -0.13 | 0.27 | 0.17 | -0.03 | 1.86 | 0.41 | 0.01 | 1.68 | 0.19 | 0.6 | 2.68 | 2.18 | -0.4 | 0.88 | 0.42 | -0.0775 | 0.6475 | 0.13 | 0.903235 | 0.373157 | 0.851578 | | 1 | 0.996511 | 0.976225 |
| Dnpep | Q3TVK3 | 0.41 | 0.08 | -0.27 | -0.22 | 0 | 0.21 | 0.24 | -0.02 | -0.11 | -0.21 | -0.1 | 0.19 | -0.25 | 0.01 | -0.11 | -0.35 | 0.1075 | -0.0575 | -0.175 | 0.598888 | 0.791891 | 0.442879 | | 1 | 0.996511 | 0.883549 |
| Dnph1 | Q80VJ3 | 0.46 | 0.21 | 0.44 | 1.17 | 0.58 | 0.83 | 0.86 | 0.64 | 0.76 | 1.76 | 0.74 | 0.66 | 0.69 | 0.55 | 1.13 | 0.89 | 0.1575 | 0.41 | 0.245 | 0.510317 | 0.236519 | 0.373025 | | 1 | 0.996511 | 0.859424 |
| Dnttip1 | Q99LB0 | -0.99 | -0.11 | -0.26 | 0.24 | -2.33 | -0.8 | 0.22 | -0.06 | 1.06 | -0.73 | -0.14 | -0.67 | 2.48 | -1.98 | -0.56 | 0 | -0.4625 | 0.16 | 0.265 | 0.428815 | 0.728144 | 0.760425 | | 1 | 0.996511 | 0.963123 |
| Dnttip2 | Q8R2M2 | -0.2 | 0.52 | 0.95 | 0.91 | -0.01 | 0 | 0.61 | 0.7 | 0.42 | 1.61 | 0.57 | -0.1 | 0.5 | 1 | 0.97 | 1.22 | -0.22 | 0.08 | 0.3775 | 0.498796 | 0.850145 | 0.249978 | | 1 | 0.996511 | 0.806092 |
| Doc2a | Q7TNF0 | 2.78 | 1.26 | 1.24 | 1.56 | 2 | 3.23 | 1.87 | 1.57 | 1.39 | 0.83 | 1.6 | 2.58 | 0.55 | 1.6 | 1.46 | 1.63 | 0.4575 | -0.11 | -0.4 | 0.351356 | 0.819286 | 0.358697 | | 1 | 0.996511 | 0.856645 |
| Doc2b | P70169 | 0.11 | -0.51 | 1.89 | 0.8 | 0 | 0.63 | 2.42 | 1.81 | -0.84 | 0.99 | 1.78 | 0.11 | -1.05 | -1.02 | 1.95 | 2.46 | 0.6425 | -0.0625 | 0.0125 | 0.359158 | 0.928231 | 0.989603 | | 1 | 0.996755 | 0.996154 |
| Dock1 | Q8BUR4 | 0.7 | 0.59 | 0.62 | 0.54 | 0.3 | 0.2 | 0.74 | 0.46 | 0.2 | -0.02 | 0.43 | 0.38 | -0.16 | -0.1 | 0.48 | 0.42 | -0.1875 | -0.365 | -0.4525 | 0.295049 | 0.065139 | 0.067122 | | 1 | 0.996511 | 0.664936 |
| Dock10 | A0A087WC | 0.97 | 0.29 | -0.31 | -0.14 | -0.07 | 0.74 | 0.05 | -0.09 | 0.92 | -0.05 | -0.31 | 0.5 | 0.98 | -0.29 | 0.05 | -0.05 | -0.045 | 0.0625 | -0.03 | 0.89329 | 0.870116 | 0.938137 | | 1 | 0.996511 | 0.989844 |
| Dock11 | A2AF47 | 0.21 | 0.43 | 0.61 | 0.47 | 0.17 | 0.1 | 0.45 | 0.99 | 0.39 | 0.34 | 0.61 | 0.23 | 0.05 | 0.67 | 0.71 | 0.61 | -0.0025 | -0.0375 | 0.08 | 0.991529 | 0.835638 | 0.721673 | | 1 | 0.996511 | 0.958398 |
| Dock2 | Q8C3J5 | -0.01 | -0.78 | -1.21 | -1.48 | -0.79 | -0.75 | -1.2 | -1.23 | -0.7 | -0.59 | -0.96 | -0.42 | -0.89 | -0.93 | -1.69 | -1.14 | -0.1225 | 0.2025 | -0.2925 | 0.715537 | 0.551393 | 0.430312 | | 1 | 0.996511 | 0.879165 |
| Dock3 | F8VPQ1 | 0.5 | -0.14 | 0.2 | 0.24 | 0.22 | 0.49 | 0.34 | 0.33 | -0.02 | -0.16 | 0.28 | 0.25 | -0.28 | 0.25 | -0.01 | 0.01 | 0.145 | -0.1125 | -0.2075 | 0.440617 | 0.597483 | 0.359761 | | 1 | 0.996511 | 0.856645 |
| Dock4 | P59764 | 0.35 | 0.42 | 0.29 | 0.12 | 0.72 | 0.35 | 0.7 | 0.2 | 0.18 | -0.11 | 0.39 | 0.31 | 0.07 | 0.24 | 0.17 | 0.41 | 0.1975 | -0.1025 | -0.0725 | 0.303598 | 0.586535 | 0.692937 | | 1 | 0.996511 | 0.955729 |
| Dock5 | B2RY04 | 0.83 | 0.7 | 0.8 | 0.87 | 0.47 | 0.11 | 0.27 | 0.4 | -0.31 | 0.57 | 0.25 | 0.33 | -0.47 | 0.41 | 0.68 | 1 | -0.4875 | -0.59 | -0.395 | 0.011548 | 0.026093 | 0.241761 | | 1 | 0.996511 | 0.796892 |
| Dock6 | A0A1L1SQ | -0.12 | 0.08 | -0.39 | -0.53 | 0.23 | -0.33 | 0.11 | -0.48 | 0.48 | 1.03 | 0.19 | 0.23 | 0.77 | 0.37 | 0.45 | 0.36 | 0.1225 | 0.7225 | 0.7275 | 0.605909 | 0.019456 | 0.008537 | | 1 | 0.996511 | 0.563386 |
| Dock7 | A2A9M5 | 0.62 | 0.31 | 0.54 | 0.63 | 0.07 | 0.35 | 0.4 | 0.48 | 0.54 | 0.22 | 0.54 | 0.51 | 0.23 | 0.28 | 0.39 | 0.4 | -0.2 | -0.0725 | -0.2 | 0.257219 | 0.683546 | 0.280572 | | 1 | 0.996511 | 0.822084 |
| Dock9 | E9QMR2 | 0.67 | 0.32 | 0.36 | 0.15 | 0.55 | 0.62 | 0.31 | 0.24 | 0.01 | 0.19 | 0.35 | 0.54 | 0.07 | 0.28 | 0.18 | 0.26 | 0.055 | -0.1025 | -0.1775 | 0.765721 | 0.617182 | 0.369003 | | 1 | 0.996511 | 0.859245 |
| Dock9 | Q8BIK4-2 | -0.21 | -0.72 | -0.05 | 0.23 | -0.99 | -0.54 | -0.83 | -1.7 | 1.85 | -1.46 | 0.49 | -0.41 | 3.79 | -0.68 | 0.71 | -0.9 | -0.8275 | 0.305 | 0.9175 | 0.025209 | 0.648216 | 0.363067 | | 1 | 0.996511 | 0.858475 |
| Dohh | D3Z7J7 | -0.93 | -0.18 | 0.65 | 1.02 | -0.17 | -1.58 | 1.11 | 1.34 | -0.08 | 1.44 | 0.58 | 0.12 | -1.05 | -0.03 | 0.23 | 0.65 | 0.035 | 0.375 | -0.19 | 0.961466 | 0.470893 | 0.718404 | | 1 | 0.996511 | 0.958398 |
| Dok4 | Q99KE3 | -0.23 | -0.32 | -0.62 | -0.47 | 0.16 | -0.12 | -0.16 | -0.31 | -0.15 | -0.88 | -0.67 | -0.23 | -0.26 | 0.37 | -1.2 | -0.46 | 0.3025 | -0.0725 | 0.0225 | 0.115624 | 0.751044 | 0.946585 | | 1 | 0.996511 | 0.992376 |
| Dok6 | Q2MHE5 | -0.17 | 0.3 | -0.17 | -0.13 | 0.46 | 0.22 | -0.56 | -0.37 | 0.02 | 0.81 | -0.07 | 0.29 | 0.89 | 0.34 | 0.29 | -0.38 | -0.02 | 0.305 | 0.3275 | 0.941288 | 0.245107 | 0.288598 | | 1 | 0.996511 | 0.824705 |
| Dok7 | A0A0R4J0R | 0.19 | 0.16 | 1.36 | 0.88 | -0.51 | 0.34 | -0.3 | 1.47 | 1.07 | 1.29 | -0.53 | 0.58 | 1.28 | -0.77 | -0.2 | 1.06 | -0.3975 | -0.045 | -0.305 | 0.426971 | 0.923165 | 0.568918 | | 1 | 0.996755 | 0.922588 |
| Dolk | Q8R2Y3 | -4.15 | -2.79 | -2.17 | -3.19 | -0.42 | -3.84 | -0.67 | -2.67 | -3.89 | -4.23 | -2.99 | -4.4 | -6.55 | -2.05 | -2.85 | -1.41 | 1.175 | -0.8025 | -0.14 | 0.180508 | 0.125964 | 0.897867 | | 1 | 0.996511 | 0.983509 |
| Dolpp1 | A2AWJ3 | 4.76 | 4.12 | 1.65 | -2.03 | 5.05 | 5.07 | 2.55 | 1.33 | 2.8 | -2.94 | 2.1 | 4.33 | 3.14 | 1.51 | 3.56 | 3.56 | 1.375 | -0.5525 | 0.8175 | 0.40232 | 0.778556 | 0.572174 | | 1 | 0.996511 | 0.924918 |
| Dop1a | H7BWZ9 | 0.24 | 0.42 | -0.19 | -0.11 | 0.77 | -0.07 | -0.28 | -0.13 | 0.3 | 0.24 | -0.01 | 0.04 | 0.06 | 0.72 | 0.03 | 0.05 | -0.0175 | 0.0525 | 0.125 | 0.950136 | 0.800351 | 0.624509 | | 1 | 0.996511 | 0.939167 |
| Dop1b | Q3UHQ6 | 0.06 | -0.38 | -0.39 | -0.21 | 0.46 | 0.06 | -0.2 | -0.22 | -0.51 | -0.32 | -0.14 | 0.12 | 0.07 | -0.31 | -0.41 | -0.25 | 0.255 | 0.0175 | 0.005 | 0.256232 | 0.934201 | 0.980783 | | 1 | 0.996755 | 0.994858 |
| Dpagt1 | P42867 | 0.99 | 1.12 | 0.98 | 1.19 | 0.07 | 0.56 | 0.3 | 1.07 | 1 | 0.74 | 0.53 | 1.07 | 0.79 | 0.65 | 0.49 | 0.6 | -0.57 | -0.235 | -0.4375 | 0.037648 | 0.238417 | 0.033318 | | 1 | 0.996511 | 0.644622 |
| Dpcd | Q8BPA8 | -0.27 | 0.32 | -0.72 | -0.84 | 0.73 | -0.54 | -1.19 | -0.77 | -0.08 | -0.23 | -0.74 | -0.35 | -0.63 | -0.11 | -0.59 | -1.06 | -0.065 | 0.0275 | -0.22 | 0.886473 | 0.927708 | 0.510625 | | 1 | 0.996755 | 0.90359 |

| | | | | | | | | | | | | | | | | | | | | | | | | | | |
|---|---|---|---|---|---|---|---|---|---|---|---|---|---|---|---|---|---|---|---|---|---|---|---|---|---|---|
| Dpf1 | D3YYG6 | -4.19 | -5.59 | -1.99 | -2.48 | -4.89 | -4.99 | -2.13 | -2.31 | -4.24 | -1.93 | -2.01 | -4.41 | -3.07 | -6.73 | -2.5 | -2.42 | -1.75E-02 | 0.415 | -0.1175 | 0.986317 | 0.667486 | 0.920179 | 1 | 0.996511 | 0.98688 |
| Dpf2 | D3Z5N6 | -1.75 | -2.76 | -2.03 | -1.9 | -3.01 | -2.22 | -2.35 | -2.45 | -1.57 | -0.95 | -2.3 | -2.25 | -1.83 | -3.1 | -2.5 | -2.5 | -0.3975 | 0.3425 | -0.3725 | 0.187727 | 0.375085 | 0.293437 | 1 | 0.996511 | 0.825612 |
| Dph1 | Q5NCQ5 | -1.47 | -0.99 | -0.96 | -0.51 | -1.15 | -1.17 | -0.65 | -0.78 | -0.77 | 1.03 | -0.92 | -1.03 | -0.47 | -0.37 | -1.22 | -0.91 | 0.045 | 0.56 | 0.24 | 0.856327 | 0.270949 | 0.423027 | 1 | 0.996511 | 0.875782 |
| Dph2 | Q9CR25 | -0.88 | -0.34 | 0.01 | 0.05 | -0.31 | -0.77 | -0.56 | 0.32 | -0.5 | 1.19 | -0.18 | -0.59 | 0.03 | -0.37 | -0.12 | -0.41 | -0.04 | 0.27 | 0.0725 | 0.898521 | 0.543172 | 0.785262 | 1 | 0.996511 | 0.967642 |
| Dph5 | Q9CWQ0 | -0.77 | -0.32 | -0.2 | -0.59 | -0.28 | 0.61 | -0.73 | -0.8 | -0.54 | -0.42 | -0.68 | -0.75 | -1.25 | -0.15 | -0.67 | -0.06 | 0.17 | -0.1275 | -0.0625 | 0.617322 | 0.526353 | 0.841337 | 1 | 0.996511 | 0.976225 |
| Dph6 | Q9CQ28-2 | -0.57 | -0.96 | -1.13 | -1.43 | -0.61 | -1.28 | -1.23 | -1.63 | 0.21 | -0.99 | -1.89 | -0.81 | -1.56 | -0.89 | -1.79 | -1.14 | -0.165 | 0.1525 | -0.3225 | 0.560415 | 0.729602 | 0.280478 | 1 | 0.996511 | 0.822084 |
| Dph7 | Q9CYU6 | 0.19 | -0.2 | -0.02 | 0.51 | -0.54 | 0.03 | -0.54 | 0.28 | 0.81 | 0.98 | 0.4 | -0.34 | 1.03 | -0.9 | 0.39 | 0.58 | -0.3125 | 0.3425 | 0.155 | 0.256911 | 0.313611 | 0.713868 | 1 | 0.996511 | 0.958106 |
| Dpm1 | O70152 | 0.07 | 0.12 | -0.99 | -0.79 | 0.43 | -0.08 | 0.36 | -0.66 | -0.93 | -1.76 | -1.07 | -0.04 | -0.36 | -0.58 | -0.13 | -0.1 | 0.41 | -0.5525 | 0.105 | 0.278641 | 0.221682 | 0.740426 | 1 | 0.996511 | 0.958796 |
| Dpm3 | Q9D1Q4 | 0.99 | 0.25 | -0.14 | 0.39 | 0.34 | 0.8 | 0.38 | -0.34 | 0.41 | -0.25 | -0.33 | 0.58 | 0.87 | 0.96 | 0.3 | 0.64 | -0.0775 | -0.27 | 0.32 | 0.81125 | 0.41828 | 0.290756 | 1 | 0.996511 | 0.825497 |
| Dpp10 | Q6NXK7 | 0.34 | 0.62 | 0.63 | 0.45 | 0.78 | 0.75 | 0.55 | 0.38 | 0.26 | -0.63 | 0.6 | 0.54 | 0.08 | 0.02 | 0.22 | 0.32 | 0.105 | -0.3175 | -0.35 | 0.540956 | 0.3041 | 0.083483 | 1 | 0.996511 | 0.689531 |
| Dpp10 | A0A5F8MP | -0.94 | -0.18 | -0.02 | -0.39 | -0.43 | -0.26 | -0.73 | -0.47 | 0.94 | 0.72 | 0.18 | -0.54 | 1.73 | 0.86 | -0.18 | -0.52 | -0.09 | 0.7075 | 0.855 | 0.70743 | 0.085635 | 0.121686 | 1 | 0.996511 | 0.721448 |
| Dpp3 | Q99KK7 | -0.33 | -0.56 | -0.18 | 0.01 | -0.64 | -0.58 | -0.05 | -0.05 | -0.11 | 0.26 | -0.08 | -0.26 | -0.34 | 0.17 | -0.35 | -0.26 | -0.065 | 0.2175 | 0.07 | 0.771911 | 0.310291 | 0.753055 | 1 | 0.996511 | 0.961479 |
| Dpp4 | P28843 | -0.39 | -0.25 | -0.4 | 0.33 | 0.11 | -0.25 | -0.27 | -0.67 | 0.02 | -0.01 | -0.26 | -0.1 | -0.23 | 0.23 | -0.63 | -0.14 | -0.0925 | 0.09 | -0.015 | 0.709854 | 0.68463 | 0.955826 | 1 | 0.996511 | 0.992535 |
| Dpp6 | E9PWX1 | 1.28 | 1.03 | 1.3 | 1.51 | 0.92 | 1.1 | 1.77 | 1.74 | 0.87 | 0.78 | 1.44 | 1.45 | 0.3 | 0.69 | 1.15 | 1.36 | 0.1025 | -0.145 | -0.405 | 0.684378 | 0.541444 | 0.170193 | 1 | 0.996511 | 0.762229 |
| Dpp6 | Q80VM5 | -1.76 | 0.13 | -0.37 | -1.15 | 0.32 | 0.08 | 0.11 | -0.24 | -0.84 | 2.51 | 0.68 | -0.92 | -0.14 | 0.65 | -0.29 | -1.24 | 0.855 | 1.145 | 0.5325 | 0.061916 | 0.187108 | 0.327809 | 1 | 0.996511 | 0.844684 |
| Dpp7 | Q9ET22 | -1.29 | -1.27 | -0.98 | -0.36 | -1.68 | -1.06 | -0.7 | -0.93 | -1.48 | 0.1 | -0.3 | -1.01 | -0.96 | -1.74 | -0.69 | -0.66 | -0.1175 | 0.3025 | -0.0375 | 0.695834 | 0.454011 | 0.910633 | 1 | 0.996511 | 0.986458 |
| Dpp8 | Q80YA7 | -0.25 | -0.54 | -0.58 | -0.42 | -0.52 | -0.42 | -0.32 | -0.46 | -0.18 | -0.78 | -0.51 | -0.5 | -0.1 | -0.39 | -0.66 | -0.68 | 0.0175 | -0.045 | -0.01 | 0.909695 | 0.818396 | 0.962332 | 1 | 0.996511 | 0.9927 |
| Dpp9 | Q8BVG4 | 0.45 | 0.06 | 0.28 | -0.2 | 0.16 | 0.44 | 0.38 | 0.25 | -0.76 | -0.8 | 0.24 | 0.25 | -0.99 | -0.43 | -0.09 | 0.04 | 0.16 | -0.415 | -0.515 | 0.413802 | 0.225316 | 0.100378 | 1 | 0.996511 | 0.698156 |
| Dpy19l1 | A6X919 | -1.06 | -0.28 | -1.46 | -1.67 | 0.35 | -0.7 | -0.31 | -1.26 | -1.28 | -3.07 | -1.27 | -0.51 | -0.66 | -0.81 | -0.64 | -0.21 | 0.6375 | -0.415 | 0.5375 | 0.161627 | 0.476212 | 0.134862 | 1 | 0.996511 | 0.731223 |
| Dpy19l3 | Q71B07 | -0.38 | 0.33 | -2.05 | -2.28 | 0.4 | -0.47 | -0.76 | -1.74 | 0.06 | -2.43 | -1.89 | -0.31 | 0.51 | 0.69 | -1.61 | -1.53 | 0.4525 | -0.0475 | 0.61 | 0.523782 | 0.952158 | 0.456336 | 1 | 0.996755 | 0.887169 |
| Dpy19l4 | A2AJQ3-2 | -1.71 | -0.89 | -1.17 | -1.86 | -0.88 | -1.14 | -1.93 | -2.68 | -1.6 | -1.35 | -1.76 | -1.54 | -0.49 | -1.67 | -0.6 | 0.14 | -0.25 | -0.155 | 0.7525 | 0.569802 | 0.557219 | 0.100452 | 1 | 0.996511 | 0.698156 |
| Dpy30 | Q99LT0 | -0.37 | -0.1 | -0.09 | 0.13 | -0.47 | -0.72 | 0.12 | 0.37 | -0.02 | 0.54 | 0.35 | -0.44 | -0.57 | 0.1 | -0.19 | 0.08 | -0.0675 | 0.215 | -0.0375 | 0.807489 | 0.416086 | 0.870961 | 1 | 0.996511 | 0.979293 |
| Dpyd | Q8CHR6 | -0.12 | 0.66 | -1.35 | 1.04 | 0.3 | 0.2 | 0.34 | -0.29 | 2.71 | 1.86 | -0.35 | -0.1 | 1.58 | 3.4 | 0.15 | 0.45 | 0.08 | 0.9725 | 1.3375 | 0.87399 | 0.258314 | 0.128015 | 1 | 0.996511 | 0.723655 |
| Dpys | Q9EQF5 | -2.84 | -2.76 | -1.54 | 0.32 | -2.92 | -3.73 | -0.33 | -0.15 | -2.98 | -0.59 | -1.85 | -2.91 | -1.29 | -2.33 | -1.26 | 1.22 | -0.0775 | -0.3775 | 0.79 | 0.940979 | 0.653518 | 0.412759 | 1 | 0.996511 | 0.874811 |
| Dpysl2 | O08553 | -0.37 | -0.24 | -0.34 | -0.05 | -0.21 | -0.52 | 0.1 | -0.02 | -0.66 | 0.18 | -0.3 | -0.26 | -0.89 | -0.52 | -0.47 | -0.54 | 0.0875 | -0.01 | -0.355 | 0.649058 | 0.964123 | 0.094046 | 1 | 0.996755 | 0.698156 |
| Dpysl3 | E9PWE8 | -0.01 | -0.08 | -0.26 | -0.12 | -0.23 | -0.05 | 0.03 | 0.14 | 0.71 | 0.51 | 0.23 | 0.72 | 0.19 | 0.54 | 0 | -0.27 | 0.09 | 0.66 | 0.2325 | 0.573379 | 0.006239 | 0.317629 | 1 | 0.996511 | 0.838864 |
| Dpysl4 | Q3TMU8 | 0.35 | 0.37 | 0.01 | 0.24 | 0.05 | 0.14 | 0.18 | 0.22 | 0.5 | 0.18 | 0.02 | 0.31 | -0.04 | 1.09 | -0.34 | -0.42 | -0.095 | 0.01 | -0.17 | 0.548418 | 0.957752 | 0.633957 | 1 | 0.996755 | 0.941132 |
| Dpysl5 | Q9EQF6 | 0.23 | 0.07 | 0.08 | 0.39 | -0.1 | -0.11 | 0.51 | 0.41 | 0.67 | 0.27 | 0.53 | 0.86 | -0.22 | 0.37 | -0.03 | 0.06 | -0.015 | 0.39 | -0.1475 | 0.942792 | 0.074508 | 0.480393 | 1 | 0.996511 | 0.893906 |
| Dr1 | Q91WV0 | -3.27 | -3.56 | -2.36 | -1.72 | -2.95 | -4.01 | -1.71 | -2.66 | -4.06 | -1.92 | -2.05 | -3.04 | -2.64 | -3.56 | -2.68 | -2.51 | -0.105 | -0.04 | -0.12 | 0.856212 | 0.946675 | 0.793484 | 1 | 0.996755 | 0.969365 |
| Drap1 | F6T9K1 | -2.39 | -2.9 | -3.97 | -4.23 | -1.75 | -3.72 | -4.18 | -4.75 | -3.04 | -1.58 | -4.62 | -2.23 | -2.76 | -1.78 | -5.31 | -4.72 | -0.2275 | 0.505 | -0.27 | 0.749109 | 0.486641 | 0.748467 | 1 | 0.996511 | 0.960744 |
| Drd1 | A0A1Y7VK | 3.41 | 2.13 | -1.37 | -0.65 | 0.63 | 1.83 | -1.34 | -0.86 | 2.26 | -0.34 | -1.4 | 3.37 | 2.54 | 0.4 | -0.15 | -1.26 | -0.815 | 0.0925 | -0.4975 | 0.503271 | 0.947851 | 0.687762 | 1 | 0.996755 | 0.955729 |
| Drg1 | P32233 | -1.09 | -1.71 | -1.7 | -1.86 | -1.4 | -1.17 | -1.76 | -1.87 | -1.62 | -1.53 | -1.81 | -0.87 | -1.69 | -1.14 | -2.35 | -2.37 | 0.04 | 0.1325 | -0.2975 | 0.871757 | 0.636672 | 0.393516 | 1 | 0.996511 | 0.865775 |
| Drg2 | Q9QXB9 | -0.02 | -0.07 | -0.32 | -0.32 | -0.2 | -0.1 | -0.34 | -0.09 | -0.6 | 0.11 | -0.25 | -0.06 | -0.68 | -0.18 | -0.47 | -0.59 | 1.96E-17 | -0.0175 | -0.2975 | 1 | 0.934513 | 0.164227 | 1 | 0.996755 | 0.758352 |
| Drp2 | Q05AA6 | 0.61 | 0.44 | 0.52 | 0.31 | 0.92 | 0.44 | 0.38 | 0.28 | 0.46 | 0.63 | 0.58 | 0.41 | 0.59 | 0.89 | 0.31 | 0.09 | 0.035 | 0.05 | -2.7E-16 | 0.855949 | 0.763475 | 1 | 1 | 0.996511 | 1 |
| Dscam | Q9ERC8 | -0.46 | 0.17 | 0.71 | 0.66 | 0.24 | 0.1 | 0.25 | -0.03 | 0.36 | 0.45 | 0.64 | 0.11 | 0.27 | 0.88 | 0.37 | 0.74 | -0.13 | 0.12 | 0.295 | 0.647126 | 0.689934 | 0.363279 | 1 | 0.996511 | 0.858475 |
| Dscaml1 | E9QPR7 | 0.38 | 0.72 | 0.4 | 0.39 | 1.04 | 0.51 | 0.55 | 0.78 | 0.42 | 1.28 | 0.7 | 0.61 | 0.73 | 0.81 | 0.85 | 0.87 | 0.2475 | 0.28 | 0.3425 | 0.209662 | 0.250048 | 0.08382 | 1 | 0.996511 | 0.690099 |
| Dst | A0A087WF | -3.59 | -2.97 | -2.76 | -3.32 | -2.23 | -4.09 | -2.13 | -2.96 | -3.23 | -4.02 | -2.97 | -3.8 | -5.89 | -3.62 | -2.43 | -4.12 | 0.3075 | -0.345 | -0.855 | 0.503979 | 0.281757 | 0.226128 | 1 | 0.996511 | 0.787447 |
| Dst | S4R1P5 | 0.73 | 0.83 | 0.75 | 0.68 | 0.78 | 0.54 | 0.69 | 0.65 | 0.63 | 0.77 | 0.68 | 0.73 | 0.48 | 0.65 | 0.74 | 0.6 | -0.0825 | -0.045 | -0.13 | 0.57137 | 0.770775 | 0.455413 | 1 | 0.996511 | 0.887169 |
| Dstn | Q9R0P5 | -0.05 | -1.61 | -0.56 | -1.03 | -0.68 | -0.35 | 0.23 | -0.62 | -1.62 | -0.27 | -0.44 | -0.41 | -1.26 | -1.86 | -0.59 | -0.37 | 0.4575 | 0.1275 | -0.2075 | 0.241065 | 0.768475 | 0.64549 | 1 | 0.996511 | 0.944749 |
| Dstyk | A0A087WF | 0.44 | 0.95 | -0.08 | -0.11 | -0.27 | 0.69 | -0.45 | -0.09 | 1.6 | 0.73 | -0.1 | 0.7 | 1.44 | 0.36 | 0.01 | -0.21 | -0.33 | 0.4325 | 0.1 | 0.348597 | 0.305446 | 0.813407 | 1 | 0.996511 | 0.972768 |
| Dtd1 | Q9DD18 | -1.03 | -1.31 | -0.28 | -0.52 | -1.76 | -1.12 | -1 | -0.62 | 1.88 | -0.18 | -0.69 | -1.49 | 3.34 | -1.54 | -0.8 | -1.07 | -0.34 | 0.665 | 0.7675 | 0.311375 | 0.347317 | 0.463932 | 1 | 0.996511 | 0.888267 |
| Dtd2 | Q8BHA3 | -0.51 | -0.89 | -0.99 | -1.27 | -1.18 | -0.59 | -0.51 | -1.29 | -0.3 | -0.82 | -1.07 | -0.83 | 0.37 | -1.31 | -1.11 | -0.98 | 0.0225 | 0.16 | 0.1575 | 0.931566 | 0.526147 | 0.693764 | 1 | 0.996511 | 0.955729 |
| Dtna | Q8CFR5 | 0.4 | -0.05 | 0.98 | 0.41 | 0.02 | 0.45 | 1.13 | 0.29 | 0.45 | 0.3 | 0.91 | 0.26 | 1.01 | -0.03 | 1.04 | 1.15 | 0.0375 | 0.045 | 0.3575 | 0.904166 | 0.869002 | 0.316565 | 1 | 0.996511 | 0.838625 |
| Dtnb | G3UXK0 | 0.91 | 0.65 | 0.08 | 0.5 | 0.44 | 0.89 | 0.28 | 0.15 | 0.52 | 0.89 | 0.42 | 1.28 | 0.35 | -0.41 | 0.5 | 0.26 | -0.095 | 0.2425 | -0.36 | 0.704233 | 0.389434 | 0.226976 | 1 | 0.996511 | 0.787447 |
| Dtnbp1 | A0A1Y7VIX | -0.94 | -1.76 | -3.1 | -2.47 | -1.76 | -0.53 | -3.96 | -2.83 | -0.62 | -1.21 | -2.12 | -1.22 | 0.48 | -0.48 | -1.62 | -3.37 | -0.2025 | 0.775 | 0.82 | 0.796306 | 0.160369 | 0.349648 | 1 | 0.996511 | 0.85327 |
| Dtx3 | D3YV13 | 0.34 | -0.23 | 0.87 | 0.66 | -0.05 | 0.85 | 1.02 | 0.95 | 0.02 | 3.7 | 0.99 | 0.69 | 0.62 | -0.54 | 0.47 | 0.95 | 0.2825 | 0.94 | -0.035 | 0.409594 | 0.239403 | 0.928142 | 1 | 0.996511 | 0.98946 |
| Dtymk | P97930 | 1.42 | 1.28 | 0.64 | 0.84 | 0.76 | 1.29 | 0.84 | 1.06 | -0.22 | -0.17 | 0.96 | 0.82 | 0.13 | -0.67 | 0.73 | 0.39 | -0.0575 | -0.6975 | -0.9 | 0.807359 | 0.076962 | 0.027722 | 1 | 0.996511 | 0.635993 |
| Dus2 | Q9D7B1 | -1.61 | -1.12 | -1.92 | -2.23 | -1.63 | -1.11 | -2.18 | -1.37 | -0.54 | -0.66 | -1.34 | -0.98 | -0.49 | -0.86 | 0.36 | -2.13 | 0.1475 | 0.84 | 0.94 | 0.648246 | 0.020487 | 0.102239 | 1 | 0.996511 | 0.700609 |
| Dus3l | A0A0R4IZY | -3.01 | -3.01 | -2.84 | -2.6 | -2.83 | -3.36 | -2.64 | -2.94 | -1.96 | -1.59 | -2.6 | -2.99 | -2.08 | -1.85 | -2.96 | -2.7 | -0.0775 | 0.58 | 0.4675 | 0.711782 | 0.104301 | 0.136884 | 1 | 0.996511 | 0.732318 |
| Dusp14 | Q9JLY7 | -0.03 | 0.34 | -0.17 | -0.39 | 0.79 | 0.1 | -0.47 | -0.27 | -0.18 | -0.61 | -0.38 | 0.1 | -1.68 | 0.47 | -0.77 | -1.11 | 0.1 | -0.205 | -0.71 | 0.748788 | 0.406438 | 0.143592 | 1 | 0.996511 | 0.742733 |
| Dusp15 | Q8R4V2 | 0.21 | -0.26 | 0.49 | 0.6 | -0.39 | 0.66 | 0.9 | 0.14 | 0.16 | 0.9 | 0.92 | 0.12 | -0.03 | 0.07 | 0.47 | -0.16 | 0.0675 | 0.265 | -0.1725 | 0.840145 | 0.387656 | 0.516758 | 1 | 0.996511 | 0.905676 |
| Dusp19 | Q99N12 | 0.76 | 0.57 | -0.49 | -0.19 | 0.39 | 0.15 | -1.4 | -0.29 | 0.58 | 0.46 | -0.43 | 0.68 | 0.76 | 0.59 | -0.16 | -0.6 | -0.45 | 0.16 | -0.015 | 0.34308 | 0.674337 | 0.97137 | 1 | 0.996511 | 0.99283 |
| Dusp23 | Q6NT99 | -0.42 | -0.25 | -0.42 | 0.37 | -0.71 | -0.09 | -0.08 | 0.31 | -0.47 | -0.15 | 0.98 | 0 | -0.95 | -0.32 | 0.12 | -0.03 | 0.0375 | 0.27 | -0.115 | 0.894932 | 0.455751 | 0.712692 | 1 | 0.996511 | 0.958106 |
| Dusp26 | Q9D700 | 0.13 | 0.4 | 0.36 | 0.61 | 0.42 | 0.51 | 0.44 | 0.15 | 0.63 | 0.73 | 0.44 | 0.63 | 0.58 | 1.16 | 0.44 | 0.6 | 0.005 | 0.2325 | 0.32 | 0.977184 | 0.220601 | 0.189644 | 1 | 0.996511 | 0.776843 |
| Dusp28 | Q8BTR5 | -0.1 | 0.17 | -0.14 | -0.09 | 0.6 | -0.24 | -0.56 | -0.2 | -0.21 | 0.42 | -0.56 | -0.19 | 0.08 | 0.64 | -0.33 | -0.78 | -0.06 | -0.095 | -0.0575 | 0.820521 | 0.695923 | 0.8563 | 1 | 0.996511 | 0.976225 |
| Dusp3 | B1AQF4 | -0.85 | -0.71 | -0.48 | -0.35 | -0.8 | -0.98 | -0.64 | -0.13 | -0.36 | -0.84 | -0.08 | -0.75 | -0.84 | -0.28 | -0.84 | -1 | -0.04 | 0.09 | -0.1425 | 0.863625 | 0.706035 | 0.548937 | 1 | 0.996511 | 0.912757 |
| Dusp6 | Q9DBB1 | -0.01 | -0.39 | 0.16 | -0.71 | -0.95 | 0.27 | 0.71 | -0.9 | -0.71 | -0.26 | -0.46 | 0.46 | -0.9 | 0.93 | 0.15 | -0.63 | 0.02 | -0.005 | 0.125 | 0.962992 | 0.987478 | 0.773589 | 1 | 0.999285 | 0.966062 |
| Dusp9 | Q7TNL7 | -5.15 | 1.46 | -4.08 | -2.18 | 0.31 | -5.12 | -5.89 | -4.36 | -1.97 | 0.46 | -4.76 | -4.82 | -1.69 | 1.21 | -5.09 | -6 | -1.2775 | -0.285 | -0.405 | 0.48314 | 0.868236 | 0.8354 | 1 | 0.996511 | 0.975524 |
| Dut | Q9CQ43-2 | 0.13 | -0.15 | -0.54 | -0.33 | -0.8 | -0.13 | -0.92 | -0.53 | 0.1 | -0.89 | -0.35 | -0.23 | 0.61 | -0.41 | 0.69 | -1.01 | -0.3725 | -0.12 | 0.1925 | 0.148279 | 0.654603 | 0.645926 | 1 | 0.996511 | 0.944749 |
| Dvl1 | P51141 | 0.09 | -0.08 | -0.22 | 0.03 | 0.3 | 0.26 | -0.09 | -0.12 | 0.29 | -0.24 | -0.21 | -0.03 | -0.17 | 0.2 | -0.14 | -0.19 | 0.1325 | -0.0025 | -0.03 | 0.463038 | 0.989689 | 0.875155 | 1 | 0.999285 | 0.979626 |
| Dvl2 | Q60838 | -1.71 | -1.59 | -1.29 | -1.42 | -2.53 | -1.3 | -2.73 | -1.6 | -1.11 | 0.1 | -1.55 | -1.82 | 0.56 | -2.52 | -2.09 | -1.85 | -0.5375 | 0.4075 | 0.0275 | 0.148191 | 0.338007 | 0.965543 | 1 | 0.996511 | 0.9927 |
| Dvl3 | E9Q150 | 0.35 | 0.6 | 0.74 | 1.22 | 0.56 | 0.46 | 0.43 | 0.9 | -0.17 | 1.04 | 0.94 | 0.41 | -0.15 | -1.23 | 0.7 | 0.37 | -0.14 | -0.1725 | -0.805 | 0.549131 | 0.604719 | 0.092793 | 1 | 0.996511 | 0.695968 |
| Dym | Q8CHY3 | -0.81 | -1.02 | -0.75 | -0.91 | -0.79 | -0.74 | -0.76 | -1.3 | -1.32 | -0.16 | -0.93 | -0.88 | -1.05 | -0.12 | -0.85 | -0.86 | -0.025 | 0.05 | 0.1525 | 0.893773 | 0.850886 | 0.543283 | 1 | 0.996511 | 0.912212 |
| Dync1h1 | Q9JHU4 | 0.45 | 0.03 | 0.32 | 0.37 | 0.22 | 0.38 | 0.54 | 0.51 | 0.28 | 0.09 | 0.58 | 0.61 | -0.25 | 0.34 | 0.12 | 0.19 | 0.12 | 0.0975 | -0.1925 | 0.486524 | 0.632319 | 0.375422 | 1 | 0.996511 | 0.859611 |
| Dync1i1 | O88485 | 0.25 | 0.21 | 0.33 | 0.5 | 0.01 | 0.11 | -0.19 | 0.36 | 0.84 | 1.03 | 0.33 | 0.56 | 0.72 | 0.77 | 0.53 | 0.08 | -0.25 | 0.3675 | 0.2025 | 0.184128 | 0.107632 | 0.370113 | 1 | 0.996511 | 0.859245 |
| Dync1i2 | Q3TPJ8 | -0.5 | -0.3 | -0.55 | -0.2 | -0.34 | -0.72 | -0.49 | -0.24 | -0.23 | -0.19 | -0.2 | -0.31 | -1.03 | 0.43 | -0.45 | -0.59 | -0.06 | 0.155 | -0.0225 | 0.738077 | 0.368159 | 0.94435 | 1 | 0.996511 | 0.991865 |
| Dync1li1 | Q8R1Q8 | 0 | -0.17 | 0.5 | 0.61 | -0.11 | -0.11 | 0.3 | 0.67 | -0.05 | 1.02 | 0.64 | 0.24 | -0.54 | 0.25 | 0.4 | 0.09 | -0.0475 | 0.2275 | -0.185 | 0.861267 | 0.463199 | 0.53526 | 1 | 0.996511 | 0.912212 |
| Dync1li2 | Q6PDL0 | -0.02 | -0.16 | -0.27 | -0.18 | -0.03 | -0.11 | -0.12 | -0.18 | 0.14 | -0.25 | 0 | 0.01 | 0.08 | 0.21 | -0.16 | -0.46 | 0.0475 | 0.1325 | 0.075 | 0.743604 | 0.449 | 0.724928 | 1 | 0.996511 | 0.958398 |
| Dync2h1 | Q45VK7 | -0.73 | -0.35 | -0.28 | -0.27 | 0.08 | -0.73 | -0.24 | -0.31 | -0.27 | 0.3 | -0.31 | 0.06 | -0.51 | 0.16 | -0.4 | -0.19 | 0.1075 | 0.3525 | 0.1725 | 0.63013 | 0.134258 | 0.458199 | 1 | 0.996511 | 0.887902 |
| Dync2i1 | Q8C761 | -1.07 | -1.09 | -1.02 | -0.81 | -1.53 | -1.15 | -1.91 | -1.56 | -0.95 | 0.59 | -0.71 | -1.16 | -0.72 | -0.84 | -1.18 | -0.99 | -0.54 | 0.44 | 0.065 | 0.023593 | 0.270014 | 0.736487 | 1 | 0.996511 | 0.958635 |
| Dync2i2 | Q5U4F6 | -0.48 | -0.36 | -0.07 | -0.01 | -0.43 | -0.63 | 0.49 | 0.18 | -0.48 | -0.17 | -0.01 | -0.6 | -0.04 | -0.2 | -0.18 | 0.01 | 0.1325 | -0.085 | 0.1275 | 0.644708 | 0.695485 | 0.519638 | 1 | 0.996511 | 0.907667 |
| Dync2li1 | Q8K0T2 | 3.86 | 3.65 | -2.21 | -1.72 | 2.07 | 2.74 | -1.66 | -2.2 | 4.62 | -1.4 | -2.04 | 3.58 | 2.82 | 4.45 | -2.48 | -1.78 | -0.6575 | 0.295 | -0.1425 | 0.723957 | 0.88888 | 0.945915 | 1 | 0.996511 | 0.992376 |
| Dynll1 | P63168 | 0.41 | 0.43 | -0.42 | -0.39 | -0.05 | 0.49 | -0.09 | -0.08 | -0.43 | -0.88 | -0.15 | 0.71 | -0.25 | -0.9 | -0.29 | -0.07 | 0.06 | -0.195 | -0.385 | 0.830335 | 0.622869 | 0.233503 | 1 | 0.996511 | 0.789665 |
| Dynll2 | Q9D0M5 | 1.37 | 0.68 | 0.46 | 0.68 | 0.32 | 1.51 | 1.01 | 1.1 | -0.13 | -0.1 | 0.54 | 1.29 | 0.22 | -0.57 | 0.77 | 0.66 | 0.1875 | -0.3975 | -0.5275 | 0.551075 | 0.305849 | 0.166092 | 1 | 0.996511 | 0.759345 |
| Dynlrb1 | P62627 | -0.46 | -0.14 | -0.09 | -0.14 | 0.02 | -0.57 | -0.2 | 0 | -0.35 | 0.12 | -0.08 | -0.16 | -0.8 | -0.02 | -0.96 | -0.1 | 0.02 | 0.09 | -0.2625 | 0.918735 | 0.634285 | 0.354631 | 1 | 0.996511 | 0.855029 |
| Dynlt1a | A0A338P7 | 0.53 | 0.31 | -2.71 | -1.94 | -0.21 | 0.14 | -2.3 | -2.32 | 1.73 | -2.74 | -1.85 | 1.39 | 1.33 | 1.66 | -2.43 | -2.93 | -0.22 | 0.585 | 0.36 | 0.814745 | 0.640066 | 0.781488 | 1 | 0.996511 | 0.967642 |
| Dynlt3 | P56387 | -2.03 | -2.62 | -6.21 | -5.18 | -2.54 | -2.16 | -4.15 | -4.72 | -2.28 | -6.68 | -5.25 | -1.59 | -2.5 | -1.69 | -5.41 | -6.56 | 0.6175 | 0.06 | -0.03 | 0.56243 | 0.965841 | 0.982414 | 1 | 0.996755 | 0.994858 |
| Dyrk1a | F6U6X3 | -0.04 | 0.03 | 0.5 | 0.07 | -0.06 | -0.12 | 0.6 | 0.55 | -0.19 | -0.17 | 0.64 | 0.04 | -0.4 | -0.05 | 0.58 | 0.3 | 0.1025 | -0.06 | -0.0325 | 0.674245 | 0.811694 | 0.90409 | 1 | 0.996511 | 0.985278 |
| Dyrk1b | Q9Z188-3 | 1.42 | 0.95 | 1.45 | 0.42 | 1.27 | 0.88 | 1.44 | 1.15 | -1.33 | -2.35 | 1.22 | 1.27 | -1.07 | -1.34 | 1.31 | 1.4 | 0.125 | -1.3575 | -0.985 | 0.64998 | 0.138776 | 0.187518 | 1 | 0.996511 | 0.776843 |
| Dyrk2 | Q5U4C9 | 0.73 | -0.07 | -0.92 | -0.63 | -0.52 | 1.21 | -0.77 | -0.48 | 0.37 | -1.12 | -0.17 | 0.87 | 0.16 | -1.12 | 1.61 | -0.41 | 0.0825 | 0.21 | 0.2825 | 0.877233 | 0.686875 | 0.652819 | 1 | 0.996511 | 0.947633 |
| Dzank1 | E9Q4Z6 | -0.12 | 0.25 | -0.02 | 0.04 | 0.25 | -0.05 | 0.35 | 0.07 | 0.24 | 0.23 | 0 | 0.22 | 0.83 | 0.78 | 0.29 | 0.37 | 0.1175 | 0.135 | 0.53 | 0.497831 | 0.441467 | 0.033159 | 1 | 0.996511 | 0.644622 |
| Dzip3 | E9QNZ2 | -0.81 | -0.7 | -1.16 | -1.16 | -0.6 | -0.27 | -1.22 | -1.22 | -0.35 | -0.72 | -0.98 | -0.68 | -0.19 | -0.25 | -1.23 | -1.15 | 0.13 | 0.275 | 0.2525 | 0.632823 | 0.223576 | 0.429342 | 1 | 0.996511 | 0.879165 |
| E2f3 | Q3TMJ9 | -3.24 | -2.67 | -2.52 | -2.57 | -3.88 | -3.9 | -2.15 | -2.7 | -3.44 | -1.74 | -2.93 | -1.78 | -4.15 | -2.16 | -1.84 | -1.63 | -0.4075 | 0.2775 | 0.305 | 0.363589 | 0.524766 | 0.58602 | 1 | 0.996511 | 0.928178 |
| E9Q0B3 | E9Q0B3 | 0.23 | 1.94 | -1.33 | -1.91 | 0.97 | -0.37 | -1.35 | -2.57 | 0.8 | -2.06 | -2.09 | 0.47 | -0.47 | 1.27 | -1.98 | -1.87 | -0.5625 | -0.4525 | -0.495 | 0.586603 | 0.66731 | 0.632282 | 1 | 0.996511 | 0.941132 |
| Ears2 | Q9CXJ1 | 0.37 | 0.35 | -0.04 | 0.05 | -0.06 | 0.39 | -0.07 | 0.04 | 0.15 | -0.16 | 0.04 | 0.18 | 0.05 | 0.04 | -0.5 | 0.01 | -0.1075 | -0.13 | -0.2825 | 0.573356 | 0.495728 | 0.221209 | 1 | 0.996511 | 0.787447 |
| Ebag9 | Q9D0V7 | -0.26 | -1.36 | -2.81 | -1 | -0.18 | -0.7 | -0.14 | -0.53 | -0.82 | -2.7 | -1.67 | -0.36 | -0.39 | 0.5 | -0.95 | -0.48 | 0.97 | -0.03 | 1.0275 | 0.084315 | 0.964587 | 0.09718 | 1 | 0.996755 | 0.698156 |
| Ebna1bp2 | Q9D903 | 1.02 | 0.91 | -0.83 | -0.98 | 0.56 | 0.69 | -1.24 | -1.05 | 0.23 | -1.66 | -0.97 | 0.48 | 0.67 | 0.59 | -0.77 | -1.07 | -0.29 | -0.51 | -0.175 | 0.669592 | 0.45689 | 0.785715 | 1 | 0.996511 | 0.967642 |
| Ebp | A2AC29 | -0.6 | -0.4 | -0.14 | -0.34 | 0.41 | -0.39 | 0.52 | 0.04 | -0.12 | 0.38 | -0.28 | -0.41 | 0.09 | 0.87 | 0.29 | 0.51 | 0.515 | 0.2625 | 0.81 | 0.058322 | 0.271193 | 0.006937 | 1 | 0.996511 | 0.552614 |
| Ebpl | Q9D0P0 | 0.59 | 0.04 | 0.05 | 0.51 | -1.11 | 1.62 | -0.91 | -0.11 | 2.09 | -0.07 | -0.1 | 0.14 | 2.95 | -0.66 | 0.72 | 0.46 | -0.425 | 0.2175 | 0.57 | 0.473021 | 0.67038 | 0.423637 | 1 | 0.996511 | 0.876113 |
| Ecd | Q9CS74 | -3.31 | -1.19 | -1.29 | -1.22 | -3.35 | -1.85 | -1.99 | -1.59 | -1.63 | 0.71 | -1.42 | -1.01 | -0.04 | -1.85 | -1.32 | -1.6 | -0.4425 | 0.915 | 0.55 | 0.464284 | 0.19989 | 0.372702 | 1 | 0.996511 | 0.859424 |
| Ece1 | Q4PZA2-3 | 0.38 | 0.59 | -0.17 | -0.14 | 0.54 | 0.2 | 0.48 | -0.03 | 0 | -0.71 | -0.28 | 0.51 | 0.7 | 0.05 | 0.13 | -0.06 | 0.1325 | -0.285 | 0.04 | 0.591637 | 0.38304 | 0.884893 | 1 | 0.996511 | 0.979767 |
| Ece2 | E9Q1S5 | -0.82 | 1.5 | 0.73 | 0.66 | 1.92 | 1.64 | 0.84 | 0.69 | 2.77 | 1.69 | 0.39 | -1.47 | 4.35 | -1.21 | 1.13 | 1.22 | 0.755 | 0.3275 | 0.855 | 0.174869 | 0.725414 | 0.447479 | 1 | 0.996511 | 0.884817 |
| Ecel1 | Q9JMI0 | 0.77 | 1.03 | 0.09 | 1.01 | 0.72 | 0.42 | 0.82 | 1.15 | 1.21 | 2.16 | 0.39 | 0.88 | 0.7 | 1.19 | 2.09 | 1.14 | 0.0525 | 0.435 | 0.555 | 0.846775 | 0.307439 | 0.149342 | 1 | 0.996511 | 0.748169 |
| Ech1 | O35459 | 0.79 | 0.71 | 0.69 | 0.71 | 0.61 | 0.79 | 0.88 | 0.52 | 0.58 | 0.24 | 0.74 | 0.58 | 1.07 | 0.41 | 0.51 | 0.8 | -0.025 | -0.19 | -0.0275 | 0.871305 | 0.299616 | 0.895531 | 1 | 0.996511 | 0.983509 |
| Echdc1 | Q9D9V3 | -0.01 | 0.2 | -0.79 | -0.4 | 0.63 | -0.05 | -0.47 | -0.56 | -0.3 | -0.9 | -0.84 | -0.13 | -0.62 | 0.25 | -0.79 | -0.48 | 0.1375 | -0.2925 | -0.16 | 0.6848 | 0.34012 | 0.621995 | 1 | 0.996511 | 0.938628 |
| Echdc2 | Q3TLP5 | 1 | 0.5 | 1.78 | 1.77 | 0.42 | -0.37 | 1.64 | 0.78 | 1.27 | 1.47 | 3 | 0.59 | -0.29 | -0.37 | 0.51 | 1.2 | -0.645 | 0.32 | -1 | 0.204338 | 0.565058 | 0.053191 | 1 | 0.996511 | 0.661738 |
| Echs1 | Q8BH95 | 0.89 | 0.87 | 0.79 | 1.23 | 0.57 | 0.8 | 1.2 | 0.91 | 0.61 | 0.3 | 0.93 | 0.55 | 0.83 | 0.86 | 0.61 | 1 | -0.075 | -0.3475 | -0.12 | 0.705116 | 0.120607 | 0.546048 | 1 | 0.996511 | 0.912321 |
| Eci1 | P42125 | 1.66 | 1.29 | 1.36 | 0.84 | 1.33 | 1.67 | 2 | 0.97 | 1.03 | 0.27 | 1.43 | 1.15 | 0.87 | 1.1 | 1.29 | 1.5 | 0.205 | -0.3175 | -0.0975 | 0.473257 | 0.313093 | 0.697543 | 1 | 0.996511 | 0.955729 |
| Eci2 | Q3TCD4 | 0.4 | 0.33 | 0.64 | 0.54 | 0.65 | 0.53 | 0.87 | 0.63 | 0.39 | 0.22 | 0.66 | 0.22 | 0.08 | 0.43 | 0.32 | 0.34 | 0.1925 | -0.105 | -0.185 | 0.251914 | 0.575432 | 0.332453 | 1 | 0.996511 | 0.845601 |
| Eci2 | E9PUY9 | 0.37 | 1.29 | 0.01 | -0.48 | 1.59 | -0.43 | 0.82 | -0.64 | -0.59 | -1.56 | -0.58 | 0.43 | 0.35 | 0.93 | -0.93 | 0.19 | 0.0375 | -0.8725 | -0.1625 | 0.949318 | 0.116293 | 0.747049 | 1 | 0.996511 | 0.960582 |
| Ecpas | Q6PDI5 | -0.95 | -0.54 | -1.09 | -1.14 | -0.47 | -0.83 | -1.18 | -1.19 | -0.64 | -0.75 | -1.08 | -0.84 | -0.82 | -0.43 | -1.28 | -1.39 | 0.0125 | 0.1025 | -0.05 | 0.957676 | 0.625888 | 0.858292 | 1 | 0.996511 | 0.976225 |
| Ecsit | J3KMM6 | -2.29 | -1.35 | -1.63 | -4.26 | -3.76 | -3.39 | -2.78 | -4.48 | -2.14 | -3.45 | -4.36 | -5.51 | -4.15 | -3.19 | -5.2 | -2.31 | -1.22 | -1.4825 | -1.33 | 0.099345 | 0.117377 | 0.129419 | 1 | 0.996511 | 0.723655 |
| Ecsit | A0A0R4J17 | -0.16 | -0.27 | -0.16 | 0.38 | -0.2 | 0.45 | -0.03 | -0.06 | 0.53 | -0.2 | 0.03 | -0.11 | 0.65 | 0.49 | -0.19 | -0.17 | 0.0925 | 0.115 | 0.2475 | 0.682446 | 0.640221 | 0.392224 | 1 | 0.996511 | 0.865775 |
| Edc3 | Q8K2D3 | -0.39 | -0.35 | -0.98 | -1.05 | -0.02 | -0.13 | -1 | -1.03 | -0.38 | 2.29 | -0.78 | -0.12 | -0.18 | -0.1 | -0.96 | -0.85 | 0.1475 | 0.945 | 0.17 | 0.649263 | 0.173823 | 0.577834 | 1 | 0.996511 | 0.927198 |
| Edc4 | G5E896 | -0.93 | -0.81 | -1.01 | -0.98 | -0.61 | -0.89 | -0.95 | -1.12 | -0.93 | -0.85 | -0.97 | -0.84 | -0.83 | -0.55 | -1.19 | -1.15 | 0.04 | 0.035 | 0.0025 | 0.812768 | 0.82345 | 0.990612 | 1 | 0.996511 | 0.996154 |
| Edem3 | A0A087WF | -0.63 | -0.54 | -0.55 | -0.14 | -0.42 | -0.36 | 0.09 | 0.02 | 0.14 | 0.8 | -0.18 | -0.53 | 0.43 | 0.44 | 0.27 | -0.03 | 0.2975 | 0.5225 | 0.7425 | 0.166572 | 0.116706 | 0.006517 | 1 | 0.996511 | 0.552614 |
| Edf1 | Q9JMG1 | -1.33 | 1.23 | -2.65 | -2.23 | -1.09 | -0.56 | -1.92 | -2.58 | -0.53 | -0.29 | -2.26 | -0.04 | -0.17 | 0.66 | -2.12 | -2.68 | -0.2925 | 0.465 | 0.1675 | 0.740026 | 0.610044 | 0.873211 | 1 | 0.996511 | 0.979626 |
| Edil3 | Q8C4U8 | 0.18 | -0.02 | 0.86 | 0.61 | -0.04 | -0.02 | 1.08 | 1.02 | -0.16 | 1.38 | 0.56 | -0.19 | -0.3 | -0.03 | 1.15 | 0.99 | 0.1025 | -0.01 | 0.045 | 0.773435 | 0.980143 | 0.910167 | 1 | 0.998773 | 0.986458 |
| Ednrb | P48302 | 0.15 | 0.42 | 0.14 | -0.02 | 0.41 | 0.78 | 0.56 | 0.23 | 0.74 | -0.45 | 0.04 | 0.53 | 1.13 | 0.22 | 1.04 | 0.29 | 0.3225 | 0.0425 | 0.4975 | 0.113716 | 0.883424 | 0.101441 | 1 | 0.996511 | 0.698697 |
| Edrf1 | D3YZE7 | -2.8 | -6.51 | 1.27 | 1.3 | -5.83 | -3.07 | 1.59 | 1.74 | -2.7 | 1.54 | 1.31 | -4.32 | 1.32 | -4.27 | 0.24 | 0.77 | 0.2925 | 0.6425 | 1.2 | 0.900787 | 0.762082 | 0.554665 | 1 | 0.996511 | 0.916796 |
| Eea1 | Q8BL66 | -0.46 | -1.11 | -0.68 | -0.68 | -0.79 | -0.85 | -0.35 | -0.52 | -1.09 | -0.58 | -0.66 | -0.85 | -1.15 | -0.68 | -0.33 | -0.56 | 0.105 | -0.0625 | 0.0525 | 0.616756 | 0.773128 | 0.835278 | 1 | 0.996511 | 0.975524 |
| Eed | A0A5F8MP | 0.01 | 0.46 | -0.07 | -0.79 | 0.69 | 0.53 | 0.17 | -1.42 | -0.38 | -1.04 | -0.66 | 0.27 | -0.49 | 0.85 | 0.35 | -0.47 | 0.09 | -0.355 | 0.1575 | 0.858603 | 0.347074 | 0.696534 | 1 | 0.996511 | 0.955729 |
| Eef1a1 | P10126 | -0.78 | -1.85 | -1.27 | -1.24 | -1.54 | -0.6 | -0.67 | -0.83 | -1.4 | -0.74 | -1.11 | -0.98 | -1.35 | -1.6 | -1.19 | -1.34 | 0.375 | 0.2275 | -0.085 | 0.238201 | 0.414646 | 0.746905 | 1 | 0.996511 | 0.960533 |
| Eef1a2 | P62631 | -1.73 | -1.81 | -1.14 | -1.57 | -1.61 | -1.67 | -1.15 | -0.87 | -2.36 | -2.52 | -1.04 | -1.55 | -2.28 | -2.24 | -1.22 | -1.33 | 0.2375 | -0.305 | -0.205 | 0.361284 | 0.416163 | 0.534731 | 1 | 0.996511 | 0.912212 |
| Eef1akmt1 | A0A0D2X7 | -0.77 | -0.88 | -0.71 | -0.76 | -0.88 | -1.3 | -0.68 | -0.33 | -0.84 | 0.19 | -0.82 | -1.14 | -0.75 | -0.56 | -0.73 | -0.88 | -0.0175 | 0.1275 | 0.05 | 0.938338 | 0.67097 | 0.775479 | 1 | 0.996511 | 0.966799 |
| Eef1akmt2 | Q9D853 | 0.99 | 0.11 | 0.76 | -0.15 | 0.49 | 0.33 | 0.79 | 0.66 | -0.66 | -0.19 | 0.46 | 0.74 | -0.92 | -0.74 | 0.35 | 0.15 | 0.14 | -0.34 | -0.7175 | 0.628017 | 0.401801 | 0.100289 | 1 | 0.996511 | 0.698156 |
| Eef1b | O70251 | -3.49 | -3.9 | -3.42 | -3.29 | -2.99 | -4.27 | -2.91 | -3.61 | -3.93 | -2.07 | -3.29 | -3.72 | -3.53 | -2.3 | -3.35 | -3.5 | 0.08 | 0.2725 | 0.355 | 0.810034 | 0.516624 | 0.292323 | 1 | 0.996511 | 0.825497 |
| Eef1d | Q91VK2 | 0.69 | 0.24 | -0.69 | -0.15 | -0.63 | -0.55 | 0.18 | 0.86 | -0.15 | 1.4 | 1.57 | 1.95 | -0.07 | -0.96 | -0.27 | -1.1 | -0.0575 | 1.17 | -0.6225 | 0.892801 | 0.044347 | 0.126393 | 1 | 0.996511 | 0.721505 |
| Eef1d | A0A0R4J1E | -2.56 | -2.8 | -3.37 | -3.33 | -2.5 | -2.69 | -3.23 | -3.34 | -2.35 | -1.99 | -3.21 | -2.46 | -2.03 | -1.62 | -3.57 | -3.81 | 0.075 | 0.5125 | 0.2575 | 0.793865 | 0.140815 | 0.634712 | 1 | 0.996511 | 0.941377 |
| Eef1d | Q80T06 | -1.61 | -3.26 | -1.08 | -0.55 | -1.63 | -1.41 | -1.13 | -0.55 | -1.45 | -0.62 | -0.7 | -1.78 | -1.67 | -1.36 | -1.1 | -1.47 | 0.445 | 0.4875 | 0.225 | 0.448496 | 0.42381 | 0.684873 | 1 | 0.996511 | 0.955729 |
| Eef1e1 | Q9D1M4 | -2.15 | -2.45 | -2.19 | -2.73 | -1.57 | -1.96 | -1.59 | -2.13 | -3.41 | -2.62 | -2.06 | -1.84 | -3.06 | -2 | -1.49 | -1.88 | 0.5675 | -0.1025 | 0.2725 | 0.026956 | 0.778782 | 0.45324 | 1 | 0.996511 | 0.885993 |
| Eef1g | Q9D8N0 | -1.32 | -2.34 | -2.19 | -2.07 | -2.16 | -1.66 | -2.16 | -2.1 | -1.3 | -1.3 | -2 | -1.22 | -1.41 | -1.02 | -1.94 | -2.12 | -0.04 | 0.525 | 0.3575 | 0.879735 | 0.105656 | 0.307417 | 1 | 0.996511 | 0.832533 |
| Eef2 | P58252 | 0.67 | 0.47 | -0.95 | -1.05 | 0.61 | 0.69 | -0.87 | -0.93 | -1.56 | -0.67 | -0.9 | 0.75 | -1.16 | -1.54 | -1.12 | -1.38 | 0.09 | -0.38 | -1.085 | 0.877369 | 0.538074 | 0.034311 | 1 | 0.996511 | 0.65013 |
| Eefsec | Q9JHW4 | -1.21 | -1 | -0.49 | -0.84 | -0.74 | -1.27 | -0.44 | -0.29 | -1.3 | 0.17 | -0.46 | -1 | -1.09 | -0.65 | -0.27 | -0.3 | 0.2 | 0.2375 | 0.3075 | 0.466237 | 0.503131 | 0.271503 | 1 | 0.996511 | 0.822084 |
| Eepd1 | Q3TGW2 | 0.82 | 0.98 | 1.22 | 1.15 | 0.87 | 0.73 | 1.22 | 1.28 | 1 | 1.01 | 1.23 | 0.87 | 0.91 | 0.76 | 0.65 | 1.26 | -0.0175 | -0.015 | -0.1475 | 0.928812 | 0.934075 | 0.498314 | 1 | 0.996755 | 0.898799 |
| Efcab14 | A2A8T9 | 0.69 | 0.56 | -1.15 | -0.82 | 0.79 | 0.5 | -1.07 | -0.85 | -1.42 | -0.62 | -0.81 | 0.91 | -0.73 | -0.16 | -0.39 | -0.66 | 0.0225 | -0.305 | -0.305 | 0.970332 | 0.627618 | 0.512689 | 1 | 0.996511 | 0.904116 |
| Efemp1 | Q8BPB5 | -2.04 | -1.29 | -2.93 | -2.5 | -3.04 | -1.94 | -1.42 | -3.38 | -1.01 | -0.38 | -1 | -2.16 | -1.63 | -1.71 | -1.41 | -0.33 | -0.255 | 1.0525 | 0.92 | 0.633215 | 0.051939 | 0.066431 | 1 | 0.996511 | 0.664936 |
| Efhd1 | Q9D4J1 | 1.1 | 1.53 | 0.35 | 0.2 | 1.23 | 0.69 | -0.45 | -0.29 | 0.92 | 0.47 | 0.33 | 1.02 | 1.18 | 1.27 | -0.06 | 0.44 | -0.5 | -0.11 | -0.0875 | 0.305636 | 0.753421 | 0.836993 | 1 | 0.996511 | 0.975831 |
| Efhd2 | Q8C845 | 0.63 | 0.72 | 1 | 1.02 | 0.8 | 0.37 | 0.38 | 1.13 | 0.58 | 0.82 | 0.77 | 1.03 | 0.16 | 0.76 | 0.22 | 0.42 | -0.1725 | -0.0425 | -0.4525 | 0.457187 | 0.824219 | 0.064582 | 1 | 0.996511 | 0.66433 |
| Efl1 | Q8C0D5 | -0.15 | -0.31 | -0.25 | -0.58 | -0.02 | -0.26 | -0.29 | -0.35 | -0.12 | -0.01 | -0.16 | -0.19 | -0.16 | 0.12 | 0.23 | -0.59 | 0.0925 | 0.2025 | 0.2225 | 0.589418 | 0.262204 | 0.370287 | 1 | 0.996511 | 0.859245 |
| Efna3 | O08545 | 1.41 | 1.11 | 2.25 | 1.25 | 2.41 | 0.48 | 1.72 | 1.37 | 1.34 | 0.98 | 1.56 | 1.15 | 1.12 | 2.24 | 1.29 | 2 | -0.01 | -0.2475 | 0.1575 | 0.982 | 0.407783 | 0.668434 | 1 | 0.996511 | 0.95204 |
| Efna5 | O08543 | 0.27 | 1.25 | -0.49 | -0.68 | 1.32 | 0.83 | -0.55 | -0.73 | 0.67 | 0.22 | -0.58 | 0.41 | -0.03 | 1.84 | -0.32 | -0.38 | 0.13 | 0.0925 | 0.19 | 0.831395 | 0.84739 | 0.761864 | 1 | 0.996511 | 0.963322 |
| Efnb1 | P52795 | 0.39 | 0.82 | 1.11 | 1.27 | 0.79 | 0.53 | 1.15 | 1.18 | 0.39 | 0.56 | 1.34 | 0.75 | 0.48 | 0.48 | 0.78 | 0.83 | 0.015 | -0.1375 | -0.255 | 0.953489 | 0.639125 | 0.320788 | 1 | 0.996511 | 0.839916 |
| Efnb2 | P52800 | -0.65 | -0.36 | 1.01 | 1.36 | -0.51 | -0.11 | 0.49 | 0.88 | 0.14 | 0.78 | 1.06 | 0.04 | 0.47 | -0.88 | 0.98 | 0.93 | -0.1525 | 0.165 | 0.035 | 0.776702 | 0.748987 | 0.953738 | 1 | 0.996511 | 0.992476 |
| Efnb3 | O35393 | -0.48 | -0.78 | -0.72 | -0.22 | -1.03 | -0.62 | -0.24 | -0.9 | -0.77 | 0.03 | -0.91 | -0.51 | -0.55 | -0.77 | -0.54 | -0.5 | -0.1475 | 0.01 | -0.04 | 0.533651 | 0.969523 | 0.844897 | 1 | 0.996755 | 0.976225 |
| Efr3a | A0A1D5RLI | 0.54 | 0.98 | 1.15 | 1.09 | 0.89 | 0.82 | 0.94 | 0.89 | 0.36 | 0.57 | 0.9 | 0.73 | 0.09 | 0.63 | 0.43 | 0.97 | -0.055 | -0.3 | -0.41 | 0.764316 | 0.192555 | 0.140374 | 1 | 0.996511 | 0.736961 |
| Efr3b | Q6ZQ18 | -0.05 | 0.12 | 1.03 | 0.75 | 0.44 | 0.1 | 0.84 | 0.84 | -0.16 | -0.26 | 0.88 | 0.05 | -0.3 | 0.33 | 0.91 | 1.08 | 0.0925 | -0.335 | 0.0425 | 0.764094 | 0.358746 | 0.913417 | 1 | 0.996511 | 0.986458 |
| Efs | A0A2I3BRP | -0.58 | 1.01 | 1.82 | 0.05 | -1 | -0.8 | -0.77 | -0.96 | 2.95 | 2.15 | -1.53 | -0.87 | 0.01 | 0.25 | 0.9 | 2.28 | -1.4575 | 0.1 | 0.285 | 0.015252 | 0.92729 | 0.670847 | 1 | 0.996755 | 0.952558 |
| Eftud2 | A2AH85 | 0.17 | -0.79 | -0.56 | -0.6 | -0.69 | -0.34 | -0.47 | 0.03 | -0.97 | -1.4 | -0.59 | -1.02 | -0.23 | -0.6 | 0.28 | -0.68 | 0.0775 | -0.55 | 0.1375 | 0.771434 | 0.076815 | 0.66191 | 1 | 0.996511 | 0.949645 |
| Egfr | Q01279 | 1.12 | 1.22 | 0.97 | 0.93 | 1.45 | 1.36 | 0.57 | 0.73 | -0.27 | 0.75 | 0.78 | 1.46 | -0.43 | -0.39 | 0.59 | 0.7 | -0.0325 | -0.38 | -0.9425 | 0.894404 | 0.300317 | 0.015415 | 1 | 0.996511 | 0.589387 |
| Egln1 | Q91YE3 | -1.59 | -1.73 | -1.79 | -1.51 | -1.66 | -1.56 | -2.55 | -2.19 | -1.61 | -0.5 | -1.84 | -1.48 | -0.31 | -1.87 | -0.87 | -2.06 | -0.335 | 0.2975 | 0.3775 | 0.207041 | 0.344941 | 0.362407 | 1 | 0.996511 | 0.858475 |
| Egr3 | Q3ZB14 | -0.77 | -1.1 | -0.53 | -0.72 | -1.45 | -1.87 | 0.15 | -1.21 | -1.67 | -1.25 | -0.64 | -0.86 | -1.23 | -1.71 | 0.09 | -1.77 | -0.315 | -0.325 | -0.375 | 0.464796 | 0.25027 | 0.3907 | 1 | 0.996511 | 0.865375 |
| Ehbp1 | Q69ZW3 | 0.11 | 0.4 | 0.13 | 0.34 | 0.16 | 0.29 | -0.17 | 0.06 | 0.45 | 0.37 | 0.15 | 0.4 | 0.75 | 0.34 | 0 | -0.19 | -0.16 | 0.0975 | -0.02 | 0.365936 | 0.576482 | 0.936664 | 1 | 0.996511 | 0.989767 |
| Ehbp1l1 | Q99MS7 | 0.63 | -0.3 | -2.8 | -2.63 | -0.58 | 0.01 | -2.35 | -3.19 | 0.02 | -2.08 | -2.8 | 0.17 | -0.22 | 0.01 | -1.58 | -3.4 | -0.2525 | 0.1025 | -0.0225 | 0.804181 | 0.919786 | 0.982646 | 1 | 0.996755 | 0.994982 |
| Ehd1 | Q9WVK4 | 0.45 | 0.5 | 0.45 | 0.47 | 0.78 | 0.07 | 0.64 | 0.36 | -0.05 | 0.37 | 0.49 | 0.42 | -0.58 | 0.46 | 0.15 | 0.37 | -0.005 | -0.16 | -0.3675 | 0.97942 | 0.395979 | 0.187001 | 1 | 0.996511 | 0.776843 |
| Ehd2 | Q8BH64 | -3.15 | -3.64 | -3.88 | -4.01 | -2.85 | -3.36 | -3.71 | -4.54 | -3.73 | -2.76 | -3.52 | -2.84 | -3.41 | -2.95 | -4.02 | -4.06 | 0.055 | 0.4575 | 0.06 | 0.885478 | 0.166335 | 0.855592 | 1 | 0.996511 | 0.976225 |
| Ehd3 | Q9QXY6 | 1.51 | 1.32 | 1.48 | 0.98 | 1.5 | 1.57 | 1.61 | 1.61 | 0.53 | 0.31 | 1.42 | 1.61 | 0.12 | 0.68 | 1.19 | 1.34 | 0.25 | -0.355 | -0.49 | 0.175516 | 0.311128 | 0.141718 | 1 | 0.996511 | 0.737973 |
| Ehd4 | Q9EQP2 | 0.07 | 0.05 | -0.06 | 0.26 | -0.02 | -0.1 | 0.22 | 0.33 | -0.04 | 0.24 | 0.12 | 0 | -0.62 | 0.21 | -0.44 | -0.13 | 0.0275 | 1.47E-17 | -0.325 | 0.872857 | 1 | 0.191093 | 1 | 1 | 0.777117 |
| Ehhadh | Q9DBM2 | 0.29 | 0.26 | -1.09 | -0.18 | 0.27 | 0.22 | -0.6 | -0.42 | 0.42 | -1.71 | -1.03 | 0.43 | 0.59 | 0.01 | -0.84 | -0.92 | 0.0475 | -0.2925 | -0.11 | 0.898314 | 0.613595 | 0.8096 | 1 | 0.996511 | 0.972113 |
| Ehmt1 | E9Q5A3 | -0.44 | -0.91 | 0.26 | 0.23 | -1.09 | -1.27 | 0.39 | 0.77 | -1.27 | 1.01 | 0.77 | -1.05 | -0.31 | 0.11 | 1.65 | 0.69 | -0.085 | 0.08 | 0.75 | 0.874702 | 0.894565 | 0.143088 | 1 | 0.996511 | 0.742711 |
| Ehmt2 | A2CG76 | -1.26 | -0.98 | -0.65 | -0.67 | -0.65 | 0.35 | -1.54 | -1.08 | -0.49 | -0.13 | -0.53 | -0.36 | -0.25 | 0.31 | 0.45 | -0.42 | 0.16 | 0.5125 | 0.9125 | 0.692908 | 0.036741 | 0.009035 | 1 | 0.996511 | 0.563386 |
| Ei24 | A0A0R4J25 | 0.08 | 0.08 | 0.23 | 0.78 | 0.56 | 0.01 | 0.37 | 0.37 | 0.2 | 0.62 | 0.38 | -0.1 | 0.32 | 0.57 | 0.98 | -0.08 | 0.035 | -0.0175 | 0.155 | 0.875796 | 0.943889 | 0.599821 | 1 | 0.996755 | 0.930673 |
| Eid2 | Q6X7S9 | -0.91 | -0.77 | -0.93 | -0.78 | -2.01 | -1.02 | -2.49 | -1.56 | 0.14 | 0.32 | -0.55 | -1.1 | 1.94 | -0.66 | -0.82 | -0.75 | -0.9225 | 0.55 | 0.775 | 0.015445 | 0.120952 | 0.229108 | 1 | 0.996511 | 0.787447 |
| Eif1 | P48024 | -1.84 | -1.41 | -0.05 | 0.46 | -1.05 | -1.48 | 0.61 | 0.61 | -2.39 | 0.11 | 0.2 | -1.45 | -1.73 | -1.12 | -0.02 | 0.46 | 0.3825 | -0.1725 | 0.1075 | 0.588233 | 0.819286 | 0.873083 | 1 | 0.996511 | 0.979626 |
| Eif1ad | Q3THJ3 | -2.65 | -2.16 | -2.28 | -2.43 | -2.16 | -2.64 | -1.95 | -2.67 | -2.47 | -0.64 | -1.57 | -1.73 | -2.47 | -1.91 | -2.73 | -1.9 | 0.025 | 0.7775 | 0.1275 | 0.91253 | 0.0655 | 0.627759 | 1 | 0.996511 | 0.939936 |
| Eif1ax | Q8BMJ3 | -1.77 | -2.27 | -4.15 | -5.52 | -1.63 | -1.52 | -4.38 | -4.67 | -0.92 | -2.3 | -4 | -1.73 | 1.05 | -1.7 | -1.08 | -4.72 | 0.3775 | 1.19 | 1.815 | 0.729641 | 0.242368 | 0.189737 | 1 | 0.996511 | 0.776843 |
| Eif1b | Q9CXU9 | -2.62 | -2.24 | -2.42 | -2.46 | -1.25 | -1.62 | -2.4 | -1.9 | -2.08 | -3.75 | -1.73 | -1.7 | -1.25 | -0.04 | -4.48 | -1.48 | 0.6425 | 0.12 | 0.6225 | 0.035085 | 0.795125 | 0.471522 | 1 | 0.996511 | 0.891144 |
| Eif2a | Q8BJW6 | -1.31 | -1.63 | -1.21 | -1.51 | -1.47 | -1.41 | -1.31 | -1.35 | -1.6 | -1.25 | -1.29 | -1.27 | -1.41 | -0.91 | -0.84 | -1.93 | 0.03 | 0.0625 | 0.1425 | 0.852919 | 0.737973 | 0.622998 | 1 | 0.996511 | 0.938628 |

| | | | | | | | | | | | | | | | | | | | | | | | | | | | |
|---|---|---|---|---|---|---|---|---|---|---|---|---|---|---|---|---|---|---|---|---|---|---|---|---|---|---|---|
| Eif2ak2 | Q03963 | -0.98 | -2.36 | -0.28 | 0.02 | -2.63 | -1.22 | 0.79 | 0.32 | 0.36 | -0.5 | -0.05 | -1.6 | 0.31 | -0.16 | 0.31 | 1.07 | 0.215 | 0.4525 | 1.2825 | 0.799856 | 0.472442 | 0.040092 | | 1 | 0.996511 | 0.658454 |
| Eif2ak3 | Q9Z2B5 | -0.46 | -0.04 | -0.69 | -0.18 | -0.21 | -0.1 | -0.35 | -0.22 | 0.75 | -0.7 | -1.09 | -0.05 | 1.03 | 0.67 | -0.56 | -0.3 | 0.1225 | 0.07 | 0.5525 | 0.52738 | 0.863741 | 0.184741 | | 1 | 0.996511 | 0.776843 |
| Eif2ak4 | A2AUM0 | -0.44 | -0.37 | -0.54 | -0.24 | -0.36 | -0.39 | -0.86 | -0.5 | -0.42 | -0.02 | -0.22 | -0.61 | -0.31 | -0.23 | -0.88 | -0.41 | -1.30E-01 | 0.08 | -0.06 | 0.471583 | 0.683008 | 0.779447 | | 1 | 0.996511 | 0.967289 |
| Eif2b1 | Q99LC8 | 0.15 | 0.05 | -0.13 | -0.07 | -0.06 | 0.14 | 0.06 | -0.06 | 0.01 | -0.36 | -0.37 | 0.02 | -0.09 | 0.16 | -0.63 | -0.23 | 0.02 | -0.175 | -0.1975 | 0.895226 | 0.360468 | 0.389939 | | 1 | 0.996511 | 0.865375 |
| Eif2b2 | Q99LD9 | -1.41 | -1.3 | -0.6 | -0.8 | -1.85 | -1.33 | -0.58 | -0.67 | -0.08 | -0.64 | -0.75 | -1.47 | 0.62 | -1.4 | -0.97 | -1.17 | -0.08 | 0.2925 | 0.2975 | 0.816159 | 0.401005 | 0.52997 | | 1 | 0.996511 | 0.908663 |
| Eif2b3 | B1AUN2 | -0.24 | -0.53 | -0.14 | -0.65 | -0.5 | -0.62 | -0.13 | -0.32 | -0.97 | -0.06 | -0.15 | -0.31 | -0.84 | -1.47 | -0.4 | -0.09 | -0.0025 | 0.0175 | -0.31 | 0.989807 | 0.9459 | 0.35541 | | 1 | 0.996755 | 0.855029 |
| Eif2b4 | D3Z780 | -0.67 | -0.64 | -0.47 | -0.17 | -0.83 | -0.7 | -0.59 | -0.32 | -0.62 | 0.67 | -0.2 | -0.69 | -0.36 | -0.34 | -0.46 | -0.45 | -0.1225 | 0.2775 | 0.085 | 0.532595 | 0.41115 | 0.661035 | | 1 | 0.996511 | 0.949645 |
| Eif2b5 | Q8CHW4 | 0.41 | -0.21 | 0.83 | 1.45 | 0.16 | 0.09 | 1.09 | 1.26 | -0.29 | 1.76 | 0.85 | 0.1 | -0.67 | -0.38 | 0.57 | 0.9 | 0.03 | -0.015 | -0.515 | 0.944799 | 0.977338 | 0.299636 | | 1 | 0.997923 | 0.829161 |
| Eif2d | Q61211 | -0.72 | -0.84 | -1.47 | -1.28 | -0.62 | -1.04 | -1.08 | -1.36 | -0.51 | -1.07 | -1.2 | -0.88 | -0.4 | -0.33 | -1.17 | -0.95 | 0.0525 | 0.1625 | 0.365 | 0.831952 | 0.527639 | 0.227964 | | 1 | 0.996511 | 0.787447 |
| Eif2s1 | Q6ZWX6 | -0.3 | -1.38 | -1.37 | -1.33 | -0.94 | -0.56 | -1.02 | -1.31 | -0.96 | -0.34 | -1.28 | -0.66 | -1.05 | -0.34 | -1.04 | -1.23 | 0.1375 | 0.285 | 0.18 | 0.652511 | 0.399202 | 0.591498 | | 1 | 0.996511 | 0.929755 |
| Eif2s2 | Q99L45 | -1.16 | -1.66 | -0.38 | -0.24 | -1.3 | -1.02 | -0.48 | -0.35 | -1.26 | 0.71 | -0.58 | -1.3 | -1.09 | -1.28 | 0.09 | -0.22 | 0.0725 | 0.2525 | 0.235 | 0.849549 | 0.637749 | 0.601093 | | 1 | 0.996511 | 0.930802 |
| Eif2s3x | Q9Z0N1 | -1.36 | -2.71 | -1.12 | -1.22 | -2.03 | -1.35 | -0.72 | -1.01 | -2.37 | -0.82 | -1.2 | -1.59 | -2.17 | -1.98 | -0.79 | -0.8 | 0.325 | 0.1075 | 0.1675 | 0.463597 | 0.818001 | 0.733907 | | 1 | 0.996511 | 0.958635 |
| Eif2s3y | Q9Z0N2 | -1.17 | 0.84 | -0.04 | -1.32 | -0.6 | -0.05 | -0.89 | 0.26 | -0.27 | -2.12 | -0.38 | 0.87 | 0.27 | 0.68 | 0.32 | -1.92 | 0.1025 | -0.0525 | 0.26 | 0.845283 | 0.942266 | 0.714375 | | 1 | 0.996755 | 0.958106 |
| Eif3a | P23116 | -1.3 | -2.32 | -1.46 | -1.34 | -1.59 | -1.58 | -1.04 | -1.49 | -1.36 | -0.76 | -1.47 | -1.49 | -0.89 | -1.21 | -0.51 | -0.88 | 0.18 | 0.335 | 0.7325 | 0.521606 | 0.284088 | 0.032687 | | 1 | 0.996511 | 0.644622 |
| Eif3b | Q8JZQ9 | -1.82 | -2.27 | -1.3 | -1.12 | -1.59 | -1.74 | -0.7 | -1.07 | -2.04 | -1.06 | -1.03 | -1.78 | -1.77 | -1.32 | -0.64 | -0.83 | 0.3525 | 0.15 | 0.4875 | 0.317583 | 0.675058 | 0.197934 | | 1 | 0.996511 | 0.78004 |
| Eif3c | Q8R1B4 | -0.61 | -1.41 | -0.99 | -1.04 | -0.85 | -0.78 | -0.37 | -0.91 | -1.36 | -1.08 | -0.98 | -0.91 | -1.17 | -0.76 | -0.53 | -0.56 | 0.285 | -0.07 | 0.2575 | 0.227506 | 0.75678 | 0.322833 | | 1 | 0.996511 | 0.840788 |
| Eif3d | O70194 | -1.31 | -1.49 | -1.61 | -1.69 | -0.74 | -1.53 | -0.44 | -1.58 | -1.91 | -1.61 | -1.4 | -1.27 | -2.04 | -1.11 | -1.24 | -0.96 | 0.4525 | -0.0225 | 0.1875 | 0.152569 | 0.913423 | 0.50362 | | 1 | 0.996755 | 0.901002 |
| Eif3e | P60229 | -1.38 | -1.88 | -1.36 | -1.48 | -1.03 | -1.44 | -0.82 | -1.42 | -1.8 | -1.24 | -1.24 | -1.31 | -1.6 | -1.02 | -0.82 | -1.11 | 0.3475 | 0.1275 | 0.3875 | 0.13869 | 0.564672 | 0.13693 | | 1 | 0.996511 | 0.732318 |
| Eif3f | Q9DCH4 | -1.29 | -1.84 | -1.43 | -1.23 | -0.93 | -1.47 | -0.7 | -1.31 | -1.32 | -1.02 | -1.3 | -1.36 | -1.3 | -0.7 | -0.88 | -0.9 | 0.345 | 0.1975 | 0.5025 | 0.173768 | 0.34722 | 0.053946 | | 1 | 0.996511 | 0.661738 |
| Eif3g | Q9Z1D1 | -1.96 | -2.18 | -1.54 | -1.67 | -1.57 | -2.02 | -1.41 | -1.77 | -1.64 | -0.76 | -1.61 | -1.81 | -1.23 | -1.14 | -1.14 | -1.52 | 0.145 | 0.3825 | 0.58 | 0.513428 | 0.2047 | 0.025752 | | 1 | 0.996511 | 0.633075 |
| Eif3h | Q91WK2 | -1.69 | -2.1 | -1.45 | -1.51 | -1.71 | -1.85 | -1.15 | -1.74 | -1.35 | -1.32 | -1.43 | -1.84 | -0.4 | -1.59 | -0.31 | -1.26 | 0.075 | 0.2025 | 0.7975 | 0.748196 | 0.378956 | 0.044453 | | 1 | 0.996511 | 0.658454 |
| Eif3i | Q9QZD9 | -1.54 | -1.93 | -1.52 | -1.36 | -1.57 | -1.81 | -0.81 | -1.42 | -1.37 | -1.51 | -1.27 | -1.58 | -0.92 | -1.27 | -0.66 | -0.82 | 0.185 | 0.155 | 0.67 | 0.475795 | 0.432445 | 0.014672 | | 1 | 0.996511 | 0.589387 |
| Eif3j1 | Q3UGC7 | -0.96 | -1.15 | -1.04 | -0.98 | -1.08 | -1.37 | -1.19 | -0.8 | -1.11 | -0.38 | -0.8 | -0.87 | -1.09 | -0.36 | -1.28 | -1.32 | -0.0775 | 0.2425 | 0.02 | 0.660149 | 0.254867 | 0.938017 | | 1 | 0.996511 | 0.989844 |
| Eif3k | Q9DBZ5 | -0.95 | -0.6 | -1.54 | -1.07 | -0.82 | -0.9 | -1.02 | -1.61 | 0.13 | -0.52 | -1.41 | -0.91 | -0.48 | 0.91 | -1.47 | -1.18 | -0.0475 | 0.3625 | 0.485 | 0.860297 | 0.337765 | 0.366923 | | 1 | 0.996511 | 0.858904 |
| Eif3l | Q8QZY1 | -0.71 | -1.34 | -1.3 | -1.26 | -0.08 | -0.47 | -0.64 | -1.21 | -2.1 | -1.02 | -1.11 | -0.86 | -1.65 | -1.65 | -0.67 | -0.59 | 0.5525 | -0.12 | 0.0125 | 0.076022 | 0.707097 | 0.969998 | | 1 | 0.996511 | 0.992767 |
| Eif3m | Q99JX4 | -3.9 | -4.14 | -1.87 | -2.08 | -2.97 | -3.95 | -1.14 | -1.99 | -4.04 | -1.62 | -2.12 | -3.99 | -3.98 | -2.95 | -1.85 | -1.55 | 0.485 | 0.055 | 0.415 | 0.532097 | 0.943868 | 0.575848 | | 1 | 0.996755 | 0.926728 |
| Eif4a1 | P60843 | -0.53 | -0.56 | -1.38 | -1.51 | -0.5 | -0.8 | -1.34 | -1.6 | -0.55 | -0.72 | -1.44 | -0.16 | -0.21 | 0.24 | -1.64 | -1.6 | -0.065 | 0.2775 | 0.1925 | 0.851887 | 0.453315 | 0.706363 | | 1 | 0.996511 | 0.957084 |
| Eif4a2 | P10630 | -1.48 | -1.66 | -1.38 | -1.63 | -1.26 | -1.49 | -1.01 | -1.51 | -1.99 | -1.53 | -1.31 | -1.47 | -1.68 | -0.97 | -1.78 | -1.6 | 0.22 | -0.0375 | 0.03 | 0.241192 | 0.855347 | 0.898634 | | 1 | 0.996511 | 0.983509 |
| Eif4a3 | Q91VC3 | 0.02 | -0.53 | -0.39 | -0.21 | -1 | -0.56 | -0.6 | 0.05 | -1.17 | -0.03 | -0.41 | -1.25 | -0.53 | -0.32 | -0.23 | -0.69 | -0.25 | -0.4375 | -0.165 | 0.344062 | 0.195131 | 0.447421 | | 1 | 0.996511 | 0.884817 |
| Eif4b | Q8BGD9 | 0.25 | -0.09 | -0.67 | -0.49 | -0.66 | 0.03 | -1.13 | -0.93 | 0.98 | 0.13 | -0.91 | 0.31 | 1.39 | 0 | -0.62 | -1.14 | -0.4225 | 0.3775 | 0.1575 | 0.208759 | 0.37992 | 0.770894 | | 1 | 0.996511 | 0.965933 |
| Eif4e | P63073 | -0.39 | -0.42 | -0.85 | -0.32 | -0.48 | -0.53 | -0.44 | -0.65 | -0.83 | -0.52 | -0.5 | -0.35 | -1.19 | -0.06 | -1.03 | -0.95 | -0.03 | -0.055 | -0.3125 | 0.864888 | 0.787503 | 0.306345 | | 1 | 0.996511 | 0.832533 |
| Eif4e2 | D3Z730 | -0.96 | -1.18 | -0.33 | -0.73 | -1.59 | -0.91 | -1.15 | -1.01 | 0.39 | -0.43 | -0.53 | -0.98 | 1.72 | -0.9 | -0.33 | -0.43 | -0.365 | 0.4125 | 0.815 | 0.16689 | 0.238759 | 0.171646 | | 1 | 0.996511 | 0.763858 |
| Eif4ebp2 | P70445 | -0.89 | -1.09 | -0.27 | -0.44 | -1.15 | -1.31 | -0.21 | -0.27 | -0.9 | -0.1 | -0.27 | -0.8 | -1.18 | -1.04 | -0.47 | -0.02 | -0.0625 | 0.155 | -0.005 | 0.851932 | 0.588965 | 0.987958 | | 1 | 0.996511 | 0.996154 |
| Eif4enif1 | Q9EST3 | -0.53 | -0.02 | -0.22 | 0.18 | -0.11 | -0.63 | 0.01 | -0.58 | 0.71 | -0.35 | -0.23 | 0.55 | -0.57 | 0.33 | -0.34 | -0.22 | -0.18 | 0.3175 | -0.0525 | 0.457603 | 0.322455 | 0.845187 | | 1 | 0.996511 | 0.976225 |
| Eif4g1 | Q6NZJ6 | -1.27 | -1.59 | -1.03 | -1.27 | -1.32 | -1.54 | -1.42 | -1.34 | -1.34 | -0.78 | -1.11 | -1.37 | -1.02 | -0.96 | -1.28 | -1.46 | -0.115 | 0.14 | 0.11 | 0.515185 | 0.524117 | 0.614338 | | 1 | 0.996511 | 0.935876 |
| Eif4g1 | D6RCH1 | -2.3 | -1.29 | -4.13 | -2.36 | -2.73 | -2.37 | -4.72 | -6.42 | -0.99 | -0.85 | -3.23 | -1.75 | 0.06 | -2.32 | -3.1 | -4.93 | -1.54 | 0.815 | -0.0525 | 0.148925 | 0.281424 | 0.96058 | | 1 | 0.996511 | 0.992693 |
| Eif4g2 | Q62448 | -0.78 | -1.4 | -0.73 | -0.75 | -1.03 | -0.9 | -0.74 | -0.85 | -0.74 | -0.56 | -0.87 | -0.78 | -0.72 | -0.64 | -0.7 | -0.8 | 0.035 | 0.1775 | 0.2 | 0.863651 | 0.418002 | 0.370397 | | 1 | 0.996511 | 0.859245 |
| Eif4g2 | G3XA17 | -0.93 | -1.23 | -1.68 | -0.56 | -1.27 | -1.59 | -0.66 | -1.07 | -0.91 | -0.88 | -1.42 | -0.65 | -2.02 | -0.46 | -1.28 | -0.82 | -0.0475 | 0.135 | -0.045 | 0.875572 | 0.648794 | 0.909759 | | 1 | 0.996511 | 0.986458 |
| Eif4g3 | Z4YKC4 | -1.54 | -2.08 | -1.52 | -1.49 | -2.75 | -2.28 | -2.05 | -1.76 | -0.82 | -1.43 | -1.73 | -1.18 | -1.67 | -1.59 | -1.8 | -2.55 | -0.5525 | 0.3675 | -0.245 | 0.059748 | 0.180138 | 0.394574 | | 1 | 0.996511 | 0.866125 |
| Eif4g3 | A0A0N4SV | -1.82 | -2.01 | -1.47 | -1.44 | -1.66 | -2.03 | -1.71 | -1.8 | -1.66 | -0.66 | -1.34 | -1.66 | -1.55 | -1.38 | -1.97 | -2.02 | -0.115 | 0.355 | -0.045 | 0.561369 | 0.232661 | 0.854557 | | 1 | 0.996511 | 0.976225 |
| Eif4h | Q9WUK2 | 0.92 | -0.31 | 0.72 | 1.2 | -1.15 | 0.11 | -0.15 | 1.09 | 0.67 | 1.39 | 0.88 | 1.07 | 0.77 | -0.43 | 0.46 | 0.07 | -0.6575 | 0.37 | -0.415 | 0.22809 | 0.31249 | 0.318467 | | 1 | 0.996511 | 0.838969 |
| Eif5 | P59325 | -0.36 | -0.5 | -1.35 | -1.54 | -0.21 | -0.5 | -1.53 | -1.42 | -0.57 | -1.51 | -1.52 | -0.25 | -0.54 | -0.11 | -1.47 | -1.23 | 0.0225 | -0.025 | 0.1 | 0.95685 | 0.952312 | 0.808881 | | 1 | 0.996755 | 0.972098 |
| Eif5a | P63242 | 1.7 | -0.04 | -0.24 | -0.37 | 0.19 | 1.19 | -0.1 | -0.14 | -0.35 | 0.44 | -0.25 | 1.65 | -0.59 | -0.54 | -0.13 | -0.49 | 0.0225 | 0.11 | -0.7 | 0.965996 | 0.857291 | 0.158542 | | 1 | 0.996511 | 0.753756 |
| Eif5a2 | Q8BGY2 | -4.89 | -4.97 | -5.07 | -5.62 | -4.5 | -5.26 | -6.24 | -5.22 | -5.75 | -2.34 | -5.37 | -4.34 | -5.19 | -2.91 | -4.69 | -5.65 | -0.1675 | 0.6875 | 0.5275 | 0.656315 | 0.3463 | 0.368505 | | 1 | 0.996511 | 0.859245 |
| Eif5b | Q05D44 | -0.98 | -1.07 | -0.81 | -1.37 | -0.85 | -1.05 | -0.55 | -1.1 | -1.31 | -0.99 | -0.97 | -1.18 | -1.27 | -1.17 | -0.57 | -0.68 | 0.17 | -0.055 | 0.135 | 0.411675 | 0.778698 | 0.586219 | | 1 | 0.996511 | 0.928178 |
| Eif6 | O55135 | -1.42 | -0.99 | -0.67 | -0.89 | -0.96 | -1.49 | -0.48 | -0.71 | -0.87 | -0.69 | -0.75 | -1.22 | -1.32 | -0.93 | -1.29 | -0.48 | 0.0825 | 0.11 | -0.0125 | 0.762806 | 0.633826 | 0.963616 | | 1 | 0.996511 | 0.9927 |
| Eipr1 | Q8K0G5 | -0.02 | -0.5 | 0.01 | -0.16 | -0.43 | -0.1 | -0.79 | -0.27 | -0.18 | 0.78 | -0.35 | 0.03 | -0.29 | -0.11 | -0.68 | -0.5 | -0.23 | 0.2375 | -0.2275 | 0.299054 | 0.4151 | 0.317921 | | 1 | 0.996511 | 0.838969 |
| Elac1 | Q8VEB6 | -0.68 | -0.29 | -0.99 | -1.56 | -0.41 | -0.35 | -0.94 | -1.71 | -0.85 | -1.59 | -1.44 | -0.28 | -0.4 | -0.6 | -1.06 | -1.81 | 0.0275 | -0.16 | -0.0875 | 0.94391 | 0.680185 | 0.825961 | | 1 | 0.996511 | 0.973155 |
| Elac2 | B1ATP7 | -1.09 | -1.48 | -1.1 | -1.32 | -1.23 | -0.58 | -1.27 | -1.26 | -1.17 | -0.56 | -1.06 | -1.18 | -1 | -0.66 | -1.44 | -1.12 | 0.1625 | 0.255 | 0.1925 | 0.462462 | 0.254095 | 0.413556 | | 1 | 0.996511 | 0.874945 |
| Elapor1 | A0A0A0MC | -2.57 | -3.28 | -3.42 | -3.29 | -3.13 | -2.46 | -3.11 | -3.52 | -2.51 | -2.21 | -3.6 | -3.16 | -0.9 | -2.56 | -3.24 | -3.24 | 0.085 | 0.27 | 0.655 | 0.771295 | 0.459507 | 0.245727 | | 1 | 0.996511 | 0.802442 |
| Elapor2 | Q3UZV7-2 | -1.69 | -2.47 | -1.78 | -1.88 | -1.97 | -2.04 | -1.63 | -1.93 | -2.53 | -1.43 | -2.01 | -2.05 | -1.91 | -2.59 | -2.13 | -1.95 | 0.0625 | -0.05 | -0.19 | 0.777623 | 0.864757 | 0.476404 | | 1 | 0.996511 | 0.892174 |
| Elavl1 | P70372 | 0.28 | -0.51 | -0.02 | 0.29 | -0.6 | -0.24 | -0.05 | 0.59 | -0.77 | -0.07 | -0.11 | -0.92 | -0.02 | -0.27 | 0.47 | -0.37 | -0.085 | -0.4775 | -0.0575 | 0.782691 | 0.13421 | 0.839699 | | 1 | 0.996511 | 0.976219 |
| Elavl2 | B1AXZ4 | -1.91 | -2.77 | -1.4 | -1.37 | -2.04 | -1.85 | -0.86 | -1.07 | -3.3 | -1.56 | -1.46 | -2.37 | -2.48 | -2.43 | -0.77 | -0.95 | 0.4075 | -0.31 | 0.205 | 0.334585 | 0.540143 | 0.697197 | | 1 | 0.996511 | 0.955729 |
| Elavl3 | Q60900 | 0.33 | -0.3 | -1.05 | -0.93 | -0.37 | 0.06 | -0.67 | -0.67 | -0.48 | -1.04 | -1.11 | -0.24 | -0.07 | 0.04 | -0.56 | -1.04 | 0.075 | -0.23 | 0.08 | 0.829638 | 0.539549 | 0.837823 | | 1 | 0.996511 | 0.975961 |
| Elavl4 | Q61701-4 | -1.01 | -0.91 | -0.21 | 0.02 | -1.44 | -1.18 | -0.35 | -0.18 | -1.16 | 0.13 | -0.34 | -1.08 | -0.56 | -0.82 | 0.36 | 0.03 | -0.26 | -0.085 | 0.28 | 0.499538 | 0.825407 | 0.450465 | | 1 | 0.996511 | 0.884817 |
| Elavl4 | A2A9R5 | -0.39 | 0.79 | 1.64 | -0.1 | 0.66 | -1.35 | 1.69 | 0.88 | -0.21 | 0.02 | 1.12 | -0.26 | -0.63 | 0.59 | 0.56 | 2.06 | -0.015 | -0.3175 | 0.16 | 0.983298 | 0.546328 | 0.806675 | | 1 | 0.996511 | 0.971715 |
| Elfn1 | Q8C8T7 | 1.17 | 0.96 | 0.94 | 1.24 | 0.93 | 1.31 | 0.64 | 0.73 | 0.68 | 1.18 | 1.14 | 1.02 | 0.48 | 0.86 | 1.25 | 1.01 | -0.175 | -0.0725 | -0.1775 | 0.391814 | 0.706412 | 0.439383 | | 1 | 0.996511 | 0.883205 |
| Elfn2 | Q68FM6 | 0.8 | 0.72 | 0.86 | 0.81 | 0.49 | 1.01 | 0.95 | 0.79 | 1.1 | 0.49 | 1.04 | 0.9 | 1.45 | 0.55 | 0.96 | 0.63 | 0.0125 | 0.085 | 0.1 | 0.941846 | 0.663194 | 0.682581 | | 1 | 0.996511 | 0.955729 |
| Elmo1 | Q8BPU7 | 0.15 | -0.14 | -0.12 | -0.03 | -0.25 | -0.01 | -0.36 | -0.24 | 0.12 | -0.51 | -0.25 | -0.1 | 0.42 | -0.28 | -0.15 | -0.25 | -0.18 | -0.15 | -0.03 | 0.280307 | 0.457467 | 0.893757 | | 1 | 0.996511 | 0.983068 |
| Elmo2 | Q8BHL5-2 | 1.02 | 0.48 | 0.75 | 0.41 | 0.84 | 1.05 | 0.55 | 0.61 | 0.63 | 0 | 0.77 | 0.97 | 0.65 | 0.49 | 0.39 | 0.26 | 0.0975 | -0.0725 | -0.2175 | 0.642471 | 0.786922 | 0.327721 | | 1 | 0.996511 | 0.844684 |
| Elmod1 | E9Q0K9 | 0.04 | 0.03 | 0.79 | 0.85 | 0.32 | 0.02 | 1.21 | 1.04 | 0.1 | 0.17 | 0.76 | 0.16 | -0.05 | 0.62 | 0.62 | 0.67 | 0.22 | -0.13 | 0.0375 | 0.534298 | 0.650393 | 0.899949 | | 1 | 0.996511 | 0.983509 |
| Elmod2 | A0A1B0GR | 0.27 | 0.45 | 0.12 | 0.16 | 0.34 | 0.32 | -0.34 | 0.05 | -1.49 | 0.18 | -0.12 | 0.51 | -1.34 | -1.45 | -0.03 | 0.13 | -0.1575 | -0.48 | -0.9225 | 0.450798 | 0.274061 | 0.047397 | | 1 | 0.996511 | 0.658454 |
| Eln | P54320 | -5.33 | -5.71 | -2.59 | -2.88 | -5.67 | -5.55 | -2.03 | -3.19 | -5.92 | -2.53 | -2.79 | -4.79 | -4.66 | -6.22 | -2.83 | -2.87 | 0.0175 | 0.12 | -0.0175 | 0.9871 | 0.907231 | 0.986371 | | 1 | 0.996744 | 0.996154 |
| Eloa | Q8CB77 | -1.48 | -3.24 | -2.05 | -1.6 | -2.51 | -1.39 | -1.15 | -1.13 | -0.21 | -1.22 | -1.03 | -1.59 | 0.74 | -2.23 | -1.49 | -1.61 | 0.5475 | 1.08 | 0.945 | 0.271926 | 0.043002 | 0.196251 | | 1 | 0.996511 | 0.78004 |
| Elob | P62869 | 0.92 | 0.1 | 0.4 | 0.65 | -0.16 | 0.51 | 0.58 | 0.66 | 0.82 | 0.25 | 0.42 | 1.2 | 0.45 | -0.33 | 0.41 | 0.31 | -0.12 | 0.155 | -0.3075 | 0.651916 | 0.590522 | 0.281296 | | 1 | 0.996511 | 0.822084 |
| Eloc | A0A087WC | -0.25 | -0.31 | -0.47 | -0.22 | -0.37 | -0.35 | -0.03 | -0.12 | -1.36 | -1.05 | -0.22 | -0.29 | -1.06 | -1.34 | -0.25 | -0.64 | 0.095 | -0.4175 | -0.51 | 0.560887 | 0.181228 | 0.086005 | | 1 | 0.996511 | 0.691619 |
| Elovl1 | Q9JLJ5 | -0.22 | 0.88 | -0.43 | -0.54 | -0.14 | 0.94 | -0.36 | -0.64 | 1.41 | -2.67 | 0.19 | 0.07 | 3.21 | -0.97 | 0.34 | 0.39 | 0.0275 | -0.1725 | 0.82 | 0.95044 | 0.835753 | 0.34496 | | 1 | 0.996511 | 0.84949 |
| Elovl4 | Q9EQC4 | 0.11 | 0.94 | 0 | -0.19 | 1.22 | 0.48 | 0.06 | 0.4 | 0.11 | -0.84 | 0.07 | 0.19 | 0.83 | 0.66 | 1.04 | 0.61 | 0.325 | -0.3325 | 0.57 | 0.349364 | 0.344999 | 0.072656 | | 1 | 0.996511 | 0.668066 |
| Elovl5 | D3Z0Z6 | 0.59 | 0.59 | 0.72 | 0.08 | -0.19 | 0.55 | 1.12 | 1.04 | 2.12 | 1.01 | 0.88 | 0.61 | 1.98 | 0.71 | 0.11 | 0.99 | 0.135 | 0.66 | 0.4525 | 0.678654 | 0.08943 | 0.275932 | | 1 | 0.996511 | 0.822084 |
| Elp1 | Q7TT37 | -0.91 | -1.1 | -0.89 | -0.77 | -0.77 | -1.03 | -0.54 | -0.79 | -1 | -0.64 | -0.81 | -1.01 | -0.95 | -0.64 | -0.87 | -0.78 | 0.135 | 0.0525 | 0.1075 | 0.44138 | 0.769495 | 0.559805 | | 1 | 0.996511 | 0.918629 |
| Elp2 | Q91WG4 | -0.38 | -0.34 | -1.16 | -1.53 | -0.06 | -0.34 | -0.92 | -1.33 | -0.83 | -1.16 | -1.19 | -0.39 | -0.79 | -0.6 | -1.19 | -1.37 | 0.19 | -0.04 | -0.135 | 0.626839 | 0.907067 | 0.695041 | | 1 | 0.996744 | 0.955729 |
| Elp3 | Q9CZX0-2 | -0.91 | -0.82 | -0.55 | -0.8 | -0.29 | -1.08 | -0.43 | -0.87 | -0.26 | 0.66 | -0.8 | -0.87 | -0.63 | -0.2 | -0.7 | -0.89 | 0.1025 | 0.4525 | 0.165 | 0.647195 | 0.222414 | 0.455828 | | 1 | 0.996511 | 0.887169 |
| Elp4 | Q9ER73 | 0.26 | -0.27 | -0.27 | -0.09 | -0.1 | 0.12 | -0.41 | -0.31 | 0.27 | -0.42 | -0.24 | 0.08 | 0.12 | 0.21 | -0.3 | -0.41 | -0.0825 | 0.015 | -0.0025 | 0.685766 | 0.948174 | 0.991597 | | 1 | 0.996755 | 0.996154 |
| Elp5 | Q99L85 | -0.82 | -0.69 | -0.75 | -0.8 | -0.68 | -0.9 | -0.7 | -1.09 | -0.6 | 0.01 | -1.03 | -0.82 | -0.75 | 0.27 | -0.8 | -0.87 | -0.0775 | 0.155 | 0.2275 | 0.633415 | 0.539399 | 0.439358 | | 1 | 0.996511 | 0.883205 |
| Elp6 | Q8BK75 | 0.04 | 0.74 | -0.44 | -0.84 | 0.61 | 0.14 | -0.33 | -0.77 | 0.24 | -0.77 | -0.36 | -0.01 | -0.21 | 0.38 | -1.47 | -0.82 | 0.0375 | -0.1 | -0.405 | 0.929329 | 0.797099 | 0.416986 | | 1 | 0.996511 | 0.875537 |
| Emb | P21995 | 0.65 | 1.13 | 0.77 | 0.54 | 1.62 | 1.05 | 0.42 | 0.45 | 0.15 | -0.41 | 0.95 | 0.98 | 0.28 | 0.4 | 0.73 | 0.93 | 0.1125 | -0.355 | -0.1875 | 0.715718 | 0.327958 | 0.437515 | | 1 | 0.996511 | 0.883161 |
| Emc1 | Q8C7X2 | -0.32 | 0.09 | -0.51 | -0.66 | 0.73 | -0.02 | 0.2 | -0.43 | -0.17 | -0.88 | -0.5 | -0.11 | -0.11 | 0.64 | -0.09 | 0.21 | 0.47 | -0.065 | 0.5125 | 0.133653 | 0.802546 | 0.080363 | | 1 | 0.996511 | 0.683877 |
| Emc1 | Q8C7X2-2 | -1.54 | -0.35 | -0.18 | -0.42 | 0.34 | -1.05 | 1.12 | -0.58 | -1.64 | -1.66 | -0.72 | -1.64 | -2.07 | -0.74 | -0.26 | 0.65 | 0.58 | -0.7925 | 0.0175 | 0.288836 | 0.059836 | 0.976421 | | 1 | 0.996511 | 0.993727 |
| Emc10 | A0A0X1KG | -0.1 | 0.36 | 0.13 | 0.41 | 0.18 | -0.1 | 0.1 | 0.11 | 0.13 | 0.6 | -0.04 | 0.34 | 0.17 | 0.78 | 1.03 | 0.34 | -0.1275 | 0.0575 | 0.38 | 0.480808 | 0.793078 | 0.167451 | | 1 | 0.996511 | 0.759484 |
| Emc2 | Q9CRD2 | -0.35 | -0.22 | -0.22 | -0.3 | 0.34 | -0.28 | 0.29 | -0.36 | -0.53 | 0.07 | -0.39 | -0.35 | 0.03 | 0.22 | 0.06 | 0.33 | 0.27 | -0.0275 | 0.4325 | 0.227369 | 0.884959 | 0.034004 | | 1 | 0.996511 | 0.647005 |
| Emc3 | Q99KI3 | 1.62 | 0.05 | 0.36 | 0.47 | 0.74 | 1.86 | 1.94 | 0.83 | -0.36 | 0.37 | 0.29 | 1.37 | -0.35 | 0.01 | 1.13 | 1.36 | 0.7175 | -0.2075 | -0.0875 | 0.130129 | 0.656641 | 0.862069 | | 1 | 0.996511 | 0.977204 |
| Emc4 | Q9CZX9 | -0.61 | 0.02 | -0.63 | -0.72 | 0.7 | -0.23 | -0.36 | -0.76 | -0.83 | 0.4 | -0.87 | 0.08 | -0.44 | 0.18 | -0.05 | -0.12 | 0.3225 | 0.18 | 0.3775 | 0.356594 | 0.614882 | 0.153321 | | 1 | 0.996511 | 0.750763 |
| Emc6 | Q9CQW0 | -0.8 | -1.03 | -1.3 | -1.18 | -0.34 | -0.69 | -0.46 | -1.17 | -0.69 | -0.49 | -1.29 | -0.76 | -0.7 | 0.16 | -0.36 | -0.55 | 0.4125 | 0.27 | 0.715 | 0.102438 | 0.264736 | 0.017777 | | 1 | 0.996511 | 0.602211 |
| Emc7 | Q9EP72 | -0.54 | -0.5 | -0.73 | -0.75 | 0.5 | -0.7 | -0.02 | -0.58 | -0.06 | -0.29 | -0.74 | -0.44 | 0.25 | 0.63 | 0.23 | 0.05 | 0.43 | 0.2475 | 0.92 | 0.157823 | 0.244006 | 0.001438 | | 1 | 0.996511 | 0.419079 |
| Emc8 | O70378 | 0.12 | 0.19 | -0.2 | 0.6 | 0.44 | 0.19 | 0.44 | 0.56 | 0.48 | -0.25 | -0.04 | 0.28 | 0.72 | 0.23 | 0.4 | 0.65 | 0.23 | -0.06 | 0.3225 | 0.290354 | 0.812872 | 0.200284 | | 1 | 0.996511 | 0.78004 |
| Emd | O08579 | 1.67 | 1.96 | 1.46 | 1.25 | 2.11 | 1.34 | 1.36 | 1.22 | 0.8 | 0.25 | 1.24 | 1.46 | 0.91 | 1.07 | 1.84 | 1.99 | -0.0775 | -0.6475 | -0.1325 | 0.767584 | 0.062707 | 0.67782 | | 1 | 0.996511 | 0.953944 |
| Emg1 | O35130 | 0.47 | 1.15 | -0.64 | 0.22 | 0.01 | 0.01 | -0.11 | 0.07 | 0.74 | -0.76 | -0.59 | 0.76 | 0.82 | 0.97 | -0.65 | -0.71 | -0.305 | -0.2625 | -0.1925 | 0.402421 | 0.612253 | 0.723531 | | 1 | 0.996511 | 0.958398 |
| Emilin1 | Q99K41 | -1.08 | -0.94 | -2.83 | -2.76 | 0.17 | -0.7 | -3.62 | -2.97 | -0.72 | -0.68 | -2.69 | -0.31 | -0.88 | -0.53 | -3.44 | -2.71 | 0.1225 | 0.8025 | 0.0125 | 0.895589 | 0.256592 | 0.987317 | | 1 | 0.996511 | 0.996154 |
| Eml1 | D3Z4J9 | -0.32 | -0.16 | 0.01 | 0.08 | -0.37 | -0.73 | -0.71 | -0.74 | 0.01 | 0.17 | -0.07 | -0.07 | -0.03 | 0.31 | -0.28 | -0.25 | -0.54 | 0.1075 | 0.035 | 0.012667 | 0.546334 | 0.871521 | | 1 | 0.996511 | 0.979293 |
| Eml2 | Q7TNG5-3 | -0.48 | -0.58 | -0.43 | -0.37 | -0.37 | -0.37 | -0.05 | -0.45 | -0.25 | 0.16 | -0.29 | -0.21 | 0.48 | 0.1 | -0.21 | -0.32 | 0.155 | 0.3175 | 0.4775 | 0.348422 | 0.104765 | 0.063197 | | 1 | 0.996511 | 0.66433 |
| Eml3 | Q8VC03 | 0.21 | 1.07 | 0.26 | 0.66 | 0.27 | -0.02 | 0.13 | 0.66 | 0.93 | -0.11 | 0.19 | 0.15 | 1.45 | 0.58 | -0.22 | -0.09 | -0.29 | -0.26 | -0.12 | 0.278059 | 0.403737 | 0.772149 | | 1 | 0.996511 | 0.965933 |
| Eml4 | A7ISP9 | 0.01 | 0.44 | -0.03 | 0.34 | 0.67 | 0.42 | 0.47 | 0.5 | 0.49 | 1.07 | 0.62 | 0.05 | -0.31 | 0.52 | -0.83 | -0.05 | 0.325 | 0.3675 | -0.3575 | 0.093756 | 0.181838 | 0.271521 | | 1 | 0.996511 | 0.822084 |
| Eml4 | F8WJ93 | -0.76 | -0.36 | -0.26 | -0.46 | -0.36 | -0.5 | 0.16 | -0.36 | -0.18 | -0.07 | -0.16 | -0.29 | -0.19 | -0.68 | -0.31 | -0.17 | 0.195 | 0.285 | 0.1225 | 0.364217 | 0.143343 | 0.572066 | | 1 | 0.996511 | 0.924918 |
| Eml5 | Q8BQM8 | 0.29 | 1.91 | 0.28 | 0.94 | 0.61 | 0.19 | 0.67 | 1.14 | -0.03 | 1.52 | 0.5 | 0.49 | 0.01 | 0.06 | 0.61 | 0.63 | -0.2025 | -0.235 | -0.5275 | 0.619754 | 0.61989 | 0.213755 | | 1 | 0.996511 | 0.786931 |
| Eml6 | Q5SQM0 | 0.17 | 0.38 | 0.25 | 0.61 | 0.52 | 0.05 | 0.64 | 0.13 | 0.11 | 0.7 | 0.44 | 0.14 | 0.01 | 0.11 | 0.67 | 0.46 | -0.0175 | -0.005 | -0.04 | 0.93169 | 0.981104 | 0.860268 | | 1 | 0.998773 | 0.977204 |
| Emp2 | O88662 | 0.71 | -0.23 | 0.43 | -0.98 | -1.05 | 2.2 | -1.61 | -0.07 | 3.84 | -4.23 | 0.1 | 1.67 | 4.95 | -4.88 | 0.73 | -0.5 | -0.115 | 0.3625 | 0.0925 | 0.889666 | 0.816389 | 0.959494 | | 1 | 0.996511 | 0.992636 |
| Emsy | Q8BMB0-2 | -1.66 | 0.77 | 1.08 | 0.57 | -2.1 | -1.71 | 0.05 | -0.12 | 0.7 | 1.09 | 0.47 | -0.6 | 3.62 | -1.1 | 0.6 | 0.51 | -1.16 | 0.225 | 0.7175 | 0.148858 | 0.733396 | 0.497234 | | 1 | 0.996511 | 0.897918 |
| Enah | A0A0A6YW | 1.09 | 0.94 | 1.59 | 1.91 | 0.97 | 0.71 | 1.2 | 1.87 | 1.28 | 2.03 | 1.82 | 1.06 | 1.17 | 1.16 | 1.6 | 1.74 | -0.195 | 0.165 | 0.035 | 0.553865 | 0.607946 | 0.902806 | | 1 | 0.996511 | 0.984938 |
| Enc1 | O35709 | -3.25 | -2.89 | -3.24 | -4.41 | -2.62 | -3.14 | -3.43 | -3.69 | -3.24 | -3.69 | -4.71 | -2.34 | -2.17 | -1.78 | -4.89 | -3.54 | 0.2275 | -0.0475 | 0.3525 | 0.555985 | 0.930863 | 0.621028 | | 1 | 0.996755 | 0.938628 |
| Endod1 | Q8C522 | 1.14 | 0.8 | 0.99 | 0.53 | 1.12 | 0.96 | 1.12 | 0.75 | 0.71 | 1.05 | 0.65 | 0.92 | 0.33 | 0.69 | 0.96 | 1.14 | 0.1225 | -0.0325 | -0.085 | 0.533045 | 0.874791 | 0.736964 | | 1 | 0.996511 | 0.958635 |
| Endog | O08600 | 0.81 | 0.88 | 0.34 | 0.53 | 1.07 | 0.57 | 0.46 | 0.3 | 0.63 | 0.36 | 0.33 | 0.76 | -0.01 | 0.94 | 0.01 | 0.13 | -0.04 | -0.12 | -0.3725 | 0.860888 | 0.567171 | 0.205173 | | 1 | 0.996511 | 0.783452 |
| Endou | Q3V188 | -0.35 | 0.04 | -2.92 | -3.92 | -0.22 | 0.04 | -2.6 | -3.46 | 0.54 | -1.47 | -3.07 | -0.02 | -0.4 | 1.99 | -1.03 | -1.83 | 0.2275 | 0.7825 | 1.47 | 0.845024 | 0.494741 | 0.216472 | | 1 | 0.996511 | 0.787447 |
| Eng | F7A1B4 | -0.41 | -0.96 | -1.27 | -1.16 | -0.65 | 0.01 | -1.91 | -0.96 | -0.12 | -0.75 | -0.89 | -0.65 | 0.39 | -1.36 | -1.03 | -0.66 | 0.0725 | 0.3475 | 0.285 | 0.861452 | 0.220208 | 0.491206 | | 1 | 0.996511 | 0.897531 |
| Engase | Q8BX80 | -0.42 | -0.93 | -0.25 | -1.09 | -0.72 | -0.13 | -0.42 | -0.89 | -1.15 | -0.62 | 0.03 | -0.5 | -0.99 | -0.68 | -0.74 | -0.07 | 0.1325 | 0.1125 | 0.0525 | 0.623217 | 0.722483 | 0.858666 | | 1 | 0.996511 | 0.976225 |
| Enho | Q8K1D8 | 0.46 | 1.35 | 1.46 | 1.56 | 1.85 | -0.25 | 1.5 | 1.73 | 0.7 | 2.06 | 1.46 | 0.66 | 0.65 | 0.74 | 1.6 | 0.99 | 1.57E-16 | 0.0125 | -0.2125 | 1 | 0.975112 | 0.528931 | | 1 | 0.997515 | 0.908663 |
| Eno1 | P17182 | 0.57 | 0.07 | -0.07 | 0.39 | 0.04 | 0.29 | -0.02 | 0.51 | 0.67 | 0.33 | -0.08 | 0.74 | -0.34 | 0.21 | -0.53 | -0.64 | -0.035 | 0.175 | -0.565 | 0.871068 | 0.503347 | 0.058261 | | 1 | 0.996511 | 0.661738 |
| Eno2 | P17183 | -1.13 | -0.55 | -0.26 | -0.13 | -0.63 | -1.36 | -0.21 | -0.22 | -1.05 | -0.19 | -0.5 | -1.03 | -1.44 | -0.17 | -0.63 | -0.32 | -0.0875 | -0.175 | -0.1225 | 0.796109 | 0.574993 | 0.731521 | | 1 | 0.996511 | 0.958635 |
| Eno3 | P21550 | -1.79 | -0.91 | -1.15 | -1.76 | -2.82 | -1.2 | -2.33 | -1.7 | -0.78 | -1.71 | -1.5 | -1.98 | 1.08 | -4.05 | -1.09 | -1.33 | -0.61 | -0.09 | 0.055 | 0.148803 | 0.788576 | 0.954357 | | 1 | 0.996511 | 0.992476 |
| Enoph1 | Q8BGB7 | -1.02 | -0.67 | -0.52 | -0.53 | -1.1 | -0.64 | -1.11 | -0.69 | -0.68 | -0.21 | -0.67 | -0.99 | -0.55 | -0.25 | -0.96 | -1.22 | -0.2 | 0.0475 | -0.06 | 0.340389 | 0.836827 | 0.823917 | | 1 | 0.996511 | 0.972768 |
| Enpep | P16406 | -0.21 | -0.08 | 0.1 | 0.02 | 0.33 | -0.17 | -0.19 | -0.13 | -0.01 | -0.15 | -0.09 | -0.11 | 0.27 | -0.11 | -0.03 | -0.06 | 0.0025 | -0.0475 | 0.06 | 0.989122 | 0.770685 | 0.750524 | | 1 | 0.996511 | 0.961479 |
| Enpp1 | G3X9S2 | -0.29 | 0.29 | -0.87 | -0.62 | 0.24 | -0.2 | -0.38 | -0.92 | -0.39 | -0.29 | -0.78 | 0.32 | -0.24 | -1.47 | -0.63 | -0.02 | 0.0575 | 0.0875 | -0.2175 | 0.864168 | 0.794267 | 0.583028 | | 1 | 0.996511 | 0.927734 |
| Enpp2 | Q9R1E6-3 | -1.39 | -1.13 | -0.98 | -1.23 | -0.24 | -1.24 | 0.98 | -0.72 | -1.32 | -1.24 | -1.15 | -0.83 | -1.2 | -0.5 | -0.29 | -0.2 | 0.8775 | 0.0475 | 0.635 | 0.077957 | 0.805446 | 0.03925 | | 1 | 0.996511 | 0.658454 |
| Enpp4 | Q8BTJ4-2 | -0.29 | -0.28 | -0.26 | -0.67 | -0.67 | -0.17 | -1.07 | -0.9 | -0.36 | 0.1 | -0.34 | 0.07 | -0.17 | -0.43 | -1.2 | -0.67 | -0.3275 | 0.2425 | -0.2425 | 0.18929 | 0.257416 | 0.374548 | | 1 | 0.996511 | 0.859424 |
| Enpp5 | Q9EQG7 | -1.17 | -0.73 | -1.4 | -1.39 | -0.35 | -1.03 | -0.93 | -1.69 | -1 | -2.04 | -1.32 | -1.26 | -1.35 | -0.22 | -2.1 | -1.8 | 0.1725 | -0.2325 | -0.195 | 0.582504 | 0.421332 | 0.645392 | | 1 | 0.996511 | 0.944749 |
| Enpp6 | Q8BGN3 | 2.35 | 1.13 | 1.16 | 0.44 | 1.17 | 2.04 | 1.13 | 0.76 | -0.35 | -0.1 | 0.63 | 2.01 | -0.49 | -0.25 | 0.95 | 1.06 | 0.005 | -0.7225 | -0.9525 | 0.991075 | 0.253193 | 0.096042 | | 1 | 0.996511 | 0.698156 |
| Ensa | P60840 | 0.53 | -0.21 | 0.47 | 0.47 | -0.74 | 0.11 | -0.37 | 0.56 | 0.13 | 0.46 | 0.28 | 0.78 | -0.23 | 0.08 | 0.06 | -0.09 | -0.425 | 0.0975 | -0.36 | 0.213252 | 0.695789 | 0.148143 | | 1 | 0.996511 | 0.746406 |
| Enthd1 | E9Q1Z2 | 1.25 | 1.47 | 1.92 | 1.64 | 1.64 | 1.82 | 1.84 | 1.94 | 1.28 | 2.21 | 1.78 | 1.29 | 1.78 | 1.38 | 1.43 | 1.85 | 0.24 | 0.07 | 0.04 | 0.232369 | 0.800783 | 0.861847 | | 1 | 0.996511 | 0.977204 |
| Entpd1 | Q8CDV7 | -1.31 | -1.42 | -1.54 | -1.23 | -1.32 | -1.47 | -1.86 | -1.66 | -1.28 | -0.8 | -1.14 | -1.17 | -1.32 | -1.08 | -2.24 | -1.96 | -0.2025 | 0.2775 | -0.275 | 0.278769 | 0.159498 | 0.361897 | | 1 | 0.996511 | 0.858475 |
| Entpd2 | O55026 | 0.09 | 0 | 0.48 | 0.17 | 0.32 | 0.66 | 0.36 | 0.36 | -1.04 | -0.06 | 0.37 | 0.18 | -0.87 | -0.94 | 0.44 | 0.22 | 0.24 | -0.3225 | -0.4725 | 0.199944 | 0.340124 | 0.220234 | | 1 | 0.996511 | 0.787447 |
| Entpd3 | Q8BFW6 | -0.13 | -0.51 | 0.19 | 0.6 | -0.19 | -0.31 | 0.59 | 0.08 | -0.03 | 0.79 | 0.01 | -0.22 | 0.5 | -0.06 | 1 | 0.47 | 0.005 | 0.1 | 0.44 | 0.986926 | 0.758085 | 0.198951 | | 1 | 0.996511 | 0.78004 |
| Entpd5 | Q3TQC7 | -0.36 | -0.33 | -0.13 | -0.41 | -0.1 | -0.13 | -0.36 | -0.61 | 0.1 | -0.71 | -0.32 | 0.08 | 0.31 | 0.57 | 0.98 | 0.55 | 0.0075 | 0.095 | 0.91 | 0.966585 | 0.68415 | 0.001934 | | 1 | 0.996511 | 0.419079 |
| Entpd6 | Q3U0P5 | 0.34 | -0.25 | -0.37 | 0.38 | 0.59 | 0.98 | 1.02 | 0.5 | -0.05 | 0.53 | 0.08 | 0.06 | 0.29 | 0.5 | 0.7 | 1.1 | 0.7475 | 0.13 | 0.6225 | 0.014279 | 0.612507 | 0.050432 | | 1 | 0.996511 | 0.658454 |
| Eny2 | A0A2I3BRK | -1.65 | -0.67 | 0.05 | -0.18 | -1.95 | -1.99 | -0.54 | 0.14 | -1.45 | 0.82 | -0.34 | -1.83 | -1.06 | -0.42 | 1.07 | 0.34 | -0.4725 | -0.0875 | 0.595 | 0.434398 | 0.89198 | 0.296131 | | 1 | 0.996511 | 0.825612 |
| Eogt | Q8BYW9 | -0.47 | -0.34 | -1.13 | -1.78 | -0.07 | -0.32 | -0.38 | -1.48 | -0.26 | -0.54 | -1.45 | -0.19 | -0.37 | -0.32 | -0.31 | -0.5 | 0.3675 | 0.32 | 0.555 | 0.39993 | 0.452406 | 0.125906 | | 1 | 0.996511 | 0.721505 |
| Ep300 | B2RWS6 | -2.12 | -5.51 | -1.43 | -1.35 | -2.49 | -3.21 | -0.25 | -1.18 | 0.4 | 0.3 | -2.92 | -2.78 | 1.45 | -2.79 | 0.43 | -1.18 | 0.82 | 1.3525 | 2.08 | 0.448511 | 0.281067 | 0.112094 | | 1 | 0.996511 | 0.713445 |
| Ep400 | Q8CHI8-3 | -0.94 | -1.96 | -1.42 | -0.88 | -1.73 | -1.49 | -1.31 | -1.14 | -0.96 | -0.72 | -1.06 | -1.82 | -1.29 | -0.66 | -0.78 | -1.13 | -0.1175 | 0.16 | 0.335 | 0.679462 | 0.640651 | 0.286027 | | 1 | 0.996511 | 0.823465 |
| Epb41 | A2A841 | 1.75 | 0.45 | 0.88 | 1.44 | 0.64 | 1.65 | 0.35 | 1.19 | 0.79 | 0.45 | 0.57 | 1.06 | 0.77 | 0.51 | 0.35 | 0.3 | -0.1725 | -0.4125 | -0.6475 | 0.657838 | 0.218771 | 0.066372 | | 1 | 0.996511 | 0.664936 |
| Epb41l1 | E9PV14 | 1.48 | 0.37 | 0.56 | 0.55 | 0.68 | 1.25 | 0.91 | 0.54 | 0.66 | 0.21 | 0.65 | 0.92 | -0.01 | 0.96 | 0.4 | 0.5 | 0.105 | -0.13 | -0.2775 | 0.721676 | 0.663502 | 0.403193 | | 1 | 0.996511 | 0.870698 |
| Epb41l1 | A2AUK5 | 1.53 | 0.45 | 0.65 | 0.77 | 0.44 | 1.13 | 0.37 | 0.45 | 1.42 | 0.18 | 0.58 | 0.99 | 0.72 | 1.12 | 0.38 | 0.54 | -0.2525 | -0.0575 | -0.16 | 0.402199 | 0.869159 | 0.59479 | | 1 | 0.996511 | 0.929843 |
| Epb41l2 | O70318 | 1 | 0.89 | 0.89 | 0.93 | 0.94 | 0.81 | 0.76 | 0.81 | 0.66 | 1.1 | 1 | 0.88 | 0.21 | 0.39 | 0.96 | 0.75 | -0.0975 | -0.0175 | -0.35 | 0.49488 | 0.919236 | 0.140827 | | 1 | 0.996755 | 0.737973 |
| Epb41l3 | A0A5F8MP | 3.4 | 1.1 | 1.31 | 1.65 | 0.73 | 2.82 | 1.23 | 1.45 | 1.7 | 0.71 | 1.14 | 2.56 | 0.72 | 0.64 | 0.94 | 1.07 | -0.3075 | -0.3375 | -1.0225 | 0.626004 | 0.579576 | 0.065884 | | 1 | 0.996511 | 0.664936 |
| Epb41l3 | A7YY80 | 0.08 | 0.25 | 0.77 | 0.08 | 0.02 | -0.87 | 0.53 | 0.25 | -0.09 | 3.98 | -0.28 | -0.28 | 0.28 | -0.74 | 0.02 | -0.02 | -0.3125 | 0.5375 | -0.41 | 0.361456 | 0.577321 | 0.181261 | | 1 | 0.996511 | 0.775592 |
| Epb41l3 | Q9WV92-8 | 0.63 | 0.7 | 0.89 | 1.19 | 1.05 | 0.72 | 0.73 | 1.02 | -0.23 | 1.16 | 0.71 | 0.71 | -0.2 | -0.32 | 0.74 | 1.13 | 0.0275 | -0.265 | -0.515 | 0.885873 | 0.415373 | 0.186757 | | 1 | 0.996511 | 0.776843 |
| Epb41l3 | D0VYV6 | 1.06 | 0.11 | 1.89 | 2.05 | 0.42 | 0.76 | 1.26 | 1.73 | 0.93 | 1.73 | 1.9 | 0.54 | -0.14 | 0.62 | 1.39 | 2.33 | -0.235 | -0.0025 | -0.2275 | 0.633417 | 0.996093 | 0.719368 | | 1 | 0.999603 | 0.958398 |
| Epb41l4b | A2ALK6 | 0.24 | 0.61 | -0.6 | -0.06 | 0.41 | 0.3 | -0.74 | -0.72 | 0.73 | -0.06 | -0.64 | 0.33 | 1.17 | 1.19 | -0.61 | -0.74 | -0.235 | 0.0425 | 0.205 | 0.544843 | 0.909714 | 0.709515 | | 1 | 0.996744 | 0.957718 |
| Epb41l5 | Q8BGS1 | 2.6 | 1.16 | 1.71 | 1.67 | 1.02 | 2.63 | 1.85 | 2.23 | 0.95 | 1.6 | 1.63 | 2.61 | 0.87 | 0.64 | 1.89 | 1.99 | 0.1475 | -0.0875 | -0.4375 | 0.730024 | 0.838788 | 0.328445 | | 1 | 0.996511 | 0.844943 |
| Epb42 | Q3UV95 | 0.16 | -0.63 | 1.36 | 2.26 | 0.04 | 1.41 | 0.81 | 2.32 | 1.01 | 0.96 | 0.9 | -0.36 | 1.11 | 1.66 | 0.42 | 0.64 | 0.3575 | -0.16 | 0.17 | 0.623734 | 0.807407 | 0.789082 | | 1 | 0.996511 | 0.968677 |
| Epc1 | Q8C9X6 | -5.64 | -7.43 | -11.58 | -6.58 | -10.48 | -4.12 | -5.74 | -10.02 | -1.93 | -4.78 | -6.38 | -4.72 | -2.81 | -6.49 | -4.27 | -5.97 | 0.2175 | 3.355 | 2.9225 | 0.905091 | 0.041577 | 0.059855 | | 1 | 0.996511 | 0.661738 |
| Epdr1 | Q99M71 | -1.24 | -1.08 | -1.29 | -1.21 | 0.03 | -1.2 | -0.41 | -1.35 | -2.23 | -1.43 | -0.44 | -1.11 | -1.32 | -1.22 | -1.16 | -0.9 | 0.4725 | -0.0975 | 0.055 | 0.168416 | 0.788993 | 0.766139 | | 1 | 0.996511 | 0.964134 |
| Epg5 | Q80TA9 | 0.18 | 0.47 | 0.11 | 0.18 | 0.32 | 0.18 | 0.13 | -0.02 | 0.32 | -0.13 | 0.2 | 0.27 | 0.12 | 0.75 | -0.23 | 0.07 | -0.0825 | -0.07 | -0.0575 | 0.618256 | 0.710561 | 0.820385 | | 1 | 0.996511 | 0.972768 |
| Epha10 | F6WVB0 | 0.89 | 0.89 | 0.68 | 0.82 | 0.66 | 0.41 | 0.53 | 0.11 | 1.26 | 1.32 | 1.01 | 0.9 | 1.32 | 0.64 | 0.29 | -0.08 | -0.3925 | 0.3025 | -0.2775 | 0.049473 | 0.118465 | 0.383769 | | 1 | 0.996511 | 0.860984 |
| Epha3 | B7ZNN5 | 0.27 | 0.49 | 0.64 | 1.17 | 0.21 | 1.12 | 0.74 | 0.99 | 0.76 | 2.14 | 0.79 | 0.83 | 1.36 | -1.13 | 0.85 | 1.04 | 0.1225 | 0.4875 | -0.1125 | 0.66396 | 0.215611 | 0.837754 | | 1 | 0.996511 | 0.975961 |
| Epha4 | Q03137 | 0.23 | 0.29 | 0.2 | 0.15 | 0.78 | 0.52 | 0.3 | 0.12 | 0.53 | 0.06 | 0.35 | 0.57 | 0.63 | 0.63 | 0.25 | 0.14 | 0.2125 | 0.16 | 0.195 | 0.272419 | 0.39284 | 0.34047 | | 1 | 0.996511 | 0.84829 |
| Epha5 | E9PUP9 | 0.01 | 0.11 | 0.31 | 0.68 | 0.95 | 0.49 | 1.1 | 1.01 | 0.76 | 0.18 | 0.92 | 0.56 | 0.72 | 1.2 | 0.94 | 1.18 | 0.61 | 0.3275 | 0.7325 | 0.022023 | 0.201977 | 0.010754 | | 1 | 0.996511 | 0.563386 |
| Epha6 | G1K381 | 0.16 | 0.44 | -0.77 | -0.57 | 1 | 0.56 | -0.11 | 0 | 0.03 | 0.14 | -0.09 | 0.48 | -0.03 | -0.16 | -0.46 | -0.16 | 0.5475 | 0.325 | -0.0175 | 0.16393 | 0.31783 | 0.955199 | | 1 | 0.996511 | 0.992482 |
| Epha7 | Q61772 | -0.32 | -0.61 | 1.04 | 0.54 | -0.45 | 0.37 | 0.72 | 1 | 0.76 | 0.81 | 0.96 | 0.14 | 0.84 | -0.23 | 1.12 | 1.05 | 0.2475 | 5.05E-01 | 0.5325 | 0.592738 | 0.2312 | 0.270018 | | 1 | 0.996511 | 0.8219 |
| Ephb1 | Q8CBF3 | 0.69 | 1.21 | 1.16 | 0.7 | 0.91 | 0.97 | 0.66 | 0.78 | 1.04 | 0.39 | 1.19 | 0.93 | 0.19 | 0.79 | 1.08 | 0.97 | -0.11 | -0.0525 | -0.1825 | 0.57435 | 0.832746 | 0.50306 | | 1 | 0.996511 | 0.901002 |
| Ephb2 | P54763 | 0.06 | 0.46 | 0.8 | 0.58 | 0.44 | 0.29 | 0.46 | 0.54 | 0.23 | 0.52 | 0.96 | 0.25 | 0.38 | 0.36 | 0.29 | 0.65 | -0.0425 | 0.015 | -0.055 | 0.830025 | 0.952583 | 0.805455 | | 1 | 0.996755 | 0.971715 |
| Ephb3 | P54754 | 0.95 | 0.42 | 0.71 | 0.68 | 0.7 | 1.68 | 0.48 | 0.54 | 0.31 | 0.76 | 0.67 | 1.08 | 0.22 | 0.37 | 0.32 | 0.61 | 0.16 | 0.015 | -0.31 | 0.595251 | 0.947043 | 0.150347 | | 1 | 0.996755 | 0.749304 |
| Ephb6 | O08644 | 0.25 | 0.44 | 0.15 | 0.33 | 0.7 | 0.45 | 0.63 | 0.51 | 0.31 | 0.23 | 0.61 | 0.69 | 0.18 | 0.66 | 0.18 | 0.37 | 0.28 | 0.1675 | 0.055 | 0.096127 | 0.382401 | 0.781391 | | 1 | 0.996511 | 0.967642 |
| Ephx1 | Q9D379 | 0.59 | 0.7 | 0.51 | 0.55 | 0.59 | 0.08 | 0.97 | 0.21 | 0.35 | 0.54 | -0.02 | 0.15 | 0.06 | 0.55 | 0.27 | 0.11 | -0.125 | -0.3325 | -0.34 | 0.583404 | 0.103713 | 0.104566 | | 1 | 0.996511 | 0.702979 |
| Ephx2 | P34914 | -0.58 | -0.34 | -0.34 | -0.09 | -0.1 | -0.57 | 0.28 | -0.22 | -0.77 | 0.98 | -0.32 | -0.69 | -1.12 | -0.33 | -0.59 | -0.61 | 0.185 | 0.1375 | -0.325 | 0.417807 | 0.73126 | 0.190176 | | 1 | 0.996511 | 0.776843 |
| Ephx4 | Q6IE26 | -0.92 | -0.13 | -2.32 | -3.38 | -0.08 | -0.72 | -2.6 | -2.82 | 0.54 | -1.37 | -2.54 | -0.53 | 1.23 | 0.32 | -1.77 | -2.14 | 0.1325 | 0.7125 | 1.0975 | 0.882052 | 0.426047 | 0.277703 | | 1 | 0.996511 | 0.822084 |
| Epm2a | Q9WUA5 | 0.21 | -1.2 | 0.64 | 0.87 | -0.34 | -0.82 | -0.45 | 4.77 | 0.18 | -0.69 | 2.26 | -0.59 | 0.31 | -0.24 | 0.45 | -1.58 | 0.66 | 0.16 | -0.395 | 0.603239 | 0.831062 | 0.515955 | | 1 | 0.996511 | 0.905674 |
| Epm2aip1 | Q8VEH5 | -0.63 | -1.46 | -1.04 | -0.8 | -1.3 | -0.44 | -0.88 | -0.64 | -0.49 | -0.21 | -0.67 | -0.76 | -0.57 | -1.2 | -0.95 | -1.1 | 0.1675 | 0.45 | 0.0275 | 0.532608 | 0.091797 | 0.914799 | | 1 | 0.996511 | 0.986556 |
| Epn1 | Q80VP1 | 0.89 | 0.81 | 0.88 | 0.9 | 0.57 | 0.46 | 0.7 | 0.61 | 0.29 | 0.33 | 0.52 | 0.57 | -0.09 | 0.53 | 0.16 | 0.33 | -0.285 | -0.4425 | -0.6375 | 0.072833 | 0.022252 | 0.010567 | | 1 | 0.996511 | 0.563386 |
| Epn1 | Q80VP1-2 | 1.3 | -0.45 | 1.72 | 0.58 | 1.34 | -0.01 | 0.46 | 0.5 | -0.81 | 0.23 | 0.3 | -0.33 | 0.41 | -0.78 | 0.84 | 0.28 | -0.215 | -0.94 | -0.6 | 0.673781 | 0.089614 | 0.285596 | | 1 | 0.996511 | 0.822669 |
| Epn2 | J3QNT7 | 0.91 | 0.68 | 0.46 | 0.73 | 0.02 | 0.55 | 0.7 | 0.62 | 1.05 | 0.39 | 0.68 | 0.83 | 1.1 | 0.71 | 0.52 | 0.38 | -0.2225 | 0.0425 | -0.0175 | 0.301937 | 0.839495 | 0.938671 | | 1 | 0.996511 | 0.989956 |

| | | | | | | | | | | | | | | | | | | | | | | | | | | |
|---|---|---|---|---|---|---|---|---|---|---|---|---|---|---|---|---|---|---|---|---|---|---|---|---|---|---|
| Epn3 | Q91W69 | 1.06 | 1.93 | 1.17 | 0.7 | 1.06 | 0.71 | 1.1 | 0.63 | 1.71 | 1.34 | 0.48 | 0.87 | 0.51 | 1.69 | 0.83 | 0.93 | -0.34 | -0.115 | -0.225 | 0.254543 | 0.751766 | 0.531762 | 1 | 0.996511 | 0.910136 |
| Eprs1 | Q8CGC7 | -1.48 | -2.31 | -2.2 | -2.5 | -1.9 | -1.63 | -1.73 | -2.42 | -1.82 | -1.72 | -2.08 | -1.51 | -1.63 | -1.46 | -1.6 | -2.21 | 0.2025 | 0.34 | 0.3975 | 0.484468 | 0.226916 | 0.198491 | 1 | 0.996511 | 0.78004 |
| Eps15 | P42567 | 1.21 | 1.28 | 0.51 | 0.37 | 1.29 | 1.03 | 0.82 | 0.22 | 1.15 | -0.54 | 0.44 | 1.39 | 0.96 | 1.2 | 0.57 | 0.52 | -0.0025 | -0.2325 | -0.03 | 0.993757 | 0.617194 | 0.91996 | 1 | 0.996511 | 0.98688 |
| Eps15l1 | Q60902 | 0.81 | 0.73 | 0.39 | 0.52 | 0.53 | 0.53 | 0.38 | 0.45 | 0.27 | 0.31 | 0.33 | 0.5 | -0.02 | 0.63 | 0.03 | 0.08 | -0.14 | -0.26 | -0.4325 | 0.39941 | 0.168403 | 0.082916 | 1 | 0.996511 | 0.688875 |
| Eps15l1 | Q60902-4 | 0.52 | 0.02 | 1.54 | 1.14 | 2.13 | -0.2 | 0.88 | 0.47 | -0.17 | 1.83 | 1.12 | -0.05 | 0.2 | 1.97 | 0.45 | -0.18 | 0.015 | -0.1225 | -0.195 | 0.978004 | 0.821045 | 0.716599 | 1 | 0.996511 | 0.958334 |
| Eps8 | Q08509 | 0.56 | 0.89 | 1.38 | 0.9 | 0.62 | 0.54 | 1.48 | 1.18 | 0.9 | 0.4 | 1.29 | 0.61 | 0.8 | 0.47 | 1.32 | 1.49 | 0.0225 | -0.1325 | 0.0875 | 0.936729 | 0.627637 | 0.772149 | 1 | 0.996511 | 0.965933 |
| Eps8l2 | Q99K30 | -0.96 | 0.04 | 0.07 | 0.16 | 0.54 | -0.65 | 0.21 | 0.38 | -1.57 | -0.19 | 0.39 | -0.88 | -1.18 | -0.6 | 0.19 | 0.57 | 0.2925 | -0.39 | -0.0825 | 0.423586 | 0.414252 | 0.853791 | 1 | 0.996511 | 0.976225 |
| Eral1 | Q9CZU4 | 0.86 | 0.95 | 0.71 | 1.04 | -0.17 | 0.74 | 0.28 | 0.57 | 1.68 | 1.67 | 0.78 | 0.64 | 1.3 | 1.41 | 0.93 | 0.86 | -0.535 | 0.3025 | 0.235 | 0.042842 | 0.321608 | 0.281726 | 1 | 0.996511 | 0.822084 |
| Erap1 | Q9EQH2 | -0.09 | -0.06 | -0.54 | -0.35 | 0.09 | -0.15 | -0.05 | -0.18 | -0.45 | -1.44 | -0.56 | -0.13 | -0.76 | -0.18 | -0.21 | -0.1 | 0.1875 | -0.385 | -0.0525 | 0.30455 | 0.23034 | 0.82129 | 1 | 0.996511 | 0.972768 |
| Erbb3 | Q61526 | -0.09 | 0.04 | 0.31 | 0.55 | 0.42 | -0.35 | 1.45 | 0.68 | 0.18 | 1.23 | 0.21 | 0.36 | -0.05 | 0.65 | 0.8 | 1.94 | 0.3475 | 0.2925 | 0.6325 | 0.369902 | 0.333926 | 0.155523 | 1 | 0.996511 | 0.750763 |
| Erbb4 | Q61527-3 | 0.65 | 0.88 | 0.93 | 0.79 | 1.37 | 0.8 | 1.08 | 0.8 | 0.48 | 0.47 | 1.01 | 0.78 | 0.42 | 0.52 | 0.94 | 0.9 | 0.2 | -0.1275 | -0.1175 | 0.305481 | 0.520775 | 0.572906 | 1 | 0.996511 | 0.925347 |
| Erbin | Q80TH2-2 | 0.91 | 0.68 | 0.91 | 0.83 | 0.47 | 0.69 | 0.34 | 0.46 | 0.94 | 0.84 | 0.5 | 0.78 | 1.39 | -0.04 | 0.85 | 0.64 | -0.3425 | -0.0675 | -0.1225 | 0.053696 | 0.704915 | 0.694072 | 1 | 0.996511 | 0.955729 |
| Erc1 | V9GWT6 | -1.02 | -0.17 | 0.07 | -0.75 | -0.19 | -2.25 | 0.25 | 0.14 | -1.37 | -1.4 | -0.09 | -0.72 | -2.55 | -0.8 | -0.3 | -0.48 | -0.045 | -0.4275 | -0.565 | 0.938438 | 0.284361 | 0.304281 | 1 | 0.996511 | 0.831662 |
| Erc1 | V9GXH3 | 1.01 | 0.87 | 0.78 | 0.8 | 0.87 | 0.98 | 0.69 | 0.81 | 1.19 | 1.05 | 1.02 | 1.31 | 0.98 | 0.89 | 0.77 | 0.63 | -0.0275 | 0.2775 | -0.0475 | 0.856531 | 0.124087 | 0.793504 | 1 | 0.996511 | 0.969365 |
| Erc2 | Q3UHT7 | 1.12 | 1.6 | 1.03 | 0.87 | 1.6 | 0.87 | 1.29 | 1.21 | 0.16 | 0.75 | 1.14 | 1.3 | -0.18 | 0.19 | 0.73 | 1 | 0.0875 | -0.3175 | -0.72 | 0.710288 | 0.310877 | 0.045728 | 1 | 0.996511 | 0.658454 |
| Erc2 | Q6PH08 | 1.61 | 1.31 | 1.42 | 1.46 | 1.35 | 1.68 | 1.23 | 1.5 | 1.34 | 1.04 | 1.48 | 1.68 | 1.05 | 1.1 | 1.36 | 1.36 | -0.01 | -0.065 | -0.2325 | 0.95278 | 0.744071 | 0.232144 | 1 | 0.996511 | 0.788863 |
| Ercc2 | O08811 | -4.33 | -4.72 | -2.51 | -3.29 | -4.03 | -4.38 | -3.12 | -3.26 | -3.91 | -2.27 | -2.39 | -3.88 | -3.51 | -3.54 | -2.54 | -2.39 | 0.015 | 0.6 | 0.7175 | 0.977729 | 0.34535 | 0.209581 | 1 | 0.996511 | 0.786218 |
| Ercc3 | P49135 | -2.49 | -2.13 | -0.59 | -0.57 | -3.07 | -2.54 | -0.37 | -0.86 | -2.21 | -0.7 | -0.68 | -1.89 | -1.01 | -1.99 | -0.49 | -0.13 | -0.265 | 0.075 | 0.54 | 0.722229 | 0.898783 | 0.374298 | 1 | 0.996511 | 0.859424 |
| Ercc4 | Q9QZD4 | -2.07 | -2.36 | -2.13 | -1.93 | -2.08 | -2.22 | -1.91 | -2.25 | -1.94 | -0.73 | -2.22 | -1.87 | -2 | -1.66 | -1.62 | -1.34 | 0.0075 | 0.4325 | 0.4675 | 0.96494 | 0.221145 | 0.055039 | 1 | 0.996511 | 0.661738 |
| Erf | P70459 | -0.33 | 0.13 | -0.83 | -0.65 | -0.55 | -0.44 | -0.8 | -0.53 | -0.45 | -0.31 | -0.67 | -0.28 | 0.27 | 0.21 | -0.91 | -1.02 | -0.16 | -0.0075 | 0.0575 | 0.510223 | 0.976161 | 0.883902 | 1 | 0.997588 | 0.979767 |
| Erg28 | Q9ERY9 | 0.84 | 0.78 | 0.76 | 0.81 | 1.24 | 0.88 | 1.52 | 1.37 | -0.35 | -0.56 | 0.6 | 0.77 | 0.64 | -1.37 | 1.76 | 1.67 | 0.455 | -0.6825 | -0.1225 | 0.031915 | 0.065858 | 0.853242 | 1 | 0.996511 | 0.976225 |
| Ergic1 | F6SLP4 | -0.47 | -0.36 | -0.73 | -0.97 | 0.41 | -0.41 | 0.17 | -0.94 | -0.52 | -0.84 | -0.73 | -0.41 | -0.06 | 0.39 | -0.12 | 0.15 | 0.44 | 0.0075 | 0.7225 | 0.198141 | 0.971565 | 0.010421 | 1 | 0.997515 | 0.563386 |
| Ergic2 | Q9CR89 | -1.22 | -0.77 | -2.25 | -2.25 | 0.02 | -1.06 | -2.23 | -2.11 | -1.04 | -1.44 | -2.29 | -0.55 | -0.59 | -0.78 | -1.05 | -1.31 | 0.2775 | 0.2925 | 0.69 | 0.639353 | 0.553793 | 0.106223 | 1 | 0.996511 | 0.703417 |
| Ergic3 | Q9CQE7 | -0.52 | -0.75 | -0.71 | -1.66 | 0.14 | -0.2 | -0.43 | -0.86 | -0.23 | -1.79 | -1.01 | -0.31 | 0.72 | -0.52 | 0.61 | 0.17 | 0.5725 | 0.075 | 1.155 | 0.10372 | 0.858638 | 0.012053 | 1 | 0.996511 | 0.566767 |
| Erh | P84089 | 0.13 | -1.28 | 0.1 | 0.03 | -1.7 | -0.26 | -0.73 | 1.17 | -0.28 | 0.7 | 0.23 | -1.18 | 1.31 | -1.14 | 2 | -0.76 | -0.125 | 0.1225 | 0.6075 | 0.841885 | 0.803588 | 0.432929 | 1 | 0.996511 | 0.879365 |
| Eri3 | Q8C460 | -1.2 | -1.09 | -0.97 | -0.97 | -1.13 | -1.15 | -0.71 | -0.99 | -1.1 | -0.63 | -0.97 | -1.19 | -0.99 | -0.09 | -0.94 | -1.15 | 0.0625 | 0.085 | 0.265 | 0.712661 | 0.657947 | 0.339187 | 1 | 0.996511 | 0.847641 |
| Erlec1 | Q8VEH8 | -1.22 | -0.99 | -1.15 | -1.39 | -0.28 | -0.81 | 0.18 | -0.96 | -1.31 | -1.52 | -1.06 | -0.96 | -0.68 | -0.35 | 0.36 | 0.68 | 0.72 | -0.025 | 1.19 | 0.027536 | 0.900618 | 0.005381 | 1 | 0.996511 | 0.552614 |
| Erlin1 | Q91X78 | 1.31 | 0.7 | -0.08 | -0.3 | 1.43 | 1.09 | 0.23 | 0.03 | 1.34 | 0.41 | -0.51 | 0.87 | 1.62 | 2.18 | 1.01 | 0.84 | 0.2875 | 0.12 | 1.005 | 0.539719 | 0.811376 | 0.050487 | 1 | 0.996511 | 0.658454 |
| Erlin2 | Q8BFZ9 | 0.01 | 0.17 | -0.27 | -0.15 | 0.5 | 0.03 | 0.49 | 0.01 | -0.07 | 0.05 | -0.26 | 0.12 | -0.06 | 0.61 | 0.83 | 0.43 | 0.3175 | 0.02 | 0.5125 | 0.139577 | 0.91465 | 0.0639 | 1 | 0.996755 | 0.66433 |
| Ermn | Q5EBJ4 | 1.18 | 1.5 | 0.87 | 0.92 | 1.08 | 0.45 | 0.83 | 0.54 | 0.81 | 0.3 | 0.58 | 0.82 | 0.74 | 1.19 | 0.66 | 0.97 | -0.3925 | -0.49 | -0.2275 | 0.108761 | 0.053558 | 0.339146 | 1 | 0.996511 | 0.847641 |
| Ermp1 | Q3UVK0 | 0.15 | -0.09 | 0.22 | 0.25 | -0.03 | 0.16 | 0.25 | 0.15 | -0.26 | 0.08 | 0.06 | -0.16 | 0.16 | 0.21 | 0.47 | 0.55 | 1.96E-17 | -0.2025 | 0.215 | 1 | 0.278303 | 0.287071 | 1 | 0.996511 | 0.824333 |
| Ero1a | Q8R180 | -1.01 | -0.61 | -0.88 | -1.09 | -0.27 | -0.94 | -0.23 | -1.04 | -0.71 | -1.07 | -1.19 | -0.96 | -0.51 | 0.12 | -0.29 | -0.64 | 0.2775 | -0.085 | 0.5675 | 0.286065 | 0.668875 | 0.037973 | 1 | 0.996511 | 0.658454 |
| Ero1a | A0A2I3BPN | -2.67 | -1.65 | -0.69 | -2.6 | -0.49 | -2.4 | -1.71 | -2.93 | -1.87 | -2.11 | -2.52 | -2.86 | -2.22 | -1.06 | -1.93 | 0.13 | 0.02 | -0.4375 | 0.6325 | 0.975035 | 0.374912 | 0.338463 | 1 | 0.996511 | 0.847578 |
| Ero1b | Q8R2E9 | -1.36 | -0.38 | -1.68 | -1.49 | -0.24 | -1.03 | -1.51 | -2.24 | -1.09 | -2.71 | -2.53 | -0.89 | -1.41 | -0.31 | -1.39 | -1.32 | -0.0275 | -0.5775 | 0.12 | 0.953586 | 0.280127 | 0.754531 | 1 | 0.996511 | 0.962333 |
| Erp29 | P57759 | -0.96 | -0.63 | -0.95 | -1.12 | 0.06 | -0.74 | -0.18 | -0.77 | -0.91 | -1.24 | -0.91 | -0.87 | -0.31 | -0.33 | 0.42 | 0.07 | 0.5075 | -0.0675 | 0.8775 | 0.06404 | 0.72416 | 0.005495 | 1 | 0.996511 | 0.552614 |
| Erp44 | Q9D1Q6 | -0.53 | -0.65 | -1.06 | -1.07 | 0.18 | -0.45 | 0.23 | -0.73 | -1.26 | -0.74 | -1.06 | -0.37 | -0.72 | -0.45 | 0 | 0.08 | 0.635 | -0.03 | 0.555 | 0.045821 | 0.907738 | 0.059928 | 1 | 0.996744 | 0.661738 |
| Esam | Q925F2 | -0.9 | -0.34 | 0.23 | -0.11 | 0.1 | -1.09 | 0.31 | -0.4 | -0.66 | -0.05 | 0.42 | -0.64 | -0.5 | 0.04 | -0.04 | 0.24 | 0.01 | 0.0475 | 0.215 | 0.978574 | 0.89064 | 0.477531 | 1 | 0.996511 | 0.892234 |
| Esd | H3BLJ9 | -2.06 | -1.93 | -2.1 | -1.72 | -2.32 | -1.71 | -1.68 | -2.14 | -2.7 | -2.48 | -2.3 | -1.91 | -2.81 | -2.22 | -2.25 | -2.48 | -0.01 | -0.395 | -0.4875 | 0.961794 | 0.10462 | 0.046736 | 1 | 0.996511 | 0.658454 |
| Ess2 | Q3UFM6 | -1.32 | -0.83 | -2.2 | -1.95 | -1.27 | -0.81 | -3.41 | -2.21 | -1.07 | 0.37 | -1.96 | -0.76 | -1.24 | -1.47 | -2.09 | -3.01 | -0.35 | 0.72 | -0.3775 | 0.560512 | 0.197435 | 0.43229 | 1 | 0.996511 | 0.879165 |
| Esyt1 | Q3U7R1 | -0.07 | 0.16 | 0.79 | 0.61 | 0.35 | -0.44 | 0.64 | 0.23 | -0.13 | 0.03 | 0.68 | -0.29 | -0.46 | 0.55 | 0.24 | 0.79 | -0.1775 | -0.3 | -0.0925 | 0.55826 | 0.32783 | 0.784078 | 1 | 0.996511 | 0.967642 |
| Esyt2 | Q3TZZ7 | 0.81 | 0.95 | 0.47 | 0.22 | 1.15 | 0.71 | 1.04 | 0.53 | -0.04 | -0.79 | 0.73 | 0.7 | -0.19 | -0.16 | 0.76 | 0.91 | 0.245 | -0.4625 | -0.2825 | 0.31467 | 0.246111 | 0.412385 | 1 | 0.996511 | 0.874811 |
| Etf1 | Q8BWY3 | -0.33 | -0.49 | -0.01 | -0.21 | -0.24 | -0.3 | 0.05 | 0.33 | -0.6 | -0.44 | 0.03 | -0.11 | -0.91 | -0.3 | -0.43 | -0.34 | 0.22 | -0.02 | -0.235 | 0.303007 | 0.926272 | 0.307046 | 1 | 0.996755 | 0.832533 |
| Etfa | Q99LC5 | 0 | -0.04 | 0.34 | 0.4 | -0.39 | 0.02 | -0.18 | -0.1 | 0.75 | -0.39 | 0.13 | -0.36 | 0.99 | 0.1 | 0.08 | 0.06 | -0.3375 | -0.1425 | 0.1325 | 0.095173 | 0.633633 | 0.633724 | 1 | 0.996511 | 0.941132 |
| Etfb | Q9DCW4 | 1.93 | 2.21 | 0.84 | 0.82 | 2.04 | 1.89 | 0.79 | 0.71 | 1.22 | 1.25 | 0.77 | 1.61 | 0.85 | 1.69 | 0.59 | 0.47 | -0.0925 | -0.2375 | -0.55 | 0.843639 | 0.542686 | 0.225109 | 1 | 0.996511 | 0.787447 |
| Etfdh | Q921G7 | 0.7 | 0.86 | 0.15 | 0.4 | 0.38 | 0.69 | 0.4 | 0.1 | 0.56 | -0.06 | 0.25 | 0.43 | 0.29 | 0.75 | -0.06 | 0.05 | -0.135 | -0.2325 | -0.27 | 0.546506 | 0.339238 | 0.324065 | 1 | 0.996511 | 0.84194 |
| Ethe1 | Q9DCM0 | 0.61 | -0.09 | 0.27 | 0.21 | -0.1 | 0.78 | 0.59 | 0.63 | -2.22 | -0.61 | 0.57 | 0.41 | -1.16 | -2.48 | 0.37 | 0.59 | 0.225 | -0.7125 | -0.92 | 0.386377 | 0.256437 | 0.190954 | 1 | 0.996511 | 0.777117 |
| Etl4 | E9QAU4 | 1.35 | 1.21 | 1.38 | 1.17 | 1.15 | 1.33 | 0.92 | 1.08 | 1.02 | 0.99 | 1.29 | 1.4 | 0.89 | 0.96 | 0.64 | 0.91 | -0.1575 | -0.1025 | -0.4275 | 0.341346 | 0.572183 | 0.03883 | 1 | 0.996511 | 0.658454 |
| Etl4 | D3Z781 | -1.42 | -0.76 | 0.54 | -0.31 | -1.09 | -0.71 | -0.29 | -1.26 | -2.2 | 0.82 | -0.13 | -1.91 | -1.06 | -1.38 | -0.64 | -0.43 | -0.35 | -0.3675 | -0.39 | 0.428942 | 0.627737 | 0.385721 | 1 | 0.996511 | 0.861533 |
| Etl4 | Q8BHR1 | 0.71 | -0.21 | 1.56 | 3.41 | -1.43 | 0.36 | -0.96 | 2.82 | 3.78 | 2.87 | 1.22 | -0.34 | 4.05 | 0.08 | 3.41 | -0.15 | -1.17 | 0.515 | 0.48 | 0.302847 | 0.632607 | 0.688766 | 1 | 0.996511 | 0.955729 |
| Etnk1 | Q9D4V0 | -0.79 | -1.1 | -0.08 | 0.3 | -0.71 | -1.39 | 0.5 | 0.12 | -1.36 | -0.47 | -0.25 | -1.01 | -1.31 | -0.6 | 0.2 | 0.29 | 0.0475 | -0.355 | 0.0625 | 0.922834 | 0.376331 | 0.892993 | 1 | 0.996511 | 0.982821 |
| Etnppl | E9PYH3 | -2.68 | -1.16 | 0.5 | 0.24 | -2.71 | -2.37 | -0.01 | -0.15 | -0.78 | -0.27 | -0.08 | -1.15 | 0.44 | -2.25 | -0.29 | -0.4 | -0.535 | 0.205 | 0.15 | 0.564319 | 0.770419 | 0.857646 | 1 | 0.996511 | 0.976225 |
| Eva1a | D3Z511 | 1.12 | 1.32 | 0.42 | 0.74 | 1.53 | 0.82 | 1.27 | 1.13 | -0.22 | 1.77 | 0.77 | 1.51 | 0.6 | 2.59 | 0.67 | 0.6 | 0.2875 | 0.0575 | 0.215 | 0.283181 | 0.900199 | 0.665999 | 1 | 0.996511 | 0.950205 |
| Evi2a | P20934 | 0.63 | 0.79 | 1.74 | 2.06 | 0.36 | 0.58 | -0.17 | 0.54 | 0.93 | -0.91 | 0.85 | 0.71 | 0.72 | -0.2 | 1.98 | 1.62 | -0.9775 | -0.91 | -0.275 | 0.026441 | 0.107865 | 0.622401 | 1 | 0.996511 | 0.938628 |
| Evi5 | E9PWR7 | 0.93 | 1.11 | 0.38 | 0.4 | 0.14 | 1.11 | -0.29 | 0.05 | 0.92 | 0.23 | 0.03 | 1.01 | 1.56 | 0.7 | -0.22 | -0.02 | -0.4525 | -0.1575 | -0.2 | 0.207604 | 0.614399 | 0.638875 | 1 | 0.996511 | 0.942804 |
| Evi5l | H3BKQ3 | 0.15 | -0.05 | 1.04 | 0.88 | -0.11 | -0.14 | 0.82 | 0.81 | 0.68 | 1.03 | 0.85 | 0.57 | 0.5 | 0.35 | 0.77 | 0.78 | -0.16 | 0.2775 | 0.095 | 0.66214 | 0.357771 | 0.752584 | 1 | 0.996511 | 0.961479 |
| Evl | P70429 | 0.86 | 0.7 | 0.41 | 0.67 | 0.57 | 0.86 | 0.47 | 0.57 | 1.13 | 0.7 | 0.67 | 0.96 | 1.28 | 0.8 | 0.64 | 0.54 | -0.0425 | 0.205 | 0.155 | 0.807921 | 0.311712 | 0.509548 | 1 | 0.996511 | 0.902804 |
| Ewsr1 | Q5SUT0 | -4.11 | -4.47 | -4.87 | -5.79 | -4.37 | -4.69 | -5.08 | -5.56 | -4.17 | -3.58 | -4.97 | -3.93 | -3.8 | -2.88 | -5.44 | -5.76 | -0.115 | 0.6475 | 0.34 | 0.782612 | 0.166946 | 0.629477 | 1 | 0.996511 | 0.940703 |
| Exd2 | Q8VEG4 | 0.4 | 0.3 | 0.79 | 0.51 | 0.36 | 0.75 | 1.41 | 1.04 | 0.73 | 0.47 | 1.54 | 0.4 | 0.67 | 0.54 | 1.01 | 1.43 | 0.39 | 0.285 | 0.4125 | 0.154221 | 0.340771 | 0.133915 | 1 | 0.996511 | 0.730839 |
| Exoc1 | Q5PPR2 | -3.58 | -2.98 | -1.75 | -0.73 | -2.59 | -2.4 | -0.91 | -0.39 | -1.83 | 0.19 | -0.36 | -3.44 | -0.74 | -1.38 | -0.37 | -1.49 | 0.6875 | 0.9 | 1.265 | 0.375332 | 0.346625 | 0.071089 | 1 | 0.996511 | 0.668066 |
| Exoc1 | Q6P1Y9 | 0.52 | -0.1 | 0.78 | 1.01 | 0.31 | 0.39 | 0.57 | 1.08 | -0.22 | 1.1 | 0.86 | 0.31 | -0.37 | -0.05 | 0.57 | 0.69 | 0.035 | -0.04 | -0.3425 | 0.905324 | 0.913464 | 0.336691 | 1 | 0.996755 | 0.847578 |
| Exoc2 | Q9D4H1 | 0.17 | -0.06 | -0.11 | -0.43 | 0.34 | 0.23 | -0.31 | -0.27 | 0.08 | -0.53 | -0.2 | 0.05 | -0.08 | 0.32 | -0.56 | -0.63 | 0.105 | -0.0425 | -0.13 | 0.647557 | 0.849076 | 0.640104 | 1 | 0.996511 | 0.942804 |
| Exoc3 | Q6KAR6 | -0.42 | -0.42 | 0.18 | 0.27 | 0.12 | -0.68 | 0.25 | 0.07 | -0.83 | 0.29 | 0.16 | -0.59 | -1.02 | 0 | -0.15 | -0.02 | 0.0375 | -0.145 | -0.2 | 0.894668 | 0.662682 | 0.530069 | 1 | 0.996511 | 0.908663 |
| Exoc4 | O35382 | -0.01 | -0.33 | 0.29 | -0.13 | -0.03 | 0.13 | 0.18 | 0.3 | -0.65 | -0.17 | 0.25 | -0.2 | -0.65 | -0.6 | -0.2 | 0.16 | 0.19 | -0.1475 | -0.2775 | 0.32458 | 0.557239 | 0.300468 | 1 | 0.996511 | 0.830276 |
| Exoc5 | Q3TPX4 | 0.14 | -0.3 | 0.32 | 0.19 | -0.04 | 0.53 | 0.41 | 0.31 | -0.12 | -0.2 | 0.31 | -0.11 | -0.22 | -0.15 | -0.04 | 0.27 | 0.215 | -0.1175 | -0.1225 | 0.321188 | 0.590244 | 0.584667 | 1 | 0.996511 | 0.927734 |
| Exoc6 | Q3U9D6 | 0.21 | -0.11 | -0.04 | -0.16 | 0.49 | 0.39 | -0.03 | -0.05 | -0.18 | -0.1 | 0.08 | 0.17 | -0.36 | 0.21 | -0.35 | -0.31 | 0.225 | 0.0175 | -0.1775 | 0.271754 | 0.922746 | 0.418415 | 1 | 0.996755 | 0.87558 |
| Exoc6b | A6H5Z3 | 0 | 0.02 | 0.45 | 0.48 | 0.3 | -0.06 | 0.18 | 0.51 | -0.05 | 0.51 | 0.39 | -0.18 | -0.23 | 0.38 | 0.01 | 0.15 | -0.005 | -0.07 | -0.16 | 0.980699 | 0.77102 | 0.49042 | 1 | 0.996511 | 0.897531 |
| Exoc7 | O35250 | 0.51 | 0.28 | 0.51 | 0.5 | 0.29 | 0.38 | 0.54 | 0.7 | 0.46 | 0.06 | 0.66 | 0.31 | 0.02 | 0.42 | 0.1 | 0.42 | 0.0275 | -0.0775 | -0.21 | 0.867249 | 0.689474 | 0.293824 | 1 | 0.996511 | 0.825612 |
| Exoc8 | Q6PGF7 | 0.62 | 0.44 | 0.26 | 0.38 | 0.57 | 0.61 | 0.19 | 0.5 | 0.74 | 0.45 | 0.39 | 0.39 | 0.82 | 0.63 | 0.19 | 0.07 | 0.0425 | 0.0675 | 0.0025 | 0.805277 | 0.70764 | 0.991494 | 1 | 0.996511 | 0.996154 |
| Exog | Q8C163 | -1.13 | -1.31 | -0.14 | 0.58 | -1.28 | -0.95 | 0.14 | 0.62 | -1.23 | 0.13 | 0.47 | -1.05 | -1.22 | -1.15 | -0.01 | 0.03 | 0.1325 | 0.08 | -0.0875 | 0.817952 | 0.887073 | 0.866581 | 1 | 0.996511 | 0.978305 |
| Exosc1 | Q9DAA6 | -0.67 | -0.64 | -1.21 | -0.72 | -0.6 | 0.42 | -0.42 | -1.87 | -0.74 | -0.82 | -1.02 | -0.25 | -0.35 | -1.02 | 0.53 | -0.59 | 0.1925 | 0.1025 | 0.4525 | 0.675035 | 0.670027 | 0.218691 | 1 | 0.996511 | 0.787447 |
| Exosc10 | P56960 | -0.98 | -1.3 | -0.79 | -0.68 | -1.3 | -1.55 | -0.07 | -0.45 | -1.78 | 0.08 | -0.65 | -1.34 | -0.81 | -0.62 | 0.3 | 0.07 | 0.095 | 0.015 | 0.6725 | 0.791283 | 0.970705 | 0.054169 | 1 | 0.997408 | 0.661738 |
| Exosc2 | Q8VBV3 | -1.38 | -1.61 | -1.42 | -1.26 | -1.06 | -1.74 | -0.6 | -1.26 | -2.3 | -1.33 | -1.03 | -1.94 | -2.71 | -0.79 | -1.23 | -0.67 | 0.2525 | -0.2325 | 0.0675 | 0.33967 | 0.449415 | 0.880402 | 1 | 0.996511 | 0.979729 |
| Exosc3 | Q7TQK4 | -1.57 | -0.6 | -2.52 | -1.83 | -0.56 | -1.55 | -0.57 | -0.75 | -1.25 | -2.97 | -3.03 | -1.36 | -1.88 | -1.27 | -0.6 | -2.02 | 0.7725 | -0.5225 | 0.1875 | 0.101436 | 0.379042 | 0.697621 | 1 | 0.996511 | 0.955729 |
| Exosc4 | Q921I9 | 1.46 | 0.77 | 0.1 | -3.3 | 1.15 | 1.48 | 2.33 | 0.06 | -1.3 | -4.48 | 1.91 | 1.21 | -0.91 | -1.37 | 1.47 | 2.86 | 1.4975 | -0.4225 | 0.755 | 0.173704 | 0.791756 | 0.563366 | 1 | 0.996511 | 0.920024 |
| Exosc5 | Q9CRA8 | -0.69 | -1.87 | -1.66 | -1.36 | -1.08 | -0.69 | -1.29 | -1.02 | -0.92 | -0.93 | -1.5 | -1.74 | -1.46 | -1.78 | -0.36 | -0.48 | 0.375 | 0.1225 | 0.375 | 0.21357 | 0.709877 | 0.380979 | 1 | 0.996511 | 0.860984 |
| Exosc6 | Q8BTW3 | -1.54 | -2.8 | -2.15 | -2.25 | -2.6 | -1.43 | -2.37 | -2.43 | -1.12 | -0.62 | -2.57 | -1.75 | 0.16 | -2.41 | -0.28 | -1.95 | -0.0225 | 0.67 | 1.065 | 0.949291 | 0.172954 | 0.112672 | 1 | 0.996511 | 0.714111 |
| Exosc7 | Q9D0M0 | -1.45 | -1.48 | -1.32 | -1.99 | -1.46 | -1.91 | -1.85 | -1.37 | -1.49 | -0.58 | -1.16 | -1.42 | -0.83 | -0.48 | -0.69 | -0.29 | -0.0875 | 0.3975 | 0.9875 | 0.695804 | 0.165931 | 0.002164 | 1 | 0.996511 | 0.419079 |
| Exosc8 | Q9D753 | -0.45 | -0.79 | -1.55 | -1.36 | -0.36 | -1.29 | -0.95 | -1.22 | -1.38 | -1.43 | -1.63 | -1.14 | -1.31 | -0.52 | -0.56 | -0.39 | 0.0825 | -0.3575 | 0.3425 | 0.798312 | 0.229306 | 0.316307 | 1 | 0.996511 | 0.838273 |
| Exosc9 | Q9JHI7 | -1.45 | -2.15 | -1 | -1.24 | -1.46 | -1.31 | -0.77 | -1.54 | -1.33 | -0.56 | -0.64 | -1.83 | -1.31 | -0.82 | -0.14 | -0.66 | 0.19 | 0.37 | 0.7275 | 0.53134 | 0.33978 | 0.060142 | 1 | 0.996511 | 0.661738 |
| Exph5 | Q0VAV2 | -0.33 | 1.05 | -1.56 | -0.27 | 0.7 | -0.77 | -2.43 | -1.06 | 1.77 | 0.51 | -1.65 | -0.26 | 1.44 | 2.41 | -1.57 | -2.01 | -0.6125 | 0.37 | 0.345 | 0.425564 | 0.648335 | 0.751355 | 1 | 0.996511 | 0.961479 |
| Ext2 | P70428 | -2.51 | -1.19 | -2.19 | -2.49 | -1.28 | -1.22 | -1.49 | -1.23 | -0.7 | -1.05 | -1.7 | -1.59 | -1.14 | -0.21 | -1.6 | -2.1 | 0.79 | 0.835 | 0.8325 | 0.030463 | 0.050604 | 0.107978 | 1 | 0.996511 | 0.706427 |
| Extl1 | Q9JKV7 | -2.51 | -2.81 | 0.69 | 0.27 | -0.44 | -1.67 | 0.65 | -0.05 | -1.05 | 2.53 | 0.71 | -2.47 | -2.7 | -0.8 | 0.63 | -0.1 | 0.7125 | 1.02 | 0.3475 | 0.451063 | 0.429863 | 0.737914 | 1 | 0.996511 | 0.958796 |
| Extl2 | Q3TSR0 | -0.9 | -3.11 | -2.36 | -2.12 | -1.25 | -0.69 | -1.24 | -2.02 | -2.91 | -3.1 | -2.04 | -1.01 | -2.51 | -2.99 | -2.26 | -2.05 | 0.8225 | -0.1425 | -0.33 | 0.122997 | 0.814458 | 0.489721 | 1 | 0.996511 | 0.897531 |
| Extl3 | Q6P1H4 | -0.79 | -0.97 | -0.67 | -0.63 | -0.12 | -0.75 | -0.13 | -0.55 | -1.43 | -0.56 | -0.55 | -0.89 | -1.1 | -0.68 | -0.37 | -0.32 | 0.3775 | -0.0925 | 0.1475 | 0.09316 | 0.707175 | 0.536649 | 1 | 0.996511 | 0.912212 |
| Ezr | P26040 | 1.47 | 0.86 | 0.93 | 0.9 | 1.23 | 1.19 | 0.5 | 0.88 | 1.42 | 0.62 | 0.72 | 1.03 | 0.81 | 1.45 | 0.27 | 0.33 | -0.09 | -0.0925 | -0.325 | 0.706772 | 0.715262 | 0.319297 | 1 | 0.996511 | 0.838969 |
| F11r | O88792 | 0.2 | -1.06 | -0.5 | -0.44 | -0.31 | -0.7 | -1.35 | -0.86 | 0.05 | 0.7 | -0.78 | 0.34 | 0.54 | 0.28 | -0.67 | -0.8 | -0.355 | 0.5275 | 0.2875 | 0.294468 | 0.20045 | 0.48612 | 1 | 0.996511 | 0.895384 |
| F13a1 | Q8BH61 | -1.02 | -1.51 | -0.92 | -0.42 | -2.34 | -1.69 | -1.49 | -0.86 | -1.19 | -0.45 | -0.61 | -2.02 | -1.08 | -1.75 | -1.53 | -1.03 | -0.6275 | -0.1 | -0.38 | 0.110477 | 0.803202 | 0.223252 | 1 | 0.996511 | 0.787447 |
| F2 | H7BX99 | -3.84 | -4.45 | -5.37 | -4.94 | -3.64 | -3.47 | -5.45 | -5.58 | -3.32 | -3.44 | -4.75 | -3.82 | -2.91 | -2.07 | -6.15 | -5.24 | 0.115 | 0.8175 | 0.5575 | 0.847583 | 0.088435 | 0.54422 | 1 | 0.996511 | 0.912212 |
| F3 | P20352 | 1.18 | 1.48 | 2.36 | 1.77 | 2.07 | 1.04 | 3.23 | 2.24 | 0.32 | 1.59 | 2.33 | 1.31 | -0.06 | 0.64 | 2.06 | 2.14 | 0.4475 | -0.31 | -0.5025 | 0.360027 | 0.502749 | 0.373851 | 1 | 0.996511 | 0.859424 |
| F8a1 | Q00558 | -0.25 | -0.17 | -0.7 | -0.11 | -0.23 | -0.31 | -0.53 | -0.48 | -0.15 | -0.9 | -0.62 | -0.32 | -0.47 | 0.32 | -0.68 | -0.77 | -0.08 | -0.19 | -0.0925 | 0.675151 | 0.437772 | 0.755399 | 1 | 0.996511 | 0.962333 |
| Fa2h | Q5MPP0-2 | 1.18 | 0.93 | -0.26 | -0.62 | 2 | 0.83 | 0.4 | -0.2 | 1.47 | 0.55 | 0.29 | 0.61 | 1.23 | 1.94 | 0.94 | 0.85 | 0.45 | 0.4225 | 0.9325 | 0.44975 | 0.386366 | 0.076029 | 1 | 0.996511 | 0.674054 |
| Faah | O08914 | -0.88 | -0.79 | -0.22 | -0.24 | -0.25 | -0.61 | -0.07 | -0.11 | -0.74 | -0.18 | -0.31 | -0.84 | -0.94 | -0.45 | -0.2 | 0.01 | 0.2725 | 0.015 | 0.1375 | 0.26071 | 0.953628 | 0.63434 | 1 | 0.996755 | 0.941132 |
| Faap100 | A2ACJ2 | -2 | -1.25 | -0.95 | 0.07 | 0.54 | 0.63 | 0.95 | 0.41 | 0.8 | 1.04 | 0.2 | -1.21 | -1.31 | -0.28 | 1.44 | 0.99 | 1.665 | 1.24 | 1.2425 | 0.003236 | 0.067365 | 0.098208 | 1 | 0.996511 | 0.698156 |
| Fabp3 | P11404 | 0.64 | 0.55 | 1.49 | 0.4 | 0.5 | 0.34 | 0.8 | 1.33 | -0.55 | -0.43 | 1.43 | 0.59 | -0.44 | -0.27 | 1 | 1 | -0.0275 | -0.51 | -0.4475 | 0.931565 | 0.314315 | 0.322939 | 1 | 0.996511 | 0.840788 |
| Fabp5 | Q05816 | -0.29 | -0.81 | -0.61 | -0.49 | -0.63 | -0.08 | 0.4 | -0.31 | -1.61 | -1.34 | -0.28 | -0.03 | -1.24 | -2.54 | -0.44 | -0.19 | 0.395 | -0.265 | -0.5525 | 0.145705 | 0.500079 | 0.28912 | 1 | 0.996511 | 0.824705 |
| Fabp7 | P51880 | -1.27 | -1.93 | -1.63 | -1.74 | -1.56 | -1.01 | -1.03 | -1.2 | -0.5 | -1.19 | -1.03 | -0.11 | -0.89 | -1.57 | -1.61 | -1.2 | 0.4425 | 0.935 | 0.325 | 0.065067 | 0.010566 | 0.218885 | 1 | 0.996511 | 0.787447 |
| Fadd | Q61160 | 0.12 | -0.09 | 8.36 | 10.71 | -0.19 | -0.39 | 9.76 | 11.13 | -1.04 | 13.64 | 10.54 | -0.36 | -0.64 | 0.43 | 9.89 | 12.91 | 0.3025 | 0.92 | 0.8725 | 0.935109 | 0.82485 | 0.821991 | 1 | 0.996511 | 0.972768 |
| Fads1 | Q920L1 | 0.05 | -0.09 | 0.09 | -0.03 | 0.51 | 0.38 | 0.92 | 0.19 | -0.23 | -0.71 | 0.26 | 0.39 | -0.35 | 0.2 | 0.2 | 0.32 | 0.495 | -0.0775 | 0.0875 | 0.030379 | 0.774295 | 0.681339 | 1 | 0.996511 | 0.955729 |
| Fads2 | Q9Z0R9 | 0.11 | 0.41 | 1.36 | 0.38 | 0.96 | 0.52 | 2.84 | 1.25 | 0.81 | 1.24 | 1.72 | 0.5 | 1.32 | 0.78 | 2.53 | 3.26 | 0.8275 | 0.5025 | 1.4075 | 0.143856 | 0.197181 | 0.035081 | 1 | 0.996511 | 0.651434 |
| Fads3 | Q9JJE7 | -1.19 | -0.93 | -0.62 | -0.32 | -0.27 | -0.9 | 0.09 | -0.78 | -0.4 | -1 | -0.91 | -0.84 | -0.66 | 0.31 | -0.58 | -0.04 | 0.3 | -0.0225 | 0.5225 | 0.325777 | 0.929076 | 0.116836 | 1 | 0.996755 | 0.718608 |
| Fads6 | Q80UG1 | 0.11 | 1.35 | 1.83 | 1 | 0.95 | 1.45 | 2.63 | 1.4 | -1.18 | 0.61 | 1.09 | 1.05 | -0.08 | 1.13 | 3.06 | 3.52 | 0.535 | -0.68 | 0.835 | 0.276277 | 0.270371 | 0.325711 | 1 | 0.996511 | 0.843092 |
| Faf1 | P54731 | -0.04 | -0.13 | -0.31 | -0.39 | -0.04 | 0 | -0.28 | -0.3 | 0.32 | -0.23 | -0.45 | -0.04 | 0.55 | 0.18 | -0.81 | -0.43 | 0.0625 | 0.1175 | 0.09 | 0.709888 | 0.595328 | 0.77941 | 1 | 0.996511 | 0.967289 |
| Faf2 | Q3TDN2 | 0.29 | 0.42 | -0.32 | -0.19 | 0.14 | 0.4 | -0.25 | -0.09 | 0.15 | -0.4 | -0.42 | 0.18 | 0.4 | 0.25 | -0.33 | -0.41 | 0 | -0.1725 | -0.0725 | 1 | 0.517341 | 0.802031 | 1 | 0.996511 | 0.971091 |
| Fah | P35505 | -0.55 | -1.02 | 0.35 | 0.36 | -1.05 | -0.71 | 0.14 | 0.59 | -0.99 | 0.13 | -0.01 | -1.03 | -1.1 | -0.9 | 0.16 | 0.41 | -0.0425 | -0.26 | -0.1425 | 0.928269 | 0.555514 | 0.765769 | 1 | 0.996511 | 0.964134 |
| Fahd1 | Q8R0F8 | 1.59 | 1.06 | 1 | 1.59 | 0.27 | 1.22 | 0.31 | 1.18 | 0.9 | 0.8 | 1.04 | 1.14 | 0.71 | -0.57 | 0.4 | 0.38 | -0.565 | -0.34 | -1.08 | 0.091997 | 0.144349 | 0.008686 | 1 | 0.996511 | 0.563386 |
| Fahd2a | A0A0R4J09 | -0.27 | 0.16 | 0.43 | 0.7 | -0.22 | -0.03 | 0.93 | 0.65 | -0.42 | 0.19 | 0.57 | -0.26 | -0.7 | -0.3 | 0.2 | 0.44 | 0.0775 | -0.235 | -0.345 | 0.815765 | 0.453983 | 0.313182 | 1 | 0.996511 | 0.835247 |
| Faim | Q9WUD8 | -0.54 | -0.42 | -0.39 | -0.01 | -0.21 | -0.56 | -0.32 | -0.15 | -0.27 | 0.94 | -0.21 | -0.32 | -0.47 | -0.21 | -0.45 | -0.51 | 0.03 | 0.375 | -0.07 | 0.872331 | 0.264659 | 0.727204 | 1 | 0.996511 | 0.958635 |
| Faim2 | Q8K097-2 | 1.14 | 0.82 | 0.72 | 1.01 | 1.03 | 1.39 | 0.67 | 0.79 | 3.64 | 1.33 | 1.07 | 1.6 | 3.23 | 1.34 | 1.24 | 1.17 | 0.0475 | 0.9875 | 0.8225 | 0.822629 | 0.099551 | 0.110193 | 1 | 0.996511 | 0.710755 |
| Fam102a | Q78T81 | -0.3 | -0.34 | 0.03 | -0.22 | -0.37 | -0.29 | -0.05 | -0.56 | -0.36 | -0.39 | -0.45 | -0.26 | -0.74 | -0.17 | -1.24 | -0.78 | -0.11 | -0.1575 | -0.525 | 0.544804 | 0.3665 | 0.071684 | 1 | 0.996511 | 0.668066 |
| Fam107a | A0A5S8E87 | 0.85 | 1.07 | 1.16 | 1.11 | 1.53 | 1.29 | 1.6 | 1.61 | 1.48 | 0.71 | 1.82 | 1.22 | 1.93 | 0.78 | 1.41 | 1.57 | 0.46 | 0.26 | 0.375 | 0.018476 | 0.335526 | 0.193093 | 1 | 0.996511 | 0.777764 |
| Fam107b | Q3TGF2 | 0.46 | 0.48 | 0.15 | -0.3 | 0.2 | 0.53 | -0.34 | -0.31 | 0.53 | 0.49 | -0.27 | 0.21 | 0.63 | 0.46 | -0.2 | -0.16 | -0.1775 | 0.0425 | -0.015 | 0.532383 | 0.876495 | 0.959371 | 1 | 0.996511 | 0.992636 |
| Fam110b | Q8C739 | 0.79 | 0.53 | 0.17 | 0.05 | -0.23 | 0.81 | -0.13 | -0.22 | 0.75 | 0.67 | 0.17 | 0.49 | 1.48 | 0.66 | -0.01 | -0.3 | -0.3275 | 0.135 | 0.0725 | 0.29331 | 0.577328 | 0.860479 | 1 | 0.996511 | 0.977204 |
| Fam114a1 | Q9D281 | -1.48 | -1.26 | -0.72 | -1 | -2.05 | -0.11 | -0.05 | -1.28 | 0.16 | -0.56 | -1.3 | -1.36 | 0.84 | -1.14 | -1.22 | -1.02 | 0.2425 | 0.35 | 0.48 | 0.611189 | 0.368913 | 0.337565 | 1 | 0.996511 | 0.847578 |
| Fam114a2 | Q8VE88-2 | 0.49 | 0.58 | -0.13 | -0.08 | 0.53 | 0.19 | -0.44 | -0.4 | -0.1 | -0.07 | -0.12 | 0.53 | -0.22 | -0.03 | -0.08 | -0.28 | -0.245 | -0.155 | -0.3675 | 0.420933 | 0.55798 | 0.145652 | 1 | 0.996511 | 0.745321 |
| Fam117b | Q3U3E2 | -1 | -1.06 | -0.88 | -0.8 | -1.4 | -1.08 | -1.41 | -0.59 | -0.13 | 0.37 | -1.4 | -1.11 | 0.16 | -0.44 | -0.24 | -0.95 | -0.185 | 0.3675 | 0.5675 | 0.417034 | 0.370104 | 0.057014 | 1 | 0.996511 | 0.661738 |
| Fam118a | Q91YN1 | -2.57 | -2.13 | -1.88 | -2.59 | -2.42 | -2.73 | -1.73 | -2.3 | -2.31 | -0.95 | -1.61 | -2.64 | -1.75 | -1.65 | -3.08 | -1.41 | -0.0025 | 0.415 | 0.32 | 0.992762 | 0.31137 | 0.431749 | 1 | 0.996511 | 0.879165 |
| FAM120A | Q6A0A9 | -1.33 | -1.24 | -1.53 | -1.53 | -1.16 | -1.3 | -1.38 | -1.71 | -1.3 | -1.02 | -1.57 | -1.26 | -0.88 | -0.86 | -1.05 | -1.58 | 0.02 | 0.12 | 0.315 | 0.912287 | 0.534021 | 0.191822 | 1 | 0.996511 | 0.777117 |
| Fam120b | Q6RI63 | 0.37 | -0.41 | -1.42 | -1.15 | -0.89 | 0.36 | -2.27 | -1.42 | -0.44 | -0.87 | -1.29 | -0.43 | -0.22 | -1.13 | -1.33 | -1.5 | -0.4025 | -0.105 | -0.3925 | 0.52188 | 0.806089 | 0.407505 | 1 | 0.996511 | 0.874084 |
| Fam120c | Q8C3F2 | -1.24 | -1.4 | -0.77 | -0.81 | -1.34 | -1.42 | -0.32 | -0.69 | -1.32 | -0.46 | -0.83 | -1.31 | -1.06 | -1.33 | -0.79 | -0.77 | 0.1125 | 0.075 | 0.0675 | 0.71209 | 0.784761 | 0.78096 | 1 | 0.996511 | 0.9676 |
| Fam124a | D3Z5V4 | -1.69 | -1.77 | -2.53 | -2.39 | -0.34 | -0.6 | -1.47 | -1.87 | -0.88 | -3.43 | -0.19 | -0.53 | -3.65 | -0.07 | -3.47 | -2.45 | 1.025 | 0.8375 | -0.315 | 0.027711 | 0.250242 | 0.682983 | 1 | 0.996511 | 0.955729 |
| Fam126a | Q6P9N1 | 0.61 | 0.41 | 0.66 | 0.82 | 0.69 | 0.62 | 1.15 | 1.06 | 1.15 | 1.56 | 0.76 | 1 | 1.31 | 1.37 | 1.28 | 1.18 | 0.255 | 0.4925 | 0.66 | 0.209905 | 0.051848 | 0.005348 | 1 | 0.996511 | 0.552614 |
| Fam126b | Q8C729-2 | 0.84 | 0.95 | 1.36 | 1.26 | 1.05 | 0.92 | 0.98 | 1.33 | 0.65 | 1.32 | 1.17 | 0.84 | 0.73 | 1.22 | 0.93 | 1.21 | -0.0325 | -0.1075 | -0.08 | 0.865008 | 0.64076 | 0.718551 | 1 | 0.996511 | 0.958398 |
| Fam131a | D3YXU9 | -1.44 | -0.4 | -0.04 | 0 | -0.08 | -1.68 | -0.33 | -1.39 | -1.14 | -1.15 | -0.76 | -0.8 | -1.03 | -0.14 | -1.03 | -1.43 | -0.4 | -0.4925 | -0.4375 | 0.41218 | 0.179107 | 0.30668 | 1 | 0.996511 | 0.832533 |
| Fam131b | E9Q8P8 | 0.64 | 0.9 | 1.23 | 1.28 | 0.95 | 0.65 | 0.74 | 1.12 | 0.47 | 1.21 | 1.14 | 0.58 | 0.38 | 0.45 | 1.15 | 0.8 | -0.1475 | -0.1625 | -0.3175 | 0.490919 | 0.537744 | 0.243598 | 1 | 0.996511 | 0.79948 |
| Fam135a | A0A087WF | 1.41 | 1.82 | 0.5 | -0.26 | 1.33 | 0.85 | 0.02 | 0.17 | 0.47 | 0.29 | -0.18 | 1.13 | 0.66 | 1.45 | -0.13 | 1.63 | -0.275 | -0.44 | 0.035 | 0.594885 | 0.391172 | 0.950779 | 1 | 0.996511 | 0.992476 |
| Fam135b | Q9DAI6 | 1.28 | 1.2 | 0.41 | 0.8 | 1.04 | 1.02 | -0.09 | 0.11 | 1.19 | 1.31 | 0.44 | 1.06 | 1.44 | 1.45 | 0.77 | 0.51 | -0.4025 | 0.0775 | 0.12 | 0.263152 | 0.788244 | 0.707256 | 1 | 0.996511 | 0.957157 |
| Fam136a | Q9CR98 | -0.86 | -1.05 | -1.27 | -1.23 | -1.51 | -0.81 | -1.46 | -1.5 | -0.18 | -0.99 | -1.18 | -0.67 | 0.56 | -1.16 | -0.57 | -1.6 | -0.2175 | 0.3475 | 0.41 | 0.334009 | 0.20126 | 0.3748 | 1 | 0.996511 | 0.859611 |
| Fam13c | G3X9S1 | 0.91 | 1.62 | 0.43 | 0.58 | 1.23 | 1.18 | 0.69 | 0.52 | 1.67 | 0.61 | 0.45 | 1.02 | 1.48 | 1.58 | 0.44 | 0.2 | 0.02 | 0.0525 | 0.04 | 0.948983 | 0.886555 | 0.924351 | 1 | 0.996511 | 0.988215 |
| Fam151b | D3YUE4 | -1.85 | -1.49 | -2.35 | -2.11 | -1.78 | -1.62 | -1.66 | -2.35 | -2.13 | -2.05 | -2.23 | -1.55 | -2.43 | -2.08 | -2.65 | -2.58 | 0.0975 | -0.04 | -0.485 | 0.707819 | 0.876897 | 0.084222 | 1 | 0.996511 | 0.690427 |
| Fam162a | Q9D6U8 | 1.39 | 1.09 | 1.51 | 1.29 | 1.03 | 1.5 | 1.46 | 1.33 | 0.45 | 1.71 | 1.56 | 1.16 | 0.56 | 0.28 | 1.62 | 1.37 | 0.01 | -0.1 | -0.3625 | 0.956196 | 0.74033 | 0.294286 | 1 | 0.996511 | 0.825612 |
| Fam163b | Q8BUM6 | -0.07 | 0.61 | 0.02 | -0.02 | 0.91 | 0.18 | 0.28 | 0.2 | -0.1 | 0.13 | 0.07 | 0.26 | -0.21 | 0.22 | -0.06 | 0.41 | 0.2575 | -0.045 | -0.045 | 0.315435 | 0.834808 | 0.855319 | 1 | 0.996511 | 0.976225 |
| Fam169a | Q5XG69 | -2.79 | -4.4 | -3.88 | -4.11 | -4.06 | -3.24 | -4.12 | -4.36 | -3.45 | -1.79 | -3.82 | -3.43 | -3.85 | -2.95 | -4.52 | -4.72 | -0.15 | 0.6725 | -2.15E-01 | 0.710665 | 0.226535 | 0.666449 | 1 | 0.996511 | 0.95055 |
| Fam171a1 | A2ATK9 | 1.41 | 1.81 | 0.94 | 0.84 | 2.11 | 0.98 | 1.07 | 0.96 | 1.06 | 0.88 | 0.91 | 1.68 | 1.15 | 1.65 | 1.29 | 1.13 | 0.03 | -0.1175 | 0.055 | 0.930484 | 0.694713 | 0.842329 | 1 | 0.996511 | 0.976225 |
| Fam171a2 | A2A699 | 0.49 | 1.34 | 0.37 | 0.52 | 1.2 | 0.77 | 0.59 | 1.02 | 1.42 | 0.47 | 1.17 | 1.21 | 1.35 | 1.69 | 0.38 | 0.35 | 0.215 | 0.3875 | 0.2625 | 0.429103 | 0.22871 | 0.510128 | 1 | 0.996511 | 0.903419 |
| Fam171b | Q14CH0-2 | 0.48 | 0.67 | 0.85 | 1.13 | 0.74 | 0.62 | 0.85 | 1.22 | 1 | 1.38 | 1.19 | 0.81 | 1.08 | 1.09 | 1.16 | 0.67 | 0.075 | 0.3125 | 0.2175 | 0.728179 | 0.182494 | 0.347288 | 1 | 0.996511 | 0.851503 |
| Fam172a | E9QMA2 | -2.28 | -2.17 | -1.41 | -0.95 | -2.31 | -2.63 | -0.8 | -0.95 | -1.02 | -1.48 | -1.89 | -2.25 | -0.3 | -1.93 | -1.1 | -1.57 | 0.03 | 0.0425 | 0.4775 | 0.953811 | 0.91437 | 0.302502 | 1 | 0.996755 | 0.830923 |
| Fam174a | Q9D3L0 | -1.71 | -2.41 | -3.1 | -3.07 | -1.86 | -2.07 | -3.54 | -3.03 | -1.84 | -2.08 | -3.41 | -2.74 | -1.89 | -2.01 | -2.7 | -2.73 | -0.0525 | 0.055 | 0.24 | 0.91238 | 0.903375 | 0.537935 | 1 | 0.996609 | 0.912212 |
| Fam177a1 | Q8BR63 | 0.39 | 0.44 | 0.45 | -0.04 | 0.33 | 0.2 | 0.21 | -0.64 | 0.92 | -0.19 | -0.83 | 0.11 | 1.07 | 0.14 | 0.7 | 0.36 | -0.285 | -0.3075 | 0.2575 | 0.291954 | 0.415448 | 0.340341 | 1 | 0.996511 | 0.84829 |
| Fam184a | E9PW83 | 0.05 | 0.14 | 0.27 | 0.53 | 0.06 | -0.13 | 0.5 | 0.46 | 0.1 | 0.91 | 0.36 | 0.35 | -0.2 | -0.03 | 0.34 | 0.33 | -0.025 | 0.1825 | -0.1375 | 0.906383 | 0.439258 | 0.537896 | 1 | 0.996511 | 0.912212 |
| Fam184b | Q0KK56 | -0.51 | 0.42 | -0.35 | -0.62 | -0.26 | 0.44 | -0.27 | -1.94 | 0.86 | -0.25 | -0.49 | 0.33 | 0.15 | 0.57 | -1.88 | 0.51 | -0.2425 | 0.3775 | 0.1025 | 0.638676 | 0.324025 | 0.85859 | 1 | 0.996511 | 0.976225 |
| Fam185a | Q7TPD2 | 0.2 | 0.5 | 0.14 | 0.34 | 0.33 | 0.3 | 0.24 | 0.14 | -0.04 | 0.4 | 0.32 | 0.01 | 0.54 | 0.32 | 0.14 | 0.3 | -0.0425 | -0.1225 | 0.03 | 0.786521 | 0.527839 | 0.875199 | 1 | 0.996511 | 0.979626 |
| Fam189a1 | Q6A044-2 | 0.32 | 1.08 | 1.26 | 1.71 | 0.19 | 0.39 | 0.63 | 1.07 | 1.49 | 2.23 | 1.21 | 0.98 | 2.31 | -0.07 | 1.22 | 1.21 | -0.5225 | 3.85E-01 | 0.075 | 0.14585 | 0.328651 | 0.886375 | 1 | 0.996511 | 0.979767 |
| Fam193a | M0QWZ1 | -1.66 | -1.98 | -0.92 | -2.64 | -2.23 | -1.7 | -2.77 | -2.08 | -0.93 | -4.44 | -1.75 | -2.17 | -0.3 | -3.18 | -0.75 | -1.48 | -0.395 | -0.5225 | 0.3725 | 0.333966 | 0.49437 | 0.576678 | 1 | 0.996511 | 0.927198 |
| Fam20b | Q8VCS3 | 0.35 | -0.4 | -1.6 | -0.97 | 0.06 | 0.13 | -0.87 | -1.17 | -0.5 | -1.49 | -1.27 | 0.3 | -0.65 | -1.02 | -0.73 | -0.72 | 0.1925 | -0.085 | -0.125 | 0.694885 | 0.874169 | 0.758981 | 1 | 0.996511 | 0.963123 |
| Fam20c | Q5MJS3 | -0.37 | -0.44 | 1.81 | -0.42 | 0.04 | -1.72 | 1.82 | 0.62 | 1.97 | 0.1 | 1.3 | -0.99 | 2.61 | 0.33 | 1 | 0.52 | 0.045 | 0.45 | 0.97 | 0.95677 | 0.566223 | 0.183586 | 1 | 0.996511 | 0.776843 |
| Fam210a | Q8BGY7 | 0.24 | 0.41 | 0.59 | 0.64 | -0.03 | 0.36 | 0.15 | 0.74 | -0.05 | 0.67 | 1.04 | 0.33 | -0.06 | -0.26 | 0.4 | 0.54 | -0.165 | 0.0275 | -0.315 | 0.450215 | 0.917899 | 0.220968 | 1 | 0.996755 | 0.787447 |
| Fam210b | Q9D8B6 | 0.2 | 0.48 | -0.32 | -0.24 | -0.04 | 0.46 | 0.03 | -0.72 | 0.51 | -1.62 | -0.23 | 0.11 | 1.22 | 0.36 | -0.99 | -0.39 | -0.0975 | -0.3375 | 0.02 | 0.749616 | 0.477615 | 0.966811 | 1 | 0.996511 | 0.9927 |
| Fam217b | A2AJW5 | -0.75 | -0.65 | -1.37 | -0.59 | -0.47 | 1.14 | -1.14 | -1.17 | -0.5 | -0.18 | -1.03 | -0.67 | -0.35 | 0.1 | -1.15 | -1.26 | 0.43 | 0.245 | 0.175 | 0.421546 | 0.373238 | 0.634271 | 1 | 0.996511 | 0.941132 |
| Fam219a | A2ANP1 | -1.76 | -0.99 | -0.28 | -0.41 | -1.12 | -1.54 | -0.23 | -0.62 | -0.99 | -0.58 | -0.41 | -0.89 | -0.97 | -0.36 | -0.05 | -0.36 | -0.0175 | 0.1425 | 0.425 | 0.966357 | 0.689228 | 0.27936 | 1 | 0.996511 | 0.822084 |
| Fam234a | Q8C0Z1 | 1.17 | 1.57 | 1.31 | 0.67 | 1.57 | 1.16 | 1.6 | 0.99 | 0.16 | 0.95 | 1.19 | 1.2 | 0.42 | -0.43 | 0.91 | 1.58 | 0.15 | -0.305 | -0.56 | 0.557632 | 0.342202 | 0.22419 | 1 | 0.996511 | 0.787447 |
| Fam234b | Q8BYI8 | -1.51 | -1.21 | -0.74 | -0.99 | -0.72 | -1.22 | -0.52 | -1.03 | -1.1 | -0.89 | -0.84 | -1.21 | -0.85 | -0.23 | -1 | -0.72 | 0.24 | 0.1025 | 0.4125 | 0.334261 | 0.645872 | 0.141506 | 1 | 0.996511 | 0.737973 |
| Fam241b | D3Z7G3 | 0.59 | 0.85 | 0.44 | 0.5 | 0.73 | 0.96 | 0.01 | 0.51 | -0.3 | -0.82 | 0.52 | 0.63 | -0.08 | -1.81 | 1.23 | 0.43 | -0.0425 | -0.5875 | -0.6525 | 0.858454 | 0.120578 | 0.290255 | 1 | 0.996511 | 0.825134 |
| Fam3c | G3X9I1 | -2.11 | -3.06 | -2.56 | -2.06 | -1.39 | -2.06 | -1.72 | -2.05 | -3.17 | -1.64 | -2.44 | -2.21 | -2.16 | -2.45 | -2.09 | -1.92 | 0.6425 | 0.0825 | 0.2925 | 0.047452 | 0.827865 | 0.309796 | 1 | 0.996511 | 0.834523 |
| Fam50a | Q9WV03 | -2.7 | -1.79 | -2.54 | -2.54 | -2.37 | -2.54 | -3.59 | -2.57 | -0.59 | -0.13 | -2.84 | -2.45 | -1.06 | 0.94 | -2.78 | -2.79 | -0.375 | 0.89 | 0.97 | 0.279774 | 0.188883 | 0.255508 | 1 | 0.996511 | 0.809524 |
| Fam76b | Q80XP8 | -3.38 | -5.58 | -4.86 | -7.72 | -4.21 | -4.16 | -3.75 | -5.42 | -6.57 | -5.08 | -4.91 | -4.48 | -3.41 | -3.95 | -0.96 | -2.32 | 1 | 0.125 | 2.725 | 0.271381 | 0.890739 | 0.022279 | 1 | 0.996511 | 0.621926 |
| Fam78b | F6ZXE6 | -1.1 | -0.53 | 0.84 | -0.42 | -0.54 | -1.04 | 0.76 | 0.3 | -1 | 1.63 | 0.06 | -1.46 | -0.56 | -0.43 | -0.33 | 1.43 | 0.1725 | 0.11 | 0.33 | 0.744995 | 0.879313 | 0.568264 | 1 | 0.996511 | 0.922462 |
| Fam81a | Q3UXZ6 | 1.87 | 1.73 | 2.27 | 1.85 | 1.96 | 1.94 | 2.55 | 2.64 | 0.9 | -0.38 | 2.3 | 1.53 | 0.87 | 0.68 | 2.05 | 2.07 | 0.3425 | -0.8425 | -0.5125 | 0.174246 | 0.142228 | 0.201268 | 1 | 0.996511 | 0.780919 |
| Fam89b | Q05DI7 | 0 | -0.53 | 7.29 | 1.01 | -0.12 | 0.87 | 6.7 | 6.87 | -0.47 | 0.46 | 7.47 | 0.14 | -0.52 | 0.4 | 8.04 | 7.79 | 1.6375 | -0.0425 | 1.985 | 0.485496 | 0.985341 | 0.452347 | 1 | 0.998892 | 0.885993 |

| | | | | | | | | | | | | | | | | | | | | | | | | | | |
|---|---|---|---|---|---|---|---|---|---|---|---|---|---|---|---|---|---|---|---|---|---|---|---|---|---|---|
| Fam8a1 | Q3URQ4 | 1.03 | -0.88 | -0.52 | 0 | -0.58 | 0.92 | -0.8 | 0.21 | 3.98 | -0.46 | -0.45 | 0.5 | 4.37 | 0.16 | 1.1 | 0.79 | 0.03 | 0.985 | 1.6975 | 0.954433 | 0.346313 | 0.093419 | 1 | 0.996511 | 0.698156 |
| Fam91a1 | Q3UVG3 | -0.01 | -0.06 | 0.13 | 0.03 | 0.06 | -0.07 | 0.36 | 0 | 0.33 | -0.44 | 0.14 | -0.02 | 0.67 | -0.89 | 0.12 | 0.16 | 0.065 | -0.02 | -0.0075 | 0.690537 | 0.924476 | 0.981952 | 1 | 0.996755 | 0.994858 |
| Fam98a | Q3TJZ6 | -1.12 | -0.64 | -1.13 | -1.43 | -0.45 | -0.8 | -1.18 | -1.41 | -0.24 | -0.77 | -1.12 | -0.38 | -0.22 | -0.02 | -0.89 | -1.28 | 0.12 | 0.4525 | 0.4775 | 0.661122 | 0.123751 | 0.180252 | 1 | 0.996511 | 0.775542 |
| Fam98b | Q80VD1 | -2.02 | -1.52 | -1.5 | -1.33 | -1.49 | -2.11 | -1.02 | -1.41 | -0.83 | -1.06 | -1.41 | -1.73 | -1.31 | 0.27 | -1.75 | -1.25 | 0.085 | 0.335 | 0.5825 | 0.757422 | 0.227205 | 0.209263 | 1 | 0.996511 | 0.786218 |
| Fam98c | E9PYD1 | -1.29 | -1.08 | -1.62 | -2.07 | -1.31 | -1.37 | -1.68 | -2.05 | -0.52 | -0.98 | -2.08 | -1.1 | 0.02 | -0.73 | -1.73 | -1.55 | -0.0875 | 0.345 | 0.5175 | 0.753537 | 0.374318 | 0.253017 | 1 | 0.996511 | 0.809524 |
| Far1 | Q922J9 | 0.03 | 0.2 | -0.41 | -0.46 | 1.25 | 0.44 | 1.05 | -0.19 | 0.14 | 0.07 | 0.04 | 0.78 | 0.51 | 0.85 | 1 | 0.92 | 7.98E-01 | 0.4175 | 0.98 | 0.046038 | 0.134004 | 0.0025 | 1 | 0.996511 | 0.42982 |
| Farp1 | F8VPU2 | 1.23 | 0.45 | 0.53 | 0.74 | 0.82 | 1.38 | 1.08 | 0.97 | 0.32 | 0.06 | 0.91 | 0.83 | 0 | 0.32 | 0.1 | 0.42 | 0.325 | -0.2075 | -0.5275 | 0.183225 | 0.467122 | 0.0516 | 1 | 0.996511 | 0.658454 |
| Farp2 | Q91VS8 | 0.74 | 0.5 | -0.05 | -0.1 | 0.53 | 0.57 | 0.55 | 0.41 | -0.21 | -1.25 | 0.19 | 0.68 | -0.38 | 0.24 | -0.09 | 0.55 | 0.2425 | -0.42 | -0.1925 | 0.306597 | 0.346601 | 0.527738 | 1 | 0.996511 | 0.908538 |
| Fars2 | Q99M01 | 0.06 | 0.25 | 1.07 | 0.92 | 0.33 | -0.17 | 1.04 | 0.88 | 0.78 | 0.5 | 1.25 | -0.17 | 0.91 | 2.15 | 0.97 | 0.98 | -0.055 | 0.015 | 0.6775 | 0.876952 | 0.967953 | 0.101115 | 1 | 0.996755 | 0.698156 |
| Farsa | Q8C0C7 | -2.2 | -2.57 | -2.73 | -3.13 | -2.47 | -2.21 | -2.59 | -3.12 | -2.39 | -2.15 | -2.9 | -2.12 | -2.4 | -2.17 | -3.04 | -2.89 | 0.06 | 0.2675 | 0.0325 | 0.827813 | 0.3477 | 0.912339 | 1 | 0.996511 | 0.986458 |
| Farsb | Q9WUA2 | -2.48 | -2.87 | -2.5 | -2.74 | -2.46 | -2.83 | -2.47 | -2.85 | -2.99 | -1.73 | -2.63 | -2.23 | -2.74 | -2.19 | -2.66 | -2.63 | -0.005 | 0.2525 | 0.0925 | 0.978469 | 0.401288 | 0.664812 | 1 | 0.996511 | 0.949825 |
| Fasn | P19096 | 0.15 | -0.09 | 0.14 | 0.12 | 0.02 | 0.32 | -0.15 | 0.16 | 0.23 | -0.1 | 0.24 | 0.23 | -0.02 | -0.1 | 0.03 | 0.12 | 0.0075 | 0.07 | -0.0725 | 0.964561 | 0.688648 | 0.67919 | 1 | 0.996511 | 0.955269 |
| Fastkd2 | Q922E6 | 0.2 | 0.05 | 0.24 | 0.51 | -1.22 | 0.2 | -0.2 | 0.13 | 0.38 | 0.64 | 0.45 | -0.19 | 0.98 | -0.68 | 0.37 | 0.54 | -0.5225 | 0.07 | 0.0525 | 0.141887 | 0.764583 | 0.8839 | 1 | 0.996511 | 0.979767 |
| Fastkd5 | Q7TMV3 | 0.04 | 0.63 | 0.19 | -0.16 | 0.28 | -0.16 | -0.2 | -0.32 | -0.31 | -0.36 | -0.11 | 0.2 | -0.71 | -0.35 | -0.32 | -0.11 | -0.275 | -0.32 | -0.5475 | 0.254743 | 0.202925 | 0.049133 | 1 | 0.996511 | 0.658454 |
| Fat1 | F2Z4A3 | 0.1 | 0.63 | -0.2 | -0.33 | 0.55 | 0.19 | -0.4 | -0.32 | 0.55 | -0.31 | -0.46 | 0.32 | 0.74 | 0.31 | 0.1 | 0.09 | -0.045 | -0.025 | 0.26 | 0.882825 | 0.938264 | 0.368073 | 1 | 0.996755 | 0.859245 |
| Fat3 | E9QK16 | 0.56 | 0.79 | 0.65 | 0.78 | 0.57 | 0.54 | 0.54 | 0.47 | 0.9 | 0.46 | 0.57 | 0.59 | 1.2 | 0.81 | 0.61 | 0.29 | -0.165 | -0.065 | 0.0325 | 0.271868 | 0.715692 | 0.891408 | 1 | 0.996511 | 0.982143 |
| Fat4 | Q2PZL6 | 0.46 | 1.58 | 0.13 | 0.48 | 1.44 | 0.52 | 0.08 | 1.91 | 0.02 | 0.65 | 2.31 | 0.86 | 0.26 | 0.57 | 0.74 | 2.6 | 0.325 | 0.2975 | 0.38 | 0.50821 | 0.581182 | 0.508286 | 1 | 0.996511 | 0.902804 |
| Faxc | Q3UMF9 | 1.11 | 1.04 | 0.5 | 0.46 | 0.75 | 0.88 | -0.07 | -0.01 | 1.13 | 0.79 | -0.09 | 1.06 | 0.68 | 1.74 | -0.16 | -0.19 | -0.39 | -0.055 | -0.26 | 0.216262 | 0.866712 | 0.574081 | 1 | 0.996511 | 0.925754 |
| Fbh1 | Q8K2I9 | -3.69 | -2.73 | -3.14 | -0.72 | -3.63 | -3.76 | -0.89 | -2.68 | -2.42 | -1.73 | -1.01 | -3.51 | -1.62 | -4.84 | -1.58 | -1.72 | -0.17 | 0.4025 | 0.13 | 0.838509 | 0.598813 | 0.887936 | 1 | 0.996511 | 0.98057 |
| Fbl | P35550 | 2.11 | 1.1 | 1.54 | 1.66 | 1.38 | 2.01 | 1.71 | 1.74 | 0.75 | 0.51 | 1.77 | 1.36 | 0.78 | 0.64 | 1.9 | 1.98 | 0.1075 | -0.505 | -0.2775 | 0.674654 | 0.171447 | 0.491975 | 1 | 0.996511 | 0.897531 |
| Fbll1 | Q80WS3 | 1.97 | 1.15 | 0.87 | 0.94 | 1.37 | 1.92 | 1.06 | 0.9 | 0.73 | 0.76 | 1.2 | 1.47 | 0.57 | 0.81 | 1.58 | 1.04 | 0.08 | -0.1925 | -2.33E-01 | 0.808055 | 0.542104 | 0.4931 | 1 | 0.996511 | 0.897531 |
| Fbln1 | Q08879-2 | 0.06 | -1.47 | -1.09 | -0.97 | -1.52 | -0.65 | -0.92 | -0.43 | -1.2 | -0.37 | -0.6 | 0.19 | -1.95 | -0.61 | -0.83 | -1.02 | -0.0125 | 0.3725 | -0.235 | 0.973894 | 0.379966 | 0.579629 | 1 | 0.996511 | 0.927734 |
| Fbln5 | A0A1Y7VJV | -3.16 | -3.82 | -3.49 | -3.21 | -3.21 | -3.27 | -3.31 | -3.55 | -3.39 | -3.16 | -3.01 | -3.01 | -2.91 | -3.57 | -3.04 | -3.8 | 0.085 | 0.2775 | 0.09 | 0.67444 | 0.220571 | 0.749178 | 1 | 0.996511 | 0.961014 |
| Fbn1 | Q61554 | -2.81 | -3.32 | -1.52 | -1.48 | -2.99 | -3.14 | -2.04 | -2.39 | -2.05 | -0.62 | -1.77 | -2.69 | -1.7 | -3.25 | -1.64 | -1.75 | -0.3575 | 0.5 | 0.1975 | 0.472655 | 0.401103 | 0.72414 | 1 | 0.996511 | 0.958398 |
| Fbn2 | Q61555 | -8.69 | -11.31 | -1.73 | -2.63 | -14.07 | -10.07 | -2.32 | -1.85 | -7.47 | -1.63 | -1.49 | -6.78 | -5.68 | -13.45 | -2.13 | -2.23 | -0.9875 | 1.7475 | 0.2175 | 0.77033 | 0.494222 | 0.944415 | 1 | 0.996511 | 0.991865 |
| Fbxl15 | Q91W61 | -0.12 | -0.59 | -2.92 | -2.64 | -0.88 | -0.12 | -2.48 | -2.75 | -0.42 | -2.34 | -2.89 | -0.34 | -0.91 | -0.39 | -3.53 | -3.26 | 0.01 | 0.07 | -0.455 | 0.990659 | 0.935836 | 0.636922 | 1 | 0.996755 | 0.942665 |
| Fbxl16 | A2RT62 | -0.88 | -0.75 | -0.19 | -0.01 | -0.84 | -0.56 | 0.18 | 0.1 | -1.26 | -0.68 | -0.09 | -0.67 | -1.26 | -0.77 | -0.85 | -0.27 | 0.1775 | -0.2175 | -0.33 | 0.582294 | 0.501719 | 0.295162 | 1 | 0.996511 | 0.825612 |
| Fbxl17 | Q9QZN1 | -2.92 | -2.84 | -0.68 | -0.13 | -2.25 | -2.02 | -1.87 | 0.12 | -1.62 | 0.21 | 0.19 | -2.62 | -1.81 | -1.65 | 0.01 | -0.36 | 0.1375 | 0.6825 | 0.69 | 0.866407 | 0.459174 | 0.382687 | 1 | 0.996511 | 0.860984 |
| Fbxl18 | F8VQK6 | -0.04 | -0.48 | -0.35 | -0.33 | -0.3 | -0.08 | -0.42 | -0.49 | 0.78 | -0.1 | -0.34 | 0.13 | 1.53 | 0.03 | -0.83 | -0.38 | -0.0225 | 0.4175 | 0.3875 | 0.89863 | 0.151997 | 0.43442 | 1 | 0.996511 | 0.880558 |
| Fbxl2 | Q8BH16 | -0.26 | -1.46 | 0.31 | -0.68 | -1.25 | -1.04 | 1.23 | -0.24 | -2.49 | -1.85 | 1.51 | -1.64 | -0.86 | -3.35 | 0.35 | 0.73 | 0.1975 | -0.595 | -0.26 | 0.748367 | 0.501579 | 0.770875 | 1 | 0.996511 | 0.965933 |
| Fbxl20 | Q9CZV8 | -1.24 | -2.4 | -0.56 | 0.32 | -2.01 | 0.09 | -0.44 | 0.06 | -1.65 | 0.8 | -0.36 | -1.28 | -0.52 | -2.64 | -0.5 | -0.11 | 0.395 | 0.3475 | 0.0275 | 0.568432 | 0.63117 | 0.970085 | 1 | 0.996511 | 0.992767 |
| Fbxl4 | Q8BH70 | 0.71 | 0.91 | 0.7 | 0.64 | 0.58 | 1.43 | -0.34 | 0.37 | 1.67 | -0.09 | 0.5 | 0.39 | 0.72 | 1.01 | 0.19 | 0.11 | -0.23 | -0.1225 | -0.2325 | 0.521256 | 0.73911 | 0.372952 | 1 | 0.996511 | 0.859424 |
| Fbxo10 | Q7TQF2 | -1.48 | -1.24 | -0.21 | -0.09 | -1.32 | -1.25 | -0.28 | 0.03 | -0.62 | 0.17 | 0.53 | -1 | 0.25 | -1.18 | 0.18 | 0.08 | 0.05 | 0.525 | 0.5875 | 0.912756 | 0.278791 | 0.224499 | 1 | 0.996511 | 0.787447 |
| Fbxo11 | Q7TPD1-3 | -1.91 | -2.24 | -1.46 | -1.15 | -1.15 | -1.81 | -0.48 | -0.44 | -1.14 | -0.45 | -1.5 | -1.83 | -0.04 | -0.16 | 0.71 | 0.26 | 0.72 | 0.46 | 1.8825 | 0.087662 | 0.23429 | 0.000247 | 1 | 0.996511 | 0.270112 |
| Fbxo2 | Q80UW2 | 0.13 | -0.35 | 0.88 | 0.96 | -0.24 | 0.15 | 1.52 | 0.67 | -1.19 | 0.61 | 0.69 | 0 | -1.3 | -0.3 | 0.45 | 1.02 | 0.12 | -0.3775 | -0.4375 | 0.793597 | 0.456264 | 0.431959 | 1 | 0.996511 | 0.879165 |
| Fbxo21 | Q8VDH1 | -0.57 | -0.5 | -0.66 | -0.98 | -0.34 | -0.76 | -0.36 | -0.84 | -0.89 | -1.34 | -0.94 | -0.75 | -1.18 | -1.02 | -1.07 | -0.98 | 0.1025 | -0.3025 | -0.385 | 0.612669 | 0.172492 | 0.069975 | 1 | 0.996511 | 0.668066 |
| Fbxo22 | Q78JE5 | -1.76 | -2.29 | -1.46 | -1.54 | -2.1 | -1.66 | -1.86 | -1.79 | -2.08 | -0.67 | -1.42 | -1.51 | -2.11 | -1.98 | -1.44 | -1.7 | -0.09 | 0.3425 | -0.045 | 0.694798 | 0.328302 | 0.865503 | 1 | 0.996511 | 0.977956 |
| Fbxo3 | Q9DC63-3 | -0.4 | -0.59 | -0.31 | -0.96 | -0.27 | -0.5 | -0.3 | -0.53 | -1.19 | -1 | -0.6 | -0.59 | -1.1 | -1.03 | -0.38 | -0.49 | 0.165 | -0.28 | -0.185 | 0.406808 | 0.256041 | 0.485965 | 1 | 0.996511 | 0.895384 |
| Fbxo30 | Q8BJL1 | -0.86 | -0.79 | -1.29 | -0.83 | -1.19 | -0.8 | -0.81 | -1.07 | -0.1 | -0.37 | -1.31 | -0.68 | 0.13 | -0.23 | -1.11 | -1.04 | -0.025 | 0.3275 | 0.38 | 0.895403 | 0.281587 | 0.268031 | 1 | 0.996511 | 0.819742 |
| Fbxo31 | Q3TQF0 | -1.39 | -1 | 0.89 | 0.67 | -0.91 | -0.91 | 0.72 | 0.55 | -0.08 | 0.93 | 0.36 | -1.3 | 0.73 | 0.42 | 0.5 | 0.44 | 0.07 | 0.185 | 0.73 | 0.915854 | 0.785747 | 0.198245 | 1 | 0.996511 | 0.78004 |
| Fbxo38 | Q8BMI0 | -0.68 | -0.31 | -0.25 | 0.09 | -1.19 | -0.63 | -0.48 | 0.41 | 0.41 | 1.2 | 0.52 | -0.43 | 1.24 | -1.35 | 0.46 | -0.13 | -0.185 | 0.7125 | 0.3425 | 0.602549 | 0.074884 | 0.520689 | 1 | 0.996511 | 0.907667 |
| Fbxo4 | Q8CHQ0 | 0.08 | -0.14 | 0.33 | 0.11 | 0.07 | 0.2 | 0.52 | 0.21 | 0.34 | 0.08 | 0.28 | 0.1 | -0.18 | 0.31 | -0.11 | 0.37 | 0.155 | 0.105 | 0.0025 | 0.400511 | 0.565838 | 0.99097 | 1 | 0.996511 | 0.996154 |
| Fbxo41 | Q6NS60 | 1.03 | 0.92 | 0.73 | 0.49 | 0.97 | 1.21 | 0.46 | 0.41 | 1.3 | 0.64 | 0.8 | 1.24 | 1.36 | 1.21 | 0.57 | 0.57 | -0.03 | 0.2025 | 0.135 | 0.901735 | 0.393828 | 0.61447 | 1 | 0.996511 | 0.935876 |
| Fbxo42 | Q6PDJ6 | -0.28 | -0.5 | -1.14 | -1.01 | -0.54 | 0.58 | -0.94 | -0.53 | -0.18 | -1.13 | -0.4 | -0.18 | -0.6 | -0.23 | -1.21 | -1.34 | 0.375 | 0.26 | -0.1125 | 0.322838 | 0.40831 | 0.736671 | 1 | 0.996511 | 0.958635 |
| Fbxo44 | A2A7H5 | -1.19 | -1.99 | -0.45 | -0.09 | -0.12 | -2.25 | 0.4 | -0.03 | -1.82 | -0.81 | 0.08 | -2.06 | -2.72 | -1.3 | -0.13 | -0.21 | 0.43 | -0.2225 | -0.16 | 0.520621 | 0.708959 | 0.81153 | 1 | 0.996511 | 0.972642 |
| Fbxo45 | Q8K3B1 | 0.25 | 0.18 | 0.72 | 0.41 | 1.02 | -0.09 | 0.55 | 0.34 | 0.61 | 0.82 | 0.58 | 0.68 | 0.81 | 1.41 | 0.37 | 0.42 | 0.065 | 0.2825 | 0.3625 | 0.807676 | 0.161118 | 0.227005 | 1 | 0.996511 | 0.787447 |
| Fbxo6 | Q9QZN4 | -0.67 | -0.55 | -0.88 | -0.83 | -0.13 | -0.49 | -0.57 | -0.7 | -0.27 | -0.13 | -0.91 | -0.73 | -0.24 | -0.07 | -0.41 | -0.54 | 0.26 | 0.2225 | 0.4175 | 0.184683 | 0.349308 | 0.059186 | 1 | 0.996511 | 0.661738 |
| Fbxo7 | Q3U7U3 | 0.23 | 0.26 | 0.42 | 0.36 | -0.05 | 0.28 | -0.69 | 0.54 | 0.53 | 1.28 | 0.06 | 0.66 | 0.27 | 1.08 | 0.02 | -0.57 | -0.2975 | 0.315 | -0.1175 | 0.294376 | 0.263276 | 0.733477 | 1 | 0.996511 | 0.958635 |
| Fbxo9 | Q8BK06-3 | -0.6 | -0.61 | -0.54 | -0.24 | -3.53 | 0.39 | -0.59 | -0.38 | 1.04 | -1.87 | -0.98 | -1.11 | 2.06 | -4.33 | -1.8 | 0.16 | -0.53 | -0.2325 | -0.48 | 0.502854 | 0.688262 | 0.69516 | 1 | 0.996511 | 0.955729 |
| Fbxw11 | Q5SRY7-4 | -0.79 | -0.59 | -1.66 | -2.06 | -0.88 | -0.28 | -1.79 | -1.89 | -0.22 | -1.32 | -1.96 | -0.71 | 0.48 | -1.92 | -1.84 | -1.52 | 0.065 | 0.2225 | 0.075 | 0.89244 | 0.645069 | 0.902005 | 1 | 0.996511 | 0.98447 |
| Fbxw2 | Q8BS25 | -0.68 | -1.96 | -0.98 | -1.21 | -2.56 | -1.78 | -1.18 | -0.77 | -3.78 | 0.22 | -0.72 | -1.97 | -3.97 | -1.83 | 1 | -1.72 | -0.365 | -0.355 | -0.4225 | 0.419665 | 0.666296 | 0.656331 | 1 | 0.996511 | 0.948332 |
| Fbxw5 | Q9QXW2-2 | -0.08 | -0.36 | -0.78 | -0.75 | -0.34 | -0.01 | -0.81 | -0.86 | 0.29 | -1.15 | -0.78 | 0.06 | 0.16 | 1.23 | -0.71 | -0.79 | -0.0125 | 0.0975 | 0.465 | 0.962875 | 0.791476 | 0.336218 | 1 | 0.996511 | 0.847578 |
| Fbxw8 | Q8BIA4-4 | -1.03 | -0.58 | -1.42 | -0.8 | -1.08 | 0.25 | -1.23 | -0.48 | 0.35 | -1.72 | -0.8 | -0.22 | -1.17 | -0.05 | -1.7 | 0.08 | 0.3225 | 0.36 | 0.2475 | 0.38724 | 0.431128 | 0.580571 | 1 | 0.996511 | 0.927734 |
| Fcer1g | P20491 | 0.47 | 0.17 | 0.33 | 0.43 | -0.7 | -1.62 | -0.61 | -0.24 | 0.67 | 0.87 | 0.08 | 0.12 | -0.12 | 0.7 | 0.21 | 0.1 | -1.1425 | 0.085 | -0.1275 | 0.004154 | 0.721222 | 0.582946 | 1 | 0.996511 | 0.927734 |
| Fcf1 | E9Q186 | -1.46 | 0.48 | -0.38 | -0.96 | -0.67 | -0.7 | 0.23 | 0.48 | 0.05 | 2.09 | 0.03 | -1.13 | 0.39 | 0.09 | 2.09 | 1.43 | 0.415 | 0.84 | 1.58 | 0.39576 | 0.259975 | 0.021073 | 1 | 0.996511 | 0.621926 |
| Fcgr1 | P26151 | -1.61 | -2.11 | -0.7 | -0.87 | -1.79 | -1.94 | -1.21 | -0.89 | -2.79 | 0.44 | -1.22 | -1.73 | -2.11 | -1.94 | -1.51 | -1.53 | -0.135 | -0.0025 | -0.45 | 0.728966 | 0.997071 | 0.228474 | 1 | 1 | 0.787447 |
| Fcgr2b | A0A0B4J1G | -1.62 | -2.17 | -2.62 | -2.41 | -2.04 | -1.68 | -3.18 | -1.69 | -1.83 | -0.29 | -2.37 | -1.81 | -1.24 | -1.29 | -3.34 | -3.01 | 0.0575 | 0.63 | -0.015 | 0.883339 | 0.199356 | 0.978223 | 1 | 0.996511 | 0.993727 |
| Fcgrt | Q61559 | 2.06 | 2.21 | 0.15 | 0.46 | 2.02 | 1.64 | 0.78 | 0.27 | 1.83 | 0.41 | 0.45 | 2.24 | 1.43 | 0.8 | 0.59 | 0.65 | -0.0425 | 0.0125 | -0.3525 | 0.944081 | 0.984615 | 0.507874 | 1 | 0.998892 | 0.902804 |
| Fcho1 | Q8K285 | 0.45 | -0.08 | 0.18 | 0.23 | 0.12 | 0.34 | 0.3 | 0.13 | 0.31 | -0.2 | 0.11 | 0.29 | 0.01 | 0.5 | -0.15 | -0.09 | 0.0275 | -0.0675 | -0.1275 | 0.873751 | 0.743811 | 0.580793 | 1 | 0.996511 | 0.927734 |
| Fcho2 | Q3UQN2 | 0.37 | 0.12 | 0.01 | -0.04 | -0.05 | 0.36 | 0.17 | -0.12 | 0.18 | -0.44 | 0.34 | 0.54 | 0.74 | -1.02 | 0.5 | 0.49 | -0.025 | 0.04 | 0.0625 | 0.892141 | 0.874164 | 0.875459 | 1 | 0.996511 | 0.979626 |
| Fchsd2 | Q3USJ8 | -0.06 | -0.28 | -0.72 | -0.13 | 0.09 | -0.36 | 0.07 | 0.48 | -0.48 | -0.32 | 0.13 | -0.32 | -0.06 | 0.03 | -0.28 | -0.04 | 0.3675 | 0.05 | 0.21 | 0.154485 | 0.82806 | 0.34478 | 1 | 0.996511 | 0.84949 |
| Fcsk | Q7TMC8 | 0.2 | -0.05 | 0.12 | -0.08 | 0.07 | -0.04 | -0.07 | 0.15 | -0.39 | -0.12 | -0.09 | 0.13 | -0.41 | -0.32 | -0.27 | -0.04 | -0.02 | -0.165 | -0.3075 | 0.896604 | 0.386876 | 0.125369 | 1 | 0.996511 | 0.721448 |
| Fdft1 | P53798 | -0.38 | 0.11 | 0.4 | -0.27 | -0.12 | 0.17 | 0.66 | 0.27 | 0.05 | 0.43 | 0.5 | -0.12 | 1 | -0.92 | 0 | 0.44 | 0.28 | 0.25 | 0.165 | 0.284395 | 0.339667 | 0.697554 | 1 | 0.996511 | 0.955729 |
| Fdps | Q920E5 | -0.3 | -1.06 | -0.24 | -0.13 | -1.2 | -1.13 | -0.77 | -0.06 | -0.83 | 0.07 | -0.18 | -0.31 | -0.69 | 0.06 | -0.1 | 0.19 | -0.3575 | 0.12 | 0.2975 | 0.291807 | 0.683078 | 0.336073 | 1 | 0.996511 | 0.847578 |
| Fdx1 | P46656 | 1 | 0.94 | 1.35 | 1.62 | -0.67 | 1.84 | -0.48 | 0.65 | 0.92 | -0.56 | 1.3 | 0.71 | 1.7 | -0.99 | 1.17 | 0.83 | -0.8925 | -0.635 | -0.55 | 0.132703 | 0.150068 | 0.338085 | 1 | 0.996511 | 0.847578 |
| Fdx2 | Q9CPW2 | 1 | 1.27 | 0.46 | 0.59 | 0.91 | 0.89 | 0.56 | 0.56 | 1 | -0.42 | 0.48 | 0.87 | 0.69 | 0.97 | 0.28 | 0.35 | -0.1 | -0.3475 | -0.2575 | 0.665415 | 0.349036 | 0.353314 | 1 | 0.996511 | 0.855029 |
| Fdxr | Q61578 | 0.44 | 0.37 | 0.6 | 1.03 | -0.21 | -0.01 | 0.53 | 0.74 | -0.17 | 0.82 | 0.89 | 0.12 | -0.38 | 0.39 | 0.18 | 0.24 | -0.3475 | -0.195 | -0.5025 | 0.224366 | 0.527233 | 0.075713 | 1 | 0.996511 | 0.674038 |
| Fech | P22315 | 0.76 | 0.57 | 0.9 | 0.85 | 0.26 | 0.78 | 0.71 | 1.07 | 0.18 | -0.13 | 0.95 | 0.41 | -0.02 | 0.03 | 0.46 | 0.31 | -0.065 | -0.4175 | -0.575 | 0.7579 | 0.133807 | 0.017867 | 1 | 0.996511 | 0.602211 |
| Fem1aa | Q9Z2G1 | 0.73 | 0.81 | 0.19 | 0.13 | 0.81 | 1.22 | 0.47 | 0.3 | 0.43 | 0.58 | 0.13 | 1.48 | -0.63 | 1.72 | 0.05 | 0.37 | 0.235 | 0.19 | -0.0875 | 0.401816 | 0.575743 | 0.858318 | 1 | 0.996511 | 0.976225 |
| Fem1b | Q9Z2G0 | -1.73 | -1.8 | -1.05 | -0.96 | -1.24 | -0.96 | -1.1 | -1.02 | -0.9 | 0.06 | -0.95 | -1.82 | -0.28 | -1.7 | -1.78 | -1.2 | 0.305 | 0.4825 | 0.145 | 0.231032 | 0.2696 | 0.714768 | 1 | 0.996511 | 0.958106 |
| Fem1c | Q8CEF1 | 0.21 | 0.43 | -0.92 | -0.36 | 0.48 | 0.69 | -0.82 | -1.43 | 1.1 | 0.52 | -1.11 | 1 | 0.39 | 0.59 | -0.08 | -1.01 | -0.11 | 0.5375 | 0.1325 | 0.84001 | 0.341127 | 0.765362 | 1 | 0.996511 | 0.964134 |
| Fer | P70451-2 | -0.42 | -0.04 | 0.24 | 0.13 | -0.2 | -0.17 | -0.02 | -0.03 | -0.25 | 0.56 | 0.05 | 0.11 | -0.34 | 0 | 0.08 | -0.03 | -0.0825 | 0.14 | -0.05 | 0.666174 | 0.572728 | 0.821437 | 1 | 0.996511 | 0.972768 |
| Fermt2 | Q8CIB5 | -0.16 | -0.5 | -0.1 | -0.08 | -0.11 | -0.16 | 0.01 | -0.18 | -0.66 | -0.39 | -0.02 | -0.23 | -0.73 | -0.51 | -0.46 | -0.21 | 0.1 | -0.115 | -0.2675 | 0.547978 | 0.586992 | 0.216683 | 1 | 0.996511 | 0.787447 |
| Fermt2 | A6X940 | 0.48 | 0.78 | 0.14 | 1.25 | 0.04 | 0.52 | -0.16 | 1.06 | 1.41 | 1.94 | 1.53 | 0.77 | 0.49 | 1.47 | 0.33 | -1.14 | -0.2975 | 0.75 | -0.375 | 0.401032 | 0.048956 | 0.495905 | 1 | 0.996511 | 0.89758 |
| Fermt3 | Q8K1B8 | -0.01 | -0.52 | -0.4 | -0.44 | -0.37 | -0.02 | -0.38 | -0.1 | -0.53 | -0.55 | 0.08 | -0.55 | -0.61 | -0.37 | -0.61 | -0.58 | 0.125 | -0.045 | -0.2 | 0.509306 | 0.842693 | 0.325038 | 1 | 0.996511 | 0.842881 |
| Fez1 | Q8K0X8 | 0.68 | 0.65 | 0.66 | 0.39 | 0.35 | 0.26 | 0.28 | 0.44 | 0.48 | 0.46 | 0.6 | 0.77 | 0.39 | 0.74 | 0.3 | 0.48 | -0.2625 | -0.0175 | -0.1175 | 0.112024 | 0.919385 | 0.545129 | 1 | 0.996755 | 0.912212 |
| Fga | E9PV24 | -1.88 | -3.79 | -3.73 | -1.98 | -3.44 | -1.92 | -3.69 | -3.04 | -3.12 | -0.98 | -3.67 | -3.1 | -2.03 | -3.1 | -3.84 | -4.31 | -0.1775 | 0.1275 | -0.475 | 0.767682 | 0.859805 | 0.478687 | 1 | 0.996511 | 0.892849 |
| Fgb | Q8K0E8 | -1.88 | -2.68 | -2.68 | -1.89 | -2.45 | -1.78 | -2.74 | -2.35 | -2.58 | -0.58 | -2.64 | -2.32 | -2.34 | -2.17 | -3.22 | -3.21 | -0.0475 | 0.2525 | -0.4525 | 0.875093 | 0.617185 | 0.226495 | 1 | 0.996511 | 0.787447 |
| Fgd1 | P52734 | 0.11 | 0.69 | 0.3 | 0.25 | 0.56 | 0.46 | 0.28 | 0.16 | 0.56 | 0.63 | 0.08 | 0.59 | 0.34 | 0.69 | -0.12 | 0.36 | 0.0275 | 0.1275 | -0.02 | 0.88554 | 0.56196 | 0.934693 | 1 | 0.996511 | 0.989767 |
| Fgd2 | Q8BY35 | -1.49 | -1.62 | -0.11 | 0.18 | -0.63 | -2.26 | 0.41 | 0.32 | -1.45 | -0.14 | 0.08 | -1.91 | -2.06 | -1.23 | -1.03 | -0.36 | 0.22 | -0.095 | -0.41 | 0.754115 | 0.877344 | 0.453292 | 1 | 0.996511 | 0.885993 |
| Fgd4 | Q91ZT5 | 0.07 | 0.26 | 0.04 | 0.28 | 0.47 | 0.14 | 0.27 | 0.34 | -0.06 | 0.23 | 0.26 | 0.03 | -0.27 | 0.09 | 0.06 | 0.37 | 0.1425 | -0.0475 | -0.1 | 0.3783 | 0.784346 | 0.630208 | 1 | 0.996511 | 0.941132 |
| Fgd6 | Q69ZL1 | 0.72 | 1.45 | 0.41 | -0.17 | 1.3 | 1.14 | 0.35 | -0.19 | 1.09 | -0.12 | -0.21 | 1.08 | 1.96 | 0.67 | -0.02 | 0.6 | 0.0475 | -0.1425 | 0.2 | 0.916143 | 0.758902 | 0.689932 | 1 | 0.996511 | 0.955729 |
| Fgf1 | P61148 | 0.24 | 0.57 | 0.56 | 0.39 | 0.95 | 0.01 | 0.26 | 0.51 | 0.04 | -0.19 | 0.22 | 0.24 | -0.11 | 0.7 | 0.18 | 0.32 | -0.0075 | -0.3625 | -0.1675 | 0.974333 | 0.079444 | 0.473602 | 1 | 0.996511 | 0.891635 |
| Fgf12 | P61329 | -0.66 | -0.57 | -0.77 | -0.8 | -0.98 | -0.48 | -0.77 | -0.8 | -0.53 | -0.62 | -0.57 | -0.58 | -0.66 | -0.96 | -0.73 | -1.38 | -0.0575 | 0.125 | -0.2325 | 0.735064 | 0.435329 | 0.308443 | 1 | 0.996511 | 0.834018 |
| Fgf13 | P70377 | 0.2 | 0.01 | 0.12 | -0.54 | 0.63 | 0.2 | 0.15 | -0.28 | -0.06 | -0.38 | -0.02 | -0.11 | -0.36 | 0.03 | -0.49 | -0.26 | 0.2275 | -0.09 | -0.2175 | 0.390917 | 0.686471 | 0.375367 | 1 | 0.996511 | 0.859611 |
| Fgf13 | P70377-3 | 1.22 | 1.27 | 0.22 | 0.6 | 0.65 | 1.35 | 0.09 | 1.2 | 1.29 | 2.11 | 0.76 | 1.36 | 0.87 | 0.7 | 1.34 | 1.57 | -0.005 | 0.5525 | 0.2925 | 0.989072 | 0.1565 | 0.383713 | 1 | 0.996511 | 0.860984 |
| Fgf14 | O89096 | -0.18 | 0.09 | -0.17 | -0.16 | 0.29 | -0.12 | -0.06 | 0.26 | -0.6 | -0.55 | 0.24 | -0.82 | -0.27 | 0.8 | -0.17 | -0.34 | 0.1975 | -0.3275 | 0.11 | 0.274792 | 0.229073 | 0.706691 | 1 | 0.996511 | 0.957157 |
| Fgfr1 | P16092 | 0.44 | 1.04 | -0.65 | 0.27 | 0.71 | 0.54 | -0.25 | -0.26 | 0.33 | 0.63 | -0.37 | 0.45 | 0.4 | 0.78 | 0.08 | 0.18 | -0.09 | -0.015 | 0.085 | 0.825486 | 0.969691 | 0.820301 | 1 | 0.996811 | 0.972768 |
| Fgfr1op2 | Q9CRA9-2 | -0.28 | 0.23 | -0.08 | 0.18 | 0.22 | -0.32 | -0.11 | -0.09 | 0.06 | -0.24 | -0.24 | -0.01 | -0.12 | 0.06 | -0.11 | -0.35 | -0.0875 | -0.12 | -0.1425 | 0.659033 | 0.542975 | 0.49644 | 1 | 0.996511 | 0.897918 |
| Fgfr2 | E9Q7T0 | 2.83 | 2.54 | 4.17 | 3.49 | 2.29 | 2.87 | 2.95 | 3.97 | 1.25 | 1.28 | 3.36 | 3.09 | 1.68 | 0.47 | 4.1 | 5.04 | -0.2375 | -1.0125 | -0.435 | 0.613951 | 0.128447 | 0.664854 | 1 | 0.996511 | 0.949825 |
| Fgfr3 | Q61563 | 0.65 | 1.33 | 0.91 | 1.3 | 1.65 | 0.81 | 0.51 | 1.51 | 0.58 | 0.61 | 0.76 | 1.01 | 0.14 | 1.02 | 0.65 | 1.07 | 0.0725 | -0.3075 | -0.3275 | 0.81718 | 0.195711 | 0.272699 | 1 | 0.996511 | 0.822084 |
| Fgg | Q3UER8 | -1.96 | -4.49 | -4.01 | -2.31 | -3.89 | -2.1 | -4.14 | -3.24 | -3.85 | -0.39 | -3.92 | -3.41 | -2.71 | -3.52 | -4.45 | -4.24 | -0.15 | 0.3 | -0.5375 | 0.830076 | 0.750634 | 0.431117 | 1 | 0.996511 | 0.879165 |
| Fggy | A2AJL3 | 0.54 | 0.4 | -0.4 | 0.13 | 0.69 | -0.11 | 0.14 | -0.56 | -1.2 | -0.38 | -0.36 | 0.33 | -1.05 | -0.46 | -0.49 | -0.22 | -0.1275 | -0.57 | -0.7225 | 0.695717 | 0.144863 | 0.031793 | 1 | 0.996511 | 0.641974 |
| Fh | P97807-2 | 0.33 | 0.27 | 0.71 | 0.71 | -0.41 | 0.45 | 0.45 | 0.71 | 0.07 | 0.14 | 0.73 | 0.01 | 0.24 | -0.94 | 0.13 | 0.27 | -0.205 | -0.2675 | -0.58 | 0.464932 | 0.272248 | 0.097191 | 1 | 0.996511 | 0.698156 |
| Fhdc1 | Q3ULZ2 | -0.66 | 0.59 | -0.77 | 0.11 | -0.19 | 0.01 | -1.02 | -0.43 | 1.61 | 1.06 | -0.59 | 0.35 | 2.08 | 1.95 | 0.3 | -1.08 | -0.225 | 0.79 | 0.995 | 0.551525 | 0.163122 | 0.20099 | 1 | 0.996511 | 0.780827 |
| Fhip1b | Q3U2I3 | -0.61 | -0.81 | -0.99 | -0.98 | -0.55 | -0.76 | -1.31 | -1.23 | -0.78 | -0.87 | -0.87 | -0.82 | -1.25 | -0.57 | -1.13 | -1.17 | -0.115 | 0.0125 | -0.1825 | 0.612279 | 0.941332 | 0.428497 | 1 | 0.996755 | 0.879165 |
| Fhip2a | Q8CDM8 | -0.28 | -0.54 | -0.95 | -1.28 | -0.2 | -0.78 | -1.05 | -1.18 | -1.2 | -0.77 | -0.96 | -0.3 | -1.48 | -1.41 | -1.2 | -1.5 | -0.04 | -0.045 | -0.635 | 0.895963 | 0.880012 | 0.035046 | 1 | 0.996511 | 0.651434 |
| Fhip2b | Q80YR2 | -1.29 | -0.77 | -0.55 | -1.48 | -0.32 | -1.55 | -0.86 | -1.33 | -1.7 | -1.28 | -0.81 | -1.26 | -1.82 | -1.22 | -1.49 | -0.78 | 0.0075 | -0.24 | -0.305 | 0.982255 | 0.420237 | 0.348769 | 1 | 0.996511 | 0.85278 |
| Fhit | E9PZ91 | 0.62 | 0.29 | 0.38 | 0.34 | -0.02 | -0.16 | 1.68 | 0.23 | -0.38 | 0.4 | 0.36 | 0.17 | 0.05 | 0.59 | 0.44 | 0.69 | 0.025 | -0.27 | 0.035 | 0.950702 | 0.254614 | 0.869972 | 1 | 0.996511 | 0.979293 |
| Fhl1 | A2AEX8 | -0.13 | -1.42 | -1.46 | -2.29 | -0.69 | 0.36 | -0.43 | -1.3 | -1.35 | -0.96 | -1.07 | 0.31 | -1.28 | -0.87 | -0.55 | -0.54 | 0.81 | 0.5575 | 0.515 | 0.144807 | 0.312808 | 0.272925 | 1 | 0.996511 | 0.822084 |
| Fhl1 | A2AEY2 | 0.72 | -0.55 | -0.02 | 0.28 | -1.14 | 0.26 | -1.61 | -1.23 | 2.38 | 0.94 | 0.38 | -0.33 | 2.15 | -0.38 | -0.08 | -0.38 | -1.0375 | 0.735 | 0.22 | 0.045856 | 0.228073 | 0.719431 | 1 | 0.996511 | 0.958398 |
| Fhl2 | O70433 | -0.75 | -1.46 | -1.89 | -1.48 | -1.11 | -0.64 | -1.59 | -1.56 | -0.95 | -0.85 | -1.84 | -0.61 | -1.22 | -1.01 | -2.02 | -1.93 | 0.17 | 0.3325 | -0.15 | 0.596823 | 0.356129 | 0.665393 | 1 | 0.996511 | 0.949825 |
| Fhod1 | Q6P9Q4 | -2 | -3.91 | -2.05 | -1.62 | -5.35 | -1.33 | -1.03 | -2.08 | -2.05 | -1.42 | -1.22 | -2.02 | -0.72 | -5.34 | -1.59 | -2.11 | -0.0525 | 0.7175 | -0.045 | 0.958198 | 0.187956 | 0.964496 | 1 | 0.996511 | 0.9927 |
| Fhod3 | Q76LL6-3 | -0.15 | 0.37 | -0.19 | 0.03 | 0.08 | 0.06 | -0.35 | -0.23 | -0.43 | 0.46 | -0.11 | -0.13 | -0.22 | -0.59 | -0.88 | 0.06 | -1.25E-01 | -0.0675 | -0.4225 | 0.535905 | 0.786679 | 0.141277 | 1 | 0.996511 | 0.737973 |
| Fibcd1 | A2AV25 | 0.29 | 0.5 | 0.84 | 1.13 | -0.06 | 0.19 | 1.26 | 0.39 | 0.95 | 1.91 | 0.54 | 0.49 | 0.53 | 0.78 | 1.2 | 1.11 | -0.245 | 0.2825 | 0.215 | 0.467212 | 0.448936 | 0.429408 | 1 | 0.996511 | 0.879165 |
| Fibp | Q8K2D8 | 0.79 | 1.25 | 0.96 | 0.81 | 1.76 | 0.43 | 0.79 | 0.8 | 0.22 | 0.64 | 0.66 | 0.8 | -0.08 | 1.2 | 0.75 | 0.64 | -0.0075 | -0.3725 | -0.325 | 0.980128 | 0.10129 | 0.293669 | 1 | 0.996511 | 0.825612 |
| Fig4 | Q91WF7 | 0.24 | 0.32 | 0.12 | -0.04 | 0.46 | 0.3 | 0.49 | 0.18 | -0.04 | 0.17 | 0 | 0.21 | 0.06 | 0.24 | 0.42 | 0.37 | 0.1975 | -0.075 | 0.1125 | 0.249756 | 0.666899 | 0.557411 | 1 | 0.996511 | 0.917092 |
| Filip1 | A0A5F8MP | -1.73 | -0.39 | 0.61 | 0.54 | -2.39 | -1.24 | 0.43 | 0.83 | -0.45 | 0.65 | 0.61 | -0.25 | 0.26 | -2.61 | 0.35 | -0.12 | -0.35 | 0.3825 | -0.2875 | 0.676209 | 0.505614 | 0.720267 | 1 | 0.996511 | 0.958398 |
| Fip1l1 | Q9D824-3 | -0.67 | -1.75 | -2.42 | -2.28 | -0.97 | -0.99 | -1 | -1.35 | -1.05 | -1.89 | -1.95 | -1.29 | 0.01 | -0.07 | 0.23 | -0.99 | 0.7025 | 0.235 | 1.575 | 0.096644 | 0.588299 | 0.006825 | 1 | 0.996511 | 0.552614 |
| Fis1 | Q9CQ92 | 1.91 | 1.1 | 0.48 | 1.23 | 0.77 | 1.51 | 0.09 | 0.92 | 0.89 | 0.03 | 0.86 | 2.05 | 1.04 | -0.18 | 0.75 | 0.68 | -0.3575 | -0.2225 | -0.6075 | 0.372724 | 0.641543 | 0.139162 | 1 | 0.996511 | 0.73493 |
| Fitm2 | P59266 | -0.44 | 0.4 | 0.3 | 0.89 | 0.84 | 0.03 | 1.03 | 0.54 | -0.33 | 0.87 | 0.63 | -0.45 | -0.08 | 0.81 | 0.59 | 0.33 | 0.3225 | -0.1075 | 0.125 | 0.354555 | 0.794052 | 0.710938 | 1 | 0.996511 | 0.957819 |
| Fkbp15 | Q6P9Q6 | -0.29 | -0.26 | -0.39 | -0.26 | -0.11 | -0.34 | -0.27 | -0.51 | 0.15 | -0.38 | -0.21 | -0.1 | -0.16 | 0.14 | -0.53 | -0.47 | -0.0075 | 0.165 | 0.045 | 0.961573 | 0.373252 | 0.833044 | 1 | 0.996511 | 0.97513 |
| Fkbp1a | P26883 | -0.99 | -1.51 | -1 | -0.46 | -1.36 | -1.02 | -0.69 | -0.37 | -1.23 | -0.46 | -0.52 | -0.85 | -0.86 | -0.7 | -0.71 | -0.94 | 0.13 | 0.225 | 0.1875 | 0.666244 | 0.442501 | 0.467482 | 1 | 0.996511 | 0.889213 |
| Fkbp1b | Q9Z2I2 | 0 | 0.4 | 0.17 | -0.07 | 0.4 | -0.29 | 0.01 | 0.57 | 0.19 | 0.06 | 0.44 | 0.3 | 0.13 | 1.18 | -0.22 | -0.29 | 0.0475 | 0.1225 | 0.075 | 0.841112 | 0.523282 | 0.831385 | 1 | 0.996511 | 0.973982 |
| Fkbp2 | P45878 | 1.24 | -0.13 | -2.65 | -2.67 | 0.33 | 1.25 | -2.04 | -2.45 | 0.48 | -4.18 | -2.82 | 1.68 | 0.19 | 0.34 | -1.73 | -2.23 | 0.325 | -0.1575 | 0.195 | 0.78392 | 0.916354 | 0.852129 | 1 | 0.996755 | 0.976225 |
| Fkbp3 | Q62446 | -2.72 | -3.46 | -2.84 | -3.16 | -2.67 | -3.11 | -2.04 | -2.9 | -4.07 | -2.86 | -2.83 | -2.9 | -4.51 | -3.05 | -3 | -3.13 | 0.365 | -0.12 | -0.3775 | 0.224305 | 0.725013 | 0.343863 | 1 | 0.996511 | 0.848473 |
| Fkbp4 | P30416 | 0.2 | -0.27 | -0.34 | -0.03 | -0.24 | 0.1 | -0.21 | -0.16 | -0.58 | -0.25 | -0.35 | -0.05 | -0.53 | -0.48 | -0.33 | -0.23 | -0.0175 | -0.1975 | -0.2825 | 0.925083 | 0.356895 | 0.189915 | 1 | 0.996511 | 0.776843 |
| Fkbp5 | Q64378 | -0.96 | -1.55 | -1.8 | -1.83 | -1.24 | -0.9 | -1.57 | -1.58 | -0.81 | -0.96 | -1.45 | -0.69 | -0.62 | -0.92 | -1.88 | -1.89 | 0.2125 | 0.5575 | 0.2075 | 0.432279 | 0.069484 | 0.583924 | 1 | 0.996511 | 0.927734 |
| Fkbp8 | O35465-2 | 1.21 | 0 | -0.03 | -0.38 | 0.76 | 1.37 | 0.28 | -0.05 | -0.47 | -0.96 | 0.01 | 1.29 | -0.17 | -0.32 | 0.78 | 0.55 | 0.39 | -0.2325 | 0.01 | 0.380602 | 0.673444 | 0.980974 | 1 | 0.996511 | 0.994858 |
| Fkbp9 | Q9Z247 | 1.25 | 1.63 | -0.97 | -0.58 | 1.8 | 1.29 | -0.6 | -1.24 | -0.34 | 0.07 | -1.02 | 1.7 | 0.22 | 0.7 | -0.28 | -0.71 | -0.02 | -0.23 | -0.35 | 0.981834 | 0.770247 | 0.595103 | 1 | 0.996511 | 0.929843 |
| Fkbpl | O35450 | -1.13 | -1.49 | -0.43 | -1.33 | -1.66 | -1.49 | -0.68 | -1.01 | -1.17 | -0.57 | -0.12 | -1.14 | -0.57 | -1.71 | -0.19 | -1.29 | -0.115 | 0.345 | 0.155 | 0.717675 | 0.323309 | 0.699663 | 1 | 0.996511 | 0.955745 |
| Flad1 | Q8R123 | 0.34 | 0.53 | -0.16 | -0.3 | 0.34 | 0.68 | -0.35 | -0.12 | -0.17 | -0.22 | 0.01 | 0.31 | -0.21 | 0.04 | -0.69 | -0.55 | 0.035 | -0.12 | -0.455 | 0.907487 | 0.637732 | 0.129507 | 1 | 0.996511 | 0.723655 |
| Flcn | Q8QZS3 | -0.88 | -0.65 | -1.15 | -1 | -0.56 | -1.05 | -1.34 | -1.45 | 2.26 | -1.2 | -0.94 | -0.88 | 1.52 | 5.54 | -1.32 | -1.27 | -0.18 | 0.73 | 2.0375 | 0.461664 | 0.343424 | 0.181256 | 1 | 0.996511 | 0.775592 |
| Flii | Q9JJ28 | 0.13 | 0.37 | 0.53 | 0.46 | 0.26 | 0.3 | 0.26 | 0.37 | 0.26 | 0.78 | 0.44 | 0.28 | 0.76 | 0.17 | 0.5 | 0.62 | -0.075 | 0.0675 | 0.14 | 0.634892 | 0.734989 | 0.512076 | 1 | 0.996511 | 0.903993 |
| Flna | B7FAU9 | -0.97 | -1.62 | -1.33 | -0.99 | -1.36 | -0.96 | -1.26 | -1.21 | -1.06 | -1.22 | -1.15 | -0.92 | -0.92 | -1.24 | -1.09 | -1.29 | 0.03 | 0.14 | 0.0925 | 0.884367 | 0.511373 | 0.681019 | 1 | 0.996511 | 0.955729 |
| Flnb | Q80X90 | -0.03 | -0.19 | -0.08 | 0.02 | -0.21 | -0.37 | -0.26 | -0.25 | -0.37 | 0.29 | 0.03 | -0.03 | -0.22 | -0.26 | -0.23 | -0.01 | -0.2025 | 0.05 | -0.11 | 0.183184 | 0.798517 | 0.532826 | 1 | 0.996511 | 0.911273 |
| Flnc | Q8VHX6-2 | 0.09 | -0.74 | 0.57 | 0.45 | -0.97 | -0.48 | 0.31 | 0.18 | 5.03 | 2.32 | 0.5 | 0.1 | 6 | -0.07 | 0.21 | 0.12 | -0.3325 | 1.895 | 1.4725 | 0.411677 | 0.097004 | 0.289379 | 1 | 0.996511 | 0.824705 |
| Flot1 | O08917 | 1.2 | 0.03 | 0.56 | 0.49 | 0.36 | 1.53 | 1.3 | 0.57 | 0.44 | -0.26 | 0.79 | 1.25 | 0.42 | 0.42 | 0.85 | 0.93 | 0.37 | -0.015 | 0.085 | 0.313234 | 0.968811 | 0.771729 | 1 | 0.996755 | 0.965933 |
| Flot2 | Q60634 | 0.49 | 0.83 | 0.85 | 1.17 | 1.22 | 0.69 | 1.66 | 1.2 | 1.04 | 1.13 | 1.09 | 0.96 | 0.54 | 1.55 | 0.58 | 0.95 | 0.3575 | 0.22 | 0.07 | 0.182396 | 0.279493 | 0.808836 | 1 | 0.996511 | 0.972098 |
| Flrt1 | Q6RKD8 | -0.26 | 0.18 | 1.51 | 1.34 | 0.06 | -0.14 | 1.19 | 1.09 | -0.4 | 0.52 | 1.66 | 0.07 | -0.34 | -0.19 | 0.83 | 1.17 | -0.1425 | -0.23 | -0.325 | 0.78014 | 0.687383 | 0.544064 | 1 | 0.996511 | 0.912212 |
| Flrt2 | Q8BLU0 | 0.26 | 0.35 | 0.69 | 0.33 | 0.76 | 0.56 | 1.04 | 0.75 | 0.08 | -0.01 | 0.98 | 0.49 | -0.24 | 0.82 | 0.09 | 0.47 | 0.37 | -0.0225 | -0.1225 | 0.0682 | 0.931947 | 0.654935 | 1 | 0.996755 | 0.94818 |
| Flrt3 | Q8BGT1 | 0.3 | 0.44 | 0.57 | 0.19 | 0.27 | 0.65 | 0.72 | 0.13 | 1.01 | 0.33 | 0.61 | 0.51 | 0.92 | 0.45 | 0.64 | 0.62 | 0.0675 | 0.24 | 0.2825 | 0.73536 | 0.270013 | 0.176698 | 1 | 0.996511 | 0.771503 |
| Flt1 | P35969 | -0.14 | 0 | 0.18 | 0.11 | 0.03 | -0.65 | -1.54 | -1.15 | -0.06 | 0.75 | -0.33 | -0.19 | -0.12 | 0.13 | -0.21 | -0.32 | -0.865 | 0.005 | -0.1675 | 0.028368 | 0.985115 | 0.395494 | 1 | 0.998892 | 0.866946 |
| Flywch1 | Q8CI03 | 1.06 | 1.03 | 1.07 | 1.04 | -0.27 | 0.88 | -0.29 | 0.65 | 0.82 | 1.11 | 0.87 | 1.28 | 1.5 | 2.27 | 0.8 | 0.47 | -0.8075 | -0.03 | 0.21 | 0.02457 | 0.865467 | 0.59091 | 1 | 0.996511 | 0.929458 |
| Fmn1 | E9Q7P6 | -0.04 | 0.23 | 0 | -0.2 | -0.05 | -0.08 | 0.45 | -0.31 | -0.12 | -0.85 | 0.07 | -0.1 | -0.68 | 0.19 | -0.37 | 0.19 | 0.005 | -0.2475 | -0.165 | 0.981049 | 0.331242 | 0.532948 | 1 | 0.996511 | 0.911312 |
| Fmn2 | Q9JL04 | -0.22 | 0.11 | 1.11 | 1.04 | 0.34 | -0.23 | 1.06 | 0.93 | 0.43 | 1.24 | 1.46 | 0.44 | 0.19 | 0.34 | 1.18 | 0.74 | 0.015 | 0.3825 | 0.1025 | 0.971374 | 0.360766 | 0.792071 | 1 | 0.996511 | 0.96933 |
| Fmnl1 | A2AB60 | 0.18 | 0.6 | 0.26 | 0.17 | 0.39 | 0.22 | -0.09 | -0.08 | 0.25 | -0.31 | 0.14 | 0.14 | 0.31 | 0.45 | -0.27 | -0.17 | -0.1925 | -0.2475 | -0.2225 | 0.331617 | 0.249239 | 0.369084 | 1 | 0.996511 | 0.859245 |
| Fmnl2 | F8VPR2 | -0.37 | -0.31 | -0.06 | 1.22 | -0.17 | -0.4 | 0.9 | 0.04 | -0.68 | 0 | 0.43 | -0.3 | -0.85 | 0.92 | -1.38 | -0.46 | -0.0275 | -0.2575 | -0.5625 | 0.949813 | 0.542659 | 0.335956 | 1 | 0.996511 | 0.847578 |
| Fmnl2 | A2APV2-3 | 0.9 | 0.86 | 0.74 | 0.47 | 1.21 | 0.98 | 0.5 | 0.45 | 0.84 | 0.5 | 0.67 | 0.93 | 0.58 | 0.86 | 0.55 | 0.88 | 0.0425 | -0.0075 | -0.025 | 0.852761 | 0.968748 | 0.899998 | 1 | 0.996755 | 0.983509 |
| Fmnl3 | Q6ZPF4 | 2 | 1.47 | 2.01 | 0.42 | 1.29 | 1.5 | 0.7 | 1.39 | 1.16 | -0.1 | 1.43 | 2.12 | 0.53 | 1.22 | 1.95 | 1.67 | -0.255 | -0.3225 | -0.1325 | 0.520064 | 0.561106 | 0.772514 | 1 | 0.996511 | 0.965933 |
| Fmo1 | P50285 | -0.29 | -0.16 | -0.9 | -0.6 | 0.65 | 0.12 | 1.22 | -0.23 | -0.17 | -1.08 | -0.14 | 0.61 | -0.18 | 0.01 | -0.37 | 0.01 | 0.9275 | 0.2925 | 0.355 | 0.023706 | 0.439243 | 0.152206 | 1 | 0.996511 | 0.750422 |
| Fmo5 | P97872 | -0.65 | -0.25 | 0.38 | 0.03 | 1.2 | -0.26 | 0.81 | 1.47 | -0.35 | 1.06 | 2.28 | -0.36 | -0.71 | -0.1 | 0.04 | 0.78 | 0.9275 | 0.78 | 0.125 | 0.048108 | 0.226245 | 0.735118 | 1 | 0.996511 | 0.958635 |
| Fmr1 | P35922-9 | -0.51 | -0.49 | -0.32 | -0.53 | -0.58 | -0.28 | -0.14 | -0.41 | 0.32 | -0.26 | -0.29 | -0.37 | 0.87 | -0.93 | 0.26 | -0.07 | 0.11 | 0.3125 | 0.495 | 0.507689 | 0.160772 | 0.204461 | 1 | 0.996511 | 0.783294 |
| Fn1 | B9EHT6 | -1.29 | -1.96 | -1.95 | -1.65 | -1.36 | -1.19 | -1.55 | -1.76 | -1.76 | -1.04 | -1.81 | -1.26 | -1.42 | -1.53 | -1.43 | -1.57 | 0.2475 | 0.245 | 0.225 | 0.283713 | 0.365932 | 0.313177 | 1 | 0.996511 | 0.835247 |
| Fn3k | Q9ER35 | 0.05 | 0.09 | -0.08 | -0.18 | -0.04 | -0.24 | -0.39 | -0.29 | -0.35 | -0.78 | -0.31 | -0.13 | -0.47 | -0.72 | -0.48 | -0.66 | -0.21 | -0.3625 | -0.5525 | 0.210104 | 0.097946 | 0.012693 | 1 | 0.996511 | 0.573604 |
| Fn3krp | Q8K274 | -0.53 | -1.25 | 0.24 | 0.37 | -1.15 | -0.55 | 0.22 | 0.4 | 0.8 | 0.07 | -0.05 | -1 | 2.21 | -0.71 | 0.35 | -0.23 | 0.0225 | 0.2475 | 0.6975 | 0.962627 | 0.61679 | 0.316171 | 1 | 0.996511 | 0.838265 |
| Fnbp1 | A2AQ42 | 0.69 | 0.6 | 0.7 | 0.57 | 0.36 | 0.27 | 0.2 | 0.16 | 0.58 | 0.67 | 0.3 | 0.5 | 0.31 | 0.51 | 0.45 | 0.3 | -0.3925 | -0.1275 | -0.2475 | 0.021694 | 0.453676 | 0.172355 | 1 | 0.996511 | 0.76392 |
| Fnbp1l | E9PUI5 | 1.06 | 0.31 | 0.81 | 0.66 | 0.01 | 1.16 | 1.11 | 1.03 | 1.01 | 0.71 | 1.39 | 1.32 | 1.21 | 0.28 | 1.1 | 0.93 | 0.1175 | 0.3975 | 0.17 | 0.706167 | 0.133328 | 0.549053 | 1 | 0.996511 | 0.912757 |
| Fndc10 | A2A9Q0 | 0.64 | 0.72 | 1.12 | 1.02 | 1.19 | 0.69 | 0.08 | 0.4 | 1.05 | 1.72 | 0.79 | 0.67 | 2.17 | 0.77 | 1.31 | 1.38 | -0.285 | 0.1825 | 0.5325 | 0.304874 | 0.512823 | 0.121023 | 1 | 0.996511 | 0.721448 |
| Fndc3a | Q8BX90 | -1.33 | -1.32 | -0.66 | -1.11 | -1.69 | -0.51 | -1.04 | -1.02 | 0.29 | -1.56 | -1.01 | -1.4 | 1.66 | -2.58 | 0.99 | -0.19 | 0.04 | 0.185 | 1.075 | 0.88976 | 0.664515 | 0.227132 | 1 | 0.996511 | 0.787447 |
| Fndc3b | A0A0R4J0H | -0.47 | 0.6 | -1.75 | -1.91 | 0.22 | -0.04 | -1.47 | -1.36 | 0.18 | -3.18 | -2.39 | -0.3 | -0.24 | 0.64 | -1.79 | -1.04 | 0.22 | -0.54 | 0.275 | 0.742093 | 0.553434 | 0.701187 | 1 | 0.996511 | 0.956536 |
| Fndc4 | A0A0J9YUI | 2.79 | 2.38 | 3.42 | 3.68 | 0.24 | 3.8 | 0 | 1.74 | 7.17 | -0.26 | 2.88 | 2.84 | 7.51 | -0.54 | 2.66 | 1.86 | -1.6225 | 0.09 | -0.195 | 0.078299 | 0.948302 | 0.897999 | 1 | 0.996755 | 0.983509 |
| Fnta | Q61239 | 0.08 | -0.25 | -1.49 | -1.12 | 0.3 | 0.08 | -1.21 | -1.32 | -0.38 | -1.64 | -1.18 | -0.08 | -0.95 | 0.26 | -1.69 | -1.71 | 0.1575 | -0.125 | -0.3275 | 0.760255 | 0.794296 | 0.552176 | 1 | 0.996511 | 0.915338 |
| Fntb | Q8K2I1 | 0.39 | -0.72 | 0.13 | -0.18 | -0.24 | 0.2 | 1.04 | 0.23 | -1.76 | -0.89 | 0.15 | 0.01 | -1.74 | -1.75 | 0.09 | 0.36 | 0.4025 | -0.5275 | -0.665 | 0.262482 | 0.280892 | 0.262036 | 1 | 0.996511 | 0.815083 |
| Focad | A2AKG8 | -0.53 | -0.25 | 0.09 | -0.13 | -0.48 | -0.39 | -0.27 | 0.09 | -0.28 | -0.36 | 0.19 | -0.35 | -0.06 | -0.52 | -0.11 | 0.04 | -0.0575 | 0.005 | 0.0425 | 0.783074 | 0.981926 | 0.850676 | 1 | 0.998773 | 0.976225 |
| Folh1 | O35409 | 0.81 | 0.95 | 0.75 | 0.7 | 0.84 | 0.86 | 0.74 | 0.92 | 0.75 | -0.6 | 0.75 | 0.81 | 0.8 | 0.29 | 1.51 | 1.59 | 0.0375 | -0.375 | 0.245 | 0.798523 | 0.289806 | 0.451473 | 1 | 0.996511 | 0.885538 |
| Folr1 | E9Q543 | -2.98 | -2.87 | -3.3 | -3.81 | -2.32 | -3.39 | -3.39 | -4.27 | -3.77 | -2.13 | -6.39 | -2.9 | -2.69 | -2.32 | -3.24 | -3.49 | -0.1025 | -0.5575 | 0.305 | 0.808576 | 0.520708 | 0.379499 | 1 | 0.996511 | 0.860984 |
| Foxg1 | Q60987 | -2.28 | -0.49 | 2.71 | 2.81 | -2.29 | -1.25 | 2.27 | 2.38 | -1.4 | 4.11 | 0.44 | -1.73 | -0.93 | -0.19 | 2.67 | 7.56 | -0.41 | -0.3325 | 1.59 | 0.791369 | 0.838943 | 0.442322 | 1 | 0.996511 | 0.883549 |

| | | | | | | | | | | | | | | | | | | | | | | | | | | | |
|---|---|---|---|---|---|---|---|---|---|---|---|---|---|---|---|---|---|---|---|---|---|---|---|---|---|---|---|
| Foxj3 | Q8BUR3-2 | -2.03 | -2.18 | -1.75 | -1.23 | -0.86 | -3.15 | -0.22 | 0.82 | 0.31 | 0.08 | 0.59 | -1.72 | 1.62 | 0.34 | -2.83 | 1.57 | 0.945 | 1.6125 | 1.9725 | 0.246797 | 0.012516 | 0.064587 | | 1 | 0.996511 | 0.66433 |
| Foxk1 | P42128 | -1.9 | -2.77 | -2.48 | -3.47 | -2.42 | -2.25 | -3.78 | -2.52 | -2.35 | -0.22 | -2.59 | -1.9 | -1.41 | -1.99 | -2.35 | -2.75 | -0.0875 | 0.89 | 0.53 | 0.844505 | 0.148686 | 0.222732 | | 1 | 0.996511 | 0.787447 |
| Foxk2 | Q3UCQ1-2 | -1.21 | -1.16 | -1.61 | -2.05 | -0.4 | -1.48 | -1.42 | -2.25 | -1.62 | -0.63 | -1.96 | -1.02 | -1.67 | -0.8 | -1.73 | -2.1 | 0.12 | 0.2 | -0.0675 | 0.768407 | 0.576107 | 0.844348 | | 1 | 0.996511 | 0.976225 |
| Foxo1 | Q9R1E0 | -0.31 | -0.11 | 0.06 | -0.7 | -1.16 | -1.41 | -1.67 | 0.06 | 0.64 | -1.01 | -0.7 | 0.12 | 0.14 | -0.1 | -0.08 | -0.79 | -0.78 | 0.0275 | 0.0575 | 0.074429 | 0.944313 | 0.83739 | | 1 | 0.996755 | 0.975831 |
| Foxo3 | Q9WVH4 | 0.21 | -0.69 | -0.6 | -0.39 | -2.2 | 0.18 | -0.32 | 0 | 1.91 | -0.86 | -0.51 | -0.75 | 2.55 | 0.61 | -0.66 | -0.37 | -0.2175 | 0.315 | 0.9 | 0.686385 | 0.619227 | 0.210659 | | 1 | 0.996511 | 0.786218 |
| Foxp1 | A0A0R4J28 | -1.74 | -2.14 | -3.35 | -2.67 | -1.46 | -2.12 | -2.75 | -4.09 | -2.35 | -2.23 | -3.02 | -1.99 | -1.6 | -2.37 | -2.91 | -3.08 | -0.13 | 0.0775 | -0.015 | 0.82906 | 0.844458 | 0.973662 | | 1 | 0.996511 | 0.992946 |
| Foxred1 | Q3TQB2 | 0.7 | 0.59 | 0.81 | 0.81 | 0.21 | 0.43 | 0.09 | 0.63 | 0.68 | 0.95 | 0.86 | 0.62 | 0.79 | 1.13 | 0.23 | 0.23 | -0.3875 | 0.05 | -0.1325 | 0.053739 | 0.769745 | 0.610073 | | 1 | 0.996511 | 0.93496 |
| Fras1 | Q80T14 | -0.52 | -0.03 | 0.04 | -0.16 | 0.42 | 0.08 | -0.23 | -0.46 | 0.05 | -0.03 | -0.54 | -0.4 | 0.63 | 0.73 | -0.37 | 0.17 | 0.12 | -0.0625 | 0.4575 | 0.623431 | 0.780797 | 0.147562 | | 1 | 0.996511 | 0.746406 |
| Frg1 | P97376 | -2.19 | -3.25 | -0.52 | -0.4 | -3.24 | -3.21 | 0.1 | -0.17 | -1.98 | -0.54 | -0.56 | -3.26 | -2.28 | -0.71 | -0.68 | -0.25 | -0.04 | 0.005 | 0.61 | 0.969044 | 0.995329 | 0.419843 | | 1 | 0.999285 | 0.87558 |
| Frmd3 | Q8BHD4 | 1.54 | 2.96 | -0.11 | 0.88 | 2.96 | 2.27 | 0.93 | 0.4 | 1.79 | 1.19 | -0.11 | 1.65 | -0.14 | 3.21 | 0.52 | 0.51 | 0.3225 | -0.1875 | -0.2925 | 0.683675 | 0.790528 | 0.741273 | | 1 | 0.996511 | 0.958796 |
| Frmd4a | H3BIZ7 | -0.02 | 0.11 | 0.02 | -0.05 | 0.07 | -0.1 | 0 | 0.25 | 0.69 | 0 | 0.35 | 0.26 | 0.98 | 0.35 | 0.48 | 0.02 | 0.04 | 0.31 | 0.4425 | 0.793779 | 0.143619 | 0.093356 | | 1 | 0.996511 | 0.698156 |
| Frmd4b | Q6ZQ04 | -0.28 | 0.27 | -1.76 | -2.04 | -0.58 | 0.23 | -0.34 | -2.25 | 1.55 | -3.79 | -0.9 | -0.26 | 1.98 | -0.06 | -1.51 | -1.85 | 0.2175 | 0.1025 | 0.5925 | 0.756815 | 0.926642 | 0.528447 | | 1 | 0.996755 | 0.908663 |
| Frmd5 | A2AR56 | 0.44 | 0.35 | -0.35 | -0.01 | 0.98 | 0.14 | -0.14 | -0.4 | 0.84 | -0.01 | -0.26 | 0.51 | 0.81 | 0.96 | 0 | -0.16 | 0.0375 | 0.1625 | 0.295 | 0.91168 | 0.603358 | 0.391431 | | 1 | 0.996511 | 0.865375 |
| Frmd8 | Q3UFK8 | 0.78 | 0.67 | 0.4 | 0.49 | 0 | 0.58 | 0.26 | 0.35 | 2.44 | 0.4 | 0.24 | 0.41 | 2.95 | 0.35 | 0.28 | 0.09 | -0.2875 | 0.2875 | 0.3325 | 0.151625 | 0.564621 | 0.5981 | | 1 | 0.996511 | 0.930093 |
| Frmpd1 | A2AKB4 | -2.79 | -2.74 | -0.36 | 0.15 | -2.17 | -2.67 | -1.12 | -0.35 | -2.41 | 0.78 | -0.01 | -1.9 | -2.34 | -2.21 | -0.12 | -0.54 | -0.1425 | 0.55 | 0.1325 | 0.865441 | 0.575957 | 0.878028 | | 1 | 0.996511 | 0.979626 |
| Frmpd3 | A0A140LIW | 0.95 | 1.23 | 0.74 | 0.78 | 1.07 | 1.17 | 0.45 | 0.86 | 1.23 | 0.47 | 0.97 | 0.8 | 1.27 | 1.31 | 0.81 | 0.86 | -0.0375 | -0.0575 | 0.1375 | 0.863811 | 0.800819 | 0.540525 | | 1 | 0.996511 | 0.912212 |
| Frmpd4 | A0A589MR | 1.14 | 0.66 | 0.39 | 0.47 | 1.13 | 1.06 | 0.41 | 0.41 | 0.57 | 0.49 | 0.53 | 0.5 | 0.89 | 0.81 | 0.13 | 0.23 | 0.0875 | -0.1425 | -0.15 | 0.743473 | 0.505776 | 0.59336 | | 1 | 0.996511 | 0.929843 |
| Frrs1l | B1AXV0 | 1.55 | 0.83 | 1.86 | 0.63 | 1.22 | 2.21 | 2.92 | 1.58 | 1.15 | 1.63 | 1.79 | 1.85 | 1.07 | 0.63 | 2.58 | 2.57 | 0.765 | 0.3875 | 0.495 | 0.111188 | 0.259077 | 0.371147 | | 1 | 0.996511 | 0.859424 |
| Frs2 | Q8C180 | 1.06 | 1.74 | 1.08 | 0.9 | 1.59 | 1.01 | 1.22 | 0.91 | 1.32 | 1.31 | 0.93 | 1.36 | 1.37 | 1.91 | 1.2 | 1.32 | -0.0125 | 0.035 | 0.255 | 0.960244 | 0.883696 | 0.355541 | | 1 | 0.996511 | 0.855029 |
| Frs3 | Q91WJ0 | 0.55 | 1.08 | 1.26 | 0.93 | 1.29 | 0.62 | 1.06 | 1.1 | -0.36 | 1.12 | 1.12 | 0.56 | 0.17 | -0.4 | 1.24 | 1.16 | 0.0625 | -0.345 | -0.4125 | 0.783755 | 0.36216 | 0.326815 | | 1 | 0.996511 | 0.843799 |
| Fry | A0A5F8MP | -0.26 | -0.28 | -0.25 | -0.24 | -0.02 | -0.26 | -0.02 | -0.14 | -0.19 | 0.08 | 0.01 | -0.15 | -0.02 | 0.12 | -0.33 | -0.07 | 0.1475 | 0.195 | 0.1825 | 0.321336 | 0.242974 | 0.33075 | | 1 | 0.996511 | 0.845601 |
| Fry | A0A0J9YUF | 0.03 | 0.17 | -0.22 | 0.43 | 0.79 | -0.35 | 0.53 | -0.21 | 0.4 | 1.06 | 0.79 | 0.11 | -0.03 | 1.52 | -0.01 | 0.37 | 0.0875 | 0.4875 | 0.36 | 0.774977 | 0.09508 | 0.354815 | | 1 | 0.996511 | 0.855029 |
| Fryl | F8VQ05 | -0.3 | -0.26 | -0.37 | -0.76 | -0.3 | -0.21 | -0.5 | -0.83 | 0.08 | -0.55 | -0.64 | -0.31 | -0.04 | -0.09 | -1 | -0.75 | -0.0375 | 0.0675 | -0.0475 | 0.856934 | 0.769398 | 0.867342 | | 1 | 0.996511 | 0.978457 |
| Fscn1 | Q61553 | 0.69 | -0.15 | 0.61 | 0.84 | -0.08 | 0.87 | 1.52 | 1.14 | 0.19 | 0.82 | 0.66 | 0.72 | -0.04 | -0.18 | 0.82 | 0.92 | 0.365 | 0.1 | -0.1175 | 0.356389 | 0.717272 | 0.74235 | | 1 | 0.996511 | 0.958976 |
| Fsd1 | Q7TPM6 | 0.61 | 0.74 | 0.32 | 0.38 | 0.73 | 0.26 | 0.33 | 0.53 | 0.51 | 0.44 | 0.45 | 0.83 | 0.06 | 0.9 | 0.2 | 0.19 | -0.05 | 0.045 | -0.175 | 0.788079 | 0.813883 | 0.487601 | | 1 | 0.996511 | 0.897232 |
| Fsd1l | E0CXZ1 | -0.38 | -0.4 | 0 | -0.12 | -0.42 | -0.37 | -0.04 | -0.21 | -0.34 | -0.08 | -0.03 | 0 | -0.27 | -0.53 | -0.31 | -0.38 | -0.035 | 0.1125 | -0.1475 | 0.843773 | 0.549421 | 0.447014 | | 1 | 0.996511 | 0.884817 |
| Fstl4 | Q5STE3 | -0.7 | 0.61 | -0.67 | -0.52 | 0.4 | -0.65 | -0.46 | -0.72 | -0.03 | -0.74 | -0.52 | -0.49 | -0.15 | 0.43 | -0.63 | -0.04 | -0.0375 | -0.125 | 0.2225 | 0.922218 | 0.714993 | 0.554308 | | 1 | 0.996511 | 0.916796 |
| Fth1 | P09528 | -0.66 | -1.9 | -0.86 | -0.81 | -0.9 | -0.95 | -0.59 | -0.83 | -1.71 | -0.64 | -0.95 | -1.2 | -1.56 | -0.98 | -0.94 | -0.6 | 0.24 | -0.0675 | 3.75E-02 | 0.423838 | 0.849101 | 0.913556 | | 1 | 0.996511 | 0.986458 |
| Ftl1 | Q9CPX4 | -3.29 | -3.5 | -1.43 | -1.34 | -2.42 | -3.31 | -0.77 | -1.2 | -3.9 | -1.42 | -0.57 | -3.28 | -3.13 | -2.37 | -1.18 | -0.67 | 0.465 | 0.0975 | 0.5525 | 0.535598 | 0.9114 | 0.456941 | | 1 | 0.996744 | 0.887902 |
| Fto | Q8BGW1 | -1.28 | -1.64 | -2.74 | -2.74 | -1.32 | -1.2 | -2.34 | -2.86 | -1.55 | -1.41 | -2.73 | -1.3 | -1.49 | -1.11 | -2.91 | -2.7 | 0.17 | 0.3525 | 0.0475 | 0.738327 | 0.460448 | 0.929617 | | 1 | 0.996511 | 0.989687 |
| Ftsj1 | F6Q475 | -1.42 | -1.28 | 0.16 | 1.62 | -2.32 | -1.37 | -0.15 | 0.5 | -0.84 | 0.33 | 0.35 | -0.74 | -0.35 | -2.5 | 1.25 | 0.81 | -0.605 | 0.005 | 0.0325 | 0.485862 | 0.994391 | 0.97364 | | 1 | 0.999285 | 0.992946 |
| Ftsj3 | Q9DBE9 | 0.66 | 0.7 | 0.27 | 0.45 | 0.76 | 0.35 | 0.31 | 0.34 | 0.29 | 0.37 | 0.11 | 0.41 | 0.66 | 0.75 | 0.49 | 0.74 | -0.08 | -0.225 | 0.14 | 0.670064 | 0.239705 | 0.472397 | | 1 | 0.996511 | 0.891144 |
| Fubp1 | Q3TUE1 | -4.1 | -4.52 | -4.06 | -3.93 | -4.4 | -4.34 | -4.44 | -4.25 | -4.4 | -1.84 | -4.31 | -3.84 | -3.41 | -3.46 | -4.42 | -4.99 | -0.205 | 0.555 | 0.0825 | 0.272128 | 0.338652 | 0.833513 | | 1 | 0.996511 | 0.975304 |
| Fubp3 | A2AJ72 | -0.22 | -0.81 | -2.02 | -1.22 | -1.13 | -0.57 | -1.99 | -2.11 | -0.29 | -1.01 | -1.53 | -0.6 | -0.09 | -0.29 | -1.23 | -2.24 | -0.3825 | 0.21 | 0.105 | 0.439933 | 0.633417 | 0.854243 | | 1 | 0.996511 | 0.976225 |
| Fuca1 | Q99LJ1 | 0.08 | 0.42 | 0.22 | 0.04 | 0.37 | 0.6 | 0.31 | 0.28 | 0.88 | 0.63 | 0.07 | 0.22 | 0.06 | 3.27 | 0.07 | 0.02 | 0.2 | 0.26 | 0.665 | 0.251984 | 0.28526 | 0.375651 | | 1 | 0.996511 | 0.859611 |
| Fundc1 | H3BJZ1 | -0.33 | 0.49 | 0.37 | 0.34 | -0.54 | -0.1 | 0.74 | 0.21 | 0.27 | 0.22 | 0.12 | -0.28 | 0.72 | -0.05 | 0.68 | 0.76 | -0.14 | -0.135 | 0.31 | 0.663012 | 0.588838 | 0.294813 | | 1 | 0.996511 | 0.825612 |
| Fundc2 | Q9D6K8 | 0.12 | 0.33 | -0.01 | -0.08 | 0.62 | 0.12 | -0.32 | -0.3 | 0.55 | 0.57 | -0.26 | -0.22 | 0.69 | 1.43 | -0.28 | -0.23 | -0.06 | 0.07 | 0.3125 | 0.811035 | 0.792169 | 0.446032 | | 1 | 0.996511 | 0.884817 |
| Fuom | Q8R2K1 | -1 | -0.61 | -0.9 | -1.01 | -1 | -1.14 | -1.2 | -1.5 | -0.98 | -0.94 | -1.32 | -0.8 | -1.37 | -0.36 | -1.37 | -1.08 | -0.33 | -0.13 | -0.165 | 0.099836 | 0.513692 | 0.556069 | | 1 | 0.996511 | 0.917092 |
| Fus | Q8CFQ9 | -3.02 | -4.38 | -4.19 | -4.23 | -3.49 | -4.12 | -3.15 | -4.6 | -3.5 | -2.58 | -5.03 | -3.36 | -2.48 | -2.28 | -1.43 | -3.84 | 0.115 | 0.3375 | 1.4475 | 0.785759 | 0.546867 | 0.024827 | | 1 | 0.996511 | 0.632054 |
| Fut11 | Q8BHC9 | -0.75 | -0.77 | -0.8 | -0.85 | -0.02 | -0.42 | -0.61 | -1.01 | -0.3 | -1.02 | -1.22 | -0.81 | 1.75 | 0.02 | -0.28 | -0.35 | 0.2775 | -0.045 | 1.0775 | 0.242636 | 0.84541 | 0.043359 | | 1 | 0.996511 | 0.658454 |
| Fut8 | Q9WTS2 | -1.17 | -2.15 | -1.33 | -1.08 | -1.6 | -0.67 | -0.29 | -1.32 | -0.77 | -1.21 | -1.22 | -1.53 | -0.65 | -1.55 | -0.83 | -0.57 | 0.4625 | 0.25 | 0.5325 | 0.230638 | 0.409969 | 0.137081 | | 1 | 0.996511 | 0.73233 |
| Fut9 | O88819 | -1.42 | -1.48 | -2.05 | -0.4 | -0.84 | -1.8 | -0.27 | -0.99 | -1.25 | -1.74 | -0.88 | -1.21 | -1.4 | -0.53 | -0.68 | -0.48 | 0.3625 | 0.0675 | 0.565 | 0.414904 | 0.856622 | 0.173674 | | 1 | 0.996511 | 0.764376 |
| Fv1 | P70213 | -1.51 | -1.6 | -0.18 | 0.52 | -2.84 | -2.04 | 0.53 | 0.81 | -1.36 | 0.37 | 0.19 | -2.03 | -0.54 | -3.87 | 0.58 | 0.6 | -0.1925 | -0.015 | -0.115 | 0.838457 | 0.983165 | 0.912736 | | 1 | 0.998773 | 0.986458 |
| Fxn | O35943 | 1.04 | 1.18 | 0.24 | 0.81 | 1.19 | 1.02 | 0.22 | 0.66 | 0.94 | -2.06 | 1.18 | 1.21 | 0.89 | 1.44 | 0.42 | 0.15 | -0.045 | -0.5 | -0.0925 | 0.879455 | 0.507531 | 0.791057 | | 1 | 0.996511 | 0.969105 |
| Fxr1 | Q61584-3 | -0.57 | -0.79 | -0.59 | -0.69 | -0.35 | -0.88 | -0.64 | -0.57 | -0.59 | -0.51 | -0.73 | -0.36 | -0.18 | 0.15 | 0.05 | -0.3 | 0.05 | 0.1125 | 0.59 | 0.771334 | 0.514006 | 0.012387 | | 1 | 0.996511 | 0.573077 |
| Fxr2 | Q6P5B5 | -0.44 | -0.24 | -0.32 | -0.29 | -0.29 | -0.59 | -0.09 | -0.47 | -0.01 | 0.46 | -0.41 | -0.33 | 0.25 | 0.26 | 0.06 | 0.12 | -0.0375 | 0.25 | 0.495 | 0.825472 | 0.299236 | 0.017266 | | 1 | 0.996511 | 0.602211 |
| Fxyd1 | A0A0U1RP | 0.61 | 0.98 | 0.94 | 0.94 | 1.08 | 0.03 | 1.13 | 0.43 | 0.78 | 2.02 | 0.36 | 0.63 | 0.42 | 1.67 | 0.79 | 0.67 | -0.2 | 0.08 | 0.02 | 0.484404 | 0.827413 | 0.946592 | | 1 | 0.996511 | 0.992376 |
| Fxyd6 | Q9D164-2 | 0.19 | 1.09 | 0.73 | 0.58 | 1.31 | 0.82 | 1.44 | 0.97 | 0.99 | 1.29 | 1.3 | 0.57 | 0.52 | 1.56 | 0.74 | 0.22 | 0.4875 | 0.39 | 0.1125 | 0.076685 | 0.172472 | 0.742867 | | 1 | 0.996511 | 0.959128 |
| Fxyd7 | A0A0U1RP | -1.45 | -1.29 | -1.75 | -2.77 | -1.38 | -0.83 | -1.41 | -1.77 | 2.8 | -2.73 | -0.74 | -1.27 | 4.9 | -1.33 | 2.74 | -1.64 | 0.4675 | 1.33 | 2.9825 | 0.224648 | 0.244197 | 0.065289 | | 1 | 0.996511 | 0.664857 |
| Fyco1 | Q8VDC1 | 0.88 | 1.1 | -0.56 | -0.96 | 1.53 | 0.62 | 0.4 | 0.1 | 0.55 | -0.26 | -0.53 | 1 | 0.12 | 1.58 | -0.65 | -0.3 | 0.5475 | 0.075 | 0.0725 | 0.333042 | 0.895953 | 0.910731 | | 1 | 0.996511 | 0.986458 |
| Fyn | P39688 | 1.02 | 1.18 | 0.82 | 0.77 | 0.92 | 0.75 | 0.92 | 0.68 | 0.21 | 0.32 | 0.92 | 1.32 | -0.15 | 0.47 | 0.87 | 0.9 | -0.13 | -0.255 | -0.425 | 0.445723 | 0.385911 | 0.15739 | | 1 | 0.996511 | 0.751678 |
| Fyttd1 | Q91Z49 | 0.17 | -0.98 | -1.14 | -1.07 | -1.36 | -0.75 | -1.36 | -0.47 | -0.98 | -1.19 | -1.32 | -1.58 | -0.05 | -0.16 | -0.46 | -1.81 | -0.23 | -0.5125 | 0.135 | 0.533613 | 0.14903 | 0.778292 | | 1 | 0.996511 | 0.967289 |
| Fzd2 | Q9JIP6 | 0.59 | 0.46 | -1.05 | -0.93 | 0.97 | 0.02 | -0.34 | -0.39 | -0.08 | -0.86 | -1.85 | 0.31 | -0.89 | 0.43 | -1.41 | 0.53 | 0.2975 | -0.3875 | -0.1025 | 0.554762 | 0.519627 | 0.863883 | | 1 | 0.996511 | 0.977235 |
| Fzd3 | Q61086 | -0.24 | -0.35 | 0.22 | 0.43 | -0.17 | 0.25 | -0.69 | 0.26 | 0.48 | 0.82 | 0.1 | 0.04 | 0.77 | 0.19 | -0.45 | 0.78 | -0.1025 | 0.345 | 0.3075 | 0.725528 | 0.229 | 0.383841 | | 1 | 0.996511 | 0.860984 |
| G0s2 | Q61585 | 1.09 | 0.17 | 1.29 | 0.28 | 1.97 | 2.02 | 0.12 | -0.27 | 0.67 | 1.51 | 0.12 | 1.04 | 1.97 | 0.13 | 0.81 | -0.57 | 0.2525 | 0.1275 | -0.1225 | 0.678696 | 0.74497 | 0.827559 | | 1 | 0.996511 | 0.973757 |
| G3bp1 | P97855 | -2.85 | -2.94 | -1.99 | -2.12 | -2.46 | -2.92 | -1.01 | -1.99 | -2.44 | -1.55 | -1.67 | -2.45 | -2.69 | -2.07 | -1.4 | -1.69 | 0.38 | 0.4475 | 0.5125 | 0.402202 | 0.210411 | 0.183949 | | 1 | 0.996511 | 0.776843 |
| G3bp2 | P97379 | -2.45 | -2.88 | -2.76 | -2.79 | -2.11 | -2.77 | -2.53 | -3.19 | -2.39 | -1.69 | -2.78 | -2.18 | -1.7 | -1.94 | -1.77 | -2.42 | 0.07 | 0.46 | 0.7625 | 0.783897 | 0.108535 | 0.008886 | | 1 | 0.996511 | 0.563386 |
| G6pc3 | Q6NSQ9 | -3.79 | -1.83 | -1.37 | -1.56 | -2.39 | -2.34 | 0.33 | -1.06 | -4 | -2.17 | -1.27 | -3.44 | -3.5 | -2.22 | -0.59 | 0.04 | 0.7725 | -0.5825 | 0.57 | 0.329906 | 0.447876 | 0.520883 | | 1 | 0.996511 | 0.907667 |
| G6pdx | Q00612 | 0.85 | 0.29 | 0.01 | 0.1 | -0.26 | 0.53 | -0.13 | 0.05 | 0.89 | 0.19 | -0.22 | 0.52 | 0.55 | 0.51 | -0.37 | -0.57 | -0.265 | 0.0325 | -0.2825 | 0.329123 | 0.915465 | 0.423249 | | 1 | 0.996755 | 0.875782 |
| Gaa | P70699 | -0.86 | -1.42 | -0.97 | -0.72 | -1.57 | -0.69 | -0.14 | -0.91 | -1.48 | -1.35 | -0.42 | -0.72 | -0.95 | -1.47 | -0.79 | -0.8 | 0.165 | 2.36E-16 | -0.01 | 0.613221 | 1 | 0.968404 | | 1 | 1 | 0.992733 |
| Gab1 | A0A1B0GS | 0.13 | 0.02 | -0.06 | 0.33 | -0.18 | -0.22 | -0.09 | -0.15 | 0.06 | 0.11 | -0.21 | -0.36 | -0.29 | 0.26 | -0.26 | -0.18 | -0.265 | -0.205 | -0.2225 | 0.116902 | 0.306436 | 0.307223 | | 1 | 0.996511 | 0.832533 |
| Gab2 | Q3ZB57 | 0.19 | 0.38 | 0.57 | 0.68 | -0.22 | 0.59 | 0.24 | 0.37 | 0.5 | 0.74 | 0.55 | 0.09 | 1.04 | -0.05 | 0.52 | 0.46 | -0.21 | 0.015 | 0.0375 | 0.361089 | 0.944215 | 0.890109 | | 1 | 0.996755 | 0.981598 |
| Gabarapl1 | Q8R3R8 | 6.17 | 2.92 | 3.06 | 3.92 | 2.5 | 5.75 | 3.87 | 4.11 | 4.14 | 3.96 | 3.35 | 5.46 | 2.65 | 2.87 | 3.43 | 2.9 | 0.04 | 0.21 | -1.055 | 0.964611 | 0.790387 | 0.15624 | | 1 | 0.996511 | 0.751493 |
| Gabarapl2 | P60521 | 0.58 | 0.54 | 0.68 | 0.54 | 0.55 | 0.29 | 0.92 | 1.07 | 0.52 | 0.55 | 0.57 | 1.06 | 0.09 | 1.51 | 0.25 | 0.41 | 0.1225 | 0.09 | -0.02 | 0.561022 | 0.637821 | 0.95126 | | 1 | 0.996511 | 0.992476 |
| Gabbr1 | Q9WV18-2 | -0.02 | 0.92 | 0.77 | 1.27 | 0.34 | 0.21 | -0.94 | -0.26 | 2.19 | 0.3 | 0.41 | -0.21 | 3.36 | 0.13 | 1.14 | 0.61 | -0.8975 | -0.0625 | 0.575 | 0.039449 | 0.908615 | 0.416434 | | 1 | 0.996744 | 0.874945 |
| Gabbr1 | Q9WV18 | 0.89 | 0.58 | 0.55 | 0.73 | 0.6 | 0.93 | 0.41 | 0.35 | 0.92 | 0.19 | 0.61 | 0.8 | 1.15 | 0.74 | 0.34 | 0.54 | -0.115 | -0.0575 | 0.005 | 0.550239 | 0.791247 | 0.982844 | | 1 | 0.996511 | 0.995073 |
| Gabbr2 | Q80T41 | 1.08 | 1.06 | 1.55 | 1.74 | 1.07 | 1.27 | 1.33 | 1.38 | 0.86 | 1.12 | 1.59 | 1.14 | 1.16 | 0.6 | 1.36 | 1.55 | -0.095 | -0.18 | -0.19 | 0.654414 | 0.479936 | 0.510828 | | 1 | 0.996511 | 0.90359 |
| Gabpa | Q00422 | -2.99 | -4.8 | -2.62 | -2.2 | -3.99 | -4.06 | -1.56 | -2.01 | -3.45 | -1.65 | -1.92 | -3.92 | -1.95 | -3.28 | -1.84 | -2.12 | 0.2475 | 0.4175 | 0.855 | 0.752661 | 0.569127 | 0.182641 | | 1 | 0.996511 | 0.776002 |
| Gabra1 | P62812 | 1.13 | 1.05 | 1.67 | 1.42 | 1.33 | 1.31 | 0.68 | 1.04 | 0.66 | 0.09 | 1.45 | 0.91 | 0.74 | 0.73 | 1.11 | 1.19 | -0.2275 | -0.54 | -0.375 | 0.332623 | 0.116178 | 0.130764 | | 1 | 0.996511 | 0.725078 |
| Gabra2 | P26048 | -0.99 | -1.33 | -2.28 | -2.14 | 1.73 | 1.61 | 1.96 | 1.58 | 1.02 | -0.07 | 1.52 | 1.63 | 0.85 | 1.07 | 1.72 | 1.66 | 3.405 | 2.71 | 3.01 | 3.61E-06 | 3.10E-04 | 2.72E-05 | 0.010958 | 0.706619 | 0.082536 | |
| Gabra3 | B1AVY2 | 0.83 | 1.18 | 1.1 | 1.62 | 0.79 | 0.38 | 0.58 | 1.15 | 0.04 | 1.16 | 1.25 | 0.51 | 0.13 | 0.69 | 0.8 | 0.72 | -0.4575 | -0.4425 | -0.5975 | 0.090292 | 0.200417 | 0.041446 | | 1 | 0.996511 | 0.658454 |
| Gabra4 | Q9D6F4 | 1.29 | 1.33 | 1.45 | 1.43 | 0.9 | 0.89 | 0.7 | 0.73 | 0.61 | 0.46 | 0.77 | 0.96 | 0.56 | 0.3 | 0.93 | 0.67 | -0.57 | -0.675 | -0.76 | 0.003715 | 0.004577 | 0.004341 | | 1 | 0.996511 | 0.548272 |
| Gabra5 | Q8BHJ7 | -0.15 | -0.26 | 1.52 | 0.98 | 0.09 | -0.64 | 1.22 | 0.79 | -0.74 | -1.27 | 1.04 | -0.27 | -0.49 | 0.15 | 0.78 | 1.32 | -0.1575 | -0.8325 | -0.0825 | 0.773607 | 0.191593 | 0.878746 | | 1 | 0.996511 | 0.979626 |
| Gabrb1 | P50571 | 0.76 | 0.46 | 0.85 | 1.39 | 1.16 | 1.57 | 1.69 | 1.94 | 1.76 | 2.01 | 1.79 | 1.6 | 1.11 | 1.94 | 1.33 | 1.35 | 0.725 | 0.925 | 0.5675 | 0.020742 | 0.003837 | 0.070852 | | 1 | 0.996511 | 0.668066 |
| Gabrb2 | P63137-2 | 0.12 | 0.53 | 0.34 | 0.23 | 1.38 | 0.66 | 0.77 | 0.36 | 0.26 | -0.13 | 0.65 | 0.48 | 0.38 | 0.73 | 0.39 | 0.76 | 0.4875 | 0.01 | 0.26 | 0.073502 | 0.964458 | 0.218696 | | 1 | 0.996755 | 0.787447 |
| Gabrb3 | Q8C446 | 0.58 | 0.97 | 0.83 | 1.02 | 1.18 | 0.57 | 1.02 | 0.92 | 0.95 | 1.18 | 1.01 | 0.61 | 0.53 | 1.48 | 0.31 | 0.64 | 0.0725 | 0.0875 | -0.11 | 0.713829 | 0.667439 | 0.706163 | | 1 | 0.996511 | 0.957084 |
| Gabrd | P22933 | 0.56 | 0.46 | 0.66 | 0.84 | 0.35 | 0.34 | 0.48 | 0.14 | -0.07 | -0.34 | 0.56 | 0.51 | -0.25 | 0.49 | -0.23 | 0 | -0.3025 | -0.465 | -0.6275 | 0.094024 | 0.097018 | 0.023276 | | 1 | 0.996511 | 0.623925 |
| Gabrg1 | Q9R0Y8 | -2.86 | -2.01 | -0.03 | 0.32 | -0.96 | -1.12 | 0.96 | 1.01 | -1.69 | 1.4 | 1.28 | -2.23 | -0.91 | -0.31 | 1.14 | 0.99 | 1.1175 | 0.835 | 1.3725 | 0.220552 | 0.456282 | 0.123323 | | 1 | 0.996511 | 0.721448 |
| Gabrg2 | P22723-2 | -0.27 | 0.53 | 0.62 | 0.4 | 1.15 | 0.53 | 0.61 | 0.52 | 0.22 | -0.02 | 0.96 | 0.01 | 0.52 | 0.68 | 0.55 | 0.58 | 0.3825 | -0.0275 | 0.2625 | 0.167376 | 0.928916 | 0.295254 | | 1 | 0.996755 | 0.825612 |
| Gabrg2 | P22723 | 1.47 | 1.99 | 1.32 | 1.39 | 1.83 | 1.51 | 0.96 | 0.92 | 1.53 | 1.57 | 0.9 | 1.16 | 1.2 | 2.29 | 0.56 | 1.01 | -0.2375 | -0.2525 | -0.2775 | 0.395187 | 0.31897 | 0.478696 | | 1 | 0.996511 | 0.892849 |
| Gabrg3 | P27681 | 7.77 | 5.46 | -0.29 | 0.06 | 4.73 | 9.56 | 0.01 | -0.26 | 3.77 | 0.34 | 0.07 | 8.09 | 5.05 | -0.89 | 0.39 | 0.08 | 0.26 | -0.1825 | -2.0925 | 0.923974 | 0.940329 | 0.338528 | | 1 | 0.996755 | 0.847578 |
| Gad1 | P48318 | 0.16 | 0.25 | 0.49 | 0.79 | 0.34 | 0.12 | 0.67 | 0.68 | 0.33 | 0.33 | 0.37 | 0 | 0.42 | 0.26 | 0.56 | 0.86 | 0.03 | -0.165 | 0.1025 | 0.891288 | 0.438152 | 0.661409 | | 1 | 0.996511 | 0.949645 |
| **Gad2** | **P48320** | **0.36** | **-0.23** | **-0.7** | **-0.2** | **0.08** | **0.63** | **0.06** | **-0.1** | **-0.04** | **0.05** | **-0.6** | **0.18** | **-0.04** | **-0.66** | **0.43** | **0.11** | 0.36 | 0.09 | 0.1525 | 0.211251 | 0.754072 | 0.637028 | | 1 | 0.996511 | 0.942665 |
| Gadd45gip | Q9CR59 | 0.26 | 0 | 0.74 | 0.71 | -0.95 | 0.49 | -0.29 | 0.05 | 1.62 | 1.19 | 0.3 | -0.26 | 2.5 | -0.96 | 0.59 | 0.38 | -0.6025 | 0.285 | 0.2 | 0.105108 | 0.518388 | 0.765117 | | 1 | 0.996511 | 0.964134 |
| Gak | A0A0R4J0F | 0.34 | 0.23 | 0.54 | 0.55 | 0.36 | 0.39 | 0.36 | 0.56 | 0.57 | 0.46 | 0.57 | 0.32 | 0.49 | 0.43 | 0.42 | 0.24 | 0.0025 | 0.065 | -0.02 | 0.987313 | 0.707703 | 0.912678 | | 1 | 0.996511 | 0.986458 |
| Gal3st3 | P61315 | 0.26 | 0.05 | -0.19 | 0.09 | 0.46 | 0.05 | 0.45 | 0.36 | 0.4 | -0.28 | -0.06 | 0.28 | -0.49 | 0.46 | 0.17 | 0.57 | 0.2775 | 0.0325 | 0.125 | 0.147934 | 0.88206 | 0.653632 | | 1 | 0.996511 | 0.947633 |
| Galc | P54818 | -0.77 | -0.77 | 0.22 | 0.19 | -0.5 | -0.78 | 0.66 | 0.2 | -1.06 | 0.65 | 0.39 | -1.32 | -0.87 | 0.04 | -0.03 | 0.17 | 0.1775 | -0.0525 | 0.11 | 0.664097 | 0.921073 | 0.761601 | | 1 | 0.996755 | 0.963123 |
| Gale | Q8R059 | 1.13 | 0.96 | 0.05 | -0.47 | 0.85 | 1.3 | -0.07 | -0.02 | 0.07 | 0.17 | -0.37 | 0.82 | 0.01 | 0.24 | 0.22 | 0 | 0.0975 | -0.245 | -0.3 | 0.835664 | 0.57035 | 0.433074 | | 1 | 0.996511 | 0.879439 |
| Galk1 | Q9R0N0 | -0.27 | -0.3 | -0.88 | -0.91 | -0.39 | -0.46 | -1.07 | -0.86 | -0.35 | -0.57 | -0.89 | -0.31 | -0.25 | -0.05 | -0.88 | -0.86 | -0.105 | 0.06 | 0.08 | 0.677443 | 0.807321 | 0.784159 | | 1 | 0.996511 | 0.967642 |
| Galk2 | Q68FH4 | -2 | -2.28 | -2.23 | -1.85 | -2.6 | -2.03 | -1.94 | -1.82 | -1.71 | -1.58 | -2.09 | -2.05 | -1.05 | -2.31 | -1.96 | -2.49 | -0.0075 | 0.2325 | 0.1375 | 0.973084 | 0.277715 | 0.684539 | | 1 | 0.996511 | 0.955729 |
| Galm | Q8K157 | 0.52 | 0.83 | -0.22 | -0.1 | 0.81 | 0.37 | -0.18 | -0.2 | -0.02 | -0.74 | -0.34 | 0.66 | -0.46 | 0.86 | -0.45 | -0.47 | -0.0575 | -0.3675 | -0.3875 | 0.864632 | 0.340582 | 0.34544 | | 1 | 0.996511 | 0.849669 |
| Galns | Q571E4 | -0.75 | -0.84 | -0.36 | -0.32 | -0.92 | -0.67 | -0.07 | -0.53 | -0.48 | 0 | -0.04 | -0.87 | -0.26 | -0.73 | -0.52 | -0.55 | 0.02 | 0.22 | 0.0525 | 0.933146 | 0.412257 | 0.809238 | | 1 | 0.996511 | 0.972098 |
| Galnt1 | O08912 | -0.84 | -1 | -0.27 | -0.03 | -0.22 | -0.7 | 0.12 | -0.57 | -1.14 | -0.04 | -0.11 | -0.82 | -1.02 | -0.14 | 0.09 | -0.01 | 0.1925 | 0.0075 | 0.265 | 0.517942 | 0.982775 | 0.447185 | | 1 | 0.998773 | 0.884817 |
| Galnt13 | Q8CF93 | -1.21 | -0.23 | -0.68 | -0.65 | 0.13 | -1.07 | 0.07 | -0.17 | -0.59 | 0.09 | -0.69 | -0.84 | -1.29 | 0.75 | -0.06 | -0.18 | 0.4325 | 0.185 | 0.4975 | 0.215764 | 0.53507 | 0.276908 | | 1 | 0.996511 | 0.822084 |
| Galnt14 | Q08EC9 | 1.98 | 1.07 | 0.33 | 0.42 | 1.03 | 1.9 | 0.07 | 0.95 | 1.65 | 2.08 | -0.13 | 1.51 | 1.2 | 2.28 | 0.61 | 0.05 | 0.0375 | 0.3275 | 0.085 | 0.939301 | 0.567383 | 0.879782 | | 1 | 0.996511 | 0.979626 |
| Galnt16 | Q9JJ61 | -0.58 | -0.52 | -0.83 | -0.75 | -0.3 | -0.37 | -0.01 | -0.56 | -0.88 | -0.41 | -0.83 | -0.5 | -0.86 | -0.41 | 0.33 | -0.14 | 0.36 | 0.015 | 0.4 | 0.072339 | 0.937911 | 0.17745 | | 1 | 0.996755 | 0.772068 |
| Galnt17 | Q7TT15 | -0.8 | -1.09 | -1.46 | -1.16 | -0.36 | -0.78 | -0.57 | -1.22 | -0.9 | -1.09 | -1.45 | -0.96 | -0.93 | -0.63 | -0.92 | -0.79 | 0.395 | 0.0275 | 0.31 | 0.13101 | 0.900596 | 0.166 | | 1 | 0.996511 | 0.759345 |
| Galnt2 | Q6PB93 | -1.35 | -2.01 | -1.73 | -1.67 | -1.35 | -0.91 | -1.81 | -1.68 | -0.88 | -2 | -1.55 | -1.6 | -0.47 | -1.31 | -0.58 | -1.18 | 0.2525 | 0.1825 | 0.805 | 0.33285 | 0.520708 | 0.015582 | | 1 | 0.996511 | 0.589387 |
| Galt | Q03249 | -0.13 | -0.37 | -1.56 | -0.88 | -0.36 | 0.03 | -1.99 | -1.61 | -0.03 | -0.83 | -1.03 | 0.03 | -0.56 | 0.18 | -1.04 | -1.54 | -0.2475 | 0.27 | -0.005 | 0.643569 | 0.504252 | 0.99125 | | 1 | 0.996511 | 0.996154 |
| Gamt | O35969 | -0.67 | -0.41 | -0.34 | -0.4 | -0.02 | 0.27 | 0.29 | -0.24 | -0.45 | -1.07 | 0.13 | -0.21 | -0.19 | -0.94 | 0.06 | 0.11 | 0.53 | 0.055 | 0.215 | 0.018694 | 0.842487 | 0.443335 | | 1 | 0.996511 | 0.883549 |
| Gan | Q8CA72 | 0.64 | 0.27 | 0.67 | 0.41 | 0.58 | 0.76 | 1.06 | 0.47 | 0.49 | 0.39 | 1.17 | 0.72 | 0.22 | 0.65 | 0.71 | 1.1 | 0.22 | 0.195 | 0.1725 | 0.278836 | 0.406862 | 0.482094 | | 1 | 0.996511 | 0.893925 |
| Ganab | Q8BHN3 | 2.71 | 2.81 | 2.14 | 2.23 | 2.13 | 2.42 | 1.5 | 2.48 | 3.82 | 3.78 | 1.95 | 2.89 | 3.54 | 4.3 | 2.83 | 2.9 | -0.34 | 0.6375 | 0.92 | 0.248966 | 0.177451 | 0.03367 | | 1 | 0.996511 | 0.644622 |
| Ganab | Q8BHN3-2 | -0.1 | -0.37 | -0.21 | -0.15 | 0.17 | -0.11 | 0.87 | 0.11 | -0.86 | -0.28 | -0.1 | -0.02 | -0.58 | -0.01 | 0.67 | 0.65 | 0.4675 | -0.1075 | 0.39 | 0.075284 | 0.64283 | 0.234172 | | 1 | 0.996511 | 0.790641 |
| Ganc | A2AQJ8 | -4.9 | -5.12 | -2.32 | -3.61 | -2.56 | -5.31 | -2.76 | -4.11 | -3.98 | -2.11 | -1.81 | -4.47 | -6.82 | -3.38 | -3.2 | -2.72 | 0.3025 | 0.895 | -0.0425 | 0.713701 | 0.301703 | 0.96676 | | 1 | 0.996511 | 0.9927 |
| Gap43 | P06837 | 2.92 | 2.55 | 1.94 | 1.89 | 2.07 | 2.92 | 0.91 | 2.28 | 2.08 | 1.35 | 1.97 | 3.31 | 2.56 | -0.48 | 3.01 | 1.9 | -0.28 | -0.1475 | -0.5775 | 0.541423 | 0.744271 | 0.441957 | | 1 | 0.996511 | 0.883549 |
| Gapdh | P16858 | 0.52 | 0.06 | 0.02 | 0.41 | -0.24 | 0.45 | 0.62 | 0.71 | 0.18 | -0.81 | 0.29 | 0.7 | -0.46 | 0.1 | -0.45 | -0.18 | 0.1325 | -0.1625 | -0.5 | 0.610521 | 0.634003 | 0.05259 | | 1 | 0.996511 | 0.659923 |
| Gapdh | S4R1W8 | -1.78 | -1.12 | 0.27 | -0.5 | -1.4 | -1.31 | -0.46 | -0.42 | -1.23 | -0.23 | -0.6 | -1.13 | -1.1 | -1.85 | -1.65 | -1.48 | -0.115 | -0.015 | -0.7375 | 0.808394 | 0.974238 | 0.121931 | | 1 | 0.997515 | 0.721448 |
| Gapdhs | A0A0R4J0X | -0.15 | 0.14 | 0.12 | 0.04 | 0.39 | 0.31 | -0.15 | 0.21 | 1.24 | 1.24 | -0.05 | 0.19 | 1.52 | 0.07 | 1.21 | 0.29 | 0.1525 | 0.6175 | 0.735 | 0.408977 | 0.099119 | 0.063338 | | 1 | 0.996511 | 0.66433 |
| Gapvd1 | Q6PAR5 | -0.07 | -0.02 | 0.08 | -0.07 | -0.1 | -0.22 | -0.15 | -0.04 | 0.14 | -0.01 | 0.03 | 0.01 | 0.02 | 0.44 | -0.18 | -0.16 | -0.1075 | 0.0625 | 0.05 | 0.457806 | 0.688749 | 0.810229 | | 1 | 0.996511 | 0.972222 |
| Gar1 | D3YZ09 | 0.79 | 0.41 | 0.88 | -0.17 | 0.91 | 0.48 | 0.34 | 0.16 | -0.19 | -1.25 | 0.49 | 0.47 | -0.41 | 0.09 | 0.35 | 0.91 | -0.005 | -0.5975 | -0.2425 | 0.986135 | 0.203103 | 0.505134 | | 1 | 0.996511 | 0.901114 |
| Garem1 | Q3UFT3 | -0.05 | 0.3 | 0.2 | 0.14 | 0.28 | 0.09 | 0.61 | 0.45 | 0.26 | 0 | 0.48 | 0.16 | 0.45 | 0.64 | 0.03 | 0.35 | 0.21 | 0.0775 | 0.22 | 0.260848 | 0.677548 | 0.305064 | | 1 | 0.996511 | 0.831662 |
| Garnl3 | Q3V0G7 | -0.1 | 0.02 | 0.32 | 0.33 | 0.41 | 0 | 0.5 | 0.54 | 0.12 | 0.67 | 0.42 | 0.26 | 0.08 | 0.65 | 0.33 | 0.48 | 0.22 | 0.225 | 0.2425 | 0.284776 | 0.291117 | 0.279175 | | 1 | 0.996511 | 0.822084 |
| Garre1 | Q8C5X1 | -0.54 | -1.06 | -1.11 | -2.31 | -0.53 | -0.49 | -0.35 | -1.3 | 0.22 | -6.5 | -1.36 | -0.04 | -0.23 | 0.22 | -0.43 | -0.84 | 0.5875 | -0.665 | 0.935 | 0.173276 | 0.645633 | 0.04816 | | 1 | 0.996511 | 0.658454 |
| Gars1 | Q9CZD3 | -0.36 | -0.56 | -0.84 | -1.22 | -0.57 | -0.55 | -0.98 | -1.05 | -0.67 | -1.38 | -1.12 | -0.47 | -0.68 | -0.51 | -1.52 | -1.56 | -0.0425 | -0.165 | -0.3225 | 0.861185 | 0.570767 | 0.34799 | | 1 | 0.996511 | 0.851888 |
| Gart | Q64737 | -0.17 | -0.03 | 0.02 | 0.17 | -0.17 | -0.2 | 0.25 | 0.31 | 0.22 | 0.14 | -0.02 | -0.15 | 0.05 | 0.43 | 0.29 | 0.14 | 0.05 | 0.05 | 0.23 | 0.793151 | 0.778408 | 0.242509 | | 1 | 0.996511 | 0.798545 |
| Gas2 | P11862 | 0.21 | 1.36 | -0.3 | -0.1 | 1.68 | -0.02 | 0.8 | -0.13 | 0.1 | -0.51 | -0.46 | 0 | -0.86 | -0.69 | -0.43 | -0.78 | 0.29 | -0.51 | -0.9825 | 0.577644 | 0.21018 | 0.025964 | | 1 | 0.996511 | 0.633075 |
| Gas2l1 | Q8JZP9 | 0.52 | 0.56 | 0.88 | 0.79 | 0.39 | 0.9 | 0.76 | 0.81 | 1.78 | 0.99 | 0.55 | 0.63 | 2.56 | 1.01 | 1.16 | 0.88 | 0.0275 | 0.3 | 0.715 | 0.881409 | 0.331874 | 0.092642 | | 1 | 0.996511 | 0.695968 |
| Gas6 | Q61592 | 0.6 | 0.03 | 1.11 | -0.03 | -2 | 3.92 | 0.43 | 0.06 | 1.4 | -0.3 | 0.72 | -1.05 | 3.17 | -6.96 | 0.22 | 0.79 | 0.175 | -0.235 | -1.1225 | 0.876428 | 0.674562 | 0.568315 | | 1 | 0.996511 | 0.922462 |
| Gas7 | F6QB26 | 4.87 | 3.48 | 3.84 | 2.21 | 4.23 | 4.6 | 4.24 | 4.13 | 0.87 | 0.31 | 3.84 | 4.44 | 0.78 | 0.6 | 3.89 | 3.92 | 0.7 | -1.235 | -1.3025 | 0.19713 | 0.260572 | 0.200664 | | 1 | 0.996511 | 0.78037 |
| Gatad1 | Q920S3 | -0.86 | -0.51 | 0.24 | -0.45 | -1.16 | -1.32 | 0.55 | 0.09 | 0.14 | 0.31 | 0.05 | -2.44 | -0.32 | -1.2 | 0.52 | 1.61 | -0.065 | -0.09 | 0.5475 | 0.891615 | 0.88696 | 0.365006 | | 1 | 0.996511 | 0.858475 |
| Gatad2a | A0A1I7Q4C | -0.6 | -2.01 | -0.56 | -1.6 | -1.52 | -0.99 | -2.77 | -1.18 | -1.34 | -1.02 | -1.04 | -1.25 | -0.1 | -2.18 | -0.92 | -0.9 | -0.4225 | 0.03 | 0.1675 | 0.407137 | 0.93388 | 0.748955 | | 1 | 0.996755 | 0.961014 |
| Gatad2b | A1L3S7 | -1.7 | -2.48 | -2.29 | -2.25 | -2.48 | -1.94 | -2.45 | -2.3 | -1.02 | -0.42 | -2.04 | -1.84 | -0.49 | -1.21 | -1.21 | -1.56 | -0.1125 | 0.85 | 1.0625 | 0.624805 | 0.056026 | 0.005611 | | 1 | 0.996511 | 0.552614 |
| Gatb | Q99JT1 | 0.58 | 0.62 | 0.51 | 0.64 | 0.4 | 0.57 | 0.51 | 0.82 | 0.96 | 0.2 | 0.77 | 0.53 | 0.51 | 0.9 | 0.49 | 0.38 | -0.0125 | 0.0275 | -0.0175 | 0.936923 | 0.895591 | 0.92739 | | 1 | 0.996511 | 0.989331 |
| Gatc | Q8CBY0 | 0.68 | 0.52 | 0.65 | 0.65 | -0.08 | 0.93 | 0.16 | 0.2 | 0.86 | 0.23 | 0.51 | 0.86 | 0.86 | 0.66 | -0.13 | 0.21 | -0.3225 | -0.01 | -0.225 | 0.198015 | 0.960808 | 0.390875 | | 1 | 0.996755 | 0.865375 |
| Gatd1 | Q8BFQ8 | 0.31 | -0.03 | 0.06 | 0.43 | 0.2 | 0.1 | -0.27 | 0.13 | 0.22 | 0.27 | 0.05 | 0.33 | -0.44 | 0.29 | -0.45 | -0.29 | -0.1525 | 0.025 | -0.415 | 0.429308 | 0.892216 | 0.114186 | | 1 | 0.996511 | 0.717287 |
| Gatd3 | Q9D172 | 0.14 | 0.14 | -0.03 | 0.19 | -0.03 | -0.06 | 0.11 | -0.06 | 0.2 | 0.4 | -0.02 | -0.16 | 0.22 | 0.23 | -0.23 | -0.01 | -0.12 | -0.005 | -0.0575 | 0.419844 | 0.978943 | 0.76642 | | 1 | 0.998749 | 0.964134 |
| Gatm | Q9D964 | 0 | -0.01 | -0.21 | 0.25 | -0.02 | 0.06 | 1.68 | 0.41 | 0.35 | 0.1 | 0.78 | 0.11 | 0.06 | 0.94 | 0.66 | 0.06 | 0.525 | 0.3275 | 0.4225 | 0.195189 | 0.164706 | 0.139322 | | 1 | 0.996511 | 0.73493 |
| Gba | P17439 | -0.7 | -0.1 | -0.39 | -0.01 | -0.15 | -0.1 | 0.19 | -0.11 | -0.48 | -0.26 | 0.26 | 0.05 | -0.1 | -0.47 | -0.61 | 0.08 | 0.2575 | 0.1925 | 0.025 | 0.230309 | 0.44861 | 0.921787 | | 1 | 0.996511 | 0.987443 |
| Gba2 | Q69ZF3 | -0.35 | -0.21 | -0.77 | -0.46 | -0.29 | -0.32 | -0.08 | -0.27 | 0.96 | -0.4 | -0.54 | -0.34 | 0.31 | 1.74 | -0.47 | -0.58 | 0.2075 | 0.3675 | 0.6975 | 0.261848 | 0.321694 | 0.194629 | | 1 | 0.996511 | 0.779027 |
| Gbe1 | Q9D6Y9 | -0.05 | -0.27 | -0.02 | -0.2 | -0.61 | -0.06 | -0.19 | 0.1 | 0.25 | -1.21 | 0.25 | 0.07 | 0.06 | -0.77 | -0.47 | -0.44 | -0.055 | -0.025 | -0.27 | 0.781112 | 0.943057 | 0.255384 | | 1 | 0.996755 | 0.809524 |
| Gbf1 | Q6DFZ1 | -0.55 | -0.76 | -1.17 | -0.95 | -0.47 | -0.66 | -0.78 | -1.11 | -0.42 | -0.29 | -1.34 | -0.46 | -0.45 | -0.21 | -1.38 | -1.13 | 0.1025 | 0.23 | 0.065 | 0.634992 | 0.428402 | 0.836316 | | 1 | 0.996511 | 0.975831 |
| Gbp11 | A9YVJ5 | -0.82 | -1 | -1.48 | -1.64 | -1.49 | -1.54 | -1.7 | -1.72 | -1.57 | -0.68 | -1.69 | -1.55 | -1.78 | -1.07 | -2.36 | -2.11 | -0.3775 | -0.1375 | -0.595 | 0.119535 | 0.655937 | 0.108135 | | 1 | 0.996511 | 0.706427 |
| Gbp4 | Q61107 | -0.77 | -0.81 | -0.66 | -0.52 | -1.44 | -1.76 | -1.22 | -0.91 | -0.68 | 0.3 | -0.38 | -1.46 | -0.56 | -1.39 | -1.17 | -0.28 | -0.6425 | 0.135 | -0.16 | 0.014969 | 0.709176 | 0.578155 | | 1 | 0.996511 | 0.927198 |
| Gbp5 | Q8BMN7 | 0.58 | 0.5 | -0.74 | -0.78 | 0.37 | 0.1 | -2.04 | 0.28 | -0.04 | -0.85 | -0.88 | 0.32 | -0.82 | -0.1 | -1.42 | 1.16 | -0.2125 | -0.2525 | -0.185 | 0.734648 | 0.579498 | 0.763276 | | 1 | 0.996511 | 0.964134 |
| Gbp7 | Q91Z40 | -1.05 | -1.53 | -1.16 | -1.06 | -2.03 | -1.34 | -1.55 | -1.1 | -0.6 | -0.3 | -1.37 | -1.46 | -0.86 | -0.98 | -1.39 | -1.45 | -0.305 | 0.2675 | 0.03 | 0.229477 | 0.398972 | 0.896114 | | 1 | 0.996511 | 0.983509 |
| Gbp9 | Q8BTS3 | 5.26 | 4.3 | -0.09 | -0.08 | 3.51 | 5.38 | -0.04 | -0.41 | -2.08 | 0.11 | 0.27 | 5.74 | -3.04 | -4.73 | -0.7 | 0.71 | -0.2375 | -1.3375 | -4.2875 | 0.893731 | 0.49856 | 0.027477 | | 1 | 0.996511 | 0.635993 |
| Gc | P21614 | -3.78 | -4.06 | -3.67 | -4.35 | -3.53 | -4.29 | -3.73 | -3.72 | -4.45 | -1.69 | -3.74 | -3.32 | -3.73 | -3.26 | -3.98 | -4.31 | 0.1475 | 0.665 | 0.145 | 0.542669 | 0.25244 | 0.615371 | | 1 | 0.996511 | 0.936621 |
| Gca | Q8VC88 | -0.56 | -0.13 | -0.03 | -0.72 | 0.58 | -0.31 | 0.87 | -0.47 | -0.34 | -0.51 | -0.03 | -0.36 | -0.57 | 0.35 | -0.03 | -0.02 | 0.5275 | 0.05 | 0.2925 | 0.161763 | 0.82625 | 0.30284 | | 1 | 0.996511 | 0.831227 |
| Gcat | O88986 | 0.17 | 0.34 | 0.23 | 0.43 | 0.17 | 0.09 | 0 | 0.22 | 0.03 | -0.1 | 0.4 | 0.04 | -0.07 | -0.09 | -0.42 | -0.32 | -0.1725 | -0.2 | -0.5175 | 0.269468 | 0.294218 | 0.02021 | | 1 | 0.996511 | 0.618919 |
| Gcc1 | Q9D4H2 | -1.24 | -1.95 | -2.21 | -2.69 | -2.25 | -1.48 | -3.01 | -2.85 | -1.23 | -0.89 | -2.16 | -1.54 | -0.65 | -1.13 | -2.6 | -2.76 | -0.375 | 0.5675 | 0.2375 | 0.394304 | 0.170272 | 0.672909 | | 1 | 0.996511 | 0.952944 |
| Gcc2 | E9PWI6 | -0.49 | -0.54 | -0.96 | -0.82 | -0.45 | -0.72 | -0.92 | -1.05 | -0.51 | -0.29 | -0.89 | -0.54 | -0.8 | -0.09 | -1.13 | -1.07 | -0.0825 | 0.145 | -0.07 | 0.686509 | 0.496463 | 0.804835 | | 1 | 0.996511 | 0.971715 |
| Gcdh | Q60759 | 1.3 | 0.96 | 0.29 | 0.44 | 0.83 | 1.32 | 1.21 | 0.63 | 0.4 | -0.03 | 1.01 | 1.27 | 0.07 | 0.75 | 0.21 | 0.43 | 0.25 | -0.085 | -0.3825 | 0.39178 | 0.815912 | 0.21291 | | 1 | 0.996511 | 0.786931 |
| Gclc | P97494 | -0.25 | -0.02 | -0.41 | -0.27 | 0.24 | -0.14 | -0.45 | -0.35 | -0.31 | 0.05 | -0.47 | -0.24 | -0.22 | 0.28 | -0.48 | -0.49 | 0.0625 | -0.005 | 0.01 | 0.759527 | 0.979139 | 0.966413 | | 1 | 0.998755 | 0.9927 |
| Gclm | O09172 | -0.61 | -0.85 | -0.8 | -0.28 | -0.5 | -0.95 | -0.32 | -0.48 | -0.82 | -0.2 | -0.82 | -0.56 | -1.18 | -0.45 | -0.91 | -0.92 | 0.0725 | 0.035 | -0.23 | 0.734792 | 0.878184 | 0.348614 | | 1 | 0.996511 | 0.85278 |
| Gcn1 | E9PVA8 | -1.61 | -1.8 | -1.71 | -1.98 | -1.61 | -1.86 | -1.82 | -2.1 | -1.77 | -1.3 | -1.82 | -1.6 | -1.66 | -1.45 | -2.05 | -1.85 | -0.0725 | 0.1525 | 0.0225 | 0.681318 | 0.441685 | 0.914378 | | 1 | 0.996511 | 0.986458 |
| Gcsh | Q91WK5 | 0.41 | 0.83 | 0.45 | 0.76 | 0.43 | 0.09 | 0.76 | 0.5 | 0.5 | 0.16 | 0.63 | 0.6 | -0.33 | 1.15 | 0.21 | 0.33 | -0.1675 | -0.14 | -0.2725 | 0.422861 | 0.491627 | 0.415351 | | 1 | 0.996511 | 0.874945 |
| Gda | Q9R111 | 0.38 | -2.59 | -0.55 | -0.42 | 0.08 | 0.28 | -0.17 | -0.3 | -0.29 | -1 | -0.52 | 0.28 | -0.86 | -0.03 | -0.89 | -0.8 | 0.7675 | 0.4125 | 0.15 | 0.215321 | 0.516184 | 0.805595 | | 1 | 0.996511 | 0.971715 |
| Gdap1 | O88741 | 0.72 | 0.69 | 0.53 | 0.35 | 0.77 | 0.77 | 0.42 | 0.61 | 0.49 | 0.61 | 0.57 | 0.64 | 0.76 | 0.63 | 0.58 | 0.47 | 0.07 | 0.005 | 0.0375 | 0.683603 | 0.976463 | 0.840503 | | 1 | 0.997588 | 0.976225 |
| Gdap1l1 | Q8VE33 | 0.78 | -0.17 | -1.2 | -1.43 | -0.01 | 1.13 | -0.37 | -1.2 | -0.23 | -1.13 | -1.12 | 0.54 | -0.12 | -0.09 | -1.02 | -1.04 | 0.3925 | 0.02 | -0.0625 | 0.542762 | 0.973126 | 0.906402 | | 1 | 0.997515 | 0.986458 |
| Gdap2 | Q9DBL2-2 | -0.61 | -0.27 | -0.04 | -0.54 | -0.45 | -0.75 | 0.34 | -0.4 | -0.42 | -0.76 | 0 | -0.46 | -0.53 | 0.03 | -0.39 | -0.13 | 0.05 | -0.045 | 0.11 | 0.853831 | 0.847559 | 0.631152 | | 1 | 0.996511 | 0.941132 |
| Gde1 | Q9JL56 | 0.42 | 0.1 | -0.04 | -0.07 | 0.53 | 0.84 | 0.54 | 0 | -0.03 | -0.41 | -0.09 | 0.51 | 1.13 | -0.71 | 0.17 | 0.57 | 0.375 | -0.1075 | 0.1875 | 0.127049 | 0.663967 | 0.633182 | | 1 | 0.996511 | 0.941132 |
| Gdi1 | P50396 | -0.08 | -0.51 | -0.37 | -0.12 | -0.61 | -0.08 | -0.3 | -0.09 | -0.14 | -0.26 | -0.33 | 0.01 | -0.4 | -0.35 | -0.7 | -0.82 | 1.18E-16 | 0.09 | -0.2975 | 1 | 0.631938 | 0.182784 | | 1 | 0.996511 | 0.776002 |
| Gdi2 | Q61598 | 1.17 | 0.57 | 1.06 | 0.94 | 0.78 | 0.95 | 1.45 | 1.29 | 0.59 | 0.61 | 1.16 | 1.19 | -0.73 | 1.15 | 0.79 | 0.98 | 0.1825 | -0.0475 | -0.3875 | 0.42318 | 0.842585 | 0.378608 | | 1 | 0.996511 | 0.860984 |
| Gdpd1 | Q9CRY7 | 0.02 | 0.52 | 0.93 | 0.36 | 0.53 | 0.26 | 0.81 | 0.73 | -1.26 | -0.83 | 0.61 | 0.05 | -1.11 | -1.06 | 0.95 | 1.17 | 0.125 | -0.815 | -0.47 | 0.606071 | 0.089744 | 0.438055 | | 1 | 0.996511 | 0.883161 |
| Gdpd2 | Q9ESM6 | 0.24 | 1.43 | 1.11 | 1.12 | 0.98 | 0.14 | 1.48 | 1.59 | 0.39 | 0.33 | 1.04 | 0.91 | 0.67 | 1.69 | 1.42 | 1.32 | 0.0725 | -0.3075 | 0.3 | 0.854283 | 0.342152 | 0.383828 | | 1 | 0.996511 | 0.860984 |
| Gdpd5 | A0A1L1STK | 0.23 | 0.5 | 0.64 | 0.95 | 0.94 | 0.07 | 0.82 | 1.27 | 0.47 | 0.85 | 0.95 | 0.64 | 0.39 | 0.79 | 1.07 | 0.54 | 0.195 | 0.1475 | 0.1175 | 0.512312 | 0.510161 | 0.635952 | | 1 | 0.996511 | 0.942144 |
| Gdpgp1 | Q3TLS3 | -0.93 | -1.02 | -0.62 | -0.68 | -0.44 | -1.17 | -0.5 | -0.41 | -1.17 | 0.04 | -0.88 | -0.81 | -1.21 | -0.64 | -0.83 | -0.72 | 0.1825 | 0.1075 | -0.0375 | 0.42838 | 0.709252 | 0.860902 | | 1 | 0.996511 | 0.977204 |
| Gemin4 | Q6P6L6 | -2.08 | -2.54 | -1.77 | -2.4 | -0.83 | -2.08 | -1.23 | -2.24 | -3.05 | -1.55 | -1.6 | -1.14 | -0.88 | -1.19 | -2.3 | -1.77 | 0.6025 | 0.3625 | 0.6625 | 0.124034 | 0.406861 | 0.089233 | | 1 | 0.996511 | 0.69217 |

| | | | | | | | | | | | | | | | | | | | | | | | | | | |
|---|---|---|---|---|---|---|---|---|---|---|---|---|---|---|---|---|---|---|---|---|---|---|---|---|---|---|
| Gemin5 | A2AFQ9 | -0.94 | -1.44 | -2.55 | -2.5 | -1.02 | -1.06 | -2.53 | -2.04 | -1.35 | -1.95 | -2.44 | -1.22 | -1.29 | -1.11 | -2.55 | -2.41 | 0.195 | 0.1175 | 0.0175 | 0.699648 | 0.797974 | 0.972481 | 1 | 0.996511 | 0.992865 |
| Gemin7 | D3Z684 | 0.29 | 0.81 | -1.18 | -1.17 | 0.02 | 1.28 | -1.54 | -0.81 | 0.75 | 0.5 | -0.76 | 0.04 | 0.94 | -0.12 | 0.05 | -0.84 | 0.05 | 0.445 | 0.32 | 0.94424 | 0.435338 | 0.582023 | 1 | 0.996511 | 0.927734 |
| Get1 | Q8K0D7 | -0.75 | -0.84 | -0.33 | -0.66 | -0.28 | -0.22 | -0.42 | -0.61 | 0.19 | -0.22 | -1.16 | -0.87 | 0.99 | -0.2 | 0.16 | 0.11 | 0.2625 | 0.13 | 0.91 | 0.17767 | 0.690332 | 0.012064 | 1 | 0.996511 | 0.566767 |
| Get3 | O54984 | -0.68 | -0.41 | -0.44 | -0.69 | 0.05 | -0.56 | -0.34 | -0.78 | -1.01 | -0.45 | -0.73 | -0.65 | -0.87 | -0.54 | -0.71 | -0.45 | 0.1475 | -0.155 | -0.0875 | 0.502705 | 0.430805 | 0.652671 | 1 | 0.996511 | 0.947633 |
| Get4 | Q9D1H7 | -0.3 | -0.21 | -0.18 | -0.14 | -0.7 | -0.51 | -0.78 | -0.63 | -0.56 | 0.13 | -0.3 | -0.37 | -0.73 | -0.5 | -0.36 | -0.66 | -0.4475 | -0.0675 | -0.355 | 0.01345 | 0.735799 | 0.075623 | 1 | 0.996511 | 0.674038 |
| Gfap | P03995 | 1.36 | 0.87 | 0.51 | 0.54 | 0.97 | 1.05 | 0.46 | 0.37 | 0.97 | 0.27 | 0.62 | 1.07 | 0.98 | 0.93 | 1.16 | 1.36 | -0.1075 | -0.0875 | 0.2875 | 0.69068 | 0.755608 | 0.273305 | 1 | 0.996511 | 0.822084 |
| Gfap | P03995-2 | -0.23 | 0.01 | -1.17 | 0.73 | -2.97 | -1.25 | 0.16 | 0.41 | -0.28 | -0.42 | 0.43 | 0.13 | -0.56 | -1.04 | -0.1 | -0.75 | -0.7475 | 0.13 | -0.4475 | 0.354251 | 0.755072 | 0.302314 | 1 | 0.996511 | 0.830768 |
| Gfer | P56213 | 0.5 | -0.22 | -0.23 | -0.76 | -0.49 | 1.17 | -0.68 | -0.7 | 0.89 | -0.25 | -0.55 | 0.28 | 2.22 | -0.76 | 0.09 | -0.03 | 0.0025 | 0.27 | 0.5575 | 0.995848 | 0.496948 | 0.388179 | 1 | 0.996511 | 0.86437 |
| Gfm1 | Q8K0D5 | 1.36 | 0.45 | 0.78 | 0.63 | 0.14 | 1.54 | 1.17 | 0.69 | 0.05 | -0.04 | 0.89 | 1.15 | 0.35 | -0.3 | 0.8 | 0.81 | 0.08 | -0.2925 | -0.39 | 0.81828 | 0.413351 | 0.257053 | 1 | 0.996511 | 0.811777 |
| Gfm2 | Q8R2Q4-3 | 0.68 | 0.52 | 0.41 | 0.32 | 0.57 | 0.74 | 0.89 | 0.31 | 0.5 | 0.29 | 0.38 | 0.55 | 0.46 | 0.71 | 0.13 | 0.67 | 0.145 | -0.0525 | 0.01 | 0.446283 | 0.761041 | 0.962185 | 1 | 0.996511 | 0.9927 |
| Gfod1 | Q3UHD2 | -0.1 | -0.06 | 0.05 | 0.14 | 0.15 | -0.02 | -0.06 | 0.38 | -0.09 | -0.2 | 0.54 | -0.24 | -0.61 | 0.39 | -0.5 | -0.37 | 0.105 | -0.005 | -0.28 | 0.537268 | 0.98223 | 0.299768 | 1 | 0.998773 | 0.829161 |
| Gfod2 | Q9CYH5 | 0.42 | 0.46 | 0.41 | 0.61 | 0.64 | 0.22 | 0.77 | 0.36 | 0.45 | 0.49 | 0.61 | 0.36 | -0.04 | 0.9 | -0.11 | -0.09 | 0.0225 | 0.0025 | -0.31 | 0.900264 | 0.987633 | 0.274185 | 1 | 0.999285 | 0.822084 |
| Gfpt1 | P47856-2 | -0.08 | -0.11 | -0.28 | -0.5 | -0.06 | -0.17 | -0.11 | -0.12 | -0.97 | 0.52 | -0.37 | 0.07 | -0.93 | -0.5 | -0.43 | -0.13 | 0.1275 | 0.055 | -0.255 | 0.436252 | 0.867599 | 0.291714 | 1 | 0.996511 | 0.825497 |
| Gfra1 | D6RDD4 | -0.61 | -1.82 | -1.75 | -1.53 | -2.43 | 0.45 | -1.37 | -1.42 | -1.31 | -0.55 | -2.18 | -0.46 | 0.17 | -2.69 | -0.74 | -1.54 | 0.235 | 0.3025 | 0.2275 | 0.697839 | 0.514323 | 0.710821 | 1 | 0.996511 | 0.957819 |
| Gfra2 | O08842 | -0.21 | -0.5 | 0.19 | 0.23 | -0.03 | -0.03 | -0.54 | -0.15 | -0.32 | 0.62 | -0.01 | 0.08 | 0.22 | 0.28 | 0.24 | -0.15 | -0.115 | 0.165 | 0.22 | 0.620603 | 0.553101 | 0.370241 | 1 | 0.996511 | 0.859245 |
| Gfus | P23591 | 0.66 | 0.66 | -0.45 | -0.59 | 0.95 | 0.32 | -0.54 | -0.45 | -0.22 | -1.88 | -0.4 | 0.77 | -0.3 | 0.31 | -0.65 | -0.79 | -1.6E-16 | -0.5025 | -0.4275 | 1 | 0.405851 | 0.306624 | 1 | 0.996511 | 0.832533 |
| Gga1 | Q8R0H9 | 0.33 | 0.09 | 0.18 | -0.02 | 0.1 | 0.39 | 0.3 | 0.04 | 0.05 | 0.29 | 0.21 | 0.45 | -0.09 | 0.18 | 0.1 | 0.13 | 0.0625 | 0.105 | -0.065 | 0.708536 | 0.561075 | 0.722229 | 1 | 0.996511 | 0.958398 |
| Gga2 | Q6P5E6 | -0.94 | -0.25 | -1.17 | -0.99 | -0.59 | -0.88 | -0.88 | -0.92 | -1.02 | -0.15 | -0.74 | -0.58 | -0.71 | 0.04 | -1.43 | -1.48 | 0.02 | 0.215 | -0.0575 | 0.93179 | 0.455082 | 0.884734 | 1 | 0.996511 | 0.979767 |
| Gga3 | A2A9W7 | 0.16 | 0.52 | 0.15 | 0.22 | 0.57 | 0.11 | -0.03 | 0.11 | -0.25 | 0.15 | 0.14 | 0.36 | -0.02 | 0.22 | -0.03 | -0.16 | -0.0725 | -0.1625 | -0.26 | 0.709203 | 0.430492 | 0.199756 | 1 | 0.996511 | 0.78004 |
| Ggact | E0CZ04 | -1 | -2.1 | -1.85 | -0.8 | -1.45 | -1.47 | -1.12 | -0.98 | -1.96 | -0.46 | -1.37 | -1.33 | -2.49 | -2.54 | -1.75 | -1.74 | 0.1825 | 0.1575 | -0.6925 | 0.583681 | 0.708728 | 0.094237 | 1 | 0.996511 | 0.698156 |
| Ggct | Q9D7X8 | -0.54 | -0.67 | -0.57 | -0.63 | -0.93 | -0.46 | -0.89 | -0.99 | -0.56 | 0.14 | -0.69 | -0.76 | -0.3 | -1.15 | -1.25 | -1.03 | -0.215 | 0.135 | -0.33 | 0.237335 | 0.573185 | 0.210616 | 1 | 0.996511 | 0.786218 |
| Ggh | Q9Z0L8-2 | -1.99 | -3.48 | -0.82 | -0.01 | -4.35 | -1.26 | 0.43 | -0.49 | -2.06 | -1.54 | -0.46 | -1.85 | -0.51 | -5.5 | -0.93 | -0.66 | 0.1575 | 0.0975 | -0.325 | 0.890784 | 0.897224 | 0.797247 | 1 | 0.996511 | 0.969727 |
| Ggps1 | Q9WTN0 | -0.73 | -1.03 | -0.43 | -0.17 | -0.67 | -0.84 | -0.43 | -0.45 | -1.32 | -0.69 | -0.34 | -0.7 | -1.23 | -0.96 | -1.05 | -1.02 | -0.0075 | -0.1725 | -0.475 | 0.97391 | 0.550742 | 0.070392 | 1 | 0.996511 | 0.668066 |
| Ggt1 | Q60928 | 1.11 | 1.31 | 1.68 | 1.69 | 1.25 | 0.32 | 1.97 | 1.88 | 1.07 | -0.76 | 1.29 | 0.78 | 0.91 | 0.73 | 1.03 | 1.19 | -0.0925 | -0.8525 | -0.4825 | 0.810327 | 0.089057 | 0.054512 | 1 | 0.996511 | 0.661738 |
| Ggt5 | Q9Z2A9 | 0.81 | 0.1 | 1.1 | 0.05 | 0.56 | 0.22 | 0.54 | 0.32 | 0.32 | 0.18 | 0.67 | 0.85 | 0.11 | -0.17 | 1.25 | 1.62 | -0.105 | -0.01 | 0.1875 | 0.706322 | 0.973998 | 0.693976 | 1 | 0.997515 | 0.955729 |
| Ggt7 | Q99JP7 | 0.37 | 0.57 | 0.52 | 0.43 | 1.04 | 0.41 | 0.97 | 0.52 | -0.19 | -0.11 | 0.54 | 0.86 | -0.13 | 0.3 | 0.52 | 0.86 | 0.2625 | -0.1975 | -0.085 | 0.208301 | 0.476109 | 0.731729 | 1 | 0.996511 | 0.958635 |
| Ghdc | Q99J23 | -0.65 | -0.35 | -0.05 | 0 | -0.5 | -0.36 | -0.54 | 0 | 0.68 | 0.7 | -1.1 | -0.89 | 1.17 | 0.58 | -0.07 | -0.01 | -0.0875 | 0.11 | 0.68 | 0.690001 | 0.817891 | 0.065506 | 1 | 0.996511 | 0.664936 |
| Ghitm | Q91VC9 | 0.69 | 0.52 | 0.65 | 0.98 | -0.09 | 1.02 | -0.4 | 0.29 | 1.64 | 0.44 | 0.95 | 0.82 | 1.93 | 0.5 | 0.14 | 0.41 | -0.505 | 0.2525 | 0.035 | 0.136147 | 0.379585 | 0.930071 | 1 | 0.996511 | 0.989687 |
| Gid4 | Q9CPY6 | -0.88 | -0.61 | -2.5 | -2.04 | 0.05 | -0.66 | -0.82 | -1.49 | 0.22 | -2.77 | -0.87 | -0.69 | 0.19 | -0.94 | -2.85 | -1.5 | 0.7775 | 0.48 | 0.2325 | 0.153197 | 0.50116 | 0.742199 | 1 | 0.996511 | 0.958976 |
| Gid8 | Q9D7M1 | -0.63 | -0.58 | 0.16 | 0.31 | -0.12 | -0.61 | 0.33 | 0.26 | -0.34 | 0.23 | 0.16 | -0.35 | -0.49 | -0.28 | 0.02 | 0.31 | 0.15 | 0.11 | 0.075 | 0.640316 | 0.712188 | 0.809123 | 1 | 0.996511 | 0.972098 |
| Gigyf1 | Q99MR1 | -0.69 | 0 | -0.06 | -0.09 | -0.6 | -0.54 | -0.64 | -0.36 | 0.26 | -0.07 | -0.25 | -0.27 | 0.5 | 0.6 | -0.71 | -0.47 | -0.325 | 0.1275 | 0.19 | 0.137448 | 0.587655 | 0.60454 | 1 | 0.996511 | 0.932185 |
| Gigyf2 | Q6Y7W8 | -1.57 | -1.86 | -1.41 | -1.38 | -0.77 | -1.6 | -1.66 | -1.41 | 0.02 | -0.89 | -1.27 | -1.4 | 2.13 | -1.4 | 1.43 | -1.71 | 0.195 | 0.67 | 1.6675 | 0.435152 | 0.07256 | 0.08575 | 1 | 0.996511 | 0.691619 |
| Gipc1 | Q9Z0G0 | 0.22 | 0 | 0.12 | 0.02 | 0.23 | 0.1 | 0.3 | 0.35 | 0.23 | 0.71 | 0.13 | 0.59 | 0.14 | 0.43 | -0.06 | -0.14 | 0.155 | 0.325 | 0.0025 | 0.316386 | 0.129122 | 0.99011 | 1 | 0.996511 | 0.996154 |
| Gipc3 | Q8R5M0 | 0.29 | 0.31 | -1.99 | -1.62 | 1.64 | 1.28 | -1.96 | -1.54 | 0.5 | -0.52 | -1.19 | 1.53 | 0.96 | 1.56 | 1.17 | -0.8 | 0.6075 | 0.8325 | 1.475 | 0.548438 | 0.298139 | 0.069262 | 1 | 0.996511 | 0.667373 |
| Git1 | Q68FF6 | 0.65 | 0.33 | 0.85 | 0.52 | 0.24 | 0.83 | 0.7 | 0.62 | 0.51 | -0.07 | 0.67 | 0.52 | 0.52 | -0.16 | 0.55 | 0.66 | 0.01 | -0.18 | -0.195 | 0.960147 | 0.440541 | 0.443546 | 1 | 0.996511 | 0.883549 |
| Git2 | E9PVA6 | -0.42 | -0.94 | -0.19 | -0.47 | -0.95 | -0.38 | -0.49 | -0.22 | 0.01 | -0.72 | -0.05 | -0.38 | 0.25 | -0.55 | -0.08 | -0.67 | -0.005 | 0.22 | 0.2425 | 0.983245 | 0.394539 | 0.402987 | 1 | 0.996511 | 0.870698 |
| Gja1 | P23242 | -0.06 | 0.24 | 0.37 | 1.19 | 0.57 | -0.12 | 1.01 | 1.08 | 0.24 | 1.08 | 0.67 | 0.29 | 0.22 | 0.53 | 0.76 | 0.51 | 0.2 | 0.135 | 0.07 | 0.588498 | 0.682896 | 0.816669 | 1 | 0.996511 | 0.972768 |
| Gjb1 | P28230 | -0.08 | 0.59 | -4.07 | -3.74 | 0.18 | -0.66 | -3.26 | -4.36 | 0.88 | -4.98 | -3.78 | 0.25 | -0.19 | 0.23 | -3.42 | -4.26 | -0.2 | -0.0825 | -0.085 | 0.889633 | 0.960973 | 0.95388 | 1 | 0.996755 | 0.992476 |
| Gjb6 | P70689 | 0.58 | 0.92 | 1.41 | 0.16 | 1.06 | 1.17 | 0.32 | -0.17 | 0.39 | 0.06 | 0.33 | 1.15 | -0.04 | 0.66 | -0.55 | 0.77 | -0.1725 | -0.285 | -0.5575 | 0.660521 | 0.420446 | 0.181784 | 1 | 0.996511 | 0.776002 |
| Gjc2 | Q8BQU6 | 1.52 | 1.16 | 0.39 | -0.49 | 1.25 | 0.92 | 0.25 | -0.56 | -0.64 | 1.66 | -0.06 | 1.21 | 0.88 | 1.06 | 0.93 | -0.04 | -0.18 | -0.1025 | 0.0625 | 0.743572 | 0.872171 | 0.896298 | 1 | 0.996511 | 0.983509 |
| Gjc3 | Q921C1 | 2.49 | 2.36 | 2.78 | 1.26 | 2.13 | 2.12 | 1.32 | 1.66 | 1.23 | 0.59 | 1.6 | 1.5 | 1.9 | 0.18 | 2.11 | 2.01 | -0.415 | -0.9925 | -0.6725 | 0.277767 | 0.029093 | 0.222187 | 1 | 0.996511 | 0.787447 |
| Gjd2 | O54851 | -0.23 | 0.21 | 0.19 | 0.96 | -1.46 | 0.77 | -0.41 | 0.71 | 0.9 | 1.32 | 0.41 | 0.62 | 1.28 | 0.91 | 0.49 | 0.5 | -0.38 | 0.53 | 0.5125 | 0.485002 | 0.122317 | 0.134234 | 1 | 0.996511 | 0.730883 |
| Gk | Q64516-1 | -0.1 | 0.27 | 0.11 | 0.32 | 0.65 | 0.28 | 0 | 0.13 | -0.06 | 0.19 | 0.36 | -0.22 | 0.07 | 0.21 | -0.08 | 0.2 | 0.115 | -0.0825 | -0.05 | 0.571913 | 0.689507 | 0.794594 | 1 | 0.996511 | 0.969367 |
| Gkap1 | Q9JMB0 | -2.61 | -1.9 | -2.41 | -2.07 | -1.33 | -2.67 | -3.32 | -2.77 | -0.69 | -1.08 | -2.33 | -2.1 | 0.11 | -0.38 | -1.9 | -2.32 | -0.275 | 0.6975 | 1.125 | 0.521138 | 0.114025 | 0.070743 | 1 | 0.996511 | 0.668066 |
| Gla | Q8BGZ6 | -1.62 | -1.47 | -2.37 | -2.03 | -1.6 | -1.38 | -1.76 | -2.65 | -1.14 | -1.84 | -1.99 | -1.29 | -1.45 | -0.66 | -2.76 | -2.2 | 0.025 | 0.3075 | 0.105 | 0.940378 | 0.315872 | 0.823161 | 1 | 0.996511 | 0.972768 |
| Glb1 | P23780 | -0.77 | -0.98 | -1.12 | -0.67 | -0.59 | -0.58 | -0.47 | -0.81 | -1.13 | -0.75 | -0.42 | -0.79 | -1.13 | -0.4 | -1.07 | -0.95 | 0.2725 | 0.1125 | -0.0025 | 0.143516 | 0.605904 | 0.991547 | 1 | 0.996511 | 0.996154 |
| Glb1l | A0A087WF | -0.41 | 0.06 | 1.9 | 1.53 | -0.98 | -0.42 | 1.41 | 0.86 | 0.26 | 1.25 | 0.99 | -0.49 | -0.34 | -1.16 | 0.92 | 1.6 | -0.5525 | -0.2675 | -0.515 | 0.445972 | 0.669769 | 0.500481 | 1 | 0.996511 | 0.900792 |
| Glcci1 | E9QKK4 | -0.32 | -0.89 | -0.07 | -0.19 | -0.95 | -0.36 | -0.6 | -0.5 | -0.43 | -0.3 | -0.17 | 0.26 | -0.73 | -0.3 | -0.36 | -0.4 | -0.235 | 0.2075 | -0.08 | 0.336326 | 0.426811 | 0.742433 | 1 | 0.996511 | 0.958976 |
| Glce | Q9EPS3 | -0.94 | -1.17 | -0.53 | -0.53 | -0.87 | -0.91 | -0.23 | -0.46 | -0.82 | -0.35 | -0.35 | -1.24 | -1.03 | -1.46 | 0.01 | -0.16 | 0.175 | 0.1025 | 0.1325 | 0.478164 | 0.714272 | 0.725155 | 1 | 0.996511 | 0.958398 |
| Gldc | Q91W43 | 1.71 | 0.77 | 1.37 | 1.41 | 0.5 | 1.83 | 1.99 | 1.48 | 1.05 | 1.71 | 1.47 | 1.09 | 0.6 | 0.48 | 0.53 | 0.6 | 0.135 | 0.015 | -0.7625 | 0.71582 | 0.955424 | 0.010449 | 1 | 0.996755 | 0.563386 |
| Gle1 | Q8R322 | -0.54 | -0.17 | 1.07 | 0.64 | 0.11 | -0.29 | 0.83 | 1.28 | -0.98 | 1.01 | 0.78 | -1.06 | -0.5 | -0.35 | 1.69 | 1.4 | 0.2325 | -0.3125 | 0.31 | 0.624896 | 0.610988 | 0.620591 | 1 | 0.996511 | 0.938143 |
| Glg1 | Q61543 | -1.94 | -2.99 | -3.11 | -2.6 | -1.78 | -2.26 | -2.43 | -2.87 | -2.46 | -1.68 | -2.97 | -2.34 | -2.02 | -2.08 | -2.56 | -2.61 | 0.325 | 0.2975 | 0.3425 | 0.346639 | 0.422491 | 0.292347 | 1 | 0.996511 | 0.825497 |
| Glipr2 | Q9CYL5 | 0.95 | 1.32 | 1.52 | 0.78 | 0.02 | 0.76 | 1.95 | 2.87 | 0.52 | -0.76 | 2.76 | 0.81 | 0.54 | -0.12 | 1.7 | 3.87 | 0.2575 | -0.31 | 0.355 | 0.667187 | 0.650197 | 0.660298 | 1 | 0.996511 | 0.949645 |
| Glmn | Q8BZM1 | -1.41 | -0.67 | -0.68 | -1.29 | -0.42 | -1.69 | -0.85 | -1.02 | -1.06 | -1.67 | -0.98 | -1.45 | -1.14 | -0.08 | -1.27 | -0.91 | 0.0175 | -0.2775 | 0.1625 | 0.956528 | 0.320243 | 0.628649 | 1 | 0.996511 | 0.940342 |
| Glo1 | Q9CPU0 | -1.04 | -1.18 | -0.33 | -0.3 | -1.09 | -0.75 | -0.28 | -0.38 | -1.38 | -0.51 | -0.42 | -1.06 | -1.45 | -0.69 | -0.9 | -0.84 | 0.0875 | -0.13 | -0.2575 | 0.767294 | 0.689687 | 0.398457 | 1 | 0.996511 | 0.867334 |
| Glod4 | Q9CPV4 | 0.49 | -0.23 | 0.54 | -0.46 | -0.06 | 0.48 | 0.99 | 0.18 | -1.16 | -1.4 | 0.45 | 0.16 | -1.12 | -0.84 | 0.2 | 0.52 | 0.3125 | -0.5725 | -0.395 | 0.356303 | 0.263714 | 0.385949 | 1 | 0.996511 | 0.861533 |
| Glra3 | G5E811 | -0.18 | -0.75 | -0.4 | 0.62 | -4.04 | 0.75 | -0.13 | 0.08 | 3.63 | 1.14 | 0.18 | -0.78 | 4.84 | -3.92 | 1.26 | 0.47 | -0.6575 | 1.22 | 0.84 | 0.519172 | 0.195555 | 0.60611 | 1 | 0.996511 | 0.932425 |
| Glrb | E9Q3E3 | 0.15 | 0.66 | 0.26 | 1 | 0.78 | 0.18 | 1 | 0.9 | 0.39 | 0.46 | 0.48 | 0.15 | 0.26 | 1.33 | 0.97 | 1.09 | 0.1975 | -0.1475 | 0.395 | 0.475887 | 0.539114 | 0.224971 | 1 | 0.996511 | 0.787447 |
| Glrx | Q9QUH0 | -0.46 | -0.57 | -1.77 | -1.89 | 0.03 | -0.12 | 0.12 | -1.79 | -0.46 | -2 | -1.91 | 0.05 | -0.35 | 0.06 | -1.23 | 0.22 | 0.7325 | 0.0925 | 0.8475 | 0.200601 | 0.875142 | 0.09906 | 1 | 0.996511 | 0.698156 |
| Glrx2 | Q923X4 | 1.23 | 1.4 | -0.19 | 0.23 | 0.99 | 0.94 | -0.01 | -0.39 | 1.12 | 0.2 | -0.1 | 0.9 | 1.09 | 1.1 | -0.56 | -0.21 | -0.285 | -0.1375 | -0.3125 | 0.555748 | 0.761273 | 0.563895 | 1 | 0.996511 | 0.920203 |
| Glrx3 | Q9CQM9 | -0.98 | -0.43 | 0.36 | 1.05 | -0.67 | -0.93 | 0.55 | 1.2 | -0.78 | 1.02 | 0.41 | -0.49 | -0.76 | -0.51 | -0.11 | 0.13 | 0.0375 | 0.04 | -0.3125 | 0.951147 | 0.94296 | 0.502228 | 1 | 0.996755 | 0.901002 |
| Glrx5 | Q80Y14 | 1.21 | 1.01 | 1.78 | 2.12 | 0.23 | 1.69 | 0.83 | 2.03 | 1.64 | 2.65 | 1.66 | 1.29 | 1.58 | 1.24 | 1.31 | 1.66 | -0.335 | 0.28 | -0.0825 | 0.462985 | 0.463119 | 0.777333 | 1 | 0.996511 | 0.967289 |
| Gls | D3Z7P3 | 0.36 | 0.03 | 0.31 | 0.54 | 0.03 | 0.59 | 0.69 | 0.46 | 0.23 | -0.14 | 0.43 | 0.21 | 0.66 | 0.18 | 0.23 | 0.38 | 0.1325 | -0.1275 | 0.0525 | 0.529791 | 0.538313 | 0.802411 | 1 | 0.996511 | 0.971091 |
| Gls | D3Z7P3-2 | 2.02 | 1.78 | 2.42 | 0.94 | 1.58 | 1.8 | 2.67 | 2.12 | 1.26 | 1.03 | 2.26 | 1.95 | 1.29 | 1.25 | 2.01 | 2.97 | 0.2525 | -0.165 | 0.09 | 0.504448 | 0.685358 | 0.850708 | 1 | 0.996511 | 0.976225 |
| Gls2 | Q571F8 | -1.08 | -0.59 | 0.4 | 0.04 | 0.2 | -1.2 | -0.43 | -0.12 | 1.25 | -0.54 | 0 | -1.36 | 0.68 | -0.49 | -0.75 | -0.17 | -0.08 | 0.145 | 0.125 | 0.848063 | 0.804882 | 0.772008 | 1 | 0.996511 | 0.965933 |
| Gltp | Q9JL62 | 0.86 | 0.99 | 1.28 | 1.24 | 1.24 | 0.53 | 1.6 | 1.08 | 0.59 | 0.73 | 1.03 | 0.62 | 0.69 | 0.66 | 1.12 | 1.64 | 0.02 | -0.35 | -0.065 | 0.937366 | 0.100605 | 0.812436 | 1 | 0.996511 | 0.972642 |
| Glud1 | P26443 | 0.55 | 0.1 | 0.67 | 0.98 | 0.09 | 0.95 | 0.75 | 1.12 | 0.51 | -0.07 | 1.12 | 0.47 | 0.6 | -0.25 | 0.77 | 0.65 | 0.1525 | -0.0675 | -0.1325 | 0.60217 | 0.826454 | 0.667571 | 1 | 0.996511 | 0.951339 |
| Glul | P15105 | 0.24 | -0.87 | -0.68 | -0.31 | -0.6 | 0.57 | -0.58 | -0.21 | -0.25 | -0.78 | -0.53 | -0.13 | -0.64 | -0.79 | -1.09 | -0.67 | 0.2 | -0.0175 | -0.3925 | 0.57375 | 0.952272 | 0.190679 | 1 | 0.996755 | 0.776865 |
| Glyctk | A0A0R4J1H | 0.25 | 0.92 | -0.45 | -0.3 | 0.99 | 0.38 | -0.03 | -0.02 | 0.13 | -1.36 | -0.25 | -0.02 | 0.13 | 0.41 | -0.94 | -0.44 | 0.225 | -0.48 | -0.315 | 0.551043 | 0.285558 | 0.454572 | 1 | 0.996511 | 0.887169 |
| Glyr1 | D3YYT1 | 0.87 | 0.21 | 0.19 | 0.19 | 0.03 | 0.51 | -0.14 | 0.04 | 0.78 | -0.81 | 0.37 | 0.19 | 1.54 | -0.14 | 1.89 | -0.13 | -2.55E-01 | -0.2325 | 0.425 | 0.296297 | 0.530361 | 0.425999 | 1 | 0.996511 | 0.878405 |
| Gm11938 | Q9D3H4 | -7.5 | -8.49 | -10.15 | -8.16 | -9.16 | -7.78 | -12.92 | -12.51 | -5.44 | -13.28 | -9.43 | -7.18 | -5.16 | -9.97 | -8.76 | -13.24 | -2.0175 | -0.2575 | -0.7075 | 0.129692 | 0.871485 | 0.652569 | 1 | 0.996511 | 0.947633 |
| Gm13889 | A0A087WS | -0.93 | -1.23 | -2.16 | -1.58 | -1.27 | -0.94 | -1.22 | -1.87 | -0.43 | -2.37 | -1.91 | -0.92 | 0.61 | -0.51 | -1.68 | -1.81 | 0.15 | 0.0675 | 0.6275 | 0.642471 | 0.888722 | 0.292045 | 1 | 0.996511 | 0.825497 |
| Gm1673 | Q3UR78 | -1.26 | -1.39 | -0.96 | -1.04 | -0.88 | -1.62 | -0.47 | -1.15 | -1.68 | -0.71 | -0.92 | -1.3 | -0.77 | -0.96 | -0.4 | -0.82 | 0.1325 | 0.01 | 0.425 | 0.622526 | 0.968866 | 0.071242 | 1 | 0.996755 | 0.668066 |
| Gm19410 | A0A140LJC | 0.13 | 0.17 | 0.14 | 0.09 | 1.02 | -0.26 | 0.65 | -0.2 | -0.5 | -0.05 | 0.2 | 0.16 | -1.2 | 0.82 | -0.16 | 0 | 0.17 | -0.18 | -0.2675 | 0.58893 | 0.394097 | 0.509381 | 1 | 0.996511 | 0.902804 |
| Gm20390 | E9PZF0 | 0.17 | -0.33 | -0.38 | -0.78 | -0.03 | 0.19 | -0.07 | -0.4 | -0.88 | -0.89 | -0.4 | 0.26 | -0.73 | -0.64 | -0.55 | -0.49 | 0.2525 | -0.1475 | -0.2725 | 0.315842 | 0.657829 | 0.274775 | 1 | 0.996511 | 0.822084 |
| Gm20431 | E9PY39 | 0.25 | 0.32 | -0.01 | -0.02 | -0.05 | 0.15 | 0.13 | 0.2 | -0.04 | -0.11 | 0.1 | 0.47 | -0.13 | 0.22 | -0.21 | -0.42 | -0.0275 | -0.03 | -0.27 | 0.865882 | 0.882726 | 0.229073 | 1 | 0.996511 | 0.787447 |
| Gm20441 | G3UYJ7 | -0.38 | -0.37 | 1 | 1.25 | -0.12 | -0.15 | 0.57 | 1.42 | -0.84 | 1.49 | 0.86 | -0.37 | -1.27 | -0.2 | 2.02 | 1.19 | 0.055 | -0.09 | 0.06 | 0.916561 | 0.886887 | 0.937639 | 1 | 0.996511 | 0.989767 |
| Gm20517 | G3UVU6 | -4.1 | -3.01 | -1.79 | -1.65 | -2.62 | -2.91 | -2.01 | -2.59 | -3.46 | -1.54 | -1.52 | -2.26 | -3.36 | -1.85 | -1.62 | -2.15 | 0.105 | 0.4425 | 0.3925 | 0.849823 | 0.513417 | 0.539646 | 1 | 0.996511 | 0.912212 |
| Gm20521 | D3Z5F7 | -1.3 | -1.97 | -1.78 | -1.66 | -1.43 | -2.16 | -2.01 | -1.78 | -1.4 | -0.35 | -1.86 | -2.38 | -0.76 | -1.54 | -1.09 | -1.83 | -0.1675 | 0.18 | 0.3725 | 0.475905 | 0.676681 | 0.223239 | 1 | 0.996511 | 0.787447 |
| Gm20604 | F8WIA3 | -0.49 | -0.42 | -1.23 | -0.22 | 5.74 | -1.79 | -0.78 | 0.05 | -1.39 | -1.32 | -0.63 | -1.36 | -1 | -0.49 | -1.11 | -0.07 | 1.395 | -0.585 | -0.0775 | 0.370544 | 0.073493 | 0.814451 | 1 | 0.996511 | 0.972768 |
| Gm20671 | F6TVX7 | -0.14 | -0.09 | 0.49 | 0.23 | -0.2 | 0.09 | 0.03 | 0.09 | -0.04 | -0.38 | 0.38 | -0.15 | -0.19 | -0.15 | -0.66 | -0.45 | -0.12 | -0.17 | -0.485 | 0.547191 | 0.490561 | 0.062675 | 1 | 0.996511 | 0.66433 |
| Gm21985 | Q6P6P5 | -1.47 | -0.89 | -1.25 | -0.6 | -2.84 | -0.61 | -2.9 | -1.24 | -0.5 | 0.63 | -1.45 | 0.17 | 0.55 | -1.22 | -0.89 | -1.44 | -0.845 | 0.765 | 0.3025 | 0.155047 | 0.124216 | 0.515478 | 1 | 0.996511 | 0.905674 |
| Gm21992 | F7BGR7 | 0.38 | -0.58 | -1.17 | -0.98 | -0.39 | -0.12 | -0.78 | -0.75 | -0.86 | -0.48 | -1.21 | -0.74 | -0.18 | -0.3 | 0.62 | -0.77 | 0.0775 | -0.235 | 0.43 | 0.830826 | 0.525821 | 0.330012 | 1 | 0.996511 | 0.845493 |
| Gm2a | Q60648 | -0.25 | -0.33 | -0.38 | -0.57 | -0.25 | -0.06 | -0.17 | -0.6 | -1.14 | -0.1 | -0.3 | 0.14 | -0.51 | 0.04 | -0.64 | -0.64 | 0.1125 | 0.0325 | -0.055 | 0.536852 | 0.911955 | 0.806267 | 1 | 0.996744 | 0.971715 |
| Gm3839 | S4R1W1 | -0.28 | -1.34 | -1.35 | -1.36 | -1.5 | 0.31 | -0.71 | -0.66 | -1.35 | -0.67 | -1.49 | -0.72 | -0.68 | -1.05 | -1.29 | -0.79 | 0.4425 | 0.025 | 0.13 | 0.31621 | 0.940814 | 0.676387 | 1 | 0.996755 | 0.952944 |
| Gm38393 | A0A087WF | 0.42 | 2.47 | -0.84 | -0.97 | 0.97 | 0.19 | 0.3 | 0.57 | 0.12 | 2.46 | -0.76 | 0.57 | -0.11 | 1.17 | -1.95 | 0.09 | 0.2375 | 0.3275 | -0.47 | 0.748095 | 0.728995 | 0.612402 | 1 | 0.996511 | 0.935166 |
| Gm42517 | A0A1Y7VL( | 0.49 | 0.19 | 1.75 | 1.29 | -0.41 | 1.21 | -0.25 | 0.45 | 2.32 | -0.47 | 0.98 | 0.02 | 3.21 | 0.09 | 1.46 | 0.82 | -0.68 | -0.2175 | 0.465 | 0.179226 | 0.738287 | 0.504357 | 1 | 0.996511 | 0.901002 |
| Gm42715 | A0A087WF | 0.25 | -0.74 | -0.7 | 0.04 | -1.2 | -0.96 | -3.24 | -1.76 | 0.18 | 0.47 | 0.01 | -0.92 | 0.84 | -1.75 | 0.29 | -0.29 | -1.5025 | 0.2225 | 0.06 | 0.018045 | 0.561197 | 0.915221 | 1 | 0.996511 | 0.986556 |
| Gm43738 | A0A0G2JE/ | -0.74 | -0.97 | -0.3 | -0.51 | -0.89 | -0.37 | -0.36 | -0.55 | -0.69 | -0.05 | -0.24 | -0.31 | -0.23 | -0.83 | -0.13 | 1.34 | 0.0875 | 0.3075 | 0.6675 | 0.686315 | 0.203626 | 0.168169 | 1 | 0.996511 | 0.759484 |
| Gm44505 | E0CY81 | -1.02 | -0.7 | 0.5 | 0.67 | -0.06 | -0.79 | 0.27 | 1.39 | -0.57 | 1.99 | 0.99 | -0.58 | -0.8 | 0.25 | 1.66 | 0.72 | 0.34 | 0.595 | 0.595 | 0.553366 | 0.399408 | 0.341359 | 1 | 0.996511 | 0.848473 |
| Gm45208 | A0A140LJ3 | 0.25 | 0.78 | 0.74 | 0.28 | 0.81 | 0.27 | 0.22 | 0.51 | 0.7 | 1.12 | 0.31 | 0.63 | 0.53 | 0.96 | 0.7 | 1.42 | -0.06 | 0.1775 | 0.39 | 0.785555 | 0.475672 | 0.16897 | 1 | 0.996511 | 0.759484 |
| Gm45785 | A0A1B0GT( | -1.2 | -1.49 | -1.16 | -1.26 | -0.2 | -1.14 | -0.38 | -0.56 | -1.32 | -1.54 | -1.12 | -1.25 | -1.07 | -1.36 | -0.85 | -0.69 | 0.7075 | -0.03 | 0.285 | 0.013983 | 0.867986 | 0.21011 | 1 | 0.996511 | 0.786218 |
| Gm45808 | A0A1B0GSI | -0.51 | -0.95 | -0.08 | 0 | -0.52 | -1.03 | 0.28 | 0 | -0.54 | 0.25 | -0.24 | -0.64 | -1 | 0.19 | -0.51 | -0.32 | 0.0675 | 0.0925 | -0.025 | 0.846785 | 0.759742 | 0.939949 | 1 | 0.996511 | 0.990645 |
| Gm45837 | F7D3W5 | 1.55 | 0.93 | 1 | 1 | 1.11 | 1.3 | 1.05 | 1.06 | 0.93 | 0.62 | 0.96 | 1.34 | 0.89 | 0.83 | 1.18 | 1.05 | 0.01 | -0.1575 | -0.1325 | 0.958562 | 0.509601 | 0.546467 | 1 | 0.996511 | 0.912406 |
| Gm49325 | G3UZ60 | 0.93 | -0.13 | 1.02 | -0.4 | 0.94 | 0.23 | -0.53 | -0.51 | -3.76 | -1.14 | 0.64 | 0.3 | -1.96 | -4.32 | 0.32 | 0.13 | -0.3225 | -1.345 | -1.8125 | 0.496572 | 0.184485 | 0.103877 | 1 | 0.996511 | 0.702846 |
| Gm49416 | Q9D937 | -1.7 | -0.12 | -2.85 | -3.29 | -3.01 | 0.19 | -0.56 | 0.76 | -2.2 | -0.87 | 1.15 | -2.03 | -1.47 | -0.76 | -3.07 | 2.12 | 1.335 | 1.0025 | 1.195 | 0.196479 | 0.303403 | 0.316042 | 1 | 0.996511 | 0.838265 |
| Gm49486 | E9QAC6 | 0.13 | 0.62 | -1.15 | -1.6 | 0.31 | 0.07 | -1.86 | -1.27 | -0.27 | -0.83 | -1.29 | -0.16 | -0.76 | 0.29 | -1.45 | -1.45 | -0.1875 | -0.1375 | -0.3425 | 0.780257 | 0.7994 | 0.577202 | 1 | 0.996511 | 0.927198 |
| Gm49510 | A0A2R8W6 | 0.18 | -0.05 | 0.02 | 0.08 | 0.03 | 0.26 | -0.12 | 0.1 | 0.52 | 6.4 | 0.06 | -0.23 | 0.48 | 0.38 | -0.15 | -0.17 | 0.01 | 1.63 | 0.0775 | 0.949409 | 0.266768 | 0.731955 | 1 | 0.996511 | 0.958635 |
| Gm49804 | A0A3B2W8 | -0.52 | -0.56 | -1.08 | -0.93 | -0.72 | -0.41 | -0.7 | -1.15 | -0.42 | -0.92 | -1.03 | -0.43 | -0.35 | -0.81 | -0.74 | -0.85 | 0.0275 | 0.0725 | 0.085 | 0.903277 | 0.762392 | 0.708168 | 1 | 0.996511 | 0.957562 |
| Gm49948 | D3YXH0 | -0.96 | -0.86 | -0.85 | -0.53 | -1.08 | -1.63 | -0.77 | -0.73 | -0.8 | -0.42 | -0.9 | -0.79 | -0.99 | -0.48 | -0.55 | -1.53 | -0.2525 | 0.0725 | -0.0875 | 0.313288 | 0.710196 | 0.755133 | 1 | 0.996511 | 0.962333 |
| Gm50364 | D6RI20 | 0.2 | 0.53 | 0.31 | -0.3 | 1.32 | 0.18 | 0.13 | -0.27 | -0.26 | -0.24 | -0.28 | 0.45 | -0.35 | 1.14 | 0.21 | 0.36 | 0.155 | -0.2675 | 0.155 | 0.674005 | 0.330169 | 0.660607 | 1 | 0.996511 | 0.949645 |
| Gm55978 | A0A1B0GS | 0.84 | -0.94 | 2.59 | 0.23 | 0.11 | 0.93 | 2.49 | 0.39 | 2.3 | 0.91 | 2.29 | 1.09 | 1.69 | 0.49 | 3.36 | 4.01 | 0.3 | 0.9675 | 1.7075 | 0.71472 | 0.218599 | 0.106677 | 1 | 0.996511 | 0.703417 |
| Gm56451 | A0A338P6I | 1.82 | 2.83 | 1.02 | 1.2 | 2.79 | 1.99 | 1.14 | 0.66 | 0.9 | 1.36 | 0.93 | 1.9 | 0.9 | 2 | 0.82 | 1.52 | -0.0725 | -0.445 | -0.4075 | 0.898688 | 0.330295 | 0.391551 | 1 | 0.996511 | 0.865375 |
| Gm7324 | A0A2I3BRL | 1.47 | -0.03 | 0.39 | 0.58 | -0.52 | 0.66 | -0.36 | 0.98 | 0.88 | 0.73 | 0.31 | -0.51 | 2.09 | 0.24 | 1.38 | -0.21 | -0.4125 | -0.25 | 0.2725 | 0.374699 | 0.556782 | 0.631493 | 1 | 0.996511 | 0.941132 |
| Gmds | Q8K0C9 | -0.14 | -0.12 | -0.31 | -0.21 | 0 | -0.11 | -0.38 | -0.31 | 0.07 | 0.09 | -0.42 | -0.12 | 0.28 | 0.01 | -0.37 | -0.55 | -0.005 | 0.1 | 0.0375 | 0.975064 | 0.594183 | 0.872612 | 1 | 0.996511 | 0.979626 |
| Gmeb1 | Q9JL60 | -3.29 | -0.58 | -1.54 | -1.78 | -1.04 | -3 | -1.81 | -1.63 | -0.79 | 0.41 | -1.45 | -3.68 | -1.11 | -2.15 | -1.97 | -2.14 | -0.0725 | 0.42 | -0.045 | 0.908688 | 0.650989 | 0.936431 | 1 | 0.996511 | 0.989767 |
| Gmeb2 | P58929 | -0.37 | -0.56 | -1.14 | -1.75 | -0.57 | 0.14 | -2.03 | -1.4 | -0.64 | -1.2 | -1.67 | -0.81 | -0.86 | -1.2 | -1.23 | -2.44 | -0.01 | -0.125 | -0.4775 | 0.984697 | 0.73875 | 0.295919 | 1 | 0.996511 | 0.825612 |
| Gmfb | Q9CQI3 | 0.94 | -0.81 | 0.03 | -1.06 | -0.44 | 1.21 | 1.3 | -0.64 | -0.57 | -1.24 | 0.07 | 0.63 | -1.2 | -0.36 | 0.45 | 0.42 | 0.5825 | -0.0525 | 0.0525 | 0.365655 | 0.925112 | 0.924148 | 1 | 0.996755 | 0.988152 |
| Gmppa | Q922H4 | -0.19 | -0.41 | -0.2 | -0.24 | -0.85 | -0.09 | -0.82 | -0.5 | 1.76 | 0.01 | -0.29 | -0.3 | 0.94 | 1.22 | -0.58 | -0.3 | -0.305 | 0.555 | 0.58 | 0.174365 | 0.252419 | 0.19845 | 1 | 0.996511 | 0.78004 |
| Gmppb | Q8BTZ7 | 0.92 | 1.08 | 0.22 | 0.41 | 1.06 | 0.82 | 0.2 | 0.31 | 0.88 | 0.69 | 0.2 | 0.87 | 0.61 | 1.61 | 0.18 | 0.38 | -0.06 | 0.0025 | 0.0375 | 0.835935 | 0.992697 | 0.919167 | 1 | 0.999285 | 0.986868 |
| Gmpr | Q9DCZ1 | -0.91 | -1.28 | -0.13 | -0.33 | -1.06 | -0.95 | -0.26 | -0.4 | -1.45 | 0.13 | -0.5 | -1.2 | -1.14 | -0.81 | 0.08 | -0.13 | -0.005 | -0.0925 | 0.1625 | 0.987612 | 0.826269 | 0.669608 | 1 | 0.996511 | 0.952558 |
| Gmpr2 | Q99L27 | -0.01 | -0.69 | -0.85 | -1.21 | -0.25 | -0.5 | -0.57 | -1.2 | -0.19 | -1.29 | -1.15 | 0.01 | -0.35 | 0.02 | -0.24 | -0.84 | 0.06 | 0.035 | 0.3375 | 0.84951 | 0.929697 | 0.302617 | 1 | 0.996755 | 0.830988 |
| Gmps | Q3THK7 | -1.14 | -1.3 | -1.24 | -0.91 | -1.15 | -1.06 | -0.98 | -1.21 | -1.44 | -1.34 | -1.15 | -0.97 | -1.33 | -1.53 | -1.25 | -1.5 | 0.0475 | -0.0775 | -0.255 | 0.768326 | 0.686348 | 0.198979 | 1 | 0.996511 | 0.78004 |
| Gna11 | P21278 | 4.34 | 3.38 | 3.43 | 1.26 | 4.31 | 4.44 | 3.56 | 3.34 | 0.25 | 0.52 | 3.47 | 4.26 | 0.05 | 0.3 | 3.53 | 3.9 | 0.81 | -0.9775 | -1.1575 | 0.228985 | 0.37932 | 0.302354 | 1 | 0.996511 | 0.830768 |
| Gna12 | P27600 | 1.78 | 1 | 0.19 | 0.16 | 1.22 | -0.93 | 1.11 | 0.17 | 1.54 | 0.23 | 0.21 | 0.75 | 1.26 | 1.79 | 0.31 | 0.76 | -0.39 | -0.1 | 0.2475 | 0.504641 | 0.829624 | 0.601243 | 1 | 0.996511 | 0.930802 |
| Gna13 | P27601 | 1.95 | 1.18 | 1.63 | 1.14 | 1.65 | 1.81 | 1.57 | 1.52 | 0.54 | 1.04 | 1.57 | 1.96 | 0.38 | 0.7 | 1.6 | 1.54 | 0.1625 | -0.1975 | -0.42 | 0.476901 | 0.582856 | 0.258083 | 1 | 0.996511 | 0.812074 |
| Gna14 | P30677 | 6.24 | 4.59 | 7.33 | 1.47 | 5.25 | 6.89 | 6.28 | 6.73 | -1.54 | -1.59 | 7.35 | 6.67 | 0.62 | -1.3 | 7.7 | 8.5 | 1.38 | -2.185 | -1.0275 | 0.264824 | 0.38975 | 0.67783 | 1 | 0.996511 | 0.953944 |
| Gnai1 | B2RSH2 | 2.58 | 1.36 | 2.09 | 1.76 | 1.48 | 2.77 | 2.26 | 2.33 | 1.82 | 1.54 | 2.1 | 2.63 | 0.94 | 1.4 | 1.87 | 2.01 | 0.2625 | 0.075 | -0.3925 | 0.469187 | 0.826624 | 0.281903 | 1 | 0.996511 | 0.822084 |
| Gnai2 | P08752 | 0.66 | 0.79 | 0.34 | 0.37 | 0.73 | 0.87 | 0.93 | 0.52 | 0.33 | -0.79 | 0.39 | 0.69 | 0.03 | -0.15 | 0.14 | 0.49 | 0.2225 | -0.385 | -0.4125 | 0.248613 | 0.2737 | 0.091709 | 1 | 0.996511 | 0.693639 |
| Gnai3 | Q9DC51 | 0.9 | 0.81 | 1.37 | 1.24 | 1.13 | 1.23 | 1.69 | 1.41 | 0.37 | 0.74 | 1.44 | 1.22 | 0.21 | -0.23 | 1.29 | 1.32 | 0.285 | -0.1375 | -0.4325 | 0.197775 | 0.632232 | 0.293891 | 1 | 0.996511 | 0.825612 |
| Gnal | Q8CGK7 | 5.54 | 1.34 | 1.3 | 0.83 | 1.99 | 3.62 | 1.9 | 1.09 | 1.77 | 1.06 | 0.84 | 5.28 | 1.51 | 1.07 | 1.99 | 1.58 | -0.1025 | -0.015 | -0.715 | 0.925391 | 0.991128 | 0.481138 | 1 | 0.999285 | 0.893906 |
| Gnal | Q66L47 | 1.22 | 1.27 | 2 | 1.47 | 1.64 | 1.37 | 0.43 | 1.65 | 2.94 | 1.32 | 1.83 | 2.11 | 2.62 | 0.88 | 0.87 | 0.14 | -0.2175 | 0.56 | -0.3625 | 0.514594 | 0.157105 | 0.489153 | 1 | 0.996511 | 0.897531 |
| Gnao1 | P18872-2 | -1.2 | -2.89 | -1.95 | -2.61 | -1.54 | 0.75 | -0.72 | -1.55 | -5.79 | -3.83 | -1.62 | -0.82 | -5.25 | -5.34 | -2.18 | -0.69 | 1.3975 | -0.8525 | -1.2025 | 0.042834 | 0.431259 | 0.284851 | 1 | 0.996511 | 0.822669 |
| Gnao1 | P18872 | 1.31 | 0.84 | 2.88 | 1.23 | 1.02 | 1.52 | 3.23 | 3.09 | 0.09 | -0.59 | 3.11 | 1.33 | -0.08 | -0.13 | 2.93 | 3.56 | 0.65 | -0.58 | 0.005 | 0.331592 | 0.495098 | 0.995849 | 1 | 0.996511 | 0.998588 |
| Gnaq | P21279 | 1.51 | 1.42 | 1.08 | 0.94 | 1.89 | 1.41 | 1.13 | 1.15 | 0.75 | 0.6 | 1.14 | 1.82 | 0.57 | 0.47 | 0.86 | 0.92 | 0.1575 | -0.16 | -0.5325 | 0.515135 | 0.606069 | 0.038329 | 1 | 0.996511 | 0.658454 |
| Gnas | A0A571BE: | -0.53 | 3.31 | -0.1 | 0.25 | -0.28 | 4.44 | -0.37 | 0.24 | 0.68 | 0.87 | -0.7 | 1.55 | 0.69 | 1.89 | 1.27 | 0.39 | 0.275 | -0.1325 | 0.3275 | 0.831584 | 0.882225 | 0.697913 | 1 | 0.996511 | 0.955729 |
| Gnas | P63094 | 0.69 | 0.85 | 1.48 | 0.94 | 1.12 | 0.88 | 1.63 | 1.29 | 0.71 | -0.12 | 1.51 | 0.67 | 0.72 | 0.34 | 1.34 | 1.24 | 0.24 | -0.2975 | -0.08 | 0.343282 | 0.422845 | 0.791356 | 1 | 0.996511 | 0.969105 |
| Gnaz | O70443 | 0.5 | 0.56 | 1.22 | 0.95 | 0.85 | 0.94 | 1.34 | 1.11 | -0.22 | 0.6 | 1.03 | 0.65 | 0.02 | -0.09 | 0.82 | 1.01 | 0.2525 | -0.2925 | -0.3675 | 0.276161 | 0.36526 | 0.281802 | 1 | 0.996511 | 0.822084 |
| Gnb1 | P62874 | -1 | -1.93 | 0 | 1.02 | -2.23 | -0.38 | 0.42 | 0.94 | -0.39 | 0.63 | 0.06 | -0.77 | -0.56 | -1.35 | -0.13 | 0.02 | 0.165 | 0.36 | -0.0275 | 0.845711 | 0.578695 | 0.965901 | 1 | 0.996511 | 0.9927 |
| Gnb2 | P62880 | -0.36 | -0.44 | -0.23 | -0.07 | -0.22 | 0.16 | -0.06 | 0.01 | -0.13 | -0.13 | -0.35 | -0.13 | -0.18 | -0.17 | -0.34 | 0.07 | 0.2475 | 0.09 | 0.12 | 0.165665 | 0.604708 | 0.53668 | 1 | 0.996511 | 0.912212 |
| Gnb2 | E9PWM7 | -4.26 | -4.19 | -3.43 | -1.95 | -4.29 | -3.78 | -2.75 | -2.77 | -3.5 | -2.51 | -2.44 | -4.22 | -4.03 | -5.54 | -3.18 | -2.63 | 0.06 | 0.29 | -0.3875 | 0.920392 | 0.645346 | 0.608377 | 1 | 0.996511 | 0.933931 |
| Gnb4 | P29387 | 0.83 | -0.2 | -0.39 | -0.19 | -0.23 | 1.52 | 0.35 | -0.02 | -0.21 | -0.73 | -0.09 | 0.72 | 0.12 | -1.1 | -0.28 | 0.04 | 0.3925 | -0.09 | -0.3175 | 0.389587 | 0.818187 | 0.415221 | 1 | 0.996511 | 0.874945 |
| Gnb5 | P62881-2 | 0.23 | 0.01 | 0.4 | 0.6 | 0.07 | 0.01 | 0.43 | 0.36 | 0.26 | 0 | 0.32 | 0 | -0.01 | 0.08 | 0.15 | 0 | -0.0925 | -0.165 | -0.255 | 0.641376 | 0.419783 | 0.221344 | 1 | 0.996511 | 0.787447 |
| Gne | Q3UW64 | -0.34 | -0.68 | -1.18 | -1.06 | -0.65 | -0.48 | -0.68 | -1.17 | -1.14 | -0.55 | -1.07 | -0.62 | -0.76 | -0.82 | -1.13 | -1.14 | 0.07 | -0.03 | -0.1475 | 0.782207 | 0.908832 | 0.558394 | 1 | 0.996744 | 0.91747 |
| Gng10 | Q9CXP8 | 2.44 | 1.42 | 1.24 | 2.85 | -0.02 | 1.89 | 2.21 | 2.73 | 0.9 | 2.52 | 1.61 | 1.27 | 0.37 | 0.34 | 0.89 | 1.36 | -0.285 | -0.4125 | -1.2475 | 0.662711 | 0.406964 | 0.017915 | 1 | 0.996511 | 0.602211 |
| Gng11 | P61953 | 1.52 | 2.32 | 1.48 | 0.93 | 1.6 | 0.03 | -0.21 | 0.13 | 1.34 | 1.05 | 0.88 | 1.69 | 1.4 | 1.49 | 1.96 | 1.04 | -1.175 | -0.3225 | -0.09 | 0.03073 | 0.34714 | 0.793098 | 1 | 0.996511 | 0.96933 |
| Gng12 | Q9DAS9 | 1.57 | 1.3 | 1.06 | 1.75 | 1.15 | 1.37 | 0.82 | 1.64 | 1.37 | 1.39 | 0.93 | 1.36 | 1.43 | 1.08 | 0.9 | 1.05 | -0.175 | -0.1575 | -0.305 | 0.480582 | 0.487048 | 0.21077 | 1 | 0.996511 | 0.786269 |
| Gng13 | Q9JMF3 | 2.05 | 2.06 | 1.62 | 2.13 | 2.45 | 1.78 | 2.03 | 1.74 | 1.43 | 1.44 | 1.71 | 1.84 | 1.4 | 2.71 | 1.3 | 1.72 | 0.035 | -0.36 | -0.1825 | 0.875114 | 0.102967 | 0.596601 | 1 | 0.996511 | 0.929911 |
| Gng2 | P63213 | 1 | 1.39 | 0.29 | 0.4 | 1.63 | 1.36 | 0.52 | 0.14 | 0.56 | 0.47 | 0.06 | 0.58 | 0.61 | 2.06 | -0.27 | 0.77 | 0.1425 | -0.3525 | 0.0225 | 0.728311 | 0.250095 | 0.964628 | 1 | 0.996511 | 0.9927 |
| Gng3 | P63216 | -0.13 | -0.66 | -0.34 | 0.46 | -0.45 | -0.07 | 0.42 | 0.4 | -0.18 | -0.04 | 0.14 | 0.35 | -0.17 | 0.17 | 0.04 | 0 | 0.2425 | 0.235 | 0.1775 | 0.442314 | 0.404371 | 0.516831 | 1 | 0.996511 | 0.905676 |
| Gng4 | P50153 | 0.76 | 0.78 | -0.02 | 0.55 | 1.01 | 0.77 | 0.42 | 0.6 | 0.78 | 1.58 | 0.27 | 1.14 | 0.58 | 1.2 | 0.22 | 0.55 | 0.1825 | 0.425 | 0.12 | 0.455454 | 0.222071 | 0.682939 | 1 | 0.996511 | 0.955729 |
| Gng5 | Q80SZ7 | 1.26 | 1.39 | 1.94 | 2.34 | 2.02 | 1.09 | 2.19 | 2.12 | 0.87 | 2.33 | 1.95 | 1.3 | 1.55 | 0.42 | 2.32 | 1.99 | 0.1225 | -0.12 | -0.1625 | 0.724074 | 0.761226 | 0.722802 | 1 | 0.996511 | 0.958398 |
| Gng7 | Q61016 | 1.84 | 0.61 | 1.12 | 1.91 | 0.51 | 0.98 | 2.01 | 1.56 | 1.26 | 1.08 | 0.86 | 1.52 | 1.18 | 0.82 | 1.34 | 1.6 | -0.105 | -0.19 | -0.135 | 0.804112 | 0.57546 | 0.699385 | 1 | 0.996511 | 0.955729 |
| Gnl1 | P36916 | -0.28 | -0.34 | 0.22 | 0.49 | -0.35 | -0.46 | 0.21 | 0.34 | -0.86 | 0.09 | 0.1 | -0.25 | -0.61 | -1.1 | 0.2 | 0.12 | -0.0875 | -0.2525 | -0.37 | 0.758676 | 0.41806 | 0.322995 | 1 | 0.996511 | 0.840788 |
| Gnl3 | Q8CI11 | 1.06 | 0.24 | 0.22 | 0.02 | 0.35 | 0.83 | 0.4 | 0.01 | 1.25 | 0.48 | 0.21 | 0.89 | 0.72 | 1.16 | 0.18 | 0.24 | 0.0125 | 0.3225 | 0.19 | 0.965159 | 0.333681 | 0.567084 | 1 | 0.996511 | 0.921806 |
| Gnl3l | Q6PGG6 | -1.09 | -0.69 | -0.83 | -0.74 | -1.08 | -0.82 | -0.77 | -1.31 | -0.81 | -1.03 | -0.96 | -0.81 | -0.46 | -1.15 | -1.18 | -0.17 | -0.1575 | -0.065 | 0.0975 | 0.419813 | 0.712372 | 0.733786 | 1 | 0.996511 | 0.958635 |
| Gnpat | P98192 | -0.15 | 0.15 | -0.11 | -0.18 | 0.86 | 0.3 | 0.73 | -0.13 | 0.16 | 0.16 | -0.31 | 0.08 | 0.1 | 0.46 | 0.34 | 0.73 | 0.5125 | 0.095 | 0.48 | 0.065976 | 0.622053 | 0.044936 | 1 | 0.996511 | 0.658454 |
| Gnpda1 | O88958 | -0.04 | -0.37 | 0.44 | -0.47 | 0.08 | 0.01 | 0.31 | 0.28 | -1.61 | -2.19 | 0.21 | -0.27 | -1.75 | -1.26 | -0.31 | 0.07 | 0.28 | -0.855 | -0.7025 | 0.253647 | 0.147423 | 0.138947 | 1 | 0.996511 | 0.73493 |
| Gnpda2 | Q9CRC9 | -0.61 | -0.72 | -0.57 | -0.41 | -0.21 | -0.67 | -0.06 | -0.51 | -0.74 | -0.66 | -0.7 | -1.02 | -0.75 | -0.13 | -0.71 | -0.84 | 0.215 | -0.2025 | -0.03 | 0.278104 | 0.266649 | 0.892867 | 1 | 0.996511 | 0.982801 |
| Gnpnat1 | Q9JK38 | -0.55 | -0.43 | 0.66 | 0.27 | -0.24 | -1.07 | 0.8 | 0.42 | -1.31 | 0.72 | 0.06 | -0.91 | -1.27 | 0.05 | 0.02 | 0.45 | -0.01 | -0.3475 | -0.175 | 0.982823 | 0.497284 | 0.696261 | 1 | 0.996511 | 0.955729 |
| Gnptg | A0A0R4J0H | -0.05 | 0.06 | 0.37 | 0.04 | 0.7 | -0.16 | 0.46 | 0.4 | 0.02 | -0.16 | -0.3 | -0.3 | -0.51 | 0.57 | 0.2 | 0.66 | 0.245 | -0.29 | 0.125 | 0.29316 | 0.139044 | 0.674056 | 1 | 0.996511 | 0.952944 |
| Gns | Q8BFR4 | -0.06 | -0.16 | 0.35 | 0.06 | 0.69 | -0.52 | 0.76 | 0 | -0.85 | -0.14 | 0.26 | 0.12 | -0.52 | -0.03 | -0.15 | 0.53 | 0.185 | -0.2 | -0.09 | 0.563101 | 0.48455 | 0.738512 | 1 | 0.996511 | 0.958796 |
| Golga1 | Q9CW79-2 | -1.25 | -1.61 | -1.85 | -1.63 | -1.44 | -1.09 | -1.66 | -1.51 | -1.22 | -1 | -1.69 | -1.25 | -1.08 | -0.84 | -1.68 | -1.99 | 0.16 | 0.295 | 0.1875 | 0.441413 | 0.212513 | 0.542086 | 1 | 0.996511 | 0.912212 |
| Golga2 | E9PUQ5 | -1.69 | -2.53 | -2.32 | -1.6 | -1.74 | -0.76 | -1.86 | -0.82 | -2.11 | -1.31 | -1.05 | -2.04 | -1.79 | -1.86 | -1.24 | -1.28 | 0.74 | 0.4075 | 0.4925 | 0.06431 | 0.256219 | 0.123918 | 1 | 0.996511 | 0.721448 |
| Golga3 | E9QP99 | 0.07 | -0.28 | 0.04 | 0.04 | -0.06 | -0.24 | -0.03 | -0.01 | -0.24 | -0.06 | -0.05 | -0.13 | -0.52 | -0.13 | -0.16 | -0.24 | -0.0525 | -0.0875 | -0.23 | 0.743877 | 0.610932 | 0.255541 | 1 | 0.996511 | 0.809524 |
| Golga4 | Q91VW5 | -1.2 | -1.45 | -1.42 | -1.55 | -1.13 | -1.45 | -1.42 | -1.53 | -1.16 | -0.72 | -1.29 | -1.09 | -1.03 | -0.86 | -1.66 | -1.65 | 0.0225 | 0.34 | 0.105 | 0.893713 | 0.109678 | 0.679324 | 1 | 0.996511 | 0.955309 |
| Golga5 | Q9QYE6 | -1.02 | -1.63 | -1.37 | -1.02 | -0.78 | -1.04 | -1.11 | -1.31 | -1.05 | -0.81 | -1.33 | -1.22 | -0.69 | -0.83 | -0.32 | -0.99 | 0.2 | 0.1575 | 0.5525 | 0.357418 | 0.485608 | 0.046186 | 1 | 0.996511 | 0.658454 |
| Golga7 | Q91W53 | 0.49 | 1.21 | 1.28 | 1.15 | 1.29 | 0.81 | 0.64 | 0.87 | 1.23 | 0.77 | 1.25 | 0.77 | 0.12 | 0.83 | 1 | 1.05 | -0.13 | -0.0275 | -0.2825 | 0.596276 | 0.912582 | 0.353707 | 1 | 0.996744 | 0.855029 |

| | | | | | | | | | | | | | | | | | | | | | | | | | | | |
|---|---|---|---|---|---|---|---|---|---|---|---|---|---|---|---|---|---|---|---|---|---|---|---|---|---|---|---|
| GOLGA7B | Q9D428 | 1 | 1.28 | 1.14 | 1.17 | 1.1 | 1.33 | 1.11 | 1.03 | 1.41 | 1.26 | 1.08 | 1.26 | 1.35 | 0.66 | 0.91 | 1.01 | -0.005 | 0.105 | -0.165 | 0.97426 | 0.538193 | 0.443158 | | 1 | 0.996511 | 0.883549 |
| Golgb1 | E9PVZ8 | -1.26 | -1.77 | -1.79 | -1.36 | -0.87 | -1.41 | -1.35 | -1.49 | -1.5 | -0.84 | -1.49 | -1.65 | -1.15 | -1 | -0.91 | -1.17 | 0.265 | 0.175 | 0.4875 | 0.247396 | 0.490131 | 0.042156 | | 1 | 0.996511 | 0.658454 |
| Golim4 | D3YVW2 | -0.49 | 0.22 | -0.16 | -0.4 | 0.07 | -0.04 | 0.17 | -0.05 | -0.16 | 0.75 | -0.05 | -0.2 | 0.94 | -0.3 | 0.2 | -0.09 | 0.245 | 0.2925 | 0.395 | 0.240868 | 0.319757 | 0.237973 | | 1 | 0.996511 | 0.792722 |
| Golm2 | E9PZJ6 | 0.23 | -0.14 | -2.65 | -2.53 | 0.38 | 0.12 | -1.9 | -2.05 | -0.63 | -1.62 | -2.37 | 0.11 | -0.04 | -0.26 | -1.64 | -2 | 0.41 | 0.145 | 0.2875 | 0.650406 | 0.864127 | 0.72589 | | 1 | 0.996511 | 0.958398 |
| Golph3 | Q9CRA5 | -1.44 | -1.96 | -0.71 | -1.13 | -1.49 | -1.23 | -1.12 | -1.26 | -0.77 | -0.6 | -1.19 | -1.5 | 0.41 | -1.92 | -0.93 | -1.15 | 0.035 | 0.295 | 0.4125 | 0.899767 | 0.383487 | 0.429995 | | 1 | 0.996511 | 0.879165 |
| Golph3l | H3BJ07 | -2.06 | -2.25 | -2.91 | -2.56 | -2.26 | -2.54 | -1.95 | -3.2 | -2.43 | -2.97 | -3.02 | -1.94 | -2.71 | -1.8 | -3.95 | -3.09 | -0.0425 | -0.145 | -0.4425 | 0.893667 | 0.648745 | 0.344959 | | 1 | 0.996511 | 0.84949 |
| Golt1b | Q9CR60 | -1.93 | -1.38 | -1.38 | -1.3 | -0.71 | -1.51 | -0.77 | -1.03 | -1.41 | -1.3 | -1.71 | -1.96 | -1.44 | -0.45 | -1.63 | -0.7 | 0.4925 | -0.0975 | 0.4425 | 0.072339 | 0.682374 | 0.195998 | | 1 | 0.996511 | 0.78004 |
| Gopc | Q8BH60 | -0.8 | -0.96 | -0.79 | -0.94 | -0.51 | -0.83 | -0.99 | -0.89 | -0.86 | -0.36 | -0.76 | -0.48 | -0.46 | -0.56 | -1.1 | -1.05 | 0.0675 | 0.2575 | 0.08 | 0.688752 | 0.19007 | 0.718233 | | 1 | 0.996511 | 0.958398 |
| Gorab | Q8BRM2 | -0.55 | 0.08 | -0.2 | -0.38 | 0.28 | -0.16 | -0.07 | -0.28 | 0 | 1.33 | -0.31 | -0.43 | -0.01 | 0.67 | 0.08 | -0.34 | 0.205 | 0.41 | 0.3625 | 0.340704 | 0.32782 | 0.206911 | | 1 | 0.996511 | 0.785338 |
| Gorasp1 | Q91X51 | 0.01 | 0.2 | -0.02 | 0.15 | 0.28 | -0.22 | 0.17 | 0.07 | 0.08 | 0.69 | -0.37 | -0.05 | -0.43 | 0.41 | 0.64 | 0.73 | -0.01 | 0.0025 | 0.2525 | 0.953445 | 0.992048 | 0.390529 | | 1 | 0.999285 | 0.865375 |
| Gorasp2 | A2ATI9 | -1.69 | -1.63 | 0.15 | 0.25 | -1.38 | -1.99 | -0.39 | -0.18 | -1.71 | 0.64 | -0.22 | -1.44 | -0.88 | -1.42 | -0.14 | -0.45 | -0.255 | 0.0475 | 0.0075 | 0.683929 | 0.945639 | 0.989259 | | 1 | 0.996755 | 0.996154 |
| Gosr1 | O88630 | -0.64 | -1.44 | -2.04 | -2.15 | -0.59 | -0.75 | -0.82 | -1.3 | -0.68 | -2.21 | -1.82 | -0.81 | -0.22 | -0.8 | -0.48 | -1.43 | 0.7025 | 0.1875 | 0.835 | 0.079647 | 0.69577 | 0.070779 | | 1 | 0.996511 | 0.668066 |
| Gosr2 | A2A9I0 | 1.04 | 0.94 | 1.48 | 1.93 | 1.51 | 0.32 | 1.36 | 1.8 | 1.38 | 2.66 | 1.2 | 1.24 | 1.89 | 1.6 | 2.47 | 2.29 | -0.1 | 0.2725 | 0.715 | 0.789947 | 0.499553 | 0.043469 | | 1 | 0.996511 | 0.658454 |
| Got1 | P05201 | -0.68 | -0.74 | -0.51 | -0.04 | -0.79 | -0.96 | -0.28 | 0.06 | -1.34 | -0.45 | -0.51 | -0.79 | -1.45 | -0.96 | -0.81 | -0.83 | 1.96E-16 | -0.28 | -0.52 | 1 | 0.319161 | 0.064066 | | 1 | 0.996511 | 0.66433 |
| Got2 | P05202 | 0.78 | 0.57 | -0.52 | 0.23 | 0.29 | 0.86 | -0.35 | 0.05 | 0.35 | -0.14 | -0.32 | 0.46 | -0.06 | 0.16 | -0.81 | -0.67 | -0.0525 | -0.1775 | -0.61 | 0.885328 | 0.601849 | 0.12024 | | 1 | 0.996511 | 0.721448 |
| Gpaa1 | Q9WTK3 | -0.77 | -0.15 | -0.54 | -0.33 | 0.48 | -0.43 | 0.58 | 0.05 | -0.58 | -0.59 | -0.22 | -0.69 | -0.33 | 0.45 | 0.52 | 0.46 | 0.6175 | -0.0725 | 0.7225 | 0.0467 | 0.732078 | 0.023147 | | 1 | 0.996511 | 0.623925 |
| Gpalpp1 | Q69ZC8 | -3.8 | -3.96 | -3.09 | -2.64 | -2.69 | -6.48 | -3.52 | -4 | -3.97 | -1.66 | -2.87 | -3 | -3.84 | -2.76 | -3.28 | -2.12 | -0.8 | 0.4975 | 0.3725 | 0.324072 | 0.354978 | 0.420074 | | 1 | 0.996511 | 0.87558 |
| Gpam | A0A494BA: | 0.09 | 0.76 | 0.83 | 0.9 | 0.34 | 0.27 | 0.98 | 1.26 | 0.66 | 1.55 | 1.64 | 0.33 | 0.17 | 0.68 | 0.37 | 2.14 | 0.0675 | 0.4 | 0.195 | 0.823935 | 0.289716 | 0.670491 | | 1 | 0.996511 | 0.952558 |
| Gpat3 | Q8C0N2 | 0.2 | 0.38 | 0.2 | 1.04 | 0.21 | 0.13 | 0.35 | 1.16 | 0.03 | 2.01 | -0.15 | 0.37 | -0.21 | 0.67 | 0.7 | 0.96 | 0.0075 | 0.11 | 0.075 | 0.980421 | 0.824421 | 0.819184 | | 1 | 0.996511 | 0.972768 |
| Gpat4 | Q8K2C8 | -0.66 | 0.03 | -0.78 | -0.8 | -0.13 | -0.34 | -0.24 | -0.64 | 0.54 | -0.69 | -0.62 | -0.78 | 1.1 | 0.29 | 0.43 | -0.58 | 0.215 | 0.165 | 0.8625 | 0.382682 | 0.64732 | 0.049156 | | 1 | 0.996511 | 0.658454 |
| Gpatch2 | M0QWJ1 | -0.02 | 0.04 | 0.71 | -0.31 | -0.54 | -0.85 | -1.24 | -0.35 | 0.46 | 1.88 | 0.17 | -0.26 | 1.15 | 0.73 | 0.35 | -0.43 | -0.85 | 0.4575 | 0.345 | 0.016343 | 0.351247 | 0.384071 | | 1 | 0.996511 | 0.860984 |
| Gpatch2l | Q6PE65 | -0.14 | -0.7 | 2.51 | 1.22 | -0.77 | -0.55 | 1.16 | 1.92 | 0.18 | 1.6 | 1.13 | -0.22 | 0.43 | 0.94 | 1.17 | 3.98 | -0.2825 | -0.05 | 0.9075 | 0.747486 | 0.947107 | 0.358273 | | 1 | 0.996755 | 0.856645 |
| Gpatch8 | A2A6A1 | 0.94 | 0.74 | -0.28 | 0.02 | -0.5 | 0.48 | -1.36 | 0.12 | 1 | 0.67 | -0.9 | 0.07 | 1.89 | 0.38 | -0.2 | -1.06 | -0.67 | -0.145 | -0.1025 | 0.171291 | 0.760445 | 0.869856 | | 1 | 0.996511 | 0.979293 |
| Gpbp1 | Q6NXH3 | -1.88 | -1.72 | 0.45 | -0.18 | -2.41 | -3.04 | 0.75 | 0.49 | 0.34 | -0.21 | 1.6 | -2.41 | 1.24 | -0.47 | -0.11 | -0.1 | -0.22 | 0.6625 | 0.9725 | 0.828529 | 0.475476 | 0.147272 | | 1 | 0.996511 | 0.746406 |
| Gpc1 | Q9QZF2 | 0.07 | 0.15 | -0.15 | 0.09 | 0.04 | 0.09 | -0.2 | -0.18 | 0.47 | 0.21 | 0.09 | 0.29 | 0.33 | 0.64 | -0.42 | -0.33 | -0.1025 | 0.225 | 0.015 | 0.528743 | 0.220422 | 0.957842 | | 1 | 0.996511 | 0.992636 |
| Gpc4 | P51655 | 0.73 | 1.47 | 0.52 | 0.51 | 1.11 | 0.96 | 0.56 | 0.75 | 0.75 | 0.32 | 0.56 | 1 | 0.26 | 0.62 | 0.35 | 0.06 | 0.0375 | -0.15 | -0.485 | 0.887041 | 0.595816 | 0.106817 | | 1 | 0.996511 | 0.703417 |
| Gpc5 | Q8CAL5 | 0.54 | 0.71 | 0.85 | 0.66 | 0.65 | 0.91 | 0.62 | 0.3 | 0.98 | 0.79 | 1.08 | 1.11 | 1.11 | 0.22 | 0.63 | 0.85 | -0.07 | 0.3 | 0.0125 | 0.703627 | 0.108784 | 0.958185 | | 1 | 0.996511 | 0.992636 |
| Gpc6 | Q8R3X6 | 0.24 | 0.14 | 0.54 | 0.57 | 1 | 0.51 | -0.3 | 0.87 | 0.27 | 1.02 | 0.68 | 0.87 | 0.03 | 0.01 | 0.61 | 0.4 | 0.1475 | 0.3375 | -0.11 | 0.63339 | 0.164153 | 0.631183 | | 1 | 0.996511 | 0.941132 |
| Gpcpd1 | Q8C0L9 | -1.15 | -0.94 | -0.19 | -0.61 | -1.09 | -1.22 | -0.15 | -0.23 | -1.17 | -0.33 | -0.4 | -1.16 | -0.74 | -0.85 | -0.61 | -0.63 | 0.05 | -0.0425 | 0.015 | 0.882502 | 0.892282 | 0.952154 | | 1 | 0.996511 | 0.992476 |
| Gpd1 | P13707 | 0.56 | -0.01 | 0.23 | -0.02 | 0.01 | 0.32 | -0.5 | -0.06 | -0.04 | 0.25 | -0.22 | 0.19 | -0.09 | -0.01 | -0.32 | -0.42 | -0.2475 | -0.145 | -0.4 | 0.307484 | 0.50451 | 0.094596 | | 1 | 0.996511 | 0.698156 |
| Gpd1l | Q3ULJ0 | -0.35 | -0.5 | -0.32 | -0.46 | -0.52 | -0.24 | -0.35 | -0.15 | -0.76 | -0.77 | -0.44 | -0.3 | -0.86 | -0.35 | -0.75 | -0.81 | 0.0925 | -0.16 | -0.285 | 0.559078 | 0.399753 | 0.169795 | | 1 | 0.996511 | 0.761572 |
| Gpd2 | Q64521 | 0.33 | 0.58 | 0.85 | 0.85 | 0.26 | 0.61 | 0.45 | 0.73 | 0.06 | 0.33 | 0.79 | 0.13 | -0.05 | 0.04 | 0.04 | 0.25 | -0.14 | -0.325 | -0.5825 | 0.482147 | 0.192077 | 0.017809 | | 1 | 0.996511 | 0.602211 |
| Gphn | A0JNY3 | 0.29 | 0.21 | 0.22 | 0.39 | 0.42 | 0.35 | 0.45 | 0.48 | 0.28 | 0.27 | 0.54 | 0.26 | 0.21 | 0.24 | 0.22 | 0.29 | 0.1475 | 0.06 | -0.0375 | 0.313723 | 0.716945 | 0.822407 | | 1 | 0.996511 | 0.972768 |
| Gpi | P06745 | -0.1 | 0.26 | -0.4 | -0.09 | 0.23 | -0.24 | -0.73 | -0.08 | 0.4 | -0.4 | -0.41 | 0.19 | -0.01 | 0.6 | -0.42 | -0.59 | -0.1225 | 0.0275 | -0.0225 | 0.630092 | 0.9171 | 0.941628 | | 1 | 0.996755 | 0.991085 |
| Gpkow | Q56A08 | -3.23 | -4.47 | -3.79 | -4.18 | -3.84 | -3.34 | -3.74 | -3.74 | -4.54 | -2.3 | -3.32 | -3.33 | -3.23 | -3.69 | -4.22 | -4.37 | 0.2525 | 0.545 | 0.04 | 0.395242 | 0.28745 | 0.913152 | | 1 | 0.996511 | 0.986458 |
| Gpld1 | Q8VCU2 | -0.47 | -0.9 | -1.13 | -0.63 | -0.18 | -0.97 | -0.7 | -1.24 | -0.05 | -0.93 | -0.82 | -1.15 | 0.53 | -0.4 | -0.83 | -0.93 | 0.01 | 0.045 | 0.375 | 0.97084 | 0.876372 | 0.311434 | | 1 | 0.996511 | 0.834523 |
| Gpm6a | P35802 | -0.64 | -0.35 | -0.58 | 0.27 | -0.11 | -0.37 | -0.17 | 0.15 | -0.19 | 0.54 | -0.44 | -0.3 | -0.71 | 0.86 | -0.71 | -0.53 | 0.2 | 0.2275 | 0.0525 | 0.427208 | 0.464352 | 0.899211 | | 1 | 0.996511 | 0.983509 |
| Gpm6b | A2AEG6 | 0.7 | 0.83 | 1.71 | 1 | 1.05 | 0.75 | 1.4 | 0.99 | 0.22 | 1.26 | 1.67 | 0.71 | -0.26 | 0.89 | 1.2 | 1.34 | -0.0125 | -0.095 | -0.2675 | 0.962819 | 0.800486 | 0.519367 | | 1 | 0.996511 | 0.907667 |
| Gpm6b | P35803-8 | -13.52 | -14.11 | -10.22 | -13.27 | -11.93 | -20.25 | -12.71 | -13.11 | -14.32 | -2.47 | -9.34 | -11.07 | -15.04 | -13.07 | -11.85 | -12.65 | -1.72 | 3.48 | -0.3725 | 0.374505 | 0.165477 | 0.70709 | | 1 | 0.996511 | 0.957157 |
| Gpn1 | Q8VCE2 | -0.95 | -1.03 | -0.71 | -0.79 | -1.1 | -0.97 | -0.75 | -0.75 | 0.7 | -0.88 | -0.72 | -0.52 | 0.82 | -0.49 | -1.11 | -0.74 | -0.0225 | 0.515 | 0.49 | 0.893335 | 0.174954 | 0.250625 | | 1 | 0.996511 | 0.807201 |
| Gpr151 | Q7TSN6 | -0.38 | -0.3 | 0.63 | 1.33 | 1.87 | -1 | 1.2 | 1.76 | 1.11 | 1.48 | 1.67 | -1.12 | -0.58 | 1.18 | 1.56 | 1.52 | 0.6375 | 0.465 | 0.6 | 0.380037 | 0.508514 | 0.328767 | | 1 | 0.996511 | 0.844943 |
| Gpr155 | A2AWR2 | -2.17 | -1.37 | -0.32 | -1.09 | -1.57 | -0.32 | -1.72 | -1.38 | 1.65 | -3.12 | -1.52 | -1.19 | 2.24 | -1.59 | -0.45 | -0.87 | -0.01 | 0.1925 | 1.07 | 0.982673 | 0.840493 | 0.217895 | | 1 | 0.996511 | 0.787447 |
| Gpr158 | Q8C419 | 0.75 | 0.92 | 0.89 | 0.98 | 0.67 | 0.53 | 0.83 | 0.86 | 0.86 | 0.82 | 0.81 | 0.74 | 0.58 | 0.54 | 0.75 | 0.5 | -0.1625 | -0.0775 | -0.2925 | 0.31553 | 0.623807 | 0.120981 | | 1 | 0.996511 | 0.721448 |
| Gpr162 | Q3UN16 | 0.02 | 0.44 | 1.03 | 0.88 | 0.33 | -0.02 | 0.21 | 0.12 | 0 | 0.91 | 0.79 | 0.11 | 0.61 | 0.22 | 0.44 | 0.17 | -0.4325 | -0.14 | -0.2325 | 0.113124 | 0.667729 | 0.404749 | | 1 | 0.996511 | 0.872135 |
| Gpr17 | Q6NS65 | -0.45 | 0.27 | -0.15 | -0.57 | 0.29 | -0.12 | -0.93 | -0.36 | 1.12 | 0.92 | -0.53 | -0.01 | 1.14 | 0.47 | 1.11 | -0.25 | -0.055 | 0.6 | 0.8425 | 0.85966 | 0.16997 | 0.045862 | | 1 | 0.996511 | 0.658454 |
| Gpr176 | Q80WT4 | -0.32 | 0.62 | 0.31 | 0.41 | 0.62 | 0.04 | 0.63 | 0.48 | 0.07 | 0.76 | 0.25 | 0.66 | -0.29 | 0.72 | 0.48 | 0.9 | 0.1875 | 0.18 | 0.1975 | 0.469966 | 0.517338 | 0.55748 | | 1 | 0.996511 | 0.917092 |
| Gpr180 | Q8BPS4 | -0.65 | -1.11 | -1.11 | -0.65 | -0.29 | -0.42 | 0.42 | -0.84 | -0.31 | -1.21 | -1.42 | -0.78 | 0.76 | -0.6 | -1.69 | -0.08 | 0.5975 | -0.05 | 0.4775 | 0.068077 | 0.862516 | 0.347277 | | 1 | 0.996511 | 0.851503 |
| Gpr22 | G3X9C3 | 1.83 | 2.2 | 1.32 | 1.27 | 1.44 | 0.87 | 0.35 | 1.19 | 1.3 | 2.1 | 1.33 | 1.46 | 0.96 | 1.02 | 1.15 | 0.88 | -0.6925 | -0.1075 | -0.6525 | 0.053481 | 0.718133 | 0.03051 | | 1 | 0.996511 | 0.641481 |
| Gpr37 | Q9QY42 | -0.05 | 0.07 | 0.42 | 0.45 | 0.11 | -0.52 | 0.03 | 0.06 | -0.19 | 0.46 | 0.12 | -0.47 | -0.77 | 0.48 | -0.15 | 0.26 | -0.3025 | -0.2425 | -0.2675 | 0.189228 | 0.357702 | 0.400193 | | 1 | 0.996511 | 0.86904 |
| Gpr37l1 | Q99JG2 | -0.17 | -0.53 | -1.04 | -1.39 | -0.28 | 0.01 | -1.04 | -1.26 | -1.15 | -1.86 | -1.29 | -0.28 | -0.98 | -1.17 | -1.97 | -1.45 | 0.14 | -0.3625 | -0.61 | 0.71641 | 0.381493 | 0.104 | | 1 | 0.996511 | 0.702979 |
| Gpr68 | Q8BFQ3 | 1.64 | 1.72 | 0.43 | 1.09 | 0.94 | 0.97 | 0.71 | 1.13 | 2.11 | 0.96 | 1.05 | 1.13 | 1.33 | 1.85 | 0.88 | 0.87 | -0.2825 | 0.0925 | 0.0125 | 0.368973 | 0.810597 | 0.973038 | | 1 | 0.996511 | 0.992865 |
| Gpr88 | Q9EPB7 | -0.3 | 0.95 | 1.91 | 1.64 | -0.1 | 0.03 | 1.86 | 1.79 | 0.25 | 0.38 | 0.63 | 0.95 | -1.05 | 0.33 | 0.66 | 0.72 | -0.155 | -0.4975 | -0.885 | 0.815116 | 0.317685 | 0.159808 | | 1 | 0.996511 | 0.755194 |
| Gpr89 | Q8BS95 | -0.43 | 0.34 | -0.34 | -0.38 | 0.47 | -0.46 | 0.38 | -0.08 | -0.47 | -0.37 | -0.31 | -0.47 | -0.66 | 0.42 | 0 | 0.33 | 0.28 | -0.2025 | 0.225 | 0.338287 | 0.373798 | 0.479429 | | 1 | 0.996511 | 0.893468 |
| Gprasp1 | Q5U4C1 | -1.41 | -0.69 | -0.36 | -0.37 | -0.77 | -1.59 | -0.48 | -0.49 | -0.58 | 0.83 | -0.52 | -0.75 | 0.38 | -0.49 | -0.15 | -0.19 | -0.125 | 0.4525 | 0.595 | 0.718737 | 0.295946 | 0.085237 | | 1 | 0.996511 | 0.691619 |
| Gprasp2 | Q8BUY8 | -0.98 | -0.94 | -1.23 | -0.98 | -1.41 | -0.75 | -1.47 | -1.56 | 0.29 | -0.35 | -1.16 | -1.07 | 1.65 | -0.83 | -1.17 | -0.85 | -0.265 | 0.46 | 0.7325 | 0.248585 | 0.201059 | 0.244353 | | 1 | 0.996511 | 0.799652 |
| Gprc5b | Q923Z0 | -0.88 | -0.02 | 0.34 | 0.78 | -1.37 | -0.63 | 0.78 | 0.56 | 0.08 | 0.87 | 0.4 | -0.08 | 0.01 | 0.28 | 0.62 | 0.62 | -0.22 | 0.2625 | 0.3275 | 0.698023 | 0.509431 | 0.391036 | | 1 | 0.996511 | 0.865375 |
| Gprin1 | Q3UNH4 | 1.2 | 1.17 | 1.14 | 1.11 | 1.04 | 0.92 | 0.55 | 0.78 | 1.75 | 0.9 | 1.21 | 1.41 | 2.08 | 1.62 | 1.15 | 0.75 | -0.3325 | 0.1625 | 0.245 | 0.070018 | 0.463016 | 0.425158 | | 1 | 0.996511 | 0.877372 |
| Gprin2 | A0A2I3BRN | 0.04 | 0.16 | -0.16 | 0.17 | -1.12 | 0.78 | -0.25 | -0.08 | 1.51 | 0.08 | -0.36 | 0.47 | 2.82 | -0.65 | 1.55 | 0.01 | -0.22 | 0.3725 | 0.88 | 0.56355 | 0.353058 | 0.236041 | | 1 | 0.996511 | 0.791493 |
| Gprin3 | Q8BWS5 | 1.15 | 0.54 | 1.03 | 1.2 | 0.12 | 0.47 | 1.39 | 1.13 | 1.15 | 1.38 | 0.95 | 1.2 | 1.35 | 1.16 | 1.74 | 1.38 | -0.2025 | 0.19 | 0.4275 | 0.533546 | 0.38785 | 0.096342 | | 1 | 0.996511 | 0.698156 |
| Gps1 | A0A140LJB | -2.87 | -2.01 | -0.53 | -0.98 | -3.46 | -2.76 | -1.91 | -0.82 | -1.49 | 0.39 | -0.98 | -1.96 | -0.27 | -1.2 | -0.57 | -1.25 | -0.64 | 0.5875 | 0.775 | 0.373344 | 0.38854 | 0.17218 | | 1 | 0.996511 | 0.76392 |
| Gps1 | G3UXW9 | -0.2 | -0.28 | -0.59 | -0.66 | 0.11 | -0.01 | -0.74 | -0.56 | -0.39 | -0.71 | -0.63 | -0.33 | -0.45 | -0.05 | -0.79 | -0.69 | 0.1325 | -0.0825 | -0.0625 | 0.596742 | 0.677158 | 0.794043 | | 1 | 0.996511 | 0.969367 |
| Gps2 | Q921N8 | -2.01 | -2.52 | 3.1 | 0.98 | -1.3 | -3.64 | 3.34 | 2.89 | -1.35 | -1.03 | 3.14 | -1.97 | -0.76 | -0.93 | 2.96 | 3 | 0.435 | -0.19 | 1.18 | 0.819693 | 0.903666 | 0.447964 | | 1 | 0.996609 | 0.884817 |
| Gpsm1 | Q6IR34 | 0.18 | -0.11 | 0.28 | 0.57 | -0.16 | 0.08 | 0.21 | 0.63 | 0.38 | 0.9 | 0.35 | 0.46 | 0.39 | 0.06 | 0.57 | 0.57 | -0.04 | 0.2925 | 0.1675 | 0.864592 | 0.215239 | 0.472556 | | 1 | 0.996511 | 0.891144 |
| Gpt | Q8QZR5 | 0.11 | 0 | 0.22 | 0.42 | 0.5 | 0.36 | -0.9 | 0.4 | -0.04 | 0.47 | 0.38 | -0.11 | -0.34 | 0.09 | -0.05 | 0.32 | -0.0975 | -0.0125 | -0.1825 | 0.770035 | 0.953115 | 0.410799 | | 1 | 0.996755 | 0.874811 |
| Gpt2 | Q8BGT5 | -0.6 | -0.43 | 0.09 | 0.44 | -0.16 | -0.43 | 1.03 | 0.29 | 0.94 | 0.64 | 0.68 | -0.23 | 1.05 | 0.84 | 0.12 | 0.35 | 0.3075 | 0.6325 | 0.715 | 0.425812 | 0.093363 | 0.052372 | | 1 | 0.996511 | 0.659923 |
| Gpx1 | P11352 | 0.04 | -0.39 | -0.43 | 0.09 | -0.12 | -0.27 | -0.15 | -0.23 | -0.37 | -0.36 | -0.6 | -0.47 | -0.67 | 0.19 | -0.71 | -0.91 | -0.02 | -0.2775 | -0.3525 | 0.913521 | 0.187116 | 0.251675 | | 1 | 0.996511 | 0.80892 |
| Gpx3 | P46412 | -1.06 | -0.85 | 0.36 | 1.01 | 0.14 | -0.92 | 1.77 | 1.8 | -0.68 | 2.8 | 0.57 | -0.76 | -1.18 | -0.11 | 1.39 | 1 | 0.8325 | 0.6175 | 0.41 | 0.283185 | 0.484813 | 0.556994 | | 1 | 0.996511 | 0.917092 |
| Gpx4 | O70325-2 | 1.89 | 0.41 | 1.17 | 0.61 | 0.08 | 1.58 | 1.86 | 0.48 | 0.58 | 1 | 0.82 | 1.21 | 0.57 | 0.38 | 1.5 | 1.68 | -0.02 | -0.1175 | 0.0125 | 0.968006 | 0.737963 | 0.977365 | | 1 | 0.996511 | 0.993727 |
| Gpx7 | Q99LJ6 | 0 | 0.44 | 0.58 | 0.74 | 1 | 0.16 | 1.76 | 1.25 | 0.56 | 0.6 | 0.87 | 0.33 | 0.9 | 1.39 | 1.04 | 0.91 | 0.6025 | 0.15 | 0.62 | 0.117263 | 0.51453 | 0.026189 | | 1 | 0.996511 | 0.633075 |
| Gpx8 | Q9D7B7 | 1.08 | 0.11 | -0.1 | -0.6 | 0.36 | 1.87 | 1.01 | -0.85 | 0.87 | -0.74 | -0.01 | 1.47 | -0.1 | -0.65 | 0.12 | 0.38 | 0.475 | 0.275 | -0.185 | 0.446386 | 0.621568 | 0.645508 | | 1 | 0.996511 | 0.944749 |
| Gramd1a | F8WJ74 | 0.18 | 0.73 | 0.26 | 0.4 | 0.44 | 0.17 | 0.75 | 0.28 | 0.67 | 0.47 | 0.37 | 0.25 | 1.25 | 1.73 | 0.69 | 0.77 | 0.0175 | 0.0475 | 0.7175 | 0.932023 | 0.812376 | 0.031984 | | 1 | 0.996511 | 0.641974 |
| Gramd1b | A0A1L1SQ4 | -0.32 | 0.13 | 0.26 | 0.15 | 0.53 | 0.26 | 0.37 | 0.13 | -0.46 | -0.31 | 0.27 | -0.23 | 0.23 | -0.04 | 0.78 | 0.72 | 0.2675 | -0.2375 | 0.3675 | 0.186615 | 0.324719 | 0.186174 | | 1 | 0.996511 | 0.776843 |
| Gramd3 | Q9D2W5 | 1.4 | 1.21 | 0.31 | 0 | 0.94 | 0.63 | 0.94 | 1.83 | -0.96 | 2.3 | 0.76 | 1.62 | -0.12 | -2.22 | 1.61 | 0.78 | 0.355 | 0.2 | -0.7175 | 0.389116 | 0.778671 | 0.38282 | | 1 | 0.996511 | 0.860984 |
| Gramd4 | Q8CB44 | -0.31 | -0.25 | 0.04 | -0.04 | -0.26 | -0.21 | 0.09 | -0.21 | -0.35 | -0.11 | -0.13 | -0.36 | -0.17 | 0.16 | 0.11 | 0.22 | -0.0075 | -0.0975 | 0.22 | 0.964601 | 0.585048 | 0.27353 | | 1 | 0.996511 | 0.822084 |
| Grb10 | Q5SUW3 | -0.72 | -0.97 | -0.23 | -0.51 | -0.88 | -0.23 | -0.6 | -0.53 | 0.04 | 0.12 | -0.16 | -0.47 | -0.31 | -1.22 | -0.22 | -0.62 | 0.0475 | 0.49 | 0.015 | 0.834177 | 0.06263 | 0.958909 | | 1 | 0.996511 | 0.992636 |
| Grb14 | Q9JLM9 | -0.81 | -1.39 | -0.73 | 0.67 | -2.22 | 0.31 | -0.45 | -0.27 | 2.49 | 0.65 | -0.51 | -0.73 | 2.47 | -1.74 | -0.31 | -0.5 | -0.0925 | 1.04 | 0.545 | 0.884258 | 0.203083 | 0.544112 | | 1 | 0.996511 | 0.912212 |
| Grb2 | Q60631 | 0.5 | -0.39 | 0 | 0.18 | -0.2 | 0.47 | 0.59 | 0.21 | 0.01 | 0.36 | 0.39 | 0.25 | -0.29 | 0.12 | 0.15 | 0.29 | 0.195 | 0.18 | -0.005 | 0.465581 | 0.4509 | 0.984338 | | 1 | 0.996511 | 0.996154 |
| Grcc10 | O35127 | 0.2 | -0.56 | 0.35 | 0.72 | -0.48 | 0.14 | 0.33 | 0.77 | -0.41 | 3.93 | 0.54 | 0.38 | -0.26 | -0.93 | 0.44 | 0.47 | 0.0125 | 0.9325 | -0.2475 | 0.972116 | 0.315785 | 0.55064 | | 1 | 0.996511 | 0.913801 |
| Grhpr | Q91Z53 | -0.44 | -0.44 | -0.26 | -0.23 | -0.48 | -0.87 | -0.08 | -0.56 | -0.96 | -0.88 | -0.56 | -0.89 | -1.02 | -0.84 | -1.09 | -0.57 | -0.155 | -0.48 | -0.5375 | 0.453845 | 0.022299 | 0.022451 | | 1 | 0.996511 | 0.621926 |
| Gria1 | Q7TNB5 | 0.63 | 0.88 | 1.19 | 0.99 | 1.17 | 0.92 | 1.06 | 0.96 | 0.31 | 0.45 | 1.01 | 0.61 | 0.15 | 0.89 | 0.91 | 1.07 | 0.105 | -0.3275 | -0.1675 | 0.556678 | 0.171502 | 0.530149 | | 1 | 0.996511 | 0.908663 |
| Gria2 | G5E8H1 | 1.61 | 1.76 | 2.44 | 1.3 | 2.27 | 1.37 | 1.51 | 1.91 | 1.24 | 0.41 | 2.36 | 0.75 | 0.74 | 1.83 | 0.75 | 1.48 | -0.0125 | -0.5875 | -0.5775 | 0.967836 | 0.222304 | 0.134183 | | 1 | 0.996511 | 0.730883 |
| Gria2 | P23819-4 | 1.47 | 1.49 | 0.95 | 0.95 | 1.76 | 1.76 | 0.88 | 0.85 | 1.39 | 0.4 | 1.03 | 1.45 | 0.79 | 1.82 | 0.66 | 0.65 | 0.0975 | -0.1475 | -0.235 | 0.744476 | 0.617649 | 0.476937 | | 1 | 0.996511 | 0.892234 |
| Gria3 | B0QZW1 | 0.48 | 0.5 | 0.81 | 0.75 | 0.99 | 0.92 | 0.62 | 0.84 | -0.05 | 0.33 | 0.85 | 0.32 | -0.31 | 0.1 | 0.12 | 0.53 | 0.2075 | -0.2725 | -0.525 | 0.241975 | 0.262738 | 0.048125 | | 1 | 0.996511 | 0.658454 |
| Gria3 | Q9Z2W9 | 1.53 | 1.77 | 1.49 | 1.2 | 1.98 | 1.26 | 1.49 | 1.52 | 1.25 | 0.68 | 1.31 | 1.16 | 0.92 | 1.22 | 0.85 | 0.86 | 0.065 | -0.3975 | -0.535 | 0.76393 | 0.100318 | 0.027611 | | 1 | 0.996511 | 0.635993 |
| Gria4 | G5E863 | 0.49 | 0.68 | 0.74 | 0.37 | 1.1 | 0.71 | 0.74 | 0.6 | 0.66 | -0.5 | 0.83 | 0.44 | 0.56 | 1.49 | 0.4 | 0.85 | 0.2175 | -0.2125 | 0.255 | 0.251405 | 0.500884 | 0.368993 | | 1 | 0.996511 | 0.859245 |
| Grid1 | Q61627 | 0.76 | 0.87 | 1.06 | 1.01 | 0.9 | 0.86 | 1.05 | 0.92 | 0.71 | 0.52 | 1.09 | 0.99 | 0.54 | 0.89 | 0.94 | 0.88 | 0.0075 | -0.0975 | -0.1125 | 0.960614 | 0.624703 | 0.559364 | | 1 | 0.996511 | 0.9184 |
| Grid2 | Q61625 | 0.27 | 0.27 | 0.32 | 0.38 | 1.04 | 0.46 | 0.56 | 0.5 | 0.4 | 0.19 | 0.43 | 0.51 | 0.51 | 0.56 | 0.26 | 0.4 | 0.33 | 0.0725 | 0.1225 | 0.096723 | 0.657556 | 0.487546 | | 1 | 0.996511 | 0.897232 |
| Grik1 | E9Q2Y1 | -1.39 | -2.54 | -3.52 | -3.88 | -2.21 | -0.72 | -3.96 | -4.17 | -1.44 | -1.28 | -4.4 | -1.15 | -2.28 | -1.35 | -4.33 | -3.68 | 0.0675 | 0.765 | -0.0775 | 0.939174 | 0.387433 | 0.921648 | | 1 | 0.996511 | 0.987433 |
| Grik2 | P39087 | 1.74 | 1.25 | 0.78 | 1.04 | 1.61 | 1.56 | 1.07 | 0.98 | 0.78 | 0.26 | 0.65 | 1.67 | 0.21 | 0.82 | 0.48 | 1.13 | 0.1025 | -0.3625 | -0.5425 | 0.702336 | 0.31941 | 0.096853 | | 1 | 0.996511 | 0.698156 |
| Grik3 | B1AS29 | 0.62 | 0.69 | 0.5 | 0.42 | 1.17 | 0.71 | 0.92 | 0.49 | 0.75 | -0.01 | 0.44 | 0.64 | 0.81 | 1.44 | 0.39 | 0.49 | 0.265 | -0.1025 | 0.225 | 0.195109 | 0.638924 | 0.41291 | | 1 | 0.996511 | 0.874811 |
| Grik5 | Q61626 | 1.43 | 0.81 | 0.65 | 0.73 | 1.04 | 1.38 | 1.04 | 0.69 | 0.84 | -0.55 | 0.71 | 1.27 | 1.05 | 0.9 | 0.24 | 0.65 | 0.1325 | -0.3375 | -0.195 | 0.587092 | 0.419225 | 0.48312 | | 1 | 0.996511 | 0.893925 |
| Grin1 | A2AI14 | -0.25 | 0.87 | -0.27 | 0.41 | 1.02 | 0.39 | 0.7 | 0.2 | 0.97 | -0.71 | 0.13 | 0.31 | -0.1 | 1.57 | 0.19 | 0.33 | 0.3875 | -0.015 | 0.3075 | 0.248881 | 0.971487 | 0.488176 | | 1 | 0.997515 | 0.897531 |
| Grin1 | A2AI21 | 1.15 | 1.12 | 1.41 | 1.03 | 1.44 | 1.32 | 1.27 | 1.06 | 1.12 | 0.28 | 1.38 | 1.03 | 1.34 | 1.12 | 1.21 | 1.07 | 0.095 | -0.225 | 0.0075 | 0.575703 | 0.408674 | 0.967603 | | 1 | 0.996511 | 0.9927 |
| Grin2a | P35436 | 1.14 | 1.08 | 1.56 | 1.13 | 1.31 | 1.18 | 1.19 | 1.42 | 1.2 | 0.62 | 1.68 | 1.09 | 0.93 | 1.15 | 1.09 | 1.31 | 0.0475 | -0.08 | -0.1075 | 0.785654 | 0.761223 | 0.597086 | | 1 | 0.996511 | 0.929911 |
| Grin2b | G3X9V4 | 0.96 | 0.95 | 0.81 | 0.75 | 1.22 | 1.3 | 1.01 | 0.98 | 0.53 | 0.04 | 1.11 | 1.23 | 0.54 | 0.16 | 0.66 | 0.88 | 0.26 | -0.14 | -0.3075 | 0.127985 | 0.630788 | 0.176133 | | 1 | 0.996511 | 0.771503 |
| Grin2d | Q03391 | 0.59 | 0.97 | 1.14 | 0.69 | 0.99 | 0.74 | 0.78 | 0.93 | 1.22 | 2.2 | 0.98 | 0.48 | 1.5 | 1.61 | 1.21 | 1.42 | 0.0125 | 0.3725 | 0.5875 | 0.94546 | 0.329469 | 0.019741 | | 1 | 0.996511 | 0.615793 |
| Grin3a | A2AIR5-2 | 0.73 | 1.2 | 0.82 | 0.86 | 1.77 | 1.36 | 1.21 | 1.29 | 1.73 | 1.51 | 1.39 | 1.13 | 1.63 | 2.23 | 0.98 | 1.16 | 0.505 | 0.5375 | 0.5975 | 0.028653 | 0.02733 | 0.079196 | | 1 | 0.996511 | 0.681114 |
| Grip1 | D3YZL8 | 1.28 | 0.99 | 0.63 | 0.84 | 1.11 | 0.84 | 0.74 | 0.65 | 1.3 | 0.28 | 0.86 | 1.38 | 1.24 | 1.21 | 0.5 | 0.92 | -0.1 | 0.02 | 0.0325 | 0.621842 | 0.945813 | 0.897382 | | 1 | 0.996755 | 0.983509 |
| Grip2 | G3XA20 | 0.77 | 0.68 | 0.98 | 1.25 | 0.67 | 0.56 | 1.05 | 0.95 | 0.74 | 0.36 | 0.95 | 0.86 | 0.61 | 0.39 | 0.85 | 0.89 | -0.1125 | -0.1925 | -0.235 | 0.582375 | 0.390595 | 0.305036 | | 1 | 0.996511 | 0.831662 |
| Gripap1 | A2AEW8 | 0.85 | 0.58 | 0.28 | 0.28 | 0.25 | 0.69 | 0.03 | 0.25 | -0.01 | 0.29 | 0.36 | 0.76 | -0.1 | 0.12 | 0.04 | 0.1 | -0.1925 | -0.1475 | -0.4575 | 0.390002 | 0.539936 | 0.05088 | | 1 | 0.996511 | 0.658454 |
| Gripap1 | Q8VD04 | -0.57 | -0.77 | 0.79 | 0.09 | -0.52 | -0.82 | 1.48 | 0.65 | 0 | 0.37 | 0.11 | -0.19 | 0.07 | -0.22 | 1.07 | 1.29 | 0.3125 | 0.1875 | 0.6675 | 0.595201 | 0.607222 | 0.187645 | | 1 | 0.996511 | 0.776843 |
| Gripap1 | A2AEW6 | 0.74 | 0.55 | 2.01 | 0.29 | 0.8 | 0.63 | 2.89 | 0.82 | 0.52 | -0.57 | 1.93 | 1.21 | -1.58 | -0.46 | 0.48 | 2.36 | 0.3875 | -0.125 | -0.6975 | 0.52399 | 0.835155 | 0.408149 | | 1 | 0.996511 | 0.874356 |
| Grk2 | Q99MK8 | 0.01 | -0.44 | -0.26 | 0.03 | -0.39 | 0.01 | 0.04 | 0.21 | -0.38 | -0.09 | -0.09 | -0.32 | -0.55 | -0.12 | -0.5 | -0.58 | 0.1325 | -0.055 | -0.2725 | 0.517148 | 0.774704 | 0.220857 | | 1 | 0.996511 | 0.787447 |
| Grk5 | Q8VEB1 | -0.34 | -0.08 | -0.3 | 0.03 | -0.55 | 0.16 | 0.02 | -0.3 | -0.05 | 0.16 | -0.56 | 0.33 | 0.02 | 1.09 | -1.2 | -0.47 | 0.005 | 0.1425 | 0.0325 | 0.981023 | 0.556495 | 0.943663 | | 1 | 0.996511 | 0.991609 |
| Grk6 | O70293-2 | 0.32 | -0.09 | 0.62 | 0.66 | 0.02 | 0.52 | 0.66 | 0.81 | 0.38 | 0.84 | 0.37 | 0.47 | 0.59 | 0.52 | 0.52 | 0.57 | 0.125 | 0.1375 | 0.1725 | 0.625453 | 0.562786 | 0.44743 | | 1 | 0.996511 | 0.884817 |
| Grm1 | P97772 | 0.25 | 0.53 | 0.82 | 0.66 | 0.84 | 0.38 | 1 | 0.72 | -0.35 | 0.48 | 0.69 | 0.36 | -0.46 | 0.34 | 0.7 | 0.88 | 0.17 | -0.27 | -0.2 | 0.422083 | 0.333342 | 0.5431 | | 1 | 0.996511 | 0.912212 |
| Grm2 | Q14BI2 | 1.57 | 1.32 | 1.5 | 1.3 | 1.38 | 1.6 | 1.01 | 0.92 | 0.86 | 0.9 | 1.16 | 1.32 | 0.45 | 1.06 | 0.65 | 0.97 | -0.195 | -0.3625 | -0.64 | 0.351407 | 0.08014 | 0.014009 | | 1 | 0.996511 | 0.589387 |
| Grm3 | Q9QYS2 | 0.64 | 0.83 | 1.16 | 0.99 | 1.08 | 0.73 | 0.52 | 0.65 | 0.44 | 0.78 | 0.88 | 0.49 | 0.41 | 0.57 | 0.65 | 0.58 | -0.16 | -0.2575 | -0.3525 | 0.42718 | 0.2252 | 0.097332 | | 1 | 0.996511 | 0.698156 |
| Grm4 | A0A338P7I | 0.81 | 0.56 | 0.41 | 0.56 | 0.45 | 0.32 | 0.56 | 0.48 | 0.52 | 0.29 | 0.39 | 0.65 | -0.1 | 0.56 | 0.07 | 0.01 | -0.1325 | -0.1225 | -0.45 | 0.416114 | 0.502322 | 0.066105 | | 1 | 0.996511 | 0.664936 |
| Grm5 | Q3UVX5 | 1.07 | 0.77 | 0.74 | 0.75 | 0.77 | 0.96 | 0.64 | 0.58 | 0.63 | 0.8 | 0.73 | 1.05 | 0.46 | 0.57 | 0.92 | 0.51 | -0.095 | -0.03 | -0.2175 | 0.577931 | 0.869777 | 0.291 | | 1 | 0.996511 | 0.825497 |
| Grm7 | Q68ED2 | 0.62 | 0.75 | 0.53 | 0.78 | 0.69 | 0.57 | 0.65 | 0.73 | 0.87 | 0.69 | 0.6 | 0.74 | 1.01 | 1.04 | 0.51 | 0.46 | -0.01 | 0.055 | 0.085 | 0.945848 | 0.740404 | 0.698607 | | 1 | 0.996511 | 0.955729 |
| Grm8 | Q05BD6 | 0.93 | 0.74 | 0.78 | 1.3 | 0.91 | 0.99 | 0.78 | 1.09 | 1.46 | 0.61 | 0.96 | 0.84 | 1.03 | 0.89 | 1.07 | 0.84 | 0.005 | 0.03 | 0.02 | 0.978428 | 0.902647 | 0.921483 | | 1 | 0.996609 | 0.987433 |
| Grm8 | P47743 | 2.17 | 5.19 | 0.43 | 2.59 | 2.65 | 2.2 | 3.08 | 2.22 | 2.9 | 3.3 | 2.32 | 4.24 | 1.77 | 4.05 | 2.1 | 2.65 | -0.0575 | 0.595 | 0.0475 | 0.949213 | 0.538258 | 0.961625 | | 1 | 0.996511 | 0.9927 |
| Grn | P28798 | -3.97 | -4.71 | -7.09 | -6.37 | -4 | -3.69 | -6.49 | -7.7 | -3.45 | -3.01 | -6.6 | -3.08 | -3.87 | -3.61 | -6.49 | -7.53 | 0.065 | 1.5 | 0.16 | 0.952168 | 0.162777 | 0.882023 | | 1 | 0.996511 | 0.979767 |
| Grp | Q8R1I2 | -0.83 | 0.72 | 0.06 | 0.21 | 0.13 | 0.25 | 0.76 | 0.58 | -0.82 | 1.11 | 0.35 | 0.36 | 0.95 | 0.46 | 1.59 | 1.67 | 0.39 | 0.21 | 1.1275 | 0.272083 | 0.662655 | 0.022022 | | 1 | 0.996511 | 0.621926 |
| Grpel1 | Q99LP6 | 0.26 | 0.28 | 0.41 | 0.55 | -0.46 | 0.57 | -0.07 | 0.46 | 1.31 | -0.17 | 0.55 | 0.12 | 2.23 | -0.28 | 0.4 | 0.2 | -0.25 | 0.0775 | 0.2625 | 0.346778 | 0.812933 | 0.61254 | | 1 | 0.996511 | 0.935166 |
| Grpel2 | O88396 | 0.11 | -0.01 | -0.41 | 0 | 0.81 | 0 | 0.45 | 0.25 | 0.31 | -0.16 | 0.46 | -0.63 | 0.67 | 0.31 | -0.9 | 1.82 | 0.455 | 0.0725 | 0.5525 | 0.071131 | 0.797994 | 0.312328 | | 1 | 0.996511 | 0.834523 |
| Grsf1 | E9Q179 | 0.14 | 0.39 | -0.18 | 0.13 | 0.28 | 0.18 | 0.1 | -0.1 | 0.59 | 0.27 | 0.09 | 0.2 | 0.33 | 0.66 | -0.25 | -0.3 | -0.005 | 0.1675 | -0.01 | 0.978315 | 0.422762 | 0.971528 | | 1 | 0.996511 | 0.992865 |
| Grtp1 | Q9D3N8-4 | 0.25 | 0.74 | 0.91 | 0.56 | 1.08 | 0.25 | 0.36 | 0.12 | 1.09 | 0.53 | -0.14 | 0.56 | 0.62 | 1.27 | 0.34 | 0.5 | -0.1625 | -0.105 | 0.0675 | 0.54336 | 0.723498 | 0.804105 | | 1 | 0.996511 | 0.971404 |
| Grwd1 | Q810D6 | -1.24 | -1.15 | 0.12 | -0.23 | -0.59 | -0.63 | -0.45 | -0.24 | -0.69 | -0.08 | -0.33 | -1.05 | -0.34 | -0.21 | -0.3 | -0.27 | 0.1475 | 0.0875 | 0.345 | 0.663539 | 0.819575 | 0.323581 | | 1 | 0.996511 | 0.841404 |
| Gsdme | Q9Z2D3 | -0.05 | 0.1 | -0.36 | -0.08 | 0.24 | -0.23 | 0.05 | -0.28 | -0.52 | -0.41 | -0.29 | 0.31 | -0.68 | -1.12 | -0.53 | -0.1 | 0.0425 | -0.13 | -0.51 | 0.82539 | 0.587029 | 0.076669 | | 1 | 0.996511 | 0.675786 |
| Gse1 | Q3U3C9-4 | -3.4 | -3.82 | 0.11 | -0.66 | -3.26 | -2.93 | -0.62 | 0.05 | -3.24 | 0.32 | -0.44 | -4.83 | -1.21 | -3.74 | -0.76 | 0.11 | 0.2525 | -0.105 | 0.5425 | 0.825774 | 0.939554 | 0.637023 | | 1 | 0.996755 | 0.942665 |
| Gsg1l | D3Z7H4 | 0.82 | 1.3 | 1.81 | 1.9 | 1.44 | 1.06 | 1.56 | 1.68 | 0.84 | 1.34 | 1.44 | 0.83 | 0.78 | 1.04 | 1.21 | 1.3 | -0.0225 | -0.345 | -0.375 | 0.937113 | 0.272941 | 0.219655 | | 1 | 0.996511 | 0.787447 |
| Gsk3a | Q2NL51 | -1.11 | -1.09 | -0.84 | -0.33 | -0.63 | -0.71 | -0.39 | -0.33 | -1.32 | -0.2 | -0.48 | -1.04 | -1.32 | -1.11 | -0.7 | -0.74 | 0.3275 | 0.0825 | -0.125 | 0.170515 | 0.793699 | 0.636404 | | 1 | 0.996511 | 0.942536 |
| Gsk3b | E9QAQ5 | -0.27 | -0.55 | -0.55 | -0.64 | -0.44 | -0.15 | -0.71 | -0.39 | -0.48 | -0.35 | -0.49 | -0.31 | -0.97 | -0.5 | -0.97 | -0.8 | 0.08 | 0.095 | -0.3075 | 0.663853 | 0.580047 | 0.153746 | | 1 | 0.996511 | 0.750763 |
| Gskip | A0A1Y7VN: | -1.68 | -1.07 | -0.91 | -0.85 | -1.48 | -0.74 | -0.92 | -1.13 | -1.3 | -0.87 | -0.93 | -1.36 | -1.12 | -0.58 | -1.02 | -0.97 | 0.06 | 0.0125 | 0.205 | 0.815754 | 0.960177 | 0.430265 | | 1 | 0.996755 | 0.879165 |
| Gsn | P13020-2 | 0.07 | -0.05 | -0.04 | -0.31 | -0.22 | -0.31 | -0.5 | -0.46 | 0.35 | 0.16 | -0.07 | 0.47 | 0.3 | -0.06 | 0.38 | -0.29 | -0.29 | 0.31 | 0.165 | 0.102922 | 0.139678 | 0.469093 | | 1 | 0.996511 | 0.890107 |
| Gsn | P13020 | -0.3 | -0.4 | 0.07 | -0.16 | -0.22 | -0.5 | 0.65 | -0.18 | 0.28 | 0.93 | 0.11 | 0.05 | -0.22 | -0.16 | 0.12 | -0.15 | 0.135 | 0.54 | 0.095 | 0.623279 | 0.055298 | 0.63162 | | 1 | 0.996511 | 0.941132 |
| Gspt1 | Q8R050-2 | 0.76 | 0.44 | 0.05 | -0.24 | 0.7 | 0.43 | 0.02 | -0.01 | 0.35 | 0.04 | 0.07 | 0.85 | 0.01 | 0.43 | 0.03 | -0.17 | 0.0325 | 0.075 | -0.1775 | 0.90776 | 0.799967 | 0.524233 | | 1 | 0.996511 | 0.908107 |
| Gspt2 | Q149F3 | 0.03 | 0.13 | 0.24 | 0.42 | 0.05 | -0.26 | 0.45 | 0.37 | -0.3 | 0.62 | 0.57 | 0.18 | 0.07 | -0.6 | 0.46 | 0.17 | -0.0525 | 0.0625 | -0.18 | 0.803167 | 0.803993 | 0.503714 | | 1 | 0.996511 | 0.901002 |
| Gsr | P47791-2 | 0.65 | 0.19 | -0.01 | -0.43 | 0.18 | 1.06 | 0.16 | -0.13 | -0.54 | -0.69 | -0.12 | 0.37 | -0.06 | 0.01 | -0.11 | -0.14 | 0.2175 | -0.345 | -1.75E-01 | 0.517082 | 0.305703 | 0.501671 | | 1 | 0.996511 | 0.901002 |
| Gss | P51855 | 0.47 | 0.18 | 1.25 | 0.88 | 0.31 | 0.11 | 1.21 | 1.11 | 0.09 | 0.99 | 1.06 | 0.26 | -0.18 | -0.14 | 0.95 | 1.01 | -0.01 | -0.095 | -0.285 | 0.977155 | 0.778477 | 0.473336 | | 1 | 0.996511 | 0.891467 |
| Gsta4 | P24472 | -0.03 | -0.5 | 0.01 | -0.17 | -0.42 | -0.63 | 0.56 | 0.17 | -0.57 | 0.06 | -0.36 | -0.53 | -0.93 | -0.37 | -0.42 | -0.12 | 0.0925 | -0.1775 | -0.2875 | 0.754305 | 0.43144 | 0.255728 | | 1 | 0.996511 | 0.809552 |
| Gsta4 | A0A1L1SS6 | -0.7 | 0.71 | -0.46 | -0.05 | 2.59 | 0.78 | -0.67 | -1.75 | 0.78 | -1.33 | -0.69 | 1.29 | -0.71 | 0.25 | -0.92 | 0.59 | 0.3625 | 0.1375 | -0.0725 | 0.684997 | 0.826763 | 0.872593 | | 1 | 0.996511 | 0.979626 |
| Gstk1 | Q9DCM2 | 1.49 | 1.86 | 1.24 | 1.37 | 1.75 | 1.72 | 2.37 | 1.79 | 0.41 | 0.62 | 1.46 | 1.34 | 0.47 | 0.48 | 1.14 | 1.4 | 0.4175 | -0.5325 | -0.6175 | 0.091817 | 0.102422 | 0.056961 | | 1 | 0.996511 | 0.661738 |
| Gstm1 | P10649 | 1.3 | 0.07 | 0.33 | 1.08 | 0.15 | 1.22 | 1.53 | 1.15 | 0.7 | -0.05 | 0.34 | 0.39 | -0.2 | 0.1 | -0.19 | 0.6 | 0.3175 | -0.35 | -0.6175 | 0.431623 | 0.304727 | 0.102772 | | 1 | 0.996511 | 0.700625 |
| Gstm2 | P15626 | -1.32 | -1.92 | -1.5 | -1.2 | -2.13 | -1.71 | -1.65 | -1.3 | -2.5 | 0.06 | -1.8 | -1.75 | -2.01 | -1.81 | -1.81 | -1.88 | -0.2125 | -0.0125 | -0.3925 | 0.397318 | 0.981042 | 0.098269 | | 1 | 0.998773 | 0.698156 |
| Gstm4 | Q8R5I6 | 0.94 | 1.15 | 1.09 | 0.27 | 0.51 | 0.96 | 1.37 | 1.31 | 0.21 | 0.05 | 1.14 | 0.87 | 0.7 | 1.06 | 1.59 | 1.47 | 0.175 | -0.295 | 0.3425 | 0.543084 | 0.380251 | 0.271362 | | 1 | 0.996511 | 0.822084 |
| Gstm5 | P48774 | -0.13 | -0.73 | -0.61 | -0.92 | -0.24 | 0.24 | -0.48 | -0.57 | -0.83 | -1.25 | -0.83 | -0.71 | -1.43 | -0.39 | -1.17 | -0.99 | 0.335 | -0.3075 | -0.3975 | 0.215124 | 0.213877 | 0.19843 | | 1 | 0.996511 | 0.78004 |
| Gstm7 | Q80W21 | 0.19 | 0.23 | 0.17 | -0.38 | 0.26 | -0.05 | -0.2 | -0.16 | -0.11 | 0.75 | -0.31 | -0.25 | -0.34 | 0.17 | 0.42 | 0.1 | -0.09 | -0.0325 | 0.035 | 0.666026 | 0.911895 | 0.888047 | | 1 | 0.996744 | 0.980575 |
| Gsto1 | O09131 | 0.19 | -0.07 | -0.36 | -0.74 | -0.18 | -0.14 | -0.5 | -0.57 | -0.12 | 0.33 | -0.44 | 0.25 | -0.41 | 0.12 | -0.39 | -0.75 | -0.1025 | 0.25 | -0.1125 | 0.673857 | 0.38257 | 0.695542 | | 1 | 0.996511 | 0.955729 |
| Gstp1 | P19157 | -1.9 | -1.88 | -0.51 | -0.38 | -2 | -2.28 | 0.33 | -0.44 | -2.22 | -1.21 | -0.68 | -2.03 | -2.35 | -2.04 | -1.13 | -1.13 | 0.07 | -0.3675 | -0.495 | 0.918126 | 0.479304 | 0.325688 | | 1 | 0.996511 | 0.843092 |
| Gstt1 | Q64471 | 0.28 | -2.82 | -0.59 | -0.51 | -2.66 | 1.58 | -0.51 | -0.78 | -2.81 | -1.4 | -1.54 | -0.62 | -2.75 | -2.32 | -1.33 | -1.55 | 0.3175 | -0.6825 | -1.0775 | 0.746981 | 0.363107 | 0.137781 | | 1 | 0.996511 | 0.733489 |
| Gstt2 | A0A1W2P7 | -1.5 | -0.27 | -3.51 | -1.72 | -1.18 | -2.7 | -2.36 | -2.65 | -0.61 | -1.32 | -2.15 | -2.21 | -0.96 | 1.28 | -2.49 | -1.31 | -0.4725 | 0.1775 | 0.88 | 0.496843 | 0.799566 | 0.354689 | | 1 | 0.996511 | 0.855029 |
| Gstz1 | Q9WVL0 | 0.16 | 0.05 | -0.37 | 0.52 | 0.24 | 0.43 | 0.06 | 0.56 | 0.22 | 1.51 | -0.13 | -0.23 | 0.47 | 0.43 | -0.01 | -0.1 | 0.2325 | 0.2525 | 0.1075 | 0.33076 | 0.550561 | 0.683723 | | 1 | 0.996511 | 0.955729 |
| Gtdc1 | G8JL42 | -1.18 | -1.15 | -0.64 | -0.98 | -0.15 | -1.32 | 0.28 | -0.86 | -0.94 | -0.38 | -1.49 | -0.51 | -1.02 | 0.03 | -1.27 | -0.52 | 0.475 | 0.1575 | 0.2925 | 0.211798 | 0.588577 | 0.372004 | | 1 | 0.996511 | 0.859424 |
| Gtf2a1 | Q99PM3 | -2.34 | -1.84 | -2.5 | -2.36 | -2.06 | -2.53 | -2.61 | -2.25 | -2.34 | -0.06 | -3.01 | -2.1 | -2.52 | -1.7 | -2.43 | -2.59 | -0.1025 | 0.3825 | -0.05 | 0.638879 | 0.527196 | 0.855387 | | 1 | 0.996511 | 0.976225 |
| Gtf2a2 | D3Z793 | -3.28 | -2.99 | -3.15 | -3.86 | -2.6 | -4.23 | -4.29 | -4.32 | -2.21 | -1.26 | -3.54 | -3.1 | -1.81 | -1.63 | -4.03 | -3.86 | -0.54 | 0.7925 | 0.4875 | 0.231673 | 0.140965 | 0.435944 | | 1 | 0.996511 | 0.88254 |
| Gtf2b | P62915 | -1.66 | -2.28 | -1.82 | -2.07 | -2.04 | -1.48 | -1.95 | -2.34 | -1.05 | -0.58 | -2.02 | -1.74 | -0.47 | -1.22 | -1.82 | -1.92 | 0.005 | 0.61 | 0.6 | 0.983371 | 0.107337 | 0.119647 | | 1 | 0.996511 | 0.721448 |
| Gtf2e1 | Q9D0D5 | -3.02 | -2.19 | -2.53 | -3.05 | -2.51 | -2.66 | -2.34 | -3.21 | -1.91 | -1.41 | -2.42 | -2.13 | -2.4 | -2.25 | -1.92 | -2.76 | 0.0175 | 0.73 | 0.365 | 0.950504 | 0.036908 | 0.22619 | | 1 | 0.996511 | 0.787447 |
| Gtf2e2 | Q9D902 | -3.38 | -3.59 | -1.42 | -2.2 | -2.9 | -2.27 | -1.97 | -1.51 | -2.78 | -0.95 | -1.42 | -3.22 | -3.26 | -2 | -1.69 | -2.7 | 0.485 | 0.555 | 0.235 | 0.380287 | 0.421611 | 0.68204 | | 1 | 0.996511 | 0.955729 |
| Gtf2f1 | Q3THK3 | -2.14 | -2.03 | -1.27 | -1.44 | -2.34 | -2.01 | -1.71 | -2 | -1.77 | -0.64 | -1.62 | -1.79 | -1.85 | -1.61 | -1.26 | -1.76 | -0.295 | 0.265 | 0.1 | 0.273671 | 0.447715 | 0.715902 | | 1 | 0.996511 | 0.958106 |
| Gtf2f2 | Q8R0A0 | -2.27 | -2.69 | -2.59 | -2.75 | -2.69 | -2.79 | -3.27 | -2.84 | -2.56 | -1.46 | -2.41 | -2.03 | -2.23 | -2.55 | -3.12 | -3.23 | -0.3225 | 0.46 | -0.2075 | 0.13366 | 0.127361 | 0.466254 | | 1 | 0.996511 | 0.888632 |
| Gtf2h1 | G3X8R4 | -4.94 | -5.68 | -4.79 | -7.02 | -4.07 | -5.1 | -5.61 | -6.65 | -4.21 | -3.19 | -3.97 | -3.54 | -3.92 | -3.49 | -3.46 | -5.26 | 0.25 | 1.88 | 1.575 | 0.710567 | 0.005462 | 0.027223 | | 1 | 0.996511 | 0.635993 |
| Gtf2h4 | O70422 | -2.71 | -2.26 | -1.6 | -2.29 | -2.07 | -3.24 | -2.31 | -2.54 | -2.89 | -1.98 | -2.05 | -2.33 | -2.33 | -2.25 | -1.93 | -1.92 | -0.325 | -0.0975 | 0.1075 | 0.340216 | 0.754708 | 0.697248 | | 1 | 0.996511 | 0.955729 |
| Gtf2h5 | Q8K2X8 | -0.44 | -0.39 | -1.87 | -2.54 | 0.32 | -0.63 | -2.25 | -1.12 | -1.09 | -1.93 | -2.12 | -0.38 | -0.78 | -0.03 | -2.57 | -0.93 | 0.39 | -0.07 | 0.2325 | 0.572908 | 0.908897 | 0.735622 | | 1 | 0.996744 | 0.958635 |
| Gtf2i | Q3UHU8 | 0.42 | -0.07 | -0.22 | 0.02 | -0.01 | 0.11 | -0.04 | 0.03 | 0.11 | 0.25 | -0.2 | 0.35 | -0.29 | 0.41 | -0.14 | -0.06 | -0.015 | 0.09 | -0.0575 | 0.934761 | 0.682956 | 0.812116 | | 1 | 0.996511 | 0.972642 |
| Gtf3c1 | A0A0U1RP | 0.21 | 0.06 | 0.1 | 0.29 | -0.06 | 0.22 | 0.28 | 0.26 | 0.4 | -0.13 | 0.26 | 0.44 | 0.61 | 0.26 | 0.35 | 0.24 | 0.01 | 0.0775 | 0.2 | 0.949753 | 0.690814 | 0.294794 | | 1 | 0.996511 | 0.825612 |

| | | | | | | | | | | | | | | | | | | | | | | | | | | |
|---|---|---|---|---|---|---|---|---|---|---|---|---|---|---|---|---|---|---|---|---|---|---|---|---|---|---|
| Gtf3c2 | D3Z7I0 | 2.03 | -0.04 | -0.05 | 0.52 | -0.09 | 1.9 | 0.24 | 0.03 | 0.61 | 1.08 | 0.15 | 1.49 | 1 | 1.74 | 0.18 | 0.01 | -0.095 | 0.2175 | 0.1175 | 0.877265 | 0.681188 | 0.839717 | 1 | 0.996511 | 0.976219 |
| Gtf3c3 | Q3TMP1 | -0.15 | -0.06 | -0.07 | -0.08 | -0.1 | -0.14 | 0.1 | -0.17 | 0.19 | -0.64 | -0.33 | 0.13 | 0.79 | 0.38 | -0.51 | -0.34 | 0.0125 | -0.0725 | 0.17 | 0.931818 | 0.753292 | 0.591579 | 1 | 0.996511 | 0.929755 |
| Gtf3c4 | Q8BMQ2 | -1.84 | -2.26 | 0.51 | 0.19 | -1.78 | -1.45 | 0.82 | 0.39 | -2.15 | -0.22 | 0.47 | -1.48 | -1.78 | -2.44 | 0.35 | 0.69 | 0.345 | 0.005 | 0.055 | 0.689751 | 0.995185 | 0.953098 | 1 | 0.999285 | 0.992476 |
| Gtf3c5 | Q8R2T8 | 1.05 | 0.36 | 0.36 | 0.23 | 0.47 | 0.74 | 0.21 | 0.15 | 0.34 | 0.28 | 0.16 | 0.92 | 1.17 | 0.48 | 0.86 | 0.42 | -0.1075 | -0.075 | 0.2325 | 0.661106 | 0.779711 | 0.411247 | 1 | 0.996511 | 0.874811 |
| Gtpbp1 | O08582 | -1.35 | -1 | -0.93 | -0.98 | -1.01 | -1.46 | -0.83 | -0.85 | 0.04 | 1.4 | -1.18 | -1.47 | 0.38 | -0.35 | -1.02 | -1.48 | 0.0275 | 0.7625 | 0.4475 | 0.893463 | 0.230054 | 0.282409 | 1 | 0.996511 | 0.822325 |
| Gtpbp10 | Q8K013 | 0.03 | 0.32 | 0.09 | 0.01 | -0.55 | -0.18 | -1.23 | 0.35 | 0.25 | 0.75 | 0.63 | -0.27 | 0.24 | -0.71 | -0.17 | -0.01 | -0.515 | 0.2275 | -0.275 | 0.145329 | 0.391928 | 0.284677 | 1 | 0.996511 | 0.822669 |
| Gtpbp2 | Q3UJK4 | -0.24 | 0.48 | 0.78 | 0.68 | 0.55 | 0.06 | -0.48 | 1.46 | 1.04 | 1.95 | 1.56 | 0.28 | 1.04 | 1.66 | 1.17 | 0.45 | -0.0275 | 0.7825 | 0.655 | 0.950144 | 0.082656 | 0.08072 | 1 | 0.996511 | 0.684354 |
| Gtpbp3 | E9PWV7 | 0.16 | 0.97 | 0.32 | 0.25 | 0.96 | 0.36 | 0.57 | 0.53 | 0.11 | -0.05 | 0.54 | 0.4 | 0.01 | 0.82 | 0.18 | 0.84 | 0.18 | -0.175 | 0.0375 | 0.460144 | 0.491965 | 0.899602 | 1 | 0.996511 | 0.983509 |
| Gtpbp4 | Q99ME9 | 0.72 | 1.07 | 0.33 | 0.66 | 0.94 | 0.48 | 0.32 | 0.19 | 0.34 | 0.13 | 0.55 | 0.68 | 0.11 | 0.96 | 0.77 | 0.51 | -0.2125 | -0.27 | -0.1075 | 0.385157 | 0.254352 | 0.686304 | 1 | 0.996511 | 0.955729 |
| Gtpbp6 | Q3U6U5 | -0.87 | 0.54 | -0.22 | 0.16 | 0.48 | 0.59 | 0.35 | 0.53 | 0.6 | 0.4 | 0.37 | -0.61 | 0.01 | 0.56 | 0.28 | 0.3 | 0.585 | 0.2875 | 0.385 | 0.080196 | 0.467095 | 0.256245 | 1 | 0.996511 | 0.810628 |
| Gtpbp8 | F6XXP5 | 0.43 | 0.79 | 0.18 | 0.11 | 0.31 | 0.22 | -0.2 | 0.1 | -0.03 | -0.42 | 0.19 | 0.46 | 0.06 | 1.34 | -0.4 | 0.35 | -0.27 | -0.3275 | -0.04 | 0.230931 | 0.229744 | 0.917635 | 1 | 0.996511 | 0.986868 |
| Gucy1a1 | Q9ERL9 | 0.31 | -0.21 | -0.6 | -0.51 | 0.21 | -0.44 | -0.02 | -0.31 | -0.72 | -0.53 | -0.57 | 0.24 | -0.75 | -0.37 | -0.3 | -0.06 | 0.1125 | -0.1425 | -0.1175 | 0.667388 | 0.639858 | 0.668762 | 1 | 0.996511 | 0.952359 |
| Gucy1a2 | F8VQK3 | 1.07 | 0.15 | 0.3 | 0.36 | 0.52 | 1.18 | 0.77 | 0.35 | 0.2 | -0.37 | 0.31 | 0.7 | -0.39 | 0.08 | -0.12 | 0.46 | 0.235 | -0.26 | -0.4625 | 0.405934 | 0.405097 | 0.136171 | 1 | 0.996511 | 0.731656 |
| Gucy1b1 | O54865 | 0.39 | 0.02 | 0.43 | 0.51 | 0.18 | 0.38 | 0.35 | 0.44 | 0.36 | 0.64 | 0.28 | 0.25 | -0.05 | -0.01 | 0.19 | 0.35 | 1.96E-17 | 0.045 | -0.2175 | 1 | 0.816898 | 0.305143 | 1 | 0.996511 | 0.831662 |
| Guf1 | Q8C3X4 | 0.17 | 0.3 | 0.33 | 0.62 | 0.01 | 0.31 | 0.36 | 0.51 | 0.32 | 0.21 | 0.5 | 0.17 | 0.31 | -0.21 | -0.01 | -0.03 | -0.0575 | -0.055 | -0.34 | 0.75503 | 0.764558 | 0.125379 | 1 | 0.996511 | 0.721448 |
| Guk1 | Q64520-2 | -0.31 | -0.17 | 0.16 | 0.31 | -0.37 | -0.3 | 0.23 | 0.27 | -0.13 | 0.14 | 0.15 | -0.27 | -0.42 | 0.36 | 0.1 | -0.21 | -0.04 | -0.025 | -0.04 | 0.866713 | 0.907912 | 0.875258 | 1 | 0.996744 | 0.979626 |
| Gulp1 | E0CXC5 | 1.68 | 1.1 | 0.04 | -0.06 | 0.96 | 1.45 | 0.48 | 0.9 | 0.16 | 0.28 | -0.05 | 0.96 | 1.13 | -0.24 | 0.12 | 0.21 | 0.2575 | -0.3525 | -0.385 | 0.558138 | 0.438826 | 0.431519 | 1 | 0.996511 | 0.879165 |
| Gusb | P12265 | -0.17 | -0.05 | -0.06 | 0.25 | 0.87 | 0.19 | 1.16 | 0.05 | -1.53 | -0.64 | 0.44 | 0.07 | -1.47 | -0.9 | -0.34 | 0.1 | 0.575 | -0.4075 | -0.645 | 0.069105 | 0.346096 | 0.092746 | 1 | 0.996511 | 0.695968 |
| Gvin1 | L7N451 | -1.55 | -2.53 | 0.98 | 0.95 | -2.4 | -3.23 | 0.38 | 0.52 | -1.16 | 2.18 | 0.46 | -1.57 | 0.63 | -2.69 | 0.97 | 1.94 | -0.645 | 0.515 | 0.75 | 0.584856 | 0.643636 | 0.535596 | 1 | 0.996511 | 0.912212 |
| Gyg1 | Q9R062 | 1.13 | 1.76 | 0.45 | 0.66 | 2.18 | 1.24 | 0.54 | 0.89 | -0.09 | 0.27 | 0.7 | 2.42 | 0.17 | 0.81 | 0.8 | 0.42 | 0.2125 | -0.175 | -0.45 | 0.621598 | 0.76184 | 0.198829 | 1 | 0.996511 | 0.78004 |
| Gypa | P14220 | 0.77 | -0.71 | -0.27 | 1.92 | -0.57 | 1.98 | 1.68 | 2.64 | 0.96 | -0.08 | 1.28 | -2.38 | 0.45 | 1.3 | -0.15 | -0.2 | 1.005 | -0.4825 | -0.0775 | 0.241354 | 0.600134 | 0.901041 | 1 | 0.996511 | 0.983757 |
| Gys1 | A0A1B0GT | -0.08 | -0.19 | 0.46 | 0.52 | -0.05 | 0.06 | 0.81 | 0.64 | -1.13 | -0.44 | 0.33 | -0.32 | -0.72 | -1.07 | 0.2 | 0.56 | 0.1875 | -0.5675 | -0.435 | 0.511501 | 0.126857 | 0.301121 | 1 | 0.996511 | 0.830276 |
| H1-0 | P10922 | 2.88 | 1.99 | 3.07 | 2.05 | 1.81 | 2.48 | 2.38 | 2.49 | 0.87 | -0.13 | 2.61 | 1.84 | 1.57 | -0.88 | 2.26 | 2.54 | -0.2075 | -1.2 | -1.125 | 0.51674 | 0.073005 | 0.157857 | 1 | 0.996511 | 0.751678 |
| H1-1 | P43275 | 1 | 0.16 | 0.54 | 0.76 | -0.13 | 0.59 | 0.65 | 0.37 | 2.04 | 0.07 | 0.62 | 0.68 | 2.38 | 0.97 | 1.26 | 0.27 | -0.245 | 0.2375 | 0.605 | 0.359115 | 0.584574 | 0.200143 | 1 | 0.996511 | 0.78004 |
| H1-2 | P15864 | 2.68 | 0.65 | 2.15 | 1.74 | -0.84 | 2.32 | 0.79 | 1.12 | 2.71 | -0.9 | 1.65 | 2.21 | 3.4 | -1.78 | 2.7 | 1.17 | -0.9575 | -0.3875 | -0.4325 | 0.199162 | 0.639513 | 0.694081 | 1 | 0.996511 | 0.955729 |
| H13 | A3KGR9 | -0.31 | -0.2 | -0.58 | -0.79 | 0.65 | -0.19 | -0.25 | -0.48 | -0.54 | -1.02 | -0.51 | 0.06 | 0.02 | -0.08 | -0.26 | -0.06 | 0.4025 | -0.0325 | 0.375 | 0.1797 | 0.905062 | 0.097034 | 1 | 0.996744 | 0.698156 |
| H1-3 | P43277 | 2.1 | 1.41 | 2.14 | 2.04 | 1.43 | 1.38 | 1.72 | 2.04 | 2.01 | 0.36 | 1.95 | 1.5 | 1.94 | 1.46 | 2.49 | 1.76 | -0.28 | -0.4675 | -0.01 | 0.269506 | 0.262402 | 0.972685 | 1 | 0.996511 | 0.992865 |
| H1-4 | P43274 | 1.31 | 0.15 | 1.56 | 2.18 | -0.01 | 0.6 | 1.3 | 1.84 | 1.93 | 0.41 | 1.65 | 0.92 | 3.15 | 0.38 | 2.42 | 1.65 | -0.3675 | -0.0725 | 0.6 | 0.500375 | 0.886583 | 0.377025 | 1 | 0.996511 | 0.860984 |
| H1-5 | P43276 | 1.86 | 1.07 | 1.81 | 1.89 | 1.13 | 0.87 | 1.21 | 1.33 | 1.26 | 0.96 | 1.59 | 1.56 | 1.16 | 1.43 | 1.81 | 1.7 | -0.5225 | -0.315 | -0.1325 | 0.051306 | 0.251967 | 0.625098 | 1 | 0.996511 | 0.939193 |
| H1f10 | Q80ZM5 | 1.43 | -0.22 | -0.3 | 0.4 | -1.5 | 1.34 | -0.89 | 0.19 | 2.33 | -1.39 | 0.64 | -0.04 | 3.23 | -0.92 | 0.49 | -0.03 | -0.5425 | 0.0575 | 0.365 | 0.429093 | 0.941738 | 0.679523 | 1 | 0.996755 | 0.955402 |
| H2ac21 | Q64522 | 12.49 | 11.3 | 9.96 | 4.84 | 10.53 | 11.7 | 9.39 | 9.4 | -0.91 | 0.97 | 9.81 | 12.56 | -1.29 | 1.2 | 9.97 | 11.87 | 0.6075 | -4.04 | -4.21 | 0.701821 | 0.239903 | 0.213201 | 1 | 0.996511 | 0.786931 |
| H2ax | P27661 | 2.49 | 0.46 | 1.15 | 1.26 | -0.48 | 2.18 | 0.8 | 1.05 | 3.61 | -2.04 | 0.98 | 2.15 | 3.98 | -2.05 | 0.93 | 1.06 | -0.4525 | -0.165 | -0.36 | 0.477889 | 0.884795 | 0.756818 | 1 | 0.996511 | 0.962333 |
| H2az2 | Q3THW5 | 1.96 | 0.65 | 0.97 | 1.02 | 0.79 | 1.27 | 1.35 | 0.82 | 0.9 | 0.8 | 0.74 | 1.85 | 0.61 | 1.06 | 1.35 | 1.12 | -0.0925 | -0.0775 | -0.115 | 0.767708 | 0.835099 | 0.725297 | 1 | 0.996511 | 0.958398 |
| H2bc15 | P10853 | 2.54 | 1.23 | 1.72 | 1.45 | 1.02 | 1.89 | 1.12 | 1.58 | 1.32 | 1.73 | 1.53 | 1.89 | 0.94 | 0.54 | 1.42 | 1.72 | -0.3325 | -0.1175 | -0.58 | 0.34118 | 0.709094 | 0.150588 | 1 | 0.996511 | 0.749304 |
| H2bu2 | Q9D2U9 | 1.97 | 1.41 | 1.26 | 1.4 | 0.72 | 1.33 | 0.82 | 0.59 | 1.21 | -1.44 | 0.56 | 1.22 | 1.47 | -0.19 | 0.96 | 1.47 | -0.645 | -1.1225 | -0.5825 | 0.024113 | 0.086512 | 0.175654 | 1 | 0.996511 | 0.771152 |
| H2-D1 | P01899 | -0.3 | -0.41 | -0.3 | 0.15 | -0.74 | -1.52 | -0.49 | -0.41 | -0.5 | 0.63 | -0.54 | -0.95 | 0.33 | -0.35 | -0.5 | 0.47 | -0.575 | -0.125 | 0.2025 | 0.069601 | 0.724126 | 0.489678 | 1 | 0.996511 | 0.897531 |
| H2-K1 | A0A494BA | -0.92 | -0.3 | -1.07 | -0.49 | 0.4 | -1.05 | -1.84 | -1.5 | -0.15 | 1.28 | -1.16 | -0.18 | 0.58 | 1.77 | -1.25 | -0.72 | -0.3025 | 0.6425 | 0.79 | 0.537796 | 0.217956 | 0.237444 | 1 | 0.996511 | 0.792722 |
| H2-Q10 | P01898 | -2.57 | -2.4 | -2.11 | -2.83 | -2.01 | -2.7 | -2.84 | -2.64 | -2.22 | -1.02 | -2.57 | -2.88 | -1.59 | -1.82 | -2.21 | -2.5 | -0.07 | 0.305 | 0.4475 | 0.779895 | 0.467543 | 0.130495 | 1 | 0.996511 | 0.724318 |
| H3c14 | A0A1W2P7 | 1.38 | 0.86 | 1.57 | 1.63 | 0.72 | 0.51 | 1.13 | 1.07 | 1.64 | 0.94 | 1.17 | 0.67 | 1.63 | 1 | 1.32 | 1.49 | -0.5025 | -0.255 | -3.5E-16 | 0.064714 | 0.376775 | 1 | 1 | 0.996511 | 1 |
| H3f3a | F8WI35 | 1.62 | 0.55 | 1.25 | 0.98 | -1.32 | 1.72 | -1.6 | -0.56 | 4.1 | -2.51 | 0.34 | -0.3 | 4.92 | -1.85 | 0.43 | 1.06 | -1.54 | -0.6925 | 0.04 | 0.055477 | 0.581675 | 0.974803 | 1 | 0.996511 | 0.992977 |
| H4f16 | P62806 | 2.24 | 1.74 | 1.2 | 2.05 | 1.05 | 1.71 | 1.68 | 2.19 | 1.83 | 0.74 | 1.72 | 2.14 | 1.19 | 1.26 | 0.87 | 0.99 | -0.15 | -0.2 | -0.73 | 0.640225 | 0.589058 | 0.022578 | 1 | 0.996511 | 0.621926 |
| H6pd | Q8CFX1 | 0.08 | 0.55 | -0.33 | -0.59 | 0.53 | -0.09 | 0.12 | -0.22 | 0.45 | 0.1 | -0.41 | 0.26 | 0.71 | 1.44 | 0.02 | -0.24 | 0.1575 | 0.1725 | 0.555 | 0.598565 | 0.584033 | 0.218713 | 1 | 0.996511 | 0.787447 |
| Habp4 | E9QKB2 | -2.24 | -2.42 | -1.33 | -1.52 | -1.99 | -2.19 | -0.93 | -1.53 | -2.11 | -0.89 | -1.35 | -1.83 | -0.9 | -2.37 | -0.66 | -0.92 | 0.2175 | 0.3325 | 0.665 | 0.559099 | 0.377153 | 0.162522 | 1 | 0.996511 | 0.757516 |
| Hacd2 | Q9D3B1 | 0.18 | -0.28 | -0.26 | -0.72 | -1.32 | 1.27 | -0.77 | -1.08 | 3.22 | -2.42 | -0.67 | 0.21 | 4.67 | -1.92 | 1.02 | -0.13 | -0.205 | 0.355 | 1.18 | 0.718727 | 0.740856 | 0.357676 | 1 | 0.996511 | 0.856624 |
| Hacd3 | Q8K2C9 | 0.78 | 0.1 | 0.66 | 0.82 | 0.6 | 1.05 | 2.15 | 0.9 | 0.54 | 0.56 | 0.56 | 0.28 | 0.89 | 0.39 | 1.26 | 1.23 | 0.585 | -0.105 | 0.3525 | 0.132245 | 0.632944 | 0.232118 | 1 | 0.996511 | 0.788863 |
| Hace1 | Q3U0D9 | 0.25 | 0.33 | 0.32 | -0.15 | 0.43 | 0.16 | 0.08 | 0.15 | -0.09 | -0.58 | 0.04 | 0.18 | -0.56 | 0.36 | -0.43 | 0.12 | 0.0175 | -0.3 | -0.315 | 0.923164 | 0.217945 | 0.264108 | 1 | 0.996511 | 0.816045 |
| Hacl1 | D3YZ54 | 0.12 | -0.41 | 0.07 | 0.68 | -0.47 | 0.73 | 1.43 | 0.28 | 0.34 | 1.08 | 0.68 | 0.73 | -0.4 | 0.32 | 0.16 | 0.13 | 0.3775 | 0.5925 | -0.0625 | 0.38823 | 0.060828 | 0.829073 | 1 | 0.996511 | 0.973939 |
| Hadh | Q61425 | 2.39 | 1.71 | 1.28 | 1.83 | 1.59 | 2.18 | 1.43 | 1.58 | 1.29 | 1.72 | 1.58 | 1.99 | 0.98 | 0.14 | 0.86 | 0.85 | -0.1075 | -0.1575 | -1.095 | 0.706257 | 0.580526 | 0.006047 | 1 | 0.996511 | 0.552614 |
| Hadha | Q8BMS1 | 0.34 | 0.27 | 0.89 | 1.11 | 0.19 | 0.16 | 1.08 | 0.85 | 0.29 | 1.16 | 1.01 | 0.19 | -0.2 | 0.29 | 0.47 | 0.66 | -0.0825 | 0.01 | -0.3475 | 0.788354 | 0.975154 | 0.253968 | 1 | 0.997515 | 0.809524 |
| Hadhb | Q99JY0 | 1.23 | 1.16 | 1.1 | 1.41 | 0.65 | 1.02 | 1.36 | 1.21 | 1.14 | 1.29 | 1.38 | 0.98 | 0.72 | 1.09 | 0.72 | 0.93 | -0.165 | -0.0275 | -0.36 | 0.419396 | 0.877493 | 0.085001 | 1 | 0.996511 | 0.691619 |
| Hagh | G5E8T9 | -0.18 | -0.51 | -0.11 | -0.43 | -0.46 | -0.16 | 0.01 | -0.09 | -1.52 | -1.35 | -0.16 | -0.33 | -1.15 | -1.22 | -0.32 | 0.11 | 0.1325 | -0.5325 | -0.3375 | 0.475563 | 0.156878 | 0.33342 | 1 | 0.996511 | 0.845601 |
| Haghl | Q9DB32 | -1.64 | -0.94 | -0.31 | -0.52 | -1.56 | -0.97 | -0.73 | -0.66 | -1.19 | 0.77 | -0.85 | -0.82 | -0.88 | -1.1 | -0.43 | -0.25 | -0.1275 | 0.33 | 0.1875 | 0.712377 | 0.507812 | 0.595605 | 1 | 0.996511 | 0.929911 |
| Hap1 | O35668 | 0.1 | 0.22 | -1.33 | -0.06 | -0.84 | -0.13 | 0.15 | -0.26 | 0.26 | 1.52 | -1.34 | -0.35 | 0.75 | -0.35 | 1.66 | 0.63 | -0.0025 | 0.29 | 0.94 | 0.994909 | 0.650816 | 0.091613 | 1 | 0.996511 | 0.693639 |
| Hapln1 | Q9QUP5 | -0.11 | -0.78 | -1.6 | -0.61 | -0.67 | -0.23 | -1.01 | -0.28 | -1.26 | -1.4 | -1.38 | -0.56 | -1.4 | -0.97 | -2.1 | -2.12 | 0.2275 | -0.375 | -0.8725 | 0.516588 | 0.31165 | 0.054961 | 1 | 0.996511 | 0.661738 |
| Hapln2 | Q9ESM3 | 1.79 | 1.4 | 2.74 | 3.3 | 1.06 | 0 | 2.75 | 2.36 | 0.69 | 2.21 | 2.35 | 1.02 | -0.52 | 0.31 | 1.99 | 3.11 | -0.765 | -0.74 | -1.085 | 0.286482 | 0.205851 | 0.214851 | 1 | 0.996511 | 0.787313 |
| Hapln3 | Q80WM5 | -0.36 | -1.83 | -0.72 | -0.24 | -1.39 | 0.07 | -0.57 | 0.07 | -1.35 | -0.79 | -0.62 | -1.02 | -1.45 | -0.04 | -0.14 | -0.4 | 0.3325 | -0.1575 | 0.28 | 0.481231 | 0.680506 | 0.544996 | 1 | 0.996511 | 0.912212 |
| Hapln4 | Q80WM4 | 1.09 | 0.96 | 0.33 | 0.58 | 0.84 | 0.84 | 0.72 | 0.66 | 0.13 | 2.18 | 0.28 | 0.7 | -0.32 | 0.34 | -0.1 | -0.46 | 0.025 | 0.0825 | -0.875 | 0.904583 | 0.860026 | 0.009916 | 1 | 0.996511 | 0.563386 |
| Hars1 | Q61035 | -1.03 | -1.34 | -0.59 | -1.06 | -1.08 | -1.06 | -0.55 | -0.77 | -1.29 | -1.18 | -0.78 | -0.85 | -0.62 | -1.02 | -0.62 | -0.55 | 0.14 | -0.02 | 0.3025 | 0.534024 | 0.930794 | 0.214054 | 1 | 0.996755 | 0.786931 |
| Hars2 | Q99KK9 | 0.34 | 0.29 | 0.35 | 0.15 | 0.03 | 0.43 | 0.49 | 0.36 | -0.03 | 0.22 | 0.57 | 0.41 | 0.06 | 0.44 | 0.06 | 0.51 | 0.045 | 0.01 | -0.015 | 0.788514 | 0.958494 | 0.939564 | 1 | 0.996755 | 0.990645 |
| Hax1 | O35387 | 0.03 | 0.07 | -0.82 | -0.76 | -0.45 | 0.17 | -1.05 | -1.1 | 1.42 | -1.18 | -0.47 | -0.13 | 2.5 | -0.67 | -0.85 | -0.81 | -0.2375 | 0.28 | 0.4125 | 0.522736 | 0.615361 | 0.596044 | 1 | 0.996511 | 0.929911 |
| Hba-a1 | Q91VB8 | -0.8 | -3.07 | -0.5 | 1.86 | -2.07 | 0.7 | -1.4 | 1.12 | -1.1 | -3.97 | -1.35 | -2.88 | -0.16 | -1.64 | -2.05 | -1.84 | 0.215 | -1.6975 | -0.795 | 0.850615 | 0.145492 | 0.426504 | 1 | 0.996511 | 0.878976 |
| Hbb-b1 | P02088 | 1.26 | -1.46 | 1.22 | 3.42 | -0.59 | 2.86 | -0.05 | 3.25 | 0.69 | -0.18 | 0.71 | -1.25 | 1.1 | 0.64 | -0.4 | -0.08 | 0.2575 | -1.1175 | -0.795 | 0.837316 | 0.276838 | 0.410185 | 1 | 0.996511 | 0.874811 |
| Hbb-bs | A8DUK4 | -1.29 | -3.84 | -0.75 | 1.86 | -2.69 | 0.66 | -0.49 | 1.9 | -1.91 | -2.9 | -0.93 | -3.5 | -1.37 | -1.79 | -2.25 | -1.87 | 0.85 | -1.305 | -0.815 | 0.536719 | 0.279645 | 0.448334 | 1 | 0.996511 | 0.884817 |
| Hbs1l | Q3TGM7 | -0.41 | -0.27 | 0.17 | -0.23 | -0.04 | -0.28 | 0.18 | -0.06 | -0.18 | -0.27 | 0.34 | 0.02 | -0.46 | -0.12 | -0.06 | -0.1 | 0.135 | 0.1625 | 9.81E-17 | 0.490281 | 0.468627 | 1 | 1 | 0.996511 | 1 |
| Hccs | P53702 | 0.69 | 0.53 | 1.87 | 0.88 | 0.92 | 0.99 | 2.42 | 2.06 | -0.62 | -0.81 | 2.54 | 1.25 | -0.74 | -0.35 | 2.34 | 2.43 | 0.605 | -0.4025 | -0.0725 | 0.202305 | 0.605327 | 0.928792 | 1 | 0.996511 | 0.989597 |
| Hcfc1 | Q61191 | -1.76 | -1.96 | -1.98 | -1.51 | -2.24 | -2.15 | -2.38 | -1.73 | -2.05 | -0.89 | -1.86 | -2.03 | -1.67 | -1.78 | -2.09 | -2.7 | -0.3225 | 0.095 | -0.2575 | 0.145859 | 0.753613 | 0.365235 | 1 | 0.996511 | 0.858475 |
| Hcls1 | P49710 | 0.15 | -0.42 | -0.21 | -0.06 | -1 | -0.7 | -0.79 | -1.28 | -0.79 | -0.57 | -0.51 | -0.63 | -0.61 | -1.14 | -0.79 | -0.37 | -0.8075 | -0.49 | -0.5925 | 0.003575 | 0.028774 | 0.033789 | 1 | 0.996511 | 0.644622 |
| Hcn1 | O88704 | 0.24 | 0.8 | 0.69 | 0.65 | 0.79 | 0.57 | -0.07 | 0.54 | 0.58 | 0.91 | 0.73 | 0.4 | 0.6 | 0.9 | 0.42 | 0.49 | -0.1375 | 0.06 | 0.0075 | 0.566909 | 0.773289 | 0.972253 | 1 | 0.996511 | 0.992865 |
| Hcn2 | O88703 | 0.95 | 0.88 | 1.03 | 1.02 | 0.47 | 0.99 | 0.39 | 0.77 | 1.39 | 0.62 | 0.86 | 1.06 | 1.51 | 0.68 | 1.46 | 0.6 | -0.315 | 0.0125 | 0.0925 | 0.115683 | 0.952437 | 0.733756 | 1 | 0.996755 | 0.958635 |
| Hcn3 | O88705 | 0.3 | -0.1 | -0.05 | -0.04 | 0.09 | 0.25 | -0.07 | 0.12 | 1.35 | 0.31 | -0.16 | 0.52 | 1.83 | 0.41 | 0.08 | -0.06 | 0.07 | 0.4775 | 0.5375 | 0.67783 | 0.169778 | 0.224809 | 1 | 0.996511 | 0.787447 |
| Hcn4 | B2RY58 | 0.51 | 0.71 | 0.79 | 0.7 | 0.99 | 0.7 | 0.56 | 0.81 | 0.76 | 0.85 | 0.51 | 0.42 | 0.57 | 0.75 | 0.47 | 0.92 | 0.0875 | -0.0425 | -6.9E-17 | 0.599739 | 0.815447 | 1 | 1 | 0.996511 | 1 |
| Hdac1 | O09106 | -0.29 | -1.46 | -1.29 | -1.21 | -1.12 | -0.39 | -1.29 | -1.9 | -0.89 | -1.4 | -1.14 | -0.76 | 0.34 | -2.98 | -0.95 | -0.71 | -0.1125 | 0.015 | -0.0125 | 0.770791 | 0.960521 | 0.9852 | 1 | 0.996755 | 0.996154 |
| Hdac10 | Q6P3E7 | -1.38 | -1.05 | -0.09 | 0.05 | -1.3 | -0.89 | 0 | -0.09 | -0.04 | -0.03 | -0.11 | -1.23 | -0.57 | -0.61 | -0.05 | -0.03 | 0.0475 | 0.265 | 0.3025 | 0.91417 | 0.54519 | 0.431449 | 1 | 0.996511 | 0.879165 |
| Hdac11 | Q91WA3 | 1.42 | 1.29 | 1.18 | 1.06 | 1.51 | 1.22 | 1.21 | 1.28 | 0.68 | 0.03 | 1.01 | 0.76 | 0.68 | 0.36 | 1.02 | 1 | 0.0675 | -0.6175 | -0.4725 | 0.680445 | 0.032069 | 0.059134 | 1 | 0.996511 | 0.661738 |
| Hdac2 | P70288 | -1.21 | -2.06 | -1.74 | -1.93 | -1.76 | -1.77 | -1.53 | -2.01 | -1.32 | -1.53 | -1.62 | -1.59 | -1.14 | -1.53 | -1.2 | -1.27 | -0.0325 | 0.22 | 0.45 | 0.88775 | 0.353156 | 0.091294 | 1 | 0.996511 | 0.693316 |
| Hdac3 | O88895 | 0.1 | 0.06 | -1.54 | -1.71 | 0.06 | -0.11 | -1.25 | -1.42 | -0.23 | -1.87 | -1.35 | 0.34 | 0.82 | -0.82 | -1.42 | -1.05 | 0.0925 | -0.005 | 0.155 | 0.871123 | 0.993808 | 0.808068 | 1 | 0.999285 | 0.971953 |
| Hdac4 | Q6NZM9 | 0.62 | 0.72 | 0.4 | -0.02 | 0.38 | 0.42 | 0.62 | 0.21 | 0.26 | -0.11 | 0.49 | 0.59 | 0.07 | 0.39 | 0.21 | 0.17 | -0.0225 | -0.1225 | -0.22 | 0.915319 | 0.62577 | 0.341706 | 1 | 0.996511 | 0.848473 |
| Hdac5 | Q9Z2V6 | 0.03 | 0.33 | 0.26 | -0.24 | 0.04 | -0.29 | -0.69 | -0.65 | 0 | 0.29 | -0.21 | -0.05 | 0.35 | 0.16 | -0.15 | -0.35 | -0.4925 | -0.0875 | -0.0925 | 0.059787 | 0.677124 | 0.702002 | 1 | 0.996511 | 0.956791 |
| Hdac6 | Q9Z2V5 | 0.18 | 0.01 | -0.09 | 0.02 | -0.24 | 0.2 | -0.55 | -0.08 | 0.23 | 0.38 | 0.04 | 0.26 | 0.46 | 0.31 | -0.11 | -0.18 | -0.1975 | 0.1975 | 0.09 | 0.336274 | 0.261871 | 0.681613 | 1 | 0.996511 | 0.955729 |
| Hdac8 | Q8VH37 | -0.09 | 0.79 | -2.47 | -2.3 | 0.76 | 0.34 | -2.03 | -2.2 | -0.33 | -0.84 | -1.77 | -0.32 | -0.75 | -0.28 | -1.91 | -1.61 | 0.235 | 0.2025 | -0.12 | 0.815395 | 0.799143 | 0.8815 | 1 | 0.996511 | 0.979767 |
| Hdac9 | A0A1B0GS | -2.29 | -0.05 | 0.22 | -0.24 | -0.39 | -2.61 | -1.62 | -1.05 | 0.1 | -2.45 | -0.92 | -2.51 | 0.27 | 1.01 | -0.22 | -1.13 | -0.8275 | -0.855 | 0.5725 | 0.239861 | 0.286539 | 0.398487 | 1 | 0.996511 | 0.867334 |
| Hddc2 | Q3SXD3 | 0.95 | 0.39 | 0.28 | 0.22 | 0.49 | 0.58 | 0.37 | 0.5 | 0.24 | -0.54 | 0.36 | 0.9 | 0.18 | 0.35 | 0.35 | 0.27 | 0.025 | -0.22 | -0.1725 | 0.902267 | 0.518874 | 0.444689 | 1 | 0.996511 | 0.884524 |
| Hddc3 | Q9D114 | -0.23 | -0.57 | 0.25 | -0.09 | -0.6 | -1.68 | 0.32 | -0.43 | 0.21 | 0.13 | -0.84 | -0.98 | 0.41 | 0.04 | 0.24 | 0.34 | -0.4375 | -0.21 | 0.4175 | 0.311462 | 0.552061 | 0.09875 | 1 | 0.996511 | 0.698156 |
| Hdgf | P51859 | -4.45 | -4.96 | -3.79 | -3.96 | -4.62 | -4.87 | -3.6 | -4.2 | -5.15 | -2.23 | -3.72 | -3.78 | -4.66 | -4.27 | -4.16 | -4.29 | -0.0325 | 0.57 | -0.055 | 0.929092 | 0.353483 | 0.853896 | 1 | 0.996511 | 0.976225 |
| Hdgfl2 | Q3UMU9 | -1.26 | -2.75 | -2.05 | -2.44 | -4.74 | -1.74 | -4.39 | -3.74 | -0.85 | -2.24 | -2.36 | -1.87 | -0.35 | -3.46 | -3.83 | -3.96 | -1.5275 | 0.295 | -0.775 | 0.047214 | 0.511176 | 0.359313 | 1 | 0.996511 | 0.856645 |
| Hdgfl3 | Q9JMG7 | -2.54 | -3.16 | -1.61 | -1.61 | -2.63 | -3.48 | -1.87 | -2.49 | -2.92 | -1.04 | -2.17 | -2.43 | -3.37 | -2.72 | -2.12 | -2.77 | -0.3875 | 0.09 | -0.515 | 0.416818 | 0.860105 | 0.255491 | 1 | 0.996511 | 0.809524 |
| Hdhd2 | Q3UGR5 | 0.13 | 0.25 | 0.23 | 0.32 | 0.11 | -0.13 | -0.07 | 0.29 | 0.12 | 0.61 | 0.09 | -0.14 | -0.33 | 0.56 | -0.06 | -0.27 | -0.1825 | -0.0625 | -0.2575 | 0.279216 | 0.763424 | 0.306877 | 1 | 0.996511 | 0.832533 |
| Hdhd3 | Q9CYW4 | 1.1 | 1.24 | 0.76 | 0.48 | 0.93 | 0.95 | 0.43 | 0.53 | 0.41 | -0.26 | 0.76 | 0.77 | 0.36 | 0.66 | 0.24 | 0.41 | -0.185 | -0.475 | -0.4775 | 0.439696 | 0.141976 | 0.067809 | 1 | 0.996511 | 0.664936 |
| Hdhd5 | Q91WM2 | 0.35 | 0.55 | 0.72 | 1.14 | 0.32 | 0.15 | 0.64 | 0.97 | 0.49 | 0.37 | 1.03 | 0.07 | 0.27 | 0.58 | 0.47 | 0.59 | -0.17 | -0.2 | -0.2125 | 0.5135 | 0.474452 | 0.365413 | 1 | 0.996511 | 0.858475 |
| Hdlbp | Q8VDJ3 | -1.18 | -1.53 | -1.07 | -0.83 | -1.06 | -1.33 | -0.44 | -0.99 | -1.29 | -0.17 | -0.73 | -1.14 | -1.04 | -0.74 | -0.53 | -0.49 | 0.1975 | 0.32 | 0.4525 | 0.436392 | 0.297303 | 0.080455 | 1 | 0.996511 | 0.684021 |
| Heatr1 | G3X9B1 | 0.24 | 0.25 | 0.47 | 0.37 | 0.1 | -0.15 | 0.23 | 0.13 | 0.07 | 0.52 | 0.42 | 0.21 | 0.07 | 0.72 | 0.53 | 0.74 | -0.255 | -0.0275 | 0.1825 | 0.137809 | 0.879052 | 0.411349 | 1 | 0.996511 | 0.874811 |
| Heatr3 | Q8BQM4 | -1.71 | -1.43 | -1.6 | -1.95 | -1.82 | -2.07 | -2.08 | -2.52 | -1.74 | -2.23 | -1.51 | -1.56 | -2.15 | -1.82 | -3.65 | -1.9 | -0.45 | -0.0875 | -0.7075 | 0.05717 | 0.704057 | 0.121404 | 1 | 0.996511 | 0.721448 |
| Heatr5a | Q5PRF0 | -1.42 | -1.46 | -1.26 | -1.39 | -1.18 | -0.49 | -0.4 | -1.7 | -1 | -0.55 | -0.98 | -0.77 | -2.07 | -0.23 | -1.4 | -0.46 | 0.44 | 0.5575 | 0.3425 | 0.175523 | 0.012373 | 0.414432 | 1 | 0.996511 | 0.874945 |
| Heatr5b | Q8C547 | -0.21 | -0.45 | -0.15 | -0.1 | -0.51 | -0.24 | -0.67 | -0.16 | -0.25 | 0.02 | -0.24 | -0.32 | -0.08 | -0.57 | -0.57 | -0.42 | -0.1675 | 0.03 | -0.1825 | 0.374269 | 0.865433 | 0.380348 | 1 | 0.996511 | 0.860984 |
| Heatr6 | Q6P1G0 | -0.36 | 0.81 | -0.45 | 0.07 | 0.28 | -1 | -0.23 | 0.27 | 1.72 | 1.26 | -0.37 | -0.39 | 1.99 | 1.29 | 0.15 | -0.22 | -0.1875 | 0.5375 | 0.785 | 0.635795 | 0.357899 | 0.171844 | 1 | 0.996511 | 0.76392 |
| Hebp1 | A0A140T8J | -2.71 | -2.7 | -2.11 | -1.94 | 1.29 | 1.23 | 1.1 | 1.01 | 1.42 | 0.32 | 1.06 | 1.02 | 1.27 | 1.85 | 0.99 | 0.6 | 3.5225 | 3.32 | 3.5425 | 2.02E-07 | 3.35E-06 | 3.32E-06 | 0.001837 | 0.015262 | 0.015132 |
| Hebp2 | Q9WU63 | 0.04 | 0.39 | 0.67 | -0.15 | -0.23 | -0.22 | 0.37 | -1.51 | -0.41 | 0.24 | 0.28 | 0.78 | -0.69 | 0.77 | -0.69 | -0.11 | -0.635 | -0.015 | -0.4175 | 0.148319 | 0.961144 | 0.288907 | 1 | 0.996755 | 0.824705 |
| Heca | Q3V1N5 | -0.35 | -0.64 | -0.32 | 0.09 | -0.48 | -0.82 | 0 | -0.47 | -0.17 | 0.7 | -0.11 | -0.12 | -0.05 | 0 | -0.13 | -0.48 | -0.1375 | 0.38 | 0.14 | 0.571779 | 0.186617 | 0.546703 | 1 | 0.996511 | 0.912406 |
| Hectd1 | F8WIE5 | -0.34 | -0.43 | -0.36 | -0.63 | -0.54 | -0.25 | -0.5 | -0.47 | -0.13 | -0.43 | -0.39 | -0.36 | 0.01 | -0.38 | -0.52 | -0.53 | -2E-17 | 0.1125 | 0.085 | 1 | 0.516111 | 0.679848 | 1 | 0.996511 | 0.955484 |
| Hectd2 | B2RPZ5 | 0.14 | 0.54 | 1.25 | -0.28 | 0.13 | 0.27 | -0.33 | -0.05 | 0.9 | 0.44 | 0.74 | 0.58 | 0.33 | 0.63 | 1.14 | 0.49 | -0.4075 | 0.2525 | 0.235 | 0.249063 | 0.460541 | 0.523182 | 1 | 0.996511 | 0.907797 |
| Hectd3 | Q3U487 | -0.2 | -0.03 | 0.48 | 0.75 | 0.03 | -0.29 | 0.34 | 0.47 | 0.12 | 1.06 | 0.44 | -0.2 | -0.02 | 0.66 | 0.41 | 0.37 | -0.1125 | 0.105 | 0.105 | 0.690121 | 0.759573 | 0.710035 | 1 | 0.996511 | 0.957819 |
| Hectd4 | E9Q2E4 | -0.32 | -0.34 | -0.25 | -0.35 | -0.16 | -0.2 | -0.01 | -0.27 | -0.27 | -0.27 | -0.11 | -0.01 | -0.36 | -0.17 | -0.38 | -0.08 | 0.155 | 0.15 | 0.0675 | 0.300157 | 0.363576 | 0.701748 | 1 | 0.996511 | 0.956791 |
| Hecw1 | Q8K4P8 | 0.67 | -0.12 | -0.02 | -0.42 | -0.47 | 1.22 | -0.62 | -0.47 | -0.02 | -0.04 | 0.1 | 0.13 | 1.01 | 0.11 | -0.43 | -0.71 | -0.1125 | 0.015 | -0.0325 | 0.806376 | 0.953122 | 0.938714 | 1 | 0.996755 | 0.989956 |
| Hecw2 | Q6I6G8 | 0.57 | 0.4 | 0.75 | 0.43 | 0.3 | 0.36 | 0.67 | 0.5 | 0.52 | 0.28 | 0.67 | 0.51 | 0.54 | 0.02 | 0.5 | 0.72 | -0.08 | -0.0425 | -0.0925 | 0.637437 | 0.81335 | 0.676001 | 1 | 0.996511 | 0.952944 |
| Helz | Q6DFV5-2 | -0.07 | 0.1 | -0.23 | -0.13 | -0.1 | -0.07 | -0.6 | -0.25 | 0.31 | 0.64 | -0.07 | 0.17 | 0.38 | 0.53 | -0.37 | -0.28 | -0.1725 | 0.345 | 0.1475 | 0.358505 | 0.124389 | 0.58108 | 1 | 0.996511 | 0.927734 |
| Hepacam | Q640R3 | 0.48 | 0.66 | 0.22 | 0.13 | 0.76 | 0.41 | 0.37 | -0.02 | 0.48 | -0.01 | 0.16 | 0.78 | 0.11 | 0.32 | 0.08 | 0.04 | 0.0075 | -0.02 | -0.235 | 0.973109 | 0.933608 | 0.262723 | 1 | 0.996755 | 0.81524 |
| Heph | Q9Z0Z4 | -0.14 | -0.07 | -0.2 | 0.02 | 0.14 | -0.28 | 0.41 | -0.25 | -0.16 | 0.24 | -0.72 | 0.06 | 0.2 | 0.14 | 0.4 | 0.02 | 0.1025 | -0.0475 | 0.2875 | 0.616729 | 0.843686 | 0.139503 | 1 | 0.996511 | 0.73493 |
| Herc1 | E9PZP8 | -0.55 | -0.21 | -0.02 | 0.05 | -0.16 | -0.62 | 0.13 | -0.12 | -0.08 | 0.2 | 0.02 | -0.44 | -0.11 | 0.47 | -0.38 | -0.22 | -0.01 | 0.1075 | 0.1225 | 0.964787 | 0.634804 | 0.63749 | 1 | 0.996511 | 0.942804 |
| Herc2 | Q4U2R1 | 0.17 | 0.51 | 0.15 | 0.01 | 0.33 | 0.37 | -0.13 | -0.13 | 0.33 | 0.45 | 0.1 | 0.36 | 0.36 | 0.65 | 0.19 | 0.09 | -0.1 | 0.1 | 0.1125 | 0.628145 | 0.598209 | 0.605146 | 1 | 0.996511 | 0.932185 |
| Herc3 | A6H6S0 | -1.23 | -0.88 | -0.65 | -1.15 | -0.81 | -0.74 | -0.66 | -1.13 | -1.06 | -0.91 | -0.67 | -0.86 | -0.96 | -1.13 | -1.14 | -1 | 0.1425 | 0.1025 | -0.08 | 0.48373 | 0.615482 | 0.695529 | 1 | 0.996511 | 0.955729 |
| Herc4 | Q6PAV2-2 | -0.66 | -0.15 | -0.21 | -0.24 | 0.02 | -0.34 | -0.08 | -0.24 | -0.34 | -0.07 | -0.25 | -0.34 | -0.65 | 0.4 | -0.57 | -0.34 | 0.155 | 0.065 | 0.025 | 0.40944 | 0.732882 | 0.929979 | 1 | 0.996511 | 0.989687 |
| Herpud1 | Q8C4N0 | 0.46 | 0.35 | -0.4 | -0.63 | 1.03 | 0.08 | -0.97 | -1.06 | 1.22 | 0.02 | -0.67 | 0.11 | 1.89 | 0.58 | 0.56 | 0.14 | -0.175 | 0.225 | 0.8475 | 0.736017 | 0.617606 | 0.082255 | 1 | 0.996511 | 0.68865 |
| Herpud2 | Q9JJC9 | -0.03 | -1.08 | -0.32 | -0.4 | 1.42 | 0.24 | 1.05 | 0.08 | -0.73 | -0.22 | -0.61 | 0.29 | 0.73 | 1.52 | 0.75 | 0.93 | 1.155 | 0.14 | 1.44 | 0.012302 | 0.663396 | 0.001053 | 1 | 0.996511 | 0.399727 |
| Hexa | P29416 | -1 | -0.99 | -0.64 | -0.6 | -0.68 | -1.05 | 0.4 | -0.56 | -1.22 | -0.93 | 0.01 | -1.13 | -1.28 | -0.09 | -0.95 | -0.34 | 0.335 | -0.01 | 0.1425 | 0.311226 | 0.973951 | 0.642563 | 1 | 0.997515 | 0.94365 |
| Hexb | P20060 | 0.18 | -1.15 | -0.9 | -0.44 | -0.9 | 0.41 | 0.13 | -0.01 | -0.36 | -0.15 | -0.22 | 0.14 | -0.54 | -0.58 | -1.18 | -0.82 | 0.485 | 0.43 | -0.2025 | 0.22968 | 0.193113 | 0.544189 | 1 | 0.996511 | 0.912212 |
| Hexdc | F6UV46 | 6.21 | 4.34 | 0.64 | 0.48 | 4.66 | 5.63 | 0.39 | 0.71 | -0.81 | 0.5 | 0.38 | 5.97 | -0.49 | -0.08 | 1.17 | 0.08 | -0.07 | -1.4075 | -2.7475 | 0.967857 | 0.454047 | 0.059458 | 1 | 0.996511 | 0.661738 |
| Hexim1 | Q8R409 | -2.78 | -1.93 | -2.3 | -2.73 | -0.95 | -2.43 | -1.81 | -2.78 | -1.98 | -0.98 | -2.1 | -2.26 | -2.26 | -1.22 | -2.07 | -1.82 | 0.4425 | 0.605 | 0.5925 | 0.308293 | 0.107681 | 0.083938 | 1 | 0.996511 | 0.690249 |
| Hgh1 | Q8C3I8 | 0.49 | 0.36 | 0.2 | -0.22 | 1.4 | 0.17 | 0.67 | 0.3 | 0.44 | 0.37 | 0.34 | 0.13 | -0.29 | 0.83 | 0.16 | 0.62 | 0.4275 | 0.1125 | 0.1225 | 0.19348 | 0.596376 | 0.688991 | 1 | 0.996511 | 0.955729 |
| Hgs | Q3UMA3 | -0.05 | -0.16 | 0.07 | -0.19 | -0.27 | -0.3 | -0.5 | -0.39 | 0.72 | 0.23 | -0.24 | -0.15 | 1.36 | -0.35 | 0.32 | -0.53 | -2.83E-01 | 0.2225 | 0.2825 | 0.089933 | 0.384167 | 0.500506 | 1 | 0.996511 | 0.900792 |
| Hhatl | Q9D1G3 | -0.48 | -0.39 | -1.09 | -0.24 | 0.36 | -0.25 | -0.78 | -1.52 | -0.67 | -1.73 | -0.62 | -0.59 | 0.2 | -0.62 | -2.48 | -1.49 | 0.0025 | -0.3525 | -0.5475 | 0.995175 | 0.303183 | 0.340148 | 1 | 0.996511 | 0.848183 |
| Hibadh | Q99L13 | 0.6 | 0.79 | 1.29 | 1.82 | 0.65 | 0.55 | 1.45 | 1.76 | 0.58 | 0.94 | 1.55 | 0.57 | 0.25 | 0.73 | 0.96 | 1.18 | -0.0225 | -0.215 | -0.345 | 0.95315 | 0.543669 | 0.323581 | 1 | 0.996511 | 0.841404 |
| Hibch | Q8QZS1 | 0.46 | 0.26 | 0.68 | 1.11 | -0.08 | 0.38 | 0.86 | 0.66 | 0.61 | 0.78 | 0.62 | -0.02 | 0.34 | 0.29 | 0.2 | 0.6 | -0.1725 | -0.13 | -0.27 | 0.538224 | 0.63194 | 0.278655 | 1 | 0.996511 | 0.822084 |
| Hid1 | Q8R1F6 | -0.78 | -1.12 | -0.59 | -0.44 | -0.72 | -0.85 | -0.88 | -0.65 | -0.56 | 0.12 | -0.25 | -0.73 | -0.44 | -0.07 | -0.64 | -0.4 | -0.0425 | 0.3775 | 0.345 | 0.82596 | 0.168366 | 0.162103 | 1 | 0.996511 | 0.757516 |
| Hif1an | Q8BLR9 | -0.37 | 0.23 | -0.25 | -0.44 | -0.8 | 0.26 | -0.52 | -0.46 | -0.06 | -0.65 | -0.14 | 0.14 | -0.44 | 0.52 | -1.41 | -0.58 | -0.1725 | 0.03 | -0.27 | 0.536348 | 0.904231 | 0.512263 | 1 | 0.996744 | 0.903993 |
| Higd1a | A0A1L1SR6 | -0.43 | -0.12 | 0.85 | 0.86 | -0.69 | -0.2 | 0.68 | 0.75 | 1.34 | 1.63 | 0.56 | -0.11 | 0.38 | 1.64 | 0.63 | 0.43 | -0.155 | 0.565 | 0.48 | 0.730756 | 0.259035 | 0.275449 | 1 | 0.996511 | 0.822084 |
| Hikeshi | Q9DD02 | -0.69 | -0.18 | 0.08 | -0.1 | 0.3 | -0.8 | 0.08 | -0.36 | -0.6 | -0.65 | -0.88 | -0.64 | -0.67 | -0.02 | -0.31 | -0.56 | 0.0275 | -0.47 | -0.1675 | 0.925362 | 0.054436 | 0.511748 | 1 | 0.996511 | 0.903993 |
| Hint1 | P70349 | -0.63 | -1.2 | -0.61 | -0.66 | -1.16 | -0.98 | -0.55 | -0.54 | -1.25 | -0.82 | -0.85 | -0.87 | -1.63 | -0.75 | -0.96 | -0.46 | -0.0325 | -0.1725 | -0.175 | 0.887709 | 0.430585 | 0.562988 | 1 | 0.996511 | 0.920024 |
| Hint2 | Q9D0S9 | -0.07 | 0.11 | 0.06 | 0.25 | -0.42 | 0.16 | -0.16 | 0.03 | -0.19 | -0.41 | 0.06 | -0.06 | -0.54 | -0.35 | -0.54 | -0.48 | -0.185 | -0.2375 | -0.565 | 0.330235 | 0.217326 | 0.010423 | 1 | 0.996511 | 0.563386 |
| Hint3 | Q9CPS6 | -0.89 | -0.82 | -0.13 | -0.29 | -0.68 | -0.38 | 0.27 | -0.47 | -1.07 | -0.65 | -0.28 | -0.83 | -1.38 | -1.65 | -0.14 | 0.12 | 0.2175 | -0.175 | -0.23 | 0.448038 | 0.519641 | 0.614423 | 1 | 0.996511 | 0.935876 |
| Hip1 | Q8VD75 | 0.4 | 0.21 | 0.28 | 0.55 | 0.44 | 0.3 | 0.45 | 0.5 | 0.45 | 0.62 | 0.51 | 0.49 | 0.05 | 0.79 | 0.42 | 0.09 | 0.0625 | 0.1575 | -0.0225 | 0.687494 | 0.35648 | 0.922462 | 1 | 0.996511 | 0.987467 |
| Hip1r | Q9JKY5 | 1.23 | 0.68 | 0.92 | 0.86 | 0.84 | 1.09 | 0.81 | 0.91 | 0.55 | 0.69 | 0.98 | 0.96 | 0.4 | 0.79 | 0.58 | 0.78 | -0.01 | -0.1275 | -0.285 | 0.95503 | 0.53448 | 0.193286 | 1 | 0.996511 | 0.777764 |
| Hirip3 | Q8BLH7 | -2.15 | -2.33 | -2.11 | -1.77 | -4.17 | -1.61 | -2.39 | -1.99 | -1.15 | 0.52 | -1.96 | -1.88 | -0.09 | -3.49 | -1.35 | -2.15 | -0.45 | 0.9725 | 0.32 | 0.406858 | 0.100772 | 0.628871 | 1 | 0.996511 | 0.940521 |
| Hist1h2ap | C0HKE9 | 8.67 | 7.88 | 10.71 | 7.27 | 8.69 | 8.18 | 14.76 | 13.45 | 3.85 | -0.93 | 13.72 | 8.77 | 5.26 | -2.45 | 10.5 | 13.16 | 2.6375 | -2.28 | -2.015 | 0.132188 | 0.437895 | 0.522312 | 1 | 0.996511 | 0.907667 |
| Hist2h2aa2 | Q6GSS7 | 1.45 | 1.18 | 2.69 | 2 | 0.69 | 1.01 | 2.5 | 2.41 | 1 | 1.05 | 2.34 | 1.16 | 1.25 | 1.14 | 2.33 | 2.99 | -0.1775 | -0.4425 | 0.0975 | 0.737526 | 0.325007 | 0.850205 | 1 | 0.996511 | 0.976225 |
| Hivep2 | Q3UHF7 | -0.04 | 0.71 | 0.04 | -0.72 | -0.21 | -0.37 | -1.05 | -0.92 | 0.71 | 0.72 | -0.17 | 0.59 | 1.45 | 0.36 | 0.14 | -0.05 | -0.635 | 0.465 | 0.4775 | 0.092963 | 0.211192 | 0.278434 | 1 | 0.996511 | 0.822084 |
| Hk1 | G3UVV4 | 0.52 | 0.24 | 0.41 | 0.67 | -0.07 | 0.48 | -0.14 | 0.52 | 0.74 | 0.31 | 0.47 | 0.33 | 0.62 | 0.14 | -0.13 | -0.2 | -0.2625 | 0.0025 | -0.3525 | 0.253809 | 0.989534 | 0.172818 | 1 | 0.999285 | 0.76428 |
| Hk2 | O08528 | -0.04 | 0.28 | -0.25 | 0.41 | -0.41 | -0.13 | -0.45 | -0.15 | -0.09 | 0.48 | 0.09 | -0.68 | -0.39 | 0.14 | -0.32 | 0.03 | -0.385 | -0.15 | -0.235 | 0.085825 | 0.610625 | 0.337799 | 1 | 0.996511 | 0.847578 |
| Hkdc1 | Q91W97 | -0.32 | 0.41 | 0.62 | 0.25 | 0.32 | -0.62 | 0.31 | -0.02 | 0.4 | -0.08 | 0.5 | -0.55 | 0.73 | 1.31 | -0.13 | -0.14 | -0.2425 | -0.1725 | 0.2025 | 0.423575 | 0.588263 | 0.609663 | 1 | 0.996511 | 0.934646 |
| Hlcs | A0A2I3BPG | 0.75 | 1.36 | 2.39 | 1.3 | 1.45 | 0.67 | 2.52 | 2.83 | 0.5 | 0.85 | 2.55 | 0.93 | 0.16 | 0.12 | 3.56 | 3.41 | 0.4175 | -0.2425 | 0.3625 | 0.45755 | 0.648124 | 0.694367 | 1 | 0.996511 | 0.955729 |
| Hltf | Q6PCN7 | -3.23 | -3.29 | -2.19 | -3.01 | -1.96 | -3.72 | -1.79 | -1.72 | -2.78 | -0.16 | -2.64 | -2.69 | -1.97 | -1.95 | -0.8 | 0.21 | 0.6325 | 0.8625 | 1.8025 | 0.226163 | 0.192593 | 0.008267 | 1 | 0.996511 | 0.563386 |
| Hmbs | P22907 | -1.02 | -0.98 | -1.16 | -0.91 | -0.38 | -0.82 | -1.25 | -0.07 | -1.01 | -0.78 | -1.4 | -1.02 | -0.61 | -0.4 | -1.32 | -1.96 | 0.3875 | -0.035 | -0.055 | 0.17486 | 0.856302 | 0.876964 | 1 | 0.996511 | 0.979626 |
| Hmcn1 | D3YXG0-2 | -0.49 | -0.08 | 0.87 | -0.27 | -0.33 | -0.21 | -1.33 | 0.47 | 0.42 | 1.49 | 0.75 | -0.12 | 1.54 | 0.43 | 0.59 | 0.14 | -0.3575 | 0.6275 | 0.6675 | 0.430105 | 0.166991 | 0.129705 | 1 | 0.996511 | 0.723659 |
| Hmg20a | Q9DC33 | 0.15 | 0.49 | 0.56 | 0.45 | -0.16 | 0.06 | 0.16 | 0.78 | 0.33 | 0.65 | 0.49 | -0.28 | 0.63 | 0.29 | 0.78 | 0.45 | -0.2025 | -0.115 | 0.125 | 0.40377 | 0.643447 | 0.543406 | 1 | 0.996511 | 0.912212 |
| Hmga1 | A0A338P6 | 3.19 | 3.21 | 3.15 | 2.46 | 1.47 | 2.05 | 2.53 | 2.02 | 2.72 | 2.1 | 1.81 | 1.47 | 2.72 | 3.29 | 4.41 | 3.74 | -0.985 | -0.9775 | 0.5375 | 0.007078 | 0.013522 | 0.19027 | 1 | 0.996511 | 0.776843 |
| Hmga1 | Q3TE85 | 0.5 | -1.3 | -0.37 | -0.39 | -1.37 | -0.22 | -1.38 | -1.58 | 1.03 | -0.53 | -1.11 | 0.09 | 0.87 | -0.79 | -0.2 | -0.4 | -0.7475 | 0.26 | 0.26 | 0.122521 | 0.633627 | 0.590291 | 1 | 0.996511 | 0.929142 |
| Hmga2 | Q6NSP9 | 0.51 | 1.75 | -0.48 | 0.91 | -0.89 | -0.52 | 0.82 | 0.13 | 1.06 | 0.42 | 0.2 | 0.3 | 2.01 | 1.46 | 1.27 | 0.96 | -0.7875 | -0.1775 | 0.7525 | 0.175035 | 0.706482 | 0.143734 | 1 | 0.996511 | 0.742777 |
| Hmgb1 | A0A0J9YUD | -3.88 | -4.57 | -1.68 | -1.44 | -4.46 | -4.21 | -1.23 | -2.06 | -4.41 | -0.51 | -1.21 | -3.48 | -4.14 | -3.57 | -2.03 | -1.69 | -0.0975 | 0.49 | 0.035 | 0.922453 | 0.653356 | 0.9683 | 1 | 0.996511 | 0.992733 |
| Hmgb2 | P30681 | -1.36 | -2.57 | -3.95 | -4.33 | -1.72 | -2.49 | -3.71 | -4.87 | -2.47 | -4.16 | -3.43 | -1.89 | -2.47 | -1.6 | -4.94 | -4.97 | -0.145 | 0.065 | -0.4425 | 0.867946 | 0.932284 | 0.653905 | 1 | 0.996755 | 0.947657 |
| Hmgb3 | A2AP78 | -3.29 | -4.13 | -4.84 | -4.26 | -3.52 | -3.29 | -4.48 | -5.26 | -2.96 | -3.51 | -4.84 | -3.23 | -3.53 | -2.83 | -5.59 | -6.01 | -0.0075 | 0.495 | -0.36 | 0.988291 | 0.326836 | 0.636424 | 1 | 0.996511 | 0.942536 |
| Hmgcl | P38060 | 0.96 | 0.54 | -0.15 | 0.22 | 0.26 | 0.64 | 0.06 | 0.03 | 0.94 | -0.52 | 0.07 | 0.72 | 0.42 | 0.8 | -0.66 | -0.32 | -0.145 | -0.09 | -0.3325 | 0.604811 | 0.817541 | 0.410026 | 1 | 0.996511 | 0.874811 |
| Hmgcll1 | D3YVU9 | 0.83 | 1.41 | 2.84 | 2.08 | 1.63 | 0.6 | 2.97 | 2.27 | 0.89 | 2.03 | 2.37 | 0.07 | 0.88 | 1.63 | 1.87 | 3.61 | 0.0775 | -0.45 | 0.2075 | 0.898077 | 0.478259 | 0.752717 | 1 | 0.996511 | 0.961479 |
| Hmgcr | F2Z470 | -0.78 | 0.1 | 0.14 | -0.08 | 0.14 | -0.75 | 0.26 | 0.51 | -0.7 | 0.46 | -0.34 | -0.41 | 0.25 | 0.81 | 0.85 | 0.97 | 0.195 | -0.0925 | 0.875 | 0.566272 | 0.7772 | 0.012839 | 1 | 0.996511 | 0.573604 |
| Hmgcs1 | Q8JZK9 | -0.7 | -0.64 | -0.37 | -0.79 | -0.73 | -0.29 | 0.65 | -0.38 | -1.23 | -1.04 | -0.3 | -0.57 | -0.72 | -0.96 | -0.26 | -0.2 | 0.4375 | -0.16 | 0.09 | 0.176305 | 0.532402 | 0.710955 | 1 | 0.996511 | 0.957819 |
| Hmgcs2 | P54869 | 0.35 | -0.21 | -0.34 | 0.69 | 0.21 | 0.54 | 1.07 | 0.48 | 0.54 | 0.76 | 0.27 | 0.85 | 0.8 | -0.39 | 0.65 | 0.42 | 0.4525 | 0.4825 | 0.2475 | 0.156268 | 0.117349 | 0.491504 | 1 | 0.996511 | 0.897531 |
| Hmgn1 | A0A3B2W3 | -3.08 | -5.86 | -5.44 | -7.94 | -7.68 | -3.61 | -6.1 | -6.43 | -1.87 | -8.47 | -5.25 | -3.75 | -0.17 | -8.03 | -2.15 | -5.47 | -0.375 | 0.745 | 1.625 | 0.749881 | 0.628681 | 0.373127 | 1 | 0.996511 | 0.859424 |
| Hmgn2 | A3KGL9 | -4.1 | -7.99 | -7.33 | -9.04 | -8.4 | -4.55 | -8.13 | -8.76 | -1.19 | -6.12 | -6.45 | -5.35 | 0.83 | -8.66 | -1.69 | -7.72 | -0.345 | 2.3375 | 2.805 | 0.789828 | 0.133011 | 0.233088 | 1 | 0.996511 | 0.788994 |
| Hmgn3 | A0A087WS | -0.11 | -0.9 | -4.97 | -9.49 | -1.98 | 0.25 | -6.28 | -4.45 | 1.41 | -4.15 | -6.75 | 0.35 | 1.58 | 0.59 | -4.37 | -6.74 | 0.7525 | 1.5825 | 1.6325 | 0.744342 | 0.541951 | 0.533672 | 1 | 0.996511 | 0.911825 |
| Hmgn5 | Q9JL35 | -5.07 | -5.87 | -5.07 | -5.75 | -5.08 | -5.67 | -5.86 | -5.84 | -5.36 | -2.24 | -5.07 | -4.76 | -3.96 | -4.49 | -5.1 | -5.87 | -0.1725 | 1.0825 | 0.585 | 0.547792 | 0.138849 | 0.20614 | 1 | 0.996511 | 0.78517 |

| | | | | | | | | | | | | | | | | | | | | | | | | | | |
|---|---|---|---|---|---|---|---|---|---|---|---|---|---|---|---|---|---|---|---|---|---|---|---|---|---|---|
| Hmox1 | P14901 | -0.13 | 0.53 | 0.14 | -0.29 | 0.42 | 0.5 | 1.13 | 0.24 | 0.23 | -1.2 | 0.21 | 0.29 | -0.19 | 0.77 | 0.11 | 0.19 | 0.51 | -0.18 | 0.1575 | 0.08629 | 0.644248 | 0.586298 | 1 | 0.996511 | 0.928178 |
| Hmox2 | O70252 | 0.95 | 0.93 | 0.68 | 0.42 | 0.81 | 0.73 | 0.67 | 0.55 | 0.74 | 0.11 | 0.23 | 1.16 | 0.8 | 0.05 | 0.74 | 0.6 | -0.055 | -0.185 | -0.1975 | 0.761292 | 0.519061 | 0.432314 | 1 | 0.996511 | 0.879165 |
| Hnmt | Q91VF2 | -0.12 | -0.73 | 0.01 | -0.87 | -1.19 | -0.27 | -0.6 | -0.2 | -0.48 | -0.97 | -0.45 | -0.5 | 0.53 | -2.18 | -0.35 | -0.69 | -0.1375 | -0.1725 | -0.245 | 0.658618 | 0.524245 | 0.66221 | 1 | 0.996511 | 0.949645 |
| Hnrnpa0 | Q9CX86 | 0.21 | -1.28 | -1.66 | -1.45 | -1.41 | -0.48 | -1.4 | -1.13 | -1.09 | -1.62 | -1.71 | -1.02 | -0.28 | -1.72 | -0.69 | -1.8 | -0.06 | -0.315 | -0.0775 | 0.892824 | 0.476291 | 0.883135 | 1 | 0.996511 | 0.979767 |
| Hnrnpa1 | Q5EBP8 | 2.4 | -0.06 | -1.25 | -1.01 | -0.14 | 1.34 | -0.85 | -0.63 | 1.15 | -0.99 | -1.29 | 1.01 | 1.3 | 1.11 | -0.25 | -1.09 | -0.09 | -0.05 | 0.2475 | 0.917692 | 0.957895 | 0.78491 | 1 | 0.996755 | 0.967642 |
| Hnrnpa2b1 | O88569 | 0.46 | -1.22 | -1.82 | -1.39 | -1.06 | -0.42 | -1.44 | -0.62 | -0.94 | -2.16 | -1.72 | -1.19 | -0.63 | -0.89 | -0.41 | -1.71 | 0.1075 | -0.51 | 0.0825 | 0.832084 | 0.347081 | 0.876908 | 1 | 0.996511 | 0.979626 |
| Hnrnpa3 | A2AL12 | 0.11 | -1.27 | -0.91 | -0.76 | -1.47 | -0.38 | -0.52 | -0.2 | -0.88 | -0.54 | -0.86 | -1.71 | 0.21 | -0.38 | 0.15 | -0.63 | 0.065 | -0.29 | 0.545 | 0.866273 | 0.444497 | 0.149503 | 1 | 0.996511 | 0.748169 |
| Hnrnpa3 | Q8BG05-2 | 2.84 | 0.52 | 1.31 | 2.73 | -0.21 | 1.68 | 1.33 | 3.39 | 2.71 | 2.38 | 1.55 | 1.17 | 3.3 | 1.9 | 3.47 | 1.53 | -0.3025 | 0.1025 | 0.7 | 0.718635 | 0.866785 | 0.317242 | 1 | 0.996511 | 0.838864 |
| Hnrnpab | Q99020 | 0.68 | -2.64 | -2.79 | -2.53 | -2.56 | -0.46 | -2.29 | -2.57 | 0.4 | -3.23 | -2.71 | -1.1 | 1.66 | -2.5 | -2.18 | -2.79 | -0.15 | 0.16 | 0.3675 | 0.864483 | 0.879087 | 0.758261 | 1 | 0.996511 | 0.963042 |
| Hnrnpc | Q9Z204-2 | 0.39 | 0.52 | 0.12 | 0.56 | 0.24 | -0.45 | 0.71 | 0.4 | 0.1 | -0.61 | 0.37 | -0.1 | -0.04 | 0.64 | 0.08 | 0.39 | -0.1725 | -0.4575 | -0.13 | 0.528293 | 0.097187 | 0.572935 | 1 | 0.996511 | 0.925347 |
| Hnrnpc | Q9Z204 | 2.98 | 1.46 | -0.2 | -0.25 | 0.74 | 2.03 | 0.02 | 0.35 | 1.31 | -0.41 | 0.04 | 1.2 | 1.33 | 1.31 | 1.08 | -0.23 | -0.2125 | -0.4625 | -0.125 | 0.790946 | 0.565334 | 0.871445 | 1 | 0.996511 | 0.979293 |
| Hnrnpd | Q60668-3 | 0.74 | -1.53 | -0.38 | 0.3 | -1.14 | -0.18 | 0.23 | 1.25 | -1.46 | -1.37 | 0.06 | -0.79 | -1.37 | -2.32 | -0.06 | -0.4 | 0.2575 | -0.6725 | -0.82 | 0.686928 | 0.24822 | 0.226481 | 1 | 0.996511 | 0.787447 |
| Hnrnpdl | Q9Z130 | 1.01 | 1.05 | 0.36 | 0.46 | 1.22 | 0.61 | 0.49 | 0.79 | 0.35 | 0.69 | 0.33 | 0.31 | -0.26 | 1.33 | 0.28 | 0.22 | 0.0575 | -0.3 | -0.3275 | 0.819829 | 0.21912 | 0.389823 | 1 | 0.996511 | 0.865375 |
| Hnrnpdl | D3YTQ3 | 0.08 | -0.49 | -0.31 | -0.22 | -0.43 | -0.46 | -0.29 | 0.29 | -0.39 | -0.74 | -0.45 | -0.89 | 0.58 | -1.03 | 0.35 | -0.5 | 0.0125 | -0.3825 | 0.085 | 0.956739 | 0.097102 | 0.823982 | 1 | 0.996511 | 0.972768 |
| Hnrnpf | Q9Z2X1 | -0.68 | -1.7 | -0.87 | -0.72 | -1.51 | -1.03 | -0.91 | -0.61 | -2.05 | -1.96 | -0.54 | -1.75 | -1.64 | -1.8 | -0.44 | -1.17 | -0.0225 | -0.5825 | -0.27 | 0.940399 | 0.175505 | 0.481619 | 1 | 0.996511 | 0.893906 |
| Hnrnph1 | Q8C2Q7 | -1.11 | -2.28 | -1.87 | -1.34 | -1.93 | -1.9 | -1.24 | -1.32 | -2.31 | -1.11 | -1.5 | -2.15 | -1.93 | -1.22 | -1.04 | -1.55 | 0.0525 | -0.1175 | 0.215 | 0.867741 | 0.752904 | 0.521412 | 1 | 0.996511 | 0.907667 |
| Hnrnph2 | P70333 | -0.05 | -2.11 | -0.7 | -0.71 | -2 | -0.65 | -0.15 | 0.08 | -1.73 | -0.94 | -0.49 | -1.55 | -1.17 | -1.51 | 0.44 | -0.42 | 0.2125 | -0.285 | 0.2275 | 0.7157 | 0.560226 | 0.688268 | 1 | 0.996511 | 0.955729 |
| Hnrnph3 | D3Z3N4 | 1.59 | 0.46 | 0.65 | 0.72 | -0.03 | 1.22 | 0.51 | 1.37 | 0.12 | 0.55 | 0.43 | 0.15 | 0.3 | 0.45 | 1.28 | 0.79 | -0.0875 | -0.5425 | -0.15 | 0.822091 | 0.082945 | 0.655261 | 1 | 0.996511 | 0.94818 |
| Hnrnpk | B2M1R6 | -0.24 | -1.89 | -1.43 | -1.17 | -2.02 | -1.35 | -1.58 | -0.95 | -1.27 | -0.45 | -1.19 | -1.55 | -1.08 | -0.84 | -0.9 | -1.74 | -0.2925 | 0.0675 | 0.0425 | 0.461728 | 0.866254 | 0.913481 | 1 | 0.996511 | 0.986458 |
| Hnrnpl | G5E924 | 0.13 | -1.44 | -0.74 | -0.47 | -1 | -0.63 | -0.01 | 0 | -1.09 | -0.55 | -0.63 | -1.61 | 0.04 | -0.12 | 0.85 | -0.23 | 0.22 | -0.34 | 0.765 | 0.572616 | 0.394708 | 0.079055 | 1 | 0.996511 | 0.681114 |
| Hnrnpll | Q921F4 | -1.46 | -1.86 | -2.41 | -2.39 | -1.88 | -1.75 | -2.55 | -2.49 | -1.56 | -1.37 | -2.11 | -1.64 | -1.41 | -1.58 | -2.15 | -2.57 | -0.1375 | 0.36 | 0.1025 | 0.653024 | 0.231394 | 0.769317 | 1 | 0.996511 | 0.965912 |
| Hnrnpm | Q9D0E1-2 | 0.32 | -1.56 | -1.26 | -0.7 | -1.59 | -0.31 | -1.07 | -0.4 | -1.08 | -0.85 | -0.67 | -1.42 | -0.11 | -1.05 | -0.12 | -1.06 | -0.0425 | -0.205 | 0.215 | 0.928524 | 0.628847 | 0.645994 | 1 | 0.996511 | 0.944749 |
| Hnrnpr | Q8VHM5 | 0.35 | -1.16 | -0.83 | -0.59 | -1.47 | -0.19 | -0.69 | -0.08 | -1.47 | -1.48 | -0.61 | -1.32 | -0.72 | -1.89 | -0.26 | -0.69 | -0.05 | -0.6625 | -0.3325 | 0.906167 | 0.102923 | 0.467632 | 1 | 0.996511 | 0.889213 |
| Hnrnpu | Q8VEK3 | 1.33 | -0.27 | -0.67 | -0.29 | -0.72 | 0.67 | -0.48 | -0.18 | 0.1 | -0.47 | -0.65 | -0.23 | 0.78 | -0.08 | -0.02 | -0.85 | -0.2025 | -0.3375 | -0.0675 | 0.684821 | 0.45657 | 0.89588 | 1 | 0.996511 | 0.983509 |
| Hnrnpul1 | Q8VDM6 | -1.15 | -1.99 | -2.5 | -2.31 | -1.9 | -1.54 | -2.27 | -1.95 | -1.45 | -1.02 | -2.39 | -1.47 | -1.27 | -1.25 | -1.8 | -2.42 | 0.0725 | 0.405 | 0.3025 | 0.823688 | 0.322856 | 0.449619 | 1 | 0.996511 | 0.884817 |
| Hnrnpul2 | Q00PI9 | 1.24 | 0.18 | 0.17 | 0.32 | -0.21 | 0.56 | -0.26 | 0.73 | 0.07 | -0.1 | -0.01 | -0.34 | 0.67 | -0.11 | 0.44 | -0.58 | -0.2725 | -0.5725 | -0.3725 | 0.443212 | 0.069342 | 0.331128 | 1 | 0.996511 | 0.845601 |
| Homer1 | Q9Z2Y3 | 1.77 | 1.16 | 2.08 | 2.05 | 1.39 | 1.48 | 1.59 | 2.04 | 0.81 | 1.9 | 2.15 | 1.42 | 0.27 | 1.19 | 1.29 | 1.63 | -0.14 | -0.195 | -0.67 | 0.600028 | 0.586719 | 0.087436 | 1 | 0.996511 | 0.691619 |
| Homer1 | Q9Z2Y3-5 | 1.08 | 1.05 | -0.65 | -1.25 | 1.57 | 0.92 | -0.55 | -0.82 | 0.91 | -1.4 | -0.56 | 0.99 | 0.54 | 1.07 | -0.72 | -0.93 | 0.2225 | -0.0725 | -0.0675 | 0.766393 | 0.923392 | 0.922676 | 1 | 0.996755 | 0.987467 |
| Homer2 | Q9QWW1- | 3.58 | 2.54 | 1.5 | 1.25 | 2.69 | 3.47 | 1.06 | 1.12 | 1.41 | 0.56 | 1.42 | 3.69 | 0.96 | 1.16 | 1.07 | 0.94 | -0.1325 | -0.4475 | -1.185 | 0.854706 | 0.567717 | 0.039322 | 1 | 0.996511 | 0.658454 |
| Homer3 | Q99JP6-2 | 1.09 | 1.69 | 1.45 | 0.96 | 1.7 | 1.05 | 0.92 | 0.92 | 0.16 | 0.76 | 1.17 | 0.91 | 0.3 | 0.49 | 0.98 | 1.41 | -0.15 | -0.5475 | -0.5025 | 0.566862 | 0.080044 | 0.132096 | 1 | 0.996511 | 0.726871 |
| Hook1 | Q8BIL5 | -0.55 | -0.9 | -0.88 | -1.01 | -0.05 | -0.53 | -0.48 | -0.78 | -0.25 | -0.55 | -0.79 | -0.48 | -0.32 | 0.22 | -1.33 | -0.85 | 0.375 | 0.3175 | 0.265 | 0.101093 | 0.13768 | 0.453096 | 1 | 0.996511 | 0.885993 |
| Hook2 | A0A1B0GS | -1.95 | -4.82 | -1.62 | -1.52 | -4.23 | -1.92 | -2.69 | -1.68 | -2.73 | 0.24 | -1.24 | -2.41 | -0.55 | -5.24 | -1.29 | -1.47 | -0.1525 | 0.9425 | 0.34 | 0.861842 | 0.326263 | 0.771968 | 1 | 0.996511 | 0.965933 |
| Hook3 | Q8BUK6 | -0.05 | -0.72 | -0.38 | -0.48 | -0.58 | -0.42 | -0.44 | -0.48 | -0.78 | -0.33 | -0.36 | -0.21 | -0.24 | -0.79 | -0.92 | -0.67 | -0.0725 | -0.0125 | -0.2475 | 0.696392 | 0.955136 | 0.319228 | 1 | 0.996755 | 0.838969 |
| Hopx | Q8R1H0 | 0.23 | 0.11 | 0.29 | -0.87 | -0.25 | 0.71 | 1.04 | -0.06 | -1.21 | -0.24 | -0.6 | 0.42 | -0.55 | -1.48 | 0.03 | 0.48 | 0.42 | -0.3475 | -0.32 | 0.29675 | 0.412088 | 0.503852 | 1 | 0.996511 | 0.901002 |
| Hp1bp3 | Q3TEA8-2 | 2.37 | 0.96 | 1.69 | 1.92 | -1.06 | 0.41 | -0.52 | 1.04 | 0.33 | 0.44 | 0.83 | -0.14 | 1.02 | -0.29 | 0.75 | 0.27 | -1.7675 | -1.37 | -1.2975 | 0.007083 | 0.003565 | 0.009455 | 1 | 0.996511 | 0.563386 |
| Hpca | P84075 | -0.86 | -0.5 | -0.12 | -0.26 | -0.02 | -0.59 | 0.3 | -0.56 | -1.12 | 0.01 | -0.49 | -0.94 | -1.09 | 0.08 | -0.55 | -0.39 | 0.2175 | -0.2 | -0.0525 | 0.437296 | 0.516698 | 0.862295 | 1 | 0.996511 | 0.977204 |
| Hpcal1 | P62748 | 0 | 0.26 | 0.82 | 0.42 | 0.13 | 0.11 | 0.85 | 0.62 | 0.05 | 0.3 | 0.69 | 0.11 | 0.62 | -1.08 | 1.08 | 0.97 | 0.0525 | -0.0875 | 2.25E-02 | 0.840147 | 0.725324 | 0.963733 | 1 | 0.996511 | 0.9927 |
| Hpcal4 | Q8BGZ1 | 0.49 | 0.84 | 0.57 | -0.01 | 1.48 | 1.11 | 0.8 | 0.48 | 0.4 | 0 | 0.75 | 1.13 | 0.25 | 0.52 | 0.34 | 0.68 | 0.495 | 0.0975 | -0.025 | 0.105639 | 0.749203 | 0.917078 | 1 | 0.996511 | 0.986641 |
| Hpgds | Q9JHF7 | -2.47 | -2.08 | -2.31 | -2.29 | -2.71 | -2.43 | -1.84 | -2.41 | -4.22 | -3.65 | -1.79 | -2.88 | -3.98 | -3.05 | -3.96 | -2.43 | -0.06 | -0.8475 | -1.0675 | 0.78719 | 0.11466 | 0.018509 | 1 | 0.996511 | 0.602211 |
| Hprt1 | P00493 | -0.49 | -0.66 | -0.67 | -0.64 | -0.55 | -0.3 | -0.29 | -0.59 | -1.69 | -0.41 | -0.53 | -0.27 | -1.82 | -1.36 | -0.72 | -0.8 | 0.1825 | -0.11 | -0.56 | 0.262887 | 0.737411 | 0.072954 | 1 | 0.996511 | 0.668066 |
| Hpx | Q91X72 | -3.05 | -3.82 | -4.4 | -5.12 | -3.01 | -3.72 | -4.33 | -4.58 | -3.67 | -0.75 | -4.26 | -3.41 | -3.2 | -2.78 | -4.55 | -4.42 | 0.1875 | 1.075 | 0.36 | 0.717545 | 0.206091 | 0.533899 | 1 | 0.996511 | 0.911825 |
| Hras | Q61411 | 0.06 | 0.01 | 0.46 | 0.36 | 0.5 | 0.41 | 0.93 | 0.58 | -0.49 | -1.14 | 0.7 | 0.17 | -1.16 | -0.69 | 0.08 | 0.78 | 0.3825 | -0.4125 | -0.47 | 0.07637 | 0.313988 | 0.282344 | 1 | 0.996511 | 0.822325 |
| Hrg | Q9ESB3 | -4.64 | -6.1 | -4.46 | -4.51 | -6.72 | -5.1 | -5.52 | -4.73 | -3.93 | -1.92 | -5.26 | -4.05 | -2.32 | -4.1 | -4.14 | -5.2 | -0.59 | 1.1375 | 0.9875 | 0.288287 | 0.14211 | 0.155209 | 1 | 0.996511 | 0.750763 |
| Hrh1 | P70174 | 1.06 | 1.3 | 1.12 | 0.98 | 0.48 | 1.13 | -0.47 | 0.77 | 0.49 | 1.14 | 1.08 | 0.91 | 1.04 | 0.95 | 0.92 | 1.09 | -0.6375 | -0.21 | -0.115 | 0.086947 | 0.32444 | 0.519338 | 1 | 0.996511 | 0.907667 |
| Hrh3 | E9Q7T5 | 0.91 | 1.08 | 1.08 | 0.96 | 1.29 | 1.06 | 0.66 | 0.53 | 2.19 | 1.59 | 0.69 | 0.78 | 2.73 | 1.59 | 1.43 | 0.82 | -0.1225 | 0.305 | 0.635 | 0.562396 | 0.395388 | 0.129409 | 1 | 0.996511 | 0.723655 |
| Hs1bp3 | Q3TC93 | -0.52 | -0.11 | -0.59 | -0.74 | -0.18 | -0.44 | -0.66 | -0.88 | 0.18 | -1.17 | -0.75 | -0.31 | -0.18 | 0.3 | -0.91 | -1.28 | -0.05 | -0.0225 | -0.0275 | 0.823121 | 0.943907 | 0.941035 | 1 | 0.996755 | 0.991085 |
| Hs2st1 | Q8R3H7 | -2.34 | -4.71 | -0.93 | -1.31 | -2.44 | -2.51 | -0.52 | -0.58 | -4.65 | -0.55 | -0.7 | -2.55 | -3.4 | -5.06 | -0.96 | -0.03 | 0.81 | 0.21 | -0.04 | 0.386039 | 0.854974 | 0.974842 | 1 | 0.996511 | 0.992977 |
| Hs3st4 | D3YVV6 | 0.16 | 0.56 | 1.2 | 1 | 0.51 | -0.13 | 0.24 | 1.12 | 0.82 | 1.17 | 1.05 | -0.19 | 0.45 | 1.44 | 0.37 | 0.21 | -0.295 | -0.0175 | -0.1125 | 0.395515 | 0.962657 | 0.753965 | 1 | 0.996755 | 0.962145 |
| Hs6st1 | Q9QYK5 | 0.42 | -0.42 | -1.87 | -1.72 | -0.01 | -0.3 | -2.08 | -1.92 | 0.01 | 0.02 | -1.44 | 0.4 | -0.37 | 0.61 | -0.53 | -1.83 | -0.18 | 0.645 | 0.3675 | 0.795627 | 0.315847 | 0.588809 | 1 | 0.996511 | 0.929131 |
| Hsbp1 | Q9CQZ1 | 0.85 | 1.59 | 1.48 | 0.88 | 0.81 | 0.44 | 0.38 | 0.66 | 1.94 | 0.85 | 1.06 | 1.31 | 1.85 | 1.35 | 0.51 | 0.93 | -0.6275 | 0.09 | -0.04 | 0.024768 | 0.771476 | 0.907796 | 1 | 0.996511 | 0.986458 |
| Hscb | Q8K3A0 | 1.63 | 1.27 | 1.1 | 1.01 | 1.14 | 1.34 | 0.78 | 0.92 | 0.19 | -0.69 | 0.73 | 1.21 | -0.07 | -0.02 | 0.51 | 0.45 | -0.2075 | -0.8925 | -1.035 | 0.340236 | 0.054085 | 0.002105 | 1 | 0.996511 | 0.419079 |
| Hsd11b1 | P50172 | -0.08 | 0.52 | 0.09 | -0.35 | 0.94 | 0.86 | -0.74 | -0.32 | 0.82 | -0.77 | 0.1 | 0.18 | 1.54 | -0.29 | -0.12 | -0.4 | 0.14 | 0.0375 | 0.1375 | 0.745251 | 0.91782 | 0.76665 | 1 | 0.996755 | 0.964195 |
| Hsd17b10 | Q99N15 | 0.54 | 0.71 | 0.44 | 0.62 | 0.42 | 0.61 | 0.61 | 0.6 | 0.47 | 0.46 | 0.73 | 0.54 | 0.07 | 0.77 | -0.11 | 0.46 | -0.0175 | -0.0275 | -0.28 | 0.906967 | 0.869299 | 0.267394 | 1 | 0.996511 | 0.818895 |
| Hsd17b11 | Q9EQ06 | 0.01 | 0.26 | 0.14 | -0.25 | 0.53 | -0.05 | 1.32 | 0.1 | 0.02 | 0.26 | 0.26 | -0.03 | -0.28 | 0.46 | 0.22 | 0.55 | 0.435 | 0.0875 | 0.1975 | 0.196604 | 0.647901 | 0.437284 | 1 | 0.996511 | 0.883161 |
| Hsd17b12 | O70503 | -0.43 | -0.15 | -0.95 | -0.92 | 0.09 | -0.22 | -0.64 | -0.96 | -0.27 | -0.88 | -1.12 | -0.43 | -0.61 | 0.17 | -1.14 | -0.91 | 0.18 | -0.0625 | -0.01 | 0.552425 | 0.82757 | 0.976837 | 1 | 0.996511 | 0.993727 |
| Hsd17b4 | P51660 | 3.92 | 0.77 | 0.38 | 0.19 | 2.85 | 4.39 | 1.14 | 0.43 | 0.48 | 0.78 | 0.57 | 4.58 | 0.91 | 0.69 | 1.31 | 0.67 | 0.8875 | 0.2875 | -0.42 | 0.435896 | 0.80881 | 0.602937 | 1 | 0.996511 | 0.931069 |
| Hsd17b7 | O88736 | 0.25 | 0.44 | 0.21 | 0.18 | 0.7 | 0.32 | 0.45 | 0.3 | 0.83 | 0.2 | 0.46 | 0.18 | 1.23 | 0.33 | 0.46 | 0.64 | 0.1725 | 0.1475 | 0.395 | 0.31344 | 0.483068 | 0.132187 | 1 | 0.996511 | 0.726871 |
| Hsd17b8 | P50171 | 0.65 | 0.64 | 0.27 | 0.82 | 0.29 | 0.62 | 0.54 | 0.58 | 0.39 | 0.57 | 0.49 | 0.37 | 0.2 | 0.64 | 0.35 | 0.46 | -0.0875 | -0.14 | -0.1825 | 0.63226 | 0.458505 | 0.391429 | 1 | 0.996511 | 0.865375 |
| Hsdl1 | Q8BTX9 | -0.23 | 0.02 | 0.18 | 0.21 | 0.27 | 0.03 | 0.39 | 0.26 | -0.17 | -0.25 | 0.41 | -0.04 | -0.57 | 0.21 | 0.04 | 0.52 | 0.1925 | -0.0575 | 0.005 | 0.287784 | 0.791514 | 0.985402 | 1 | 0.996511 | 0.996154 |
| Hsdl2 | Q2TPA8 | 0.21 | -0.01 | 0.63 | 0.79 | 0.11 | 0.4 | 1.59 | 0.78 | 0.3 | 0.9 | 0.91 | 0.51 | 0.43 | 0.35 | 0.72 | 0.79 | 0.315 | 0.25 | 0.1675 | 0.386173 | 0.346403 | 0.505482 | 1 | 0.996511 | 0.901382 |
| Hsf1 | A0A2R8W6 | -4.42 | -5.53 | -4.46 | -5.27 | -3.85 | -3.78 | -5.91 | -5.34 | -4.56 | -3.1 | -5.68 | -5.11 | -3.38 | -2.26 | -5.86 | -6.66 | 0.2 | 0.3075 | 0.38 | 0.719167 | 0.593591 | 0.692806 | 1 | 0.996511 | 0.955729 |
| Hsp90aa1 | P07901 | 0.5 | -0.1 | -0.02 | -0.03 | 0.25 | 0.73 | 0.26 | 0.22 | -0.51 | -0.36 | 0.18 | 0.54 | -0.64 | -0.31 | -0.1 | 0.24 | 0.2775 | -0.125 | -0.29 | 0.21262 | 0.666857 | 0.281936 | 1 | 0.996511 | 0.822084 |
| Hsp90ab1 | P11499 | 0.95 | 0.03 | -0.43 | -0.25 | 0.23 | 0.86 | 0.22 | -0.02 | 0.2 | -0.05 | -0.12 | 1.03 | -0.12 | 0.18 | -0.03 | -0.2 | 0.2475 | 0.19 | -0.1175 | 0.481748 | 0.627227 | 0.716805 | 1 | 0.996511 | 0.958334 |
| Hsp90b1 | P08113 | -0.94 | -1.05 | -0.95 | -0.19 | -0.38 | -0.89 | 0.11 | -0.02 | -0.51 | -0.1 | -0.75 | -0.82 | -0.11 | 0.27 | 0.67 | 0.21 | 0.4875 | 0.2375 | 1.0425 | 0.129006 | 0.393134 | 0.004391 | 1 | 0.996511 | 0.548272 |
| Hspa12a | Q8K0U4 | 0.58 | 0.01 | 0.03 | 0.06 | 0.38 | 0.39 | 0.62 | 0.4 | -0.23 | -0.39 | 0.17 | 0.4 | -0.74 | -0.02 | -0.15 | 0.02 | 0.2775 | -0.1825 | -0.3925 | 0.166098 | 0.472746 | 0.149355 | 1 | 0.996511 | 0.748169 |
| Hspa12a | A0A494B9 | 0.25 | -0.03 | 0.17 | 0.4 | 0.14 | 0.54 | 1.11 | 0.58 | 0.2 | 0.73 | 0.54 | 0.16 | -0.53 | -0.26 | 0.33 | 1.18 | 0.395 | 0.21 | -0.0175 | 0.121011 | 0.327388 | 0.963144 | 1 | 0.996511 | 0.9927 |
| Hspa12b | Q9CZJ2 | -0.39 | -0.63 | -0.37 | -0.68 | -0.48 | -0.76 | -0.51 | -0.96 | -0.17 | -0.79 | 0.01 | -0.55 | -0.35 | -0.53 | -0.68 | -0.3 | -0.16 | 0.1425 | 0.0525 | 0.390869 | 0.540185 | 0.785559 | 1 | 0.996511 | 0.967642 |
| Hspa13 | Q8BM72 | -0.41 | 0.1 | 0.06 | -0.36 | 0.72 | -0.31 | 1 | 0.32 | -0.46 | -0.87 | -0.08 | -0.29 | -0.04 | 0.77 | 1.26 | 1.48 | 0.585 | -0.2725 | 1.02 | 0.086604 | 0.277562 | 0.018232 | 1 | 0.996511 | 0.602211 |
| Hspa14 | Q99M31 | -1.83 | -1.87 | -1.3 | -1.14 | -1.86 | -1.95 | -1.27 | -1.63 | -2.09 | -0.99 | -1.33 | -1.55 | -1.85 | -1.22 | -1.43 | -1.6 | -0.1425 | 0.045 | 0.01 | 0.573653 | 0.881227 | 0.969042 | 1 | 0.996511 | 0.992767 |
| Hspa1a | Q61696 | 0.28 | 0.2 | -0.07 | 0.2 | 0.19 | -0.56 | -0.34 | -0.61 | -0.01 | 0.02 | -0.24 | -0.22 | -1.02 | -0.4 | -1.13 | -0.13 | -0.4825 | -0.265 | -0.8225 | 0.053923 | 0.155261 | 0.014538 | 1 | 0.996511 | 0.589387 |
| Hspa2 | P17156 | 1.06 | 0.7 | 0.67 | 0.87 | -0.01 | 1.05 | 0.54 | 0.83 | 1.61 | 1.05 | 0.75 | 0.78 | 1.76 | 0.4 | 0.61 | 0.22 | -0.2225 | 0.2225 | -0.0775 | 0.395953 | 0.374109 | 0.827064 | 1 | 0.996511 | 0.973757 |
| Hspa4 | Q3U2G2 | 0.28 | 0.13 | 0.5 | 0.39 | -0.24 | 0.36 | 0.01 | 0.4 | 0.09 | 0.47 | 0.39 | 0.4 | -0.19 | -0.23 | 0.15 | 0.08 | -0.1925 | 0.0125 | -0.3725 | 0.355423 | 0.944878 | 0.083543 | 1 | 0.996755 | 0.689531 |
| Hspa4l | P48722 | 0.21 | -0.2 | 0.53 | 0.59 | -0.13 | 0.24 | 0.03 | 0.46 | 0.36 | 0.5 | 0.33 | 0.06 | 0.63 | -0.02 | 0.35 | 0.23 | -0.1325 | 0.03 | 0.015 | 0.581347 | 0.897768 | 0.953325 | 1 | 0.996511 | 0.992476 |
| Hspa5 | P20029 | 0.68 | 0.33 | -0.13 | -0.08 | 0.98 | 0.69 | 0.39 | 0.2 | 0.38 | -0.48 | -0.36 | 0.97 | 0.93 | 0.57 | 1.27 | 0.95 | 0.365 | -0.0725 | 0.73 | 0.190327 | 0.847011 | 0.021016 | 1 | 0.996511 | 0.621926 |
| Hspa8 | P63017 | 1.4 | 0.55 | 0.97 | 0.88 | 0.74 | 1.27 | 1.08 | 1.37 | 0.61 | 0.54 | 1.26 | 1.63 | 0.34 | 0.24 | 1.03 | 1.16 | 0.165 | 0.06 | -0.2575 | 0.496447 | 0.849794 | 0.407476 | 1 | 0.996511 | 0.874084 |
| Hspa9 | P38647 | 0.97 | 0.8 | 0.48 | 0.91 | 0.21 | 0.84 | 0.41 | 0.77 | 0.76 | 0.59 | 0.68 | 0.82 | 0.83 | 0.35 | 0.21 | 0.53 | -0.2325 | -0.0775 | -0.31 | 0.289888 | 0.673691 | 0.186846 | 1 | 0.996511 | 0.776843 |
| Hspb1 | P14602 | -3.03 | -3.56 | -4.27 | -4 | -3.85 | -3.84 | -4.68 | -4.66 | -2.2 | -2.19 | -3.75 | -3.74 | -0.93 | -3.45 | -3.56 | -3.09 | -0.5425 | 0.745 | 0.9575 | 0.145237 | 0.153021 | 0.14679 | 1 | 0.996511 | 0.746406 |
| Hspb11 | Q9D6H2 | 1.57 | 1.7 | 0.83 | 0.58 | 1.69 | 1.31 | 0.4 | 0.68 | 1.54 | 0.64 | 0.38 | 1.08 | 1.17 | 1.09 | 0.08 | 0.64 | -0.15 | -0.26 | -0.425 | 0.695173 | 0.481809 | 0.263338 | 1 | 0.996511 | 0.81524 |
| Hspb8 | Q9JK92 | -0.19 | -0.55 | -0.92 | -1.19 | -1.66 | -0.55 | -1.31 | -1.21 | -2.09 | -1.46 | -0.71 | -0.33 | -0.33 | -4.85 | -0.85 | -1.12 | -0.47 | -0.435 | -1.075 | 0.158982 | 0.321296 | 0.27412 | 1 | 0.996511 | 0.822084 |
| Hspbp1 | Q99P31 | -0.69 | -0.65 | -1.3 | -1.59 | -1.06 | -0.88 | -1.48 | -1.82 | -0.74 | -0.53 | -1.77 | -0.71 | -0.61 | -0.52 | -1.89 | -1.88 | -0.2525 | 0.12 | -0.1675 | 0.423092 | 0.736578 | 0.694962 | 1 | 0.996511 | 0.955729 |
| Hspd1 | P63038 | 0.78 | 0.73 | 0.19 | 0.61 | 0.28 | 0.65 | 0.37 | 0.5 | 1.12 | 0.63 | 0.42 | 0.74 | 0.48 | 0.71 | -0.11 | 0.08 | -0.1275 | 0.15 | -0.2875 | 0.51451 | 0.520502 | 0.284125 | 1 | 0.996511 | 0.822669 |
| Hspe1 | Q64433 | 2.28 | 2.52 | 1.98 | 1.98 | 1.91 | 2.09 | 1.84 | 2.12 | 2.03 | 0.72 | 2.05 | 2.13 | 1.04 | 2.11 | 1.37 | 2.12 | -0.2 | -0.4575 | -0.53 | 0.302584 | 0.21884 | 0.115168 | 1 | 0.996511 | 0.717287 |
| Hspg2 | E9PZ16 | -0.7 | -0.89 | -0.67 | -0.63 | -0.69 | -0.61 | -0.73 | -0.69 | -0.33 | -0.43 | -0.62 | -0.64 | 0.07 | -0.68 | -0.34 | -0.62 | 0.0425 | 0.2175 | 0.33 | 0.771006 | 0.225389 | 0.170806 | 1 | 0.996511 | 0.762322 |
| Hsph1 | Q61699 | 0.27 | 0.07 | -0.14 | -0.04 | 0.06 | 0.66 | -0.26 | -0.07 | 0.29 | -0.12 | -0.07 | 0.56 | 0.34 | 0.28 | -0.37 | -0.23 | 0.0575 | 0.125 | -0.035 | 0.806703 | 0.573474 | 0.88342 | 1 | 0.996511 | 0.979767 |
| Htatip2 | Q3U816 | -0.62 | -0.38 | 0.21 | 0.44 | -0.1 | 0.2 | 0.75 | 0.3 | -0.07 | 0.11 | 0.58 | -0.23 | -0.61 | -0.1 | -0.11 | 0.55 | 0.375 | 0.185 | 0.02 | 0.234232 | 0.551159 | 0.953444 | 1 | 0.996511 | 0.992476 |
| Htatsf1 | Q8BGC0 | -2.39 | -2.95 | -1.5 | -1.87 | -2.68 | -2.56 | -2.35 | -1.94 | -2.16 | -1.03 | -1.74 | -2.24 | -1.78 | -2.41 | -2.1 | -2.09 | -0.205 | 0.385 | 0.0825 | 0.553724 | 0.350442 | 0.808962 | 1 | 0.996511 | 0.972098 |
| Htr1a | Q64264 | 1.6 | 1.16 | 0.84 | 0.4 | 1.46 | 2.28 | 0.41 | 0.8 | 1.99 | 0.72 | 0.78 | 1.91 | 1.8 | 1.77 | 0.64 | 1.26 | 0.2375 | 0.35 | 0.3675 | 0.599317 | 0.402851 | 0.329134 | 1 | 0.996511 | 0.844943 |
| Htr1b | P28334 | 0.68 | -0.87 | -0.75 | 0.2 | -0.56 | 2.35 | 0.54 | 0.33 | 1.67 | 1.16 | -0.41 | 0.7 | 0.36 | 0.53 | -0.67 | 0.15 | 0.85 | 0.965 | 0.2775 | 0.21384 | 0.099951 | 0.529525 | 1 | 0.996511 | 0.908663 |
| Htr1f | Q02284 | 0.61 | 0.4 | 1.58 | 1.69 | 0.95 | 0.57 | 0.57 | 1.55 | 0.49 | 1.92 | 1.76 | 0.42 | 0.27 | 0.86 | 1.26 | 0.99 | -0.16 | 0.0775 | -0.225 | 0.676848 | 0.87291 | 0.5579 | 1 | 0.996511 | 0.917135 |
| Htr2a | P35363 | 0.5 | 0.36 | 0.81 | 1.29 | 0.54 | 1.02 | -0.54 | 1.12 | 1.61 | 1.24 | 1.1 | 0.56 | 1.9 | -0.33 | 1 | 0.04 | -0.205 | 0.3875 | -0.0875 | 0.616807 | 0.224587 | 0.862391 | 1 | 0.996511 | 0.977204 |
| Htr5a | P30966 | 1.73 | 2.64 | 1.7 | 1.83 | 2.09 | 2.17 | 2.29 | 2.34 | 1.02 | 2.36 | 2.51 | 1.92 | 1.03 | 2.55 | 1.55 | 2.71 | 0.2475 | -0.0225 | -0.015 | 0.326101 | 0.953601 | 0.972589 | 1 | 0.996755 | 0.992865 |
| Htra1 | Q9R118 | -0.02 | -0.25 | -0.32 | -0.17 | 0.02 | -0.06 | -0.3 | 0.37 | -0.12 | -0.76 | -0.2 | -0.35 | -0.23 | 0.42 | -0.52 | -0.44 | 0.1975 | -0.1675 | -0.0025 | 0.316322 | 0.419696 | 0.992147 | 1 | 0.996511 | 0.996154 |
| Htra2 | Q9JIY5 | 0.45 | 0.69 | 0.31 | 0.65 | 0.45 | 0.37 | 0.49 | 0.57 | 0.31 | 0.2 | 0.59 | 0.39 | -0.47 | 0.17 | -0.02 | -0.04 | -0.055 | -0.1525 | -0.615 | 0.7322 | 0.416084 | 0.018058 | 1 | 0.996511 | 0.602211 |
| Htt | G3X9H5 | 0.1 | 0.11 | 0.35 | 0.15 | 0.29 | -0.03 | 0.04 | 0.11 | -0.2 | 0.06 | 0.07 | -0.11 | -0.34 | 0.16 | -0.25 | -0.06 | -0.075 | -0.2225 | -0.3 | 0.634183 | 0.209291 | 0.151927 | 1 | 0.996511 | 0.750422 |
| Huwe1 | A2AFQ0 | -0.6 | -0.93 | -0.57 | -0.65 | -0.73 | -0.49 | -0.6 | -0.65 | -0.83 | -0.57 | -0.57 | -0.73 | -0.61 | -0.74 | -0.92 | -0.74 | 0.07 | 0.0125 | -0.065 | 0.662496 | 0.943182 | 0.727904 | 1 | 0.996755 | 0.958635 |
| Hyi | Q8R1F5 | -2.64 | -2.57 | -3.42 | -4.18 | 2.01 | 2.13 | 1.53 | 2.4 | -1.16 | -5.06 | -4.63 | -2.34 | 2.31 | 2.67 | 3.27 | 3.17 | 5.22 | -0.095 | 6.0575 | 7.43E-07 | 0.916238 | 3.21E-07 | 0.003385 | 0.996755 | 0.002926 |
| Hykk | Q5U5V2 | -0.4 | -0.83 | -0.23 | 0.23 | -0.26 | -0.81 | 0.2 | 0.08 | 0.05 | 1.38 | -0.3 | -0.7 | -0.43 | -0.16 | -0.49 | -0.39 | 0.11 | 0.415 | -0.06 | 0.723609 | 0.387685 | 0.817477 | 1 | 0.996511 | 0.972768 |
| Hyou1 | Q9JKR6 | -0.29 | -0.12 | -0.73 | -0.45 | 0.24 | -0.44 | 0.13 | -0.29 | -0.52 | -0.06 | -0.75 | -0.32 | 0.01 | 0.35 | 0.52 | 0.27 | 0.3075 | -0.015 | 0.685 | 0.201598 | 0.947648 | 0.011661 | 1 | 0.996755 | 0.566767 |
| Hypk | Q9CR41 | -0.63 | -1.68 | -0.37 | 0.22 | -1.12 | -0.58 | 0.67 | 0.38 | -3.38 | -2.46 | 0.26 | -0.67 | -2.6 | -1.87 | 0.89 | 0.73 | 0.4525 | -0.9475 | -0.0975 | 0.402226 | 0.271611 | 0.911412 | 1 | 0.996511 | 0.986458 |
| Iah1 | Q9DB29 | -0.76 | -0.49 | -0.39 | -0.38 | -0.99 | -0.37 | -0.39 | -0.91 | -0.44 | -1.75 | -0.66 | -0.24 | -0.75 | -0.61 | -0.74 | -1.44 | -0.16 | -0.2675 | -0.38 | 0.461769 | 0.44361 | 0.145892 | 1 | 0.996511 | 0.745741 |
| Iap | P03975 | -0.73 | 1.88 | 0.03 | -0.63 | 0.89 | 0.28 | -0.14 | -0.11 | -0.5 | 0.15 | 0.16 | -0.65 | 1.11 | -0.32 | 1.14 | 0.38 | 0.0925 | -0.3475 | 0.44 | 0.876146 | 0.558862 | 0.495194 | 1 | 0.996511 | 0.89758 |
| Iars1 | Q8BU30 | -1.97 | -2.24 | -2 | -2.31 | -1.7 | -1.83 | -1.65 | -2.22 | -2.48 | -1.72 | -2.04 | -1.81 | -2.24 | -1.76 | -2.03 | -1.89 | 0.28 | 0.1175 | 0.15 | 0.170045 | 0.603856 | 0.461037 | 1 | 0.996511 | 0.887902 |
| Iars2 | Q8BIJ6 | 0.59 | 0.42 | 0.59 | 0.65 | -0.26 | 0.65 | 0.3 | 0.53 | -0.16 | 0.09 | 0.5 | 0.29 | -0.45 | -0.44 | -0.28 | 0.02 | -0.2575 | -0.3825 | -0.85 | 0.276766 | 0.082253 | 0.001781 | 1 | 0.996511 | 0.419079 |
| Iba57 | Q8CAK1 | -0.07 | -0.01 | 0.24 | 0.38 | 0.05 | 0.11 | 0.19 | 0.04 | -0.06 | 0.28 | 0.75 | -0.08 | -0.13 | 0.06 | -0.05 | 0.14 | -0.0375 | 0.0875 | -0.13 | 0.822271 | 0.722944 | 0.5086 | 1 | 0.996511 | 0.902804 |
| Ica1 | D3Z118 | -0.34 | 0.03 | 0.14 | 0 | -0.14 | -0.22 | 0.21 | -0.03 | 0.07 | 0.26 | 0.27 | -0.26 | -0.02 | 0.01 | -0.1 | -0.06 | -0.0025 | 0.1275 | 1.96E-17 | 0.989077 | 0.541633 | 1 | 1 | 0.996511 | 1 |
| Ica1l | A0A087WS | -0.51 | -0.76 | -0.86 | -0.71 | -0.19 | 0.35 | -0.43 | -0.63 | -0.41 | -0.29 | -0.57 | -0.28 | -0.87 | -0.38 | -0.9 | -0.7 | 0.485 | 0.3225 | -0.0025 | 0.069655 | 0.090584 | 0.990198 | 1 | 0.996511 | 0.996154 |
| Icam2 | G5E8M9 | -0.02 | -0.23 | -0.13 | -0.85 | 0.53 | -0.61 | -0.03 | -1.49 | -0.06 | -0.29 | -0.1 | 0.34 | 0.11 | -0.15 | -0.67 | 0 | -0.0925 | 0.28 | 0.13 | 0.832968 | 0.282213 | 0.639406 | 1 | 0.996511 | 0.942804 |
| Icam5 | Q60625 | -0.03 | -0.18 | 0.44 | 0.57 | -0.14 | -0.02 | 0.48 | 0.49 | -0.78 | 0.16 | 0.61 | 0.1 | -0.87 | -0.83 | 0.11 | 0.11 | 0.0025 | -0.1775 | -0.57 | 0.992171 | 0.602332 | 0.114555 | 1 | 0.996511 | 0.717287 |
| Icmt | Q3U4N2 | -1.51 | -0.95 | 1.96 | 1.77 | -0.42 | -1.56 | 1.4 | 1.49 | -0.85 | 2.49 | 2.24 | -0.98 | -0.86 | -0.48 | 2.57 | 1.43 | -0.09 | 0.4075 | 0.3475 | 0.931375 | 0.72885 | 0.748469 | 1 | 0.996511 | 0.960744 |
| Ide | Q8CGB9 | -0.01 | -0.1 | 0.08 | -0.14 | 0.17 | 0.03 | -0.09 | -0.03 | -0.02 | -0.15 | 0.2 | 0.36 | -0.05 | 0.21 | -0.26 | -0.12 | 0.0625 | 0.14 | -0.0125 | 0.678054 | 0.45692 | 0.947188 | 1 | 0.996511 | 0.992376 |
| Idh1 | O88844 | 0.63 | 0.63 | -0.37 | -0.27 | 0.65 | 0.79 | -0.53 | -0.32 | 0.47 | -0.69 | -0.27 | 0.84 | 0.32 | 0.44 | -0.3 | -0.3 | -0.0075 | -0.0675 | -0.115 | 0.985298 | 0.872338 | 0.735367 | 1 | 0.996511 | 0.958635 |
| Idh2 | P54071 | 1.18 | 0.74 | 0.91 | 1.1 | 0.5 | 1.14 | 1.12 | 1.15 | 0.15 | 0.33 | 1.02 | 0.72 | -0.15 | 0.14 | 0.49 | 0.7 | -0.005 | -0.4275 | -0.6875 | 0.981313 | 0.107979 | 0.020237 | 1 | 0.996511 | 0.618919 |
| Idh3a | Q9D6R2 | 1.22 | 0.92 | 0.69 | 0.43 | 0.74 | 1.23 | 0.46 | 0.58 | 0.11 | 0.1 | 0.66 | 0.96 | -0.2 | 0.28 | 0.13 | 0.25 | -0.0625 | -0.3575 | -0.7 | 0.803088 | 0.226919 | 0.016169 | 1 | 0.996511 | 0.589387 |
| Idh3b | Q91VA7 | 0.29 | 0.69 | 0.51 | 0.59 | 0.5 | 0.25 | 0.49 | 0.54 | 0.14 | 0.11 | 0.58 | 0.22 | 0.03 | 0.51 | 0.02 | 0.31 | -0.075 | -0.2575 | -0.3025 | 0.651217 | 0.204409 | 0.168456 | 1 | 0.996511 | 0.759484 |
| Idh3g | P70404 | -0.36 | -0.26 | -0.04 | -0.18 | -0.62 | -0.22 | 0.26 | 0.04 | -0.25 | -0.85 | 0.01 | -0.41 | -0.07 | -0.28 | -0.42 | 0.03 | 0.075 | -0.165 | 0.025 | 0.73733 | 0.473353 | 0.89765 | 1 | 0.996511 | 0.983509 |
| Idi1 | P58044 | -0.99 | -0.14 | -0.45 | -0.21 | -0.11 | -0.47 | -0.07 | -0.41 | -0.7 | -0.93 | -0.2 | -0.51 | -1.07 | -0.43 | -0.28 | 0.35 | 0.1825 | -0.1375 | 0.09 | 0.446209 | 0.605875 | 0.795929 | 1 | 0.996511 | 0.969727 |
| Ids | Q08890 | -0.04 | -0.52 | 0.27 | 0.24 | -0.85 | -0.15 | 0.75 | 0.47 | -1.13 | -1.45 | 0.53 | -0.15 | -0.9 | -0.79 | 0.34 | 0.3 | 0.0675 | -0.5375 | -0.25 | 0.858867 | 0.261026 | 0.510171 | 1 | 0.996511 | 0.903419 |
| Iffo1 | D6RH90 | 0.58 | 1.76 | 0.92 | 1.44 | 1.73 | 0.79 | 1.07 | 1.48 | 1.07 | 3.11 | 1.02 | 1.28 | 0.9 | 1.28 | 0.43 | 0.87 | 0.0925 | 0.445 | -0.305 | 0.777854 | 0.405202 | 0.35543 | 1 | 0.996511 | 0.855029 |
| Ifi47 | A0A1L1SQI | -0.06 | -1.3 | -2.12 | -1.4 | -2.39 | -1.93 | -1.91 | -3.58 | -0.6 | -0.87 | -2.65 | -2.5 | -1.96 | -3.28 | -2.02 | -0.83 | -1.2325 | -0.435 | -0.8025 | 0.043597 | 0.493395 | 0.206034 | 1 | 0.996511 | 0.78517 |
| Ifngr1 | P15261 | 0.03 | 0.63 | 1.09 | 1.21 | 0.77 | 0.32 | 0.64 | 0.8 | 0.48 | 1.45 | 1.1 | 0.36 | 1.02 | 0.31 | 1.22 | 0.84 | -0.1075 | 0.1075 | 0.1075 | 0.71146 | 0.766793 | 0.747801 | 1 | 0.996511 | 0.960744 |
| Ifngr2 | Q63953 | 0.71 | 0.88 | -3.2 | -3.88 | 1.92 | 0.82 | -3.04 | -3.05 | 0.89 | -2.89 | -3.06 | 0.96 | 0.93 | 0.78 | -4.15 | -2.65 | 0.535 | 0.3475 | 0.1 | 0.740484 | 0.817956 | 0.949643 | 1 | 0.996511 | 0.992476 |
| Ift122 | E9Q9G8 | -0.86 | -0.76 | -0.32 | -0.34 | -0.55 | -0.94 | -0.16 | -0.15 | -0.52 | 0.19 | -0.39 | -0.74 | -0.43 | 0.03 | -0.5 | -0.18 | 0.12 | 0.205 | 0.3 | 0.629925 | 0.441744 | 0.214092 | 1 | 0.996511 | 0.786931 |
| Ift140 | E9PY46 | -0.77 | -0.71 | -0.95 | -0.96 | -0.39 | -0.89 | -0.74 | -1.16 | -0.14 | -0.84 | -1.13 | -0.31 | -0.01 | -0.39 | -0.93 | -0.79 | 0.0525 | 0.2425 | 0.3175 | 0.796812 | 0.360498 | 0.226298 | 1 | 0.996511 | 0.787447 |
| Ift172 | Q6VH22 | -0.83 | -0.75 | 0.17 | -0.31 | -0.49 | -1.05 | -0.74 | -0.87 | -0.85 | 0.49 | -0.38 | -0.89 | -0.96 | 0 | -0.26 | -0.44 | -0.3575 | 0.0225 | 0.015 | 0.201894 | 0.952822 | 0.961982 | 1 | 0.996755 | 0.9927 |
| Ift22 | Q9DAI2 | 2.67 | 0.38 | 1.01 | -0.78 | 1.74 | 2.67 | 1.89 | 0.92 | -0.66 | -0.35 | 1.34 | 2.95 | -0.39 | 0.27 | 1.75 | 2.04 | 0.985 | 3.14E-16 | 0.0975 | 0.19908 | 1 | 0.906798 | 1 | 1 | 0.986458 |
| Ift27 | Q9D0P8 | -1.08 | -0.89 | -0.93 | 0.05 | -1.94 | -0.99 | -0.61 | -0.77 | 0.43 | -0.16 | -0.18 | -0.91 | 1.54 | -1.39 | -0.6 | -0.3 | -0.365 | 0.5075 | 0.525 | 0.343498 | 0.189033 | 0.402742 | 1 | 0.996511 | 0.870698 |
| Ift46 | A0A1L1SR1 | -1.04 | -1.04 | -0.87 | -0.68 | -0.11 | -1.4 | -0.38 | -1.08 | -1.18 | 0.8 | -0.61 | -0.53 | -1.28 | -0.72 | -0.88 | -0.5 | 0.165 | 0.5275 | 0.0625 | 0.594567 | 0.217995 | 0.786204 | 1 | 0.996511 | 0.967714 |
| Ift52 | Q62559 | -1.33 | -0.23 | 0.57 | 0.34 | -0.56 | -1.53 | 0.03 | 0.02 | -1.31 | 0.44 | 0.17 | -1.12 | -1.14 | -0.81 | -0.24 | -0.28 | -0.3475 | -0.2925 | -0.455 | 0.506358 | 0.607401 | 0.326741 | 1 | 0.996511 | 0.843799 |
| Ift57 | F7B458 | -0.35 | -0.3 | -0.42 | -1 | -0.6 | -0.32 | -1.37 | -0.97 | -0.48 | -0.83 | -0.78 | -0.26 | 0.01 | -1.23 | -0.29 | -0.84 | -0.2975 | -0.07 | -0.07 | 0.307991 | 0.769733 | 0.830051 | 1 | 0.996511 | 0.973939 |
| Ift74 | Q8BKE9 | -0.04 | 0 | 0.18 | -0.04 | -0.75 | -0.17 | -0.29 | -0.41 | 0.79 | 1.11 | 0.41 | -0.01 | 1.86 | -0.53 | 0.01 | 0.3 | -0.43 | 0.55 | 0.385 | 0.038781 | 0.063835 | 0.433824 | 1 | 0.996511 | 0.879859 |
| Ift80 | Q8K057-2 | -0.16 | 0.32 | -1.71 | -1.08 | 0.93 | -0.82 | -0.56 | -1.55 | 0.34 | -1.33 | -1.52 | 0.24 | -0.16 | 0.94 | -1.7 | -1.21 | 0.1575 | 0.09 | 0.125 | 0.802575 | 0.883967 | 0.853095 | 1 | 0.996511 | 0.976225 |
| Ift81 | O35594 | -0.77 | -0.65 | 1.16 | -0.28 | -0.62 | -1 | 1.47 | 0.67 | -0.34 | 0.54 | 1.18 | -0.91 | -0.5 | -0.08 | 1.46 | 1.37 | 0.265 | 0.2525 | 0.6975 | 0.688168 | 0.669572 | 0.272415 | 1 | 0.996511 | 0.822084 |
| Ift88 | G3X9Z7 | -1.09 | -1.53 | -0.3 | -0.73 | -1.24 | -1.23 | -0.87 | -1.15 | -0.16 | 0.03 | -0.61 | -0.77 | 0.29 | -0.67 | -0.77 | -0.64 | -0.21 | 0.535 | 0.465 | 0.458972 | 0.123849 | 0.213715 | 1 | 0.996511 | 0.786931 |
| Igbp1 | Q61249 | 0.05 | 0.03 | 0.41 | 0.2 | -0.03 | 0.33 | 0.01 | 0.37 | 0.1 | 0.37 | 0.58 | 0.02 | -0.37 | -0.26 | -0.03 | 0.36 | -0.0025 | 0.095 | -0.2475 | 0.988967 | 0.641701 | 0.2961 | 1 | 0.996511 | 0.825612 |
| Igbp1b | A0A0B4J1F | -0.43 | -0.96 | -0.57 | -0.39 | -2.6 | -0.95 | -1.8 | -0.48 | 0.35 | 1.9 | 0.14 | -0.01 | 1.38 | -1.71 | 0.09 | 0.53 | -0.87 | 1.1825 | 0.66 | 0.082013 | 0.022586 | 0.294555 | 1 | 0.996511 | 0.825612 |
| Igdcc4 | Q9EQS9 | 0.85 | 1.44 | 0.34 | 0.03 | 0.93 | 0.95 | -0.49 | -0.15 | 0.82 | -0.37 | -0.3 | 1.09 | 0.81 | 1.54 | -0.27 | -0.06 | -0.355 | -0.355 | -0.16 | 0.437615 | 0.445693 | 0.742204 | 1 | 0.996511 | 0.958976 |
| Igf1r | E9QNX9 | 0.86 | 0.7 | 0.94 | 0.96 | 0.54 | 1.14 | 0.59 | 0.78 | 0.99 | 1.01 | 0.77 | 0.96 | 1.34 | 0.43 | 0.56 | 0.75 | -0.1025 | 0.0675 | -0.095 | 0.588971 | 0.684704 | 0.69985 | 1 | 0.996511 | 0.955776 |
| Igf2bp1 | O88477 | -2.69 | -1.32 | -2.63 | -2.49 | -2.24 | -2.16 | -1.41 | -2.07 | -0.44 | 5.09 | -2.42 | -1.41 | -0.41 | -1.29 | -2.68 | -2.43 | 0.3125 | 2.4875 | 0.58 | 0.395 | 0.130317 | 0.322587 | 1 | 0.996511 | 0.840788 |
| Igf2r | Q07113 | -1.11 | -0.9 | -0.89 | -1.12 | -0.47 | -1.1 | -1.18 | -1.17 | -0.78 | -1.07 | -1.15 | -0.94 | -0.22 | -0.27 | -0.97 | -1.01 | 0.025 | 0.02 | 0.3875 | 0.9053 | 0.908871 | 0.155288 | 1 | 0.996744 | 0.750763 |
| Igfbp4 | P47879 | -0.46 | -0.79 | -1.38 | 0.3 | -0.51 | -0.26 | 0.87 | 0.2 | 1.47 | 0.43 | -0.7 | -0.23 | 2.62 | -0.8 | 0.3 | 0.6 | 0.6575 | 0.825 | 1.2625 | 0.155074 | 0.153956 | 0.106933 | 1 | 0.996511 | 0.703671 |
| Igfbp5 | Q07079 | 0.14 | 0.23 | -0.9 | -1.75 | 0.29 | 0.27 | -0.15 | -1.42 | 0.14 | 0.33 | -1.28 | 0.47 | 0.8 | 0.99 | 0.02 | 0.56 | 0.3175 | 0.485 | 1.1625 | 0.577997 | 0.404886 | 0.036815 | 1 | 0.996511 | 0.658454 |
| Igfbp6 | P47880 | -0.54 | -0.82 | -1.07 | -0.28 | 0.06 | -0.82 | -1.08 | -0.65 | -0.15 | -0.14 | -0.37 | -0.63 | 0.89 | 0.46 | -0.65 | -0.2 | 0.055 | 0.355 | 0.8025 | 0.8529 | 0.15807 | 0.056746 | 1 | 0.996511 | 0.661738 |
| Igfbp7 | E9Q5D9 | -0.09 | -0.16 | -1.05 | -0.65 | -0.42 | -0.8 | -0.32 | -1.39 | -0.2 | 0.01 | -0.49 | 0.13 | -0.19 | 0.47 | -0.45 | -0.03 | -0.245 | 0.35 | 0.4375 | 0.455665 | 0.226754 | 0.179175 | 1 | 0.996511 | 0.775542 |
| Ighg1 | A0A075B5I | -0.99 | -1.38 | -2.69 | -2.92 | -0.81 | -1.92 | -1.89 | -3.29 | -2.06 | 3.18 | -3.08 | -1.33 | -2.74 | -1.9 | -3.55 | -3.1 | 0.0175 | 1.1725 | -0.8275 | 0.977941 | 0.38115 | 0.155464 | 1 | 0.996511 | 0.750763 |
| Ighg2b | A0A075B5I | -2.6 | -2.81 | 0.06 | -0.73 | -2.8 | -2.49 | -0.86 | -0.37 | -2.27 | 1.17 | -0.85 | -2.46 | -2.53 | -2.32 | -0.48 | -1.25 | -0.11 | 0.4175 | -0.125 | 0.894777 | 0.672814 | 0.870813 | 1 | 0.996511 | 0.979293 |
| Ighg2c | F6TQW2 | -2.73 | -1.93 | -1.03 | -1.1 | -1.87 | -2.58 | -1.58 | -1.47 | -2.06 | 0.17 | -1.2 | -1.83 | -1.19 | -3.17 | -1.49 | -1.23 | -0.1775 | 0.4675 | -0.0725 | 0.68796 | 0.435603 | 0.898573 | 1 | 0.996511 | 0.983509 |
| Ighm | A0A075B5I | -2.8 | -4.17 | -0.84 | -1.01 | -4.43 | -2.61 | -1.49 | -0.92 | -2.72 | 0.07 | -1.11 | -3.58 | -1.6 | -3.06 | -0.89 | -0.17 | -0.1575 | 0.37 | 0.775 | 0.87389 | 0.717723 | 0.39946 | 1 | 0.996511 | 0.868275 |
| Ighmbp2 | D3Z5P5 | -0.58 | -0.61 | -3.47 | -2.93 | -0.7 | -0.68 | -5.83 | -5.44 | 0.21 | -2.1 | -3.55 | -0.43 | 0.92 | -0.26 | -1.53 | -4.61 | -1.265 | 0.43 | 0.5275 | 0.392489 | 0.675516 | 0.676133 | 1 | 0.996511 | 0.952944 |
| Ighv6-7 | A0A0A6YW | 2.23 | 2.03 | 2.56 | 3.18 | 2.19 | 1.95 | 1.81 | 2.42 | 2.42 | 3.27 | 2.05 | 1.87 | 1.99 | 3.67 | 2.15 | 2.23 | -0.4075 | -0.0975 | 0.01 | 0.179698 | 0.800205 | 0.981851 | 1 | 0.996511 | 0.994858 |
| Igkc | A0A5H1ZRI | -2.77 | -4.61 | -3.32 | -4.45 | -3.76 | -4.34 | -3.14 | -3.66 | -4.74 | 1.76 | -3.25 | -3.59 | -3.24 | -4.27 | -3.21 | -2.7 | 0.0625 | 1.3325 | 0.4325 | 0.894338 | 0.336928 | 0.409876 | 1 | 0.996511 | 0.874811 |
| Iglon5 | Q8HW98 | 0.47 | 1.09 | 0.85 | 1 | 0.85 | 0.62 | 0.62 | 0.72 | 0.49 | 0.71 | 0.91 | 0.92 | 0.14 | 0.67 | 0.69 | 0.63 | -0.15 | -0.095 | -0.32 | 0.432339 | 0.655892 | 0.192551 | 1 | 0.996511 | 0.777539 |
| Igsf1 | Q7TQA1-5 | 0.32 | 0.45 | -2.08 | -0.82 | -0.32 | 0.14 | -0.25 | -0.74 | 0.81 | 0.23 | -0.93 | 0.59 | 0.72 | 0.66 | 1.62 | -0.23 | 0.24 | 0.7075 | 1.225 | 0.671939 | 0.289047 | 0.082248 | 1 | 0.996511 | 0.68865 |
| Igsf11 | D3Z6S7 | -0.76 | 0.51 | 1.14 | 1.3 | -0.65 | -0.16 | 1.51 | 0.95 | -0.64 | 1.24 | 1.28 | -0.7 | -0.64 | 0.26 | 1.23 | 0.9 | -0.135 | -0.2525 | -0.11 | 0.827898 | 0.704459 | 0.847857 | 1 | 0.996511 | 0.976225 |
| Igsf21 | Q7TNR6 | 0.87 | 1.04 | 1.4 | 0.76 | 1.04 | 1.01 | 1.35 | 0.97 | 0.47 | 0.38 | 1.59 | 1.4 | 0.32 | 0.79 | 0.69 | 0.88 | 0.075 | -0.0575 | -0.3475 | 0.706609 | 0.864754 | 0.157328 | 1 | 0.996511 | 0.751678 |
| Igsf3 | Q6ZQA6 | 0.55 | 0.59 | 0.65 | 0.88 | 1 | 0.68 | 0.93 | 0.98 | 1.31 | 0.66 | 1.11 | 1.08 | 1.18 | 1.46 | 1.09 | 0.71 | 0.23 | 0.3725 | 0.4425 | 0.183643 | 0.093839 | 0.07216 | 1 | 0.996511 | 0.668066 |
| Igsf8 | G3UYZ1 | 0.65 | 0.78 | -0.06 | -0.52 | 0.57 | 0.66 | -0.27 | -0.5 | 0.34 | 0.99 | -0.53 | 0.68 | 0.19 | 0.36 | -0.22 | -0.74 | -0.0975 | 0.1575 | -0.315 | 0.807411 | 0.711929 | 0.418273 | 1 | 0.996511 | 0.87558 |
| Igsf9b | A0A1L1SR8 | 1.29 | 0.94 | 0.72 | 0.89 | 0.81 | 1.28 | 0.69 | 0.57 | 1.89 | 0.74 | 0.96 | 1.06 | 2.23 | 0.61 | 0.18 | 0.58 | -0.1225 | 0.2025 | -0.06 | 0.580594 | 0.48844 | 0.892594 | 1 | 0.996511 | 0.982737 |
| Igtp | Q9DCE9 | -0.14 | -0.45 | 0.25 | 0.33 | -0.17 | -0.6 | -0.27 | -0.09 | -0.89 | 0.81 | 0.01 | -1 | 0.05 | 0.17 | 0.3 | 0.8 | -0.28 | -0.265 | 0.3325 | 0.246904 | 0.547429 | 0.237858 | 1 | 0.996511 | 0.792722 |

| | | | | | | | | | | | | | | | | | | | | | | | | | | | |
|---|---|---|---|---|---|---|---|---|---|---|---|---|---|---|---|---|---|---|---|---|---|---|---|---|---|---|---|
| Ik | Q9Z1M8 | 1.54 | 0.29 | -0.43 | -0.51 | 0.17 | 0.61 | -0.9 | 0.02 | -0.05 | -0.12 | -0.66 | -0.51 | 0.63 | 0.74 | -0.15 | -1.33 | -0.2475 | -0.5575 | -0.25 | 0.63927 | 0.251206 | 0.685865 | | 1 | 0.996511 | 0.955729 |
| Ikbkb | A0A0R4J0T | -0.49 | -0.36 | 0.2 | -0.18 | -0.69 | -0.55 | 0.14 | 0.11 | -0.49 | 0.34 | 0.21 | -0.47 | -0.74 | -0.94 | -0.2 | 0.11 | -0.04 | 0.105 | -0.235 | 0.882054 | 0.707199 | 0.438784 | | 1 | 0.996511 | 0.883205 |
| Ikbkg | E9Q2Y3 | 0.1 | 0.13 | -0.01 | 0.06 | -0.51 | 0.02 | 0.02 | 0.14 | -0.84 | 0.23 | 0.19 | 0.11 | 0.12 | -1.55 | -0.14 | -0.16 | -0.1525 | -0.1475 | -0.5025 | 0.426704 | 0.590789 | 0.199004 | | 1 | 0.996511 | 0.78004 |
| Il16 | O54824-2 | 0.45 | 0.3 | 2.65 | -0.89 | -0.27 | 0.74 | 2.44 | 2.12 | 0.63 | -0.08 | 2.71 | 0.74 | 0.39 | 0.57 | 4.22 | 2.97 | 0.63 | 0.3725 | 1.41 | 0.476905 | 0.665328 | 0.207743 | | 1 | 0.996511 | 0.785338 |
| Il17ra | Q60943 | 1.03 | 1.64 | 0.66 | 0.73 | 1.83 | 1 | -0.55 | -0.4 | 1.21 | 0.2 | -0.19 | 1.08 | 0.81 | 1.28 | 0.42 | 0.04 | -0.545 | -0.44 | -0.3775 | 0.345797 | 0.277402 | 0.2917 | | 1 | 0.996511 | 0.825497 |
| Il18 | P70380 | 0.53 | 0.42 | 1.08 | 1.27 | -0.01 | 0.61 | 0.8 | 1.33 | -0.05 | 0.46 | 1.44 | 0.61 | 0 | -0.97 | 0.53 | 0.81 | -0.1425 | -0.21 | -0.7325 | 0.671843 | 0.565618 | 0.110857 | | 1 | 0.996511 | 0.712422 |
| Il19 | A0A087WS | 0.31 | 3.29 | 1.33 | 1.56 | 4.76 | -0.34 | 5.48 | 4.98 | -0.57 | -5.89 | 3.73 | -0.74 | -2.47 | 0.08 | -0.01 | 0.96 | 2.0975 | -2.49 | -1.9825 | 0.14342 | 0.199229 | 0.044707 | | 1 | 0.996511 | 0.658454 |
| Il1rap | Q61730 | 0.54 | 0.76 | 0.17 | -0.07 | 0.86 | 0.75 | 0.2 | -0.23 | 0.69 | 0.57 | -0.04 | 0.55 | 0.7 | 0.79 | 0.02 | 0.1 | 0.045 | 0.0925 | 0.0525 | 0.884322 | 0.727709 | 0.855961 | | 1 | 0.996511 | 0.976225 |
| Il1rapl1 | B1ASU0 | 1.1 | 1.29 | 1.32 | 1.74 | 1.55 | 1.07 | 1.49 | 1.71 | 1.03 | 1.25 | 1.78 | 1.09 | 0.23 | 1.57 | 0.89 | 1.03 | 0.0925 | -0.075 | -0.4325 | 0.671134 | 0.758551 | 0.193111 | | 1 | 0.996511 | 0.777764 |
| Il1rapl2 | Q9ERS6 | -0.61 | 0.03 | 0.14 | -0.36 | 0.52 | -0.42 | 0.38 | 0.34 | -0.45 | 0.46 | 0.44 | -0.64 | -0.59 | 0.64 | -0.03 | 0.13 | 0.405 | 0.1525 | 0.2375 | 0.170035 | 0.650353 | 0.458538 | | 1 | 0.996511 | 0.887902 |
| Il6st | V9GX00 | 0.11 | 0.3 | -0.01 | 0.28 | 0.03 | -0.52 | -0.18 | -0.32 | 0.91 | 0.65 | 0.48 | 0.48 | 1.04 | 0.77 | 0.63 | -0.02 | -0.4175 | 0.46 | 0.435 | 0.044297 | 0.033481 | 0.126327 | | 1 | 0.996511 | 0.721505 |
| Ildr2 | A0A0A6YV' | 0.31 | 0.82 | 0.97 | 1.05 | 0.88 | 0.58 | 0.54 | 0.93 | 1.09 | 0.8 | 1.15 | 0.69 | 0.79 | 0.66 | 0.55 | 0.67 | -0.055 | 0.145 | -0.12 | 0.801343 | 0.535609 | 0.592731 | | 1 | 0.996511 | 0.929843 |
| Ilf2 | Q9CXY6 | -0.08 | -0.89 | -0.75 | -0.66 | -1.29 | -0.36 | -0.55 | -0.25 | -1.3 | -1.14 | -0.86 | -1.71 | -0.37 | -1.19 | -0.55 | -1.14 | -0.0175 | -0.6575 | -0.2175 | 0.952414 | 0.034519 | 0.460383 | | 1 | 0.996511 | 0.887902 |
| Ilf3 | Q9Z1X4-3 | 0.01 | -0.96 | -0.07 | -0.08 | -1 | -0.48 | -0.24 | 0.27 | -0.67 | 0.65 | -0.2 | -1.13 | -0.22 | -0.43 | 0.5 | -0.26 | -0.0875 | -0.0625 | 0.1725 | 0.796006 | 0.881943 | 0.588124 | | 1 | 0.996511 | 0.929131 |
| Ilk | O55222 | -0.08 | -0.59 | -0.57 | -0.58 | -0.09 | -0.17 | -0.57 | -0.55 | -0.51 | -0.69 | -0.39 | -0.16 | -0.63 | -0.33 | -0.8 | -0.54 | 0.11 | 0.0175 | -0.12 | 0.597437 | 0.933599 | 0.578405 | | 1 | 0.996755 | 0.927198 |
| Ilkap | Q8R0F6 | -4.1 | -4.36 | -2.65 | -2.75 | -4.31 | -4.74 | -3.26 | -2.86 | -2.86 | -1.55 | -3.11 | -3.72 | -1.74 | -2.09 | -2.96 | -3.7 | -0.3275 | 0.655 | 0.8425 | 0.570647 | 0.281422 | 0.168945 | | 1 | 0.996511 | 0.759484 |
| Ilrun | A0A3B2W3 | -3.4 | -3.23 | -2.79 | -2.79 | -1.59 | -3.9 | -2.05 | -3.42 | -3.41 | -0.94 | -1.73 | -3.57 | -3.31 | -2.29 | -1.87 | -2.47 | 0.3125 | 0.64 | 0.5675 | 0.554934 | 0.307291 | 0.122557 | | 1 | 0.996511 | 0.721448 |
| Ilvbl | Q8BU33 | 0.02 | 0.59 | -0.26 | 0.02 | 0.7 | 0.16 | 0.63 | -0.09 | 0.12 | -0.41 | -0.27 | 0.14 | 0.18 | 0.26 | 0.16 | 0.57 | 0.2575 | -0.1975 | 0.2 | 0.348152 | 0.437139 | 0.415459 | | 1 | 0.996511 | 0.874945 |
| Immp1l | Q9CQU8 | 1.77 | 1.26 | 1.15 | 1.07 | 1.08 | 1.51 | -0.04 | 0.84 | 0.86 | 3.8 | 0.86 | 1.46 | 1.02 | 1.27 | 0.13 | 2.89 | -0.465 | 0.4325 | 0.015 | 0.205667 | 0.513773 | 0.978244 | | 1 | 0.996511 | 0.993727 |
| Immp2l | Q8BPT6 | -4.23 | -3.25 | -2.14 | -1.64 | -2.41 | -3.12 | -1.24 | -2.07 | -3.46 | -2.7 | -1.66 | -4.21 | -4.35 | -4.29 | -2.43 | -1.15 | 0.605 | -0.1925 | -0.24 | 0.353991 | 0.789974 | 0.78288 | | 1 | 0.996511 | 0.967642 |
| Immt | E9Q800 | 1.85 | 1.74 | 1.67 | 2.26 | 0.92 | 1.8 | 1.03 | 2.13 | -0.3 | 2.68 | 2.27 | 1.58 | -0.01 | -0.11 | 1.86 | 1.36 | -0.41 | -0.3225 | -1.105 | 0.217249 | 0.603127 | 0.043604 | | 1 | 0.996511 | 0.658454 |
| Immt | Q8CAQ8-3 | 0.94 | 0.95 | 0.5 | 0.6 | 0.19 | 0.58 | -0.81 | 0.57 | 0.72 | 0.97 | -0.47 | 0.48 | 0.67 | -0.06 | 0.19 | 0.19 | -0.615 | -0.3225 | -0.5 | 0.095066 | 0.345586 | 0.057491 | | 1 | 0.996511 | 0.661738 |
| Immt | Q8CAQ8-2 | 1.74 | 1.48 | 1.53 | 1.17 | 1.21 | 1.78 | 1.51 | 1.58 | 0.76 | 0.8 | 1.59 | 1.46 | 1.14 | 0.41 | 1.42 | 1.51 | 0.04 | -0.3275 | -0.36 | 0.841389 | 0.236207 | 0.237752 | | 1 | 0.996511 | 0.792722 |
| Immt | Q8CAQ8 | -3.5 | -2.17 | -2.52 | -1.78 | -2.95 | -4.61 | -1.41 | -1.69 | -3.15 | -2.02 | -1.98 | -4.05 | -2.72 | -1.09 | -1.52 | -2.38 | -0.1725 | -0.3075 | 0.565 | 0.815531 | 0.592108 | 0.269089 | | 1 | 0.996511 | 0.820534 |
| Immt | E9PVS5 | -2.74 | -1.57 | -2.81 | -1.52 | -3.03 | 5.5 | -0.8 | 4.66 | -2.77 | -2.67 | 4.51 | -2.58 | -2.58 | -2.08 | -2.42 | 2.94 | 3.7425 | 1.2825 | 1.125 | 0.073321 | 0.440859 | 0.370416 | | 1 | 0.996511 | 0.859245 |
| Imp3 | Q921Y2 | 0.4 | 0.2 | 1.03 | 0.77 | 1.18 | 0.35 | 2.72 | 1.35 | 0.53 | 1.43 | 1.15 | -0.79 | 0.39 | 0.76 | 2.21 | 2.32 | 8.00E-01 | -0.02 | 0.82 | 0.126958 | 0.967508 | 0.123483 | | 1 | 0.996755 | 0.721448 |
| Impa1 | Q924B0 | 0.21 | -0.14 | 0.1 | 0.28 | -0.41 | 0.03 | -0.13 | -0.01 | 0 | 0.59 | -0.21 | 0.27 | -0.42 | 0.25 | -0.36 | -0.34 | -0.2425 | 0.05 | -0.33 | 0.201811 | 0.827075 | 0.17209 | | 1 | 0.996511 | 0.76392 |
| Impa2 | Q91UZ5 | 0.4 | 0.54 | -2.11 | -3.29 | 0.19 | 0.97 | -2.64 | -3.51 | 1.98 | -2.53 | -3.1 | 0.14 | 2.02 | 1 | -3.19 | -3.09 | -0.1325 | 0.2375 | 0.3 | 0.917819 | 0.860981 | 0.838399 | | 1 | 0.996511 | 0.976133 |
| Impact | O55091 | -0.5 | -0.38 | -0.67 | -0.38 | -0.92 | -0.59 | -0.58 | -0.16 | -0.38 | 0.01 | -0.52 | -0.24 | -0.22 | -0.49 | -0.22 | -0.76 | -0.08 | 0.2 | 0.06 | 0.693332 | 0.307447 | 0.772343 | | 1 | 0.996511 | 0.965933 |
| Impdh1 | F7DEU6 | -0.76 | -1.09 | -0.61 | -0.49 | -0.95 | -1.06 | -0.65 | -0.22 | -1.12 | -0.13 | -0.44 | -0.96 | -1.31 | -0.91 | -0.69 | -0.95 | 0.0175 | 0.075 | -0.2275 | 0.942517 | 0.786661 | 0.332997 | | 1 | 0.996511 | 0.845601 |
| Impdh2 | P24547 | 0.32 | 0.14 | -0.21 | -0.22 | 0.12 | -0.06 | -0.15 | -0.09 | -0.11 | -0.28 | -0.3 | 0.2 | -0.65 | -0.01 | -0.32 | -0.47 | -0.0525 | -0.13 | -0.37 | 0.778662 | 0.551682 | 0.138904 | | 1 | 0.996511 | 0.73493 |
| Ina | P46660 | 1.22 | 0.94 | 1.44 | 1.29 | 1.01 | 0.36 | 1.05 | 1.3 | 0.77 | 0.21 | 1.13 | 0.63 | 0.52 | 0.39 | 1.56 | 1.48 | -0.2925 | -0.5375 | -0.235 | 0.245713 | 0.051339 | 0.482511 | | 1 | 0.996511 | 0.893925 |
| Inafm2 | A0A1L1STD | 1.78 | 1.96 | 1.16 | 0.67 | 1.78 | 1.92 | 0.02 | 0.28 | 2.01 | 1.3 | 0.62 | 1.82 | 2.06 | 2.35 | 1.59 | 0.84 | -0.3925 | 0.045 | 0.3175 | 0.465612 | 0.912294 | 0.459889 | | 1 | 0.996744 | 0.887902 |
| Inf2 | E9QLA5 | 0.14 | -0.49 | 0.23 | 0.61 | -0.71 | -0.54 | 0.25 | 0.32 | -0.15 | 0.27 | -0.04 | -0.1 | -0.1 | -0.81 | -0.05 | 0.27 | -0.2925 | -0.1275 | -0.295 | 0.398055 | 0.631959 | 0.378029 | | 1 | 0.996511 | 0.860984 |
| Ing2 | Q9ESK4 | -2.19 | -5 | -1.45 | -1.22 | -2.42 | -3.16 | -1.37 | -0.68 | -4.35 | -2.13 | -1.71 | -2.48 | -0.66 | -4.8 | 1.48 | 0.51 | 0.5575 | -0.2025 | 1.5975 | 0.55079 | 0.829853 | 0.287844 | | 1 | 0.996511 | 0.824705 |
| Inip | Q3TXT3 | -0.26 | -0.35 | -0.71 | -0.83 | -0.87 | -1.71 | -2.8 | 0.15 | -0.54 | 0.7 | -1.8 | -0.66 | -1.29 | 0.17 | -0.58 | -1.36 | -0.77 | -0.0375 | -0.2275 | 0.211154 | 0.939305 | 0.547999 | | 1 | 0.996755 | 0.912571 |
| Inpp1 | P49442 | 0.58 | 0.19 | 0.12 | 0.27 | -0.41 | 0.29 | 0.06 | 0.41 | 0.05 | -1.33 | -0.01 | 0.64 | -0.38 | -0.4 | -0.47 | -0.1 | -0.2025 | -0.4525 | -0.6275 | 0.385188 | 0.284888 | 0.011016 | | 1 | 0.996511 | 0.563386 |
| Inpp4a | F6V2U0 | -0.34 | -0.32 | 0.01 | 0.1 | -0.18 | -0.27 | -0.05 | -0.01 | -0.36 | 0.06 | -0.01 | -0.15 | -0.53 | -0.05 | -0.15 | -0.18 | 0.01 | 0.0225 | -0.09 | 0.954596 | 0.90916 | 0.67144 | | 1 | 0.996744 | 0.952631 |
| Inpp4b | E9PVM1 | -0.05 | 0.05 | 0.11 | 0.53 | -0.15 | -0.11 | 0.24 | 0.43 | 0.01 | 0.99 | 0.06 | -0.09 | -0.38 | 0.57 | 0.02 | 0.06 | -0.0575 | 0.0825 | -0.0925 | 0.789492 | 0.776737 | 0.724072 | | 1 | 0.996511 | 0.958398 |
| Inpp5a | Q7TNC9 | 0.29 | 0.34 | 0.73 | 0.6 | 0.58 | 0.22 | 0.56 | 0.34 | -0.16 | 0.53 | 0.48 | 0.33 | -0.17 | 0.13 | 0.76 | 0.68 | -0.065 | -0.195 | -0.14 | 0.720321 | 0.395254 | 0.607593 | | 1 | 0.996511 | 0.933262 |
| Inpp5b | Q8K337 | -0.13 | -0.05 | 0.34 | 0.4 | -0.11 | -0.08 | 0.1 | 0.28 | -0.1 | 0.69 | 0.64 | 0.04 | -0.15 | 0.25 | 0.11 | 0.07 | -0.0925 | 0.1775 | -0.07 | 0.640158 | 0.503727 | 0.743596 | | 1 | 0.996511 | 0.959321 |
| Inpp5d | Q9ES52-3 | -0.32 | -0.38 | -1.2 | -0.74 | -0.52 | -0.06 | -0.7 | -0.23 | 2.04 | -0.19 | -0.7 | -0.63 | 0.67 | -0.02 | -1.55 | -0.17 | 0.2825 | 0.79 | 0.3925 | 0.290066 | 0.224287 | 0.418668 | | 1 | 0.996511 | 0.87558 |
| Inpp5e | Q9JII1 | -0.33 | 0.14 | -0.35 | -0.4 | 0.07 | 0.06 | -0.58 | -0.29 | 0.57 | -1.02 | -0.12 | -0.02 | 0.91 | 0.31 | -0.15 | -0.08 | 0.05 | 0.0875 | 0.4825 | 0.822529 | 0.800247 | 0.12343 | | 1 | 0.996511 | 0.721448 |
| Inpp5f | Q8CDA1 | 0.11 | 0.2 | 0.2 | 0.04 | 0.19 | 0.2 | -0.12 | -0.25 | 0.35 | 0.18 | -0.13 | 0.11 | 0.45 | 0.1 | 0.08 | 0.03 | -0.1325 | -0.01 | 0.0275 | 0.447067 | 0.954823 | 0.882461 | | 1 | 0.996755 | 0.979767 |
| Inpp5j | P59644 | -0.16 | -0.03 | 0.28 | 0.23 | -0.17 | -0.1 | -0.46 | -0.09 | 0.61 | 0.29 | 0.21 | -0.04 | 0.81 | 0.14 | -0.02 | -0.12 | -0.285 | 0.1875 | 0.1225 | 0.141685 | 0.387593 | 0.642356 | | 1 | 0.996511 | 0.94365 |
| Inpp5k | Q8C5L6 | 0.24 | 0.71 | 0.21 | -0.03 | 0.8 | 0.1 | 0.58 | -0.01 | 0.39 | 0.13 | 0.26 | 0.4 | 0.47 | 0.97 | 0.36 | 0.67 | 0.085 | 0.0125 | 0.335 | 0.741495 | 0.952539 | 0.19042 | | 1 | 0.996755 | 0.776843 |
| Inppl1 | Q6P549 | 0.1 | 0.26 | 0.57 | 0.2 | 0.28 | 0.06 | -0.19 | 0.01 | 0.57 | 1.19 | 0.5 | 0.32 | 0.59 | 0.95 | 0.11 | 0.38 | -0.2425 | 0.3625 | 0.225 | 0.207963 | 0.159099 | 0.364619 | | 1 | 0.996511 | 0.858475 |
| Insc | A0A0U1RN | -1.33 | -0.71 | -1.53 | -0.71 | -1.44 | -0.53 | -0.72 | -0.6 | 0.56 | 0.51 | -0.9 | -0.13 | -0.76 | 0.71 | -1.1 | -1.26 | 0.2475 | 1.08 | 0.4675 | 0.413637 | 0.02008 | 0.331438 | | 1 | 0.996511 | 0.845601 |
| Insr | P15208 | 0.21 | 0.42 | 0.31 | 0.21 | 0.61 | 0.43 | 0.55 | 0.22 | 0.29 | -0.26 | 0.28 | 0.29 | 0.37 | 0.34 | 0.15 | 0.39 | 0.165 | -0.1375 | 0.025 | 0.320442 | 0.491696 | 0.886304 | | 1 | 0.996511 | 0.979767 |
| Insyn1 | Q8CD60 | 0.48 | 0.47 | 0.81 | 0.42 | 0.84 | 0.06 | 0.99 | 0.81 | 0.61 | 0.28 | 0.69 | 0.82 | -0.19 | 0.87 | 0.55 | 1.5 | 0.13 | 0.055 | 0.1375 | 0.594082 | 0.780639 | 0.702004 | | 1 | 0.996511 | 0.956791 |
| Insyn2a | E9PZW7 | 0.55 | 0.71 | 0.5 | 1.15 | 0.57 | 1.62 | 0.86 | 0.69 | 1.14 | 1.4 | 0.2 | 0.78 | 0.95 | 1.09 | 1.43 | 0.85 | 0.2075 | 0.1525 | 0.3525 | 0.467595 | 0.618101 | 0.160409 | | 1 | 0.996511 | 0.75577 |
| Ints1 | K3W4P2 | 0.04 | -0.78 | -0.73 | -1.47 | -1 | -0.47 | -0.41 | -0.54 | -1.19 | -2.65 | -0.86 | -0.78 | -1.12 | -1.52 | -0.48 | -0.89 | 0.13 | -0.635 | -0.2675 | 0.692638 | 0.223145 | 0.476323 | | 1 | 0.996511 | 0.892174 |
| Ints10 | E9Q360 | -0.76 | -1.11 | -0.98 | -1.16 | -1.32 | -1.37 | -1.07 | -0.99 | -1.58 | -0.93 | -1.19 | -1.39 | -1.69 | -0.97 | -1.31 | -1.19 | -0.185 | -0.27 | -0.2875 | 0.310377 | 0.218399 | 0.220433 | | 1 | 0.996511 | 0.787447 |
| Ints11 | Q9CWS4 | -0.53 | -0.9 | -0.61 | -0.99 | -0.41 | -0.96 | -0.77 | -0.6 | -0.83 | -0.55 | -0.84 | -0.97 | -1.28 | -1.09 | -0.9 | -0.25 | 0.0725 | -0.04 | -0.1225 | 0.713119 | 0.837569 | 0.655528 | | 1 | 0.996511 | 0.94818 |
| Ints12 | Q9D168 | -0.41 | -2.63 | -1.51 | -1.51 | -2.65 | -1.32 | -2.75 | -1.18 | -2.05 | -1.33 | -1.81 | -1.97 | -1.81 | -1.53 | -0.54 | -2.96 | -0.46 | -0.275 | -0.195 | 0.425001 | 0.547355 | 0.751952 | | 1 | 0.996511 | 0.961479 |
| IntS13 | Q8QZV7 | -1.19 | -1.89 | -1.49 | -1.53 | -1.8 | -1.58 | -1.99 | -1.84 | -2.26 | 0.54 | -1.76 | -1.69 | -2.09 | -1.66 | -1.22 | -1.91 | -0.2775 | 0.2325 | -0.195 | 0.188709 | 0.693517 | 0.467276 | | 1 | 0.996511 | 0.889213 |
| Ints14 | Q8R3P6 | 0.35 | -0.1 | -0.94 | -1.36 | -0.9 | 1.12 | -1.07 | -0.83 | 0.06 | -1 | -0.66 | -0.34 | 0.12 | -0.93 | -0.29 | -0.11 | 0.0925 | 0.0275 | 0.21 | 0.875307 | 0.948495 | 0.626052 | | 1 | 0.996755 | 0.939602 |
| Ints15 | Q8BGA7 | -0.67 | -1.14 | -0.71 | -0.93 | -1.28 | -0.53 | -1.91 | -1.55 | -0.56 | -0.86 | -0.23 | -0.55 | -1.53 | -0.15 | -2.44 | -0.48 | -0.455 | 0.3125 | -0.2875 | 0.165043 | 0.163957 | 0.566328 | | 1 | 0.996511 | 0.921535 |
| Ints2 | Q80UK8 | -1.09 | -1.06 | -1.08 | -0.65 | -1.03 | -1.43 | -0.92 | -1.05 | -0.42 | -0.7 | -1.08 | -0.78 | -1.73 | -0.21 | -0.52 | -0.72 | -0.1375 | 0.225 | 0.175 | 0.48012 | 0.308027 | 0.614291 | | 1 | 0.996511 | 0.935876 |
| Ints3 | A0A0G2JFJ | -1.27 | -1.98 | -1.75 | -1.39 | -1.62 | -2.11 | -1.91 | -1.94 | -2.39 | -1.38 | -1.71 | -2 | -1.94 | -1.53 | -1.79 | -1.89 | -0.2975 | -0.2725 | -0.19 | 0.194308 | 0.3463 | 0.420094 | | 1 | 0.996511 | 0.87558 |
| Ints4 | Q8CIM8 | 0.03 | -0.68 | -1.01 | -1.66 | -1.02 | -0.1 | -2.61 | -1.23 | -0.32 | -0.24 | -1.47 | 0 | -1.1 | -0.81 | -0.96 | -1.23 | -0.41 | 0.3225 | -0.195 | 0.480706 | 0.482136 | 0.589458 | | 1 | 0.996511 | 0.929142 |
| Ints5 | Q8CHT3 | -0.5 | -0.53 | -0.08 | -0.75 | -0.7 | -0.9 | -0.57 | -0.66 | -0.51 | 0.51 | -0.4 | -0.77 | -0.58 | -0.27 | -0.32 | -0.24 | -0.2425 | 0.1725 | 0.1125 | 0.230421 | 0.5856 | 0.603095 | | 1 | 0.996511 | 0.931155 |
| Ints6 | Q6PCM2-2 | 0.45 | -0.15 | -0.75 | -0.41 | -0.17 | -0.17 | -0.52 | -0.27 | -0.12 | -1.02 | -0.59 | 0.53 | -0.71 | -0.41 | -0.44 | -0.27 | -0.0675 | -0.085 | -0.2425 | 0.804061 | 0.831559 | 0.408424 | | 1 | 0.996511 | 0.874356 |
| Ints7 | A0A0R4J0E | -0.44 | -0.3 | -0.28 | -0.12 | -0.69 | -0.24 | -0.24 | -0.14 | -0.12 | -0.14 | -0.28 | -0.32 | -0.07 | -0.08 | -0.52 | -0.14 | -0.0425 | 0.07 | 0.0825 | 0.816206 | 0.674899 | 0.675446 | | 1 | 0.996511 | 0.952944 |
| Ints9 | A0A0R4J0J | -1.6 | -1.04 | -0.03 | -0.06 | -0.2 | -2.06 | 0.34 | 0.08 | -1.18 | -0.12 | -0.24 | -1.82 | 0.12 | -1.03 | -0.73 | -0.48 | 0.2225 | -0.1575 | 0.1525 | 0.715299 | 0.761246 | 0.725853 | | 1 | 0.996511 | 0.958398 |
| Intu | Q059U7-2 | -0.51 | 0 | 1.01 | -0.32 | -1.2 | -0.87 | -0.98 | 0.94 | 0.33 | 1.73 | 0.09 | -0.14 | 2.22 | -0.46 | 1.23 | 0.33 | -0.5725 | 0.4575 | 0.785 | 0.312557 | 0.37367 | 0.221733 | | 1 | 0.996511 | 0.787447 |
| Ip6k1 | Q6PD10 | 0.05 | -0.09 | -0.42 | -0.3 | 0.05 | 0.12 | -0.57 | -0.38 | 0.07 | 0.33 | -0.6 | 0.15 | -0.15 | 0.44 | 0.06 | -0.49 | -0.005 | 0.1775 | 0.155 | 0.981891 | 0.48668 | 0.544958 | | 1 | 0.996511 | 0.912212 |
| Ipcef1 | Q5DU31 | 0.62 | -0.42 | 0.33 | -0.19 | -0.42 | 0.71 | -0.16 | 0 | -0.72 | -0.29 | -0.11 | 0.07 | -0.33 | -0.81 | -0.21 | -0.4 | -0.0525 | -0.3475 | -0.5225 | 0.873379 | 0.262738 | 0.097192 | | 1 | 0.996511 | 0.698156 |
| Ipo11 | Q8K2V6 | -1.13 | -0.68 | -0.49 | -0.55 | -1 | -0.72 | -0.78 | -0.66 | -0.48 | 0.68 | -0.5 | -0.61 | 0.04 | -0.62 | -0.57 | -0.34 | -0.0775 | 0.485 | 0.34 | 0.695373 | 0.17114 | 0.188891 | | 1 | 0.996511 | 0.776843 |
| Ipo13 | Q8K0C1 | -0.55 | -0.44 | -0.79 | -1.37 | -0.31 | -0.21 | -1.69 | -1.03 | 0.58 | -1.21 | -0.82 | -0.28 | 1.46 | -0.57 | -0.39 | -0.83 | -0.0225 | 0.355 | 0.705 | 0.953025 | 0.405415 | 0.199468 | | 1 | 0.996511 | 0.78004 |
| Ipo4 | Q8VI75 | -0.41 | -0.34 | -0.65 | -0.56 | -0.3 | -0.2 | -0.77 | -0.41 | -0.29 | 0.33 | -0.36 | -0.45 | -0.31 | -0.05 | -0.74 | -0.46 | 0.07 | 0.2975 | 0.1 | 0.705777 | 0.209177 | 0.643728 | | 1 | 0.996511 | 0.944148 |
| Ipo5 | Q8BKC5 | -0.45 | -0.57 | -0.32 | -0.51 | -0.38 | -0.15 | -0.52 | -0.41 | -0.68 | -0.33 | -0.57 | -0.37 | -0.81 | -0.72 | -0.63 | -0.45 | 0.0975 | -0.025 | -0.19 | 0.542883 | 0.885002 | 0.311627 | | 1 | 0.996511 | 0.834523 |
| Ipo7 | Q9EPL8 | -0.89 | -0.86 | -0.28 | -0.44 | -0.46 | -0.74 | -0.21 | -0.28 | -0.91 | -0.42 | -0.3 | -1 | -1.4 | -0.64 | -0.7 | -0.51 | 0.195 | -0.04 | -0.195 | 0.381232 | 0.874579 | 0.483311 | | 1 | 0.996511 | 0.893925 |
| Ipo8 | Q7TMY7 | -1.01 | -0.48 | 0.07 | -0.65 | -0.71 | -1.01 | -0.54 | -0.28 | -1.01 | -0.65 | -0.48 | -0.66 | -1.02 | -0.64 | -1.11 | -0.01 | -0.1175 | -0.1825 | -0.1775 | 0.67188 | 0.500794 | 0.601638 | | 1 | 0.996511 | 0.930802 |
| Ipo9 | E9QKZ2 | 0.33 | -0.17 | -0.02 | -0.57 | 0.15 | 0.58 | 0.12 | -0.38 | -0.25 | -0.06 | 0.03 | 0.52 | -0.18 | 0.04 | -0.05 | 0.26 | 0.225 | 0.1675 | 0.125 | 0.422626 | 0.532894 | 0.61184 | | 1 | 0.996511 | 0.935047 |
| Iqcb1 | Q8BP00-2 | 0.09 | 0.51 | 1.58 | 0 | 0.31 | 0.13 | 1.01 | 0.93 | 0.2 | 0.18 | 1.47 | 0.49 | 0.02 | 0.69 | 1.34 | 1.59 | 0.05 | 0.04 | 0.365 | 0.900449 | 0.928343 | 0.44936 | | 1 | 0.996755 | 0.884817 |
| Iqgap1 | Q9JKF1 | -0.21 | -0.59 | -1.12 | -1.05 | -0.41 | -0.44 | -0.94 | -1.34 | -0.64 | -0.76 | -0.98 | -0.25 | -0.82 | -0.47 | -1.38 | -1.05 | -0.04 | 0.085 | -0.1875 | 0.89534 | 0.758367 | 0.53604 | | 1 | 0.996511 | 0.912212 |
| Iqgap2 | Q3UQ44 | 0 | -0.3 | -0.07 | -0.09 | -0.57 | -0.18 | -0.23 | -0.02 | -0.36 | -0.42 | -0.35 | -0.69 | 0.07 | -0.42 | -0.82 | -0.84 | -0.135 | -0.34 | -0.3875 | 0.457404 | 0.078794 | 0.154172 | | 1 | 0.996511 | 0.750763 |
| Iqsec1 | E9PUA3 | 1.26 | 1.22 | 0.21 | 0.7 | 0.61 | 1.34 | 1.1 | 0.28 | 1.1 | 1.81 | 0.79 | 1.52 | 1.13 | 1.69 | 1.16 | 1.45 | -0.015 | 0.4575 | 0.51 | 0.96411 | 0.192319 | 0.111736 | | 1 | 0.996511 | 0.713445 |
| Iqsec1 | A0A1D5RN | 0.49 | 1.11 | 1.66 | 1.08 | 0.55 | 0.46 | 1.16 | 1.27 | 0.9 | 1.4 | 1.55 | 0.93 | 1.62 | 0.39 | 1.68 | 1.63 | -0.225 | 0.11 | 0.245 | 0.476351 | 0.712026 | 0.527574 | | 1 | 0.996511 | 0.908538 |
| Iqsec2 | A4GZ26 | 0.88 | 0.76 | 1.64 | 1.58 | 0.53 | 0.83 | 0.67 | 1.23 | 1.15 | 1.39 | 1.61 | 0.87 | 1.41 | 0.65 | 1.14 | 1.05 | -0.4 | 0.04 | -0.1525 | 0.175899 | 0.88976 | 0.606311 | | 1 | 0.996511 | 0.932425 |
| Iqsec3 | Q3TES0 | 0.55 | 0.47 | 0.17 | 0.27 | 0.44 | 0.45 | 0.33 | 0.23 | 0.72 | 0.15 | 0.2 | 0.57 | 0.99 | 0.49 | 0.71 | 0.54 | -0.0025 | 0.045 | 0.3175 | 0.987684 | 0.82929 | 0.147865 | | 1 | 0.996511 | 0.746406 |
| Irak1 | B1AUW9 | 1.71 | 0.95 | 1.99 | 1.75 | 0.81 | 0.74 | 1.84 | 0.94 | 0.28 | 1.79 | 0.94 | 0.7 | 4.3 | 1.19 | 1.16 | 2.25 | -0.5175 | -0.6725 | 0.625 | 0.144869 | 0.101995 | 0.38236 | | 1 | 0.996511 | 0.860984 |
| Ireb2 | Q811J3 | 0.35 | -0.16 | -0.52 | -0.68 | -0.53 | 0.85 | -1 | -0.91 | -0.67 | -0.58 | -0.75 | 0.46 | -0.28 | -2.07 | -0.75 | -0.65 | -0.145 | -0.1325 | -0.685 | 0.748664 | 0.710469 | 0.138448 | | 1 | 0.996511 | 0.73493 |
| Irf2bp1 | Q8R3Y8-2 | 2 | -3.26 | 0.17 | -1.98 | 1.21 | -3.2 | -2.68 | -2.67 | -1.23 | -0.41 | -2.25 | -1.75 | 2.38 | -2.07 | 0.3 | -2.52 | -1.0675 | -0.6425 | 0.29 | 0.448308 | 0.563167 | 0.840842 | | 1 | 0.996511 | 0.976225 |
| Irf2bp2 | E9Q1P8 | -0.02 | -0.29 | -1.08 | -0.42 | 1.15 | 0.3 | -0.91 | -0.94 | 0.22 | 0.03 | -1.15 | 0.47 | -0.27 | 0.34 | -0.69 | -1.16 | 0.3525 | 0.345 | 0.0075 | 0.496287 | 0.40383 | 0.984234 | | 1 | 0.996511 | 0.996154 |
| Irf2bpl | Q8K3X4 | -3.13 | -3.04 | -3.39 | -3.46 | -2.85 | -3.35 | -3.97 | -4 | -2.69 | -1.72 | -3.36 | -2.65 | -2.92 | -2.13 | -3.59 | -4.07 | -0.2875 | 0.65 | 0.0775 | 0.339848 | 0.087673 | 0.851821 | | 1 | 0.996511 | 0.976225 |
| Irf3 | P70671 | 0.19 | -0.11 | -0.05 | -0.31 | -0.31 | 0.28 | -0.58 | -0.52 | 0.7 | -0.18 | -0.21 | -0.11 | 0.94 | -0.63 | 0.07 | -0.2 | -0.2125 | 0.12 | 0.115 | 0.383155 | 0.646411 | 0.740026 | | 1 | 0.996511 | 0.958796 |
| Irgm1 | Q60766-2 | 1.29 | 1.1 | 1.11 | 0.87 | 1.12 | 0.61 | 1.07 | 0.85 | -1.23 | 0.08 | 0.29 | 0.61 | -0.99 | -1.03 | 0.2 | 0.56 | -0.18 | -1.155 | -1.4075 | 0.346706 | 0.016487 | 0.006545 | | 1 | 0.996511 | 0.552614 |
| Irgq | Q8VIM9 | -0.45 | -0.7 | -0.16 | -0.49 | -0.58 | -0.5 | -0.23 | -0.32 | -0.3 | 0.25 | -0.03 | -0.21 | -0.4 | 0.08 | -0.2 | -0.12 | 0.0425 | 0.3775 | 0.29 | 0.814837 | 0.097779 | 0.188149 | | 1 | 0.996511 | 0.776843 |
| Irs1 | Q543V3 | 1 | 0.56 | 0.9 | 0.65 | 1.57 | 0.39 | -0.04 | 0.46 | 0.02 | 0.47 | 0.91 | 1.41 | -0.41 | -0.8 | -0.02 | 0.52 | -0.1825 | -0.075 | -0.955 | 0.600184 | 0.812743 | 0.012494 | | 1 | 0.996511 | 0.573604 |
| Irs2 | P81122 | -1.07 | -0.76 | -0.64 | -0.76 | -1.07 | -1.43 | -0.76 | -0.66 | -0.88 | -0.13 | -0.71 | -0.95 | -0.45 | -0.36 | -0.84 | -0.58 | -0.1725 | 0.14 | 0.25 | 0.441206 | 0.558068 | 0.238894 | | 1 | 0.996511 | 0.793481 |
| Isca1 | Q9D924 | 0.25 | 0.45 | -0.38 | -0.34 | 0.75 | 0.38 | -0.27 | -0.26 | 0.12 | -0.33 | -0.28 | 0.05 | 0.61 | 0.9 | -0.25 | -0.34 | 0.155 | -0.105 | 0.235 | 0.630042 | 0.6855 | 0.526491 | | 1 | 0.996511 | 0.908107 |
| Isca2 | A0A1Y7VK: | 0.14 | 0.44 | 0.14 | 0.27 | 0.01 | -0.05 | -0.03 | -0.03 | -0.07 | -0.06 | 0.11 | 0.28 | 0.26 | 0.29 | -0.31 | 0.16 | -0.2725 | -0.1825 | -0.1475 | 0.095286 | 0.319631 | 0.494155 | | 1 | 0.996511 | 0.89758 |
| Iscu | Q9D7P6 | 0.09 | -0.28 | 0.08 | 0.27 | -0.53 | 0.25 | -0.14 | 0.06 | -0.08 | 0.21 | 0.01 | 0.21 | 0 | -0.2 | 0.02 | 0 | -0.13 | 0.0475 | -0.085 | 0.56621 | 0.804759 | 0.667061 | | 1 | 0.996511 | 0.951125 |
| Isg15 | Q64339 | -1.11 | -2.47 | -0.86 | -0.14 | -1.77 | 0.14 | -0.58 | 0.18 | -1.14 | -1.42 | 0.23 | -1.89 | -1.48 | -1.77 | -1.77 | -0.96 | 0.6375 | 0.09 | -0.35 | 0.310138 | 0.88279 | 0.480833 | | 1 | 0.996511 | 0.893906 |
| Isg20l2 | A0A0R4J0R | 0.98 | -0.14 | -0.22 | 0.33 | -1.85 | 0.45 | -0.79 | -0.57 | 0.66 | -0.41 | -0.01 | -0.33 | 1.3 | -0.61 | -0.49 | -0.92 | -0.9275 | -0.26 | -0.4175 | 0.093394 | 0.475171 | 0.439391 | | 1 | 0.996511 | 0.883205 |
| Islr2 | E9Q9N7 | 0.95 | 1.07 | 0.84 | 0.56 | 1.18 | 0.82 | 1.12 | 0.99 | 0.73 | 1.02 | 1.07 | 1.17 | 0.4 | 0.98 | 0.76 | 0.56 | 0.1725 | 0.1425 | -0.18 | 0.351283 | 0.474878 | 0.419044 | | 1 | 0.996511 | 0.87558 |
| Isoc1 | Q91V64 | 1.3 | 0.06 | 0.07 | 0.68 | 0.07 | 0.88 | 0.65 | 0.96 | 1.13 | 1.15 | 0.27 | 1.45 | 0.52 | 0.89 | 0.56 | 0.17 | 0.1125 | 0.4725 | 0.0075 | 0.744638 | 0.230711 | 0.981996 | | 1 | 0.996511 | 0.994858 |
| Isoc2a | P85094 | 2.19 | 1.8 | 2.58 | 2.75 | 1.04 | 2.03 | 2.14 | 2.55 | 0.56 | 1.52 | 2.61 | 1.63 | 0.2 | -0.12 | 2.09 | 2.4 | -0.39 | -0.75 | -1.1875 | 0.303561 | 0.117893 | 0.082178 | | 1 | 0.996511 | 0.68865 |
| Isoc2b | Q9DCC7 | -0.96 | 0.08 | -0.29 | -0.11 | -0.44 | -0.83 | 0.53 | -0.62 | -0.09 | -0.21 | -0.09 | -0.77 | -0.41 | 0.05 | -1.11 | -0.39 | -0.02 | 0.03 | -0.145 | 0.955677 | 0.917013 | 0.664205 | | 1 | 0.996755 | 0.949825 |
| Ist1 | Q9CX00 | 0.18 | 0.44 | -0.52 | -0.56 | 0.34 | 0.13 | -0.55 | -0.7 | 0.51 | -0.19 | -0.48 | 0.53 | 0.89 | 1.03 | -0.67 | -0.51 | -0.08 | 0.2075 | 0.3 | 0.816382 | 0.557214 | 0.538365 | | 1 | 0.996511 | 0.912212 |
| Isy1 | Q69ZQ2 | -0.27 | -0.39 | -0.88 | 0.11 | -0.74 | -0.34 | -1.41 | -0.05 | -0.53 | 1.58 | -0.79 | -0.74 | 0.33 | -0.21 | 0.04 | -0.7 | -0.2775 | 0.2375 | 0.2225 | 0.430591 | 0.671537 | 0.47801 | | 1 | 0.996511 | 0.892279 |
| Isyna1 | Q9JHU9 | -1.27 | -1.29 | -0.42 | -0.55 | -1.17 | -1.01 | -0.1 | -0.09 | -0.86 | -0.29 | -0.06 | -0.85 | -0.9 | -1.11 | 0.07 | 0.32 | 0.29 | 0.3675 | 0.4775 | 0.423242 | 0.255146 | 0.25811 | | 1 | 0.996511 | 0.812074 |
| Itch | Q8C863 | 0.99 | 1.04 | 0.14 | 0.19 | 1.03 | 1.39 | 0.9 | 0.64 | 0.31 | 0.03 | 0.4 | 1.09 | 0.63 | -0.27 | 0.43 | 0.91 | 0.4 | -0.1325 | -0.165 | 0.193396 | 0.69 | 0.638898 | | 1 | 0.996511 | 0.942804 |
| Itfg1 | Q99KW9 | -1.25 | -0.72 | -0.45 | 0.36 | 0.06 | -0.99 | 0.2 | 0.5 | -0.74 | 0.02 | 0.27 | -0.69 | -1.16 | -0.63 | -0.04 | 0.06 | 0.4575 | 0.23 | 0.0725 | 0.309618 | 0.570763 | 0.862892 | | 1 | 0.996511 | 0.977235 |
| Itfg2 | Q91WI7-2 | -0.05 | 0.37 | 0.34 | -0.05 | -0.34 | -0.24 | -0.76 | -0.13 | 0.21 | 0.22 | -0.07 | 0.02 | 0.99 | -0.21 | 0.18 | -0.61 | -0.52 | -0.0575 | -0.065 | 0.032812 | 0.76551 | 0.855391 | | 1 | 0.996511 | 0.976225 |
| Itga1 | Q3V3R4 | -0.46 | -0.24 | 0.1 | -0.52 | 0.12 | -0.4 | -0.69 | -0.5 | 1.18 | -0.74 | 0.31 | -0.05 | 0.56 | -1.22 | 0.19 | 0.29 | -0.0875 | 0.455 | 0.235 | 0.715088 | 0.278258 | 0.567148 | | 1 | 0.996511 | 0.921806 |
| Itga3 | Q62470 | 0.13 | 0.36 | 0.15 | -0.07 | 0.5 | 0.13 | 0.21 | 0.2 | 0.95 | 0.85 | 0.32 | 0.33 | 1.04 | 0.77 | 0.25 | 0.52 | 0.1175 | 0.47 | 0.5025 | 0.499548 | 0.061623 | 0.055942 | | 1 | 0.996511 | 0.661738 |
| Itga4 | Q792F9 | -1.02 | -0.65 | -0.66 | -0.62 | -0.88 | 0.39 | -0.61 | -0.66 | 0.01 | -1.56 | -0.08 | -0.7 | 0.13 | 0.97 | -0.49 | -0.39 | 0.2975 | 0.155 | 0.7925 | 0.330551 | 0.671494 | 0.044756 | | 1 | 0.996511 | 0.658454 |
| Itga5 | P11688 | 0.3 | 0.7 | 1.23 | 1.18 | 0.2 | -0.14 | 1.52 | 1.38 | 0.39 | 1.01 | 1.58 | 0.74 | 0.48 | 0.79 | 1.62 | 1.17 | -0.1125 | 0.0775 | 0.1625 | 0.798169 | 0.815027 | 0.627199 | | 1 | 0.996511 | 0.939737 |
| Itga6 | Q61739-2 | -0.18 | 0.14 | 0.09 | 0.22 | 0.29 | 0.04 | -0.17 | -0.31 | 0.16 | 0.31 | 0.34 | 0.41 | 0.15 | -0.02 | 0.18 | 0.23 | -0.105 | 0.2375 | 0.0675 | 0.590447 | 0.196275 | 0.716764 | | 1 | 0.996511 | 0.958334 |
| Itga7 | G3X9Q1 | -0.94 | -0.56 | -0.1 | -0.08 | -0.24 | -0.91 | 0.02 | -0.28 | -0.82 | -0.28 | -0.07 | -0.51 | -0.68 | -0.51 | -0.59 | 0.16 | 0.0675 | 1.96E-16 | 0.015 | 0.813768 | 1 | 0.959502 | | 1 | 1 | 0.992636 |
| Itgam | E9Q604 | -0.78 | -0.67 | 0.1 | 0.01 | -1 | -1.08 | -0.04 | 0.22 | -1.25 | 0.07 | 0.02 | -1.13 | -1.31 | -0.91 | -0.84 | -0.49 | -0.14 | -0.2375 | -0.5525 | 0.714079 | 0.560713 | 0.089844 | | 1 | 0.996511 | 0.69217 |
| Itgav | P43406 | 0.91 | 0.82 | 0.3 | 0.39 | 1.02 | 1.02 | 0.34 | 0.18 | 0.63 | -0.13 | 0.34 | 1.01 | 0.3 | 0.54 | 0.21 | 0.34 | 0.035 | -0.1425 | -0.2575 | 0.898226 | 0.628664 | 0.258907 | | 1 | 0.996511 | 0.812074 |
| Itgb1 | P09055 | -0.48 | -0.64 | 0 | 0.08 | -0.61 | -0.48 | 0.16 | -0.4 | -0.53 | 0.46 | 0.24 | 0.01 | -0.27 | -0.25 | 0.75 | 0.34 | -0.0725 | 0.305 | 0.4025 | 0.776949 | 0.303427 | 0.219713 | | 1 | 0.996511 | 0.787447 |
| Itgb1bp1 | O35671 | -0.52 | -0.88 | -0.3 | -0.25 | -0.35 | -0.81 | -0.23 | -0.48 | -0.5 | -0.48 | -0.71 | -1.12 | -0.43 | -0.06 | -0.74 | -0.59 | 0.02 | -0.215 | 0.0325 | 0.926057 | 0.373802 | 0.893258 | | 1 | 0.996511 | 0.982875 |
| Itgb2 | Q542I8 | -0.91 | -1.22 | -1.78 | -1.21 | -1.08 | -1.07 | -1.81 | -1.24 | -1.44 | -1.88 | -1.63 | -1.32 | -1.52 | -1.34 | -2.06 | -1.87 | -0.02 | -0.2875 | -0.4175 | 0.93873 | 0.258651 | 0.147046 | | 1 | 0.996511 | 0.746406 |
| Itgb3 | O54890 | 1.97 | 0.58 | 0.64 | 0.45 | 1.48 | 2.61 | 0.75 | 0.76 | 2.05 | 1.3 | 0.89 | 2.07 | 1.79 | 0.88 | 1.34 | 1.02 | 0.49 | 0.6675 | 0.3475 | 0.357961 | 0.150559 | 0.390001 | | 1 | 0.996511 | 0.865375 |
| Itgb4 | A2A864 | -1.36 | -1.73 | 0.22 | -0.18 | -1.38 | -1.16 | -1.33 | -1.2 | -0.37 | -0.22 | -1.34 | -0.47 | -0.83 | -1.72 | -1.79 | -0.31 | -0.505 | 0.1625 | -0.4 | 0.266736 | 0.742173 | 0.468301 | | 1 | 0.996511 | 0.889423 |
| Itgb5 | Q6PE70 | 0.59 | 0.79 | 0.48 | 0.25 | 0.88 | 0.17 | -0.13 | -0.24 | -0.72 | -0.58 | 0.21 | 0.16 | -0.48 | -1.77 | 0.36 | 0.48 | -0.3575 | -0.76 | -0.88 | 0.222336 | 0.022708 | 0.103122 | | 1 | 0.996511 | 0.700625 |
| Itgb8 | Q0VBD0 | 0.91 | 1.07 | -0.45 | -0.21 | 1.37 | 0.98 | -0.17 | -0.33 | 0.79 | 0.5 | -0.4 | 1 | 0.47 | 1.09 | -0.09 | -0.14 | 0.1325 | 0.1425 | 0.0025 | 0.800689 | 0.758639 | 0.995603 | | 1 | 0.996511 | 0.998452 |
| Itih1 | F8WJ05 | -1.1 | -0.68 | 0.13 | -0.3 | 0.25 | -1.11 | 0.1 | -0.16 | -0.76 | 3.66 | 0.23 | -0.87 | -1.5 | -0.56 | -0.36 | -0.2 | 0.2575 | 1.0525 | -0.1675 | 0.50612 | 0.30044 | 0.662307 | | 1 | 0.996511 | 0.949645 |
| Itih2 | Q61703 | -1.53 | -0.83 | -1.12 | -1.01 | -0.77 | -1.35 | -0.99 | -0.93 | -1.74 | 0.01 | -0.78 | -0.98 | -1.45 | -1.32 | -1.09 | -1.02 | 0.1125 | 0.25 | -0.0975 | 0.607133 | 0.511403 | 0.667652 | | 1 | 0.996511 | 0.951339 |
| Itih3 | Q61704 | -0.71 | -1.53 | -0.46 | -0.27 | -1.24 | -1.92 | -0.26 | -0.52 | -1.72 | -0.46 | -0.5 | -1.83 | -1.24 | -1.38 | -0.37 | 0.05 | -0.2425 | -0.385 | 0.0075 | 0.580739 | 0.392153 | 0.985807 | | 1 | 0.996511 | 0.996154 |
| Itih4 | E9Q5L2 | -2.35 | -3.27 | -2.11 | -1.84 | -1.97 | -2.53 | -2.26 | -2.36 | -3.61 | 0.14 | -2.53 | -2.57 | -2.68 | -3.95 | -2.33 | -3.06 | 0.1125 | 0.25 | -0.6125 | 0.72901 | 0.748305 | 0.189932 | | 1 | 0.996511 | 0.776843 |
| Itih5 | Q8BJD1 | -0.59 | -0.28 | -0.49 | -0.59 | 0.34 | -0.34 | 0.1 | -0.39 | -0.44 | -0.12 | -0.35 | -0.33 | -0.6 | -0.37 | -1.12 | -0.59 | 0.415 | 0.1775 | -0.1825 | 0.081936 | 0.320866 | 0.423567 | | 1 | 0.996511 | 0.876113 |
| Itm2a | Q61500 | -0.22 | -0.22 | 0.03 | 0.2 | 0.25 | -0.33 | 0.05 | -0.35 | 0.05 | 0.2 | 0.6 | -0.06 | -0.03 | 0.2 | 0.36 | 0.16 | -0.0425 | 0.25 | 0.225 | 0.83843 | 0.266695 | 0.275082 | | 1 | 0.996511 | 0.822084 |
| Itm2b | O89051 | 0.09 | 0.29 | 1.11 | 1.12 | 0.7 | -0.14 | 1.28 | 1.06 | -0.08 | 1.15 | 0.93 | 0.08 | -0.26 | 0.48 | 0.9 | 1.11 | 0.0725 | -0.1325 | -0.095 | 0.852702 | 0.735455 | 0.808834 | | 1 | 0.996511 | 0.972098 |
| Itm2c | Q91VK4 | -0.21 | -0.21 | -0.07 | -0.06 | 0.38 | -0.6 | 0.48 | 0.02 | -0.25 | -0.5 | -0.13 | -0.42 | -0.2 | -0.06 | 0.35 | 0.15 | 0.2075 | -0.1875 | 0.1975 | 0.429119 | 0.288762 | 0.330814 | | 1 | 0.996511 | 0.845601 |
| Itpa | Q9D892 | -0.79 | -0.37 | -0.62 | -0.72 | -0.4 | -0.93 | -0.77 | -0.74 | -1.02 | -0.09 | -0.98 | -0.7 | -1.37 | -0.05 | -0.95 | -1.42 | -0.085 | -0.0725 | -0.3225 | 0.649045 | 0.776348 | 0.345765 | | 1 | 0.996511 | 0.850095 |
| Itpk1 | Q8BYN3 | -1.36 | -1.52 | -0.55 | -0.7 | -1.59 | -1.36 | -0.36 | -1.32 | -1.63 | -0.47 | -0.61 | -1.18 | -1.23 | -1.7 | -1.12 | -0.61 | -0.125 | 0.06 | -0.1325 | 0.720988 | 0.864461 | 0.689964 | | 1 | 0.996511 | 0.955729 |
| Itpka | Q8R071 | 0.34 | -0.58 | 0.48 | 0.36 | -0.77 | 0.29 | -0.26 | 0.41 | -0.55 | -0.08 | 0.26 | -0.57 | -1.4 | -0.9 | -0.98 | -1.04 | -0.2325 | -0.385 | -1.23 | 0.513904 | 0.245549 | 0.002052 | | 1 | 0.996511 | 0.419079 |
| Itpkc | Q7TS72 | -0.27 | -0.3 | -1.14 | -0.74 | -2.13 | -0.71 | -2.02 | -2.08 | -0.07 | -0.8 | -0.88 | -1.25 | -0.17 | -0.42 | -0.59 | -2.4 | -1.1225 | -0.1375 | -0.2825 | 0.01552 | 0.67043 | 0.583432 | | 1 | 0.996511 | 0.927734 |
| Itpr1 | P11881-4 | 1.26 | 2.45 | 0.99 | 1.07 | 0.16 | 0.14 | 0.26 | 1 | 0.12 | -0.44 | 0.14 | 0.79 | -0.62 | 1.63 | -0.08 | 1.9 | -1.0525 | -1.29 | -0.735 | 0.020376 | 0.010738 | 0.273979 | | 1 | 0.996511 | 0.822084 |
| Itpr1 | P11881-2 | 0.5 | 0.32 | 0.53 | 0.33 | 0.37 | 0.39 | 0.48 | 0.54 | 0.5 | -0.16 | 0.14 | 0.2 | 0.75 | 0.39 | 0.63 | 0.35 | 0.025 | -0.25 | 0.11 | 0.865447 | 0.226892 | 0.564625 | | 1 | 0.996511 | 0.920315 |
| Itpr2 | Q9Z329-3 | 0.62 | 0.87 | 0.61 | 0.53 | 1.29 | 0.82 | 0.73 | 0.6 | 0.48 | 0.78 | 0.57 | 0.77 | 0.76 | 1.1 | 0.55 | 0.71 | 0.2025 | -0.0075 | 0.1225 | 0.326529 | 0.965919 | 0.548665 | | 1 | 0.996755 | 0.912757 |
| Itpr3 | P70227 | -3.57 | -4.43 | -0.09 | -0.53 | -4.96 | -2.82 | -1.08 | -1.1 | -2.39 | -1.12 | -0.72 | -2.98 | -2 | -6.36 | -0.99 | -1.34 | -0.335 | 0.3525 | -0.5175 | 0.792195 | 0.745098 | 0.725034 | | 1 | 0.996511 | 0.958398 |
| Itprid2 | A2AQD6 | -1.03 | -0.14 | -1 | -1.2 | -1.25 | 0.18 | -0.76 | -0.38 | 0 | 0.17 | 0.14 | 0.34 | -1.01 | -1.28 | -1.16 | 0.61 | 0.29 | 1.005 | 0.1325 | 0.43819 | 0.004331 | 0.779504 | | 1 | 0.996511 | 0.967289 |
| Itsn1 | E9Q0N0 | 0.93 | 0.29 | 0.2 | 0.3 | 0.36 | 0.84 | 0.39 | 0.22 | 0.4 | 0.28 | 0.12 | 0.45 | 0.25 | 1 | 0.03 | 0.03 | 0.0225 | -0.1175 | -0.1025 | 0.923135 | 0.597077 | 0.732065 | | 1 | 0.996511 | 0.958635 |
| Itsn2 | E9QNG1 | 0.79 | 0.35 | 1.08 | 1.05 | 0.54 | 0.69 | 0.96 | 1.38 | 0.28 | 0.54 | 0.91 | 0.15 | 0.2 | 0.77 | 0.72 | 0.72 | 0.075 | -0.3475 | -0.215 | 0.772568 | 0.2012 | 0.399688 | | 1 | 0.996511 | 0.868507 |
| Ivd | Q9JHI5 | 1.08 | 0.91 | 0.61 | 0.64 | 0.6 | 0.64 | 0.78 | 0.64 | 1.14 | 0.45 | 1.17 | 1.1 | 0.87 | 0.29 | 0.27 | 0.48 | -0.145 | 0.155 | -0.3325 | 0.406823 | 0.515497 | 0.166316 | | 1 | 0.996511 | 0.759484 |
| Ivns1abp | Q920Q8-4 | -2.26 | -3.32 | -2.38 | -2.9 | -2.83 | -2.24 | -1.56 | -2.12 | -1.37 | -1.25 | -2.1 | -2.02 | 0.79 | -2.21 | -0.39 | -1.67 | 0.5275 | 1.03 | 1.845 | 0.152278 | 0.011436 | 0.018738 | | 1 | 0.996511 | 0.602211 |
| Iws1 | A0A1D5RL' | -1.09 | -2.32 | -0.72 | -0.53 | -2.96 | -1.75 | -1.32 | -0.24 | -1.32 | -1.05 | -1.02 | -2.04 | -0.68 | -1.7 | -0.71 | -1.32 | -0.4025 | -0.1925 | 0.0625 | 0.52723 | 0.663619 | 0.888735 | | 1 | 0.996511 | 0.98074 |
| Jagn1 | Q5XKN4 | -0.57 | -0.18 | -0.39 | -0.27 | -0.41 | -0.03 | -2.56 | -0.67 | 0.41 | 0.36 | -1.25 | -0.47 | 0.68 | -0.91 | 0.51 | -0.28 | -0.565 | 0.115 | 0.3525 | 0.29713 | 0.766513 | 0.353935 | | 1 | 0.996511 | 0.855029 |
| Jak1 | B1ASP2 | 0.56 | 0.71 | 0.66 | 0.53 | 0.72 | 0.26 | 0.46 | 0.09 | 0.2 | 1.05 | 0.39 | 0.55 | -0.01 | 0.5 | 0.6 | 0.33 | -0.2325 | -6.75E-02 | -0.26 | 0.228278 | 0.762323 | 0.223449 | | 1 | 0.996511 | 0.787447 |
| Jak2 | Q62120 | -0.3 | 0.29 | 0.86 | 0.7 | 0.36 | -0.31 | 0.45 | 0.34 | 0.24 | 0.3 | 0.69 | -0.14 | 0.62 | 0.53 | 0.26 | 0.49 | -0.1775 | -0.115 | 0.0875 | 0.568099 | 0.713304 | 0.76126 | | 1 | 0.996511 | 0.963123 |
| Jakmip1 | D3Z4U5 | -0.06 | 0.08 | 0.37 | 0.18 | 0.09 | -0.54 | 0.35 | 0.2 | -0.06 | 0.48 | 0.18 | 0.13 | -0.12 | 0.15 | 0.51 | 0.25 | -0.1175 | 0.04 | 0.055 | 0.617451 | 0.838438 | 0.797274 | | 1 | 0.996511 | 0.969727 |
| Jakmip2 | D3YXK0 | 0.14 | 0.51 | 0.68 | 0.79 | 0.87 | 0.08 | 0.83 | 0.81 | 0.27 | 1.08 | 0.62 | 0.36 | 0.05 | 0.35 | 0.82 | 0.66 | 0.1175 | 0.0525 | -0.06 | 0.639441 | 0.835553 | 0.813334 | | 1 | 0.996511 | 0.972768 |
| Jakmip3 | Q5DTN8 | 1.17 | 1.19 | 0.92 | 1.11 | 1.17 | 1.02 | 1.03 | 1.07 | 1.19 | 1.02 | 0.93 | 1.32 | 0.92 | 1.45 | 1.16 | 0.9 | -0.025 | 0.0175 | 0.01 | 0.866248 | 0.920996 | 0.961054 | | 1 | 0.996755 | 0.9927 |
| Jam2 | Q9JI59 | 0.62 | 0.68 | 0.64 | 0.7 | 0.76 | 0.73 | 0.96 | 0.41 | 0.1 | 0.71 | 0.2 | 0.75 | -0.21 | 0.39 | 0.62 | 0.48 | 0.055 | -0.22 | -0.34 | 0.745385 | 0.314329 | 0.162938 | | 1 | 0.996511 | 0.758224 |
| Jam3 | Q9D8B7 | 1.83 | 1.51 | 0.59 | 0.24 | 0.82 | 1.59 | -0.12 | -0.24 | 0.66 | 0.2 | 0.25 | 1.61 | 1.01 | -0.79 | 0.86 | 0.95 | -0.53 | -0.3625 | -0.535 | 0.327174 | 0.444745 | 0.328131 | | 1 | 0.996511 | 0.844932 |
| Jcad | Q5DTX6 | 0.55 | 0.48 | 0.74 | 0.81 | 0.21 | 0.32 | 0.36 | 0.52 | 0.52 | 0.4 | 0.79 | 0.71 | 0.71 | -0.2 | 0.59 | 0.52 | -0.2925 | -0.04 | -0.24 | 0.097972 | 0.826262 | 0.35418 | | 1 | 0.996511 | 0.855029 |
| Jmjd6 | G3UVU9 | -1.61 | -1.68 | -1.04 | -0.91 | -2.53 | -2.69 | -1.71 | -1.67 | -1.03 | -2.03 | -1.22 | -1.48 | -3.12 | -1.57 | -1.83 | -2.06 | -0.84 | -0.13 | -0.835 | 0.02764 | 0.664972 | 0.053132 | | 1 | 0.996511 | 0.661738 |
| Jmy | A0A0R4J0\ | 0.86 | 0.53 | 0.47 | 0.39 | 1.1 | 1 | 0.33 | 0.24 | 0.51 | 0.57 | 0.47 | 1.23 | 0.94 | 0.48 | 0.63 | 0.39 | 0.105 | 0.1325 | 0.0475 | 0.682482 | 0.578359 | 0.824452 | | 1 | 0.996511 | 0.972852 |
| Josd1 | Q9DBJ6 | 0.07 | 0.67 | 2.47 | 0.52 | -0.18 | 0.55 | 1.79 | 2.04 | 0.58 | 1.51 | 1.58 | -0.61 | -0.77 | -0.6 | 1.92 | -0.06 | 0.1175 | -0.1675 | -0.81 | 0.861707 | 0.80276 | 0.287813 | | 1 | 0.996511 | 0.824705 |
| Josd2 | D3YTZ7 | -0.3 | 0.21 | 0.32 | -0.43 | 0.51 | -0.34 | -0.16 | -1.01 | 0.14 | 1.14 | -0.19 | -0.11 | -0.16 | 0.15 | 0.67 | 0.27 | -0.2 | 0.295 | 0.2825 | 0.570586 | 0.410179 | 0.319237 | | 1 | 0.996511 | 0.838969 |
| Jph1 | Q9ET80 | -0.18 | -0.11 | 0.23 | 0.43 | -0.03 | -0.18 | 0.06 | 0.57 | 0.06 | 0.38 | -0.08 | -0.62 | 0.87 | 0.47 | 0.33 | -0.33 | 0.0125 | -0.1575 | 0.2425 | 0.957489 | 0.562343 | 0.428841 | | 1 | 0.996511 | 0.879165 |

| | | | | | | | | | | | | | | | | | | | | | | | | | | |
|---|---|---|---|---|---|---|---|---|---|---|---|---|---|---|---|---|---|---|---|---|---|---|---|---|---|---|
| Jph3 | Q9ET77 | 1.27 | 0.57 | 0.83 | 1.14 | 0.95 | 1.17 | 0.65 | 0.94 | 1.15 | 0.5 | 0.46 | 0.6 | 1.46 | 1.1 | 1.09 | 0.8 | -0.025 | -0.275 | 0.16 | 0.907621 | 0.285742 | 0.517394 | 1 | 0.996511 | 0.905961 |
| Jph4 | Q80WT0 | 0.51 | 0.08 | -0.11 | 0.01 | 0.73 | 0.19 | 0.44 | 0.12 | -0.03 | -0.13 | -0.12 | -0.35 | -0.3 | 0.82 | 0.54 | 0.18 | 0.2475 | -0.28 | 0.1875 | 0.273827 | 0.186504 | 0.526647 | 1 | 0.996511 | 0.908204 |
| Jpt1 | P97825 | -0.84 | -0.96 | -0.68 | -0.47 | -1 | -0.71 | -0.15 | -0.66 | 0.62 | -0.88 | -0.56 | -0.25 | 1.16 | -0.49 | -0.35 | -0.54 | 0.1075 | 0.47 | 0.6825 | 0.637509 | 0.186073 | 0.118969 | 1 | 0.996511 | 0.721448 |
| Jpt2 | Q6PGH2 | 0.25 | 0.18 | -0.12 | -0.28 | -0.15 | 0.44 | -0.01 | -0.09 | 1.84 | 0.16 | 0.23 | 0.83 | 2.33 | 0.26 | 1.42 | 0.64 | 0.04 | 0.7575 | 1.155 | 0.84954 | 0.080432 | 0.028394 | 1 | 0.996511 | 0.637553 |
| Jtb | D3Z4D1 | -3.15 | -4.87 | -3.16 | -1.43 | -4 | -4.26 | -2.51 | -1.77 | -3.65 | 0.2 | -2.49 | -3.93 | -1.94 | -2.02 | -1.8 | -1.14 | 0.0175 | 0.685 | 1.4275 | 0.983159 | 0.52113 | 0.056615 | 1 | 0.996511 | 0.661738 |
| Junb | P09450 | -3.78 | -3.36 | -2.12 | -2.63 | -3.98 | -2.67 | -2.71 | -2.16 | -2.2 | 2.47 | -2.74 | -3.48 | -0.72 | -1.66 | -1.46 | -3.27 | 0.0925 | 1.485 | 1.195 | 0.85173 | 0.253914 | 0.072221 | 1 | 0.996511 | 0.668066 |
| Jund | P15066 | -1.47 | -1.62 | -2.79 | -2.66 | -1.21 | -1.58 | -3.17 | -3.14 | -2.31 | -1.67 | -2.61 | -1.54 | -1.92 | -1.62 | -2.97 | -3.17 | -0.14 | 0.1025 | -0.285 | 0.804525 | 0.801725 | 0.557596 | 1 | 0.996511 | 0.917092 |
| Jup | Q02257 | 1.1 | 1.2 | 1.5 | 1.11 | 1.1 | 0.89 | 1.24 | 1.23 | 0.81 | 0.84 | 1.44 | 1.05 | 0.26 | 0.46 | 0.75 | 1.24 | -0.1125 | -0.1925 | -0.55 | 0.522586 | 0.37896 | 0.060361 | 1 | 0.996511 | 0.661738 |
| Kalrn | D3Z534 | 2.33 | 2.4 | 1.93 | 0.51 | 2.81 | 2.85 | 1.88 | 1.39 | -0.5 | -1.71 | 1.83 | 2.72 | -0.47 | -0.76 | 1.99 | 1.6 | 0.44 | -1.2075 | -1.2025 | 0.409466 | 0.247488 | 0.135377 | 1 | 0.996511 | 0.731223 |
| Kalrn | A2CG49 | 0.63 | 0.67 | 0.68 | 0.25 | 0.78 | 0.76 | 0.52 | 0.45 | 0.33 | -0.19 | 0.84 | 0.74 | 0.26 | 0.38 | 0.41 | 0.2 | 0.07 | -0.1275 | -0.245 | 0.69646 | 0.640201 | 0.220832 | 1 | 0.996511 | 0.787447 |
| Kalrn | A2CG49-9 | 7.28 | 2.45 | 4.61 | 1.83 | 3.01 | 7.43 | 7.05 | 5.08 | -0.17 | 0.97 | 4.59 | 5.67 | -0.74 | -0.41 | 4.75 | 6.66 | 1.6 | -1.2775 | -1.4775 | 0.280207 | 0.451073 | 0.461368 | 1 | 0.996511 | 0.887902 |
| Kank1 | E9Q238 | -1.02 | -0.47 | -0.33 | -0.91 | -1.66 | 0.01 | -2.25 | -1.08 | 3.19 | 0.27 | -1.16 | -0.55 | 3.44 | -0.7 | 0.12 | -0.69 | -0.5625 | 1.12 | 1.225 | 0.253608 | 0.225361 | 0.195202 | 1 | 0.996511 | 0.78004 |
| Kank2 | Q8BX02 | -0.33 | -0.27 | -0.6 | -0.83 | -0.64 | -0.84 | -0.61 | -0.97 | -0.36 | -0.25 | -0.48 | 0 | -0.11 | 0.1 | -0.3 | -0.46 | -0.2575 | 0.235 | 0.315 | 0.203405 | 0.277899 | 0.185154 | 1 | 0.996511 | 0.776843 |
| Kank3 | Q9Z1P7 | 0.16 | 0.44 | -0.01 | 0.56 | 0.16 | 0.1 | -0.32 | -0.29 | 0.38 | 1.27 | 0.47 | 0.56 | 0.42 | -0.09 | 0.15 | 0.27 | -0.375 | 0.3825 | -0.1 | 0.101061 | 0.167228 | 0.650974 | 1 | 0.996511 | 0.947407 |
| Kank4 | Q6P9J5 | 1.24 | 0.74 | 0.74 | 1 | 0.19 | 0.3 | -0.28 | -0.16 | 0.84 | 1.85 | 0.64 | 0.69 | 1.15 | 2.57 | 0.89 | 0.58 | -0.9175 | 0.075 | 0.3675 | 0.001927 | 0.810003 | 0.406492 | 1 | 0.996511 | 0.87356 |
| Kars | Q8R2P8 | -1.61 | -1.71 | -1.69 | -1.47 | -1.26 | -1.88 | -1.37 | -1.53 | -1.95 | -1.03 | -1.55 | -1.7 | -1.84 | -1.21 | -1.5 | -1.7 | 0.11 | 0.0625 | 0.0575 | 0.559354 | 0.788448 | 0.78197 | 1 | 0.996511 | 0.967642 |
| Kars1 | Q99MN1 | -0.53 | -0.04 | -0.61 | -1.86 | -0.68 | -0.13 | -1.91 | -0.33 | -0.04 | 0.36 | -0.89 | -0.36 | 0.44 | 0.81 | -0.2 | -0.22 | -0.0025 | 0.5275 | 0.9675 | 0.996102 | 0.252633 | 0.05176 | 1 | 0.996511 | 0.658454 |
| Kat2a | Q6P3Z8 | -0.82 | -0.51 | -2.79 | -2.71 | 0.05 | -0.48 | -1.56 | -2.22 | -0.76 | -2.85 | -2.8 | -0.88 | -0.26 | 0.3 | -0.86 | -0.92 | 0.655 | -0.115 | 1.2725 | 0.373752 | 0.879546 | 0.063673 | 1 | 0.996511 | 0.66433 |
| Kat5 | A0A494BAI | -1.78 | -5.91 | -0.89 | -0.11 | -4.38 | -3.06 | -0.83 | -0.5 | -0.72 | -1.29 | -0.75 | -3.71 | 1.34 | -6.16 | -0.65 | -0.95 | -0.02 | 0.555 | 0.5675 | 0.98874 | 0.675085 | 0.756076 | 1 | 0.996511 | 0.962333 |
| Kat7 | Q5SVQ0-3 | 0.97 | 0.41 | -0.29 | -0.8 | 0.33 | 0.93 | -0.78 | -1.05 | 0.75 | -1.73 | -0.94 | 0.85 | 1.23 | 0.26 | 0.05 | -0.1 | -0.215 | -0.34 | 0.2875 | 0.699838 | 0.619281 | 0.53816 | 1 | 0.996511 | 0.912212 |
| Katnal1 | Q8K0T4 | 0.21 | 0.02 | -0.08 | 0.13 | 0.12 | 0.18 | 0.08 | 0.1 | 0.48 | 0.56 | 0.03 | 0.21 | 0.2 | 0.98 | -0.19 | -0.03 | 0.05 | 0.25 | 0.17 | 0.73475 | 0.216592 | 0.55537 | 1 | 0.996511 | 0.917092 |
| Katnb1 | Q8BG40 | -0.06 | -0.28 | -0.41 | -0.62 | -0.07 | -0.08 | -0.21 | -0.31 | -0.62 | -1.24 | -0.23 | 0.02 | -0.6 | -0.25 | -0.57 | -0.55 | 0.175 | -0.175 | -0.15 | 0.338105 | 0.567798 | 0.471641 | 1 | 0.996511 | 0.891144 |
| Kazn | G3UY74 | 0.46 | 0.33 | 1.46 | 1.03 | 0.11 | -0.21 | 0.66 | 0.63 | 0.46 | 1.9 | 0.96 | 0.28 | 0.25 | 0.42 | 0.85 | 0.55 | -0.5225 | 0.08 | -0.3025 | 0.137658 | 0.850448 | 0.332241 | 1 | 0.996511 | 0.845601 |
| Kazn | Q69ZS8-2 | 0.55 | 0.55 | -0.04 | 0.19 | 0.9 | 0.12 | 0.35 | -0.06 | 0.47 | 0.49 | 0.22 | 0.75 | 0.58 | 0.81 | 0.52 | 0.06 | 0.015 | 0.17 | 0.18 | 0.954313 | 0.445388 | 0.475277 | 1 | 0.996511 | 0.892134 |
| Kbtbd11 | Q8BNW9 | 0.34 | 0.23 | 0.76 | 1.17 | -0.16 | 0.08 | 0.68 | 0.92 | -0.15 | 0.98 | 0.77 | -0.08 | -0.73 | 0.18 | 0.6 | 0.54 | -0.245 | -0.245 | -0.4775 | 0.45607 | 0.493137 | 0.215831 | 1 | 0.996511 | 0.787447 |
| Kbtbd2 | G3X9X1 | -0.48 | -0.96 | -1.22 | -1.08 | -0.97 | -0.47 | -0.76 | -0.86 | -1.45 | -0.32 | -0.86 | -0.44 | -1.27 | -0.98 | -0.83 | -0.89 | 0.17 | 0.1675 | -0.0575 | 0.443165 | 0.5871 | 0.804101 | 1 | 0.996511 | 0.971404 |
| Kbtbd3 | Q8BHI4 | -0.27 | 0.14 | -0.23 | -0.14 | 0.69 | -0.18 | 0.45 | 0.05 | -0.5 | -0.42 | 0.04 | 0.12 | -0.43 | -0.06 | -0.07 | 0.06 | 0.3775 | -0.065 | 7.85E-17 | 0.133813 | 0.768031 | 1 | 1 | 0.996511 | 1 |
| Kbtbd4 | Q8R179-2 | 0.92 | 1.24 | 0.83 | 0.82 | 0.22 | 1.69 | 0.63 | 0.89 | 1.88 | 2.92 | 0.9 | 0.93 | 2.48 | 1.28 | 0.58 | 0.22 | -0.095 | 0.705 | 0.1875 | 0.765763 | 0.150533 | 0.694798 | 1 | 0.996511 | 0.955729 |
| Kbtbd7 | G5E8C2 | -1.44 | -1.5 | -0.04 | 0.56 | -0.39 | -1.63 | 0.46 | 0.62 | -1.4 | -0.41 | 0.32 | -1.19 | -0.68 | -0.81 | 0.31 | 0.36 | 0.37 | -0.065 | 0.4 | 0.578955 | 0.912755 | 0.478044 | 1 | 0.996747 | 0.892279 |
| Kcmf1 | Q80UY2 | -0.45 | -0.25 | -0.21 | -0.57 | -0.38 | -0.23 | -1.57 | -0.58 | 0.97 | 0.11 | -0.44 | 0.08 | 0.58 | -0.8 | -0.32 | -0.88 | -0.32 | 0.55 | 0.015 | 0.315948 | 0.101711 | 0.965274 | 1 | 0.996511 | 0.9927 |
| Kcna1 | P16388 | 0.5 | 1.38 | 1.71 | 1.46 | 1.07 | 0.36 | 1.37 | 1.16 | 0.09 | 0.69 | 1.09 | 0.37 | -0.42 | 0.83 | 1.14 | 1.21 | -0.2725 | -0.7025 | -0.5725 | 0.422108 | 0.062782 | 0.21365 | 1 | 0.996511 | 0.786931 |
| Kcna2 | P63141 | 0.92 | 1.51 | 1.02 | 1.17 | 1.5 | 0.6 | 1.11 | 0.92 | 0.85 | 1.01 | 0.84 | 0.93 | 0.51 | 1.87 | 0.58 | 1.07 | -0.1225 | -0.2475 | -0.1475 | 0.616218 | 0.217454 | 0.665433 | 1 | 0.996511 | 0.949825 |
| Kcna3 | P16390 | 0.46 | 0.75 | 1.24 | 1.04 | 0.86 | 0.68 | 0.7 | 1.23 | 0.64 | 1.09 | 0.75 | 0.54 | -0.11 | 0.63 | 0.83 | 0.29 | -0.005 | -0.1175 | -0.4625 | 0.982753 | 0.622453 | 0.132231 | 1 | 0.996511 | 0.726871 |
| Kcna4 | Q61423 | 0.53 | 0.46 | 2.32 | 2.67 | 0.36 | 0.52 | 1.92 | 2.51 | 0.64 | 2.78 | 2.06 | 0.6 | 0.52 | 0.03 | 2.58 | 2.09 | -0.1675 | 0.025 | -0.19 | 0.813859 | 0.9723 | 0.803323 | 1 | 0.997515 | 0.971149 |
| Kcna6 | Q61923 | -0.13 | 0.32 | 0.52 | 0.79 | 0.42 | 0.05 | 0.97 | 0.8 | 0.07 | 0.75 | 0.71 | 0.24 | -0.4 | 0.37 | 0.45 | 0.44 | 0.185 | 0.0675 | -0.16 | 0.520226 | 0.80452 | 0.59297 | 1 | 0.996511 | 0.929843 |
| Kcnab1 | P63143 | 0.77 | 0.53 | 0.18 | 0.52 | 0.43 | 0.83 | 0.29 | 0.37 | 0.24 | -0.3 | -0.06 | 0.49 | 0.18 | -0.18 | -0.16 | -0.13 | -0.02 | -0.4075 | -0.5725 | 0.921253 | 0.115587 | 0.021126 | 1 | 0.996511 | 0.621926 |
| Kcnab2 | P62482 | 0.29 | 0.54 | 0.62 | 0.53 | 0.51 | 0.12 | 0.4 | 0.41 | 0.06 | -0.21 | 0.23 | 0.25 | -0.17 | 0.05 | 0.14 | 0.43 | -0.135 | -0.4125 | -0.3825 | 0.425724 | 0.052071 | 0.089756 | 1 | 0.996511 | 0.69217 |
| Kcnab3 | Q8VD73 | 0.02 | 0.54 | 0.74 | 0.5 | 0.4 | 0.03 | 0.84 | 0.65 | -0.41 | -0.67 | 0.42 | -0.25 | 0.45 | 0.13 | 0.36 | 0.7 | 0.03 | -0.6775 | -0.04 | 0.902786 | 0.040688 | 0.864641 | 1 | 0.996511 | 0.977686 |
| Kcnb1 | Q03717 | 0.81 | 0.98 | 0.77 | 0.66 | 1.11 | 1.17 | 0.08 | 0.32 | 0.76 | -0.25 | 0.54 | 0.96 | 1 | 0.25 | 0.69 | 0.3 | -0.135 | -0.3025 | -0.245 | 0.638222 | 0.303167 | 0.306757 | 1 | 0.996511 | 0.832533 |
| Kcnb2 | A6H8H5 | 0.77 | 0.94 | 1.08 | 1.57 | 1.69 | 0.33 | 0.48 | 0.51 | 0.88 | 1.44 | 0.77 | 0.78 | 0.17 | 1.44 | 0.83 | 0.69 | -0.3375 | -0.1225 | -0.3075 | 0.34283 | 0.633904 | 0.348777 | 1 | 0.996511 | 0.85278 |
| Kcnc1 | P15388-3 | 0.46 | 0.82 | 1.14 | 0.72 | 1.26 | 0.27 | 0.54 | 0.31 | 0.35 | 0.43 | 0.54 | -0.03 | 0.85 | 0.79 | 0.77 | 0.71 | -0.19 | -0.4625 | -0.005 | 0.493918 | 0.064017 | 0.98057 | 1 | 0.996511 | 0.994858 |
| Kcnc2 | Q14B80 | -0.09 | 0.61 | 1.11 | 0.85 | 1.21 | 0.18 | 1.41 | 1.03 | -0.28 | 0.02 | 0.95 | 0.27 | -0.01 | 0.95 | 0.5 | 1.18 | 0.3375 | -0.38 | 0.035 | 0.358536 | 0.305111 | 0.923006 | 1 | 0.996511 | 0.987595 |
| Kcnc3 | E9QKI2 | 0.78 | 1.33 | 0.99 | 1.05 | 1.58 | 0.72 | 0.68 | 0.41 | 0.73 | 0.56 | 0.45 | 0.6 | 0.04 | 1.3 | 0.31 | 0.72 | -0.19 | -0.4525 | -0.445 | 0.504181 | 0.036488 | 0.172321 | 1 | 0.996511 | 0.76392 |
| Kcnc4 | A0A571BG! | -0.18 | 0.25 | 0.25 | 0.39 | 0.71 | 0.54 | 0.6 | 0.12 | 0.06 | 1.01 | 0.24 | 0.28 | 0.96 | 0.76 | 0.47 | -0.02 | 0.315 | 0.22 | 0.365 | 0.155594 | 0.410858 | 0.200193 | 1 | 0.996511 | 0.78004 |
| Kcnd1 | Q03719 | 0.59 | 0.54 | 0.52 | 0.96 | 0.8 | 0.75 | 0.99 | 0.7 | -0.06 | 1.67 | 0.6 | 1.04 | -0.31 | -0.79 | 0.25 | 0.46 | 0.1575 | 0.16 | -0.75 | 0.372635 | 0.664893 | 0.03644 | 1 | 0.996511 | 0.658454 |
| Kcnd2 | Q9Z0V2 | 1.67 | 1.42 | 1.16 | 0.99 | 1.56 | 1.77 | 1.21 | 1 | 0.77 | 0.78 | 1.09 | 1.58 | -0.07 | 1.06 | 0.57 | 1.02 | 0.075 | -0.255 | -0.665 | 0.757804 | 0.341584 | 0.057721 | 1 | 0.996511 | 0.661738 |
| Kcnd3 | Q9Z0V1 | 1.39 | 1.53 | 0.76 | 0.98 | 1.65 | 1.37 | 0.91 | 1.01 | 0.89 | -0.48 | 0.61 | 1.26 | 0.5 | 1.04 | 0.21 | 0.62 | 0.07 | -0.595 | -0.5725 | 0.785134 | 0.159927 | 0.06049 | 1 | 0.996511 | 0.661738 |
| Kcne5 | Q9QZ26 | -0.56 | -0.43 | -0.33 | -0.58 | 0.4 | -0.07 | 1.27 | 0.43 | 0.1 | -1.12 | 0.85 | 0.52 | 0.63 | 0.68 | 0.69 | 1.03 | 0.9825 | 0.5625 | 1.2325 | 0.00742 | 0.197375 | 0.000125 | 1 | 0.996511 | 0.228506 |
| Kcnf1 | E9PYC0 | -0.67 | 0.05 | -0.21 | -0.13 | -0.46 | -1.1 | -0.77 | -0.45 | -0.29 | -0.38 | -0.72 | -0.97 | -2.31 | -0.48 | -0.25 | -0.47 | -0.455 | -0.35 | -0.6375 | 0.079904 | 0.17751 | 0.201126 | 1 | 0.996511 | 0.780919 |
| Kcnh1 | A0A1L1M1 | 0.47 | 0.68 | 1.04 | 0.64 | 0.72 | -0.07 | 0.47 | 0.33 | 0.59 | 1.49 | 0.73 | 0.46 | 0.93 | 0.96 | 0.59 | 0.84 | -0.345 | 0.11 | 0.1225 | 0.152008 | 0.689522 | 0.557989 | 1 | 0.996511 | 0.917135 |
| Kcnh2 | Q53Z09 | -0.43 | 0.04 | 0.3 | -0.3 | -0.11 | -0.02 | 0.29 | 0.68 | 1.87 | 0.04 | 0.47 | 0.11 | 3.13 | -0.43 | 0.93 | 1.05 | 0.3075 | 0.72 | 1.2675 | 0.246423 | 0.123409 | 0.091235 | 1 | 0.996511 | 0.693316 |
| Kcnh3 | Q9WVJ0 | 0.45 | 1.84 | 1.32 | 1.95 | 0.8 | 0.81 | 1.7 | 1.47 | 1.1 | 1.93 | 1.47 | 0.46 | -0.02 | 2.31 | 1.44 | 1.36 | -0.195 | -0.15 | -0.1175 | 0.619528 | 0.73167 | 0.829586 | 1 | 0.996511 | 0.973939 |
| Kcnh5 | Q920E3 | 0.61 | 0.47 | 1.27 | 0.76 | 0.73 | 1.03 | 0.49 | 0.53 | -0.38 | -0.8 | 1.43 | 0.62 | -1 | -0.67 | 1.11 | 0.98 | -0.0825 | -0.56 | -0.6725 | 0.723234 | 0.275862 | 0.227609 | 1 | 0.996511 | 0.787447 |
| Kcnh7 | Q9ER47 | 0.84 | 1.1 | 0.77 | 0.83 | 0.82 | 1.06 | 0.58 | 0.65 | 0.76 | 0.83 | 0.76 | 0.9 | 0.6 | 0.84 | 0.65 | 0.53 | -0.1075 | -0.0725 | -0.23 | 0.547707 | 0.662849 | 0.232227 | 1 | 0.996511 | 0.788863 |
| Kcnip1 | Q3YAB5 | 1.08 | 0.52 | -0.03 | 1.06 | 0.82 | 0.7 | 0.82 | 0.41 | 0.15 | 0.51 | 0.44 | 0.81 | -1.06 | -0.01 | 0.33 | 0.9 | 0.03 | -0.18 | -0.6175 | 0.915438 | 0.554661 | 0.202971 | 1 | 0.996511 | 0.782817 |
| Kcnip2 | Q3YAB3 | 1.29 | 1.12 | 1.44 | 1.2 | 1.34 | 0.86 | 2.08 | 1.32 | 0.28 | 0.83 | 0.99 | 0.96 | 0.28 | 0.99 | 1.72 | 1.92 | 0.1375 | -0.4975 | -0.035 | 0.610411 | 0.045361 | 0.924956 | 1 | 0.996511 | 0.988301 |
| Kcnip3 | Q9QXT8 | 1.31 | 1.06 | 1.25 | 0.7 | 1.68 | 1.29 | 1.15 | 1.26 | 0.47 | -0.2 | 1.09 | 1.3 | -0.12 | 1.3 | 0.6 | 1.08 | 0.265 | -0.415 | -0.365 | 0.225078 | 0.262926 | 0.302892 | 1 | 0.996511 | 0.831227 |
| Kcnip4 | Q6PHZ8 | 0.72 | 1.23 | 0.39 | 0.39 | 1.35 | 0.93 | 0.52 | 0.26 | 0.21 | 0.18 | 0.24 | 0.92 | -0.18 | 1.04 | -0.12 | 0.3 | 0.0825 | -0.295 | -0.4225 | 0.788073 | 0.3063 | 0.239026 | 1 | 0.996511 | 0.79362 |
| Kcnj10 | Q9JM63 | 2.27 | 0.24 | 1.04 | 1.46 | 0.13 | 1.89 | 0.78 | 0.88 | 0.29 | 1.63 | 0.61 | 1.27 | 0.48 | 0.15 | 1.87 | 2.39 | -0.3325 | -0.3025 | -0.03 | 0.52252 | 0.538567 | 0.961728 | 1 | 0.996511 | 0.9927 |
| Kcnj11 | Q61743 | 0.84 | 1.32 | 1.28 | 1 | 1.18 | 0.63 | 1.06 | 1.09 | 0.7 | 1.01 | 1.19 | 0.48 | 0.57 | 1.31 | 0.97 | 0.91 | -0.12 | -0.265 | -0.17 | 0.553891 | 0.264748 | 0.472387 | 1 | 0.996511 | 0.891144 |
| Kcnj12 | B1ATI1 | 1.37 | 1.48 | 1.64 | 1.87 | 1.23 | 1.44 | 0.27 | 0.6 | 1.17 | 0.97 | 1.42 | 1.67 | 1.24 | 0.97 | 0.91 | 0.79 | -0.705 | -0.2825 | -0.6125 | 0.037223 | 0.225902 | 0.014827 | 1 | 0.996511 | 0.589387 |
| Kcnj16 | Q9Z307 | 2.92 | 1.17 | 0.12 | 0.44 | 2.19 | 2.5 | -0.09 | 0.25 | 0.59 | 0.36 | 0.09 | 2.21 | 0.49 | 1.23 | 1.33 | 0.73 | 0.05 | -0.35 | -0.2175 | 0.951326 | 0.626506 | 0.718667 | 1 | 0.996511 | 0.958398 |
| Kcnj3 | P63250 | 0.83 | 0.99 | 1.35 | 1.17 | 0.96 | 0.93 | 1.3 | 1.21 | 0.47 | 0.59 | 1.34 | 0.83 | 0.48 | 0.61 | 0.48 | 0.84 | 0.015 | -0.2775 | -0.4825 | 0.935685 | 0.27966 | 0.039707 | 1 | 0.996511 | 0.658454 |
| Kcnj4 | Q8R435 | 1.53 | 0.93 | 1 | 1.15 | 0.97 | 1.32 | 0.81 | 1.03 | 1.48 | 0.89 | 0.85 | 1.41 | 0.96 | 1.46 | 0.62 | 1.03 | -0.12 | 0.005 | -0.135 | 0.557846 | 0.983431 | 0.594284 | 1 | 0.998773 | 0.929843 |
| Kcnj6 | Q0VB45 | -0.29 | 0.42 | 0.94 | 0.71 | 0.19 | -0.1 | 0.59 | 0.81 | 0.12 | 0.08 | 0.7 | 0.05 | -0.36 | -0.13 | 0.47 | 0.17 | -0.0725 | -0.2075 | -0.4075 | 0.8245 | 0.510661 | 0.233066 | 1 | 0.996511 | 0.788994 |
| Kcnj9 | P48543 | -1.4 | -1.65 | -1.23 | -2.08 | -2.79 | -0.35 | -1.58 | -1.71 | -1.22 | -1.19 | -1.54 | -1.95 | 0.29 | -2.58 | -2.3 | -1.68 | -0.0175 | 0.115 | 0.0225 | 0.971579 | 0.672843 | 0.970806 | 1 | 0.996511 | 0.992803 |
| Kcnk1 | O08581 | -0.48 | -0.74 | -0.49 | -0.81 | -1.08 | -0.46 | -0.04 | 1.91 | 0.86 | -1.77 | -0.43 | -0.4 | 0.04 | 1.02 | -1.06 | -1 | 0.7125 | 0.195 | 0.38 | 0.250343 | 0.700529 | 0.427626 | 1 | 0.996511 | 0.879165 |
| Kcnk10 | Q8BUW1 | 0.43 | 1.09 | 0.58 | 1.26 | 0.42 | 0.34 | 1.34 | 2.03 | 1.33 | 1.55 | 1.61 | 0.6 | 0.26 | 2.39 | 1.19 | 1.89 | 0.1925 | 0.4325 | 0.5925 | 0.649225 | 0.185876 | 0.230555 | 1 | 0.996511 | 0.788863 |
| Kcnk2 | Q6P6P9 | 0.04 | 0.76 | 0.73 | 0.45 | 0.33 | -0.16 | 0.5 | 0.39 | 0.79 | 0.74 | 0.48 | 0.42 | 0.96 | 1.31 | 0.78 | 0.78 | -0.23 | 0.1125 | 0.4625 | 0.347403 | 0.619957 | 0.086006 | 1 | 0.996511 | 0.691619 |
| Kcnk4 | O88454 | -0.5 | -0.12 | -0.1 | -0.82 | -0.06 | -0.07 | -3.05 | -1.34 | 0.75 | -3.4 | -0.7 | -0.37 | 1.01 | -0.65 | -0.92 | -1.78 | -0.745 | -0.545 | -0.2 | 0.27743 | 0.506785 | 0.722102 | 1 | 0.996511 | 0.958398 |
| Kcnma1 | E3VRY9 | 0.17 | 0.49 | 0.98 | 0.75 | 0.6 | 0.48 | 1.12 | 0.87 | -0.48 | 0.21 | 1.13 | 0.12 | -0.54 | -0.15 | 0.42 | 0.57 | 0.17 | -0.3525 | -0.5225 | 0.486437 | 0.347095 | 0.127008 | 1 | 0.996511 | 0.722414 |
| Kcnmb4 | Q9JIN6 | 0.58 | 0.77 | -0.49 | -0.69 | 1.61 | 0.55 | -0.37 | -0.96 | 0.51 | -1.54 | -0.71 | 0.36 | 0.62 | 1.26 | -1.52 | -0.96 | 0.165 | -0.3875 | -0.1925 | 0.787791 | 0.494724 | 0.77789 | 1 | 0.996511 | 0.967289 |
| Kcnn2 | P58390 | 0.94 | 1.25 | 1.6 | 1.27 | 1.6 | 1.12 | 1.15 | 1.63 | 0.28 | 0.09 | 1.82 | 1.42 | 0.47 | 0.54 | 1.43 | 1.3 | 0.11 | -0.3625 | -0.33 | 0.616241 | 0.400701 | 0.285156 | 1 | 0.996511 | 0.822669 |
| Kcnn3 | P58391 | -0.38 | 0.48 | 0.78 | 0.24 | 0.09 | -0.5 | 0.85 | 0.86 | 1 | 0.47 | 0.55 | 0.39 | 1.2 | 0.08 | 0.71 | 0.79 | 0.045 | 0.3225 | 0.415 | 0.907793 | 0.284087 | 0.240678 | 1 | 0.996511 | 0.795658 |
| Kcnq2 | B7ZBW2 | 0.55 | 0.74 | 0.41 | 0.53 | 0.99 | 0.87 | 0.34 | 0.51 | 1.14 | 0.84 | 0.27 | 0.66 | 1.52 | 0.93 | 0.47 | 0.54 | 0.12 | 0.17 | 0.3075 | 0.551978 | 0.462426 | 0.278213 | 1 | 0.996511 | 0.822084 |
| Kcnq2 | B7ZBW1 | 0.41 | 0.54 | 1 | 1.48 | 0.24 | 0.31 | 0.56 | 1.29 | 1.02 | 1.88 | 0.74 | 0.67 | 1.66 | 0.68 | 0.79 | 2.17 | -0.2575 | 0.22 | 0.4675 | 0.445698 | 0.545209 | 0.276331 | 1 | 0.996511 | 0.822084 |
| Kcnq3 | Q8K3F6 | -0.01 | 0.49 | 0.49 | 0.24 | 0.65 | 0.15 | 0.28 | 0.09 | 0.01 | 0.12 | 0.38 | 0.04 | 0.08 | 0.84 | 0.27 | 0.07 | -0.01 | -0.165 | 0.0125 | 0.96092 | 0.413256 | 0.960153 | 1 | 0.996511 | 0.992636 |
| Kcnq5 | G3UYG5 | 0.59 | 0.78 | 0.95 | 0.58 | 0.8 | 0.84 | -0.14 | 0.38 | 1.58 | 0.25 | 0.97 | 0.55 | 1.51 | 1.27 | 1.36 | 0.34 | -0.255 | 0.1125 | 0.395 | 0.331934 | 0.71368 | 0.202105 | 1 | 0.996511 | 0.78218 |
| Kcns2 | O35174 | -0.68 | 0.82 | -0.11 | -0.25 | 0.29 | -0.72 | -0.18 | -0.23 | 1.04 | 0.43 | -0.01 | -0.22 | 0.83 | 0.22 | -0.21 | 0.59 | -0.155 | 0.365 | 0.4125 | 0.669169 | 0.37591 | 0.293159 | 1 | 0.996511 | 0.825612 |
| Kcnt1 | A2AHB9 | 0.69 | 0.69 | 0.71 | 0.66 | 0.65 | 0.25 | 0.25 | 0.4 | 0.59 | 0.59 | 0.12 | 0.51 | 0.56 | 1.16 | 0.48 | 0.52 | -0.3 | -0.235 | -0.0075 | 0.086284 | 0.213112 | 0.972197 | 1 | 0.996511 | 0.992865 |
| Kcnt2 | D3Z592 | -0.41 | 0.51 | 0.06 | 0.04 | 0.54 | 0.16 | -0.11 | -0.46 | 0.18 | -0.33 | -0.13 | -0.07 | -0.32 | 0.24 | -0.62 | -0.48 | -0.0175 | -0.1375 | -0.345 | 0.950915 | 0.573309 | 0.245632 | 1 | 0.996511 | 0.802442 |
| Kcnv1 | Q8BZN2 | -1.07 | -0.23 | 0 | -0.29 | -1.36 | -0.7 | -0.14 | 0.44 | -0.85 | -2.02 | -0.42 | -0.37 | -0.84 | 0.12 | -1.43 | -0.04 | -0.0425 | -0.5175 | -0.15 | 0.919639 | 0.243977 | 0.716736 | 1 | 0.996511 | 0.958334 |
| Kctd1 | A0A3Q4EG | -1.21 | -1.13 | -1.88 | -2.32 | -0.56 | -1.18 | -1.86 | -2.03 | -1.98 | -0.91 | -1.46 | -0.99 | -2.41 | -1.21 | -1.89 | -2.18 | 0.2275 | 0.3 | -0.2875 | 0.585525 | 0.420742 | 0.45198 | 1 | 0.996511 | 0.885993 |
| Kctd10 | Q922M3 | -0.43 | -0.09 | -0.21 | -0.05 | 0.09 | -0.2 | -0.43 | 0.05 | -0.57 | 0.29 | 0.74 | -0.36 | -0.59 | -0.03 | 1.15 | 0.38 | 0.0725 | 0.22 | 0.4225 | 0.700736 | 0.490154 | 0.270089 | 1 | 0.996511 | 0.8219 |
| Kctd12 | A0A0R4J2E | 0.12 | -0.03 | -0.26 | -0.08 | -0.01 | -0.12 | 0.01 | -0.08 | 0.39 | 1.52 | -0.05 | 0.27 | 0.41 | 0.08 | -0.3 | -0.21 | 0.0125 | 0.595 | 0.0575 | 0.935203 | 0.112329 | 0.799114 | 1 | 0.996511 | 0.969791 |
| Kctd12b | Q8C7J6 | 0.17 | 0.78 | -0.24 | -0.21 | 0.61 | 0.49 | -0.47 | 0.04 | 0.8 | 0.55 | -0.66 | 0.15 | 1.17 | 0.15 | 0.28 | -0.2 | 0.0425 | 0.085 | 0.225 | 0.897939 | 0.824533 | 0.54596 | 1 | 0.996511 | 0.912321 |
| Kctd13 | Q8BGV7 | 0.94 | 0.77 | -0.1 | -0.4 | 1.19 | 0.3 | -0.56 | -0.02 | 0.71 | 0.54 | -0.17 | 0.76 | 0.51 | 0.85 | -0.58 | -0.01 | -0.075 | 0.1575 | -0.11 | 0.869469 | 0.677977 | 0.798371 | 1 | 0.996511 | 0.969727 |
| Kctd16 | Q5DTY9 | 1.47 | 1.13 | 1.71 | 1.71 | 1.33 | 1.27 | 1.47 | 1.7 | 1.2 | 0.66 | 1.75 | 1.5 | 0.75 | 0.86 | 1.22 | 1.23 | -0.0625 | -0.2275 | -0.49 | 0.755302 | 0.430194 | 0.057646 | 1 | 0.996511 | 0.661738 |
| Kctd17 | E0CYQ0 | -1.61 | -0.97 | -0.85 | -0.52 | -1.19 | -1.7 | -1.25 | -1.28 | -1.18 | -1.13 | -0.59 | -1.38 | -1.68 | -1.38 | -0.67 | -1.11 | -0.3675 | -0.0825 | -0.2225 | 0.187971 | 0.778007 | 0.492527 | 1 | 0.996511 | 0.897531 |
| Kctd2 | Q4VBE7 | -0.23 | 0.35 | 0.39 | 0.01 | -0.04 | 0.37 | 0.23 | 0.66 | -0.99 | -0.87 | 0 | 0.41 | -0.44 | -1.68 | -0.45 | 0.42 | 0.175 | -0.4925 | -0.6675 | 0.449775 | 0.195144 | 0.150017 | 1 | 0.996511 | 0.749179 |
| Kctd21 | B2RTJ2 | -0.31 | -0.77 | -0.05 | -0.86 | -0.22 | 0.25 | -0.04 | -0.03 | -0.58 | 0.09 | 0.1 | -0.27 | -0.37 | -0.49 | -0.04 | 0.15 | 0.4875 | 0.3325 | 0.31 | 0.062274 | 0.235456 | 0.264973 | 1 | 0.996511 | 0.817248 |
| Kctd3 | Q8BFX3 | -0.01 | -0.3 | -0.8 | -1.07 | -0.34 | 0.03 | -0.96 | -1.04 | -0.12 | -0.14 | -0.79 | 0.28 | 0.44 | -0.43 | -0.68 | -0.95 | -0.0325 | 0.3525 | 0.14 | 0.92351 | 0.294964 | 0.710176 | 1 | 0.996511 | 0.957819 |
| Kctd4 | Q9D7X1 | -2.02 | -1.29 | 0.12 | -0.4 | -0.82 | -2.04 | 0.53 | -0.39 | -2.01 | -0.57 | 0.11 | -1.77 | -0.92 | -1.08 | 0.23 | 1.05 | 0.2175 | -0.1625 | 0.7175 | 0.737986 | 0.796852 | 0.273666 | 1 | 0.996511 | 0.822084 |
| Kctd5 | A0A0R4J00 | -0.12 | -0.29 | 1.33 | 0.72 | 0.59 | -0.61 | 0.94 | 0.8 | -0.35 | 1.02 | 0.88 | -0.15 | -0.34 | -0.77 | 1.87 | 1.1 | 0.02 | -0.06 | 0.055 | 0.966682 | 0.90064 | 0.933305 | 1 | 0.996511 | 0.989767 |
| Kctd6 | Q8BNL5 | -0.61 | 0.05 | -0.06 | -0.01 | -0.25 | -0.21 | 0.2 | -0.45 | -0.1 | 0.78 | -0.53 | -0.07 | -0.27 | 0.03 | -0.33 | -0.26 | -0.02 | 0.1775 | -0.05 | 0.929333 | 0.576969 | 0.821987 | 1 | 0.996511 | 0.972768 |
| Kctd7 | F6QXA3 | 3.97 | 3.64 | 3.47 | 2.31 | 3.98 | 3.8 | 3.99 | 3.73 | 0.1 | -0.16 | 3.38 | 4.11 | -1.77 | -0.19 | 4.15 | 4.68 | 0.5275 | -1.49 | -1.63 | 0.160385 | 0.177019 | 0.27925 | 1 | 0.996511 | 0.822084 |
| Kctd8 | Q50H33 | 0.51 | 0.38 | 0.18 | 0.48 | 0.7 | 0.48 | 0.47 | 0.43 | 0.3 | 0.74 | 0.29 | 0.7 | 0.1 | 0.97 | 0.85 | 0.6 | 0.1325 | 0.12 | 0.2425 | 0.415483 | 0.544985 | 0.333546 | 1 | 0.996511 | 0.845601 |
| Kdelr1 | Q99JH8 | -2.22 | -3.32 | -4.36 | -14.86 | -2.17 | -3.26 | -5.22 | -6.96 | -1.4 | -14.63 | -5.75 | -3.69 | 0.22 | -1.93 | -4.94 | -4.27 | 1.7875 | -0.1775 | 3.46 | 0.523501 | 0.961258 | 0.235018 | 1 | 0.996755 | 0.791437 |
| Kdm1a | A3KG93 | -0.73 | -2.33 | -1.88 | -1.93 | -2.22 | -1.68 | -1.63 | -1.78 | -2.67 | -1.27 | -1.37 | -1.49 | -1.55 | -2.09 | -0.92 | -1.32 | -0.11 | 0.0175 | 0.2475 | 0.756862 | 0.968711 | 0.546205 | 1 | 0.996755 | 0.912406 |
| Kdm2a | F6YRW4 | -0.84 | -0.46 | -2.4 | -3.85 | -0.11 | -1.4 | -3 | -4.09 | -1.03 | -2.2 | -3.21 | -1.11 | -1.46 | -0.18 | -3.7 | -2.77 | -0.2625 | -1.6E-16 | -0.14 | 0.802741 | 1 | 0.886138 | 1 | 1 | 0.979767 |
| Kdm3b | Q6ZPY7 | -0.83 | -1.16 | -1.26 | -0.95 | -1.57 | -0.39 | -1.12 | -1.62 | -0.04 | -0.4 | -1.46 | -0.42 | -0.45 | -1.05 | -1.42 | -0.91 | -0.125 | 0.47 | 0.0925 | 0.677314 | 0.168927 | 0.717148 | 1 | 0.996511 | 0.958398 |
| Kdm4a | A2A8L9 | -3.94 | -4.3 | -2.01 | -2.69 | -6.57 | -4.96 | -3.75 | -3.05 | -1.71 | -2.26 | -3.35 | -4.87 | 0.12 | -5.65 | -3.2 | -7.45 | -1.3475 | 0.1875 | -0.81 | 0.137806 | 0.813456 | 0.599446 | 1 | 0.996511 | 0.930565 |
| Kdm5a | Q3UXZ9 | -0.3 | -0.11 | 0.42 | 1.51 | 1.55 | -0.68 | 0.4 | 1.93 | -0.32 | 2.05 | 1.37 | -0.54 | -0.4 | -0.33 | 0.22 | 1.48 | 0.42 | 0.26 | -0.1375 | 0.524135 | 0.705576 | 0.802634 | 1 | 0.996511 | 0.971091 |
| Kdm5b | Q80Y84-2 | -3.89 | 1.42 | 1.27 | 1.85 | -0.22 | -3.42 | 0.75 | 1.62 | 3.59 | 3.07 | 1.07 | -2.13 | 2.2 | 2.65 | 1.13 | 1.54 | -0.48 | 1.2375 | 1.7175 | 0.758589 | 0.466872 | 0.191328 | 1 | 0.996511 | 0.777117 |
| Kdsr | Q6GV12 | -0.76 | -0.53 | -1.18 | -1.52 | 0.51 | -0.56 | -0.67 | -1.34 | -0.53 | -0.98 | -1.3 | -0.8 | -0.58 | 0.33 | -1.02 | -0.61 | 0.4825 | 0.095 | 0.5275 | 0.263447 | 0.738556 | 0.163795 | 1 | 0.996511 | 0.758352 |
| Keap1 | Q9Z2X8 | -1.26 | -0.85 | -0.82 | -0.85 | -0.99 | -1.34 | -0.3 | -0.95 | -0.41 | -1.06 | -0.85 | -0.97 | 0.14 | -0.71 | -0.1 | -0.47 | 0.05 | 0.1225 | 0.66 | 0.842948 | 0.576261 | 0.025419 | 1 | 0.996511 | 0.633075 |
| Khdc4 | Q3TCX3-2 | -4.75 | -5.78 | -1.57 | -0.95 | -7.43 | -5.92 | -1.6 | -1.47 | -2.53 | 0.69 | -0.83 | -4.62 | 1.01 | -4.76 | -0.87 | -1.68 | -0.8425 | 1.44 | 1.6875 | 0.625403 | 0.340264 | 0.277368 | 1 | 0.996511 | 0.822084 |
| Khdrbs1 | Q60749 | -0.96 | -2.55 | -2.01 | -1.65 | -2.64 | -1.91 | -1.51 | -1.57 | -1.51 | -1.16 | -1.58 | -2.1 | -0.82 | -1.77 | -1.54 | -1.99 | -0.115 | 0.205 | 0.2625 | 0.773343 | 0.58601 | 0.520351 | 1 | 0.996511 | 0.907667 |
| Khdrbs2 | Q9WU01 | 0.75 | -0.53 | -0.73 | -0.34 | -1.35 | -0.22 | -0.59 | 0.16 | -2.47 | -0.43 | -0.85 | -1.6 | -1.87 | -2.39 | 0.2 | -0.57 | -0.2875 | -1.125 | -0.945 | 0.510698 | 0.054293 | 0.153613 | 1 | 0.996511 | 0.750763 |
| Khdrbs3 | Q9R226 | 1.64 | -0.91 | -0.52 | -0.48 | -1.07 | 1.16 | -0.43 | -0.03 | 0.58 | -0.74 | -0.21 | -0.89 | 2.7 | -0.66 | -0.24 | -0.68 | -0.025 | -0.2475 | 0.3475 | 0.970417 | 0.686171 | 0.698846 | 1 | 0.996511 | 0.955729 |
| Khk | A0A0J9YTU | 0.06 | 0.45 | 1.41 | 0.38 | 0.49 | -0.15 | 0.96 | 0.55 | -0.33 | -0.03 | 0.77 | -0.29 | 0.06 | 0.59 | 0.96 | 1.15 | -0.1125 | -0.545 | 0.115 | 0.752063 | 0.170801 | 0.756206 | 1 | 0.996511 | 0.962333 |
| Khnyn | Q80U38 | -0.59 | -0.76 | 0.17 | -0.02 | -0.61 | -0.14 | 0.18 | -0.28 | 0.22 | 1.03 | 0.16 | 0.16 | -0.2 | -0.49 | -0.77 | 0.46 | 0.0875 | 0.6925 | 0.05 | 0.754748 | 0.05003 | 0.8845 | 1 | 0.996511 | 0.979767 |
| Khsrp | Q3U0V1 | -4.2 | -4.89 | -4.47 | -4.66 | -4.37 | -4.72 | -4.44 | -4.92 | -5.23 | -2.55 | -4.36 | -3.98 | -4.54 | -3.94 | -4.84 | -5.16 | -0.0575 | 0.525 | -0.065 | 0.792391 | 0.339219 | 0.833029 | 1 | 0.996511 | 0.97513 |
| Kiaa0100 | Q5SYL3 | 0.45 | 0.54 | 0.66 | 0.58 | 0.62 | 0.6 | 0.84 | 0.42 | 1.03 | 0.99 | 0.32 | 0.43 | 1.3 | 1.19 | 0.65 | 0.68 | 0.0625 | 0.135 | 0.3975 | 0.696297 | 0.552185 | 0.102499 | 1 | 0.996511 | 0.700625 |
| Kiaa0319 | Q5SZV5 | -6.99 | -6.88 | -0.66 | -0.29 | -6.01 | -6.95 | -0.19 | -0.71 | -5.97 | 0.2 | 0.12 | -8.35 | -4.52 | -4.32 | 0.53 | -0.07 | 0.24 | 0.205 | 1.61 | 0.916166 | 0.935782 | 0.438217 | 1 | 0.996755 | 0.883161 |
| Kiaa0319l | Q8K135 | 0.76 | -0.4 | -0.95 | -0.73 | 0.13 | 0.17 | -0.67 | -0.8 | -1.05 | -0.64 | -1 | -0.08 | -0.31 | -0.12 | -0.97 | -1.07 | 0.0375 | -0.3625 | -0.2875 | 0.930357 | 0.397542 | 0.506283 | 1 | 0.996511 | 0.902457 |
| Kiaa0513 | Q8R0A7 | 0.43 | 0.41 | 0.57 | 0.46 | 0.08 | 0.56 | -0.13 | 0.37 | -0.34 | -0.95 | 0.12 | 0.3 | -0.15 | -0.39 | 0.25 | 0.07 | -0.2475 | -0.685 | -0.5225 | 0.222632 | 0.040201 | 0.029525 | 1 | 0.996511 | 0.637651 |
| Kiaa1107 | Q80TK0 | -0.64 | -0.46 | -0.24 | -0.56 | -0.65 | -0.96 | -1.31 | -1.12 | -0.2 | -0.23 | -0.66 | -0.47 | 0.34 | 0.14 | -0.4 | -1.2 | -0.535 | 0.085 | 0.195 | 0.023301 | 0.661659 | 0.582795 | 1 | 0.996511 | 0.927734 |
| Kiaa1109 | A2AAE1-6 | 0.53 | 0.97 | 1.15 | 1.12 | 1.13 | 0.54 | 0.9 | 0.99 | 0.83 | 1.23 | 1.06 | 0.71 | -0.09 | -0.2 | -0.22 | 1.11 | -0.0525 | 0.015 | -0.7925 | 0.808009 | 0.945853 | 0.046536 | 1 | 0.996755 | 0.658454 |
| Kiaa1522 | A2A7S8-4 | 0.19 | 0.27 | 1.31 | 0.84 | 0.71 | 0.75 | 0.01 | 0.49 | 1.03 | 0.7 | 0.69 | 0.36 | 1.98 | 0.55 | 1.18 | 1.12 | -0.1625 | 0.0425 | 0.555 | 0.601074 | 0.887871 | 0.172498 | 1 | 0.996511 | 0.76392 |
| Kidins220 | E9Q9B7 | 0.35 | 0.6 | 0.56 | 0.58 | 0.79 | 0.47 | 1.35 | 0.77 | 0.46 | 0.44 | 0.73 | 0.56 | 0.26 | 0.66 | 0.51 | 0.71 | 0.3225 | 0.025 | 0.0125 | 0.163284 | 0.881951 | 0.948007 | 1 | 0.996511 | 0.992435 |
| Kif11 | Q6P9P6 | -2.56 | -1.5 | -2.99 | -3.38 | -1.11 | -2.39 | -2.84 | -3.69 | -2.27 | -3.59 | -3.04 | -2.46 | -1.99 | -1.32 | -3.01 | -3.44 | 0.1 | -0.2325 | 0.1675 | 0.870537 | 0.624056 | 0.772492 | 1 | 0.996511 | 0.965933 |
| Kif13a | Q9EQW7 | -1.84 | -3.41 | -1.27 | -0.43 | -1.32 | -0.96 | -1.41 | -0.62 | -1.95 | -2.56 | -1.75 | -1.08 | -0.2 | -0.33 | -0.57 | 0.68 | 0.66 | -0.0975 | 1.6325 | 0.285878 | 0.878466 | 0.026771 | 1 | 0.996511 | 0.635993 |
| Kif13b | A0A286YC\ | 0.57 | 0.19 | 0.42 | 0.47 | -0.07 | 0.12 | 0.22 | 0.49 | 0.55 | 0.83 | 0.28 | 0.53 | 0.15 | 0.58 | 0.45 | 0.29 | -0.2225 | 0.135 | -0.045 | 0.246863 | 0.490434 | 0.817786 | 1 | 0.996511 | 0.972768 |
| Kif1a | E9Q9G6 | -0.93 | -0.6 | -0.62 | -0.32 | -0.39 | -0.79 | -0.31 | -0.71 | -0.89 | -0.53 | -0.7 | -0.75 | -0.92 | -0.01 | -1.03 | -0.67 | 0.0675 | -0.1 | -0.04 | 0.740079 | 0.614063 | 0.886614 | 1 | 0.996511 | 0.979823 |
| Kif1a | Q6TA13 | 0.14 | 0.21 | 0.2 | 0.08 | 0.24 | 0.28 | 0.54 | 0.29 | 0.18 | -0.1 | 0.36 | 0.26 | 0.12 | 0.27 | 0.22 | 0.08 | 0.18 | 0.0175 | 0.015 | 0.250175 | 0.920296 | 0.929135 | 1 | 0.996755 | 0.989687 |
| Kif1b | A0A5F8MP | -0.01 | 0.25 | 0.39 | 0.26 | 0.17 | 0 | 0.83 | 0.17 | -0.29 | -0.14 | 0.28 | 0.2 | -0.44 | -0.11 | 0.14 | 0.4 | 0.07 | -0.21 | -0.225 | 0.754895 | 0.321701 | 0.356811 | 1 | 0.996511 | 0.855416 |
| Kif1c | O35071 | -0.37 | -0.78 | 0.42 | 0.34 | 0.37 | -0.39 | 0.79 | 0.68 | 1.26 | -0.69 | 0.61 | -0.52 | 1.52 | 0.59 | 0.24 | 0.56 | 0.46 | 0.2625 | 0.825 | 0.238982 | 0.60827 | 0.058334 | 1 | 0.996511 | 0.661738 |
| Kif21a | F8WGN6 | -2.54 | -1.68 | -1.55 | -1.08 | -1.12 | -0.99 | -1.37 | -0.69 | 0.2 | -0.49 | -0.5 | -2.92 | 0.94 | -2.19 | -0.4 | -0.06 | 0.67 | 0.785 | 1.285 | 0.06685 | 0.268061 | 0.076948 | 1 | 0.996511 | 0.67628 |
| Kif21a | Q9QXL2-2 | 0.47 | -0.47 | -0.37 | -0.05 | -0.74 | 0.48 | -0.37 | 0.16 | 0.2 | 1.02 | 0.32 | 1.28 | 0.34 | 0.04 | 0.87 | 0.4 | -0.0125 | 0.81 | 0.5175 | 0.970101 | 0.036618 | 0.101064 | 1 | 0.996511 | 0.698156 |
| Kif21a | Q9QXL2-3 | -1.84 | -1.36 | -1.83 | -2.34 | -0.94 | -0.74 | -0.86 | -1.49 | -1.24 | -1.38 | -2.12 | -2.62 | -1.45 | -0.78 | -2.33 | -1.88 | 0.835 | 0.0025 | 0.2325 | 0.011807 | 0.994589 | 0.541324 | 1 | 0.999285 | 0.912212 |
| Kif21a | E9Q0J5 | -0.08 | -0.03 | -0.45 | -0.21 | -0.15 | -0.05 | -0.47 | -0.37 | 0.43 | -0.31 | -0.49 | -0.11 | 0.75 | -0.17 | -0.28 | -0.63 | -0.0675 | 0.0725 | 0.11 | 0.70858 | 0.767976 | 0.728882 | 1 | 0.996511 | 0.958635 |
| Kif21b | Q9QXL1 | -0.03 | 0.13 | 0.22 | 0.16 | 0.33 | 0.03 | 0.1 | 0.07 | 0.54 | -0.45 | 0.48 | 0.66 | 0.48 | 0.8 | 0.12 | 0.06 | 0.0125 | 0.1875 | 0.245 | 0.935557 | 0.500785 | 0.296402 | 1 | 0.996511 | 0.825612 |
| Kif2a | P28740-2 | 0.61 | 0.78 | 0.31 | 1.26 | 0.46 | -0.12 | -0.4 | 1.68 | 1.53 | 3.37 | 0.71 | 0.7 | 2.06 | 1.94 | 1.23 | 0.84 | -0.335 | 0.8375 | 0.7775 | 0.478921 | 0.189649 | 0.05011 | 1 | 0.996511 | 0.658454 |
| Kif2a | E0CZ72 | 0.75 | 0.13 | 0.78 | 0.59 | 0.57 | 0.8 | 0.79 | 0.74 | 0 | 1.25 | 0.74 | 0.69 | -0.08 | 0.08 | 0.87 | 0.66 | 0.1625 | 0.1075 | -0.18 | 0.414296 | 0.723104 | 0.538354 | 1 | 0.996511 | 0.912212 |
| Kif3a | Q3TET1 | -0.29 | -0.77 | -0.39 | -0.46 | -1.17 | -0.45 | -0.87 | -0.5 | 0.47 | 0.45 | -0.56 | -0.19 | 0.74 | -0.27 | -0.29 | -1.01 | -0.27 | 0.52 | 0.27 | 0.242749 | 0.094694 | 0.469372 | 1 | 0.996511 | 0.890107 |
| Kif3b | Q61771 | -0.42 | -0.59 | 0.18 | 0.2 | -0.76 | -0.84 | -0.54 | 0.05 | -0.54 | 0.38 | -0.07 | -0.76 | -0.41 | -0.74 | -0.28 | -0.71 | -0.365 | -0.09 | -0.3775 | 0.225464 | 0.781891 | 0.17325 | 1 | 0.996511 | 0.764376 |
| Kif3c | O35066 | -0.7 | -0.41 | -0.09 | -0.12 | 0.79 | -0.63 | -0.06 | -0.2 | 0.12 | 0.49 | -0.08 | -0.53 | 0.53 | 0.38 | -0.11 | -0.1 | 0.305 | 0.33 | 0.505 | 0.356827 | 0.24397 | 0.070587 | 1 | 0.996511 | 0.668066 |
| Kif5a | P33175 | 0.33 | 0.11 | 0.44 | 0.6 | -0.13 | 0.15 | 0.51 | 0.52 | 0.32 | 0.85 | 0.44 | 0.43 | 0.11 | 0.21 | 0.42 | 0.31 | -0.1075 | 0.14 | -0.1075 | 0.617375 | 0.496339 | 0.584104 | 1 | 0.996511 | 0.927734 |
| Kif5b | Q61768 | 1.37 | 0.86 | 0.88 | 0.87 | 0.56 | 1.37 | 0.58 | 0.9 | 1.23 | 1.21 | 0.86 | 1.53 | 1.28 | 0.39 | 0.97 | 0.67 | -0.1425 | 0.2125 | -0.1675 | 0.559598 | 0.351195 | 0.522143 | 1 | 0.996511 | 0.907667 |
| Kif5c | P28738 | 0.76 | 0.77 | 0.9 | 0.89 | 0.65 | 0.81 | 0.68 | 0.84 | 0.91 | 1.07 | 0.92 | 0.88 | 0.89 | 0.88 | 0.8 | 0.68 | -0.085 | 0.115 | -0.0175 | 0.561285 | 0.472318 | 0.918406 | 1 | 0.996511 | 0.986868 |
| Kifap3 | P70188-2 | -0.58 | -0.6 | -0.44 | -0.54 | -0.52 | -0.53 | -0.28 | -0.61 | -0.62 | -0.64 | -0.62 | -0.63 | -0.49 | -0.2 | -0.92 | -0.81 | 0.055 | -0.0875 | -0.065 | 0.718407 | 0.570105 | 0.76668 | 1 | 0.996511 | 0.964195 |
| Kifbp | Q6ZPU9 | -0.28 | -0.38 | -1.21 | -1.07 | -0.91 | 0.01 | -1.09 | -1.17 | 0.49 | -0.52 | -1.1 | -0.17 | 0.39 | -0.36 | -1.46 | -1.43 | -0.055 | 0.41 | 0.02 | 0.874075 | 0.310667 | 0.966419 | 1 | 0.996511 | 0.9927 |
| Kifc2 | A0A2R8W6 | -1.56 | -1.4 | -1.38 | -1.54 | -1.49 | -1.76 | -1.33 | -1.97 | -0.41 | -1.13 | -1.51 | -1.39 | 0.28 | -1.05 | -2.03 | -1.66 | -0.1675 | 0.36 | 0.355 | 0.386079 | 0.201336 | 0.46431 | 1 | 0.996511 | 0.888267 |
| Kifc3 | O35231 | -0.37 | -0.09 | 0.05 | -0.26 | -0.16 | 0.28 | 0.12 | -0.04 | 0.71 | -0.61 | 0.3 | -0.07 | 0.52 | 0.41 | -0.28 | 0.02 | 0.2175 | 0.25 | 0.335 | 0.244607 | 0.415742 | 0.191599 | 1 | 0.996511 | 0.777117 |
| Kirrel2 | Q7TSU7-2 | 1.12 | 1.38 | 0 | 0.3 | 1.24 | 0.32 | 0.11 | -0.91 | 0.89 | 0.24 | -0.5 | 0.81 | 0.32 | 2.67 | -0.64 | -0.53 | -0.51 | -0.34 | -0.245 | 0.328619 | 0.441308 | 0.746166 | 1 | 0.996511 | 0.960124 |
| Kirrel3 | Q8BR86-2 | -0.81 | -0.59 | -0.33 | 0.02 | -0.83 | -0.59 | -0.26 | -0.18 | -0.07 | -0.4 | -0.07 | -0.26 | -0.14 | -1.12 | 0.05 | -0.39 | -0.0375 | 0.2275 | 0.0275 | 0.879173 | 0.333844 | 0.931271 | 1 | 0.996511 | 0.989713 |
| Kit | P05532 | 0.11 | 0.66 | 0.73 | 0.72 | 0.89 | 0.32 | 1.11 | 0.48 | 0.62 | 0.68 | 0.51 | 0.29 | 0.99 | 1.05 | 0.69 | 0.63 | 0.145 | -0.03 | 0.285 | 0.562587 | 0.888053 | 0.232592 | 1 | 0.996511 | 0.788994 |
| Kitlg | P20826-2 | -0.24 | 1.25 | 2.59 | 2.06 | 1.46 | -0.42 | 2.73 | 2.48 | 1.03 | 2.77 | 2.41 | 0.51 | -0.41 | 2.01 | 2.15 | 2.66 | 0.1475 | 0.265 | 0.1875 | 0.862294 | 0.722404 | 0.821274 | 1 | 0.996511 | 0.972768 |
| Kl | O35082-2 | 0.62 | 0.52 | -0.68 | -0.67 | -0.08 | 0.56 | -2.31 | -1.78 | 3.29 | 0.68 | 0.23 | 1.56 | 2.34 | 4.91 | -1.83 | -0.59 | -0.85 | 1.4925 | 1.26 | 0.24363 | 0.057363 | 0.373602 | 1 | 0.996511 | 0.859424 |
| Klc1 | Q8CD76 | 0.65 | 0.62 | 1.03 | 1.07 | 0.57 | 0.66 | 0.96 | 1.29 | 0.32 | 1.58 | 1.12 | 0.85 | 0.11 | 0.78 | 1.12 | 1.16 | 0.0275 | 0.125 | -0.05 | 0.902339 | 0.674623 | 0.862331 | 1 | 0.996511 | 0.977204 |
| Klc2 | D3YXZ3 | 0.73 | 1.12 | 0.78 | 0.77 | 0.86 | 0.49 | 0.44 | 0.58 | 0.68 | 0.68 | 0.61 | 1.01 | 0.96 | 0.64 | 0.71 | 0.57 | -0.2575 | -0.105 | -0.13 | 0.171785 | 0.578215 | 0.511605 | 1 | 0.996511 | 0.903991 |
| Klc4 | Q9DBS5 | 0.82 | 0.75 | 0.54 | 0.97 | 0.27 | 0.64 | -0.08 | 0.62 | 1.12 | 0.58 | 0.65 | 0.85 | 1.08 | 0.32 | 0.29 | 0.22 | -0.4075 | 3.00E-02 | -0.2925 | 0.087946 | 0.880717 | 0.266148 | 1 | 0.996511 | 0.818107 |
| Klhdc10 | Q6PAR0 | -0.68 | -0.08 | -1.79 | -2.11 | 0.12 | 4.81 | -2.45 | -2.98 | 0.03 | -1.29 | -2.46 | -0.1 | 1.61 | -0.27 | -2.65 | -2.7 | 1.04 | 0.21 | 0.1625 | 0.531384 | 0.759206 | 0.873514 | 1 | 0.996511 | 0.979626 |
| Klhdc2 | Q4G5Y1 | 0.08 | 0 | -1.71 | -1.45 | 0.04 | -0.32 | -0.99 | -1.73 | -0.84 | -1.65 | -1.64 | 0.31 | -1.3 | -0.53 | -2.41 | -1.47 | 0.02 | -0.185 | -0.6575 | 0.971474 | 0.760116 | 0.260285 | 1 | 0.996511 | 0.813803 |
| Klhdc3 | A0A3B2WC | 0.03 | 0.34 | 0.24 | -1.12 | 0.97 | 0.23 | -0.57 | -0.16 | 1.09 | 0.86 | -0.81 | 0.3 | 0.26 | 0.97 | 0.25 | -0.8 | 0.245 | 0.4875 | 0.2975 | 0.579953 | 0.346243 | 0.528849 | 1 | 0.996511 | 0.908663 |
| Klhdc4 | G3UZG5 | -1.08 | -0.83 | -1.71 | -1.29 | 0.01 | -0.58 | -1.86 | -1.59 | -0.33 | -1.31 | -1.99 | -0.66 | -0.17 | -0.28 | -2.92 | -1.7 | 0.2225 | 0.155 | -0.04 | 0.617446 | 0.695616 | 0.948481 | 1 | 0.996511 | 0.992476 |
| Klhdc7a | A2APT9 | 0.24 | 1.07 | 0.22 | 0.13 | 1.21 | 0.07 | 0.93 | 0.23 | 1.44 | 0.29 | 0.52 | 0.31 | 1.12 | 1.28 | 0.29 | 0.58 | 0.195 | 0.225 | 0.4025 | 0.569072 | 0.517974 | 0.234716 | 1 | 0.996511 | 0.791172 |

| | | | | | | | | | | | | | | | | | | | | | | | | | | | |
|---|---|---|---|---|---|---|---|---|---|---|---|---|---|---|---|---|---|---|---|---|---|---|---|---|---|---|---|
| Klhdc8b | Q9D2D9 | 0.17 | -1.17 | -0.67 | -0.38 | 0.51 | 0.69 | -0.8 | -0.75 | -0.26 | -0.32 | -0.97 | 0.07 | 0.07 | 1.3 | -1.5 | -0.92 | 0.425 | 0.1425 | 0.25 | 0.361032 | 0.683386 | 0.686384 | | 1 | 0.996511 | 0.955729 |
| Klhdc9 | Q3USL1 | 1.34 | 1.35 | 0.77 | 0.31 | 1.14 | 0.83 | 1.34 | 0.81 | 0.28 | 0.15 | -0.01 | 0.71 | 1.08 | 0.56 | 0.89 | 1.22 | 0.0875 | -0.66 | -0.005 | 0.75802 | 0.052412 | 0.98674 | | 1 | 0.996511 | 0.996154 |
| Klhl11 | Q8CE33 | -0.83 | -0.38 | -0.62 | -0.91 | -0.29 | -0.78 | -0.82 | -0.82 | -0.33 | -0.9 | -0.72 | -0.44 | -0.05 | -0.96 | -0.83 | -1.11 | 0.0075 | 0.0875 | -0.0525 | 0.970889 | 0.686237 | 0.853077 | | 1 | 0.996511 | 0.976225 |
| Klhl13 | Q80TF4-5 | -2.06 | -0.71 | -0.94 | -0.09 | -0.09 | -0.51 | 0.38 | -0.81 | -0.17 | -2.74 | -0.89 | -0.35 | -0.67 | -2.59 | -1.5 | -1.05 | 0.6925 | -0.0875 | -0.5025 | 0.152014 | 0.893556 | 0.364748 | | 1 | 0.996511 | 0.858475 |
| Klhl2 | Q8JZP3 | 0.57 | 1.4 | 0.28 | 0.11 | 1.37 | 0.46 | 0.92 | 0.3 | -1 | -1.44 | 0.53 | 0.84 | -1.05 | -3.28 | -0.02 | 0.3 | 0.1725 | -0.8575 | -1.6025 | 0.632634 | 0.164069 | 0.064377 | | 1 | 0.996511 | 0.66433 |
| Klhl20 | Q8VCK5 | -0.49 | -1.99 | 0.57 | 0.1 | -3.9 | -0.14 | -1.46 | -0.62 | 2.67 | 0.06 | 0.08 | -0.99 | 4.1 | -2.9 | 0.79 | -0.5 | -1.0775 | 0.9075 | 0.825 | 0.252423 | 0.309275 | 0.556175 | | 1 | 0.996511 | 0.917092 |
| Klhl22 | Q99JN2 | -0.93 | -0.35 | 0.32 | 0.26 | -0.44 | -0.57 | 0.26 | 0.31 | -0.3 | 0.66 | 0.64 | -0.5 | -0.14 | 0.21 | 0.27 | -0.02 | 0.065 | 0.3 | 0.255 | 0.855654 | 0.465732 | 0.429352 | | 1 | 0.996511 | 0.879165 |
| Klhl26 | Q8BGY4 | 0.54 | 0.65 | 0.72 | -0.06 | 0.61 | 0.11 | 0.63 | 0.49 | 0.77 | 0.41 | 0.84 | 0.64 | 1.4 | 0.9 | 0.52 | 0.55 | -0.0025 | 0.2025 | 0.38 | 0.99144 | 0.394537 | 0.210305 | | 1 | 0.996511 | 0.786218 |
| Klhl3 | E0CZ16 | -0.57 | -0.85 | -0.21 | -0.12 | 0.14 | -0.68 | -0.02 | 0.02 | -1.26 | -0.83 | 0.32 | -0.44 | -0.91 | -0.23 | 0.04 | 0.38 | 0.3025 | -0.115 | 0.2575 | 0.262105 | 0.753082 | 0.437709 | | 1 | 0.996511 | 0.883161 |
| Klhl36 | Q8R124 | -0.21 | 0.27 | -0.93 | 0.06 | 0.71 | -0.99 | -0.29 | -0.55 | 0.46 | 0.52 | -0.68 | -0.41 | 0.28 | 0.76 | -0.27 | -0.05 | -0.0775 | 0.175 | 0.3825 | 0.852804 | 0.651886 | 0.283145 | | 1 | 0.996511 | 0.822669 |
| Klhl4 | Q8BLJ6 | 0.23 | -0.38 | -0.17 | -0.17 | 0.44 | -0.8 | 0.81 | -0.35 | 0.59 | 3.65 | -0.3 | 0.11 | 0.15 | 1.15 | -0.6 | -0.61 | 0.1475 | 1.135 | 0.145 | 0.69107 | 0.190012 | 0.728317 | | 1 | 0.996511 | 0.958635 |
| Klhl41 | A2AUC9 | 0.64 | 0.05 | -1.04 | -0.6 | 0.21 | -1.19 | 1.2 | -0.62 | 1.77 | 0.77 | 0.99 | -0.36 | -0.04 | 2.57 | 0.45 | 2.18 | 0.1375 | 1.03 | 1.5275 | 0.813383 | 0.079669 | 0.045971 | | 1 | 0.996511 | 0.658454 |
| Klhl42 | Q8BFQ9 | 1.03 | 1.12 | 1.16 | 1.24 | 0.54 | 1.05 | 0.25 | 1.35 | -0.26 | 1.44 | 0.42 | 1.41 | -0.34 | 0.91 | 0.71 | 0.57 | -0.34 | -0.385 | -0.675 | 0.213622 | 0.345604 | 0.044848 | | 1 | 0.996511 | 0.658454 |
| Klhl5 | E9Q1L2 | -1.19 | -0.02 | 1.13 | 0.3 | -1.75 | -0.78 | 1.91 | 0.64 | 1.12 | 0.41 | 1.49 | -0.32 | 0.51 | -0.55 | 0.99 | 1.89 | -0.05 | 0.62 | 0.655 | 0.95262 | 0.297561 | 0.319592 | | 1 | 0.996511 | 0.838969 |
| Klhl8 | A2AH60 | -1.81 | -0.89 | -0.28 | -1.02 | -1.3 | -1.27 | -1.2 | -1.65 | -1.1 | -1.82 | -1.5 | -0.84 | -0.57 | -1.93 | -2.64 | -1.61 | -0.355 | -0.315 | -0.6875 | 0.28815 | 0.404566 | 0.190503 | | 1 | 0.996511 | 0.776843 |
| Klhl9 | Q6ZPT1 | -0.27 | -0.01 | -0.74 | -0.86 | 0.15 | -0.16 | -0.84 | -0.86 | -1.23 | -0.06 | -0.57 | 0.03 | -0.88 | -0.84 | -0.85 | -0.79 | 0.0425 | 0.0125 | -0.37 | 0.892695 | 0.971095 | 0.148272 | | 1 | 0.997515 | 0.746406 |
| Kmo | Q91WN4-2 | -0.35 | 0.33 | -0.14 | -0.05 | -0.22 | -0.97 | -0.86 | -0.09 | -0.82 | -0.67 | -0.22 | -0.07 | -2.09 | 0.85 | -0.78 | -1.13 | -0.4825 | -0.3925 | -0.735 | 0.101389 | 0.144493 | 0.224614 | | 1 | 0.996511 | 0.787447 |
| Kmt2a | P55200-2 | 0.6 | 1.37 | 0.8 | 0.93 | 1.1 | 0.61 | 0.46 | 1.09 | -0.11 | 1.58 | 0.07 | 0.09 | -0.99 | 0.57 | 0.25 | 0.62 | -0.11 | -0.5175 | -0.8125 | 0.655686 | 0.224403 | 0.06644 | | 1 | 0.996511 | 0.664936 |
| Kndc1 | Q0KK55 | -0.49 | -0.47 | -0.9 | -0.75 | -0.6 | -0.65 | -1.02 | -1.46 | 0.35 | -0.83 | -1.12 | -0.79 | 0.75 | 0.43 | -1.13 | -1.1 | -0.28 | 0.055 | 0.39 | 0.262959 | 0.870218 | 0.421688 | | 1 | 0.996511 | 0.875736 |
| Kng1 | O08677-2 | -2.5 | -3.37 | -2.92 | -2.75 | -3.17 | -2.6 | -2.53 | -2.56 | -2.62 | -0.98 | -2.43 | -2.76 | -2.04 | -1.89 | -2.75 | -2.6 | 0.17 | 0.6875 | 0.565 | 0.502785 | 0.134204 | 0.081283 | | 1 | 0.996511 | 0.687007 |
| Kng2 | Q6S9I3 | -0.44 | 0.2 | -0.33 | -0.44 | 1.36 | -0.42 | 0.61 | -0.6 | -0.65 | -2.44 | -0.01 | -1.13 | -1.98 | 1.3 | -1.52 | -0.8 | 0.49 | -0.805 | -0.4975 | 0.293855 | 0.134247 | 0.467403 | | 1 | 0.996511 | 0.889213 |
| Kpna1 | Q60960 | -1.92 | -2.29 | 0.08 | 0.46 | -1.76 | -2.09 | -0.99 | 0.37 | -1.11 | 1.38 | 0.05 | -2.1 | -0.94 | -1.07 | -0.25 | -0.5 | -0.2 | 0.4725 | 0.2275 | 0.802014 | 0.610258 | 0.72926 | | 1 | 0.996511 | 0.958635 |
| Kpna2 | P52293 | -0.83 | -0.08 | -1.31 | -1.15 | 0.11 | -0.48 | -0.88 | -0.31 | -0.19 | -1.46 | -1.34 | -0.37 | -0.73 | 0.48 | -1.76 | -0.14 | 0.4525 | 0.0025 | 0.305 | 0.196085 | 0.99508 | 0.554474 | | 1 | 0.999285 | 0.916796 |
| Kpna3 | O35344 | 0.86 | 0.16 | 0.22 | 0.1 | -0.15 | 0.7 | -0.4 | 0.03 | 1.32 | -0.01 | 0.2 | 1 | 1.15 | 0.59 | 0.22 | 0.13 | -0.29 | 0.2925 | 0.1875 | 0.337515 | 0.419123 | 0.540841 | | 1 | 0.996511 | 0.912212 |
| Kpna4 | O35343 | -0.66 | -0.29 | -0.86 | -0.62 | -0.64 | -0.16 | -0.39 | -0.43 | -0.9 | -0.58 | -0.71 | -0.41 | -0.96 | -0.04 | -0.85 | -0.48 | 0.2025 | -0.0425 | 0.025 | 0.306244 | 0.834972 | 0.924958 | | 1 | 0.996511 | 0.988301 |
| Kpna6 | O35345 | -0.95 | -1.07 | -0.54 | -0.77 | -0.8 | -1.09 | -0.73 | -0.68 | -0.68 | -0.04 | -0.67 | -1.2 | -0.27 | -0.05 | -0.4 | -0.69 | 0.0075 | 0.185 | 0.48 | 0.968065 | 0.509796 | 0.057483 | | 1 | 0.996511 | 0.661738 |
| Kpnb1 | P70168 | -0.17 | -0.4 | -0.02 | 0.02 | -0.32 | -0.21 | 0.01 | -0.05 | -0.1 | -0.42 | -0.05 | -0.34 | -0.14 | -0.15 | -0.45 | -0.24 | 5.89E-17 | -0.085 | -0.1025 | 1 | 0.654621 | 0.598483 | | 1 | 0.996511 | 0.930093 |
| Kptn | G3X8R1 | 0.15 | -0.18 | 0.12 | 0.13 | -0.27 | -0.09 | 0.08 | 0.04 | 0.26 | 0.17 | 0.22 | 0.01 | 0.91 | 0.64 | 0.61 | 0.02 | -0.115 | 0.11 | 0.49 | 0.496838 | 0.526635 | 0.068529 | | 1 | 0.996511 | 0.665034 |
| Kras | P32883-2 | 1.17 | 1.26 | 1.27 | 1.17 | 1.48 | 1.46 | 1.48 | 1.42 | 0.83 | 0.59 | 1.31 | 1.7 | 0.57 | 0.56 | 1.13 | 1.42 | 0.2425 | -0.11 | -0.2975 | 0.105429 | 0.680853 | 0.251997 | | 1 | 0.996511 | 0.80892 |
| Kri1 | V9GXJ9 | 0.21 | -0.59 | -0.18 | 0.38 | 0.04 | -0.35 | 0.58 | -0.16 | -1.28 | -0.57 | 0.08 | -0.12 | -0.94 | 0.73 | 0.55 | 0.09 | 0.0725 | -0.4275 | 0.1525 | 0.80548 | 0.256101 | 0.713527 | | 1 | 0.996511 | 0.958106 |
| Krit1 | A0A0G2JG( | 0.58 | 0.39 | -0.2 | 0.04 | 0.1 | 0.48 | -0.25 | -0.2 | 1.27 | 0.22 | -0.64 | 0.15 | 1.12 | 0.75 | -0.8 | -0.7 | -0.17 | 0.0475 | -0.11 | 0.507774 | 0.907445 | 0.82199 | | 1 | 0.996744 | 0.972768 |
| Krr1 | Q8BGA5 | 0.58 | -0.4 | 1.89 | 0.39 | 0.1 | 0.46 | 1.69 | 1.37 | 1.54 | 0.02 | 1.72 | 0.24 | 2.52 | 0.42 | 1.76 | 2.76 | 0.29 | 0.265 | 1.25 | 0.60334 | 0.656221 | 0.080213 | | 1 | 0.996511 | 0.683237 |
| Krt12 | Z4YJD9 | -1.66 | -0.62 | 0.32 | -0.39 | 0.06 | -0.69 | 0.33 | -0.02 | 0.04 | 0.18 | 0.73 | -0.54 | -0.66 | -0.3 | -1.05 | 0 | 0.5075 | 0.69 | 0.085 | 0.260488 | 0.156463 | 0.848056 | | 1 | 0.996511 | 0.976225 |
| Krt33a | Q8K0Y2 | -0.71 | -0.01 | -0.39 | -0.34 | -0.23 | -0.18 | -0.83 | -0.34 | -0.07 | -0.97 | -0.42 | -0.49 | 0 | 0.05 | -0.58 | -0.85 | -0.0325 | -0.125 | 0.0175 | 0.886168 | 0.626671 | 0.950594 | | 1 | 0.996511 | 0.992476 |
| Krt76 | Q3UV17 | -1.52 | -0.28 | 0.35 | -1.84 | 1.1 | 2.09 | 0.14 | -1.43 | -1.64 | 0.42 | 1.31 | -1.22 | -0.32 | -2.3 | -1.7 | 0.01 | 1.2975 | 0.54 | -0.255 | 0.140471 | 0.495457 | 0.710667 | | 1 | 0.996511 | 0.957819 |
| Krt81 | Q9ERE2 | -6.98 | -7.05 | -4.72 | -4.47 | -6.84 | -7.11 | -4.82 | -5.31 | -6.96 | -6.19 | -4.83 | -5.89 | -3.89 | -8.77 | -5.51 | -5.31 | -0.215 | -0.1625 | -0.065 | 0.790981 | 0.828606 | 0.95337 | | 1 | 0.996511 | 0.992476 |
| Krtap3-1 | A2A591 | -3.59 | -4.8 | -4.89 | -2.69 | -4.54 | -3.22 | -3.78 | -5.01 | -1.95 | -4.99 | -4.91 | -1.66 | -3.22 | -4.86 | -4.34 | -3.24 | -0.145 | 0.615 | 0.0775 | 0.809431 | 0.520532 | 0.898818 | | 1 | 0.996511 | 0.983509 |
| Krtcap2 | A0A0A0M( | 0.06 | 0.77 | -1.08 | -0.47 | 0.76 | 0.49 | 0.51 | -0.73 | 0.52 | -1.06 | -0.12 | 0.24 | -0.01 | -0.29 | 0.65 | -0.07 | 0.4375 | 0.075 | 0.25 | 0.372114 | 0.877607 | 0.557661 | | 1 | 0.996511 | 0.917092 |
| Ksr1 | Q61097-2 | 0.16 | 1.63 | 0.27 | 0.09 | 0.32 | 0.31 | 0.24 | 0.07 | 0.27 | 0.39 | 0.11 | 0.42 | 0.77 | -0.21 | 0.05 | 0.08 | -0.3025 | -0.24 | -0.365 | 0.40489 | 0.513453 | 0.379535 | | 1 | 0.996511 | 0.860984 |
| Ksr2 | M0QW59 | -0.36 | -0.19 | 0.37 | -0.08 | -0.11 | 0.17 | 0.34 | 0.12 | -0.12 | -0.37 | 0.17 | 0.31 | 0.22 | -0.07 | 0.35 | 0.47 | 0.195 | 0.0625 | 0.3075 | 0.364759 | 0.797717 | 0.214056 | | 1 | 0.996511 | 0.786931 |
| Kti12 | Q9D1R2 | 0.27 | -0.65 | 0.27 | 0.61 | -1.06 | 0.15 | 0.89 | 0.28 | 0.74 | 0.98 | 0.76 | 0.11 | 1.62 | -0.98 | -0.1 | 0.43 | -0.06 | 0.5225 | 0.1175 | 0.895111 | 0.137605 | 0.83372 | | 1 | 0.996511 | 0.975371 |
| Ktn1 | A0A087WC | -0.79 | -0.64 | -0.3 | -0.36 | -0.18 | -0.4 | 0.53 | 0.1 | -0.48 | -0.45 | -0.34 | -0.49 | 0.4 | -0.37 | 1.24 | 1.04 | 0.535 | 0.0825 | 1.1 | 0.053913 | 0.655016 | 0.015459 | | 1 | 0.996511 | 0.589387 |
| Ktn1 | A0A087WF | -1.43 | -0.91 | -1.98 | -0.75 | -0.43 | -1.02 | -0.56 | -1.93 | -1.03 | -1.12 | -1.54 | -0.8 | -1.36 | -0.71 | -0.8 | -0.52 | 0.2825 | 0.145 | 0.42 | 0.499324 | 0.651237 | 0.229532 | | 1 | 0.996511 | 0.787447 |
| Kxd1 | E9Q4P0 | -0.12 | 0.34 | 0.25 | 0.39 | 0.43 | -0.28 | -0.49 | 1.03 | -0.75 | 0.56 | -0.04 | -0.17 | 0.15 | -0.2 | -0.11 | 0.29 | -0.0425 | -0.315 | -0.1825 | 0.903676 | 0.30758 | 0.40753 | | 1 | 0.996511 | 0.874084 |
| Kyat1 | Q8BTY1 | 0.01 | -0.03 | 0.41 | 0.37 | 0.08 | -0.49 | 0.48 | 0.5 | -0.44 | 0.18 | -0.05 | -0.6 | -0.59 | -0.12 | -0.16 | 0.18 | -0.0475 | -0.4175 | -0.3625 | 0.858414 | 0.109777 | 0.152002 | | 1 | 0.996511 | 0.750422 |
| Kyat3 | Q71RI9-2 | 0.36 | 0.68 | 0.52 | 0.89 | 0.56 | 0.09 | 0.95 | 0.08 | -0.09 | 0.13 | 0.55 | -0.07 | -0.47 | 0.3 | 0.18 | -0.11 | -0.1925 | -0.4825 | -0.6375 | 0.4492 | 0.055056 | 0.025974 | | 1 | 0.996511 | 0.633075 |
| L1cam | Q6PGJ3 | 1.63 | 1.23 | 1.44 | 1.86 | 0.94 | 1.55 | 1.79 | 2.16 | 1.53 | 1.73 | 1.83 | 2.11 | 1.01 | 1.26 | 1.79 | 1.54 | 0.07 | 0.26 | -0.14 | 0.809164 | 0.253883 | 0.578558 | | 1 | 0.996511 | 0.927198 |
| L2hgdh | Q91YP0 | 1.15 | 1.48 | 0.55 | 0.8 | 1.13 | 1.35 | 0.47 | 0.52 | 0.66 | 0.15 | 0.53 | 0.87 | 0.51 | 1.01 | -0.01 | 0.24 | -0.1275 | -0.4425 | -0.5575 | 0.66993 | 0.127509 | 0.098075 | | 1 | 0.996511 | 0.698156 |
| L3hypdh | Q9CXA2 | 0.99 | 1.05 | -0.03 | -0.1 | 1.17 | 1 | -1.47 | -0.09 | 0.03 | -0.25 | -0.61 | 1.08 | -0.24 | 0.22 | -0.77 | -0.79 | -0.325 | -0.415 | -0.8725 | 0.604811 | 0.370131 | 0.046545 | | 1 | 0.996511 | 0.658454 |
| Lactb | Q9EP89 | 0.47 | 0.5 | 0.87 | 0.91 | 0.21 | 0.49 | 0.91 | 1.03 | -0.38 | -0.52 | 1.01 | 0.19 | -0.43 | -0.78 | 0.16 | 0.53 | -0.0275 | -0.6125 | -0.8175 | 0.908301 | 0.114604 | 0.029842 | | 1 | 0.996511 | 0.640893 |
| Lactb2 | Q99KR3 | 0.12 | -0.01 | 0.2 | -0.08 | 0.26 | -0.11 | 0.08 | 0.05 | -0.19 | 0.65 | 0.11 | -0.14 | 0.03 | 0.12 | 0.12 | 0.03 | 0.0125 | 0.05 | 0.0175 | 0.937879 | 0.83024 | 0.919331 | | 1 | 0.996511 | 0.986868 |
| Lama1 | P19137 | -1.18 | -1.8 | -1.19 | -0.81 | -1.31 | -1.21 | -1 | -1.43 | -1.26 | -0.61 | -0.82 | -1.42 | -1.13 | -1.79 | -0.89 | -1.02 | 0.0075 | 0.2175 | 0.0375 | 0.975042 | 0.457666 | 0.900392 | | 1 | 0.996511 | 0.983718 |
| Lama2 | A0A1W2P7 | 0.12 | -0.26 | 1.05 | 0.61 | -0.46 | 0.15 | 0.17 | -0.15 | -0.21 | 0.55 | 0.57 | 0.36 | 0.22 | 0.3 | 1.11 | 0.8 | -0.4525 | -0.0625 | 0.2275 | 0.176779 | 0.85209 | 0.523223 | | 1 | 0.996511 | 0.907797 |
| Lama2 | Q60675 | -0.08 | -0.11 | -0.28 | -0.39 | 0.18 | 0.07 | -0.43 | -0.61 | 0.09 | 0.18 | -0.1 | 0.04 | 0.41 | -0.54 | -0.06 | -0.07 | 0.0175 | 0.2675 | 0.15 | 0.938153 | 0.143722 | 0.543069 | | 1 | 0.996511 | 0.912212 |
| Lama4 | P97927 | -0.26 | 0.05 | -0.74 | -0.97 | 0.18 | -0.18 | -1.02 | -1.15 | 0.02 | -0.89 | -0.49 | 0.01 | 0.22 | 0.18 | -0.29 | -0.27 | -0.0625 | 0.1425 | 0.44 | 0.868287 | 0.656536 | 0.151025 | | 1 | 0.996511 | 0.749304 |
| Lama5 | Q61001 | -1.46 | -1.15 | -1.16 | -1.33 | -1.36 | -1.37 | -1.42 | -1.77 | -1.15 | -0.97 | -1.19 | -1.36 | -0.84 | -1.19 | -1.23 | -1.63 | -0.205 | 0.1075 | 0.0525 | 0.255428 | 0.549628 | 0.815897 | | 1 | 0.996511 | 0.972768 |
| Lamb1 | E9QN70 | -0.49 | -0.62 | -0.06 | -0.5 | -0.02 | -0.3 | -0.2 | -0.44 | 0.11 | 0.15 | 0.27 | 0.01 | 0.39 | -0.31 | 0.44 | 0.31 | 0.1775 | 0.5525 | 0.625 | 0.362474 | 0.016894 | 0.030839 | | 1 | 0.996511 | 0.641481 |
| Lamb2 | Q61292 | -0.74 | -1.3 | -0.85 | -1.02 | -0.91 | -0.7 | -1.17 | -1.42 | -0.79 | -0.5 | -0.92 | -0.77 | -0.19 | -1.32 | -0.87 | -0.88 | -0.0725 | 0.2325 | 0.1625 | 0.743773 | 0.26388 | 0.568993 | | 1 | 0.996511 | 0.922588 |
| Lamc1 | F8VQJ3 | 0.41 | -0.42 | -0.31 | -0.77 | 0.25 | 0.37 | -0.22 | -0.9 | -0.37 | -0.74 | -0.29 | 0.53 | -0.18 | -0.41 | -0.22 | -0.16 | 0.1475 | 0.055 | 0.03 | 0.685836 | 0.877223 | 0.913467 | | 1 | 0.996511 | 0.986458 |
| Lamp1 | P11438 | -0.79 | -0.52 | -0.01 | -0.05 | -0.43 | -0.66 | 0.18 | 0.04 | -0.61 | 0.42 | 0.19 | -0.51 | -0.33 | 0.08 | -0.18 | -0.14 | 0.125 | 0.215 | 0.2 | 0.65324 | 0.504776 | 0.421199 | | 1 | 0.996511 | 0.875639 |
| Lamp2 | P17047-3 | -0.54 | -0.44 | -0.51 | -0.06 | 0.33 | -0.42 | 0.61 | -0.23 | -0.62 | -0.48 | -0.03 | -0.38 | -0.4 | 0.29 | -0.61 | -0.33 | 0.46 | 0.01 | 0.125 | 0.116147 | 0.962088 | 0.625775 | | 1 | 0.996755 | 0.939447 |
| Lamp5 | Q9D387 | 2.2 | 2.13 | 2.12 | 2.18 | 2.53 | 1.59 | 2.77 | 3 | -0.1 | 0.78 | 2.59 | 1.89 | 0.39 | -0.44 | 3.69 | 2.85 | 0.315 | -0.8675 | -0.535 | 0.318789 | 0.140881 | 0.548003 | | 1 | 0.996511 | 0.912571 |
| Lamtor1 | Q9CQ22 | -0.18 | -0.16 | -0.08 | 0.45 | -0.39 | -0.35 | 0.43 | 0.4 | -0.43 | 0.27 | 0.33 | 0.14 | -0.03 | -0.46 | -0.01 | 0.25 | 0.015 | 0.07 | -0.07 | 0.956557 | 0.781122 | 0.7758 | | 1 | 0.996511 | 0.966799 |
| Lamtor2 | Q9JHS3 | -0.57 | -0.23 | -0.37 | -0.44 | -0.01 | -0.15 | -0.61 | -0.35 | -0.92 | -0.16 | -0.46 | -0.59 | -0.94 | -0.45 | -0.6 | -0.62 | 0.1225 | -0.13 | -0.25 | 0.52005 | 0.546998 | 0.223502 | | 1 | 0.996511 | 0.787447 |
| Lamtor3 | O88653 | -1.5 | -1.45 | -1.51 | -1.39 | -1.27 | -1.67 | -1.08 | -1.93 | -1.88 | -1.53 | -1.4 | -1.42 | -1.73 | -1.89 | -1.33 | -1.92 | -0.025 | -0.095 | -0.255 | 0.90875 | 0.601383 | 0.229679 | | 1 | 0.996511 | 0.787447 |
| Lamtor4 | Q8CF66 | 5.83 | 5.24 | 0.69 | -1.79 | 5.95 | 4.26 | -1.4 | -0.39 | 0.92 | -1.99 | -0.21 | 5.12 | 3.05 | 2.18 | 0.03 | -1.49 | -0.3875 | -1.5325 | -1.55 | 0.864671 | 0.475595 | 0.414768 | | 1 | 0.996511 | 0.874945 |
| Lamtor5 | Q9D1L9 | 1.45 | 2.02 | 0.41 | 0.25 | 1.94 | 1.36 | 0.7 | 0.63 | 1.07 | 0.11 | 0.65 | 1.78 | 0.34 | 1.15 | 0.51 | 0.97 | 0.125 | -0.13 | -0.29 | 0.796525 | 0.799417 | 0.515138 | | 1 | 0.996511 | 0.905674 |
| Lancl1 | O89112 | -0.62 | -1.1 | -0.67 | -0.61 | -1.07 | -0.37 | -0.36 | -0.58 | -1.45 | 0.05 | -0.79 | -0.84 | -1.25 | -0.81 | -0.65 | -0.69 | 0.155 | -0.0075 | -0.1 | 0.496835 | 0.981741 | 0.661438 | | 1 | 0.998773 | 0.949645 |
| Lancl2 | F6RJV6 | 0.51 | 0.47 | 0.29 | 0.65 | 0.42 | 0.57 | 0.62 | 0.73 | -0.18 | 0.23 | 0.43 | 0.75 | -0.42 | 0.28 | -0.07 | 0.34 | 0.105 | -0.1725 | -0.4475 | 0.517774 | 0.47379 | 0.082191 | | 1 | 0.996511 | 0.68865 |
| Lancl3 | Q8CD19 | -1.52 | -1.02 | -1.78 | -1.32 | -1.25 | -0.82 | -1.59 | -1.23 | -1.39 | 0.43 | -1.86 | -1.37 | -1.67 | -0.99 | -1.4 | -1.94 | 0.1875 | 0.3625 | -0.09 | 0.443643 | 0.469913 | 0.747809 | | 1 | 0.996511 | 0.960744 |
| Lap3 | Q9CPY7 | 0.15 | 0.12 | 0.47 | 0.43 | 0.31 | 0.36 | 0.84 | 0.46 | -0.15 | 0.09 | 0.45 | 0 | -0.24 | -0.23 | 0.24 | 0.46 | 0.2 | -0.195 | -0.235 | 0.307626 | 0.35184 | 0.336306 | | 1 | 0.996511 | 0.847578 |
| Laptm4a | E9QLR3 | -2.56 | -2.01 | -0.08 | 0.7 | -1.77 | -1.95 | 0.73 | -0.01 | -1.49 | -0.36 | 0.28 | -0.88 | -1.68 | -1.12 | -1.23 | 0.99 | 0.2375 | 0.375 | 0.2275 | 0.795185 | 0.632402 | 0.795572 | | 1 | 0.996511 | 0.969727 |
| Large1 | Q9Z1M7 | -1.67 | -2.46 | -2.17 | -2.71 | -1.61 | -2.1 | -1.76 | -2.19 | -1.37 | -4.47 | -1.97 | -2.67 | -0.94 | -1.88 | -1.01 | -2.32 | 0.3375 | -0.3675 | 0.715 | 0.230098 | 0.572907 | 0.095377 | | 1 | 0.996511 | 0.698156 |
| Larp1 | Z4YJT3 | 0.39 | 0.12 | -0.23 | -0.25 | 0.09 | 0.29 | -0.4 | -0.25 | -0.27 | 0.22 | -0.41 | 0.31 | 0.05 | -0.17 | -0.06 | -0.47 | -0.075 | -0.045 | -0.17 | 0.75161 | 0.860326 | 0.471966 | | 1 | 0.996511 | 0.891144 |
| Larp1b | F6U5V1 | -1.89 | -1.9 | -2.2 | -3.32 | -3.98 | -0.72 | -1.5 | -0.49 | 1.25 | -3.97 | -0.3 | -1.13 | 3.19 | -2.44 | -0.44 | -0.05 | 0.655 | 1.29 | 2.3925 | 0.412634 | 0.231246 | 0.051432 | | 1 | 0.996511 | 0.658454 |
| Larp4 | E9Q066 | -1.37 | -1.24 | -1.06 | -1.06 | -1.09 | -1.25 | -0.76 | -1.29 | -0.52 | -0.27 | -1.28 | -1.37 | 0.1 | -0.56 | -0.38 | -1.11 | 0.085 | 0.3225 | 0.695 | 0.646713 | 0.287225 | 0.033308 | | 1 | 0.996511 | 0.644622 |
| Larp4b | Q6A0A2 | -1.83 | -2.09 | -0.35 | -0.74 | -1.44 | -1.95 | -0.19 | -0.51 | -1.02 | 0.71 | -0.52 | -1.43 | -0.16 | -1.07 | 0.18 | 1.11 | 0.23 | 0.6875 | 1.2675 | 0.669748 | 0.251714 | 0.049055 | | 1 | 0.996511 | 0.658454 |
| Larp6 | Q8BN59 | -0.28 | -0.11 | 0.06 | -0.05 | -0.23 | -0.3 | 0.56 | 0.02 | 0.1 | 0.52 | 0.22 | -0.02 | 0.33 | 0.68 | 0.16 | 0.5 | 0.1075 | 0.3 | 0.5125 | 0.639037 | 0.144085 | 0.028534 | | 1 | 0.996511 | 0.637553 |
| Larp7 | Q05CL8 | -1.63 | -2.8 | -0.94 | -0.7 | -3.33 | -1.33 | -1.13 | -1.11 | -0.23 | -0.03 | -0.68 | -3.18 | 1.65 | -2.3 | -0.56 | -0.87 | -0.2075 | 0.4875 | 0.9975 | 0.749715 | 0.539356 | 0.258307 | | 1 | 0.996511 | 0.812074 |
| Lars1 | Q8BMJ2 | -1.74 | -1.93 | -1.89 | -1.78 | -1.9 | -1.79 | -1.42 | -1.74 | -1.84 | -1.65 | -1.6 | -1.69 | -2.04 | -1.87 | -1.82 | -1.88 | 0.1225 | 0.14 | -0.0675 | 0.471507 | 0.395436 | 0.695999 | | 1 | 0.996511 | 0.955729 |
| Lars2 | Q8VDC0 | 0.47 | 0.49 | 0.45 | 0.2 | 0.2 | 0.41 | 0.68 | 0.33 | 0.32 | -0.13 | 0.39 | 0.33 | 0.17 | 0.19 | 0.2 | -0.01 | 0.0025 | -0.175 | -0.265 | 0.988438 | 0.375902 | 0.162494 | | 1 | 0.996511 | 0.757516 |
| Las1l | A2BE28-2 | -0.86 | -1.27 | -1.14 | -1.7 | -1.47 | -0.71 | -1.65 | -1.37 | -0.24 | -0.53 | -1.25 | -0.87 | 0.73 | -1.72 | -1.04 | -0.94 | -0.0575 | 0.52 | 0.5 | 0.833887 | 0.09943 | 0.341209 | | 1 | 0.996511 | 0.848473 |
| Lasp1 | A2A6H1 | -1.5 | -0.72 | -0.46 | -2.15 | -0.17 | -0.68 | -1.7 | -1.2 | -4.42 | -3.13 | -0.95 | -2.31 | -1.48 | -4.88 | -0.7 | -1.38 | 0.27 | -1.495 | -0.9025 | 0.568892 | 0.071937 | 0.335473 | | 1 | 0.996511 | 0.847442 |
| Lasp1 | A2A6G9 | -2.78 | -0.79 | -1.66 | -3.69 | -0.68 | -0.21 | -4.26 | -0.07 | 1.44 | -1.02 | -1.49 | -2.86 | 4.55 | -4.41 | 3.18 | -4.96 | 0.925 | 1.2475 | 1.82 | 0.392257 | 0.227942 | 0.431652 | | 1 | 0.996511 | 0.879165 |
| Lasp1 | Q61792 | -0.75 | -1.03 | -0.71 | -0.51 | -1.05 | -0.72 | -1.3 | -0.48 | -0.46 | -0.35 | -0.48 | -0.71 | 0 | -1.41 | -0.91 | -1.46 | -0.1375 | 0.25 | -0.195 | 0.553109 | 0.208654 | 0.582969 | | 1 | 0.996511 | 0.927734 |
| Lbh | Q9CX60 | 0.27 | 0.48 | 0.32 | 0.79 | 0.51 | 0.2 | 0.68 | 0.46 | 0.18 | 0.08 | 0.33 | 0.42 | -0.3 | 0.9 | 0.12 | 0.59 | -0.0025 | -0.2125 | -0.1375 | 0.989571 | 0.289791 | 0.64975 | | 1 | 0.996511 | 0.946733 |
| Lbhd2 | G3UW55 | -0.49 | -0.93 | -0.09 | 0.89 | -0.81 | -0.52 | 0.6 | 0.58 | -1.02 | 0.51 | 0.56 | -0.31 | -1.32 | -0.81 | 0.72 | 0.82 | 0.1175 | 0.09 | 0.0075 | 0.811925 | 0.857768 | 0.990162 | | 1 | 0.996511 | 0.996154 |
| Lbr | Q3U9G9 | 0.77 | 0.64 | 0.79 | 0.74 | 0.6 | 0.65 | 0.29 | 0.18 | 1.18 | -0.38 | 0.61 | 0.57 | 1.4 | 0.48 | 0.65 | 0.61 | -0.305 | -0.24 | 0.05 | 0.103106 | 0.466386 | 0.839214 | | 1 | 0.996511 | 0.976218 |
| Lcat | P16301 | -0.53 | -0.37 | -1.44 | -1.93 | 0.97 | -2.24 | -1.73 | -2.06 | 0.52 | -0.49 | -1.19 | -0.14 | 0.74 | 1.1 | -1.71 | -1.28 | -0.1975 | 0.7425 | 0.78 | 0.79467 | 0.149028 | 0.297038 | | 1 | 0.996511 | 0.826122 |
| Lclat1 | Q3UN02 | 0.94 | 0.65 | 0.52 | 1.01 | 0.96 | 0.85 | 0.3 | 0.95 | 0.22 | 1.18 | 0.43 | 0.91 | 0.51 | 0.74 | 1.21 | 0.76 | -0.015 | -0.095 | 0.025 | 0.945425 | 0.721327 | 0.913793 | | 1 | 0.996511 | 0.986458 |
| Lcmt1 | A2RTH5 | -0.6 | -0.62 | -0.3 | -0.51 | -0.77 | -0.65 | -0.4 | -0.48 | -1.39 | -0.44 | -0.27 | -0.52 | -1.62 | -1.25 | -0.81 | -0.66 | -0.0675 | -0.1475 | -0.5775 | 0.686771 | 0.594515 | 0.04959 | | 1 | 0.996511 | 0.658454 |
| Lcp1 | Q61233 | -0.54 | -0.63 | -1.14 | -1 | 0.06 | -0.93 | -0.95 | -1 | -1.31 | -1.14 | -0.84 | -1.28 | -1.19 | -0.86 | -1.69 | -1.17 | 0.1225 | -0.315 | -0.4 | 0.677062 | 0.173553 | 0.14344 | | 1 | 0.996511 | 0.742733 |
| Ldah | A0A0F6AIX | -3.21 | -3.94 | -0.9 | -0.92 | -3.56 | -3.17 | 0 | -0.85 | -1.77 | -1.2 | -0.64 | -3.22 | -0.2 | -4.64 | 0.05 | -0.05 | 0.3475 | 0.535 | 1.0325 | 0.741187 | 0.541054 | 0.412787 | | 1 | 0.996511 | 0.874811 |
| Ldb1 | P70662 | -2.6 | -3.45 | -1.12 | -1.13 | -2.87 | -2.83 | -3.11 | -1.11 | -1.74 | -0.19 | -0.8 | -2.5 | -1.04 | -2.23 | -1.54 | -0.79 | -0.405 | 0.7675 | 0.675 | 0.549066 | 0.289169 | 0.279785 | | 1 | 0.996511 | 0.822084 |
| Ldha | P06151 | 0.22 | 0.12 | 0.25 | 0.59 | 0.22 | 0.08 | 0.27 | 1.06 | -0.12 | 0.43 | 0.58 | 0.45 | -0.69 | 0.24 | -0.28 | -0.16 | 0.1125 | 0.04 | -0.5175 | 0.659873 | 0.857054 | 0.064557 | | 1 | 0.996511 | 0.66433 |
| Ldhb | A0A6I8MX: | 0.03 | -0.45 | -0.01 | 0.38 | -0.36 | -0.31 | -0.32 | 0.41 | -0.13 | 0.24 | -0.21 | -0.41 | -0.74 | 0.16 | -0.68 | -0.48 | -0.1325 | -0.115 | -0.4225 | 0.613288 | 0.638994 | 0.164174 | | 1 | 0.996511 | 0.758352 |
| Ldhd | Q7TNG8 | 0.04 | -0.03 | -0.08 | -0.02 | -0.02 | 0.09 | -0.11 | 0.33 | -0.07 | -0.08 | -0.15 | -0.07 | 0.37 | -0.38 | 0.41 | 0.32 | 0.095 | -0.07 | 0.2025 | 0.5575 | 0.646689 | 0.394049 | | 1 | 0.996511 | 0.865806 |
| Ldlr | A0A1L1SRE | 0.66 | 0.62 | 0.54 | 0.22 | 0.8 | 0.54 | 0.36 | 0.04 | -0.16 | 0.71 | 0.22 | 1.21 | 0.28 | 0.78 | 0.67 | 0.46 | -0.075 | -0.015 | 0.0375 | 0.727141 | 0.962024 | 0.85743 | | 1 | 0.996755 | 0.976225 |
| Ldlrap1 | Q8C142 | -0.58 | -1.57 | 1 | 0.54 | -2.18 | -1.36 | -2.04 | -0.62 | -0.19 | 0.62 | 0.62 | -0.49 | 0.16 | -3.56 | -0.18 | 0.43 | -1.3975 | 0.2925 | -0.635 | 0.047628 | 0.622657 | 0.5227 | | 1 | 0.996511 | 0.907667 |
| Lemd2 | Q6DVA0 | -0.11 | 0.19 | -0.23 | -0.19 | 0.41 | 0.11 | -0.46 | -0.33 | -1.05 | 0.16 | -0.39 | -0.08 | -0.64 | 0.03 | 0.57 | 0.44 | 0.0175 | -0.255 | 0.185 | 0.941377 | 0.387403 | 0.54317 | | 1 | 0.996511 | 0.912212 |
| Lemd3 | E9QP59 | 0.57 | 0.19 | 0.26 | 0.12 | 0.2 | 0.3 | 0.1 | 0.28 | 0.31 | -0.28 | 0.04 | -0.23 | 1.13 | 0.95 | 0.7 | 0.8 | -0.065 | -0.325 | 0.61 | 0.696105 | 0.149685 | 0.013699 | | 1 | 0.996511 | 0.589387 |
| Leng1 | Q9DB98 | -3.4 | -3.66 | -4.43 | -3.4 | -3.05 | -1.75 | -6.95 | -6.74 | 1.12 | -3.53 | -4.54 | -1.97 | 1.97 | -3.78 | -4.41 | -6.74 | -0.9 | 1.4925 | 0.4825 | 0.458286 | 0.208898 | 0.770981 | | 1 | 0.996511 | 0.965933 |
| Leo1 | Q5XJE5 | -1.79 | -1.04 | -0.47 | -0.4 | -3.7 | -0.97 | -0.07 | 0.96 | 0 | 0.48 | 0.29 | -1.28 | 0.07 | -1.53 | -0.41 | 0.12 | -0.02 | 0.7975 | 0.4875 | 0.983062 | 0.12237 | 0.315148 | | 1 | 0.996511 | 0.837638 |
| Leprotl1 | Q9CQ74 | 3.79 | -1.21 | -5.94 | -3.95 | -0.67 | 3.79 | -2.35 | -4.87 | -1.41 | -4.16 | -5.7 | 2.14 | -0.91 | -1.44 | -4.28 | -4.43 | 0.8025 | -4.55E-01 | -0.9375 | 0.746976 | 0.851043 | 0.647535 | | 1 | 0.996511 | 0.945623 |
| Letm1 | Q9Z2I0 | -0.43 | 0.18 | 0.86 | 1.21 | 0.12 | -0.28 | 0.66 | 1.17 | -0.48 | 1.17 | 0.92 | -0.54 | -0.44 | 0.13 | 0.29 | 0.45 | -0.0375 | -0.1875 | -0.3475 | 0.933407 | 0.72748 | 0.393498 | | 1 | 0.996511 | 0.865775 |
| Letmd1 | Q924L1 | 0.29 | 0.06 | 0.62 | 0.57 | 0.33 | 0.68 | 0.28 | 0.65 | -0.17 | 0.25 | 0.64 | 0.2 | -0.19 | -0.21 | 0.11 | 0.29 | 0.1 | -1.55E-01 | -0.385 | 0.619725 | 0.521539 | 0.114972 | | 1 | 0.996511 | 0.717287 |
| Lgals1 | P16045 | -1.4 | -1.35 | -0.65 | -0.89 | -1.96 | -1.47 | -1.75 | -1.33 | -2.43 | -0.53 | -0.92 | -1.56 | -2.09 | -2.18 | -1.73 | -1.99 | -0.555 | -0.2875 | -0.925 | 0.046375 | 0.508152 | 0.004131 | | 1 | 0.996511 | 0.548272 |
| Lgals3bp | Q07797 | 0.59 | 1.37 | 0.14 | 0.29 | 0.92 | 0.1 | 0.68 | 0.55 | 0.07 | 0.62 | 0.08 | 0.19 | -0.03 | 0.31 | -0.08 | 0.48 | -0.035 | -0.3575 | -0.4275 | 0.912034 | 0.263098 | 0.196361 | | 1 | 0.996511 | 0.78004 |
| Lgals8 | Q9JL15 | 1.2 | 1.52 | 0.57 | 0.49 | 2.12 | 1.76 | 0.83 | 0.73 | 0.85 | 0.57 | 0.54 | 1 | 0.36 | 0.92 | 0.65 | 0.64 | 0.415 | -0.205 | -0.3025 | 0.314969 | 0.476583 | 0.312922 | | 1 | 0.996511 | 0.835247 |
| Lgals9 | B1AQR8 | 0.4 | -0.43 | 0.26 | 0.44 | -1.27 | -0.92 | -1.62 | -1.15 | -0.74 | -0.24 | -0.33 | -0.47 | -0.1 | -1.05 | -0.73 | -0.06 | -1.4075 | -0.6125 | -0.6525 | 0.000478 | 0.036198 | 0.06909 | 0.544245 | | 0.996511 | 0.667373 |
| Lgalsl | Q8VED9 | 0.4 | -0.51 | -0.93 | -1.06 | -0.65 | 0.22 | -1.06 | -1.25 | 0.06 | -1.12 | -1.29 | 0.41 | -1.05 | 0.2 | -2.14 | -1.9 | -0.16 | 0.04 | -0.6975 | 0.713124 | 0.936069 | 0.243758 | | 1 | 0.996755 | 0.799567 |
| Lgi1 | Q9JIA1 | -0.21 | 0.15 | 0.24 | -0.01 | 0.47 | 0.05 | 0.43 | 0.05 | -0.83 | -0.85 | 0.04 | -0.41 | -1.23 | -0.59 | -0.25 | 0.15 | 0.2075 | -0.555 | -0.5225 | 0.292936 | 0.053434 | 0.125387 | | 1 | 0.996511 | 0.721448 |
| Lgi2 | Q8K4Z0-2 | 0.55 | 1.18 | 1.03 | 0.76 | 0.96 | 0.48 | 1.57 | 0.48 | 0.03 | -0.81 | 0.71 | 0.43 | -0.28 | 0.37 | 0.29 | 1.01 | -0.0075 | -0.79 | -0.5325 | 0.97959 | 0.048626 | 0.112097 | | 1 | 0.996511 | 0.713445 |
| Lgi3 | Q8K406 | 1.06 | 1.1 | 1.03 | 0.57 | 1.41 | 0.69 | 0.91 | 0.52 | -0.05 | 0.22 | 0.7 | 0.87 | 0.23 | 0.19 | 1.04 | 1.32 | -0.0575 | -0.505 | -0.245 | 0.813702 | 0.08288 | 0.449748 | | 1 | 0.996511 | 0.884817 |
| Lgi4 | Q8K1S1 | 0.35 | 0.03 | 0.94 | 0.79 | 0.95 | 0.78 | 0.29 | 0.44 | 0.63 | 1.06 | 0.22 | 0.86 | 0.19 | 1.22 | 0.72 | 0.67 | 0.0875 | 0.165 | 0.1725 | 0.741493 | 0.567016 | 0.577076 | | 1 | 0.996511 | 0.927198 |
| Lgmn | O89017 | 0.03 | -0.71 | -1.36 | -1.13 | -0.71 | 0.14 | -0.92 | -1.23 | 0.04 | -1.29 | -0.51 | 0.22 | 0.49 | -0.63 | -0.61 | -1.29 | 0.1125 | 0.4075 | 0.2825 | 0.778547 | 0.357916 | 0.536818 | | 1 | 0.996511 | 0.912212 |
| Lhfpl2 | Q8BGA2 | 0.47 | 1.03 | 1.94 | 0.65 | 0.01 | 0.45 | -1.45 | -0.31 | -0.46 | 2.38 | 0.31 | -0.15 | 0.94 | -0.03 | 1.47 | 0.49 | -1.3475 | -0.5025 | -0.305 | 0.020176 | 0.451231 | 0.489189 | | 1 | 0.996511 | 0.897531 |
| Lhfpl3 | Q9CTN8 | 4.11 | 0.74 | 1.27 | 1.5 | 1.43 | 3.57 | 2.89 | 2.57 | 0.52 | 0.89 | 1.9 | 3.64 | 0.17 | 1.35 | 2.5 | 2.55 | 0.71 | -0.1675 | -0.2625 | 0.379776 | 0.856083 | 0.756571 | | 1 | 0.996511 | 0.962333 |
| Lhfpl4 | Q5U4E0 | 0.07 | -1.23 | -0.65 | -1.74 | -1.01 | 0.78 | -0.53 | -1.18 | -0.48 | -0.04 | -0.49 | 0.17 | -0.11 | -0.04 | -1.03 | -0.81 | 0.4025 | 0.6775 | 0.39 | 0.464496 | 0.120666 | 0.384528 | | 1 | 0.996511 | 0.860984 |
| Lhpp | Q9D7I5 | -0.27 | -0.49 | -0.49 | -0.41 | -0.35 | -0.15 | -0.17 | -0.39 | -1.26 | -1.15 | -0.21 | -0.32 | -1.3 | -1.56 | -0.66 | -0.57 | 0.15 | -0.32 | -0.6075 | 0.338656 | 0.283595 | 0.048757 | | 1 | 0.996511 | 0.658454 |
| Lias | Q99M04 | 0.55 | 0.56 | 0.4 | 0.41 | 1.07 | 0.49 | 0.17 | 0.33 | 0.82 | 0.69 | 0.75 | 0.85 | 0.81 | 1.22 | 0.3 | 0.72 | 0.035 | 0.2975 | 0.2825 | 0.875325 | 0.086127 | 0.249291 | | 1 | 0.996511 | 0.804842 |
| Lifr | P42703 | 0.3 | 0.07 | 0.24 | 0.22 | 0.19 | 0.37 | 0.34 | 0.37 | 0.17 | 0.46 | 0.23 | 0.55 | 0.36 | 0.33 | 0.46 | 0.28 | 0.11 | 0.145 | 0.15 | 0.460372 | 0.416689 | 0.391293 | | 1 | 0.996511 | 0.865375 |
| Lig3 | Q3UC82 | -2.58 | -3 | -2.38 | -2.27 | -1.92 | -3.58 | -1.53 | -3.16 | -2.01 | -1.23 | -2.98 | -2.56 | -2.71 | -2.16 | -1.9 | -0.27 | 0.01 | 0.3625 | 0.7975 | 0.983248 | 0.369358 | 0.144575 | | 1 | 0.996511 | 0.743156 |
| Lima1 | Q9ERG0 | -0.06 | -0.03 | 0.18 | 0.07 | 0.12 | -0.27 | -0.06 | 0.36 | 0.48 | 0.55 | 0.47 | 0.29 | 1.07 | 0.25 | 0.42 | 0.07 | -0.0025 | 0.4075 | 0.4125 | 0.989249 | 0.033114 | 0.133249 | | 1 | 0.996511 | 0.728417 |
| Limch1 | Q3UH68 | 0.43 | 0.38 | 0.5 | 0.36 | 0.09 | 0 | 0.49 | 0.01 | 0.63 | 0.45 | 0.41 | 0.3 | 1.04 | -0.03 | 0.57 | 0.52 | -0.27 | 0.03 | 0.1075 | 0.142289 | 0.854423 | 0.672681 | | 1 | 0.996511 | 0.952944 |
| Limd1 | Q9QXD8 | -1.34 | -1.84 | -1.26 | -0.25 | -1.4 | -1.16 | -0.86 | -0.68 | -0.43 | -0.51 | -0.36 | -0.88 | 0.53 | -1.46 | -0.06 | -0.08 | 0.1475 | 0.6275 | 0.905 | 0.678656 | 0.097946 | 0.098452 | | 1 | 0.996511 | 0.698156 |
| Limd2 | Q8BGB5 | -1.31 | -1.87 | -1.77 | 1.42 | -1.62 | -0.48 | -1.52 | -1.95 | -0.08 | -2.43 | -1.65 | -1.29 | 0.88 | -1.37 | -1.06 | -2.06 | -0.51 | -0.48 | -0.02 | 0.506906 | 0.566432 | 0.982204 | | 1 | 0.996511 | 0.994858 |
| Limk1 | P53668 | -0.35 | -0.19 | -0.57 | 0.07 | -0.02 | -0.38 | -0.3 | -0.13 | 0.16 | 0.28 | -0.47 | -0.22 | -0.09 | -0.07 | -0.54 | -0.12 | 0.0525 | 0.1975 | 0.055 | 0.786788 | 0.42828 | 0.806042 | | 1 | 0.996511 | 0.971715 |
| Limk2 | Q5NC03 | -0.28 | 0.19 | 0.11 | -0.32 | -0.16 | -0.18 | -0.17 | -0.31 | 0.67 | 0.19 | 0.31 | 0.26 | 0.62 | 0.3 | 0.38 | 0.1 | -0.13 | 0.4325 | 0.425 | 0.4783 | 0.067049 | 0.0807 | | 1 | 0.996511 | 0.684354 |
| Lims1 | A0A5F8MP | -1.42 | -2.3 | -0.81 | -0.89 | -1.81 | -1.45 | -1.33 | -1.12 | -2.01 | 0.11 | -0.44 | -1.61 | -1 | -2.17 | -0.43 | -0.79 | -0.0725 | 0.3675 | 0.2575 | 0.838956 | 0.513176 | 0.592587 | | 1 | 0.996511 | 0.929843 |
| Lin7a | Q8JZS0 | 0.45 | 0.21 | 0.81 | 1.04 | 0.42 | 0.52 | 0.47 | 0.83 | 0.35 | 0.65 | 0.8 | 0.28 | 0.23 | 0.65 | 0.23 | 0.3 | -0.0675 | -0.1075 | -0.275 | 0.766644 | 0.664485 | 0.281034 | | 1 | 0.996511 | 0.822084 |
| Lin7b | O88951 | 0.89 | 0.92 | 0.79 | 0.88 | 0.78 | 1.06 | 0.57 | 0.86 | 1.58 | 1.56 | 1.05 | 0.97 | 1.57 | 1.42 | 1.27 | 0.4 | -0.0525 | 0.42 | 0.295 | 0.749049 | 0.070745 | 0.312664 | | 1 | 0.996511 | 0.835177 |
| Lin7c | O88952 | 1.04 | 0.62 | 0.86 | 1.07 | 0.61 | 1.15 | 0.06 | 0.55 | 1.65 | 0.61 | 0.48 | 0.61 | 1.76 | 0.33 | 0.9 | 0.75 | -0.305 | -0.06 | 0.0375 | 0.253017 | 0.840483 | 0.907543 | | 1 | 0.996511 | 0.986458 |
| Lin9 | A0A0A6YW | -2.08 | -0.31 | 0.66 | -0.21 | -0.28 | 0.59 | -0.43 | -0.15 | 1.74 | 0.76 | -0.34 | -0.59 | -1.02 | -0.82 | -0.44 | 0.21 | 0.4175 | 0.8775 | -0.0325 | 0.467231 | 0.239676 | 0.955522 | | 1 | 0.996511 | 0.992482 |
| Lingo1 | A9DA50 | -0.08 | 0.19 | 1.08 | 1.04 | 0.69 | 0.15 | 0.91 | 1.07 | 0.22 | 0.74 | 1.24 | 0.33 | 0.27 | 0.31 | 1.02 | 1.08 | 0.1475 | 0.075 | 0.1125 | 0.669926 | 0.837065 | 0.756796 | | 1 | 0.996511 | 0.962333 |
| Lingo2 | Q3URE9 | 1.48 | 1.66 | 0.99 | 0.88 | 1.96 | 1.52 | 1.39 | 1.07 | 0.64 | 0.93 | 1.31 | 1.74 | -0.06 | 0.63 | 0.86 | 0.86 | 0.2325 | -0.0975 | -0.68 | 0.398804 | 0.751729 | 0.046959 | | 1 | 0.996511 | 0.658454 |
| Lingo3 | Q6GQU6 | 0.97 | 0.83 | 0.28 | 0.45 | 0.37 | 0.21 | 0.35 | 0.31 | 0.37 | -0.02 | -0.16 | 0.83 | 0.58 | -0.41 | 0.46 | 0.21 | -0.3225 | -0.3775 | -0.4225 | 0.131421 | 0.207793 | 0.169738 | | 1 | 0.996511 | 0.761572 |
| Lipa | Q9Z0M5 | -0.57 | -0.56 | -1.24 | -1.01 | 0.79 | -0.85 | -0.48 | -0.88 | -0.06 | -1.02 | -0.46 | -0.51 | -1.44 | -0.09 | -1.46 | -1.34 | 0.49 | 0.3325 | -0.2375 | 0.243676 | 0.244205 | 0.521063 | | 1 | 0.996511 | 0.907667 |
| Lipe | E9Q4M2 | -0.57 | 0.33 | -0.25 | -0.21 | -0.07 | -0.59 | -0.2 | -0.25 | 0.03 | -1.04 | -0.22 | -0.08 | -0.64 | 0.45 | -0.65 | -0.11 | -0.1025 | -0.1525 | -0.0625 | 0.663861 | 0.62453 | 0.847944 | | 1 | 0.996511 | 0.976225 |
| Lipt1 | Q8VCM4 | -0.61 | -1.12 | 0.4 | 1.01 | -1.6 | 0.25 | 0.56 | 0.93 | 1.51 | 0.79 | 0.32 | -0.67 | -1.48 | 0.4 | 1 | 0.67 | 0.115 | 0.5675 | 0.2275 | 0.864027 | 0.363061 | 0.734478 | | 1 | 0.996511 | 0.958635 |
| Lipt2 | Q9D009 | -0.36 | 0.08 | -0.26 | 0.15 | -0.29 | -0.48 | -1.05 | -0.51 | 0.36 | 1.16 | -0.66 | -0.44 | 0.9 | 0.84 | -0.52 | -0.74 | -0.485 | 0.2025 | 0.2175 | 0.057475 | 0.625337 | 0.616386 | | 1 | 0.996511 | 0.936924 |
| Lix1 | Q6P566 | 0.64 | 1.26 | 2.07 | 0.94 | 1.97 | 1.21 | 0.51 | 0.85 | 1.41 | 3.21 | 0.69 | 1.12 | 1.14 | 2.02 | 0.86 | 1.39 | -0.0925 | 0.38 | 0.125 | 0.822664 | 0.519354 | 0.745332 | | 1 | 0.996511 | 0.959729 |
| Llgl1 | A0A0R4J0S | 0.53 | 0.52 | 0.33 | 0.2 | 0.75 | 0.37 | 0.34 | 0.06 | 0.23 | -0.37 | 0.29 | 0.57 | 0.09 | -0.26 | 0.24 | 0.29 | -0.015 | -0.215 | -0.305 | 0.939291 | 0.382601 | 0.167056 | | 1 | 0.996511 | 0.759484 |
| Lman1 | Q9D0F3 | -0.99 | -0.5 | -1.81 | -2.06 | 0.25 | -1.18 | -1.37 | -2.11 | -0.57 | -1.19 | -1.83 | -0.49 | -0.36 | 0.2 | -0.76 | -1.31 | 0.2375 | 0.32 | 0.7825 | 0.672903 | 0.482806 | 0.112825 | | 1 | 0.996511 | 0.714581 |
| Lman2 | Q9DBH5 | -1.03 | -0.29 | -0.53 | -0.37 | 0.81 | -0.98 | 0.79 | -0.3 | -1.68 | -0.71 | -0.49 | -0.52 | -1.67 | -0.32 | 0.34 | 0.49 | 0.635 | -0.295 | 0.265 | 0.17196 | 0.376214 | 0.588799 | | 1 | 0.996511 | 0.929131 |
| Lman2l | D3Z4P2 | -1.38 | -1.97 | -2.22 | -2.01 | -0.93 | -1.03 | -0.71 | -1.93 | -1.14 | -1.27 | -2.15 | -1.36 | -0.22 | -0.51 | -0.9 | -0.96 | 0.745 | 0.415 | 1.2475 | 0.041037 | 0.186883 | 0.001407 | | 1 | 0.996511 | 0.419079 |
| Lmbr1 | J3QM78 | -0.01 | 1.12 | -0.52 | -0.76 | -1.42 | 2.36 | -0.25 | -0.68 | 2.54 | -0.22 | -0.54 | 3.39 | 2.68 | 3.33 | -0.31 | -0.54 | 0.045 | 1.335 | 1.3325 | 0.95676 | 0.18887 | 0.193269 | | 1 | 0.996511 | 0.777764 |
| Lmbrd1 | Q8K0B2-3 | -1.16 | -0.82 | -0.66 | -0.83 | -0.44 | -0.76 | 0.03 | -0.61 | -0.11 | -0.38 | -0.59 | -0.3 | 0.28 | 0.49 | -0.15 | -0.3 | 0.4225 | 0.5225 | 0.9475 | 0.085693 | 0.024818 | 0.003928 | | 1 | 0.996511 | 0.542407 |
| Lmbrd2 | Q8C561 | 2.01 | 0.61 | -0.39 | -0.6 | 0.49 | 1.98 | -0.63 | -0.58 | 0.32 | -0.42 | -0.17 | 1.72 | -0.02 | -0.33 | 1.08 | -0.45 | -0.0925 | -0.045 | -0.3375 | 0.904465 | 0.948292 | 0.595268 | | 1 | 0.996755 | 0.929911 |
| Lmcd1 | Q8VEE1 | -0.41 | -0.75 | -0.61 | -0.57 | -1.03 | -1.06 | -1.37 | -0.58 | -0.77 | -0.17 | -0.48 | -0.38 | -0.58 | -0.41 | -0.72 | -0.82 | -0.425 | 0.135 | -0.0475 | 0.065901 | 0.496209 | 0.803086 | | 1 | 0.996511 | 0.971091 |
| Lmf1 | Z4YJR1 | -0.05 | 0.34 | -0.18 | -0.01 | 0.39 | 0.16 | -0.4 | -0.6 | 0.83 | 0.84 | -0.83 | 0.04 | 1.1 | -0.26 | 0.78 | 0.04 | -0.1375 | 0.195 | 0.39 | 0.606274 | 0.623806 | 0.264964 | | 1 | 0.996511 | 0.817248 |
| Lmf2 | Q8C3X8 | -0.26 | 0.06 | -0.77 | -0.23 | 0.58 | -0.02 | 0.19 | -0.08 | 0.22 | -1.28 | -0.2 | -0.14 | 0.82 | -0.88 | -0.32 | -0.46 | 0.4675 | -0.05 | 0.09 | 0.081317 | 0.888769 | 0.817503 | | 1 | 0.996511 | 0.972768 |
| Lmln | Q8BMN4 | -0.63 | -0.78 | -0.02 | 0.38 | -0.67 | -0.25 | 0.41 | 0.09 | -0.71 | 1.16 | 0.41 | -0.19 | -0.21 | -0.6 | 0.18 | 0.36 | 0.1575 | 0.43 | 0.195 | 0.648118 | 0.357504 | 0.573896 | | 1 | 0.996511 | 0.92575 |
| Lmna | P48678 | 2.44 | 0.93 | 1.28 | 1 | 0.26 | 1.6 | 1.07 | 1.41 | 0.47 | 0.55 | 0.83 | 1.01 | 1.1 | 0.4 | 1.73 | 1.38 | -0.3275 | -0.6975 | -0.26 | 0.453378 | 0.081616 | 0.548314 | | 1 | 0.996511 | 0.912754 |
| Lmnb1 | P14733 | 1.8 | 0.85 | 1.33 | 1.36 | 0.57 | 1.17 | 1.2 | 1.63 | 0.77 | 1.13 | 1.05 | 0.94 | 0.98 | 0.88 | 1.96 | 1.57 | -0.1925 | -0.3625 | 0.0125 | 0.514586 | 0.153381 | 0.969295 | | 1 | 0.996511 | 0.992767 |
| Lmnb2 | P21619 | 2.26 | 1.05 | 0.78 | 1.13 | 0.34 | 1.39 | 0.6 | 1.42 | 1.04 | 0.76 | 0.51 | 0.95 | 1.53 | 0.78 | 1.62 | 1.01 | -0.3675 | -0.49 | -0.07 | 0.373348 | 0.177045 | 0.852178 | | 1 | 0.996511 | 0.976225 |
| Lmo4 | P61969 | 0.57 | 0.85 | 0.6 | 0.61 | 0.21 | -0.04 | 0.14 | 0.71 | 0.86 | 1.08 | 0.92 | 0.71 | 0.05 | 1.7 | 0.65 | -0.16 | -0.4025 | 0.235 | -0.0975 | 0.075456 | 0.199232 | 0.810222 | | 1 | 0.996511 | 0.972222 |
| Lmo7 | E9PYI7 | 1.24 | 0.85 | 0.99 | 1.12 | 0.53 | 0.86 | 0.48 | 0.86 | 1.15 | 0.86 | 0.8 | 0.8 | 1.4 | 0.92 | 0.98 | 0.8 | -0.3675 | -0.1475 | -0.025 | 0.066076 | 0.427771 | 0.906058 | | 1 | 0.996511 | 0.986321 |

| | | | | | | | | | | | | | | | | | | | | | | | | | | |
|---|---|---|---|---|---|---|---|---|---|---|---|---|---|---|---|---|---|---|---|---|---|---|---|---|---|---|
| Lmtk2 | Q3TYD6 | 0.77 | 0.98 | 0.74 | 0.69 | 1.12 | 0.49 | 0.61 | 0.46 | 0.45 | 0.69 | 0.77 | 0.97 | 0.68 | 1 | 0.63 | 0.62 | -0.125 | -0.075 | -0.0625 | 0.535579 | 0.68771 | 0.741557 | 1 | 0.996511 | 0.958796 |
| Lmtk3 | A0A3B2WC | 0.29 | 0.11 | 0.4 | 0.32 | 0.11 | 0.43 | 0.31 | 0.36 | 0.54 | 0.6 | 0.37 | 0.29 | 1.21 | 0.85 | 0.52 | 0.31 | 0.0225 | 0.17 | 0.4425 | 0.88632 | 0.334779 | 0.096279 | 1 | 0.996511 | 0.698156 |
| Lmtk3 | A0A1B0GSI | 0.36 | 0.25 | 0.19 | 0.21 | 0.41 | 0.02 | -0.3 | -0.29 | 0.59 | -0.23 | 0.03 | 0.42 | 0.61 | 1.02 | 0.23 | -0.13 | -0.2925 | -0.05 | 0.18 | 0.174671 | 0.823882 | 0.515548 | 1 | 0.996511 | 0.905674 |
| Lnpep | Q8C129 | 0.06 | -0.02 | -0.7 | -0.75 | 0.18 | 0.16 | -0.48 | -0.97 | 0.38 | -0.58 | -0.86 | 0.35 | 0.28 | 0.77 | -0.8 | -0.75 | 0.075 | 0.175 | 0.2275 | 0.825273 | 0.640373 | 0.594577 | 1 | 0.996511 | 0.929843 |
| Lnpk | Q7TQ95 | -0.04 | 0.5 | 0.51 | 0.51 | 0.7 | 0.06 | 0.66 | 0.65 | 0.1 | 0.87 | 0.48 | 0.29 | 0.09 | 1.33 | 0.71 | 0.8 | 0.1475 | 0.065 | 0.3625 | 0.519294 | 0.787604 | 0.248454 | 1 | 0.996511 | 0.803985 |
| Lonp1 | Q8CGK3 | 1.06 | 0.83 | 0.95 | 0.94 | 0.16 | 0.95 | 0.64 | 1.07 | 0.43 | 1.07 | 1.15 | 0.87 | -0.08 | 0.24 | 0.21 | 0.24 | -0.24 | -0.065 | -0.7925 | 0.309153 | 0.758001 | 0.001796 | 1 | 0.996511 | 0.419079 |
| Lonp2 | Q9DBN5 | -0.41 | 0.03 | 0.36 | 0.46 | 0.34 | -0.06 | 1.41 | 0.36 | 0.55 | 0.87 | 0.33 | 0.31 | 0.63 | 0.16 | 0.9 | 1.02 | 0.4025 | 0.405 | 0.5675 | 0.275621 | 0.140682 | 0.077289 | 1 | 0.996511 | 0.676667 |
| Lonrf2 | F6ZE64 | -2.09 | -2.26 | -3.03 | -2.19 | -2.38 | -2.52 | -3.61 | -3.03 | -2.11 | -0.77 | 0.05 | -1.43 | -1.12 | -1.58 | -1.45 | -3.21 | -0.4925 | 1.3275 | 0.5525 | 0.173391 | 0.020264 | 0.269509 | 1 | 0.996511 | 0.821231 |
| Lpar1 | P61793-2 | 0.61 | 0.37 | 1.02 | 0.67 | -0.16 | 0.73 | 0.75 | 0.45 | 0.72 | 0.39 | 0.7 | 0.31 | 0.73 | -0.29 | 0.08 | 0.54 | -0.225 | -0.1375 | -0.4025 | 0.397217 | 0.522158 | 0.181785 | 1 | 0.996511 | 0.776002 |
| Lpcat1 | Q3TFD2 | -0.36 | 0.01 | -0.54 | -0.7 | 0.61 | 0.26 | -0.27 | -0.58 | -0.14 | -0.31 | -0.83 | -0.15 | -0.07 | 0.11 | -0.55 | -0.23 | 0.4025 | 0.04 | 0.2125 | 0.207643 | 0.871401 | 0.394182 | 1 | 0.996511 | 0.86589 |
| Lpcat2 | Q8BYI6 | 0.03 | -0.36 | -0.64 | 0.19 | -0.45 | -0.01 | 0.1 | 0.15 | 0.14 | 0.53 | 0.03 | -0.17 | 0.49 | -0.39 | 0.23 | 0.27 | 0.1425 | 0.3275 | 0.345 | 0.56601 | 0.226811 | 0.245977 | 1 | 0.996511 | 0.802442 |
| Lpcat3 | Q91V01 | -0.08 | 0.28 | -0.39 | 0.26 | 0.44 | -0.01 | 0.61 | 0.18 | 0.39 | 0.34 | -0.02 | -0.09 | 0.48 | 0.4 | 0.26 | -0.07 | 0.2875 | 0.1375 | 0.25 | 0.23163 | 0.557284 | 0.311508 | 1 | 0.996511 | 0.834523 |
| Lpcat4 | Q6NVG1 | 0.4 | 0.32 | 0.2 | 0.33 | 0.86 | 0.51 | 0.4 | 0.52 | -0.05 | 0.08 | 0.18 | 0.24 | -0.15 | 0.2 | 0.21 | 0.36 | 0.26 | -0.2 | -0.1575 | 0.142115 | 0.242041 | 0.420034 | 1 | 0.996511 | 0.87558 |
| Lpgat1 | E9QL80 | 0.76 | -0.37 | -0.23 | -0.2 | 0.1 | 1.18 | 0.15 | -0.19 | 0.3 | -0.45 | -0.39 | 0.7 | 0.88 | -0.51 | 0.64 | 0.23 | 0.32 | 0.05 | 0.32 | 0.405662 | 0.892073 | 0.418771 | 1 | 0.996511 | 0.87558 |
| Lpin1 | A0A1Y7VLI | -1.23 | -0.37 | -0.4 | 0.29 | -1.71 | -0.49 | -0.43 | 1.09 | -1.04 | 1.02 | 0.14 | -1.42 | -0.44 | 0.18 | 0.26 | 0.78 | 0.0425 | 0.1025 | 0.6225 | 0.943021 | 0.861268 | 0.13488 | 1 | 0.996511 | 0.731223 |
| Lpin2 | E9PWN0 | 0.35 | -0.46 | -0.78 | -0.87 | 0.5 | 0.17 | -0.85 | -0.85 | -1.18 | -0.82 | -0.72 | 0.54 | -1.43 | -0.29 | -1.07 | -0.63 | 0.1825 | -0.105 | -0.415 | 0.664055 | 0.811531 | 0.275419 | 1 | 0.996511 | 0.822084 |
| Lpp | Q8BFW7 | -0.12 | -0.19 | -0.88 | -0.62 | -0.29 | -0.25 | 0.05 | -0.38 | -0.18 | -0.52 | -0.47 | 0.48 | -0.31 | 0.31 | -0.99 | -0.82 | 0.235 | 0.28 | 7.85E-17 | 0.311065 | 0.361001 | 1 | 1 | 0.996511 | 1 |
| Lratd1 | A0A0R4J01 | 4.9 | 3.71 | 7.62 | 4.03 | 3.78 | 5.15 | 7.81 | 7.46 | 0.75 | 4.7 | 7.18 | 4.91 | 1.86 | 0 | 8.78 | 8.7 | 0.985 | -0.68 | -0.23 | 0.409832 | 0.63712 | 0.915579 | 1 | 0.996511 | 0.986556 |
| Lratd2 | D3YXJ5 | 0.52 | 1.04 | 0.72 | 1.37 | 1.1 | 0.46 | 0.59 | 1.58 | 1.74 | 3.29 | 1.28 | 0.7 | 0.44 | 2.62 | 1.83 | 1.32 | 0.02 | 0.84 | 0.64 | 0.948776 | 0.146921 | 0.191996 | 1 | 0.996511 | 0.777117 |
| Lrba | E9Q3Y4 | -0.36 | -0.25 | -0.49 | -0.39 | 0.07 | -0.58 | -0.51 | -0.74 | -0.28 | -0.03 | -0.53 | -0.22 | -0.21 | 0 | -0.71 | -0.57 | -0.0675 | 0.1075 | -7.9E-17 | 0.749587 | 0.555155 | 1 | 1 | 0.996511 | 1 |
| Lrch1 | P62046 | 0.48 | 0.62 | 0.39 | 0.39 | 0.73 | 0.25 | 0.74 | -0.04 | 0.12 | 0.29 | 0.08 | 0.29 | -0.17 | 0.35 | 0.29 | 0.65 | -0.05 | -0.275 | -0.19 | 0.821629 | 0.121618 | 0.408973 | 1 | 0.996511 | 0.874356 |
| Lrch2 | F6VQ19 | -0.26 | -0.8 | -0.12 | -0.31 | -0.39 | -0.3 | 0.07 | -0.31 | -0.01 | 0.06 | -0.2 | -0.47 | -0.63 | 0.35 | -0.28 | -0.51 | 0.14 | 0.2175 | 0.105 | 0.508205 | 0.346142 | 0.711592 | 1 | 0.996511 | 0.957969 |
| Lrch3 | B2RXP1 | 0.32 | 0.75 | 1.2 | 0.83 | 0.23 | 0.19 | 1.09 | 0.76 | 0.29 | 1.75 | 1.4 | -0.04 | 0.03 | 0.43 | 0.66 | 0.97 | -0.2075 | 0.075 | -0.2525 | 0.472474 | 0.864585 | 0.388329 | 1 | 0.996511 | 0.864493 |
| Lrfn1 | Q2WF71 | 1.8 | 1.49 | 1.41 | 0.97 | 1.87 | 1.7 | 1.53 | 1.22 | 0.26 | 0.75 | 1.36 | 2.15 | -0.09 | 0.77 | 1.12 | 1.49 | 0.1625 | -0.2875 | -0.595 | 0.498921 | 0.499288 | 0.135429 | 1 | 0.996511 | 0.731223 |
| Lrfn2 | Q80TG9 | 0.83 | 0.72 | 0.76 | 0.58 | 0.58 | 0.8 | 0.81 | 1.27 | 0.38 | 0 | 1.03 | 0.92 | -0.21 | 0.36 | 0.85 | 1.19 | 0.1425 | -0.14 | -0.175 | 0.464933 | 0.598844 | 0.583685 | 1 | 0.996511 | 0.927734 |
| Lrfn3 | Q8BLY3 | 1.14 | 1.48 | 1.48 | 1.06 | 1.35 | 1.17 | 1.32 | 1.02 | 0.5 | 0.41 | 1.39 | 1.34 | 0.68 | 0.93 | 1.18 | 1.18 | -0.075 | -0.38 | -0.2975 | 0.67789 | 0.217868 | 0.193009 | 1 | 0.996511 | 0.777764 |
| Lrfn4 | Q80XU8 | 0.56 | 0.93 | 0.68 | 0.84 | 0.87 | 0.6 | 1.15 | 0.83 | 1.45 | 0.91 | 0.66 | 0.95 | 1.85 | 0.93 | 0.58 | 0.65 | 0.11 | 0.24 | 0.25 | 0.55164 | 0.295012 | 0.432541 | 1 | 0.996511 | 0.879165 |
| Lrfn5 | Q8BXA0 | 0.18 | 0.11 | 0.33 | 0.38 | 0.36 | 0.37 | 0.85 | 0.58 | -0.24 | 0.2 | 0.66 | 0.39 | 0.05 | -0.25 | 0.27 | 0.25 | 0.29 | 0.0025 | -0.17 | 0.130273 | 0.991363 | 0.408946 | 1 | 0.999285 | 0.874356 |
| Lrif1 | G5E8M8 | 1.19 | 1.19 | 1.15 | 0.88 | 2.33 | 0.87 | 1.14 | 0.75 | -0.04 | -0.33 | 1.3 | 1.35 | -0.19 | 0.45 | 1.62 | 1.54 | 0.17 | -0.5325 | -0.2475 | 0.633676 | 0.229635 | 0.563696 | 1 | 0.996511 | 0.920143 |
| Lrig1 | P70193 | 0.1 | 0.42 | 0.79 | 1.08 | -0.1 | -0.17 | 0.74 | 0.77 | 0.98 | 1.23 | 0.42 | -0.05 | 0.67 | 0.29 | 1.08 | 1.18 | -0.2875 | 0.0475 | 0.2075 | 0.38889 | 0.892165 | 0.504008 | 1 | 0.996511 | 0.901002 |
| Lrp1 | A0A0R4J0I | 0 | -0.08 | -0.13 | -0.19 | 0.02 | 0.11 | -0.01 | -0.14 | -0.33 | -0.23 | -0.09 | 0.28 | -0.1 | -0.01 | -0.12 | -0.1 | 0.095 | 0.0075 | 0.0175 | 0.52227 | 0.969078 | 0.916686 | 1 | 0.996755 | 0.986556 |
| Lrp11 | Q8CB67 | -1.02 | -1.4 | -0.78 | -1.03 | -1.21 | -1.01 | -0.74 | -0.52 | -1.55 | 0.54 | -0.67 | -0.59 | -0.24 | -0.27 | -0.74 | -0.07 | 0.1875 | 0.49 | 0.7275 | 0.406575 | 0.266434 | 0.012053 | 1 | 0.996511 | 0.566767 |
| Lrp1b | A2API5 | 0.13 | 0.04 | -0.06 | 0 | 0.45 | 0.31 | -0.27 | -0.06 | 0.61 | -0.02 | 0.14 | 0.53 | 0.81 | 0.69 | 0.19 | -0.03 | 0.08 | 0.2875 | 0.3875 | 0.69347 | 0.18271 | 0.135767 | 1 | 0.996511 | 0.731656 |
| Lrp4 | Q8VI56-2 | 0.23 | 0.45 | 0.72 | 0.45 | 0.15 | 0.89 | 0.31 | 0.5 | 1.67 | -0.16 | 0.62 | 0.43 | 1.97 | 0.11 | 0.43 | 0.32 | -1.6E-16 | 0.1775 | 0.245 | 1 | 0.642565 | 0.56207 | 1 | 0.996511 | 0.920024 |
| Lrp6 | O88572 | -0.16 | -0.11 | 0.66 | 0.73 | 0.12 | -0.04 | 0.84 | 0.79 | 0.64 | 1.73 | 0.39 | 0.12 | 0.24 | 0.72 | 0.37 | 0.61 | 0.1475 | 0.44 | 0.205 | 0.649048 | 0.296501 | 0.476008 | 1 | 0.996511 | 0.892174 |
| Lrp8 | B1AXJ4 | 0.55 | 0.68 | 0.81 | 0.54 | 0.52 | 0.36 | -0.13 | 0.28 | 0.67 | 0.72 | 0.56 | 0.51 | 1.21 | 0.92 | 0.47 | 0.29 | -0.3875 | -0.03 | 0.0775 | 0.068487 | 0.855881 | 0.758772 | 1 | 0.996511 | 0.963116 |
| Lrpap1 | P55302 | 1.7 | 0.79 | 0.59 | 1.08 | 0.82 | 1.39 | 1.18 | 1.1 | 1.2 | 0.71 | 0.23 | 1.97 | 1.8 | 1.15 | 2.02 | 2.02 | 0.0825 | -0.0125 | 0.7075 | 0.763614 | 0.976262 | 0.05284 | 1 | 0.997588 | 0.660608 |
| Lrpprc | Q6PB66 | 0.33 | 0.01 | 0.32 | 0.31 | -0.17 | 0.62 | 0.5 | 0.42 | 0.2 | 0.52 | 0.3 | 0.14 | 0.23 | -0.04 | 0.29 | 0.56 | 0.1 | 0.0475 | 0.0175 | 0.646716 | 0.792265 | 0.932443 | 1 | 0.996511 | 0.989713 |
| Lrrc1 | Q80VQ1 | 1.11 | 1.29 | 0.91 | 1.08 | 0.7 | 0.77 | 1.27 | 0.33 | 1.52 | 1.11 | 0.62 | 1.08 | 0.65 | 0.34 | 1.02 | 1.39 | -0.33 | -0.015 | -0.2475 | 0.17357 | 0.948244 | 0.365119 | 1 | 0.996755 | 0.858475 |
| Lrrc10b | G3XA50 | 0.13 | 0.65 | 0.87 | 1.08 | -0.88 | -0.63 | 0.95 | 0.65 | 0.82 | 0.25 | 0.05 | 0.25 | 1.52 | 0.08 | 0.63 | 0.46 | -0.66 | -0.34 | -0.01 | 0.179455 | 0.239239 | 0.977907 | 1 | 0.996511 | 0.993727 |
| Lrrc14 | Q8VC16 | 0.48 | 0.03 | -0.74 | -0.84 | 0.54 | 0.55 | -0.33 | -0.56 | 0.51 | -0.58 | -0.64 | 0.16 | 0.33 | 0.51 | -1.07 | -0.42 | 0.3175 | 0.13 | 0.105 | 0.440086 | 0.748388 | 0.817414 | 1 | 0.996511 | 0.972768 |
| Lrrc20 | Q8CI70 | -1.6 | -0.95 | -0.66 | -0.86 | -0.62 | -1.29 | -0.9 | -0.98 | -1.34 | -0.98 | -0.95 | -1.4 | -1.68 | -1.64 | -1.51 | -1.42 | 0.07 | -0.15 | -0.545 | 0.78467 | 0.561633 | 0.051476 | 1 | 0.996511 | 0.658454 |
| Lrrc24 | Q8BHA1 | -0.58 | -0.36 | -1.21 | -1.28 | -0.42 | -0.12 | -0.88 | -0.59 | -0.15 | 0.82 | -0.62 | 0 | -0.15 | -0.06 | -0.68 | -0.59 | 0.355 | 0.87 | 0.4875 | 0.22818 | 0.039363 | 0.122322 | 1 | 0.996511 | 0.721448 |
| Lrrc4 | Q99PH1 | 0.36 | 1.56 | 0.14 | 0.69 | 0.93 | 0.27 | 0.53 | 0.29 | 0.08 | -1 | 0.16 | 0.62 | 0 | -0.19 | -0.83 | 0.24 | -0.1825 | -0.7225 | -0.8825 | 0.590854 | 0.126228 | 0.041621 | 1 | 0.996511 | 0.658454 |
| Lrrc40 | A0A0R4J0V | -0.4 | -0.43 | -0.27 | -0.26 | -0.31 | -0.37 | 0.01 | -0.06 | -0.08 | 0 | -0.14 | -0.25 | -0.1 | -0.17 | -0.37 | -0.02 | 0.1575 | 0.2225 | 0.175 | 0.345899 | 0.191138 | 0.342378 | 1 | 0.996511 | 0.848473 |
| Lrrc47 | Q505F5 | 0.38 | -0.19 | 0.52 | -0.3 | -0.35 | 0.17 | 1.04 | 0.11 | -0.59 | -0.37 | 0.59 | 0.24 | -1.01 | -0.37 | 0.55 | 0.76 | 0.14 | -0.135 | -0.12 | 0.683799 | 0.689079 | 0.783625 | 1 | 0.996511 | 0.967642 |
| Lrrc49 | D3YWN5 | 0.23 | 0.64 | 0.2 | -0.05 | 0.61 | 0.26 | 0.39 | -0.2 | 0.27 | -0.5 | -0.05 | 0.45 | 0.67 | 0.19 | -0.12 | 0.09 | 0.01 | -0.2125 | -0.0475 | 0.966573 | 0.437479 | 0.850742 | 1 | 0.996511 | 0.976225 |
| Lrrc4b | P0C192 | 1.6 | 1.25 | 0.73 | 0.76 | 1.31 | 1.7 | 1.01 | 0.69 | 0.45 | -0.09 | 1.09 | 1.82 | 0.07 | 0.68 | 0.66 | 0.67 | 0.0925 | -0.2675 | -0.565 | 0.756885 | 0.544183 | 0.068541 | 1 | 0.996511 | 0.665034 |
| Lrrc4c | Q8C031 | 0.49 | 0.12 | 1.32 | 0.93 | 0.38 | 1.04 | 1.14 | 1.07 | -0.58 | 0.53 | 1.24 | 0.32 | -0.42 | -0.69 | 0.44 | 1.2 | 0.1925 | -0.3375 | -0.5825 | 0.539704 | 0.44285 | 0.237744 | 1 | 0.996511 | 0.792722 |
| Lrrc57 | Q8JZX5 | 1.48 | 0.76 | 0.81 | 0.98 | 0.94 | 1.4 | 0.96 | 1.16 | 0.95 | 1.1 | 0.75 | 1.19 | 0.65 | 0.89 | 0.71 | 0.38 | 0.1075 | -0.01 | -0.35 | 0.627118 | 0.96453 | 0.163845 | 1 | 0.996755 | 0.758352 |
| Lrrc58 | Q3UGP9 | -0.45 | -0.36 | -0.96 | -1.44 | -0.22 | 0.07 | -0.93 | -1.17 | -0.86 | -0.02 | -1.58 | -0.36 | -0.44 | -1.45 | -1.52 | -0.75 | 0.24 | 0.0975 | -0.2375 | 0.518173 | 0.808663 | 0.514183 | 1 | 0.996511 | 0.90567 |
| Lrrc59 | Q922Q8 | -1.08 | -0.76 | -0.65 | -0.77 | -0.19 | -0.96 | 0.07 | -0.45 | -1.39 | 0.02 | -0.84 | -0.84 | -0.75 | -0.82 | 0.49 | 0.39 | 0.4325 | 0.0525 | 0.6425 | 0.111823 | 0.865028 | 0.102613 | 1 | 0.996511 | 0.700625 |
| Lrrc7 | A0A571BE/ | 2 | 0.5 | 0.91 | 1.17 | 0.25 | 0.43 | -0.26 | 0.59 | 4.93 | 1.06 | 0.84 | 1.66 | 6.14 | 1.48 | 0.22 | 0.16 | -0.8925 | 0.9775 | 0.855 | 0.03046 | 0.296003 | 0.512378 | 1 | 0.996511 | 0.903993 |
| Lrrc7 | A0A0G2JD | 0.96 | 0.88 | 0.88 | 1.03 | 0.69 | 0.95 | 0.78 | 0.91 | 1.14 | 0.83 | 1.09 | 1.15 | 0.92 | 1.05 | 0.67 | 0.62 | -0.105 | 0.115 | -0.1225 | 0.485185 | 0.496 | 0.520059 | 1 | 0.996511 | 0.907667 |
| Lrrc73 | B2RWC4 | 1.35 | 1.21 | 1.29 | 0.83 | 1.46 | 1.73 | 1.55 | 1.81 | 0.58 | -0.17 | 1.35 | 1.8 | -0.07 | 0.88 | 1.45 | 2.06 | 0.4675 | -0.28 | -0.09 | 0.030154 | 0.516035 | 0.839046 | 1 | 0.996511 | 0.976218 |
| Lrrc75a | Q7TSF4 | 1.08 | 1.37 | 1.92 | 1.71 | 1.35 | 0.85 | 1.26 | 1.64 | 0.3 | 0.85 | 1.53 | 0.92 | 0.1 | 1.23 | 1.37 | 1.45 | -0.245 | -0.62 | -0.4825 | 0.35312 | 0.075592 | 0.204042 | 1 | 0.996511 | 0.782817 |
| Lrrc8a | Q80WG5 | 0.42 | 0.65 | 0.9 | 1.03 | 0.86 | 0.35 | 0.82 | 1.06 | 0.42 | 0.62 | 0.84 | 0.43 | 0.1 | 0.91 | 0.69 | 0.82 | 0.0225 | -0.1725 | -0.12 | 0.919954 | 0.421515 | 0.642871 | 1 | 0.996511 | 0.94365 |
| Lrrc8b | Q5DU41 | 0.33 | 0.46 | 0.48 | 0.78 | 0.53 | 0.39 | 0.53 | 0.52 | 0.16 | 0.37 | 0.43 | 0.5 | -0.17 | 0.21 | -0.05 | 0.22 | -0.02 | -0.1475 | -0.46 | 0.901544 | 0.429716 | 0.045322 | 1 | 0.996511 | 0.658454 |
| Lrrc8c | Q8R502 | -0.28 | 0.15 | 0.29 | 0.13 | 0.31 | 0.38 | 0.03 | 0.2 | -0.05 | -0.2 | 0.25 | -0.16 | 0.2 | -0.22 | 0.06 | 0 | 0.1575 | -0.1125 | -0.0625 | 0.406621 | 0.587006 | 0.76581 | 1 | 0.996511 | 0.964134 |
| Lrrc8d | Q8BGR2 | 0.18 | 0.56 | 0.45 | 0.14 | 0.7 | 0.25 | 1.17 | 0.36 | 0.79 | 0.61 | 0.23 | 0.35 | 0.6 | 1.09 | 0.79 | 0.98 | 0.2875 | 0.1625 | 0.5325 | 0.259271 | 0.441437 | 0.02909 | 1 | 0.996511 | 0.637553 |
| Lrrc9 | Q8CDN9 | -0.5 | -0.61 | -13.03 | -9.65 | 0.7 | -0.38 | -11.82 | -14.52 | -0.38 | -16.94 | -11.36 | -0.1 | -0.06 | 1.01 | -14.91 | -14.3 | -0.5575 | -1.2475 | -1.1175 | 0.900843 | 0.789626 | 0.814962 | 1 | 0.996511 | 0.972768 |
| Lrrfip1 | A0A087WN | -0.72 | 0.42 | 1.45 | 0.8 | 0.54 | -0.13 | 0.79 | 0.36 | 0.83 | 2.58 | 0.74 | 0.23 | 0.87 | 1.55 | 0.85 | 0.63 | -0.0975 | 0.6075 | 0.4875 | 0.832153 | 0.345828 | 0.31221 | 1 | 0.996511 | 0.834523 |
| Lrrfip1 | Q3UZ39-2 | 1.09 | 1.13 | 0.77 | 0.59 | 1.33 | 0.69 | 0.38 | 0.42 | 1.23 | 1.29 | 0.75 | 1.31 | 1.29 | 1.8 | 0.8 | 0.58 | -0.19 | 0.25 | 0.2225 | 0.476569 | 0.277739 | 0.480188 | 1 | 0.996511 | 0.893906 |
| Lrrfip1 | Q3UZ39 | -1.33 | -0.06 | -0.7 | -1.06 | -0.62 | -1.51 | -2.97 | -0.88 | -0.33 | -1.59 | -2 | -1.49 | 0.17 | 0.22 | -0.99 | -1.33 | -0.7075 | -0.565 | 0.305 | 0.215944 | 0.209036 | 0.508883 | 1 | 0.996511 | 0.902804 |
| Lrrfip2 | A0A1L1SSC | -0.16 | 0.29 | 0.29 | 0.44 | 0.36 | -0.04 | 0.22 | 0.58 | 0.33 | 1.02 | 0.46 | 0.19 | 0.84 | 0.32 | 1 | 0.22 | 0.065 | 0.285 | 0.38 | 0.758686 | 0.268445 | 0.169259 | 1 | 0.996511 | 0.760288 |
| Lrrk2 | Q5S006 | -1.05 | -0.79 | -0.6 | -0.79 | -0.68 | -1.21 | -1.13 | -0.72 | -0.79 | -0.56 | -0.63 | -0.78 | -0.68 | -0.63 | -1.13 | -0.6 | -0.1275 | 0.1175 | 0.0475 | 0.526286 | 0.513249 | 0.822706 | 1 | 0.996511 | 0.972768 |
| Lrrn1 | Q61809 | 1.62 | 0.57 | 0.24 | -0.04 | 0.36 | 1.51 | 0.43 | 0.23 | 1.73 | -0.06 | -0.07 | 1.79 | 1.23 | 1.09 | 0.25 | 0.08 | 0.035 | 0.25 | 0.065 | 0.936092 | 0.671861 | 0.882741 | 1 | 0.996511 | 0.979767 |
| Lrrtm1 | Q8K377 | -0.14 | 0.89 | 0.31 | 0.51 | 1.13 | 0.48 | 0.8 | 0.59 | -0.14 | -0.77 | 0.7 | 0.54 | 0.13 | 0.67 | 0.09 | 0.75 | 0.3575 | -0.31 | 0.0175 | 0.200709 | 0.4293 | 0.952187 | 1 | 0.996511 | 0.992476 |
| Lrrtm2 | Q8BGA3 | 0.49 | 0.54 | 0.75 | 0.7 | 0.74 | 0.8 | 0.68 | 0.65 | -0.06 | 0.23 | 0.68 | 0.55 | -0.01 | 0.32 | 0.25 | 0.72 | 0.0975 | -0.27 | -0.3 | 0.516811 | 0.233574 | 0.189922 | 1 | 0.996511 | 0.776843 |
| Lrrtm3 | Q8BZ81 | 1.26 | 0.39 | 0.46 | 0.46 | 0.88 | 1.6 | 0.21 | 0.44 | 0.05 | 0.42 | 0.68 | 1.13 | -0.25 | -0.08 | -0.28 | 0.26 | 0.14 | -0.0725 | -0.73 | 0.693775 | 0.815174 | 0.021695 | 1 | 0.996511 | 0.621926 |
| Lrrtm4 | A0A0R4J1T | 0.91 | 1.06 | 1.12 | 1.12 | 1.29 | 1.14 | 0.95 | 1.11 | -0.1 | -0.15 | 1.35 | 0.95 | -0.24 | 0 | 0.58 | 1.01 | 0.07 | -0.54 | -0.715 | 0.652351 | 0.168611 | 0.038503 | 1 | 0.996511 | 0.658454 |
| Lrsam1 | Q80ZI6 | 0.12 | 0 | 0.17 | 0.4 | 0.22 | -0.04 | 0.56 | 0.57 | 0.11 | 0.24 | 0.09 | 0.13 | -0.14 | 0.34 | 0.08 | 0.11 | 0.155 | -0.03 | -0.075 | 0.449675 | 0.859345 | 0.705799 | 1 | 0.996511 | 0.957084 |
| Lrtm2 | Q8BGX3 | 0.85 | 0.52 | 0.75 | 1.03 | 0.42 | 0.68 | 0.97 | 1.03 | 1.21 | 1.96 | 0.89 | 0.83 | 1.25 | 0.91 | 0.77 | 1.15 | -0.0125 | 0.435 | 0.2325 | 0.951638 | 0.158954 | 0.286706 | 1 | 0.996511 | 0.824333 |
| Lsamp | Q3TYE5 | 0.21 | 0.59 | 1.22 | 1.23 | 0.56 | 0.15 | 1.68 | 1.54 | 0.18 | 0.82 | 1.39 | 0.63 | -0.52 | 0.14 | 1.37 | 1.21 | 0.17 | -0.0575 | -0.2625 | 0.687644 | 0.868621 | 0.588846 | 1 | 0.996511 | 0.929131 |
| Lsg1 | Q3UM18-2 | -0.74 | -0.69 | -0.31 | -0.37 | -0.44 | -0.6 | -0.41 | -0.33 | -0.3 | -0.02 | -0.23 | -0.02 | 0.6 | -0.41 | 0.74 | 0.18 | 0.0825 | 0.385 | 0.805 | 0.636199 | 0.067984 | 0.022029 | 1 | 0.996511 | 0.621926 |
| Lsm1 | Q8VC85 | 0.08 | 0.3 | -0.74 | -0.5 | 0.75 | -0.16 | -0.05 | -0.31 | -0.19 | -1.26 | -0.32 | 0.11 | -0.63 | 0.42 | -1 | 0.05 | 0.2725 | -0.2 | -0.075 | 0.418296 | 0.592638 | 0.847658 | 1 | 0.996511 | 0.976225 |
| Lsm11 | Q8BUV6 | 0.74 | 1.23 | 1.13 | 0.8 | 0.62 | 1.15 | 0.43 | 1.21 | 1.36 | 1.39 | 1.78 | 1.1 | 1.57 | 1.11 | 0.75 | 0.76 | -0.1225 | 0.4325 | 0.0725 | 0.616179 | 0.077902 | 0.778867 | 1 | 0.996511 | 0.967289 |
| Lsm12 | Q9D0R8 | -2.33 | -2.28 | -2.74 | -2.86 | -1.74 | -2.82 | -2.13 | -3.34 | -2.22 | -2.32 | -2.7 | -1.98 | -1.55 | -1.37 | -2.64 | -3.3 | 0.045 | 0.2475 | 0.3375 | 0.902369 | 0.312649 | 0.463931 | 1 | 0.996511 | 0.888267 |
| Lsm14a | Q8K2F8 | -0.8 | -1.43 | -0.36 | -0.11 | -0.94 | -0.73 | -0.6 | -0.38 | -0.71 | 0.28 | -0.5 | -0.54 | -0.34 | -0.13 | -0.24 | -0.56 | 0.0125 | 0.3075 | 0.3575 | 0.96758 | 0.397374 | 0.270497 | 1 | 0.996511 | 0.822084 |
| Lsm14b | Q8BM41 | -1.98 | 0.1 | -0.55 | -1.25 | -0.19 | -0.77 | -1.2 | -1.23 | 0.16 | -1.09 | -0.67 | -1.04 | -0.69 | 0.45 | 0.46 | -0.3 | 0.0725 | 0.26 | 0.9 | 0.878002 | 0.603113 | 0.097056 | 1 | 0.996511 | 0.698156 |
| Lsm14b | Q8CGC4 | -1.91 | -2.42 | -1.61 | -1.7 | -1.91 | -1.98 | -1.74 | -1.67 | -1.79 | -1.56 | -1.58 | -2.08 | -0.6 | -1.92 | -1.21 | -1.29 | 0.085 | 0.1575 | 0.655 | 0.699037 | 0.523258 | 0.072913 | 1 | 0.996511 | 0.668066 |
| Lsm2 | O35900 | -2.34 | -2.42 | -0.99 | -0.82 | -2.16 | -2.55 | -0.48 | 0.12 | -2.67 | -2.03 | -0.85 | -3 | -1.71 | -1.98 | -0.48 | -0.64 | 0.375 | -0.495 | 0.44 | 0.595447 | 0.408354 | 0.414179 | 1 | 0.996511 | 0.874945 |
| Lsm3 | P62311 | -0.43 | -0.75 | -0.74 | -0.47 | -0.32 | -1.07 | -0.57 | -0.56 | -1.42 | -0.48 | -0.85 | -1.04 | -1.37 | -0.65 | -0.94 | -1.33 | -0.0325 | -0.35 | -0.475 | 0.87594 | 0.171223 | 0.068139 | 1 | 0.996511 | 0.664936 |
| Lsm4 | Q9CY46 | -0.6 | -1.14 | -1.29 | -1.27 | -1.15 | -1.24 | -1.14 | -0.74 | -1.71 | -0.96 | -1.33 | -1.74 | -1.24 | -0.38 | -0.95 | -1.63 | 0.0075 | -0.36 | 0.025 | 0.972776 | 0.194705 | 0.93699 | 1 | 0.996511 | 0.989767 |
| Lsm5 | P62322 | -0.92 | -1.13 | -1.27 | -1.13 | -1.49 | -0.9 | -1.63 | -1.66 | -0.63 | -0.84 | -1.82 | -1.94 | -0.84 | 0.34 | -1.55 | -0.61 | -0.3075 | -0.195 | 0.4475 | 0.180015 | 0.567392 | 0.264256 | 1 | 0.996511 | 0.816139 |
| Lsm6 | P62313 | -0.1 | -1.09 | -0.9 | -0.97 | -1.32 | -0.48 | -1.05 | -0.64 | -1.7 | -1.11 | -1.01 | -1.19 | -0.92 | -0.76 | -0.95 | -1.17 | -0.1075 | -0.4875 | -0.185 | 0.716052 | 0.113207 | 0.493596 | 1 | 0.996511 | 0.89758 |
| Lsm7 | Q9CQQ8 | -0.49 | -0.08 | -0.04 | -0.96 | -0.09 | -0.73 | 0.16 | -0.51 | -0.26 | 0.45 | -0.28 | -0.31 | -0.42 | -0.16 | -0.87 | 0.45 | 0.1 | 0.2925 | 0.1425 | 0.733974 | 0.329541 | 0.683154 | 1 | 0.996511 | 0.955729 |
| Lsm8 | Q6ZWM4 | -0.77 | -1.01 | -0.85 | -0.69 | 0.67 | -1.47 | -0.97 | -0.32 | -1.58 | -0.84 | -1.15 | -1.07 | -1.59 | -1.02 | -0.83 | -0.53 | 0.3075 | -0.33 | -0.1625 | 0.48714 | 0.144868 | 0.539085 | 1 | 0.996511 | 0.912212 |
| Lsp1 | A2A6J4 | 0.91 | 1.94 | 0.21 | 0.18 | 2.76 | 1.77 | 0.65 | 0.26 | -0.2 | 2.41 | 0.42 | 1.97 | 1.39 | 2.45 | 0.32 | -0.2 | 0.55 | 0.34 | 0.18 | 0.398314 | 0.618588 | 0.783625 | 1 | 0.996511 | 0.967642 |
| Lsr | Q99KG5-3 | 0.78 | -2.11 | -1.23 | -0.8 | -4.34 | -2.39 | -1.05 | -1.28 | 2.79 | -0.41 | -1.33 | 0.55 | 3.53 | -2.43 | -1.82 | -1.31 | -1.425 | 1.24 | 0.3325 | 0.128009 | 0.220126 | 0.802471 | 1 | 0.996511 | 0.971091 |
| Lss | Q8BLN5 | -0.17 | -0.03 | 0.76 | 0.19 | 0 | 0.02 | 1.74 | 0.8 | -0.25 | -0.23 | 1.11 | 0.36 | -0.07 | -0.22 | 1.36 | 1.51 | 0.4525 | 0.06 | 0.4575 | 0.308212 | 0.870688 | 0.345062 | 1 | 0.996511 | 0.849512 |
| Lta4h | P24527 | -0.37 | -0.4 | 0.09 | 0.03 | -0.41 | -0.7 | -0.25 | -0.01 | -1.34 | 0.04 | -0.43 | -0.82 | -0.87 | -0.57 | -0.39 | -0.58 | -0.18 | -0.475 | -0.44 | 0.418997 | 0.163787 | 0.068137 | 1 | 0.996511 | 0.664936 |
| Ltbp4 | E9QPS5 | -5.74 | -7.29 | -5.59 | -5.6 | -7.09 | -5.81 | -6.01 | -5.61 | -5.71 | -4.65 | -5.38 | -5.84 | -2.12 | -9.46 | -5.45 | -5.93 | -0.075 | 0.66 | 0.315 | 0.878004 | 0.177259 | 0.820106 | 1 | 0.996511 | 0.972768 |
| Ltn1 | Q6A009 | -1.02 | -1.6 | -2.27 | -2.39 | -0.89 | -1.71 | -1.58 | -2.22 | -1.44 | -1.49 | -2.16 | -1.54 | -1.07 | -0.97 | -1.79 | -1.86 | 0.22 | 0.1625 | 0.3975 | 0.582306 | 0.646677 | 0.31557 | 1 | 0.996511 | 0.838071 |
| Luc7l | Q9CYI4 | -2.02 | -2.66 | -2.91 | -4.07 | -1.94 | -2.14 | -2.68 | -3.34 | -2.11 | -2.01 | -3.47 | -2.02 | -1.31 | -1.07 | -1.43 | -2.31 | 0.39 | 0.5125 | 1.385 | 0.435011 | 0.335376 | 0.016394 | 1 | 0.996511 | 0.589387 |
| Luc7l2 | Q7TNC4-2 | -0.97 | -2.85 | -3.34 | -3.31 | -2.19 | -1.29 | -3.1 | -3.4 | -3.2 | -1.68 | -3.44 | -1.59 | -2.51 | -2.16 | -2.05 | -3.35 | 0.1225 | 0.14 | 0.1 | 0.854461 | 0.836301 | 0.863453 | 1 | 0.996511 | 0.977235 |
| Luc7l3 | Q5SUF2 | -0.73 | -1.75 | -1.61 | -2.09 | -1.74 | -1.61 | -1.9 | -1.99 | -0.99 | -0.74 | -1.8 | -1.79 | -0.15 | -0.32 | -0.95 | -1.42 | -0.265 | 0.215 | 0.835 | 0.387504 | 0.578404 | 0.061208 | 1 | 0.996511 | 0.661738 |
| Lum | P51885 | -2.2 | -1.92 | -4.76 | -3.58 | -1.37 | -1.04 | -5.64 | -5.88 | -0.83 | -3.2 | -7.18 | -1.49 | -1.88 | -0.84 | -7.89 | -3.73 | -0.3675 | -0.06 | -0.47 | 0.780367 | 0.965843 | 0.754304 | 1 | 0.996755 | 0.962308 |
| Lurap1 | Q9D6I9 | 0.2 | 0.15 | 0.24 | 0.56 | 0.27 | 0.17 | -0.19 | 0.23 | 0.59 | 1.42 | 0.39 | 0.63 | 0.33 | 1.17 | 0.09 | -0.19 | -0.1675 | 0.47 | 0.0625 | 0.372552 | 0.101871 | 0.842972 | 1 | 0.996511 | 0.976225 |
| Lurap1l | Q8K2P1 | 0.11 | 0.91 | 1.99 | 0.74 | 1.66 | 0.25 | 0.1 | 1.1 | 0.27 | -1.7 | 0.79 | 0.12 | 0.16 | 0.41 | 0.47 | 1.87 | -0.16 | -1.0675 | -0.21 | 0.747167 | 0.109085 | 0.682544 | 1 | 0.996511 | 0.955729 |
| Luzp1 | Q8R4U7 | 0.32 | 0.45 | 0.07 | -0.16 | 0.18 | 0.01 | -0.38 | -0.22 | 0.18 | 0.17 | -0.07 | 0.36 | 0.74 | 0.23 | -0.08 | -0.04 | -0.2725 | -0.01 | 0.0425 | 0.218335 | 0.961367 | 0.870567 | 1 | 0.996755 | 0.979293 |
| Luzp2 | Q8BGY3 | 2.3 | 2.42 | 0.29 | 1 | 1.96 | 2.49 | 1.12 | 0.52 | 2.11 | 0.78 | 0.45 | 2.63 | 1.65 | 3.31 | 1.06 | 0.69 | 0.02 | -0.01 | 0.175 | 0.974083 | 0.98806 | 0.803727 | 1 | 0.999285 | 0.97131 |
| Lxn | P70202 | -1 | -0.75 | -0.08 | -0.19 | -0.78 | -0.46 | -0.02 | -0.09 | -0.62 | 0.08 | 0 | -0.82 | -0.8 | -0.15 | -0.41 | -0.16 | 0.1675 | 0.165 | 0.125 | 0.559682 | 0.604067 | 0.663987 | 1 | 0.996511 | 0.949825 |
| Ly6a | P05533 | -4.64 | -3.63 | -2.09 | -2.42 | -5.57 | -5.37 | -1.58 | -2.62 | -3.56 | -0.63 | -2.32 | -3.03 | -3.59 | -5.49 | -2.57 | -1.81 | -0.59 | 0.81 | -0.17 | 0.572548 | 0.316149 | 0.848019 | 1 | 0.996511 | 0.976225 |
| Ly6c1 | A0A087WF | -1.41 | -2.84 | -2.33 | -3.72 | -2.24 | -1.31 | -0.69 | -2.57 | -2.18 | -1.88 | -1.66 | -1.09 | -2.08 | -3.38 | -2.21 | -1.51 | 0.8725 | 0.8725 | 0.28 | 0.165099 | 0.107502 | 0.626919 | 1 | 0.996511 | 0.939737 |
| Ly6e | A0A087WC | -7.67 | -8.44 | -4.96 | -3.81 | -4.58 | -7.1 | -2.43 | -4.57 | -7.85 | -2.33 | -3.64 | -6.7 | -8.01 | -5.22 | -3.87 | -4.27 | 1.55 | 1.09 | 0.8775 | 0.252802 | 0.477372 | 0.499266 | 1 | 0.996511 | 0.899626 |
| Ly6g6e | Q8K1T6-2 | 4.83 | 1.1 | -2.77 | -2.73 | 1.17 | 1.81 | -5.07 | -3.45 | 1.63 | -0.73 | -2.79 | 2.95 | 1.53 | 1.76 | -2.87 | -4.08 | -1.4925 | 0.1575 | -1.0225 | 0.506133 | 0.936417 | 0.626249 | 1 | 0.996755 | 0.939683 |
| Ly6h | Q8K356 | -3.05 | -4.41 | -1.74 | -2.21 | -3.46 | -2.03 | -1.92 | -2.28 | -3.76 | -1.01 | -2.63 | -2.65 | -2.83 | -5.58 | -1.02 | -0.99 | 0.43 | 0.34 | 0.2475 | 0.496023 | 0.6472 | 0.821624 | 1 | 0.996511 | 0.972768 |
| Ly75 | A2AW86 | -0.69 | 0.29 | -0.12 | -0.38 | -0.62 | 4.58 | -0.82 | -1.39 | 0.66 | -1.61 | -0.12 | -0.17 | 1.06 | 1.34 | -0.56 | -0.34 | 0.6625 | -0.085 | 0.6 | 0.601045 | 0.860068 | 0.241287 | 1 | 0.996511 | 0.796473 |
| Lyar | Q08288 | 0.09 | -0.78 | -0.58 | -0.5 | -0.23 | 0.18 | -0.49 | -0.13 | -0.14 | -0.37 | 0.52 | -1.06 | 0.58 | -0.26 | 0.83 | 0.42 | 0.275 | 0.18 | 0.835 | 0.281498 | 0.624284 | 0.022936 | 1 | 0.996511 | 0.623147 |
| Lyn | P25911-2 | -0.11 | -0.44 | 0.7 | 0.59 | -0.33 | -0.4 | 0.55 | 0.78 | -0.57 | 1.26 | 0.69 | 0.24 | -0.6 | -0.99 | 0.78 | 0.69 | -0.035 | 0.22 | -0.215 | 0.927764 | 0.624417 | 0.663638 | 1 | 0.996511 | 0.949825 |
| Lynx1 | P0DP60 | -1.15 | -1.74 | -1.8 | -1.31 | -1.42 | -0.79 | -0.92 | -1.64 | -1.52 | -0.88 | -1.49 | -0.86 | -1.16 | -1.44 | -1.6 | -1.08 | 0.3075 | 0.3125 | 0.18 | 0.264252 | 0.252344 | 0.459847 | 1 | 0.996511 | 0.887902 |
| Lypd1 | Q8BLC3 | 0.04 | -0.13 | -0.08 | 0.63 | -0.41 | 0.26 | 0.61 | 0.16 | 0.13 | 0.96 | 0.31 | 0 | 0.08 | 0.28 | 0.13 | 0.28 | 0.04 | 0.235 | 0.0775 | 0.885651 | 0.420722 | 0.734981 | 1 | 0.996511 | 0.958635 |
| Lypd6 | Q8BPP5 | -1.21 | -3.63 | -0.82 | -1.53 | -5.69 | 0.68 | -1.04 | -1 | 1.45 | -1.15 | -0.87 | -1.49 | 4.25 | -8.47 | 0.68 | -0.91 | 0.035 | 1.2825 | 0.685 | 0.979231 | 0.147714 | 0.778757 | 1 | 0.996511 | 0.967289 |
| Lypla1 | P97823 | -0.37 | -1.08 | 0.36 | 0.32 | -0.92 | -0.23 | -0.48 | 0.25 | -0.42 | 0.11 | 0.05 | -0.82 | -0.26 | -1.01 | -0.1 | -0.19 | -0.1525 | -0.0775 | -0.1975 | 0.700759 | 0.841769 | 0.61267 | 1 | 0.996511 | 0.935166 |
| Lypla2 | Q9WTL7 | -0.06 | 0 | -0.74 | -0.58 | -0.05 | -0.03 | -0.67 | -0.8 | 0.36 | -0.09 | -0.83 | -0.03 | 0.39 | 0.67 | -0.59 | -0.93 | -0.0425 | 0.1975 | 0.23 | 0.878413 | 0.531825 | 0.576901 | 1 | 0.996511 | 0.927198 |
| Lyplal1 | E9QLB2 | -0.01 | 0.08 | -1.11 | -0.54 | -0.02 | -0.12 | -0.04 | -0.66 | -0.59 | -0.42 | -0.68 | -0.09 | -0.44 | -0.06 | -1.2 | -1.22 | 0.185 | -0.05 | -0.335 | 0.553726 | 0.870878 | 0.396312 | 1 | 0.996511 | 0.867334 |
| Lyrm1 | Q9CQB7 | -0.93 | -0.98 | 1 | 0.53 | -1.5 | -1.11 | 1.1 | 0.64 | -0.04 | 0.45 | 0.9 | -1.08 | 0.64 | -0.73 | 0.39 | 1.07 | -0.1225 | 0.1525 | 0.4375 | 0.868024 | 0.801272 | 0.460618 | 1 | 0.996511 | 0.887902 |
| Lyrm2 | Q8R033 | 0.61 | 0.6 | -1.28 | -0.22 | -0.85 | 1.36 | -1.63 | -0.84 | 3.65 | 0.95 | 0.2 | 1.27 | 5.03 | 0.63 | 0.16 | 0.66 | -0.4175 | 1.59 | 1.6925 | 0.56019 | 0.069841 | 0.148392 | 1 | 0.996511 | 0.746406 |
| Lyrm4 | Q8K215 | 1.32 | 0.83 | 1.73 | -0.24 | 0.94 | 1.26 | 1.83 | 1.62 | -2.05 | -1.42 | 2.17 | 1.24 | -1.63 | -0.82 | 1.78 | 1.99 | 0.5025 | -0.925 | -0.58 | 0.269154 | 0.363712 | 0.526168 | 1 | 0.996511 | 0.908107 |
| Lyrm7 | Q9DA03 | 0.43 | 0.91 | -0.48 | -0.46 | 0.98 | 0.15 | -0.02 | -0.44 | 0.11 | -0.37 | -0.08 | 0.1 | 0.47 | 1.39 | -0.18 | 0.3 | 0.0675 | -0.16 | 0.395 | 0.873943 | 0.652406 | 0.390137 | 1 | 0.996511 | 0.865375 |
| Lyrm9 | E9PX24 | 1.01 | 0.74 | 0.03 | 0.6 | 0.33 | 0.61 | 1.09 | 0.64 | 0.91 | 0.53 | 0.68 | 0.89 | 2.11 | 0.95 | 0.96 | 0.67 | 0.0725 | 0.1575 | 5.78E-01 | 0.785852 | 0.532172 | 0.147208 | 1 | 0.996511 | 0.746406 |
| Lysmd1 | Q9D0E3 | 1.33 | 1.08 | 2.74 | 2.47 | 0.85 | 1.35 | 3.27 | 3.2 | 0.27 | 1.31 | 3.19 | 1.53 | 0.07 | 0.31 | 2.94 | 3.25 | 0.2625 | -0.33 | -0.2625 | 0.69982 | 0.622642 | 0.755977 | 1 | 0.996511 | 0.962333 |
| Lysmd2 | G3X915 | -1.9 | -1.24 | -0.91 | -1.42 | -0.92 | -1.69 | -1.73 | -1.4 | -1.31 | -1.07 | -0.96 | -1.6 | -1.2 | -0.88 | -1.24 | -1.28 | -0.0675 | 0.1325 | 0.2175 | 0.81038 | 0.622347 | 0.406331 | 1 | 0.996511 | 0.87356 |
| Lyst | G5E8Q0 | 0.05 | 0.36 | 0.76 | 0.78 | 0.23 | 0.02 | 0.61 | 0.83 | 0.14 | 0.47 | 0.61 | 0.01 | -0.05 | 1.09 | 0.45 | 0.63 | -0.065 | -0.18 | 0.0425 | 0.803761 | 0.474891 | 0.889095 | 1 | 0.996511 | 0.981018 |
| Lzic | Q8K3C3 | 0.85 | 0.73 | 0.48 | 0.44 | 0.81 | 0.8 | 0.41 | 0.36 | 0.76 | 0.42 | 0.46 | 1.49 | 0.09 | 1.48 | -0.02 | -0.01 | -0.03 | 0.1575 | -0.24 | 0.876667 | 0.576045 | 0.52275 | 1 | 0.996511 | 0.907667 |
| Lztfl1 | Q9JHQ5 | -0.55 | -0.01 | -0.35 | -0.52 | -0.51 | -0.49 | -0.5 | -0.59 | -0.12 | -0.28 | -0.43 | -0.19 | -0.24 | -0.32 | -0.51 | -0.5 | -0.165 | 0.1025 | -0.035 | 0.358208 | 0.600902 | 0.863831 | 1 | 0.996511 | 0.977235 |
| Lztr1 | Q9CQ33 | -0.55 | 0.06 | -0.01 | -0.11 | 0.1 | -0.94 | 0.12 | -0.34 | -0.19 | -0.4 | 0 | -0.23 | 0.06 | 0.46 | -0.02 | -0.03 | -0.1125 | -0.0525 | 0.27 | 0.69466 | 0.798603 | 0.252308 | 1 | 0.996511 | 0.80892 |
| Lzts1 | P60853 | 1.17 | 0.85 | 1.18 | 0.63 | 1.26 | 0.98 | 1.46 | 1.19 | 0.52 | 0.44 | 1.07 | 1.38 | 0.75 | 1.29 | 1.36 | 1.36 | 0.265 | -0.105 | 0.2325 | 0.206772 | 0.703788 | 0.343238 | 1 | 0.996511 | 0.848473 |
| Lzts3 | A2AHG0 | 0.58 | 0.4 | 1.03 | 1.23 | 0.65 | 0.32 | 0.67 | 0.98 | 0.68 | 0.63 | 1.13 | 0.4 | 1.11 | 0.68 | 1.36 | 0.73 | -0.155 | -0.1 | 0.16 | 0.536578 | 0.705603 | 0.563482 | 1 | 0.996511 | 0.920024 |
| M6pr | P24668 | -0.15 | -0.42 | -0.02 | -0.39 | -0.42 | -0.27 | -0.72 | -0.67 | -0.37 | -0.41 | -0.57 | 0.13 | 0.35 | -1.37 | -0.03 | -0.25 | -0.275 | -0.06 | -0.08 | 0.162837 | 0.783269 | 0.83047 | 1 | 0.996511 | 0.973939 |
| Macf1 | A0A0A0MC | -1.34 | -0.65 | -0.62 | -1.61 | 0.55 | -0.22 | -0.72 | -0.67 | -1.42 | -1.18 | 0.12 | -0.95 | -0.8 | -0.48 | -0.92 | 0.83 | 0.79 | 0.1975 | 0.7125 | 0.056837 | 0.625641 | 0.137695 | 1 | 0.996511 | 0.733463 |
| Macf1 | F6Q750 | 1.21 | -0.1 | 0.32 | 0.62 | 0.56 | 0.04 | 0.99 | 1.14 | 1.19 | 2.15 | 1.18 | 0.76 | 1.32 | 0.59 | 0.96 | 0.8 | 0.17 | 0.8075 | 0.405 | 0.634026 | 0.063129 | 0.22892 | 1 | 0.996511 | 0.787447 |
| Macf1 | F7ACR9 | 1.8 | 1.1 | 0.55 | 0.96 | 1.75 | 1.54 | 1.09 | 0.38 | 1.39 | 0.11 | 0.63 | 1.61 | 1.01 | 0.85 | 0.86 | 0.64 | 0.0875 | -0.1675 | -0.2625 | 0.817667 | 0.685562 | 0.374237 | 1 | 0.996511 | 0.859424 |
| Macf1 | E9PVY8 | 0.92 | 0.57 | 0.71 | 0.81 | 0.63 | 0.85 | 0.79 | 0.88 | 0.6 | 0.7 | 0.89 | 0.82 | 0.51 | 0.59 | 0.77 | 0.67 | 0.035 | -3.9E-17 | -0.1175 | 0.824644 | 1 | 0.522653 | 1 | 1 | 0.907667 |
| Macf1 | Q9QXZ0-3 | 0.6 | -0.2 | -0.51 | -0.13 | 0.89 | -0.27 | -0.21 | -0.47 | -0.09 | 0.11 | -0.9 | -0.06 | -0.49 | 0.84 | -0.56 | -1.05 | 0.045 | -0.175 | -0.255 | 0.902746 | 0.59337 | 0.568737 | 1 | 0.996511 | 0.922588 |
| Maco1 | Q7TQE6 | -0.69 | 0.14 | -1.24 | -1.24 | 0.58 | -0.3 | -0.74 | -0.97 | -0.16 | -1.13 | -1.65 | -0.12 | 0.23 | 0 | 0.43 | 0.23 | 0.4 | -0.0075 | 0.98 | 0.375907 | 0.987154 | 0.016855 | 1 | 0.999285 | 0.595413 |
| Macrod1 | Q922B1 | 0.53 | -0.8 | -1.38 | -0.41 | -1.09 | 0.94 | -0.61 | -0.73 | 0.4 | -0.18 | -1.01 | 0.14 | 0.88 | -0.95 | -0.81 | -0.24 | 0.1425 | 0.3525 | 0.235 | 0.796475 | 0.462842 | 0.662093 | 1 | 0.996511 | 0.949645 |
| Macrod2 | Q3UYG8 | -0.84 | -0.82 | 1.11 | 0.29 | -1.04 | -0.52 | 1.4 | 1.35 | -0.91 | 0.42 | 1.21 | -0.47 | -0.05 | -0.66 | 1.4 | 1.6 | 0.3625 | 0.1275 | 0.6375 | 0.613979 | 0.834873 | 0.347874 | 1 | 0.996511 | 0.851888 |
| Macroh2a1 | Q9QZQ8 | 0.79 | 0.23 | 0.89 | 1.1 | 0.08 | 0.49 | 1.03 | 0.86 | 0.2 | 0.44 | 0.66 | 0.11 | 0.22 | 0.54 | 0.58 | 0.92 | -0.1375 | -0.4 | -0.1875 | 0.629512 | 0.133176 | 0.481404 | 1 | 0.996511 | 0.893906 |
| Macroh2a2 | Q8CCK0 | 1 | 0.08 | 1.71 | 1.74 | 0.04 | 0.39 | 1.8 | 1.8 | 0.03 | 1.17 | 1.69 | 0.5 | 0.04 | 0.4 | 1.56 | 1.74 | -0.125 | -0.285 | -0.1975 | 0.821817 | 0.570368 | 0.711179 | 1 | 0.996511 | 0.957874 |
| Mad1l1 | Q9WTX8 | 0.47 | -0.8 | -0.49 | -0.47 | -1.54 | -0.54 | -2.11 | -0.08 | -0.28 | -0.28 | -0.74 | -1.13 | 0.7 | -0.68 | -0.34 | -0.72 | -0.745 | -0.285 | 0.0625 | 0.159376 | 0.409773 | 0.879523 | 1 | 0.996511 | 0.979626 |
| Mad2l1 | Q9Z1B5 | 0.66 | -0.14 | -0.12 | -0.53 | -0.47 | 0.38 | -1.53 | -0.73 | 0.14 | -0.96 | -0.52 | 0.24 | 0.98 | -0.62 | -1.05 | -0.41 | -0.555 | -0.2425 | -0.2425 | 0.223988 | 0.513833 | 0.611383 | 1 | 0.996511 | 0.935047 |
| Mad2l1bp | Q9DCX1 | -0.95 | -1.33 | -1.39 | -1.6 | -2.2 | -0.66 | -3.41 | -2.02 | -0.86 | -1.71 | -1.99 | -1.83 | 0.24 | -1.85 | -2.58 | -1.8 | -0.755 | -0.28 | -0.18 | 0.180483 | 0.35413 | 0.753188 | 1 | 0.996511 | 0.961479 |
| Madd | A2AGQ8 | -1.74 | -2.96 | -1.24 | -4.03 | -2.26 | -0.62 | -3.16 | -2.31 | 2.95 | -2.39 | -0.59 | -2.04 | 6.87 | -4.69 | 1.8 | -3.04 | 0.405 | 1.975 | 2.7275 | 0.588649 | 0.134828 | 0.268446 | 1 | 0.996511 | 0.820082 |
| Madd | E9QN47 | 0.17 | 0.25 | 0.41 | 0.43 | 0.16 | 0.31 | 0.5 | 0.44 | 0.46 | 0.14 | 0.64 | 0.24 | 0.49 | 0.4 | 0.24 | 0.39 | 0.0375 | 0.055 | 0.065 | 0.815212 | 0.76984 | 0.714821 | 1 | 0.996511 | 0.958106 |
| Madd | A2AGR0 | 6.8 | 1.54 | 6.82 | 1.55 | 1.88 | 7.03 | 10.98 | 3.68 | 1.79 | 1.28 | 8.18 | 6.52 | -0.05 | 1.11 | 10.19 | 9.61 | 1.715 | 0.265 | 1.0375 | 0.451828 | 0.896691 | 0.707313 | 1 | 0.996511 | 0.957157 |
| Maea | Q4VC33 | -0.96 | -0.96 | -0.07 | -0.37 | -0.65 | -0.72 | -0.5 | -0.2 | -0.81 | 0.93 | -0.49 | -0.77 | -0.53 | -0.43 | -0.42 | -0.19 | 0.0725 | 0.305 | 0.1975 | 0.780175 | 0.495045 | 0.45801 | 1 | 0.996511 | 0.887902 |
| Mafg | O54790 | -0.06 | -0.27 | -0.91 | -1.5 | 0.32 | -0.53 | -1.06 | -0.91 | 0.84 | 0.7 | -0.84 | -0.59 | 1.64 | 0 | 0.32 | -0.88 | 0.14 | 0.7125 | 0.955 | 0.740276 | 0.181808 | 0.119468 | 1 | 0.996511 | 0.721448 |
| Mag | P20917-2 | 2.85 | 2.89 | 4.6 | 3.8 | 1.06 | 1.55 | 1.1 | 3.4 | 2.14 | 1.96 | 3.37 | 1.67 | 2.24 | 1.56 | 4.21 | 3.96 | -1.7575 | -1.25 | -0.5425 | 0.020218 | 0.037539 | 0.445785 | 1 | 0.996511 | 0.884817 |
| Mag | P20917 | 0.4 | 0.59 | 0.52 | 0.39 | 0.44 | 0.1 | -0.06 | -0.22 | 1.01 | -0.7 | 0.35 | 0.2 | 1.09 | -0.04 | 0.33 | 0.71 | -0.41 | -0.26 | 0.0475 | 0.054485 | 0.463119 | 0.861313 | 1 | 0.996511 | 0.977204 |
| Maged1 | Q9QYH6 | -0.84 | 0.14 | -0.24 | -0.44 | -0.47 | -0.67 | -0.39 | -0.44 | -0.37 | 0.47 | -0.23 | -0.16 | 0.35 | -0.38 | 0.04 | -0.34 | -0.1475 | 0.2725 | 0.2625 | 0.529586 | 0.354197 | 0.369912 | 1 | 0.996511 | 0.859245 |
| Maged2 | Q9ER67 | 0.39 | -0.06 | 0.16 | -0.36 | -0.04 | 0.48 | 0.4 | 0 | 0.58 | -0.28 | 0.3 | 0.41 | 0.71 | 0.35 | 0.01 | 0.76 | 0.1775 | 0.22 | 0.425 | 0.445593 | 0.413681 | 0.134244 | 1 | 0.996511 | 0.730883 |
| Magee1 | Q6PCZ4 | -0.45 | -0.38 | -0.35 | -0.21 | 0.1 | -0.59 | -0.28 | -0.66 | -0.78 | 0.11 | -0.24 | -0.32 | -0.15 | -0.46 | 0 | 0.04 | -0.01 | 0.04 | 0.205 | 0.961811 | 0.858717 | 0.309942 | 1 | 0.996511 | 0.834523 |
| Magee2 | Q52KG3 | 0.54 | 0.47 | -2.87 | -3.43 | 0.32 | 0.06 | -4.55 | -2.25 | 0.01 | 0.24 | -3.29 | 0.41 | -0.34 | 0.92 | -2.75 | -4.39 | -0.2825 | 0.665 | -0.3175 | 0.839069 | 0.593218 | 0.822907 | 1 | 0.996511 | 0.972768 |
| Magi1 | A0A0N4SU | 0.46 | 0.57 | 1.07 | 1.34 | 0.55 | 0.63 | 1.27 | 1.35 | 0.41 | 1.3 | 1.3 | 0.69 | 0.15 | 0.43 | 1.06 | 1.08 | 0.09 | 0.065 | -0.18 | 0.760027 | 0.833384 | 0.575584 | 1 | 0.996511 | 0.926728 |
| Magi2 | Q9WVQ1 | 2.68 | 0.44 | 1.7 | 1.31 | -0.08 | 4.7 | 1.07 | 1.83 | -0.49 | -0.51 | 2.8 | 3.4 | -1.67 | -0.18 | 0.89 | 2.18 | 0.3475 | -0.2325 | -1.2275 | 0.730067 | 0.821162 | 0.171063 | 1 | 0.996511 | 0.762752 |
| Magi2 | A0A0G2JEC | 0.52 | 0.85 | 1.28 | 1.04 | 0.84 | 0.8 | 1.02 | 0.93 | 0.74 | 0.52 | 1.33 | 0.73 | 0.87 | 0.58 | 0.8 | 0.87 | -0.025 | -0.0925 | -0.1425 | 0.900722 | 0.719965 | 0.528223 | 1 | 0.996511 | 0.908663 |

| | | | | | | | | | | | | | | | | | | | | | | | | | |
|---|---|---|---|---|---|---|---|---|---|---|---|---|---|---|---|---|---|---|---|---|---|---|---|---|---|
| Magi3 | G5E8T6 | 1 | 1.08 | 0.82 | 0.97 | 1.05 | 1.02 | 0.83 | 0.82 | 0.86 | 0.86 | 0.88 | 0.94 | 0.52 | 0.76 | 0.67 | 0.86 | -0.0375 | -0.0825 | -0.265 | 0.806425 | 0.60375 | 0.167371 | 1 | 0.996511 | 0.759484 |
| Magoh | P61327 | 1.3 | -0.3 | -0.85 | -1.23 | -0.57 | 0.58 | -1.52 | -0.68 | -1.6 | -2.34 | -1.12 | 0.08 | -1.08 | -1.64 | -1 | -1.97 | -0.2775 | -0.975 | -1.1525 | 0.666428 | 0.179984 | 0.064143 | 1 | 0.996511 | 0.66433 |
| Magt1 | A2ADH1 | -0.42 | 0.08 | 0.58 | -0.24 | 1.07 | 0.05 | 0.76 | 1.15 | -0.2 | -0.79 | 0.8 | -0.08 | 0.19 | 0.39 | 1.08 | 1.72 | 0.7575 | -0.0675 | 0.845 | 0.042181 | 0.859168 | 0.058707 | 1 | 0.996511 | 0.661738 |
| Maip1 | Q8BHE8 | 0.53 | 0.62 | -0.29 | -0.24 | 0.58 | 0.68 | -0.25 | -0.3 | 0.29 | -0.27 | 0.02 | 0.37 | 0.59 | 0.16 | -0.42 | -0.41 | 0.0225 | -0.0525 | -0.175 | 0.947935 | 0.857174 | 0.613607 | 1 | 0.996511 | 0.935813 |
| Mal2 | Q8BI08 | 0.09 | 0.82 | 1.02 | 0.77 | 0.78 | 0.23 | 0.98 | 0.65 | 0.73 | 0.61 | 0.82 | 0.26 | 0.34 | 0.77 | 0.5 | 0.84 | -0.015 | -0.07 | -0.0625 | 0.954735 | 0.785701 | 0.811429 | 1 | 0.996511 | 0.972642 |
| Malsu1 | Q9CWV0 | 0.02 | 0.6 | -0.13 | 0.31 | -0.12 | 0.1 | -0.21 | 0.09 | 0.79 | 1.48 | -0.51 | 0.13 | 1.3 | 0.73 | 0.16 | 0.52 | -0.235 | 0.2725 | 0.4775 | 0.276701 | 0.533384 | 0.139039 | 1 | 0.996511 | 0.73493 |
| Man1a1 | P45700 | -1.13 | -1.5 | -1 | -1.04 | -0.92 | -1.27 | -1.36 | -1.14 | -1.57 | 0.33 | -1.32 | -0.92 | -0.94 | -0.87 | -0.51 | -0.71 | -0.005 | 0.2975 | 0.41 | 0.978808 | 0.480514 | 0.075805 | 1 | 0.996511 | 0.674038 |
| Man1a2 | P39098-2 | -1.06 | -1.22 | -1.38 | -1.05 | -0.11 | -1.23 | -0.04 | -0.79 | -1.98 | -1.1 | -0.81 | -1.08 | -1.03 | -1.11 | -0.04 | 0.3 | 0.635 | -0.065 | 0.7075 | 0.056219 | 0.815774 | 0.074608 | 1 | 0.996511 | 0.671911 |
| Man1b1 | A2AJ15 | -1.24 | -2.34 | -1.23 | -0.65 | -1.58 | -1.4 | -0.68 | -0.41 | -1.07 | -0.31 | -0.77 | -1.87 | -0.8 | -0.98 | -0.42 | -0.22 | 0.3475 | 0.36 | 0.76 | 0.420315 | 0.435327 | 0.073598 | 1 | 0.996511 | 0.669435 |
| Man2a1 | P27046 | -1.8 | -1.13 | -5.86 | -1.71 | -0.99 | -0.97 | -6.86 | -5.55 | -0.28 | -0.18 | -4.98 | -1.53 | -0.53 | -1.81 | -4.79 | -8.25 | -0.9675 | 0.8825 | -1.22 | 0.567536 | 0.53233 | 0.504268 | 1 | 0.996511 | 0.901002 |
| Man2a2 | Q8BRK9 | -0.72 | -1.85 | -1.69 | -1.12 | -0.36 | -0.42 | -1.05 | -1.23 | -0.91 | -0.82 | -1.52 | -0.94 | -1.46 | -0.63 | -1.17 | -0.9 | 0.58 | 0.2975 | 0.305 | 0.107819 | 0.348855 | 0.355744 | 1 | 0.996511 | 0.855029 |
| Man2b1 | O09159 | -0.2 | 0.35 | -0.64 | -0.65 | 0.17 | 0.16 | 0.27 | -0.58 | -0.31 | 0.98 | -0.11 | 0.26 | 0.02 | 0.12 | -0.64 | -0.49 | 0.29 | 0.49 | 0.0375 | 0.353209 | 0.19702 | 0.903876 | 1 | 0.996511 | 0.985278 |
| Man2b2 | O54782 | -0.26 | -0.21 | -0.02 | -0.24 | -0.11 | -0.36 | 0.36 | 0.04 | 0.25 | 0.19 | 0.05 | -0.08 | 0.56 | 0.21 | 0.14 | 0.32 | 0.165 | 0.285 | 0.49 | 0.408904 | 0.121247 | 0.02654 | 1 | 0.996511 | 0.634876 |
| Man2c1 | F8WIE1 | -0.23 | -0.21 | -0.01 | 0.02 | -0.4 | -0.07 | -0.58 | -0.07 | -0.12 | 0.29 | -0.37 | -0.35 | 0.01 | -0.11 | -0.59 | -0.76 | -0.1725 | -0.03 | -0.255 | 0.36251 | 0.886271 | 0.298326 | 1 | 0.996511 | 0.828189 |
| Manba | A0A0R4J09 | -1.09 | -1.37 | -1.33 | -0.95 | -1.25 | -0.87 | -1.15 | -1.38 | -1.19 | 0.17 | -1.05 | -1.26 | -1.29 | -1.09 | -1.48 | -1.01 | 0.0225 | 0.3525 | -0.0325 | 0.904404 | 0.322466 | 0.87487 | 1 | 0.996511 | 0.979626 |
| Manbal | A2BGN7 | 2.44 | 0.83 | 0.65 | -0.62 | 2.02 | 2.54 | 2.18 | 0.83 | 3.56 | -0.14 | 0.5 | 1.75 | 2.82 | 1.89 | 2.78 | 3.22 | 1.0675 | 0.5925 | 1.8525 | 0.133883 | 0.527137 | 0.015491 | 1 | 0.996511 | 0.589387 |
| Manea | Q6NXH2 | 0.05 | -0.17 | -0.09 | -1.5 | -0.23 | -0.93 | -1.01 | -0.25 | 0.02 | 0.26 | -1.21 | 0.85 | 0.36 | -0.61 | 0.47 | -0.27 | -0.1775 | 0.4075 | 0.415 | 0.654642 | 0.44335 | 0.339933 | 1 | 0.996511 | 0.848092 |
| Maneal | Q6P1J0 | -2.96 | -3.91 | -1.18 | -1.07 | -3.16 | -2.19 | -1.59 | -0.95 | -1.86 | -1.03 | -1.21 | -2.78 | -0.15 | -3.99 | -0.79 | -0.99 | 0.3075 | 0.56 | 0.8 | 0.686094 | 0.448361 | 0.426253 | 1 | 0.996511 | 0.87873 |
| Manf | Q80ZP8 | 0.04 | 0.02 | -0.7 | -1.14 | 0.8 | -0.1 | -0.08 | -0.44 | -0.73 | -1.3 | -0.98 | 0.25 | -0.03 | -0.23 | 0.55 | 0.37 | 0.49 | -0.245 | 0.61 | 0.211549 | 0.562834 | 0.100233 | 1 | 0.996511 | 0.698156 |
| Maoa | Q64133 | 1.72 | 1.24 | 1.21 | 0.94 | 1.71 | 1.83 | 1.58 | 1.32 | 0.19 | 0.36 | 1.25 | 1.46 | -0.07 | 0.62 | 0.97 | 1.22 | 0.3325 | -0.4625 | -0.5925 | 0.155645 | 0.207811 | 0.098046 | 1 | 0.996511 | 0.698156 |
| Maob | Q8BW75 | 1.01 | 0.69 | 0.74 | 0.74 | 1.07 | 1.15 | 0.91 | 0.84 | 0.16 | 0.35 | 0.8 | 0.71 | -0.14 | 0.7 | 0.44 | 0.51 | 0.1975 | -0.29 | -0.4175 | 0.243221 | 0.19138 | 0.10437 | 1 | 0.996511 | 0.702979 |
| Map1a | A2ARP8 | -0.18 | -0.64 | -0.44 | -0.49 | -0.98 | -0.99 | -0.75 | -0.73 | -0.72 | -0.33 | -0.72 | -0.37 | -0.62 | -0.53 | -0.64 | -0.9 | -0.425 | -0.0975 | -0.235 | 0.032932 | 0.62137 | 0.249405 | 1 | 0.996511 | 0.804913 |
| Map1b | P14873 | -0.72 | -0.52 | 0.2 | 0.22 | -1.07 | -1.15 | -0.06 | 0.17 | -0.38 | 0.24 | 0.02 | -0.43 | -0.16 | -0.35 | 0.21 | -0.22 | -0.3225 | 0.0675 | 0.075 | 0.423212 | 0.820978 | 0.795044 | 1 | 0.996511 | 0.969498 |
| Map1lc3a | Q91VR7 | -0.77 | -0.72 | -1.14 | -0.75 | -0.53 | -1.07 | -0.54 | -0.96 | -1.17 | -1.03 | -0.8 | -0.84 | -0.99 | -0.06 | -0.77 | -0.99 | 0.07 | -0.115 | 0.1425 | 0.731076 | 0.547252 | 0.596821 | 1 | 0.996511 | 0.929911 |
| Map1lc3b | Q9CQV6 | -0.86 | -0.27 | -0.93 | -0.87 | -0.39 | -0.74 | -0.4 | -0.71 | 0.56 | -0.89 | -0.9 | -0.83 | 0.41 | -0.06 | -1.17 | -1.21 | 0.1725 | 0.2175 | 0.225 | 0.420742 | 0.567913 | 0.59135 | 1 | 0.996511 | 0.929717 |
| Map1s | Q8C052 | 0.13 | -0.17 | -0.01 | -0.17 | -0.6 | 0.06 | -0.8 | -0.29 | 0.23 | 0.65 | -0.23 | 0.2 | 0.84 | -0.66 | -0.08 | -0.57 | -0.3525 | 0.2675 | -0.0625 | 0.141287 | 0.258367 | 0.857884 | 1 | 0.996511 | 0.976225 |
| Map2 | G3UZJ2 | -2.61 | -2.67 | -0.9 | -1.15 | -3.58 | -3.12 | -0.79 | -1.72 | -2.61 | -1.4 | -1.04 | -2.75 | -2.51 | -2.49 | -1.91 | -2.26 | -0.47 | -0.1175 | -0.46 | 0.518347 | 0.840501 | 0.332578 | 1 | 0.996511 | 0.845601 |
| Map2 | A0A668KL( | -1.64 | -1.65 | -1.48 | -1.71 | -2.24 | -2.19 | -1.8 | -1.58 | -2.02 | -1.47 | -1.78 | -1.63 | -1.45 | -1.14 | -1.93 | -2.22 | -0.3325 | -0.105 | -0.065 | 0.122587 | 0.57738 | 0.810297 | 1 | 0.996511 | 0.972222 |
| Map2k1 | P31938 | 1.21 | 0.43 | 1.08 | 1.53 | 0.41 | 1.13 | 1.22 | 1.45 | 0.01 | 1.03 | 0.96 | 0.95 | -0.37 | 0.75 | 1.06 | 1.18 | -0.01 | -0.325 | -0.4075 | 0.974721 | 0.3422 | 0.329851 | 1 | 0.996511 | 0.845407 |
| Map2k2 | Q91YS7 | 0.3 | 0.25 | 0.38 | 0.09 | 0.5 | 0.12 | 0.18 | 0.29 | -0.17 | -0.17 | 0.25 | 0.28 | -0.1 | 0.09 | 0.02 | 0.22 | 0.0175 | -0.2075 | -0.1975 | 0.914427 | 0.300751 | 0.291217 | 1 | 0.996511 | 0.825497 |
| Map2k3 | O09110 | 1.08 | 3 | 1.27 | 1.22 | 2.38 | 1.22 | 1.32 | 1.48 | 2.12 | 1.91 | 1.24 | 2.02 | 2.77 | 4.1 | 1.88 | 1.46 | -0.0425 | 0.18 | 0.91 | 0.930318 | 0.699563 | 0.199184 | 1 | 0.996511 | 0.78004 |
| Map2k4 | P47809 | 0.35 | -0.2 | 0.39 | 0.25 | -0.07 | 0.44 | 0.21 | 0.34 | -0.45 | -0.14 | 0.44 | 0.32 | -0.32 | -0.52 | 0.19 | 0.11 | 0.0325 | -0.155 | -0.3325 | 0.874202 | 0.561807 | 0.20776 | 1 | 0.996511 | 0.785338 |
| Map2k5 | Q9WVS7 | -0.61 | 0.04 | -1.34 | -1.01 | -0.13 | -0.72 | -1.21 | -1.07 | 0.08 | 0.01 | -0.95 | -0.53 | 0 | 0.29 | -1.9 | -1.2 | -0.0525 | 0.3825 | 0.0275 | 0.885584 | 0.318232 | 0.959902 | 1 | 0.996511 | 0.992636 |
| Map2k6 | P70236 | -0.38 | -0.29 | -0.05 | -0.56 | -0.39 | -0.23 | -0.63 | -0.13 | -0.64 | 0.12 | -0.16 | -0.18 | -0.58 | -0.26 | -0.29 | -0.48 | -0.025 | 0.105 | -0.0825 | 0.895529 | 0.642149 | 0.679599 | 1 | 0.996511 | 0.955402 |
| Map2k7 | Q8CE90-2 | 1.36 | 1.19 | 0.02 | -0.07 | 1.09 | 1.5 | -0.67 | -0.08 | 0.78 | 0.61 | -0.13 | 1.28 | 0.2 | 1.03 | -0.8 | -0.32 | -0.165 | 0.01 | -0.5975 | 0.774636 | 0.982207 | 0.257132 | 1 | 0.998773 | 0.811777 |
| Map3k10 | D3YXM8 | 0.4 | 0.21 | 0.49 | 0.58 | 0.36 | 0.27 | 0.19 | 0.13 | 0.74 | 0.28 | 0.24 | 0.39 | 0.67 | 0.48 | 0.03 | 0.15 | -0.1825 | -0.0075 | -0.0875 | 0.26702 | 0.968987 | 0.690586 | 1 | 0.996755 | 0.955729 |
| Map3k11 | Q80XI6 | -0.18 | -0.14 | 0.61 | 0.7 | 0.27 | -0.6 | -0.11 | 0.32 | 0.56 | 0.84 | 0.35 | -0.2 | 1.04 | 0.39 | 0.33 | 0.73 | -0.2775 | 0.14 | 0.375 | 0.385747 | 0.665754 | 0.233132 | 1 | 0.996511 | 0.788994 |
| Map3k12 | Q60700 | -0.08 | 0.28 | 0.77 | 0.71 | -0.27 | 1.27 | 0.56 | 0.45 | 1.43 | 0.17 | 0.97 | 0.19 | 2 | 0.36 | 1.2 | 1.04 | 0.0825 | 0.27 | 0.73 | 0.817493 | 0.458953 | 0.082407 | 1 | 0.996511 | 0.68865 |
| Map3k13 | Q1HKZ5 | 0.67 | -1 | 0.35 | 0.42 | -2.5 | 0.82 | -0.16 | 0.13 | 0.48 | 0.18 | 0.46 | 0.51 | 0.87 | -2.59 | 0.38 | -0.3 | -0.5375 | 0.2975 | -0.52 | 0.471732 | 0.432363 | 0.505013 | 1 | 0.996511 | 0.901074 |
| Map3k2 | G5E8L8 | -0.41 | 0.05 | -0.52 | -0.24 | -1.34 | -0.75 | -0.87 | -0.5 | 0.95 | 1.64 | -0.09 | 0.05 | 2.57 | -1.93 | 0.38 | 0.88 | -0.585 | 0.9175 | 0.755 | 0.03182 | 0.047125 | 0.381472 | 1 | 0.996511 | 0.860984 |
| Map3k20 | Q9ESL4-2 | -0.75 | -0.94 | -1.25 | -0.58 | -0.73 | -1.31 | 0.39 | -0.55 | -0.86 | -1.09 | -0.77 | -0.82 | -0.95 | 1.07 | -1.43 | -0.59 | 0.33 | -0.005 | 0.405 | 0.376285 | 0.980592 | 0.445244 | 1 | 0.998773 | 0.884817 |
| Map3k4 | O08648 | -0.46 | -0.44 | -0.11 | -0.06 | -0.61 | -0.92 | -0.38 | 0.04 | -0.61 | -0.09 | 0.02 | -0.65 | -0.74 | -0.25 | -0.17 | -0.34 | -0.2 | -0.065 | -0.1075 | 0.419501 | 0.781166 | 0.623682 | 1 | 0.996511 | 0.938628 |
| Map3k5 | E9PWG9 | -0.01 | 0.3 | 0.63 | 0.36 | 0.4 | 0.04 | 0.07 | 0.05 | 0.24 | 0.57 | 0.38 | 0.21 | 0.52 | 0.63 | -0.02 | 0.17 | -0.18 | 0.03 | 0.005 | 0.36544 | 0.882266 | 0.983321 | 1 | 0.996511 | 0.995445 |
| Map3k7 | Q62073 | 0.88 | 0.4 | 0.34 | 0.33 | 0.26 | 0.93 | 1.09 | 0.09 | 1.26 | 0.82 | 0.36 | 0.95 | 1.2 | 0.36 | 0.31 | 0.69 | 0.105 | 0.36 | 0.1525 | 0.709977 | 0.176846 | 0.573792 | 1 | 0.996511 | 0.925747 |
| Map3k9 | A0A1Y7VK) | 0.33 | 0.57 | 0.69 | 0.71 | 0.96 | 0.01 | 0.53 | 0.59 | 0.3 | 0.98 | 0.74 | 0.82 | 0.09 | 0.05 | 0.18 | 0.92 | -0.0525 | 0.135 | -0.265 | 0.821488 | 0.528517 | 0.313049 | 1 | 0.996511 | 0.835247 |
| Map4 | E9PZ43 | -0.85 | -0.66 | -1.21 | -1.1 | -0.93 | -1.15 | -1.41 | -1.17 | -0.18 | -0.67 | -1.15 | -0.53 | 0.13 | -0.31 | -1.3 | -1.38 | -0.21 | 0.3225 | 0.24 | 0.296175 | 0.230091 | 0.534129 | 1 | 0.996511 | 0.911961 |
| Map4 | P27546-2 | -0.91 | -0.77 | -0.39 | -0.48 | -1.03 | -1.33 | -0.89 | -0.88 | -0.46 | 0.11 | -0.53 | -0.69 | -0.21 | -0.54 | -0.51 | -0.81 | -0.395 | 0.245 | 0.12 | 0.070362 | 0.322395 | 0.592352 | 1 | 0.996511 | 0.929843 |
| Map4 | P27546 | 1.21 | 1.1 | -0.91 | -0.86 | 2.17 | 1.63 | 0.8 | 0.16 | 0.43 | 1.31 | 0.08 | 1.87 | 0.05 | 1.95 | -0.72 | -0.34 | 1.055 | 0.7875 | 0.1 | 0.142043 | 0.248585 | 0.894622 | 1 | 0.996511 | 0.983453 |
| Map4k2 | Q61161 | -0.61 | -0.24 | -0.14 | -0.24 | -0.1 | -0.64 | 0.96 | -0.05 | -0.35 | -0.59 | 0.33 | -0.2 | -0.66 | -0.55 | 0.3 | 0.61 | 0.35 | 0.105 | 0.2325 | 0.315944 | 0.670602 | 0.490621 | 1 | 0.996511 | 0.897531 |
| Map4k3 | A0A3Q4EG | -0.12 | -0.08 | 0.11 | 0.22 | 0.29 | -0.15 | 0.28 | -0.05 | 0.22 | -0.49 | 0.16 | -0.06 | 0.12 | 0.38 | -0.49 | -0.08 | 0.06 | -0.075 | -0.05 | 0.742507 | 0.731601 | 0.834636 | 1 | 0.996511 | 0.975487 |
| Map4k4 | A0A0A6YW | -0.05 | 0.08 | -0.29 | -0.1 | -0.02 | 0.29 | -0.18 | -0.18 | 1.39 | 0.05 | 0.24 | 0.58 | 1.7 | 0.47 | 0 | -0.41 | 0.0675 | 0.655 | 0.53 | 0.708525 | 0.059632 | 0.24763 | 1 | 0.996511 | 0.803677 |
| Map4k4 | B2RUE8 | 1.65 | 0.69 | 1.48 | 2.05 | -0.02 | 0.87 | 1.18 | 0.98 | 0.46 | 2.61 | 1.17 | 1.36 | 0.52 | 2.44 | 1.81 | 0.57 | -0.715 | -0.0675 | -0.1325 | 0.081562 | 0.891283 | 0.796864 | 1 | 0.996511 | 0.969727 |
| Map4k5 | E9PX30 | 0.17 | -0.17 | -0.16 | 0.03 | -0.09 | -0.02 | -0.12 | 0.21 | -0.8 | 0.06 | 0.28 | 0.24 | -0.46 | -0.83 | -0.14 | -0.28 | 0.0275 | -0.0225 | -0.395 | 0.86885 | 0.935542 | 0.100872 | 1 | 0.996755 | 0.698156 |
| Map6 | Q7TSJ2 | 0.45 | 0.48 | 0.62 | 0.92 | -0.13 | 0.29 | 0.03 | 0.54 | 0.63 | 0.87 | 0.3 | 0.42 | 0.66 | 0.31 | 0.51 | -0.07 | -0.435 | -0.0625 | -0.265 | 0.064544 | 0.765052 | 0.274439 | 1 | 0.996511 | 0.822084 |
| Map6d1 | Q14BB9 | -1.53 | -1.38 | -1.57 | -1.6 | -1.85 | -2.03 | -2.05 | -2.06 | -0.78 | -1.86 | -2.04 | -1.53 | -0.46 | -1.73 | -2.13 | -2.14 | -0.4775 | -0.0325 | -0.095 | 0.010145 | 0.911058 | 8.07E-01 | 1 | 0.996744 | 0.971715 |
| Map7 | E9QMU3 | 1.15 | 1.44 | -0.33 | -0.89 | 1.45 | 1.07 | -0.67 | -1.02 | 1.21 | -0.05 | -0.54 | 1.75 | 1.44 | 1.19 | -0.77 | -1.21 | -0.135 | 0.25 | -0.18 | 0.858162 | 0.724043 | 0.820392 | 1 | 0.996511 | 0.972768 |
| Map7 | A0A1L1SUI | -0.6 | 0.02 | -1.24 | -1.47 | 0.21 | -1.64 | -0.92 | -2.33 | -0.48 | -1.8 | -1.56 | -0.43 | -1.31 | 0.33 | -1.51 | -1.08 | -0.3475 | -0.245 | -0.07 | 0.555092 | 0.597157 | 0.888418 | 1 | 0.996511 | 0.980661 |
| Map7d1 | A2AJI0 | -0.73 | -0.34 | -0.6 | -0.62 | -0.33 | -0.96 | -0.91 | -1.03 | 0.05 | 0.28 | -0.98 | -0.53 | 0.64 | 0.45 | -0.15 | -1.13 | -0.235 | 0.2775 | 0.525 | 0.279392 | 0.370145 | 0.204805 | 1 | 0.996511 | 0.783452 |
| Map7d1 | A8Y5P4 | -2.61 | -2.15 | -3.55 | -3.37 | -3.09 | -1.86 | -3.99 | -4.58 | -1.92 | -0.73 | -4.06 | -2.33 | -0.65 | -1.69 | -3.1 | -3.57 | -0.46 | 0.66 | 0.6675 | 0.462831 | 0.35463 | 0.336945 | 1 | 0.996511 | 0.847578 |
| Map7d2 | A2AG50 | -0.68 | -1.02 | 0.54 | -0.14 | -1.18 | -0.87 | 0.29 | 0.06 | -0.66 | 0.58 | 0.62 | -0.87 | -0.72 | -0.61 | 0.8 | 0.22 | -0.1 | 0.2425 | 0.2475 | 0.827014 | 0.62111 | 0.59825 | 1 | 0.996511 | 0.930093 |
| Map9 | Q3TRR0 | -0.31 | -0.27 | -0.18 | -0.47 | -0.98 | -0.76 | -0.65 | -0.2 | -0.02 | -0.27 | -0.5 | 0.04 | 0.53 | -0.09 | 0.12 | -0.37 | -0.34 | 0.12 | 0.355 | 0.125049 | 0.53825 | 0.162222 | 1 | 0.996511 | 0.757516 |
| Mapk1 | P63085 | 0.28 | -0.14 | -0.23 | 0.04 | -0.07 | 0.2 | 0.36 | 0.11 | -0.08 | 0.44 | -0.19 | 0.34 | -0.42 | 0.05 | -0.22 | -0.18 | 0.1625 | 0.14 | -0.18 | 0.391602 | 0.53933 | 0.397314 | 1 | 0.996511 | 0.867334 |
| Mapk10 | Q3TQZ7 | 1.05 | 0.16 | -0.1 | 0.29 | 0.4 | 0.87 | 0.06 | 0.27 | 0.33 | 0.03 | 0.13 | 1.06 | 0.28 | 0.04 | 0.56 | -0.29 | 0.05 | 0.0375 | -0.2025 | 0.867069 | 0.910991 | 0.524213 | 1 | 0.996744 | 0.908107 |
| Mapk11 | Q9WUI1 | 0.49 | 1.51 | -0.42 | -0.25 | 0.4 | 0.15 | -0.31 | -0.44 | 0.55 | -0.13 | 0.12 | -0.25 | 1.41 | 0.94 | 0.02 | -1.13 | -0.3825 | -0.26 | -0.0225 | 0.403638 | 0.563727 | 0.972302 | 1 | 0.996511 | 0.992865 |
| Mapk14 | P47811 | -1.42 | -1.13 | -1.54 | -1.01 | -0.38 | -0.94 | -1.23 | -1.19 | -0.73 | -1.02 | -1.5 | -0.57 | -0.62 | 0.34 | -1.57 | -1.7 | 0.34 | 0.32 | 0.3875 | 0.189522 | 0.236359 | 0.411203 | 1 | 0.996511 | 0.874811 |
| Mapk1ip1l | A0A2I3BRS | -0.45 | -0.48 | -0.8 | 0.1 | -0.7 | -1.97 | 0.08 | 0.12 | -1.3 | 0.23 | -0.34 | 0.22 | -1.17 | 0.73 | -0.59 | -0.14 | -0.21 | 0.11 | 0.115 | 0.665891 | 0.77779 | 0.785567 | 1 | 0.996511 | 0.967642 |
| Mapk3 | Q63844 | 0.33 | 0.19 | -0.08 | 0.22 | 0.27 | 0.17 | 0.86 | 0.42 | 0.29 | 0.46 | 0.37 | 0.46 | 0.39 | 0.43 | 0.53 | 0.34 | 0.265 | 0.23 | 0.2575 | 0.219306 | 0.203736 | 0.183083 | 1 | 0.996511 | 0.776002 |
| Mapk4 | Q6P5G0 | -0.31 | 0.36 | -0.02 | 0.28 | 0.33 | -0.53 | 0.29 | 0.14 | -0.11 | 0.96 | 0.02 | -0.14 | -0.01 | 0.56 | -0.54 | 0.06 | -0.02 | 0.105 | -0.06 | 0.93874 | 0.732879 | 0.835628 | 1 | 0.996511 | 0.975524 |
| Mapk7 | Q9WVS8-4 | -0.56 | -1.14 | -0.99 | -0.52 | -2.83 | -2.54 | -0.51 | -1.12 | -1.95 | -0.81 | -1.11 | -0.78 | -1.29 | -2.22 | -1.35 | -1.42 | -0.9475 | -0.36 | -0.7675 | 0.102363 | 0.271898 | 0.023576 | 1 | 0.996511 | 0.624626 |
| Mapk8 | Q91Y86 | 0.21 | -0.07 | -0.43 | -0.11 | -0.32 | 0.22 | -0.51 | -0.17 | 0.68 | -0.22 | -0.13 | 0.66 | -0.06 | 0.35 | -1.07 | -0.73 | -0.095 | 0.3475 | -0.2775 | 0.673582 | 0.246569 | 0.43034 | 1 | 0.996511 | 0.879165 |
| Mapk8ip1 | Q9WVI9-3 | 0.62 | 0.86 | 0.95 | 0.77 | 0.62 | 0.7 | 0.68 | 0.76 | 0.85 | 1.51 | 1.14 | 0.88 | 1.28 | 0.75 | 1.54 | 1.11 | -0.11 | 0.295 | 0.37 | 0.472131 | 0.185917 | 0.129369 | 1 | 0.996511 | 0.723655 |
| Mapk8ip2 | Q9ERE9 | 0.35 | 0.65 | 0.62 | 0.31 | 0.45 | -0.13 | 0.11 | -0.01 | 0.45 | 1.42 | 0.77 | 0.82 | 0.72 | 0.45 | 0.97 | 0.53 | -0.3775 | 0.3825 | 0.185 | 0.075228 | 0.146258 | 0.382001 | 1 | 0.996511 | 0.860984 |
| Mapk8ip3 | J3QNR6 | 0.45 | 0.77 | 0.48 | 0.41 | 0.22 | 0.77 | 0.39 | 0.38 | 0.58 | 0.46 | 0.42 | 0.54 | 0.26 | 0.46 | 0.22 | 0.11 | -0.0875 | -0.0275 | -0.265 | 0.63768 | 0.870811 | 0.184487 | 1 | 0.996511 | 0.776843 |
| Mapk8ip3 | Q9ESN9-5 | 2.43 | 2.68 | 1.1 | 2.15 | 2.12 | 1.38 | 2.25 | 2.56 | 1.66 | 3.27 | 1.97 | 2.29 | -0.08 | 2.22 | 2.53 | 1.82 | -0.0125 | 0.2075 | -0.4675 | 0.975242 | 0.654772 | 0.459968 | 1 | 0.996511 | 0.887902 |
| Mapk9 | Q9WTU6-4 | -1.64 | 0.25 | -0.43 | -0.36 | -3.97 | 0.68 | -6.12 | -6.54 | 1.76 | 1.06 | -0.85 | 0.13 | 2.74 | -1.7 | -1.48 | -3.52 | -3.4425 | 1.07 | -0.445 | 0.047033 | 0.118075 | 0.718575 | 1 | 0.996511 | 0.958398 |
| Mapk9 | Q9WTU6-3 | 0.43 | 0.03 | 1.28 | 0.62 | -0.74 | 0.39 | 0 | -0.43 | -0.33 | 0.31 | 1.1 | 0.3 | -0.82 | 0.25 | 1.41 | 1.23 | -0.785 | -0.245 | -0.0725 | 0.046945 | 0.522414 | 0.891807 | 1 | 0.996511 | 0.982213 |
| Mapkap1 | Q8BKH7-2 | 0.56 | 0.36 | -0.21 | -0.28 | 0.24 | 0.36 | 0.18 | -0.37 | 0.01 | -0.2 | 0 | 0.64 | -0.21 | -0.44 | -0.27 | 0.02 | -0.005 | 0.005 | -0.3325 | 0.985185 | 0.986069 | 0.219908 | 1 | 0.998892 | 0.787447 |
| Mapkapk2 | P49138 | -0.43 | -0.55 | -0.38 | -1.2 | -0.48 | -0.71 | -1.6 | -0.89 | -0.11 | 0.53 | -0.87 | -0.23 | -0.17 | -0.13 | -1.04 | -1.2 | -0.28 | 0.47 | 0.005 | 0.369339 | 0.190303 | 0.988268 | 1 | 0.996511 | 0.996154 |
| Mapkbp1 | A0A0A0M( | -0.76 | -0.47 | 0.87 | 1.12 | -0.66 | -1.02 | 0.34 | 0.66 | 0.28 | 0.96 | 0.85 | -0.12 | 1.25 | 0.37 | 0.89 | 0.29 | -0.36 | 0.3025 | 0.51 | 0.528963 | 0.547527 | 0.31225 | 1 | 0.996511 | 0.834523 |
| Mapre1 | Q61166 | 0.73 | 0.84 | 1.24 | 0.93 | 0.66 | 0.62 | 1.29 | 1.32 | 0.9 | 1.27 | 1.73 | 1.35 | 0.78 | 1.45 | 1.21 | 1.15 | 0.0375 | 0.3775 | 0.2125 | 0.874632 | 0.132419 | 0.356759 | 1 | 0.996511 | 0.855416 |
| Mapre2 | Q8R001 | 2.69 | 1.73 | 1.27 | 0.64 | 1.65 | 2.65 | 1.54 | 1.22 | 1.04 | -1 | 1.29 | 2.98 | 0.77 | -0.43 | 1.15 | 1.18 | 0.1825 | -0.505 | -0.915 | 0.710627 | 0.54865 | 0.112373 | 1 | 0.996511 | 0.714111 |
| Mapre2 | E9Q6X0 | 1.21 | 0.93 | -0.47 | -0.24 | 0.72 | 1.41 | 0.22 | -0.3 | 0.33 | 0.55 | -0.21 | 1.62 | -0.48 | -0.04 | -0.28 | 2.22 | 0.155 | 0.215 | -0.0025 | 0.761978 | 0.683697 | 0.997086 | 1 | 0.996511 | 0.999279 |
| Mapre3 | D3Z6G3 | -0.29 | 0.19 | 0.11 | -0.29 | -0.73 | -0.44 | -0.74 | -0.21 | 0.44 | 1.26 | 0.18 | -0.15 | 1.06 | -0.16 | 0.81 | 0.2 | -0.46 | 0.5025 | 0.5475 | 0.052198 | 0.149724 | 0.10951 | 1 | 0.996511 | 0.709362 |
| Mapre3 | Q6PER3 | 0.41 | 0.65 | 1.01 | 0.83 | 0.81 | 0.06 | 1.26 | 1.42 | -0.14 | 0.38 | 1.16 | 0.83 | -0.44 | 0.43 | 0.85 | 1.02 | 0.1625 | -0.1675 | -0.26 | 0.616934 | 0.594693 | 0.462863 | 1 | 0.996511 | 0.888267 |
| Mapt | P10637-5 | -1.49 | -0.8 | -2.3 | -2.13 | -1.43 | -1.76 | -2.23 | -1.93 | -0.36 | -1.84 | -1.33 | -1.45 | 0.25 | 0.14 | -1.66 | -1.46 | -0.1575 | 0.435 | 0.9975 | 0.666042 | 0.334778 | 0.104608 | 1 | 0.996511 | 0.702979 |
| Mapt | P10637-4 | 0.99 | 0.95 | 1.59 | 1.17 | 1.47 | 0.6 | 1.93 | 1.65 | 1.79 | 2.73 | 2.12 | 1.89 | 1.01 | 2.82 | 1.99 | 1.08 | 0.2375 | 0.9575 | 0.55 | 0.459628 | 0.006394 | 0.222827 | 1 | 0.996511 | 0.787447 |
| Mapt | A2A5Y6 | 0.31 | 0.64 | 0.59 | 0.72 | 0.2 | 0.18 | 0.64 | 0.63 | 0.65 | 1.34 | 0.65 | 0.67 | 0.64 | 0.46 | 0.89 | 0.29 | -0.1525 | 0.2625 | 0.005 | 0.438346 | 0.265395 | 0.981218 | 1 | 0.996511 | 0.994858 |
| Mapt | P10637-2 | -1.71 | 0.9 | -1.48 | -0.66 | -1.79 | -0.34 | -0.05 | 1.27 | -2.09 | -0.54 | 0.27 | -0.71 | -1.5 | 0.14 | -1.11 | -0.51 | 0.51 | -0.03 | -0.0075 | 0.51731 | 0.965662 | 0.990501 | 1 | 0.996755 | 0.996154 |
| Marchf1 | D3YVY0 | 0.55 | 1.54 | 0.6 | 0.39 | 1 | 0.55 | 1.12 | 0.56 | 0.72 | 2.2 | 0.15 | 1.48 | 0.53 | -0.89 | 0.27 | 1.33 | 0.0375 | 0.3675 | -0.46 | 0.899737 | 0.454238 | 0.363157 | 1 | 0.996511 | 0.858475 |
| Marchf5 | Q3KNM2 | -0.71 | -0.74 | -1.1 | -0.99 | -0.5 | -0.26 | -1.01 | -1.07 | -0.82 | -0.68 | -1.08 | -0.58 | -0.61 | -0.61 | -1.19 | -1.02 | 0.175 | 0.095 | 0.0275 | 0.465659 | 0.63087 | 0.902118 | 1 | 0.996511 | 0.98447 |
| Marchf6 | Q6ZQ89 | -0.57 | -0.61 | 0.48 | -1.37 | 0.54 | -0.73 | 1.01 | -1.11 | -1.45 | -0.94 | -1.05 | -0.75 | -1.06 | -0.14 | 0.49 | 0.45 | 0.445 | -0.53 | 0.4525 | 0.448836 | 0.199012 | 0.368008 | 1 | 0.996511 | 0.859245 |
| Marchf7 | Q9WV66 | -1.31 | -0.81 | 0.49 | -0.57 | -1.44 | -1.49 | -0.62 | -1.3 | 0.36 | 1.58 | 0.14 | -1.07 | 1.67 | -0.79 | 0.53 | -0.05 | -0.6625 | 0.8025 | 0.89 | 0.12893 | 0.208764 | 0.157206 | 1 | 0.996511 | 0.751678 |
| Marcks | P26645 | -0.67 | 0.23 | 0.51 | 0.36 | -0.07 | -0.4 | 0.18 | 0.59 | -0.46 | 1.17 | 0.83 | -0.41 | 0.2 | 0.34 | 1 | 0.22 | -0.0325 | 0.175 | 0.3325 | 0.921118 | 0.707974 | 0.327319 | 1 | 0.996511 | 0.844633 |
| Marcksl1 | P28667 | 1.01 | 0.82 | 0.52 | 0.49 | 1.13 | 1.29 | 0.57 | 1.11 | 2.37 | 1.54 | 1.24 | 1.98 | 2.51 | 1.91 | 2.03 | 0.66 | 0.315 | 1.0725 | 1.0675 | 0.18187 | 0.00469 | 0.024064 | 1 | 0.996511 | 0.626254 |
| Mark1 | Q8VHJ5 | -0.35 | -0.16 | -0.08 | -0.31 | -0.32 | -0.11 | -0.67 | -0.36 | 0.51 | 0.28 | -0.15 | 0.02 | 1.04 | -0.43 | 0.14 | -0.54 | -0.14 | 0.39 | 0.2775 | 0.440383 | 0.084012 | 0.450415 | 1 | 0.996511 | 0.884817 |
| Mark2 | E9QMP6 | -0.13 | -0.38 | 0.45 | 0.3 | -0.63 | 0.27 | 0.08 | 0.09 | -0.27 | 0.25 | 0.26 | 0 | 0.02 | -0.68 | 0 | 0.05 | -0.1075 | 9.08E-17 | -0.2125 | 0.701331 | 1 | 0.456139 | 1 | 1 | 0.887169 |
| Mark3 | Q03141 | 0.06 | 1.31 | 0.88 | 0.07 | 0.18 | -0.69 | -0.79 | 0.08 | -0.55 | 0.16 | 0.52 | -0.05 | 0.88 | -1.29 | 0.2 | -0.11 | -0.885 | -0.56 | -0.66 | 0.04247 | 0.15658 | 0.218184 | 1 | 0.996511 | 0.787447 |
| Mark3 | A0A1Y7VN | -0.17 | -0.12 | 0.79 | 0.69 | -0.58 | -0.05 | 0.26 | 0.9 | 0.79 | 0.37 | 0.9 | 0.34 | 1.59 | -0.34 | 1.01 | 0.13 | -0.165 | 0.3025 | 0.3 | 0.666643 | 0.327585 | 0.530409 | 1 | 0.996511 | 0.908674 |
| Mark4 | Q8CIP4-2 | -0.38 | 0.18 | 0.57 | 0.3 | 0.06 | -0.2 | 0.53 | 0.58 | 0.52 | 0.14 | 0.6 | 0.22 | 1.08 | 0.6 | 0.39 | 0.35 | 0.075 | 0.2025 | 0.4375 | 0.787691 | 0.430008 | 0.145641 | 1 | 0.996511 | 0.745321 |
| Mars1 | E9QB02 | -1.42 | -1.74 | -2.03 | -1.98 | -1.02 | -1.43 | -1.74 | -1.91 | -1.74 | -1.03 | -1.69 | -1.17 | -1.61 | -1.06 | -1.36 | -1.79 | 0.2675 | 0.385 | 0.3375 | 0.303872 | 0.150861 | 0.194317 | 1 | 0.996511 | 0.778959 |
| Mars2 | Q499X9 | 0.33 | 0.49 | 0.83 | 0.74 | 0.3 | 0.07 | 0.57 | 0.92 | -0.04 | 0.41 | 1.03 | 0.53 | -0.16 | -0.4 | 0.29 | 0.64 | -0.1325 | -0.115 | -0.505 | 0.574328 | 0.666188 | 0.098016 | 1 | 0.996511 | 0.698156 |
| Mas1 | P30554 | 1.68 | 1.56 | 2.41 | 1.69 | -0.03 | 0.51 | 1.5 | 2.77 | -0.71 | 2.1 | 2.73 | 0.83 | -0.56 | -1.88 | 2.26 | 2.34 | -0.6475 | -0.5975 | -1.295 | 0.288824 | 0.411447 | 0.199392 | 1 | 0.996511 | 0.78004 |
| Masp1 | A0A2R8VH | -4.26 | -4.31 | -3.11 | -3.98 | -6.3 | -4.33 | -5.05 | -4.24 | -2.12 | -1.02 | -3.34 | -4.12 | -1.07 | -4.38 | -2.64 | -2.29 | -1.065 | 1.265 | 1.32 | 0.061908 | 0.086856 | 0.075972 | 1 | 0.996511 | 0.674054 |
| Mast1 | Q9R1L5 | 0.62 | 0.65 | 0.6 | 0.47 | 0.44 | 1 | 0.2 | 0.62 | 0.7 | -0.08 | 0.47 | 1.02 | 0.89 | -0.37 | 0.58 | 0.43 | -0.02 | -0.0575 | -0.2025 | 0.921882 | 0.822392 | 0.489124 | 1 | 0.996511 | 0.897531 |
| Mast2 | B1AST8 | -0.21 | 0.1 | 1.09 | 1.55 | -0.38 | 1.03 | 1.53 | 1.23 | 1.87 | 1.81 | 1.2 | 0.08 | 1.13 | 0.65 | 0.43 | 1.85 | 0.22 | 0.6075 | 0.3825 | 0.686307 | 0.279174 | 0.439089 | 1 | 0.996511 | 0.883205 |
| Mast3 | A0A1D5RN | -0.19 | 0.31 | 0.46 | 0.17 | 0 | -0.41 | -0.09 | -0.19 | 0.51 | 0.26 | 0.23 | 0.17 | 0.68 | 0.54 | 0.36 | -0.08 | -0.36 | 0.105 | 0.1875 | 0.097388 | 0.610189 | 0.460216 | 1 | 0.996511 | 0.887902 |
| Mast4 | A0A0A6YX( | 0.25 | 0.51 | 1.14 | 0.89 | 0.18 | 0.36 | -0.15 | 1.24 | 0.61 | 1.28 | 1.13 | 0.59 | 1.27 | 0.24 | 0.84 | 0.92 | -0.29 | 0.205 | 0.12 | 0.409097 | 0.468388 | 0.693377 | 1 | 0.996511 | 0.955729 |
| Mat2a | Q3THS6 | -0.51 | -0.84 | -0.37 | -0.48 | -0.78 | -0.89 | 0.02 | -0.2 | -1.76 | -0.53 | -0.42 | -0.68 | -1.52 | -1.54 | -0.75 | -0.78 | 0.0875 | -0.2975 | -0.5975 | 0.730963 | 0.370887 | 0.04971 | 1 | 0.996511 | 0.658454 |
| Mat2b | Q99LB6-2 | -0.59 | -0.73 | -0.62 | -0.13 | -0.76 | -0.55 | -0.04 | 0.11 | -0.51 | -0.42 | -0.25 | -0.51 | -0.87 | -0.23 | -0.71 | -0.93 | 0.2075 | 0.095 | -0.1675 | 0.425535 | 0.632578 | 0.497729 | 1 | 0.996511 | 0.898105 |
| Matk | A0A0R4J1P | -0.46 | -0.68 | -0.41 | -0.65 | -0.61 | -0.09 | -0.9 | -0.66 | 0.01 | -0.01 | -0.35 | -0.4 | 0.1 | -0.03 | -0.42 | -0.63 | -0.015 | 0.3625 | 0.305 | 0.943261 | 0.080186 | 0.203851 | 1 | 0.996511 | 0.782817 |
| Matn2 | A0A0A0M( | 0.48 | 0.37 | -0.5 | -1 | 0.71 | 1.06 | -0.05 | 0.34 | 0.5 | -0.18 | 0.21 | 0.69 | 0 | 0.62 | 0.07 | 0.04 | 0.6775 | 0.4675 | 0.345 | 0.120473 | 0.246886 | 0.369583 | 1 | 0.996511 | 0.859245 |
| Matn4 | F2Z468 | -0.46 | -0.91 | -0.75 | -0.58 | 0.41 | -0.18 | 0.28 | 0.12 | -0.18 | -0.08 | -0.02 | 0.34 | -0.34 | 0.21 | 0.74 | 0.45 | 0.8325 | 0.69 | 0.94 | 0.002228 | 0.007091 | 0.007062 | 1 | 0.996511 | 0.552614 |
| Matr3 | A0A494B9( | -3.29 | -2.91 | -3.78 | -7.17 | -2.03 | -3.49 | -4.67 | -6.11 | -3.74 | -4.48 | -5.01 | -4.16 | -3.6 | -2.46 | -4.89 | -4.33 | 0.2125 | -0.06 | 0.4675 | 0.855776 | 0.947516 | 0.639538 | 1 | 0.996755 | 0.942804 |
| Matr3 | Q8K310 | 0.26 | -1.42 | -0.71 | -0.73 | -1.74 | -0.29 | -0.97 | -0.1 | -1.1 | -1.09 | -0.72 | -1.55 | -0.12 | -1.37 | 0.19 | -1.19 | -0.125 | -0.465 | 0.0275 | 0.790748 | 0.23297 | 0.954742 | 1 | 0.996511 | 0.992482 |
| Mau2 | Z4YNA3 | -0.67 | -0.85 | -0.57 | -0.99 | -0.28 | -0.68 | -0.66 | -1.11 | -0.86 | -1.37 | -0.95 | -1.09 | -0.99 | -0.41 | -0.67 | -0.44 | 0.0875 | -0.2975 | 0.1425 | 0.689403 | 0.157001 | 0.516072 | 1 | 0.996511 | 0.905674 |
| Mavs | Q8VCF0 | -0.48 | -0.36 | -0.42 | -0.05 | -0.85 | -0.34 | -0.22 | -0.56 | 0.5 | -0.88 | -0.21 | -0.43 | 0.66 | -0.79 | -0.79 | -0.27 | -0.165 | 0.0725 | 0.03 | 0.420645 | 0.813478 | 0.932034 | 1 | 0.996511 | 0.989713 |
| Max | P28574 | -0.15 | -0.88 | -0.57 | -1 | -0.43 | -0.32 | -0.98 | -1.3 | -0.26 | -0.25 | -0.97 | -0.6 | -0.22 | -0.36 | 0.13 | -0.41 | -0.1075 | 0.13 | 0.435 | 0.718703 | 0.633026 | 0.116716 | 1 | 0.996511 | 0.718608 |
| Mb21d2 | D3Z742 | 0.17 | 0.45 | 0.36 | 0.39 | 0.51 | 0 | 0.07 | 0.42 | 0.58 | 0.48 | 0.6 | 0.13 | -0.06 | 0.77 | -0.02 | 0.18 | -0.0925 | 0.105 | -0.125 | 0.615867 | 0.575078 | 0.604902 | 1 | 0.996511 | 0.932185 |
| Mbd1 | A0A0R4J15 | 1.84 | 0.34 | 1.06 | 1.05 | 0.25 | 1.67 | 0.25 | 0.78 | 1.5 | -0.43 | 0.64 | 0.65 | 2.7 | -1.42 | 1 | 1.31 | -0.335 | -0.4825 | -0.175 | 0.439111 | 0.318055 | 0.830838 | 1 | 0.996511 | 0.973939 |
| Mbd2 | Q9Z2E1 | -0.67 | -2.48 | -0.91 | -1.13 | -1.71 | -1.76 | -0.76 | -1.26 | -1.7 | -0.01 | -1.1 | -1.3 | -1.62 | -1.09 | -0.86 | -0.46 | -0.075 | 0.27 | 0.29 | 0.863549 | 0.595503 | 0.521237 | 1 | 0.996511 | 0.907667 |
| Mbd3 | Q9Z2D8 | -2.87 | -3.83 | -2.14 | -1.97 | -4.04 | -3.52 | -1.81 | -2.32 | -1.95 | -2.09 | -2.18 | -2.71 | -2.06 | -2.94 | -1.66 | -1.69 | -0.22 | 0.47 | 0.615 | 0.718731 | 0.291754 | 0.225952 | 1 | 0.996511 | 0.787447 |
| Mblac1 | Q8BWY4 | -0.36 | -0.34 | -0.3 | -0.33 | -0.2 | -0.24 | -0.44 | -0.33 | 0.21 | 0.08 | -0.21 | -0.16 | 0.51 | 0.06 | -0.11 | -0.15 | 0.03 | 0.3125 | 0.41 | 0.834084 | 0.099516 | 0.077669 | 1 | 0.996511 | 0.676942 |
| Mblac2 | Q8BL86 | 0.47 | 0.38 | 0.77 | 0.93 | 0.76 | 0.59 | 0.62 | 0.81 | 0.13 | 0.32 | 0.78 | 0.38 | 0.14 | 0.25 | 0.64 | 0.88 | 0.0575 | -0.235 | -0.16 | 0.753097 | 0.307979 | 0.52498 | 1 | 0.996511 | 0.908107 |
| Mbnl1 | A0A0A6YXI | 10.41 | 1.59 | -2.79 | -6.8 | 5.53 | 9.93 | -1.23 | -3.81 | -3.1 | -4.11 | -1.63 | 9.79 | -3.71 | -4.17 | -0.89 | -1.02 | 2.0025 | -0.365 | -3.05 | 0.644074 | 0.933188 | 0.374411 | 1 | 0.996755 | 0.859424 |
| Mbnl2 | Q8C181-4 | -1.42 | -2.93 | -1.51 | -3.11 | -2.68 | -1.95 | -0.01 | -2.54 | -2.95 | -2.17 | -0.91 | -1.84 | -2.58 | -3.09 | -0.7 | -0.95 | 0.4475 | 0.275 | 0.4125 | 0.522608 | 0.630714 | 0.546051 | 1 | 0.996511 | 0.912321 |
| Mboat2 | A0A217FL6 | 0.09 | 0.93 | 0.85 | 0.56 | 0.95 | 0.58 | 1.27 | 0.57 | 0.57 | 1.08 | 0.87 | 0.63 | 0.91 | 0.44 | 1.1 | 1.49 | 0.235 | 0.18 | 0.3775 | 0.38061 | 0.473846 | 0.231697 | 1 | 0.996511 | 0.788863 |
| Mboat7 | Q8CHK3 | -0.35 | -0.19 | 0.08 | 0.71 | -0.66 | 0.04 | 0.01 | 1.04 | 2.49 | 0.84 | 0.22 | -0.32 | 3.6 | -0.73 | 1.46 | 0.47 | 0.045 | 0.745 | 1.1375 | 0.909753 | 0.233839 | 0.203958 | 1 | 0.996511 | 0.782817 |
| Mbp | P04370-11 | 5.42 | 1.81 | 3.47 | 4.7 | 1.03 | 3 | 2.73 | 4.51 | 2.3 | 0.2 | 2.51 | 4.2 | 0.7 | -1.05 | 2.96 | 2.64 | -1.0325 | -1.5475 | -2.5375 | 0.296726 | 0.156331 | 0.042426 | 1 | 0.996511 | 0.658454 |
| Mbp | P04370-4 | 2.82 | 2.06 | 1.65 | 1 | 2.36 | 2.23 | 1.24 | 1.26 | 0.13 | 0.51 | 1.34 | 2.91 | 1.06 | -2.01 | 2.16 | 1.94 | -0.11 | -0.66 | -1.095 | 0.808252 | 0.332296 | 0.257108 | 1 | 0.996511 | 0.811777 |
| Mbp | P04370 | 1.75 | 1.45 | 1.24 | 1.45 | 1.31 | 1.64 | 2.33 | 1.59 | 0.78 | 1.76 | 1.85 | 1.99 | 0.73 | 1.93 | 2.05 | 2.09 | 0.245 | 0.1225 | 0.2275 | 0.344731 | 0.685021 | 0.510829 | 1 | 0.996511 | 0.90359 |
| Mbp | P04370-10 | -0.13 | -2.16 | 0.61 | 0.97 | -2.68 | -1.84 | 1.42 | 1.07 | 5.97 | -1.31 | 1.44 | -1.12 | 7.67 | -5.18 | 0.31 | 0.76 | -0.33 | 1.4225 | 1.0675 | 0.767465 | 0.395313 | 0.659464 | 1 | 0.996511 | 0.949645 |
| Mbtps2 | Q8CHX6 | -4.46 | -5.29 | -5.7 | -5.19 | -4.92 | -4.05 | -4.21 | -4.61 | -4.14 | -1.53 | -6.93 | -4.54 | -2.85 | -4.03 | -3.1 | -3.59 | 0.7125 | 0.875 | 1.7675 | 0.048971 | 0.401211 | 0.00093 | 1 | 0.996511 | 0.3852 |
| Mcam | Q8R2Y2 | 0.25 | 0.44 | 0.76 | -0.16 | 0.39 | 0.13 | 0.97 | 0.12 | -0.32 | -0.94 | 0.8 | 0.43 | -0.12 | -0.5 | 1.08 | 1.18 | 0.08 | -0.33 | 0.0875 | 0.775309 | 0.43239 | 0.842684 | 1 | 0.996511 | 0.976225 |
| Mcat | Q8R3F5 | 0.51 | 0.08 | 0.2 | 0.33 | -0.22 | 0.29 | 0.75 | 0.13 | -0.04 | 0.3 | 0.43 | 0.35 | 0.27 | 0.12 | 0.24 | 0.43 | -0.0425 | -0.02 | -0.015 | 0.858156 | 0.917473 | 0.936952 | 1 | 0.996755 | 0.989767 |
| Mcc | G3UW40 | -0.33 | 0.25 | 0.19 | -0.01 | 0.05 | -0.05 | 0.27 | 0.01 | 0.19 | -0.04 | 0.47 | 0.18 | 0.97 | -0.67 | 0.42 | 0.04 | 0.045 | 0.175 | 0.165 | 0.811004 | 0.414771 | 0.650898 | 1 | 0.996511 | 0.947407 |
| Mccc1 | Q99MR8 | 0.54 | 0.54 | 0.72 | 0.76 | 0.53 | 0.46 | 0.99 | 0.94 | 0.29 | 0.03 | 1.13 | 0.35 | 0.07 | 0.44 | 0.52 | 0.9 | 0.09 | -0.19 | -0.1575 | 0.634905 | 0.475985 | 0.493099 | 1 | 0.996511 | 0.897531 |
| Mccc2 | Q3ULD5 | 0.78 | 0.63 | 0.7 | 1 | 0.35 | 0.7 | 0.73 | 0.97 | 0.45 | 0.69 | 0.94 | 0.45 | 0.31 | 0.14 | 0.79 | 1.04 | -0.09 | -0.145 | -0.2075 | 0.637201 | 0.464547 | 0.424469 | 1 | 0.996511 | 0.876965 |
| Mcee | Q9D1I5 | -1.38 | -0.86 | -1.53 | -0.82 | -1.86 | -1 | -1.1 | -1.13 | -0.88 | -2.38 | -1.03 | -1.45 | -0.84 | -1.44 | -2.09 | -1.75 | -0.125 | -0.2875 | -0.3825 | 0.648532 | 0.445934 | 0.259176 | 1 | 0.996511 | 0.812074 |
| Mcf2l | E9PXE2 | 1.53 | 1.01 | 0.86 | 0.55 | 1.09 | 1.03 | 0.45 | 0.42 | 0.78 | 0.42 | 0.69 | 1.42 | 0.93 | 1.21 | 1.06 | 0.59 | -0.24 | -0.16 | -0.04 | 0.396823 | 0.597117 | 0.881636 | 1 | 0.996511 | 0.979767 |
| Mcfd2 | Q8K5B2 | -1.77 | -0.39 | 0.06 | -0.47 | 1.03 | -0.85 | 0.24 | -0.77 | -3.43 | -0.33 | -0.83 | -1.39 | -2.19 | -1.98 | 0.03 | 0.39 | 0.555 | -0.8525 | -0.295 | 0.324614 | 0.252171 | 0.676151 | 1 | 0.996511 | 0.952944 |
| Mchr1 | Q8JZL2 | 1.23 | 0.74 | 0.69 | 1.37 | -0.01 | 1 | -2.26 | -0.63 | 1.49 | -0.55 | -0.21 | 0.52 | 3.07 | -0.77 | 0.05 | 0.04 | -1.4825 | -0.695 | -0.41 | 0.043524 | 0.152085 | 0.601498 | 1 | 0.996511 | 0.930802 |
| Mcl1 | P97287 | 1.3 | 1.37 | -0.6 | -0.25 | 0.45 | -0.08 | -0.63 | -0.39 | 0.84 | -0.17 | -0.52 | 0.34 | 1.07 | 0.94 | -0.36 | -0.79 | -0.6175 | -0.3325 | -0.24 | 0.25355 | 0.547801 | 0.705268 | 1 | 0.996511 | 0.957084 |
| Mcm3ap | Q9WUU9 | 0.61 | 0.41 | 1.64 | 2.09 | 0.13 | 0.37 | 0.67 | 1.88 | 0.79 | 2.39 | 1.04 | 0.15 | 0.74 | 0.41 | 1.92 | 2.03 | -0.425 | -0.095 | 0.0875 | 0.419407 | 0.867823 | 0.869594 | 1 | 0.996511 | 0.979293 |
| Mcm4 | P49717 | -0.64 | -0.93 | -1.45 | -1.89 | -0.03 | -0.7 | -2.11 | -0.85 | -0.45 | -2.15 | -1.45 | -1.22 | -0.08 | 0.09 | -0.94 | 1.23 | 0.305 | -0.09 | 1.3025 | 0.527468 | 0.831758 | 0.024914 | 1 | 0.996511 | 0.63249 |
| Mcmbp | H9KV04 | -1.74 | -2.38 | -1.64 | -0.74 | -2.13 | -1.94 | -1.59 | -1.24 | -3.07 | 0.65 | -1.49 | -1.91 | -2.59 | -2.77 | -1.27 | -1.95 | -0.1 | 0.17 | -0.52 | 0.788294 | 0.82472 | 0.269164 | 1 | 0.996511 | 0.820534 |
| Mcoln1 | Q99J21 | 0.98 | 0.55 | 0.83 | -0.78 | 0.4 | 1.38 | 1.84 | 0.2 | 1.13 | -0.14 | 0.71 | 1.02 | 1.35 | 0.69 | 1.25 | 1.29 | 0.56 | 0.285 | 0.75 | 0.295012 | 0.542659 | 0.09629 | 1 | 0.996511 | 0.698156 |
| Mcrip1 | Q3UGS4 | -1.76 | -1.71 | -1.52 | -2.03 | -1.42 | -2.33 | -1.74 | -2.56 | -1.36 | -1.6 | -1.67 | -1.82 | -1.11 | -0.88 | -2.46 | -2.47 | -0.2575 | 0.1425 | 0.025 | 0.376968 | 0.472551 | 0.952507 | 1 | 0.996511 | 0.992476 |
| Mcrip2 | A0A494BB( | -3.68 | -5.31 | 0.12 | 0.59 | -3.14 | -2.61 | 0 | 0.52 | -3.62 | 0.81 | -0.33 | -3.03 | -1.73 | -3.29 | 0.43 | 0.84 | 0.7625 | 0.5275 | 1.1325 | 0.619694 | 0.741984 | 0.46936 | 1 | 0.996511 | 0.890107 |
| Mcrs1 | Q3TJY1 | 0.08 | -0.11 | -0.08 | -0.69 | 0.95 | 0.01 | -0.74 | -0.17 | -0.49 | 0.42 | -0.42 | -0.31 | -1.14 | 0.13 | -0.2 | -0.29 | 0.2125 | 7.85E-17 | -0.175 | 0.5704 | 1 | 0.592068 | 1 | 1 | 0.929843 |
| Mctp1 | E9PV86 | 0.67 | 0.3 | 0.44 | 0.49 | 0.49 | 0.69 | 0.49 | 0.55 | 0.62 | 0.32 | 0.28 | 0.45 | 0.79 | 0.21 | 0.8 | 0.41 | 0.08 | -0.0575 | 0.0775 | 0.611665 | 0.746344 | 0.722327 | 1 | 0.996511 | 0.958398 |
| Mcts1 | Q9DB27 | 0.01 | -0.09 | 0.09 | -0.04 | 0.2 | 0.29 | 0.63 | 0.48 | -0.42 | -0.48 | 0.13 | 0.19 | -0.75 | -0.23 | 0.09 | 0.38 | 0.4075 | -0.1375 | -0.12 | 0.032112 | 0.534646 | 0.658082 | 1 | 0.996511 | 0.949013 |
| Mcu | Q3UMR5 | 0.9 | 0.72 | 0.81 | 1.35 | 0.25 | 0.74 | 0.76 | 1.17 | 0.94 | 1.45 | 1 | 0.97 | 0.17 | 0.91 | 0.33 | 0.37 | -0.215 | 0.145 | -0.5 | 0.395565 | 0.517606 | 0.069983 | 1 | 0.996511 | 0.668066 |
| Mcub | Q810S1 | -1.15 | -0.87 | 1.54 | 1.28 | -0.98 | -0.14 | 1.44 | 1.9 | -0.2 | -0.42 | 1.8 | -0.68 | -1.45 | -1.76 | 0.52 | 0.89 | 0.355 | -0.075 | -0.65 | 0.686333 | 0.926698 | 0.464644 | 1 | 0.996755 | 0.888267 |
| Mcur1 | Q9CXD6 | 0.38 | 0.48 | 0.57 | 1 | 0.31 | 0.51 | 0.81 | 0.52 | 0.78 | 0.71 | 0.94 | 0.42 | 1.12 | 0.61 | 0.58 | 0.59 | -0.07 | 0.105 | 0.1175 | 0.730414 | 0.6271 | 0.616087 | 1 | 0.996511 | 0.93691 |
| Mdc1 | Q5PSV9 | 10.55 | 7.82 | -0.5 | -1 | 8.62 | 10.15 | -3.33 | -2.45 | 2.98 | 1.44 | -3.79 | 9.86 | 1.95 | 2.47 | -2.33 | 1.45 | -0.97 | -1.595 | -3.3325 | 0.813084 | 0.660159 | 0.247104 | 1 | 0.996511 | 0.803237 |
| Mdga1 | D3Z499 | 0.39 | 0.26 | 0.12 | 0.38 | -0.16 | 0.56 | -0.4 | 0.2 | 0.37 | 0.67 | 0.53 | 0.81 | 0.15 | -0.58 | 0.49 | -0.23 | -0.2375 | 0.3075 | -0.33 | 0.329314 | 0.114944 | 0.235947 | 1 | 0.996511 | 0.791493 |
| Mdga2 | Q3UV20 | 0.52 | 1.01 | 0.8 | 0.97 | 0.64 | 0.87 | 0.59 | 0.91 | 0.49 | 0.83 | 1 | 0.67 | 0.55 | 0.77 | 1.34 | 0.22 | -0.0725 | -0.0775 | -0.105 | 0.690767 | 0.703989 | 0.709363 | 1 | 0.996511 | 0.957656 |
| Mdh1 | P14152 | -0.48 | -0.53 | 0.66 | 1.06 | -0.61 | -0.88 | 0.3 | 0.93 | -0.55 | 0.93 | 0.46 | -0.59 | -0.81 | -0.25 | 0.29 | 0.11 | -0.2425 | -0.115 | -0.3425 | 0.650442 | 0.822437 | 0.449343 | 1 | 0.996511 | 0.884817 |

| | | | | | | | | | | | | | | | | | | | | | | | | | | |
|---|---|---|---|---|---|---|---|---|---|---|---|---|---|---|---|---|---|---|---|---|---|---|---|---|---|---|
| Mdh2 | P08249 | -0.46 | -0.82 | 0.01 | 0.23 | -0.86 | -0.27 | 0.03 | 0.14 | -0.85 | 0.6 | -0.06 | -0.6 | -0.86 | -0.64 | -0.4 | -0.24 | 0.02 | 0.0325 | -0.275 | 0.949862 | 0.932424 | 0.354292 | 1 | 0.996755 | 0.855029 |
| Mdk | P12025 | -2.01 | -1.95 | -2.3 | -2.9 | -2.3 | -0.85 | -0.32 | -2.89 | -0.76 | -1.08 | -2.63 | -1.12 | 1.15 | -3.77 | -1.56 | -1.86 | 0.7 | 0.8925 | 0.78 | 0.251184 | 0.071376 | 0.410593 | 1 | 0.996511 | 0.874811 |
| Mdm1 | E9Q0B1 | -0.68 | -0.37 | -0.29 | -0.32 | -1.66 | -0.5 | 0.16 | -0.71 | 0.12 | 0.35 | -0.27 | 0.13 | 1.5 | 0.49 | -0.12 | -0.23 | -0.2625 | 0.4975 | 0.825 | 0.48329 | 0.035388 | 0.061424 | 1 | 0.996511 | 0.662517 |
| Mdm4 | A0A087WF | -1.71 | -0.82 | -1.03 | -0.12 | -1.4 | -1.67 | -1.28 | -1.39 | 0.17 | 0.69 | -0.35 | -0.85 | 1.3 | 0.66 | -0.6 | -0.74 | -0.515 | 0.835 | 1.075 | 0.143788 | 0.085132 | 0.07659 | 1 | 0.996511 | 0.675786 |
| Mdn1 | J3QMC5 | -2.04 | -2.89 | -0.85 | -0.53 | -3.17 | -2.72 | -0.5 | -0.87 | 0.64 | -0.41 | -0.63 | -2.49 | 1.21 | -2.91 | -0.81 | -0.25 | -0.2375 | 0.855 | 0.8875 | 0.759634 | 0.283632 | 0.342188 | 1 | 0.996511 | 0.848473 |
| Mdp1 | Q9D967 | -0.49 | -0.11 | -0.34 | -0.77 | -0.54 | -0.7 | -0.34 | 0.3 | -0.14 | -0.56 | 0.11 | -0.81 | 0.14 | -0.28 | 0.95 | -0.68 | 0.1075 | 0.0775 | 0.46 | 0.687658 | 0.770882 | 0.232192 | 1 | 0.996511 | 0.788863 |
| Me1 | P06801 | -0.49 | -0.41 | -0.36 | -0.73 | -0.28 | -0.46 | -0.89 | -0.66 | -0.31 | -0.28 | -0.58 | -0.37 | -0.91 | -0.24 | -0.92 | -0.62 | -0.075 | 0.1125 | -0.175 | 0.697373 | 0.528577 | 0.447796 | 1 | 0.996511 | 0.884817 |
| Me2 | Q99KE1 | 0.29 | 0.33 | -0.06 | 0.32 | -0.26 | -0.13 | -0.03 | -0.24 | 0.31 | 0.22 | -0.15 | 0.25 | 0.16 | 0.09 | -0.46 | -0.05 | -0.385 | -0.0625 | -0.285 | 0.042152 | 0.747935 | 0.215233 | 1 | 0.996511 | 0.787313 |
| Me3 | Q8BMF3 | 0.96 | 0.89 | 1.23 | 0.99 | 0.19 | 0.23 | 0.42 | 0.77 | 0.16 | 0.75 | 0.49 | 0.18 | 0.09 | 0.14 | 0.18 | -0.03 | -0.615 | -0.6225 | -0.9225 | 0.009977 | 0.013527 | 0.000649 | 1 | 0.996511 | 0.29579 |
| Mea1 | Q64327 | 1.74 | 1.89 | -0.78 | -0.06 | 1.23 | 1.31 | 0.14 | -0.51 | 1.42 | -1.8 | -0.57 | 2.46 | 1.54 | 0.82 | -0.63 | -0.59 | -0.155 | -0.32 | -0.4125 | 0.829602 | 0.760329 | 0.595082 | 1 | 0.996511 | 0.929843 |
| Meaf6 | V9GX42 | -0.62 | -1.16 | 0.03 | 0.03 | -0.84 | -1.24 | -0.25 | -0.54 | -0.92 | 0.36 | -0.38 | -0.87 | -0.5 | -0.05 | 0.61 | 0.03 | -0.2875 | -0.0225 | 0.4525 | 0.413488 | 0.954625 | 0.231581 | 1 | 0.996755 | 0.788863 |
| Meak7 | Q8K0P3 | -0.64 | -0.24 | -0.18 | -0.19 | -0.03 | -0.46 | 0.22 | -0.29 | 0.09 | -0.62 | -0.16 | -0.1 | -0.11 | 0.06 | 0.03 | -0.06 | 0.1725 | 0.115 | 0.2925 | 0.425373 | 0.607844 | 0.153441 | 1 | 0.996511 | 0.750763 |
| Mecp2 | Q9Z2D6-2 | 1.57 | 0.91 | 0.58 | 0.94 | 0.85 | 1.03 | 0.41 | 0.53 | 0.7 | 0.6 | 0.52 | 0.71 | 0.84 | 1.18 | 1.06 | 0.71 | -0.295 | -0.3675 | -0.0525 | 0.274752 | 0.150983 | 0.840552 | 1 | 0.996511 | 0.976225 |
| Mecr | Q9DCS3 | 1.96 | 1.16 | 0.8 | 0.72 | 0.66 | 2.11 | 1.02 | 1.04 | 0.31 | 0.29 | 0.93 | 1.8 | -0.02 | 0.27 | 0.2 | 0.73 | 0.0475 | -0.3275 | -0.865 | 0.905053 | 0.453824 | 0.025554 | 1 | 0.996511 | 0.633075 |
| Med1 | Q925J9 | -4.51 | -2.72 | -2.08 | -2.63 | -3.8 | -4.72 | -2.39 | -2.01 | -2.49 | -1.66 | -2.26 | -3.39 | -0.43 | -2.08 | -1.87 | -1.66 | -0.245 | 0.535 | 1.475 | 0.74176 | 0.373096 | 0.032243 | 1 | 0.996511 | 0.641974 |
| Med12 | A2AGH8 | -3.31 | -3.93 | -1.94 | -2.79 | -3.97 | -4.45 | -4.53 | -3.6 | -4 | -0.71 | -2.73 | -4.07 | -1.41 | -3.54 | -2.07 | -2.7 | -1.145 | 0.115 | 0.5625 | 0.027913 | 0.886427 | 0.337529 | 1 | 0.996511 | 0.847578 |
| Med13l | A0A0J9YUA | -2.16 | -10.36 | 0.72 | 0.5 | -3.15 | -1.86 | -2.09 | -0.84 | -1.66 | 0 | -0.6 | -2.66 | -3.89 | -2.76 | -2.86 | 0.7 | 0.84 | 1.595 | 0.6225 | 0.721549 | 0.506497 | 0.8005 | 1 | 0.996511 | 0.970568 |
| Med14 | A2ABV5 | -1.21 | -1.36 | -2.52 | 0.31 | -0.88 | -0.65 | 0.14 | -1.05 | -1.17 | 1.1 | 0 | -0.77 | -1.28 | -1.1 | -0.29 | -1.44 | 0.585 | 0.985 | 0.1675 | 0.329711 | 0.182 | 0.774129 | 1 | 0.996511 | 0.966062 |
| Med15 | A0A338P73 | 0.32 | 0.03 | 1.03 | 0.98 | 0.85 | -0.24 | 0.41 | 0.5 | 1.05 | 4.75 | 0.61 | -0.15 | 0.99 | 1.52 | 0.8 | 0.39 | -0.21 | 0.975 | 0.335 | 0.525635 | 0.345168 | 0.339333 | 1 | 0.996511 | 0.84777 |
| Med16 | E9QP84 | 0.27 | 0.99 | -1.23 | -0.83 | 1.5 | 0.35 | -0.5 | -0.48 | -1.29 | -1 | -1.1 | -0.2 | -0.69 | -0.51 | -0.91 | -0.81 | 0.4175 | -0.6975 | -0.53 | 0.513099 | 0.203467 | 0.288824 | 1 | 0.996511 | 0.824705 |
| Med17 | Q8VCD5 | -2.38 | -1.96 | -1.82 | -2.21 | -1.48 | -3.22 | -2.69 | -2.74 | -1.45 | -0.11 | -1.6 | -3.37 | -2.15 | -1.44 | -2.02 | -1.43 | -0.44 | 0.46 | 0.3325 | 0.256985 | 0.467202 | 0.21814 | 1 | 0.996511 | 0.787447 |
| Med19 | Q8C1S0 | -3.43 | -3.03 | -4.2 | -3.62 | -2.57 | -4.87 | -3.18 | -2.55 | -4.54 | -2.82 | -3.71 | -2.49 | -5.04 | -3.59 | -3.31 | -3.5 | 0.2775 | 0.18 | -0.29 | 0.614981 | 0.712461 | 0.516834 | 1 | 0.996511 | 0.905676 |
| Med22 | A2ALA4 | -1.97 | -1.27 | -1.13 | -0.7 | -1.59 | -1.55 | -1.72 | -2 | -0.16 | 0.38 | -1.28 | -1.69 | 0 | -0.25 | -1.34 | -0.81 | -0.4475 | 0.58 | 0.6675 | 0.142461 | 0.272941 | 0.11204 | 1 | 0.996511 | 0.713445 |
| Med23 | F8WJB0 | -1.38 | -0.47 | -0.67 | -1.61 | -0.74 | -1.83 | -1.52 | -1.21 | -2.17 | -0.11 | -1.34 | -0.84 | -2.12 | -1.95 | -0.61 | -0.44 | -0.2925 | -0.0825 | -0.2475 | 0.408277 | 0.863141 | 0.611766 | 1 | 0.996511 | 0.935047 |
| Med24 | A6PW47 | -2.82 | -1.85 | -0.79 | -1.5 | -1.8 | -2.61 | -0.29 | -1.32 | -1.72 | -2.73 | -0.92 | -2.67 | -0.22 | -0.93 | -1.19 | -1.05 | 0.235 | -0.27 | 0.8925 | 0.690012 | 0.629375 | 0.073165 | 1 | 0.996511 | 0.668161 |
| Med28 | Q920D3 | -4.57 | -4.17 | -2.57 | -3.09 | -3.13 | -3.86 | -3.72 | -3.17 | -4.3 | -0.28 | -2.66 | -3.44 | -3.56 | -2.34 | -3.1 | -3.56 | 0.13 | 0.93 | 0.46 | 0.779936 | 0.308755 | 0.377071 | 1 | 0.996511 | 0.860984 |
| Med29 | Q9DB91 | 0.59 | 1.6 | -0.36 | 0.31 | 0.26 | -0.93 | 0 | 0.32 | 0.45 | 0.32 | 1.03 | 0.44 | 0.74 | 0.34 | 0.23 | 0.08 | -0.6225 | 0.025 | -0.1875 | 0.202687 | 0.951982 | 0.65161 | 1 | 0.996755 | 0.947633 |
| Med30 | Q9CQI9 | -4.2 | -3.71 | -1.81 | -1.5 | -3.1 | -3.63 | -1.92 | -2.05 | 0.37 | -0.4 | -1.87 | -3.99 | -0.89 | 2.85 | -2.68 | -2.16 | 0.13 | 1.3325 | 2.085 | 0.855919 | 0.227014 | 0.125824 | 1 | 0.996511 | 0.721505 |
| Med4 | Q9CQA5-2 | -3.25 | -3.22 | -1.48 | -1.87 | -3.15 | -3.08 | -1.88 | -1.93 | -2.94 | -0.34 | -1.71 | -2.91 | -2.23 | -2.07 | -1.61 | -1.06 | -0.055 | 0.48 | 0.7125 | 0.91713 | 0.497246 | 0.172961 | 1 | 0.996511 | 0.764376 |
| Med6 | Q921D4 | -5.84 | -5.4 | -4.5 | -7.06 | -2.83 | -6.98 | -4.7 | -6.36 | -5.37 | -3.77 | -5.88 | -5.27 | -3.97 | -4 | -5.13 | -4.1 | 0.4825 | 0.6275 | 1.4 | 0.618464 | 0.340014 | 0.030587 | 1 | 0.996511 | 0.641481 |
| Med8 | A2A9C6 | -2.03 | -2.27 | -3.5 | -2.79 | -4.33 | -0.8 | -5.91 | -3.04 | 1.92 | 0.2 | -2.23 | -2.02 | 4.06 | -2.96 | -2.32 | -3.29 | -0.8725 | 2.115 | 1.52 | 0.399 | 0.047132 | 0.346799 | 1 | 0.996511 | 0.851134 |
| Med9 | Q8VCS6 | -0.42 | -0.2 | -0.78 | -1.15 | -0.18 | -1.03 | -0.71 | -1.12 | -1.55 | -2.38 | -1.01 | -0.67 | -1.37 | -1.48 | -1.31 | -1.53 | -0.1225 | -0.765 | -0.785 | 0.680603 | 0.087978 | 0.011119 | 1 | 0.996511 | 0.563386 |
| Mef2c | Q8CFN5 | -2.31 | -2.46 | -4.76 | -5.19 | -3.63 | -2.12 | -4.64 | -5.19 | -1.64 | -2.8 | -3.81 | -1.22 | -1.1 | -1.78 | -4.13 | -4.78 | -0.215 | 1.3125 | 0.7325 | 0.813108 | 0.153566 | 0.488661 | 1 | 0.996511 | 0.897531 |
| Mef2d | E9QKT0 | -6.84 | -7.32 | -6.1 | -6.05 | -6.59 | -7.63 | -7.06 | -7.15 | -4.64 | -3.33 | -5.83 | -5.93 | -2.44 | -4.87 | -5.72 | -7.38 | -0.53 | 1.645 | 1.475 | 0.16406 | 0.02583 | 0.151145 | 1 | 0.996511 | 0.749304 |
| Megf10 | A0A140T8S | -1.65 | -1.53 | -0.99 | -1.07 | -2.03 | -2.2 | -0.34 | -1.26 | -0.36 | 0.39 | 0.18 | -0.96 | -0.22 | -1.64 | -0.59 | -0.18 | -0.1475 | 1.1225 | 0.6525 | 0.73015 | 0.008983 | 0.10545 | 1 | 0.996511 | 0.703417 |
| Megf6 | B1ASH4 | -2.09 | -1.48 | -3.03 | -2.07 | -2.06 | -1.76 | -1.18 | -4.2 | 0.01 | -2.3 | -1.57 | -1.17 | -0.11 | -0.94 | -1.23 | -2.95 | -0.1325 | 0.91 | 0.86 | 0.842134 | 0.117105 | 0.188751 | 1 | 0.996511 | 0.776843 |
| Megf8 | P60882 | -0.17 | -0.21 | -0.49 | -0.3 | 0.06 | -0.48 | -0.33 | -0.6 | 0.53 | -0.13 | -1.1 | 0.03 | 0.4 | 0.28 | 0.2 | 0.28 | -0.045 | 0.125 | 0.5825 | 0.81819 | 0.716407 | 0.009155 | 1 | 0.996511 | 0.563386 |
| Megf9 | Q8BH27 | -0.05 | 0.4 | 0.39 | 0.14 | 0.69 | 0.13 | 0.37 | 0.34 | 0.08 | -0.32 | 0.46 | 1.02 | 0.46 | -0.94 | 0.79 | 0.8 | 0.1625 | 0.09 | 0.0575 | 0.412947 | 0.770883 | 0.888448 | 1 | 0.996511 | 0.980661 |
| Meis2 | Q3TYM2 | -1.16 | -0.82 | -2.07 | -1.72 | -0.63 | -0.86 | -1.7 | -2.21 | -1.09 | -0.81 | -2.02 | -0.82 | -0.96 | -1.4 | -2.15 | -1.81 | 0.0925 | 0.2575 | -0.1375 | 0.830384 | 0.508831 | 0.712466 | 1 | 0.996511 | 0.958014 |
| Memo1 | Q91VH6 | -0.12 | 0.04 | -0.31 | -0.04 | 0.48 | -0.36 | 0.01 | -0.08 | -0.54 | -0.62 | -0.29 | -0.01 | -1.19 | -0.08 | -0.44 | -0.5 | 0.12 | -0.2575 | -0.445 | 0.581 | 0.227607 | 0.124816 | 1 | 0.996511 | 0.721448 |
| Men1 | F8WHD9 | 0.17 | -0.12 | -0.47 | -0.7 | -0.19 | -0.03 | -0.37 | -0.69 | -0.55 | -0.89 | -0.57 | -0.53 | 0.17 | -0.27 | -0.07 | -0.26 | -0.04 | -0.355 | 0.1725 | 0.872968 | 0.161698 | 0.497707 | 1 | 0.996511 | 0.898105 |
| Mepce | Q8K3A9 | -2.88 | -4.08 | -2.74 | -2.35 | -3.78 | -3.36 | -3.02 | -2.47 | -3.57 | -0.6 | -2.23 | -3.38 | -1.97 | -2.43 | -2.5 | -3.16 | -0.145 | 0.5675 | 0.4975 | 0.738972 | 0.430863 | 0.261308 | 1 | 0.996511 | 0.81412 |
| Mertk | Q60805 | 1.04 | 1.05 | 0.7 | 0.56 | 1.48 | 1.13 | 0.3 | 0.01 | 0.52 | 0.26 | 0.72 | 1 | 0.42 | -0.05 | 0.16 | 0.42 | -0.1075 | -0.2125 | -0.6 | 0.760298 | 0.369442 | 0.021668 | 1 | 0.996511 | 0.621926 |
| Mesd | Q9ERE7 | -0.3 | 0.03 | -0.93 | -1.2 | 0.49 | -0.29 | -0.2 | -0.97 | -1.61 | -0.75 | -1.04 | -0.25 | -0.23 | -0.78 | 0.67 | -0.06 | 0.3575 | -0.3125 | 0.5 | 0.370008 | 0.426261 | 0.229097 | 1 | 0.996511 | 0.787447 |
| Met | F8VQL0 | 1.76 | 0.99 | -0.89 | -1.24 | 1.91 | 2.26 | 0.47 | -0.65 | 2.6 | -1.96 | 0.02 | 2.39 | 3 | -0.43 | -2.06 | 0.32 | 0.8425 | 0.6075 | 0.0525 | 0.356295 | 0.604844 | 0.963284 | 1 | 0.996511 | 0.9927 |
| Metap1 | Q8BP48 | -1.09 | -1.85 | -1.58 | -2.11 | -1.56 | -1.27 | -0.7 | -1.95 | -1.65 | -1.68 | -1.31 | -1.49 | -1.44 | -1.56 | -1.4 | -1.6 | 0.2875 | 0.125 | 0.1575 | 0.396783 | 0.626467 | 0.540538 | 1 | 0.996511 | 0.912212 |
| Metap1d | Q9CPW9-2 | -0.48 | -0.5 | -0.38 | -0.57 | -0.6 | -0.74 | 0.44 | 0.06 | 0.89 | 1.53 | 0.16 | -0.2 | 0.16 | 2.24 | -0.05 | -0.74 | 0.2725 | 1.0775 | 0.885 | 0.349 | 0.01762 | 0.159363 | 1 | 0.996511 | 0.754511 |
| Metap2 | Q3UI33 | -0.21 | -0.57 | -1.59 | -1.64 | -0.49 | -0.09 | -1.19 | -1.82 | -0.62 | -1.49 | -1.81 | -0.08 | -0.72 | -0.27 | -1.59 | -1.76 | 0.105 | 0.0025 | -0.0825 | 0.829153 | 0.995998 | 0.862071 | 1 | 0.999603 | 0.977204 |
| Metap2 | M0QW48 | -2.44 | -2.63 | -2.58 | -3 | -3.77 | -2.55 | -2.53 | -2.99 | -3.44 | -0.5 | -2.1 | -2.3 | -3.29 | -0.5 | -2.52 | -3.91 | -0.2975 | 0.5775 | 0.1075 | 0.349223 | 0.323883 | 0.874787 | 1 | 0.996511 | 0.979626 |
| Mettl1 | Q9Z120 | -0.54 | -1.95 | -1.81 | 0.1 | -0.99 | -0.3 | -3.28 | -1.37 | -1.79 | -1.73 | -1.47 | -1.3 | -0.37 | -2.19 | -1.6 | -2.63 | -0.435 | -0.5225 | -0.6475 | 0.556171 | 0.290472 | 0.324261 | 1 | 0.996511 | 0.84221 |
| Mettl13 | Q91YR5 | 0.03 | -0.14 | -0.04 | -0.11 | -0.41 | 0.06 | -0.04 | -0.12 | 0.1 | -0.09 | -0.46 | 0.05 | -0.31 | -0.23 | -0.55 | -0.49 | -0.0625 | -0.035 | -0.33 | 0.706173 | 0.853546 | 0.091378 | 1 | 0.996511 | 0.693316 |
| Mettl14 | Q3UIK4-2 | -1.32 | -1.56 | -3.47 | -4.38 | -1.51 | -0.48 | -4.88 | -3.9 | -2.87 | -2.55 | -3.61 | -0.99 | -2.14 | -2.18 | -3.54 | -4.17 | -0.01 | 0.1775 | -0.325 | 0.992939 | 0.831753 | 0.68924 | 1 | 0.996511 | 0.955729 |
| Mettl15 | Q9DCL4 | 0.36 | 0.78 | 0.29 | 0.69 | 0.53 | 0.44 | 0.35 | 0.58 | 0.2 | 0.58 | 0.28 | 0.07 | 0.31 | -0.04 | -0.14 | 0.22 | -0.055 | -0.2475 | -0.4425 | 0.757379 | 0.251404 | 0.065516 | 1 | 0.996511 | 0.664936 |
| Mettl16 | Q5SW15 | -1.28 | -0.44 | 0.83 | -0.43 | -0.45 | -1.61 | -0.15 | -0.88 | -0.82 | 0 | -0.28 | -1.52 | -1.52 | -0.57 | -0.83 | -0.66 | -0.4425 | -0.325 | -0.565 | 0.384261 | 0.528503 | 0.237488 | 1 | 0.996511 | 0.792722 |
| Mettl17 | E9Q7K9 | 0.13 | 0.13 | 0.3 | 0.49 | 0.28 | 0.47 | 0.12 | 0.17 | -0.08 | 0.33 | 0.23 | -0.37 | -0.18 | 0.87 | 0.02 | -0.11 | -0.0025 | -0.235 | -0.1125 | 0.988199 | 0.29855 | 0.688083 | 1 | 0.996511 | 0.955729 |
| Mettl2 | Q8BMK1 | -0.68 | 0.38 | 0.03 | -0.08 | -1.18 | 0.07 | 0.47 | 0.66 | 0.62 | 0.14 | 1.13 | 0.43 | 0.69 | -1.32 | 0.24 | 1.45 | 0.0925 | 0.6675 | 0.3525 | 0.832649 | 0.054603 | 0.54374 | 1 | 0.996511 | 0.912212 |
| Mettl21A | Q9CQL0 | 0.19 | 0.36 | 0.16 | -0.65 | 0.09 | 0.1 | 0.68 | -0.21 | -0.82 | -0.33 | -0.15 | 0.34 | -0.38 | -0.8 | 0.26 | 0.41 | 0.15 | -0.255 | -0.1425 | 0.61061 | 0.444773 | 0.691081 | 1 | 0.996511 | 0.955729 |
| Mettl22 | Q8R1C6 | -6.46 | -1.46 | -4.1 | 0.33 | -1.82 | -6.03 | -9.29 | -3 | 0.3 | 0.8 | -5.33 | -6.01 | -0.63 | 0.38 | -5.29 | -7.62 | -2.1125 | 0.3625 | -0.3675 | 0.305968 | 0.861884 | 0.863438 | 1 | 0.996511 | 0.977235 |
| Mettl26 | Q9DCS2 | -0.06 | -0.17 | -0.54 | -0.65 | -0.02 | -0.15 | -0.64 | -0.47 | 0.4 | -0.28 | -0.38 | 0.09 | 0.76 | 0.97 | 0.09 | -0.92 | 0.035 | 0.3125 | 0.58 | 0.87547 | 0.234386 | 0.197933 | 1 | 0.996511 | 0.78004 |
| Mettl3 | A0A0R4J04 | -1.71 | -1.09 | -3.02 | -2.76 | -2.03 | -1.33 | -2.92 | -2.95 | -0.96 | -1.57 | -3 | -0.97 | -0.28 | -1.01 | -3.17 | -3.28 | -0.1625 | 0.52 | 0.21 | 0.766674 | 0.399621 | 0.792328 | 1 | 0.996511 | 0.96933 |
| Mettl7a1 | Q8C6B0 | -0.44 | -0.06 | -1.31 | -1.29 | 0.35 | -0.08 | -1.36 | -1.3 | -0.58 | 0.91 | -1.43 | -0.42 | -1.27 | -0.14 | -0.89 | -1.26 | 0.1775 | 0.395 | -0.115 | 0.719495 | 0.465737 | 0.771957 | 1 | 0.996511 | 0.965933 |
| Mettl9 | Q9EPL4 | 1.65 | 1.98 | 0.2 | 0.94 | 1.59 | 1.46 | 0.76 | 1.06 | 0.57 | 0.7 | 0.46 | 1.38 | 0.66 | 1.2 | -0.39 | 0.04 | 0.025 | -0.415 | -0.815 | 0.951594 | 0.340633 | 0.125173 | 1 | 0.996511 | 0.721448 |
| Mex3c | Q05A36 | -0.37 | -0.02 | -0.15 | -0.93 | -2.51 | -2.16 | -1.05 | -1.17 | -1.56 | 0.07 | -0.16 | -0.51 | 0.02 | -2.76 | -0.91 | -1.08 | -1.355 | -0.1725 | -0.815 | 0.007099 | 0.664033 | 0.173473 | 1 | 0.996511 | 0.764376 |
| Mfap1b | C0HKD9 | 1.73 | 0.62 | 0.39 | 0.75 | -0.97 | 0.94 | -0.71 | 0.9 | 1.13 | 1.03 | 0.41 | 0.18 | 1.65 | 0.92 | 1.09 | -0.3 | -0.8325 | -0.185 | -0.0325 | 0.150614 | 0.614842 | 0.945363 | 1 | 0.996511 | 0.992376 |
| Mfap3l | Q9D3X9-2 | -1.22 | 0.83 | 0.84 | 0.56 | -1.17 | -0.44 | 1.53 | 0.5 | 0.62 | 0.03 | -0.23 | 0.8 | 0.61 | -0.67 | 0.43 | 0.24 | -0.1475 | 0.0525 | -0.1 | 0.831863 | 0.918209 | 0.850092 | 1 | 0.996755 | 0.976225 |
| Mff | Q6PCP5 | 0.37 | 0.46 | 0.23 | 0.27 | 0.43 | 0.27 | 0.39 | 0.29 | -0.18 | -0.08 | 0.51 | 0.4 | -0.07 | -0.22 | -0.1 | 0.13 | 0.0125 | -0.17 | -0.3975 | 0.93187 | 0.443167 | 0.051369 | 1 | 0.996511 | 0.658454 |
| Mff | Q6PCP5-2 | -1.16 | -0.7 | 0.77 | 1.31 | -1.16 | -1.3 | 1.17 | 1.25 | -1.71 | 1.85 | 1.55 | -0.98 | -1.08 | -0.81 | 1.64 | 1.22 | -0.065 | 0.1225 | 0.1875 | 0.937261 | 0.898551 | 0.818747 | 1 | 0.996511 | 0.972768 |
| Mfge8 | P21956-2 | -0.47 | -0.02 | -0.07 | 0.35 | 0.82 | -0.19 | -0.2 | 0.49 | -0.31 | -0.39 | 0.16 | -0.13 | -0.36 | 0.18 | 0.4 | 0.31 | 0.2825 | -0.115 | 0.185 | 0.362078 | 0.629015 | 0.492076 | 1 | 0.996511 | 0.897531 |
| Mfhas1 | Q3V1N1 | -0.34 | 0.35 | -0.25 | 0.36 | -0.63 | -0.06 | -1 | 0.23 | 0.61 | 0.76 | -0.05 | 0.28 | 0.54 | 0.03 | -0.4 | -0.51 | -0.395 | 0.37 | -0.115 | 0.24636 | 0.201467 | 0.714103 | 1 | 0.996511 | 0.958106 |
| Mfn1 | Q811U4 | 0.7 | 0.7 | 0.89 | 0.98 | 0.83 | 0.64 | 1.25 | 1.27 | 0.49 | 0.92 | 1.18 | 0.6 | 0.35 | 0.62 | 1.32 | 1.69 | 0.18 | -0.02 | 0.1775 | 0.386356 | 0.925138 | 0.585233 | 1 | 0.996755 | 0.928105 |
| Mfn2 | Q80U63 | 0.27 | 0.42 | 0.49 | 0.7 | 0.03 | 0.65 | 0.58 | 0.75 | 0.25 | 0.69 | 0.76 | 0.37 | -0.3 | 0.37 | 0.16 | 0.21 | 0.0325 | 0.0475 | -0.36 | 0.877927 | 0.813297 | 0.128942 | 1 | 0.996511 | 0.723655 |
| Mfsd1 | Q9DC37 | -0.76 | -2.88 | -2.44 | -2.29 | -3.2 | -0.95 | -2.31 | -1.34 | -0.16 | -0.19 | -1.2 | -0.83 | 2.66 | -4.53 | 4.99 | -2.67 | 0.1425 | 1.4975 | 2.205 | 0.819155 | 0.013386 | 0.290767 | 1 | 0.996511 | 0.825497 |
| Mfsd10 | Q9D2V8 | -1.16 | -1.02 | 0.23 | 0.73 | -0.44 | -1.1 | -0.11 | 0.69 | -1.11 | 0.51 | 0.37 | -1.13 | -0.67 | -0.99 | 0.88 | 0.54 | 0.065 | -0.035 | 0.245 | 0.905237 | 0.953019 | 0.682551 | 1 | 0.996755 | 0.955729 |
| Mfsd11 | Q8BJ51 | -0.19 | -1.59 | -0.88 | -1.81 | -2.63 | -1.01 | -2.32 | -2.22 | 2.68 | -1.75 | -3.17 | -1.2 | 3.15 | -2.15 | -1.94 | -1.79 | -0.9275 | 0.2575 | 0.435 | 0.077582 | 0.825564 | 0.714718 | 1 | 0.996511 | 0.958106 |
| Mfsd2a | Q9DA75 | -4.04 | -4.13 | -1.01 | -0.78 | -2.17 | -3.15 | 0.13 | -0.41 | -1.65 | -0.56 | -0.61 | -2.56 | -2.91 | -2.99 | -2.62 | -1.35 | 1.09 | 1.145 | 0.0225 | 0.324307 | 0.240927 | 0.979932 | 1 | 0.996511 | 0.994776 |
| Mfsd3 | A0A0R4J00 | 0.76 | 0.53 | -0.54 | -0.2 | 1.05 | 1.28 | 0.33 | -0.09 | 1.26 | -0.07 | 0.11 | 0.37 | 0.35 | 1.02 | 0.33 | -0.29 | 0.505 | 0.28 | 0.215 | 0.242102 | 0.495146 | 0.586935 | 1 | 0.996511 | 0.92838 |
| Mfsd4a | Q6PDC8 | -0.04 | 0.42 | 0.57 | 0.17 | 0.76 | 0.06 | 0.07 | 0.01 | -0.13 | 0.93 | 0.25 | 0 | 0.2 | 1.18 | -0.22 | 0.16 | -0.055 | -0.0175 | 0.05 | 0.818628 | 0.95065 | 0.879836 | 1 | 0.996755 | 0.979626 |
| Mfsd6 | F6U3S2 | 0.73 | 0.54 | 1.62 | 1.78 | 0.79 | -0.26 | 1.19 | 1.25 | 0.93 | 1.55 | 1.2 | 0.92 | 0.72 | 2.2 | 1.13 | 1.38 | -0.425 | -0.0175 | 0.19 | 0.344495 | 0.958941 | 0.653656 | 1 | 0.996755 | 0.947633 |
| Mfsd6 | D3Z183 | -2.01 | 0.17 | 0.97 | 0.7 | -0.71 | -1.64 | 0.32 | 0.41 | 0.03 | 1.09 | 0.72 | -1.11 | -0.29 | -0.86 | 0.99 | 1.01 | -0.3625 | 0.225 | 0.255 | 0.631698 | 0.765663 | 0.732643 | 1 | 0.996511 | 0.958635 |
| Mfsd9 | Q8C0T7 | 0.35 | -0.58 | -0.13 | 1.37 | 0.77 | 0.06 | 1.17 | 1.06 | 1.48 | 2.4 | 1.48 | -0.66 | 1.98 | -0.15 | 1.56 | 0.79 | 0.5125 | 0.9225 | 0.7925 | 0.276089 | 0.21129 | 0.193289 | 1 | 0.996511 | 0.777764 |
| Mgat1 | J3JS24 | -3.81 | -4.02 | -1.51 | -2.57 | -2.96 | -1.15 | -3.4 | -0.76 | -1.63 | -0.94 | 0.06 | -1.41 | -0.07 | -2.88 | -2.85 | -3.29 | 0.91 | 1.9975 | 0.705 | 0.268535 | 0.011516 | 0.415566 | 1 | 0.996511 | 0.874945 |
| Mgat3 | Q5RKT9 | -1.39 | -0.16 | -0.55 | -0.09 | -0.52 | 0.06 | -0.18 | -0.38 | -0.8 | -0.44 | -0.76 | 0.46 | -0.59 | -1.56 | -0.09 | 0.13 | 0.2925 | 0.1625 | 0.02 | 0.368961 | 0.685712 | 0.964755 | 1 | 0.996511 | 0.9927 |
| Mgat5 | Q8R4G6 | -1.62 | -0.61 | -0.82 | -0.72 | -0.18 | -1.52 | -0.1 | -0.41 | -1.22 | -0.43 | -0.64 | -1.52 | -1.56 | -0.67 | -0.28 | -0.46 | 0.39 | -0.01 | 0.2 | 0.321125 | 0.976323 | 0.581991 | 1 | 0.997588 | 0.927734 |
| Mgll | E9Q3B9 | 0.25 | 0.58 | 0.91 | 0.93 | 0.74 | 0.13 | 0.52 | 0.85 | 0.31 | 0.68 | 0.88 | 0.33 | -0.64 | 0.24 | 0.23 | 0.5 | -0.1075 | -0.1175 | -0.585 | 0.65747 | 0.626656 | 0.083665 | 1 | 0.996511 | 0.689531 |
| Mgrn1 | Q9D074 | 0.51 | 0.46 | 0.8 | 0.69 | 0.69 | 0.74 | 0.63 | 0.79 | 0.12 | 0.71 | 0.77 | 0.85 | 0.42 | 0.36 | 1.13 | 1.09 | 0.0975 | -0.0025 | 0.135 | 0.533489 | 0.99097 | 0.598529 | 1 | 0.999285 | 0.930093 |
| Mgst1 | D3YU60 | 0.8 | 0.46 | -0.65 | 1.34 | 1.33 | 0.71 | 2.13 | 1.45 | -0.42 | 0.74 | -0.73 | 0.93 | -0.21 | -1.03 | 1.59 | 1.12 | 0.9175 | -0.3575 | -0.12 | 0.080118 | 0.516686 | 0.857362 | 1 | 0.996511 | 0.976225 |
| Mgst3 | Q9CPU4 | -1.4 | -1.37 | -2.06 | -1.37 | -1.61 | 0.02 | -1.24 | -1.03 | -0.85 | -1.57 | -1.36 | -1.32 | -0.28 | -1.57 | -2.13 | -1.92 | 0.585 | 0.275 | 0.075 | 0.141238 | 0.290186 | 0.860609 | 1 | 0.996511 | 0.977204 |
| Mia2 | E9PY90 | 0.17 | 0.35 | -0.6 | -0.44 | 0.87 | 0.17 | 0.22 | -0.3 | 0.4 | -0.11 | -0.65 | 0.53 | 0.97 | 0.46 | 0.77 | 0.77 | 0.37 | 0.1725 | 0.8725 | 0.272895 | 0.622098 | 0.010948 | 1 | 0.996511 | 0.563386 |
| Mia3 | A0A0R4J2D | -0.66 | -0.51 | -0.93 | -1.01 | 0.06 | -0.69 | -0.38 | -0.88 | -0.34 | -0.85 | -1.03 | -0.42 | 0.52 | -0.13 | 0.57 | 0.17 | 0.305 | 0.1175 | 1.06 | 0.240313 | 0.617529 | 0.001708 | 1 | 0.996511 | 0.419079 |
| Mib1 | Q80SY4 | 0.9 | 0.64 | 0.67 | 0.28 | 0.73 | 1.1 | 0.6 | 0.62 | 0.46 | -0.21 | 0.73 | 0.84 | 0.86 | 0.35 | 0.66 | 0.67 | 0.14 | -0.1675 | 0.0125 | 0.498217 | 0.554135 | 0.954277 | 1 | 0.996511 | 0.992476 |
| Mib2 | Q8R516-2 | 0.5 | 0.54 | 0.62 | 0.47 | 0.91 | 0.43 | 0.7 | 0.52 | 0.27 | 0.26 | 0.76 | 0.69 | 0.59 | 0.6 | 0.76 | 1.06 | 0.1075 | -0.0375 | 0.22 | 0.524073 | 0.845604 | 0.267388 | 1 | 0.996511 | 0.818895 |
| Mical1 | Q8VDP3-3 | 0.14 | 0.3 | 0.11 | -0.01 | 0.45 | -0.19 | 0.3 | 0.34 | -0.33 | -0.53 | 0.39 | -0.1 | -0.52 | 0.48 | 0.28 | 0.5 | 0.09 | -0.2775 | 0.05 | 0.642404 | 0.260636 | 0.854261 | 1 | 0.996511 | 0.976225 |
| Mical2 | Q8BML1-2 | 0.33 | -0.07 | 0.96 | 0.4 | -0.49 | 0.67 | 0.09 | -0.15 | 0.04 | -0.87 | 0.61 | 0 | 0.44 | -1.51 | -0.24 | 0.48 | -0.375 | -0.46 | -0.6125 | 0.256261 | 0.225193 | 0.222249 | 1 | 0.996511 | 0.787447 |
| Mical3 | A0A5F8MP | 0.47 | 0.71 | 0.74 | 0.79 | 0.48 | 0.3 | 0.66 | 0.68 | 0.53 | 0.93 | 0.81 | 0.41 | 0.39 | 0.77 | 0.61 | 0.46 | -0.1475 | -0.0075 | -0.12 | 0.391445 | 0.969121 | 0.53028 | 1 | 0.996755 | 0.908663 |
| Micall1 | Q8BGT6 | -0.12 | 0.29 | 0.5 | 0.15 | -0.02 | -0.07 | 0.59 | -0.05 | 0.23 | 3.65 | 0.58 | 0.25 | -0.09 | 0.28 | 0.67 | 0 | -0.0925 | 0.9725 | 0.01 | 0.684362 | 0.222194 | 0.967929 | 1 | 0.996511 | 0.992733 |
| Micos10 | Q7TNS2 | -2.21 | -1.99 | -3 | -2.77 | -2.35 | -1.9 | -3.31 | -3.33 | -1.48 | -3.33 | -3.05 | -2.32 | -1.13 | -2.97 | -2.98 | -2.48 | -0.23 | -0.0525 | 0.1025 | 0.571835 | 0.906704 | 0.826018 | 1 | 0.996744 | 0.973155 |
| Micos13 | Q8R404 | 0.17 | 0.57 | 0.47 | 0.72 | 0.29 | 0.13 | 0.88 | 0.59 | 0.11 | -0.28 | 0.74 | -0.03 | -0.5 | 0.66 | -0.05 | 0.23 | -0.01 | -0.3475 | -0.3975 | 0.964401 | 0.210741 | 0.190416 | 1 | 0.996511 | 0.776843 |
| Micu1 | Q8VCX5 | 1.3 | 0.21 | 0.74 | 0.53 | -0.04 | 1.03 | 0.32 | 0.27 | 0.43 | 0.62 | 0.71 | 0.87 | 0.47 | -0.25 | 0.49 | 0.4 | -0.3 | -0.0375 | -0.4175 | 0.355082 | 0.887215 | 0.190396 | 1 | 0.996511 | 0.776843 |
| Micu2 | Q8CD10 | -0.01 | 0.4 | 1.28 | 0.96 | 0.72 | -0.25 | 1.59 | 0.84 | -0.24 | 0.52 | 1.15 | 0.12 | 0.07 | 0.32 | 1.03 | 1.26 | 0.0675 | -0.27 | 0.0125 | 0.878725 | 0.50145 | 0.974442 | 1 | 0.996511 | 0.992977 |
| Micu3 | Q9CTY5 | 0.85 | 1.19 | 0.41 | 0.39 | 0.83 | 0.76 | 0.23 | 0.25 | -0.06 | -0.11 | 0.4 | 0.9 | -0.24 | 0.19 | -0.25 | -0.16 | -0.1925 | -0.4275 | -0.825 | 0.465478 | 0.188808 | 0.009104 | 1 | 0.996511 | 0.563386 |
| Mid1ip1 | Q9CQ20 | 0.75 | 0.81 | 0.68 | 0.8 | 0.39 | 0.71 | 0.97 | 0.91 | 0.2 | 0.13 | 0.99 | 0.46 | -0.34 | -0.01 | 0.72 | 0.71 | -0.015 | -0.315 | -0.49 | 0.933362 | 0.193834 | 0.113502 | 1 | 0.996511 | 0.716416 |
| Mid2 | B1AVF4 | 0.45 | 0.33 | 0.23 | 0.75 | 0.09 | 0.36 | 0.11 | 0.66 | 0.07 | 0.56 | 0.22 | 0.23 | 0.2 | 0.32 | 0.76 | 0.32 | -0.135 | -0.17 | -0.04 | 0.515544 | 0.408631 | 0.854971 | 1 | 0.996511 | 0.976225 |
| Mief1 | Q8BGV8 | -0.07 | 0.05 | -0.2 | -0.16 | 0.02 | 0.12 | -0.28 | -0.09 | 0.06 | 0.05 | -0.03 | -0.12 | -0.28 | 0.52 | -0.37 | -0.71 | 0.0375 | 0.085 | -0.115 | 0.817289 | 0.601375 | 0.687611 | 1 | 0.996511 | 0.955729 |
| Mien1 | Q9CQ86 | 0.1 | 0.22 | -0.02 | -0.3 | 0.08 | 0.25 | -0.6 | -0.17 | -0.15 | 0.33 | -0.29 | 0.09 | 0.62 | -0.03 | -0.39 | -0.43 | -0.11 | -0.005 | -0.0575 | 0.639835 | 0.981493 | 0.840524 | 1 | 0.998773 | 0.976225 |
| Mif | P34884 | -0.28 | -0.51 | 1.12 | 0.67 | -0.56 | 0.44 | 1.28 | 1.13 | -1.1 | -0.61 | 1.32 | -0.09 | -1.22 | -1.35 | 0.91 | 0.64 | 0.3225 | -0.37 | -0.505 | 0.542947 | 0.53925 | 0.444373 | 1 | 0.996511 | 0.884411 |
| Mif4gd | A2A9W1 | 1.12 | -0.3 | 0.59 | 0.88 | -0.09 | 0.75 | 1.57 | 1.88 | -3.06 | -1.89 | 1.73 | 1.69 | -3.26 | -4.76 | 1.05 | 1.01 | 0.455 | -0.955 | -2.0625 | 0.373322 | 0.411265 | 0.152736 | 1 | 0.996511 | 0.750422 |
| Miga1 | Q4QQM5 | -0.84 | -0.78 | -0.08 | -0.04 | -0.81 | -0.6 | 0.18 | -0.26 | -0.91 | -0.78 | 0.06 | -1.04 | -0.58 | -1.24 | -0.26 | -0.06 | 0.0625 | -2.33E-01 | -0.1 | 0.836738 | 0.484949 | 0.767888 | 1 | 0.996511 | 0.965084 |
| Miga2 | Q8BK03 | -0.01 | 0.18 | 0.09 | 0.2 | -0.17 | 0.14 | 0.07 | 0.08 | -0.13 | 0.22 | 0.42 | 0.31 | -0.2 | -0.53 | -0.57 | -0.19 | -0.085 | 0.09 | -0.4875 | 0.583554 | 0.633373 | 0.028879 | 1 | 0.996511 | 0.637553 |
| Minar2 | Q8C4X7-2 | -4.89 | -6 | -0.28 | -0.34 | -5.55 | -4.83 | -0.28 | -0.54 | -6.26 | 0.52 | -0.25 | -5.18 | -1.87 | -6.67 | -0.77 | -0.68 | 0.0775 | 0.085 | 0.38 | 0.966041 | 0.966526 | 0.835412 | 1 | 0.996755 | 0.975524 |
| Mindy1 | Q76LS9-2 | -0.95 | -0.79 | 0.04 | 0.33 | -1.32 | -1.22 | 0.49 | 0.08 | -0.92 | 0.37 | -0.05 | -0.85 | -0.52 | -0.68 | -0.16 | -0.06 | -0.15 | -0.02 | -0.0125 | 0.768953 | 0.961984 | 0.970981 | 1 | 0.996755 | 0.992803 |
| Mindy2 | A0A1L1STF | 0.23 | 0.57 | 1.12 | 0.94 | 0.62 | 0.2 | 0.17 | 0.62 | 0.58 | 1.41 | 0.84 | 0.28 | 0.64 | 0.44 | 1.11 | 0.83 | -0.3125 | 0.0625 | 0.04 | 0.228204 | 0.841874 | 0.881785 | 1 | 0.996511 | 0.979767 |
| Mindy3 | Q9CV28-2 | -1.93 | -2.86 | 1.94 | 0.02 | -2.75 | -2.44 | 1.15 | 1.38 | -2.25 | -0.38 | 0.97 | -1.71 | -2 | -1.6 | 1.64 | 1.46 | 0.0425 | -0.135 | 0.5825 | 0.975408 | 0.906791 | 0.651841 | 1 | 0.996744 | 0.947633 |
| Mindy4 | A0A0N4SV | -1.31 | -0.54 | 2.09 | 2.03 | -1.22 | 2.07 | 2.14 | 2.26 | 1.23 | 2.38 | 1.89 | -0.22 | 1.08 | 0.37 | 2.24 | 2.2 | 0.745 | 0.7525 | 0.905 | 0.500389 | 0.431201 | 0.322034 | 1 | 0.996511 | 0.840788 |
| Mink1 | J3QM71 | 11.53 | 6.86 | 13.29 | 7.39 | 5.66 | 12.75 | 11.36 | 12.31 | 5.66 | 2.27 | 12.54 | 11.1 | 5.27 | -0.46 | 13.68 | 15.67 | 0.7525 | -1.875 | -1.2275 | 0.711457 | 0.468456 | 0.733429 | 1 | 0.996511 | 0.958635 |
| Mink1 | Q9JM52 | 0.39 | 0.25 | 0.37 | 0.46 | 0.11 | 0.65 | 0.65 | 0.56 | 0.42 | 0.14 | 0.64 | 0.41 | 0.63 | -0.03 | 0.53 | 0.2 | 0.125 | 0.035 | -0.035 | 0.495684 | 0.844844 | 0.86979 | 1 | 0.996511 | 0.979293 |
| Minpp1 | Q9Z2L6 | -0.23 | 0.05 | 0.16 | -0.38 | 0.12 | -0.33 | 0.23 | -0.14 | -0.18 | 0.12 | -0.04 | -0.29 | -0.03 | -0.02 | 0.35 | 0.14 | 0.07 | 0.0025 | 0.21 | 0.735782 | 0.990082 | 0.332923 | 1 | 0.999285 | 0.845601 |
| Mios | Q8VE19 | -0.35 | -0.33 | -0.67 | -0.71 | -0.1 | -0.63 | -1.04 | -1.02 | -1.08 | -1.17 | -0.89 | -0.36 | -1.06 | -0.99 | -0.57 | -0.46 | -0.1825 | -0.36 | -0.255 | 0.478364 | 0.154425 | 0.279115 | 1 | 0.996511 | 0.822084 |
| Mipep | A6H611 | 2.36 | 0.84 | 0.67 | 0.36 | 0.83 | 2.37 | 0.62 | 0.15 | 0.11 | -0.54 | 0.68 | 1.77 | 0.19 | -0.43 | 0.44 | 0.52 | -0.065 | -0.5525 | -0.8775 | 0.91334 | 0.374163 | 0.086625 | 1 | 0.996511 | 0.691619 |
| Mipol1 | E9Q8Z7 | 0.43 | -1.15 | 0.8 | 0.5 | -1.85 | 0.08 | -0.55 | 0.88 | 3.04 | 0.77 | 0.39 | 0.54 | 3.2 | 2.62 | 2.22 | -0.11 | -0.505 | 1.04 | 1.8375 | 0.450707 | 0.159603 | 0.037689 | 1 | 0.996511 | 0.658454 |
| Mix23 | Q8R3Q6 | 0.72 | -0.77 | 0.88 | -0.03 | -0.79 | 1.7 | 0.73 | 0.84 | -1.35 | 0.2 | 1.08 | 0.86 | -0.22 | -2.47 | 0.17 | 0.91 | 0.42 | -0.0025 | -0.6025 | 0.480194 | 0.996731 | 0.425941 | 1 | 1 | 0.878405 |
| Mkln1 | O89050 | -0.27 | 0.1 | -0.54 | -0.44 | 0.61 | -0.33 | -0.31 | -0.3 | -0.62 | -0.54 | -0.42 | -0.27 | -0.69 | 0.09 | -0.46 | -0.53 | 0.205 | -0.175 | -0.11 | 0.463095 | 0.40499 | 0.664854 | 1 | 0.996511 | 0.949825 |
| Mkrn2 | Q9ERV1 | -1.57 | -1.12 | -1.03 | -1.35 | -1.08 | -1.47 | -0.53 | -1.06 | -1.16 | -0.83 | -0.91 | -1.3 | 0.17 | -1.81 | -0.73 | -0.87 | 0.2325 | 0.2175 | 0.4575 | 0.351021 | 0.310394 | 0.278181 | 1 | 0.996511 | 0.822084 |
| Mlc1 | E9QP87 | -0.13 | 0.16 | 1.46 | 1.33 | 0.21 | 0.13 | 1.53 | 1.41 | 0.65 | 2.23 | 1.54 | 0.4 | 0.6 | 0.51 | 1.67 | 1.45 | 0.115 | 0.5 | 0.3525 | 0.821181 | 0.365373 | 0.460011 | 1 | 0.996511 | 0.887902 |
| Mlec | Q6ZQI3 | 0.05 | -0.15 | -0.04 | -0.43 | 0.44 | 0.15 | 0.63 | 0.15 | -0.07 | -0.26 | -0.45 | 0.26 | -0.31 | 0.29 | 1.26 | 1.34 | 0.485 | 0.0125 | 0.7875 | 0.03188 | 0.95479 | 0.074023 | 1 | 0.996755 | 0.671115 |
| Mlf2 | Q99KX1 | -0.3 | 0.05 | 0.77 | 0.45 | -0.33 | -0.42 | 0.88 | 0.67 | -0.22 | 0.57 | 0.43 | -0.7 | 0.1 | -1.02 | 0.73 | 0.73 | -0.0425 | -0.2225 | -0.1075 | 0.912441 | 0.546972 | 0.810612 | 1 | 0.996511 | 0.972222 |
| Mlip | V9GXA0 | 1.96 | 1.96 | 0.87 | 0.24 | 2.56 | 2.2 | 2.97 | 1.21 | -0.48 | 0.31 | 0.75 | 2.51 | -0.31 | 0.2 | -0.47 | 0.64 | 0.9775 | -0.485 | -1.2425 | 0.088978 | 0.490495 | 0.024305 | 1 | 0.996511 | 0.626254 |
| Mlip | V9GWW6 | 0.34 | 0.06 | -0.15 | -0.09 | -0.1 | 0.33 | -0.86 | -0.5 | 0.83 | -0.99 | 0.15 | 0.02 | 1.08 | 0.04 | -0.26 | -0.55 | -0.3225 | -0.0375 | 0.0375 | 0.270267 | 0.920813 | 0.918166 | 1 | 0.996755 | 0.986868 |
| Mllt11 | P97783 | -1.84 | -2.02 | -1.96 | -2.08 | -2.2 | -2.19 | -3.13 | -1.4 | 0.2 | -1.43 | -1.09 | -1.31 | 1.39 | -0.98 | -0.46 | -1.72 | -0.255 | 1.0675 | 1.5325 | 0.466544 | 0.016948 | 0.03111 | 1 | 0.996511 | 0.641481 |
| Mllt6 | B1AR09 | -2.8 | -1.95 | -1.81 | -2.68 | -2.06 | -2.17 | -2.15 | -2.64 | -2.96 | -1.28 | -1.73 | -1.55 | -0.72 | -2.05 | -1.27 | -1.32 | 0.055 | 0.43 | 0.97 | 0.846821 | 0.326406 | 0.024632 | 1 | 0.996511 | 0.628846 |
| Mlst8 | Q9DCJ1 | 0.71 | 0.36 | -0.62 | -1.04 | 0.58 | 0.91 | -0.68 | -0.9 | 0.57 | -1.83 | -0.74 | 0.78 | 0.92 | -0.45 | -0.72 | -0.86 | 1.25E-01 | -0.1575 | -0.13 | 0.822674 | 0.813997 | 0.808214 | 1 | 0.996511 | 0.971953 |
| Mlycd | Q99J39-2 | 0.05 | 0.31 | 0.4 | 0.61 | 0.51 | 0.11 | 0.53 | 0.52 | -0.16 | 0.03 | 0.6 | -0.13 | -0.18 | 0.37 | 0.36 | 0.36 | 0.075 | -0.2575 | -0.115 | 0.697473 | 0.29821 | 0.613105 | 1 | 0.996511 | 0.935516 |
| Mmaa | Q8C7H1 | 0.77 | 0 | 0.67 | 0.83 | 0.17 | 0.75 | 0.69 | 0.58 | 0.09 | 0.51 | 1.02 | 0.63 | 0.59 | 0.22 | 0.85 | 0.7 | -0.02 | -0.005 | 0.0225 | 0.935096 | 0.985864 | 0.931584 | 1 | 0.998892 | 0.989713 |
| Mmab | Q9D273 | 1.3 | 1.28 | 1.17 | 1.19 | 1.13 | 1.27 | 1.04 | 1.3 | 0.62 | 0.67 | 1.37 | 1.23 | 0.83 | 0.45 | 1.22 | 1.07 | -0.05 | -0.2625 | -0.3425 | 0.735964 | 0.267295 | 0.147659 | 1 | 0.996511 | 0.746406 |
| Mme | Q61391 | 0.6 | 0.35 | 0.44 | 0.46 | 0.58 | 0.12 | 1.07 | 0.81 | 0.23 | 0.41 | 0.46 | 0.8 | 0.16 | -0.49 | 0.94 | 0.87 | 0.1825 | 0.0125 | -0.0925 | 0.43267 | 0.947033 | 0.786247 | 1 | 0.996755 | 0.967714 |
| Mmgt1 | Q8K273 | 0.66 | 1.27 | 1.12 | 0.6 | 1.33 | 0.6 | 0.96 | 0.92 | 0.46 | 0.22 | 0.77 | 0.84 | 0.57 | 0.83 | 1.83 | 1.55 | 0.04 | -0.34 | 0.2825 | 0.867315 | 0.189368 | 0.415709 | 1 | 0.996511 | 0.874945 |
| Mmp17 | Q9R0S3 | -0.43 | -0.4 | 0.8 | 0.7 | 0.03 | -0.43 | 1.2 | 1.1 | -0.35 | 0.66 | 1.15 | -0.13 | -0.56 | -0.13 | 0.45 | 0.65 | 0.3075 | 0.165 | -0.065 | 0.530623 | 0.717829 | 0.876015 | 1 | 0.996511 | 0.979626 |
| Mms19 | Q9D071 | -0.35 | -0.57 | -0.14 | -0.67 | -0.33 | -0.43 | -0.12 | -0.34 | -0.49 | -0.95 | -0.11 | -0.2 | -0.6 | -0.54 | -0.6 | -0.31 | 0.1275 | -0.005 | -0.08 | 0.48568 | 0.983856 | 0.693186 | 1 | 0.998876 | 0.955729 |
| Mmtag2 | Q99LX5 | -0.49 | -1.34 | -0.49 | -0.63 | -0.43 | -1.5 | -1.47 | -1.49 | -0.4 | 0.06 | -1.03 | -1.02 | -0.29 | 0.16 | -0.54 | -1.44 | -0.485 | 0.14 | 0.21 | 0.161659 | 0.673722 | 0.5864 | 1 | 0.996511 | 0.928178 |
| Mmut | P16332 | 0.61 | 0.7 | 0.25 | 0.42 | 0.5 | 0.64 | 0.59 | 0.42 | 0.58 | 0.17 | 0.63 | 0.45 | 0.49 | 0.87 | 0.05 | 0.26 | 0.0425 | -0.0375 | -0.0775 | 0.799371 | 0.848413 | 0.748037 | 1 | 0.996511 | 0.960744 |
| Mnat1 | P51949 | -4.14 | -4.72 | -2.68 | -3.49 | -3.91 | -4.71 | -2.45 | -2.81 | -2.17 | -1.28 | -2.5 | -3.73 | -2.71 | -2.77 | -3.1 | -2.88 | 0.2875 | 1.3375 | 0.8925 | 0.64331 | 0.054118 | 0.061669 | 1 | 0.996511 | 0.662517 |
| Moap1 | Q9ERH6 | 0.31 | 0.33 | -1.69 | -0.79 | 0.44 | 0.04 | -0.75 | -1.54 | 0.66 | -3.19 | -1.45 | -0.18 | 1.18 | 0.9 | -2.28 | -1.59 | 0.0075 | -0.58 | 0.0125 | 0.989974 | 0.512057 | 0.988859 | 1 | 0.996511 | 0.996154 |
| Mob1b | Q3UDM0 | -0.98 | -0.69 | -0.08 | 0.51 | -0.55 | -1.16 | 0.25 | 0.63 | -0.61 | 0.38 | 0.2 | -0.9 | -1.09 | 0.06 | -0.24 | 0.06 | 0.1025 | 0.0775 | 0.0075 | 0.831587 | 0.8567 | 0.985432 | 1 | 0.996511 | 0.996154 |
| Mob2 | A0A1B0GR | 1.11 | 1.27 | 1.32 | 1.47 | 1.52 | 0.31 | 1 | 1.52 | 0.27 | 1 | 1.42 | 1.23 | -0.25 | 0.33 | 1.11 | 1.08 | -0.205 | -0.3125 | -0.725 | 0.493403 | 0.27555 | 0.055992 | 1 | 0.996511 | 0.661738 |
| Mob4 | E0CXA9 | -0.17 | -0.58 | -1.16 | -0.91 | -0.16 | 0.23 | -0.42 | -0.87 | -0.94 | -1.13 | -0.96 | -0.35 | -1.24 | -0.51 | -1.05 | -0.87 | 0.4 | -0.14 | -0.2125 | 0.219289 | 0.62493 | 0.461255 | 1 | 0.996511 | 0.887902 |
| Mobp | Q9D2P8 | 2.42 | 2 | 2.39 | 1.95 | 1.27 | 1.65 | 0.77 | 1.14 | 2.04 | 1.6 | 1.52 | 1.63 | 1.74 | 1.69 | 2.43 | 1.54 | -0.9825 | -0.4925 | -0.34 | 0.002575 | 0.04294 | 0.216315 | 1 | 0.996511 | 0.787447 |
| Mocs1 | Q5RKZ7 | 0.96 | 1.29 | 3.02 | 2.51 | 1.01 | 0.95 | 2.35 | 2.1 | 0.76 | 2.07 | 2.4 | 0.89 | 0.08 | 1.39 | 1.62 | 1.68 | -0.3425 | -0.415 | -0.7525 | 0.543796 | 0.486802 | 0.207924 | 1 | 0.996511 | 0.785338 |
| Mocs2 | F6ZGI7 | -2.25 | -2.42 | -2.36 | -1.94 | -2.13 | -2.08 | -1.77 | -2.26 | -2.82 | -1.69 | -2.1 | -1.99 | -2.91 | -2.25 | -2.56 | -2.56 | 0.1825 | 0.0925 | -0.3275 | 0.346412 | 0.738018 | 0.165002 | 1 | 0.996511 | 0.758837 |
| Mocs2 | E9PVP5 | 2.61 | 2.81 | 4.11 | 3.38 | 0.85 | 2.86 | 2.24 | 3.33 | 2.19 | 4.18 | 3.18 | 2.16 | 2.56 | 0.47 | 3.6 | 3.93 | -0.9075 | -0.3 | -0.5875 | 0.144475 | 0.582767 | 0.451467 | 1 | 0.996511 | 0.885538 |
| Mocs3 | A2BDX3 | -0.18 | -0.09 | -0.18 | 0.05 | 0.04 | -0.26 | -0.39 | 0 | -0.22 | 0.73 | -0.1 | -0.3 | -0.25 | 0.27 | -0.25 | -0.35 | -0.0525 | 0.1275 | -0.045 | 0.757677 | 0.629138 | 0.829866 | 1 | 0.996511 | 0.973939 |
| Mog | Q61885 | 4.62 | 1.98 | 1.61 | 0.83 | 1.92 | 3.71 | 1.86 | 0.96 | 1.56 | 0.48 | 0.88 | 2.75 | 1.45 | 0.78 | 1.72 | 1.89 | -0.1475 | -0.8425 | -0.8 | 0.870165 | 0.344501 | 0.316624 | 1 | 0.996511 | 0.838625 |
| Mogs | Q80UM7 | -0.85 | -0.92 | -0.39 | -0.38 | -0.04 | -0.49 | 0.33 | -0.21 | -0.82 | 0.02 | -0.15 | -0.7 | -0.59 | 0.19 | 1.09 | 1 | 0.5325 | 0.2225 | 1.0575 | 0.050762 | 0.415363 | 0.026326 | 1 | 0.996511 | 0.633075 |
| Mon1a | A0A0R4J0D | 0.21 | 0.24 | 0.32 | 0.25 | 0.89 | -0.32 | 0.05 | 0.18 | -1.51 | -0.51 | 0.22 | 0.23 | -1.36 | -1.29 | 0.03 | 0.05 | -0.055 | -0.6475 | -0.8975 | 0.834015 | 0.127966 | 0.041581 | 1 | 0.996511 | 0.658454 |
| Mon1b | Q8BMQ8 | -0.35 | 0.17 | 0.19 | -0.42 | 0.66 | -0.14 | -0.35 | -0.12 | 0.38 | 0.26 | 0 | -0.08 | 0.86 | -0.18 | 0.43 | -0.12 | 0.115 | 0.2425 | 0.35 | 0.681639 | 0.305406 | 0.271452 | 1 | 0.996511 | 0.822084 |
| Mon2 | Q80TL7 | -0.53 | -0.52 | -0.5 | -0.62 | -0.33 | -0.54 | -0.56 | -0.73 | -0.5 | 0.29 | -0.58 | -0.51 | -0.42 | -0.23 | -0.79 | -0.42 | 0.0025 | 0.2175 | 0.0775 | 0.987102 | 0.369881 | 0.690119 | 1 | 0.996511 | 0.955729 |
| Mon2 | B9EKJ3 | -0.56 | 1.11 | -1.98 | -0.39 | -0.64 | -0.1 | -0.82 | -0.97 | 0.42 | -1.64 | -2.22 | 0.6 | 0.62 | 2.02 | -2.7 | -2.81 | -0.1775 | -0.255 | -0.2625 | 0.768325 | 0.767423 | 0.829203 | 1 | 0.996511 | 0.973939 |
| Mon2 | Q80TL7-2 | -0.96 | 0.52 | 0.82 | 0.68 | 0 | -0.31 | -1.3 | 1.14 | 0.21 | 0 | -0.17 | 0.23 | -0.74 | -0.52 | 0.85 | 1.54 | -0.3825 | -0.1975 | 0.0175 | 0.524295 | 0.627469 | 0.977679 | 1 | 0.996511 | 0.993727 |
| Morc2a | Q69ZX6 | 0.45 | -0.63 | -1.12 | -0.89 | -1.3 | 0.07 | -0.72 | -0.89 | -0.26 | 0.78 | -0.17 | 0.55 | -0.24 | -0.52 | -0.45 | -0.05 | -0.1625 | 0.7725 | 0.2325 | 0.701353 | 0.088625 | 0.521723 | 1 | 0.996511 | 0.907667 |
| Morf4l1 | P60762-2 | 3.09 | -1.67 | -2.85 | -3.49 | -4.03 | 3.45 | -2.95 | -3.36 | -2.54 | -2.19 | -2.85 | 6.02 | -4.89 | -4.12 | -2.49 | -2.54 | -0.4925 | 0.84 | -2.28 | 0.809321 | 0.718349 | 0.136264 | 1 | 0.996511 | 0.731656 |

| | | | | | | | | | | | | | | | | | | | | | | | | | | |
|---|---|---|---|---|---|---|---|---|---|---|---|---|---|---|---|---|---|---|---|---|---|---|---|---|---|---|
| Morf4l2 | Q9R0Q4 | -1.01 | -2.26 | -1.29 | -1.85 | -1.48 | -1.73 | -2.53 | -2.12 | -1.95 | -0.95 | -2.23 | -1.63 | -0.86 | -2.05 | -1.04 | -1.42 | -0.3625 | -0.0875 | 0.26 | 0.314075 | 0.817543 | 0.494572 | 1 | 0.996511 | 0.89758 |
| Morn4 | Q6PGF2 | -0.48 | -0.65 | -0.63 | -0.79 | -0.25 | -0.46 | -1.22 | -1.09 | -0.62 | 0.14 | -0.52 | -0.39 | -0.77 | -0.25 | -1.09 | -0.9 | -0.1175 | 0.29 | -0.115 | 0.647006 | 0.207562 | 0.624466 | 1 | 0.996511 | 0.939167 |
| Mospd1 | Q8VEL0 | -0.49 | 0.16 | 0.68 | 0.06 | 0.25 | 0.81 | 0.14 | -0.02 | 0.39 | 1.73 | -0.75 | 0.37 | -0.35 | 0.42 | 0.91 | 0.37 | 0.1925 | 0.3325 | 0.235 | 0.523616 | 0.527412 | 0.508328 | 1 | 0.996511 | 0.902804 |
| Mospd2 | B1AU74 | 0.26 | -0.17 | 1.76 | 0.65 | 0.24 | -0.21 | 0.56 | 0.57 | 0.32 | 0.88 | 0.66 | -0.11 | 0.78 | 0.07 | 2.51 | 1.52 | -0.335 | -0.1875 | 0.595 | 0.437899 | 0.671555 | 0.342611 | 1 | 0.996511 | 0.848473 |
| Mov10 | D3Z3E8 | 1.68 | 0.73 | 0.44 | -0.15 | 0.57 | 1.34 | 0.09 | -1.08 | 0.66 | -0.67 | 0.83 | 2.07 | 1.31 | -0.65 | 0 | 0.47 | -0.445 | 0.0475 | -0.3925 | 0.450708 | 0.938918 | 0.45912 | 1 | 0.996755 | 0.887902 |
| Moxd1 | Q9CXI3 | 0.76 | 0.67 | -0.5 | -0.51 | 0.2 | 1.45 | -0.43 | -0.2 | 0.4 | 0.96 | -0.55 | 1.33 | -0.03 | 0.45 | 0.09 | -0.55 | 0.15 | 0.43 | -0.115 | 0.766796 | 0.402093 | 0.77138 | 1 | 0.996511 | 0.965933 |
| Mpc1 | **D3YWY6** | **0.88** | 0.97 | 0.82 | 0.53 | **0.59** | 0.83 | 0.12 | 0.6 | 0.26 | -0.13 | 0.85 | 0.53 | **0.32** | -0.12 | 0 | 0.21 | -0.265 | -0.4225 | -0.6975 | 0.221242 | 0.120068 | 0.007081 | 1 | 0.996511 | 0.552614 |
| Mpc2 | Q9D023 | -0.92 | -0.85 | -0.55 | -0.5 | -0.56 | -0.43 | 0.05 | -0.22 | -1.92 | -3.18 | 0.16 | -0.93 | -2.19 | -1.32 | -1.58 | -0.96 | 0.415 | -0.7625 | -0.8075 | 0.06601 | 0.264027 | 0.021567 | 1 | 0.996511 | 0.621926 |
| Mpdu1 | Q8R0J2 | -0.38 | -0.57 | 0.08 | -0.38 | -0.5 | -1.28 | -0.7 | -1.15 | -0.53 | -0.03 | -0.38 | -1.65 | -0.07 | -0.07 | -0.53 | 0.33 | -0.595 | -0.335 | 0.2275 | 0.035638 | 0.37157 | 0.383342 | 1 | 0.996511 | 0.860984 |
| Mpdz | D3YUX2 | 0.89 | 0.71 | 0.23 | 0.3 | 0.17 | 0.35 | 0.11 | -0.37 | 0.96 | -0.26 | 0.42 | 1.04 | 1.18 | 0.25 | 0.54 | 0.65 | -0.4675 | 0.0075 | 0.1225 | 0.076331 | 0.982175 | 0.656691 | 1 | 0.998773 | 0.948657 |
| Mphosph6 | Q9D1Q1 | 3.56 | 4.33 | 1.28 | 0.55 | 5.1 | 2.41 | 1.01 | 1.33 | 3.37 | 2.66 | 1.15 | 4.23 | 3.56 | 4.82 | 1.64 | 0.75 | 0.0325 | 0.4225 | 0.2625 | 0.977592 | 0.673873 | 0.819281 | 1 | 0.996511 | 0.972768 |
| Mphosph8 | Q3TYA6 | 1.03 | 1.9 | 1.04 | 0.83 | 1.87 | 1.17 | 1.29 | 1.26 | 1.02 | 2.19 | 1.6 | 1.24 | 0.07 | 2.56 | 1.06 | 0.43 | 0.1975 | 0.3125 | -0.17 | 0.498311 | 0.375073 | 0.758834 | 1 | 0.996511 | 0.963116 |
| Mpi | Q924M7 | -0.98 | -1.27 | -0.79 | -0.43 | -1.12 | -0.69 | -0.27 | -0.38 | -1.14 | -0.28 | -0.67 | -0.83 | -1.28 | -0.99 | -0.94 | -0.62 | 0.2525 | 0.1375 | -0.09 | 0.355842 | 0.609918 | 0.723695 | 1 | 0.996511 | 0.958398 |
| Mpp1 | P70290 | 0.35 | -0.24 | 0.21 | 0.53 | -0.32 | 0.25 | 0.65 | 0.64 | 0.5 | -0.51 | 0.3 | -0.32 | 0.23 | -0.23 | 0.11 | 0.24 | 0.0925 | -0.22 | -0.125 | 0.74464 | 0.468662 | 0.601964 | 1 | 0.996511 | 0.930802 |
| Mpp2 | Q9WV34 | 2.09 | 0.83 | 1.45 | 1.13 | 1.18 | 2.17 | 1.11 | 1.14 | 1.2 | 0.64 | 1.42 | 1.7 | 0.87 | 0.52 | 1.07 | 0.9 | 0.025 | -0.135 | -0.535 | 0.944144 | 0.696783 | 0.106678 | 1 | 0.996511 | 0.703417 |
| Mpp3 | Q6XE40 | 1.41 | 1.35 | 0.86 | 0.52 | 1.37 | 1.44 | 0.64 | 0.4 | 0.73 | 0.22 | 0.56 | 1.29 | 0.87 | 0.72 | 0.7 | 0.79 | -0.0725 | -0.335 | -0.265 | 0.824437 | 0.297111 | 0.303136 | 1 | 0.996511 | 0.831227 |
| Mpp7 | G5E8S8 | 1.02 | 1.07 | 0.2 | 0.43 | 1 | 1.11 | 1.17 | 0.76 | 0.99 | -0.71 | 0.37 | 1.5 | 0.42 | 1.99 | 0.28 | 0.67 | 0.33 | -0.1425 | 0.16 | 0.205089 | 0.77027 | 0.708453 | 1 | 0.996511 | 0.957562 |
| Mpped1 | E9PYE1 | -1.8 | -2.08 | -1.36 | -1.13 | -0.68 | -1.63 | -1.28 | -1.19 | -0.41 | -0.56 | -1.36 | -0.8 | -0.29 | 1.04 | 0.17 | -0.48 | 0.3975 | 0.81 | 1.7025 | 0.194833 | 0.024519 | 0.001822 | 1 | 0.996511 | 0.419079 |
| Mpped2 | Q9CZJ0 | -0.3 | -0.85 | 0.15 | -0.27 | -0.27 | -0.62 | 0.57 | -0.06 | -1.76 | -0.94 | -0.09 | -0.36 | -1.53 | -1.35 | -0.12 | 0.19 | 0.2225 | -0.47 | -0.385 | 0.488868 | 0.263062 | 0.404173 | 1 | 0.996511 | 0.871454 |
| Mprip | Q5SWZ5 | 1.05 | 0.86 | 1.29 | 0.89 | 0.86 | 0.88 | 0.87 | 0.87 | 0.68 | 1.07 | 1.33 | 1.14 | 0.73 | 0.94 | 0.99 | 0.93 | -0.1525 | 0.0325 | -0.125 | 0.354895 | 0.877456 | 0.517239 | 1 | 0.996511 | 0.905864 |
| Mprip | P97434 | 1.07 | 0.96 | 0.09 | -0.2 | 0.85 | 1.1 | -0.14 | 0.16 | 0.96 | 0.07 | 0.1 | 0.69 | 0.54 | 1.37 | -0.84 | -0.32 | 0.0125 | -0.025 | -0.2925 | 0.975253 | 0.946433 | 0.588514 | 1 | 0.996755 | 0.929131 |
| Mpst | Q99J99 | 1.49 | 1.28 | 0.89 | 0.56 | 1 | 1.48 | 1.11 | 0.76 | 0.44 | 0.12 | 0.9 | 1.26 | 0.07 | 0.92 | 0.43 | 0.55 | 0.0325 | -0.375 | -0.5625 | 0.901626 | 0.266355 | 0.077748 | 1 | 0.996511 | 0.676942 |
| Mpv17 | D3YWI4 | 0.41 | 0.83 | 1.27 | 0.15 | 0.87 | 0.73 | 0.43 | 0.15 | 0.28 | 0.62 | 0.66 | 0.42 | 0.47 | -0.38 | -0.02 | 1.31 | -0.12 | -0.17 | -0.32 | 0.683294 | 0.540913 | 0.4547 | 1 | 0.996511 | 0.887169 |
| Mpv17l | D3Z6M3 | -0.36 | 0.59 | -0.03 | -1.14 | 1.28 | -0.44 | -0.61 | -0.64 | -0.76 | 0.75 | -0.4 | -0.03 | 0.08 | 0.65 | -0.88 | -0.57 | 0.1325 | 0.125 | 0.055 | 0.805741 | 0.783622 | 0.906129 | 1 | 0.996511 | 0.986321 |
| Mpv17l2 | Q8VIK2 | -0.23 | -0.01 | -1.67 | -1.81 | 0.52 | -0.06 | -0.24 | -1.21 | -0.17 | -0.66 | -0.88 | -0.37 | 0.28 | 0.9 | -1.7 | -1.29 | 0.6825 | 0.41 | 0.4775 | 0.23033 | 0.388765 | 0.504635 | 1 | 0.996511 | 0.901002 |
| Mpzl1 | A0A0A6YW | 0.51 | 0.13 | 0.98 | 0.61 | 0.57 | 0.74 | -1.05 | -0.07 | 1.37 | 2.5 | 0.59 | 1.03 | 1.04 | 1.19 | 1.42 | 1.09 | -0.51 | 0.815 | 0.6275 | 0.239185 | 0.08045 | 0.024325 | 1 | 0.996511 | 0.626254 |
| Mras | O08989 | 0.53 | 0.81 | 1.18 | 1.04 | 1.14 | 0.53 | 1.19 | 0.94 | 0.19 | 0.68 | 1.05 | 0.39 | -0.07 | 0.87 | 0.54 | 0.9 | 0.06 | -0.3125 | -0.33 | 0.791988 | 0.242125 | 0.265371 | 1 | 0.996511 | 0.817264 |
| Mrc1 | Q61830 | -0.69 | -1.54 | -0.82 | -0.82 | -0.96 | -0.14 | -1.15 | -1.02 | -1.56 | 0.1 | -1.15 | -1.28 | -1.21 | -1.22 | -1.12 | -0.53 | 0.15 | -0.005 | -0.0525 | 0.61727 | 0.989951 | 0.849214 | 1 | 0.999285 | 0.976225 |
| Mrc2 | A2AAA9 | -1.82 | -1.49 | 1.9 | 0.14 | -1.81 | -1.78 | -0.2 | 1.24 | -1.96 | 1.93 | -0.33 | -1.31 | -2.19 | -1.32 | -0.37 | 2.04 | -0.32 | -0.1 | -0.1425 | 0.751442 | 0.926286 | 0.898262 | 1 | 0.996755 | 0.983509 |
| Mre11 | Q61216-2 | -5.99 | -6 | -2.78 | -3.6 | -5.91 | -7.15 | -3.08 | -3.23 | -3.89 | -2.93 | -3.08 | -5.43 | -3.17 | -6.4 | -3.44 | -3.34 | -0.25 | 0.76 | 0.505 | 0.830334 | 0.411328 | 0.620167 | 1 | 0.996511 | 0.93807 |
| Mri1 | Q9CQT1 | 0.32 | -0.03 | 0.59 | -0.15 | -0.24 | 0.09 | 0.85 | 0.3 | 0.12 | 0.17 | 0.63 | 0.02 | -0.01 | 0.02 | 0.56 | 0.54 | 0.0675 | 0.0525 | 0.095 | 0.813283 | 0.828693 | 0.715413 | 1 | 0.996511 | 0.958106 |
| Mrm1 | Q99J25 | 0.49 | 0.2 | 0.47 | 0.9 | 0.15 | 0.47 | 0.1 | 0.58 | 0.69 | 0.44 | 0.45 | 0.27 | 0.86 | 0.93 | -0.24 | 0 | -0.19 | -0.0525 | -0.1275 | 0.383823 | 0.803476 | 0.702863 | 1 | 0.996511 | 0.956859 |
| Mrm3 | Q5ND52 | 0.4 | 1.06 | 0.55 | 0.38 | 0.67 | 0.59 | -0.14 | 0.14 | 0.03 | 0.62 | 0.16 | 0.31 | -0.4 | 0.64 | 0.34 | 0.09 | -0.2825 | -0.3175 | -0.43 | 0.290714 | 0.197683 | 0.16158 | 1 | 0.996511 | 0.757516 |
| Mroh1 | E0CZ22 | 0.25 | 0.73 | 0.39 | -0.06 | 1.51 | 0.38 | 0.4 | 0.18 | 0.2 | 0.14 | 0.4 | 0.39 | 0.24 | 0.99 | -0.08 | 0.35 | 0.29 | -0.045 | 0.0475 | 0.394556 | 0.835108 | 0.871313 | 1 | 0.996511 | 0.979293 |
| Mrpl1 | Q99N96 | 0.62 | 0.66 | -0.1 | 0.05 | 0.55 | 0.93 | 0.23 | 0.17 | 0.77 | -0.39 | -0.12 | 0.34 | 0.65 | 1.4 | -0.25 | -0.36 | 0.1625 | -0.1575 | 0.0525 | 0.5483 | 0.62634 | 0.903792 | 1 | 0.996511 | 0.985278 |
| Mrpl10 | Q3TBW2 | 0.42 | 0.46 | 0.39 | 0.62 | 0.34 | 0.51 | 0.57 | 0.64 | 0.95 | 0.32 | 0.7 | 0.42 | 0.81 | 1.12 | 0.16 | 0.27 | 0.0425 | 0.125 | 0.1175 | 0.781557 | 0.53757 | 0.654227 | 1 | 0.996511 | 0.947657 |
| Mrpl11 | Q9CQF0 | 0.68 | 0.63 | 1.19 | 0.61 | 0.66 | 0.74 | 1.53 | 1.01 | 0.62 | 0.21 | 0.93 | 0.67 | 0.71 | 0.29 | 1.14 | 1.53 | 0.2075 | -0.17 | 0.14 | 0.419246 | 0.473762 | 0.654208 | 1 | 0.996511 | 0.947657 |
| Mrpl12 | Q9DB15 | 0.51 | 0.82 | 0.69 | 1.13 | 0.74 | 0.66 | 1.01 | 1.09 | 0.01 | 0.46 | 1.02 | 0.88 | -0.06 | 0.43 | 0.69 | 0.57 | 0.0875 | -0.195 | -0.38 | 0.663139 | 0.486338 | 0.148397 | 1 | 0.996511 | 0.746406 |
| Mrpl13 | Q9D1P0 | 0.62 | 0.51 | 0.82 | 0.4 | 0.64 | 0.57 | 1.15 | 1.02 | 0.5 | 0.47 | 1.14 | 0.46 | 0.17 | 0.76 | 0.88 | 0.96 | 0.2575 | 0.055 | 0.105 | 0.220873 | 0.806128 | 0.66248 | 1 | 0.996511 | 0.949645 |
| Mrpl14 | Q9D1I6 | 0.38 | 0.67 | 0.46 | -0.16 | 0.57 | 0.52 | 0.55 | 0.37 | 0.58 | 0.26 | 0.37 | 0.16 | 0.05 | 0.23 | 0.47 | 0.31 | 0.165 | 0.005 | -0.0725 | 0.441107 | 0.982665 | 0.760542 | 1 | 0.998773 | 0.963123 |
| Mrpl15 | Q9CPR5 | 0.57 | 0.21 | 0.44 | 0.73 | 0.13 | 0.48 | 0.62 | 0.57 | 0.82 | 0.82 | 0.52 | 0.51 | 0.52 | 0.85 | 0.23 | 0.27 | -0.0375 | 0.18 | -0.02 | 0.845831 | 0.366731 | 0.929654 | 1 | 0.996511 | 0.989687 |
| Mrpl16 | Q99N93 | -0.56 | -0.27 | 0.43 | 0.35 | -0.14 | -0.43 | 0.84 | 0.81 | -0.17 | 0.89 | 0.56 | -0.59 | -0.17 | 0.4 | 0.15 | -0.04 | 0.2825 | 0.185 | 0.0975 | 0.468993 | 0.642655 | 0.735504 | 1 | 0.996511 | 0.958635 |
| Mrpl17 | Q9D8P4 | 0.7 | 0.52 | 0.72 | 0.6 | 0.3 | 0.44 | 0.48 | 0.96 | 0.3 | 0.82 | 0.7 | 0.45 | 0.44 | 0.3 | 0.49 | 0.24 | -0.09 | -0.0675 | -0.2675 | 0.637337 | 0.718486 | 0.152244 | 1 | 0.996511 | 0.750422 |
| Mrpl18 | Q9CQL5 | -0.59 | -0.34 | -0.34 | 0.3 | -0.27 | -0.26 | -0.07 | 0.02 | -0.25 | 0.14 | 0.11 | -0.81 | -0.12 | 0.26 | -0.32 | -1.03 | 0.0975 | 0.04 | -0.06 | 0.666019 | 0.893196 | 0.8571 | 1 | 0.996511 | 0.976225 |
| Mrpl19 | Q9D338 | -0.23 | -0.16 | -0.32 | -0.12 | -0.2 | -0.38 | 0.08 | -0.3 | 0.03 | -0.57 | -0.27 | -0.52 | -0.51 | 0.55 | -0.84 | -0.69 | 0.0075 | -0.125 | -0.165 | 0.96392 | 0.529377 | 0.612317 | 1 | 0.996511 | 0.935166 |
| Mrpl2 | Q9D773 | -0.45 | -0.72 | 0 | 0.18 | -0.66 | -0.34 | -0.11 | 0.24 | -0.08 | 0.45 | 0.28 | -0.81 | 0.38 | -0.46 | 0.58 | -0.09 | 0.03 | 0.2075 | 0.35 | 0.915285 | 0.548129 | 0.289019 | 1 | 0.996511 | 0.824705 |
| Mrpl20 | Q9CQL4 | 0.32 | 0.47 | 0.31 | 0.7 | 0.08 | 0.53 | 0.62 | 0.73 | 0.1 | -0.84 | 0.65 | 0.4 | -0.02 | -0.77 | 0.07 | 0.24 | 0.04 | -0.3725 | -0.57 | 0.84265 | 0.283573 | 0.057724 | 1 | 0.996511 | 0.661738 |
| Mrpl21 | Q9D1N9 | 0.46 | 0.59 | 0.66 | 1.23 | 0.62 | 0.37 | 1.09 | 1.55 | 0.87 | 0.82 | 1.11 | 0.34 | 0.19 | 1.04 | 0.09 | 0.3 | 0.1725 | 0.05 | -0.33 | 0.578117 | 0.844865 | 0.274922 | 1 | 0.996511 | 0.822084 |
| Mrpl22 | Q8BU88 | -0.24 | 0.17 | 0.71 | 1.25 | 0.17 | 0.22 | 0.93 | 1.22 | 0.44 | 1.23 | 1.03 | -0.25 | 0.34 | 1.3 | 0.64 | 0.78 | 0.1625 | 0.14 | 0.2925 | 0.680768 | 0.749964 | 0.440896 | 1 | 0.996511 | 0.883549 |
| Mrpl23 | O35972 | -0.42 | 0.09 | -0.53 | -0.69 | 0.14 | -0.29 | -0.29 | -0.6 | -0.27 | -0.18 | -0.29 | -0.27 | -0.51 | 0.44 | -0.9 | -0.74 | 0.1275 | 0.135 | -0.04 | 0.601006 | 0.52929 | 0.907248 | 1 | 0.996511 | 0.986458 |
| Mrpl24 | Q9CQ06 | -0.29 | 0.32 | 0.37 | 0.42 | -0.05 | -0.11 | 0.21 | 0.34 | 0.79 | 0.29 | 0.36 | -0.13 | 0.56 | 0.66 | 0.13 | -0.06 | -0.1075 | 0.1225 | 0.1175 | 0.62844 | 0.649046 | 0.65973 | 1 | 0.996511 | 0.949645 |
| Mrpl27 | Q99N92 | 0.19 | 0.39 | 0.19 | 0.09 | -0.4 | 0.3 | -0.1 | -0.02 | 0.35 | 0.49 | -0.46 | 0.25 | 0.39 | 1.08 | 0.04 | -0.07 | -0.27 | -0.0575 | 0.145 | 0.187194 | 0.815171 | 0.61375 | 1 | 0.996511 | 0.935874 |
| Mrpl28 | Q9D1B9 | 2.02 | 1.69 | 0.57 | 0.6 | 1.8 | 1.98 | 0.72 | 0.84 | 2.42 | 0.79 | 0.74 | 1.85 | 2.14 | 2.43 | 0.53 | 0.63 | 0.115 | 0.23 | 0.2125 | 0.802195 | 0.657203 | 0.71069 | 1 | 0.996511 | 0.957819 |
| Mrpl3 | Q99N95 | 0.69 | 0.54 | 0.55 | 0.55 | 0.37 | 0.6 | 0.71 | 0.45 | 0.06 | 0.85 | 0.57 | 0.26 | 0.57 | 0.17 | 0.88 | 0.28 | -0.05 | -0.1475 | -0.1075 | 0.745835 | 0.501517 | 0.621996 | 1 | 0.996511 | 0.938628 |
| Mrpl30 | Q9D7N6 | 0.35 | 0.2 | 0.72 | 0.64 | -0.47 | 0.5 | 0.25 | 0.84 | 0.95 | 1.87 | 0.56 | -0.03 | 1.88 | 0.37 | 1.08 | 0.46 | -0.1975 | 0.36 | 0.47 | 0.516736 | 0.378339 | 0.217794 | 1 | 0.996511 | 0.787447 |
| Mrpl32 | Q9DCI9 | 0.05 | 0.15 | -0.07 | -0.18 | 0.16 | 0.59 | 0.28 | 0.12 | 1.07 | 1.25 | -0.15 | 0.3 | 0.12 | 1.12 | 0.39 | 0.34 | 0.3 | 0.63 | 0.505 | 0.116217 | 0.086259 | 0.077435 | 1 | 0.996511 | 0.676942 |
| Mrpl33 | Q9CQP0 | -0.23 | -0.18 | 1.08 | 0.93 | -0.63 | -0.19 | 0.88 | 0.7 | -0.52 | 1.01 | 0.78 | -0.31 | -0.44 | 0.19 | 0.37 | 0.15 | -0.21 | -0.16 | -0.3325 | 0.654154 | 0.741979 | 0.394341 | 1 | 0.996511 | 0.86603 |
| Mrpl34 | Q99N91 | 0.03 | 0.04 | 1.35 | -0.12 | -0.07 | 0 | -1.26 | -0.93 | 0.16 | 0.69 | -1.36 | -0.44 | 2.08 | 1.51 | 0.49 | -1.36 | -0.89 | -0.5625 | 0.355 | 0.067041 | 0.293779 | 0.638015 | 1 | 0.996511 | 0.942804 |
| Mrpl35 | Q9CQL6 | 0.31 | 0.75 | 2.31 | 0.47 | 1.18 | 1.1 | 3.47 | 2.96 | -0.81 | -1.95 | 3.44 | 0.92 | -0.25 | -0.37 | 3.15 | 3.59 | 1.2175 | -0.56 | 0.57 | 0.10579 | 0.622147 | 0.585911 | 1 | 0.996511 | 0.928178 |
| Mrpl37 | Q921S7 | 0.44 | 0.55 | 0.31 | 0.61 | 0.22 | 0.68 | 0.79 | 0.64 | 0.63 | 0.55 | 0.57 | 0.53 | 0.62 | 0.61 | 0.4 | 0.31 | 0.105 | 0.0925 | 0.0075 | 0.571462 | 0.570285 | 0.967664 | 1 | 0.996511 | 0.9927 |
| Mrpl38 | Q8K2M0 | 0.26 | 0.09 | 0.02 | -0.1 | -0.18 | 0.15 | -0.11 | -0.2 | 0.43 | 0.3 | 0.25 | -0.04 | 0.41 | 0.11 | 0.1 | -0.06 | -0.1525 | 0.1675 | 0.0725 | 0.371309 | 0.378104 | 0.711311 | 1 | 0.996511 | 0.957874 |
| Mrpl39 | Q9JKF7 | 3.66 | 0.27 | 0.5 | 0.65 | 0.54 | 3.81 | 1 | 0.84 | -0.05 | -0.02 | 0.84 | 2.77 | 0.2 | -0.14 | 0.51 | 0.64 | 0.2775 | -0.385 | -0.9675 | 0.779682 | 0.681634 | 0.213792 | 1 | 0.996511 | 0.786931 |
| Mrpl4 | Q9DCU6 | -0.36 | -0.32 | -0.26 | -0.09 | -0.46 | 0.1 | 0.23 | -0.11 | 2.02 | -0.11 | -0.06 | -0.54 | 3.02 | -0.08 | -0.09 | -0.61 | 0.1975 | 0.585 | 0.8175 | 0.329834 | 0.290337 | 0.29332 | 1 | 0.996511 | 0.825612 |
| Mrpl40 | D3Z7C0 | -0.49 | -0.7 | -0.32 | -0.13 | 2.48 | -0.67 | -0.31 | -0.16 | 0.98 | 0.02 | 0.54 | -0.31 | 0.93 | 0.18 | -0.3 | -0.16 | 0.745 | 0.7175 | 0.5725 | 0.280646 | 0.04437 | 0.092248 | 1 | 0.996511 | 0.695968 |
| Mrpl41 | Q9CQN7 | 0.34 | 0.44 | 0.64 | 0.99 | 0.46 | 0.14 | 0.5 | 0.76 | 0.42 | 0.73 | 0.66 | 0.27 | 0.29 | 0.69 | 0.67 | 0.18 | -0.1375 | -0.0825 | -0.145 | 0.530447 | 0.705319 | 0.542218 | 1 | 0.996511 | 0.912212 |
| Mrpl42 | Q9CPV3 | 1.46 | 1.26 | -1.11 | -1.5 | 1.02 | 1.71 | -1.34 | -1.27 | -0.57 | -1.5 | -0.92 | 1.43 | -0.68 | -0.33 | -1.36 | -2.77 | 0.0025 | -0.4175 | -1.3125 | 0.99798 | 0.645674 | 0.148159 | 1 | 0.996511 | 0.746406 |
| Mrpl43 | Q5RL20 | -0.3 | -0.36 | 0.11 | 0.15 | -0.44 | -0.3 | -0.09 | 0.13 | 1.74 | 0.66 | -0.05 | -0.42 | 1.66 | 0.15 | 0.34 | 0.32 | -0.075 | 0.5825 | 0.7175 | 0.722363 | 0.229609 | 0.077908 | 1 | 0.996511 | 0.676942 |
| Mrpl44 | Q9CY73 | -0.25 | 0.06 | 0.04 | 1.05 | -0.03 | 0.06 | 0.23 | 1.01 | 0.72 | 1.2 | 0.55 | -0.3 | 1.26 | -0.25 | 0.51 | 0.23 | 0.0925 | 0.3175 | 0.2125 | 0.79457 | 0.439604 | 0.605194 | 1 | 0.996511 | 0.932185 |
| Mrpl45 | Q9D0Q7 | 0.67 | 0.61 | 0.79 | 0.59 | 0.64 | 0.67 | 0.54 | 0.76 | -0.67 | 0.09 | 0.75 | 0.51 | -0.56 | -0.48 | 0.57 | 0.49 | -0.0125 | -0.495 | -0.66 | 0.931766 | 0.143995 | 0.061796 | 1 | 0.996511 | 0.662517 |
| Mrpl46 | Q9EQI8 | -0.39 | -0.33 | -0.21 | -0.02 | -0.25 | -0.22 | -0.24 | -0.12 | -0.32 | 0.46 | -0.24 | -0.32 | -0.44 | -0.48 | -0.35 | -0.09 | 0.03 | 0.1325 | -0.1025 | 0.846671 | 0.577311 | 0.598948 | 1 | 0.996511 | 0.930565 |
| Mrpl47 | Q8K2Y7 | 1.02 | -0.21 | -0.52 | -0.86 | -0.07 | 0.84 | -0.48 | -0.96 | -0.24 | -0.46 | -0.53 | 0.06 | -0.27 | -0.21 | -0.57 | -0.92 | -0.025 | -0.15 | -0.35 | 0.961279 | 0.715392 | 0.415303 | 1 | 0.996511 | 0.874945 |
| Mrpl48 | A0A140LHT | 0.34 | 0.48 | -0.33 | 0.09 | 0.17 | 0.5 | -0.12 | -0.02 | 0.68 | -0.28 | -0.49 | 0.19 | 0.77 | 0.79 | -0.58 | -0.46 | -0.0125 | -0.12 | -0.015 | 0.958381 | 0.705632 | 0.970119 | 1 | 0.996511 | 0.992767 |
| Mrpl49 | Q9CQ40 | 1.58 | 1.14 | 0.32 | 0.89 | 1.27 | 1.29 | 0.55 | 0.44 | 1.06 | 0.6 | 0.21 | 1.34 | 0.52 | 1.34 | 0.48 | 0.32 | -0.095 | -0.18 | -0.3175 | 0.77817 | 0.61393 | 0.372129 | 1 | 0.996511 | 0.859424 |
| Mrpl49 | A0A494B9( | -0.22 | 1.43 | -0.19 | -1.07 | 0.7 | 0.04 | 0.69 | -0.12 | 0.6 | -1.16 | -0.66 | -0.21 | -0.33 | -0.57 | -0.18 | -0.27 | 0.34 | -0.345 | -0.325 | 0.518067 | 0.562144 | 0.51574 | 1 | 0.996511 | 0.905674 |
| Mrpl50 | Q8VDT9 | 0.47 | 0.6 | 0.37 | -0.29 | 0.49 | 0.3 | 0.44 | 0.17 | -0.06 | -0.03 | -0.03 | 0.38 | -0.39 | 1.13 | 0.03 | 0.07 | 0.0625 | -0.2225 | -0.0775 | 0.786318 | 0.381619 | 0.834848 | 1 | 0.996511 | 0.975487 |
| Mrpl51 | Q9CPY1 | -0.12 | 0.37 | -0.13 | 0.09 | 0.26 | 0.19 | 0.23 | -0.18 | 0.15 | -0.12 | -0.03 | 0.2 | 0.2 | 0.57 | -0.37 | 0.16 | 0.0725 | -0.0025 | 0.0875 | 0.707975 | 0.989686 | 0.733908 | 1 | 0.999285 | 0.958635 |
| Mrpl52 | Q9D0Y8 | -0.25 | 0.14 | 4.54 | 1.47 | -0.16 | 0.01 | 3.61 | 3.14 | 0.3 | 0.75 | 3.59 | 0.15 | 0.13 | -1.07 | 4.68 | 4.8 | 0.175 | -0.2775 | 6.60E-01 | 0.894481 | 0.818951 | 0.692065 | 1 | 0.996511 | 0.955729 |
| Mrpl53 | Q9D1H8 | 0.36 | -0.68 | -0.13 | 0.41 | -0.45 | 0.64 | -0.37 | -0.06 | 1.33 | -0.3 | -0.3 | 0.25 | 1.43 | -0.87 | -0.38 | -0.5 | -0.05 | 0.255 | -0.07 | 0.884002 | 0.561804 | 0.89511 | 1 | 0.996511 | 0.983509 |
| Mrpl54 | Q9CPW3 | -0.4 | -0.06 | -0.05 | 0.52 | -0.24 | 0.02 | 0.84 | 0.51 | 0.37 | 0.9 | 0.27 | -0.54 | 0 | 0.36 | 1.04 | 0.66 | 0.28 | 0.2475 | 0.5125 | 0.370397 | 0.482129 | 0.118673 | 1 | 0.996511 | 0.721448 |
| Mrpl55 | Q9CZ83 | -0.12 | -0.15 | -0.45 | -0.07 | -1.27 | 0.55 | -0.49 | -0.36 | 1.08 | -0.52 | -0.3 | -0.45 | 0.97 | 0.07 | -0.07 | -0.46 | -0.195 | 0.15 | 0.325 | 0.596528 | 0.690493 | 0.325259 | 1 | 0.996511 | 0.842881 |
| Mrpl57 | Q9CQF8 | -0.16 | 0.48 | 1.01 | 1.04 | 0.06 | 0.33 | 0.41 | 0.83 | 0.86 | 1.52 | 0.81 | -0.11 | 0.85 | 0.83 | 0.81 | 0.04 | -0.185 | 0.1775 | 0.04 | 0.563677 | 0.671132 | 0.907279 | 1 | 0.996511 | 0.986458 |
| Mrpl58 | Q8R035 | 0.19 | 0 | -0.31 | 0.01 | -0.76 | 0.4 | -0.54 | -0.34 | 1.76 | 1.81 | -0.21 | 0.07 | 2.33 | -0.28 | -0.6 | -0.2 | -0.2825 | 0.885 | 0.34 | 0.321627 | 0.110084 | 0.590417 | 1 | 0.996511 | 0.929142 |
| Mrpl9 | Q99N94 | 0.18 | 0.38 | 0.62 | 0.83 | 0.04 | 0.13 | 0.31 | 0.76 | 0.49 | 1.23 | 0.42 | 0.28 | 0.27 | 0.6 | 0.67 | 0.53 | -0.1925 | 0.1025 | 0.015 | 0.416647 | 0.706113 | 0.945184 | 1 | 0.996511 | 0.992376 |
| Mrps10 | E9QJS0 | -0.14 | 0.34 | -0.23 | -0.43 | 0.66 | 0.1 | -0.09 | -0.33 | 0.22 | -0.44 | -0.25 | -0.46 | 0 | 0.81 | -0.51 | -0.24 | 0.2 | -0.1175 | 0.13 | 0.469252 | 0.640889 | 0.695821 | 1 | 0.996511 | 0.955729 |
| Mrps11 | Q3U8Y1 | 0.49 | 0.64 | 0.12 | -0.07 | 0.34 | 0.87 | 0.33 | 0.83 | 0.64 | -0.87 | -0.06 | 0.55 | 0.28 | 0.8 | 0.07 | 0.3 | 0.2975 | -0.23 | 0.0675 | 0.231744 | 0.541287 | 0.792277 | 1 | 0.996511 | 0.96933 |
| Mrps12 | O35680 | 0.48 | -0.53 | 0.23 | 0.96 | -0.67 | 0.98 | 0.64 | 1.05 | 0.4 | 1.45 | 0.29 | 0.03 | 1.54 | -1.2 | 0.48 | 0.59 | 0.215 | 0.2575 | 0.0675 | 0.649144 | 0.542814 | 0.909624 | 1 | 0.996511 | 0.986458 |
| Mrps14 | D3Z2K2 | 0.68 | 0.49 | 1.21 | 0.7 | 0.4 | -0.33 | 1.82 | 1.17 | 0.42 | 0.53 | 1.62 | 0.21 | 0.18 | 0.78 | 1.42 | 3.15 | -0.005 | -0.075 | 0.6125 | 0.991252 | 0.828 | 0.325195 | 1 | 0.996511 | 0.842881 |
| Mrps15 | Q9DC71 | 0.88 | 0.67 | 1.52 | 1.42 | 0.41 | 0.59 | 0.96 | 1.35 | 1.1 | 1.42 | 1.15 | 0.36 | 1.78 | 0.3 | 1.35 | 1.31 | -0.295 | -0.115 | 0.0625 | 0.328226 | 0.711351 | 0.86517 | 1 | 0.996511 | 0.977944 |
| Mrps16 | Q9CPX7 | -0.16 | -0.02 | -0.2 | 0.38 | -0.37 | 0.24 | -0.32 | 0.21 | 1.48 | 0.75 | 0.11 | -0.69 | 1.49 | 0.38 | -0.78 | -0.34 | -0.06 | 0.4125 | 0.1875 | 0.795017 | 0.373105 | 0.697509 | 1 | 0.996511 | 0.955729 |
| Mrps17 | Q9CQE3 | 2.95 | 1.06 | -0.6 | -0.67 | 1.87 | 3.25 | 0.14 | -0.43 | 1.1 | 0.26 | -0.57 | 2.4 | 0.71 | 1.13 | -0.46 | -0.11 | 0.5225 | 0.1125 | -0.3675 | 0.628392 | 0.906322 | 0.661788 | 1 | 0.996744 | 0.949645 |
| Mrps18a | Q99N85 | -0.3 | -0.31 | -0.53 | -0.12 | -0.12 | -0.04 | -0.23 | -0.91 | 0.23 | 0.22 | -0.27 | 0.09 | 0.33 | 0.86 | -0.96 | -0.19 | -0.01 | 0.3825 | 0.325 | 0.965847 | 0.078275 | 0.408968 | 1 | 0.996511 | 0.874356 |
| Mrps18b | Q99N84 | 0.88 | 0.76 | -0.86 | -1.15 | 0.31 | 0.47 | -0.75 | -0.81 | 0.48 | -0.8 | -1.19 | 0.71 | 0.47 | -0.02 | -0.7 | -0.81 | -0.1025 | -0.1075 | -0.1725 | 0.858711 | 0.867799 | 0.759366 | 1 | 0.996511 | 0.963123 |
| Mrps18c | Q8R2L5 | -0.59 | 0.79 | -0.54 | -0.75 | -0.3 | 0.27 | -0.01 | -0.07 | 1.02 | 0.26 | -0.26 | -0.09 | -1.13 | 0.12 | -1.5 | 1.22 | 0.245 | 0.505 | -0.05 | 0.501615 | 0.260869 | 0.938796 | 1 | 0.996511 | 0.989956 |
| Mrps2 | Q924T2 | 0.38 | 0.28 | 0.1 | -0.08 | 0.11 | 0.02 | -0.23 | 0.07 | -0.66 | -1.87 | 0.14 | -0.1 | -0.23 | -1.33 | -0.1 | -0.18 | -0.1775 | -0.7925 | -0.63 | 0.326456 | 0.095868 | 0.072741 | 1 | 0.996511 | 0.668066 |
| Mrps21 | P58059 | 0.18 | 0.06 | -1.41 | -1.26 | -0.31 | 0.39 | -1.1 | -1.5 | -0.33 | -0.97 | -1.61 | -0.32 | -0.35 | -0.56 | -1.19 | -1.48 | -0.0225 | -0.2 | -0.2875 | 0.967098 | 0.682526 | 0.543343 | 1 | 0.996511 | 0.912212 |
| Mrps22 | Q9CXW2 | 0.98 | 0.96 | 1.18 | 1.68 | 0.82 | 0.85 | 1.29 | 1.66 | 0.92 | 0.64 | 1.19 | 0.42 | 0.92 | 1.42 | 1.21 | 1.05 | -4.50E-02 | -0.4075 | -0.05 | 0.86627 | 0.139883 | 0.834182 | 1 | 0.996511 | 0.97546 |
| Mrps23 | Q8VE22 | -0.05 | 0.26 | 1.01 | 0.73 | 0.1 | 0.06 | 0.98 | 0.9 | -0.53 | 0.27 | 0.66 | -0.36 | -0.71 | -0.06 | 0.56 | 0.65 | 0.0225 | -0.4775 | -0.3775 | 0.946115 | 0.20262 | 0.338724 | 1 | 0.996511 | 0.847578 |
| Mrps24 | Q9CQV5 | 3.24 | 2.99 | -0.72 | -0.48 | 2.08 | 3.52 | -0.44 | -0.73 | 0.2 | -1.71 | -0.85 | 3.07 | 0.54 | 0.93 | -1.37 | -0.43 | -0.15 | -1.08 | -1.34 | 0.909841 | 0.428308 | 0.229286 | 1 | 0.996511 | 0.787447 |
| Mrps25 | Q9D125 | 0.29 | 0.41 | 0.52 | 1.24 | -0.03 | 0.53 | 0.67 | 1.1 | 1.1 | 1.44 | 0.66 | 0.02 | 0.55 | 0.4 | 0.49 | 0.45 | -0.0475 | 0.19 | -0.1425 | 0.878609 | 0.603735 | 0.571806 | 1 | 0.996511 | 0.92467 |
| Mrps26 | Q80ZS3 | 0.39 | 0.19 | 0.45 | 0.21 | 0.39 | 0.16 | 0.11 | 0.43 | 0.87 | -1.39 | 0.39 | -0.2 | 1.63 | 0.12 | 0.83 | 0.43 | -0.0375 | -0.3925 | 0.4425 | 0.817973 | 0.405085 | 0.20884 | 1 | 0.996511 | 0.786218 |
| Mrps27 | Q8BK72 | 0.82 | 0.63 | 0.65 | 0.91 | 0.37 | 0.77 | 0.62 | 0.82 | 0.58 | 0.75 | 0.57 | 0.46 | 0.5 | 0.39 | 0.32 | 0.42 | -0.1075 | -0.1625 | -0.345 | 0.537586 | 0.350358 | 0.076436 | 1 | 0.996511 | 0.675786 |
| Mrps28 | Q9CY16 | 1.84 | 1.67 | 2.12 | 1.79 | 1.49 | 1.92 | 2.16 | 2.39 | 2.01 | 1.33 | 2.25 | 1.78 | 1.42 | 1.32 | 2.16 | 2.33 | 0.135 | -0.0125 | -0.0475 | 0.565892 | 0.959062 | 0.869748 | 1 | 0.996755 | 0.979293 |
| Mrps30 | Q9D0G0 | -0.23 | -0.35 | 0.38 | 0.69 | -0.43 | -0.34 | 0.56 | 0.74 | -0.14 | 0.53 | 0.6 | -0.2 | -0.45 | -0.12 | 0.48 | 0.45 | 0.01 | 0.075 | -0.0325 | 0.978512 | 0.817917 | 0.92313 | 1 | 0.996511 | 0.987595 |
| Mrps31 | Q61733 | 0.65 | 0.7 | -0.24 | 0.17 | 0.58 | 0.45 | -0.31 | -0.09 | 0.29 | -0.05 | -0.12 | 0.29 | 0.7 | 0.59 | -0.28 | -0.43 | -0.1625 | -0.2175 | -0.175 | 0.595923 | 0.420436 | 0.629721 | 1 | 0.996511 | 0.940865 |
| Mrps34 | Q9JIK9 | 0.73 | 0.26 | -0.05 | -0.15 | 0.62 | 0.76 | -0.11 | -0.04 | 0.43 | 0.06 | 0.66 | 0.09 | 0.29 | 0.84 | -0.36 | -0.51 | 0.11 | 0.1125 | -0.1325 | 0.711758 | 0.670027 | 0.715926 | 1 | 0.996511 | 0.958106 |
| Mrps35 | Q8BJZ4 | 0.24 | 0.05 | 0.02 | 0.11 | -0.67 | 0.36 | -0.65 | -0.3 | 0.49 | 1.25 | -0.45 | -0.4 | 1.03 | -1.09 | -0.52 | -0.21 | -0.42 | 0.1175 | -0.3025 | 0.127706 | 0.764344 | 0.487066 | 1 | 0.996511 | 0.896644 |
| Mrps36 | Q9CQX8 | 0.55 | 0.99 | 0.37 | 0.52 | 0.56 | 0.48 | 0.26 | 0.5 | 0.5 | 0.19 | 0.48 | 0.62 | 0.38 | 1.27 | -0.19 | -0.29 | -0.1575 | -0.16 | -0.315 | 0.412729 | 0.446964 | 0.408586 | 1 | 0.996511 | 0.874356 |
| Mrps5 | Q99N87 | 0.81 | 0.25 | -0.09 | -0.03 | 0.54 | 0.92 | 0.33 | 0.07 | 0.41 | -0.48 | -0.08 | 0.22 | 0.5 | 0.6 | -0.25 | -0.27 | 0.23 | -0.2175 | -0.09 | 0.415587 | 0.462542 | 0.778031 | 1 | 0.996511 | 0.967289 |
| Mrps6 | P58064 | 1.09 | 0.78 | 1.31 | 0.94 | 0.84 | 0.9 | 0.99 | 1.39 | 0.25 | 1.48 | 1.12 | 0.54 | 0.12 | 0.52 | 1.23 | 0.99 | -9.8E-17 | -0.1825 | -0.315 | 1 | 0.55295 | 0.292064 | 1 | 0.996511 | 0.825497 |
| Mrps7 | Q80X85 | 0.25 | -0.03 | 1.48 | 0.89 | -0.5 | -0.02 | 1.91 | 0.95 | 0.05 | 1.6 | 1.43 | -0.57 | 0.31 | 0.28 | 1.54 | 1.96 | -0.0625 | -0.02 | 0.375 | 0.913827 | 0.972312 | 0.46746 | 1 | 0.997515 | 0.889213 |
| Mrps9 | Q9D7N3 | -0.21 | 0.21 | 0.28 | -0.15 | -0.05 | 0.19 | -0.22 | 0.18 | 0.31 | 0.15 | 0.02 | -0.16 | -0.1 | -0.07 | -0.08 | 0.2 | -0.0075 | 0.0475 | -0.045 | 0.96922 | 0.817041 | 0.826559 | 1 | 0.996511 | 0.973289 |
| Mrrf | Q9D6S7 | 0.11 | 0.14 | 0.62 | 0.65 | -0.05 | 0.22 | 0.44 | 0.33 | 0.64 | 0.27 | 0.8 | -0.02 | 0.53 | 0.62 | 0.23 | 0.11 | -0.145 | 0.0425 | -0.0075 | 0.494721 | 0.868536 | 0.974329 | 1 | 0.996511 | 0.992977 |
| Mrs2 | Q5NCE8 | 2 | 2.31 | 0.89 | 0.85 | 2.44 | 1.13 | 0.75 | 0.89 | 0.94 | 0.5 | 1.41 | 1.87 | 0.07 | 1.46 | 0.86 | 1.23 | -0.21 | -0.3325 | -0.6075 | 0.674852 | 0.467357 | 0.206157 | 1 | 0.996511 | 0.78517 |
| Mrtfa | A0A5F8MC | -0.3 | 0.03 | -0.89 | -0.94 | -0.48 | -0.31 | -1.51 | -1.17 | -0.03 | -0.44 | -1.02 | 0 | 0.48 | 1.02 | -1.43 | -1.58 | -0.3425 | 0.1525 | 0.1475 | 0.346375 | 0.647817 | 0.817801 | 1 | 0.996511 | 0.972768 |
| Mrtfb | P59759 | -0.29 | 0.11 | -0.19 | -0.33 | -0.35 | -0.57 | -0.76 | -0.47 | 0.1 | 0.47 | -0.26 | 0.07 | 0.74 | 0.61 | -0.37 | -0.86 | -0.3625 | 0.27 | 0.205 | 0.068462 | 0.235938 | 0.59938 | 1 | 0.996511 | 0.930565 |
| Mrto4 | Q9D0I8 | -0.58 | -0.52 | 0.27 | 0.47 | -0.46 | -0.87 | -0.01 | 0.02 | -0.69 | -0.54 | 0.27 | -0.7 | -0.66 | -0.81 | 0.23 | 0.06 | -0.24 | -0.325 | -0.205 | 0.476319 | 0.362874 | 0.57898 | 1 | 0.996511 | 0.927466 |
| Msh2 | P43247 | -2.92 | -3.03 | -2.88 | -2.93 | -2.35 | -3.53 | -2.47 | -3.51 | -3.02 | 0.33 | -2.72 | -2.65 | -2.91 | -2.49 | -3.43 | -3.26 | -0.025 | 0.925 | -0.0825 | 0.936996 | 0.217526 | 0.738018 | 1 | 0.996511 | 0.958796 |
| Msh6 | P54276 | -3.43 | -3.75 | -2.56 | -2.63 | -3.93 | -3.71 | -3.39 | -2.44 | -3.56 | -1.48 | -2.84 | -3.4 | -2.88 | -1.56 | -3.27 | -3.41 | -0.275 | 0.2725 | 0.3125 | 0.512484 | 0.600647 | 0.521817 | 1 | 0.996511 | 0.907667 |
| Msi1 | F8WJA5 | 0.12 | 0.27 | -0.25 | -0.65 | 0.16 | 0.26 | -0.48 | -0.51 | -0.43 | -0.06 | -0.18 | 0.41 | 0.89 | -0.26 | -0.3 | 0.22 | -0.015 | 0.0625 | 0.265 | 0.958747 | 0.824786 | 0.449422 | 1 | 0.996511 | 0.884817 |
| Msi2 | Q920Q6 | -0.68 | -0.53 | -0.91 | -0.67 | -0.51 | -0.9 | -0.83 | -0.8 | -0.95 | -1.33 | -0.93 | -0.94 | -1.04 | -0.76 | -0.6 | -1.04 | -0.0625 | -0.34 | -0.1625 | 0.713496 | 0.095095 | 0.426988 | 1 | 0.996511 | 0.879165 |
| Msl1 | Q6PDM1-2 | -0.17 | -0.6 | -0.62 | -0.29 | -0.46 | -0.07 | -0.72 | -0.43 | -0.27 | -0.27 | -0.29 | -0.54 | 0.91 | 0.2 | -0.32 | -0.05 | 0 | 0.0775 | 0.605 | 1 | 0.68253 | 0.069972 | 1 | 0.996511 | 0.668066 |
| Msl3 | Q9WVG9-2 | 0.01 | -0.7 | -0.17 | -1 | -0.08 | -0.49 | -0.44 | -0.71 | -0.59 | -2.39 | -0.64 | -1.4 | -0.59 | -0.14 | -0.62 | -0.26 | 0.035 | -0.79 | 0.0625 | 0.897769 | 0.10882 | 0.824713 | 1 | 0.996511 | 0.972852 |
| Msn | P26041 | -0.27 | -1.04 | -1.57 | -1.15 | -0.96 | -0.85 | -1.39 | -1.39 | -0.47 | -1.64 | -1.36 | -0.72 | -0.09 | -1.16 | -1.35 | -1.87 | -0.14 | -0.04 | -0.11 | 0.64581 | 0.914284 | 0.802001 | 1 | 0.996755 | 0.971091 |
| Msra | Q9D6Y7-2 | 1.6 | 0.89 | 0.14 | 0.16 | 0.81 | 1.63 | 0.38 | 0.19 | 0.8 | -0.08 | 1.14 | 1.37 | 0.53 | 0.58 | 0.18 | -0.19 | 0.055 | 0.11 | -0.4225 | 0.900643 | 0.804522 | 0.284321 | 1 | 0.996511 | 0.822669 |
| Msrb2 | Q78J03 | 0.58 | 0.55 | 0.28 | 0.46 | 0.37 | 0.46 | -0.05 | 0.13 | 0.68 | 0.59 | 0.43 | 0.69 | 0.43 | 0.61 | -0.08 | -0.03 | -0.24 | 0.13 | -0.235 | 0.204257 | 0.450662 | 0.318955 | 1 | 0.996511 | 0.838969 |
| Msto1 | E9PUB7 | -0.96 | -1.16 | -0.64 | -0.27 | -0.96 | -1.38 | -0.54 | -0.35 | -1.01 | -0.03 | -0.54 | -1.06 | -1.05 | -0.61 | -0.57 | -0.52 | -0.05 | 0.0975 | 0.07 | 0.867105 | 0.755037 | 0.787346 | 1 | 0.996511 | 0.968142 |
| Mt1 | P02802 | -0.36 | -2.87 | -2 | -1.13 | -3.16 | -0.31 | -0.18 | -1.34 | -0.1 | -0.31 | -1.88 | -0.11 | -0.13 | -0.77 | -1.33 | -0.57 | 0.3425 | 0.99 | 0.89 | 0.666386 | 0.14415 | 0.13309 | 1 | 0.996511 | 0.728417 |
| Mt2 | P02798 | -1.36 | -4.56 | -2.96 | -2.96 | -3.67 | -1.45 | -1.72 | -2.68 | -3.47 | -3.19 | -3.31 | -0.94 | -2.07 | -7.16 | -2.84 | -3.25 | 0.58 | 0.2325 | -0.87 | 0.445019 | 0.771974 | 0.463798 | 1 | 0.996511 | 0.888267 |
| Mt3 | P28184 | 1.53 | -3.82 | -1.8 | 1.28 | -5.09 | 0.98 | -2.02 | 0.35 | 1.85 | -1.23 | -1.41 | 1.57 | 1.79 | -6.91 | -1.32 | -2.22 | -0.7425 | 0.8975 | -1.4625 | 0.660136 | 0.523845 | 0.462731 | 1 | 0.996511 | 0.888267 |
| Mta1 | Q8K4B0 | -0.42 | -1.12 | -0.8 | -0.87 | -0.84 | -0.58 | -0.6 | -1.33 | -0.55 | -0.61 | -1.04 | -0.59 | 0.19 | -0.81 | -0.36 | -0.3 | -0.035 | 0.105 | 0.4825 | 0.884837 | 0.637358 | 0.10528 | 1 | 0.996511 | 0.703417 |
| Mta2 | Q9R190 | -1.19 | -2.17 | -1.91 | -1.66 | -1.24 | -1.84 | -1.95 | -2.31 | -1.65 | -1.01 | -1.82 | -1.62 | -1.21 | -1.3 | -1.75 | -1.8 | -0.1025 | 0.2075 | 0.2175 | 0.735008 | 0.472696 | 0.443541 | 1 | 0.996511 | 0.883549 |
| Mta3 | Q3UII8 | -1.04 | -1.22 | -1.53 | -1.36 | -0.96 | -1.01 | -1.59 | -1.73 | -0.89 | -0.92 | -0.97 | -1.16 | -0.69 | -0.94 | -1.39 | -1.42 | -0.035 | 0.3025 | 0.1775 | 0.883513 | 0.126294 | 0.472429 | 1 | 0.996511 | 0.891144 |
| Mtap | Q9CQ65 | 0.42 | 0.05 | 0.1 | -0.05 | -0.26 | -0.26 | -0.57 | -0.55 | -0.4 | 0.22 | -0.15 | 0.37 | -0.68 | -0.33 | 0.15 | 0.05 | -0.54 | -0.12 | -0.3325 | 0.014111 | 0.609246 | 0.204989 | 1 | 0.996511 | 0.783452 |
| Mtarc2 | Q922Q1 | 0.56 | 0.48 | 0.17 | 0.46 | 0.25 | 0.62 | 0.35 | 0.3 | 0.46 | 0.38 | 0.3 | 0.28 | 0.5 | 0.55 | -0.18 | 0.06 | -0.0375 | -0.0625 | -0.185 | 0.826355 | 0.715903 | 0.442256 | 1 | 0.996511 | 0.883549 |
| Mtatp6 | P00848 | -1.75 | -0.53 | -0.26 | -1.65 | 0.76 | -0.69 | 0.74 | -0.36 | -2.92 | -1.53 | 0.07 | -1.86 | -1.92 | -1.99 | -0.58 | -0.07 | 1.16 | -0.5125 | -0.0925 | 0.041035 | 0.447354 | 0.870141 | 1 | 0.996511 | 0.979293 |
| Mtatp8 | P03930 | 1.66 | 1.51 | 1.15 | 1.37 | 1.35 | 2.09 | 1.52 | 1.92 | 1.23 | 1.97 | 1.32 | 1.47 | 1.17 | 1.26 | 1.09 | 1.17 | 0.2975 | 0.075 | -0.25 | 0.20815 | 0.744719 | 0.21188 | 1 | 0.996511 | 0.786269 |
| Mtch1 | Q791T5-2 | -0.14 | 0.26 | -0.6 | -0.37 | 0.31 | 0.05 | 0.1 | -0.28 | 0.01 | -1.09 | -0.05 | -0.03 | 0.08 | 0.45 | -0.58 | -0.11 | 0.2575 | -0.0775 | 0.1725 | 0.29408 | 0.810726 | 0.564292 | 1 | 0.996511 | 0.920315 |
| Mtch2 | Q791V5 | -0.28 | -0.29 | 0.12 | 0.31 | -0.44 | 0.06 | 0.29 | 0.19 | -1.15 | -0.09 | 0.29 | -0.44 | -1.34 | -1.08 | -0.47 | 0.26 | 0.06 | -0.3125 | -0.6225 | 0.800427 | 0.36584 | 0.125625 | 1 | 0.996511 | 0.721451 |
| Mtcl1 | Q3UHU5-3 | -1.2 | -0.88 | -0.56 | -0.8 | -1.01 | -1.48 | -1.57 | -1.43 | -0.77 | -0.37 | -0.86 | -0.72 | 0.18 | 0.02 | -0.48 | -0.87 | -0.5125 | 0.18 | 0.5725 | 0.035078 | 0.405978 | 0.075201 | 1 | 0.996511 | 0.673156 |
| Mtcl1 | H3BL83 | 2.36 | 2.09 | 1.15 | 1.39 | 0.57 | 1.62 | -0.18 | 1.03 | 0.41 | 2.69 | 0.76 | 0.9 | 0.71 | 1.29 | 2.78 | 2.57 | -0.9875 | -0.5575 | 0.09 | 0.050016 | 0.317421 | 0.86584 | 1 | 0.996511 | 0.978094 |
| Mtco1 | P00397 | -4.26 | -4.52 | -3.18 | -2.72 | -3.97 | -3.44 | -2.37 | -2.84 | -3.64 | -4.27 | -2.79 | -4.23 | -3.6 | -4.28 | -4.12 | -3.3 | 0.515 | -0.0625 | -0.155 | 0.327054 | 0.902702 | 0.735757 | 1 | 0.996609 | 0.958635 |
| Mtco2 | P00405 | -1.12 | -0.19 | 0.21 | -0.04 | -0.11 | -0.64 | -0.42 | -0.31 | -0.43 | 0.1 | -0.11 | -1.21 | -0.67 | -0.03 | -0.77 | -0.46 | -0.085 | -0.1275 | -0.1975 | 0.781925 | 0.74541 | 0.559661 | 1 | 0.996511 | 0.918558 |
| mt-Co3 | P00416 | -2.84 | -2.53 | -2.08 | -1.82 | -2.2 | -2.35 | -1.99 | -1.93 | -0.83 | -2.29 | -2.13 | -3.19 | 0.12 | -1.2 | -1.38 | -2.04 | 0.2 | 0.2075 | 1.1925 | 0.444847 | 0.679137 | 0.0315 | 1 | 0.996511 | 0.641974 |
| Mt-Cyb | P00158 | -2.43 | -1.83 | -2.15 | -2.27 | -0.87 | -2.14 | -1.41 | -2.48 | -3.34 | -3.46 | -1.74 | -2.81 | -3.8 | -2.2 | -2.68 | -2.64 | 0.445 | -0.6675 | -0.66 | 0.244655 | 0.118633 | 0.093532 | 1 | 0.996511 | 0.698156 |
| Mtdh | Q80WJ7 | -0.87 | -0.45 | -1.09 | -1.06 | 0.33 | -0.89 | -0.22 | -0.73 | -0.38 | -0.46 | -1.03 | -0.68 | 0.77 | 0.08 | 0.84 | 0.08 | 0.49 | 0.23 | 1.31 | 0.138997 | 0.343892 | 0.00114 | 1 | 0.996511 | 0.412464 |
| Mtdh | F6ZSG0 | -0.94 | -0.05 | -0.93 | -1.39 | 0.2 | -0.57 | 0.44 | -0.2 | -0.79 | -1.37 | -0.72 | -0.15 | 0.01 | 0.3 | 0.37 | 0.24 | 0.795 | 0.07 | 1.0575 | 0.044325 | 0.847807 | 0.006637 | 1 | 0.996511 | 0.552614 |
| Mtdh | F6ZQL0 | -1.4 | -1.1 | -0.3 | -1.25 | -0.49 | -0.66 | 1.24 | 0.37 | -1.58 | -0.31 | -0.76 | -1.18 | -0.91 | -0.49 | 0.2 | 0.61 | 1.1275 | 0.055 | 0.865 | 0.036584 | 0.877862 | 0.057316 | 1 | 0.996511 | 0.661738 |
| Mterf2 | Q8BKY8 | 0.13 | 0.72 | -0.47 | -0.27 | 0.37 | 0.72 | -0.01 | -0.36 | 1.23 | 0.21 | -0.37 | 0.17 | 1.66 | 0.76 | -0.91 | -0.31 | 0.1525 | 0.2825 | 0.2725 | 0.655163 | 0.491901 | 0.638594 | 1 | 0.996511 | 0.942804 |
| Mtfmt | Q9D799 | -0.37 | 0.22 | 0.05 | 0.02 | -0.48 | -0.37 | -0.3 | 0.03 | -0.75 | -1.37 | 0 | -0.21 | -0.42 | -0.48 | -0.44 | 0.15 | -0.26 | -0.5625 | -0.2775 | 0.21466 | 0.114944 | 0.258529 | 1 | 0.996511 | 0.812074 |
| Mtfp1 | Q9CRB8 | 0.04 | 0.39 | 0.83 | 1.44 | 0.82 | 0.12 | 0.96 | 1.42 | 0 | 0.19 | 1.02 | 0.04 | 0.24 | 0.76 | 0.21 | 0.79 | 0.155 | -0.3625 | -0.175 | 0.687321 | 0.344451 | 0.610678 | 1 | 0.996511 | 0.935047 |
| Mtfr1l | Q9CWE0 | 1.12 | 0.9 | 0.43 | 1.02 | 0.43 | 1.16 | 0.56 | 0.79 | 1.21 | 0.47 | 0.58 | 0.9 | 0.94 | 1.08 | 0.34 | 0.5 | -0.1325 | -0.0775 | -0.1525 | 0.580469 | 0.756744 | 0.563091 | 1 | 0.996511 | 0.920024 |
| Mtg1 | Q8R2R6 | -0.7 | -0.87 | 0.05 | 0.14 | -0.78 | -0.32 | -0.44 | 0.28 | -0.82 | 0.95 | -0.04 | -0.63 | -1 | -0.43 | -0.39 | -0.23 | 0.03 | 0.21 | -0.1675 | 0.927399 | 0.639827 | 0.597953 | 1 | 0.996511 | 0.930093 |
| Mthfd1 | Q922D8 | 0.42 | 0.17 | 0.19 | 0.4 | 0.29 | 0.31 | 0.43 | 0.35 | 0.22 | 0.42 | 0.2 | 0.21 | -0.29 | 0.47 | -0.02 | -0.01 | 0.05 | -0.0325 | -0.2575 | 0.738854 | 0.845845 | 0.263705 | 1 | 0.996511 | 0.816045 |
| Mthfd1l | Q3V3R1 | 0.92 | 0.87 | 0.68 | 0.38 | 0.6 | 0.99 | 0.39 | 0.66 | 0.25 | -0.05 | 0.63 | 0.77 | 0.17 | 0.39 | 0.14 | 0.24 | -0.0525 | -0.3125 | -0.4775 | 0.79797 | 0.227252 | 0.038756 | 1 | 0.996511 | 0.658454 |
| Mthfd2l | D3YZG8 | 0.62 | 0.8 | 1.33 | 0.9 | 0.2 | 0.43 | 0.58 | 0.45 | 0.68 | 1.34 | 0.87 | 0.6 | 0.46 | 0.53 | 1.45 | 0.69 | -0.4975 | -0.04 | -0.13 | 0.03415 | 0.8719 | 0.655434 | 1 | 0.996511 | 0.94818 |
| Mthfsl | L7N466 | 0.17 | 0.96 | 0.27 | 0.08 | -0.72 | 0.43 | -1.14 | -0.07 | 0.74 | 0.17 | -0.06 | 0.28 | -2 | -0.2 | -0.78 | -0.51 | -0.745 | -0.0875 | -1.2425 | 0.077452 | 0.751199 | 0.015662 | 1 | 0.996511 | 0.589387 |
| Mtif2 | Q91YJ5 | -0.14 | 0.37 | -0.91 | -0.77 | 0.09 | -0.23 | -0.78 | -0.74 | 0.2 | -0.18 | -0.77 | -0.23 | 0.12 | 0.24 | -1.11 | -1.27 | -0.0525 | 0.1175 | -0.1425 | 0.8804 | 0.73766 | 0.760309 | 1 | 0.996511 | 0.963123 |
| Mtif3 | Q9CZD5 | 3.43 | 2.18 | 2.34 | 0.53 | 2.7 | 3.9 | 3.79 | 2.93 | 1.09 | -0.32 | 2.74 | 3.04 | 1.69 | 0.49 | 2.56 | 3.57 | 1.21 | -0.4825 | -0.0425 | 0.074595 | 0.589348 | 0.957527 | 1 | 0.996511 | 0.992636 |

| | | | | | | | | | | | | | | | | | | | | | | | | | | | |
|---|---|---|---|---|---|---|---|---|---|---|---|---|---|---|---|---|---|---|---|---|---|---|---|---|---|---|---|
| Mtm1 | Q9Z2C5 | 0.4 | 0.11 | 0.23 | 0.05 | 0 | 0.59 | 0.31 | 0.07 | 0.55 | 0.22 | 0.49 | 0.32 | 0.63 | 0.03 | 0.27 | 0.63 | 0.045 | 0.1975 | 0.1925 | 0.814625 | 0.283042 | 0.388567 | | 1 | 0.996511 | 0.86481 |
| Mtmr1 | F6T9H9 | -1.19 | -0.28 | 0.57 | 0.01 | 0.28 | 0.36 | -0.33 | 0.02 | 0.22 | 1.06 | 1.22 | -1.43 | -0.93 | 2.37 | -0.03 | 1.45 | 0.305 | 0.49 | 0.9375 | 0.429961 | 0.456854 | 0.229737 | | 1 | 0.996511 | 0.787447 |
| Mtmr1 | Q9Z2C4 | 0.41 | 0.31 | 0.63 | 0.69 | 0.1 | 0.51 | 0.51 | 0.78 | 0.9 | 0.16 | 0.59 | 0.39 | 1.08 | 0.1 | 0.29 | 0.15 | -0.035 | -1.2E-16 | -0.105 | 0.860879 | 1 | 0.698556 | | 1 | 1 | 0.955729 |
| Mtmr10 | Q7TPM9 | -0.53 | 0.17 | 0.27 | 0.22 | 0.07 | -0.35 | 0.32 | 0.31 | -0.08 | 0.33 | 0.38 | 0.28 | -0.44 | 0.09 | -0.12 | 0.55 | 0.055 | 0.195 | -0.0125 | 0.829672 | 0.429605 | 0.966194 | | 1 | 0.996511 | 0.9927 |
| Mtmr12 | Q80TA6 | -0.33 | -0.45 | 0.27 | -0.14 | -0.7 | -0.52 | 0.07 | 0.31 | -0.13 | -0.19 | -0.02 | -0.33 | 0.02 | -0.96 | -0.3 | -0.07 | -0.0475 | -0.005 | -0.165 | 0.868626 | 0.981175 | 0.571394 | | 1 | 0.998773 | 0.924332 |
| Mtmr14 | Q8VEL2 | -1.59 | -0.86 | -1.31 | -1.7 | -0.97 | -0.91 | -1.16 | -1.71 | -0.77 | -2.44 | -1.01 | -1.55 | 0.03 | -1.03 | -1.73 | -1.77 | 0.1775 | -0.0775 | 0.24 | 0.512766 | 0.845316 | 0.58626 | | 1 | 0.996511 | 0.928178 |
| Mtmr2 | Q9Z2D1 | 0.57 | 0.43 | 0.26 | 0.33 | 0.69 | 0.51 | 0.48 | 0.15 | 0.17 | 0.19 | 0.19 | 0.45 | -0.07 | 0.55 | 0.15 | 0.07 | 0.06 | -0.1475 | -0.2225 | 0.736427 | 0.399414 | 0.302049 | | 1 | 0.996511 | 0.830681 |
| Mtmr3 | B1ATD5 | 0.26 | 0.01 | 0.38 | 0.05 | 0.04 | 0.35 | 0.24 | 0.34 | 0.48 | 0.04 | 0.4 | 0.34 | 0.55 | -0.06 | 0.58 | 0.69 | 0.0675 | 0.14 | 0.265 | 0.689023 | 0.464791 | 0.273952 | | 1 | 0.996511 | 0.822084 |
| Mtmr6 | Q8VE11 | 0.4 | -0.12 | 0.03 | -0.05 | -0.05 | 0.11 | 0.46 | -0.02 | -0.15 | -0.43 | 0.19 | 0.29 | -0.29 | -0.36 | 0.06 | -0.13 | 0.06 | -0.09 | -0.245 | 0.762795 | 0.699524 | 0.258671 | | 1 | 0.996511 | 0.812074 |
| Mtmr7 | Q9Z2C9 | -0.37 | -0.34 | -0.24 | -0.28 | 0.17 | -0.31 | 0.24 | -0.09 | -0.47 | -0.72 | -0.12 | -0.2 | -0.8 | 0 | -0.4 | -0.18 | 0.31 | -0.07 | -0.0375 | 0.107281 | 0.718728 | 0.866715 | | 1 | 0.996511 | 0.978305 |
| Mtmr9 | Q9Z2D0 | -0.47 | -0.33 | -0.31 | -0.1 | -0.13 | -0.18 | -0.14 | -0.16 | -0.37 | -0.29 | -0.03 | -0.09 | -0.49 | -0.32 | -0.14 | -0.39 | 0.15 | 0.1075 | -0.0325 | 0.334745 | 0.551546 | 0.862113 | | 1 | 0.996511 | 0.977204 |
| Mtnd1 | P03888 | -1.27 | -0.67 | -1.25 | -1.37 | -0.37 | -0.97 | -0.95 | -1.69 | -1.24 | -1.41 | -1.13 | -1.37 | -1.57 | -0.58 | -2.43 | -1.49 | 0.145 | -0.1475 | -0.3775 | 0.640795 | 0.493983 | 0.353467 | | 1 | 0.996511 | 0.855029 |
| mt-Nd2 | P03893 | -2.07 | -1.46 | -2.38 | -1.4 | -1.49 | -1.61 | -0.95 | -1.49 | 0.17 | -1.61 | -1.03 | -2.18 | 1.02 | -1.15 | -0.28 | -1.78 | 0.4425 | 0.665 | 1.28 | 0.144271 | 0.219321 | 0.057093 | | 1 | 0.996511 | 0.661738 |
| mt-Nd3 | P03899 | -0.09 | 0.04 | -1.42 | -1.31 | -0.35 | -0.23 | -1.08 | -1.51 | -0.63 | -0.52 | -1.43 | -0.6 | 0.02 | 0.08 | -1.71 | -1.95 | -0.0975 | -0.1 | -0.195 | 0.831619 | 0.812082 | 0.750656 | | 1 | 0.996511 | 0.961479 |
| Mtnd4 | P03911 | -1 | -0.6 | 0.05 | -0.71 | -1.31 | -0.73 | -1.12 | -1.08 | -0.26 | -1.12 | -0.64 | -1.21 | -0.55 | -1.54 | -1.42 | 0.27 | -0.495 | -0.2425 | -0.245 | 0.085411 | 0.449209 | 0.589735 | | 1 | 0.996511 | 0.929142 |
| Mtnd4l | P03903 | -2.54 | -4.18 | -4.38 | -1.69 | -3.76 | -1.29 | -2.52 | -2.89 | -6.06 | -5.99 | -4.09 | -1.92 | -8.57 | -6.46 | -3.73 | -3.42 | 0.5825 | -1.3175 | -2.3475 | 0.444303 | 0.232414 | 0.083353 | | 1 | 0.996511 | 0.689531 |
| Mtnd5 | P03921 | -0.73 | -0.4 | -2.34 | -2 | -0.09 | -0.51 | -1.94 | -2.47 | -0.7 | -2.12 | -1.95 | -0.94 | -0.67 | -0.37 | -2.95 | -2.27 | 0.115 | -0.06 | -0.1975 | 0.86372 | 0.91251 | 0.780751 | | 1 | 0.996744 | 0.9676 |
| Mto1 | G5E889 | 0.11 | 0.3 | 0.43 | 0.44 | -0.27 | 0.16 | 0.37 | 0.32 | 0.68 | 0.33 | 0.18 | -0.27 | 0.72 | 0.34 | 0.19 | 0.51 | -0.175 | -0.09 | 0.12 | 0.388666 | 0.707725 | 0.557015 | | 1 | 0.996511 | 0.917092 |
| Mtor | Q9JLN9 | 0.5 | 0.29 | 0.78 | 0.32 | 0.4 | 0.43 | 0.68 | 0.68 | 0.17 | 0.32 | 0.77 | 0.4 | -0.17 | 0.28 | 0.31 | 0.38 | 0.075 | -0.0575 | -0.2725 | 0.679593 | 0.786434 | 0.235156 | | 1 | 0.996511 | 0.791437 |
| Mtpap | Q9D0D3 | 0.4 | 0.11 | 0.76 | 0.86 | -0.07 | 0.45 | 0.23 | 0.53 | 0.32 | 0.76 | 0.42 | 0.06 | 0.16 | -0.15 | 0.18 | 0.08 | -0.2475 | -0.1425 | -0.465 | 0.309062 | 0.57022 | 0.070953 | | 1 | 0.996511 | 0.668066 |
| Mtpn | P62774 | -0.61 | -0.44 | 1.06 | 2.12 | -0.34 | -0.7 | 0.91 | 2.72 | -2.01 | 1.63 | 1.27 | -0.4 | -1.68 | -2.17 | 0.25 | 0.89 | 0.115 | -0.41 | -1.21 | 0.899 | 0.668417 | 0.194008 | | 1 | 0.996511 | 0.778959 |
| Mtr | A6H5Y3 | -0.73 | -0.4 | 0.04 | -0.05 | -0.58 | -0.58 | 0.27 | 0.36 | -0.05 | 0.01 | -0.02 | -0.36 | -0.3 | -0.21 | -0.25 | -0.03 | 0.1525 | 0.18 | 0.0875 | 0.623654 | 0.439679 | 0.704801 | | 1 | 0.996511 | 0.957084 |
| Mtres1 | Q9CQF4 | -0.16 | 0.07 | 1.99 | 2.55 | 0.23 | -0.01 | 1.4 | 2.67 | -0.45 | 2.71 | 2.4 | -0.14 | -0.13 | -0.57 | 2.18 | 1.98 | -0.04 | 0.0175 | -0.2475 | 0.961332 | 0.985485 | 0.779176 | | 1 | 0.998892 | 0.967289 |
| Mtrex | Q9CZU3 | -0.9 | -1.79 | -1.64 | -1.94 | -1.36 | -1.58 | -1.39 | -1.77 | -1.68 | -0.3 | -2 | -1.64 | -1.65 | -1.15 | -1.16 | -1.53 | 0.0425 | 0.1625 | 0.195 | 0.869533 | 0.699621 | 0.496948 | | 1 | 0.996511 | 0.897918 |
| Mtrf1l | Q8BJU9 | 2.09 | 1.65 | 0.57 | 2.05 | 1.75 | 1.9 | 1.24 | 2.81 | 1.19 | 2.6 | 1.09 | 2.21 | 0.99 | 0.66 | 1.08 | 1.23 | 0.335 | 0.1825 | -0.6 | 0.462929 | 0.705434 | 0.129327 | | 1 | 0.996511 | 0.723655 |
| Mtrfr | Q80VP5 | 0.49 | 0.49 | 1.26 | 0.66 | -0.19 | 0.49 | 0.82 | 0.97 | 1.15 | 0.59 | 1.25 | 0.28 | 1.63 | -0.31 | 1.43 | 1.99 | -0.2025 | 0.0925 | 0.46 | 0.520106 | 0.7584 | 0.374823 | | 1 | 0.996511 | 0.859611 |
| Mtrr | Q8C1A3 | -0.96 | -0.78 | -0.73 | -0.97 | -1.45 | -0.74 | -1.51 | -2.41 | -0.37 | 0.19 | -0.21 | -0.7 | 0.31 | -0.63 | 0.13 | -0.44 | -0.6675 | 0.5875 | 0.7025 | 0.074442 | 0.0283 | 0.023373 | | 1 | 0.996511 | 0.624626 |
| Mtss1 | G3X9H7 | 0.84 | 0.64 | -0.77 | -0.95 | 0.28 | 0.33 | 0.04 | -1.11 | 0.98 | -1.22 | -0.86 | 0.81 | 0.95 | 0.66 | -0.97 | 0.07 | -0.055 | -0.0125 | 0.2375 | 0.916998 | 0.98502 | 0.682667 | | 1 | 0.998892 | 0.955729 |
| Mtss2 | F7D291 | -0.01 | 0.01 | -0.65 | -0.95 | -0.02 | 0.14 | -1.12 | -0.74 | 1.39 | 0.53 | 0.03 | 0 | 2.91 | 0.84 | 2.15 | -1.36 | -0.035 | 0.8875 | 1.535 | 0.923499 | 0.045158 | 0.10422 | | 1 | 0.996511 | 0.702979 |
| Mtss2 | Q6P9S0 | 0.53 | 0.8 | -0.03 | 0.02 | 0.2 | 0.67 | 0.11 | -0.34 | 0.24 | -0.15 | 0.21 | 0.71 | 0.52 | 0.03 | 0.53 | 0.43 | -0.17 | -0.0775 | 0.0475 | 0.560632 | 0.782158 | 0.855932 | | 1 | 0.996511 | 0.976225 |
| Mturn | A0A087WC | -0.36 | 0.3 | 2.76 | 0.36 | 0.12 | -0.58 | 3.45 | 0.44 | -0.77 | 2.35 | 2.62 | 0.17 | -1.02 | -0.17 | 3.24 | 4.38 | 0.0925 | 0.3275 | 0.8425 | 0.926776 | 0.735343 | 0.525216 | | 1 | 0.996511 | 0.908107 |
| Mtus2 | Q3UHD3 | 0.58 | 0.99 | 0.67 | 0.55 | 0.62 | 0.48 | -0.24 | 0.31 | 1.03 | 1.55 | 0.49 | 0.59 | 1 | 0.58 | 0.87 | -0.24 | -0.405 | 0.2175 | -0.145 | 0.108871 | 0.438685 | 0.638488 | | 1 | 0.996511 | 0.942804 |
| Mtx1 | Q8R5C0 | 0.84 | 0.51 | -0.12 | 0.83 | -0.35 | 1.01 | -0.16 | 0.68 | 1.13 | 0.39 | 0.3 | 0.73 | 0.86 | 0.91 | -0.57 | 0.08 | -0.22 | 0.1225 | -0.195 | 0.56359 | 0.683925 | 0.630293 | | 1 | 0.996511 | 0.941132 |
| Mtx2 | O88441 | 2.25 | 1.1 | 0.37 | -0.34 | 1.29 | 2.71 | 0.9 | 0.09 | -0.77 | -1.34 | 0.57 | 2.02 | -0.08 | -1.07 | 0.41 | 0.52 | 0.4025 | -0.725 | -0.9 | 0.571551 | 0.398856 | 0.162657 | | 1 | 0.996511 | 0.757516 |
| Mtx3 | D3YTP3 | 0.36 | 0.15 | -0.35 | 0.34 | 0.08 | 0.5 | -0.04 | 0.08 | 0.22 | 0.22 | -0.19 | -0.06 | -0.11 | 0.24 | -0.53 | -0.08 | 0.03 | -0.0775 | -0.245 | 0.893789 | 0.73454 | 0.355627 | | 1 | 0.996511 | 0.855029 |
| Muc5b | E9Q5I3 | 1.31 | 2.01 | 1.43 | 1.02 | 1.44 | 0.68 | -0.06 | 0.7 | -0.77 | 0.87 | 0.6 | -0.12 | -0.55 | 0.79 | -0.15 | 0.01 | -0.7525 | -1.2975 | -1.4175 | 0.059694 | 0.010515 | 0.002785 | | 1 | 0.996511 | 0.453261 |
| Mug1 | P28665 | -3.15 | -3.16 | -2.67 | -3.42 | -3.04 | -3.21 | -3.01 | -3.27 | -2.84 | -2.06 | -2.97 | -3 | -2.23 | -1.88 | -3.09 | -3.03 | -0.0325 | 0.3825 | 0.5425 | 0.871637 | 0.199457 | 0.135559 | | 1 | 0.996511 | 0.731487 |
| Mul1 | Q8VCM5 | 1.07 | 1.08 | 1.15 | 1 | 1.09 | 1.1 | 1.62 | 1.45 | 0.3 | 0.78 | 1.57 | 1.23 | -0.01 | 0.54 | 1.21 | 1.62 | 0.24 | -0.105 | -0.235 | 0.205847 | 0.71561 | 0.51594 | | 1 | 0.996511 | 0.905674 |
| Mvb12a | Q78HU3 | 0.03 | -0.27 | 0.58 | 0.69 | -0.06 | -0.9 | 0.39 | -0.01 | 0.37 | 1.16 | 0.92 | -0.36 | 0.01 | 0.93 | 0.66 | -0.21 | -0.4025 | 0.265 | 0.09 | 0.258208 | 0.503243 | 0.796879 | | 1 | 0.996511 | 0.969727 |
| Mvb12b | Q6KAU4 | -0.08 | -0.05 | 0.19 | 0.05 | -0.08 | 0.25 | -0.12 | -0.06 | 1.01 | 0.69 | 0.18 | 0.27 | 1.1 | 0.32 | 0.15 | -0.14 | -0.03 | 0.51 | 0.33 | 0.854407 | 0.053066 | 0.271642 | | 1 | 0.996511 | 0.822084 |
| Mvd | Q99JF5 | -1.52 | -1.06 | -0.77 | -1.26 | -1.46 | -1.27 | -1.23 | -1.17 | -0.99 | -0.47 | -0.72 | -0.74 | -0.11 | -0.93 | -0.8 | -1.03 | -0.13 | 0.4225 | 0.435 | 0.525388 | 0.088227 | 0.148219 | | 1 | 0.996511 | 0.746406 |
| Mvk | Q9R008 | -0.3 | 0.05 | -0.31 | -0.34 | 0.05 | -0.23 | -0.24 | -0.42 | -0.41 | -0.8 | -0.32 | -0.03 | -0.77 | -0.01 | -0.31 | -0.42 | 0.015 | -0.165 | -0.1525 | 0.933295 | 0.461905 | 0.50878 | | 1 | 0.996511 | 0.902804 |
| Mvp | Q9EQK5 | 0.06 | -0.09 | 0.31 | 0.09 | -0.03 | 0 | 0.32 | 0.06 | -0.1 | -0.14 | 0.09 | 0.03 | -0.3 | 0.19 | 0.14 | 0.24 | -0.005 | -0.1225 | -0.025 | 0.976333 | 0.485089 | 0.904537 | | 1 | 0.996511 | 0.985525 |
| Mx1 | A0A338P72 | 0.37 | -0.2 | 0.92 | 1.13 | 0.1 | -0.22 | -0.11 | -0.01 | 0.27 | 0.56 | -0.04 | -0.24 | -0.16 | 0.58 | 0.22 | 0.27 | -0.615 | -0.4175 | -0.3275 | 0.069043 | 0.241593 | 0.343465 | | 1 | 0.996511 | 0.848473 |
| Mxra7 | Q9D1T2 | -0.63 | -0.13 | 0.43 | 0.64 | -0.09 | -0.36 | 0.68 | 0.12 | 0.26 | 0.7 | -0.04 | -0.37 | 0.68 | 0.14 | 0.06 | 0.33 | 0.01 | 0.06 | 0.225 | 0.977047 | 0.866456 | 0.492572 | | 1 | 0.996511 | 0.897531 |
| Myadm | O35682 | 0.46 | 0.44 | 0.59 | 1.45 | 0.25 | 0.77 | 1.38 | 1.49 | 1.12 | 1.7 | 0.76 | 0.57 | 0.65 | 0.51 | 0.23 | 0.9 | 0.2375 | 0.3025 | -0.1625 | 0.514188 | 0.386164 | 0.583225 | | 1 | 0.996511 | 0.927734 |
| Mybbp1a | Q7TPV4 | 1.04 | 0.57 | 0.17 | 0.28 | 0.29 | 0.54 | 0.08 | 0.25 | 0.63 | -0.6 | 0.79 | 0.57 | 0.95 | 1 | 0.59 | 0.5 | -0.225 | -0.1675 | 0.245 | 0.34965 | 0.646703 | 0.359122 | | 1 | 0.996511 | 0.856645 |
| Mycbp | Q9EQS3 | -1.19 | -1.66 | -0.97 | -1.1 | -1.36 | -1.47 | -0.86 | -1.49 | -1.68 | -0.57 | -1.3 | -1.03 | -0.97 | -0.92 | -0.99 | -0.73 | -0.065 | 0.085 | 0.3275 | 0.77793 | 0.767818 | 0.154603 | | 1 | 0.996511 | 0.750763 |
| Mycbp2 | Q7TPH6 | 0.94 | 0.76 | 0.67 | 0.44 | 0.77 | 0.51 | 0.76 | 0.55 | 0.81 | 0.86 | 0.56 | 0.92 | 0.6 | 0.68 | 1 | 0.67 | -0.055 | 0.085 | 0.035 | 0.752463 | 0.654271 | 0.862479 | | 1 | 0.996511 | 0.977204 |
| Mydgf | Q9CPT4 | -0.87 | -0.75 | 0.23 | -0.68 | -0.1 | -0.74 | 0.7 | 0.23 | -1.57 | -1.05 | -0.12 | -0.67 | -0.64 | -0.72 | 0.96 | 0.57 | 0.54 | -0.335 | 0.56 | 0.174706 | 0.391417 | 0.248248 | | 1 | 0.996511 | 0.803985 |
| Myef2 | Q8C854 | 1.33 | -0.77 | -1.08 | -0.39 | -0.85 | 0.48 | -1.01 | 0.06 | -0.37 | -0.23 | -0.56 | -0.79 | 0.47 | -0.23 | 0.27 | -1.16 | -0.1025 | -0.26 | 0.065 | 0.862065 | 0.613999 | 0.913003 | | 1 | 0.996511 | 0.986458 |
| Myf6 | P15375 | 3.27 | 1.33 | 2.07 | 0.82 | 2.17 | 3.09 | 2.36 | 2.07 | -3.44 | 0.05 | 3.58 | 3.42 | -0.6 | -3.62 | 3.45 | 1.92 | 0.55 | -0.97 | -1.585 | 0.316893 | 0.537745 | 0.292196 | | 1 | 0.996511 | 0.825497 |
| Myg1 | Q9JK81 | -0.18 | -0.49 | -1.35 | -1.52 | -0.37 | -0.52 | -1.41 | -0.6 | -0.46 | -1.04 | -0.79 | 0 | -0.67 | -0.11 | -1.64 | -1.46 | 0.16 | 0.3125 | -0.085 | 0.675341 | 0.421768 | 0.851843 | | 1 | 0.996511 | 0.976225 |
| Myh10 | Q5SV64 | 0.98 | 0.59 | 0.49 | 0.6 | 0.52 | 0.76 | 0.29 | 0.48 | 0.66 | 0.71 | 0.49 | 1.02 | 0.45 | 0.95 | 0.34 | 0.11 | -0.1525 | 0.055 | -0.2025 | 0.421433 | 0.786063 | 0.416336 | | 1 | 0.996511 | 0.874945 |
| Myh10 | Q61879 | 10.04 | 0.08 | 3.15 | 0.32 | -0.27 | 10.16 | 4.49 | 3.53 | 1.29 | 1.42 | 1.2 | 4.66 | 1.4 | 0.6 | 2.74 | 4.79 | 1.08 | -1.255 | -1.015 | 0.702723 | 0.571732 | 0.647721 | | 1 | 0.996511 | 0.945744 |
| Myh11 | A0A338P6I | -3.47 | -3.84 | -3.62 | -3.62 | -3.49 | -3.73 | -3.62 | -3.87 | -3.46 | -2.52 | -3.59 | -3.35 | -3.11 | -3.95 | -4.15 | -4.22 | -0.04 | 0.4075 | -0.22 | 0.81059 | 0.156206 | 0.44844 | | 1 | 0.996511 | 0.884817 |
| Myh14 | Q6URW6-2 | 0.85 | 0.82 | 0.59 | 0.64 | 0.79 | 0.71 | 0.57 | 0.46 | 0.84 | 0.56 | 0.3 | 0.58 | 0.61 | 0.54 | 0.55 | 0.39 | -0.0925 | -0.155 | -0.2025 | 0.567129 | 0.417334 | 0.270025 | | 1 | 0.996511 | 0.8219 |
| Myh9 | Q8VDD5 | 0.7 | -0.06 | 0.46 | 0.49 | -0.01 | 0.34 | 0.33 | 0.26 | 0.36 | 0.52 | 0.57 | 0.66 | 0.21 | 0.24 | 0.27 | 0.33 | -0.1675 | 0.13 | -0.135 | 0.432618 | 0.547835 | 0.538035 | | 1 | 0.996511 | 0.912212 |
| Myl1 | A0A087WF | -0.24 | -0.07 | 0.41 | -0.07 | 0.02 | -0.15 | -0.41 | 0.15 | -1.27 | -2.36 | 0.35 | 0.29 | -1.57 | -1.9 | 0.16 | 0.33 | -0.105 | -0.755 | -0.7525 | 0.622866 | 0.239179 | 0.193245 | | 1 | 0.996511 | 0.777764 |
| Myl12a | Q6ZWQ9 | -1.92 | -2.16 | 2.96 | 2.83 | -0.63 | -0.63 | 4.11 | 3.16 | -0.68 | 1.92 | 3.22 | -1.05 | -0.74 | -1.36 | 3.46 | 3.82 | 1.075 | 0.425 | 0.8675 | 0.529377 | 0.786818 | 0.622032 | | 1 | 0.996511 | 0.938628 |
| Myl12b | Q3THE2 | -0.17 | 0.02 | -0.11 | -0.09 | -0.43 | -0.25 | -0.15 | -0.11 | -0.05 | -0.02 | -0.06 | 0.2 | -0.79 | -0.73 | -0.65 | -0.64 | -0.1475 | 0.105 | -0.615 | 0.347961 | 0.523736 | 0.005245 | | 1 | 0.996511 | 0.552614 |
| Myl4 | A2A6Q8 | 0.06 | 0.42 | 1.64 | 1.03 | 0.16 | -0.03 | 0.43 | 0.7 | -0.76 | 0.43 | 1.16 | -0.16 | -0.41 | -0.33 | 0.07 | 0.29 | -0.4725 | -0.62 | -0.8825 | 0.217913 | 0.236079 | 0.040938 | | 1 | 0.996511 | 0.658454 |
| Myl6 | Q60605-2 | 0.4 | 1.24 | -0.34 | -1.2 | 0.94 | 0.32 | 0.07 | -0.55 | -0.22 | -2.71 | -0.31 | 0.77 | -0.14 | -0.39 | -1.06 | -0.48 | 0.17 | -0.6425 | -0.5425 | 0.759481 | 0.43992 | 0.308979 | | 1 | 0.996511 | 0.834523 |
| Myl6b | Q8CI43 | 0.39 | 0.74 | 0.54 | 0.33 | 0.46 | 0.28 | -0.08 | 0.11 | 0.27 | 0.48 | 0.32 | 0.41 | 0.67 | 1.11 | -0.21 | 0.31 | -0.3075 | -0.13 | -0.03 | 0.128256 | 0.464169 | 0.921669 | | 1 | 0.996511 | 0.987433 |
| Myl9 | Q9CQ19 | -2.15 | -2.14 | -2.46 | -2.04 | -1.79 | -2.39 | -2.25 | -2.66 | -2.02 | -1.92 | -2.04 | -1.92 | -1.99 | -2.21 | -2.64 | -2.52 | -0.075 | 0.2225 | -0.1425 | 0.739391 | 0.217697 | 0.527075 | | 1 | 0.996511 | 0.90845 |
| Mylk | B1B1A8 | -0.26 | -0.28 | -4.13 | -4.36 | 0.31 | -0.59 | -4.03 | -4.49 | -3.06 | -2.93 | -3.96 | -0.42 | -2.88 | -3.16 | -4.42 | -4.1 | 0.0575 | -0.335 | -1.3825 | 0.969132 | 0.786018 | 0.220913 | | 1 | 0.996511 | 0.787447 |
| Myo16 | F8WGF8 | 0.18 | 0.47 | 0.7 | 0.58 | 0.69 | 0.1 | 0.57 | 0.63 | 0.64 | 1 | 0.78 | 0.33 | 0.43 | 1.07 | 0.25 | 0.41 | 0.015 | 0.205 | 0.0575 | 0.941652 | 0.358886 | 0.816442 | | 1 | 0.996511 | 0.972768 |
| Myo18a | B2RRE2 | 0.96 | 0.85 | 1.02 | 1.16 | 0.72 | 0.93 | 0.71 | 0.96 | 1.09 | 0.65 | 1 | 1.11 | 0.8 | 0.72 | 0.64 | 0.47 | -0.1675 | -0.035 | -0.34 | 0.304608 | 0.850527 | 0.089982 | | 1 | 0.996511 | 0.692261 |
| Myo1b | E9QNH6 | 0.59 | 0.36 | 0.81 | 0.44 | 0.35 | 0.37 | 0.39 | 0.28 | 0.5 | -0.03 | 0.66 | 0.36 | 0.86 | 0.17 | 0.54 | 0.54 | -0.2025 | -0.1775 | -0.0225 | 0.232535 | 0.421967 | 0.919196 | | 1 | 0.996511 | 0.986868 |
| Myo1c | Q9WTI7-3 | -0.7 | -0.62 | 0.59 | 1.12 | -0.42 | -0.7 | 0.66 | 1.04 | -0.28 | -0.17 | 0.65 | -0.53 | -0.39 | -0.2 | -0.02 | 0.2 | 0.0475 | -0.18 | -0.2 | 0.932682 | 0.710626 | 0.653694 | | 1 | 0.996511 | 0.947633 |
| Myo1d | Q5SYD0 | 1.79 | 1.58 | 1.62 | 0.69 | 1.51 | 1.01 | 1.27 | 0.38 | 0.16 | 0.07 | 0.77 | 1.25 | -0.17 | -0.01 | 0.78 | 1.27 | -0.3775 | -0.8575 | -0.9525 | 0.279338 | 0.039662 | 0.040179 | | 1 | 0.996511 | 0.658454 |
| Myo1e | E9Q634 | -0.43 | 0.04 | 1.01 | 0.23 | -0.23 | -0.63 | 0.66 | 0.54 | 0.26 | -0.89 | 0.6 | -0.1 | 0.06 | -0.03 | 0.8 | 1.2 | -0.1275 | -0.245 | 0.295 | 0.753726 | 0.560196 | 0.473714 | | 1 | 0.996511 | 0.891644 |
| Myo1f | Q8CG29 | -0.29 | -1.45 | -1.21 | -0.57 | -1.53 | -1.96 | -2.04 | -2.27 | 0.2 | 0.93 | -1.25 | -0.96 | 0.51 | -0.24 | -1.77 | -2.35 | -1.07 | 0.61 | -0.0825 | 0.006918 | 0.270331 | 0.899237 | | 1 | 0.996511 | 0.983509 |
| Myo5a | D3YZ62 | 1.25 | 1.07 | 1.23 | 0.99 | 1.3 | 1.21 | 1.06 | 1.19 | 1.17 | 0.7 | 1.18 | 1.43 | 0.64 | 1.25 | 0.93 | 0.96 | 0.055 | -0.015 | -0.19 | 0.718708 | 0.942646 | 0.362645 | | 1 | 0.996755 | 0.858475 |
| Myo5b | G5E8G6 | 1.32 | 0.71 | 0.85 | 1.02 | 1.75 | 0.31 | 1.24 | 0.51 | 1.06 | 1.26 | 0.54 | 0.45 | 1.32 | 1.05 | 1.05 | 0.73 | -0.0225 | -0.1475 | 0.0625 | 0.947906 | 0.569989 | 0.782068 | | 1 | 0.996511 | 0.967642 |
| Myo6 | E9Q3L1 | 1.16 | 0.79 | 0.95 | 0.56 | 0.91 | 1.03 | 0.95 | 0.52 | 0.87 | 0.26 | 0.71 | 1.12 | 0.86 | 0.34 | 0.84 | 0.73 | -0.0125 | -0.125 | -0.1725 | 0.950645 | 0.613727 | 0.448589 | | 1 | 0.996511 | 0.884817 |
| Myo9a | Q8C170 | -0.01 | 0.29 | 0.45 | 0.18 | 0.26 | 0.22 | 0.16 | 0.17 | 0.05 | -0.04 | 0.38 | 0.29 | 0.42 | 0.44 | 0.21 | 0.31 | -0.025 | -0.0575 | 0.1175 | 0.876497 | 0.766085 | 0.538684 | | 1 | 0.996511 | 0.912212 |
| Myo9b | E9QKV6 | 0.13 | 0.46 | 0.54 | 0.35 | -0.48 | 0.25 | 0.03 | 0.33 | 1.03 | 0.23 | 0.69 | 0.29 | 1.33 | 0.54 | 0.48 | 0.77 | -0.3375 | 0.19 | 0.41 | 0.158528 | 0.430177 | 0.125156 | | 1 | 0.996511 | 0.721448 |
| Myoc | O70624 | -0.55 | -0.58 | 1 | 0.31 | 0.27 | -0.99 | 0.92 | 0.99 | -0.95 | 2.3 | 1.06 | -0.51 | -0.78 | 0.49 | 1.01 | 1.47 | 0.2525 | 0.43 | 0.5025 | 0.645561 | 0.574042 | 0.386743 | | 1 | 0.996511 | 0.863071 |
| Myof | A0A286YDI | 0.71 | 0.65 | -0.77 | -0.52 | 0.74 | 0.29 | -0.64 | -0.93 | 0.54 | 0.15 | -0.9 | 0.84 | 0.28 | 0.65 | -0.66 | -0.48 | -0.1525 | 0.14 | -0.07 | 0.763699 | 0.781191 | 0.880531 | | 1 | 0.996511 | 0.979754 |
| Myorg | Q69ZQ1 | -0.47 | -0.13 | -0.31 | 0.06 | 0.44 | 0.11 | 0.72 | 0.12 | -0.21 | 0.21 | -0.1 | 0.21 | -0.08 | -0.37 | -0.22 | 0.23 | 0.56 | 0.24 | 0.1025 | 0.02569 | 0.258476 | 0.64545 | | 1 | 0.996511 | 0.944749 |
| Mypop | Q8R4U1 | 1.67 | 0.93 | 1.03 | 1.44 | -0.62 | 0.89 | -0.69 | 0.73 | 0.8 | 0.84 | 0.87 | 0.31 | 2.48 | -2.04 | 1.95 | 1.04 | -1.19 | -0.5625 | -0.41 | 0.020778 | 0.044071 | 0.657318 | | 1 | 0.996511 | 0.948659 |
| Myrip | Q8K3I4 | 0.02 | 0.45 | 1.22 | 1.19 | 0.2 | 0.33 | 1.38 | 1.41 | 0.79 | 1.34 | 1.42 | 0.32 | 1.3 | 0.78 | 0.97 | 0.92 | 0.11 | 0.2475 | 0.2725 | 0.790222 | 0.516689 | 0.404409 | | 1 | 0.996511 | 0.871757 |
| Myt1l | P97500-2 | -0.84 | -3.3 | -0.91 | -0.95 | -2.46 | -0.7 | -0.97 | -0.82 | -1.34 | 0.09 | -0.79 | -1.68 | 0.48 | -4.78 | 0.08 | 0.09 | 0.2625 | 0.57 | 0.4675 | 0.692298 | 0.392356 | 0.706414 | | 1 | 0.996511 | 0.957084 |
| Mzt1 | Q8BUR9 | 0.09 | -0.34 | 0.38 | -0.25 | 0.47 | 0.07 | 0.37 | 0.28 | -0.88 | -0.38 | -0.01 | -0.34 | -0.88 | -1.68 | -0.76 | 0.04 | 0.3275 | -0.3725 | -0.79 | 0.149746 | 0.180108 | 0.062712 | | 1 | 0.996511 | 0.66433 |
| Mzt2 | Q9CQ25 | 0.7 | 1.47 | 0.05 | 1 | 1.1 | -0.67 | 0.49 | 0.6 | 1.19 | 1.04 | 1.16 | 0.96 | 0.35 | 1.77 | 0.22 | 0.95 | -0.425 | 0.2825 | 0.0175 | 0.353169 | 0.368435 | 0.968127 | | 1 | 0.996511 | 0.992733 |
| N4bp1 | Q6A037 | -0.06 | 0.52 | 0.12 | 0.33 | 0.01 | -0.32 | 0.25 | -0.29 | 0.07 | 0.71 | -0.12 | 0.53 | 0.17 | 0.98 | -0.1 | 0.14 | -0.315 | 0.07 | 0.07 | 0.162527 | 0.782379 | 0.806395 | | 1 | 0.996511 | 0.971715 |
| N4bp2l2 | Q8JZS6 | 2.44 | 1.64 | 0.93 | 1.52 | 2.71 | 1.68 | 1.29 | 1.77 | 2.85 | 2.05 | 1.66 | 2.3 | 2.32 | 2.87 | 1.42 | 1.06 | 0.23 | 0.5825 | 0.285 | 0.575588 | 0.15499 | 0.559916 | | 1 | 0.996511 | 0.918645 |
| N4bp3 | Q8C7U1 | 0.43 | 0.77 | 1.3 | 1.24 | 0.2 | 0.43 | 0.68 | 0.86 | 1.11 | 1.01 | 0.93 | 0.45 | 1.71 | 0.6 | 1.05 | 0.79 | -0.3925 | -0.06 | 0.1025 | 0.157383 | 0.823379 | 0.751794 | | 1 | 0.996511 | 0.961479 |
| N6amt1 | Q6SKR2 | -0.66 | -0.37 | -0.5 | -0.23 | -0.69 | -0.9 | -0.49 | -0.13 | -1.1 | -0.87 | -0.65 | -0.66 | -1.63 | -0.42 | -1.32 | -0.81 | -0.1125 | -3.80E-01 | -0.605 | 0.601841 | 0.07733 | 0.068066 | | 1 | 0.996511 | 0.664936 |
| Naa10 | Q3V4D5 | -2.65 | -3.92 | -1.4 | -0.94 | -4.29 | -2.65 | -5.36 | -1.92 | -1.14 | 1.58 | -2.31 | -1.8 | -0.34 | -3.52 | -1.99 | -1.83 | -1.3275 | 1.31 | 0.3075 | 0.175684 | 0.206218 | 0.715039 | | 1 | 0.996511 | 0.958106 |
| Naa10 | B1AUY7 | -1.65 | -1.93 | -1.69 | -1.88 | -1.62 | -1.77 | -2.05 | -1.89 | -1.95 | -2.13 | -1.47 | -1.97 | -1.67 | -1.43 | -1.75 | -2.29 | -0.045 | -0.0925 | 0.0025 | 0.789699 | 0.653128 | 0.991524 | | 1 | 0.996511 | 0.996154 |
| Naa12 | Q9CQX6 | -0.37 | -0.9 | -0.06 | 0.05 | -0.96 | -1.1 | 0.18 | 0.24 | -1.06 | 0.12 | 0.56 | -0.7 | -1.15 | -0.93 | 0.43 | 0.73 | -0.09 | 0.05 | 0.09 | 0.819635 | 0.902294 | 0.853048 | | 1 | 0.996609 | 0.976225 |
| Naa15 | G3X8Y3 | -1.09 | -1.35 | -1.83 | -1.8 | -1.39 | -1.26 | -1.99 | -1.96 | -1.39 | -1.24 | -1.89 | -1.14 | -1.36 | -1.15 | -1.95 | -2.23 | -0.1325 | 0.1025 | -0.155 | 0.623123 | 0.698025 | 0.626724 | | 1 | 0.996511 | 0.939737 |
| Naa20 | P61600 | -0.64 | -0.69 | -0.9 | -1.69 | -1.13 | -0.87 | -1.91 | -1.43 | -0.62 | -0.3 | -1.32 | -0.94 | -0.55 | -0.11 | -1.96 | -1.66 | -0.355 | 0.185 | -0.09 | 0.288122 | 0.573669 | 0.849007 | | 1 | 0.996511 | 0.976225 |
| Naa25 | Q8BWZ3 | -1.48 | -1.65 | -2.47 | -2.61 | -1.51 | -1.5 | -2.35 | -2.85 | -1.76 | -2.36 | -2.65 | -1.43 | -1.15 | -1.81 | -2.6 | -2.78 | -1.2E-16 | 0.0025 | -0.0325 | 1 | 0.994787 | 0.941933 | | 1 | 0.999285 | 0.991085 |
| Naa30 | E9QMB7 | -0.52 | -0.51 | -0.62 | -0.44 | -0.59 | -0.26 | -0.13 | -0.3 | 0.06 | -0.06 | -0.62 | -0.28 | 0.03 | -0.16 | -0.56 | -0.87 | 0.2025 | 0.2975 | 0.1325 | 0.23528 | 0.166196 | 0.588108 | | 1 | 0.996511 | 0.929131 |
| Naa35 | Q6PHQ8 | -1 | -1.19 | -1.28 | -1.17 | -0.86 | -1.12 | -1.21 | -1.13 | -1.29 | -0.68 | -0.94 | -0.88 | -1.19 | -1.44 | -1.16 | -1.62 | 0.08 | 0.2125 | -0.1925 | 0.617869 | 0.289986 | 0.338077 | | 1 | 0.996511 | 0.847578 |
| Naa38 | Q9D2U5 | -0.84 | -0.45 | -2.02 | -2.25 | -0.39 | -1.18 | -1.86 | -2.07 | -0.56 | -1.4 | -2.06 | -0.15 | -1.34 | 0.34 | -2.41 | -2.05 | 0.015 | 0.3475 | 0.025 | 0.977541 | 0.542735 | 0.970785 | | 1 | 0.996511 | 0.992803 |
| Naa40 | A0A494B9I | -1.27 | 0.05 | -1.3 | -1.63 | 0.14 | -0.88 | -1.26 | -2.15 | 0.63 | -1.13 | -1.43 | -0.79 | 0.16 | -0.29 | -0.61 | -0.92 | -1.2E-16 | 0.3575 | 0.6225 | 1 | 0.515622 | 0.162208 | | 1 | 0.996511 | 0.757516 |
| Naa50 | Q6PGB6-2 | -1.42 | -1.36 | -1.46 | -1.84 | -1.16 | -1.76 | -1.19 | -1.77 | -1.48 | -0.67 | -1.14 | -1.49 | -1.67 | -0.83 | -1.84 | -1.92 | 0.05 | 0.325 | -0.045 | 0.823437 | 0.209553 | 0.876561 | | 1 | 0.996511 | 0.979626 |
| Naaa | Q9D7V9 | -0.26 | 0.08 | -0.75 | -0.4 | 0.46 | 0.12 | -0.27 | -0.25 | -0.24 | -0.75 | 0.03 | -0.2 | -0.01 | -0.26 | -1.38 | -1.01 | 0.3475 | 0.0425 | -0.3325 | 0.195958 | 0.869111 | 0.365236 | | 1 | 0.996511 | 0.858475 |
| Nab2 | Q3TYF1 | -4.42 | -3.66 | -1.87 | -2.25 | -3.84 | -4.78 | -2.07 | -3.22 | -3.24 | -2.28 | -2.67 | -3.65 | -0.21 | -4.58 | -2.76 | -2.8 | -0.4275 | 0.09 | 0.4625 | 0.569787 | 0.883253 | 0.634303 | | 1 | 0.996511 | 0.941132 |
| Nabp2 | E9Q199 | -2.98 | -3.04 | -2.58 | -2.08 | -3.29 | -3.46 | -3.74 | -2.98 | -3.57 | -2.07 | -3.34 | -3.05 | -3.08 | -3.74 | -4.31 | -3.05 | -0.6975 | -0.3375 | -0.875 | 0.03116 | 0.389045 | 0.03843 | | 1 | 0.996511 | 0.658454 |
| Naca | Q60817 | -1.62 | -1.5 | -1.49 | -1.4 | -1.64 | -1.45 | -1.37 | -1.72 | -0.78 | -1.21 | -1.57 | -1.14 | -0.66 | -1 | -1.94 | -1.89 | -0.0425 | 0.3275 | 0.13 | 0.788047 | 0.146511 | 0.693062 | | 1 | 0.996511 | 0.955729 |
| Nacad | Q5SWP3 | -1.8 | -1.91 | -1.32 | -0.97 | -1.56 | -1.44 | -1.39 | -1.3 | -0.78 | -0.72 | -1.29 | -1.28 | 0.22 | -0.94 | -0.32 | -0.48 | 0.0775 | 0.4825 | 1.12 | 0.747668 | 0.112325 | 0.007437 | | 1 | 0.996511 | 0.552614 |
| Nacc1 | Q7TSZ8 | -3.21 | -0.2 | -1.15 | -2.86 | -1.67 | -3.77 | -2.21 | -3.31 | -1.76 | -0.5 | -1.86 | -1.26 | 0.96 | 0.1 | -1.11 | -1.75 | -0.885 | 0.51 | 1.405 | 0.273059 | 0.476785 | 0.122153 | | 1 | 0.996511 | 0.721448 |
| Nacc2 | Q9DCM7 | -4.09 | -2.83 | -4.07 | -2.65 | -4.11 | -3.46 | -2.94 | -3.75 | -1.28 | -2.01 | -2.46 | -2.72 | 1.71 | -3.35 | -2.77 | -3.57 | -0.155 | 1.2925 | 1.415 | 0.719705 | 0.021274 | 0.244151 | | 1 | 0.996511 | 0.799567 |
| Nadk | P58058 | 0.59 | -0.07 | 0.23 | -0.12 | 0.88 | 0.62 | -0.08 | -0.06 | 0.49 | 0.27 | 0.19 | -0.13 | 0.11 | 0.24 | -0.04 | -0.17 | 0.1825 | 0.0475 | -0.1225 | 0.537015 | 0.841117 | 0.598045 | | 1 | 0.996511 | 0.930093 |
| Nadk2 | Q8C5H8-2 | 0.98 | 0.82 | 0.86 | 1.06 | 0.62 | 0.84 | 0.92 | 0.96 | -0.09 | 0.3 | 1.09 | 0.68 | -0.23 | -0.18 | 0.7 | 0.83 | -0.095 | -0.435 | -0.65 | 0.552396 | 0.138444 | 0.054761 | | 1 | 0.996511 | 0.661738 |
| Nae1 | E0CZE0 | 0.35 | 0.27 | -0.01 | -0.35 | -0.26 | 0.37 | -0.29 | -0.14 | -0.59 | -0.24 | -0.14 | 0.37 | -0.35 | -0.45 | -0.56 | -0.54 | -0.145 | -0.215 | -0.54 | 0.545355 | 0.433852 | 0.03301 | | 1 | 0.996511 | 0.644622 |
| Naf1 | E9QJT2 | 0 | 0.04 | -1.23 | -1.37 | -0.21 | 0.18 | -0.95 | -1.6 | 0.62 | -1.39 | -1.27 | 0.16 | -0.75 | 0.63 | -2.02 | -1.54 | -0.005 | 0.17 | -0.28 | 0.992119 | 0.770743 | 0.660068 | | 1 | 0.996511 | 0.949645 |
| Naga | Q9QWR8 | 0.34 | 0.03 | -0.58 | -0.36 | 0.47 | 0.37 | 0.39 | -0.45 | -0.48 | -0.88 | -0.36 | 0.22 | -0.87 | -0.38 | -0.89 | -0.65 | 0.3375 | -0.2325 | -0.555 | 0.272759 | 0.459357 | 0.060698 | | 1 | 0.996511 | 0.661738 |
| Nagk | Q9QZ08 | -0.08 | -0.11 | 0.15 | -0.08 | -0.05 | 0.2 | 0.3 | 0.04 | 0.46 | 0.4 | -0.16 | -0.18 | 0.16 | 1.19 | -0.22 | 0 | 0.1525 | 0.16 | 0.3125 | 0.355702 | 0.475452 | 0.34727 | | 1 | 0.996511 | 0.851503 |
| Naglu | O88325 | -0.38 | -0.4 | -0.55 | 0.11 | -0.19 | -0.87 | -0.49 | -0.48 | 0.08 | 2.85 | -0.17 | -0.36 | 0.34 | 0.83 | 0.1 | -0.25 | -0.2025 | 0.905 | 0.56 | 0.374524 | 0.21765 | 0.077234 | | 1 | 0.996511 | 0.676667 |
| Nalcn | E9QLE4 | 0.65 | 0.63 | 0.29 | 0.74 | 0.47 | 0.95 | 0.74 | 0.53 | 2.29 | 0.45 | 0.58 | 0.44 | 3.08 | 0.37 | 0.38 | 0.61 | 0.095 | 0.3625 | 0.5325 | 0.614759 | 0.415447 | 0.393224 | | 1 | 0.996511 | 0.865775 |
| Nampt | Q99KQ4 | -0.89 | -1.15 | -1.39 | -1.69 | -1.08 | -1.45 | -1.55 | -1.73 | -1.29 | -1.09 | -1.48 | -0.88 | -1.47 | -0.9 | -1.81 | -1.84 | -0.1725 | 0.095 | -0.225 | 0.471688 | 0.694611 | 0.450047 | | 1 | 0.996511 | 0.884817 |
| Nanp | Q9CPT3 | -0.36 | -0.16 | -3.09 | -3.48 | 0.05 | 0.14 | -2.98 | -3.47 | -0.67 | -1.02 | -3.03 | -0.38 | -0.88 | 0.48 | -3.26 | -3.3 | 0.2075 | 0.4975 | 0.0325 | 0.85884 | 0.605531 | 0.977259 | | 1 | 0.996511 | 0.993727 |
| Nans | Q99J77 | 0.14 | -0.34 | -0.65 | -0.37 | -0.37 | 0.45 | -0.27 | -0.48 | -0.36 | -0.23 | -0.41 | 0.26 | -0.48 | -0.11 | -0.69 | -0.77 | 0.1375 | 0.12 | -0.2075 | 0.615552 | 0.63106 | 0.42097 | | 1 | 0.996511 | 0.875639 |
| Nap1l1 | Q3TF41 | 0.75 | 0.71 | -0.11 | 0.03 | 0.24 | 1.07 | -0.01 | 0.06 | 0.65 | -0.66 | 0 | 1.16 | 0.17 | 0.43 | -0.66 | -0.13 | -0.005 | -0.0575 | -0.3925 | 0.987763 | 0.893517 | 0.251091 | | 1 | 0.996511 | 0.807507 |
| Nap1l4 | B7ZNL2 | 0.27 | -0.35 | -0.58 | -0.43 | -0.05 | 0.36 | -0.18 | -0.23 | -0.3 | -1.31 | -0.4 | 0.45 | 0.06 | -0.49 | 0.24 | -0.61 | 0.2475 | -0.1175 | 0.0725 | 0.325354 | 0.763573 | 0.805417 | | 1 | 0.996511 | 0.971715 |
| Nap1l5 | Q9JJF0 | 0.86 | 1.52 | 1.19 | 1.2 | -0.41 | 0.46 | 1.21 | 1.35 | -0.96 | 1.42 | 0.73 | 0.95 | -1.73 | -2.29 | 1.09 | 0.44 | -0.54 | -0.6575 | -1.815 | 0.202305 | 0.209987 | 0.036604 | | 1 | 0.996511 | 0.658454 |
| Napa | Q9DB05 | -0.14 | 0.01 | 0.39 | 0.29 | -0.08 | -0.27 | 0.48 | 0.09 | -0.74 | -0.07 | 0.24 | -0.15 | -0.85 | -0.66 | 0.17 | 0.3 | -0.0825 | -0.3175 | -0.3975 | 0.712758 | 0.239871 | 0.235794 | | 1 | 0.996511 | 0.791493 |
| Napb | P28663 | 0.67 | 0.03 | 0.68 | 1 | 0.16 | 1.23 | 1.09 | 1.23 | 0.27 | 0.91 | 0.86 | 0.77 | 0.19 | 0.3 | 0.75 | 0.83 | 0.3325 | 0.1075 | -0.0775 | 0.31573 | 0.688937 | 0.781747 | | 1 | 0.996511 | 0.967642 |
| Napepld | Q8BH82 | 2.22 | 1.61 | 1.33 | 1.47 | 1.65 | 2.28 | 0.83 | 1.28 | 0.78 | 1.46 | 1.58 | 1.93 | 1.04 | 0.3 | 1.62 | 1.86 | -0.1475 | -0.22 | -0.4525 | 0.674771 | 0.488121 | 0.261219 | | 1 | 0.996511 | 0.81412 |
| Napg | Q9CWZ7 | 0.8 | 1.13 | 1.25 | 0.86 | 1.14 | 0.8 | 1.26 | 1.37 | 0.07 | 0.14 | 1.07 | 0.91 | 0.11 | 0.22 | 1.15 | 1 | 0.1325 | -0.4625 | -0.39 | 0.508607 | 0.136218 | 0.214978 | | 1 | 0.996511 | 0.787313 |
| Naprt | Q8CC86 | -0.14 | -1.03 | -0.38 | -0.12 | -0.8 | 0.21 | -1.95 | -0.78 | -1.25 | -0.33 | -0.89 | -0.6 | -1.05 | -1.64 | -0.9 | -0.64 | -0.4125 | -0.35 | -0.64 | 0.376987 | 0.257728 | 0.064621 | | 1 | 0.996511 | 0.66433 |
| NARS1 | Q8BP47 | -0.05 | -0.34 | -0.99 | -0.99 | -0.12 | -0.17 | -0.76 | -0.86 | -0.76 | -1.31 | -0.86 | -0.07 | -0.86 | -0.99 | -1.19 | -1.35 | 0.115 | -0.1575 | -0.505 | 0.705123 | 0.648422 | 0.099398 | | 1 | 0.996511 | 0.698156 |
| Nars2 | Q8BGV0 | 0.46 | 0.31 | 0.15 | -0.14 | 0.25 | 0.19 | 0.71 | 0.22 | -0.32 | -0.19 | 0.59 | 0.41 | -0.29 | -0.04 | 0.24 | 0.45 | 0.1475 | -0.0725 | -0.105 | 0.483101 | 0.789997 | 0.667732 | | 1 | 0.996511 | 0.951339 |
| Nasp | B1AU75 | -3.83 | -3.76 | -4.22 | -4.73 | -3.67 | -4.04 | -4.19 | -4.33 | -3.77 | -2.57 | -4.27 | -3.32 | -3.42 | -2.52 | -4.53 | -4.55 | 0.0775 | 0.6525 | 0.38 | 0.774391 | 0.133997 | 0.455676 | | 1 | 0.996511 | 0.887169 |
| Nat1 | A0A1D5RL( | -1.22 | -0.49 | 0.32 | -0.27 | -1.17 | -1.93 | 0.12 | -0.12 | 0.26 | 0.13 | -0.87 | -0.82 | 0.24 | -0.41 | -0.5 | 0 | -0.36 | 0.09 | 0.2475 | 0.499897 | 0.829008 | 0.495474 | | 1 | 0.996511 | 0.89758 |
| Nat10 | Q8K224 | 0.85 | 0.79 | 1.12 | 0.87 | 0.34 | 0.46 | 0.89 | 0.82 | 0.06 | 0.79 | 1.44 | 0.43 | 0.09 | 0.44 | 1.2 | 1.51 | -0.28 | -0.2275 | -0.0975 | 0.167873 | 0.465572 | 0.773231 | | 1 | 0.996511 | 0.966062 |
| Nat14 | Q8BVG8 | 0.3 | 0.79 | 0.2 | 0.34 | 0.94 | 0.32 | 0.58 | 0.16 | -0.07 | -0.37 | 0.16 | 0.26 | 0.16 | 0.3 | 0.59 | 0.29 | 0.0925 | -0.4125 | -0.0725 | 0.692546 | 0.095352 | 0.735798 | | 1 | 0.996511 | 0.958635 |
| Nat2 | P50295 | -0.09 | 0.6 | -0.16 | 0.48 | -0.26 | 0.55 | -0.19 | -0.05 | 1.04 | 0.41 | -0.84 | 0.44 | 0.17 | 1.02 | -0.95 | -0.13 | -0.195 | 0.055 | -0.18 | 0.481232 | 0.895271 | 0.676264 | | 1 | 0.996511 | 0.952944 |
| Nat8f1 | E9PUL7 | 0.34 | 0.33 | 1.61 | 1.24 | 0.76 | 0.72 | 3.02 | 1.51 | -1.53 | -0.19 | 1.15 | 0.35 | -0.91 | -2.59 | 1.4 | 1.84 | 0.6225 | -0.935 | -0.945 | 0.294618 | 0.143472 | 0.343955 | | 1 | 0.996511 | 0.848473 |
| Nat8l | A0A0R4J0F | 0.51 | 0.47 | 1.07 | 0.14 | 0.38 | 0.54 | 1.67 | 1.26 | 0.17 | 2.1 | 0.84 | 0.3 | 1.1 | -0.58 | 1.68 | 1.89 | 0.415 | 0.305 | 0.475 | 0.25099 | 0.505507 | 0.395529 | | 1 | 0.996511 | 0.866946 |
| Natd1 | Q9DBW3 | 1.41 | 0.75 | 1.19 | 0.48 | 1.6 | 1.23 | 1.38 | 1.28 | 1.61 | 2.94 | 0.84 | 1.29 | 0.37 | 1.69 | 1.21 | 1.28 | 0.415 | 0.7125 | 0.18 | 0.112527 | 0.153193 | 0.605831 | | 1 | 0.996511 | 0.932425 |
| Nav1 | Q8CH77 | 0.14 | 0.27 | 0.59 | 0.73 | 0.2 | 0.3 | 0.55 | 0.61 | 1.04 | 0.94 | 0.94 | 0.89 | 1.38 | 0.77 | 0.77 | 0.41 | -0.0175 | 0.52 | 0.4 | 0.930577 | 0.0241 | 0.161867 | | 1 | 0.996511 | 0.757516 |
| Nav2 | V9GXT3 | -1.08 | -0.34 | 0 | 0.24 | -0.2 | -0.25 | -0.27 | -0.46 | 0.23 | 0.97 | -0.14 | -0.34 | 0.88 | -0.14 | 0.21 | 0.41 | 1.96E-17 | 0.475 | 0.635 | 1 | 0.244586 | 0.100429 | | 1 | 0.996511 | 0.698156 |
| Nav3 | A0A668KM | 0.35 | 0.62 | 0.74 | 0.66 | 0.48 | 0.49 | 0.12 | 0.41 | 0.81 | 0.84 | 1.2 | 0.88 | 0.97 | 1.03 | 0.42 | 0.18 | -0.2175 | 0.34 | 0.0575 | 0.228043 | 0.093837 | 0.822208 | | 1 | 0.996511 | 0.972768 |
| Naxd | J3QMM7 | 0.52 | 0.59 | 0.41 | 0.51 | 0.67 | 0.33 | 0.11 | 0.43 | 0.42 | 1.17 | 0.45 | 0.39 | -0.32 | 0.86 | 0.27 | 0.14 | -0.1225 | 0.1 | -0.27 | 0.484404 | 0.659506 | 0.331997 | | 1 | 0.996511 | 0.845601 |
| Naxd | Q9CZ42-2 | -0.18 | -1.24 | -0.98 | -0.25 | 0.38 | -0.46 | -2.01 | -1.49 | 0.17 | 1.27 | -1.75 | -1.14 | 0.59 | 1.46 | -2.51 | -1.36 | -0.2325 | 0.3 | 0.2075 | 0.671503 | 0.651883 | 0.806547 | | 1 | 0.996511 | 0.971715 |
| Naxe | Q8K4Z3 | 0.21 | 0 | -0.1 | 0.19 | -0.3 | -0.2 | -0.23 | 0.06 | -0.35 | 0.32 | -0.24 | -0.18 | -0.38 | -0.39 | -0.34 | -0.48 | -0.2425 | -0.1875 | -0.4725 | 0.168392 | 0.382312 | 0.024196 | | 1 | 0.996511 | 0.626254 |
| Nbas | E9Q411 | 0.35 | 0.48 | 0.92 | 0.87 | 0.73 | 0.35 | 1.16 | 1.17 | 0.16 | 0.86 | 0.92 | 0.43 | 0.14 | 0.54 | 1.48 | 1.32 | 0.1975 | -0.0625 | 0.215 | 0.443431 | 0.804307 | 0.539826 | | 1 | 0.996511 | 0.912212 |
| Nbdy | A0A0N4SU | -0.51 | -0.73 | -0.54 | -0.49 | -1.43 | -0.7 | -1.35 | -0.49 | -0.3 | 0.94 | -0.42 | -0.53 | 0.68 | -0.33 | 0.07 | -0.78 | -0.425 | 0.49 | 0.4775 | 0.119117 | 0.176518 | 0.16216 | | 1 | 0.996511 | 0.757516 |
| Nbea | Q9EPN1-4 | -0.62 | -0.5 | -0.25 | -0.37 | -0.24 | -0.38 | -0.1 | -0.35 | -0.42 | -0.23 | -0.05 | -0.33 | -0.34 | -0.11 | -0.37 | -0.42 | 0.1675 | 0.1775 | 0.125 | 0.317146 | 0.336736 | 0.508982 | | 1 | 0.996511 | 0.902804 |
| Nbeal1 | E9PYP2 | 0.58 | 0.96 | 0.81 | 1.05 | 1.79 | 0.81 | 1.82 | 0.95 | 0.68 | 0.67 | 0.94 | 1.29 | -0.96 | -0.2 | 0.75 | 0.84 | 0.4925 | 0.045 | -0.7425 | 0.115343 | 0.835889 | 0.104186 | | 1 | 0.996511 | 0.702979 |
| Nbr1 | Q05BC8 | -0.59 | -1.01 | 0.22 | 0.7 | -0.08 | -0.58 | 1.01 | 0.78 | 0.32 | 1.13 | 0.71 | 0.01 | 0.67 | 0.79 | 1.56 | 1.2 | 0.4525 | 0.7125 | 1.225 | 0.372203 | 0.126084 | 0.015891 | | 1 | 0.996511 | 0.589387 |
| Ncald | Q91X97 | 0.1 | 1.03 | 0.7 | 0.24 | 0.92 | 0.53 | 0.99 | 0.64 | -0.34 | 0.28 | 0.69 | 0.13 | -0.75 | -0.47 | 0.22 | 0.59 | 0.2525 | -0.3275 | -0.62 | 0.330321 | 0.300336 | 0.119067 | | 1 | 0.996511 | 0.721448 |
| Ncam1 | E9QB01 | 1.47 | 1.63 | 1.3 | 1.85 | 2.37 | 1.29 | 2.23 | 1.82 | 0.3 | 0.42 | 2.1 | 1.6 | -0.25 | 0.98 | 1.2 | 1.49 | 0.365 | -0.4575 | -0.7075 | 0.205757 | 0.306603 | 0.09592 | | 1 | 0.996511 | 0.698156 |
| Ncam1 | A0A0A6YY! | 1.35 | 1.5 | 1.21 | 1.43 | 1.33 | 1.25 | 1.2 | 1.49 | 1.32 | 1.02 | 1.51 | 1.87 | 0.89 | 1.25 | 1.09 | 1.05 | -0.055 | 0.0575 | -0.3025 | 0.725195 | 0.797478 | 0.125531 | | 1 | 0.996511 | 0.721449 |
| Ncam1 | A0A0A6YY4 | 12.52 | 7.96 | 9.3 | 5.22 | 9.25 | 12.84 | 10.34 | 9.66 | 2.12 | 2.42 | 9.93 | 12.6 | -0.13 | 0.62 | 9.47 | 11.54 | 1.7725 | -1.9825 | -3.375 | 0.266216 | 0.473093 | 0.273641 | | 1 | 0.996511 | 0.822084 |
| Ncam1 | P13595 | 1.76 | 1.19 | 1.87 | 3.26 | 0.76 | 1.36 | 1.55 | 3.28 | 2.66 | 3.89 | 2.49 | 2.69 | 2.59 | 2.08 | 2.87 | 1.31 | -0.2825 | 0.9125 | 0.1925 | 0.65754 | 0.099271 | 0.710757 | | 1 | 0.996511 | 0.957819 |
| Ncam2 | O35136 | 0.27 | 0.53 | 0.9 | 1.19 | 0.92 | 0.84 | 1.45 | 1.17 | 0.69 | 0.49 | 1.34 | 0.82 | 0.68 | 0.83 | 1.38 | 1.31 | 0.3725 | 0.1125 | 0.3275 | 0.169676 | 0.692211 | 0.269103 | | 1 | 0.996511 | 0.820534 |
| Ncan | A0A0R4IZX | -1.56 | -2.37 | -1.63 | -1.56 | -2.12 | -1.82 | -1.71 | -1.61 | -1.97 | -0.52 | -1.53 | -1.72 | -1.63 | -1.74 | -1.56 | -2.17 | -0.035 | 0.345 | 0.005 | 0.884616 | 0.355933 | 0.985004 | | 1 | 0.996511 | 0.996154 |
| Ncbp1 | Q3UYV9 | -0.95 | -1.88 | -1.54 | -1.48 | -1.9 | -1.68 | -1.27 | -1.24 | -2.96 | -0.99 | -1.45 | -2.06 | -2.33 | -2.16 | -1.34 | -1.72 | -0.06 | -0.4025 | -0.425 | 0.817358 | 0.372786 | 0.188219 | | 1 | 0.996511 | 0.776843 |
| Ncbp2 | Q9CQ49 | -1.61 | -2.51 | -2.05 | -1.92 | -2.67 | -2.5 | -2.28 | -1.82 | -1.96 | -0.92 | -1.38 | -2.71 | -1.09 | -1.93 | -0.92 | -1.86 | -0.295 | 0.28 | 0.5725 | 0.289142 | 0.499429 | 0.105422 | | 1 | 0.996511 | 0.703417 |
| Ncbp3 | Q8BZR9 | 0.26 | -0.86 | -1.28 | -0.9 | -1.65 | 0.02 | -1.37 | -0.58 | -0.56 | -1.06 | -0.98 | -1.04 | -0.29 | -0.13 | -0.61 | -1.66 | -0.2 | -0.215 | 0.0225 | 0.67064 | 0.539084 | 0.960229 | | 1 | 0.996511 | 0.992636 |

| | | | | | | | | | | | | | | | | | | | | | | | | | | |
|---|---|---|---|---|---|---|---|---|---|---|---|---|---|---|---|---|---|---|---|---|---|---|---|---|---|---|
| Ncdn | Q9Z0E0-2 | -1.07 | 0.46 | -1.32 | -0.4 | -1.1 | 0.28 | -2.42 | -1.94 | 2.37 | 1.53 | -0.84 | 0.88 | 1.79 | 0.23 | -0.29 | -1.92 | -0.7125 | 1.5675 | 0.535 | 0.288737 | 0.053293 | 0.497557 | 1 | 0.996511 | 0.898105 |
| Ncdn | Q9Z0E0 | -1.09 | -1.12 | -1.08 | -1.16 | -0.9 | -1.07 | -0.61 | -0.93 | -0.97 | -1.02 | -0.94 | -1.07 | -1.21 | -0.98 | -1.14 | -1.13 | 0.235 | 0.1125 | -0.0025 | 0.16808 | 0.467155 | 0.988145 | 1 | 0.996511 | 0.996154 |
| Nceh1 | Q8BLF1 | -0.14 | 0.27 | -0.35 | -0.76 | 0.73 | 0.13 | 0.1 | -0.48 | -0.28 | -1.77 | -0.61 | -0.01 | -0.07 | 0.13 | -0.13 | -0.02 | 0.365 | -0.4225 | 0.2225 | 0.272448 | 0.329331 | 0.391012 | 1 | 0.996511 | 0.865375 |
| Ncf1 | Q09014 | -0.88 | -3.15 | -1.7 | -1.19 | -2.63 | -1.68 | -1.17 | -1.43 | -1.48 | -0.79 | -1.12 | -2.12 | -0.67 | -1.43 | -1.98 | -1.96 | 0.0025 | 0.3525 | 0.22 | 0.996339 | 0.514646 | 0.687393 | 1 | 0.996511 | 0.955729 |
| Ncf2 | O70145 | -0.11 | -1.5 | -0.72 | -0.68 | -2.1 | 1.11 | -1.15 | -0.9 | 2.42 | -1.64 | -0.84 | -1.23 | 3.44 | -0.71 | -2.34 | -1.1 | -0.0075 | 0.43 | 0.575 | 0.990991 | 0.625747 | 0.618716 | 1 | 0.996511 | 0.937856 |
| Nck1 | Q99M51 | 0.09 | -0.03 | 0.07 | 0.35 | 0.2 | -0.12 | 0.18 | 0.11 | -0.53 | 0.03 | 0.21 | 0.13 | -1.12 | -0.12 | -0.13 | -0.33 | -0.0275 | -0.16 | -0.545 | 0.868312 | 0.478117 | 0.072606 | 1 | 0.996511 | 0.668066 |
| Nck2 | O55033 | 1.1 | 0.81 | 0.94 | 0.79 | 1.11 | 0.99 | 0.82 | 0.7 | 0.48 | 0.83 | 0.89 | 0.9 | -0.2 | 1.08 | 0.71 | 0.56 | -0.005 | -0.135 | -0.3725 | 0.976386 | 0.471364 | 0.225926 | 1 | 0.996511 | 0.787447 |
| Nckap1 | A2AS98 | 0.59 | 0.38 | 0.88 | 0.78 | 0.63 | 0.63 | 0.6 | 0.71 | 0.1 | 0.61 | 0.82 | 0.52 | -0.18 | 0.36 | 0.44 | 0.59 | -0.015 | -0.145 | -0.355 | 0.928673 | 0.520522 | 0.163987 | 1 | 0.996511 | 0.758352 |
| Nckap1l | Q8K1X4 | -1.82 | -1.63 | -1.89 | -1.26 | -1.75 | -1.51 | -1.96 | -1.46 | -1.26 | -0.71 | -1.29 | -1.96 | -0.63 | -1.05 | -1.76 | -2.02 | -0.02 | 0.345 | 0.285 | 0.924169 | 0.266497 | 0.421337 | 1 | 0.996511 | 0.875728 |
| Nckap5l | Q6GQX2 | 0.27 | 1.15 | -1.39 | -1.7 | 1.3 | 0.42 | -1.31 | -1.71 | 0.78 | -4.11 | -0.82 | 0.54 | 1.06 | 1.94 | -1.04 | -2.35 | 0.0925 | -0.485 | 0.32 | 0.916647 | 0.681641 | 0.763628 | 1 | 0.996511 | 0.964134 |
| Nckipsd | Q9ESJ4 | 0.9 | 0.32 | 0.64 | 0.36 | 0.62 | 0.56 | 0.33 | 0.33 | 0.36 | 0.1 | 0.38 | 0.45 | -0.09 | 0.17 | 0.35 | 0.5 | -0.095 | -0.2325 | -0.3225 | 0.623798 | 0.270344 | 0.184926 | 1 | 0.996511 | 0.776843 |
| Ncl | P09405 | -1.83 | -2.87 | -2.16 | -2.16 | -2.48 | -2.77 | -2.05 | -2.45 | -2.4 | -1.52 | -1.98 | -2.5 | -2.02 | -2.35 | -2.06 | -2.27 | -0.1825 | 0.155 | 0.08 | 0.505194 | 0.624915 | 0.760234 | 1 | 0.996511 | 0.963123 |
| Ncln | Q8VCM8 | -0.61 | -0.37 | -1.21 | -1.17 | 0.22 | -0.22 | -0.02 | -0.86 | -0.91 | -0.85 | -1.22 | -0.17 | -0.29 | 0.13 | 0.07 | 0.16 | 0.62 | 0.0525 | 0.8575 | 0.0703 | 0.864396 | 0.009102 | 1 | 0.996511 | 0.563386 |
| Ncoa1 | P70365-4 | -2.11 | -1.53 | -0.89 | -1.22 | -1.17 | -1.49 | -1.02 | -1.21 | -1.44 | 0.82 | -0.86 | -1.52 | -1.25 | -0.96 | -1.34 | -1.38 | 0.215 | 0.6875 | 0.205 | 0.450971 | 0.236519 | 0.489073 | 1 | 0.996511 | 0.897531 |
| Ncoa2 | Q61026 | -5.21 | -7.36 | -3.42 | -5.24 | -5.02 | -3.31 | -5.82 | -7.09 | -3.54 | -1.74 | -3.77 | -5.07 | -1.52 | -5.45 | -4.57 | -4.93 | -0.0025 | 1.7775 | 1.19 | 0.998029 | 0.089412 | 0.282782 | 1 | 0.996511 | 0.822595 |
| Ncoa5 | B7ZC24 | -2.57 | -3.83 | -2.66 | -2.2 | -4.64 | -2.56 | -2.93 | -1.12 | -0.69 | -2.63 | -0.9 | -3.42 | 0.75 | -2.72 | 0.01 | -1.58 | 0.0025 | 0.905 | 1.93 | 0.997257 | 0.209249 | 0.032244 | 1 | 0.996511 | 0.641974 |
| Ncoa5 | Q91W39 | 2.07 | -0.05 | 0.77 | 0.72 | -0.85 | 1.52 | 0.24 | 1.28 | 0.16 | 0.34 | 0.5 | 0.08 | 0.65 | -0.16 | 1.23 | 0.33 | -0.33 | -0.6075 | -0.365 | 0.605957 | 0.178527 | 0.465876 | 1 | 0.996511 | 0.888469 |
| Ncoa7 | A0A1L1SSS | 0.11 | 0.64 | 0.71 | 0.71 | 0.52 | 0.33 | 0.4 | 0.76 | 0.28 | 1.01 | 0.74 | 0.5 | 0.09 | 0.71 | 0.59 | 0.55 | -0.04 | 0.09 | -0.0575 | 0.8446 | 0.70968 | 0.809667 | 1 | 0.996511 | 0.972113 |
| Ncor1 | Q5RIM6 | -1.96 | -2.24 | -1.8 | -1.95 | -2.79 | -2.22 | -2.75 | -2.16 | -1.12 | -1.34 | -1.97 | -1.79 | 0.14 | -1.54 | -2.03 | -2.23 | -0.4925 | 0.4325 | 0.5725 | 0.046271 | 0.102758 | 0.277633 | 1 | 0.996511 | 0.822084 |
| Ncor2 | F8VQL9 | -2.24 | -2.49 | -1.67 | -1.6 | -2.88 | -2.37 | -1.69 | -1.83 | -1.62 | -1.32 | -1.39 | -2.22 | 0.01 | -2.27 | -1.7 | -1.68 | -0.1925 | 0.3625 | 0.59 | 0.571482 | 0.251776 | 0.258796 | 1 | 0.996511 | 0.812074 |
| Ncs1 | Q8BNY6 | -0.74 | -0.11 | 0.76 | 0.16 | 0.57 | -0.53 | 1.23 | 0.01 | -1.06 | -1.77 | 0.63 | -0.71 | -0.86 | 0.29 | 0.36 | 1.3 | 0.3025 | -0.745 | 0.255 | 0.511518 | 0.196242 | 0.615142 | 1 | 0.996511 | 0.936534 |
| Ncstn | P57716 | -1.09 | -0.64 | -0.25 | -0.32 | -0.3 | -0.95 | -0.3 | -0.68 | -0.58 | -1.32 | -0.48 | -1.01 | -0.46 | -0.83 | -0.77 | -0.3 | 0.0175 | -0.2725 | -0.015 | 0.945885 | 0.350681 | 0.953743 | 1 | 0.996511 | 0.992476 |
| Ndc1 | Q8VCB1 | 0.44 | 0.5 | 0.36 | 0.11 | 0.44 | 0.43 | 0.17 | 0.51 | 0.76 | -0.79 | 0.12 | 0.46 | 1.41 | -0.45 | 1.26 | 0.7 | 0.035 | -0.215 | 0.3775 | 0.835189 | 0.533554 | 0.372182 | 1 | 0.996511 | 0.859424 |
| Nde1 | Q9CZA6-3 | -0.08 | 0.7 | 1.02 | 1 | 0.89 | -0.19 | 0.38 | 0.64 | 0.39 | 1.54 | 0.88 | 0.19 | 0.21 | 1.4 | 0.8 | 0.61 | -0.23 | 0.09 | 0.095 | 0.498697 | 0.813524 | 0.788509 | 1 | 0.996511 | 0.968677 |
| Ndel1 | Q9ERR1 | 0.7 | 0.83 | 0.62 | 0.46 | 0.79 | 0.6 | 0.45 | 0.33 | 0.74 | 0.44 | 0.59 | 0.85 | 0.45 | 1.16 | 0.49 | 0.08 | -0.11 | 0.0025 | -0.1075 | 0.533943 | 0.989044 | 0.685844 | 1 | 0.999285 | 0.955729 |
| Ndfip1 | A0A494BA | -1.3 | -1.4 | -0.38 | -0.52 | -1.55 | -1.32 | -0.31 | -0.86 | -2.25 | -0.77 | -0.8 | -0.73 | -2.12 | -1.52 | -0.9 | -1 | -0.11 | -0.2375 | -0.485 | 0.761973 | 0.583873 | 0.21761 | 1 | 0.996511 | 0.787447 |
| Ndfip2 | F6RT43 | -0.05 | 0.68 | 0.49 | 0.44 | 0.16 | 0.46 | 0.17 | 0.38 | 1.94 | 0.42 | 0.77 | -0.11 | 2.72 | 0.36 | 0.24 | 0.4 | -0.0975 | 0.365 | 0.54 | 0.634576 | 0.412049 | 0.35489 | 1 | 0.996511 | 0.855029 |
| Ndn | P25233 | -0.45 | 0.03 | -0.83 | -1.6 | -0.05 | -0.64 | -0.41 | -0.74 | -0.9 | 1.22 | 0.02 | -0.39 | -1.4 | -1.12 | -0.17 | 0.43 | 0.2525 | 0.7 | 0.1475 | 0.489954 | 0.206184 | 0.77183 | 1 | 0.996511 | 0.965933 |
| Ndor1 | A2AI05 | -0.23 | -0.16 | 0.67 | 0.36 | 0.2 | -0.72 | 0.23 | 0.02 | -0.17 | -0.12 | -0.39 | -0.51 | -0.99 | 0.01 | -0.09 | 0.99 | -0.2275 | -0.4575 | -0.18 | 0.46396 | 0.100505 | 0.680638 | 1 | 0.996511 | 0.955729 |
| Ndrg1 | Q62433 | 1.16 | 0.53 | 0.47 | 0.28 | 0.2 | 0.79 | 0.46 | -0.04 | -0.02 | -0.57 | 0.5 | 1 | -0.15 | -0.34 | 0.32 | 0.38 | -0.2575 | -0.3825 | -0.5575 | 0.349159 | 0.322633 | 0.073245 | 1 | 0.996511 | 0.668219 |
| Ndrg2 | Q9QYG0 | 0.02 | 0.32 | 0.12 | 0.08 | 0.63 | 0 | 1 | 0.33 | -0.61 | -1.29 | 0.14 | 0.14 | -0.85 | -0.98 | -0.16 | -0.01 | 0.355 | -0.54 | -0.635 | 0.162927 | 0.143512 | 0.043055 | 1 | 0.996511 | 0.658454 |
| Ndrg2 | A0A2I3BQI | -0.94 | 0.3 | 1.93 | 0.96 | 0.1 | -0.18 | 1.19 | 1.19 | -0.23 | 1.29 | 1.01 | -0.33 | -0.33 | -0.05 | 1.3 | 2.59 | 0.0125 | -0.1275 | 0.315 | 0.984349 | 0.848562 | 0.699546 | 1 | 0.996511 | 0.955729 |
| Ndrg2 | Q9QYG0-2 | 2.42 | 3.83 | 4.04 | 1.75 | 3.82 | 2.02 | 6.54 | 4.23 | 1.84 | 1.65 | 4.05 | 3.32 | 0.95 | 1.64 | 4.67 | 5.76 | 1.1425 | -0.295 | 0.245 | 0.258598 | 0.686444 | 0.830958 | 1 | 0.996511 | 0.973939 |
| Ndrg3 | Q9QYF9 | 0.55 | 1.21 | 0.68 | 0.61 | 1.08 | 0.28 | 0.9 | 0.58 | 0.73 | 0.81 | 0.81 | 1.26 | 0.04 | 1.27 | 0.47 | 0.37 | -0.0525 | 0.14 | -0.225 | 0.831199 | 0.542245 | 0.476134 | 1 | 0.996511 | 0.892174 |
| Ndrg4 | Q8BTG7-3 | -0.42 | 0.11 | 0 | -0.04 | 0.14 | -0.31 | 0.55 | -0.08 | -0.33 | -0.36 | 0.25 | 0.37 | -0.39 | 0.22 | 0.49 | 0.37 | 0.1625 | 0.07 | 0.26 | 0.493781 | 0.777446 | 0.326513 | 1 | 0.996511 | 0.843799 |
| Ndufa1 | O35683 | -0.39 | -0.27 | -0.58 | 0.18 | -0.26 | -0.29 | 0.39 | -0.13 | -0.34 | -2.78 | 0.47 | -0.44 | -0.99 | -0.05 | -1.67 | -0.94 | 0.1925 | -0.5075 | -0.6475 | 0.433412 | 0.446417 | 0.102323 | 1 | 0.996511 | 0.700625 |
| Ndufa10 | Q99LC3 | 0.89 | 0.24 | -0.4 | 0.11 | -0.05 | 1.13 | -0.15 | -0.04 | 0.72 | -0.34 | -0.01 | 0.8 | 0.43 | 0.16 | -0.35 | -0.05 | 0.0125 | 0.0825 | -0.1625 | 0.973899 | 0.824774 | 0.613223 | 1 | 0.996511 | 0.93554 |
| Ndufa11 | G5E814 | -0.52 | -0.33 | -0.92 | -0.43 | -0.06 | 0.52 | -0.5 | -0.98 | -0.04 | -0.53 | -0.6 | -0.35 | 0.15 | 0.25 | 0.21 | -0.96 | 0.295 | 0.17 | 0.4625 | 0.388326 | 0.443966 | 0.178052 | 1 | 0.996511 | 0.772747 |
| Ndufa12 | Q7TMF3 | 2.29 | 1.47 | 2.62 | 1.35 | 1.85 | 2.29 | 3.03 | 2.8 | 0.31 | 0.36 | 2.62 | 2.15 | -0.38 | 0.59 | 2.41 | 2.81 | 0.56 | -0.5725 | -0.575 | 0.172038 | 0.366324 | 0.44381 | 1 | 0.996511 | 0.883549 |
| Ndufa13 | Q9ERS2 | 0.09 | 0.52 | 0.39 | 0.68 | 0.11 | 0.08 | 0.25 | 0.33 | 0.29 | 0.02 | 0.42 | -0.07 | -0.1 | 0.72 | -0.03 | 0.21 | -0.2275 | -0.255 | -0.22 | 0.232264 | 0.24788 | 0.398641 | 1 | 0.996511 | 0.867334 |
| Ndufa2 | Q9CQ75 | 0.76 | 0.81 | 0.06 | 0.73 | 0.08 | 0.86 | 0.19 | 0.49 | -0.04 | -0.51 | 0.47 | 0.6 | 0.34 | -1.04 | 0.07 | -0.06 | -0.185 | -0.46 | -0.7625 | 0.480569 | 0.166331 | 0.052537 | 1 | 0.996511 | 0.659923 |
| Ndufa3 | Q9CQ91 | 1.23 | 1.05 | 1.32 | 1.93 | 0.87 | 1.1 | 1.15 | 1.26 | 1.15 | 1 | 1.51 | 1.01 | 0.99 | 0.86 | 1.24 | 0.81 | -0.2875 | -0.215 | -0.4075 | 0.228712 | 0.398024 | 0.127709 | 1 | 0.996511 | 0.723391 |
| Ndufa4 | Q62425 | 0.78 | 0.83 | 0.62 | 1.09 | 0.89 | 0.69 | 0.4 | 0.79 | 0.84 | 1.21 | 0.68 | 0.47 | 0.6 | 1.52 | 0.31 | -0.04 | -0.1375 | -0.03 | -0.2325 | 0.465493 | 0.891963 | 0.507944 | 1 | 0.996511 | 0.902804 |
| Ndufa4l2 | Q4FZG9 | -0.38 | 0.32 | -0.02 | 0.85 | 0.14 | -0.28 | 0.06 | 0.83 | -0.26 | 1.76 | 0.66 | -0.27 | 0.55 | 0.64 | 0.44 | 0.68 | -0.005 | 0.28 | 0.385 | 0.988207 | 0.585271 | 0.200359 | 1 | 0.996511 | 0.78004 |
| Ndufa5 | Q9CPP6 | 0 | -0.02 | 0.58 | 0.99 | -0.25 | 0 | 0.4 | 0.95 | 0.03 | 0.76 | 0.86 | 0.06 | -0.09 | 0.4 | 0.7 | 0.49 | -0.1125 | 0.04 | -0.0125 | 0.745037 | 0.902955 | 0.967469 | 1 | 0.996609 | 0.9927 |
| Ndufa6 | Q9CQZ5 | 2.17 | 0.99 | 1.13 | 0.63 | 0.78 | 1.94 | 1.05 | 1.13 | 0.97 | 0.1 | 1.24 | 1.85 | 0.81 | 0.67 | 0.95 | 1.28 | -0.005 | -0.19 | -0.3025 | 0.989806 | 0.681399 | 0.400864 | 1 | 0.996511 | 0.869875 |
| Ndufa7 | Q9Z1P6 | 1.15 | 0.7 | 1.05 | 1.49 | 0.29 | 0.95 | 0.12 | 1.32 | 0.59 | 1.39 | 1.17 | 1.04 | 0.67 | 0.4 | 0.63 | 0.43 | -0.4275 | -0.05 | -0.565 | 0.202087 | 0.844816 | 0.030959 | 1 | 0.996511 | 0.641481 |
| Ndufa8 | Q9DCJ5 | -1.08 | -1.15 | -1.54 | -1.16 | -1.04 | -0.93 | -1.25 | -1.56 | -0.68 | -1.1 | -1.39 | -1.39 | 0.81 | -0.53 | -1.35 | -2 | 0.0375 | 0.0925 | 0.465 | 0.854272 | 0.688496 | 0.420512 | 1 | 0.996511 | 0.87558 |
| Ndufa9 | A0A0R3P9 | -0.13 | 0.22 | -0.29 | -0.36 | -0.3 | 0.17 | -0.16 | -0.15 | -0.34 | -1.3 | -0.09 | -0.14 | -0.86 | -0.94 | -1.02 | -0.7 | 0.03 | -0.3275 | -0.74 | 0.87921 | 0.311176 | 0.005817 | 1 | 0.996511 | 0.552614 |
| Ndufab1 | F6ZFT1 | 4.39 | 1.34 | 1.21 | 1.63 | -0.08 | 3.48 | 0.92 | 1.63 | -0.81 | 1.09 | 1.04 | 4.91 | -0.56 | -1.81 | 0.91 | 0.88 | -0.655 | -0.585 | -2.2875 | 0.499128 | 0.646801 | 0.029202 | 1 | 0.996511 | 0.637553 |
| Ndufaf1 | A0A0R4J08 | 0.53 | 0.65 | 0.76 | 0.69 | 0.4 | 0.94 | 0.78 | 1.05 | 0.89 | 0.75 | 1.05 | 0.47 | 1.25 | 0.87 | 0.64 | 0.63 | 0.135 | 0.1325 | 0.19 | 0.482058 | 0.490851 | 0.378631 | 1 | 0.996511 | 0.860984 |
| Ndufaf2 | Q59J78 | 1.68 | 1.46 | 1.4 | 1.03 | 1.52 | 1.75 | 1.11 | 1.27 | 0.4 | 0.51 | 1.69 | 1.64 | 0.58 | 1.47 | 1.24 | 1.42 | 0.02 | -0.3325 | -0.215 | 0.92715 | 0.373731 | 0.434643 | 1 | 0.996511 | 0.880687 |
| Ndufaf3 | Q9JKL4 | -0.04 | 0.36 | 0.9 | 0.89 | -0.12 | 0.07 | 0.63 | 0.75 | 0.66 | 0.96 | 0.96 | 0.02 | 0.05 | 0.67 | 0.54 | 0.93 | -0.195 | 0.1225 | 0.02 | 0.529536 | 0.701351 | 0.947567 | 1 | 0.996511 | 0.992414 |
| Ndufaf4 | Q9D1H6 | 0.67 | 0.67 | 0.95 | 0.94 | 0.47 | 0.66 | 0.54 | 0.62 | 0.71 | 0.97 | 0.89 | 0.61 | 0.44 | 0.94 | 0.3 | 0.3 | -0.235 | -0.0125 | -0.3125 | 0.159062 | 0.94468 | 0.182392 | 1 | 0.996755 | 0.776002 |
| Ndufaf5 | A2APY7 | 0.92 | 0.22 | -0.11 | -0.1 | 0.62 | 1.1 | -0.05 | -0.26 | 1 | -0.38 | 0.06 | 0.46 | 1.27 | -0.47 | -0.07 | 0.13 | 0.12 | 0.0525 | -0.0175 | 0.749677 | 0.886724 | 0.967051 | 1 | 0.996511 | 0.9927 |
| Ndufaf6 | A2AIL4 | -0.66 | -0.22 | 0.26 | 0.77 | -0.29 | 0.08 | 0.61 | 0.09 | 1.15 | 0.05 | 0.08 | -0.47 | 1.48 | -0.16 | -0.12 | 0.53 | 0.085 | 0.165 | 0.395 | 0.806198 | 0.704585 | 0.403631 | 1 | 0.996511 | 0.870698 |
| Ndufaf7 | Q9CWG8 | 0.63 | 0.92 | 0.22 | 0.44 | 0.58 | 0.77 | 0.52 | 0.46 | 0.51 | 1.06 | 0.44 | 0.69 | 0.67 | 1.08 | 0.28 | 0.39 | 0.03 | 0.1225 | 0.0525 | 0.879125 | 0.602786 | 0.840425 | 1 | 0.996511 | 0.976225 |
| Ndufaf8 | A0A087WC | 0.24 | -3.15 | -1.26 | -1.7 | -3.22 | 0.32 | -2.66 | -2.2 | 1.74 | 1.68 | -1.8 | 0.91 | 1.32 | 0.75 | -0.64 | -1.48 | -0.4725 | 2.1 | 1.455 | 0.618116 | 0.056766 | 0.114674 | 1 | 0.996511 | 0.717287 |
| Ndufb1 | P0DN34 | -0.22 | 0.13 | 0.42 | 0.31 | 0.53 | -0.15 | 0.58 | 0.33 | -0.61 | -0.34 | 0.77 | -0.53 | -0.97 | 0.62 | -0.39 | 1.31 | 0.1625 | -0.3375 | -0.0175 | 0.495396 | 0.340807 | 0.971666 | 1 | 0.996511 | 0.992865 |
| Ndufb10 | Q9DCS9 | 1.41 | 0.43 | -0.22 | 0.75 | -0.25 | 1.36 | -0.56 | 0.23 | 1.13 | 0.53 | -0.27 | 1.19 | 0.54 | 0.64 | -0.51 | -0.46 | -0.3975 | 0.0525 | -0.54 | 0.434178 | 0.907296 | 0.237095 | 1 | 0.996744 | 0.792722 |
| Ndufb11 | O09111 | 0.42 | 0.92 | 0.45 | 0.62 | 0.56 | 0.3 | 0.45 | 0.52 | 0.4 | 0 | 0.54 | 0.22 | -0.3 | 0.74 | 0.22 | 0.27 | -0.145 | -0.3125 | -0.37 | 0.419646 | 0.158672 | 0.189989 | 1 | 0.996511 | 0.776843 |
| Ndufb2 | Q9CPU2 | 0.91 | 1.26 | 0.84 | 0.81 | 0.95 | 0.48 | 0.5 | 0.47 | 0.63 | 1.23 | 0.68 | 0.55 | 0.68 | 1.72 | 0.35 | 0.31 | -0.355 | -0.1825 | -0.19 | 0.093631 | 0.421188 | 0.584117 | 1 | 0.996511 | 0.927734 |
| Ndufb3 | Q9CQZ6 | 0.81 | 0.74 | 0.06 | 0.54 | 0.13 | 0.48 | 0.25 | 0.59 | 0.73 | -0.39 | 0.27 | 0.7 | 0.2 | 0.6 | -0.46 | -0.6 | -0.175 | -0.21 | -0.6025 | 0.438606 | 0.508278 | 0.096732 | 1 | 0.996511 | 0.698156 |
| Ndufb4 | Q9CQC7 | 2.2 | 1.23 | 1.52 | 1.86 | 0.73 | 1.88 | 2.24 | 2.13 | 1.62 | 1.14 | 1.75 | 2.07 | 0.7 | 0.55 | 1.28 | 1.35 | 0.0425 | -0.0575 | -0.7325 | 0.911846 | 0.844536 | 0.036557 | 1 | 0.996511 | 0.658454 |
| Ndufb5 | Q9CQH3 | -1.34 | -0.89 | -0.45 | 0.08 | -1.06 | -1.28 | -0.31 | -0.01 | -1.71 | -0.41 | -0.22 | -1.53 | -1.36 | -0.5 | -0.9 | -1.01 | -0.015 | -0.3175 | -0.2925 | 0.970279 | 0.493416 | 0.41206 | 1 | 0.996511 | 0.874811 |
| Ndufb6 | Q3UIU2 | 1.15 | 0.91 | 1.12 | 1.09 | 0.08 | 1.26 | 1 | 1.4 | 0.57 | 0.59 | 1.2 | 1.18 | 0.52 | 0.33 | 1.08 | 1.57 | -0.1325 | -0.1825 | -0.1925 | 0.659834 | 0.418349 | 0.525274 | 1 | 0.996511 | 0.908107 |
| Ndufb7 | Q9CR61 | -0.37 | -0.07 | -0.61 | -0.57 | 0.11 | 0.02 | -0.3 | -0.82 | -0.77 | -1.03 | -0.53 | -0.33 | -0.89 | 0.05 | -0.82 | -0.49 | 0.1575 | -0.26 | -0.1325 | 0.540249 | 0.272881 | 0.627586 | 1 | 0.996511 | 0.939833 |
| Ndufb8 | Q9D6J5 | 0.49 | 0.54 | 0.82 | 0.75 | 0.06 | 0.46 | 0.62 | 0.73 | 0.4 | 0.51 | 0.81 | 0.29 | 0.34 | 0.21 | 0.38 | 0.55 | -0.1825 | -0.1475 | -0.28 | 0.373435 | 0.451527 | 0.160211 | 1 | 0.996511 | 0.755385 |
| Ndufb9 | Q9CQJ8 | 1.41 | 0.94 | -0.73 | -0.24 | 0.08 | 1.82 | -0.63 | -0.66 | 2.15 | -1.36 | -0.58 | 1.24 | 2.39 | 0.54 | -0.98 | -1.09 | -0.1925 | 0.0175 | -0.13 | 0.78184 | 0.983673 | 0.879614 | 1 | 0.998876 | 0.979626 |
| Ndufc2 | Q9CQ54 | 2.05 | 0.98 | 1.41 | 1.57 | 0.75 | 2.35 | 1.52 | 1.65 | 0.87 | 1.24 | 1.4 | 1.38 | 0.71 | 0.69 | 1.24 | 1.53 | 0.065 | -2.80E-01 | -0.46 | 0.862704 | 0.313091 | 0.165032 | 1 | 0.996511 | 0.758837 |
| Ndufs1 | Q91VD9 | 0.41 | 0.06 | 0.22 | 0.62 | -0.22 | 0.71 | 0.68 | 0.58 | -0.16 | -0.12 | 0.42 | 0.03 | -0.35 | -0.29 | 0.06 | 0.3 | 0.11 | -0.285 | -0.3975 | 0.674281 | 0.212992 | 0.119318 | 1 | 0.996511 | 0.721448 |
| Ndufs1 | A0A087WS | -1.66 | -0.88 | -0.63 | -2.44 | -1.18 | -1.72 | -1.81 | -2.62 | -1.85 | -0.99 | -1.75 | -2.56 | -2.05 | -0.09 | -1.82 | -2.07 | -0.43 | -0.385 | -0.105 | 0.371421 | 0.436327 | 0.855624 | 1 | 0.996511 | 0.976225 |
| Ndufs2 | Q91WD5 | 0.35 | 0.4 | 0.32 | 0.41 | 0.27 | 0.41 | 0.4 | 0.28 | 0.06 | -0.17 | 0.42 | 0.27 | -0.16 | 0.02 | -0.14 | 0.22 | -0.03 | -0.225 | -0.385 | 0.830448 | 0.252231 | 0.058143 | 1 | 0.996511 | 0.661738 |
| Ndufs2 | A0A0A6YXI | 0.27 | 0.41 | 0.04 | -1.79 | 0.91 | -0.37 | -0.92 | -1.39 | 4.32 | 5.81 | -0.3 | -0.97 | 3.59 | -0.46 | -0.91 | -1.14 | -0.175 | 2.4825 | 0.5375 | 0.787649 | 0.1401 | 0.626092 | 1 | 0.996511 | 0.939602 |
| Ndufs3 | Q9DCT2 | 1.68 | 1.05 | -0.01 | 0.08 | 0.82 | 1.81 | 0.17 | -0.02 | 0.84 | -1.1 | 0.12 | 1.34 | 0.41 | 0.25 | -0.67 | -0.19 | -0.005 | -0.4 | -0.75 | 0.992485 | 0.517739 | 0.120594 | 1 | 0.996511 | 0.721448 |
| Ndufs4 | Q9CXZ1 | 0.7 | 0.79 | 0.66 | 1.03 | 0.44 | 0.52 | -0.03 | 0.12 | 0.55 | -0.28 | 0.6 | 0.25 | 1.07 | 0.53 | 0.12 | -0.09 | -0.5325 | -0.515 | -0.3875 | 0.020619 | 0.06042 | 0.199495 | 1 | 0.996511 | 0.78004 |
| Ndufs5 | Q99LY9 | 2.01 | 1.16 | 1.23 | 0.79 | 1.36 | 2.28 | 0.93 | 1.17 | 0.14 | -0.34 | 1.5 | 1.88 | 0.11 | -0.23 | 0.64 | 1.02 | 0.1375 | -0.5025 | -0.9125 | 0.712736 | 0.368531 | 0.033722 | 1 | 0.996511 | 0.644622 |
| Ndufs6 | P52503 | 2.91 | 0.66 | 1.14 | 1.47 | 0.28 | 2.45 | 1.59 | 1.77 | 1.3 | 0.73 | 1.43 | 2.23 | 0.72 | 0.62 | 1 | 1.6 | -0.0225 | -0.1225 | -0.56 | 0.970302 | 0.817837 | 0.277193 | 1 | 0.996511 | 0.822084 |
| Ndufs7 | Q9DC70 | 2.2 | 1.48 | 2.03 | 1.92 | 1.14 | 2.2 | 1.36 | 2.13 | -1.7 | -0.86 | 2.15 | 1.84 | -1.5 | -1.7 | 1.66 | 2.01 | -0.2 | -1.55 | -1.79 | 0.517962 | 0.105695 | 0.074216 | 1 | 0.996511 | 0.671707 |
| Ndufs8 | Q8K3J1 | 0.1 | 0.22 | 0.27 | 0.65 | -0.39 | 0.02 | -0.08 | 0.16 | 0.29 | 0.41 | 0.34 | -0.2 | 0.62 | -0.01 | 0.11 | -0.1 | -0.3825 | -0.1 | -0.155 | 0.082775 | 0.651293 | 0.522125 | 1 | 0.996511 | 0.907667 |
| Ndufv1 | D3YUM1 | 0.59 | 0.45 | 0.41 | 0.57 | 0.2 | 0.56 | 0.46 | 0.48 | -0.39 | -0.43 | 0.57 | 0.35 | -0.61 | -0.02 | 0.24 | 0.14 | -0.08 | -0.48 | -0.5675 | 0.611572 | 0.106891 | 0.037321 | 1 | 0.996511 | 0.658454 |
| Ndufv2 | Q9D6J6 | 1.23 | 1.33 | 0.58 | 0.7 | 1 | 1.31 | 0.78 | 0.61 | 1.05 | 0.56 | 0.68 | 1.25 | 0.55 | 1.07 | 0.31 | 0.52 | -0.035 | -0.075 | -0.3475 | 0.889565 | 0.776924 | 0.222037 | 1 | 0.996511 | 0.787447 |
| Ndufv3 | Q3U422 | 1.81 | 1.59 | 0.89 | 1.4 | 0.8 | 0.79 | 0.37 | 0.77 | 1.73 | 1.92 | 0.91 | 1.38 | 1.67 | 1.06 | 0.96 | 0.47 | -0.74 | 0.0625 | -0.3825 | 0.012244 | 0.835914 | 0.252999 | 1 | 0.996511 | 0.809524 |
| Ndufv3 | Q8BK30 | 4.19 | 1.06 | 2.41 | 4.66 | -0.06 | 2.48 | 2.19 | 4.29 | 3.3 | 3.29 | 2.81 | 4 | 2.21 | 2.07 | 2.31 | 2.17 | -0.855 | 0.27 | -0.89 | 0.441761 | 0.730816 | 0.255343 | 1 | 0.996511 | 0.809524 |
| Nebl | Q9DC07 | 0.89 | 0.16 | 0.03 | 0.31 | 0.09 | 0.46 | -0.19 | 0.25 | 0.23 | -0.09 | 0 | 0.36 | 0.25 | 0.06 | -0.22 | -0.24 | -0.195 | -0.2225 | -0.385 | 0.438442 | 0.370707 | 0.156308 | 1 | 0.996511 | 0.751493 |
| Necab1 | Q8BG18 | 0.04 | -0.18 | 0.32 | 0.39 | -0.08 | 0.08 | 0.54 | 1.15 | -0.06 | 0.7 | 0.49 | 0.25 | -0.65 | 0.44 | -0.11 | 0.13 | 0.28 | 0.2025 | -0.19 | 0.36702 | 0.405037 | 0.509027 | 1 | 0.996511 | 0.902804 |
| Necab2 | Q91ZP9 | 1.27 | 0.55 | 0.49 | 0.47 | 0.6 | 0.67 | 0.97 | 0.24 | 0.26 | 0.63 | 0.19 | 1.17 | 0.21 | 0.78 | 0.9 | 0.75 | -0.075 | -0.1325 | -0.035 | 0.768638 | 0.661548 | 0.896996 | 1 | 0.996511 | 0.983509 |
| Necab3 | Q9D6J4 | 0.01 | -0.07 | 0.52 | 0.43 | 0.07 | -0.31 | -0.14 | 0.13 | -0.1 | 1.66 | 0.34 | -0.18 | 0.05 | 0.03 | -0.36 | -0.45 | -0.285 | 0.2075 | -0.405 | 0.194177 | 0.628991 | 0.114972 | 1 | 0.996511 | 0.717287 |
| Necap1 | Q9CR95 | 0.06 | 0.28 | 0.13 | 0.13 | 0.3 | -0.02 | 0.02 | -0.16 | 0.17 | -0.74 | 0.13 | 0.03 | -0.04 | 0.07 | -0.1 | -0.38 | -0.115 | -0.2525 | -0.2625 | 0.491162 | 0.31769 | 0.185033 | 1 | 0.996511 | 0.776843 |
| Necap2 | Q9D1J1 | -0.66 | -0.27 | -0.27 | -0.24 | 0.08 | -0.43 | -0.07 | -0.38 | -0.19 | -0.55 | -0.34 | -0.55 | -0.42 | 0.12 | -0.74 | -0.34 | 0.16 | -0.0475 | 0.015 | 0.419999 | 0.802983 | 0.95048 | 1 | 0.996511 | 0.992476 |
| Nectin1 | Q9JKF6 | 1.03 | 1.53 | 1.75 | 2.11 | 1.31 | 0.89 | 1.47 | 1.83 | 0.94 | 1.49 | 1.74 | 1.1 | 0.59 | 1.4 | 2.18 | 1.66 | -0.23 | -0.2875 | -0.1475 | 0.44653 | 0.346442 | 0.705468 | 1 | 0.996511 | 0.957084 |
| Nectin2 | P32507-2 | 0.97 | 1.16 | -0.6 | 0.19 | 1.29 | 1.47 | -0.03 | -0.17 | 1.46 | 0.85 | -0.04 | 1.52 | 1.44 | 1.72 | -0.25 | -0.17 | 0.21 | 0.5175 | 0.255 | 0.698539 | 0.31902 | 0.673493 | 1 | 0.996511 | 0.952944 |
| Nectin3 | Q9JLB9 | 0.64 | 0.99 | 0.87 | 0.93 | 0.84 | 0.8 | 0.94 | 0.8 | 1.29 | 0.9 | 1.41 | 1.13 | 1.5 | 1.47 | 0.89 | 1 | -0.0125 | 0.325 | 0.3575 | 0.935087 | 0.115909 | 0.136662 | 1 | 0.996511 | 0.732318 |
| Nedd4 | P46935 | -0.46 | -0.55 | -0.24 | -0.19 | -0.81 | -0.51 | 0.02 | -0.21 | -0.27 | 0.15 | -0.23 | -0.46 | -0.13 | -0.29 | -0.21 | -0.44 | -0.0175 | 0.1575 | 0.0925 | 0.937252 | 0.444204 | 0.625505 | 1 | 0.996511 | 0.93934 |
| Nedd4l | A0A494B9 | 0.53 | 0.81 | 1.04 | 0.87 | 0.91 | 0.71 | 0.92 | 0.74 | 0.65 | 0.67 | 1.08 | 0.57 | 0.34 | 0.94 | 0.68 | 0.66 | 0.0075 | -0.07 | -0.1575 | 0.965045 | 0.731365 | 0.472858 | 1 | 0.996511 | 0.891155 |
| Nedd8 | P29595 | -0.39 | -0.72 | -0.08 | 0.21 | -0.98 | -0.91 | -0.57 | 0.18 | -0.25 | 1.19 | -0.22 | -0.54 | 0.76 | 0.24 | 0.36 | -0.35 | -0.325 | 0.29 | 0.4975 | 0.330774 | 0.491316 | 0.13874 | 1 | 0.996511 | 0.73493 |
| Nefh | P19246 | 1.1 | 0.43 | 1 | 1.35 | 0.53 | 0.35 | 0.68 | 1.15 | 1.48 | 0.74 | 0.71 | 0.5 | 2.29 | -0.14 | 2.38 | 1.37 | -0.2925 | -0.1125 | 0.505 | 0.288941 | 0.705226 | 0.384447 | 1 | 0.996511 | 0.860984 |
| Nefl | P08551 | 1.09 | 0.94 | 0.9 | 0.58 | 1.34 | 0.69 | 1.12 | 0.76 | 0.99 | -0.12 | 1.03 | 1.02 | 1.05 | 0.26 | 1.65 | 1.64 | 0.1 | -0.1475 | 0.2725 | 0.641416 | 0.633013 | 0.437227 | 1 | 0.996511 | 0.883161 |
| Nefm | A0A0R4J03 | 1.38 | 0.99 | 0.92 | 0.62 | 1.23 | 0.93 | 0.71 | 0.75 | 0.75 | 0.83 | 0.89 | 1.07 | 1.05 | 0.34 | 1.69 | 1.45 | -0.0725 | -0.0925 | 0.155 | 0.74207 | 0.664468 | 0.646225 | 1 | 0.996511 | 0.944769 |
| Negr1 | D3Z4T6 | 0.98 | 1.06 | 0.71 | 1.04 | 1.2 | 0.92 | 1.11 | 1.18 | 0.69 | -0.13 | 0.92 | 1.14 | 0.09 | 1.1 | 0.52 | 0.36 | 0.155 | -0.2925 | -0.43 | 0.353795 | 0.337092 | 0.12283 | 1 | 0.996511 | 0.721448 |
| Nek1 | B2RXQ4 | -0.49 | -0.28 | 0.46 | -0.13 | 0.02 | -0.7 | 0.08 | 0.06 | -0.32 | 0.52 | -0.16 | -0.23 | -0.04 | -0.05 | -1.06 | -0.18 | -0.025 | 0.0625 | -0.2225 | 0.928949 | 0.828972 | 0.497732 | 1 | 0.996511 | 0.898105 |
| Nek4 | A0A2I3BPN | -0.36 | 0.06 | -0.22 | 0.08 | 0.05 | -0.2 | 0.24 | 0.14 | -0.12 | -0.06 | -0.11 | -0.23 | -0.02 | 0.19 | -0.16 | 0.37 | 0.1675 | -0.02 | 0.205 | 0.37684 | 0.911462 | 0.351116 | 1 | 0.996744 | 0.853984 |
| Nek6 | Q9ES70 | 0.67 | 0.69 | 0.64 | 0.45 | 0.52 | 0.98 | 0.28 | 0.55 | 0.74 | 0.93 | 0.65 | 0.72 | 0.93 | 0.87 | 0.17 | 0.79 | -0.03 | 0.1475 | 0.0775 | 0.876276 | 0.384683 | 0.73593 | 1 | 0.996511 | 0.958635 |
| Nek7 | Q9ES74 | 0.07 | -0.12 | 0.4 | 0.33 | -0.28 | 0.3 | 0.43 | 0.22 | -0.02 | 0.1 | 0.23 | -0.18 | -0.67 | 0.39 | -0.38 | 0.34 | -0.0025 | -0.1375 | -0.25 | 0.990917 | 0.495115 | 0.41737 | 1 | 0.996511 | 0.87558 |
| Nek9 | Q8K1R7 | 1.44 | 0.57 | -0.32 | -0.31 | 0.59 | 1.44 | -0.05 | -0.33 | 0.98 | -0.08 | -0.2 | 0.5 | 1.14 | -0.07 | -0.28 | -0.01 | 0.0675 | -0.045 | -0.15 | 0.898353 | 0.923563 | 0.761591 | 1 | 0.996755 | 0.963123 |
| Nelfa | Q8BG30 | -1.58 | -2.25 | -0.98 | -0.8 | -1.76 | -2.69 | -0.91 | -1.08 | -2.18 | -0.46 | -0.55 | -2.16 | -1.94 | -1.93 | -1.33 | -0.77 | -0.2075 | 0.065 | -0.09 | 0.668105 | 0.903772 | 0.827661 | 1 | 0.996609 | 0.973757 |
| Nelfb | A0A5H1ZR | -0.91 | -1.75 | -0.44 | -0.95 | -0.91 | -1.46 | -0.7 | -0.67 | -0.87 | -0.6 | -0.72 | -1.21 | -0.56 | -1.56 | -0.7 | -0.31 | 0.0775 | 0.1625 | 0.23 | 0.808955 | 0.598572 | 0.544126 | 1 | 0.996511 | 0.912212 |
| Nelfcd | Q3TW27 | -1.26 | -1.12 | -1.88 | -2.32 | -0.56 | -1.35 | -2.7 | -2.33 | -1.12 | -2.5 | -1.67 | -1.13 | -1.74 | -0.9 | -1.97 | -1.61 | -0.09 | 0.04 | 0.09 | 0.861129 | 0.921865 | 0.801496 | 1 | 0.996755 | 0.971091 |
| Nelfe | P19426 | -1.1 | -1.13 | -0.29 | -0.91 | -1.41 | -2.84 | -0.06 | -1.64 | -0.52 | -0.04 | -0.57 | -1.3 | -3.56 | -0.92 | -0.89 | -0.53 | -0.63 | 0.25 | -0.6175 | 0.272103 | 0.44888 | 0.363211 | 1 | 0.996511 | 0.858475 |
| Nell2 | A0A2R8VH | -2.21 | -2.14 | -1.93 | -2.43 | -1.72 | -2.15 | -1.7 | -2.08 | -2.17 | -1.53 | -1.8 | -1.82 | -1.79 | -1.55 | -1.22 | -1.6 | 0.265 | 0.3475 | 0.6375 | 0.19406 | 0.125131 | 0.014396 | 1 | 0.996511 | 0.589387 |
| Nemf | Q8CCP0 | -0.66 | -1.02 | -0.44 | -0.32 | -0.57 | -0.96 | -0.55 | -0.47 | -1.15 | 0.32 | -0.14 | -0.88 | -0.65 | -0.56 | -0.13 | -0.39 | -0.0275 | 0.1475 | 0.1775 | 0.898022 | 0.683901 | 0.455496 | 1 | 0.996511 | 0.887169 |
| Nenf | Q9CQ45 | -1.24 | -0.92 | -0.16 | -0.39 | -0.84 | -1.7 | 0.34 | 0 | -1.75 | -0.41 | -0.8 | -1.18 | -0.92 | -0.18 | 0.58 | 0.25 | 0.1275 | -0.3575 | 0.61 | 0.790748 | 0.342274 | 0.147308 | 1 | 0.996511 | 0.746406 |
| Neo1 | E9QK04 | 1.38 | 0.78 | 0.94 | 0.68 | 0.68 | 1.34 | 1.11 | 0.83 | 0.94 | 0.89 | 1.21 | 1.79 | 0.62 | 0.88 | 0.75 | 0.77 | 0.045 | 0.2625 | -0.19 | 0.846134 | 0.348871 | 0.391437 | 1 | 0.996511 | 0.865375 |
| Nes | Q6P5H2-2 | -2.24 | -2.6 | -1.33 | -1.03 | -3.26 | -2.54 | -1.87 | -1.83 | -1.7 | -0.26 | -0.93 | -2.68 | -0.24 | -2.88 | -0.9 | -0.76 | -0.575 | 0.4075 | 0.605 | 0.236973 | 0.493312 | 0.348568 | 1 | 0.996511 | 0.85278 |
| Neto1 | Q8R4I7 | 0.77 | 0.76 | 0.7 | 0.69 | 0.63 | 1.16 | 0.34 | 0.35 | 0.95 | 0.41 | 0.4 | 0.65 | 0.54 | 0.9 | 0.38 | 0.61 | -0.11 | -0.1275 | -0.1225 | 0.615907 | 0.505608 | 0.522228 | 1 | 0.996511 | 0.907667 |
| Neto2 | B2RXQ6 | 0.71 | 0.71 | 0.74 | 0.81 | 1.25 | 0.73 | 1.11 | 0.87 | 0.52 | 0.4 | 0.78 | 0.9 | 0.07 | 0.92 | 0.79 | 0.86 | 0.2475 | -0.0925 | -0.0825 | 0.173198 | 0.61358 | 0.730849 | 1 | 0.996511 | 0.958635 |
| Neu1 | O35657 | 2.45 | 1.9 | -0.4 | -0.5 | 1.75 | 2.16 | 0.48 | 0.44 | 0.1 | -0.6 | -0.24 | 2.19 | -0.2 | 0.83 | -2.22 | -0.03 | 0.345 | -0.5 | -1.2675 | 0.666581 | 0.578273 | 0.182899 | 1 | 0.996511 | 0.776002 |
| Neu2 | E9Q0P5 | -1.09 | -1.6 | 0.39 | -0.31 | -1.98 | -0.27 | -0.95 | -0.94 | 1.28 | -0.66 | -0.15 | -3.24 | 1.64 | -5.24 | -0.61 | -0.47 | -0.3825 | -0.04 | -0.5175 | 0.466663 | 0.965859 | 0.701965 | 1 | 0.996755 | 0.956791 |
| Neu3 | Q9JMH7 | -0.17 | 0.35 | 0.28 | -0.16 | 1.37 | -0.94 | 0.59 | -0.18 | -0.04 | -0.75 | 0.28 | -0.12 | -0.57 | 0.8 | -1.23 | 0.35 | 0.135 | -0.2325 | -0.2375 | 0.777905 | 0.402367 | 0.600907 | 1 | 0.996511 | 0.930802 |
| Neu4 | Q8BZL1 | 0.8 | 0.74 | 0.03 | 0.12 | 1.11 | 0.89 | 0.1 | 0.21 | 0.86 | 0.78 | 0.05 | 1.07 | 0.94 | 0.85 | -0.03 | 0.17 | 0.155 | 0.2675 | 0.06 | 0.624584 | 0.390674 | 0.851911 | 1 | 0.996511 | 0.976225 |
| Neurl1 | Q923S6 | 4.82 | 5.66 | 0.77 | 0.18 | 5.32 | 4.71 | 0.83 | 0.77 | -0.52 | 0.17 | 0.38 | 4.12 | 0.57 | 0.88 | 0.99 | 0.88 | 0.05 | -1.82 | -2.0275 | 0.975839 | 0.260578 | 0.129463 | 1 | 0.996511 | 0.723655 |
| Neurl4 | H3BK72 | 0.12 | 0.65 | 0.61 | 0.87 | 0.17 | -0.03 | 0.17 | 0.61 | 0.94 | 1.02 | 0.65 | 0.37 | 0.91 | 0.46 | 0.77 | 0.79 | -0.3325 | 0.1825 | 0.17 | 0.170294 | 0.460022 | 0.466471 | 1 | 0.996511 | 0.888757 |
| Nexn | A0A0G2JE | 1.1 | -0.29 | 1.11 | 1.39 | -0.98 | -0.81 | 1.46 | 0.37 | 0.68 | 1.07 | -0.01 | 0.31 | 0.97 | 0.31 | 1.04 | 1.31 | -0.8175 | -0.315 | 0.08 | 0.211489 | 0.462523 | 0.847319 | 1 | 0.996511 | 0.976225 |
| Nf1 | Q04690-2 | 0.58 | 0.45 | 0.37 | 0.18 | 0.79 | 0.46 | 0.45 | 0.2 | -0.3 | 0.21 | 0.35 | 0.44 | -0.41 | 0.08 | 0.35 | 0.4 | 0.08 | -0.22 | -0.29 | 0.670921 | 0.335033 | 0.249524 | 1 | 0.996511 | 0.805014 |
| Nf1 | Q5SYH9 | 0.14 | 0.48 | 0.51 | 0.6 | -0.06 | 0.11 | -0.36 | 0.63 | 0.86 | 0.56 | 0.14 | 0.57 | 0.74 | 0.87 | 0.42 | 0.53 | -0.3525 | 0.1 | 0.2075 | 0.174546 | 0.647902 | 0.32645 | 1 | 0.996511 | 0.843799 |
| Nf2 | P46662-2 | 0.64 | 0.67 | 0.54 | 0.8 | 0.3 | 0.62 | 0.82 | 0.51 | 0.12 | 0.84 | 0.54 | 0.67 | 0.38 | -0.36 | 0.7 | 0.65 | -0.1 | -0.12 | -0.32 | 0.565022 | 0.566525 | 0.260727 | 1 | 0.996511 | 0.81412 |
| Nfasc | A0A087WF | 1.04 | 0.86 | 0.59 | 0.74 | 0.7 | 0.8 | 0.41 | 0.51 | 0.76 | 0.18 | 0.47 | 1.04 | 0.46 | 0.56 | 0.12 | 0.2 | -0.2025 | -0.195 | -0.4725 | 0.271394 | 0.420168 | 0.043189 | 1 | 0.996511 | 0.658454 |
| Nfasc | Q810U3 | 2.68 | 1.98 | 1.79 | 2.59 | 1.93 | 1.56 | 2.22 | 2.68 | 1.84 | 2.46 | 1.55 | 2.11 | 1.62 | 2.12 | 1.92 | 1.51 | -0.1625 | -0.27 | -0.4675 | 0.610835 | 0.379303 | 0.123358 | 1 | 0.996511 | 0.721448 |
| Nfasc | E9Q171 | -2.05 | -1.01 | 0.6 | 0.48 | 0.09 | -2.08 | 1.55 | -0.13 | -2.65 | -0.75 | 0.63 | -2.96 | -1.57 | -1.85 | -0.41 | 0.87 | 0.3525 | -0.9375 | -0.245 | 0.690205 | 0.337477 | 0.759844 | 1 | 0.996511 | 0.963123 |
| Nfat5 | A0A1U6UR | 0.89 | 1.01 | 0.6 | -0.04 | 1.7 | 0.61 | 0.33 | 0.5 | 0.54 | 0.13 | 0.56 | 0.6 | -0.83 | 0.75 | 0.47 | 0.59 | 0.17 | -0.1575 | -0.37 | 0.648435 | 0.568382 | 0.382701 | 1 | 0.996511 | 0.860984 |
| Nfia | Q02780-3 | -1.32 | -1.58 | -2.44 | -3.06 | -1.08 | -0.64 | -3.03 | -3.19 | -0.91 | -1.36 | -2.46 | -1.32 | -0.76 | -0.86 | -2.5 | -2.33 | 0.115 | 0.5875 | 0.4875 | 0.868781 | 0.245575 | 0.398334 | 1 | 0.996511 | 0.867334 |
| Nfib | A2BG76 | -4.24 | -4.37 | -3.24 | -4.02 | -4.54 | -4.3 | -3.11 | -3.71 | -4.66 | -2.26 | -3.33 | -3.91 | -3.71 | -2.9 | -3.16 | -3.43 | 0.0525 | 0.4275 | 0.6675 | 0.891878 | 0.423801 | 0.059429 | 1 | 0.996511 | 0.661738 |
| Nfic | P70255-3 | -1.5 | 0.4 | -0.09 | -0.9 | 1.17 | -0.98 | -0.75 | -2.46 | 0.14 | 0.58 | 0.16 | -0.57 | -0.31 | 0.83 | -1.14 | -0.77 | -0.2325 | 0.6 | 0.175 | 0.764001 | 0.210243 | 0.752664 | 1 | 0.996511 | 0.961479 |
| Nfix | P70257-1 | -2.27 | -2.3 | -2.14 | -1.86 | -2.3 | -2.44 | -1.91 | -2.03 | -3.15 | -0.22 | -1.7 | -2.39 | -2.25 | -1.51 | -1.65 | -1.43 | -0.0275 | 0.2775 | 0.4325 | 0.887178 | 0.634433 | 0.106694 | 1 | 0.996511 | 0.703417 |
| Nfkb1 | P25799-4 | -1.13 | -1.25 | -0.21 | -0.07 | -0.91 | -1.26 | -0.33 | -0.37 | -1.46 | -1.21 | -0.03 | -1.29 | -0.81 | -1.54 | -1.5 | -0.52 | -0.0525 | -0.3325 | -0.4275 | 0.884701 | 0.441197 | 0.284989 | 1 | 0.996511 | 0.822669 |
| Nfrkb | Q6PIJ4 | -0.65 | -0.26 | -0.76 | 0.21 | -1.27 | -1.02 | -0.56 | -0.24 | 0.01 | -0.71 | -0.16 | -1.98 | 1.69 | -1.25 | 0.47 | -0.56 | -0.4075 | -0.345 | 0.4525 | 0.215189 | 0.469147 | 0.471698 | 1 | 0.996511 | 0.891144 |
| Nfs1 | Q9Z1J3 | 0.43 | 0.62 | 0.72 | 0.7 | 0.37 | 0.35 | 0.47 | 0.58 | 0.36 | 0.98 | 0.7 | 0.19 | 0.27 | 0.98 | 0.5 | 0.34 | -0.175 | -0.06 | -0.095 | 0.27439 | 0.788738 | 0.670764 | 1 | 0.996511 | 0.952558 |
| Nfu1 | Q9QZ23 | 0.61 | 0.8 | 1.16 | 0.24 | 0.52 | 0.53 | 0.93 | 0.89 | -0.06 | -0.5 | 1.02 | 0.88 | -0.14 | 0.37 | 0.77 | 1.18 | 0.015 | -0.3675 | -0.1575 | 0.949725 | 0.36652 | 0.646107 | 1 | 0.996511 | 0.944749 |
| Nfx1 | B1AY10 | -1.08 | -1.29 | -0.23 | -0.45 | -1.7 | -1.34 | -0.21 | -0.36 | -0.79 | -0.04 | -0.26 | -1.08 | -0.77 | -1.14 | 0.1 | 0.01 | -0.14 | 0.22 | 0.3125 | 0.737317 | 0.524595 | 0.422722 | 1 | 0.996511 | 0.875782 |
| Nfxl1 | E9Q8I7 | 1.39 | -0.11 | -0.36 | -0.06 | 0.37 | 1.47 | -0.53 | 0.6 | 0.12 | 1.26 | -0.13 | 0.87 | 1.25 | 0.27 | 0.23 | 0.26 | 0.2625 | 0.315 | 0.2875 | 0.619722 | 0.515434 | 0.522399 | 1 | 0.996511 | 0.907667 |
| Nfyc | P70353 | -1.56 | -1.68 | -3.07 | -1.31 | -1.95 | -1.46 | -1.85 | -2.66 | -2.03 | -0.76 | -2.29 | -1.15 | -1.99 | -1.96 | -2.99 | -2.89 | -0.075 | 0.3475 | -0.5525 | 0.863767 | 0.49179 | 0.246592 | 1 | 0.996511 | 0.802442 |
| Ngb | D3YW07 | -3.13 | -1.77 | -3.28 | -2.13 | -1.47 | -0.73 | 1.25 | -1.31 | -1.28 | 0.56 | -4.5 | -0.25 | -4.06 | -1.39 | -2.7 | 0.25 | 2.0125 | 1.21 | 0.6025 | 0.013468 | 0.268006 | 0.504429 | 1 | 0.996511 | 0.901002 |
| Ngef | Q8CHT1-2 | 0.45 | 0.23 | 0.23 | 0.54 | 0.25 | 0.29 | 0.45 | 0.33 | 0.7 | -0.2 | 0.31 | 0.55 | 0.28 | 0.43 | -0.22 | 0.05 | -0.0325 | -0.0225 | -0.2275 | 0.835445 | 0.925198 | 0.30786 | 1 | 0.996755 | 0.833087 |
| Ngly1 | Q9JI78 | 0.11 | -0.02 | -0.02 | -0.12 | 0.18 | 0 | -0.21 | 0.17 | 0.29 | 0.18 | -0.15 | 0.2 | 0.2 | 0.34 | 0.02 | -0.38 | 0.0475 | 0.1425 | 0.0575 | 0.770631 | 0.429363 | 0.790795 | 1 | 0.996511 | 0.969105 |
| Ngp | O08692 | 3.2 | 0.77 | 3.51 | 2.42 | -1.36 | 0.64 | 0.22 | 0.56 | -1.37 | 1.95 | -0.21 | -0.12 | -1.39 | -0.64 | 0.63 | -0.06 | -2.46 | -2.4125 | -2.84 | 0.006483 | 0.017148 | 0.002431 | 1 | 0.996511 | 0.427013 |
| Nhlrc2 | Q8BZW8 | 0.21 | 0.23 | -0.89 | -1.02 | 0.15 | 0.19 | -1.15 | -1.06 | -0.36 | -0.98 | -0.94 | 0.04 | -0.37 | -0.47 | -1.05 | -1.38 | -0.1 | -0.1925 | -0.45 | 0.829977 | 0.633133 | 0.279683 | 1 | 0.996511 | 0.822084 |
| Nhlrc3 | Q8CCH2 | -1.18 | -0.75 | -0.32 | -0.02 | -0.84 | -0.78 | 0.1 | 0.07 | -1.04 | 0.19 | -0.47 | -0.95 | -0.75 | -1.16 | -0.74 | -0.55 | 0.205 | 2.75E-16 | -0.2325 | 0.560154 | 1 | 0.443127 | 1 | 1 | 0.883549 |
| Nhp2 | Q9CRB2 | 0.69 | 0.66 | 1.27 | 0.81 | -0.03 | 0.73 | -0.13 | 0.22 | 0.51 | 1.05 | 0.85 | 0.05 | 1.17 | 0.77 | 0.87 | 0.93 | -0.66 | -0.2425 | 0.0775 | 0.025819 | 0.388027 | 0.72247 | 1 | 0.996511 | 0.958398 |
| Nhsl1 | A0A140LIZ | -0.39 | -0.12 | 0 | 0.2 | -0.34 | 0.19 | 0.33 | -0.24 | 0.6 | -0.05 | 0.18 | 0.09 | 0.77 | 0 | 0.17 | -0.18 | 0.0625 | 0.2825 | 0.2675 | 0.781822 | 0.225651 | 0.329488 | 1 | 0.996511 | 0.845234 |
| Nhsl2 | B1AXH1 | 0.16 | 0.23 | 0.68 | 0.57 | 0.14 | -0.4 | 0.96 | 0.82 | 0.26 | 0.85 | 0.76 | 0.13 | 0.81 | 0.8 | 0.48 | 0.49 | -0.03 | 0.09 | 0.235 | 0.927631 | 0.714118 | 0.287009 | 1 | 0.996511 | 0.824333 |
| Niban1 | E9PYV4 | 1.18 | 1.45 | -0.3 | -0.24 | 1.33 | 0.85 | -0.5 | -0.3 | 0.88 | -0.42 | 0.02 | 1.29 | 0.82 | 1.08 | -0.81 | 0.13 | -0.1775 | -0.08 | -0.2175 | 0.761583 | 0.885726 | 0.70614 | 1 | 0.996511 | 0.957084 |
| Niban2 | Q8R1F1 | 0.36 | -0.26 | -0.52 | -0.68 | -0.07 | 0.02 | -0.91 | -0.54 | -0.44 | 0.11 | -0.45 | 0.19 | -0.34 | 0.27 | -0.87 | -0.98 | -0.1 | 0.1275 | -0.205 | 0.747622 | 0.666224 | 0.573744 | 1 | 0.996511 | 0.925747 |
| Nicn1 | Q9CQM0 | 1.4 | 0.78 | 1.64 | 1.59 | 0.79 | 1.57 | 1.69 | 1.16 | 0.85 | 1.58 | 1.35 | 1.06 | 0.25 | 1.05 | 1.04 | 1.19 | -0.05 | -0.1425 | -0.47 | 0.86127 | 0.600679 | 0.147094 | 1 | 0.996511 | 0.746406 |
| Nid1 | P10493 | -0.82 | -1.02 | -1.06 | -1.16 | -0.67 | -0.68 | -0.86 | -1.36 | -0.46 | -1.27 | -0.63 | -0.62 | -0.06 | -0.66 | -0.82 | -0.86 | 0.1225 | 0.27 | 0.415 | 0.557119 | 0.253302 | 0.108848 | 1 | 0.996511 | 0.707444 |
| Nid2 | O88322 | -0.62 | -0.38 | -0.01 | -0.34 | -0.26 | -0.62 | -0.39 | -0.39 | -0.15 | 0.48 | 0.02 | -0.29 | 0.57 | -0.13 | 0.22 | 0.01 | -0.0775 | 0.3525 | 0.505 | 0.679509 | 0.163743 | 0.058548 | 1 | 0.996511 | 0.661738 |
| Nif3l1 | Q9EQ80 | -0.85 | -0.51 | -1.27 | -1.22 | -0.81 | -0.76 | -1.43 | -1.14 | -0.6 | -1.44 | -1.28 | -0.83 | -0.43 | -0.3 | -1.57 | -1.98 | -0.0725 | -0.075 | -0.1075 | 0.771295 | 0.786697 | 0.803094 | 1 | 0.996511 | 0.971091 |
| Nifk | Q91VE6-2 | 0.63 | 0.39 | 2.1 | 1.43 | -0.26 | 0.42 | 1.36 | 1.6 | 1.2 | 2.29 | 1.74 | 0.03 | 1.57 | 0.68 | 2.43 | 2.15 | -0.3575 | 0.1775 | 0.57 | 0.508292 | 0.75638 | 0.282022 | 1 | 0.996511 | 0.822084 |
| Nin | A0A1Y7VN | 0.02 | -0.5 | -0.51 | 0.04 | 0.25 | 0.58 | -1.25 | -0.42 | -0.3 | 1.05 | -1.07 | -0.24 | 0.28 | -0.68 | 1.09 | 0.51 | 0.0275 | 0.0975 | 0.5375 | 0.94611 | 0.82447 | 0.189361 | 1 | 0.996511 | 0.776843 |

| | | | | | | | | | | | | | | | | | | | | | | | | | | |
|---|---|---|---|---|---|---|---|---|---|---|---|---|---|---|---|---|---|---|---|---|---|---|---|---|---|---|
| Nip7 | E9Q109 | 0.21 | -0.1 | -0.84 | -0.41 | 0.25 | -0.28 | -0.26 | -0.44 | 1.3 | 0.07 | -0.22 | -0.39 | 1.64 | 1.83 | 0.17 | 0.28 | 0.1025 | 0.475 | 1.265 | 0.709436 | 0.276501 | 0.022026 | 1 | 0.996511 | 0.621926 |
| Nipa1 | Q8BHK1 | -0.59 | -0.15 | -0.52 | -0.67 | -0.32 | 0.28 | -0.92 | -0.25 | -0.07 | -0.3 | -0.41 | -0.62 | 0.35 | 0.24 | -0.67 | -0.1 | 0.18 | 0.1325 | 0.4375 | 0.518645 | 0.527238 | 0.141759 | 1 | 0.996511 | 0.737973 |
| Nipal3 | Q8BGN5 | 0.76 | 0.71 | 0.86 | 0.61 | 1.64 | 0.48 | -0.17 | 0.7 | 1.29 | 1.82 | 0.65 | 0.75 | 2.4 | 1.92 | 0.98 | 0.57 | -0.0725 | 0.3925 | 0.7275 | 0.841142 | 0.193096 | 0.099286 | 1 | 0.996511 | 0.698156 |
| Nipbl | Q6KCD5 | 0.6 | -0.13 | -0.26 | -0.13 | -0.4 | 0.55 | -0.49 | -0.38 | 0.97 | -0.73 | -0.46 | 0.01 | 1.46 | -0.03 | 0.17 | 0.47 | -0.2 | -0.0725 | 0.4975 | 0.522198 | 0.857131 | 0.207456 | 1 | 0.996511 | 0.785338 |
| Nipsnap1 | O55125 | 0.78 | 0.34 | 0.55 | 0.5 | 0.58 | 0.97 | -0.54 | 0.2 | -0.8 | -0.54 | 0.2 | 0.47 | -0.89 | -1.92 | 0.35 | 0.27 | -0.24 | -0.71 | -1.09 | 0.46856 | 0.048005 | 0.056857 | 1 | 0.996511 | 0.661738 |
| Nipsnap2 | O55126 | 0.72 | 0.56 | 0.58 | 0.94 | -0.09 | 0.84 | 0.09 | 0.61 | -0.41 | 0.24 | 0.64 | 0.3 | -0.55 | -1.38 | 0.02 | -0.55 | -0.3375 | -0.5075 | -1.315 | 0.196288 | 0.074897 | 0.002222 | 1 | 0.996511 | 0.419079 |
| Nipsnap3b | Q9CQE1 | -0.4 | 0.03 | 0.02 | -0.24 | 0.08 | -0.1 | 0.29 | -0.16 | -0.46 | -0.49 | 0.29 | -0.31 | -0.82 | 0.19 | -0.3 | -0.27 | 0.175 | -0.095 | -0.1525 | 0.36074 | 0.691212 | 0.56184 | 1 | 0.996511 | 0.91998 |
| Nisch | Q80TM9 | -0.92 | -0.74 | -0.38 | -0.5 | -0.87 | -0.77 | -0.31 | -0.29 | -0.46 | -0.29 | -0.36 | -0.85 | -0.29 | -0.33 | -0.56 | -0.67 | 0.075 | 0.145 | 0.1725 | 0.731751 | 0.504451 | 0.420394 | 1 | 0.996511 | 0.87558 |
| Nit1 | Q8VDK1-2 | 0.31 | 0.32 | 0.04 | -0.19 | 0.31 | 0.4 | 0.17 | -0.04 | -0.6 | 0.17 | 0.17 | 0.35 | -0.74 | -0.58 | 0.04 | 0.22 | 0.09 | -0.0975 | -0.385 | 0.641884 | 0.711457 | 0.196503 | 1 | 0.996511 | 0.78004 |
| Nit2 | Q9JHW2 | 0.43 | 0.38 | 0.46 | 0.06 | 0.48 | 0.47 | 0.59 | 0.56 | 0.17 | -0.02 | 0.56 | 0.44 | -0.07 | 0.63 | 0.19 | 0.49 | 0.1925 | -0.045 | -0.0225 | 0.248102 | 0.827044 | 0.921213 | 1 | 0.996511 | 0.987433 |
| Nkain4 | Q9JMG4 | -0.33 | 0.89 | -4.16 | -3.91 | 1.27 | 0.14 | -4.3 | -4.5 | 1.86 | -2.66 | -3.87 | -0.03 | 0.35 | 4.07 | -4.18 | -4.87 | 0.03 | 0.7025 | 0.72 | 0.986293 | 0.665453 | 0.74032 | 1 | 0.996511 | 0.958796 |
| Nkiras1 | Q8CEC5 | -0.22 | -0.26 | -0.4 | -0.57 | -0.41 | -0.55 | -0.77 | -0.9 | 0.17 | -1.62 | -0.86 | -0.35 | 0.57 | -0.03 | -1.22 | -1.16 | -0.295 | -0.3025 | -0.0975 | 0.128908 | 0.430408 | 0.818828 | 1 | 0.996511 | 0.972768 |
| Nkiras2 | Q9CR56 | 0.82 | -0.22 | 0.22 | 0.8 | -0.21 | 0.33 | 0.38 | 0.91 | 0.71 | 0.53 | 0.61 | 0.65 | -0.09 | 1.28 | 0 | 0.41 | -0.0525 | 0.22 | -0.005 | 0.873485 | 0.423035 | 0.989723 | 1 | 0.996511 | 0.996154 |
| Nkrf | A0A668KL6 | 0.46 | 0.12 | -0.15 | 0.2 | 0.07 | 0.14 | 0.25 | 0.12 | 0.15 | -0.16 | 0.44 | -0.13 | 0.74 | 0.43 | 0.84 | 0.37 | -0.0125 | -0.0825 | 0.4375 | 0.943925 | 0.713117 | 0.074077 | 1 | 0.996511 | 0.671115 |
| Nlgn1 | Q8VHQ9 | 0.28 | 0.55 | 0.25 | 0.49 | 0.76 | 0.13 | 0.58 | 0.69 | 0.09 | 0.31 | 0.45 | 0.42 | -0.41 | 0.75 | -0.04 | 0.33 | 0.1475 | -0.075 | -0.235 | 0.458331 | 0.675862 | 0.411193 | 1 | 0.996511 | 0.874811 |
| Nlgn2 | Q69ZK9 | 0.74 | 0.49 | 0.75 | 0.64 | 0.94 | 0.91 | 0.94 | 0.77 | 0.04 | 0.44 | 0.95 | 0.81 | -0.36 | 0.39 | 0.6 | 1 | 0.235 | -0.095 | -0.2475 | 0.142224 | 0.693607 | 0.423919 | 1 | 0.996511 | 0.876297 |
| Nlgn3 | A2AGI2 | 0.72 | 0.89 | 1.29 | 1.44 | 0.95 | 0.78 | 1.14 | 1.22 | 0.64 | 0.84 | 1.51 | 0.79 | -0.16 | 1.03 | 0.79 | 1.2 | -0.0625 | -0.14 | -0.37 | 0.775637 | 0.608377 | 0.300751 | 1 | 0.996511 | 0.830276 |
| Nlgn4l | B0F2B4 | 1.05 | 1.22 | 1.18 | 1.08 | 0.95 | 1.2 | 0.37 | 0.95 | 1.6 | 1.21 | 1.35 | 1.12 | 1.53 | 1.42 | 0.73 | 0.43 | -0.265 | 0.1875 | -0.105 | 0.226362 | 0.311124 | 0.715553 | 1 | 0.996511 | 0.958106 |
| Nlk | O54949 | 0.56 | 0.47 | 0.31 | -0.97 | 0.89 | 0.51 | -0.4 | -1.1 | 0.05 | -1.29 | -0.54 | 0.29 | -0.14 | 0.42 | -1.14 | -1 | -0.1175 | -0.465 | -0.5575 | 0.823881 | 0.336466 | 0.265304 | 1 | 0.996511 | 0.817264 |
| Nln | Q91YP2 | 0.82 | 0.58 | -0.12 | -0.14 | 0.36 | 0.92 | 0.16 | -0.03 | 0.59 | -0.12 | 0.18 | 0.64 | 0.49 | 0.45 | -0.32 | -0.31 | 0.0675 | 0.0375 | -0.2075 | 0.829644 | 0.902848 | 0.540473 | 1 | 0.996609 | 0.912212 |
| Nlrx1 | Q3TL44 | 0.4 | 0.36 | 0.42 | 0.47 | -0.04 | 0.41 | 0.44 | 0.23 | 0.34 | 0.19 | 0.39 | 0.09 | 0.13 | -0.17 | -0.25 | -0.01 | -0.1525 | -0.16 | -0.4875 | 0.374555 | 0.338685 | 0.022464 | 1 | 0.996511 | 0.621926 |
| Nmd3 | Q99L48 | -1.13 | -2.26 | -1.86 | -1.58 | -3.17 | -0.61 | -1.78 | -1.62 | 0.28 | -1.48 | -1.58 | -1.6 | 0.06 | -2.47 | -1.9 | -1.51 | -0.0875 | 0.6125 | 0.2525 | 0.868992 | 0.225947 | 0.646086 | 1 | 0.996511 | 0.944749 |
| Nme1 | P15532 | 0.69 | 0.03 | -0.17 | 0.14 | -0.37 | 0.38 | 0.15 | 0.2 | 0.31 | -0.1 | 0.2 | 0.71 | -0.03 | -0.36 | 0.07 | -0.5 | -0.0825 | 0.1075 | -0.3775 | 0.746868 | 0.687261 | 0.168299 | 1 | 0.996511 | 0.759484 |
| Nme3 | Q9WV85 | 0.37 | 0.04 | 0.38 | -0.01 | 0.02 | 0.97 | 0.52 | 0.34 | 0.58 | 0.97 | 0.5 | 0.6 | 0.73 | 0.68 | 0.31 | 0.53 | 0.2675 | 0.4675 | 0.3675 | 0.282192 | 0.040552 | 0.099009 | 1 | 0.996511 | 0.698156 |
| Nme7 | Q3UMG6 | -0.75 | -0.72 | -0.57 | -0.25 | -0.3 | -0.98 | -0.69 | -0.53 | -0.82 | -0.3 | -0.25 | -0.89 | -0.47 | -0.13 | -0.15 | -0.33 | -0.0525 | 0.0075 | 0.3025 | 0.80345 | 0.974394 | 0.162651 | 1 | 0.997515 | 0.757516 |
| Nmnat1 | Q9EPA7 | -1.81 | -1.68 | 0.03 | 0.04 | -1.15 | -1.91 | 0.52 | -0.36 | -1.99 | -0.89 | -0.65 | -2.08 | -1.26 | -1.18 | -0.46 | 0.24 | 0.13 | -0.5475 | 0.19 | 0.845059 | 0.359242 | 0.7408 | 1 | 0.996511 | 0.958796 |
| Nmnat2 | Q8BNJ3 | -0.5 | 0.04 | 0.7 | 0.33 | 0.01 | -0.12 | 0.82 | 0.69 | 0.73 | 1.4 | 1.05 | -0.01 | 0.73 | 0.54 | 1.28 | 1.27 | 0.2075 | 0.65 | 0.8125 | 0.541131 | 0.114137 | 0.030585 | 1 | 0.996511 | 0.641481 |
| Nmnat3 | Q99JR6 | 0.12 | 0.37 | 0.33 | 0.63 | -0.08 | 0.14 | 0.31 | 0.05 | 0.61 | 0.52 | 0.05 | -0.13 | 0.6 | 0.21 | 0.2 | 1.13 | -0.2575 | -0.1 | 0.1725 | 0.174894 | 0.674138 | 0.525023 | 1 | 0.996511 | 0.908107 |
| Nmral1 | D3YU12 | 0.03 | 0.61 | 0.72 | 0.54 | 0.18 | -0.5 | 0.53 | -0.1 | -0.1 | 0.15 | 0.38 | 0.13 | -0.17 | 0.21 | 0.15 | 0.26 | -0.4475 | -0.335 | -0.3625 | 0.126881 | 0.152764 | 0.137246 | 1 | 0.996511 | 0.732462 |
| Nmt1 | O70310 | -3.23 | -3.81 | -2.49 | -2.39 | -3.61 | -3.44 | -2.38 | -2.35 | -3.08 | -1.73 | -2.63 | -3.21 | -2.28 | -2.6 | -2.83 | -2.68 | 0.035 | 0.3175 | 0.3825 | 0.936873 | 0.482539 | 0.292556 | 1 | 0.996511 | 0.825497 |
| Nmt2 | O70311 | -1.24 | -1.49 | -0.96 | -1.12 | -1.52 | -0.99 | -0.97 | -1 | -1.29 | -1.63 | -0.86 | -1.21 | -0.83 | -1.56 | -1.53 | -1.4 | 0.0825 | -0.045 | -0.1275 | 0.688094 | 0.842968 | 0.600525 | 1 | 0.996511 | 0.930802 |
| Nnt | Q61941 | -4.94 | -6.32 | -3.48 | -3.57 | 0.98 | 1.23 | 1.03 | -3.56 | 0.91 | 2.88 | 1.26 | 0.49 | -5.3 | 4.61 | 4.08 | 1.53 | 4.4975 | 5.9625 | 5.8075 | 0.004635 | 4.07E-05 | 0.020929 | 1 | 0.1236 | 0.621926 |
| Noa1 | Q9JJG9 | 0.26 | 0.43 | 0.45 | 0.37 | -0.92 | 0.61 | 0.3 | 0.25 | 1.84 | 0.8 | 0.28 | -0.02 | 2.2 | -0.67 | -0.08 | 0.16 | -0.3175 | 0.3475 | 0.025 | 0.348619 | 0.388368 | 0.965291 | 1 | 0.996511 | 0.9927 |
| Nob1 | Q8BW10 | 0.58 | 0.84 | 2.55 | 2.12 | 0.48 | 0.51 | 1.05 | 1.94 | 0.14 | 1.94 | 2.3 | 0.61 | -1.33 | 3.85 | -0.65 | 1.74 | -0.5275 | -0.275 | -0.62 | 0.342759 | 0.671211 | 0.589059 | 1 | 0.996511 | 0.929131 |
| Noc2l | J3QK52 | 2.37 | 0.66 | 2.42 | 1.1 | 1.31 | 1.69 | 3.42 | 1.54 | 0.63 | 1.09 | 2.28 | 1.97 | 0.96 | 0.14 | 3.2 | 2.82 | 0.3525 | -0.145 | 0.1425 | 0.560721 | 0.789625 | 0.85414 | 1 | 0.996511 | 0.976225 |
| Noc3l | Q8VI84 | 1.28 | 0.94 | 1.42 | 0.9 | 1.85 | 1.19 | 1.21 | 1.63 | 1.69 | 1.23 | 1.2 | 0.64 | 1.47 | 1.8 | 1.44 | 1.34 | 0.335 | 0.055 | 0.3775 | 0.165573 | 0.836684 | 0.107724 | 1 | 0.996511 | 0.706141 |
| Noc4l | Q8BHY2 | -0.5 | 0.42 | -2.32 | -1.98 | 0.48 | -0.48 | -1.74 | -2.06 | 0.05 | -3.19 | -1.7 | 0.08 | -0.01 | 0.39 | -1.89 | -1.78 | 0.145 | -0.095 | 0.2725 | 0.853235 | 0.917087 | 0.729611 | 1 | 0.996755 | 0.958635 |
| Noct | O35710 | 0.17 | 0.43 | 0.74 | 0.8 | -0.33 | 0.22 | -0.15 | 0.84 | -0.08 | 0.29 | 0.51 | 0.04 | -0.4 | 0.18 | 0.46 | 1.17 | -0.39 | -0.345 | -0.1825 | 0.209993 | 0.158192 | 0.608694 | 1 | 0.996511 | 0.934103 |
| Nol10 | Q5RJG1 | 0.31 | 0.74 | 0.07 | 0.82 | 0.67 | 0.36 | -0.8 | 0.5 | 0.73 | 0.97 | 0.68 | 0.46 | 1.32 | 1.22 | 0.65 | 0.4 | -0.3025 | 0.225 | 0.4125 | 0.412636 | 0.353097 | 0.190253 | 1 | 0.996511 | 0.776843 |
| Nol11 | Q8BJW5-2 | -0.23 | 0.58 | 1.17 | 1.08 | -0.52 | 0.41 | 0.78 | 1.43 | 1.29 | 4.51 | 1.83 | -1.13 | 1.83 | 0.29 | 1.13 | 0.93 | -0.125 | 0.975 | 0.395 | 0.794656 | 0.377069 | 0.369604 | 1 | 0.996511 | 0.859245 |
| Nol3 | Q9D1X0 | -0.82 | -1.46 | -2.32 | -1.88 | -1.24 | -1.03 | -2.17 | -2.32 | -0.64 | -1.08 | -2.19 | -1.12 | -0.84 | -0.16 | -2.36 | -2.16 | -0.07 | 0.3625 | 0.24 | 0.869533 | 0.412738 | 0.674121 | 1 | 0.996511 | 0.952944 |
| Nol4 | F7BIN0 | -7.03 | -9.28 | -2.22 | -3.05 | -5.98 | -6.06 | -2.64 | -2.81 | -5.95 | -0.66 | -2.13 | -5.71 | -3.75 | -6.23 | -2.11 | -2.24 | 1.0225 | 1.7825 | 1.8125 | 0.554457 | 0.359102 | 0.303744 | 1 | 0.996511 | 0.831662 |
| Nol6 | Q8R5K4-2 | 1.15 | 1.11 | 0.26 | 0.18 | 1.08 | 0.61 | -0.02 | 0.15 | 1.18 | -0.06 | 0.39 | 0.87 | 0.71 | 1.24 | 0.24 | 0.56 | -0.22 | -0.08 | 0.0125 | 0.531686 | 0.827713 | 0.970397 | 1 | 0.996511 | 0.992767 |
| Nol7 | Q9D7Z3-2 | 1.09 | 2.01 | -0.29 | -1.2 | 0.11 | 0.53 | -1.54 | -0.42 | 1.24 | 0.04 | -1.18 | 0.93 | 0.65 | 1.43 | -1.37 | 0.64 | -0.7325 | -0.145 | -0.065 | 0.348848 | 0.857886 | 0.938084 | 1 | 0.996511 | 0.989844 |
| Nol8 | E9QKD1 | 0.43 | -1.23 | 0.53 | 0.61 | -1.91 | 1.14 | 0.43 | 0.34 | 0.22 | 0.54 | 0.81 | -0.42 | 0.54 | -2.3 | 0.03 | 0.55 | -0.085 | 0.2025 | -0.38 | 0.905792 | 0.674237 | 0.606502 | 1 | 0.996511 | 0.932425 |
| Nol9 | Q3TZX8-2 | -0.76 | -1.1 | -1.94 | -1.62 | -1.01 | -0.7 | -1.9 | -2.04 | -0.65 | -1.7 | -1.55 | -1.09 | 0.01 | 0.14 | -2.57 | -2.23 | -0.0575 | 0.1075 | 0.1925 | 0.885023 | 0.757844 | 0.781861 | 1 | 0.996511 | 0.967642 |
| Nolc1 | A0A286YDA | 0.99 | 0.64 | 1.02 | 0.44 | 0.06 | 0.7 | -0.17 | 0.08 | 1.91 | -0.03 | 0.89 | 0.41 | 2.63 | -0.74 | 1.56 | 0.9 | -0.605 | 0.0225 | 0.315 | 0.034633 | 0.956973 | 0.632867 | 1 | 0.996755 | 0.941132 |
| Nomo1 | Q6GQT9 | -1.45 | -1.52 | -0.49 | -0.38 | -0.78 | -1.04 | 0.34 | 0.07 | -1.17 | -0.18 | -0.45 | -1.11 | -0.41 | -0.88 | 0.9 | 0.8 | 0.6075 | 0.2325 | 1.0625 | 0.174453 | 0.541947 | 0.059102 | 1 | 0.996511 | 0.661738 |
| Nono | Q99K48 | -1.62 | -4.02 | -2.97 | -3.04 | -3.18 | -2.75 | -2.6 | -2.96 | -2.78 | -1.46 | -2.36 | -3.21 | -2.17 | -2.43 | -1.7 | -2.81 | 0.04 | 0.46 | 0.635 | 0.932366 | 0.427933 | 0.231862 | 1 | 0.996511 | 0.788863 |
| Nop10 | Q9CQS2 | 0.78 | 0.44 | 0.97 | 0.96 | 0.8 | 0.24 | 0.61 | 1.02 | 1.13 | -0.21 | 1.18 | -0.01 | 1.45 | 0.95 | 1.29 | 1.27 | -0.12 | -0.265 | 0.4525 | 0.600059 | 0.486107 | 0.061586 | 1 | 0.996511 | 0.662517 |
| Nop14 | Q8R3N1 | 2.25 | 2.23 | 0.51 | 0.92 | 2.12 | 1.47 | 0.98 | 0.58 | 1.3 | 0.2 | 0.73 | 1.92 | 0.32 | 2.04 | 1.36 | 0.65 | -0.19 | -0.44 | -0.385 | 0.712242 | 0.421976 | 0.486111 | 1 | 0.996511 | 0.895384 |
| Nop16 | Q9CPT5 | -0.47 | -0.94 | -1.3 | -0.95 | -0.47 | -0.88 | -1.68 | -1.26 | -0.62 | -0.23 | -0.81 | -0.7 | 0.46 | -0.51 | -0.35 | -0.62 | -0.1575 | 0.325 | 0.66 | 0.60947 | 0.198412 | 0.057234 | 1 | 0.996511 | 0.661738 |
| Nop2 | E9QN31 | 1.63 | 0.73 | 1.32 | 1.07 | 0.93 | 1.06 | 1.12 | 1.6 | -0.3 | -0.45 | 1.72 | 0.92 | -0.21 | -0.28 | 1.56 | 1.78 | -0.01 | -0.715 | -0.475 | 0.968294 | 0.186285 | 0.39182 | 1 | 0.996511 | 0.865498 |
| Nop53 | Q8BK35 | 0.41 | 0.33 | 0.64 | 0.03 | 0.37 | 0.6 | 0.52 | 0.02 | 0.5 | -0.04 | -0.41 | 0.3 | 0.48 | -0.09 | 0.17 | 0.67 | 0.025 | -0.265 | -0.045 | 0.904529 | 0.31727 | 0.854699 | 1 | 0.996511 | 0.976225 |
| Nop56 | Q9D6Z1 | 0.86 | 0.4 | 1.14 | 1.02 | 0.51 | 0.64 | 0.89 | 0.97 | 0.22 | 0.67 | 1 | 0.19 | 0.47 | 0.57 | 1.35 | 1.39 | -0.1025 | -0.335 | 0.09 | 0.64099 | 0.233748 | 0.769013 | 1 | 0.996511 | 0.965796 |
| Nop58 | Q6DFW4 | 1.22 | 0.67 | 0.35 | -0.09 | 0.8 | 0.98 | 0.94 | 0.5 | 0.24 | -1.45 | 0.7 | 0.69 | 0.42 | 0.5 | 1.15 | 1.07 | 0.2675 | -0.4925 | 0.2475 | 0.376716 | 0.369992 | 0.468297 | 1 | 0.996511 | 0.889423 |
| Nop9 | Q8BMC4 | -1.55 | -0.81 | 0.08 | -0.39 | -0.46 | -1.7 | -0.62 | -0.64 | -1.35 | -0.99 | -0.54 | -0.97 | -1.73 | -1.88 | -1.21 | -0.54 | -0.1875 | -0.295 | -0.6725 | 0.656899 | 0.435531 | 0.149568 | 1 | 0.996511 | 0.748169 |
| Nopchap1 | Q9CX66 | -2.3 | -1.91 | -1.68 | -2.35 | -2.54 | -3.7 | -2.26 | -2.32 | -2.09 | -0.01 | -1.69 | -2.93 | -1.3 | -0.93 | -2.11 | -2.54 | -0.645 | 0.38 | 0.34 | 0.099218 | 0.519474 | 0.392895 | 1 | 0.996511 | 0.865775 |
| Nos1 | F8WGF2 | 1.11 | 0.76 | 0.81 | 0.85 | 0.55 | 1 | 0.91 | 1.21 | 0.57 | 0.65 | 0.88 | 0.73 | 0.32 | 0.84 | 0.78 | 0.81 | 0.035 | -0.175 | -0.195 | 0.85709 | 0.330492 | 0.358125 | 1 | 0.996511 | 0.856645 |
| Nos1ap | Q9D3A8 | 0.84 | 0.9 | 1.84 | 1.02 | 0.47 | 1.24 | 2.01 | 1.97 | 0.45 | 0.72 | 2.16 | 0.71 | 0.02 | 0.36 | 1.41 | 1.68 | 0.2725 | -0.14 | -0.2825 | 0.50811 | 0.74455 | 0.525696 | 1 | 0.996511 | 0.908107 |
| Nos3 | P70313 | 4.1 | 3.21 | 0.41 | -0.27 | 3.02 | 4.08 | 1.59 | 0.87 | 0.33 | -0.9 | 1.04 | 4.3 | 0.4 | 0.68 | -0.03 | -0.02 | 0.5275 | -0.67 | -1.595 | 0.647348 | 0.627605 | 0.124125 | 1 | 0.996511 | 0.721448 |
| Nosip | A0A1B0GT | -1.35 | -1.55 | -6.36 | -7.97 | -1.8 | -1.99 | -7.58 | -6.32 | -1.19 | -5.41 | -5.74 | -1.68 | 0.92 | -6.52 | -4.87 | -6.71 | -0.115 | 0.8025 | 1.5125 | 0.954064 | 0.664835 | 0.49333 | 1 | 0.996511 | 0.89758 |
| Nostrin | Q6WKZ7 | -0.08 | -0.29 | 1.42 | 0.82 | 0.3 | -0.56 | -0.27 | 1.36 | -0.27 | 1.64 | 1.62 | -0.33 | -0.67 | 0.25 | 1.38 | 1.25 | -0.26 | 0.1975 | 0.085 | 0.628591 | 0.753036 | 0.882085 | 1 | 0.996511 | 0.979767 |
| Notch1 | Q01705-4 | 0.6 | 1.57 | 0.02 | 0.77 | 0.44 | 0.13 | 0.32 | 0.38 | 1.46 | 2.52 | 0.11 | 1.21 | 0.42 | 1.83 | 0.8 | 0.12 | -0.4225 | 0.585 | 0.0525 | 0.209157 | 0.298615 | 0.909557 | 1 | 0.996511 | 0.986458 |
| Notch2 | O35516 | 0.07 | -0.23 | -0.53 | -0.9 | -1.5 | 0.15 | -1.33 | -1.82 | 0.59 | -2.3 | -1.45 | -0.09 | 1.66 | -1.4 | -0.18 | -0.65 | -0.7275 | -0.415 | 0.255 | 0.133435 | 0.512813 | 0.684271 | 1 | 0.996511 | 0.955729 |
| Nova1 | Q9JKN6 | -1.77 | -2.26 | -2.17 | -1.91 | -2.61 | -2.11 | -1.81 | -2.49 | -2.04 | -1 | -1.85 | -2.22 | -1.85 | -2.09 | -1.5 | -2.1 | -0.2275 | 0.25 | 0.1425 | 0.34289 | 0.412842 | 0.535016 | 1 | 0.996511 | 0.912212 |
| Nova2 | A0A1W2P8 | -1.23 | -1.84 | -1.52 | -1.58 | -1.86 | -2.09 | -1.88 | -1.89 | -2.04 | -0.86 | -1.52 | -1.95 | -1.54 | -1.42 | -1.59 | -1.83 | -0.3875 | -0.05 | -0.0525 | 0.057264 | 0.868553 | 0.802905 | 1 | 0.996511 | 0.971091 |
| Npc1 | O35604 | 0.14 | 0.21 | 0.1 | -0.02 | -0.05 | 0.68 | 0.75 | 0.08 | -0.51 | 0.53 | 0.28 | 0.71 | 0.02 | -0.51 | 0.03 | 0.43 | 0.2575 | 0.145 | -0.115 | 0.279659 | 0.612387 | 0.631762 | 1 | 0.996511 | 0.941132 |
| Npc1l1 | Z4YJC9 | 0.74 | 0.25 | 0.67 | 0.15 | -0.31 | 0.26 | -0.07 | 0.56 | 0.61 | 0.92 | 0.91 | -0.24 | 0.43 | 0.38 | 0.36 | 0.68 | -0.3425 | 0.0975 | 0.01 | 0.197978 | 0.755702 | 0.963353 | 1 | 0.996511 | 0.9927 |
| Npc2 | Q9Z0J0 | -0.91 | -0.66 | -2.42 | -1.79 | -0.74 | -0.85 | -1.7 | -2.33 | -0.5 | -0.93 | -2.12 | -0.32 | -0.36 | -0.94 | -1.77 | -1.79 | 0.04 | 0.4775 | 0.23 | 0.937322 | 0.379259 | 0.64593 | 1 | 0.996511 | 0.944749 |
| Npdc1 | Q64322 | -0.53 | -0.12 | -0.01 | 0.03 | -0.39 | -0.2 | -0.58 | -0.21 | 0.31 | 0.25 | -0.16 | -0.45 | 1.01 | -0.47 | 0.27 | -0.13 | -0.1875 | 0.145 | 0.3275 | 0.345247 | 0.558805 | 0.352179 | 1 | 0.996511 | 0.854088 |
| Npepl1 | Q6NSR8 | -1.08 | -0.95 | -0.66 | -0.4 | -1.18 | -1.07 | -0.76 | -0.38 | -0.21 | -0.31 | -0.83 | -0.73 | 0.23 | -0.95 | -0.83 | -1.01 | -0.075 | 0.2525 | 0.1325 | 0.762737 | 0.312653 | 0.69122 | 1 | 0.996511 | 0.955729 |
| Npepps | Q11011 | 0.29 | 0.11 | 0.48 | 0.35 | 0.4 | 0.24 | 0.55 | 0.58 | -0.34 | 0.31 | 0.58 | 0.33 | -0.4 | 0.07 | 0.41 | 0.35 | 0.135 | -0.0875 | -0.2 | 0.425224 | 0.715153 | 0.412853 | 1 | 0.996511 | 0.874811 |
| Nphp1 | A2APS1 | -0.09 | 0.38 | 0.58 | 0.88 | 0.73 | -0.22 | 0.8 | 0.57 | 0.05 | 0.77 | 0.52 | 0.08 | -0.16 | 1.27 | 0.3 | 0.91 | 0.0325 | -0.0825 | 0.1425 | 0.915601 | 0.769115 | 0.700999 | 1 | 0.996511 | 0.956468 |
| Nphp4 | P59240 | 0.98 | 0.62 | 0.93 | 0.61 | 0.06 | 1.32 | 1.23 | 1.13 | 1.4 | 0.78 | 1.11 | 0.66 | 1.16 | 0.35 | 0.78 | 1.48 | 0.15 | 0.2025 | 0.1575 | 0.626749 | 0.384678 | 0.580588 | 1 | 0.996511 | 0.927734 |
| Npl | Q9DCJ9 | -0.78 | -0.83 | -2.59 | -2.52 | -0.7 | -0.86 | -2.19 | -3.07 | -1.43 | -3.08 | -2.72 | -1.13 | -1.07 | -0.89 | -3.41 | -2.79 | -0.025 | -0.41 | -0.36 | 0.97093 | 0.523668 | 0.623411 | 1 | 0.996511 | 0.938628 |
| Nploc4 | P60670 | 0.98 | 0.61 | -0.81 | -0.91 | 0.6 | 0.75 | -0.68 | -0.96 | 0.15 | -0.55 | -0.73 | 1.17 | -0.21 | 0.07 | -1.05 | -0.91 | -0.04 | 0.0425 | -0.4925 | 0.94632 | 0.942947 | 0.352116 | 1 | 0.996755 | 0.854088 |
| Npm1 | Q61937 | -0.4 | -1.35 | -0.58 | 0.04 | -1.01 | -1.1 | -0.58 | -0.19 | -1.09 | 0.17 | -0.13 | -1.11 | -0.5 | -1.22 | -0.17 | 0.28 | -0.1475 | 0.0325 | 0.17 | 0.670581 | 0.937787 | 0.680701 | 1 | 0.996755 | 0.955729 |
| Npm3 | Q9CPP0 | -0.88 | -1.09 | -1.51 | -0.34 | -1.16 | -0.46 | -1.57 | -1.5 | -0.18 | -2.93 | -1.51 | -0.98 | -0.88 | -1.52 | -2.31 | -1.32 | -0.2175 | -0.445 | -0.5525 | 0.527071 | 0.448564 | 0.167179 | 1 | 0.996511 | 0.759484 |
| Npr2 | B1AWI7 | 0.25 | 0.63 | 0.76 | 0.68 | 0.16 | 0.38 | 0.62 | 0.59 | 0.48 | 1.14 | 0.66 | 0.54 | 0.68 | 0.5 | 0.79 | 0.63 | -0.1425 | 0.125 | 0.07 | 0.466798 | 0.578981 | 0.723837 | 1 | 0.996511 | 0.958398 |
| Nprl2 | Q9WUE4 | 0.67 | 0.07 | -0.23 | -0.22 | 0.29 | -0.77 | -0.07 | 0.22 | -0.58 | 0.27 | -0.36 | 0.47 | -0.31 | -0.43 | -0.65 | 0.32 | -0.155 | -0.1225 | -0.34 | 0.625038 | 0.707989 | 0.285249 | 1 | 0.996511 | 0.822669 |
| Nprl3 | A7M7S2 | -0.86 | -0.91 | -1.55 | -1.42 | -1.71 | -0.8 | -1.78 | -1.89 | 0.1 | -1.93 | -1.83 | -1.23 | 0.38 | -0.93 | -1.91 | -1.4 | -0.36 | 0.0375 | 0.22 | 0.254764 | 0.935987 | 0.65357 | 1 | 0.996755 | 0.947633 |
| Nptn | P97300-3 | -0.51 | 0.28 | 1.64 | 1.17 | 0.23 | 0.23 | 0.77 | 1.14 | 0.46 | 1.79 | 1.24 | -0.68 | 1.15 | 0.54 | 1.33 | 1.05 | -0.1525 | 0.0575 | 0.3725 | 0.91405 | 0.929908 | 0.441255 | 1 | 0.996755 | 0.883549 |
| Nptn | P97300 | 0.46 | 0.01 | 0.6 | 0.63 | -0.2 | 0.75 | -0.11 | 0.19 | 1.1 | 0.17 | 0.55 | 0.46 | 1.68 | -0.18 | 1.13 | -0.43 | -0.2675 | 0.145 | 0.125 | 0.327557 | 0.581171 | 0.800249 | 1 | 0.996511 | 0.970393 |
| Nptx1 | Q62443 | 0.43 | 0.75 | 0.73 | 0.63 | 0.61 | 0.27 | 0.21 | 0.54 | 0.24 | 0.99 | 0.51 | 0.42 | 0.59 | 0.15 | 0.73 | 0.39 | -0.2275 | -0.095 | -0.17 | 0.211227 | 0.661324 | 0.420155 | 1 | 0.996511 | 0.87558 |
| Nptx2 | O70340 | 0.7 | 1.01 | 0.52 | 1.05 | 0.82 | 0.45 | 0.37 | 0.33 | 0.64 | -0.23 | 0.68 | 0.98 | 0.43 | 0.52 | 0.58 | 0.36 | -0.3275 | -0.3025 | -0.3475 | 0.131243 | 0.322795 | 0.111886 | 1 | 0.996511 | 0.713445 |
| Nptxr | E9PZM8 | -0.24 | 0.21 | -0.23 | -0.02 | 0.39 | -0.14 | -0.07 | 0.11 | -0.03 | 0.07 | 0.09 | 0.1 | 0.15 | 0.29 | 0.16 | -0.03 | 0.1425 | 0.1275 | 0.2125 | 0.471434 | 0.481298 | 0.294581 | 1 | 0.996511 | 0.825612 |
| Npy | P57774 | 0.98 | 0.28 | -0.34 | 0.57 | 1.25 | -0.26 | 1.39 | -0.14 | 0.55 | -0.38 | 0.17 | 0.44 | 0.88 | 1.31 | 1.56 | 1.58 | 0.1875 | -0.1775 | 0.96 | 0.698153 | 0.605948 | 0.015621 | 1 | 0.996511 | 0.589387 |
| Npy1r | Q04573 | -0.95 | -1.16 | 0.77 | 0.98 | -0.38 | -0.76 | 0.65 | 0.78 | -1.06 | 0.22 | 0.82 | -1.18 | -0.06 | -1.39 | 1.53 | 1.66 | 0.1625 | -0.21 | 0.525 | 0.792362 | 0.757076 | 0.528633 | 1 | 0.996511 | 0.908663 |
| Nqo1 | Q64669 | -0.38 | -0.18 | -0.5 | -0.1 | 0.62 | -0.83 | -0.1 | -0.09 | -1.05 | 0.01 | -0.29 | -0.68 | -0.94 | -0.02 | -0.41 | -0.68 | 0.19 | -0.2125 | -0.2225 | 0.539198 | 0.432206 | 0.385197 | 1 | 0.996511 | 0.861519 |
| Nqo2 | Q9JI75 | 0.12 | 0.13 | 0.41 | 0.17 | -0.18 | -0.38 | 0.22 | 0.61 | -0.2 | -0.4 | 0.38 | -0.3 | -0.72 | 0.23 | 0.31 | 0.14 | -0.14 | -0.3375 | -0.2575 | 0.57004 | 0.156584 | 0.433158 | 1 | 0.996511 | 0.879439 |
| Nr1d2 | A0A286YD | 0.96 | -2.1 | -0.29 | 0.12 | 2.37 | -1.55 | -0.8 | 1.44 | 0.17 | 1.39 | -0.8 | 0.52 | -0.41 | -1.35 | -1.42 | -1.13 | 0.2825 | 0.6475 | -0.75 | 0.783462 | 0.377562 | 0.250727 | 1 | 0.996511 | 0.807201 |
| Nr2c2 | G3X9W4 | -0.41 | 0.67 | -0.51 | -1.18 | 0.65 | -0.25 | -0.39 | -1.25 | 0.02 | -0.98 | -0.67 | -0.15 | -0.26 | -0.26 | -1.2 | -0.6 | 0.0475 | -0.0875 | -0.2225 | 0.92473 | 0.836458 | 0.601078 | 1 | 0.996511 | 0.930802 |
| Nr2c2ap | Q3TV70 | -3.09 | -1.97 | -2.96 | -2.76 | -3.22 | -3.02 | -4.15 | -3.06 | -2 | -4.75 | -2.27 | -2.66 | -1.32 | -1.8 | -4.21 | -5.16 | -0.6675 | -0.225 | -0.4275 | 0.085136 | 0.715919 | 0.622957 | 1 | 0.996511 | 0.938628 |
| Nr2f1 | Q32NY6 | -1.92 | -3.08 | -2.15 | -2.06 | -2.45 | -1.8 | -1.6 | -1.86 | -2.43 | -1.74 | -1.56 | -2.41 | 0.19 | -1.61 | 0.03 | 0.02 | 0.375 | 0.2675 | 1.96 | 0.253178 | 0.441858 | 0.002436 | 1 | 0.996511 | 0.427013 |
| Nr3c1 | P06537-5 | -2.18 | -1.58 | -1.67 | -1.83 | -2.03 | -1.63 | -2.44 | -2.32 | -1.17 | -1.62 | -2.13 | -1.54 | 0.17 | -2.22 | -1.89 | -2.24 | -0.29 | 0.2 | 0.27 | 0.24579 | 0.446173 | 0.623878 | 1 | 0.996511 | 0.938628 |
| Nr4a3 | F6R7A2 | -2.1 | -2.03 | -1.07 | -1.78 | -0.93 | -0.92 | -1.35 | -1.6 | -1.75 | -0.96 | -1.62 | -1.1 | -0.49 | -1.87 | -1.82 | -0.67 | 0.545 | 0.3875 | 0.5325 | 0.086903 | 0.229578 | 0.223358 | 1 | 0.996511 | 0.787447 |
| Nras | A0A0G2JG | 3.37 | 0.53 | 0.78 | 0.87 | 0.79 | 4.34 | 1.66 | 0.47 | -0.41 | -2.43 | 0.83 | 3.53 | -0.35 | -2.4 | 0.69 | 0.65 | 0.4275 | -1.0075 | -1.74 | 0.666951 | 0.434102 | 0.075364 | 1 | 0.996511 | 0.673156 |
| Nrbf2 | Q8VCQ3 | 0.13 | 0.41 | 0.12 | -0.08 | 0.64 | 0.13 | 0.07 | -0.31 | 0.07 | 0.4 | -0.1 | 0.29 | -0.69 | 0.37 | -0.2 | 0.26 | -0.0125 | 0.02 | -0.21 | 0.957817 | 0.920293 | 0.463695 | 1 | 0.996755 | 0.888267 |
| Nrbp1 | D3YUV1 | -0.95 | -0.92 | -0.78 | -0.84 | -0.9 | -1.07 | -1.04 | -0.66 | -1 | -0.29 | -0.97 | -1.07 | -0.92 | -0.74 | -1.1 | -1.06 | -0.045 | 0.04 | -0.0825 | 0.781169 | 0.857142 | 0.649455 | 1 | 0.996511 | 0.946607 |
| Nrbp2 | Q91V36 | -0.15 | 0.09 | 0.63 | 0.23 | 0.28 | 0.07 | 0.32 | 0.22 | -0.07 | 0.47 | 0.53 | 0.26 | 0.11 | 0.2 | 0.51 | 0.52 | 0.0225 | 0.0975 | 0.135 | 0.91192 | 0.685643 | 0.570471 | 1 | 0.996511 | 0.923986 |
| Nrcam | Q810U4-2 | -0.58 | -0.01 | 0.24 | 0.23 | 0.39 | -0.28 | 0.71 | 0.36 | -1.3 | -1.4 | -0.01 | 0.2 | -1.16 | -3.27 | -0.06 | 0.54 | 0.325 | -0.9575 | -0.9775 | 0.271732 | 0.19441 | 0.237856 | 1 | 0.996511 | 0.792722 |
| Nrcam | Q810U4 | -4.57 | -4.02 | -1.67 | -1.77 | -4.22 | -3.74 | -1.42 | -2.15 | -3.39 | -0.54 | -2.04 | -4.12 | -3.04 | -1.7 | -1.96 | -1.98 | 0.125 | 0.485 | 0.8375 | 0.889317 | 0.62291 | 0.270623 | 1 | 0.996511 | 0.822084 |
| Nrcam | Q810U4-3 | -0.43 | -0.18 | -0.77 | -0.7 | -0.12 | -0.34 | -0.66 | -0.87 | -0.54 | -0.39 | -0.49 | -0.01 | -0.53 | 0 | -1.07 | -1.06 | 0.0225 | 0.1625 | -0.145 | 0.922904 | 0.464582 | 0.632359 | 1 | 0.996511 | 0.941132 |
| Nrcam | A0A1W2P7 | 7.97 | 2.8 | 5.13 | 0.26 | 5.09 | 7.93 | 6.11 | 3.64 | 0.98 | 0.51 | 5.63 | 7.85 | 0.58 | 1.27 | 5.53 | 6.67 | 1.6525 | -0.2975 | -0.5275 | 0.337378 | 0.890706 | 0.790438 | 1 | 0.996511 | 0.969105 |
| Nrd1 | A2A9Q2 | -0.7 | -0.97 | -0.87 | -1.27 | -1.33 | -1.14 | -1.19 | -1.31 | -1.21 | -0.94 | -0.69 | -0.29 | -1.34 | -1 | -1.34 | -1.56 | -0.29 | 0.17 | -0.1575 | 0.126949 | 0.505183 | 0.128603 | 1 | 0.996511 | 0.723655 |
| Nrf1 | Q99K73 | -0.85 | -1.64 | -1.38 | -1.4 | -1.37 | -1.56 | -1.4 | -1.16 | -1.37 | -0.75 | -1.03 | -1.3 | -1.3 | -1.08 | -1.99 | -1.65 | -0.055 | 0.205 | -0.1875 | 0.796103 | 0.410995 | 0.509287 | 1 | 0.996511 | 0.902804 |
| Nrg3 | G3V023 | -0.03 | 0.59 | 0.82 | 0.62 | 0.32 | -0.37 | -0.46 | -0.21 | 0.69 | 0.86 | 0.97 | 0.31 | 0.35 | 1.38 | 0.79 | 0.43 | -0.68 | 0.2075 | 0.2375 | 0.027481 | 0.425038 | 0.448961 | 1 | 0.996511 | 0.884817 |
| Nrgn | P60761 | -0.14 | -0.82 | -0.35 | 0.36 | -1.38 | -0.57 | -0.97 | -0.21 | 0.06 | 0.03 | -0.3 | 0.05 | 0.48 | -0.32 | -0.48 | -0.74 | -0.545 | 0.1975 | -0.0275 | 0.135945 | 0.477618 | 0.938263 | 1 | 0.996511 | 0.989851 |
| Nrip2 | Q9JHR9-2 | 0.71 | 0.64 | -0.06 | 0.27 | -0.15 | 0.76 | -0.55 | -0.33 | 1.37 | -0.09 | -0.5 | 0.19 | 2.08 | 0.06 | 0.52 | 0.07 | -0.4575 | -0.1475 | 0.2925 | 0.188784 | 0.724612 | 0.545116 | 1 | 0.996511 | 0.912212 |
| Nrip3 | Q9JJR9 | -1.14 | -0.68 | -0.3 | -0.18 | -0.97 | -0.73 | 1.08 | -0.64 | -0.61 | 0.94 | -0.21 | -0.52 | -0.19 | -0.46 | -0.55 | -0.16 | 0.26 | 0.475 | 0.235 | 0.590469 | 0.253862 | 0.384928 | 1 | 0.996511 | 0.861128 |
| Nrm | Q8VC65 | -0.77 | -0.24 | 0.47 | 0.21 | -0.39 | 0.14 | 1.18 | 1.01 | -0.12 | -0.51 | 0.76 | -0.52 | 0.76 | 0.52 | 1.01 | 0.8 | 0.3625 | -0.215 | 0.27 | 0.493837 | 0.654018 | 0.497233 | 1 | 0.996511 | 0.897918 |
| Nrn1 | A0A286YC | -0.92 | -0.76 | 0.57 | 0.88 | -0.64 | -0.74 | 1.11 | 1.19 | -1.24 | 0.68 | 0.65 | -1.09 | -1.04 | -0.84 | 0.36 | 0.54 | 0.2875 | -0.1925 | -0.1875 | 0.6539 | 0.76357 | 0.739616 | 1 | 0.996511 | 0.958796 |
| Nrp1 | P97333 | 0.51 | 0.91 | 0.37 | 0.27 | 1 | 0.37 | 0.85 | 0 | -0.01 | -0.13 | 0.25 | 0.64 | -0.37 | 0.33 | 0.25 | 0.22 | 0.04 | -0.3275 | -0.4075 | 0.883572 | 0.205755 | 0.127571 | 1 | 0.996511 | 0.723391 |
| Nrp2 | O35375-5 | 0.75 | 1.43 | 0.32 | 0.88 | 1.05 | 1.09 | 0.89 | 1.01 | 0.49 | 0.39 | 0.38 | 1.18 | 0 | 0.12 | 0.97 | 0.56 | 0.165 | -0.235 | -0.4325 | 0.508356 | 0.446631 | 0.204252 | 1 | 0.996511 | 0.783092 |
| Nrsn1 | P97799 | 1.11 | 0.72 | 2.83 | 1.68 | -1.11 | 1.65 | 1.54 | 2.14 | 3.48 | 1.4 | 2.51 | 1.35 | 4.41 | -1.54 | 4.34 | 4.14 | -0.53 | 0.6 | 1.2525 | 0.503143 | 0.351205 | 0.369398 | 1 | 0.996511 | 0.859245 |
| Nrxn1 | Q9CS84-4 | 0.32 | 0.71 | 0.48 | 0.44 | 0.83 | 0.59 | 0.42 | 0.36 | 0.45 | 0.1 | 0.58 | 0.66 | 0.35 | 0.68 | 0.27 | 0.26 | 0.0625 | -0.04 | -0.0975 | 0.727755 | 0.840446 | 0.623807 | 1 | 0.996511 | 0.938628 |
| Nrxn1 | H3BK41 | 1.94 | 0.5 | -0.24 | 0.32 | 0.23 | 1.86 | -0.11 | 0.25 | 2.3 | 0.89 | 0.21 | 1.99 | 1.31 | 1.4 | 0.54 | -0.61 | -0.0725 | 0.7175 | 0.03 | 0.901358 | 0.262114 | 0.960219 | 1 | 0.996511 | 0.992636 |
| Nrxn2 | E9PUM9 | 0.55 | 0.78 | 0.84 | 0.98 | 0.89 | 0.53 | 0.98 | 0.93 | 0.53 | 0.77 | 1.02 | 0.8 | 0.37 | 0.69 | 0.77 | 0.87 | 0.045 | -0.0075 | -0.1125 | 0.803567 | 0.968623 | 0.584254 | 1 | 0.996755 | 0.927734 |
| Nrxn3 | Q6P9K9 | 0.35 | 0.58 | 0.37 | 0.55 | 0.82 | 0.53 | 0.73 | 0.59 | 0.14 | 0.05 | 0.61 | 0.61 | -0.12 | 0.42 | 0.3 | 0.43 | 0.205 | -0.11 | -0.205 | 0.211351 | 0.597099 | 0.331229 | 1 | 0.996511 | 0.845601 |
| Nsd1 | E9QAE4 | 0.25 | -0.55 | -0.82 | -0.83 | -0.76 | 0.2 | -1.11 | -0.85 | 1.33 | -1.22 | -0.76 | -0.2 | 1.87 | 0.84 | 1.18 | -0.95 | -0.1425 | 0.275 | 1.2225 | 0.69786 | 0.626912 | 0.067147 | 1 | 0.996511 | 0.664936 |
| Nsd2 | Q8BVE8 | -0.59 | -0.35 | -0.94 | -0.97 | 0.02 | -0.22 | -0.73 | -1.71 | 0.19 | -1.71 | -1.1 | -0.57 | -0.08 | -0.76 | 0.09 | -0.56 | 0.0525 | -0.085 | 0.385 | 0.892598 | 0.835542 | 0.181446 | 1 | 0.996511 | 0.776002 |
| Nsd3 | D3Z197 | 0.13 | -0.36 | -0.5 | -0.22 | -1.63 | -0.94 | 0.12 | -0.13 | -0.05 | 0.48 | -0.3 | -0.68 | 0.19 | 0.16 | 0.26 | -0.15 | -0.4075 | 0.1 | 0.3525 | 0.320261 | 0.729017 | 0.130133 | 1 | 0.996511 | 0.723659 |
| Nsdhl | Q9R1J0 | -0.7 | -0.17 | -0.41 | -0.82 | -0.14 | -0.4 | -0.36 | -0.6 | 0.1 | 0.13 | -0.51 | -0.07 | 0.69 | 0.61 | 0.19 | 0.18 | 0.15 | 0.4375 | 0.9425 | 0.470685 | 0.092198 | 0.003343 | 1 | 0.996511 | 0.483611 |
| Nsf | P46460 | 0.43 | 0.48 | 0.93 | 0.95 | 0.7 | 0.46 | 1.07 | 1.35 | 0.53 | 0.25 | 1.04 | 0.54 | 0.28 | 0.57 | 1.02 | 0.86 | 0.1975 | -1.08E-01 | -0.015 | 0.443116 | 0.659332 | 0.951996 | 1 | 0.996511 | 0.992476 |
| Nsfl1c | Q9CZ44 | 0.9 | 0.51 | 0.26 | 0.02 | 0.12 | 1.08 | 0.1 | 0.19 | 0.54 | -1.14 | 0.11 | 1.08 | 1.07 | -1.09 | 0.3 | 0 | -0.05 | -0.275 | -0.3525 | 0.867568 | 0.566839 | 0.449106 | 1 | 0.996511 | 0.884817 |
| Nsg1 | Q62092 | 0.77 | -0.27 | 0.81 | 1.06 | -1.34 | 0.57 | -0.25 | 0.65 | 0.61 | 2.13 | 0.63 | 0.58 | 1.74 | -0.8 | 0.88 | 0.8 | -0.685 | 0.395 | 0.0625 | 0.199132 | 0.393619 | 0.910858 | 1 | 0.996511 | 0.986458 |
| Nsg2 | P47759 | -0.36 | -0.7 | -0.33 | -0.55 | -1.09 | -0.04 | -1.41 | -0.62 | -0.39 | -1.34 | -0.54 | -0.16 | 0.42 | -2.11 | -0.14 | -0.88 | -0.305 | -0.1225 | -0.1925 | 0.333865 | 0.665663 | 0.709186 | 1 | 0.996511 | 0.957656 |
| Nsmaf | O35242 | -1.53 | -0.81 | -0.65 | -1.37 | -1.31 | -1.17 | -1.44 | -1.78 | -0.16 | -1.08 | -3.3 | -1.06 | -0.01 | -0.09 | -2.13 | -0.49 | -0.335 | -0.31 | 0.41 | 0.217876 | 0.630745 | 0.42305 | 1 | 0.996511 | 0.875782 |
| Nsmce3 | Q9CPR8 | 3.29 | 1.75 | -3.98 | -3.03 | 1.48 | 2.99 | -5.61 | -6.52 | 1.39 | -1.57 | -2.96 | 3.1 | -0.37 | -0.16 | -2.88 | -0.91 | -1.4225 | 0.4825 | -0.5875 | 0.597889 | 0.809886 | 0.725789 | 1 | 0.996511 | 0.958398 |
| Nsmf | Q99NF2-2 | 0.21 | 0.27 | -0.85 | -2 | 1.54 | 0.57 | -1.61 | -0.97 | -0.86 | -2.28 | -0.76 | 1.02 | 1.3 | -0.78 | -1.15 | -2.13 | 0.475 | -0.1275 | -0.0975 | 0.559952 | 0.870141 | 0.904057 | 1 | 0.996511 | 0.985278 |
| Nsmf | Q99NF2-3 | 0.82 | 0.51 | 0.36 | 0.15 | 0.7 | 0.81 | 0.21 | 0.33 | 0.63 | 0.08 | 0.34 | 0.88 | 0.64 | 0.87 | -0.09 | 0.31 | 0.0525 | 0.0225 | -0.0275 | 0.814378 | 0.927603 | 0.920141 | 1 | 0.996755 | 0.98688 |
| Nsrp1 | Q5NCR9 | -1.54 | -3.13 | 2.99 | 0.71 | -4.41 | -2.3 | 2.23 | 2.62 | -2.01 | 0.93 | 2.85 | -2.85 | -0.7 | -4.54 | 2.4 | 3.37 | -0.2225 | -0.0275 | 0.375 | 0.908816 | 0.986876 | 0.849253 | 1 | 0.999285 | 0.976225 |
| Nsun2 | Q1HFZ0 | -2.01 | -2.36 | -1.93 | -1.73 | -2.21 | -2.09 | -1.43 | -1.71 | -2.31 | -0.81 | -1.67 | -2.02 | -2.08 | -1.68 | -1.73 | -1.68 | 0.1475 | 0.305 | 0.215 | 0.540071 | 0.387326 | 0.334679 | 1 | 0.996511 | 0.846824 |
| Nsun4 | Q9CZ57 | 0.39 | 0.62 | 1.18 | 1.1 | 0.75 | 0.67 | 0.32 | 0.83 | 0.58 | 1 | 0.62 | 0.79 | 0.83 | 1.15 | 0.63 | 0.66 | -0.18 | -0.075 | -0.005 | 0.455796 | 0.755292 | 0.984363 | 1 | 0.996511 | 0.996154 |
| Nt5c | Q9JM14 | -1.92 | -2.19 | -3.01 | -2.63 | -1.73 | -2.12 | -2.22 | -3.1 | -2.42 | -1.55 | -3.01 | -2.15 | -2.3 | -0.96 | -2.74 | -2.52 | 0.145 | 0.155 | 0.3075 | 0.688304 | 0.679972 | 0.49312 | 1 | 0.996511 | 0.897531 |
| Nt5c1a | A3KFX0 | 2.02 | 1.2 | 1.2 | 1.28 | 1.25 | 1.8 | 0.79 | 1.56 | 0.1 | 1.6 | 1.02 | 1.93 | -0.33 | 0.21 | 1.21 | 0.85 | -0.075 | -0.2625 | -0.94 | 0.79913 | 0.540694 | 0.034807 | 1 | 0.996511 | 0.651434 |
| Nt5c2 | A0A494B9 | -0.19 | -0.09 | -0.93 | -0.9 | -0.32 | -0.48 | -0.31 | -1.2 | -0.72 | -1.14 | -1.37 | -0.44 | -1.25 | -0.19 | -1.94 | -1.52 | -0.05 | -0.39 | -6.98E-01 | 0.869675 | 0.229179 | 0.121469 | 1 | 0.996511 | 0.721448 |
| Nt5c3a | Q9D020-1 | -0.42 | -0.56 | -0.03 | -0.13 | -0.4 | -0.35 | 0.06 | -0.02 | -0.92 | -0.33 | 0 | -0.55 | -0.98 | -0.23 | -0.71 | -0.52 | 0.1075 | -0.165 | -0.325 | 0.596501 | 0.517521 | 0.198072 | 1 | 0.996511 | 0.78004 |
| Nt5c3b | Q3UFY7 | 0.39 | 0.52 | -0.76 | -0.98 | 0.31 | 0.14 | -0.81 | -0.69 | 0.16 | -0.33 | -0.89 | 0.49 | 0.02 | 0.58 | -0.88 | -0.87 | -0.055 | 0.065 | -0.08 | 0.901969 | 0.887382 | 0.870742 | 1 | 0.996511 | 0.979293 |
| Nt5dc2 | A0A2I3BRL | 0.67 | 0.48 | -0.24 | 0.55 | 0.21 | 0.87 | 0.58 | 0.52 | 0.49 | 0.02 | 0.7 | 0.74 | 0.22 | 0.19 | -0.44 | 0.46 | 0.18 | 0.1225 | -0.2575 | 0.488452 | 0.659803 | 0.394681 | 1 | 0.996511 | 0.866151 |
| Nt5dc3 | Q3UHB1 | 0.34 | 0.65 | 0.68 | 1 | 0.47 | 0.61 | 0.75 | 0.88 | 0.3 | 0.54 | 0.76 | 0.06 | -0.23 | 1.02 | 0.25 | 0.38 | 0.01 | -0.2525 | -0.3125 | 0.959332 | 0.295514 | 0.317168 | 1 | 0.996511 | 0.838864 |
| Nt5e | Q61503 | 1.34 | 1.06 | 0.01 | 0.51 | 1.02 | 0.24 | 1.08 | -0.09 | 0.76 | 0.17 | -0.3 | 0.91 | 0.3 | 0.92 | 0.07 | 0.46 | -0.1675 | -0.345 | -0.2925 | 0.670103 | 0.387341 | 0.406258 | 1 | 0.996511 | 0.87356 |
| Nt5m | Q8VCE6 | 1.45 | 1.43 | 0.49 | 0.76 | 1.52 | 1.39 | 0.5 | 0.49 | 0.21 | 0.46 | 0.57 | 1.03 | 0.02 | 0.95 | 0.24 | 0.21 | -0.0575 | -0.465 | -0.6775 | 0.870877 | 0.14974 | 0.060872 | 1 | 0.996511 | 0.661738 |
| Ntan1 | Q64311 | 1.66 | 0.15 | 1.58 | -0.47 | 0.18 | 1.22 | -0.43 | 0.19 | 0.11 | 2.3 | -0.03 | 0.21 | -0.68 | 0.08 | 0.86 | 0.72 | -0.44 | -0.0825 | -0.525 | 0.452922 | 0.905601 | 0.374096 | 1 | 0.996744 | 0.859424 |
| Ntaq1 | Q80WB5 | -0.04 | -0.19 | 0.37 | 0.21 | -0.5 | -0.75 | 0.16 | -0.35 | 0.91 | -0.13 | 0.36 | 0.07 | 1.42 | 0.97 | -0.03 | -0.32 | -0.4475 | 0.215 | 0.4225 | 0.093756 | 0.439378 | 0.319447 | 1 | 0.996511 | 0.838969 |
| Ntm | D3Z396 | 1.03 | 1.26 | 0.83 | 1.03 | 1.37 | 0.95 | 0.98 | 1.1 | 0.7 | 0.67 | 0.93 | 1.23 | 0.15 | 1.59 | 0.2 | 0.29 | 0.0625 | -0.155 | -0.48 | 0.724964 | 0.454269 | 0.197321 | 1 | 0.996511 | 0.78004 |
| Ntmt1 | Q8R2U4 | -0.13 | -0.51 | -0.47 | -0.3 | 0.01 | -0.26 | -0.41 | -0.73 | -0.89 | 0.07 | -0.41 | 0.26 | -0.92 | -1.61 | -1.23 | -1.35 | 0.005 | 0.11 | -0.925 | 0.980684 | 0.698785 | 0.002253 | 1 | 0.996511 | 0.419079 |
| Ntng1 | Q8R4G0-5 | 2.76 | 1.52 | 0.6 | 0.72 | 2.46 | 3.01 | 0.42 | 0.46 | -0.99 | -0.3 | 0.58 | 2.64 | -1.4 | -1.39 | -0.34 | 0.05 | 0.1875 | -0.9175 | -2.17 | 0.804077 | 0.292154 | 0.004158 | 1 | 0.996511 | 0.548272 |
| Ntng2 | J3QNA0 | -0.03 | 0.12 | 0.42 | 0.41 | 0 | 0.64 | 0.24 | 0.24 | -0.38 | 0.42 | 0.41 | 0.47 | -0.35 | -0.92 | -0.11 | 0.08 | 0.05 | 1.57E-16 | -0.555 | 0.806554 | 1 | 0.064751 | 1 | 1 | 0.66433 |
| Ntpcr | Q9CQA9 | -0.79 | -0.33 | 0.55 | -0.56 | -0.32 | -0.55 | 1.09 | 0.85 | 0.55 | -0.69 | 0.59 | -0.25 | -0.46 | -0.04 | 0.65 | 1.52 | 0.55 | 0.3325 | 0.7 | 0.259511 | 0.425161 | 0.176671 | 1 | 0.996511 | 0.771503 |
| Ntrk2 | P15209 | -0.05 | 0.1 | 0.05 | 0.08 | 0.29 | 0.31 | 0.11 | -0.03 | 0.14 | -0.11 | 0.11 | 0.14 | -0.08 | 0.71 | -0.25 | -0.19 | 0.125 | 0.025 | 0.0025 | 0.429456 | 0.876783 | 0.992219 | 1 | 0.996511 | 0.996154 |
| Ntrk3 | Q6VNS1-3 | 0.48 | 1.47 | 1.38 | 0.67 | 1.2 | 0.65 | 1.13 | 0.81 | 0.21 | -0.15 | 0.86 | 0.79 | -0.08 | 0.59 | -0.14 | 0.69 | -0.0525 | -0.5725 | -0.735 | 0.852978 | 0.121479 | 0.051519 | 1 | 0.996511 | 0.658454 |
| Ntrk3 | Q6VNS1 | 3.04 | 0.97 | 1.9 | 0.84 | 1.95 | 3.27 | 2.06 | 1.2 | 0.61 | 0.45 | 1.77 | 2.64 | 0.28 | 0.88 | 1.58 | 1.99 | 0.4325 | -0.32 | -0.505 | 0.482202 | 0.630381 | 0.396325 | 1 | 0.996511 | 0.867334 |
| Nts | Q9D3P9 | 1.36 | 0.77 | -1.61 | 0.47 | 0.93 | 0.71 | 0.03 | 0.22 | 1.2 | 1.6 | -1.08 | 0.63 | 1.4 | 1.57 | 1.98 | 0.43 | 0.225 | 0.34 | 1.0975 | 0.716859 | 0.669075 | 0.123454 | 1 | 0.996511 | 0.721448 |
| Ntsr2 | P70310 | 0.26 | 0.83 | 1.02 | 1.3 | 0.3 | 0.15 | 0.42 | 0.54 | 0.95 | 0.63 | 1.36 | 1.35 | 0.27 | 1.97 | 0.97 | 1.13 | -0.71 | 0.22 | 0.2325 | 0.039042 | 0.456165 | 0.56301 | 1 | 0.996511 | 0.920024 |
| Nuak1 | Q641K5 | -0.77 | -0.43 | 0.8 | -0.47 | 0.17 | -1.09 | 0.59 | 0.01 | -0.49 | -1.74 | 1.09 | -0.62 | 1.18 | -2.11 | 1.09 | 1.26 | 0.1375 | -0.2225 | 0.5725 | 0.767046 | 0.720606 | 0.483139 | 1 | 0.996511 | 0.893925 |
| Nub1 | A0A0G2JG | -1.14 | -1.21 | -1.05 | -0.84 | -1.24 | -1.19 | -1 | -1.16 | -0.94 | -0.56 | -0.79 | -1.04 | -1.02 | -0.82 | -1.38 | -1.28 | -0.0875 | 0.2275 | -0.065 | 0.585551 | 0.249395 | 0.756446 | 1 | 0.996511 | 0.962333 |

| | | | | | | | | | | | | | | | | | | | | | | | | | | | |
|---|---|---|---|---|---|---|---|---|---|---|---|---|---|---|---|---|---|---|---|---|---|---|---|---|---|---|---|
| Nubp1 | Q9R060 | -0.74 | -0.87 | -1.26 | -0.82 | -1.41 | -1.18 | -1.33 | -1.1 | -0.69 | -0.7 | -1.01 | -0.83 | -0.58 | -1.3 | -1.31 | -1.25 | -0.3325 | 0.115 | -0.1875 | 0.092742 | 0.554586 | 0.455106 | | 1 | 0.996511 | 0.887169 |
| Nubp2 | A0A3Q4L2 | -0.23 | 0.42 | -0.3 | -0.38 | -0.12 | -0.36 | -0.6 | -0.34 | 0.32 | -0.41 | -0.45 | -0.37 | 0.45 | -0.3 | -1.21 | -0.86 | -0.2325 | -0.105 | -0.3575 | 0.323044 | 0.702184 | 0.374483 | | 1 | 0.996511 | 0.859424 |
| Nubpl | Q9CWD8 | -0.33 | 0.01 | 0.24 | 0.7 | -0.13 | -0.77 | 0.83 | 0.43 | 0.29 | 0.24 | 0.36 | -0.28 | -0.07 | 0.03 | -0.03 | 0.68 | -0.065 | -0.0025 | -0.0025 | 0.866818 | 0.992735 | 0.993218 | | 1 | 0.999285 | 0.996845 |
| Nucb1 | Q02819 | -1.04 | -0.78 | -1.54 | -1.24 | -0.13 | -1.05 | -0.69 | -1.12 | -1.76 | -0.86 | -1.16 | -1.05 | -0.75 | -0.91 | -0.12 | -0.58 | 0.4025 | -0.0575 | 0.56 | 0.176141 | 0.830352 | 0.057634 | | 1 | 0.996511 | 0.661738 |
| Nucb2 | P81117 | 1.03 | -0.95 | -1.68 | -1.37 | -0.25 | 1.01 | -0.63 | -1.18 | -0.6 | -1.24 | -1.7 | 0.71 | -0.32 | -0.63 | -0.28 | -0.6 | 0.48 | 0.035 | 0.285 | 0.495634 | 0.961671 | 0.617856 | | 1 | 0.996755 | 0.937274 |
| Nucks1 | A0A087WF | -3.06 | -4.49 | -2.24 | -2.77 | -4.52 | -4.24 | -4.78 | -5.44 | -2.63 | -0.86 | -2.13 | -2.78 | -3.51 | -3.28 | -4.99 | -4.06 | -1.605 | 1.04 | -0.82 | 0.010747 | 0.108562 | 0.171442 | | 1 | 0.996511 | 0.763325 |
| Nudc | O35685 | 0.23 | 0.26 | -0.01 | -0.16 | 0.11 | 0.31 | -0.07 | -0.1 | -0.59 | -0.41 | -0.05 | 0.43 | -0.14 | -0.33 | 0.18 | 0.09 | -0.0175 | -0.235 | -0.13 | 0.923247 | 0.384717 | 0.541362 | | 1 | 0.996511 | 0.912212 |
| Nudcd1 | Q6PIP5 | 1.98 | 1.5 | -0.3 | -0.32 | 0.8 | 1.6 | -0.4 | -0.46 | -0.7 | -0.61 | -0.73 | 1.85 | -1.24 | -1.04 | -0.94 | -0.74 | -0.33 | -0.7625 | -1.705 | 0.642196 | 0.347196 | 0.013457 | | 1 | 0.996511 | 0.586837 |
| Nudcd2 | Q9CQ48 | 0.77 | 1.02 | 0.75 | 0.68 | 0.73 | 0.24 | 0.78 | 0.71 | -1.67 | 0.14 | 0.87 | 0.89 | -1.71 | -1.23 | 0.69 | 0.61 | -0.19 | -0.7475 | -1.215 | 0.323815 | 0.203984 | 0.059709 | | 1 | 0.996511 | 0.661738 |
| Nudcd3 | Q8R1N4 | -0.48 | -0.62 | -0.5 | -0.68 | -0.2 | -0.41 | -0.86 | -0.5 | -1.27 | -0.38 | -0.63 | -0.37 | -0.91 | -0.86 | -0.49 | -0.48 | 0.0775 | -0.0925 | -0.115 | 0.679201 | 0.704333 | 0.561544 | | 1 | 0.996511 | 0.91997 |
| Nudt11 | P0C028 | 0.16 | 0.02 | -0.86 | -0.14 | -0.48 | 0.24 | -0.23 | -0.01 | 0.44 | -0.08 | -0.03 | 0.44 | -0.05 | -0.13 | -0.33 | -0.58 | 0.085 | 0.3975 | -0.0675 | 0.759897 | 0.180739 | 0.807846 | | 1 | 0.996511 | 0.971953 |
| Nudt12 | Q9DCN1 | 0.36 | 0.36 | 0.58 | -0.25 | 0 | 0.41 | -0.23 | 0.22 | 2.88 | -0.08 | -0.07 | -0.11 | 1.62 | 0.04 | 0.58 | 0.31 | -0.1625 | 0.3925 | 0.375 | 0.505896 | 0.575071 | 0.336371 | | 1 | 0.996511 | 0.847578 |
| Nudt13 | Q8JZU0 | 1.94 | -0.72 | -1.36 | -0.83 | -1.37 | -2.09 | -2 | -1.8 | -0.46 | -3.02 | -1.36 | -1.56 | -2.5 | -1.11 | -0.24 | -1.61 | -1.5725 | -1.3575 | -1.1225 | 0.045387 | 0.126493 | 0.18089 | | 1 | 0.996511 | 0.775542 |
| Nudt14 | Q9D142 | 1.06 | 1.07 | -0.85 | -0.32 | 1.34 | 1.48 | -0.7 | -0.64 | 1.52 | -0.39 | -0.53 | 0.98 | 0.68 | 1.69 | -0.32 | -0.91 | 0.13 | 0.155 | 0.045 | 0.852996 | 0.80951 | 0.947428 | | 1 | 0.996511 | 0.992414 |
| Nudt15 | Q8BG93 | -0.63 | -0.03 | -1.14 | -1.23 | -1.05 | -1.12 | -0.97 | -1.73 | 0.01 | -0.68 | -1.12 | -0.82 | -0.08 | 0.65 | -1.28 | -0.91 | -0.46 | 0.105 | 0.3525 | 0.174617 | 0.768769 | 0.47003 | | 1 | 0.996511 | 0.890428 |
| Nudt16 | Q6P3D0 | -0.41 | -0.54 | -0.1 | -0.45 | -0.34 | -0.52 | -0.03 | -0.28 | -0.75 | -0.86 | -0.48 | -0.77 | -0.96 | -0.2 | -0.23 | -0.38 | 0.0825 | -0.34 | -0.0675 | 0.653219 | 0.095528 | 0.778831 | | 1 | 0.996511 | 0.967289 |
| Nudt16l1 | Q8VHN8 | -0.71 | -1.88 | -0.93 | -1.29 | -1.44 | -0.71 | -0.65 | -0.86 | -1.62 | -1.3 | -0.77 | -1.5 | -1.66 | -1.4 | -1.18 | -0.94 | 0.2875 | -0.095 | -0.0925 | 0.363704 | 0.765526 | 0.764629 | | 1 | 0.996511 | 0.964134 |
| Nudt17 | Q9CWD3-2 | 0.72 | 0.8 | 0.85 | 0.76 | -0.41 | 1.04 | -0.52 | 0.19 | 1.1 | 1.27 | 0.52 | -0.03 | 0.97 | 0.05 | -0.13 | 0.15 | -0.7075 | -0.0675 | -0.5225 | 0.067082 | 0.823373 | 0.079718 | | 1 | 0.996511 | 0.681818 |
| Nudt18 | Q3U2V3 | 1.08 | 1.05 | -0.35 | -0.29 | 1.43 | 1.01 | -0.16 | -0.61 | 0.6 | -0.61 | -0.2 | 1.18 | -0.06 | 0.83 | -0.56 | -0.77 | 0.045 | -0.13 | -0.5125 | 0.937171 | 0.804409 | 0.319699 | | 1 | 0.996511 | 0.838969 |
| Nudt19 | P11930 | -0.18 | 0.41 | 0.15 | 0.32 | 0.78 | 0.15 | 0.53 | 0.29 | -0.57 | -0.94 | 0.1 | 0.09 | -1.02 | -0.63 | 0.06 | 0.11 | 0.2625 | -0.505 | -0.545 | 0.243452 | 0.11474 | 0.108717 | | 1 | 0.996511 | 0.707444 |
| Nudt2 | P56380 | -2.04 | -2.47 | -0.5 | -0.67 | -1.95 | -1.67 | -0.02 | -0.23 | -2.62 | -1.78 | -0.5 | -2.26 | -2.77 | -2.93 | -0.43 | -0.18 | 0.4525 | -0.37 | -0.1575 | 0.480792 | 0.55336 | 0.843641 | | 1 | 0.996511 | 0.976225 |
| Nudt21 | Q9CQF3 | -4.14 | -5.65 | -4.53 | -4.96 | -4.94 | -5.03 | -4.04 | -4.62 | -4.62 | -2.51 | -4.27 | -4.61 | -3.95 | -3.95 | -2.92 | -4.37 | 0.1625 | 0.8175 | 1.0225 | 0.665687 | 0.164519 | 0.037675 | | 1 | 0.996511 | 0.658454 |
| Nudt3 | I1E4X7 | 2.48 | 0.92 | 2.24 | -0.11 | -1.37 | 3.08 | 0.43 | 1.39 | -3.22 | -2.02 | 1.6 | 1.98 | -2.31 | -6.01 | 1.44 | 0.75 | -0.5 | -1.7975 | -2.915 | 0.61766 | 0.185125 | 0.096199 | | 1 | 0.996511 | 0.698156 |
| Nudt4 | Q8R2U6 | -0.54 | -0.62 | 0.09 | -0.41 | -0.49 | -0.95 | -0.5 | -0.44 | -0.47 | 0.25 | -0.27 | -0.55 | -1.26 | -0.18 | -0.73 | -0.14 | -0.225 | 0.11 | -0.2075 | 0.324901 | 0.673362 | 0.516974 | | 1 | 0.996511 | 0.905748 |
| Nudt5 | Q9JKX6 | -0.71 | -0.9 | -2.03 | -2.16 | -0.98 | -0.79 | -1.81 | -2.1 | -1.89 | -2.32 | -1.66 | -0.9 | -1.53 | -1.05 | -2.57 | -2.85 | 0.03 | -0.2425 | -0.55 | 0.947569 | 0.59334 | 0.311225 | | 1 | 0.996511 | 0.834523 |
| Nudt7 | Q99P30-3 | 0.36 | 0.45 | -0.4 | -0.37 | 0.75 | 0.71 | 0.13 | -0.26 | 0.62 | -0.76 | -0.34 | 0.57 | 0.62 | 0.67 | -0.31 | -0.27 | 0.3225 | 0.0125 | 1.68E-01 | 0.335764 | 0.974667 | 0.63562 | | 1 | 0.997515 | 0.9419 |
| Nudt8 | Q9CR24 | 0.05 | 0.11 | -1.12 | -0.36 | -0.29 | -0.18 | -0.07 | -0.4 | 0.01 | -0.61 | -0.5 | 0.06 | 0.07 | 0.17 | -0.91 | -0.55 | 0.095 | 0.07 | 0.025 | 0.745365 | 0.831776 | 0.946632 | | 1 | 0.996511 | 0.992376 |
| Nudt9 | Q8BVU5 | -0.37 | -0.24 | 0.37 | 0.62 | -0.62 | -0.63 | 0.59 | 0.77 | -0.1 | 0.49 | 0.9 | -0.43 | 0.26 | -1.61 | 0.6 | 0.36 | -0.0675 | 0.12 | -0.1925 | 0.87207 | 0.745667 | 0.712973 | | 1 | 0.996511 | 0.958106 |
| Nufip2 | Q5F2E7 | -2.13 | -2.19 | -2.28 | -2.11 | -1.67 | -2.21 | -1.45 | -2.44 | -1.72 | -1.52 | -1.88 | -1.79 | -1.22 | -1.44 | -1.22 | -1.79 | 0.235 | 0.45 | 0.76 | 0.353599 | 0.023625 | 0.004583 | | 1 | 0.996511 | 0.550241 |
| Numa1 | E9Q7G0 | 0.96 | 0.59 | 0.78 | 0.95 | 0.07 | 0.46 | 0.07 | 0.5 | 0.63 | 0.74 | 0.59 | 0.44 | 0.91 | 0.36 | 0.8 | 0.59 | -0.545 | -0.22 | -0.155 | 0.016658 | 0.234097 | 0.464964 | | 1 | 0.996511 | 0.888267 |
| Numb | Q9QZS3-4 | 1.82 | 0.48 | 1.74 | 1.64 | 0.21 | 1.87 | 1.1 | 1.54 | 0.63 | 1.31 | 1.44 | 1.72 | 0.65 | 0.08 | 1.24 | 1.28 | -0.24 | -0.145 | -0.6075 | 0.593385 | 0.702061 | 0.161553 | | 1 | 0.996511 | 0.757516 |
| Numbl | O08919 | -0.02 | 0.19 | 0.35 | 0.73 | 0.25 | -0.05 | 0.6 | 0.74 | -0.01 | 0.82 | 0.34 | 0.14 | -0.11 | 0.57 | 0.51 | 0.31 | 0.0725 | 0.01 | 0.0075 | 0.772383 | 0.969256 | 0.976327 | | 1 | 0.996755 | 0.993727 |
| Nup107 | Q8BH74 | 0.57 | 0.37 | 0.18 | 0.22 | 0.27 | 0.05 | 0.45 | 0.5 | -0.33 | -0.06 | 0.18 | -0.22 | 0.15 | 0.21 | 0.91 | 0.63 | -0.0175 | -0.4425 | 0.14 | 0.922504 | 0.046533 | 0.562874 | | 1 | 0.996511 | 0.920024 |
| Nup133 | Q8R0G9 | 0.87 | 1.04 | 0.48 | 0.61 | 1.04 | 0.79 | 0.51 | 0.92 | 0.88 | 0.71 | 0.25 | 0.41 | 1.09 | 1.61 | 1.07 | 0.93 | 0.065 | -0.1875 | 0.425 | 0.748051 | 0.414209 | 0.099522 | | 1 | 0.996511 | 0.698156 |
| Nup153 | E9Q3G8 | 0.78 | 0.31 | 0.57 | 0.82 | -0.21 | 0.34 | 0.16 | 1 | 0.5 | 0.99 | 0.65 | 0.05 | 1 | 1.2 | 1.11 | 0.66 | -0.2975 | -0.0725 | 0.3725 | 0.306586 | 0.772569 | 0.115297 | | 1 | 0.996511 | 0.717287 |
| Nup155 | A0A2R8VH | 0.25 | 0.17 | -0.24 | -0.41 | 0.26 | -0.12 | 0.39 | 0.19 | -0.33 | -1.01 | -0.22 | -0.43 | 0.07 | 0.27 | 0.63 | 0.44 | 0.2375 | -0.44 | 0.41 | 0.291158 | 0.115164 | 0.11304 | | 1 | 0.996511 | 0.714953 |
| Nup160 | Q9Z0W3 | 1.18 | 1.04 | 0.43 | 0.24 | 0.88 | 0.93 | 0.85 | 0.8 | -0.06 | -0.77 | 0.17 | 0.59 | 0.12 | 0.26 | 0.94 | 0.89 | 0.1425 | -0.74 | -0.17 | 0.563781 | 0.064057 | 0.596448 | | 1 | 0.996511 | 0.929911 |
| Nup188 | Q6ZQH8 | 0.29 | -0.08 | 0.77 | 0.79 | -0.38 | 0.24 | 0.67 | 1.03 | 0.06 | 0.53 | 0.62 | -0.43 | 0.72 | 0.19 | 1.47 | 1.21 | -0.0525 | -0.2475 | 0.455 | 0.8819 | 0.447303 | 0.214675 | | 1 | 0.996511 | 0.787313 |
| Nup205 | B9EJ54 | 0.63 | 0.41 | 0.33 | -0.03 | 0.61 | 0.46 | 0.4 | 0.59 | -0.73 | -0.39 | 0.13 | -0.07 | -0.46 | -0.3 | 0.83 | 0.85 | 0.18 | -0.6 | -0.105 | 0.348696 | 0.040394 | 0.777715 | | 1 | 0.996511 | 0.967289 |
| Nup210 | Q9QY81 | 0.71 | 0.71 | 0.52 | 0.47 | 0.75 | 0.64 | 0.73 | 0.91 | 0.2 | 0.34 | 0.48 | 0.05 | 0.83 | 0.8 | 1.37 | 1.19 | 0.155 | -0.335 | 0.445 | 0.328761 | 0.08879 | 0.059887 | | 1 | 0.996511 | 0.661738 |
| Nup214 | Q80U93 | 1.17 | 0.81 | 0.44 | 0.62 | 0.71 | 0.69 | 0.19 | 0.6 | 0.21 | 0.29 | 0.52 | 0.43 | 0.69 | 0.57 | 1.01 | 0.81 | -0.2125 | -0.3975 | 0.01 | 0.349471 | 0.088813 | 0.96498 | | 1 | 0.996511 | 0.9927 |
| Nup35 | Q8R4R6 | 0.71 | 0.98 | 0.87 | 0.72 | 0.54 | 0.43 | 0.58 | 0.85 | 0.75 | 0.34 | 0.54 | 0.19 | 1.46 | 0.9 | 1.55 | 0.84 | -0.22 | -0.365 | 0.3675 | 0.208599 | 0.085622 | 0.146326 | | 1 | 0.996511 | 0.746406 |
| Nup37 | Q9CWU9 | -0.18 | -0.3 | -0.03 | -0.41 | -0.98 | 0 | -0.75 | -0.28 | -0.25 | -0.3 | -0.58 | -1.16 | 0.97 | -0.85 | 0.56 | 0.34 | -0.2725 | -0.3425 | 0.485 | 0.290693 | 0.191467 | 0.231484 | | 1 | 0.996511 | 0.788863 |
| Nup43 | P59235 | 0.24 | 0.01 | 0.26 | 0.48 | -0.56 | 0.28 | 0.37 | 0.58 | 0.21 | -0.27 | -0.13 | -0.24 | 1.21 | -0.31 | 0.13 | 0.39 | -0.08 | -0.355 | 0.1075 | 0.770011 | 0.099786 | 0.749105 | | 1 | 0.996511 | 0.961014 |
| Nup50 | Q9JIH2 | 0.53 | -0.35 | 0.39 | 0.46 | -0.85 | 0.16 | 0.07 | 0.66 | -0.12 | 0.4 | 0.47 | -0.05 | 0.8 | -0.67 | 0.95 | 1.3 | -0.2475 | -0.0825 | 0.3375 | 0.496953 | 0.760455 | 0.463904 | | 1 | 0.996511 | 0.888267 |
| Nup54 | Q8BTS4 | 0.74 | 0.31 | -0.21 | -0.11 | 0.78 | 0.44 | 0.24 | 0.07 | -0.09 | -0.09 | -0.08 | 0.14 | 0.53 | 0.06 | 0.5 | 0.15 | 0.2 | -0.2125 | 0.1275 | 0.467747 | 0.402656 | 0.641178 | | 1 | 0.996511 | 0.943421 |
| Nup58 | Q8R332-4 | 1.85 | 1.58 | 0.92 | 0.92 | 0.86 | 1.41 | 1.12 | 1.47 | 1.36 | 1.1 | 0.69 | 1.05 | 1.68 | 2.1 | 1.68 | 1.07 | -0.1025 | -0.2675 | 0.315 | 0.713552 | 0.359172 | 0.342687 | | 1 | 0.996511 | 0.848473 |
| Nup62 | Q63850 | 0.31 | 0.3 | 0.39 | 1.12 | 0.26 | -0.04 | 0.82 | 1.43 | -0.29 | -0.05 | 0.92 | -0.18 | 0.08 | 0.88 | 1.38 | 1.01 | 0.0875 | -0.43 | 0.3075 | 0.809646 | 0.223671 | 0.375329 | | 1 | 0.996511 | 0.859611 |
| Nup85 | Q8R480 | 0.69 | 0.54 | 0.31 | -0.27 | 0.25 | 0.45 | 0.49 | 0.56 | -0.02 | -1.34 | -0.04 | -0.04 | 0.74 | 0.19 | 0.97 | 0.81 | 0.12 | -0.6775 | 0.36 | 0.61602 | 0.0992 | 0.232057 | | 1 | 0.996511 | 0.788863 |
| Nup88 | Q8CEC0-3 | 1.42 | 0.63 | 0.58 | 0.59 | 0.89 | 0.63 | 1.19 | 1.29 | 0.4 | 0.18 | 0.8 | 0.37 | 0.99 | 1.06 | 1.6 | 1.36 | 0.195 | -0.3675 | 0.4475 | 0.464457 | 0.18463 | 0.127513 | | 1 | 0.996511 | 0.723391 |
| Nup93 | Q8BJ71 | 1.18 | 0.55 | 0.51 | 0.25 | 0.43 | 0.77 | 0.46 | 0.58 | 0.45 | -0.46 | 0.37 | 0.32 | 0.99 | 0.34 | 1.06 | 0.68 | -0.0625 | -0.4525 | 0.145 | 0.786955 | 0.153097 | 0.605012 | | 1 | 0.996511 | 0.932185 |
| Nup98 | A0A1B0GS | 1.15 | 0.81 | 0.18 | 0.05 | 0.58 | 0.77 | 0.4 | 0.57 | 0 | -0.58 | -0.06 | 0.26 | 0.62 | 0.18 | 0.99 | 0.6 | 0.0325 | -0.6425 | 0.05 | 0.906036 | 0.068385 | 0.87436 | | 1 | 0.996511 | 0.979626 |
| Nupl2 | E9QL43 | 1.65 | 0.59 | 2.32 | 1.12 | 1.35 | 1.34 | 0.67 | 2.23 | 2.43 | 1.07 | 1.84 | 1.99 | 2.11 | 1.03 | 2.07 | 2.51 | -0.0225 | 0.4125 | 0.51 | 0.9605 | 0.360614 | 0.283473 | | 1 | 0.996511 | 0.822669 |
| Nutf2 | P61971 | -0.85 | -0.5 | -0.49 | -0.51 | -0.31 | -0.94 | -0.28 | -0.4 | -0.75 | -0.58 | -0.53 | -0.83 | -1.34 | -0.41 | -0.84 | -0.8 | 0.105 | -0.085 | -0.26 | 0.614706 | 0.637685 | 0.305759 | | 1 | 0.996511 | 0.832533 |
| Nvl | Q9DBY8 | 1.9 | -0.59 | -0.74 | -0.95 | -0.46 | 1.27 | -0.64 | -0.97 | 1.66 | -0.35 | -1.24 | 0.65 | 2.97 | -0.09 | -1.09 | -0.79 | -0.105 | 0.275 | 0.345 | 0.889371 | 0.740322 | 0.736952 | | 1 | 0.996511 | 0.958635 |
| Nwd1 | E0CX99 | 1.48 | 1.69 | 0.78 | 0.56 | 1.71 | 1.46 | 0.3 | 0.68 | 1.8 | 0.18 | 0.52 | 1.15 | 2.44 | 1.97 | 0.26 | 0.64 | -0.09 | -0.215 | 0.2 | 0.823028 | 0.615699 | 0.713645 | | 1 | 0.996511 | 0.958106 |
| Nwd2 | Q6P5U7 | 0.52 | 0.47 | 0.56 | 0.47 | 0.81 | 0.77 | 0.76 | 0.54 | 0.26 | 0.02 | 0.81 | 0.55 | 0.05 | 0.53 | 0.43 | 0.78 | 0.215 | -0.095 | -0.0575 | 0.16734 | 0.658612 | 0.785327 | | 1 | 0.996511 | 0.967642 |
| Nxf1 | Q99JX7 | -0.28 | -0.58 | -0.12 | 0.17 | -0.1 | -1 | 0.09 | 0.27 | -0.88 | -0.07 | -0.39 | -1.13 | -0.14 | 0.18 | 0.84 | 0.14 | 0.0175 | -0.415 | 0.4575 | 0.955818 | 0.181627 | 0.129581 | | 1 | 0.996511 | 0.723655 |
| Nxn | P97346 | -0.2 | -0.8 | -0.09 | -0.05 | -0.34 | -0.17 | 0.21 | -0.12 | 0.14 | -0.7 | -0.43 | 0.15 | -0.52 | -0.48 | -0.05 | -0.02 | 0.18 | 0.075 | 0.0175 | 0.440087 | 0.792849 | 0.94476 | | 1 | 0.996511 | 0.992112 |
| Nxph1 | Q61200 | -0.87 | -0.1 | 0.85 | 0.45 | 0.89 | -0.41 | 2.3 | 1.08 | -1.71 | -1.78 | 1.39 | -1.06 | -2.09 | -0.9 | 1.07 | 1.25 | 0.8825 | -0.8725 | -0.25 | 0.172466 | 0.266216 | 0.754924 | | 1 | 0.996511 | 0.962333 |
| Nxph3 | Q91VX5 | -2.14 | -1.52 | 0.39 | -1.06 | -1.13 | -0.84 | 0.58 | 0.2 | -1.45 | -1.57 | -0.49 | -1.68 | -0.55 | -1.62 | -1.48 | -0.78 | 0.785 | -0.215 | -0.025 | 0.224099 | 0.70026 | 0.963856 | | 1 | 0.996511 | 0.9927 |
| Nxt1 | Q9QZV9 | -0.48 | -1.42 | -1.31 | -1.2 | -1.09 | -0.4 | -0.59 | -0.89 | -1.75 | -2.33 | -0.86 | -1.09 | -0.81 | -2.38 | -0.06 | -1.47 | 0.36 | -0.405 | -0.0775 | 0.202939 | 0.303424 | 0.876859 | | 1 | 0.996511 | 0.979626 |
| Nyap1 | E9PXJ0 | -0.41 | 0.32 | 0.36 | -0.14 | 0.12 | 2.31 | -0.18 | 0.04 | 0.73 | 0.6 | 0.17 | 0.06 | 1.14 | 0.67 | 0.61 | 0.01 | 0.54 | 0.3575 | 0.575 | 0.348233 | 0.199986 | 0.088847 | | 1 | 0.996511 | 0.691619 |
| Nyap2 | A0A0R4J1X | 0.32 | 0.25 | -0.63 | -0.7 | -0.23 | 0.76 | -0.73 | -0.9 | 2.09 | -1.11 | -0.27 | 0.65 | 2.53 | 0.06 | -0.81 | -0.49 | -0.085 | 0.53 | 0.5125 | 0.844447 | 0.439056 | 0.488511 | | 1 | 0.996511 | 0.897531 |
| Oard1 | Q8R5F3 | -0.25 | -0.79 | -3.62 | -3.52 | -0.42 | -1.27 | -4.06 | -3.34 | -0.69 | -0.51 | -4.05 | -0.99 | -1.54 | -1.25 | -3.74 | -4.53 | -0.2275 | 0.485 | -0.72 | 0.836763 | 0.659026 | 0.507599 | | 1 | 0.996511 | 0.902804 |
| Oat | P29758 | 0.66 | 0.2 | 0.7 | 1.34 | 0.53 | 0.79 | 1.52 | 1.17 | -0.06 | 0.29 | 1.21 | 0.47 | -0.32 | 0.21 | 0.81 | 0.87 | 0.2775 | -0.2475 | -0.3325 | 0.387284 | 0.484606 | 0.368045 | | 1 | 0.996511 | 0.859245 |
| Obp1a | Q9D3H2 | -4.48 | -4.35 | -3.9 | -5.56 | -4.19 | -13.25 | -11.13 | -11.25 | -4.18 | -5.64 | -7.48 | -5.74 | 0.83 | -7.65 | -5.91 | -2.39 | -5.3825 | -1.1875 | 0.7925 | 0.014561 | 0.114115 | 0.643334 | | 1 | 0.996511 | 0.943874 |
| Obscn | H7BX05 | -0.82 | -0.66 | 0.38 | 0.43 | -0.56 | -1.51 | -0.22 | 0.53 | -1.65 | 1.11 | 0.4 | -0.58 | -1.28 | -4.2 | 0.87 | -0.35 | -0.2725 | -0.0125 | -1.0725 | 0.5864 | 0.98408 | 0.304622 | | 1 | 0.998876 | 0.831662 |
| Ociad1 | Q9CRD0 | 0.18 | 0.16 | 0.14 | 0.57 | 0.05 | 0.12 | 0.5 | 0.4 | 0.11 | -0.11 | 0.42 | 0.19 | 0.04 | -0.04 | 0.09 | 0.19 | 0.005 | -0.11 | -0.1925 | 0.978843 | 0.58519 | 0.326626 | | 1 | 0.996511 | 0.843799 |
| Ociad1 | Q9CRD0-3 | 1.58 | 0.6 | 1 | 1.76 | -0.31 | 1.05 | 1.61 | 1.47 | 1.29 | 1.64 | 1.11 | 1.27 | 2.11 | 0.6 | 1.94 | 1.45 | -0.28 | 0.0925 | 0.29 | 0.559568 | 0.75526 | 0.488855 | | 1 | 0.996511 | 0.897531 |
| Ociad2 | Q9D8W7 | 0.28 | -0.65 | 1.75 | 0.59 | -0.75 | 0.94 | 1.77 | 1.64 | -0.73 | 1.26 | 1.68 | 0.31 | -0.11 | -1.5 | 1.71 | 2.22 | 0.4075 | 0.1375 | 0.0875 | 0.559117 | 0.836271 | 0.921339 | | 1 | 0.996511 | 0.987433 |
| Ocln | Q61146 | -0.82 | -0.39 | 0.62 | 0.59 | -0.23 | -0.96 | 0.32 | 0.38 | -0.26 | 1.48 | 0.98 | -0.18 | -0.01 | -0.09 | 0.44 | 1.18 | -0.1225 | 0.505 | 0.38 | 0.782755 | 0.345552 | 0.397465 | | 1 | 0.996511 | 0.867334 |
| Ocrl | Q6NVF0 | 0.23 | 0.21 | 0.56 | 0.79 | 0.57 | 0.32 | 0.96 | 0.87 | -0.22 | -0.14 | 0.48 | 0.14 | -0.29 | -0.06 | 0.41 | 0.64 | 0.2325 | -0.3825 | -0.2725 | 0.314539 | 0.136476 | 0.337871 | | 1 | 0.996511 | 0.847578 |
| Odr4 | Q4PJX1 | 1.34 | 0.96 | 0.04 | -0.06 | 1.24 | 1.36 | 0.35 | 0.12 | 1.13 | -0.15 | -0.06 | 1.03 | 1.7 | 0.64 | 1.03 | 0.86 | 0.1975 | -0.0825 | 0.4875 | 0.650943 | 0.856574 | 0.241708 | | 1 | 0.996511 | 0.796892 |
| Oga | Q9EQQ9 | -1.09 | -1.51 | -0.53 | -0.33 | -1.42 | -0.95 | -0.3 | -0.48 | -0.8 | -0.23 | -0.39 | -0.87 | -1 | -0.64 | -0.61 | -0.89 | 0.0775 | 0.2925 | 0.08 | 0.826382 | 0.36204 | 0.788947 | | 1 | 0.996511 | 0.968677 |
| Ogdh | Q60597-3 | 0.57 | 0.24 | 0.3 | 0.71 | -0.05 | 0.58 | 0.17 | 0.59 | 0.27 | 0.39 | 0.52 | 0.59 | -0.15 | 0.21 | 0.08 | 0.15 | -0.1325 | -0.0125 | -0.3825 | 0.547207 | 0.947351 | 0.084984 | | 1 | 0.996755 | 0.691619 |
| Ogdh | Q60597 | 0.26 | 0.53 | -0.1 | -0.02 | 0.41 | 0.45 | 0.01 | -0.02 | 0.21 | -0.67 | 0.08 | -0.05 | -0.08 | 0.9 | -0.69 | -1.06 | 0.045 | -0.275 | -0.4 | 0.835094 | 0.307699 | 0.364301 | | 1 | 0.996511 | 0.858475 |
| Ogdhl | B2RXT3 | 1.21 | 0.89 | 0.81 | 0.82 | 0.44 | 1.14 | 0.88 | 0.9 | 0.45 | 0.34 | 0.96 | 1.05 | -0.07 | 0.24 | 0.32 | 0.66 | -0.0925 | -0.2325 | -0.645 | 0.652616 | 0.331978 | 0.017545 | | 1 | 0.996511 | 0.602211 |
| Ogfod1 | Q3U0K8-2 | -3.65 | -3.77 | -1.69 | -1.78 | -3.33 | -4.24 | -1.55 | -1.88 | -4.22 | -0.92 | -1.96 | -3.47 | -3.17 | -2.59 | -1.74 | -1.75 | -0.0275 | 0.08 | 0.41 | 0.971378 | 0.924503 | 0.508405 | | 1 | 0.996755 | 0.902804 |
| Ogfod3 | Q9D136 | -0.15 | -0.34 | -0.78 | 0.88 | -0.35 | -0.02 | -0.11 | 0.15 | 0.95 | 1.3 | 0.27 | -0.87 | 2.34 | 0.64 | 2.15 | 1.28 | 0.015 | 0.51 | 1.7 | 0.965991 | 0.36402 | 0.007188 | | 1 | 0.996511 | 0.552614 |
| Ogfr | Q99PG2 | -0.5 | -0.6 | -0.79 | -0.57 | -1.19 | -0.73 | -0.85 | -0.7 | -0.55 | -0.55 | -0.59 | -0.36 | -0.21 | -0.89 | -1.23 | -1.17 | -0.2525 | 0.1025 | -0.26 | 0.176618 | 0.540166 | 0.344376 | | 1 | 0.996511 | 0.84897 |
| Ogfrl1 | Q8VE52 | 0.16 | 0.3 | 0.38 | 0.04 | 0.15 | 0.21 | 0.06 | 0.22 | 1.46 | -0.99 | 0.41 | -0.08 | 1.78 | 0.71 | 0.58 | 0.48 | -0.06 | -0.02 | 0.6675 | 0.697512 | 0.96684 | 0.061023 | | 1 | 0.996755 | 0.661738 |
| Ogt | Q8CGY8-2 | -0.68 | -0.46 | -0.22 | 0.51 | 0.53 | -0.02 | 1.03 | 0.94 | 0.38 | 0.38 | 0.92 | 0.25 | -0.17 | 0.94 | 0.67 | 0.49 | 0.8325 | 0.695 | 0.695 | 0.035253 | 0.045093 | 0.07251 | | 1 | 0.996511 | 0.668066 |
| Ogt | Q8CGY8 | 0.41 | 0.45 | 0.56 | 0.54 | 0.53 | 0.64 | 0.5 | 0.55 | 0.37 | 0.45 | 0.75 | 0.56 | 0.13 | 0.52 | 0.3 | 0.41 | 0.065 | 0.0425 | -0.15 | 0.646582 | 0.80174 | 0.415885 | | 1 | 0.996511 | 0.874945 |
| Ola1 | Q9CZ30 | 1.25 | -0.93 | 0.63 | -0.59 | -0.35 | 1.29 | 1.14 | 0.22 | -1.54 | -0.41 | 0.68 | 1.07 | -2.04 | -0.94 | 0.51 | 0.49 | 0.485 | -0.14 | -0.585 | 0.417821 | 0.843171 | 0.428866 | | 1 | 0.996511 | 0.879165 |
| Olfm1 | Q8R357 | 0.39 | 1.08 | 0.75 | 0.82 | 1.3 | 0.5 | 0.7 | 0.8 | 0.49 | 0.21 | 0.88 | 0.46 | 0.5 | 1.11 | 0.49 | 0.62 | 0.065 | -0.25 | -0.08 | 0.785173 | 0.294704 | 0.741518 | | 1 | 0.996511 | 0.958796 |
| Olfm2 | A0A1L1SV4 | 0.26 | 0.79 | 0.5 | 0.52 | 1.11 | 0.42 | 1.07 | 0.43 | 0.58 | -0.37 | 0.64 | 0.63 | 0.76 | 0.58 | 0.35 | 0.34 | 0.24 | -0.1475 | -0.01 | 0.326371 | 0.602993 | 0.961609 | | 1 | 0.996511 | 0.9927 |
| Olfm3 | P63056 | 0.63 | 1.19 | 1.07 | 1.19 | 1.15 | 0.52 | 0.95 | 1.87 | 0.05 | 1.74 | 1.33 | 0.29 | -0.27 | 0.32 | 0.68 | 1.21 | 0.1025 | -0.1675 | -0.535 | 0.739403 | 0.682085 | 0.141007 | | 1 | 0.996511 | 0.737973 |
| Olfml3 | A0A0R4J08 | -1.55 | -1.92 | -6.52 | -14.17 | -1.93 | -1.3 | -10.12 | -7.38 | -0.05 | -5.57 | -5.4 | -1.25 | 0.56 | -1.59 | -6.43 | -11.05 | 0.8575 | 2.9725 | 1.4125 | 0.791041 | 0.320665 | 0.684496 | | 1 | 0.996511 | 0.955729 |
| Olig1 | Q9JKN5 | -1.19 | -0.4 | -1.61 | -1.34 | -0.38 | -0.87 | -1.23 | -2.11 | -0.5 | -1.61 | -1.97 | -0.73 | -0.03 | -0.12 | -0.59 | -0.88 | -0.0125 | -0.0675 | 0.73 | 0.976241 | 0.870667 | 0.051628 | | 1 | 0.996511 | 0.658454 |
| Oma1 | Q9D8H7 | 0.81 | 1.43 | -0.4 | -0.37 | 1.64 | 1.14 | 0.85 | 1.03 | 1.64 | -0.97 | -0.06 | 1.51 | 1.16 | 1.73 | -2.36 | -0.5 | 0.7975 | 0.1625 | -0.36 | 0.104136 | 0.818118 | 0.696463 | | 1 | 0.996511 | 0.955729 |
| Omg | G3XA53 | 0.41 | 0.82 | 0.25 | 0.19 | 0.76 | 0.3 | 0.43 | 0.21 | 0.1 | -0.76 | 0.03 | 0.47 | -0.24 | 0.51 | -0.04 | 0.14 | 0.0075 | -0.4575 | -0.325 | 0.971865 | 0.154388 | 0.212348 | | 1 | 0.996511 | 0.786404 |
| Omp | Q64288 | 3.69 | -1.53 | -5.92 | -5.07 | -0.06 | -1 | -5.25 | -5.36 | 3.01 | 3.69 | -5.39 | 6.68 | -2.57 | -2.15 | -3.24 | -6.45 | -0.71 | 4.205 | -1.395 | 0.758663 | 0.18711 | 0.515291 | | 1 | 0.996511 | 0.905674 |
| Opa1 | P58281-2 | 0.54 | 0.54 | 1.94 | 1.55 | 0.39 | 0.55 | 1.8 | 2.48 | 0.14 | -0.13 | 2.15 | 0.45 | -0.1 | 0.44 | 1.69 | 1.75 | 0.1625 | -0.49 | -0.1975 | 0.773658 | 0.403991 | 0.71501 | | 1 | 0.996511 | 0.958106 |
| Opa3 | Q505D7 | 0.97 | 1.16 | 1.78 | 1.82 | 0.68 | 1.14 | 1.02 | 1.74 | 1.12 | 1.87 | 1.73 | 0.51 | 1.3 | 1.26 | 1.15 | 0.9 | -0.2875 | -0.125 | -0.28 | 0.359022 | 0.734634 | 0.301173 | | 1 | 0.996511 | 0.830276 |
| Opalin | Q7M750 | 1.02 | 1.75 | 1.08 | 0.9 | 0.87 | 0.7 | 0.58 | 0.5 | 2.29 | 0.85 | 0.44 | 1.09 | 2.06 | 1.09 | 1.55 | 0.87 | -0.525 | -0.02 | 0.205 | 0.044017 | 0.961871 | 0.538154 | | 1 | 0.996755 | 0.912212 |
| Opcml | Q6DFY2 | 0.95 | 1.26 | 1.5 | 1.43 | 1.29 | 0.95 | 0.84 | 1.37 | 0.82 | 0.92 | 1.28 | 1.1 | 0.64 | 1.02 | 0.84 | 0.88 | -0.1725 | -0.255 | -0.44 | 0.414406 | 0.234728 | 0.058211 | | 1 | 0.996511 | 0.661738 |
| Opcml | G5E8G3 | 0.7 | 1.39 | 2.21 | 2.22 | 1.18 | 0.86 | 1.88 | 2.52 | 0.28 | 0.42 | 2.32 | 1.16 | -0.32 | -0.29 | 1.43 | 1.94 | -0.02 | -0.585 | -0.94 | 0.966861 | 0.301517 | 0.160784 | | 1 | 0.996511 | 0.756132 |
| Ophn1 | Q99J31 | 0.28 | 0.37 | 0.69 | 0.68 | 0.31 | 0.18 | 0 | 0.53 | 0.46 | 0.44 | 0.72 | 0.37 | 0.09 | 0.24 | 0.41 | 0.33 | -0.25 | -0.0075 | -0.2375 | 0.21383 | 0.96826 | 0.246343 | | 1 | 0.996755 | 0.802442 |
| Oplah | Q8K010 | 0.03 | -0.14 | 0.29 | 0.38 | -0.22 | 0.18 | 0.44 | 0.41 | -0.12 | -0.1 | 0.22 | -0.36 | -0.24 | -0.41 | 0.04 | 0.12 | 0.0625 | -0.23 | -0.2625 | 0.774579 | 0.292392 | 0.254986 | | 1 | 0.996511 | 0.809524 |
| Oprk1 | A0A5F8MP | -0.59 | -1.19 | 2.62 | 3.17 | -0.31 | -1.15 | 1.21 | 0.47 | 0.45 | 5.63 | -0.12 | 0.25 | 0.93 | 0.69 | 3.18 | 3.84 | -0.9475 | 0.55 | 1.1575 | 0.395219 | 0.726378 | 0.352262 | | 1 | 0.996511 | 0.854088 |
| Oprl1 | B0R0C0 | 0.2 | 0.71 | 0.43 | 1.39 | -0.6 | 0.79 | -0.91 | 0.29 | 1.36 | 1.53 | 1.8 | 0.97 | 2.29 | -0.16 | 0.67 | 1.02 | -0.79 | 0.7325 | 0.2725 | 0.099458 | 0.041732 | 0.609008 | | 1 | 0.996511 | 0.934202 |
| Oprm1 | P42866-14 | 1.12 | -0.18 | 0.47 | 0.5 | -1.15 | 0.2 | -1.62 | -0.19 | 1.95 | -0.39 | -0.57 | 0.79 | 1.85 | -0.26 | 1.1 | -0.02 | -1.1675 | -0.0325 | 0.19 | 0.030772 | 0.956017 | 0.715637 | | 1 | 0.996755 | 0.958106 |
| Optn | Q8K3K8 | 0.4 | 0.51 | -0.02 | 1.45 | 0.3 | 0.52 | -0.27 | 0.67 | 0.98 | 0.17 | 0.01 | 0.75 | 1.16 | 1.21 | -0.41 | 0.15 | -0.28 | -0.1075 | -0.0575 | 0.44114 | 0.77407 | 0.903107 | | 1 | 0.996511 | 0.985031 |
| Orai1 | Q8BWG9 | 1.21 | 1.28 | 1.31 | 0.86 | 1.21 | 0.71 | 0.98 | 1.13 | 0.02 | -0.55 | 1.04 | 1.19 | -0.81 | -1.52 | 0.68 | 1.34 | -0.1575 | -0.74 | -1.2425 | 0.416525 | 0.098038 | 0.067848 | | 1 | 0.996511 | 0.664936 |
| Orai2 | Q8BH10 | -0.37 | 0.09 | 0.64 | 0.47 | 0.71 | -0.84 | 0.65 | 0.88 | 0.47 | 1.58 | 0.2 | 0.34 | -0.39 | 1.62 | 0.92 | 0.43 | 0.1425 | 0.44 | 0.4375 | 0.740196 | 0.259163 | 0.346344 | | 1 | 0.996511 | 0.851096 |
| Ormdl3 | Q9CPZ6 | 0.47 | 0.81 | 0.1 | 0.46 | -0.12 | 0.53 | -0.14 | 0.18 | 0.89 | 0.96 | 0.04 | 1.1 | 1.27 | 0.59 | 0.63 | 0.81 | -0.3475 | 0.2875 | 0.365 | 0.160547 | 0.33462 | 0.165767 | | 1 | 0.996511 | 0.758837 |
| Os9 | Q8K2C7 | -0.35 | -0.2 | -0.69 | -0.78 | 0.21 | 0 | 0.02 | -0.54 | 0.24 | -0.01 | -0.92 | 0.08 | 1.26 | 0.66 | 0.89 | 0.78 | 0.4275 | 0.3525 | 1.4025 | 0.09177 | 0.260249 | 0.00022 | | 1 | 0.996511 | 0.270112 |
| Osbp | Q3B7Z2 | 0.1 | 0.1 | 0.22 | 0.57 | -0.02 | 0.37 | 0.17 | 0.49 | 0.5 | 0.7 | 0.36 | 0.42 | 0.59 | 0.31 | 0.32 | -0.02 | 0.005 | 0.2475 | 0.0525 | 0.979437 | 0.21581 | 0.81061 | | 1 | 0.996511 | 0.972222 |
| Osbp2 | Q8K0C7 | -0.44 | -0.05 | 1.38 | 1.38 | 0.88 | -0.58 | 1.04 | 0.66 | 0.28 | 0.1 | 0.83 | 0.07 | -0.81 | 0.64 | 0.07 | 0.28 | -0.0675 | -0.2475 | -0.5225 | 0.902503 | 0.604619 | 0.334344 | | 1 | 0.996511 | 0.846677 |
| Osbp2 | Q5QNQ6 | -0.13 | 0.02 | 0.73 | 0.32 | 0.1 | -0.1 | 1.14 | 0.29 | -0.01 | 1.15 | 0.65 | -0.02 | -0.23 | -0.04 | 0.77 | 0.73 | 0.1225 | 0.2075 | 0.0725 | 0.706365 | 0.542376 | 0.823783 | | 1 | 0.996511 | 0.972768 |
| Osbpl10 | S4R1M9 | 0.14 | -0.01 | 0.01 | 0.27 | 0.36 | 0.49 | 0.41 | 0.14 | 0.19 | 0.1 | 0.18 | 0.31 | -0.11 | 0.67 | 0.28 | 0.14 | 0.2475 | 0.0925 | 0.1425 | 0.150104 | 0.577333 | 0.529285 | | 1 | 0.996511 | 0.908663 |
| Osbpl11 | G5E8A0 | -0.61 | -0.65 | -0.81 | -0.84 | -0.56 | -0.55 | -0.59 | -0.69 | -0.42 | -1 | -0.63 | -0.45 | 0.18 | -1.15 | -0.68 | -0.84 | 0.13 | 0.1025 | 0.105 | 0.386569 | 0.605329 | 0.729311 | | 1 | 0.996511 | 0.958635 |
| Osbpl1a | Q91XL9 | -0.8 | -0.75 | -0.52 | -0.72 | -0.63 | -0.2 | -0.62 | -0.59 | -1.11 | -0.8 | -0.49 | -0.66 | -1.22 | -1.16 | -1.27 | -0.94 | 0.1875 | -0.0675 | -0.45 | 0.292132 | 0.7323 | 0.033673 | | 1 | 0.996511 | 0.644622 |
| Osbpl2 | Q8BX94 | 0.78 | 0.95 | 0.17 | 0.41 | 1.02 | 0.8 | 0.53 | 0.44 | 0.88 | 0.13 | 0.82 | 0.75 | 0.31 | 1.36 | 0.41 | 0.44 | 0.12 | 0.0675 | 0.0525 | 0.615254 | 0.799181 | 0.865998 | | 1 | 0.996511 | 0.978151 |
| Osbpl3 | F8WH20 | 0.13 | 0.06 | 0.58 | 0.77 | -0.17 | -0.04 | 0.6 | 0.6 | 0.56 | 0.19 | 0.57 | 0.03 | 0.8 | 0.26 | 0.4 | 0.49 | -0.1375 | -0.0475 | 0.1025 | 0.616997 | 0.846119 | 0.675574 | | 1 | 0.996511 | 0.952944 |
| Osbpl5 | Q9ER64 | -0.36 | -0.32 | 1.1 | 0.23 | 0.71 | 0.5 | 0.66 | 0.53 | -0.06 | 1.87 | 0.8 | -0.79 | 1.13 | -0.43 | 0.78 | 1.6 | 0.4375 | 0.2925 | 0.6075 | 0.212348 | 0.633762 | 0.255292 | | 1 | 0.996511 | 0.809524 |
| Osbpl6 | F8WIZ7 | 0.34 | 0.51 | 0.4 | -0.08 | 0.38 | 0.47 | 0.17 | 0.15 | 0.21 | -0.51 | 0.19 | 0.41 | 0.27 | 0.41 | 0.09 | 0.23 | -3.9E-17 | -0.2175 | -0.0425 | 1 | 0.410526 | 0.836802 | | 1 | 0.996511 | 0.975831 |
| Osbpl7 | F8VQM4 | -0.05 | 0.27 | 0.39 | 0.59 | 0.11 | -0.17 | 0.31 | 0.53 | 0.7 | 0.4 | 0.54 | 0.05 | 0.66 | 0 | 0.47 | 0.36 | -0.105 | 0.1225 | 0.0725 | 0.639311 | 0.591813 | 0.757899 | | 1 | 0.996511 | 0.962717 |
| Osbpl8 | B9EJ86 | 0.43 | 0.04 | 0.14 | 0 | 0.49 | 0.23 | 0.89 | 0.13 | 0.14 | -0.31 | 0.23 | 0.32 | 0.32 | 0.71 | 1.15 | 0.6 | 0.2825 | -0.0575 | 0.5425 | 0.219533 | 0.786555 | 0.045549 | | 1 | 0.996511 | 0.658454 |
| Osbpl9 | A0A0A6YXI | -0.2 | 0.14 | 0.63 | 0.54 | -1.17 | -0.78 | 0.16 | 0.38 | 0.25 | -1.61 | -0.32 | -0.47 | -2.74 | 1.18 | 0.74 | -0.47 | -0.63 | -0.815 | -0.6 | 0.137209 | 0.075597 | 0.465782 | | 1 | 0.996511 | 0.888469 |
| Osbpl9 | E9PXZ2 | 0.49 | 0.24 | 0.35 | 0.72 | 0.38 | 0.34 | 0.68 | 0.82 | 0.53 | 0.96 | 0.45 | 0.41 | 0.5 | 0.67 | 0.38 | 0.51 | 0.105 | 0.1375 | 0.065 | 0.589153 | 0.513731 | 0.737217 | | 1 | 0.996511 | 0.958635 |
| Oscp1 | A0ZV96 | 0.6 | 0.34 | 0.39 | -0.01 | 0.2 | 0.35 | 0.25 | 0.09 | -0.7 | -0.76 | 0.34 | 0.71 | -0.85 | -0.9 | 0.4 | 0.21 | -0.1075 | -0.4325 | -0.615 | 0.557922 | 0.270684 | 0.115305 | | 1 | 0.996511 | 0.717287 |
| Osgep | A0A0R4J1Y | -2.05 | -2.3 | -1.38 | -1.55 | -1.75 | -2.02 | -1.31 | -1.15 | -2.89 | -0.98 | -1.43 | -1.78 | -2.56 | -1.91 | -1.18 | -1.5 | 0.2625 | 0.05 | 0.0325 | 0.380756 | 0.908413 | 0.928259 | | 1 | 0.996744 | 0.98946 |
| Ostc | Q78XF5 | -0.87 | -1.32 | -1.84 | -1.95 | 0.07 | -0.12 | -0.39 | -1.3 | -1.17 | -1.29 | -1.41 | -1.87 | -0.89 | -0.12 | -0.65 | -0.4 | 1.06 | 0.06 | 0.98 | 0.018843 | 0.84097 | 0.010901 | | 1 | 0.996511 | 0.563386 |
| Ostf1 | Q62422 | -1.41 | -1.29 | -0.96 | -1.05 | -1.1 | -1.35 | -0.82 | -0.92 | -0.69 | -0.31 | -0.98 | -1.42 | -0.3 | -0.95 | -1.05 | -1.1 | 0.13 | 0.3275 | 0.3275 | 0.506262 | 0.247743 | 0.208902 | | 1 | 0.996511 | 0.786218 |
| Ostm1 | Q8BGT0 | -1.64 | -2.58 | 0.74 | 0.28 | -2.35 | -1.8 | 0.15 | 0.4 | -2.25 | 1.31 | 0.48 | -1.63 | -2.18 | -3 | 0.49 | -0.06 | -0.1 | 0.2775 | -0.3875 | 0.915139 | 0.789096 | 0.706272 | | 1 | 0.996511 | 0.957084 |
| Otof | A0A0R4J1K | -2.16 | -1.86 | -0.53 | 0.15 | -2.18 | -2.27 | -0.4 | -0.78 | 1.72 | 1.97 | -0.86 | -1.34 | 0.87 | 4.89 | -0.52 | -1.04 | -0.3075 | 1.4725 | 2.15 | 0.643142 | 0.134942 | 0.122672 | | 1 | 0.996511 | 0.721448 |
| Otub1 | Q7TQI3 | -0.96 | -0.82 | -0.77 | -0.79 | -1.08 | -1.06 | 1.11 | -0.96 | -1.25 | -0.82 | -0.91 | -0.98 | -1.44 | -1.04 | -0.89 | -0.56 | 0.3375 | -0.155 | -0.1475 | 0.502684 | 0.385496 | 0.528482 | | 1 | 0.996511 | 0.908663 |
| Otub2 | Q9CQX0 | -0.54 | -0.19 | -0.49 | -0.44 | 0.34 | -1.05 | -0.53 | -0.62 | -0.67 | -0.3 | -0.63 | -0.22 | -1.13 | 1.05 | -0.88 | -0.73 | -0.05 | -0.04 | -0.0075 | 0.86733 | 0.835739 | 0.987332 | | 1 | 0.996511 | 0.996154 |
| Otud4 | A0A0R4J26 | -0.99 | -1.17 | -1.22 | -1.63 | -1.39 | -1.23 | -1.74 | -1.34 | -0.16 | -0.63 | -1.71 | -0.46 | 0.48 | -0.92 | -1.28 | -1.63 | -0.1725 | 0.5125 | 0.415 | 0.409719 | 0.174235 | 0.373179 | | 1 | 0.996511 | 0.859424 |
| Otud5 | A2AES3 | 3.79 | 4.46 | 3.05 | 3.12 | 5.03 | 3.78 | 2.72 | 2.63 | 2.5 | 3.81 | 2.77 | 3.45 | 1.84 | 4.02 | 2.84 | 2.83 | -0.065 | -0.4725 | -0.7225 | 0.912808 | 0.282635 | 0.185184 | | 1 | 0.996511 | 0.776843 |
| Otud6b | Q8K2H2 | 0.91 | 0.28 | -0.68 | -1.19 | 0.51 | 0.86 | 0.1 | -0.46 | -0.52 | -1.42 | -0.28 | 0.68 | -0.95 | 0.23 | -0.7 | -0.48 | 0.4225 | -0.215 | -0.305 | 0.41456 | 0.715001 | 0.545699 | | 1 | 0.996511 | 0.912321 |
| Otud7a | Q8R554 | -0.26 | -0.23 | 0.49 | 0.52 | -0.13 | -0.23 | -0.1 | 0.28 | -0.28 | 0.3 | 0.46 | -0.37 | -0.14 | -0.37 | 0.13 | -0.11 | -0.175 | -0.1025 | -0.2525 | 0.497592 | 0.736877 | 0.355094 | | 1 | 0.996511 | 0.855029 |
| Otud7b | B2RUR8 | 0.26 | -0.1 | 1.05 | 0.95 | -0.64 | 0.04 | 0.69 | 0.79 | 0.05 | 1.24 | 0.72 | 0.13 | 0.43 | -0.02 | 0.92 | 0.98 | -0.32 | -0.005 | 0.0375 | 0.43921 | 0.989449 | 0.916454 | | 1 | 0.999285 | 0.986556 |
| Otulin | Q3UCV8 | 0.13 | -0.01 | -0.39 | -0.64 | -0.44 | -0.18 | -1.2 | -0.86 | -0.47 | -0.11 | -1 | -0.01 | -0.34 | -0.33 | -0.57 | -0.86 | -0.4425 | -0.17 | -0.2975 | 0.149711 | 0.565263 | 0.252654 | | 1 | 0.996511 | 0.809096 |
| Otulinl | A0A2I3BPS | -0.01 | -0.05 | -0.05 | -0.02 | 0.64 | -0.1 | 0.64 | -0.39 | -0.18 | -1.01 | -0.25 | -0.44 | -0.46 | 0.77 | 0.23 | 0.01 | 0.23 | -0.4375 | 0.17 | 0.402221 | 0.077583 | 0.543913 | | 1 | 0.996511 | 0.912212 |
| Ovos | Q3UU35 | 0.63 | 0.74 | 1.19 | 0.95 | 0.96 | 0.27 | -0.01 | 0.45 | -0.01 | 0.53 | 0.55 | -0.03 | -0.35 | 0.83 | 0.23 | -0.33 | -0.46 | -0.6175 | -0.7825 | 0.093976 | 0.025537 | 0.032246 | | 1 | 0.996511 | 0.641974 |
| Oxa1l | Q8BGA9 | 0.86 | 0.02 | -0.44 | -0.11 | 0.75 | 1.16 | -0.89 | -0.82 | 0.19 | -0.08 | -0.42 | 0.66 | 0.97 | 0.35 | -0.64 | -0.88 | -0.0325 | 0.005 | -0.1325 | 0.952622 | 0.988624 | 0.782987 | | 1 | 0.999285 | 0.967642 |
| Oxct1 | Q9D0K2 | 0.77 | 0.58 | 0.42 | 0.29 | 0.25 | 0.99 | 0.22 | 0.42 | -0.29 | 0.45 | 0.69 | 0.77 | -0.36 | -0.36 | -0.16 | -0.29 | -0.045 | -0.11 | -0.8075 | 0.843042 | 0.691606 | 0.002206 | | 1 | 0.996511 | 0.419079 |
| Oxld1 | Q9CR10 | -0.68 | -0.22 | -0.4 | -0.53 | -0.14 | -1.58 | -0.5 | -1.11 | -1.07 | -0.85 | -0.6 | -0.94 | -0.09 | -0.36 | -0.64 | -0.87 | -0.375 | -0.4075 | -0.0325 | 0.268209 | 0.060806 | 0.890571 | | 1 | 0.996511 | 0.981669 |
| Oxnad1 | Q8VE38-2 | 1.2 | 1.05 | 0.49 | 0.33 | 0.86 | 1.05 | 0.5 | 0.31 | 0.44 | 0.48 | 0.69 | 1.16 | 0.14 | 0.68 | 0.06 | 0.22 | -0.0875 | -0.075 | -0.4925 | 0.750349 | 0.78923 | 0.100897 | | 1 | 0.996511 | 0.698156 |
| Oxr1 | Q4KMM3 | 0.77 | 0.61 | 1.48 | 1.15 | 0.79 | 0.86 | 1.04 | 1.67 | 0.17 | 0.68 | 1.5 | 0.71 | -0.04 | 0.37 | 1.14 | 1.24 | 0.0875 | -0.2375 | -0.325 | 0.756983 | 0.483294 | 0.377912 | | 1 | 0.996511 | 0.860984 |
| Oxr1 | E9Q0A7 | 1.24 | 0.17 | 0.58 | 0.98 | 1.05 | 0.37 | 0.76 | 0.18 | 0.01 | -0.8 | 0.32 | -0.16 | -0.55 | 2.33 | -0.21 | 0.34 | -0.1525 | -0.9 | -0.265 | 0.616097 | 0.022447 | 0.673514 | | 1 | 0.996511 | 0.952944 |
| Oxsm | Q9D404 | 0.29 | 0.38 | 0.04 | 0.64 | -0.26 | 0.28 | 0.17 | 0.33 | 0 | -0.17 | 0 | 0.03 | -0.38 | -0.64 | -0.26 | 0.25 | -0.2075 | -0.3725 | -0.595 | 0.338063 | 0.076272 | 0.041953 | | 1 | 0.996511 | 0.658454 |
| Oxsr1 | Q6P9R2 | -0.62 | -0.49 | -0.43 | -0.39 | -0.05 | -0.47 | -0.49 | -0.5 | -0.4 | -0.33 | -0.41 | -0.37 | -0.75 | 0.24 | -0.72 | -0.44 | 0.105 | 0.105 | 0.065 | 0.545297 | 0.508062 | 0.804992 | | 1 | 0.996511 | 0.971715 |
| P10404 | P10404 | -0.93 | 0.02 | 0.36 | -0.54 | -0.05 | -0.54 | 0.44 | 0.71 | -0.02 | -1.39 | 0.15 | 0.07 | -1.04 | -0.78 | -1.34 | -0.46 | 0.4125 | -0.025 | -0.6325 | 0.292551 | 0.95452 | 0.092165 | | 1 | 0.996755 | 0.695932 |
| P2ry12 | Q9CPV9 | -2 | -2.36 | -1.78 | -2.01 | -2.17 | -2.04 | -2.02 | -1.85 | -2.22 | -1.32 | -1.85 | -2.4 | -2.01 | -2 | -2.81 | -2.45 | 0.0175 | 0.09 | -0.28 | 0.922959 | 0.74782 | 0.295112 | | 1 | 0.996511 | 0.825612 |
| P33monox | Q9DBN4 | 0.7 | 0.25 | 0.34 | 0.15 | -0.22 | 0.88 | -0.26 | -0.29 | 1.04 | 0.39 | 0.17 | 0.47 | 1.55 | -0.21 | -0.34 | -0.42 | -0.3325 | 0.1575 | -0.215 | 0.292807 | 0.52651 | 0.640472 | | 1 | 0.996511 | 0.942906 |
| P4ha1 | Q60715-2 | 0.27 | 0.82 | -0.26 | 0.36 | 0.88 | 0.25 | 0.22 | 0.07 | 0.27 | 0.59 | -0.3 | 0.53 | 1.15 | 1.14 | 1.46 | 0.24 | 0.0575 | -0.025 | 0.7 | 0.840823 | 0.934603 | 0.067608 | | 1 | 0.996755 | 0.664936 |
| P4hb | P09103 | 0.68 | 0.4 | -0.13 | -0.05 | 0.47 | 0.62 | 0.51 | 0.18 | 0.34 | -0.36 | -0.21 | 0.77 | 0.74 | 0.6 | 0.69 | 0.42 | 0.22 | -0.09 | 0.3875 | 0.355476 | 0.780379 | 0.135655 | | 1 | 0.996511 | 0.731572 |
| P4htm | Q8BG58 | -1.18 | -1.17 | -0.43 | -0.98 | 0.23 | -1 | -0.27 | -0.52 | -1.11 | -0.73 | -0.77 | -0.67 | -1.26 | 0.12 | -0.02 | -0.13 | 0.55 | 0.12 | 0.6175 | 0.100996 | 0.608865 | 0.111827 | | 1 | 0.996511 | 0.713445 |
| Pa2g4 | P50580 | -1.86 | -2.2 | -1.19 | -1.65 | -1.65 | -2 | -1.02 | -1.42 | -2.42 | -1.32 | -1.26 | -1.73 | -2.08 | -1.24 | -0.99 | -1.1 | 0.2025 | 0.0425 | 0.3725 | 0.497048 | 0.899483 | 0.275755 | | 1 | 0.996511 | 0.822084 |
| Paat | Q9D2Q3 | -2.38 | -2.94 | 1.62 | 1.44 | -0.91 | -2.98 | 0.54 | 1.12 | -2.63 | 2.4 | 1.57 | -2.31 | -5.13 | -1.94 | 0.71 | 0.74 | 0.0075 | 0.3225 | -0.84 | 0.99559 | 0.839062 | 0.610891 | | 1 | 0.996511 | 0.935047 |
| Pabpc1 | P29341 | -0.61 | -0.86 | -1.11 | -1.04 | -0.75 | -0.66 | -0.98 | -1.1 | -0.31 | -1.21 | -1.07 | -0.51 | -0.06 | -0.25 | -1.01 | -1.25 | 0.0325 | 0.13 | 0.2625 | 0.86391 | 0.62226 | 0.416513 | | 1 | 0.996511 | 0.874945 |
| Pabpc4 | Q6PHQ9 | -1.63 | -1.84 | -0.72 | -0.66 | -1.19 | -1.74 | -0.37 | -0.84 | -1.52 | -0.97 | -0.68 | -1.43 | -1.35 | -1.68 | -0.81 | -0.76 | 0.1775 | 0.0625 | 0.0625 | 0.656142 | 0.860233 | 0.865624 | | 1 | 0.996511 | 0.977971 |
| Pabpn1 | G3UY42 | 4.3 | -0.1 | 3.55 | 3.26 | 1.02 | 4.43 | 4.03 | 4.52 | 0.07 | 4.16 | 2.62 | 3.41 | -0.36 | 0.68 | 4.13 | 4.11 | 0.7475 | -0.1875 | -0.6125 | 0.520446 | 0.873953 | 0.651731 | | 1 | 0.996511 | 0.947633 |
| Pacs1 | Q8K212 | -0.04 | -0.14 | 0.45 | 0.59 | 0.21 | -0.02 | 0.53 | 0.64 | -0.32 | 0.18 | 0.58 | 0.01 | -0.5 | -0.02 | 0.29 | 0.26 | 0.125 | -0.1025 | -0.2075 | 0.615899 | 0.709009 | 0.462356 | | 1 | 0.996511 | 0.888262 |

| | | | | | | | | | | | | | | | | | | | | | | | | | | | |
|---|---|---|---|---|---|---|---|---|---|---|---|---|---|---|---|---|---|---|---|---|---|---|---|---|---|---|---|
| Pacs2 | A0A1Y7VLZ | 0.28 | 0.26 | -0.22 | -0.47 | 0.15 | 0.3 | 0.01 | -0.15 | 1.21 | -0.69 | -0.02 | 0.32 | 1.13 | -0.53 | 0.17 | 0.13 | 0.115 | 0.2425 | 0.2625 | 0.617052 | 0.562681 | 0.494159 | | 1 | 0.996511 | 0.89758 |
| Pacsin1 | Q61644 | 0.67 | 0.45 | 0.53 | 0.8 | 0.25 | 0.47 | 0.47 | 0.64 | 0.31 | 0.15 | 0.39 | 0.57 | 0.19 | 0.47 | 0.21 | 0.19 | -0.155 | -0.2575 | -0.3475 | 0.365184 | 0.18042 | 0.08877 | | 1 | 0.996511 | 0.691619 |
| Pacsin2 | A0A2R8W6 | 1.46 | 1.96 | 1.55 | 1.52 | 1.28 | 0.57 | 0.92 | 1.26 | -0.35 | -0.78 | 1.22 | 0.89 | 0.23 | -0.78 | 0.36 | 0.87 | -0.615 | -1.3775 | -1.4525 | 0.021764 | 0.015089 | 0.002921 | | 1 | 0.996511 | 0.45967 |
| Pacsin2 | Q9WVE8 | 0.69 | 0.53 | 0.6 | 0.41 | 0.47 | 0.58 | 0.43 | 0.44 | 0.65 | 1.19 | 0.36 | 0.7 | 0.5 | 0.98 | 0.24 | 0.4 | -0.0775 | 0.1675 | -0.0275 | 0.602556 | 0.453294 | 0.900704 | | 1 | 0.996511 | 0.983757 |
| Pacsin3 | Q99JB8 | -0.34 | -0.24 | 0.35 | 0.28 | 0.02 | -0.51 | 0.45 | 0.38 | -0.57 | 0.33 | 0.55 | -0.24 | -0.76 | -0.04 | 0.13 | -0.02 | 0.0725 | 0.005 | -0.185 | 0.798447 | 0.987271 | 0.521174 | | 1 | 0.999285 | 0.907667 |
| Padi2 | Q08642 | 0.9 | 0.97 | 0.59 | 0.23 | 1.5 | 0.45 | 0.43 | 0.07 | 0.57 | 0.3 | 0.41 | 0.81 | 0.37 | 0.63 | 0.76 | 0.66 | -0.06 | -0.15 | -0.0675 | 0.859883 | 0.524485 | 0.771315 | | 1 | 0.996511 | 0.965933 |
| Paf1 | Q8K2T8 | -0.01 | -1.4 | -0.8 | -1 | -1.32 | -1.08 | -0.83 | -0.77 | -0.94 | -0.56 | -0.71 | -1.38 | -0.14 | -0.18 | -0.05 | -1.25 | -0.1975 | -0.095 | 0.3975 | 0.532408 | 0.779111 | 0.326817 | | 1 | 0.996511 | 0.843799 |
| Pafah1b1 | P63005 | 0.11 | -0.02 | 0.25 | 0.28 | 0.19 | 0.16 | 0.4 | 0.24 | -0.03 | 0.17 | 0.32 | 0.22 | -0.17 | 0.63 | 0.23 | 0.08 | 0.0925 | 0.015 | 0.0375 | 0.555563 | 0.931231 | 0.869003 | | 1 | 0.996755 | 0.979293 |
| Pafah1b2 | A0A1L1SVK | -1.24 | -1.08 | -1.04 | -1.35 | -0.94 | -1.74 | -1.41 | -1.76 | -0.68 | -2.36 | -1.45 | -1.16 | -1 | -0.76 | -2 | -1.58 | -0.285 | -0.235 | -0.1575 | 0.228377 | 0.51093 | 0.604793 | | 1 | 0.996511 | 0.932185 |
| Pafah1b3 | Q61205 | 0.51 | -0.6 | 0.39 | -0.54 | -0.77 | 0.56 | 0.72 | -0.1 | -0.79 | -0.37 | 0.5 | 0.76 | -1.03 | -0.79 | 0.38 | 0.6 | 0.1625 | 0.085 | -0.15 | 0.701685 | 0.847444 | 0.752194 | | 1 | 0.996511 | 0.961479 |
| Pafah2 | E9QNW6 | -0.41 | 0.17 | 0.45 | 0.35 | 0.02 | 0.22 | 0.36 | 0.07 | 0.37 | -0.99 | -0.12 | -0.03 | -0.37 | -0.31 | -0.02 | 0.17 | 0.0275 | -0.3325 | -0.2725 | 0.903733 | 0.342078 | 0.309066 | | 1 | 0.996511 | 0.834523 |
| Pag1 | Q3U1F9 | 0.45 | 0.93 | 0.67 | 0.53 | 1.13 | 0.08 | 0.51 | 0.48 | 0.19 | 0.82 | 0.73 | 0.79 | 0.59 | -0.03 | 0.25 | 0.43 | -0.095 | -0.0125 | -0.335 | 0.707497 | 0.95458 | 0.15422 | | 1 | 0.996755 | 0.750763 |
| Paics | Q9DCL9 | 0.09 | -0.46 | 0.81 | 0.91 | -0.63 | -0.15 | 0.46 | 0.99 | -0.24 | 1.03 | 0.64 | 0.02 | -0.58 | -0.43 | 0.69 | 0.38 | -0.17 | 0.025 | -0.3225 | 0.703972 | 0.951587 | 0.456046 | | 1 | 0.996755 | 0.887169 |
| Paip1 | F6Y616 | 0.25 | 0.2 | -0.08 | -0.11 | -0.08 | 0.06 | -0.36 | 0.04 | 0.48 | 0.52 | 0.07 | 0.31 | 0.67 | 0.6 | -0.02 | -0.31 | -0.15 | 0.28 | 0.17 | 0.413467 | 0.17201 | 0.54481 | | 1 | 0.996511 | 0.912212 |
| Paip2b | Q91W45 | -0.76 | -0.73 | 0.65 | 0.12 | -1.82 | -0.96 | -1.23 | -0.79 | -0.96 | 0.85 | -0.11 | -1.17 | -0.38 | -0.2 | 0.07 | 0.14 | -1.02 | -0.1675 | 0.0875 | 0.02654 | 0.753085 | 0.807798 | | 1 | 0.996511 | 0.971953 |
| Pak1 | G5E884 | 0.2 | 0.37 | 0.06 | 0.2 | 0.23 | 0.26 | -0.09 | 0.14 | -0.08 | 0 | 0.01 | 0.28 | -0.43 | 0.21 | -0.21 | -0.13 | -0.0725 | -0.155 | -0.3475 | 0.656065 | 0.384061 | 0.121122 | | 1 | 0.996511 | 0.721448 |
| Pak1ip1 | Q9DCE5 | -1.41 | -1.57 | -2.6 | -2.39 | -1.47 | -2.24 | -2.63 | -2.89 | -1.88 | -1.83 | -2.44 | -1.5 | -1.32 | -1.08 | -2.05 | -2.39 | -0.315 | 0.08 | 0.2825 | 0.442606 | 0.817871 | 0.49526 | | 1 | 0.996511 | 0.89758 |
| Pak2 | Q8CIN4 | 0.17 | 0.32 | 0.72 | 1.24 | 0.23 | 0.02 | 0.66 | 1.23 | 0.14 | 0.9 | 0.66 | 0.27 | -0.21 | 0.51 | 0.23 | 0.53 | -0.0775 | -0.12 | -0.3475 | 0.822658 | 0.692095 | 0.27397 | | 1 | 0.996511 | 0.822084 |
| Pak3 | Q61036 | 0.74 | -0.47 | 0.01 | 0.09 | -0.53 | 0.58 | 0 | 0.19 | -0.1 | 0.12 | 0.1 | 0.61 | 0.54 | -0.2 | 0.19 | -0.11 | -3.25E-02 | 0.09 | 0.0125 | 0.921435 | 0.762739 | 0.967758 | | 1 | 0.996511 | 0.9927 |
| Pak4 | Q8BTW9 | -0.95 | 0.12 | -0.22 | -0.02 | -0.61 | 0.04 | -0.35 | -0.62 | 0.98 | 0.42 | -0.19 | 0.16 | 0.87 | 2.99 | 1.08 | -0.2 | -0.1175 | 0.61 | 1.4525 | 0.681856 | 0.099248 | 0.04729 | | 1 | 0.996511 | 0.658454 |
| Pak5 | Q8C015 | 1.22 | 1.01 | 0.85 | 0.73 | 0.99 | 1.09 | 0.61 | 0.83 | 0.47 | 0.34 | 0.97 | 1.12 | 0.28 | 0.61 | 0.27 | 0.57 | -0.0725 | -0.2275 | -0.52 | 0.701644 | 0.361703 | 0.029149 | | 1 | 0.996511 | 0.637553 |
| Pak6 | Q3ULB5 | 0.44 | -0.18 | 0.39 | 1.04 | 0.24 | 0.69 | 0.67 | 1.03 | 1.23 | 0.9 | 0.96 | 0.21 | 2.2 | 0.64 | 0.82 | 1.12 | 0.235 | 0.4025 | 0.7725 | 0.435781 | 0.241385 | 0.087853 | | 1 | 0.996511 | 0.691619 |
| Pakap | F7AA26 | 1.04 | 0.81 | 1.3 | 1.21 | 1.08 | 0.95 | 1.01 | 0.92 | 1.09 | 0.97 | 1.21 | 1.32 | 0.87 | 0.81 | 1.45 | 1.52 | -0.1 | 0.0575 | 0.0725 | 0.555775 | 0.761778 | 0.77198 | | 1 | 0.996511 | 0.965933 |
| Pald1 | A0A0R4J00 | -0.59 | -0.39 | -0.91 | -0.84 | -0.26 | -0.45 | -0.6 | -0.67 | -0.46 | -0.36 | -0.89 | -0.33 | -0.52 | -0.58 | -0.81 | -0.91 | 0.1875 | 0.1725 | -0.0225 | 0.335473 | 0.432796 | 0.914466 | | 1 | 0.996511 | 0.986458 |
| Palld | Q9ET54-3 | -1.34 | -1.32 | 0.13 | -0.01 | -1.2 | -1.47 | -0.15 | 0.15 | -1.12 | -0.31 | -0.05 | -1.18 | -1.2 | -1.09 | -0.36 | -0.31 | -0.0325 | -0.03 | -0.105 | 0.949939 | 0.94822 | 0.812214 | | 1 | 0.996755 | 0.972642 |
| Palm | Q9Z0P4-2 | -0.16 | 0.58 | -0.1 | 0.44 | 1.17 | -0.3 | 0.86 | 0.19 | 0.61 | 0.25 | 0.64 | 0.4 | 0.37 | 2.09 | 0.42 | 0.76 | 0.29 | 0.285 | 0.72 | 0.433509 | 0.249805 | 0.116741 | | 1 | 0.996511 | 0.718608 |
| Palm | Q9Z0P4 | 1.88 | 1.53 | 1.54 | 2 | 1.63 | 1.52 | 1.51 | 1.37 | 1.49 | 1.36 | 1.41 | 1.86 | 1.75 | 1.58 | 1.57 | 1.26 | -0.23 | -0.2075 | -0.1975 | 0.217531 | 0.333841 | 0.366095 | | 1 | 0.996511 | 0.858839 |
| Palm2 | Q8BR92 | 1.22 | 1.73 | 0.98 | 0.83 | 1.42 | 1.15 | 0.49 | 0.27 | 1.27 | -0.04 | 0.56 | 1.27 | 0.71 | 1.82 | 0.6 | 0.04 | -0.3575 | -0.425 | -0.3975 | 0.290283 | 0.260253 | 0.340462 | | 1 | 0.996511 | 0.84829 |
| Palm3 | A2TJV2 | 0.78 | 1.37 | 0.84 | 1.07 | 1.3 | 1.13 | 1.13 | 1.05 | 1.49 | 1.27 | 1.3 | 1.73 | 1.47 | 1.96 | 1.08 | 1.23 | 0.1375 | 0.4325 | 0.42 | 0.465452 | 0.067988 | 0.136061 | | 1 | 0.996511 | 0.731656 |
| Palmd | A0A0R4J0J | 0.15 | 0.3 | -0.43 | -0.43 | 0.17 | -0.28 | -0.46 | -0.45 | 0.3 | -0.08 | -0.06 | 0.29 | 0.55 | 0.36 | -0.45 | -0.56 | -0.1525 | 0.215 | 0.0775 | 0.550709 | 0.389746 | 0.819905 | | 1 | 0.996511 | 0.972768 |
| Pals1 | Q9JLB2 | 0.67 | 0.84 | 0.84 | 1.06 | 0.78 | 0.38 | 0.34 | 0.53 | 0.71 | 1.14 | 0.84 | 0.68 | 0.75 | 0.95 | 0.76 | 0.69 | -0.345 | -0.01 | -0.065 | 0.076818 | 0.957882 | 0.724068 | | 1 | 0.996755 | 0.958398 |
| Pals2 | Q9JLB0 | 0.66 | -0.1 | 1.17 | 0.94 | 0.12 | 0.08 | 0.35 | 0.71 | 0.91 | 1.9 | 1.05 | 0.82 | 0.64 | 2.3 | 0.61 | 1.58 | -0.3525 | 0.5025 | 0.615 | 0.270282 | 0.188522 | 0.206904 | | 1 | 0.996511 | 0.785338 |
| Pals2 | Q9JLB0-2 | 0.99 | 0.31 | 1.14 | 1.29 | 0.84 | 0.76 | 1.49 | 1.32 | 0.5 | 0.81 | 1.25 | 0.49 | -0.61 | 0.55 | 0.73 | 1.35 | 0.17 | -0.17 | -0.4275 | 0.551994 | 0.56112 | 0.343804 | | 1 | 0.996511 | 0.848473 |
| Pam | P97467 | 0.37 | 0.42 | -0.38 | 0.02 | 0.49 | 0.21 | -0.09 | 0.16 | 0.66 | 0.97 | -0.13 | 0.45 | 0.44 | 0.48 | 0.78 | 0.53 | 0.085 | 0.38 | 0.45 | 0.720577 | 0.229733 | 0.087456 | | 1 | 0.996511 | 0.691619 |
| Pam16 | Q9CQV1 | 0.69 | 0.57 | -0.2 | 0.14 | -0.23 | 1.13 | -0.64 | -0.11 | 2.09 | 0.78 | -0.29 | 0.92 | 2.6 | 0.15 | -0.52 | -0.73 | -0.2625 | 0.575 | 0.075 | 0.524139 | 0.261479 | 0.916663 | | 1 | 0.996511 | 0.986556 |
| Pan2 | Q8BGF7-2 | -0.36 | 0.47 | 0.71 | 0.03 | 0.13 | -0.44 | 0.12 | 0 | 0.05 | 1.09 | 0.53 | -0.23 | -0.18 | -0.52 | 0.24 | 0.62 | -0.26 | 0.1475 | -0.1725 | 0.359628 | 0.686414 | 0.617254 | | 1 | 0.996511 | 0.937141 |
| Pan3 | H3BKF3 | 0.27 | 0.56 | 0.16 | -0.07 | 0.57 | -0.49 | -0.33 | -0.1 | 0.24 | 1.36 | -0.01 | 0.28 | 0.54 | 1.07 | -0.26 | -0.34 | -0.3175 | 0.2375 | 0.0225 | 0.263089 | 0.477504 | 0.949749 | | 1 | 0.996511 | 0.992476 |
| Pank1 | Q8K4K6 | 2.45 | 2.81 | 2.12 | 1.91 | 2.18 | 1.14 | 1.65 | 2.5 | 1.04 | 2.38 | 2.02 | 2.24 | 1.03 | 1.2 | 2.88 | 3.2 | -0.455 | -0.4025 | -0.245 | 0.210262 | 0.272607 | 0.656869 | | 1 | 0.996511 | 0.948659 |
| Pank2 | Q7M753 | 0.48 | 0.18 | 0.59 | -0.15 | 0.24 | 0.31 | 0.43 | 0.43 | 0.08 | 0 | 0.17 | 0.57 | -0.01 | -0.09 | 0.1 | 0.01 | 0.0775 | -0.07 | -0.2725 | 0.70459 | 0.768533 | 0.237866 | | 1 | 0.996511 | 0.792722 |
| Pank4 | Q80YV4-2 | -0.31 | -0.19 | 0.16 | 0 | 0.04 | -0.22 | 0.27 | 0.19 | -1.11 | -0.56 | 0.06 | -0.34 | -0.93 | -0.4 | -0.07 | 0.11 | 0.155 | -0.4025 | -0.2375 | 0.421116 | 0.172639 | 0.396153 | | 1 | 0.996511 | 0.867334 |
| Panx1 | Q9JIP4 | 0.28 | 0.83 | 0.67 | 0.65 | 1.02 | 0.64 | 0.81 | 1.1 | 1.14 | 0.34 | 0.89 | 0.84 | 0.86 | 1.2 | 0.4 | 1.14 | 0.285 | 0.195 | 0.2925 | 0.165218 | 0.414612 | 0.260885 | | 1 | 0.996511 | 0.81412 |
| Panx2 | Q6IMP4-2 | 1.19 | 1.22 | 0.78 | 1.09 | 1.13 | 1.24 | 0.8 | 1.06 | 1.07 | 1.57 | 0.88 | 1.36 | 0.74 | 1.25 | 0.72 | 0.82 | -0.0125 | 0.15 | -0.1875 | 0.945029 | 0.501473 | 0.393127 | | 1 | 0.996511 | 0.865775 |
| Paox | Q8C0L6 | -0.29 | -0.59 | -0.88 | -1.13 | 0.1 | -0.24 | -0.48 | -0.78 | -1.17 | -0.73 | -0.63 | -0.5 | -0.74 | -0.2 | -1.05 | -0.99 | 0.3725 | -0.035 | -0.0225 | 0.187019 | 0.890416 | 0.936865 | | 1 | 0.996511 | 0.989767 |
| Papola | Q61183 | -1.26 | -0.72 | -2.47 | -2.61 | -0.74 | -1.31 | -1.85 | -3 | -1.17 | -1.11 | -2.18 | -1.2 | -1.65 | -0.86 | -2.72 | -2.52 | 0.04 | 0.35 | -0.1725 | 0.947497 | 0.483088 | 0.765926 | | 1 | 0.996511 | 0.964134 |
| Papss1 | Q6NZM8 | -1.06 | -1.15 | -0.95 | -1.3 | -0.91 | -1.14 | -0.8 | -1.24 | -1.29 | -0.28 | -1.1 | -0.88 | -1.08 | -0.88 | -1.09 | -1.12 | 0.0925 | 0.2275 | 0.0725 | 0.599176 | 0.380423 | 0.690403 | | 1 | 0.996511 | 0.955729 |
| Papss2 | A0A494B9Z | 0.47 | 0.21 | 0.73 | 0.49 | 0.51 | 0.4 | 1.27 | 0.8 | 0.13 | 0.26 | 0.41 | 0.35 | 0.17 | 0.24 | 0.81 | 0.73 | 0.27 | -0.1875 | 0.0125 | 0.274424 | 0.32415 | 0.957828 | | 1 | 0.996511 | 0.992636 |
| Paqr4 | Q9JJE4 | -1.05 | -1.24 | -0.59 | -0.65 | -1.24 | -0.58 | -0.33 | -1.04 | -3.56 | -1.75 | -0.87 | -0.68 | -3.25 | -3.72 | -1.21 | -0.86 | 0.085 | -0.8325 | -1.3775 | 0.751057 | 0.201758 | 0.065111 | | 1 | 0.996511 | 0.66433 |
| Paqr7 | Q80ZE4 | 0.05 | 0.42 | 1.28 | -0.56 | -0.32 | 0.46 | -0.51 | 0.04 | 0.53 | -0.09 | -0.01 | -0.68 | 0.99 | 0.59 | 0.05 | 3.18 | -0.38 | -0.36 | 0.905 | 0.370159 | 0.414913 | 0.225909 | | 1 | 0.996511 | 0.787447 |
| Paqr9 | Q6TCG2 | -1.27 | -1.11 | -0.98 | -1.82 | -2.32 | 0.71 | -1.71 | -1.55 | 2.48 | -3.61 | -1.44 | -0.38 | 4.49 | -4.02 | -0.09 | -0.88 | 0.0775 | 0.5575 | 1.17 | 0.901538 | 0.628922 | 0.462833 | | 1 | 0.996511 | 0.888267 |
| Pard3 | Q99NH2 | 0.79 | 0.83 | 0.61 | 0.48 | 0.96 | 0.61 | 0.45 | 0.35 | 0.49 | 0.97 | 0.24 | 0.85 | 1.2 | 1.27 | 0.89 | 0.64 | -0.085 | -0.04 | 0.3225 | 0.661678 | 0.857086 | 0.165539 | | 1 | 0.996511 | 0.758837 |
| Pard6a | Q9Z101-3 | 0.4 | 0.11 | 0.59 | 0.61 | 0.25 | 0.22 | 0.35 | 0.6 | -0.13 | 0.3 | 0.94 | 0.56 | 0.01 | -0.58 | 0.35 | 0.4 | -0.0725 | -0.01 | -0.3825 | 0.697449 | 0.970313 | 0.189261 | | 1 | 0.99723 | 0.776843 |
| Pard6b | Q9JK83 | -0.93 | -0.49 | -1.38 | -0.46 | -0.65 | -1.02 | -0.89 | -1.28 | -0.45 | -2.19 | -1.36 | -0.81 | -1.31 | 0.31 | -2.59 | -1.23 | -0.145 | -0.3875 | -0.39 | 0.582921 | 0.362744 | 0.507202 | | 1 | 0.996511 | 0.902614 |
| Parg | G3X8U8 | -0.89 | -0.95 | -0.85 | -1.2 | -1.1 | -1.06 | -1.24 | -1.29 | -1.25 | -1.13 | -0.99 | -0.63 | -1.28 | -1.37 | -1.42 | -1.43 | -0.2 | -0.0275 | -0.4025 | 0.230321 | 0.892212 | 0.047818 | | 1 | 0.996511 | 0.658454 |
| Park7 | Q99LX0 | 0.05 | -0.31 | -0.15 | -0.21 | -0.16 | -0.18 | 0.05 | 0.01 | -0.35 | -0.51 | -0.22 | -0.16 | -0.27 | -0.07 | -0.18 | -0.33 | 0.085 | -0.155 | -0.0575 | 0.596312 | 0.393206 | 0.752978 | | 1 | 0.996511 | 0.961479 |
| Parl | A0A338P6L | 0.36 | 0.04 | 0.91 | 1.2 | -0.52 | 0.46 | 0.84 | 1.17 | 1.96 | 0.77 | 1.28 | 0.09 | 2.64 | -0.61 | 0.8 | 0.61 | -0.14 | 0.3975 | 0.2325 | 0.740532 | 0.38487 | 0.723632 | | 1 | 0.996511 | 0.958398 |
| Parm1 | Q923D3 | 0.16 | -0.2 | -0.72 | -0.74 | -2.02 | 0.8 | -1.83 | -1.5 | 4.7 | -1.25 | -0.88 | 0.02 | 5.27 | -1.72 | -0.46 | -0.38 | -0.7625 | 1.0225 | 1.0525 | 0.244882 | 0.42173 | 0.459919 | | 1 | 0.996511 | 0.887902 |
| Parn | Q8VDG3 | -1.7 | -2.15 | -1.59 | -1.51 | -1.8 | -2.14 | -1.73 | -2.28 | -2.05 | -0.73 | -1.33 | -1.8 | -1.39 | -1.16 | -0.91 | -2.14 | -0.25 | 0.26 | 0.3375 | 0.271577 | 0.43007 | 0.294148 | | 1 | 0.996511 | 0.825612 |
| Parp1 | Q921K2 | -2.4 | -2.81 | -2.34 | -2.26 | -2.63 | -2.8 | -2.05 | -2.99 | -2.14 | -1.15 | -2.24 | -2.04 | -2.12 | -2.44 | -2.2 | -2.02 | -0.165 | 0.56 | 0.2575 | 0.513615 | 0.0795 | 0.240546 | | 1 | 0.996511 | 0.795658 |
| Parp12 | Q8BZ20 | -0.49 | -0.34 | -0.33 | -0.14 | -0.56 | -0.84 | -0.34 | -0.2 | -0.73 | -0.72 | -0.59 | -0.57 | -0.76 | -0.87 | -0.79 | -0.53 | -0.16 | -0.3275 | -0.4125 | 0.417831 | 0.07601 | 0.049922 | | 1 | 0.996511 | 0.658454 |
| Pars2 | Q8CFI5 | -0.65 | -0.16 | -0.51 | -0.46 | 0.22 | 0.31 | 0.23 | 0.18 | -0.21 | -0.79 | -0.6 | -0.58 | 0.41 | 0.78 | -0.19 | -0.31 | 0.68 | -0.1 | 0.6175 | 0.002621 | 0.628313 | 0.059881 | | 1 | 0.996511 | 0.661738 |
| Parva | Q9EPC1 | -0.6 | -0.53 | 0.12 | -0.06 | 0 | -0.44 | 0.37 | 0.2 | -0.71 | 0.47 | 0.42 | -0.64 | -0.62 | -0.11 | -0.11 | 0.16 | 0.3 | 0.1525 | 0.0975 | 0.263322 | 0.672663 | 0.715061 | | 1 | 0.996511 | 0.958106 |
| Parvb | Q9ES46 | -0.6 | -0.7 | -0.52 | -0.17 | -1.01 | -0.6 | 0.02 | -0.39 | -0.4 | 0.33 | 0.03 | -0.29 | -0.8 | -0.63 | -0.31 | -0.43 | 0.0025 | 0.415 | -0.045 | 0.992146 | 0.101676 | 0.833235 | | 1 | 0.996511 | 0.975228 |
| Patl1 | Q3TC46 | -0.77 | -0.89 | -0.35 | -0.3 | -0.72 | -1.04 | -0.72 | -0.49 | 2.14 | 0.05 | -0.53 | -0.93 | 1.8 | 1.92 | -0.5 | -0.63 | -0.165 | 0.76 | 1.225 | 0.446609 | 0.253389 | 0.087852 | | 1 | 0.996511 | 0.691619 |
| Patz1 | Q9JMG9 | -1.74 | 0.06 | -0.19 | -0.19 | -0.14 | -1.82 | -2.05 | -0.55 | -0.14 | 0.33 | -0.95 | -0.97 | -0.24 | -1.11 | 0.28 | 0.14 | -0.625 | 0.0825 | 0.2825 | 0.290919 | 0.865318 | 0.563482 | | 1 | 0.996511 | 0.920024 |
| Pawr | Q925B0-2 | -0.86 | -1.47 | -0.07 | 0.22 | -0.57 | -0.79 | -0.6 | 0.32 | -0.23 | 1 | 0.23 | -0.57 | -0.33 | -0.05 | -0.14 | -0.92 | 0.135 | 0.6525 | 0.185 | 0.752679 | 0.196458 | 0.655676 | | 1 | 0.996511 | 0.948243 |
| Paxbp1 | P58501 | 0.04 | -0.31 | -0.95 | -0.63 | 0.01 | -0.36 | -0.57 | -0.49 | -0.8 | -0.73 | -0.9 | -0.69 | -0.82 | -0.21 | -0.3 | -1.21 | 0.11 | -0.3175 | -0.1725 | 0.671004 | 0.214353 | 0.595047 | | 1 | 0.996511 | 0.929843 |
| Paxx | A2AJ30 | 0.48 | 1.6 | 0.72 | -1.24 | 0.7 | 0.25 | -0.83 | -0.46 | 0.57 | -0.4 | 0.01 | 0.74 | 0.11 | 1.42 | 0.15 | -0.13 | -0.475 | -0.16 | -0.0025 | 0.455115 | 0.788438 | 0.996828 | | 1 | 0.996511 | 0.99924 |
| Pbdc1 | Q9D0B6 | 0.95 | 0.28 | 0.48 | 1.41 | 0.33 | 0.24 | 0.68 | 0.8 | 0.71 | 1.42 | 1.03 | 0.93 | -0.22 | 0.81 | 0.59 | 0.85 | -0.2675 | 0.2425 | -0.2725 | 0.364288 | 0.426606 | 0.446683 | | 1 | 0.996511 | 0.884817 |
| Pbrm1 | D3Z1N4 | 1.21 | 0.86 | 1.69 | 1.5 | 1 | 0.84 | 1.63 | 1.56 | 0.42 | 0.42 | 1.5 | 0.79 | 0.35 | 0.94 | 1.89 | 1.77 | -0.0575 | -0.5325 | -0.0775 | 0.833345 | 0.117732 | 0.843635 | | 1 | 0.996511 | 0.976225 |
| Pbx1 | P41778-2 | -1.57 | -1.48 | -1.87 | -1.22 | -1.34 | -0.9 | -0.76 | -1.09 | -1.36 | -0.55 | -1.4 | -0.89 | -2.01 | -0.93 | -2.09 | -2.42 | 0.5125 | 0.485 | -0.3275 | 0.036126 | 0.09153 | 0.359281 | | 1 | 0.996511 | 0.856645 |
| Pbxip1 | Q3TVI8 | -0.33 | 0.14 | 0.13 | 0.18 | 0.43 | -0.76 | 1.93 | -0.45 | 1.29 | 0.36 | -0.06 | 0.21 | 0.69 | 0.87 | -0.3 | 0.27 | 0.2575 | 0.42 | 0.3525 | 0.648982 | 0.209161 | 0.25847 | | 1 | 0.996511 | 0.812074 |
| Pcbd1 | P61458 | -0.03 | -0.01 | -0.06 | 0.45 | 0.02 | -0.38 | 0.63 | 0.39 | 0.14 | 1.58 | 0.11 | -0.1 | 0.25 | 1.32 | 0.49 | 0.02 | 0.0775 | 0.345 | 0.4325 | 0.766523 | 0.386049 | 0.194491 | | 1 | 0.996511 | 0.778959 |
| Pcbd2 | A0A0R4J1R | -0.97 | -0.63 | -1.8 | -1.42 | -0.56 | -0.4 | -0.68 | -1.47 | -1.22 | -0.92 | -1.55 | -1.05 | -0.97 | -0.89 | -1.45 | -1.31 | 0.4275 | 0.02 | 0.05 | 0.227553 | 0.946229 | 0.868247 | | 1 | 0.996755 | 0.978872 |
| Pcbp1 | P60335 | -0.72 | -1.15 | -1.52 | -1.3 | -1.06 | -0.69 | -1.41 | -1.39 | -0.96 | -0.83 | -1.35 | -0.59 | -1.03 | -0.96 | -1.59 | -1.36 | 0.035 | 0.24 | -0.0625 | 0.889034 | 0.356833 | 0.806639 | | 1 | 0.996511 | 0.971715 |
| Pcbp2 | Q61990-2 | -1.95 | -1.78 | -1.12 | -1.2 | -2.05 | -1.55 | -1.4 | -1.21 | -2.61 | -3.63 | -2.43 | -2.12 | -3.06 | -3.36 | -3.07 | -2.68 | -0.04 | -1.185 | -1.53 | 0.885484 | 0.011172 | 0.000368 | | 1 | 0.996511 | 0.270112 |
| Pcbp2 | B2M1R7 | -1.92 | -2.24 | -1.13 | -1.71 | -2.22 | -2.02 | -2.15 | -1.54 | -2.93 | -2.5 | -1.22 | -1.93 | -2.56 | -2.82 | -1.42 | -1.82 | -0.2325 | -0.395 | -0.405 | 0.418878 | 0.355296 | 0.311387 | | 1 | 0.996511 | 0.834523 |
| Pcbp3 | P57722 | -1.03 | -1.27 | -2.14 | -1.2 | -1.26 | -1.2 | -2.43 | -2.42 | -1.05 | 3.59 | -1.87 | -0.95 | -1.56 | -1.08 | -2.37 | -2.46 | -0.4175 | 1.34 | -0.4575 | 0.31338 | 0.255681 | 0.270387 | | 1 | 0.996511 | 0.822084 |
| Pcbp4 | A0A0R4J04 | -0.99 | -1.12 | -2.21 | -2.44 | -1.76 | -1.28 | -2.6 | -3.02 | -1.15 | -1.35 | -2.25 | -0.93 | -0.58 | -1.2 | -2.89 | -2.06 | -0.475 | 0.27 | 0.0075 | 0.35451 | 0.546717 | 0.989586 | | 1 | 0.996511 | 0.996154 |
| Pcca | Q91ZA3 | 0.74 | 0.84 | 0.89 | 0.78 | 0.31 | 0.74 | 0.96 | 0.91 | 0.63 | 0.64 | 1 | 0.62 | 0.69 | 0.23 | 0.75 | 0.7 | -0.0825 | -0.09 | -0.22 | 0.665697 | 0.605155 | 0.280168 | | 1 | 0.996511 | 0.822084 |
| Pccb | Q99MN9 | 0.64 | 0.77 | 0.99 | 0.77 | 0.42 | 0.72 | 0.94 | 0.86 | 0.03 | -0.2 | 0.97 | 0.63 | -0.17 | -0.17 | 0.53 | 0.91 | -0.0575 | -0.435 | -0.5175 | 0.750446 | 0.156999 | 0.105172 | | 1 | 0.996511 | 0.703417 |
| Pcdh1 | E0CXD4 | 0 | 1.03 | 0 | 0.49 | 0.66 | -0.17 | 0.11 | 0.25 | 1.21 | 1.13 | 0.57 | 0.5 | 1.84 | 1.86 | 0.51 | 0.23 | -0.1675 | 0.4725 | 0.73 | 0.577957 | 0.153859 | 0.144087 | | 1 | 0.996511 | 0.743156 |
| Pcdh1 | F7BJK1 | 1.11 | 1.01 | 1.13 | 1.26 | 0.96 | 0.84 | 0.71 | 1.19 | 1.15 | 1.31 | 1.29 | 1.54 | 1.17 | 0.37 | 1.31 | 0.78 | -0.2025 | 0.195 | -0.22 | 0.249182 | 0.274008 | 0.39112 | | 1 | 0.996511 | 0.865375 |
| Pcdh1 | Q8CFX3 | 0.65 | 1.16 | 0.85 | 0.48 | 1.14 | 0.43 | 0.78 | 0.49 | 1.05 | 0.45 | 0.75 | 1.23 | 0.21 | 1.4 | 0.35 | 0.24 | -0.075 | 0.085 | -0.235 | 0.750847 | 0.733471 | 0.476442 | | 1 | 0.996511 | 0.892174 |
| Pcdh10 | Q925I8 | 1.56 | 1.47 | 0.64 | 0.59 | 1.62 | 1.69 | 0.51 | 0.73 | 1.94 | 1.01 | 0.65 | 1.98 | 1.6 | 1.96 | 0.53 | 0.35 | 0.0725 | 0.33 | 0.045 | 0.848388 | 0.423982 | 0.919922 | | 1 | 0.996511 | 0.98688 |
| Pcdh11x | F6ZNL5 | -1.36 | -1.26 | 0.66 | 0.47 | -1.63 | -0.88 | 0.16 | 0.79 | -0.53 | 0.72 | 0.69 | -0.76 | -0.55 | -0.42 | 0.95 | -0.19 | -0.0175 | 0.4025 | 0.32 | 0.97982 | 0.517028 | 0.590052 | | 1 | 0.996511 | 0.929142 |
| Pcdh15 | Q99PJ1 | 0.46 | -1.32 | 0.63 | 1.84 | -0.2 | 2.6 | 0.77 | 2.2 | 4.43 | 3.7 | 0.44 | -3.25 | 8.31 | 3.68 | 1.39 | 0.61 | 0.94 | 0.9275 | 3.095 | 0.274006 | 0.582085 | 0.086585 | | 1 | 0.996511 | 0.691619 |
| Pcdh17 | E9PXF0 | 0.62 | 0.9 | 0.55 | 0.71 | 0.94 | 0.77 | 0.86 | 0.73 | 0.84 | 0.6 | 0.69 | 1.14 | 0.57 | 0.69 | 1.04 | 0.85 | 0.13 | 0.1225 | 0.0925 | 0.414691 | 0.53303 | 0.640502 | | 1 | 0.996511 | 0.942906 |
| Pcdh19 | Q80TF3 | 0 | 0.49 | 0.3 | 0.74 | 0.61 | 0.15 | 0.98 | 0.88 | 0.02 | -0.49 | -0.41 | -0.58 | 0.89 | 0.64 | 1.05 | 1.07 | 0.2725 | -0.7475 | 0.53 | 0.298807 | 0.010797 | 0.043633 | | 1 | 0.996511 | 0.658454 |
| Pcdh7 | E9Q2S0 | 0.76 | 0.96 | 1.03 | 1.07 | 0.96 | 0.61 | 0.93 | 0.88 | 0.96 | 0.72 | 1.2 | 1.11 | 0.79 | 0.77 | 0.91 | 0.93 | -0.11 | 0.0425 | -0.105 | 0.507915 | 0.819405 | 0.556571 | | 1 | 0.996511 | 0.917092 |
| Pcdh8 | Q7TSK3 | 0.35 | 0.55 | 0.64 | 0.41 | 0.79 | 0.63 | 0.32 | 0.13 | 0.88 | 0.83 | 0.11 | 0.71 | 0.97 | 1.16 | 1.14 | 0.54 | -0.02 | 0.145 | 0.465 | 0.919198 | 0.523426 | 0.05405 | | 1 | 0.996511 | 0.661738 |
| Pcdh9 | A0A0A6YY( | 0.84 | 1.27 | 1.03 | 1.03 | 1.51 | 1.03 | 0.88 | 0.87 | 1.42 | 0.64 | 1.19 | 1.19 | 1.13 | 1.05 | 0.74 | 0.84 | 0.03 | 0.0675 | -0.1025 | 0.883496 | 0.762848 | 0.604762 | | 1 | 0.996511 | 0.932185 |
| Pcdh9 | A0A0A6YY3 | 1.29 | 3.09 | 1.38 | 1.12 | 0.11 | 1.1 | 1.05 | 1.67 | 1.8 | 1.05 | 1.58 | 2.59 | 2.5 | 2.9 | 0.98 | 1.06 | -0.7375 | 0.035 | 0.14 | 0.178723 | 0.946192 | 0.820287 | | 1 | 0.996755 | 0.972768 |
| Pcdha12 | Q91Y18 | 0.46 | 0.24 | 0.38 | 0.72 | 0.09 | 0.96 | 0.27 | 0.31 | 1.16 | 1.2 | -0.08 | 1.63 | 1.47 | 0.02 | 0.18 | -0.18 | -0.0425 | 0.5275 | -0.0775 | 0.85553 | 0.178723 | 0.837212 | | 1 | 0.996511 | 0.975831 |
| Pcdhac2 | Q91Y09 | 0.43 | 0.67 | 0.24 | 0.69 | 0.58 | 0.3 | 0.33 | 0.49 | 0.94 | 0.59 | 0.37 | 0.55 | 0.65 | 0.33 | 0.04 | 0.02 | -0.0825 | 0.105 | -0.2475 | 0.639056 | 0.612469 | 0.294793 | | 1 | 0.996511 | 0.825612 |
| Pcdhb12 | Q91Y07 | 0.42 | 0.23 | -0.7 | 0.79 | 1.09 | 0.12 | -0.55 | -1.55 | -0.15 | 1.72 | -1.46 | 0.51 | -0.77 | 2.14 | -0.72 | -0.85 | -0.4075 | -0.03 | -0.235 | 0.49192 | 0.964227 | 0.745098 | | 1 | 0.996755 | 0.959729 |
| Pcdhb14 | Q6PB90 | 0.29 | 0.44 | 0.9 | 1.21 | 0.94 | -0.12 | 1.12 | 0.87 | 0.1 | 1.15 | 0.58 | 0.26 | -0.16 | 0.47 | 0.2 | 0.76 | -0.0075 | -0.1875 | -0.3925 | 0.982308 | 0.555922 | 0.214146 | | 1 | 0.996511 | 0.786931 |
| Pcdhb17 | Q91VD8 | 0.87 | 1.76 | -0.28 | -0.71 | 1.49 | 4.36 | -0.14 | -2.89 | 1.23 | -2.77 | -0.38 | 0.73 | 0.53 | 0.93 | -0.83 | 0.08 | 0.295 | -0.7075 | -0.2325 | 0.838166 | 0.462041 | 0.707645 | | 1 | 0.996511 | 0.957213 |
| Pcdhb18 | Q91Y02 | 0.66 | 1.39 | 0.53 | 0.87 | 0.48 | 0.64 | 0.22 | 0.52 | 1.3 | 1.2 | 0.59 | 1.28 | 1 | 1.62 | 0.68 | 1.3 | -0.3975 | 0.23 | 0.2875 | 0.109994 | 0.403536 | 0.338923 | | 1 | 0.996511 | 0.847578 |
| Pcdhb20 | Q91XZ9 | 0.54 | 0.21 | 0.51 | 0.03 | -0.56 | -0.1 | -0.14 | 0.23 | 0.04 | -0.92 | 0.47 | 0.21 | 0.57 | 0.65 | 0.08 | 0 | -0.465 | -0.3725 | 0.0025 | 0.064578 | 0.271481 | 0.991801 | | 1 | 0.996511 | 0.996154 |
| Pcdhb21 | Q91V48 | 0.49 | 0.22 | -0.7 | -0.1 | 0.05 | -0.07 | -0.04 | -0.86 | 0.49 | -0.73 | -0.97 | -0.12 | -1.14 | -0.5 | -1.38 | -0.6 | -0.2075 | -0.31 | -0.8825 | 0.526785 | 0.443905 | 0.0254 | | 1 | 0.996511 | 0.633075 |
| Pcdhb9 | E9Q5G2 | 1.05 | 0.41 | 0.39 | 0.37 | 1.04 | 0.15 | 0.47 | 0.83 | 1.09 | -0.55 | -0.47 | 0.73 | -0.53 | 1.7 | -0.42 | 0.12 | 0.0675 | -0.355 | -0.3375 | 0.798627 | 0.413093 | 0.507907 | | 1 | 0.996511 | 0.902804 |
| Pcdhga10 | Q91XY9 | 0.55 | 0.21 | 0.72 | 1.44 | 1.28 | 0.6 | 1.19 | 1.15 | 0.7 | 0.36 | 0.78 | 0.65 | 0.42 | 1.21 | 1.02 | 1.14 | 0.325 | -0.1075 | 0.2175 | 0.294673 | 0.707393 | 0.504603 | | 1 | 0.996511 | 0.901002 |
| Pcdhga11 | Q91XY8 | 0.83 | 1.83 | 1.08 | 1.11 | 1.38 | 1.06 | 0.92 | 1.17 | 1 | 0.57 | 1.13 | 1.42 | 0.72 | 1.48 | 0.82 | 0.72 | -0.08 | -0.1825 | -0.2775 | 0.748182 | 0.531085 | 0.362129 | | 1 | 0.996511 | 0.858475 |
| Pcdhga12 | Q91XY7 | 0.02 | 0.69 | 0.89 | 0.55 | 0.98 | 0.41 | 0.89 | 1.23 | 0.09 | -0.24 | 0.66 | 0.4 | 0.39 | 0.35 | 0.75 | 0.72 | 3.40E-01 | -0.31 | 0.015 | 0.215834 | 0.287264 | 0.951826 | | 1 | 0.996511 | 0.992476 |
| Pcdhga3 | Q91XY5 | 0.22 | 0.67 | 0.47 | 0.26 | 0.58 | 0.42 | 0.92 | 0.52 | 0.68 | 0.18 | 0.6 | 0.69 | 0.58 | 1.22 | 0.85 | 0.36 | 0.205 | 0.1325 | 0.3475 | 0.29555 | 0.523806 | 0.184227 | | 1 | 0.996511 | 0.776843 |
| Pcdhga5 | Q91XY3 | 1.02 | -0.51 | 0.47 | -0.08 | 0.34 | 0.27 | 0.52 | 0.48 | 0.53 | 2.4 | 0.14 | 0.27 | 0.74 | 0.4 | 0.14 | 1.03 | 0.1775 | 0.61 | 0.3525 | 0.591809 | 0.303818 | 0.360843 | | 1 | 0.996511 | 0.857677 |
| Pcdhga6 | Q91XY2 | -0.3 | 0.02 | 0.5 | 0.2 | -0.17 | -0.66 | 0.28 | 0.31 | 0.06 | 0.63 | 0.64 | -0.2 | -0.53 | 0.47 | -0.15 | 0.47 | -0.165 | 0.1775 | -0.04 | 0.565915 | 0.531989 | 0.896662 | | 1 | 0.996511 | 0.983509 |
| Pcdhga7 | Q6DD96 | 1.28 | 0.39 | 1.07 | 0.78 | 1.65 | 0.98 | 0.39 | 0.21 | 2 | 0.44 | 0.45 | 0.44 | 2 | -0.38 | -0.31 | 0.4 | -0.0725 | -0.0475 | -0.4525 | 0.841193 | 0.908412 | 0.412925 | | 1 | 0.996744 | 0.874811 |
| Pcdhga8 | Q91XY0 | 0.42 | 0.18 | 0.96 | 0.67 | 0.68 | 0.18 | 0.75 | 0.18 | -0.17 | 0.17 | 0.97 | 0.56 | -0.64 | -0.07 | 0.76 | 0.36 | -0.11 | -0.175 | -0.455 | 0.65229 | 0.566953 | 0.207279 | | 1 | 0.996511 | 0.785338 |
| Pcdhgb1 | Q91XX8 | -0.18 | 0.93 | 1.82 | 1.46 | 0.46 | 0.69 | 3.28 | 2.24 | 0.14 | 2.06 | 2.64 | 0.03 | 0.09 | 0.73 | 3.13 | 2.22 | 0.66 | 0.21 | 0.535 | 0.372037 | 0.771442 | 0.476359 | | 1 | 0.996511 | 0.892174 |
| Pcdhgb2 | Q91XX7 | -0.01 | 1.01 | 0.57 | 1.07 | 1.46 | 0.58 | 0.91 | 0.46 | 0.49 | 0.99 | 1.09 | 0.54 | 0.22 | 1.47 | 1.01 | -0.39 | 0.1925 | 0.1175 | -0.0825 | 0.559361 | 0.695534 | 0.856231 | | 1 | 0.996511 | 0.976225 |
| Pcdhgb5 | Q91XX5 | 1.34 | 1.42 | 1.23 | 1.5 | 1.79 | 0.59 | 1.35 | 0.81 | 1.16 | 1.45 | 0.98 | 1.11 | 0.18 | 1.89 | 0.65 | -0.2 | -0.2375 | -0.1975 | -0.7425 | 0.405822 | 0.290892 | 0.115455 | | 1 | 0.996511 | 0.717287 |
| Pcdhgb6 | Q91XX4 | 1.18 | 1.12 | 1.32 | 1.21 | 1.05 | 0.57 | 1.07 | 1.14 | 0.44 | 1.02 | 1.13 | 1.03 | 0.15 | 0.18 | 0.74 | 0.88 | -0.25 | -0.3025 | -0.72 | 0.191901 | 0.169542 | 0.012804 | | 1 | 0.996511 | 0.573604 |
| Pcdhgc3 | Q91XX1 | 0.82 | 1.18 | 0.95 | 0.67 | 0.88 | 0.85 | 0.19 | 0.6 | 1.47 | 0.89 | 0.6 | 1.07 | 1.77 | 1.28 | 0.73 | 0.24 | -0.275 | 0.1025 | 0.1 | 0.227212 | 0.670115 | 0.773777 | | 1 | 0.996511 | 0.966062 |
| Pcdhgc5 | Q91XW9 | 0.1 | 0.24 | 0.5 | 0.49 | 0.21 | 0.06 | 0.38 | 0.45 | 0.42 | 0.49 | 0.57 | 0.09 | 0.35 | 0.3 | 0.38 | 0.25 | -0.0575 | 0.06 | -0.0125 | 0.74731 | 0.760232 | 0.946204 | | 1 | 0.996511 | 0.992376 |
| Pcf11 | G3X9Z4 | -2.83 | -2.88 | -2.08 | -1.41 | -2.87 | -3.38 | -2.3 | -2.42 | -2.82 | -0.43 | -1.85 | -2.54 | -2.46 | -1.86 | -2.54 | -2.57 | -0.4425 | 0.39 | -0.0575 | 0.288013 | 0.510772 | 0.8788 | | 1 | 0.996511 | 0.979626 |
| Pcgf2 | P23798 | 0.07 | -0.39 | 0.13 | -0.11 | -0.71 | -0.41 | -0.09 | -0.02 | -0.56 | -0.36 | -0.01 | -0.7 | -0.02 | 0.22 | 0.48 | -0.16 | -0.2325 | -0.3325 | 0.205 | 0.307373 | 0.164427 | 0.380577 | | 1 | 0.996511 | 0.860984 |
| Pcid2 | Q8BFV2 | -1.37 | -1.17 | -0.44 | -0.18 | -2.16 | -0.88 | -0.83 | -0.23 | -0.16 | -0.54 | -0.46 | -1.51 | 0.87 | -2.35 | -0.26 | -0.68 | -0.235 | 0.1225 | 0.185 | 0.612714 | 0.755021 | 0.779736 | | 1 | 0.996511 | 0.967289 |
| Pcif1 | P59114 | -7.16 | -6.62 | -3.98 | -4.84 | -7.55 | -7.89 | -3.92 | -4.27 | -7.58 | -1.4 | -3.76 | -6.22 | -6.04 | -7.43 | -3.55 | -4.16 | -0.2575 | 0.91 | 0.355 | 0.823446 | 0.517982 | 0.732419 | | 1 | 0.996511 | 0.958635 |
| Pck2 | A0A0R4J0G | -0.12 | -0.26 | -0.2 | 0.01 | -0.21 | -0.26 | -0.01 | -0.16 | -0.01 | 0.03 | 0.13 | -0.25 | -0.09 | 0.39 | -0.02 | 0.15 | -0.0175 | 0.1175 | 0.25 | 0.908285 | 0.503506 | 0.21994 | | 1 | 0.996511 | 0.787447 |
| Pclo | Q9QYX7 | 0.84 | 0.91 | 1.19 | 1.15 | 0.57 | 0.7 | 0.5 | 0.93 | 1.04 | 0.96 | 1.18 | 0.82 | 1.03 | 0.94 | 0.86 | 0.64 | -0.3475 | -0.0225 | -0.155 | 0.075538 | 0.900784 | 0.432158 | | 1 | 0.996511 | 0.879165 |
| Pcm1 | Q9R0L6-2 | -0.61 | -0.33 | -0.41 | -0.48 | -0.43 | -0.6 | -0.52 | -0.96 | -0.15 | -0.37 | -0.61 | -0.56 | 0.46 | -0.03 | -0.03 | -0.31 | -0.17 | 0.035 | 0.48 | 0.351126 | 0.848568 | 0.054372 | | 1 | 0.996511 | 0.661738 |
| Pcmt1 | F7D432 | 0.93 | 0.85 | 0.63 | 0.72 | 0.32 | 0.63 | 0.14 | 0.71 | 0.96 | -0.45 | 0.37 | 1.09 | 0.43 | 0.95 | -0.07 | 0.07 | -0.3325 | -0.29 | -0.4375 | 0.105513 | 0.417065 | 0.124737 | | 1 | 0.996511 | 0.721448 |
| Pcmtd1 | P59913 | 0.95 | 0.29 | 0.13 | -0.64 | 0.45 | 0.77 | 0.04 | -0.72 | 0.04 | -1.6 | -0.09 | 0.91 | -0.39 | -0.34 | 0.12 | 0.41 | -0.0475 | -0.3675 | -0.2325 | 0.911748 | 0.521417 | 0.535227 | | 1 | 0.996511 | 0.912212 |
| Pcmtd2 | Q8BHD8 | -1.01 | -0.52 | -0.63 | 0.37 | -0.76 | -1.13 | 0.03 | -0.41 | -0.24 | 1.03 | -0.46 | -0.56 | -0.64 | 0.28 | -0.37 | -0.65 | -0.12 | 0.39 | 0.1025 | 0.743277 | 0.389801 | 0.775935 | | 1 | 0.996511 | 0.966799 |
| Pcna | P17918 | -1.57 | -1.97 | -1.93 | -1.55 | -1.76 | -1.97 | -2.01 | -2.25 | -2.29 | -1.08 | -2.09 | -1.4 | -1.69 | -2.52 | -2.07 | -2.36 | -0.2425 | 0.04 | -0.405 | 0.223257 | 0.897135 | 0.130326 | | 1 | 0.996511 | 0.724262 |
| Pcnp | Q6P8I4 | -2.05 | -2.32 | -2.21 | -2.62 | -2.29 | -2.57 | -2.57 | -2.59 | -2.38 | -0.45 | -2.28 | -1.81 | -1.55 | -1.65 | -1.89 | -2.36 | -0.205 | 0.57 | 0.4375 | 0.280506 | 0.212536 | 0.107943 | | 1 | 0.996511 | 0.706427 |
| Pcnt | A0A1W2P7 | 0.72 | 0.19 | -0.47 | 0.34 | -0.29 | -0.2 | -0.13 | 0.08 | 0.33 | 0.79 | 0.28 | 0.56 | 0.37 | 1.14 | 0.79 | 0.55 | -0.33 | 0.295 | 0.5175 | 0.237352 | 0.315879 | 0.120963 | | 1 | 0.996511 | 0.721448 |
| Pcnx | A0A1Y7VKI | 0.26 | 0.81 | 0.18 | 0.57 | 0.44 | 0.51 | -0.86 | -0.2 | 0.9 | 0.48 | 0.05 | 0.96 | 1.26 | 0.46 | 0.11 | -0.17 | -0.4825 | 0.1425 | -0.04 | 0.180399 | 0.602536 | 0.906817 | | 1 | 0.996511 | 0.986458 |
| Pcnx2 | Q5DU28 | 0.81 | 0.47 | 1.04 | 0.45 | 0.46 | 0.81 | 0.34 | 1.03 | 0.51 | 0.3 | 0.94 | 0.71 | 1.74 | 0.25 | 0.86 | 0.79 | -0.0325 | -0.0775 | 0.2175 | 0.888436 | 0.736521 | 0.527288 | | 1 | 0.996511 | 0.90845 |
| Pcnx3 | Q8VI59 | 0.24 | 0.38 | -0.3 | -0.36 | 0.53 | 0.3 | -0.13 | -0.37 | 0.38 | -0.57 | -0.29 | 0.66 | 0.34 | 0.73 | -0.7 | -0.1 | 0.0925 | 0.055 | 0.0775 | 0.741503 | 0.870579 | 0.827576 | | 1 | 0.996511 | 0.973757 |
| Pcolce | D3YWH0 | -2.13 | -2.15 | -4.04 | -3.41 | -2.77 | -1.78 | -3.44 | -3.87 | -0.48 | -1.02 | -2.08 | -0.96 | 0.91 | -2.51 | -1.69 | -3.03 | -0.0325 | 1.7975 | 1.3525 | 0.956771 | 0.008584 | 0.155907 | | 1 | 0.996511 | 0.751421 |
| Pcp4l1 | Q6W8Q3 | -0.12 | -0.92 | -0.56 | 0.16 | -0.85 | -1.64 | -0.82 | -0.21 | -0.59 | 1.11 | -0.99 | -0.1 | -0.72 | -0.05 | -0.19 | -0.85 | -0.52 | 0.2175 | -0.0925 | 0.175248 | 0.652246 | 0.770222 | | 1 | 0.996511 | 0.965933 |
| Pcsk1 | P63239 | -0.14 | -0.27 | 0.34 | 0.2 | -0.24 | 0.32 | 0.55 | 0.72 | 0.75 | 0.21 | 0.68 | 0.22 | 1.6 | -0.14 | 1.65 | 1.33 | 0.305 | 0.4325 | 1.0775 | 0.262529 | 0.091833 | 0.030263 | | 1 | 0.996511 | 0.641481 |
| Pcsk1n | Q9QXV0 | 0.26 | 0.42 | -0.07 | 0.44 | 0.03 | 0.16 | 0.86 | 0.6 | 0.6 | 0.92 | -0.14 | 0.15 | 0.49 | 0.83 | 1.54 | 0.92 | 0.15 | 0.12 | 0.6825 | 0.538432 | 0.665737 | 0.03148 | | 1 | 0.996511 | 0.641974 |
| Pcsk2 | P21661 | 0.45 | 0.23 | -0.06 | 0.07 | 0.59 | 0.37 | 0.67 | 0.29 | 0.44 | -0.21 | 0.11 | 0.39 | 0.45 | 0.93 | 1.01 | 0.65 | 0.3075 | 0.01 | 0.5875 | 0.122338 | 0.964022 | 0.024254 | | 1 | 0.996755 | 0.626254 |
| Pcx | G5E8R3 | 1.28 | 0.85 | 0.77 | 0.82 | 0.6 | 0.79 | 1.23 | 0.77 | 0.18 | 0.54 | 0.94 | 0.55 | -0.35 | 0.49 | 0.49 | 1.03 | -0.0825 | -0.3775 | -0.515 | 0.692556 | 0.125096 | 0.130138 | | 1 | 0.996511 | 0.723659 |
| Pcyox1 | Q9CQF9 | 0.12 | 0.2 | 0.15 | 0.09 | 0.61 | 0.1 | 1.06 | 0.3 | -0.21 | -0.43 | 0.2 | 0.04 | 0.05 | 0.34 | 0.89 | 1.05 | 0.3775 | -0.24 | 0.4425 | 0.12776 | 0.237421 | 0.119212 | | 1 | 0.996511 | 0.721448 |
| Pcyox1l | Q8C7K6 | -0.3 | 0.08 | 0.39 | 0.34 | 0.47 | -0.27 | 0.74 | 0.4 | -0.44 | -1.02 | 0.31 | -0.32 | -0.34 | 0.48 | 0.69 | 0.64 | 0.2075 | -0.495 | 0.24 | 0.452893 | 0.144346 | 0.432764 | | 1 | 0.996511 | 0.879365 |
| Pcyt1a | P49586 | -2.47 | -3.13 | -2.74 | -2.99 | -2.56 | -3.14 | -2.69 | -3.2 | -2.98 | -0.86 | -2.43 | -2.15 | -1.98 | -2.37 | -2.7 | -2.66 | -0.065 | 0.7275 | 0.405 | 0.781483 | 0.129343 | 0.136105 | | 1 | 0.996511 | 0.731656 |
| Pcyt1b | Q811Q9 | 0.41 | 0.24 | 0.08 | -0.42 | -0.08 | 0.17 | -0.79 | -0.47 | -0.38 | -0.63 | 0.02 | 1.09 | 0.55 | -0.49 | -0.02 | -0.28 | -0.37 | -0.0525 | -0.1375 | 0.209476 | 0.895752 | 0.648955 | | 1 | 0.996511 | 0.946117 |
| Pcyt2 | A2ABY3 | -0.3 | -0.06 | -0.36 | -0.41 | 0.15 | -0.34 | -0.13 | -0.36 | -0.11 | -0.57 | -0.26 | -0.05 | 0.03 | 0.04 | -0.12 | 0.02 | 0.1125 | 0.035 | 0.275 | 0.545484 | 0.856672 | 0.150504 | | 1 | 0.996511 | 0.749304 |
| Pdap1 | Q3UHX2 | 0.07 | -0.25 | 0.03 | -0.12 | -0.79 | 0.02 | -0.14 | 0 | 0.79 | -0.72 | 0.01 | 0.34 | 1.22 | -0.13 | -0.04 | -0.39 | -0.16 | 0.1725 | 0.2325 | 0.484686 | 0.598913 | 0.524936 | | 1 | 0.996511 | 0.908107 |
| Pdcd10 | Q8VE70 | -1.29 | -0.97 | -0.4 | -0.28 | -1.05 | -1.32 | -0.44 | -0.4 | -1.2 | -0.37 | -0.38 | -1.47 | -1.71 | -0.39 | -0.98 | -1.02 | -0.0675 | -0.12 | -0.29 | 0.83404 | 0.739616 | 0.424544 | | 1 | 0.996511 | 0.876965 |
| Pdcd11 | Q6NS46 | 0.36 | 0.28 | 0.69 | 0.45 | 0.11 | 0.17 | 0.37 | 0.26 | 0.48 | 0.8 | 0.32 | 0.04 | 0.11 | 0.45 | -0.03 | 0.65 | -0.2175 | -0.035 | -0.15 | 0.207077 | 0.873223 | 0.51229 | | 1 | 0.996511 | 0.903993 |
| Pdcd2 | A2T4N6 | -1.02 | 0.89 | -0.85 | -0.03 | 1.25 | -0.76 | 0.08 | -0.46 | 0.05 | -0.24 | -0.28 | 0.62 | 0.41 | 1.18 | 0.52 | 0.96 | 0.28 | 0.29 | 1.02 | 0.625367 | 0.528949 | 0.046095 | | 1 | 0.996511 | 0.658454 |
| Pdcd4 | Q61823 | -0.77 | -0.87 | -1.07 | -1.1 | -0.17 | -0.96 | -0.83 | -1.11 | -0.55 | -0.58 | -0.91 | -0.78 | -0.65 | -0.32 | -1.12 | -0.45 | 0.185 | 0.2475 | 0.3175 | 0.445498 | 0.196024 | 0.198759 | | 1 | 0.996511 | 0.78004 |
| Pdcd5 | P56812 | 1.7 | 0.62 | 1.09 | 0.44 | 0.66 | 2.14 | 1.09 | 0.95 | 0.41 | -0.84 | 1.13 | 1.95 | 0.98 | -0.21 | 1.05 | 1.47 | 0.2475 | -0.3 | -0.14 | 0.543876 | 0.619746 | 0.747978 | | 1 | 0.996511 | 0.960744 |
| Pdcd6 | P12815-2 | -0.57 | -0.29 | -0.04 | -0.54 | -0.01 | -0.68 | -0.51 | -0.54 | -1.18 | -0.81 | -0.7 | -0.89 | -0.86 | -0.51 | -0.14 | -0.41 | -0.075 | -0.535 | -0.12 | 0.729978 | 0.027499 | 0.612368 | | 1 | 0.996511 | 0.935166 |
| Pdcd6ip | Q9WU78 | 0.69 | 0.36 | 0.62 | 0.37 | 0.02 | 0.45 | 0.38 | 0.44 | 0.08 | 0.22 | 0.3 | 0.61 | 0.1 | -0.23 | 0.18 | 0.18 | -0.1875 | -0.2075 | -0.4525 | 0.309979 | 0.301884 | 0.045306 | | 1 | 0.996511 | 0.658454 |
| Pdcd6ip | Q9WU78-3 | -1.5 | -0.54 | -0.78 | -1.52 | -1.37 | -2.45 | -0.77 | -1.15 | -2.55 | -2.28 | -1.35 | -0.79 | -2.66 | -2.87 | -2.52 | -0.94 | -0.35 | -0.6575 | -1.1625 | 0.405943 | 0.169843 | 0.035161 | | 1 | 0.996511 | 0.651434 |
| Pdcl | Q9DBX2 | -0.23 | -0.27 | 0.01 | -0.25 | 0.02 | -0.48 | -0.03 | -0.75 | 0.2 | -0.37 | -0.08 | -0.3 | 0.68 | 0.63 | -0.14 | -0.37 | -0.125 | 0.0475 | 0.385 | 0.572953 | 0.80895 | 0.208573 | | 1 | 0.996511 | 0.786218 |
| Pdcl3 | Q8BVF2 | -0.38 | 0.2 | -0.75 | -0.9 | 0.31 | -0.64 | -0.85 | -1.17 | -0.31 | -0.05 | -0.94 | -0.04 | -0.55 | 0.42 | -0.85 | -0.9 | -0.13 | 0.1225 | -0.0125 | 0.733759 | 0.70544 | 0.973851 | | 1 | 0.996511 | 0.992977 |
| Pde10a | Q8CA95-3 | 0.87 | -0.13 | 0.95 | 1.67 | -0.75 | -0.64 | 2.06 | 1.39 | 0.39 | 0.69 | 0.4 | 0.39 | 0.3 | 0.07 | 1.56 | 1.52 | -0.325 | -0.3725 | 0.0225 | 0.656113 | 0.324136 | 0.964323 | | 1 | 0.996511 | 0.9927 |
| Pde11a | P0C1Q2 | 1.04 | 0.92 | 1.05 | 1.71 | 0.94 | 0.73 | 1.59 | 1.76 | 0.78 | 0.94 | 1.44 | 1.16 | 0.38 | 1.14 | 1.27 | 1.24 | 0.075 | -0.1 | -0.1725 | 0.80487 | 0.692408 | 0.559281 | | 1 | 0.996511 | 0.9184 |
| Pde12 | Q3TIU4 | -0.19 | -0.17 | -0.41 | -0.21 | -0.24 | -0.53 | -0.24 | -0.2 | -0.19 | -0.5 | -0.68 | -0.22 | -0.05 | -0.31 | -0.58 | -0.63 | -0.0575 | -0.1525 | -0.1475 | 0.717768 | 0.426993 | 0.48136 | | 1 | 0.996511 | 0.893906 |
| Pde1a | Q61481 | -1.23 | -0.91 | -0.67 | -0.98 | -0.9 | -0.91 | -0.44 | -0.86 | -0.96 | -1.33 | -0.56 | -0.94 | -0.93 | -0.89 | -1.06 | -0.66 | 0.17 | 1.23E-16 | 0.0625 | 0.397159 | 1 | 0.760679 | | 1 | 1 | 0.963123 |
| Pde1b | Q01065 | -0.96 | -1.48 | -1.43 | -1.23 | -1.37 | -1.82 | -0.96 | -1.48 | -0.63 | -0.91 | -1.8 | -0.91 | -0.87 | -0.45 | -1.36 | -1.4 | -0.1325 | 0.2125 | 0.255 | 0.571064 | 0.470213 | 0.367765 | | 1 | 0.996511 | 0.859245 |

| | | | | | | | | | | | | | | | | | | | | | | | | | | | |
|---|---|---|---|---|---|---|---|---|---|---|---|---|---|---|---|---|---|---|---|---|---|---|---|---|---|---|---|
| Pde1c | Q8CDV2 | 1.41 | 0.29 | -4.62 | -3.02 | -0.76 | 0.49 | -4.08 | -3.91 | 0.79 | -2.38 | -3.5 | 1.36 | 0.89 | 1.14 | -3.02 | -5.43 | -0.58 | 0.5525 | -0.12 | 0.720637 | 0.737104 | 0.949098 | | 1 | 0.996511 | 0.992476 |
| Pde2a | Q5PR72 | 1.39 | 0.91 | 1.67 | 2.05 | -1.14 | 1.68 | -0.44 | 1.04 | 2.28 | 0.59 | 1.16 | 0.83 | 2.76 | -2.42 | 1.48 | 0.88 | -1.22 | -0.29 | -0.83 | 0.080394 | 0.497316 | 0.420447 | | 1 | 0.996511 | 0.87558 |
| Pde3b | E9QLQ3 | -0.52 | 1.09 | -0.75 | 0.14 | -0.37 | 0.14 | -0.49 | 0 | 0.74 | 1.29 | 0.25 | 1.28 | 0.47 | 0.84 | 0.42 | 0.37 | -0.17 | 0.9 | 0.535 | 0.681219 | 0.073813 | 0.213095 | | 1 | 0.996511 | 0.786931 |
| Pde4a | O89084 | 0.59 | 0.39 | -0.03 | 0.11 | 0.16 | 0.38 | 0.28 | 0.01 | 0.77 | 0.38 | 0.16 | 0.79 | 0.61 | 0.47 | -0.09 | 0.17 | -0.0575 | 0.26 | 0.025 | 0.769288 | 0.288892 | 0.918792 | | 1 | 0.996511 | 0.986868 |
| Pde4b | Q8VBU5 | 0.6 | 0.43 | 0.25 | 0.5 | 0.14 | 0.29 | 0.06 | 0.34 | 0.36 | 0.3 | 0.28 | 0.76 | 0.35 | 0.04 | 0 | -0.07 | -0.2375 | -0.02 | -0.365 | 0.164215 | 0.916598 | 0.085468 | | 1 | 0.996755 | 0.691619 |
| Pde4d | E9PVU7 | 2.66 | 2.01 | 0.56 | 0.47 | 2.3 | 3.09 | 0.46 | 0.6 | 0.55 | 0.61 | 0.66 | 2.87 | 1.1 | 0.46 | 0.48 | 0.41 | 0.1875 | -0.2525 | -0.8125 | 0.805727 | 0.723689 | 0.148033 | | 1 | 0.996511 | 0.746406 |
| Pde4dip | Q80YT7-2 | -1.51 | -1.14 | -1.58 | -1.6 | -2.17 | -2.14 | -2.29 | -2.07 | -0.49 | -0.75 | -2.08 | -1.48 | -0.11 | 0.05 | -2.13 | -1.92 | -0.71 | 0.2575 | 0.43 | 0.002456 | 0.487372 | 0.437463 | | 1 | 0.996511 | 0.883161 |
| Pde5a | A0A0G2JF6 | -1.46 | -0.97 | 0.64 | 0.29 | -0.9 | -0.5 | -0.29 | 0 | -0.43 | 1.07 | -0.02 | -0.33 | -1.18 | -0.3 | 0.2 | 0.94 | -0.0475 | 0.4475 | 0.29 | 0.923329 | 0.432372 | 0.637977 | | 1 | 0.996511 | 0.942804 |
| Pde6d | O55057 | 0.99 | 1.03 | 0.32 | 0.6 | 1.16 | 0.92 | 0.95 | 0.59 | -0.79 | 0.04 | 0.58 | 1.2 | -0.54 | -2.04 | 0.86 | 0.75 | 0.17 | -0.4775 | -0.9775 | 0.46076 | 0.284353 | 0.151384 | | 1 | 0.996511 | 0.749435 |
| Pde7a | P70453 | -1.95 | 0.09 | -2.45 | -1.99 | -1.58 | 0.86 | -2.21 | -2.38 | -0.91 | -0.48 | -2.06 | -1.03 | -2.19 | -0.34 | -2.25 | -2.55 | 0.2475 | 0.455 | -0.2575 | 0.770163 | 0.458163 | 0.709234 | | 1 | 0.996511 | 0.957656 |
| Pde7b | E9PW23 | -1.15 | -1.29 | -0.46 | -1.55 | -1.36 | -1.82 | -0.09 | -0.76 | -0.96 | -0.39 | -1.69 | -1.44 | -1.37 | -0.7 | -1.39 | -1.87 | 0.105 | -0.0075 | -0.22 | 0.800183 | 0.983348 | 0.517993 | | 1 | 0.998773 | 0.90668 |
| Pde8a | O88502 | 0.74 | 2.59 | -1.02 | -1.74 | 4.01 | 1.6 | -1.23 | -1.64 | 0.44 | -1.27 | -1.3 | 0.17 | 0.99 | 1.58 | -1.02 | -1.41 | 0.5425 | -0.6325 | -0.1075 | 0.711667 | 0.517264 | 0.921047 | | 1 | 0.996511 | 0.987433 |
| Pde8b | E9Q8J6 | 0.27 | -0.05 | 0.61 | 1.19 | -0.48 | 0.44 | 0.14 | 0.48 | 1.58 | 2.4 | 0.49 | -0.05 | 1.75 | 0.46 | 2.31 | -0.26 | -0.36 | 0.6 | 0.56 | 0.299021 | 0.301529 | 0.356288 | | 1 | 0.996511 | 0.855416 |
| Pde8b | E9PX38 | 0.64 | 0.38 | 1.61 | 1.61 | 0.85 | 1 | 1.84 | 1.76 | 0.49 | 0.53 | 1.5 | 0.35 | 0.5 | 0.51 | 1.38 | 1.71 | 0.3025 | -0.3425 | -0.035 | 0.444659 | 0.400053 | 0.934269 | | 1 | 0.996511 | 0.989767 |
| Pde9a | A0A494BAI | 0.8 | 1.63 | 0.51 | 0.78 | 0.06 | 0.38 | 1.07 | 0.71 | 0.69 | 0.74 | 0.61 | 0.47 | 1.02 | 1.35 | 0.5 | 1.45 | -0.375 | -0.3025 | 0.15 | 0.258463 | 0.274379 | 0.648837 | | 1 | 0.996511 | 0.946117 |
| Pdf | S4R2K0 | 0.21 | 0.25 | 0.96 | 1.04 | 0.48 | 0.66 | 1 | 1.14 | 0.87 | 1.04 | 1.14 | 0.52 | 1.04 | 1.48 | 0.74 | 1.2 | 0.205 | 0.2775 | 0.5 | 0.462252 | 0.327681 | 0.110472 | | 1 | 0.996511 | 0.711551 |
| Pdgfra | P26618 | 0.51 | 1.23 | -0.15 | 0.29 | 0.59 | 0.13 | 0.09 | -0.02 | 1.07 | 1.08 | 0.31 | 0.88 | 0.38 | 1.04 | 0.14 | 0.37 | -0.2725 | 0.365 | 0.0125 | 0.394056 | 0.294721 | 0.97115 | | 1 | 0.996511 | 0.992803 |
| Pdgfrb | E9QN12 | -0.35 | -0.17 | 0.72 | 0.77 | 0.45 | -0.61 | 0.58 | 0.76 | -0.62 | 0.02 | 0.52 | -0.46 | -0.46 | -0.26 | 0.77 | 0.68 | 0.0525 | -0.3775 | -0.06 | 0.895709 | 0.330282 | 0.884451 | | 1 | 0.996511 | 0.979767 |
| Pdha1 | P35486 | 0.93 | 0.62 | 0.46 | 1.01 | 0.3 | 0.65 | 0.27 | 0.83 | 0.38 | 0.53 | 0.6 | 0.76 | 0.3 | -0.45 | 0.08 | 0.15 | -0.2425 | -0.1875 | -0.735 | 0.275587 | 0.362456 | 0.014133 | | 1 | 0.996511 | 0.589387 |
| Pdhb | Q9D051 | 0.31 | 0.48 | 0.67 | 0.95 | 0.11 | 0.35 | 0.44 | 0.74 | 0.2 | 0.54 | 0.77 | 0.23 | -0.14 | 0.53 | 0.05 | 0.26 | -0.1925 | -0.1675 | -0.4275 | 0.381736 | 0.467135 | 0.100379 | | 1 | 0.996511 | 0.698156 |
| Pdhx | Q8BKZ9 | 1.36 | 1.25 | 0.66 | 0.93 | 0.86 | 1.1 | 0.46 | 0.67 | 1.65 | 0.98 | 0.69 | 1.19 | 1.54 | 1.54 | 0.42 | 0.44 | -0.2775 | 0.0775 | -0.065 | 0.245865 | 0.775988 | 0.854364 | | 1 | 0.996511 | 0.976225 |
| Pdia3 | P27773 | 0.71 | 0.04 | -0.04 | 0.16 | 0.58 | 0.68 | 0.7 | 0.37 | -0.19 | 0.18 | -0.08 | 0.96 | 0.17 | 0.1 | 1.01 | 0.81 | 0.365 | 7.85E-17 | 0.305 | 0.113151 | 1 | 0.320463 | | 1 | 1 | 0.839916 |
| Pdia4 | A0A0R4J0Z | 0.2 | 0.22 | -0.23 | -0.04 | 0.75 | 0.28 | 0.11 | 0.02 | -0.16 | 0.33 | -0.31 | 0.5 | 0.48 | 0.6 | 1.24 | 0.81 | 0.2525 | 0.0525 | 0.745 | 0.266843 | 0.830985 | 0.01162 | | 1 | 0.996511 | 0.566767 |
| Pdia5 | Q921X9 | -0.55 | -0.42 | -1.14 | -0.72 | -0.1 | -0.32 | 0.63 | -0.57 | 0.06 | -2.55 | -0.3 | -0.16 | -0.68 | 0.47 | 0.12 | -0.14 | 0.6175 | -0.03 | 0.65 | 0.065897 | 0.95854 | 0.055577 | | 1 | 0.996755 | 0.661738 |
| Pdia6 | Q3TML0 | -0.97 | -0.88 | -0.76 | -0.84 | -0.27 | -0.72 | -0.28 | -0.9 | 0.52 | -1.14 | -1.01 | -0.81 | 1.63 | -0.27 | 0.58 | 0.13 | 0.32 | 0.2525 | 1.38 | 0.133946 | 0.503899 | 0.006873 | | 1 | 0.996511 | 0.552614 |
| Pdk1 | Q8BFP9 | -0.42 | 0.04 | 0.12 | -0.13 | -0.32 | 0.27 | -0.25 | -0.14 | 0.07 | -0.55 | 0.14 | -0.28 | 0.65 | -0.89 | -0.21 | -0.22 | -0.0125 | -0.0575 | -0.07 | 0.951877 | 0.804059 | 0.83613 | | 1 | 0.996511 | 0.975831 |
| Pdk2 | Q9JK42 | 1 | 0.6 | 0.38 | 0.16 | 0.64 | 1.13 | 0.75 | 0.29 | 0.28 | 0.1 | 0.53 | 0.63 | 0.41 | 0.48 | 0.28 | 0.41 | 0.1675 | -0.15 | -0.14 | 0.521911 | 0.540291 | 0.545406 | | 1 | 0.996511 | 0.912246 |
| Pdk3 | Q922H2 | 0.73 | 0.97 | 0.01 | 0.24 | 0.29 | 0.76 | 0.2 | 0.42 | 0.34 | -0.71 | 0.2 | 0.49 | 0.18 | 0.16 | -0.66 | -0.49 | -7.00E-02 | -0.4075 | -0.69 | 0.788482 | 0.254183 | 0.054 | | 1 | 0.996511 | 0.661738 |
| Pdk4 | O70571 | 1.33 | 1.09 | 1.37 | 0.51 | 1.5 | -0.14 | 2.08 | 0.88 | 0.73 | 0.06 | 0.77 | -0.35 | -0.39 | 1.07 | 0.16 | 2.55 | 0.005 | -0.7725 | -0.2275 | 0.991618 | 0.043286 | 0.712295 | | 1 | 0.996511 | 0.958014 |
| Pdlim1 | O70400 | -1.49 | -1.16 | -0.38 | -0.57 | -0.41 | -1.54 | -0.09 | -0.18 | -0.99 | -1.28 | -0.03 | -0.77 | -1.07 | -0.41 | -0.67 | -0.24 | 0.345 | 0.1325 | 0.3025 | 0.397421 | 0.714563 | 0.358092 | | 1 | 0.996511 | 0.856645 |
| Pdlim3 | O70209 | -0.72 | -0.54 | -1.48 | -0.81 | -0.52 | -0.92 | -1.11 | -1.09 | -0.75 | -2.65 | -1.47 | -0.45 | -1.04 | -0.61 | -1.61 | -0.96 | -0.0225 | -0.4425 | -0.1675 | 0.930076 | 0.38185 | 0.584295 | | 1 | 0.996511 | 0.927734 |
| Pdlim4 | P70271 | -0.19 | -0.52 | -0.61 | -0.3 | -0.44 | -0.89 | -0.41 | -0.38 | 0.01 | 1.2 | -0.25 | -0.34 | 0.1 | 0.56 | -0.1 | -0.43 | -0.125 | 0.56 | 0.4375 | 0.520368 | 0.1461 | 0.1175 | | 1 | 0.996511 | 0.718609 |
| Pdlim5 | Q8CI51 | -1.29 | -1.71 | -0.95 | -0.56 | -1.75 | -0.99 | -0.52 | -0.37 | -0.93 | -0.51 | -0.54 | -1.15 | -0.47 | -1.46 | -0.62 | -0.84 | 0.22 | 0.345 | 0.28 | 0.5622 | 0.264025 | 0.407231 | | 1 | 0.996511 | 0.874084 |
| Pdlim7 | Q3TJD7 | -2.04 | -2.37 | -2.56 | -3.08 | -2.15 | -2.22 | -2.66 | -2.93 | -2.02 | -4.43 | -2.72 | -1.59 | -2.36 | -2.27 | -2.84 | -2.3 | 0.0225 | -0.1775 | 0.07 | 0.937397 | 0.770069 | 0.800739 | | 1 | 0.996511 | 0.970599 |
| Pdp1 | A2AJP9 | 0.2 | 0.43 | 0.46 | 0.72 | 0.25 | 0.21 | 0.7 | 0.9 | 0.05 | 0.61 | 0.52 | 0 | 0.01 | 0.22 | 0.29 | 0.42 | 0.0625 | -0.1575 | -0.2175 | 0.779303 | 0.48965 | 0.297583 | | 1 | 0.996511 | 0.826881 |
| Pdp2 | Q504M2 | -0.19 | -0.18 | 0.81 | 0.94 | -0.75 | -0.08 | 0.16 | 1.16 | -0.41 | 1.31 | 0.57 | 0.24 | 0.03 | -0.41 | 0.72 | 0.87 | -0.2225 | 0.0825 | -0.0425 | 0.63441 | 0.852657 | 0.917695 | | 1 | 0.996511 | 0.986868 |
| Pdpk1 | Q9Z2A0 | 0.54 | 0.27 | -0.42 | -0.31 | 0.36 | 0.26 | -0.08 | -0.29 | -0.08 | -0.86 | -0.52 | 0.34 | -0.26 | 0.61 | -0.52 | -0.5 | 0.0425 | -0.3 | -0.1875 | 0.878633 | 0.39216 | 0.594427 | | 1 | 0.996511 | 0.929843 |
| Pdpr | Q7TSQ8 | 0.17 | 0.21 | 0.45 | 0.86 | 0.06 | 0.37 | 0.15 | 0.85 | -0.14 | 0.07 | 0.66 | 0.03 | -0.33 | 0.02 | 0.12 | 0.15 | -0.065 | -0.2675 | -0.4325 | 0.794683 | 0.312205 | 0.092726 | | 1 | 0.996511 | 0.695968 |
| Pds5a | Q6A026 | -1.47 | -2.45 | -2.22 | -1.95 | -2.06 | -2.11 | -3.06 | -3.12 | -1.84 | -0.61 | -2.28 | -1.99 | -1.64 | -2.03 | -2.73 | -2.81 | -0.565 | 0.3425 | -0.28 | 0.12966 | 0.407468 | 0.431833 | | 1 | 0.996511 | 0.879165 |
| Pds5b | F8WHU5 | -0.81 | -1.64 | -1.23 | -1.07 | -0.8 | -1.43 | -0.79 | -1 | -1.76 | -1.48 | -1.07 | -1.34 | -1.29 | -1.21 | -0.41 | -0.52 | 0.1825 | -0.225 | 0.33 | 0.461423 | 0.378121 | 0.288354 | | 1 | 0.996511 | 0.824705 |
| Pdxdc1 | A0A0R4J03 | -2.01 | -1.47 | -1.2 | -1.6 | -0.96 | -1.86 | -1.36 | -1.53 | -1.8 | -0.9 | -1.27 | -1.27 | -1.48 | -1.49 | -1.69 | -1.31 | 0.1425 | 0.26 | 0.0775 | 0.587722 | 0.343923 | 0.737008 | | 1 | 0.996511 | 0.958635 |
| Pdxk | Q8K183 | -1.18 | -1.23 | -0.89 | -0.8 | -1.1 | -1.49 | -0.59 | -0.7 | -1.43 | -1.22 | -1.02 | -1.37 | -1.32 | -0.99 | -1 | -1.25 | 0.055 | -0.235 | -0.115 | 0.822305 | 0.245722 | 0.571647 | | 1 | 0.996511 | 0.924577 |
| Pdxp | P60487 | -0.25 | 0.06 | -0.44 | -0.26 | 0.01 | -0.37 | -0.5 | -0.38 | -0.2 | 0.53 | -0.69 | -0.11 | -0.24 | 0.65 | -0.65 | -0.88 | -0.0875 | 0.105 | -0.0575 | 0.647431 | 0.711152 | 0.869561 | | 1 | 0.996511 | 0.979293 |
| Pdyn | O35852 | 1.61 | 0.63 | -0.22 | 0.85 | 0.89 | 0.65 | 1.1 | 1.09 | 1.51 | 1.33 | 0.31 | 1.3 | 1.34 | 0.98 | 1.75 | 0.57 | 0.215 | 0.395 | 0.4425 | 0.567389 | 0.378768 | 0.319513 | | 1 | 0.996511 | 0.838969 |
| Pdzd11 | Q9CZG9 | 1.08 | 1.39 | 0.44 | 0.62 | 1.23 | 1.28 | 1.07 | 0.67 | 1.37 | 0.61 | 0.81 | 2.28 | 0.96 | 2.06 | 0.34 | 0.79 | 0.18 | 0.385 | 0.155 | 0.50157 | 0.361667 | 0.704753 | | 1 | 0.996511 | 0.957084 |
| Pdzd2 | E9Q1M1 | 0.22 | 0.29 | 0.63 | 0.8 | 0.13 | 0.09 | 1.38 | 0.71 | 0.79 | 0.96 | 0.89 | 0.58 | 0.69 | 0.15 | 0.55 | 0.61 | 0.0925 | 0.32 | 0.015 | 0.77587 | 0.147179 | 0.947608 | | 1 | 0.996511 | 0.992414 |
| Pdzd4 | A2AFG3 | 1.13 | 1.55 | 1.07 | 1.1 | 0.77 | 1.18 | 0.18 | 1.16 | 1.68 | 0.32 | 0.57 | 1.27 | 2.24 | 1.12 | 1.85 | 1.22 | -0.39 | -0.2525 | 0.395 | 0.169107 | 0.452794 | 0.212111 | | 1 | 0.996511 | 0.786366 |
| Pdzd8 | B9EJ80 | 0.71 | 1.03 | 0.73 | 0.36 | 0.84 | 0.54 | 0.9 | 0.64 | 0.88 | 0.66 | 0.75 | 1.04 | 1.09 | 1.34 | 0.69 | 0.83 | 0.0225 | 0.125 | 0.28 | 0.908559 | 0.547659 | 0.25909 | | 1 | 0.996511 | 0.812074 |
| Pdzrn3 | Q69ZS0-2 | 0.24 | 1.18 | 0.41 | -0.2 | 1.06 | 0.46 | 0.33 | -0.02 | 0.15 | 0.16 | 0.17 | 1.15 | 0.22 | 1.04 | 0.32 | 0.59 | 0.05 | 7.85E-17 | 0.135 | 0.88669 | 1 | 0.693549 | | 1 | 1 | 0.955729 |
| Pea15 | Q62048 | -0.74 | -1.06 | -0.63 | 0.06 | -1.36 | -0.91 | -0.4 | 0.12 | -0.95 | -0.54 | -0.5 | -0.57 | -1.76 | -1.1 | -1.16 | -0.5 | -0.045 | -0.0475 | -0.5375 | 0.905059 | 0.861651 | 0.148062 | | 1 | 0.996511 | 0.746406 |
| Peak1 | Q69Z38 | 0.95 | 0.95 | 1.26 | 0.86 | 0.44 | 1.05 | 0.67 | 0.95 | 1.03 | 0.99 | 1.21 | 0.94 | 1.14 | 0.26 | 0.92 | 1.17 | -0.2275 | 0.0375 | -0.1325 | 0.268739 | 0.831399 | 0.611023 | | 1 | 0.996511 | 0.935047 |
| Pebp1 | P70296 | 0.18 | -0.22 | -1.15 | -0.73 | -0.18 | -0.08 | -0.53 | -0.92 | -0.75 | -0.98 | -0.91 | 0.43 | -1.23 | -0.4 | -1.29 | -1.53 | 0.0525 | -0.0725 | -0.6325 | 0.875992 | 0.862408 | 0.117001 | | 1 | 0.996511 | 0.718608 |
| Pecam1 | B1ARB3 | -0.63 | -0.81 | -0.36 | -0.22 | -0.57 | -0.82 | -0.72 | -0.71 | -0.66 | -0.64 | -0.2 | -0.57 | 0.01 | -1.5 | -0.4 | -0.5 | -0.2 | -0.0125 | -0.0925 | 0.294749 | 0.953014 | 0.789209 | | 1 | 0.996755 | 0.968677 |
| Pecr | Q3UXD9 | 0.32 | 0.8 | 0.72 | 0.56 | 1.12 | 0.16 | 0.5 | 0.87 | 0.59 | 0.39 | 0.39 | 0.46 | 0.43 | 0.6 | 0.39 | 0.67 | 0.0625 | -0.1425 | -0.0775 | 0.801394 | 0.439702 | 0.69378 | | 1 | 0.996511 | 0.955729 |
| PEDS1 | Q99LQ7 | 1.19 | 1.81 | 2.52 | 2.24 | 3.03 | 1.57 | 3 | 2.66 | 2.28 | 3.17 | 2.45 | 1.58 | 2.48 | 2.94 | 2.82 | 3.78 | 0.625 | 0.43 | 1.065 | 0.16274 | 0.315387 | 0.021516 | | 1 | 0.996511 | 0.621926 |
| Pef1 | Q8BFY6 | 0.43 | 0.38 | -0.17 | -0.09 | 0.46 | 0.8 | -0.14 | -0.02 | -0.56 | 0.55 | -0.77 | -0.1 | 0.7 | 0.1 | -0.35 | -0.29 | 0.1375 | -0.3575 | -0.0975 | 0.616599 | 0.294327 | 0.746003 | | 1 | 0.996511 | 0.960124 |
| Peg3 | A0A5F8MP | -1.92 | -4.14 | -1.28 | -0.67 | -4.11 | -2.46 | -2.05 | -1.84 | 4.74 | 0.64 | -1.84 | -3.13 | 6.37 | 1.22 | -0.35 | -1.65 | -0.6125 | 2.105 | 3.4 | 0.464969 | 0.23311 | 0.071722 | | 1 | 0.996511 | 0.668066 |
| Peli1 | Q8C669 | -0.02 | 0.16 | -0.24 | -0.45 | -1.56 | -1.15 | -0.1 | -0.4 | -1.73 | -0.22 | -0.32 | -0.83 | -1.68 | -1.5 | -0.51 | -0.83 | -0.665 | -0.6375 | -0.9925 | 0.08299 | 0.104591 | 0.011109 | | 1 | 0.996511 | 0.563386 |
| Pelo | Q80X73 | 0.49 | -0.33 | 0.74 | -0.54 | -0.53 | 0.7 | -0.06 | 0 | 0.1 | -1.95 | 0.73 | 0.98 | 1.21 | -2.67 | 1.32 | 0.34 | -0.0625 | -0.125 | -0.04 | 0.869473 | 0.851632 | 0.963816 | | 1 | 0.996511 | 0.9927 |
| Pelp1 | Q9DBD5 | -1.88 | -3.23 | -2.06 | -2.25 | -2.87 | -1.98 | -2.12 | -2.1 | -2.49 | -1.47 | -1.79 | -2.36 | -1.86 | -2.87 | -2.59 | -2.31 | 0.0875 | 0.3275 | -0.0525 | 0.802495 | 0.390564 | 0.885338 | | 1 | 0.996511 | 0.979767 |
| Penk | P22005 | 1.8 | -0.03 | -1.3 | 0.78 | 0.14 | -0.84 | 1.6 | 0.65 | 0.06 | 1.05 | -1.04 | 0.59 | -0.82 | 0.02 | 2.18 | 0.97 | 0.075 | -0.1475 | 0.275 | 0.920225 | 0.838005 | 0.740122 | | 1 | 0.996511 | 0.958796 |
| Pepd | Q11136 | 0.02 | -0.3 | -1.05 | -1.16 | -0.31 | -0.2 | -1.15 | -1.16 | -0.39 | -0.55 | -1.12 | 0.06 | -0.53 | -0.04 | -1.48 | -1.68 | -0.0825 | 0.1225 | -0.31 | 0.823479 | 0.73824 | 0.50248 | | 1 | 0.996511 | 0.901002 |
| Pes1 | Q5SQ20 | -0.48 | -0.16 | -0.18 | -0.28 | -0.58 | -0.14 | 0.17 | -0.33 | -0.57 | -0.1 | -0.15 | -1.01 | -0.13 | -0.6 | -0.16 | -0.15 | 0.055 | -0.1825 | 0.015 | 0.788555 | 0.469575 | 0.940472 | | 1 | 0.996511 | 0.990894 |
| Pex1 | Q5BL07 | 0.07 | 0.19 | 0.13 | 0.32 | 0.46 | 0.02 | 0.46 | 0.34 | 0.48 | 0.55 | 0.34 | 0.27 | 0.4 | 0.48 | 0.52 | 0.38 | 0.1425 | 0.2325 | 0.2675 | 0.411286 | 0.186709 | 0.144733 | | 1 | 0.996511 | 0.743156 |
| Pex10 | B1AUE5 | -0.73 | -0.2 | 0.3 | 0.12 | 0.22 | -0.56 | 0.58 | 0.19 | 0.34 | 0.24 | -0.32 | -0.52 | 0.51 | 0.38 | 0.71 | 0.28 | 0.235 | 0.0625 | 0.5975 | 0.471983 | 0.84082 | 0.05026 | | 1 | 0.996511 | 0.658454 |
| Pex11b | Q9Z210 | 0.68 | 0.28 | 1.99 | 0.54 | 0.83 | 0.88 | 2.19 | 1.92 | -0.03 | 0.76 | 1.85 | 0.73 | 0.65 | -0.03 | 2.64 | 2.87 | 0.5825 | -0.045 | 0.66 | 0.245088 | 0.928911 | 0.382267 | | 1 | 0.996755 | 0.860984 |
| Pex12 | Q8VC48 | 0.79 | 0.82 | 0.13 | -0.56 | 1.22 | 0.37 | 0.86 | -0.57 | -0.01 | -1.59 | -0.01 | 0.89 | 0.07 | 0.96 | 0.71 | -0.12 | 0.175 | -0.475 | 0.11 | 0.710642 | 0.408032 | 0.783687 | | 1 | 0.996511 | 0.967642 |
| Pex13 | Q9D0K1 | -0.88 | -0.2 | -0.28 | 0.2 | 0.39 | -0.83 | 0.66 | 0.37 | -0.11 | -0.21 | -0.13 | -0.06 | 0.41 | 0.65 | -0.09 | 0.75 | 0.4375 | 0.1625 | 0.72 | 0.26834 | 0.518947 | 0.039062 | | 1 | 0.996511 | 0.658454 |
| Pex14 | Q9R0A0 | -0.07 | 0.18 | 0.17 | 0.37 | 0.4 | 0.16 | 0.32 | 0.15 | 0.43 | 0.38 | 0.26 | 0.31 | 0.67 | 0.9 | 0.52 | 0.05 | 0.095 | 0.1825 | 0.3725 | 0.570155 | 0.305942 | 0.146592 | | 1 | 0.996511 | 0.746406 |
| Pex16 | Q91XC9 | 0.17 | 0.89 | 0.03 | 0 | -0.19 | 0.4 | 0.21 | -0.21 | 0.1 | -0.07 | 0.01 | 1.06 | 0.85 | 1.04 | 0.45 | -0.02 | -0.22 | 0.0025 | 0.3075 | 0.416268 | 0.994015 | 0.35059 | | 1 | 0.999285 | 0.853439 |
| Pex19 | Q8VCI5 | -0.46 | -0.31 | -0.43 | -0.25 | -0.33 | -0.25 | -0.32 | -0.42 | -0.43 | 0.04 | -0.33 | 0.42 | -0.13 | -0.14 | -0.09 | -0.45 | 0.0325 | 0.2875 | 0.16 | 0.82276 | 0.234728 | 0.393061 | | 1 | 0.996511 | 0.865775 |
| Pex2 | Q91YZ5 | 0.92 | 0.57 | 0.64 | 0.68 | 0.85 | 0.29 | 0.44 | -0.53 | 0.78 | 1.14 | 0.36 | 0.89 | 1.17 | 0.36 | 0.27 | 0.45 | -0.44 | 0.09 | -0.14 | 0.164964 | 0.68097 | 0.581894 | | 1 | 0.996511 | 0.927734 |
| Pex26 | Q8BGI5-2 | 1.24 | 0.6 | -0.05 | -0.13 | 0.66 | 0.83 | 0.28 | 0 | -0.03 | -0.02 | 0.07 | 1.32 | 0.35 | 0.17 | 0.31 | 0.23 | 0.0275 | -0.08 | -0.15 | 0.938266 | 0.853652 | 0.648055 | | 1 | 0.996511 | 0.945928 |
| Pex3 | Q9QXY9 | 0.3 | 0.14 | 0.55 | 0.37 | 0.2 | 0.48 | 0.71 | 0.55 | -0.06 | 0.87 | 0.6 | 0.53 | 0.57 | -0.15 | 0.58 | 0.87 | 0.145 | 0.145 | 0.1275 | 0.43138 | 0.551115 | 0.628142 | | 1 | 0.996511 | 0.940342 |
| Pex5 | O09012-2 | -0.12 | 1.24 | 0.29 | 0.91 | -0.45 | -0.22 | 1.27 | 0.39 | -0.47 | -0.82 | 1.26 | 1.41 | -0.14 | 0.45 | 0.08 | 0.45 | -0.3325 | -0.235 | -0.37 | 0.473081 | 0.695661 | 0.291813 | | 1 | 0.996511 | 0.825497 |
| Pex5 | O09012 | -0.06 | 0.44 | 0.05 | -0.24 | 0.3 | 0.36 | 0.22 | 0 | -0.16 | 0.35 | 0.06 | 0.59 | -0.11 | 0.14 | -0.37 | 0.04 | 0.1725 | 0.1625 | -0.1225 | 0.394408 | 0.510957 | 0.593519 | | 1 | 0.996511 | 0.929843 |
| Pex5l | F8SLQ1 | 0.51 | 1.16 | 0.32 | 0.25 | 1.34 | 0.44 | 0.36 | 0.09 | 0.47 | 0.08 | 0.28 | 0.89 | 0.47 | 1.09 | 0.15 | 0.28 | -0.0025 | -0.13 | -0.0625 | 0.993978 | 0.64621 | 0.837752 | | 1 | 0.996511 | 0.975961 |
| Pex5l | D3YYH0 | 0.21 | 0.93 | 0.46 | 0.15 | 1.14 | -0.23 | 0.09 | 0.14 | 0.41 | 0.95 | -0.04 | 0.39 | 0.68 | 1.03 | -0.18 | 0.27 | -0.1525 | -0.01 | 0.0125 | 0.650648 | 0.971598 | 0.968968 | | 1 | 0.997515 | 0.992767 |
| Pex6 | Q99LC9 | 0.99 | 0.98 | 0.91 | 0.62 | 0.83 | 1.13 | 1.04 | 0.83 | 0.53 | 1.02 | 0.82 | 1.18 | 0.78 | 0.35 | 0.95 | 0.97 | 0.0825 | 0.0125 | -0.1125 | 0.627744 | 0.952185 | 0.610833 | | 1 | 0.996755 | 0.935047 |
| Pex7 | P97865 | 0.27 | 1.72 | -0.75 | -0.37 | 0.94 | 0.79 | 0.49 | 0.45 | 0.4 | 0.05 | 1.13 | 0.96 | 0.8 | 0.9 | 0.67 | 1.6 | 0.45 | 0.4175 | 0.775 | 0.390421 | 0.456615 | 0.174582 | | 1 | 0.996511 | 0.767185 |
| Pfas | Q5SUR0 | -1.73 | -2.13 | -1.58 | -1.57 | -1.76 | -1.81 | -1.42 | -1.57 | -2.11 | -0.91 | -1.77 | -1.65 | -1.81 | -1.57 | -1.55 | -1.92 | 0.1125 | 0.1425 | 0.04 | 0.567133 | 0.62878 | 0.852047 | | 1 | 0.996511 | 0.976225 |
| Pfdn1 | Q9CQF7 | 0.25 | 0.03 | 0.61 | 0.04 | 1.32 | -0.07 | 0.42 | 0.37 | -0.68 | -0.75 | 0.26 | 0.24 | -0.39 | 0.46 | 0.29 | 0.69 | 0.2775 | -0.465 | 0.03 | 0.390009 | 0.162538 | 0.916677 | | 1 | 0.996511 | 0.986556 |
| Pfdn2 | O70591 | -0.85 | -0.47 | -0.72 | -0.43 | -0.23 | -0.88 | -0.19 | -0.59 | -0.79 | 0.1 | -0.59 | -0.58 | -0.29 | 0.2 | -0.45 | -0.51 | 0.145 | 0.1525 | 0.355 | 0.508619 | 0.537398 | 0.153111 | | 1 | 0.996511 | 0.750649 |
| Pfdn4 | E9Q6U4 | 1.58 | 2.19 | 0.68 | 0.82 | 1.99 | 1.55 | 0.8 | 0.8 | 1.49 | 0.47 | 0.41 | 2.13 | 1.12 | 2.52 | 0.34 | 0.6 | -0.0325 | -0.1925 | -0.1725 | 0.939532 | 0.704887 | 0.75581 | | 1 | 0.996511 | 0.962333 |
| Pfdn5 | Q9WU28 | 0.56 | 0.27 | 0.16 | 0.22 | 0.44 | 0.68 | 0.47 | 0.49 | 0.13 | 0.44 | 0.13 | 0.4 | 0.19 | 0.93 | -0.05 | 0.33 | 0.2175 | -0.0275 | 0.0475 | 0.205524 | 0.881374 | 0.853402 | | 1 | 0.996511 | 0.976225 |
| Pfdn6 | Q03958 | -0.5 | -0.4 | -0.41 | -0.46 | -0.58 | -0.47 | -0.35 | -0.51 | -0.27 | -0.08 | -0.24 | -0.31 | 0.2 | -0.28 | -0.43 | -0.48 | -0.035 | 0.2175 | 0.195 | 0.806552 | 0.190298 | 0.371528 | | 1 | 0.996511 | 0.859424 |
| Pfkfb2 | A0A087WF | -0.06 | 0.15 | -0.1 | 0.21 | -0.03 | -0.07 | -0.01 | 0.34 | -0.33 | 0.18 | 0.19 | 0.03 | -0.22 | -0.33 | -0.01 | -0.07 | 0.0075 | -0.0325 | -0.2075 | 0.965314 | 0.868267 | 0.283914 | | 1 | 0.996511 | 0.822669 |
| Pfkfb2 | B2Z893 | 1.14 | 2.06 | -0.47 | 0.81 | 0.96 | 0.49 | -0.07 | -0.07 | 1.83 | 2.67 | 0.07 | 1.05 | 1.06 | 1.26 | -0.48 | -0.64 | -0.5575 | 0.52 | -0.585 | 0.310659 | 0.460801 | 0.385407 | | 1 | 0.996511 | 0.861533 |
| Pfkfb4 | Q6DTY7-5 | -0.21 | -0.95 | 0.13 | -2.13 | -0.96 | 1.12 | -1.1 | -0.61 | 0.04 | -6.72 | 0.2 | -0.49 | 1.31 | -1.91 | 0.24 | 0.68 | 0.4025 | -0.9525 | 0.87 | 0.540854 | 0.543937 | 0.279589 | | 1 | 0.996511 | 0.822084 |
| Pfkl | P12382 | 0.2 | 0.18 | -0.24 | -0.26 | 0.35 | 0.16 | -0.35 | -0.2 | 0.17 | -1.08 | -0.28 | 0.02 | -0.24 | 0.15 | -0.89 | -0.73 | 0.02 | -0.2625 | -0.3975 | 0.929425 | 0.406382 | 0.189498 | | 1 | 0.996511 | 0.776843 |
| Pfkm | P47857 | -0.2 | -0.34 | 0.17 | 0.31 | -0.13 | -0.13 | 0.08 | 0.4 | -0.63 | -0.49 | 0.1 | -0.48 | -0.63 | -0.33 | -0.28 | -0.15 | 0.07 | -0.36 | -0.3325 | 0.751609 | 0.170268 | 0.17087 | | 1 | 0.996511 | 0.762322 |
| Pfkp | Q8C605 | 0.3 | 0.15 | 0.07 | 0.06 | 0.1 | 0.25 | -0.18 | 0.25 | 0.73 | -0.32 | 0.03 | 0.45 | 0.85 | 0.47 | -0.07 | -0.44 | -0.04 | 0.0775 | 0.0575 | 0.813197 | 0.764117 | 0.849442 | | 1 | 0.996511 | 0.976225 |
| Pfkp | D3YUA3 | 1.12 | 0.71 | -0.81 | -2.44 | -2.14 | -0.13 | -1.03 | -3.04 | 1.44 | 0.6 | -1.41 | -1.7 | -0.96 | 0.8 | -1.33 | -0.25 | -1.23 | 0.0875 | -0.08 | 0.206602 | 0.930267 | 0.924183 | | 1 | 0.996755 | 0.988152 |
| Pfn2 | Q9JJV2 | -0.24 | -0.31 | 0.38 | 0.5 | -0.38 | -0.15 | 0.67 | 0.76 | -0.13 | -0.08 | 0.62 | -0.13 | -0.42 | -0.48 | 0.1 | 0.38 | 0.1425 | -0.0125 | -0.1875 | 0.67878 | 0.965285 | 0.54351 | | 1 | 0.996755 | 0.912212 |
| Pgam1 | Q9DBJ1 | 1.52 | 1.59 | 0.2 | 0.44 | 1.46 | 1.55 | 0.59 | 1.04 | 0.51 | -0.61 | 0.51 | 1.55 | 0.34 | 1.29 | 0.42 | 0.34 | 0.2225 | -0.4475 | -0.34 | 0.579747 | 0.405964 | 0.417249 | | 1 | 0.996511 | 0.87558 |
| Pgam2 | O70250 | 1.28 | -0.26 | -0.8 | -1.23 | 0.69 | 0 | -0.11 | -1.12 | 0.23 | -0.25 | -0.83 | 0.29 | 0.31 | 0.82 | -0.86 | -0.44 | 0.1175 | 0.1125 | 0.21 | 0.845856 | 0.840464 | 0.730604 | | 1 | 0.996511 | 0.958635 |
| Pgam5 | Q8BX10-2 | 1.25 | 1.39 | 0.11 | 0.41 | 1.08 | 0.84 | 1.06 | 0.95 | 0.93 | -1.27 | 0.29 | 1.57 | 0.29 | 1.1 | -0.38 | -0.06 | 0.1925 | -0.41 | -0.5525 | 0.542665 | 0.517865 | 0.216562 | | 1 | 0.996511 | 0.787447 |
| Pgam5 | Q8BX10 | 0.47 | 0.77 | 0.8 | 0.8 | 0.47 | 0.47 | 0.98 | 0.99 | 0.68 | 0.67 | 1.1 | 0.25 | 0.88 | 0.99 | 0.65 | 0.44 | 0.0175 | -0.035 | 0.03 | 0.930671 | 0.876664 | 0.88481 | | 1 | 0.996511 | 0.979767 |
| Pgap1 | Q3UUQ7 | -1.29 | -1.19 | -1.17 | -1.27 | -0.49 | -1.05 | -0.48 | -0.88 | -1 | -1.36 | -1.13 | -1.04 | -0.65 | -0.6 | -0.42 | -0.51 | 0.505 | 0.0975 | 0.685 | 0.023002 | 0.56421 | 0.002986 | | 1 | 0.996511 | 0.461278 |
| Pgap3 | A2A559-2 | -1.84 | -2.17 | 1.35 | 1.34 | 0.45 | -1.46 | 1.4 | 0.65 | -0.7 | 0.25 | 0.1 | -1.47 | -0.81 | -0.22 | 0.72 | 3.5 | 0.59 | -0.125 | 1.1275 | 0.569119 | 0.894446 | 0.364344 | | 1 | 0.996511 | 0.858475 |
| Pgbd5 | D3YZI9 | 0.59 | 0.28 | 0.25 | 0.06 | -0.09 | 0.11 | 0.14 | -0.09 | 0.28 | -0.22 | 0.11 | 0.4 | -0.18 | 0.59 | 0.03 | 0.06 | -0.2775 | -0.1525 | -0.17 | 0.139658 | 0.483094 | 0.480695 | | 1 | 0.996511 | 0.893906 |
| Pgd | Q9DCD0 | -0.1 | 0.17 | -0.59 | -0.56 | 0.33 | -0.25 | -0.27 | -0.74 | -0.16 | -0.09 | -0.73 | 0.26 | -0.26 | 0.46 | -0.87 | -0.77 | 0.0375 | 0.09 | -0.09 | 0.896129 | 0.753586 | 0.798793 | | 1 | 0.996511 | 0.969753 |
| Pgghg | Q8BP56 | -0.47 | 0.25 | -0.62 | -0.06 | 0.15 | -0.28 | -0.24 | 0.23 | -0.53 | -0.21 | -0.28 | -0.04 | -0.76 | -1 | -1.49 | -0.9 | 0.19 | -0.04 | -0.8125 | 0.454108 | 0.871255 | 0.015469 | | 1 | 0.996511 | 0.589387 |
| Pggt1b | Q8BUY9 | -0.56 | -0.84 | -0.08 | -0.13 | -1.47 | -1.23 | -0.23 | -0.3 | -1.03 | -0.32 | 0.06 | -1.18 | -0.46 | -1.33 | -0.22 | -0.05 | -0.405 | -0.215 | -0.1125 | 0.267592 | 0.532219 | 0.74002 | | 1 | 0.996511 | 0.958796 |
| Pgk1 | P09411 | 0.28 | -0.03 | 0.35 | 0.17 | -0.08 | 0.18 | 0.61 | 0.56 | -0.76 | -0.09 | 0.31 | 0.23 | -0.86 | -0.55 | -0.07 | 0.07 | 0.125 | -0.27 | -0.545 | 0.557526 | 0.334986 | 0.061172 | | 1 | 0.996511 | 0.661738 |
| Pgls | Q9CQ60 | 0.02 | -0.46 | -0.18 | 0.19 | -0.55 | -0.12 | -0.05 | 0.27 | 0.29 | 0.76 | -0.25 | 0.3 | -0.02 | 0.72 | -0.58 | -0.66 | -0.005 | 0.3825 | -0.0275 | 0.983089 | 0.17573 | 0.936679 | | 1 | 0.996511 | 0.989767 |
| Pgm1 | Q9D0F9 | 0.32 | 0.28 | 0.18 | 0.46 | -0.63 | -0.65 | -1.06 | -0.45 | 0.11 | 0.65 | 0.07 | 0.1 | -0.16 | 0.06 | 0.32 | 0.09 | -1.0075 | -0.0775 | -0.2325 | 0.000446 | 0.699916 | 0.242846 | 0.544245 | | 0.996511 | 0.798789 |
| Pgm2 | Q7TSV4 | -1.25 | -1.22 | -0.71 | -0.85 | -1.62 | -1.91 | -0.65 | -0.86 | -1.85 | -0.05 | -0.74 | -1.22 | -1.08 | -1.99 | -1.17 | -1.25 | -0.2525 | 0.0425 | -0.365 | 0.440916 | 0.912643 | 0.202582 | | 1 | 0.996744 | 0.782817 |
| Pgm2l1 | Q8CAA7 | -0.99 | -1.16 | -0.42 | -0.18 | -1.09 | -0.8 | -0.36 | 0.03 | -0.77 | 0.29 | 0.01 | -0.65 | -0.86 | -0.93 | -0.7 | -1.12 | 0.1325 | 0.4075 | -0.215 | 0.688079 | 0.251082 | 0.43674 | | 1 | 0.996511 | 0.883161 |
| Pgm3 | Q9CYR6 | -0.8 | -0.58 | -0.39 | -0.32 | -0.78 | -0.82 | -0.67 | -0.42 | -0.81 | -0.2 | -0.32 | -0.86 | -0.26 | -1.06 | -0.18 | -0.63 | -0.15 | -0.025 | -0.01 | 0.422125 | 0.913826 | 0.969062 | | 1 | 0.996755 | 0.992767 |
| Pgm5 | Q8BZF8 | -2.11 | -0.62 | -1.88 | -0.9 | -1.35 | -1.4 | -2.47 | -1.28 | -1.47 | -0.36 | -1.53 | -1.91 | -1.87 | -1.58 | -2.07 | -0.94 | -0.2475 | 0.06 | -0.2375 | 0.569465 | 0.896834 | 0.575312 | | 1 | 0.996511 | 0.926723 |
| Pgp | Q8CHP8 | -1.03 | -0.85 | -0.7 | -0.36 | -0.75 | -0.93 | -0.62 | -0.52 | -1.39 | -0.56 | -0.77 | -1.04 | -1.04 | -1 | -0.61 | -0.54 | 0.03 | -0.205 | -0.0625 | 0.880879 | 0.424016 | 0.789898 | | 1 | 0.996511 | 0.969105 |
| Pgpep1 | Q9ESW8 | -0.03 | 0.24 | -0.51 | -0.96 | 0.19 | -0.07 | 1.44 | -0.87 | -4.12 | -0.27 | 0.18 | 0.07 | -2.93 | -2.05 | -1.04 | -0.91 | 0.4875 | -0.72 | -1.4175 | 0.347471 | 0.460258 | 0.019539 | | 1 | 0.996511 | 0.612607 |
| Pgrmc1 | O55022 | 0.73 | 0.28 | 0.18 | 0.49 | 0.38 | 0.5 | 1.06 | 0.55 | 0.27 | 0.39 | 0.12 | 0.87 | 0.22 | 0.32 | 1.15 | 0.65 | 0.2025 | -0.0075 | 0.165 | 0.364876 | 0.974341 | 0.542324 | | 1 | 0.997515 | 0.912212 |
| Pgrmc2 | Q80UU9 | 0.96 | 1.14 | 0.79 | 0.68 | 0.7 | 0.81 | 0.98 | 0.56 | 1.31 | 0.84 | 0.68 | 0.95 | 0.82 | 1.7 | 1.18 | 1.2 | -0.13 | 0.0525 | 0.3325 | 0.47596 | 0.802968 | 0.195875 | | 1 | 0.996511 | 0.78004 |
| Pgs1 | Q8BHF7 | 0.19 | 0.58 | 0.54 | 0.7 | 0.69 | 0.83 | 0.79 | 0.65 | 0.07 | 0.5 | 0.53 | 0.36 | 0.02 | 0.35 | -0.02 | 0.46 | 0.2375 | -0.1375 | -0.3 | 0.186875 | 0.499378 | 0.18843 | | 1 | 0.996511 | 0.776843 |
| Phactr1 | Q2M3X8 | 0.39 | 0.33 | 0.15 | 0.02 | 0.04 | 0.07 | -0.17 | 0.02 | 0.06 | 0.01 | 0.03 | 0.37 | 0.41 | -0.08 | -0.08 | -0.19 | -0.2325 | -0.105 | -0.2075 | 0.174803 | 0.569881 | 0.345419 | | 1 | 0.996511 | 0.849669 |
| Phactr2 | F7D4H5 | -0.05 | -0.44 | 0.14 | 0.57 | -0.96 | -0.04 | -0.48 | 0.34 | 0.38 | 0.13 | -0.14 | -0.65 | 0.87 | -0.26 | 0.03 | 0.32 | -0.34 | -0.125 | 0.185 | 0.330115 | 0.686141 | 0.572276 | | 1 | 0.996511 | 0.924918 |
| Phactr3 | Q8BYK5 | -0.37 | -0.5 | -0.19 | -0.21 | -0.84 | -0.4 | -0.78 | -0.17 | -0.8 | -0.64 | -0.29 | -0.63 | -0.35 | -1 | -0.18 | -0.68 | -0.23 | -0.2725 | -0.235 | 0.28038 | 0.172888 | 0.334527 | | 1 | 0.996511 | 0.846677 |
| Phactr4 | Q501J7-2 | -0.32 | -0.59 | 0.42 | 0.17 | -0.73 | -0.24 | 0.14 | 0.55 | 0.57 | 0.17 | 0.64 | -0.17 | 0.59 | 0.25 | 0.93 | 0.84 | 0.01 | 0.3825 | 0.7325 | 0.976761 | 0.226848 | 0.031086 | | 1 | 0.996511 | 0.641481 |
| Phaf1 | Q922R1 | 0.51 | 0.48 | 0.47 | 0.62 | 0.62 | 0.59 | 0.9 | 0.7 | 0.17 | -0.03 | 0.4 | 0.55 | -0.02 | 0.01 | 0.29 | 0.52 | 0.1825 | -0.2475 | -0.32 | 0.247755 | 0.214045 | 0.135329 | | 1 | 0.996511 | 0.731223 |
| Phax | Q9JJT9 | -3.28 | -4.02 | -1.42 | -2.21 | -3.43 | -3.96 | -1.2 | -1.48 | -4.67 | -1.34 | -0.87 | -3.36 | -3.84 | -3.63 | -1.34 | -1.16 | 0.215 | 0.1725 | 0.24 | 0.790818 | 0.855934 | 0.77237 | | 1 | 0.996511 | 0.965933 |
| Phb1 | P67778 | -0.56 | -0.27 | 0.44 | 0.32 | -0.36 | -0.28 | 0.17 | 0.12 | -0.02 | 0.54 | 0.43 | -0.55 | 0.13 | 0.07 | 0.17 | -0.08 | -0.07 | 0.1175 | 0.09 | 0.801216 | 0.729967 | 0.738912 | | 1 | 0.996511 | 0.958796 |
| Phb2 | O35129 | 0.57 | 0.65 | 0.45 | 0.42 | 0.45 | 0.81 | 0.02 | 0.35 | 1.87 | 0.5 | 0.54 | 0.6 | 1.81 | 0.55 | 0.22 | 0.03 | -0.115 | 0.355 | 0.13 | 0.57409 | 0.302456 | 0.7403 | | 1 | 0.996511 | 0.958796 |
| Phc1 | Q3V116 | -1.83 | -1.15 | -0.2 | 0.41 | 0.77 | -1.28 | -0.24 | -0.43 | -0.6 | -0.24 | -1.15 | -0.14 | -0.1 | -0.9 | -2.43 | 0.4 | 0.3975 | 0.16 | -0.065 | 0.509004 | 0.753161 | 0.927858 | | 1 | 0.996511 | 0.98946 |
| Phc2 | Q9QWH1- | -0.09 | -1.2 | -1.28 | -0.78 | -1.47 | -0.83 | -0.64 | -1.22 | 0.07 | 0.16 | -0.89 | -1.18 | 0.19 | -0.07 | -0.85 | -0.68 | -0.2025 | 0.3775 | 0.485 | 0.534939 | 0.372316 | 0.202882 | | 1 | 0.996511 | 0.782817 |
| Phf10 | K4DI61 | -0.56 | 0.41 | 0.12 | 0.58 | -0.72 | -0.62 | 0.26 | 0.26 | -0.59 | -0.49 | 0.42 | -0.52 | -0.43 | -1.8 | 0.82 | -0.19 | -0.3425 | -0.4325 | -0.5375 | 0.346063 | 0.227182 | 0.341559 | | 1 | 0.996511 | 0.848473 |
| Phf14 | Q9D4H9-2 | 0.64 | 0.59 | 0.35 | 0.87 | 0.51 | 0.62 | 0.44 | 0.59 | 0.77 | 1.04 | 0.59 | 0.65 | 0.73 | 0.24 | 1.09 | 0.56 | -0.0725 | 0.15 | 0.0425 | 0.668317 | 0.455063 | 0.861501 | | 1 | 0.996511 | 0.977204 |
| Phf2 | Q9WTU0 | 0.41 | -0.31 | -0.1 | 0.36 | -0.48 | 0.37 | -0.61 | -0.34 | 2.15 | -0.19 | -0.15 | 0.2 | 2.62 | -0.26 | 0.59 | -0.18 | -0.355 | 0.4125 | 0.6025 | 0.230808 | 0.451879 | 0.353837 | | 1 | 0.996511 | 0.855029 |
| Phf20l1 | A0A2R8VH | -1.97 | -1.72 | -1.12 | -1.63 | -1.9 | -1.37 | -1.16 | -1.55 | -1.51 | -0.97 | -1.87 | -0.96 | -1.15 | -0.55 | -0.93 | -1.31 | 0.115 | 0.2825 | 0.625 | 0.647037 | 0.347955 | 0.041941 | | 1 | 0.996511 | 0.658454 |
| Phf24 | Q80TL4 | 0.59 | 0.29 | 0.86 | 0.99 | 0.46 | 0.53 | 0.51 | 0.93 | -0.12 | 0.81 | 0.61 | 0.18 | -0.35 | 0.05 | 0.5 | 0.56 | -0.075 | -0.3125 | -0.4925 | 0.727959 | 0.273518 | 0.108981 | | 1 | 0.996511 | 0.707444 |
| Phf24 | Q80TL4-2 | 3.71 | -0.68 | 1.89 | 6.37 | -1.97 | 2.56 | 2.36 | 6.45 | 2.51 | 5.54 | 2.33 | 3.62 | 0.79 | 0.51 | 0.01 | 1.44 | -0.4725 | 0.6775 | -2.135 | 0.815708 | 0.649343 | 0.139852 | | 1 | 0.996511 | 0.735494 |
| Phf3 | B2RQG2 | -1.23 | -1.98 | -1.64 | -1.56 | -2.22 | -2.09 | -1.59 | -1.34 | -0.64 | -0.25 | -1.73 | -1.93 | 1.15 | -0.31 | 0.13 | -0.94 | -0.2075 | 0.465 | 1.61 | 0.442632 | 0.281361 | 0.005277 | | 1 | 0.996511 | 0.552614 |
| Phf5a | P83870 | 0.05 | -0.78 | -3.13 | -2.92 | -0.82 | -0.69 | -2.49 | -2.57 | -1.09 | -2.2 | -2.98 | -1.09 | -0.31 | -0.86 | -2.37 | -3.81 | 0.0525 | -0.145 | -0.1425 | 0.950494 | 0.86036 | 0.886291 | | 1 | 0.996511 | 0.979767 |
| Phf6 | Q9D4J7 | -1.8 | -1.86 | -0.45 | -1.14 | -0.5 | -2.45 | 0.27 | -0.73 | -2.62 | 0.36 | -0.43 | -2.36 | -1.89 | -0.7 | 0.28 | 0.49 | 0.46 | 0.05 | 0.8575 | 0.453896 | 0.945056 | 0.167616 | | 1 | 0.996755 | 0.759484 |
| Phf8 | Q80TJ7 | -2.55 | -3.17 | -4.64 | -5.08 | -3.5 | -3.1 | -7.32 | -6.49 | -2.36 | -1.67 | -3.73 | -3.01 | -1.69 | -3.62 | -3.61 | -7.69 | -1.2425 | 1.1675 | -0.2925 | 0.272046 | 0.112149 | 0.814291 | | 1 | 0.996511 | 0.972768 |
| Phgdh | Q61753 | 0.53 | 0.05 | 0.27 | 0.25 | 0.46 | 0.7 | 0.44 | 0.53 | -0.04 | -0.5 | 0.13 | 0.51 | -0.45 | -0.3 | 0.09 | 0.26 | 0.2575 | -0.25 | -0.375 | 0.153203 | 0.337594 | 0.136313 | | 1 | 0.996511 | 0.731656 |
| Phip | F8VQ93 | 0.63 | 0.61 | 1.19 | 1.09 | 0.56 | 0.4 | 0.89 | 1.15 | 0.16 | -0.49 | 1.29 | 0.33 | 0.33 | 0.52 | 1.58 | 1.49 | -0.13 | -0.5575 | 0.1 | 0.593109 | 0.170814 | 0.777678 | | 1 | 0.996511 | 0.967289 |
| Phka1 | A2AI89 | 0.68 | 0.57 | 0.27 | 0.51 | 1.07 | 0.82 | 0.6 | 0.73 | 0.37 | 0.33 | 0.42 | 0.61 | 0.11 | 0.38 | 0.32 | 0.52 | 0.2975 | -0.075 | -0.175 | 0.122378 | 0.672409 | 0.378963 | | 1 | 0.996511 | 0.860984 |
| Phka2 | A2AHQ7 | -0.25 | -0.26 | 0.15 | 0.13 | -0.18 | -0.49 | 0.61 | 0.43 | -1.07 | 0.21 | 0.52 | -0.49 | -1.24 | -0.1 | -0.4 | 0.08 | 0.15 | -0.15 | -0.3575 | 0.599715 | 0.68164 | 0.281591 | | 1 | 0.996511 | 0.822084 |
| Phkb | E0CX43 | -1.11 | -0.04 | -0.2 | -0.34 | -1.23 | -0.39 | 0.36 | 0.73 | 0.4 | -1.29 | -0.59 | -1.21 | -1.58 | -0.4 | -1.37 | -0.08 | 0.29 | -0.25 | -0.435 | 0.532559 | 0.566329 | 0.311842 | | 1 | 0.996511 | 0.834523 |
| Phkb | Q7TSH2 | 0.5 | 0.21 | -0.21 | -0.2 | 0.83 | 0.56 | -0.14 | -0.05 | 0.04 | 0.09 | -0.12 | 0.25 | -0.07 | 0.12 | -0.34 | -0.23 | 0.225 | -0.01 | -0.205 | 0.448233 | 0.964301 | 0.400683 | | 1 | 0.996755 | 0.86969 |
| Phkg1 | P07934 | 0.53 | 0.4 | 0.3 | 0.6 | 1.01 | 0.84 | 0.55 | 0.77 | -0.27 | -0.19 | 0.5 | 0.35 | -0.4 | 0.06 | 0.2 | 0.68 | 0.335 | -0.36 | -0.3225 | 0.075149 | 0.150003 | 0.236697 | | 1 | 0.996511 | 0.792722 |
| Phkg2 | Q9DB30 | 0.36 | -0.13 | -0.66 | -0.36 | 0.65 | 0.25 | -0.16 | -0.54 | -1.13 | -0.69 | -0.66 | 0.12 | -1.45 | -0.22 | -1.19 | -0.8 | 0.2475 | -0.3925 | -0.7175 | 0.456025 | 0.259441 | 0.061893 | | 1 | 0.996511 | 0.662517 |
| Phlda1 | Q62392 | -0.12 | 0.1 | -0.18 | -0.23 | -0.53 | -0.13 | -0.37 | -0.4 | -0.28 | -0.75 | -0.08 | -0.53 | 0.18 | 0.42 | -0.52 | -0.32 | -0.25 | -0.3025 | 0.0475 | 0.15923 | 0.17026 | 0.854366 | | 1 | 0.996511 | 0.976225 |
| Phlda3 | Q9WV95 | 0.83 | 0.89 | 0.88 | 0.76 | 1.4 | 0.17 | 0.81 | 0.61 | -0.42 | 0.45 | 0.43 | 0.21 | 0.57 | -1.38 | 0.36 | 1.14 | -0.0925 | -0.6725 | -0.6675 | 0.727268 | 0.017953 | 0.209921 | | 1 | 0.996511 | 0.786218 |
| Phldb1 | Q6PDH0-2 | 0.32 | 0.49 | 0.22 | 0.45 | 0.02 | -0.03 | 0.45 | -0.11 | 0.32 | 0.5 | 0.12 | 0.41 | 0.16 | 0.2 | 0.39 | 0.24 | -0.2875 | -0.0325 | -0.1225 | 0.142692 | 0.852244 | 0.494078 | | 1 | 0.996511 | 0.89758 |
| Phldb1 | E9PWW4 | 0.49 | 1.04 | 2.04 | 1.1 | 0.65 | 0.46 | -0.4 | 0.35 | 1.56 | -2.13 | 0.87 | 0.32 | 1.47 | 1.06 | 1.64 | 0.87 | -0.9025 | -1.0125 | 0.0925 | 0.037919 | 0.217915 | 0.798529 | | 1 | 0.996511 | 0.969727 |
| Phldb2 | F6QU68 | -0.15 | -1.68 | -0.99 | -2.88 | -1.54 | -0.98 | -1.42 | -3.23 | -0.13 | -1.97 | -1.78 | -0.86 | 0.86 | -1.62 | -0.59 | -1.36 | -0.3675 | 0.24 | 0.7475 | 0.596478 | 0.714868 | 0.318411 | | 1 | 0.996511 | 0.838969 |
| Phlpp1 | Q8CHE4 | 0.16 | -0.18 | 0.36 | 0.35 | -0.83 | 0.11 | 0.05 | 0.3 | -0.12 | 0.34 | -0.14 | -0.64 | -0.1 | 0.19 | 0.21 | 0.48 | -0.265 | -0.3125 | 0.0225 | 0.362357 | 0.244311 | 0.919379 | | 1 | 0.996511 | 0.986868 |
| Phlpp2 | J3QM82 | 0.17 | -0.49 | -1.01 | -1.29 | -0.16 | 0.33 | -1.19 | -1.29 | -0.39 | 1.05 | -1.05 | -0.12 | -0.39 | -0.18 | -1.37 | -0.69 | 0.0775 | 0.5275 | -0.0025 | 0.869658 | 0.311836 | 0.994987 | | 1 | 0.996511 | 0.998224 |
| Phospho1 | Q8R2H9 | 1.57 | 0.63 | 0.96 | 0.51 | 0.65 | 1.19 | -0.08 | 0.64 | -0.06 | 0.65 | 0.6 | 1.22 | -0.2 | 0.31 | 0.58 | 0.78 | -0.3175 | -0.315 | -0.55 | 0.363641 | 0.375698 | 0.116608 | | 1 | 0.996511 | 0.718608 |
| Phospho2 | Q9D9M5 | -0.63 | -0.64 | -1.32 | -0.57 | -0.35 | -0.36 | -0.69 | -1.09 | 0.45 | -0.25 | -0.92 | -0.9 | 0.36 | -0.74 | -0.87 | -1.28 | 0.1675 | 0.385 | 0.1575 | 0.521812 | 0.301703 | 0.681372 | | 1 | 0.996511 | 0.955729 |
| Phpt1 | Q9DAK9 | 0.21 | -0.44 | 0.2 | -0.15 | -0.59 | 0.28 | 0.77 | 0.19 | -0.97 | -0.37 | 0.55 | 0.33 | -1.58 | 0.68 | 0.74 | 0.65 | 0.2075 | -0.07 | 0.1675 | 0.516134 | 0.849185 | 0.758668 | | 1 | 0.996511 | 0.963116 |
| Phrf1 | A6H619-2 | -0.69 | -2.15 | -1.82 | -0.73 | -0.42 | -1.12 | -1.15 | -0.71 | 1.66 | 1.39 | -1.76 | -0.32 | 3.86 | -0.79 | 1.67 | -1.23 | 0.4975 | 1.59 | 2.225 | 0.224734 | 0.073103 | 0.070638 | | 1 | 0.996511 | 0.668066 |
| Phyh | O35386 | 0.38 | 1.06 | 0.45 | 0.69 | 0.48 | 0.46 | 0.46 | 0.15 | 1.29 | 1.24 | 0.86 | 0.92 | 1.99 | 2 | 0.83 | 0.68 | -0.2575 | 0.4325 | 0.73 | 0.227753 | 0.080411 | 0.082062 | | 1 | 0.996511 | 0.68865 |
| Phyhd1 | A2AQZ2 | -0.28 | -1.24 | -0.68 | -0.4 | -1.2 | -1.55 | -0.25 | -0.19 | -0.55 | -0.26 | -0.28 | -1.02 | -2.45 | -0.76 | -0.62 | -1.38 | -0.1475 | 0.1225 | -0.6525 | 0.700461 | 0.671318 | 0.16654 | | 1 | 0.996511 | 0.759484 |
| Phyhip | Q8K0S0 | 3.11 | -1.13 | -0.08 | -1.37 | 0.64 | 3.15 | 1.51 | -0.14 | -1.54 | -1.46 | 0.08 | 2.34 | -1.72 | -1.15 | -0.19 | 0.4 | 1.1575 | -0.2775 | -0.7975 | 0.316394 | 0.821609 | 0.440424 | | 1 | 0.996511 | 0.883549 |

| | | | | | | | | | | | | | | | | | | | | | | | | | | | |
|---|---|---|---|---|---|---|---|---|---|---|---|---|---|---|---|---|---|---|---|---|---|---|---|---|---|---|---|
| Phyhipl | Q8BGT8 | 2.93 | 1.39 | 1.11 | 0.13 | 1.7 | 3.02 | 1.05 | 1.27 | -0.48 | 0.42 | 1.01 | 2.87 | -0.26 | -0.57 | 1.08 | 1.18 | 0.37 | -0.435 | -1.0325 | 0.579353 | 0.601552 | 0.148563 | | 1 | 0.996511 | 0.746831 |
| Pi4k2a | Q2TBE6 | 0.28 | 0.49 | 0.35 | -0.15 | 0.8 | 0.24 | 0.67 | 0.22 | -0.21 | 0.13 | 0.43 | 0.48 | -0.13 | 0.24 | 0.49 | 0.46 | 0.24 | -0.035 | 0.0225 | 0.300777 | 0.883396 | 0.924753 | | 1 | 0.996511 | 0.988301 |
| Pi4ka | E9Q3L2 | 0.39 | 0.52 | 0.93 | 0.57 | 0.78 | 0.56 | 1.09 | 0.84 | -0.06 | 0.05 | 0.91 | 0.39 | -0.06 | 0.31 | 0.8 | 1.08 | 0.215 | -0.28 | -0.07 | 0.28682 | 0.305567 | 0.811912 | | 1 | 0.996511 | 0.972642 |
| Pi4kb | Q8BKC8-2 | -0.73 | -0.69 | -0.08 | -0.37 | -0.31 | -0.63 | -0.34 | -0.5 | -0.19 | 0.17 | -0.09 | -0.56 | -0.28 | -0.06 | -0.4 | -0.56 | 0.0225 | 0.3 | 0.1425 | 0.911086 | 0.234932 | 0.539677 | | 1 | 0.996511 | 0.912212 |
| Pianp | A0A0N4SV | 1.83 | 1.3 | 1.23 | 2.03 | 0.1 | 1.75 | 1.01 | 1.45 | 2.88 | 2.78 | 0.95 | 1.6 | 2.95 | 1.08 | 2.57 | 1.19 | -0.52 | 0.455 | 0.35 | 0.203445 | 0.351363 | 0.474832 | | 1 | 0.996511 | 0.891741 |
| Pias4 | Q9JM05 | -1.17 | -0.75 | -2.48 | -3.75 | -2.04 | -1.59 | -7.28 | -6.26 | 0.05 | -2.49 | -4.91 | -0.17 | -0.52 | -1.39 | -5.23 | -3.58 | -2.255 | 0.1575 | -0.6425 | 0.141434 | 0.895963 | 0.572365 | | 1 | 0.996511 | 0.924918 |
| Picalm | Q7M6Y3-6 | 1.14 | 0.2 | 0.36 | 0.73 | -0.07 | -0.21 | 0.9 | 0.65 | -0.07 | -0.21 | 0.17 | 0.2 | -0.99 | 0.26 | -0.71 | -0.03 | -0.29 | -0.585 | -0.975 | 0.393265 | 0.043616 | 0.021253 | | 1 | 0.996511 | 0.621926 |
| Picalm | A0A1L1SU | -0.76 | 1.98 | 2.01 | 2.99 | 2.26 | -0.25 | 1.38 | 3.56 | 1.38 | 2.43 | 2.36 | -0.1 | -1.44 | 1.02 | 1.41 | 1.96 | 0.1825 | -0.0375 | -0.8175 | 0.857801 | 0.966736 | 0.416544 | | 1 | 0.996755 | 0.874945 |
| Picalm | Q7M6Y3-5 | -0.2 | -0.67 | -0.37 | -0.2 | -0.43 | -0.26 | -0.34 | -0.05 | -0.26 | -0.62 | 0.01 | -0.44 | -0.19 | 0.08 | -0.87 | -0.82 | 0.09 | 0.0325 | -0.09 | 0.622311 | 0.879546 | 0.748546 | | 1 | 0.996511 | 0.960744 |
| Pick1 | E9PUZ5 | 0.45 | 0.49 | 0.43 | 0.24 | 0.23 | 0.6 | 0.32 | 0.35 | 1.25 | 0.15 | 0.43 | 0.53 | 1.87 | 0.26 | 0.26 | 0.29 | -0.0275 | 0.1875 | 0.2675 | 0.863226 | 0.477629 | 0.499871 | | 1 | 0.996511 | 0.90036 |
| Pid1 | Q3UBG2-2 | 0.97 | 1.21 | -2.86 | -2.4 | 1.28 | 0.76 | -2.42 | -2.41 | 1.68 | -2.39 | -2.13 | 1.15 | 0.8 | 1.5 | -2.95 | -2.72 | 0.0725 | 0.3475 | -0.0725 | 0.955931 | 0.798087 | 0.958892 | | 1 | 0.996511 | 0.992636 |
| Pigb | Q9JJQ0 | -1.59 | -0.78 | -0.62 | 1.54 | -2.37 | -0.73 | -2.83 | 1.87 | 0.42 | 0.06 | 1.28 | -1.64 | 1.14 | -1.75 | 1.34 | 1.02 | -0.6525 | 0.3925 | 0.8 | 0.565305 | 0.634036 | 0.380112 | | 1 | 0.996511 | 0.860984 |
| Pigg | D3Z3Y1 | -0.16 | -0.17 | -1.65 | -0.96 | 0.41 | -0.53 | 0.35 | -1.05 | -0.69 | -1.39 | -0.97 | -0.38 | -0.62 | 0.09 | -1.04 | -1.01 | 0.53 | -0.1225 | 0.09 | 0.274848 | 0.759343 | 0.831889 | | 1 | 0.996511 | 0.974403 |
| Pigk | Q9CXY9 | -1.6 | -1.28 | -0.38 | -0.4 | -0.63 | -1.17 | 0.04 | -0.14 | -0.4 | -0.89 | -0.34 | -1.37 | 0.06 | -0.82 | 0.46 | 0.05 | 0.44 | 0.165 | 0.8525 | 0.277211 | 0.664611 | 0.056717 | | 1 | 0.996511 | 0.661738 |
| Pign | A0A087WI | -0.54 | -0.28 | 0.14 | -0.5 | -1.06 | -0.22 | 0.53 | -0.24 | 0.11 | -2.28 | -0.8 | 0.01 | -0.86 | 0.33 | -1.2 | -0.94 | 0.0475 | -0.445 | -0.3725 | 0.891028 | 0.411683 | 0.325213 | | 1 | 0.996511 | 0.842881 |
| Pigo | A2AG36 | -0.33 | -0.75 | -0.45 | -0.98 | -0.15 | 0.29 | 0.11 | -0.17 | 2.19 | 0.21 | -0.33 | -0.1 | 3.7 | -0.1 | 0.86 | 0.71 | 0.6475 | 1.12 | 1.92 | 0.012988 | 0.067817 | 0.030813 | | 1 | 0.996511 | 0.641481 |
| Pigp | B8JK67 | 0.3 | 0.09 | 0.58 | 0.49 | 0.32 | 0.06 | 0.6 | 0.32 | 0.14 | 1.04 | -0.15 | 0.2 | 0.16 | 0.87 | 0.85 | 0.54 | -0.04 | -0.0575 | 0.24 | 0.83517 | 0.841594 | 0.328592 | | 1 | 0.996511 | 0.844943 |
| Pigs | Q6PD26 | -0.54 | 0.04 | 0.35 | 0.52 | 0.76 | -0.31 | 0.91 | 0.53 | -0.61 | 0.59 | 0.26 | -0.28 | -0.64 | -0.1 | 1.01 | 0.93 | 0.38 | -0.1025 | 0.2075 | 0.288552 | 0.769206 | 0.639967 | | 1 | 0.996511 | 0.942804 |
| Pigt | Q8BXQ2 | -0.67 | -0.42 | -0.6 | -0.5 | 0.8 | -0.35 | -0.24 | -0.53 | -0.48 | -0.91 | -0.56 | -0.56 | -0.06 | -0.01 | 0.54 | 0.3 | 0.4675 | -0.08 | 0.74 | 0.147811 | 0.655787 | 0.006221 | | 1 | 0.996511 | 0.552614 |
| Pigu | Q3TAA8 | -1.35 | -1.09 | -0.77 | 0.32 | -0.58 | -0.38 | -0.34 | -0.01 | -1.04 | -0.58 | -0.74 | -1.53 | -0.73 | 0.57 | 0.04 | 0.11 | 0.395 | -0.25 | 0.72 | 0.299547 | 0.53988 | 0.123597 | | 1 | 0.996511 | 0.721448 |
| Pih1d1 | Q9CQJ2 | -0.44 | -0.35 | -0.26 | 0.08 | -0.52 | 0.02 | -0.58 | -0.33 | 0.38 | -0.45 | -0.4 | -0.39 | 0.38 | -0.49 | -0.62 | -0.63 | -0.11 | 0.0275 | -0.0975 | 0.596872 | 0.912896 | 0.733606 | | 1 | 0.996755 | 0.958635 |
| Pik3ap1 | Q9EQ32 | -3.46 | -3.43 | -4.11 | -4.12 | -3.74 | -2.96 | -2.29 | -3.78 | -3.82 | -5.03 | -3.18 | -4.06 | -2.41 | -2.64 | -2.94 | -3.85 | 0.5875 | -0.2425 | 0.82 | 0.151965 | 0.557634 | 0.047906 | | 1 | 0.996511 | 0.658454 |
| Pik3c2a | F8VPL2 | -0.53 | -0.09 | 0.1 | 0.07 | 0.18 | -0.44 | 0.07 | -0.04 | -0.29 | -0.21 | 0.25 | -0.25 | 0.01 | -0.1 | 0.27 | 0.19 | 0.055 | -0.0125 | 0.205 | 0.803969 | 0.955916 | 0.362273 | | 1 | 0.996755 | 0.858475 |
| Pik3c3 | E9QLS6 | 0.31 | 0.41 | 0.39 | 0.13 | 0.43 | 0.22 | 0.08 | 0.05 | 0.22 | -0.2 | 0.25 | 0.33 | -0.03 | 0.34 | 0 | 0.05 | -0.115 | -0.16 | -0.22 | 0.49197 | 0.412295 | 0.258807 | | 1 | 0.996511 | 0.812074 |
| Pik3ca | P42337 | -0.92 | -0.58 | -0.71 | -0.24 | -0.41 | -0.96 | -0.66 | -0.45 | -0.63 | -0.23 | -0.38 | -0.94 | -0.84 | -0.2 | -0.91 | -1.01 | -0.0075 | 0.0675 | -0.1275 | 0.972225 | 0.777845 | 0.627213 | | 1 | 0.996511 | 0.939737 |
| Pik3cb | Q8BTI9 | 0.73 | 0.17 | 0.28 | 0.19 | 0.58 | 0.94 | 0.14 | 0.25 | -0.21 | -0.6 | 0.33 | 0.62 | -0.43 | -0.6 | -0.16 | 0.03 | 0.135 | -0.3075 | -0.6325 | 0.575726 | 0.330455 | 0.02294 | | 1 | 0.996511 | 0.623147 |
| Pik3cd | B0QZL5 | 0.7 | 1.18 | 0.28 | 0.15 | 1.52 | 0.59 | 0.15 | -0.02 | 0.98 | 2.21 | 0.88 | 1.43 | 0.91 | 1.08 | 0.83 | -0.22 | -0.0175 | 0.7975 | 0.0725 | 0.964456 | 0.056351 | 0.843938 | | 1 | 0.996511 | 0.976225 |
| Pik3ip1 | Z4YN57 | -0.47 | 0.11 | -1.15 | -0.41 | 0.17 | -0.2 | -0.8 | -0.73 | 0.78 | 0.64 | -1.02 | 0.03 | 1.05 | 1.08 | 0.06 | -1.19 | 0.09 | 0.5875 | 0.73 | 0.78853 | 0.218761 | 0.206254 | | 1 | 0.996511 | 0.785211 |
| Pik3r1 | P26450 | 0.25 | 0.19 | 0.21 | 0.33 | 0.4 | 0.18 | 0.33 | 0.57 | -0.4 | -0.43 | 0.4 | 0.27 | -0.56 | -0.55 | -0.06 | -0.2 | 0.125 | -0.285 | -0.5875 | 0.428975 | 0.264512 | 0.015091 | | 1 | 0.996511 | 0.589387 |
| Pik3r2 | O08908 | 0.71 | 0.51 | -0.24 | -0.34 | 0.66 | 0.4 | -0.12 | -0.26 | 0.55 | -0.28 | -0.14 | 0.65 | 0.06 | 0.44 | -0.59 | -0.31 | 0.01 | 0.035 | -0.26 | 0.975897 | 0.919742 | 0.458508 | | 1 | 0.996755 | 0.887902 |
| Pik3r4 | Q8VD65 | 0.02 | 0.12 | 0.25 | 0.25 | -0.02 | 0.05 | 0.06 | 0.42 | 0.03 | -0.08 | 0.2 | 0.06 | -0.09 | 0.13 | -0.06 | 0.21 | -0.0325 | -0.1075 | -0.1125 | 0.846802 | 0.520644 | 0.539097 | | 1 | 0.996511 | 0.912212 |
| Pikfyve | Q9Z1T6 | 0.61 | 0.82 | 0.07 | 0.14 | 0.63 | 0.41 | 0.31 | 0.2 | 0.81 | 0.04 | 0.26 | 0.58 | 0.48 | 0.66 | 0.31 | 0.08 | -0.0225 | 0.0125 | -0.0275 | 0.920274 | 0.962491 | 0.913171 | | 1 | 0.996755 | 0.986458 |
| Pin1 | Q9QUR7 | -0.2 | -0.49 | 0.08 | 0.39 | -0.19 | -0.34 | 0.4 | 0.49 | -0.42 | 0.14 | 0.15 | -0.13 | -0.84 | 0.19 | 0.02 | -0.24 | 0.145 | -0.01 | -0.1625 | 0.610914 | 0.968519 | 0.597349 | | 1 | 0.996755 | 0.930093 |
| Pin4 | Q9CWW6 | -0.43 | -1.12 | -0.86 | -0.47 | -1.21 | 1.52 | -0.83 | -0.74 | -0.59 | -0.14 | -0.89 | -0.28 | -0.76 | 0.16 | -1.07 | -1.04 | 0.405 | 0.245 | 0.0425 | 0.495481 | 0.351271 | 0.898646 | | 1 | 0.996511 | 0.983509 |
| Pip4k2a | O70172 | 0.18 | -0.16 | 0.25 | 0.3 | 0.11 | -0.24 | 0.29 | 0.37 | -0.48 | 0.41 | 0.23 | -0.53 | -0.8 | 0.5 | 0.16 | 0.17 | -0.01 | -0.235 | -0.135 | 0.960567 | 0.404756 | 0.663837 | | 1 | 0.996511 | 0.949825 |
| Pip4k2b | Q80XI4 | 1.05 | 0.31 | 1.09 | 0.78 | 0.63 | 1.09 | 0.89 | 1.54 | -0.69 | 0.84 | 0.92 | 0.7 | -0.83 | -0.38 | 0.81 | 1.03 | 0.23 | -0.365 | -0.65 | 0.402916 | 0.375342 | 0.180645 | | 1 | 0.996511 | 0.775542 |
| Pip4k2c | Q91XU3 | 0.59 | -0.1 | -0.36 | -0.98 | 0.15 | 0.47 | -0.54 | -0.79 | -0.18 | -1.47 | -0.77 | 0.1 | 0.1 | -0.46 | -0.83 | -0.92 | 0.035 | -0.3675 | -0.315 | 0.932156 | 0.421268 | 0.424993 | | 1 | 0.996511 | 0.877323 |
| Pip4p1 | Q3TWL2 | -0.44 | -0.52 | -0.52 | -0.51 | -0.54 | -0.14 | -0.32 | -0.54 | -1.09 | -0.84 | -0.66 | -0.35 | -0.82 | -1.04 | -0.25 | 0.06 | 0.1125 | -0.2375 | -0.015 | 0.489745 | 0.262816 | 0.956625 | | 1 | 0.996511 | 0.992636 |
| Pip4p2 | Q9CZX7 | -0.54 | -0.92 | -0.62 | -0.59 | -0.82 | -0.26 | -0.23 | -0.54 | -0.61 | -0.77 | -0.37 | -0.37 | -0.54 | -0.54 | -1.1 | -0.22 | 0.205 | 0.1375 | 0.0675 | 0.313841 | 0.472804 | 0.779369 | | 1 | 0.996511 | 0.967289 |
| Pip5k1a | P70182-3 | 1.57 | -0.3 | 0.13 | 0.95 | -0.87 | 1.77 | 0.12 | 0.83 | 2.05 | 0.33 | 0.49 | 0.14 | 2.1 | -0.61 | 0.46 | 0.24 | -0.125 | 0.165 | -0.04 | 0.843723 | 0.767816 | 0.950263 | | 1 | 0.996511 | 0.992476 |
| Pip5k1b | P70181 | 0.64 | 0.68 | 0.52 | 0.71 | 0.98 | 0.77 | 0.57 | 1.05 | -0.14 | 0.83 | 0.38 | -0.6 | 1.53 | 1.52 | 0.35 | 0.47 | 0.205 | -0.52 | 0.33 | 0.246718 | 0.126211 | 0.330278 | | 1 | 0.996511 | 0.845524 |
| Pip5k1c | F8WHW6 | 0.46 | 0.28 | 1.02 | 1.21 | 0.19 | 0.47 | 0.62 | 1.3 | 0.4 | 1.02 | 1.01 | 0.28 | 0.31 | 0.22 | 0.76 | 0.42 | -0.0975 | -0.065 | -0.315 | 0.758916 | 0.829407 | 0.268637 | | 1 | 0.996511 | 0.820219 |
| Pir | A2AIH8 | 1.54 | 0.48 | 0.03 | 0.17 | 1.01 | 1.59 | -0.42 | 0.19 | 0.6 | -0.4 | -0.4 | 1.4 | 0.46 | 0.74 | -0.4 | -0.52 | 0.0375 | -0.255 | -0.485 | 0.941899 | 0.621992 | 0.28645 | | 1 | 0.996511 | 0.824295 |
| Pithd1 | Q8BWR2 | 0.32 | 0.08 | -0.42 | -0.12 | -0.19 | 0.42 | -0.89 | -0.55 | 0.09 | 0.37 | -0.45 | 0.27 | -0.06 | -0.33 | -0.41 | -0.71 | -0.2675 | 0.105 | -0.3425 | 0.405754 | 0.68761 | 0.183145 | | 1 | 0.996511 | 0.776002 |
| Pitpna | P53810 | 1.03 | -0.03 | 0.23 | 0.22 | -0.02 | 0.89 | -0.09 | 0.15 | -0.13 | -0.29 | 0.06 | 1.11 | -0.46 | -0.16 | -0.16 | -0.17 | -0.13 | -0.175 | -0.6 | 0.681903 | 0.644764 | 0.048534 | | 1 | 0.996511 | 0.658454 |
| Pitpnb | Q8JZZ5 | 0.07 | -0.19 | -0.05 | 0.26 | -0.29 | -0.06 | 0.07 | 0.47 | -0.34 | 0.34 | 0.32 | 0.35 | -0.61 | -0.45 | -0.01 | -0.23 | 0.025 | 0.145 | -0.3475 | 0.906289 | 0.52898 | 0.134004 | | 1 | 0.996511 | 0.730883 |
| Pitpnc1 | Q8K4R4-2 | 8.78 | 1.1 | -0.99 | -1.26 | 1.23 | 8.36 | -0.73 | -0.36 | 1.79 | -2.6 | -2.26 | 7.81 | 2.9 | 0.42 | -0.12 | 1.26 | 0.2175 | -0.7225 | -0.7925 | 0.938438 | 0.810046 | 0.714739 | | 1 | 0.996511 | 0.958106 |
| Pitpnc1 | Q8K4R4 | 0.77 | 0.37 | -0.18 | -0.01 | 0.22 | 0.54 | 0.2 | -0.13 | 0.8 | 0.09 | -0.23 | 0.84 | 0.96 | 0.64 | -0.39 | -0.1 | -0.03 | 0.1375 | 0.04 | 0.908271 | 0.683292 | 0.914065 | | 1 | 0.996511 | 0.986458 |
| Pitpnm1 | O35954 | 0.27 | 0.12 | 0.41 | 0.33 | 0.37 | 0.36 | 0.67 | 0.4 | 0.19 | -0.1 | 0.29 | 0.4 | 0.29 | 0.24 | 0.41 | 0.46 | 0.1675 | -0.0875 | 0.0675 | 0.308723 | 0.638303 | 0.703525 | | 1 | 0.996511 | 0.957004 |
| Pitpnm2 | E9PYJ7 | 0.14 | -0.16 | 0.32 | 0.14 | 0.06 | 0.26 | 0.5 | 0.39 | -0.25 | -0.18 | 0.04 | -0.01 | -0.27 | -0.36 | 0.12 | 0.18 | 0.1925 | -0.21 | -0.1925 | 0.304719 | 0.271127 | 0.391669 | | 1 | 0.996511 | 0.865375 |
| Pitpnm3 | Q3UHE1 | 0.62 | -0.1 | -0.18 | -0.06 | 0.25 | 0.69 | 0.11 | 0.02 | -0.24 | -0.04 | -0.14 | 0.65 | 0.01 | -0.29 | 0.2 | -0.02 | 0.1975 | -0.0125 | -0.095 | 0.437204 | 0.964908 | 0.700754 | | 1 | 0.996755 | 0.956416 |
| Pitrm1 | Q8K411 | 1.6 | 0.76 | 0.99 | 0.76 | 0.8 | 1.58 | 0.88 | 0.87 | 0.5 | 0.73 | 1.13 | 1.43 | 0.3 | 0.52 | 0.6 | 0.7 | 0.005 | -0.08 | -0.4975 | 0.985458 | 0.786 | 0.072773 | | 1 | 0.996511 | 0.668066 |
| Pja1 | O55176 | -1.14 | 0 | -0.98 | -2.1 | -0.79 | -0.71 | -2.04 | -1.99 | 0.29 | -0.91 | -1.52 | 0.19 | 2.12 | 0.05 | -0.72 | -1.29 | -0.3275 | 0.5675 | 1.095 | 0.532971 | 0.329962 | 0.182884 | | 1 | 0.996511 | 0.776002 |
| Pja2 | Q80U04-2 | -2.2 | -1.9 | -1.91 | -1.98 | -2.83 | -2.13 | -2.74 | -2.44 | 0.08 | -0.4 | -1.98 | -1.76 | 1.15 | -0.56 | -1.86 | -2.39 | -0.5375 | 0.9825 | 1.0825 | 0.025912 | 0.065572 | 0.156912 | | 1 | 0.996511 | 0.751493 |
| Pkd2 | O35245-5 | -0.24 | -0.07 | 1.62 | 0.54 | -0.02 | -0.67 | 1.57 | 0.82 | -0.46 | 0.5 | 1.24 | -0.09 | -0.67 | -0.56 | 1.52 | 1.42 | -0.0375 | -0.165 | -0.035 | 0.949142 | 0.751455 | 0.958198 | | 1 | 0.996511 | 0.992636 |
| Pkia | P63248 | -0.99 | -1.01 | -0.78 | -0.67 | -1.5 | -1.23 | -0.51 | -0.54 | 0.89 | -0.58 | -0.83 | -0.49 | 2.17 | -0.5 | -0.36 | -0.72 | -0.0825 | 0.61 | 1.01 | 0.759 | 0.137174 | 0.132593 | | 1 | 0.996511 | 0.728417 |
| Pkib | Q04758-2 | -0.46 | 0.3 | 0.22 | -0.67 | -0.7 | -0.97 | -0.73 | -0.12 | -0.17 | -0.78 | -0.37 | -0.56 | 0.7 | -0.23 | -0.15 | -0.27 | -0.4775 | -0.3175 | 0.165 | 0.138491 | 0.283914 | 0.625783 | | 1 | 0.996511 | 0.939447 |
| Pkig | O70139 | -0.48 | -0.47 | -0.34 | -0.59 | -0.84 | -0.79 | -0.08 | -0.53 | 0.17 | 0.26 | -0.14 | -0.37 | -0.84 | 0.72 | -0.63 | -0.3 | -0.09 | 0.45 | 0.2075 | 0.668961 | 0.049701 | 0.554794 | | 1 | 0.996511 | 0.916796 |
| Pkm | P52480 | 0.01 | -0.14 | 0.22 | 0.45 | -0.22 | -0.15 | 0.16 | 0.44 | -0.68 | -0.7 | 0.06 | -0.22 | -1.22 | -0.5 | -0.24 | -0.07 | -0.0775 | -0.52 | -0.6425 | 0.727382 | 0.062538 | 0.05571 | | 1 | 0.996511 | 0.661738 |
| Pkm | P52480-2 | -0.77 | -0.76 | -0.63 | -0.47 | -0.74 | -0.74 | -0.84 | -0.73 | -0.74 | -0.85 | -0.83 | -1.03 | -1.05 | -0.54 | -1.2 | -0.95 | -0.105 | -0.205 | -0.2775 | 0.490509 | 0.24876 | 0.215568 | | 1 | 0.996511 | 0.787447 |
| Pkn1 | P70268 | -0.14 | -0.11 | 0.18 | 0.46 | -0.36 | -0.18 | -0.2 | 0.22 | 0.15 | 0.76 | 0.34 | -0.1 | 0.13 | 0.18 | -0.14 | -0.16 | -0.2275 | 0.19 | -0.095 | 0.302933 | 0.458518 | 0.665196 | | 1 | 0.996511 | 0.949825 |
| Pkn2 | G3UXH4 | 0.22 | 0.61 | 0.8 | 0.82 | 0.4 | -0.31 | 0.44 | 0.19 | -0.15 | 1.4 | -0.08 | -0.31 | 0.19 | 0 | 0.32 | 1.1 | -0.4325 | -0.3975 | -0.21 | 0.096686 | 0.337505 | 0.481575 | | 1 | 0.996511 | 0.893906 |
| Pkp2 | Q9CQ73 | 0.93 | 0.53 | 1.78 | 0.43 | 0.97 | 1.17 | 2.1 | 1.01 | 1.1 | 0.22 | 1.89 | 1 | 1.14 | -0.16 | 2.7 | 2.26 | 0.395 | 0.135 | 0.5675 | 0.320153 | 0.756215 | 0.390702 | | 1 | 0.996511 | 0.865375 |
| Pkp4 | Q68FH0 | 1.26 | 1.08 | 1.05 | 1.04 | 0.85 | 1.16 | 1.11 | 0.96 | 0.9 | 1.01 | 1.25 | 1.22 | 0.93 | 0.85 | 1.08 | 1.02 | -0.0875 | -0.0125 | -0.1375 | 0.576939 | 0.942323 | 0.43819 | | 1 | 0.996755 | 0.883161 |
| Pla2g15 | Q8VEB4 | 0.13 | -0.07 | -0.1 | -0.02 | 0.32 | 0.33 | -0.11 | -0.43 | -0.6 | 0.15 | 0.09 | 0.23 | -0.9 | -0.58 | -0.73 | -0.42 | 0.0425 | -0.0175 | -0.6425 | 0.84405 | 0.939334 | 0.008032 | | 1 | 0.996755 | 0.563386 |
| Pla2g4e | Q50L42 | 0.77 | 1.27 | 0.31 | 0.38 | 1.61 | 0.6 | 0.89 | 0.58 | 0.88 | 0.16 | 0.42 | 1.11 | 0.35 | 1.28 | 0.27 | 0.52 | 0.2375 | -0.04 | -0.0775 | 0.464332 | 0.897607 | 0.811294 | | 1 | 0.996511 | 0.972642 |
| Pla2g6 | P97819-2 | 0.62 | 0.49 | 0.31 | -0.22 | 0.66 | 0.75 | 0.21 | -0.03 | -0.52 | -0.21 | 0.14 | 0.73 | -0.46 | -0.06 | -0.33 | -0.2 | 0.0975 | -0.265 | -0.5625 | 0.71672 | 0.423531 | 0.042125 | | 1 | 0.996511 | 0.658454 |
| Pla2g7 | Q60963 | -1.75 | -1.35 | -1.4 | -1.14 | -0.62 | -1.79 | 0.28 | -1.3 | -1.16 | -1.22 | -1.38 | -1.53 | -1.42 | -0.58 | -1.5 | -1.44 | 0.5525 | 0.0875 | 0.175 | 0.226123 | 0.664307 | 0.529797 | | 1 | 0.996511 | 0.908663 |
| Plaa | P27612 | -0.43 | -0.51 | -0.04 | -0.12 | -0.22 | -0.44 | -0.24 | -0.11 | -0.69 | 0.23 | 0.08 | -0.35 | -0.83 | -0.12 | -0.15 | -0.44 | 0.0225 | 0.0925 | -0.11 | 0.900164 | 0.721279 | 0.648473 | | 1 | 0.996511 | 0.946083 |
| Plaat3 | Q8R3U1-2 | 0.74 | 0.65 | 0.04 | 0.03 | 0.56 | 0.94 | -0.56 | -0.29 | 0.45 | 1.86 | 0.08 | 0.46 | 0.04 | 1.06 | 0.13 | 0.56 | -0.2025 | 0.3475 | 0.0825 | 0.597592 | 0.412405 | 0.791197 | | 1 | 0.996511 | 0.969105 |
| Plat | P11214 | -0.5 | 0.06 | -0.57 | -0.37 | 0.07 | -0.29 | 0.2 | -0.01 | -0.49 | 0.99 | -0.27 | -0.22 | -0.82 | -0.85 | 0.57 | 0.56 | 0.3375 | 0.3475 | 0.21 | 0.127624 | 0.340993 | 0.611731 | | 1 | 0.996511 | 0.935047 |
| Plbd2 | Q3TCN2 | 0.02 | -0.17 | -0.99 | -1.04 | 0.06 | -0.05 | -0.31 | -1.04 | -1 | -0.87 | -0.52 | -0.41 | -0.95 | -0.3 | -1.24 | -1.17 | 0.21 | -0.155 | -0.37 | 0.55756 | 0.62038 | 0.301743 | | 1 | 0.996511 | 0.830589 |
| Plcb1 | Q9Z1B3 | 0.17 | -0.3 | 0.14 | 0.37 | -0.52 | 0.17 | 0.33 | 0.56 | -0.14 | -0.05 | 0.18 | -0.39 | -0.57 | -0.18 | -0.39 | -0.39 | 0.04 | -0.195 | -0.4775 | 0.88477 | 0.388118 | 0.051012 | | 1 | 0.996511 | 0.658454 |
| Plcb1 | Q9Z1B3-2 | -0.6 | -0.1 | -0.39 | -0.91 | 0.03 | -0.03 | -0.3 | -0.72 | -0.06 | -0.08 | -1.01 | -0.84 | -1.05 | 0.37 | -0.89 | -1.79 | 0.245 | 0.0025 | -0.34 | 0.345845 | 0.99348 | 0.460454 | | 1 | 0.999285 | 0.887902 |
| Plcb3 | P51432 | 0.18 | 0.55 | 0.59 | 0.98 | 0.5 | -0.06 | 0.33 | 0.68 | 0.14 | 1.17 | 0.57 | -0.11 | -0.02 | 0.25 | 0.52 | 0.46 | -0.2125 | -0.1325 | -0.2725 | 0.391078 | 0.683693 | 0.277667 | | 1 | 0.996511 | 0.822084 |
| Plcb4 | Q91UZ1 | 0.05 | -0.54 | -0.52 | -0.45 | -0.06 | -0.34 | -0.53 | -0.65 | -0.57 | -0.28 | -0.85 | -0.65 | -1.23 | -0.02 | -1.17 | -1.1 | -0.03 | -0.2225 | -0.515 | 0.889187 | 0.32684 | 0.139485 | | 1 | 0.996511 | 0.73493 |
| Plcd1 | Q8R3B1 | 0.86 | 0.66 | 0.69 | 0.87 | 0.8 | 0.76 | 0.79 | 0.82 | 0.37 | 0.24 | 0.58 | 0.39 | 0.11 | 0.42 | 0.28 | 0.28 | 0.0225 | -0.375 | -0.4975 | 0.875584 | 0.050451 | 0.019868 | | 1 | 0.996511 | 0.615793 |
| Plcd3 | Q8K2J0 | 1.78 | 1.97 | 0.9 | 1.03 | 2.06 | 1.54 | 0.24 | 0.16 | 0.92 | 0.36 | 0.42 | 1.06 | 0.14 | 1.72 | -0.32 | 0.3 | -0.42 | -0.73 | -0.96 | 0.412429 | 0.045882 | 0.073008 | | 1 | 0.996511 | 0.668066 |
| Plcd4 | Q8K3R3-2 | -1.55 | -1.39 | 0.68 | 0.38 | -0.87 | -1.74 | 2.02 | 0.65 | -1.09 | 0.96 | 0.68 | -0.89 | -1.28 | -0.69 | 1.02 | 1.14 | 0.485 | 0.385 | 0.5175 | 0.59765 | 0.592228 | 0.501877 | | 1 | 0.996511 | 0.901002 |
| Plcg1 | A2A4A6 | 0.51 | 0.43 | 0.42 | 0.46 | 0.44 | 0.28 | 0.14 | 0.48 | 0.44 | 0.43 | 0.43 | 0.37 | 0.2 | 0.93 | 0.24 | 0.21 | -0.12 | -0.0375 | -0.06 | 0.439628 | 0.804088 | 0.791286 | | 1 | 0.996511 | 0.969105 |
| Plcg2 | Q8CIH5 | 0.1 | -0.93 | -0.67 | -0.64 | -0.16 | -0.33 | -1.21 | -0.73 | 0.39 | -0.03 | -0.71 | 0.01 | 0.02 | 0.63 | -0.83 | -1.17 | -0.0725 | 0.45 | 0.1975 | 0.818492 | 0.18291 | 0.655511 | | 1 | 0.996511 | 0.94818 |
| Plch2 | H3BIW6 | 0.97 | 0.76 | 0.88 | 0.91 | 0.52 | 0.69 | 0.63 | 0.85 | 0.44 | 0.44 | 0.49 | 0.57 | 0.37 | 0.4 | 0.64 | 0.73 | -0.2075 | -0.395 | -0.345 | 0.198293 | 0.030986 | 0.088166 | | 1 | 0.996511 | 0.691619 |
| Plch2 | A2AP18-4 | 1.75 | 0.32 | -0.21 | 1.72 | 0.76 | 1.32 | -1.15 | 0.31 | 5.6 | 4.14 | 0.03 | 0.76 | 3.35 | 6.98 | 0.34 | 0.84 | -0.585 | 1.7375 | 1.9825 | 0.385753 | 0.196217 | 0.186451 | | 1 | 0.996511 | 0.776843 |
| Plcl1 | Q3USB7 | -0.16 | -0.02 | 1.21 | 1.2 | 0.08 | -0.2 | 1.22 | 1.28 | 0.04 | 0.88 | 0.95 | -0.22 | -0.22 | 0.04 | 0.92 | 0.79 | 0.0375 | -0.145 | -0.175 | 0.939511 | 0.747297 | 0.693869 | | 1 | 0.996511 | 0.955729 |
| Plcl2 | Q8K394 | 0.4 | 0.62 | 0.48 | 0.24 | 0.47 | 0.51 | -0.06 | 0.2 | 0.59 | 0.78 | 0.25 | 0.38 | 0.4 | 0.56 | 0.05 | -0.08 | -0.155 | 0.065 | -0.2025 | 0.428402 | 0.738979 | 0.368963 | | 1 | 0.996511 | 0.859245 |
| Plcxd2 | B2RXA1 | -0.87 | -0.61 | -0.58 | -0.27 | -0.22 | -1.01 | -0.38 | -0.13 | -0.52 | -0.32 | -0.6 | -0.85 | -0.2 | 0.21 | -0.64 | -0.69 | 0.1475 | 0.01 | 0.2525 | 0.553967 | 0.96172 | 0.360514 | | 1 | 0.996755 | 0.85741 |
| Plcxd3 | G3X9A7 | -0.05 | 0.65 | -0.37 | -0.19 | 0.54 | 0.27 | -0.08 | 0.16 | -0.41 | -1.13 | -0.23 | 0.2 | -1.22 | -0.72 | -0.99 | -0.09 | 0.2125 | -0.4025 | -0.765 | 0.431261 | 0.267376 | 0.044419 | | 1 | 0.996511 | 0.658454 |
| Pld1 | Q6NVF2 | 1.22 | 1.49 | 0.66 | 0.79 | 1.6 | 0.73 | 0.47 | 0.29 | 1.46 | 0.68 | 0.57 | 1.07 | 1.4 | 1.53 | 1.14 | 0.82 | -0.2675 | -0.095 | 0.1825 | 0.435738 | 0.742421 | 0.507262 | | 1 | 0.996511 | 0.902614 |
| Pld2 | Q6NV49 | 1.15 | 1.43 | 0.86 | 0.49 | 1.47 | 1.35 | 0.62 | 0.61 | 0.49 | -0.28 | 0.95 | 1.37 | 0.27 | 0.63 | 0.89 | 1.14 | 0.03 | -0.35 | -0.25 | 0.921095 | 0.381222 | 0.399757 | | 1 | 0.996511 | 0.868507 |
| Pld3 | O35405 | 0.33 | 0.5 | 0.17 | 0.38 | 0.32 | 0.76 | 1.19 | 0.21 | 0.46 | 0.07 | 0.77 | 0.56 | 0.43 | 0.9 | -0.21 | 0.54 | 0.275 | 0.12 | 0.07 | 0.283649 | 0.565499 | 0.793425 | | 1 | 0.996511 | 0.969365 |
| Plec | Q9QXS1-6 | 1.18 | 0.85 | 1.2 | 1.21 | 0.75 | 1.06 | 0.65 | 1.02 | 1.07 | 1.03 | 1.23 | 0.93 | 1.08 | 0.87 | 0.96 | 1.06 | -0.24 | -0.045 | -0.1175 | 0.202784 | 0.799236 | 0.5286 | | 1 | 0.996511 | 0.908663 |
| Plek | Q9JHK5 | -2.04 | -3.16 | -0.6 | -0.56 | -2.77 | -1.86 | -1.08 | -0.8 | -2.22 | 0.48 | -0.29 | -3.37 | -1.9 | -2.03 | -1.63 | -0.45 | -0.0375 | 0.24 | 0.0875 | 0.956898 | 0.805081 | 0.894157 | | 1 | 0.996511 | 0.98327 |
| Plek2 | Q9WV52 | -0.69 | 0.31 | -0.65 | -1.29 | 1.15 | -0.62 | -1.48 | -2.08 | -0.5 | 0.4 | -0.9 | -1.01 | 0.5 | 0.78 | -0.42 | -0.42 | -0.1775 | 0.0775 | 0.69 | 0.800982 | 0.858544 | 0.136998 | | 1 | 0.996511 | 0.732318 |
| Plekha1 | Q8BUL6 | -0.17 | -0.31 | 0.45 | 0.36 | -0.3 | -0.25 | -0.09 | 0.15 | -0.71 | 1.08 | -0.03 | -0.22 | -0.07 | -0.74 | -0.16 | 0.06 | -0.205 | -0.0525 | -0.31 | 0.389736 | 0.896472 | 0.284737 | | 1 | 0.996511 | 0.822669 |
| Plekha2 | Q9ERS5 | -0.42 | -0.31 | 0.78 | 0.35 | -0.89 | -0.46 | -0.1 | 0.37 | 0.26 | 0.79 | -0.01 | -0.17 | 0.18 | 0.2 | 0.05 | -0.39 | -0.37 | 0.1175 | -0.09 | 0.333251 | 0.735261 | 0.77952 | | 1 | 0.996511 | 0.967289 |
| Plekha5 | E9Q6H8 | 1.53 | 1.03 | 1.01 | 0.77 | 1.17 | 1.3 | 0.98 | 0.89 | 0.91 | 0.58 | 1.2 | 1.58 | 0.95 | 1.17 | 1.01 | 1.03 | -2.4E-16 | -0.0175 | -0.045 | 1 | 0.949862 | 0.836814 | | 1 | 0.996755 | 0.975831 |
| Plekha5 | A0A0N4SW | -0.89 | 1.02 | 0.52 | 0.62 | -0.29 | 0.04 | 0.82 | 2.21 | 0.95 | -0.59 | 1.3 | 0.39 | 0.19 | 0.35 | -0.63 | 1.5 | 0.3775 | 0.195 | 0.035 | 0.554433 | 0.719614 | 0.949919 | | 1 | 0.996511 | 0.992476 |
| Plekha6 | Q7TQG1 | 1.04 | 1.09 | 1.54 | 1.26 | 1 | 0.95 | 0.98 | 0.85 | 1.56 | 0.51 | 1.3 | 0.79 | 2.03 | 0.75 | 1.47 | 1.37 | -0.2875 | -0.1925 | 0.1725 | 0.118798 | 0.493018 | 0.567518 | | 1 | 0.996511 | 0.921806 |
| Plekha7 | A0A0U1RP | 0.7 | 0.84 | 0.76 | 0.82 | 0.89 | 1.04 | 0.8 | 0.65 | 1.07 | 0.44 | 1.1 | 0.55 | 0.97 | 0.17 | 0.5 | 0.76 | 0.065 | 0.01 | -0.18 | 0.677225 | 0.96285 | 0.430131 | | 1 | 0.996755 | 0.879165 |
| Plekhb1 | Q9QYE9-4 | 1.11 | 0.76 | 1.01 | 0.96 | 1.26 | 0.54 | 0.44 | 0.8 | 0.32 | 1.3 | 0.77 | 1.13 | -0.4 | 0.6 | 0.89 | 1.12 | -0.2 | -0.08 | -0.4075 | 0.376306 | 0.751088 | 0.253547 | | 1 | 0.996511 | 0.809524 |
| Plekhd1 | B2RPU2 | 0.37 | 0.07 | 0.63 | 0.75 | 0.27 | 0.03 | 0.39 | 0.58 | 0.73 | -0.54 | 0.79 | 0.04 | 1.03 | 0.85 | 0.46 | 0.56 | -0.1375 | -0.2 | 0.27 | 0.52803 | 0.564281 | 0.276665 | | 1 | 0.996511 | 0.822084 |
| Plekhf2 | Q91WB4 | 0.54 | 0.15 | 0.01 | 0 | 0.02 | 0.55 | -0.39 | 0.14 | 0.31 | 1.74 | -0.11 | 0.52 | 0.02 | 0.43 | -0.55 | -0.17 | -0.095 | 0.44 | -0.2425 | 0.699184 | 0.28712 | 0.373763 | | 1 | 0.996511 | 0.859424 |
| Plekhg1 | Q3UX37 | -0.34 | 0.13 | 0.67 | 0.06 | -0.55 | 0.39 | -1.08 | -0.49 | 1.65 | -1.08 | 0.24 | 0.33 | 1.99 | -0.22 | 0.81 | 0.09 | -0.5625 | 0.155 | 0.5375 | 0.138882 | 0.777965 | 0.296165 | | 1 | 0.996511 | 0.825612 |
| Plekhg3 | Q4VAC9-2 | 0.36 | 1.18 | -0.23 | 0.98 | 1.28 | -0.13 | -0.55 | 0.37 | 0.77 | -0.53 | 0.38 | 0.5 | 0.26 | 1.04 | 0.84 | -0.41 | -0.33 | -0.2925 | -0.14 | 0.488962 | 0.478404 | 0.747419 | | 1 | 0.996511 | 0.960744 |
| Plekhg5 | B1AS67 | -0.13 | -0.37 | -0.36 | -0.35 | -0.07 | -0.23 | -0.51 | -0.43 | -0.51 | -0.22 | -0.42 | -0.24 | -0.19 | -0.07 | -1.11 | -0.69 | -0.0075 | -0.045 | -0.2125 | 0.96452 | 0.791011 | 0.440623 | | 1 | 0.996511 | 0.883549 |
| Plekhh2 | Q8C115-2 | 0.96 | 0.59 | 0.47 | 0.34 | 0.69 | 0.77 | 0.17 | 0.64 | 0.62 | 0.63 | 0.56 | 0.65 | 0.49 | 1.52 | 0.35 | -0.54 | -0.0225 | 0.025 | -0.135 | 0.916778 | 0.895865 | 0.749839 | | 1 | 0.996511 | 0.961456 |
| Plekhm2 | A2ADE0 | 0.86 | 0.79 | 1.72 | 0.31 | 0.95 | 0.85 | 1.31 | 1.03 | -0.15 | -0.08 | 1.5 | 0.83 | 0.18 | -0.47 | 1.89 | 2.05 | 0.115 | -0.395 | -0.0075 | 0.707802 | 0.401788 | 0.990504 | | 1 | 0.996511 | 0.996154 |
| Plekho1 | F6XQM2 | 0.06 | -0.25 | 1.94 | 1.78 | -0.01 | 0.14 | 1.64 | 2.46 | 1.09 | 1.13 | 1.86 | -0.12 | 1.39 | -0.99 | 1.72 | 1.98 | 0.175 | 0.1075 | 0.1425 | 0.814585 | 0.866661 | 0.858722 | | 1 | 0.996511 | 0.976225 |
| Plekho2 | Q8K124 | 1.55 | 1.39 | 0.55 | 0.73 | 1.02 | 0.87 | 0.64 | 0.64 | 0.95 | 2.32 | 0.7 | 0.87 | 0.85 | 0.71 | 0.21 | 0.59 | -0.2625 | 0.155 | -0.465 | 0.341337 | 0.715162 | 0.141687 | | 1 | 0.996511 | 0.737973 |
| Plg | P20918 | -3.99 | -4.67 | -5.07 | -5.38 | -3.9 | -4.39 | -5.25 | -5.11 | -3.82 | -1.53 | -4.38 | -3.8 | -3.31 | -3.18 | -4.18 | -4.83 | 0.115 | 1.395 | 0.9025 | 0.779384 | 0.053818 | 0.077646 | | 1 | 0.996511 | 0.676942 |
| Plgrkt | Q9D3P8 | 0.88 | 0.86 | 1.44 | 1.97 | 0.41 | 0.53 | 1.16 | 1.56 | 1.25 | 1.19 | 1.53 | 0.58 | 1.41 | -0.41 | 0.98 | 0.89 | -0.3725 | -0.15 | -0.57 | 0.318456 | 0.650645 | 0.224378 | | 1 | 0.996511 | 0.787447 |
| Plin3 | Q9DBG5 | -0.82 | -0.57 | 0.24 | 0.33 | -0.47 | -1.05 | 0.65 | 0.28 | -0.68 | 0.09 | 0.05 | -0.87 | -0.36 | -0.78 | 0.36 | 0.71 | 0.0575 | -0.1475 | 0.1875 | 0.897075 | 0.68984 | 0.659741 | | 1 | 0.996511 | 0.949645 |
| Pllp | Q9DCU2 | 0.16 | 0.49 | 1.04 | 0.91 | -0.95 | 0.13 | 0.64 | 0.4 | 2.3 | 0.72 | 0.28 | -0.32 | 2.55 | -0.7 | 0.6 | 0.61 | -0.595 | 0.095 | 0.115 | 0.147391 | 0.86267 | 0.856946 | | 1 | 0.996511 | 0.976225 |
| Plod1 | Q9R0E2 | -0.94 | -0.35 | -0.68 | -0.78 | 0.11 | -0.55 | -0.6 | -0.71 | -1.08 | -0.19 | -0.64 | -0.54 | -0.58 | -1.23 | -0.86 | -0.31 | 0.25 | 0.075 | -0.0575 | 0.311851 | 0.761265 | 0.825927 | | 1 | 0.996511 | 0.973155 |
| Plod3 | Q9R0E1 | -1.6 | -2.22 | -1.35 | -1.86 | -1.69 | -1.49 | -1.52 | -1.43 | -1.47 | -0.18 | -1.3 | -1.11 | 0.14 | -2.4 | -0.35 | -0.48 | 0.225 | 0.7425 | 0.985 | 0.318113 | 0.052899 | 0.097783 | | 1 | 0.996511 | 0.698156 |
| Plp1 | P60202 | -2.72 | -3 | -2.66 | -2.95 | -2.12 | -3.3 | -2.29 | -3.84 | -3.72 | -2.89 | -3.95 | -2.84 | -3.79 | -4.1 | -3.67 | -3.84 | -0.055 | -0.5175 | -1.0175 | 0.889415 | 0.113891 | 0.000596 | | 1 | 0.996511 | 0.294873 |
| Plp2 | Q9R1Q7 | 5.46 | 3.08 | 1.7 | -1.27 | 4.48 | 5.84 | 1.54 | 2.09 | 1.12 | -4.99 | 3.21 | 6.07 | 2.48 | -0.94 | 2.99 | 2.28 | 1.245 | -0.89 | -0.54 | 0.429247 | 0.715455 | 0.715818 | | 1 | 0.996511 | 0.958106 |
| Plpbp | A0A1B0GR | 0.7 | 0.34 | 0.16 | 0.36 | 0.28 | 0.47 | 0.06 | 0.13 | 0.34 | 0.84 | 0.17 | 0.65 | 0.31 | 0.72 | 0.26 | 0.13 | -0.155 | 0.11 | -0.035 | 0.414058 | 0.625046 | 0.873948 | | 1 | 0.996511 | 0.979626 |
| Plpp1 | Q61469-2 | -0.18 | 0.41 | 0.28 | 0.68 | 0.2 | 0.56 | -0.92 | 0.63 | 0.47 | 1.14 | 0.27 | 0.45 | 0.85 | 0.86 | 1.03 | 0.69 | -0.18 | 0.285 | 0.56 | 0.638282 | 0.316884 | 0.038217 | | 1 | 0.996511 | 0.658454 |
| Plpp3 | Q99JY8 | -0.01 | 0.52 | 0.11 | 0.61 | 0.6 | 0.27 | 0.83 | 0.14 | 0.08 | 1.02 | -0.1 | 0.03 | -0.09 | 0.33 | 0.15 | 0.04 | 0.1525 | -0.05 | -0.2 | 0.522386 | 0.869067 | 0.383394 | | 1 | 0.996511 | 0.860984 |
| Plpp4 | Q0VBU9 | 0.16 | 0.66 | -0.03 | 0.35 | 0.72 | 0.39 | 0.29 | 0.01 | 0.05 | -0.23 | 0.13 | 0.11 | -0.39 | 0.41 | -0.54 | -1.27 | 0.0675 | -0.27 | -0.7325 | 0.767482 | 0.223061 | 0.072841 | | 1 | 0.996511 | 0.668066 |
| Plpp6 | Q9D4F2 | -1.61 | -2.03 | -1.76 | -2.19 | -1.02 | -1.27 | -1.38 | -2.15 | -1.03 | -2.29 | -2.05 | -2 | -0.84 | -1.95 | -2.14 | -1.68 | 0.4425 | 0.055 | 0.245 | 0.140356 | 0.85927 | 0.453311 | | 1 | 0.996511 | 0.885993 |
| Plpp7 | Q91WB2 | -0.27 | -0.05 | -0.1 | -0.46 | 0.07 | -0.81 | -0.57 | -0.7 | 0.45 | 0.41 | -0.8 | -0.55 | 0.8 | 1.12 | -0.1 | -0.65 | -0.2825 | 0.0975 | 0.5125 | 0.249583 | 0.770182 | 0.223681 | | 1 | 0.996511 | 0.787447 |
| Plppr1 | Q8BFZ2-2 | 2.12 | 1.66 | 0.67 | 0.6 | 1.42 | 1.74 | 0.36 | 0.4 | 2.33 | 2.41 | 0.81 | 2.47 | 2.28 | 1.46 | 0.69 | 0.13 | -0.2825 | 0.7425 | -0.1225 | 0.557193 | 0.169571 | 0.82472 | | 1 | 0.996511 | 0.972852 |
| Plppr2 | A0A087WC | -0.01 | 0.27 | 0.48 | 0.54 | 0.74 | 0.59 | 0.31 | 0.3 | 0.56 | 1.37 | 0.41 | 0.43 | 0.63 | 0.96 | 0.54 | 0.51 | 0.165 | 0.3725 | 0.34 | 0.415732 | 0.197593 | 0.141259 | | 1 | 0.996511 | 0.737973 |
| Plppr3 | A0A0R3P9 | 1.04 | 0.96 | 0.95 | 0.91 | 0.4 | 0.79 | 0.9 | 1 | 1.5 | 1.16 | 1.3 | 1.53 | 1.51 | 1.34 | 1.52 | 0.62 | -0.1925 | 0.4075 | 0.2825 | 0.300146 | 0.037708 | 0.274619 | | 1 | 0.996511 | 0.822084 |
| Plppr4 | A0A0G2JG | 0.61 | 0.41 | 1.01 | 0.89 | 0.75 | 0.7 | 0.93 | 0.85 | 1.36 | 1.38 | 1.29 | 0.88 | 1.86 | 1.1 | 1.17 | 0.86 | 0.0775 | 0.4975 | 0.5175 | 0.678467 | 0.045559 | 0.088009 | | 1 | 0.996511 | 0.691619 |
| Plppr5 | A0A668KM | 0.57 | 0.37 | 1.25 | 0.97 | -0.34 | 0.8 | -0.14 | 0.6 | 1.95 | 1.84 | 1.18 | 0.65 | 1.88 | 0.85 | 0.61 | 0.03 | -0.56 | 0.615 | 0.0525 | 0.118289 | 0.110324 | 0.899378 | | 1 | 0.996511 | 0.983509 |
| Plrg1 | Q922V4 | 0.73 | -0.96 | -1.29 | -0.75 | -1.1 | -0.13 | -1.2 | -0.16 | -0.35 | -1.24 | -0.76 | -0.86 | 0.62 | -0.42 | -0.03 | -0.81 | -0.08 | -0.235 | 0.4075 | 0.870865 | 0.606993 | 0.427408 | | 1 | 0.996511 | 0.879165 |
| Pls1 | Q3V0K9 | 0.32 | 0.33 | 0.19 | -0.06 | 0.44 | 0.04 | -0.48 | -0.42 | -0.14 | -0.32 | -0.11 | -0.45 | -0.17 | -0.11 | 0.06 | -0.24 | -0.3 | -0.45 | -0.31 | 0.244839 | 0.034733 | 0.128521 | | 1 | 0.996511 | 0.723655 |
| Pls3 | B1AX58 | 0.61 | 0.25 | 0.38 | 0.06 | 0.17 | 0.61 | 0.54 | 0.53 | 0.2 | -0.48 | 0.53 | 0.34 | 0.25 | -0.25 | 0.05 | 0.26 | 0.1375 | -0.1775 | -0.2475 | 0.477266 | 0.509385 | 0.274768 | | 1 | 0.996511 | 0.822084 |
| Plscr3 | Q9JIZ9 | 2.18 | 1.94 | 1.47 | 2 | 0.95 | 1.81 | 1.5 | 2.12 | 2.82 | 2.09 | 1.57 | 1.94 | 2.36 | 3.1 | 1.39 | 1.77 | -0.3025 | 0.2075 | 0.2575 | 0.315143 | 0.504414 | 0.514347 | | 1 | 0.996511 | 0.90567 |
| Pltp | P55065 | -2.57 | -2.58 | -1.52 | -2.3 | -1.97 | -2.4 | -2.54 | -2.97 | -2.22 | -0.12 | -2.15 | -2.39 | -2.22 | -1.58 | -2.31 | -2.29 | -0.2275 | 0.5225 | 0.1425 | 0.479723 | 0.351276 | 0.650726 | | 1 | 0.996511 | 0.947407 |
| Plxdc1 | Q91ZV7 | 0.28 | 0.8 | 0.34 | 0.08 | 0.07 | 0.14 | 0.01 | 0.17 | 0.76 | 0.63 | 0.37 | 0.34 | 1.27 | 1.26 | -0.4 | 0.25 | -0.2775 | 0.15 | 0.22 | 0.175683 | 0.501696 | 0.600963 | | 1 | 0.996511 | 0.930802 |
| Plxdc2 | Q9DC11 | -0.67 | -0.55 | 0.2 | 0.94 | -0.79 | -0.18 | 0.39 | 0.28 | -0.94 | 0.33 | 0.48 | -0.15 | -1.28 | -0.93 | 1.18 | 1.06 | -0.055 | -0.05 | 0.0275 | 0.89818 | 0.913571 | 0.967641 | | 1 | 0.996755 | 0.9927 |
| Plxna1 | P70206 | 0.53 | 0.72 | 0.79 | 0.79 | 0.94 | 0.56 | 0.99 | 0.81 | 0.78 | 1.04 | 0.99 | 0.92 | 0.61 | 1.21 | 0.9 | 0.78 | 0.1175 | 0.225 | 0.1675 | 0.491465 | 0.199961 | 0.420754 | | 1 | 0.996511 | 0.875639 |
| Plxna2 | P70207 | 0.11 | 0.51 | 0.65 | 0.49 | 0.74 | 0.36 | 0.8 | 0.45 | 0.26 | -0.03 | 0.74 | 0.4 | 0.07 | 0.7 | 0.17 | 0.38 | 0.1475 | -0.0975 | -0.11 | 0.455193 | 0.672637 | 0.630522 | | 1 | 0.996511 | 0.941132 |
| Plxna3 | P70208 | 0.65 | 0.62 | 0.69 | 0.72 | 0.71 | 0.91 | 0.73 | 0.84 | 1.08 | 1.17 | 0.9 | 1.16 | 0.99 | 0.98 | 1.26 | 0.78 | 0.1275 | 0.4075 | 0.3325 | 0.381858 | 0.030005 | 0.099498 | | 1 | 0.996511 | 0.698156 |
| Plxna4 | Q80UG2 | 0.24 | 0.7 | 0.98 | 0.95 | 0.71 | 0.41 | 1.02 | 0.96 | 0.42 | 0.82 | 0.86 | 0.33 | 0.22 | 0.43 | 0.77 | 0.63 | 0.0575 | -0.11 | -0.205 | 0.808543 | 0.654043 | 0.412526 | | 1 | 0.996511 | 0.874811 |
| Plxnb1 | Q8CJH3 | 0.45 | 0.82 | 0.82 | 0.61 | 0.95 | 0.73 | 0.72 | 0.67 | 0.55 | 0.45 | 0.74 | 0.73 | 0.3 | 0.72 | 0.6 | 0.64 | 0.0925 | -0.0575 | -0.11 | 0.580419 | 0.750428 | 0.581154 | | 1 | 0.996511 | 0.927734 |
| Plxnb2 | B2RXS4 | 1.18 | 0.69 | 0.5 | 0.45 | 0.69 | 1.04 | 0.49 | 0.35 | 0.56 | 0.5 | 0.29 | 1.25 | 0.43 | 0.44 | 0.37 | 0.35 | -0.0625 | -0.055 | -0.3075 | 0.794524 | 0.843806 | 0.185078 | | 1 | 0.996511 | 0.776843 |
| Plxnb3 | Q9QY40 | -0.42 | 0.31 | 0.37 | 0.39 | 0.45 | -0.11 | 0.11 | 0.19 | -0.31 | -0.06 | 0.45 | 0.01 | -0.55 | 0.05 | 0.31 | 0.16 | -0.0025 | -0.14 | -0.17 | 0.991737 | 0.603525 | 0.55947 | | 1 | 0.996511 | 0.918409 |
| Plxnc1 | Q9QZC2 | 0.32 | 0.25 | 0.36 | 0.44 | 0.1 | 0.47 | 1.53 | 0.71 | 0.59 | 0.24 | 0.6 | 0.49 | 0.31 | -0.07 | 0.96 | 0.9 | 0.36 | 0.1375 | 0.1825 | 0.253909 | 0.428418 | 0.509466 | | 1 | 0.996511 | 0.902804 |
| Plxnd1 | Q3UH93 | 0.83 | 0.69 | 0.82 | 0.69 | 0.91 | 0.98 | 1.16 | 0.83 | 0.7 | 0.5 | 0.98 | 1.14 | 0.39 | 0.67 | 0.73 | 0.99 | 0.2125 | 0.0725 | -0.0625 | 0.187319 | 0.715843 | 0.75279 | | 1 | 0.996511 | 0.961479 |
| Pm20d1 | Q8C165 | -0.25 | -0.38 | 0.74 | 0.99 | 0.5 | 0.05 | 1.23 | 0.33 | 0.41 | 1.74 | 0.41 | 0.29 | 0.89 | 0.63 | 0.66 | 1.28 | 0.2525 | 0.4375 | 0.59 | 0.53544 | 0.352021 | 0.136761 | | 1 | 0.996511 | 0.732318 |
| Pmch | P56942 | -0.09 | 0.31 | -0.02 | 0.42 | 0.71 | -0.34 | 0.22 | -0.04 | -0.59 | -0.02 | -0.49 | -0.36 | -0.78 | 0.26 | 0.71 | 0.3 | -0.0175 | -0.52 | -0.0325 | 0.946849 | 0.03688 | 0.924025 | | 1 | 0.996511 | 0.988152 |
| Pml | D3Z3A6 | 1.47 | 0.68 | -0.06 | 0.43 | 0.15 | 0.43 | -0.69 | -0.16 | 0.06 | 0.46 | 0.17 | 0.45 | -0.63 | 0.64 | -0.28 | 0.09 | -0.6975 | -0.345 | -0.675 | 0.093816 | 0.314466 | 0.121639 | | 1 | 0.996511 | 0.721448 |
| Pmm1 | O35621 | 0 | 0.35 | -0.18 | -0.15 | 0.19 | -0.08 | 0.16 | -0.16 | 0.25 | -0.02 | 0.05 | 0.18 | -0.49 | 0.48 | -0.27 | -0.64 | 0.0225 | 0.11 | -0.235 | 0.905086 | 0.569854 | 0.430406 | | 1 | 0.996511 | 0.879165 |
| Pmm2 | Q9Z2M7 | 0.25 | 0.08 | 0.97 | 0.95 | 0.09 | -0.06 | 0.3 | 0.7 | -0.06 | 0.91 | 0.78 | 0.28 | -0.64 | 0.14 | 0.42 | 0.54 | -0.305 | -0.085 | -0.4475 | 0.302661 | 0.792532 | 0.222659 | | 1 | 0.996511 | 0.787447 |
| Pmpca | Q9DC61 | -0.01 | -0.19 | 0.29 | 0.6 | -0.45 | 0.14 | 0.28 | 0.3 | 0.16 | 0.3 | 0.51 | 0.05 | 0.26 | 0.28 | 0.13 | 0.34 | -0.105 | 0.0825 | 0.08 | 0.684166 | 0.722229 | 0.724216 | | 1 | 0.996511 | 0.958398 |
| Pmpcb | Q9CXT8 | 0.6 | 0.38 | 0.55 | 0.78 | 0.19 | 0.61 | 0.6 | 0.8 | 0.29 | 0.35 | 0.93 | 0.68 | 0.28 | 0.37 | 0.5 | 0.55 | -0.0275 | -0.015 | -0.1525 | 0.885346 | 0.943624 | 0.420593 | | 1 | 0.996755 | 0.87558 |
| Pmvk | D3YXP6 | -1.85 | -2.17 | -2.11 | -2.03 | -0.88 | -1.4 | -1.16 | -2.3 | -0.25 | -2.64 | -1.5 | -1.32 | 0.26 | -1.35 | -1.75 | -1.81 | 0.605 | 0.6125 | 0.8775 | 0.078431 | 0.209005 | 0.084466 | | 1 | 0.996511 | 0.690427 |
| Pnck | Q9QYK9 | -0.35 | 0.08 | -1.09 | -0.96 | -0.67 | -0.46 | -1 | -0.76 | -0.8 | -0.07 | -0.45 | -0.33 | -0.33 | -1.22 | -0.56 | -0.96 | -0.1425 | 0.1675 | -0.1875 | 0.630161 | 0.596663 | 0.583007 | | 1 | 0.996511 | 0.927734 |
| Pnisr | A2AJT4 | 1.47 | 0.94 | 0.37 | -0.67 | -1.16 | 0.59 | -1.83 | -0.02 | 0.87 | -1.55 | -0.4 | -0.24 | 1.65 | 0.19 | -0.05 | -1.34 | -1.1325 | -0.8575 | -0.415 | 0.108229 | 0.188936 | 0.554044 | | 1 | 0.996511 | 0.916796 |
| Pnkd | Q69ZP3 | -0.08 | 0.21 | -0.33 | -0.26 | 0.66 | -0.33 | 0.27 | -0.29 | 0.29 | 0.46 | -0.31 | -0.09 | 0.27 | 1.05 | 0.2 | -0.01 | 0.1925 | 0.2025 | 0.4925 | 0.485127 | 0.40913 | 0.10736 | | 1 | 0.996511 | 0.704959 |
| Pnkd | A0A0R4J1J | 2.01 | 0.87 | 0.8 | -0.07 | -1.06 | 1.39 | -0.99 | -0.04 | 0.79 | -0.19 | 0.76 | 1.38 | 1.97 | -0.62 | 0.57 | 1.16 | -1.0775 | -0.2175 | -0.1325 | 0.123812 | 0.664337 | 0.833939 | | 1 | 0.996511 | 0.975371 |
| Pnkp | G5E8N7 | -2.12 | -2.83 | -1.91 | -1.79 | -2.19 | -3.19 | -1.6 | -1.87 | -2.44 | -1.24 | -2.05 | -2.31 | -2.87 | -2.09 | -2.11 | -1.6 | -0.05 | 0.1525 | -0.005 | 0.899203 | 0.663641 | 0.988543 | | 1 | 0.996511 | 0.996154 |
| Pnma2 | Q8BHK0 | -0.63 | 0.13 | -0.55 | -1.13 | 0.36 | -0.7 | -0.9 | -0.85 | 0.4 | -0.35 | -0.15 | 0.05 | 2.35 | 1.85 | 0.59 | 0.34 | 0.0225 | 0.5325 | 1.8275 | 0.952109 | 0.111549 | 0.005913 | | 1 | 0.996511 | 0.552614 |

| | | | | | | | | | | | | | | | | | | | | | | | | | | | |
|---|---|---|---|---|---|---|---|---|---|---|---|---|---|---|---|---|---|---|---|---|---|---|---|---|---|---|---|
| Pnma8a | Q80VM8 | 0.47 | 0.95 | 0.58 | 1.33 | 0.64 | 1.22 | 0.39 | 0.91 | 0.69 | 0.89 | 0.99 | 0.18 | 0.77 | 1.02 | 1.41 | 1.19 | -0.0425 | -0.145 | 0.265 | 0.874928 | 0.605133 | 0.330361 | | 1 | 0.996511 | 0.845524 |
| Pnma8b | G3X9N3 | -0.83 | -1.22 | -1.2 | -1.12 | -1.72 | -1.01 | -1.8 | -1.69 | -0.48 | -0.85 | -1.36 | -1.07 | 0.04 | -1.23 | -1.73 | -2.36 | -0.4625 | 0.1525 | -0.2275 | 0.066378 | 0.522728 | 0.640122 | | 1 | 0.996511 | 0.942804 |
| Pnn | O35691 | 0.97 | -0.19 | 0.43 | 0.95 | -1.19 | 0.4 | -0.64 | 1.07 | 0.81 | 0.95 | 0.18 | -1.01 | 2.23 | 0.31 | 1.03 | -0.27 | -0.63 | -0.3075 | 0.285 | 0.254353 | 0.5336 | 0.611428 | | 1 | 0.996511 | 0.935047 |
| Pno1 | Q9CPS7 | 2.78 | 1.6 | 1.72 | 2.09 | -0.14 | 1.7 | -0.71 | 0.74 | 2.73 | 2.04 | 1.22 | 2.45 | 2.98 | 0.42 | 1.66 | 1.41 | -1.65 | 0.0625 | -0.43 | 0.013448 | 0.876846 | 0.438786 | | 1 | 0.996511 | 0.883205 |
| Pnoc | Q64387 | -0.07 | 1.22 | 0.4 | 0.49 | 0.59 | 0.42 | 0.23 | 1.03 | 0.71 | 0.76 | -0.92 | 0.49 | 0.7 | 0.76 | 1.76 | 1.66 | 0.0575 | -0.25 | 0.71 | 0.853531 | 0.58208 | 0.088634 | | 1 | 0.996511 | 0.691619 |
| Pnp | Q543K9 | -0.48 | -0.49 | -0.29 | -0.37 | -0.14 | 0 | -0.02 | 0.02 | -0.56 | -0.24 | -0.3 | -0.66 | -0.65 | -0.74 | -0.87 | -0.22 | 0.3725 | -0.0325 | -0.2125 | 0.028469 | 0.855939 | 0.322408 | | 1 | 0.996511 | 0.840788 |
| Pnpla2 | Q8BJ56-3 | 2.73 | 2.11 | -0.15 | -0.14 | 1.47 | 2.49 | -1.02 | -0.28 | 2.29 | 0.82 | -0.08 | 1.84 | 1.08 | 0.45 | 1.44 | -0.78 | -0.4725 | 0.08 | -0.59 | 0.633886 | 0.923197 | 0.47153 | | 1 | 0.996755 | 0.891144 |
| Pnpla6 | A0A140LHU | -0.61 | -0.05 | 0.01 | -0.09 | 0.2 | -0.46 | -0.03 | 0 | -0.19 | 0.15 | -0.07 | -0.6 | -0.18 | 0.45 | 0.63 | 0.55 | 0.1125 | 0.0075 | 0.5475 | 0.61499 | 0.975068 | 0.061926 | | 1 | 0.997515 | 0.662517 |
| Pnpla7 | A2AJ88-2 | -0.49 | 0.01 | 0.78 | 0.43 | 0.63 | 0.12 | 1.4 | 0.93 | -0.59 | 0.96 | 1.07 | -0.08 | 0.67 | -0.46 | 0.68 | 1.63 | 0.5875 | 0.1575 | 0.4475 | 0.135719 | 0.731703 | 0.359798 | | 1 | 0.996511 | 0.856645 |
| Pnpla8 | Q8K1N1 | 0.37 | 0.76 | -0.19 | -0.02 | 0.71 | 0.45 | 0.05 | -0.11 | 0.58 | 0.15 | 0.14 | 0.75 | 0.8 | 1.08 | -0.26 | -0.22 | 0.045 | 0.175 | 0.12 | 0.874213 | 0.530421 | 0.760134 | | 1 | 0.996511 | 0.963123 |
| Pnpo | Q91XF0 | 1.18 | 0.39 | 0.6 | 1.03 | 0.56 | 0.87 | 1 | 1.06 | 1.52 | 1.15 | 0.5 | 1.04 | 1.29 | 1.11 | 1.35 | 0.93 | 0.0725 | 0.2525 | 0.37 | 0.756313 | 0.392989 | 0.154756 | | 1 | 0.996511 | 0.750763 |
| Pnpt1 | Q8K1R3 | 1.02 | 0.78 | 1.05 | 1.04 | 0.68 | 1.58 | 1.31 | 0.82 | 0.96 | 0.72 | 1.23 | 0.86 | 0.68 | 0.93 | 1.02 | 0.97 | 0.125 | -0.03 | -0.0725 | 0.600105 | 0.871881 | 0.695057 | | 1 | 0.996511 | 0.955729 |
| Podxl | Q9R0M4 | 2.13 | 1.68 | 0.5 | -0.04 | 2.99 | 1.07 | 1.01 | 0.06 | 0.7 | 0.01 | 0.4 | 1.75 | 0.27 | 1.1 | 0.59 | 1.12 | 0.215 | -0.3525 | -0.2975 | 0.765406 | 0.545276 | 0.561503 | | 1 | 0.996511 | 0.91997 |
| Podxl2 | G3X9D3 | 1.38 | 1.19 | 0.83 | 0.83 | 1.55 | 1.18 | 1.03 | 0.68 | 0.32 | 0.71 | 0.92 | 1.28 | 0.56 | 1.09 | 1.39 | 0.79 | 0.0525 | -0.25 | -0.1 | 0.827953 | 0.352317 | 0.698177 | | 1 | 0.996511 | 0.955729 |
| Pofut1 | A2AMC3 | -0.75 | -0.45 | -1.4 | -1.63 | -0.03 | -0.4 | -0.33 | -1.01 | -0.53 | -2.78 | -1.34 | -0.51 | -0.76 | -1.08 | -0.59 | -1.47 | 0.615 | -0.2325 | 0.0825 | 0.092716 | 0.675593 | 0.807126 | | 1 | 0.996511 | 0.97178 |
| Pofut2 | Q8VHI3 | -0.94 | -0.98 | -0.9 | -0.59 | -0.19 | -1.27 | -0.35 | -0.68 | -0.12 | 0.21 | 0.2 | -0.19 | 0.02 | 0.54 | 1.02 | 0.94 | 0.23 | 0.8775 | 1.4825 | 0.392213 | 0.001452 | 0.000423 | | 1 | 0.996511 | 0.270112 |
| Poglut1 | Q8BYB9 | -0.42 | -0.15 | -1.58 | -2.07 | 0.55 | -0.88 | -1.2 | -1.55 | -1.13 | -0.82 | -2.4 | -0.76 | -0.84 | 0.53 | -0.61 | -0.91 | 0.285 | -0.2225 | 0.5975 | 0.633205 | 0.687703 | 0.274819 | | 1 | 0.996511 | 0.822084 |
| Poglut3 | G5E897 | -0.13 | -0.38 | 0.57 | 0.33 | 0.17 | 0.18 | 0.58 | 0.43 | 0 | 0.15 | -0.14 | 0.11 | 0.41 | -0.3 | 0.48 | 0.62 | 0.2425 | -0.0675 | 0.205 | 0.345073 | 0.786428 | 0.511182 | | 1 | 0.996511 | 0.90359 |
| Pogz | D3YUW8 | 0.38 | -0.13 | 0.07 | -0.06 | -0.32 | 0.21 | -0.04 | 0.03 | 0.49 | -0.13 | -0.35 | -0.06 | 0.53 | 0.14 | 0.18 | 0.06 | -0.095 | -0.0775 | 0.1625 | 0.6266 | 0.746444 | 0.449035 | | 1 | 0.996511 | 0.884817 |
| Polb | Q8K409 | -3.24 | -3.69 | -3.94 | -3.65 | -3.15 | -3.17 | -3.78 | -3.98 | -3.47 | -2.11 | -3.6 | -2.63 | -2.98 | -2.95 | -3.7 | -3.54 | 0.11 | 0.6775 | 0.3375 | 0.678557 | 0.095129 | 0.226735 | | 1 | 0.996511 | 0.787447 |
| Pold2 | A8Y5G7 | 0.66 | 0.81 | -2.75 | -0.33 | 0.46 | 0.54 | -0.82 | -0.67 | 1.1 | 1.73 | -1.68 | 0.55 | 0.44 | 1.5 | -0.45 | -1.89 | 0.28 | 0.8275 | 0.3025 | 0.730184 | 0.415323 | 0.757268 | | 1 | 0.996511 | 0.962481 |
| Poldip2 | Q91VA6 | -0.01 | 0.23 | -0.19 | -0.13 | -0.17 | 0.11 | -0.08 | -0.24 | 0.41 | -0.58 | 0.13 | -0.02 | 0.82 | -0.4 | -0.6 | -0.32 | -0.07 | 0.01 | -0.1 | 0.68441 | 0.968185 | 0.765982 | | 1 | 0.996755 | 0.964134 |
| Poldip3 | Q8BG81 | 0.18 | -1.1 | -0.39 | 0.29 | -1.5 | -0.29 | -0.78 | -0.1 | -0.08 | 0.26 | 0.14 | -0.83 | 0.16 | -0.31 | -0.23 | -0.32 | -0.4125 | 0.1275 | 0.08 | 0.338298 | 0.741896 | 0.813555 | | 1 | 0.996511 | 0.972768 |
| Polg | Q75WC0 | -1.06 | -0.89 | -2.52 | -1.97 | -0.53 | -0.69 | -3.15 | -3.27 | -0.46 | 0.61 | -3.25 | -1.64 | -1.48 | -1.39 | -0.39 | -3.04 | -0.3 | 0.425 | 0.035 | 0.695183 | 0.608845 | 0.954362 | | 1 | 0.996511 | 0.992476 |
| Poll | Q9QXE2 | -5.51 | -6.53 | -7.09 | -8.33 | -9.67 | -7.84 | -11.42 | -10.79 | -5.31 | -4.55 | -6.07 | -6.66 | -0.96 | -15.02 | -5.66 | -6.59 | -3.065 | 1.2175 | -0.1925 | 0.006983 | 0.10066 | 0.941815 | | 1 | 0.996511 | 0.991085 |
| Polr1c | P52432 | -0.37 | 0.07 | 0.08 | -0.7 | -0.06 | -0.71 | -0.78 | -1.08 | -0.04 | 0.04 | -0.68 | 0.28 | -0.41 | -0.28 | -0.99 | -0.31 | -0.4275 | 0.13 | -0.2675 | 0.162 | 0.653515 | 0.342961 | | 1 | 0.996511 | 0.848473 |
| Polr1d | P97304 | -0.98 | -0.6 | -1.12 | -2.01 | -2.42 | -1.55 | -1.84 | -1.85 | 0.97 | -0.85 | -1.79 | -2.04 | 2.08 | 0.49 | -1.52 | -1.32 | -0.7375 | 0.25 | 1.11 | 0.054104 | 0.713264 | 0.193445 | | 1 | 0.996511 | 0.777907 |
| Polr1h | Q791N7 | -0.02 | 0.03 | 0.89 | 1.28 | 1 | -0.71 | 1.05 | 0.76 | 0.39 | 0.41 | 1.42 | -0.39 | -0.47 | 0.52 | 0.21 | 1.27 | -0.02 | -0.0875 | -0.1625 | 0.967185 | 0.849383 | 0.722485 | | 1 | 0.996511 | 0.958398 |
| Polr2a | A0A0R4J0V | 0.22 | -0.23 | -0.48 | -0.16 | -0.35 | -0.28 | -0.4 | -0.17 | -0.17 | -0.26 | -0.49 | -0.6 | 0.32 | 0.25 | 0.12 | -0.54 | -0.1375 | -0.2175 | 0.2 | 0.478604 | 0.327464 | 0.464892 | | 1 | 0.996511 | 0.888267 |
| Polr2b | Q8CFI7 | 0.03 | -0.79 | -0.29 | -0.21 | -1.14 | -0.42 | -0.54 | -0.12 | -0.9 | -0.08 | -0.45 | -0.97 | -0.12 | -0.89 | 0.01 | -0.35 | -0.24 | -0.285 | -0.0225 | 0.398911 | 0.326479 | 0.936464 | | 1 | 0.996511 | 0.989767 |
| Polr2c | P97760 | 0.35 | 1.07 | 0.06 | -0.26 | 0.53 | 0.21 | -0.38 | -0.43 | -0.36 | -0.01 | -0.61 | 0.43 | -0.21 | 1.65 | -0.15 | 0.06 | -0.3225 | -0.4425 | 0.0325 | 0.372579 | 0.232916 | 0.947101 | | 1 | 0.996511 | 0.992376 |
| Polr2d | Q9D7M8 | -0.75 | -1 | -0.08 | -0.42 | -1.34 | -0.78 | -0.04 | 0.04 | 0.49 | -0.38 | -0.68 | -1.02 | 1.6 | -0.88 | -0.03 | 0.08 | 0.0325 | 0.165 | 0.755 | 0.929172 | 0.6562 | 0.167256 | | 1 | 0.996511 | 0.759484 |
| Polr2e | Q80UW8 | 0.25 | -0.28 | -1 | -0.83 | -0.1 | -0.62 | -1.41 | -0.97 | -0.66 | -0.39 | -1.13 | -0.52 | -0.07 | -0.18 | -0.57 | -1.32 | -0.31 | -0.21 | -0.07 | 0.420096 | 0.523592 | 0.856949 | | 1 | 0.996511 | 0.976225 |
| Polr2g | P62488 | 1.06 | 0.57 | -0.27 | -0.06 | 0.45 | 0.47 | 0.04 | 0.21 | 0.1 | -0.49 | -0.28 | 0.26 | -0.32 | 0.7 | -0.15 | -0.54 | -0.0325 | -0.4275 | -0.4025 | 0.917561 | 0.233655 | 0.321131 | | 1 | 0.996511 | 0.839916 |
| Polr2h | Q923G2 | 0.22 | 0.43 | -0.68 | -1.09 | 0.55 | 0.14 | -1.32 | -1.1 | -0.02 | -1.21 | -1.1 | -0.16 | 0.34 | 0.19 | -1.14 | -0.73 | -0.1525 | -0.3425 | -0.055 | 0.776479 | 0.452487 | 0.908145 | | 1 | 0.996511 | 0.986458 |
| Polr2j | D3YZ20 | -0.68 | -0.77 | -0.6 | -1.08 | -0.91 | -0.57 | -1.57 | -0.78 | -0.36 | 0.99 | -0.93 | -1.24 | -0.19 | 0.13 | -0.1 | -0.96 | -0.175 | 0.3975 | 0.5025 | 0.492889 | 0.409409 | 0.099539 | | 1 | 0.996511 | 0.698156 |
| Polr2l | P62876 | 0.44 | 0.97 | -0.7 | -0.6 | 0.63 | 0.32 | -0.43 | -0.48 | -0.77 | -2.14 | -0.88 | -0.28 | 0.14 | -0.92 | -0.71 | -0.79 | -0.0175 | -1.045 | -0.5975 | 0.969398 | 0.073716 | 0.204032 | | 1 | 0.996511 | 0.782817 |
| Polr2m | Q6P6I6 | -0.04 | 0.02 | 0.01 | -0.42 | -0.09 | 0.29 | -0.63 | -0.14 | 1.23 | 0.18 | -0.02 | 0.23 | 1.9 | -0.69 | 0.04 | -0.08 | -0.035 | 0.5125 | 0.4 | 0.881004 | 0.119415 | 0.45577 | | 1 | 0.996511 | 0.887169 |
| Polr2m | A0A3B2WE | 1.04 | 0.32 | 0.79 | -0.03 | 0.85 | 1.05 | 0.98 | 0.57 | -0.02 | 0.22 | 0.69 | 1.23 | -0.08 | -0.03 | 0.96 | 0.42 | 0.3325 | 7.85E-17 | -0.2125 | 0.235431 | 1 | 0.536949 | | 1 | 1 | 0.912212 |
| Polr3a | B2RXC6 | -1.47 | -0.63 | -3.21 | -2.61 | -0.22 | -1.6 | -2.2 | -2.83 | -1.04 | -1.97 | -2.84 | -1.02 | -1.81 | -0.7 | -1.79 | -2.56 | 0.2675 | 0.2625 | 0.265 | 0.713242 | 0.691398 | 0.676353 | | 1 | 0.996511 | 0.952944 |
| Polr3b | P59470 | 1.16 | -0.53 | -0.24 | -1.78 | 0.29 | 1.01 | 0.03 | -1.61 | -0.17 | -0.3 | -0.3 | 1.07 | -0.49 | -0.02 | 0.03 | -0.33 | 0.2775 | 0.4225 | 0.145 | 0.708706 | 0.508056 | 0.798428 | | 1 | 0.996511 | 0.969727 |
| Polr3c | Q9D483 | -0.15 | 2.14 | -0.66 | -1.13 | 0.31 | 0.57 | -1.34 | -1.42 | -1.24 | 0.01 | -1.27 | 1.09 | 0.67 | 0.84 | -1.93 | -0.05 | -0.52 | -0.4025 | -0.1675 | 0.525185 | 0.630056 | 0.846531 | | 1 | 0.996511 | 0.976225 |
| Polr3d | Q91WD1 | 1.01 | 1.02 | -0.17 | -1.26 | 1.65 | 0.84 | -0.61 | -0.67 | 0.12 | -0.91 | -0.2 | 1.68 | -0.28 | 0.37 | -0.31 | -0.14 | 0.1525 | 0.0225 | -0.24 | 0.830955 | 0.974428 | 0.650886 | | 1 | 0.997515 | 0.947407 |
| Polr3e | Q9CZT4-2 | 0.25 | -0.24 | -0.88 | -0.87 | 0.6 | 0.18 | -0.45 | -0.75 | 0.43 | -0.37 | -0.87 | 0.46 | 0.63 | 0.86 | -0.41 | -1.06 | 0.33 | 0.3475 | 0.44 | 0.404437 | 0.40053 | 0.381995 | | 1 | 0.996511 | 0.860984 |
| Polr3f | Q9CZ23 | -1.22 | -1.51 | -0.98 | -1.04 | -1.21 | -1.11 | -1.02 | -0.13 | -3.04 | -2.44 | -1.3 | -1.61 | -3.33 | -2.62 | -1.2 | -0.38 | 0.32 | -0.91 | -0.695 | 0.270089 | 0.044587 | 0.280363 | | 1 | 0.996511 | 0.822084 |
| Polrmt | Q8BKF1 | -0.26 | 0.39 | 0.34 | 1.16 | -0.76 | -0.28 | -0.18 | 0.51 | 0.27 | 2.01 | -0.76 | -0.29 | 0.08 | -0.19 | 0.25 | 0.45 | -0.585 | -0.1 | -0.26 | 0.143368 | 0.870755 | 0.433371 | | 1 | 0.996511 | 0.879675 |
| Pom121 | Q8K3Z9 | 0.63 | 0.38 | -0.74 | -0.63 | -0.44 | 0.95 | -1.05 | -0.76 | 1.67 | -1.35 | -1.13 | -0.19 | 2.99 | 0.63 | -0.02 | -0.23 | -0.235 | -0.16 | 0.9325 | 0.652022 | 0.819448 | 0.228068 | | 1 | 0.996511 | 0.787447 |
| Pomgnt2 | Q8BW41-2 | 0.1 | -0.17 | -0.25 | -0.48 | 0.22 | -0.37 | 0.6 | 0.3 | -0.41 | -0.82 | -0.05 | 0.02 | -0.29 | -0.17 | 0.58 | 0.49 | 0.3875 | -0.115 | 0.3525 | 0.145251 | 0.647629 | 0.22205 | | 1 | 0.996511 | 0.787447 |
| Pomk | Q3TUA9 | -0.78 | -0.75 | -0.33 | -0.09 | -0.4 | -0.17 | 0.66 | -0.41 | -0.49 | -0.14 | -0.47 | -0.42 | 0.09 | -0.31 | 0.33 | 0.61 | 0.4075 | 0.1075 | 0.6675 | 0.199095 | 0.630983 | 0.037527 | | 1 | 0.996511 | 0.658454 |
| Pomp | Q9CQT5 | -1.11 | -1.04 | -3.4 | -3.51 | -1.11 | -1.48 | 0.06 | -2.52 | -1 | -0.81 | -2.19 | -1.64 | -1.07 | -0.44 | -2.71 | -2.55 | 1.0025 | 0.855 | 0.5725 | 0.223223 | 0.234969 | 0.479575 | | 1 | 0.996511 | 0.893468 |
| Pomt2 | Q8BGQ4-2 | 0.84 | 0.86 | 4.03 | 4.02 | 0.98 | 0.96 | 2.62 | 4.12 | 1.06 | 5.17 | 3.73 | 0.88 | 1.23 | 1.53 | 3.46 | 3.02 | -0.2675 | 0.2725 | -0.1275 | 0.80198 | 0.826892 | 0.893879 | | 1 | 0.996511 | 0.983083 |
| Pon2 | Q62086 | -0.45 | 0.33 | -0.09 | 0.05 | 0.59 | -0.42 | 0.84 | 0.05 | -0.01 | -0.17 | 0.04 | 0 | -0.35 | 0.55 | -0.07 | 0.41 | 0.305 | 0.005 | 0.175 | 0.351004 | 0.981078 | 0.541786 | | 1 | 0.998773 | 0.912212 |
| Pon3 | Q62087 | 0.9 | 0.97 | 0.3 | 0.32 | 0.6 | 0.59 | 0.83 | 0.52 | 0.64 | -0.48 | -0.02 | 1.7 | 0.56 | 0.4 | 0.93 | 0.94 | 0.0125 | -0.1625 | 0.085 | 0.954167 | 0.73218 | 0.741046 | | 1 | 0.996511 | 0.958796 |
| Pop7 | Q9DCH2 | -1.23 | -1.4 | -1.16 | -0.81 | -1.32 | -1.44 | -0.15 | -0.66 | -1.69 | -0.22 | -1.04 | -1.49 | -1.17 | -0.25 | -0.46 | -0.84 | 0.2575 | 0.04 | 0.47 | 0.42814 | 0.907234 | 0.103821 | | 1 | 0.996744 | 0.702846 |
| Por | P37040 | 0.74 | 0.71 | 0.69 | 0.69 | 0.73 | 0.69 | 0.85 | 0.95 | 1.23 | 0.61 | 0.65 | 0.89 | 1.14 | 0.74 | 0.97 | 0.74 | 0.0975 | 0.1375 | 0.19 | 0.506657 | 0.488459 | 0.316095 | | 1 | 0.996511 | 0.838265 |
| Porcn | Q9JJJ7-4 | -1.42 | -0.58 | -0.22 | -1.2 | -0.09 | -1.07 | -0.08 | -1.02 | -1.11 | -0.79 | -0.9 | -1.35 | -0.48 | -0.1 | -0.42 | -0.25 | 0.29 | -0.1825 | 0.5425 | 0.444838 | 0.555899 | 0.099734 | | 1 | 0.996511 | 0.698156 |
| Postn | Q62009-5 | -4.7 | -3.78 | -0.36 | -0.64 | -2.91 | -3.91 | -1.37 | -0.94 | -3.46 | -1.64 | -1.03 | -4.25 | -3.55 | -2.72 | -1.67 | -1.09 | 0.0875 | -0.225 | 0.1125 | 0.939779 | 0.850413 | 0.918062 | | 1 | 0.996511 | 0.986868 |
| Pou3f1 | P21952 | -3.8 | -4.19 | -2.11 | -1.56 | -4.46 | -4.27 | -1.78 | -1.3 | -2.82 | -0.11 | -2.16 | -3.71 | -0.67 | -4.25 | -1.9 | -1.55 | -0.0375 | 0.715 | 0.8225 | 0.967983 | 0.434646 | 0.369426 | | 1 | 0.996511 | 0.859245 |
| Pou3f3 | P31361 | -6.15 | -6.15 | -2.68 | -4.05 | -6.31 | -6.45 | -4.07 | -3.17 | -3.22 | -2.77 | -4.36 | -5.19 | -1.39 | -3.96 | -3.43 | -4.29 | -0.2425 | 0.8725 | 1.49 | 0.818855 | 0.351875 | 0.146699 | | 1 | 0.996511 | 0.746406 |
| Pou6f1 | Q5U4D4 | -2.02 | -2.12 | 0.57 | 0.97 | -2.09 | -1.96 | 0.69 | 0.73 | -0.82 | 0.36 | 0.19 | -1.38 | -1.26 | 0.33 | -0.06 | 0.3 | -0.0075 | 0.2375 | 0.4775 | 0.994149 | 0.77587 | 0.561632 | | 1 | 0.996511 | 0.91997 |
| Pou6f1 | H3BKJ6 | -0.06 | -0.77 | -1.52 | -0.9 | -2.15 | -2.38 | -2.34 | -1.49 | 0.06 | -2.02 | -2.84 | -0.77 | -0.88 | -0.66 | -0.79 | -1.19 | -1.2775 | -0.58 | -0.0675 | 0.005341 | 0.382577 | 0.83548 | | 1 | 0.996511 | 0.975524 |
| Ppa1 | Q9D819 | -0.13 | -0.55 | -0.22 | -0.22 | -0.35 | -0.21 | 0.27 | -0.13 | -0.22 | 0.11 | -0.03 | 0.29 | -0.78 | 0.43 | -0.38 | -0.32 | 0.175 | 0.3175 | 0.0175 | 0.384956 | 0.1309 | 0.951286 | | 1 | 0.996511 | 0.992476 |
| Ppa2 | Q91VM9 | 0.6 | 0.36 | 0.28 | 0.02 | 0.03 | 0.74 | 1.33 | 0.24 | 0.75 | -0.66 | 0.48 | 0.48 | 0.87 | 0.26 | 0.04 | 0.44 | 0.27 | -0.0525 | 0.0875 | 0.392688 | 0.874698 | 0.724605 | | 1 | 0.996511 | 0.958398 |
| Ppan | Q91YU8 | 0.64 | 0.7 | 0.4 | -0.08 | 0.2 | 0.37 | -0.38 | -0.23 | 0.37 | -0.08 | 0.05 | -0.01 | 1.21 | -0.22 | 0.9 | 0.38 | -0.425 | -0.3325 | 0.1525 | 0.128433 | 0.179372 | 0.669899 | | 1 | 0.996511 | 0.952558 |
| Ppat | Q8CIH9 | 0.16 | 0.01 | 0.25 | 0.63 | 0.03 | 0.42 | 0.42 | 0.7 | -0.01 | 1.23 | 0.76 | 0.32 | -0.12 | 0.85 | 0.46 | 0.23 | 0.13 | 0.3125 | 0.0925 | 0.551546 | 0.319992 | 0.730776 | | 1 | 0.996511 | 0.958635 |
| Ppbp | Q9EQI5 | -0.38 | -3.51 | -0.48 | -0.16 | -0.86 | -0.99 | -2.07 | -0.12 | -3.48 | -1.18 | -0.48 | -3.42 | -3.22 | -0.47 | 1.69 | 0.13 | 0.1225 | -1.0075 | 0.665 | 0.878781 | 0.325839 | 0.568751 | | 1 | 0.996511 | 0.922588 |
| Ppcdc | Q8BZB2 | -0.05 | 0.67 | -0.63 | -0.37 | 0.02 | -0.19 | -0.46 | -0.41 | -0.68 | -0.46 | -0.61 | -0.08 | -0.73 | -0.49 | -0.59 | -0.77 | -0.165 | -0.3625 | -0.55 | 0.585303 | 0.266362 | 0.095168 | | 1 | 0.996511 | 0.698156 |
| Ppcs | Q8VDG5 | -0.34 | -0.39 | -0.94 | -0.94 | -0.55 | -0.49 | -0.54 | -0.56 | -0.51 | -1.45 | -0.72 | -0.3 | -0.51 | 0.2 | -1.37 | -1.22 | 0.1175 | -0.0925 | -0.0725 | 0.560808 | 0.762018 | 0.850957 | | 1 | 0.996511 | 0.976225 |
| Ppdpf | Q9CR37 | 0.67 | 0.68 | 1.37 | 1.43 | -1.12 | 0.68 | 0.19 | 1.3 | 1.13 | 2.05 | 0.03 | 0.15 | 2.45 | -1.05 | 1.15 | 0.97 | -0.775 | -0.1975 | -0.1575 | 0.155642 | 0.683611 | 0.817904 | | 1 | 0.996511 | 0.972768 |
| Ppfia1 | B2RXW8 | 0.9 | 0.69 | 0.86 | 0.96 | 0.41 | 0.73 | 0.41 | 0.62 | 0.67 | 0.54 | 0.64 | 0.95 | 0.6 | 0.09 | 0.48 | 0.31 | -0.31 | -0.1525 | -0.4825 | 0.080641 | 0.396577 | 0.033808 | | 1 | 0.996511 | 0.644622 |
| Ppfia2 | B8QI34 | 1.16 | 0.62 | 0.44 | 0.46 | 0.77 | 1.24 | 0.37 | 0.28 | 0.69 | 0.12 | 0.55 | 1.13 | 0.74 | 0.53 | 0.4 | 0.33 | -0.005 | -0.0475 | -0.17 | 0.985711 | 0.865197 | 0.473251 | | 1 | 0.996511 | 0.891467 |
| Ppfia3 | P60469 | 1.11 | 0.74 | 0.68 | 0.91 | 0.7 | 1.23 | 0.61 | 0.83 | 0.95 | 0.65 | 0.84 | 1.12 | 0.97 | 0.45 | 0.47 | 0.45 | -0.0175 | 0.03 | -0.275 | 0.930324 | 0.876324 | 0.222577 | | 1 | 0.996511 | 0.787447 |
| Ppfia4 | B8QI36 | 1.48 | 1.02 | 0.65 | 0.62 | 0.67 | 1.39 | 0.71 | 0.69 | 1.03 | 0.7 | 0.78 | 1.55 | 1.17 | 0.07 | 0.74 | 0.64 | -0.0775 | 0.0725 | -0.2875 | 0.775916 | 0.801057 | 0.368401 | | 1 | 0.996511 | 0.859245 |
| Ppfibp1 | Q8C8U0 | -0.37 | -0.28 | -0.43 | -0.17 | -0.6 | -0.47 | -0.19 | -0.44 | 0.21 | -0.52 | -0.17 | -0.21 | 0.44 | -0.47 | -0.8 | -0.33 | -0.1125 | 0.14 | 0.0225 | 0.494894 | 0.501854 | 0.937359 | | 1 | 0.996511 | 0.989767 |
| Ppfibp2 | G3X957 | 2.77 | 2.58 | -0.59 | -3.37 | 1.99 | 2.59 | -1.25 | 0.05 | 0.7 | 0.24 | 1.46 | 1.85 | 0.53 | -0.18 | 4.89 | -1.05 | 0.4975 | 0.715 | 0.7 | 0.744485 | 0.597768 | 0.689515 | | 1 | 0.996511 | 0.955729 |
| Pphln1 | G3UWD4 | 3.61 | 2.37 | 0.02 | -0.55 | 1.08 | 3.57 | -0.09 | 0.48 | 2.05 | -0.33 | -0.47 | 2.07 | 3.83 | -1.48 | 0.09 | -0.38 | -0.1025 | -0.5325 | -0.8475 | 0.928061 | 0.626143 | 0.53346 | | 1 | 0.996511 | 0.911674 |
| Ppia | P17742 | 2.1 | 0.56 | 0.92 | 2.08 | 0.5 | 1.43 | 1.17 | 2.16 | 0.11 | 0.58 | 1.18 | 2.35 | -0.46 | -1.5 | 0.84 | 0.9 | -0.1 | -0.36 | -1.47 | 0.836671 | 0.536111 | 0.043746 | | 1 | 0.996511 | 0.658454 |
| Ppib | P24369 | 0.61 | 0.07 | -0.31 | -0.2 | 0.56 | 0.22 | 0.85 | 0.14 | 0.14 | -0.7 | -0.24 | 0.91 | 0.24 | 0.11 | 0.67 | 0.5 | 0.4 | -0.015 | 0.3375 | 0.162339 | 0.968723 | 0.227196 | | 1 | 0.996755 | 0.787447 |
| Ppic | P30412 | 0.07 | 0.17 | -1.17 | 0.67 | 1.12 | -0.26 | 0.8 | -1.04 | -0.45 | -0.75 | -0.81 | 0.25 | -0.54 | 0.61 | -0.77 | -1.03 | 0.22 | -0.375 | -0.3675 | 0.703967 | 0.399038 | 0.466709 | | 1 | 0.996511 | 0.888757 |
| Ppid | Q9CR16 | 1.2 | 1.09 | 1.57 | 1.23 | 1.12 | 1.25 | 1.23 | 1.74 | 0.19 | -0.05 | 1.58 | 1.22 | -0.16 | 0.06 | 1.3 | 1.44 | 0.0625 | -0.5375 | -0.6125 | 0.759418 | 0.193037 | 0.160005 | | 1 | 0.996511 | 0.755194 |
| Ppie | Q9QZH3 | 2.14 | 0.69 | -4.63 | -6.78 | -0.55 | 1.42 | -6.33 | -5.87 | 0.74 | -4.97 | -5.69 | 1.53 | 0.39 | 1.05 | -4.49 | -3.22 | -0.6875 | 0.0475 | 0.5775 | 0.788127 | 0.984964 | 0.795666 | | 1 | 0.998892 | 0.969727 |
| Ppif | Q99KR7 | 1.77 | 1.72 | 0.79 | 0.87 | 2.03 | 2.21 | 0.87 | 0.81 | 0.8 | 0.54 | 0.86 | 1.58 | 0.98 | 0.82 | 0.75 | 0.7 | 0.1925 | -0.3425 | -0.475 | 0.653887 | 0.329813 | 0.12706 | | 1 | 0.996511 | 0.722414 |
| Ppig | A2AR01 | 0.39 | 0.66 | 0.73 | -0.25 | 0.22 | 1.11 | 0.64 | 1.03 | 1.48 | 1.5 | 1.15 | 0.48 | 0.31 | 1.15 | 0.1 | 0.92 | 0.3675 | 0.77 | 0.2375 | 0.241516 | 0.039984 | 0.485176 | | 1 | 0.996511 | 0.894759 |
| Ppig | A2AR02 | -0.01 | -1.12 | -0.43 | -0.22 | -1.22 | -0.69 | -0.45 | 0.16 | -0.87 | -0.86 | -0.65 | -1.67 | 0.37 | -1.1 | 0.17 | -0.5 | -0.105 | -0.5675 | 0.18 | 0.769807 | 0.111293 | 0.652805 | | 1 | 0.996511 | 0.947633 |
| Ppih | Q9D868-2 | -0.25 | -0.46 | 0.03 | -0.12 | -0.25 | -0.62 | -0.15 | 0.01 | -1.67 | 0.72 | -0.19 | -0.44 | -1.16 | -1.39 | -0.12 | -0.16 | -0.0525 | -0.195 | -0.5075 | 0.794677 | 0.680274 | 0.167299 | | 1 | 0.996511 | 0.759484 |
| Ppil1 | Q9D0W5 | 0.44 | -0.41 | -0.13 | -0.01 | -0.83 | 0.12 | -0.66 | 0.07 | -1.73 | -0.78 | -0.25 | -0.47 | -1.03 | -1.68 | -0.18 | -0.38 | -0.2975 | -0.78 | -0.79 | 0.335587 | 0.055952 | 0.060278 | | 1 | 0.996511 | 0.661738 |
| Ppil2 | Q9D787 | -3.14 | -3.02 | -3.05 | -3.45 | -3.04 | -3.43 | -2.68 | -3.31 | -3.35 | -2.4 | -3.12 | -2.8 | -2.14 | -2.2 | -3.36 | -3.1 | 0.05 | 0.2475 | 0.465 | 0.818506 | 0.338293 | 0.182395 | | 1 | 0.996511 | 0.776002 |
| Ppil3 | Q9D6L8 | 0.75 | 0.67 | -0.84 | -0.6 | 0.04 | 0.25 | -0.91 | -0.82 | -0.18 | -0.58 | -0.63 | 0.49 | -0.58 | -0.06 | -1.14 | -1.35 | -0.355 | -0.22 | -0.7775 | 0.460249 | 0.634333 | 0.12938 | | 1 | 0.996511 | 0.723655 |
| Ppil4 | Q9CXG3 | -2.75 | -2.96 | -2.92 | -3.26 | -3.44 | -3.06 | -2.93 | -3.04 | -2.04 | -1.5 | -3 | -2.27 | -1.5 | -2.15 | -3.13 | -3.36 | -0.145 | 0.77 | 0.4375 | 0.456546 | 0.040834 | 0.318842 | | 1 | 0.996511 | 0.838969 |
| Ppip5k1 | A2ARP1 | -0.52 | -0.47 | 0.04 | 0.04 | -0.91 | -0.5 | -0.44 | -0.1 | 0.02 | 0.11 | -0.27 | -0.67 | 0.36 | -0.04 | -0.32 | -0.46 | -0.26 | 0.025 | 0.1125 | 0.299007 | 0.921926 | 0.672414 | | 1 | 0.996755 | 0.952944 |
| Ppip5k2 | Q6ZQB6-3 | 1.14 | 0.56 | 1.49 | -0.24 | 0.42 | 0.72 | 1.11 | 0.94 | 0.38 | -1.69 | 1.2 | 1.38 | 0.23 | 0.14 | 1.4 | 1.76 | 0.06 | -0.42 | 0.145 | 0.876153 | 0.566119 | 0.779753 | | 1 | 0.996511 | 0.967289 |
| Ppm1a | P49443 | 0.43 | 0.69 | -0.05 | 0 | 0.48 | 0.19 | -0.08 | -0.05 | 0.26 | -0.19 | 0.04 | 0.68 | -0.05 | 0.78 | -0.3 | -0.42 | -0.1325 | -0.07 | -0.265 | 0.578959 | 0.796747 | 0.427264 | | 1 | 0.996511 | 0.879165 |
| Ppm1b | P36993 | 0.42 | 1.13 | 1.9 | 1.98 | 0.92 | 0.53 | 1.81 | 1.74 | 1.06 | 0.93 | 1.73 | 0.75 | -0.59 | -0.41 | 0.48 | 1.46 | -0.1075 | -0.24 | -1.1225 | 0.811008 | 0.557381 | 0.067678 | | 1 | 0.996511 | 0.664936 |
| Ppm1b | Q99NF7 | 0.08 | -0.47 | -0.06 | 0.33 | -0.37 | 0.02 | 0.01 | 0.33 | -0.06 | 0.63 | 0.31 | 0.12 | -0.1 | 0.02 | 0.05 | 0.14 | 0.0275 | 0.28 | 0.0575 | 0.907582 | 0.275762 | 0.797038 | | 1 | 0.996511 | 0.969727 |
| Ppm1e | Q80TL0 | -0.65 | -0.43 | -0.61 | -1.02 | -0.69 | -0.27 | -0.99 | -0.81 | -0.5 | -0.87 | -0.69 | -0.71 | -0.59 | -0.23 | -0.73 | -1.27 | -0.0125 | -0.015 | -0.0275 | 0.954691 | 0.939219 | 0.919517 | | 1 | 0.996755 | 0.986868 |
| Ppm1f | Q8CGA0 | -0.49 | -0.44 | 0.04 | 0.3 | 0.09 | -0.4 | 0.35 | 0.21 | -0.8 | 0.24 | -0.06 | -0.55 | -0.76 | -0.25 | -0.15 | -0.02 | 0.21 | -0.145 | -0.1475 | 0.428305 | 0.638412 | 0.592827 | | 1 | 0.996511 | 0.929843 |
| Ppm1g | Q61074 | -4.5 | -5.19 | -4.44 | -5.65 | -4.64 | -5.19 | -5.15 | -5.58 | -4.56 | -2.62 | -4.5 | -4.28 | -3.57 | -3.63 | -4.93 | -5.26 | -0.195 | 0.955 | 0.5975 | 0.566779 | 0.086604 | 0.243031 | | 1 | 0.996511 | 0.798789 |
| Ppm1h | Q3UYC0 | 0.15 | 0.41 | 0.95 | 0.99 | 0.78 | 0.32 | 1.08 | 1.05 | -0.13 | 1.02 | 1.03 | 0.31 | -0.27 | 1.01 | 0.6 | 0.67 | 0.1825 | -0.0675 | -0.1225 | 0.512979 | 0.845055 | 0.72084 | | 1 | 0.996511 | 0.958398 |
| Ppm1k | Q8BXN7 | -0.25 | -0.64 | -1.09 | -0.75 | -0.45 | -0.27 | -0.48 | -0.68 | -0.29 | -0.02 | -0.7 | -0.42 | 0.02 | 0.62 | -1.24 | 0.19 | 0.2125 | 0.325 | 0.58 | 0.341471 | 0.211904 | 0.187934 | | 1 | 0.996511 | 0.776843 |
| Ppm1l | Q8BHN0 | -0.14 | -0.37 | -0.08 | 0.43 | 0.26 | -0.16 | -0.32 | -0.04 | -0.25 | 0.15 | -0.33 | -0.54 | -0.18 | 0.73 | -0.14 | -0.06 | -0.025 | -0.2025 | 0.1275 | 0.912824 | 0.421005 | 0.662135 | | 1 | 0.996511 | 0.949645 |
| Ppm1m | A0A8I4Q8Y | -1.26 | 0.16 | 5.45 | 3.96 | -3 | 0.15 | 1.86 | 3.73 | 4.12 | 1.24 | 4.15 | -1.86 | 4.75 | -2.18 | 3.96 | 4.77 | -1.3925 | -0.165 | 0.7475 | 0.469649 | 0.93059 | 0.714674 | | 1 | 0.996755 | 0.958106 |
| Ppme1 | Q8BVQ5 | -1.01 | -1.09 | -0.82 | -0.71 | -1.03 | -1.29 | -0.39 | -0.74 | -0.9 | -0.78 | -0.73 | -1 | -1.35 | 0.02 | -1.01 | -1.24 | 0.045 | 0.055 | 0.0125 | 0.845602 | 0.755526 | 0.969633 | | 1 | 0.996511 | 0.992767 |
| Ppox | P51175 | 0.1 | 0.26 | -0.1 | 0.17 | -0.09 | 0.31 | -0.1 | -0.05 | 0.39 | 0.23 | 0.12 | 0.12 | 0.33 | 0.47 | -0.14 | -0.24 | -0.09 | 0.1075 | -0.0025 | 0.607695 | 0.539741 | 0.991426 | | 1 | 0.996511 | 0.996154 |
| Ppp1ca | P62137 | 0.95 | -0.72 | -1.75 | -1.75 | -0.14 | 1.28 | -0.63 | -1.3 | -0.43 | -1.15 | -1.59 | 0.78 | -0.82 | -0.36 | -1.56 | -1.42 | 0.62 | 0.22 | -0.2225 | 0.422938 | 0.767944 | 0.726528 | | 1 | 0.996511 | 0.958398 |
| Ppp1cb | P62141 | -0.16 | -0.27 | -1.42 | -1.2 | -0.48 | -0.45 | -1.31 | -1.45 | -0.55 | -1.26 | -1.27 | -0.08 | -1.11 | -0.76 | -1.56 | -1.6 | -0.16 | -0.0275 | -0.495 | 0.6853 | 0.946662 | 0.203436 | | 1 | 0.996755 | 0.782817 |
| Ppp1cc | P63087-2 | 0.96 | 1.23 | 0.7 | 0.68 | 1.56 | 1.35 | 0.91 | 0.63 | 1.17 | 0.09 | 0.64 | 1.06 | 0.57 | 1.07 | 0.24 | 0.28 | 0.22 | -0.1525 | -0.3525 | 0.401756 | 0.597932 | 0.198214 | | 1 | 0.996511 | 0.78004 |
| Ppp1r10 | Q80W00 | -1.32 | -2.81 | -3.15 | -3.5 | -2.02 | -2.23 | -2.6 | -2.65 | -2.12 | -2 | -2.94 | -2.57 | -1.1 | -1.81 | -0.44 | -1.88 | 0.32 | 0.2875 | 1.3875 | 0.498612 | 0.561237 | 0.029135 | | 1 | 0.996511 | 0.637553 |
| Ppp1r11 | G3UZ30 | 0.01 | -0.31 | -0.48 | 0.4 | -0.76 | -0.56 | -0.47 | 0.21 | 0.1 | -0.14 | -0.19 | 0.59 | -0.6 | 0.3 | -0.46 | -1.05 | -0.3 | 0.185 | -0.3575 | 0.311854 | 0.509156 | 0.310174 | | 1 | 0.996511 | 0.834523 |
| Ppp1r12a | Q9DBR7 | -0.28 | -0.3 | 0.27 | -0.07 | -0.16 | -0.62 | -0.23 | -0.14 | -0.26 | 0.42 | 0.24 | -0.16 | 0.25 | -0.17 | 0.38 | -0.12 | -0.1925 | 0.155 | 0.18 | 0.35901 | 0.519064 | 0.447042 | | 1 | 0.996511 | 0.884817 |
| Ppp1r12a | A0A1W2P7 | -0.27 | -1.84 | -0.13 | -0.69 | -3.42 | 0.56 | -3.61 | -2.24 | 3.13 | -2.72 | -0.59 | -0.8 | 4.4 | -3.3 | -1.57 | -3.95 | -1.445 | 0.4875 | -0.3725 | 0.147483 | 0.672932 | 0.828773 | | 1 | 0.996511 | 0.973757 |
| Ppp1r12b | Q8BG95 | -0.62 | -0.56 | -0.53 | -0.84 | -0.57 | -1.06 | -0.65 | -1.12 | -0.37 | -0.64 | -0.4 | -0.08 | 0.34 | -0.33 | 0.16 | -0.51 | -0.2125 | 0.265 | 0.5525 | 0.288769 | 0.188943 | 0.049067 | | 1 | 0.996511 | 0.658454 |
| Ppp1r12c | F6XWD4 | -0.1 | 0.05 | -0.2 | 0.71 | -1.05 | -0.5 | -2.69 | 0.4 | 0.04 | 1.95 | 0.01 | 0.05 | 1.44 | -0.27 | -0.91 | -1.32 | -1.075 | 0.3975 | -0.38 | 0.11083 | 0.422738 | 0.523671 | | 1 | 0.996511 | 0.90791 |
| Ppp1r12c | Q3UMT1 | -0.12 | -0.01 | 0.53 | 0.78 | -0.05 | -0.09 | -0.11 | 0.58 | 0.03 | 1.21 | 0.68 | 0.03 | 0.1 | 0.04 | 0.58 | 0.22 | -0.2125 | 0.1925 | -0.06 | 0.449421 | 0.585781 | 0.824875 | | 1 | 0.996511 | 0.972852 |
| Ppp1r13b | Q62415 | 0.57 | 0.21 | 0.92 | 1.26 | -0.4 | 0.37 | 0.67 | 1.01 | 1.13 | 1.05 | 1.08 | 0.62 | 1.18 | -0.28 | 0.91 | 0.75 | -0.3275 | 0.23 | -0.1 | 0.374815 | 0.404957 | 0.793049 | | 1 | 0.996511 | 0.96933 |
| Ppp1r14a | Q91VC7 | 0.5 | 0.58 | 1.17 | 1.28 | 0.47 | 0.27 | -0.24 | 0.56 | 0.43 | 0.77 | 0.82 | -0.12 | 0.8 | 0.74 | 1.23 | 1.33 | -0.6175 | -0.4075 | 0.1425 | 0.047396 | 0.198256 | 0.60401 | | 1 | 0.996511 | 0.931779 |
| Ppp1r14b | Q62084 | 0.03 | 0.54 | -2.03 | -0.83 | -0.04 | -0.12 | -1.31 | -0.71 | 0.81 | -0.23 | -1.55 | -0.05 | 0.97 | 0.81 | -1.57 | -1.72 | 0.0275 | 0.3175 | 0.195 | 0.962155 | 0.641328 | 0.814656 | | 1 | 0.996511 | 0.972768 |
| Ppp1r14c | Q8R4S0 | -1.7 | -1.8 | 0.14 | -0.67 | -0.71 | 0.34 | -1.67 | -0.03 | -1.77 | 0.04 | -0.12 | -1.21 | -1.38 | -1.52 | 0.88 | -0.28 | 0.49 | 0.2425 | 0.4325 | 0.409975 | 0.678096 | 0.5167 | | 1 | 0.996511 | 0.905676 |
| Ppp1r16a | Q923M0 | 1.33 | 1.45 | 0.7 | 1.28 | 1.74 | 1.12 | 0.78 | 0.79 | 0.06 | 1.16 | 1.01 | 0.96 | -0.11 | -0.33 | 0.97 | 0.42 | -0.0825 | -0.3925 | -0.9525 | 0.770757 | 0.219878 | 0.018696 | | 1 | 0.996511 | 0.602211 |
| Ppp1r16b | Q8VHQ3 | 1.91 | 2.06 | 1.36 | 1.67 | 1.53 | 1.62 | 0.55 | 1.28 | 1.96 | 1.25 | 1.27 | 1.87 | 1.59 | 2.08 | 1.14 | 0.94 | -0.505 | -0.1625 | -0.3125 | 0.10696 | 0.540047 | 0.32316 | | 1 | 0.996511 | 0.840788 |
| Ppp1r18 | Q8BQ30-2 | 0.6 | 0.56 | 0.8 | 0.35 | 0.55 | 0.17 | -0.47 | -0.19 | -0.03 | 1.54 | 0.63 | 0.87 | -0.12 | 0.54 | 0.23 | -0.96 | -0.5625 | 0.175 | -0.655 | 0.049158 | 0.602969 | 0.079248 | | 1 | 0.996511 | 0.681114 |
| Ppp1r1a | Q9ERT9 | -0.67 | -0.48 | -0.24 | 0.08 | -0.9 | -0.73 | -0.54 | -0.18 | 0.1 | 0 | -0.16 | -0.23 | 0.37 | 0.27 | -0.05 | -0.39 | -0.26 | 0.255 | 0.3775 | 0.294436 | 0.259492 | 0.176437 | | 1 | 0.996511 | 0.771503 |
| Ppp1r1b | Q60829 | -0.71 | -2.29 | -1.94 | -1.14 | -2.81 | -1.88 | -0.75 | -1.21 | -1.35 | -0.64 | -2.09 | -0.78 | -0.5 | -2.27 | -0.48 | -1.66 | -0.1425 | 0.305 | 0.2925 | 0.787736 | 0.512009 | 0.584437 | | 1 | 0.996511 | 0.927734 |
| Ppp1r2 | D3Z3A0 | -0.15 | -0.57 | -0.06 | 0.08 | -0.46 | -0.72 | 0.07 | 0.2 | -2.01 | -0.15 | -0.29 | -0.1 | -1.66 | -0.86 | -0.04 | -0.33 | -0.0525 | -0.4625 | -0.5475 | 0.843406 | 0.320677 | 0.167946 | | 1 | 0.996511 | 0.759484 |
| Ppp1r21 | Q3TDD9 | -0.41 | -0.47 | -0.24 | -0.08 | -0.44 | -0.56 | -0.41 | -0.1 | -0.37 | 0.16 | -0.19 | -0.35 | -0.63 | -0.17 | -0.42 | -0.52 | -0.0775 | 0.1125 | -0.135 | 0.66518 | 0.577745 | 0.504046 | | 1 | 0.996511 | 0.901002 |
| Ppp1r37 | Q8BKR5 | -1.03 | -1.45 | -1.04 | -0.75 | -1 | -1.14 | -1.01 | -0.98 | -2.72 | -0.29 | -0.81 | -0.77 | -2.33 | -2.33 | -1.03 | -1 | 0.035 | -8.00E-02 | -0.605 | 0.852568 | 0.876722 | 0.149776 | | 1 | 0.996511 | 0.748798 |
| Ppp1r3d | A2AJW4 | -0.17 | 0.27 | -0.3 | -0.1 | 0.1 | 0.82 | -0.02 | 0.1 | -0.01 | -0.86 | 0.04 | 0.36 | 0.14 | -0.47 | -0.45 | -0.11 | 0.325 | -0.0425 | -0.1475 | 0.203147 | 0.885506 | 0.53198 | | 1 | 0.996511 | 0.910339 |
| Ppp1r3f | Q9JIG4-2 | 0.73 | 0.67 | 0.72 | 0.66 | 0.7 | 0.15 | -0.04 | 0.45 | 0.31 | 0.21 | 0.19 | 0.51 | 0.27 | 0.34 | 0.55 | 0.18 | -0.38 | -0.39 | -0.36 | 0.082455 | 0.038582 | 0.068098 | | 1 | 0.996511 | 0.664936 |
| Ppp1r3g | Q9CW07 | 0.43 | -0.99 | -0.4 | 0.14 | -0.6 | 0.89 | -1.45 | -0.16 | -0.25 | -1.09 | -0.54 | 0.12 | 0.02 | -0.73 | -1.1 | -0.59 | -0.125 | -0.235 | -0.395 | 0.814253 | 0.549113 | 0.314017 | | 1 | 0.996511 | 0.836827 |
| Ppp1r7 | Q3UM45 | -0.26 | -0.22 | -0.33 | -0.08 | -0.32 | -0.33 | -0.28 | -0.04 | -0.06 | 0.35 | -0.29 | -0.21 | -0.29 | 0.31 | -0.59 | -0.85 | -0.02 | 0.17 | -0.1325 | 0.897193 | 0.407166 | 0.634203 | | 1 | 0.996511 | 0.941132 |
| Ppp1r8 | A2ADR8 | 0.11 | -1.22 | -2.49 | -2.02 | -0.86 | 0.1 | -1.89 | -1.58 | -1.16 | -1.53 | -1.82 | -0.99 | -0.88 | -0.38 | -1.16 | -1.99 | 0.3475 | 0.03 | 0.3025 | 0.598306 | 0.956621 | 0.619767 | | 1 | 0.996755 | 0.93807 |
| Ppp1r9a | H3BJD6 | 1.26 | 0.77 | 1.23 | 1.28 | 0.62 | 1.07 | 0.58 | 1.03 | 1.28 | 0.71 | 1.11 | 0.97 | 1.45 | 0.52 | 0.95 | 0.75 | -0.31 | -0.1175 | -0.2175 | 0.161373 | 0.584198 | 0.414382 | | 1 | 0.996511 | 0.874945 |
| Ppp1r9b | Q6R891 | 0.63 | 0.56 | 0.64 | 0.6 | 0.76 | 0.78 | 0.74 | 0.63 | 0.97 | 0.74 | 0.72 | 0.82 | 1.05 | 0.73 | 0.49 | 0.45 | 0.12 | 0.205 | 0.0725 | 0.397976 | 0.217435 | 0.722013 | | 1 | 0.996511 | 0.958398 |
| Ppp2ca | P63330 | -1.84 | -2.73 | -0.57 | -0.92 | -3.28 | -1.41 | -1.63 | -0.52 | -2.06 | 0.46 | -1.04 | -1.82 | -1.49 | -2.85 | -0.15 | -0.28 | -0.195 | 0.4 | 0.3225 | 0.775524 | 0.560526 | 0.655924 | | 1 | 0.996511 | 0.948301 |
| Ppp2cb | P62715 | -0.57 | -0.99 | -1.16 | -0.89 | -1.14 | -0.22 | -0.89 | -0.95 | -0.67 | -0.99 | -1.19 | -0.43 | -0.77 | -1.31 | -1.2 | -1.07 | 0.1025 | 0.0825 | -0.185 | 0.681237 | 0.729297 | 0.411679 | | 1 | 0.996511 | 0.874811 |
| Ppp2r1a | Q76MZ3 | -0.22 | -0.34 | -0.18 | 0.16 | -0.13 | -0.34 | -0.14 | 0.12 | -0.2 | 0.54 | -0.01 | -0.28 | -0.78 | 0.05 | -0.34 | -0.34 | 0.0225 | 0.1575 | -0.2075 | 0.902978 | 0.518231 | 0.397016 | | 1 | 0.996511 | 0.867334 |
| Ppp2r1a | G3UWL2 | -2.15 | -1.21 | -4.01 | -3.34 | -1.03 | -1 | -6.94 | -2.52 | -0.92 | -2.9 | -1.64 | -1.73 | -2.39 | -1.67 | -2.67 | -3.13 | -0.195 | 0.88 | 0.2125 | 0.886763 | 0.216799 | 0.737661 | | 1 | 0.996511 | 0.958796 |
| Ppp2r1b | H3BK50 | -2.44 | -4.17 | -3.75 | -3.12 | -2.03 | -1.69 | -2.46 | -3.28 | -2.01 | -2.65 | -3.76 | -2.07 | -3 | -1.51 | -4.08 | -3.65 | 1.005 | 0.7475 | 0.31 | 0.059199 | 0.171308 | 0.619057 | | 1 | 0.996511 | 0.93807 |
| Ppp2r1b | Q3TTF6 | -1.21 | -0.51 | -0.22 | -0.33 | 0.01 | -1.86 | 0.27 | -0.08 | 0 | -0.02 | 0.08 | -0.66 | -1.29 | -1.32 | -0.51 | 0.41 | 0.1525 | 0.4175 | -0.11 | 0.758288 | 0.174579 | 0.803051 | | 1 | 0.996511 | 0.971091 |
| Ppp2r2a | Q6P1F6 | 0.29 | 0.34 | 0.45 | 0.32 | 0.18 | 0.19 | 0.2 | 0.26 | 0.21 | 0.37 | 0.32 | 0.34 | -0.38 | 0.72 | -0.02 | -0.2 | -0.1425 | -0.04 | -0.32 | 0.32074 | 0.797268 | 0.253504 | | 1 | 0.996511 | 0.809524 |
| Ppp2r2b | A0A668KLA | -0.05 | -0.08 | 0.81 | 0.95 | 0.39 | 0.29 | 0.91 | 0.95 | 0.17 | 0.53 | 1.01 | 0 | -0.52 | 0.63 | 0.94 | 1.2 | 0.2275 | 0.02 | 0.155 | 0.479849 | 0.954029 | 0.727366 | | 1 | 0.996755 | 0.958635 |
| Ppp2r2c | Q8BG02 | 0.01 | -0.41 | -0.77 | -0.92 | -0.08 | -0.06 | -0.22 | -1.12 | -0.14 | -0.55 | -0.81 | -0.01 | -0.44 | -0.48 | -0.57 | -0.43 | 0.1525 | 0.145 | 0.0425 | 0.635317 | 0.616212 | 0.86261 | | 1 | 0.996511 | 0.977231 |
| Ppp2r2d | Q925E7 | 0.98 | 1.27 | 1.4 | 1.51 | 1.11 | 1.02 | 1.14 | 1.83 | 1.28 | 1.7 | 1.24 | 1.33 | 1.06 | 1.56 | 1.44 | 1.05 | -0.015 | 0.0975 | -0.0125 | 0.94929 | 0.633434 | 0.955226 | | 1 | 0.996511 | 0.992482 |
| Ppp2r5a | Q6PD03 | 0.14 | 0.29 | 0.38 | 0.16 | 0.3 | 0.38 | 0.21 | 0.21 | -0.08 | 0.07 | 0.37 | 0.31 | 0.09 | -0.26 | -0.09 | 0.39 | 0.0325 | -0.075 | -0.21 | 0.826547 | 0.68286 | 0.329506 | | 1 | 0.996511 | 0.845234 |
| Ppp2r5b | Q6PD28 | -0.36 | -0.69 | 0.62 | -0.42 | -0.54 | -0.33 | 0.25 | 0.39 | -0.43 | -1.9 | 0.66 | -0.24 | -0.72 | -0.46 | -0.08 | 0.32 | 0.155 | -0.265 | -0.0225 | 0.659346 | 0.635542 | 0.950174 | | 1 | 0.996511 | 0.992476 |
| Ppp2r5c | Q60996-3 | 0.14 | 0.45 | 0.44 | 0.27 | 0.43 | 0.22 | 0.43 | 0.23 | 0.2 | -0.13 | 0.32 | 0.31 | -0.07 | -0.09 | 0.17 | 0.32 | 0.0025 | -0.15 | -0.2425 | 0.987425 | 0.433104 | 0.234817 | | 1 | 0.996511 | 0.791172 |
| Ppp2r5c | A0A1Y7VIR | 1.54 | 1.27 | 1.29 | 1.17 | 0.31 | 1.67 | -0.1 | 0.51 | 2.7 | -1.05 | 1.01 | 1.26 | 3.06 | -0.15 | 1.03 | 0.85 | -0.72 | -0.3375 | -0.12 | 0.07857 | 0.634488 | 0.84638 | | 1 | 0.996511 | 0.976225 |
| Ppp2r5d | Q91V89 | 0.39 | 0.33 | 0.04 | -0.05 | 0.66 | 0.39 | 0.36 | 0.17 | -0.08 | -0.43 | 0.23 | 0.65 | -0.41 | 0.04 | -0.16 | -0.15 | 0.2175 | -0.085 | -0.3475 | 0.265137 | 0.752873 | 0.116641 | | 1 | 0.996511 | 0.718608 |
| Ppp2r5e | Q61151 | 0.59 | 0.52 | 0.55 | 0.76 | 0.59 | 0.46 | 0.74 | 0.71 | 0.25 | 0.54 | 0.58 | 0.55 | 0.03 | 0.01 | 0.23 | 0.58 | 0.02 | -0.125 | -0.3925 | 0.896356 | 0.471417 | 0.082594 | | 1 | 0.996511 | 0.68865 |
| Ppp3ca | P63328 | 0.55 | 1.1 | 0.64 | 0.68 | 1.03 | 0.2 | 0.49 | 0.69 | 0.17 | 0.51 | 0.62 | 0.5 | -0.33 | 0.72 | 0.23 | 0.2 | -0.14 | -0.2925 | -0.5375 | 0.54994 | 0.17526 | 0.073871 | | 1 | 0.996511 | 0.670574 |
| Ppp3ca | P63328-2 | 1.43 | 0.84 | 1.25 | 1.3 | 0.27 | 0.86 | -0.25 | 0.8 | 1.01 | 1.49 | 0.67 | 0.74 | 0.51 | 1.48 | 0.38 | 0.04 | -0.785 | -0.2275 | -0.6025 | 0.023007 | 0.372823 | 0.100148 | | 1 | 0.996511 | 0.698156 |
| Ppp3cb | E0CZ78 | 0.22 | 0.27 | -0.16 | 0.23 | 0.31 | 0.27 | 0 | 0.06 | 0.47 | 0.68 | 0.04 | 0.4 | -0.1 | 0.6 | -0.07 | -0.07 | 0.02 | 0.2575 | -0.05 | 0.909008 | 0.240473 | 0.833496 | | 1 | 0.996511 | 0.975304 |
| Ppp3cc | A0A2I3BPC | 0.89 | 1.09 | 0.93 | 0.96 | 1.16 | 0.72 | 0.64 | 1.03 | 0.67 | 1.29 | 1.04 | 0.89 | 0.59 | 1.06 | 0.47 | 0.52 | -0.08 | 0.005 | -0.3075 | 0.654563 | 0.97925 | 0.158926 | | 1 | 0.998756 | 0.754492 |
| Ppp3r1 | Q63810-2 | -0.48 | 1.31 | 1.3 | -0.28 | 2.45 | 0.44 | 1.48 | 0.1 | -2.89 | -1.16 | 1 | -0.16 | -1.88 | -2.11 | 1.51 | 1.64 | 0.655 | -1.265 | -0.6725 | 0.33238 | 0.168512 | 0.515221 | | 1 | 0.996511 | 0.905674 |
| Ppp4c | P97470 | -0.79 | -1.04 | -1.16 | -0.68 | -0.76 | -0.87 | -0.7 | -1.05 | -1.02 | -0.59 | -0.72 | -0.92 | -0.97 | -0.62 | -1.16 | -0.77 | 0.0725 | 0.105 | 0.0375 | 0.688086 | 0.598691 | 0.861878 | | 1 | 0.996511 | 0.977204 |
| Ppp4r1 | E9PX53 | -0.12 | -0.34 | -1.25 | -1.12 | -0.49 | -0.16 | -1.27 | -1.13 | 0.37 | -0.25 | -1.03 | -0.4 | 0.49 | 0.22 | -1.55 | -1.57 | -0.055 | 0.38 | 0.105 | 0.880815 | 0.339084 | 0.854477 | | 1 | 0.996511 | 0.976225 |
| Ppp4r2 | A0A0R4J0U | -4.7 | -4.56 | -2.52 | -3.18 | -4.68 | -4.9 | -2.63 | -3.08 | -4.39 | -1.1 | -2.59 | -4.32 | -3.45 | -4.42 | -2.92 | -2.92 | -0.0825 | 0.64 | 0.3125 | 0.906729 | 0.462176 | 0.597 | | 1 | 0.996511 | 0.929911 |
| Ppp4r3a | E9Q481 | -3.66 | -4.02 | -3.92 | -4.74 | -3.15 | -3.9 | -3.78 | -4.14 | -4.23 | -2.39 | -4.21 | -3.53 | -3.3 | -3.36 | -4.34 | -4.39 | 0.3425 | 0.495 | 0.2375 | 0.284647 | 0.298028 | 0.526255 | | 1 | 0.996511 | 0.908107 |
| Ppp4r4 | Q8C0Y0 | -0.31 | -0.05 | 0.07 | -0.1 | -0.11 | -0.45 | -0.14 | -0.16 | -0.08 | 0.44 | 0.16 | -0.08 | -0.26 | -0.25 | -0.06 | -0.32 | -0.1175 | 0.2075 | -0.125 | 0.488634 | 0.310164 | 0.501211 | | 1 | 0.996511 | 0.901002 |

| | | | | | | | | | | | | | | | | | | | | | | | | | | |
|---|---|---|---|---|---|---|---|---|---|---|---|---|---|---|---|---|---|---|---|---|---|---|---|---|---|---|
| Ppp5c | Q60676 | 0.39 | 0.39 | 0.61 | 0.66 | 0.46 | 0.36 | 0.21 | 0.68 | -0.47 | -0.69 | -0.24 | 0.44 | -0.57 | -0.74 | -0.75 | -0.65 | -0.085 | -0.7525 | -1.19 | 0.624588 | 0.020302 | 0.000107 | 1 | 0.996511 | 0.228506 |
| Ppp6c | Q9CQR6 | 1.51 | -0.26 | 0.19 | 0.62 | -0.63 | 1.25 | 0.14 | 0.62 | 1.06 | 0.51 | 0.24 | 1.31 | 0.27 | 0.08 | -0.11 | -0.11 | -0.17 | 0.265 | -0.4825 | 0.736823 | 0.538751 | 0.223609 | 1 | 0.996511 | 0.787447 |
| Ppp6r1 | Q7TSI3 | -0.34 | -0.68 | -0.4 | -0.31 | -0.5 | -0.72 | -0.44 | -0.54 | -1.48 | -0.26 | -0.09 | -0.35 | -1.43 | -1.23 | -0.59 | -0.65 | -0.1175 | -0.1125 | -0.5425 | 0.478031 | 0.730877 | 0.059559 | 1 | 0.996511 | 0.661738 |
| Ppp6r2 | Q8R3Q2 | -0.3 | -0.17 | -0.18 | -0.16 | -0.01 | -0.25 | 0.11 | -0.16 | -0.09 | -0.38 | -0.26 | -0.16 | -0.42 | 0.04 | -0.57 | -0.28 | 0.125 | -0.02 | -0.105 | 0.428402 | 0.901679 | 0.603367 | 1 | 0.996609 | 0.931296 |
| Ppp6r3 | G5E8R4 | -0.22 | 0.2 | -0.17 | -0.15 | -0.21 | -0.14 | -0.07 | -0.28 | -0.2 | -0.35 | -0.26 | -0.2 | -0.41 | -0.24 | -0.74 | -0.42 | -0.09 | -0.1675 | -0.3675 | 0.587005 | 0.350658 | 0.099893 | 1 | 0.996511 | 0.698156 |
| Ppp6r3 | A0A286YC( | -0.56 | 1.12 | -0.27 | 0.31 | 0.47 | -0.68 | 0.86 | 0.78 | 0.07 | 1.12 | 0.37 | -1.09 | -0.99 | 0.3 | 0.06 | 0.97 | 0.2075 | -0.0325 | -0.065 | 0.664363 | 0.952363 | 0.898849 | 1 | 0.996755 | 0.983509 |
| Ppt1 | O88531 | -1.11 | -1.57 | -1.58 | -0.45 | -0.16 | -0.48 | 0.14 | -0.31 | -2.02 | -1.56 | -0.7 | -1.45 | -0.79 | -0.69 | -0.74 | -0.24 | 0.975 | -0.255 | 0.5625 | 0.009144 | 0.496568 | 0.093171 | 1 | 0.996511 | 0.697749 |
| Pptc7 | Q6NVE9 | 1.06 | 0.84 | -0.6 | -0.77 | 0.94 | 0.75 | -0.32 | -0.63 | -0.84 | -1.1 | -0.73 | 1.08 | -0.44 | -0.65 | -0.59 | -0.16 | 0.0525 | -0.53 | -0.5925 | 0.92548 | 0.409864 | 0.21924 | 1 | 0.996511 | 0.787447 |
| Ppwd1 | Q8CEC6 | -0.83 | -0.99 | -2.05 | -0.05 | -1.2 | -1.06 | -1.81 | -1.39 | -1.45 | 0.98 | -1.59 | -0.89 | -1.31 | -1 | -1.7 | -2.31 | -0.385 | 0.2425 | -0.6 | 0.366479 | 0.713179 | 0.222417 | 1 | 0.996511 | 0.787447 |
| Pqbp1 | Q91VJ5 | -0.18 | -2.13 | -1.42 | -1.28 | -2.2 | -1.58 | -1.45 | -1.2 | -1.75 | 0.19 | -1.28 | -1.91 | -1.14 | -0.81 | -0.56 | -1.71 | -0.355 | 0.065 | 0.1975 | 0.414894 | 0.909931 | 0.660841 | 1 | 0.996744 | 0.949645 |
| Pradc1 | A0A0N4SV: | 5.67 | 0.8 | 13.15 | -0.86 | 4.1 | 5.07 | 13.42 | 9.73 | -2.75 | 0.3 | 12.26 | 5.62 | -1.99 | -2.15 | 14.52 | 15.36 | 3.39 | -0.8325 | 1.745 | 0.332334 | 0.837036 | 0.735455 | 1 | 0.996511 | 0.958635 |
| Praf2 | Q9JIG8 | -0.2 | -0.23 | 0.45 | 0.34 | 0.71 | -0.35 | 0.8 | 0.28 | -0.07 | 1.91 | 0.4 | -0.17 | 0.69 | 0.01 | 0.86 | 1.36 | 0.27 | 0.4275 | 0.64 | 0.396951 | 0.383013 | 0.081719 | 1 | 0.996511 | 0.68865 |
| Prag1 | Q571I4 | 0.48 | 0.88 | 0.3 | 0.34 | 0.52 | 0.85 | -0.1 | 0.17 | 0.91 | -0.22 | 0.28 | 0.8 | 1.47 | 1.32 | 0.37 | 0.31 | -0.14 | -0.0575 | 0.3675 | 0.587309 | 0.847148 | 0.290147 | 1 | 0.996511 | 0.825134 |
| Prcc | Q9EQC8 | -4.19 | -3.63 | -5.13 | -4.98 | -3.2 | -3.52 | -5.21 | -5.75 | -3.29 | -2.08 | -4.88 | -3.64 | -3.06 | -3.57 | -4.53 | -5.91 | 0.0625 | 1.01 | 0.215 | 0.923499 | 0.1292 | 0.742984 | 1 | 0.996511 | 0.959128 |
| Prcp | Q7TMR0 | -0.81 | -1.11 | -0.3 | -0.79 | -1.33 | -0.49 | 0.16 | -0.74 | -0.77 | -0.13 | 0.05 | -1.61 | -0.49 | -0.56 | -0.66 | -0.53 | 0.1525 | 0.1375 | 0.1925 | 0.654981 | 0.727857 | 0.39532 | 1 | 0.996511 | 0.866926 |
| Prdm2 | A2A7B5 | 0.58 | 0.01 | 0.66 | 0.5 | 1.12 | 0.23 | 1.03 | 1.07 | 1.78 | 0.82 | 1.23 | 0.26 | 1.44 | 4.27 | 0.8 | 0.91 | 0.425 | 0.585 | 1.4175 | 0.135365 | 0.119188 | 0.086107 | 1 | 0.996511 | 0.691619 |
| Prdx1 | P35700 | 0.64 | 0.1 | -0.16 | -0.21 | 0.53 | 0.46 | 0.35 | 0.04 | 0.16 | -0.18 | -0.05 | 0.5 | -0.33 | 0.47 | -0.58 | -0.43 | 0.2525 | 0.015 | -0.31 | 0.306367 | 0.954476 | 0.336726 | 1 | 0.996755 | 0.847578 |
| Prdx2 | Q61171 | -1.46 | -1.18 | -0.71 | -0.1 | -1.11 | -1.34 | -0.61 | -0.33 | -1.39 | -0.4 | -0.63 | -1.4 | -1.36 | -0.31 | -1.06 | -1.03 | 0.015 | -0.0925 | -0.0775 | 0.966733 | 0.807814 | 0.832058 | 1 | 0.996511 | 0.974476 |
| Prdx3 | P20108 | 0.13 | 0.43 | 0.82 | 0.66 | 0.64 | 0.15 | 1.01 | 0.8 | 0.09 | 0.52 | 0.85 | -0.05 | 0.02 | 0.36 | 0.69 | 1.12 | 0.14 | -0.1575 | 0.0375 | 0.577384 | 0.563356 | 0.898384 | 1 | 0.996511 | 0.983509 |
| Prdx4 | O08807 | -0.32 | -0.29 | -0.3 | -0.04 | 0.11 | -0.05 | 0.79 | -0.01 | -0.18 | -0.37 | -0.09 | -0.28 | -0.04 | 1.11 | 0.53 | 0.16 | 0.4475 | 0.0075 | 0.6775 | 0.076184 | 0.964587 | 0.036885 | 1 | 0.996755 | 0.658454 |
| Prdx5 | P99029-2 | 1.39 | 0.95 | 1.46 | 0.81 | 0.98 | 1.27 | 3.41 | 1.42 | 0.33 | 0.35 | 1.52 | 0.86 | 0.25 | 0.35 | 1.74 | 2.13 | 0.6175 | -0.3875 | -0.035 | 0.263111 | 0.249433 | 0.941102 | 1 | 0.996511 | 0.991085 |
| Prdx6 | Q6GT24 | 0.66 | -0.12 | -0.02 | 0.14 | -0.03 | 0.54 | 0.17 | 0.18 | 0.31 | -0.54 | -0.17 | 0.32 | -0.83 | -0.61 | -0.59 | -0.21 | 0.05 | -0.185 | -0.725 | 0.827757 | 0.51716 | 0.016616 | 1 | 0.996511 | 0.591538 |
| Preb | Q9WUQ2 | -0.4 | -0.26 | -0.03 | -0.44 | 0.24 | -0.35 | -0.1 | -0.45 | -0.1 | -0.06 | -0.39 | -0.36 | 0.3 | 0.16 | 0.65 | 0.13 | 0.1175 | 0.055 | 0.5925 | 0.576175 | 0.768354 | 0.018756 | 1 | 0.996511 | 0.602211 |
| Prelid3a | A0A494B9I | -0.2 | -0.16 | -0.09 | 0.15 | -0.29 | 0.47 | 0.05 | -0.04 | -0.08 | 0.17 | -0.01 | -0.48 | 0.02 | 0.56 | -0.09 | -0.41 | 0.1225 | -0.025 | 0.095 | 0.556646 | 0.902637 | 0.705067 | 1 | 0.996609 | 0.957084 |
| Prelp | Q9JK53 | -2.96 | -2.9 | -2 | -1.99 | -2.87 | -3.37 | -0.83 | -1.88 | -3.18 | -3.28 | -1.47 | -2.99 | -3.14 | -2.29 | -2.47 | -1.88 | 0.225 | -0.2675 | 0.0175 | 0.694345 | 0.57337 | 0.962164 | 1 | 0.996511 | 0.9927 |
| Prep | Q9QUR6 | -0.74 | -0.5 | -0.52 | -0.25 | -0.45 | -0.8 | -0.22 | -0.37 | -0.41 | -0.03 | -0.34 | -0.47 | -0.78 | -0.13 | -0.66 | -0.7 | 0.0425 | 0.19 | -0.065 | 0.827187 | 0.341076 | 0.773846 | 1 | 0.996511 | 0.966062 |
| Prepl | Q8C167-2 | -1.46 | -1.29 | -2.09 | -2.07 | -1.21 | -1.65 | -1.67 | -1.94 | -1.7 | -2.07 | -2.01 | -1.35 | -1.56 | -1.26 | -1.85 | -1.97 | 0.11 | -0.055 | 0.0675 | 0.677814 | 0.842951 | 0.810096 | 1 | 0.996511 | 0.972222 |
| Prex1 | Q69ZK0 | 1.02 | 0.53 | 0.33 | 0.41 | 0.68 | 1.17 | 0.12 | 0.47 | 0.35 | -0.24 | 0.26 | 0.84 | 0.1 | 0.38 | -0.03 | 0.17 | 0.0375 | -0.27 | -0.4175 | 0.890844 | 0.350815 | 0.090446 | 1 | 0.996511 | 0.692714 |
| Prex2 | Q3LAC4 | 0.45 | 1.13 | 0.81 | 0.23 | 1.96 | 1.02 | 1.38 | 0.58 | 0.64 | 0.66 | 0.86 | 1.19 | 0.7 | 0.8 | 0.86 | 0.99 | 0.58 | 0.1825 | 0.1825 | 0.116605 | 0.483325 | 0.462031 | 1 | 0.996511 | 0.888262 |
| Prickle1 | Q3U5C7 | 0.45 | 0.82 | 1.24 | 0.97 | 1.33 | 0.55 | 1.66 | 0.87 | 1.32 | 0.59 | 1.37 | 1.16 | 1.42 | 2.16 | 1.39 | 1.63 | 0.2325 | 0.24 | 0.78 | 0.438451 | 0.371388 | 0.016426 | 1 | 0.996511 | 0.589387 |
| Prickle2 | A7YQ68 | 0.08 | 0.61 | 0.28 | 0.25 | 0.67 | 0.23 | 0.43 | 0.15 | 0.66 | 0.62 | 0.57 | 0.73 | 0.91 | 0.82 | 0.61 | 0.42 | 0.065 | 0.34 | 0.385 | 0.740479 | 0.087859 | 0.097271 | 1 | 0.996511 | 0.698156 |
| Prkaa1 | Q3TUQ7 | 0.85 | 0.75 | 0.91 | 0.94 | 0.69 | 0.8 | 0.59 | 0.73 | 0.05 | 1.32 | 0.8 | 0.79 | -0.26 | 0.3 | 0.9 | 0.82 | -0.16 | -0.1225 | -0.4225 | 0.287167 | 0.660565 | 0.168628 | 1 | 0.996511 | 0.759484 |
| Prkaa2 | Q8BRK8 | 0.98 | 0.53 | 0.54 | -0.03 | 0.83 | 1.13 | 0.35 | 0.03 | 0.13 | -0.43 | 0.42 | 1.15 | 0.08 | 0.41 | 0.11 | 0.14 | 0.08 | -0.1875 | -0.32 | 0.799535 | 0.619886 | 0.226711 | 1 | 0.996511 | 0.787447 |
| Prkab1 | Q9R078 | -0.54 | 0.01 | 0 | -0.36 | -0.05 | -0.23 | 0.58 | -0.2 | -0.45 | -1.57 | -0.07 | -0.34 | -1.1 | -0.51 | -0.23 | 0.18 | 0.2475 | -0.385 | -0.1925 | 0.329799 | 0.289065 | 0.540077 | 1 | 0.996511 | 0.912212 |
| Prkab2 | Q6PAM0 | 2.62 | 2.01 | 2.44 | 0.71 | 3.04 | 2.44 | 2.7 | 2.44 | 0.09 | -0.51 | 2.87 | 3.15 | 0.19 | -0.44 | 2.74 | 2.75 | 0.71 | -0.545 | -0.635 | 0.122469 | 0.561798 | 0.461173 | 1 | 0.996511 | 0.887902 |
| Prkaca | P05132 | 0.48 | 0.31 | 0.8 | 0.79 | 0 | 0.44 | 0.39 | 0.67 | 1.43 | 0.33 | 0.78 | 0.83 | 1.56 | 0.07 | 1.11 | 0.5 | -0.22 | 0.2475 | 0.215 | 0.313569 | 0.373244 | 0.541329 | 1 | 0.996511 | 0.912212 |
| Prkacb | P68181 | 0.24 | -0.33 | -0.02 | 0.24 | -0.13 | 0.02 | 0.19 | 0.01 | 0.59 | 0.21 | 0.1 | 0.39 | -0.11 | 0.2 | -0.05 | 0.11 | -0.01 | 0.29 | 0.005 | 0.957865 | 0.198125 | 0.981059 | 1 | 0.996511 | 0.994858 |
| Prkag1 | O54950 | -0.47 | 0.45 | 1.21 | 0.5 | 0.63 | -0.23 | 1.33 | 0.55 | -0.02 | 0.26 | 0.71 | -0.2 | -0.37 | 0.52 | 0.83 | 0.83 | 0.1475 | -0.235 | 0.03 | 0.737064 | 0.543579 | 0.943737 | 1 | 0.996511 | 0.991609 |
| Prkag2 | Q91WG5 | -0.17 | 0.34 | 0.45 | -0.07 | 0.24 | -0.2 | 0.21 | 0.05 | -0.15 | -0.32 | 0.29 | 0.28 | -0.28 | -0.04 | 0.04 | -0.03 | -0.0625 | -0.1125 | -0.215 | 0.766497 | 0.644805 | 0.339517 | 1 | 0.996511 | 0.84777 |
| Prkar1a | Q9DBC7 | 1.03 | 0.8 | 0.25 | 0.35 | 0.82 | 1.16 | 0.18 | 0.21 | 0.94 | 0.58 | 0.1 | 1.16 | 0.19 | 0.83 | 0.07 | 0.12 | -0.015 | 0.0875 | -0.305 | 0.960128 | 0.772132 | 0.288941 | 1 | 0.996511 | 0.824705 |
| Prkar1b | P12849 | -0.34 | -0.3 | -0.5 | -0.6 | 0.01 | -0.25 | -0.59 | -0.54 | -0.19 | -0.49 | -0.57 | -0.03 | -0.74 | -0.06 | -0.59 | -0.54 | 0.0925 | 0.115 | -0.0475 | 0.63367 | 0.56245 | 0.826347 | 1 | 0.996511 | 0.973235 |
| Prkar2a | Q8K1M3 | 0.55 | 0.62 | 0.68 | 0.75 | 0.61 | 0.59 | 0.47 | 0.63 | -0.2 | -0.78 | 0.53 | 0.45 | -0.18 | -0.74 | 0.35 | 0.51 | -0.075 | -0.65 | -0.665 | 0.603932 | 0.063477 | 0.050243 | 1 | 0.996511 | 0.658454 |
| Prkar2b | P31324 | 0.29 | -0.15 | -0.16 | 0.09 | 0.08 | 0.31 | 0.07 | -0.18 | -0.4 | 0.23 | -0.3 | 0.15 | -0.59 | -0.77 | -0.46 | -0.04 | 0.0525 | -0.0975 | -0.4825 | 0.779562 | 0.667153 | 0.063348 | 1 | 0.996511 | 0.66433 |
| Prkca | Q4VA93 | 1.45 | 0.58 | -0.18 | 0.15 | 0.53 | 1.68 | -0.43 | -0.23 | 1.32 | 0.01 | -0.35 | 0.87 | 1.16 | 0.28 | -0.45 | -0.5 | -0.1125 | -0.0375 | -0.3775 | 0.836426 | 0.93845 | 0.449643 | 1 | 0.996755 | 0.884817 |
| Prkcb | P68404-2 | 0.34 | 0 | 0.71 | 0.49 | 0.03 | 0.27 | 0.49 | 0.69 | 0.03 | -0.83 | 0.48 | -0.04 | -0.45 | 0.26 | -0.31 | 0.17 | -0.015 | -0.475 | -0.4675 | 0.947204 | 0.154913 | 0.098745 | 1 | 0.996511 | 0.698156 |
| Prkcb | P68404 | 0.61 | -0.71 | 1.54 | 0.84 | -1.07 | 0.8 | 0.29 | 1.14 | 2.17 | 0.02 | 0.58 | -0.33 | 3.08 | -1.29 | 1.67 | 1.5 | -0.28 | 0.04 | 0.67 | 0.65114 | 0.951737 | 0.47429 | 1 | 0.996755 | 0.891644 |
| Prkcd | P28867 | -0.92 | -0.04 | -0.09 | -0.27 | 0.19 | -0.81 | 0.04 | -0.28 | -0.39 | -0.76 | -0.21 | -0.42 | -0.75 | 0.06 | -0.84 | -0.41 | 0.115 | -0.115 | -0.155 | 0.700845 | 0.653102 | 0.608583 | 1 | 0.996511 | 0.93409 |
| Prkce | P16054 | -0.39 | -0.8 | -0.39 | -0.28 | -0.71 | -0.39 | -0.18 | -0.15 | -0.83 | -0.24 | -0.51 | -0.42 | -0.95 | -0.73 | -0.61 | -0.67 | 0.1075 | -0.035 | -0.275 | 0.60056 | 0.868198 | 0.196248 | 1 | 0.996511 | 0.78004 |
| Prkcg | P63318 | 1 | 0.29 | -0.29 | 0.09 | 0.37 | 1.02 | 0.21 | 0.24 | 0.69 | -0.05 | 0.1 | 0.94 | -0.19 | 0.58 | -0.45 | -0.42 | 0.1875 | 0.1475 | -0.3925 | 0.565074 | 0.676065 | 0.289361 | 1 | 0.996511 | 0.824705 |
| Prkch | P23298 | 2.59 | 1.77 | 0.56 | 0.2 | 2.44 | 2.65 | 0.61 | 0.56 | 1.41 | 0.98 | 0.27 | 2.05 | 1.12 | 0.91 | 0.49 | 0.35 | 0.285 | -0.1025 | -0.5625 | 0.691669 | 0.866471 | 0.309677 | 1 | 0.996511 | 0.834523 |
| Prkci | Q62074 | 0.07 | -0.23 | -0.61 | -0.67 | -0.35 | 0.11 | -0.7 | -0.33 | -0.54 | -0.14 | -0.42 | 0.03 | -0.27 | -0.95 | 0.01 | -0.32 | 0.0425 | 0.0925 | -0.0225 | 0.865785 | 0.704271 | 0.937052 | 1 | 0.996511 | 0.989767 |
| Prkcq | Q02111 | -0.66 | -0.6 | 0.01 | 0.1 | -0.67 | -0.48 | -0.34 | -0.14 | 0.08 | 0.37 | 0.11 | -0.2 | 0.34 | -0.08 | 0.42 | 0.43 | -0.12 | 0.3775 | 0.565 | 0.623582 | 0.160979 | 0.05515 | 1 | 0.996511 | 0.661738 |
| Prkcsh | O08795-2 | -1.02 | -1.18 | -0.74 | -0.38 | -1.11 | -1.34 | -0.24 | -0.3 | -1.1 | -0.45 | -0.66 | -0.83 | -0.89 | -0.67 | 0.18 | -0.36 | 0.0825 | 0.07 | 0.395 | 0.798305 | 0.777406 | 0.21431 | 1 | 0.996511 | 0.786931 |
| Prkcz | Q02956-2 | -0.67 | -0.35 | -1.85 | -1.8 | 0.02 | -0.62 | -1.8 | -1.96 | -0.83 | -1.33 | -1.69 | -0.44 | -0.78 | 0.19 | -2.18 | -2.31 | 0.0775 | 0.095 | -0.1025 | 0.889938 | 0.831271 | 0.874238 | 1 | 0.996511 | 0.979626 |
| Prkd1 | Q62101 | -0.59 | -0.4 | -0.88 | -0.26 | 0.49 | -0.12 | -0.54 | -0.87 | -0.81 | -0.29 | -0.8 | 0.08 | 0.3 | -0.68 | 0.21 | -0.52 | 0.2725 | 0.0775 | 0.36 | 0.399315 | 0.774217 | 0.245931 | 1 | 0.996511 | 0.802442 |
| Prkdc | P97313 | -1.09 | -0.63 | -0.4 | -2.03 | -1.51 | -3.16 | -1.26 | -2 | -0.85 | -2.7 | -0.99 | -1.13 | -0.1 | -3.69 | -1 | -0.75 | -0.945 | -0.38 | -0.3475 | 0.092172 | 0.472665 | 0.659408 | 1 | 0.996511 | 0.949645 |
| Prkg1 | P0C605-2 | 0.12 | 0.37 | -0.1 | -0.18 | 0.82 | 0.19 | 0.12 | -0.01 | 0.01 | 0.22 | -0.03 | 0.05 | -0.07 | -0.29 | -0.08 | 0.14 | 0.2275 | 0.01 | -0.1275 | 0.352875 | 0.958251 | 0.547727 | 1 | 0.996755 | 0.912571 |
| Prkg2 | E9QPH0 | 0.93 | 1.25 | 0.9 | 0.95 | 1.56 | 1.31 | 0.67 | 1.1 | 0.52 | 0.47 | 0.96 | 1.03 | 0.4 | 0.46 | 1.06 | 0.9 | 0.1525 | -0.2625 | -0.3025 | 0.505934 | 0.231942 | 0.206735 | 1 | 0.996511 | 0.785338 |
| Prkn | B8YP90 | -0.11 | 1.12 | -0.62 | -0.32 | 0.9 | -2.34 | -0.78 | -0.91 | -1.68 | -1.52 | -0.46 | 0.99 | -2.4 | -0.65 | -0.49 | 0.28 | -0.8 | -0.685 | -0.8325 | 0.267268 | 0.315002 | 0.204944 | 1 | 0.996511 | 0.783452 |
| Prkra | Q9WTX2 | -0.51 | -0.51 | -2.01 | -2.24 | -0.19 | -0.9 | -1.36 | -2.24 | -0.7 | -0.71 | -2.02 | -0.54 | -0.31 | -0.26 | -1.69 | -1.86 | 0.145 | 0.325 | 0.2875 | 0.803002 | 0.54967 | 0.625354 | 1 | 0.996511 | 0.939288 |
| Prmt1 | Q9JIF0-2 | -0.19 | -0.93 | -1.1 | -0.74 | -0.68 | -0.41 | -0.67 | -0.71 | -0.39 | -0.67 | -0.84 | -0.11 | -0.98 | -0.44 | -1.23 | -1.13 | 0.1225 | 0.2375 | -0.205 | 0.595966 | 0.390022 | 0.476556 | 1 | 0.996511 | 0.892174 |
| Prmt2 | M0QW88 | 4.23 | 2.7 | 1.78 | 0.72 | 3.17 | 4.29 | 2.64 | 1.21 | 1.4 | 0.55 | 2.34 | 4.55 | 1.42 | 0.67 | 1.91 | 2.17 | 0.47 | -0.1475 | -0.815 | 0.597194 | 0.885196 | 0.285425 | 1 | 0.996511 | 0.822669 |
| Prmt3 | Q922H1 | 0.04 | 0.18 | -0.42 | -0.44 | 0.73 | -0.2 | -0.79 | -0.82 | -0.05 | -0.22 | -0.19 | 0.17 | -0.16 | 0.56 | -0.66 | -0.81 | -0.11 | 0.0875 | -0.1075 | 0.770388 | 0.691203 | 0.756434 | 1 | 0.996511 | 0.962333 |
| Prmt5 | A0A0R4J04 | 0.65 | -0.07 | -0.12 | 0.38 | -0.41 | 0.58 | 0.23 | 0.51 | 0.95 | 0.31 | 0.14 | 1.07 | 0.55 | 0.42 | -0.12 | -0.25 | 0.0175 | 0.4075 | -0.06 | 0.952078 | 0.199296 | 0.83484 | 1 | 0.996511 | 0.975487 |
| Prmt7 | Q922X9 | 0.22 | 0.18 | -1.1 | -1.13 | -0.26 | -0.21 | -0.69 | -0.87 | 0.36 | -0.01 | -0.64 | 0.22 | 0.31 | 0.44 | -0.59 | -1.78 | -0.05 | 4.40E-01 | 0.0525 | 0.898054 | 0.307773 | 0.928494 | 1 | 0.996511 | 0.98946 |
| Prmt8 | Q6PAK3 | 0.83 | 0.72 | 1.38 | 1.19 | 1.08 | 1.03 | 1.98 | 1.47 | -0.2 | 0.33 | 1.74 | 0.92 | -0.46 | 0.75 | 1.14 | 1.33 | 0.36 | -0.3325 | -0.34 | 0.210413 | 0.438023 | 0.419153 | 1 | 0.996511 | 0.87558 |
| Prmt9 | F8WIU7 | -0.01 | -0.3 | -0.85 | -0.13 | -0.23 | 0.28 | 0.2 | -0.4 | -0.08 | -0.09 | -0.41 | -0.55 | -0.73 | 0.39 | -0.68 | -0.08 | 0.285 | 0.04 | 0.0475 | 0.286175 | 0.870314 | 0.884974 | 1 | 0.996511 | 0.979767 |
| Prnp | P04925 | 1.11 | 0.92 | 0.48 | 0.35 | 0.25 | 1.39 | 0.22 | 0.23 | 0.4 | 0.38 | 0.62 | 0.91 | 0.65 | -0.09 | 0.35 | -0.22 | -0.1925 | -0.1375 | -0.5425 | 0.564278 | 0.576106 | 0.085774 | 1 | 0.996511 | 0.691619 |
| Prodh | Q9WU79 | 1.34 | 0.75 | 0.81 | 1.3 | 0.48 | 1.11 | 0.4 | 0.75 | 0.71 | 0.49 | 0.96 | 0.75 | 0.63 | 0.31 | 0.25 | 0.33 | -0.365 | -0.3225 | -0.67 | 0.154023 | 0.169215 | 0.014897 | 1 | 0.996511 | 0.589387 |
| Prom1 | Q8CDK8 | 1.14 | 1.61 | 1.67 | 1.2 | 1.59 | 0.93 | 1.4 | 1.48 | 0.76 | 0.1 | 1.5 | 1.22 | 0.27 | 1.06 | 0.99 | 1.66 | -0.055 | -0.51 | -0.41 | 0.804647 | 0.151165 | 0.223913 | 1 | 0.996511 | 0.787447 |
| Prorp | V9GXP4 | 0.57 | 0.77 | -0.32 | 0.34 | 0.47 | 0.71 | 0.44 | 0.33 | 0.62 | -1.07 | 0.36 | 0.28 | 0.22 | 0.63 | -0.48 | -0.01 | 0.1475 | -0.2925 | -0.25 | 0.572793 | 0.496351 | 0.460923 | 1 | 0.996511 | 0.887902 |
| Prorsd1 | Q9D820 | 2.08 | -0.93 | -0.53 | -0.96 | -2.76 | 1.75 | -4.15 | 3.48 | 1.67 | 5.82 | -2.81 | 4.01 | -0.81 | -1.36 | -5.92 | -5.91 | -0.335 | 2.2575 | -3.415 | 0.847328 | 0.227665 | 0.035436 | 1 | 0.996511 | 0.652696 |
| Prpf18 | Q8BM39 | -1.27 | -1.06 | -1.7 | -1.55 | -1.16 | -1.92 | -1.34 | -0.9 | -1.07 | -0.38 | -2.36 | -1.55 | -0.65 | -0.56 | -1.73 | -1.29 | 0.065 | 0.055 | 0.3375 | 0.807276 | 0.895246 | 0.305139 | 1 | 0.996511 | 0.831662 |
| Prpf19 | Q99KP6-2 | -0.64 | -1.93 | -1.07 | -0.64 | -2.02 | -1.02 | -1 | -0.33 | -1.43 | -0.02 | -0.77 | -1.75 | -0.84 | -0.63 | -0.32 | -1.08 | -0.0225 | 0.0775 | 0.3525 | 0.958432 | 0.865869 | 0.319814 | 1 | 0.996511 | 0.839028 |
| Prpf3 | Q922U1 | 0.2 | -0.8 | -1 | -0.75 | -1.34 | -0.69 | -0.7 | -0.13 | -0.35 | -0.66 | -1.14 | -1.63 | 0.44 | -0.37 | 0.07 | -0.78 | -0.1275 | -0.3575 | 0.4275 | 0.716913 | 0.353267 | 0.265606 | 1 | 0.996511 | 0.817264 |
| Prpf31 | Q8CCF0 | -1.22 | -2.24 | -1.3 | -1.19 | -2.67 | -1.74 | -1.28 | -0.81 | -1.32 | -1.15 | -1.39 | -2.06 | -0.63 | -1.85 | -0.71 | -1.4 | -0.1375 | 0.0075 | 0.34 | 0.754379 | 0.981367 | 0.377117 | 1 | 0.998773 | 0.860984 |
| Prpf38a | Q4FK66 | -0.29 | -1.22 | -0.68 | -0.99 | -1.1 | -0.89 | -0.29 | -0.65 | -1.94 | -1.56 | -0.92 | -1.72 | -1.53 | -1.35 | -0.71 | -0.42 | 0.0625 | -0.74 | -0.2075 | 0.818006 | 0.03523 | 0.537697 | 1 | 0.996511 | 0.912212 |
| Prpf38b | Q80SY5 | -2.23 | -2.9 | -2.74 | -2.66 | -1.9 | -2.61 | -1.37 | -2.37 | -2.54 | -2.02 | -2.5 | -2.71 | -1.07 | -1.72 | -1.27 | -1.87 | 0.57 | 0.19 | 1.15 | 0.08972 | 0.428658 | 0.001837 | 1 | 0.996511 | 0.419079 |
| Prpf39 | E9QJV4 | -1.41 | -0.18 | -2.17 | -2.51 | -0.03 | -1.61 | -1.48 | -2.08 | -1.89 | -0.97 | -2.59 | -0.9 | -1.13 | -0.75 | -2.18 | -2.18 | 0.2675 | -0.02 | 0.0075 | 0.667307 | 0.973408 | 0.989701 | 1 | 0.997515 | 0.996154 |
| Prpf4 | Q9DAW6 | -0.72 | -1.04 | -0.86 | -0.73 | -1.28 | -1.09 | -0.7 | -0.15 | -0.9 | -0.63 | -1.05 | -1.46 | -0.41 | -0.45 | 0.09 | -0.75 | 0.0325 | -0.1725 | 0.4575 | 0.903089 | 0.449106 | 0.075743 | 1 | 0.996511 | 0.674038 |
| Prpf40a | A0A1B0GR | 0.51 | -0.76 | -1.12 | -1.33 | -0.83 | -0.16 | -0.82 | -0.2 | -0.87 | -0.03 | -1.28 | -1.12 | 0.07 | -0.15 | 0.93 | -0.72 | 0.1725 | -0.15 | 0.7075 | 0.685179 | 0.747979 | 0.181114 | 1 | 0.996511 | 0.775592 |
| Prpf40b | D3Z4N6 | -5.14 | -6.54 | -2.51 | -1.56 | -6.25 | -7.75 | -2 | -2.47 | -5.82 | -0.88 | -1.78 | -5.7 | -5.23 | -3.6 | -1 | -2.58 | -0.68 | 0.3925 | 0.835 | 0.677155 | 0.799657 | 0.523687 | 1 | 0.996511 | 0.90791 |
| Prpf4b | Q61136 | 1.37 | 0.34 | -0.11 | -0.38 | 0.14 | 0.55 | -0.45 | 0.25 | 0.52 | -0.33 | -0.62 | -0.53 | 1.13 | 1.1 | 0.31 | -0.62 | -0.1825 | -0.545 | 0.175 | 0.659276 | 0.234907 | 0.738815 | 1 | 0.996511 | 0.958796 |
| Prpf6 | Q91YR7 | 0.03 | -1.21 | -1.77 | -1.68 | -1.23 | -0.4 | -1.2 | -1.07 | -1.06 | -1.99 | -1.7 | -1.55 | -0.34 | -0.84 | -0.36 | -1.4 | 0.1825 | -0.4175 | 0.4225 | 0.671585 | 0.348014 | 0.366828 | 1 | 0.996511 | 0.858904 |
| Prpf8 | Q99PV0 | 0.32 | -0.56 | -0.41 | -0.35 | -0.86 | -0.27 | -0.42 | 0.12 | -1.1 | -0.63 | -0.5 | -1.24 | -0.32 | -0.44 | 0.22 | -0.36 | -0.1075 | -0.6175 | 0.025 | 0.705292 | 0.050241 | 0.926345 | 1 | 0.996511 | 0.989128 |
| Prph | G5E846 | 1.53 | 1.52 | 1.46 | 0.62 | 1.35 | 0.92 | 0.98 | 0.79 | 1.5 | 0.65 | 1.48 | 1.82 | 1.04 | 1.53 | 0.95 | 0.96 | -0.2725 | 0.08 | -0.1625 | 0.311233 | 0.809356 | 0.566793 | 1 | 0.996511 | 0.921806 |
| Prps1l1 | Q8C5R8 | -1.38 | -0.82 | -2.68 | -1.04 | -1.54 | -1.98 | -2.45 | -1.66 | 4.07 | 2.09 | -2.44 | -1.9 | 2.83 | 6.07 | -1.84 | -3.27 | -0.4275 | 1.935 | 2.4275 | 0.337457 | 0.205961 | 0.231324 | 1 | 0.996511 | 0.788863 |
| Prps1l3 | G3UXL2 | -1.76 | -2.43 | -0.5 | -0.67 | -2 | -1.27 | -0.32 | -0.6 | -2.27 | -0.1 | -0.49 | -1.86 | -1.97 | -2.27 | -0.3 | -0.33 | 0.2925 | 0.16 | 0.1225 | 0.593132 | 0.801485 | 0.847186 | 1 | 0.996511 | 0.976225 |
| Prps2 | Q9CS42 | -1.39 | -1.55 | -1.08 | -0.75 | -1.1 | -1.57 | -0.79 | -0.74 | -1.65 | -1.25 | -1.25 | -1.62 | -1.38 | -1 | -1.42 | -1.11 | 0.1425 | -0.25 | -0.035 | 0.596291 | 0.308454 | 0.885011 | 1 | 0.996511 | 0.979767 |
| Prpsap1 | B1AT82 | -0.66 | -0.4 | -0.3 | -0.52 | 0.63 | -0.62 | 0.03 | -0.38 | 0.95 | -0.38 | -0.36 | -0.34 | -0.61 | -0.08 | -0.6 | -0.28 | 0.385 | 0.4375 | 0.0775 | 0.201214 | 0.212275 | 0.712404 | 1 | 0.996511 | 0.958014 |
| Prpsap2 | Q8R574 | -0.59 | -0.59 | -0.41 | -0.34 | -0.29 | -0.87 | -0.51 | -0.34 | -0.74 | 0.13 | -0.57 | -0.83 | -1.16 | -0.69 | -0.55 | -0.53 | -0.02 | -0.02 | -0.25 | 0.914593 | 0.936185 | 0.263111 | 1 | 0.996755 | 0.81524 |
| Prr12 | E9PYL2 | 0.22 | -3.11 | -1.73 | -1.39 | -2.94 | -0.83 | -1.6 | -2.17 | -0.67 | 0.52 | -1.1 | -0.2 | -0.67 | -2.2 | -1.2 | -1.7 | -0.3825 | 1.14 | 0.06 | 0.607116 | 0.129466 | 0.930546 | 1 | 0.996511 | 0.989713 |
| Prr14 | Q7TPN9 | 0.26 | 0.62 | 0.6 | 0.6 | -0.26 | 0.07 | 0.48 | 0.53 | 0.22 | 1.37 | 0.98 | 0.77 | -0.39 | 0.19 | 0.8 | 0.24 | -0.315 | 0.315 | -0.31 | 0.187981 | 0.262991 | 0.283858 | 1 | 0.996511 | 0.822669 |
| Prr18 | Q6PAN7 | -0.08 | -0.58 | -0.25 | -0.45 | -1.71 | 0.61 | -2.75 | -0.8 | 1.96 | -0.36 | -0.42 | -0.27 | 2.83 | -1.07 | -0.33 | -0.13 | -0.8225 | 0.5675 | 0.665 | 0.230452 | 0.312112 | 0.404776 | 1 | 0.996511 | 0.872135 |
| Prr3 | Q811B5 | 0.84 | -0.49 | -0.67 | -0.04 | -0.31 | -0.4 | -0.51 | -1.05 | -1.36 | -1.73 | -2.95 | -0.64 | -0.54 | 0.06 | 0.39 | -0.08 | -0.4775 | -1.58 | 0.0475 | 0.207311 | 0.016837 | 0.900093 | 1 | 0.996511 | 0.983509 |
| Prr36 | E9PV26 | 0.38 | 0.52 | 0.58 | 0.42 | -0.01 | 0.59 | -0.48 | 0.12 | 1.54 | 0.67 | 0.58 | 0.65 | 1.88 | 0.69 | 0.69 | -0.43 | -0.42 | 0.385 | 0.2325 | 0.108112 | 0.155256 | 0.606577 | 1 | 0.996511 | 0.932425 |
| Prr7 | Q3V0I2 | 0.82 | 1.07 | 1.47 | 1.54 | 1.6 | 0.55 | 1.71 | 1.59 | 0.88 | 1.65 | 1.5 | 0.89 | 0.87 | 1.55 | 1.48 | 1.39 | 0.1375 | 0.005 | 0.0975 | 0.664022 | 0.98558 | 0.70764 | 1 | 0.998892 | 0.957213 |
| Prrc1 | Q3UPH1 | 0.43 | -0.24 | -0.39 | -0.23 | 0.24 | -0.35 | -0.63 | -0.48 | -0.19 | 0.52 | -0.16 | 0.1 | 0 | 0.3 | -0.31 | -0.7 | -0.1975 | 0.175 | -0.07 | 0.471329 | 0.512386 | 0.813215 | 1 | 0.996511 | 0.972768 |
| Prrc2a | Q7TSC1 | -0.77 | -1.05 | -0.82 | -0.91 | -0.78 | -1.11 | -0.86 | -1.14 | -0.29 | 0.12 | -0.8 | -0.65 | 0.48 | -0.03 | -0.22 | -0.6 | -0.085 | 0.4825 | 0.795 | 0.610352 | 0.072057 | 0.013282 | 1 | 0.996511 | 0.584789 |
| Prrc2b | F8WHT3 | -1.71 | -2.45 | -1.84 | -2.05 | -2 | -1.92 | -1.47 | -1.82 | -1.35 | -1.18 | -1.62 | -1.81 | -0.55 | -1.41 | -0.75 | -1.33 | 0.21 | 0.5225 | 1.0025 | 0.35725 | 0.054953 | 0.006398 | 1 | 0.996511 | 0.552614 |
| Prrc2c | A0A0A0M( | -0.92 | -1.33 | -1.37 | -1.19 | -1.25 | -1.02 | -1.46 | -1.44 | -0.97 | -0.25 | -1.08 | -0.7 | -0.17 | -0.44 | -0.43 | -1.21 | -0.09 | 0.4525 | 0.64 | 0.630132 | 0.08565 | 0.039817 | 1 | 0.996511 | 0.658454 |
| Prrt1 | O35449 | 1 | 0.81 | 0.88 | 0.84 | 0.3 | 0.78 | 0.94 | 0.92 | 1.32 | 1.17 | 0.91 | 1.11 | 0.93 | 1.44 | 0.57 | 0.29 | -0.1475 | 0.245 | -0.075 | 0.450964 | 0.177012 | 0.784909 | 1 | 0.996511 | 0.967642 |
| Prrt2 | E9PUL5 | 2.63 | 2.3 | 2.33 | 1.95 | 1.61 | 2.19 | 1.73 | 1.89 | 2.15 | 2.39 | 1.93 | 2.45 | 2.97 | 0.87 | 1.66 | 1.68 | -0.4475 | -0.0725 | -0.5075 | 0.062008 | 0.742697 | 0.261002 | 1 | 0.996511 | 0.81412 |
| Prrt3 | Q6PE13 | 0.32 | 0.71 | 0.58 | 0.55 | 0.04 | -0.01 | -0.72 | 0.16 | 0.69 | 0.86 | 0.32 | 0.45 | 0.74 | 0.56 | -0.74 | -0.79 | -0.6725 | 0.04 | -0.5975 | 0.017297 | 0.83922 | 0.163309 | 1 | 0.996511 | 0.758224 |
| Prune1 | Q8BIW1 | -0.44 | -0.3 | 0.43 | 0.63 | -0.26 | -0.42 | 0.08 | 0.5 | -0.4 | 0.77 | 0.31 | -0.12 | -0.68 | -0.12 | -0.06 | 0.21 | -0.105 | 0.06 | -0.2425 | 0.747481 | 0.867135 | 0.466656 | 1 | 0.996511 | 0.888757 |
| Prune2 | Q52KR3 | 0.35 | 0.05 | 0.13 | 0.59 | 0.05 | 0.13 | 0.6 | 0.42 | 1.26 | 1.75 | 0.03 | 0.34 | 1.56 | 0.38 | 0.17 | 0.08 | 0.02 | 0.565 | 0.2675 | 0.922616 | 0.181299 | 0.463741 | 1 | 0.996511 | 0.888267 |
| Prxl2a | Q3U125 | 0.92 | 0.62 | 1.02 | 1.37 | 0.89 | 0.81 | 1.96 | 1.5 | 0.79 | 1.37 | 0.84 | 0.65 | 0.19 | 1.12 | 1.01 | 0.86 | 0.3075 | -0.07 | -0.1875 | 0.332433 | 0.776195 | 0.508746 | 1 | 0.996511 | 0.902804 |
| Prxl2b | Q9DB60 | -0.68 | -1.02 | -0.06 | -0.1 | -0.77 | -0.64 | 0.21 | -0.13 | -0.72 | -0.98 | -0.14 | -0.64 | -0.57 | -1.05 | -0.1 | 0.18 | 0.1325 | -0.155 | 0.08 | 0.678993 | 0.605993 | 0.821072 | 1 | 0.996511 | 0.972768 |
| Psap | E9PZ00 | -3.81 | -3.93 | -4.28 | -5.05 | -3.27 | -3.98 | -4.11 | -4.69 | -4.22 | -2.52 | -4.07 | -3.29 | -3.68 | -3.23 | -4.27 | -4.41 | 0.255 | 0.7425 | 0.37 | 0.511001 | 0.128237 | 0.343988 | 1 | 0.996511 | 0.848473 |
| Psat1 | Q99K85 | 0.25 | -0.04 | 0.08 | -0.02 | 0.49 | 0.37 | 0.05 | -0.03 | -0.31 | -0.54 | -0.03 | 0.2 | -0.48 | -0.28 | -0.5 | -0.06 | 0.1525 | -0.2375 | -0.3975 | 0.416276 | 0.286083 | 0.067095 | 1 | 0.996511 | 0.664936 |
| Psd | Q5DTT2 | 1.82 | 0.6 | 0.43 | 0.66 | 0.17 | 1.72 | 0.78 | 0.71 | 0.69 | -0.61 | 0.54 | 0.55 | 0.82 | 0.23 | -0.19 | -0.24 | -0.0325 | -0.585 | -0.7225 | 0.938815 | 0.185176 | 0.090885 | 1 | 0.996511 | 0.693316 |
| Psd | Q5DTT2-2 | 2.1 | 1.12 | 0.84 | 1.14 | 1.8 | 1.03 | 0.77 | 1.02 | 1.15 | -0.34 | 0.84 | 0.51 | 1.47 | 1.56 | -0.32 | -0.17 | -0.145 | -0.76 | -0.665 | 0.673179 | 0.087461 | 0.23529 | 1 | 0.996511 | 0.791493 |
| Psd2 | Q6P1I6-2 | 0.37 | 0.49 | 0.78 | 0.58 | 0.45 | 0.56 | 0.24 | 0.6 | 0.22 | 0.81 | 0.43 | 0.26 | 0.21 | 0.26 | 0.62 | 0.52 | -0.0925 | -0.125 | -0.1525 | 0.590774 | 0.547644 | 0.451945 | 1 | 0.996511 | 0.885993 |
| Psd3 | E9PUC5 | 0.68 | 0.31 | -0.11 | 0.05 | 0.11 | 0.38 | -0.55 | 0.11 | -0.63 | -2.08 | -0.35 | 0.29 | -0.51 | -0.11 | -0.76 | -0.9 | -0.22 | -0.925 | -0.8025 | 0.422281 | 0.088501 | 0.014685 | 1 | 0.996511 | 0.589387 |
| Psd3 | F6Z9E6 | 1.21 | 0.4 | 1.09 | 0.9 | 0.28 | 1.09 | 1.05 | 1.51 | -0.15 | 0.33 | 1.05 | 0.6 | -0.18 | 0.15 | 0.69 | 0.52 | 0.0825 | -0.4425 | -0.605 | 0.789026 | 0.179325 | 0.057559 | 1 | 0.996511 | 0.661738 |
| Psen1 | P49769 | 0.8 | 0.86 | 0.37 | 0.77 | 0.81 | 0.46 | 0.48 | 0.56 | 1.46 | 1.28 | 0.52 | 0.88 | 1.06 | 1.77 | 0.48 | 0.57 | -0.1225 | 0.335 | 0.27 | 0.505629 | 0.216293 | 0.41113 | 1 | 0.996511 | 0.874811 |
| Psip1 | A2BI12 | -2.26 | -3.86 | -3.47 | -2.82 | -4.51 | -3.92 | -3.23 | -3.66 | -3.76 | -1.17 | -3.11 | -0.85 | -5.1 | -3.9 | -3.4 | -3.57 | -0.7275 | 0.88 | -0.89 | 0.108249 | 0.244697 | 0.095473 | 1 | 0.996511 | 0.698156 |
| Psip1 | Q99JF8 | -2.13 | -3.83 | -2.29 | -1.66 | -3.71 | -2.78 | -3.09 | -2.77 | -2.34 | -1.4 | -2.48 | -2.58 | -2.72 | -3.72 | -2.26 | -2.58 | -0.61 | 0.2775 | -0.3425 | 0.226031 | 0.585444 | 0.519242 | 1 | 0.996511 | 0.907667 |
| Psma1 | Q9R1P4 | 2.18 | 0.83 | 0.64 | 0.61 | 1.21 | 2.14 | 0.62 | 0.91 | -0.1 | 0.2 | 0.74 | 1.9 | -0.21 | -0.17 | 0.59 | 0.61 | 0.155 | -0.38 | -0.86 | 0.738601 | 0.483943 | 0.066423 | 1 | 0.996511 | 0.664936 |
| Psma2 | P49722 | -0.21 | -0.27 | 0.02 | -0.31 | -0.31 | -0.24 | -0.14 | -0.11 | -0.47 | -0.55 | 0.07 | -0.09 | -0.68 | -0.49 | -0.67 | -0.46 | -0.0075 | -0.0675 | -0.3825 | 0.961407 | 0.747841 | 0.060941 | 1 | 0.996511 | 0.661738 |
| Psma3 | O70435 | 1.17 | 0.49 | 0.58 | 0.51 | 0.01 | 1.13 | 0.15 | 0.58 | 0.81 | 0.22 | 0.5 | 0.92 | 0.56 | 0.01 | 0.32 | 0.39 | -0.22 | -0.075 | -0.3675 | 0.467248 | 0.76439 | 0.149214 | 1 | 0.996511 | 0.748169 |
| Psma4 | Q9R1P0 | -0.68 | -0.77 | -0.18 | -0.6 | -0.5 | -0.74 | -0.62 | -0.52 | -0.67 | 0.55 | -0.43 | -0.93 | -0.53 | -0.16 | -0.24 | -0.2 | -0.0375 | 0.1875 | 0.275 | 0.83859 | 0.587873 | 0.216543 | 1 | 0.996511 | 0.787447 |
| Psma5 | Q9Z2U1 | -1.06 | -1.06 | -0.75 | -0.71 | -1.32 | -1.28 | -0.48 | -0.89 | -1.51 | -1.18 | -0.8 | -1.24 | -2.09 | -0.95 | -1.12 | -1.17 | -0.0975 | -0.2875 | -0.4375 | 0.680454 | 0.203957 | 0.157727 | 1 | 0.996511 | 0.751678 |
| Psma6 | Q9QUM9 | -1.87 | -1.38 | -1.23 | -0.73 | -1.67 | -0.99 | -0.97 | -0.78 | -1.2 | 0.09 | -0.84 | -1.13 | -1.98 | -0.02 | -1.26 | -1.02 | 0.2 | 0.5325 | 0.2325 | 0.51359 | 0.172418 | 0.602858 | 1 | 0.996511 | 0.931069 |
| Psma7 | Q9Z2U0 | -0.97 | -1.07 | -0.35 | 0.12 | -1.08 | -1.07 | -0.24 | 0.34 | 0.34 | -0.71 | -0.24 | -1.09 | 0.87 | -1.2 | -0.63 | -0.9 | 0.055 | 0.1425 | 0.1025 | 0.89498 | 0.721167 | 0.838067 | 1 | 0.996511 | 0.975961 |
| Psmb1 | O09061 | -1.18 | -1.15 | -1.13 | -1.07 | -1.28 | -1.32 | -0.95 | -0.95 | -1.39 | -0.75 | -1.06 | -1.12 | -1.63 | -1.13 | -1.28 | -1.16 | 0.0075 | 0.0525 | -0.1675 | 0.963302 | 0.782909 | 0.393765 | 1 | 0.996511 | 0.865775 |
| Psmb10 | O35955 | 0 | -0.12 | -0.38 | -0.39 | -0.54 | -0.59 | -1.11 | -0.76 | -0.48 | 0.84 | -0.79 | -0.48 | -0.74 | -0.51 | -1.31 | -1 | -0.5275 | -0.005 | -0.6675 | 0.023728 | 0.989065 | 0.019318 | 1 | 0.999285 | 0.611343 |
| Psmb2 | Q9R1P3 | 0.25 | -0.62 | -0.39 | -0.42 | -0.63 | 1.01 | 0.42 | 0.08 | -0.97 | -1.32 | -0.04 | 0.06 | -0.7 | -1.2 | -0.6 | -0.23 | 0.515 | -0.2725 | -0.3875 | 0.190598 | 0.476453 | 0.20586 | 1 | 0.996511 | 0.785026 |
| Psmb3 | Q9R1P1 | -0.67 | -0.5 | -0.22 | -0.35 | -0.7 | -0.56 | -0.07 | -0.03 | -0.36 | -0.03 | -0.16 | -0.67 | -0.99 | -0.38 | -0.9 | -0.38 | 0.095 | 0.13 | -0.2275 | 0.666712 | 0.543616 | 0.342932 | 1 | 0.996511 | 0.848473 |
| Psmb4 | P99026 | 0.48 | 0.25 | -0.34 | -0.34 | 0.4 | 0.04 | 0.09 | -0.13 | -0.58 | -0.8 | -0.29 | 0.43 | -0.87 | -0.55 | -0.68 | -0.35 | 0.0875 | -0.3225 | -0.625 | 0.725841 | 0.350322 | 0.039092 | 1 | 0.996511 | 0.658454 |
| Psmb5 | O55234 | -0.37 | -0.82 | -0.41 | -0.4 | -0.45 | -0.34 | -0.41 | -0.5 | -0.73 | -0.14 | -0.3 | -0.5 | -0.71 | -0.82 | -0.46 | -0.62 | 0.075 | 0.0825 | -0.1525 | 0.655827 | 0.695142 | 0.451484 | 1 | 0.996511 | 0.885538 |
| Psmb6 | Q60692 | -1.05 | -1.37 | -0.81 | -0.86 | -1.6 | -1.26 | -1.54 | -1.05 | -1.04 | -0.5 | -1.07 | -1.02 | -1.09 | -1.29 | -1.27 | -1.49 | -0.34 | 0.115 | -0.2625 | 0.13047 | 0.607518 | 0.23129 | 1 | 0.996511 | 0.788863 |
| Psmb7 | P70195 | -0.07 | -0.64 | -1 | -0.91 | 0.02 | -0.67 | -1.51 | -1.3 | -0.22 | 0.08 | -1.15 | -0.17 | -0.39 | -0.35 | -1.36 | -1.83 | -0.21 | 0.29 | -0.3275 | 0.586346 | 0.400518 | 0.428568 | 1 | 0.996511 | 0.879165 |
| Psmb8 | P28063 | 0.96 | -0.73 | -0.05 | 0.36 | -1.03 | -0.1 | -0.67 | -0.61 | -0.01 | -0.23 | -0.42 | -0.48 | -0.17 | 0.95 | -1.26 | -0.56 | -0.7375 | -0.42 | -0.395 | 0.081422 | 0.263503 | 0.47067 | 1 | 0.996511 | 0.891144 |
| Psmc1 | P62192 | -0.61 | -0.67 | 0.13 | 0.02 | -0.41 | -0.52 | 0.08 | 0.35 | -0.6 | -0.1 | 0.16 | -0.45 | -0.65 | -0.37 | 0.01 | 0.03 | 0.1575 | 0.035 | 0.0375 | 0.59255 | 0.900932 | 0.894748 | 1 | 0.996511 | 0.983453 |
| Psmc2 | P46471 | -0.61 | -0.88 | -0.32 | -0.09 | -0.6 | -0.77 | 0.17 | -0.11 | -0.51 | 0.11 | -0.04 | -0.6 | -0.68 | -0.09 | -0.3 | -0.28 | 0.1475 | 0.215 | 0.1375 | 0.600955 | 0.422554 | 0.580938 | 1 | 0.996511 | 0.927734 |
| Psmc3 | A0A087WF | 0.38 | -0.18 | 0.1 | 0.56 | 1.15 | 0.05 | -0.05 | 0.11 | 0.11 | 0.11 | 0.07 | -0.17 | -1.29 | 0.55 | -0.68 | 0.21 | 0.1 | -0.185 | -0.5175 | 0.753097 | 0.400747 | 0.246551 | 1 | 0.996511 | 0.802442 |
| Psmc3 | O88685 | 0.32 | -0.46 | -0.09 | -0.33 | -0.1 | 0.2 | 0.16 | -0.12 | -0.16 | -0.21 | 0.07 | 0.36 | -0.41 | 0.29 | -0.03 | -0.17 | 0.175 | 0.155 | 0.06 | 0.427209 | 0.526646 | 0.814866 | 1 | 0.996511 | 0.972768 |
| Psmc4 | P54775 | -0.49 | -0.6 | 0.02 | 0.18 | -0.49 | -0.57 | -0.11 | 0.32 | -0.53 | -0.09 | 0.23 | -0.32 | -0.81 | -0.01 | 0.54 | 0.12 | 0.01 | 0.045 | 0.1825 | 0.971632 | 0.866215 | 0.595088 | 1 | 0.996511 | 0.929843 |
| Psmc5 | P62196 | 0.7 | -0.01 | -0.34 | -0.07 | 0.23 | 0.62 | -0.05 | -0.08 | -0.28 | 0.24 | -0.06 | 0.75 | -0.6 | 0.14 | -0.29 | -0.4 | 0.11 | 0.0925 | -0.3575 | 0.693935 | 0.770076 | 0.236023 | 1 | 0.996511 | 0.791493 |
| Psmc6 | P62334 | 0.1 | -0.35 | 0.2 | 0.13 | -0.35 | -0.01 | 0.35 | 0.5 | -0.76 | 0.16 | 0.3 | 0.1 | -0.87 | -0.75 | -0.06 | 0.14 | 0.1025 | -0.07 | -0.405 | 0.674009 | 0.804369 | 0.192608 | 1 | 0.996511 | 0.777539 |
| Psmd1 | Q3TXS7 | -0.17 | -0.41 | -0.54 | -0.34 | -0.36 | -0.02 | -0.27 | -0.34 | -0.09 | -0.03 | -0.33 | -0.14 | -0.23 | -0.1 | -0.68 | -0.67 | 0.1175 | 0.2175 | -0.055 | 0.485856 | 0.231992 | 0.802409 | 1 | 0.996511 | 0.971091 |
| Psmd10 | Q9Z2X2 | 0.4 | -0.83 | -1.16 | -0.78 | 0.09 | 0.61 | -0.72 | 0.26 | -1.05 | 0.21 | -0.45 | 0.17 | -0.59 | 0.14 | 0.03 | -0.62 | 0.6525 | 0.3125 | 0.3325 | 0.143149 | 0.473067 | 0.397764 | 1 | 0.996511 | 0.867334 |
| Psmd11 | Q8BG32 | -0.68 | -1 | -0.12 | -0.12 | -0.67 | -1.09 | -0.28 | -0.03 | -0.52 | -0.05 | -0.04 | -0.69 | -1.04 | 0.05 | -0.25 | -0.47 | -0.0375 | 0.155 | 0.0525 | 0.904478 | 0.588721 | 0.870713 | 1 | 0.996511 | 0.979293 |
| Psmd12 | Q9D8W5 | 0.45 | -0.08 | -0.1 | -0.22 | 0.16 | 0.33 | 0.38 | -0.11 | 0.01 | -0.36 | 0.09 | 0.46 | -0.55 | 0.03 | -0.21 | -0.04 | 0.1775 | 0.0375 | -0.205 | 0.413144 | 0.880053 | 0.398544 | 1 | 0.996511 | 0.867334 |
| Psmd13 | Q9WVJ2 | -1.23 | -1.4 | -1.39 | -1.18 | -0.99 | -1.03 | -1.1 | -1.33 | -1.5 | -1.59 | -1.27 | -1.21 | -1.42 | -0.87 | -1.8 | -1.57 | 0.1875 | -0.0925 | -0.115 | 0.256153 | 0.603134 | 0.637846 | 1 | 0.996511 | 0.942804 |
| Psmd14 | O35593 | -1.07 | -1.17 | -0.46 | -0.28 | -0.66 | -1.22 | 0.04 | -0.15 | -1.52 | -0.74 | -0.26 | -1.03 | -1.45 | -1.2 | -0.54 | -0.54 | 0.2475 | -0.1425 | -0.1875 | 0.480369 | 0.675958 | 0.567814 | 1 | 0.996511 | 0.921978 |
| Psmd2 | Q8VDM4 | -0.83 | -1.12 | -0.38 | -0.25 | -0.9 | -0.83 | -0.26 | -0.18 | -1.25 | -0.24 | -0.25 | -0.78 | -1.58 | -0.82 | -0.64 | -0.57 | 0.1025 | 0.015 | -0.2575 | 0.713691 | 0.962121 | 0.422868 | 1 | 0.996755 | 0.875782 |
| Psmd3 | P14685 | 0.92 | -0.09 | -0.25 | -0.27 | -0.09 | 0.81 | 0.12 | -0.08 | 0.68 | -0.5 | -0.06 | 0.87 | 0.39 | 0.63 | -0.18 | -0.57 | 0.1125 | 0.17 | -0.01 | 0.742826 | 0.677969 | 0.97911 | 1 | 0.996511 | 0.994274 |
| Psmd4 | O35226 | 1.5 | 1.35 | 1.59 | 1.68 | 1.92 | 0.9 | 2.24 | 1.76 | -0.21 | 1.07 | 0.95 | 1.7 | -0.61 | 0.42 | 0.54 | 1.22 | 0.175 | -0.6525 | -1.1375 | 0.555774 | 0.120534 | 0.013603 | 1 | 0.996511 | 0.589387 |
| Psmd4 | O35226-2 | 1.99 | -0.67 | -0.59 | -0.63 | 0.03 | 2 | -0.55 | -0.51 | -0.92 | -0.43 | -0.46 | 1.85 | -1.12 | -0.32 | -0.79 | -0.72 | 0.2175 | -0.015 | -0.7625 | 0.786368 | 0.985338 | 0.237547 | 1 | 0.998892 | 0.792722 |
| Psmd5 | Q8BJY1 | -0.75 | -0.47 | -0.97 | -0.78 | -0.27 | -0.68 | -0.72 | -0.76 | -0.44 | -0.13 | -1.09 | -0.53 | -0.65 | 0.2 | -1.25 | -1.14 | 0.135 | 0.195 | 0.0325 | 0.486794 | 0.441542 | 0.924868 | 1 | 0.996511 | 0.988301 |
| Psmd6 | Q99JI4 | -0.03 | -0.59 | -0.32 | -0.44 | -0.84 | 0.14 | -0.59 | -0.49 | 0.16 | -0.74 | -0.44 | -0.1 | 0.15 | -0.42 | -0.53 | -0.84 | -0.1 | 0.065 | -0.065 | 0.69201 | 0.796652 | 0.806461 | 1 | 0.996511 | 0.971715 |
| Psmd7 | P26516 | -0.75 | -1.38 | -1.26 | -1.17 | -1.09 | -1.16 | -1.1 | -1.18 | -0.57 | -0.65 | -1.06 | -0.54 | -1.17 | -0.63 | -1.48 | -1.48 | 0.0075 | 0.435 | -0.05 | 0.967147 | 0.074683 | 0.852086 | 1 | 0.996511 | 0.976225 |
| Psmd8 | Q9CX56 | -0.83 | -1.17 | -1.11 | -1.43 | -0.83 | -0.69 | -0.43 | -0.75 | -0.41 | -1.45 | -0.6 | -0.91 | -0.13 | -0.79 | -1.24 | -1.06 | 0.46 | 0.2925 | 0.33 | 0.036071 | 0.300297 | 0.27237 | 1 | 0.996511 | 0.822084 |
| Psmd9 | Q9CR00 | -0.45 | -0.49 | -0.04 | -0.07 | -0.95 | -0.37 | -0.15 | -0.01 | -1.05 | 0.65 | -0.09 | -0.4 | -1.36 | -1.11 | -0.23 | -0.73 | -0.1075 | 0.04 | -0.595 | 0.670099 | 0.912122 | 0.066374 | 1 | 0.996744 | 0.664936 |
| Psme1 | G3UXZ5 | -1.63 | -1.39 | -1.67 | -1.49 | -1.49 | -1.74 | -1.4 | -1.65 | -2.2 | -2.28 | -1.44 | -1.62 | -2.25 | -1.82 | -2.19 | -2.17 | -0.025 | -0.34 | -0.5625 | 0.876796 | 0.187998 | 0.015889 | 1 | 0.996511 | 0.589387 |
| Psme2 | P97372 | -0.44 | -1.11 | -0.72 | -0.83 | -1.05 | -0.67 | -0.44 | -0.55 | -1.17 | -0.58 | -0.61 | -0.63 | -1.37 | -1.02 | -0.83 | -0.84 | 0.0975 | 0.0275 | -0.24 | 0.65469 | 0.904781 | 0.314556 | 1 | 0.996744 | 0.836924 |

| | | | | | | | | | | | | | | | | | | | | | | | | | | |
|---|---|---|---|---|---|---|---|---|---|---|---|---|---|---|---|---|---|---|---|---|---|---|---|---|---|---|
| Psme3 | P61290 | -3.73 | -4.08 | -3.17 | -3.26 | -3.36 | -4.26 | -3.22 | -3.78 | -4.41 | -1.66 | -3.32 | -3.31 | -3.93 | -3.95 | -3.82 | -3.47 | -0.095 | 0.385 | -0.2325 | 0.7605 | 0.496265 | 0.392793 | | 1 | 0.996511 | 0.865775 |
| Psme3ip1 | Q91WE2 | -1.56 | -1.51 | -1.79 | -1.89 | -1.55 | -1.54 | -2.02 | -2.23 | -1.86 | -0.86 | -1.78 | -1.34 | -1.84 | -0.92 | -2.34 | -2.42 | -0.1475 | 0.2275 | -0.1925 | 0.506625 | 0.401041 | 0.58891 | | 1 | 0.996511 | 0.929131 |
| Psme4 | Q5SSW2 | -0.68 | -0.45 | 0.4 | 0.77 | -0.56 | -0.61 | 0.39 | 0.82 | -0.55 | 0.3 | 0.76 | -0.21 | -0.59 | -0.78 | 0.23 | 0.27 | 1.96E-16 | 0.065 | -0.2275 | 1 | 0.878288 | 0.590577 | | 1 | 0.996511 | 0.929142 |
| Psmf1 | Q8BHL8 | -0.06 | -0.16 | -0.39 | 0.18 | -0.37 | -0.1 | -0.74 | -0.52 | 0.04 | -0.36 | -0.58 | -0.13 | -0.27 | 0.14 | -0.54 | -0.96 | -0.325 | -0.15 | -0.3 | 0.144629 | 0.49727 | 0.300312 | | 1 | 0.996511 | 0.830161 |
| Psmg1 | Q9JK23 | -1.3 | -1.5 | 0.55 | 0.73 | -1.61 | -1.63 | 0.73 | 0.71 | -1.14 | 0.58 | 0.19 | -1.41 | -1.27 | -1.01 | 0.33 | 0.75 | -0.07 | -0.065 | 0.08 | 0.931022 | 0.925707 | 0.908924 | | 1 | 0.996755 | 0.986458 |
| Psmg2 | Q9EST4-3 | -1.42 | -1.53 | -1.28 | -0.91 | -1.3 | -1.35 | -1.13 | -1.23 | -1.66 | -0.57 | -1.54 | -1.37 | -1.3 | -0.88 | -1.21 | -1.37 | 0.0325 | 3.73E-16 | 0.095 | 0.860541 | 1 | 0.670874 | | 1 | 1 | 0.952558 |
| Psmg3 | Q9CZH3 | -0.37 | -0.36 | -0.32 | 0.13 | -0.34 | -0.44 | 0.6 | 0.37 | 0 | -1.33 | -0.64 | 0.41 | -1.36 | -0.44 | 0.8 | 0.35 | 0.2775 | -0.16 | 0.0675 | 0.345238 | 0.678432 | 0.883541 | | 1 | 0.996511 | 0.979767 |
| Pspc1 | Q8R326 | -1.7 | -4.06 | -3.92 | -3.6 | -3.39 | -3.28 | -3.12 | -3.68 | -3.95 | -2.19 | -3.09 | -3.54 | -3.14 | -2.85 | -2.33 | -3.28 | -0.0475 | 0.1275 | 0.42 | 0.926645 | 0.834433 | 0.447131 | | 1 | 0.996511 | 0.884817 |
| Psph | Q99LS3 | 0.31 | 0.3 | -0.16 | 0.15 | -0.01 | 0.76 | -0.27 | 0.09 | -0.1 | 0.33 | -0.24 | 0.46 | -0.49 | -0.17 | -0.62 | -0.66 | -0.0075 | -0.0375 | -0.635 | 0.976558 | 0.871401 | 0.014481 | | 1 | 0.996511 | 0.589387 |
| Psrc1 | A2AEF3 | 0.23 | 0 | -0.09 | -0.2 | -2.57 | -0.79 | -2.46 | -1.04 | -0.13 | -1.98 | -1.12 | -0.37 | 1.89 | -2.6 | -2.02 | -0.64 | -1.7 | -0.885 | -0.8275 | 0.004029 | 0.054681 | 0.36998 | | 1 | 0.996511 | 0.859245 |
| Ptar1 | A0A494B9\ | 0.08 | 0.27 | 0 | -0.3 | 0.29 | 0.15 | 0 | -0.36 | -0.1 | -0.24 | -0.3 | -0.04 | -0.35 | 0 | -0.35 | -0.64 | 0.0075 | -0.1825 | -0.3475 | 0.971605 | 0.350137 | 0.147905 | | 1 | 0.996511 | 0.746406 |
| Ptbp1 | Q8BGJ5 | 0.08 | -0.88 | -1.07 | -1.06 | -0.88 | -0.4 | -1.33 | -0.97 | 0.16 | -1.79 | -0.44 | -0.75 | 0.01 | -0.29 | -0.42 | -0.87 | -0.1625 | 0.0275 | 0.34 | 0.620504 | 0.952398 | 0.321128 | | 1 | 0.996755 | 0.839916 |
| Ptbp2 | Q91Z31 | -1.44 | -2.7 | -2.21 | -2.49 | -2.47 | -1.94 | -1.94 | -2.35 | -1.82 | -1.86 | -1.94 | -2.51 | -1.31 | -2.32 | -1.75 | -2.09 | 0.035 | 0.1775 | 0.3425 | 0.908477 | 0.582006 | 0.34137 | | 1 | 0.996511 | 0.848473 |
| Ptbp2 | A0A0G2JF\ | -0.48 | -2.71 | -0.3 | -0.31 | -5.02 | -0.63 | 0.26 | -0.46 | -1.15 | 0.81 | -1.34 | -0.58 | -1.13 | -1.87 | -0.19 | 2.63 | -0.5125 | 0.385 | 0.81 | 0.669997 | 0.582257 | 0.438091 | | 1 | 0.996511 | 0.883161 |
| Ptbp3 | G8JL74 | -0.66 | -2.14 | -0.53 | -0.79 | -2.24 | -0.81 | -0.6 | -0.26 | -0.71 | -0.15 | -0.74 | -1.47 | 0.65 | -1.73 | -0.33 | -0.58 | 0.0525 | 0.2625 | 0.5325 | 0.920677 | 0.551123 | 0.359934 | | 1 | 0.996511 | 0.856735 |
| Ptcd1 | Q8C2E4 | 0.76 | 0.44 | -0.02 | 0.18 | 0.41 | 0.95 | -0.23 | -0.07 | 0.62 | -0.07 | 0 | 0.87 | 0.2 | 0.74 | -0.75 | -0.16 | -0.075 | 0.015 | -0.3325 | 0.808965 | 0.959271 | 0.358098 | | 1 | 0.996755 | 0.856645 |
| Ptcd2 | Q8R3K3 | -0.56 | -0.21 | 0.22 | 0.08 | -0.25 | -0.3 | 0.25 | 0.45 | -0.37 | 0.69 | 0.39 | -0.34 | 0.3 | 0.41 | -0.26 | 0.29 | 0.155 | 0.21 | 0.3025 | 0.557388 | 0.513953 | 0.26111 | | 1 | 0.996511 | 0.81412 |
| Ptcd3 | Q14C51 | 0.2 | 0.25 | 0.49 | 0.42 | 0.03 | 0.27 | 0.29 | 0.59 | 0.12 | 0.19 | 0.37 | -0.3 | 0.22 | -0.13 | 0.77 | 0.42 | -0.045 | -0.245 | -0.02 | 0.802046 | 0.250057 | 0.93339 | | 1 | 0.996511 | 0.989767 |
| Ptdss1 | Q99LH2 | -2.06 | -1.35 | -0.53 | 0.64 | -0.69 | -1.64 | 0.37 | 0.8 | -2.27 | 0.2 | -0.31 | -1.4 | -2.42 | -1.78 | 0.94 | 1.01 | 0.535 | -0.12 | 0.2625 | 0.46592 | 0.868944 | 0.784088 | | 1 | 0.996511 | 0.967642 |
| Ptdss2 | Q9Z1X2 | -0.99 | -0.33 | -0.59 | -0.68 | 0.26 | -0.6 | 0.02 | -0.61 | -1.1 | -1.05 | -0.4 | -0.75 | -0.73 | -0.75 | -0.27 | -0.37 | 0.415 | -0.1775 | 0.1175 | 0.145623 | 0.465095 | 0.610188 | | 1 | 0.996511 | 0.934978 |
| Pten | O08586 | 0.13 | -0.23 | -0.29 | 0.32 | -0.47 | 0 | -0.73 | 0.15 | -0.14 | 0.38 | -0.48 | -0.03 | -0.54 | -0.4 | -0.98 | -0.78 | -0.245 | -0.05 | -0.6575 | 0.358464 | 0.842744 | 0.019873 | | 1 | 0.996511 | 0.615793 |
| Pter | A2AUR3 | -0.42 | -0.65 | -0.23 | -0.11 | -0.45 | -0.48 | -0.62 | -0.08 | -0.44 | 5.72 | -0.04 | -0.15 | -0.47 | 0.13 | -0.34 | -0.74 | -0.055 | 1.625 | -0.0025 | 0.782177 | 0.242878 | 0.992023 | | 1 | 0.996511 | 0.996154 |
| Ptgds | O09114 | -4.44 | -5.37 | -4.93 | -5.1 | -4.81 | -5.23 | -5.13 | -5.39 | -5.83 | -2.2 | -4.65 | -4.64 | -5.07 | -3.75 | -5.37 | -5.62 | -0.18 | 0.63 | 0.0075 | 0.468983 | 0.389434 | 0.986283 | | 1 | 0.996511 | 0.996154 |
| Ptges2 | Q8BWM0 | 0.08 | 0.29 | -0.42 | -0.15 | -0.23 | 0.16 | -0.34 | -0.47 | 0.45 | -0.84 | -0.2 | 0.02 | 0.26 | -0.01 | -0.7 | -0.67 | -0.17 | -0.0925 | -0.23 | 0.45849 | 0.767035 | 0.448619 | | 1 | 0.996511 | 0.884817 |
| Ptges3 | Q9R0Q7 | -2.07 | -2.51 | -2.66 | -2.58 | -1.42 | -2.26 | -1.8 | -2.84 | -2.95 | -2.8 | -2.46 | -2.4 | -3.17 | -2.06 | -2.52 | -2.94 | 0.375 | -0.1975 | -0.2175 | 0.267183 | 0.386571 | 0.465531 | | 1 | 0.996511 | 0.888369 |
| Ptgfrn | Q9WV91 | 0.64 | 0.99 | 1.23 | 1.03 | 1.05 | 0.65 | 0.97 | 0.85 | 0.64 | 1.22 | 1.33 | 1.01 | 0.89 | 0.79 | 1.13 | 1.26 | -0.0925 | 0.0775 | 0.045 | 0.627499 | 0.734905 | 0.835502 | | 1 | 0.996511 | 0.975524 |
| Ptgis | Q8BXC0 | -0.57 | 0.6 | -1.06 | 0.33 | 0.16 | 0.55 | -0.2 | -0.52 | -0.67 | -1.24 | -1.18 | -0.58 | -0.78 | -0.32 | -1.82 | -1.07 | 0.1725 | -0.7425 | -0.8225 | 0.683982 | 0.093513 | 0.106744 | | 1 | 0.996511 | 0.703417 |
| Ptgr1 | Q91YR9 | -0.89 | 0.71 | 0.64 | 0.66 | 0.66 | 0.07 | 1.52 | 0.53 | 0.11 | 0.12 | 0.24 | 0.4 | -0.26 | -0.05 | 0.93 | 1.05 | 0.415 | -0.0625 | 0.1375 | 0.377535 | 0.869881 | 0.775428 | | 1 | 0.996511 | 0.966799 |
| Ptgr2 | Q8VDQ1 | 0.35 | 0.09 | -0.54 | -0.15 | 0.34 | -0.09 | 0.26 | -0.07 | -0.04 | -0.64 | -0.74 | 0.18 | -0.3 | 0.54 | -0.86 | -0.4 | 0.1725 | -0.2475 | -0.1925 | 0.472291 | 0.418465 | 0.581536 | | 1 | 0.996511 | 0.927734 |
| Ptgr3 | Q8BGC4 | 0.31 | 0.26 | 0.35 | 0.76 | 0.31 | 0.3 | 0.53 | 0.75 | 0.09 | 0.61 | 0.3 | 0.06 | 0.06 | 0.59 | -0.07 | 0.25 | 0.0525 | -0.155 | -0.2125 | 0.786282 | 0.472786 | 0.365469 | | 1 | 0.996511 | 0.858475 |
| Ptgs1 | P22437 | -0.51 | -1.02 | 0.51 | 0.76 | -0.65 | 0.03 | 0.18 | 0.73 | -0.44 | 0.95 | 0.74 | -0.29 | 0.01 | -0.31 | 0.99 | 0.96 | 0.1375 | 0.305 | 0.4775 | 0.77042 | 0.553748 | 0.351771 | | 1 | 0.996511 | 0.854088 |
| Ptgs2 | Q05769 | -2.6 | -1.31 | -0.64 | 0.3 | -0.05 | -2.53 | 0.45 | 0.55 | -1.35 | -0.35 | -0.53 | -1.75 | -1.79 | -1.01 | 0.32 | 0.21 | 0.6675 | 0.0675 | 0.495 | 0.441655 | 0.915222 | 0.496603 | | 1 | 0.996755 | 0.897918 |
| Ptk2 | A0A5F8MP | 0.41 | -0.22 | 0.39 | 0.1 | -0.09 | 0.34 | 0.34 | 0.3 | -0.05 | -0.05 | 0.3 | 0.23 | -0.2 | -0.44 | -0.18 | 0.09 | 0.0525 | -0.0625 | -0.3525 | 0.802326 | 0.771257 | 0.14941 | | 1 | 0.996511 | 0.748169 |
| Ptk2b | Q9QVP9 | -0.26 | -0.15 | -0.44 | -0.39 | -0.24 | -0.23 | -0.36 | -0.39 | -0.61 | -0.33 | -0.5 | -0.35 | -0.63 | -0.25 | -0.65 | -0.59 | 0.005 | -0.1375 | -0.22 | 0.973568 | 0.428816 | 0.267721 | | 1 | 0.996511 | 0.819502 |
| Ptk7 | Q8BKG3 | 0.26 | 0.41 | 0.85 | 0.7 | 0.41 | 0.14 | 0.33 | 0.41 | 0.76 | 0.64 | 0.77 | 0.38 | 0.62 | 0.6 | 0.79 | 0.66 | -0.2325 | 0.0825 | 0.1125 | 0.237521 | 0.69131 | 0.584654 | | 1 | 0.996511 | 0.927734 |
| Ptma | P26350 | -4.57 | -5.01 | -2.95 | -3.44 | -5.04 | -5.45 | -4.26 | -3.88 | -4.85 | -0.51 | -3.38 | -3.94 | -3.11 | -3.22 | -3.22 | -3.78 | -0.665 | 0.8225 | 0.66 | 0.245689 | 0.39572 | 0.186302 | | 1 | 0.996511 | 0.776843 |
| Ptms | Q9D0J8 | -2.86 | -4.23 | -4.3 | -4.41 | -5.6 | -2.74 | -5.92 | -6.04 | -1.77 | -1.62 | -5.24 | -2.59 | -0.91 | -5.15 | -5.17 | -6.1 | -1.125 | 1.145 | -0.3825 | 0.174563 | 0.190086 | 0.725773 | | 1 | 0.996511 | 0.958398 |
| Ptn | P63089 | 2.81 | -0.4 | -1.38 | -1.77 | 1.49 | 4.53 | -0.69 | -1.85 | 0.22 | -0.76 | -1.56 | 2.77 | 1.14 | -0.64 | -0.41 | -0.65 | 1.055 | 0.3525 | 0.045 | 0.504061 | 0.778991 | 0.964277 | | 1 | 0.996511 | 0.9927 |
| Ptov1 | Q91VU8 | -1.37 | -2.35 | -1.26 | -1.72 | -2.61 | -1.95 | -1.62 | -0.97 | -1.38 | -1.04 | -1.4 | -1.88 | 0.3 | -3.02 | -0.77 | -1.69 | -0.1125 | 0.25 | 0.38 | 0.777184 | 0.421198 | 0.578374 | | 1 | 0.996511 | 0.927198 |
| Ptp4a1 | Q63739 | 0.82 | 0.71 | 1.42 | 1.27 | -0.01 | 0.75 | 0.24 | 1.1 | 1.25 | 1.07 | 1.29 | 0.91 | 1.17 | 1.07 | 0.28 | -0.05 | -0.535 | 0.075 | -0.4375 | 0.103037 | 0.74126 | 0.224482 | | 1 | 0.996511 | 0.787447 |
| Ptp4a2 | O70274 | -0.14 | -0.37 | 0.04 | -0.47 | -0.33 | 0.45 | -0.43 | -0.48 | -0.41 | -0.33 | -0.59 | -0.06 | -0.12 | -0.55 | -0.35 | -0.54 | 0.0375 | -0.1125 | -0.155 | 0.883679 | 0.586637 | 0.469005 | | 1 | 0.996511 | 0.890107 |
| Ptp4a3 | Q9D658 | -0.16 | 0.44 | -0.13 | -0.35 | 0.04 | 0.35 | -0.69 | -0.54 | 1.48 | -1.01 | -0.23 | 0.26 | 1.65 | 1.46 | -0.77 | -0.07 | -0.16 | 0.175 | 0.6175 | 0.592365 | 0.732029 | 0.291055 | | 1 | 0.996511 | 0.825497 |
| Ptpa | P58389 | 0.09 | 0.14 | 0.65 | 0.37 | -0.06 | 0.09 | 0.38 | 0.6 | -0.45 | -0.58 | 0.44 | 0.06 | -0.38 | -0.19 | 0.57 | 0.37 | -0.06 | -0.445 | -0.22 | 0.784538 | 0.139078 | 0.439674 | | 1 | 0.996511 | 0.883419 |
| Ptpdc1 | A0A5F8MP | 0.68 | 1.06 | -0.01 | 0.28 | 0.29 | 0.01 | 0.02 | 0 | 1.15 | 1.2 | 0.43 | 1.05 | 0.58 | 0.63 | 0.92 | 0.52 | -0.4225 | 0.455 | 0.16 | 0.124264 | 0.155357 | 0.561795 | | 1 | 0.996511 | 0.91998 |
| Ptpmt1 | Q66GT5 | 0.24 | 0.51 | -0.51 | -0.89 | 0.04 | 0.11 | 0.12 | -0.79 | 0.76 | -1.01 | -0.15 | 0.29 | 0.54 | 1.15 | -0.77 | -1.18 | 0.0325 | 0.135 | 0.0975 | 0.930614 | 0.772272 | 0.867299 | | 1 | 0.996511 | 0.978457 |
| Ptpn1 | P35821 | 0.08 | 0.33 | 0.17 | 0.09 | 0.56 | 0.2 | 0.35 | 0.23 | 0.36 | 0.25 | 0.43 | 0.22 | 0.32 | 0.76 | 0.34 | 0.6 | 0.1675 | 0.1475 | 0.3375 | 0.314314 | 0.378743 | 0.109201 | | 1 | 0.996511 | 0.708369 |
| Ptpn11 | P35235-1 | 1.19 | 0.31 | 1.01 | 0.4 | 0.12 | 1.17 | 1.26 | 0.88 | -0.2 | 0.32 | 0.93 | 0.7 | -0.47 | -0.3 | 0.71 | 0.65 | 0.13 | -0.29 | -0.58 | 0.694392 | 0.388075 | 0.14462 | | 1 | 0.996511 | 0.743156 |
| Ptpn11 | P35235 | -1.06 | -1.24 | 0.32 | -2.4 | -4 | -0.09 | -1.51 | 0.03 | 2.3 | -3.61 | 0.56 | -1.84 | 3.96 | -6.53 | 1.65 | -1.14 | -0.2975 | 0.4475 | 0.58 | 0.761522 | 0.724751 | 0.778714 | | 1 | 0.996511 | 0.967289 |
| Ptpn12 | P35831 | -0.7 | -0.77 | 0.18 | 0.21 | -1.04 | -0.52 | -0.36 | 0.01 | 0.2 | 0.55 | 0.22 | -0.62 | 0.15 | -0.46 | 0.25 | 0.11 | -0.2075 | 0.3575 | 0.2825 | 0.540577 | 0.33182 | 0.38766 | | 1 | 0.996511 | 0.864251 |
| Ptpn13 | G5E8B1 | 1.32 | 0.21 | 0.58 | 0.77 | -0.18 | 1.3 | 0.97 | 0.54 | 0.97 | 1.33 | 0.82 | 1.43 | 0.88 | -0.63 | 0.9 | 1.13 | -0.0625 | 0.4175 | -0.15 | 0.867691 | 0.166726 | 0.734598 | | 1 | 0.996511 | 0.958635 |
| Ptpn2 | Q06180 | 0.7 | 1.07 | 0.31 | 0.24 | 0.33 | 0.72 | -0.14 | 0.1 | 0.05 | 0.12 | 0.13 | 0.64 | 0.1 | -0.74 | 0.03 | 0.19 | -0.3275 | -0.345 | -0.685 | 0.246841 | 0.201375 | 0.045663 | | 1 | 0.996511 | 0.658454 |
| Ptpn23 | Q6PB44 | 0.59 | 0.53 | 0.39 | 0.37 | 0.32 | 0.44 | 0.14 | 0.18 | 0.06 | 0.34 | 0.24 | 0.28 | 0.02 | 0.29 | 0.12 | 0.06 | -0.2 | -0.24 | -0.3475 | 0.219413 | 0.171965 | 0.076025 | | 1 | 0.996511 | 0.674054 |
| Ptpn3 | A2ALK8 | 1.38 | 0.4 | 0.33 | 0.4 | 0.64 | 1.05 | 0.16 | 0.35 | 0.05 | 0.02 | 0.53 | 0.54 | -0.38 | -0.04 | -0.28 | 0.12 | -0.0775 | -0.3425 | -0.7725 | 0.804017 | 0.267176 | 0.025064 | | 1 | 0.996511 | 0.633075 |
| Ptpn4 | Q9WU22 | 0.17 | 0.04 | 1.16 | 1 | 0.43 | 0.27 | 1.52 | 1.1 | -0.17 | 0.46 | 1.35 | 0.32 | -0.05 | -0.03 | 0.73 | 1.13 | 0.2375 | -0.1025 | -0.1475 | 0.543016 | 0.800967 | 0.709109 | | 1 | 0.996511 | 0.957656 |
| Ptpn5 | P54830 | -0.19 | -0.51 | -0.13 | -0.39 | -0.03 | -0.36 | 0.53 | -0.41 | -0.67 | -0.94 | -0.45 | -0.09 | -0.03 | -0.92 | 0.19 | 0.33 | 0.2375 | -0.2325 | 0.1975 | 0.349028 | 0.329748 | 0.522923 | | 1 | 0.996511 | 0.907794 |
| Ptpn6 | P29351 | -0.54 | -0.83 | -0.76 | -0.38 | -1.27 | -0.55 | -1.14 | -0.78 | -0.04 | -0.07 | -0.89 | -0.87 | 0.25 | -1.39 | -0.86 | -0.84 | -0.3075 | 0.16 | -0.0825 | 0.183923 | 0.562884 | 0.816652 | | 1 | 0.996511 | 0.972768 |
| Ptpn9 | O35239 | 0.29 | 0.54 | 0.41 | 0.17 | 0.59 | 0.32 | 0.43 | 0.17 | 0.19 | -0.36 | 0.4 | 0.19 | -0.12 | 0.4 | 0.11 | 0.3 | 0.025 | -0.2475 | -0.18 | 0.883735 | 0.276231 | 0.386911 | | 1 | 0.996511 | 0.863233 |
| Ptpra | Q91V35 | 0.76 | 1.24 | 0.68 | 0.68 | 1.34 | 1 | 0.65 | 0.62 | 0.85 | 3.54 | 0.68 | 0.8 | 0.73 | 1.28 | 0.56 | 0.6 | 0.0625 | 0.6275 | -0.0475 | 0.789834 | 0.343097 | 0.848465 | | 1 | 0.996511 | 0.976225 |
| Ptprc | S4R1M0 | -0.08 | -0.45 | -1.09 | -0.66 | -1.71 | -0.46 | -1.37 | -0.48 | -0.88 | 1.52 | -1.41 | -1.13 | 0.64 | -0.51 | -0.91 | -1.76 | -0.435 | 0.095 | -0.065 | 0.251278 | 0.882754 | 0.897355 | | 1 | 0.996511 | 0.983509 |
| Ptprd | Q64487-5 | 0.72 | 0.86 | 0.85 | 1.05 | 0.85 | 0.76 | 0.81 | 0.85 | 0.94 | 0.84 | 1.02 | 0.96 | 0.19 | 1.09 | 0.49 | 0.62 | -0.0525 | 0.07 | -0.2725 | 0.7249 | 0.671555 | 0.271776 | | 1 | 0.996511 | 0.822084 |
| Ptprd | Q64487-1 | -2.38 | -2.72 | 1.43 | 1.87 | -3.38 | -2.63 | 2 | 1.26 | -1.54 | 2.4 | 1.86 | -2.63 | -2.52 | 0.51 | 2.71 | 2.23 | -0.2375 | 0.4725 | 1.1825 | 0.883788 | 0.761309 | 0.440971 | | 1 | 0.996511 | 0.883549 |
| Ptprd | G3UY75 | -0.46 | -0.02 | -0.72 | -0.44 | -0.89 | -0.98 | -0.9 | -1.14 | -0.41 | -0.01 | -0.65 | -0.48 | 0.07 | 0.79 | -0.1 | -1.42 | -0.5675 | 0.0225 | 0.245 | 0.015773 | 0.92213 | 0.593729 | | 1 | 0.996755 | 0.929843 |
| Ptpre | A0A1B0GR | 0.81 | 0.92 | 0.67 | 0.81 | 1.03 | 0.54 | 1.32 | 1.18 | 0.78 | 0.88 | 1.11 | 0.83 | 0.11 | 1.16 | 0.95 | 1.6 | 0.215 | 0.0975 | 0.1525 | 0.314162 | 0.56728 | 0.638218 | | 1 | 0.996511 | 0.942804 |
| Ptprf | F6Y3V0 | 0.13 | 0.42 | 0.31 | 0.4 | 0.38 | 0.46 | -0.75 | 0.28 | 0.38 | -0.46 | 0.61 | 0.52 | 1.1 | 0.23 | 0.91 | 0.54 | -0.2225 | -0.0525 | 0.38 | 0.452242 | 0.84577 | 0.143209 | | 1 | 0.996511 | 0.742711 |
| Ptprf | A0A1Y7VN | 1.7 | 1.23 | 1.03 | 1.18 | 1.07 | 1.29 | 2.32 | 1.95 | 1.24 | 3.35 | 0.61 | 1.33 | 0.82 | 1.11 | 1.84 | 0.42 | 0.3725 | 0.3475 | -0.2375 | 0.259291 | 0.541801 | 0.484384 | | 1 | 0.996511 | 0.894266 |
| Ptprf | A2A8L5 | 0.45 | 0.65 | 0.69 | 0.93 | 0.8 | 0.37 | 1.11 | 0.8 | 0.57 | 0.59 | 0.74 | 0.86 | 0.28 | 0.68 | 0.89 | 0.91 | 0.09 | 0.01 | 0.01 | 0.668915 | 0.956397 | 0.96442 | | 1 | 0.996755 | 0.9927 |
| Ptprg | Q05909 | 0.59 | 0.64 | 0.87 | 1.06 | 0.3 | 1.02 | 0.77 | 1.05 | 1.16 | 0.91 | 0.97 | 0.68 | 0.99 | -0.25 | 0.68 | 0.54 | -0.005 | 0.14 | -0.3 | 0.982314 | 0.485835 | 0.329842 | | 1 | 0.996511 | 0.845407 |
| Ptprj | A2AWF8 | 0.56 | 0.85 | 0.84 | 0.68 | 1.63 | 0.44 | 0.88 | 0.78 | 0.03 | -0.15 | 0.95 | 0.61 | -0.04 | 0.33 | 0.64 | 0.61 | 0.2 | -0.3725 | -0.3475 | 0.461252 | 0.202267 | 0.143976 | | 1 | 0.996511 | 0.743032 |
| Ptprk | A0A1W2P7 | 0.11 | 0.27 | 0.47 | 0.15 | 0.53 | 0.44 | 0.5 | 0.46 | 0.34 | 0.47 | 0.69 | 0.33 | 0.55 | 0.32 | 0.61 | 0.53 | 0.2325 | 0.2075 | 0.2525 | 0.156076 | 0.270916 | 0.196896 | | 1 | 0.996511 | 0.78004 |
| Ptprm | Q68FM4 | 0.97 | 1.75 | 1.1 | 1.24 | -1.46 | -0.38 | 0.96 | -0.81 | -0.4 | 1.04 | -0.73 | 1.28 | 0.73 | 1.25 | 1.41 | 1.22 | -1.6875 | -0.9675 | -0.1125 | 0.007929 | 0.078404 | 0.662012 | | 1 | 0.996511 | 0.949645 |
| Ptprn | A6MDD2 | 0.25 | 0.15 | -0.33 | -0.05 | -0.18 | -0.21 | 0.07 | -0.08 | 0.55 | 0.73 | -0.05 | 0.26 | 0.54 | 0.37 | 0.59 | 0.1 | -0.105 | 0.3675 | 0.395 | 0.573109 | 0.152918 | 0.099984 | | 1 | 0.996511 | 0.698156 |
| Ptprn2 | P80560 | 0.58 | 0.69 | 0.45 | 0.6 | 0.64 | 0.42 | 0.53 | 0.71 | 1.08 | 0.95 | 0.55 | 0.72 | 0.94 | 1.02 | 0.96 | 0.92 | -0.005 | 0.245 | 0.38 | 0.973807 | 0.213394 | 0.047966 | | 1 | 0.996511 | 0.658454 |
| Ptpro | E9Q4I1 | 0.72 | 0.79 | 0.97 | 1.26 | 0.96 | 0.67 | 1.09 | 0.94 | 0.85 | 0.33 | 0.9 | 0.57 | -0.27 | 0.75 | 1.32 | 1.36 | -0.02 | -0.2725 | -0.145 | 0.915538 | 0.231638 | 0.70864 | | 1 | 0.996511 | 0.957656 |
| Ptprr | Q62132 | 1.19 | 0.98 | 0.53 | 0.68 | 1.18 | 1.32 | 0.97 | 0.47 | 1.05 | 0.5 | 0.88 | 1.29 | 0.5 | 1.45 | 0.93 | 0.62 | 0.14 | 0.085 | 0.03 | 0.57891 | 0.731605 | 0.914442 | | 1 | 0.996511 | 0.986458 |
| Ptprs | B0V2N1-2 | 0.57 | 0.78 | 0.69 | 0.89 | 0.95 | 0.65 | 0.89 | 0.84 | 0.4 | 0.5 | 0.89 | 0.79 | 0.14 | 0.54 | 0.59 | 0.64 | 0.1 | -0.0875 | -0.255 | 0.532304 | 0.648626 | 0.223615 | | 1 | 0.996511 | 0.787447 |
| Ptprt | Q99M80-3 | 0.89 | 0.83 | 0.68 | 0.9 | 0.75 | 1.07 | 0.97 | 0.97 | 1.11 | 0.56 | 0.85 | 1.32 | 0.76 | 1.28 | 0.98 | 0.78 | 0.115 | 0.135 | 0.125 | 0.462868 | 0.531299 | 0.534365 | | 1 | 0.996511 | 0.912193 |
| Ptpru | B1AUH1 | -0.04 | -0.11 | 0.96 | 0.9 | 0 | 0.26 | 1.03 | 1.13 | -0.1 | 0.78 | 0.74 | 0.03 | -0.55 | 0.13 | 0.9 | 0.41 | 0.1775 | -0.065 | -0.205 | 0.644857 | 0.857304 | 0.614265 | | 1 | 0.996511 | 0.935876 |
| Ptprz1 | B9EKR1 | -1.97 | -2.36 | -2.22 | -2.04 | -1.92 | -2.28 | -1.98 | -2.23 | -2.06 | -1.66 | -2.1 | -1.88 | -1.94 | -0.88 | -1.75 | -2.19 | 0.045 | 0.2225 | 0.4575 | 0.796966 | 0.261698 | 0.16309 | | 1 | 0.996511 | 0.758224 |
| Ptrh1 | Q8BW00 | -0.01 | 0.14 | -0.12 | -0.43 | 0.08 | -0.26 | -0.17 | -0.98 | -0.21 | -1.16 | -0.38 | -0.14 | -0.18 | 1.3 | -0.84 | -0.6 | -0.2275 | -0.3675 | 0.025 | 0.400775 | 0.207465 | 0.957119 | | 1 | 0.996511 | 0.992636 |
| Ptrh2 | Q8R2Y8 | 0.09 | 0.4 | 0.39 | 0.47 | 0.49 | 0.34 | 0.79 | 0.25 | -0.03 | 0.46 | 0.19 | -0.03 | -0.32 | 0.53 | 0.67 | 0.76 | 0.13 | -0.19 | 0.0725 | 0.491503 | 0.345996 | 0.79711 | | 1 | 0.996511 | 0.969727 |
| Ptrhd1 | D3Z4S3 | -0.42 | -0.45 | -0.47 | -0.56 | -0.72 | -0.17 | -0.64 | -0.69 | -0.57 | -0.39 | -0.65 | -0.51 | -0.35 | -0.23 | -0.86 | -0.67 | -0.08 | -0.055 | -0.0525 | 0.656343 | 0.730471 | 0.800736 | | 1 | 0.996511 | 0.970599 |
| Pts | Q9R1Z7 | 0.73 | 1.29 | 0.55 | -0.06 | 0.9 | 0.77 | 0.51 | 0.78 | 0.54 | 0.14 | 0.77 | 1.05 | 0.82 | 0.01 | 0.81 | 1.29 | 0.1125 | -0.0025 | 0.105 | 0.699348 | 0.994034 | 0.780068 | | 1 | 0.999285 | 0.967548 |
| Pttg1ip | Q8R143 | 3.05 | 1.93 | -0.16 | -0.76 | 0.97 | 2.35 | -0.51 | -0.44 | 3.08 | -0.42 | -0.49 | 3.49 | 2.32 | 3.1 | 0.06 | -0.52 | -0.4225 | 0.4 | 0.225 | 0.675233 | 0.750355 | 0.839624 | | 1 | 0.996511 | 0.976219 |
| Puf60 | Q3UEB3-2 | -1.14 | -1.84 | -1.68 | -1.62 | -1.37 | -1.71 | -0.91 | -1.3 | -1.74 | -0.66 | -1.54 | -1.58 | -1.72 | -1.03 | -1.05 | -1.31 | 0.2475 | 0.19 | 0.2925 | 0.315406 | 0.523551 | 0.265523 | | 1 | 0.996511 | 0.817264 |
| Pum1 | Q80U78-3 | -0.91 | -0.72 | -0.74 | -0.68 | -0.41 | -1.1 | -0.6 | -1.06 | -1.28 | -0.47 | -0.72 | -0.93 | -1.21 | -0.68 | -0.49 | -0.57 | -0.03 | -0.0875 | 0.025 | 0.884908 | 0.688619 | 0.90971 | | 1 | 0.996511 | 0.986458 |
| Pum1 | Q80U78-2 | 0.51 | 0.75 | -2.73 | -1.59 | 1 | 0.11 | -2.59 | -0.69 | 0.67 | -2.07 | -1.39 | 0.92 | -0.1 | 0.32 | -2.26 | -1.23 | 0.2225 | 0.2975 | -0.0525 | 0.827185 | 0.768203 | 0.954252 | | 1 | 0.996511 | 0.992476 |
| Pum2 | Q3UR91 | -1.66 | -2.31 | -1.89 | -1.46 | -1.47 | -2.25 | -1.58 | -2.31 | -1.54 | -1.03 | -1.62 | -1.84 | -1.12 | -1.37 | -1.33 | -1.18 | -0.0725 | 0.3225 | 0.58 | 0.80048 | 0.247151 | 0.033041 | | 1 | 0.996511 | 0.644622 |
| Pum3 | Q8BKS9 | 1.26 | 1.3 | 0.9 | 0.76 | 1.02 | 1.53 | 0.44 | 0.77 | 0.18 | 0.84 | 0.88 | 1.08 | 0.01 | 1.06 | 1.15 | 0.73 | -0.115 | -0.31 | -0.3175 | 0.672863 | 0.247778 | 0.310313 | | 1 | 0.996511 | 0.834523 |
| Pura | P42669 | -0.91 | -0.75 | -1.04 | -1.05 | -0.62 | -1.17 | -1.08 | -1.09 | -1.07 | -0.98 | -1.31 | -0.82 | -0.94 | -0.5 | -0.76 | -0.97 | -0.0525 | -0.1075 | 0.145 | 0.776972 | 0.565558 | 0.470514 | | 1 | 0.996511 | 0.891144 |
| Purb | O35295 | -1.38 | -2.07 | -1.25 | -0.06 | -1.68 | -1.34 | -0.97 | -0.18 | -1.35 | -0.69 | -1.14 | -1.18 | -1.21 | -1.41 | -0.95 | -1.32 | 0.1475 | 0.1 | -0.0325 | 0.762566 | 0.811221 | 0.937002 | | 1 | 0.996511 | 0.989767 |
| Purg | Q8R4E6 | -1.65 | -1.84 | -1.24 | -1.58 | -0.83 | -1.83 | -1.24 | -1.62 | -1.58 | -0.7 | -1.58 | -1.37 | -1.05 | -0.9 | -0.91 | -1.05 | 0.1975 | 0.27 | 0.6 | 0.457982 | 0.317578 | 0.014039 | | 1 | 0.996511 | 0.589387 |
| Pus7 | Q91VU7-2 | -2.41 | -2.2 | -2.71 | -2.91 | -1.96 | -2.88 | -2.63 | -3.3 | -1.42 | -1.81 | -3.29 | -2.08 | -1.19 | -1.37 | -3.74 | -2.94 | -0.135 | 0.4075 | 0.2475 | 0.670244 | 0.336819 | 0.673816 | | 1 | 0.996511 | 0.952944 |
| Pusl1 | A2ADA5 | 0.35 | 0.85 | 1.02 | 1.14 | 0.82 | 0.41 | 0.79 | 0.88 | 0.72 | 1.44 | 1.16 | 0.25 | 0.48 | 0.89 | 0.39 | 0.33 | -0.115 | 0.0525 | -0.3175 | 0.612268 | 0.867527 | 0.223764 | | 1 | 0.996511 | 0.787447 |
| Pvalb | P32848 | 2.97 | -0.02 | -0.37 | -1.67 | -1.29 | 3.47 | -0.64 | -1.63 | -1.38 | -2.53 | -0.54 | 2.08 | -0.52 | -0.56 | -2.33 | -1.62 | -0.25 | -0.82 | -1.485 | 0.855433 | 0.514032 | 0.148907 | | 1 | 0.996511 | 0.748146 |
| Pvr | Q8K094 | -0.56 | -0.05 | 1.15 | 0.54 | -0.87 | -0.14 | -0.01 | 0.82 | 0.42 | 0.59 | 0.56 | -0.32 | 1.28 | -0.61 | 0.32 | 0.16 | -0.32 | 0.0425 | 0.0175 | 0.501454 | 0.9168 | 0.972032 | | 1 | 0.996755 | 0.992865 |
| Pwp1 | Q99LL5 | -0.08 | -0.17 | -0.23 | -0.37 | -0.34 | -0.12 | -0.74 | -0.48 | 0.24 | -0.15 | -0.62 | -0.26 | 0.09 | 0.41 | -0.41 | -0.46 | -0.2075 | 0.015 | 0.12 | 0.280073 | 0.946308 | 0.63416 | | 1 | 0.996755 | 0.941132 |
| Pwp2 | Q8BU03 | 0.58 | 0.03 | 1.7 | 1.5 | -2.41 | 0.69 | 0.34 | 1.37 | 3.87 | 0.97 | 1.46 | -0.28 | 4.59 | -0.58 | 1.98 | 1.84 | -0.955 | 0.5525 | 1.005 | 0.267262 | 0.525709 | 0.332688 | | 1 | 0.996511 | 0.845601 |
| Pwwp3a | Q6DID5 | -2.69 | -4.64 | -2.45 | -2.3 | -3.73 | -3.3 | -1.45 | -2.38 | -3.14 | -0.81 | -1.6 | -3.48 | -1.07 | -2.42 | -3.06 | -2.18 | 0.305 | 0.7625 | 0.8375 | 0.653514 | 0.328413 | 0.204007 | | 1 | 0.996511 | 0.782817 |
| Pxdn | Q3UQ28 | 0.21 | 0.33 | -0.41 | -0.66 | 0.04 | -0.12 | 0.16 | -0.36 | -0.69 | -0.49 | -0.26 | 0.59 | -0.06 | -0.47 | -0.13 | -0.21 | 0.0625 | -0.08 | -0.085 | 0.817098 | 0.824177 | 0.759467 | | 1 | 0.996511 | 0.963123 |
| Pxk | A0A2C9F2E | -0.58 | -1.44 | -0.64 | -0.36 | -0.95 | -0.19 | -0.18 | -0.28 | -1.12 | -0.02 | -0.35 | 0.9 | -1.91 | -1.06 | -0.96 | -0.76 | 0.355 | 0.6075 | -0.4175 | 0.253277 | 0.201931 | 0.246614 | | 1 | 0.996511 | 0.802442 |
| Pxmp2 | D6REH1 | -0.16 | 0.6 | 0.02 | 0.02 | 0.78 | 0.35 | 2.19 | 0.18 | 0.44 | 0.73 | 0.21 | -0.35 | 0.6 | 0.89 | -0.06 | 0.38 | 0.755 | 0.1375 | 0.3325 | 0.122174 | 0.638586 | 0.254637 | | 1 | 0.996511 | 0.809524 |
| Pxmp4 | Q9JJW0 | -1.69 | -1.62 | -0.45 | -1.31 | -1.06 | -0.43 | 0.31 | -1.13 | -0.36 | -0.89 | -2.56 | -1.04 | -0.11 | -0.32 | -1.6 | 0.16 | 0.69 | 0.055 | 0.8 | 0.12246 | 0.914349 | 0.106661 | | 1 | 0.996755 | 0.703417 |
| Pxn | F8VQ28 | -1.43 | -1.48 | -1.43 | -1.82 | -1.75 | -1.35 | -2 | -1.91 | -1.64 | -0.6 | -1.45 | -1.56 | -0.33 | -1.93 | -1.48 | -2.11 | -0.2125 | 0.2275 | 0.0775 | 0.311268 | 0.414234 | 0.845183 | | 1 | 0.996511 | 0.976225 |
| Pycard | Q9EPB4 | -0.96 | -0.96 | -1.63 | -1.03 | -1.62 | -1.01 | -1.74 | -1.24 | -0.76 | -0.99 | -1.28 | -1.05 | -1.5 | -0.83 | -2.43 | -1.44 | -0.2575 | 0.125 | -0.405 | 0.313262 | 0.586326 | 0.281081 | | 1 | 0.996511 | 0.822084 |
| Pycr1 | Q922W5 | 0.18 | 0.28 | 1.45 | 0.81 | -1.25 | 0 | 0.83 | 0.7 | 2.38 | 1.61 | 0.91 | -0.35 | 1.41 | 0.32 | 0.48 | 0.07 | -0.61 | 0.4575 | -0.11 | 0.255004 | 0.449998 | 0.782753 | | 1 | 0.996511 | 0.967642 |
| Pycr2 | Q922Q4 | -0.12 | -0.15 | 0.29 | 0.53 | -0.33 | -0.06 | 0.33 | 0.58 | 0.05 | 0.52 | 0.67 | -0.1 | 0.36 | -0.13 | 0.43 | 0.09 | -0.0075 | 0.1475 | 0.05 | 0.977601 | 0.580716 | 0.838516 | | 1 | 0.996511 | 0.976145 |
| Pycr3 | Q9DCC4 | -0.69 | -1.3 | -0.58 | -0.36 | -1.3 | -1.08 | -1.08 | -0.85 | -0.27 | 0.06 | -0.85 | -1.08 | 0.24 | -0.81 | -0.47 | -0.92 | -0.345 | 0.1975 | 0.2425 | 0.171748 | 0.55151 | 0.472749 | | 1 | 0.996511 | 0.891144 |
| Pygb | Q8CI94 | 0.64 | 0.23 | 0.4 | 0.46 | 0.48 | 0.72 | 0.47 | 0.77 | 0.21 | -0.33 | 0.28 | 0.28 | -0.18 | 0.32 | 0.07 | 0.3 | 0.1775 | -0.3225 | -0.305 | 0.309675 | 0.154191 | 0.164074 | | 1 | 0.996511 | 0.758352 |
| Pygl | Q9ET01 | 0.71 | 0.25 | -0.27 | 0.1 | 0.45 | 0.62 | 0.45 | 0.16 | 0.08 | 0.23 | -0.19 | 0.31 | -0.35 | 0.65 | -0.08 | -0.03 | 0.2225 | -0.09 | -0.15 | 0.365804 | 0.722666 | 0.625367 | | 1 | 0.996511 | 0.939288 |
| Pygm | Q9WUB3 | 0.92 | 0.57 | 0.89 | 1.32 | 0.75 | 0.43 | 0.68 | 0.84 | -0.19 | 0.27 | 0.38 | -0.24 | -0.71 | 0.03 | 0.15 | 0.67 | -0.25 | -0.87 | -0.89 | 0.246498 | 0.006187 | 0.0224 | | 1 | 0.996511 | 0.621926 |
| Pygo2 | Q3V113 | -2.63 | -2.56 | -1.19 | -0.88 | -1.36 | -2.34 | -1.16 | -0.75 | -2.26 | -0.23 | -1.51 | -2.93 | -1.39 | -1.65 | -0.58 | -0.91 | 0.4125 | 0.0825 | 0.6825 | 0.43703 | 0.902027 | 0.180829 | | 1 | 0.996609 | 0.775542 |
| Pym1 | Q8CHP5-2 | -1.72 | -1.67 | -1.66 | -1.89 | -1.9 | -1.48 | -1.6 | -1.7 | -2.25 | -1.74 | -1.26 | -1.77 | -1.64 | -0.84 | -1.55 | -2.01 | 0.065 | -0.02 | 0.225 | 0.691234 | 0.93288 | 0.419791 | | 1 | 0.996755 | 0.87558 |
| Pyroxd1 | Q3TMV7 | -0.96 | -1.31 | -0.42 | -0.21 | 0.24 | -1.35 | -1.75 | -0.5 | -0.95 | 1.04 | -0.84 | -2.06 | -2.11 | -1.8 | -0.78 | -1.18 | -0.115 | 0.0225 | -0.7425 | 0.808046 | 0.971513 | 0.077989 | | 1 | 0.997515 | 0.676942 |
| Pyroxd2 | A0A494BA! | -1.69 | -3.19 | -0.76 | -1.64 | -3.3 | -1.02 | 0.03 | -1.15 | 3.83 | -1.08 | -1.15 | -1.54 | 6.61 | -2.26 | -0.61 | -1.29 | 0.46 | 1.835 | 2.4325 | 0.55807 | 0.160947 | 0.211313 | | 1 | 0.996511 | 0.786269 |
| Pyurf | Q9D1C3 | -1.52 | -0.13 | 0.37 | 0.47 | -1.13 | 0.28 | -0.19 | 0.34 | 2.14 | 1.45 | 0.3 | -1.62 | 2.43 | -1.32 | 0.45 | 0.62 | 0.0275 | 0.77 | 0.7475 | 0.958141 | 0.376295 | 0.365029 | | 1 | 0.996511 | 0.858475 |
| Pzp | Q61838 | -3.38 | -3.93 | -4.07 | -4.66 | -3.22 | -3.91 | -3.78 | -4.24 | -4.02 | -1.32 | -3.84 | -3.2 | -3.65 | -2.48 | -4.3 | -4.48 | 0.2225 | 0.915 | 0.2825 | 0.504311 | 0.162122 | 0.567224 | | 1 | 0.996511 | 0.921806 |
| Q0VG49 | Q0VG49 | 2.11 | 1.95 | -1.51 | -0.39 | 2.99 | 2.18 | -0.75 | -0.61 | 0.86 | -2.27 | -1.23 | 1.77 | 0.48 | 1.53 | -2.07 | -1.04 | 0.4125 | -0.7575 | -0.815 | 0.725258 | 0.516474 | 0.453026 | | 1 | 0.996511 | 0.885993 |
| Q3TQI7 | Q3TQI7 | -2.62 | -2.85 | -4.52 | -4.25 | -2.56 | -2.81 | -4.1 | -4.88 | -2.61 | -1.51 | -4.15 | -2.76 | -2.02 | -1.89 | -4.41 | -4.34 | -0.0275 | 0.8025 | 0.395 | 0.966822 | 0.244804 | 0.60904 | | 1 | 0.996511 | 0.934202 |
| Q3TTL0 | Q3TTL0 | -2.39 | -1.69 | -1.36 | -0.9 | -1.44 | -1.73 | -0.47 | 0.18 | -0.73 | -0.25 | -1.57 | -0.71 | -1.29 | -1.61 | -1.4 | -0.64 | 0.72 | 0.77 | 0.35 | 0.174012 | 0.080914 | 0.356047 | | 1 | 0.996511 | 0.8553 |
| Q5EBG8 | Q5EBG8 | -1.76 | -1.8 | -1.92 | -1.38 | -0.97 | -2.28 | -1.78 | -2.01 | -1.97 | -1.24 | -1.69 | -1.42 | -1.82 | 0.75 | -2.26 | -2.14 | -0.045 | 0.135 | 0.3475 | 0.881752 | 0.560937 | 0.599625 | | 1 | 0.996511 | 0.930673 |
| Q80X32 | Q80X32 | 0.96 | 0.14 | -0.15 | -0.2 | 0 | 0.59 | -0.2 | -0.29 | -1.31 | 1.01 | -0.08 | 0.43 | -0.54 | -0.42 | 0.88 | -0.03 | -0.1625 | -0.175 | -0.215 | 0.619389 | 0.737458 | 0.596644 | | 1 | 0.996511 | 0.929911 |
| Q8BGK9 | Q8BGK9 | 0.39 | 0.13 | -0.22 | 1.55 | -0.15 | -0.65 | -0.18 | 1.76 | 0.88 | 1.18 | 0.1 | 0.04 | 0.26 | 0.66 | -0.34 | -0.44 | -0.2675 | 0.0875 | -0.4275 | 0.657207 | 0.845498 | 0.343819 | | 1 | 0.996511 | 0.848473 |
| Q8BH50 | Q8BH50 | 0.35 | 0.69 | 0.74 | 0.25 | 0.41 | -0.09 | -0.07 | 0.35 | 0.72 | -0.21 | 0.24 | 0.57 | 0.1 | 0.84 | -0.79 | -0.25 | -0.3575 | -0.1775 | -0.5325 | 0.115008 | 0.499188 | 0.162443 | | 1 | 0.996511 | 0.757516 |
| Q8BN57 | Q8BN57 | 0.72 | 0.9 | 0.56 | 0.76 | 0.08 | 1.23 | -0.26 | 0.08 | 0.99 | 1.2 | 0.79 | 0.98 | 1.42 | -0.11 | 0.63 | 0.56 | -0.4525 | 0.255 | -0.11 | 0.187976 | 0.176266 | 0.736098 | | 1 | 0.996511 | 0.958635 |
| Q8BP99 | Q8BP99 | 0.52 | 0.19 | 2.15 | 0.26 | 0.7 | 0.28 | 1.21 | 1.01 | 0.03 | 1.05 | 1.07 | 0.17 | 0.87 | 0.03 | 1.87 | 1.07 | 0.02 | -0.2 | 0.18 | 0.96588 | 0.691338 | 0.74447 | | 1 | 0.996511 | 0.959432 |
| Q8C3W1 | Q8C3W1 | 0.09 | 0.65 | 0.48 | 0.65 | 0.56 | 0.13 | 0.56 | 0.31 | 0.3 | 0.51 | 0.62 | 0.36 | 0.14 | 1.02 | 0.53 | 0.74 | -0.0775 | -0.02 | 0.14 | 0.700945 | 0.920329 | 0.589925 | | 1 | 0.996755 | 0.929142 |
| Q8C5K5 | Q8C5K5 | -0.29 | -0.32 | -0.13 | -0.19 | 0.35 | -0.31 | -0.13 | 0.08 | -1.43 | 0.02 | 0.5 | -0.12 | -1.37 | -0.78 | 0.1 | -0.16 | 0.23 | -0.025 | -0.32 | 0.242325 | 0.949773 | 0.351245 | | 1 | 0.996755 | 0.853984 |
| Q91WE4 | Q91WE4 | -0.82 | -0.64 | -0.36 | -0.69 | -0.6 | -0.48 | -1.16 | -0.43 | 2.94 | -0.15 | -0.22 | -0.85 | 5.04 | -0.36 | 2.18 | -0.61 | -0.04 | 1.0575 | 2.19 | 0.854571 | 0.196412 | 0.090645 | | 1 | 0.996511 | 0.693068 |
| Q922M7 | Q922M7 | -4.36 | -5.5 | -2.73 | -2.24 | -4.59 | -6.48 | -2.87 | -2.51 | -5.86 | -1.76 | -2.92 | -4.42 | -5.39 | -4.66 | -1.28 | -2.6 | -0.405 | -0.0325 | 0.225 | 0.702677 | 0.975187 | 0.834298 | | 1 | 0.997515 | 0.975471 |
| Q99M08 | Q99M08 | -1.12 | 0.07 | 1.74 | 1.29 | -0.64 | -0.16 | 1.68 | 1.64 | 0.2 | 3.03 | 2.11 | -0.78 | 0.93 | -0.47 | 1.14 | 1.17 | 0.135 | 0.645 | 0.1975 | 0.865187 | 0.512795 | 0.773146 | | 1 | 0.996511 | 0.966062 |
| Q9CPS8 | Q9CPS8 | -0.41 | 0.96 | 0.66 | 0.13 | 1.13 | -0.17 | 0.4 | 0.02 | -0.75 | 0.02 | 0.1 | -1.15 | -0.68 | 1.37 | -0.07 | 0.33 | 0.01 | -0.78 | -0.0975 | 0.979714 | 0.083291 | 0.842762 | | 1 | 0.996511 | 0.976225 |
| Q9CQ90 | Q9CQ90 | -4.62 | -5.18 | -5.01 | -5.3 | -5.72 | -5.03 | -3.52 | -4.57 | -2.57 | 0.52 | -4.25 | -6.5 | -1.67 | -1.42 | -2.27 | -5.26 | 0.3175 | 1.8275 | 2.3725 | 0.488113 | 0.192886 | 0.015988 | | 1 | 0.996511 | 0.589387 |
| Q9CRC3 | Q9CRC3 | -1.45 | -0.9 | -1.39 | -1.59 | -1.49 | -1.15 | -1.3 | -1.61 | -0.8 | -2.48 | -1.32 | -1.47 | -1.6 | -1.63 | -2.18 | -2.01 | -0.055 | -0.185 | -0.5225 | 0.793203 | 0.619763 | 0.057158 | | 1 | 0.996511 | 0.661738 |
| Q9CYS6 | Q9CYS6 | 0.2 | 0.9 | 0.64 | 0.41 | 0.85 | -0.1 | 0.83 | -0.08 | -0.15 | -0.51 | 0.44 | 0.24 | -0.65 | 0.33 | -0.36 | 0.24 | -0.1625 | -0.5325 | -0.6475 | 0.59645 | 0.078651 | 0.052382 | | 1 | 0.996511 | 0.659923 |
| Q9D7E4 | Q9D7E4 | 0.01 | 0.48 | 0.52 | 0.07 | 0.26 | 0.53 | 0.95 | 0.19 | 0.59 | -0.94 | 0.58 | 0.2 | 0.23 | 0.73 | 0.3 | 0.54 | 0.2125 | -0.1625 | 0.18 | 0.378079 | 0.664103 | 0.430758 | | 1 | 0.996511 | 0.879165 |
| Q9D9H8 | Q9D9H8 | -1.42 | -1.32 | -0.91 | -0.84 | -1.09 | -1.16 | -0.81 | -0.95 | -1.03 | -0.98 | -0.93 | -1.54 | -1.01 | -1.44 | -1.15 | -1.03 | 0.12 | 0.0025 | -0.035 | 0.549474 | 0.991439 | 0.875811 | | 1 | 0.999285 | 0.979626 |
| Qars1 | Q8BML9 | -1.1 | -1.03 | -1.67 | -1.8 | -0.76 | -1.18 | -1.39 | -1.73 | -1.22 | -1.27 | -1.78 | -1.05 | -0.87 | -0.66 | -1.69 | -1.66 | 0.135 | 0.07 | 0.18 | 0.636443 | 0.793834 | 0.591775 | | 1 | 0.996511 | 0.929843 |
| Qdpr | Q8BVI4 | 0.37 | 0.39 | 0.38 | 0.36 | 0.4 | -0.04 | 0.05 | 0.15 | -0.31 | 0.33 | 0.02 | 0.01 | -0.77 | 0.4 | 0.06 | 0.22 | -0.235 | -0.3625 | -0.3975 | 0.164686 | 0.082204 | 0.17927 | | 1 | 0.996511 | 0.775542 |
| Qki | Q9QYS9-8 | -0.36 | -0.42 | -0.22 | -0.25 | -0.25 | -0.43 | -0.05 | -0.23 | -0.18 | -0.15 | -0.08 | -0.46 | -0.39 | -0.15 | -0.49 | -0.26 | 0.0725 | 0.095 | -0.01 | 0.64577 | 0.583188 | 0.955566 | | 1 | 0.996511 | 0.992482 |
| Qpct | A0A3B2W7 | -0.35 | -0.52 | -0.34 | 1.04 | 0.13 | -1.24 | 0.53 | 0.54 | -0.58 | 2.19 | -1.08 | -0.68 | -0.39 | -0.4 | 0.49 | 0.15 | 0.0325 | 0.005 | 0.005 | 0.949338 | 0.994712 | 0.990186 | | 1 | 0.999285 | 0.996154 |
| Qpctl | Q8BH73 | -0.96 | -0.66 | -0.04 | -1.32 | -0.24 | -0.22 | -1.35 | -0.6 | -0.62 | -0.24 | -1.25 | -0.9 | -0.75 | 0.34 | -0.33 | -0.74 | 0.1425 | -0.0075 | 0.375 | 0.694823 | 0.982404 | 0.320715 | | 1 | 0.998773 | 0.839916 |
| Qrich1 | G3X8R5 | -4.28 | -3.94 | -4.18 | -4.07 | -3.24 | -4.38 | -3.65 | -4.97 | -4.28 | -3.3 | -4.21 | -4.05 | -4.24 | -3.17 | -4.78 | -4.59 | 0.0575 | 0.1575 | -0.0775 | 0.877236 | 0.544609 | 0.830274 | | 1 | 0.996511 | 0.973939 |
| Qrsl1 | Q9CZN8 | -0.22 | 0 | 0.26 | 0.46 | 0.02 | 0.32 | 0.34 | 0.58 | -0.06 | 0.72 | 0.56 | 0.06 | 0.16 | -0.02 | 0.27 | 0.29 | 0.19 | 0.195 | 0.05 | 0.38564 | 0.460613 | 0.818277 | | 1 | 0.996511 | 0.972768 |
| Qsox1 | Q8BND5-3 | -3.53 | -4.43 | -2.66 | -3.89 | -3.29 | -4.14 | -3.59 | -2.89 | -2.62 | -2.67 | -4.68 | -3.17 | -3.64 | -3.51 | -4.73 | -2.91 | 0.15 | 0.3425 | -0.07 | 0.725769 | 0.545289 | 0.887616 | | 1 | 0.996511 | 0.980447 |
| Qsox2 | Q3TMX7 | -1.09 | -1.41 | -2.82 | -3.07 | -1.01 | -1.12 | -1.83 | -2.94 | -1.89 | -1.97 | -2.99 | -1 | -1.86 | -1.9 | -2.86 | -2.95 | 0.3725 | 0.135 | -0.295 | 0.543745 | 0.818842 | 0.584644 | | 1 | 0.996511 | 0.927734 |
| Qtrt1 | Q9JMA2 | 0.26 | -0.01 | 1.07 | 0.84 | 0.39 | 0.73 | 0.41 | 0.56 | 0.65 | 1.56 | 0.79 | -0.2 | 0.07 | -0.03 | 0.24 | 0.21 | -0.0175 | 0.16 | -0.4175 | 0.948011 | 0.702244 | 0.160011 | | 1 | 0.996511 | 0.755194 |
| Qtrt2 | A0A5F8MP | -3.2 | -0.05 | -0.34 | -1.04 | -1.01 | -3.4 | -0.97 | -1.48 | -0.89 | -0.7 | -1.17 | 0.83 | -0.37 | 0.2 | -1.19 | -1.24 | -0.5575 | 0.675 | 0.5075 | 0.504251 | 0.386742 | 0.485546 | | 1 | 0.996511 | 0.895078 |
| R3hcc1 | E9PUE1 | -1.01 | -0.74 | 0.4 | -0.07 | -1.72 | -1.02 | 0.33 | 0.34 | -0.42 | 0.63 | 0.18 | -0.77 | 0.34 | -0.21 | -0.28 | 0.15 | -0.1625 | 0.26 | 0.355 | 0.769454 | 0.543386 | 0.324897 | | 1 | 0.996511 | 0.842881 |
| R3hdm1 | E9Q9Q2 | -1.02 | -0.98 | -0.54 | -0.49 | -0.98 | -0.95 | -0.33 | -0.26 | -0.97 | -0.82 | -0.21 | -0.97 | -1.11 | -0.96 | -0.88 | -0.42 | 0.1275 | 0.015 | -0.085 | 0.614421 | 0.952443 | 0.726477 | | 1 | 0.996755 | 0.958398 |
| R3hdm2 | Q80TM6-2 | -0.01 | 0.14 | 0.26 | 0.22 | -0.33 | -0.13 | -0.43 | -0.04 | 0.68 | 0.68 | 0.42 | 0.36 | 1.77 | 0.59 | 0.09 | -0.4 | -0.385 | 0.3825 | 0.36 | 0.042009 | 0.053545 | 0.426709 | | 1 | 0.996511 | 0.879074 |
| Rab10 | P61027 | 0.42 | 0.91 | 1.25 | 0.83 | 1.1 | 0.67 | 1.11 | 1.02 | -1.25 | -1.03 | 1.27 | 0.45 | -1.14 | -1.72 | 0.76 | 0.85 | 0.1225 | -0.9925 | -1.165 | 0.584576 | 0.112611 | 0.087478 | | 1 | 0.996511 | 0.691619 |
| Rab11b | P46638 | 0.49 | 0.77 | 0.8 | 0.61 | 1.05 | 0.65 | 0.55 | 0.7 | 0.43 | 0.51 | 0.77 | 0.47 | 0.25 | 0.87 | 0.61 | 0.79 | 0.07 | -0.1225 | -0.0375 | 0.694515 | 0.492714 | 0.859766 | | 1 | 0.996511 | 0.977046 |
| Rab11fip2 | G3XA57-2 | 0.45 | 0.82 | 0.92 | 0.79 | 0.45 | 0.39 | 0.55 | 0.53 | 0.75 | 1.15 | 0.68 | 0.78 | 0.94 | 0.99 | 0.79 | 0.63 | -0.265 | 0.095 | 0.0925 | 0.136353 | 0.63329 | 0.643478 | | 1 | 0.996511 | 0.943933 |
| Rab11fip3 | Q8CHD8-3 | 0.23 | -0.23 | -0.16 | -0.16 | 0.5 | -0.31 | -0.19 | -0.08 | 0.04 | 2.48 | -0.29 | 0.2 | 0.06 | 1.22 | -0.19 | 0.6 | 0.06 | 0.6875 | 0.5025 | 0.792701 | 0.261778 | 0.156395 | | 1 | 0.996511 | 0.751493 |
| Rab11fip4 | Q8BQP8 | 1.3 | 0.89 | 1.48 | 1.46 | 0.93 | 0.95 | 1.44 | 1.2 | 1.12 | 0.94 | 1.18 | 1.05 | 1.7 | 2.18 | 1.68 | 2.12 | -0.1525 | -0.21 | 0.6375 | 0.474878 | 0.303865 | 0.021956 | | 1 | 0.996511 | 0.621926 |
| Rab11fip5 | Q8R361 | 0.97 | 2.35 | 2.03 | 1.42 | 0.99 | 0.87 | 1.93 | 2.01 | 2.24 | 2.43 | 2.19 | 1.17 | 3.5 | 3.2 | 2.19 | 1.05 | -0.2425 | 0.315 | 0.7925 | 0.554555 | 0.440249 | 0.19717 | | 1 | 0.996511 | 0.78004 |
| Rab11fip5 | A0A0N4SW | 0.44 | 0.68 | 0.75 | 0.7 | 0.39 | 0.39 | 0.66 | 0.74 | 0.37 | 0.49 | 0.59 | 0.71 | 0.9 | 0.62 | 0.63 | 0.74 | -0.0975 | -0.1025 | 0.08 | 0.566439 | 0.55871 | 0.662585 | | 1 | 0.996511 | 0.949646 |
| Rab12 | P35283 | 0.5 | 2.27 | 2.61 | 1.5 | -0.2 | -0.77 | -0.03 | -0.18 | -0.88 | -0.38 | -0.1 | -0.87 | -0.84 | -0.59 | -0.9 | -0.64 | -2.015 | -2.2775 | -2.4625 | 0.001902 | 0.001057 | 0.000432 | | 1 | 0.996511 | 0.270112 |

| | | | | | | | | | | | | | | | | | | | | | | | | | | |
|---|---|---|---|---|---|---|---|---|---|---|---|---|---|---|---|---|---|---|---|---|---|---|---|---|---|---|
| Rab13 | Q9DD03 | 2.16 | 1.49 | 1.11 | -0.9 | 1.32 | 2.55 | 1.43 | 0.46 | -0.43 | -3.09 | 0.54 | 2.15 | 1.25 | -2.96 | 0.55 | 1.18 | 0.475 | -1.1725 | -0.96 | 0.50983 | 0.321892 | 0.38083 | 1 | 0.996511 | 0.860984 |
| Rab14 | Q91V41 | -0.34 | -0.07 | -0.39 | -0.23 | 0.33 | -0.25 | 0.06 | -0.23 | -0.25 | -0.81 | -0.21 | -0.08 | -0.31 | -0.32 | -0.36 | -0.21 | 0.235 | -0.08 | -0.0425 | 0.24123 | 0.712014 | 0.809341 | 1 | 0.996511 | 0.972098 |
| Rab15 | Q3TYH2 | -0.27 | 0.41 | 0.48 | 0.34 | 0.83 | 0.34 | 0.72 | 0.4 | 0.65 | 0.04 | 0.64 | -0.03 | 0.53 | 1.31 | 0.25 | 0.57 | 0.3325 | 0.085 | 0.425 | 0.172429 | 0.752568 | 0.17862 | 1 | 0.996511 | 0.77484 |
| Rab18 | P35293 | 0.19 | 0.32 | 1.03 | 0.76 | 0.32 | 0.18 | 0.87 | 0.88 | 0.13 | 3.36 | 0.79 | 0 | 0.61 | 0.33 | 1.03 | 0.74 | -0.0125 | 0.495 | 0.1025 | 0.963324 | 0.50446 | 0.703378 | 1 | 0.996511 | 0.956947 |
| Rab1a | Q5SW88 | 0.83 | 0.88 | 0.77 | 0.57 | 0.89 | 0.83 | 0.76 | 0.56 | 0.45 | 0.02 | 0.8 | 0.86 | 0.4 | 0.35 | 0.34 | 0.6 | -0.0025 | -0.23 | -0.34 | 0.987571 | 0.341001 | 0.087329 | 1 | 0.996511 | 0.691619 |
| Rab1A | P62821 | 0.19 | 0.67 | 0.97 | 0.35 | 1.27 | 0.22 | 0.94 | 0.59 | 0.25 | 1.11 | 0.86 | 0.45 | -0.45 | 1.44 | 0.33 | 0.88 | 0.21 | 0.1225 | 0.005 | 0.470015 | 0.6572 | 0.990476 | 1 | 0.996511 | 0.996154 |
| Rab1b | Q9D1G1 | 0.61 | 0.48 | 0.72 | 0.59 | 0.39 | 0.82 | 0.47 | 0.54 | 1.21 | 0.22 | 0.78 | 0.59 | 1.78 | -0.14 | 0.64 | 0.46 | -0.045 | 0.1 | 0.085 | 0.784063 | 0.677656 | 0.828162 | 1 | 0.996511 | 0.973757 |
| Rab21 | P35282 | -0.09 | -0.85 | -0.45 | -0.91 | -0.62 | 0.78 | -0.21 | -0.57 | 0.75 | -1.63 | -0.54 | -0.26 | 2.06 | -1.19 | -0.3 | -0.41 | 0.42 | 0.155 | 0.615 | 0.263186 | 0.752055 | 0.365116 | 1 | 0.996511 | 0.858475 |
| Rab22a | P35285 | -0.39 | -0.19 | 0.01 | -0.01 | 0.03 | -0.4 | -0.1 | -0.01 | 0.12 | 0.06 | -0.14 | -0.23 | 0.69 | 0.63 | -0.2 | 0.06 | 0.025 | 0.0975 | 0.44 | 0.889432 | 0.601921 | 0.122624 | 1 | 0.996511 | 0.721448 |
| Rab23 | Q9D4I9 | 1.34 | 1.36 | 1.12 | 1.43 | 1.61 | 1.42 | 1.24 | 1.47 | 1.14 | 1.54 | 1.11 | 1.36 | 0.78 | 1.69 | 1.07 | 1.09 | 0.1225 | -0.025 | -0.155 | 0.457174 | 0.891743 | 0.524898 | 1 | 0.996511 | 0.908107 |
| Rab24 | P35290 | -0.39 | -0.38 | -0.45 | -0.26 | -0.23 | -0.04 | -0.31 | -0.47 | -0.28 | -0.46 | -0.41 | -0.25 | 0.2 | 0.4 | -0.08 | -0.2 | 0.1075 | 0.02 | 0.45 | 0.507693 | 0.900322 | 0.051753 | 1 | 0.996511 | 0.658454 |
| Rab26 | Q504M8 | 0.97 | 1.03 | 0.53 | 0.47 | 1.38 | 0.63 | 0.75 | 0.59 | -0.18 | -0.13 | 0.78 | 1.03 | -0.08 | 0.25 | 2.06 | 1.09 | 0.0875 | -0.375 | 0.08 | 0.724424 | 0.285286 | 0.864728 | 1 | 0.996511 | 0.977686 |
| Rab27a | Q9ERI2 | 1 | 1.22 | -0.57 | 0.27 | 1.18 | 0.38 | -0.38 | -0.16 | -0.35 | 1.16 | -0.73 | 0.6 | 0.16 | 0.3 | 0.21 | 0.47 | -0.225 | -0.31 | -0.195 | 0.649582 | 0.574152 | 0.624967 | 1 | 0.996511 | 0.939167 |
| Rab27b | Q99P58 | 0.2 | 0.89 | 0.68 | 1.01 | 0.7 | 0.43 | 0.77 | 0.97 | 0.48 | 0.17 | 0.61 | -0.08 | 0 | 0.27 | 0.28 | 0.32 | 0.0225 | -0.4 | -0.4775 | 0.922004 | 0.146224 | 0.068046 | 1 | 0.996511 | 0.664936 |
| Rab28 | Q99KL7 | -0.61 | -0.65 | -0.98 | -0.83 | -0.29 | -0.83 | -0.81 | -1.13 | -0.12 | 0.45 | -0.66 | -0.38 | 0.2 | 0.46 | -0.56 | -1.51 | 0.0025 | 0.59 | 0.415 | 0.990859 | 0.051955 | 0.346676 | 1 | 0.996511 | 0.851134 |
| Rab29 | E9QLQ7 | 0.07 | 0.95 | -0.17 | 0.29 | 0.09 | -0.33 | 0.95 | -0.79 | -0.56 | -0.56 | -0.22 | -0.24 | -0.09 | 1.1 | 0.53 | -0.25 | -0.305 | -0.68 | 0.0375 | 0.468918 | 0.032742 | 0.921575 | 1 | 0.996511 | 0.987433 |
| Rab2a | P53994 | -0.02 | 0.38 | 2.18 | 1.36 | 0.87 | 0.28 | 2.55 | 2.52 | -0.83 | 0.1 | 2.19 | 0.32 | -0.26 | -0.79 | 2.25 | 2.95 | 0.58 | -0.53 | 0.0625 | 0.410595 | 0.474245 | 0.94672 | 1 | 0.996511 | 0.992376 |
| Rab2b | P59279 | 0.14 | 0.55 | -0.62 | -0.87 | 0.53 | 1.44 | -0.49 | -0.27 | 0.25 | -1.65 | -0.47 | 1.03 | 0.18 | -0.06 | -0.44 | -0.12 | 0.5025 | -0.01 | 0.09 | 0.334205 | 0.986753 | 0.797727 | 1 | 0.999285 | 0.969727 |
| Rab30 | Q923S9 | 0.57 | 0.93 | 1.21 | 1.13 | 0.83 | 0.24 | 0.58 | 0.95 | 0.48 | 2.08 | 1.13 | 0.45 | 0.8 | 1.41 | 1.22 | 1.2 | -0.31 | 0.075 | 0.1975 | 0.202864 | 0.847997 | 0.408565 | 1 | 0.996511 | 0.874356 |
| Rab31 | Q3TXV4 | -0.5 | -0.2 | 0.78 | 0.37 | 0.15 | -0.6 | 0.43 | 0.48 | -0.32 | 1.39 | 0.68 | -0.14 | -0.06 | -0.34 | 1.08 | 1.03 | 0.0025 | 0.29 | 0.315 | 0.994497 | 0.531165 | 0.481844 | 1 | 0.996511 | 0.893925 |
| Rab33a | P97950 | -0.28 | 0.17 | 0.63 | 0.37 | 0.16 | -0.06 | 0.99 | 0.5 | -0.05 | -0.72 | 0.59 | -0.14 | 0.11 | 0.07 | 0.33 | 0.54 | 0.175 | -0.3025 | 0.04 | 0.559762 | 0.368814 | 0.874354 | 1 | 0.996511 | 0.979626 |
| Rab33b | O35963 | 1.04 | 1.13 | 1.02 | 0.66 | 1.48 | 1.6 | 1.04 | 0.93 | -0.07 | -0.49 | 0.9 | 1.44 | 0.43 | 1.1 | 1.26 | 0.94 | 0.3 | -0.5175 | -0.03 | 0.192663 | 0.247745 | 0.902161 | 1 | 0.996511 | 0.98447 |
| Rab34 | Q0PD20 | 0.1 | 0.62 | -0.23 | -0.8 | 0.42 | -0.06 | -0.27 | -0.73 | 0.23 | -0.05 | -0.22 | -0.09 | -0.02 | 0.55 | -0.31 | -0.13 | -0.0825 | 0.045 | 0.1 | 0.821013 | 0.886078 | 0.774673 | 1 | 0.996511 | 0.966468 |
| Rab35 | Q6PHN9 | 1.4 | 0.86 | 1.91 | 1.87 | 1.17 | 1.41 | 1.5 | 1.91 | 0.68 | 1.76 | 1.85 | 1.27 | 0.77 | 0.7 | 1.67 | 1.89 | -0.0125 | -0.12 | -0.2525 | 0.965611 | 0.736437 | 0.513256 | 1 | 0.996511 | 0.904284 |
| Rab36 | Q8CAM5 | -0.11 | -0.14 | 0.43 | 0.41 | -0.53 | 0.14 | 0.82 | 0.12 | 1.51 | 0.34 | 0.22 | 0.23 | 2.23 | -0.2 | 0.22 | 0.19 | -0.01 | 0.4275 | 0.4625 | 0.974414 | 0.235513 | 0.392425 | 1 | 0.996511 | 0.865775 |
| Rab37 | Q8BQX0 | 0.35 | 0.91 | 0.73 | 0.72 | 0.78 | 0.56 | -0.27 | 0.33 | 1.13 | 1 | 0.62 | 0.64 | 1.97 | 1.42 | 0.61 | 0.52 | -0.3275 | 0.17 | 0.4525 | 0.233819 | 0.436531 | 0.229714 | 1 | 0.996511 | 0.787447 |
| Rab37 | Q544E8 | -3.01 | 0.55 | 0.28 | -0.43 | -2.31 | -1.34 | -1.31 | -0.35 | 3.26 | 1.63 | 0.02 | -3.56 | 0.5 | 0.5 | -1.4 | -3.03 | -0.675 | 0.99 | -0.205 | 0.417817 | 0.511996 | 0.84543 | 1 | 0.996511 | 0.976225 |
| Rab39b | Q8BHC1 | 0.25 | 0.54 | 0.99 | 1.22 | 0.77 | 0.26 | 0.93 | 1.15 | 0.74 | 1.14 | 0.93 | 0.5 | 0.79 | 1.06 | 0.94 | 1.25 | 0.0275 | 0.0775 | 0.26 | 0.924247 | 0.776275 | 0.341147 | 1 | 0.996511 | 0.848473 |
| Rab3a | P63011 | 0.04 | 0.77 | 0.64 | 0.53 | 1.02 | 0.67 | 0.74 | 0.56 | -0.52 | -0.94 | 0.67 | 0.31 | -0.37 | -1.18 | 0.11 | 0.39 | 0.2525 | -0.615 | -0.7575 | 0.256408 | 0.139104 | 0.067304 | 1 | 0.996511 | 0.664936 |
| Rab3b | Q9CZT8 | 1.62 | 1.77 | 1.28 | 1.38 | -0.33 | 0.48 | -0.44 | -0.42 | 1.36 | 1.31 | 1.24 | 1.81 | 0.36 | -1.5 | -0.33 | -0.56 | -1.69 | -0.0825 | -2.02 | 0.000129 | 0.69861 | 0.000615 | 0.235326 | 0.996511 | 0.294873 |
| Rab3c | P62823 | 1.25 | 1.03 | 0.98 | 0.67 | 1.33 | 1.95 | 1.69 | 1.13 | 0.41 | -0.88 | 1.2 | 1.51 | 0.13 | 0.21 | 1.06 | 1.15 | 0.5425 | -0.4225 | -0.345 | 0.044327 | 0.414566 | 0.278594 | 1 | 0.996511 | 0.822084 |
| Rab3d | P35276 | 0.52 | 0.81 | 1.18 | 0.86 | 1.47 | 0.82 | 1.26 | 1.03 | 0.54 | -0.61 | 0.95 | 0.43 | -0.19 | 0.91 | 1.61 | 1.1 | 0.3025 | -0.515 | 0.015 | 0.191213 | 0.167803 | 0.96929 | 1 | 0.996511 | 0.992767 |
| Rab3gap1 | Q80UJ7 | 1.19 | 0.62 | -0.11 | 0.57 | 1.43 | 0.79 | 0.79 | 0.41 | 0.84 | 0.77 | 0.1 | 1.21 | 1.06 | 1.84 | 0.34 | 0.24 | 0.2875 | 0.1625 | 0.3025 | 0.394763 | 0.640751 | 0.492067 | 1 | 0.996511 | 0.897531 |
| Rab3gap1 | A0A1D5RL( | 0.01 | -0.26 | 0.57 | 0.58 | 0.02 | 0.08 | 0.9 | 0.74 | -0.31 | 0.81 | 0.57 | 0.13 | -0.34 | 0.16 | 0.72 | 0.79 | 0.21 | 0.075 | 0.1075 | 0.495864 | 0.816803 | 0.751313 | 1 | 0.996511 | 0.961479 |
| Rab3gap2 | A0A0A6YW | 0.73 | 0.17 | -0.06 | -0.26 | 0.35 | 0.71 | 0.73 | -0.13 | 0.18 | -0.25 | 0 | 0.55 | -0.23 | 0.53 | 0 | 0.16 | 0.27 | -0.025 | -0.03 | 0.369421 | 0.929608 | 0.916227 | 1 | 0.996755 | 0.986556 |
| Rab3il1 | D3YZ86 | 0.15 | -0.06 | -0.66 | -0.42 | 0.66 | 0.59 | -0.32 | 0.26 | 0.02 | -0.25 | -0.29 | -0.29 | -0.84 | 0.49 | -0.05 | -0.07 | 0.545 | 0.045 | 0.13 | 0.086359 | 0.844012 | 0.695648 | 1 | 0.996511 | 0.955729 |
| Rab3ip | Q68EF0 | -0.71 | -0.98 | -0.86 | -1.41 | -0.71 | 0.21 | -1.77 | 0.9 | -0.68 | -1.35 | -1.11 | -0.52 | -0.34 | -1.84 | -0.86 | -1.16 | 0.6475 | 0.075 | -0.06 | 0.257114 | 0.774929 | 0.862344 | 1 | 0.996511 | 0.977204 |
| Rab40c | E9PXR4 | -3.23 | -3.64 | 0.51 | 1.37 | -0.38 | -3.22 | 0.43 | 1.68 | -3.4 | -0.26 | 0.97 | -2.31 | -2.22 | -1.82 | 0.91 | 0.7 | 0.875 | -0.0025 | 0.64 | 0.555928 | 0.998615 | 0.637569 | 1 | 1 | 0.942804 |
| Rab43 | Q8CG50 | 0.35 | 0.74 | 0.81 | 0.92 | 0.58 | 1.16 | 0.54 | 1.3 | 0.43 | 0.61 | 1.29 | 0.91 | 2.06 | -0.96 | 1.04 | 1.45 | 0.19 | 0.105 | 0.1925 | 0.44679 | 0.674685 | 0.75242 | 1 | 0.996511 | 0.961479 |
| Rab4a | P56371 | 1.37 | 0.44 | -0.02 | 0.06 | 1.06 | 1.59 | 0.56 | 0.3 | -0.15 | -0.25 | 0.39 | 0.8 | 0.13 | -0.57 | 0.13 | -0.25 | 0.415 | -0.265 | -0.6025 | 0.318258 | 0.498931 | 0.118387 | 1 | 0.996511 | 0.721448 |
| Rab4b | Q91ZR1 | -1.52 | -1.45 | -0.18 | -0.1 | -0.8 | -1.29 | 0.48 | -0.23 | -2.07 | -0.79 | -0.06 | -1.48 | -2 | -2.47 | -0.32 | 0.21 | 0.3525 | -0.2875 | -0.3325 | 0.488913 | 0.596434 | 0.629378 | 1 | 0.996511 | 0.940703 |
| Rab5a | Q9CQD1 | 0.55 | 0.49 | 1.02 | 0.53 | 0.64 | 0.7 | 0.93 | 0.79 | 0.56 | -1.59 | 0.77 | 0.62 | 0.31 | 0.37 | 0.44 | 0.73 | 0.1175 | -0.5575 | -0.185 | 0.525668 | 0.310613 | 0.393974 | 1 | 0.996511 | 0.865806 |
| Rab5b | P61021 | -0.09 | 0.13 | 0.48 | 0.26 | 0.28 | 0.06 | 0.33 | 0.33 | -0.12 | -0.5 | 0.23 | -0.02 | 0.01 | -0.06 | -0.33 | 0.1 | 0.055 | -0.2975 | -0.265 | 0.760969 | 0.211893 | 0.227422 | 1 | 0.996511 | 0.787447 |
| Rab5c | P35278 | 0.23 | 0.55 | 0.54 | 0.64 | 0.27 | 0.34 | 0.71 | 0.76 | 0.56 | -0.37 | 0.72 | 0.41 | 0.26 | 0.11 | -0.25 | 0.48 | 0.03 | -0.16 | -0.34 | 0.875608 | 0.561017 | 0.156798 | 1 | 0.996511 | 0.751493 |
| Rab5if | Q9CQT9 | 1.12 | 1.49 | -1.01 | -1.27 | 1.44 | 1.77 | -0.21 | -1.03 | 1.83 | -0.15 | -1.2 | 1.93 | 1.62 | 2.73 | -0.62 | -0.5 | 0.41 | 0.52 | 0.725 | 0.642365 | 0.584333 | 0.463558 | 1 | 0.996511 | 0.888267 |
| Rab6a | P35279 | 0.08 | 0.26 | 0.79 | 0.75 | 0.6 | 0 | 0.91 | 0.88 | 0.18 | 0.53 | 0.86 | 0.2 | -0.16 | 0.47 | 0.15 | 0.3 | 0.1275 | -0.0275 | -0.28 | 0.649391 | 0.915363 | 0.287078 | 1 | 0.996755 | 0.824333 |
| Rab6a | P35279-2 | 1.04 | 2 | 0.22 | 0.01 | 1.54 | 0.96 | 0.79 | 0.43 | 0.93 | -0.69 | 0.54 | 1.49 | -0.38 | 1.34 | 0.23 | 0.48 | 0.1125 | -0.25 | -0.4 | 0.811406 | 0.674973 | 0.46072 | 1 | 0.996511 | 0.887902 |
| Rab6b | P61294 | 0.01 | 0.28 | 0.74 | 1.06 | 0.48 | 0.11 | 0.85 | 1.34 | 0.97 | 0.4 | 1.08 | 0.24 | 0.7 | 1.42 | 0.24 | 0.4 | 0.1725 | 0.15 | 0.1675 | 0.61454 | 0.635501 | 0.633185 | 1 | 0.996511 | 0.941132 |
| Rab7a | P51150 | 0.5 | 0.8 | 0.74 | 1.24 | 0.66 | 0.37 | 1.06 | 1.23 | 0.57 | 0.86 | 0.8 | 0.61 | 0.05 | 1.04 | 0.36 | 0.21 | 0.01 | -0.11 | -0.405 | 0.969016 | 0.606112 | 0.180039 | 1 | 0.996511 | 0.775542 |
| Rab8a | P55258 | 0.31 | 0.36 | 0.82 | 0.5 | 0.35 | 0.02 | 0.38 | 0.68 | 0.1 | 0.07 | 0.54 | 0 | -0.17 | 0.22 | 0.17 | 0.31 | -0.14 | -0.32 | -0.365 | 0.504015 | 0.154704 | 0.112648 | 1 | 0.996511 | 0.714111 |
| Rab8b | P61028 | 0.16 | 0.22 | 0.09 | 0.04 | 1 | -0.25 | -0.1 | -0.01 | -0.21 | -0.54 | -0.01 | -0.19 | 0.35 | 0.28 | 1.57 | -0.26 | 0.0325 | -0.365 | 0.3575 | 0.910052 | 0.071234 | 0.357726 | 1 | 0.996511 | 0.856624 |
| Rab9a | Q9R0M6 | 1.3 | 0.85 | 0.41 | 0.65 | 0.86 | 1.3 | 0.97 | 0.64 | 0.58 | 0.15 | 0.64 | 1.24 | 0.45 | 0.88 | 0.5 | 0.77 | 0.14 | -0.15 | -0.1525 | 0.57387 | 0.618522 | 0.545208 | 1 | 0.996511 | 0.912212 |
| Rab9b | Q8BHH2 | 0.59 | 0.99 | 0.86 | 0.88 | 1.44 | 0.43 | 1.34 | 0.98 | 0.49 | 0.49 | 0.73 | 0.73 | 0.05 | 0.87 | 1.05 | 1.4 | 0.2175 | -0.22 | 0.0125 | 0.40259 | 0.237204 | 0.967661 | 1 | 0.996511 | 0.9927 |
| Rabac1 | Q9Z0S9 | -0.56 | -0.9 | -3.82 | -3.83 | -0.34 | -0.29 | -3.79 | -3.11 | -0.99 | -1.54 | -3.46 | -0.46 | -0.83 | -0.64 | -3.65 | -3.28 | 0.395 | 0.665 | 0.1775 | 0.731267 | 0.510246 | 0.868019 | 1 | 0.996511 | 0.978872 |
| Rabep1 | O35551 | 0.26 | 0.1 | 0.06 | -0.01 | -0.13 | 0.15 | -0.02 | 0.06 | 0.53 | 0.12 | 0.15 | 0.34 | 0.41 | 0.26 | 0.28 | -0.11 | -0.0875 | 0.1825 | 0.1075 | 0.571644 | 0.321963 | 0.586899 | 1 | 0.996511 | 0.92838 |
| Rabep2 | Q91WG2 | 0.33 | 0.65 | -0.14 | 0.17 | -0.03 | -0.05 | -0.92 | -0.58 | 0.41 | 0.19 | -0.2 | 0.21 | 0.32 | 0.44 | -0.11 | -0.46 | -0.6475 | -0.1 | -0.205 | 0.041341 | 0.674561 | 0.47581 | 1 | 0.996511 | 0.892174 |
| Rabepk | B0R0S4 | -1.02 | -0.65 | -0.36 | -0.39 | -0.24 | -0.54 | -0.6 | -0.4 | -1.34 | -0.35 | -0.51 | -0.61 | -0.9 | 0.17 | -0.4 | -0.76 | 0.16 | -0.0975 | 0.1325 | 0.440584 | 0.727979 | 0.657629 | 1 | 0.996511 | 0.948659 |
| Rabgap1 | A2AWA9 | 0.09 | -0.19 | 0.2 | 0.17 | -0.15 | 0.44 | 0.14 | 0.39 | 0.1 | 0.5 | 0.16 | 0.18 | 0.32 | -0.61 | 0.05 | 0.13 | 0.1375 | 0.1675 | -0.095 | 0.49077 | 0.380667 | 0.708376 | 1 | 0.996511 | 0.957562 |
| Rabgap1l | A6H6A9 | -0.33 | -0.23 | -0.03 | 0.04 | 0.13 | -0.25 | 0.11 | 0.09 | -0.21 | 0.17 | -0.04 | -0.17 | -0.55 | -0.17 | -0.21 | -0.03 | 0.1575 | 0.075 | -0.1025 | 0.377476 | 0.684582 | 0.617027 | 1 | 0.996511 | 0.937141 |
| Rabgap1l | A6H6A9-3 | 1.02 | 0.85 | 0.78 | 0.55 | 0.92 | 0.81 | 0.31 | 0.42 | 1.11 | 1.46 | 0.59 | 1.11 | 0.88 | 1.54 | 0.58 | 0.49 | -0.185 | 0.2675 | 0.0725 | 0.381965 | 0.271642 | 0.794525 | 1 | 0.996511 | 0.969367 |
| Rabgef1 | Q9JM13 | 0.97 | 1.84 | 1.34 | 0.19 | 1.18 | 0.98 | 1.59 | 1.38 | 1.03 | 0.06 | 1.61 | 1.58 | 0.97 | 0.95 | 0.87 | 1.39 | 0.1975 | -0.015 | -0.04 | 0.582466 | 0.974499 | 0.911784 | 1 | 0.997515 | 0.986458 |
| Rabggta | Q9JHK4 | -0.57 | -0.57 | -0.57 | -0.83 | -0.55 | -0.51 | -0.81 | -0.79 | -0.67 | -0.28 | -0.76 | -0.63 | -0.69 | -0.26 | -0.7 | -1.06 | -0.03 | 0.05 | -0.0425 | 0.853208 | 0.787321 | 0.849698 | 1 | 0.996511 | 0.976225 |
| Rabggtb | Q3TVF4 | -1.01 | -1 | -0.96 | -1 | -0.61 | -1.21 | -1.68 | -1.74 | -1.35 | -0.97 | -1.38 | -1.03 | -1.43 | -1.2 | -1.59 | -1.64 | -0.3175 | -0.19 | -0.4725 | 0.256063 | 0.298652 | 0.02873 | 1 | 0.996511 | 0.637553 |
| Rabif | Q91X96 | -1.26 | -0.42 | -0.99 | -1.79 | -0.08 | -1.65 | -1.36 | -1.83 | -1.48 | -1.78 | -1.23 | -0.94 | -1.28 | -0.52 | -1.78 | -1.3 | -0.115 | -0.2425 | -0.105 | 0.799976 | 0.474046 | 0.780768 | 1 | 0.996511 | 0.9676 |
| Rabl2 | E9Q9D5 | -1.41 | -0.94 | 0.46 | -0.35 | -1.32 | -1.17 | 0.44 | 0.43 | -0.51 | -2.05 | 0.38 | -0.94 | 0.06 | -0.65 | 0.48 | 0.25 | 0.155 | -0.22 | 0.595 | 0.788497 | 0.711435 | 0.205042 | 1 | 0.996511 | 0.783452 |
| Rabl3 | Q9D4V7 | -0.42 | -0.37 | -0.02 | -0.1 | -0.38 | -0.49 | 0.18 | 0.17 | -1.12 | -1.26 | -0.32 | -0.65 | -0.88 | -1.63 | 0.44 | 0.24 | 0.0975 | -0.61 | -0.23 | 0.665838 | 0.039849 | 0.62475 | 1 | 0.996511 | 0.939167 |
| Rabl6 | Q5U3K5 | 0.46 | 0.74 | 0.69 | 1.11 | 0.7 | 0.15 | 0.81 | 0.97 | 0.59 | 0.84 | 0.87 | 0.42 | 0.55 | 0.58 | 0.82 | 0.69 | -0.0925 | -0.07 | -0.09 | 0.699948 | 0.743157 | 0.666616 | 1 | 0.996511 | 0.95064 |
| Rac1 | Q3TLP8 | 1.46 | 1 | 1.22 | 1.75 | 0.83 | 1.25 | 1.27 | 1.65 | 1.31 | 1.59 | 1.18 | 1.32 | 0.52 | 1.57 | 0.95 | 0.75 | -0.1075 | -0.0075 | -0.41 | 0.663586 | 0.972772 | 0.185219 | 1 | 0.997515 | 0.776843 |
| Rac2 | Q05144 | -0.25 | 0.33 | -0.74 | -0.41 | -0.9 | 0.52 | -0.83 | -0.02 | -0.3 | 0.53 | -0.25 | -1.8 | -0.51 | 0.28 | -1.55 | -0.73 | -0.04 | -0.1875 | -0.36 | 0.917407 | 0.7078 | 0.400044 | 1 | 0.996511 | 0.868923 |
| Rac3 | P60764 | 0.58 | 0.61 | 3.4 | 4.56 | 1.15 | 0.58 | 3.1 | 5.17 | 1.24 | 6.81 | 4.44 | 1.19 | 0.56 | 1.78 | 4.46 | 4.09 | 0.2125 | 1.1325 | 0.435 | 0.869491 | 0.461184 | 0.722595 | 1 | 0.996511 | 0.958398 |
| Rack1 | P68040 | -2.32 | -2.41 | -1.26 | -1.15 | -1.38 | -2.1 | -0.49 | -1.18 | -2.93 | -1.31 | -1.26 | -2.1 | -2.42 | -1.88 | -0.73 | -0.66 | 0.4975 | -0.115 | 0.3625 | 0.276534 | 0.811933 | 0.484419 | 1 | 0.996511 | 0.894266 |
| Rad21 | Q61550 | 0.33 | -0.1 | 0.33 | 0.93 | -0.52 | 0 | 0.34 | 0.61 | 0.06 | 0.72 | 0.16 | -0.18 | 0.33 | -0.33 | 0.63 | 0.61 | -0.265 | -0.1825 | -0.0625 | 0.412007 | 0.537752 | 0.843222 | 1 | 0.996511 | 0.976225 |
| Rad23a | Q8CAP3 | 0.82 | 0.03 | 0.88 | -0.33 | -0.2 | 0.67 | 1.99 | -0.61 | -0.1 | -0.16 | 0.74 | 0.73 | -0.03 | 0.52 | 1.04 | 1.88 | 0.1125 | -0.0475 | 0.5025 | 0.849044 | 0.899385 | 0.304068 | 1 | 0.996511 | 0.831662 |
| Rad23b | P54728 | 0.04 | 0.51 | 0.48 | 0.42 | -0.54 | 0.03 | -0.01 | 0.45 | 0.66 | 0.06 | 0.45 | 0.49 | -0.44 | 0.08 | -0.88 | -0.25 | -0.38 | 0.0525 | -0.735 | 0.14648 | 0.802939 | 0.018113 | 1 | 0.996511 | 0.602211 |
| Rad50 | Q5SV02 | -1.01 | -0.36 | -2.03 | -2.43 | -0.03 | -1.74 | -2.16 | -2.67 | 0.36 | -1.21 | -2.15 | -1.27 | 0.19 | -0.83 | -2.5 | -2.46 | -0.1925 | 0.39 | 0.0575 | 0.775291 | 0.548322 | 0.937381 | 1 | 0.996511 | 0.989767 |
| Radil | D3YWJ9 | 1.47 | 0.68 | 0.35 | 0.56 | 0.48 | 1.39 | 0.2 | 0.18 | 0.86 | 1.02 | 0.04 | 0.8 | 0.75 | 0.44 | 0.49 | 0.47 | -0.2025 | -0.085 | -0.2275 | 0.576274 | 0.794882 | 0.420273 | 1 | 0.996511 | 0.87558 |
| Rae1 | Q8C570 | 1.04 | 0.62 | 0.33 | 0.4 | 0.63 | 0.6 | 1.01 | 0.89 | -0.22 | -0.63 | 0.57 | 1.01 | -0.09 | -0.36 | 1.34 | 0.74 | 0.185 | -0.415 | -0.19 | 0.398893 | 0.301874 | 0.640677 | 1 | 0.996511 | 0.943012 |
| Raf1 | Q99N57 | -0.19 | -0.1 | -0.63 | -1.06 | -0.03 | 0.34 | -0.63 | -1.21 | 0.76 | -0.64 | -0.82 | 0.28 | 1.08 | -0.39 | -0.94 | -1.27 | 0.1125 | 0.39 | 0.115 | 0.770417 | 0.359471 | 0.826246 | 1 | 0.996511 | 0.973235 |
| Rai1 | Q61818 | 1.23 | 0.45 | 1.1 | 0.54 | 0.45 | 0.8 | 0.94 | 1.19 | 0.33 | 0.38 | 1.14 | 0.74 | 1.1 | 0.45 | 1.78 | 1.27 | 0.015 | -0.1825 | 0.32 | 0.953735 | 0.523748 | 0.356746 | 1 | 0.996511 | 0.855416 |
| Rai14 | A0A2I3BPY | 0.76 | 0.57 | 0.21 | -0.08 | 0.79 | 0.62 | 0.25 | -0.18 | 0.48 | -0.27 | 0.37 | 0.69 | 0.29 | 1.1 | 0.25 | 0.19 | 0.005 | -0.0475 | 0.0925 | 0.986055 | 0.86912 | 0.757252 | 1 | 0.996511 | 0.962481 |
| Rala | P63321 | 0.47 | 0.41 | 0.69 | 0.84 | 0.68 | 0.32 | 0.66 | 0.64 | 0.75 | 1.52 | 0.78 | 0.34 | 0.39 | 1.11 | 0.32 | 0.12 | -0.0275 | 0.245 | -0.1175 | 0.877049 | 0.388885 | 0.658775 | 1 | 0.996511 | 0.949645 |
| Ralb | Q9JIW9 | 0 | 0.38 | 1 | 0.84 | 0.47 | -0.16 | 0.83 | 0.54 | 0.56 | 0.6 | 0.92 | 0.29 | 0.17 | 1.33 | 0.63 | 0.6 | -0.135 | 0.0375 | 0.1275 | 0.659329 | 0.891458 | 0.702684 | 1 | 0.996511 | 0.956859 |
| Ralbp1 | Q62172 | 1.29 | 0.68 | 0.95 | 1.31 | 0.51 | 0.94 | 0.42 | 1.01 | 1.08 | 0.77 | 0.67 | 0.47 | 0.77 | 0.73 | 0.61 | 0.61 | -0.3375 | -0.31 | -0.3775 | 0.169099 | 0.199998 | 0.103001 | 1 | 0.996511 | 0.700625 |
| Ralgapa1 | A0A1W2P8 | -0.03 | -0.19 | 0.05 | 0.05 | -0.11 | 0.11 | -0.04 | -0.12 | 0.43 | 0.15 | 0.09 | -0.23 | 0.3 | 0 | -0.23 | -0.15 | -0.01 | 0.14 | 0.01 | 0.947224 | 0.485553 | 0.959751 | 1 | 0.996511 | 0.992636 |
| Ralgapa2 | A0A0A0M( | 0.02 | -0.14 | 0.42 | 0.33 | 0.12 | -0.11 | -0.09 | 0.3 | -0.14 | -0.14 | 0.23 | -0.21 | -0.28 | 0.3 | 0.11 | 0.45 | -0.1025 | -0.2225 | -0.0125 | 0.606242 | 0.302254 | 0.95887 | 1 | 0.996511 | 0.992636 |
| Ralgapb | A2ACC6 | 0.05 | -0.19 | -0.09 | -0.28 | -0.03 | 0.03 | -0.28 | -0.05 | -0.36 | -0.55 | -0.08 | -0.11 | -0.73 | -0.26 | -0.55 | -0.33 | 0.045 | -0.1475 | -0.34 | 0.779237 | 0.444213 | 0.112639 | 1 | 0.996511 | 0.714111 |
| Ralgapb | Q8BQZ4 | 0.48 | 0.26 | 0.89 | 0.57 | 0.63 | 0.46 | 0 | 0.76 | 0.34 | 1.51 | 0.37 | 0.43 | 0.03 | 0.43 | 0.31 | 0.46 | -0.0875 | 0.1125 | -0.2425 | 0.705203 | 0.720782 | 0.28056 | 1 | 0.996511 | 0.822084 |
| Ralgps1 | A2AR50-3 | 0.63 | 0.57 | 0.12 | 0.14 | 0.67 | 0.47 | -0.58 | 0 | -0.09 | 0.74 | 0.21 | 0.51 | 0.13 | 0.67 | 0.82 | -0.23 | -0.225 | -0.0225 | -0.0175 | 0.469117 | 0.928107 | 0.952461 | 1 | 0.996755 | 0.992476 |
| Ralgps2 | A0A087WC | 0.09 | -0.02 | 0.01 | -0.78 | -0.15 | 0.3 | -0.51 | -0.2 | -0.33 | -0.29 | -0.27 | 0.11 | 0.01 | -0.21 | -0.57 | -0.2 | 0.035 | -0.02 | -0.0675 | 0.896388 | 0.936299 | 0.797977 | 1 | 0.996755 | 0.969727 |
| Raly | Q64012-2 | 1.2 | 0.4 | 0.64 | 0.57 | 0.02 | 0.46 | 0.14 | 0.93 | 0.23 | 0.61 | 0.31 | -0.14 | 1.07 | 0.19 | 0.78 | 0.03 | -0.315 | -0.45 | -0.185 | 0.266123 | 0.103853 | 0.554295 | 1 | 0.996511 | 0.916796 |
| Ralyl | Q8BTF8 | 1.17 | -0.1 | 0.28 | -0.72 | -0.47 | 0.92 | -0.4 | 0.33 | -0.17 | -1.52 | 0.33 | -0.55 | 1.2 | -1.35 | 0.68 | -0.39 | -0.0625 | -0.635 | -0.1225 | 0.895585 | 0.239157 | 0.846284 | 1 | 0.996511 | 0.976225 |
| Ralyl | A0A0A6YX( | 1 | -0.45 | 0.52 | -0.54 | -0.06 | 0.27 | -1.41 | -1.24 | -0.41 | -2.42 | 0.23 | -0.4 | 0.14 | -0.5 | 2.1 | -1.28 | -0.7425 | -0.8825 | -0.0175 | 0.176986 | 0.185053 | 0.981029 | 1 | 0.996511 | 0.994858 |
| Ramac | Q9CQY2 | -4.24 | -4.65 | -4.47 | -4.57 | -4.87 | -5.26 | -4.51 | -4.85 | -4.66 | -2.77 | -4.07 | -4.45 | -3.86 | -4 | -4.4 | -5.16 | -0.39 | 0.495 | 0.1275 | 0.086725 | 0.249641 | 0.685441 | 1 | 0.996511 | 0.955729 |
| Ran | P62827 | 1.36 | 0.12 | 0.19 | 0.61 | 0.03 | 1.44 | 0.5 | 1.56 | -0.26 | -1.11 | 0.48 | 1.39 | -1.06 | -0.58 | -0.22 | 0.4 | 0.3125 | -0.445 | -0.935 | 0.48019 | 0.432535 | 0.043007 | 1 | 0.996511 | 0.658454 |
| Ranbp1 | P34022 | -0.79 | -1.22 | -1.3 | -0.64 | -1.02 | -0.4 | -0.75 | -0.82 | -1.33 | -0.74 | -1.3 | -0.78 | -1.97 | -1.09 | -1.15 | -1.29 | 0.24 | -0.05 | -0.3875 | 0.306517 | 0.84183 | 0.189482 | 1 | 0.996511 | 0.776843 |
| Ranbp10 | Q6VN19 | -0.81 | -0.85 | 0.69 | 1.17 | -0.31 | -0.86 | 0.97 | 1.21 | -1.2 | 1.53 | 0.93 | -0.8 | -0.85 | -0.71 | 0.5 | 0.98 | 0.2025 | 0.065 | -0.07 | 0.756895 | 0.931781 | 0.911001 | 1 | 0.996755 | 0.986458 |
| Ranbp17 | Q99NF8 | -2.28 | -1.06 | -0.34 | 0.4 | -1.7 | -1.34 | -0.42 | -0.36 | -1.23 | 0.2 | 0.55 | -0.64 | -2.57 | -0.68 | 0.39 | 0.1 | -0.135 | 0.54 | 0.13 | 0.822975 | 0.407526 | 0.869409 | 1 | 0.996511 | 0.979293 |
| Ranbp2 | Q9ERU9 | 0.64 | 0.05 | 0.31 | 0.3 | -0.03 | 0.48 | 0.19 | 0.72 | 0.19 | 0.22 | 0.22 | -0.09 | 0.85 | 0.17 | 1.09 | 0.63 | 0.015 | -0.19 | 0.36 | 0.94686 | 0.344388 | 0.18884 | 1 | 0.996511 | 0.776843 |
| Ranbp3 | A0A3B2W( | -2.65 | -3.05 | -3.2 | -3.48 | -2.68 | -2.87 | -3.56 | -3.39 | -2.69 | -2.06 | -3.11 | -2.44 | -2.16 | -2.18 | -3.34 | -4.06 | -0.03 | 5.20E-01 | 0.16 | 0.91318 | 0.10004 | 0.732556 | 1 | 0.996511 | 0.958635 |
| Ranbp6 | Q8BIV3 | 0.24 | -0.31 | -0.64 | -0.65 | -0.12 | 0.3 | -0.11 | -0.5 | -0.11 | -1.1 | -0.71 | 0.14 | 0.25 | -0.32 | -0.11 | 0.16 | 0.2325 | -0.105 | 0.335 | 0.400864 | 0.761232 | 0.235899 | 1 | 0.996511 | 0.791493 |
| Ranbp9 | E9Q5D6 | -0.18 | 0.27 | -0.41 | -0.32 | 0.92 | -0.13 | -0.06 | -0.24 | -0.09 | -0.39 | -0.14 | 0.11 | -0.12 | 0.61 | -0.49 | -0.29 | 0.2825 | 0.0325 | 0.0875 | 0.36586 | 0.882494 | 0.768911 | 1 | 0.996511 | 0.965796 |
| Rangap1 | P46061 | 0.02 | -0.1 | -0.6 | -0.76 | -0.12 | 0.18 | -0.51 | -0.17 | -0.32 | -0.54 | -0.39 | -0.45 | -0.2 | 0.27 | -0.46 | -0.7 | 0.205 | -0.065 | 0.0875 | 0.41918 | 0.775936 | 0.767598 | 1 | 0.996511 | 0.965083 |
| Rap1a | P62835 | 2.3 | 2.14 | 1.74 | 0.68 | 1.87 | 2.11 | 1.53 | 1.69 | 0.61 | -0.2 | 1.47 | 2.33 | 0.33 | 0.06 | 1.34 | 1.78 | 0.085 | -0.6625 | -0.8375 | 0.817207 | 0.287927 | 0.12701 | 1 | 0.996511 | 0.722414 |
| Rap1b | Q99JI6 | 1.67 | 1.36 | 1.25 | 0.83 | 1.4 | 1.51 | 0.93 | 0.99 | 0.22 | 0.57 | 1.06 | 1.81 | -0.18 | -0.52 | 0.47 | 0.74 | -0.07 | -0.3625 | -1.15 | 0.772237 | 0.344969 | 0.007659 | 1 | 0.996511 | 0.554014 |
| Rap1gap | A2ALS5 | -0.09 | -0.67 | -0.33 | -0.24 | -0.59 | -0.56 | -0.4 | -0.36 | 0.56 | -0.05 | -0.27 | -0.08 | 0.29 | 0.3 | -0.45 | -0.66 | -0.145 | 0.3725 | 0.2025 | 0.430427 | 0.15275 | 0.495827 | 1 | 0.996511 | 0.89758 |
| Rap1gap | A2ALS7 | -0.43 | -0.39 | -1.06 | -1.3 | -0.02 | -1.12 | -0.91 | -1 | -0.56 | -0.17 | -0.41 | -0.43 | -0.93 | 0.17 | -1.49 | -0.62 | 0.0325 | 0.4025 | 0.0775 | 0.921395 | 0.149113 | 0.846262 | 1 | 0.996511 | 0.976225 |
| Rap1gap2 | A0A571BEI | -0.12 | -0.41 | 0.18 | 0.17 | -0.24 | 0.01 | 0.52 | 0.5 | -0.19 | -0.43 | 0.35 | -0.02 | -0.27 | -0.13 | 0.06 | -0.18 | 0.2425 | -0.0275 | -0.085 | 0.340365 | 0.909572 | 0.690649 | 1 | 0.996744 | 0.955729 |
| Rap1gds1 | E9Q912 | 0.17 | -0.09 | 0.18 | 0 | 0.29 | 0.43 | 0.32 | 0.23 | -0.21 | -0.06 | 0.19 | 0.26 | -0.54 | 0.35 | -0.12 | -0.02 | 0.2525 | -0.02 | -0.1475 | 0.122992 | 0.914871 | 0.536142 | 1 | 0.996755 | 0.912212 |
| Rap1gds1 | E9Q912-3 | -3.78 | -3.94 | -3.9 | -2.99 | -4.36 | -3.13 | -2.73 | -3.44 | -2.66 | -3.08 | -3.48 | -3.43 | -4.12 | -3.15 | -4.14 | -3.47 | 0.2375 | 0.49 | -0.0675 | 0.547195 | 0.128789 | 0.839947 | 1 | 0.996511 | 0.976225 |
| Rap1gds1 | E9Q912-2 | -0.02 | 0.31 | 0 | 0.43 | 0.47 | -0.13 | 0.51 | 0.44 | -0.17 | 0.52 | 0.37 | 0.5 | -1 | 1.2 | 0.03 | 0.51 | 0.1425 | 0.125 | 0.005 | 0.512109 | 0.589068 | 0.991119 | 1 | 0.996511 | 0.996154 |
| Rap2a | Q80ZJ1 | 0.49 | 0.86 | 0.77 | 1.29 | 0.95 | 0.78 | 1.19 | 1.65 | 1.27 | 1 | 1.07 | 0.76 | 0.43 | 1.28 | 0.66 | 0.58 | 0.29 | 0.1725 | -0.115 | 0.282516 | 0.458818 | 0.675155 | 1 | 0.996511 | 0.952944 |
| Rap2b | P61226 | 1.65 | 0.73 | 1.24 | 1.67 | 1.49 | 1.49 | 1.26 | 1.52 | 2.19 | 1.98 | 1.03 | 1.72 | 1.46 | 1.21 | 1.3 | 1.07 | 0.1175 | 0.4075 | -0.0625 | 0.631528 | 0.241032 | 0.812402 | 1 | 0.996511 | 0.972642 |
| Rap2c | Q8BU31 | 0.24 | 0.18 | 0.47 | 0.38 | 0.45 | 0.77 | -0.08 | 0.34 | 1.5 | 0.58 | 0.24 | -0.09 | 1.83 | -0.21 | 0.55 | 0.42 | 0.0525 | 0.24 | 0.33 | 0.806169 | 0.490819 | 0.434482 | 1 | 0.996511 | 0.880558 |
| Rapgef1 | A1L338 | -0.53 | -0.37 | 0.01 | -0.37 | -0.62 | -0.34 | -0.58 | -0.8 | 0.61 | 0.27 | -0.19 | -0.33 | 1.27 | -0.44 | -0.17 | -0.6 | -0.27 | 0.405 | 0.33 | 0.176567 | 0.149236 | 0.443096 | 1 | 0.996511 | 0.883549 |
| Rapgef2 | A0A0A6YW | 0.07 | 0 | 0.55 | 0.76 | 0.12 | 0 | 0.53 | 0.71 | 0.21 | 0.86 | 0.47 | 0 | -0.11 | 0.45 | 0.39 | 0.41 | -0.005 | 0.04 | -0.06 | 0.984557 | 0.884421 | 0.815733 | 1 | 0.996511 | 0.972768 |
| Rapgef3 | H3BK39 | 0.54 | 0.91 | 0.62 | 1.01 | 0.82 | 0.53 | 0.63 | 1.04 | -0.39 | 0.03 | 0.76 | 0.55 | -1.28 | -0.16 | -0.64 | 0.29 | -0.015 | -0.5325 | -1.2175 | 0.938628 | 0.095311 | 0.006948 | 1 | 0.996511 | 0.552614 |
| Rapgef4 | Q9EQZ6 | 0.84 | 1.14 | 0.98 | 0.88 | 1.11 | 0.89 | 0.6 | 0.85 | 0.9 | 0.6 | 0.93 | 0.66 | 0.73 | 0.8 | 0.87 | 0.77 | -0.0975 | -0.1875 | -0.1675 | 0.57823 | 0.304695 | 0.350026 | 1 | 0.996511 | 0.85327 |
| Rapgef5 | Q8C0Q9 | 1.83 | 0.54 | 1.66 | 0.53 | 0.54 | 2.15 | 2.52 | 0.9 | 0.1 | -0.07 | 1.67 | 1.67 | -0.3 | 0.5 | 1.21 | 1.97 | 0.3875 | -0.2975 | -0.295 | 0.482749 | 0.590276 | 0.596511 | 1 | 0.996511 | 0.929911 |
| Rapgef6 | Q5NCJ4 | -0.21 | -0.06 | 0.11 | 0.19 | -0.61 | -0.32 | 0.08 | -0.21 | 0.48 | 0.17 | -0.41 | -0.53 | -0.59 | 0.58 | -0.29 | 0.4 | -0.2725 | -0.08 | 0.0175 | 0.198268 | 0.768039 | 0.953993 | 1 | 0.996511 | 0.992476 |
| Rapgefl1 | Q68EF8 | -0.48 | -1.46 | -0.84 | -0.95 | -1.34 | -0.47 | -0.85 | -0.78 | -1.04 | -2.01 | -0.84 | -1.07 | -0.89 | -1.71 | -1.95 | -1.39 | 0.0725 | -0.3075 | -0.5525 | 0.792473 | 0.362411 | 0.105967 | 1 | 0.996511 | 0.703417 |
| Raph1 | F2Z3U3 | 0.34 | 0.23 | 0.57 | 0.65 | 0.24 | 0.41 | 0.42 | 0.53 | 0.77 | 0.13 | 0.73 | 0.46 | 0.94 | 0.36 | 0.54 | 0.21 | -0.0475 | 0.075 | 0.065 | 0.778598 | 0.729756 | 0.777884 | 1 | 0.996511 | 0.967289 |
| Rars1 | Q9D0I9 | -1.32 | -1.68 | -0.29 | -0.68 | -1.36 | -1.47 | -0.28 | -0.01 | -2.27 | -1.09 | -0.22 | -1.17 | -2.27 | -1.78 | -0.43 | -0.47 | 0.2125 | -0.195 | -0.245 | 0.640004 | 0.690659 | 0.63981 | 1 | 0.996511 | 0.942804 |
| Rars2 | Q3U186 | 1.14 | 0.88 | 0.33 | 0.45 | 0.95 | 1.11 | 0.55 | 0.37 | 0.82 | 0.01 | 0.18 | 0.59 | 0.89 | 1.09 | 0.14 | 0.37 | 0.045 | -0.3 | -0.0775 | 0.864022 | 0.296082 | 0.798058 | 1 | 0.996511 | 0.969727 |
| Rasa1 | E9PYG6 | 0.23 | 0.04 | 0.33 | 0.27 | 0.2 | -0.19 | 0.08 | 0.21 | 0.21 | 0.49 | 0.16 | 0.06 | -0.41 | 0.57 | -0.11 | 0.1 | -0.1425 | 0.0125 | -0.18 | 0.404742 | 0.944237 | 0.4771 | 1 | 0.996755 | 0.892234 |
| Rasa2 | P58069 | 0.72 | 0.29 | 0.2 | -0.14 | 0.2 | 0.3 | 0.66 | 0.22 | 0.14 | -0.79 | 0.12 | -0.06 | -0.49 | 0.62 | -0.29 | -0.24 | 0.0775 | -0.415 | -0.3675 | 0.733905 | 0.177623 | 0.257946 | 1 | 0.996511 | 0.812074 |
| Rasa3 | Q60790 | 2.25 | 0.91 | 1.32 | 1.25 | 1.11 | 2.04 | 1.58 | 1.18 | 0.28 | 0.33 | 1.14 | 1.57 | 0.04 | 0.38 | 1.22 | 1.34 | 0.045 | -0.6025 | -0.6875 | 0.896179 | 0.164255 | 0.121291 | 1 | 0.996511 | 0.721448 |
| Rasal1 | Q9Z268 | 0.41 | 0.01 | 0.39 | 0.62 | 0.24 | 0.61 | 0.96 | 0.83 | 0.13 | -0.23 | 0.79 | 0.24 | -0.05 | -0.32 | 0.61 | 0.48 | 0.3025 | -0.125 | -0.1775 | 0.200385 | 0.638123 | 0.524696 | 1 | 0.996511 | 0.908107 |
| Rasal2 | D3YUS5 | 0.77 | 0.6 | 1.14 | 0.66 | 0.61 | 0.63 | 0.58 | 0.61 | 0.32 | 0.37 | 1.16 | 1.06 | 0.43 | 0.33 | 0.98 | 0.85 | -0.185 | -0.065 | -0.145 | 0.299607 | 0.809134 | 0.54772 | 1 | 0.996511 | 0.912571 |
| Rasd2 | P63032 | 1.86 | 0.24 | 0.68 | 0.81 | 0.72 | -0.35 | 0.98 | 0.77 | 0.38 | 1.27 | 0.52 | 1.32 | 1.27 | 2.03 | 1.15 | 0.79 | -0.3675 | -0.025 | 0.4125 | 0.398755 | 0.950499 | 0.332207 | 1 | 0.996755 | 0.845601 |
| Rasgef1a | A6H650 | 0.03 | 0.26 | 0.4 | 0.54 | 0.18 | 0.25 | 0.63 | 0.46 | -0.09 | 0.58 | 0.6 | -0.09 | 0.23 | 0.61 | 0.74 | 0.66 | 0.0725 | -0.0575 | 0.2525 | 0.701954 | 0.817044 | 0.255056 | 1 | 0.996511 | 0.809524 |
| Rasgef1c | B1ATE5 | -1.12 | -3.05 | -1.05 | 0.57 | -0.95 | -0.38 | -0.25 | -0.23 | -1.03 | 1.64 | -0.76 | -1.26 | -0.03 | -0.11 | -0.71 | 1.43 | 0.71 | 0.81 | 1.3075 | 0.317657 | 0.38 | 0.122461 | 1 | 0.996511 | 0.721448 |
| Rasgrf1 | P27671 | 0.27 | 0.11 | -0.25 | -0.71 | -0.07 | 0.23 | -0.31 | -0.53 | 0.22 | -0.15 | -0.37 | 0.31 | 0.25 | -0.18 | -0.46 | -0.69 | -0.025 | 0.1475 | -0.125 | 0.927751 | 0.602742 | 0.685615 | 1 | 0.996511 | 0.955729 |
| Rasgrf2 | P70392 | 0.1 | 0.3 | 0.07 | -0.14 | 0.41 | 0.34 | 0.01 | -0.01 | -0.18 | 0 | -0.02 | 0.04 | 0.03 | 0.23 | -0.18 | -0.17 | 0.105 | -0.1225 | -0.105 | 0.571826 | 0.489011 | 0.601759 | 1 | 0.996511 | 0.930802 |
| Rasgrp1 | Q9Z1S3 | -4.77 | -0.32 | -1.47 | 0.12 | -0.52 | -4.93 | -2.48 | -0.34 | 0.05 | 0.04 | -1.52 | -3.91 | -0.66 | 0.23 | -1.87 | -2.15 | -0.4575 | 0.275 | 0.4975 | 0.739876 | 0.831827 | 0.653577 | 1 | 0.996511 | 0.947633 |
| Rasgrp2 | Q9QUG9 | -0.91 | -1.01 | -0.76 | -0.44 | -1.11 | -1.71 | -0.24 | -0.88 | -0.47 | 0.14 | -0.8 | -0.36 | 0.25 | -0.97 | 0.26 | -0.24 | -0.205 | 0.4075 | 6.05E-01 | 0.527789 | 0.134371 | 0.086841 | 1 | 0.996511 | 0.691619 |
| Rasip1 | Q3U0S6 | -2.11 | -2.73 | 0.56 | 0.54 | -1.84 | -1.59 | -0.13 | 0.33 | -2.33 | 0.7 | 0.83 | -1.91 | -1.91 | -2.39 | -0.72 | -0.33 | 0.1275 | 0.2575 | -0.4025 | 0.889156 | 0.811884 | 0.653466 | 1 | 0.996511 | 0.947633 |
| Rassf2 | Q8BMS9 | 0 | -0.37 | -0.28 | 0.33 | -0.29 | 0.06 | -0.31 | 0.03 | 0.03 | 0.08 | -0.01 | -0.25 | 0.41 | -0.33 | -0.44 | 0.12 | -0.0475 | 0.0425 | 0.02 | 0.823806 | 0.842859 | 0.942076 | 1 | 0.996511 | 0.991114 |
| Rassf5 | Q5EBH1 | 1.38 | 0.14 | 0.2 | -0.71 | 0.81 | 0.04 | -0.64 | -0.87 | 0.31 | -0.53 | -0.63 | 0.06 | 1.74 | 0.08 | 0.23 | -0.34 | -0.4175 | -0.45 | 0.175 | 0.435906 | 0.338124 | 0.76111 | 1 | 0.996511 | 0.963123 |
| Rassf8 | Q8CJ96 | -0.86 | -0.57 | 1.1 | 0.54 | 1.4 | -0.5 | -0.43 | 1.62 | -0.31 | 0.14 | 0.23 | 0.26 | -0.47 | -0.66 | -0.37 | 0.85 | 0.47 | 0.0275 | -0.215 | 0.487412 | 0.951372 | 0.687517 | 1 | 0.996755 | 0.955729 |
| Raver1 | Q9CW46 | -3.12 | -2.82 | -1.49 | -0.98 | -3.84 | -3.99 | -2.09 | -1.85 | -2.71 | -0.61 | -1.85 | -3.46 | -2.67 | -2.82 | -2.49 | -1.39 | -0.84 | -0.055 | -0.24 | 0.245411 | 0.939545 | 0.670315 | 1 | 0.996755 | 0.952558 |
| Rb1cc1 | Q9ESK9 | 0.62 | 0.53 | 0.71 | 0.58 | 0.71 | 0.41 | 0.3 | 0.55 | 0.27 | 1.17 | 0.61 | 0.6 | 0.56 | 0.35 | 0.71 | 0.57 | -0.1175 | 0.0525 | -0.0625 | 0.468579 | 0.815742 | 0.726259 | 1 | 0.996511 | 0.958398 |
| Rbbp4 | Q60972 | -0.31 | -1.19 | -1.45 | -1.64 | -0.85 | -0.74 | -0.97 | -1.42 | -1.82 | -1.16 | -1.43 | -0.59 | -1.42 | -0.85 | -1.09 | -0.88 | 0.1525 | -0.1025 | 0.0875 | 0.63786 | 0.786412 | 0.788924 | 1 | 0.996511 | 0.968677 |
| Rbbp5 | Q8BX09-2 | 0.04 | -1.66 | 0.63 | -0.58 | -1.43 | -1.5 | -0.92 | -0.33 | -1.4 | -0.13 | -0.63 | -0.82 | -1.98 | -1.03 | 0.81 | -0.25 | -0.6525 | -0.3525 | -0.22 | 0.226795 | 0.499626 | 0.752642 | 1 | 0.996511 | 0.961479 |
| Rbbp5 | A0A0R4J2E | -2.28 | -2.25 | -2 | -2.14 | -2.44 | -2.91 | -2.31 | -2.26 | -2.22 | -1.48 | -2.01 | -2.46 | -1.7 | -1.78 | -2.33 | -2.52 | -0.3125 | 0.125 | 8.50E-02 | 0.138235 | 0.611076 | 0.731513 | 1 | 0.996511 | 0.958635 |
| Rbbp6 | P97868-2 | -1.16 | -2.22 | -0.33 | -0.64 | -1.94 | -1.91 | -0.46 | -0.72 | -1.94 | -0.34 | -0.79 | -2.22 | -0.24 | -1.49 | 0.26 | -0.51 | -0.17 | -0.235 | 0.5925 | 0.745601 | 0.678104 | 0.268436 | 1 | 0.996511 | 0.820082 |
| Rbbp7 | Q60973 | -0.22 | -1.54 | -0.75 | -0.75 | -1.28 | -0.93 | -0.8 | -0.47 | -1.18 | 0.04 | -0.8 | -0.65 | -1.04 | -0.78 | -0.06 | -0.47 | -0.055 | 0.1675 | 0.2275 | 0.860825 | 0.646037 | 0.513261 | 1 | 0.996511 | 0.904284 |
| Rbbp9 | O88851 | -0.19 | -0.66 | -0.5 | -0.2 | -0.49 | 0.13 | -0.16 | -0.1 | -0.79 | -0.62 | -0.37 | -0.49 | -1.01 | -0.47 | -0.72 | -0.93 | 0.2325 | -0.18 | -0.395 | 0.272029 | 0.374589 | 0.098692 | 1 | 0.996511 | 0.698156 |
| Rbck1 | Q9WUB0 | -0.63 | -0.58 | -0.03 | 0.06 | -0.38 | -0.67 | 0.3 | 0 | -0.49 | 0.28 | 0.01 | -0.46 | -0.4 | 0.39 | 0.16 | -0.04 | 0.1075 | 0.13 | 0.3225 | 0.703413 | 0.637398 | 0.252383 | 1 | 0.996511 | 0.80892 |
| Rbfa | Q6P3B9 | 2.03 | 2.42 | 1 | 0.68 | 2.47 | 1.79 | 0.7 | 0.62 | 1.87 | -1.04 | 0.41 | 0.94 | 1.22 | 1.88 | 0.61 | 1.44 | -0.1375 | -0.9875 | -0.245 | 0.80521 | 0.166031 | 0.59848 | 1 | 0.996511 | 0.930093 |
| Rbfox1 | A0A2R8VK | -0.46 | -1.06 | -1.03 | -0.44 | -1.41 | -0.93 | -0.93 | -0.51 | -1.71 | -0.96 | -0.71 | -1.31 | -0.8 | -0.89 | -0.77 | -1.14 | -0.1975 | -0.425 | -0.1525 | 0.455951 | 0.164248 | 0.519218 | 1 | 0.996511 | 0.907667 |
| Rbfox1 | Q9JJ43-6 | -2.23 | -0.97 | -3.63 | -3.45 | -4.67 | -4.87 | -4.47 | -4.37 | -3.53 | -1.57 | -3.59 | -3.68 | -1.73 | -0.53 | -4.13 | -5.16 | -2.025 | -0.5225 | -0.3175 | 0.006409 | 0.478366 | 0.772894 | 1 | 0.996511 | 0.966062 |
| Rbfox2 | Q8BP71-7 | -1.31 | -1.1 | -0.62 | -0.3 | -0.86 | -1.98 | -0.75 | -1.49 | -1.2 | -0.34 | -1.31 | -1.68 | -1.49 | -1.11 | -1.63 | -1.73 | -0.4375 | -0.3 | -0.6575 | 0.235857 | 0.408964 | 0.04347 | 1 | 0.996511 | 0.658454 |
| Rbfox3 | Q8BIF2-2 | -0.22 | -0.64 | -1.06 | -0.79 | -0.81 | -0.77 | -1.1 | -0.31 | -2.14 | -1.04 | -0.99 | -1.21 | -1.25 | -1.4 | -0.35 | -1.49 | -0.07 | -0.6675 | -0.445 | 0.780958 | 0.06386 | 0.190043 | 1 | 0.996511 | 0.776843 |
| Rbks | Q8R1Q9 | -0.78 | -2.01 | -3.98 | -3.78 | -3.14 | -0.45 | -4.55 | -3.83 | 0.37 | -2.61 | -4.31 | -1.32 | -0.18 | -3.58 | -4.12 | -3.95 | -0.355 | 0.67 | -0.32 | 0.736712 | 0.553826 | 0.766316 | 1 | 0.996511 | 0.964134 |
| Rbl2 | A0A1B0GR | -1.55 | -1.84 | 0.08 | 0.07 | -1.89 | -1.52 | -2.66 | -1.13 | -0.27 | -0.18 | -2.03 | -2 | 1 | -1.37 | -0.19 | -0.5 | -0.99 | -0.31 | 0.545 | 0.103745 | 0.642902 | 0.409738 | 1 | 0.996511 | 0.87476 |
| Rbm10 | Q99KG3-3 | -0.18 | -1.43 | -1.93 | -1.93 | -1.42 | -1.07 | -1.43 | -1.52 | -1.34 | -1.76 | -1.77 | -1.29 | -1.01 | -1.19 | -0.47 | -1.38 | 0.0075 | -0.1725 | 0.355 | 0.985042 | 0.676931 | 0.422947 | 1 | 0.996511 | 0.875782 |
| Rbm12 | Q8R4X3 | -2.67 | -3.15 | -2.9 | -3.22 | -2.67 | -3.26 | -3.37 | -3.82 | -2.89 | -1.94 | -3.29 | -2.6 | -3.08 | -2.21 | -3.48 | -3.65 | -0.295 | 0.305 | -0.12 | 0.296347 | 0.343456 | 0.728093 | 1 | 0.996511 | 0.958635 |
| Rbm12b2 | Q66JV4 | -0.03 | -0.6 | -0.19 | -0.34 | -0.73 | -0.37 | -0.81 | -0.07 | -0.72 | -0.04 | -0.81 | -1.35 | 0.1 | -0.79 | 1 | -0.4 | -0.205 | -0.44 | 0.2675 | 0.382525 | 0.168605 | 0.500805 | 1 | 0.996511 | 0.900975 |

| | | | | | | | | | | | | | | | | | | | | | | | | | | | |
|---|---|---|---|---|---|---|---|---|---|---|---|---|---|---|---|---|---|---|---|---|---|---|---|---|---|---|---|
| Rbm14 | Q8C2Q3 | 0.03 | -2.78 | -1.8 | -1.23 | -2.85 | -1.14 | -2.1 | -1.09 | -1.81 | -1.1 | -1.13 | -2.29 | -0.58 | -1.2 | -0.67 | -1.82 | -0.35 | -0.1375 | 0.3775 | 0.596621 | 0.818515 | 0.532621 | | 1 | 0.996511 | 0.911094 |
| Rbm15 | Q0VBL3-2 | 0.59 | -1.11 | -0.96 | -0.77 | -0.53 | -0.09 | -0.46 | -0.07 | -0.06 | -0.74 | -0.97 | -0.66 | 0.7 | -0.31 | 0.37 | -0.42 | 0.275 | -0.045 | 0.6475 | 0.483886 | 0.913489 | 0.173691 | | 1 | 0.996755 | 0.764376 |
| Rbm15b | Q6PHZ5 | 0.43 | 0.09 | -0.3 | -0.47 | -0.1 | 0.1 | -0.17 | 0.38 | 0.04 | -0.03 | -0.47 | 0 | 0.54 | -0.51 | 0.64 | -0.35 | 0.115 | -0.0525 | 0.1425 | 0.645917 | 0.837185 | 0.689466 | | 1 | 0.996511 | 0.955729 |
| Rbm17 | Q8JZX4 | -0.84 | -1.72 | -0.94 | -1.17 | -1.69 | -1.31 | -0.83 | -0.82 | -1.66 | -0.47 | -1.08 | -1.43 | -1.14 | -1.03 | -0.14 | -0.74 | 0.005 | 0.0075 | 0.405 | 0.986146 | 0.981539 | 0.211264 | | 1 | 0.998773 | 0.786269 |
| Rbm19 | Q8R3C6 | -0.36 | -0.89 | -0.5 | -0.76 | -1.17 | -0.16 | -0.93 | -1.37 | 0.55 | -2.58 | -0.08 | -0.84 | 0.9 | -0.54 | 0.61 | -0.8 | -0.28 | -0.11 | 0.67 | 0.349347 | 0.861016 | 0.135033 | | 1 | 0.996511 | 0.731223 |
| Rbm22 | Q8BHS3 | 0.61 | -0.65 | -0.69 | -0.65 | -1.6 | 0.01 | -1.44 | -0.3 | -0.15 | -0.31 | -0.78 | -0.9 | 0.99 | -0.03 | 0.05 | -1.33 | -0.4875 | -0.19 | 0.265 | 0.320818 | 0.597786 | 0.62014 | | 1 | 0.996511 | 0.93807 |
| Rbm25 | B2RY56 | -0.59 | -2 | -2.36 | -2.95 | -1.8 | -1.42 | -2.13 | -1.89 | -1.41 | -1.55 | -2.48 | -2.25 | -0.16 | -0.94 | -0.64 | -2.33 | 0.165 | 0.0525 | 0.9575 | 0.733587 | 0.920084 | 0.151837 | | 1 | 0.996755 | 0.750422 |
| Rbm26 | E9Q640 | -1.77 | -3.86 | -3.38 | -3.43 | -3.41 | -2.64 | -3.39 | -3.37 | -2.78 | -2.3 | -2.83 | -2.59 | -2.91 | -1.96 | -3.86 | -3.84 | -0.0925 | 0.485 | -0.0325 | 0.840999 | 0.294786 | 0.95613 | | 1 | 0.996511 | 0.992636 |
| Rbm27 | Q5SFM8 | 0.13 | -1.43 | -2.11 | -2.11 | -1.41 | -0.56 | -1.52 | -2 | -1.59 | -0.92 | -1.78 | -0.06 | -1.44 | -2.27 | -1.86 | -1.47 | 0.0075 | 0.2925 | -0.38 | 0.989229 | 0.628893 | 0.473877 | | 1 | 0.996511 | 0.891644 |
| Rbm28 | Q8CGC6 | 0.97 | 0.5 | 0.68 | 0.68 | 0.16 | 0.31 | 0.03 | 0.28 | 0.48 | 1.06 | 0.47 | 0.05 | 1 | 0.77 | 0.58 | 0.32 | -0.5125 | -0.1925 | -0.04 | 0.0139 | 0.452211 | 0.857609 | | 1 | 0.996511 | 0.976225 |
| Rbm3 | O89086 | -3.1 | -3.8 | -1.91 | -2.26 | -4.27 | -3.55 | -2.28 | -1.92 | -3.57 | -1.72 | -1.78 | -3.37 | -2.79 | -3.23 | -1.5 | -2.51 | -0.2375 | 0.1575 | 0.26 | 0.707504 | 0.793336 | 0.620336 | | 1 | 0.996511 | 0.93807 |
| Rbm34 | A0A1D5RLI | 0.98 | 0.99 | 0.89 | 0.84 | -0.15 | 0.86 | 0.47 | 0.61 | 0.71 | 0.71 | 0.82 | 0.71 | 0.48 | 0.32 | 0.54 | 0.32 | -0.4775 | -1.88E-01 | -0.51 | 0.068843 | 0.245362 | 0.015241 | | 1 | 0.996511 | 0.589387 |
| Rbm39 | Q8VH51-2 | 0.1 | -1.16 | -0.58 | -1 | -1.03 | 0.01 | -0.62 | -0.58 | -0.94 | -0.83 | -0.74 | -0.89 | -0.2 | -0.15 | -0.71 | -0.39 | 0.105 | -0.19 | 0.2975 | 0.758848 | 0.521364 | 0.359314 | | 1 | 0.996511 | 0.856645 |
| Rbm42 | D3YYW8 | -0.47 | -1.63 | -1.19 | -1.2 | -1.54 | -1.39 | -1.38 | -0.64 | -2.18 | -1.58 | -1.39 | -2.06 | -1.06 | -1.11 | 0.01 | -1.25 | -0.115 | -0.68 | 0.27 | 0.711268 | 0.052309 | 0.471833 | | 1 | 0.996511 | 0.891144 |
| Rbm45 | Q8BHN5 | -0.49 | -2.66 | -0.72 | -0.56 | -1.81 | -2.01 | -1.4 | -0.08 | -1.92 | -0.87 | -0.24 | -2.24 | -0.05 | -0.9 | 0.89 | -0.18 | -0.2175 | -0.21 | 1.0475 | 0.725065 | 0.741866 | 0.100701 | | 1 | 0.996511 | 0.698156 |
| Rbm4b | Q8VE92 | -1.7 | -0.9 | -0.49 | 1.12 | -0.96 | -1.02 | 1.23 | -0.09 | 0.17 | -0.58 | 0.1 | -1.19 | -0.05 | 1.2 | -1.4 | 0.05 | 0.2825 | 0.1175 | 0.4425 | 0.694671 | 0.848957 | 0.544071 | | 1 | 0.996511 | 0.912212 |
| Rbm5 | Q91YE7-2 | -0.66 | -0.65 | -1.37 | -1.61 | -0.73 | -1.43 | -1.22 | -1.21 | -1.33 | -0.6 | -1.66 | -1.44 | -0.4 | -0.55 | 0.24 | -0.33 | -0.075 | -0.185 | 0.8125 | 0.794715 | 0.582321 | 0.02625 | | 1 | 0.996511 | 0.633075 |
| Rbm6 | S4R1W5 | 1.25 | -0.42 | -0.6 | -0.7 | -0.72 | 0.23 | -0.43 | -0.34 | 0.83 | -0.14 | -0.48 | 0.18 | 1.77 | -0.51 | 1.2 | -0.22 | -0.1975 | 0.215 | 0.6775 | 0.672308 | 0.668642 | 0.314558 | | 1 | 0.996511 | 0.836924 |
| Rbm7 | Q9CQT2 | -0.17 | -2.6 | -1.29 | -1.45 | -3.75 | -0.6 | -2.16 | -1.41 | 0.08 | -1.27 | -1.4 | -2.45 | 1.1 | -3.44 | -1.23 | -1.97 | -0.6025 | 0.1175 | -0.0075 | 0.433045 | 0.857705 | 0.993746 | | 1 | 0.996511 | 0.997248 |
| Rbm8a | Q9CWZ3 | 1.18 | -0.35 | -0.34 | -1.14 | -0.5 | 0.45 | -0.38 | -0.24 | -1.64 | -1.08 | -0.2 | -0.51 | -0.44 | -1.48 | 0.15 | -0.34 | -0.005 | -0.695 | -0.365 | 0.99184 | 0.217225 | 0.511155 | | 1 | 0.996511 | 0.90359 |
| Rbms3 | G3UZJ9 | -1.41 | -2.01 | -3.34 | -2.37 | -1.28 | -1.13 | -1.52 | -2.79 | -1.65 | -1.14 | -2.6 | -1.27 | 0.01 | -1.01 | -1.08 | -2.15 | 0.6025 | 0.6175 | 1.225 | 0.257125 | 0.226076 | 0.050551 | | 1 | 0.996511 | 0.658454 |
| Rbmx | Q9WV02 | 1.72 | -0.16 | 0.33 | 0.23 | -1.13 | 1.03 | -0.33 | 0.65 | 0.28 | -1.34 | -0.01 | -0.02 | 1.34 | -1.3 | 0.48 | -0.4 | -0.475 | -0.8025 | -0.5 | 0.423975 | 0.141466 | 0.442357 | | 1 | 0.996511 | 0.883549 |
| Rbmx2 | Q8R0F5 | -0.99 | -1.05 | -0.07 | -1.14 | -0.7 | -1.24 | -0.39 | -0.43 | 0.32 | -2.2 | -0.65 | -1.49 | 1.06 | 2.27 | 1.11 | 0.32 | 0.1225 | -0.1925 | 2.0025 | 0.695568 | 0.727789 | 0.001619 | | 1 | 0.996511 | 0.419079 |
| Rbp1 | Q00915 | -0.54 | -0.95 | -0.15 | -0.09 | -0.69 | -0.42 | 0.7 | 0.25 | -1.82 | -0.99 | -0.42 | -0.66 | -1.99 | -1.2 | -0.42 | -0.81 | 0.3925 | -0.54 | -0.6725 | 0.289992 | 0.155761 | 0.104141 | | 1 | 0.996511 | 0.702979 |
| Rbsn | Q80Y56 | 0.5 | 0.6 | 0.03 | 0.21 | 0.31 | 0.47 | 0.07 | 0.03 | 0.65 | 0.22 | 0.14 | 0.58 | 0.76 | 0.57 | -0.06 | -0.15 | -0.115 | 0.0625 | -0.055 | 0.569163 | 0.776735 | 0.845121 | | 1 | 0.996511 | 0.976225 |
| Rbx1 | P62878 | 0.56 | -0.54 | -0.21 | 1.39 | -0.51 | 0.29 | -0.76 | 1.08 | 1.1 | 2.17 | 0.62 | 0.77 | 0.45 | 0.61 | 0.26 | -0.28 | -0.275 | 0.865 | -0.04 | 0.618952 | 0.120226 | 0.928556 | | 1 | 0.996511 | 0.98946 |
| Rc3h1 | Q4VGL6 | 0.11 | -0.46 | -0.54 | -0.8 | -0.8 | 0.1 | -0.65 | -0.87 | 0.39 | -0.35 | -0.63 | 0.31 | 0.58 | 0.07 | -0.41 | -0.82 | -0.1325 | 0.3525 | 0.2775 | 0.653486 | 0.282469 | 0.44195 | | 1 | 0.996511 | 0.883549 |
| Rc3h2 | A2AVP4 | -1.52 | -1.37 | -3.43 | -3.43 | -2.03 | -1.87 | -2.85 | -2.73 | -0.79 | -0.55 | -3.41 | -1.01 | -0.66 | -0.61 | -2.06 | -3.01 | 0.0675 | 0.9975 | 0.8525 | 0.905817 | 0.229948 | 0.267256 | | 1 | 0.996511 | 0.818895 |
| Rcan1 | Q9JHG6 | -0.62 | 0.2 | 0.1 | 0.01 | 0.07 | -0.72 | 0.06 | -0.2 | 0.04 | 0.2 | 0.1 | -0.25 | 0.05 | 0.68 | -0.15 | -0.38 | -0.12 | 0.1 | 0.1275 | 0.656065 | 0.674982 | 0.677023 | | 1 | 0.996511 | 0.953398 |
| Rcan2 | A0A2R8VKI | -3.38 | -4.65 | 0.46 | -0.1 | -4.07 | -3.5 | 1.15 | 0.85 | -4.31 | -0.17 | 0.46 | -3.05 | -4.1 | -2.58 | 0.21 | 0.1 | 0.525 | 0.15 | 0.325 | 0.752631 | 0.920653 | 0.822642 | | 1 | 0.996755 | 0.972768 |
| Rcan3 | Q9JKK0 | -0.47 | 0.12 | -1.17 | -0.34 | 0.45 | -0.13 | -0.4 | -0.97 | -0.13 | 0.52 | -0.48 | 0.18 | -0.42 | -0.23 | -0.14 | -1.28 | 0.2025 | 0.4875 | -0.0525 | 0.595164 | 0.174062 | 0.886032 | | 1 | 0.996511 | 0.979767 |
| Rcbtb1 | A0A0R4J02 | 0.86 | 0.64 | 0.69 | 0.22 | 0.88 | 0.63 | 0.68 | 0.8 | 0.25 | -0.95 | 0.71 | 1.2 | -0.37 | 0.3 | -0.12 | 1.02 | 0.145 | -0.3 | -0.395 | 0.447343 | 0.511707 | 0.257376 | | 1 | 0.996511 | 0.812003 |
| Rcbtb2 | E9Q3B1 | -0.39 | -0.02 | 0.06 | -0.2 | -0.51 | -0.42 | -0.17 | -0.36 | 0.65 | 0.17 | -0.4 | 0.12 | -0.05 | -0.04 | -0.16 | 0.3 | -0.2275 | 0.2725 | 0.15 | 0.212037 | 0.305474 | 0.469859 | | 1 | 0.996511 | 0.890289 |
| Rcc1 | Q8VE37 | 0.77 | -0.86 | -0.62 | -0.11 | -0.8 | -0.05 | -1.09 | -0.13 | -0.61 | 0.08 | -0.1 | -0.78 | -0.59 | -0.77 | -0.49 | -0.39 | -0.3125 | -0.1475 | -0.355 | 0.458402 | 0.710949 | 0.34251 | | 1 | 0.996511 | 0.848473 |
| Rcc2 | Q8BK67 | -4.72 | -5.68 | -3.71 | -4.66 | -5.5 | -5.35 | -3.75 | -4.62 | -4.53 | -3.92 | -3.67 | -4.77 | -4.16 | -4.08 | -3.62 | -4.37 | -0.1125 | 0.47 | 0.635 | 0.829497 | 0.310977 | 0.151266 | | 1 | 0.996511 | 0.749304 |
| Rchy1 | G3UWD8 | -2.11 | -2.18 | -1.69 | -1.79 | -1.89 | -1.46 | -1.98 | -2.03 | -1.26 | -0.13 | -1.44 | -1.45 | 0.15 | -1.8 | -1.49 | -1.23 | 0.1025 | 0.8725 | 0.85 | 0.621747 | 0.027163 | 0.072454 | | 1 | 0.996511 | 0.668066 |
| Rcl1 | Q9JJT0 | 1.52 | 1.55 | -0.26 | -0.6 | 1.59 | 0.97 | -0.34 | -1.57 | 0.89 | -1.24 | -0.22 | 1.02 | 0.35 | 1.28 | -0.33 | -0.13 | -0.39 | -0.44 | -0.26 | 0.635252 | 0.538862 | 0.674877 | | 1 | 0.996511 | 0.952944 |
| Rcn1 | Q05186 | 0.45 | 0.01 | -0.15 | -0.1 | 0.35 | 0.12 | 0.88 | 0.44 | -0.59 | -0.87 | 0.14 | 1.04 | -0.36 | -0.38 | 1.58 | 1.04 | 0.395 | -0.1225 | 0.4175 | 0.112713 | 0.77337 | 0.397683 | | 1 | 0.996511 | 0.867334 |
| Rcn2 | Q8BP92 | 1.1 | 1.27 | 1.31 | 0.41 | 1.44 | 1.13 | 1.98 | 1.14 | -1.07 | -0.48 | 1.09 | 1.43 | -0.17 | -0.38 | 2.56 | 2.33 | 0.4 | -0.78 | 0.0625 | 0.190921 | 0.206604 | 0.932387 | | 1 | 0.996511 | 0.989713 |
| Rcor1 | A0A140T8I | -0.77 | -1.28 | -0.81 | -1.09 | -0.78 | -1.31 | -0.22 | -1.41 | -1.18 | -1.18 | -0.72 | -0.79 | -1.28 | -0.69 | -1.06 | -0.98 | 0.0575 | 0.02 | -0.015 | 0.846882 | 0.925341 | 0.945973 | | 1 | 0.996755 | 0.992376 |
| Rcsd1 | Q3UZA1-2 | -1.15 | -1.87 | -1.16 | -1.6 | -2.54 | -1.37 | -2.9 | -2.16 | 0.3 | -0.64 | -1.49 | -1.78 | 0.23 | 1.03 | -2.31 | -2.12 | -0.7975 | 0.5425 | 0.6525 | 0.049471 | 0.265593 | 0.40853 | | 1 | 0.996511 | 0.874356 |
| Rdh11 | Q9R1R8 | 1.11 | 1.02 | 1.15 | 1.12 | 1.34 | 0.87 | 1.42 | 0.92 | -0.02 | 0.03 | 1.12 | 1.27 | 0.2 | 0.18 | 1.58 | 1.39 | 0.0375 | -0.5 | -0.2625 | 0.839846 | 0.168241 | 0.483795 | | 1 | 0.996511 | 0.894091 |
| Rdh13 | Q8CEE7 | 0.5 | 0.67 | 0.37 | 0.57 | 0.13 | 0.49 | -0.09 | 0.27 | 0.41 | 0.76 | 0.39 | 0.51 | -0.01 | 0.57 | -0.35 | -0.21 | -0.3275 | -0.01 | -0.5275 | 0.098612 | 0.9547 | 0.057766 | | 1 | 0.996755 | 0.661738 |
| Rdh14 | Q9ERI6 | 0.85 | 0.82 | 0.36 | 0.4 | 1 | 0.99 | 0.08 | 0.26 | 0.75 | 0.83 | 0.43 | 0.84 | 1.14 | 0.75 | 0.41 | 0.16 | -0.025 | 0.105 | 0.0075 | 0.928321 | 0.615606 | 0.978091 | | 1 | 0.996511 | 0.993727 |
| Rdx | P26043 | 0.53 | 0.44 | 0.18 | -0.14 | 0.59 | 0.4 | -0.36 | -0.29 | 0.53 | -0.45 | -0.11 | 0.52 | 0.6 | -0.28 | 0.35 | -0.31 | -0.1675 | -0.13 | -0.1625 | 0.560711 | 0.659275 | 0.578548 | | 1 | 0.996511 | 0.927198 |
| Recql5 | Q8VID5 | 13.46 | 2.69 | 16.26 | -0.27 | 9.94 | 13.92 | 16.94 | 11.65 | 1.22 | -0.13 | 15.76 | 13.44 | 1.59 | 1.16 | 20.09 | 21.54 | 5.0775 | -0.4625 | 3.06 | 0.207666 | 0.927756 | 0.618695 | | 1 | 0.996755 | 0.937856 |
| Reep1 | Q8BGH4 | 0.5 | 0.28 | 0.83 | 0.92 | 0.47 | 0.05 | 0.8 | 1.07 | 0.91 | 1.2 | 0.87 | 0.52 | 1.03 | 0.92 | 1.17 | 1.16 | -0.035 | 0.2425 | 0.4375 | 0.897089 | 0.314928 | 0.067411 | | 1 | 0.996511 | 0.664936 |
| Reep2 | Q8VCD6 | 0.83 | 0.63 | 0.34 | 0.42 | 0.53 | 0.79 | -0.04 | 0.74 | 1.35 | 0.59 | 0.58 | 0.95 | 1.8 | 0.65 | 0.44 | 0.51 | -0.05 | 0.3125 | 0.295 | 0.832742 | 0.215883 | 0.394542 | | 1 | 0.996511 | 0.866125 |
| Reep3 | A0A1W2P8 | -0.29 | -0.24 | 0.09 | 0.01 | -0.72 | -0.25 | -0.08 | -0.58 | 1.35 | -0.85 | 0.2 | -0.87 | 1.5 | -1.29 | 0.3 | 0.53 | -0.3 | 0.065 | 0.3675 | 0.167931 | 0.896097 | 0.503132 | | 1 | 0.996511 | 0.901002 |
| Reep5 | G3X8R0 | 2.95 | 1.43 | 0.84 | 0.21 | 1.17 | 2.65 | 0.51 | 0.52 | 2.77 | -0.19 | 0.72 | 3.32 | 2 | 1.58 | 0.76 | 0.92 | -0.145 | 0.2975 | -0.0425 | 0.83594 | 0.745904 | 0.943398 | | 1 | 0.996511 | 0.991596 |
| Reep6 | Q9JM62 | 0.92 | 0.84 | 0.51 | 0.71 | 1.07 | 0.95 | 0.92 | 0.76 | 0.95 | 0.65 | 0.66 | 0.98 | 0.16 | 0.68 | 0.36 | 0.49 | 0.18 | 0.065 | -0.3225 | 0.295444 | 0.728444 | 0.140965 | | 1 | 0.996511 | 0.737973 |
| Rel | A4QPD3 | -0.06 | 1.02 | 0.97 | 1.57 | -0.05 | 0.19 | 0.57 | 0.19 | 0.21 | 2.16 | 0.25 | 0.29 | 0.4 | 1.57 | 0.23 | -0.14 | -0.65 | -0.1475 | -0.36 | 0.090671 | 0.785619 | 0.452814 | | 1 | 0.996511 | 0.885993 |
| Rela | A0A494BA | -0.35 | -0.11 | -0.48 | -0.55 | -0.37 | 0.44 | -0.86 | -0.55 | -0.21 | -0.25 | -0.59 | -0.32 | 0.05 | -0.03 | -0.63 | -0.91 | 0.0375 | 0.03 | -0.0075 | 0.898196 | 0.873197 | 0.978204 | | 1 | 0.996511 | 0.993727 |
| Relch | E9QM90 | 0.33 | 0.23 | 0.6 | 0.12 | 0.24 | 0.49 | 0.73 | 0.28 | 0.31 | 0.15 | 0.61 | 0.53 | 0 | 0.2 | 0.2 | 0.49 | 0.115 | 0.08 | -0.0975 | 0.550572 | 0.687106 | 0.637639 | | 1 | 0.996511 | 0.942804 |
| Relch | A0A087WS | -1.09 | -0.79 | -0.81 | 0.5 | -1.13 | -0.97 | -1.45 | 0.01 | 0.16 | 1.03 | -0.6 | -0.59 | 0.19 | 0.02 | -1.47 | -1.52 | -0.3375 | 0.5475 | -0.1475 | 0.453772 | 0.281321 | 0.785083 | | 1 | 0.996511 | 0.967642 |
| Rell1 | Q8K2J7-2 | -0.72 | -1.92 | 0.35 | 0.57 | -1.64 | -0.86 | 0.93 | 1.23 | -2.4 | 1.42 | 0.99 | -0.42 | -0.43 | -2.54 | 0.97 | 1.13 | 0.345 | 0.3275 | 0.2125 | 0.671665 | 0.725393 | 0.817055 | | 1 | 0.996511 | 0.972768 |
| Rell2 | Q8BRJ3 | 0.75 | 0.63 | 0.38 | 0.6 | -0.21 | 0.74 | -0.32 | 0.19 | 1.33 | 0.69 | 0.42 | 0.93 | 1.41 | 0.79 | -0.01 | -0.2 | -0.49 | 0.2525 | -0.0925 | 0.087621 | 0.302518 | 0.803777 | | 1 | 0.996511 | 0.97131 |
| Reln | Q60841-3 | 0.31 | 0.15 | -0.38 | 0.03 | 0.12 | 0.18 | -0.23 | 0.04 | 0.22 | 0.17 | -0.17 | 0.35 | 0.06 | 0.42 | -0.15 | -0.09 | -3.9E-17 | 0.115 | 0.0325 | 1 | 0.605782 | 0.890496 | | 1 | 0.996511 | 0.981669 |
| Rem2 | E9Q4D5 | 0.06 | -0.29 | -0.05 | 0.15 | -0.9 | -1.01 | 0.31 | 0.17 | -0.05 | 0.75 | -0.02 | 0.11 | 0.14 | 0.12 | 0.78 | -0.09 | -0.325 | 0.23 | 0.27 | 0.360828 | 0.347953 | 0.28993 | | 1 | 0.996511 | 0.825134 |
| Renbp | D6RHA2 | -1.03 | -1.95 | -3.64 | -3.75 | -1.65 | -1.34 | -5.38 | -4.27 | -1.77 | -2.52 | -3.32 | -1.63 | -1.1 | -0.96 | -2.57 | -3.71 | -0.5675 | 0.2825 | 0.5075 | 0.598106 | 0.68763 | 0.548761 | | 1 | 0.996511 | 0.912757 |
| Reps1 | O54916 | 0.35 | 0.4 | 0 | 0.1 | 0.01 | 0.34 | -0.38 | -0.12 | 0.16 | 0.45 | -0.07 | 0.18 | -0.01 | 0.01 | -0.29 | -0.38 | -0.25 | -0.0325 | -0.38 | 0.248003 | 0.868311 | 0.087819 | | 1 | 0.996511 | 0.691619 |
| Reps2 | B9EI38 | 0.03 | 0.14 | -0.17 | 0.08 | 0.01 | -0.21 | -0.58 | -0.2 | 0.1 | 0.46 | -0.15 | -0.1 | -0.15 | 0.45 | -0.15 | -0.66 | -0.265 | 0.0575 | -0.1475 | 0.172581 | 0.77629 | 0.579556 | | 1 | 0.996511 | 0.927734 |
| Rer1 | Q9CQU3 | -2.24 | -1.93 | -2.41 | -2 | -1.14 | -1.65 | -0.76 | -2.26 | -1.99 | -1.73 | -2.16 | -2.04 | -0.89 | -0.71 | -1.14 | -1.69 | 0.6925 | 0.165 | 1.0375 | 0.063759 | 0.408854 | 0.003605 | | 1 | 0.996511 | 0.513346 |
| Retreg1 | Q8VE91 | -0.08 | 0.09 | 0.02 | 0.47 | -0.01 | -0.26 | 0.86 | 0.49 | 0.16 | 0.16 | 0.11 | -0.01 | 1 | 0.6 | 0.18 | 0.72 | 0.145 | -0.02 | 0.5 | 0.608469 | 0.914618 | 0.067106 | | 1 | 0.996755 | 0.664936 |
| Retreg2 | Q6NS82-3 | 0.3 | 0.55 | -0.68 | -0.82 | 0.54 | 1.99 | -0.96 | -1.09 | 0.69 | -0.08 | -1.01 | 0.2 | 0.77 | 0.75 | -0.9 | -0.83 | 0.2825 | 0.1125 | 0.11 | 0.698191 | 0.808948 | 0.838144 | | 1 | 0.996511 | 0.975961 |
| Retreg3 | Q9CQV4 | -0.38 | -0.56 | 0.28 | 0.62 | -0.98 | -0.04 | 0.39 | -0.06 | 1.47 | -2.96 | 0.41 | -0.52 | 1.59 | 0.6 | -0.01 | 0.33 | -0.1625 | -0.39 | 0.6375 | 0.670178 | 0.661995 | 0.156587 | | 1 | 0.996511 | 0.751493 |
| Retsat | Q64FW2 | -0.22 | -0.46 | -1.88 | -2.11 | 0.49 | -0.83 | -1.6 | -1.78 | 1.45 | -1.16 | -2.07 | -0.36 | 2.14 | 1.81 | -2.43 | -2.12 | 0.2375 | 0.6325 | 1.0175 | 0.712469 | 0.440326 | 0.398025 | | 1 | 0.996511 | 0.867334 |
| Rexo2 | Q9D8S4 | 0.48 | 0.55 | 0.99 | 0.72 | 0.36 | 0.76 | 1.43 | 1.2 | -0.07 | 0.54 | 1.18 | 0.55 | -0.23 | 0.02 | 1.21 | 1.39 | 0.2525 | -0.135 | -0.0875 | 0.360857 | 0.641963 | 0.830601 | | 1 | 0.996511 | 0.973939 |
| Rexo4 | A2ALB1 | 0.86 | 0.29 | 0.98 | 0.82 | -0.42 | -0.11 | 0.58 | 0.59 | -0.07 | 0.46 | 1.08 | 0.43 | 0.2 | -0.36 | 0.55 | 0.54 | -0.5775 | -0.2625 | -0.505 | 0.077518 | 0.376691 | 0.101035 | | 1 | 0.996511 | 0.698156 |
| Rfc2 | Q9WUK4 | -1.92 | -1.07 | -1.31 | -1.18 | -0.33 | -2.45 | -1.07 | -1.49 | -2.66 | -1.08 | -1.64 | -1.69 | -2.19 | -1 | -1.75 | -1.47 | 0.035 | -0.3975 | -0.2325 | 0.937701 | 0.296362 | 0.474291 | | 1 | 0.996511 | 0.891644 |
| Rfc3 | Q8R323 | -0.93 | -0.94 | -0.38 | -0.51 | -0.57 | -1.32 | 0.14 | -0.91 | 0.61 | -1.96 | -0.11 | -0.01 | -0.97 | -1.67 | 0.08 | -0.54 | 0.025 | 0.3225 | -0.085 | 0.939758 | 0.546987 | 0.824907 | | 1 | 0.996511 | 0.972852 |
| Rfc4 | D6RGM1 | 0.01 | -0.03 | 0.31 | 0.58 | -1.9 | 1.43 | -0.83 | -0.26 | 1.83 | 1.68 | 0.07 | 0.69 | 0.96 | 0.07 | -0.44 | -0.42 | -0.6075 | 0.85 | -0.175 | 0.359103 | 0.069161 | 0.62337 | | 1 | 0.996511 | 0.938628 |
| Rfc5 | Q9D0F6 | 1.99 | 1.3 | -0.07 | 0.81 | 0.69 | 0.54 | -1.83 | -2.78 | 1.64 | 0.23 | -7.13 | 0.95 | 1.44 | 1.4 | -0.05 | 1.39 | -1.8525 | -2.085 | 0.0375 | 0.058965 | 0.279475 | 0.943038 | | 1 | 0.996511 | 0.991519 |
| Rfesd | Q8K2P6 | -0.31 | -0.34 | 2.26 | 0.89 | -0.59 | -0.48 | 0.15 | 0.38 | 0.68 | 1.6 | 0.5 | -0.19 | 0.68 | 0.25 | -0.18 | 0.05 | -0.76 | 0.0225 | -0.425 | 0.227929 | 0.972549 | 0.477478 | | 1 | 0.997515 | 0.892234 |
| Rfk | Q8CFV9 | 1 | 0.52 | 0.5 | 0.11 | 1.03 | 0.7 | 1.01 | 0.7 | -2.39 | -1.47 | 0.33 | 0.81 | -1.89 | -2.23 | 0.25 | 0.62 | 0.3275 | -1.2125 | -1.345 | 0.171568 | 0.112518 | 0.075139 | | 1 | 0.996511 | 0.673156 |
| Rft1 | A0A2I3BRR | -1.25 | -1.03 | -3.64 | -4.87 | -2.4 | 0.65 | -6.39 | -5.12 | 0.13 | -2.75 | -4.67 | -1.61 | 1.75 | -2.48 | -3.31 | -2.91 | -0.6175 | 0.4725 | 0.96 | 0.704767 | 0.701915 | 0.478753 | | 1 | 0.996511 | 0.892849 |
| Rftn1 | Q6A0D4 | 0.48 | 0.5 | 1.07 | 0.77 | 0.91 | 0.68 | 0.84 | 1.22 | 1.45 | 1.31 | 0.85 | 1.02 | 1.1 | 0.99 | 1.42 | 0.88 | 0.2075 | 0.4525 | 0.3925 | 0.332522 | 0.073496 | 0.111439 | | 1 | 0.996511 | 0.713445 |
| Rftn2 | Q8CHX7 | 1.07 | 1.37 | 0.82 | 0.5 | 1.07 | 1.58 | 0.35 | 0.16 | 0.9 | -0.25 | 0.5 | 1.12 | 1.29 | -0.05 | 1.18 | 0.7 | -0.15 | -0.3725 | -0.16 | 0.678675 | 0.299473 | 0.652337 | | 1 | 0.996511 | 0.947633 |
| Rfx3 | G3UWP0 | -2.75 | -2.34 | -1.33 | -1.84 | -2.36 | -3.08 | -1.28 | -1.5 | -2.43 | -2.23 | -2.01 | -2.46 | -1.27 | -1.29 | -1.42 | -0.97 | 0.01 | -0.2175 | 0.8275 | 0.983265 | 0.507785 | 0.030122 | | 1 | 0.996511 | 0.641481 |
| Rfxap | F7DBT7 | -0.94 | -0.12 | -1.1 | -1.85 | -1.01 | -0.54 | -2.73 | -2.3 | -0.8 | -2.07 | -2.24 | -0.4 | -1.75 | 0.16 | -2.86 | -1.46 | -0.6425 | -0.375 | -0.475 | 0.281242 | 0.489835 | 0.47416 | | 1 | 0.996511 | 0.891644 |
| Rgcc | Q9DBX1 | -0.02 | 1.22 | -0.4 | -1.04 | -0.32 | -0.96 | -0.84 | -1.95 | -0.06 | -0.92 | -0.14 | -0.38 | 0.24 | 1.19 | -0.64 | -0.92 | -0.9575 | -0.315 | 0.0275 | 0.102473 | 0.516769 | 0.964346 | | 1 | 0.996511 | 0.9927 |
| Rgl1 | E9Q8M9 | -0.78 | 0.01 | 0.05 | -0.21 | -0.56 | -0.57 | 0.28 | 0.3 | -0.15 | -0.88 | 0.36 | -0.46 | -0.02 | -0.1 | 0.14 | -0.03 | 0.095 | -0.05 | 0.23 | 0.758177 | 0.87704 | 0.346841 | | 1 | 0.996511 | 0.851134 |
| Rgl2 | Q61193 | 0.81 | 0.21 | -0.1 | 0.39 | 0.12 | 0.52 | -0.42 | -0.23 | 0.9 | 0.39 | -0.41 | 0.66 | 0.59 | 0.83 | -0.28 | -0.05 | -0.33 | 0.0575 | -0.055 | 0.262788 | 0.86492 | 0.866579 | | 1 | 0.996511 | 0.978305 |
| Rgma | Q8C4C4 | 0.68 | 0.32 | -0.75 | -0.79 | -0.16 | 0.98 | -0.75 | -0.52 | 1.7 | -0.48 | -0.7 | 1.12 | 2.4 | -0.25 | 0.26 | -0.77 | 0.0225 | 0.545 | 0.545 | 0.963694 | 0.403804 | 0.454062 | | 1 | 0.996511 | 0.88691 |
| Rgmb | Q7TQ33 | -0.73 | 1.97 | -1.08 | -0.1 | 0.43 | -0.38 | -0.2 | -0.09 | -0.11 | -0.09 | -0.02 | 1.61 | -1.19 | -0.89 | -0.81 | -0.15 | -0.075 | 0.3325 | -0.775 | 0.906687 | 0.649609 | 0.254603 | | 1 | 0.996511 | 0.809524 |
| Rgp1 | Q8BHT7 | -0.48 | -0.26 | 0.67 | 1.37 | -1.53 | -1.93 | 0.34 | 1.36 | -0.68 | 1.15 | 0.72 | -1.19 | -1.91 | 0.27 | 1.62 | 0.6 | -0.765 | -0.325 | -0.18 | 0.352433 | 0.614875 | 0.816187 | | 1 | 0.996511 | 0.972768 |
| Rgs10 | Q9CQE5 | -0.86 | -0.65 | -0.71 | -0.71 | -0.79 | -1.06 | -0.72 | -0.58 | -0.85 | 0.05 | -0.66 | -0.76 | -1.02 | -0.48 | -0.85 | -0.35 | -0.055 | 0.1775 | 0.0575 | 0.741574 | 0.46391 | 0.790571 | | 1 | 0.996511 | 0.969105 |
| Rgs12 | E9Q652 | 1.04 | 1.38 | 1.13 | 0.99 | 1.17 | 1.34 | 0.71 | 1.19 | 1.07 | 0.47 | 1.28 | 1.57 | 1.44 | 0.31 | 1.14 | 0.94 | -0.0325 | -0.0375 | -0.1775 | 0.868547 | 0.88771 | 0.525465 | | 1 | 0.996511 | 0.908107 |
| Rgs14 | P97492 | 0.67 | 0.74 | 0.64 | 0.54 | 0.59 | 0.35 | 1 | 0.37 | 0.37 | -0.17 | 0.63 | 0.96 | 0.09 | 0.27 | 0.82 | 0.84 | -0.07 | -0.2 | -0.1425 | 0.718081 | 0.451525 | 0.550947 | | 1 | 0.996511 | 0.913801 |
| Rgs17 | G5E8E0 | 0.34 | 0.49 | 0.22 | 0.59 | 1.08 | 0.36 | 0.92 | 0.62 | -0.04 | -0.15 | 0.57 | 0.43 | -0.41 | 0.09 | 0.05 | 0.64 | 0.335 | -0.2075 | -0.3175 | 0.13478 | 0.372838 | 0.240775 | | 1 | 0.996511 | 0.795658 |
| Rgs19 | Q9CX84 | 0.3 | 0.7 | 0.65 | 0.86 | 0.74 | 0.51 | 0.31 | 0.15 | 0.94 | 0.79 | 0.71 | 0.79 | 0.7 | 1.06 | 0.56 | 0.33 | -0.2 | 0.18 | 0.035 | 0.341727 | 0.349102 | 0.881494 | | 1 | 0.996511 | 0.979767 |
| Rgs20 | Q9QZB1 | 0.84 | 0.98 | 0.94 | 0.99 | 0.86 | 0.99 | 1.16 | 1.25 | 1.12 | -0.48 | 0.87 | 1.04 | 1.21 | -0.02 | 0.88 | 0.81 | 0.1275 | -0.3 | -0.2175 | 0.428978 | 0.422444 | 0.450213 | | 1 | 0.996511 | 0.884817 |
| Rgs3 | Q9DC04-4 | 0.26 | 0.16 | 0.61 | 0.58 | 0.67 | -0.08 | 0.08 | -0.33 | 0.85 | 1.01 | 0.4 | 0.2 | 0.7 | 0.56 | 0.19 | 0.53 | -0.3175 | 0.2125 | 0.0925 | 0.22922 | 0.397388 | 0.6653 | | 1 | 0.996511 | 0.949825 |
| Rgs4 | O08899 | 0.23 | 0.01 | 0.27 | 0.6 | 0.11 | 0.24 | -1.7 | 0.62 | 0.05 | 2.17 | -0.26 | 0.41 | -0.24 | 0.43 | 1.27 | 0.54 | -0.46 | 0.315 | 0.2225 | 0.361377 | 0.546144 | 0.511147 | | 1 | 0.996511 | 0.90359 |
| Rgs6 | A0A087WS | 0.4 | 0.39 | 0.59 | 0.63 | 0.61 | 0.31 | 0.49 | 0.48 | 0.27 | -0.02 | 0.48 | 0.19 | -0.45 | 0.25 | -0.42 | 0.07 | -0.03 | -0.2725 | -0.64 | 0.84712 | 0.163356 | 0.020423 | | 1 | 0.996511 | 0.621926 |
| Rgs6 | A0A0J9YTS | -1.26 | -0.71 | 1.22 | 1.44 | -0.31 | -0.36 | 0.54 | 1.67 | -2.35 | 2.27 | 0.91 | -0.85 | -1.38 | -2.18 | 0.45 | -0.2 | 0.2125 | -0.1775 | -1 | 0.776993 | 0.870736 | 0.237365 | | 1 | 0.996511 | 0.792722 |
| Rgs7 | A0A0A6YW | 0.65 | 0.65 | 0.61 | 0.73 | 1.06 | 0.67 | 0.96 | 0.74 | 0.34 | 0.33 | 0.65 | 0.53 | -0.2 | 0.47 | 0.14 | 0.24 | 0.1975 | -0.1975 | -0.4975 | 0.234434 | 0.253205 | 0.035449 | | 1 | 0.996511 | 0.652696 |
| Rgs7 | O54829 | 0.91 | 0.5 | 0.77 | 0.49 | -1.12 | -0.16 | 0.23 | 0.38 | -0.04 | 0.85 | 0.93 | -0.38 | -0.52 | 0.65 | -0.4 | 0.65 | -0.835 | -0.3275 | -0.5725 | 0.03499 | 0.34535 | 0.118047 | | 1 | 0.996511 | 0.720612 |
| Rgs7bp | Q8BQP9 | 0.56 | 1.01 | 1.2 | 0.8 | 1.3 | 0.57 | 1.29 | 0.36 | 0.46 | 0.64 | 0.47 | 0.72 | 1.27 | 0.3 | 0.72 | 1.68 | -0.0125 | -0.32 | 0.1 | 0.964615 | 0.137945 | 0.765667 | | 1 | 0.996511 | 0.964134 |
| Rgs8 | D3Z289 | 0.33 | -0.03 | 0.74 | 0.9 | -0.62 | -0.58 | 1.13 | 0.34 | -0.34 | 0.14 | -0.09 | 0.2 | -0.19 | -0.61 | 0.94 | 0.57 | -0.4175 | -0.5075 | -0.3075 | 0.351931 | 0.079477 | 0.445564 | | 1 | 0.996511 | 0.884817 |
| Rgs9 | O54828 | 2.07 | 0.66 | 1.57 | 1.59 | -0.74 | -0.02 | 1.63 | 1.15 | 1.36 | 0.55 | 0.5 | 1.52 | 1.46 | 1.5 | 1.56 | 1.58 | -0.9675 | -0.49 | 0.0525 | 0.113735 | 0.222007 | 0.864007 | | 1 | 0.996511 | 0.977235 |
| Rhbdd2 | Q8VEK2 | -0.05 | -0.85 | -1.27 | -1.21 | -1.5 | -0.24 | -0.7 | -0.5 | 0.46 | -1.38 | -1.02 | -1.12 | 1.3 | -1.5 | -0.65 | -0.52 | 0.11 | 0.08 | 0.5025 | 0.767932 | 0.86461 | 0.410551 | | 1 | 0.996511 | 0.874811 |
| Rhbdf1 | Q6PIX5 | -0.32 | 0.22 | 0.32 | 0.18 | -0.25 | -0.38 | -0.41 | 0.07 | 0.17 | 0.63 | 0.45 | -0.01 | 0.15 | 0.56 | -0.22 | 0.09 | -0.3425 | 0.21 | 0.045 | 0.12832 | 0.378571 | 0.856616 | | 1 | 0.996511 | 0.976225 |
| Rhbdl3 | P58873 | 0.54 | 0.52 | 1.58 | 1.55 | 0.22 | 0 | 1.05 | 0.32 | 0.8 | 3.08 | 0.92 | 0.05 | 0.42 | 1.37 | 1.91 | 0.69 | -0.65 | 0.165 | 0.05 | 0.098556 | 0.800959 | 0.907083 | | 1 | 0.996511 | 0.986458 |
| Rheb | Q921J2 | 1.02 | 0.56 | 1.54 | 2.6 | 0.61 | 0.79 | 2.04 | 2.93 | 0.64 | 2.02 | 2.06 | 1.28 | 0.24 | -0.31 | 1.52 | 1.04 | 0.1625 | 0.07 | -0.8075 | 0.799085 | 0.891534 | 0.168978 | | 1 | 0.996511 | 0.759484 |
| Rhoa | Q9QUI0 | 2.04 | 1.04 | 1.74 | 1.38 | 1.58 | 2 | 2.59 | 2.23 | 0.75 | 1.13 | 2 | 2.08 | 0.64 | 0.41 | 1.89 | 3.22 | 0.55 | -0.06 | -0.01 | 0.0952 | 0.874127 | 0.9872 | | 1 | 0.996511 | 0.996154 |
| Rhob | P62746 | -0.24 | -2.17 | 0.57 | 0.77 | -0.93 | -1.63 | 1.3 | 0.92 | -0.38 | 1.01 | 0.58 | -1.59 | 0.1 | -0.3 | 0.8 | 1.04 | 0.1825 | 0.1725 | 0.6775 | 0.835365 | 0.829309 | 0.327576 | | 1 | 0.996511 | 0.844684 |
| Rhobtb2 | Q91V93 | 1.21 | 0.67 | 0.47 | 1.15 | 0.93 | 1.54 | 0.83 | 0.85 | 0.94 | 1.13 | 0.85 | 1.29 | 0.77 | 1.49 | 0.94 | 1.01 | 0.1625 | 0.1775 | 0.1775 | 0.53252 | 0.458839 | 0.508168 | | 1 | 0.996511 | 0.902804 |
| Rhoc | Q62159 | -0.53 | -0.62 | -0.99 | -0.36 | -1.53 | 0.21 | -0.57 | -1.12 | -0.38 | -1.2 | -1.31 | -0.1 | -0.63 | -0.37 | -1.11 | -0.42 | -0.1275 | -0.1225 | -0.0075 | 0.737261 | 0.708341 | 0.975958 | | 1 | 0.996511 | 0.993727 |
| Rhof | Q8BYP3 | 0.47 | 0.42 | 0.32 | 0.39 | 0.31 | 0.52 | 0.52 | 0.68 | 0.89 | 0.37 | 0.59 | 0.55 | 0.42 | 0.89 | 0.06 | 0.39 | 0.1075 | 0.2 | 0.04 | 0.488812 | 0.282284 | 0.857485 | | 1 | 0.996511 | 0.976225 |
| Rhog | P84096 | 1.36 | 0.67 | 0.76 | 0.84 | 0.53 | 0.58 | -0.17 | -0.04 | 0.86 | 0.81 | 0.1 | 0.94 | 0.35 | 0.45 | 0.21 | 0.32 | -0.6825 | -0.23 | -0.575 | 0.024983 | 0.397092 | 0.024542 | | 1 | 0.996511 | 0.628313 |
| Rhoq | Q8R527 | 1.49 | -0.14 | -0.35 | -1.03 | -0.13 | 1.29 | 0.17 | -0.66 | 0.52 | -2 | -0.52 | 0.88 | 1.36 | 0.25 | -2.86 | -0.65 | 0.175 | -0.2725 | -0.4675 | 0.776034 | 0.720403 | 0.619117 | | 1 | 0.996511 | 0.93807 |
| Rhot1 | Q8BG51-3 | 1.07 | 0.13 | -0.21 | -0.35 | 0.1 | 1.65 | -0.13 | -0.07 | 0.23 | -0.63 | -0.08 | 0.6 | 0.54 | -0.22 | -0.46 | -0.28 | 0.2275 | -0.13 | -0.265 | 0.644596 | 0.742568 | 0.490284 | | 1 | 0.996511 | 0.897531 |
| Rhot2 | Q8JZN7 | -0.09 | -0.2 | -0.91 | -0.38 | -0.25 | 0.24 | -0.61 | -0.65 | 0.07 | -0.69 | -0.5 | 0.19 | -0.04 | -0.36 | -0.73 | -0.51 | 0.0775 | 0.1625 | -0.015 | 0.781081 | 0.578554 | 0.95412 | | 1 | 0.996511 | 0.992476 |
| Rhou | Q9EQT3 | 1.76 | 2.36 | 1.11 | 0.44 | 2.7 | 1.9 | 1.59 | 1.04 | 1.42 | 2.09 | 1.16 | 2.01 | 0.99 | 1.24 | 0.91 | 1.13 | 0.39 | 0.2525 | -0.35 | 0.442043 | 0.573434 | 0.397214 | | 1 | 0.996511 | 0.867334 |
| Rhpn2 | Q8BWR8 | -0.1 | -0.22 | 0.64 | 0.6 | 0.12 | -0.38 | 0.44 | 0.41 | 0.18 | 1 | 0.48 | 0.43 | 0.29 | 0.92 | 0.27 | 0.71 | -0.0825 | 0.2925 | 0.3175 | 0.779906 | 0.332003 | 0.295257 | | 1 | 0.996511 | 0.825612 |
| Ric1 | E9QPA1 | -0.93 | -1.57 | -0.82 | -0.83 | -0.84 | -2.01 | -1.17 | -1.27 | -0.56 | -0.07 | -1.17 | -1.48 | -0.01 | -0.72 | -1.12 | -0.67 | -0.285 | 0.2175 | 0.4075 | 0.358442 | 0.543357 | 0.20211 | | 1 | 0.996511 | 0.78218 |
| Ric3 | D3YTZ9 | -0.29 | 0.11 | 0.94 | -0.06 | -0.4 | -0.44 | 0.59 | 0.54 | -0.59 | -2 | 0.72 | -0.35 | 0.57 | -2 | 0.85 | 1.33 | -0.1025 | -0.73 | 0.0125 | 0.783295 | 0.222682 | 0.986092 | | 1 | 0.996511 | 0.996154 |
| Ric8a | Q3TIR3 | -0.04 | 0.08 | 0.27 | 0.75 | 0.24 | 0.17 | 1.28 | 0.45 | 0.32 | 0.4 | 1.23 | 0.43 | 0.08 | 0.48 | 0.65 | 0.54 | 0.27 | 0.33 | 0.1725 | 0.387539 | 0.266115 | 0.493179 | | 1 | 0.996511 | 0.897531 |
| Ric8b | Q80XE1-1 | -0.79 | -0.85 | -0.49 | -0.19 | -0.53 | -1.09 | -0.69 | -0.66 | -1.19 | 0.74 | -0.15 | -0.57 | -1.34 | -0.58 | 0.01 | -0.24 | -0.1625 | 0.2875 | 0.0425 | 0.464213 | 0.49129 | 0.898394 | | 1 | 0.996511 | 0.983509 |
| Rictor | Q6QI06 | 0.28 | 0.46 | 0.29 | 0.03 | 0.53 | 0.07 | 0.43 | -0.11 | 0.61 | 0.27 | 0.26 | 0.64 | 0.26 | 1.3 | 0.18 | 0.27 | -0.035 | 0.18 | 0.2375 | 0.864386 | 0.36127 | 0.430389 | | 1 | 0.996511 | 0.879165 |
| Rida | P52760 | -0.15 | -0.44 | 0.09 | 0.43 | -0.06 | -0.52 | 1.37 | 0.15 | -1.02 | -0.15 | -0.25 | -0.95 | -1.92 | -0.29 | -0.57 | -0.26 | 0.2525 | -0.575 | -0.7425 | 0.54838 | 0.08206 | 0.10149 | | 1 | 0.996511 | 0.698697 |
| Rif1 | Q6PR54-2 | 0.24 | -0.69 | -0.25 | 0.11 | -1.56 | -0.42 | -0.86 | -0.16 | 0.27 | 1.41 | -0.41 | -0.6 | 1.37 | 0.29 | 0.74 | 0.1 | -0.6025 | 0.315 | 0.7725 | 0.117657 | 0.506583 | 0.050759 | | 1 | 0.996511 | 0.658454 |
| Rilpl1 | Q9JJC6 | -1.02 | -1.8 | -1.19 | -1.29 | -0.14 | -0.49 | -0.02 | -0.08 | -0.65 | -0.35 | -0.18 | -0.48 | -0.2 | 0.04 | -0.31 | -0.36 | 1.1425 | 0.91 | 1.1175 | 0.000658 | 0.003216 | 0.00102 | 0.599778 | 0.996511 | 0.399727 | |
| Rimbp2 | A0A0G2JFE | 0.73 | 0.91 | 0.14 | 0.88 | 1.02 | 0.97 | 0.98 | 1.09 | 0.68 | -0.64 | 0.95 | 0.83 | 0.61 | 1.05 | 0.9 | 1.5 | 0.35 | -0.21 | 0.35 | 0.122405 | 0.596438 | 0.230668 | | 1 | 0.996511 | 0.788863 |
| Rimbp2 | A0A571BD | 1.32 | 0.84 | 1.27 | 1.18 | 1 | 1.36 | 1.37 | 1.24 | 0.72 | 1.13 | 1.33 | 0.88 | 0.73 | 0.97 | 1.11 | 1.22 | 0.09 | -0.1375 | -0.145 | 0.623242 | 0.524747 | 0.495348 | | 1 | 0.996511 | 0.89758 |
| Rimoc1 | Q8BR90 | -0.81 | -0.86 | -1.09 | -1.6 | -0.5 | -0.94 | -1.5 | -1.8 | -1.1 | -1 | -1.45 | -0.75 | -1.78 | -0.21 | -2.3 | -1.69 | -0.095 | 0.015 | -0.405 | 0.774743 | 0.95277 | 0.38596 | | 1 | 0.996755 | 0.861533 |
| Rims1 | F6TZK4 | 11.32 | 7.56 | 6.89 | 1.79 | 2.35 | 12.03 | 5.55 | 6.63 | 4.69 | 0.62 | 7.48 | 10.07 | 5 | 1.47 | 6.51 | 8.65 | -0.25 | -1.175 | -1.4825 | 0.920341 | 0.641269 | 0.504228 | | 1 | 0.996511 | 0.901002 |
| Rims1 | E9PWP6 | 0.78 | 0.67 | 0.81 | 0.8 | 0.79 | 0.68 | 0.6 | 0.63 | 0.93 | 0.72 | 0.85 | 0.87 | 0.94 | 0.87 | 0.6 | 0.5 | -0.09 | 0.0775 | -3.75E-02 | 0.532659 | 0.623807 | 0.842744 | | 1 | 0.996511 | 0.976225 |
| Rims2 | Q0VF51 | 0.44 | 0.58 | 0.65 | 0.55 | 0.33 | 0.41 | 0.71 | 0.42 | 0.63 | 0.75 | 0.64 | 0.35 | 0.88 | 0.64 | 0.75 | 0.7 | -0.0875 | 0.0375 | 0.1875 | 0.58388 | 0.827171 | 0.293961 | | 1 | 0.996511 | 0.825612 |
| Rims3 | Q80U57 | 0.33 | 0.18 | 0.5 | 0.46 | 0.3 | 0.27 | 0.26 | 0.34 | -0.53 | 0.21 | 0.09 | 0.1 | -0.98 | -0.23 | -0.37 | 0.15 | -0.075 | -0.4 | -0.725 | 0.619468 | 0.096841 | 0.023628 | | 1 | 0.996511 | 0.624626 |
| Rims4 | P60191 | -0.08 | 0.02 | 0.53 | 0.35 | -0.14 | -0.15 | 0.7 | 0.24 | -0.14 | 0.26 | 0.34 | -0.25 | 0.06 | 0.83 | -0.29 | -0.19 | -0.0425 | -0.1525 | -0.1025 | 0.86816 | 0.519343 | 0.735522 | | 1 | 0.996511 | 0.958635 |
| Rin1 | A0A2C9F2 | 0.29 | 0.22 | -0.28 | -0.37 | 0.14 | 0.14 | -0.64 | -0.5 | 0.58 | 0.15 | -0.54 | 0.38 | 0.5 | 0.54 | -0.82 | -0.57 | -0.18 | 0.1775 | -0.0525 | 0.514046 | 0.560635 | 0.890654 | | 1 | 0.996511 | 0.981669 |
| Ring1 | O35730 | -1.29 | -2.68 | -0.5 | -0.37 | -3.5 | -0.67 | -0.96 | -0.4 | -0.06 | -0.62 | -0.89 | -1.27 | 2.45 | -0.13 | -0.49 | -0.25 | -0.1725 | 0.5 | 1.605 | 0.829932 | 0.369455 | 0.066859 | | 1 | 0.996511 | 0.664936 |
| Rint1 | Q8BZ36-2 | 0.3 | 0.47 | -0.33 | -0.02 | 0.49 | 0.19 | 0.31 | 0.12 | 0.26 | 0.01 | -0.32 | 0.17 | 0.22 | 0.84 | 0.3 | 0.26 | 0.1725 | -0.075 | 0.3 | 0.438787 | 0.759029 | 0.265075 | | 1 | 0.996511 | 0.817264 |
| Riok3 | A0A3Q4EG | 0.74 | -0.19 | 0.34 | 0.53 | 0.32 | 0.31 | 0.72 | -0.16 | -1.62 | 0.35 | 0.44 | 0.7 | -1.71 | -0.92 | 1.11 | 0.42 | -0.0575 | -0.3875 | -0.63 | 0.833403 | 0.46992 | 0.317711 | | 1 | 0.996511 | 0.838864 |
| Riox1 | Q9JJF3 | 0.47 | -0.05 | 0.22 | 0.31 | 0.17 | 0.42 | 0.27 | 0.51 | 0.47 | -0.09 | 0.59 | 0 | 1.32 | 0.7 | 1.83 | 0.86 | 0.105 | 0.005 | 0.94 | 0.560811 | 0.982792 | 0.010113 | | 1 | 0.998773 | 0.563386 |
| Ripk1 | F7D1J2 | -0.55 | -0.21 | 0.21 | -0.03 | 0.67 | -0.21 | 0.55 | -0.35 | 0.79 | 0.1 | -0.17 | -0.2 | -0.43 | -0.54 | -0.43 | -0.4 | 0.31 | 0.275 | -0.305 | 0.320112 | 0.354949 | 0.182986 | | 1 | 0.996511 | 0.776002 |
| Ripor1 | Q68FE6 | -0.21 | -0.49 | 0.17 | 0.03 | 0 | -0.24 | 0.05 | 0.15 | -0.25 | 0.51 | -0.02 | -0.31 | -0.49 | 0.04 | -0.18 | -0.25 | 0.115 | 0.1075 | -0.095 | 0.568668 | 0.676828 | 0.677564 | | 1 | 0.996511 | 0.953864 |
| Ripor2 | A6PW28 | -0.2 | -0.6 | -0.05 | -0.03 | 0.2 | -0.38 | 0.29 | 0.19 | -0.25 | 0.11 | -0.31 | -0.26 | -0.56 | -0.12 | -0.1 | 0.25 | 0.295 | 0.0425 | 0.0875 | 0.210834 | 0.838181 | 0.724177 | | 1 | 0.996511 | 0.958398 |
| Ripor2 | A0A5F8MP | -0.64 | 0.19 | -1.31 | -1.05 | -0.17 | -0.87 | 0.21 | -0.51 | 0.41 | -1.77 | -0.58 | 0.19 | -1.53 | -0.02 | -0.2 | -1.46 | 0.3675 | 0.265 | -0.1 | 0.347904 | 0.629586 | 0.836761 | | 1 | 0.996511 | 0.975831 |
| Rit1 | P70426 | 0.02 | 0.18 | -0.35 | -0.8 | 0.71 | -0.01 | 0.22 | -0.11 | -0.08 | -1.4 | -0.48 | 0.05 | -1.73 | 0.89 | -2.34 | -0.45 | 0.44 | -0.24 | -0.67 | 0.15011 | 0.531808 | 0.338835 | | 1 | 0.996511 | 0.847578 |
| Rit2 | P70425 | -0.55 | -0.27 | 1.19 | 0.16 | -0.41 | -0.69 | 1.5 | 1.03 | -0.97 | -0.45 | 1.05 | -0.28 | -0.49 | -0.42 | 0.69 | 1.41 | 0.225 | -0.295 | 0.165 | 0.708522 | 0.582321 | 0.764922 | | 1 | 0.996511 | 0.964134 |
| Rlbp1 | Q9Z275 | 0.29 | -0.28 | 1.77 | 1.56 | -0.06 | 0.03 | 0.28 | 1.1 | 0.47 | 1.43 | 1.26 | -0.02 | 0.18 | 1.09 | 0.87 | 1.2 | -0.4975 | -0.05 | 1.57E-16 | 0.34868 | 0.927991 | 1 | | 1 | 0.996755 | 1 |
| Rlf | E9Q532 | -4.21 | -2.66 | -0.87 | -1.43 | -1.53 | -2.65 | -0.03 | -0.26 | -1.24 | 0.03 | -0.5 | -0.95 | -1.64 | -0.73 | -0.29 | -0.05 | 1.175 | 1.6275 | 1.615 | 0.19671 | 0.047 | 0.053804 | | 1 | 0.996511 | 0.661738 |
| Rlim | Q9WTV7 | -4.83 | -5.31 | -3.52 | -2.93 | -8 | -4.19 | -5.55 | -5.88 | -3.79 | -0.73 | -4.17 | -3.98 | 1.98 | -6.02 | -2.11 | -1.72 | -1.7575 | 0.98 | 2.18 | 0.069432 | 0.287766 | 0.179474 | | 1 | 0.996511 | 0.775542 |
| Rln3 | Q8CHK2 | 0.32 | 1.94 | 1.37 | 1.05 | 0.43 | 1.07 | 1.22 | 0.97 | 0.86 | -0.45 | 1.34 | 0.19 | 1 | 0.44 | 1.5 | 1.85 | -0.2475 | -0.685 | 0.0275 | 0.500622 | 0.180191 | 0.949407 | | 1 | 0.996511 | 0.992476 |
| Rmc1 | Q8VC42 | 0.59 | 0.44 | 0.45 | 0.15 | 0.56 | 0.52 | 0.67 | 0.53 | 0.25 | -0.16 | 0.42 | 0.86 | -0.09 | 0.43 | 0.35 | 0.84 | 0.1625 | -0.065 | -0.025 | 0.326077 | 0.797183 | 0.919325 | | 1 | 0.996511 | 0.986868 |
| Rmdn1 | Q9DCV4 | -0.04 | 0.23 | 0.12 | 0.38 | 0.11 | 0.25 | 0.44 | 0.3 | 0.07 | -0.17 | 0.48 | -0.16 | -0.07 | -0.33 | -0.35 | -0.11 | 0.1025 | -0.1175 | -0.3875 | 0.544143 | 0.589097 | 0.068589 | | 1 | 0.996511 | 0.665034 |
| Rmdn2 | Q8BSE0 | -1.5 | -1.5 | -1.71 | -1.44 | -0.97 | -1.47 | -1.02 | -1.51 | -1.94 | -3.69 | -1.45 | -1.33 | -1.95 | -0.57 | -1.13 | -0.4 | 0.295 | -0.565 | 0.525 | 0.151527 | 0.284783 | 0.160713 | | 1 | 0.996511 | 0.756132 |
| Rmdn3 | Q3UJU9 | 0.08 | 0.11 | 0.39 | 0.34 | 0.08 | 0.08 | 0.6 | 0.17 | -0.28 | 0.04 | 0.35 | -0.02 | -0.23 | -0.29 | 0 | 0.49 | 0.0025 | -0.2075 | -0.2375 | 0.989347 | 0.316269 | 0.327418 | | 1 | 0.996511 | 0.844633 |
| Rmnd1 | Q8CI78 | 0.2 | 0.65 | 0.44 | 0.35 | 0.74 | 0.07 | 0.58 | 0.27 | 0.24 | 0 | 0.59 | 0.33 | -0.02 | 0.96 | -0.12 | 0.41 | 0.005 | -0.12 | -0.1025 | 0.980722 | 0.556592 | 0.717862 | | 1 | 0.996511 | 0.958398 |
| Rmnd5a | Q80YQ8 | -0.74 | -0.6 | -0.02 | -0.67 | -0.08 | -0.64 | 0 | -0.49 | -0.32 | 0.13 | -0.05 | -0.6 | -0.75 | 0.42 | 0.15 | 0.41 | 0.205 | 0.2975 | 0.565 | 0.406407 | 0.257021 | 0.110747 | | 1 | 0.996511 | 0.712422 |
| Rnase4 | Q8C7E4 | -2.58 | -3.19 | -6.52 | -7.39 | -2.71 | -2.17 | -6.52 | -7.69 | -2.25 | -0.6 | -6.13 | -2.2 | -2.13 | -1.97 | -7.2 | -5.73 | 0.1475 | 2.125 | 0.6625 | 0.927538 | 0.180465 | 0.675092 | | 1 | 0.996511 | 0.952944 |
| Rnaseh2a | Q05C51 | -2.1 | -1.18 | -2.05 | -1.73 | -1.14 | -0.66 | -1.17 | -1.45 | -0.92 | -1.14 | -1.56 | -1.14 | -2.29 | -0.9 | -2.12 | -0.98 | 0.66 | 0.575 | 0.1925 | 0.036631 | 0.056282 | 0.638393 | | 1 | 0.996511 | 0.942804 |
| Rnaseh2b | Q80ZV0 | -2.95 | -4.23 | -3.02 | -5.45 | -5.99 | -3.29 | -6.51 | -5.45 | -2.33 | -3.28 | -4.34 | -3.2 | -0.23 | -5.25 | -4 | -4.96 | -1.3975 | 0.625 | 0.3025 | 0.119983 | 0.354143 | 0.794271 | | 1 | 0.996511 | 0.969367 |
| Rnasek | H3BLR9 | -7.12 | -9.59 | -5.46 | -5.85 | -8.65 | -6.12 | -5.56 | -5.91 | -8.85 | -3.25 | -4.89 | -7.65 | -7.93 | -5.02 | -5.18 | -4.88 | 0.445 | 0.845 | 1.2525 | 0.672578 | 0.55356 | 0.255092 | | 1 | 0.996511 | 0.809524 |
| Rnasel | S4R1J5 | -0.3 | -1.05 | 0.42 | 0.18 | 0.48 | -0.39 | -0.33 | 0.06 | -0.02 | 0.29 | 0.02 | -0.97 | -0.83 | 0.45 | -0.47 | -1.09 | 0.1425 | 0.0175 | -0.2975 | 0.696966 | 0.965559 | 0.507181 | | 1 | 0.996755 | 0.902614 |

| | | | | | | | | | | | | | | | | | | | | | | | | | | | |
|---|---|---|---|---|---|---|---|---|---|---|---|---|---|---|---|---|---|---|---|---|---|---|---|---|---|---|---|
| Rnaset2b | A0A338P7E | 1.08 | 0.93 | 0.41 | 1.72 | 0.45 | 1.04 | 2.2 | 1.43 | 0.96 | -0.63 | 1.27 | 1.13 | 0.59 | 0.17 | 0.15 | 1.18 | 0.245 | -0.3525 | -0.5125 | 0.569108 | 0.472702 | 0.176432 | | 1 | 0.996511 | 0.771503 |
| Rnd2 | Q9QYM5 | -0.06 | 0.42 | 0.27 | 0.08 | 0.08 | -0.26 | 0.23 | -0.08 | 0.09 | -0.33 | 0.06 | 0.33 | 0.03 | 0.23 | 0.43 | 0.45 | -0.185 | -0.14 | 0.1075 | 0.340243 | 0.517193 | 0.604332 | | 1 | 0.996511 | 0.932118 |
| Rnd3 | E9Q8D7 | -1.96 | -0.71 | -2.52 | -1.33 | -0.89 | -1.84 | -0.03 | -2.07 | -2.15 | -3.03 | -1.58 | -2.49 | -1.75 | -0.65 | -0.67 | -0.51 | 0.4225 | -0.6825 | 0.735 | 0.4572 | 0.167206 | 0.135119 | | 1 | 0.996511 | 0.731223 |
| Rnf10 | F7AN16 | -1.42 | -1.54 | -0.92 | -1.47 | -3.36 | -0.45 | -2.84 | -1.62 | -2.86 | -1.29 | -0.97 | -1.23 | -1.54 | -7.16 | 0.51 | -2.01 | -0.73 | -0.25 | -1.2125 | 0.249184 | 0.56321 | 0.414013 | | 1 | 0.996511 | 0.874945 |
| Rnf11 | Q9QYK7 | -0.1 | 0.5 | 0.1 | 0.37 | 1.14 | 0.69 | 0.18 | -0.02 | 0.67 | 0.73 | -0.13 | 0.2 | 0.2 | 1 | -0.73 | -0.32 | 0.28 | 0.15 | -0.18 | 0.352828 | 0.571459 | 0.642245 | | 1 | 0.996511 | 0.94365 |
| Rnf112 | Q96DY5-2 | 1.34 | 0.87 | -0.45 | -0.21 | 0.78 | 1.32 | -0.11 | -0.31 | 0.44 | -1.25 | -0.19 | 1.54 | 0.46 | 0.56 | -0.06 | 0.4 | 0.0325 | -0.2525 | -0.0475 | 0.950812 | 0.703063 | 0.911605 | | 1 | 0.996511 | 0.986458 |
| Rnf113a2 | Q14B01 | -3.83 | -4.71 | -4.26 | -4.39 | -4.45 | -4.32 | -4.3 | -5.28 | -4.58 | -2.27 | -4.23 | -3.73 | -4.45 | -3.05 | -3.22 | -4.56 | -0.29 | 0.595 | 0.4775 | 0.339481 | 0.255518 | 0.271815 | | 1 | 0.996511 | 0.822084 |
| Rnf114 | Q9ET26 | -0.17 | -1.42 | -0.93 | -0.17 | -1.3 | -0.41 | -0.89 | -0.63 | -0.83 | -0.03 | -0.98 | -0.23 | -0.12 | -0.83 | -0.54 | -0.6 | -0.135 | 0.155 | 0.15 | 0.699155 | 0.677541 | 0.661512 | | 1 | 0.996511 | 0.949645 |
| Rnf115 | Q9D0C1 | 0 | 0.96 | 3.43 | 2.9 | 0.62 | 0.1 | 1.09 | 1.71 | 1.88 | 3.63 | 1.94 | 1.14 | 3.45 | 1.15 | 0.6 | 2.15 | -0.9425 | 0.325 | 0.015 | 0.253028 | 0.709266 | 0.986916 | | 1 | 0.996511 | 0.996154 |
| Rnf121 | Q8R1Z9 | -1.45 | -1.7 | -2.14 | -1.56 | -0.63 | -1.28 | -0.69 | -1.59 | -1.18 | -2.21 | -1.18 | -1.44 | -0.49 | -0.94 | -0.94 | -1.43 | 0.665 | 0.21 | 0.7625 | 0.039762 | 0.482291 | 0.018629 | | 1 | 0.996511 | 0.602211 |
| Rnf123 | Q5XPI3 | 0.42 | 0.39 | 0.14 | -0.21 | 0.5 | 0.29 | -0.31 | -0.14 | 0.5 | -0.21 | -0.15 | 0.42 | 0.52 | 0.66 | -0.3 | -0.35 | -0.1 | -0.045 | -0.0525 | 0.689894 | 0.860896 | 0.866434 | | 1 | 0.996511 | 0.978305 |
| Rnf126 | Q91YL2 | -0.97 | 0.08 | -0.05 | -0.41 | -0.64 | -0.73 | -0.75 | -0.65 | -0.36 | -0.87 | -0.15 | -0.22 | 0.4 | -0.49 | -0.28 | 0.07 | -0.355 | -0.0625 | 0.2625 | 0.178679 | 0.83147 | 0.413083 | | 1 | 0.996511 | 0.874811 |
| Rnf13 | Q8C4F9 | -1.61 | -1.97 | -0.97 | -1.07 | -1.94 | -1.19 | -0.51 | -0.76 | -2.36 | -0.52 | -1.7 | -2.25 | -1.83 | -2.07 | -0.72 | -0.77 | 0.305 | -0.3025 | 0.0575 | 0.422564 | 0.510162 | 0.887428 | | 1 | 0.996511 | 0.980365 |
| Rnf130 | Q5SVR5 | 1.8 | 0.61 | -1.99 | -3.07 | 0.93 | 1.38 | 0.17 | -3.09 | 0.47 | -2.8 | -1.63 | 1.64 | -0.36 | 0.84 | -2.68 | -2.33 | 0.51 | 0.0825 | -0.47 | 0.707096 | 0.951168 | 0.707283 | | 1 | 0.996755 | 0.957157 |
| Rnf14 | Q9JI90 | -2.7 | -2.66 | -0.52 | -0.49 | -2.28 | -2.45 | -0.89 | -0.86 | -2.22 | 0.01 | -0.61 | -2.33 | -2.35 | -1.83 | -0.18 | -0.89 | -0.0275 | 0.305 | 0.28 | 0.968198 | 0.695571 | 0.697594 | | 1 | 0.996511 | 0.955729 |
| Rnf141 | H3BJB4 | 2.11 | 1.75 | 2.21 | 1.14 | 2.4 | 2.21 | 2.16 | 1.86 | 0.8 | 1.45 | 2.08 | 1.89 | 1.33 | 1.43 | 1.73 | 2.38 | 0.355 | -0.2475 | -0.085 | 0.213995 | 0.501148 | 0.802243 | | 1 | 0.996511 | 0.971091 |
| Rnf146 | Q9CZW6 | 1.37 | 1.16 | 1.13 | -0.58 | 1.58 | 0.99 | 0.9 | 0.54 | -0.39 | 0.88 | 1.15 | 1.36 | 0.2 | -0.17 | 2.02 | 1.45 | 0.2325 | -0.02 | 0.105 | 0.620022 | 0.971107 | 0.866962 | | 1 | 0.997515 | 0.978392 |
| Rnf149 | Q3U2C5 | 0.16 | -0.43 | 1.7 | 1.6 | -0.7 | -0.49 | 0.58 | 1.1 | 0.24 | 0.75 | 1.28 | 0.27 | 0.86 | -0.81 | 1.22 | 1.08 | -0.635 | -0.1225 | -0.17 | 0.321524 | 0.8203 | 0.792776 | | 1 | 0.996511 | 0.96933 |
| Rnf150 | Q5DTZ6 | 3.3 | 1.37 | 0.37 | 0.66 | 2.1 | 3.01 | 0.02 | 0.25 | 0.58 | 1.27 | 0.5 | 2.78 | 0.92 | 0.98 | 1.42 | 0.05 | -0.08 | -0.1425 | -0.5825 | 0.927596 | 0.852252 | 0.384784 | | 1 | 0.996511 | 0.861128 |
| Rnf157 | A2AAP0 | -0.46 | -0.01 | 0.86 | -0.52 | -0.84 | 0 | 0.48 | 0.25 | 0.21 | -0.48 | 0.43 | -0.31 | 2.3 | -1.25 | 1.96 | 0.97 | 0.005 | -0.005 | 1.0275 | 0.990125 | 0.989251 | 0.208496 | | 1 | 0.999285 | 0.786218 |
| Rnf168 | E9PYW4 | -3.93 | -7.26 | 0.69 | -0.04 | -5.67 | -3.22 | -0.43 | -0.18 | -2.63 | -0.05 | 1.85 | -4.3 | 1.95 | -4 | 2.54 | 0.88 | 0.26 | 1.3525 | 2.9775 | 0.897005 | 0.513416 | 0.180407 | | 1 | 0.996511 | 0.775542 |
| Rnf170 | A0A1B0GR | 0.3 | 0.14 | 0.29 | 0.13 | -0.36 | 0.66 | -0.13 | -0.14 | 0.33 | 1.2 | 0.1 | -0.68 | 1.43 | -0.82 | 0.51 | 0.31 | -0.2075 | 0.0225 | 0.1425 | 0.403563 | 0.952275 | 0.746797 | | 1 | 0.996755 | 0.960533 |
| Rnf181 | Q9CY62 | -1.06 | -1.68 | -14.27 | -12.88 | -0.86 | -0.96 | -12.29 | -11.07 | -0.99 | -18.46 | -13.02 | -0.62 | -0.28 | -1.24 | -13.3 | -13.87 | 1.1775 | -0.8 | 0.3 | 0.779276 | 0.874222 | 0.94712 | | 1 | 0.996511 | 0.992376 |
| Rnf185 | Q91YT2 | -1.05 | -0.6 | -1.06 | -1.56 | -1.11 | -0.21 | -2.06 | -2.23 | 0.61 | -1.86 | -1.76 | -0.85 | 1.39 | -0.65 | 0.52 | -0.8 | -0.335 | 0.1025 | 1.1825 | 0.482757 | 0.854059 | 0.04429 | | 1 | 0.996511 | 0.658454 |
| Rnf2 | Q9CQJ4 | -0.02 | 0.49 | -0.5 | -0.1 | -0.5 | -0.56 | -0.39 | -0.15 | -1.23 | -0.35 | -0.16 | -0.76 | 0.66 | 1 | -0.12 | -0.17 | -0.3675 | -0.5925 | 0.375 | 0.149958 | 0.087036 | 0.302042 | | 1 | 0.996511 | 0.830681 |
| Rnf20 | Q5DTM8 | -1.69 | -1.8 | -2.23 | -2.04 | -1.75 | -1.85 | -1.51 | -1.92 | -2.2 | -0.31 | -1.93 | -1.47 | -1.96 | -1.56 | -2.16 | -2.18 | 0.1825 | 0.4625 | -0.025 | 0.348993 | 0.280977 | 0.91401 | | 1 | 0.996511 | 0.986458 |
| Rnf213 | A0A8I4P9C | -0.53 | -1.16 | 0.29 | 0.68 | -1.28 | -1.15 | 0.21 | 0.33 | 0.44 | 0.55 | 0.3 | -0.86 | 0.12 | -0.61 | 0.5 | 0.52 | -0.2925 | 0.2875 | 0.3125 | 0.595503 | 0.561515 | 0.503918 | | 1 | 0.996511 | 0.901002 |
| Rnf214 | Q8BFU3 | -0.46 | -0.46 | -0.5 | -0.46 | -0.33 | -0.51 | -0.59 | -0.56 | -0.83 | -0.5 | -0.45 | -0.34 | -0.32 | -0.87 | -0.57 | -0.75 | -0.0275 | -0.06 | -0.1575 | 0.849096 | 0.734218 | 0.424723 | | 1 | 0.996511 | 0.876965 |
| Rnf214 | Q8BFU3-2 | -0.72 | -0.99 | -0.35 | -0.03 | -0.28 | -0.49 | -0.95 | 0.6 | -2.14 | 1.05 | -0.43 | -0.38 | -0.1 | -3.15 | 0.9 | -0.4 | 0.2425 | 0.0475 | -0.165 | 0.516382 | 0.939531 | 0.837333 | | 1 | 0.996755 | 0.975831 |
| Rnf216 | P58283-2 | -1.52 | -1.87 | -1.1 | -0.98 | -2.11 | -2.23 | -1.8 | -1.65 | -0.45 | -0.36 | -2.02 | -1.59 | 0.56 | -0.39 | -1.02 | -1.05 | -0.58 | 0.2625 | 0.8925 | 0.046014 | 0.551022 | 0.05457 | | 1 | 0.996511 | 0.661738 |
| Rnf220 | Q9D4L6 | -2.61 | -2.99 | -2.82 | -4.05 | -3.23 | -2.36 | -4.5 | -3.18 | -2.07 | -1.34 | -2.99 | -1.75 | -2.06 | -1.6 | -4.57 | -4.89 | -0.2 | 1.08 | -0.1625 | 0.69219 | 0.037004 | 0.841758 | | 1 | 0.996511 | 0.976225 |
| Rnf25 | E9Q446 | -0.29 | -0.15 | -0.57 | -0.48 | 0.02 | -0.68 | -0.82 | -0.71 | -0.36 | -0.11 | -0.66 | -0.38 | -0.39 | 0.08 | -0.96 | -0.98 | -0.175 | -0.005 | -0.19 | 0.458455 | 0.979817 | 0.515232 | | 1 | 0.998773 | 0.905674 |
| Rnf26 | A0A1L1SQ( | 0.43 | 0.38 | 0.46 | 0.09 | 0.35 | 0.23 | -0.49 | 0.63 | -0.58 | 0.1 | 0.51 | -0.18 | 2 | 1.09 | 1.14 | 0.5 | -0.16 | -0.3775 | 0.8425 | 0.545082 | 0.176117 | 0.027486 | | 1 | 0.996511 | 0.635993 |
| Rnf31 | Q924T7-2 | 0.01 | 0.14 | 0.16 | 0.24 | 0.26 | 0.14 | 0.32 | 0.05 | 0.22 | 0.58 | -0.08 | 0.15 | 0.63 | 0.55 | 0.12 | 0.19 | 0.055 | 0.08 | 0.235 | 0.716519 | 0.685518 | 0.262373 | | 1 | 0.996511 | 0.81524 |
| Rnf34 | Q99KR6 | -0.49 | -1.47 | -0.41 | -0.27 | -1.31 | -0.13 | -0.66 | -0.35 | -0.62 | 0.78 | -0.96 | -0.01 | -0.51 | -1 | -0.78 | -0.78 | 0.0475 | 0.4575 | -0.1075 | 0.894768 | 0.316953 | 0.723927 | | 1 | 0.996511 | 0.958398 |
| Rnf40 | Q3U319 | 0.03 | -1.68 | -2.67 | -2.58 | -1.8 | -1.26 | -2.89 | -2.87 | -0.79 | -0.85 | -2.47 | -0.09 | -0.88 | -1.26 | -2.76 | -2.76 | -0.48 | 0.675 | -0.19 | 0.484448 | 0.366933 | 0.792734 | | 1 | 0.996511 | 0.96933 |
| Rnf41 | Q8BH75 | 0.08 | 0.2 | -0.48 | -1.33 | -1.03 | 1.49 | -2.53 | -2.27 | 2.79 | -2.48 | -1.16 | 0.08 | 3.88 | -1.47 | -0.97 | -1.77 | -0.7025 | 0.19 | 0.3 | 0.435369 | 0.857132 | 0.806755 | | 1 | 0.996511 | 0.971715 |
| Rnf5 | O35445 | -1.5 | -2.32 | 0.42 | 1.74 | -3.74 | -0.49 | 1.46 | 2.63 | 0.84 | 2.98 | 0.93 | -1.24 | 2.38 | -1.59 | 2.73 | 1.95 | 0.38 | 1.2925 | 1.7825 | 0.798875 | 0.270955 | 0.165772 | | 1 | 0.996511 | 0.758837 |
| Rnf7 | Q9WTZ1 | -2.79 | -4.28 | -1.81 | -2.03 | -3.4 | -2.72 | -1.92 | -1.43 | -4.39 | -1.11 | -1.45 | -3.04 | -2.24 | -2.94 | -1.92 | -2.27 | 0.36 | 0.23 | 0.385 | 0.579149 | 0.786891 | 0.491664 | | 1 | 0.996511 | 0.897531 |
| Rnft2 | Q3UF64 | -0.57 | -1.74 | -2.73 | -2.71 | -1.03 | -1.03 | -1.6 | -2.81 | -1.24 | -2.08 | -2.44 | -1.03 | -0.96 | -0.89 | -1.81 | -2.08 | 0.32 | 0.24 | 0.5025 | 0.598458 | 0.67137 | 0.370407 | | 1 | 0.996511 | 0.859245 |
| Rngtt | Q9DCC1 | -3.9 | -3.88 | -2.75 | -3.86 | -4.05 | -4.03 | -4.4 | -4.27 | -3.88 | -2.86 | -3.51 | -3.41 | -2.89 | -3.28 | -3.86 | -3.46 | -0.59 | 0.1825 | 0.225 | 0.072645 | 0.601033 | 0.520345 | | 1 | 0.996511 | 0.907667 |
| Rnh1 | Q91VI7 | 0.13 | -0.33 | -0.61 | -0.54 | -0.52 | 0.06 | -0.51 | -0.71 | -0.26 | -0.02 | -0.75 | 0.13 | -0.44 | -0.1 | -0.96 | -0.78 | -0.0825 | 0.1125 | -0.2325 | 0.740345 | 0.678584 | 0.408253 | | 1 | 0.996511 | 0.874356 |
| Rnmt | Q9D0L8 | -3.73 | -4.34 | -4.28 | -4.73 | -3.63 | -4.32 | -3.9 | -4.27 | -4.38 | -2.55 | -3.84 | -3.51 | -4.22 | -3.64 | -4.29 | -4.86 | 0.24 | 0.7 | 0.0175 | 0.383055 | 0.118994 | 0.957343 | | 1 | 0.996511 | 0.992636 |
| Rnpep | Q8VCT3 | -0.53 | -0.56 | -0.47 | -0.26 | -0.56 | -0.54 | -0.76 | -0.25 | -0.39 | 0.42 | -0.29 | -0.39 | -0.79 | -0.35 | -0.53 | -0.78 | -0.0725 | 0.2925 | -0.1575 | 0.678769 | 0.236336 | 0.431237 | | 1 | 0.996511 | 0.879165 |
| Rnps1 | Q99M28 | 1.32 | 0.74 | 0.01 | 0.14 | 0.41 | 1.22 | -0.4 | 0.35 | 0.44 | -0.6 | -0.37 | 0.45 | 1.04 | -0.4 | 0.77 | -0.19 | -0.1575 | -0.5725 | -0.2475 | 0.708388 | 0.167588 | 0.577437 | | 1 | 0.996511 | 0.927198 |
| RO60 | O08848 | 0.02 | -0.01 | -0.15 | -0.38 | 0.12 | 0.19 | -0.27 | -0.14 | -0.31 | -1.17 | -0.46 | 0 | -0.38 | -0.12 | -0.49 | -0.29 | 0.105 | -0.355 | -0.19 | 0.571735 | 0.22184 | 0.339965 | | 1 | 0.996511 | 0.848092 |
| Robo1 | G5E843 | 1.1 | 1.04 | 0.8 | 1.15 | 0.91 | 1.22 | 1.17 | 1.34 | 1.56 | 1.38 | 0.96 | 1.32 | 1.47 | 0.69 | 1.37 | 0.99 | 0.1375 | 0.2825 | 0.1075 | 0.430604 | 0.17912 | 0.651026 | | 1 | 0.996511 | 0.947407 |
| Robo2 | A0A2I3BR7 | 0.45 | 0.61 | 0.33 | 0.67 | 0.79 | 0.57 | 0.78 | 0.73 | 0.82 | 0.62 | 0.71 | 1.01 | 0.56 | 0.69 | 0.58 | 0.47 | 0.2025 | 0.275 | 0.06 | 0.220795 | 0.153073 | 0.739867 | | 1 | 0.996511 | 0.958796 |
| Rock1 | P70335-2 | 0.74 | 0.45 | 0.6 | 0.63 | 0.41 | 0.45 | 0.33 | 0.27 | 0.78 | 0.82 | 0.5 | 0.66 | 1.01 | 0.34 | 0.47 | 0.15 | -0.24 | 0.085 | -0.1125 | 0.134439 | 0.620853 | 0.634858 | | 1 | 0.996511 | 0.941407 |
| Rock2 | F8VPK5 | 0.92 | 0.52 | 0.74 | 0.76 | 0.19 | 0.74 | 0.23 | 1.69 | 0.5 | 0.25 | 0.59 | 0.55 | 0.14 | 0.29 | 0.5 | 0.36 | -0.0225 | -0.2625 | -0.4125 | 0.948036 | 0.168433 | 0.053815 | | 1 | 0.996511 | 0.661738 |
| Rogdi | Q3TDK6 | 0.39 | 0.43 | 0.75 | 0.83 | 0.49 | 0.31 | 0.64 | 1.04 | 0.05 | 0.69 | 0.83 | 0.31 | -0.03 | 0.42 | 0.41 | 0.47 | 0.02 | -0.13 | -0.2825 | 0.926255 | 0.58835 | 0.211845 | | 1 | 0.996511 | 0.786269 |
| Romo1 | P60603 | 0.54 | 1.5 | 1.99 | 1.08 | 1.42 | 1.65 | 0.01 | 0.68 | -0.74 | -2.01 | 1.13 | 0.98 | -0.28 | -2.96 | 0.71 | 0.99 | -0.3375 | -1.4375 | -1.6625 | 0.460966 | 0.077479 | 0.078972 | | 1 | 0.996511 | 0.681114 |
| Rp2 | Q9EPK2 | 1.1 | 0.8 | 0.89 | 0.78 | 1.08 | 0.83 | 0.85 | 0.79 | 0.86 | 0.37 | 1.11 | 1.32 | 0.65 | 0.2 | 0.99 | 0.72 | -0.005 | 0.0225 | -0.2525 | 0.975091 | 0.926269 | 0.282056 | | 1 | 0.996755 | 0.822084 |
| Rpa1 | Q5SWN2 | -2.78 | -2.51 | -0.25 | -0.42 | -2.33 | -2.96 | -0.34 | -0.4 | -3.63 | -0.76 | -0.32 | -2.28 | -3.42 | -3.34 | -0.39 | 0.2 | -0.0175 | -0.2575 | -0.2475 | 0.983606 | 0.777295 | 0.812579 | | 1 | 0.996511 | 0.972662 |
| Rpa2 | Q62193 | -1.15 | -1.05 | -2.25 | -2.26 | -1.46 | -1.14 | -2 | -2.52 | -0.59 | -0.78 | -1.65 | -1.11 | -0.03 | -0.49 | -2.51 | -2.8 | -0.1025 | 0.645 | 0.22 | 0.808707 | 0.126022 | 0.754985 | | 1 | 0.996511 | 0.962333 |
| Rpa3 | Q9CQ71 | -1.29 | -0.84 | -2.36 | -2.41 | -0.84 | -1.02 | -2.4 | -2.75 | -1.2 | -2.23 | -2.13 | -0.94 | -1.27 | -0.19 | -3 | -2.35 | -0.0275 | 0.1 | 0.0225 | 0.96135 | 0.83373 | 0.973003 | | 1 | 0.996511 | 0.992865 |
| Rpap1 | Q80TE0 | 0.24 | 0.65 | -0.26 | -0.3 | 0.13 | 0.18 | -0.64 | -0.53 | 0.8 | 0.12 | -0.32 | 0.14 | 0.95 | 0.5 | -0.56 | -0.53 | -0.2975 | 0.1025 | 0.0075 | 0.346712 | 0.75111 | 0.985755 | | 1 | 0.996511 | 0.996154 |
| Rpap3 | Q9D706 | 0.71 | 0.46 | 0.28 | 0.55 | -0.05 | 0.98 | 0.14 | 0.66 | 0.41 | 1.54 | 0.49 | 0.82 | -0.11 | 0.29 | 0.27 | 0.29 | -0.0675 | 0.315 | -0.315 | 0.796286 | 0.284169 | 0.141781 | | 1 | 0.996511 | 0.737973 |
| Rpe | B2KGF0 | -0.94 | -1.33 | -1.18 | -0.78 | -1.27 | -0.23 | -0.84 | -1.21 | -1.22 | -0.64 | -1.54 | -0.96 | -1.38 | -1.25 | -1.42 | -1.04 | 0.17 | -0.0325 | -0.215 | 0.538529 | 0.89662 | 0.318058 | | 1 | 0.996511 | 0.838969 |
| Rpf2 | G3X926 | 0.7 | 0.56 | 0.99 | 0.95 | 0.33 | 0.13 | 0.48 | 0.49 | 0.72 | 1 | 0.96 | 0.26 | 0.7 | 0.93 | 1.06 | 0.56 | -0.4425 | -0.065 | 0.0125 | 0.033376 | 0.778239 | 0.952548 | | 1 | 0.996511 | 0.992476 |
| Rpgr | Q9R0X5-2 | 0.19 | 0.22 | -0.25 | -0.51 | -0.06 | 0.34 | -0.26 | -0.36 | 0.05 | 0.25 | -0.6 | 0.27 | -0.11 | -0.12 | -0.02 | -0.36 | 0.0025 | 0.08 | -0.065 | 0.991948 | 0.776636 | 0.781315 | | 1 | 0.996511 | 0.967642 |
| Rpgrip1l | Q8CG73 | 0.7 | 0.36 | -0.73 | 0.19 | 0.51 | 0.67 | -0.05 | -0.37 | 0.89 | 0.46 | 0.36 | 0.62 | 0.6 | 0.7 | 0.59 | 0.73 | 0.06 | 0.4525 | 0.525 | 0.871648 | 0.188903 | 0.123248 | | 1 | 0.996511 | 0.721448 |
| Rph3a | P47708 | 1.42 | 1.14 | 1.58 | 2.19 | 0.77 | 1.67 | 1.51 | 2.02 | 1.32 | 1.17 | 1.74 | 1.47 | 1.16 | 1.06 | 1.32 | 1.28 | -0.09 | -0.1575 | -0.3775 | 0.7878 | 0.562101 | 0.170018 | | 1 | 0.996511 | 0.762194 |
| Rpia | P47968 | -0.32 | -0.34 | -0.11 | 0.31 | -0.61 | 0.72 | -0.74 | 0.19 | -0.01 | 0.32 | -0.25 | -0.39 | 0.46 | -0.81 | -0.78 | 0.08 | 0.005 | 0.0325 | -0.1475 | 0.98891 | 0.893509 | 0.674028 | | 1 | 0.996511 | 0.952944 |
| Rpl10 | I7HLV2 | -0.83 | -1.32 | -1.54 | -1.9 | -0.53 | -0.86 | -1.24 | -1.56 | -1.41 | -1.1 | -1.7 | -0.5 | -1.18 | -1.08 | -0.64 | -0.71 | 0.35 | 0.22 | 0.495 | 0.27913 | 0.518821 | 0.10534 | | 1 | 0.996511 | 0.703417 |
| Rpl10a | A0A3B2WD | -2.44 | -2.43 | -0.11 | 0.01 | -1.52 | -2.38 | 0.87 | 0.34 | -2.97 | -1.57 | -0.09 | -2.19 | -2.38 | -1.98 | 0.82 | 0.73 | 0.57 | -0.4625 | 0.54 | 0.542474 | 0.581055 | 0.586338 | | 1 | 0.996511 | 0.928178 |
| Rpl11 | Q9CXW4 | -2.28 | -2.42 | -0.82 | -0.13 | -1.69 | -2.33 | -0.65 | -0.17 | -2.07 | 0.06 | -0.95 | -2.13 | -1.61 | -1.3 | -0.17 | -0.59 | 0.2025 | 0.14 | 0.495 | 0.764552 | 0.840436 | 0.415069 | | 1 | 0.996511 | 0.874945 |
| Rpl12 | P35979 | -0.15 | -0.47 | -1.66 | -1.67 | 0.27 | 0.29 | -1.09 | -1.53 | -1.54 | -0.81 | -1.38 | 0.39 | -1.13 | -1.42 | -1.29 | -1.04 | 0.4725 | 0.1525 | -0.2325 | 0.409826 | 0.779909 | 0.557433 | | 1 | 0.996511 | 0.917092 |
| Rpl13 | P47963 | -1.87 | -1.93 | -1.6 | -1.57 | -1.56 | -1.44 | -0.9 | -1.48 | -1.79 | -1.39 | -1.4 | -1.74 | -0.75 | -1.89 | -0.77 | -1.17 | 0.3975 | 0.1625 | 0.5975 | 0.081085 | 0.413615 | 0.069984 | | 1 | 0.996511 | 0.668066 |
| Rpl13a | P19253 | -0.83 | -1.94 | -1.77 | -2.17 | -1.33 | -0.9 | -0.54 | -2.24 | -2.16 | -1.5 | -1.85 | -0.99 | -1.51 | -1.34 | -1.12 | -1.01 | 0.425 | 0.0525 | 0.4325 | 0.346155 | 0.888284 | 0.2007 | | 1 | 0.996511 | 0.78037 |
| Rpl14 | Q9CR57 | -3.39 | -4.28 | -3.09 | -3.74 | -3.27 | -3.59 | -2.5 | -3.23 | -4.25 | -1.35 | -3.32 | -3.5 | -4.31 | -3.45 | -2.31 | -2.85 | 0.4775 | 0.52 | 0.395 | 0.177389 | 0.406257 | 0.409499 | | 1 | 0.996511 | 0.87457 |
| Rpl15 | Q9CZM2 | -1.91 | -2.39 | -1.12 | -1.64 | -1.14 | -1.99 | -0.94 | -1.35 | -2.31 | -1.53 | -1.13 | -1.64 | -1.71 | -2.39 | -0.03 | -0.44 | 0.41 | 0.1125 | 0.6225 | 0.245464 | 0.750529 | 0.284699 | | 1 | 0.996511 | 0.822669 |
| Rpl17 | Q6ZWZ7 | -2.27 | -2.12 | -2.36 | -2.3 | -1.79 | -2.13 | -1.52 | -2.2 | -1.96 | -2.02 | -2.33 | -1.96 | -1.61 | -1.54 | -1.47 | -1.87 | 0.3525 | 0.195 | 0.64 | 0.104627 | 0.281574 | 0.006932 | | 1 | 0.996511 | 0.552614 |
| Rpl18 | P35980 | -2.52 | -2.43 | -1.89 | -2.37 | -1.83 | -2.28 | -1.22 | -2.12 | -2.56 | -1.63 | -2.17 | -2.25 | -1.79 | -1.44 | -1.3 | -1.38 | 0.44 | 0.15 | 0.825 | 0.138779 | 0.56621 | 0.005084 | | 1 | 0.996511 | 0.552614 |
| Rpl18a | A0A1D5RL\ | -1.6 | -1.65 | -1.96 | -2.07 | -1.17 | -1.48 | -1.3 | -2.18 | -1.84 | -0.8 | -1.89 | -1.38 | -0.98 | -1.37 | -1.49 | -1.54 | 0.2875 | 0.3425 | 0.475 | 0.288328 | 0.253776 | 0.056747 | | 1 | 0.996511 | 0.661738 |
| Rpl19 | A2A547 | -0.73 | -2.17 | -2.19 | -2.4 | -1.45 | -0.79 | -1.07 | -1.75 | -2.53 | -1.88 | -1.86 | -0.72 | -1.42 | -2.38 | -0.96 | -0.49 | 0.6075 | 0.125 | 0.56 | 0.165928 | 0.802829 | 0.295706 | | 1 | 0.996511 | 0.825612 |
| Rpl21 | Q9CQM8 | -3.22 | -3.49 | -1.74 | -1.92 | -2.83 | -3.13 | -1.42 | -1.97 | -3.4 | -1.52 | -1.9 | -3.06 | -2.22 | -3.99 | -0.92 | -1.24 | 0.255 | 0.1225 | 0.5 | 0.641371 | 0.833374 | 0.506443 | | 1 | 0.996511 | 0.902565 |
| Rpl22 | P67984 | -1.6 | -1.65 | -1.94 | -1.6 | -0.72 | -2.05 | -1.12 | -1.74 | -1.82 | -1.22 | -1.7 | -1.85 | -1.01 | -0.74 | -1.49 | -1.23 | 0.29 | 0.05 | 0.58 | 0.357287 | 0.812187 | 0.028671 | | 1 | 0.996511 | 0.637553 |
| Rpl22l1 | Q9D7S7-2 | -3.12 | -3.15 | -1.66 | -2.91 | -2.66 | -2.96 | -0.83 | -2.39 | -3.04 | -0.94 | -2.75 | -2.72 | -1.18 | -1.57 | -0.89 | -1.63 | 0.5 | 0.3475 | 1.3925 | 0.36931 | 0.532205 | 0.005388 | | 1 | 0.996511 | 0.552614 |
| Rpl23 | P62830 | -2.42 | -2.75 | -2.24 | -2.26 | -2.16 | -2.16 | -1.44 | -2.2 | -1.23 | -1.63 | -2.39 | -2.49 | 0.53 | -2.5 | -1.83 | -1.93 | 0.4275 | 0.4825 | 0.985 | 0.097136 | 0.163079 | 0.140047 | | 1 | 0.996511 | 0.736096 |
| Rpl23a | P62751 | -0.39 | -1.49 | -0.98 | -1.12 | -0.75 | -0.49 | 0.94 | -1.12 | -1.33 | -0.38 | -1.03 | -0.75 | -1.01 | -0.16 | -0.5 | -0.47 | 0.64 | 0.1225 | 0.46 | 0.19578 | 0.692657 | 0.152517 | | 1 | 0.996511 | 0.750422 |
| Rpl24 | Q8BP67 | -3.27 | -3.67 | -2.54 | -1.98 | -2.74 | -3.06 | -2.34 | -2.96 | -2.36 | -2.71 | -2.53 | -2.86 | -1.44 | -3.49 | -1.66 | -2.13 | 0.09 | 0.25 | 0.685 | 0.816338 | 0.513914 | 0.232773 | | 1 | 0.996511 | 0.788994 |
| Rpl26 | P61255 | -1.48 | -1.74 | -1.14 | -1.72 | -1.05 | -1.63 | -1.19 | -1.55 | -2.72 | -2.02 | -1.43 | -1.29 | -1.69 | -1.97 | -0.49 | -0.88 | 0.165 | -0.345 | 0.2625 | 0.462188 | 0.335603 | 0.479308 | | 1 | 0.996511 | 0.893468 |
| Rpl27 | P61358 | -1.29 | -1.28 | -1.25 | -1.45 | -0.57 | -1.42 | -0.73 | -1.41 | -1.85 | -1.18 | -1.38 | -1.12 | -1.24 | -0.93 | -0.74 | -0.79 | 0.285 | -0.065 | 0.3925 | 0.259016 | 0.760472 | 0.069271 | | 1 | 0.996511 | 0.667373 |
| Rpl27a | P14115 | -2.51 | -2.91 | -4.09 | -4.7 | -1.53 | -2.76 | -3.42 | -4.38 | -3.21 | -3.44 | -4.57 | -2.49 | -2.63 | -2.4 | -3.42 | -3.75 | 0.53 | 0.125 | 0.5025 | 0.463388 | 0.837857 | 0.376144 | | 1 | 0.996511 | 0.860376 |
| Rpl28 | P41105 | -1.78 | -2.23 | -2.19 | -2.25 | -1.15 | -1.55 | -1.86 | -2.17 | -1.74 | -1.18 | -2.15 | -1.53 | -0.88 | -1.21 | -1.31 | -1.55 | 0.43 | 0.4625 | 0.875 | 0.1196 | 0.095513 | 0.003687 | | 1 | 0.996511 | 0.517003 |
| Rpl29 | P47915 | -1.48 | -2.32 | -1.11 | -1.39 | -1.56 | -1.71 | -1.19 | -1.32 | -1.34 | -1.47 | -1.32 | -1.71 | 0.33 | -1.92 | 0.96 | -0.65 | 0.13 | 0.115 | 1.255 | 0.652032 | 0.688635 | 0.069741 | | 1 | 0.996511 | 0.668066 |
| Rpl3 | P27659 | -1.86 | -1.86 | -1.91 | -2.32 | -0.96 | -1.92 | -1.69 | -2.19 | -3.17 | -1.7 | -2.18 | -1.8 | -1.94 | -2.43 | -1.11 | -1.6 | 0.2975 | -0.225 | 0.2175 | 0.31606 | 0.521902 | 0.488521 | | 1 | 0.996511 | 0.897531 |
| Rpl30 | P62889 | -1.75 | -2.62 | -1.89 | -1.98 | -2.12 | -1.7 | -1.95 | -2.32 | -1.19 | -1.49 | -2.11 | -1.62 | -0.39 | -1.56 | -1.49 | -1.6 | 0.0375 | 0.4575 | 0.8 | 0.879024 | 0.132932 | 0.043887 | | 1 | 0.996511 | 0.658454 |
| Rpl31 | P62900 | -1.37 | -1.69 | -2.3 | -3.13 | -1.29 | -1.55 | -1.85 | -2.31 | -0.12 | -2.18 | -2.35 | -1.25 | 2.34 | -2.19 | -1.06 | -1.8 | 0.3725 | 0.6475 | 1.445 | 0.38351 | 0.289191 | 0.168327 | | 1 | 0.996511 | 0.759484 |
| Rpl32 | P62911 | -2.44 | -2.22 | -3.03 | -3.01 | -1.63 | -2.24 | -2.32 | -2.79 | -2.54 | -2 | -2.83 | -2.21 | -2.14 | -1.68 | -2.14 | -2.43 | 0.43 | 0.28 | 0.5775 | 0.18852 | 0.339161 | 0.063243 | | 1 | 0.996511 | 0.66433 |
| Rpl34 | Q9D1R9 | -2.68 | -2.94 | -1.54 | -2.28 | -1.72 | -2.45 | -1.56 | -2.2 | -3.24 | -2.14 | -1.71 | -2.42 | -2.15 | -2.62 | -1.03 | -1.44 | 0.3775 | -0.0175 | 0.55 | 0.30236 | 0.966823 | 0.235909 | | 1 | 0.996755 | 0.791493 |
| Rpl35 | Q6ZWV7 | -0.13 | -1.03 | -1.08 | -1.28 | -0.61 | -0.27 | -0.43 | -0.49 | -0.36 | -1.61 | -0.64 | 0.2 | 0.31 | -0.29 | 0.99 | -0.7 | 0.43 | 0.2775 | 0.9575 | 0.139429 | 0.52555 | 0.048667 | | 1 | 0.996511 | 0.658454 |
| Rpl35a | O55142 | -0.22 | -1.24 | -1.38 | -2.18 | 0.01 | 0.07 | -0.83 | -1.42 | -1.22 | -1.11 | -1.39 | 0.29 | -0.81 | -1.7 | -0.18 | -0.67 | 0.7125 | 0.3975 | 0.415 | 0.176499 | 0.450319 | 0.397562 | | 1 | 0.996511 | 0.867334 |
| Rpl36 | Q6ZWZ4 | -1.72 | -1.59 | -1.45 | -1.4 | -0.52 | -1.84 | -0.66 | -1.35 | -1.76 | -0.73 | -1.32 | -1.46 | -1.28 | -0.69 | -0.67 | -1.05 | 0.4475 | 0.2225 | 0.6175 | 0.175767 | 0.386287 | 0.018314 | | 1 | 0.996511 | 0.602211 |
| Rpl36a | P83882 | 0.74 | -0.72 | -0.82 | -0.49 | 0.1 | 1.01 | -0.61 | -0.16 | -1.32 | 0.48 | -0.81 | 0.88 | -0.25 | -0.97 | 0.47 | 0.28 | 0.4075 | 0.13 | 0.205 | 0.388184 | 0.823348 | 0.65506 | | 1 | 0.996511 | 0.94818 |
| Rpl37 | Q9D823 | -0.25 | -0.31 | -3.89 | -3.06 | -1.18 | 1.27 | -2.74 | -2.97 | 4.45 | -6.11 | -2.67 | -0.35 | 6.25 | -1.51 | -0.04 | -2.4 | 0.4725 | 0.7075 | 2.4525 | 0.697529 | 0.7419 | 0.222968 | | 1 | 0.996511 | 0.787447 |
| Rpl37a | P61514 | -3.43 | -3.42 | -5.46 | -6.33 | -2.26 | -3.2 | -4.53 | -5.74 | -3.21 | -5.22 | -5.51 | -2.99 | -2.3 | -1.76 | -4.93 | -4.66 | 0.7275 | 0.4275 | 1.2475 | 0.451854 | 0.632704 | 0.223332 | | 1 | 0.996511 | 0.787447 |
| Rpl38 | Q9JJI8 | 1.31 | 0.1 | 0.16 | 0.67 | -0.14 | 1.59 | 1.1 | 0.82 | 0.9 | 0.34 | -0.06 | 1.26 | -0.02 | 1.05 | 0.26 | 1.05 | 0.2825 | 0.05 | 0.025 | 0.516516 | 0.897792 | 0.947769 | | 1 | 0.996511 | 0.992414 |
| Rpl39 | P62892 | -1.47 | -1.71 | -1.07 | -1.03 | -1.35 | -2.36 | -1.4 | -1.52 | -1.82 | -0.7 | -1.07 | -1.9 | -1.56 | -0.59 | -1.03 | -1.43 | -0.3375 | -0.0525 | 0.1675 | 0.261268 | 0.87428 | 0.567092 | | 1 | 0.996511 | 0.921806 |
| Rpl4 | Q9D8E6 | -2.61 | -2.85 | -2.44 | -2.37 | -1.99 | -2.54 | -1.78 | -2.48 | -2.97 | -2.07 | -2.39 | -2.41 | -2.42 | -2.09 | -1.59 | -1.95 | 0.37 | 0.1075 | 0.555 | 0.138686 | 0.658562 | 0.043753 | | 1 | 0.996511 | 0.658454 |
| Rpl5 | P47962 | -1.54 | -1.79 | -1.88 | -2.01 | -1.34 | -1.58 | -1.43 | -2.09 | -1.8 | -1.63 | -2.07 | -1.27 | -1.23 | -0.94 | -1.28 | -1.4 | 0.195 | 0.1125 | 0.5925 | 0.383892 | 0.623833 | 0.016255 | | 1 | 0.996511 | 0.589387 |
| Rpl6 | P47911 | -1.84 | -1.86 | -1.46 | -1.82 | -0.86 | -2.09 | -1.22 | -1.92 | -2.43 | -1.42 | -1.91 | -2 | -1.21 | -1.57 | -0.93 | -1.35 | 0.2225 | -0.195 | 0.48 | 0.469156 | 0.445243 | 0.051192 | | 1 | 0.996511 | 0.658454 |
| Rpl7 | P14148 | 0.16 | -0.76 | -1.43 | -1.89 | 0.29 | 0.03 | -0.32 | -1.4 | -0.43 | -1.65 | -1.43 | 0.46 | -0.34 | -0.38 | -0.44 | -0.75 | 0.63 | 0.2175 | 0.5025 | 0.256385 | 0.719684 | 0.264047 | | 1 | 0.996511 | 0.816045 |
| Rpl7a | P12970 | 0.24 | -0.98 | -1.37 | -1.33 | -0.36 | 0.27 | -0.69 | -1.11 | -1.07 | -0.56 | -1.33 | 0.23 | -0.91 | -0.11 | -0.72 | -0.84 | 0.3875 | 0.1775 | 0.215 | 0.393612 | 0.710552 | 0.596799 | | 1 | 0.996511 | 0.929911 |
| Rpl8 | P62918 | -2.11 | -1.93 | -2.44 | -2.34 | -1.62 | -1.91 | -2.02 | -2.49 | -1.13 | -2.52 | -2.42 | -1.89 | -0.46 | -1.29 | -0.86 | -1.9 | 0.195 | 0.215 | 1.0775 | 0.411639 | 0.525836 | 0.00965 | | 1 | 0.996511 | 0.563386 |
| Rpl9 | P51410 | -0.9 | -0.88 | -1.97 | -2.12 | 0.17 | -0.77 | -0.96 | -2.09 | -1.72 | -2.73 | -2.06 | -0.69 | -1.64 | -1.36 | -1.72 | -1.06 | 0.555 | -0.3325 | 0.0225 | 0.307182 | 0.514104 | 0.950308 | | 1 | 0.996511 | 0.992476 |
| Rplp0 | P14869 | -1.76 | -2.1 | -1.54 | -1.72 | -1.63 | -1.77 | -0.96 | -1.53 | -1.69 | -0.68 | -1.5 | -1.53 | -1.77 | -0.97 | -0.72 | -0.94 | 0.3075 | 0.43 | 0.68 | 0.207045 | 0.139158 | 0.035278 | | 1 | 0.996511 | 0.652172 |
| Rplp1 | P47955 | -4.81 | -4.05 | -5.31 | -4.38 | -3.15 | -3.67 | -3.22 | -4.45 | -3.08 | -3.09 | -4.42 | -3.94 | -3.91 | -1.61 | -4.15 | -3.84 | 1.015 | 1.005 | 1.26 | 0.025549 | 0.034445 | 0.060647 | | 1 | 0.996511 | 0.661738 |
| Rplp2 | P99027 | -1.44 | -1.17 | -1.03 | -0.97 | -0.63 | -1.56 | -0.4 | -1.07 | -1.46 | -0.71 | -0.86 | -0.96 | -0.98 | -0.67 | -0.56 | -0.63 | 0.2375 | 0.155 | 0.4425 | 0.405822 | 0.500462 | 0.054178 | | 1 | 0.996511 | 0.661738 |
| Rpn1 | Q91YQ5 | -1.61 | -1.8 | -1.87 | -1.74 | -1.06 | -1.29 | -0.77 | -1.46 | -1.43 | -2.15 | -1.72 | -1.19 | -0.92 | -0.91 | -0.63 | -0.6 | 0.61 | 0.1325 | 0.99 | 0.011783 | 0.585523 | 0.000515 | | 1 | 0.996511 | 0.286246 |
| Rpn2 | A2ACG7 | -1.02 | -0.76 | -0.99 | -1.04 | 0.05 | -0.53 | -0.01 | -0.8 | -0.27 | -1.29 | -0.92 | -0.74 | 0.52 | 0.1 | -0.04 | 0.3 | 0.63 | 0.1475 | 1.1725 | 0.023377 | 0.553539 | 0.000296 | | 1 | 0.996511 | 0.270112 |
| Rpp14 | J3QMX0 | -0.79 | -1.09 | 0.05 | 0.03 | -0.18 | -2.11 | -0.2 | -0.21 | 0.04 | -0.62 | -0.27 | -0.72 | -1.78 | -0.68 | -0.89 | 0.45 | -0.225 | 0.0575 | -0.275 | 0.663612 | 0.863375 | 0.589143 | | 1 | 0.996511 | 0.929131 |
| Rpp25 | Q91WE3 | -3.08 | -3.44 | -5.17 | -5.58 | -8.1 | -3.95 | -9.27 | -3.81 | -5.36 | -0.84 | -5.1 | -3.39 | -2.39 | -7.3 | -4.79 | -6.08 | -1.965 | 0.645 | -0.8225 | 0.177777 | 0.556776 | 0.457554 | | 1 | 0.996511 | 0.887902 |
| Rpp30 | O88796 | -1.39 | -1.87 | -2.57 | -2.57 | -2.65 | -1.36 | -3.4 | -3.56 | -1.87 | -2.62 | -2.9 | -1.91 | -1.67 | -1.93 | -3.12 | -3.06 | -0.6425 | -0.225 | -0.345 | 0.246973 | 0.551666 | 0.449587 | | 1 | 0.996511 | 0.884817 |
| Rprd1a | Q8VDS4 | -2.38 | -2.37 | -1.91 | -1.61 | -2.72 | -2.74 | -1.76 | -1.86 | -2.93 | -0.04 | -2.59 | -2.23 | -2.39 | -1.79 | -2.19 | -2.8 | -0.2025 | 0.12 | -0.225 | 0.53036 | 0.846658 | 0.455177 | | 1 | 0.996511 | 0.887169 |
| Rprd1b | A0A0R4J19 | -0.97 | -1.11 | -4.34 | -4.36 | -1.74 | -1.88 | -3.93 | -4.64 | -1.63 | -2.96 | -3.86 | -0.99 | -0.81 | -0.63 | -3.83 | -4.26 | -0.3525 | 0.335 | 0.3125 | 0.744155 | 0.747022 | 0.796348 | | 1 | 0.996511 | 0.969727 |
| Rprd2 | A0A0G2JFV | -0.98 | -1.56 | -0.67 | -0.71 | -2.14 | -1.16 | -1.12 | -1.1 | -1.33 | -0.2 | -0.8 | -1.41 | 0.17 | -1.12 | -0.59 | -0.85 | -0.4 | 0.045 | 0.3825 | 0.231483 | 0.895221 | 0.284123 | | 1 | 0.996511 | 0.822669 |
| Rps10 | P63325 | 2.02 | -0.13 | 0.13 | 1.94 | -0.17 | 1.19 | 1.26 | 2 | 1.77 | 2.1 | 0.88 | 2.94 | 1.52 | 0.37 | 1.57 | 0.66 | 0.08 | 0.9325 | 0.04 | 0.903956 | 0.17772 | 0.946392 | | 1 | 0.996511 | 0.992376 |
| Rps11 | P62281 | -0.2 | -0.65 | -0.1 | -0.13 | 0.23 | -0.33 | 0.19 | -0.14 | -0.17 | 0.8 | -0.41 | -0.12 | 0.03 | 0.53 | 0.87 | 0.81 | 0.2575 | 0.295 | 0.83 | 0.246072 | 0.340775 | 0.010436 | | 1 | 0.996511 | 0.563386 |
| Rps12 | Q6ZWZ6 | -2.56 | -3.3 | -2.57 | -2.42 | -2.43 | -2.66 | -2.13 | -2.57 | -3.21 | -1.27 | -2.54 | -2.57 | -2.57 | -1.87 | -1.93 | -2.19 | 0.265 | 0.315 | 0.5725 | 0.294654 | 0.469087 | 0.064183 | | 1 | 0.996511 | 0.66433 |
| Rps13 | P62301 | -0.28 | -2.2 | -1.89 | -1.74 | -1.41 | -0.46 | -0.76 | -1.71 | -1.92 | -1.21 | -1.77 | -1.31 | -1.82 | -0.91 | -1.07 | -1.37 | 0.4425 | -0.025 | 0.235 | 0.365496 | 0.954112 | 0.601084 | | 1 | 0.996755 | 0.930802 |
| Rps14 | P62264 | -1.92 | -1.77 | -2.46 | -2.4 | -1.2 | -1.83 | -1.22 | -2.12 | -2.33 | -1.73 | -2.14 | -1.53 | -1.68 | -1.24 | -1.21 | -1.42 | 0.545 | 0.205 | 0.75 | 0.085317 | 0.452001 | 0.01198 | | 1 | 0.996511 | 0.566767 |
| Rps15 | D3YTQ9 | -1.39 | -1.47 | -2.04 | -1.83 | -0.98 | -1.24 | -1.4 | -1.89 | -1.46 | -0.79 | -1.9 | -1.32 | -1.43 | -0.69 | -0.75 | -1.15 | 0.305 | 0.315 | 0.6775 | 0.25201 | 0.28711 | 0.027106 | | 1 | 0.996511 | 0.635993 |
| Rps15a | P62245 | -1.91 | -1.91 | -1.56 | -0.83 | -1.15 | -1.62 | -1.06 | -0.91 | -2.25 | -2.2 | -1.79 | -1.7 | -1.71 | -1.7 | -1.34 | -1.64 | 0.3675 | -0.4325 | -0.045 | 0.235225 | 0.170974 | 0.875344 | | 1 | 0.996511 | 0.979626 |
| Rps16 | P14131 | -2.52 | -2.94 | -1.2 | -0.62 | -1.91 | -2.59 | 0 | -0.06 | -2.38 | -0.35 | -0.76 | -2.49 | -1.69 | -1.9 | -0.25 | -0.3 | 0.68 | 0.325 | 0.785 | 0.388273 | 0.646062 | 0.240145 | | 1 | 0.996511 | 0.795595 |
| Rps17 | P63276 | -1.59 | -1.59 | -1.02 | -0.96 | -1.14 | -1.47 | -0.41 | -0.77 | -1.79 | -0.7 | -1.04 | -1.3 | -1.14 | -2.04 | -0.37 | -0.46 | 0.3425 | 0.0825 | 0.2875 | 0.254433 | 0.780269 | 0.486203 | | 1 | 0.996511 | 0.895384 |
| Rps18-ps5 | A0A1Y7VK\ | -1.84 | -2.44 | -1.51 | -2.03 | -1.39 | -2.05 | -1.46 | -2.09 | -2.03 | -1.01 | -1.86 | -1.79 | -1.25 | -1.14 | -0.62 | -1.25 | 0.2075 | 0.2825 | 0.89 | 0.456707 | 0.36396 | 0.008952 | | 1 | 0.996511 | 0.563386 |
| Rps19 | Q9CZX8 | -0.78 | -1.3 | -2.69 | -2.85 | -0.27 | -1.06 | -2.13 | -2.93 | -0.9 | -2.09 | -2.83 | -0.65 | -0.33 | 0.16 | -1.46 | -2 | 0.3075 | 0.2875 | 0.9975 | 0.663825 | 0.664607 | 0.151146 | | 1 | 0.996511 | 0.749304 |
| Rps2 | P25444 | -2.44 | -2.71 | -2.59 | -2.61 | -1.75 | -2.55 | -1.48 | -2.56 | -2.69 | -2.33 | -2.56 | -2.32 | -2.29 | -1.78 | -1.65 | -2.14 | 0.5025 | 0.1125 | 0.6225 | 0.104979 | 0.528661 | 0.016692 | | 1 | 0.996511 | 0.591942 |
| Rps20 | P60867 | -2.59 | -3.09 | -5.02 | -5.97 | -2.68 | -1.92 | -3.57 | -4.25 | -4.39 | -4.23 | -4.64 | -2.25 | -2.62 | -3.96 | -2.09 | -3.54 | 1.0625 | 0.29 | 1.115 | 0.233316 | 0.740267 | 0.194215 | | 1 | 0.996511 | 0.778959 |
| Rps21 | Q9CQR2 | -0.62 | -1.11 | -1.33 | -1.4 | -1.07 | -0.66 | -1.33 | -1.32 | -0.18 | -1.2 | -1.43 | -0.62 | 0.37 | 0.03 | -0.79 | -0.73 | 0.02 | 0.2575 | 0.835 | 0.935813 | 0.444629 | 0.0331 | | 1 | 0.996511 | 0.644622 |
| Rps23 | P62267 | 0.21 | 0 | 0.17 | -1.38 | 1.18 | 0.46 | 0.2 | -0.2 | -1.32 | -0.25 | -0.14 | 0.69 | -0.5 | -0.47 | 1.26 | 0.65 | 0.66 | -0.005 | 0.485 | 0.163993 | 0.99231 | 0.375114 | | 1 | 0.999285 | 0.859611 |
| Rps24 | P62849-2 | -0.4 | -1.13 | -2.5 | -2.46 | -0.34 | -0.66 | -0.87 | -2.44 | -1.15 | -2.99 | -2.55 | -0.49 | -1.27 | -0.81 | -1.76 | -1.48 | 0.545 | -0.1725 | 0.2925 | 0.400227 | 0.808134 | 0.573745 | | 1 | 0.996511 | 0.925747 |
| Rps25 | A0A1L1SQ/ | -1.13 | -1.42 | -1.65 | -1.92 | -0.16 | -1.34 | -1.57 | -2.08 | -1.48 | -1.26 | -1.8 | -1.24 | -0.75 | -0.09 | -0.67 | -1.4 | 0.2425 | 0.085 | 0.8025 | 0.561059 | 0.724057 | 0.032338 | | 1 | 0.996511 | 0.641974 |
| Rps26 | P62855 | -3.5 | -3.78 | -3.78 | -3.8 | -2.49 | -3.47 | -3.96 | -4.28 | -3.48 | -1.74 | -3.67 | -3.26 | -2.67 | -3.13 | -2.94 | -3.75 | 0.165 | 0.6775 | 0.5925 | 0.663595 | 0.136492 | 0.049999 | | 1 | 0.996511 | 0.658454 |
| Rps27 | A0A0G2JD\ | 0.59 | 0.71 | 0.69 | 0.05 | 0.67 | 0.4 | 0.33 | 0.6 | -0.48 | 0.52 | 0.42 | 0.74 | -0.59 | -0.16 | 1.46 | 1.33 | -0.01 | -0.21 | 3.53E-16 | 0.961134 | 0.507318 | 1 | | 1 | 0.996511 | 1 |
| Rps27a | P62983 | 1.1 | -0.71 | 0.07 | 0.59 | -0.87 | 0.94 | 0.11 | 1.92 | -1.14 | -1.11 | 0.93 | 0.97 | -0.88 | -1.74 | 0.3 | 0.14 | 0.2625 | -0.35 | -0.8075 | 0.684896 | 0.593523 | 0.17747 | | 1 | 0.996511 | 0.772068 |
| Rps28 | P62858 | -1.94 | -1.88 | -1.74 | -1.94 | -1.61 | -2.31 | -1.66 | -1.96 | -1.22 | -0.6 | -1.7 | -1.11 | -0.16 | -0.38 | -1.21 | -1.29 | -0.01 | 0.7175 | 1.115 | 0.960232 | 0.018567 | 0.004956 | | 1 | 0.996511 | 0.552614 |
| Rps29 | P62274 | -3.24 | -4.82 | -5.66 | -6.08 | -2.81 | -3.54 | -5.28 | -3.64 | -6.47 | -4.09 | -5.47 | -2.24 | -3.8 | -5.07 | -5.43 | -4.63 | 1.1325 | 0.3825 | 0.2175 | 0.151087 | 0.702513 | 0.740351 | | 1 | 0.996511 | 0.958796 |
| Rps3 | P62908 | -2.4 | -2.36 | -1.88 | -2.21 | -1.34 | -2.42 | -1.34 | -2.08 | -2.54 | -1.27 | -2.11 | -2.23 | -1.91 | -1.35 | -1.35 | -1.43 | 0.4175 | 0.175 | 0.7025 | 0.181957 | 0.565337 | 0.012006 | | 1 | 0.996511 | 0.566767 |
| Rps3a | P97351 | -1.39 | -2.36 | -1.76 | -1.66 | -1.76 | -1.84 | -1.25 | -1.45 | -1.11 | -0.84 | -1.77 | -1.13 | -0.55 | -1.2 | -0.6 | -1.18 | 0.2175 | 0.58 | 0.91 | 0.406272 | 0.074231 | 0.011127 | | 1 | 0.996511 | 0.563386 |
| Rps4x | P62702 | -1.86 | -2.05 | -0.94 | -1.45 | -0.78 | -2.03 | -0.13 | -1.12 | -2.56 | -1.02 | -0.94 | -1.81 | -1.68 | -1.42 | 0.11 | -0.08 | 0.56 | -0.0075 | 0.8075 | 0.220266 | 0.986026 | 0.123287 | | 1 | 0.998892 | 0.721448 |
| Rps5 | Q91V55 | -2.85 | -4.02 | -2.42 | -2.79 | -3.19 | -2.91 | -2.55 | -2.52 | -3.28 | -2.07 | -2.36 | -2.91 | -1.99 | -3.87 | -1.27 | -1.8 | 0.2275 | 0.365 | 0.7875 | 0.537104 | 0.393512 | 0.217285 | | 1 | 0.996511 | 0.787447 |
| Rps6 | P62754 | -2.37 | -2.41 | -1.43 | -2.32 | -2.18 | -1.97 | -1.08 | -1.76 | -2.32 | -2.96 | -1.39 | -1.94 | -0.95 | -3.03 | -0.51 | -0.99 | 0.385 | -0.02 | 0.7625 | 0.257166 | 0.958952 | 0.198677 | | 1 | 0.996755 | 0.78004 |
| Rps6ka1 | E9PWV3 | -1.21 | -1.03 | -0.84 | -1.13 | -0.98 | -1.31 | -1 | -1 | -1.22 | 1.75 | -0.83 | -1.18 | -1.13 | -0.92 | -1.27 | -0.7 | -0.02 | 0.6825 | 0.0475 | 0.904927 | 0.312053 | 0.819409 | | 1 | 0.996511 | 0.972768 |
| Rps6ka2 | Q9WUT3 | 0.23 | -0.07 | -0.04 | 0.45 | 0.45 | -0.13 | 0.83 | 0.7 | -0.14 | 0.84 | -0.1 | -0.04 | 0.36 | 0.64 | 1.17 | 1.15 | 0.32 | -0.0025 | 0.6875 | 0.231947 | 0.992803 | 0.025963 | | 1 | 0.999285 | 0.633075 |
| Rps6ka3 | B1AXN9 | 0.14 | -0.12 | 0.11 | 0.25 | -0.02 | 0.33 | -0.04 | 0.36 | 0 | 0.96 | 0.3 | 0.02 | 0.25 | 0.08 | 0.09 | -0.15 | 0.0625 | 0.225 | -0.0275 | 0.727992 | 0.392875 | 0.885063 | | 1 | 0.996511 | 0.979767 |
| Rps6ka4 | Q9Z2B9 | -2.79 | -2.59 | -1.96 | -1.68 | -2.21 | -3.22 | -1.2 | -2.19 | -2.34 | -1.18 | -1.46 | -2.42 | -1.33 | -2.62 | -1.86 | -1.66 | 0.05 | 0.405 | 0.3875 | 0.912208 | 0.314696 | 0.31085 | | 1 | 0.996511 | 0.834523 |
| Rps6ka5 | Q8C050 | -3.67 | -3.75 | -2.54 | -3.04 | -3.3 | -4.4 | -2.46 | -3.26 | -3.65 | -1.15 | -2.81 | -3.19 | -3.1 | -2.95 | -3.15 | -2.6 | -0.105 | 0.55 | 0.3 | 0.8178 | 0.344498 | 0.357991 | | 1 | 0.996511 | 0.856645 |
| Rps6kb1 | Q8BSK8-2 | 0.5 | 0.04 | 0.22 | -0.41 | 0.29 | 0.26 | 0.43 | -0.07 | -1.14 | -0.8 | 0.05 | 0.33 | -0.97 | -0.49 | 0.07 | 0.12 | 0.14 | -0.4775 | -0.405 | 0.556344 | 0.231829 | 0.234109 | | 1 | 0.996511 | 0.790641 |
| Rps6kc1 | E9QMX4 | 0.61 | -0.49 | -0.21 | -0.04 | -0.56 | 0.48 | -0.22 | -0.14 | 0.26 | 0.01 | -0.27 | 0.03 | 0 | -0.25 | -0.48 | -0.5 | -0.0775 | 0.04 | -0.275 | 0.804584 | 0.883133 | 0.341609 | | 1 | 0.996511 | 0.848473 |
| Rps7 | P62082 | -0.63 | -0.6 | -1.11 | -0.91 | -0.13 | -0.78 | -0.48 | -0.98 | -0.85 | -0.71 | -0.99 | -0.6 | -0.47 | -0.34 | -0.47 | -0.41 | 0.22 | 0.025 | 0.39 | 0.367162 | 0.899724 | 0.0731 | | 1 | 0.996511 | 0.668161 |
| Rps8 | P62242 | -2.53 | -2.43 | -1.76 | -1.55 | -1.56 | -2.47 | -0.68 | -1.43 | -2.71 | -0.99 | -1.55 | -2.2 | -2.4 | -1.78 | -0.82 | -0.84 | 0.5325 | 0.205 | 0.6075 | 0.219828 | 0.630776 | 0.18469 | | 1 | 0.996511 | 0.776843 |
| Rps9 | Q6ZWN5 | -0.91 | -1.34 | -0.52 | -0.58 | -0.51 | -1.25 | 0.12 | -0.3 | -1.14 | -0.3 | -0.61 | -0.63 | -0.53 | -0.58 | 0.41 | 0.3 | 0.3525 | 0.1675 | 0.7375 | 0.305533 | 0.541524 | 0.048038 | | 1 | 0.996511 | 0.658454 |
| Rpsa | P14206 | -0.12 | -0.19 | -0.69 | -0.65 | 0.72 | -0.2 | 0.01 | -0.51 | -1.16 | -0.74 | -0.68 | 0.06 | -1.21 | -0.53 | -0.16 | -0.02 | 0.4175 | -0.2175 | -0.0675 | 0.186878 | 0.475647 | 0.829691 | | 1 | 0.996511 | 0.973939 |
| Rptor | A2ACM0 | 0.15 | 0 | 0.09 | 0.31 | 0.06 | 0.09 | -0.23 | 0.15 | 0.17 | -0.02 | 0.05 | 0.08 | -0.35 | -0.24 | -0.43 | -0.39 | -0.12 | -0.0675 | -0.49 | 0.472921 | 0.681604 | 0.019964 | | 1 | 0.996511 | 0.615793 |
| Rpusd3 | Q14AI6 | -0.06 | 0.39 | 0.78 | 0.06 | 0.29 | 0.2 | -0.08 | -0.03 | 0.45 | 0.23 | 0.08 | -0.18 | 0.59 | 1.07 | 0.3 | 0.47 | -0.1975 | -0.1475 | 0.315 | 0.397649 | 0.562818 | 0.267951 | | 1 | 0.996511 | 0.819742 |

| | | | | | | | | | | | | | | | | | | | | | | | | | | |
|---|---|---|---|---|---|---|---|---|---|---|---|---|---|---|---|---|---|---|---|---|---|---|---|---|---|---|
| Rpusd4 | Q9CWX4 | 0 | 0.37 | 0.29 | 0.94 | -0.02 | 0.01 | -0.16 | 0.77 | 0.53 | 0.81 | 0.49 | -0.04 | 0.22 | 0.49 | 1.03 | 0.11 | -0.25 | 0.0475 | 0.0625 | 0.396673 | 0.864231 | 0.833999 | 1 | 0.996511 | 0.975371 |
| Rraga | Q80X95 | -0.26 | -0.91 | -1.27 | -0.94 | -2 | 0.66 | -1.56 | -0.81 | 0.67 | 0.2 | -1.36 | 1.63 | -0.22 | -0.36 | -1.56 | -2.74 | -0.0825 | 1.13 | -0.375 | 0.884387 | 0.088903 | 0.519789 | 1 | 0.996511 | 0.907667 |
| Rragb | Q6NTA4 | -1.98 | -2.09 | -1.83 | -1.77 | -2.07 | -1.99 | -1.57 | -2.15 | -1.91 | -1.84 | -1.9 | -1.81 | -1.32 | -1.86 | -2.39 | -2.23 | -0.0275 | 0.0525 | -0.0325 | 0.883668 | 0.74922 | 0.904858 | 1 | 0.996511 | 0.985525 |
| Rragc | Q99K70 | -1.15 | -0.73 | 0.06 | 0.05 | -0.44 | -0.83 | -0.24 | -0.19 | -0.89 | -0.04 | 0.05 | -0.86 | -0.74 | -0.65 | -0.52 | -0.33 | 0.0175 | 0.0075 | -0.1175 | 0.956989 | 0.984212 | 0.713801 | 1 | 0.998892 | 0.958106 |
| Rragd | B1AWT2 | -1.32 | -0.6 | -1.34 | -2.07 | -1.44 | -1.65 | -2.22 | -2.69 | -1.85 | -1.97 | -1.84 | -1.57 | -0.77 | -2.46 | -2.41 | -2 | -0.6675 | -0.475 | -0.5775 | 0.114921 | 0.156138 | 0.235617 | 1 | 0.996511 | 0.791493 |
| Rras | P10833 | 0.36 | 1.04 | 0.9 | 0.93 | 1.41 | 0.54 | 1.39 | 0.85 | -0.6 | 0.19 | 0.72 | 0.47 | -0.13 | -0.81 | 0.44 | 0.87 | 0.24 | -0.6125 | -0.715 | 0.382302 | 0.085991 | 0.089587 | 1 | 0.996511 | 0.69217 |
| Rras2 | P62071 | 1.26 | 1.43 | 1.5 | 1.18 | 1.7 | 1.23 | 1.5 | 1.23 | 0.87 | 0.92 | 1.65 | 1.48 | 0.7 | 0.64 | 1.55 | 1.59 | 0.0725 | -0.1125 | -0.2225 | 0.68961 | 0.639089 | 0.447673 | 1 | 0.996511 | 0.884817 |
| Rrbp1 | A2AVJ7 | -0.2 | -0.3 | -0.35 | -0.37 | 0.12 | 0.03 | 0.31 | -0.21 | 0.92 | 0.18 | -0.21 | 0.15 | 2.32 | 0.19 | 2.21 | 0.9 | 0.3675 | 0.565 | 1.71 | 0.053926 | 0.054246 | 0.006378 | 1 | 0.996511 | 0.552614 |
| Rrm1 | P07742 | -1.85 | -1.29 | -0.03 | -0.41 | -1.27 | -1.25 | -0.86 | -0.12 | -1.57 | 0.04 | -0.25 | -1.27 | -2.08 | -0.17 | -0.95 | -1.42 | 0.02 | 0.1325 | -0.26 | 0.965154 | 0.801159 | 0.628429 | 1 | 0.996511 | 0.940342 |
| Rrm2b | Q6PEE3 | -0.02 | 0.05 | -0.79 | -0.73 | -0.5 | -0.11 | -0.67 | -0.84 | -0.7 | -1.12 | -1.08 | -0.17 | -0.5 | -0.7 | -1.45 | -1.45 | -0.1575 | -0.395 | -0.6525 | 0.573565 | 0.232492 | 0.079395 | 1 | 0.996511 | 0.681213 |
| Rrp1 | P56183-2 | -1.52 | -3.54 | -1.95 | -2.46 | -3.27 | -2.94 | -2.92 | -2.95 | -1.21 | -2.42 | -2.47 | -2.57 | 0.89 | -5.02 | -1.38 | -2.25 | -0.6525 | 0.2 | 0.4275 | 0.143762 | 0.691918 | 0.712446 | 1 | 0.996511 | 0.958014 |
| Rrp12 | Q6P5B0 | 0.6 | 0.69 | 1.13 | 0.75 | 0.33 | 0.62 | 0.55 | 0.78 | 0.4 | 0.8 | 0.73 | 0.53 | 0.77 | -0.16 | 1.16 | 1.05 | -0.2225 | -0.1775 | -0.0875 | 0.25785 | 0.383126 | 0.788768 | 1 | 0.996511 | 0.968677 |
| Rrp15 | Q9CYX7 | 0.2 | 0.78 | -0.5 | -0.71 | 0.13 | 0.31 | -0.03 | -0.8 | 0.25 | -1.27 | -0.55 | 0.89 | -0.19 | 0.49 | -0.17 | -0.77 | -0.04 | -0.1125 | -0.1025 | 0.919385 | 0.833962 | 0.802558 | 1 | 0.996511 | 0.971091 |
| Rrp9 | Q91WM3 | 0.53 | 0.2 | 1.37 | 1.17 | 0.68 | 0.24 | 0.68 | 0.83 | 0.54 | 1.19 | 1.17 | -0.12 | 1.02 | 0.67 | 1.38 | 1.42 | -0.21 | -0.1225 | 0.305 | 0.488954 | 0.757523 | 0.365117 | 1 | 0.996511 | 0.858475 |
| Rrs1 | Q9CYH6 | 0.8 | 0.69 | 0.24 | 0.5 | 0.59 | -0.14 | 0.12 | -0.02 | 0.25 | 0.09 | 0.57 | -0.05 | -0.24 | 0.99 | 0.62 | 0.59 | -0.42 | -0.3425 | -0.0675 | 0.087762 | 0.144319 | 0.822223 | 1 | 0.996511 | 0.972768 |
| Rsf1 | E9PWW9 | 0.74 | 0.51 | 0.55 | 0.47 | 0.3 | 0.33 | 0.48 | 0.04 | 0.96 | 0.71 | 0.27 | 0.79 | 0.98 | 1.16 | 0.29 | 0.5 | -0.28 | 0.115 | 0.165 | 0.118811 | 0.578098 | 0.509253 | 1 | 0.996511 | 0.902804 |
| Rsl1d1 | Q8BVY0 | -0.27 | -0.3 | 3.22 | 1.81 | -0.66 | -0.36 | 2.45 | 1.88 | 0.82 | 1.25 | 2.33 | -0.81 | 1.67 | -0.93 | 2.29 | 2.6 | -0.2875 | -0.2175 | 0.2925 | 0.782828 | 0.822454 | 0.780815 | 1 | 0.996511 | 0.9676 |
| Rsl24d1 | E9PY85 | 1.55 | 1.79 | 0.01 | -0.89 | 1.15 | 1.53 | -0.22 | -0.71 | 0.73 | -2.92 | -1.21 | 1.61 | 0.93 | 1.2 | -0.92 | 0.39 | -0.1775 | -1.0625 | -0.215 | 0.813772 | 0.336642 | 0.765146 | 1 | 0.996511 | 0.964134 |
| Rspry1 | Q8BVR6 | 0.14 | 0.42 | 0.31 | 0.23 | 0.7 | 0.1 | 0.45 | 0.16 | 0.72 | 0.41 | 0.51 | -0.07 | 0.43 | 0.7 | 0.05 | 0.43 | 0.0775 | 1.18E-01 | 0.1275 | 0.684277 | 0.590574 | 0.542364 | 1 | 0.996511 | 0.912212 |
| Rsrc1 | Q3TGY1 | -0.07 | -0.47 | -0.86 | -1.29 | 0.23 | -1.23 | -0.02 | -0.85 | -0.45 | -1.09 | -1.23 | -0.52 | -0.18 | 0.14 | -0.49 | -1.85 | 0.205 | -0.15 | 0.0775 | 0.616285 | 0.647683 | 0.871148 | 1 | 0.996511 | 0.979293 |
| Rsrc2 | S4R2L4 | -2.14 | -2.96 | -2.57 | -2.37 | -2.74 | -2.29 | -2.62 | -2.75 | -2.3 | -1.17 | -2.52 | -2.53 | -0.93 | -2.14 | 0.24 | -1.43 | -0.09 | 0.38 | 1.445 | 0.691078 | 0.305597 | 0.016259 | 1 | 0.996511 | 0.589387 |
| Rsu1 | Q9D031 | 0.16 | -0.13 | 0.6 | 0.6 | 0.28 | 0.4 | 0.62 | 0.81 | 0.02 | 0.26 | 0.72 | 0.4 | -0.19 | 0.51 | 0.53 | 0.58 | 0.22 | 0.0425 | 0.05 | 0.356897 | 0.866619 | 0.857018 | 1 | 0.996511 | 0.976225 |
| RtcA | Q9D7H3 | -1.34 | -1.45 | -1.51 | -1.42 | -1.55 | -1.57 | -1.09 | -1.49 | -0.76 | -0.69 | -1.29 | -1.34 | 0.1 | -0.8 | -1.39 | -1.41 | 0.005 | 0.41 | 0.555 | 0.976604 | 0.083759 | 0.142334 | 1 | 0.996511 | 0.740431 |
| Rtcb | Q99LF4 | -1.6 | -1.79 | -1.5 | -1.87 | -1.2 | -1.69 | -1.26 | -1.58 | -2.31 | -1.99 | -1.52 | -1.22 | -1.98 | -1.99 | -1.45 | -1.54 | 0.2575 | -0.07 | -0.05 | 0.192665 | 0.796944 | 0.818905 | 1 | 0.996511 | 0.972768 |
| Rtf1 | A2AQ19 | -2.04 | -2.48 | -1.78 | -1.46 | -3.23 | -2.82 | -2.77 | -2.54 | -1.86 | -0.48 | -1.73 | -1.88 | -2.34 | -1.78 | -3.01 | -2.89 | -0.9 | 0.4525 | -0.565 | 0.007985 | 0.259396 | 0.135356 | 1 | 0.996511 | 0.731223 |
| Rtkn | A0A1D5RLJ | 0.56 | -1.29 | -0.63 | -0.32 | -1.12 | -1.06 | -0.48 | -0.44 | 0.59 | -0.3 | -0.23 | -0.92 | 0.77 | -0.27 | -0.74 | -0.16 | -0.355 | 0.205 | 0.32 | 0.387105 | 0.659444 | 0.499711 | 1 | 0.996511 | 0.90025 |
| Rtkn | Q8C6B2-2 | 0.31 | 0.18 | 0.92 | 1.1 | 0.38 | -0.02 | 0.8 | 0.9 | 0.11 | 1.26 | 1.04 | 0.31 | 0.57 | 0.06 | 0.99 | 1.41 | -0.1125 | 0.0525 | 0.13 | 0.71325 | 0.880932 | 0.719652 | 1 | 0.996511 | 0.958398 |
| Rtkn | D3Z4N1 | 0.55 | -0.22 | 0.72 | 0.02 | -2.64 | 1.04 | -1.92 | -0.68 | 1.09 | 0.26 | -0.54 | 0.06 | 2.51 | -1.06 | -0.43 | -0.72 | -1.3175 | -0.05 | -0.1925 | 0.108712 | 0.897137 | 0.802309 | 1 | 0.996511 | 0.971091 |
| Rtl8b | Q9D1F0 | -1.68 | -1.77 | -0.9 | -2.14 | -1.18 | -1.92 | -1.7 | -1.52 | -1.08 | -1.6 | -1.06 | -1.6 | -1.72 | -1.59 | -1.94 | -1.41 | 0.0425 | 0.2875 | -0.0425 | 0.88777 | 0.360516 | 0.886333 | 1 | 0.996511 | 0.979767 |
| Rtn1 | Q8K0T0 | 1.39 | 1.24 | 0.51 | 0.5 | 0.87 | 0.78 | 0.46 | 0.44 | 0.59 | 0.57 | 0.26 | 1.24 | 0.37 | 1 | 0.57 | 0.45 | -0.2725 | -0.245 | -0.3125 | 0.32225 | 0.44465 | 0.298801 | 1 | 0.996511 | 0.828257 |
| Rtn1 | Q7M6W1 | -0.88 | -0.65 | -0.81 | -0.44 | -0.26 | -0.42 | -0.12 | -0.72 | -0.81 | -1.18 | -0.52 | -0.84 | -0.4 | -0.96 | -0.46 | -0.76 | 0.315 | -0.1425 | 0.05 | 0.13598 | 0.502792 | 0.817741 | 1 | 0.996511 | 0.972768 |
| Rtn2 | O70622 | 0.79 | 0.68 | 0.75 | 0.74 | 1.23 | 0.49 | 0.9 | 1.14 | 1.09 | 1.66 | 1.32 | 0.89 | 1 | 1.9 | 1.55 | 1.41 | 0.2 | 0.5 | 0.725 | 0.331561 | 0.038816 | 0.011496 | 1 | 0.996511 | 0.566767 |
| Rtn3 | Q9ES97-2 | 0.83 | 0.57 | 0.36 | 0.45 | 0.21 | 0.35 | 0.19 | 0.19 | 0.55 | 0.24 | 0.22 | 0.82 | 0.34 | 0.77 | 0.16 | 0.05 | -0.3175 | -0.095 | -0.2225 | 0.082732 | 0.660541 | 0.350267 | 1 | 0.996511 | 0.853281 |
| Rtn3 | Q9ES97-3 | -7.51 | -8.18 | -4.94 | -4.83 | -6.49 | -5.93 | -3.12 | -5.04 | -7.61 | -7 | -5.19 | -7.89 | -7.02 | -6.97 | -4.46 | -4.03 | 1.22 | -0.5575 | 0.745 | 0.251644 | 0.561528 | 0.485313 | 1 | 0.996511 | 0.894829 |
| Rtn4 | Q99P72-1 | -0.93 | -0.66 | 0.06 | -0.26 | 0.64 | -0.93 | 0.67 | 0.27 | -3.27 | -4.88 | 0.52 | -0.96 | -4.19 | -5.15 | -1.62 | -0.42 | 0.61 | -1.7 | -2.3975 | 0.161145 | 0.146517 | 0.038124 | 1 | 0.996511 | 0.658454 |
| Rtn4 | Q99P72 | 0.35 | 0.7 | 0.65 | 0.59 | 0.76 | -0.07 | 1.07 | 0.57 | -0.07 | 0.12 | 0.6 | 0.45 | -0.09 | 0.86 | 0.67 | 0.59 | 0.01 | -0.2975 | -0.065 | 0.969438 | 0.185879 | 0.797624 | 1 | 0.996511 | 0.969727 |
| Rtn4ip1 | A0A0R4J0S | 0.99 | 1.09 | 0.84 | 1.38 | 0.58 | 0.69 | 1.27 | 0.82 | 0.73 | 0.92 | 0.75 | 0.73 | 0.45 | 1.02 | 0.48 | 0.49 | -0.235 | -0.2925 | -0.465 | 0.291714 | 0.139943 | 0.064989 | 1 | 0.996511 | 0.66433 |
| Rtn4r | Q99PI8 | 0.08 | 0.19 | 0.31 | 0.41 | 0.48 | 0.46 | 0.52 | 0.51 | 0.24 | 0.42 | 0.88 | 0.41 | 0.14 | 0.89 | -0.36 | 0.01 | 0.245 | 0.24 | -0.0775 | 0.128304 | 0.255501 | 0.788829 | 1 | 0.996511 | 0.968677 |
| Rtn4rl1 | Q8K0S5 | 0.93 | 0.71 | -0.01 | -0.35 | 1.07 | 1.44 | 0.39 | -0.04 | -0.69 | -1.06 | 0.57 | 0.76 | -0.95 | -0.31 | -0.67 | -0.11 | 0.395 | -0.425 | -0.83 | 0.359792 | 0.409648 | 0.039757 | 1 | 0.996511 | 0.658454 |
| Rtn4rl2 | A2ATS6 | -0.11 | -0.32 | 0.41 | 0.25 | 0.16 | 0.45 | 0.37 | 0.59 | 0.04 | 0.3 | 0.61 | 0.16 | 0.15 | -0.19 | 0.37 | 0.28 | 0.335 | 0.22 | 0.095 | 0.145434 | 0.363772 | 0.697686 | 1 | 0.996511 | 0.955729 |
| RTRAF | Q9CQE8 | -0.36 | -1.6 | -1.36 | -2.04 | -0.95 | -0.23 | -1.07 | -1.57 | -1.42 | -0.83 | -1.22 | -0.05 | -0.84 | -1.31 | -1.42 | -1.4 | 0.385 | 0.46 | 0.0975 | 0.373664 | 0.311951 | 0.793805 | 1 | 0.996511 | 0.969367 |
| Rubcn | D3Z7B1 | -0.72 | -0.83 | -1.46 | -0.57 | -1.59 | -1.26 | -1.28 | -0.57 | -0.54 | -0.36 | -0.49 | -0.98 | 0.65 | -2.44 | -0.84 | -1.08 | -0.28 | 0.3025 | -0.0325 | 0.34932 | 0.259639 | 0.957226 | 1 | 0.996511 | 0.992636 |
| Rufy1 | Q8BIJ7 | -0.84 | -1.76 | -1.49 | -1.85 | -1.36 | -0.74 | -1.24 | -1.61 | -1.12 | -1.08 | -1.19 | -1.05 | -0.65 | -1.74 | -1.64 | -1.46 | 0.2475 | 0.375 | 0.1125 | 0.406976 | 0.163033 | 0.739752 | 1 | 0.996511 | 0.958796 |
| Rufy2 | Q8R4C2 | 0 | -0.13 | 0.22 | -0.12 | 0.48 | -0.17 | 0.53 | 0.18 | 0.23 | 0.3 | 0.42 | 0.4 | 0.5 | 0.76 | 0.49 | 0.63 | 0.2625 | 0.345 | 0.6025 | 0.230646 | 0.069253 | 0.009722 | 1 | 0.996511 | 0.563386 |
| Rufy2 | D3Z4D2 | 0.71 | -0.75 | 0.72 | -0.3 | 0.65 | 0.79 | -0.14 | -0.15 | 0.44 | 0.56 | -0.09 | 0.92 | -0.08 | 1.82 | -0.06 | 0.34 | 0.1925 | 0.3625 | 0.41 | 0.647902 | 0.382276 | 0.453625 | 1 | 0.996511 | 0.886245 |
| Rufy3 | Q9D394-3 | 2.17 | 2.49 | 2.01 | 2.13 | -0.32 | 2.56 | 1.02 | 2.57 | 1.24 | 3.15 | 2.11 | 1.55 | -0.91 | 0.61 | 1.77 | 2.11 | -0.7425 | -0.1875 | -1.305 | 0.264296 | 0.6503 | 0.064074 | 1 | 0.996511 | 0.66433 |
| Rufy3 | Q9D394-4 | -1.08 | -0.65 | 0.64 | 0.28 | -0.15 | -0.72 | -0.41 | -0.43 | 0.66 | -0.6 | -0.2 | 0.28 | 0.12 | -0.09 | 0.21 | -0.46 | -0.225 | 0.2375 | 0.1475 | 0.570668 | 0.604815 | 0.719257 | 1 | 0.996511 | 0.958398 |
| Rufy3 | Q9D394 | 0.46 | 0.36 | 1.1 | 1.03 | 0.44 | 0.45 | 0.87 | 1.19 | 0.48 | 0.96 | 1.38 | 0.86 | 0.4 | 0.01 | 0.93 | 1.18 | -2E-17 | 0.1825 | -0.1075 | 1 | 0.517336 | 0.74415 | 1 | 0.996511 | 0.959371 |
| Rundc1 | Q0VDN7 | -0.72 | -1.3 | -0.72 | -0.61 | -1.2 | -0.73 | -0.86 | -0.76 | -1.49 | -0.53 | -1.11 | -0.39 | -0.71 | -1.35 | -1.91 | -1.58 | -0.05 | -0.0425 | -0.55 | 0.816585 | 0.889009 | 0.101586 | 1 | 0.996511 | 0.698697 |
| Rundc3a | O08576-4 | -0.91 | -0.34 | -0.16 | 1.45 | 0.13 | -0.72 | 1.39 | 1.09 | -0.31 | 1.7 | 0.94 | -0.06 | -0.33 | 0.47 | 1.6 | 2.05 | 0.4625 | 0.5575 | 0.9375 | 0.472786 | 0.385636 | 0.187503 | 1 | 0.996511 | 0.776843 |
| Rundc3a | O08576 | 0.35 | 0.43 | 0.17 | 0.35 | 0.8 | 0.58 | 0.87 | 0.34 | 0.09 | 0.05 | 0.53 | 0.53 | 0.01 | 0.78 | 0.44 | 0.43 | 0.3225 | -0.025 | 0.09 | 0.09796 | 0.898563 | 0.682094 | 1 | 0.996511 | 0.955729 |
| Rundc3b | D3YUQ5 | -0.2 | 0.34 | 0.31 | 0.06 | 0.14 | -0.46 | 0.22 | -0.3 | 0.25 | 0.15 | -0.22 | 0.16 | 0.03 | 0.41 | 0.34 | -0.06 | -0.2275 | -0.0425 | 0.0525 | 0.333137 | 0.837977 | 0.812282 | 1 | 0.996511 | 0.972642 |
| Rusc1 | Q8BG26 | 0.15 | 0.61 | 0.15 | 0.85 | 0.64 | -0.06 | 0.31 | -0.47 | 1.24 | 2.16 | -0.05 | 0.37 | 0.86 | 1.15 | 0.44 | -0.19 | -0.335 | 0.49 | 0.125 | 0.27434 | 0.327316 | 0.71374 | 1 | 0.996511 | 0.958106 |
| Rusc2 | G8JL85 | 0.03 | 0.07 | 0.67 | 0.74 | 0.49 | 0.23 | 0.82 | 0.77 | 0.22 | 0.86 | 0.56 | 0.06 | 0.38 | 0.75 | 0.78 | 0.53 | 0.2 | 0.0475 | 0.2325 | 0.427149 | 0.862655 | 0.359482 | 1 | 0.996511 | 0.856645 |
| Rusf1 | Q91W34 | 0.06 | 0.67 | -0.07 | 0.02 | 1.28 | 0.17 | 1.98 | 1.38 | 1.25 | 0.59 | 0.67 | 0.56 | 0.64 | 0.63 | -0.37 | 1.02 | 1.0325 | 0.5975 | 0.31 | 0.025708 | 0.041618 | 0.37619 | 1 | 0.996511 | 0.860376 |
| Rusf1 | A0A0R4J0E | -0.62 | -0.14 | -0.76 | -0.19 | 0.2 | -0.46 | -0.55 | -0.85 | 0.09 | -0.08 | -0.88 | -0.53 | 0.12 | 0.42 | -0.57 | -0.62 | 0.0125 | 0.0775 | 0.265 | 0.963636 | 0.782738 | 0.402549 | 1 | 0.996511 | 0.870698 |
| Ruvbl1 | P60122 | -0.65 | -0.8 | -0.64 | -0.21 | -0.71 | -1.13 | -0.01 | -0.24 | -0.98 | -0.66 | -0.37 | -0.86 | -0.87 | -0.27 | -0.43 | -0.23 | 0.0525 | -0.1425 | 0.125 | 0.852458 | 0.523984 | 0.59824 | 1 | 0.996511 | 0.930093 |
| Ruvbl2 | Q9WTM5 | 0.1 | -0.1 | 0.07 | 0.23 | -0.26 | -0.26 | 0.09 | 0.19 | -0.21 | -0.13 | -0.04 | -0.07 | -0.43 | -0.24 | -0.09 | 0.17 | -0.135 | -0.1875 | -0.2225 | 0.461621 | 0.272043 | 0.295668 | 1 | 0.996511 | 0.825612 |
| Rwdd1 | Q9CQK7 | -0.41 | 0.17 | -0.24 | -0.1 | 0.14 | -0.52 | -0.13 | -0.17 | 0.15 | -0.02 | -0.22 | -0.15 | -0.04 | 0.29 | -0.4 | -0.43 | -0.025 | 0.085 | 0 | 0.905411 | 0.669323 | 1 | 1 | 0.996511 | 1 |
| Rwdd2a | Q14BN8 | 0.12 | 0.3 | -0.39 | -1.33 | 0.18 | 0.65 | 0.22 | 0.06 | -1.59 | -0.92 | -0.19 | 0.66 | -0.46 | -1.92 | 0.6 | 0.11 | 0.6025 | -0.185 | -0.0925 | 0.130267 | 0.741045 | 0.877774 | 1 | 0.996511 | 0.979626 |
| Rwdd4 | Q9CPR1 | -0.24 | 0 | 0.24 | -0.35 | 0.12 | -0.07 | -1.31 | -0.83 | 0.13 | -1.5 | -0.42 | -0.25 | 0.47 | -1.68 | -0.7 | -1.21 | -0.435 | -0.4225 | -0.6925 | 0.229082 | 0.26353 | 0.152907 | 1 | 0.996511 | 0.750456 |
| Rxrb | Q8VCR0 | -1.19 | -1.49 | -2.92 | -2.14 | -1.91 | -1.18 | -2.33 | -3.76 | 0.38 | -3.37 | -3.09 | -1.28 | -0.53 | -1.36 | -4.53 | -1.21 | -0.36 | 0.095 | 0.0275 | 0.556495 | 0.911985 | 0.974833 | 1 | 0.996744 | 0.992977 |
| Ryr2 | E9Q401 | 0.53 | 0.68 | 0.77 | 0.62 | 0.64 | 0.51 | 0.55 | 0.49 | 0.29 | 0.18 | 0.45 | 0.15 | 0.21 | 0.47 | 0.32 | 0.4 | -0.1025 | -0.3825 | -0.3 | 0.485947 | 0.045043 | 0.114071 | 1 | 0.996511 | 0.717287 |
| Ryr2 | A0A1Y7VK( | -0.95 | -0.85 | -1.63 | -2.03 | -0.89 | -1.31 | -0.82 | -1.27 | -0.41 | -3.49 | -1.48 | -1.26 | -3.01 | -1.52 | -2.67 | -1.38 | 0.2925 | -0.295 | -0.78 | 0.350752 | 0.650826 | 0.121635 | 1 | 0.996511 | 0.721448 |
| Ryr2 | A0A1Y7VJP | 1.34 | 1.66 | 1.44 | 1.08 | 1.39 | 0.72 | -0.09 | 0.84 | 1.28 | 1.71 | 1.15 | -0.3 | 0.63 | 1.84 | 0.72 | 0.72 | -0.665 | -0.42 | -0.4025 | 0.064395 | 0.341046 | 0.228302 | 1 | 0.996511 | 0.787447 |
| Ryr3 | A0A140LJK | 0.71 | 0.4 | 0.7 | 0.8 | 0.73 | 0.23 | 1 | 0.69 | 0.48 | 0.43 | 0.38 | 0.47 | 0.12 | 1.01 | 0.87 | 0.86 | 0.01 | -0.2125 | 0.0625 | 0.961981 | 0.230338 | 0.804316 | 1 | 0.996511 | 0.971446 |
| S100a1 | Q91V77 | -1.57 | -1.62 | -0.82 | -0.61 | -1.32 | -1.22 | -0.33 | -0.73 | -1.38 | -1.51 | -0.96 | -1.87 | -1.49 | -1.12 | -1.44 | -1.07 | 0.255 | -0.275 | -0.125 | 0.453852 | 0.399764 | 0.67266 | 1 | 0.996511 | 0.952944 |
| S100a10 | P08207 | 1.1 | 0.65 | -0.37 | 1.65 | 0.76 | 1.14 | 0.4 | 0.47 | 1.05 | 0.95 | -0.17 | 0.77 | 0.21 | 1.41 | -0.31 | 0.56 | -0.065 | -0.1075 | -0.29 | 0.879793 | 0.821949 | 0.580321 | 1 | 0.996511 | 0.927734 |
| S100a11 | P50543 | -4.5 | -4.63 | -3.2 | -3.22 | -3.85 | -4.07 | -3 | -3.53 | -3.87 | -3.33 | -3.62 | -3.53 | -4.23 | -3.94 | -4.5 | -3.4 | 0.275 | 0.3 | -0.13 | 0.524182 | 0.451091 | 0.765234 | 1 | 0.996511 | 0.964134 |
| S100a13 | A0A0A0MC | 1.04 | 1.46 | 1.8 | 1.82 | 2.02 | 0.92 | 2.3 | 1.41 | 0.82 | 1.79 | 1.54 | 0.98 | 0.81 | 1.64 | 1.89 | 1.75 | 0.1325 | -0.2475 | -0.0075 | 0.703061 | 0.417714 | 0.980857 | 1 | 0.996511 | 0.994858 |
| S100a16 | Q9D708 | -0.77 | -1.01 | 0.23 | 0.07 | -0.25 | -0.86 | -0.2 | -0.3 | -1.16 | 0.33 | -0.27 | -0.79 | -1.44 | -0.81 | 0.69 | 0.39 | -0.0325 | -0.1025 | 0.0775 | 0.92204 | 0.808448 | 0.88642 | 1 | 0.996511 | 0.979767 |
| S100a4 | A0A0G2JGI | -0.33 | -0.55 | -0.2 | 0.13 | -0.96 | -0.66 | -1.82 | -0.35 | -0.75 | -0.26 | -0.85 | -0.5 | -0.02 | -1.73 | 0.05 | -0.65 | -0.71 | -0.3525 | -0.35 | 0.060309 | 0.147846 | 0.411081 | 1 | 0.996511 | 0.874811 |
| S100a6 | P14069 | 0.78 | 0.12 | 1.19 | 0.69 | 0.59 | 0.34 | 0.42 | 0.92 | 0.69 | 0.87 | 1.16 | 0.47 | 0.34 | -0.07 | 1.47 | 0.96 | -0.1275 | 0.1025 | -0.02 | 0.630078 | 0.712634 | 0.959228 | 1 | 0.996511 | 0.992636 |
| S100a8 | P27005 | 0.53 | -1.97 | 0.25 | 0.58 | -3.85 | -0.93 | -0.86 | 0.51 | 1.06 | 0.24 | -0.13 | -1.98 | 0.97 | -2.99 | 0.16 | 0.41 | -1.13 | -0.05 | -0.21 | 0.271664 | 0.950184 | 0.82825 | 1 | 0.996755 | 0.973757 |
| S100a9 | P31725 | 1.68 | 0.64 | 4.58 | 2.31 | -1.36 | 0.65 | 0.29 | 2.14 | -0.26 | -0.2 | 1.39 | -0.59 | -0.69 | -2.02 | -0.5 | 1.08 | -1.8725 | -2.2175 | -2.835 | 0.085593 | 0.027302 | 0.013889 | 1 | 0.996511 | 0.589387 |
| S100b | P50114 | -1.55 | -1.43 | -3.55 | -2.5 | -1.24 | -2.05 | -2.48 | -3.77 | -1.57 | -3.52 | -3.55 | -1.64 | -1.82 | -2.31 | -4.23 | -3.23 | -0.1275 | -0.3125 | -0.64 | 0.845949 | 0.646454 | 0.345499 | 1 | 0.996511 | 0.849669 |
| S1pr1 | O08530 | 0.76 | 1.15 | 1.01 | 0.72 | 1.64 | 1.16 | 0.94 | 0.75 | 1.3 | 1.2 | 1.18 | 0.47 | 1.57 | 1.31 | 1.3 | 0.28 | 0.2125 | 0.1275 | 0.205 | 0.377382 | 0.602582 | 0.516515 | 1 | 0.996511 | 0.905676 |
| S1pr5 | Q91X56 | 0.81 | 1.32 | -0.95 | 1.48 | -0.1 | 0.35 | 0.52 | 0.19 | 3.3 | -0.61 | 0.63 | 0.43 | 3.56 | 0.25 | 2.02 | 1.04 | -0.425 | 0.2725 | 1.0525 | 0.428172 | 0.763154 | 0.218301 | 1 | 0.996511 | 0.787447 |
| Saal1 | Q9D2C2-2 | -3.02 | -2.39 | -1.75 | -2.15 | -3.01 | -3.5 | -1.21 | -3.78 | -3.4 | -1.84 | -2.4 | -3.34 | -2.89 | -2.2 | -3.52 | -2.1 | -0.5475 | -0.4175 | -0.35 | 0.358542 | 0.352359 | 0.401188 | 1 | 0.996511 | 0.870023 |
| Sacm1l | Q9EP69 | 0.29 | 0.11 | 0.57 | 0.64 | 0.43 | 0.1 | 0.87 | 0.86 | -0.39 | -0.03 | 0.43 | 0.33 | -0.08 | -0.45 | 1.11 | 0.89 | 0.1625 | -0.3175 | -0.035 | 0.502445 | 0.221966 | 0.927347 | 1 | 0.996511 | 0.989331 |
| Sacs | E9QNY8 | -0.21 | 0 | 0.04 | -0.13 | 0.16 | -0.4 | -0.04 | -0.31 | -0.24 | -0.31 | 0.01 | -0.12 | -0.68 | 0.02 | -0.42 | 0.01 | -0.0725 | -0.09 | -0.1925 | 0.693548 | 0.598584 | 0.406883 | 1 | 0.996511 | 0.874084 |
| Sae1 | Q9R1T2 | -4.91 | -5.21 | -3.91 | -4.49 | -4.42 | -5.55 | -3.86 | -4.45 | -5.21 | -2.27 | -3.8 | -4.39 | -4.7 | -3.81 | -4.26 | -4.18 | 0.06 | 0.7125 | 0.3925 | 0.887312 | 0.271482 | 0.262965 | 1 | 0.996511 | 0.81524 |
| Safb | S4R1M2 | 1.13 | -0.2 | -0.11 | -0.19 | -0.55 | 0.51 | -0.39 | 0.29 | 0.16 | -0.3 | -0.24 | -0.37 | 1.02 | -0.1 | 1.03 | -0.17 | -0.1925 | -0.345 | 0.2875 | 0.625446 | 0.326269 | 0.520763 | 1 | 0.996511 | 0.907667 |
| Safb2 | Q80YR5 | -0.37 | -0.74 | -0.66 | -0.09 | -0.98 | -0.98 | -0.55 | 0.39 | -0.82 | -0.31 | -0.39 | -1.52 | -0.21 | -0.34 | 0.34 | -0.62 | -0.065 | -0.295 | 0.2575 | 0.849654 | 0.362399 | 0.358668 | 1 | 0.996511 | 0.856645 |
| Sall2 | A0A2I3BQ\ | -0.29 | -0.75 | -0.7 | -0.77 | -1.13 | -0.78 | -1.6 | -1.44 | 0 | 0.74 | -1.19 | -0.87 | 0.83 | -0.23 | 0.36 | -0.64 | -0.61 | 0.2975 | 0.7075 | 0.026133 | 0.492128 | 0.064663 | 1 | 0.996511 | 0.66433 |
| Samd12 | Q0VE29 | 0.11 | 0.29 | 1.71 | 1.44 | -0.16 | -0.04 | 0.71 | 1.97 | 0.1 | 1.57 | 1.67 | 0.02 | 0.88 | 0.27 | 1.64 | 1.09 | -0.2675 | -0.0475 | 0.0825 | 0.645956 | 0.931987 | 0.858607 | 1 | 0.996755 | 0.976225 |
| Samd14 | Q8K070 | -0.97 | -1.11 | -0.56 | -0.38 | -2.06 | -0.39 | -0.25 | -0.64 | 2.68 | -0.88 | -0.55 | -0.72 | 3.6 | -1.54 | -0.52 | -0.34 | -0.08 | 0.8875 | 1.055 | 0.849548 | 0.276911 | 0.317436 | 1 | 0.996511 | 0.838864 |
| Samd4a | Q8CBY1-3 | -1.6 | -1.45 | -1.29 | 0.48 | -1.09 | -1.24 | -2.65 | -1.76 | 0.26 | -5.91 | -1.41 | -0.12 | 0.42 | 0.5 | -1.66 | -0.29 | -0.72 | -0.83 | 0.7075 | 0.213051 | 0.540065 | 0.283449 | 1 | 0.996511 | 0.822669 |
| Samd4b | G5E8A7 | -0.3 | 0.31 | 0.02 | 0.15 | 0.5 | -0.02 | 0.41 | 0.4 | 0.34 | 0.43 | 0.32 | -0.01 | 0.46 | 0.65 | -0.12 | -0.06 | 0.2775 | 0.225 | 0.1875 | 0.198496 | 0.292755 | 0.476727 | 1 | 0.996511 | 0.892174 |
| Samd5 | A0A0R4J18 | -0.81 | -1.18 | 0.56 | 0.54 | -1.32 | -0.21 | -1.31 | -0.62 | 2.35 | 1.34 | -0.08 | -0.79 | 3.07 | -0.75 | 0.1 | 1.38 | -0.6425 | 0.9275 | 1.1725 | 0.21099 | 0.242016 | 0.19187 | 1 | 0.996511 | 0.777117 |
| Samd8 | Q9DA37 | -0.69 | -0.37 | -0.2 | -0.57 | 0.63 | -0.59 | -0.03 | 0.2 | 0.5 | 0.11 | -0.92 | -0.4 | 0.47 | 0.35 | -0.55 | 0.64 | 0.51 | 0.28 | 0.685 | 0.095762 | 0.400575 | 0.045399 | 1 | 0.996511 | 0.658454 |
| Samd9l | E9PX59 | 4.53 | 5.46 | -2.72 | -0.74 | 4.67 | 4.97 | -0.67 | 0.18 | 5.74 | -0.04 | 1.26 | 5.6 | 6.21 | 4.32 | -0.76 | 0.41 | 0.655 | 1.5075 | 0.9125 | 0.767062 | 0.501543 | 0.69025 | 1 | 0.996511 | 0.955729 |
| Samhd1 | E0CXZ5 | -0.37 | 1.18 | 0.66 | 0.16 | 0.77 | -0.16 | 0.5 | -0.15 | 0.28 | 1.17 | 0.24 | 0.48 | 0.31 | 1.03 | 0.51 | 0.48 | -0.1675 | 0.135 | 0.175 | 0.665535 | 0.724407 | 0.630547 | 1 | 0.996511 | 0.941132 |
| Samhd1 | Q60710 | -1.26 | -1.2 | -2.02 | -2.15 | -1.86 | -1.55 | -1.89 | -2.37 | -0.96 | -0.57 | -1.79 | -0.88 | -0.51 | -1.23 | -2.28 | -1.9 | -0.26 | 0.6075 | 0.1775 | 0.394821 | 0.112504 | 0.686776 | 1 | 0.996511 | 0.955729 |
| Samm50 | Q8BGH2 | -0.35 | -0.2 | -0.13 | -0.01 | -0.34 | -0.24 | -0.03 | -0.12 | -0.6 | -0.34 | 0.12 | -0.46 | -0.76 | -0.62 | -0.54 | -0.3 | -0.01 | -0.1475 | -0.3825 | 0.950223 | 0.494224 | 0.074483 | 1 | 0.996511 | 0.671911 |
| Sanbr | M0QWF3 | -0.1 | 0.45 | 0.69 | 0.22 | 0.74 | -0.36 | 0.35 | 0.28 | -0.08 | 0.24 | 0.54 | 0.19 | 0.18 | -0.14 | 1.14 | 0.69 | -0.0625 | -0.0925 | 0.1525 | 0.82645 | 0.699708 | 0.647149 | 1 | 0.996511 | 0.945362 |
| Sap130 | A0A0R4J06 | -1.64 | -1.63 | -0.84 | -0.44 | -4.36 | -0.16 | -1.76 | -0.6 | 1.23 | 0.07 | -0.96 | -2.24 | 2 | -1.98 | 0.62 | -0.62 | -0.5825 | 0.6625 | 1.1425 | 0.517662 | 0.371826 | 0.183752 | 1 | 0.996511 | 0.776843 |
| Sap18 | E9Q317 | 1.68 | 0.63 | 0.45 | 0.47 | -0.73 | 1.13 | -0.73 | 0.89 | 0.82 | 0.64 | 0.16 | -0.4 | 2.3 | 0.4 | 0.95 | -0.43 | -0.6675 | -0.5025 | -0.0025 | 0.234102 | 0.21504 | 0.996621 | 1 | 0.996511 | 0.999143 |
| Sap30bp | Q02614 | -0.87 | -1.59 | -1.04 | -0.89 | -0.94 | -1.35 | -0.58 | -0.93 | -1.56 | -0.27 | -0.76 | -0.81 | -1 | -0.35 | -0.43 | -0.5 | 0.1475 | 0.2475 | 0.5275 | 0.550631 | 0.442244 | 0.064078 | 1 | 0.996511 | 0.66433 |
| Sap30l | Q5SQF8 | -0.58 | -1.79 | -1.19 | -1.55 | -1.1 | -1.87 | -1.33 | -0.61 | -1.09 | 0.2 | -1.13 | -1.74 | 0.1 | -0.66 | 1.16 | -0.62 | 0.05 | 0.3375 | 1.2725 | 0.888186 | 0.466465 | 0.022903 | 1 | 0.996511 | 0.623147 |
| Sar1a | Q99JZ4 | -0.47 | -0.92 | -0.5 | -0.46 | -1.84 | -0.29 | -1.23 | -0.86 | 0.84 | -0.25 | -0.56 | -0.61 | 0.65 | -0.4 | -1.34 | -1.36 | -0.4675 | 0.4425 | -0.025 | 0.18538 | 0.226177 | 0.956716 | 1 | 0.996511 | 0.992636 |
| Sar1b | Q9CQC9 | -0.21 | -2.18 | -0.9 | -0.94 | -2.16 | -0.67 | -0.58 | -0.96 | -0.46 | -1.82 | -1.15 | 0 | -1.04 | -1.45 | -1.28 | -1.29 | -0.035 | 0.2 | -0.2075 | 0.944791 | 0.706363 | 0.609239 | 1 | 0.996511 | 0.934202 |
| Saraf | A0A0R4J0D | 1.34 | 0 | 0.07 | -0.3 | -0.48 | 1.74 | 0.79 | 0.23 | 0.28 | -0.73 | 0 | 1.25 | 1.05 | -1.89 | 1.25 | 1.27 | 0.2925 | -0.0775 | 0.1425 | 0.593642 | 0.878782 | 0.853147 | 1 | 0.996511 | 0.976225 |
| Sardh | Q99LB7 | 0.54 | 0.73 | 1.16 | 1.24 | 0.44 | 0.17 | 1.24 | 0.7 | 0.4 | 1.03 | 1.15 | 0.04 | 0.3 | 0.8 | 0.72 | 0.9 | -0.28 | -0.2625 | -0.2375 | 0.33999 | 0.412984 | 0.351983 | 1 | 0.996511 | 0.854088 |
| Sarm1 | Q6PDS3 | 0.01 | -0.04 | -0.14 | -0.01 | 0.26 | 0.24 | 0.53 | 0.08 | 0.32 | 0.22 | 0.21 | 0.48 | -0.2 | 0.51 | -0.17 | 0.3 | 0.3225 | 0.3525 | 0.155 | 0.071482 | 0.054192 | 0.498589 | 1 | 0.996511 | 0.899062 |
| Sarnp | Q9D1J3 | 0.82 | -1.99 | -1.47 | -1.29 | -1.82 | -0.34 | -1.7 | -1.06 | -1.3 | -1.24 | -1.24 | -1.62 | -2.3 | -1.22 | -1.37 | -1.99 | -0.2475 | -0.3675 | -0.7375 | 0.700911 | 0.527748 | 0.248356 | 1 | 0.996511 | 0.803985 |
| Sars | Q8C483 | -1.03 | -1.04 | -0.93 | -0.84 | -1.06 | -1.02 | -1.1 | -0.8 | -1.12 | -0.77 | -0.92 | -0.88 | -1.27 | -0.95 | -1.42 | -1.47 | -0.035 | 3.75E-02 | -0.3175 | 0.819159 | 0.823394 | 0.132995 | 1 | 0.996511 | 0.728417 |
| Sars2 | Q9JJL8 | 1.24 | 1.02 | 0.33 | 0.26 | 0.43 | 1.03 | 0.25 | 0.56 | 0.7 | -0.19 | 0.35 | 1.02 | 0.16 | 0.59 | -0.01 | 0.18 | -0.145 | -0.2425 | -0.4825 | 0.626131 | 0.494324 | 0.125942 | 1 | 0.996511 | 0.721505 |
| Sart1 | Q9Z315 | 0.12 | -0.84 | -0.87 | -0.75 | -0.88 | -0.3 | -0.59 | -0.28 | -0.75 | -0.3 | -0.78 | -1.25 | -0.1 | -0.85 | 0.17 | -0.34 | 0.0725 | -0.185 | 0.305 | 0.795053 | 0.553593 | 0.361033 | 1 | 0.996511 | 0.857783 |
| Sart3 | Q9JLI8 | -2.94 | -3.63 | -2.65 | -2.84 | -3.02 | -3.15 | -2.59 | -3.02 | -2.48 | -1.41 | -2.56 | -2.79 | -1.94 | -2.82 | -2.83 | -2.87 | 0.07 | 0.705 | 0.4 | 0.784946 | 0.07992 | 0.229008 | 1 | 0.996511 | 0.787447 |
| Sash1 | F8VQK5 | 0.59 | 0.76 | 0.97 | 0.72 | 0.61 | 0.52 | 0.94 | 0.84 | 0.77 | 0.85 | 0.83 | 0.86 | 1.2 | 0.53 | 0.69 | 0.82 | -0.0325 | 0.0675 | 0.05 | 0.852403 | 0.68509 | 0.817545 | 1 | 0.996511 | 0.972768 |
| Sat2 | Q6P8J2 | -0.5 | -0.71 | -1.44 | -1.65 | 0.35 | -0.69 | -0.95 | -1.17 | -0.5 | -1.28 | -1.5 | -0.59 | -0.28 | 0.35 | -0.54 | -1.01 | 0.46 | 0.1075 | 0.705 | 0.280009 | 0.767407 | 0.095025 | 1 | 0.996511 | 0.698156 |
| Satb1 | E9PVB7 | -3.31 | -2.97 | -2.79 | -3.18 | -3.27 | -3.25 | -2.32 | -3.26 | -2.39 | -2.07 | -2.39 | -2.5 | -2.34 | -2.21 | -0.73 | -2.77 | 0.0375 | 0.725 | 1.05 | 0.888644 | 0.005235 | 0.036954 | 1 | 0.996511 | 0.658454 |
| Satb2 | H3BKH3 | -4.16 | -3.41 | -4.71 | -6.48 | -3.4 | -3.47 | -4.87 | -6.96 | -2.67 | -4.87 | -5.24 | -3.6 | -3.04 | -2.51 | -4.35 | -4.66 | 0.015 | 0.595 | 1.05 | 0.987403 | 0.462488 | 0.186579 | 1 | 0.996511 | 0.776843 |
| Saysd1 | Q8K190 | -0.16 | 0.09 | -1.15 | -1.93 | -0.71 | 0.36 | 0.64 | -0.18 | 0.66 | 2.91 | -0.57 | -0.51 | 0.31 | 1.34 | 2.3 | 1.33 | 0.815 | 1.41 | 2.1075 | 0.13693 | 0.123052 | 0.004829 | 1 | 0.996511 | 0.552614 |
| Sbds | P70122 | -0.69 | -1.24 | -0.56 | -0.92 | -1.08 | -1.02 | -0.4 | -0.84 | -0.83 | -1.39 | -0.64 | -0.94 | -1.13 | -0.77 | -0.93 | -0.88 | 0.0175 | -0.0975 | -0.075 | 0.93996 | 0.690563 | 0.732491 | 1 | 0.996511 | 0.958635 |
| Sbf1 | Q6ZPE2 | 0.33 | 0.26 | 0.6 | 0.53 | 0.5 | 0.42 | 0.67 | 0.44 | 0.19 | 0.08 | 0.5 | 0.16 | -0.22 | 0.55 | 0.26 | 0.46 | 0.0775 | -0.1975 | -0.1675 | 0.631344 | 0.300462 | 0.478382 | 1 | 0.996511 | 0.892706 |
| Sbf2 | E9Q0D4 | 0.71 | 0.65 | 0.88 | 0.8 | 0.47 | 0.64 | 0.63 | 0.79 | 0.55 | 0.95 | 1.07 | 0.74 | 0.38 | 0.34 | 0.77 | 0.67 | -0.1275 | 0.0675 | -0.22 | 0.415124 | 0.717607 | 0.270912 | 1 | 0.996511 | 0.822084 |
| Sbno1 | F7CVW0 | -2.86 | -2.69 | -1.91 | -1.71 | -2.39 | -2.94 | -1.75 | -1.92 | -3.09 | -0.97 | -1.97 | -2.46 | -2.87 | -2.25 | -2.29 | -1.82 | 0.0425 | 0.17 | -0.015 | 0.908746 | 0.731078 | 0.966077 | 1 | 0.996511 | 0.9927 |
| Sc5d | O88822 | 0.5 | 0.07 | 0.89 | -0.49 | -0.04 | 0.77 | 0.24 | 0.41 | 0.9 | -0.06 | 0.26 | 0.72 | 1.82 | -0.71 | 1.43 | 1.48 | 0.1025 | 0.2125 | 0.7625 | 0.75745 | 0.557435 | 0.222095 | 1 | 0.996511 | 0.787447 |
| Scaf1 | Q5U4C3 | 0.83 | 0.14 | 0.08 | 0.02 | 0 | 0.3 | -0.75 | 0.3 | 0.05 | -0.29 | -0.15 | -0.47 | 0.21 | 0.17 | 0.04 | -0.29 | -0.305 | -0.4825 | -0.235 | 0.335464 | 0.075193 | 0.364822 | 1 | 0.996511 | 0.858475 |
| Scaf11 | A0A2I3BQL | 0.65 | 0.07 | 0.75 | 0.43 | -0.92 | 0.5 | -0.91 | 0.09 | 1.56 | -0.81 | -0.84 | -1.15 | 2.58 | -0.19 | 0.39 | -0.13 | -0.785 | -0.785 | 0.1875 | 0.060099 | 0.20816 | 0.75995 | 1 | 0.996511 | 0.963123 |
| Scaf4 | Q7TSH6 | -2.89 | -2.94 | 0 | -0.17 | -2.97 | -3.27 | -0.43 | -0.37 | -2.87 | 2.33 | 0.18 | -3.52 | -1.9 | -2.76 | 1.45 | 0.67 | -0.26 | 0.53 | 0.865 | 0.798677 | 0.710076 | 0.461798 | 1 | 0.996511 | 0.888262 |
| Scaf8 | Q6DID3 | -1.52 | -2.41 | -2.57 | -2.64 | -2.1 | -2.69 | -2.79 | -2.72 | -1.08 | -1.69 | -2.45 | -1.65 | -1.17 | -0.46 | -2.68 | -2.94 | -0.29 | 0.5675 | 0.4725 | 0.35007 | 0.152071 | 0.436499 | 1 | 0.996511 | 0.882882 |
| Scai | Q8C8N2 | 1.03 | 0.84 | 1.04 | 1.01 | 0.85 | 1.21 | 1.01 | 0.96 | 0.59 | 0.35 | 1.14 | 0.59 | 0.34 | 0.36 | 0.45 | 0.79 | 0.0275 | -0.3125 | -0.495 | 0.860054 | 0.169979 | 0.027387 | 1 | 0.996511 | 0.635993 |
| Scai | B0R0E3 | 3.12 | 0.88 | 1.43 | 3.87 | 0.76 | 2.42 | 0.89 | 3.77 | 3.8 | 4.03 | 2.22 | 3.16 | 3.44 | 2.15 | 2.41 | 1.23 | -0.365 | 0.9775 | -0.0175 | 0.685798 | 0.206496 | 0.981496 | 1 | 0.996511 | 0.994858 |
| Scamp1 | Q8K021 | 1.14 | 0.88 | 1.25 | 1.05 | 0.69 | 0.93 | 1.11 | 1.3 | 0.93 | 0.87 | 1.14 | 1.28 | 1.06 | 1.05 | 1.38 | 1.3 | -0.0725 | -0.025 | 0.1175 | 0.704145 | 0.892252 | 0.542814 | 1 | 0.996511 | 0.912212 |
| Scamp2 | Q9ERN0 | -0.24 | -0.79 | 0.08 | -0.03 | -1.7 | 0.95 | 0.3 | 0.54 | 2.68 | -0.39 | 0.21 | 0.01 | 4.43 | -2.44 | 0.1 | 0.28 | 0.2675 | 0.8725 | 0.8375 | 0.639487 | 0.207878 | 0.517163 | 1 | 0.996511 | 0.905864 |
| Scamp3 | E9Q855 | -3.14 | -2.36 | 0.91 | 0.74 | -1.31 | -0.04 | 1.23 | 0.52 | -4.89 | 0.93 | 0.49 | -2.14 | -3.06 | -3.94 | 1.23 | 0.65 | 1.0625 | -0.44 | -0.3175 | 0.327665 | 0.772723 | 0.830301 | 1 | 0.996511 | 0.973939 |
| Scamp3 | Q3UXS0 | 1.09 | 1.26 | 1.1 | 1.24 | 0.61 | 1.23 | 0.38 | 0.78 | 1.77 | 1.12 | 1.08 | 0.95 | 2.22 | 0.37 | 1.15 | 1.03 | -0.4225 | 0.0575 | 0.02 | 0.073333 | 0.797759 | 0.95758 | 1 | 0.996511 | 0.992636 |
| Scamp4 | Q9JKV5 | 0.35 | 0.03 | 0.28 | 0.53 | -0.9 | 1.05 | 0.02 | 0.4 | 3.46 | -2.17 | 0.59 | 0.03 | 3.59 | -2.78 | 0.24 | -0.6 | -0.155 | 0.18 | -0.185 | 0.697171 | 0.863146 | 0.875586 | 1 | 0.996511 | 0.979626 |
| Scamp5 | Q9JKD3 | 1.04 | 1 | 0.61 | 0.83 | 1.16 | 1.22 | 1.15 | 0.91 | -0.6 | -1.16 | 1.07 | 1.27 | -0.4 | -1.45 | 0.88 | 0.63 | 0.24 | -0.725 | -0.955 | 0.185365 | 0.220261 | 0.0856 | 1 | 0.996511 | 0.691619 |
| Scap | Q6GQT6 | -0.49 | -0.68 | -0.13 | -0.62 | 0.21 | -0.78 | 0.05 | -0.26 | 0.17 | 0.53 | 0 | -0.43 | 0.58 | -0.38 | 0.98 | 0.56 | 0.285 | 0.5475 | 0.915 | 0.289165 | 0.057163 | 0.017843 | 1 | 0.996511 | 0.602211 |
| Scaper | B2RXS3 | -1.06 | -1.11 | -1.47 | -1.57 | -1.13 | -1.14 | -1.66 | -1.69 | -0.93 | -0.99 | -1.63 | -0.96 | -0.64 | -0.64 | -1.32 | -1.55 | -0.1025 | 0.175 | 0.265 | 0.648491 | 0.470946 | 0.365017 | 1 | 0.996511 | 0.858475 |
| Scara3 | Q8C850 | -1.19 | -0.49 | 1.1 | 0.15 | -0.07 | -0.71 | 0.09 | 0.35 | 0.27 | 0.89 | 0.75 | 0.03 | 0.97 | 1.12 | 1.08 | 0.59 | 0.0225 | 0.5925 | 1.0475 | 0.963725 | 0.247474 | 0.050089 | 1 | 0.996511 | 0.658454 |
| Scarb2 | O35114 | -1.31 | -1 | -0.59 | -0.72 | -0.72 | -1.16 | -0.61 | -0.55 | -1.35 | -1.05 | -0.46 | -1.22 | -0.91 | -1.58 | -0.88 | -0.7 | 0.145 | -0.115 | -0.1125 | 0.533767 | 0.670712 | 0.682373 | 1 | 0.996511 | 0.955729 |
| Sccpdh | Q8R127 | -0.08 | -0.12 | -0.43 | 0.13 | -0.15 | 0.34 | 0.07 | -0.09 | 0.37 | -0.09 | -0.48 | 0.19 | 0.31 | -0.04 | -0.25 | -0.42 | 0.1675 | 0.1225 | 0.025 | 0.400539 | 0.618116 | 0.915619 | 1 | 0.996511 | 0.986556 |
| Scd2 | A0A494BA' | 0.33 | 0.74 | 0.35 | 0.59 | 0.3 | 0.36 | 0.73 | 0.19 | -0.16 | 1 | -0.03 | -0.18 | 0.13 | -1.17 | 1.08 | 0.3 | -0.1075 | -0.345 | -0.4175 | 0.577279 | 0.274923 | 0.366457 | 1 | 0.996511 | 0.858904 |
| Scfd1 | Q8BRF7 | 0.37 | 0.35 | 0.45 | 0.21 | 0.32 | 0.43 | 0.6 | 0.7 | -0.43 | -1.15 | 0.44 | 0.55 | -0.37 | -1.13 | 0.75 | 0.7 | 0.1675 | -0.4925 | -0.3575 | 0.312254 | 0.224977 | 0.418883 | 1 | 0.996511 | 0.87558 |
| Scfd2 | Q8BTY8 | -0.36 | -0.03 | 0.99 | 0.38 | -0.08 | -0.24 | 1.66 | 1.14 | -0.46 | -0.37 | 0.86 | -0.27 | -0.42 | -0.04 | 1.07 | 1.49 | 0.375 | -0.305 | 0.28 | 0.464791 | 0.458925 | 0.57903 | 1 | 0.996511 | 0.927466 |
| Scg2 | Q03517 | 0.94 | 0.64 | -0.01 | 0.65 | 0.64 | 0.33 | 0.71 | 0.64 | 0.8 | 0.79 | 0.09 | 0.98 | 0.78 | 1.03 | 1.92 | 1.65 | 0.025 | 0.11 | 0.79 | 0.915347 | 0.705393 | 0.039893 | 1 | 0.996511 | 0.658454 |
| Scg3 | P47867 | 1.16 | 1.08 | 0.93 | 0.75 | 0.58 | 1.21 | 0.69 | 1.23 | 1.66 | 0 | 1.27 | 1.94 | 1.84 | 0.66 | 1.52 | 1.32 | -0.0525 | 0.2375 | 0.355 | 0.809386 | 0.571751 | 0.232095 | 1 | 0.996511 | 0.788863 |
| Scg5 | P12961 | 1.84 | 0.81 | 0.15 | 0.14 | 1.11 | 1.41 | 1.42 | 0.24 | 0.6 | 0.63 | 0.57 | 1.66 | 0.54 | 0.65 | 1.44 | 1.64 | 0.31 | 0.13 | 0.3325 | 0.499946 | 0.773752 | 0.475335 | 1 | 0.996511 | 0.892134 |
| Scgn | Q91WD9 | 3.5 | -0.87 | -2.46 | -1.11 | -1.83 | 0.21 | -1.21 | -1.82 | 5.21 | 0.61 | -2.84 | 4.59 | 3.25 | -4.02 | -1.45 | -1.61 | -0.9275 | 2.1275 | -0.7225 | 0.459749 | 0.31077 | 0.684335 | 1 | 0.996511 | 0.955729 |
| Schip1 | A0A0A6YX( | 1.45 | 1.08 | -0.2 | -0.04 | 1.67 | 2.27 | 0.13 | -0.55 | 0.86 | -0.43 | -0.68 | 2.45 | 0.48 | -0.59 | -0.22 | 0.23 | 0.3075 | -0.0225 | -0.5975 | 0.661984 | 0.975987 | 0.204079 | 1 | 0.997588 | 0.782817 |
| Scly | A0A0R4J06 | -1.46 | -2.32 | -2.38 | -3.2 | -1.79 | -1.31 | -2.89 | -2.31 | -1.63 | -2.45 | -2.61 | -1.16 | -1.32 | -1.4 | -2.6 | -2.14 | 0.265 | 0.3775 | 0.475 | 0.565848 | 0.424102 | 0.301478 | 1 | 0.996511 | 0.830308 |
| Scn1a | A2APX8-3 | 0.97 | 1.62 | 1.24 | 0.92 | 1.24 | 1.07 | 0.5 | 0.73 | 0.63 | -0.01 | 0.86 | 0.67 | 0.83 | 0.19 | 0.36 | 0.82 | -0.3025 | -0.65 | -0.6375 | 0.237617 | 0.034817 | 0.033451 | 1 | 0.996511 | 0.644622 |
| Scn1b | P97952 | 1.75 | 1.84 | 1.49 | 0.58 | 2.25 | 1.64 | 0.99 | 0.97 | -0.01 | 0.68 | 1.1 | 1.85 | -0.29 | 0.85 | 0.98 | 1.36 | 0.0475 | -5.10E-01 | -0.69 | 0.904509 | 0.280109 | 0.139006 | 1 | 0.996511 | 0.73493 |
| Scn2a | A0A5H1ZRI | 0.76 | 1.27 | 0.55 | 0.65 | 0.92 | 0.78 | 0.49 | 0.58 | 0.54 | 0.42 | 0.55 | 0.75 | 0.7 | 0.22 | 0.25 | 0.31 | -0.115 | -0.2425 | -0.4375 | 0.593229 | 0.276046 | 0.091145 | 1 | 0.996511 | 0.693316 |
| Scn2b | Q56A07 | 2.21 | 0.89 | 0.93 | 0.71 | 0.23 | 1.87 | 0.07 | 0.36 | 2.86 | 1.44 | 0.46 | 2.23 | 2.61 | 0.86 | 0.68 | 0.28 | -0.5525 | 0.5625 | -0.0775 | 0.285826 | 0.339776 | 0.891902 | 1 | 0.996511 | 0.982213 |
| Scn3a | A2ASI5 | -0.7 | -0.54 | 1.34 | 1.21 | -0.75 | -0.85 | 1.47 | 1.54 | 2.23 | 1.57 | 1.24 | -0.78 | 2.26 | 0.71 | 1.44 | 1.86 | 0.025 | 0.7375 | 1.24 | 0.974359 | 0.350842 | 0.060466 | 1 | 0.996511 | 0.661738 |
| Scn3b | H3BJR6 | 0.6 | 0.97 | 1.57 | 1.94 | 0.87 | 0.57 | 1.89 | 2.43 | 1.18 | 1.81 | 1.89 | 1.11 | -0.22 | 1.4 | 1.77 | 1.73 | 0.17 | 0.2275 | -0.1 | 0.728201 | 0.526159 | 0.846988 | 1 | 0.996511 | 0.976225 |
| Scn4b | Q7M729 | 1.56 | 0.44 | 1.49 | 1.16 | -0.46 | -0.11 | 1.69 | 0.69 | 2.16 | 2.24 | 0.66 | 0.93 | 2.67 | -0.39 | 1.7 | 1.39 | -0.71 | 0.335 | 0.18 | 0.180342 | 0.467728 | 0.774857 | 1 | 0.996511 | 0.966477 |
| Scn7a | B1AYL1 | 0.8 | 0.9 | 0.48 | 0.78 | 1.29 | 0.58 | 0.12 | 0.84 | 0.9 | 0.98 | 0.86 | 0.89 | 0.68 | 1.73 | 0.56 | 0.58 | -0.0325 | 0.1675 | 0.1475 | 0.903217 | 0.340857 | 0.632946 | 1 | 0.996511 | 0.941132 |
| Scn8a | A0A0J9YU\ | 0.41 | 0.81 | 1.07 | 0.93 | 0.88 | 0.48 | 0.86 | 1 | 0.29 | 0.35 | 0.82 | 0.35 | 0.23 | 0.38 | 0.72 | 0.79 | -1.6E-16 | -0.3525 | -0.275 | 1 | 0.141431 | 0.263183 | 1 | 0.996511 | 0.81524 |
| Scnm1 | A0A0R4J26 | -0.91 | -2.47 | -1.44 | -1.93 | -2.28 | -1.21 | -2.45 | -2.37 | -0.92 | 0.01 | -1.2 | -0.49 | -2.06 | -0.1 | -2.04 | -2.29 | -0.39 | 1.0375 | 0.065 | 0.360908 | 0.02946 | 0.907872 | 1 | 0.996511 | 0.986458 |
| Sco1 | Q5SUD5 | 0.77 | 0.22 | 1.17 | 1.89 | 0 | 0.45 | 1.18 | 1.97 | 1.14 | 1.77 | 1.51 | 0.51 | 0.89 | 0.57 | 0.88 | 0.88 | -0.1125 | 0.22 | -0.2075 | 0.826336 | 0.605055 | 0.563757 | 1 | 0.996511 | 0.920143 |
| Sco2 | Q8VCL2 | 0.9 | 0.9 | -0.17 | -0.22 | 0.69 | 1.05 | -0.11 | -0.11 | 0.41 | 0.06 | -0.43 | 1.15 | 0.2 | 0.72 | -0.52 | -0.15 | 0.0275 | -0.055 | -0.29 | 0.945869 | 0.898984 | 0.472649 | 1 | 0.996511 | 0.891144 |

| | | | | | | | | | | | | | | | | | | | | | | | | | | |
|---|---|---|---|---|---|---|---|---|---|---|---|---|---|---|---|---|---|---|---|---|---|---|---|---|---|---|
| Scoc | Q78YZ6-2 | 2.56 | 1.34 | 0.16 | -0.09 | 0.33 | 1.43 | 0.15 | 0.07 | 2.87 | -0.74 | -0.26 | 2.99 | 2.09 | 2.08 | -0.38 | 0.25 | -0.4975 | 0.2225 | 0.0175 | 0.434144 | 0.831775 | 0.982356 | 1 | 0.996511 | 0.994858 |
| Scp2 | P32020 | 0.06 | 0.26 | 0.11 | 0.04 | 0.84 | 0.35 | 0.87 | 0.31 | 0.2 | 0.18 | 0.16 | 0.62 | 0.44 | 0.65 | 0.58 | 0.48 | 0.475 | 0.1725 | 0.42 | 0.035867 | 0.36025 | 0.035916 | 1 | 0.996511 | 0.658454 |
| Scpep1 | Q920A5 | -1.07 | -1.09 | -1.21 | -1.11 | -0.88 | -1.14 | -0.75 | -1.33 | -1.08 | -1.12 | -0.87 | -0.98 | -0.85 | -1.26 | -1.59 | -1.36 | 0.095 | 0.1075 | -0.145 | 0.599156 | 0.50638 | 0.502599 | 1 | 0.996511 | 0.901002 |
| Scrg1 | O88745 | -1.5 | -1.28 | -1.3 | -1.46 | -0.79 | -1.12 | -0.42 | -1.17 | -1.66 | -0.53 | -1.31 | -1.13 | -2 | -1.01 | -0.62 | -1.17 | 0.51 | 0.2275 | 0.185 | 0.036134 | 0.394972 | 0.55046 | 1 | 0.996511 | 0.913801 |
| Scrib | Q80U72 | 0.39 | 0.34 | 1.21 | 1.26 | 0.4 | 0.25 | 1.16 | 1.3 | 0.71 | 1.43 | 1.29 | 0.51 | 0.48 | 0.4 | 1.18 | 1.2 | -0.0225 | 0.185 | 0.015 | 0.948823 | 0.581525 | 0.964183 | 1 | 0.996511 | 0.9927 |
| Scrn1 | Q9CZC8 | -0.21 | -0.55 | -0.4 | -0.23 | -0.94 | -0.4 | -0.37 | -0.25 | -0.25 | -0.33 | -0.51 | -0.08 | -0.45 | -0.31 | -0.77 | -0.97 | -0.1425 | 0.055 | -0.2775 | 0.490976 | 0.76426 | 0.229296 | 1 | 0.996511 | 0.787447 |
| Scrn2 | Q8VCA8 | -0.5 | -0.7 | -1.05 | -1.15 | -0.97 | -0.57 | -0.25 | -0.43 | -0.27 | -0.27 | -1.08 | -0.7 | -0.18 | -1.78 | -0.48 | -0.51 | 0.295 | 0.27 | 0.1125 | 0.227605 | 0.322286 | 0.765727 | 1 | 0.996511 | 0.964134 |
| Scrn3 | Q3TMH2 | -0.2 | -0.25 | -0.38 | -0.19 | -0.37 | -0.29 | -0.22 | -0.3 | -0.34 | -0.83 | -0.54 | -0.21 | -0.61 | -0.61 | -0.94 | -0.52 | -0.04 | -0.225 | -0.415 | 0.779661 | 0.267193 | 0.048823 | 1 | 0.996511 | 0.658454 |
| Scrt1 | Q99M85 | -1.61 | -1.11 | -1.61 | -0.41 | -1.27 | -2.64 | -0.84 | -1.85 | -1.71 | -0.92 | -1.43 | -2.03 | 0.13 | -2.86 | -0.77 | -0.92 | -0.465 | -0.3375 | 0.08 | 0.315646 | 0.360288 | 0.898945 | 1 | 0.996511 | 0.983509 |
| Scube1 | A2A442 | -1.36 | -2.34 | -1.36 | -0.79 | -2.49 | -1.01 | -1.55 | -1.22 | 0.02 | -1.75 | -0.86 | -1.28 | 0.66 | -1.99 | -1.29 | -1.22 | -0.105 | 0.495 | 0.5025 | 0.806877 | 0.302293 | 0.410368 | 1 | 0.996511 | 0.874811 |
| Scyl1 | R4H4V1 | -0.48 | -0.12 | 0.69 | -0.08 | 0.07 | -0.23 | 0.57 | 0.17 | 0.14 | -1.19 | 0.53 | -0.27 | 0.1 | -0.04 | 0.5 | 1.4 | 0.1425 | -0.2 | 0.4875 | 0.631655 | 0.636689 | 0.236259 | 1 | 0.996511 | 0.791642 |
| Scyl2 | Q8CFE4 | 0.83 | 0.86 | 0.72 | 0.49 | 0.92 | 0.82 | 0.6 | 0.6 | 0.98 | 1.14 | 0.91 | 1.11 | 0.09 | 1.02 | 0.44 | 0.61 | 0.01 | 0.31 | -0.185 | 0.95293 | 0.101616 | 0.459837 | 1 | 0.996511 | 0.887902 |
| Scyl3 | Q9DBQ7-3 | -1.66 | -1.07 | -1.79 | -1.43 | -0.76 | -0.55 | -1.19 | -1.7 | -0.73 | -2.45 | -1.4 | -0.99 | -0.71 | -0.77 | -2.49 | -2.47 | 0.4375 | 0.095 | -0.1225 | 0.167146 | 0.809201 | 0.803527 | 1 | 0.996511 | 0.971265 |
| Sdad1 | A0A0J9YV2 | -0.17 | -0.12 | 0.43 | -0.13 | -0.77 | -1.2 | -0.13 | -2.68 | 0.35 | -0.02 | -0.91 | -0.15 | 0.7 | 0.93 | -0.63 | -0.29 | -1.1975 | -0.185 | 0.175 | 0.042983 | 0.548596 | 0.656088 | 1 | 0.996511 | 0.948332 |
| Sdc2 | P43407 | 1.76 | 1.23 | 0.96 | 0.9 | 1.29 | 1.16 | 0.61 | 0.35 | 0.85 | 1.78 | 0.72 | 1.41 | 0.57 | 1.02 | 1.76 | 0.89 | -0.36 | -0.0225 | -0.1525 | 0.243992 | 0.943306 | 0.640109 | 1 | 0.996755 | 0.942804 |
| Sdc3 | Q64519 | 0.48 | 1.12 | 0.52 | 0.11 | 0.64 | 0.8 | 0.52 | 0.7 | 1.24 | 1.17 | 1.07 | 1.4 | 1.18 | 1.5 | 1.14 | 0.28 | 0.1075 | 0.6625 | 0.4675 | 0.648701 | 0.023211 | 0.187948 | 1 | 0.996511 | 0.776843 |
| Sdc4 | O35988 | 0.72 | 1.27 | 1.01 | 1.37 | 1.13 | 0.41 | 1.3 | 0.7 | 1.34 | 1.68 | 0.95 | 1.01 | 1.41 | 2.09 | 0.05 | 0.51 | -0.2075 | 0.1525 | -0.0775 | 0.431199 | 0.541542 | 0.863931 | 1 | 0.996511 | 0.977235 |
| Sdcbp | Q3TMX0 | -0.24 | 0.16 | -0.29 | 0.38 | 0.24 | -0.19 | 0.43 | 0.77 | 0.18 | 0.84 | -0.19 | 0.36 | 0.1 | 0.07 | -0.38 | -0.28 | 0.31 | 0.295 | -0.125 | 0.260287 | 0.308044 | 0.605972 | 1 | 0.996511 | 0.932425 |
| Sde2 | Q8K1J5 | -1.2 | -1.39 | -2.81 | -3.17 | -1.54 | -0.95 | -3.15 | -2.75 | -0.7 | -1.25 | -2.61 | -1.24 | 0.33 | -0.11 | -2.87 | -3.09 | 0.045 | 0.6925 | 0.7075 | 0.944641 | 0.259293 | 0.44961 | 1 | 0.996511 | 0.884817 |
| Sdf2 | A0A0R4J2V | 2.15 | 0.53 | 1.17 | -1.15 | 1.9 | 2.79 | 3.12 | 0.6 | -1.33 | -2.12 | 1.51 | 2.51 | 0.18 | -3.39 | 3.65 | 3.03 | 1.4275 | -0.5325 | 0.1925 | 0.104284 | 0.651311 | 0.901364 | 1 | 0.996511 | 0.983954 |
| Sdf2l1 | Q9ESP1 | -0.96 | -0.93 | -1.77 | -1.87 | -0.53 | -0.73 | -1.28 | -1.85 | -0.98 | -2.59 | -2 | -0.68 | 1.22 | -2.67 | -0.25 | -1.1 | 0.285 | -0.18 | 0.6825 | 0.451171 | 0.707254 | 0.385693 | 1 | 0.996511 | 0.861533 |
| Sdf4 | Q61112 | -0.17 | -0.73 | -1.49 | -1.3 | -0.74 | -0.35 | -1.09 | -0.9 | -0.83 | -0.4 | -0.88 | -0.6 | -0.7 | -0.52 | 0.07 | -0.74 | 0.1525 | 0.245 | 0.45 | 0.644006 | 0.450208 | 0.219958 | 1 | 0.996511 | 0.787447 |
| Sdha | Q8K2B3 | 0.78 | 0.42 | 0.76 | 1.12 | 0.2 | 0.93 | 0.57 | 0.92 | 0.37 | 0.85 | 0.81 | 0.59 | 0.45 | 0.16 | 0.52 | 0.85 | -0.115 | -0.115 | -0.275 | 0.634173 | 0.602034 | 0.270224 | 1 | 0.996511 | 0.822038 |
| Sdhaf1 | Q3U276 | 0.16 | 0 | 0.48 | 0.91 | -0.45 | -0.16 | 0.22 | 0.66 | 0.66 | 1.04 | 0.39 | 0.24 | 1.9 | 1.12 | 0.72 | 0.41 | -0.32 | 0.195 | 0.65 | 0.315439 | 0.490987 | 0.10768 | 1 | 0.996511 | 0.706141 |
| Sdhaf2 | A0A494B9I | 0.69 | 0.88 | 0.12 | 0.14 | 0.22 | 0.92 | -1.06 | -0.53 | 1.16 | 0 | 0.17 | 0.54 | 1.18 | 0.91 | -0.45 | -0.43 | -0.57 | 0.01 | -0.155 | 0.219348 | 0.975148 | 0.72993 | 1 | 0.997515 | 0.958635 |
| Sdhaf3 | Q8BQU3 | -0.32 | 0.49 | 0.24 | 1.03 | 0.23 | -0.2 | -1.93 | 0.12 | 0.29 | 1.52 | 1.05 | -0.48 | 1.85 | 0.38 | 1.23 | 2.13 | -0.805 | 0.235 | 1.0375 | 0.154583 | 0.630675 | 0.04458 | 1 | 0.996511 | 0.658454 |
| Sdhaf4 | Q8BTE0 | 0.58 | 0.46 | 0.24 | 0.19 | -0.02 | 0.47 | 0.53 | -0.31 | 0.73 | 0.23 | -0.05 | 0.78 | 0.6 | 1.42 | 0.34 | 0.58 | -0.2 | 0.055 | 0.3675 | 0.410043 | 0.822967 | 0.204859 | 1 | 0.996511 | 0.783452 |
| Sdhb | Q9CQA3 | 1.11 | 0.75 | 1.33 | 1.41 | 0.13 | 0.93 | 1.4 | 1.62 | 0.3 | 0.13 | 1.56 | 1.06 | 0.29 | -0.36 | 1.52 | 1.37 | -0.13 | -0.3875 | -0.445 | 0.709479 | 0.293865 | 0.333284 | 1 | 0.996511 | 0.845601 |
| Sdhc | Q9CZB0 | -1.15 | -2.17 | 1.71 | -0.4 | -1.36 | -0.36 | 2.1 | 1.2 | -4.44 | -1.48 | 2.29 | -1.4 | -3.54 | -4.46 | -0.01 | 0.93 | 0.8975 | -0.755 | -1.2675 | 0.386865 | 0.601198 | 0.369808 | 1 | 0.996511 | 0.859245 |
| Sdhd | Q9CXV1 | -3.8 | -3.21 | -2.03 | -1.65 | -2.24 | -3.26 | -0.31 | -1.69 | -4.18 | -2.82 | -0.94 | -3.58 | -2.63 | -3.36 | -1.87 | -1.86 | 0.7975 | -0.2075 | 0.2425 | 0.284218 | 0.791016 | 0.670649 | 1 | 0.996511 | 0.952558 |
| Sdk1 | Q3UH53 | 0.54 | 0.69 | 0.46 | -0.28 | 0.33 | 0.95 | 0 | -0.1 | 1.97 | 1.25 | 0.58 | 0.65 | 2.01 | 0.96 | -0.41 | 0.34 | -0.0575 | 0.76 | 0.3725 | 0.855024 | 0.069653 | 0.477176 | 1 | 0.996511 | 0.892234 |
| Sdk2 | Q6V4S5 | 0.43 | 0.84 | 0.86 | 0.68 | 1.19 | 0.76 | 0.93 | 0.7 | 0.93 | 0.83 | 1.03 | 1 | 0.59 | 1.16 | 0.35 | 0.49 | 0.1925 | -0.055 | -0.4725 | 0.320738 | 0.191654 | 0.820064 | 1 | 0.996511 | 0.972768 |
| Sdr39u1 | Q5M8N4 | 1.21 | 0.85 | 0.67 | 0.5 | 0.59 | 0.93 | 1.15 | 0.84 | -0.39 | -0.68 | 0.81 | 0.72 | -0.29 | -0.7 | 0.48 | 0.78 | 0.07 | -0.6925 | -0.74 | 0.746758 | 0.105728 | 0.069844 | 1 | 0.996511 | 0.668066 |
| Sdsl | Q8R238 | -0.41 | -0.14 | 0.04 | 0.64 | 0.16 | -0.13 | 0.02 | 0.5 | -0.5 | 0.37 | 0.41 | -0.67 | -1.14 | 0.25 | -0.07 | -0.25 | 1.05E-01 | -0.13 | -0.335 | 0.695102 | 0.713389 | 0.370996 | 1 | 0.996511 | 0.859424 |
| Sec11a | D3YWT0 | -1.23 | -0.89 | -0.51 | -1.09 | -0.38 | -0.97 | -0.14 | -0.66 | -0.3 | -0.6 | -0.97 | -1.04 | -0.65 | 0.38 | -0.24 | -0.23 | 0.3925 | 0.2025 | 0.745 | 0.142977 | 0.434246 | 0.025656 | 1 | 0.996511 | 0.633075 |
| Sec11c | D3Z5H3 | -0.75 | -0.62 | 0.18 | 0.54 | -0.05 | -0.62 | 0.88 | 0.39 | -0.3 | 0.6 | 0.1 | -0.5 | -0.21 | -0.03 | 1.22 | 0.86 | 0.3125 | 0.1375 | 0.6225 | 0.462638 | 0.718858 | 0.182082 | 1 | 0.996511 | 0.776002 |
| Sec13 | Q9D1M0 | -1.01 | -1.38 | -0.13 | 0.55 | -1.61 | -1 | 0.21 | 0.67 | -1.1 | 0.06 | 0.11 | -1.21 | -0.7 | -1.61 | -0.12 | -0.49 | 0.06 | -0.0425 | -0.2375 | 0.923143 | 0.935117 | 0.638009 | 1 | 0.996755 | 0.942804 |
| Sec14l1 | A8Y5H7 | -0.32 | -0.25 | -0.77 | -0.52 | -0.26 | -0.6 | -0.45 | -1.01 | -0.57 | -1.17 | -0.78 | -0.19 | -0.68 | 0.37 | -1.12 | -1.03 | -0.115 | -0.2125 | -0.15 | 0.605049 | 0.416492 | 0.675661 | 1 | 0.996511 | 0.952944 |
| Sec14l2 | Q99J08 | -0.03 | -0.19 | 0.17 | 0 | -0.14 | 0.13 | -0.05 | 0.27 | -0.42 | 0.17 | -0.22 | -0.35 | -0.58 | -1.05 | -0.68 | -0.49 | 0.065 | -0.1925 | -0.6875 | 0.703708 | 0.349654 | 0.008255 | 1 | 0.996511 | 0.563386 |
| Sec16a | E9QAT4-2 | -1.17 | -1.1 | -1.45 | -1.46 | -1 | -1.42 | -1.33 | -1.57 | -0.99 | -0.94 | -1.43 | -1.02 | -0.87 | -0.55 | -1.57 | -1.62 | -0.035 | 0.2 | 0.1425 | 0.854452 | 0.326402 | 0.631064 | 1 | 0.996511 | 0.941132 |
| Sec22b | O08547 | -0.49 | -0.71 | -0.02 | -0.02 | -0.29 | -0.52 | 0.02 | -0.04 | -0.46 | -0.14 | -0.11 | -0.62 | 0.12 | -0.57 | 0.42 | 0.33 | 0.1025 | -0.0225 | 0.385 | 0.660007 | 0.925433 | 0.217141 | 1 | 0.996755 | 0.787447 |
| Sec23a | E9Q1S3 | -0.47 | -0.88 | -0.59 | -0.49 | -0.67 | -0.47 | -0.77 | -0.44 | -0.88 | -0.01 | -0.51 | -0.24 | -1.19 | -0.72 | -0.76 | -0.89 | 0.02 | 0.7575 | -0.2825 | 0.908162 | 0.422379 | 0.191513 | 1 | 0.996511 | 0.777117 |
| Sec23b | Q9D662 | -1.53 | -1.86 | -1.36 | -1.18 | -1.3 | -1.49 | -1.82 | -1.41 | -1.29 | -0.43 | -1.54 | -1.26 | -0.26 | -1.59 | -1.58 | -1.17 | -0.0225 | 0.3525 | 0.3325 | 0.914892 | 0.24507 | 0.346522 | 1 | 0.996511 | 0.851096 |
| Sec23ip | G3X928 | -1.66 | -1.67 | -1.53 | -1.48 | -1.7 | -1.61 | -1.45 | -1.68 | -1.65 | -1.04 | -1.61 | -1.42 | -1.25 | -1.52 | -1.72 | -1.58 | -0.025 | 0.155 | 0.0675 | 0.867399 | 0.442163 | 0.721144 | 1 | 0.996511 | 0.958398 |
| Sec24a | A2AA71 | -0.7 | -0.84 | -0.45 | -0.48 | -0.47 | -1.02 | -0.37 | -0.57 | -0.35 | -0.09 | -0.49 | -1.04 | -0.62 | -0.15 | -1.1 | -0.54 | 0.01 | 0.125 | 0.015 | 0.96058 | 0.613698 | 0.95207 | 1 | 0.996511 | 0.992476 |
| Sec24b | Q80ZX0 | -1.06 | -1.19 | -0.92 | -0.98 | -1.18 | -1.07 | -0.87 | -0.74 | -0.58 | -0.63 | -0.97 | -0.89 | -0.68 | -0.84 | -1.11 | -1.33 | 0.0725 | 0.27 | 0.0475 | 0.668848 | 0.15786 | 0.822856 | 1 | 0.996511 | 0.972768 |
| Sec24c | A0A286YDI | 0.11 | 0.26 | 0.27 | 0.73 | 1.52 | -0.45 | 0.86 | 0.76 | -0.18 | 1.48 | 1.07 | -0.21 | -0.31 | 0.01 | 0.06 | 1.13 | 0.33 | 0.1975 | -0.12 | 0.425896 | 0.646867 | 0.725711 | 1 | 0.996511 | 0.958398 |
| Sec24c | G3X972 | -0.86 | -0.23 | 0.08 | -0.27 | -0.57 | -0.83 | 0.05 | -0.2 | -0.69 | -0.08 | 0.09 | -0.24 | -0.78 | -0.26 | -0.09 | -0.03 | -0.0675 | 0.09 | 0.03 | 0.809408 | 0.741956 | 0.914715 | 1 | 0.996511 | 0.986556 |
| Sec24d | Q6NXL1 | -0.45 | 0.01 | 0.11 | 0.18 | 0.04 | -0.49 | -0.32 | 0 | 0.31 | 1.24 | 0.2 | -0.29 | 0.64 | 1.09 | -0.4 | -0.61 | -0.155 | 0.4025 | 0.2175 | 0.479922 | 0.260979 | 0.602151 | 1 | 0.996511 | 0.930802 |
| Sec31a | Q3UPL0 | -1.25 | -1.37 | -0.9 | -0.91 | -1.25 | -1.39 | -0.94 | -0.88 | -1.14 | -0.18 | -0.84 | -1.17 | -1.09 | -1.23 | -1.22 | -1.28 | -0.0075 | 0.275 | -0.0975 | 0.970448 | 0.329109 | 0.622129 | 1 | 0.996511 | 0.938628 |
| Sec61a1 | P61620 | -1.65 | -1.28 | -0.42 | -0.7 | -0.46 | -1.56 | -0.23 | -0.61 | -1.06 | -0.01 | -0.75 | -1.35 | -0.46 | 0.05 | 0.1 | 0.14 | 0.2975 | 0.22 | 0.97 | 0.444983 | 0.571414 | 0.013222 | 1 | 0.996511 | 0.584789 |
| Sec61a2 | Q9JLR1 | -1.5 | -2.08 | -1.96 | -2.05 | -0.67 | -0.42 | -1.37 | -1.73 | -1.26 | -0.58 | -2.07 | -1.37 | -0.84 | -0.9 | -0.46 | -0.52 | 0.5775 | 1.2175 | 1.2175 | 0.026467 | 0.108913 | 0.000445 | 1 | 0.996511 | 0.270112 |
| Sec61b | Q9CQS8 | -1.48 | -1.21 | -1.01 | -1.23 | -0.14 | -1.51 | -0.38 | -0.86 | -1.56 | -0.95 | -0.54 | -1.46 | -0.65 | -0.39 | 1.1 | -0.29 | 0.51 | 0.105 | 1.175 | 0.129399 | 0.699578 | 0.014357 | 1 | 0.996511 | 0.589387 |
| Sec61g | P60060 | 0.74 | 0.72 | 0.2 | -1.2 | 1.41 | 0.87 | 1.67 | 0.02 | 1.52 | -1.3 | 0.27 | 1.02 | -0.39 | 1.52 | 1.25 | 1.72 | 0.8775 | 0.2625 | 0.91 | 0.128881 | 0.707142 | 0.159462 | 1 | 0.996511 | 0.754589 |
| Sec62 | Q8BU14 | 0.63 | 0.43 | 0.27 | 0.37 | 0.44 | 1.24 | 0.22 | 0.28 | 1.08 | 0.69 | 0.2 | 0.52 | 2.08 | 0.85 | 0.91 | 0.4 | 0.12 | 0.1975 | 0.635 | 0.643428 | 0.401474 | 0.106151 | 1 | 0.996511 | 0.703417 |
| Sec63 | Q8VHE0 | -0.32 | 0.32 | -0.21 | -0.16 | 0.71 | -0.22 | 0.38 | -0.07 | 0.05 | -0.36 | -0.35 | 0.13 | 0.03 | 0.69 | 0.4 | 0.4 | 0.2925 | -0.04 | 0.4725 | 0.283872 | 0.859503 | 0.072147 | 1 | 0.996511 | 0.668066 |
| Secisbp2l | Q6A098 | 0.19 | 0.75 | -0.81 | -0.97 | -0.07 | -0.22 | -0.7 | -1.07 | 0.48 | 0.27 | -0.98 | 0.55 | 0.71 | 0.22 | -0.39 | -0.1 | -0.305 | 0.29 | 0.32 | 0.492711 | 0.57017 | 0.481109 | 1 | 0.996511 | 0.893906 |
| Seh1l | Q8R2U0-2 | -0.51 | -0.47 | -0.49 | -0.93 | -0.91 | -0.7 | -0.77 | -0.84 | -0.59 | -1.14 | -0.73 | -1.34 | 0.28 | -0.9 | -0.47 | -0.59 | -0.205 | -0.35 | 0.18 | 0.250706 | 0.163544 | 0.539488 | 1 | 0.996511 | 0.912212 |
| Sel1l | Q9Z2G6 | -1.59 | -1.68 | -1.02 | -1.05 | -1.11 | -1.4 | -0.22 | -0.82 | -1.46 | -1.24 | -1.09 | -1.57 | -0.28 | -1.32 | 0.67 | 0.24 | 0.4475 | -0.005 | 1.1625 | 0.165746 | 0.982998 | 0.025213 | 1 | 0.998773 | 0.633075 |
| Selenbp1 | P17563 | 0.53 | 0.27 | -0.25 | 0.21 | 0.5 | 0.39 | 0.12 | -0.05 | -0.6 | -0.35 | -0.3 | -0.21 | -1.27 | -0.01 | -0.63 | -0.53 | 0.05 | -0.555 | -0.8 | 0.825037 | 0.030818 | 0.029381 | 1 | 0.996511 | 0.637553 |
| Selenof | A0A0R4J0K | -0.82 | -0.36 | -0.3 | -0.15 | 0.23 | -0.8 | 0.3 | -0.07 | -0.65 | -0.2 | -0.28 | -0.76 | -0.48 | 0.02 | -0.09 | 0.44 | 0.3225 | -0.065 | 0.38 | 0.284061 | 0.778614 | 0.175865 | 1 | 0.996511 | 0.771503 |
| Selenoh | A0A087WF | -0.79 | -1.84 | -1.33 | -0.51 | -1.08 | -2.38 | -3.71 | 0.54 | -2.56 | 1.83 | -0.03 | -0.7 | -2.67 | 0.47 | 0.18 | -0.36 | -0.54 | 0.7525 | 0.1025 | 0.534499 | 0.392308 | 0.869435 | 1 | 0.996511 | 0.979293 |
| Selenoi | D6RE20 | -0.21 | -0.06 | -0.65 | 0.38 | 0.9 | -0.44 | 0.62 | -0.84 | 0.06 | -0.11 | 0.1 | -0.45 | -0.27 | 1.29 | -0.12 | 1.23 | 0.195 | 0.035 | 0.6675 | 0.656691 | 0.894598 | 0.160541 | 1 | 0.996511 | 0.75577 |
| Selenom | Q8VHC3 | -0.37 | -0.09 | -0.3 | -0.05 | 0.27 | -0.29 | 0.49 | 0.05 | -0.89 | -0.57 | -0.37 | -0.23 | -0.92 | 0.26 | 0.19 | 0.45 | 0.3325 | -0.3125 | 0.1975 | 0.142192 | 0.158145 | 0.547173 | 1 | 0.996511 | 0.912571 |
| Selenon | D3Z2R5 | 1.01 | 0.54 | 0.7 | 0.53 | 0.76 | 0.11 | 0.9 | -0.25 | -0.29 | -1.37 | 0.42 | -2.07 | 1.6 | -0.51 | 0.82 | 0.69 | -0.315 | -1.5225 | -0.045 | 0.29915 | 0.016764 | 0.916298 | 1 | 0.996511 | 0.986556 |
| Selenoo | Q9DBC0 | -0.48 | -0.71 | -0.09 | -0.02 | -1.06 | -0.43 | -0.54 | -0.19 | -0.63 | 0.01 | -0.13 | -0.85 | -0.62 | -0.62 | -0.57 | -0.55 | -0.23 | -0.075 | -0.265 | 0.380194 | 0.785213 | 0.242699 | 1 | 0.996511 | 0.798789 |
| Selenop | P70274 | -1.27 | -1.64 | -1.79 | -1.67 | -1.24 | -1.74 | -2.19 | -2.08 | -0.72 | -1.24 | -1.69 | -0.91 | -0.28 | -0.76 | -2.37 | -1.62 | -0.22 | 0.4525 | 0.335 | 0.394278 | 0.109105 | 0.464104 | 1 | 0.996511 | 0.888267 |
| Selenos | Q9BCZ4 | 1.58 | 1.82 | -0.19 | -0.36 | 1.08 | 1.79 | -0.25 | -0.7 | 0.21 | 0.25 | -0.45 | 1.79 | 0.69 | 0.2 | 0.11 | 0.05 | -0.2325 | -0.2625 | -0.45 | 0.75249 | 0.699535 | 0.419486 | 1 | 0.996511 | 0.87558 |
| Selenot | P62342 | -1.1 | -0.53 | -0.25 | -0.44 | -0.04 | -0.29 | -0.04 | -0.71 | -0.51 | -0.68 | -0.7 | -0.72 | -0.07 | 0.5 | -0.17 | -0.13 | 0.31 | -0.0725 | 0.6125 | 0.240733 | 0.746989 | 0.044545 | 1 | 0.996511 | 0.658454 |
| Selenow | P63300 | -1.48 | -3 | -0.43 | -0.72 | -3.78 | -0.9 | -2.86 | -0.77 | -0.68 | 1.03 | -0.5 | -2.23 | 0.59 | -3.56 | -0.82 | -0.57 | -0.67 | 0.8125 | 0.3175 | 0.436625 | 0.322416 | 0.736459 | 1 | 0.996511 | 0.958635 |
| Selplg | Q3TA56 | 1.56 | 1.17 | 0.27 | 0.33 | 1.4 | 0.67 | 1.7 | 0.87 | -0.27 | 1.29 | -0.07 | 0.62 | 1.25 | 0.38 | 1.62 | -0.12 | 0.22 | -0.1575 | -0.06 | -0.3075 | 0.721203 | 0.888216 | 0.514367 | 1 | 0.996511 | 0.90567 |
| Sema4a | Q62178 | 1.49 | 0.46 | 0.42 | 0.87 | 0.48 | 1.8 | 0.88 | 0.52 | 0.96 | 0.86 | 0.81 | 1.29 | 1.38 | 0.73 | 1.04 | 0.71 | 0.11 | 0.17 | 0.155 | 0.769911 | 0.55118 | 0.614303 | 1 | 0.996511 | 0.935876 |
| Sema4b | Q62179 | 0.61 | 0.74 | 0.4 | 0.63 | 0.65 | 0.62 | 0.41 | 0.29 | 0.97 | 0.38 | 0.36 | 0.61 | 0.9 | 1.09 | 0.3 | 0.32 | -0.1025 | -0.015 | 0.0575 | 0.543867 | 0.94158 | 0.81696 | 1 | 0.996755 | 0.972768 |
| Sema4c | Q64151 | 0.55 | 0.66 | 0.88 | 0.81 | 0.18 | 0.74 | 0.78 | 0.51 | 0.9 | 0.33 | 0.77 | 0.57 | 0.96 | -0.22 | 0.77 | 0.84 | -0.1725 | -0.0825 | -0.1375 | 0.383176 | 0.676297 | 0.644543 | 1 | 0.996511 | 0.944749 |
| Sema4d | O09126 | 0.87 | 0.99 | -0.24 | -0.41 | 1.15 | 0.9 | -0.16 | -0.39 | 0.54 | -0.24 | -0.27 | 0.94 | 0.79 | 1.11 | -0.18 | -0.3 | 0.0725 | -0.06 | 0.0525 | 0.881838 | 0.892484 | 0.911868 | 1 | 0.996511 | 0.986458 |
| Sema4f | Q9Z123 | -0.52 | -0.12 | 0.45 | 0.78 | 0.24 | -0.27 | 0.87 | 0.3 | 0.54 | -0.7 | 0.64 | -0.29 | 0.09 | 0.53 | 0.16 | 0.87 | 0.1375 | -0.1 | 0.265 | 0.700887 | 0.809311 | 0.445786 | 1 | 0.996511 | 0.884817 |
| Sema4g | Q9WUH7 | 0.66 | 0.61 | -0.84 | 0.52 | 1.17 | 0.5 | 0.62 | 0.02 | 0.32 | 0.77 | 0.26 | 0.59 | 0.63 | 0.88 | 0.94 | 0.72 | 0.34 | 0.2475 | 0.555 | 0.411727 | 0.506964 | 0.151085 | 1 | 0.996511 | 0.749304 |
| Sema5a | Q3UPZ0 | -0.02 | -0.28 | -0.81 | -0.5 | -0.74 | 0.44 | -1.35 | -1.02 | -0.35 | -1.42 | -0.76 | 0.4 | 0.27 | -1.79 | -0.19 | -0.69 | -0.265 | -0.13 | -0.1975 | 0.513058 | 0.744652 | 0.660622 | 1 | 0.996511 | 0.949645 |
| Sema5b | Q60519-2 | -0.55 | -1.27 | -1.06 | -0.94 | -0.13 | 0.05 | -0.88 | -1.37 | 0.5 | -1.34 | -0.66 | -0.33 | 0.94 | -0.6 | -0.76 | -1.65 | 0.3725 | 0.4975 | 0.4375 | 0.303225 | 0.228098 | 0.408936 | 1 | 0.996511 | 0.874356 |
| Sema6a | K3W4S5 | 0.51 | 0.64 | -0.04 | 0.25 | 0.51 | 0.24 | 0.24 | 0.4 | 0.86 | 0.14 | 0.19 | 0.33 | 0.56 | 0.72 | 0.47 | 0.03 | 0.0075 | 0.04 | 0.105 | 0.969785 | 0.871594 | 0.671367 | 1 | 0.996511 | 0.952631 |
| Sema6b | O54951 | 0.1 | 0.31 | 0.74 | 0.69 | 0.38 | 0.05 | 1.34 | 0.84 | 0.47 | 0.85 | 0.97 | 0.84 | -0.18 | 0.8 | 0.14 | 0.72 | 0.1925 | 0.3225 | -0.09 | 0.544407 | 0.173896 | 0.76112 | 1 | 0.996511 | 0.963123 |
| Sema6d | A2AW73 | 0.16 | 0.21 | 0.83 | 0.71 | 0.38 | -0.15 | 0.5 | 0.42 | 0.42 | 0.94 | 0.65 | 0.05 | 0.98 | 0.03 | 0.85 | 0.44 | -0.19 | 0.0975 | 0.0975 | 0.439372 | 0.889401 | 0.738233 | 1 | 0.996511 | 0.958796 |
| Sema7a | Q9QUR8 | 0.24 | 0.46 | 0.5 | 0.37 | 0.81 | 0.13 | 0.75 | 0.29 | 0.47 | 0.62 | 0.61 | 0.42 | 0.45 | 0.72 | 0.34 | 0.83 | 0.1025 | 0.1375 | 0.1925 | 0.623134 | 0.411241 | 0.342919 | 1 | 0.996511 | 0.848473 |
| Senp2 | A0A338P6/ | -0.95 | -0.34 | -0.7 | -0.7 | 0.25 | -0.48 | -1.46 | -0.94 | 0.42 | -2 | -1.03 | -0.84 | 0.42 | 0.23 | -0.62 | -0.16 | 0.015 | -0.19 | 0.64 | 0.967307 | 0.692785 | 0.04626 | 1 | 0.996511 | 0.658454 |
| Senp3 | Q9EP97 | -4.56 | -3.49 | -2.99 | -2.6 | -4.14 | -4.98 | -1.97 | -3.14 | -0.76 | 3.38 | -2.72 | -5.37 | -0.35 | -3.85 | -3.63 | -2.66 | -0.1475 | 2.0425 | 0.7875 | 0.833725 | 0.246213 | 0.347629 | 1 | 0.996511 | 0.851691 |
| Senp7 | Q8BUH8 | 1.49 | 0.92 | 4.87 | 2.05 | 0.39 | 1.14 | 4.35 | 3.95 | 0.45 | 0.42 | 4.66 | 1.02 | 1.01 | 0.41 | 4.69 | 5.56 | 0.125 | -0.695 | 0.585 | 0.915929 | 0.566913 | 0.674862 | 1 | 0.996511 | 0.952944 |
| Senp8 | Q9D2Z4 | -1.18 | -1.7 | 0.79 | 0.66 | -0.79 | -1.56 | 0.83 | 0.56 | -1.43 | 0.65 | 0.48 | -1.66 | -1.53 | 0.17 | 0.4 | 1.09 | 0.1175 | -0.1325 | 0.39 | 0.878213 | 0.867942 | 0.611403 | 1 | 0.996511 | 0.935047 |
| Sephs1 | Q8BH69 | -1.47 | -1.24 | -2.93 | -2.6 | -0.94 | -1.55 | -2.59 | -2.61 | -0.87 | -1.38 | -2.9 | -0.96 | -1.43 | -0.46 | -3.44 | -3.09 | 0.1375 | 0.5325 | -0.045 | 0.797749 | 0.368263 | 0.951317 | 1 | 0.996511 | 0.992476 |
| Sephs2 | P97364 | -2.02 | -1.47 | -1.53 | -1.69 | -1.39 | -1.46 | -1.48 | -1.28 | -1.56 | -1.46 | -1.75 | -1.76 | -2.16 | -1.42 | -1.68 | -1.61 | 0.275 | 0.045 | -0.04 | 0.148558 | 0.818517 | 0.86718 | 1 | 0.996511 | 0.978457 |
| Sepsecs | Q6P6M7 | -1 | -0.57 | -0.49 | -0.8 | -0.4 | -1.16 | -1.13 | -0.88 | -1.64 | -0.75 | -0.57 | -1.45 | -0.86 | -0.31 | -0.63 | -0.56 | -0.1775 | -0.3875 | 0.125 | 0.447907 | 0.209217 | 0.566006 | 1 | 0.996511 | 0.921174 |
| Septin10 | A0A0R4J23 | 0.54 | 0.29 | 0.21 | 0 | 0.67 | 0.47 | 0.62 | -0.09 | -0.5 | -0.5 | -0.22 | 0.47 | -0.95 | -0.58 | -0.07 | 0.27 | 0.1575 | -0.4475 | -0.5925 | 0.494779 | 0.124991 | 0.077958 | 1 | 0.996511 | 0.676942 |
| Septin11 | A0A0J9YUL | 0.67 | 0.43 | 0.54 | 0.52 | 0.43 | 0.46 | 0.47 | 0.35 | 0.42 | 0.41 | 0.57 | -0.09 | 0.69 | 0.63 | -0.27 | 0.27 | 0.1125 | -0.0775 | -0.3125 | 0.441989 | 0.628116 | 0.189597 | 1 | 0.996511 | 0.776843 |
| Septin2 | E9Q3V6 | -2.77 | -1.76 | -1.98 | -3.6 | -0.52 | -2.64 | -2.96 | -3.78 | -3.02 | -0.74 | -2.64 | -3.04 | -1.57 | -0.64 | -2.45 | -2.88 | 0.0525 | 0.1675 | 0.6425 | 0.942848 | 0.790539 | 0.296844 | 1 | 0.996511 | 0.825836 |
| Septin2 | P42208 | 0.94 | 0.67 | 1.34 | 1.22 | 0.74 | 0.79 | 1.7 | 1.09 | 0.43 | 0.88 | 1.19 | 0.76 | -0.16 | 0.56 | 0.92 | 1.45 | 0.0375 | -0.2275 | -0.35 | 0.890267 | 0.362005 | 0.348916 | 1 | 0.996511 | 0.85278 |
| Septin3 | Q9Z1S5-2 | 1.58 | 1.08 | 1.69 | 1.15 | 1.24 | 1.35 | 1.81 | 1.66 | 0.59 | 0.89 | 1.67 | 1.47 | -0.05 | 0.62 | 1.36 | 1.47 | 0.14 | -0.22 | -0.525 | 0.536425 | 0.469614 | 0.187175 | 1 | 0.996511 | 0.776843 |
| Septin3 | A0A5F8MP | 0.42 | 0.57 | 0.99 | 0.63 | 0.72 | 0.48 | 0.3 | 0.52 | 1.23 | 1.17 | 0.6 | 0.44 | 0.9 | 1.82 | 0.13 | 0.23 | -0.1475 | 0.2075 | 0.1175 | 0.441478 | 0.424145 | 0.766098 | 1 | 0.996511 | 0.964134 |
| Septin3 | A0A2R8VH | 0.35 | 0.44 | 0.91 | 0.37 | 1.08 | 1 | 1.4 | 1.62 | -0.15 | 0.88 | 1.42 | 1.16 | -0.89 | -1.1 | 1.96 | 1.94 | 0.7575 | 0.31 | -0.04 | 0.007107 | 0.398578 | 0.958918 | 1 | 0.996511 | 0.992636 |
| Septin4 | A0A5F8MP | 1.54 | 1.18 | 1.33 | 1.04 | 0.95 | 1.05 | 0.47 | 0.84 | 0.17 | 0.64 | 0.88 | 0.99 | 0.38 | -0.24 | 1.11 | 1.25 | -0.4475 | -0.6025 | -0.4675 | 0.049454 | 0.031019 | 0.097562 | 1 | 0.996511 | 0.698156 |
| Septin5 | Q9Z2Q6 | 2.36 | 1.17 | 1.53 | 1.23 | 1.2 | 2.32 | 1.2 | 1.58 | 0.73 | 0.74 | 1.38 | 2.2 | 0.42 | 0.18 | 1.15 | 1.29 | 0.0025 | -0.31 | -0.8125 | 0.994508 | 0.467693 | 0.055727 | 1 | 0.996511 | 0.661738 |
| Septin5 | A0A338P7: | -0.36 | -0.92 | -0.83 | -2.73 | -0.43 | -2.36 | -3.04 | -3.03 | -2.33 | -1.04 | -1.89 | -2.74 | 1.59 | -1.4 | -1.33 | -2.2 | -1.005 | -0.79 | 0.375 | 0.192461 | 0.199592 | 0.672024 | 1 | 0.996511 | 0.952835 |
| Septin5 | A0A338P7( | 0.26 | 0.59 | 0.33 | 1.03 | 0.7 | 0.39 | 0.44 | 0.62 | 0.59 | 1.16 | 0.94 | 0.51 | 0.39 | 1.27 | 0.27 | 0.13 | -0.015 | 0.2475 | -0.0375 | 0.94429 | 0.342052 | 0.906019 | 1 | 0.996511 | 0.986321 |
| Septin6 | Q9R1T4-2 | -0.56 | -0.07 | 0.22 | 0.06 | -0.05 | -0.33 | 0.28 | -0.01 | -0.6 | -0.26 | 0.23 | -0.61 | -0.21 | 0.2 | 0.72 | 0.3 | 0.06 | -0.2225 | 0.1125 | 0.79375 | 0.425468 | 0.768692 | 1 | 0.996511 | 0.965659 |
| Septin7 | O55131 | 1.84 | 0.8 | 1.09 | 1.09 | 0.77 | 1.6 | 1.27 | 1.32 | 0.53 | 0.37 | 1.01 | 1.44 | -0.21 | 0.54 | 0.66 | 0.92 | 0.035 | -0.3675 | -0.7275 | 0.901711 | 0.279707 | 0.05264 | 1 | 0.996511 | 0.659923 |
| Septin7 | E9Q1G8 | 0.54 | -0.03 | 0.09 | 0.33 | -0.81 | 1.2 | -0.84 | 0.49 | -0.34 | 0.26 | -0.03 | 0.84 | 0.09 | -2.33 | 0.06 | -0.04 | -0.2225 | -0.05 | -0.7875 | 0.645232 | 0.863191 | 0.182757 | 1 | 0.996511 | 0.776002 |
| Septin8 | B7ZC46 | 0.19 | 0.36 | 1.46 | 1.35 | 0.47 | -0.05 | 0.84 | 1.14 | -0.55 | 0.85 | 1.14 | 0.17 | -1.56 | -0.39 | 0.13 | 1.01 | -0.24 | -0.4375 | -1.0425 | 0.546687 | 0.362752 | 0.0991 | 1 | 0.996511 | 0.698156 |
| Septin8 | B1AQZ0 | 0.42 | 0.46 | 1.72 | 1.61 | 0.54 | 0.37 | 1.24 | 1.45 | 0.49 | 1.43 | 1.42 | 0.31 | -0.12 | 0.69 | 1.09 | 1.33 | -0.1525 | -0.14 | -0.305 | 0.713714 | 0.749345 | 0.503608 | 1 | 0.996511 | 0.901002 |
| Septin9 | Q80UG5-3 | 1.42 | 0.87 | 1.51 | 0.75 | 0.65 | 1.3 | 1.23 | 1.11 | 0.21 | 0.24 | 1.3 | 1.25 | 0.4 | -0.07 | 0.93 | 0.79 | -0.065 | -0.3875 | -0.625 | 0.796893 | 0.287304 | 0.067064 | 1 | 0.996511 | 0.664936 |
| Serac1 | Q3U213 | -0.01 | 0.29 | 0 | -0.13 | -0.05 | 0.33 | -0.64 | -0.32 | 0.06 | 0.19 | -0.1 | 0.09 | -0.12 | 0.06 | -0.85 | -0.75 | -0.2075 | 0.0225 | -0.4525 | 0.397539 | 0.898815 | 0.119012 | 1 | 0.996511 | 0.721448 |
| Serbp1 | Q9CY58 | -2.4 | -2.41 | -1.74 | -1.94 | -1.78 | -2.64 | -1.59 | -1.97 | -2.53 | -0.66 | -1.73 | -2.3 | -1.67 | -1.53 | -1.42 | -1.23 | 0.1275 | 0.3175 | 0.66 | 0.656631 | 0.463414 | 0.019016 | 1 | 0.996511 | 0.604671 |
| Serbp1 | Q9CY58-2 | -5.09 | -6.51 | -0.9 | -3.1 | -4.66 | -7.84 | -3.04 | -3.6 | -8.44 | -0.52 | -3.95 | -6.75 | -6.45 | -4.58 | -1.09 | -4.58 | -0.885 | -1.015 | -0.275 | 0.546136 | 0.594096 | 0.851642 | 1 | 0.996511 | 0.976225 |
| Serf2 | P84102 | -3.22 | -3.72 | -2.15 | -2.48 | -3.43 | -2.54 | -3.43 | -3.11 | -2.8 | -0.71 | -2.35 | -2.91 | -2.11 | -2.17 | -1.67 | -2.65 | -0.235 | 0.7 | 0.7425 | 0.551221 | 0.240645 | 0.087002 | 1 | 0.996511 | 0.691619 |
| Sergef | Q80YD6 | 0.17 | 0.15 | -0.04 | 0.49 | 0.29 | 0.25 | 0.19 | 0.28 | 0.52 | 0.37 | 0.36 | 0.16 | -0.2 | -0.91 | 0.58 | -0.11 | 0.06 | -0.035 | -0.3525 | 0.720773 | 0.943371 | 0.29867 | 1 | 0.996755 | 0.828257 |
| Serhl | Q9EPB5 | 0.24 | 0.23 | 1.48 | 0.67 | 0.65 | -0.3 | 2.54 | 0.64 | 0.19 | 0.37 | 1.5 | 0.16 | -0.42 | 0.56 | 1.67 | 1.57 | 0.2275 | -0.1 | 0.19 | 0.708013 | 0.80842 | 0.720605 | 1 | 0.996511 | 0.958398 |
| Serinc1 | Q9QZI8 | -1.59 | -1.56 | -0.76 | -0.5 | -1.29 | -1.01 | -0.73 | -0.69 | -1.24 | 0.36 | -0.64 | -1.27 | -1.13 | -0.84 | -0.62 | -1 | 0.1725 | 0.405 | 0.205 | 0.577583 | 0.374002 | 0.512324 | 1 | 0.996511 | 0.903993 |
| Serinc3 | Q9QZI9 | 0.86 | -0.56 | 0.42 | -0.79 | 0.04 | 1.13 | -1.22 | -0.57 | -0.94 | -1.97 | -0.06 | 0.72 | -1.2 | -1.73 | 0.56 | -0.62 | -0.1375 | -0.545 | -0.73 | 0.812946 | 0.403893 | 0.22794 | 1 | 0.996511 | 0.787447 |
| Serinc5 | Q8BHJ6 | -0.64 | -0.63 | -0.61 | -2.61 | -0.73 | -1.05 | -3.38 | -3.27 | -1.29 | -5.77 | -2.27 | -1.3 | -0.55 | -2.44 | -1.27 | -1.89 | -0.985 | -1.535 | -0.415 | 0.22755 | 0.170077 | 0.487084 | 1 | 0.996511 | 0.896644 |
| Serpina1b | P22599 | -5.77 | -6.11 | -5.17 | -6.35 | -3.96 | -6.36 | -6.54 | -6.35 | -5.96 | -3.6 | -5.31 | -4.96 | -4.65 | -3.51 | -6.02 | -6.46 | 0.0475 | 0.8925 | 0.69 | 0.937641 | 0.113124 | 0.307576 | 1 | 0.996511 | 0.832565 |
| Serpina1c | A0A0R4J0X | -3.77 | -4.51 | -3.52 | -4.94 | -3.35 | -3.72 | -3.54 | -4.04 | -4.37 | -3.01 | -3.49 | -3.35 | -3.62 | -2.61 | -4.26 | -4.43 | 0.5225 | 0.63 | 0.455 | 0.157256 | 0.155984 | 0.367065 | 1 | 0.996511 | 0.858904 |
| Serpina1d | Q00897 | -3.99 | -4.8 | -1.9 | -2.67 | -3.2 | -4.58 | -2.42 | -2.74 | -3.99 | -1.45 | -2.28 | -4.06 | -3.58 | -2.21 | -2.52 | -1.95 | 0.105 | 0.395 | 0.775 | 0.885422 | 0.635443 | 0.269128 | 1 | 0.996511 | 0.820534 |
| Serpina1e | Q00898 | -2.09 | -3.27 | -1.41 | -1.08 | -2.55 | -1.58 | -1.02 | -1.19 | -1.42 | -1.81 | -0.9 | -1.88 | 0.12 | -2.39 | -1.46 | -1.22 | 0.3775 | 0.46 | 0.725 | 0.4936 | 0.366558 | 0.27951 | 1 | 0.996511 | 0.822084 |
| Serpina3b | Q8BYY9 | -1.52 | -0.67 | -1.31 | -1.79 | 0.15 | -1.63 | -1.66 | -1.72 | -1.97 | -0.4 | -1.67 | -1.27 | -1.39 | -0.27 | -1.77 | -2 | 0.1075 | -0.005 | -0.035 | 0.821483 | 0.989967 | 0.935083 | 1 | 0.999285 | 0.989767 |
| Serpina3k | A0A0R4J0I | -3.46 | -3.9 | -3.37 | -3.61 | -3.18 | -3.26 | -3.27 | -3.44 | -3.79 | 0.01 | -2.93 | -2.95 | -3.33 | -1.9 | -3.5 | -3.85 | 0.2975 | 1.17 | 0.44 | 0.118859 | 0.150937 | 0.314604 | 1 | 0.996511 | 0.836924 |
| Serpina3m | Q03734 | -8.62 | -10.63 | -4.56 | -5.71 | -6.5 | -7.63 | -3.9 | -4.76 | -8.99 | -2.43 | -4.91 | -7.27 | -6.59 | -4.32 | -5.01 | -5.1 | 1.6825 | 1.48 | 2.125 | 0.26268 | 0.412861 | 0.128919 | 1 | 0.996511 | 0.723655 |
| Serpina3n | G3X8T9 | -0.68 | -0.72 | -1.29 | -1.61 | -1.37 | -0.58 | -1.05 | -0.69 | -1.12 | -0.79 | -1.52 | -0.72 | -1.71 | 0 | -2.01 | -1.74 | 0.1525 | 0.0375 | -0.29 | 0.601118 | 0.899674 | 0.550108 | 1 | 0.996511 | 0.913572 |
| Serpina6 | Q06770 | -2.85 | -3.68 | -5.86 | -4.62 | -3.98 | -3.12 | -5.64 | -5.7 | -3.36 | -1.34 | -5.16 | -3.09 | -3.89 | -3.88 | -6.61 | -7.16 | -0.3575 | 1.015 | -1.1325 | 0.663582 | 0.284018 | 0.263021 | 1 | 0.996511 | 0.81524 |
| Serpinb1a | Q9D154 | 1.8 | 1.91 | 0.96 | 0.53 | 1.55 | 0.98 | 0.34 | 0.62 | -0.18 | -0.13 | -0.06 | 0.53 | -0.35 | -0.18 | 0.63 | 0.11 | -0.4275 | -1.26 | -1.2475 | 0.300588 | 0.00688 | 0.009789 | 1 | 0.996511 | 0.563386 |
| Serpinb1b | Q8VHP7 | -0.25 | -0.31 | -0.9 | -0.23 | 0.08 | -0.8 | -1.04 | -0.37 | -0.11 | 1.66 | -1.09 | -0.58 | -0.55 | 0.21 | 0.45 | -0.18 | -0.11 | 0.3925 | 0.405 | 0.708771 | 0.496639 | 0.185129 | 1 | 0.996511 | 0.776843 |
| Serpinb6 | Q60854 | 0.58 | 0.79 | 0.28 | 0.23 | 0.66 | 0.56 | 0.23 | 0.3 | 0.86 | 0.76 | 0.25 | 0.63 | 0.38 | 1.16 | -0.33 | -0.37 | -0.0325 | 0.155 | -0.26 | 0.870893 | 0.493514 | 0.494037 | 1 | 0.996511 | 0.89758 |
| Serpinb6a | F8WIV2 | -2.94 | -3.31 | -4.65 | -2.65 | -4.39 | -1.08 | -4.21 | -3.83 | 0.26 | -2.72 | -5.93 | -2.26 | 0.72 | -7.3 | -2.9 | -1.61 | 0.01 | 0.725 | 0.615 | 0.990067 | 0.551493 | 0.691954 | 1 | 0.996511 | 0.955729 |
| Serpinb8 | O08800 | -1.57 | -1.43 | -0.76 | -0.82 | -1.21 | -1.37 | -1.08 | -1.11 | -1.44 | -0.86 | -0.99 | -1.59 | -1.29 | -0.95 | -1.63 | -1.52 | -0.0475 | -0.075 | -0.2025 | 0.839785 | 0.790978 | 0.472464 | 1 | 0.996511 | 0.891144 |
| Serpinb9 | O08797 | -0.7 | -0.48 | -1.28 | -1.35 | -0.46 | -0.69 | -0.79 | -1.08 | -1.14 | -0.91 | -1.49 | -0.72 | -1.54 | -0.66 | -1.67 | -1.3 | 0.1975 | -0.1125 | -0.34 | 0.453795 | 0.691548 | 0.300548 | 1 | 0.996511 | 0.830276 |
| Serpinc1 | P32261 | -2.91 | -3.41 | -2.81 | -2.51 | -1.45 | -3 | -2.46 | -2.67 | -3.43 | -1.78 | -3.01 | -2.68 | -3.25 | -2.24 | -2.94 | -2.53 | 0.515 | 0.185 | 0.17 | 0.183472 | 0.630862 | 0.577893 | 1 | 0.996511 | 0.927198 |
| Serpine2 | Q07235 | -0.9 | -1.35 | -0.85 | -0.54 | -1.2 | -1.1 | -0.4 | -0.76 | -0.99 | 0.07 | -0.74 | -0.55 | -1.18 | -0.65 | -0.47 | -0.57 | 0.045 | 0.3575 | 0.1925 | 0.860684 | 0.238385 | 0.465205 | 1 | 0.996511 | 0.888369 |
| Serpinf1 | P97298 | -1.8 | -1.12 | -1.76 | -2.16 | -1.69 | -1.7 | -1.26 | -2.36 | -1.23 | -1.41 | -1.76 | -1.42 | -1.8 | -1.03 | -2.31 | -2.01 | -0.0425 | 0.255 | -0.0775 | 0.890533 | 0.343281 | 0.823345 | 1 | 0.996511 | 0.972768 |
| Serpinf2 | Q61247 | -0.91 | -1.4 | -2.24 | -3.26 | -1 | -1.47 | -2.41 | -3.1 | -2.16 | -0.85 | -2.03 | -1.29 | -2.12 | -1.29 | -2.23 | -1.48 | -0.0425 | 0.37 | 0.1725 | 0.946554 | 0.510058 | 0.742673 | 1 | 0.996511 | 0.959115 |
| Serping1 | A2ATR8 | -3.41 | -4.8 | -5.12 | -4.11 | -2.87 | -4.27 | -3.41 | -4.32 | -3.44 | -2.46 | -3.27 | -3.29 | -3.07 | -1.26 | -4.41 | -4.86 | 0.6425 | 1.245 | 0.96 | 0.203337 | 0.015019 | 0.252491 | 1 | 0.996511 | 0.80892 |
| Serpinh1 | P19324 | -0.38 | 0.09 | 0.02 | 0.31 | 0.52 | 0.07 | 0.58 | 0.15 | 0.16 | 0.18 | 0.35 | 0.19 | 0.62 | 0.55 | 0.82 | 0.8 | 0.32 | 0.21 | 0.6875 | 0.166823 | 0.30941 | 0.010149 | 1 | 0.996511 | 0.563386 |
| Serpini1 | O35684 | -0.05 | -0.22 | -1.22 | -1.55 | 0.77 | 0.25 | -0.51 | -1 | -0.76 | -1.51 | -0.74 | 0.5 | -0.91 | 0.42 | -0.19 | -0.7 | 0.6125 | 0.1325 | 0.415 | 0.249286 | 0.797454 | 0.365932 | 1 | 0.996511 | 0.858761 |
| Sesn3 | Q9CYP7 | -0.25 | -0.74 | 2.35 | 0.36 | -0.1 | -0.98 | 1.8 | 1.77 | -0.1 | 2.99 | 0.51 | 0.06 | 0.53 | -0.18 | 0.84 | 3.05 | 0.1925 | 0.435 | 0.63 | 0.825694 | 0.62779 | 0.477462 | 1 | 0.996511 | 0.892234 |
| Sestd1 | Q80UK0 | 0.06 | 0.05 | 0.07 | 0 | 0.11 | 0.28 | -0.09 | 0.06 | 0.56 | -0.27 | 0.26 | 0.58 | 0.49 | -0.17 | -0.2 | 0.01 | 0.045 | 0.2375 | -0.0125 | 0.766577 | 0.318295 | 0.953555 | 1 | 0.996511 | 0.992476 |
| Set | A2BE93 | -1.49 | -4.91 | -1.82 | -2.46 | -4.21 | -2.87 | -1.14 | -2.59 | -4.43 | -1.22 | -2.15 | -1.18 | -4.27 | -3.45 | -2.23 | -1.68 | -0.0325 | 0.425 | -0.2375 | 0.971053 | 0.664802 | 0.785464 | 1 | 0.996511 | 0.967642 |
| Set | Q9EQU5-2 | -8.7 | -8.71 | -8.4 | -9.66 | -8.31 | -8.27 | -11.25 | -11.99 | -7.38 | -4.05 | -9.71 | -7.27 | -4.9 | -9.73 | -10.15 | -11.61 | -1.0875 | 1.765 | -0.23 | 0.251283 | 0.127011 | 0.861427 | 1 | 0.996511 | 0.977204 |
| Setd2 | E9Q5F9 | 0.16 | 0.1 | 0.15 | -0.33 | -1 | -0.98 | -0.54 | -0.15 | -0.92 | -0.05 | -0.17 | -0.75 | 0.54 | 1.44 | 0.61 | -0.58 | -0.7125 | -0.4925 | 0.4825 | 0.020665 | 0.08777 | 0.261637 | 1 | 0.996511 | 0.81412 |
| Setd3 | Q91WC0 | -2.67 | -2.8 | -2.73 | -3.16 | -2.27 | -2.73 | -2.7 | -3.41 | -2.54 | -2.13 | -2.78 | -2.28 | -2.42 | -2.23 | -2.89 | -2.9 | 0.0625 | 0.4075 | 0.23 | 0.814858 | 0.089971 | 0.351533 | 1 | 0.996511 | 0.853984 |
| Setd7 | Q8VHL1 | -0.28 | -0.18 | -0.58 | -0.55 | -0.43 | -0.44 | -0.63 | -0.92 | -0.91 | -0.56 | -0.62 | -0.19 | -0.75 | -0.59 | -0.85 | -0.98 | -0.2075 | -0.1725 | -0.395 | 0.291922 | 0.435195 | 0.072256 | 1 | 0.996511 | 0.668066 |
| Setdb1 | G5E8N3 | -2.62 | -1.49 | 0.35 | 0.53 | -2.72 | -3.06 | -0.98 | -0.45 | 6.69 | 1.44 | -0.12 | -4.13 | 5.93 | 7.07 | -0.16 | -0.24 | -0.995 | 1.7775 | 3.9575 | 0.282391 | 0.408454 | 0.057732 | 1 | 0.996511 | 0.661738 |
| Sez6 | Q7TSK2 | -0.37 | -1.43 | -0.37 | -0.42 | -1.43 | -1.14 | -0.31 | -0.47 | -2.2 | -0.56 | -0.3 | -0.15 | -1.35 | -1.94 | -0.67 | -0.12 | -0.19 | -0.155 | -0.3725 | 0.598332 | 0.758241 | 0.416305 | 1 | 0.996511 | 0.874945 |
| Sez6l | A0A1D5RN | -0.47 | -0.02 | 0.24 | -0.54 | 0.3 | -0.26 | -0.5 | -0.23 | -0.93 | -0.79 | -0.26 | -0.04 | -0.88 | -0.45 | -0.26 | -0.16 | 0.025 | -0.3075 | -0.24 | 0.923298 | 0.305862 | 0.384338 | 1 | 0.996511 | 0.860984 |
| Sez6l2 | Q4V9Z5-2 | 0.04 | 0.13 | 0.06 | 0.08 | 0.24 | 0.14 | -0.1 | -0.07 | 0.11 | 0.49 | -0.01 | 0.56 | 0.08 | 0.49 | 0.11 | 0.24 | -0.025 | 0.21 | 0.1525 | 0.871068 | 0.297363 | 0.413156 | 1 | 0.996511 | 0.874811 |
| Sez6l2 | Q4V9Z5 | -0.57 | -0.44 | -0.12 | -0.47 | -2.03 | 0.4 | -2.68 | -1.71 | 3.58 | -1.92 | -0.8 | -0.6 | 4.8 | -1.74 | -0.52 | -0.4 | -1.105 | 0.465 | 0.935 | 0.099533 | 0.670076 | 0.477375 | 1 | 0.996511 | 0.892234 |
| Sf1 | F6SRV1 | -1.56 | -2.39 | -2.83 | -3.53 | -3.89 | -1.48 | -3.71 | -3.18 | 1.18 | -3.98 | -2.56 | -2.89 | 2.83 | -4.89 | -0.74 | -3.17 | -0.4875 | 0.515 | 1.085 | 0.443821 | 0.633381 | 0.484675 | 1 | 0.996511 | 0.894377 |
| Sf1 | E9Q4Q2 | -3.03 | -3.31 | -2.84 | -3.01 | -3.19 | -3.32 | -2.86 | -2.98 | -3.17 | -1.6 | -2.58 | -3.03 | -2.7 | -2.57 | -2.8 | -3.49 | -0.04 | 0.4525 | 0.1575 | 0.82815 | 0.228498 | 0.545035 | 1 | 0.996511 | 0.912212 |
| Sf3a1 | Q8K4Z5 | 0.67 | -0.31 | -0.98 | -0.85 | -0.16 | 0.06 | -0.73 | -0.35 | -0.81 | -0.9 | -0.78 | -0.85 | -0.19 | -0.05 | -0.16 | -1.35 | 0.0725 | -0.4675 | -0.07 | 0.852051 | 0.222364 | 0.878051 | 1 | 0.996511 | 0.979626 |
| Sf3a2 | G3UVU2 | 0.35 | -0.83 | -0.41 | -0.7 | -1.61 | -0.13 | -1.66 | -0.1 | -0.84 | -1.22 | -0.65 | -1.3 | 0.12 | -1.51 | -0.21 | -1.45 | -0.4775 | -0.605 | -0.365 | 0.32898 | 0.077576 | 0.442195 | 1 | 0.996511 | 0.883549 |
| Sf3a3 | Q9D554 | 0.28 | -0.94 | -0.82 | -0.52 | -1.2 | -0.63 | -0.55 | -0.08 | -1.35 | -0.91 | -0.66 | -1.51 | -0.75 | -1 | -0.49 | -1.08 | -0.115 | -0.6075 | -0.33 | 0.739539 | 0.096797 | 0.308479 | 1 | 0.996511 | 0.834018 |
| Sf3b1 | G5E866 | 0.53 | -0.75 | -0.76 | -0.73 | -1.23 | -0.27 | -0.83 | -0.14 | -1.16 | -1.07 | -0.84 | -1.22 | -0.53 | -0.67 | -0.49 | -1.19 | -0.19 | -0.645 | -0.2925 | 0.624865 | 0.076552 | 0.418194 | 1 | 0.996511 | 0.87558 |
| Sf3b2 | Q3UJB0 | -0.04 | -0.94 | -1.05 | -0.88 | -1.34 | -0.75 | -1.14 | -0.49 | -0.87 | -0.67 | -1.03 | -1.41 | -0.37 | -0.56 | -0.67 | -1.42 | -0.2025 | -0.2675 | -0.0275 | 0.503342 | 0.366981 | 0.93354 | 1 | 0.996511 | 0.989767 |
| Sf3b3 | Q921M3 | -0.66 | -1.27 | -1.15 | -1.19 | -1.34 | -1.24 | -1 | -0.83 | -1.84 | -0.36 | -1.13 | -1.82 | -1.39 | -1.03 | -0.88 | -1.44 | -0.035 | -0.22 | -0.1175 | 0.866785 | 0.551714 | 0.620355 | 1 | 0.996511 | 0.93807 |

| | | | | | | | | | | | | | | | | | | | | | | | | | | |
|---|---|---|---|---|---|---|---|---|---|---|---|---|---|---|---|---|---|---|---|---|---|---|---|---|---|---|
| Sf3b4 | Q8QZY9 | 1.25 | 0.42 | -1.82 | -1.14 | -0.6 | 0.89 | -0.85 | -1.19 | -0.74 | -3.66 | -1.42 | 0.18 | -0.82 | -1.48 | -1.67 | -1.64 | -0.115 | -1.0875 | -1.08 | 0.87943 | 0.28017 | 0.131266 | 1 | 0.996511 | 0.726389 |
| Sf3b5 | Q923D4 | -3.42 | -3.17 | -3.73 | 0.95 | -3.43 | -3.99 | -2.43 | -3.24 | -3.45 | -0.88 | -3.99 | -4.24 | -2.54 | -3.81 | -2.71 | -3.72 | -0.93 | -0.7975 | -0.8525 | 0.378629 | 0.51361 | 0.417174 | 1 | 0.996511 | 0.87558 |
| Sf3b6 | P59708 | 0 | -1.46 | -0.14 | 0.27 | -1.59 | -1.15 | -0.51 | 0.16 | -1.05 | 0.03 | -0.13 | -1.8 | -1.45 | -0.48 | 0.02 | -0.32 | -0.44 | -0.405 | -0.225 | 0.390211 | 0.454074 | 0.632803 | 1 | 0.996511 | 0.941132 |
| Sfn | O70456 | -1 | -0.98 | -0.69 | -0.73 | -0.69 | -1.14 | -0.97 | -0.5 | -1.14 | 0.1 | -0.52 | -0.97 | -0.62 | -0.83 | -0.54 | -0.43 | 0.025 | 0.2175 | 0.245 | 0.89956 | 0.469327 | 0.223849 | 1 | 0.996511 | 0.787447 |
| Sfpq | Q8VIJ6 | -1.71 | -3.89 | -2.83 | -2.25 | -3.35 | -2.81 | -2.25 | -2.39 | -2.61 | -1.48 | -1.9 | -3.23 | -1.39 | -2.73 | -0.7 | -2.01 | -0.03 | 0.365 | 0.9625 | 0.950933 | 0.518461 | 0.126826 | 1 | 0.996511 | 0.722414 |
| Sfr1 | Q8BP27 | -2.36 | -2.21 | -6.3 | -6.82 | -1.98 | -2.81 | -6.46 | -7.37 | -2.23 | -3.3 | -5.52 | -2.11 | -1.89 | -1.97 | -6.87 | -7.31 | -0.2325 | 1.1325 | -0.0875 | 0.885901 | 0.39925 | 0.959385 | 1 | 0.996511 | 0.992636 |
| Sfrp1 | Q8C4U3 | -1.29 | -1.64 | -0.32 | 0.13 | -1.02 | -1.69 | 0.12 | 0.67 | -2.4 | -0.1 | -0.26 | -1.17 | -1.06 | -1.75 | -0.42 | -0.34 | 0.3 | -0.2025 | -0.1125 | 0.628134 | 0.741848 | 0.819287 | 1 | 0.996511 | 0.972768 |
| Sfswap | Q3USH5 | 0.21 | -0.99 | -3 | -3.8 | -0.03 | -1.3 | -3.3 | -3.74 | 0.12 | -1.89 | -3.34 | -1.09 | 0.47 | 0.89 | -2.02 | -1.81 | -0.1975 | 0.345 | 1.2775 | 0.861393 | 0.743044 | 0.2485 | 1 | 0.996511 | 0.803985 |
| Sft2d2 | A0A0A6YX2 | 1.47 | 0.33 | 0.38 | 0.63 | -2.06 | 1.19 | 0.01 | -0.12 | 0.36 | -0.52 | 0.42 | 0.83 | 1.82 | -5.34 | -0.14 | 0.21 | -0.9475 | -0.43 | -1.565 | 0.174096 | 0.269946 | 0.280968 | 1 | 0.996511 | 0.822084 |
| Sfxn1 | Q99JR1 | -0.01 | -0.03 | 0.52 | 0.7 | -0.23 | 0.23 | 0.22 | 0.39 | 0.97 | 0.03 | 0.51 | -0.15 | 0.83 | 0.08 | -0.03 | 0.08 | -0.1425 | 0.045 | -0.055 | 0.561516 | 0.886242 | 0.848969 | 1 | 0.996511 | 0.976225 |
| Sfxn2 | Q925N2 | -0.17 | -1.61 | 0.21 | -0.45 | -1.33 | -0.87 | -3.4 | -0.49 | -0.34 | -2.01 | -0.02 | -0.96 | 1.41 | -3.24 | 0.22 | -0.46 | -1.0175 | -0.3275 | -0.0125 | 0.164537 | 0.550742 | 0.9895 | 1 | 0.996511 | 0.996154 |
| Sfxn3 | Q91V61 | 0.8 | -0.05 | 0.66 | 0.71 | 0.04 | 1.11 | 1.57 | 0.63 | -0.16 | 0.68 | 0.77 | 0.37 | -0.43 | 0.29 | 0.44 | 0.68 | 0.3075 | -0.115 | -0.285 | 0.408997 | 0.697214 | 0.378962 | 1 | 0.996511 | 0.860984 |
| Sfxn4 | Q925N1 | 6.79 | 4.91 | 2.32 | 1.63 | 5.75 | 5.73 | 2.19 | 2.23 | -0.44 | 0.78 | 2.51 | 6.01 | -0.25 | -0.8 | 2.1 | 2.83 | 0.0625 | -1.6975 | -2.9425 | 0.964358 | 0.316473 | 0.049497 | 1 | 0.996511 | 0.658454 |
| Sfxn5 | Q925N0 | 1.02 | 1.36 | 1.51 | 1.53 | 1.13 | 1.09 | 1.76 | 0.97 | 1.17 | 1.04 | 1.57 | 0.91 | 0.48 | 0.79 | 0.98 | 1.79 | -0.1175 | -0.1825 | -0.345 | 0.614729 | 0.419537 | 0.286523 | 1 | 0.996511 | 0.824295 |
| Sgcd | P82347 | 0.67 | 0.24 | 0.34 | 0.87 | -0.42 | 0.86 | -0.19 | 0.63 | 1.66 | 1.16 | 0.73 | 1.01 | 2.04 | 0.98 | 1.13 | 0.72 | -0.31 | 0.61 | 0.6875 | 0.363925 | 0.042367 | 0.060516 | 1 | 0.996511 | 0.661738 |
| Sgce | F6WKE4 | 0.73 | 0.7 | 1.3 | 1.47 | 0.92 | 0.61 | 0.1 | 1.11 | 1.3 | 1.58 | 0.7 | 1.14 | 0.59 | 1.82 | 0.81 | 0.9 | -0.365 | 0.13 | -0.02 | 0.236225 | 0.645645 | 0.952553 | 1 | 0.996511 | 0.992476 |
| Sgcz | Q8BX51 | 0.86 | 0.48 | 0.53 | 0.46 | 1.84 | 0.33 | -0.46 | 0.26 | 1.28 | 1.39 | 0.4 | 0.9 | 0.21 | 2.52 | 0.49 | -0.03 | -0.09 | 0.41 | 0.215 | 0.8441 | 0.14323 | 0.695007 | 1 | 0.996511 | 0.955729 |
| Sgip1 | Q8VD37 | 2.07 | 1.33 | 2.21 | 2.13 | 1.91 | 0.15 | 1.2 | 2.27 | 0.91 | 2.58 | 0.32 | 2.31 | 1.57 | 0.9 | 1.07 | 0.83 | -0.5525 | -0.405 | -0.8425 | 0.261251 | 0.458614 | 0.014694 | 1 | 0.996511 | 0.589387 |
| Sgip1 | Q8VD37-3 | 0.17 | 1.15 | 1.1 | 1.3 | 0.17 | 0.32 | 0.67 | 0.56 | 1.62 | 2.17 | 1.11 | 1.07 | 1.12 | 2.05 | 0.85 | 0.48 | -0.5 | 0.5625 | 0.195 | 0.105077 | 0.140703 | 0.63307 | 1 | 0.996511 | 0.941132 |
| Sgip1 | Q8VD37-8 | 1.14 | 1.37 | 2.18 | 1.43 | 0.9 | 0.9 | 2.23 | 1.89 | 0.89 | 0.61 | 2.02 | 1.05 | 0.55 | 0.82 | 1.88 | 2.06 | -0.05 | -0.3875 | -0.2025 | 0.897361 | 0.309228 | 0.631597 | 1 | 0.996511 | 0.941132 |
| Sgk2 | Q9QZS5-2 | 2.97 | 2.63 | 8.05 | 3.41 | 3.64 | 2.93 | 8.87 | 6.17 | 2.42 | 1.15 | 8.57 | 3.2 | 2.48 | 2.84 | 7.55 | 8.56 | 1.1375 | -0.43 | 1.0925 | 0.497575 | 0.815905 | 0.546535 | 1 | 0.996511 | 0.912406 |
| Sgpl1 | Q8R0X7 | 0.81 | -0.05 | 0.8 | 0.4 | 0.92 | 0.83 | 0.99 | 0.74 | 0 | 0.92 | 0.69 | 1 | 0.29 | 0.74 | 1.31 | 1.6 | 0.38 | 0.1625 | 0.495 | 0.126105 | 0.60133 | 0.185148 | 1 | 0.996511 | 0.776843 |
| Sgpp1 | Q9JI99 | 0.06 | -0.28 | -0.31 | -0.23 | -0.48 | 0.02 | -0.65 | -0.84 | 0.6 | -0.2 | -0.22 | 0.04 | 1.5 | 0.39 | 0.18 | -0.03 | -0.2975 | 0.245 | 0.7 | 0.20815 | 0.317761 | 0.071424 | 1 | 0.996511 | 0.668066 |
| Sgsh | Q9EQ08 | -1.13 | -0.23 | -1.09 | -3.44 | -0.15 | -0.98 | 1.07 | -0.88 | -0.62 | -0.58 | -2.13 | -1.08 | -1.59 | -0.26 | -1.58 | -0.46 | 1.2375 | 0.37 | 0.5 | 0.128211 | 0.602939 | 0.483063 | 1 | 0.996511 | 0.893925 |
| Sgsm1 | B2RQR5 | 0.94 | 1.01 | 0.63 | 0.85 | 1 | 0.74 | 0.38 | 0.61 | 1.12 | 1.28 | 0.61 | 1.19 | 1.14 | 1.42 | 1.01 | 0.75 | -0.175 | 0.1925 | 0.2225 | 0.372307 | 0.377301 | 0.317726 | 1 | 0.996511 | 0.838864 |
| Sgsm2 | Q80U12 | 0.17 | 0.26 | 0.63 | 0.51 | 0.48 | -0.07 | 0.57 | 0.5 | 0.87 | 1.9 | 0.55 | 0.12 | 1.47 | 0.77 | 0.82 | 0.66 | -0.0225 | 0.4675 | 0.5375 | 0.914773 | 0.239197 | 0.0541 | 1 | 0.996511 | 0.661738 |
| Sgsm3 | Q8VCZ6 | -2.25 | -2.53 | -0.47 | -0.78 | -2.53 | -2.24 | -1.04 | -1.25 | -2.09 | -0.32 | -0.59 | -1.91 | -1.1 | -2.21 | -1.17 | -0.61 | -0.2575 | 0.28 | 0.235 | 0.657748 | 0.656852 | 0.680117 | 1 | 0.996511 | 0.955686 |
| Sgta | Q8BJU0 | 0.88 | 0.39 | 0.05 | -0.66 | 0.47 | 0.87 | 0.08 | 0.12 | -0.56 | -0.67 | 0.22 | 1.11 | -0.08 | -1.86 | 0.2 | 0.04 | 0.22 | -0.14 | -0.59 | 0.541819 | 0.774776 | 0.288845 | 1 | 0.996511 | 0.824705 |
| Sgtb | Q8VD33 | -1.35 | -1.14 | -0.58 | -0.64 | -1.13 | -1.23 | -0.61 | -0.79 | -0.84 | -0.68 | -0.72 | -1.06 | -0.6 | -0.62 | -0.83 | -0.61 | -0.0125 | 0.1025 | 0.2625 | 0.960072 | 0.665993 | 0.285289 | 1 | 0.996511 | 0.822669 |
| Sh2b1 | Q91ZM2-2 | 0.5 | -1 | -2.07 | -0.02 | -3.73 | -0.81 | -2 | -0.73 | 2.35 | -0.52 | -0.86 | -0.77 | 1.61 | -5.72 | 0.1 | -1.89 | -1.17 | 0.6975 | -0.8275 | 0.174928 | 0.427693 | 0.583341 | 1 | 0.996511 | 0.927734 |
| Sh2b1 | Q91ZM2-4 | -0.1 | -0.18 | -0.05 | -0.27 | -0.18 | -0.16 | -0.62 | -0.4 | 0.07 | -0.12 | -0.32 | -0.04 | 0.12 | 0.25 | -0.87 | -0.6 | -0.19 | 0.0475 | -0.125 | 0.282957 | 0.782062 | 0.670601 | 1 | 0.996511 | 0.952558 |
| Sh2b2 | A0A0G2JED | 0.23 | 1.58 | 0.11 | 0.4 | 0.83 | -0.4 | 0.3 | 0.29 | 0.13 | -0.09 | 0.28 | -0.56 | 1.41 | 1.56 | -0.86 | 1.01 | -0.325 | -0.64 | 0.2 | 0.423759 | 0.113589 | 0.739017 | 1 | 0.996511 | 0.958796 |
| Sh2d3c | Q9QZS8-2 | -0.15 | -0.55 | -0.19 | -0.26 | 0.01 | 0.01 | 0.07 | -0.43 | 0.07 | -0.04 | -0.17 | 0.18 | 0.21 | 0.43 | -0.15 | -0.17 | 0.2025 | 0.2975 | 0.3675 | 0.296213 | 0.129048 | 0.124099 | 1 | 0.996511 | 0.721448 |
| Sh2d5 | Q8JZW5 | -0.31 | -0.97 | 0.3 | -0.78 | -0.66 | -0.46 | -0.69 | -0.84 | -0.4 | -0.59 | -1.21 | -0.44 | -0.29 | -0.66 | 0.41 | -1.16 | -0.2225 | -0.22 | 0.015 | 0.455456 | 0.517809 | 0.971234 | 1 | 0.996511 | 0.992803 |
| Sh3bgrl | Q9JJU8 | -0.29 | -0.46 | -0.59 | -0.31 | -0.98 | -0.27 | -0.16 | -0.01 | 0.69 | -0.23 | -0.16 | -0.13 | 0.15 | -0.47 | -0.61 | -0.58 | 0.0575 | 0.455 | 0.035 | 0.811845 | 0.097348 | 0.881053 | 1 | 0.996511 | 0.979767 |
| Sh3bgrl2 | Q8BG73 | 0.44 | -0.63 | -0.11 | -0.1 | -1.12 | 0.45 | -0.72 | -0.03 | -0.98 | -0.69 | -0.16 | -0.35 | -1.01 | -1.74 | -1.33 | -0.66 | -0.255 | -0.445 | -1.085 | 0.518942 | 0.154184 | 0.008096 | 1 | 0.996511 | 0.563386 |
| Sh3bgrl3 | Q91VW3 | -0.36 | -0.27 | -0.4 | -0.65 | -0.41 | -0.3 | -1.08 | -0.58 | -0.33 | -2.12 | -0.54 | -0.38 | 0.13 | -1.5 | -0.69 | -0.99 | -0.1725 | -0.4225 | -0.3425 | 0.433081 | 0.322949 | 0.339751 | 1 | 0.996511 | 0.848092 |
| Sh3bp1 | P55194 | 0.27 | 0.71 | 0.4 | 0.19 | 0.4 | -0.07 | 0.12 | 0.03 | 0.65 | -0.77 | 0.39 | 0.22 | 0.95 | 0.51 | 0.05 | 0.61 | -0.2725 | -0.27 | 0.1375 | 0.177946 | 0.420966 | 0.587208 | 1 | 0.996511 | 0.92849 |
| Sh3bp4 | Q921I6 | -0.3 | 0.24 | -0.08 | -0.1 | 0.1 | -0.21 | -0.07 | 0.04 | 0.08 | 0.4 | 0.1 | -0.12 | 0.2 | 0.39 | 0.13 | -0.16 | 0.025 | 0.175 | 0.2 | 0.888034 | 0.398072 | 0.363426 | 1 | 0.996511 | 0.858475 |
| Sh3bp5 | Q9Z131 | -1.2 | -0.48 | 0.27 | -0.78 | -0.14 | -0.98 | 0.18 | -0.19 | -1.43 | -0.93 | 0.16 | -1 | -0.74 | -1.67 | -0.34 | 0.13 | 0.265 | -0.2525 | -0.1075 | 0.487885 | 0.564248 | 0.816587 | 1 | 0.996511 | 0.972768 |
| Sh3bp5l | Q99LH9 | -0.44 | -0.52 | -0.04 | -0.63 | -1.36 | -0.13 | -2.12 | -0.48 | 1.31 | 1.18 | -0.14 | 0.02 | 2.17 | -0.12 | 0.13 | -0.71 | -0.615 | 1 | 0.775 | 0.182426 | 0.02757 | 0.209077 | 1 | 0.996511 | 0.786218 |
| Sh3d19 | A0A571BEI | -0.78 | -0.43 | 0.33 | -0.15 | -0.92 | -0.36 | -0.42 | 0.41 | 1.17 | 1.38 | 0.26 | -0.51 | 0.97 | 0.33 | 0.64 | -0.01 | -0.065 | 0.8325 | 0.74 | 0.851642 | 0.100518 | 0.043673 | 1 | 0.996511 | 0.658454 |
| Sh3gl1 | Q62419 | 1.24 | 1.25 | 0.86 | 0.75 | 0.8 | 1.51 | 0.39 | 0.59 | 1.22 | 0.78 | 0.75 | 1.25 | 0.96 | 0.75 | 0.07 | 0.29 | -0.2025 | -0.025 | -0.5075 | 0.475141 | 0.910712 | 0.084845 | 1 | 0.996744 | 0.691619 |
| Sh3gl2 | A2ALV3 | 1.71 | 1.26 | 1.85 | 1.78 | 0.85 | 1.38 | 0.77 | 1.49 | 1.66 | 1.05 | 1.48 | 1.7 | 1.04 | 1.01 | 1 | 0.87 | -0.5275 | -0.1775 | -0.67 | 0.053282 | 0.450535 | 0.008808 | 1 | 0.996511 | 0.563386 |
| Sh3gl3 | A0A0R4J0E | 0.93 | 1.17 | 0.85 | 1.08 | 0.8 | 0.95 | 1.06 | 0.75 | 0.91 | 1.13 | 0.54 | 0.7 | 0.89 | 1.02 | 0.69 | 0.76 | -0.1175 | -0.1875 | -0.1675 | 0.474439 | 0.356441 | 0.378202 | 1 | 0.996511 | 0.860984 |
| Sh3glb1 | Q9JK48-2 | 0.49 | 0.62 | 1.56 | 1.65 | 0.73 | 0.59 | 1.58 | 1.8 | 0.85 | 1.77 | 2.01 | 0.58 | 0.47 | 1.09 | 1.48 | 1.57 | 0.095 | 0.2225 | 0.0725 | 0.814291 | 0.61257 | 0.850014 | 1 | 0.996511 | 0.976225 |
| Sh3glb2 | A2AWI9 | 1.16 | 1.45 | 1.34 | 2.08 | 1.08 | 0.61 | 1.71 | 1.88 | 0.03 | 1.32 | 1.77 | 0.84 | -0.51 | 1.68 | 0.96 | 0.54 | -0.1875 | -0.5175 | -0.84 | 0.586858 | 0.221419 | 0.101018 | 1 | 0.996511 | 0.698156 |
| Sh3glb2 | A2AWI7 | 0.86 | 0.32 | -0.74 | -0.83 | -0.11 | 0.72 | -1.31 | -0.69 | 0.24 | -0.74 | -0.24 | 0.42 | -0.92 | 0.28 | -0.88 | -0.67 | -0.25 | 0.0175 | -0.45 | 0.649778 | 0.969474 | 0.350439 | 1 | 0.996755 | 0.853299 |
| Sh3glb2 | Q8R3V5-1 | 0.87 | 0.23 | 1.76 | 0.9 | 0.55 | 1.02 | 1.87 | 1.45 | -0.4 | 0.57 | 1.82 | 0.86 | -0.74 | -0.52 | 1.57 | 2.04 | 0.2825 | -0.2275 | -0.3525 | 0.485437 | 0.659905 | 0.619296 | 1 | 0.996511 | 0.93807 |
| Sh3kbp1 | B0R0Y8 | 0.74 | 0.42 | 0.98 | 0.84 | 0.34 | 0.57 | 0.67 | 1.21 | 1.35 | 1.81 | 1.16 | 0.2 | 1.23 | 1.29 | 2.06 | 1.48 | -0.0475 | 0.385 | 0.77 | 0.840746 | 0.290451 | 0.013762 | 1 | 0.996511 | 0.589387 |
| Sh3kbp1 | Q8R550 | 0.98 | 0.77 | 0.51 | 0.73 | 0.94 | 0.68 | 0.65 | 0.64 | 1.01 | -0.24 | 0.42 | 0.82 | 0.88 | 1.39 | 0.48 | 0.45 | -0.02 | -0.245 | 0.0525 | 0.907093 | 0.421666 | 0.843655 | 1 | 0.996511 | 0.976225 |
| Sh3pxd2a | O89032-2 | 0.52 | 0.95 | 0.56 | 0.67 | 0.75 | 0.4 | 0.3 | 0.44 | 0.52 | 1.43 | 0.57 | 0.57 | 0.56 | 0.28 | 0.49 | 0.24 | -0.2025 | 0.0975 | -0.2825 | 0.281892 | 0.708066 | 0.173679 | 1 | 0.996511 | 0.764376 |
| Sh3pxd2b | A0A5F8MP | 0.28 | 0.57 | 0.42 | 0.61 | 0.29 | 0.29 | 0.15 | 0.21 | 0.82 | 0.71 | 0.76 | 0.62 | 0.67 | 1.36 | 0.47 | 0.04 | -0.235 | 0.2575 | 0.165 | 0.151613 | 0.150979 | 0.583954 | 1 | 0.996511 | 0.927734 |
| Sh3rf1 | E9QQ33 | -0.01 | 0.93 | -0.19 | -0.48 | 0.95 | 0.03 | -0.42 | -0.48 | 0.47 | -0.6 | -0.41 | 0.03 | -0.26 | 0.79 | -0.86 | -0.47 | -0.0425 | -0.19 | -0.2625 | 0.919582 | 0.614454 | 0.556421 | 1 | 0.996511 | 0.917092 |
| Sh3rf2 | Q8BZT2 | -1.55 | 0.67 | -0.96 | -0.22 | -0.77 | -1.16 | -0.05 | -0.95 | 0.4 | 0.04 | -0.39 | 0.21 | -0.87 | -0.63 | -0.63 | 0.2 | -0.2175 | 0.58 | 0.0325 | 0.662347 | 0.242748 | 0.947912 | 1 | 0.996511 | 0.992435 |
| Sh3rf3 | Q8C120 | -0.77 | -0.22 | 0.39 | 0.09 | 0.45 | -0.55 | 0.19 | 0.41 | -0.05 | 0.23 | 0.91 | -0.65 | -0.58 | 0.19 | 0 | 0.65 | 0.2525 | 0.2375 | 0.1925 | 0.452018 | 0.547974 | 0.587842 | 1 | 0.996511 | 0.929131 |
| Sh3yl1 | O08641-3 | -0.61 | 0.15 | 0.36 | 0.63 | 1.2 | 0.01 | 0.23 | 0.84 | 0.39 | 0.49 | 0.79 | 0.66 | 0.01 | 0.3 | 0.48 | 0.86 | 0.4375 | 0.45 | 0.28 | 0.250295 | 0.146887 | 0.39933 | 1 | 0.996511 | 0.8682 |
| Shank1 | D3YZU5 | 0.73 | 1.03 | 1.37 | 1.21 | 1.04 | 1.07 | 0.92 | 1.16 | 1.02 | 0.94 | 1.47 | 0.74 | 0.75 | 1.01 | 0.71 | 0.77 | -0.0375 | -0.0425 | -0.275 | 0.840779 | 0.856975 | 0.214063 | 1 | 0.996511 | 0.786931 |
| Shank1 | D3YZU4 | 0.95 | 1.14 | 1.16 | 1.1 | 0.87 | 0.98 | 1.26 | 1.46 | 0.31 | 0.27 | 1.56 | 0.79 | -0.01 | 1.43 | 1.32 | 0.44 | 0.055 | -0.355 | -0.2925 | 0.765819 | 0.267035 | 0.411692 | 1 | 0.996511 | 0.874811 |
| Shank2 | Q80Z38-2 | 1.28 | 1.35 | 1.4 | 1.38 | 1.21 | 1.26 | 0.92 | 1.18 | 1.25 | 1.28 | 1.59 | 1.32 | 0.71 | 1.43 | 0.8 | 0.69 | -0.21 | 0.0075 | -0.445 | 0.191949 | 0.963949 | 0.074811 | 1 | 0.996755 | 0.672844 |
| Shank2 | D3Z5K8 | 8.61 | 8.15 | 5.27 | 3.23 | 8.47 | 9.25 | 5.05 | 4.39 | 3.03 | 2.84 | 4.94 | 8.31 | 3.27 | 2.97 | 4.65 | 4.85 | 0.475 | -1.535 | -2.38 | 0.761938 | 0.350506 | 0.07505 | 1 | 0.996511 | 0.673156 |
| Shank2 | A0A1C7ZM | 1.7 | 2.6 | 2.26 | 2.6 | 2.25 | 2.77 | 2.35 | 1.65 | 1.08 | 1.48 | 2.31 | 1.88 | 0.97 | 2.38 | 2 | 2.34 | -0.035 | -0.6025 | -0.3675 | 0.909804 | 0.100224 | 0.347912 | 1 | 0.996511 | 0.851888 |
| Shank3 | Q4ACU6 | 1.17 | 0.69 | 0.94 | 1.16 | 0.75 | 0.97 | 1.02 | 1.01 | 1.17 | 0.41 | 1.04 | 1 | 1.36 | 0.9 | 0.78 | 0.78 | -0.0525 | -0.085 | -0.035 | 0.767224 | 0.716994 | 0.876729 | 1 | 0.996511 | 0.979626 |
| Shank3 | Q4ACU6-9 | 1.78 | 1.32 | 1.49 | 1.44 | 1.69 | 0.95 | 0.4 | 0.49 | 1.31 | 1.23 | 0.82 | 1.41 | 1.92 | 1.52 | 1.18 | 1.28 | -0.625 | -0.315 | -0.0325 | 0.068083 | 0.15418 | 0.889367 | 1 | 0.996511 | 0.981199 |
| Sharpin | Q91WA6 | -0.24 | 0.33 | 0.46 | 0.04 | -0.1 | -0.09 | 0.15 | -0.02 | -0.04 | 0.15 | 0.11 | 0.51 | 0.25 | -0.42 | 0.31 | 0.28 | -0.1625 | 0.035 | -0.0425 | 0.424914 | 0.877976 | 0.871426 | 1 | 0.996511 | 0.979293 |
| Shb | Q6PD21 | 0.68 | 0.53 | 0.76 | 0.86 | 0.05 | -0.1 | 0.82 | 1.12 | 0.93 | 0.61 | 0.34 | 0.71 | 1.03 | 1.27 | 1.56 | 1.44 | -0.235 | -0.06 | 0.6175 | 0.443045 | 0.758092 | 0.012589 | 1 | 0.996511 | 0.573604 |
| Shc2 | Q8BMC3 | 0.91 | 0.74 | 1.52 | 1.61 | 0.17 | 0.06 | 1.71 | 1.25 | 0.96 | 1.45 | 0.84 | 0.84 | 1.17 | 1.48 | 1.32 | 1.27 | -0.3975 | -0.1725 | 0.115 | 0.367854 | 0.535323 | 0.656211 | 1 | 0.996511 | 0.948332 |
| Shc3 | A0A5F8MP | 0.4 | 0.49 | 0.45 | 0.55 | 0.7 | 0.3 | 0.41 | 0.4 | -0.12 | 0.4 | 0.41 | 0.38 | -0.56 | 0.57 | 0.34 | 0.35 | -0.02 | -0.205 | -0.2975 | 0.898795 | 0.297514 | 0.296013 | 1 | 0.996511 | 0.825612 |
| Shd | O88834-2 | -0.09 | 0.02 | -0.07 | 0.01 | -0.3 | 0.13 | -0.22 | -0.08 | 0.92 | 0.27 | 0.18 | 0.19 | 1.26 | 1.13 | 0.24 | -0.09 | -0.085 | 0.4225 | 0.6675 | 0.598913 | 0.080003 | 0.072466 | 1 | 0.996511 | 0.668066 |
| Shf | A2AQ87 | -0.44 | -0.4 | 0.34 | 0.33 | -0.56 | -0.83 | 0.72 | 0.24 | 0.93 | -0.25 | 0.51 | -0.53 | 1.69 | 0.42 | 0.37 | 0.59 | -0.065 | 0.2075 | 0.81 | 0.869382 | 0.593632 | 0.0539 | 1 | 0.996511 | 0.661738 |
| Shfl | Q8CAK3 | -1.52 | -1.31 | -0.8 | -0.9 | -1.19 | -1.96 | -0.82 | -1.02 | -1.91 | -0.39 | -0.93 | -1.64 | -0.8 | -1.37 | -0.36 | -0.89 | -0.115 | -0.085 | 0.2775 | 0.701993 | 0.819339 | 0.345236 | 1 | 0.996511 | 0.849669 |
| Shisa4 | Q8CA71 | -0.33 | 0.34 | 0.69 | 0.4 | 0.79 | -0.51 | 1.48 | 1.24 | 1.03 | 1.27 | 1.47 | -0.77 | -0.1 | 1.73 | 0.83 | 2.27 | 0.475 | 0.475 | 0.9075 | 0.315596 | 0.370537 | 0.110059 | 1 | 0.996511 | 0.710755 |
| Shisa6 | Q3UH99-2 | 0.83 | 1.56 | 1.31 | 1.06 | 1.86 | 0.9 | 1.31 | 1.38 | 1.02 | 0.5 | 1.24 | 1.06 | 0.97 | 1.14 | 1.29 | 1.46 | 0.1725 | -0.235 | 0.025 | 0.513516 | 0.356074 | 0.914245 | 1 | 0.996511 | 0.986458 |
| Shisa7 | Q8C3Q5-2 | 0.35 | 0.48 | 0.22 | 0.72 | 0.55 | 0.73 | 0.08 | 0.6 | 1.07 | 0.66 | 1.03 | 0.49 | 1.51 | 0.38 | 0.88 | 0.41 | 0.0475 | 0.37 | 0.3525 | 0.81857 | 0.115031 | 0.256966 | 1 | 0.996511 | 0.811777 |
| Shisa9 | Q9CZN4-3 | 1.68 | 1.26 | 1.09 | 1.05 | 1.46 | 1.73 | 1.2 | 1.29 | 0.97 | 1.45 | 1.29 | 1.95 | 0.85 | 0.22 | 0.8 | 1.03 | 0.15 | 0.145 | -0.545 | 0.486175 | 0.589878 | 0.059271 | 1 | 0.996511 | 0.661738 |
| Shmt1 | G3UZ26 | 0.9 | -0.64 | -0.42 | -0.36 | -0.94 | 0.05 | -0.45 | -0.62 | -0.89 | 0.9 | -0.72 | 0.9 | -0.33 | -0.27 | -0.25 | -0.51 | -0.36 | 0.1775 | -0.21 | 0.361315 | 0.750381 | 0.553578 | 1 | 0.996511 | 0.916662 |
| Shmt2 | Q9CZN7-2 | 1.28 | 0.01 | 0.45 | 0.91 | -0.69 | 1.01 | 0.45 | 0.58 | 0.56 | 0.61 | 0.5 | 1.19 | 0.28 | -0.05 | -0.14 | 0.02 | -0.325 | 0.0525 | -0.635 | 0.453972 | 0.869407 | 0.061206 | 1 | 0.996511 | 0.661738 |
| Shoc2 | O88520 | 0.04 | -0.44 | -0.12 | -0.13 | -0.47 | 0.24 | 1.01 | -0.18 | -0.24 | -1.39 | -0.22 | -0.22 | -0.3 | -1.42 | -0.25 | -0.3 | 0.3125 | -0.355 | -0.405 | 0.353542 | 0.271388 | 0.214786 | 1 | 0.996511 | 0.787313 |
| Shpk | Q9D5J6 | -0.48 | -0.56 | -0.46 | -0.21 | 0.15 | -0.44 | -0.52 | -0.27 | -0.51 | 0.42 | -0.5 | -0.44 | -0.21 | 0.22 | -0.29 | -0.68 | 0.1575 | 0.17 | 0.1875 | 0.440076 | 0.516134 | 0.4408 | 1 | 0.996511 | 0.883549 |
| Shprh | Q7TPQ3-2 | -1.94 | -2.26 | -0.42 | -0.78 | -1.08 | -1.35 | -0.76 | -1.59 | -1.23 | 0.18 | -0.64 | -1.28 | -1.73 | -1.52 | -0.16 | 0.53 | 0.155 | 0.6075 | 0.63 | 0.728867 | 0.260497 | 0.338485 | 1 | 0.996511 | 0.847578 |
| Shq1 | Q7TMX5-2 | -0.63 | 0.12 | 0.58 | -0.23 | -0.08 | 0.09 | 2.24 | 0.86 | 0.9 | -1.58 | 1.08 | 0.19 | -0.51 | 0.05 | 1.47 | 1.16 | 0.8175 | 0.1875 | 0.5825 | 0.155908 | 0.756763 | 0.259053 | 1 | 0.996511 | 0.812074 |
| Shroom1 | D3Z3D5 | -0.37 | -2.86 | 1.21 | 0.17 | -0.03 | -0.76 | -0.38 | 0.06 | 1.08 | -1.67 | 0.58 | 0.71 | -0.14 | 0.35 | 0.35 | -1.38 | 0.185 | 0.6375 | 0.2575 | 0.816569 | 0.511416 | 0.764678 | 1 | 0.996511 | 0.964134 |
| Shroom2 | A7TU71 | 0.14 | -0.05 | -0.04 | 0.08 | 0.15 | 0 | 0.09 | 0.21 | -0.74 | -0.06 | 0.09 | 0.11 | -0.46 | -0.92 | -0.15 | 0.13 | 0.08 | -0.1825 | -0.3825 | 0.587734 | 0.446035 | 0.165071 | 1 | 0.996511 | 0.758837 |
| Shroom4 | Q1W617-2 | 0.97 | -3.71 | -0.19 | 1.53 | 0.63 | -0.64 | -0.19 | -0.76 | -1.79 | 0.21 | -2.05 | -0.26 | -0.04 | -0.38 | 1.03 | 0.78 | 0.11 | -0.6225 | 0.6975 | 0.919596 | 0.596355 | 0.527437 | 1 | 0.996511 | 0.908536 |
| Shtn1 | Q8K2Q9-2 | 0.52 | 0.21 | 0.33 | 0.43 | 0.41 | 0.53 | 0.17 | 0.61 | 0.89 | 1.16 | 0.53 | 1.1 | 0.52 | 1.32 | 0.5 | 0.29 | 0.0575 | 0.5475 | 0.285 | 0.735581 | 0.023797 | 0.2963 | 1 | 0.996511 | 0.825612 |
| Siae | A0A1N9M4 | -2.1 | -2.45 | -1.94 | -2.23 | -2.62 | -2.53 | -1.11 | -2.13 | -2.29 | -0.79 | -1.56 | -1.98 | -1.74 | -1.74 | -2.08 | -1.45 | 0.0825 | 0.525 | 0.4275 | 0.813076 | 0.147006 | 0.07811 | 1 | 0.996511 | 0.677244 |
| Sidt1 | E9Q247 | -0.06 | 0.34 | 0.24 | 0.4 | 1.12 | 0.29 | 0.23 | 0.46 | 0.63 | 0.87 | 0.42 | 0.5 | 0.58 | 1.35 | 0.67 | 0.36 | 0.295 | 0.375 | 0.51 | 0.244782 | 0.081668 | 0.080173 | 1 | 0.996511 | 0.683237 |
| Sidt2 | E0CXC7 | -0.33 | -0.79 | 1.73 | 1.7 | -0.4 | -1.44 | 1.9 | 1.77 | -0.47 | 0.76 | 1.83 | -0.18 | -1.28 | -2.87 | 2.13 | 2.11 | -0.12 | -0.0925 | -0.555 | 0.899398 | 0.90347 | 0.661835 | 1 | 0.996609 | 0.949645 |
| Sigmar1 | O55242 | -0.17 | 0.19 | 0.33 | 0.25 | 1.22 | -0.05 | 0.64 | 0.39 | -0.33 | -2.08 | 0.3 | -0.65 | 0.44 | 0.86 | 0.44 | 0.89 | 0.4 | -0.84 | 0.5075 | 0.188916 | 0.10879 | 0.042888 | 1 | 0.996511 | 0.658454 |
| Sik2 | H3BJC9 | -0.44 | -0.53 | 0.11 | -0.35 | -0.84 | -0.04 | 0.1 | -0.05 | -0.77 | -1.68 | -0.08 | -0.56 | -0.29 | -2.08 | -0.25 | 0 | 0.095 | -0.47 | -0.3525 | 0.719995 | 0.209133 | 0.460445 | 1 | 0.996511 | 0.887902 |
| Sik3 | Q6P4S6 | 1.82 | 1.09 | 0.77 | 0.3 | 1 | 1.88 | 0.44 | 0.45 | -0.96 | 0.61 | 0.71 | 1.88 | -0.83 | -1 | 0.58 | 0.56 | -0.0525 | -0.435 | -1.1675 | 0.903841 | 0.481501 | 0.041638 | 1 | 0.996511 | 0.658454 |
| Sike1 | Q9CPR7 | 0.35 | 0.01 | -0.12 | -0.42 | -0.05 | 0.49 | -0.06 | -0.1 | -0.27 | -0.05 | -0.41 | 0.2 | -0.43 | -0.24 | 0.21 | -0.41 | 0.115 | -0.0875 | -0.1725 | 0.621559 | 0.712702 | 0.498155 | 1 | 0.996511 | 0.898691 |
| Sil1 | A0A494B9I | -0.99 | -1.68 | 1.56 | 1.64 | -0.47 | -1.11 | 1.4 | 1.79 | -2.17 | 2.11 | 1.54 | -0.85 | -1.43 | -1.11 | 0.94 | 1.07 | 0.27 | 0.025 | -0.265 | 0.786852 | 0.983151 | 0.784987 | 1 | 0.998773 | 0.967642 |
| Sin3a | Q60520-1 | 1.18 | -0.39 | -0.46 | -0.56 | 0.28 | 0.44 | -0.08 | -0.11 | 0.27 | -0.82 | -0.3 | 0.02 | 0.24 | -0.01 | 0.88 | 0.54 | 0.19 | -0.15 | 0.47 | 0.644651 | 0.738415 | 0.295196 | 1 | 0.996511 | 0.825612 |
| Sin3b | Q62141-1 | -1.75 | -1.8 | -2.5 | -2.54 | -1.78 | -2.07 | -1.56 | -2.29 | -1.77 | -2.8 | -2.39 | -2.07 | -0.8 | -1.06 | -2.22 | -1.96 | 0.2225 | -0.11 | 0.6375 | 0.42489 | 0.724476 | 0.131033 | 1 | 0.996511 | 0.725981 |
| Sipa1 | P46062 | 0.8 | -0.2 | 0.66 | 0.54 | -0.63 | 1.25 | -0.14 | 0.28 | 6.47 | 1.29 | 0.82 | 1.25 | 2.86 | -2.64 | 0.19 | 0.97 | -0.26 | 2.0075 | -0.105 | 0.547768 | 0.126673 | 0.9195 | 1 | 0.996511 | 0.986868 |
| Sipa1l1 | Q8C0T5 | 1.17 | 0.75 | 0.87 | 1.02 | 0.9 | 0.94 | 0.76 | 0.99 | 0.73 | 0.86 | 0.93 | 1.1 | 0.89 | 0.54 | 0.76 | 0.63 | -0.055 | -0.0475 | -0.2475 | 0.736465 | 0.795263 | 0.221283 | 1 | 0.996511 | 0.787447 |
| Sipa1l2 | A0A1D5RL' | -0.37 | 0.21 | 1.09 | 0.77 | 0.07 | -0.2 | 1.29 | 1.33 | 0.17 | 0.34 | 1.29 | -0.24 | 0.62 | 0.26 | 1.1 | 1.37 | 0.1975 | -0.035 | 0.4125 | 0.679457 | 0.935276 | 0.308826 | 1 | 0.996755 | 0.834463 |
| Sipa1l3 | G3X9J0 | 0.54 | 0.77 | 0.95 | 0.56 | 0.87 | 0.44 | 0.77 | 0.62 | 0.4 | 0.37 | 0.84 | 0.46 | 0.3 | 0.72 | 0.61 | 0.7 | -0.03 | -0.1875 | -0.1225 | 0.867555 | 0.354883 | 0.548695 | 1 | 0.996511 | 0.912757 |
| Sirpa | P97797-2 | 1.24 | 1.36 | 1.01 | 0.57 | 1.4 | 1.1 | 1.23 | 0.99 | 0.55 | 0.01 | 1.07 | 1.39 | 0.21 | 1.11 | 1.23 | 0.94 | 0.135 | -0.29 | -0.1725 | 0.542315 | 0.408652 | 0.569423 | 1 | 0.996511 | 0.922945 |
| Sirpa | P97797 | 0.59 | 1.24 | 0.23 | 1.21 | 0.98 | -0.11 | 0.76 | 0.7 | 0.7 | 0.38 | 0.13 | 1.17 | 0.14 | 2.12 | 0.08 | 0.13 | -0.235 | -0.2225 | -0.2 | 0.486619 | 0.507003 | 0.700741 | 1 | 0.996511 | 0.956416 |
| Sirt2 | Q8VDQ8 | 1.18 | 1.1 | 1.39 | 1.23 | 1.15 | 0.66 | 1.65 | 0.95 | 0.56 | 0.47 | 1.07 | 0.78 | 0.18 | 0.4 | 1.13 | 1.52 | -0.1225 | -0.505 | -0.4175 | 0.604549 | 0.029495 | 0.21887 | 1 | 0.996511 | 0.787447 |
| Sirt3 | Q8R104 | -0.51 | -0.48 | -1.65 | -1.64 | -0.4 | -0.48 | -1.06 | -1.88 | -0.6 | -1.14 | -1.79 | -0.66 | -1.67 | -0.82 | -2.06 | -1.74 | 0.115 | 0.0225 | -0.5025 | 0.79586 | 0.956271 | 0.238525 | 1 | 0.996755 | 0.793481 |
| Sirt4 | Q8R216 | -0.81 | -1.26 | -0.31 | -0.24 | -0.54 | -0.95 | -0.06 | -0.6 | -0.53 | 0.43 | -0.35 | -0.73 | -0.7 | -0.95 | -0.08 | 0 | 0.1175 | 0.36 | 0.2225 | 0.69501 | 0.309388 | 0.511988 | 1 | 0.996511 | 0.903993 |
| Sirt5 | A0A1Y7VJY | 0.95 | 0.36 | 0.18 | 0.27 | -0.34 | 0.67 | 0.09 | 0.24 | 1.15 | 0.53 | -0.05 | 0.4 | 0.82 | 0 | -0.04 | 0.02 | -0.275 | 0.0675 | -0.24 | 0.332804 | 0.82582 | 0.413982 | 1 | 0.996511 | 0.874945 |
| Skiv2l | Q6NZR5 | -0.47 | -0.3 | -0.22 | -0.1 | -0.36 | -0.54 | -0.32 | -0.23 | -0.26 | -0.09 | -0.16 | -0.37 | -0.58 | -0.25 | -0.46 | -0.5 | -0.09 | 0.0525 | -0.175 | 0.581459 | 0.761896 | 0.360594 | 1 | 0.996511 | 0.85741 |
| Skp1 | Q9WTX5 | 1.41 | 1.23 | 1.01 | 1.38 | 0.57 | 1.12 | 0.71 | 1.02 | -0.11 | -1.04 | 1.03 | 1.35 | -0.03 | -0.71 | 0.97 | 0.7 | -0.4025 | -0.95 | -1.025 | 0.064275 | 0.092849 | 0.023042 | 1 | 0.996511 | 0.623147 |
| Skp2 | A0A087WC | 0.32 | 0.07 | -0.6 | 0.15 | -0.09 | 0.37 | -1.39 | 0.03 | 1.33 | 0.89 | -0.92 | 0.4 | 2.06 | 0.5 | 0.2 | -0.89 | -0.255 | 0.44 | 0.4825 | 0.538027 | 0.381659 | 0.42173 | 1 | 0.996511 | 0.875736 |
| Slain1 | Q68FF7 | -3 | -3.16 | 0.04 | -0.4 | -1.89 | -4.92 | -1.27 | -1.29 | -3.53 | 0.36 | -0.42 | -3.52 | -1.94 | -1.87 | 0.28 | 0 | -0.7125 | -0.1475 | 0.7475 | 0.517019 | 0.90095 | 0.427368 | 1 | 0.996511 | 0.879165 |
| Slain2 | Q8CI08-2 | 0.34 | -0.14 | 0.49 | 0.04 | -0.62 | 0.46 | -0.43 | -0.04 | -0.16 | -0.24 | 0.29 | 0 | 0.17 | -0.87 | 0.73 | -0.03 | -0.34 | -0.21 | -0.1825 | 0.246049 | 0.355066 | 0.61151 | 1 | 0.996511 | 0.935047 |
| Slc10a4 | Q3UEZ8 | 1.1 | 0.14 | -0.78 | -0.42 | -0.27 | -0.15 | -0.73 | -0.79 | 1.44 | 0.41 | -1.06 | 1.04 | 0.37 | 0.96 | -0.44 | -1.15 | -0.495 | 0.4475 | -0.075 | 0.251137 | 0.481297 | 0.894918 | 1 | 0.996511 | 0.983509 |
| Slc11a2 | P49282-2 | 0.6 | 0.01 | 1.92 | 1.69 | 0.38 | 0.52 | 2.27 | 1.48 | -0.04 | 1.22 | 2.61 | 0.45 | -1.02 | 0.65 | 0.17 | 0.82 | 0.1075 | 0.005 | -0.9 | 0.852483 | 0.994026 | 0.137946 | 1 | 0.999285 | 0.733943 |
| Slc11a2 | P49282 | -0.8 | -1.3 | 0.08 | -0.08 | -1.04 | -0.84 | -1.22 | -0.3 | 1.22 | 3.42 | -0.79 | -0.66 | 0.1 | 0.03 | -0.6 | -0.11 | -0.325 | 1.3225 | 0.38 | 0.38036 | 0.181679 | 0.299987 | 1 | 0.996511 | 0.829516 |
| Slc12a2 | E9QM38 | 2.21 | 1.83 | 1.23 | 1.09 | 1.72 | 1.76 | 1.03 | 0.84 | 0.75 | 0.63 | 0.88 | 2.06 | 0.42 | 0.96 | 1.01 | 0.92 | -0.2525 | -0.51 | -0.7625 | 0.465412 | 0.226902 | 0.032402 | 1 | 0.996511 | 0.641974 |
| Slc12a4 | Q9JIS8 | -0.46 | 0.23 | 0.05 | 0.43 | -0.87 | -0.06 | 0.3 | 0.24 | 0.28 | 0.19 | 0.38 | -0.23 | 0.92 | -0.29 | 0.35 | 0.2 | -0.16 | 0.0925 | 0.2325 | 0.621861 | 0.717158 | 0.474493 | 1 | 0.996511 | 0.891711 |
| Slc12a5 | Q91V14 | 0.68 | 0.74 | 1.02 | 1.22 | 0.9 | 0.73 | 0.66 | 0.85 | 0.9 | 0.86 | 0.98 | 0.58 | 0.74 | 1.02 | 0.55 | 0.57 | -0.13 | -0.085 | -0.195 | 0.480252 | 0.674511 | 0.383035 | 1 | 0.996511 | 0.860984 |
| Slc12a5 | Q91V14-2 | 0.45 | 0.6 | 0.07 | 0.5 | 0.36 | 0.72 | 0.04 | 0.05 | 0.58 | -0.19 | 0.16 | 0.17 | 0.28 | 1.16 | -0.26 | -0.12 | -0.1125 | -0.225 | -0.14 | 0.613554 | 0.338545 | 0.682028 | 1 | 0.996511 | 0.955729 |
| Slc12a6 | Q924N4 | 2.03 | 1.62 | 1.8 | 1.92 | 1.74 | 1.63 | 1.22 | 1.71 | 1.67 | 1.99 | 1.62 | 1.97 | 1.49 | 1.41 | 1.65 | 1.32 | -0.2675 | -0.03 | -0.375 | 0.180733 | 0.873863 | 0.074889 | 1 | 0.996511 | 0.672844 |
| Slc12a9 | Q99MR3 | 1.49 | 1.4 | 0.52 | 0.34 | 1.44 | 1.59 | 1.05 | 0.36 | 0.91 | 0.19 | 0.39 | 1.44 | 0.77 | 0.59 | 0.31 | 0.04 | 0.1725 | -0.205 | -0.51 | 0.654099 | 0.602986 | 0.156541 | 1 | 0.996511 | 0.751493 |
| Slc13a3 | A2A4U9 | -0.14 | 0.45 | 0.36 | 0.66 | 0.19 | 0.3 | 0.32 | -0.1 | 1.22 | 1.68 | 0.53 | 0.43 | 0.88 | 0.14 | 0.32 | 0.35 | -0.155 | 0.6325 | 0.09 | 0.485827 | 0.088333 | 0.731094 | 1 | 0.996511 | 0.958635 |
| Slc13a4 | Q8BZ82 | -0.36 | -0.68 | -1.05 | 1.05 | -2.9 | -0.34 | -1.99 | -1.63 | 1.25 | -2.52 | -1.83 | -0.4 | 1.8 | -2.67 | -0.66 | -1.33 | -1.455 | -0.615 | -0.455 | 0.045955 | 0.48074 | 0.628636 | 1 | 0.996511 | 0.940342 |
| Slc13a5 | Q5NBV0 | 0.26 | 1.06 | 0.42 | 0.51 | 0.9 | 0.67 | 0.52 | 0.32 | 0.06 | -0.43 | 0.4 | 0.09 | -0.01 | 0.11 | 0.1 | 0.37 | 0.04 | -0.5325 | -0.42 | 0.862724 | 0.068893 | 0.099813 | 1 | 0.996511 | 0.698156 |
| Slc14a1 | Q8VHL0 | -0.2 | 0.43 | 0.39 | 0.41 | -0.1 | 0.56 | 0.24 | 0.18 | 0.59 | 1.4 | 0.51 | 0.01 | 1.1 | -0.11 | 0.37 | 0.12 | -0.0375 | 0.37 | 0.1125 | 0.867827 | 0.273007 | 0.719595 | 1 | 0.996511 | 0.958398 |
| Slc15a2 | Q9JM03 | 1.68 | 0.58 | 0.66 | 0.42 | 1.9 | 2.13 | 0.88 | 0.86 | 0.59 | -0.44 | 1.08 | 2.14 | 0.41 | 0.59 | 1.48 | 1.6 | 0.6075 | 0.0075 | 0.185 | 0.166916 | 0.989297 | 0.647004 | 1 | 0.999285 | 0.945362 |
| Slc15a4 | Q91W98 | -0.32 | -0.34 | -0.57 | -0.92 | -0.17 | 0.17 | -0.37 | -0.64 | 1.07 | -0.52 | -0.39 | -0.15 | 1.26 | -0.24 | -0.99 | -0.25 | 0.285 | 0.54 | 0.4825 | 0.249233 | 0.176491 | 0.31324 | 1 | 0.996511 | 0.835247 |
| Slc16a1 | P53986 | -0.23 | 0.23 | 0.43 | 0.37 | 0.17 | 0.61 | 0.09 | -0.03 | 0.33 | 0.66 | 0.45 | 0.11 | 0.78 | 0.43 | 0.03 | -0.65 | 0.01 | 0.1875 | -0.0525 | 0.964603 | 0.411774 | 0.877704 | 1 | 0.996511 | 0.979626 |
| Slc16a10 | Q3U9N9 | -0.38 | -0.57 | -0.29 | 1.17 | -0.02 | 0.39 | 0.71 | 1.58 | 0.25 | -0.37 | 1.4 | -0.34 | -0.14 | 0.63 | 0.21 | 1.12 | 0.6825 | 0.2525 | 0.4725 | 0.184157 | 0.637307 | 0.315679 | 1 | 0.996511 | 0.838071 |
| Slc16a2 | O70324 | 0.65 | 0.9 | 0.59 | 0.97 | 1.51 | 0.33 | 1.01 | 0.38 | 0.73 | 0.64 | 0.98 | 1.18 | 0.42 | 2 | 0.52 | 1.56 | 0.03 | 0.105 | 0.3475 | 0.91885 | 0.605788 | 0.381824 | 1 | 0.996511 | 0.860984 |
| Slc16a3 | P57787 | 2.21 | 1.96 | 4.39 | 3.01 | 1.47 | 1.46 | 4.76 | 4.51 | 1.37 | 2.37 | 4.84 | 2.16 | 1.48 | 1.76 | 5.16 | 5.83 | 0.1575 | -0.2075 | 0.665 | 0.869439 | 0.804588 | 0.556198 | 1 | 0.996511 | 0.917092 |
| Slc16a7 | O70451 | -0.64 | -0.49 | 0.47 | 0.91 | 0.22 | 0.21 | -0.1 | 0.34 | 0.52 | 0.6 | 0.96 | 0.58 | 0.89 | -0.46 | 0.6 | 0.61 | 0.105 | 0.6025 | 0.3475 | 0.775877 | 0.134983 | 0.449792 | 1 | 0.996511 | 0.884817 |
| Slc17a5 | Q8BN82 | -1.44 | -1.44 | -1.11 | -1.32 | -0.88 | -0.82 | 0.01 | -0.89 | -1.17 | -1.21 | -0.69 | -1.13 | -0.47 | -1.05 | -1.38 | -0.34 | 0.6825 | 0.2775 | 0.5175 | 0.020574 | 0.181737 | 0.090008 | 1 | 0.996511 | 0.692261 |
| Slc17a6 | A0A0R4J0A | 0.77 | 0.9 | 1.48 | 1.48 | 0.64 | 0.78 | 1.22 | 1.4 | 0.7 | 1.37 | 1.55 | 0.95 | 0.1 | 1.3 | 0.88 | 1.24 | -0.1475 | -0.015 | -0.2775 | 0.583468 | 0.957488 | 0.417782 | 1 | 0.996755 | 0.87558 |
| Slc17a7 | Q3TXX4 | 2.23 | 1.5 | 1.26 | 1.15 | 1.86 | 2.16 | 1.93 | 1.15 | 1.14 | 0.34 | 1.36 | 1.87 | 0.98 | 1.66 | 1 | 1.45 | 0.24 | -0.3575 | -0.2625 | 0.459822 | 0.365652 | 0.401987 | 1 | 0.996511 | 0.870698 |
| Slc17a8 | Q8BFU8 | 2.26 | 2.07 | 1.54 | 1.68 | 1.64 | 2.36 | 1.65 | 1.51 | -1.01 | -0.21 | -0.67 | 1.12 | 0.81 | -0.3 | 1.27 | 1.08 | -0.0975 | -2.08 | -1.1725 | 0.711813 | 0.00166 | 0.012343 | 1 | 0.996511 | 0.573077 |
| Slc18a2 | Q8BRU6 | 0.22 | -0.99 | 0.09 | 0.59 | -1.51 | -0.47 | 0.57 | 0.68 | 2.54 | -0.33 | -0.06 | -0.32 | 2.58 | -1.42 | 0.31 | -0.45 | -0.16 | 0.48 | 0.2775 | 0.776509 | 0.501513 | 0.737383 | 1 | 0.996511 | 0.958703 |
| Slc18a3 | Q3TYJ1 | 0.58 | 0.48 | 0.38 | 0.6 | 0.47 | 0.15 | 0.71 | 0.44 | 0.58 | 0 | 0.03 | 0.39 | 0.1 | -0.11 | 0.2 | 0.38 | -0.0675 | -0.26 | -0.3675 | 0.700053 | 0.215276 | 0.080883 | 1 | 0.996511 | 0.685104 |
| Slc19a1 | P41438-2 | 1.38 | 0.02 | -0.2 | -0.08 | -0.07 | -0.31 | -0.64 | -0.58 | 0.22 | 0.08 | -0.61 | 1.39 | -0.27 | -1.65 | -1.05 | -0.07 | -0.68 | -0.01 | -1.04 | 0.096317 | 0.984501 | 0.057184 | 1 | 0.998892 | 0.661738 |
| Slc19a2 | Q9EQN9 | -0.62 | -0.49 | 1.24 | 0.98 | 0.03 | -0.35 | 1.8 | 2.11 | 1.2 | 0.33 | 2.26 | -0.25 | 0.42 | 0.92 | 1.71 | 1.92 | 0.62 | 0.6075 | 0.965 | 0.394109 | 0.373249 | 0.107277 | 1 | 0.996511 | 0.704919 |
| Slc1a1 | P51906 | 0.13 | 0.61 | -0.47 | -0.23 | 0.77 | 0.41 | -0.15 | -0.25 | 0.52 | -0.3 | -0.29 | 0.34 | 0.25 | 0.95 | -0.62 | -0.23 | 0.185 | 0.0575 | 0.0775 | 0.575373 | 0.856379 | 0.845408 | 1 | 0.996511 | 0.976225 |
| Slc1a2 | P43006 | 1.1 | 1.81 | 1.03 | 1.24 | 1.67 | 1.11 | 0.2 | 0.85 | 0.75 | 1.03 | 0.91 | 1.21 | 0.48 | 0.42 | 0.32 | 0.65 | -0.3375 | -0.32 | -0.8275 | 0.336069 | 0.19302 | 0.006025 | 1 | 0.996511 | 0.552614 |
| Slc1a2 | A2APL5 | -0.94 | -0.43 | -1.14 | 0.11 | -0.11 | 1.46 | -1.1 | -0.74 | 0.29 | -0.53 | -2.13 | 0.5 | -2.13 | 1.41 | -0.52 | 0.26 | 0.4775 | 0.1325 | 0.355 | 0.417363 | 0.826506 | 0.623827 | 1 | 0.996511 | 0.938628 |
| Slc1a3 | P56564 | -0.5 | -0.29 | -0.01 | 0.07 | -0.41 | 0.19 | 0.71 | -0.22 | -0.63 | -0.81 | -0.2 | -0.45 | -0.99 | -0.58 | -0.9 | -0.34 | 0.25 | -0.34 | -0.52 | 0.387686 | 0.150683 | 0.053577 | 1 | 0.996511 | 0.661738 |
| Slc1a4 | O35874 | 0.33 | 0.89 | 0.97 | 1.15 | 0.47 | 0.21 | 0.45 | 0.48 | 0.7 | 2.18 | 0.88 | 0.51 | 0.51 | 1.39 | 0.55 | 0.77 | -0.4325 | 0.2325 | -0.03 | 0.070118 | 0.564329 | 0.916737 | 1 | 0.996511 | 0.986556 |
| Slc1a5 | Q9ESU7 | 0.39 | 1.38 | 1.65 | 0.89 | 2.26 | -1.26 | 2.34 | 0.56 | 0.71 | 2.22 | -0.07 | 0.78 | -0.34 | 0.45 | 0.52 | 2.29 | -0.1025 | -0.1675 | -0.3475 | 0.898841 | 0.744363 | 0.547284 | 1 | 0.996511 | 0.912571 |
| Slc1a6 | O35544 | 0.41 | 0.48 | 0.67 | 0.84 | 0.32 | 0.33 | 0.47 | 0.84 | 1.02 | 0.37 | 0.61 | 0.61 | 1.16 | 1.25 | 0.57 | 1.01 | -0.11 | 0.0525 | 0.3975 | 0.571592 | 0.802327 | 0.106313 | 1 | 0.996511 | 0.703417 |
| Slc20a1 | Q61609 | 0.61 | 1.17 | 1.25 | 1.06 | 1.38 | 0.62 | -0.11 | 0.88 | 0.8 | 1.1 | 1.07 | 0.78 | 1.33 | 1.24 | 1.1 | 1.32 | -0.33 | -0.085 | 0.225 | 0.334281 | 0.686494 | 0.301107 | 1 | 0.996511 | 0.830276 |
| Slc20a2 | Q80UP8 | 1.24 | 1.56 | 1.1 | 1.29 | 1.21 | 1.38 | -0.63 | 0.69 | 1.23 | 2.33 | 0.87 | 1.04 | 1.93 | 0.79 | 1.07 | 1.02 | -0.635 | 0.07 | -0.095 | 0.169425 | 0.836247 | 0.740461 | 1 | 0.996511 | 0.958796 |
| Slc22a17 | A0A2I3BQ( | -0.5 | -0.16 | 0 | -0.32 | 0.15 | -0.38 | -0.07 | -0.05 | 0.19 | 0.59 | -0.41 | -0.25 | 0.01 | 0.85 | 0.15 | -0.15 | 0.1575 | 0.275 | 0.46 | 0.419335 | 0.317621 | 0.115382 | 1 | 0.996511 | 0.717287 |
| Slc22a23 | Q3UHH2 | 0.27 | 0.62 | 0.4 | 0.68 | 0.63 | 0.34 | 0.11 | 0.34 | 0.65 | 0.92 | 0.46 | 0.27 | 0.82 | 1.05 | 0.13 | 0.21 | -0.1375 | 0.0825 | 0.06 | 0.464383 | 0.696765 | 0.824356 | 1 | 0.996511 | 0.972852 |
| Slc22a4 | Q9Z306 | -0.11 | 0.39 | 0.23 | -0.29 | 0.89 | -0.2 | 0.43 | -0.11 | -0.36 | -2.5 | -1.03 | -0.71 | -0.87 | -0.33 | -0.89 | 0.12 | 0.1975 | -1.205 | -0.5475 | 0.510574 | 0.027617 | 0.095661 | 1 | 0.996511 | 0.698156 |
| Slc22a5 | Q9Z0E8 | 0.57 | 0.34 | -1.4 | 1.1 | -0.67 | -1.1 | -0.15 | 0.37 | -1.12 | -1.06 | 0.46 | -0.82 | -1.74 | -0.05 | 0.47 | 0.25 | -0.54 | -0.7875 | -0.42 | 0.35972 | 0.212978 | 0.536252 | 1 | 0.996511 | 0.912212 |
| Slc23a2 | Q9EPR4 | 0.22 | 0.81 | 0.82 | 0.3 | 1.04 | 0.72 | 0.72 | 0.54 | 0.35 | -0.11 | 0.5 | -0.03 | -0.18 | -0.06 | 0.22 | 1.06 | 0.2175 | -0.36 | -0.2775 | 0.329961 | 0.165404 | 0.405902 | 1 | 0.996511 | 0.873384 |
| Slc24a2 | Q8BUN9-2 | -0.69 | -0.26 | 0.46 | 0.21 | -0.27 | -1.08 | -0.15 | -0.1 | -0.74 | 0.51 | 0.18 | -0.53 | -0.4 | -0.34 | -0.22 | -0.11 | -0.33 | -0.075 | -0.1975 | 0.335564 | 0.841884 | 0.490612 | 1 | 0.996511 | 0.897531 |
| Slc24a2 | Q8BUN9 | 1.51 | 2.31 | 4.66 | 2.33 | 1.46 | 1.72 | 3.54 | 3.55 | -0.32 | -10.97 | 4.35 | 2.89 | 1.56 | -5.92 | 4.39 | 4.96 | -0.135 | -3.715 | -1.455 | 0.865684 | 0.255769 | 0.530797 | 1 | 0.996511 | 0.908996 |
| Slc24a3 | Q8VD29 | 1.11 | 1.02 | 1.33 | 1.25 | -0.1 | 1.35 | 0.67 | 1.29 | -0.47 | 1.37 | 1.46 | 1.63 | -0.71 | -2.07 | 1.44 | 1.38 | -0.375 | -0.18 | -1.1675 | 0.279906 | 0.700131 | 0.157215 | 1 | 0.996511 | 0.751678 |
| Slc24a4 | E9PUL6 | 0.24 | 0.7 | 1.07 | 0.87 | 0.83 | 0.49 | 1.2 | 0.86 | -0.03 | 0.2 | 0.78 | 0.3 | -0.36 | 0.18 | 0.68 | 0.69 | 0.125 | -0.4075 | -0.4225 | 0.609998 | 0.148549 | 0.201578 | 1 | 0.996511 | 0.781447 |
| Slc25a1 | Q8JZU2 | 0.12 | -0.6 | 0.07 | -0.43 | -0.09 | 0.49 | 0.11 | 0.08 | -0.93 | 0.65 | 0.42 | 0.15 | -0.41 | -0.79 | 0.23 | 0.47 | 0.3575 | 0.2825 | 0.085 | 0.154233 | 0.463502 | 0.80317 | 1 | 0.996511 | 0.971091 |
| Slc25a10 | Q9QZD8 | 1.32 | 1.26 | 2.1 | 1.27 | 1.14 | 1.94 | 1.83 | 1.94 | 0.71 | 0.85 | 2.03 | 1.69 | 0.89 | -0.15 | 1.98 | 2.79 | 0.225 | -0.1675 | -0.11 | 0.435384 | 0.6515 | 0.858106 | 1 | 0.996511 | 0.976225 |

| | | | | | | | | | | | | | | | | | | | | | | | | | | | |
|---|---|---|---|---|---|---|---|---|---|---|---|---|---|---|---|---|---|---|---|---|---|---|---|---|---|---|---|
| Slc25a11 | Q9CR62 | -0.34 | 0.16 | 0.24 | 0.15 | -0.07 | 0.03 | -0.28 | 0.12 | -0.22 | -0.23 | 0.21 | -0.69 | -0.18 | -0.07 | -0.43 | -0.13 | -0.1025 | -0.285 | -0.255 | 0.600263 | 0.272193 | 0.24753 | | 1 | 0.996511 | 0.803677 |
| Slc25a12 | Q8BH59 | 0.29 | 0.12 | 0.46 | 0.41 | -0.46 | 0.07 | 0.24 | 0.28 | -0.04 | -0.04 | 0.4 | 0.12 | -0.41 | -0.12 | -0.34 | 0.01 | -0.2875 | -0.21 | -0.535 | 0.200384 | 0.280563 | 0.02145 | | 1 | 0.996511 | 0.621926 |
| Slc25a13 | Q9QXX4 | -0.28 | 0.2 | 0.06 | 0.32 | -0.1 | -0.31 | 0.21 | -0.1 | 0.06 | 0.01 | -0.22 | -0.44 | -0.49 | -0.32 | -0.12 | 0.22 | -0.15 | -0.2225 | -0.2525 | 0.462473 | 0.313892 | 0.307309 | | 1 | 0.996511 | 0.832533 |
| Slc25a14 | A0A0A0M( | -0.92 | -0.48 | -0.27 | -0.59 | -0.3 | -0.57 | -0.17 | -0.46 | -1.12 | -0.43 | -0.36 | -0.78 | -0.74 | -0.68 | -0.33 | -0.19 | 0.19 | -0.1075 | 0.08 | 0.347135 | 0.664168 | 0.732175 | | 1 | 0.996511 | 0.958635 |
| Slc25a15 | Q9WVD5 | 0.21 | 0.48 | 0.61 | 1.15 | 0.69 | 0.08 | 1 | 1.23 | -0.23 | 0.39 | 0.99 | -0.04 | -0.16 | 0.43 | 0.34 | 0.88 | 0.1375 | -0.335 | -0.24 | 0.661669 | 0.327996 | 0.436435 | | 1 | 0.996511 | 0.882882 |
| Slc25a16 | D3YVC9 | -0.67 | -0.15 | -0.46 | -0.47 | -0.2 | -0.15 | -0.3 | -0.59 | -0.42 | -1.03 | -0.25 | -0.12 | 0.21 | -0.61 | -0.57 | -0.4 | 0.1275 | -0.0175 | 0.095 | 0.49999 | 0.94436 | 0.706187 | | 1 | 0.996755 | 0.957084 |
| Slc25a17 | O70579 | 0.52 | 0.3 | 0.22 | 0.6 | 1.08 | 0.88 | 1.04 | 0.46 | 0.42 | -0.39 | 0.49 | 0.44 | -0.37 | 0.79 | 0.04 | 0.68 | 0.455 | -0.17 | -0.125 | 0.046635 | 0.504724 | 0.679191 | | 1 | 0.996511 | 0.955269 |
| Slc25a18 | Q9DB41 | 1.22 | 0.85 | 0.55 | 0.9 | 0.57 | 0.8 | -0.17 | -0.04 | 0.68 | 0.27 | 0.88 | 0.62 | 0.99 | 0.32 | 0.76 | 0.74 | -0.59 | -0.2675 | -0.1775 | 0.057745 | 0.249613 | 0.460305 | | 1 | 0.996511 | 0.887902 |
| Slc25a19 | Q9DAM5 | 1.05 | 0.63 | 0.76 | 0.98 | 0.09 | 0.85 | -0.42 | 0.51 | 0.14 | 0.95 | 0.58 | 0.65 | 0 | -0.27 | 0.02 | 0.2 | -0.5975 | -0.275 | -0.8675 | 0.066388 | 0.245838 | 0.002042 | | 1 | 0.996511 | 0.419079 |
| Slc25a20 | Q9Z2Z6 | 0.87 | 0.28 | -0.12 | 0.02 | 0.4 | 0.57 | 0.04 | -0.38 | 1.4 | -0.5 | -0.15 | 0.79 | 0.63 | 1.53 | -0.64 | -0.26 | -0.105 | 0.1225 | 0.0525 | 0.72798 | 0.788747 | 0.91567 | | 1 | 0.996511 | 0.986556 |
| Slc25a21 | B6CI26 | -0.56 | 0.37 | 0.56 | 0.74 | 1 | 0.21 | 1.31 | 0.54 | -0.16 | 0.93 | 0.66 | -0.78 | -0.06 | -0.1 | 0.78 | 0.07 | 0.4875 | -0.115 | -0.105 | 0.20095 | 0.801176 | 0.765853 | | 1 | 0.996511 | 0.964134 |
| Slc25a22 | Q9D6M3 | 0.07 | -0.38 | 0.1 | -0.03 | -0.12 | 0.73 | 0.73 | 0.21 | -0.6 | -0.43 | 0.5 | 0.29 | -0.89 | -0.35 | -0.18 | 0.23 | 0.4475 | 0 | -0.2375 | 0.099721 | 1 | 0.403193 | | 1 | 1 | 0.870698 |
| Slc25a23 | Q6GQS1 | 0.53 | 0.7 | 0.68 | 0.58 | 0.36 | 0.68 | 0.35 | 0.63 | 0.5 | 0.31 | 0.69 | 0.44 | 0.67 | 0.59 | 0.19 | 0.26 | -0.1175 | -0.1375 | -0.195 | 0.468082 | 0.424134 | 0.33437 | | 1 | 0.996511 | 0.846677 |
| Slc25a24 | Q8BMD8 | 1.05 | 0.44 | 1.04 | 1.27 | 0.53 | 0.92 | 1.26 | 2.17 | 0.86 | 0.08 | 1.24 | 0.79 | 1.5 | 0.84 | 0.53 | 1.09 | 0.27 | -0.2075 | 0.04 | 0.476862 | 0.501987 | 0.889758 | | 1 | 0.996511 | 0.98149 |
| Slc25a25 | A2ASZ8-2 | -0.16 | 0.91 | 0.41 | 0.18 | 1.19 | 0.7 | 1.25 | 0.51 | 0.4 | 0.24 | 0.84 | 0.14 | 0.06 | 1.28 | 0.17 | 0.8 | 0.5775 | 0.07 | 0.2425 | 0.073053 | 0.805225 | 0.504374 | | 1 | 0.996511 | 0.901002 |
| Slc25a25 | A2ASZ8-4 | 0.12 | 0.32 | 0.32 | 0.36 | 0.13 | 0.35 | 0.1 | -0.02 | 0.47 | -0.31 | 0.22 | 0.07 | 0.48 | 0.25 | 0.16 | 0.3 | -0.14 | -0.1675 | 0.0175 | 0.387992 | 0.440272 | 0.922116 | | 1 | 0.996511 | 0.987447 |
| Slc25a26 | Q5U680 | -0.83 | 0.45 | 0.01 | -0.22 | 0.78 | -0.3 | -0.11 | -0.59 | -0.55 | -0.37 | -0.16 | -0.68 | -0.49 | 0.06 | -0.68 | -0.53 | 0.0925 | -0.2925 | -0.2625 | 0.806841 | 0.337823 | 0.419484 | | 1 | 0.996511 | 0.87558 |
| Slc25a27 | Q9D6D0 | 0.15 | 0.26 | 0.09 | 0.17 | 0.11 | 0.4 | -0.34 | -0.19 | -0.22 | -0.57 | 0.37 | 0.09 | 0.08 | -0.47 | -0.23 | -0.52 | -0.1725 | -0.25 | -0.4525 | 0.400298 | 0.303701 | 0.05084 | | 1 | 0.996511 | 0.658454 |
| Slc25a28 | A0A494BA( | 2.7 | -0.24 | 2.2 | 0.7 | 1.48 | 1.19 | -1.25 | 0.5 | 1.23 | -1.13 | 0.31 | 0.23 | 4.3 | -2.35 | 2.46 | 0.08 | -0.86 | -1.18 | -0.2175 | 0.311364 | 0.144814 | 0.877985 | | 1 | 0.996511 | 0.979626 |
| Slc25a29 | Q8BL03 | -0.06 | 0.12 | 0.11 | 0.24 | 0.2 | -0.12 | -0.21 | -0.27 | -0.11 | 0.48 | -0.07 | -0.09 | 0.94 | 0.17 | 0.02 | 0.42 | -0.2025 | -0.05 | 0.285 | 0.259261 | 0.805603 | 0.266042 | | 1 | 0.996511 | 0.818107 |
| Slc25a3 | Q8VEM8 | -0.02 | 0.2 | 0.22 | 0.35 | -0.14 | 0.08 | 0.06 | 0.04 | -0.07 | -0.06 | 0.22 | -0.22 | -0.28 | 0.28 | -0.51 | -0.4 | -0.1775 | -0.22 | -0.415 | 0.277375 | 0.249117 | 0.103387 | | 1 | 0.996511 | 0.701092 |
| Slc25a31 | Q3V132 | -0.54 | -0.66 | 0.01 | -0.49 | -0.66 | -0.3 | -0.88 | -0.86 | 1.43 | 2.5 | -0.5 | -0.84 | 1.95 | -0.61 | -1 | -0.68 | -0.255 | 1.0675 | 0.335 | 0.269804 | 0.170667 | 0.602323 | | 1 | 0.996511 | 0.93091 |
| Slc25a32 | Q8BMG8 | -1.11 | 0.51 | -0.04 | 0.37 | -0.21 | -0.61 | -0.04 | -0.06 | 0.57 | -0.61 | 0.1 | -0.77 | 0.6 | 1.96 | -0.56 | -0.43 | -0.1625 | -0.11 | 0.46 | 0.663177 | 0.808245 | 0.471621 | | 1 | 0.996511 | 0.891144 |
| Slc25a33 | Q3TZX3 | -0.14 | 0.57 | 0.58 | 0.3 | -0.15 | -0.2 | 0.5 | 0.32 | -0.03 | -0.15 | 0.48 | -0.18 | 0.62 | -0.23 | 0.13 | 0.11 | -0.21 | -0.2975 | -0.17 | 0.415915 | 0.255305 | 0.531143 | | 1 | 0.996511 | 0.909419 |
| Slc25a35 | Q5SWT3 | -0.88 | -0.28 | -1.02 | -0.97 | -0.31 | -0.95 | -0.64 | -0.91 | -0.26 | -0.79 | -0.27 | -0.61 | 0.08 | -0.26 | -1.17 | -0.64 | 0.085 | 3.05E-01 | 0.29 | 0.725714 | 0.229401 | 0.38146 | | 1 | 0.996511 | 0.860984 |
| Slc25a36 | Q922G0 | -1.45 | -1.23 | -0.38 | -1.33 | -1.43 | -0.41 | -1.03 | -0.64 | -1.16 | -0.73 | -0.05 | -1.13 | -0.58 | -1.11 | -0.19 | -0.47 | 0.22 | 0.33 | 0.51 | 0.501647 | 0.356227 | 0.134971 | | 1 | 0.996511 | 0.731223 |
| Slc25a4 | P48962 | -0.2 | 0.04 | -0.42 | -0.23 | -0.04 | 0.23 | -0.12 | -0.17 | -0.27 | -1.08 | -0.01 | -0.35 | -0.73 | 0.02 | -1.2 | -0.69 | 0.1775 | -0.225 | -0.4475 | 0.330521 | 0.406358 | 0.144715 | | 1 | 0.996511 | 0.743156 |
| Slc25a40 | Q8BGP6 | -0.82 | -0.32 | -0.5 | -0.62 | -0.37 | -0.35 | -0.34 | -0.76 | -0.6 | -1.01 | -0.46 | -0.52 | -1.23 | -0.5 | -1.69 | -0.89 | 0.11 | -0.0825 | -0.5125 | 0.560002 | 0.691838 | 0.104887 | | 1 | 0.996511 | 0.703417 |
| Slc25a42 | Q8R0Y8 | 0.33 | 0.43 | 0.29 | 0.29 | 0.17 | 0.66 | 0.11 | -0.05 | 0.05 | -0.32 | 0.21 | 0.43 | 0.33 | -1.08 | -0.22 | -0.15 | -0.1125 | -0.2425 | -0.615 | 0.564328 | 0.259573 | 0.070888 | | 1 | 0.996511 | 0.668066 |
| Slc25a44 | Q3TMC4 | 1.39 | 1.52 | 0.06 | -0.03 | 0.41 | 1.68 | -0.41 | -0.82 | 2.45 | -1.1 | -0.31 | 1.43 | 3.13 | 1.26 | -1.23 | -0.71 | -0.52 | -0.1175 | -0.1225 | 0.417796 | 0.886156 | 0.899092 | | 1 | 0.996511 | 0.983509 |
| Slc25a46 | Q9CQS4 | 0.38 | 0.67 | 0.94 | 0.34 | 0.58 | 0.79 | 1.62 | 0.57 | -0.14 | 0.74 | 1.05 | 0.35 | -0.05 | 0.34 | 1.2 | 1.36 | 0.3075 | -0.0825 | 0.13 | 0.299615 | 0.7828 | 0.719718 | | 1 | 0.996511 | 0.958398 |
| Slc25a5 | P51881 | -0.8 | -0.74 | -1.4 | -0.73 | -0.38 | -0.31 | -0.67 | -1.26 | -0.98 | -1.62 | -1.09 | -1.04 | -1.47 | -0.38 | -1.79 | -1.34 | 0.2625 | -0.265 | -0.3275 | 0.35197 | 0.29496 | 0.352918 | | 1 | 0.996511 | 0.855029 |
| Slc25a51 | Q5HZI9 | 0.26 | 0.27 | 0.17 | 0.16 | -0.11 | 0.35 | -0.02 | 0.01 | 0.32 | -0.06 | 0.07 | -0.11 | 0.69 | -0.09 | -0.12 | -0.07 | -0.1575 | -0.16 | -0.1125 | 0.348977 | 0.369084 | 0.638641 | | 1 | 0.996511 | 0.942804 |
| Slc25a53 | A2AF34 | -1.55 | -0.75 | -0.01 | -0.49 | -0.74 | -0.98 | -0.21 | -0.43 | -0.92 | 0.25 | -0.17 | -1.34 | -0.96 | -1.19 | -1.04 | -0.3 | 0.11 | 0.155 | -0.1725 | 0.753791 | 0.733022 | 0.642854 | | 1 | 0.996511 | 0.94365 |
| Slc26a2 | E9PZ06 | -0.05 | -0.24 | 0.34 | 1.2 | 0.01 | 0.79 | -0.36 | 0.1 | -0.24 | -0.11 | -0.86 | 0.44 | -0.7 | 0.14 | -0.53 | 1.22 | -0.1775 | -0.505 | -0.28 | 0.641918 | 0.226413 | 0.581156 | | 1 | 0.996511 | 0.927734 |
| Slc26a6 | Q8CIW6-3 | 2.89 | 2.5 | 1.16 | 0.59 | 2.14 | 3.18 | 0.67 | 0.74 | 1.2 | 0.27 | 1.08 | 3.46 | 1.14 | 0.72 | 1.58 | 0.7 | -0.1025 | -0.2825 | -0.75 | 0.888946 | 0.721705 | 0.187817 | | 1 | 0.996511 | 0.776843 |
| Slc27a1 | Q60714 | 0.46 | 0.38 | 0.56 | 0.2 | 0.84 | 0.6 | 1.1 | 0.62 | 0.22 | 0.33 | 0.77 | 0.51 | -0.15 | 0.52 | 0.92 | 1.1 | 0.39 | 0.0575 | 0.1975 | 0.058059 | 0.768728 | 0.516008 | | 1 | 0.996511 | 0.905674 |
| Slc27a2 | O35488 | 0.13 | 0.3 | -0.31 | -0.19 | 0.43 | 0.25 | -0.24 | -0.15 | 0.9 | 0.2 | -0.96 | 0.2 | 0.98 | 0.51 | 0.36 | -0.2 | 0.09 | 0.1025 | 0.43 | 0.69881 | 0.794629 | 0.170641 | | 1 | 0.996511 | 0.762322 |
| Slc27a3 | G3X8Y7 | 0.12 | 0.72 | 0.32 | 0.24 | 1.85 | 0.85 | -0.05 | 0.59 | 1.4 | 1.14 | 0.46 | 0.92 | 2.62 | 2.31 | 1.61 | 0.96 | 0.46 | 0.63 | 1.525 | 0.260922 | 0.03519 | 0.00318 | | 1 | 0.996511 | 0.467521 |
| Slc27a4 | Q91VE0 | 0.65 | 0.57 | 0.31 | 0.1 | 0.7 | 0.67 | 0.23 | 0.26 | 0.1 | -0.16 | 0.23 | 0.78 | 0.16 | 0.21 | 0.46 | 0.45 | 0.0575 | -0.17 | -0.0875 | 0.782376 | 0.511489 | 0.675485 | | 1 | 0.996511 | 0.952944 |
| Slc29a1 | Q9JIM1-2 | 1.93 | 1.44 | 0.92 | 0.3 | 1.51 | 2.27 | 1.86 | 1.46 | 0.06 | -0.65 | 1.46 | 2.11 | 0.93 | -0.7 | 1.4 | 1.41 | 0.6275 | -0.4025 | -0.3875 | 0.123742 | 0.545355 | 0.496271 | | 1 | 0.996511 | 0.897918 |
| Slc29a2 | A0A494BA( | 1.28 | 1.37 | 1.39 | 1.33 | 1.57 | 1.04 | 0.61 | 1.19 | 1.51 | 2.01 | 1.45 | 1.51 | 1.32 | 1.64 | 1.12 | 0.79 | -0.24 | 0.2775 | -0.125 | 0.297174 | 0.169503 | 0.584388 | | 1 | 0.996511 | 0.927734 |
| Slc2a1 | P17809 | 1.4 | 0.97 | 1.42 | 0.88 | 0.85 | 0.83 | 1.07 | 0.99 | 1.2 | -0.4 | 1.54 | 1.14 | 0.97 | 0.74 | 0.94 | 1.13 | -0.2325 | -0.2975 | -0.2225 | 0.243084 | 0.495401 | 0.318197 | | 1 | 0.996511 | 0.838969 |
| Slc2a13 | Q3UHK1 | -0.07 | 0.13 | -0.51 | -0.51 | 0.29 | -0.07 | -1.19 | -0.77 | -0.06 | 0.25 | -0.72 | -0.18 | 0.14 | 0.51 | -0.33 | -0.63 | -0.195 | 0.0625 | 0.1625 | 0.586903 | 0.819101 | 0.601779 | | 1 | 0.996511 | 0.930802 |
| Slc2a3 | P32037 | 0.82 | 1.05 | 1.26 | 1.38 | 1.13 | 0.72 | 0.79 | 1.39 | 0.46 | 0.36 | 1.32 | 0.76 | -0.04 | 1.34 | 1.08 | 0.7 | -0.12 | -0.4025 | -0.3575 | 0.591007 | 0.155698 | 0.29297 | | 1 | 0.996511 | 0.825612 |
| Slc2a6 | Q3UDF0 | -1.14 | -0.88 | -0.68 | -0.16 | -0.42 | -1.9 | -0.2 | -0.56 | -0.57 | -0.88 | -0.46 | -0.66 | -1 | -0.09 | -1.06 | -0.27 | -0.055 | 0.0725 | 0.11 | 0.893303 | 0.77166 | 0.737804 | | 1 | 0.996511 | 0.958796 |
| Slc30a1 | Q60738 | 0.44 | 0.47 | -0.4 | -0.18 | 1.08 | 0.9 | 0.12 | -0.23 | 1.14 | -0.75 | -0.4 | 0.7 | 1.17 | 0.29 | 0 | -0.23 | 0.385 | 0.09 | 0.225 | 0.307115 | 0.846945 | 0.54699 | | 1 | 0.996511 | 0.912571 |
| Slc30a10 | Q3UVU3 | 0.95 | 1.18 | -0.3 | 0.3 | 0.81 | 1.7 | -0.36 | 0.2 | 2.16 | -1.08 | 0.41 | 1.34 | 2.44 | 0.19 | -0.06 | 0.07 | 0.055 | 0.175 | 0.1275 | 0.914029 | 0.803 | 0.838126 | | 1 | 0.996511 | 0.975961 |
| Slc30a3 | P97441 | -0.34 | 0.32 | 0.82 | 0.72 | 0.83 | 0.13 | 0.62 | 0.72 | -0.1 | 0.51 | 0.7 | -0.2 | 0.06 | 0.69 | 0.03 | 0.46 | 0.195 | -0.1525 | -0.07 | 0.523886 | 0.655337 | 0.824693 | | 1 | 0.996511 | 0.972852 |
| Slc30a6 | J3QMX8 | -2.04 | -2.27 | -0.41 | -0.44 | -1.48 | -1.41 | 0.59 | 0.09 | -1.6 | -0.73 | -0.67 | -2.19 | -0.73 | -0.79 | 0.16 | 0.42 | 0.7375 | -0.0075 | 1.055 | 0.281288 | 0.989508 | 0.079452 | | 1 | 0.999285 | 0.681213 |
| Slc30a7 | Q9JKN1 | -0.48 | 0.82 | -1.75 | 5.35 | 0.08 | -0.87 | -2.03 | 2.41 | -0.62 | 4.58 | -0.07 | -0.33 | -0.02 | -0.41 | -2.35 | -1.48 | -1.0875 | -0.095 | -2.05 | 0.506604 | 0.957032 | 0.182239 | | 1 | 0.996755 | 0.776002 |
| Slc30a9 | Q5IRJ6 | 2.16 | 1.04 | 0.77 | 0.86 | 1.15 | 1.47 | 0.53 | 0.71 | 0.76 | 0.54 | 0.86 | 2.1 | 0.44 | 0.81 | 0.24 | 0.39 | -0.2425 | -0.1425 | -0.7375 | 0.515758 | 0.751334 | 0.056805 | | 1 | 0.996511 | 0.661738 |
| Slc31a1 | Q8K211 | -0.23 | 0.56 | -0.54 | 0.08 | 0.3 | 0.42 | 0.44 | 0.15 | -0.74 | 0.71 | 1.64 | 0.11 | 0.46 | 0.46 | 0.49 | 0.48 | 0.36 | 0.4625 | 0.505 | 0.181749 | 0.379178 | 0.081334 | | 1 | 0.996511 | 0.687007 |
| Slc32a1 | O35633-2 | 0.99 | 0.74 | 0.59 | 0.91 | 0.8 | 0.89 | 0.56 | 0.84 | 0.88 | 1.33 | 0.47 | 0.97 | 0.51 | 0.93 | 1.05 | 0.5 | -0.035 | 0.105 | -0.06 | 0.836079 | 0.649954 | 0.784703 | | 1 | 0.996511 | 0.967642 |
| Slc33a1 | Q99J27 | -1.08 | -0.87 | -0.74 | -0.83 | -0.67 | -0.89 | -0.52 | -1.04 | -1.09 | -1 | -1.19 | -1.07 | -0.22 | -2.18 | -0.57 | -0.27 | 0.1 | -0.2075 | 0.07 | 0.583083 | 0.232277 | 0.874668 | | 1 | 0.996511 | 0.979626 |
| Slc35a2 | A0A140LHI | -0.23 | -1.28 | -0.92 | -1.5 | 6.16 | -0.78 | 0.7 | 1.81 | 0.04 | -0.59 | -2.72 | -1.55 | 0.21 | 0.4 | -0.75 | 0.96 | 2.955 | -0.2225 | 1.1875 | 0.054377 | 0.71484 | 0.020907 | | 1 | 0.996511 | 0.621926 |
| Slc35a3 | E9Q3V2 | -0.25 | 0.44 | -0.52 | 0.08 | -0.99 | 2.19 | -1.48 | -0.11 | 1.96 | -0.22 | -1.07 | 0.76 | 3.69 | -1.7 | 0.39 | 0.91 | -0.035 | 0.42 | 0.885 | 0.963148 | 0.507673 | 0.391482 | | 1 | 0.996511 | 0.865375 |
| Slc35a4 | A0A087WC | -2 | -0.72 | -0.91 | -0.96 | -0.56 | -1.26 | 0.15 | -0.61 | -1.82 | -0.54 | -0.53 | -1.72 | -1.41 | -0.63 | -1.34 | -0.06 | 0.5775 | -0.005 | 0.2875 | 0.161014 | 0.990797 | 0.492344 | | 1 | 0.999285 | 0.897531 |
| Slc35a5 | M0QWE5 | -1.69 | -3.19 | -2.09 | -1.89 | -4.31 | -0.26 | -4.99 | -2 | -2.1 | -0.42 | -2.48 | -2.57 | 0.77 | -4.87 | -1.34 | -2.52 | -0.675 | 0.3225 | 0.225 | 0.513049 | 0.565476 | 0.836802 | | 1 | 0.996511 | 0.975831 |
| Slc35b2 | Q91ZN5-2 | 0.12 | -0.52 | -3.08 | -4.52 | -0.03 | -0.85 | -3.4 | -4.1 | 0.2 | -2.04 | -3.37 | 0.13 | -1.1 | -0.29 | -2.69 | -2.93 | -0.095 | 0.73 | 0.2475 | 0.942093 | 0.562334 | 0.825699 | | 1 | 0.996511 | 0.973155 |
| Slc35d3 | Q8BGF8 | -0.18 | -1.99 | -0.71 | 1.12 | -2.25 | 0.12 | 1.22 | 1.04 | 3.05 | 1.61 | 0.27 | 0.21 | 2.5 | -1.8 | 2.16 | 0.6 | 0.4725 | 1.725 | 1.305 | 0.610759 | 0.0663 | 0.233705 | | 1 | 0.996511 | 0.790054 |
| Slc35f1 | Q8BGK5 | -0.1 | 0.26 | -0.03 | -1.94 | 0.64 | 0.14 | 2.61 | 1.94 | 0.16 | -0.37 | 3.09 | 0.87 | 0 | 0.76 | 8.57 | 0.93 | 1.785 | 1.39 | 3.0175 | 0.027798 | 0.119101 | 0.127671 | | 1 | 0.996511 | 0.723391 |
| Slc35f3 | Q1LZI2 | -1.51 | -3.63 | -2.62 | -3.53 | -3.26 | -2.08 | -3.57 | -2.8 | -0.17 | -0.95 | -3.33 | -2.85 | 1.38 | -2.4 | -2.42 | -3.25 | -0.105 | 0.9975 | 1.15 | 0.846149 | 0.240954 | 0.279975 | | 1 | 0.996511 | 0.822084 |
| Slc35f6 | A0A0G2JE( | -1.89 | -2.19 | -1.75 | -2 | -1.66 | -1.23 | -2.3 | -2.33 | -0.56 | -0.49 | -2 | -1.56 | 0.35 | -0.69 | -1.54 | -1.52 | 0.0775 | 0.805 | 1.1075 | 0.785051 | 0.055788 | 0.02922 | | 1 | 0.996511 | 0.637553 |
| Slc35g2 | D3YVE8 | 1.11 | 0.82 | 0.28 | 0.67 | 0.62 | 1.24 | 0.55 | 2.03 | -0.6 | -0.42 | 1.61 | 0.31 | -0.74 | -1.3 | 1.24 | 1.24 | 0.39 | -0.495 | -0.61 | 0.30418 | 0.330943 | 0.342844 | | 1 | 0.996511 | 0.848473 |
| Slc36a1 | Q8K4D3 | -1.59 | -0.96 | -0.81 | -1.61 | -0.36 | -0.51 | -0.43 | -0.7 | -1.06 | 0.1 | -0.29 | -1.3 | -0.41 | -0.22 | -0.85 | -0.16 | 0.7425 | 0.605 | 0.8325 | 0.011854 | 0.133756 | 0.015035 | | 1 | 0.996511 | 0.589387 |
| Slc36a4 | D3Z0L1 | -1.48 | -2.1 | -0.52 | -1.2 | -2.47 | -1.52 | -0.65 | -0.09 | -0.62 | -0.83 | -0.55 | -2.14 | -1.69 | -2.25 | -1.32 | -1.34 | 0.1425 | 0.29 | -0.325 | 0.800283 | 0.537672 | 0.405374 | | 1 | 0.996511 | 0.872803 |
| Slc37a4 | A0A1L1SUI | 0.12 | 0.78 | -0.37 | 0.13 | 0.72 | -0.01 | 0.35 | -0.08 | 0.27 | 0.21 | -0.48 | 0.15 | 0.45 | 1.37 | 0.15 | -0.25 | 0.08 | -0.1275 | 0.265 | 0.788329 | 0.671631 | 0.515233 | | 1 | 0.996511 | 0.905674 |
| Slc38a1 | Q8K2P7 | -0.32 | -0.03 | 0 | -0.22 | 0.22 | -0.12 | -0.76 | -0.16 | -0.31 | 0.57 | -0.26 | -0.1 | -0.23 | -0.03 | -0.12 | -0.52 | -0.0625 | 0.1175 | -0.0825 | 0.790596 | 0.631389 | 0.679867 | | 1 | 0.996511 | 0.955484 |
| Slc38a10 | J3QNE8 | 0.9 | 1.32 | -3.62 | -4.45 | 1.44 | 1.38 | -4.08 | -4.41 | 1.27 | -2.13 | -3.37 | 1.24 | 1.48 | 1.21 | -5.23 | -4.72 | 0.045 | 0.715 | -0.3525 | 0.981803 | 0.675891 | 0.866193 | | 1 | 0.996511 | 0.978251 |
| Slc38a2 | Q8CFE6 | 0.97 | 0.19 | 0.74 | -0.31 | 0.45 | 1.04 | -0.37 | -0.1 | 0 | -0.75 | 0.32 | 0.69 | -1.2 | -0.9 | 0.57 | 1.09 | -0.1425 | -0.3325 | -0.5075 | 0.722476 | 0.41655 | 0.388829 | | 1 | 0.996511 | 0.864915 |
| Slc38a3 | Q9DCP2 | 1.94 | 1.07 | 0.48 | -0.05 | 0.83 | 1.55 | 0.27 | -0.52 | 0.31 | -0.44 | -0.24 | 1.87 | 1.51 | -0.52 | 1 | -0.34 | -0.3275 | -0.485 | -0.4475 | 0.562246 | 0.440633 | 0.463981 | | 1 | 0.996511 | 0.888267 |
| Slc38a7 | A0A1D5RL) | -0.55 | -0.47 | -0.35 | -0.05 | -0.66 | -0.53 | 0.19 | -0.3 | 0 | 0.78 | 0.43 | 0.06 | 0.3 | 0.37 | -0.47 | 0.28 | 0.03 | 0.6725 | 0.475 | 0.898164 | 0.019508 | 0.091363 | | 1 | 0.996511 | 0.693316 |
| Slc39a10 | Q6P5F6 | 0.5 | 0.64 | 0.25 | 0.14 | 0.8 | 0.55 | -0.03 | -0.1 | 0.85 | -0.16 | 0.37 | 0.98 | 0.99 | 0.93 | 0.58 | -0.02 | -0.0775 | 0.1275 | 0.2375 | 0.76391 | 0.663455 | 0.404874 | | 1 | 0.996511 | 0.872139 |
| Slc39a11 | Q8BWY7 | -0.08 | -0.92 | -0.79 | -0.47 | -3.75 | 0.19 | -1.01 | -0.54 | 1.16 | 0.46 | -0.82 | -1.24 | 2.69 | -1.78 | -0.2 | -0.57 | -0.7125 | 0.455 | 0.6 | 0.381108 | 0.411351 | 0.495108 | | 1 | 0.996511 | 0.89758 |
| Slc39a12 | Q5FWH7 | 0.63 | 0.35 | 0.78 | 0.7 | 0.34 | 0.94 | -0.17 | 0.82 | 0.82 | 0.8 | 0.93 | 1.06 | 1.31 | -0.64 | 1.06 | 1.11 | -0.1325 | 0.2875 | 0.095 | 0.631417 | 0.134556 | 0.828756 | | 1 | 0.996511 | 0.973757 |
| Slc39a14 | Q75N73-3 | 0.79 | 0.15 | 1.41 | 1.25 | -0.78 | 1.41 | -0.61 | 0.35 | 1.8 | 0.33 | 1.11 | 0.39 | 3.24 | -2.5 | 0.93 | 0.33 | -0.8075 | 0.0075 | -0.4 | 0.154792 | 0.985858 | 0.7129 | | 1 | 0.998892 | 0.958106 |
| Slc39a3 | A0A452J89 | -0.56 | -2.61 | -0.77 | -0.79 | -2.86 | 0.14 | -1.05 | -0.69 | 4.85 | -0.03 | -0.64 | -1.47 | 8.06 | -2.3 | -0.42 | -0.54 | 0.0675 | 1.86 | 2.3825 | 0.925036 | 0.189658 | 0.275249 | | 1 | 0.996511 | 0.822084 |
| Slc39a6 | Q8C145 | 1.01 | 1.15 | 0.39 | 0.66 | 1.08 | 1.22 | -0.02 | 0.43 | 0.99 | 1.7 | 0.33 | 1.32 | 1.34 | 1.05 | 0.82 | 0.44 | -0.125 | 0.2825 | 0.11 | 0.703693 | 0.407963 | 0.693121 | | 1 | 0.996511 | 0.955729 |
| Slc39a7 | Q31125 | 1.17 | 1.41 | 0.77 | -0.23 | 1.48 | 0.54 | -0.57 | -0.07 | 1.13 | -0.3 | -0.03 | 1.11 | 1.83 | 2.06 | 3.07 | 0.13 | -0.435 | -0.3025 | 0.9925 | 0.416428 | 0.538147 | 0.150834 | | 1 | 0.996511 | 0.749304 |
| Slc3a2 | P10852-2 | 1.85 | 1.76 | 1.41 | 1.81 | 1.35 | 1.79 | 1.25 | 2.13 | 1.29 | 1.17 | 1.51 | 1.94 | 0.61 | 1.14 | 1.05 | 1.24 | -0.0775 | -0.23 | -0.6975 | 0.749535 | 0.330811 | 0.011251 | | 1 | 0.996511 | 0.566517 |
| Slc3a2 | P10852 | -0.62 | -0.23 | 0.76 | 0.75 | -0.41 | -0.56 | -0.45 | 0.08 | 2.27 | 4.67 | 0.78 | -0.72 | 2.03 | 4.14 | 0.76 | 0.58 | -0.5 | 1.585 | 1.7125 | 0.190338 | 0.166804 | 0.058241 | | 1 | 0.996511 | 0.661738 |
| Slc43a2 | Q8CGA3 | 0.31 | 0.23 | 0.1 | -0.3 | -0.99 | 0.05 | -0.32 | -0.54 | 3.16 | 0.49 | -0.36 | -0.4 | 4.43 | -0.92 | 1.86 | -0.78 | -0.535 | 0.6375 | 1.0625 | 0.069061 | 0.41645 | 0.362265 | | 1 | 0.996511 | 0.858475 |
| Slc44a1 | A2AMH5 | 0.29 | 0.31 | 0.51 | 0.08 | 0.69 | 0.25 | 0.09 | -0.33 | -0.14 | -0.09 | -0.16 | -0.16 | 0.1 | -0.89 | -0.01 | 0.32 | -0.1225 | -0.435 | -0.4175 | 0.616564 | 0.0284 | 0.180914 | | 1 | 0.996511 | 0.775542 |
| Slc44a2 | A0A1L1SU/ | 0.27 | 0.29 | 1.26 | 1.13 | -0.66 | 0.66 | 0.71 | 0.8 | 1.71 | 1.82 | 1.24 | 0.23 | 2.82 | 0.45 | 1.24 | 0.64 | -0.36 | 0.5125 | 0.55 | 0.391655 | 0.248635 | 0.334346 | | 1 | 0.996511 | 0.846677 |
| Slc44a2 | Q8BY89-2 | -0.16 | 1 | 0.85 | 0.49 | 1.06 | 0.5 | 0.99 | 0.5 | 0.92 | 0.38 | 0.92 | -0.14 | 0.63 | 0.63 | 0.48 | 0.81 | 0.2175 | -0.025 | 0.0925 | 0.472415 | 0.943668 | 0.74636 | | 1 | 0.996755 | 0.960238 |
| Slc45a1 | Q8BIV7 | 0.73 | 1.29 | 0.33 | 0.28 | 0.68 | 1.13 | -0.52 | -0.36 | 1.83 | 0.71 | 0.29 | 1.1 | 2.85 | 0.94 | -0.2 | 0.37 | -0.425 | 0.325 | 0.3325 | 0.340374 | 0.410447 | 0.606334 | | 1 | 0.996511 | 0.932425 |
| Slc45a4 | F7BEQ1 | 0.86 | 2.11 | 2.09 | 1.28 | 1.29 | 1.5 | 0.91 | 0.91 | 2.23 | 1.38 | 0.65 | 1.83 | 2.52 | 2.13 | 3.45 | 0.43 | -0.4325 | -0.0625 | 0.5475 | 0.215479 | 0.885328 | 0.40335 | | 1 | 0.996511 | 0.870698 |
| Slc47a1 | D6RGQ3 | -0.76 | -2.56 | -2.8 | 0.52 | -3.98 | 0.27 | -1.84 | -1.01 | 1.28 | -0.03 | -2.11 | -1.6 | 3.49 | 0.09 | -1.19 | -2.42 | -0.24 | 0.785 | 1.3925 | 0.821887 | 0.436067 | 0.310875 | | 1 | 0.996511 | 0.834523 |
| Slc49a4 | D6RI31 | 0.35 | 0.74 | -0.09 | -0.08 | -0.8 | 0.63 | -0.61 | -0.27 | 1.02 | 0.43 | 0.14 | 1.06 | 0.65 | 0.09 | -0.23 | 0.01 | -0.4925 | 0.4325 | -0.1 | 0.193224 | 0.181084 | 0.730195 | | 1 | 0.996511 | 0.958635 |
| Slc4a1 | P04919-2 | 0.52 | -1.5 | 2.38 | 4.63 | -0.57 | 2.08 | 1.92 | 5 | 0.54 | 0.95 | 1.9 | -0.98 | 1.07 | 0.15 | 0.7 | 1.07 | 0.6 | -0.905 | -0.76 | 0.6994 | 0.488321 | 0.525049 | | 1 | 0.996511 | 0.908107 |
| Slc4a10 | Q5DTL9 | 1.25 | 1.06 | 1.16 | 0.73 | 1.2 | 1.49 | 1.18 | 0.77 | 1.22 | -0.22 | 1.3 | 1.29 | 1.31 | 0.55 | 0.55 | 0.7 | 0.11 | -0.1525 | -0.2725 | 0.608423 | 0.686681 | 0.289181 | | 1 | 0.996511 | 0.824705 |
| Slc4a10 | Q5DTL9-2 | 0.45 | 0.84 | 1.5 | 1.11 | 1.39 | 0.17 | 1.21 | 0.93 | 0.22 | 1.16 | 1.32 | 0.65 | 0.46 | 1.67 | 0.35 | 0.5 | -0.05 | -0.1375 | -0.23 | 0.882032 | 0.67979 | 0.541425 | | 1 | 0.996511 | 0.912212 |
| Slc4a1ap | E9Q585 | -2.18 | -2.48 | -2.55 | -2.68 | -2.3 | -2.11 | -2.82 | -2.15 | -1.67 | -1.75 | -2.19 | -2.07 | -1.48 | -2.16 | -2.84 | -2.23 | 0.1275 | 0.5525 | 0.295 | 0.563981 | 0.024872 | 0.351117 | | 1 | 0.996511 | 0.853984 |
| Slc4a2 | A0A0R4J1K | -0.96 | -0.2 | 0.47 | 0.14 | 0.12 | -0.53 | -0.08 | -0.55 | -0.46 | -1.2 | -0.58 | -0.49 | -0.29 | 0.07 | 0.05 | 0.25 | -0.1225 | -0.545 | 0.1575 | 0.717542 | 0.142353 | 0.635104 | | 1 | 0.996511 | 0.941407 |
| Slc4a3 | P16283 | 0.32 | 0.19 | 0.8 | 0.97 | 0.34 | 0.38 | 0.26 | 0.76 | 1.36 | 1.23 | 0.65 | 0.27 | 1.04 | 0.47 | 1.1 | 0.68 | -0.135 | 0.3075 | 0.2525 | 0.569375 | 0.345575 | 0.355052 | | 1 | 0.996511 | 0.855029 |
| Slc4a4 | O88343 | 1.52 | 1.02 | 0.89 | 1.63 | 0.79 | 0.19 | 1.2 | 0.71 | 0.42 | 0.87 | 0.94 | 1.16 | 0.24 | 0.19 | 0.26 | 0.88 | -0.5425 | -0.4175 | -0.8725 | 0.079449 | 0.134152 | 0.009774 | | 1 | 0.996511 | 0.563386 |
| Slc4a4 | A7E1Z5 | -0.6 | -0.54 | -2.58 | -1.3 | -2.14 | -0.54 | -1.71 | -1.17 | 1.17 | -0.66 | -0.62 | -0.33 | 2.47 | -1.89 | 1.36 | -1.79 | -0.135 | 1.145 | 1.2925 | 0.802269 | 0.07935 | 0.249294 | | 1 | 0.996511 | 0.804842 |
| Slc4a4 | A0A571BE! | 0.49 | 0.55 | 0.39 | 1.01 | 0.72 | 0.11 | 0.66 | 0.89 | 0.59 | 0.99 | 0.54 | 0.82 | 0.69 | 0.55 | 0.67 | 0.14 | -0.015 | 0.125 | -0.0975 | 0.949081 | 0.562168 | 0.674966 | | 1 | 0.996511 | 0.952944 |
| Slc4a4 | E9Q8N8 | 0.36 | 0.62 | 1.37 | 1.65 | 0.62 | 0.39 | 1.61 | 1.33 | 0.67 | 1.04 | 1.3 | 0.5 | 0.64 | 0.53 | 1.42 | 1.43 | -0.0125 | -0.1225 | 0.005 | 0.974788 | 0.725174 | 0.989486 | | 1 | 0.996511 | 0.996154 |
| Slc4a7 | F8VQC9 | 0.25 | 0.54 | 0.82 | 0.83 | 0.67 | 0.26 | 0.83 | 0.77 | 0.58 | 1.83 | 1.17 | 0.49 | 0.53 | 0.98 | 0.76 | 0.7 | 0.0225 | 0.4075 | 0.1325 | 0.916308 | 0.244852 | 0.547957 | | 1 | 0.996511 | 0.912571 |
| Slc4a8 | Q8JZR6 | 1.04 | 1.32 | 0.87 | 1.25 | 1.53 | 1.22 | 1.25 | 1.13 | 0.53 | 0.54 | 1.17 | 1.33 | 0.56 | 0.51 | 0.89 | 0.68 | 0.1625 | -0.2275 | -0.46 | 0.376848 | 0.382236 | 0.043991 | | 1 | 0.996511 | 0.658454 |
| Slc52a2 | Q9D8F3-2 | -0.05 | 1.07 | 0.53 | -0.42 | 0.91 | 0.4 | -0.54 | -0.28 | 0.97 | 1.42 | -0.01 | -0.41 | 2.38 | 1.08 | 0.62 | -0.24 | -0.16 | 0.21 | 0.6775 | 0.712998 | 0.674774 | 0.265457 | | 1 | 0.996511 | 0.817264 |
| Slc5a3 | Q9JKZ2 | -0.15 | 0.03 | -0.06 | 0.8 | -0.12 | 0.43 | -0.19 | 0.46 | 0.31 | 0.19 | 0.36 | -0.23 | 0.38 | 0.11 | 0.41 | 0.39 | -0.01 | 0.0025 | 0.1675 | 0.971633 | 0.992637 | 0.52265 | | 1 | 0.999285 | 0.907667 |
| Slc5a6 | Q5U4D8 | -0.89 | -1.62 | 0.92 | 0.66 | -0.07 | 1.91 | -1.52 | -2.15 | 1.39 | -1.34 | 0.9 | -1.25 | 4.39 | -2.54 | 0.15 | 0.4 | -0.225 | 0.1575 | 0.8325 | 0.818052 | 0.852407 | 0.550648 | | 1 | 0.996511 | 0.913801 |
| Slc5a7 | Q8BGY9 | 0.12 | 0.04 | 0.54 | 0.8 | -0.4 | -0.32 | 0.09 | 0.48 | 0.67 | 0.41 | -0.06 | -0.25 | 0.67 | -0.06 | 0.42 | 0.08 | -0.4125 | -0.1825 | -0.0975 | 0.159851 | 0.52945 | 0.717619 | | 1 | 0.996511 | 0.958398 |
| Slc66a2 | Q80XM9-2 | -3.09 | -2.97 | -1.09 | -1.46 | -1.6 | -3.14 | -1.11 | -1.55 | -3.34 | -3.3 | -1.37 | -1.9 | -1.89 | -3.08 | -1.18 | -1.04 | 0.3025 | -0.325 | 0.355 | 0.626563 | 0.620735 | 0.578068 | | 1 | 0.996511 | 0.927198 |
| Slc6a1 | P31648 | 2.67 | 0.6 | 1.34 | 0.58 | 0.98 | 3.22 | 3.63 | 0.65 | 0.17 | 0.15 | 1.49 | 2.28 | -0.16 | 0.55 | 1.69 | 2.26 | 0.8225 | -0.275 | -0.2125 | 0.328149 | 0.675661 | 0.750865 | | 1 | 0.996511 | 0.961479 |
| Slc6a11 | P31650 | 1.11 | 0.99 | 1.37 | 1.63 | 0.76 | 0.58 | 1.28 | 1.09 | 0.92 | 1.79 | 0.91 | 0.92 | 0.61 | 0.47 | 1.87 | 1.59 | -0.3475 | -0.14 | -0.14 | 0.159226 | 0.612836 | 0.705948 | | 1 | 0.996511 | 0.957084 |
| Slc6a12 | E9PXG1 | -2.22 | -1.52 | -0.99 | 0.9 | 0.66 | -1.28 | -3.91 | -0.69 | 4.31 | 1.92 | -2.38 | -0.66 | 4.57 | -2.1 | -2.83 | -0.12 | -0.3475 | 1.755 | 0.8375 | 0.740727 | 0.243349 | 0.601625 | | 1 | 0.996511 | 0.930802 |
| Slc6a13 | D3YWZ5 | -2.25 | -1.29 | -0.72 | 0.83 | -1.85 | -1.96 | -1.39 | -1.21 | 0.37 | 1.91 | -1.19 | -2.19 | -0.15 | 1.69 | -1.2 | 0.24 | -0.745 | 0.5825 | 1.0025 | 0.242585 | 0.561772 | 0.227445 | | 1 | 0.996511 | 0.787447 |
| Slc6a15 | Q8BG16 | -0.02 | 0.72 | -0.6 | -0.57 | 1.34 | 0.53 | -0.75 | -0.82 | 0.24 | -0.48 | -0.67 | 0.23 | 0.28 | 0.82 | -0.47 | -0.28 | 0.1925 | -0.0525 | 0.205 | 0.7303 | 0.889026 | 0.617971 | | 1 | 0.996511 | 0.937292 |
| Slc6a17 | A0A0R4J08 | -0.85 | -0.1 | -0.32 | -0.15 | -0.14 | -0.33 | -0.07 | -0.23 | 0.13 | -0.09 | -0.01 | -0.21 | 0.2 | 0.07 | -0.32 | -0.26 | 0.1625 | 0.31 | 0.2775 | 0.445146 | 0.186803 | 0.28021 | | 1 | 0.996511 | 0.822084 |
| Slc6a20a | E9Q0H0 | -0.45 | 0 | -0.42 | -1.2 | 0.32 | 0.54 | -0.08 | -1.6 | 0.78 | 0.08 | -0.23 | 0.78 | 0.08 | -0.4 | -1.7 | -0.18 | 0.3125 | 0.87 | -0.0325 | 0.537392 | 0.032841 | 0.941489 | | 1 | 0.996511 | 0.991085 |
| Slc6a3 | Q61327 | 0.34 | -0.71 | 0.67 | 1.03 | -0.24 | -0.88 | 1.33 | 0.89 | 0.22 | 0.81 | -0.04 | 0.35 | -0.35 | -0.55 | 0.96 | 0.64 | -0.0575 | 0.0025 | -0.1575 | 0.920511 | 0.994973 | 0.749139 | | 1 | 0.999285 | 0.961014 |
| Slc6a4 | Q60857 | -0.13 | -0.36 | -0.47 | 0.03 | -0.76 | -0.66 | 0.31 | -0.22 | 0.38 | 1.46 | -0.15 | 0.17 | 0.25 | 0.56 | 0.13 | -0.21 | -0.1 | 0.6975 | 0.415 | 0.715653 | 0.078505 | 0.10604 | | 1 | 0.996511 | 0.703417 |
| Slc6a6 | O35316 | -0.01 | 0.38 | 0.74 | 1.29 | 0.72 | -0.07 | -0.34 | 0.57 | 0.77 | 2.5 | 0.55 | -0.04 | 1.51 | 1.84 | 1.12 | 0.63 | -0.38 | 0.345 | 0.675 | 0.305885 | 0.5446 | 0.096846 | | 1 | 0.996511 | 0.698156 |
| Slc6a7 | Q6PGE7 | -0.29 | 0.02 | -0.09 | 0.07 | 0.28 | -0.05 | 0.31 | 0.44 | 0.18 | 0.18 | 0.52 | 0.28 | 0.35 | 0.44 | 0.46 | -0.19 | 0.3175 | 0.3625 | 0.3375 | 0.102084 | 0.070228 | 0.15493 | | 1 | 0.996511 | 0.750763 |
| Slc6a8 | E9Q151 | 1.27 | 1.49 | 1.52 | 1.21 | 1.71 | 1.5 | 1.38 | 0.92 | 0.52 | 0.22 | 1.3 | 1.35 | 0.13 | 1.1 | 0.28 | 1.18 | 0.005 | -0.525 | -0.7 | 0.981123 | 0.10708 | 0.04025 | | 1 | 0.996511 | 0.658454 |
| Slc6a9 | P28571-1 | 1.26 | 1.25 | 0.15 | 0.15 | 1.49 | 0.79 | -0.2 | -0.02 | 1.28 | -0.03 | 0.27 | 1.6 | 0.92 | 1.08 | 0.63 | 0.54 | -0.1875 | 0.0775 | 0.09 | 0.689439 | 0.869637 | 0.792954 | | 1 | 0.996511 | 0.96933 |
| Slc7a1 | E9Q3N1 | -0.68 | 0.37 | 0.03 | 0.3 | 0.24 | -0.4 | 0 | 0.12 | -0.07 | -0.19 | 0.47 | -0.04 | -1.05 | 0.53 | 0.21 | -0.24 | -0.015 | 0.0375 | -0.1425 | 0.957248 | 0.896994 | 0.724239 | | 1 | 0.996511 | 0.958398 |
| Slc7a10 | P63115 | 0.14 | -0.14 | 0.65 | 0.6 | 0.06 | 0.37 | 0.06 | 0.48 | -1.16 | 0.76 | 0.46 | 0.28 | -0.73 | -2.19 | 0.61 | 0.54 | -0.07 | -0.2275 | -0.755 | 0.766554 | 0.608045 | 0.24782 | | 1 | 0.996511 | 0.803677 |
| Slc7a11 | Q9WTR6 | 0.53 | 0.86 | 0.54 | 0.72 | 0.5 | 0.69 | -0.28 | 0.51 | 0.74 | 0.68 | 0.41 | 0.72 | 0.78 | 1.03 | 0.33 | 0.54 | -0.3075 | -0.025 | 0.0075 | 0.229261 | 0.888445 | 0.972892 | | 1 | 0.996511 | 0.992865 |
| Slc7a14 | Q8BXR1 | 0.2 | 0.55 | -0.2 | -0.15 | 0.44 | 0.23 | -0.04 | -0.07 | 0.27 | -0.5 | -0.2 | 0.01 | -0.05 | 0.6 | -0.65 | -0.26 | 0.04 | -0.205 | -0.19 | 0.862545 | 0.436453 | 0.557297 | | 1 | 0.996511 | 0.917092 |
| Slc7a2 | P18581-2 | 0.76 | 0.94 | 0.7 | 0.38 | 1.69 | 1.14 | -0.62 | 0.94 | 0.56 | -0.14 | 0.83 | 0.95 | 0.35 | 0.88 | 0.16 | 0.74 | 0.0925 | -0.145 | -0.1625 | 0.844528 | 0.609775 | 0.507226 | | 1 | 0.996511 | 0.902614 |
| Slc7a3 | P70423 | -0.26 | 0.46 | -0.18 | 0.27 | 0.88 | -0.51 | 0.66 | 0.32 | -0.16 | 0.49 | 0.45 | 0.25 | -0.81 | -0.2 | -0.02 | 0.11 | 0.265 | 0.185 | -0.3025 | 0.44338 | 0.469191 | 0.305936 | | 1 | 0.996511 | 0.832533 |
| Slc7a4 | A0A338P6\ | -0.72 | 0.08 | 0.74 | 0.73 | 0.52 | -0.14 | 0.5 | 0.35 | -0.15 | 1.98 | 0.93 | -0.3 | -0.16 | 1.09 | 0.66 | 0.84 | 0.1 | 0.4075 | 0.4 | 0.782529 | 0.49015 | 0.353242 | | 1 | 0.996511 | 0.855029 |
| Slc7a5 | Q9Z127 | -0.09 | 0.52 | 0.75 | 0.33 | 0.49 | 0.23 | -0.01 | 0.23 | 0.63 | 0.66 | 0.58 | 0.33 | 0.45 | 0.85 | 0.04 | -0.05 | -0.1425 | 0.1725 | -0.055 | 0.533272 | 0.454757 | 0.849396 | | 1 | 0.996511 | 0.976225 |
| Slc7a6 | Q8BGK6 | -0.14 | 0.81 | 0.24 | 0.38 | 0.95 | 0.82 | 0.36 | 0.66 | 0.66 | 0.87 | 0.39 | 0.61 | 1.21 | 1.3 | 0.67 | 1.13 | 0.375 | 0.31 | 0.755 | 0.15493 | 0.226975 | 0.018718 | | 1 | 0.996511 | 0.602211 |
| Slc7a6os | Q7TPE5 | -0.64 | -0.73 | -0.24 | 2.6 | -1.65 | -1.03 | 0.95 | 0.62 | -0.47 | 3.36 | 0.74 | -0.91 | 0.26 | 0.34 | 0.17 | 0.74 | -0.525 | 0.4325 | 0.13 | 0.567373 | 0.699088 | 0.857769 | | 1 | 0.996511 | 0.976225 |
| Slc7a8 | Q9QXW9 | -0.77 | 0.12 | 0.68 | 0.23 | 0.11 | -0.28 | -0.28 | 0.3 | -0.78 | 0.3 | 0.48 | -0.7 | -0.56 | -0.8 | 0.12 | 0.05 | -0.1025 | -0.24 | -0.3625 | 0.754814 | 0.574915 | 0.342032 | | 1 | 0.996511 | 0.848473 |
| Slc8a1 | Q68FL0 | 0.65 | 1.03 | 1.23 | 1.23 | 1.22 | 0.83 | 0.96 | 1.16 | 0.75 | 1.05 | 1.23 | 0.84 | 0.56 | 1.01 | 1.25 | 0.93 | 0.0075 | -0.0675 | -0.0975 | 0.969733 | 0.753452 | 0.683195 | | 1 | 0.996511 | 0.955729 |
| Slc8a1 | G3X9J1 | 0.21 | -0.27 | 1.88 | 1.04 | 1.28 | -1.21 | 0.47 | 0.83 | -0.44 | -1.31 | 0.56 | -0.11 | -0.48 | 0.78 | -0.31 | 2.06 | -0.3725 | -1.04 | -0.2025 | 0.5728 | 0.09291 | 0.767737 | | 1 | 0.996511 | 0.965084 |
| Slc8a2 | Q8K596 | 1.49 | 1.39 | 2.49 | 2.26 | 1.42 | 2.31 | 1.91 | 2.55 | 0.59 | 1.48 | 2.6 | 1.87 | 0.27 | 0.74 | 1.88 | 1.99 | 0.14 | -0.2725 | -0.6875 | 0.694168 | 0.564779 | 0.172207 | | 1 | 0.996511 | 0.76392 |
| Slc8a3 | S4R2P9-2 | 0.54 | 0.92 | 1.31 | 1.43 | 0.51 | 0.06 | 1.96 | 1.21 | 0.88 | 1.24 | 1.09 | 0.47 | 0.28 | 1 | 1.47 | 1.3 | -0.115 | -0.13 | -0.0375 | 0.790265 | 0.639314 | 0.91045 | | 1 | 0.996511 | 0.986458 |
| Slc9a1 | Q61165 | 0.64 | 0.7 | 0.51 | 0.62 | 0.23 | 0.48 | 0.69 | 0.52 | 0.22 | 0.99 | 0.09 | 0.59 | -0.12 | 0.22 | -0.15 | -0.05 | -0.1375 | -0.145 | -0.6425 | 0.407804 | 0.542895 | 0.006173 | | 1 | 0.996511 | 0.552614 |
| Slc9a3r1 | P70441 | 1.57 | 1.7 | 1.58 | 1.56 | 1.79 | 1.26 | 0.65 | 1.42 | 0.83 | 1.7 | 1.35 | 1.64 | 0.88 | 1.81 | 1.15 | 1.04 | -0.3225 | -0.2225 | -0.3825 | 0.221418 | 0.351353 | 0.14299 | | 1 | 0.996511 | 0.742711 |
| Slc9a3r2 | A0A0R4IZX | -0.18 | -0.24 | -0.29 | -0.65 | -0.41 | -0.21 | -0.87 | -1.05 | 0.69 | 0.46 | -0.44 | 0.16 | 1.1 | 0.09 | 0.01 | -0.65 | -0.295 | 0.5575 | 0.4775 | 0.236813 | 0.072396 | 0.217518 | | 1 | 0.996511 | 0.787447 |
| Slc9a5 | B2RXE2 | -0.1 | 1.46 | 0.13 | 0.14 | 1.4 | -0.46 | -0.93 | -0.67 | 0.5 | 1.34 | 0.52 | -0.25 | -0.63 | 0.91 | 0.41 | 0.07 | -0.5725 | 0.12 | -0.2175 | 0.340059 | 0.791045 | 0.633555 | | 1 | 0.996511 | 0.941132 |
| Slc9a6 | D3Z0Q9 | -0.24 | 0.1 | 0.28 | 0.38 | 0.35 | -1.01 | 0.1 | 0.07 | 1.11 | 1.13 | -0.31 | -0.67 | 1.36 | 1.95 | 0.11 | 0.18 | -0.2525 | 0.185 | 0.77 | 0.443529 | 0.688537 | 0.112469 | | 1 | 0.996511 | 0.714111 |
| Slc9a6 | A1L3P4 | 0.62 | 0.91 | 0.96 | 1.11 | 1.15 | 0.29 | 0.79 | 0.72 | 0.18 | 0.78 | 0.81 | 0.77 | 0.32 | 0.63 | 0.92 | 1.08 | -0.1625 | -0.265 | -0.1625 | 0.477588 | 0.249026 | 0.498657 | | 1 | 0.996511 | 0.899062 |
| Slc9a7 | Q8BLV3 | -0.02 | 0.07 | 0.76 | 0.67 | 0.11 | -0.04 | -0.02 | 0.35 | 0.29 | 0.89 | 0.4 | -0.02 | 1.13 | -0.19 | 1.24 | 0.77 | -0.27 | 0.02 | 0.3675 | 0.272465 | 0.944191 | 0.338975 | | 1 | 0.996755 | 0.847578 |
| Slc9a9 | Q8BZ00 | -0.49 | 0.47 | 0.15 | -0.65 | 0.63 | 0.59 | 0.27 | -0.74 | 0.23 | -0.01 | -0.3 | -0.05 | 0.32 | 0.47 | -0.03 | 0.02 | 0.3175 | 0.0975 | 0.325 | 0.427063 | 0.740417 | 0.298843 | | 1 | 0.996511 | 0.828257 |
| Slc9b2 | Q5BKR2 | -1.29 | 6.98 | -0.77 | -0.03 | 0.94 | 1.57 | -1.14 | 0.29 | 5.02 | 1.74 | -0.22 | 8.87 | -1.4 | 6.37 | -1.88 | 0.79 | -0.8075 | 2.63 | -0.2525 | 0.655898 | 0.303733 | 0.915981 | | 1 | 0.996511 | 0.986556 |
| Slco1a4 | Q9EP96 | -0.41 | -0.43 | 0.52 | 1.34 | -1.18 | 0.15 | -0.22 | 0.86 | 0.4 | 1.29 | 1.13 | -0.26 | 0.98 | -1.47 | 1.08 | 1.76 | -0.3525 | 0.385 | 0.3325 | 0.526638 | 0.461569 | 0.657352 | | 1 | 0.996511 | 0.948659 |
| Slco1a6 | Q99J94 | -1.99 | -0.81 | -0.46 | -2.96 | 0.03 | -1.32 | -0.4 | -1.89 | -2.43 | -1.64 | -0.12 | -1.34 | -0.89 | -0.86 | -1.37 | -1.25 | 0.66 | 0.1725 | 0.4625 | 0.3264 | 0.799661 | 0.403039 | | 1 | 0.996511 | 0.870698 |
| Slco1c1 | A0A0N4SU | 0.61 | 1.26 | 0.61 | 0.09 | 0.95 | 0.97 | -0.47 | -0.12 | 1.04 | 0.08 | 0.53 | 1.18 | 1.13 | 0.44 | -0.41 | -0.23 | -0.31 | 0.065 | -0.41 | 0.460881 | 0.84931 | 0.329207 | | 1 | 0.996511 | 0.844943 |
| Slco2b1 | Q3V1K7 | -1.64 | -2.54 | 0.16 | 0.36 | -2.33 | -2.33 | -0.99 | 0.35 | 0.87 | 2.12 | 0.13 | -1.74 | 0.69 | -0.13 | 0.62 | 0.79 | -0.41 | 1.26 | 1.4075 | 0.634598 | 0.213346 | 0.060671 | | 1 | 0.996511 | 0.661738 |
| Slco3a1 | Q8R3L5-2 | 1.32 | 0.83 | 0.97 | 0.76 | -0.18 | 1.74 | 0.43 | 0.68 | 2.01 | 1.06 | 0.92 | 1.14 | 2.47 | 0.54 | 0.44 | 0.7 | -0.3025 | 0.3125 | 0.0675 | 0.452835 | 0.292218 | 0.884969 | | 1 | 0.996511 | 0.979767 |
| Slirp | Q9D8T7 | 2.52 | 1.29 | 3.66 | 2.35 | 1.17 | 2.5 | 3.46 | 3.99 | 0.47 | -0.26 | 3.68 | 2.34 | 0.44 | 0.16 | 4.49 | 4.45 | 0.325 | -0.8975 | -0.07 | 0.649826 | 0.341141 | 0.951756 | | 1 | 0.996511 | 0.992476 |
| Slit1 | Q80TR4 | -0.18 | 0.16 | -0.26 | -0.3 | 0.3 | -0.4 | 0.01 | -0.29 | -0.03 | -0.25 | -0.41 | 0.24 | 0.02 | 0.47 | -0.44 | -0.04 | 0.05 | 0.0325 | 0.1475 | 0.816186 | 0.880042 | 0.556388 | | 1 | 0.996511 | 0.917092 |
| Slit2 | A0A140T81 | 0.89 | 1.04 | 0.46 | 0.36 | 1.14 | 0.96 | 0.4 | 0.3 | 0.79 | 0.63 | 0.63 | 1.09 | 1.14 | 0.93 | 0.62 | 0.73 | 0.0125 | 0.0975 | 0.1675 | 0.963002 | 0.672278 | 0.490552 | | 1 | 0.996511 | 0.897531 |
| Slit3 | Q9WVB4 | -0.23 | 0.32 | 0.53 | -0.02 | 0.84 | 0.29 | 1.55 | 0.34 | 0.1 | 1.38 | 0.77 | 0.11 | -0.04 | 0.56 | 0.91 | 1.81 | 0.605 | 0.44 | 0.66 | 0.093829 | 0.223441 | 0.131113 | | 1 | 0.996511 | 0.725983 |
| Slitrk1 | Q810C1 | 1.02 | 0.3 | 0.47 | 0.27 | 0.65 | 1.29 | 0.59 | 0.34 | -0.48 | -0.19 | 0.58 | 0.84 | -0.31 | -0.13 | 0.08 | 0.18 | 0.2025 | -0.3275 | -0.56 | 0.464012 | 0.362002 | 0.043697 | | 1 | 0.996511 | 0.658454 |
| Slitrk2 | Q810C0 | 0.08 | 0.44 | 1.2 | 0.96 | 0.18 | 0.5 | 1.06 | 1.34 | 0.64 | 0.27 | 1.33 | 0.36 | 0.14 | 0.31 | 0.6 | 1.17 | 0.1 | -0.02 | -0.115 | 0.775784 | 0.953453 | 0.735466 | | 1 | 0.996755 | 0.958635 |
| Slitrk3 | Q810B9 | 0.57 | 0.45 | 0.38 | 0.28 | 0.39 | 0.74 | 0.72 | 0.37 | 0.81 | 0.4 | 0.54 | 1.08 | 0.52 | 0.57 | -0.05 | 0.23 | 0.135 | 0.2875 | -0.1025 | 0.437145 | 0.189189 | 0.631626 | | 1 | 0.996511 | 0.941132 |
| Slitrk4 | Q810B8 | 0.38 | 0.44 | 0.5 | 0.35 | 0.84 | 0.5 | 0.83 | 0.2 | -0.51 | 0.26 | 0.49 | 0.43 | 0.18 | -0.37 | 0.29 | 0.56 | 0.175 | -0.25 | -0.2525 | 0.37664 | 0.340206 | 0.3048 | | 1 | 0.996511 | 0.831662 |
| Slitrk5 | Q810B7 | 1.14 | 1.18 | 1.34 | 0.98 | 1.37 | 1.46 | 1.19 | 1.27 | 0.56 | 0.65 | 1.45 | 1.36 | 0.2 | 0.95 | 1.15 | 1.16 | 0.1625 | -0.155 | -0.295 | 0.320478 | 0.558529 | 0.283357 | | 1 | 0.996511 | 0.822669 |

| | | | | | | | | | | | | | | | | | | | | | | | | | | | |
|---|---|---|---|---|---|---|---|---|---|---|---|---|---|---|---|---|---|---|---|---|---|---|---|---|---|---|---|
| Slk | O54988-2 | 0.94 | 0.32 | 0.14 | 0.37 | 0.04 | 0.58 | -0.19 | 0.2 | 0.4 | -0.02 | -0.19 | 0.61 | 0.29 | -0.3 | -0.11 | -0.56 | -0.285 | -0.2425 | -0.6125 | 0.271383 | 0.378752 | 0.048008 | | 1 | 0.996511 | 0.658454 |
| Slmap | F6WY34 | 0.07 | 0.26 | -0.02 | 0.45 | 1 | -0.2 | 0.81 | -1.19 | 0.5 | 0.52 | 0.32 | 0.37 | -0.28 | 1.05 | -0.37 | 1.14 | -0.085 | 0.2375 | 0.195 | 0.858861 | 0.211523 | 0.634148 | | 1 | 0.996511 | 0.941132 |
| Slmap | D3Z7V3 | 0.72 | 0.42 | 0.61 | 0.37 | 0.24 | 0.65 | 0.63 | 0.48 | 0.4 | 0.39 | 0.32 | 0.56 | 0.79 | -0.09 | 0.93 | 0.72 | -0.03 | -0.1125 | 0.0575 | 0.863376 | 0.51792 | 0.83086 | | 1 | 0.996511 | 0.973939 |
| Sltm | Q8CH25-2 | 1.71 | 0.5 | 0.67 | 0.74 | -0.67 | 1.05 | -0.52 | 1.2 | 0.69 | -0.78 | 0.71 | -0.04 | 1.77 | -0.12 | 1.13 | 0.08 | -0.64 | -0.76 | -0.19 | 0.241281 | 0.100894 | 0.698536 | | 1 | 0.996511 | 0.955729 |
| Slu7 | Q8BHJ9 | 0.56 | 0.32 | 1.3 | 0.66 | -0.71 | 1.12 | 1.11 | 1.39 | 1.19 | -1.88 | 1.56 | 1.21 | 2.62 | -0.69 | 0.98 | 0.64 | 0.0175 | -0.19 | 0.1775 | 0.971306 | 0.80104 | 0.784708 | | 1 | 0.996511 | 0.967642 |
| Smad2 | Q62432-2 | -1.12 | -1.32 | -1.59 | 0.12 | -1.39 | -1.32 | -1.37 | -1.68 | -0.47 | 2.42 | -2.02 | -0.9 | -0.04 | 0.41 | -2.14 | -2.04 | -0.4625 | 0.735 | 0.025 | 0.231298 | 0.431679 | 0.971197 | | 1 | 0.996511 | 0.992803 |
| Smad3 | Q8BUN5 | -1.41 | -1.82 | -1.27 | -1 | -1.3 | -0.81 | -1.54 | -1.57 | -1.25 | 0.03 | -2.12 | -1.57 | -0.63 | -0.78 | -0.95 | -2.11 | 0.07 | 0.1475 | 0.2575 | 0.78436 | 0.747559 | 0.492995 | | 1 | 0.996511 | 0.897531 |
| Smad4 | P97471 | -1.17 | -1.43 | -0.47 | -0.41 | -1.33 | -1.24 | -0.71 | -0.82 | -0.6 | -0.53 | -0.86 | -1.15 | -0.91 | -0.79 | -0.75 | -0.5 | -0.155 | 0.085 | 0.1325 | 0.602667 | 0.77529 | 0.645514 | | 1 | 0.996511 | 0.944749 |
| Smad5 | P97454 | -1.55 | -0.93 | 0.53 | 0.19 | -0.88 | -1.76 | 0.1 | 0.01 | 1.3 | 1.03 | -0.08 | -1.95 | 1.45 | 3.93 | 0.51 | 0.23 | -0.1925 | 0.515 | 1.97 | 0.746581 | 0.523762 | 0.047279 | | 1 | 0.996511 | 0.658454 |
| Smap | Q9R0P4 | -4.87 | -5.3 | -5.51 | -6.09 | -5.15 | -6.04 | -6.27 | -7.04 | -4.85 | -2.92 | -6.01 | -4.4 | -5.03 | -3.73 | -5.93 | -6.76 | -0.6825 | 0.8975 | 0.08 | 0.142605 | 0.178193 | 0.899782 | | 1 | 0.996511 | 0.983509 |
| Smap1 | Q91VZ6 | -0.08 | 0.06 | -0.03 | 0.25 | 0.05 | -0.05 | 0.21 | 0.36 | 0.05 | 0.04 | 0.11 | 0.15 | 0.34 | 0.15 | -0.01 | 0.04 | 0.0925 | 0.0375 | 0.08 | 0.587923 | 0.819423 | 0.670606 | | 1 | 0.996511 | 0.952558 |
| Smap2 | Q7TN29 | -0.35 | -0.5 | -0.52 | -0.45 | -0.8 | -0.68 | -0.66 | -0.91 | -0.52 | -0.93 | -0.52 | -0.39 | -0.45 | -0.86 | -0.79 | -1.02 | -0.3075 | -0.135 | -0.325 | 0.063768 | 0.472347 | 0.124981 | | 1 | 0.996511 | 0.721448 |
| Smarca1 | Q6PGB8 | 1.92 | 0.98 | 1 | 1.27 | 0.72 | 1.01 | 0.78 | 1.25 | 1.93 | 2.05 | 1 | 1.11 | 1.92 | 1.98 | 1.74 | 0.92 | -0.3525 | 0.23 | 0.3475 | 0.197751 | 0.508999 | 0.310623 | | 1 | 0.996511 | 0.834523 |
| Smarca2 | Q6DIC0 | -0.74 | -1.28 | -0.78 | -0.77 | -1.33 | -1.48 | -1 | -0.91 | -1.65 | -0.48 | -0.87 | -1.37 | -2.12 | -1.22 | -1.22 | -1.41 | -0.2875 | -0.2 | -0.6 | 0.200967 | 0.506805 | 0.052278 | | 1 | 0.996511 | 0.659923 |
| Smarca4 | A0A0R4J17 | -0.15 | -1.15 | -1.21 | -1.17 | -1.04 | -0.65 | -1.34 | -1.36 | -0.67 | -0.81 | -1.37 | -0.97 | -0.53 | -0.87 | -1.46 | -1.15 | -0.1775 | -0.035 | -0.0825 | 0.561806 | 0.908034 | 0.801843 | | 1 | 0.996744 | 0.971091 |
| Smarca5 | Q91ZW3 | 0.5 | 0.08 | 0.66 | 0.63 | 0.08 | 0.43 | 0.69 | 0.59 | -0.24 | -0.15 | 0.71 | 0.1 | 0.13 | -0.38 | 0.72 | 0.84 | -0.02 | -0.3625 | -0.14 | 0.925787 | 0.200367 | 0.662457 | | 1 | 0.996511 | 0.949645 |
| Smarcal1 | Q8BJL0 | -2.28 | -1.9 | -1.87 | -2.19 | -1.93 | -2.4 | -1.99 | -2.51 | -2.01 | -0.77 | -1.97 | -1.78 | -1.21 | -1.72 | -2.3 | -1.92 | -0.1475 | 0.4275 | 0.2725 | 0.483198 | 0.194307 | 0.331768 | | 1 | 0.996511 | 0.845601 |
| Smarcb1 | Q9Z0H3 | -1.44 | -1.63 | -1.85 | -1.69 | -1.42 | -1.47 | -1.84 | -2.02 | -1.85 | -1.35 | -1.97 | -1.67 | -1.53 | -1.24 | -2.44 | -2.41 | -0.035 | -0.0575 | -0.2525 | 0.861532 | 0.779704 | 0.442625 | | 1 | 0.996511 | 0.883549 |
| Smarcc1 | Q3UNN4 | -0.03 | -0.9 | 0.22 | 0.12 | -0.78 | -0.7 | 0.65 | 0.26 | -0.16 | 0.76 | 0.02 | -0.87 | -0.73 | -0.38 | 0.34 | 0.81 | 0.005 | 0.085 | 0.1575 | 0.990282 | 0.832809 | 0.704305 | | 1 | 0.996511 | 0.957084 |
| Smarcc2 | Q3UID0 | -1.14 | -2.22 | -1.49 | -1.72 | -2.23 | -1.74 | -2.01 | -1.81 | -1.69 | -1.46 | -1.52 | -1.82 | -1.34 | -1.98 | -1.45 | -1.69 | -0.305 | 0.02 | 0.0275 | 0.260785 | 0.93865 | 0.923229 | | 1 | 0.996755 | 0.987595 |
| Smarcd1 | Q61466 | -0.82 | -1.25 | -1.03 | -1.08 | -1.23 | -1.24 | -1.25 | -1.16 | -1.4 | -1.04 | -1.12 | -1.39 | -1.43 | -1.06 | -1.26 | -1.17 | -0.175 | -0.1925 | -0.185 | 0.282307 | 0.319302 | 0.349879 | | 1 | 0.996511 | 0.85327 |
| Smarcd3 | Q6P9Z1 | -1.47 | -1.62 | -1.94 | -1.43 | -1.99 | -2.2 | -1.78 | -1.94 | -1.14 | -0.42 | -1.98 | -1.69 | -1.08 | -1.22 | -2.14 | -1.81 | -0.3625 | 0.3075 | 0.0525 | 0.0782 | 0.395865 | 0.856845 | | 1 | 0.996511 | 0.976225 |
| Smarce1 | O54941 | -1.93 | -2.63 | -2.59 | -2.81 | -2.46 | -2.2 | -3.47 | -3.66 | -1.97 | -2.15 | -2.87 | -2.48 | -1.85 | -2.25 | -3.76 | -3.68 | -0.4575 | 0.1225 | -0.395 | 0.258452 | 0.67032 | 0.430134 | | 1 | 0.996511 | 0.879165 |
| Smc1a | Q9CU62 | 0.4 | -0.06 | -0.12 | 0.06 | -0.39 | 0.16 | -0.19 | 0.02 | 0.49 | 0.18 | -0.33 | -0.02 | 0.86 | 0.22 | 0.55 | 0.12 | -0.17 | 0.01 | 0.3675 | 0.405996 | 0.96628 | 0.154813 | | 1 | 0.996755 | 0.750763 |
| Smc1b | Q920F6 | 0.48 | -0.31 | 0.7 | 0.74 | 0.03 | -0.47 | -0.49 | -0.06 | -0.38 | -0.37 | 0.17 | -0.12 | 0.12 | 0.41 | 0.64 | 0.16 | -0.65 | -0.5775 | -0.07 | 0.044108 | 0.070527 | 0.809063 | | 1 | 0.996511 | 0.972098 |
| Smc2 | Q8CG48 | -0.27 | -0.02 | -1.39 | -1.6 | 1.01 | -0.02 | -0.3 | -0.79 | 0.09 | 0.55 | -1.33 | 0.07 | 1.52 | 0.64 | -0.78 | -0.84 | 0.795 | 0.665 | 0.955 | 0.142621 | 0.227171 | 0.15911 | | 1 | 0.996511 | 0.754492 |
| Smc3 | Q9CW03 | -0.11 | -0.62 | 0.07 | -0.06 | -0.64 | -0.33 | -0.31 | -0.15 | -0.41 | 0 | -0.31 | -0.73 | -0.14 | -0.29 | 0.57 | -0.06 | -0.1775 | -0.1825 | 0.2 | 0.409221 | 0.455982 | 0.462779 | | 1 | 0.996511 | 0.888267 |
| Smc5 | A0A286YD( | 1.32 | 2.5 | 0.63 | 0.82 | 3.04 | 1.99 | -0.55 | -0.99 | 2.61 | 9.36 | -1.36 | 1.19 | 2.49 | 3.57 | -0.98 | 0.02 | -0.445 | 1.6325 | -0.0425 | 0.643019 | 0.439512 | 0.966627 | | 1 | 0.996511 | 0.9927 |
| Smchd1 | Q6P5D8 | 1 | 0.14 | 0.69 | 0.28 | 0.28 | 0.41 | 1.2 | 0.15 | 0.24 | -0.27 | 0.85 | -0.04 | 0 | 0.3 | 1.08 | 1.5 | -0.0175 | -0.3325 | 0.1925 | 0.953987 | 0.30491 | 0.620152 | | 1 | 0.996511 | 0.93807 |
| Smcr8 | Q3UMB5 | 0.41 | 0.9 | 0.68 | 0.95 | 0.66 | 0.5 | 0.56 | 0.9 | 0.24 | 0.54 | 0.53 | 0.41 | 0.06 | 0.64 | 0.55 | 0.63 | -0.08 | -0.305 | -0.265 | 0.67522 | 0.144132 | 0.267225 | | 1 | 0.996511 | 0.818895 |
| Smdt1 | Q9DB10 | -2.11 | -1.26 | -0.53 | -0.95 | -0.29 | -1.25 | -0.26 | -0.74 | -3.01 | -1.37 | -0.19 | -2.41 | -2.8 | -2.36 | -0.9 | -0.97 | 0.5775 | -0.5325 | -0.545 | 0.160532 | 0.41681 | 0.330163 | | 1 | 0.996511 | 0.845493 |
| Smg1 | Q8BKX6 | -0.09 | -0.46 | 0.24 | 0.4 | -0.61 | -0.06 | -0.1 | 0.25 | -0.2 | 0.1 | 0.26 | -0.15 | 0.23 | -0.99 | 0.12 | 0.18 | -0.1525 | -0.02 | -0.1375 | 0.571967 | 0.934758 | 0.693384 | | 1 | 0.996755 | 0.955729 |
| Smg5 | Q6ZPY2 | -2.13 | -0.55 | -1.31 | -1.53 | -0.15 | -1.67 | -1.03 | -1.54 | -1.31 | -0.63 | -1.16 | -1.35 | -0.92 | -0.59 | -1.71 | -1.22 | 0.2825 | 0.2675 | 0.27 | 0.528265 | 0.461499 | 0.495963 | | 1 | 0.996511 | 0.89758 |
| Smg6 | P61406 | 0.01 | 0.47 | 0.43 | 0.02 | -0.04 | -0.15 | 0.03 | 0.02 | 0.04 | 0.44 | -0.01 | 0.07 | 0.32 | 0.45 | -0.29 | 0.18 | -0.2675 | -0.0975 | -0.0675 | 0.160008 | 0.640143 | 0.780866 | | 1 | 0.996511 | 0.9676 |
| Smg7 | Q5RJH6-3 | -0.01 | 0.55 | 0.64 | 0.02 | 0.21 | 0.28 | 0.63 | -0.01 | 0.1 | 0.5 | 0.09 | 0.27 | 0.54 | 0.05 | 0.63 | 0.81 | -0.0225 | -0.06 | 0.2075 | 0.923499 | 0.793627 | 0.439429 | | 1 | 0.996511 | 0.883205 |
| Smg8 | Q8VE18 | -0.44 | -0.19 | -0.27 | -0.34 | -0.14 | -0.35 | -0.57 | -0.44 | -0.02 | -0.37 | -0.37 | -0.07 | 0.12 | 0.03 | -0.21 | -0.81 | -0.065 | 0.1025 | 0.0925 | 0.692298 | 0.567045 | 0.712101 | | 1 | 0.996511 | 0.958014 |
| Smg9 | Q9DB90 | -0.86 | -0.68 | -0.55 | -0.43 | -0.55 | -0.83 | -0.68 | 0.56 | 0.54 | 1.03 | 2 | -0.63 | 0.21 | -0.09 | -0.41 | 2.78 | 0.255 | 1.365 | 1.2525 | 0.437309 | 0.025171 | 0.088683 | | 1 | 0.996511 | 0.691619 |
| Smim12 | Q78RX3 | 1.65 | 1.53 | 1.33 | 0.79 | 1.04 | 1.58 | 0.09 | 0.38 | 0.67 | 0.85 | 0.95 | 1.14 | 0.84 | 1.17 | 0.71 | 0.64 | -0.5525 | -0.4225 | -0.485 | 0.158519 | 0.107263 | 0.084119 | | 1 | 0.996511 | 0.690427 |
| Smim13 | E9Q942 | -0.24 | -0.55 | 0.1 | 0.26 | -1.11 | 0.63 | -1.51 | -0.95 | 2.42 | -2.09 | -0.32 | -0.65 | 3.76 | -2.24 | 0.32 | 0.42 | -0.6275 | -0.0525 | 0.6725 | 0.202465 | 0.951552 | 0.548563 | | 1 | 0.996755 | 0.912757 |
| Smim18 | J3QNP2 | 0.94 | 0.88 | 1.16 | 1.14 | 1.05 | 0.55 | 0.14 | 0.87 | -0.17 | 0.88 | 0.97 | 0.88 | -0.43 | 0.28 | 1.49 | 0.73 | -0.3775 | -0.39 | -0.5125 | 0.129184 | 0.200591 | 0.216521 | | 1 | 0.996511 | 0.787447 |
| Smim20 | D3Z7Q2 | 0.14 | 0.85 | 0.43 | 1.16 | 0.61 | 0.86 | 1.04 | 0.63 | 0.43 | -0.02 | 1.4 | 0.91 | 0.56 | 0.8 | 0.62 | 0.31 | 0.14 | 0.035 | -0.0725 | 0.589133 | 0.924257 | 0.789358 | | 1 | 0.996755 | 0.968677 |
| Smim26 | Q3V460 | 0.72 | 0.91 | 1.2 | 0.13 | 0.36 | 1.41 | -0.13 | 0.28 | -0.13 | -1.58 | 0.83 | 0.6 | 1.87 | -3.96 | 0.82 | 0.63 | -0.26 | -0.81 | -0.9 | 0.498053 | 0.161876 | 0.450466 | | 1 | 0.996511 | 0.884817 |
| Smim4 | A0A0R3P9( | 0.95 | 1.64 | 1.14 | 1.21 | 1.09 | 0.73 | 1.13 | 0.25 | 0.33 | -0.12 | 1.24 | -1.09 | -0.25 | 1.41 | 0.9 | 0.97 | -0.435 | -1.145 | -0.4775 | 0.121563 | 0.036692 | 0.223619 | | 1 | 0.996511 | 0.787447 |
| Smim7 | F8WIU9 | -4.07 | -3.11 | -3.75 | -4.72 | -3.83 | -3.01 | -5.47 | -4.24 | -3.73 | -1.52 | -4.09 | -4.75 | -0.89 | -3.04 | -3.59 | -4.2 | -0.225 | 0.39 | 0.9825 | 0.689323 | 0.583572 | 0.195459 | | 1 | 0.996511 | 0.78004 |
| Smim8 | Q9CQQ0 | 0.27 | 0.4 | -0.34 | -0.15 | 0.71 | 0.03 | -0.24 | 0.01 | -0.23 | 0.05 | -0.05 | 0.4 | 0.02 | 0.35 | -0.11 | 0 | 0.0825 | -0.0025 | 0.02 | 0.762756 | 0.991826 | 0.933368 | | 1 | 0.999285 | 0.989767 |
| Smn1 | P97801 | 0.01 | 0.39 | -0.48 | -0.59 | 0.39 | -0.19 | 0.62 | 0.35 | -0.94 | -0.25 | -0.75 | 0.04 | -0.12 | 0.11 | -0.28 | 0.14 | 0.46 | -0.3075 | 0.13 | 0.134775 | 0.349831 | 0.634463 | | 1 | 0.996511 | 0.941161 |
| Smndc1 | Q8BGT7 | -0.71 | -0.77 | -0.81 | 0.02 | -1.72 | -1.5 | -1.29 | 0.48 | -1.59 | -0.35 | -0.98 | -1.63 | -1.08 | -1.17 | -0.36 | -0.64 | -0.44 | -0.57 | -0.245 | 0.388827 | 0.133108 | 0.4085 | | 1 | 0.996511 | 0.874356 |
| Smoc1 | D3Z3A3 | -0.28 | 0.19 | 0.52 | 0.76 | -1.37 | -1.5 | 0.25 | 1.17 | -1.29 | 1.57 | 1.57 | -0.49 | 0.52 | -0.31 | 1.31 | 1.01 | -0.66 | 0.0425 | 0.335 | 0.306308 | 0.951092 | 0.416063 | | 1 | 0.996755 | 0.874945 |
| Smpd1 | Q04519 | -1.26 | -0.92 | -1.63 | -1.65 | -0.49 | -1.18 | -0.93 | -1.71 | -0.81 | -1.21 | -1.17 | -1.16 | -0.9 | -0.63 | -2 | -1.4 | 0.2875 | 0.2775 | 0.1325 | 0.357893 | 0.246224 | 0.703065 | | 1 | 0.996511 | 0.956859 |
| Smpd2 | O70572 | -0.14 | 0.49 | -0.24 | -0.06 | 0.66 | 0.19 | 0.66 | 0.07 | -0.29 | 0.01 | 0.2 | 0.26 | -0.04 | -0.23 | 0.4 | 0.61 | 0.3825 | 0.0325 | 0.1725 | 0.138694 | 0.889873 | 0.535934 | | 1 | 0.996511 | 0.912212 |
| Smpd3 | Q9JJY3 | 1.16 | 0.76 | 1.22 | 1.37 | -0.1 | 0.94 | 0.51 | 1.13 | 1.19 | 1.51 | 0.85 | 1.19 | 1.56 | -0.01 | 1.2 | 0.88 | -0.5075 | 0.0575 | -0.22 | 0.118407 | 0.796794 | 0.541234 | | 1 | 0.996511 | 0.912212 |
| Smpd4 | Q6ZPR5 | 0.51 | 0.07 | -0.05 | -0.07 | -0.03 | 0.54 | 0.31 | 0.14 | 0.3 | -0.63 | -0.2 | -0.14 | 0.76 | 0.53 | 0.74 | 0.71 | 0.125 | -0.2825 | 0.57 | 0.555691 | 0.284855 | 0.020829 | | 1 | 0.996511 | 0.621926 |
| Smpdl3a | P70158 | -1.09 | 0.22 | -1.27 | -1.13 | 1.1 | -0.07 | -0.7 | -1.11 | -0.46 | -1.14 | -0.55 | -0.65 | -0.07 | -0.58 | -1.54 | -1.02 | 0.6225 | 0.1175 | 0.015 | 0.271771 | 0.750144 | 0.972989 | | 1 | 0.996511 | 0.992865 |
| Smpdl3b | P58242 | 0.94 | 0.82 | 0.23 | 1.2 | 0.11 | 0.35 | 1.16 | 1.07 | 0.81 | 1.45 | 0.11 | 0.61 | 0.35 | 0.19 | 1.26 | 0.94 | -0.125 | -0.0525 | -0.1125 | 0.700426 | 0.877295 | 0.732127 | | 1 | 0.996511 | 0.958635 |
| Sms | P97355 | -0.51 | -0.53 | -0.47 | -0.62 | -0.55 | -0.65 | -0.6 | -0.45 | -0.28 | -0.35 | -0.74 | -0.63 | -0.45 | -0.13 | -0.74 | -0.9 | -0.03 | 0.0325 | -0.0225 | 0.833562 | 0.857054 | 0.919243 | | 1 | 0.996511 | 0.986868 |
| Smu1 | Q3UKJ7 | 0.4 | -0.68 | -0.34 | -0.45 | -0.96 | 0.16 | -0.13 | 0.29 | -0.96 | -0.98 | -0.16 | -1.08 | -0.03 | -1.16 | 0.18 | -0.35 | 0.1075 | -0.5275 | -0.0725 | 0.759817 | 0.123887 | 0.844019 | | 1 | 0.996511 | 0.976225 |
| Smug1 | Q6P5C5 | -0.49 | 0.5 | -0.34 | -0.68 | -0.03 | -0.87 | -0.32 | -0.77 | -0.29 | -0.78 | 0.11 | -0.42 | -0.68 | 0.12 | -0.96 | -0.07 | -0.245 | -0.0925 | -0.145 | 0.450566 | 0.772298 | 0.687217 | | 1 | 0.996511 | 0.955729 |
| Smurf2 | A2A5Z6 | 0.17 | 0.12 | 0.21 | 0.45 | 0.3 | 0.14 | -0.15 | 0.09 | 0.07 | 0.79 | 0.03 | 0.22 | 0.45 | -0.25 | 0.31 | 0.06 | -0.1425 | 0.04 | -0.095 | 0.413554 | 0.858879 | 0.668424 | | 1 | 0.996511 | 0.95204 |
| Smyd2 | Q8R5A0 | -1.48 | -0.8 | -0.86 | -0.36 | -0.5 | -1.48 | -0.37 | -1.41 | -1.51 | -0.67 | -0.3 | -1.65 | -0.65 | -0.76 | -3.04 | -2.58 | -0.065 | -0.1575 | -0.8825 | 0.855632 | 0.683261 | 0.16779 | | 1 | 0.996511 | 0.759484 |
| Smyd3 | Q9CWR2 | -0.13 | -0.71 | -0.58 | -0.16 | -0.45 | -0.79 | -0.88 | -0.55 | -1.59 | -0.62 | -0.89 | -0.43 | -0.47 | -1 | -0.77 | -0.73 | -0.2725 | -0.4875 | -0.3475 | 0.211221 | 0.130782 | 0.153777 | | 1 | 0.996511 | 0.750763 |
| Smyd5 | Q3TYX3 | 0.32 | -0.26 | -1.84 | -2.29 | -0.14 | 0.15 | -2.21 | -1.94 | -0.77 | -1.64 | -1.72 | 0.48 | 0.44 | -1.86 | -0.26 | -2.5 | -0.0175 | 0.105 | -0.0275 | 0.982176 | 0.8858 | 0.973606 | | 1 | 0.996511 | 0.992946 |
| Snap23 | Q9D3L3 | 1.5 | 0.84 | 0.29 | 0.45 | 0.73 | 1.51 | 0.57 | 0.5 | 0.58 | 0.19 | 0.26 | 1.47 | 0.11 | 0.95 | 0.23 | 0.06 | 0.0575 | -0.145 | -0.4325 | 0.866957 | 0.706633 | 0.225005 | | 1 | 0.996511 | 0.787447 |
| Snap25 | P60879 | 2.31 | 1.61 | 1.35 | 1.72 | 2.6 | 2.13 | 2.24 | 2.42 | 1.22 | 0.71 | 1.86 | 2.68 | 0.52 | -0.17 | 1.05 | 1.04 | 0.6 | -0.13 | -1.1375 | 0.033408 | 0.769984 | 0.00961 | | 1 | 0.996511 | 0.563386 |
| Snap25 | P60879-2 | 0.72 | 0.12 | 0.72 | 1.44 | 0.55 | 0.42 | 1.41 | 1.47 | 1.42 | 1.95 | 1.59 | 1.74 | 0.64 | 1.51 | 1.5 | 1.14 | 0.2125 | 0.925 | 0.4475 | 0.568362 | 0.012423 | 0.209818 | | 1 | 0.996511 | 0.786218 |
| Snap29 | Q9ERB0 | -1.24 | -0.69 | -1.58 | -1.3 | -0.64 | -0.84 | -0.8 | -0.87 | -0.42 | -1.84 | -0.56 | -0.37 | -0.2 | -0.24 | -1.15 | -1.45 | 0.415 | 0.405 | 0.4425 | 0.083921 | 0.304938 | 0.242525 | | 1 | 0.996511 | 0.798545 |
| Snap47 | Q8R570 | -0.62 | -0.27 | -0.26 | -0.13 | 0.13 | -0.44 | -0.26 | -0.19 | -0.28 | 0.23 | -0.02 | -0.24 | 0.63 | 0.5 | 0.01 | 0.06 | 0.13 | 0.2425 | 0.62 | 0.50965 | 0.254389 | 0.023804 | | 1 | 0.996511 | 0.625216 |
| Snap91 | Q61548-3 | 0.2 | 0.44 | 0.17 | 0.38 | 0.6 | -0.19 | 0.31 | 0.17 | 0.13 | 0.04 | 0.25 | 0.17 | -0.33 | 0.78 | -0.03 | -0.08 | -0.075 | -0.15 | -0.2125 | 0.717499 | 0.375045 | 0.444534 | | 1 | 0.996511 | 0.884411 |
| Snapin | Q9Z266 | -0.39 | -0.2 | -0.14 | 0.3 | -0.08 | -0.16 | -0.14 | -0.02 | -0.4 | 0.96 | -0.45 | -0.38 | -0.64 | 0.44 | -0.04 | 0.6 | 0.0075 | 0.04 | 0.1975 | 0.968268 | 0.912061 | 0.542527 | | 1 | 0.996744 | 0.912212 |
| Snca | O55042 | 1.61 | 0.86 | 0.96 | 0.71 | 0.28 | 1.56 | 0.63 | 0.86 | 1.36 | -0.09 | 0.85 | 1.89 | 0.83 | 0.18 | 0.26 | 0.34 | -0.2025 | -0.0325 | -0.6325 | 0.539669 | 0.940997 | 0.041575 | | 1 | 0.996755 | 0.658454 |
| Sncb | Q91ZZ3 | -0.32 | 0.14 | 1.33 | 2.4 | -0.29 | -0.43 | 1.3 | 2.9 | -0.73 | 0.22 | 1.81 | -0.16 | -0.44 | -0.17 | 0.37 | 0.57 | -0.0175 | -0.6025 | -0.805 | 0.984353 | 0.426488 | 0.203136 | | 1 | 0.996511 | 0.782817 |
| Sncg | Q9Z0F7 | 0.32 | 0.29 | 0.1 | 0.43 | -0.02 | -0.8 | 0.09 | -0.22 | 0.13 | 0.98 | 0.02 | 0.08 | 0.49 | 0.8 | 0.85 | 0.56 | -0.5225 | 0.0175 | 0.39 | 0.046013 | 0.945749 | 0.065657 | | 1 | 0.996755 | 0.664936 |
| Snd1 | Q78PY7 | -1 | -0.97 | -0.98 | -0.9 | 0.12 | -0.83 | 0.09 | -0.55 | -0.91 | -0.72 | -1.04 | -0.51 | -0.42 | -0.01 | 0.56 | 0.43 | 0.67 | 0.1675 | 1.1025 | 0.023926 | 0.369242 | 0.001961 | | 1 | 0.996511 | 0.419079 |
| Snf8 | Q9CZ28 | -0.27 | -0.27 | 0.19 | 0.13 | -0.28 | -0.47 | 0.61 | 0.38 | -0.08 | -0.08 | 0.14 | -0.16 | -0.27 | 0.32 | -0.21 | 0.29 | 0.115 | 0.01 | 0.0875 | 0.690643 | 0.958892 | 0.716369 | | 1 | 0.996755 | 0.958312 |
| Snip1 | Q8BIZ6 | -0.41 | -2.4 | 0.28 | -1.09 | -4.38 | 0.56 | -0.26 | 0 | 1.44 | -0.1 | -0.93 | -2.51 | 4.12 | -4.45 | 0 | 0.23 | -0.115 | 0.38 | 0.88 | 0.919315 | 0.67534 | 0.593343 | | 1 | 0.996511 | 0.929843 |
| Snn | P61807 | -0.2 | -0.07 | 0.19 | 0.27 | 0.34 | -0.35 | 0.51 | 0.51 | -0.26 | 0.65 | 0.38 | 0.84 | -1.09 | 1.13 | -0.22 | 0.21 | 0.205 | 0.355 | -0.04 | 0.414132 | 0.2221 | 0.928907 | | 1 | 0.996511 | 0.989604 |
| Snph | Q80U23-2 | 2.9 | 2.34 | 0.73 | 0.78 | 2.5 | 3.05 | 0.28 | 0.71 | 1.06 | 0.98 | 0.61 | 3.16 | 1.75 | 0.05 | 0.76 | 1.05 | -0.0525 | -0.235 | -0.785 | 0.946613 | 0.746116 | 0.212221 | | 1 | 0.996511 | 0.786366 |
| Snrk | Q8VDU5 | 0.02 | 0.69 | 0.91 | 0.85 | 0.75 | 0.2 | 0.57 | 0.65 | 0.56 | 0.94 | 0.9 | 0.55 | 0.2 | 1.23 | 0.85 | 0.88 | -0.075 | 0.12 | 0.1725 | 0.764041 | 0.636649 | 0.577766 | | 1 | 0.996511 | 0.927198 |
| Snrnp200 | Q6P4T2 | 0.6 | -0.56 | -0.69 | -0.75 | -0.8 | 0.03 | -0.71 | -0.21 | -0.96 | -1.29 | -0.87 | -0.96 | 0.01 | -0.65 | -0.19 | -0.84 | -0.0725 | -0.67 | -0.0675 | 0.840409 | 0.068967 | 0.854491 | | 1 | 0.996511 | 0.976225 |
| Snrnp27 | Q8K194 | 0.06 | 0.13 | -0.09 | -0.28 | -0.16 | 0.25 | 0 | 0.06 | 2.52 | -1.3 | -0.57 | -0.34 | 3.63 | -1.55 | -0.02 | -0.26 | 0.0825 | 0.1225 | 0.495 | 0.637206 | 0.872692 | 0.622229 | | 1 | 0.996511 | 0.938628 |
| Snrnp40 | Q6PE01 | 0.89 | -0.13 | -0.02 | 0 | -0.2 | 0.28 | 0.28 | 1.16 | -0.79 | -1.05 | 0.04 | -0.84 | 0.19 | -0.6 | 1.03 | 1.15 | 0.195 | -0.845 | 0.2575 | 0.585425 | 0.03069 | 0.567608 | | 1 | 0.996511 | 0.921806 |
| Snrnp70 | Q62376 | 1.15 | -0.55 | -0.51 | -0.51 | -0.09 | 0.18 | 0.12 | 0.4 | -0.59 | -0.1 | -0.62 | -0.4 | 0.29 | -0.07 | 1.5 | 0.13 | 0.2575 | -0.3225 | 0.5675 | 0.529993 | 0.4441 | 0.282907 | | 1 | 0.996511 | 0.822595 |
| Snrpa | Q62189 | 1.48 | -1.81 | -0.71 | -1.21 | -1.55 | 0.89 | 0.22 | -0.78 | -1.41 | -1.41 | -1 | -0.53 | -0.02 | -1.94 | 1.03 | -0.1 | 0.2575 | -0.525 | 0.305 | 0.750599 | 0.448109 | 0.720683 | | 1 | 0.996511 | 0.958398 |
| Snrpa1 | P57784 | 0.23 | -0.97 | -1.04 | -1.16 | -1.46 | -0.71 | -1.16 | -0.51 | -1.57 | -1.27 | -1.44 | -1.51 | -0.51 | -0.08 | -0.65 | -1.51 | -0.225 | -0.7125 | 0.0475 | 0.548034 | 0.055169 | 0.910229 | | 1 | 0.996511 | 0.986458 |
| Snrpb | P27048 | -0.38 | -1.05 | -0.79 | -0.85 | -0.22 | -0.6 | -0.49 | -0.45 | -0.92 | -0.54 | -0.78 | -1.31 | -0.69 | -0.03 | -0.55 | -0.73 | 0.3275 | -0.12 | 0.2675 | 0.123119 | 0.62094 | 0.297993 | | 1 | 0.996511 | 0.827516 |
| Snrpb2 | Q9CQI7 | 0.22 | -0.97 | -0.99 | -0.92 | -1.22 | -0.69 | -1.75 | -0.91 | -0.72 | -0.34 | -1.2 | -1.09 | 0.82 | -0.76 | -0.23 | -1.63 | -0.4775 | -0.1725 | 0.215 | 0.205896 | 0.621883 | 0.695006 | | 1 | 0.996511 | 0.955729 |
| Snrpc | Q62241 | -0.82 | -1.74 | -0.94 | -1.38 | -1.5 | -1.55 | -1.12 | -0.72 | -1.6 | 0.37 | -1.14 | -1.76 | -0.33 | -0.82 | -0.32 | -1.18 | -0.0025 | 0.1875 | 0.5575 | 0.993044 | 0.704758 | 0.096852 | | 1 | 0.996511 | 0.698156 |
| Snrpd1 | P62315 | -2.08 | -3.42 | -1.1 | -0.62 | -1.63 | -3.26 | -0.54 | -0.53 | -3.46 | -0.58 | -0.87 | -4.26 | -2.2 | -1.17 | -0.36 | -0.82 | 0.315 | -0.4875 | 0.6675 | 0.696619 | 0.627136 | 0.330149 | | 1 | 0.996511 | 0.845493 |
| Snrpd2 | P62317 | -0.04 | -0.81 | -1.71 | -1.36 | -0.59 | -0.29 | -0.89 | -1.11 | -1.46 | -2.21 | -1.28 | -1.16 | -0.96 | -0.33 | -1.06 | -1.62 | 0.26 | -0.5475 | -0.0125 | 0.504473 | 0.207472 | 0.976698 | | 1 | 0.996511 | 0.993727 |
| Snrpd3 | P62320 | -0.98 | -1.16 | -1.51 | -1.45 | -0.77 | -1.12 | -1.16 | -1.2 | -1.5 | -1.2 | -1.49 | -1.78 | -1.32 | -0.25 | -1.18 | -1.39 | 0.2125 | -0.2175 | 0.24 | 0.292314 | 0.322202 | 0.439009 | | 1 | 0.996511 | 0.883205 |
| Snrpe | P62305 | -0.55 | -1.18 | -0.27 | -0.54 | -0.95 | -1.01 | -0.05 | 0.18 | -2.07 | 2.86 | -0.01 | -1.65 | -2.05 | -1.3 | 0.32 | -0.14 | 0.1775 | 0.4175 | -0.1575 | 0.612402 | 0.68326 | 0.76729 | | 1 | 0.996511 | 0.96483 |
| Snrpf | P62307 | -1.27 | -2.57 | -3.46 | -3.4 | -2.1 | -1.81 | -2.93 | -2.99 | -3.1 | -2.99 | -3.43 | -2.65 | -2.22 | -1.91 | -2.78 | -3.6 | 2.18E-01 | -0.3675 | 0.0475 | 0.689134 | 0.467678 | 0.934641 | | 1 | 0.996511 | 0.989767 |
| Snrpg | P62309 | 0.8 | -0.78 | -1.27 | -1.34 | -0.51 | -0.01 | -0.84 | -0.68 | -0.51 | -0.9 | -1.21 | -0.47 | -0.06 | 0.13 | -0.19 | -0.96 | 0.1375 | -0.125 | 0.3775 | 0.779109 | 0.799637 | 0.469522 | | 1 | 0.996511 | 0.890206 |
| Snta1 | A2AKD7 | 0.82 | 0.37 | 0.54 | 0.52 | 0.65 | 0.74 | 0.56 | 0.39 | 0.5 | 0.42 | 0.65 | 0.83 | 0.43 | 0.48 | 0.65 | 0.53 | 0.0225 | 0.0375 | -0.04 | 0.895581 | 0.842391 | 0.830378 | | 1 | 0.996511 | 0.973939 |
| Sntb1 | Q99L88 | 1.51 | 1.03 | 0.91 | 0.81 | 0.71 | 0.99 | 0.68 | 0.71 | 0.99 | 0.91 | 0.7 | 1.22 | 0.57 | 0.79 | 0.84 | 0.56 | -0.2925 | -0.11 | -0.375 | 0.174215 | 0.625445 | 0.11747 | | 1 | 0.996511 | 0.718609 |
| Sntb2 | Q61235 | 0.27 | 0.38 | 1.2 | 1.15 | 0.78 | -0.01 | 0.9 | 1 | -0.24 | 1.28 | 1.02 | 0.28 | -0.22 | 0.27 | 0.91 | 1 | -0.0825 | -0.165 | -0.26 | 0.801738 | 0.685781 | 0.489252 | | 1 | 0.996511 | 0.897531 |
| Snu13 | Q9D0T1 | -0.78 | -1.55 | -2.4 | -3.16 | -1.96 | -0.33 | -1.67 | -1.43 | 0.77 | -0.38 | -1.22 | -1.31 | 3.08 | -3.08 | 2.45 | -2.47 | 0.625 | 1.4375 | 1.9675 | 0.292441 | 0.049996 | 0.212126 | | 1 | 0.996511 | 0.786366 |
| Snupn | Q80W37 | -0.96 | -0.52 | -0.89 | -0.95 | -0.16 | -0.78 | -0.88 | -1.04 | -0.68 | -0.64 | -0.35 | -0.77 | -0.46 | -0.03 | -0.68 | -1.09 | 0.115 | 0.22 | 0.265 | 0.628522 | 0.272744 | 0.338853 | | 1 | 0.996511 | 0.847578 |
| Snw1 | A0A0B4J1E | 0.44 | -0.7 | -0.32 | -0.32 | -1.23 | -0.58 | -0.48 | -0.05 | -0.85 | -0.15 | -0.45 | -1.15 | -0.45 | -0.17 | -0.02 | -0.65 | -0.36 | -0.425 | -0.0975 | 0.294451 | 0.212335 | 0.739988 | | 1 | 0.996511 | 0.958796 |
| Snx1 | Q6NZD2 | 0.41 | 0.32 | 0.9 | 0.9 | 0.37 | -0.01 | 0.86 | 1.29 | -0.39 | 0.62 | 0.89 | 0.16 | -0.57 | 0.09 | 0.88 | 0.77 | -0.005 | -0.3125 | -0.34 | 0.987402 | 0.343779 | 0.363169 | | 1 | 0.996511 | 0.858475 |
| Snx10 | D3YW62 | -0.62 | -1.19 | -1.21 | -1.13 | -0.89 | -1.03 | -0.72 | -0.96 | -0.69 | 2.14 | -0.99 | -0.25 | -0.21 | -0.47 | -1.26 | -0.92 | 0.1375 | 1.09 | 0.3225 | 0.481398 | 0.126958 | 0.282278 | | 1 | 0.996511 | 0.822325 |
| Snx11 | A2A880 | -0.46 | -0.48 | -0.1 | -0.46 | -0.37 | -0.62 | -0.08 | -0.49 | 0.46 | -0.08 | -0.03 | -0.75 | 0.78 | -0.61 | -0.3 | -0.75 | -0.015 | 0.275 | 0.155 | 0.936187 | 0.336343 | 0.663632 | | 1 | 0.996511 | 0.949825 |
| Snx12 | Q6ZWQ5 | 0.66 | 0.1 | 0.15 | -0.26 | 0.51 | 0.48 | -0.59 | -0.07 | -0.1 | -0.28 | -0.06 | 0.27 | -0.12 | 0.36 | -0.23 | -0.46 | -0.08 | -0.205 | -0.275 | 0.800333 | 0.414587 | 0.335886 | | 1 | 0.996511 | 0.847578 |
| Snx13 | Q6PHS6 | 0.24 | 0.47 | 0.47 | 0.21 | 0.6 | 0.42 | 0.6 | 0.36 | 0.11 | 0.19 | 0.54 | 0.27 | 0.2 | -0.03 | 0.75 | 0.57 | 0.1475 | -0.07 | 0.025 | 0.364169 | 0.701278 | 0.914481 | | 1 | 0.996511 | 0.986458 |
| Snx14 | Q8BHY8 | 0.29 | 0.49 | 0.14 | -0.13 | 1.07 | 0.4 | 0.58 | 0.29 | -0.28 | -1.14 | 0.2 | 0.51 | -0.35 | 0.32 | 0.17 | 0.56 | 0.3875 | -0.375 | -0.0225 | 0.125179 | 0.326369 | 0.931278 | | 1 | 0.996511 | 0.989713 |
| Snx15 | Q91WE1 | -0.25 | -0.12 | -0.24 | -0.05 | -0.29 | -0.29 | -0.72 | -0.72 | 0.03 | 0.51 | -0.26 | -0.16 | -0.34 | 0.03 | -0.26 | -0.15 | -0.34 | 0.195 | -0.015 | 0.084972 | 0.379725 | 0.934218 | | 1 | 0.996511 | 0.989767 |
| Snx16 | Q8C080 | -0.02 | 0.18 | 0.6 | 0.57 | -0.08 | 0.04 | 0.34 | 0.54 | -0.11 | 0.41 | 0.37 | 0.18 | -0.3 | 0.44 | 0.12 | 0.53 | -0.1225 | -0.12 | -0.135 | 0.593201 | 0.59896 | 0.615701 | | 1 | 0.996511 | 0.93691 |
| Snx17 | Q8BVL3 | -1.4 | -1.16 | -1.62 | -1.3 | -1.02 | -1.26 | -0.82 | -1.2 | -0.86 | -1.29 | -1.3 | -1.2 | -1.54 | -0.08 | -1.18 | -1.3 | 0.295 | 0.2075 | 0.345 | 0.132472 | 0.30305 | 0.32312 | | 1 | 0.996511 | 0.840788 |
| Snx18 | Q8C788 | -0.67 | -0.72 | -0.46 | -0.05 | -0.5 | -0.82 | -0.37 | -0.22 | -0.28 | 0 | -0.05 | -0.58 | -0.1 | -0.25 | -0.65 | -0.63 | -0.0025 | 0.2475 | 0.0675 | 0.990992 | 0.303728 | 0.781332 | | 1 | 0.996511 | 0.967642 |
| Snx19 | Q6P4T1 | -0.09 | 0.41 | 0.34 | 0.34 | 0.49 | -0.24 | 0.02 | 0.36 | 0.27 | 0.46 | 0.22 | -0.02 | -0.03 | 1.16 | 0.56 | 0.48 | -0.0925 | -0.0175 | 0.2925 | 0.680086 | 0.930375 | 0.323017 | | 1 | 0.996755 | 0.840788 |
| Snx2 | Q9CWK8 | 0.79 | 0.76 | 0.95 | 1.21 | 0.74 | 0.59 | 0.37 | 1.17 | 1.14 | 1.27 | 1 | 0.86 | 0.91 | 0.84 | 0.56 | 0.58 | -0.21 | 0.14 | -0.205 | 0.354765 | 0.471915 | 0.324054 | | 1 | 0.996511 | 0.84194 |
| Snx21 | G3UYX2 | -1.89 | -1.35 | -0.98 | -1.02 | -1.54 | -2.83 | -1.1 | -1.95 | -1.19 | -1 | -1.39 | -2.38 | -1.98 | -0.83 | -1.63 | -1.52 | -0.545 | -0.18 | -0.18 | 0.196812 | 0.621615 | 0.582649 | | 1 | 0.996511 | 0.927734 |
| Snx24 | Q9CRB0 | -1.27 | -0.94 | 0.1 | -1.16 | -0.78 | -1.61 | -0.15 | -0.53 | -1.44 | -2.21 | -0.1 | -1.03 | -1.91 | -2.34 | -0.04 | 0.01 | 0.05 | -0.3775 | -0.2525 | 0.903708 | 0.459394 | 0.690031 | | 1 | 0.996511 | 0.955729 |
| Snx25 | E9Q1K0 | 0.14 | 0.23 | 0.02 | -0.17 | 0.26 | -0.1 | 0.02 | -0.12 | 0.09 | -0.22 | -0.15 | 0.03 | -0.05 | -0.09 | 0.05 | 0.25 | -0.04 | -0.1175 | -0.015 | 0.817427 | 0.518633 | 0.937409 | | 1 | 0.996511 | 0.989767 |
| Snx27 | Q3UHD6-2 | -0.25 | -0.07 | 0.09 | 0.39 | -0.19 | -0.28 | 0.35 | 0.26 | 0.15 | 0.14 | 0.23 | -0.09 | -0.32 | 0.34 | -0.13 | 0.16 | -0.005 | 0.0675 | -0.0275 | 0.982516 | 0.7377 | 0.908498 | | 1 | 0.996511 | 0.986458 |
| Snx29 | Q9D3S3 | 0.05 | 0.16 | 0 | 0.05 | 0.44 | 0.27 | 0.12 | 0.36 | 0.74 | 0 | 0.23 | -0.31 | 1.09 | 0.19 | 0.16 | 0.68 | 0.2325 | 0.1 | 0.465 | 0.149411 | 0.687976 | 0.095792 | | 1 | 0.996511 | 0.698156 |
| Snx3 | Q78ZM0 | 1 | 0.56 | 0.77 | -0.18 | 0.76 | 0.89 | 1.31 | 0.49 | 0.74 | -0.18 | 0.58 | 0.86 | 0.46 | 0.99 | 0.72 | 0.91 | 0.325 | -0.0375 | 0.2325 | 0.301465 | 0.912439 | 0.440262 | | 1 | 0.996744 | 0.883549 |
| Snx30 | Q8CE50 | 0.47 | 0.7 | 0.76 | 0.48 | 0.66 | 0.36 | 0.69 | 0.64 | 0.35 | 0.4 | 0.8 | 0.43 | -0.16 | 1.09 | 0.6 | 0.38 | -0.015 | -0.1075 | -0.125 | 0.927349 | 0.568914 | 0.665139 | | 1 | 0.996511 | 0.949825 |
| Snx32 | Q80ZJ7 | 0.22 | -0.02 | -1.19 | -1.28 | -0.56 | -0.47 | -1.24 | -1.39 | -0.35 | -0.76 | -1.48 | -0.19 | -0.34 | -0.49 | -1.54 | -1.64 | -0.3475 | -0.1275 | -0.435 | 0.422685 | 0.779416 | 0.380332 | | 1 | 0.996511 | 0.860984 |
| Snx33 | Q4VAA7 | -0.02 | -0.76 | 1 | 0.8 | -0.69 | 0.55 | 0.9 | 0.67 | 0.77 | 1.41 | 0.69 | 0.46 | 0.06 | 1.23 | 0.23 | 0.61 | 0.1025 | 0.5775 | 0.2775 | 0.836746 | 0.20074 | 0.544696 | | 1 | 0.996511 | 0.912212 |
| Snx4 | Q91YJ2 | 0.35 | 0.67 | 0.48 | 0.4 | 0.48 | 0.26 | 0.41 | 0.45 | 0.35 | 0.06 | 0.46 | 0.32 | -0.15 | 0.71 | 0.22 | 0.24 | -0.075 | -0.1775 | -0.22 | 0.629723 | 0.333488 | 0.357033 | | 1 | 0.996511 | 0.855416 |
| Snx5 | Q9D8U8 | 0.07 | 0.18 | 0.48 | 0.44 | 0.21 | -0.05 | 0.33 | 0.36 | 0.02 | 0.18 | 0.18 | -0.17 | -0.43 | -0.16 | 0.22 | 0.63 | -0.08 | -0.24 | -0.2275 | 0.66009 | 0.224132 | 0.416099 | | 1 | 0.996511 | 0.874945 |
| Snx6 | Q6P8X1 | 0.52 | -0.1 | 0.23 | 0.44 | 0.04 | 0.29 | 0.09 | 0.3 | 0.51 | 0.41 | 0.21 | 0.52 | 0.12 | 0.44 | -0.13 | -0.07 | -0.0925 | 0.14 | -0.1825 | 0.631388 | 0.496983 | 0.439902 | | 1 | 0.996511 | 0.883445 |
| Snx7 | F8WI30 | 0.84 | 1.19 | 1.2 | 0.92 | 1.28 | 0.76 | 0.88 | 0.97 | 0.1 | 0.48 | 1.24 | 1.05 | 0.17 | -0.37 | 0.97 | 1 | -0.065 | -0.32 | -0.595 | 0.727222 | 0.282843 | 0.111064 | | 1 | 0.996511 | 0.712422 |
| Snx9 | Q91VH2 | -0.83 | -1.95 | 0.19 | 0.05 | -2.18 | -0.67 | 0.21 | -0.14 | 0.78 | -0.38 | -0.11 | -0.59 | 2.11 | -2 | 0.14 | -0.08 | -0.06 | 0.56 | 0.6775 | 0.926996 | 0.310375 | 0.446733 | | 1 | 0.996511 | 0.884817 |
| Soat1 | Q61263 | 0.38 | 0.67 | 0.22 | 0.29 | 0.51 | 0.59 | 0.6 | 0.49 | -1.21 | 0.52 | 0.38 | 0.47 | -1.26 | -1.22 | 0.71 | 1.53 | 0.1575 | -0.35 | -0.45 | 0.346011 | 0.402374 | 0.492069 | | 1 | 0.996511 | 0.897531 |
| Sobp | Q0P5V2 | -1.56 | -2.25 | -0.91 | -0.34 | -2.1 | -2.41 | -0.69 | -0.86 | -2.56 | -0.24 | -0.92 | -2.01 | -1.07 | -1.85 | -1.12 | -0.75 | -0.25 | -0.1675 | 0.0675 | 0.65019 | 0.783991 | 0.880093 | | 1 | 0.996511 | 0.979626 |
| Sod1 | P08228 | -0.1 | -0.53 | -0.2 | -0.55 | -0.97 | 0.12 | 0.1 | 1.2 | 1.14 | -1.77 | 0.34 | -0.18 | 2.56 | -0.85 | 1.84 | -0.94 | 0.4575 | 0.2275 | 0.9975 | 0.301585 | 0.693187 | 0.244859 | | 1 | 0.996511 | 0.800734 |
| Sod2 | P09671 | 2.66 | 2.03 | 1.51 | 1.04 | 1.78 | 2.42 | 1.19 | 1.46 | 0.59 | 0.35 | 1.43 | 1.96 | 0.06 | 1.2 | 0.88 | 1 | -0.0975 | -0.7275 | -1.025 | 0.812655 | 0.153358 | 0.032385 | | 1 | 0.996511 | 0.641974 |
| Sod3 | O09164 | -3.15 | -3.32 | -1.88 | -1.72 | -2.89 | -3.1 | -1.73 | -1.86 | -2.38 | -0.7 | -1.96 | -2.42 | -1.91 | -1.33 | -1.65 | -1.63 | 0.1225 | 0.6525 | 0.8875 | 0.807405 | 0.245391 | 0.057606 | | 1 | 0.996511 | 0.661738 |
| Soga1 | A2ACV6 | -0.65 | -0.33 | -0.14 | -0.33 | -0.76 | -0.73 | -0.19 | -0.6 | -0.49 | 0.56 | -0.15 | -0.1 | -0.19 | -0.11 | 0.49 | 0.17 | -0.2075 | 0.3175 | 0.4525 | 0.317358 | 0.245913 | 0.076747 | | 1 | 0.996511 | 0.675822 |
| Soga3 | Q6NZL0 | 0.79 | 0.59 | 0.68 | 0.74 | 0.75 | 0.77 | 0.63 | 0.68 | 1.24 | 0.66 | 0.69 | 0.96 | 1.24 | 1.64 | 0.71 | 0.71 | 0.0075 | 0.1875 | 0.125 | 0.958096 | 0.349908 | 0.549638 | | 1 | 0.996511 | 0.913178 |
| Son | H9KV00 | 1.56 | 0.42 | 0.18 | 0.48 | -0.25 | 0.35 | -0.15 | 0.6 | 0.05 | 0.37 | 0.07 | -0.28 | 0.64 | 0.48 | 0.59 | -0.62 | -0.5225 | -0.6075 | -0.3875 | 0.164599 | 0.094909 | 0.358628 | | 1 | 0.996511 | 0.856645 |
| Son | Q9QX47-2 | 0.46 | 0.53 | -0.29 | -0.51 | 0.46 | 0.38 | 0.44 | 0.8 | 0.8 | -0.81 | -0.31 | -0.24 | -0.18 | 0.5 | 0.01 | -0.04 | 0.4725 | -0.1875 | 0.025 | 0.121151 | 0.647703 | 0.935848 | | 1 | 0.996511 | 0.989767 |
| Sorbs1 | A0A286YCI | 0.98 | 0.9 | 0.75 | 0.78 | 0.78 | 0.79 | 0.79 | 0.58 | 1.05 | 0.68 | 0.79 | 1.03 | 1.49 | 1.64 | 1.17 | 0.81 | -0.1175 | 0.035 | 0.175 | 0.441586 | 0.842453 | 0.468037 | | 1 | 0.996511 | 0.889423 |
| Sorbs1 | A0A668KL6 | -0.67 | -0.13 | 0.96 | 1.23 | -1.44 | 0.38 | 0.68 | 0.38 | -1.53 | 1.79 | 0.22 | -0.18 | -0.5 | -0.99 | 0.99 | 0.29 | -0.3475 | -0.2725 | -0.4 | 0.567869 | 0.714066 | 0.493073 | | 1 | 0.996511 | 0.897531 |
| Sorbs1 | A0A5F8MP | 0.06 | 0.2 | 0.22 | 0.2 | -0.09 | 0.05 | -0.44 | -0.06 | 0.18 | 0.15 | 0.1 | -0.01 | 0.46 | -0.65 | 0.91 | 0.03 | -0.305 | -0.065 | 0.0175 | 0.096152 | 0.680646 | 0.958383 | | 1 | 0.996511 | 0.992636 |
| Sorbs2 | Q3UTJ2-2 | 1.24 | 0.72 | 1.1 | 0.85 | 0.6 | 1.31 | 0.55 | 0.77 | 0.7 | 0.31 | 1.26 | 1.23 | 0.97 | 0.29 | 0.66 | 0.61 | -0.17 | -0.1025 | -0.345 | 0.46638 | 0.706949 | 0.156084 | | 1 | 0.996511 | 0.751493 |
| Sorbs2 | D3Z080 | -0.6 | 0.73 | 0.73 | -0.21 | -1.4 | -0.11 | -0.75 | -1.16 | 0.08 | -0.85 | 0.56 | -0.59 | 0.35 | 1.58 | -0.02 | -2.19 | -1.0175 | -0.3625 | -0.2325 | 0.034237 | 0.41818 | 0.763474 | | 1 | 0.996511 | 0.964134 |
| Sorbs2 | B7ZWM6 | -0.01 | -0.2 | -1.55 | -2.47 | -0.29 | 0.23 | -2.11 | -2.35 | 1.69 | -1.47 | -1.49 | -0.32 | 1.98 | -0.07 | -3.18 | -3.31 | -0.0725 | 0.66 | -0.0875 | 0.926318 | 0.448191 | 0.944326 | | 1 | 0.996511 | 0.991865 |
| Sorbs3 | Q8K0M3 | -0.07 | 0.19 | 0.02 | 0.04 | 0.32 | -0.14 | 0.28 | -0.04 | 0.02 | 0.09 | 0.5 | 0.42 | -0.28 | -0.26 | 0.32 | 0.75 | 0.06 | 0.2125 | 0.0875 | 0.732883 | 0.278064 | 0.751943 | | 1 | 0.996511 | 0.961479 |
| Sorcs1 | Q9JLC4-2 | 0.66 | 0.41 | 0.08 | -0.05 | 0.22 | 0.89 | -0.25 | -0.06 | 0.54 | -0.37 | -0.03 | 1.03 | 0.75 | -1.35 | -0.01 | -0.19 | -0.075 | 0.0175 | -0.475 | 0.799884 | 0.959278 | 0.295939 | | 1 | 0.996755 | 0.825612 |
| Sorcs2 | Q9EPR5 | 0.16 | 0.45 | 0.84 | 0.69 | 0.55 | 0.39 | 0.72 | 0.54 | -0.26 | 0.28 | 0.3 | 0.29 | 0.15 | 0.33 | 0.72 | 0.94 | 0.015 | -0.3825 | -0.075 | 0.939412 | 0.128078 | 0.800238 | | 1 | 0.996755 | 0.970393 |
| Sorcs3 | Q8VI51 | 1.03 | 0.87 | 0.69 | 0.35 | 1.02 | 1.15 | 0.14 | 0.43 | 0.35 | -1.16 | 0.75 | 1.06 | -0.01 | 0.32 | 0.61 | 0.22 | -0.05 | -0.485 | -4.50E-01 | 0.859716 | 0.326287 | 0.083241 | | 1 | 0.996511 | 0.689531 |
| Sord | Q64442 | -0.01 | 0.09 | 0.43 | 0.47 | 0.05 | 0.23 | 0.55 | 0.67 | -0.33 | -0.49 | 0.11 | 0.09 | -0.18 | -0.17 | 0.08 | 0.13 | 0.13 | -0.4 | -0.28 | 0.546357 | 0.105398 | 0.198309 | | 1 | 0.996511 | 0.78004 |
| Sorl1 | O88307 | -1.32 | -0.89 | -1.07 | -1.04 | -0.67 | -1.18 | -1.05 | -1.15 | -1.34 | -0.9 | -1.21 | -1.12 | -0.64 | -0.55 | -1.14 | -0.97 | 0.0675 | -0.0625 | 0.255 | 0.71984 | 0.73972 | 0.259843 | | 1 | 0.996511 | 0.812701 |
| Sort1 | Q6PHU5 | -0.23 | -0.39 | -0.63 | -1.16 | -0.02 | 0 | -0.67 | -0.92 | -0.69 | -1.4 | -0.65 | -0.18 | -0.1 | -0.66 | -0.86 | -0.69 | 0.2 | -0.1275 | 0.025 | 0.517357 | 0.694151 | 0.929225 | | 1 | 0.996511 | 0.989687 |
| Sos1 | Q62245 | 0.27 | 0.37 | 0.31 | 0.2 | 0.26 | 0.39 | 0.04 | 0.33 | 0.04 | 0.11 | 0.29 | 0.27 | -0.17 | 0.18 | 0.15 | -0.04 | -0.0325 | -0.11 | -0.2575 | 0.83295 | 0.502537 | 0.178018 | | 1 | 0.996511 | 0.772747 |
| Sos2 | S4R2C4 | 0.6 | 0.63 | 0.04 | -0.04 | 0.8 | 0.3 | -0.84 | -0.27 | 0.54 | 0.4 | -0.18 | 0.24 | 0.25 | 0.33 | -0.15 | 0.09 | -0.31 | -0.0575 | -0.1775 | 0.420565 | 0.823242 | 0.473963 | | 1 | 0.996511 | 0.891644 |
| Sowaha | Q8BLS7 | -0.02 | 0.18 | 0.56 | 0.62 | 0.78 | -0.52 | 0.62 | 0.31 | 0.09 | 1.16 | 0.23 | -0.02 | 0.34 | -0.4 | 0.42 | 0.32 | -0.0375 | 0.03 | -0.165 | 0.906717 | 0.9235 | 0.545063 | | 1 | 0.996755 | 0.912212 |
| Sowahb | Q8BZW2 | 0.69 | 1.43 | 1 | 1.31 | 1.44 | 1.2 | 0.81 | 1.04 | 0.73 | 0.71 | 1 | 1 | 0.41 | 1.19 | 0.34 | 0.59 | 0.015 | -0.2475 | -0.475 | 0.948271 | 0.281063 | 0.113194 | | 1 | 0.996511 | 0.715427 |
| Sowahc | Q8C0J6 | 1.15 | 1 | 0.64 | 0.48 | 0.8 | 0.88 | -0.73 | 0.46 | 1.49 | 0.79 | 0.16 | 0.87 | 1.13 | 0.81 | 0.55 | 0.43 | -0.465 | 0.01 | -0.0875 | 0.244459 | 0.974636 | 0.729291 | | 1 | 0.997515 | 0.958635 |
| Sox1 | P53783 | -1.03 | -0.64 | -1.73 | -1.71 | -1.59 | -0.62 | -2.22 | -0.45 | 0.43 | 0.84 | -0.87 | -0.96 | 0.55 | -0.21 | 1.69 | -3.41 | 0.0575 | 1.1375 | 0.9325 | 0.900572 | 0.043746 | 0.366552 | | 1 | 0.996511 | 0.858904 |
| Sox13 | D3Z7I3 | -2.89 | -4.39 | -4.82 | -2.88 | -8.36 | -1.82 | -6.08 | -5.27 | 1.04 | -1.23 | -3.17 | -2.44 | 3.29 | -4.93 | -3.94 | -4.89 | -1.6375 | 2.295 | 1.1275 | 0.226988 | 0.03617 | 0.538323 | | 1 | 0.996511 | 0.912212 |
| Sox5 | B2KFM4 | -1.45 | -1.92 | -2.82 | -4.21 | -1.97 | -2.37 | -3.46 | -3.22 | -0.93 | -1.11 | -3.33 | -1.4 | -0.11 | -1.59 | -3.32 | -3.26 | -0.155 | 0.9075 | 0.53 | 0.808116 | 0.243815 | 0.550314 | | 1 | 0.996511 | 0.913747 |
| Sox9 | Q04887 | 0.67 | -0.13 | -2.13 | -2.71 | -0.79 | 0.03 | -2.7 | -2.61 | 2.48 | -2.23 | -1.07 | 0.27 | 4.48 | -1.33 | 3.07 | -2.98 | -0.4425 | 0.9375 | 1.885 | 0.641055 | 0.426586 | 0.290501 | | 1 | 0.996511 | 0.825497 |
| Sp3 | O70494-3 | -5.05 | -5.8 | -3.34 | -2.48 | -5.04 | -5.74 | -3.88 | -2.96 | -5.15 | -0.74 | -3.59 | -4.47 | -4.34 | -3.63 | -3.29 | -4.52 | -0.2175 | 0.7 | 0.2425 | 0.804251 | 0.530275 | 0.741204 | | 1 | 0.996511 | 0.958796 |
| Spacdr | D3YZZ4 | 0.68 | 0.08 | 0.83 | 0.81 | 0.5 | 0.7 | 3.22 | 1.1 | 0.31 | 1.51 | 0.73 | 0.67 | 0.92 | 0.25 | 1.07 | 1.23 | 0.78 | 0.205 | 0.2675 | 0.21138 | 0.515058 | 0.374103 | | 1 | 0.996511 | 0.859424 |
| Spag1 | Q80ZX8-2 | 0.12 | 0.13 | -0.11 | 0.48 | 0.14 | 0.4 | -0.02 | 0.29 | 0.3 | 0.25 | 0.32 | 0.12 | 0.07 | 0.15 | 0.27 | 0.2 | 0.0475 | 0.0925 | 0.0175 | 0.803273 | 0.62555 | 0.929452 | | 1 | 0.996511 | 0.989687 |

| | | | | | | | | | | | | | | | | | | | | | | | | | | |
|---|---|---|---|---|---|---|---|---|---|---|---|---|---|---|---|---|---|---|---|---|---|---|---|---|---|---|
| Spag7 | Q7TNE3 | -1.67 | -1.83 | -2.5 | -2.41 | -2.14 | -2.04 | -2.64 | -2.84 | -1.45 | -1.13 | -3.02 | -1.74 | -1 | -1.43 | -2.92 | -2.45 | -0.3125 | 0.2675 | 0.1525 | 0.28958 | 0.544888 | 0.74152 | 1 | 0.996511 | 0.958796 |
| Spag9 | Q58A65-2 | -0.19 | -0.35 | -0.63 | -0.64 | -0.33 | -0.58 | -0.74 | -0.8 | -0.91 | -0.48 | -0.7 | -0.48 | -0.98 | -0.28 | -1.18 | -1.23 | -0.16 | -0.19 | -0.465 | 0.411002 | 0.35575 | 0.112356 | 1 | 0.996511 | 0.714111 |
| Sparc | Q5NCU4 | -0.43 | -0.13 | 0 | -0.53 | 0.29 | -0.27 | 0.83 | 0.25 | -1.25 | 2.8 | 0.05 | -0.54 | -1.22 | -0.77 | 0.71 | 1.45 | 0.5475 | 0.5375 | 0.315 | 0.064464 | 0.511616 | 0.59308 | 1 | 0.996511 | 0.929843 |
| Sparcl1 | P70663 | 0.54 | -0.05 | -0.32 | -0.54 | -0.06 | 0.45 | -0.36 | -0.35 | -1.05 | 0 | -0.39 | 0.85 | -0.29 | -1.11 | 0.03 | -0.12 | 0.0125 | -0.055 | -0.28 | 0.966604 | 0.899309 | 0.425209 | 1 | 0.996511 | 0.877372 |
| Spart | Q8R1X6 | 0.6 | 0.4 | 0.66 | 0.65 | 0.29 | 0.34 | 0.39 | 0.47 | 0.29 | 1 | 0.37 | 0.31 | 0.71 | 0.36 | 0.74 | 0.51 | -0.205 | -0.085 | 0.0025 | 0.192495 | 0.698444 | 0.989376 | 1 | 0.996511 | 0.996154 |
| Spast | A0A286YE2 | 0.48 | 0.43 | -0.11 | 0.04 | 0.05 | 0.43 | -0.49 | 0.14 | 0.42 | 0.49 | -0.07 | 0.83 | 0.57 | -0.26 | 0.06 | -0.42 | -0.1775 | 0.2075 | -0.2225 | 0.487773 | 0.42712 | 0.438286 | 1 | 0.996511 | 0.883161 |
| Spata1 | Q9D5R4 | 4.31 | 3.81 | -0.21 | -0.29 | 4.61 | 4.3 | 0.33 | -0.02 | 0.43 | -1.41 | -0.29 | 3.78 | 1 | 2.01 | 0.27 | 0.25 | 0.4 | -1.2775 | -1.0225 | 0.799516 | 0.403348 | 0.393746 | 1 | 0.996511 | 0.865775 |
| Spata2 | Q8K004 | -0.14 | -0.05 | 0.49 | 0.33 | -0.59 | -0.03 | 0.21 | 0.09 | 0.36 | 0.64 | 0.43 | -0.2 | 0.98 | -0.51 | 0 | -0.1 | -0.2375 | 0.15 | -0.065 | 0.347334 | 0.560677 | 0.851816 | 1 | 0.996511 | 0.976225 |
| Spata2l | Q8BNN1 | 0.39 | 0.2 | 0.14 | 0.4 | -0.07 | 0.22 | 0.54 | 0.09 | 0.38 | 0.89 | 0.2 | 0.69 | 0.29 | 0.31 | 0.4 | 0.27 | -0.0875 | 0.2575 | 0.035 | 0.640535 | 0.241872 | 0.840746 | 1 | 0.996511 | 0.976225 |
| Spata5 | A0A0G2JFY | -0.77 | -0.36 | -0.28 | -0.44 | -0.49 | -1.25 | 0.15 | -0.44 | -0.08 | -0.19 | -0.26 | -0.56 | -0.93 | 0.27 | -0.3 | -0.52 | -0.045 | 0.19 | 0.0925 | 0.882023 | 0.352695 | 0.749638 | 1 | 0.996511 | 0.961334 |
| Spats2 | Q8K1N4 | -1.42 | -1.31 | -1.89 | -1.64 | -1.08 | -1.67 | -1.46 | -1.96 | -0.93 | -0.81 | -1.34 | -1.29 | -0.43 | -1.05 | -0.9 | -1.09 | 0.0225 | 0.4725 | 0.6975 | 0.925437 | 0.057227 | 0.015962 | 1 | 0.996511 | 0.589387 |
| Spats2l | E9PVQ3 | -1.08 | -1.45 | -1.47 | -1.38 | -1.13 | -0.96 | -1.74 | -1.75 | -1.05 | -0.94 | -1.29 | -1.01 | 0.19 | -1.16 | -0.87 | -0.61 | -0.05 | 0.2725 | 0.7325 | 0.834884 | 0.160501 | 0.041165 | 1 | 0.996511 | 0.658454 |
| Spcs1 | A0A2I3BRV | -0.15 | 0.35 | -0.32 | -0.14 | 0.61 | -0.18 | 0.62 | 0.31 | -0.38 | -0.27 | -0.01 | -0.08 | -0.29 | 1.12 | 0.25 | 0.1 | 0.405 | -0.12 | 0.36 | 0.131223 | 0.571965 | 0.296374 | 1 | 0.996511 | 0.825612 |
| Spcs2 | A0A140LH( | 2.19 | 1.98 | 2.34 | 2.38 | 2.19 | 2.14 | 2.96 | 2.78 | 2.05 | 2.52 | 2.21 | 2.01 | 1.96 | 2.48 | 3.53 | 2.85 | 0.295 | -0.025 | 0.4825 | 0.24171 | 0.899476 | 0.182606 | 1 | 0.996511 | 0.776002 |
| Spcs3 | Q6ZWQ7 | -0.19 | -0.15 | 0.56 | -0.19 | 0.38 | -0.28 | 0.82 | -0.03 | 0 | 0.69 | 0.3 | -0.56 | -0.35 | 0.59 | 0.65 | 1.06 | 0.215 | 0.1 | 0.48 | 0.481726 | 0.756314 | 0.19202 | 1 | 0.996511 | 0.777117 |
| Specc1 | A0A0J9YUF | 0.48 | 1.46 | 1.2 | 1.58 | 1.12 | 0.86 | 1.63 | 1.45 | 0.9 | 1.78 | 1.57 | 1.11 | 1.5 | 2.25 | 2.21 | 2.31 | 0.085 | 0.16 | 0.8875 | 0.776017 | 0.619004 | 0.019999 | 1 | 0.996511 | 0.615793 |
| Specc1 | A0A0J9YTU | -0.09 | 0.01 | 0.13 | 0.07 | -0.02 | -0.4 | 0.14 | -0.14 | -0.18 | 0.54 | -0.04 | 0.16 | 0.33 | 0.31 | 0.04 | 0.27 | -0.135 | 0.09 | 0.2075 | 0.444144 | 0.66637 | 0.261013 | 1 | 0.996511 | 0.81412 |
| Specc1l | Q2KN98 | 0.43 | 0.67 | 0.89 | 0.54 | 0.56 | 0.1 | 0.49 | 0.26 | 0.59 | 0.79 | 0.66 | 0.83 | 0.27 | 0.96 | 0.52 | 0.28 | -0.28 | 0.085 | -0.125 | 0.158022 | 0.638689 | 0.593784 | 1 | 0.996511 | 0.929843 |
| Speg | E9QQ25 | 0.27 | 0.42 | 0.6 | 0.54 | 0.03 | 0.23 | 0.11 | 0.45 | 0.9 | 1.02 | 0.63 | 0.45 | 1.18 | 0.79 | 0.62 | 0.33 | -0.2525 | 0.2925 | 0.2725 | 0.162816 | 0.166309 | 0.26304 | 1 | 0.996511 | 0.81524 |
| Spen | A2ADB1 | -0.84 | -0.71 | -0.03 | 0.28 | -1.2 | -1.49 | -0.35 | 0.07 | 0.13 | 0.34 | -0.54 | -1.18 | 0.3 | -0.66 | 0.42 | 0.25 | -0.4175 | 0.0125 | 0.4025 | 0.337894 | 0.975971 | 0.281918 | 1 | 0.997588 | 0.822084 |
| Spg11 | Q3UHA3 | -1 | -1.5 | 0.67 | 0.59 | -1 | -0.78 | 0.42 | 0.13 | 2.28 | 0.4 | 0.97 | -0.81 | 2.32 | -0.1 | 0.53 | 0.7 | 0.0025 | 1.02 | 1.1725 | 0.996638 | 0.206391 | 0.114897 | 1 | 0.996511 | 0.717287 |
| Spg21 | Q9CQC8 | -0.4 | 0.14 | -0.64 | 0.16 | 0.78 | -0.18 | -0.33 | 0.12 | -0.33 | -0.54 | -0.42 | -0.37 | -0.33 | 1.03 | -0.6 | -0.32 | 0.2825 | -0.23 | 0.13 | 0.376288 | 0.340745 | 0.747326 | 1 | 0.996511 | 0.960744 |
| Spg7 | D3YZN4 | 0.39 | 0.36 | 0.08 | 0.37 | 0.41 | 0.45 | 0.05 | 0.19 | 0.49 | 0.01 | 0.43 | 0.22 | 0.61 | 0.24 | -0.11 | -0.03 | -0.025 | -0.0125 | -0.1225 | 0.88396 | 0.947308 | 0.590255 | 1 | 0.996755 | 0.929142 |
| Sphk2 | Q9JIA7 | -0.03 | 0.3 | -0.59 | -0.43 | 0.34 | 0.16 | -0.41 | -0.35 | 0.47 | -1.27 | -0.23 | 0.34 | 0.27 | 0.56 | -1.05 | -0.52 | 0.1225 | 0.015 | 0.0025 | 0.660333 | 0.971567 | 0.995048 | 1 | 0.997515 | 0.998224 |
| Sphkap | E9PUC4 | -0.68 | -0.65 | -0.65 | -0.71 | 0.01 | -0.35 | -0.49 | -0.71 | 0.01 | -0.13 | -0.39 | -0.11 | 0.74 | 0.55 | 0.18 | -0.41 | 0.2875 | 0.5175 | 0.9375 | 0.156732 | 0.01242 | 0.007614 | 1 | 0.996511 | 0.554014 |
| Spin1 | Q61142-2 | -1.83 | -1.56 | -1.48 | -1.81 | -1.85 | -1.98 | -1.33 | -2 | -1.66 | -0.56 | -1.97 | -1.07 | -1.43 | -0.86 | -1.22 | -1.53 | -0.12 | 0.355 | 0.41 | 0.568884 | 0.291266 | 0.092111 | 1 | 0.996511 | 0.695932 |
| Spire1 | A0A5H1ZRI | 0.97 | 1.1 | 0.64 | 0.64 | 1.04 | 0.89 | 0.13 | 0.45 | 0.65 | -0.19 | -0.65 | 0.21 | 1.55 | 1.36 | -0.2 | 0.26 | -0.21 | -0.8325 | -0.095 | 0.411519 | 0.022416 | 0.821174 | 1 | 0.996511 | 0.972768 |
| Spire2 | Q8K1S6 | -0.65 | -0.03 | -0.04 | -0.21 | 0.18 | -0.8 | 0.12 | -0.44 | -0.08 | 0.57 | -0.27 | -0.45 | 0.59 | 0.98 | 0.08 | -0.19 | -0.0025 | 0.175 | 0.5975 | 0.992843 | 0.534173 | 0.079897 | 1 | 0.996511 | 0.681818 |
| Spns1 | D3YTR6 | -0.9 | -0.63 | -0.7 | -0.37 | -1.07 | -0.31 | -0.32 | -0.64 | 0.11 | -0.33 | -0.29 | -0.17 | -0.37 | -0.67 | -0.63 | -0.65 | 0.065 | 0.48 | 0.07 | 0.777237 | 0.036565 | 0.725173 | 1 | 0.996511 | 0.958398 |
| Spns2 | Q91VM4 | 0.76 | 1.68 | 0.83 | 0.26 | 0.72 | 1.04 | 0.69 | 0.96 | 2.1 | 0.76 | 0.14 | 1.22 | 2.27 | -0.05 | 0.35 | 1.38 | -0.03 | 0.1725 | 0.105 | 0.921286 | 0.716815 | 0.849611 | 1 | 0.996511 | 0.976225 |
| Spock1 | Q6DIC9 | 2.25 | -0.55 | 1.24 | -0.03 | 1.29 | -0.67 | 0.51 | -0.32 | -0.66 | -0.34 | -0.33 | -0.24 | 2.55 | -0.82 | 1.91 | 0.22 | -0.525 | -1.12 | 0.2375 | 0.458724 | 0.083485 | 0.790947 | 1 | 0.996511 | 0.969105 |
| Spock2 | Q9ER58 | -1.13 | -1.14 | -1.45 | -0.71 | -0.67 | -0.67 | -0.28 | -0.38 | -0.97 | 0.59 | -0.87 | -0.78 | -0.72 | -0.7 | -0.3 | -0.43 | 0.6075 | 0.6 | 0.57 | 0.017071 | 0.144186 | 0.032331 | 1 | 0.996511 | 0.641974 |
| Spock3 | Q8BKV0 | 1.17 | 0.48 | -0.32 | -0.23 | 0.73 | 1.31 | 0 | 0.03 | -0.06 | 0.08 | -0.35 | 1.06 | 0.22 | -0.42 | 1.02 | 0.48 | 0.2425 | -0.0925 | 0.05 | 0.582368 | 0.832519 | 0.908579 | 1 | 0.996511 | 0.986458 |
| Spon1 | Q8VCC9 | -1.91 | -2.09 | -1.19 | -1.56 | -1.66 | -1.88 | -1.15 | -1.3 | -1.64 | -1.3 | -1.19 | -1.85 | -1.25 | -1.38 | -1.62 | -1.68 | 0.19 | 0.1925 | 0.205 | 0.482207 | 0.477961 | 0.429561 | 1 | 0.996511 | 0.879165 |
| Spout1 | Q3UHX9+C | 0.49 | -0.63 | 1.26 | 1.83 | -1.57 | -0.66 | -1.58 | -0.38 | 0.51 | 1.89 | 0.93 | -0.56 | 1.58 | -1.74 | 1.06 | -0.47 | -1.785 | -0.045 | -0.63 | 0.011177 | 0.946396 | 0.455689 | 1 | 0.996755 | 0.887169 |
| Sppl2a | Q9JJF9 | -0.08 | 0.12 | 0.74 | 0.66 | -0.16 | 0.64 | 0.38 | 0.3 | 1.41 | 0.87 | 0.43 | 0 | 2.11 | 0.43 | 0.45 | 0.8 | -0.07 | 0.3175 | 0.5875 | 0.793942 | 0.381905 | 0.189986 | 1 | 0.996511 | 0.776843 |
| Sppl2b | Q3TD49 | 0.18 | 0.35 | 0.77 | 0.34 | 0.61 | 0.05 | -0.04 | 0.22 | 1.69 | -0.62 | 0.87 | 0.89 | 1.73 | 1.06 | 0.24 | 0.21 | -0.2 | 0.2975 | 0.4 | 0.367542 | 0.528655 | 0.30389 | 1 | 0.996511 | 0.831662 |
| Spr | Q91XH5 | 0.2 | 0.13 | -0.26 | -0.12 | 0.23 | 0.14 | -0.1 | -0.2 | 0.08 | -0.14 | -0.35 | 0.08 | -0.15 | 0.43 | -0.63 | -0.31 | 0.03 | -0.07 | -0.1525 | 0.872749 | 0.725867 | 0.576796 | 1 | 0.996511 | 0.927198 |
| Spred1 | Q924S8 | 1.49 | 1.94 | 0.95 | 1.12 | 1.55 | 1.32 | 0.79 | 1.08 | 1.65 | 1.08 | 1.08 | 1.66 | 1.95 | 1.98 | 1.03 | 1.06 | -0.19 | -0.0075 | 0.13 | 0.498916 | 0.979024 | 0.706419 | 1 | 0.998749 | 0.957084 |
| Spred2 | Q924S7 | 0.04 | 0.62 | 0.93 | 0.74 | 0.25 | 0.36 | 0.66 | 0.94 | 0.04 | 0.12 | 0.91 | 0.49 | 0.12 | -0.87 | 0.77 | 0.88 | -0.03 | -0.1925 | -0.3575 | 0.906929 | 0.507258 | 0.409553 | 1 | 0.996511 | 0.87457 |
| Spred3 | A0A0R4J0K | 1.05 | 0.05 | 0.72 | 1.26 | -0.37 | 1.17 | -0.07 | 0.65 | 1.83 | 1.59 | 0.66 | 1.6 | 1.13 | 0.46 | 0.37 | 0.62 | -0.425 | 0.65 | -0.125 | 0.316689 | 0.099396 | 0.697641 | 1 | 0.996511 | 0.955729 |
| Sprn | A0A1B0GSI | -0.13 | 0.25 | -0.22 | -0.11 | 0.21 | -0.94 | 0.1 | 0 | 0.51 | 0.14 | -0.21 | -1.32 | -0.34 | -0.46 | 0.13 | 0.82 | -0.105 | -0.1675 | 0.09 | 0.713845 | 0.670443 | 0.77607 | 1 | 0.996511 | 0.966799 |
| Spry2 | Q9QXV8 | 1.24 | 1.19 | 1.65 | 1.27 | 1.06 | 1.47 | 1.59 | 1.84 | 1.61 | -0.09 | 2.03 | 1.62 | 1.55 | 1.85 | 2.52 | 1.26 | 0.1525 | -0.045 | 0.4575 | 0.490877 | 0.920919 | 0.155759 | 1 | 0.996755 | 0.751103 |
| Spryd3 | E9Q9B3 | 1.13 | 1.06 | 0.97 | 0.43 | 1.22 | 0.71 | 0.43 | 0.66 | -0.1 | 0.02 | 0.9 | 1.34 | -0.38 | 0.38 | 1.3 | 1.23 | -0.1425 | -0.3575 | -0.265 | 0.563739 | 0.347815 | 0.523496 | 1 | 0.996511 | 0.90791 |
| Spryd4 | Q91WK1 | 0.61 | 0.4 | 0.69 | 0.4 | 0.49 | 0.52 | 0.76 | 0.48 | 0.19 | -0.14 | 0.53 | 0.34 | -0.05 | 0.65 | -0.04 | 0.13 | 0.0375 | -0.295 | -0.3525 | 0.815851 | 0.176071 | 0.146382 | 1 | 0.996511 | 0.746406 |
| Spryd7 | Q3TFQ1 | -1.27 | -1.46 | -0.12 | -0.71 | -1.09 | -0.87 | -0.16 | -0.42 | -1.38 | -0.85 | -0.15 | -1.05 | -1.17 | -1.23 | -0.72 | -0.28 | 0.255 | 0.0325 | 0.04 | 0.47764 | 0.932418 | 0.913273 | 1 | 0.996755 | 0.986458 |
| Spta1 | P08032 | 0.88 | -0.78 | 1.39 | 2.59 | -0.42 | 2.06 | 1.04 | 2.46 | 0.57 | 0.12 | 0.83 | -0.3 | 0.97 | 0.38 | 0.44 | 1.25 | 0.265 | -0.715 | -0.26 | 0.756909 | 0.306047 | 0.696513 | 1 | 0.996511 | 0.955729 |
| Sptan1 | A3KGU9 | 1.82 | 1.14 | 1.22 | 1.43 | 1.21 | 1.55 | 1.52 | 1.61 | 0.81 | 0.84 | 1.34 | 1.39 | 0.36 | 0.96 | 1.06 | 1.1 | 0.07 | -0.3075 | -0.5325 | 0.73496 | 0.228656 | 0.06585 | 1 | 0.996511 | 0.664936 |
| Sptan1 | E9Q447 | 1.6 | 1.25 | 1.4 | 2.73 | 1.24 | 1.64 | 2.1 | 2.99 | 1.58 | 3.04 | 2.63 | 1.8 | 0.26 | 2.13 | 1.17 | 0.89 | 0.2475 | 0.5175 | -0.6325 | 0.599652 | 0.270766 | 0.211272 | 1 | 0.996511 | 0.786269 |
| Sptan1 | A3KGU7 | 1.23 | 0.27 | -0.84 | 0.3 | 0.23 | 1.12 | 0.48 | 0.1 | 0.71 | 0.41 | -0.79 | 1.25 | -0.36 | 0.36 | -0.52 | -0.38 | 0.2425 | 0.155 | -0.465 | 0.590823 | 0.781133 | 0.30942 | 1 | 0.996511 | 0.834523 |
| Sptan1 | P16546 | -5.32 | -7.05 | 1.26 | 1.8 | -7.38 | -9.55 | 0.69 | 1.09 | -5.32 | 2.31 | 1.55 | -7.17 | -3.98 | -9.24 | 1.65 | 1.26 | -1.46 | 0.17 | -0.25 | 0.645788 | 0.953661 | 0.933989 | 1 | 0.996755 | 0.989767 |
| Sptb | Q3UGX2 | 1.92 | 1.09 | 1.21 | 1.46 | 1.18 | 1.47 | 1.19 | 1.39 | 0.45 | 0.34 | 0.94 | 0.79 | 0.11 | 0.5 | 0.68 | 0.94 | -0.1125 | -0.79 | -0.8625 | 0.612783 | 0.011934 | 0.011567 | 1 | 0.996511 | 0.566767 |
| Sptbn1 | Q62261 | 1.32 | 0.9 | 1.06 | 1.13 | 1.04 | 1.29 | 1.46 | 1.38 | 0.72 | 0.65 | 1.23 | 1.16 | 0.38 | 0.59 | 1 | 1.04 | 0.19 | -0.1625 | -0.35 | 0.294031 | 0.453855 | 0.151094 | 1 | 0.996511 | 0.749304 |
| Sptbn2 | Q68FG2 | 1.76 | 0.82 | 1.67 | 1.93 | 1.4 | 1.61 | 1.69 | 1.98 | 0.56 | 0.92 | 1.91 | 0.99 | 0.24 | 1 | 1.29 | 1.33 | 0.125 | -0.45 | -0.58 | 0.655361 | 0.241962 | 0.125545 | 1 | 0.996511 | 0.721449 |
| Sptbn4 | E9PX29 | 1.14 | 0.99 | 0.87 | 0.94 | 0.83 | 0.93 | 0.8 | 0.81 | 0.83 | 0.8 | 0.84 | 0.87 | 0.62 | 0.67 | 0.77 | 0.62 | -0.1425 | -0.15 | -0.315 | 0.343729 | 0.355761 | 0.095393 | 1 | 0.996511 | 0.698156 |
| Sptlc1 | O35704 | 1.92 | 0.43 | 0.69 | 0.64 | 0.77 | 2.03 | 0.55 | 0.47 | 1.03 | 0.33 | 0.3 | 1.76 | 1.46 | 0.41 | 1.03 | 0.96 | 0.035 | -0.065 | 0.045 | 0.939465 | 0.886159 | 0.907772 | 1 | 0.996511 | 0.986458 |
| Sptlc2 | P97363 | 0.17 | 0.44 | 0.59 | 0.73 | 0.6 | 0.1 | 0.48 | 0.63 | 0.38 | 1.3 | 0.72 | 0.42 | 0.24 | 0.52 | 1.39 | 1.42 | -0.03 | 0.2225 | 0.41 | 0.882235 | 0.406262 | 0.232484 | 1 | 0.996511 | 0.788994 |
| Sqle | P52019 | 0.25 | 0.68 | 0.25 | 0.1 | 0.74 | 0.32 | 1.09 | 0.11 | 0.37 | 0.41 | -0.79 | 0.11 | 0.74 | 0.32 | 0.32 | 0.71 | 0.245 | -0.295 | 0.2025 | 0.359644 | 0.353593 | 0.372126 | 1 | 0.996511 | 0.859424 |
| Sqor | Q9R112 | 1.39 | 1.19 | 0.73 | 1.05 | 0.89 | 1.45 | 1.33 | 0.96 | 0.25 | 0.84 | 0.84 | 1.14 | -0.4 | 0.76 | 0.34 | 1.25 | 0.0675 | -0.3225 | -0.6025 | 0.758471 | 0.225787 | 0.129224 | 1 | 0.996511 | 0.723655 |
| Sqstm1 | Q64337-2 | -0.58 | 0.22 | 0.19 | -0.65 | -0.53 | -0.77 | 0.9 | 0.47 | 0.31 | 3.11 | 1.65 | -0.28 | 2.03 | 2.39 | 3.36 | 0.96 | 0.2225 | 1.4025 | 2.39 | 0.610145 | 0.078102 | 0.001222 | 1 | 0.996511 | 0.412464 |
| Sqstm1 | Q64337 | 0.69 | 0.41 | 0.11 | -0.03 | 0.81 | 0.67 | 0.73 | 0.29 | -0.04 | -0.68 | 0.13 | 0.94 | -0.13 | 0.39 | 0.79 | 0.5 | 0.33 | -0.2075 | 0.0925 | 0.160771 | 0.568438 | 0.735793 | 1 | 0.996511 | 0.958635 |
| Sra1 | Q80VJ2 | -2.32 | -1.59 | -0.62 | -0.65 | -2.71 | -2.06 | -1.74 | -0.95 | -1.19 | 0.27 | -0.87 | -2.2 | -0.4 | -2.08 | -0.87 | -1.64 | -0.57 | 0.2975 | 0.0475 | 0.27783 | 0.621592 | 0.926733 | 1 | 0.996511 | 0.989128 |
| Srbd1 | F8WGW3 | 0.21 | -0.65 | 0.22 | 0.04 | 0.03 | -0.33 | 0.25 | -0.18 | -0.74 | -0.4 | -0.1 | -0.96 | -0.77 | 0.4 | -0.35 | 0.03 | -0.0125 | -0.505 | -0.1275 | 0.960509 | 0.108108 | 0.699047 | 1 | 0.996511 | 0.955729 |
| Src | P05480-1 | 0.99 | 1.22 | 0.78 | 0.81 | 1.45 | 1.15 | 0.96 | 0.76 | 0.25 | 0.33 | 0.75 | 1.22 | -0.08 | 0.72 | 0.53 | 0.94 | 0.13 | -0.3125 | -0.4225 | 0.535691 | 0.254327 | 0.140261 | 1 | 0.996511 | 0.736797 |
| Srcin1 | A0A571BD | 1.02 | 1.3 | 1.39 | 1.28 | 1.1 | 1.07 | 0.86 | 1.2 | 1.19 | 1.27 | 1.42 | 1.3 | 1.22 | 1.03 | 0.98 | 0.97 | -0.19 | 0.0475 | -0.1975 | 0.267108 | 0.780667 | 0.299043 | 1 | 0.996511 | 0.828257 |
| Srcin1 | B1AQX7 | 0.8 | 0.25 | 0.74 | 0.49 | -1.08 | 0.97 | 0.77 | 0.44 | 2.5 | -1.72 | 0.9 | 0.43 | 2.68 | -1.65 | -0.55 | 0.04 | -0.295 | -0.0425 | -0.44 | 0.515914 | 0.957338 | 0.600953 | 1 | 0.996755 | 0.930802 |
| Srcin1 | B1AQX9 | 0.45 | 0.85 | 0.92 | 0.68 | 1.49 | 0.24 | 1.21 | 0.81 | 0.56 | 0.62 | 0.99 | 0.7 | 0.35 | 1.71 | 0.48 | 0.86 | 0.2125 | -0.0075 | 0.125 | 0.471854 | 0.969257 | 0.703956 | 1 | 0.996755 | 0.957084 |
| Srd5a3 | Q9WUP4 | -1.44 | -1.76 | 0.09 | 0.36 | -0.69 | -1.37 | 0.02 | 0.35 | -1.74 | 1.35 | -0.02 | -1.71 | -1.65 | -0.89 | -0.28 | 0.18 | 0.265 | 0.1575 | 0.0275 | 0.659968 | 0.849014 | 0.96391 | 1 | 0.996511 | 0.9927 |
| Srebf2 | Q3U1N2 | -1.34 | -1.2 | -1.83 | -1.99 | -1.41 | -2.24 | -1.93 | -2.44 | 3.67 | -0.55 | -2.27 | -0.68 | 3.89 | 2.82 | -1.09 | -1.7 | -0.415 | 1.6325 | 2.57 | 0.181766 | 0.180906 | 0.066234 | 1 | 0.996511 | 0.664936 |
| Srek1 | Q8BZX4-2 | -1.34 | -2.35 | -1.97 | -1.44 | -2.23 | -2.1 | -1.82 | -1.4 | -1.37 | -1.37 | -1.81 | -2.49 | -0.69 | -1.93 | -1.01 | -1.35 | -0.1125 | 0.015 | 0.53 | 0.706378 | 0.965582 | 0.157975 | 1 | 0.996755 | 0.751845 |
| Srgap1 | Q91Z69 | 0.17 | -0.23 | 0.43 | 0.57 | -0.09 | 0.08 | 0.41 | 0.32 | 0.77 | 0.58 | 0.23 | 0.16 | 0.99 | 0.08 | 0.59 | 0.33 | -0.055 | 0.2 | 0.2625 | 0.810456 | 0.433523 | 0.363897 | 1 | 0.996511 | 0.858475 |
| Srgap2 | Q91Z67 | 0.49 | 0.12 | 0.74 | 0.33 | 0.23 | 0.79 | 1.03 | 0.74 | 0.41 | 0.55 | 1.2 | 0.75 | 0.19 | 0.41 | 0.6 | 0.87 | 0.2775 | 0.3075 | 0.0975 | 0.250997 | 0.226532 | 0.680718 | 1 | 0.996511 | 0.955729 |
| Srgap3 | E9QN14 | 2.26 | 1.53 | -1.19 | -3.52 | 1.44 | 4.4 | -0.77 | 0.63 | -1.58 | -2.7 | -1.3 | 2.6 | -1.52 | -3.15 | -1.92 | 0.85 | 1.655 | -0.515 | -1.205 | 0.296691 | 0.743255 | 0.397215 | 1 | 0.996511 | 0.867334 |
| Srgap3 | F8VPQ4 | 0.05 | 0.12 | 0.47 | 0.41 | 0.35 | 0.17 | 0.44 | 0.54 | 0.05 | 0.05 | 0.55 | -0.06 | -0.02 | 0.41 | 0.24 | 0.21 | 0.1125 | -0.115 | -0.0525 | 0.530943 | 0.59238 | 0.795032 | 1 | 0.996511 | 0.969498 |
| Sri | Q6P069-2 | -0.42 | -0.21 | 0.1 | 0.06 | 0.15 | -0.49 | 0.38 | 0.12 | -0.72 | -0.45 | 0.08 | -0.38 | -0.87 | -0.22 | 0.19 | 0.18 | 0.1575 | -0.25 | -0.0625 | 0.514549 | 0.304872 | 0.831122 | 1 | 0.996511 | 0.973982 |
| Srm | Q64674 | -1.18 | -1.07 | -1.33 | -0.98 | -0.84 | -1.33 | -0.79 | -0.99 | -1.18 | -1.99 | -1.11 | -0.96 | -1.58 | -0.68 | -1.33 | -1.1 | 0.1525 | -0.17 | -0.0325 | 0.419191 | 0.521306 | 0.893496 | 1 | 0.996511 | 0.983001 |
| Srp14 | P16254 | -2.19 | -1.87 | -1.49 | -1 | -1.19 | -1.95 | -0.83 | -1.2 | -2.22 | -1.62 | -1.55 | -1.95 | -1.43 | -1.6 | -1.64 | -0.97 | 0.345 | -0.1975 | 0.2275 | 0.320847 | 0.521295 | 0.468274 | 1 | 0.996511 | 0.889423 |
| Srp19 | Q9D104 | -3.61 | -3.59 | -3.06 | -3.65 | -3.32 | -3.34 | -3.29 | -3.92 | -3.54 | -3.2 | -2.97 | -3.19 | -3.65 | -3.63 | -2.99 | -3.64 | 0.01 | 0.2525 | -9.8E-17 | 0.964769 | 0.269449 | 1 | 1 | 0.996511 | 1 |
| Srp54 | P14576 | -1.5 | -1.6 | -1.02 | -0.97 | -1.25 | -1.57 | -0.73 | -1.05 | -1.11 | -1.07 | -1.04 | -1.43 | -1.09 | -1.28 | -1.24 | -0.99 | 0.1225 | 0.11 | 0.1225 | 0.627808 | 0.622159 | 0.587171 | 1 | 0.996511 | 0.92849 |
| Srp68 | Q8BMA6 | -1.46 | -1.75 | -0.42 | -0.33 | -1.66 | -1.59 | -0.8 | -0.82 | -0.88 | -0.17 | -0.67 | -1.57 | -0.52 | -0.91 | -0.93 | -0.55 | -0.2275 | 0.1675 | 0.2625 | 0.578158 | 0.702463 | 0.484116 | 1 | 0.996511 | 0.894252 |
| Srp72 | F8VQC1 | -1.36 | -1.51 | -1.56 | -1.9 | -1.45 | -1.52 | -1.53 | -2.01 | -1.58 | -1.03 | -1.72 | -1.37 | -1.23 | -1.38 | -1.77 | -1.58 | -0.045 | 0.1575 | 0.0925 | 0.824906 | 0.487847 | 0.671348 | 1 | 0.996511 | 0.952631 |
| Srp9 | P49962 | 0.19 | -1.37 | 0.13 | -1.61 | -0.32 | -0.44 | 0.07 | 0.12 | -2.76 | -1.17 | 0.07 | 0.12 | -1.93 | -2.23 | 0.94 | 0.84 | 0.5225 | -0.27 | 0.07 | 0.277368 | 0.720363 | 0.936802 | 1 | 0.996511 | 0.989767 |
| Srpk1 | O70551 | -0.8 | -1.07 | -0.04 | -0.98 | -0.54 | -0.84 | -0.1 | -0.31 | -1.62 | -0.77 | -0.08 | -0.65 | -1.61 | -0.95 | -0.38 | 0.4 | 0.275 | -0.0575 | 0.0875 | 0.347496 | 0.879891 | 0.848801 | 1 | 0.996511 | 0.976225 |
| Srpk2 | A0A0R4J12 | -0.93 | -1.18 | -0.43 | -0.67 | -0.63 | -0.93 | -0.79 | -0.87 | -1.2 | -0.18 | -0.37 | -0.66 | -1.08 | -0.47 | -0.49 | -1.15 | -0.0025 | 0.2 | 0.005 | 0.990261 | 0.489799 | 0.985198 | 1 | 0.996511 | 0.996154 |
| Srpra | Q9DBG7 | -0.28 | -0.51 | -0.15 | -0.05 | 0.2 | -0.71 | 0.41 | 0.14 | -0.01 | 0.63 | -0.1 | -0.33 | 0.39 | 0.32 | 0.74 | 0.57 | 0.2575 | 0.295 | 0.7525 | 0.355828 | 0.259205 | 0.004649 | 1 | 0.996511 | 0.550241 |
| Srprb | P47758 | -0.35 | 0.02 | -0.29 | -0.3 | 0.89 | 0.15 | 0.31 | 0.92 | -0.01 | 0.3 | 0.49 | -0.05 | 0.09 | 0.93 | 0.2 | 0.69 | 0.7975 | 0.4125 | 0.7075 | 0.007485 | 0.067561 | 0.018703 | 1 | 0.996511 | 0.602211 |
| Srr | Q9QZX7-2 | -0.9 | -0.68 | 0.76 | 1.03 | -0.78 | -0.97 | 0.89 | 0.99 | -0.76 | 1.05 | 0.42 | -0.84 | -1.02 | -0.36 | 0.35 | 0.08 | -0.02 | -0.085 | -0.29 | 0.975568 | 0.890403 | 0.589337 | 1 | 0.996511 | 0.929142 |
| Srrm1 | E9QKA4 | 0.63 | -0.52 | -1.13 | -1.43 | -0.48 | -0.16 | -1.37 | -0.8 | -0.26 | -0.51 | -0.91 | -1.02 | 0.74 | 0.53 | 0.9 | -1.59 | -0.09 | -0.0625 | 0.7575 | 0.852511 | 0.891486 | 0.277945 | 1 | 0.996511 | 0.822084 |
| Srrm2 | Q8BTI8 | 1.3 | 0.38 | 0.43 | 0.36 | -0.26 | 0.44 | 0 | 0.93 | 0.38 | 0.58 | 0.37 | -0.14 | 0.93 | 0.83 | 0.93 | -0.06 | -0.34 | -0.32 | 0.04 | 0.326025 | 0.279848 | 0.904487 | 1 | 0.996511 | 0.985525 |
| Srrm3 | D3Z3C2 | 0.12 | -0.08 | -0.6 | -2.9 | -0.65 | -0.44 | -2.11 | -1.73 | -0.16 | 0.86 | -1.97 | -1.85 | 1.08 | -0.79 | -0.93 | -0.5 | -0.3675 | 0.085 | 0.58 | 0.616378 | 0.922973 | 0.450214 | 1 | 0.996755 | 0.884817 |
| Srrt | Q99MR6-3 | -1.52 | -3.31 | -3.53 | -3.42 | -3.35 | -2.12 | -3.69 | -3.67 | -2.88 | -2.24 | -3.3 | -2.84 | -1.48 | -2.81 | -2.98 | -3.89 | -0.2625 | 0.13 | 0.155 | 0.637418 | 0.790573 | 0.805525 | 1 | 0.996511 | 0.971715 |
| Srsf1 | H7BX95 | 0.86 | -0.02 | 0.01 | -0.11 | -0.24 | 0.03 | -0.01 | 0.43 | -0.7 | -0.4 | -0.26 | -0.27 | -0.15 | -0.41 | 0.43 | -0.11 | -0.1325 | -0.5925 | -0.245 | 0.627471 | 0.049754 | 0.423285 | 1 | 0.996511 | 0.875782 |
| Srsf10 | Q3TFP0 | 2.55 | 1.03 | 0.34 | 0.63 | 0.53 | 1.51 | 0.52 | 1.25 | 1.13 | -0.43 | 0.34 | 0.84 | 0.85 | 0.88 | 0.64 | 0 | -0.185 | -0.6675 | -0.545 | 0.717198 | 0.24674 | 0.289614 | 1 | 0.996511 | 0.825111 |
| Srsf11 | Q3UIX4 | -1.47 | -2.19 | -2.58 | -2.65 | -1.79 | -2.21 | -1.9 | -2.36 | -2.03 | -1.73 | -2.65 | -2.01 | -2.72 | -1.38 | -1.72 | -2.43 | 0.1575 | 0.1175 | 0.16 | 0.601357 | 0.723058 | 0.688108 | 1 | 0.996511 | 0.955729 |
| Srsf12 | A3KG57 | 0.78 | -0.32 | 0.06 | 0.18 | -0.12 | -0.14 | -0.17 | 0.62 | -0.74 | 0.19 | -0.3 | -0.59 | -0.22 | 0.31 | 0.19 | 0.87 | -0.1275 | -0.535 | 0.1125 | 0.668414 | 0.111769 | 0.730058 | 1 | 0.996511 | 0.958635 |
| Srsf2 | Q62093 | 1.32 | 0.52 | 0.27 | 0.17 | -0.44 | 0.5 | -0.13 | 0.52 | -0.19 | -0.14 | 0.01 | 0.21 | 0.5 | -0.49 | 0.58 | -0.36 | -0.4575 | -0.5975 | -0.5125 | 0.202958 | 0.062467 | 0.194167 | 1 | 0.996511 | 0.778959 |
| Srsf3 | P84104-2 | 1.11 | -0.61 | -0.4 | -0.01 | -0.95 | 0.31 | -0.53 | 0.51 | -0.09 | 0.23 | -0.65 | -0.29 | 0.32 | -0.4 | 0.23 | -1.16 | -0.1875 | -0.2225 | -0.275 | 0.695755 | 0.586535 | 0.57127 | 1 | 0.996511 | 0.924332 |
| Srsf4 | Q8VE97 | 0.82 | -0.15 | 0.33 | 0.49 | -0.71 | 0.03 | -0.21 | 0.88 | 0.17 | 0.66 | 0.12 | -1.04 | 1.19 | 0.16 | 0.68 | -0.28 | -0.375 | -0.395 | 0.065 | 0.326049 | 0.33238 | 0.860355 | 1 | 0.996511 | 0.977204 |
| Srsf5 | O35326 | 1.48 | 0.67 | 0.45 | 0.26 | -0.11 | 0.5 | -0.19 | 0.72 | 0.25 | -0.09 | 0.15 | 0.21 | 0.13 | 0.04 | 1.07 | -0.2 | -0.485 | -0.585 | -0.455 | 0.176882 | 0.069022 | 0.247875 | 1 | 0.996511 | 0.803677 |
| Srsf6 | Q3TWW8 | 2.35 | 0.43 | 1.09 | 0.66 | -0.71 | 1.35 | -0.07 | 1.96 | -0.38 | 0.13 | 0.77 | 0.59 | 0.79 | -1 | 2.08 | 0.59 | -0.5 | -0.855 | -0.5175 | 0.470066 | 0.095431 | 0.462247 | 1 | 0.996511 | 0.888262 |
| Srsf7 | A0A3Q4L3 | 1.09 | 0.24 | -0.08 | -0.13 | -0.39 | 0.65 | -0.15 | 0.57 | 0.23 | -0.25 | -0.11 | -0.26 | 0.51 | 0.47 | 0.42 | -0.38 | -0.11 | -0.3775 | -0.025 | 0.764062 | 0.240931 | 0.943107 | 1 | 0.996511 | 0.991519 |
| Srsf9 | Q9D0B0 | 2.07 | 0.56 | 1.18 | 0.53 | -0.43 | 1.61 | 0.46 | 0.68 | 1.14 | -0.23 | 0.58 | 0.63 | 1.78 | -1.01 | 1.67 | 1.27 | -0.505 | -0.555 | -0.1575 | 0.335642 | 0.222348 | 0.816828 | 1 | 0.996511 | 0.972768 |
| Srxn1 | A2AQU8 | -0.86 | 0.13 | -0.82 | -0.3 | -0.79 | -0.29 | -1.57 | -0.96 | -0.58 | -0.25 | -1 | -0.73 | -0.36 | -0.4 | -0.11 | 0.2 | -0.44 | -0.1775 | 0.295 | 0.219619 | 0.546184 | 0.323039 | 1 | 0.996511 | 0.840788 |
| Ss18 | A0A3Q4EI( | -0.97 | -0.33 | 0.47 | 0.83 | -0.98 | -1.23 | -0.82 | -0.09 | -0.34 | 0.36 | 0.41 | -1.41 | 0.97 | -0.41 | -0.2 | -0.56 | -0.78 | -0.245 | -0.05 | 0.10472 | 0.652565 | 0.919876 | 1 | 0.996511 | 0.98688 |
| Ss18l1 | Q8BW22 | -0.67 | -0.54 | 0.41 | 0.18 | -0.78 | -0.84 | -0.18 | -0.13 | -0.82 | 1.63 | -0.11 | -0.75 | -0.78 | -1.05 | -0.2 | -0.52 | -0.3275 | 0.1425 | -0.4825 | 0.320038 | 0.805462 | 0.164251 | 1 | 0.996511 | 0.758352 |
| Ssb | P32067 | -0.75 | -1.3 | -1 | -1.2 | -1.54 | -1.55 | -0.86 | -0.84 | -2.22 | -1.53 | -1.06 | -1.73 | -1.41 | -1.45 | -0.5 | -1.34 | -0.135 | -0.5725 | -0.1125 | 0.588578 | 0.06804 | 0.686926 | 1 | 0.996511 | 0.955729 |
| Ssbp1 | Q8R2K3 | 1.18 | 0.83 | 0.45 | 0.73 | 0.8 | 0.35 | 0.46 | 0.57 | 0.42 | 0.41 | 0.54 | 0.48 | 0.42 | 1.06 | 0.51 | 0.59 | -0.2525 | -0.335 | -0.1525 | 0.244615 | 0.124414 | 0.536822 | 1 | 0.996511 | 0.912212 |
| Ssbp2 | A0A338P6( | -9.06 | -11.43 | -3.25 | -3.52 | -9.73 | -12.49 | -2.72 | -3.53 | -20.9 | -3.23 | -2.96 | -8.78 | -10.69 | -11.11 | -3.58 | -3.53 | -0.3025 | -2.1525 | -0.4125 | 0.913572 | 0.606656 | 0.874061 | 1 | 0.996511 | 0.979626 |
| Ssbp3 | B1AS34 | -8.09 | -7.88 | -5.39 | -5.06 | -9.37 | -8.2 | -6.36 | -4.86 | -5.87 | -0.77 | -3.73 | -6.4 | -3.03 | -10.14 | -5.77 | -5.67 | -0.5925 | 2.4125 | 0.4525 | 0.607135 | 0.101511 | 0.761586 | 1 | 0.996511 | 0.963123 |
| Ssh1 | Q76I79-2 | -0.44 | -0.19 | 0.52 | 0.55 | 0.22 | -0.02 | 0.41 | 0.57 | -0.16 | 0.82 | 0.8 | -0.52 | -0.34 | -0.34 | 0.34 | 1.14 | 0.185 | 0.125 | 0.09 | 0.518763 | 0.757101 | 0.827958 | 1 | 0.996511 | 0.973757 |
| Ssh2 | Q5SW75 | -0.57 | -0.25 | -0.14 | -0.17 | -0.37 | -0.54 | -0.21 | -0.14 | -0.11 | 0.21 | -0.14 | -0.11 | 0 | 0.39 | -0.32 | -0.04 | -0.0325 | 0.245 | 0.29 | 0.855817 | 0.21426 | 0.2179 | 1 | 0.996511 | 0.787447 |
| Ssh3 | A0A498WG | 0.19 | -0.94 | -0.41 | -0.25 | 0.28 | 0.32 | -0.79 | -0.15 | 0.21 | 0.68 | 0.04 | -1.52 | 0.32 | -0.45 | 0.06 | -0.07 | 0.2675 | 0.205 | 0.3175 | 0.435622 | 0.679778 | 0.301423 | 1 | 0.996511 | 0.830308 |
| Ssna1 | Q9JJ94 | -1.4 | -1.12 | -1.66 | -1.88 | 1.39 | -0.5 | -0.96 | -1.35 | -0.11 | 0.07 | -1.67 | 1.02 | -0.91 | -0.24 | -2.22 | -0.81 | 1.16 | 1.3425 | 0.47 | 0.070346 | 0.032225 | 0.28938 | 1 | 0.996511 | 0.824705 |
| Sspn | Q62147 | -1.92 | -0.7 | 0.1 | -0.16 | -2.06 | -1.34 | -0.25 | -0.43 | 0.09 | 0.56 | 0.03 | -1.01 | 0.55 | -1.24 | -0.05 | 0.01 | -0.35 | 0.5875 | 0.4875 | 0.53873 | 0.274092 | 0.379832 | 1 | 0.996511 | 0.860984 |
| Ssr1 | Q9CY50 | -0.67 | -0.36 | 0.26 | 0.05 | 0.28 | -0.31 | 1.04 | 0.56 | -0.98 | -0.24 | 0.42 | -0.21 | -0.47 | -0.35 | 0.83 | 1.3 | 0.5725 | -0.0725 | 0.5075 | 0.118749 | 0.834695 | 0.283734 | 1 | 0.996511 | 0.822669 |
| Ssr3 | Q9DCF9-2 | -1.57 | -0.04 | -1.02 | -1.3 | 0.02 | -1.19 | -0.06 | -0.92 | -0.38 | -1.88 | -0.8 | -0.88 | 0.37 | -0.56 | 0.82 | 0.23 | 0.445 | -0.0025 | 1.1975 | 0.308728 | 0.995412 | 0.018274 | 1 | 0.999285 | 0.602211 |
| Ssr4 | Q9D8L3 | 0.54 | -0.44 | 1.45 | -0.75 | 2.08 | 0.85 | 4.59 | 0.02 | -2.39 | -2.54 | 1.22 | 0.51 | -1.53 | -3.73 | 3.81 | 4.75 | 1.685 | -1 | 0.625 | 0.119549 | 0.323741 | 0.739197 | 1 | 0.996511 | 0.958796 |
| Ssrp1 | Q08943-2 | 0.07 | -0.75 | -0.19 | -0.18 | 0.25 | -0.2 | 0.69 | -0.85 | -0.61 | -1.66 | 0.29 | 0.28 | -1.02 | -0.57 | 0.37 | 0.85 | 0.235 | -0.1625 | 0.17 | 0.515 | 0.726399 | 0.698734 | 1 | 0.996511 | 0.955729 |
| Sst | P60041 | 1.08 | 0.91 | 0.76 | 0.67 | 0.34 | 1.33 | -0.66 | 0.63 | 0.88 | 0.14 | 0.6 | 1 | 1.43 | -0.4 | 1.09 | 0.61 | -0.445 | -0.2 | -0.1725 | 0.28192 | 0.412517 | 0.663524 | 1 | 0.996511 | 0.949825 |
| Sstr1 | P30873 | -3.35 | 0.8 | -0.57 | -0.86 | 0.35 | 0.67 | 0.51 | -0.86 | 0.71 | 0.97 | 0.6 | -0.98 | 2.3 | 1.15 | -0.92 | -0.04 | 1.1625 | 1.32 | 1.6175 | 0.190246 | 0.157597 | 0.131365 | 1 | 0.996511 | 0.726398 |
| Sstr2 | P30875 | -0.09 | 0.55 | -0.38 | -1 | 0.53 | 0.44 | -0.38 | -1.24 | 0.48 | -1.69 | -0.26 | 0.07 | 0.71 | -0.19 | -1.58 | -0.77 | 0.0675 | -0.12 | -0.2275 | 0.889158 | 0.820444 | 0.673024 | 1 | 0.996511 | 0.952944 |
| Ssu72 | Q9CY97-2 | -3.06 | -3.3 | -2.26 | -2.32 | -3.88 | -3.11 | -2.69 | -2.26 | -0.47 | -1.6 | -2.79 | -3.19 | 0.18 | -1.8 | -2.22 | -3 | -0.25 | 0.7225 | 1.025 | 0.544031 | 0.257292 | 0.147219 | 1 | 0.996511 | 0.746406 |
| St13 | Q99L47 | -0.03 | 0.07 | -1.2 | -0.95 | -0.02 | 0.06 | -0.97 | -1.1 | -0.18 | -1.06 | -1.22 | 0.28 | 0.23 | -0.17 | -1.21 | -1.36 | 0.02 | -0.0175 | -0.1 | 0.961635 | 0.969063 | 0.832829 | 1 | 0.996755 | 0.975129 |
| St3gal5 | O88829 | 0.09 | -0.7 | -0.86 | -0.82 | -0.49 | 0.31 | -0.77 | -1.11 | 0.3 | -0.57 | -0.69 | -0.2 | 0.87 | -0.78 | -0.27 | -0.21 | 0.0575 | 0.2825 | 0.475 | 0.873117 | 0.383525 | 0.252427 | 1 | 0.996511 | 0.80892 |
| St6galnac5 | Q6GTI0 | -0.36 | -0.09 | -0.89 | -1.66 | -0.34 | 0.16 | -1.4 | -1.48 | 1.96 | -2.75 | -1.1 | -0.88 | 3.11 | -1.01 | 0.41 | -0.63 | -0.015 | 0.0575 | 1.22 | 0.97559 | 0.950824 | 0.194182 | 1 | 0.996755 | 0.778959 |
| St7 | Q99M96-9 | -1.07 | -0.93 | -0.94 | -0.97 | -0.38 | -0.87 | -0.11 | -0.38 | -0.81 | -1.3 | -1.03 | -0.82 | -0.11 | 0.47 | -0.2 | 0.06 | 0.5425 | -0.0125 | 1.0325 | 0.021059 | 0.945465 | 0.000876 | 1 | 0.996755 | 0.3804 |
| St8sia3 | Q64689 | -1.01 | -1.12 | -1.24 | -1 | -0.67 | -1.18 | -0.6 | -1.16 | -0.72 | -1.13 | -1.33 | -0.92 | -0.77 | -0.95 | -0.66 | -1.16 | 0.19 | 0.0675 | 0.2075 | 0.352068 | 0.730957 | 0.30316 | 1 | 0.996511 | 0.831227 |
| St8sia5 | P70126-3 | -1.54 | -0.83 | -1.27 | -1.33 | -0.23 | -0.43 | -1.04 | -1.34 | -0.09 | -1.42 | -0.36 | -0.63 | 0.6 | -0.17 | -0.76 | -0.77 | 0.4825 | 0.6175 | 0.9675 | 0.131854 | 0.083432 | 0.02163 | 1 | 0.996511 | 0.621926 |
| Stab1 | G3X973 | -1.01 | -1 | -2.05 | -1.79 | -0.93 | -1.26 | -2.76 | -1.93 | -0.22 | -0.75 | -1.65 | -0.66 | -0.93 | -0.54 | -1.58 | -1.91 | -0.2575 | 0.6425 | 0.2225 | 0.572521 | 0.124773 | 0.577982 | 1 | 0.996511 | 0.927198 |
| Stac2 | Q8R1B0 | 0.24 | 1.03 | 3.25 | 3.11 | 0.91 | 0.25 | 3.16 | 3.01 | 0.51 | 3.5 | 2.97 | 0.41 | 0.41 | 0.63 | 2.99 | 3.43 | -0.075 | -0.06 | -0.0425 | 0.936728 | 0.951756 | 0.965097 | 1 | 0.996755 | 0.9927 |
| Stag1 | F8WHU7 | 0.05 | -0.12 | 0.1 | 0.08 | -0.26 | -0.59 | 0.19 | 0.41 | -0.71 | 0.35 | -0.13 | -0.59 | -0.45 | -0.36 | 1.18 | -0.13 | -0.09 | -0.2975 | 0.0325 | 0.713258 | 0.278258 | 0.930696 | 1 | 0.996511 | 0.989713 |
| Stag2 | A2AFF6 | 0.01 | -0.35 | -0.67 | -0.57 | -0.2 | -0.47 | -0.55 | -0.57 | -0.31 | -0.14 | -0.61 | -0.3 | -0.14 | -0.07 | 0.28 | -0.42 | -0.0525 | 0.055 | 0.3075 | 0.797154 | 0.801084 | 0.22991 | 1 | 0.996511 | 0.787745 |
| Stam | Q3UGN9 | -0.03 | 0.16 | 0.03 | 0.04 | 0.29 | -0.07 | 0.13 | -0.12 | -0.07 | -0.4 | 0.15 | 0.19 | -0.22 | 0.66 | -0.37 | -0.5 | 0.0075 | -0.0825 | -0.1575 | 0.963135 | 0.67286 | 0.582537 | 1 | 0.996511 | 0.927734 |
| Stam2 | O88811 | 1.36 | 1.88 | 0.45 | 0.23 | 1.48 | 0.82 | 0.18 | -0.12 | 1.27 | 1.33 | 0.01 | 1.04 | 1.49 | 2.46 | 0.17 | 0.22 | -0.39 | -0.0675 | 0.105 | 0.431111 | 0.884103 | 0.864554 | 1 | 0.996511 | 0.977686 |
| Stambp | Q9CQ26 | -1.2 | -0.99 | -0.46 | -0.36 | -1.29 | -0.78 | -0.14 | -0.46 | -0.89 | -0.11 | -0.37 | -0.92 | -0.97 | -0.22 | -0.62 | -0.72 | 0.085 | 0.18 | 0.12 | 0.786229 | 0.543126 | 0.667177 | 1 | 0.996511 | 0.951142 |
| Stambpl1 | Q76N33 | -0.06 | 0.04 | 0.8 | 0.79 | 0.07 | 0.12 | 0.67 | 0.78 | -0.05 | 0.32 | 0.67 | 0.05 | -0.13 | -0.28 | 0.52 | 0.63 | 0.0175 | -0.145 | -0.2075 | 0.95275 | 0.62169 | 0.533949 | 1 | 0.996511 | 0.911825 |
| Stard10 | Q9JMD3 | -0.01 | -0.4 | -0.63 | 0.13 | -0.71 | -0.45 | -0.01 | -0.14 | -0.3 | -0.52 | -0.46 | -0.7 | -0.53 | -0.09 | -0.61 | -0.58 | -0.1 | -0.2675 | -0.225 | 0.688037 | 0.257923 | 0.375667 | 1 | 0.996511 | 0.859611 |
| Stard3 | Q61542 | 0.44 | -1.49 | -0.47 | -0.49 | -1.33 | 0.34 | -1.18 | -0.46 | 1.06 | -0.68 | -0.26 | 1.19 | 0.6 | -0.32 | -0.35 | -0.56 | -0.155 | 0.83 | 0.345 | 0.760024 | 0.16678 | 0.447133 | 1 | 0.996511 | 0.884817 |
| Stard3nl | A0A1Y7VJA | -0.14 | 0.15 | -0.22 | -0.27 | 0.28 | -0.45 | -0.1 | -0.07 | 0.7 | 0.68 | 0 | -0.35 | 0.23 | 0.87 | -0.29 | 0.22 | 0.035 | 0.3775 | 0.3775 | 0.865151 | 0.20999 | 0.196334 | 1 | 0.996511 | 0.78004 |
| Stard5 | D3YU00 | -1.75 | -2.06 | -0.1 | 0.26 | -1.05 | -0.81 | -0.09 | 0.15 | -0.55 | 0.5 | -0.06 | -0.59 | -1.15 | -1 | -0.41 | -0.21 | 0.4625 | 0.7375 | 0.22 | 0.442236 | 0.227425 | 0.702382 | 1 | 0.996511 | 0.956859 |
| Stard7 | Q8R1R3 | 1.74 | 1.43 | 1.02 | 0.11 | 1.27 | 1.96 | 0.8 | 0.87 | 0.32 | 0 | 1.25 | 2.18 | 0.43 | -0.36 | 0.61 | 0.92 | 0.15 | -0.1375 | -0.675 | 0.718951 | 0.805585 | 0.13967 | 1 | 0.996511 | 0.735387 |
| Stat1 | Q8C3V4 | 0.06 | -0.15 | -0.29 | -0.29 | -0.27 | -0.4 | -0.02 | -0.28 | -0.48 | 0.17 | -0.25 | -0.22 | -0.99 | -0.12 | -0.44 | -0.29 | -0.075 | -0.0275 | -0.2925 | 0.658555 | 0.893023 | 0.248387 | 1 | 0.996511 | 0.803985 |
| Stat2 | Q9QXJ2 | 1.23 | 0.46 | 0.07 | -0.54 | 0.16 | 0.59 | -0.72 | -0.4 | -0.61 | 0.01 | -0.32 | 0.76 | -1.27 | -0.96 | -0.26 | -0.16 | -0.3975 | -0.345 | -0.9675 | 0.377782 | 0.447472 | 0.050241 | 1 | 0.996511 | 0.658454 |
| Stat3 | B7ZC18 | -0.4 | -0.37 | -0.59 | -0.25 | 0.05 | -0.64 | -0.15 | -0.42 | -0.45 | -0.61 | -0.3 | -0.42 | -0.69 | -0.18 | -0.79 | -0.36 | 0.1125 | -0.0425 | -0.1025 | 0.577048 | 0.804269 | 0.633885 | 1 | 0.996511 | 0.941132 |
| Stat5b | P42232 | 0.15 | -0.12 | 0.74 | 0.31 | 0.31 | -0.04 | 1.35 | 0.78 | -0.25 | 0.11 | 0.77 | 0.13 | -0.05 | -0.16 | 0.98 | 0.93 | 0.33 | -0.08 | 0.155 | 0.344335 | 0.781376 | 0.661961 | 1 | 0.996511 | 0.949645 |
| Stat6 | P52633 | -1.78 | -1.67 | -1.12 | -1.11 | -2.25 | -1.58 | -1.04 | -1.09 | -1.31 | -2.2 | -1.64 | -1.09 | -2.8 | -1.7 | -2.63 | -1.25 | -0.07 | -0.14 | -0.675 | 0.829143 | 0.647361 | 0.11665 | 1 | 0.996511 | 0.718608 |
| Stau1 | A2A5S3 | -1.65 | -1.92 | -1.15 | -1.54 | -1.97 | -1.94 | -1.69 | -1.56 | -0.92 | -0.8 | -1.43 | -1.69 | 0.03 | -1.94 | -1.14 | -1.35 | -0.225 | 0.355 | 0.465 | 0.309039 | 0.222127 | 0.288309 | 1 | 0.996511 | 0.824705 |
| Stau2 | Q8CJ67-2 | -0.86 | -1.13 | -1.05 | -0.91 | -0.66 | -0.98 | -0.47 | -0.8 | -1.25 | -0.25 | -0.92 | -0.65 | -0.63 | -0.51 | -0.43 | -0.56 | 0.26 | 0.22 | 0.455 | 0.161119 | 0.381893 | 0.027141 | 1 | 0.996511 | 0.635993 |
| Steap2 | D3YUM6 | -0.1 | 0.64 | 0.61 | 1.5 | 0.02 | 0.42 | 0.89 | 1.32 | 0.12 | 0.63 | 1.47 | -0.29 | 0.4 | -0.91 | 1.91 | 2.1 | 7.85E-17 | -0.18 | 0.2125 | 1 | 0.702126 | 0.763735 | 1 | 0.996511 | 0.964134 |
| Steap3 | E9QN92 | 0.1 | 0.82 | 0.82 | 1.04 | 0.79 | 0.76 | 0.73 | 0.88 | 0.25 | 1.01 | 0.93 | 0.39 | 0.43 | 0.11 | 0.99 | 1.12 | 0.095 | -0.05 | -0.0325 | 0.678059 | 0.862665 | 0.919055 | 1 | 0.996511 | 0.986868 |
| Steep1 | Q8VDP2 | -2.57 | -3.54 | -1.15 | -0.79 | -5.11 | -2.88 | -1.95 | -1.25 | -1.33 | 0.05 | -1.24 | -3.55 | 0.56 | -2.95 | -1.37 | -1.69 | -0.785 | 0.495 | 0.65 | 0.416853 | 0.579497 | 0.461389 | 1 | 0.996511 | 0.887902 |
| Stim1 | A0A1B0GR | -0.1 | -0.01 | -0.18 | -0.01 | 0.25 | -0.41 | 0 | -0.07 | 0.47 | -0.2 | -0.28 | 0.02 | 0.85 | 0.54 | 0.47 | 0.08 | 0.0175 | 0.0775 | 0.56 | 0.924306 | 0.71793 | 0.027773 | 1 | 0.996511 | 0.635993 |
| Stim2 | P83093 | -0.46 | -0.09 | 0.05 | 0 | -0.04 | -0.62 | 0.04 | 0.05 | -0.19 | -0.16 | -0.14 | -0.53 | 0.5 | 0.44 | 0.55 | 0.31 | -0.0175 | -0.13 | 0.575 | 0.936771 | 0.517518 | 0.016054 | 1 | 0.996511 | 0.589387 |
| Sting1 | Q3TBT3-3 | -0.35 | 0.28 | -0.46 | 0.09 | 0.06 | 0.35 | -0.6 | -0.11 | 0.51 | -0.37 | -0.51 | 0.2 | 0.26 | 0.6 | -1.02 | -0.01 | 0.035 | 0.0675 | 0.0675 | 0.89728 | 0.823592 | 0.859193 | 1 | 0.996511 | 0.976627 |
| Stip1 | Q60864 | 1.08 | 0.23 | 0.16 | 0.39 | -0.01 | 0.56 | -0.17 | 0.46 | -0.63 | 0.71 | 0.11 | 0.74 | -0.44 | -1.01 | 0.03 | -0.05 | -0.255 | -0.2325 | -0.8325 | 0.372186 | 0.537003 | 0.028084 | 1 | 0.996511 | 0.637553 |
| Stk11 | Q9WTK7-2 | -1.19 | -0.15 | 0.53 | 0.34 | -0.87 | -1.17 | 0.47 | 0.47 | 1.63 | -1.11 | 0.29 | -0.93 | 1.29 | 1.92 | 0.19 | 0.31 | -0.1575 | 0.0875 | 1.045 | 0.768068 | 0.896955 | 0.072497 | 1 | 0.996511 | 0.668066 |
| Stk11ip | Q3TAA7 | -1.14 | -0.1 | -1.88 | -0.98 | -0.43 | -1.16 | -1.09 | -1.53 | -0.85 | -0.27 | -0.09 | -0.58 | -0.55 | -0.51 | -0.25 | -1.25 | -0.0275 | 0.5775 | 0.385 | 0.945922 | 0.161143 | 0.356733 | 1 | 0.996511 | 0.855416 |
| Stk16 | A0A087WC | 0.15 | -0.26 | 0.47 | 0.87 | 0.09 | 0.41 | 1.32 | 1.05 | 0.25 | 1.04 | 1.46 | 0.15 | 0.73 | 0.02 | 1.46 | 1.37 | 0.41 | 0.4175 | 0.5875 | 0.267833 | 0.291322 | 0.1642 | 1 | 0.996511 | 0.758352 |

| | | | | | | | | | | | | | | | | | | | | | | | | | | |
|---|---|---|---|---|---|---|---|---|---|---|---|---|---|---|---|---|---|---|---|---|---|---|---|---|---|---|
| Stk24 | Q99KH8 | 0.57 | 0.46 | 0.64 | 0.74 | 0.09 | 0.25 | 0.78 | 1.1 | -0.89 | -0.36 | 0.65 | 0.36 | -1.1 | -1.21 | 0.21 | 0.26 | -0.0475 | -0.6625 | -1.0625 | 0.850807 | 0.08389 | 0.023017 | 1 | 0.996511 | 0.623147 |
| Stk25 | Q9Z2W1 | -0.81 | -1.46 | -0.37 | -0.02 | -1.38 | -1.27 | -0.55 | -0.18 | -0.69 | -0.12 | -0.08 | -1.21 | -0.12 | -1.13 | -0.27 | -0.65 | -0.18 | 0.14 | 0.1225 | 0.654008 | 0.722355 | 0.744314 | 1 | 0.996511 | 0.959371 |
| Stk26 | A2AD84 | -0.44 | 0.18 | 0.33 | -0.5 | -0.66 | -0.49 | -0.33 | 0 | 0.8 | 0.47 | -0.34 | 0.08 | 0.36 | 0.54 | -0.09 | 0.58 | -0.2625 | 0.36 | 0.455 | 0.331419 | 0.284296 | 0.132806 | 1 | 0.996511 | 0.728417 |
| Stk3 | A0A2I3BQE | -0.88 | -0.61 | -0.28 | -0.09 | -0.35 | -1.09 | -0.43 | 0 | -0.09 | 0.21 | -0.4 | -0.66 | -1.12 | 0.66 | 0.07 | -0.33 | -0.0025 | 0.23 | 0.285 | 0.993065 | 0.408103 | 0.480628 | 1 | 0.996511 | 0.893906 |
| Stk32c | Q8QZV4 | 0.36 | -0.24 | 0.67 | 0.66 | -0.52 | 0.36 | 0.79 | 0.47 | -0.16 | -0.36 | 0.21 | -0.07 | -0.34 | -0.13 | 0.05 | 0.28 | -0.0875 | -0.4575 | -0.3975 | 0.797299 | 0.108835 | 0.171357 | 1 | 0.996511 | 0.763316 |
| Stk38 | Q91VJ4 | 0.45 | -0.11 | 0.41 | 0.34 | 0.02 | 0.23 | 0.23 | 0.25 | 0.42 | 0.82 | 0.46 | 0.62 | -0.14 | 0.5 | -0.14 | 0.1 | -0.09 | 0.3075 | -0.1925 | 0.62578 | 0.158745 | 0.428568 | 1 | 0.996511 | 0.879165 |
| Stk38l | Q7TSE6 | 0.13 | 0.2 | -0.52 | -0.45 | -0.01 | 0.32 | -1.2 | -0.58 | -0.5 | -0.84 | -0.73 | 0.17 | -0.7 | -0.37 | -1.17 | -0.97 | -0.2075 | -0.315 | -0.6425 | 0.574486 | 0.310227 | 0.043126 | 1 | 0.996511 | 0.658454 |
| Stk39 | Q9Z1W9 | -1.05 | -1.06 | -0.59 | -0.7 | -0.52 | -0.86 | -0.37 | -0.77 | -1.29 | -0.74 | -0.68 | -1.19 | -1 | -1.02 | -1.18 | -0.74 | 0.22 | -0.125 | -0.135 | 0.285462 | 0.58816 | 0.52456 | 1 | 0.996511 | 0.908107 |
| Stk4 | Q9JI11 | -0.23 | -0.52 | -0.42 | -0.26 | 0.16 | -0.4 | 0.07 | -0.3 | -0.42 | -0.38 | -0.21 | -0.11 | -0.66 | -0.1 | -0.61 | -0.22 | 0.24 | 0.0775 | -0.04 | 0.230379 | 0.656686 | 0.850468 | 1 | 0.996511 | 0.976225 |
| Stmn1 | P54227 | 2.6 | -0.15 | -0.47 | -0.38 | -0.5 | 2.94 | 0.51 | -0.13 | 1.58 | -1.31 | 0.07 | 2.26 | 1.78 | -0.24 | -0.13 | -0.06 | 0.305 | 0.25 | -0.0625 | 0.75099 | 0.798019 | 0.93717 | 1 | 0.996511 | 0.989767 |
| Stmn2 | P55821 | 0.62 | 0.73 | 0.92 | 0.65 | 0.78 | 0.63 | 0.55 | 0.96 | 0.25 | 1.18 | 0.8 | 1.01 | 1.1 | 0.14 | 0.87 | 0.87 | -9.8E-18 | 0.08 | 0.015 | 1 | 0.740003 | 0.952513 | 1 | 0.996511 | 0.992476 |
| Stmn3 | O70166 | 4.21 | 1.09 | 2.26 | 0.82 | 1.71 | 4.24 | 2.08 | 1.03 | 1.44 | 1.22 | 1.88 | 4.26 | 1.58 | 1.03 | 2.41 | 2.47 | 0.17 | 0.105 | -0.2225 | 0.855209 | 0.910741 | 0.77109 | 1 | 0.996744 | 0.965933 |
| Stmn4 | Q05DI3 | 1.57 | 1.74 | 0.76 | 0.55 | 0.85 | 1.38 | 1.08 | 0.38 | 0.58 | 0.57 | 0.72 | 1.75 | 0.62 | -0.06 | 0.55 | 0.64 | -0.2325 | -0.25 | -0.7175 | 0.509784 | 0.528362 | 0.059413 | 1 | 0.996511 | 0.661738 |
| Stn1 | Q3UXU8 | -1.67 | -2.08 | -2.86 | -3.33 | -1.28 | -2.61 | -2.93 | -2.91 | -2.13 | -1.54 | -2.66 | -1.73 | -2.29 | -1.2 | -2.61 | -3.24 | 0.0525 | 0.47 | 0.15 | 0.916237 | 0.28658 | 0.775987 | 1 | 0.996511 | 0.966799 |
| Stom | P54116 | -0.52 | -1.09 | -0.41 | 0.02 | -0.58 | -0.31 | -0.57 | -0.5 | -0.72 | -0.37 | -0.79 | -0.85 | -0.78 | -0.52 | -0.72 | -0.93 | 0.01 | -0.1825 | -0.2375 | 0.967973 | 0.502757 | 0.387979 | 1 | 0.996511 | 0.864251 |
| Stoml2 | Q99JB2 | 0.69 | 1.02 | 0.79 | 0.71 | 0.99 | -0.33 | 0.39 | 0.55 | 0.64 | 0.92 | 0.87 | 0.41 | 0.79 | 1.51 | 0.55 | 0.75 | -0.4025 | -0.0925 | 0.0975 | 0.183824 | 0.634976 | 0.702839 | 1 | 0.996511 | 0.956859 |
| Ston2 | Q8BZ60 | 1.29 | 1.43 | 0.87 | 0.31 | 1.42 | 1.1 | 0.47 | 0.28 | 1.24 | 0.59 | -0.17 | 1.24 | 1.13 | 1.56 | 0.79 | 0.3 | -0.1575 | -0.25 | -0.03 | 0.656335 | 0.537286 | 0.933741 | 1 | 0.996511 | 0.989767 |
| Stox2 | Q499E5 | 0.61 | 0.57 | 0.48 | 0.15 | 0.8 | 0.86 | -0.22 | 0.1 | 0.29 | 3.73 | -0.29 | 0.97 | -0.05 | 0.72 | -0.06 | 0.22 | -0.0675 | 0.7225 | -0.245 | 0.813804 | 0.383498 | 0.333545 | 1 | 0.996511 | 0.845601 |
| Strada | Q3UUJ4 | 0.47 | -0.45 | 1.07 | 0.57 | -0.28 | 0.46 | 0.82 | 0.71 | -0.94 | 0.69 | 0.9 | 0.04 | -0.97 | -0.59 | 0.58 | 0.52 | 0.0125 | -0.2425 | -0.53 | 0.973834 | 0.620106 | 0.280127 | 1 | 0.996511 | 0.822084 |
| Stradb | Q8K4T3 | 0.96 | 0.93 | -0.4 | -0.09 | 0.87 | 0.17 | -0.02 | -0.23 | 0.66 | 0.63 | 0 | 0.99 | 1.45 | 1.06 | 0.08 | 0.29 | -0.1525 | 0.22 | 0.37 | 0.70337 | 0.575896 | 0.419326 | 1 | 0.996511 | 0.87558 |
| Strap | Q9Z1Z2 | -1.75 | -1.68 | -0.31 | -0.45 | -1.51 | -1.68 | -0.16 | -0.36 | -2.17 | -1.63 | -0.23 | -1.39 | -1.82 | -1.87 | -0.41 | -0.55 | 0.12 | -0.3075 | -0.115 | 0.812539 | 0.559313 | 0.823026 | 1 | 0.996511 | 0.972768 |
| Strbp | Q91WM1 | -0.32 | -1.12 | -1.15 | -1.54 | -1.2 | -0.61 | -1.85 | -0.91 | -1.63 | -1.76 | -1.28 | -1.58 | -0.49 | -1.27 | -0.68 | -1.71 | -0.11 | -0.53 | -0.005 | 0.756197 | 0.09122 | 0.98923 | 1 | 0.996511 | 0.996154 |
| Strip1 | Q8C079 | 0.58 | 0.6 | 0.59 | 0.61 | 0.58 | 0.63 | 0.53 | 0.7 | -0.06 | 0.09 | 0.73 | 0.57 | -0.49 | -0.04 | 0.35 | 0.23 | 0.015 | -0.2625 | -0.5825 | 0.913768 | 0.260301 | 0.030522 | 1 | 0.996511 | 0.641481 |
| Strip2 | Q8C9H6 | 0.03 | -0.3 | 0.26 | 0.42 | -0.19 | -0.32 | 0.31 | 0.29 | 0.24 | 0.55 | 0.06 | -0.09 | 0.22 | 0.24 | 0.48 | 0.17 | -0.08 | 0.0875 | 0.175 | 0.740276 | 0.71297 | 0.437047 | 1 | 0.996511 | 0.883161 |
| Strn | O55106 | 0.35 | 0 | 0.5 | 0.5 | 0.23 | 0.05 | 0.68 | 0.56 | -0.32 | 0.18 | 0.51 | 0.35 | -0.71 | -0.01 | 0.5 | 0.3 | 0.0425 | -0.1575 | -0.3175 | 0.842312 | 0.519894 | 0.309179 | 1 | 0.996511 | 0.834523 |
| Strn3 | Q9ERG2 | -0.66 | -0.47 | -0.19 | -0.34 | -0.23 | -0.83 | -0.09 | -0.22 | -1.16 | -0.34 | -0.11 | -0.94 | -1.43 | -0.51 | -0.51 | -0.83 | 0.0725 | -0.2225 | -0.405 | 0.739837 | 0.433532 | 0.152731 | 1 | 0.996511 | 0.750422 |
| Strn3 | B2RQS1 | 0.02 | 0.23 | 0.38 | 0.73 | 0.08 | -0.13 | 0.98 | 1 | -1.3 | -0.09 | 0.86 | -0.13 | -1.47 | -0.07 | 0.38 | 0.28 | 0.1425 | -0.505 | -0.56 | 0.661372 | 0.269829 | 0.215979 | 1 | 0.996511 | 0.787447 |
| Strn4 | P58404-2 | 0.93 | 0.5 | 0.52 | 0.6 | 0.73 | 0.83 | 0.61 | 0.63 | 0.53 | 0.69 | 0.66 | 0.81 | -0.01 | 0.96 | 0.19 | 0.25 | 0.0625 | 0.035 | -0.29 | 0.7096 | 0.846499 | 0.285541 | 1 | 0.996511 | 0.822669 |
| Sts | P50427 | -0.35 | 0.46 | 0.48 | -0.14 | -0.11 | -0.33 | 0 | 0.04 | -0.02 | -0.81 | -0.05 | 0.47 | 0.87 | 0.28 | 0.81 | -0.19 | -0.2125 | -0.215 | 0.33 | 0.390043 | 0.525581 | 0.331146 | 1 | 0.996511 | 0.845601 |
| Stt3a | P46978 | -0.38 | -0.61 | -0.64 | -1.15 | 0.56 | 0.08 | 0.43 | -0.26 | -1.58 | -0.62 | -0.61 | -0.25 | -0.32 | -1.23 | 1.27 | 1.13 | 0.8975 | -0.07 | 0.9075 | 0.0066 | 0.830379 | 0.139321 | 1 | 0.996511 | 0.73493 |
| Stt3b | A0A0R4J0D | -0.5 | -0.45 | -0.53 | -0.56 | 0.19 | -0.04 | -0.34 | -0.74 | 0.35 | -1.5 | -0.69 | -0.34 | 1.14 | 0.03 | 0.68 | 0.5 | 0.2775 | -0.035 | 1.0975 | 0.236432 | 0.925162 | 0.002243 | 1 | 0.996755 | 0.419079 |
| Stub1 | Q9WUD1 | 0.29 | -0.08 | -0.53 | -0.29 | -0.02 | 0.53 | -0.47 | -0.46 | 0.26 | 0.03 | -0.44 | 0.38 | 0.15 | -0.35 | -0.31 | -0.45 | 0.0475 | 0.21 | -0.0875 | 0.871106 | 0.440726 | 0.728976 | 1 | 0.996511 | 0.958635 |
| Stum | A0A1B0GS | 0.39 | 0.55 | 0.46 | 0.55 | 0.5 | 0.22 | 0.28 | 0.61 | 0.28 | 0.05 | 0.5 | 0.43 | 0.43 | -0.17 | 0.28 | -0.11 | -0.085 | -0.1725 | -0.38 | 0.600364 | 0.342546 | 0.09775 | 1 | 0.996511 | 0.698156 |
| Stx12 | Q9ER00 | 0.52 | 0.53 | 0.49 | 0.32 | 0.83 | 0.55 | 0.64 | 0.71 | 0.13 | -0.29 | 0.7 | 0.84 | 0.47 | 0.18 | 0.6 | 0.56 | 0.2175 | -0.12 | -0.0125 | 0.174446 | 0.668261 | 0.946774 | 1 | 0.996511 | 0.992376 |
| Stx16 | Q8BVI5 | -0.16 | 0.16 | 0.26 | 0.11 | 0.46 | -0.12 | 0.28 | 0.21 | -0.4 | -0.31 | 0.19 | 0.03 | -0.4 | -0.17 | 0.22 | 0.28 | 0.115 | -0.215 | -0.11 | 0.548547 | 0.319275 | 0.634281 | 1 | 0.996511 | 0.941132 |
| Stx17 | Q9D0I4 | 1.14 | 0.87 | -0.4 | -0.75 | 1.84 | 1.39 | -0.1 | -0.73 | 1.29 | 0.09 | -0.51 | 1.46 | 1.24 | 1.32 | 0.24 | 0.08 | 0.385 | 0.3675 | 0.505 | 0.581717 | 0.550248 | 0.350242 | 1 | 0.996511 | 0.853281 |
| Stx18 | Q8VDS8-3 | -0.45 | 0.45 | 1.01 | 0.41 | 0.42 | 0.12 | 0.76 | 0.62 | 0.48 | 0.79 | 0.49 | 0.25 | 0.86 | 0.5 | 1.55 | 1.39 | 0.125 | 0.1475 | 0.72 | 0.700369 | 0.647781 | 0.083237 | 1 | 0.996511 | 0.689531 |
| Stx1a | O35526 | 1.66 | 2.13 | 2.12 | 1.23 | 2.19 | 1.43 | 1.45 | 1.78 | 0.58 | 1.04 | 1.84 | 1.64 | -0.18 | 0.79 | 1.22 | 1.75 | -0.0725 | -0.51 | -0.89 | 0.797165 | 0.171928 | 0.068507 | 1 | 0.996511 | 0.665034 |
| Stx1b | P61264 | 1.59 | 1.49 | 1.84 | 2.03 | 1.47 | 1.35 | 1.62 | 1.74 | 1.33 | 1.3 | 1.6 | 1.44 | 0.83 | 1.47 | 1.38 | 1.31 | -0.1925 | -0.32 | -0.49 | 0.322162 | 0.125996 | 0.059609 | 1 | 0.996511 | 0.661738 |
| Stx2 | Q3TJ55 | 0.99 | 1.08 | 0.92 | 0.57 | 0.63 | 0.53 | 1.47 | 0.98 | 0.65 | 1.44 | 1.15 | 1.34 | 0.58 | 0.73 | 1.28 | 1.25 | 0.0125 | 0.255 | 0.07 | 0.960316 | 0.298338 | 0.776615 | 1 | 0.996511 | 0.967045 |
| Stx3 | A0A1B0GR | -0.18 | -0.42 | 0.6 | 0.75 | -0.23 | -0.09 | 0.61 | 0.85 | -0.57 | 0.54 | 0.25 | -0.03 | -0.39 | 0.11 | 1.19 | 0.81 | 0.0975 | -0.14 | 0.2425 | 0.793008 | 0.700063 | 0.578713 | 1 | 0.996511 | 0.927283 |
| Stx4 | P70452 | 0.52 | 1.06 | 1.16 | 0.89 | 0.76 | 0.15 | 0.31 | 1.07 | 0.88 | 1.41 | 1.04 | 0.64 | 0.25 | 0.94 | 0.99 | 1.98 | -0.335 | 0.085 | 0.1325 | 0.221751 | 0.725362 | 0.723934 | 1 | 0.996511 | 0.958398 |
| Stx5 | Q8K1E0 | -0.07 | 0.28 | 0.11 | -0.02 | 0.5 | -0.28 | 0.41 | -0.18 | -0.13 | 0.06 | 0.13 | -0.05 | 0.09 | 0.4 | 0.22 | 0.17 | 0.0375 | -0.0725 | 0.145 | 0.871968 | 0.675229 | 0.442317 | 1 | 0.996511 | 0.883549 |
| Stx6 | Q9JKK1-2 | 0.23 | 0.55 | 0.7 | 0.98 | 0.74 | -0.03 | 0.84 | 0.91 | 0.55 | 0.86 | 0.58 | 0.43 | 0.4 | 0.98 | 0.52 | 0.62 | 0 | -0.01 | 0.015 | 1 | 0.963503 | 0.949982 | 1 | 0.996755 | 0.992476 |
| Stx7 | Q8BH40 | 0.54 | 0.54 | 0.4 | 0.42 | 0.59 | 1.01 | 0.29 | 0.53 | 0.13 | -0.11 | 0.65 | 0.73 | -0.07 | -0.4 | 0.04 | 0.36 | 0.13 | -0.125 | -0.4925 | 0.503945 | 0.599014 | 0.045126 | 1 | 0.996511 | 0.658454 |
| Stx8 | O88983 | -1.12 | -0.76 | -1.41 | -1.1 | -0.19 | -0.93 | -1.27 | -1.45 | -0.68 | -0.87 | -1.38 | -0.88 | -0.26 | 0.47 | -1.01 | -1.11 | 0.1375 | 0.145 | 0.62 | 0.653819 | 0.535948 | 0.130791 | 1 | 0.996511 | 0.725078 |
| Stxbp1 | O08599 | 1.41 | 0.31 | 1.17 | 1 | 0.55 | 1.73 | 1.43 | 1.12 | 0.11 | 0.5 | 1.1 | 1.27 | -0.4 | -0.29 | 0.71 | 1.11 | 0.235 | -0.2275 | -0.69 | 0.489137 | 0.521286 | 0.128303 | 1 | 0.996511 | 0.723655 |
| Stxbp1 | O08599-2 | -0.53 | -0.5 | -1.43 | -1.2 | -0.39 | -0.98 | -1.06 | -2.19 | -0.59 | -3.73 | -2.21 | -1.16 | -0.58 | -0.69 | -2.97 | -1.5 | -0.24 | -1.0075 | -0.52 | 0.568111 | 0.155512 | 0.359719 | 1 | 0.996511 | 0.856645 |
| Stxbp2 | Q64324 | 0.86 | 0.86 | -1.05 | 0.62 | 1.02 | 1.08 | -2.03 | -2.93 | -0.07 | 7.42 | -3.75 | 0.95 | -0.05 | 0.88 | -1.26 | -2.89 | -1.0375 | 0.815 | -1.1525 | 0.32251 | 0.700518 | 0.194466 | 1 | 0.996511 | 0.778959 |
| Stxbp3 | Q60770 | 0.7 | 0.58 | 0.51 | 0.13 | 0.62 | 0.68 | 0.46 | 0.19 | 0.74 | -0.35 | 0.23 | 0.61 | 0.62 | -0.33 | 0.48 | 0.47 | 0.0075 | -0.1725 | -0.17 | 0.969835 | 0.54839 | 0.536961 | 1 | 0.996511 | 0.912212 |
| Stxbp4 | Q9WV89-4 | -0.93 | -0.46 | 0.12 | -0.27 | -0.28 | -1.02 | -0.19 | -0.52 | -0.36 | -0.61 | -0.06 | -0.57 | -0.55 | -1.19 | -0.76 | -1.27 | -0.1175 | -0.015 | -0.5575 | 0.684117 | 0.955343 | 0.084438 | 1 | 0.996755 | 0.690427 |
| Stxbp5 | D3Z079 | -0.21 | -0.21 | 0.17 | 0.19 | -0.12 | -0.02 | 0.23 | 0.24 | -0.1 | 0.42 | 0.16 | 0.13 | -0.09 | -0.14 | -0.02 | 0.16 | 9.75E-02 | 0.1675 | -0.0075 | 0.602442 | 0.417968 | 0.969851 | 1 | 0.996511 | 0.992767 |
| Stxbp5l | Q5DQR4 | -0.07 | 0.18 | 0.26 | -0.14 | 0.26 | 0.38 | -0.12 | -0.08 | -0.09 | -0.46 | -0.06 | 0.08 | 0.02 | -0.31 | -0.27 | -0.17 | 0.0525 | -0.19 | -0.24 | 0.786463 | 0.354637 | 0.236771 | 1 | 0.996511 | 0.792722 |
| Stxbp5l | Q5DQR4-3 | 0.1 | -0.7 | 0.52 | 0.55 | 0.93 | 0.22 | 0.25 | 0.98 | 0.06 | 0.71 | 0.47 | -0.77 | -1.37 | 1.2 | 0.12 | 1.19 | 0.4775 | -7.9E-17 | 0.1675 | 0.190526 | 1 | 0.785874 | 1 | 1 | 0.967642 |
| Stxbp6 | Q3TYA4 | 0.16 | -0.03 | 0.11 | -0.01 | 0 | 0.09 | -0.11 | 0.16 | -0.6 | -1.04 | -0.03 | -0.11 | -0.6 | -1.01 | -0.13 | -0.39 | -0.0225 | -0.5025 | -0.59 | 0.880684 | 0.079569 | 0.030669 | 1 | 0.996511 | 0.641481 |
| Sub1 | P11031 | -1.68 | -1.87 | -0.6 | -0.98 | -2.88 | -3.34 | -1.81 | -2.96 | -1.51 | -0.43 | -1.35 | -1.63 | -2.86 | -2.57 | -3.55 | -2.64 | -1.465 | 0.0525 | -1.6225 | 0.006477 | 0.892221 | 0.001683 | 1 | 0.996511 | 0.419079 |
| Sucla2 | Q9Z2I9 | 1.43 | 0.96 | 1.43 | 1.32 | 0.94 | 1.43 | 1.32 | 1.51 | 0.02 | 0.65 | 1.36 | 0.99 | -0.23 | -0.09 | 0.97 | 0.92 | 0.015 | -0.53 | -0.8925 | 0.940419 | 0.113992 | 0.025944 | 1 | 0.996511 | 0.633075 |
| Suclg1 | Q9WUM5 | 3.01 | 1.89 | 0.6 | 0.71 | 1.68 | 2.81 | 1.14 | 0.8 | 0.21 | -0.06 | 0.83 | 2.34 | 0.04 | -0.05 | 0.88 | 0.48 | 0.055 | -0.7225 | -1.215 | 0.932686 | 0.323239 | 0.054276 | 1 | 0.996511 | 0.661738 |
| Suclg2 | Q9Z2I8 | 1.04 | 1.28 | 0.87 | 0.89 | 0.73 | 1.01 | 0.78 | 0.7 | 0.87 | 0.51 | 0.91 | 0.65 | 0.72 | 0.78 | 0.42 | 0.61 | -0.215 | -0.285 | -0.3875 | 0.228381 | 0.160143 | 0.073529 | 1 | 0.996511 | 0.669435 |
| Suco | Q8C341 | 1.1 | 1.49 | 0.72 | 0.41 | 1.69 | 0.99 | 0.48 | 0.02 | 0.31 | -1.42 | 0.6 | 1.29 | 0.2 | 0.14 | 0.34 | 0.21 | -0.135 | -0.735 | -0.7075 | 0.738282 | 0.220818 | 0.024016 | 1 | 0.996511 | 0.626254 |
| Suds3 | A0A0R4J24 | -0.28 | -1.42 | -1.54 | -1.33 | -0.37 | -0.82 | -1.06 | -1.35 | -0.55 | -0.46 | -1.25 | -0.69 | -0.55 | -0.37 | -0.62 | -0.87 | 0.2425 | 0.405 | 5.40E-01 | 0.487569 | 0.248921 | 0.115008 | 1 | 0.996511 | 0.717287 |
| Sugct | Q7TNE1 | 1.2 | 0.73 | 0.73 | 0.88 | -0.59 | 0.94 | -1 | 0.03 | 1.43 | 1.45 | 0.76 | 0.48 | 1.98 | -0.26 | 0.55 | 0.34 | -1.04 | 0.145 | -0.2325 | 0.03046 | 0.606999 | 0.61521 | 1 | 0.996511 | 0.936534 |
| Sugp1 | Q8CH02 | -3.38 | -3.61 | -2.36 | -2.04 | -3.81 | -3.25 | -3.29 | -2.21 | -2.36 | -1.04 | -1.79 | -2.83 | -0.28 | -3.85 | -2.5 | -2.7 | -0.2925 | 0.8425 | 0.515 | 0.539464 | 0.122578 | 0.502251 | 1 | 0.996511 | 0.901002 |
| Sugp2 | Q8CH09 | 0.93 | -0.07 | -0.1 | -0.14 | -0.49 | 0.38 | -0.34 | 0.31 | -0.15 | -0.2 | -0.1 | -0.66 | 0.38 | -0.16 | 0.29 | -0.25 | -0.19 | -0.4325 | -0.09 | 0.569722 | 0.169563 | 0.77344 | 1 | 0.996511 | 0.966062 |
| Sugt1 | Q9CX34 | -0.52 | -0.82 | -0.22 | -0.08 | -0.5 | -0.53 | -0.25 | -0.04 | -0.84 | 0.2 | -0.05 | -0.59 | -1.07 | -0.59 | -0.32 | -0.48 | 0.08 | 0.09 | -0.205 | 0.720785 | 0.762404 | 0.438259 | 1 | 0.996511 | 0.883161 |
| Sulf2 | Q8CFG0 | 0.44 | 1.12 | 0.64 | 0.31 | 1.13 | 0.92 | 0.77 | 0.67 | 0.78 | 0.17 | 0.68 | 0.46 | 1.1 | 1.76 | 1.25 | 0.71 | 0.245 | -0.105 | 0.5775 | 0.293465 | 0.672861 | 0.077257 | 1 | 0.996511 | 0.676667 |
| Sult1a1 | Q91W19 | -0.75 | -1.31 | -3.32 | -2.18 | 0.07 | -0.78 | -0.92 | -0.98 | 0.17 | -0.28 | -1.02 | 0.09 | -0.9 | 0.43 | -2.22 | -2.5 | 1.2375 | 1.63 | 0.5925 | 0.051437 | 0.018477 | 0.4605 | 1 | 0.996511 | 0.887902 |
| Sult4a1 | P63046 | -1.39 | -1.04 | 0.16 | -0.22 | -1 | -0.97 | 0.41 | 0.23 | -1.49 | -0.98 | 0.21 | -1.12 | -1.74 | -1.03 | -0.21 | -0.19 | 0.29 | -0.2225 | -0.17 | 0.551607 | 0.644425 | 0.725619 | 1 | 0.996511 | 0.958398 |
| Sumf2 | Q8BPG6 | -1.47 | -1.37 | 0.03 | -0.28 | -0.3 | -0.8 | 0.12 | 0.03 | -0.79 | 0.57 | 0.13 | -1.68 | 0.42 | 1.02 | 0.41 | 0.53 | 0.535 | 0.33 | 1.3675 | 0.211786 | 0.571219 | 0.006788 | 1 | 0.996511 | 0.552614 |
| Sumo1 | P63166 | 0.57 | -0.25 | -0.06 | 0.29 | -0.72 | 0.12 | -0.49 | 0.36 | 0.1 | 0.66 | 0.23 | -0.05 | 0.86 | 0.71 | 0.7 | -0.46 | -0.32 | 0.0975 | 0.315 | 0.314087 | 0.706531 | 0.383713 | 1 | 0.996511 | 0.860984 |
| Sumo2 | H7BWX9 | -0.83 | -1.88 | -2.43 | -2.87 | -2.09 | -1.64 | -2.49 | -2.56 | -1.56 | -2.47 | -2.06 | -1.03 | -1.3 | -1.62 | -2.25 | -2.74 | -0.1925 | 0.2225 | 0.025 | 0.673267 | 0.658917 | 0.96067 | 1 | 0.996511 | 0.992693 |
| Sun1 | Q9D666-3 | 0.8 | 0.54 | -0.33 | -0.1 | 0.43 | 0.56 | -0.07 | 0.13 | -0.33 | -0.87 | -0.16 | -0.02 | 0.63 | -0.02 | 0.44 | 0.14 | 0.035 | -0.5725 | 0.07 | 0.906771 | 0.104554 | 0.822473 | 1 | 0.996511 | 0.972768 |
| Sun2 | Q8BJS4-3 | 0.66 | 0.32 | 0.78 | 1.31 | 0.42 | 0.28 | 1.16 | 1.27 | 0.37 | 1.21 | 0.79 | 0.24 | 0.51 | 0.79 | 1.17 | 1.36 | 0.015 | -0.115 | 0.19 | 0.962397 | 0.707309 | 0.524476 | 1 | 0.996511 | 0.908107 |
| Suox | Q8R086 | 1.49 | 0.96 | 0.3 | 0.65 | 0.65 | 1.36 | 1.12 | 0.54 | 0.47 | 0.23 | 0.69 | 1.01 | 0.36 | 0.56 | 0.24 | 0.54 | 0.0675 | -0.25 | -0.425 | 0.829177 | 0.423294 | 0.158688 | 1 | 0.996511 | 0.754059 |
| Supt16 | G3X956 | -0.21 | -0.97 | -0.17 | -0.32 | -0.1 | -0.33 | 0.26 | -1.26 | 0.01 | -0.69 | 0 | -0.19 | -0.29 | -0.2 | 0.07 | 0.92 | 0.06 | 0.2 | 0.5425 | 0.867076 | 0.459313 | 0.132212 | 1 | 0.996511 | 0.726871 |
| Supt20 | A0A0G2JEY | -0.45 | -0.38 | 0.03 | 0.54 | -0.24 | -1.13 | 0.08 | -0.65 | -3.37 | -2.01 | -0.05 | -0.07 | -2.42 | -3.93 | 0.24 | 1.22 | -0.42 | -1.31 | -1.1575 | 0.232614 | 0.11287 | 0.29922 | 1 | 0.996511 | 0.8284 |
| Supt4h1b | Q9Z199 | -0.89 | -2.27 | -1.56 | -1.79 | -2.05 | -1.66 | -1.13 | -0.68 | -4.09 | -3.55 | -0.97 | -1.61 | -3.57 | -3.73 | -1.06 | -1.7 | 0.2475 | -0.9275 | -0.8875 | 0.532953 | 0.224823 | 0.203252 | 1 | 0.996511 | 0.782817 |
| Supt5h | O55201 | -0.71 | -1.08 | -1.03 | -0.92 | -1.27 | -0.61 | -1.27 | -0.95 | -1.34 | -0.5 | -0.9 | -1.19 | -1.1 | -0.69 | -1 | -1.38 | -0.09 | -0.0475 | -0.1075 | 0.665425 | 0.83855 | 0.623075 | 1 | 0.996511 | 0.938628 |
| Supt6h | Q62383 | -0.54 | -0.73 | -0.13 | -0.04 | -0.59 | -0.82 | -0.3 | 0.25 | -0.7 | -0.08 | -0.22 | -0.99 | -0.95 | -0.58 | -0.12 | -0.5 | -0.005 | -0.1375 | -0.1775 | 0.985992 | 0.625481 | 0.506778 | 1 | 0.996511 | 0.902614 |
| Supv3l1 | Q80YD1 | 0.08 | 0.14 | 0.58 | 0.56 | -0.09 | -0.01 | 0.42 | 0.41 | -0.2 | 0.61 | 0.54 | -0.18 | -0.07 | -0.17 | 0.28 | 0.44 | -0.1575 | -0.1475 | -0.22 | 0.471809 | 0.592037 | 0.365029 | 1 | 0.996511 | 0.858475 |
| Surf1 | A0A0A6YVI | 0.25 | 0.45 | 0.33 | 0.65 | 0.3 | 0.27 | 0.04 | 0.12 | 0.62 | 0.61 | 0.64 | 0.2 | 0.57 | 0.76 | 0.11 | -0.01 | -0.2375 | 0.0975 | -0.0625 | 0.173809 | 0.614767 | 0.79575 | 1 | 0.996511 | 0.969727 |
| Surf2 | P09926 | -5.47 | -8.92 | -5.38 | -6.98 | -9.02 | -11.04 | -10.47 | -6.84 | -4.38 | -2.06 | -7.53 | -7.06 | -5.32 | -4.58 | -5 | -6.52 | -2.655 | 1.43 | 1.3325 | 0.039824 | 0.30805 | 0.137427 | 1 | 0.996511 | 0.732462 |
| Surf4 | Q64310 | -1 | -0.33 | -2.25 | -2.25 | 0.33 | -0.81 | -1.59 | -2.36 | -0.93 | -2.38 | -2.27 | -0.41 | -0.99 | -1.28 | -2.09 | -1.69 | 0.35 | -0.04 | -0.055 | 0.608014 | 0.948969 | 0.91209 | 1 | 0.996755 | 0.986458 |
| Surf6 | P70279 | 0.53 | 0.76 | 0.77 | 0.35 | 0.37 | 0.55 | 0.47 | 0.61 | 0.81 | 0.56 | 0.51 | -0.01 | 1.12 | 0.6 | 0.29 | 0.4 | -0.1025 | -0.135 | -2E-16 | 0.546106 | 0.562968 | 1 | 1 | 0.996511 | 1 |
| Susd2 | Q9DBX3 | 0.94 | 0.4 | -0.24 | 0.8 | 0.48 | 0.7 | 0.5 | 0.16 | 0.93 | -0.08 | 0.07 | 0.63 | 1.2 | 0.39 | 0.89 | 0.34 | -0.015 | -0.0875 | 0.23 | 0.958393 | 0.801733 | 0.500469 | 1 | 0.996511 | 0.900792 |
| Susd5 | G3UW60 | -0.6 | -1.11 | -0.93 | -0.47 | -0.5 | -0.42 | -0.43 | -1.09 | -0.39 | 0.53 | 0.35 | 0.02 | -0.05 | 0.51 | -1.48 | -1.37 | 0.1675 | 0.905 | 0.18 | 0.483327 | 0.007829 | 0.709075 | 1 | 0.996511 | 0.957656 |
| Sv2a | Q9JIS5 | 1.58 | 1.15 | -0.01 | -0.29 | 1.11 | 1.73 | -0.21 | -0.4 | 0.95 | -0.45 | 0.01 | 1.68 | 0.46 | 0.78 | -0.31 | -0.31 | -0.05 | -0.06 | -0.4525 | 0.935928 | 0.920391 | 0.370514 | 1 | 0.996755 | 0.859253 |
| Sv2b | Q8BG39 | 1.24 | 0.56 | 0.03 | -0.4 | 1.07 | 1.43 | 0.49 | -0.33 | -0.18 | -0.51 | 0.05 | 1.14 | -0.43 | -0.02 | -0.4 | -0.01 | 0.3075 | -0.2325 | -0.5725 | 0.529201 | 0.623513 | 0.143792 | 1 | 0.996511 | 0.742777 |
| Sv2c | Q69ZS6 | 0.42 | -0.49 | -2.39 | -1.23 | -0.94 | 0.25 | -1.76 | -2.12 | 1.8 | -1.32 | -2.38 | 0.38 | 2.14 | -1.53 | -1.32 | -1.84 | -0.22 | 0.5425 | 0.285 | 0.759737 | 0.5855 | 0.773549 | 1 | 0.996511 | 0.966062 |
| Svbp | A2A7P9 | 0.12 | -0.34 | -0.02 | 0.02 | -0.42 | 0.37 | -0.14 | 0.04 | 1.11 | -0.38 | -0.4 | 0.11 | 1.09 | -1.38 | 0.22 | -0.29 | 0.0175 | 0.165 | -0.035 | 0.935865 | 0.646917 | 0.943099 | 1 | 0.996511 | 0.991519 |
| Svil | Q8K4L2 | 0.16 | 0.04 | -0.2 | 0.27 | -0.04 | 0.17 | 0.05 | -0.01 | 0.7 | 0.36 | -0.06 | 0.47 | 0.95 | 0.1 | 0.12 | -0.1 | -0.025 | 0.3 | 0.2 | 0.880769 | 0.202955 | 0.475254 | 1 | 0.996511 | 0.892134 |
| Svip | Q3UZP4 | 0.48 | 0.92 | 2.04 | 1.41 | 0.27 | 0.78 | 1.02 | 0.87 | 0.82 | 0.84 | 1.02 | 0.39 | 0.53 | 0.81 | 1.1 | 0.58 | -0.4775 | -0.445 | -0.4575 | 0.204085 | 0.228568 | 0.219788 | 1 | 0.996511 | 0.787447 |
| Svop | Q8BFT9 | 0 | 0.09 | 0.45 | 0.62 | 0.23 | 0.26 | 0.01 | 0.32 | 0.26 | 0.75 | 0.47 | 0.33 | 0.42 | 0.46 | 0.46 | 0.7 | -0.085 | 1.63E-01 | 0.22 | 0.66691 | 0.46698 | 0.320628 | 1 | 0.996511 | 0.839916 |
| Swap70 | Q6A028 | -0.08 | -0.23 | 0.46 | 0.44 | -0.01 | -0.13 | -0.44 | 0.32 | -0.03 | 0.8 | 0.37 | -0.13 | -0.13 | 0.44 | 0.22 | 0.21 | -0.2125 | 0.105 | 0.0375 | 0.404098 | 0.714886 | 0.879624 | 1 | 0.996511 | 0.979626 |
| Swi5 | A0A0A0MC | -2.5 | -3.05 | -3.08 | -3.78 | -2.68 | -2.56 | -2.99 | -3.18 | -3.02 | -1.83 | -3.59 | -2.23 | -2.02 | -2.38 | -3.61 | -3.48 | 0.25 | 0.435 | 0.23 | 0.409384 | 0.343173 | 0.611134 | 1 | 0.996511 | 0.935047 |
| Syap1 | Q9D5V6 | -0.09 | 0.18 | 0.59 | 0.66 | -0.24 | -0.13 | 0.24 | 0.34 | -0.17 | 0.79 | 0.67 | 0.01 | -0.38 | 0.15 | 0.14 | 0.01 | -0.2825 | -0.01 | -0.355 | 0.260341 | 0.973543 | 0.179695 | 1 | 0.997515 | 0.775542 |
| Sybu | Q8BHS8-2 | -0.32 | -0.42 | -0.04 | 0.4 | 0.3 | -0.31 | -0.63 | -0.4 | 1.12 | 0.08 | -0.01 | -0.34 | 1.05 | 0.77 | 0.17 | 0.01 | -0.165 | 0.3075 | 0.595 | 0.552312 | 0.398802 | 0.085901 | 1 | 0.996511 | 0.691619 |
| Syf2 | Q9D198 | 0.41 | -0.01 | -1.05 | -0.99 | -0.47 | -0.09 | -1.18 | -0.72 | 0.24 | -1.31 | -1.45 | 0.05 | 0.19 | 1.07 | -1.72 | -1.65 | -0.205 | -0.2075 | -0.1175 | 0.614214 | 0.696318 | 0.868222 | 1 | 0.996511 | 0.978872 |
| Syk | E9PWE9 | -0.09 | 0.1 | -2.98 | -2.63 | -0.37 | -0.46 | -2.67 | -2.21 | 0.82 | 0.76 | -3.33 | -0.35 | 0.4 | -0.09 | -0.79 | -4.58 | -0.0275 | 0.875 | 0.135 | 0.975756 | 0.449757 | 0.913365 | 1 | 0.996511 | 0.986458 |
| Sympk | Q80X82 | -1.57 | -2.51 | -1.81 | -1.9 | -2.19 | -2.12 | -0.95 | -1.45 | -2.12 | -2.07 | -1.39 | -2.65 | -0.68 | -1.99 | -0.76 | -1.38 | 0.27 | -0.11 | 0.745 | 0.440034 | 0.736149 | 0.064401 | 1 | 0.996511 | 0.66433 |
| **Syn1** | **O88935** | **1.12** | **1.12** | **1.06** | **1.01** | **0.59** | **1.03** | **0.75** | **0.92** | **0.9** | **0.05** | **1.16** | **0.94** | **0.65** | **0.89** | **0.69** | **0.36** | -0.255 | -0.315 | -0.43 | 0.1406 | 0.251845 | 0.046806 | 1 | 0.996511 | 0.658454 |
| Syn1 | O88935-1 | -1.55 | -0.88 | -1.95 | -1.95 | -1.02 | -1.37 | -1.31 | -2.23 | -2.15 | -2.83 | -1.57 | -1.49 | -1.64 | -1.4 | -2.45 | -3.38 | 0.1 | -0.4275 | -0.635 | 0.774833 | 0.284252 | 0.209745 | 1 | 0.996511 | 0.786218 |
| Syn2 | Q64332 | 0.87 | 0.76 | 0.98 | 1.25 | 0.53 | 0.93 | 1.68 | 1.58 | 0.58 | 0.9 | 1.06 | 0.66 | -0.01 | 0.87 | 0.87 | 1.1 | 0.215 | -0.165 | -0.2575 | 0.469151 | 0.421081 | 0.378114 | 1 | 0.996511 | 0.860984 |
| Syn2 | Q64332-2 | -1.03 | -2.33 | -4.1 | -2.72 | -2.78 | -0.12 | -2.98 | -3.19 | -1.76 | -1.95 | -3.58 | -0.64 | -1.87 | -1.92 | -4 | -2.83 | 0.2775 | 0.5625 | -0.11 | 0.748219 | 0.483934 | 0.880029 | 1 | 0.996511 | 0.979626 |
| Syn3 | Q8JZP2 | 1 | 0.84 | 0.97 | 0.9 | 0.14 | 1.09 | 0.72 | 0.75 | 0.79 | 0.35 | 0.98 | 0.69 | 0.67 | 0.1 | 0.67 | 0.52 | -0.2525 | -0.225 | -0.4375 | 0.274963 | 0.260689 | 0.056176 | 1 | 0.996511 | 0.661738 |
| Syncrip | A0A0R4J25 | -0.88 | 0.34 | -0.53 | 0.15 | 0.03 | -0.74 | 0.29 | -0.98 | -0.69 | 0.34 | -0.09 | 0.13 | -0.77 | -0.93 | -1.09 | -0.81 | -0.12 | 0.1525 | -0.67 | 0.761036 | 0.668199 | 0.052804 | 1 | 0.996511 | 0.660608 |
| Syncrip | Q7TMK9 | 0.32 | -0.54 | -0.34 | -0.3 | -0.09 | 0.18 | 0 | 0 | -0.96 | -0.76 | -0.3 | 0.33 | -0.85 | -1.19 | -0.04 | -0.23 | 0.2375 | -0.2075 | -0.3625 | 0.29385 | 0.542923 | 0.288062 | 1 | 0.996511 | 0.824705 |
| Syndig1 | A2ANU3 | 0.08 | 0.27 | 0.91 | 0.57 | 0.88 | -0.29 | 0.04 | 0.61 | 0.19 | -0.15 | 0.69 | 0.33 | 0.68 | -0.09 | 0.99 | 0.65 | -0.1475 | -0.1925 | 0.1 | 0.642258 | 0.478816 | 0.742751 | 1 | 0.996511 | 0.959115 |
| Syne1 | Q6ZWR6-4 | 0.88 | 0.76 | 0.8 | 0.83 | 0.58 | 0.63 | 0.66 | 0.73 | 0.63 | 0.56 | 0.73 | 0.61 | 0.55 | 0.55 | 0.67 | 0.76 | -0.1675 | -0.185 | -0.185 | 0.249271 | 0.25011 | 0.292463 | 1 | 0.996511 | 0.825497 |
| Syne2 | Q6ZWQ0 | 0.82 | 0.93 | 0.45 | -0.08 | 0.57 | 0.51 | -0.52 | -0.06 | 1.04 | 0.5 | -0.28 | 0.34 | 1.6 | 0.53 | 0.92 | 0.22 | -0.405 | -0.13 | 0.2875 | 0.24497 | 0.709555 | 0.442477 | 1 | 0.996511 | 0.883549 |
| Syngap1 | F6SEU4 | 1.3 | 0.97 | 1.41 | 1.35 | 0.96 | 1.28 | 1.58 | 1.59 | 0.64 | 0.79 | 1.52 | 1.22 | 0.35 | 0.71 | 1.07 | 1.23 | 0.095 | -0.215 | -0.4175 | 0.649496 | 0.396499 | 0.124806 | 1 | 0.996511 | 0.721448 |
| Syngap1 | A0A0J9YUN | 1.86 | 1.65 | 2.85 | 2.75 | 1 | 1.25 | 2.48 | 2.4 | 1.91 | 2.58 | 2.49 | 1.65 | 1.05 | 1.43 | 2.12 | 1.62 | -0.495 | -0.12 | -0.7225 | 0.294281 | 0.744907 | 0.077846 | 1 | 0.996511 | 0.676942 |
| Syngap1 | J3QQ18 | 1.26 | 1.8 | 1.59 | 1.77 | 1.65 | 1.26 | 1.89 | 1.73 | 0.66 | 1.18 | 1.75 | 1.31 | 0.61 | 1.82 | 1.57 | 1.27 | 0.0275 | -0.38 | -0.2875 | 0.895912 | 0.18578 | 0.353418 | 1 | 0.996511 | 0.855029 |
| Syngap1 | A0A0J9YVH | 0.86 | 0.12 | 1.07 | 1.46 | 0.65 | 0.57 | 0.74 | 1.16 | 0.41 | 1.08 | 1.2 | 0.8 | 0.72 | 1.42 | 0.5 | 0.76 | -0.0975 | -0.005 | -0.0275 | 0.751003 | 0.987882 | 0.936173 | 1 | 0.999285 | 0.989767 |
| Syngr1 | O55100-2 | 0.35 | 0.78 | 0.76 | 0.48 | 1.22 | 0.94 | 0.41 | 0.54 | 1.63 | 0.62 | 0.54 | 0.39 | 1.67 | 1.38 | 1.04 | 0.63 | 0.185 | 0.2025 | 0.5875 | 0.434435 | 0.513284 | 0.055197 | 1 | 0.996511 | 0.661738 |
| Syngr2 | H3BJY1 | -1.45 | -1.18 | -1.34 | -1.61 | -1.23 | -1.94 | -1.55 | -1.07 | -1.5 | -0.77 | -0.76 | -1.5 | -1.28 | -1.02 | -1.51 | -2.29 | -0.0525 | 0.2625 | -0.13 | 0.820551 | 0.31456 | 0.667528 | 1 | 0.996511 | 0.951339 |
| Syngr3 | Q8R191 | -1.1 | -0.78 | -0.75 | -1.16 | -0.38 | -0.74 | -1.67 | -1.47 | -0.22 | -0.17 | -0.93 | -1.09 | 0.22 | -0.58 | -0.48 | -1.33 | -0.1175 | 0.345 | 0.405 | 0.710407 | 0.229969 | 0.247857 | 1 | 0.996511 | 0.803677 |
| Synj1 | F7BQW7 | 0.35 | 0.35 | 0.95 | 0.97 | 0.06 | 0.44 | 0.48 | 1.04 | -0.03 | 0.65 | 0.85 | 0.27 | -0.21 | 0.1 | 0.52 | 0.17 | -0.15 | -0.22 | -0.51 | 0.58577 | 0.43546 | 0.076629 | 1 | 0.996511 | 0.675786 |
| Synj1 | A0A338P6 | -0.66 | -1.04 | 0.11 | 0.79 | -1.1 | -0.82 | -1.21 | 0.01 | 0.29 | -1.03 | 0.04 | -1.11 | 0.81 | 0.93 | 0.14 | -1.34 | -0.58 | -0.2525 | 0.335 | 0.226456 | 0.619864 | 0.583639 | 1 | 0.996511 | 0.927734 |
| Synj1 | F6XIX0 | -2.64 | -2.78 | -0.05 | -0.3 | -1.9 | -3.73 | -0.85 | -0.3 | -3.57 | 0.15 | 0.07 | -1.62 | -3.28 | -2.79 | -0.49 | -0.48 | -0.2525 | 0.2 | -0.3175 | 0.789765 | 0.84557 | 0.735098 | 1 | 0.996511 | 0.958635 |
| Synj2 | D3YZB2 | 0.54 | 0.31 | 0.32 | 0.17 | 0.19 | -0.64 | -0.19 | -0.1 | 0.27 | 0.61 | 0.25 | -0.02 | 1.32 | 0.43 | 1.2 | 0.44 | -0.52 | -0.0575 | 0.5125 | 0.036115 | 0.773775 | 0.088791 | 1 | 0.996511 | 0.691619 |
| Synj2bp | Q9D6K5 | 0.28 | 0.13 | 0.49 | 0.61 | -0.15 | 0.27 | 0.08 | 0.07 | 0.81 | 0.27 | 0.24 | -0.07 | 1.08 | 0.04 | -0.23 | -0.16 | -0.31 | -0.065 | -0.195 | 0.115037 | 0.786395 | 0.555067 | 1 | 0.996511 | 0.916796 |
| Synm | Q70IV5-2 | -0.14 | -0.1 | -0.2 | -0.63 | -0.2 | -0.54 | -0.87 | -0.85 | 0.12 | 0.41 | -0.04 | -0.23 | 0.32 | -0.24 | -0.13 | -0.09 | -0.3475 | 0.3325 | 0.2325 | 0.144484 | 0.156582 | 0.311622 | 1 | 0.996511 | 0.834523 |
| Synpo | Q8CC35-3 | 1.39 | 0.63 | 1.12 | 1.59 | 0.55 | 1.17 | 0.79 | 1.23 | 1.03 | 1.01 | 1.12 | 0.9 | 0.91 | 1.18 | 0.85 | 0.49 | -0.2475 | -0.1675 | -0.325 | 0.369703 | 0.491679 | 0.255501 | 1 | 0.996511 | 0.809524 |
| Synpo2 | D3YVV9 | -0.06 | -0.04 | -0.24 | 0.13 | -1.29 | -0.6 | 0.03 | 0.14 | 0.6 | 1.06 | -0.43 | 0.87 | 1.2 | -0.4 | -0.02 | 0.14 | -0.3775 | 0.5775 | 0.2825 | 0.269804 | 0.114037 | 0.42687 | 1 | 0.996511 | 0.879165 |
| Synpr | Q8BGN8-2 | 0.79 | 1.08 | 1.01 | 0.75 | 0.9 | 0.78 | 0.34 | 0.52 | 0.49 | 1.07 | 0.8 | 0.56 | 0.49 | 1.07 | 0.56 | 0.6 | -0.2725 | -0.1775 | -0.2275 | 0.174775 | 0.391086 | 0.299031 | 1 | 0.996511 | 0.828257 |
| Synrg | Q5SV85-2 | 0.13 | -0.08 | -0.15 | -0.07 | -0.34 | -0.13 | -0.45 | -0.21 | -0.1 | 0.11 | -0.3 | 0.15 | -0.3 | -0.03 | -0.34 | -0.58 | -0.24 | 0.0075 | -0.27 | 0.154365 | 0.96732 | 0.194526 | 1 | 0.996755 | 0.778959 |
| Syp | Q62277 | 1.15 | 0.33 | 2.13 | 0.2 | 0.28 | 1.5 | 2.9 | 2.89 | 2.82 | -0.49 | 2.63 | 0.99 | 4.4 | -0.69 | 5.86 | 2.76 | 0.94 | 0.535 | 2.13 | 0.201969 | 0.513873 | 0.131837 | 1 | 0.996511 | 0.726871 |
| Sypl1 | O09117-2 | -0.94 | -0.92 | -1.97 | -1.83 | -0.25 | -1.16 | -1.81 | -2.38 | -1.03 | -0.22 | -2.13 | -1.12 | -1.09 | -0.71 | -1.68 | -2.1 | 0.015 | 0.29 | 0.02 | 0.975832 | 0.527079 | 0.960289 | 1 | 0.996511 | 0.992636 |
| Sys1 | Q78S06 | -1.39 | -1.32 | -1.25 | -0.86 | -1.52 | -2.51 | -0.5 | -0.04 | -2.55 | 0.73 | -0.24 | -1.26 | 0.07 | -1.09 | 0.06 | -0.2 | 0.0625 | 0.375 | 0.915 | 0.903975 | 0.567795 | 0.015039 | 1 | 0.996511 | 0.589387 |
| Syt1 | P46096 | 0.28 | 0.26 | 1.38 | 1.18 | 0.43 | 0.39 | 1.52 | 1.41 | -0.26 | 0.72 | 1.45 | 0.42 | -0.39 | 0.23 | 0.75 | 1.06 | 0.1625 | -0.1925 | -0.3625 | 0.68588 | 0.65993 | 0.392789 | 1 | 0.996511 | 0.865775 |
| Syt10 | Q9R0N4 | 0.13 | 0.67 | -0.37 | 0.26 | 0.93 | 0.87 | 1.45 | 0.9 | 0.25 | 1.07 | 0.28 | 0.77 | 0.12 | 0.14 | 1.35 | 1.42 | 0.865 | 0.42 | 0.585 | 0.009198 | 0.184064 | 0.173575 | 1 | 0.996511 | 0.764376 |
| Syt11 | Q9R0N3 | 0.42 | 0.61 | 0.66 | 0.44 | 0.78 | 0.24 | 0.6 | 0.24 | 0.25 | -0.17 | 0.31 | 0.65 | 0.03 | 0.84 | 0.29 | 0.63 | -0.0675 | -0.2725 | -0.085 | 0.720111 | 0.231139 | 0.71546 | 1 | 0.996511 | 0.958106 |
| Syt12 | Q920N7 | 0.76 | 1.05 | 1.24 | 1.49 | 0.96 | 0.72 | 0.93 | 1.15 | 0.99 | 1.15 | 1.1 | 0.71 | 0.36 | 1.05 | 0.8 | 0.87 | -0.195 | -0.1475 | -0.365 | 0.358731 | 0.508257 | 0.165491 | 1 | 0.996511 | 0.758837 |
| Syt13 | Q9EQT6 | 0.34 | 0.37 | 0 | -0.2 | 1 | 0.02 | -0.16 | -0.17 | 0.82 | 0.41 | 0.48 | 0.56 | 1 | 1.08 | 0.64 | 0.15 | 0.045 | 0.44 | 0.59 | 0.883502 | 0.060166 | 0.056894 | 1 | 0.996511 | 0.661738 |
| Syt16 | Q7TN83-2 | 0.86 | 0.44 | 0.66 | 0.33 | 0.89 | 1.21 | 1 | 0.83 | 0.92 | 0.63 | 1.03 | 0.66 | 0.89 | 1.55 | -0.13 | 1.23 | 0.41 | 0.2375 | 0.3125 | 0.051628 | 0.258178 | 0.413118 | 1 | 0.996511 | 0.874811 |
| Syt17 | Q920M7 | 0.3 | 0.19 | 0.25 | 0.33 | 0.47 | 0.57 | 0.59 | 0.68 | 1.34 | 0.32 | 0.44 | 0.58 | 1.23 | 1.22 | 0.76 | 0.2 | 0.31 | 0.4025 | 0.585 | 0.054507 | 0.139072 | 0.054854 | 1 | 0.996511 | 0.661738 |
| Syt2 | A0A0R4J2C | 1.4 | 0.89 | 1.15 | 1.37 | 1.14 | 1.25 | 0.86 | 0.65 | 0.41 | 0.61 | 0.73 | 0.66 | 0.37 | 0.56 | 0.69 | 1.11 | -0.2275 | -0.6 | -0.52 | 0.292596 | 0.011835 | 0.052391 | 1 | 0.996511 | 0.659923 |
| Syt3 | G3X9Y1 | 1.02 | 0.94 | 1.13 | 1.03 | 0.96 | 0.94 | 0.98 | 1.01 | 0.64 | 0.47 | 1.22 | 0.9 | 0.84 | 0.52 | 1.59 | 0.82 | -0.0575 | -0.2225 | -0.0875 | 0.681927 | 0.306783 | 0.737226 | 1 | 0.996511 | 0.958635 |
| Syt4 | P40749 | 0.43 | 0.9 | -0.35 | -0.38 | 1.2 | 0.71 | -0.28 | -0.92 | 0.7 | -0.23 | -0.5 | 1.02 | 0.48 | 0.81 | -0.11 | 0 | 0.0275 | 0.0975 | 0.145 | 0.958186 | 0.828902 | 0.696981 | 1 | 0.996511 | 0.955729 |
| Syt5 | Q9R0N5 | 0.98 | 0.88 | 0.27 | 0.37 | 0.89 | 1.15 | 0.46 | 0.39 | 0.6 | 0.34 | 0.21 | 1.33 | 0.62 | 0.77 | 0.6 | 0.57 | 0.0975 | -0.005 | 0.015 | 0.710649 | 0.987128 | 0.947743 | 1 | 0.999285 | 0.992414 |
| Syt6 | Q8C8S6 | -1.04 | -1.83 | 0.32 | -0.07 | -2.09 | -1.34 | 0.75 | -0.12 | 0.67 | -1.11 | 0.13 | -0.85 | 0.38 | -1.47 | 1.11 | 0.85 | -0.045 | 0.365 | 0.8725 | 0.950127 | 0.538638 | 0.226169 | 1 | 0.996511 | 0.787447 |
| Syt7 | A0A2C9F2 | 1.07 | 1.05 | 0.44 | 0.75 | 0.5 | 1.1 | 0.55 | 1.01 | 1.04 | 1.28 | 1 | 1.6 | 0.21 | 0.59 | 0.4 | 0.71 | -0.0375 | 0.4025 | -0.35 | 0.871899 | 0.112029 | 0.153047 | 1 | 0.996511 | 0.750649 |
| Syt7 | Q9R0N7-4 | 0.31 | 0.98 | 1.53 | 1.47 | 1.11 | 0.6 | 0.4 | 1.01 | 0.75 | 0.83 | 1.11 | 0.84 | -0.06 | 0.2 | 1.26 | 1.46 | -0.2925 | -0.19 | -0.3575 | 0.374232 | 0.530321 | 0.432267 | 1 | 0.996511 | 0.879165 |
| Syt7 | A0A498WG | 0.53 | 0.77 | 0.91 | 0.89 | 1.21 | 0.5 | 1.12 | 0.66 | 0.44 | 0.05 | 1.15 | 0.64 | 0.29 | 0.68 | 1.02 | 1.1 | 0.0975 | -0.205 | -0.0025 | 0.656578 | 0.44407 | 0.991754 | 1 | 0.996511 | 0.996154 |
| Syt9 | Q9R0N9 | -0.04 | 0.12 | 0.47 | 1.61 | -0.98 | 1.15 | 0.31 | 0.34 | 1.82 | -0.78 | 1.52 | -0.06 | 2.17 | 0.27 | 1.74 | 1.77 | -0.335 | 0.085 | 0.9475 | 0.531893 | 0.897818 | 0.095567 | 1 | 0.996511 | 0.698156 |
| Sytl2 | A0A5F8MP | 0.12 | -0.04 | -0.95 | -0.83 | 0.09 | -0.07 | -0.21 | -0.84 | -1.07 | -0.69 | -0.97 | -0.08 | -1.29 | -0.9 | -1.31 | -0.94 | 0.1675 | -0.2775 | -0.685 | 0.61417 | 0.430221 | 0.047917 | 1 | 0.996511 | 0.658454 |
| Sytl5 | Q80T23-2 | 0.25 | 0.2 | -0.27 | 0.59 | 0.09 | -0.2 | 0.4 | 0.28 | 0.51 | 0 | 0.07 | 0.02 | 0.5 | 0.27 | 0.32 | 0.42 | -0.05 | -0.0425 | 0.185 | 0.832813 | 0.860135 | 0.42844 | 1 | 0.996511 | 0.879165 |
| Syvn1 | E9QQ99 | -0.67 | -0.77 | -0.44 | -0.88 | 0.26 | -0.38 | -0.02 | -0.44 | -1.24 | -0.84 | -0.47 | -0.92 | -0.18 | -1.69 | 0.38 | 0.48 | 0.545 | -0.1775 | 0.4375 | 0.030096 | 0.4302 | 0.371461 | 1 | 0.996511 | 0.859424 |
| Szrd1 | A2AA85 | -0.28 | -0.55 | 0.18 | -0.09 | -2.09 | -0.27 | -1.58 | -0.62 | 0.91 | 0.22 | -0.41 | -0.63 | 1.46 | -1.54 | -0.26 | -0.73 | -0.955 | 0.2075 | -0.0825 | 0.046515 | 0.576007 | 0.890159 | 1 | 0.996511 | 0.981598 |
| Szt2 | A2A9C3 | 0.93 | 1.24 | -0.04 | 0.22 | -0.06 | -0.34 | -0.89 | -0.54 | 1.29 | -0.02 | 0.34 | 1.04 | 0.26 | -1.44 | -0.72 | -0.64 | -1.045 | 0.075 | -1.2225 | 0.012109 | 0.853545 | 0.01956 | 1 | 0.996511 | 0.612607 |
| Tab1 | Q8CF89 | 0.01 | 0.1 | 0.33 | 0.39 | 0.07 | 0.13 | 0.86 | 0.49 | -0.3 | 0.45 | 0.62 | 0.22 | -0.64 | 0.24 | 0.23 | 0.63 | 0.18 | 0.04 | -0.0925 | 0.433148 | 0.870301 | 0.756181 | 1 | 0.996511 | 0.962333 |
| Tab2 | D3Z564 | -0.72 | -0.68 | -0.31 | -0.63 | -0.8 | -1.14 | -0.93 | -0.79 | -0.96 | -0.04 | -0.09 | -0.16 | 0.67 | -1.06 | -0.6 | -1.35 | -0.33 | 0.2725 | -3.9E-16 | 0.084987 | 0.304983 | 1 | 1 | 0.996511 | 1 |
| Tab3 | Q571K4 | -0.05 | 0.05 | 0.86 | 0.44 | -0.18 | 0.12 | 1.49 | 0.72 | 0.29 | 0.4 | 0.87 | 0.01 | 0.66 | -0.28 | 0.86 | 0.88 | 0.2125 | 0.0675 | 0.205 | 0.597141 | 0.812661 | 0.554588 | 1 | 0.996511 | 0.916796 |
| Tacc1 | F8VQ95 | -0.28 | -0.27 | 0.22 | 0.4 | -0.01 | -0.36 | 0.35 | 0.18 | -0.44 | 0.07 | 0.17 | -0.18 | -0.39 | -0.29 | 0.09 | 0.11 | 0.0225 | -0.1125 | -0.1375 | 0.926587 | 0.648538 | 0.584711 | 1 | 0.996511 | 0.927734 |
| Tacc2 | E9QL08 | 0.31 | -0.07 | 0.69 | 0.48 | -0.38 | 0.37 | 0.01 | -0.27 | 0.8 | 0.25 | 0.29 | 0.39 | 1.18 | 1.72 | 0.21 | 0.16 | -0.42 | 0.08 | 0.465 | 0.114976 | 0.733567 | 0.263057 | 1 | 0.996511 | 0.81524 |
| Tacc2 | E9Q8T1 | -0.1 | 0.6 | 0.25 | 0 | 0.45 | -0.19 | -0.16 | -0.22 | 0.58 | 0.57 | 0.2 | 0.11 | 0.52 | 1.58 | 0.23 | -0.27 | -0.2175 | 0.1775 | 0.3275 | 0.375619 | 0.449011 | 0.427269 | 1 | 0.996511 | 0.879165 |
| Taco1 | Q8K0Z7 | -0.04 | 0.03 | 0.28 | 0.31 | 0.08 | 0.06 | 0.3 | 0.3 | 0.39 | -0.11 | 0.3 | -0.28 | 0.18 | 0.55 | -0.27 | -0.03 | 0.04 | -0.07 | -0.0375 | 0.810165 | 0.751267 | 0.873683 | 1 | 0.996511 | 0.979626 |

| | | | | | | | | | | | | | | | | | | | | | | | | | | | |
|---|---|---|---|---|---|---|---|---|---|---|---|---|---|---|---|---|---|---|---|---|---|---|---|---|---|---|---|
| Tacr1 | P30548 | 1.66 | 0.56 | 0.12 | 0.62 | -0.34 | -0.42 | 0.43 | 0.19 | 0.54 | 0.93 | -1.17 | 1.46 | 1.38 | 0.83 | 1.4 | -0.02 | -0.775 | -0.3 | 0.1575 | 0.061013 | 0.620131 | 0.722339 | | 1 | 0.996511 | 0.958398 |
| Tada1 | Q99LM9 | -1.34 | -2.12 | -1.24 | -1.37 | 0.19 | -0.41 | -0.62 | -1.42 | -0.44 | -0.52 | -1.35 | 0.01 | -2.42 | -0.07 | -0.24 | -0.44 | 0.9525 | 0.9425 | 0.725 | 0.02899 | 0.021649 | 0.202477 | | 1 | 0.996511 | 0.782817 |
| Taf1 | S4R1B9 | 3.07 | 4.43 | -0.28 | 0.64 | 2.36 | 3.05 | -1.36 | -0.62 | 1.03 | -0.36 | 0.01 | 3.6 | 0.09 | 1.03 | 0.82 | -0.69 | -1.1075 | -0.895 | -1.6525 | 0.428206 | 0.48287 | 0.135277 | | 1 | 0.996511 | 0.731223 |
| Taf10 | Q8K0H5 | -1.41 | -1.19 | -0.68 | -0.65 | -0.9 | -1.35 | -0.16 | -0.62 | -0.59 | 0.2 | -0.82 | -0.69 | -0.47 | 0.42 | -0.83 | -0.06 | 0.225 | 0.5075 | 0.7475 | 0.472778 | 0.121063 | 0.047582 | | 1 | 0.996511 | 0.658454 |
| Taf11 | Q99JX1 | -0.08 | -0.23 | -2.14 | -2.67 | -0.98 | 0.17 | -2.76 | -1.71 | -1.18 | -0.56 | -1.53 | -0.18 | 0.39 | -3.42 | -1.33 | -1.29 | -0.04 | 0.4175 | -0.1325 | 0.960745 | 0.532668 | 0.884913 | | 1 | 0.996511 | 0.979767 |
| Taf15 | Q8BQ46 | 2.54 | -0.29 | 2.75 | -1.62 | -0.04 | 2.56 | 3.57 | 0.97 | 0.69 | -0.83 | 2.18 | 2.13 | 1.77 | -0.2 | 5.53 | 4.58 | 0.92 | 0.1975 | 2.075 | 0.451815 | 0.864309 | 0.191772 | | 1 | 0.996511 | 0.777117 |
| Taf2 | B9EJX5 | 0.71 | -0.22 | -0.44 | -0.08 | -0.26 | 0.7 | -0.33 | -0.22 | 1.76 | -1.03 | -1.01 | 0.05 | 0.73 | 1.39 | -0.47 | 0.14 | -0.02 | -0.05 | 0.455 | 0.952761 | 0.937802 | 0.321849 | | 1 | 0.996755 | 0.840788 |
| Taf5 | F8VPY2 | -0.34 | -1.34 | -0.26 | 0.27 | -1.84 | -0.14 | -0.67 | 0.28 | 0.38 | -1.32 | 0.38 | -0.89 | 1.35 | -2.04 | 0.1 | 0.03 | -0.175 | 0.055 | 0.2775 | 0.738573 | 0.914381 | 0.695284 | | 1 | 0.996755 | 0.955729 |
| Taf5l | Q91WQ5 | -0.98 | -0.74 | -1.05 | -0.78 | -1.8 | 0.49 | -0.7 | -0.06 | 1.37 | -0.67 | -0.44 | 0.28 | 3.28 | -2.33 | 1.17 | 0.95 | 0.37 | 1.0225 | 1.655 | 0.432065 | 0.042366 | 0.137365 | | 1 | 0.996511 | 0.732462 |
| Taf6 | D3Z0T0 | 1.24 | 1.62 | 0.23 | -0.31 | 1.21 | 1.78 | -1.22 | -0.21 | 0.81 | 0.57 | 0.64 | 0.9 | 0.93 | 1.16 | 0.81 | 0.34 | -0.305 | 0.035 | 0.115 | 0.679619 | 0.934705 | 0.799013 | | 1 | 0.996755 | 0.969791 |
| Taf6l | K3W4L1 | 0.3 | 0.31 | -0.26 | -0.22 | 0.93 | -0.45 | 0.48 | 0.31 | 0.05 | 0.36 | -0.19 | 0.55 | 0.11 | -0.22 | 1.33 | 1.75 | 0.285 | 0.16 | 0.71 | 0.384946 | 0.527818 | 0.155691 | | 1 | 0.996511 | 0.751103 |
| Taf8 | Q9EQH4-5 | 0.04 | -1.32 | 3.61 | 2.93 | -0.19 | 0.13 | 2.5 | 1.94 | -0.67 | 4.27 | 2.52 | 0.61 | 0.79 | 1.42 | 3.22 | 3.37 | -0.22 | 0.3675 | 0.885 | 0.854955 | 0.796685 | 0.464747 | | 1 | 0.996511 | 0.888267 |
| Taf9 | Q8VI33 | -0.56 | -1.5 | -1.99 | -1.99 | -1.19 | -1.25 | -2.33 | -1.69 | -1.07 | -1.31 | -1.71 | -1.11 | -0.01 | -0.32 | -1.13 | -1.09 | -0.105 | 0.21 | 0.8725 | 0.794598 | 0.561707 | 0.063056 | | 1 | 0.996511 | 0.66433 |
| Tafa1 | Q7TPG8 | -2.62 | -3.19 | -2.81 | -2.59 | -2.69 | -1.49 | -1.86 | -2.77 | -3.09 | -1.88 | -2.16 | -1.87 | -2.23 | -3.13 | -2.64 | -1.83 | 0.6 | 0.5525 | 0.345 | 0.099445 | 0.111753 | 0.294487 | | 1 | 0.996511 | 0.825612 |
| Tafa5 | Q91WE9-2 | -0.16 | -0.69 | -0.43 | -0.29 | -1.95 | 0.24 | -2.28 | -1.53 | 2.93 | -2.17 | -1.22 | -0.36 | 3.55 | -1.78 | -0.97 | -0.32 | -0.9875 | 0.1875 | 0.5125 | 0.088378 | 0.851427 | 0.631001 | | 1 | 0.996511 | 0.941132 |
| Tafazzin | J3JS92 | 0.42 | 0.42 | 0.48 | 0.38 | -0.33 | 0.55 | 0.56 | 0.27 | -0.35 | 0.18 | 0.91 | 0.73 | -1.19 | -0.55 | -0.2 | 0.15 | -0.1625 | -0.0575 | -0.8725 | 0.485021 | 0.844803 | 0.016385 | | 1 | 0.996511 | 0.589387 |
| Tagln | P37804 | -4.11 | -4.91 | -5.23 | -5.15 | -4.63 | -4.46 | -4.7 | -5.53 | -4.91 | -4.25 | -5.16 | -4.05 | -4.03 | -5.32 | -5.04 | -5.94 | 0.02 | 0.2575 | -0.2325 | 0.952832 | 0.479042 | 0.606147 | | 1 | 0.996511 | 0.932425 |
| Tagln2 | Q9WVA4 | -2.39 | -3.17 | -2.28 | -2 | -2.61 | -2.59 | -2.13 | -2.2 | -3.26 | -1.52 | -1.93 | -2.48 | -2.68 | -2.75 | -1.97 | -2.35 | 0.0775 | 0.1625 | 0.0225 | 0.784331 | 0.70514 | 0.943097 | | 1 | 0.996511 | 0.991519 |
| Tagln3 | Q9R1Q8 | 0.74 | 0.03 | -0.5 | 0.03 | 0.39 | 0.46 | -0.38 | 0.05 | 0.14 | -0.78 | -0.18 | 0.71 | -0.08 | -0.06 | -0.62 | -0.52 | 0.055 | -0.1025 | -0.395 | 0.860603 | 0.791043 | 0.216767 | | 1 | 0.996511 | 0.787447 |
| Taldo1 | A0A1B0GR | -2.34 | -2.42 | -1.94 | -1.98 | -2.22 | -2.6 | -1.71 | -2.21 | -2.12 | -0.74 | -1.71 | -1.96 | -1.4 | -2 | -1.79 | -1.98 | -0.015 | 0.5375 | 0.3775 | 0.949426 | 0.131052 | 0.127695 | | 1 | 0.996511 | 0.723391 |
| Tamalin | Q9JJA9 | 0.23 | -0.2 | 0.92 | 0.73 | -0.34 | 0.18 | 0.28 | 0.38 | 0.82 | 1.33 | 1.04 | 0.14 | 1.47 | 0.07 | 0.17 | 0.31 | -0.295 | 0.4125 | 0.085 | 0.335721 | 0.25865 | 0.830816 | | 1 | 0.996511 | 0.973939 |
| Tamm41 | G5E881 | 0.8 | 1.23 | 0.34 | 0.23 | 0.78 | 0.61 | 0.26 | -0.01 | 0.54 | 0.52 | 0.23 | 0.39 | 1.1 | 0.58 | 0.12 | -0.27 | -0.24 | -0.23 | -0.2675 | 0.417432 | 0.386881 | 0.473231 | | 1 | 0.996511 | 0.891467 |
| Tanc1 | E9QAF9 | 0.46 | -0.62 | -0.32 | -0.34 | -0.35 | 0.46 | -0.63 | -0.35 | 0.05 | -0.21 | -0.27 | 0.42 | 0.15 | -0.6 | -0.77 | -0.34 | -0.0125 | 0.2025 | -0.185 | 0.969058 | 0.490208 | 0.560563 | | 1 | 0.996511 | 0.919045 |
| Tanc2 | A2A690-2 | -0.07 | 0.18 | 0.77 | 0.63 | 0.29 | 0.1 | 0.63 | 0.68 | 0.4 | 0.72 | 0.96 | 0.37 | 0.66 | 0.38 | 0.83 | 0.77 | 0.0475 | 0.235 | 0.2825 | 0.85 | 0.377252 | 0.279683 | | 1 | 0.996511 | 0.822084 |
| Tanc2 | F6VCI2 | 4.51 | 4.68 | 5.16 | 2.26 | 2.7 | 5.12 | 2.91 | 3.54 | 1.87 | 0.38 | 6.96 | 4.49 | 2.76 | -3.23 | 7.83 | 5.56 | -0.585 | -0.7275 | -0.9225 | 0.451975 | 0.6111 | 0.675183 | | 1 | 0.996511 | 0.952944 |
| Tanc2 | A0A140LJD | -0.22 | 0.77 | -0.77 | -0.62 | 0.72 | -0.42 | -0.23 | -0.3 | 0.25 | 0.57 | 0.08 | 0.15 | 0.61 | 0.86 | -1.04 | -0.56 | 0.1525 | 0.4725 | 0.1775 | 0.710031 | 0.206046 | 0.738836 | | 1 | 0.996511 | 0.958796 |
| Tanc2 | E9Q2K2 | 0.19 | 2.4 | -1.07 | -0.83 | 0.03 | 0.84 | -0.45 | -1.05 | 1.43 | -1.42 | -0.96 | -0.18 | 2.86 | -1.4 | -0.78 | -0.83 | -0.33 | -0.455 | -0.21 | 0.680974 | 0.618859 | 0.851373 | | 1 | 0.996511 | 0.976225 |
| Tango2 | Q8BTN3 | -1.15 | -1.06 | -1.19 | -1.05 | -0.8 | -1.27 | -0.92 | -1.05 | -1.05 | -0.9 | -1.11 | -1.03 | -1.43 | -0.74 | -1.09 | -1.03 | 0.1025 | 0.09 | 0.04 | 0.537331 | 0.570781 | 0.846561 | | 1 | 0.996511 | 0.976225 |
| Tango6 | Q8C3S2-2 | 0.76 | 0.55 | 0.43 | 0.6 | 0.08 | 0.76 | 0.35 | 0.37 | 0.92 | 1.19 | 0.1 | 0.43 | 0.06 | 0.45 | -0.1 | 0.33 | -0.195 | 0.075 | -0.4 | 0.326395 | 0.780964 | 0.07786 | | 1 | 0.996511 | 0.676942 |
| Tank | A2AR78 | 0.05 | -0.66 | 0.23 | 0.66 | 0.26 | -1.43 | -0.26 | -0.02 | 0.32 | 0.93 | 0.23 | -0.46 | 0.18 | 1.14 | 0.38 | -0.26 | -0.4325 | 0.185 | 0.29 | 0.331105 | 0.629752 | 0.463285 | | 1 | 0.996511 | 0.888267 |
| Taok1 | Q5F2E8 | 0.89 | 0.65 | 0.53 | 0.63 | 0.78 | 0.8 | 0.94 | 0.89 | 0.7 | 0.77 | 0.62 | 0.7 | 0.21 | 0.8 | 0.76 | 0.46 | 0.1775 | 0.0225 | -0.1175 | 0.268718 | 0.892184 | 0.585515 | | 1 | 0.996511 | 0.928178 |
| Taok2 | Q6ZQ29-2 | 1.11 | 0.6 | 1.37 | 1.29 | 0.29 | 0.91 | 1.62 | 1.57 | 0.97 | 1.24 | 1.48 | 0.81 | 0.64 | 1.5 | 1.12 | 1.01 | 0.005 | 0.0325 | -0.025 | 0.98844 | 0.896754 | 0.926559 | | 1 | 0.996511 | 0.989128 |
| Taok3 | Q8BYC6 | -0.57 | -0.58 | -0.57 | -0.57 | -0.41 | -0.89 | -1.07 | -0.89 | -0.53 | 0.55 | -0.45 | -0.76 | -0.84 | 0.05 | -0.26 | -0.55 | -0.2425 | 0.275 | 0.1725 | 0.21008 | 0.366545 | 0.467304 | | 1 | 0.996511 | 0.889213 |
| Tapbp | Q3TCU5 | 0.29 | -0.41 | -0.17 | 0.14 | 0.38 | -1.32 | -0.55 | 0.06 | 1.15 | 0.09 | -0.34 | -0.48 | 0.92 | 0.99 | -0.66 | 1.07 | -0.32 | 0.1425 | 0.6175 | 0.41515 | 0.71254 | 0.169401 | | 1 | 0.996511 | 0.760553 |
| Tapbpl | Q8VD31 | 0.22 | 0.47 | -0.35 | 0.31 | -0.57 | 0.71 | -1.48 | -0.03 | 1.07 | -0.24 | -0.88 | -0.15 | 2.77 | -0.93 | 0.76 | 0.29 | -0.505 | -0.2125 | 0.56 | 0.288873 | 0.616799 | 0.442311 | | 1 | 0.996511 | 0.883549 |
| Tapt1 | Q4VBD2 | 0.37 | 0.52 | 0.66 | 0.36 | 1.09 | 0.36 | 0.54 | 0.52 | 0.47 | 1.04 | 0.56 | 0.35 | 0.38 | 0.98 | 0.92 | 1.04 | 0.15 | 0.1275 | 0.3525 | 0.471203 | 0.548122 | 0.134588 | | 1 | 0.996511 | 0.731223 |
| Tarbp2 | D3Z282 | -0.46 | -0.65 | -0.95 | -0.97 | -0.42 | 0.15 | -0.82 | -1.33 | 0.59 | 0.44 | -0.6 | -0.63 | 0.94 | -0.18 | -0.33 | -1.1 | 0.1525 | 0.7075 | 0.59 | 0.643395 | 0.066254 | 0.183324 | | 1 | 0.996511 | 0.7764 |
| Tardbp | Q921F2 | -1.14 | -3.4 | -1.87 | -1.88 | -2.92 | -2.1 | -2.06 | -1.83 | -3.21 | -1.32 | -1.78 | -2.97 | -1.99 | -3.43 | -1.09 | -2.16 | -0.155 | -0.2475 | -0.095 | 0.752818 | 0.683338 | 0.87793 | | 1 | 0.996511 | 0.979626 |
| Tars1 | Q9D0R2 | -1.07 | -1.07 | -1.19 | -1.48 | -0.93 | -1.13 | -0.99 | -1.45 | -1.18 | -1.36 | -1.26 | -1.05 | -1.12 | -0.71 | -1.35 | -1.49 | 0.0775 | -0.01 | 0.035 | 0.684438 | 0.956034 | 0.882417 | | 1 | 0.996755 | 0.979767 |
| Tars2 | Q3UQ84 | 0.93 | 0.91 | 1.28 | 0.92 | 0.92 | 0.92 | 1.66 | 1.39 | 0.61 | 0.28 | 1.41 | 0.55 | 0.92 | 1.01 | 1.69 | 1.69 | 0.2125 | -0.2975 | 0.3175 | 0.358432 | 0.293936 | 0.239364 | | 1 | 0.996511 | 0.794419 |
| Tars3 | Q8BLY2 | -1.36 | -2.06 | -1.44 | -1.23 | -1.78 | -1.53 | -1.43 | -1.3 | -1.33 | -0.62 | -1.22 | -1.65 | -1.02 | -1.44 | -1.19 | -1.56 | 0.0125 | 0.3175 | 0.22 | 0.956598 | 0.293396 | 0.395159 | | 1 | 0.996511 | 0.866783 |
| Tasor | Q69ZR9-2 | 0.68 | 0.57 | -0.07 | -0.34 | 0.48 | 0.81 | -0.01 | -0.04 | 0.04 | -0.45 | -0.59 | 0.52 | 0.01 | 0.21 | 0.47 | -0.25 | 0.1 | -0.33 | -0.1 | 0.751291 | 0.354248 | 0.741506 | | 1 | 0.996511 | 0.958796 |
| Tasor2 | Q5DTT3 | -1.9 | -2.24 | -0.41 | 0.11 | -1.99 | -1.98 | 0.61 | 0.22 | -2.02 | 0.13 | 0.52 | -2.43 | -1.53 | -1.43 | -0.39 | 0.01 | 0.325 | 0.16 | 0.275 | 0.689848 | 0.849975 | 0.661858 | | 1 | 0.996511 | 0.949645 |
| Tatdn1 | Q6P8M1-2 | -0.41 | -0.13 | -0.25 | -0.43 | -0.02 | -0.36 | -0.13 | -0.22 | -0.37 | -0.08 | -0.49 | -0.4 | -0.33 | -0.06 | -0.36 | -0.21 | 0.1225 | -0.03 | 0.065 | 0.456308 | 0.867573 | 0.724406 | | 1 | 0.996511 | 0.958398 |
| Tax1bp1 | Q3UKC1 | 0.16 | 0.49 | 0.72 | 0.83 | 0.49 | 0.36 | 0.42 | 0.9 | 0.42 | 0.81 | 0.51 | 0.33 | 0.46 | 0.52 | 0.65 | 0.66 | -0.0075 | -0.0325 | 0.0225 | 0.972376 | 0.881941 | 0.915599 | | 1 | 0.996511 | 0.986556 |
| Tax1bp3 | Q9DBG9 | 0.2 | 0.1 | 0.86 | 0.72 | -0.92 | 0.36 | 0.04 | 0.46 | 0.68 | -0.24 | 0.95 | 0.41 | 1.54 | -2.4 | 0.49 | 0.79 | -0.485 | -0.02 | -0.365 | 0.19241 | 0.94977 | 0.648701 | | 1 | 0.996755 | 0.946113 |
| Tbc1d1 | Q60949 | 1.14 | 1.85 | 0.94 | 0.87 | 1.14 | 1.29 | 0.43 | 0.72 | 1 | 0.41 | 0.68 | 0.95 | 0.88 | 1.07 | 0.07 | 0.5 | -0.305 | -0.44 | -0.57 | 0.318966 | 0.137541 | 0.103164 | | 1 | 0.996511 | 0.700625 |
| Tbc1d10a | Q5SPX8 | 1.89 | 0.2 | 0 | -0.12 | 0.31 | 1.97 | 0.35 | 0.11 | -0.63 | -1.53 | 0.25 | 1.03 | -0.58 | -0.37 | -0.31 | -0.22 | 0.1925 | -0.7125 | -0.8625 | 0.741901 | 0.298205 | 0.082693 | | 1 | 0.996511 | 0.68865 |
| Tbc1d10b | Q8BHL3 | 1.05 | 0.64 | 0.48 | 0.73 | 0.59 | 1 | 0.71 | 1.04 | 0.26 | -0.03 | 0.56 | 0.67 | 0.09 | 0.45 | 0.46 | 0.35 | 0.11 | -0.36 | -0.3875 | 0.580397 | 0.144392 | 0.089473 | | 1 | 0.996511 | 0.69217 |
| Tbc1d12 | A0A494BA | 0.18 | 0.73 | 0.64 | 0.31 | 0.73 | -0.44 | 0.08 | 0.24 | 0.75 | 1.3 | -0.03 | 0.7 | -0.29 | 0.36 | -0.22 | 0.09 | -0.3125 | 0.215 | -0.48 | 0.278827 | 0.489602 | 0.070578 | | 1 | 0.996511 | 0.668066 |
| Tbc1d13 | Q8R3D1 | 0.34 | 0.44 | -0.09 | -0.39 | 0.68 | 0.26 | 0.02 | -0.19 | -1.26 | -0.23 | 0.21 | 0.59 | -0.86 | -0.9 | -0.28 | -0.06 | 0.1175 | -0.2475 | -0.6 | 0.668814 | 0.560351 | 0.070894 | | 1 | 0.996511 | 0.668066 |
| Tbc1d14 | E9PYB1 | -0.78 | -0.47 | -0.39 | 0 | -0.52 | -0.76 | -0.05 | -0.33 | -1.27 | -0.53 | 0.24 | -0.45 | -1.17 | -0.47 | -0.91 | 0.4 | -0.005 | -0.0925 | -0.1275 | 0.983124 | 0.787412 | 0.732305 | | 1 | 0.996511 | 0.958635 |
| Tbc1d15 | Q9CXF4 | -0.73 | -0.82 | -0.99 | -0.86 | -0.64 | -1.05 | -0.78 | -1.02 | -1.15 | -0.31 | -0.93 | -0.77 | -1.17 | -0.48 | -1.1 | -0.98 | -0.0225 | 0.06 | -0.0825 | 0.89299 | 0.78734 | 0.705583 | | 1 | 0.996511 | 0.957084 |
| Tbc1d17 | Q8BYH7 | 0.28 | 0.16 | 0.04 | -0.51 | 0.17 | 0 | -0.38 | -0.11 | -0.35 | -1.01 | -0.09 | 0.06 | -0.26 | -0.38 | -0.36 | -0.29 | -0.0725 | -0.34 | -0.315 | 0.752093 | 0.274993 | 0.184953 | | 1 | 0.996511 | 0.776843 |
| Tbc1d19 | Q8VDV7 | 0.39 | 0.26 | 0.48 | 0.27 | 0.69 | 0.32 | 1.39 | 0.42 | 0.34 | 0.02 | 0.7 | 0.6 | 0.42 | 0.93 | 0.54 | 0.9 | 0.355 | 0.065 | 0.3475 | 0.190848 | 0.753057 | 0.113879 | | 1 | 0.996511 | 0.717275 |
| Tbc1d20 | Q9D9I4 | 0.24 | 0.02 | 0 | 0 | 0.25 | 0.01 | 0.26 | -0.2 | 0.53 | 0.2 | -0.29 | 0.23 | -0.03 | 0.44 | 0.19 | 0.34 | 0.015 | 0.1025 | 0.17 | 0.931246 | 0.640152 | 0.387902 | | 1 | 0.996511 | 0.864251 |
| Tbc1d22a | Q8R5A6-2 | 0.07 | -0.15 | -0.34 | -0.56 | 0 | -0.18 | -0.45 | -0.64 | -0.17 | -0.56 | -0.24 | -0.18 | -0.46 | -0.17 | -0.81 | -0.73 | -0.0725 | -4.25E-02 | -0.2975 | 0.741433 | 0.837736 | 0.231677 | | 1 | 0.996511 | 0.788863 |
| Tbc1d22b | Q80VE5 | -0.06 | 0.05 | 0.23 | -0.06 | 0.45 | -0.23 | 0.15 | 0.22 | -0.3 | -0.33 | 0.36 | 0.11 | -0.2 | -0.24 | -0.01 | 0.01 | 0.1075 | -0.08 | -0.15 | 0.58184 | 0.715135 | 0.419039 | | 1 | 0.996511 | 0.87558 |
| Tbc1d23 | A0A2I3BRD | -1.53 | -1.4 | -1.74 | -1.84 | -1.09 | -1.08 | -1.69 | -1.7 | -1.14 | -1.07 | -1.63 | -1.29 | -1.65 | -0.5 | -2.14 | -1.73 | 0.2375 | 0.345 | 0.1225 | 0.305311 | 0.120459 | 0.734618 | | 1 | 0.996511 | 0.958635 |
| Tbc1d24 | Q3UUG6-2 | -0.02 | 0.13 | 0.03 | -0.11 | 0.15 | 0.14 | 0.23 | 0.09 | -0.24 | -1.02 | 0.04 | 0.01 | -0.83 | 0.28 | -0.62 | -0.28 | 0.145 | -0.31 | -0.37 | 0.328879 | 0.26681 | 0.196074 | | 1 | 0.996511 | 0.78004 |
| Tbc1d25 | A0A6I8MX | 0.23 | 0.72 | -0.4 | -0.37 | 0.58 | 0.17 | 0.06 | 0.02 | 0.34 | -0.45 | -0.39 | 0.63 | 0.16 | 0.65 | -0.36 | -0.37 | 0.1625 | -0.0125 | -0.025 | 0.58576 | 0.972845 | 0.944218 | | 1 | 0.997515 | 0.991865 |
| Tbc1d30 | Q69ZT9 | 1.46 | 0.82 | -0.64 | -1.06 | 1.36 | -0.14 | -2.06 | -1.11 | 1.09 | 0.7 | -0.52 | -0.05 | 1.84 | 0.52 | -0.71 | -0.65 | -0.6325 | 0.16 | 0.105 | 0.4632 | 0.801799 | 0.890613 | | 1 | 0.996511 | 0.981669 |
| Tbc1d32 | Q3URV1 | 0.38 | 0.83 | -1.5 | -1.06 | 1.11 | 1.1 | -1.5 | -0.85 | 0.18 | -0.89 | -1.21 | 0.33 | -0.5 | 0.66 | -1.09 | -1.03 | 0.3025 | -0.06 | -0.1525 | 0.702193 | 0.922881 | 0.809066 | | 1 | 0.996755 | 0.972098 |
| Tbc1d4 | E9Q6Q8 | -0.27 | -0.33 | -0.56 | -0.18 | -0.98 | -0.06 | -0.72 | -0.58 | -0.3 | -0.53 | -0.75 | -0.24 | -0.31 | -0.32 | -0.6 | -0.1 | -0.25 | -0.12 | 0.0025 | 0.292915 | 0.542857 | 0.989958 | | 1 | 0.996511 | 0.996154 |
| Tbc1d5 | A0A286YDI | 0.1 | -0.26 | 0.01 | -0.21 | -0.82 | 0.57 | -0.26 | -0.22 | 0.72 | -0.36 | 0.17 | 0.1 | 0.86 | -0.76 | -0.46 | -0.53 | -0.0925 | 0.2475 | -0.1325 | 0.755935 | 0.350063 | 0.720423 | | 1 | 0.996511 | 0.958398 |
| Tbc1d7 | Q9D0K0 | 0.59 | 1.05 | 0.7 | 0.4 | 1.22 | 0.38 | 0.54 | 0.28 | -0.01 | -0.06 | 0.44 | 0.47 | -0.25 | 0.46 | 0.31 | 0.78 | -0.08 | -0.475 | -0.36 | 0.759379 | 0.064382 | 0.215272 | | 1 | 0.996511 | 0.787313 |
| Tbc1d8 | Q9Z1A9 | -2.06 | -1 | 0.07 | -0.72 | -0.75 | -1.48 | -0.16 | -0.09 | -0.15 | -1.01 | -1.05 | -1.53 | -1.03 | -0.96 | -1.24 | -0.31 | 0.3075 | -0.0075 | 0.0425 | 0.545557 | 0.987769 | 0.925746 | | 1 | 0.999285 | 0.988894 |
| Tbc1d8b | A3KGB4 | -0.12 | 0.02 | -0.36 | -0.08 | 0.15 | -0.83 | -0.39 | -0.36 | 0.13 | 0.23 | -0.51 | -0.4 | -0.31 | 0.53 | -0.4 | 0.4 | -0.2225 | -0.0025 | 0.19 | 0.354382 | 0.991437 | 0.495205 | | 1 | 0.999285 | 0.89758 |
| Tbc1d9 | Q3UYK3 | 2.16 | 1.71 | 1.24 | 1.92 | 1.72 | 1.64 | 1.64 | 1.88 | 2.39 | 2.45 | 1.57 | 2.07 | 2.07 | 2.26 | 1.79 | 1.83 | -0.0375 | 0.3625 | 0.23 | 0.867574 | 0.231939 | 0.379097 | | 1 | 0.996511 | 0.860984 |
| Tbc1d9b | Q5SVR0-2 | -0.22 | -0.02 | 0.27 | 0.19 | 0.33 | 0.21 | 0.55 | 0.51 | 0.45 | -0.02 | 0.37 | 0.28 | 0.3 | 0.23 | 0.23 | 0.34 | 0.345 | 0.215 | 0.22 | 0.083371 | 0.299027 | 0.266466 | | 1 | 0.996511 | 0.818107 |
| Tbca | A0A1Y7VJ4 | -1.84 | -1.79 | -0.53 | 0.08 | -1.99 | -1.65 | -0.83 | 0.13 | -2.15 | 0.75 | -0.73 | -1.74 | -1.4 | -1.16 | -0.94 | -1.19 | -0.065 | 0.0525 | -0.1525 | 0.915202 | 0.942432 | 0.739378 | | 1 | 0.996755 | 0.958796 |
| Tbcb | Q9D1E6 | -0.11 | -0.51 | -0.72 | -0.76 | -0.83 | -0.27 | -1.19 | -0.73 | 0.46 | -0.29 | -0.8 | -0.06 | 0.73 | -0.11 | -0.15 | -0.92 | -0.23 | 0.3525 | 0.4125 | 0.37369 | 0.266829 | 0.273412 | | 1 | 0.996511 | 0.822084 |
| Tbcc | A0A0R4J0M | -0.32 | -0.44 | -0.94 | -0.74 | -0.52 | -0.23 | -0.83 | -0.55 | 0.5 | -0.55 | -0.83 | -0.31 | 0.86 | -0.41 | -1.53 | -1.3 | 0.0775 | 0.3125 | 0.015 | 0.717308 | 0.341909 | 0.976951 | | 1 | 0.996511 | 0.993727 |
| Tbcd | Q8BYA0 | 0.09 | -0.57 | -0.48 | -0.52 | -0.25 | 0.25 | -0.5 | -0.54 | -0.78 | -0.06 | -0.57 | -0.04 | -0.54 | -0.4 | -0.66 | -0.66 | 0.11 | 0.0075 | -0.195 | 0.662091 | 0.977007 | 0.383158 | | 1 | 0.997808 | 0.860984 |
| Tbce | Q8CIV8 | 0.5 | 0.26 | -0.12 | 0.09 | 0.14 | 0.15 | -0.14 | -0.04 | 0.39 | 0.17 | -0.04 | 0.51 | 0.25 | 0.44 | 0.11 | -0.29 | -0.155 | 0.075 | -0.055 | 0.420646 | 0.731037 | 0.819713 | | 1 | 0.996511 | 0.972768 |
| Tbcel | Q8C5W3-2 | 0.3 | -0.05 | 1.02 | 0.14 | 0.12 | 0.25 | 0.7 | 0.67 | -0.16 | 0.48 | 0.89 | 0.36 | -0.33 | 0.24 | 0.66 | 0.96 | 0.0825 | 0.04 | 0.03 | 0.768035 | 0.900166 | 0.933443 | | 1 | 0.996511 | 0.989767 |
| Tbck | E9Q1W7 | 0.67 | 0.49 | -0.03 | 0.14 | 0.33 | 0.52 | -0.02 | 0.09 | -0.32 | 0.33 | 0.1 | 0.31 | -0.14 | -0.16 | -0.34 | -0.19 | -0.0875 | -0.2125 | -0.525 | 0.695691 | 0.39646 | 0.03714 | | 1 | 0.996511 | 0.658454 |
| Tbk1 | Q9WUN2 | 0.41 | 0.49 | 0.13 | -0.08 | 0.5 | 0.64 | -0.02 | 0 | 0.78 | -0.05 | 0.16 | 0.82 | 0.8 | 0.41 | 0 | -0.21 | 0.0425 | 0.19 | 0.0125 | 0.85517 | 0.489304 | 0.964358 | | 1 | 0.996511 | 0.9927 |
| Tbkbp1 | Z4YKN2 | 0.71 | 0.98 | 0.64 | 0.35 | 0.43 | 0.71 | 0.64 | 0.8 | 0.9 | 1.37 | 0.6 | 0.84 | 1.61 | 0.91 | 0.54 | 0.5 | -0.025 | 0.2575 | 0.22 | 0.895323 | 0.2921 | 0.471313 | | 1 | 0.996511 | 0.891144 |
| Tbl1x | Q9QXE7 | -1.24 | -0.47 | -0.26 | -0.26 | -0.48 | -0.46 | -0.01 | -0.39 | -0.17 | 0.18 | -0.18 | -0.55 | -0.54 | -0.83 | -0.31 | -0.33 | 0.2225 | 0.3775 | 0.055 | 0.410713 | 0.210253 | 0.845421 | | 1 | 0.996511 | 0.976225 |
| Tbl1xr1 | Q8BHJ5 | -2.26 | -2.64 | -2.6 | -2.49 | -2.26 | -2.46 | -2.34 | -2.96 | -2.4 | -1.63 | -2.26 | -2.3 | -1.47 | -2.18 | -2.17 | -2.72 | -0.0075 | 0.35 | 0.3625 | 0.971185 | 0.150808 | 0.228582 | | 1 | 0.996511 | 0.787447 |
| Tbl2 | Q9R099 | -0.49 | -0.37 | -0.85 | -0.79 | 0.07 | -0.53 | -0.1 | -0.54 | -0.16 | -0.67 | -0.74 | -0.35 | 0.15 | 0.33 | 0.16 | 0.04 | 0.35 | 0.145 | 0.795 | 0.134468 | 0.510566 | 0.003086 | | 1 | 0.996511 | 0.461628 |
| Tbl3 | Q8C4J7 | 0.01 | 1.18 | 1.22 | 0.73 | 1.41 | 0.33 | 0.78 | 0.66 | 2.15 | 0.32 | 0.98 | 0.26 | -0.22 | 1.69 | 0.75 | 1.16 | 0.01 | 0.1425 | 0.06 | 0.977007 | 0.769798 | 0.896875 | | 1 | 0.996511 | 0.983509 |
| Tbpl1 | P62340 | -2.79 | -3.43 | -2.79 | -2.92 | -3.02 | -3.22 | -3.02 | -3.38 | -2.17 | -1.84 | -2.68 | -2.87 | -2.33 | -2.08 | -3.41 | -3.74 | -0.1775 | 0.5925 | 0.0925 | 0.398442 | 0.067008 | 0.82328 | | 1 | 0.996511 | 0.972768 |
| Tbr1 | Q64336 | -1.8 | -1.56 | -0.7 | -0.86 | -2.15 | -1.67 | -1.58 | -1.01 | 0.05 | -0.29 | -0.72 | -1.53 | -0.44 | -1.21 | -1.28 | -1.55 | -0.3725 | 0.6075 | 0.11 | 0.293239 | 0.165865 | 0.756414 | | 1 | 0.996511 | 0.962333 |
| Tbrg4 | Q91YM4 | 1.18 | 0.31 | 0.42 | 0.43 | 0.28 | 1.21 | 0.25 | 0.64 | 0.69 | 0.62 | 0.55 | 0.85 | 1.17 | -0.26 | 0.13 | 0.09 | 0.01 | 0.0925 | -0.3025 | 0.97321 | 0.699157 | 0.413166 | | 1 | 0.996511 | 0.874811 |
| Tcaf1 | Q8BNE1 | -1.27 | -1.05 | -1.11 | -1.31 | -1.16 | -1.3 | -0.92 | -1.36 | -0.87 | -0.55 | -1.18 | -1.18 | -1.45 | -0.28 | -1.32 | -1.13 | 1.96E-17 | 0.24 | 0.14 | 1 | 0.26616 | 0.629127 | | 1 | 0.996511 | 0.940595 |
| TCAIM | Q66JZ4 | 0.69 | 1.07 | -0.09 | 0.94 | 0.87 | 0.55 | 0.74 | 0.47 | -0.15 | 1.24 | 0.8 | 0.68 | 0.19 | -1.22 | 0.93 | 0.34 | 0.005 | -0.01 | -0.5925 | 0.985661 | 0.978774 | 0.246607 | | 1 | 0.998749 | 0.802442 |
| Tcea1 | P10711 | -2.02 | -2.13 | -2.17 | -2.51 | -2.03 | -2.08 | -2.96 | -2.97 | -1.29 | -1.81 | -2.4 | -2.02 | -0.63 | -1.96 | -2.21 | -2.95 | -0.3025 | 0.3275 | 0.27 | 0.304615 | 0.245386 | 0.5659 | | 1 | 0.996511 | 0.921174 |
| Tceal1 | Q921P9 | -3.71 | -5.33 | -1.47 | -1.2 | -4.84 | -3.95 | -1.23 | -1.21 | -5.14 | 0.02 | -1.27 | -2.98 | -3.73 | -3.12 | -0.88 | -0.67 | 0.12 | 0.585 | 0.8275 | 0.92097 | 0.661123 | 0.464976 | | 1 | 0.996511 | 0.888267 |
| Tceal3 | Q8R0A5 | -5 | -5.55 | -1.53 | -0.66 | -4 | -4.8 | -2.16 | -1.69 | -5.46 | 0.25 | -1.9 | -4.79 | -5.19 | -2.73 | -1.5 | -2.32 | 0.0225 | 0.21 | 0.25 | 0.985949 | 0.896121 | 0.847353 | | 1 | 0.996511 | 0.976225 |
| Tceal5 | Q8CCT4 | -3.32 | -4.92 | -5.25 | -7.23 | -4.84 | -3.17 | -6.4 | -5.92 | -2.69 | -3.11 | -5.63 | -3.41 | -1.45 | -3.67 | -5.37 | -6.77 | 0.0975 | 1.47 | 0.865 | 0.919528 | 0.142179 | 0.493554 | | 1 | 0.996511 | 0.89758 |
| Tcerg1 | Q8CGF7-2 | -3.59 | -5.28 | -2.84 | -3.01 | -4.38 | -4.49 | -3.2 | -2.95 | -4.86 | -1.56 | -2.6 | -4.39 | -3.63 | -3.37 | -2.3 | -2.98 | -0.075 | 0.3275 | 0.61 | 0.90402 | 0.703625 | 0.306234 | | 1 | 0.996511 | 0.832533 |
| Tcf20 | Q9EPQ8 | 0.38 | 0.04 | 0.47 | 0.61 | -0.14 | 0.07 | 0.14 | 0.34 | 0.06 | 0.47 | 0.15 | 0.06 | 0.27 | 0.05 | 0.88 | 0.41 | -0.2725 | -0.19 | 0.0275 | 0.183011 | 0.361754 | 0.911668 | | 1 | 0.996511 | 0.986458 |
| Tcf25 | B2ZAC8 | -0.72 | -1.16 | -0.36 | -0.86 | -0.37 | -0.83 | -0.38 | -0.61 | -1.34 | 0.09 | -0.54 | -0.65 | -0.61 | -0.77 | -0.32 | 0.15 | 0.2275 | 0.165 | 0.3875 | 0.319659 | 0.623178 | 0.191997 | | 1 | 0.996511 | 0.777117 |
| Tcf4 | A0A0J9YTV | -1.76 | -1.69 | -0.48 | -0.85 | -1.58 | -1.84 | -1.58 | -0.67 | -2.26 | 0.1 | -0.84 | -1.85 | -0.12 | -0.82 | 0.19 | -0.37 | -0.2225 | -0.0175 | 0.915 | 0.567393 | 0.975363 | 0.034308 | | 1 | 0.997582 | 0.65013 |
| Tcirg1 | Q9JHF5 | -1.25 | -1.09 | 0.57 | 0.4 | -0.5 | -0.64 | 1.04 | 0.24 | 1.01 | 0.35 | 0.71 | -0.4 | 1.57 | -0.16 | 0.36 | 0.97 | 0.3775 | 0.76 | 1.0275 | 0.508946 | 0.173744 | 0.094577 | | 1 | 0.996511 | 0.698156 |
| Tcn2 | O88968 | 1.04 | 0.71 | -0.76 | -1.45 | 1.22 | 0.95 | -0.52 | -1.48 | 0.12 | -0.21 | -0.97 | 0.63 | -0.63 | 1.36 | -1.04 | -1.28 | 0.1575 | 0.0075 | -0.2825 | 0.840796 | 0.99038 | 0.711821 | | 1 | 0.999285 | 0.958014 |
| Tcof1 | H3BL37 | 0.25 | -1.55 | -1.34 | -0.8 | -1.39 | -0.91 | -2.49 | -2.11 | -0.39 | -0.69 | -0.65 | -1.08 | -0.31 | -1.56 | -1.1 | -0.94 | -0.865 | 0.1575 | -0.1175 | 0.108545 | 0.699757 | 0.794975 | | 1 | 0.996511 | 0.969498 |
| Tcp1 | P11983 | -0.57 | -0.62 | -0.64 | -0.88 | -0.53 | -0.67 | -0.54 | -0.81 | -1.24 | -1.22 | -0.76 | -0.52 | -1.41 | -1 | -0.78 | -0.57 | 0.04 | -0.2575 | -0.2625 | 0.801549 | 0.270683 | 0.280277 | | 1 | 0.996511 | 0.822084 |
| Tcp11l1 | Q8BTG3 | -0.18 | -0.14 | -0.56 | -0.51 | 0.49 | -0.23 | -0.09 | -0.66 | -0.38 | -0.71 | -0.36 | 0 | -0.86 | 0.32 | -1.14 | -0.47 | 0.225 | -0.015 | -0.19 | 0.410611 | 0.945393 | 0.575994 | | 1 | 0.996755 | 0.926728 |
| Tctn3 | Q8R2Q6 | -0.2 | -0.33 | 0.56 | 0.98 | -0.68 | 0.23 | -1.17 | 0.11 | 2.02 | 1.02 | 0.07 | -0.32 | 2.84 | -0.57 | 1.07 | 0.44 | -0.63 | 0.445 | 0.6925 | 0.165514 | 0.436152 | 0.341772 | | 1 | 0.996511 | 0.848473 |
| Tdg | P56581-2 | -1.88 | -2.44 | -3.96 | -5.67 | -2.33 | -2.51 | -4.21 | -4.7 | -2.09 | -1.74 | -3.99 | -1.36 | -1.85 | -1.57 | -4.1 | -4.55 | 0.05 | 1.1925 | 0.47 | 0.957223 | 0.220937 | 0.647076 | | 1 | 0.996511 | 0.945362 |
| Tdp1 | B8JJC1 | 0.3 | 0.37 | -0.73 | 0.09 | 1.15 | 0.45 | 0.62 | 0.29 | 0.15 | -1.26 | 0.27 | 0.56 | -0.79 | 0.18 | -1.26 | -0.02 | 0.62 | -0.0775 | -0.48 | 0.072166 | 0.863198 | 0.254533 | | 1 | 0.996511 | 0.809524 |
| Tdp2 | Q9JJX7 | -1.61 | -1.39 | 0.16 | 0.48 | -0.8 | -2.27 | 0.06 | 0.82 | -0.13 | 1.46 | 0.09 | -1.11 | -0.64 | -1.1 | -0.05 | -0.04 | 0.0425 | 0.6675 | 0.1325 | 0.955763 | 0.341977 | 0.80804 | | 1 | 0.996511 | 0.971953 |
| Tdrd1 | Q99MV1 | -1.72 | -1.48 | -1.26 | -2.9 | -0.54 | -2.23 | -2.31 | -2.55 | -2.77 | -1.4 | -2.24 | -1.91 | -1.84 | -1.29 | -2.18 | -2.38 | -0.0675 | -0.24 | -0.0825 | 0.900424 | 0.587686 | 0.843394 | | 1 | 0.996511 | 0.976225 |
| Tdrd3 | Q91W18-2 | -0.75 | -0.45 | -1.6 | -1.72 | -0.48 | -0.58 | -1.72 | -2.07 | -0.13 | -1.67 | -1.8 | -0.65 | 0.41 | -0.54 | -1.02 | -0.74 | -0.0825 | 0.0675 | 0.6575 | 0.861094 | 0.887438 | 0.144722 | | 1 | 0.996511 | 0.743156 |
| Tdrd7 | A0A0A0MC | 0.31 | -0.3 | -0.54 | -0.84 | -0.23 | 0.45 | -0.38 | -0.71 | -0.77 | -0.9 | -0.59 | 0.22 | -0.41 | -0.99 | -0.59 | -0.23 | 0.125 | -0.1675 | -0.2125 | 0.709562 | 0.62906 | 0.491554 | | 1 | 0.996511 | 0.897531 |
| Tdrkh | A0A0G2JFE | 0.9 | 0.58 | 0.57 | 0.77 | 0.62 | 1.08 | 0.85 | 0.78 | 0.62 | 0.45 | 0.86 | 1.08 | 0.45 | 0.96 | 0.77 | 0.53 | 0.1275 | 0.0475 | -0.0275 | 0.470686 | 0.817055 | 0.892758 | | 1 | 0.996511 | 0.982799 |
| Tdrp | Q8C5P7 | -1.57 | -1.76 | 0.66 | 0.71 | -1.29 | -2.21 | 0.66 | 0.87 | -1.87 | 0.3 | 0.72 | -2.06 | -1.56 | -1.17 | 0.7 | 0.93 | -0.0025 | -0.2375 | 0.215 | 0.997807 | 0.790392 | 0.797555 | | 1 | 0.996511 | 0.969727 |
| Tecpr1 | Q80VP0 | 0.42 | 0.09 | 0.32 | 0.51 | 0.32 | 0.66 | 0.27 | 0.19 | 0.59 | -0.13 | 0.5 | 0.35 | 0.41 | 0.58 | 0.12 | 0.03 | 0.025 | -0.0075 | -0.05 | 0.890415 | 0.972886 | 0.814189 | | 1 | 0.997515 | 0.972768 |
| Tecpr2 | Q3UH45 | -0.49 | -0.09 | -0.35 | -0.31 | -0.11 | -0.31 | -0.3 | -0.62 | -0.48 | 0.03 | -0.38 | -0.35 | -0.44 | -0.3 | -0.74 | -0.52 | -0.025 | 0.015 | -0.19 | 0.889255 | 0.93816 | 0.344163 | | 1 | 0.996755 | 0.848675 |
| Tecr | Q9CY27 | 1.1 | 0.21 | 1.54 | 2.06 | 0.66 | 1.32 | 1.91 | 2.59 | 0.23 | 1.45 | 1.66 | 0.97 | -0.23 | -0.4 | 2.5 | 2.77 | 0.3925 | -0.15 | -0.0675 | 0.460313 | 0.750988 | 0.936274 | | 1 | 0.996511 | 0.989767 |
| Tef | Q9JLC6 | -1.66 | -1.73 | 0.19 | 0.58 | -1.34 | -2.19 | -0.98 | 0.45 | 6.07 | 0.61 | 0.35 | -2.66 | 7.32 | 0.68 | 0.3 | -0.04 | -0.36 | 1.7475 | 2.72 | 0.628795 | 0.321542 | 0.126255 | | 1 | 0.996511 | 0.721505 |
| Tefm | Q5SSK3 | 0.95 | 1 | -0.47 | -0.81 | 0.65 | 0.94 | -0.19 | -0.97 | 0.52 | -1.81 | -0.29 | 1.11 | 0.41 | 0.45 | -1.39 | -0.74 | -0.06 | -0.285 | -0.485 | 0.918084 | 0.69204 | 0.426565 | | 1 | 0.996511 | 0.878976 |
| Telo2 | Q9DC40 | -1.38 | -1.45 | -0.86 | -0.82 | -1.44 | -1.6 | -0.61 | -0.96 | -0.88 | -0.49 | -1.38 | -1.55 | -0.73 | -0.5 | -1.39 | -0.4 | -0.025 | 0.0525 | 3.73E-01 | 0.929593 | 0.860809 | 0.225752 | | 1 | 0.996511 | 0.787447 |
| Ten1 | Q9D7K2 | -0.24 | -0.42 | -1.26 | -1.03 | 2.14 | -0.65 | -0.64 | -1.73 | -4.53 | -1.71 | -1.48 | -0.55 | 0.24 | 0.8 | -2.86 | -2.17 | 0.5175 | -1.33 | -0.26 | 0.511455 | 0.126473 | 0.755722 | | 1 | 0.996511 | 0.962333 |
| Tenm1 | Q9WTS4 | 0.87 | 0.83 | 0.57 | 0.61 | 0.71 | 1.08 | 0.39 | 0.57 | 0.83 | 0.32 | 0.77 | 0.94 | 0.82 | 0.48 | 0.63 | 0.32 | -0.0325 | -0.005 | -0.1575 | 0.869987 | 0.980358 | 0.436894 | | 1 | 0.998773 | 0.883161 |
| Tenm2 | A0A0A0MC | 0.83 | 0.95 | 0.91 | 0.72 | 1.15 | 1.14 | 1.19 | 0.74 | 0.59 | 0.53 | 1.05 | 1 | 0.39 | 0.95 | 0.58 | 1.05 | 0.2025 | -0.06 | -0.11 | 0.253158 | 0.760506 | 0.613943 | | 1 | 0.996511 | 0.935876 |
| Tenm3 | G3X907 | 0.86 | 0.87 | 0.86 | 0.72 | 0.5 | 1.33 | 1.09 | 0.87 | 0.64 | 0.44 | 0.95 | 1.23 | 0.39 | 0.87 | 0.68 | 0.66 | 0.12 | -0.0125 | -0.1775 | 0.568756 | 0.953916 | 0.354434 | | 1 | 0.996755 | 0.855029 |
| Tenm4 | Q3UHK6-4 | 0.75 | 0.94 | 0.83 | 1.27 | 0.67 | 1.01 | 0.61 | 0.84 | 1.57 | 0.76 | 0.87 | 1.11 | 0.77 | 0.5 | 0.77 | 0.62 | -0.165 | 0.13 | -0.2825 | 0.387207 | 0.591664 | 0.180642 | | 1 | 0.996511 | 0.775542 |
| Tent4b | F8WI26 | -0.35 | 0.29 | 0.58 | -0.29 | -1.23 | -0.37 | -0.54 | 0.05 | 0.07 | 1.08 | 0.11 | 0.08 | 0.97 | 0.34 | -0.46 | -0.11 | -0.58 | 0.2775 | 0.1275 | 0.114077 | 0.414372 | 0.733844 | | 1 | 0.996511 | 0.958635 |
| Tepsin | Q3U3N6 | 0.47 | 0.38 | -0.05 | -0.48 | 0.94 | 0.84 | -0.63 | 0.04 | 1.06 | 0.75 | -0.86 | 0.86 | 0.37 | 0.71 | -0.82 | 0.13 | 0.2175 | 0.3725 | 0.0175 | 0.593236 | 0.429434 | 0.963543 | | 1 | 0.996511 | 0.9927 |
| Terf2 | D3YZ08 | -0.59 | -2.45 | -2.33 | -1.53 | -2 | -0.32 | -1.29 | -1.76 | -1.37 | -2.97 | -1.05 | -2.16 | -0.05 | -1.95 | 0.26 | -0.79 | 0.3825 | -0.1625 | 1.0925 | 0.471119 | 0.771994 | 0.095366 | | 1 | 0.996511 | 0.698156 |
| Terf2ip | Q91VL8 | -0.47 | -1.43 | -0.7 | 0.27 | -0.81 | -0.95 | -0.21 | 0.02 | -1.71 | 0.42 | -0.03 | -1.34 | -0.08 | -1.08 | -0.16 | -0.17 | 0.095 | -0.0825 | 0.21 | 0.811059 | 0.884785 | 0.607896 | | 1 | 0.996511 | 0.933507 |
| Tes | Q3ULB1 | 0.55 | 0.23 | 0.42 | -0.46 | 0.44 | 0.68 | 1.51 | -0.12 | 0.36 | -0.48 | 1.6 | 0.85 | 0.3 | 0.58 | 2.25 | 0.86 | 0.4425 | 0.3975 | 0.8125 | 0.269198 | 0.39852 | 0.10465 | | 1 | 0.996511 | 0.702979 |
| Tesc | Q9JKL5 | -0.09 | 0.66 | 1.21 | 1.12 | 0.24 | -0.06 | 1.29 | 0.97 | -0.43 | -1.77 | 0.48 | -0.06 | -0.56 | -0.42 | 0.76 | 0.87 | -0.115 | -1.17 | -0.5625 | 0.777522 | 0.049192 | 0.236217 | | 1 | 0.996511 | 0.791642 |
| Tesk1 | O70146 | 0.33 | 0.76 | 0.22 | -0.13 | 0.28 | 0.22 | -0.56 | -0.51 | 1 | 0.3 | -0.45 | 0.61 | 0.85 | 0.47 | -1.02 | -1.24 | -0.4375 | 0.07 | -0.53 | 0.15979 | 0.841695 | 0.319504 | | 1 | 0.996511 | 0.838969 |
| Tex10 | Q3URQ0 | -0.3 | -0.72 | -1.31 | -1.36 | -0.43 | -0.6 | -1.36 | -1.48 | -0.18 | -1.5 | -1.24 | -0.47 | -0.08 | -0.38 | -1.67 | -1.61 | -0.045 | 0.075 | -0.0125 | 0.898202 | 0.846103 | 0.978112 | | 1 | 0.996511 | 0.993727 |
| Tex2 | Q6ZPJ0 | 0.47 | 0.4 | 0.24 | 0.36 | 0.46 | 0.25 | 0.22 | 0.11 | 0.58 | -0.43 | 0.2 | 0.52 | 0.63 | 0.29 | 0.35 | 0.05 | -0.1075 | -0.15 | -0.0375 | 0.495267 | 0.563074 | 0.849579 | | 1 | 0.996511 | 0.976225 |
| Tex264 | E9Q137 | 0.23 | 0.54 | 0.06 | -0.16 | 0.73 | 0.26 | 0.4 | -0.03 | 0.15 | 0.3 | -0.1 | 0.34 | 1.2 | 0.72 | 1.06 | 0.52 | 0.1725 | 0.005 | 0.7075 | 0.468132 | 0.981599 | 0.018219 | | 1 | 0.998773 | 0.602211 |
| Tf | Q921I1 | -2.13 | -2.51 | -2.3 | -2.37 | -2.06 | -2.28 | -1.81 | -1.93 | -2.59 | -1.75 | -2.02 | -1.81 | -2.29 | -1.81 | -2.16 | -2.22 | 0.3075 | 0.285 | 0.2075 | 0.108237 | 0.247746 | 0.314371 | | 1 | 0.996511 | 0.836924 |
| Tfam | A0A0R4J1N | 0.63 | 0.73 | 1.52 | 1.7 | 0 | 0.62 | 0.79 | 1.43 | 0.71 | 1.77 | 1.69 | 0.36 | 0.58 | 0.38 | 1.13 | 1.02 | -0.435 | -0.0125 | -0.3675 | 0.270901 | 0.976356 | 0.280902 | | 1 | 0.997588 | 0.822084 |
| Tfb1m | Q8JZM0 | 1.2 | 1.27 | 2.08 | 0.91 | 0.92 | 1.26 | 2.44 | 2.07 | 0.62 | 0.95 | 2.12 | 1.7 | -0.43 | 0.51 | 1.75 | 2.11 | 0.3075 | -0.0175 | -0.38 | 0.456982 | 0.965503 | 0.519579 | | 1 | 0.996755 | 0.907667 |
| Tfcp2 | A0A2R8VKI | -0.91 | -0.7 | -0.8 | -1 | -0.71 | -0.53 | -0.98 | -0.58 | -0.55 | -0.94 | -0.85 | -0.93 | -0.27 | -0.4 | -0.92 | -0.42 | 0.1525 | 0.035 | 0.35 | 0.385268 | 0.845278 | 0.127528 | | 1 | 0.996511 | 0.723391 |
| Tfdp1 | Q08639 | -2.59 | -3.32 | -0.14 | 0.38 | -1.63 | -3.25 | 0.37 | 0.24 | -2.91 | 0.63 | 0.2 | -1.7 | -2.04 | -1.83 | -0.15 | 0.5 | 0.35 | 0.4725 | 0.5375 | 0.754801 | 0.669223 | 0.588422 | | 1 | 0.996511 | 0.929131 |
| Tfe3 | Q64092 | 0.43 | -0.09 | -0.36 | 0.08 | -0.12 | 0.71 | -0.83 | -0.25 | 1.13 | 0.56 | -0.59 | 0.23 | 2.99 | -1.23 | 0.16 | -0.64 | -0.1375 | 0.3175 | 0.305 | 0.691462 | 0.414544 | 0.72149 | | 1 | 0.996511 | 0.958398 |
| Tfeb | Q3UKG7 | 0.1 | -0.23 | -0.98 | -2.47 | -0.7 | 0.19 | -0.05 | -1.43 | -0.69 | -0.5 | 0.12 | 0.2 | 1.39 | -1.93 | 0.19 | -1.5 | 0.3975 | 0.6775 | 0.4325 | 0.524285 | 0.249991 | 0.617744 | | 1 | 0.996511 | 0.937259 |
| Tfg | Q9Z1A1 | -0.05 | 0.31 | 0.05 | 0.13 | -0.01 | -0.04 | 0.29 | 0.16 | 0.11 | 0.07 | 0.13 | 0.19 | 0.38 | 0.17 | -0.12 | 0.11 | -0.01 | 0.015 | 0.025 | 0.951745 | 0.927629 | 0.899176 | | 1 | 0.996755 | 0.983509 |
| Tfip11 | Q9ERA6 | -0.73 | -1.31 | -1.01 | -1.25 | -1.66 | -1.26 | -1.17 | -1 | -0.64 | -0.44 | -1.39 | -1.86 | 0.97 | -1.43 | -0.46 | -1.36 | -0.1975 | -0.0075 | 0.505 | 0.374831 | 0.982819 | 0.355109 | | 1 | 0.998773 | 0.855029 |
| Tfrc | Q62351 | -0.04 | 0.23 | 0.55 | 0.26 | 0.01 | -0.02 | 0.15 | -0.05 | 0.41 | 0.33 | 0.42 | 0.16 | -0.13 | -0.29 | -0.64 | -0.39 | -0.2275 | 0.08 | -0.6125 | 0.21893 | 0.676797 | 0.018257 | | 1 | 0.996511 | 0.602211 |
| Tgfb1i1 | Q62219-2 | 1.25 | 2.17 | -1.68 | -1.33 | -0.03 | 0.69 | -1.12 | -1.57 | 1.24 | -1.91 | -1.33 | 2.04 | 1.51 | 0.8 | -1.9 | -1.63 | -0.61 | -0.0925 | -0.4075 | 0.53382 | 0.939032 | 0.721705 | | 1 | 0.996755 | 0.958398 |
| Tgfbi | Q3UXJ2 | 1.36 | 1.15 | 0.28 | 0.08 | 1.05 | 1.04 | 0.26 | 0.47 | -1.66 | 3.05 | 1.05 | 1.6 | -0.22 | -0.2 | 1.03 | 0.44 | -0.0125 | 0.2925 | -0.455 | 0.972161 | 0.75385 | 0.290793 | | 1 | 0.996511 | 0.825497 |
| Tgfbrap1 | Q3UR70 | -0.08 | -0.01 | 0.02 | 0.11 | 0.15 | -0.08 | -0.04 | -0.19 | -0.04 | 0.23 | -0.01 | -0.2 | 0.03 | 0.42 | -0.16 | -0.1 | -0.05 | -0.015 | 0.0375 | 0.743811 | 0.930541 | 0.852454 | | 1 | 0.996755 | 0.976225 |
| Tgm2 | P21981 | -3.35 | -4.01 | -2.42 | -1.84 | -3.92 | -3.3 | -2.96 | -2.21 | -3.39 | -1.13 | -2.13 | -3.47 | -3.14 | -3.29 | -2.56 | -3.31 | -0.1925 | 0.375 | -0.17 | 0.726824 | 0.580175 | 0.724347 | | 1 | 0.996511 | 0.958398 |
| Tgoln1 | Q62313 | -1.62 | -2.18 | -1.59 | -1.09 | -1.19 | -2.1 | -1.35 | -1.36 | -1.41 | -0.36 | -0.98 | -2.42 | -0.44 | -2.55 | -0.61 | -0.62 | 0.12 | 0.3275 | 0.565 | 0.690017 | 0.480577 | 0.284578 | | 1 | 0.996511 | 0.822669 |
| Tgs1 | Q923W1 | -1.17 | -1 | -0.16 | 0.14 | -0.66 | -1.59 | -0.7 | -0.14 | -3.88 | 0.68 | 0.05 | -1.21 | -2.99 | -3.98 | -0.15 | -0.37 | -0.225 | -0.5425 | -1.325 | 0.587747 | 0.572401 | 0.167099 | | 1 | 0.996511 | 0.759484 |
| Th | P24529 | 0.92 | -0.54 | 0.37 | 1.2 | -0.87 | -0.18 | 1.35 | 0.87 | 1.35 | 0.52 | -0.2 | 1.13 | 0.62 | 0.34 | 1.4 | 0.63 | -0.195 | 0.2125 | 0.26 | 0.736232 | 0.661572 | 0.545214 | | 1 | 0.996511 | 0.912212 |
| Tha1 | Q6XPS7 | 0.99 | 0.95 | 1.01 | 1.33 | 0.59 | 0.61 | 1.2 | 0.84 | 1.24 | 0.95 | 1.01 | 0.65 | 1.68 | 1.62 | 0.91 | 1.04 | -0.26 | -0.1075 | 0.2425 | 0.21541 | 0.593152 | 0.344522 | | 1 | 0.996511 | 0.8491 |
| Thada | A8C756-3 | -0.28 | -0.08 | -0.11 | 0.85 | 0.16 | -0.18 | 0.36 | 1.62 | 0.45 | 1.62 | 0.17 | 0.22 | 0.41 | 0.05 | 0.07 | -0.66 | 0.395 | 0.52 | -0.1275 | 0.378168 | 0.222208 | 0.709098 | | 1 | 0.996511 | 0.957656 |
| Thbd | P15306 | 0.99 | 1.64 | 0.69 | 0.51 | 0.94 | 0.48 | -0.18 | 0.43 | 0.8 | 1.61 | 0.5 | 0.26 | 0.28 | 0.59 | 0.75 | 0.58 | -0.54 | -0.165 | -0.4075 | 0.128099 | 0.659325 | 0.177585 | | 1 | 0.996511 | 0.772068 |
| Thbs1 | Q80YQ1 | 0.86 | -1.13 | -0.15 | -0.13 | -0.35 | 0.69 | -1.63 | -0.11 | -0.71 | 0.19 | -0.37 | -0.67 | -0.52 | 0.07 | 1.01 | 0.16 | -0.2125 | -0.2525 | 0.3175 | 0.712864 | 0.56222 | 0.51493 | | 1 | 0.996511 | 0.905674 |
| Thbs4 | Q9Z1T2 | 0.39 | -2.89 | -1.49 | -2.16 | -2.05 | -0.87 | -0.71 | -1.81 | -2.81 | -1.37 | -1.06 | 0.23 | 1.82 | -3.08 | -1.02 | -1.22 | 0.1775 | 0.285 | 0.6625 | 0.801418 | 0.737579 | 0.550779 | | 1 | 0.996511 | 0.913801 |
| Them4 | Q3UUI3 | -0.06 | 0.4 | -0.1 | -0.17 | 0.19 | 0.11 | 0.04 | -0.26 | 0.03 | 0.15 | -0.08 | -0.1 | -0.72 | 0.44 | -0.87 | -0.6 | 0.0025 | -0.0175 | -0.455 | 0.98987 | 0.928415 | 0.1897 | | 1 | 0.996755 | 0.776843 |
| Them6 | Q80ZW2 | -0.49 | -0.35 | 0.04 | 0.01 | -0.3 | -0.35 | 0.23 | 0.07 | -0.52 | -0.27 | -0.19 | -0.4 | -0.41 | 0.03 | -0.03 | -0.05 | 0.11 | -0.1475 | 0.0825 | 0.615895 | 0.467918 | 0.705829 | | 1 | 0.996511 | 0.957084 |
| Thg1l | Q9CQT0 | -0.66 | -0.81 | -0.83 | -0.3 | -0.66 | -1.15 | -1.37 | -0.79 | -1.82 | -0.24 | -0.82 | -1.11 | -1.65 | -1.07 | -0.59 | -1.22 | -0.3425 | -0.3475 | -0.4825 | 0.154573 | 0.328044 | 0.105331 | | 1 | 0.996511 | 0.703417 |
| Thnsl1 | Q8BH55 | 2.18 | 0.82 | 0.34 | 0.04 | 0.65 | 2.07 | 1.11 | 0.65 | 0.12 | -0.58 | 0.53 | 1.49 | -0.26 | 0.54 | 0.1 | 0.34 | 0.275 | -0.455 | -0.665 | 0.608068 | 0.447382 | 0.183628 | | 1 | 0.996511 | 0.776843 |
| Thnsl2 | Q80W22 | -0.58 | 0.15 | 0.1 | 0.23 | 1.01 | -0.31 | 1.06 | 0.31 | -0.85 | 0.96 | 0.29 | -0.11 | -1.42 | -0.18 | 0.51 | 0.63 | 0.5425 | 0.0975 | -0.09 | 0.156657 | 0.80902 | 0.849647 | | 1 | 0.996511 | 0.976225 |

| | | | | | | | | | | | | | | | | | | | | | | | | | | | |
|---|---|---|---|---|---|---|---|---|---|---|---|---|---|---|---|---|---|---|---|---|---|---|---|---|---|---|---|
| Thoc1 | Q8R3N6 | 0.36 | -0.97 | -1.75 | -2.22 | -0.97 | 0.66 | -1.16 | -1.36 | -0.56 | -1.53 | -1.92 | -0.81 | 0.64 | -0.3 | -0.07 | -0.76 | 0.4375 | -0.06 | 1.0225 | 0.51424 | 0.919291 | 0.107097 | | 1 | 0.996755 | 0.704241 |
| Thoc2 | B1AZI6 | -2.09 | -0.86 | -1.32 | -1.31 | -1.13 | -2.27 | -1.45 | -1.15 | 0.59 | -1.08 | -1.65 | -2.54 | 1.55 | 0.18 | -0.36 | -0.78 | -0.105 | 0.225 | 1.5425 | 0.767405 | 0.728021 | 0.015989 | | 1 | 0.996511 | 0.589387 |
| Thoc3 | Q8VE80 | -1.46 | -1.29 | -1.6 | -1.84 | -0.91 | -1.75 | -1.09 | -1.24 | -1.98 | -1.2 | -1.87 | -1.54 | -1.21 | -1.57 | -0.99 | -1.07 | 0.3 | -0.1 | 0.3375 | 0.219873 | 0.676996 | 0.155049 | | 1 | 0.996511 | 0.750763 |
| Thoc5 | A0A0R4J0J | -1.16 | -1.78 | -1.26 | -1.01 | -1.87 | -1.29 | -0.6 | -0.56 | -1.5 | -0.69 | -1.41 | -1.92 | -0.06 | -2.5 | 0.17 | -0.61 | 0.2225 | -0.0775 | 0.5525 | 0.520346 | 0.801964 | 0.351856 | | 1 | 0.996511 | 0.854088 |
| Thoc6 | D3Z132 | -1.22 | -1.93 | -1.37 | -1.25 | -1.92 | -1.97 | 0.13 | -0.58 | -1.71 | -0.77 | -1.42 | -1.93 | -0.85 | -0.57 | 0.26 | -0.28 | 0.3575 | -0.015 | 1.0825 | 0.482155 | 0.960827 | 0.005714 | | 1 | 0.996755 | 0.552614 |
| Thoc7 | A0A286YD | -1.26 | -0.98 | -1.49 | -1.16 | -2.48 | -2.14 | -1.6 | -1.29 | -0.11 | -0.71 | -1.88 | -1.25 | 0.7 | -1.78 | -0.34 | -0.7 | -0.655 | 0.235 | 0.6925 | 0.046445 | 0.539385 | 0.180707 | | 1 | 0.996511 | 0.775542 |
| Thop1 | A0A0R4IZY | -1.17 | -1.11 | -0.85 | -0.98 | -0.96 | -1.53 | -0.73 | -1.03 | -1.5 | -0.83 | -0.94 | -1.04 | -1.7 | -0.95 | -1.34 | -1.72 | -0.035 | -0.05 | -0.4 | 0.868012 | 0.810067 | 0.117384 | | 1 | 0.996511 | 0.718609 |
| Thra | A0A0R4J1R | -1.02 | -0.82 | -1.37 | -1.15 | -0.79 | -0.88 | -1.12 | -1.19 | -0.59 | -1.19 | -1.58 | -1.07 | 0.05 | -0.62 | -1.41 | -0.4 | 0.095 | -0.0175 | 0.495 | 0.617559 | 0.945331 | 0.159017 | | 1 | 0.996755 | 0.754492 |
| Thrap3 | Q569Z6 | 1.59 | 0.69 | 0.6 | 0.49 | -0.76 | 0.7 | -0.34 | 0.74 | 0.05 | 0.43 | 0.38 | 0.09 | 0.82 | -0.52 | 1.01 | -0.14 | -0.7575 | -0.605 | -0.55 | 0.10233 | 0.057204 | 0.219241 | | 1 | 0.996511 | 0.787447 |
| Thsd7a | E9PWD2 | 0.28 | -0.23 | -0.21 | 0.08 | -0.14 | 0.24 | 0.03 | -0.1 | 0.61 | 0.51 | -0.02 | 0.57 | 0.47 | 0.15 | 0.21 | 0.17 | 0.0275 | 0.4375 | 0.27 | 0.884166 | 0.079843 | 0.21102 | | 1 | 0.996511 | 0.786269 |
| Thsd7b | Q6P4U0 | 2.29 | 1.14 | 0.49 | 0.37 | 1.9 | 2.12 | 1.11 | 0.5 | 0.86 | -0.35 | 0.92 | 2.84 | 0.41 | 0.7 | 0.77 | 0.92 | 0.335 | -0.005 | -0.3725 | 0.531557 | 0.994449 | 0.397434 | | 1 | 0.999285 | 0.867334 |
| Thtpa | Q8JZL3 | -0.83 | -0.67 | -0.75 | -0.47 | -0.96 | -1.14 | -0.79 | -0.58 | -1.11 | 0.26 | -0.57 | -0.77 | -1.1 | -0.82 | -0.9 | -1.08 | -0.1875 | 0.1325 | -0.295 | 0.324182 | 0.666213 | 0.139002 | | 1 | 0.996511 | 0.73493 |
| Thumpd1 | Q99J36 | -3.99 | -4.11 | -3.92 | -4.33 | -3.91 | -4.08 | -3.9 | -4.62 | -4.29 | -1.78 | -4.05 | -3.74 | -3.54 | -3.14 | -4.25 | -4.7 | -0.04 | 0.6225 | 0.18 | 0.853517 | 0.264379 | 0.616597 | | 1 | 0.996511 | 0.937079 |
| Thumpd3 | P97770 | -0.65 | -0.6 | -0.54 | -0.78 | -0.77 | -0.46 | -0.64 | -0.68 | -0.87 | -0.5 | -1.08 | -0.74 | -0.17 | -1.74 | -0.64 | -0.49 | 0.005 | -0.155 | -0.1175 | 0.973962 | 0.422272 | 0.733422 | | 1 | 0.996511 | 0.958635 |
| Thy1 | A0A1L1SU | 0.81 | 0.76 | 1.05 | 1.26 | 0.84 | 0.61 | 0.93 | 1.3 | 0.85 | 1.45 | 0.89 | 0.73 | 0.35 | 1.37 | 0.83 | 0.96 | -0.05 | 0.01 | -0.0925 | 0.81327 | 0.965264 | 0.728832 | | 1 | 0.996755 | 0.958635 |
| Thyn1 | Q91YJ3 | -4.7 | -5.9 | -5.03 | -5.17 | -4.3 | -5.57 | -5.49 | -5.9 | -4.89 | -2.86 | -5.24 | -4.49 | -3.34 | -3.8 | -4.73 | -5.98 | -0.115 | 0.83 | 0.7375 | 0.77754 | 0.150373 | 0.227473 | | 1 | 0.996511 | 0.787447 |
| Tia1 | P52912 | -2.32 | -4.16 | -4.51 | -3.23 | -4.57 | -3.28 | -3.15 | -4.16 | 0.12 | -1.65 | -2.45 | -3.92 | 0.01 | -1.18 | -4.48 | -2.99 | -0.235 | 1.58 | 1.395 | 0.670733 | 0.100143 | 0.182516 | | 1 | 0.996511 | 0.776002 |
| Tia1 | Q8CII5 | -1.57 | -0.55 | -2.23 | -2.68 | -0.96 | -1.66 | -1.81 | -1.91 | -0.82 | -1.91 | -2.1 | -0.69 | -0.97 | -0.93 | -1.18 | -2.28 | 0.1725 | 0.3775 | 0.4175 | 0.715955 | 0.493389 | 0.431656 | | 1 | 0.996511 | 0.879165 |
| Tial1 | Q921W2 | -1.44 | -2 | -1.52 | -1.79 | -1.62 | -1.28 | -1.5 | -1.55 | -1.75 | -1.52 | -1.55 | -1.73 | -1.94 | -1.72 | -1.39 | -2.06 | 0.2 | 0.05 | -0.09 | 0.303066 | 0.797634 | 0.704151 | | 1 | 0.996511 | 0.957084 |
| Tiam1 | G3UWG2 | 0.03 | -0.14 | -0.2 | -0.11 | -0.65 | -0.01 | -0.11 | -0.11 | 0.63 | 0.35 | -0.24 | 0.04 | 1.08 | -0.13 | -0.05 | -0.15 | -0.115 | 0.3 | 0.2925 | 0.551618 | 0.21063 | 0.360062 | | 1 | 0.996511 | 0.856815 |
| Tiam2 | Q6ZPF3-3 | 1.19 | 1.15 | -2.37 | 0.07 | -0.32 | 0.88 | -0.45 | 0.46 | 2.33 | 0.99 | 0.53 | 1.93 | 3.12 | 0.92 | 0.95 | 0.11 | 0.1325 | 1.435 | 1.265 | 0.869257 | 0.115772 | 0.203624 | | 1 | 0.996511 | 0.782817 |
| Tiam2 | Q6ZPF3-2 | -0.19 | -0.5 | -0.25 | -0.17 | -0.45 | -0.53 | -0.24 | -0.14 | 0.48 | -0.14 | -0.1 | 0.25 | 0.03 | 0.3 | -0.61 | -0.31 | -0.0625 | 0.4 | 0.13 | 0.714963 | 0.083626 | 0.602078 | | 1 | 0.996511 | 0.930802 |
| Tigar | Q8BZA9 | 0.63 | 0.19 | -1.2 | -1.31 | 0.5 | 0.11 | -0.86 | -1.19 | 0.26 | -0.77 | -1.09 | 0.57 | -0.15 | 0.96 | -1.69 | -1.32 | 0.0625 | 0.165 | -0.1275 | 0.913602 | 0.776407 | 0.85604 | | 1 | 0.996511 | 0.976225 |
| Timm10 | P62073 | 0.15 | 0.54 | 0.5 | 0.76 | 0.65 | -0.03 | 0.14 | 0.57 | -0.04 | 1.07 | 0.51 | -0.04 | -0.16 | 1.11 | -0.03 | 0.31 | -0.155 | -0.1125 | -0.18 | 0.502051 | 0.708503 | 0.575721 | | 1 | 0.996511 | 0.926728 |
| Timm10b | Q9WV96 | 1.08 | 0.69 | -1.43 | -1.32 | 0.61 | 1.3 | -1.24 | -1.41 | -0.35 | -0.56 | -1.2 | 1.12 | -1.04 | -1.31 | -1.72 | -1.28 | 0.06 | -0.0025 | -1.0925 | 0.943529 | 0.997315 | 0.103281 | | 1 | 1 | 0.700896 |
| Timm13 | P62075 | 0.42 | 0.16 | 0.35 | 0.1 | 0.09 | 0.51 | 0.18 | 0.16 | -0.46 | 0.31 | 0.39 | 0.25 | -0.43 | -0.15 | 0.12 | 0.03 | -0.0225 | -0.135 | -0.365 | 0.89581 | 0.575041 | 0.102796 | | 1 | 0.996511 | 0.700625 |
| Timm17a | Q9Z0V8 | -0.76 | -0.12 | -0.78 | -0.31 | -0.09 | -0.49 | -0.4 | -0.28 | 0.1 | -0.95 | 0.07 | -0.41 | -0.09 | 0.26 | -0.22 | -1.36 | 0.1775 | 0.195 | 0.14 | 0.414016 | 0.523743 | 0.711973 | | 1 | 0.996511 | 0.958014 |
| Timm17b | Q9Z0V7 | 0.09 | 0.66 | 0.83 | 0.68 | 0.44 | 0.1 | 0.3 | 0.1 | 0.93 | 0.38 | 0.28 | -0.07 | 0.76 | 1.21 | 0 | 0.13 | -0.33 | -0.185 | -0.04 | 0.143629 | 0.509367 | 0.903291 | | 1 | 0.996511 | 0.985087 |
| Timm21 | Q8CCM6 | 0.43 | 0.58 | -0.03 | 0.69 | 0.72 | 0.45 | 0.11 | -0.41 | 0.57 | -0.46 | 0.42 | 0.46 | 0.39 | 0.59 | -0.44 | 0.21 | -0.2 | -0.17 | -0.23 | 0.497403 | 0.566393 | 0.436501 | | 1 | 0.996511 | 0.882882 |
| Timm22 | Q9CQ85 | 0.16 | 0.32 | -1.54 | -1.36 | 0.26 | 0.28 | -1.59 | -1.59 | 0.47 | -0.98 | -1.4 | 0.47 | -0.15 | 0.65 | -1.45 | -1.62 | -0.055 | 0.245 | -0.0375 | 0.933543 | 0.699096 | 0.954968 | | 1 | 0.996511 | 0.992482 |
| Timm23 | A0A2I3BS4 | 0.01 | 0.77 | 0.61 | 0.55 | 0.59 | 0.2 | 0.41 | 0.45 | 1.31 | 1.27 | 0.21 | 0 | 0.79 | 1.11 | -0.03 | 0.74 | -0.0725 | 0.2125 | 0.1675 | 0.732026 | 0.569763 | 0.585224 | | 1 | 0.996511 | 0.928105 |
| Timm29 | Q8BGX2 | -0.09 | -0.13 | -0.27 | -0.23 | 0.38 | 0.63 | 0.08 | -0.01 | -1.05 | -0.24 | -0.08 | -0.13 | -1.47 | 0.01 | -0.64 | -0.26 | 0.45 | -0.195 | -0.41 | 0.04007 | 0.449284 | 0.233054 | | 1 | 0.996511 | 0.788994 |
| Timm44 | O35857 | 0.14 | 0.45 | 0.15 | 0.03 | 0.06 | 0.34 | -0.33 | -0.05 | 0.57 | 0.16 | 0.49 | 0.24 | 0.85 | -0.14 | 0.03 | -0.03 | -0.1875 | 0.1725 | -0.015 | 0.358488 | 0.375908 | 0.955505 | | 1 | 0.996511 | 0.992482 |
| Timm50 | Q9D880 | 0.74 | 0 | 0.28 | 0.25 | -0.49 | 1.17 | 0 | 0.2 | 1.01 | 0.42 | 0.22 | 0.62 | 1.18 | -0.01 | -0.26 | -0.36 | -0.0975 | 0.25 | -0.18 | 0.789008 | 0.333849 | 0.635684 | | 1 | 0.996511 | 0.9419 |
| Timm8a1 | Q9WVA2 | 0.07 | 0.6 | -0.73 | -0.41 | 0.51 | -0.04 | -0.53 | -0.74 | -0.52 | -0.78 | -0.56 | 0.1 | -0.97 | -0.34 | -1.53 | -0.7 | -0.0825 | -0.3225 | -0.7675 | 0.828905 | 0.357407 | 0.066588 | | 1 | 0.996511 | 0.664936 |
| Timm8b | P62077 | -0.52 | -0.62 | -1.1 | -0.8 | -0.94 | -0.05 | -1.86 | -1.62 | -0.74 | -2 | -1.48 | -0.41 | 0.2 | -1.55 | -1.87 | -1.74 | -0.3575 | -0.3975 | -0.48 | 0.383777 | 0.298284 | 0.322475 | | 1 | 0.996511 | 0.840788 |
| Timm9 | Q9WV98 | -0.61 | -0.22 | 0.21 | 0.3 | -0.37 | -0.44 | 0.33 | 0.09 | -1.07 | 0.09 | 0.32 | -0.33 | -0.83 | -0.68 | -0.07 | 0 | -0.0175 | -0.1675 | -0.315 | 0.950479 | 0.644456 | 0.320907 | | 1 | 0.996511 | 0.839916 |
| Timmdc1 | Q8BUY5 | 0.14 | 0.07 | 0.12 | 0.27 | -0.42 | 0.48 | -0.1 | 0.27 | 1.4 | 0.46 | 0.13 | -0.1 | 1.57 | 0.98 | -0.23 | 0.36 | -0.0925 | 0.3225 | 0.52 | 0.682485 | 0.342597 | 0.195295 | | 1 | 0.996511 | 0.78004 |
| Timp2 | P25785 | 0.71 | 0.4 | 0.94 | 1.26 | -0.1 | 1 | 1.15 | 1.57 | 0.11 | 1.43 | 0.97 | 0.65 | 0.33 | 0.27 | 2.1 | 1.54 | 0.0775 | -0.0375 | 0.2325 | 0.838259 | 0.909354 | 0.616006 | | 1 | 0.996744 | 0.93691 |
| Tinagl1 | Q99JR5 | -1.22 | -2.15 | -2.7 | -2.76 | -1.7 | -1.29 | -1.84 | -3.16 | -0.92 | -2.38 | -3.3 | -1.56 | -0.31 | -2.02 | -2.44 | -2.72 | 0.21 | 0.1675 | 0.335 | 0.674546 | 0.771632 | 0.576027 | | 1 | 0.996511 | 0.926728 |
| Tiprl | Q8BH58 | 0.25 | 0.38 | -0.15 | 0.04 | 0.34 | -0.05 | 0.06 | 0.17 | 0.8 | 0.18 | -0.03 | 0.29 | 1.02 | 1 | -0.35 | -0.38 | -5.89E-17 | 0.18 | 0.1925 | 1 | 0.459397 | 0.631722 | | 1 | 0.996511 | 0.941132 |
| Tjp1 | B9EHJ3 | 0.68 | 0.82 | 0.91 | 0.77 | 0.6 | 0.48 | 0.71 | 0.54 | 0.98 | 1.03 | 0.98 | 0.91 | 1.16 | 0.71 | 0.84 | 0.78 | -0.2125 | 0.18 | 0.0775 | 0.175395 | 0.269292 | 0.683136 | | 1 | 0.996511 | 0.955729 |
| Tjp2 | A0A3B2W( | 1.68 | 1.57 | 1.62 | 1.3 | 1.21 | 1.22 | 1.16 | 0.97 | 1.07 | 1.35 | 1.41 | 1.66 | 0.98 | 0.51 | 1.5 | 1.35 | -0.4025 | -0.17 | -0.4575 | 0.033086 | 0.401275 | 0.109798 | | 1 | 0.996511 | 0.709835 |
| Tk2 | Q9R088 | 0.47 | 0.38 | 0.06 | 0.25 | 0.56 | 0.02 | 0.63 | 0.32 | 0.46 | 0.03 | 0.24 | 0.29 | -0.36 | 0.71 | -0.38 | -0.11 | 0.0925 | -0.035 | -0.325 | 0.642527 | 0.850798 | 0.277156 | | 1 | 0.996511 | 0.822084 |
| Tkfc | Q8VC30 | -0.45 | -0.26 | -0.22 | -0.04 | -0.05 | -0.17 | -0.1 | -0.01 | -0.32 | 0 | -0.08 | -0.25 | -0.51 | -0.56 | -0.27 | -0.44 | 0.16 | 0.08 | -0.2025 | 0.322799 | 0.656951 | 0.294751 | | 1 | 0.996511 | 0.825612 |
| Tkt | P40142 | 0.84 | 0.45 | -0.33 | 0.01 | 0.38 | 0.67 | -1.15 | -0.07 | 0.67 | -0.01 | -0.41 | 0.93 | 0.25 | 0.74 | -0.71 | -0.65 | -0.285 | 0.0525 | -0.335 | 0.52408 | 0.891332 | 0.430935 | | 1 | 0.996511 | 0.879165 |
| Tlcd1 | Q99JT6 | -0.97 | 0.2 | -0.37 | -1.19 | 1.08 | -0.46 | 0.85 | -0.14 | 0.1 | -0.94 | -0.28 | -0.82 | 0.45 | 2.09 | -0.88 | 0.11 | 0.915 | 0.0975 | 1.025 | 0.070477 | 0.798438 | 0.131756 | | 1 | 0.996511 | 0.726871 |
| Tlcd3b | Q7TNV1 | -0.73 | 0.23 | 0.44 | 0.09 | 0.14 | -0.38 | 0.01 | 0.1 | 0.47 | -0.29 | 0.49 | -0.47 | 0.02 | 0.33 | 0.14 | 0.45 | -0.04 | 0.0425 | 0.2275 | 0.88803 | 0.903607 | 0.44122 | | 1 | 0.996609 | 0.883549 |
| Tlcd4 | Q8CGF5 | -0.55 | -0.56 | -0.78 | -0.64 | 0.11 | 0.03 | -0.74 | -1.53 | 0.3 | -1.29 | -0.63 | 0.32 | -0.04 | -0.83 | -1.78 | -1.46 | 0.1 | 0.3075 | -0.395 | 0.786846 | 0.427525 | 0.311296 | | 1 | 0.996511 | 0.834523 |
| Tle1 | Q5SQA2 | -1.48 | -0.34 | 0.17 | 0.54 | -0.08 | 0.7 | -0.36 | -0.15 | -1.04 | 1.33 | 1.13 | -2.14 | 0.34 | -0.41 | -0.73 | 0.62 | 0.305 | 0.0975 | 0.2325 | 0.514132 | 0.909621 | 0.646371 | | 1 | 0.996744 | 0.944831 |
| Tle3 | Q08122-4 | -2.46 | -2.26 | -1.35 | -2.77 | -3.03 | -2.29 | -2.62 | -1.94 | -1.08 | -0.18 | -2.21 | -1.12 | -0.79 | -1.93 | -2.01 | -2.46 | -0.26 | 1.0625 | 0.4125 | 0.484393 | 0.051722 | 0.364885 | | 1 | 0.996511 | 0.858475 |
| Tle4 | F6ZBY9 | -1.91 | -2.02 | -1.56 | -2.11 | -1.64 | -2.32 | -1.64 | -1.65 | -2.53 | -1.46 | -1.35 | -1.74 | -2.01 | -1.86 | -1.55 | -2.25 | 0.0875 | 0.13 | -0.0175 | 0.702108 | 0.664881 | 0.939993 | | 1 | 0.996511 | 0.990645 |
| Tle5 | P63002 | -2.23 | -2.89 | -2.06 | -3.3 | -3.2 | -2.36 | -4.31 | -3.27 | -1.98 | -1.88 | -3.26 | -2.14 | -0.91 | -3.6 | -2.23 | -2.5 | -0.665 | 0.305 | 0.31 | 0.171843 | 0.46446 | 0.591902 | | 1 | 0.996511 | 0.929843 |
| Tln1 | P26039 | -0.42 | -0.97 | -0.54 | -0.46 | -0.99 | -0.93 | -0.73 | -0.71 | -0.53 | -0.06 | -0.33 | -0.67 | -0.27 | -0.65 | -0.53 | -0.71 | -0.2425 | 0.2 | 0.0575 | 0.214705 | 0.375683 | 0.789024 | | 1 | 0.996511 | 0.968677 |
| Tln2 | E9PUM4 | 0.8 | 0.51 | 0.57 | 0.6 | 0.3 | 0.66 | 0.37 | 0.55 | 0.7 | 0.27 | 0.61 | 0.6 | 0.55 | 0.4 | 0.3 | 0.27 | -0.15 | -0.075 | -0.24 | 0.369423 | 0.678049 | 0.205362 | | 1 | 0.996511 | 0.783452 |
| Tln2 | A0A1L1SRI | -1.42 | -1.6 | 0.42 | 0.96 | -1.38 | -1.31 | 0.39 | 0.65 | -0.99 | 0.78 | 0.52 | -1.24 | -1.71 | -0.85 | 0.12 | 0.22 | -0.0025 | 0.1775 | -0.145 | 0.997375 | 0.812666 | 0.839388 | | 1 | 0.996511 | 0.976219 |
| Tlnrd1 | Q9ERE8 | 0.21 | -0.29 | -0.17 | -0.25 | -1.19 | 0.39 | -1.14 | -0.06 | 0.93 | 0.39 | -0.45 | -0.81 | 1.74 | -1.25 | -0.26 | -0.48 | -0.375 | 0.14 | 0.0625 | 0.348538 | 0.723318 | 0.916101 | | 1 | 0.996511 | 0.986556 |
| Tm2d1 | Q99MB3-2 | -3.3 | -2.03 | -2.94 | -3.18 | -1.2 | -3.28 | -2.63 | -3.56 | -3.53 | -4.24 | -3 | -3.21 | -2.86 | -1.84 | -4.46 | -3.15 | 0.195 | -0.6325 | -0.215 | 0.723603 | 0.124314 | 0.70367 | | 1 | 0.996511 | 0.957059 |
| Tm7sf2 | Q71KT5 | -0.75 | -0.1 | -0.51 | -0.36 | -0.12 | -0.3 | -0.29 | -0.37 | 0.4 | 0.22 | -0.3 | 0 | 0.67 | 0.96 | -0.06 | 0.12 | 0.16 | 0.51 | 0.8525 | 0.401963 | 0.053639 | 0.01567 | | 1 | 0.996511 | 0.589387 |
| Tm9sf1 | D6RGM8 | 0.12 | -1.13 | -2.31 | -1.66 | -0.77 | -1.31 | 1.07 | -0.52 | -0.24 | -1.09 | -0.86 | -0.55 | -0.6 | -0.42 | -1.14 | 0.01 | 0.8625 | 0.56 | 0.7075 | 0.213183 | 0.287777 | 0.201715 | | 1 | 0.996511 | 0.781647 |
| Tm9sf2 | P58021 | -3.11 | -4.11 | -2.1 | -1.9 | -2.33 | -3.12 | -1.12 | -2.02 | -4.78 | -1.75 | -1.93 | -3.61 | -4.48 | -3.56 | -1.91 | -1.57 | 0.6575 | -0.2125 | -0.075 | 0.28902 | 0.790385 | 0.922601 | | 1 | 0.996511 | 0.987467 |
| Tm9sf3 | A0A498WG | -2.77 | -3.83 | -2.54 | -3.11 | -2.7 | -2.94 | -1.92 | -2.35 | -3.94 | -1.18 | -2.44 | -3.27 | -3.25 | -2.84 | -2.43 | -2.06 | 0.585 | 0.355 | 0.4175 | 0.117966 | 0.559797 | 0.280221 | | 1 | 0.996511 | 0.822084 |
| Tm9sf4 | Q8BH24 | -2.08 | -2.59 | -2.41 | -2.04 | -1.93 | -1.88 | -1.28 | -1.97 | -1.95 | -2.1 | -2.17 | -2.09 | -1.25 | -1.85 | -1.73 | -1.57 | 0.515 | 0.2025 | 0.68 | 0.048581 | 0.312378 | 0.015275 | | 1 | 0.996511 | 0.589387 |
| Tma16 | Q9CR02 | -2.43 | -2.69 | -1.97 | -1.29 | -1.56 | -3.44 | -1.14 | -2.17 | 0.14 | -0.37 | -2.16 | -2.42 | 0.28 | -0.6 | -0.6 | -1.05 | 0.0175 | 0.8925 | 1.6025 | 0.974076 | 0.191562 | 0.00309 | | 1 | 0.996511 | 0.461628 |
| Tma7 | Q8K003 | -2.01 | -2.69 | -1.82 | -2.59 | -3.93 | -2.89 | -2.36 | -2.36 | -1.03 | -1.29 | -1.73 | -2.16 | 1.51 | -3.23 | -0.82 | -2.2 | -0.6075 | 0.725 | 1.0925 | 0.157543 | 0.050763 | 0.263063 | | 1 | 0.996511 | 0.81524 |
| Tmbim1 | D3Z0A5 | 1.53 | 0.63 | 1.14 | 1.04 | -0.98 | 1.73 | 1.62 | 0.59 | 0.66 | 0.88 | 0.73 | -0.18 | 1.04 | -2.44 | 1.35 | 1.39 | -0.345 | -0.5625 | -0.75 | 0.567255 | 0.093752 | 0.388994 | | 1 | 0.996511 | 0.864915 |
| Tmcc1 | F8WJ98 | 0.59 | 0.82 | 0.72 | 0.55 | 1.06 | 0.45 | 1.19 | 0.7 | 0.3 | -0.1 | 0.51 | 0.36 | 0.24 | 0.74 | 0.63 | 1 | 0.18 | -0.4025 | -0.0175 | 0.398496 | 0.066519 | 0.936637 | | 1 | 0.996511 | 0.989767 |
| Tmcc2 | Q80W04 | 1.07 | 1.26 | 0.27 | 0.22 | 1.04 | 0.95 | 0.59 | 0.5 | 1.06 | 0.67 | 0.17 | 1.12 | 1.04 | 0.82 | 0.9 | 0.63 | 0.065 | 0.05 | 0.1425 | 0.827266 | 0.883539 | 0.632676 | | 1 | 0.996511 | 0.941132 |
| Tmcc3 | Q8R310-2 | -0.24 | 0.58 | 0.66 | 0.34 | 0.06 | -0.08 | 0.44 | 0.14 | 0.45 | 0.97 | 0.15 | 0.39 | 0.24 | 0.24 | 0.84 | 0.16 | -0.195 | 0.155 | 0.035 | 0.434673 | 0.58208 | 0.89994 | | 1 | 0.996511 | 0.983509 |
| Tmco1 | A0A0A6YV | -1.29 | -0.59 | -1.09 | -1.22 | -0.29 | -1.08 | -0.65 | -0.77 | 0.17 | -1.03 | -1.24 | -1.06 | 0.26 | -0.02 | -0.46 | -0.41 | 0.35 | 0.2575 | 0.89 | 0.173273 | 0.471714 | 0.007475 | | 1 | 0.996511 | 0.552614 |
| Tmco6 | Q8BQX5 | 0.86 | 0.82 | 0.01 | 0.13 | 1.69 | 1.58 | -0.43 | -0.45 | -0.21 | 0.38 | -0.44 | 0.74 | 1.2 | 1.56 | -1.14 | -0.11 | 0.1425 | -3.38E-01 | -0.0775 | 0.807485 | 0.341375 | 0.897818 | | 1 | 0.996511 | 0.983509 |
| Tmed10 | Q9D1D4 | -1.89 | -1.94 | -1.26 | -1.26 | -1.05 | -1.9 | -0.16 | -1.12 | -1.57 | -1.42 | -1.2 | -1.89 | -0.71 | -0.96 | -0.32 | -0.48 | 0.53 | 0.0675 | 0.97 | 0.189864 | 0.794117 | 0.004887 | | 1 | 0.996511 | 0.552614 |
| Tmed2 | F6V6T4 | -0.43 | -1.07 | -2.03 | -2.07 | 0.15 | -0.35 | -0.97 | -2.12 | -1.28 | -1.79 | -1.94 | -0.58 | -0.11 | -0.74 | -0.84 | -0.91 | 0.5775 | 0.0025 | 0.75 | 0.330721 | 0.995751 | 0.100666 | | 1 | 0.99948 | 0.698156 |
| Tmed3 | Q78IS1 | 1.9 | -1.93 | 0.58 | -0.88 | -1.53 | 0.32 | -1.3 | -1.52 | -0.85 | -1.18 | -0.33 | -0.52 | -0.57 | -1.54 | -0.33 | -0.13 | -0.925 | -0.6375 | -0.56 | 0.295597 | 0.421162 | 0.49296 | | 1 | 0.996511 | 0.897531 |
| Tmed4 | Q8R1V4 | -1.39 | -1.39 | -2.2 | -2.49 | -0.66 | -1.4 | -1.82 | -2.76 | -1.18 | -1.66 | -2.57 | -1.48 | -0.34 | -1.26 | -1.4 | -1.79 | 0.2075 | 0.145 | 0.67 | 0.668376 | 0.713804 | 0.1225 | | 1 | 0.996511 | 0.721448 |
| Tmed5 | Q9CXE7 | -0.65 | -0.2 | -0.44 | -0.84 | 1.34 | -0.54 | 0.57 | -0.33 | -2.4 | -1.14 | -0.17 | -0.14 | -1.34 | -1.68 | 0.09 | 0.44 | 0.7925 | -0.43 | -0.09 | 0.090272 | 0.409549 | 0.857976 | | 1 | 0.996511 | 0.976225 |
| Tmed7 | E9Q7G1 | -3.92 | -4.13 | -1.13 | -1.49 | -2.91 | -3.42 | -0.31 | -1.13 | -4.14 | -1.48 | -1.3 | -3.6 | -3.87 | -4.28 | -0.68 | -0.43 | 0.725 | 0.0375 | 0.3525 | 0.460599 | 0.968913 | 0.759649 | | 1 | 0.996755 | 0.963123 |
| Tmed8 | Q3UHI4 | 0.06 | 0.57 | 0.1 | -0.06 | 0.94 | -0.13 | 0.2 | 0.03 | 0.03 | 0.26 | -0.33 | 0.58 | -0.23 | 0.72 | 0.3 | -0.02 | 0.0925 | -0.0325 | 0.025 | 0.73963 | 0.899287 | 0.926833 | | 1 | 0.996511 | 0.989128 |
| Tmed9 | Q99KF1 | -1.23 | -1.08 | -1.86 | -1.71 | -0.02 | -1 | -0.67 | -1.69 | -1.54 | -1.94 | -1.74 | -0.93 | -1.07 | -0.89 | -0.67 | -0.89 | 0.625 | -0.0675 | 0.59 | 0.123462 | 0.819052 | 0.035166 | | 1 | 0.996511 | 0.651434 |
| Tmeff1 | A0A0A0M( | 0.7 | 0.82 | -0.62 | -0.63 | 0.69 | 0.89 | -0.03 | -1.07 | 1.58 | -1.36 | -0.77 | 1.15 | 1.23 | 1.59 | -0.88 | -0.72 | 0.0525 | 0.0825 | 0.2375 | 0.92356 | 0.911724 | 0.730591 | | 1 | 0.996744 | 0.958635 |
| Tmeff2 | Q9QYM9 | 1.07 | 0.76 | 0.32 | -0.42 | 0.26 | 1.21 | 0.57 | 0.48 | 1.06 | -0.33 | 0.33 | 1.49 | 2.15 | -1.35 | 2.13 | 1.43 | 0.1975 | 0.205 | 0.6575 | 0.591495 | 0.671593 | 0.422014 | | 1 | 0.996511 | 0.875736 |
| Tmem106b | Q80X71 | -1.62 | 0.09 | -2.62 | -3.39 | -0.25 | -1.02 | -2.71 | -3.57 | -0.34 | -0.82 | -2.45 | -0.93 | -0.07 | 0.83 | -2.75 | -2.57 | -0.0025 | 0.75 | 0.745 | 0.99792 | 0.358361 | 0.482694 | | 1 | 0.996511 | 0.893925 |
| Tmem109 | Q3UBX0 | 0.27 | 0.12 | 0.22 | 0.22 | 0.68 | -0.18 | 0.08 | 0.39 | -0.56 | 0.03 | -0.04 | -0.3 | 0.87 | -0.91 | 1.21 | 0.91 | 0.035 | -0.425 | 0.3125 | 0.870659 | 0.051881 | 0.499263 | | 1 | 0.996511 | 0.899626 |
| Tmem11 | Q8BK08 | -0.31 | -0.4 | 0.3 | 0.34 | -1.12 | -0.04 | -1.32 | -0.56 | 0.64 | -1.29 | -0.26 | -0.59 | 1.18 | -0.83 | -0.37 | -0.49 | -0.7425 | -0.3575 | -0.11 | 0.05256 | 0.407936 | 0.81096 | | 1 | 0.996511 | 0.972384 |
| Tmem114 | Q9D563 | 0.12 | 0.61 | 1.04 | 0.28 | 0.11 | 0.58 | -0.3 | 0.33 | 2.25 | 2.18 | 0.95 | 0.68 | 3.6 | 1.8 | 0.87 | 1.22 | -0.3325 | 1.0025 | 1.36 | 0.253327 | 0.042789 | 0.043194 | | 1 | 0.996511 | 0.658454 |
| Tmem115 | Q9WUH1 | 0.04 | -0.21 | -0.55 | -0.31 | -0.13 | 0.22 | -0.11 | -0.46 | 0.26 | 0.56 | -0.27 | -0.37 | 0.51 | -0.15 | -0.42 | 0.21 | 0.1375 | 0.3025 | 0.295 | 0.521763 | 0.277327 | 0.281968 | | 1 | 0.996511 | 0.822084 |
| Tmem117 | Q8BH18 | -8.04 | -9.88 | -6.17 | -4.09 | -5.48 | -6.23 | -5.71 | -6.1 | -5.85 | -10.47 | -5.94 | -5.49 | -3.76 | -4.34 | -5.97 | -7.36 | 1.165 | 0.1075 | 1.6875 | 0.315214 | 0.943958 | 0.223241 | | 1 | 0.996755 | 0.787447 |
| Tmem120a | Q8C1E7 | -0.14 | 0.13 | 0.99 | 0.16 | 0.82 | 0.25 | 1.34 | 0.97 | -0.68 | 0.87 | 0.68 | 0.18 | -0.46 | -0.31 | 1.27 | 1.35 | 0.56 | -0.0225 | 0.1775 | 0.112588 | 0.955595 | 0.728466 | | 1 | 0.996755 | 0.958635 |
| Tmem121b | Q99MX7 | 0.37 | 0.02 | -0.67 | -0.82 | 0.45 | 0.6 | -0.53 | -1.12 | -0.37 | -1.28 | -0.33 | 0.56 | -0.9 | -0.67 | -0.9 | -0.62 | 0.125 | -0.08 | -0.4975 | 0.786788 | 0.856639 | 0.128294 | | 1 | 0.996511 | 0.723655 |
| Tmem126a | Q9D8Y1 | -0.26 | 0.5 | 0.15 | 0.41 | 0.62 | 0.38 | 0.08 | 0.28 | 0.8 | 0.55 | 0.49 | -0.08 | 0.71 | 0.26 | -0.24 | -0.08 | 0.14 | 0.24 | -0.0375 | 0.538891 | 0.382411 | 0.896341 | | 1 | 0.996511 | 0.983509 |
| Tmem126b | Q9D1R1 | -0.95 | -0.49 | -1.37 | -1.84 | 0.07 | -0.82 | -1.07 | -2.35 | -1.27 | -2.37 | -1.74 | -1.03 | -1.37 | -0.13 | -2.19 | -1.22 | 0.12 | -0.44 | -0.065 | 0.821036 | 0.283452 | 0.892246 | | 1 | 0.996511 | 0.982473 |
| Tmem127 | G3UXI2 | -0.75 | -0.51 | -1.27 | -0.76 | 0.34 | -0.44 | -1.49 | -1.01 | 0.11 | 1.03 | -0.81 | -0.16 | 1.19 | 1.67 | -1.07 | -1.36 | 0.1725 | 0.865 | 0.93 | 0.668282 | 0.05382 | 0.215121 | | 1 | 0.996511 | 0.787313 |
| Tmem130 | Q6NXM3 | -3.17 | 1.07 | -1.35 | -1.02 | -2.58 | -1.11 | -0.48 | -1.19 | -1.81 | -1.28 | -1.1 | -0.25 | -1.16 | -2.36 | -1.72 | -1.2 | -0.2225 | 0.0075 | -0.4925 | 0.8 | 0.992844 | 0.553165 | | 1 | 0.999285 | 0.916262 |
| Tmem131 | O70472 | -0.39 | -0.28 | -0.69 | -0.81 | 0.06 | -0.47 | -0.07 | -1.63 | 0.67 | 0.09 | -1.61 | 0.23 | 1.47 | 1.79 | 0.03 | -1.13 | 0.015 | 0.3875 | 1.0825 | 0.968738 | 0.429782 | 0.111911 | | 1 | 0.996511 | 0.713445 |
| Tmem132a | Q922P8 | -0.75 | -0.19 | -0.03 | 0.18 | 0.27 | -0.81 | 0.6 | 0.14 | -0.05 | 0.71 | 0.45 | -0.2 | -0.27 | 0.83 | 0.06 | -0.22 | 0.2475 | 0.425 | 0.2975 | 0.484165 | 0.178333 | 0.374543 | | 1 | 0.996511 | 0.859424 |
| Tmem132b | F7BAB2 | 1.27 | 0.78 | 0.97 | 0.64 | 0.81 | 1.25 | 0.85 | 0.52 | 0.51 | 0.14 | 0.75 | 1.04 | 0.6 | 0.39 | 0.19 | 0.65 | -0.0575 | -0.305 | -0.4575 | 0.797974 | 0.25194 | 0.065119 | | 1 | 0.996511 | 0.66433 |
| Tmem132c | Q8CEF9 | 0.35 | 0.77 | 0.68 | 0.56 | 0.64 | 0.84 | 0.16 | 0.34 | -0.09 | 1.78 | 0.7 | 0.71 | 0.05 | 0.22 | -0.56 | 0.06 | -0.095 | 0.185 | -0.6475 | 0.647417 | 0.628864 | 0.02133 | | 1 | 0.996511 | 0.621926 |
| Tmem132c | A0A0R4J0I | 0.43 | 0.85 | 0.53 | 0.87 | 0.4 | 0.54 | -0.05 | 0.35 | 0.59 | 0.58 | 0.37 | 0.55 | 0.56 | 0.67 | 0.05 | 0.15 | -0.36 | -0.1475 | -0.3125 | 0.101226 | 0.433255 | 0.200701 | | 1 | 0.996511 | 0.78037 |
| Tmem132e | Q6IEE6 | 0.89 | 0.76 | 0.75 | 1.45 | 0.19 | 0.83 | 1.04 | 1.33 | 1.28 | 1.8 | 0.86 | 0.99 | 0.53 | 0.92 | 1.23 | 1.09 | -0.115 | 0.27 | -0.02 | 0.695575 | 0.346331 | 0.937542 | | 1 | 0.996511 | 0.989767 |
| Tmem135 | Q9CYV5 | 0.14 | 0.88 | 0.34 | -0.34 | 1.25 | 0.14 | 1.05 | 0.07 | -0.45 | -0.01 | 0.52 | 0.71 | -0.08 | -0.33 | 0.41 | 0.88 | 0.3725 | -0.0625 | -0.035 | 0.336324 | 0.860574 | 0.923066 | | 1 | 0.996511 | 0.987595 |
| Tmem141 | A2AJB2 | 1.04 | 0.44 | 1.22 | 1.72 | -0.17 | 1.72 | 0.94 | 1.86 | 1.47 | 0.66 | 1.61 | 0.51 | 3.13 | -0.76 | 2.19 | 1.44 | -0.0175 | -0.0425 | 0.395 | 0.971723 | 0.90895 | 0.617135 | | 1 | 0.996744 | 0.937141 |
| Tmem143 | A0A1B0GT | 0.82 | 1.42 | 0.1 | 0.21 | 0.65 | 0.93 | 0.24 | -0.3 | 1.94 | -0.21 | 0.07 | 0.55 | 2.1 | 0.44 | -0.31 | 0.19 | -0.2575 | -0.05 | -0.0325 | 0.508655 | 0.924007 | 0.953474 | | 1 | 0.996755 | 0.992476 |
| Tmem144 | Q8VEH0 | -1 | -0.11 | -1.01 | -0.45 | 0.51 | -0.72 | -0.08 | -0.78 | -0.86 | -1.12 | 0.16 | -1.6 | 0.38 | -0.38 | -0.62 | -0.46 | 0.375 | -0.2125 | 0.3725 | 0.312361 | 0.607995 | 0.262669 | | 1 | 0.996511 | 0.81524 |
| Tmem145 | E9PUE6 | -0.38 | -0.83 | -0.77 | -0.91 | -1.03 | -0.26 | -1.02 | -0.85 | -0.69 | -1.19 | -1.22 | -0.38 | 0.6 | -0.72 | -1.1 | -0.33 | -0.0675 | -0.1475 | 0.335 | 0.773442 | 0.568211 | 0.382272 | | 1 | 0.996511 | 0.860984 |
| Tmem147 | A0A140LIQ | -2.08 | -1.38 | -1.02 | -1.23 | -1.5 | -1.3 | -0.54 | -1.58 | 0.64 | -1.36 | -1.48 | -0.43 | 0.16 | -0.88 | -1.24 | -0.16 | 0.1975 | 0.77 | 0.8975 | 0.544679 | 0.153325 | 0.041592 | | 1 | 0.996511 | 0.658454 |
| Tmem14c | Q9CQN6 | -0.11 | 1.18 | 0.37 | -0.77 | 0.25 | 2.49 | -0.21 | -0.24 | 1.53 | -0.34 | -0.19 | 0.14 | 1.04 | 0.15 | -0.78 | 0.62 | 0.405 | 0.1175 | 0.09 | 0.564242 | 0.829853 | 0.86399 | | 1 | 0.996511 | 0.977235 |
| Tmem151a | Q6GQT5 | 0.18 | 0.69 | 0.15 | 0.28 | 0.34 | 0.19 | 0.7 | 0.36 | 0.03 | 0.04 | 0.19 | 0.09 | -0.15 | -0.02 | 0.14 | 0.54 | 0.0725 | -0.2375 | -0.1975 | 0.715971 | 0.228886 | 0.412307 | | 1 | 0.996511 | 0.874811 |
| Tmem151b | Q68FE7 | 0.02 | 0.64 | -0.43 | -0.07 | 0.65 | 0.26 | -0.4 | -0.48 | 0.54 | -0.25 | -0.23 | 0.48 | 1.33 | 0.95 | 0.19 | -0.14 | -0.0325 | 0.095 | 0.5425 | 0.923534 | 0.761832 | 0.189781 | | 1 | 0.996511 | 0.776843 |
| Tmem160 | Q9D938 | -1.32 | -1.2 | -0.49 | -0.47 | -0.79 | -0.5 | -1.11 | -0.38 | 0.22 | -0.15 | -0.29 | -1.62 | 2.11 | -1.68 | 0.91 | -0.73 | 0.175 | 0.41 | 1.0225 | 0.538827 | 0.358976 | 0.216368 | | 1 | 0.996511 | 0.787447 |
| Tmem161b | Q8C2L6 | -3.02 | -3.02 | 0.24 | 0.19 | -2.88 | -3.57 | 1.09 | 0.2 | -2.41 | 0.23 | -0.15 | -1.97 | -1.86 | -2.25 | 0.59 | 0.8 | 0.1125 | 0.3275 | 0.7225 | 0.931896 | 0.749664 | 0.515383 | | 1 | 0.996511 | 0.905674 |
| Tmem163 | Q8C996 | 0.91 | 0.93 | 0.84 | 1.07 | 0.51 | 1.17 | 0.26 | 0.95 | 1.15 | 1.71 | 0.97 | 1.12 | 1.45 | 0.74 | 1.23 | 0.44 | -0.215 | 0.3 | 0.0275 | 0.364265 | 0.18055 | 0.91646 | | 1 | 0.996511 | 0.986556 |
| Tmem164 | Q6PHN7 | -1.3 | -0.89 | -1.47 | -0.92 | -0.44 | 0.05 | -1.02 | -0.93 | 1.29 | 0.13 | -0.78 | 0.91 | 3.13 | 0 | -0.08 | -0.26 | 0.56 | 1.5325 | 1.8425 | 0.077969 | 0.007841 | 0.033571 | | 1 | 0.996511 | 0.644622 |
| Tmem165 | P52875 | -0.81 | -0.51 | -0.49 | -0.23 | -0.2 | -0.97 | -0.31 | -0.96 | -0.73 | -0.76 | -0.71 | -0.91 | -0.82 | 0.15 | -0.72 | -0.28 | -0.1 | -0.2675 | 0.0925 | 0.69032 | 0.176941 | 0.737058 | | 1 | 0.996511 | 0.958635 |
| Tmem167 | E0CY07 | -0.57 | -1.06 | -2.69 | -2.48 | 0.22 | -0.14 | -0.88 | -1.95 | -0.57 | -6.34 | -1.91 | -0.61 | -1.26 | 0.3 | -3.35 | -2.45 | 1.0125 | -0.6575 | 0.01 | 0.142891 | 0.617095 | 0.990614 | | 1 | 0.996511 | 0.996154 |
| Tmem168 | A0A0N4SW | -0.06 | 0.55 | -1.16 | -0.65 | 0.05 | 0.16 | -1.01 | -0.98 | 0.57 | 1.7 | -0.8 | -0.05 | -0.62 | -1.24 | -0.11 | -0.68 | -0.115 | 0.685 | -0.3325 | 0.800101 | 0.265909 | 0.433848 | | 1 | 0.996511 | 0.879859 |
| Tmem169 | Q8BG50 | 1.04 | 0.81 | 0.8 | 0.37 | 0.24 | 0.89 | 0.17 | 0.13 | 1.49 | 0.92 | 0.23 | 1.26 | 0.86 | 0.8 | -0.13 | -0.09 | -0.3975 | 0.22 | -0.395 | 0.127706 | 0.48558 | 0.229401 | | 1 | 0.996511 | 0.787447 |
| Tmem175 | Q9CXY1 | -1.11 | -0.48 | -0.13 | -0.55 | -0.06 | -0.62 | -1.42 | -0.59 | -0.3 | -1.17 | 0.18 | -1.12 | -0.48 | 0.41 | 0.95 | -0.74 | -0.105 | -0.035 | 0.6025 | 0.75466 | 0.925244 | 0.177483 | | 1 | 0.996755 | 0.772068 |
| Tmem177 | Q8BPE4 | -0.51 | 0.29 | -0.08 | 0.52 | 0.32 | -0.09 | 0.31 | 0.03 | 0.3 | 1.54 | 0.01 | -0.11 | 0.72 | 0.17 | 0.38 | -0.14 | 0.0875 | 0.38 | 0.2275 | 0.734408 | 0.3752 | 0.457851 | | 1 | 0.996511 | 0.887902 |
| Tmem178a | Q9CZ16 | -0.07 | 0.3 | 1.1 | 0.76 | 0.13 | 0.33 | 0.45 | 0.48 | 0.93 | -0.94 | 0.88 | 0.25 | 0.99 | -0.07 | 0.08 | 0.44 | -0.175 | -0.2425 | -0.1625 | 0.527161 | 0.610426 | 0.64145 | | 1 | 0.996511 | 0.943421 |
| Tmem178b | M0QWJ9 | 0.07 | 3.03 | -0.1 | 0.19 | 3.98 | 2.93 | 0.37 | 0.12 | 2.54 | -0.6 | 0.46 | -0.09 | 1.02 | 3.74 | -0.06 | -0.25 | 1.0525 | -0.22 | 0.315 | 0.345291 | 0.810079 | 0.766268 | | 1 | 0.996511 | 0.964134 |
| Tmem181a | Q3U3W2 | 0.95 | 1.27 | -1.73 | -1.65 | 1.35 | 0.78 | -1.18 | -1.87 | 0.11 | -1.41 | -1.27 | 1.12 | 0.42 | 1.02 | -1.72 | -1.2 | 0.06 | -0.0725 | -0.08 | 0.952257 | 0.936398 | 0.931501 | | 1 | 0.996755 | 0.989713 |
| Tmem184b | A0A2R8VI0 | 1.4 | 2.26 | 0.73 | 0.99 | 2.95 | 1.11 | 1.06 | 1.73 | 1.34 | 0.33 | 0.4 | 0.9 | 0.8 | 1.59 | 1.2 | 1.12 | 0.3675 | -0.6025 | -0.1675 | 0.477221 | 0.151461 | 0.648644 | | 1 | 0.996511 | 0.946113 |
| Tmem185a | Q3TYW3 | 1.27 | 1.16 | 0.54 | 0.8 | 0.73 | 1.2 | 0.63 | 1.07 | 1.11 | 3.12 | 0.37 | 1.13 | 1.92 | 2.47 | 1.41 | 0.67 | -0.035 | 0.49 | 0.675 | 0.880835 | 0.395878 | 0.120797 | | 1 | 0.996511 | 0.721448 |
| Tmem186 | Q9CR76 | -0.78 | 0.34 | -0.56 | -0.61 | -0.02 | -0.74 | -0.22 | -0.77 | -0.2 | -1.55 | -0.35 | -0.74 | -0.44 | -0.29 | -1.41 | -1.12 | -0.035 | -0.3075 | -0.4125 | 0.909923 | 0.426827 | 0.272974 | | 1 | 0.996511 | 0.822084 |
| Tmem19 | A0A1W2P7 | 0.59 | 0.62 | 0.11 | 0.35 | 0.66 | 0.64 | 0.84 | 0.25 | 0.56 | 1.11 | 0.69 | 0.6 | 1.08 | 1.03 | 1.02 | 0.36 | 0.18 | 0.3225 | 0.455 | 0.387072 | 0.157857 | 0.090407 | | 1 | 0.996511 | 0.692714 |
| Tmem198 | Q8BG75 | 0.78 | 0.96 | 0.15 | 0.09 | 0.94 | 0.53 | 0.54 | 0.37 | 1.08 | -0.23 | 0.14 | 1.29 | 0.66 | 1.79 | 0.18 | 0.55 | 1.00E-01 | 0.075 | 0.3 | 0.701618 | 0.853716 | 0.457227 | | 1 | 0.996511 | 0.887902 |
| Tmem198b | Q8CFU0 | -0.04 | 0.32 | -0.4 | -0.16 | 0.42 | 0.37 | -1.94 | -1.53 | 0.9 | -4.53 | -2.05 | 0.47 | 0.66 | 1.18 | -0.95 | -0.31 | -0.6 | -1.2325 | 0.215 | 0.318798 | 0.294289 | 0.649606 | | 1 | 0.996511 | 0.946676 |
| Tmem199 | Q5SYH2 | 1.13 | 0.85 | -1.33 | -2 | 1.24 | 1.14 | -0.08 | -0.32 | 0.55 | 0.1 | -1.38 | 0.67 | 1.65 | 1.77 | 0.02 | -0.51 | 0.8325 | 0.3225 | 1.07 | 0.309751 | 0.696481 | 0.239678 | | 1 | 0.996511 | 0.794626 |
| Tmem200a | Q8C817 | -0.28 | 0.27 | 1.4 | 0.83 | 0.22 | -0.05 | 0.43 | 1.33 | 0.77 | -2.56 | 1.41 | 0.01 | 0.35 | 0.48 | 1.3 | 1.03 | -0.0725 | -0.6475 | 0.235 | 0.868483 | 0.454379 | 0.569028 | | 1 | 0.996511 | 0.922588 |
| Tmem200c | J3QK46 | 0.34 | 1.43 | 0.85 | 1.55 | 1.35 | 0.38 | 1.53 | 1.57 | 1.93 | 1.46 | 1.37 | 1 | 1.28 | 1.18 | 1.22 | 1.25 | 0.165 | 0.3975 | 0.19 | 0.662569 | 0.255308 | 0.524776 | | 1 | 0.996511 | 0.908107 |
| Tmem201 | A2A8U2-2 | -0.12 | 0.09 | -0.84 | -0.85 | 0.05 | -1.41 | -0.43 | -0.02 | -0.57 | -2.34 | -0.99 | -0.93 | -0.57 | -0.33 | 0.13 | 0.06 | -0.0225 | -0.7775 | 0.2525 | 0.95423 | 0.100715 | 0.417814 | | 1 | 0.996511 | 0.87558 |
| Tmem205 | A0A1L1SSA | 0.08 | 0.82 | 1.55 | 1.09 | 0.57 | 0.15 | 2.08 | 1.87 | -0.03 | 1.69 | 1.53 | 0.21 | 0.28 | 0.66 | 2.28 | 2.11 | 0.2825 | -0.035 | 0.4475 | 0.59096 | 0.944286 | 0.421961 | | 1 | 0.996755 | 0.875736 |
| Tmem209 | Q8BRG8-2 | 0.9 | 0.77 | 0.84 | 0.59 | 0.43 | 0.41 | 1.08 | 1.3 | 0.51 | 1.02 | 0.69 | 0.52 | 0.19 | 1.02 | 1.5 | 1.26 | 0.03 | -0.09 | 0.2175 | 0.904036 | 0.641417 | 0.481253 | | 1 | 0.996511 | 0.893906 |
| Tmem214 | Q8BM55 | -0.16 | -0.21 | -0.77 | -0.67 | 0.54 | -0.19 | 0.24 | -0.3 | -0.76 | -0.99 | -0.64 | -0.15 | 0.44 | -0.85 | 1.3 | 0.9 | 0.525 | -0.1825 | 0.9 | 0.068949 | 0.483703 | 0.079076 | | 1 | 0.996511 | 0.681114 |
| Tmem222 | Q8BVA2 | -0.01 | 0.27 | 0.78 | 0.43 | 0.42 | -0.37 | 1.49 | 0.79 | -1.04 | -0.13 | 0.75 | -0.14 | -0.82 | 0.96 | 0.74 | 1.1 | 0.215 | -0.5075 | 0.1275 | 0.591475 | 0.210781 | 0.77646 | | 1 | 0.996511 | 0.967021 |
| Tmem223 | Q9CQE2 | -0.93 | 0.13 | -0.14 | -0.65 | 0.56 | -0.26 | 0.46 | -0.14 | -0.85 | -2.12 | -0.32 | -0.14 | -0.81 | -0.16 | -0.76 | 0.06 | 0.5525 | -0.46 | -0.02 | 0.104498 | 0.34576 | 0.951297 | | 1 | 0.996511 | 0.992476 |
| Tmem229a | B9EJI9 | -0.36 | 0.36 | 0.83 | 0.47 | 0.37 | 0.08 | 0.26 | 0.3 | 1.02 | 1.23 | 0.28 | 0.46 | 2.32 | 0.57 | 1.02 | 0.31 | -0.0725 | 0.4225 | 0.73 | 0.784233 | 0.226194 | 0.153669 | | 1 | 0.996511 | 0.750763 |
| Tmem230 | Q8CIB6 | 0.1 | -0.66 | 0.07 | -0.73 | 0.17 | -0.04 | 0.29 | 0.04 | 0.57 | 0.87 | 0.16 | 0.65 | 0.62 | 0.95 | 0.88 | 0.47 | 0.42 | 0.8675 | 1.035 | 0.119693 | 0.012729 | 0.004349 | | 1 | 0.996511 | 0.548272 |
| Tmem231 | Q3U284 | -0.53 | -1.2 | -0.51 | -0.64 | -0.87 | 0.35 | 0.58 | 0.29 | -1.12 | 0.49 | -0.55 | 0.7 | -2.04 | -0.57 | 1.41 | 0.37 | 0.8075 | 0.6 | 0.5125 | 0.04391 | 0.191904 | 0.459591 | | 1 | 0.996511 | 0.887902 |
| Tmem237 | Q3V0J1-3 | 1.13 | 1.81 | 1.43 | 1.21 | 1.72 | 1.45 | 0.51 | 1.05 | 3.55 | 2.16 | 1.21 | 2.02 | 3.38 | 4.3 | 1.46 | 1.1 | -0.2125 | 0.84 | 1.165 | 0.486989 | 0.105356 | 0.126981 | | 1 | 0.996511 | 0.722414 |
| Tmem240 | B2RWJ3 | -0.38 | -0.7 | 0.88 | 1.02 | -1.64 | -0.43 | 1.3 | 1.67 | 0.38 | 0.35 | 1.45 | -0.3 | 0.29 | -0.59 | 0.76 | 0.71 | 0.02 | 0.265 | 0.0875 | 0.980021 | 0.61618 | 0.861089 | | 1 | 0.996511 | 0.977204 |
| Tmem243 | A0A0G2JEI | -0.79 | 0.23 | 0.95 | -0.37 | -0.3 | -0.54 | 0.78 | 0.48 | -1.29 | 0.14 | 0.97 | -0.51 | -0.57 | -1.55 | 0.56 | 0.55 | 0.1 | -0.1775 | -0.2575 | 0.826655 | 0.75295 | 0.659427 | | 1 | 0.996511 | 0.949645 |
| Tmem245 | B1AZA5 | -0.26 | 0.22 | -0.19 | -0.41 | 0.6 | -0.29 | -0.54 | 0.29 | 0.19 | 0.11 | -0.12 | -0.08 | 0.79 | 0.77 | 0.83 | -0.13 | 0.175 | 0.185 | 0.725 | 0.554858 | 0.371754 | 0.030214 | | 1 | 0.996511 | 0.641481 |
| Tmem25 | A0A1N9M | -0.55 | 0.17 | 0.29 | 0.12 | -2.01 | 0.46 | -0.93 | -0.47 | 1.09 | -2.87 | 0.05 | -1.04 | 2.1 | -3.71 | -0.65 | 0.06 | -0.745 | -0.7 | -0.5575 | 0.165343 | 0.383521 | 0.610871 | | 1 | 0.996511 | 0.935047 |
| Tmem254 | V9GX22 | 0.53 | 0.44 | -2.63 | -2 | 0.92 | 0.69 | -2.35 | -1.59 | 0.05 | -3.95 | -1.69 | 0.32 | 0.92 | 0.85 | -1.87 | -1.73 | 0.3325 | -0.4025 | 0.4575 | 0.749074 | 0.726408 | 0.651882 | | 1 | 0.996511 | 0.947633 |
| Tmem256 | F6YWB5 | -0.59 | 0.13 | 0.04 | -0.52 | 0.13 | -0.52 | -1.1 | -1.12 | 0.55 | -0.39 | -0.44 | -0.33 | 0.43 | 0.42 | -0.09 | -0.51 | -0.4175 | 0.0825 | 0.2975 | 0.237621 | 0.786773 | 0.342061 | | 1 | 0.996511 | 0.848473 |
| Tmem256 | Q5F285 | -1.93 | -1.71 | -1.83 | -0.8 | -3.17 | -2.22 | -0.96 | -1.06 | -1.74 | -2.19 | -1.81 | -1.96 | -1.02 | -1.43 | -1.63 | -1.18 | -0.285 | -0.3575 | 0.2525 | 0.598425 | 0.235075 | 0.415966 | | 1 | 0.996511 | 0.874945 |
| Tmem258 | P61166 | 0.38 | 1.07 | 0.22 | 1.06 | 0.5 | 1.22 | 0.62 | 0.69 | 2.28 | 1.87 | 0.84 | 0.51 | 3.05 | 1.09 | 3.14 | 1.47 | 0.075 | 0.6925 | 1.505 | 0.787392 | 0.147388 | 0.018888 | | 1 | 0.996511 | 0.604026 |
| Tmem259 | Q8CIV2-2 | -0.78 | -0.91 | -0.83 | -1.43 | -0.09 | -0.23 | -1.55 | -1.4 | -1.18 | -1.57 | -1.87 | -0.81 | -0.27 | -1 | -0.33 | -0.38 | 0.17 | -0.37 | 0.4925 | 0.663188 | 0.217204 | 0.082215 | | 1 | 0.996511 | 0.68865 |
| Tmem260 | A0A2I3BQ | 0.07 | -0.07 | -1.11 | -1.44 | 0.91 | 0.14 | 0.28 | -1.15 | -0.44 | -2.58 | -0.91 | -0.53 | -0.01 | 1.24 | 0.85 | -0.88 | 0.6825 | -0.4775 | 0.9375 | 0.2168 | 0.415101 | 0.119192 | | 1 | 0.996511 | 0.721448 |
| Tmem263 | Q9DAM7 | 0.6 | 0.15 | 0.1 | -0.01 | 0.92 | 0.49 | 0.89 | 0.96 | -0.22 | -0.11 | 2.95 | 0.83 | -0.06 | 0.29 | 0.18 | 1.22 | 0.605 | 0.6525 | 0.1975 | 0.015212 | 0.350393 | 0.539098 | | 1 | 0.996511 | 0.912212 |
| Tmem266 | Q8BZB3 | 1.47 | 1.37 | 0.83 | 1.18 | 0.84 | 1.25 | 1.16 | 1.14 | 1.57 | 0.81 | 1.07 | 1.31 | 1.06 | 2 | 1.08 | 1.27 | -0.115 | -0.0225 | 0.14 | 0.568684 | 0.925848 | 0.621765 | | 1 | 0.996755 | 0.938628 |
| Tmem29 | E9PZH0 | -4.67 | -5.12 | -3.84 | -7.55 | -6.08 | -4.03 | -5.41 | -5.02 | -3.63 | -3.31 | -5.7 | -4.57 | -1.23 | -5.36 | -5.14 | -5.4 | 0.16 | 0.9925 | 1.0125 | 0.844492 | 0.270651 | 0.390826 | | 1 | 0.996511 | 0.865375 |
| Tmem30a | Q8VEK0 | 1.11 | 1.32 | 1.11 | 1.48 | 1.35 | 0.72 | 1.15 | 1.39 | 0.77 | 1.25 | 1.23 | 1.26 | 0.73 | 0.4 | 1.63 | 1.33 | -0.1025 | -0.1275 | -0.2325 | 0.623134 | 0.52738 | 0.453378 | | 1 | 0.996511 | 0.885993 |
| Tmem33 | A0A0R4J1Z | -1.28 | -0.89 | -0.33 | -0.3 | 0.04 | -1.39 | 0.4 | -0.68 | -1.32 | -0.53 | -0.1 | -1.22 | -0.66 | -0.31 | 0.06 | -0.22 | 0.2925 | -0.0925 | 0.4175 | 0.503976 | 0.7994 | 0.180074 | | 1 | 0.996511 | 0.775542 |
| Tmem35a | Q9D328 | 0.02 | -0.31 | -0.67 | -0.26 | -0.27 | 0.29 | 0 | -0.21 | 0.15 | -0.01 | -0.86 | -0.04 | 0.88 | 0.22 | 0.84 | 0.84 | 0.2575 | 0.115 | 1 | 0.251744 | 0.682521 | 0.002925 | | 1 | 0.996511 | 0.45967 |
| Tmem38a | Q3TMP8 | -4.36 | -3.73 | -1.74 | -1.94 | -4.11 | -3.16 | -1.91 | -1.74 | -3 | -2.66 | -1.95 | -4.1 | -0.32 | -5.08 | -1.46 | -0.9 | 0.2125 | 0.015 | 1.0025 | 0.783937 | 0.983297 | 0.381538 | | 1 | 0.998773 | 0.860984 |
| Tmem38b | Q9DAV9 | -1.13 | 3.25 | 0.45 | 0.42 | -1.42 | 0.51 | 0.62 | 0.34 | 2.47 | 0.19 | 0.23 | -0.08 | 4.7 | -1.74 | 0.52 | 1.04 | -0.735 | -0.045 | 0.3825 | 0.436225 | 0.963263 | 0.790179 | | 1 | 0.996755 | 0.969105 |

| | | | | | | | | | | | | | | | | | | | | | | | | | | |
|---|---|---|---|---|---|---|---|---|---|---|---|---|---|---|---|---|---|---|---|---|---|---|---|---|---|---|
| Tmem41a | Q9D8U2 | -0.7 | -0.74 | -0.7 | -0.56 | -0.6 | -0.28 | 0.51 | -0.33 | -0.51 | -1.72 | 0.27 | -0.06 | -0.52 | -0.13 | -0.66 | 0.82 | 0.5 | 0.17 | 0.5525 | 0.07525 | 0.683754 | 0.12823 | 1 | 0.996511 | 0.723655 |
| Tmem43 | Q9DBS1 | 0.42 | 0.68 | 0.36 | 0.44 | 0.58 | 0.27 | 1.04 | 0.31 | -0.42 | -0.01 | 0.36 | 0.57 | -0.26 | 0.28 | 0.53 | 0.73 | 0.075 | -0.35 | -0.155 | 0.729151 | 0.18799 | 0.549277 | 1 | 0.996511 | 0.912858 |
| Tmem47 | Q9JJG6 | 0.28 | 0.77 | 0.64 | 0.44 | 1.38 | 0.68 | 0.19 | 0.21 | -1.15 | -1.56 | 0.55 | 0.67 | -0.22 | -2.97 | 0.96 | 0.37 | 0.0825 | -0.905 | -0.9975 | 0.781861 | 0.121084 | 0.227771 | 1 | 0.996511 | 0.787447 |
| Tmem50b | Q9D1X9 | -2.46 | -3.18 | -2.77 | -2.08 | -1.6 | -1.72 | -3.22 | -2.92 | -2.25 | -0.6 | -2.44 | -2.66 | -2.12 | -2 | -2.29 | -2.43 | 0.2575 | 0.635 | 0.4125 | 0.563842 | 0.216479 | 0.1584 | 1 | 0.996511 | 0.753473 |
| Tmem51 | Q99LG1 | 0.17 | 1.12 | 1.15 | 0.55 | 0.29 | 0.65 | -0.99 | -0.08 | 0.58 | 0.01 | 0.07 | 0.52 | 0.91 | -0.85 | 0.63 | 0.41 | -0.78 | -0.4525 | -0.4725 | 0.078637 | 0.143756 | 0.291782 | 1 | 0.996511 | 0.825497 |
| Tmem59l | Q7TNI2 | -1.49 | -1.9 | -0.77 | -0.71 | -1.6 | -1.51 | 0.61 | -0.08 | -1.44 | -2.25 | -0.14 | -1.37 | -1.41 | -1.21 | -0.19 | -0.43 | 0.5725 | -0.0825 | 0.4075 | 0.324642 | 0.865397 | 0.321329 | 1 | 0.996511 | 0.840101 |
| Tmem62 | Q8BXJ9 | -0.94 | -0.64 | -0.16 | -0.39 | -0.62 | -0.48 | -0.01 | -0.32 | -0.55 | 0.55 | -0.39 | -0.17 | -0.9 | -0.02 | -0.58 | -0.42 | 0.175 | 0.3925 | 0.0525 | 0.457294 | 0.214176 | 0.846886 | 1 | 0.996511 | 0.976225 |
| Tmem63b | Q3TWI9 | -0.21 | 0.04 | 0.02 | -0.31 | 0.48 | 0.13 | 0.43 | -0.46 | -0.23 | -0.12 | 0.06 | -0.08 | -0.12 | 0.11 | 0.26 | 0.21 | 0.26 | 0.0225 | 0.23 | 0.306553 | 0.898129 | 0.254584 | 1 | 0.996511 | 0.809524 |
| Tmem63b | D3Z2D8 | 1.53 | 0.85 | 0.19 | 0.45 | 1.89 | 2.99 | 0.41 | 1.05 | 1.09 | 0.43 | 2.27 | 1.87 | 1.66 | 0.61 | -0.25 | 0.32 | 0.83 | 0.66 | -0.17 | 0.174333 | 0.185266 | 0.716109 | 1 | 0.996511 | 0.958106 |
| Tmem63c | Q8CBX0 | 0.2 | 0.32 | 0.46 | 0.43 | 0.19 | 0.67 | 0.3 | 0.23 | 0.33 | 0.71 | 0.49 | 0.33 | 0.11 | 0.24 | 0.45 | 0.57 | -0.005 | 0.1125 | -0.01 | 0.977078 | 0.530002 | 0.958638 | 1 | 0.996511 | 0.992636 |
| Tmem65 | Q4VAE3 | 0.7 | 0.84 | 0.86 | 0.96 | 0.78 | 0.65 | 0.68 | 0.76 | 0.4 | 0.91 | 0.92 | 0.46 | 0.46 | 1.31 | 0.69 | 0.81 | -0.1225 | -0.1675 | -0.0225 | 0.409574 | 0.411963 | 0.922415 | 1 | 0.996511 | 0.987467 |
| Tmem67 | A2AJP5 | -4.5 | -3.06 | -1.59 | -0.39 | -3.4 | -4.1 | -1.75 | -1.51 | -2.78 | -0.28 | -2.1 | -3.93 | -1.77 | -2.26 | -1.03 | -1.11 | -0.305 | 0.1125 | 0.8425 | 0.756128 | 0.91492 | 0.332092 | 1 | 0.996755 | 0.845601 |
| Tmem68 | Q9D850 | 0.52 | 0.51 | 0.1 | 0.2 | 0.8 | 0.29 | -0.25 | 0.53 | 0.4 | 1.16 | 0.22 | 0.18 | 0.19 | 1.32 | 0.24 | -0.48 | 0.01 | 0.1575 | -0.015 | 0.968978 | 0.561357 | 0.968299 | 1 | 0.996511 | 0.992733 |
| Tmem69 | Q3KQJ0 | -1.55 | -0.86 | -1.7 | -1.44 | -1.06 | -1.77 | -1.06 | -0.43 | -1.81 | 0.57 | -0.48 | -2.07 | -2.52 | -1.71 | -0.27 | -0.89 | 0.3075 | 0.44 | 0.04 | 0.35273 | 0.46201 | 0.934728 | 1 | 0.996511 | 0.989767 |
| Tmem74b | A2AU34 | 0.44 | 0.76 | 0.24 | 1.81 | 0.49 | 0.86 | 0.33 | 1.22 | 1.56 | 1.38 | 0.78 | 0.93 | 1.48 | 1.97 | 1.61 | 0.68 | -0.0875 | 0.35 | 0.6225 | 0.81866 | 0.369798 | 0.166133 | 1 | 0.996511 | 0.759345 |
| Tmem87a | A2AQJ5 | -1.95 | -2.42 | -2.13 | -2.53 | -1.12 | -1.71 | -2.1 | -2.46 | -1.45 | -2.58 | -2.41 | -2.34 | -1.53 | -1.32 | -2.19 | -2.28 | 0.41 | 0.0625 | 0.4275 | 0.210156 | 0.831615 | 0.165212 | 1 | 0.996511 | 0.758837 |
| Tmem87b | Z4YML9 | -2.26 | -3.58 | -5.34 | -3.89 | -2.62 | -1.8 | -2.76 | -4.37 | -3.82 | -3.55 | -3.86 | -2.51 | -3.52 | -3.96 | -4.53 | -3.91 | 0.88 | 0.3325 | -0.2125 | 0.259829 | 0.608906 | 0.727923 | 1 | 0.996511 | 0.958635 |
| Tmem88b | Q3TYP4 | -1.32 | -1.46 | -1.4 | -1.86 | -0.9 | -0.97 | -1.09 | -2.01 | -1.02 | -0.85 | -1.3 | -0.97 | -0.11 | -0.97 | -1.47 | -1.58 | 0.2675 | 0.475 | 0.4775 | 0.363012 | 0.040549 | 0.19892 | 1 | 0.996511 | 0.78004 |
| Tmem9 | Q9CR23 | 0.32 | 0.5 | 0.35 | 0.26 | 0.64 | 0.73 | -0.2 | 0.61 | 0.82 | 0.99 | 0.44 | 0.94 | 1.27 | 1.53 | 0.39 | 0.38 | 0.0875 | 0.44 | 0.535 | 0.714701 | 0.043974 | 0.113038 | 1 | 0.996511 | 0.714953 |
| Tmem94 | A0A668KL5 | 0.68 | 1.08 | 0.92 | 1.08 | 1.01 | 0.62 | 0.89 | 0.93 | 0.07 | 0.89 | 0.84 | 0.66 | 0 | 0.36 | 0.85 | 0.86 | -0.0775 | -0.325 | -0.4225 | 0.660216 | 0.196894 | 0.128926 | 1 | 0.996511 | 0.723655 |
| Tmem97 | Q8VD00 | -3.16 | -2.17 | -0.7 | -0.15 | -1.74 | -2.34 | 0.59 | -0.48 | -1.74 | -0.82 | -0.56 | -3.25 | -1.64 | -1.36 | -0.57 | -0.31 | 0.5525 | -0.0475 | 0.575 | 0.522775 | 0.954213 | 0.411855 | 1 | 0.996755 | 0.874811 |
| Tmem9b | Q9JJR8 | 0.24 | -0.53 | -0.06 | 0.28 | -0.31 | 0.1 | 0.67 | 0.59 | -0.28 | -0.07 | 0.29 | 0.22 | 0.31 | -0.17 | 0.1 | 0.04 | 0.28 | 0.0575 | 0.0875 | 0.354669 | 0.819291 | 0.723649 | 1 | 0.996511 | 0.958398 |
| Tmf1 | B9EKI3 | 0.1 | 0.06 | -0.01 | -0.17 | 0.23 | -0.04 | -0.12 | -0.01 | -0.04 | 0.11 | -0.3 | 0.01 | -0.11 | 0.27 | -0.02 | 0.03 | 0.02 | -0.05 | 0.0475 | 0.900026 | 0.776892 | 0.797338 | 1 | 0.996511 | 0.969727 |
| Tmlhe | Q91ZE0 | 1.06 | 1.3 | 0.99 | 0.52 | -0.65 | 1.54 | 1.42 | 1.1 | 0.26 | 0.02 | 1.98 | 1.39 | 0.23 | 0.24 | 0.27 | 1.42 | -0.115 | -0.055 | -0.4275 | 0.81601 | 0.905128 | 0.225991 | 1 | 0.996744 | 0.787447 |
| Tmod1 | P49813 | 1.05 | 0.64 | 0.92 | 1.64 | 0.64 | 0.6 | 0.75 | 1.44 | 0.99 | 1.14 | 0.72 | 0.61 | 0.53 | 1.55 | 0.53 | 0.43 | -0.205 | -0.1975 | -0.3025 | 0.48434 | 0.458208 | 0.383102 | 1 | 0.996511 | 0.860984 |
| Tmod2 | Q9JKK7 | 1.5 | 1 | 1.47 | 1.37 | 1.22 | 1.29 | 1.97 | 1.86 | 0.65 | 0.88 | 1.55 | 1.52 | 0.15 | 0.66 | 1.23 | 1.66 | 0.25 | -0.185 | -0.41 | 0.312792 | 0.499534 | 0.257143 | 1 | 0.996511 | 0.811777 |
| Tmod3 | Q9JHJ0 | 0.25 | 0.1 | 0.59 | 0.45 | 0.22 | -0.29 | 0.48 | 0.28 | -0.13 | 0.24 | 0.48 | 0.11 | 0.04 | 0.03 | 0.6 | 0.69 | -0.175 | -0.1725 | -0.0075 | 0.434708 | 0.420693 | 0.975465 | 1 | 0.996511 | 0.993339 |
| Tmpo | Q61033 | 1.37 | 1.47 | 0.27 | 0.21 | 1.26 | 1.29 | 0.37 | 0.63 | 2.16 | 0.42 | 1.17 | 1.07 | 1.71 | 1.37 | 0.84 | 1.33 | 0.0575 | 0.375 | 0.4825 | 0.882611 | 0.428372 | 0.221451 | 1 | 0.996511 | 0.787447 |
| Tmpo | Q61029 | 1.22 | 0.69 | 0.73 | 0.84 | 0.33 | 0.7 | 0.71 | 0.91 | 1.07 | 0.39 | 0.69 | 0.52 | 1.85 | 0.19 | 1.14 | 0.73 | -0.2075 | -0.2025 | 0.1075 | 0.320852 | 0.380154 | 0.768187 | 1 | 0.996511 | 0.965206 |
| Tmppe | D3Z286 | 0.48 | 0.35 | 0.2 | 0.23 | 0.42 | 0.18 | -0.59 | 0.33 | 0.39 | 1.49 | -0.38 | -0.1 | 1 | 0.65 | 1.08 | 0.59 | -0.23 | 0.035 | 0.515 | 0.371198 | 0.929993 | 0.029886 | 1 | 0.996755 | 0.640893 |
| Tmsb10 | Q6ZWY8 | -0.2 | -0.08 | 0.66 | 1.48 | -0.83 | 0.58 | 0.34 | 0.78 | 1.01 | 0.92 | 0.6 | 0.54 | 1.41 | 1.56 | 0.72 | 0.75 | -0.2475 | 0.3025 | 0.645 | 0.61565 | 0.445024 | 0.154914 | 1 | 0.996511 | 0.750763 |
| Tmsb15l | Q8C0W0 | 0.28 | 0.54 | 0.07 | 0.23 | 0.17 | 0.27 | 0.31 | 0.21 | -0.7 | -0.12 | 0.14 | 0.91 | -0.3 | -0.7 | -0.15 | -0.32 | -0.04 | -0.2225 | -0.6475 | 0.805584 | 0.521196 | 0.012585 | 1 | 0.996511 | 0.573604 |
| Tmsb4x | P20065-2 | 1.44 | 1.16 | 0.58 | 1.46 | 0.19 | 0.69 | 0.64 | 1.46 | 1.49 | 1.55 | 0.6 | 1.33 | 0.97 | 1.97 | 0.63 | 0.07 | -0.415 | 0.0825 | -0.25 | 0.224041 | 0.786839 | 0.562483 | 1 | 0.996511 | 0.920024 |
| Tmtc1 | Q3UV71 | -0.56 | -0.66 | -1.92 | -1.24 | -0.54 | -0.95 | -1.09 | -0.94 | 7.69 | -0.93 | -2.71 | -1.36 | 5.91 | 8.92 | 0.21 | 0.19 | 0.215 | 1.7675 | 4.9025 | 0.515138 | 0.416058 | 0.030829 | 1 | 0.996511 | 0.641481 |
| Tmtc3 | G5E8C4 | -0.58 | -0.3 | -0.36 | -0.09 | 0.08 | -0.45 | 0.4 | -0.07 | 0.26 | -0.11 | -0.11 | -0.2 | 1.33 | 0.81 | 0.56 | 0.9 | 0.3225 | 0.2925 | 1.2325 | 0.176078 | 0.161251 | 0.000534 | 1 | 0.996511 | 0.286246 |
| Tmub1 | Q9JMG3-2 | 0.27 | 0.13 | 0.94 | 0.77 | 0.75 | 0.09 | 0.47 | 0.26 | 1.66 | 0.94 | -0.12 | 0.08 | 0.77 | 1.27 | 2.27 | 1.22 | -0.135 | 0.1125 | 0.855 | 0.595436 | 0.793675 | 0.041287 | 1 | 0.996511 | 0.658454 |
| Tmx1 | Q8VBT0 | 0.37 | 0.38 | 0.41 | 0.47 | 0.43 | 0.51 | 0.52 | 0.54 | 1.22 | 0.12 | 0.44 | 0.22 | 1.17 | 1.09 | 1.24 | 0.68 | 0.0925 | 0.0925 | 0.6375 | 0.507817 | 0.729614 | 0.009842 | 1 | 0.996511 | 0.563386 |
| Tmx2 | Q9D710 | -0.67 | -0.25 | 0.08 | 0.15 | -0.57 | -0.73 | 0.46 | 0.14 | -0.86 | -0.11 | -0.07 | 0.29 | -0.67 | 0.45 | 0.78 | 0.39 | -0.0025 | -0.015 | 0.41 | 0.993956 | 0.961243 | 0.273746 | 1 | 0.996755 | 0.822084 |
| Tmx3 | Q8BXZ1 | -0.47 | -0.16 | -0.34 | -0.32 | 0.36 | -0.48 | 0.33 | -0.38 | -0.56 | -0.39 | -0.37 | -0.41 | -0.61 | 0.09 | 0.09 | 0.15 | 0.28 | -0.11 | 0.2525 | 0.274616 | 0.508124 | 0.296505 | 1 | 0.996511 | 0.825612 |
| Tmx4 | Q8C0L0 | -0.54 | -0.53 | -0.84 | -1.12 | -0.69 | -0.69 | -0.66 | -0.7 | -0.65 | -1.21 | -0.91 | -0.51 | -0.67 | -0.27 | -0.05 | -0.78 | 0.0725 | -0.0625 | 0.315 | 0.694051 | 0.792558 | 0.234225 | 1 | 0.996511 | 0.790641 |
| Tnc | Q80YX1 | -0.96 | -1.34 | -1.57 | -1.47 | -0.54 | -0.89 | -0.56 | -0.89 | -1 | -0.83 | -0.98 | -0.34 | -0.5 | -0.84 | 0.25 | -0.61 | 0.615 | 0.5475 | 0.91 | 0.012559 | 0.041602 | 0.011115 | 1 | 0.996511 | 0.563386 |
| Tnfaip1 | O70479 | -1.77 | -2 | -0.65 | 0.14 | -1.16 | -0.77 | -1.06 | -0.17 | -2.5 | 0.75 | -0.49 | -0.82 | -1.62 | -0.86 | -0.41 | -0.53 | 0.28 | 0.305 | 0.215 | 0.582032 | 0.688264 | 0.68476 | 1 | 0.996511 | 0.955729 |
| Tnfaip3 | Q60769 | -0.26 | -0.52 | -1.16 | 0.17 | -0.78 | 0.55 | -2.2 | -1.65 | 1.01 | 2.01 | -0.12 | 0.09 | 1.23 | 0.18 | -0.42 | -0.52 | -0.5775 | 1.19 | 0.56 | 0.351075 | 0.045364 | 0.243997 | 1 | 0.996511 | 0.799567 |
| Tnfaip8 | D3Z325 | -1.33 | -0.77 | -0.84 | -0.26 | -0.94 | -1.52 | -0.08 | -0.7 | -1.27 | 0.41 | -0.47 | -1.31 | -0.81 | -0.65 | -0.21 | -0.02 | -0.01 | 0.14 | 0.3775 | 0.977488 | 0.748356 | 0.228701 | 1 | 0.996511 | 0.787447 |
| Tnfaip8l3 | Q3TBL6 | -0.46 | 0.23 | 0.19 | 0.24 | 0.21 | -0.45 | 0.22 | 0.12 | 0.74 | 0.71 | 0.25 | -0.22 | 0.4 | 0.87 | 0.48 | 0.69 | -0.025 | 0.32 | 0.56 | 0.919265 | 0.289771 | 0.041569 | 1 | 0.996511 | 0.658454 |
| Tnfrsf21 | Q9EPU5 | 0.23 | 0.31 | -0.24 | -0.29 | 0.32 | 0.38 | -0.44 | -0.43 | 0.44 | 0.85 | -0.38 | 0.45 | 0.47 | 0.27 | -0.47 | -0.25 | -0.045 | 0.3375 | 0.0025 | 0.871635 | 0.287025 | 0.993038 | 1 | 0.996511 | 0.996845 |
| Tnik | B2RQ80 | 1.2 | 0.07 | 1.77 | 0.22 | 0.03 | 1.26 | 2.54 | 0.79 | 0.64 | -0.29 | 2.12 | 1.1 | 0.49 | 0.05 | 2.03 | 2.32 | 0.34 | 0.0775 | 0.4075 | 0.577948 | 0.89565 | 0.523567 | 1 | 0.996511 | 0.90791 |
| Tnip1 | D3Z2W0 | 0.82 | 1.6 | -0.03 | -0.03 | 2.09 | 0.57 | -0.29 | -0.28 | 1.07 | 0.71 | 0.07 | 0.84 | 0.92 | 1.34 | -0.33 | -0.16 | -0.0675 | 0.0825 | -0.1475 | 0.91354 | 0.845489 | 0.778781 | 1 | 0.996511 | 0.967289 |
| Tnk2 | O54967-2 | 0.2 | 0.88 | 0.53 | 0.4 | 0.12 | 0.46 | 0.26 | 0.1 | 0.13 | 0.67 | 0.13 | 0.21 | 0.52 | -0.01 | 0.49 | 0.15 | -0.2675 | -0.2175 | -0.215 | 0.201867 | 0.350692 | 0.371785 | 1 | 0.996511 | 0.859424 |
| Tnks | Q6PFX9 | -0.03 | 0.48 | -0.13 | 0.12 | -0.21 | 0.07 | -0.35 | -0.01 | 0.18 | 0.94 | -0.14 | -0.13 | 0.39 | 1.03 | -0.24 | -0.12 | -0.235 | 0.1025 | 0.155 | 0.255088 | 0.728347 | 0.634325 | 1 | 0.996511 | 0.941132 |
| Tnks1bp1 | P58871 | 0.03 | -0.16 | -0.12 | -0.06 | -0.29 | -0.07 | -0.6 | -0.23 | 0.59 | 0.07 | -0.07 | -0.11 | 0.58 | 0.45 | -0.44 | -0.67 | -0.22 | 0.1975 | 0.0575 | 0.219467 | 0.358997 | 0.858295 | 1 | 0.996511 | 0.976225 |
| Tnpo1 | Q8BFY9-2 | -0.35 | -0.09 | -0.01 | 0.01 | 0.37 | -0.17 | 0.35 | 0.24 | -1.05 | -0.6 | 0.08 | -0.37 | -1.27 | -0.77 | 0.09 | 0.11 | 0.3075 | -0.375 | -0.35 | 0.131861 | 0.183659 | 0.328071 | 1 | 0.996511 | 0.844932 |
| Tnpo2 | Q99LG2 | -1.82 | -1.51 | -0.36 | -0.31 | -1.14 | -1.64 | -0.53 | -0.65 | -1.6 | -0.79 | -0.53 | -1.65 | -1.92 | -1.47 | -0.99 | -0.5 | 0.01 | -0.1425 | -0.22 | 0.9816 | 0.753458 | 0.638682 | 1 | 0.996511 | 0.942804 |
| Tnpo3 | Q6P2B1 | -0.26 | 0.12 | -0.7 | -0.92 | 0.52 | -0.22 | -0.42 | -0.76 | -0.91 | -0.76 | -0.66 | -0.35 | -0.92 | -0.13 | -0.63 | -0.57 | 0.22 | -0.23 | -0.1225 | 0.527016 | 0.411475 | 0.68167 | 1 | 0.996511 | 0.955729 |
| Tnr | Q8BYI9 | 0.06 | -0.4 | -0.05 | 0.25 | -0.05 | -0.37 | 0.35 | 0.33 | -0.98 | -0.43 | -0.07 | -0.31 | -1.03 | -0.49 | 0.09 | -0.13 | 0.1 | -0.4125 | -0.355 | 0.673677 | 0.135014 | 0.248421 | 1 | 0.996511 | 0.803985 |
| Tnrc18 | Q80WC3 | -0.25 | 0.73 | -0.13 | -0.93 | 0.24 | -0.26 | -1.63 | -1.69 | -0.95 | -0.14 | -0.52 | -0.1 | -1.15 | 0.24 | 0.2 | -0.53 | -0.69 | -0.2825 | -0.165 | 0.227266 | 0.464635 | 0.714291 | 1 | 0.996511 | 0.958106 |
| Tnrc6b | Q8BKI2 | -0.79 | -1.29 | -1.45 | -1.83 | -1.26 | -1.04 | -2.1 | -2.01 | 0.14 | -1.34 | -1.51 | -0.74 | 0.99 | -0.5 | -1.16 | -1.98 | -0.2625 | 0.4775 | 0.6775 | 0.437798 | 0.263231 | 0.282739 | 1 | 0.996511 | 0.822595 |
| Tnrc6c | B1ATC3 | -0.75 | -0.57 | -0.79 | -0.58 | -1.98 | -0.66 | -0.9 | -0.91 | 0.06 | -1.25 | -0.73 | -0.24 | 1.06 | -1.25 | -0.94 | -0.71 | -0.44 | 0.1325 | 0.2125 | 0.16602 | 0.659363 | 0.664683 | 1 | 0.996511 | 0.949825 |
| Tns1 | A0A087WC | -0.09 | -0.83 | 1.89 | 0.56 | -0.77 | -0.9 | 0.23 | -0.14 | -0.3 | 0.38 | 0.37 | -0.39 | -0.29 | -0.47 | 1.84 | 0.59 | -0.7775 | -0.3675 | 0.035 | 0.203833 | 0.520549 | 0.960581 | 1 | 0.996511 | 0.992693 |
| Tns2 | A0A2R8VH | -0.97 | -0.98 | -0.28 | 0.22 | -0.78 | -1.92 | -0.48 | -0.31 | -1.02 | -1.25 | -1.1 | -1.17 | -1.19 | -1.16 | -0.98 | -1.45 | -0.37 | -0.6325 | -0.6925 | 0.404523 | 0.061347 | 0.052089 | 1 | 0.996511 | 0.659923 |
| Tns3 | A0A5F8MP | -0.18 | 0.24 | -0.05 | -0.03 | 0.07 | -0.04 | -0.08 | -0.25 | 0.49 | -0.06 | -0.28 | 0.12 | 0.59 | 0.28 | -0.29 | 0.1 | -0.07 | 0.0725 | 0.175 | 0.675449 | 0.744157 | 0.47583 | 1 | 0.996511 | 0.892174 |
| Toe1 | Q9D2E2 | -0.39 | -0.66 | -0.99 | -1.66 | -0.16 | -0.78 | -0.81 | -1.38 | -1.78 | -0.85 | -1.26 | -1 | -1.72 | -1.45 | -1.69 | -0.06 | 0.1425 | -0.2975 | -0.305 | 0.689532 | 0.389204 | 0.506897 | 1 | 0.996511 | 0.902614 |
| Togaram1 | A0A1Y7VLS | -0.75 | -0.07 | 0.66 | 0.87 | -0.43 | -0.78 | -1.02 | 0.88 | -0.67 | 0.91 | 0.16 | 0.02 | 0.1 | -1.38 | 1.26 | 0.4 | -0.515 | -0.0725 | -0.0825 | 0.333439 | 0.874877 | 0.89169 | 1 | 0.996511 | 0.982213 |
| Tollip | Q9QZ06 | -0.4 | 0 | 0.4 | 0.55 | 0.12 | -0.48 | 0.44 | 0.51 | -0.19 | 0.33 | 0.45 | -0.28 | -0.44 | 0.27 | 0.16 | 0.37 | 1.00E-02 | -0.06 | -0.0475 | 0.973961 | 0.836189 | 0.872131 | 1 | 0.996511 | 0.979626 |
| Tom1 | O88746 | 0.13 | 0.37 | -0.14 | 0.1 | 0.35 | -0.05 | 0.07 | -0.06 | 0.46 | 0.31 | 0.1 | 0.45 | 0.25 | 0.66 | 0.12 | 0.12 | -0.0375 | 0.215 | 0.1725 | 0.838468 | 0.277102 | 0.436486 | 1 | 0.996511 | 0.882882 |
| Tom1l1 | Q8BZR6 | 1.29 | 1.3 | 1.05 | 0.86 | 1.56 | 1.34 | 1.72 | 1.14 | 1 | -0.23 | 1.47 | 1.24 | 0.99 | 0.91 | 1.14 | 1.21 | 0.315 | -0.255 | -0.0625 | 0.139649 | 0.507215 | 0.750772 | 1 | 0.996511 | 0.961479 |
| Tom1l2 | Q5SRX1 | 0.85 | 0.58 | 0.28 | 0.67 | 0.38 | 0.52 | 0.35 | 0.58 | 0.84 | 0.19 | 0.32 | 1 | 0.22 | 0.98 | -0.3 | -0.05 | -0.1375 | -0.0075 | -0.3825 | 0.447556 | 0.976248 | 0.238898 | 1 | 0.997588 | 0.793481 |
| Tom1l2 | Q5SRX1-2 | 1.82 | 1.75 | 3.42 | 2.67 | 0.62 | 2.45 | 3.4 | 2.85 | 1.88 | -0.85 | 3.43 | 3.16 | 2.67 | 0.92 | 2.9 | 3.53 | -0.085 | -0.51 | 0.09 | 0.896555 | 0.59409 | 0.88546 | 1 | 0.996511 | 0.979767 |
| Tomm20 | Q9DCC8 | -1.25 | -0.77 | -0.88 | -1.42 | -1.47 | -1.18 | -1.64 | -1.62 | -0.56 | -1.44 | -0.58 | -1.59 | 0.26 | -1.04 | -1.11 | -1.5 | -0.3975 | 0.0375 | 0.2325 | 0.089368 | 0.905388 | 0.56272 | 1 | 0.996744 | 0.920024 |
| Tomm22 | Q9CPQ3 | 0.32 | 0.37 | 0.4 | 0.68 | 0.37 | 0.14 | 0.39 | 0.56 | 0.57 | 1.16 | 0.81 | 0.24 | 0.22 | 0.65 | -0.24 | -0.24 | -0.0775 | 0.2525 | -0.345 | 0.651441 | 0.305675 | 0.204322 | 1 | 0.996511 | 0.783092 |
| Tomm34 | Q9CYG7 | -1.42 | -1.76 | -1.5 | -1.49 | -1.44 | -1.51 | -1.48 | -1.23 | -1.75 | -0.83 | -1.42 | -1.44 | -1.45 | -1.71 | -1.53 | -1.65 | 0.1275 | 0.1825 | -0.0425 | 0.434684 | 0.446704 | 0.815872 | 1 | 0.996511 | 0.972768 |
| Tomm34 | Q9CYG7-2 | -1.23 | -1.48 | -0.4 | -1.08 | -1.73 | -1.1 | -0.85 | -0.95 | -1.25 | -0.6 | -0.83 | -1.21 | -0.29 | -2.69 | -0.93 | -0.63 | -0.11 | 0.075 | -0.0875 | 0.715993 | 0.79499 | 0.87093 | 1 | 0.996511 | 0.979293 |
| Tomm40 | Q9QYA2 | 0.05 | 0.62 | -0.01 | 0 | 0.24 | 0.6 | 0.24 | 0.05 | 0.29 | -1 | 0.37 | 0.27 | 0.48 | 0.19 | -0.11 | 0.16 | 0.1175 | -0.1825 | 0.015 | 0.589581 | 0.608564 | 0.949181 | 1 | 0.996511 | 0.992476 |
| Tomm40l | Q9CZR3 | -0.7 | -0.24 | 0.04 | -0.22 | 0.08 | -0.22 | -0.3 | -0.3 | -0.32 | 0.97 | 0.03 | -0.62 | -0.26 | 0.57 | -0.23 | 0.1 | 0.095 | 0.295 | 0.325 | 0.649044 | 0.429903 | 0.250035 | 1 | 0.996511 | 0.806092 |
| Tomm5 | B1AXP6 | 2.09 | 1.79 | 1.29 | 1.32 | 1.42 | 2.45 | 1.45 | 1.63 | 1.75 | 1.2 | 1.38 | 1.44 | 2.06 | 1.96 | 1.12 | 1.69 | 0.115 | -0.18 | 0.085 | 0.707963 | 0.475794 | 0.776798 | 1 | 0.996511 | 0.967045 |
| Tomm6 | Q9CQN3 | 0.75 | 1.5 | 0.81 | 1.43 | 1.53 | 0.44 | 0.75 | 1.25 | 1.25 | 2.16 | 1.12 | 1.41 | 0.98 | 1.87 | 0.8 | 0.71 | -0.13 | 0.3625 | -0.0325 | 0.676709 | 0.260192 | 0.922449 | 1 | 0.996511 | 0.987467 |
| Tomm7 | Q9D173 | 0.61 | 1.4 | 0.69 | 0.93 | 1.6 | 0.95 | 0.94 | 0.85 | 1.42 | 1.83 | 1.39 | 0.18 | -0.66 | 2.19 | 1.12 | 1.03 | 0.1775 | 0.2975 | 0.0125 | 0.496901 | 0.445673 | 0.982371 | 1 | 0.996511 | 0.994858 |
| Tomm70 | Q9CZW5 | 0.1 | 0.1 | 0.83 | 0.85 | -0.45 | -0.07 | 0.38 | 0.82 | -0.23 | 0.02 | 0.62 | -0.11 | -0.29 | -0.57 | 0.4 | 0.28 | -0.3 | -0.395 | -0.515 | 0.384889 | 0.200988 | 0.135066 | 1 | 0.996511 | 0.731223 |
| Top1 | Q04750 | 0.93 | 0.02 | 0.17 | 0.62 | 0.23 | 0.73 | 0.54 | 0.77 | 0.66 | 0.78 | 0.49 | 0.29 | 1.16 | 0.22 | 0.73 | 0.52 | 0.1325 | 0.12 | 0.2225 | 0.60354 | 0.640306 | 0.465741 | 1 | 0.996511 | 0.888469 |
| Top2b | Q64511 | 1.38 | 0.99 | 1.36 | 1.07 | 1.25 | 1.23 | 1.29 | 0.76 | 0.81 | 0.26 | 1.28 | 1.29 | 0.72 | 0.72 | 1.35 | 1.63 | -0.0675 | -0.29 | -0.095 | 0.730285 | 0.310326 | 0.728928 | 1 | 0.996511 | 0.958635 |
| Top3b | Q9Z321 | -1.47 | -0.74 | -1.31 | -1.46 | -0.55 | -1.46 | -0.41 | -1.19 | -0.66 | -0.97 | -1.35 | -1.55 | -0.75 | 0.39 | -0.54 | -0.4 | 0.3425 | 0.1125 | 0.92 | 0.27519 | 0.684908 | 0.015398 | 1 | 0.996511 | 0.589387 |
| Tor1a | Q9ER39 | 0.2 | 0.24 | -0.21 | -0.71 | 0.9 | 0.06 | 0.09 | -0.52 | 0.1 | -1.46 | -0.61 | 0.23 | 0.36 | 0.59 | -0.45 | -0.12 | 0.2525 | -0.315 | 0.215 | 0.479735 | 0.46457 | 0.515716 | 1 | 0.996511 | 0.905674 |
| Tor1aip1 | Q921T2-2 | 2.97 | 1.46 | 1.18 | 0.39 | 1.32 | 2.87 | 1.69 | 0.62 | 1.17 | 0.45 | 1.27 | 2.39 | 1.08 | 0.34 | 1.78 | 1.78 | 0.125 | -0.18 | -0.255 | 0.847646 | 0.770264 | 0.665776 | 1 | 0.996511 | 0.950036 |
| Tor1aip2 | Q8BYU6 | 1.21 | -0.65 | 0.26 | 0.49 | 1.12 | 0.65 | 0.88 | 0.52 | 1.4 | 0.94 | 0.43 | 1.11 | 1.89 | 1.54 | 0.79 | 0.35 | 0.465 | 0.6425 | 0.815 | 0.246749 | 0.146508 | 0.120651 | 1 | 0.996511 | 0.721448 |
| Tor1aip2 | Q9ER81 | -1.35 | -1.13 | 0.19 | 0.56 | -1.62 | -2.12 | 0.43 | 0.6 | -1.72 | 1.86 | 1.14 | -1.53 | -2.55 | -3.58 | 0.36 | 0.79 | -0.245 | 0.37 | -0.8125 | 0.748158 | 0.691144 | 0.446998 | 1 | 0.996511 | 0.884817 |
| Tor1b | Q9ER41 | 0.34 | 0.45 | 0.31 | 0.49 | 0.91 | 0.15 | 0.53 | 0.54 | -0.1 | 0.03 | 0.1 | 0.46 | 0.31 | 0.26 | 0.82 | 0.74 | 0.135 | -0.275 | 0.135 | 0.497246 | 0.166855 | 0.523981 | 1 | 0.996511 | 0.908107 |
| Tor2a | Q8R1J9 | -1.06 | -0.84 | -0.89 | -0.34 | -0.23 | -0.47 | -0.76 | -0.22 | -0.11 | -0.49 | -1.11 | 0.07 | 0.36 | 0.22 | -1.46 | 0.17 | 0.3625 | 0.3725 | 0.605 | 0.130802 | 0.245496 | 0.187166 | 1 | 0.996511 | 0.776843 |
| Tor3a | Q9ER38 | 0.93 | 1.2 | 0.81 | 0.33 | 1.44 | 1.4 | 1.22 | -0.26 | 1 | 0.4 | -0.18 | 0.19 | 0.2 | 1.83 | 0.45 | 0.02 | 0.1325 | -0.465 | -0.1925 | 0.751486 | 0.159181 | 0.654795 | 1 | 0.996511 | 0.948172 |
| Tox2 | E9Q0B9 | -1.88 | -2.44 | -2.46 | -1.96 | -2.02 | -2.77 | -1.26 | -1.05 | -0.75 | -4.45 | -0.97 | -2.24 | -0.28 | -3.09 | -2.15 | -1.53 | 0.41 | 0.0825 | 0.4225 | 0.317317 | 0.915751 | 0.458891 | 1 | 0.996755 | 0.887902 |
| Tox4 | Q8BU11 | -0.61 | -1.95 | -1.32 | -1.45 | -1.98 | -0.93 | -1.91 | -0.95 | -0.83 | -3.07 | -1.73 | -1.2 | -0.52 | -1.55 | -0.71 | -1.5 | -0.11 | -0.375 | 0.2625 | 0.772955 | 0.478797 | 0.489732 | 1 | 0.996511 | 0.897531 |
| Tp53bp1 | P70399-5 | 0.38 | -0.83 | -1.18 | -0.81 | -1.46 | -0.63 | -1.83 | -0.64 | -1.02 | 0.18 | -1.13 | -1.24 | -1.35 | -1.4 | -1.86 | -1.8 | -0.53 | -0.1925 | -0.9925 | 0.234288 | 0.668185 | 0.021163 | 1 | 0.996511 | 0.621926 |
| Tpbg | Q9Z0L0 | 0.71 | 0.52 | 0.35 | 0.09 | 0.14 | 0.78 | 0.13 | 0.23 | -0.17 | -0.23 | 0.05 | 0.83 | -0.24 | 0.16 | -0.49 | 0.18 | -0.0975 | -0.2975 | -0.515 | 0.666079 | 0.315671 | 0.061185 | 1 | 0.996511 | 0.661738 |
| Tpbgl | Q8C013 | -0.16 | -0.44 | 1.26 | 1.18 | -0.52 | -0.22 | 1.56 | 1.26 | 1.2 | 2.04 | 1.16 | 0.28 | 1.07 | 1.04 | 1.49 | 1.05 | 0.06 | 0.71 | 0.7025 | 0.923147 | 0.201887 | 0.132106 | 1 | 0.996511 | 0.726871 |
| Tpcn1 | Q9EQJ0 | -0.18 | 0.17 | -0.06 | -0.82 | -0.39 | 1.32 | 0.86 | -0.45 | 1.59 | -2.97 | 0.48 | 0.37 | 2.75 | -1.22 | 0.6 | -0.45 | 0.5575 | 0.09 | 0.6425 | 0.244771 | 0.92094 | 0.43046 | 1 | 0.996755 | 0.879165 |
| Tpd52 | Q62393-3 | 1.31 | 0.63 | 0.71 | 1.19 | 0.39 | 0.91 | 0.69 | 0.96 | 1.36 | 0.83 | 0.98 | 1.83 | 0.92 | 0.1 | 0.55 | 0.27 | -0.2225 | 0.29 | -0.5 | 0.351453 | 0.331055 | 0.092445 | 1 | 0.996511 | 0.695968 |
| Tpd52l1 | B2RUC1 | 1.17 | 0.56 | 0.38 | 0.51 | 0.57 | 1.1 | 0.1 | 0.24 | -0.03 | 1.09 | -0.1 | 1.15 | -0.18 | 0.62 | -0.22 | -0.11 | -0.1525 | -0.1275 | -0.6275 | 0.5955 | 0.732909 | 0.051277 | 1 | 0.996511 | 0.658454 |
| Tpd52l2 | A2AUD5 | -1.43 | -0.61 | -0.57 | 0.05 | -1.44 | -1.14 | 0.52 | 0.14 | -0.14 | 0.4 | 1.04 | -1.18 | 0.38 | -0.67 | -0.12 | -0.14 | 0.16 | 0.67 | 0.5025 | 0.75914 | 0.218526 | 0.192538 | 1 | 0.996511 | 0.777539 |
| Tpd52l2 | Q9CYZ2 | 0.22 | 0.32 | 0.56 | 0.37 | -0.2 | 0.05 | 0.6 | 0.35 | -0.02 | 0.35 | 0.42 | 0.33 | 0.18 | 0.46 | 0.16 | 0.23 | -0.1675 | -0.0975 | -0.11 | 0.442937 | 0.599153 | 0.55444 | 1 | 0.996511 | 0.916796 |
| Tpgs1 | A0A6Q6PK | -0.19 | 0.08 | 0.64 | 0.06 | 0.45 | -0.44 | 0.63 | -0.05 | 0.09 | -0.98 | 0.24 | -0.22 | -0.44 | 0.31 | 0.3 | 0.25 | -7.9E-17 | -0.365 | -0.0425 | 1 | 0.276367 | 0.877238 | 1 | 0.996511 | 0.979626 |
| Tpgs2 | E9PUD7 | -0.11 | 0.42 | 0.66 | 0.83 | 0.48 | -0.86 | 1.93 | 1.86 | 0.8 | 1.33 | 1.24 | 0.06 | 0.43 | 1.54 | 0.93 | 1.44 | 0.4025 | 0.4075 | 0.635 | 0.527688 | 0.261048 | 0.081975 | 1 | 0.996511 | 0.68865 |
| Tph2 | Q8CGV2 | 0.16 | 0.62 | 0.17 | 0.53 | 0.05 | 0.06 | -0.18 | 0.17 | 0.13 | 0.53 | -0.15 | 0.35 | -0.14 | 0.24 | 0.19 | 0.07 | -0.345 | -0.155 | -0.28 | 0.087421 | 0.495641 | 0.199652 | 1 | 0.996511 | 0.78004 |
| Tpi1 | P17751-2 | 0.38 | 0.19 | 0.28 | 0.14 | 0.55 | 0.39 | 0.45 | 0.58 | -0.75 | -1.45 | 0.24 | 0.15 | -0.84 | -0.64 | -0.05 | 0.13 | 0.245 | -0.7 | -0.5975 | 0.124816 | 0.099227 | 0.047231 | 1 | 0.996511 | 0.658454 |
| Tpk1 | Q9R0M5 | -2.11 | -2.57 | -1.3 | -1.01 | -1.18 | -1.28 | -1.39 | -1.28 | -1.8 | -1.4 | -1.31 | -1.69 | -1.61 | -1.42 | -1.81 | -1.17 | 0.465 | 0.1975 | 0.245 | 0.205246 | 0.593329 | 0.519342 | 1 | 0.996511 | 0.907667 |
| Tpm1 | F8WID5 | -6.08 | -7.81 | -8.16 | -6.66 | -7.25 | -7.33 | -7.48 | -8.74 | -7.71 | -5.82 | -7.52 | -5.88 | -6.44 | -9.31 | -8.25 | -9.62 | -0.5225 | 0.445 | -1.2275 | 0.354363 | 0.495665 | 0.142996 | 1 | 0.996511 | 0.742711 |
| Tpm1 | Q8BP43 | -0.14 | -1.02 | -0.5 | -0.78 | -0.92 | -0.95 | -1.56 | -0.82 | -0.07 | 0.51 | -0.06 | -0.23 | 0.21 | 0.26 | -0.88 | -0.9 | -0.4525 | 0.6475 | 0.2825 | 0.111344 | 0.035832 | 0.451418 | 1 | 0.996511 | 0.885538 |
| Tpm1 | G5E8R0 | 0.64 | 0.22 | 0.7 | 1.32 | -0.19 | 0.33 | 0.03 | 1.09 | 0.61 | 2.12 | 0.9 | 0.34 | 0.59 | 0.68 | 0.51 | 0.43 | -0.405 | 0.2725 | -0.1675 | 0.261736 | 0.530321 | 0.526444 | 1 | 0.996511 | 0.908107 |
| Tpm1 | E9Q452 | 0.83 | -0.14 | 0.03 | 0.61 | -0.46 | 0.28 | 0.24 | 0.66 | 1.47 | -0.27 | 0.43 | 1.13 | 1.54 | -0.06 | 0.45 | 0.16 | -0.1525 | 0.3575 | 0.19 | 0.636621 | 0.411462 | 0.642428 | 1 | 0.996511 | 0.94365 |
| Tpm1 | P58771 | 0.64 | 0.75 | 1.02 | 1.22 | 1.16 | -0.02 | 1.34 | 0.89 | 0.42 | 1.52 | 1.05 | 0.7 | -0.49 | 0.65 | 1.06 | 1.59 | -0.065 | 0.015 | -0.205 | 0.839894 | 0.957606 | 0.641213 | 1 | 0.996755 | 0.943421 |
| Tpm3 | P21107-2 | 1.86 | 0.8 | 1.24 | 1.99 | 0.59 | 1.46 | 0.94 | 1.99 | 1.12 | 0.66 | 1.56 | 1.91 | 0.49 | 0.84 | 1.25 | 1.04 | -0.2275 | -0.16 | -0.5675 | 0.564184 | 0.671021 | 0.108949 | 1 | 0.996511 | 0.707444 |
| Tpm3 | D3Z6I8 | 0.82 | 0.5 | 0.05 | 0.06 | -0.2 | 0.61 | -1.03 | -0.2 | 0.24 | -0.03 | -0.23 | 0.93 | 0.27 | -0.08 | 0.07 | -0.07 | -0.5625 | -0.13 | -0.31 | 0.150665 | 0.681958 | 0.221034 | 1 | 0.996511 | 0.787447 |
| Tpm3 | E9Q7Q3 | -0.09 | 0.16 | 0.28 | 0.19 | -0.07 | -0.34 | -0.1 | 0.44 | -0.32 | -0.85 | 0.21 | -0.36 | -0.57 | 0.42 | -0.25 | 0.14 | -0.1525 | -0.465 | -0.2 | 0.474847 | 0.093404 | 0.449875 | 1 | 0.996511 | 0.884817 |
| Tpm4 | Q6IRU2 | 2.18 | 0.45 | 1.12 | 2.05 | 0.22 | 1.35 | 0.28 | 1.18 | 1.44 | 0.36 | 1.02 | 1.97 | 1.26 | 0.14 | 0.39 | 1.14 | -0.6925 | -0.2525 | -0.7175 | 0.164315 | 0.612434 | 0.148137 | 1 | 0.996511 | 0.746406 |
| Tpmt | A0A0R4J01 | 1.33 | 0.64 | 3 | 1.89 | 1.04 | 1.28 | 3.36 | 2.98 | -0.86 | 0.55 | 2.87 | 1.14 | -1.16 | -0.74 | 2.75 | 3.56 | 0.45 | -0.79 | -0.6125 | 0.524221 | 0.354286 | 0.599732 | 1 | 0.996511 | 0.930673 |
| Tpp1 | O89023 | -0.1 | -0.3 | -0.85 | -0.43 | -0.41 | 0.02 | -0.71 | -0.42 | -0.8 | -1.29 | -0.42 | -0.27 | -0.53 | -1.57 | -1.44 | -0.47 | 0.04 | -0.275 | -0.5825 | 0.865254 | 0.3514 | 0.108617 | 1 | 0.996511 | 0.707444 |
| Tpp2 | Q64514-2 | -0.08 | -0.43 | -0.06 | -0.1 | -0.1 | -0.07 | 0.54 | 0.35 | -0.6 | -0.22 | 0.3 | 0.07 | -0.73 | -0.01 | 0.53 | 0.4 | 0.3475 | 0.055 | 0.215 | 0.124018 | 0.819216 | 0.490459 | 1 | 0.996511 | 0.897531 |
| Tppp | Q7TQD2 | 0.4 | 0.45 | 0.44 | 0.26 | 0.42 | 0.25 | 0.56 | 0.21 | 0.17 | 0.31 | -0.02 | 0.1 | -0.45 | 0.57 | 0.26 | 0.03 | -0.0275 | -0.2475 | -0.285 | 0.861262 | 0.161324 | 0.275859 | 1 | 0.996511 | 0.822084 |
| Tppp3 | Q9CRB6 | 0.56 | -0.04 | -0.18 | 0 | 0.16 | 0.23 | 0.04 | -0.02 | 1.09 | 1.2 | 0.04 | 1 | 0.9 | 0.36 | 0.68 | 0.04 | 0.0175 | 0.7475 | 0.41 | 0.931465 | 0.039409 | 0.157769 | 1 | 0.996511 | 0.751678 |
| Tpr | Q7M739 | 0.75 | -0.41 | -0.59 | -0.47 | -1.1 | -0.15 | -0.76 | -0.18 | -0.27 | -0.87 | -0.39 | -0.62 | 0.72 | -0.56 | 0.27 | -0.52 | -0.3675 | -0.3575 | 0.1575 | 0.335364 | 0.303002 | 0.70987 | 1 | 0.996511 | 0.957772 |
| Tprg1l | Q9DBS2 | 0.67 | 0.68 | -0.32 | 0 | 0.61 | 0.73 | 0.16 | -0.02 | 0.87 | -0.41 | -0.11 | 0.87 | 0.01 | 1.1 | -0.53 | -0.33 | 0.1125 | 0.0475 | -0.195 | 0.712099 | 0.904817 | 0.645225 | 1 | 0.996744 | 0.944749 |
| Tprkb | Q8QZZ7 | -2.92 | -2.48 | -2.05 | -3.84 | -2.51 | -2.71 | -4.63 | -2.29 | -2.02 | -4.38 | -2.35 | -2.54 | -1.66 | -2.8 | -1.72 | -2.56 | -0.2125 | -2E-16 | 0.6375 | 0.725062 | 1 | 0.184063 | 1 | 1 | 0.776843 |
| Tprn | A2AI08 | 1.18 | 0.84 | 0.71 | 0.48 | 1.85 | 1.19 | 0.6 | 0.37 | 1.77 | -0.47 | 0.43 | 1.76 | 2.31 | 0.77 | 0.62 | 0.61 | 0.2 | 0.07 | 0.275 | 0.569477 | 0.893744 | 0.516293 | 1 | 0.996511 | 0.905676 |
| Tpst1 | O70281 | 0.04 | -0.06 | -0.78 | 0.4 | 0.21 | -0.32 | 0.65 | -0.2 | -0.46 | 0.24 | -0.27 | -0.06 | -1.32 | -0.54 | -0.15 | 0.98 | 0.185 | -0.0375 | -0.1575 | 0.570382 | 0.89921 | 0.754296 | 1 | 0.996511 | 0.962308 |
| Tpt1 | P63028 | 1.09 | 0.73 | -0.64 | -0.52 | 0.28 | 0.87 | -0.76 | -0.59 | 0.88 | 0.54 | -0.64 | 1.14 | -0.21 | 0.71 | -0.9 | -0.84 | -0.215 | 0.315 | -0.475 | 0.687921 | 0.565412 | 0.383738 | 1 | 0.996511 | 0.860984 |
| Tra2a | A0A0N4SV | 2.13 | 1.17 | 0.2 | 0.15 | 0.83 | 1.66 | 0.27 | 1.21 | -0.27 | -0.06 | 0.19 | 0.66 | 0 | 0.05 | 0.74 | 0.02 | 0.08 | -0.7825 | -0.71 | 0.875324 | 0.127904 | 0.157819 | 1 | 0.996511 | 0.751678 |
| Tra2b | P62996 | 1.13 | 0.25 | 0.27 | 0.34 | -0.28 | 0.38 | -0.11 | 0.7 | 0.5 | -0.39 | 0.19 | -0.46 | 1.21 | 0.36 | 0.65 | -0.36 | -0.325 | -0.5375 | -0.0325 | 0.303101 | 0.115919 | 0.93166 | 1 | 0.996511 | 0.989713 |
| Trabd | E9Q7A5 | 0.32 | 1.09 | 0.03 | 0.49 | 0.37 | 0 | -0.22 | 1.06 | 0.94 | 2.02 | 0.44 | 0.62 | 0.79 | 1.41 | 0.53 | 0.1 | -0.18 | 0.5225 | 0.225 | 0.605196 | 0.214075 | 0.526405 | 1 | 0.996511 | 0.908107 |
| Traf2 | P39429 | -0.75 | -0.35 | -0.27 | 0.27 | -0.49 | -0.33 | -0.33 | 0.22 | 0.66 | 0.47 | 0.02 | -0.55 | 0.63 | 0.14 | 0.1 | 0.34 | 0.0425 | 0.425 | 0.5775 | 0.873799 | 0.229022 | 0.055737 | 1 | 0.996511 | 0.661738 |
| Traf3 | Q3UHJ1 | -0.16 | -0.3 | 0.3 | 1.07 | -0.4 | 0.09 | 0.64 | 0.19 | 0.19 | 0.39 | 0.44 | 0.2 | 0.29 | -0.37 | 0.75 | 0.89 | -0.0975 | 0.0775 | 0.1625 | 0.786358 | 0.806747 | 0.688453 | 1 | 0.996511 | 0.955729 |
| Traf6 | P70196 | 0.53 | 0.39 | 0.23 | 0.06 | 0.56 | 0.34 | 0.8 | 0.21 | 0.19 | 0.26 | -0.12 | 0.24 | 0.08 | 0.45 | 0.32 | 0.48 | 0.175 | -0.16 | 0.03 | 0.388685 | 0.412036 | 0.881716 | 1 | 0.996511 | 0.979767 |
| Traf7 | F8WJF7 | 0.03 | 0.52 | 0.47 | -0.09 | -0.01 | -0.28 | 0.1 | 0.17 | 0.87 | -0.95 | 0.31 | 0.54 | 0.4 | 1.01 | 0.21 | 0.06 | -0.2375 | -0.04 | 0.1875 | 0.277388 | 0.921589 | 0.508288 | 1 | 0.996755 | 0.902804 |
| Trafd1 | Q3UDK1-2 | -0.24 | 0.12 | -0.3 | -0.97 | -0.02 | 0.22 | -0.88 | -0.62 | -0.21 | -1.04 | -0.21 | 0.56 | 0.58 | -0.64 | -0.24 | -0.65 | 0.0225 | 0.1225 | 0.11 | 0.945937 | 0.749738 | 0.761601 | 1 | 0.996511 | 0.963123 |
| Trak1 | A0A5F8MP | 0.25 | 0.62 | -0.03 | 0.1 | 0.75 | 0.23 | -1.23 | -0.31 | -0.31 | -0.1 | -0.25 | 0.44 | -0.04 | -0.17 | -0.14 | -0.32 | -0.375 | -0.29 | -0.4025 | 0.381149 | 0.25783 | 0.082472 | 1 | 0.996511 | 0.68865 |
| Trak2 | Q6P9N8 | 0.22 | 0.79 | 1.22 | 1.02 | 0.68 | 0.34 | 0.52 | 1.27 | 0.41 | 0.44 | 1.17 | 0.73 | 0.68 | 0.68 | 1.09 | 1.15 | -0.11 | -0.125 | 0.0875 | 0.709798 | 0.666226 | 0.749956 | 1 | 0.996511 | 0.961471 |
| Tram1 | Q91V04 | -1.22 | -0.09 | 0.5 | 0.37 | 0.11 | -0.39 | -0.66 | -0.45 | -0.69 | 0.14 | -0.02 | -0.91 | 0.52 | -0.46 | 0.1 | 0.57 | -0.2375 | -0.26 | 0.2925 | 0.556316 | 0.558552 | 0.507117 | 1 | 0.996511 | 0.902614 |
| Tram1l1 | Q8QZR0 | -1.4 | -0.94 | -1.7 | -1.1 | -0.83 | -0.78 | -2.31 | -1.91 | -0.61 | -1.38 | -1.63 | -1.74 | -0.17 | -1.28 | 1.4 | 0.59 | -0.1725 | -0.055 | 1.42 | 0.665496 | 0.858523 | 0.027766 | 1 | 0.996511 | 0.635993 |
| Trap1 | Q9CQN1 | 0.63 | 0.7 | 1.17 | 1.25 | 0.7 | 0.68 | 1.21 | 1.23 | 0.12 | 0.75 | 1.29 | 0.47 | 0.43 | 0.48 | 0.94 | 1.38 | 0.0175 | -0.28 | -0.13 | 0.941146 | 0.362668 | 0.655824 | 1 | 0.996511 | 0.948301 |
| Trappc1 | Q5NCF2 | 0.47 | 0.53 | 0.58 | 0.71 | 0.59 | 0.54 | 0.68 | 0.56 | 0.45 | 0.69 | 0.68 | 0.38 | 0.04 | 0.86 | 0.35 | 0.66 | 0.02 | -0.0225 | -0.095 | 0.89003 | 0.895271 | 0.681981 | 1 | 0.996511 | 0.955729 |
| Trappc10 | Q3TLI0 | 0.23 | 0.24 | -0.07 | -0.16 | 0.66 | 0.6 | 0.11 | -0.03 | -0.26 | -0.82 | 0 | 0.54 | -0.45 | -0.37 | -0.4 | -0.14 | 0.275 | -0.195 | -0.4 | 0.238925 | 0.527715 | 0.066405 | 1 | 0.996511 | 0.664936 |
| Trappc11 | B2RXC1 | -0.28 | 0 | -0.58 | -0.65 | -0.04 | -0.17 | -0.37 | -0.74 | -0.23 | -0.89 | -0.49 | -0.29 | -0.3 | -1.02 | -0.57 | -0.58 | 0.0475 | -0.0975 | -0.24 | 0.837857 | 0.68452 | 0.34348 | 1 | 0.996511 | 0.848473 |
| Trappc12 | Q8K2L8 | -0.34 | -0.12 | -0.08 | -0.14 | -0.15 | -0.4 | -0.13 | -0.2 | -0.27 | 0.38 | -0.01 | -0.22 | -0.29 | 0.02 | -0.46 | -0.47 | -0.05 | 0.14 | -0.13 | 0.74632 | 0.500456 | 0.515776 | 1 | 0.996511 | 0.905674 |
| Trappc13 | Q3TIR1 | -0.88 | -0.88 | -0.21 | -0.49 | -1.15 | -0.8 | -0.97 | -0.65 | -0.77 | -0.61 | -0.42 | -1 | -0.34 | -1.13 | -0.48 | -0.53 | -0.2775 | -0.085 | -0.005 | 0.227517 | 0.717823 | 0.984966 | 1 | 0.996511 | 0.996154 |
| Trappc14 | Q3UTZ3 | 0.03 | 0.26 | 0.33 | 0.22 | 0.08 | 0.2 | -0.25 | 0.26 | 0.58 | 0.89 | 0.16 | -0.02 | 0.71 | 0.62 | 0.46 | 0.13 | -0.1375 | 0.1925 | 0.27 | 0.446787 | 0.434804 | 0.21045 | 1 | 0.996511 | 0.786218 |
| Trappc2 | Q9CQP2 | -0.98 | -0.79 | -0.37 | -0.53 | 0 | -0.85 | -0.06 | -0.46 | -2.4 | -2.07 | -0.14 | -0.44 | -2.3 | -3.25 | -0.92 | -0.75 | 0.325 | -0.595 | -1.1375 | 0.217895 | 0.287922 | 0.068939 | 1 | 0.996511 | 0.666991 |
| Trappc2l | Q9JME7 | -0.26 | 0.02 | 0.34 | 0.2 | 0.19 | -0.39 | 0.39 | 0.27 | 0.07 | 1.05 | 0.18 | -0.17 | -0.2 | 0.57 | 0.17 | 0.49 | 0.04 | 0.2075 | 0.1825 | 0.864321 | 0.497137 | 0.474115 | 1 | 0.996511 | 0.891644 |
| Trappc3 | O55013 | 0.76 | 0.79 | 0.6 | 1.2 | 0.59 | 0.64 | 0.09 | 1.3 | 0.66 | 1.96 | 0.72 | 0.57 | 0.59 | 0.76 | 0.45 | 0.05 | -0.1825 | 0.14 | -0.375 | 0.522257 | 0.687465 | 0.14162 | 1 | 0.996511 | 0.737973 |
| Trappc4 | Q9ES56 | -0.43 | -0.18 | -0.21 | -0.48 | 0.08 | -0.06 | 0.78 | -0.38 | -0.52 | -1.27 | -0.2 | -0.35 | -0.9 | -0.16 | -0.4 | -0.06 | 0.43 | -0.26 | -0.055 | 0.130493 | 0.342542 | 0.819394 | 1 | 0.996511 | 0.972768 |
| Trappc5 | Q9CQA1 | -0.25 | -0.29 | -0.23 | -0.54 | -0.09 | 0.01 | -0.33 | -0.42 | -0.17 | -0.2 | -0.32 | -0.24 | -0.29 | -0.27 | -0.5 | -0.51 | 0.12 | 0.095 | -0.065 | 0.495368 | 0.568014 | 0.723615 | 1 | 0.996511 | 0.958398 |
| Trappc6b | Q9D289 | -0.62 | -0.29 | -0.12 | -0.6 | -0.01 | -0.72 | 0.07 | -0.66 | -0.76 | -1.5 | -0.19 | -0.5 | -0.75 | -0.83 | -0.21 | -0.21 | 0.0775 | -0.33 | -0.0925 | 0.759789 | 0.30112 | 0.706027 | 1 | 0.996511 | 0.957084 |
| Trappc8 | A0A286YC) | 0.5 | 0.02 | -0.01 | -0.35 | -0.01 | 0.36 | -0.25 | -0.4 | 0.2 | -0.48 | -0.15 | 0.43 | 0.11 | -0.17 | -0.36 | -0.24 | -0.115 | -0.04 | -0.205 | 0.65026 | 0.885577 | 0.403216 | 1 | 0.996511 | 0.870698 |
| Trappc9 | Q3U0M1 | 0.48 | 0.49 | 0.37 | 0.22 | 0.93 | 0.45 | 0.18 | 0.29 | 0.42 | 0.46 | 0.35 | 0.53 | 0.29 | 0.61 | 0.6 | 0.32 | 0.0725 | 0.05 | 0.065 | 0.726404 | 0.759142 | 0.729705 | 1 | 0.996511 | 0.958635 |
| Trex1 | Q91XB0 | -1.11 | -0.86 | -0.37 | 0.48 | 0.08 | -0.84 | 0.88 | 0.23 | -0.84 | -0.08 | 0.8 | -0.91 | -1.5 | 0.07 | 1.14 | 0.95 | 0.5525 | 0.2075 | 0.63 | 0.252445 | 0.676306 | 0.335573 | 1 | 0.996511 | 0.847442 |
| Trh | Q62361 | -1.14 | -1.07 | -7.04 | -10.17 | -6.68 | -2.27 | -8.41 | -7.47 | -1.6 | -1.59 | -7.56 | -1.47 | -1.53 | -6.75 | -5.68 | -6.52 | -1.3525 | 1.8 | -0.265 | 0.568109 | 0.46325 | 0.907022 | 1 | 0.996511 | 0.986458 |
| Trhde | Q8K093 | -0.02 | 0.63 | 0.79 | 0.68 | 1.08 | 0.24 | 0.65 | 0.58 | 0.53 | 0.19 | 0.66 | 0.23 | 0.42 | 0.84 | 0.27 | 0.33 | 0.1175 | -0.1175 | -0.055 | 0.653478 | 0.630106 | 0.829349 | 1 | 0.996511 | 0.973939 |
| Trhr | P21761 | 0.99 | 1.96 | -0.87 | 0.06 | 1.74 | 1.25 | 1.07 | 0.23 | 0.6 | -2.09 | 0.16 | 1.42 | 0.24 | 0.93 | -0.27 | 0.31 | 0.5375 | -0.5125 | -0.2325 | 0.39888 | 0.560293 | 0.699896 | 1 | 0.996511 | 0.955776 |
| Triap1 | Q9D8Z2 | -0.51 | -1.36 | -0.5 | -0.5 | -0.85 | -0.09 | -0.48 | -0.73 | -1.03 | -0.59 | -0.58 | -0.79 | 0.14 | -1.56 | -0.91 | -0.3 | 0.18 | -0.03 | 0.06 | 0.519772 | 0.907631 | 0.883653 | 1 | 0.996744 | 0.979767 |
| Tril | Q9DBY4 | 0.04 | 0.01 | -0.07 | -0.7 | 0.36 | 0.22 | -0.55 | -0.96 | 0.53 | -1.37 | -1.5 | 0.3 | 0.58 | -0.35 | 0.01 | 1.77 | -0.0525 | -0.33 | 0.6825 | 0.879522 | 0.53273 | 0.168388 | 1 | 0.996511 | 0.759484 |
| Trim2 | Q9ESN6 | -0.35 | -0.52 | 0.17 | 0.05 | -0.62 | -0.3 | 0.18 | 0.22 | -0.27 | -0.34 | 0.22 | -0.21 | -0.5 | -0.26 | 0.02 | -0.29 | 0.0325 | 0.0125 | -0.095 | 0.902755 | 0.957648 | 0.688436 | 1 | 0.996755 | 0.955729 |
| Trim23 | Q8BGX0-3 | 1.59 | 1.99 | 2.7 | 2.13 | 2.15 | 1.71 | 3.03 | 2.91 | 1.61 | 0.67 | 2.88 | 1.53 | 0.33 | 1.52 | 0.3 | 2.02 | 0.3475 | -0.43 | -1.06 | 0.361415 | 0.378478 | 0.044337 | 1 | 0.996511 | 0.658454 |

| | | | | | | | | | | | | | | | | | | | | | | | | | |
|---|---|---|---|---|---|---|---|---|---|---|---|---|---|---|---|---|---|---|---|---|---|---|---|---|---|
| Trim24 | Q64127 | -0.57 | -1.22 | -1.2 | -1.77 | -2.66 | -0.58 | -2.39 | -1.53 | -0.41 | -0.53 | -1.54 | -0.69 | 0.14 | -1.52 | -0.38 | -1.57 | -0.6 | 0.3975 | 0.3575 | 0.241097 | 0.27135 | 0.447069 | 1 | 0.996511 | 0.884817 |
| Trim25 | Q61510 | -0.39 | -0.65 | -0.87 | -0.71 | -1.64 | -1.32 | -0.42 | -0.79 | -0.23 | -0.4 | -0.61 | -0.49 | -0.62 | -0.23 | -1.03 | -0.91 | -0.3875 | 0.2225 | -0.0425 | 0.204233 | 0.254401 | 0.86054 | 1 | 0.996511 | 0.977204 |
| Trim26 | J3QN94 | -1.06 | -0.8 | 0.51 | -0.79 | -0.99 | -1.1 | 0.21 | -1.73 | -0.48 | -2.24 | -0.81 | 0.47 | -0.04 | 0.23 | -0.9 | -0.43 | -0.3675 | -0.23 | 0.25 | 0.466442 | 0.706198 | 0.549076 | 1 | 0.996511 | 0.912757 |
| Trim28 | Q62318 | -0.5 | -1.49 | -1.18 | -0.83 | -1.32 | -0.84 | -0.92 | -0.78 | -1.32 | -0.96 | -0.84 | -1.12 | -1.08 | -0.9 | -1.1 | -1.34 | 0.035 | -0.06 | -0.105 | 0.891403 | 0.816383 | 0.688766 | 1 | 0.996511 | 0.955729 |
| Trim3 | Q9R1R2 | 1.38 | 0.84 | -0.17 | 0.03 | 0.99 | 1.44 | 0.04 | -0.05 | 0.14 | 0.72 | 0.08 | 1.74 | 0.06 | 0.17 | 0.14 | -0.21 | 0.085 | 0.15 | -0.48 | 0.857903 | 0.760479 | 0.210209 | 1 | 0.996511 | 0.786218 |
| Trim32 | Q8CH72 | -0.25 | -0.71 | -0.46 | -0.34 | -0.26 | -0.06 | 0.19 | -0.58 | -0.87 | -0.21 | -0.3 | -0.23 | -0.69 | -0.62 | -0.39 | 0.02 | 0.2625 | 0.0375 | 0.02 | 0.244635 | 0.865943 | 0.931245 | 1 | 0.996511 | 0.989713 |
| Trim33 | E9QME5 | -2.06 | -2.62 | -2.63 | -2.7 | -2.04 | -2.35 | -2.69 | -3.24 | -2.37 | -1.43 | -2.55 | -1.96 | -1.62 | -1.33 | -3.33 | -2.99 | -0.0775 | 0.425 | 0.185 | 0.793419 | 0.176729 | 0.703103 | 1 | 0.996511 | 0.956859 |
| Trim35 | Q8C006 | 0.48 | -1.23 | -1.26 | -0.9 | -1.21 | 4.78 | -1.77 | -1.24 | -0.69 | -0.33 | -0.99 | 0.92 | 0.61 | -2.14 | 0 | -0.52 | 0.8675 | 0.455 | 0.215 | 0.549377 | 0.411284 | 0.743077 | 1 | 0.996511 | 0.959128 |
| Trim36 | Q80WG7 | 3.11 | 1.12 | 0.66 | 0.5 | 0.53 | 3.6 | 0.51 | 0.74 | 2.09 | 1.58 | 0.91 | 3.14 | 2.16 | 0.33 | 1.03 | 0.86 | -0.0025 | 0.5825 | -0.2525 | 0.997699 | 0.411522 | 0.69919 | 1 | 0.996511 | 0.955729 |
| Trim37 | F6XMH5 | 0.73 | 1.33 | 2.45 | 1.12 | 1.62 | 0.82 | 2.94 | 1.71 | -2.87 | -1.56 | 3.33 | 1.4 | -1.38 | -3.75 | 2.16 | 3.85 | 0.365 | -1.3325 | -1.1875 | 0.493985 | 0.320307 | 0.453186 | 1 | 0.996511 | 0.885993 |
| Trim44 | F7B7X2 | -1.84 | -0.16 | 1.42 | -0.02 | -1.21 | -2.23 | 0.81 | 1.79 | -0.5 | 1.46 | 0.55 | -1.03 | -1.06 | 1.39 | 2.06 | 1.68 | -0.06 | 0.27 | 1.1675 | 0.952953 | 0.731467 | 0.203686 | 1 | 0.996511 | 0.782817 |
| Trim46 | Q7TNM2 | -0.92 | -1.98 | -0.63 | -0.6 | -2.13 | -1.31 | -1.42 | -0.67 | -1.09 | -0.18 | -0.57 | -1.59 | -0.29 | -1.67 | -0.74 | -0.76 | -0.35 | 0.175 | 0.1675 | 0.407948 | 0.680596 | 0.688308 | 1 | 0.996511 | 0.955729 |
| Trim47 | Q8C0E3 | -0.09 | -1.1 | -1.68 | -1.52 | -0.54 | -1.94 | -1.91 | -1.7 | -1.1 | -1.71 | -0.69 | -0.76 | 0.32 | -2.3 | -1.16 | -1.35 | -0.425 | 0.0325 | -0.025 | 0.36144 | 0.93626 | 0.966452 | 1 | 0.996755 | 0.9927 |
| Trim56 | Q80VI1 | -1.02 | -0.17 | 0.07 | -0.49 | -0.09 | -1.66 | -0.23 | -0.44 | -0.8 | -0.95 | -0.53 | -0.51 | -0.5 | -0.34 | -0.28 | 0.04 | -0.2025 | -0.295 | 0.1325 | 0.618964 | 0.296926 | 0.639743 | 1 | 0.996511 | 0.942804 |
| Trim65 | Q8BFW4 | -0.54 | -0.53 | -0.67 | -0.82 | 0.09 | -0.4 | -0.51 | -0.64 | 0.3 | 0.99 | -0.75 | -0.66 | -0.67 | 0.47 | -0.93 | -0.61 | 0.275 | 0.61 | 0.205 | 0.20119 | 0.154994 | 0.529866 | 1 | 0.996511 | 0.908663 |
| Trim66 | E9PZP2 | 0.22 | 1.18 | 0.32 | -0.2 | 0.46 | 0.43 | 1.1 | 0.3 | -0.15 | -0.25 | 0.04 | 1.29 | -0.09 | 0.12 | 0.6 | 0.74 | 0.1925 | -0.1475 | -0.0375 | 0.564358 | 0.734998 | 0.914076 | 1 | 0.996511 | 0.986458 |
| Trim72 | Q1XH17 | -0.32 | -0.41 | 1.08 | 2.43 | -0.06 | -0.85 | -0.32 | 1.13 | 0.06 | 1.74 | 1.06 | -0.44 | -0.59 | -0.17 | 1.18 | 1.05 | -0.72 | -0.09 | -0.3275 | 0.329237 | 0.904844 | 0.654129 | 1 | 0.996744 | 0.947657 |
| Trim9 | Q8C7M3 | -1.16 | -0.96 | 0.08 | -0.28 | -0.68 | -0.95 | 0.7 | 0.13 | -1.49 | -1.54 | 0.4 | -0.16 | -1.05 | -0.94 | -1.17 | -1.33 | 0.38 | -0.1175 | -0.5425 | 0.402051 | 0.822573 | 0.108282 | 1 | 0.996511 | 0.706427 |
| Trim9 | A0A6I8MW | 0.57 | 0.49 | 1.39 | 1.65 | 0.43 | -0.03 | -0.28 | 1.07 | 0.07 | 2.32 | 0.66 | -0.11 | 0.79 | 0.24 | 0.75 | 0.73 | -0.7275 | -0.29 | -0.3975 | 0.091797 | 0.616419 | 0.240114 | 1 | 0.996511 | 0.795595 |
| Trim9 | E9QLH4 | -0.4 | -0.15 | -0.33 | -0.29 | -0.09 | -0.17 | 0.24 | -0.35 | -0.24 | -0.25 | -0.18 | -0.16 | -0.08 | 0.06 | -0.48 | -0.33 | 0.2 | 0.085 | 0.085 | 0.283045 | 0.592725 | 0.671768 | 1 | 0.996511 | 0.952718 |
| Trio | Q0KL02-3 | 0.76 | 0.41 | 0.57 | 0.4 | 0.61 | 0.73 | 0.5 | 0.69 | 0.27 | -0.03 | 0.72 | 0.79 | 0.26 | 0.05 | 0.41 | 0.35 | 0.0975 | -0.0975 | -0.2675 | 0.54782 | 0.684923 | 0.186138 | 1 | 0.996511 | 0.776843 |
| Trio | A0A2I3BQ1 | -1.36 | -0.6 | -1.29 | -2.91 | -4.19 | -2.18 | -2.23 | -0.6 | -0.01 | -0.57 | -0.99 | -1.16 | 2.04 | -2.07 | -3.87 | -5.9 | -0.76 | 0.8575 | -0.91 | 0.352567 | 0.121176 | 0.563479 | 1 | 0.996511 | 0.920024 |
| Triobp | Q99KW3-4 | -0.61 | -0.91 | -0.49 | -0.69 | -1.13 | -0.54 | -0.78 | -0.46 | 0.61 | 0.03 | 0.29 | -0.44 | 1.79 | -0.19 | -0.22 | -0.17 | -0.0525 | 0.7975 | 0.9775 | 0.798047 | 0.01261 | 0.064639 | 1 | 0.996511 | 0.66433 |
| Trip11 | E9Q512 | -0.77 | -1.48 | -1.62 | -1.56 | -1.32 | -1.11 | -1.87 | -2.03 | -0.79 | -0.71 | -1.54 | -0.9 | -0.66 | -1.28 | -1.82 | -2 | -0.225 | 0.3725 | -0.0825 | 0.4524 | 0.212764 | 0.817168 | 1 | 0.996511 | 0.972768 |
| Trip12 | G5E870 | 0.51 | -0.18 | -0.24 | -0.15 | -0.32 | 0.07 | -0.44 | -0.41 | -0.39 | -0.52 | -0.26 | 0.22 | -0.14 | -0.48 | -0.6 | -0.44 | -0.26 | -0.2225 | -0.4 | 0.27794 | 0.400051 | 0.122836 | 1 | 0.996511 | 0.721448 |
| Trip4 | Q9QXN3-2 | -0.39 | 0.07 | 0.88 | -0.01 | 0.01 | -0.38 | 1.07 | 0.75 | -0.52 | 0.97 | 0.83 | -0.4 | -0.61 | 0.42 | 1.49 | 1.15 | 0.225 | 0.0825 | 0.475 | 0.578981 | 0.853965 | 0.354625 | 1 | 0.996511 | 0.855029 |
| Trir | Q9D735 | -3.2 | -3.92 | -5.18 | -9.44 | -3.64 | -3.43 | -7.66 | -8.08 | -2.43 | -4.24 | -6.72 | -3.19 | -1.74 | -3.16 | -5.93 | -7.32 | -0.2675 | 1.29 | 0.8975 | 0.87306 | 0.400376 | 0.595036 | 1 | 0.996511 | 0.929843 |
| Trit1 | Q80UN9 | -1.44 | -1.13 | 0.02 | 0.68 | -1.44 | -0.81 | -0.63 | 0.14 | -2.18 | 1.86 | 1.05 | -1.64 | -2.44 | -1.68 | 0.66 | 0.68 | -0.2175 | 0.24 | -0.2275 | 0.690034 | 0.809673 | 0.788799 | 1 | 0.996511 | 0.968677 |
| Trmt1 | H3BKS8 | -2.19 | -2.29 | -0.8 | -1.41 | -2.38 | -2.5 | -1.89 | -1.18 | -1.47 | -0.49 | -0.93 | -2.3 | -0.56 | -2.14 | -1.28 | -1.66 | -0.315 | 0.375 | 0.2625 | 0.472462 | 0.450955 | 0.569091 | 1 | 0.996511 | 0.922588 |
| Trmt10c | Q3UFY8 | 0.3 | 0.88 | 0.33 | 0.3 | 0.74 | 0.4 | 0.19 | 0.32 | 0.46 | 1.04 | 0.36 | 0.48 | 0.63 | 0.84 | 0.18 | 0.28 | -0.04 | 0.1325 | 0.03 | 0.850297 | 0.581371 | 0.902661 | 1 | 0.996511 | 0.984898 |
| Trmt11 | E9QKG3 | 0.31 | 0.39 | 0.18 | 0.34 | 0.55 | -0.04 | 0.03 | 0.05 | 0.1 | 1.33 | 0.73 | -0.25 | -0.26 | 0.6 | -0.47 | -0.11 | -0.1575 | 0.1725 | -0.365 | 0.404014 | 0.620283 | 0.189934 | 1 | 0.996511 | 0.776843 |
| Trmt112 | Q9DCG9 | -1.69 | -1.73 | -0.49 | -0.6 | -1.58 | -1.82 | -0.21 | -0.18 | -1.72 | 0.29 | -0.46 | -1.57 | -0.64 | -1.21 | -0.48 | -0.57 | 0.18 | 0.2625 | 0.4025 | 0.724239 | 0.628712 | 0.291138 | 1 | 0.996511 | 0.825497 |
| Trmt1l | A0A0R4J0L | -1.49 | -2.09 | -0.72 | 0.39 | 3.61 | -1.85 | -1.48 | 0.42 | -1.23 | 0.02 | -0.64 | -1.47 | -1.74 | -0.8 | -1.81 | -1.81 | 1.1525 | 0.1475 | -0.5625 | 0.355788 | 0.799129 | 0.316902 | 1 | 0.996511 | 0.838864 |
| Trmt44 | Q9D2Q2 | 0.06 | -0.41 | 0.03 | -0.24 | -2.78 | -0.1 | -1.07 | -0.63 | -0.76 | -1.23 | -0.47 | -0.44 | 1.37 | -5.1 | -0.94 | -0.43 | -1.005 | -0.585 | -1.135 | 0.091458 | 0.036473 | 0.365307 | 1 | 0.996511 | 0.858475 |
| Trmt6 | Q8CE96 | -3.23 | -3.78 | -4.07 | -4.45 | -3.78 | -2.95 | -4.5 | -4.92 | -2.8 | -2.59 | -2.54 | -3.36 | -2.67 | -3.43 | -3.69 | -4.34 | -0.155 | 1.06 | 0.35 | 0.740366 | 0.00876 | 0.406375 | 1 | 0.996511 | 0.87356 |
| Trmt61a | Q80XC2 | -3.78 | -3.82 | -2.39 | -2.47 | -3.88 | -3.11 | -2.1 | -2.01 | -1.12 | -1.19 | -2.2 | -3.76 | -0.74 | -1.76 | -2.34 | -2.59 | 0.34 | 1.0475 | 1.2575 | 0.538055 | 0.143146 | 0.038998 | 1 | 0.996511 | 0.658454 |
| Trmu | Q9DAT5 | -0.49 | -0.36 | -0.26 | -0.04 | -0.74 | -0.31 | -0.17 | -0.29 | -0.37 | -0.06 | -0.11 | -0.96 | 0.05 | 0.06 | 0.19 | -0.21 | -0.09 | -0.0875 | 0.31 | 0.643653 | 0.727509 | 0.14155 | 1 | 0.996511 | 0.737973 |
| Trnt1 | Q8K1J6 | -0.73 | -0.76 | -0.62 | -0.69 | -0.06 | -0.71 | -0.47 | -0.74 | -0.74 | -0.39 | -0.35 | -0.68 | -0.78 | -0.02 | -0.84 | -0.86 | 0.205 | 0.16 | 0.075 | 0.30998 | 0.373075 | 0.757371 | 1 | 0.996511 | 0.962481 |
| Tro | Q6DIC6 | -0.79 | -0.89 | -0.34 | -0.88 | 1.57 | 1.2 | -0.6 | 0.04 | 0.73 | -1.89 | 0.17 | 0.09 | 0.7 | 0.54 | 1.41 | -0.99 | 1.2775 | 0.5 | 1.14 | 0.025211 | 0.368928 | 0.041644 | 1 | 0.996511 | 0.658454 |
| Trp53bp2 | E9QJU8 | 0.15 | 0.48 | 0.96 | 0.79 | 0.17 | 0.11 | 1 | 1.08 | 0.81 | 0.88 | 1.09 | 0.14 | 0.74 | 0.86 | 0.98 | 0.97 | -0.005 | 0.135 | 0.2925 | 0.987153 | 0.635774 | 0.22681 | 1 | 0.996511 | 0.787447 |
| Trp53i11 | A2AGS6 | 5.19 | 3.65 | 3.86 | 1.91 | 4.11 | 4.59 | 3.95 | 4.09 | 0.09 | 0.89 | 3.91 | 5 | 0.59 | 0.37 | 4.47 | 4.81 | 0.5325 | -1.18 | -1.0925 | 0.405097 | 0.344777 | 0.384863 | 1 | 0.996511 | 0.861128 |
| Trp53rkb | Q543M9 | -1.12 | -0.94 | -1.35 | -1.86 | -0.76 | -1.46 | -1.2 | -1.6 | -0.4 | -0.94 | -1.54 | -1.15 | -0.58 | -0.64 | -1.95 | -2.08 | 0.0625 | 0.31 | 0.005 | 0.820859 | 0.335323 | 0.990732 | 1 | 0.996511 | 0.996154 |
| Trpc1 | A0A087WS | 1.91 | 0.84 | 0.99 | 1.37 | 1.29 | 0.9 | 1.13 | 1.03 | 0.72 | 1.1 | 0.93 | -0.04 | 1.06 | 1.48 | 0.92 | 1.04 | -0.19 | -0.6 | -0.1525 | 0.473358 | 0.106782 | 0.596996 | 1 | 0.996511 | 0.929911 |
| Trpc4 | Q9QUQ5 | -0.04 | 0.81 | -1.12 | -1.86 | 1.23 | 0.24 | -0.67 | -1.63 | 0.53 | -1.42 | -1.06 | 0.09 | 0.44 | 1.02 | -1.41 | -0.94 | 0.345 | 0.0875 | 0.33 | 0.654521 | 0.897688 | 0.657784 | 1 | 0.996511 | 0.948732 |
| Trpc4ap | Q9JLV2-2 | -1.47 | -1.72 | -0.52 | -0.8 | -1.75 | -0.94 | -0.34 | -1.03 | -1.06 | -0.68 | -0.5 | -1.18 | -0.77 | -0.86 | -0.6 | -0.04 | 0.1125 | 0.2725 | 0.56 | 0.768801 | 0.407937 | 0.123499 | 1 | 0.996511 | 0.721448 |
| Trpc6 | Q3UZG1 | 0.99 | 0.7 | 0.72 | 0.58 | 0.47 | 2.62 | 0.59 | 1.46 | 1.74 | 1.26 | 1.61 | 0.85 | 1.48 | 1.19 | 1.17 | 1.13 | 0.5375 | 0.6175 | 0.495 | 0.26904 | 0.030177 | 0.028356 | 1 | 0.996511 | 0.637553 |
| Trpm3 | Q5F4S7 | 0.33 | 0.81 | 0.38 | 0.7 | 0.89 | 0.07 | 0.96 | 0.23 | 0.15 | 0.18 | 0.2 | 0.12 | 0.2 | 0.1 | 0.66 | 1.09 | -0.0175 | -0.3925 | -0.0425 | 0.946955 | 0.056909 | 0.878442 | 1 | 0.996511 | 0.979626 |
| Trpm7 | Q923J1 | 0.89 | 1.1 | 0.55 | 0.7 | -0.04 | 0.92 | 0.85 | 0.68 | 2.16 | 0.73 | 0.58 | -0.19 | 2.64 | -2.32 | 0.71 | 0.92 | -0.2075 | 0.01 | -0.3225 | 0.432259 | 0.98307 | 0.729507 | 1 | 0.998773 | 0.958635 |
| Trpv2 | Q9WTR1 | 0.46 | 0.71 | 0.66 | 0.7 | 1.05 | 0.13 | 0.35 | 0.55 | 0.08 | 0.92 | 0.67 | 0.43 | -0.95 | 1.03 | 0.36 | 0.31 | -0.1125 | -0.1075 | -0.445 | 0.620952 | 0.632854 | 0.285475 | 1 | 0.996511 | 0.822669 |
| Trrap | E9QLK7 | -4.83 | -8.27 | -10.64 | -5.6 | -2.46 | -4.53 | -3.55 | -3.59 | -2.06 | -1.86 | -3.89 | -5.58 | -2.02 | -3.96 | -2.42 | -2.42 | 3.8025 | 3.9875 | 4.63 | 0.013308 | 0.019544 | 0.00464 | 1 | 0.996511 | 0.550241 |
| Trrap | E9PWT1 | -0.64 | -1.44 | -1.46 | -1.44 | -0.74 | -0.97 | -0.97 | -1.05 | -1.23 | -1.11 | -1.17 | -0.93 | -1.05 | -0.72 | -0.22 | -0.38 | 0.3125 | 0.135 | 0.6525 | 0.200139 | 0.576566 | 0.04794 | 1 | 0.996511 | 0.658454 |
| Trub1 | Q8C0D0 | 0.04 | -0.11 | -2.33 | -2.04 | -1.1 | -0.28 | -2.2 | -1.4 | -0.29 | -0.25 | -1.45 | 0.31 | 0.61 | -1.1 | -2.07 | -2.33 | -0.135 | 0.69 | -0.1125 | 0.84058 | 0.312424 | 0.891025 | 1 | 0.996511 | 0.981841 |
| Try10 | Q792Z1 | -2.82 | -1.98 | -2.73 | -5.38 | -1.02 | -2.3 | -4.08 | -5.01 | -2.89 | -3.97 | -4.09 | -3.28 | -2.79 | -2.34 | -4.98 | -4.93 | 0.125 | -0.33 | -0.5325 | 0.904361 | 0.649222 | 0.563223 | 1 | 0.996511 | 0.920024 |
| Tsc1 | Q9EP53-3 | 0.7 | 0.87 | 0.47 | 0.31 | 0.4 | 0.67 | -0.31 | 0.02 | 1.05 | 0.74 | 0.34 | 0.54 | 1.2 | 0.4 | 0.31 | 0.09 | -0.3925 | 0.08 | -0.0875 | 0.153522 | 0.726896 | 0.762498 | 1 | 0.996511 | 0.963857 |
| Tsc2 | A0A2I3BPP | 0.27 | 0.45 | 0.22 | 0.18 | 0.3 | 0.2 | 0.02 | 0.05 | 0.28 | 0.49 | 0.28 | 0.32 | 0.14 | 0.3 | 0.06 | 0.08 | -0.1375 | 0.0625 | -0.135 | 0.388774 | 0.704828 | 0.453398 | 1 | 0.996511 | 0.885993 |
| Tsc22d1 | P62500-2 | -2.55 | -1.35 | -0.95 | -1.34 | -1.47 | -2.57 | -0.52 | -2.1 | -3.31 | -1.69 | -0.61 | -1.67 | -1.35 | -0.15 | -0.79 | -0.74 | -0.1175 | -0.2725 | 0.79 | 0.820613 | 0.652314 | 0.079883 | 1 | 0.996511 | 0.681818 |
| Tsc22d1 | D3Z0V7 | 1.01 | 0.21 | 0.38 | 0.57 | -0.12 | 0.52 | 1.53 | -0.07 | 0.36 | 2.02 | 0.44 | 0.71 | 0.94 | 0.23 | 0.09 | 1.06 | -0.0775 | 0.34 | 0.0375 | 0.845218 | 0.40984 | 0.903545 | 1 | 0.996511 | 0.985155 |
| Tsc22d2 | A0A0G2JFZ | 0.15 | 1.84 | 0.99 | 0.69 | 0.69 | 0.14 | 0.33 | 0.95 | -0.85 | 0.66 | 0.72 | 0.45 | -1.1 | 1.07 | 0.84 | 1.14 | -0.39 | -0.6725 | -0.43 | 0.316672 | 0.183657 | 0.47078 | 1 | 0.996511 | 0.891144 |
| Tsc22d3 | Q9Z2S7-3 | -0.17 | 0.18 | -0.14 | 0.73 | 0.4 | -0.17 | 0.67 | 0.38 | 0.06 | 1.06 | 0.44 | 0.17 | -0.06 | 1.14 | 0.32 | 0.5 | 0.17 | 0.2825 | 0.325 | 0.543959 | 0.373104 | 0.337989 | 1 | 0.996511 | 0.847578 |
| Tsc22d4 | Q9EQN3 | 0.16 | 0.68 | 0.69 | 0.7 | 0.77 | 0.92 | 1.07 | 0.51 | 0.94 | 0.89 | 1.03 | 0.8 | 1.34 | 1.1 | 0.78 | 1 | 0.26 | 0.3575 | 0.4975 | 0.231511 | 0.093674 | 0.050464 | 1 | 0.996511 | 0.658454 |
| Tsen34 | H7BX14 | -2.18 | -2.54 | -2.67 | -3.45 | -2.64 | -2.78 | -3.27 | -3.44 | -2.52 | -2.03 | -3.21 | -2.21 | -2.39 | -2.74 | -3.85 | -2.52 | -0.3225 | 0.2175 | -0.165 | 0.330287 | 0.552833 | 0.68849 | 1 | 0.996511 | 0.955729 |
| Tsfm | Q9CZR8 | 0.67 | 0.53 | -0.1 | -0.05 | 0.17 | 0.44 | -0.24 | -0.32 | 0.33 | 0.02 | -0.09 | 0.54 | 0.18 | 0.41 | -0.55 | -0.48 | -0.25 | -0.0625 | -0.3725 | 0.369009 | 0.811987 | 0.259152 | 1 | 0.996511 | 0.812074 |
| Tsg101 | Q61187 | 0.73 | 0.34 | 0.69 | 0.75 | 0.32 | 0.73 | 0.53 | 0.8 | 0.97 | 0.85 | 0.69 | 1.02 | 0.86 | 0.45 | 0.1 | 0.19 | -0.0325 | 0.255 | -0.2275 | 0.861323 | 0.189469 | 0.349351 | 1 | 0.996511 | 0.85327 |
| Tsga10 | Q6NY15-2 | 1.42 | 1.39 | 0.77 | 0.31 | 0.3 | -0.02 | -0.82 | -0.68 | 1.36 | -0.5 | 0.07 | 0.81 | -0.11 | 1.1 | -0.7 | 0.62 | -1.2775 | -0.5375 | -0.745 | 0.006329 | 0.259892 | 0.126243 | 1 | 0.996511 | 0.721505 |
| Tshz3 | A0A0R4J01 | 0.39 | 1.64 | 2.03 | -0.33 | 1.06 | 1.03 | 0.81 | 1.27 | -4.31 | -1.69 | 1.5 | 0.69 | -1.81 | -4.89 | 1.11 | 1.06 | 0.11 | -1.885 | -2.065 | 0.829748 | 0.163072 | 0.154372 | 1 | 0.996511 | 0.750763 |
| Tsn | Q62348 | -0.44 | -0.42 | 0.18 | -0.08 | -0.97 | -0.83 | 0.04 | 0.03 | -0.58 | 0.03 | -0.35 | -0.69 | -1.4 | -0.43 | -0.56 | -0.67 | -0.2425 | -0.2075 | -0.575 | 0.436144 | 0.404405 | 0.067967 | 1 | 0.996511 | 0.664936 |
| Tsnax | Q9QZE7 | -0.54 | -0.57 | -0.57 | -0.49 | -0.9 | -0.57 | -0.26 | -0.35 | -0.72 | -0.55 | -0.63 | -0.77 | -1.18 | -0.66 | -0.88 | -0.77 | 2.25E-02 | -0.125 | -0.33 | 0.903281 | 0.432546 | 0.110453 | 1 | 0.996511 | 0.711551 |
| Tspan13 | Q9D8C2 | -3.58 | -4.38 | -4.66 | -3.73 | -4.09 | -2.9 | -4.47 | -3.8 | -4.59 | -1.97 | -3.47 | -3.16 | -3.52 | -4.81 | -3.41 | -3.03 | 0.2725 | 0.79 | 0.395 | 0.499994 | 0.176501 | 0.381649 | 1 | 0.996511 | 0.860984 |
| Tspan14 | Q8QZY6 | -1.12 | -0.3 | 0.37 | 0.48 | 0.49 | -0.48 | 0.49 | 0.43 | -0.97 | 0.93 | 0.53 | -0.94 | -0.72 | -0.21 | 0.31 | 0.43 | 0.375 | 0.03 | 0.095 | 0.374805 | 0.95774 | 0.825704 | 1 | 0.996755 | 0.973155 |
| Tspan15 | F7BWT7 | 0.49 | 0.95 | 0.56 | 0 | 1.16 | 0.17 | 1.1 | -0.43 | -0.03 | -0.47 | 0.1 | 0.74 | -0.03 | 0.55 | 0.48 | 0.38 | -2.4E-16 | -0.415 | -0.155 | 1 | 0.214265 | 0.558321 | 1 | 0.996511 | 0.91747 |
| Tspan2 | Q9D1X8 | 3.81 | 2.72 | 3.39 | 1.51 | 3.16 | 3.46 | 2.81 | 2.2 | 0.22 | 0.18 | 2.92 | 3.22 | 0.59 | -0.37 | 3.15 | 3.7 | 0.05 | -1.2225 | -1.09 | 0.923979 | 0.186775 | 0.286987 | 1 | 0.996511 | 0.824333 |
| Tspan3 | Q9QY33 | -1.12 | -0.19 | -1.01 | -1.54 | 0.87 | -0.48 | -0.46 | -1 | -1.73 | -1.94 | -0.85 | -0.67 | -0.48 | -3.71 | -0.72 | -0.92 | 0.6975 | -0.3325 | -0.4925 | 0.151565 | 0.420029 | 0.505833 | 1 | 0.996511 | 0.901832 |
| Tspan31 | Q9CQ88 | -0.94 | -2.86 | -2.12 | -3.1 | -1.16 | -2.62 | -1.8 | -2.99 | -1.63 | -2.54 | -3.22 | -1.28 | 1.01 | -2.22 | -1.91 | -1.81 | 0.1125 | 0.0875 | 1.0225 | 0.846516 | 0.884113 | 0.226369 | 1 | 0.996511 | 0.787447 |
| Tspan33 | Q8R3S2-2 | -0.82 | 0.6 | -0.64 | 0.08 | 0.39 | 0.03 | 0.54 | -0.09 | -0.04 | -1.98 | 0.1 | -0.77 | -0.13 | 0.14 | -0.59 | 0.7 | 0.4125 | -0.4775 | 0.225 | 0.254394 | 0.383079 | 0.584673 | 1 | 0.996511 | 0.927734 |
| Tspan4 | A0A668KLI | -1.53 | -1.28 | -2.34 | -2.34 | 0.23 | -0.29 | -2.89 | -3.03 | -1.83 | 0.5 | -3.26 | -1.4 | -0.73 | -1.03 | -0.69 | -1.53 | 0.3775 | 0.375 | 0.8775 | 0.642262 | 0.61756 | 0.027055 | 1 | 0.996511 | 0.635993 |
| Tspan5 | D3Z641 | 0.87 | 1.31 | 0.08 | -0.08 | 2.15 | 1.12 | 0.09 | -0.13 | 0.9 | 1.28 | -0.02 | 1.11 | 0.78 | 2.75 | 0.68 | 0.03 | 0.2625 | 0.2725 | 0.515 | 0.644684 | 0.518177 | 0.412499 | 1 | 0.996511 | 0.874811 |
| Tspan6 | Q99L96 | -1.22 | -1.38 | -1.42 | -0.9 | -1.46 | -1.04 | -0.22 | -0.86 | -1.12 | -0.6 | -0.21 | -0.9 | -0.56 | -1.49 | -0.28 | -0.26 | 0.335 | 0.5225 | 0.5825 | 0.259285 | 0.064249 | 0.095263 | 1 | 0.996511 | 0.698156 |
| Tspan7 | Q62283 | 1.43 | 1.23 | 0.84 | 0.48 | 0.97 | 1.74 | 0.78 | 0.76 | 0.85 | 0.85 | 1.03 | 1.53 | 1.03 | 0.17 | 0.7 | 0.76 | 0.0675 | 0.07 | -0.33 | 0.826725 | 0.801375 | 0.277484 | 1 | 0.996511 | 0.822084 |
| Tspan8 | Q8R3G9 | 0.56 | 0.23 | -0.28 | 0.65 | -0.16 | 0.78 | -1.17 | -0.79 | -0.25 | 0.87 | 0.86 | -0.4 | 0.57 | -0.05 | 0.5 | 0.39 | -0.625 | -0.02 | 0.0625 | 0.182901 | 0.958859 | 0.820204 | 1 | 0.996755 | 0.972768 |
| Tspan9 | Q8BJU2 | -0.5 | -0.22 | 0 | 0.48 | 0.11 | -0.71 | 0.76 | 0.04 | -0.8 | 0.18 | 0.46 | -0.14 | -0.44 | -0.7 | 0.3 | 0.27 | 0.11 | -0.015 | -0.0825 | 0.754425 | 0.964463 | 0.80273 | 1 | 0.996755 | 0.971091 |
| Tspoap1 | Q5NCP6 | -1.79 | -1.14 | -0.31 | -0.53 | -0.63 | -1.76 | 0.05 | -0.32 | 0.34 | 0.06 | -0.34 | -1.23 | 0.72 | 0.04 | 0.07 | 0.38 | 0.2775 | 0.65 | 1.245 | 0.56308 | 0.172918 | 0.007289 | 1 | 0.996511 | 0.552614 |
| Tspyl1 | O88852 | -1.37 | -1.58 | -3.42 | -5.07 | -1.52 | -0.35 | -3.32 | -4.63 | -1.35 | -6.21 | -1.23 | -0.92 | -0.75 | -0.9 | -2.47 | -6.4 | 0.405 | 0.4325 | 0.23 | 0.72572 | 0.752902 | 0.869567 | 1 | 0.996511 | 0.979293 |
| Tspyl4 | Q8VD63 | -1.41 | -1.37 | -1.09 | -1.27 | -1.35 | -1.54 | -1.66 | -1.49 | -1.37 | -0.98 | -1.18 | -1.55 | -1.29 | -1.42 | -1.8 | -1.96 | -0.225 | 0.015 | -0.3325 | 0.182205 | 0.938648 | 0.160069 | 1 | 0.996755 | 0.755194 |
| Tspyl5 | Q69ZB3 | -0.97 | -0.69 | -1.57 | -0.95 | -0.07 | -0.93 | -1.83 | -1.73 | -0.69 | -0.16 | -1.59 | -0.71 | 0.3 | 0.19 | -1.72 | -1.52 | -0.095 | 0.2575 | 0.3575 | 0.821849 | 0.461816 | 0.505289 | 1 | 0.996511 | 0.901215 |
| Tsr2 | Q8C8T8 | -0.82 | -0.61 | -0.98 | -0.72 | -0.38 | -1.17 | -0.78 | -1.01 | -0.85 | -0.33 | -0.9 | -0.92 | -0.2 | 0.2 | -1.07 | -0.96 | -0.0525 | 0.0325 | 0.275 | 0.806855 | 0.874876 | 0.401868 | 1 | 0.996511 | 0.870698 |
| Tsr2 | H7BWY8 | -0.8 | -1.03 | 1.21 | -0.07 | 0.08 | 0.11 | 0.08 | -1 | -1.01 | 0.64 | -0.69 | -1.47 | -1.24 | 0.42 | 0.23 | 2.01 | -0.01 | -0.46 | 0.5275 | 0.984838 | 0.465653 | 0.490331 | 1 | 0.996511 | 0.897531 |
| Tssc4 | Q9JHE7 | -0.91 | -1.58 | -0.88 | -0.91 | -2.48 | -1.21 | -0.81 | -0.7 | 0.45 | -0.82 | -0.65 | -1.17 | 1.16 | -2.12 | -0.15 | -0.71 | -0.23 | 0.5225 | 0.615 | 0.582968 | 0.188553 | 0.34888 | 1 | 0.996511 | 0.85278 |
| Tst | P52196 | 1.39 | 0.53 | 1.4 | 1.11 | 0.66 | 1.42 | 2.45 | 1.5 | 0.68 | 1.12 | 1.85 | 1.06 | 0.33 | 0.15 | 1.39 | 1.87 | 0.4 | 0.07 | -0.1725 | 0.327389 | 0.826342 | 0.69513 | 1 | 0.996511 | 0.955729 |
| Tstd3 | Q9D0B5 | 0.51 | 0.88 | -0.06 | 0.18 | 0.96 | 0.38 | 0.41 | -0.57 | 0.57 | 0.39 | 0.32 | 0.48 | 0.05 | 1.24 | -0.3 | -0.25 | -0.0825 | 0.0625 | -0.1925 | 0.819457 | 0.794105 | 0.631825 | 1 | 0.996511 | 0.941132 |
| Ttbk1 | A0A286YDI | 1.09 | 1.59 | 0.73 | 0.11 | 1.41 | 0.96 | 0.58 | 0.06 | 1.52 | 0.64 | 0.71 | 0.97 | 1.31 | 2.39 | 0.49 | 0.84 | -0.1275 | 0.08 | 0.3775 | 0.749906 | 0.824371 | 0.443776 | 1 | 0.996511 | 0.883549 |
| Ttbk1 | Q6PCN3 | 0.5 | 0.37 | 0.44 | 0.63 | 0.12 | 0.41 | 0.02 | 0.38 | 1.06 | 0.69 | 0.49 | 0.76 | 0.96 | 0.59 | 0.43 | -0.26 | -0.2525 | 0.265 | -0.055 | 0.155855 | 0.184351 | 0.844997 | 1 | 0.996511 | 0.976225 |
| Ttbk2 | A2AW15 | -0.5 | -0.27 | -0.44 | -0.16 | -0.49 | -0.45 | -0.4 | -0.18 | 0.26 | 0.24 | 0.13 | -0.24 | 0.09 | 0.78 | -0.45 | 1.15 | -0.0375 | 0.44 | 0.735 | 0.818454 | 0.046583 | 0.067235 | 1 | 0.996511 | 0.664936 |
| Ttc1 | Q91Z38 | -0.05 | 0.1 | 0.35 | 0.36 | 0.02 | 0.08 | 0.1 | 0.33 | 0.48 | 0.52 | 0.52 | 0.21 | 0.35 | 0.33 | 0.11 | 0.34 | -0.0575 | 0.2425 | 0.0925 | 0.738948 | 0.213988 | 0.631016 | 1 | 0.996511 | 0.941132 |
| Ttc13 | A0A338P6I | 0.56 | 0.93 | -0.59 | -1.15 | 1.38 | 0.45 | 0.18 | 0.56 | 0.27 | -2.89 | 0.61 | 0.42 | 0.07 | 0.43 | 0.2 | -0.72 | 0.705 | -0.335 | 0.0575 | 0.188774 | 0.701279 | 0.909888 | 1 | 0.996511 | 0.986458 |
| Ttc14 | A0A0G2JES | 0.62 | -0.37 | -0.02 | -0.24 | -0.41 | 0.13 | -0.1 | -0.13 | 0.39 | -0.77 | -0.35 | 0.17 | -0.16 | 0.14 | 0.23 | -0.54 | -0.125 | -0.1375 | -0.08 | 0.627708 | 0.684711 | 0.786513 | 1 | 0.996511 | 0.967771 |
| Ttc19 | Q8CC21 | -0.08 | -0.02 | 0.04 | -0.21 | 0.19 | 0.58 | -0.57 | -0.31 | 0.28 | -0.66 | 0.24 | 0.15 | 0.97 | -0.2 | -0.04 | 0.19 | 0.04 | 0.07 | 0.2975 | 0.881498 | 0.781056 | 0.310033 | 1 | 0.996511 | 0.834523 |
| Ttc21b | Q0HA38 | 0.42 | 0.88 | 0.08 | -0.1 | 1.17 | -0.1 | 0.25 | 0.16 | 0.64 | 0.26 | -0.13 | 0.53 | 0.09 | 0.77 | -0.25 | 0.39 | 0.05 | 0.005 | -0.07 | 0.882764 | 0.986022 | 0.823806 | 1 | 0.998892 | 0.972768 |
| Ttc26 | Q8BS45 | -0.78 | -0.47 | -0.64 | -0.27 | 0.05 | -1.36 | -0.16 | -0.61 | -0.95 | -0.85 | -0.49 | -0.7 | -0.87 | -0.69 | -0.51 | -0.9 | 0.02 | -0.2075 | -0.2025 | 0.950569 | 0.312565 | 0.336866 | 1 | 0.996511 | 0.847578 |
| Ttc27 | Q8CD92-2 | -0.82 | -0.44 | -0.74 | -0.68 | -0.44 | -0.84 | -0.24 | -0.62 | -1.46 | -0.77 | -0.49 | -0.84 | -1.33 | -0.86 | -0.58 | -0.4 | 0.135 | -2.20E-01 | -0.1225 | 0.484575 | 0.381415 | 0.628417 | 1 | 0.996511 | 0.940342 |
| Ttc3 | G5E8T2 | -2.1 | -1.81 | -1.14 | -1.18 | -1.92 | -2 | -1.09 | -1.39 | -1.62 | -0.78 | -1.19 | -1.81 | -0.99 | -1.78 | -0.51 | -0.88 | -0.0425 | 0.2075 | 0.5175 | 0.892787 | 0.532453 | 0.16863 | 1 | 0.996511 | 0.759484 |
| Ttc30b | Q9CY00 | -0.56 | -0.44 | 0.02 | -0.24 | -1.27 | -0.43 | -0.74 | -0.74 | 0.99 | 1.13 | -0.51 | -0.48 | 0.85 | 0.11 | -0.34 | -0.23 | -0.49 | 0.5875 | 0.4025 | 0.061693 | 0.20561 | 0.212677 | 1 | 0.996511 | 0.786931 |
| Ttc33 | D3Z767 | -4.49 | -4.15 | -3.04 | -3.85 | -3.77 | -4.47 | -3.3 | -3.28 | -4.31 | -1.3 | -2.81 | -3.98 | -4.34 | -3.73 | -3.52 | -3.27 | 0.1775 | 0.7825 | 0.1675 | 0.65402 | 0.269044 | 0.657508 | 1 | 0.996511 | 0.948659 |
| Ttc37 | F8VPK0 | -0.05 | 0.05 | -0.1 | -0.2 | 0.2 | -0.21 | -0.29 | -0.28 | -0.09 | 0.2 | -0.29 | -0.14 | -0.36 | 0.3 | -0.31 | -0.01 | -0.07 | -0.005 | -0.02 | 0.691572 | 0.977897 | 0.925994 | 1 | 0.998381 | 0.988944 |
| Ttc38 | A3KMP2 | -0.64 | -0.96 | -0.58 | -0.29 | -0.77 | -0.58 | 0.19 | -0.28 | -0.64 | -0.22 | -0.53 | -0.84 | -0.85 | -1.23 | -0.62 | -0.4 | 0.2575 | 0.06 | -0.1575 | 0.334937 | 0.789058 | 0.54185 | 1 | 0.996511 | 0.912212 |
| Ttc39b | Q8BYY4 | -1.8 | -1.52 | -0.59 | -0.67 | -1.51 | -1.64 | -1.22 | -1.13 | -1.83 | -0.2 | -0.3 | -1.45 | -2.03 | -1.39 | -0.95 | -0.85 | -0.23 | 0.2 | -0.16 | 0.477778 | 0.675765 | 0.684481 | 1 | 0.996511 | 0.955729 |
| Ttc39c | A0A3Q4EG | 0.57 | 0.42 | 0.19 | 0.01 | 0.23 | 0.65 | 0.38 | 0.14 | -0.26 | -0.16 | -0.05 | 0.45 | -0.45 | 0.07 | 0.15 | 0.23 | 0.0525 | -0.3025 | -0.2975 | 0.792886 | 0.213987 | 0.231068 | 1 | 0.996511 | 0.788863 |
| Ttc5 | Q99LG4 | -0.23 | -0.33 | 0.41 | -1 | 0.01 | -0.48 | -0.69 | -0.88 | -0.67 | -0.46 | -0.71 | 0.1 | -0.49 | -0.3 | 0.83 | -0.77 | -0.2225 | -0.1475 | 0.105 | 0.512752 | 0.665164 | 0.808189 | 1 | 0.996511 | 0.971953 |
| Ttc7b | A0A1Y7VL4 | 0.84 | 1.01 | 0.8 | 0.76 | 1.4 | 0.85 | 0.8 | 0.71 | 0.66 | 0.38 | 0.75 | 0.86 | 0.37 | 1.25 | 0.56 | 0.79 | 0.0875 | -0.19 | -0.11 | 0.661778 | 0.310516 | 0.64538 | 1 | 0.996511 | 0.944749 |
| Ttc8 | Q8VD72 | -0.16 | 0.6 | 1.53 | 1.54 | -0.47 | -0.21 | 0.88 | 1.04 | -0.7 | 1.76 | 1.31 | -0.29 | -0.15 | -0.9 | 1.4 | 0.45 | -0.5675 | -0.3575 | -0.6775 | 0.287737 | 0.592095 | 0.264667 | 1 | 0.996511 | 0.816857 |
| Ttc9 | Q3V038 | -0.41 | -0.65 | -0.77 | -0.48 | -0.74 | -0.14 | -0.54 | -0.61 | 0.42 | -0.77 | -0.48 | -0.14 | 0.69 | -0.85 | -0.58 | -0.96 | 0.07 | 0.335 | 0.1525 | 0.714845 | 0.251848 | 0.688526 | 1 | 0.996511 | 0.955729 |
| Ttc9b | Q9D6E4 | -0.81 | -0.58 | -0.14 | -0.08 | -0.58 | -1.04 | -0.23 | -0.49 | 0.17 | 0.51 | 0.18 | -0.43 | 0.94 | -0.05 | -0.47 | -0.48 | -0.1825 | 0.51 | 0.3875 | 0.479708 | 0.092672 | 0.310055 | 1 | 0.996511 | 0.834523 |
| Ttc9c | Q810A3 | -0.12 | -0.16 | -2.94 | -0.67 | -0.08 | -0.05 | -0.52 | -0.95 | 0.28 | -0.45 | -0.49 | -0.44 | -0.42 | -0.14 | -1.92 | -1.93 | 0.5725 | 0.6975 | -0.13 | 0.380385 | 0.287925 | 0.861033 | 1 | 0.996511 | 0.977204 |
| Tti2 | A0A1B0GS. | 0.33 | 0.79 | -0.41 | -1.09 | 0.53 | 0.18 | -0.48 | -0.5 | -0.11 | -1.93 | -0.56 | 0.16 | -0.4 | 0.44 | -1.04 | -0.1 | 0.0275 | -0.515 | -0.18 | 0.951415 | 0.379454 | 0.710383 | 1 | 0.996511 | 0.957819 |
| Ttl | P38585 | 0.16 | -0.7 | 0.22 | 0.25 | -0.45 | 0.17 | -0.25 | 0.33 | -0.19 | 0.07 | 0.28 | -0.28 | -0.44 | -0.21 | -0.54 | -0.44 | -0.0325 | -0.0125 | -0.39 | 0.911083 | 0.963727 | 0.166702 | 1 | 0.996755 | 0.759484 |
| Ttll1 | Q91V51 | -0.52 | -0.03 | -0.28 | -0.4 | 0.23 | 0.71 | 0.12 | 0.24 | -0.59 | -0.23 | 0.46 | -0.62 | -0.51 | -0.1 | -0.19 | 0.06 | 0.6325 | 0.0625 | 0.1225 | 0.011309 | 0.825418 | 0.571142 | 1 | 0.996511 | 0.924332 |
| Ttll12 | Q3UDE2 | -0.32 | -0.58 | -0.59 | -0.57 | -0.39 | -0.26 | -0.12 | -0.36 | -0.16 | -1.11 | -0.16 | -0.14 | -0.46 | 0.15 | -0.55 | -0.2 | 0.2325 | 0.1225 | 0.25 | 0.162138 | 0.646098 | 0.274891 | 1 | 0.996511 | 0.822084 |
| Ttll7 | E9Q6C6 | 0.01 | 0.16 | 0.09 | 0.24 | -0.12 | -0.35 | -0.12 | -0.01 | 1.09 | 0.84 | 0.03 | -0.06 | 1.5 | 0.54 | 0.63 | 0.07 | -0.275 | 0.35 | 0.56 | 0.103859 | 0.260202 | 0.099464 | 1 | 0.996511 | 0.698156 |
| Ttn | A0A5K1VV | -0.05 | -0.23 | 0.39 | 0.79 | -0.69 | -0.29 | 0.06 | 0.55 | 0.01 | 1.13 | 0.57 | -0.16 | 0.28 | -0.72 | 0.57 | -0.03 | -0.3175 | 0.1625 | -0.2 | 0.359769 | 0.654857 | 0.577049 | 1 | 0.996511 | 0.927198 |
| Ttpa | Q8BWP5 | 0.99 | 1.9 | -0.15 | -1.12 | 1.21 | -0.15 | 0.05 | -1.38 | 1.54 | 0.49 | -1.04 | 0.44 | 0.99 | 0.86 | 0.15 | -1.34 | -0.4725 | -0.0475 | -0.24 | 0.540707 | 0.950507 | 0.754572 | 1 | 0.996755 | 0.962333 |
| Ttpal | Q9D3D0 | -0.07 | -0.17 | 0.29 | 0.06 | 0.37 | 0.11 | 0.17 | 0.49 | -0.22 | -0.37 | 0.13 | -0.17 | -0.6 | 0.22 | -0.63 | 0.22 | 0.2575 | -0.185 | -0.225 | 0.174737 | 0.359073 | 0.431851 | 1 | 0.996511 | 0.879165 |
| Ttr | P07309 | -3.72 | -4.68 | -3.25 | -3.05 | -3.66 | -3.69 | -3.45 | -3.41 | -5.32 | -2.12 | -3.95 | -3.67 | -4.77 | -3.8 | -3.57 | -3.47 | 0.1225 | -0.09 | -0.2275 | 0.730826 | 0.89505 | 0.61134 | 1 | 0.996511 | 0.935047 |
| Ttyh1 | Q9D3A9 | 0.73 | 1.26 | 0.89 | 1.13 | 1.43 | 0.83 | 0.77 | 1.24 | 0.97 | 1.03 | 1.01 | 1.01 | 0.46 | 1.85 | 0.49 | 0.82 | 0.065 | 0.0025 | -0.0975 | 0.769879 | 0.989084 | 0.777985 | 1 | 0.999285 | 0.967289 |
| Ttyh2 | Q3TH73 | 1.25 | 1.28 | 0.69 | 0.55 | 1.47 | 0.53 | 0.62 | 0.85 | 1.09 | 0.92 | 0.95 | 0.92 | 1.5 | 1.69 | 1.56 | 1.15 | -0.075 | 0.0275 | 0.5325 | 0.792735 | 0.903368 | 0.060756 | 1 | 0.996609 | 0.661738 |
| Ttyh3 | Q6P5F7 | 2.19 | -0.06 | 0.93 | 0.7 | 0.56 | 1.92 | 0.39 | 0.38 | 1.07 | 1.34 | 0.87 | 2.44 | 1.06 | 0.88 | 0.8 | 0.97 | -0.1275 | 0.49 | -0.0125 | 0.815893 | 0.37547 | 0.977592 | 1 | 0.996511 | 0.993727 |
| Tub | P50586 | 0.14 | -0.21 | 0.35 | 0.38 | -0.16 | 0.22 | -0.05 | 0.16 | 0.6 | 0.42 | 0.13 | -0.24 | 0.8 | -0.45 | -0.17 | -0.29 | -0.1225 | 0.0625 | -0.1925 | 0.538679 | 0.803637 | 0.550883 | 1 | 0.996511 | 0.913801 |
| Tuba1a | P68369 | 1.29 | -0.06 | 0.43 | -0.09 | 0.02 | 1.13 | 0.94 | 0.19 | 0.18 | 0.24 | 0.48 | 1.32 | -0.26 | 0.11 | 0.35 | 0.59 | 0.1775 | 0.1625 | -0.195 | 0.657774 | 0.684248 | 0.594683 | 1 | 0.996511 | 0.929843 |
| Tuba1b | P05213 | -2.85 | -4.19 | -1.63 | -2.03 | -3.75 | -2.43 | -1.34 | -1.82 | -4.24 | -2.13 | -1.45 | -2.92 | -4.26 | -3.64 | -2.23 | -1.8 | 0.34 | -0.01 | -0.3075 | 0.627862 | 0.989302 | 0.676005 | 1 | 0.999285 | 0.952944 |
| Tuba3b | P05214 | -1.22 | -0.99 | -0.64 | -1.53 | -0.9 | -2.26 | -2.31 | -1.59 | -1.76 | -0.31 | -2.17 | -1.63 | -1.51 | 0.41 | -2.72 | -2.59 | -0.67 | -0.3725 | -0.5075 | 0.09365 | 0.38782 | 0.462297 | 1 | 0.996511 | 0.888262 |
| Tuba4a | A0A0A0MC | -0.85 | -0.55 | 0.85 | 0.41 | -0.41 | -1.14 | 0.25 | 0.54 | -0.61 | 1.46 | 0.42 | -0.59 | -1.21 | 0.16 | 0.2 | 0.83 | -0.155 | 0.205 | 0.03 | 0.758953 | 0.72559 | 0.955835 | 1 | 0.996511 | 0.992535 |
| Tuba8 | Q9JJZ2 | -1.24 | -1.14 | -0.4 | -1.18 | -0.94 | -2.11 | -0.86 | -1.32 | -2.24 | -1.18 | -1.14 | -1.54 | -1.93 | -1.03 | -1.49 | -1.08 | -0.3175 | -0.535 | -0.3925 | 0.358119 | 0.124393 | 0.214146 | 1 | 0.996511 | 0.786931 |
| Tubal3 | Q3UX10 | -7.32 | -6.61 | -5.86 | -7.26 | -5.37 | -7.56 | -5.26 | -6.72 | -10.45 | -7.39 | -5.53 | -6.77 | -8.32 | -9.3 | -6.67 | -6.74 | 0.535 | -0.7725 | -0.995 | 0.379324 | 0.442829 | 0.157465 | 1 | 0.996511 | 0.751678 |
| Tubb1 | A2AQ07 | -3.38 | -2.83 | -2.96 | -1.79 | -3.64 | -2.98 | -1.7 | -1.82 | -1.11 | -2.88 | -1.84 | -3.27 | -1.82 | -0.41 | -2.91 | -2.47 | 0.205 | 0.465 | 0.8375 | 0.700044 | 0.4081 | 0.179414 | 1 | 0.996511 | 0.775542 |
| Tubb2a | Q7TMM9 | -0.96 | -0.73 | -0.85 | -0.69 | -0.87 | -0.95 | -0.66 | -0.6 | -1.67 | -1.13 | -0.58 | -0.39 | -1.99 | -1.11 | -1.14 | -1.06 | 0.0375 | -0.135 | -0.5175 | 0.818017 | 0.65539 | 0.072704 | 1 | 0.996511 | 0.668066 |
| Tubb2b | Q9CWF2 | -0.09 | 0.79 | 0.38 | 0.36 | -0.34 | -0.11 | 1.3 | 1.26 | -0.47 | 0.03 | 1.45 | 0.77 | -0.19 | -0.86 | 0.16 | -0.18 | 0.1675 | 0.085 | -0.6275 | 0.704969 | 0.844239 | 0.057976 | 1 | 0.996511 | 0.661738 |
| Tubb3 | Q9ERD7 | -1.97 | -2.14 | -1.96 | -1.69 | -2.11 | -1.67 | -1.78 | -1.61 | -1.93 | -1.67 | -1.64 | -1.61 | -1.95 | -2.12 | -2.37 | -2.07 | 0.1475 | 0.2275 | -0.1875 | 0.436841 | 0.233365 | 0.355334 | 1 | 0.996511 | 0.855029 |
| Tubb4a | Q9D6F9 | -1.35 | -0.56 | -0.19 | -0.27 | -0.96 | -1.32 | -0.78 | -0.45 | -1.03 | -0.51 | -0.49 | -1.01 | -1.09 | -1.46 | -0.93 | -0.91 | -0.285 | -0.1675 | -0.505 | 0.377132 | 0.589634 | 0.124898 | 1 | 0.996511 | 0.721448 |
| Tubb4b | P68372 | -1.22 | -1.19 | -1.26 | -0.68 | -1.49 | -1.1 | -1.01 | -0.76 | -1.29 | -0.66 | -1.03 | -0.67 | -1.69 | -1.44 | -1.4 | -1.22 | -0.0025 | 0.175 | -0.35 | 0.991137 | 0.463137 | 0.13671 | 1 | 0.996511 | 0.732318 |
| Tubb5 | P99024 | -1.48 | -0.94 | -2.63 | -2.04 | -1.63 | -1.56 | -2.16 | -2.05 | -0.44 | -3.1 | -2.21 | -0.77 | -1.39 | -0.38 | -3.07 | -3.15 | -0.0775 | 0.1425 | -0.225 | 0.835899 | 0.828481 | 0.746852 | 1 | 0.996511 | 0.960533 |
| Tubb6 | Q922F4 | -3.33 | -2.91 | -4.63 | -4.71 | -2.58 | -3.49 | -4.78 | -4.71 | -2.65 | 2.4 | -4.48 | -2.73 | -2.49 | -4.08 | -5.07 | -5.36 | 0.005 | 2.03 | -0.355 | 0.993702 | 0.168654 | 0.623005 | 1 | 0.996511 | 0.938628 |
| Tubg1 | P83887 | 0.71 | 0.17 | 0.72 | 0.3 | 0.68 | 0.62 | 0.73 | 0.65 | -0.17 | 0.25 | 0.74 | 0.9 | -0.25 | 0.54 | 0.79 | 0.63 | 0.195 | -0.045 | -0.0475 | 0.307436 | 0.876665 | 0.8692 | 1 | 0.996511 | 0.979293 |
| Tubg2 | Q8VCK3 | -0.09 | 0.28 | 0.5 | 0.77 | 0.07 | 0.08 | 0.24 | 0.66 | 0.69 | 1.21 | 0.61 | -0.4 | 0.36 | 0.37 | 0.49 | 1.06 | -0.1025 | 0.1625 | 0.205 | 0.67443 | 0.662592 | 0.456139 | 1 | 0.996511 | 0.887169 |
| Tubgcp2 | A0A1B0GR | 0.82 | 0.71 | 1.31 | 1.16 | 0.79 | 0.61 | 1.11 | 1.08 | 0.55 | 0.61 | 0.7 | 0.53 | 0.19 | 0.74 | 0.93 | 2.22 | -0.1025 | -0.4025 | 0.02 | 0.631075 | 0.067715 | 0.962894 | 1 | 0.996511 | 0.9927 |
| Tubgcp3 | P58854 | 0.38 | 0.7 | 0.74 | -0.02 | 0.58 | 0.75 | 0.92 | 0.46 | -0.44 | -0.46 | 0.78 | 0.46 | -0.84 | 0.22 | 0.2 | 0.74 | 0.2275 | -0.365 | -0.37 | 0.32542 | 0.317203 | 0.329045 | 1 | 0.996511 | 0.844943 |
| Tubgcp4 | Q9D4F8 | -0.67 | -0.18 | -0.66 | -0.61 | -0.08 | -0.23 | -0.13 | -0.43 | -0.22 | -0.99 | -0.43 | -0.59 | -1.19 | -0.2 | -0.75 | -0.19 | 0.3125 | -0.0275 | -0.0525 | 0.115997 | 0.905582 | 0.854481 | 1 | 0.996744 | 0.976225 |
| Tubgcp5 | Q8BKN5-3 | -0.34 | -0.1 | 0.07 | 0.18 | 0.01 | -0.36 | -0.24 | 0.09 | 0.1 | 0.55 | 0.66 | -0.51 | 1.05 | 0.33 | 0.99 | 0.21 | -0.0775 | 0.2475 | 0.6925 | 0.687756 | 0.41289 | 0.028783 | 1 | 0.996511 | 0.637553 |
| Tubgcp6 | E9Q6P4 | -0.65 | -0.67 | 0.53 | -0.1 | -0.7 | -1 | 0.11 | -0.21 | -0.65 | 0.87 | -0.13 | -0.78 | -0.83 | -0.28 | -0.2 | -0.69 | -0.2275 | 0.05 | -0.2775 | 0.532485 | 0.910047 | 0.405739 | 1 | 0.996744 | 0.873383 |
| Tufm | Q8BFR5 | 0.88 | 0.94 | 0.24 | 0.67 | 0.59 | 0.78 | 0.18 | 0.46 | 0.61 | 0.4 | 0.31 | 0.66 | 0.2 | 0.89 | -0.11 | 0.2 | -0.18 | -1.88E-01 | -0.3875 | 0.43086 | 0.39942 | 0.194816 | 1 | 0.996511 | 0.779437 |
| Tulp3 | O88413 | 0.55 | 0.57 | 0.54 | 0.66 | 0.73 | 0.82 | 1.04 | 0.95 | -0.38 | 0.58 | 0.78 | 0.6 | -0.73 | -0.31 | 0.6 | 0.72 | 0.305 | -0.185 | -0.51 | 0.068052 | 0.509542 | 0.170709 | 1 | 0.996511 | 0.762322 |
| Tulp4 | Q9JIL5 | 0.33 | 0.63 | 0.68 | 0.91 | 0.75 | 0.41 | 0.42 | 0.66 | 0.53 | 0.9 | 0.64 | 0.3 | 0.39 | 0.96 | 0.95 | 0.73 | -0.0775 | -0.045 | 0.12 | 0.680263 | 0.833081 | 0.59838 | 1 | 0.996511 | 0.930093 |
| Tusc2 | Q9WVF8 | 0.92 | 1 | -0.06 | 0.28 | 1.59 | 0.75 | 0.34 | 0.62 | 0.15 | 0.28 | 0.31 | 1.09 | -0.08 | 0.48 | 0.08 | 0.2 | 0.29 | -0.0775 | -0.365 | 0.424176 | 0.814837 | 0.237967 | 1 | 0.996511 | 0.792722 |
| Tusc3 | Q8BTV1 | -0.93 | 0.05 | -0.63 | -0.72 | 0.95 | -0.69 | 0.24 | -0.21 | -1.62 | -1.26 | -0.69 | -0.88 | -1.01 | -1.6 | 0.26 | 0.26 | 0.63 | -0.555 | 0.035 | 0.130532 | 0.093307 | 0.941926 | 1 | 0.996511 | 0.991085 |
| Tut1 | Q8R3F9 | -6.04 | -7.21 | -1.66 | -1.64 | -5.74 | -7.37 | -2.1 | -2.67 | -4.4 | -1.1 | -1.38 | -5.99 | -1.83 | -6.22 | -1.59 | -0.94 | -0.3325 | 0.92 | 1.4925 | 0.846196 | 0.586364 | 0.38458 | 1 | 0.996511 | 0.860984 |
| Tut4 | B2RX14 | -0.41 | -0.5 | -0.45 | -0.56 | -0.75 | 0.07 | -0.93 | -0.99 | -1.87 | -1.06 | -0.65 | -0.44 | -1.65 | -2.41 | -2.07 | -0.66 | -0.17 | -0.525 | -1.2175 | 0.514028 | 0.126399 | 0.009353 | 1 | 0.996511 | 0.563386 |
| Tut7 | E9PUA2 | -1.86 | -1.46 | -0.73 | -0.7 | -1.21 | -1.6 | -0.78 | -0.87 | -1.01 | -0.16 | -0.27 | -1.41 | -0.87 | -0.68 | -0.57 | -0.46 | 0.0725 | 0.475 | 0.5425 | 0.826515 | 0.249999 | 0.105755 | 1 | 0.996511 | 0.703417 |
| Twf1 | Q91YR1 | -0.36 | -0.56 | 0.42 | 0.49 | 0.01 | -0.34 | 0.74 | 0.68 | -0.95 | 0.19 | 0.52 | -0.7 | -1.03 | -0.18 | 0.18 | 0.38 | 0.275 | -0.2325 | -0.16 | 0.451748 | 0.582307 | 0.687855 | 1 | 0.996511 | 0.955729 |
| Twf2 | Q9Z0P5 | 0.46 | 0.74 | 0.52 | 0.12 | 1.29 | 0.66 | 0.54 | 0.6 | -0.26 | -0.41 | 0.59 | 0.34 | -0.34 | 0.2 | 0.36 | 0.49 | 0.3125 | -0.395 | -0.2825 | 0.205023 | 0.185925 | 0.284272 | 1 | 0.996511 | 0.822669 |
| Txlna | Q6PAM1 | -2.2 | -2.11 | -1.38 | -1.74 | -2.64 | -1.96 | -2.25 | -1.96 | -0.2 | -1.6 | -1.43 | -1.88 | 0.37 | -1.4 | -1.49 | -2.22 | -0.345 | 0.58 | 0.6725 | 0.202807 | 0.17021 | 0.231888 | 1 | 0.996511 | 0.788863 |
| Txlng | Q8BHN1-3 | -2.07 | -1.76 | -0.65 | -1.69 | -1.68 | -2.52 | -1.5 | -0.33 | -1.56 | 0.67 | -0.93 | -1.91 | -1.23 | -0.86 | -0.52 | -0.13 | 0.035 | 0.61 | 0.8575 | 0.944578 | 0.320891 | 0.046672 | 1 | 0.996511 | 0.658454 |

| | | | | | | | | | | | | | | | | | | | | | | | | | | | |
|---|---|---|---|---|---|---|---|---|---|---|---|---|---|---|---|---|---|---|---|---|---|---|---|---|---|---|---|
| Txn | P10639 | -1.08 | -1.44 | -0.55 | -0.56 | -0.87 | -0.74 | -0.04 | -0.69 | -1.63 | -0.25 | -0.63 | -1.01 | -1.48 | -0.94 | -0.74 | -0.28 | 0.3225 | 0.0275 | 0.0475 | 0.277982 | 0.938455 | 0.886275 | | 1 | 0.996755 | 0.979767 |
| Txn2 | G3UZY2 | 0.12 | 0.78 | 1.83 | 2.36 | 0.54 | -0.02 | 1.25 | 2.16 | -0.06 | 1.39 | 2.04 | 0.19 | -0.79 | 0.84 | 0.99 | 1.36 | -0.29 | -0.3825 | -0.6725 | 0.646323 | 0.558311 | 0.304501 | | 1 | 0.996511 | 0.831662 |
| Txndc12 | Q9CQU0 | -0.7 | -0.22 | -0.41 | -0.75 | 0.02 | -0.39 | 0.04 | 0.25 | -0.89 | -1.21 | -0.33 | -0.44 | -0.96 | -0.26 | 0.67 | -0.2 | 0.5 | -0.1975 | 0.3325 | 0.039481 | 0.453433 | 0.359029 | | 1 | 0.996511 | 0.856645 |
| Txndc15 | Q6P6J9 | -0.53 | -0.25 | -0.49 | -0.32 | 0.27 | -0.47 | 0.39 | -0.36 | -0.52 | -0.85 | -0.67 | -0.37 | -0.29 | 0.38 | 0.34 | 0.52 | 0.355 | -0.205 | 0.635 | 0.168432 | 0.284078 | 0.022587 | | 1 | 0.996511 | 0.621926 |
| Txndc16 | Q7TN22 | -1.33 | -0.38 | -1.68 | -1.12 | -0.05 | -1.32 | -0.03 | -0.3 | -0.4 | -1.62 | -1.05 | -1.43 | -0.55 | -0.63 | -0.72 | -0.84 | 0.7025 | 0.0025 | 0.4425 | 0.098273 | 0.994635 | 0.160472 | | 1 | 0.999285 | 0.75577 |
| Txndc17 | Q9CQM5 | -1.52 | -1.02 | -1.09 | -1.34 | -2.07 | -1.11 | -1.85 | -2.39 | -1.39 | -0.93 | -1.79 | -1.44 | -0.37 | -0.98 | -1.73 | -1.58 | -0.6125 | -0.145 | 0.0775 | 0.063646 | 0.548075 | 0.816397 | | 1 | 0.996511 | 0.972768 |
| Txndc5 | Q91W90 | -0.72 | -0.53 | -0.81 | -0.66 | -0.19 | -0.86 | 0.41 | -0.54 | -0.99 | -0.46 | -0.68 | -0.59 | -0.36 | -0.77 | 0.94 | 0.04 | 0.385 | 1.96E-16 | 0.6425 | 0.194367 | 1 | 0.105051 | | 1 | 1 | 0.703417 |
| Txndc9 | Q9CQ79 | -0.97 | -0.65 | -0.55 | -0.41 | -0.44 | -0.67 | 0 | -0.2 | -0.98 | -0.32 | -0.19 | -0.85 | -0.72 | -0.57 | -0.55 | -0.09 | 0.3175 | 0.06 | 0.1625 | 0.163506 | 0.811009 | 0.480981 | | 1 | 0.996511 | 0.893906 |
| Txnip | Q8BG60-2 | -0.84 | -1.43 | 0.4 | -0.94 | -1.36 | -1.28 | 0.88 | -0.52 | -1.06 | -0.23 | -0.46 | -1.6 | -1.28 | -0.88 | -1.56 | -0.71 | 0.1325 | -0.135 | -0.405 | 0.823046 | 0.772172 | 0.343557 | | 1 | 0.996511 | 0.848473 |
| Txnl1 | A0A498WG | -0.19 | -0.3 | 0.12 | -0.55 | 0.03 | -0.15 | 0.57 | -0.03 | -0.95 | -1.1 | -0.08 | -0.12 | -0.68 | -0.43 | 0.07 | 0.41 | 0.335 | -0.3325 | 0.0725 | 0.171461 | 0.294956 | 0.806777 | | 1 | 0.996511 | 0.971715 |
| Txnl4a | P83877 | 1.64 | 0.42 | 0.87 | 0.36 | 0.8 | 1.4 | 1.32 | 1.27 | -0.23 | 0.15 | 1.11 | 1.13 | -0.48 | 0.19 | 1.22 | 1.56 | 0.375 | -0.2825 | -0.2 | 0.257757 | 0.515428 | 0.698793 | | 1 | 0.996511 | 0.955729 |
| Txnl4b | Q8BUH1 | -0.78 | 0.01 | 0.51 | -0.43 | -0.53 | -1.23 | -0.67 | -0.09 | -0.93 | 0.31 | -0.69 | -0.58 | -1.04 | -0.26 | -0.72 | 0.65 | -0.4575 | -0.3 | -0.17 | 0.214564 | 0.433378 | 0.698955 | | 1 | 0.996511 | 0.955729 |
| Txnrd1 | Q9JMH6 | -0.49 | -0.47 | -0.26 | -0.76 | -0.24 | -0.59 | -0.42 | -0.75 | -0.3 | -0.07 | -0.35 | -0.38 | -0.48 | 0 | -0.54 | -0.47 | -0.005 | 0.22 | 0.1225 | 0.978904 | 0.254839 | 0.573662 | | 1 | 0.996511 | 0.925747 |
| Txnrd2 | J3QMN4 | 0.02 | 0 | -0.33 | -0.12 | -0.06 | 0.25 | -0.35 | -0.33 | 0.05 | -0.59 | -0.22 | -0.01 | 0.21 | 0.5 | -0.65 | -0.74 | -0.015 | -0.085 | -0.0625 | 0.939181 | 0.685532 | 0.847353 | | 1 | 0.996511 | 0.976225 |
| Txnrd3 | Q99MD6 | 0.25 | 0.51 | -0.2 | -0.17 | 0.81 | 0.57 | -0.26 | 0.36 | 0.69 | 0.44 | -0.02 | 0.37 | 0.56 | 0.93 | -0.02 | -0.62 | 0.2725 | 0.2725 | 0.115 | 0.355859 | 0.291618 | 0.757785 | | 1 | 0.996511 | 0.962717 |
| Tyk2 | Q9R117 | -0.71 | -0.43 | 2.21 | 1.4 | -0.03 | -0.45 | 2.72 | 2.04 | 2.47 | 2.18 | 2.21 | -0.04 | 4.06 | 0.66 | 2.84 | 2.73 | 0.4525 | 1.0875 | 1.955 | 0.633039 | 0.212307 | 0.053725 | | 1 | 0.996511 | 0.661738 |
| Tyro3 | P55144-3 | 0.24 | 0.82 | 0.06 | 0.44 | 0.63 | 1.11 | 0.22 | 0.18 | 0.4 | 0.56 | 0.3 | 0.41 | 0.4 | 0.73 | 0.14 | 0.24 | 0.145 | 0.0275 | -0.0125 | 0.601613 | 0.897161 | 0.959179 | | 1 | 0.996511 | 0.992636 |
| Tyw1 | Q8BJM7 | -0.22 | -0.07 | -0.17 | -0.03 | 0.76 | -0.03 | -0.14 | -0.16 | -0.14 | 0.46 | -0.28 | 0.31 | -0.14 | 0.66 | -0.29 | 0.18 | 0.23 | 0.21 | 0.225 | 0.350103 | 0.352069 | 0.377424 | | 1 | 0.996511 | 0.860984 |
| Tyw3 | Q8BSA9 | -1.41 | -1.94 | -0.71 | -0.88 | -3.32 | -0.49 | -1.04 | -1.09 | 1.47 | 0.23 | -0.57 | -1.69 | 1.9 | -2.2 | -0.57 | -0.15 | -0.25 | 1.095 | 0.98 | 0.68989 | 0.123562 | 0.241136 | | 1 | 0.996511 | 0.796273 |
| Tyw5 | A2RSX7 | -0.02 | 0.04 | 0.95 | 0.49 | 0.12 | 0.21 | 1.4 | 1.47 | -1.02 | -0.05 | 1.59 | 0.1 | -1.22 | -0.41 | 0.72 | 0.99 | 0.435 | -0.21 | -0.345 | 0.300322 | 0.698307 | 0.512402 | | 1 | 0.996511 | 0.903993 |
| U2af1 | Q9D883 | -3.49 | -5.71 | -2.51 | -2.57 | -5.67 | -4.4 | -2.57 | -2.2 | -5.04 | -1.03 | -2.46 | -5.07 | -3.29 | -4.21 | -1.84 | -2.47 | -0.14 | 0.17 | 0.6175 | 0.887496 | 0.879175 | 0.457223 | | 1 | 0.996511 | 0.887902 |
| U2af1l4 | Q8BGJ9 | -1.07 | -1.34 | -0.8 | -1.42 | -1.76 | -1.06 | -0.84 | -0.81 | -0.14 | -0.34 | -1.24 | -1.1 | 0.96 | 0.8 | 0.36 | -1.02 | 0.04 | 0.4525 | 1.4325 | 0.88153 | 0.1699 | 0.010278 | | 1 | 0.996511 | 0.563386 |
| U2af2 | Q80XR5 | -1.31 | -2.88 | -1.62 | -1.39 | -2.21 | -2.15 | -1.28 | -1.58 | -1.47 | -0.04 | -1.39 | -2.42 | -1.42 | -1.59 | -0.79 | -1.37 | -0.005 | 0.47 | 0.5075 | 0.990131 | 0.413364 | 0.217604 | | 1 | 0.996511 | 0.787447 |
| U2surp | Q6NV83-2 | -1.35 | -2.4 | -1.96 | -2.04 | -2.21 | -2.06 | -1.79 | -1.53 | -2.8 | -1.63 | -2.32 | -2.77 | -1.77 | -1.64 | -1.06 | -2.09 | 0.04 | -0.4425 | 0.2975 | 0.882349 | 0.220019 | 0.356827 | | 1 | 0.996511 | 0.855416 |
| Uaca | A0A1L1STY | -2.41 | 0.09 | 1 | 0.71 | 0.94 | -0.55 | 0.74 | 0.44 | -0.58 | 0.83 | 0.77 | -1.23 | 0.15 | 0.63 | 1.14 | 1.56 | 0.545 | 0.1 | 1.0225 | 0.481004 | 0.904978 | 0.198263 | | 1 | 0.996744 | 0.78004 |
| Uap1 | A0A0R4J1F | -1.02 | -0.92 | -0.52 | -0.42 | -0.44 | -0.92 | -0.6 | -0.67 | -0.08 | 0.15 | -0.54 | -0.61 | -0.05 | 0.25 | -0.8 | -0.92 | 0.0625 | 0.45 | 0.34 | 0.763385 | 0.106053 | 0.311546 | | 1 | 0.996511 | 0.834523 |
| Uap1l1 | Q3TW96 | 0.03 | 0.07 | -0.55 | -0.57 | -0.24 | 0.04 | -0.87 | -0.3 | -0.1 | -0.39 | -0.59 | -0.09 | -0.22 | 0.06 | -0.6 | -0.58 | -0.0875 | -0.0375 | -0.08 | 0.743359 | 0.876319 | 0.762254 | | 1 | 0.996511 | 0.963682 |
| Uba1 | Q02053 | -0.52 | -0.39 | -0.28 | -0.26 | -0.65 | -0.76 | -0.43 | -0.28 | -0.5 | -0.32 | -0.32 | -0.48 | -0.83 | -0.14 | -0.77 | -0.87 | -0.1675 | -0.0425 | -0.29 | 0.347495 | 0.796062 | 0.224687 | | 1 | 0.996511 | 0.787447 |
| Uba2 | Q9Z1F9 | -4.65 | -5.12 | -4.79 | -6.25 | -4.73 | -5.11 | -4.99 | -5.45 | -4.78 | -2.78 | -5.07 | -4.12 | -3.89 | -3.77 | -5.11 | -5.64 | 0.1325 | 1.015 | 0.6 | 0.723527 | 0.105433 | 0.284981 | | 1 | 0.996511 | 0.822669 |
| Uba3 | Q8C878 | -1.1 | -1.26 | -0.87 | -0.98 | -1.17 | -1.43 | -0.96 | -0.94 | -2.11 | -0.78 | -0.99 | -1.17 | -2.15 | -1.27 | -1.32 | -1.42 | -0.0725 | -0.21 | -0.4875 | 0.694662 | 0.502028 | 0.081781 | | 1 | 0.996511 | 0.68865 |
| Uba5 | Q8VE47 | -0.93 | -0.7 | -0.7 | -0.76 | -0.77 | -0.66 | -0.64 | -0.63 | -0.39 | -0.9 | -0.47 | -0.82 | -0.17 | -1.14 | -0.22 | -0.8 | 0.0975 | 0.1275 | 0.19 | 0.509461 | 0.513241 | 0.48266 | | 1 | 0.996511 | 0.893925 |
| Uba6 | Q8C7R4 | -0.83 | -0.63 | -0.69 | -0.52 | -0.42 | -0.98 | -0.39 | -0.6 | -0.81 | -0.46 | -0.54 | -0.62 | -0.92 | -0.44 | -0.59 | -0.58 | 0.07 | 0.06 | 0.035 | 0.712492 | 0.729558 | 0.856657 | | 1 | 0.996511 | 0.976225 |
| Ubac1 | Q8VDI7 | 0.8 | 1.21 | 0.37 | -0.25 | 1.39 | 1.39 | -0.51 | -0.02 | 1.17 | 1.28 | 0.26 | 0.9 | 1.35 | 1.71 | -0.13 | -0.2 | 0.03 | 0.37 | 0.15 | 0.954996 | 0.336733 | 0.781892 | | 1 | 0.996511 | 0.967642 |
| Ubac2 | Q8R1K1 | 0.34 | 0.72 | 0.67 | 0.8 | 1.01 | 0.1 | 1.43 | 1.03 | 0.08 | 0.04 | 0.55 | 0.24 | 0.48 | 0.2 | 1.28 | 1.36 | 0.26 | -0.405 | 0.1975 | 0.392678 | 0.071531 | 0.534595 | | 1 | 0.996511 | 0.912212 |
| Ubald1 | Q6P3B2 | 0.52 | 0.62 | -1.06 | 0.15 | -1.92 | 0.18 | -2.8 | -1.15 | -0.39 | 0.61 | -0.18 | 0.76 | 1.14 | -2.91 | -0.52 | -2.48 | -1.48 | 0.1425 | -1.25 | 0.052092 | 0.752593 | 0.19216 | | 1 | 0.996511 | 0.77733 |
| Ubap1 | Q8BH48 | 0.41 | 1.01 | 0.06 | 0.29 | 0.91 | 0.68 | 0.06 | 0.08 | 0.41 | -0.35 | -0.12 | 0.42 | -0.5 | 1.17 | -0.62 | 0.03 | -0.01 | -0.3525 | -0.4225 | 0.972861 | 0.243347 | 0.343233 | | 1 | 0.996511 | 0.848473 |
| Ubap2 | A2AMY5 | -3.11 | -3.59 | -1.86 | -2.44 | -3.48 | -2.78 | -2.38 | -2.78 | -2.29 | -1.32 | -2.26 | -2.63 | -1.9 | -2.57 | -1.88 | -2.22 | -0.105 | 0.625 | 0.6075 | 0.80068 | 0.184202 | 0.152843 | | 1 | 0.996511 | 0.750456 |
| Ubap2l | Q80X50 | -2.61 | -2.92 | -3.5 | -3.37 | -2.78 | -2.99 | -3.52 | -3.76 | -2.63 | -2.47 | -3.38 | -2.38 | -1.63 | -2.09 | -3.03 | -3.86 | -0.1625 | 0.385 | 0.4475 | 0.595019 | 0.234642 | 0.383016 | | 1 | 0.996511 | 0.860984 |
| Ubash3b | Q8BGG7 | 0.56 | 0.8 | 1.13 | 0.86 | 0.5 | 0.91 | 1.28 | 1.49 | 0.81 | 0.9 | 0.99 | 0.49 | 0.95 | 0.49 | 1.24 | 1.26 | 0.2075 | -0.04 | 0.1475 | 0.428306 | 0.845793 | 0.557073 | | 1 | 0.996511 | 0.917092 |
| Ube2a | A0A0J9YUN | -1.33 | -1.09 | -0.61 | -0.63 | -1.13 | -1.21 | -0.76 | -0.71 | -1.47 | -0.56 | -0.73 | -1.2 | -1.59 | -0.54 | -1.45 | -2.18 | -0.0375 | -0.075 | -0.525 | 0.87341 | 0.792671 | 0.184543 | | 1 | 0.996511 | 0.776843 |
| Ube2b | P63147 | -1.05 | -0.51 | -0.26 | -0.1 | -0.68 | -0.35 | -0.03 | -0.35 | -0.15 | 0.37 | -0.6 | -0.59 | -1.36 | -0.34 | -0.93 | -0.76 | 0.1275 | 0.2375 | -0.3675 | 0.621588 | 0.454463 | 0.25074 | | 1 | 0.996511 | 0.807201 |
| Ube2d1 | P61080 | -0.5 | -0.23 | -0.41 | -0.81 | 1.04 | -0.67 | 0.29 | -0.34 | -1.52 | -1.6 | -0.48 | -0.81 | -1.14 | 0.04 | -0.72 | -0.47 | 0.5675 | -0.615 | -0.085 | 0.157759 | 0.069384 | 0.770627 | | 1 | 0.996511 | 0.965933 |
| Ube2d3 | A0A0G2JE3 | -0.94 | -0.41 | -1.11 | -1.18 | -0.35 | -0.89 | -0.38 | -0.97 | -0.56 | -0.18 | -0.62 | -0.74 | -1.26 | 0.98 | -1.06 | -1.11 | 0.2625 | 0.385 | 0.2975 | 0.311262 | 0.135135 | 0.570878 | | 1 | 0.996511 | 0.924317 |
| Ube2e2 | D3YXD1 | -3.55 | -2.79 | -4.17 | -4.33 | -3.06 | -3.08 | -3.46 | -4.37 | -3.46 | -2.26 | -3.97 | -2.75 | -2.36 | -2.39 | -4.18 | -4.69 | 0.2175 | 0.6 | 0.305 | 0.619123 | 0.233189 | 0.633973 | | 1 | 0.996511 | 0.941132 |
| Ube2e3 | P52483 | -11.39 | -10.42 | -9.4 | -7.07 | -9.01 | -10.12 | -15.75 | -5.24 | -12.81 | -4.57 | -7.61 | -8.95 | -7.69 | -6.5 | -4.93 | -8.18 | -0.46 | 1.085 | 2.745 | 0.827149 | 0.535998 | 0.026623 | | 1 | 0.996511 | 0.635197 |
| Ube2f | Q9CY34 | 0.45 | 0.58 | 0.28 | 0.17 | 0.13 | 0.46 | 0.21 | 0.67 | 0.45 | -0.67 | 0.44 | 0.4 | 0.99 | -1.04 | 0.03 | -0.13 | -0.0025 | -0.215 | -0.4075 | 0.989538 | 0.475467 | 0.332749 | | 1 | 0.996511 | 0.845601 |
| Ube2g1 | P62254 | -1.28 | -1.07 | -0.95 | -0.76 | -1.21 | -1.24 | -1.05 | -0.88 | -1.18 | -0.38 | -0.6 | -1.3 | -1.2 | 0.17 | -0.84 | -0.7 | -0.08 | 0.15 | 0.3725 | 0.660544 | 0.575006 | 0.259731 | | 1 | 0.996511 | 0.812629 |
| Ube2g2 | P60605 | -0.12 | -0.05 | -0.3 | -0.32 | 0.83 | 0.22 | 0.43 | -0.17 | -0.11 | 0.13 | 0.09 | -0.15 | -0.84 | 0.36 | 0.06 | 0.43 | 0.525 | 0.1875 | 0.2 | 0.050217 | 0.292804 | 0.523052 | | 1 | 0.996511 | 0.907797 |
| Ube2h | P62257 | 1.51 | 1.1 | 0.73 | 0.27 | 1.56 | 1.45 | 0.65 | 0.28 | 0.75 | 1.44 | 0.5 | 1.54 | 0.32 | 1.77 | 0.01 | 0.07 | 0.0825 | 0.155 | -0.36 | 0.830888 | 0.66701 | 0.444818 | | 1 | 0.996511 | 0.884589 |
| Ube2i | P63280 | -0.74 | -1.09 | -1.55 | -1.49 | -0.67 | -1.1 | -0.81 | -1 | -1.14 | -0.68 | -1.2 | -0.85 | -1.34 | -0.28 | -1.12 | -1.25 | 0.3225 | 0.25 | 0.22 | 0.18704 | 0.329816 | 0.492329 | | 1 | 0.996511 | 0.897531 |
| Ube2j1 | Q9JJZ4 | -0.78 | -0.52 | -0.85 | -0.93 | -0.59 | -0.26 | -0.07 | -0.42 | 1.85 | -1.13 | -0.77 | -0.9 | 2.92 | 0.37 | 1.31 | 0.13 | 0.435 | 0.5325 | 1.9525 | 0.040323 | 0.417793 | 0.008553 | | 1 | 0.996511 | 0.563386 |
| Ube2j2 | B1ASK8 | 0.25 | 0.93 | 0.46 | 1.03 | 0.58 | 0.41 | -0.93 | 0.11 | 1.5 | 0.95 | -0.75 | 0.78 | 1.42 | 0.82 | 1.32 | -0.66 | -0.625 | -0.0475 | 0.0575 | 0.117636 | 0.921474 | 0.905018 | | 1 | 0.996755 | 0.985582 |
| Ube2k | P61087 | -0.68 | -0.74 | -1.27 | -0.87 | -0.7 | -0.53 | -1.34 | -1.02 | -0.64 | -0.71 | -1.13 | -0.53 | -0.74 | -0.28 | -1.57 | -1.4 | -0.0075 | 0.1375 | -0.1075 | 0.974933 | 0.541148 | 0.745319 | | 1 | 0.996511 | 0.959729 |
| Ube2l3 | P68037 | 2.39 | -0.72 | -1.38 | -1.13 | -0.47 | 2.31 | -1.41 | -1.08 | -0.67 | -0.6 | -1.79 | 2 | -1.21 | -0.41 | -1.52 | -1.75 | 0.0475 | -0.055 | -1.0125 | 0.96528 | 0.958844 | 0.243803 | | 1 | 0.996755 | 0.799567 |
| Ube2m | P61082 | -0.76 | -0.47 | -0.32 | -0.28 | -0.29 | -0.4 | -0.16 | -0.07 | 0.63 | 0.31 | -0.12 | -0.16 | 1.34 | 0.06 | 0.41 | -0.41 | 0.2275 | 0.6225 | 0.8075 | 0.221993 | 0.02915 | 0.056998 | | 1 | 0.996511 | 0.661738 |
| Ube2n | P61089 | 0.87 | -0.97 | 0.83 | -0.23 | -0.78 | 1.02 | 1.19 | 0.13 | -0.98 | 0 | 0.75 | 0.75 | -0.79 | -1.1 | 0.43 | 0.65 | 0.265 | 0.005 | -0.3275 | 0.650031 | 0.992838 | 0.57108 | | 1 | 0.999285 | 0.924332 |
| Ube2o | Q6ZPJ3 | -0.44 | -0.61 | -0.59 | -0.53 | -0.57 | -0.43 | -0.53 | -0.52 | -0.43 | -0.2 | -0.48 | -0.29 | -0.45 | -0.21 | -0.74 | -0.73 | 0.03 | 0.1925 | 0.01 | 0.831966 | 0.25758 | 0.960017 | | 1 | 0.996511 | 0.992636 |
| Ube2q1 | Q7TSS2 | -2.09 | -1.75 | -0.7 | -0.63 | -1.85 | -1.84 | -0.97 | -0.57 | -1.44 | -0.33 | -0.67 | -1.83 | -0.7 | -1.34 | -0.91 | -0.65 | -0.015 | 0.225 | 0.3925 | 0.973665 | 0.635793 | 0.327379 | | 1 | 0.996511 | 0.844633 |
| Ube2q2 | D3Z647 | -1.79 | -1.32 | -1.23 | -1.75 | -1.18 | -1.52 | -1.09 | -2.03 | -2.12 | -2.33 | -1.57 | -1.89 | -1.49 | -2.41 | -2.13 | -1.42 | 0.0675 | -0.455 | -0.34 | 0.798829 | 0.089232 | 0.269742 | | 1 | 0.996511 | 0.821668 |
| Ube2ql1 | E9QMD2 | -3.22 | -3.14 | -0.66 | -1.38 | -5.28 | -2.15 | -3.58 | -1.78 | -0.86 | -2.29 | -1.9 | -2.87 | 0.57 | -4.1 | -1.8 | -2.02 | -1.0975 | 0.12 | 0.2625 | 0.251678 | 0.863513 | 0.798675 | | 1 | 0.996511 | 0.969753 |
| Ube2r2 | Q6ZWZ2 | -0.84 | -0.99 | -1.42 | -1.19 | -0.94 | -1.52 | -1.37 | -1.66 | 0.62 | -0.5 | -1.67 | -0.95 | 1.15 | -0.47 | -1.47 | -1.72 | -0.2625 | 0.485 | 0.4825 | 0.257709 | 0.312435 | 0.435314 | | 1 | 0.996511 | 0.881656 |
| Ube2v1 | Q9CZY3 | 1.97 | -0.36 | -0.11 | 0.65 | -2.37 | 1.43 | -1.81 | -0.23 | 4.03 | 0.31 | -0.34 | -0.33 | 3.15 | 3.06 | -0.04 | -0.85 | -1.2825 | 0.38 | 0.7925 | 0.179591 | 0.718742 | 0.453344 | | 1 | 0.996511 | 0.885993 |
| Ube2v2 | Q9D2M8 | 0.3 | 0.51 | 0.99 | 0.59 | 0.12 | 0.2 | 0.54 | 0.7 | -0.48 | -0.41 | 0.83 | 0.31 | -0.4 | 0.15 | 0.67 | 0.52 | -0.2075 | -0.535 | -0.3625 | 0.362741 | 0.141663 | 0.237747 | | 1 | 0.996511 | 0.792722 |
| Ube2w | Q8VDW4 | -0.58 | -0.13 | 1.25 | 2.12 | -0.57 | -1.55 | 1.92 | 1.95 | -0.09 | 1.25 | 1.54 | -0.46 | -0.05 | -0.05 | 0.64 | 2.17 | -0.2275 | -0.105 | 0.0125 | 0.815205 | 0.883955 | 0.986424 | | 1 | 0.996511 | 0.996154 |
| Ube2z | Q3UE37 | 0.18 | 0.14 | -0.21 | -0.38 | 0.28 | 0.4 | -0.4 | -0.32 | -0.73 | 0.72 | -0.25 | 0.37 | -0.25 | -1.05 | -0.32 | -0.57 | 0.0575 | 0.095 | -0.48 | 0.822127 | 0.782843 | 0.088891 | | 1 | 0.996511 | 0.691619 |
| Ube3a | O08759 | -0.06 | -0.11 | -0.28 | -0.35 | -0.38 | 0.03 | -0.33 | -0.41 | 0.21 | -0.63 | -0.38 | -0.14 | 0.13 | -0.27 | -0.71 | -0.69 | -0.0725 | -0.035 | -0.185 | 0.677046 | 0.87682 | 0.460159 | | 1 | 0.996511 | 0.887902 |
| Ube3b | Q9ES34 | -0.07 | 0.5 | 0.41 | 0.28 | 0.47 | 0.07 | 0.62 | 0.5 | -0.17 | -0.46 | 0.69 | 0.06 | -0.28 | 0.28 | 0.1 | 0.24 | 0.135 | -0.25 | -0.195 | 0.513435 | 0.391301 | 0.398284 | | 1 | 0.996511 | 0.867334 |
| Ube3c | Q80U95 | -0.58 | -0.44 | -0.71 | -0.52 | -0.23 | -0.7 | -0.7 | -0.58 | -0.38 | -0.72 | -0.68 | -0.64 | -0.43 | -0.14 | -0.96 | -0.64 | 0.01 | -0.0425 | 0.02 | 0.954198 | 0.804523 | 0.930003 | | 1 | 0.996511 | 0.989687 |
| Ube4a | G3X9Y5 | 0 | -0.17 | 0.47 | 0.08 | -0.15 | 0.14 | 0.5 | 0.35 | -0.11 | 0.13 | 0.19 | -0.03 | 0.37 | -0.22 | 0.99 | 0.58 | 0.115 | -0.05 | 0.335 | 0.602745 | 0.803753 | 0.280642 | | 1 | 0.996511 | 0.822084 |
| Ube4b | A0A571BD! | -0.4 | -0.45 | -0.6 | -0.87 | -0.26 | -0.33 | -0.65 | -0.66 | -0.4 | -0.94 | -0.57 | -0.19 | -0.29 | -0.54 | -0.62 | -0.72 | 0.105 | 0.055 | 0.0375 | 0.580748 | 0.807196 | 0.853887 | | 1 | 0.996511 | 0.976225 |
| Ubfd1 | Q78JW9 | 0.19 | 0.23 | -0.14 | 0.19 | 0.18 | 0.26 | -0.01 | 0.03 | 0.39 | 0.58 | 0.2 | 0.44 | 0.25 | 0.63 | -0.39 | -0.44 | -0.0025 | 0.285 | -0.105 | 0.987878 | 0.143015 | 0.717885 | | 1 | 0.996511 | 0.958398 |
| Ubiad1 | Q9DC60 | 0.55 | 1 | 0.82 | 0.96 | 1.07 | 0.56 | 0.48 | 1.06 | 0.42 | 1.49 | 1.13 | 0.45 | -0.15 | 1.56 | 0.85 | 0.68 | -0.04 | 0.04 | -0.0975 | 0.85186 | 0.890549 | 0.78705 | | 1 | 0.996511 | 0.968039 |
| Ubl3 | Q9Z2M6 | 3.76 | 1.35 | 1.21 | 0.46 | 2.9 | 4.11 | 1.76 | 0.92 | 0.51 | 0.21 | 1.24 | 3.66 | 0.02 | 1.15 | 1.38 | 1.96 | 0.7275 | -0.29 | -0.5675 | 0.425952 | 0.761449 | 0.453394 | | 1 | 0.996511 | 0.885993 |
| Ubl4a | P21126 | 0.43 | 0.92 | 0.08 | -0.5 | 0.63 | 0.27 | 0.17 | 0.35 | 0.13 | -1.29 | 0.39 | 0.49 | 0.07 | 1 | 0.07 | 0.15 | 0.1225 | -0.3025 | 0.09 | 0.693797 | 0.530676 | 0.806916 | | 1 | 0.996511 | 0.971753 |
| Ubl5 | E0CY91 | -0.63 | -1.24 | -2.6 | -1.73 | -2.31 | -1.05 | -2.34 | -1.77 | -1.23 | -1.38 | -1.96 | -1.27 | -1.52 | -0.96 | -1.67 | -2.61 | -0.3175 | 0.09 | -0.14 | 0.510582 | 0.832602 | 0.780735 | | 1 | 0.996511 | 0.9676 |
| Ubl7 | Q91W67 | -1.56 | -1.33 | -1.27 | -1.18 | -1.74 | -1.2 | -1.46 | -1.72 | -0.73 | -1.27 | -1.19 | -1.44 | 0.07 | -1.68 | -1.85 | -2.02 | -0.195 | 0.1775 | -0.035 | 0.319006 | 0.414631 | 0.939676 | | 1 | 0.996511 | 0.990645 |
| Ublcp1 | Q8BGR9 | -1.77 | -2.01 | -0.73 | -1.59 | -1.61 | -2.13 | -1.76 | -2.18 | -2.61 | -1.42 | -1.83 | -1.73 | -2.16 | -2.09 | -0.36 | -2.3 | -0.395 | -0.3725 | -0.2025 | 0.220838 | 0.32312 | 0.686567 | | 1 | 0.996511 | 0.955729 |
| Ubn1 | Q4G0F8-2 | -0.07 | -0.13 | 0.01 | 0.03 | -0.6 | 0.08 | -0.45 | 0.23 | 0.82 | 0.09 | 0.27 | 0.68 | 0.73 | 0.19 | -0.29 | -0.13 | -0.145 | 0.505 | 0.165 | 0.525345 | 0.040976 | 0.527228 | | 1 | 0.996511 | 0.90845 |
| Ubp1 | Q811S7 | -0.01 | -0.01 | -0.03 | 0.27 | 0 | -0.28 | -0.03 | 0.52 | -0.97 | -0.5 | 0.03 | -0.11 | -0.89 | -0.77 | -0.11 | -0.2 | -0.0025 | -0.4425 | -0.5475 | 0.990514 | 0.111743 | 0.049614 | | 1 | 0.996511 | 0.658454 |
| Ubqln1 | Q8R317 | -0.27 | -0.18 | -0.2 | -0.13 | -0.32 | -0.35 | -0.03 | -0.35 | -0.41 | -0.19 | -0.18 | -0.1 | -0.37 | -0.17 | -0.56 | -0.31 | -0.0675 | -0.025 | -0.1575 | 0.661707 | 0.87769 | 0.389521 | | 1 | 0.996511 | 0.865375 |
| Ubqln1 | Q8R317-2 | -0.22 | 0.75 | -1 | 0.52 | 0.44 | 0.06 | -0.79 | 0.49 | -0.3 | -0.46 | -0.36 | -0.2 | -0.07 | 0.6 | -1.36 | -1.03 | 0.0375 | -0.3425 | -0.4775 | 0.934866 | 0.384353 | 0.397328 | | 1 | 0.996511 | 0.867334 |
| Ubqln2 | Q9QZM0 | -0.25 | 0.23 | 0.04 | 0.07 | 0.04 | -0.21 | 0.39 | 0.02 | -0.62 | -0.13 | 0.12 | 0.12 | -0.89 | -0.3 | -0.15 | 0.1 | 0.0375 | -0.15 | -0.3325 | 0.847433 | 0.523267 | 0.224353 | | 1 | 0.996511 | 0.787447 |
| Ubqln4 | Q99NB8 | 0.63 | 0.86 | 0.02 | -0.21 | 1.02 | 0.59 | 0 | -0.17 | -0.35 | 0.4 | -0.07 | 1.18 | -0.37 | -0.1 | -0.31 | 0.15 | 0.035 | -0.035 | -0.4825 | 0.921686 | 0.930229 | 0.126719 | | 1 | 0.996755 | 0.722414 |
| Ubr1 | O70481 | -0.16 | -0.09 | -0.12 | -0.06 | 0 | -0.18 | 0.33 | -0.11 | -0.23 | 0.19 | -0.17 | -0.11 | -0.51 | 0.19 | -0.2 | -0.06 | 0.1175 | 0.0275 | -0.0375 | 0.492575 | 0.872664 | 0.856727 | | 1 | 0.996511 | 0.976225 |
| Ubr2 | Q6WKZ8-2 | -0.2 | -0.42 | -0.15 | -0.34 | 0.1 | 0.3 | -0.2 | -0.2 | -0.54 | 0.11 | -0.25 | 0.1 | -0.34 | -0.08 | -0.1 | 0.08 | 0.2775 | 0.1325 | 0.1675 | 0.152462 | 0.533976 | 0.382493 | | 1 | 0.996511 | 0.860984 |
| Ubr3 | Q5U430 | -0.63 | -0.45 | -0.32 | -0.22 | -0.38 | -0.63 | -0.18 | 0.1 | -0.12 | -0.4 | -0.06 | -0.32 | -0.14 | -0.13 | -0.2 | -0.31 | 0.1325 | 0.18 | 0.21 | 0.527818 | 0.339232 | 0.27074 | | 1 | 0.996511 | 0.822084 |
| Ubr4 | A2AN08-3 | -0.35 | -0.26 | -0.2 | -0.22 | 0.05 | -0.19 | -0.18 | -0.35 | -0.21 | -0.12 | -0.19 | -0.23 | -0.1 | -0.01 | -0.42 | -0.19 | 0.09 | 0.07 | 0.0775 | 0.567591 | 0.650484 | 0.671836 | | 1 | 0.996511 | 0.952718 |
| Ubr5 | A0A2I3BQS | -0.5 | -0.97 | -1.11 | -1.06 | -0.83 | -1.26 | -1.68 | -0.71 | -0.67 | -0.24 | -1.4 | -1.66 | 0.22 | 0.2 | -0.58 | -1.43 | -0.21 | -0.0825 | 0.5125 | 0.4413 | 0.813048 | 0.222303 | | 1 | 0.996511 | 0.787447 |
| Ubr7 | Q8BU04 | -3.57 | -3.26 | -3.13 | -3.86 | -3.28 | -3.59 | -3.12 | -3.46 | -3.43 | -2.21 | -3.51 | -3.2 | -1.71 | -3.23 | -3.44 | -3.27 | 0.0925 | 0.3675 | 0.5425 | 0.67236 | 0.293017 | 0.214936 | | 1 | 0.996511 | 0.787313 |
| Ubtd2 | Q6PGH0 | 0.34 | 1.62 | 1.01 | 1.26 | 2.02 | 0.93 | 0.65 | 1.11 | 1.14 | 1.11 | 0.81 | 1.56 | 0.67 | 2.39 | 0.55 | 0.43 | 0.12 | 0.0975 | -0.0475 | 0.752994 | 0.755946 | 0.924167 | | 1 | 0.996511 | 0.988152 |
| Ubtf | A2AWT6 | 0.92 | 0.52 | 0.18 | 0.36 | 0.5 | 0.38 | 0.29 | -0.07 | 0.6 | -0.21 | 0.48 | 1.16 | 0.32 | 0.82 | 0.16 | 0.46 | -0.22 | 0.0125 | -0.055 | 0.336609 | 0.969032 | 0.823702 | | 1 | 0.996755 | 0.972768 |
| Ubxn1 | Q922Y1 | -1.43 | -1.04 | -0.41 | -0.32 | -1.26 | -1.63 | -0.54 | -0.64 | -1.13 | -0.48 | -0.43 | -1.35 | -1.52 | -0.53 | -0.94 | -0.9 | -0.2175 | -0.0475 | -0.1725 | 0.544276 | 0.890454 | 0.609943 | | 1 | 0.996511 | 0.934918 |
| Ubxn2a | Q99KJ0 | -0.98 | -0.35 | 0.17 | -0.33 | -0.71 | -1.1 | -0.26 | 0.03 | -0.54 | -0.08 | -0.46 | -1.26 | -1.72 | -1.01 | -0.81 | 0.14 | -0.1375 | -0.2125 | -0.4775 | 0.680475 | 0.532927 | 0.282621 | | 1 | 0.996511 | 0.822595 |
| Ubxn2b | Q0KL01 | -0.28 | -0.02 | 0.19 | 0.03 | -0.26 | -0.29 | 0.22 | -0.19 | -0.99 | 0.15 | 0.41 | -0.09 | -0.3 | -1.18 | -0.01 | 0.24 | -0.11 | -0.11 | -0.2925 | 0.56905 | 0.731664 | 0.384148 | | 1 | 0.996511 | 0.860984 |
| Ubxn4 | A0A0R4J07 | -0.24 | 0.07 | -0.43 | -0.23 | -0.08 | 0.11 | 0.46 | -0.36 | 1.15 | -0.07 | -0.36 | -0.01 | 1.92 | 0.57 | 0.24 | -0.07 | 0.24 | 0.385 | 0.8725 | 0.298302 | 0.279785 | 0.068105 | | 1 | 0.996511 | 0.664936 |
| Ubxn6 | Q99PL6 | -0.09 | -0.12 | 0.17 | 0.31 | -0.36 | 0.13 | 0.16 | 0.38 | 0.79 | 0.01 | 0.34 | -0.13 | 1.33 | -0.3 | 0.25 | -0.05 | 0.01 | 0.185 | 0.24 | 0.962649 | 0.469633 | 0.5181 | | 1 | 0.996511 | 0.90668 |
| Ubxn7 | G5E8R8 | -2.65 | -2.99 | -1.69 | -1.46 | -2.56 | -3.08 | -2.05 | -2.03 | -1.64 | -0.73 | -1.77 | -2.2 | -1.46 | -1.78 | -2.12 | -2.09 | -0.2325 | 0.6125 | 0.335 | 0.581082 | 0.199351 | 0.398368 | | 1 | 0.996511 | 0.867334 |
| Uchl1 | Q9R0P9 | -0.01 | -0.48 | 0.13 | 1.06 | -0.37 | -0.13 | 0.2 | 1.15 | 0.03 | 0.99 | 0.27 | 0.22 | 0.04 | 0.59 | 0.21 | 0.33 | 0.0375 | 0.2025 | 0.1175 | 0.931021 | 0.590055 | 0.732154 | | 1 | 0.996511 | 0.958635 |
| Uchl3 | Q9JKB1 | -0.64 | -0.59 | -0.41 | -0.42 | -0.54 | -0.29 | -0.47 | -0.17 | -0.46 | -0.57 | -0.4 | -0.29 | -0.99 | 0.08 | -0.6 | -0.62 | 0.1475 | 0.085 | -0.0175 | 0.375813 | 0.611851 | 0.946224 | | 1 | 0.996511 | 0.992376 |
| Uchl5 | Q9WUP7-2 | -0.76 | -0.67 | -0.25 | -0.05 | 0.18 | 0.26 | 0.34 | -0.31 | -0.19 | 0.19 | 0.08 | -0.33 | -0.75 | 0.57 | 0.03 | 0.47 | 0.55 | 0.37 | 0.5125 | 0.044581 | 0.144056 | 0.1618 | | 1 | 0.996511 | 0.757516 |
| Uchl5 | Q9WUP7 | -0.35 | -0.65 | 0.11 | 0.41 | -0.38 | -0.16 | 0.71 | 0.8 | -0.6 | 0.02 | 0.37 | -0.3 | -0.93 | -0.21 | 0.11 | 0.24 | 0.3625 | -0.0075 | -0.0775 | 0.334157 | 0.981056 | 0.824843 | | 1 | 0.998773 | 0.972852 |
| Uck1 | P52623 | -1.08 | -1.04 | -1.04 | -0.9 | 0.04 | -1.08 | -0.55 | -0.68 | -0.92 | -0.77 | -0.38 | -1.04 | -0.81 | -0.47 | -0.48 | -0.72 | 0.4475 | 0.2375 | 0.395 | 0.098798 | 0.252456 | 0.054702 | | 1 | 0.996511 | 0.661738 |
| Uckl1 | Q91YL3 | -0.3 | -0.6 | 0.58 | 0.06 | -0.16 | -0.3 | 0.51 | 0.5 | -1.08 | 0.75 | 0.44 | -0.33 | -1.04 | -0.81 | 0.29 | 0.34 | 0.2025 | 0.01 | -0.24 | 0.536507 | 0.982381 | 0.572664 | | 1 | 0.998773 | 0.925236 |
| Ucp2 | D3YXM5 | -4.59 | 1.4 | 0.61 | 0.49 | -2.98 | 1.73 | -0.93 | 0.84 | 1.24 | 3.12 | -0.31 | 0.19 | 1.4 | -0.28 | 0.44 | 1.22 | 0.1875 | 1.5825 | 1.2175 | 0.90279 | 0.27644 | 0.349876 | | 1 | 0.996511 | 0.85327 |
| Ucp3 | P56501 | 1.16 | 1.83 | 0.47 | 0.63 | 1.2 | 1.32 | 1.43 | 0.75 | 1.31 | -0.64 | 0.79 | 2.01 | 1.24 | 2.19 | -0.54 | 0.41 | 0.1525 | -0.155 | -0.1975 | 0.647272 | 0.792053 | 0.743858 | | 1 | 0.996511 | 0.959321 |
| Uevld | Q3U1V6 | -0.16 | 0.07 | -0.34 | -0.22 | 0.18 | -0.2 | -0.08 | -0.31 | -0.87 | -0.36 | -0.39 | -0.28 | -0.29 | 0.52 | -0.27 | -0.71 | 0.06 | -0.3125 | -0.025 | 0.740395 | 0.153682 | 0.930804 | | 1 | 0.996511 | 0.989713 |
| Ufc1 | Q9CR09 | 0.36 | 0.2 | 0.16 | 0.84 | 0.27 | 0.51 | 0.91 | 1.33 | 0.04 | 0.07 | 0.63 | 0.46 | -0.26 | 0.09 | 0.41 | 0.73 | 0.365 | -0.09 | -0.1475 | 0.218533 | 0.709503 | 0.604891 | | 1 | 0.996511 | 0.932185 |
| Ufd1 | P70362 | -0.54 | -0.29 | -0.3 | -0.35 | -0.41 | -0.48 | -0.61 | -0.51 | 0.05 | -0.59 | -0.34 | -0.24 | -0.01 | 0.28 | -0.66 | -0.81 | -0.1325 | 0.09 | 0.07 | 0.38453 | 0.6489 | 0.806069 | | 1 | 0.996511 | 0.971715 |
| Ufl1 | Q8CCJ3 | -0.16 | -0.03 | -0.29 | -0.17 | 0.21 | 0.24 | 0.37 | -0.01 | 0.11 | -0.48 | -0.36 | 0.19 | 0.82 | -0.07 | 0.6 | 0.44 | 0.365 | 0.0275 | 0.61 | 0.044569 | 0.89831 | 0.028262 | | 1 | 0.996511 | 0.637553 |
| Ufm1 | P61961 | 0.01 | 0.15 | -0.35 | -0.3 | 0.34 | -0.14 | -0.45 | -0.38 | 0 | 0.33 | -0.14 | 0.2 | -0.37 | 0.68 | -0.22 | -0.05 | -0.035 | 0.22 | 0.1325 | 0.880877 | 0.300547 | 0.64036 | | 1 | 0.996511 | 0.942906 |
| Ufsp1 | Q9CZP0 | 0.1 | -0.58 | 0.25 | -0.1 | 0.14 | -1.21 | 0.24 | 0.2 | -0.08 | 0.07 | -1.72 | -0.35 | -0.6 | -0.01 | -0.33 | -1.99 | -0.075 | -0.4375 | -0.65 | 0.841827 | 0.317544 | 0.170673 | | 1 | 0.996511 | 0.762322 |
| Ufsp2 | Q99K23 | 0.25 | 0.55 | 0.6 | 0.21 | 1.28 | 0.28 | 2.4 | 0.15 | 0.23 | 0.77 | 0.74 | 0.3 | 0.24 | 0.75 | 1.23 | 1.7 | 0.625 | 0.1075 | 0.5775 | 0.225253 | 0.618737 | 0.109496 | | 1 | 0.996511 | 0.709362 |
| Ugcg | O88693 | -0.7 | -1.34 | -1.42 | -0.91 | 0.41 | -0.34 | 0.02 | -0.33 | -0.27 | -0.75 | -1.06 | -0.78 | -0.6 | 0.29 | -0.95 | 0.28 | 1.0325 | 0.3775 | 0.8475 | 0.00312 | 0.168514 | 0.038177 | | 1 | 0.996511 | 0.658454 |
| Ugdh | O70475 | 0.34 | -0.27 | -0.93 | -0.89 | -0.05 | -0.58 | -1.39 | -1.33 | -0.32 | -1.09 | -1.26 | 0.25 | -0.2 | 0.29 | -1.4 | -1.63 | -0.4 | -0.1675 | -0.2975 | 0.345828 | 0.701798 | 0.565952 | | 1 | 0.996511 | 0.921174 |
| Uggt1 | Q6P5E4 | 0.01 | -0.46 | -0.45 | -0.44 | 0.41 | 0.1 | 0.25 | -0.28 | -0.75 | -0.8 | -0.46 | 0.13 | -0.41 | 0.02 | 0.37 | 0.22 | 0.455 | -0.135 | 0.385 | 0.058423 | 0.608193 | 0.138611 | | 1 | 0.996511 | 0.73493 |
| Uggt2 | E9Q4X2 | -2.91 | -2.65 | -0.48 | -0.46 | -2.88 | -3.03 | 0.81 | -0.08 | -0.8 | -0.68 | -0.34 | -2.23 | 0.65 | -1.79 | 0.87 | 0.35 | 0.33 | 0.6125 | 1.645 | 0.756147 | 0.4026 | 0.070448 | | 1 | 0.996511 | 0.668066 |
| Ugp2 | Q91ZJ5 | 0.4 | 0.38 | 0.55 | 0.59 | 0.79 | 0.59 | 0.47 | 0.77 | -0.48 | -0.4 | 0.53 | 0.23 | -0.65 | -0.3 | 0.26 | 0.52 | 0.175 | -0.51 | -0.5225 | 0.284785 | 0.083694 | 0.096786 | | 1 | 0.996511 | 0.698156 |
| Ugt1a7 | Q6ZQM8 | -0.28 | 0.22 | -0.33 | 0.81 | -0.38 | -0.15 | -0.29 | 0.38 | -0.22 | 0.25 | 0.13 | 0.15 | -0.11 | -0.69 | -0.4 | -0.45 | -0.215 | -0.0275 | -0.5175 | 0.494878 | 0.925146 | 0.115115 | | 1 | 0.996755 | 0.717287 |
| Ugt8 | Q64676 | -0.77 | -0.46 | -2.85 | -2.59 | 0.16 | -0.46 | -2.2 | -2.06 | 0.1 | -2.7 | -2.18 | -0.24 | 0.27 | 0.18 | -1.46 | -1.53 | 0.5275 | 0.4125 | 1.0325 | 0.497058 | 0.624567 | 0.173686 | | 1 | 0.996511 | 0.764376 |
| Uhmk1 | P97343 | -1.25 | -1.2 | -1.02 | -1.55 | -1.22 | -1.47 | -1.5 | -1.2 | -1.2 | -1.46 | -0.68 | -0.83 | -0.48 | -2.45 | -0.28 | -1.45 | -0.0925 | 0.2125 | 0.09 | 0.611146 | 0.381602 | 0.850871 | | 1 | 0.996511 | 0.976225 |
| Uhrf1bp1 | B2KF50 | 0.32 | 0.58 | 0.57 | 0.43 | 0.42 | 0.57 | 0.61 | 0.18 | 0.38 | 2.03 | 0.59 | 0.35 | 0.51 | -0.12 | 0.34 | 0.35 | -0.03 | 0.3625 | -0.205 | 0.859157 | 0.364579 | 0.339462 | | 1 | 0.996511 | 0.84777 |
| Uhrf1bp1l | A2RSJ4 | 0.22 | 0.38 | 0.31 | 0.11 | 0.08 | 0.18 | -0.27 | -0.03 | 0.65 | 0.68 | 0.22 | 0.27 | 0.26 | 0.21 | -0.3 | -0.31 | -0.265 | 0.2 | -0.29 | 0.140962 | 0.311213 | 0.207335 | | 1 | 0.996511 | 0.785338 |
| Ulk1 | A0A0R4J0E | 0.3 | 0.82 | -0.34 | -0.35 | 1.19 | 0.33 | -0.4 | -0.22 | -0.09 | -0.83 | -0.07 | 0.64 | 0.86 | -0.28 | 0.11 | 0.2 | 0.1175 | -0.195 | 0.115 | 0.782977 | 0.623492 | 0.751535 | | 1 | 0.996511 | 0.961479 |
| Ulk2 | Q9QY01 | 0.24 | 0.59 | 0.89 | 0.74 | 0.59 | 0.21 | 0.21 | 0.52 | 0.52 | 0.59 | 0.91 | 0.6 | 0.45 | 0.88 | 0.39 | 0.47 | -0.2325 | 0.04 | -0.0675 | 0.270768 | 0.847661 | 0.765477 | | 1 | 0.996511 | 0.964134 |
| Ulk3 | Q3U3Q1 | -0.6 | 0.31 | -0.38 | -0.39 | 0.11 | -0.85 | -0.26 | -0.45 | -0.33 | 0.06 | -0.4 | -0.41 | 0.1 | -0.23 | -0.42 | -0.11 | -0.0975 | -0.005 | 0.1 | 0.731602 | 0.984058 | 0.697962 | | 1 | 0.998876 | 0.955729 |
| Umad1 | E0CZ61 | 0.1 | -0.15 | -1.67 | -2.01 | -0.22 | -0.18 | -3.54 | -3.83 | 0.29 | 0.04 | -1.79 | 0.47 | 0.07 | 0.38 | -2.81 | -3.75 | -1.01 | 0.685 | -0.595 | 0.336947 | 0.326844 | 0.569709 | | 1 | 0.996511 | 0.923081 |
| Umps | P13439 | -1.13 | -1.1 | -1.26 | -1.24 | -0.89 | -1.54 | -0.75 | -1.18 | -1.95 | -1.41 | -1.03 | -1.14 | -1.79 | -1.64 | -0.99 | -1.38 | 0.0925 | -0.2 | -2.68E-01 | 0.658332 | 0.409934 | 0.256525 | | 1 | 0.996511 | 0.811232 |
| Unc119 | Q9Z2R6 | 0.09 | -0.18 | 0.06 | -0.26 | -0.07 | -0.04 | -0.23 | 0.25 | 1.01 | 1.82 | 0.14 | 0.45 | 1.12 | 0.78 | -0.1 | -0.26 | 0.05 | 0.9275 | 0.4575 | 0.779829 | 0.031004 | 0.207545 | | 1 | 0.996511 | 0.785338 |
| Unc119b | Q8C4B4 | 0.35 | 0.6 | 0.18 | 0.21 | 0.73 | -0.22 | 0.24 | 0.15 | -0.05 | 0.56 | 0.17 | 0.37 | -0.43 | 0.55 | -0.25 | 0.43 | -0.11 | -0.0725 | -0.26 | 0.641947 | 0.727104 | 0.367925 | | 1 | 0.996511 | 0.859245 |
| Unc13a | H3BJZ7 | 1.36 | 1.34 | 1.25 | 1.11 | 1.53 | 1.25 | 1 | 0.97 | 1.13 | 0.47 | 1.36 | 1.22 | 1.17 | 1.1 | 1.12 | 1.06 | -0.0775 | -0.22 | -0.1525 | 0.675629 | 0.362022 | 0.383421 | | 1 | 0.996511 | 0.860984 |
| Unc13b | F6X605 | 0.88 | 1.18 | 0.67 | 1.63 | 0.7 | 0.43 | 1.25 | 1.24 | 1.07 | 1.28 | 1.05 | 0.36 | -0.93 | 0.92 | 0.12 | 1.04 | -0.185 | -0.15 | -0.8025 | 0.530293 | 0.614809 | 0.114865 | | 1 | 0.996511 | 0.717287 |
| Unc13b | A0A140LH) | -0.32 | -0.05 | -0.09 | 1.15 | -0.57 | 0.17 | -0.83 | -0.45 | 1.35 | 0.44 | 0.18 | -0.17 | 1.37 | -0.47 | -0.36 | -0.39 | -0.5925 | 0.2775 | -0.135 | 0.14033 | 0.531115 | 0.793552 | | 1 | 0.996511 | 0.969365 |
| Unc13c | Q8K0T7 | 1.37 | 1.08 | 0.63 | 1.85 | 0.61 | 0.97 | 0.97 | 1.44 | 0.9 | 1.43 | 0.87 | 1.06 | 1.05 | 0.26 | 2.05 | 1.53 | -0.235 | -0.1675 | -0.01 | 0.447685 | 0.572628 | 0.98166 | | 1 | 0.996511 | 0.994858 |
| Unc45a | Q99KD5 | -1.34 | -1.65 | -1.25 | -1.22 | -0.89 | -1.36 | -0.25 | -1.29 | -1.56 | -0.8 | -0.83 | -1.13 | -0.38 | -0.77 | -0.74 | -0.53 | 0.4175 | 0.285 | 0.76 | 0.157506 | 0.241374 | 0.004253 | | 1 | 0.996511 | 0.548272 |
| Unc5a | Q8K1S4-2 | 0.97 | 1.24 | 0.71 | 0.44 | 1.65 | 1.31 | 0.5 | 0.49 | 0.45 | 0.38 | 0.56 | 1.01 | 0.56 | 0.76 | 0.68 | 0.64 | 0.1475 | -0.24 | -0.18 | 0.65645 | 0.344807 | 0.431609 | | 1 | 0.996511 | 0.879165 |
| Unc5b | Q8K1S3-2 | -0.1 | -0.06 | 0.33 | 0.36 | -0.37 | 0.71 | 0.24 | 0.12 | 0.46 | 1.06 | 0.49 | 0.3 | -0.05 | 0.16 | 0.83 | 0.62 | 0.0425 | 0.445 | 0.2575 | 0.871073 | 0.086861 | 0.343223 | | 1 | 0.996511 | 0.848473 |
| Unc5c | O08747-2 | 0.18 | 0.34 | 0.8 | 1.34 | 0.66 | 0.58 | 0.72 | 0.81 | 0 | 0.73 | 0.83 | 0.47 | 0.03 | 0.25 | 0.87 | 0.79 | 0.0275 | -0.1575 | -0.18 | 0.91901 | 0.624999 | 0.5926 | | 1 | 0.996511 | 0.929843 |
| Unc5d | B7ZMT2 | 0.53 | 0.71 | 1.28 | 1.19 | 0.43 | 1.03 | 0.87 | 1.33 | 0.91 | 1.2 | 1.4 | 0.72 | 1.35 | 0.43 | 0.98 | 1.32 | -0.0125 | 0.13 | 0.0925 | 0.962759 | 0.615551 | 0.75481 | | 1 | 0.996511 | 0.962333 |
| Unc79 | E5CYJ9 | 0.25 | 0.49 | 0.76 | 0.66 | 0.43 | 0.3 | 0.73 | 0.65 | -0.15 | 1.15 | 0.75 | 0.28 | 0.42 | 0.68 | 0.98 | 1.17 | -0.0125 | -0.0325 | 0.2725 | 0.947104 | 0.915702 | 0.271977 | | 1 | 0.996755 | 0.822084 |
| Unc80 | A0A1D5RL\ | 0.91 | 0.82 | 0.32 | 0.3 | 0.98 | 1.14 | 0.3 | 0.11 | 0.28 | 0.43 | 0.42 | 0.72 | 0.66 | 0.54 | 0.16 | 0.47 | 0.045 | -0.125 | -0.13 | 0.879731 | 0.576984 | 0.583404 | | 1 | 0.996511 | 0.927734 |
| Unc93b1 | E9PYK0 | 1.15 | 0.35 | 1.33 | -0.83 | -0.21 | 1.27 | 0.26 | -0.12 | -0.56 | -0.26 | 1.27 | 0.91 | -0.24 | -2.86 | 0.15 | 0.69 | -0.2 | -0.16 | -1.065 | 0.71581 | 0.792217 | 0.224167 | | 1 | 0.996511 | 0.787447 |
| Unk | Q8BL48-2 | 0.2 | -0.02 | 0.41 | 0.19 | 0.54 | -0.23 | 0.17 | 0.42 | 0.63 | 0.78 | 0.34 | 0.07 | 0.51 | 0.57 | 0.91 | 0.57 | 0.03 | 0.26 | 0.445 | 0.88974 | 0.253525 | 0.046895 | | 1 | 0.996511 | 0.658454 |
| Unkl | E0CYD7 | -1.7 | -1.37 | -1.34 | -1.08 | -2.06 | -1.81 | -1.89 | -1.36 | -1.21 | 0.08 | -0.86 | -1.7 | -0.51 | -2 | -0.92 | -1.56 | -0.4075 | 0.45 | 0.125 | 0.091284 | 0.258462 | 0.723076 | | 1 | 0.996511 | 0.958398 |
| Upf1 | Q9EPU0-2 | -1.34 | -1.18 | -1.31 | -1.5 | -0.98 | -1.42 | -0.87 | -1.41 | -1.46 | -0.88 | -1.43 | -1.27 | -1.13 | -0.85 | -1.12 | -1.25 | 0.1625 | 0.0725 | 0.245 | 0.411861 | 0.71642 | 0.213529 | | 1 | 0.996511 | 0.786931 |
| Upf2 | A2AT37 | 0.25 | 0.19 | -0.01 | -0.15 | 0.09 | 0.11 | -0.48 | -0.43 | -0.08 | -0.08 | -0.21 | 0.31 | -0.13 | -0.18 | -0.24 | -0.49 | -0.2475 | -0.085 | -0.33 | 0.26225 | 0.667728 | 0.116822 | | 1 | 0.996511 | 0.718608 |
| Upf3b | Q3ULL6 | -0.17 | -0.12 | 0.52 | 0.26 | -0.26 | -0.53 | -0.05 | 0.2 | 0.24 | 0.55 | 0.14 | -0.24 | 0.25 | 0.12 | 0.11 | 0.06 | -0.2825 | 0.05 | 0.0125 | 0.259822 | 0.843329 | 0.954595 | | 1 | 0.996511 | 0.992482 |
| Upp1 | P52624 | -0.22 | -0.09 | 0.12 | -0.28 | 0.6 | 0.29 | 0.55 | 0.37 | -0.16 | 0.39 | 0.1 | -0.33 | 0.24 | 0.01 | 0.01 | 0.24 | 0.57 | 0.1175 | 0.2425 | 0.008016 | 0.593803 | 0.2214 | | 1 | 0.996511 | 0.787447 |
| Uprt | B1AVZ0 | -1.4 | -1.34 | -1.54 | -2.22 | -0.52 | -1.54 | -1.39 | -2.22 | -0.82 | -1.12 | -1.86 | -0.95 | -1.1 | -0.74 | -2.22 | -2.14 | 0.2075 | 0.4375 | 0.075 | 0.591168 | 0.18409 | 0.853685 | | 1 | 0.996511 | 0.976225 |
| Uqcc1 | Q9CWU6 | 0.16 | 0.31 | 0.36 | 0.42 | 0.37 | 0.16 | 0.58 | 0.29 | 0.57 | -0.07 | 0.4 | 0.42 | 0.43 | 0.52 | 0 | 0.62 | 0.0375 | 0.0175 | 0.08 | 0.81841 | 0.929989 | 0.701032 | | 1 | 0.996755 | 0.956468 |
| Uqcc2 | Q9CQY6 | -0.06 | 0.36 | 0.34 | 0.31 | 0.4 | -0.14 | 0.43 | 0.44 | 0.15 | 0.01 | 0.68 | 0.22 | 0.25 | 0.87 | 0.3 | 0.66 | 0.045 | 0.0275 | 0.2825 | 0.825339 | 0.898494 | 0.2317 | | 1 | 0.996511 | 0.788863 |
| Uqcc3 | Q8K2T4 | -0.02 | -0.76 | -0.02 | 0.48 | 0.54 | 0.11 | 0.73 | 0.03 | -0.59 | -0.13 | -0.09 | -0.15 | -1.3 | 1.23 | 0.58 | 0.65 | 0.4325 | -0.16 | 0.37 | 0.178304 | 0.584446 | 0.513185 | | 1 | 0.996511 | 0.904284 |
| Uqcr10 | Q8R1I1 | 1.23 | 0.5 | 2.26 | -0.01 | 0.84 | 1.64 | 2.24 | 1.81 | -1.37 | -4.61 | 2.38 | 1.22 | -1.23 | -2.04 | 2.98 | 2.27 | 0.6375 | -1.59 | -0.5 | 0.246273 | 0.291085 | 0.67735 | | 1 | 0.996511 | 0.953711 |
| Uqcr11 | Q9CPX8 | -2.35 | -0.48 | -1.39 | -0.64 | -0.08 | -0.41 | 0.46 | -1.17 | -1.95 | -3.63 | -0.36 | -1.91 | -2.36 | -1.14 | -2.38 | -1.05 | 0.915 | -0.7475 | -0.5175 | 0.09729 | 0.314505 | 0.336682 | | 1 | 0.996511 | 0.847578 |
| Uqcrb | Q9CQB4 | 0.77 | 0.86 | 0.6 | 1.5 | 0.79 | 0.53 | 0.59 | 1.58 | 0.65 | 1.33 | 0.99 | 0.65 | 0.55 | 0.96 | 0.63 | 0.61 | -0.06 | -0.0275 | -0.245 | 0.845469 | 0.919017 | 0.34193 | | 1 | 0.996755 | 0.848473 |
| Uqcrc1 | Q9CZ13 | 0.59 | 0.63 | 0.47 | 0.42 | 0.59 | 0.54 | 0.43 | 0.41 | -0.12 | 0.3 | 0.46 | 0.38 | -0.49 | 0.02 | 0.08 | 0.14 | -0.035 | -0.2725 | -0.59 | 0.811609 | 0.18242 | 0.019544 | | 1 | 0.996511 | 0.612607 |
| Uqcrc2 | Q9DB77 | -0.1 | 0.1 | 0.51 | 0.75 | -0.15 | 0 | 0.51 | 0.73 | 0 | -0.03 | 0.65 | -0.27 | 0.06 | -0.07 | -0.05 | 0.28 | -0.0425 | -0.2275 | -0.26 | 0.881428 | 0.434738 | 0.304462 | | 1 | 0.996511 | 0.831662 |
| Uqcrfs1 | Q9CR68 | 1.12 | 1.35 | 0.99 | 1.86 | 0.96 | 1.48 | 1.05 | 1.7 | 0.85 | 1.6 | 1.31 | 0.75 | 0.98 | 1.1 | 0.7 | 1.08 | -0.0325 | -0.2025 | -0.365 | 0.902987 | 0.486098 | 0.165609 | | 1 | 0.996511 | 0.758837 |
| Uqcrh | P99028 | -0.12 | -0.16 | -1.29 | -1.4 | -0.13 | -0.13 | -1.06 | -1.31 | -0.26 | -0.48 | -1.14 | -0.14 | -0.54 | 0.07 | -1.56 | -1.33 | 0.085 | 0.2375 | -0.0975 | 0.845111 | 0.553092 | 0.838757 | | 1 | 0.996511 | 0.976176 |

| | | | | | | | | | | | | | | | | | | | | | | | | | | | |
|---|---|---|---|---|---|---|---|---|---|---|---|---|---|---|---|---|---|---|---|---|---|---|---|---|---|---|---|
| Uqcrq | Q9CQ69 | 1.45 | 1.34 | 1.48 | 1.71 | 0.5 | 1.17 | 0.56 | 1.42 | 1.56 | 2.02 | 1.44 | 1.32 | 1.92 | 0.95 | 1.11 | 1.17 | -0.5825 | 0.09 | -0.2075 | 0.043304 | 0.673963 | 0.431562 | | 1 | 0.996511 | 0.879165 |
| Urb1 | E9PU96 | 1.27 | 0.71 | 0.94 | 0.81 | 0.65 | 0.69 | 0.37 | 0.89 | 1.25 | 0.91 | 1.57 | 0.74 | 0.7 | 1.2 | 1.12 | 0.89 | -0.2825 | 0.185 | 0.045 | 0.176308 | 0.459507 | 0.836927 | | 1 | 0.996511 | 0.975831 |
| Urb2 | E9Q7L1 | 1.15 | 1.43 | 1.11 | 0.98 | 1.75 | 1.32 | 1.23 | 0.37 | 1.29 | 1.24 | 1.29 | 0.98 | 0.73 | 1.82 | 1.07 | 1.48 | -5.9E-17 | 0.0325 | 0.1075 | 1 | 0.859315 | 0.699455 | | 1 | 0.996511 | 0.955729 |
| Urgcp | Q5NCI0-2 | 0.26 | 0.63 | -0.45 | -0.54 | 0.05 | 0.01 | -0.3 | -0.39 | 0.31 | -0.39 | -0.45 | 0.13 | -0.31 | 0.5 | -0.85 | -0.53 | -0.1325 | -0.075 | -0.2725 | 0.660939 | 0.82339 | 0.49158 | | 1 | 0.996511 | 0.897531 |
| Uri1 | A0A0U1RP | 0.09 | 0.02 | -0.54 | -0.07 | -0.55 | -1.05 | -0.8 | -0.65 | 0.44 | 0.17 | -0.52 | -0.14 | -0.35 | 0.66 | -0.64 | -1.06 | -0.6375 | 0.1125 | -0.2225 | 0.01299 | 0.675261 | 0.564043 | | 1 | 0.996511 | 0.920254 |
| Urm1 | Q9D2P4 | 0.44 | 0.21 | 0.26 | 0.39 | 0.1 | -0.11 | 0.87 | 0.77 | 0.14 | -0.3 | 0.42 | 0.84 | -0.72 | -0.89 | -0.21 | -0.09 | 0.0825 | -0.05 | -0.8025 | 0.749838 | 0.849561 | 0.008293 | | 1 | 0.996511 | 0.563386 |
| Urod | P70697 | 0.38 | 0.07 | -0.34 | -0.1 | 0.01 | 0.55 | -0.15 | 0.04 | 0.05 | 0.37 | -0.31 | 0.42 | -0.03 | 0.01 | -0.51 | -0.31 | 0.11 | 0.13 | -0.2125 | 0.638396 | 0.605726 | 0.379194 | | 1 | 0.996511 | 0.860984 |
| Uros | P51163 | 0.85 | 0.42 | -0.5 | 0.38 | 0.66 | 0.82 | 0.31 | 0.52 | 0.65 | 0.62 | 0.15 | 0.78 | 0.33 | 0.85 | 0.21 | -0.18 | 0.29 | 0.2625 | 0.015 | 0.348244 | 0.416105 | 0.965909 | | 1 | 0.996511 | 0.9927 |
| Usb1 | A0A0R4J0E | 0.64 | -0.18 | -0.84 | -0.55 | -0.13 | -0.58 | -1.15 | -0.31 | 1.13 | 1.4 | -0.94 | 0.21 | 1.48 | 2.11 | -0.3 | -0.65 | -0.31 | 0.6825 | 0.8925 | 0.414085 | 0.250228 | 0.209521 | | 1 | 0.996511 | 0.786218 |
| Use1 | Q9CQ56 | 0.8 | 1.08 | 0.75 | 0.49 | 1.67 | 0.57 | 0.94 | 0.86 | 0.48 | 0.6 | 0.47 | 0.65 | 0.65 | 0.86 | 0.93 | 0.95 | 0.23 | -0.23 | 0.0675 | 0.403343 | 0.241135 | 0.740256 | | 1 | 0.996511 | 0.958796 |
| Usf1 | Q61069 | -2.59 | -5.67 | -3.4 | -3.4 | -2.07 | -3.66 | -2.94 | -2.99 | -3.7 | -1.46 | -3.34 | -2.11 | -2.82 | -5.41 | -1.37 | -2.48 | 0.85 | 1.1125 | 0.745 | 0.226463 | 0.171183 | 0.449614 | | 1 | 0.996511 | 0.884817 |
| Usf2 | F6PWU6 | -1.48 | -2.2 | -1.35 | -0.15 | -1.55 | -1.16 | -1.07 | -0.78 | -1.52 | -0.1 | -1.76 | -0.84 | -0.43 | -2.49 | 0.18 | 0.03 | 0.155 | 0.24 | 0.6175 | 0.716108 | 0.648666 | 0.376346 | | 1 | 0.996511 | 0.860517 |
| Uso1 | Q9Z1Z0 | -1.52 | -1.7 | -1.45 | -1.48 | -1.29 | -1.59 | -1.04 | -1.55 | -1.54 | -1.41 | -1.35 | -1.2 | -1.29 | -1.71 | -1.37 | -1.43 | 0.17 | 0.1625 | 0.0875 | 0.364696 | 0.350099 | 0.642793 | | 1 | 0.996511 | 0.94365 |
| Usp10 | P52479-2 | -0.89 | -1.07 | -1.24 | -1.26 | -0.73 | -1.17 | -1.6 | -1.62 | -0.99 | -0.58 | -1.41 | -0.72 | -0.63 | -0.59 | -0.85 | -1.14 | -0.165 | 0.19 | 0.3125 | 0.501889 | 0.423654 | 0.163272 | | 1 | 0.996511 | 0.758224 |
| Usp11 | Q99K46 | -0.31 | -0.11 | -1.17 | -1.38 | -0.39 | -0.62 | -0.94 | -1.36 | -0.46 | -0.39 | -1.26 | -0.34 | -0.47 | -0.15 | -1.43 | -1.35 | -0.085 | 0.13 | -0.1075 | 0.8139 | 0.725626 | 0.800983 | | 1 | 0.996511 | 0.970765 |
| Usp12 | Q9D9M2 | -0.6 | -1.16 | 0.15 | -0.47 | -0.31 | -0.41 | 0.58 | 0.3 | -3.52 | -0.63 | 0.22 | -0.33 | -2.5 | -4.81 | 1.03 | 0.46 | 0.56 | -0.545 | -0.935 | 0.133203 | 0.498974 | 0.456322 | | 1 | 0.996511 | 0.887169 |
| Usp13 | J3QSN2 | -0.1 | 0.16 | -0.24 | 0.07 | 0.14 | 0 | 0.26 | 0.22 | 0.2 | 0.08 | -0.02 | 0.13 | -0.22 | 0.09 | -0.33 | -0.23 | 0.1825 | 0.125 | -0.145 | 0.283647 | 0.478389 | 0.46888 | | 1 | 0.996511 | 0.8901 |
| Usp14 | Q9JMA1 | -0.03 | -0.11 | 0.41 | 0.59 | -0.27 | -0.01 | 0.27 | 0.54 | 0.32 | 0.98 | 0.42 | 0.16 | 0.12 | 0.43 | 0.06 | -0.01 | -0.0825 | 0.255 | -0.065 | 0.746094 | 0.34702 | 0.783453 | | 1 | 0.996511 | 0.967642 |
| Usp15 | Q8R5H1 | -0.36 | -0.57 | -0.26 | -0.45 | -0.3 | -0.44 | -0.17 | -0.18 | -1.5 | -0.61 | -0.06 | -0.48 | -1.49 | -1.08 | -0.37 | -0.34 | 0.1375 | -0.2525 | -0.41 | 0.392269 | 0.427804 | 0.197039 | | 1 | 0.996511 | 0.78004 |
| Usp19 | E9Q9M5 | -0.9 | -1.15 | -0.33 | -0.04 | -0.86 | -1.01 | -0.29 | 0.04 | -1.18 | -0.52 | 0 | -0.84 | -1.16 | -0.78 | -0.3 | -0.39 | 0.075 | -0.03 | -0.0525 | 0.826242 | 0.931697 | 0.872261 | | 1 | 0.996755 | 0.979626 |
| Usp20 | Q8C6M1 | -0.55 | -0.67 | 0.16 | 0.04 | -0.22 | -0.49 | 0.24 | -0.15 | 0.06 | 0.71 | 0 | -0.25 | -0.23 | 0.05 | 0.1 | 0.5 | 0.1 | 0.385 | 0.36 | 0.706111 | 0.218687 | 0.217021 | | 1 | 0.996511 | 0.787447 |
| Usp22 | Q5DU02 | -6.23 | -7.13 | -1.97 | -2.11 | -7.31 | -5.4 | -1.96 | -1.94 | -5.98 | -1.43 | -2.04 | -6.4 | -5.13 | -7.94 | -1.32 | -1.93 | 0.2075 | 0.3975 | 0.28 | 0.902326 | 0.812026 | 0.877194 | | 1 | 0.996511 | 0.979626 |
| Usp24 | E9PV45 | -0.46 | -0.38 | -0.47 | -0.63 | -0.11 | -0.29 | -0.24 | -0.59 | -0.47 | -0.47 | -0.36 | -0.53 | -0.45 | 0.01 | -0.63 | -0.58 | 0.1775 | 0.0275 | 0.0725 | 0.307858 | 0.86304 | 0.732879 | | 1 | 0.996511 | 0.958635 |
| Usp25 | A0A5F8MP | 0.15 | -0.05 | 0.11 | 0.11 | 0.4 | 0.13 | -0.1 | 0.16 | 0.12 | 0.27 | 0.16 | 0.18 | 0.19 | 0.24 | 0.09 | 0.06 | 0.0675 | 0.1025 | 0.065 | 0.687287 | 0.518766 | 0.704655 | | 1 | 0.996511 | 0.957084 |
| Usp30 | Q3UN04 | -0.65 | -1.66 | -0.47 | -0.1 | -1.8 | -0.37 | -0.13 | 0 | -0.51 | -0.48 | 0.19 | -0.6 | -0.67 | -0.52 | -0.58 | -1.01 | 0.145 | 0.37 | 0.025 | 0.768659 | 0.329762 | 0.942836 | | 1 | 0.996511 | 0.991519 |
| Usp31 | E9Q6Y8 | 0.25 | 0.52 | 0.1 | 0.11 | 0.18 | 0.15 | -0.15 | -0.09 | 0.47 | -0.26 | 0.14 | 0.35 | 0.83 | 0.24 | 0.29 | -0.06 | -0.2225 | -0.07 | 0.08 | 0.228431 | 0.754414 | 0.745069 | | 1 | 0.996511 | 0.959729 |
| Usp32 | F8VPZ3 | 0.3 | 0.28 | 1.24 | 0.49 | 0.44 | 0.57 | 1.27 | 1.32 | 0.67 | 0.49 | 1.42 | 0.52 | 0.96 | 0.11 | 1.59 | 1.63 | 0.3225 | 0.1975 | 0.495 | 0.322412 | 0.536263 | 0.242066 | | 1 | 0.996511 | 0.797611 |
| Usp33 | Q8R5K2-2 | -0.73 | -0.6 | 0.41 | 0.18 | 0.11 | -0.61 | 0.37 | 0.1 | -0.8 | -0.44 | -0.19 | -0.63 | -0.56 | -0.21 | 0.02 | 0.14 | 0.1775 | -0.33 | 0.0325 | 0.604613 | 0.308114 | 0.920563 | | 1 | 0.996511 | 0.98706 |
| Usp34 | F6WJB7 | -1.31 | -1.13 | -1.5 | -1.66 | -1.09 | -1.14 | -1.52 | -1.66 | -0.98 | -1.49 | -1.6 | -1.26 | -1.18 | -0.68 | -1.86 | -1.2 | 0.0475 | 0.0675 | 0.17 | 0.820882 | 0.75689 | 0.556316 | | 1 | 0.996511 | 0.917092 |
| Usp35 | M0QWN7 | -0.59 | -0.75 | -0.27 | -0.38 | -0.61 | -0.32 | -0.6 | -0.45 | -0.47 | -1.01 | -0.5 | -0.7 | -0.32 | -0.87 | -0.76 | -0.39 | 0.0025 | -0.1725 | -0.0875 | 0.988635 | 0.416935 | 0.69489 | | 1 | 0.996511 | 0.955729 |
| Usp38 | Q8BW70 | -1.81 | -0.05 | -0.72 | -0.89 | -2.29 | -0.33 | -0.56 | -1.05 | 0.35 | -1.18 | -0.38 | -0.93 | 0.83 | -1.74 | -1.31 | -0.79 | -0.19 | 0.3325 | 0.115 | 0.717006 | 0.4814 | 0.850974 | | 1 | 0.996511 | 0.976225 |
| Usp39 | Q3TIX9 | -0.35 | -1.04 | -1.28 | -1.3 | -0.74 | -0.95 | -0.74 | -0.43 | -1.09 | -0.72 | -1.43 | -1.49 | -0.29 | 0.33 | 0.15 | -0.46 | 0.2775 | -0.19 | 0.925 | 0.296115 | 0.52117 | 0.012752 | | 1 | 0.996511 | 0.573604 |
| Usp4 | P35123 | -1.04 | -0.98 | 0.17 | -0.31 | -0.96 | -0.77 | -0.06 | -0.06 | -0.77 | -0.35 | -0.08 | -0.97 | -0.58 | -0.88 | -0.31 | -0.34 | 0.0775 | -0.0025 | 0.0125 | 0.828281 | 0.994224 | 0.969103 | | 1 | 0.999285 | 0.992767 |
| Usp43 | A6PWR8 | -0.25 | -0.5 | 0.2 | -0.1 | -0.66 | -0.03 | -0.45 | 0.05 | 1.95 | 0.33 | 0.52 | 0 | 2.9 | -0.21 | 0.13 | -0.5 | -0.11 | 0.8625 | 0.7425 | 0.648059 | 0.071685 | 0.31892 | | 1 | 0.996511 | 0.838969 |
| Usp45 | Q8K387 | -1.34 | -1.23 | -0.6 | -0.53 | -3.33 | -1.27 | -1.67 | -1.16 | 1.25 | -0.16 | -1.06 | -2.1 | 1.82 | -2.72 | -1.14 | -1.17 | -0.9325 | 0.4075 | 0.1225 | 0.091244 | 0.549797 | 0.888326 | | 1 | 0.996511 | 0.980661 |
| Usp46 | P62069 | 0.47 | 0.36 | 0.17 | 0.29 | 0.74 | 0.65 | 0.2 | 0.14 | -0.17 | -0.06 | 0.11 | 0.44 | -0.44 | -0.15 | 0.01 | 0.35 | 0.11 | -0.2425 | -0.38 | 0.583829 | 0.241296 | 0.11756 | | 1 | 0.996511 | 0.718609 |
| Usp47 | A0A1L1SV7 | -0.79 | -0.96 | -0.75 | -1.01 | -0.98 | -0.81 | -0.96 | -0.86 | -0.54 | -0.46 | -0.82 | -0.81 | -0.07 | -0.28 | -0.7 | -0.98 | -0.025 | 0.22 | 0.37 | 0.867864 | 0.240082 | 0.162256 | | 1 | 0.996511 | 0.757516 |
| Usp48 | A2ALR9 | -0.82 | 1.16 | -0.68 | -0.03 | 0.16 | 0.33 | 0.95 | 0.46 | 1.61 | -0.54 | -0.25 | 0.25 | 1.86 | 2.16 | 0.73 | 0.23 | 0.5675 | 0.36 | 1.3375 | 0.227995 | 0.55368 | 0.046643 | | 1 | 0.996511 | 0.658454 |
| Usp5 | Q3U4W8 | 0.84 | 0.39 | -0.41 | -0.2 | 0.1 | 0.56 | -0.17 | -0.4 | 0.44 | -0.16 | -0.44 | 0.77 | -0.17 | 0.67 | -0.56 | -0.58 | -0.1325 | -0.0025 | -0.315 | 0.697433 | 0.994767 | 0.432928 | | 1 | 0.999285 | 0.879365 |
| Usp53 | P15975 | -0.02 | 0.22 | 0.54 | 0.01 | -1.86 | 0.39 | -1.87 | -0.36 | 1.74 | 0.45 | 0.04 | -0.45 | 2.93 | -1.71 | 0.25 | 0.33 | -1.1125 | 0.2575 | 0.2625 | 0.062251 | 0.57651 | 0.761439 | | 1 | 0.996511 | 0.963123 |
| Usp54 | Q8BL06 | 0.79 | 0.67 | 0.51 | 0.58 | 0.85 | 0.39 | 1.64 | 0.58 | 0.56 | 0.75 | 0.88 | 1.02 | 0.64 | 1.12 | 0.58 | 0.66 | 0.2275 | 0.165 | 0.1125 | 0.431265 | 0.372912 | 0.582324 | | 1 | 0.996511 | 0.927734 |
| Usp6nl | Q80XC3 | -0.31 | -0.15 | 0.59 | 0.79 | -0.25 | -0.57 | 0.33 | 0.99 | -1.01 | 0.57 | 0.83 | 0.1 | -1.08 | -0.44 | 0.41 | 0.34 | -0.105 | -0.1075 | -0.4225 | 0.798501 | 0.814721 | 0.333531 | | 1 | 0.996511 | 0.845601 |
| Usp7 | F8VPX1 | -0.19 | -0.59 | -0.56 | -0.82 | -0.39 | -0.29 | -0.27 | -0.54 | -0.9 | -0.91 | -0.47 | -0.33 | -0.96 | -0.74 | -0.63 | -0.69 | 0.1675 | -0.1125 | -0.215 | 0.378307 | 0.626821 | 0.317368 | | 1 | 0.996511 | 0.838864 |
| Usp8 | Q80U87 | -0.31 | -0.41 | 0.51 | 0.43 | -0.9 | -0.21 | 0.11 | 0.62 | 0.64 | 0.63 | 0.5 | -0.39 | 1.21 | -0.91 | 0.59 | 0.08 | -0.15 | 0.29 | 0.1875 | 0.69347 | 0.404358 | 0.695499 | | 1 | 0.996511 | 0.955729 |
| Usp9x | Q4FE56 | 0.06 | -0.11 | 0.27 | 0.09 | 0.17 | 0.16 | 0.39 | 0.33 | -0.38 | 0.26 | 0.39 | 0.2 | -0.24 | -0.16 | 0.29 | 0.25 | 0.185 | 0.04 | -0.0425 | 0.265393 | 0.857698 | 0.84209 | | 1 | 0.996511 | 0.976225 |
| Usp9x | G3UY52 | -1.75 | -3.62 | 1.91 | 2.89 | -3.17 | 1.21 | 2.58 | 2.74 | -2.57 | 3.37 | 2.05 | -1.29 | -1.61 | -2.3 | 1.62 | 2.88 | 0.9825 | 0.5325 | 0.29 | 0.595714 | 0.773024 | 0.868416 | | 1 | 0.996511 | 0.978942 |
| Ust | Q8BUB6 | 1 | 0.81 | -7.08 | -0.98 | 0.62 | 0.83 | -8.88 | -10.37 | 1.34 | 0.87 | -4.92 | 0.74 | -0.74 | 0.6 | -7.89 | -9.79 | -2.8875 | 1.07 | -2.8925 | 0.372709 | 0.619623 | 0.321012 | | 1 | 0.996511 | 0.839916 |
| Utp11 | Q9CZJ1 | 1.82 | 1.21 | 2.07 | 1.48 | 1.38 | 1.24 | 1.5 | 1.15 | 0.24 | 0.33 | 2.16 | 1.41 | 0.63 | 0.62 | 2.2 | 2.32 | -0.3275 | -0.61 | -0.2025 | 0.171103 | 0.21268 | 0.672735 | | 1 | 0.996511 | 0.952944 |
| Utp14a | Q640M1 | 0.88 | 0.94 | 1.09 | 1.28 | 0.61 | 0.02 | 0.57 | 1.45 | 1.39 | 0.89 | 1.34 | 0.6 | 1.15 | 1.48 | 0.98 | 1.37 | -0.385 | 0.0075 | 0.1975 | 0.227921 | 0.974779 | 0.349071 | | 1 | 0.997515 | 0.85293 |
| Utp15 | Q8C7V3 | 0.55 | 0.42 | 0.96 | 0.61 | -0.1 | 0.48 | 0.55 | 0.33 | 0.24 | 0.31 | 1 | 0.24 | 0.63 | 0.19 | 0.65 | 0.99 | -0.32 | -0.1875 | -0.02 | 0.158277 | 0.449792 | 0.93342 | | 1 | 0.996511 | 0.989767 |
| Utp18 | Q5SSI6 | 1.19 | 1.56 | 0.6 | 1.82 | 0.4 | 1.59 | 0.97 | 1.53 | 2.93 | 2.76 | 0.58 | 0.87 | 3.41 | 0.97 | 1.51 | 0.83 | -0.17 | 0.4925 | 0.3875 | 0.644714 | 0.431028 | 0.52131 | | 1 | 0.996511 | 0.907667 |
| Utp20 | E9QK83 | 0.36 | 0.57 | 0.91 | 0.7 | 0.12 | 0.23 | 0.64 | 0.7 | 0.68 | 0.33 | 1.05 | 0.53 | 0.26 | 0.81 | 0.66 | 1.12 | -0.2125 | 0.0125 | 0.0775 | 0.331311 | 0.95573 | 0.754754 | | 1 | 0.996755 | 0.962333 |
| Utp3 | Q9JI13-2 | 0.77 | 0.97 | 1.13 | 1.05 | 0.84 | -0.05 | 0.26 | 0.93 | 0.78 | 1.92 | 1.05 | 0.45 | 0.56 | 1.35 | 1.83 | 1.09 | -0.485 | 0.07 | 0.2275 | 0.086504 | 0.82883 | 0.443681 | | 1 | 0.996511 | 0.883549 |
| Utp4 | Q8R2N2 | 2.45 | 1.63 | 2.64 | 1.93 | 1.85 | 1.85 | 2.51 | 2.47 | -0.03 | -0.1 | 2.81 | 2.04 | 0.23 | 0.47 | 2.64 | 2.91 | 0.0075 | -0.9825 | -0.6 | 0.979749 | 0.185734 | 0.384401 | | 1 | 0.996511 | 0.860984 |
| Utp6 | Q8VCY6 | -0.64 | -0.67 | -0.58 | -0.15 | -0.49 | -0.78 | -0.57 | 0.03 | -0.8 | -0.02 | -0.36 | -0.63 | -1.1 | -0.59 | -0.15 | 0.16 | 0.0575 | 0.0575 | 0.09 | 0.802786 | 0.809013 | 0.771571 | | 1 | 0.996511 | 0.965933 |
| Utrn | E9Q6R7 | -0.16 | -0.1 | 0.1 | 0.13 | -0.28 | -0.4 | -0.18 | 0 | 0.09 | 0.08 | 0.4 | -0.08 | 0.34 | 0.09 | 0.21 | 0.26 | -0.2075 | 0.13 | 0.2325 | 0.234444 | 0.488573 | 0.218511 | | 1 | 0.996511 | 0.787447 |
| Uvrag | Q8K245 | -0.07 | 0.24 | -0.03 | -0.55 | 0.11 | -0.15 | -0.39 | -0.6 | -0.2 | -0.95 | -0.42 | -0.19 | 0.38 | 0.02 | -0.58 | -0.3 | -0.155 | -0.3375 | -0.0175 | 0.523719 | 0.218267 | 0.950711 | | 1 | 0.996511 | 0.992476 |
| Uxs1 | Q91XL3 | -1.29 | -0.89 | -0.41 | -0.31 | 0.23 | -0.6 | -0.16 | -0.47 | -0.79 | -0.23 | -0.82 | -1.27 | -1.53 | -1.38 | 0.06 | 0.33 | 0.475 | -0.0525 | 0.095 | 0.131466 | 0.866901 | 0.848083 | | 1 | 0.996511 | 0.976225 |
| Vac14 | Q80WQ2 | 0.68 | 0.71 | 0.81 | 0.63 | 0.48 | 0.62 | 0.71 | 0.66 | 0.14 | 0.67 | 0.53 | 0.66 | -0.16 | 0.65 | 0.58 | 0.71 | -0.09 | -0.2075 | -0.2625 | 0.540878 | 0.287774 | 0.297887 | | 1 | 0.996511 | 0.827473 |
| Vamp1 | D3YTU0 | 2.25 | 1.51 | 0.31 | 0.53 | 0.94 | 1.63 | 0.62 | 0.29 | 2.63 | 0.38 | 0.54 | 2.22 | 1.19 | 0.88 | 0.01 | 0.2 | -0.28 | 0.2925 | -0.58 | 0.573176 | 0.661624 | 0.259459 | | 1 | 0.996511 | 0.812337 |
| Vamp2 | B0QZN5 | 0.55 | 0.6 | 0.57 | 0.97 | 0.82 | 0.38 | 0.89 | 1.03 | 0.83 | 1.44 | 1.25 | 1.33 | -0.09 | 0.52 | 0.6 | 0.75 | 0.1075 | 0.54 | -0.2275 | 0.599967 | 0.028371 | 0.366507 | | 1 | 0.996511 | 0.858904 |
| Vamp3 | P63024 | 0.89 | 1.17 | 0.15 | -0.56 | 0.81 | 0.73 | 0.25 | 0.21 | 1.57 | 16.35 | 1.16 | 1.47 | 0.72 | 0.81 | -0.35 | -0.43 | 0.0875 | 4.725 | -0.225 | 0.825252 | 0.182207 | 0.641592 | | 1 | 0.996511 | 0.943421 |
| Vamp4 | O70480 | 0.55 | -1.29 | 1.24 | 0.99 | -1.06 | -0.37 | 2.71 | 2.01 | 0.44 | 1.14 | 1.83 | 0.3 | 0.25 | 0.55 | 1.86 | 2.45 | 0.45 | 0.555 | 0.905 | 0.642993 | 0.378556 | 0.220249 | | 1 | 0.996511 | 0.787447 |
| Vamp7 | P70280 | 0.02 | 0.35 | -0.05 | -0.23 | 0.8 | 0.33 | 0.19 | -0.27 | 0.28 | -0.89 | -0.08 | 0.39 | 0.56 | -0.04 | -0.03 | 0.03 | 0.24 | -0.0975 | 0.1075 | 0.368465 | 0.757737 | 0.644775 | | 1 | 0.996511 | 0.944749 |
| Vamp8 | O70404 | -1.29 | -1.61 | -0.17 | 1.12 | -2.28 | -1.5 | 0.11 | 0.71 | 0.09 | 2.02 | 1.09 | -0.91 | -0.14 | -1.07 | 1.04 | 0.7 | -0.2525 | 1.06 | 0.62 | 0.763321 | 0.207804 | 0.389992 | | 1 | 0.996511 | 0.865375 |
| Vangl2 | D3YY75 | 1.78 | 1.62 | 0.83 | 0.84 | 1.64 | 1.52 | 1.21 | 1.04 | 0.84 | 0.22 | 0.93 | 1.77 | 0.46 | 1.05 | 0.63 | 1.07 | 0.085 | -0.3275 | -0.465 | 0.768258 | 0.410684 | 0.153995 | | 1 | 0.996511 | 0.750763 |
| Vapa | Q9WV55 | 2.37 | 0.87 | 1.54 | 1.13 | 1.26 | 2.45 | 2.32 | 1.63 | 1.07 | 1.08 | 1.64 | 2.22 | 1.24 | 0.82 | 2.01 | 2.41 | 0.4375 | 0.025 | 0.1425 | 0.299504 | 0.951 | 0.756858 | | 1 | 0.996755 | 0.962333 |
| Vapb | Q8BH80 | 0.42 | 0.21 | 0.05 | 0.55 | 0.52 | 0.24 | 0.51 | 0.36 | 0.66 | 0.75 | 0.09 | 0.26 | 0.58 | 0.97 | 0.36 | 0.38 | 0.1 | 0.1325 | 0.265 | 0.575152 | 0.562774 | 0.260784 | | 1 | 0.996511 | 0.81412 |
| Vars1 | Q9Z1Q9 | -1.84 | -2.33 | -1.72 | -1.81 | -1.6 | -2.07 | -1.66 | -1.82 | -2.27 | -0.97 | -1.58 | -2.07 | -2.29 | -1.79 | -1.9 | -1.86 | 0.1375 | 0.2025 | -0.035 | 0.505775 | 0.532597 | 0.876403 | | 1 | 0.996511 | 0.979626 |
| Vars2 | Q3U2A8 | 0.66 | 0.87 | 0.97 | 1.06 | 0.71 | 0.74 | 0.87 | 0.98 | 0.38 | 1.19 | 0.8 | 0.68 | 0.19 | 0.67 | 0.8 | 0.8 | -0.065 | -0.1275 | -0.275 | 0.69342 | 0.572296 | 0.231543 | | 1 | 0.996511 | 0.788863 |
| Vash1 | Q8C1W1 | 0.03 | -0.32 | -0.75 | -1.12 | -0.48 | -0.25 | -1.07 | -1.06 | 0.22 | 0.21 | -0.72 | 0.01 | 0.69 | 0.84 | -0.16 | -1.19 | -0.175 | 0.47 | 0.585 | 0.585906 | 0.18164 | 0.256187 | | 1 | 0.996511 | 0.810628 |
| Vasn | Q9CZT5 | 0.28 | 0.58 | 0.57 | 0.26 | 1 | 0.06 | -0.21 | -0.59 | -0.54 | 0.1 | 0.06 | 0.12 | 0.55 | -0.32 | -0.01 | -0.06 | -0.3575 | -0.4875 | -0.3825 | 0.307254 | 0.050279 | 0.139485 | | 1 | 0.996511 | 0.73493 |
| Vasp | P70460 | -0.28 | -0.52 | 0.24 | 0.65 | -0.78 | -0.67 | -0.13 | 0.18 | -0.3 | 1.46 | 0.64 | -0.27 | -0.41 | -0.11 | 0.56 | 0.02 | -0.3725 | 0.36 | -0.0075 | 0.285009 | 0.44629 | 0.98206 | | 1 | 0.996511 | 0.994858 |
| Vat1 | Q62465 | 1.08 | 0.42 | 0.39 | 0.12 | 0.69 | 0.89 | 2.17 | 0.68 | 0.48 | 0.07 | 1.02 | 1.24 | 0.14 | 0.73 | 1.7 | 1.34 | 0.605 | 0.2 | 0.475 | 0.146977 | 0.550402 | 0.240701 | | 1 | 0.996511 | 0.795658 |
| Vat1l | Q80TB8 | 0.3 | 0.04 | -0.38 | 0.58 | -0.39 | 0.17 | 0.53 | 0.68 | 0.74 | 1.02 | -0.04 | 0.04 | 0.78 | -0.62 | 0.92 | 0.52 | 0.1125 | 0.305 | 0.265 | 0.716557 | 0.366488 | 0.504775 | | 1 | 0.996511 | 0.901002 |
| Vav1 | Q8VDU4 | -1.02 | -0.87 | 0.02 | -0.76 | -1.25 | -1.54 | -0.33 | 0.02 | -0.73 | -0.86 | -0.11 | -1.3 | -1.81 | -1.05 | -0.99 | -0.06 | -0.1175 | -0.0925 | -0.32 | 0.77498 | 0.782487 | 0.442935 | | 1 | 0.996511 | 0.883549 |
| Vav2 | Q60992 | 0.65 | 1.68 | -0.18 | -1.29 | 1.52 | 0.73 | -0.36 | -0.01 | 0.84 | 0.19 | 0.01 | 0.77 | 0.91 | 1.16 | -0.26 | -0.39 | 0.255 | 0.2375 | 0.14 | 0.710508 | 0.696794 | 0.836065 | | 1 | 0.996511 | 0.975831 |
| Vav3 | Q9R0C8 | 0.69 | 0.54 | 0.54 | 0.29 | 0.39 | 0.91 | 0.6 | 0.46 | 0.3 | -0.3 | 0.66 | 0.75 | -0.54 | 0.55 | -0.19 | 0.56 | 0.075 | -0.1625 | -0.42 | 0.685226 | 0.549148 | 0.1858 | | 1 | 0.996511 | 0.776843 |
| Vbp1 | P61759 | -0.28 | -0.38 | 0.6 | -0.22 | -0.02 | -0.47 | 0.08 | 0.11 | 0.54 | -0.53 | 0.3 | -0.52 | 2.3 | -1.37 | 0.57 | 0.49 | -0.005 | 0.0175 | 0.5675 | 0.985151 | 0.960108 | 0.432488 | | 1 | 0.996755 | 0.879165 |
| Vcam1 | Q3UPN1 | 1.03 | 1.16 | 1.38 | 1.05 | 1.22 | 1.11 | 0.61 | 1.25 | 0.46 | 0.54 | 1.18 | 1.35 | 0.51 | 0.89 | 1.32 | 1.18 | -0.1075 | -0.2725 | -0.18 | 0.596501 | 0.306984 | 0.454811 | | 1 | 0.996511 | 0.887169 |
| Vcan | E9PYH0 | -0.28 | -0.3 | -0.59 | -0.66 | -0.32 | -0.64 | -0.64 | -0.63 | -0.94 | -0.33 | -0.91 | -0.1 | -0.69 | -0.93 | -0.28 | -0.88 | -0.1 | -0.1125 | -0.2375 | 0.570944 | 0.659182 | 0.306706 | | 1 | 0.996511 | 0.832533 |
| Vcl | Q64727 | -0.41 | -0.71 | -0.13 | -0.25 | -0.44 | -0.43 | -0.03 | -0.29 | -0.43 | -0.2 | -0.2 | -0.5 | -0.43 | -0.26 | -0.62 | -0.57 | 0.0775 | 0.0425 | -0.095 | 0.69077 | 0.830432 | 0.650696 | | 1 | 0.996511 | 0.947407 |
| Vcp | Q01853 | -0.16 | -0.29 | -0.28 | -0.13 | -0.23 | -0.28 | -0.23 | -0.06 | -0.43 | 0.31 | -0.13 | 0.01 | -0.8 | -0.46 | -0.38 | -0.45 | 0.015 | 0.155 | -0.3075 | 0.918038 | 0.457508 | 0.124668 | | 1 | 0.996511 | 0.721448 |
| Vcpip1 | A0A0R4J0N | 0.13 | -0.13 | -0.21 | -0.27 | -0.17 | 0.03 | -0.24 | -0.23 | 0.07 | -0.38 | -0.15 | 0.06 | 0.01 | 0.09 | 0 | -0.55 | -0.0325 | 0.02 | 0.0075 | 0.844157 | 0.917343 | 0.972967 | | 1 | 0.996755 | 0.992865 |
| Vcpkmt | A0A571BD | -2.08 | -2.4 | -1 | -2.96 | -2.71 | -1.42 | -0.8 | 0.41 | -0.83 | -0.66 | -6.67 | -1.04 | 0.07 | -0.13 | -3.64 | -2.45 | 0.98 | -0.19 | 0.5725 | 0.184161 | 0.888439 | 0.525999 | | 1 | 0.996511 | 0.908107 |
| Vdac1 | Q60932-2 | 0.93 | 0.84 | 0.89 | 1.23 | 0.74 | 1.35 | 0.99 | 1.29 | 0.5 | 0.29 | 1.14 | 0.9 | 0.27 | 0.43 | 0.59 | 0.68 | 0.12 | -0.265 | -0.48 | 0.551978 | 0.284784 | 0.034899 | | 1 | 0.996511 | 0.651434 |
| Vdac2 | Q60930 | 0.2 | 0.35 | 0.05 | -0.14 | 0.38 | 0.43 | 0.25 | -0.15 | -0.63 | -1.38 | 0.01 | -0.04 | -0.46 | -0.7 | -0.47 | -0.09 | 0.1125 | -0.625 | -0.545 | 0.578534 | 0.091339 | 0.031264 | | 1 | 0.996511 | 0.641974 |
| Vdac2 | G3UX26 | -1.47 | -1.66 | -2.11 | -1.38 | -2.05 | -0.92 | -1.27 | -1.4 | -1.67 | -0.23 | -2.35 | -1.05 | -1.49 | -1.44 | -1.8 | -1.74 | 0.245 | 0.33 | 0.0375 | 0.404453 | 0.471443 | 0.870495 | | 1 | 0.996511 | 0.979293 |
| Vdac3 | J3QMG3 | 0.14 | 0.67 | 0.27 | 0.18 | 0.66 | 0.76 | 0.62 | 0.31 | -0.91 | -1.77 | 0.63 | 0.19 | -0.55 | -1.49 | 0.02 | 0.57 | 0.2725 | -0.78 | -0.6775 | 0.181572 | 0.156339 | 0.144734 | | 1 | 0.996511 | 0.743156 |
| Vezt | D3Z4E6 | -0.42 | -0.2 | 0.57 | 0.46 | 0.13 | -0.33 | 0.66 | 0.61 | -0.37 | 0.55 | 0.53 | -0.01 | -0.08 | -0.36 | 0.6 | 0.84 | 0.165 | 0.0725 | 0.1475 | 0.616828 | 0.82538 | 0.687911 | | 1 | 0.996511 | 0.955729 |
| Vgf | Q0VGU4 | 0.72 | 0.62 | -0.01 | 0.38 | -0.87 | 0.98 | -0.59 | 0.1 | 1.47 | 0.52 | 0.16 | 0.84 | 1.62 | -0.25 | 0.67 | 0.31 | -0.5225 | 0.32 | 0.16 | 0.230337 | 0.334293 | 0.696175 | | 1 | 0.996511 | 0.955729 |
| Vhl | P40338 | -1.33 | -1.28 | -0.66 | -0.68 | -1.36 | -1.31 | 0.25 | -0.14 | -0.21 | 0.71 | -0.19 | -1.75 | -0.17 | 0.36 | -0.4 | -0.27 | 0.3475 | 0.6275 | 0.8675 | 0.418058 | 0.23438 | 0.010519 | | 1 | 0.996511 | 0.563386 |
| Vill | G5E8H0 | 2.54 | 1.45 | 0.02 | 0.05 | 2.59 | 1.18 | 0.2 | 0.09 | 2.15 | 0.54 | -0.4 | 0.45 | 3.2 | 1.16 | 0.91 | 0.85 | -8.8E-16 | -0.33 | 0.515 | 1 | 0.654342 | 0.496883 | | 1 | 0.996511 | 0.897918 |
| Vim | P20152 | 0.27 | -0.8 | -0.31 | -0.55 | -0.83 | -0.42 | -0.62 | -0.81 | -0.15 | -0.31 | 0.03 | 0.63 | -0.08 | -1 | 0.23 | -0.01 | -0.3225 | 0.3975 | 0.1325 | 0.231878 | 0.222215 | 0.70691 | | 1 | 0.996511 | 0.957157 |
| Vip | A0A0R4J00 | 1.04 | 1.13 | 0.97 | 0.7 | 1.72 | 0.72 | 1.34 | 1.17 | 0.24 | 0.79 | 1.36 | 1.04 | 1.08 | 2.17 | 0.87 | 1.52 | 0.2775 | -0.1025 | 0.45 | 0.269883 | 0.704983 | 0.172831 | | 1 | 0.996511 | 0.76428 |
| Vipas39 | Q8BGQ1 | 1.27 | 0.84 | 0.43 | 0.15 | 0.75 | 1.35 | 0.55 | 0.44 | 0.16 | 0.46 | 0.62 | 1.36 | -0.59 | 0.48 | 0.19 | 0.61 | 0.1 | -0.0225 | -0.5 | 0.749343 | 0.948234 | 0.187242 | | 1 | 0.996755 | 0.776843 |
| Virma | E9PZY8 | -0.19 | -2.18 | -1.03 | -1.37 | -2.85 | -0.13 | -0.52 | -0.37 | 1.41 | -1.05 | -1.29 | -2.03 | 3.48 | -2.2 | -0.13 | -0.81 | 0.225 | 0.4525 | 1.2775 | 0.742948 | 0.561587 | 0.280296 | | 1 | 0.996511 | 0.822084 |
| Vkorc1 | Q9CRC0 | -1.23 | -1.98 | -1.18 | -1.05 | -2.48 | -1.23 | -1.97 | -1.75 | 1.31 | -1.81 | -1.72 | -0.92 | 3.18 | -2.49 | 0.77 | -1.15 | -0.4975 | 0.575 | 1.4375 | 0.152402 | 0.413015 | 0.219054 | | 1 | 0.996511 | 0.787447 |
| Vkorc1l1 | Q6TEK5 | -1.19 | -0.91 | -0.49 | -0.25 | 0.14 | -0.79 | 0.09 | -0.21 | -1.02 | -1.09 | -0.71 | -1.04 | -1.01 | 0.01 | 0.89 | 0.03 | 0.5175 | -0.255 | 0.69 | 0.110273 | 0.323268 | 0.128959 | | 1 | 0.996511 | 0.723655 |
| Vldlr | Q91YY0 | 0.41 | 0.47 | 0.24 | 0.4 | 1.1 | 0.63 | 0.68 | 0.35 | -0.02 | 0.89 | 0.45 | 0.5 | 0.35 | 0.16 | 0.75 | 0.62 | 0.31 | 0.075 | 0.09 | 0.142277 | 0.741291 | 0.661418 | | 1 | 0.996511 | 0.949645 |
| Vma21 | Q78T54 | 0.26 | 0.29 | 0.52 | 0.41 | 0.47 | 0.24 | 1.05 | 0.77 | -1.27 | -0.11 | 0.3 | -0.2 | -0.3 | -0.32 | 0.74 | 0.68 | 0.2625 | -0.69 | -0.17 | 0.238267 | 0.066584 | 0.586893 | | 1 | 0.996511 | 0.92838 |
| Vmp1 | Q99KU0 | -0.93 | -0.56 | 0.32 | 0.1 | -0.19 | -0.81 | 1.12 | 0.08 | -1.13 | 0.08 | 0.42 | -0.64 | -2.23 | 0.06 | 0.86 | 1.12 | 0.3175 | -0.05 | 0.22 | 0.496473 | 0.90723 | 0.765728 | | 1 | 0.996744 | 0.964134 |
| Vopp1 | A0A0N4SV | -0.07 | -0.34 | -3.09 | -3.15 | -0.13 | -0.06 | -2.96 | -3.58 | 1.06 | -2 | -2.19 | 0.04 | 0.8 | -0.45 | -2.44 | -2.74 | -0.02 | 0.89 | 0.455 | 0.985753 | 0.401923 | 0.67016 | | 1 | 0.996511 | 0.952558 |
| Vps11 | Q91W86 | -0.01 | -0.25 | 0.53 | 0.09 | 0.18 | -0.11 | 0.85 | 0.41 | -0.31 | -0.38 | 0.52 | 0.12 | -0.37 | 0.01 | 0.49 | 0.55 | 0.2425 | -0.1025 | 0.08 | 0.375293 | 0.713267 | 0.781869 | | 1 | 0.996511 | 0.967642 |
| Vps13a | A0A494B9( | 0.31 | 0.43 | 1.05 | 1.02 | 0.56 | -0.13 | 0.8 | 0.14 | 0.29 | 1.25 | 0.82 | -0.08 | -0.28 | 0.63 | 0.73 | 0.9 | -0.36 | -0.1325 | -0.2075 | 0.229231 | 0.701779 | 0.535036 | | 1 | 0.996511 | 0.912212 |
| Vps13a | Q5H8C4 | -0.19 | 0.03 | 0.17 | 0.04 | 0.27 | 0.03 | 0.01 | -0.19 | -0.09 | -0.14 | 0.19 | -0.21 | -0.16 | 0.27 | -0.27 | -0.07 | 0.0175 | -0.075 | -0.07 | 0.918738 | 0.679508 | 0.730683 | | 1 | 0.996511 | 0.958635 |
| Vps13b | Q80TY5 | -0.7 | -0.37 | -1.29 | 1.16 | 0.67 | -1.92 | -1.47 | -0.53 | -0.38 | -0.03 | -0.83 | -1.24 | -0.36 | 0.4 | 0.16 | -0.04 | -0.5125 | -0.32 | 0.34 | 0.472451 | 0.557884 | 0.509416 | | 1 | 0.996511 | 0.902804 |
| Vps13c | Q8BX70 | 0.52 | 0.72 | 0.54 | 0.46 | 1 | 0.63 | 0.42 | 0.42 | 0.69 | 0.2 | 0.53 | 0.57 | 0.36 | 0.9 | 0.33 | 0.63 | 0.0575 | -0.0625 | -0.005 | 0.759908 | 0.732816 | 0.980626 | | 1 | 0.996511 | 0.994858 |
| Vps13d | B1ART2 | 0.28 | 0.41 | 0.21 | 0.01 | 0.87 | 0.59 | 0.32 | 0.05 | 0.29 | -0.36 | 0.27 | 0.43 | 0.48 | 0.45 | 0.08 | 0.31 | 0.23 | -0.07 | 0.1025 | 0.30932 | 0.759348 | 0.60243 | | 1 | 0.996511 | 0.930918 |
| Vps16 | G3X8X7 | 0.9 | 0.55 | -0.37 | -0.38 | 0.94 | 0.91 | -0.16 | -0.54 | 0.13 | -0.86 | -0.35 | 0.97 | 0.25 | 0.34 | -0.58 | -0.44 | 0.1125 | -0.2025 | -0.2825 | 0.807747 | 0.670903 | 0.474603 | | 1 | 0.996511 | 0.891711 |
| Vps18 | Q8R307 | 0.41 | 0.06 | 0.55 | 0.66 | 0.44 | 0.19 | 0.47 | 0.55 | -0.4 | 0.54 | 0.33 | 0.25 | -0.44 | -0.15 | 0.46 | 0.41 | -0.0075 | -0.24 | -0.35 | 0.968532 | 0.368612 | 0.227225 | | 1 | 0.996511 | 0.787447 |
| Vps25 | Q9CQ80 | -0.03 | 0.04 | -0.66 | -0.34 | 0.32 | -0.21 | 0.12 | -0.38 | 0.02 | -0.7 | -0.64 | -0.03 | -0.44 | 0.57 | -0.88 | -0.47 | 0.21 | -0.09 | -0.0575 | 0.39343 | 0.736949 | 0.868093 | | 1 | 0.996511 | 0.978872 |
| Vps26a | P40336 | 1.98 | 1.18 | 2.31 | 0.5 | 1.68 | 2.11 | 2.64 | 1.84 | -0.11 | -0.06 | 2.46 | 2.14 | -0.56 | 0.41 | 1.76 | 2.37 | 0.575 | -0.385 | -0.4975 | 0.203115 | 0.599916 | 0.485898 | | 1 | 0.996511 | 0.895384 |
| Vps26b | Q8C0E2 | 0.52 | 0.09 | 0.1 | 0.44 | 0.3 | 0.61 | 0.25 | 0.18 | 0.41 | 0.22 | 0.58 | 0.54 | 0.13 | 0.51 | 0.04 | 0.01 | 0.0475 | 0.15 | -0.115 | 0.800053 | 0.445507 | 0.596207 | | 1 | 0.996511 | 0.929911 |
| Vps26c | O35075 | -0.94 | -0.73 | -1.31 | -1.2 | -0.6 | -0.59 | -1.08 | -1.24 | -0.52 | -0.56 | -1.3 | -0.71 | -0.48 | 0.4 | -1.64 | -0.95 | 0.1675 | 0.2725 | 0.3775 | 0.474546 | 0.287512 | 0.387232 | | 1 | 0.996511 | 0.863737 |
| Vps28 | Q9D1C8 | 0.56 | 0.36 | 0.28 | 0.43 | 0.39 | 0.38 | 0.53 | 0.61 | 0.59 | 0.41 | 0.46 | 0.65 | 0.54 | 0.46 | 0.34 | 0.25 | 0.07 | 0.12 | -0.01 | 0.648791 | 0.475913 | 0.955522 | | 1 | 0.996511 | 0.992482 |
| Vps29 | D3Z645 | -0.01 | -0.78 | -0.4 | -0.66 | -0.46 | 0.58 | -0.48 | -0.65 | -0.33 | -1.91 | -0.32 | 0.12 | -0.42 | -0.58 | -0.71 | -0.41 | 0.21 | -0.1475 | -0.0675 | 0.518197 | 0.743151 | 0.769177 | | 1 | 0.996511 | 0.96587 |
| Vps33a | Q9D2N9 | 0.31 | 0.54 | -0.09 | -0.02 | 0.43 | 0.35 | -0.02 | -0.03 | -0.31 | -0.04 | 0.02 | 0.36 | -0.31 | -0.21 | -0.37 | -0.35 | -0.0025 | -0.1775 | -0.495 | 0.990744 | 0.452837 | 0.040003 | | 1 | 0.996511 | 0.658454 |
| Vps33b | P59016 | 0.09 | 0.04 | -0.01 | 0.17 | 0.01 | -0.16 | 0.22 | 0.01 | 0.09 | 0.6 | 0.12 | -0.04 | -0.14 | 0.11 | 0.06 | 0.13 | -0.0525 | 0.12 | -0.0325 | 0.735602 | 0.546772 | 0.852133 | | 1 | 0.996511 | 0.976225 |
| Vps35 | Q9EQH3 | 0.26 | 0.19 | -0.04 | -0.55 | 0.3 | 0.57 | -0.16 | -0.12 | 0.44 | -1.09 | 0.14 | 0.3 | 0.48 | -0.09 | -0.24 | -0.08 | 0.1825 | -0.0175 | 0.0525 | 0.49183 | 0.963391 | 0.844592 | | 1 | 0.996755 | 0.976225 |
| Vps35l | Q8BWQ6 | -0.8 | -0.77 | -0.1 | -0.09 | -0.8 | -0.7 | 0.02 | 0.03 | -0.89 | -0.2 | 0.03 | -0.78 | -0.8 | -0.61 | -0.1 | 0.03 | 0.0775 | -0.02 | 0.07 | 0.795303 | 0.947439 | 0.813259 | | 1 | 0.996755 | 0.972768 |
| Vps36 | Q91XD6 | -0.6 | -0.54 | -0.03 | -0.11 | -0.72 | -0.74 | 0.04 | -0.07 | -0.78 | -0.46 | -0.18 | -0.51 | -0.59 | -0.44 | -0.25 | -0.19 | -0.0525 | -0.1625 | -0.0475 | 0.841181 | 0.477359 | 0.830497 | | 1 | 0.996511 | 0.973939 |
| Vps37a | Q8CHS8 | 0.08 | 0.71 | 0.94 | 0.75 | 0.5 | -0.11 | 1.21 | 0.61 | 0.16 | 0.39 | 0.93 | 0.29 | -0.42 | 1.18 | 0.38 | 1.13 | -0.0675 | -0.1775 | -0.0525 | 0.833587 | 0.513615 | 0.896665 | | 1 | 0.996511 | 0.983509 |
| Vps37b | Q8R0J7 | 0.78 | 0.64 | 0.55 | 0.52 | 1.03 | 0.52 | 0.72 | 0.55 | 0.58 | 0.3 | 0.56 | 0.79 | 0.6 | 0.91 | 0.29 | 0.54 | 0.0825 | -0.065 | -0.0375 | 0.644171 | 0.720629 | 0.853965 | | 1 | 0.996511 | 0.976225 |
| Vps37c | Q8R105 | 0.63 | 0.68 | 0.39 | 0.33 | 0.84 | 0.26 | 0.62 | 0.45 | 0.49 | 0.82 | 0.54 | 0.97 | -0.08 | 0.93 | 0.1 | 0.29 | 0.035 | 0.1975 | -0.1975 | 0.853601 | 0.327982 | 0.461331 | | 1 | 0.996511 | 0.887902 |
| Vps37d | Q810I0 | 3.11 | 1.02 | -0.15 | -0.37 | 0.49 | 1.78 | -0.01 | -0.15 | 3.02 | 1.5 | -0.26 | 3.13 | 2.82 | 0.64 | 0.1 | -0.29 | -0.375 | 0.945 | -0.085 | 0.649227 | 0.362365 | 0.928386 | | 1 | 0.996511 | 0.98946 |
| Vps39 | Q8R5L3-2 | -0.12 | 0.18 | -0.31 | -0.61 | 0.55 | 0.03 | 0.1 | -0.54 | -0.24 | -0.93 | -0.34 | -0.08 | -0.4 | 0.5 | -0.49 | -0.21 | 0.25 | -0.1825 | 0.065 | 0.385035 | 0.498367 | 0.825216 | | 1 | 0.996511 | 0.973091 |
| Vps41 | Q5KU39 | 0.28 | 0.36 | 0.12 | 0.25 | 0.79 | 0.36 | 0.6 | 0.31 | -1.01 | -0.23 | 0.21 | 0.24 | -0.84 | -0.78 | 0.22 | 0.28 | 0.2625 | -0.45 | -0.5325 | 0.154596 | 0.160928 | 0.120887 | | 1 | 0.996511 | 0.721448 |
| Vps45 | P97390 | -0.56 | -0.54 | -0.29 | -0.14 | -0.34 | -0.6 | -0.36 | -0.21 | -0.24 | 0.09 | -0.35 | -0.51 | -0.3 | -0.22 | -0.32 | -0.45 | 5.00E-03 | 0.13 | 0.06 | 0.977467 | 0.535133 | 0.752462 | | 1 | 0.996511 | 0.961479 |
| Vps4a | Q8VEJ9 | -0.5 | -0.06 | -0.59 | -0.85 | -0.03 | -0.6 | -0.5 | -0.88 | -0.13 | -0.92 | -0.71 | -0.24 | 0.48 | 0.17 | -0.46 | -0.76 | -0.0025 | -3.9E-17 | 0.3575 | 0.992096 | 1 | 0.296673 | | 1 | 1 | 0.825612 |
| Vps4b | P46467 | -0.5 | -0.34 | -0.4 | -0.91 | -0.18 | -0.5 | -0.49 | -0.7 | -0.31 | -0.85 | -0.64 | -0.36 | 0.74 | -0.52 | -0.41 | -0.56 | 0.07 | -0.0025 | 0.35 | 0.727462 | 0.990855 | 0.314885 | | 1 | 0.999285 | 0.837427 |
| Vps50 | Q8CI71 | -0.73 | -0.64 | 0.56 | 0.42 | -0.69 | -0.67 | 0.27 | 0.64 | -1.04 | -0.09 | 0.63 | -0.61 | -1.57 | -1.01 | -0.16 | 0.14 | -0.015 | -0.18 | -0.5525 | 0.973149 | 0.699312 | 0.273405 | | 1 | 0.996511 | 0.822084 |
| Vps51 | Q3UVL4 | -0.37 | -0.66 | -0.02 | 0.07 | -0.41 | -0.11 | -0.05 | 0.1 | -0.75 | 0.09 | 0.17 | -0.48 | -1.26 | -0.45 | -0.75 | -0.48 | 0.1275 | 0.0025 | -0.49 | 0.568991 | 0.993072 | 0.101507 | | 1 | 0.999285 | 0.698697 |
| Vps52 | Q8C754 | -0.26 | -0.02 | 0.31 | 0.49 | -0.06 | -0.45 | -0.12 | 0.32 | -0.06 | 0.41 | 0.3 | -0.12 | -0.49 | 0.25 | -0.47 | -0.3 | -0.2075 | 0.0025 | -0.3825 | 0.405613 | 0.991684 | 0.176033 | | 1 | 0.999285 | 0.771503 |
| Vps53 | Q8CCB4 | 0.1 | 0.29 | -0.23 | -0.16 | -0.06 | 0.11 | -0.37 | -0.17 | 0.05 | -0.65 | -0.03 | 0.15 | -0.22 | -0.56 | -0.62 | -0.6 | -0.1225 | -0.12 | -0.5 | 0.530993 | 0.623659 | 0.038971 | | 1 | 0.996511 | 0.658454 |
| Vps54 | Q5SRW8 | -1.45 | -0.88 | -0.38 | -0.99 | -1.53 | -1.6 | -1.9 | -0.83 | -0.42 | -1.64 | -0.66 | -0.85 | 0.39 | -1.13 | -0.5 | -0.32 | -0.54 | 0.0325 | 0.535 | 0.109862 | 0.923583 | 0.176704 | | 1 | 0.996755 | 0.771503 |
| Vps8 | D3YUP0 | 0.41 | 0.18 | -0.18 | -0.25 | 0.08 | 0.04 | -0.17 | -0.27 | 0.05 | -0.46 | -0.3 | 0.13 | 0.38 | -0.04 | -0.51 | -0.45 | -0.12 | -0.185 | -0.195 | 0.564387 | 0.444825 | 0.491128 | | 1 | 0.996511 | 0.897531 |
| Vps9d1 | Q8C190 | 0.22 | 0.37 | -0.13 | -0.37 | 0.32 | 0.3 | -0.6 | -0.67 | 0.47 | 0.12 | -0.42 | 0.59 | 1.13 | 1.4 | -0.51 | -0.68 | -0.185 | 0.1675 | 0.3125 | 0.560393 | 0.567914 | 0.555755 | | 1 | 0.996511 | 0.917092 |
| Vrk1 | A0A1Y7VM | -0.74 | -0.5 | -1.57 | -2.07 | -1.4 | -0.59 | -1.09 | -1.69 | -0.18 | -1.44 | -1.25 | -0.91 | 0.32 | -1.53 | -1.7 | -1.15 | 0.0275 | 0.275 | 0.205 | 0.946217 | 0.530182 | 0.706341 | | 1 | 0.996511 | 0.957084 |
| Vrk2 | Q8BN21-2 | -0.32 | -1.99 | 0.02 | 0.17 | 0.13 | -0.02 | 0.14 | 0.03 | 0.96 | 0.28 | 0.54 | -1.7 | 1.29 | 0.14 | -0.58 | 0.42 | 0.6 | 0.55 | 0.8475 | 0.217438 | 0.441399 | 0.168345 | | 1 | 0.996511 | 0.759484 |
| Vrk3 | Q8K3G5 | -0.91 | -1.37 | -0.78 | 0.16 | -0.44 | -1.74 | 0.74 | -0.18 | -0.97 | 0.25 | 0.41 | -1.52 | -1.49 | -0.59 | 0.07 | -0.23 | 0.32 | 0.2675 | 0.165 | 0.566946 | 0.613919 | 0.709758 | | 1 | 0.996511 | 0.957763 |
| Vsnl1 | P62761 | 0.13 | 0.26 | 0.51 | 0.5 | 0.56 | 0.28 | 0.44 | 0.66 | 0.03 | -0.19 | 0.78 | 0.26 | -0.14 | 0.13 | -0.02 | 0.55 | 0.135 | -0.13 | -0.22 | 0.44557 | 0.607095 | 0.341474 | | 1 | 0.996511 | 0.848473 |
| Vstm2a | A0A140LHE | -0.11 | -1.08 | -1.2 | -1 | -1.47 | 0.3 | -0.68 | -0.14 | -1.69 | -1.5 | -0.9 | 0.45 | -1.18 | -2.4 | 0.07 | -0.52 | 0.35 | -0.0625 | -0.16 | 0.420949 | 0.901528 | 0.767916 | | 1 | 0.996609 | 0.965084 |
| Vstm2b | Q9JME9 | 0.56 | -0.23 | -0.07 | -0.09 | 0.04 | 0.89 | 0.76 | 0.51 | -0.46 | 0.33 | -0.04 | 0.26 | -0.21 | -1.34 | -0.48 | -0.25 | 0.5075 | -0.02 | -0.6125 | 0.082521 | 0.940455 | 0.084713 | | 1 | 0.996755 | 0.691619 |
| Vstm2l | A2ACD2 | -1.39 | -2.52 | -0.65 | -0.72 | -2.13 | -1.38 | 0.03 | 0.1 | -0.72 | 0.16 | -0.5 | -1.62 | 0.4 | -0.89 | 0.67 | -0.45 | 0.475 | 0.65 | 1.2525 | 0.461874 | 0.237532 | 0.038376 | | 1 | 0.996511 | 0.658454 |
| Vta1 | Q9CR26 | 0.21 | 0.17 | 0.03 | -0.16 | 0.25 | 0.59 | -0.29 | -0.3 | -0.33 | -0.52 | -0.15 | 0.3 | 0.86 | -1.19 | 0.27 | -0.15 | -1.2E-16 | -0.2375 | -0.115 | 1 | 0.312379 | 0.784727 | | 1 | 0.996511 | 0.967642 |
| Vti1a | A0A286YDI | -0.35 | 0.62 | -0.99 | -1.25 | -0.1 | 0.08 | -0.44 | -0.08 | 0.95 | 0.33 | -0.88 | 0.51 | 1.79 | 0.83 | -0.2 | -0.61 | 0.3575 | 0.72 | 0.945 | 0.388688 | 0.196959 | 0.154816 | | 1 | 0.996511 | 0.750763 |
| Vti1a | Q5FWJ7 | 0.91 | 1.02 | 0.59 | 0.61 | 1.45 | 1.09 | 0.83 | 0.42 | 1.11 | 0.69 | 0.55 | 1.1 | 1.18 | 1.48 | 0.72 | 1.05 | 0.165 | 0.08 | 0.325 | 0.519916 | 0.71405 | 0.187608 | | 1 | 0.996511 | 0.776843 |
| Vti1b | Q91XH6 | -0.26 | -0.08 | -0.35 | -0.27 | 0.36 | -0.29 | -0.37 | -0.26 | -0.31 | 0.15 | -0.26 | -0.36 | 0.05 | 0.73 | 0.02 | -0.16 | 0.1 | 0.045 | 0.4 | 0.631361 | 0.811386 | 0.124927 | | 1 | 0.996511 | 0.721448 |
| Vtn | P29788 | -0.95 | -1.23 | -1.52 | -1.66 | -0.73 | -1.25 | -1.57 | -1.79 | -0.89 | 0.08 | -1.71 | -0.58 | 0.44 | -0.42 | -0.98 | -1.21 | 0.005 | 0.565 | 0.7975 | 0.985818 | 0.170552 | 0.065666 | | 1 | 0.996511 | 0.664936 |
| Vwa1 | Q8R2Z5 | -0.07 | 0.54 | 0.16 | 0.32 | 0.02 | 0.71 | 0.83 | 0.21 | 0.11 | 0.5 | 0.47 | 0.94 | 0.87 | 0.58 | 0.74 | 0.4 | 0.205 | 0.2675 | 0.41 | 0.415336 | 0.283481 | 0.086021 | | 1 | 0.996511 | 0.691619 |
| Vwa5a | Q99KC8 | -0.29 | -0.35 | -0.52 | -0.56 | -0.37 | -0.44 | -0.63 | -0.47 | -0.87 | -1.15 | -0.43 | -0.58 | -0.89 | -0.65 | -0.57 | -0.71 | -0.0475 | -0.3275 | -0.275 | 0.758866 | 0.151032 | 0.15672 | | 1 | 0.996511 | 0.751493 |
| Vwa5b2 | E0CZ01 | -2.24 | -2.71 | -0.87 | -0.69 | -2.1 | -1.91 | -0.65 | -0.82 | -0.95 | -0.71 | -0.79 | -1.44 | -1.09 | -1.98 | -1.13 | -0.17 | 0.2575 | 0.655 | 0.535 | 0.652677 | 0.204619 | 0.36202 | | 1 | 0.996511 | 0.858475 |
| Vwa8 | Q8CC88 | 0.79 | 0.91 | 0.9 | 0.94 | 0.48 | 0.69 | 0.96 | 0.81 | 0.53 | 0.64 | 0.74 | 0.4 | 0.51 | 0.95 | 0.64 | 0.61 | -0.15 | -0.3075 | -0.2075 | 0.37352 | 0.090283 | 0.278858 | | 1 | 0.996511 | 0.822084 |
| Vwc2l | Q505H4-3 | -0.76 | 0.1 | 1.33 | 0.83 | -0.38 | 0.61 | 0.95 | 0.76 | 0.21 | 0.05 | 0.6 | 0.08 | 0.01 | 0.28 | 0.38 | 0.38 | 0.11 | -0.14 | -0.1125 | 0.826254 | 0.753413 | 0.798115 | | 1 | 0.996511 | 0.969727 |
| Vwf | E9QPU1 | -2.16 | -2.83 | -3.15 | -1.92 | -2.42 | -2.04 | -3.38 | -2.59 | -2.59 | -0.33 | -2.73 | -2.19 | -1.58 | -2.23 | -1.72 | -3.06 | -0.0925 | 0.555 | 0.3675 | 0.808447 | 0.347309 | 0.392991 | | 1 | 0.996511 | 0.865775 |
| Vxn | Q8BG31 | -0.92 | -0.82 | -1 | -1.12 | -2.26 | -1.55 | -3.47 | -1.65 | -0.45 | -0.82 | -1.21 | -0.81 | 1.34 | -0.39 | -1.31 | -2.06 | -1.2675 | 0.1425 | 0.36 | 0.013845 | 0.504016 | 0.593443 | | 1 | 0.996511 | 0.929843 |
| Wac | Q924H7-3 | -3.4 | -4.25 | -3.75 | -2.87 | -4.15 | -6.04 | -4.38 | -3.51 | -4.44 | -2.03 | -3.65 | -4.13 | -3.47 | -2.35 | -4.13 | -4.05 | -0.9525 | 0.005 | 0.0675 | 0.117067 | 0.992888 | 0.886302 | | 1 | 0.999285 | 0.979767 |

| | | | | | | | | | | | | | | | | | | | | | | | | | | | |
|---|---|---|---|---|---|---|---|---|---|---|---|---|---|---|---|---|---|---|---|---|---|---|---|---|---|---|---|
| Wapl | B7ZP47 | -4.62 | -7.86 | -5.17 | -6.69 | -6.58 | -6.61 | -7.33 | -7.2 | -6.75 | -1.66 | -4.79 | -5.72 | -4.94 | -6.19 | -4.78 | -5.86 | -0.845 | 1.355 | 0.6425 | 0.241345 | 0.271089 | 0.392843 | | 1 | 0.996511 | 0.865775 |
| Wars1 | P32921-2 | -0.62 | -0.47 | -0.43 | -0.05 | -0.38 | -0.57 | -0.16 | -0.1 | -0.7 | -0.31 | -0.35 | -0.38 | -0.81 | -0.48 | -0.43 | -0.67 | 0.09 | -0.0425 | -0.205 | 0.649338 | 0.831902 | 0.340018 | | 1 | 0.996511 | 0.848092 |
| Wars2 | Q9CYK1 | 1.24 | 0.84 | 0.96 | 0.92 | 1.02 | 1.25 | 0.53 | 0.96 | 0.16 | 0.11 | 0.52 | 0.65 | -0.01 | 0.6 | 0.1 | 0.46 | -0.05 | -0.63 | -0.7025 | 0.806908 | 0.012819 | 0.01045 | | 1 | 0.996511 | 0.563386 |
| Wasf1 | Q8R5H6 | 1.88 | 1.17 | 1.07 | 1.25 | 0.74 | 1.74 | 0.83 | 1.17 | 1.97 | 0.74 | 1.16 | 2.01 | 1.33 | 1.27 | 0.72 | 0.71 | -0.2225 | 0.1275 | -0.335 | 0.452531 | 0.71943 | 0.239593 | | 1 | 0.996511 | 0.794626 |
| Wasf2 | Q8BH43 | -0.5 | -0.73 | -0.06 | 0.14 | -0.84 | -0.96 | 0.08 | 0.18 | -0.16 | 1.24 | 0.42 | -0.34 | -0.56 | 0.15 | -0.03 | -0.44 | -0.0975 | 0.5775 | 0.0675 | 0.778469 | 0.165474 | 0.809989 | | 1 | 0.996511 | 0.972222 |
| Wasf3 | A0A571BF7 | 1.19 | 0.97 | 1.26 | 1.32 | 1.14 | 1.48 | 1.06 | 1.13 | 1.32 | 2.29 | 1.38 | 0.9 | 1.7 | 1.56 | 1.06 | 0.86 | 0.0175 | 0.2875 | 0.11 | 0.919033 | 0.360834 | 0.659491 | | 1 | 0.996511 | 0.949645 |
| Wasf3 | Q8VHI6 | 0.38 | 0.91 | 0.94 | 1.32 | 0.69 | 0.06 | 0.42 | 0.92 | 0.94 | 0.97 | 0.63 | 0.08 | 0.87 | 1.36 | 0.68 | 0.43 | -0.365 | -0.2325 | -0.0525 | 0.203657 | 0.433509 | 0.85722 | | 1 | 0.996511 | 0.976225 |
| Washc1 | Q8VDD8 | -0.16 | -0.18 | -0.07 | -0.36 | -0.26 | -0.45 | -0.65 | -0.17 | -1.59 | 0.62 | 0.08 | -0.14 | -1.13 | -1.7 | 0.56 | -0.37 | -0.19 | -0.065 | -0.4675 | 0.289277 | 0.884556 | 0.328276 | | 1 | 0.996511 | 0.844932 |
| Washc2 | Q6PGL7 | -0.2 | -0.42 | -0.57 | -0.28 | -0.6 | -0.38 | -0.34 | -0.5 | 0.08 | 0.08 | -0.5 | -0.22 | -0.18 | -0.08 | -0.75 | -0.59 | -0.0875 | 0.2275 | -0.0325 | 0.590994 | 0.287782 | 0.885943 | | 1 | 0.996511 | 0.979767 |
| Washc3 | S4R287 | -0.16 | 0.17 | -0.37 | -0.21 | 0.51 | -0.24 | -0.6 | -0.41 | 0.1 | 1.09 | -0.48 | 0.02 | -0.46 | 0.77 | -0.46 | -0.39 | -0.0425 | 0.325 | 0.0075 | 0.876297 | 0.355337 | 0.981681 | | 1 | 0.996511 | 0.994858 |
| Washc4 | Q3UMB9 | -0.1 | -0.03 | -0.07 | -0.46 | -0.01 | -0.09 | -0.24 | -0.16 | -0.18 | -0.24 | -0.05 | -0.31 | 0.22 | -0.09 | 0.01 | -0.21 | 0.04 | -0.03 | 1.48E-01 | 0.81047 | 0.867599 | 0.4698 | | 1 | 0.996511 | 0.890289 |
| Washc5 | Q8C2E7 | -0.21 | -0.22 | 0.01 | -0.02 | -0.28 | -0.41 | -0.03 | -0.16 | -0.14 | 0.12 | -0.04 | -0.29 | -0.43 | 0.25 | -0.34 | -0.03 | -0.11 | 0.0225 | -0.0275 | 0.502931 | 0.898267 | 0.899815 | | 1 | 0.996511 | 0.983509 |
| Wasl | Q91YD9 | -0.6 | -0.38 | -0.19 | 0.25 | -0.61 | -0.62 | -0.48 | 0.09 | -0.31 | 0.45 | -0.31 | -0.93 | -0.4 | -0.09 | -0.31 | -0.22 | -0.175 | -0.045 | -0.025 | 0.501104 | 0.892258 | 0.91499 | | 1 | 0.996511 | 0.986556 |
| Wbp11 | Q923D5 | -0.44 | -2.2 | -1.55 | -1.45 | -1.45 | -1.43 | -2.09 | -1.51 | -1.79 | -0.85 | -1.66 | -2.27 | -1.01 | -1.05 | -1.01 | -2.13 | -0.21 | -0.2325 | 0.11 | 0.580541 | 0.601209 | 0.800223 | | 1 | 0.996511 | 0.970393 |
| Wbp2 | P97765 | -0.27 | -0.72 | -0.33 | -0.12 | -0.49 | -0.5 | -0.4 | -0.04 | -0.38 | -0.61 | -0.11 | 0.01 | -0.66 | -0.14 | -0.53 | -0.47 | 0.0025 | 0.0875 | -0.09 | 0.990033 | 0.697189 | 0.683842 | | 1 | 0.996511 | 0.955729 |
| Wbp4 | Q61048 | -2.58 | -2.45 | -2.1 | -1.82 | -4.05 | -1.68 | -1.47 | -1.75 | -0.36 | -1.04 | -2.09 | -2.24 | 0.24 | -3.37 | -1.85 | -2.35 | 1.26E-15 | 0.805 | 0.405 | 1 | 0.101398 | 0.569959 | | 1 | 0.996511 | 0.923321 |
| Wdcp | A0A1Y7VJP | -1.96 | -1.16 | -0.76 | 0.51 | -1.44 | -0.55 | -0.16 | 0.61 | -0.71 | 0.02 | 0.25 | -1.8 | 0.07 | -0.57 | -0.12 | -0.54 | 0.4575 | 0.2825 | 0.5525 | 0.460315 | 0.656565 | 0.2883 | | 1 | 0.996511 | 0.824705 |
| Wdfy1 | E9Q4P1 | -4.38 | -5.23 | -3.01 | -2.54 | -4.92 | -4.47 | -2.48 | -2.75 | 4.41 | 5.16 | 4.7 | 4.57 | -1.32 | -5.82 | -2.65 | -3.24 | 0.135 | 8.5 | 0.5325 | 0.863245 | 3.49E-07 | 0.60055 | | 1 | 0.003181 | 0.930802 |
| Wdfy2 | Q8BUB4 | 1.42 | -0.03 | 0.05 | -0.26 | -0.26 | 1.63 | -0.89 | 0.44 | 0.45 | 1 | -0.25 | 0.97 | 1.23 | 0.42 | 0.64 | 0.34 | -0.065 | 0.2475 | 0.3625 | 0.914008 | 0.586808 | 0.390542 | | 1 | 0.996511 | 0.865375 |
| Wdfy3 | Q6VNB8 | 0.42 | 0.49 | 0.37 | 0.28 | 0.6 | 0.45 | 0.39 | 0.43 | 0.23 | -0.15 | 0.39 | 0.41 | 0.03 | 0.36 | 0.19 | 0.24 | 0.0775 | -0.17 | -0.185 | 0.598547 | 0.387709 | 0.312094 | | 1 | 0.996511 | 0.834523 |
| Wdr1 | O88342 | 1.6 | -0.18 | 0.1 | 0.2 | 0.08 | 1.38 | 0.31 | 0.7 | -1.11 | -0.38 | 0.31 | 1.28 | -1.24 | -1.35 | -0.05 | -0.18 | 0.1875 | -0.405 | -1.135 | 0.681671 | 0.500437 | 0.043334 | | 1 | 0.996511 | 0.658454 |
| Wdr11 | G5E8J3 | -1.06 | -0.83 | -0.58 | -0.49 | -0.52 | -1.06 | -0.29 | -0.65 | -0.59 | -0.71 | -0.68 | -0.93 | -0.52 | -0.59 | -0.68 | -0.54 | 0.11 | 0.0125 | 0.1575 | 0.630274 | 0.949682 | 0.43982 | | 1 | 0.996755 | 0.883445 |
| Wdr12 | Q9JJA4 | -1.49 | -1.52 | -1.74 | -2.48 | -1.64 | -1.26 | -1.81 | -2.15 | -1.32 | -2.44 | -1.83 | -1.66 | -2.21 | -2.02 | -1.11 | -1.74 | 0.0925 | -0.005 | 0.0375 | 0.754314 | 0.987809 | 0.91084 | | 1 | 0.999285 | 0.986458 |
| Wdr13 | Q91V09 | 0.37 | 0.46 | 0.32 | -0.09 | 0.12 | 0.47 | 0.16 | 0.23 | -0.24 | -0.23 | 0.3 | 0.61 | -0.37 | -0.19 | -0.01 | 0.23 | -0.02 | -0.155 | -0.35 | 0.914309 | 0.556436 | 0.145551 | | 1 | 0.996511 | 0.745321 |
| Wdr17 | H3BLI8 | 0.16 | -0.09 | 0 | 0.42 | -0.54 | -0.57 | 0.04 | 0.25 | -0.21 | 1.07 | 0.18 | -0.04 | -0.44 | 0.13 | 0.26 | -0.2 | -0.3275 | 0.1275 | -0.185 | 0.20904 | 0.681378 | 0.440474 | | 1 | 0.996511 | 0.883549 |
| Wdr18 | Q4VBE8 | -0.58 | -0.58 | -1.8 | -2.15 | -0.89 | -0.04 | -2.39 | -1.95 | -0.79 | -0.93 | -1.39 | -0.75 | -0.44 | -0.67 | -1.05 | -2.08 | -0.04 | 0.3125 | 0.2175 | 0.947639 | 0.457294 | 0.670353 | | 1 | 0.996511 | 0.952558 |
| Wdr19 | Q3UGF1-3 | -0.1 | -0.32 | 0.38 | 0.35 | -0.81 | -0.14 | -0.96 | 0.08 | 0.27 | -0.2 | -0.06 | -0.01 | 0.68 | -1.31 | -0.16 | -0.22 | -0.535 | -0.0775 | -0.33 | 0.1051 | 0.737156 | 0.443124 | | 1 | 0.996511 | 0.883549 |
| Wdr20 | Q3UWE6 | -0.3 | -0.77 | -0.48 | -0.52 | -0.48 | -0.35 | -0.11 | -0.75 | -0.63 | -0.37 | -0.44 | -0.4 | -0.41 | -0.66 | -0.57 | -0.65 | 0.095 | 0.0575 | -0.055 | 0.634492 | 0.749535 | 0.772698 | | 1 | 0.996511 | 0.966032 |
| Wdr24 | Q8CFJ9 | -0.76 | -1.37 | -0.75 | -0.31 | -1.42 | -0.89 | -1.78 | -1.05 | -0.72 | 0.17 | -0.63 | -1.06 | -0.06 | -0.09 | 2.07 | -1.41 | -0.4875 | 0.2375 | 0.925 | 0.12496 | 0.485792 | 0.198986 | | 1 | 0.996511 | 0.78004 |
| Wdr25 | A0A1Y7VK | -3.55 | 0.29 | 0 | -0.55 | -4.19 | -2.66 | -2.81 | -1.73 | 2.69 | -0.18 | -0.98 | -2.96 | 3.44 | -2.24 | -0.48 | -1.76 | -1.895 | 0.595 | 0.6925 | 0.064667 | 0.652045 | 0.620244 | | 1 | 0.996511 | 0.93807 |
| Wdr26 | A0A494BB | -0.9 | -0.72 | -0.59 | -0.63 | -0.52 | -0.29 | -0.34 | -0.39 | -0.78 | -0.87 | -0.53 | -0.59 | -0.72 | -0.71 | -0.62 | -0.58 | 0.325 | 0.0175 | 0.0525 | 0.060694 | 0.920723 | 0.765558 | | 1 | 0.996755 | 0.964134 |
| Wdr3 | Q8BHB4 | 0.68 | 0.57 | 0.53 | 0.39 | 0.59 | 0.4 | 0.67 | 0.17 | 0.52 | 0.16 | 0.63 | 0.43 | 0.85 | 1.4 | 0.2 | 0.73 | -0.085 | -0.1075 | 0.2525 | 0.629647 | 0.557845 | 0.371414 | | 1 | 0.996511 | 0.859424 |
| Wdr33 | Q8K4P0 | -2.13 | -4.49 | -2.01 | -1.79 | -3.25 | -3.16 | -2.53 | -1.21 | -3.5 | -0.65 | -1.51 | -4.38 | -1.21 | -2.53 | 0.9 | -1.23 | 0.0675 | 0.095 | 1.5875 | 0.924555 | 0.921274 | 0.090602 | | 1 | 0.996755 | 0.693068 |
| Wdr35 | Q8BND3 | -0.03 | -0.35 | -0.27 | -0.83 | 0.16 | -0.4 | -0.3 | -0.86 | -0.55 | 0 | -0.49 | -0.3 | -0.59 | 0.09 | -0.99 | -0.08 | 0.02 | 0.035 | -0.0225 | 0.941559 | 0.882659 | 0.941729 | | 1 | 0.996511 | 0.991085 |
| Wdr36 | Q3TA68 | 0.56 | 0.76 | 1.52 | 0.6 | 0.72 | -0.02 | 1.04 | 1.16 | 0.45 | 0.93 | 1.46 | 0.32 | 0.25 | 1.02 | 1.03 | 1.37 | -0.135 | -0.07 | 0.0575 | 0.689264 | 0.836102 | 0.861514 | | 1 | 0.996511 | 0.977204 |
| Wdr37 | Q8CBE3 | 0.42 | 0.44 | 1.2 | 0.77 | 0.54 | 0.43 | 0.6 | 0.87 | 0.18 | -0.55 | 1.06 | 0.31 | 0.25 | -0.38 | 0.75 | 0.65 | -0.0975 | -0.4575 | -0.39 | 0.667985 | 0.232512 | 0.24434 | | 1 | 0.996511 | 0.799652 |
| Wdr41 | Q3UDP0 | -0.63 | 0.57 | 0.55 | 0.54 | 0.18 | -0.22 | 0.29 | 0.34 | -0.26 | 0.93 | 0.3 | -0.05 | -0.26 | 0.58 | 0.27 | 0.65 | -0.11 | -0.0275 | 0.0525 | 0.728313 | 0.942258 | 0.883027 | | 1 | 0.996755 | 0.979767 |
| Wdr43 | Q6ZQL4 | 1.41 | 1.18 | 0.82 | 1.28 | 1.26 | 0.68 | 0.73 | 0.69 | 0.53 | 0.69 | 1.15 | 0.75 | 0.8 | 1.46 | 0.74 | 0.84 | -0.3325 | -0.3925 | -0.2125 | 0.148117 | 0.10208 | 0.398038 | | 1 | 0.996511 | 0.867334 |
| Wdr44 | Q6NVE8 | 0.4 | 0.6 | 0.61 | 0.39 | 0.77 | 0.55 | 0.64 | 0.64 | 0.18 | 0.22 | 0.54 | 0.47 | 0.17 | 0.73 | 0.52 | 0.49 | 0.15 | -0.1475 | -0.0225 | 0.333076 | 0.415117 | 0.909793 | | 1 | 0.996511 | 0.986458 |
| Wdr45 | Q91VM3 | -0.32 | -0.17 | -0.02 | -0.55 | 0.36 | -0.4 | -0.2 | -0.58 | -1.56 | -1.41 | -0.11 | -0.18 | -1.5 | -1.54 | -0.76 | 0.05 | 0.06 | -0.55 | -0.6725 | 0.807706 | 0.181499 | 0.105096 | | 1 | 0.996511 | 0.703417 |
| Wdr45b | Q9CR39 | 0.38 | 0.33 | 0.71 | 0.46 | 0.86 | 0.52 | 0.63 | 0.56 | -0.27 | -1.69 | 0.38 | 0.77 | -1.24 | -0.45 | 0.11 | 0.76 | 0.1725 | -0.6725 | -0.675 | 0.319825 | 0.20826 | 0.130406 | | 1 | 0.996511 | 0.724265 |
| Wdr46 | Q9Z0H1 | 0.32 | 0.72 | 2.09 | 1.43 | -0.06 | 0.58 | 1.59 | 1.48 | 2.59 | 1.44 | 2.21 | 0.48 | 3.25 | 0.28 | 1.83 | 1.72 | -0.2425 | 0.54 | 0.63 | 0.636249 | 0.348611 | 0.350802 | | 1 | 0.996511 | 0.853727 |
| Wdr47 | Q8CGF6 | 0.37 | 0.4 | 0.52 | 0.34 | 0.46 | 0.48 | 0.23 | 0.59 | 0.04 | 0.02 | 0.45 | 0.84 | 0.18 | -0.4 | 0.59 | 0.37 | 0.0325 | -0.07 | -0.2225 | 0.833378 | 0.761808 | 0.384006 | | 1 | 0.996511 | 0.860984 |
| Wdr48 | Q8BH57 | 0.27 | -0.29 | 0.51 | 0.43 | -0.08 | 0.15 | 0.56 | 0.63 | -0.11 | 0.05 | 0.54 | 0.05 | -0.36 | 0.01 | 0.63 | 0.46 | 0.085 | -0.0975 | -0.045 | 0.74139 | 0.699397 | 0.880884 | | 1 | 0.996511 | 0.979767 |
| Wdr5 | P61965 | -1.7 | -2.01 | -1.7 | -2.11 | -1.19 | -2.28 | -1.77 | -2.03 | -0.9 | -1.52 | -2.01 | -2.04 | -0.58 | -0.09 | -1.87 | -2.04 | 0.0625 | 0.2625 | 0.735 | 0.813056 | 0.384587 | 0.138592 | | 1 | 0.996511 | 0.73493 |
| Wdr54 | Q9R0D8 | -0.15 | 0.02 | -0.82 | -0.27 | 0.09 | -0.47 | -0.36 | -0.49 | -0.08 | -0.33 | -0.27 | 0.25 | -0.72 | 0.3 | -0.84 | -0.61 | -0.0025 | 0.1975 | -0.1625 | 0.991738 | 0.434208 | 0.617161 | | 1 | 0.996511 | 0.937141 |
| Wdr59 | A0A1D5RL | -0.64 | -1.13 | -1.27 | -1.14 | -1.07 | -0.45 | -1.29 | -1.59 | -1.35 | -1.09 | -1.04 | -0.64 | -1.14 | -0.84 | -1.49 | -1.44 | -0.055 | 0.015 | -0.1825 | 0.844911 | 0.948566 | 0.458819 | | 1 | 0.996755 | 0.887902 |
| Wdr6 | Q99ME2 | -0.53 | -0.27 | -0.68 | -0.64 | 0.07 | -0.39 | -0.05 | -0.26 | -0.06 | -0.27 | -0.35 | -0.37 | -0.17 | 0.2 | -0.07 | 0.15 | 0.3725 | 0.2675 | 0.5575 | 0.067121 | 0.165595 | 0.018765 | | 1 | 0.996511 | 0.602211 |
| Wdr61 | Q9ERF3 | 0.64 | 0.16 | -1.16 | -1.29 | 0.28 | 0.47 | -1.41 | -1.26 | -0.78 | -1.38 | -1.35 | 0.17 | -0.76 | -0.66 | -1.99 | -1.8 | -0.0675 | -0.4225 | -0.89 | 0.914352 | 0.453778 | 0.130087 | | 1 | 0.996511 | 0.723659 |
| Wdr7 | Q920I9 | 0.64 | 0.53 | 0.42 | 0.53 | 0.56 | 0.65 | 0.6 | 0.41 | 0.78 | 0.29 | 0.49 | 0.57 | 0.65 | 0.69 | 0.23 | 0.36 | 0.025 | 0.0025 | -0.0475 | 0.865606 | 0.988821 | 0.806467 | | 1 | 0.999285 | 0.971715 |
| Wdr73 | Q9CWR1 | -1.99 | -1.14 | -0.27 | -0.54 | -1.52 | -1.6 | -1.19 | -0.17 | -1.57 | 0.1 | -0.7 | -1.24 | -1.63 | -1.99 | -0.47 | -1.28 | -0.135 | 0.1325 | -0.3575 | 0.772886 | 0.787691 | 0.454993 | | 1 | 0.996511 | 0.887169 |
| Wdr75 | Q3U821 | 1.18 | 1.43 | 0.99 | 0.81 | 0.64 | 1.31 | 0.47 | 1.26 | 0.88 | 0.52 | 1.79 | 1.23 | 0.88 | 0.17 | 0.67 | 1.36 | -0.1825 | 0.0025 | -0.3325 | 0.489662 | 0.993465 | 0.277359 | | 1 | 0.999285 | 0.822084 |
| Wdr77 | Q99J09 | -1.34 | -0.49 | -0.15 | -0.13 | -0.84 | -1.49 | 0.23 | 0 | -0.72 | -0.28 | 0.05 | -0.97 | -1.24 | -0.1 | 0.06 | -0.18 | 0.0025 | 0.0475 | 0.1625 | 0.995587 | 0.893396 | 0.68259 | | 1 | 0.996511 | 0.955729 |
| Wdr81 | K4DI77 | 0.03 | 0.2 | 0.47 | 0.49 | 0.17 | 0.04 | 0.15 | 0.35 | 0.17 | 0.7 | 0.24 | 0.2 | 0.38 | 0.24 | 0.46 | 0.42 | -0.12 | 0.03 | 0.0775 | 0.501602 | 0.886421 | 0.690618 | | 1 | 0.996511 | 0.955729 |
| Wdr82 | Q8BFQ4 | -1.78 | -1.94 | -2.1 | -2.02 | -1.64 | -1.51 | -1.35 | -2.37 | -2.28 | -0.79 | -2.12 | -1.96 | -1.78 | -1.63 | -1.64 | -1.72 | 0.2425 | 0.1725 | 0.2675 | 0.342332 | 0.615212 | 0.153799 | | 1 | 0.996511 | 0.750763 |
| Wdr89 | Q9D0R9 | -1.35 | -1.07 | -0.82 | -0.53 | -1.53 | -2.17 | 0.16 | -0.92 | -0.81 | -0.43 | -0.3 | -2.08 | -1.4 | 0.72 | -0.04 | -0.13 | -0.1725 | 0.0375 | 0.73 | 0.723454 | 0.928777 | 0.129763 | | 1 | 0.996755 | 0.723659 |
| Wdr91 | Q7TMQ7 | 0.42 | 0.63 | 0.49 | 0.05 | 0.93 | 0.6 | 0.04 | 0.18 | -1.43 | 0.48 | 0.27 | 0.45 | -0.87 | -1.14 | 0.14 | 0.33 | 0.04 | -0.455 | -0.7825 | 0.872715 | 0.324809 | 0.062927 | | 1 | 0.996511 | 0.66433 |
| Wdsub1 | Q9D0I6 | -0.48 | -0.12 | -2.26 | -2.26 | -0.35 | -1.08 | -2.07 | -2.1 | -0.07 | -1.22 | -2.14 | -0.56 | -0.26 | 0.28 | -1.29 | -2.29 | -0.12 | 0.2825 | 0.39 | 0.852472 | 0.670668 | 0.595768 | | 1 | 0.996511 | 0.929911 |
| Wdtc1 | Q80ZK9 | -0.69 | -0.74 | -0.83 | -0.62 | -1.06 | -0.69 | -0.99 | -1.13 | -0.36 | -0.39 | -0.7 | -0.73 | -0.01 | 0.22 | -0.46 | -0.77 | -0.2475 | 0.175 | 0.465 | 0.15807 | 0.337377 | 0.097729 | | 1 | 0.996511 | 0.698156 |
| Wfs1 | P56695 | 0.08 | -0.54 | -0.21 | 0.37 | -0.32 | -0.03 | 0.52 | 0.87 | 0.61 | 0.05 | 0.02 | -0.06 | 0.79 | 0.14 | 0.85 | 0.13 | 0.335 | 0.23 | 0.5525 | 0.315518 | 0.396726 | 0.086868 | | 1 | 0.996511 | 0.691619 |
| Whrn | Q80VW5-4 | 0.48 | 1.09 | 1 | 0.88 | 1.35 | 0.7 | 1.11 | 0.55 | 0.54 | 0.82 | 1.14 | 0.96 | 0.3 | 1.42 | 0.45 | 1 | 0.065 | 0.0025 | -0.07 | 0.789012 | 0.990978 | 0.817286 | | 1 | 0.999285 | 0.972768 |
| Wipf2 | Q6PEV3 | 0.47 | 0.16 | 0.26 | 0.42 | 0.13 | 0.42 | -0.2 | 0.47 | 0.43 | 0.71 | 0.37 | 0.14 | 0.68 | 0.52 | 0.02 | -0.06 | -0.1225 | 0.085 | -0.0375 | 0.54868 | 0.660407 | 0.874151 | | 1 | 0.996511 | 0.979626 |
| Wipf3 | E9Q2D0 | 0.32 | 0.23 | 0.24 | 0.36 | 0.22 | 0.23 | 0.18 | 0.59 | 0.03 | 0.37 | 0.01 | 0.01 | 0.3 | 0.41 | 0 | -0.34 | 0.0175 | -0.1825 | -0.195 | 0.913663 | 0.301641 | 0.388974 | | 1 | 0.996511 | 0.864915 |
| Wipi1 | Q8R3E3-2 | 0.13 | -0.24 | -0.08 | -0.19 | -0.05 | 0.16 | 0.1 | -0.05 | -0.6 | -1.39 | -0.08 | -0.15 | -0.38 | -0.46 | -0.01 | -0.36 | 0.135 | -0.46 | -0.2075 | 0.409303 | 0.168472 | 0.310343 | | 1 | 0.996511 | 0.834523 |
| Wipi2 | Q80W47 | -0.56 | -0.42 | -0.34 | -0.39 | -0.33 | -0.83 | -0.54 | -0.56 | 0.28 | -0.28 | -0.36 | -0.61 | 0.63 | -0.11 | -0.35 | -0.5 | -0.1375 | 0.185 | 0.345 | 0.421559 | 0.424597 | 0.233453 | | 1 | 0.996511 | 0.789665 |
| Wiz | G5E8J8 | 0.83 | 0.06 | -0.7 | -0.26 | -0.61 | 0.15 | -0.82 | -0.69 | 0.56 | 1.29 | 0.01 | 0.54 | 0.32 | 0.6 | 0.26 | -0.06 | -0.475 | 0.6175 | 0.2975 | 0.222565 | 0.147528 | 0.402753 | | 1 | 0.996511 | 0.870698 |
| Wls | Q6DID7 | -1.05 | -1.06 | -0.31 | -0.44 | 0.05 | -0.73 | 0.13 | -0.43 | -0.61 | -0.14 | -0.63 | -0.8 | -0.46 | -0.46 | -0.53 | -0.06 | 0.47 | 0.17 | 0.3375 | 0.127154 | 0.521576 | 0.209756 | | 1 | 0.996511 | 0.786218 |
| Wnk1 | P83741-4 | 0.6 | 0.69 | 0.75 | 0.54 | 0.35 | 0.27 | 0.19 | 0.45 | 0.56 | 0.58 | 0.56 | 0.44 | 0.64 | -0.46 | 0.58 | 0.45 | -0.33 | -0.11 | -0.3425 | 0.051296 | 0.491024 | 0.243567 | | 1 | 0.996511 | 0.79948 |
| Wnk1 | F6V9V5 | 0.13 | -0.13 | -1.07 | -0.12 | 0.32 | 0.71 | 0.6 | -0.33 | 0.28 | 0.45 | -0.09 | -0.33 | -1.01 | -0.87 | 0.18 | 0.1 | 0.6225 | 0.375 | -0.1025 | 0.095507 | 0.25853 | 0.796045 | | 1 | 0.996511 | 0.969727 |
| Wnk2 | E9QJQ9 | 0.01 | 0.11 | 0.67 | 0.63 | 0.07 | 0.06 | 0.47 | 0.63 | 0.43 | 0.91 | 0.65 | 0.19 | 0.27 | 0.4 | 0.55 | 0.45 | -0.0475 | 0.19 | 0.0625 | 0.842953 | 0.45955 | 0.783788 | | 1 | 0.996511 | 0.967642 |
| Wnk3 | V9GWU7 | 1.84 | 1.97 | 1.25 | 1.48 | 1.51 | 1.86 | 0.75 | 1.44 | 1.25 | 0.67 | 1.4 | 2.18 | 1.31 | 0.3 | 1.41 | 0.9 | -0.245 | -0.26 | -0.655 | 0.402222 | 0.459813 | 0.060319 | | 1 | 0.996511 | 0.661738 |
| Wrnip1 | Q91XU0 | -1.47 | -1.67 | -1.21 | -2.15 | -1.33 | -1.87 | -1.16 | -1.84 | -2.04 | -0.96 | -1.38 | -1.12 | -1.5 | -1.3 | -1.06 | -1.13 | 0.075 | 0.25 | 0.3775 | 0.783683 | 0.432321 | 0.161944 | | 1 | 0.996511 | 0.757516 |
| Wsb2 | O54929 | -0.78 | 0.02 | -1.35 | -1.1 | 0.12 | -0.54 | -0.49 | -1.41 | -0.36 | -2.3 | -0.45 | -0.58 | -1.77 | -0.32 | -1.78 | -0.54 | 0.2225 | -0.12 | -0.3 | 0.588425 | 0.814052 | 0.521811 | | 1 | 0.996511 | 0.907667 |
| Wtap | E0CYH0 | -0.38 | -2.12 | -1.71 | -2.74 | -1.32 | -1.59 | -1.61 | -1.33 | -2.19 | -3.67 | -2.53 | -1.36 | -0.46 | -0.52 | 0.96 | -1.62 | 0.275 | -0.7 | 1.3275 | 0.561612 | 0.285702 | 0.071454 | | 1 | 0.996511 | 0.668066 |
| Wwc1 | Q5SXA9 | 0.17 | 0.6 | 0.81 | 0.83 | 0.68 | 0.17 | 0.51 | 0.72 | 0.95 | 1.46 | 0.75 | 0.81 | 0.86 | 0.84 | 0.74 | 0.59 | -0.0825 | 0.39 | 0.155 | 0.709884 | 0.141106 | 0.483061 | | 1 | 0.996511 | 0.893925 |
| Wwc2 | Q6NXJ0 | -0.53 | -0.59 | 0.01 | -0.38 | -0.43 | -0.57 | 0.13 | -0.14 | -0.23 | 0.09 | -0.13 | -0.54 | 0.2 | -0.26 | -0.25 | -0.1 | 0.12 | 0.17 | 0.27 | 0.599605 | 0.454473 | 0.243936 | | 1 | 0.996511 | 0.799567 |
| Wwox | Q91WL8 | 0.41 | 0.27 | 0.09 | 0.33 | 0.65 | 0.26 | 0.91 | 0.61 | 0.47 | -0.05 | 0.62 | 0.37 | 0.42 | 1.19 | 0.62 | 0.88 | 0.3325 | 0.0775 | 0.5025 | 0.106355 | 0.70647 | 0.05049 | | 1 | 0.996511 | 0.658454 |
| Wwp1 | Q8BZZ3 | 0.06 | 0.43 | -0.23 | 0.4 | 0.31 | 0.66 | 0.23 | 0.21 | 1.07 | 0.67 | 0.44 | 0.42 | 1.2 | 0.47 | 0.98 | 0.13 | 0.1875 | 0.485 | 0.53 | 0.39201 | 0.072943 | 0.105787 | | 1 | 0.996511 | 0.703417 |
| Wwp2 | Q9DBH0 | 0.04 | 0.17 | 0.36 | 0.26 | 0.26 | 0.4 | -0.27 | -0.01 | -0.19 | -1.1 | 0.12 | 0.09 | 0.27 | -1.62 | 0.51 | -0.04 | -0.1125 | -0.4775 | -0.4275 | 0.57268 | 0.137702 | 0.362174 | | 1 | 0.996511 | 0.858475 |
| Xab2 | Q9DCD2 | -1.12 | -1.18 | -1.74 | -1.51 | -1.57 | -1.82 | -1.73 | -1.58 | -0.66 | -0.02 | -1.64 | -1.63 | -0.58 | 0.11 | -1.12 | -1.79 | -0.2875 | 0.4 | 0.5425 | 0.163788 | 0.334961 | 0.209513 | | 1 | 0.996511 | 0.786218 |
| Xdh | Q00519 | -2.59 | -3.68 | -1.43 | -1.02 | -2.41 | -3.62 | -1.26 | -1.69 | -2.29 | -0.02 | -1.11 | -2.04 | -2.69 | -2.19 | -1.78 | -1.36 | -0.065 | 0.815 | 0.175 | 0.927655 | 0.274778 | 0.773868 | | 1 | 0.996511 | 0.966062 |
| Xiap | Q60989 | -0.17 | -0.09 | -0.85 | -0.09 | -0.1 | 0.1 | 0.12 | -0.28 | 0.5 | -0.3 | -0.58 | 0.01 | 0.26 | 0.6 | -0.13 | -0.24 | 0.26 | 0.2075 | 0.4225 | 0.271244 | 0.497031 | 0.163182 | | 1 | 0.996511 | 0.758224 |
| Xk | Q9QXY7 | 1.31 | 1.37 | 1.47 | 1.32 | 1.43 | 1.45 | 0.22 | 1.18 | 1.35 | 0.52 | 0.86 | 1.55 | 1.19 | 1.04 | 1.86 | 2.01 | -0.2975 | -0.2975 | 0.1575 | 0.32322 | 0.26575 | 0.561435 | | 1 | 0.996511 | 0.91997 |
| Xkr4 | Q5GH67 | 1.66 | 1.19 | 1.63 | 1.71 | 1.16 | 1.52 | 1 | 1.57 | 2.08 | 2.13 | 1.84 | 1.99 | 1.6 | 2.07 | 2.05 | 0.85 | -0.235 | 0.4625 | 0.095 | 0.282959 | 0.036991 | 0.764981 | | 1 | 0.996511 | 0.964134 |
| Xkr6 | E9Q6C8 | 0.91 | 1.24 | 0.07 | -0.05 | 0.11 | 1.75 | 0.1 | -0.07 | 2.07 | 0.07 | 0.17 | 1.16 | 2.41 | 1.33 | 0.72 | 0.22 | -0.07 | 0.325 | 0.6275 | 0.886623 | 0.540018 | 0.25251 | | 1 | 0.996511 | 0.80892 |
| Xkr7 | Q5GH64 | 0.08 | 0.13 | 0.26 | 0.29 | -0.18 | 0.35 | -0.67 | -0.46 | 0.19 | 1.03 | 0.85 | 0.16 | 0.13 | 0.39 | -0.39 | -1.46 | -0.43 | 0.3675 | -0.5225 | 0.102752 | 0.169965 | 0.211614 | | 1 | 0.996511 | 0.786269 |
| Xpc | A0A0U1RP | 1.13 | 1.19 | -1.71 | -1.77 | 1.81 | 1.48 | -2.44 | -2.42 | 1.63 | -1.5 | -2.63 | 1.14 | 0.98 | 0.77 | -1.77 | -1.89 | -0.1025 | -0.05 | -0.1875 | 0.93672 | 0.96636 | 0.85477 | | 1 | 0.996755 | 0.976225 |
| Xpnpep1 | Q3UE92 | -0.15 | -0.27 | -0.26 | -0.44 | -0.17 | -0.44 | -0.29 | -0.31 | 0.05 | -0.35 | -0.26 | -0.08 | -0.11 | -0.32 | -0.49 | -0.77 | -0.0225 | 0.12 | -0.1425 | 0.882904 | 0.503941 | 0.503245 | | 1 | 0.996511 | 0.901002 |
| Xpnpep3 | B7ZMP1 | 0.51 | 0.44 | 0.49 | 0.5 | 0.42 | 0.41 | 0.77 | 0.61 | -0.01 | -0.5 | 0.38 | 0.27 | -0.08 | 0.03 | -0.01 | 0.3 | 0.0675 | -0.45 | -0.425 | 0.665382 | 0.07677 | 0.038598 | | 1 | 0.996511 | 0.658454 |
| Xpo1 | Q6P5F9 | -0.3 | -0.13 | -0.61 | -0.49 | 0.08 | -0.28 | -0.13 | -0.42 | -0.81 | -1.12 | -0.48 | -0.31 | -0.76 | -0.52 | -0.8 | -0.67 | 0.195 | -0.2975 | -0.305 | 0.318366 | 0.230907 | 0.143587 | | 1 | 0.996511 | 0.742733 |
| Xpo4 | A0A0R4J25 | 0.38 | 0.58 | -0.03 | -0.14 | 0.53 | 0.07 | 0.36 | 0.31 | 0.23 | 0.17 | 0.22 | 0.46 | 0.07 | 0.09 | 0.54 | 0.09 | 0.12 | 0.0725 | 7.85E-17 | 0.586169 | 0.741497 | 1 | | 1 | 0.996511 | 1 |
| Xpo5 | Q924C1 | -0.32 | -1.35 | -1.45 | -1.92 | -0.95 | -0.05 | -1.4 | -2.06 | -1.57 | -1.14 | -1.55 | -0.27 | -1.43 | -1.82 | -1.97 | -1.31 | 0.145 | 0.1275 | -0.3725 | 0.771417 | 0.766984 | 0.322586 | | 1 | 0.996511 | 0.840788 |
| Xpo6 | Q924Z6 | -1.26 | -1.23 | -1.61 | -2.17 | -0.34 | -0.93 | -1.38 | -1.66 | -1.38 | -1.39 | -1.92 | -1.06 | -1.52 | -1.28 | -2.88 | -1.31 | 0.49 | 0.13 | -0.18 | 0.183789 | 0.656448 | 0.669678 | | 1 | 0.996511 | 0.952558 |
| Xpo7 | Q9EPK7 | -0.57 | -0.31 | -0.32 | -0.16 | -0.34 | -0.45 | -0.27 | -0.22 | -0.38 | -0.57 | -0.4 | -0.57 | -0.27 | -0.67 | -0.62 | -0.62 | 0.02 | -0.14 | -0.205 | 0.900759 | 0.428318 | 0.311524 | | 1 | 0.996511 | 0.834523 |
| Xpot | Q9CRT8 | -0.88 | -0.76 | -0.98 | -1.13 | -0.73 | -0.76 | -1.01 | -1.13 | -0.84 | -0.86 | -1.08 | -0.75 | -1.01 | -0.51 | -1.36 | -1.08 | 0.03 | 0.055 | -0.0525 | 0.863281 | 0.754981 | 0.823407 | | 1 | 0.996511 | 0.972768 |
| Xpr1 | Q9Z0U0 | 0.34 | 0.27 | 0.1 | 0.23 | 0.74 | 0.75 | 0.18 | 0.14 | 0.01 | -0.07 | 0.21 | 0.58 | -0.26 | 0.27 | -0.06 | 0.17 | 0.2175 | -0.0525 | -0.205 | 0.307544 | 0.794964 | 0.315593 | | 1 | 0.996511 | 0.838071 |
| Xrcc1 | Q60596 | -0.49 | -0.81 | 0.35 | 0.12 | -0.15 | -1.11 | -0.06 | -0.14 | -0.96 | 1.45 | -0.25 | -0.14 | -1.07 | -0.1 | -0.28 | -0.6 | -0.1575 | 0.2325 | -0.305 | 0.65581 | 0.663471 | 0.383462 | | 1 | 0.996511 | 0.860984 |
| Xrcc4 | A0A0R4J1Y | -0.66 | -0.52 | -1.39 | -1.83 | -0.22 | -0.92 | -0.75 | -1.24 | -1.06 | -1.19 | -1.1 | -0.49 | -1.09 | -0.92 | -1.15 | -1.72 | 0.3175 | 0.14 | -0.12 | 0.388268 | 0.683861 | 0.733633 | | 1 | 0.996511 | 0.958635 |
| Xrcc5 | P27641 | -3.29 | -3.79 | -2.43 | -3.01 | -3.11 | -3.35 | -2.1 | 4 | -2.97 | -1.07 | 4.48 | -1.95 | -3.33 | -0.95 | -2.92 | -1.42 | 1.99 | 2.7525 | 0.975 | 0.226217 | 0.095821 | 0.124426 | | 1 | 0.996511 | 0.721448 |
| Xrcc6 | A0A0R4J18 | -0.21 | -0.8 | -0.91 | -1.4 | -0.55 | -1.14 | -0.79 | -1.28 | -0.53 | 0.46 | -1.23 | 0.15 | -0.61 | 0.51 | -1.84 | -0.19 | -0.11 | 0.5425 | 0.2975 | 0.709678 | 0.223576 | 0.564837 | | 1 | 0.996511 | 0.920315 |
| Xrn1 | A0A087WC | -0.51 | -0.73 | -0.74 | -0.71 | -1.31 | -0.23 | -1.02 | -0.94 | 0.65 | -0.04 | -0.54 | -0.13 | 1.4 | -1.12 | -0.4 | -0.8 | -0.2025 | 0.6575 | 0.4425 | 0.423908 | 0.035057 | 0.408736 | | 1 | 0.996511 | 0.874356 |
| Xrn2 | Q9DBR1-2 | -2.29 | -3.49 | -3.33 | -3.09 | -3.29 | -2.82 | -2.95 | -3.18 | -2.89 | -2.19 | -3.08 | -2.72 | -2.68 | -2.67 | -3.21 | -3.3 | -0.01 | 0.33 | 0.085 | 0.972253 | 0.327015 | 0.791531 | | 1 | 0.996511 | 0.969105 |
| Xxylt1 | Q3U4G3 | -0.09 | 0.24 | -0.09 | -0.42 | -0.15 | -0.55 | 0.01 | -0.46 | -0.08 | -0.44 | -0.82 | 0.07 | -0.39 | 0.01 | -0.04 | 0.18 | -0.1975 | -0.2275 | 0.03 | 0.368351 | 0.391622 | 0.8947 | | 1 | 0.996511 | 0.983453 |
| Xylb | Q3TNA1 | -0.48 | -0.8 | -0.26 | 0.43 | -2.62 | 0.68 | -0.24 | 0.19 | 2.01 | 1.11 | -0.56 | -0.82 | 2.72 | -0.94 | -0.34 | -0.94 | -0.22 | 0.7125 | 0.4025 | 0.755031 | 0.297944 | 0.627083 | | 1 | 0.996511 | 0.939737 |
| Yae1d1 | Q9DAY6 | -0.8 | 0.15 | 0.07 | -0.38 | -0.23 | 0.19 | -0.84 | -0.37 | -1.17 | -2.33 | 0.07 | 0.19 | -0.38 | 0.9 | 0.19 | 0.45 | -0.0725 | -0.57 | 0.53 | 0.810905 | 0.340269 | 0.150646 | | 1 | 0.996511 | 0.749304 |
| Yaf2 | Q99LW6 | -5.23 | -6.5 | -7.59 | -8.22 | -7.17 | -4.93 | -16.31 | -9.42 | -4.64 | -5.86 | -10.08 | -5.39 | -2.65 | -7.19 | -6.66 | -8.31 | -2.5725 | 0.3925 | 0.6825 | 0.27507 | 0.752354 | 0.586293 | | 1 | 0.996511 | 0.928178 |
| Yap1 | P46938 | 0.2 | 0.01 | -0.79 | -0.11 | -1.33 | -0.56 | -0.55 | -1.1 | -1.75 | -1.47 | -0.79 | -0.09 | -1.5 | -2.75 | -1 | -0.89 | -0.7125 | -0.8525 | -1.3625 | 0.035249 | 0.063125 | 0.013881 | | 1 | 0.996511 | 0.589387 |
| Yars | A2A7S7 | -0.7 | -0.98 | -0.33 | -0.41 | -1.02 | -0.77 | -0.47 | -0.26 | -1.12 | -0.11 | -0.42 | -0.71 | -0.85 | -0.78 | -0.54 | -0.82 | -0.025 | 0.015 | -0.1425 | 0.916666 | 0.95645 | 0.516828 | | 1 | 0.996755 | 0.905676 |
| Yars1 | Q91WQ3 | -0.79 | -0.32 | -1.8 | -1.4 | -2.07 | -0.11 | -1.55 | -1.98 | 0.19 | -1.63 | -0.65 | -0.02 | 1.14 | -1.76 | -0.81 | -1.11 | -0.35 | 0.55 | 0.4425 | 0.502324 | 0.277536 | 0.496847 | | 1 | 0.996511 | 0.897918 |
| Yars2 | Q8BYL4 | 0.42 | 0.48 | 0.71 | 0.49 | 0.2 | 0.63 | 0.3 | 0.67 | 0.66 | 0.24 | 0.74 | 0.62 | 0.16 | 0.31 | 0.34 | 0.62 | -0.075 | 0.04 | -0.1675 | 0.677097 | 0.831046 | 0.391662 | | 1 | 0.996511 | 0.865375 |
| Ybx1 | P62960 | 0.05 | -0.24 | -0.62 | -0.51 | -0.67 | 0.21 | -0.47 | -0.22 | 0.48 | -0.19 | -0.4 | 0.69 | 1.48 | 0.08 | 0.2 | 0.12 | 0.0425 | 0.475 | 0.8 | 0.866467 | 0.146426 | 0.051906 | | 1 | 0.996511 | 0.658874 |
| Ybx3 | Q9JKB3 | -1.14 | -1.1 | -1.04 | -0.86 | -1.19 | -0.98 | -1.42 | -1.39 | 0.01 | -1.32 | -0.85 | -0.6 | 0.46 | -0.53 | -1.07 | -1.52 | -0.21 | 0.345 | 0.37 | 0.240408 | 0.256929 | 0.381288 | | 1 | 0.996511 | 0.860984 |
| Ydjc | Q14BV6 | -0.25 | 0.02 | -0.18 | -0.15 | 0.07 | 0.07 | -0.47 | -0.42 | 0.05 | 0.74 | -0.2 | 0.35 | -0.09 | 0.69 | -0.32 | -0.24 | -0.0475 | 0.375 | 0.15 | 0.807953 | 0.142585 | 0.575161 | | 1 | 0.996511 | 0.926643 |
| Yeats4 | Q9CR11 | 0.25 | -0.3 | 0.02 | 0.09 | -0.11 | 0 | -0.49 | -0.1 | 0.05 | 0.92 | -0.08 | 0.17 | 0.39 | -0.24 | 0.13 | 0.35 | -0.19 | 0.25 | 0.1425 | 0.341679 | 0.363989 | 0.539387 | | 1 | 0.996511 | 0.912212 |
| Yes1 | Q04736 | -0.12 | 0.24 | 0.06 | 0.31 | 0.43 | 0.4 | 0.51 | 0.15 | 0.44 | 0.68 | -0.11 | 0.56 | -0.53 | -0.05 | 0.02 | 0.04 | 0.25 | 0.27 | -0.2525 | 0.175018 | 0.261583 | 0.265381 | | 1 | 0.996511 | 0.817264 |
| Yif1b | Q9CX30-2 | -0.62 | -0.28 | -0.81 | -0.89 | 0.08 | -0.62 | 0.1 | -0.67 | -0.35 | -1.02 | -0.75 | -0.6 | 0.02 | 0.72 | -1.12 | -0.46 | 0.3725 | -0.03 | 0.44 | 0.177891 | 0.895347 | 0.285053 | | 1 | 0.996511 | 0.822669 |
| Yipf1 | A2A8S2 | -0.1 | -0.62 | -0.83 | -0.89 | 0.2 | -0.82 | -0.05 | -1.36 | 0.77 | -0.79 | -0.25 | -0.3 | 0.68 | 0.82 | -1.11 | 0.15 | 0.1025 | 0.4675 | 0.745 | 0.787571 | 0.220229 | 0.12413 | | 1 | 0.996511 | 0.721448 |
| Yipf3 | Q3UDR8 | -0.82 | -0.75 | -2.21 | -1.29 | -0.44 | -0.25 | -1.22 | -1.97 | 0.37 | -1.1 | -1.82 | -0.6 | -0.54 | -0.54 | -1.49 | -1.12 | 0.2975 | 0.48 | 0.345 | 0.539425 | 0.374538 | 0.393665 | | 1 | 0.996511 | 0.865775 |
| Yipf4 | Q8C407 | -1.33 | -2.75 | -2.32 | -1.4 | -1.34 | -1.84 | -0.29 | -1.74 | -3.04 | -2.47 | -2.32 | -2.37 | -2.19 | -1.95 | -1.02 | -0.94 | 0.6475 | -0.6 | 0.425 | 0.185538 | 0.135839 | 0.35494 | | 1 | 0.996511 | 0.855029 |
| Yipf5 | Q9EQQ2 | 1.13 | 1.55 | 0.06 | -0.09 | -0.54 | 0.55 | -0.15 | 0.31 | -3.28 | -1.89 | -0.12 | 1.08 | -1.63 | -7.3 | 0.71 | 0.78 | -0.62 | -1.715 | -2.5225 | 0.181519 | 0.09507 | 0.167718 | | 1 | 0.996511 | 0.759484 |
| Yipf6 | Q8BR70 | -1.38 | -1.75 | -2.77 | -2.1 | -2.15 | -1.35 | -2.17 | -1.65 | 1.08 | -1.48 | -1.76 | -2.29 | 2.28 | -1.17 | -1.27 | -1.82 | 0.17 | 0.8875 | 1.505 | 0.623884 | 0.245221 | 0.115135 | | 1 | 0.996511 | 0.717287 |
| Yjefn3 | F6W8I0 | -0.16 | -0.48 | 0.54 | 0.38 | -0.23 | 0.55 | 0.56 | 0.27 | 0.4 | 0.75 | 0.77 | 0.1 | 0.97 | 0.89 | 0.32 | 0.46 | 0.2175 | 0.435 | 0.59 | 0.473287 | 0.163746 | 0.074933 | | 1 | 0.996511 | 0.672844 |
| Yju2 | Q9D6J3 | -0.17 | -0.52 | -2.85 | -1.8 | -0.67 | -0.06 | -0.75 | -1.83 | 0.29 | -3.45 | -1.26 | -0.29 | 0.1 | 0.24 | -2.23 | -1.77 | 0.5075 | 0.1575 | 0.42 | 0.443733 | 0.864636 | 0.600562 | | 1 | 0.996511 | 0.930802 |
| Ykt6 | Q9CQW1 | 0.03 | 0.16 | 0.17 | 0.94 | 0.41 | 0.33 | 0.36 | 1.14 | 0.31 | 1.31 | 0.72 | 0.37 | 0.12 | 0.43 | 0.11 | 0.22 | 0.235 | 0.3525 | -0.105 | 0.419875 | 0.27633 | 0.679142 | | 1 | 0.996511 | 0.955269 |
| Ylpm1 | D3YWX2 | 1.06 | -0.09 | -0.07 | -0.24 | -0.34 | 0.59 | -0.43 | 0.05 | 0.79 | -0.61 | -0.23 | -0.03 | 1.19 | 0.17 | 0.76 | -0.24 | -0.1975 | -0.185 | 0.305 | 0.589656 | 0.64781 | 0.471665 | | 1 | 0.996511 | 0.891144 |
| Yme1l1 | O88967 | 0.26 | 0.48 | 0.07 | 0.72 | 0.15 | 0.34 | -0.02 | 0.14 | 0.89 | 0.74 | 0.37 | 0.28 | 0.51 | 0.93 | 0.1 | 0.03 | -0.23 | 0.1875 | 0.01 | 0.256288 | 0.429983 | 0.970848 | | 1 | 0.996511 | 0.992803 |
| Yod1 | Q8CB27 | 0.29 | -0.01 | -0.38 | 0.29 | 0.06 | -0.01 | -0.71 | -0.33 | 0.61 | 0.88 | -0.52 | 0.03 | 0.97 | 1.07 | -0.4 | -0.33 | -0.295 | 0.2025 | 0.28 | 0.256451 | 0.560411 | 0.503573 | | 1 | 0.996511 | 0.901002 |
| Ypel1 | A0A338P7 | 0.99 | 0 | 0.56 | -0.18 | -0.49 | 1.57 | -0.44 | -0.47 | -2.81 | -8.39 | -0.06 | 0.29 | -0.92 | -8.07 | 0.39 | 0.48 | -0.3 | -3.085 | -2.3725 | 0.573781 | 0.115078 | 0.215056 | | 1 | 0.996511 | 0.787313 |
| Ypel5 | P62700 | -0.23 | 0.19 | -1.1 | -0.46 | 0.26 | 0 | -1.44 | -0.67 | -0.12 | -0.96 | -1.11 | 0.04 | 0.03 | 0.03 | -0.87 | -0.43 | -0.0625 | -0.1375 | 0.09 | 0.885752 | 0.719577 | 0.794036 | | 1 | 0.996511 | 0.969367 |
| Yrdc | Q3U5F4 | -1.5 | -0.62 | -0.11 | -0.15 | -1.49 | -1.53 | -0.81 | 0.05 | -0.82 | -0.25 | -0.24 | -1.16 | -1.46 | -0.83 | -0.49 | 0.37 | -0.35 | -0.0225 | -0.0075 | 0.451658 | 0.952759 | 0.987245 | | 1 | 0.996755 | 0.996154 |
| Ythdc1 | E9Q5K9-3 | 1.33 | 0 | 0.63 | 0.15 | -0.26 | 0.33 | -0.12 | 1.38 | 0.55 | -0.33 | 0.01 | -0.14 | 1.26 | 0.33 | 1.05 | 1.39 | -0.195 | -0.505 | 0.48 | 0.66244 | 0.171447 | 0.219955 | | 1 | 0.996511 | 0.787447 |
| Ythdc2 | B2RR83 | -0.73 | -0.61 | -0.5 | -0.62 | -0.48 | -0.97 | -0.57 | -0.44 | -1.15 | -0.17 | -0.36 | -0.66 | -1.12 | -0.44 | -0.45 | -0.76 | 1.18E-16 | 0.03 | -0.0775 | 1 | 0.902239 | 0.724286 | | 1 | 0.996609 | 0.958398 |
| Ythdf1 | A2AWN8 | -3.5 | -3.96 | -1.06 | -0.88 | -4.36 | -4.05 | -0.87 | -0.46 | -1.5 | -0.32 | -0.6 | -3.93 | -0.13 | -3.55 | -0.31 | -0.85 | -0.085 | 0.7625 | 1.14 | 0.941886 | 0.466704 | 0.277033 | | 1 | 0.996511 | 0.822084 |
| Ythdf2 | Q91YT7 | -3.57 | -3.54 | -2.57 | -3.47 | -3.79 | -3.39 | -3.03 | -3.05 | -4.71 | -0.9 | -3.05 | -3.22 | -2.04 | -6.47 | -1.73 | -2.69 | -0.0275 | 0.3175 | 0.055 | 0.926298 | 0.670888 | 0.956304 | | 1 | 0.996511 | 0.992636 |
| Ythdf3 | Q8BYK6-3 | -2.55 | -2.77 | -2.35 | -2.42 | -2.59 | -2.51 | -2.22 | -2.48 | -1.56 | -1.98 | -2.02 | -2.1 | -1.03 | -1.71 | -2.14 | -2.53 | 0.0725 | 0.6075 | 0.67 | 0.676191 | 0.01387 | 0.072546 | | 1 | 0.996511 | 0.668066 |
| Ywhab | Q9CQV8 | -0.02 | -0.64 | -0.49 | 0.19 | -0.52 | -0.62 | -0.31 | 0.11 | 0.61 | 0.21 | -0.13 | -0.01 | -0.4 | 0.18 | -0.42 | -0.74 | -0.095 | 0.41 | -0.105 | 0.717369 | 0.154426 | 0.718106 | | 1 | 0.996511 | 0.958398 |
| Ywhae | P62259 | 0.74 | 0.08 | -0.97 | -0.37 | 0.4 | 0.54 | -0.54 | -0.49 | 0.57 | -0.94 | -0.79 | 0.7 | -0.08 | 0.28 | -0.86 | -1.28 | 0.1075 | 0.015 | -0.355 | 0.803054 | 0.977116 | 0.462803 | | 1 | 0.997808 | 0.888267 |
| Ywhag | P61982 | 0.2 | 0.25 | 0.34 | 0.34 | 0.44 | 0.02 | -0.2 | 0.32 | 0.94 | 0.52 | 0.38 | 0.27 | 0.58 | 0.22 | 0.46 | -0.06 | -0.1375 | 0.245 | 0.0175 | 0.472917 | 0.241366 | 0.932463 | | 1 | 0.996511 | 0.989713 |
| Ywhah | P68510 | 0.24 | -0.15 | -0.2 | -0.31 | 0.01 | 0.09 | -0.73 | -0.56 | 0.69 | -0.43 | -0.19 | 0.06 | 0.72 | 0.19 | -0.71 | -0.66 | -0.1925 | 0.1375 | -0.01 | 0.447799 | 0.626768 | 0.977894 | | 1 | 0.996511 | 0.993727 |
| Ywhaq | P68254 | 0.04 | -0.8 | -0.25 | 0.64 | -0.88 | 0.42 | 0.21 | 0.49 | -0.01 | 0.95 | -0.02 | -0.07 | -0.49 | -0.04 | -0.09 | -0.14 | 0.1525 | 0.305 | -0.0975 | 0.711821 | 0.425203 | 0.762817 | | 1 | 0.996511 | 0.963993 |
| Ywhaz | P63101 | -0.32 | -0.5 | 0.04 | 0.53 | -0.35 | -0.7 | -0.28 | 0.55 | 0 | 1.1 | 0 | -0.66 | -0.96 | -0.11 | -0.06 | -0.38 | -0.1325 | 0.1725 | -0.315 | 0.695825 | 0.674916 | 0.331523 | | 1 | 0.996511 | 0.845601 |
| Yy1 | Q00899 | -5.8 | 2.91 | -5.01 | -4.79 | -5.68 | -4.45 | -3.12 | -6.13 | -3.81 | 0.99 | -4.55 | 3.15 | -4.13 | 5.25 | -5.7 | -5.02 | -1.6725 | 2.1175 | 0.7725 | 0.393204 | 0.399623 | 0.790247 | | 1 | 0.996511 | 0.969105 |
| Zbed3 | Q9D0L1 | -0.84 | -0.59 | -0.13 | -0.71 | -6.19 | -0.54 | -3.78 | -1.82 | 0.72 | 0.85 | -1.03 | -1.02 | 1.94 | -4.03 | -0.5 | -1.32 | -2.515 | 0.4475 | -0.41 | 0.047394 | 0.38821 | 0.711947 | | 1 | 0.996511 | 0.958014 |
| Zbtb11 | G5E8B9 | 0.67 | 1.05 | 1.65 | 1.53 | 0.18 | 0.8 | 0.82 | 1.5 | 1.2 | 2.31 | 1.27 | 0.43 | 1.22 | 0.96 | 1.87 | 1.49 | -0.4 | 0.0775 | 0.16 | 0.25868 | 0.854933 | 0.606487 | | 1 | 0.996511 | 0.932425 |
| Zbtb16 | Q3UQ17 | 1.16 | 1.63 | -0.77 | -0.81 | 1.86 | 0.99 | -0.76 | -1.11 | 0.95 | -0.55 | -0.62 | 1.46 | 0.79 | 1.37 | -0.54 | -0.71 | -0.0575 | 0.0075 | -0.075 | 0.946496 | 0.992015 | 0.91893 | | 1 | 0.999285 | 0.986868 |
| Zbtb18 | A0A0A6YY | -1.37 | -1.77 | -1.74 | -2.65 | -1.27 | -0.9 | -2.04 | -2.85 | 3.79 | -1.41 | -2.7 | -2.59 | 3.59 | -1.93 | -2.63 | -2.83 | 0.1175 | 1.155 | 0.9325 | 0.80411 | 0.416286 | 0.50409 | | 1 | 0.996511 | 0.901002 |
| Zbtb20 | Q8K0L9 | -1.39 | -1.09 | -1.84 | -2.18 | -1.17 | -0.98 | -2.01 | -2.49 | -0.41 | -1.19 | -1.55 | -1.28 | 0.31 | -0.45 | -1.18 | -1.33 | -0.0375 | 0.5175 | 0.9625 | 0.926008 | 0.153081 | 0.047397 | | 1 | 0.996511 | 0.658454 |
| Zbtb38 | F6Z595 | -0.41 | -0.74 | 0.51 | -0.42 | -2.17 | 0.17 | 0.21 | -0.33 | -0.67 | 0.32 | -0.07 | -0.61 | 1.26 | -2 | 0.3 | 0.79 | -0.265 | 0.0075 | 0.3525 | 0.643162 | 0.982901 | 0.616917 | | 1 | 0.998773 | 0.937141 |
| Zbtb45 | Q52KG4 | -0.46 | -0.53 | -0.52 | -0.94 | -0.65 | -0.84 | -1.07 | -1.37 | 0.03 | -1.45 | -0.21 | -1.21 | 0.13 | -0.02 | -0.76 | -0.51 | -0.37 | -0.0975 | 0.3225 | 0.113881 | 0.791961 | 0.240278 | | 1 | 0.996511 | 0.795658 |
| Zbtb7a | O88939-2 | -1.04 | -0.61 | -2.05 | -2.91 | -0.93 | -1 | -3.34 | -2.86 | -0.71 | -1.88 | -2.78 | -0.7 | -0.78 | 0.04 | -2.51 | -1.64 | -0.38 | 0.135 | 0.43 | 0.606703 | 0.837457 | 0.534705 | | 1 | 0.996511 | 0.912212 |
| Zbtb8os | B2KGA7 | -1.01 | -1.34 | -0.8 | -0.97 | 0.09 | -0.33 | 0.16 | 0.36 | -1.8 | -0.87 | -0.92 | -1.15 | 0.07 | -0.4 | 0.63 | 0.25 | 1.1 | -0.155 | 1.1675 | 0.000625 | 0.556362 | 0.001841 | 0.599778 | 0.996511 | 0.419079 | |
| Zc2hc1a | Q8BJH1 | -0.88 | -1.19 | -0.37 | -0.31 | -0.72 | -1.16 | -0.08 | -0.29 | -1.2 | 0.26 | -0.11 | -0.72 | -0.86 | -0.42 | 0.17 | -0.32 | 0.125 | 0.245 | 0.33 | 0.691211 | 0.516985 | 0.30102 | | 1 | 0.996511 | 0.830276 |
| Zc3h11a | Q6NZF1 | -0.58 | -1.63 | -0.56 | -0.41 | -2 | -1.32 | -0.59 | -0.2 | -1.57 | -0.07 | -0.34 | -2.06 | -0.88 | -1.19 | -0.18 | -0.97 | -0.2325 | -0.215 | -0.01 | 0.610924 | 0.677538 | 0.977352 | | 1 | 0.996511 | 0.993727 |
| Zc3h12b | G3X9I7 | -0.14 | 0.51 | -0.94 | -1.07 | 0.33 | -0.17 | -0.71 | -0.83 | 0.26 | -2.08 | -1.29 | 0.3 | 0.32 | 0.78 | -1.74 | -1.07 | 0.065 | -0.2925 | -0.0175 | 0.878804 | 0.647286 | 0.97795 | | 1 | 0.996511 | 0.993727 |
| Zc3h13 | E9Q784 | -1.23 | -0.84 | -0.35 | -0.84 | -1.35 | -0.99 | -0.75 | 1.14 | -4.15 | -0.75 | -1.26 | -0.82 | -2.42 | -6.51 | 0.94 | -0.28 | 0.3275 | -0.93 | -1.2525 | 0.546123 | 0.236848 | 0.401658 | | 1 | 0.996511 | 0.870698 |
| Zc3h14 | Q8BJ05-2 | 0.11 | -0.56 | -0.76 | -0.73 | -1.42 | -0.25 | -0.81 | -0.36 | -0.55 | -1.4 | -0.68 | -1.15 | -0.11 | -0.43 | -0.18 | -1.06 | -0.225 | -0.46 | 0.04 | 0.496108 | 0.142598 | 0.896368 | | 1 | 0.996511 | 0.983509 |
| Zc3h15 | Q3TIV5 | -1.77 | -2 | -1.81 | -1.77 | -2.15 | -2.13 | -1.96 | -1.67 | -1.68 | -1.1 | -1.81 | -1.62 | -1.04 | -1.3 | -1.94 | -2.18 | -0.14 | 0.285 | 0.2225 | 0.429176 | 0.196193 | 0.449822 | | 1 | 0.996511 | 0.884817 |
| Zc3h18 | H3BIW0 | 0.07 | -0.63 | -1.11 | -0.82 | -1.66 | -0.78 | -1.77 | -0.74 | -0.5 | -0.35 | -0.91 | -1.24 | 0.36 | -0.93 | -0.7 | -1.42 | -0.615 | -0.1275 | -0.05 | 0.113577 | 0.693193 | 0.907356 | | 1 | 0.996511 | 0.986458 |

| | | | | | | | | | | | | | | | | | | | | | | | | | | | |
|---|---|---|---|---|---|---|---|---|---|---|---|---|---|---|---|---|---|---|---|---|---|---|---|---|---|---|---|
| Zc3h3 | Q8CHP0 | -0.74 | -1.27 | -1.36 | -1.93 | -0.96 | -2.23 | -1.95 | -1.98 | -1.35 | 0.39 | -1.95 | -1.97 | 0.62 | -0.01 | -1.72 | -2.06 | -0.455 | 0.105 | 0.5325 | 0.223323 | 0.850992 | 0.41055 | | 1 | 0.996511 | 0.874811 |
| Zc3h4 | E9Q8K8 | -2.25 | -2.55 | -1.47 | -1.58 | -3.02 | -2.42 | -2.29 | -1.96 | -1.02 | -0.28 | -1.21 | -1.72 | -0.39 | -2 | -1.83 | -1.8 | -0.46 | 0.905 | 0.4575 | 0.190218 | 0.039763 | 0.307218 | | 1 | 0.996511 | 0.832533 |
| Zc3h6 | A2AP88 | 1.95 | 1.43 | 1.58 | 1.71 | 1.96 | 1.37 | 0.9 | 1.6 | 2.13 | 3.27 | 0.97 | 1.65 | 1.99 | 2.31 | 1.49 | 0.44 | -0.21 | 0.3375 | -0.11 | 0.423861 | 0.475081 | 0.787199 | | 1 | 0.996511 | 0.968092 |
| Zc3h7b | F8VPP8 | -1.84 | -2.01 | -0.58 | -1.15 | -1.61 | -1.91 | -0.7 | -1.03 | -1.99 | -1.03 | -0.64 | -1.45 | -1.07 | -1.99 | -0.94 | -0.69 | 0.0825 | 0.1175 | 0.2225 | 0.838006 | 0.778173 | 0.59565 | | 1 | 0.996511 | 0.929911 |
| Zc3hc1 | H3BKM2 | 0.17 | -1.52 | -0.95 | -0.59 | -0.72 | -0.35 | -1 | -0.98 | -0.57 | -1.16 | -0.23 | -0.67 | -1.05 | -0.67 | -0.63 | -0.87 | -0.04 | 0.065 | -0.0825 | 0.913201 | 0.866578 | 0.819542 | | 1 | 0.996511 | 0.972768 |
| Zcchc10 | F6QZB1 | 0.87 | 1.48 | -0.05 | 0.49 | 1.27 | 0.3 | 0.05 | 0.79 | 1.08 | 0.25 | -0.1 | 0.63 | 0.22 | 1.06 | -0.62 | 0.81 | -0.095 | -0.2325 | -0.33 | 0.810959 | 0.557951 | 0.483365 | | 1 | 0.996511 | 0.893925 |
| Zcchc17 | Q9ESX4-2 | -0.66 | -1.13 | 0.03 | 0.28 | -1.42 | -1.73 | 0.86 | -0.48 | -0.83 | 1.18 | 0.02 | -0.86 | -0.55 | -0.34 | 0.03 | -0.23 | -0.3225 | 0.2475 | 0.0975 | 0.597564 | 0.645039 | 0.776756 | | 1 | 0.996511 | 0.967045 |
| Zcchc18 | Q8VD24 | -1.2 | -0.05 | -0.89 | -0.66 | -0.07 | -1.36 | -1.77 | -0.99 | -0.17 | -0.74 | -1.13 | -0.21 | -1.62 | -0.57 | -0.97 | -0.65 | -0.3475 | 0.1375 | -0.2525 | 0.407349 | 0.679888 | 0.466113 | | 1 | 0.996511 | 0.888548 |
| Zcchc8 | Q9CYA6 | -2.01 | -2.61 | -0.74 | -0.77 | -1.82 | 0.22 | -0.85 | -0.79 | -3.13 | -0.3 | -1.5 | -2.99 | -2.74 | -1.02 | -0.92 | -1.28 | 0.7225 | -0.4475 | 0.0425 | 0.22744 | 0.549651 | 0.941359 | | 1 | 0.996511 | 0.991085 |
| Zcrb1 | Q9CZ96-2 | -1.11 | -1.98 | -2.45 | -2.45 | -1.26 | -1.15 | -2.71 | -2.4 | -1.45 | -1.1 | -3.13 | -1.82 | 0.24 | -1.08 | -1.3 | -2.42 | 0.1175 | 0.1225 | 0.8575 | 0.802497 | 0.80877 | 0.164229 | | 1 | 0.996511 | 0.758352 |
| Zdhhc13 | Q9CWU2 | -0.92 | -1.84 | -0.81 | -0.58 | -0.25 | -0.64 | -0.77 | -1.04 | -0.33 | -1.28 | -0.78 | -0.81 | -0.8 | -0.94 | -0.88 | -0.69 | 0.3625 | 0.2375 | 0.21 | 0.269404 | 0.484329 | 0.484902 | | 1 | 0.996511 | 0.894615 |
| Zdhhc14 | Q8BQQ1 | 0.04 | 0.54 | 0.46 | 0.83 | 0.14 | 0.41 | 0.44 | 0.96 | 1.53 | 1.97 | 0.93 | 0.05 | 0.96 | 2.66 | 1.37 | 0.31 | 0.02 | 0.6525 | 0.8575 | 0.935933 | 0.149631 | 0.106353 | | 1 | 0.996511 | 0.703417 |
| Zdhhc17 | Q80TN5 | -1.23 | -1.56 | -1.48 | -1.19 | -0.73 | -1.23 | -1.25 | -1.51 | -1.44 | -0.7 | -1.44 | -1.63 | -0.61 | -0.34 | -1.52 | -1.37 | 0.185 | 0.0625 | 0.405 | 0.396367 | 0.801892 | 0.21439 | | 1 | 0.996511 | 0.786931 |
| Zdhhc18 | Q5Y5T2 | 2.19 | 2.43 | -0.64 | -0.46 | 1.03 | 1.98 | -0.78 | -0.29 | 1.83 | 1.34 | -0.11 | 1.7 | 1.72 | 1.38 | -0.01 | -0.09 | -0.395 | 0.31 | -0.13 | 0.673645 | 0.715677 | 0.879044 | | 1 | 0.996511 | 0.979626 |
| Zdhhc2 | P59267 | 0.85 | 0.8 | 0.67 | 1.18 | 0.38 | 0.55 | 0.74 | 1.15 | 1.05 | 2.3 | 1.2 | 0.8 | 1.58 | 0.94 | 1.39 | 0.9 | -1.70E-01 | 0.4625 | 0.3275 | 0.449763 | 0.201029 | 0.19432 | | 1 | 0.996511 | 0.778959 |
| Zdhhc20 | Q5Y5T1-2 | 1.15 | -0.51 | 0.22 | 0.77 | 0.81 | -1.13 | 0.35 | 0.79 | 0.67 | 4.76 | 0.26 | 0.51 | 0.36 | 0.63 | 0.29 | -0.2 | -0.2025 | 1.1425 | -0.1375 | 0.705927 | 0.27912 | 0.723727 | | 1 | 0.996511 | 0.958398 |
| Zdhhc22 | A0PK84 | -1.41 | -1.14 | -0.94 | -1.44 | 0.02 | -1.69 | -0.13 | -1.26 | -0.29 | -1.99 | -0.77 | -1.17 | -0.52 | 0.55 | -0.35 | -1.26 | 0.4675 | 0.1775 | 0.8375 | 0.273155 | 0.631349 | 0.051735 | | 1 | 0.996511 | 0.658454 |
| Zdhhc3 | A0A571BE\ | -0.28 | -0.54 | -4.4 | -2.73 | 0.5 | -1.36 | -3.11 | -1.41 | -0.85 | -2.17 | -2.56 | -1.2 | -0.66 | -1.02 | -3.96 | -5.19 | 0.6425 | 0.2925 | -0.72 | 0.561019 | 0.758031 | 0.586657 | | 1 | 0.996511 | 0.92838 |
| Zdhhc5 | Q8VDZ4 | 1.57 | 1.52 | 1.84 | 1.69 | 0.77 | 1.55 | 0.87 | 1.48 | 1.95 | 1.35 | 1.75 | 1.82 | 1.7 | 1.41 | 1.15 | 1.36 | -0.4875 | 0.0625 | -0.25 | 0.061931 | 0.753189 | 0.233001 | | 1 | 0.996511 | 0.788994 |
| Zdhhc6 | Q9CPV7 | 0.46 | 0.91 | 2.02 | 1.53 | 1.13 | 0.35 | 2.94 | 2.03 | -0.43 | 2.56 | 0.87 | 0.75 | 0.14 | -0.6 | 2.37 | 4.05 | 0.3825 | -0.2925 | 0.26 | 0.527894 | 0.650973 | 0.794617 | | 1 | 0.996511 | 0.969367 |
| Zdhhc8 | Q5Y5T5 | 0.48 | 0.82 | 1.21 | 1.13 | 0.42 | 0.63 | 0.38 | 0.77 | 1 | 1.12 | 1.39 | 0.68 | 0.38 | 1 | 0.66 | 1.04 | -0.36 | 0.1375 | -0.14 | 0.121953 | 0.58053 | 0.589898 | | 1 | 0.996511 | 0.929142 |
| Zdhhc9 | P59268 | 0.62 | 0.94 | 1.31 | 0.68 | 0.83 | 0.41 | 0.77 | 0.77 | 0.6 | 1.97 | 0.95 | 0.56 | 0.33 | 0.49 | 1.56 | 0.89 | -0.1925 | 0.1325 | -0.07 | 0.374334 | 0.710391 | 0.827858 | | 1 | 0.996511 | 0.973757 |
| Zeb1 | Q64318 | -1.47 | -1.52 | -1.9 | -2.18 | -1.3 | -1.49 | -1.58 | -2.15 | -2.15 | -1.66 | -1.25 | -1.89 | -1.46 | -2.07 | -2.26 | -1.94 | 0.1375 | 0.03 | -0.165 | 0.596586 | 0.91139 | 0.539075 | | 1 | 0.996744 | 0.912212 |
| Zeb2 | A0A0J9YVF | 1.87 | -0.24 | -2.23 | -2.4 | -1.3 | 0.23 | -2.69 | -3.52 | 1.07 | -1.43 | -2.69 | 1.53 | -0.14 | 0.03 | -2.42 | -3.02 | -1.07 | 0.37 | -0.6375 | 0.368552 | 0.771365 | 0.576756 | | 1 | 0.996511 | 0.927198 |
| Zer1 | Q80ZJ6 | 0.41 | -0.34 | -1.06 | -1.11 | -0.35 | 0.53 | -1.14 | -0.83 | -0.3 | -1.3 | -0.84 | 0.46 | -0.59 | -0.63 | -1.57 | -1.48 | 0.0775 | 0.03 | -0.5425 | 0.869848 | 0.950698 | 0.223025 | | 1 | 0.996755 | 0.787447 |
| Zfa | P23607 | -2.39 | -1.63 | -1.59 | -0.64 | 0.21 | -2.08 | -0.8 | -0.7 | 0.68 | -0.46 | -1.21 | -2.38 | 1.56 | -0.64 | 0.42 | -0.04 | 0.72 | 0.72 | 1.8875 | 0.207648 | 0.299517 | 0.00682 | | 1 | 0.996511 | 0.552614 |
| Zfand1 | Q8BFR6 | 1.12 | 1.1 | 0.67 | 1.29 | 0.51 | 0.33 | 0.09 | 1.02 | 0.77 | -1.05 | 0.09 | 0.6 | 0.94 | 0.68 | -0.64 | -0.1 | -0.5575 | -0.9425 | -0.825 | 0.049491 | 0.044867 | 0.052582 | | 1 | 0.996511 | 0.659923 |
| Zfand2b | Q91X58 | 0.5 | 0.63 | 1.15 | 0.83 | 0.3 | 0.11 | -0.48 | 0.15 | 0.83 | 1.48 | 0.56 | 1.1 | 1.05 | 1.35 | 0.93 | 0.43 | -0.7575 | 0.215 | 0.1625 | 0.010939 | 0.419388 | 0.543968 | | 1 | 0.996511 | 0.912212 |
| Zfand5 | O88878 | -1.35 | -1.5 | -1.03 | -0.95 | -0.68 | -1.22 | -1.91 | -1.53 | -0.8 | -0.52 | -0.82 | -0.84 | -0.32 | 0.35 | -1.45 | -1.3 | -0.1275 | 0.4625 | 0.5275 | 0.662718 | 0.043494 | 0.23436 | | 1 | 0.996511 | 0.790657 |
| Zfand6 | Q9DCH6 | -0.61 | -0.71 | -1.15 | -0.98 | -0.77 | -0.7 | -1.67 | -0.92 | -0.07 | 0.04 | -1.15 | -0.3 | -0.34 | 0.69 | -1.04 | -1.58 | -0.1525 | 0.4925 | 0.295 | 0.565747 | 0.130006 | 0.538427 | | 1 | 0.996511 | 0.912212 |
| Zfc3h1 | B2RT41 | 0.27 | -0.04 | -2.5 | -1.64 | -1.8 | -0.41 | -1.51 | -0.93 | 0.34 | -1.94 | -1.83 | 0.36 | 1.73 | -0.5 | -1.99 | -1.6 | -0.185 | 0.21 | 0.3875 | 0.779516 | 0.800957 | 0.685295 | | 1 | 0.996511 | 0.955729 |
| Zfp180 | Q6NZI9 | -2.38 | -2.11 | -1.81 | -1.07 | -1.28 | -1.44 | -2.38 | -2.24 | -0.22 | -2.51 | -1.48 | -1.45 | -0.99 | -0.82 | -0.68 | -1.64 | 0.0075 | 0.4275 | 0.81 | 0.984094 | 0.409934 | 0.043512 | | 1 | 0.996511 | 0.658454 |
| Zfp219 | Q4VA98 | 0.19 | -0.47 | -0.61 | -1.08 | -1.39 | -1.12 | -3.39 | -1.9 | 1.41 | 0.32 | -1.54 | -0.73 | 1.59 | 1.52 | 0.26 | -1.62 | -1.4575 | 0.3575 | 0.93 | 0.020599 | 0.575367 | 0.218541 | | 1 | 0.996511 | 0.787447 |
| Zfp280d | V9GWW1 | 0.96 | -2.67 | 1.46 | 0.86 | -3.48 | -2.08 | 1.11 | 1.62 | -0.24 | 2.78 | 1.99 | -1.31 | -1.14 | -0.33 | 1.67 | 2.04 | -0.86 | 0.6525 | 0.4075 | 0.541064 | 0.590944 | 0.709734 | | 1 | 0.996511 | 0.957763 |
| Zfp324 | Q78F42 | -3.76 | -2.84 | -4.45 | -4.72 | -1.3 | -1.95 | -3.42 | -4 | 1.82 | -3.93 | -2.42 | -3.33 | 1.39 | 3.48 | -2.77 | -3.25 | 1.275 | 1.9775 | 3.655 | 0.090194 | 0.132907 | 0.035057 | | 1 | 0.996511 | 0.651434 |
| Zfp326 | A0A0R4J09 | 0.68 | 0.04 | -0.05 | -0.12 | -0.52 | 0.05 | -0.7 | 0.27 | -0.53 | 0.01 | -0.46 | -0.82 | 0.25 | -0.25 | 0.31 | -0.83 | -0.3625 | -0.5875 | -0.2675 | 0.237714 | 0.052896 | 0.422164 | | 1 | 0.996511 | 0.875782 |
| Zfp346 | E9PYN2 | 0.38 | -0.04 | -0.35 | 0.32 | -0.2 | -0.2 | -0.34 | -0.11 | 0.51 | 0.09 | -0.02 | -0.16 | 0.62 | 0.75 | 0.95 | -0.43 | -0.29 | 0.0275 | 0.395 | 0.184197 | 0.911431 | 0.276596 | | 1 | 0.996744 | 0.822084 |
| Zfp362 | E9QA68 | -1.19 | -1.86 | 3.92 | 1.99 | -2.73 | -0.9 | 3.22 | 3.57 | 0.12 | 1 | 4.39 | -1.37 | 0.92 | -1.16 | 4.8 | 5.77 | 0.075 | 0.32 | 1.8675 | 0.967428 | 0.844363 | 0.334519 | | 1 | 0.996511 | 0.846677 |
| Zfp36l2 | P23949 | -0.51 | -0.92 | -0.99 | -0.69 | -1.07 | -0.66 | -1.87 | -0.76 | -0.42 | 0.7 | -0.8 | -1.25 | 0.94 | 0.19 | -1.04 | -1.58 | -0.3125 | 0.335 | 0.405 | 0.304845 | 0.423685 | 0.46044 | | 1 | 0.996511 | 0.887902 |
| Zfp37 | A0A1Y7VJF | 0.29 | 0.23 | -1.17 | -0.84 | -0.02 | 0.41 | -2.39 | -1.07 | 0.51 | 2.31 | -1.74 | -0.27 | -0.57 | 0.3 | 0.52 | -3.49 | -0.395 | 0.575 | -0.4375 | 0.553155 | 0.495577 | 0.626681 | | 1 | 0.996511 | 0.939737 |
| Zfp385a | A0A2R8VH | 0.06 | -1.51 | -1.73 | -2.14 | -1.2 | -0.55 | -1.03 | -1.39 | -1.97 | -1.68 | -1.61 | -0.78 | -1.12 | -0.99 | -0.43 | -1.19 | 0.2875 | -0.18 | 0.3975 | 0.550032 | 0.722959 | 0.416285 | | 1 | 0.996511 | 0.874945 |
| Zfp385b | G5E8Y8 | 1.02 | -0.18 | -0.97 | -1.72 | -0.29 | -0.58 | -1.47 | -0.93 | -0.2 | 0.53 | -2.59 | -1.04 | 0.98 | 0.62 | -1.86 | -1.23 | -0.355 | -0.3625 | 0.09 | 0.546905 | 0.654034 | 0.912219 | | 1 | 0.996511 | 0.986458 |
| Zfp428 | H3BL05 | -0.37 | -0.27 | -1.52 | -2.18 | 0.2 | -0.51 | -2.61 | -2.08 | -1.88 | -2.1 | -0.83 | -0.24 | -1.71 | -0.51 | -0.37 | -2.6 | -0.165 | -0.1775 | -0.2125 | 0.820554 | 0.762198 | 0.740514 | | 1 | 0.996511 | 0.958796 |
| Zfp512b | Q6PHP4 | -0.24 | -1.06 | -1.8 | -0.71 | -1.7 | 0.15 | 0.11 | 0.3 | 0.05 | -1.82 | 0.04 | -0.78 | 0.38 | -1.5 | -1.51 | -1.04 | 0.6675 | 0.325 | 0.035 | 0.228917 | 0.530066 | 0.945761 | | 1 | 0.996511 | 0.992376 |
| Zfp532 | S4R2I9 | 0.69 | 0.3 | -0.15 | -0.1 | 0.25 | 0.37 | 0.35 | 0.13 | -0.11 | -0.28 | 0.2 | 0.35 | 0.23 | 0.48 | 0.35 | 0.58 | 0.09 | -0.145 | 0.225 | 0.690481 | 0.582725 | 0.372341 | | 1 | 0.996511 | 0.859424 |
| Zfp638 | A0A0N4SV: | 0.36 | -1.02 | -1.14 | -0.8 | -0.74 | -0.23 | -1.15 | -0.72 | -0.26 | -0.67 | -0.9 | -1.19 | 0.43 | -0.5 | -0.24 | -1.18 | -0.06 | -0.105 | 0.2775 | 0.872238 | 0.783469 | 0.543718 | | 1 | 0.996511 | 0.912212 |
| Zfp651 | A0A1L1STN | 0.16 | -0.56 | -4.02 | -4.92 | 1.09 | 0.13 | -6.09 | -5.8 | 0.52 | -1.23 | -3.87 | 0.22 | -0.98 | -0.24 | -2.28 | -5.73 | -0.3325 | 1.245 | 0.0275 | 0.86996 | 0.396172 | 0.98584 | | 1 | 0.996511 | 0.996154 |
| Zfp708 | A0A087WF | 9.92 | 7.43 | 11.19 | 7.1 | 8.42 | 9.89 | 9.82 | 11.96 | 5.58 | 4.36 | 11.19 | 9.23 | 5.35 | 1.64 | 11.68 | 12.13 | 1.1125 | -1.32 | -1.21 | 0.326231 | 0.436542 | 0.618755 | | 1 | 0.996511 | 0.937856 |
| Zfp947 | Q8BIQ6 | -3.6 | -2.35 | -3.53 | -4.19 | -1.18 | -3.44 | -3.99 | -4.32 | -3.26 | -1.09 | -3.6 | -2.84 | -1.81 | -0.16 | -3.56 | -3.54 | 0.185 | 0.72 | 1.15 | 0.79984 | 0.264699 | 0.180497 | | 1 | 0.996511 | 0.775542 |
| Zfpl1 | A0A494B9\ | 0.97 | -0.4 | -3.48 | -2.1 | 0.02 | 0.69 | -2.38 | -3.07 | 0.46 | -2.18 | -2.71 | -0.44 | 1.14 | -0.43 | -1.97 | -0.87 | 0.0675 | 0.035 | 0.72 | 0.954796 | 0.974493 | 0.495764 | | 1 | 0.997515 | 0.89758 |
| Zfpm1 | O35615 | -2.82 | -3.39 | -1.25 | -0.97 | -4 | -1.6 | -2.49 | -0.99 | -0.6 | 0.25 | -1.24 | -1.65 | -0.01 | -1.32 | -0.98 | -1.73 | -0.1625 | 1.2975 | 1.0975 | 0.837972 | 0.075569 | 0.111077 | | 1 | 0.996511 | 0.712422 |
| Zfr | O88532 | -0.23 | -1.26 | -0.7 | -0.61 | -1.32 | -0.81 | -0.53 | -0.02 | -1.33 | -0.8 | -0.4 | -1.63 | -0.37 | -0.75 | 0.34 | -0.66 | 0.03 | -0.34 | 0.34 | 0.928431 | 0.333767 | 0.317163 | | 1 | 0.996511 | 0.838864 |
| Zfr2 | E9Q5M4 | 1.44 | 0.14 | -0.57 | -0.68 | 0.65 | 1.17 | -0.6 | -0.19 | -2.91 | -1.71 | -3.47 | -3.23 | -2.54 | -2.47 | -3.38 | -4.28 | 0.175 | -2.9125 | -3.25 | 0.761914 | 7.58E-04 | 4.37E-04 | | 1 | 0.996511 | 0.270112 |
| Zfyve1 | Q810J8 | 0.81 | 0.52 | 1.21 | 0.56 | 0.53 | 0.87 | 1.51 | 1.3 | -0.02 | -0.13 | 1.43 | 0.8 | -0.08 | 0.05 | 1.25 | 1.63 | 0.2775 | -0.255 | -0.0625 | 0.327545 | 0.513217 | 0.885388 | | 1 | 0.996511 | 0.979767 |
| Zfyve16 | Q80U44 | -0.43 | 0.14 | 0.01 | -0.07 | -0.29 | -1.27 | -0.43 | -0.4 | -0.01 | 0.34 | -0.24 | -0.41 | 0.12 | 0.48 | 0.09 | -0.14 | -0.51 | 0.0075 | 0.225 | 0.081169 | 0.974353 | 0.33104 | | 1 | 0.997515 | 0.845601 |
| Zfyve19 | Q9DAZ9 | 0.25 | 0.46 | -0.31 | 0 | 0.06 | 0.52 | -0.76 | -0.23 | 1.14 | 0.64 | -0.33 | 0.16 | 1.28 | 0.04 | -0.13 | -0.22 | -0.2025 | 0.3025 | 0.1425 | 0.519315 | 0.397348 | 0.70694 | | 1 | 0.996511 | 0.957157 |
| Zfyve21 | A0A1Y7VN | 0.28 | -0.19 | -1.4 | -1.17 | 0.19 | -0.19 | -1.85 | -0.86 | -0.78 | -1.83 | -0.88 | -0.05 | -0.83 | -0.07 | -1.42 | -0.91 | -0.0575 | -0.265 | -0.1875 | 0.916513 | 0.600811 | 0.68371 | | 1 | 0.996511 | 0.955729 |
| Zfyve26 | Q5DU37 | 0.72 | 0.18 | 2.63 | 0.25 | 1.07 | 0.7 | 2.56 | 1.67 | -0.62 | 0.25 | 2.57 | 0.88 | -0.83 | -0.34 | 2.71 | 3.6 | 0.555 | -0.175 | 0.34 | 0.39604 | 0.827071 | 0.759137 | | 1 | 0.996511 | 0.963123 |
| Zfyve27 | Q3TXX3 | 0.65 | 0.95 | 0.54 | -0.23 | 0.75 | 0.33 | 0.34 | 0 | -0.86 | -0.12 | 0.16 | 0.73 | -0.48 | -0.54 | 0.7 | 0.54 | -0.1225 | -0.5 | -0.4225 | 0.67803 | 0.229591 | 0.305143 | | 1 | 0.996511 | 0.831662 |
| Zfyve28 | Q6ZPK7 | -1.34 | -1.2 | -0.54 | -1.02 | -0.94 | -0.96 | -0.58 | -0.97 | -0.56 | -0.24 | -1.33 | -1.05 | 0 | -0.57 | -1.14 | -0.64 | 0.1625 | 0.23 | 0.4375 | 0.469827 | 0.457706 | 0.173834 | | 1 | 0.996511 | 0.764637 |
| Zfyve9 | A2A8R0 | -1.11 | -1.41 | -1.72 | -1.97 | -1.1 | -1.43 | -1.77 | -2.14 | -2.12 | -1.02 | -1.92 | -1.3 | -1.64 | -1.58 | -2.05 | -1.8 | -0.0575 | -3.75E-02 | -0.215 | 0.843475 | 0.90638 | 0.397535 | | 1 | 0.996744 | 0.867334 |
| Zgpat | Q8VDM1 | -5.02 | -3.98 | -1.01 | -1.7 | -5.2 | -8.67 | -1.74 | -1.83 | -2.47 | -0.62 | -0.91 | -4.3 | -1.89 | -5.03 | -1.37 | -1.56 | -1.4325 | 0.8525 | 0.465 | 0.407448 | 0.460163 | 0.684504 | | 1 | 0.996511 | 0.955729 |
| Zhx2 | Q8C0C0 | -0.71 | -0.85 | -1.86 | -1.78 | -1.77 | -1.04 | -2.19 | -1.85 | -0.25 | -1.36 | -1.29 | -0.85 | 1.15 | -1.62 | -1.36 | -1 | -0.4125 | 0.3625 | 0.5925 | 0.281363 | 0.354628 | 0.365968 | | 1 | 0.996511 | 0.858761 |
| Zhx3 | Q8C0Q2 | -1.18 | -2.24 | -1.29 | -0.95 | -2.12 | -1.39 | -2.05 | -1.62 | -0.68 | 0.09 | -1.01 | -2.16 | -0.31 | -1.62 | -1.59 | -2.44 | -0.38 | 0.475 | -0.075 | 0.261356 | 0.362212 | 0.878154 | | 1 | 0.996511 | 0.979626 |
| Zkscan1 | Q8BGS3-2 | -3.54 | -3.92 | -3.68 | -3.13 | -6.96 | -4.54 | -5.1 | -3.87 | -1.02 | -3.26 | -2.63 | -3.94 | -0.19 | -4.89 | -2.23 | -3.81 | -1.55 | 0.855 | 0.7875 | 0.032815 | 0.174242 | 0.405931 | | 1 | 0.996511 | 0.873384 |
| Zkscan16 | A2ALW2 | 3.32 | 0.8 | -0.26 | -0.2 | 0.9 | 3.82 | 0.25 | -0.18 | 0.11 | -0.73 | -0.74 | 2.32 | 0.61 | 0.93 | 0.37 | -0.03 | 0.2825 | -0.675 | -0.445 | 0.797991 | 0.502632 | 0.570715 | | 1 | 0.996511 | 0.924218 |
| Zmat2 | Q9CPW7 | -1.41 | -2.39 | -1.99 | -2.09 | -2.18 | -2.04 | -2 | -1.71 | -2.8 | -0.67 | -1.93 | -2.38 | -1 | -1.28 | -1.06 | -1.77 | -0.0125 | 0.025 | 0.6925 | 0.958829 | 0.95769 | 0.0368 | | 1 | 0.996755 | 0.658454 |
| Zmat3 | B9EI21 | -1.19 | -0.8 | -1.51 | 1.39 | -2.33 | -3.62 | -0.42 | -1.78 | -0.22 | 2.49 | -0.66 | -0.75 | 0.24 | -1.21 | -0.18 | -0.95 | -1.51 | 0.7425 | 0.0025 | 0.099984 | 0.423095 | 0.997019 | | 1 | 0.996511 | 0.999279 |
| Zmiz1 | Q6P1E1-2 | -3.63 | -3.88 | -2.75 | -3.68 | -3.46 | -4.09 | -2.66 | -3.22 | -4.77 | -1.97 | -2.93 | -2.77 | -3.94 | -3.64 | -3.18 | -2.68 | 0.1275 | 0.375 | 0.125 | 0.73106 | 0.529861 | 0.733321 | | 1 | 0.996511 | 0.958635 |
| Zmpste24 | Q80W54 | 0.03 | 0.17 | 0.22 | -0.1 | 0.74 | 0.35 | 0.61 | 0.2 | -0.09 | 0.17 | -0.1 | -0.12 | 0.01 | 0.39 | 0.6 | 0.51 | 0.395 | -0.115 | 0.2975 | 0.057275 | 0.512523 | 0.177643 | | 1 | 0.996511 | 0.772068 |
| Zmym2 | Q9CU65 | -2.3 | -3.81 | -3.59 | -3.84 | -3.5 | -2.91 | -4.18 | -4.73 | -2.64 | -3.76 | -3.5 | -2.55 | -1.53 | -3.98 | -3.98 | -4.47 | -0.445 | 0.2725 | -0.105 | 0.382918 | 0.547092 | 0.878651 | | 1 | 0.996511 | 0.979626 |
| Zmym3 | B1AXS8 | -0.39 | -0.98 | -1.03 | -1.09 | -0.77 | -1.07 | -1.22 | -1.24 | -0.07 | -0.18 | -0.95 | -1.18 | 0.07 | -0.62 | -0.61 | -1.43 | -0.2025 | 0.2775 | 0.225 | 0.367708 | 0.397369 | 0.520411 | | 1 | 0.996511 | 0.907667 |
| Zmym4 | A2A791-2 | 0.35 | -0.4 | -0.51 | -0.2 | -0.93 | -0.46 | -1.33 | -0.89 | 0.29 | -0.49 | -0.07 | -0.7 | 1.41 | 0.84 | 0.13 | -0.58 | -0.7125 | -0.0525 | 0.64 | 0.024765 | 0.860269 | 0.176652 | | 1 | 0.996511 | 0.771503 |
| Zmynd11 | Q8R5C8 | 1.25 | 0.87 | 1.66 | 1.02 | -0.24 | 1.22 | 0.78 | 1.07 | 1.23 | 0.73 | 1.02 | 0.51 | 1.87 | -0.62 | 1.49 | 1.16 | -0.4925 | -0.3275 | -0.225 | 0.190463 | 0.221082 | 0.675149 | | 1 | 0.996511 | 0.952944 |
| Zmynd8 | A2A482 | 0.07 | 0.12 | 0.57 | 0.69 | 0.03 | 0.41 | 0.69 | 1.01 | 0.09 | 0.39 | 0.75 | -0.09 | 0.55 | 0.2 | 0.84 | 0.7 | 0.1725 | -0.0775 | 0.21 | 0.523656 | 0.766637 | 0.401219 | | 1 | 0.996511 | 0.870023 |
| Znf148 | Q61624 | -0.69 | -1.11 | -1.56 | -1.04 | -1.03 | -1.21 | -0.8 | -1.31 | -1.28 | -0.14 | -0.89 | -1.19 | -1.66 | -0.36 | -1.07 | -0.85 | 0.0125 | 0.225 | 0.115 | 0.956662 | 0.482885 | 0.726245 | | 1 | 0.996511 | 0.958398 |
| Znf207 | Q9JMD0-2 | -2.77 | -3.74 | -0.51 | -1.17 | -3.39 | -4.06 | -0.29 | -0.6 | -3.27 | -0.25 | -0.32 | -2.44 | -3.33 | -3.02 | -0.62 | -0.48 | -0.0375 | 0.4775 | 0.185 | 0.972364 | 0.618792 | 0.84546 | | 1 | 0.996511 | 0.976225 |
| Znf22 | Q9ERU3 | 5.58 | 4.59 | 3.43 | 2.64 | 6.29 | 5.19 | 2.61 | 2.94 | 1.6 | 1.31 | 3.27 | 5.09 | 1.17 | 2.32 | 3.96 | 3.09 | 0.1975 | -1.2425 | -1.425 | 0.841111 | 0.224489 | 0.098816 | | 1 | 0.996511 | 0.698156 |
| Znf280c | Q6P3Y5-2 | 0.98 | 0.75 | 0.73 | 1.71 | 0.52 | 1.24 | 1.15 | 1.29 | 2.07 | 1.51 | 1.07 | 0.76 | 3.08 | 0.12 | 1.13 | 1.18 | 0.0075 | 0.31 | 0.335 | 0.979426 | 0.395499 | 0.582197 | | 1 | 0.996511 | 0.927734 |
| Znf318 | Q99PP2 | -0.71 | -1.3 | -2.05 | -3.16 | -0.54 | -1.78 | -3.45 | -2.39 | -0.68 | 0.74 | -2.21 | -1.18 | -1.27 | -0.95 | -2.67 | -2.54 | -0.235 | 0.9725 | -0.0525 | 0.747457 | 0.208286 | 0.933155 | | 1 | 0.996511 | 0.989767 |
| Znf32 | Q80V23 | 0.46 | -0.77 | 0.07 | -0.8 | -0.91 | -0.31 | -0.35 | -1 | 0.78 | 1.45 | -2 | -0.17 | 2 | -0.75 | 0.8 | -0.05 | -0.3825 | 0.275 | 0.76 | 0.289653 | 0.709888 | 0.2356 | | 1 | 0.996511 | 0.791493 |
| Znf330 | Q922H9 | -0.6 | -0.08 | -0.19 | -0.63 | -0.59 | -0.96 | -0.73 | -0.39 | 0.39 | -0.62 | -0.5 | -0.46 | -0.12 | 0.79 | -0.39 | -0.91 | -0.2925 | 0.0775 | 0.2175 | 0.191267 | 0.783995 | 0.564536 | | 1 | 0.996511 | 0.920315 |
| Znf365 | Q8BG89 | 0.84 | 1 | 1.5 | 1.42 | 1.56 | 0.84 | 1.57 | 1.63 | 1.51 | 1.42 | 1.63 | 1.06 | 1.28 | 1.88 | 1.42 | 1.91 | 0.21 | 0.215 | 0.4325 | 0.422016 | 0.367579 | 0.119308 | | 1 | 0.996511 | 0.721448 |
| Znf414 | Q9DCK4 | -3.55 | -4.09 | -2.62 | -2.46 | -5.99 | -3.44 | -3.55 | -2.85 | -1.15 | -1.18 | -2.31 | -4.04 | 0.36 | -3.22 | -1.74 | -3.08 | -0.7775 | 1.01 | 1.26 | 0.296626 | 0.179764 | 0.152409 | | 1 | 0.996511 | 0.750422 |
| Znf48 | Q3US17 | 0.02 | -2.27 | -1.02 | -0.82 | -3.43 | -0.22 | -2.86 | -1.17 | -0.1 | -0.51 | -0.97 | -1.77 | 1.76 | -2.47 | -1.07 | -2.34 | -0.8975 | 0.185 | -0.0075 | 0.276475 | 0.73603 | 0.99386 | | 1 | 0.996511 | 0.997252 |
| Znf512 | Q69Z99 | 1.39 | 0.41 | 1.15 | 1.21 | 0.04 | 1 | 0.94 | 0.66 | 1.23 | 0.27 | 1.09 | 0.55 | 1.75 | 0.64 | 1.24 | 1.49 | -0.38 | -0.255 | 0.24 | 0.233337 | 0.426283 | 0.468242 | | 1 | 0.996511 | 0.889423 |
| Znf598 | Q80YR4-2 | -1.58 | -1.13 | -0.68 | -0.51 | -0.92 | -1.45 | -0.42 | -0.76 | -0.81 | -0.22 | -0.38 | -1.37 | -0.47 | -1.06 | -0.62 | -0.41 | 0.0875 | 0.28 | 0.335 | 0.782099 | 0.427201 | 0.275702 | | 1 | 0.996511 | 0.822084 |
| Znf622 | Q91VY9 | -1.26 | -0.92 | 0.68 | 0.13 | -0.67 | -0.72 | 0.2 | 0.7 | -0.86 | 1.86 | 0.76 | -0.85 | -0.59 | 0.15 | 0.18 | 1.27 | 0.22 | 0.57 | 0.595 | 0.675392 | 0.442336 | 0.293214 | | 1 | 0.996511 | 0.825612 |
| Znf668 | Q8K2R5 | 0.35 | 0.09 | -1.23 | -1.31 | 0.13 | -0.27 | -0.64 | -0.67 | -0.82 | -1.96 | -1.67 | -0.57 | -0.67 | -0.49 | -3.18 | -0.82 | 0.1625 | -0.73 | -0.765 | 0.713292 | 0.175444 | 0.289149 | | 1 | 0.996511 | 0.824705 |
| Znf687 | Q9D2D7 | 0.82 | 1.08 | -0.18 | -0.15 | 0.47 | 0.67 | -0.7 | -0.45 | -0.42 | -0.24 | -0.61 | 0.66 | 0.75 | -0.78 | 0.44 | -0.71 | -0.395 | -0.545 | -0.4675 | 0.378547 | 0.20693 | 0.341891 | | 1 | 0.996511 | 0.848473 |
| Znf704 | Q9ERQ3 | -0.33 | 0.15 | -0.49 | 0.02 | -0.09 | 0.64 | -0.63 | -0.1 | 1.09 | -0.37 | 0.42 | 0.99 | -0.26 | 0.87 | -0.08 | -1.07 | 0.1175 | 0.695 | 0.0275 | 0.694946 | 0.079326 | 0.946252 | | 1 | 0.996511 | 0.992376 |
| Znf706 | Q9D115 | -0.74 | -0.1 | -3.65 | -4.33 | -0.09 | -1.02 | -3.91 | -3.7 | -0.1 | -2.97 | -3.12 | -1.29 | -0.15 | 0.1 | -3.77 | -4.02 | 0.025 | 0.335 | 0.245 | 0.984274 | 0.769263 | 0.857376 | | 1 | 0.996511 | 0.976225 |
| Znf771 | Q8BJ90 | -0.65 | -1.77 | -1.95 | -1.25 | -1.03 | -0.23 | -1.95 | -1.08 | 0.43 | -0.85 | -1.98 | -0.71 | 0.81 | -1.05 | -1.79 | -2.41 | 0.3325 | 0.6275 | 0.295 | 0.445032 | 0.255957 | 0.66907 | | 1 | 0.996511 | 0.952558 |
| Znf787 | Q8BIF9 | -2.81 | -9.36 | -2.74 | -7.61 | -9.94 | -3.57 | -7.07 | -8.3 | 0.35 | 0.21 | -3.55 | -4.79 | 2.49 | -9.39 | -1.79 | -4.67 | -1.59 | 3.685 | 2.29 | 0.418616 | 0.080216 | 0.400344 | | 1 | 0.996511 | 0.86916 |
| Znf800 | Q0VEE6 | 0.35 | 0.23 | 0.49 | 0.54 | 0.71 | 0.07 | 0.71 | 0.68 | 0.2 | 0.02 | 0.67 | 0.15 | 0.55 | 0.77 | 0.33 | 0.91 | 0.14 | -0.1425 | 0.2375 | 0.497824 | 0.492366 | 0.268285 | | 1 | 0.996511 | 0.820082 |
| Znf830 | Q8R1N0 | 0.65 | -2.75 | -3.41 | -4.27 | -0.44 | 0.33 | -3.36 | -2.01 | 0.59 | -2.28 | -3.24 | 0.07 | 2.89 | 1.23 | -1.11 | -2.07 | 1.075 | 1.23 | 2.68 | 0.386623 | 0.345339 | 0.079852 | | 1 | 0.996511 | 0.681818 |
| Znfx1 | Q8R151 | -0.5 | -0.53 | -0.21 | -0.46 | -0.34 | -0.5 | -0.36 | -0.41 | -0.21 | -0.4 | -0.35 | -0.43 | 0.15 | -0.27 | -0.44 | -0.3 | 0.0225 | 0.0775 | 0.21 | 0.882856 | 0.647049 | 0.325458 | | 1 | 0.996511 | 0.843085 |
| Znhit2 | Q9QY66 | 4.97 | 5.53 | 0.08 | -0.36 | 4.17 | 2.85 | 0.39 | -0.56 | 5.17 | 0.59 | -0.32 | 4.61 | 6.56 | 0.42 | -0.57 | 0.27 | -0.8425 | -0.0425 | -0.885 | 0.622395 | 0.981792 | 0.661211 | | 1 | 0.998773 | 0.949645 |
| Znrd2 | P56873 | -0.35 | -0.3 | 0.1 | 1 | -0.1 | 0.4 | 0.62 | 1.09 | -0.6 | 0.95 | 0.23 | 0.16 | -0.39 | -0.43 | -0.13 | 0.61 | 0.39 | 0.0725 | -0.1975 | 0.318054 | 0.863581 | 0.609271 | | 1 | 0.996511 | 0.934202 |
| Znrf1 | E9PZB7 | 0.08 | 0.75 | 0.13 | 0.23 | 0.85 | -0.29 | 0.47 | -0.02 | 0.46 | -0.18 | 0.44 | 0.57 | 0.24 | 1.02 | 0.43 | 0.73 | -0.045 | 0.025 | 0.3075 | 0.878987 | 0.92087 | 0.255216 | | 1 | 0.996755 | 0.809524 |
| Znrf2 | Q71FD5 | -0.08 | 0.72 | 1.04 | 0.31 | -0.19 | 0.03 | -0.66 | 0.9 | 1.42 | -2.01 | 1.11 | 0.39 | 1.49 | 1.3 | 0.85 | 0.87 | -0.4775 | -0.27 | 0.63 | 0.2371 | 0.715348 | 0.063294 | | 1 | 0.996511 | 0.66433 |
| Zpr1 | Q62384 | -0.73 | -0.41 | -0.71 | -0.8 | -0.4 | -1.13 | -0.73 | -0.48 | -0.84 | -0.25 | -0.62 | -0.7 | -0.65 | -0.03 | -0.74 | -0.67 | -0.0225 | 0.06 | 0.14 | 0.915569 | 0.76611 | 0.548119 | | 1 | 0.996511 | 0.912597 |
| Zranb2 | D3Z4U0 | -3.01 | -2.79 | -0.47 | -1.18 | -4.79 | -3.6 | -2.59 | -2.14 | -2.35 | -0.03 | -1.4 | -1.75 | -2.32 | -2.79 | -3.02 | -2.97 | -1.4175 | 0.48 | -0.9125 | 0.093634 | 0.508806 | 0.145728 | | 1 | 0.996511 | 0.745321 |
| Zscan18 | E9PUD6 | 0.25 | 0.5 | -0.55 | 0.78 | 0.58 | 0.11 | 0.47 | 0.46 | 0.61 | 1.18 | 0.85 | 0.06 | 1.26 | 1.55 | 0.49 | 0.27 | 0.16 | 0.43 | 0.6475 | 0.598258 | 0.253488 | 0.134783 | | 1 | 0.996511 | 0.731223 |
| Zscan26 | F8WJ31 | 0.24 | -1.39 | 0.54 | 0.01 | 0.05 | -0.99 | -0.38 | 0.19 | 1.22 | 0.52 | -0.32 | -0.66 | 1.73 | 0.5 | 0.57 | 0.43 | -0.1325 | 0.34 | 0.9575 | 0.776768 | 0.544241 | 0.078827 | | 1 | 0.996511 | 0.680976 |
| Zswim8 | Q3UHH1-3 | -0.5 | -0.1 | 0.05 | -0.64 | -0.15 | -0.41 | 0.01 | -0.49 | -0.34 | -0.8 | -0.06 | -0.14 | -0.61 | 0.14 | -0.8 | 0.28 | 0.0375 | -0.0375 | 0.05 | 0.865991 | 0.882597 | 0.875932 | | 1 | 0.996511 | 0.979626 |
| Zw10 | O54692 | 0.57 | 0.33 | -0.16 | -0.06 | 0.64 | 0.4 | 0.07 | 0.08 | 0.27 | 0.14 | -0.22 | 0.47 | 0.01 | 0.84 | 0.23 | 0.21 | 0.1275 | -0.005 | 0.1525 | 0.591928 | 0.983887 | 0.5775 | | 1 | 0.998876 | 0.927198 |
| Zwint | Q9CQU5 | -1.39 | -1.16 | -0.95 | -1.09 | -1.47 | -1.3 | -1.19 | -0.91 | -0.01 | -0.29 | -0.83 | -1.15 | 1.16 | -0.93 | -0.42 | -0.89 | -0.07 | 0.5775 | 0.8775 | 0.711545 | 0.068619 | 0.088704 | | 1 | 0.996511 | 0.691619 |
| Zyg11b | Q3UFS0 | -0.03 | -0.08 | -0.38 | -0.44 | 0.21 | -0.05 | -0.27 | -0.49 | -0.63 | -0.79 | -0.3 | -0.01 | -0.67 | -0.17 | -0.63 | -0.64 | 0.0825 | -0.2 | -0.295 | 0.695822 | 0.401268 | 0.191058 | | 1 | 0.996511 | 0.777117 |
| Zyx | Q62523 | 1.71 | 0.57 | 0.07 | 0.26 | 0.96 | 1.55 | 0.26 | 0.28 | 0.08 | 1.09 | 0.48 | 1.57 | 0.48 | 0.96 | 0.53 | 0.46 | 0.11 | 0.1525 | -0.045 | 0.805768 | 0.742055 | 0.904816 | | 1 | 0.996511 | 0.985525 |
| Zzef1 | E9Q5W5 | 0.01 | -0.48 | -0.27 | -0.19 | -0.22 | -0.11 | -0.4 | -0.21 | -0.26 | -0.11 | -0.19 | -0.05 | -0.12 | -0.06 | -0.32 | -0.25 | -0.0025 | 0.08 | 0.045 | 0.988286 | 0.656306 | 0.815101 | | 1 | 0.996511 | 0.972768 |

**GSEA synaptoneurosomes**

** GSEA mouse MSigDB canonical pathways: m2.cp.v2023.1.Mm.symbols*

**significant pathways (FDR<0.25) are marked in yellow*

| SYP MNK1KO / WT | | | | SYP MNK2KO / WT | | | | SYP MNKDKO / WT | | | |
|---|---|---|---|---|---|---|---|---|---|---|---|
| canonical pathway | NES | NOM p-val | FDR q-val | canonical pathway | NES | NOM p-val | FDR q-val | canonical pathway | NES | NOM p-val | FDR q-val |
| REACTOME_TRANSPORT | -0.88811 | 0.667607 | 1 | REACTOME_TRANSPORT_OF_MATURE_TRANSCRIPT_T | -2.3427 | 0 | 2.89E-04 | REACTOME_TRANSPORT_ | 1.651372 | 0.003751 | 0.089592 |
| REACTOME_TRANSPORT | 0.777301 | 0.816186 | 1 | REACTOME_TRANSPORT_OF_MATURE_MRNAS_DERIV | -2.21124 | 0 | 0.002145 | REACTOME_TRANSPORT_ | 1.833126 | 9.04E-04 | 0.031142 |
| REACTOME_NUCLEAR_E | -0.51275 | 0.995295 | 1 | REACTOME_NUCLEAR_ENVELOPE_BREAKDOWN | -2.18728 | 0 | 0.002614 | REACTOME_NUCLEAR_EN | 1.407661 | 0.065347 | 0.291224 |
| REACTOME_TRANSPORT | -0.78723 | 0.783312 | 1 | REACTOME_TRANSPORT_OF_THE_SLBP_DEPENDANT_ | -2.2161 | 0 | 0.00293 | REACTOME_TRANSPORT_ | 1.46113 | 0.054284 | 0.238355 |
| REACTOME_NUCLEAR_F | -0.69559 | 0.887884 | 1 | REACTOME_NUCLEAR_PORE_COMPLEX_NPC_DISASSE | -2.16179 | 0 | 0.003003 | REACTOME_NUCLEAR_PC | 1.636719 | 0.013033 | 0.098659 |
| REACTOME_TRANSCRIP | -0.55007 | 0.986412 | 1 | REACTOME_TRANSCRIPTIONAL_REGULATION_BY_SMA | -2.15233 | 0 | 0.003062 | REACTOME_TRANSCRIPTI | 1.430609 | 0.066361 | 0.270306 |
| REACTOME_SUMOYLATI | -1.18072 | 0.226846 | 0.884357 | REACTOME_SUMOYLATION_OF_DNA_REPLICATION_PI | -2.01883 | 2.96E-04 | 0.009089 | REACTOME_SUMOYLATIC | 1.379512 | 0.080762 | 0.303196 |
| REACTOME_SUMOYLATI | -1.20708 | 0.207437 | 0.850923 | REACTOME_SUMOYLATION_OF_SUMOYLATION_PROT | -2.03173 | 5.86E-04 | 0.00921 | REACTOME_SUMOYLATIC | 1.487596 | 0.043593 | 0.206361 |
| REACTOME_COMPLEX_I | -1.26704 | 0.138691 | 0.889517 | REACTOME_COMPLEX_I_BIOGENESIS | -2.02219 | 0 | 0.009478 | REACTOME_COMPLEX_I_ | -2.03005 | 0 | 0.010675 |
| REACTOME_GLUCOSE_N | 0.742212 | 0.913124 | 1 | REACTOME_GLUCOSE_METABOLISM | -2.03181 | 0 | 0.010131 | REACTOME_GLUCOSE_MI | 0.721595 | 0.909601 | 1 |
| REACTOME_RAB_GERAN | -0.91277 | 0.596666 | 1 | REACTOME_RAB_GERANYLGERANYLATION | -2.03501 | 0 | 0.011011 | REACTOME_RAB_GERANY | -1.48063 | 0.030855 | 0.117621 |
| REACTOME_POSTMITOT | -0.59096 | 0.958891 | 1 | REACTOME_POSTMITOTIC_NUCLEAR_PORE_COMPLEX | -1.99266 | 5.54E-04 | 0.011187 | REACTOME_POSTMITOTI( | 1.048955 | 0.408874 | 0.799858 |
| WP_TCA_CYCLE | 0.983286 | 0.469392 | 1 | WP_TCA_CYCLE | -1.99699 | 0.001707 | 0.011225 | WP_TCA_CYCLE | -2.23446 | 0.00E+00 | 0.002085 |
| REACTOME_MRNA_SPLI | -0.88051 | 0.650592 | 1 | REACTOME_MRNA_SPLICING_MINOR_PATHWAY | -2.03613 | 6.04E-04 | 0.012296 | REACTOME_MRNA_SPLIC | 0.700826 | 0.897681 | 0.986234 |
| REACTOME_SUMOYLATI | -0.99988 | 0.447847 | 1 | REACTOME_SUMOYLATION_OF_RNA_BINDING_PROTE | -2.0389 | 3.01E-04 | 0.0135 | REACTOME_SUMOYLATIC | 1.71632 | 0.004045 | 0.076469 |
| REACTOME_GLYCOLYSIS | 0.679522 | 0.949098 | 1 | REACTOME_GLYCOLYSIS | -1.95653 | 0 | 0.016054 | REACTOME_GLYCOLYSIS | 0.956136 | 0.52593 | 0.903601 |
| WP_ELECTRON_TRANSP | 0.834097 | 0.784144 | 1 | WP_ELECTRON_TRANSPORT_CHAIN | -1.94647 | 0 | 0.016955 | WP_ELECTRON_TRANSPO | -2.06865 | 0 | 0.008129 |
| REACTOME_SNRNP_ASS | 0.991794 | 0.458971 | 1 | REACTOME_SNRNP_ASSEMBLY | -1.92618 | 0 | 0.020223 | REACTOME_SNRNP_ASSE | 1.352834 | 0.088353 | 0.33032 |
| REACTOME_SUMOYLATI | -1.24275 | 0.179029 | 0.855109 | REACTOME_SUMOYLATION_OF_UBIQUITINYLATION_P | -1.92209 | 8.92E-04 | 0.020231 | REACTOME_SUMOYLATIC | 1.414128 | 0.070178 | 0.289829 |
| REACTOME_PROCESSIN | -1.28226 | 0.044764 | 0.868146 | REACTOME_PROCESSING_OF_CAPPED_INTRON_CONT | -1.91038 | 0 | 0.021825 | REACTOME_PROCESSING_ | 1.692861 | 0 | 0.086389 |
| REACTOME_RESPIRATOI | -0.93888 | 0.573824 | 1 | REACTOME_RESPIRATORY_ELECTRON_TRANSPORT | -1.89568 | 0 | 0.02461 | REACTOME_RESPIRATORY | -1.84939 | 0 | 0.029073 |
| REACTOME_MITOTIC_PI | 0.72832 | 0.90833 | 1 | REACTOME_MITOTIC_PROPHASE | -1.84208 | 6.71E-04 | 0.040134 | REACTOME_MITOTIC_PR( | 1.204012 | 0.192488 | 0.51304 |
| REACTOME_MRNA_SPLI | -1.55241 | 0.002954 | 0.579474 | REACTOME_MRNA_SPLICING | -1.835 | 0 | 0.041206 | REACTOME_MRNA_SPLIC | 1.350378 | 0.030838 | 0.330305 |
| REACTOME_REGULATIO | -0.62035 | 0.981706 | 1 | REACTOME_REGULATION_OF_HSF1_MEDIATED_HEAT_ | -1.82768 | 0.001016 | 0.04225 | REACTOME_REGULATION | 1.201697 | 0.194084 | 0.5135 |
| REACTOME_NEUREXINS | 0.519758 | 0.994252 | 1 | REACTOME_NEUREXINS_AND_NEUROLIGINS | -1.79752 | 0.003818 | 0.054243 | REACTOME_NEUREXINS_ | -2.25592 | 2.81E-04 | 0.002357 |
| WP_OXIDATIVE_PHOSPI | -1.04895 | 0.382864 | 1 | WP_OXIDATIVE_PHOSPHORYLATION | -1.79167 | 0.004863 | 0.054902 | WP_OXIDATIVE_PHOSPH( | -1.73451 | 7.54E-03 | 0.043218 |
| REACTOME_GLUTATHIO | 1.255178 | 0.170213 | 0.733114 | REACTOME_GLUTATHIONE_CONJUGATION | -1.74353 | 0.00985 | 0.080299 | REACTOME_GLUTATHION | -1.55866 | 0.028271 | 0.093732 |
| REACTOME_RESPIRATOI | -0.84537 | 0.771608 | 1 | REACTOME_RESPIRATORY_ELECTRON_TRANSPORT_AT | -1.73265 | 7.61E-04 | 0.085184 | REACTOME_RESPIRATORY | -1.79349 | 0 | 0.030652 |
| REACTOME_SUMOYLATI | -0.94157 | 0.548091 | 1 | REACTOME_SUMOYLATION_OF_CHROMATIN_ORGANI | -1.72061 | 0.005761 | 0.091142 | REACTOME_SUMOYLATIC | 1.847096 | 3.02E-04 | 0.028584 |
| REACTOME_RHOBTB2_( | 0.557364 | 0.972089 | 1 | REACTOME_RHOBTB2_GTPASE_CYCLE | -1.70872 | 0.010071 | 0.09384 | REACTOME_RHOBTB2_GT | -1.2933 | 0.142743 | 0.246071 |
| REACTOME_VOLTAGE_( | -0.90878 | 0.584769 | 1 | REACTOME_VOLTAGE_GATED_POTASSIUM_CHANNELS | -1.70201 | 0.010196 | 0.095854 | REACTOME_VOLTAGE_GA | -1.31662 | 0.118224 | 0.224851 |
| REACTOME_GENE_SILEN | -0.78141 | 0.804383 | 1 | REACTOME_GENE_SILENCING_BY_RNA | -1.70913 | 0.005075 | 0.096696 | REACTOME_GENE_SILENC | 1.365919 | 0.087892 | 0.314519 |
| REACTOME_TRAFFICKIN | 1.165314 | 0.242165 | 0.85019 | REACTOME_TRAFFICKING_OF_AMPA_RECEPTORS | -1.67256 | 0.011711 | 0.117409 | REACTOME_TRAFFICKING | -1.48981 | 0.04442 | 0.116573 |
| REACTOME_REGULATIO | -0.65621 | 0.956608 | 1 | REACTOME_REGULATION_OF_INSULIN_SECRETION | -1.66055 | 0.006538 | 0.125238 | REACTOME_REGULATION | -1.65025 | 0.005825 | 0.062541 |
| REACTOME_RAB_REGUL | -0.72139 | 0.948201 | 1 | REACTOME_RAB_REGULATION_OF_TRAFFICKING | -1.65534 | 0.00198 | 0.12671 | REACTOME_RAB_REGULA | -1.44862 | 0.01916 | 0.133359 |
| REACTOME_TBC_RABGA | 0.971685 | 0.482241 | 1 | REACTOME_TBC_RABGAPS | -1.64493 | 0.020185 | 0.133213 | REACTOME_TBC_RABGAP | -1.45702 | 0.051667 | 0.128903 |
| REACTOME_TRAFFICKIN | 1.685841 | 0.016799 | 0.161513 | REACTOME_TRAFFICKING_OF_GLUR2_CONTAINING_A | -1.63804 | 0.026592 | 0.136517 | REACTOME_TRAFFICKING | -1.62217 | 0.022942 | 0.073379 |
| REACTOME_POTASSIUM | -1.03822 | 0.391105 | 1 | REACTOME_POTASSIUM_CHANNELS | -1.62591 | 0.008046 | 0.145134 | REACTOME_POTASSIUM_ | -1.19951 | 0.157843 | 0.320736 |
| REACTOME_COMPLEME | 1.159981 | 0.272902 | 0.844041 | REACTOME_COMPLEMENT_CASCADE | 1.949624 | 9.83E-04 | 0.146041 | REACTOME_COMPLEMEN | 1.264958 | 0.186331 | 0.426354 |
| REACTOME_THE_CITRIC | 0.921426 | 0.64801 | 1 | REACTOME_THE_CITRIC_ACID_TCA_CYCLE_AND_RESPI | -1.58813 | 4.73E-04 | 0.185502 | REACTOME_THE_CITRIC_ | -1.94134 | 0 | 0.017206 |
| REACTOME_CELLULAR_I | -0.7214 | 0.937947 | 1 | REACTOME_CELLULAR_RESPONSE_TO_HEAT_STRESS | -1.57444 | 0.005745 | 0.198811 | REACTOME_CELLULAR_RE | 1.05994 | 0.361254 | 0.780881 |
| WP_G_PROTEIN_SIGNAI | 0.998573 | 0.441384 | 1 | WP_G_PROTEIN_SIGNALING_PATHWAYS | -1.56746 | 0.006225 | 0.203232 | WP_G_PROTEIN_SIGNALI | -2.13382 | 0 | 0.004704 |
| REACTOME_CHAPERONI | 0.782386 | 0.78629 | 1 | REACTOME_CHAPERONIN_MEDIATED_PROTEIN_FOLD | -1.5436 | 0.031277 | 0.232583 | REACTOME_CHAPERONIN | -1.82633 | 0.002555 | 0.030379 |
| REACTOME_SIGNAL_AM | 1.161491 | 0.25852 | 0.847016 | REACTOME_SIGNAL_AMPLIFICATION | -1.53827 | 0.037486 | 0.235398 | REACTOME_SIGNAL_AMP | -1.81031 | 0.003789 | 0.02846 |
| REACTOME_RAB_GEFS_ | -0.86397 | 0.711223 | 1 | REACTOME_RAB_GEFS_EXCHANGE_GTP_FOR_GDP_OI | -1.50058 | 0.014435 | 0.258446 | REACTOME_RAB_GEFS_E) | -1.2564 | 0.103878 | 0.272787 |
| REACTOME_SIGNALING_ | -0.70359 | 0.903442 | 1 | REACTOME_SIGNALING_BY_FGFR2 | -1.48966 | 0.03596 | 0.260931 | REACTOME_SIGNALING_E | -1.12072 | 0.27064 | 0.416863 |
| REACTOME_PYRUVATE_ | -0.71755 | 0.834852 | 1 | REACTOME_PYRUVATE_METABOLISM | -1.51857 | 0.048873 | 0.261447 | REACTOME_PYRUVATE_N | -1.8249 | 0.005005 | 0.030165 |
| REACTOME_INTEGRATIC | 1.008806 | 0.425757 | 1 | REACTOME_INTEGRATION_OF_ENERGY_METABOLISM | -1.49199 | 0.024965 | 0.262254 | REACTOME_INTEGRATION | -1.67088 | 0.002931 | 0.059663 |
| REACTOME_G_PROTEIN | 1.175139 | 0.230099 | 0.831868 | REACTOME_G_PROTEIN_MEDIATED_EVENTS | -1.50459 | 0.035237 | 0.262312 | REACTOME_G_PROTEIN_ | -2.11315 | 0.00E+00 | 0.005728 |
| REACTOME_GLUTAMAT | 0.770026 | 0.796321 | 1 | REACTOME_GLUTAMATE_NEUROTRANSMITTER_RELE/ | -1.50111 | 0.048861 | 0.262869 | REACTOME_GLUTAMATE_ | -1.84188 | 0.00481 | 0.029712 |
| REACTOME_THROMBO) | 0.944508 | 0.522536 | 1 | REACTOME_THROMBOXANE_SIGNALLING_THROUGH_ | -1.50695 | 0.05281 | 0.263909 | REACTOME_THROMBOXA | -1.65622 | 0.021294 | 0.061757 |
| REACTOME_MITOCHON | -0.38833 | 0.999643 | 1 | REACTOME_MITOCHONDRIAL_FATTY_ACID_BETA_OXI | -1.48428 | 0.044126 | 0.264735 | REACTOME_MITOCHOND | -1.95184 | 0.001111 | 0.01872 |
| REACTOME_PROTEIN_P | 1.025124 | 0.399678 | 1 | REACTOME_PROTEIN_PROTEIN_INTERACTIONS_AT_SY | -1.49252 | 0.019732 | 0.266519 | REACTOME_PROTEIN_PR( | -2.02865 | 0 | 0.010233 |
| REACTOME_ADP_SIGNA | 1.038396 | 0.397871 | 1 | REACTOME_ADP_SIGNALLING_THROUGH_P2Y_PURIN( | -1.50855 | 0.053749 | 0.266805 | REACTOME_ADP_SIGNALI | -1.55536 | 0.040449 | 0.095016 |
| REACTOME_NUCLEAR_E | -0.80024 | 0.803877 | 1 | REACTOME_NUCLEAR_ENVELOPE_NE_REASSEMBLY | -1.50875 | 0.018058 | 0.272339 | REACTOME_NUCLEAR_EN | 0.542948 | 0.995712 | 0.994209 |
| REACTOME_G_PROTEIN | -0.83757 | 0.677809 | 1 | REACTOME_G_PROTEIN_ACTIVATION | -1.4682 | 0.070282 | 0.286969 | REACTOME_G_PROTEIN_/ | -1.49205 | 0.060669 | 0.117121 |
| REACTOME_CHROMOSC | -1.35193 | 0.099305 | 0.782877 | REACTOME_CHROMOSOME_MAINTENANCE | -1.46222 | 0.046787 | 0.292088 | REACTOME_CHROMOSON | -0.63535 | 0.958132 | 0.969069 |
| REACTOME_NEURONAL | 0.951952 | 0.61934 | 1 | REACTOME_NEURONAL_SYSTEM | -1.43761 | 7.13E-04 | 0.299999 | REACTOME_NEURONAL_S | -1.73981 | 0 | 0.042537 |
| REACTOME_INOSITOL_F | -1.12422 | 0.282255 | 0.985565 | REACTOME_INOSITOL_PHOSPHATE_METABOLISM | -1.44315 | 0.049629 | 0.300175 | REACTOME_INOSITOL_PH | -1.76166 | 0.003159 | 0.037477 |
| REACTOME_L1CAM_INT | 1.114467 | 0.265837 | 0.888394 | REACTOME_L1CAM_INTERACTIONS | -1.44019 | 0.034853 | 0.300412 | REACTOME_L1CAM_INTE | -2.14328 | 0.00E+00 | 0.00809 |
| REACTOME_RNA_POLYN | -0.68575 | 0.943379 | 1 | REACTOME_RNA_POLYMERASE_II_TRANSCRIPTION_TE | -1.44517 | 0.028581 | 0.301626 | REACTOME_RNA_POLYMI | 1.657974 | 0.005026 | 0.091269 |
| WP_MECP2_AND_ASSO | -1.60838 | 0.020112 | 0.562329 | WP_MECP2_AND_ASSOCIATED_RETT_SYNDROME | -1.44621 | 0.063652 | 0.304782 | WP_MECP2_AND_ASSOCI | -1.08486 | 0.323291 | 0.464488 |
| REACTOME_GABA_B_RE | -0.93255 | 0.551356 | 1 | REACTOME_GABA_B_RECEPTOR_ACTIVATION | -1.44992 | 0.052957 | 0.308679 | REACTOME_GABA_B_REC | -1.63943 | 0.012082 | 0.066475 |
| WP_GLYCOGEN_METAB | 1.330506 | 0.11479 | 0.678822 | WP_GLYCOGEN_METABOLISM | -1.44628 | 0.053877 | 0.309937 | WP_GLYCOGEN_METABO | -1.81154 | 0.001936 | 0.028592 |
| REACTOME_CA2_PATHV | 0.975278 | 0.479858 | 1 | REACTOME_CA2_PATHWAY | -1.42223 | 0.066181 | 0.322717 | REACTOME_CA2_PATHW/ | -1.83695 | 0.002488 | 0.029663 |
| WP_GLYCOLYSIS_AND_( | -0.96186 | 0.506852 | 1 | WP_GLYCOLYSIS_AND_GLUCONEOGENESIS | -1.41449 | 0.05903 | 0.332213 | WP_GLYCOLYSIS_AND_GL | -2.0012 | 8.48E-04 | 0.01362 |
| WP_OXIDATIVE_STRESS_ | 1.109517 | 0.274501 | 0.891193 | WP_OXIDATIVE_STRESS_AND_REDOX_PATHWAY | -1.40707 | 0.040208 | 0.341345 | WP_OXIDATIVE_STRESS_/ | -1.20093 | 0.160995 | 0.319787 |
| BIOCARTA_VIP_PATHW/ | 0.622929 | 0.925112 | 1 | BIOCARTA_VIP_PATHWAY | -1.40054 | 0.098535 | 0.349016 | BIOCARTA_VIP_PATHWAY | -1.82158 | 0.004969 | 0.028506 |
| REACTOME_LDL_CLEAR/ | 1.119454 | 0.314035 | 0.893653 | REACTOME_LDL_CLEARANCE | -1.3977 | 0.098839 | 0.349399 | REACTOME_LDL_CLEARAI | -1.5773 | 0.039132 | 0.086673 |
| REACTOME_PHASE_II_C | 1.372637 | 0.07758 | 0.649992 | REACTOME_PHASE_II_CONJUGATION_OF_COMPOUNI | -1.38641 | 0.058233 | 0.366606 | REACTOME_PHASE_II_CO | -1.50822 | 0.023946 | 0.112103 |
| WP_DRAVET_SYNDROM | -0.85258 | 0.728174 | 1 | WP_DRAVET_SYNDROME_SCN1AA1783V_POINT_MUT | -1.37402 | 0.046999 | 0.381446 | WP_DRAVET_SYNDROME_ | -1.24844 | 0.113304 | 0.277707 |
| BIOCARTA_TCR_PATHW | 0.819334 | 0.717031 | 1 | BIOCARTA_TCR_PATHWAY | -1.36753 | 0.098491 | 0.384293 | BIOCARTA_TCR_PATHWA | -1.91942 | 0.001575 | 0.019305 |
| REACTOME_GLUCAGON | -0.98136 | 0.478814 | 1 | REACTOME_GLUCAGON_LIKE_PEPTIDE_1_GLP1_REGU | -1.37446 | 0.098109 | 0.386004 | REACTOME_GLUCAGON_I | -1.53871 | 0.036946 | 0.101447 |
| REACTOME_SUMOYLATI | -1.50292 | 0.023615 | 0.65337 | REACTOME_SUMOYLATION_OF_DNA_DAMAGE_RESP( | -1.363 | 0.068827 | 0.388418 | REACTOME_SUMOYLATIC | 1.23112 | 0.162754 | 0.468519 |
| REACTOME_BIOLOGICAI | 1.460846 | 0.023382 | 0.479209 | REACTOME_BIOLOGICAL_OXIDATIONS | -1.36773 | 0.033449 | 0.389264 | REACTOME_BIOLOGICAL_ | -1.48924 | 0.014058 | 0.116272 |
| REACTOME_SIGNALING_ | 0.928704 | 0.573502 | 1 | REACTOME_SIGNALING_BY_NTRKS | -1.34711 | 0.069727 | 0.396265 | REACTOME_SIGNALING_E | -1.66654 | 0.003929 | 0.059666 |
| REACTOME_INTERCONV | -0.97258 | 0.495083 | 1 | REACTOME_INTERCONVERSION_OF_NUCLEOTIDE_DI_/ | -1.35534 | 0.115117 | 0.399237 | REACTOME_INTERCONVE | -1.28333 | 0.160636 | 0.251287 |
| REACTOME_EPHB_MED | 1.19078 | 0.212363 | 0.801972 | REACTOME_EPHB_MEDIATED_FORWARD_SIGNALING | -1.34746 | 0.094488 | 0.400572 | REACTOME_EPHB_MEDIA | -2.05566 | 0 | 0.008414 |
| REACTOME_DOWNSTRE | 0.696148 | 0.946281 | 1 | REACTOME_DOWNSTREAM_SIGNALING_EVENTS_OF_I | -1.34166 | 0.06678 | 0.40276 | REACTOME_DOWNSTREA | -2.09968 | 0 | 0.005959 |
| WP_METAPATHWAY_BI | 1.873613 | 7.00E-04 | 0.070935 | WP_METAPATHWAY_BIOTRANSFORMATION | -1.34878 | 0.074206 | 0.402971 | WP_METAPATHWAY_BIO | -1.0075 | 0.432662 | 0.568663 |
| REACTOME_BETA_CATE | 1.089346 | 0.283459 | 0.912221 | REACTOME_BETA_CATENIN_INDEPENDENT_WNT_SIGI | -1.34976 | 0.039279 | 0.406125 | REACTOME_BETA_CATEN | -2.13455 | 0 | 0.005645 |
| WP_FATTY_ACID_BETAC | 0.542251 | 0.985661 | 1 | WP_FATTY_ACID_BETAOXIDATION | -1.32955 | 0.110341 | 0.423836 | WP_FATTY_ACID_BETAO) | -1.82466 | 0.004974 | 0.029571 |
| REACTOME_METABOLIS | -0.82349 | 0.791066 | 1 | REACTOME_METABOLISM_OF_WATER_SOLUBLE_VITA | -1.32532 | 0.059438 | 0.427833 | REACTOME_METABOLISM | -1.36662 | 0.041595 | 0.187898 |
| REACTOME_INWARDLY_ | -1.20769 | 0.216743 | 0.859282 | REACTOME_INWARDLY_RECTIFYING_K_CHANNELS | -1.31038 | 0.126057 | 0.456941 | REACTOME_INWARDLY_R | -1.40202 | 0.088117 | 0.159407 |
| REACTOME_COLLAGEN_ | 1.112193 | 0.309182 | 0.889034 | REACTOME_COLLAGEN_BIOSYNTHESIS_AND_MODIFYI | 1.769456 | 0.005895 | 0.459878 | REACTOME_COLLAGEN_B | 1.748952 | 0.006182 | 0.057042 |
| REACTOME_REGULATIO | 1.466837 | 0.035747 | 0.475008 | REACTOME_REGULATION_OF_INSULIN_LIKE_GROWTH | 1.609199 | 0.013784 | 0.460601 | REACTOME_REGULATION | 1.66291 | 0.00363 | 0.090987 |
| BIOCARTA_NOS1_PATH\ | 1.267109 | 0.177949 | 0.741898 | BIOCARTA_NOS1_PATHWAY | -1.30604 | 0.151205 | 0.461479 | BIOCARTA_NOS1_PATHW | -1.61751 | 0.02126 | 0.074495 |
| WP_P38_MAPK_SIGNAL | -0.8539 | 0.666791 | 1 | WP_P38_MAPK_SIGNALING_PATHWAY | 1.614124 | 0.021971 | 0.479322 | WP_P38_MAPK_SIGNALIN | 1.283235 | 0.154411 | 0.41114 |
| REACTOME_DNA_REPAI | -1.48855 | 0.012133 | 0.622732 | REACTOME_DNA_REPAIR | 1.69266 | 6.48E-04 | 0.485663 | REACTOME_DNA_REPAIR | 1.095401 | 0.288255 | 0.704381 |
| REACTOME_COLLAGEN_ | 1.13776 | 0.275301 | 0.875127 | REACTOME_COLLAGEN_FORMATION | 1.709567 | 0.008426 | 0.499095 | REACTOME_COLLAGEN_F | 1.632597 | 0.015488 | 0.092223 |
| REACTOME_FORMATIOI | -0.95662 | 0.508188 | 1 | REACTOME_FORMATION_OF_INCISION_COMPLEX_IN_ | 1.616899 | 0.019742 | 0.510959 | REACTOME_FORMATION_ | 1.047815 | 0.394762 | 0.789861 |

| | | | | | | | | | | | |
|---|---|---|---|---|---|---|---|---|---|---|---|
| REACTOME_ACTIVATION | 0.679364 | 0.890411 | 1 | REACTOME_ACTIVATION_OF_KAINATE_RECEPTORS_UI | -1.27294 | 0.153131 | 0.514299 | REACTOME_ACTIVATION_ | -1.68987 | 0.011678 | 0.055743 |
| REACTOME_ACTIVATION | -1.25497 | 0.180597 | 0.88378 | REACTOME_ACTIVATION_OF_G_PROTEIN_GATED_POT | -1.27708 | 0.164578 | 0.515528 | REACTOME_ACTIVATION_ | -1.34191 | 0.120769 | 0.204872 |
| BIOCARTA_GPCR_PATHW | 0.919286 | 0.565547 | 1 | BIOCARTA_GPCR_PATHWAY | -1.2738 | 0.164885 | 0.517944 | BIOCARTA_GPCR_PATHW | -1.81751 | 0.004465 | 0.028506 |
| REACTOME_RNA_POLYM | 0.781322 | 0.744783 | 1 | REACTOME_RNA_POLYMERASE_III_TRANSCRIPTION | -1.26618 | 0.190722 | 0.519881 | REACTOME_RNA_POLYMI | -0.75166 | 0.791217 | 0.908826 |
| REACTOME_EXTRACELLU | 0.892881 | 0.682702 | 1 | REACTOME_EXTRACELLULAR_MATRIX_ORGANIZATION | 1.728427 | 0.0016 | 0.520231 | REACTOME_EXTRACELLUI | 1.278696 | 0.096616 | 0.416085 |
| REACTOME_THROMBIN_ | 1.033317 | 0.406168 | 1 | REACTOME_THROMBIN_SIGNALLING_THROUGH_PROT | -1.2772 | 0.15511 | 0.521182 | REACTOME_THROMBIN_S | -1.44748 | 0.067274 | 0.133491 |
| REACTOME_RNA_POLYM | 0.782943 | 0.752278 | 1 | REACTOME_RNA_POLYMERASE_III_TRANSCRIPTION_IN | -1.27945 | 0.177507 | 0.521465 | REACTOME_RNA_POLYMI | -0.74347 | 0.800469 | 0.912531 |
| REACTOME_METABOLIS | -0.86215 | 0.748197 | 1 | REACTOME_METABOLISM_OF_VITAMINS_AND_COFAC | -1.2671 | 0.060703 | 0.523247 | REACTOME_METABOLISM | -1.25818 | 0.078343 | 0.272231 |
| REACTOME_AMINE_LIG | -1.24806 | 0.194986 | 0.885711 | REACTOME_AMINE_LIGAND_BINDING_RECEPTORS | 1.558486 | 0.036729 | 0.525804 | REACTOME_AMINE_LIGAI | 1.248901 | 0.202562 | 0.452755 |
| REACTOME_FORMATIOI | 2.717268 | 0 | 0 | REACTOME_FORMATION_OF_A_POOL_OF_FREE_40S_ | 1.624374 | 0.006044 | 0.529141 | REACTOME_FORMATION_ | 2.660923 | 0 | 0 |
| REACTOME_TRANSPORT | -0.78373 | 0.842022 | 1 | REACTOME_TRANSPORT_OF_INORGANIC_CATIONS_AI | -1.25911 | 0.10378 | 0.532521 | REACTOME_TRANSPORT_ | -1.25679 | 0.106025 | 0.273053 |
| REACTOME_ASSEMBLY_ | 0.858363 | 0.644237 | 1 | REACTOME_ASSEMBLY_OF_COLLAGEN_FIBRILS_AND_ | 1.549106 | 0.040072 | 0.533267 | REACTOME_ASSEMBLY_O | 1.307489 | 0.153618 | 0.388813 |
| REACTOME_DEACTIVATI | -0.86407 | 0.64825 | 1 | REACTOME_DEACTIVATION_OF_THE_BETA_CATENIN_ | 1.649411 | 0.020421 | 0.535442 | REACTOME_DEACTIVATIO | 1.159415 | 0.280803 | 0.576883 |
| WP_GLUTATHIONE_MET | 0.6945 | 0.859495 | 1 | WP_GLUTATHIONE_METABOLISM | -1.25533 | 0.18329 | 0.536522 | WP_GLUTATHIONE_META | -1.02817 | 0.422136 | 0.546892 |
| REACTOME_EUKARYOTI | 2.592438 | 0 | 0 | REACTOME_EUKARYOTIC_TRANSLATION_INITIATION | 1.661947 | 0.00303 | 0.548489 | REACTOME_EUKARYOTIC_ | 2.485513 | 0 | 0 |
| REACTOME_MAPK_TAR | -1.35501 | 0.108249 | 0.788882 | REACTOME_MAPK_TARGETS_NUCLEAR_EVENTS_MED | 1.559376 | 0.031711 | 0.554826 | REACTOME_MAPK_TARGI | 0.678135 | 0.888213 | 0.97727 |
| REACTOME_ACTIVATION | 2.220735 | 0 | 5.97E-04 | REACTOME_ACTIVATION_OF_THE_MRNA_UPON_BIND | 1.536282 | 0.022018 | 0.555484 | REACTOME_ACTIVATION_ | 2.117987 | 1.46E-04 | 3.20E-04 |
| REACTOME_COLLAGEN_ | 0.756433 | 0.785606 | 1 | REACTOME_COLLAGEN_DEGRADATION | 1.575613 | 0.031655 | 0.558842 | REACTOME_COLLAGEN_D | 1.304602 | 0.159098 | 0.389919 |
| REACTOME_METABOLIS | 0.763617 | 0.963665 | 1 | REACTOME_METABOLISM_OF_CARBOHYDRATES | -1.23789 | 0.058212 | 0.559498 | REACTOME_METABOLISM | 0.762572 | 0.927348 | 1 |
| REACTOME_TNFR1_IND | -0.52572 | 0.975877 | 1 | REACTOME_TNFR1_INDUCED_PROAPOPTOTIC_SIGNAL | 1.456118 | 0.072709 | 0.559755 | REACTOME_TNFR1_INDU | 1.391445 | 0.101118 | 0.290725 |
| REACTOME_NONSENSE_ | 2.607835 | 0 | 0 | REACTOME_NONSENSE_MEDIATED_DECAY_NMD_IND | 1.464736 | 0.029525 | 0.562454 | REACTOME_NONSENSE_N | 2.493042 | 0 | 0 |
| REACTOME_RHO_GTPA | -1.55719 | 0.037736 | 0.599589 | REACTOME_RHO_GTPASES_ACTIVATE_NADPH_OXIDAS | -1.24281 | 0.195251 | 0.563962 | REACTOME_RHO_GTPASE | -1.20783 | 0.222599 | 0.315372 |
| REACTOME_RECYCLING_ | 1.015377 | 0.42977 | 1 | REACTOME_RECYCLING_PATHWAY_OF_L1 | -1.24042 | 0.164288 | 0.564459 | REACTOME_RECYCLING_F | -1.92826 | 8.64E-04 | 0.019115 |
| REACTOME_SIGNALING_ | -0.83639 | 0.725706 | 1 | REACTOME_SIGNALING_BY_FGFR | -1.2382 | 0.149443 | 0.564479 | REACTOME_SIGNALING_E | -0.92725 | 0.575092 | 0.695844 |
| REACTOME_RNA_POLYM | -0.81344 | 0.735145 | 1 | REACTOME_RNA_POLYMERASE_I_TRANSCRIPTION | 1.458889 | 0.060724 | 0.566765 | REACTOME_RNA_POLYMI | 1.618266 | 0.014954 | 0.101661 |
| REACTOME_ACETYLCHO | 0.663372 | 0.897342 | 1 | REACTOME_ACETYLCHOLINE_NEUROTRANSMITTER_RI | -1.22651 | 0.208551 | 0.567138 | REACTOME_ACETYLCHOLI | -1.58235 | 0.031652 | 0.085769 |
| REACTOME_SIGNALING_ | 0.898842 | 0.592915 | 1 | REACTOME_SIGNALING_BY_FGFR3 | -1.22792 | 0.203998 | 0.569021 | REACTOME_SIGNALING_E | -1.44425 | 0.063102 | 0.134608 |
| WP_AMINO_ACID_MET | -0.40054 | 1 | 1 | WP_AMINO_ACID_METABOLISM | -1.22963 | 0.126546 | 0.570203 | WP_AMINO_ACID_METAI | -1.96393 | 0 | 0.016956 |
| REACTOME_PLATELET_ | -0.70644 | 0.840297 | 1 | REACTOME_PLATELET_CALCIUM_HOMEOSTASIS | -1.23034 | 0.213622 | 0.574029 | REACTOME_PLATELET_CA | -1.52517 | 0.052785 | 0.107717 |
| WP_ESTROGEN_SIGNAL | -1.14457 | 0.249117 | 0.951197 | WP_ESTROGEN_SIGNALING | 1.447393 | 0.043214 | 0.575431 | WP_ESTROGEN_SIGNALIN | 1.366449 | 0.070621 | 0.318426 |
| WP_EUKARYOTIC_TRAN | -1.1281 | 0.291568 | 0.979592 | WP_EUKARYOTIC_TRANSCRIPTION_INITIATION | 1.626507 | 0.017557 | 0.577361 | WP_EUKARYOTIC_TRANS | 1.669328 | 0.011422 | 0.089052 |
| WP_MECHANISMS_ASS | -1.4188 | 0.018965 | 0.675419 | WP_MECHANISMS_ASSOCIATED_WITH_PLURIPOTENC | 1.464872 | 0.012733 | 0.580694 | WP_MECHANISMS_ASSO | 1.400718 | 0.021295 | 0.283338 |
| REACTOME_SIGNALING_ | -0.65151 | 0.961634 | 1 | REACTOME_SIGNALING_BY_TGFB_FAMILY_MEMBERS | 1.469896 | 0.039613 | 0.580915 | REACTOME_SIGNALING_E | 1.23897 | 0.165415 | 0.460291 |
| REACTOME_RETROGRAI | 0.594806 | 0.983243 | 1 | REACTOME_RETROGRADE_TRANSPORT_AT_THE_TRAN | 1.44005 | 0.051439 | 0.586182 | REACTOME_RETROGRADE | 0.996021 | 0.473369 | 0.855312 |
| REACTOME_INTEGRIN_ | 0.963672 | 0.49834 | 1 | REACTOME_INTEGRIN_CELL_SURFACE_INTERACTIONS | 1.56013 | 0.021998 | 0.588323 | REACTOME_INTEGRIN_CE | 0.873282 | 0.662777 | 0.978004 |
| REACTOME_CLASS_A_1_ | -1.30575 | 0.091192 | 0.862622 | REACTOME_CLASS_A_1_RHODOPSIN_LIKE_RECEPTORS | 1.472974 | 0.027899 | 0.589289 | REACTOME_CLASS_A_1_F | 1.568388 | 0.011886 | 0.124447 |
| REACTOME_TELOMERE_ | -1.26598 | 0.170966 | 0.878949 | REACTOME_TELOMERE_MAINTENANCE | -1.21455 | 0.201513 | 0.593879 | REACTOME_TELOMERE_N | 0.658098 | 0.908698 | 0.980504 |
| REACTOME_RHOB_GTPA | -1.59937 | 0.011408 | 0.550087 | REACTOME_RHOB_GTPASE_CYCLE | 1.429262 | 0.048667 | 0.594711 | REACTOME_RHOB_GTPAS | -0.98154 | 0.494015 | 0.61235 |
| REACTOME_NUCLEOTID | 1.259301 | 0.185543 | 0.738547 | REACTOME_NUCLEOTIDE_SALVAGE | -1.21101 | 0.228084 | 0.597966 | REACTOME_NUCLEOTIDE_ | -0.85664 | 0.651801 | 0.796233 |
| REACTOME_HDACS_DE | -0.98263 | 0.475168 | 1 | REACTOME_HDACS_DEACETYLATE_HISTONES | 1.475355 | 0.060591 | 0.601341 | REACTOME_HDACS_DEAC | 1.771666 | 0.003632 | 0.047288 |
| WP_OXIDATIVE_DAMAG | 0.867688 | 0.644329 | 1 | WP_OXIDATIVE_DAMAGE_RESPONSE | 1.392193 | 0.097938 | 0.602143 | WP_OXIDATIVE_DAMAGE | 0.733576 | 0.821331 | 1 |
| REACTOME_GLOBAL_GE | -1.48041 | 0.037323 | 0.630451 | REACTOME_GLOBAL_GENOME_NUCLEOTIDE_EXCISION | 1.430679 | 0.052094 | 0.605666 | REACTOME_GLOBAL_GEN | 0.850538 | 0.699299 | 0.973715 |
| REACTOME_SIGNALING_ | -0.65021 | 0.952849 | 1 | REACTOME_SIGNALING_BY_TGF_BETA_RECEPTOR_CO | 1.48473 | 0.039466 | 0.608491 | REACTOME_SIGNALING_E | 1.162123 | 0.247765 | 0.575401 |
| REACTOME_IMMUNORE | -0.5732 | 0.96114 | 1 | REACTOME_IMMUNOREGULATORY_INTERACTIONS_BI | 1.516324 | 0.045123 | 0.610006 | REACTOME_IMMUNOREG | 0.913392 | 0.58363 | 0.968529 |
| WP_DELTANOTCH_SIGN | -0.83068 | 0.738418 | 1 | WP_DELTANOTCH_SIGNALING_PATHWAY | 1.496186 | 0.034858 | 0.611236 | WP_DELTANOTCH_SIGNA | 1.406949 | 0.066346 | 0.287244 |
| WP_MICROGLIA_PATHO | -0.79089 | 0.765484 | 1 | WP_MICROGLIA_PATHOGEN_PHAGOCYTOSIS_PATHW | 1.39253 | 0.084066 | 0.614316 | WP_MICROGLIA_PATHOG | -0.83681 | 0.710814 | 0.823806 |
| BIOCARTA_CXCR4_PATH | 1.130935 | 0.295092 | 0.867815 | BIOCARTA_CXCR4_PATHWAY | -1.20164 | 0.223814 | 0.618623 | BIOCARTA_CXCR4_PATHV | -1.67195 | 0.0182 | 0.059876 |
| REACTOME_DEGRADATI | -0.73298 | 0.878729 | 1 | REACTOME_DEGRADATION_OF_THE_EXTRACELLULAR_ | 1.779553 | 0.003147 | 0.619553 | REACTOME_DEGRADATIO | 1.18264 | 0.228783 | 0.534175 |
| WP_CYTOPLASMIC_RIB | 2.706107 | 0 | 0 | WP_CYTOPLASMIC_RIBOSOMAL_PROTEINS | 1.413896 | 0.047885 | 0.621841 | WP_CYTOPLASMIC_RIBOS | 2.519593 | 0 | 0 |
| BIOCARTA_P38MAPK_P | -0.71446 | 0.843195 | 1 | BIOCARTA_P38MAPK_PATHWAY | 1.47592 | 0.063116 | 0.622095 | BIOCARTA_P38MAPK_PAT | 1.301463 | 0.15367 | 0.386573 |
| REACTOME_RHOQ_GTP | -0.3845 | 0.999823 | 0.999653 | REACTOME_RHOQ_GTPASE_CYCLE | 1.382841 | 0.077248 | 0.623178 | REACTOME_RHOQ_GTPAS | -1.21714 | 0.153465 | 0.306329 |
| WP_TRANSLATION_FAC | 0.923761 | 0.460915 | 1 | WP_TRANSLATION_FACTORS | 1.393487 | 0.073051 | 0.624655 | WP_TRANSLATION_FACT | 0.822741 | 0.746673 | 0.973545 |
| REACTOME_NON_INTEG | 0.827961 | 0.688706 | 1 | REACTOME_NON_INTEGRIN_MEMBRANE_ECM_INTER | 1.486411 | 0.065819 | 0.626536 | REACTOME_NON_INTEGR | 1.18213 | 0.256905 | 0.530273 |
| REACTOME_NONHOMO | -1.50837 | 0.046736 | 0.664832 | REACTOME_NONHOMOLOGOUS_END_JOINING_NHEJ | 1.40405 | 0.089874 | 0.628489 | REACTOME_NONHOMOL | -0.86212 | 0.65609 | 0.791308 |
| WP_TGFBETA_SIGNALIN | -1.42676 | 0.082725 | 0.685194 | WP_TGFBETA_SIGNALING_PATHWAY | 1.415986 | 0.08804 | 0.629911 | WP_TGFBETA_SIGNALING | 1.118781 | 0.314851 | 0.651094 |
| REACTOME_RNA_POLYM | -1.15378 | 0.237844 | 0.933654 | REACTOME_RNA_POLYMERASE_II_PRE_TRANSCRIPTIO | 1.399688 | 0.060562 | 0.630377 | REACTOME_RNA_POLYMI | 1.599164 | 0.010983 | 0.109632 |
| REACTOME_RHOD_GTP | -0.72157 | 0.895549 | 1 | REACTOME_RHOD_GTPASE_CYCLE | 1.406533 | 0.06212 | 0.634531 | REACTOME_RHOD_GTPAS | 0.846915 | 0.70025 | 0.965327 |
| REACTOME_RNA_POLYM | -1.03123 | 0.410886 | 1 | REACTOME_RNA_POLYMERASE_II_PROMOTER_ESCAP | 1.394272 | 0.085798 | 0.636186 | REACTOME_RNA_POLYMI | 1.537369 | 0.03008 | 0.15573 |
| REACTOME_ECM_PROTI | 1.475272 | 0.064367 | 0.504541 | REACTOME_ECM_PROTEOGLYCANS | 1.496754 | 0.050542 | 0.636274 | REACTOME_ECM_PROTE | 1.302503 | 0.143062 | 0.389396 |
| WP_MITOCHONDRIAL_L | 1.193304 | 0.238095 | 0.810659 | WP_MITOCHONDRIAL_LONG_CHAIN_FATTY_ACID_BET | -1.19264 | 0.248452 | 0.638812 | WP_MITOCHONDRIAL_LO | -1.35018 | 0.1301 | 0.200672 |
| REACTOME_SIGNALING_ | -0.70584 | 0.951475 | 1 | REACTOME_SIGNALING_BY_THE_B_CELL_RECEPTOR_E | -1.18811 | 0.157266 | 0.640054 | REACTOME_SIGNALING_E | -1.88875 | 0 | 0.024266 |
| REACTOME_RMTS_MET | -1.11173 | 0.309839 | 0.998103 | REACTOME_RMTS_METHYLATE_HISTONE_ARGININES | -1.18966 | 0.218197 | 0.641425 | REACTOME_RMTS_METH | -1.24532 | 0.162584 | 0.279775 |
| REACTOME_FORMATIOI | -1.20021 | 0.205893 | 0.847097 | REACTOME_FORMATION_OF_TC_NER_PRE_INCISION_ | 1.373642 | 0.084718 | 0.645232 | REACTOME_FORMATION_ | 1.391276 | 0.073064 | 0.286274 |
| BIOCARTA_TPO_PATHW | 0.961394 | 0.497722 | 1 | BIOCARTA_TPO_PATHWAY | -1.17977 | 0.242008 | 0.646923 | BIOCARTA_TPO_PATHWA | -1.4153 | 0.089364 | 0.151383 |
| REACTOME_RND1_GTP | 0.735338 | 0.847807 | 1 | REACTOME_RND1_GTPASE_CYCLE | -1.17497 | 0.223703 | 0.649171 | REACTOME_RND1_GTPAS | -1.3522 | 0.094211 | 0.199466 |
| REACTOME_RHOF_GTP | -1.04285 | 0.383687 | 1 | REACTOME_RHOF_GTPASE_CYCLE | 1.500469 | 0.040012 | 0.649732 | REACTOME_RHOF_GTPAS | 0.657283 | 0.929243 | 0.976916 |
| REACTOME_PLASMA_LI | 1.000952 | 0.451613 | 1 | REACTOME_PLASMA_LIPOPROTEIN_CLEARANCE | -1.18061 | 0.244536 | 0.650306 | REACTOME_PLASMA_LIP | -1.51523 | 0.048896 | 0.110831 |
| REACTOME_EXTRA_NU | 0.995191 | 0.445339 | 1 | REACTOME_EXTRA_NUCLEAR_ESTROGEN_SIGNALING | -1.17644 | 0.201662 | 0.650716 | REACTOME_EXTRA_NUCL | -1.45916 | 0.038392 | 0.127995 |
| REACTOME_RHOBTB_G | -0.62619 | 0.957052 | 1 | REACTOME_RHOBTB_GTPASE_CYCLE | -1.18143 | 0.210917 | 0.653915 | REACTOME_RHOBTB_GTF | -0.88682 | 0.644955 | 0.75679 |
| BIOCARTA_EGF_PATHW | 0.675787 | 0.891525 | 1 | BIOCARTA_EGF_PATHWAY | -1.17096 | 0.243366 | 0.655279 | BIOCARTA_EGF_PATHWA | -1.76427 | 0.006023 | 0.03778 |
| REACTOME_ENERGY_DE | -0.71881 | 0.867797 | 1 | REACTOME_ENERGY_DEPENDENT_REGULATION_OF_N | -1.16866 | 0.23247 | 0.656168 | REACTOME_ENERGY_DEP | -1.62461 | 0.014722 | 0.072744 |
| REACTOME_TRANSMISS | 1.117609 | 0.198969 | 0.884736 | REACTOME_TRANSMISSION_ACROSS_CHEMICAL_SYN | -1.15448 | 0.126837 | 0.66917 | REACTOME_TRANSMISSIO | -1.60898 | 4.90E-04 | 0.075578 |
| REACTOME_NOREPINEP | 0.866862 | 0.6341 | 1 | REACTOME_NOREPINEPHRINE_NEUROTRANSMITTER_ | -1.15981 | 0.26527 | 0.670475 | REACTOME_NOREPINEPH | -1.64187 | 0.024609 | 0.065875 |
| REACTOME_G_ALPHA_ | -1.21571 | 0.146835 | 0.871071 | REACTOME_G_ALPHA_Q_SIGNALLING_EVENTS | -1.15708 | 0.181676 | 0.672909 | REACTOME_G_ALPHA_Q_ | -1.32003 | 0.051215 | 0.223906 |
| REACTOME_SIGNALING_ | -0.86777 | 0.666788 | 1 | REACTOME_SIGNALING_BY_EGFR | -1.15469 | 0.232464 | 0.674192 | REACTOME_SIGNALING_E | -1.57523 | 0.016974 | 0.087231 |
| REACTOME_SIGNALING_ | 1.138549 | 0.25046 | 0.880551 | REACTOME_SIGNALING_BY_NTRK1_TRKA | -1.16028 | 0.217915 | 0.674973 | REACTOME_SIGNALING_E | -1.3975 | 0.047975 | 0.162816 |
| BIOCARTA_CASPASE_PA | 0.866829 | 0.632337 | 1 | BIOCARTA_CASPASE_PATHWAY | 1.358108 | 0.125061 | 0.677769 | BIOCARTA_CASPASE_PATI | 0.812076 | 0.708037 | 0.989287 |
| REACTOME_GLYCOSAM | 0.879811 | 0.659954 | 1 | REACTOME_GLYCOSAMINOGLYCAN_METABOLISM | 1.305819 | 0.121001 | 0.678869 | REACTOME_GLYCOSAMIN | 1.047486 | 0.387597 | 0.78432 |
| REACTOME_RHOH_GTP | -0.55378 | 0.985421 | 1 | REACTOME_RHOH_GTPASE_CYCLE | 1.350554 | 0.111698 | 0.681034 | REACTOME_RHOH_GTPAS | -1.28657 | 0.134874 | 0.250345 |
| REACTOME_NUCLEOTID | -1.54895 | 0.012276 | 0.557529 | REACTOME_NUCLEOTIDE_EXCISION_REPAIR | 1.288297 | 0.11532 | 0.684251 | REACTOME_NUCLEOTIDE_ | 1.193153 | 0.191802 | 0.518705 |
| REACTOME_CITRIC_ACI | 1.825698 | 0.005648 | 0.097376 | REACTOME_CITRIC_ACID_CYCLE_TCA_CYCLE | 1.359805 | 0.114091 | 0.684786 | REACTOME_CITRIC_ACID_ | -1.23579 | 0.181936 | 0.289785 |
| REACTOME_AURKA_ACT | -0.54472 | 0.989218 | 1 | REACTOME_AURKA_ACTIVATION_BY_TPX2 | 1.306844 | 0.132241 | 0.685349 | REACTOME_AURKA_ACTIV | 0.789073 | 0.780203 | 0.99584 |
| BIOCARTA_MAPK_PATH | -1.32294 | 0.096757 | 0.838056 | BIOCARTA_MAPK_PATHWAY | 1.291595 | 0.12073 | 0.691186 | BIOCARTA_MAPK_PATHW | 0.774956 | 0.828951 | 1 |
| REACTOME_HEPARAN_ | 1.383258 | 0.092039 | 0.641652 | REACTOME_HEPARAN_SULFATE_HEPARIN_HS_GAG_M | 1.293759 | 0.153057 | 0.693007 | REACTOME_HEPARAN_SU | 0.967292 | 0.502973 | 0.885382 |
| REACTOME_ROLE_OF_P | 0.668672 | 0.890318 | 1 | REACTOME_ROLE_OF_PHOSPHOLIPIDS_IN_PHAGOCYT | 1.350887 | 0.129425 | 0.693052 | REACTOME_ROLE_OF_PH | -1.70053 | 0.016262 | 0.052956 |
| REACTOME_REGULATIO | 0.898438 | 0.612918 | 1 | REACTOME_REGULATION_OF_TNFR1_SIGNALING | 1.288485 | 0.149485 | 0.69308 | REACTOME_REGULATION | 1.025903 | 0.422741 | 0.814471 |
| REACTOME_CHONDROI | -1.1201 | 0.308153 | 0.992773 | REACTOME_CHONDROITIN_SULFATE_DERMATAN_SUL | 1.307568 | 0.142568 | 0.693124 | REACTOME_CHONDROITI | 0.890995 | 0.620356 | 0.970327 |
| REACTOME_ESTROGEN_ | -1.01953 | 0.425474 | 1 | REACTOME_ESTROGEN_DEPENDENT_GENE_EXPRESSI | 1.321618 | 0.120502 | 0.695359 | REACTOME_ESTROGEN_D | 1.40435 | 0.071125 | 0.281732 |
| REACTOME_GPCR_LIGA | -1.36378 | 0.045586 | 0.769599 | REACTOME_GPCR_LIGAND_BINDING | 1.340256 | 0.065289 | 0.695684 | REACTOME_GPCR_LIGAN | 1.274464 | 0.098847 | 0.420404 |
| REACTOME_SIGNALING_ | 0.928183 | 0.554164 | 1 | REACTOME_SIGNALING_BY_FGFR4 | -1.14074 | 0.273397 | 0.698983 | REACTOME_SIGNALING_E | -1.40508 | 0.085684 | 0.157424 |
| WP_FATTY_ACID_BIOSY | -0.92022 | 0.566851 | 1 | WP_FATTY_ACID_BIOSYNTHESIS | -1.13686 | 0.281675 | 0.69924 | WP_FATTY_ACID_BIOSYN | -1.43213 | 0.075586 | 0.143079 |
| REACTOME_TRANSCRIP | 0.575216 | 0.964064 | 1 | REACTOME_TRANSCRIPTIONAL_ACTIVITY_OF_SMAD2_ | 1.34257 | 0.127643 | 0.699446 | REACTOME_TRANSCRIPTI | 1.033354 | 0.42352 | 0.805767 |
| REACTOME_SYNTHESIS_ | -0.96435 | 0.503517 | 1 | REACTOME_SYNTHESIS_OF_IP3_AND_IP4_IN_THE_CYT | -1.13119 | 0.291014 | 0.69954 | REACTOME_SYNTHESIS_C | -1.66164 | 0.016175 | 0.060569 |
| REACTOME_METABOLIS | 1.294835 | 0.100966 | 0.757247 | REACTOME_METABOLISM_OF_STEROIDS | 1.278848 | 0.121951 | 0.699992 | REACTOME_METABOLISM | 1.330508 | 0.086772 | 0.355311 |
| REACTOME_RAC3_GTPA | -0.89165 | 0.65552 | 1 | REACTOME_RAC3_GTPASE_CYCLE | 1.32345 | 0.092013 | 0.70011 | REACTOME_RAC3_GTPAS | 0.684817 | 0.946111 | 0.975687 |
| REACTOME_NOD1_2_SI | -0.75265 | 0.806167 | 1 | REACTOME_NOD1_2_SIGNALING_PATHWAY | 1.317341 | 0.143697 | 0.700244 | REACTOME_NOD1_2_SIG | 0.704179 | 0.856612 | 0.996121 |
| REACTOME_CLEC7A_DE | 0.683829 | 0.972619 | 1 | REACTOME_CLEC7A_DECTIN_1_SIGNALING | -1.14207 | 0.213145 | 0.700666 | REACTOME_CLEC7A_DEC | -1.87473 | 0 | 0.025437 |

| | | | | | | | | | | | |
|---|---|---|---|---|---|---|---|---|---|---|---|
| REACTOME_RHOC_GTPA | -1.31868 | 0.08988 | 0.839222 | REACTOME_RHOC_GTPASE_CYCLE | 1.314129 | 0.109676 | 0.701156 | REACTOME_RHOC_GTPAS | -0.91386 | 0.622715 | 0.712327 |
| REACTOME_TP53_REGU | -1.201 | 0.218841 | 0.854245 | REACTOME_TP53_REGULATES_TRANSCRIPTION_OF_D | 1.308058 | 0.132569 | 0.702048 | REACTOME_TP53_REGUL | 1.502405 | 0.034179 | 0.1895 |
| REACTOME_INTERLEUKI | -0.49843 | 0.983117 | 1 | REACTOME_INTERLEUKIN_2_FAMILY_SIGNALING | 1.293779 | 0.160116 | 0.702976 | REACTOME_INTERLEUKIN | -0.795 | 0.74466 | 0.870649 |
| REACTOME_ARACHIDON | 1.216713 | 0.206866 | 0.791341 | REACTOME_ARACHIDONIC_ACID_METABOLISM | -1.13744 | 0.279596 | 0.703117 | REACTOME_ARACHIDONI | -0.99509 | 0.455755 | 0.587379 |
| REACTOME_METABOLIS | 0.770193 | 0.877809 | 1 | REACTOME_METABOLISM_OF_NUCLEOTIDES | -1.13182 | 0.23513 | 0.703214 | REACTOME_METABOLISM | -1.203 | 0.151218 | 0.318984 |
| REACTOME_OPIOID_SIG | 1.034079 | 0.390542 | 1 | REACTOME_OPIOID_SIGNALLING | -1.13311 | 0.227593 | 0.704938 | REACTOME_OPIOID_SIGN | -1.85573 | 3.22E-04 | 0.028146 |
| REACTOME_DUAL_INCIS | -1.4727 | 0.041531 | 0.610841 | REACTOME_DUAL_INCISION_IN_TC_NER | 1.279496 | 0.145144 | 0.707068 | REACTOME_DUAL_INCISI | 1.448726 | 0.054393 | 0.254295 |
| REACTOME_FOXO_MED | -0.81457 | 0.713511 | 1 | REACTOME_FOXO_MEDIATED_TRANSCRIPTION | 1.295296 | 0.165444 | 0.707705 | REACTOME_FOXO_MEDIA | 1.594428 | 0.029258 | 0.111073 |
| REACTOME_NUCLEAR_E | 0.736708 | 0.828166 | 1 | REACTOME_NUCLEAR_EVENTS_KINASE_AND_TRANSCI | 1.324481 | 0.132467 | 0.708075 | REACTOME_NUCLEAR_EV | 0.866931 | 0.647802 | 0.958782 |
| BIOCARTA_VDR_PATHW | -0.84464 | 0.666237 | 1 | BIOCARTA_VDR_PATHWAY | 1.308747 | 0.154466 | 0.710594 | BIOCARTA_VDR_PATHWA | 1.711797 | 0.008622 | 0.07555 |
| WP_MRNA_PROCESSING | 0.82932 | 0.957885 | 1 | WP_MRNA_PROCESSING | -1.12514 | 0.113171 | 0.712458 | WP_MRNA_PROCESSING | 1.680201 | 0 | 0.08391 |
| REACTOME_DNA_DAMA | -1.30419 | 0.144744 | 0.835304 | REACTOME_DNA_DAMAGE_RECOGNITION_IN_GG_NE | 1.32531 | 0.132958 | 0.717136 | REACTOME_DNA_DAMAG | -1.01336 | 0.427109 | 0.565672 |
| REACTOME_LYSOSOME_ | 0.826935 | 0.719828 | 1 | REACTOME_LYSOSOME_VESICLE_BIOGENESIS | 1.270604 | 0.164483 | 0.721085 | REACTOME_LYSOSOME_V | 1.134293 | 0.287127 | 0.620745 |
| REACTOME_FORMATIOI | -1.24334 | 0.163133 | 0.865481 | REACTOME_FORMATION_OF_RNA_POL_II_ELONGATIC | 1.327467 | 0.11237 | 0.721266 | REACTOME_FORMATION_ | 1.574085 | 0.015726 | 0.121738 |
| REACTOME_TRANSPORT | -1.33994 | 0.09823 | 0.79856 | REACTOME_TRANSPORT_OF_BILE_SALTS_AND_ORGAN | 1.3299 | 0.110153 | 0.724232 | REACTOME_TRANSPORT_ | 1.15353 | 0.265132 | 0.58673 |
| WP_DYSREGULATED_MI | 0.677708 | 0.890776 | 1 | WP_DYSREGULATED_MIRNA_TARGETING_IN_INSULINI | 1.243833 | 0.193048 | 0.72918 | WP_DYSREGULATED_MIR | 0.727278 | 0.835811 | 1 |
| REACTOME_GOLGI_ASSO | 0.811301 | 0.781612 | 1 | REACTOME_GOLGI_ASSOCIATED_VESICLE_BIOGENESIS | 1.265477 | 0.147692 | 0.730525 | REACTOME_GOLGI_ASSO | 1.018951 | 0.430666 | 0.821702 |
| REACTOME_PEPTIDE_LI | -1.09582 | 0.320467 | 1 | REACTOME_PEPTIDE_LIGAND_BINDING_RECEPTORS | 1.253837 | 0.166891 | 0.736519 | REACTOME_PEPTIDE_LIG | 1.61435 | 0.011807 | 0.101943 |
| WP_ENDOCHONDRAL_C | 0.72484 | 0.850022 | 1 | WP_ENDOCHONDRAL_OSSIFICATION | 1.244075 | 0.194946 | 0.736862 | WP_ENDOCHONDRAL_OS | 1.093273 | 0.347317 | 0.704202 |
| REACTOME_INTRAFLAGI | -1.19697 | 0.20584 | 0.849784 | REACTOME_INTRAFLAGELLAR_TRANSPORT | 1.250953 | 0.161571 | 0.738147 | REACTOME_INTRAFLAGEL | -0.99466 | 0.458022 | 0.586655 |
| BIOCARTA_MITOCHOND | 1.096704 | 0.332335 | 0.907708 | BIOCARTA_MITOCHONDRIA_PATHWAY | 1.255584 | 0.195298 | 0.739194 | BIOCARTA_MITOCHONDR | 0.69864 | 0.851729 | 0.979698 |
| WP_GLYCEROLIPIDS_AN | -0.80963 | 0.727122 | 1 | WP_GLYCEROLIPIDS_AND_GLYCEROPHOSPHOLIPIDS | 1.245397 | 0.209745 | 0.740813 | WP_GLYCEROLIPIDS_AND | 1.347362 | 0.124533 | 0.331426 |
| REACTOME_ANTIGEN_P | 0.807731 | 0.811238 | 1 | REACTOME_ANTIGEN_PROCESSING_CROSS_PRESENTA | 1.235762 | 0.160517 | 0.741648 | REACTOME_ANTIGEN_PR | -1.12676 | 0.241595 | 0.408708 |
| REACTOME_SIGNALING_ | 0.814975 | 0.754679 | 1 | REACTOME_SIGNALING_BY_PDGF | 1.237454 | 0.185764 | 0.743924 | REACTOME_SIGNALING_E | -0.752 | 0.863218 | 0.910455 |
| REACTOME_DUAL_INCIS | -1.3754 | 0.106163 | 0.801903 | REACTOME_DUAL_INCISION_IN_GG_NER | 1.230158 | 0.2092 | 0.745085 | REACTOME_DUAL_INCISI | 0.75723 | 0.79621 | 1 |
| REACTOME_TRANS_GOI | 0.652827 | 0.973516 | 1 | REACTOME_TRANS_GOLGI_NETWORK_VESICLE_BUDD | 1.255839 | 0.144387 | 0.747579 | REACTOME_TRANS_GOLG | 1.019604 | 0.432962 | 0.826282 |
| REACTOME_CHROMATII | -1.67487 | 0.002785 | 0.454887 | REACTOME_CHROMATIN_MODIFYING_ENZYMES | 1.23177 | 0.148374 | 0.74764 | REACTOME_CHROMATIN_ | 1.395877 | 0.044338 | 0.287397 |
| BIOCARTA_CHEMICAL_F | 0.763778 | 0.777416 | 1 | BIOCARTA_CHEMICAL_PATHWAY | 1.258325 | 0.194987 | 0.747776 | BIOCARTA_CHEMICAL_PA | 0.430667 | 0.995346 | 0.998589 |
| WP_ADIPOGENESIS_GEI | 0.946319 | 0.536785 | 1 | WP_ADIPOGENESIS_GENES | 1.245434 | 0.170289 | 0.749611 | WP_ADIPOGENESIS_GENE | 0.861291 | 0.686465 | 0.966167 |
| WP_FOLIC_ACID_NETW | 1.150257 | 0.282189 | 0.870944 | WP_FOLIC_ACID_NETWORK | 1.226514 | 0.221059 | 0.750003 | WP_FOLIC_ACID_NETWOI | 0.957325 | 0.518562 | 0.906577 |
| WP_IL6_SIGNALING_PA | 0.702271 | 0.953207 | 1 | WP_IL6_SIGNALING_PATHWAY | 1.222963 | 0.163536 | 0.754667 | WP_IL6_SIGNALING_PATH | 0.853302 | 0.720924 | 0.978591 |
| REACTOME_TRANSCRIP | -1.47623 | 0.03355 | 0.621601 | REACTOME_TRANSCRIPTION_COUPLED_NUCLEOTIDE_ | 1.215157 | 0.189972 | 0.758315 | REACTOME_TRANSCRIPTI | 1.312445 | 0.107075 | 0.388563 |
| WP_FAS_PATHWAY_AN | -1.20623 | 0.220293 | 0.843729 | WP_FAS_PATHWAY_AND_STRESS_INDUCTION_OF_HS | 1.217329 | 0.209855 | 0.758537 | WP_FAS_PATHWAY_AND_ | -0.81567 | 0.744474 | 0.843182 |
| WP_P53_SIGNALING | -1.37222 | 0.108239 | 0.772702 | WP_P53_SIGNALING | 1.193421 | 0.24443 | 0.759826 | WP_P53_SIGNALING | -1.01891 | 0.420078 | 0.560298 |
| REACTOME_NONSENSE_ | 2.432419 | 0 | 0 | REACTOME_NONSENSE_MEDIATED_DECAY_NMD | 1.195268 | 0.178053 | 0.760719 | REACTOME_NONSENSE_N | 2.327559 | 0 | 0 |
| WP_G13_SIGNALING_PA | -0.85336 | 0.687478 | 1 | WP_G13_SIGNALING_PATHWAY | -1.10779 | 0.302271 | 0.76145 | WP_G13_SIGNALING_PAT | -1.84014 | 0.001125 | 0.02947 |
| WP_GPCRS_CLASS_A_RI | -1.54648 | 0.027938 | 0.534789 | WP_GPCRS_CLASS_A_RHODOPSINLIKE | 1.218527 | 0.203839 | 0.762361 | WP_GPCRS_CLASS_A_RH | 0.977497 | 0.48912 | 0.875714 |
| REACTOME_CYCLIN_D_ | -0.86579 | 0.652726 | 1 | REACTOME_CYCLIN_D_ASSOCIATED_EVENTS_IN_G1 | 1.188528 | 0.252815 | 0.762441 | REACTOME_CYCLIN_D_AS | 0.906866 | 0.588568 | 0.972934 |
| WP_TYROBP_CAUSAL_N | -1.37648 | 0.096588 | 0.820426 | WP_TYROBP_CAUSAL_NETWORK_IN_MICROGLIA | 1.190473 | 0.247342 | 0.762841 | WP_TYROBP_CAUSAL_NE | -1.04308 | 0.387131 | 0.521741 |
| REACTOME_CARDIAC_C | 0.709138 | 0.936096 | 1 | REACTOME_CARDIAC_CONDUCTION | -1.1052 | 0.263049 | 0.763813 | REACTOME_CARDIAC_CO | -1.02388 | 0.404404 | 0.554417 |
| WP_SIDS_SUSCEPTIBILIT | 1.261048 | 0.180893 | 0.753036 | WP_SIDS_SUSCEPTIBILITY_PATHWAYS | 1.186024 | 0.247009 | 0.764039 | WP_SIDS_SUSCEPTIBILITY | 1.067704 | 0.376349 | 0.764877 |
| REACTOME_CELL_EXTRA | -0.73418 | 0.810338 | 1 | REACTOME_CELL_EXTRACELLULAR_MATRIX_INTERACT | 1.196347 | 0.2487 | 0.764309 | REACTOME_CELL_EXTRAC | 0.768177 | 0.770324 | 1 |
| WP_PRIMARY_FOCAL_S | 0.848592 | 0.715111 | 1 | WP_PRIMARY_FOCAL_SEGMENTAL_GLOMERULOSCLEI | 1.154383 | 0.249928 | 0.766109 | WP_PRIMARY_FOCAL_SE | 0.534681 | 0.995178 | 0.991884 |
| WP_WNT_SIGNALING_P | -0.9446 | 0.538077 | 1 | WP_WNT_SIGNALING_PATHWAY_AND_PLURIPOTENCY | 1.150345 | 0.259805 | 0.767245 | WP_WNT_SIGNALING_PA | 0.63417 | 0.960831 | 0.975588 |
| REACTOME_CYTOKINE_ | -1.00244 | 0.44 | 1 | REACTOME_CYTOKINE_SIGNALING_IN_IMMUNE_SYST | 1.155846 | 0.178559 | 0.767427 | REACTOME_CYTOKINE_SI | -1.01191 | 0.410011 | 0.56554 |
| REACTOME_MYOGENES | -0.5847 | 0.945503 | 1 | REACTOME_MYOGENESIS | 1.208175 | 0.241244 | 0.767904 | REACTOME_MYOGENESIS | -0.88552 | 0.605243 | 0.75758 |
| REACTOME_MRNA_CAP | -1.23696 | 0.200514 | 0.853917 | REACTOME_MRNA_CAPPING | 1.164885 | 0.271508 | 0.768891 | REACTOME_MRNA_CAPP | 1.534822 | 0.037376 | 0.154975 |
| REACTOME_DNA_DAMA | -1.19433 | 0.229171 | 0.850158 | REACTOME_DNA_DAMAGE_BYPASS | 1.151361 | 0.281304 | 0.769909 | REACTOME_DNA_DAMAG | -1.05077 | 0.377257 | 0.51345 |
| REACTOME_PROCESSIN | 1.148401 | 0.272016 | 0.869343 | REACTOME_PROCESSING_OF_INTRONLESS_PRE_MRN | 1.156757 | 0.276 | 0.7707 | REACTOME_PROCESSING_ | 1.856646 | 0.001303 | 0.02821 |
| BIOCARTA_AGR_PATHW | -1.00023 | 0.453521 | 1 | BIOCARTA_AGR_PATHWAY | 1.133007 | 0.299624 | 0.771005 | BIOCARTA_AGR_PATHWA | 0.847941 | 0.677755 | 0.973912 |
| REACTOME_ERK_MAPK_ | 0.91143 | 0.574919 | 1 | REACTOME_ERK_MAPK_TARGETS | 1.209394 | 0.232671 | 0.771396 | REACTOME_ERK_MAPK_T | 0.571222 | 0.95672 | 0.994917 |
| REACTOME_SYNTHESIS_ | 1.24009 | 0.1958 | 0.776343 | REACTOME_SYNTHESIS_OF_PC | 1.196391 | 0.236613 | 0.771645 | REACTOME_SYNTHESIS_C | 0.759649 | 0.790148 | 1 |
| REACTOME_TRANSLATI | 2.198588 | 0 | 7.58E-04 | REACTOME_TRANSLATION | 1.134119 | 0.211078 | 0.773134 | REACTOME_TRANSLATIOI | 1.811309 | 0 | 0.037521 |
| REACTOME_GAMMA_C | 0.997411 | 0.443518 | 1 | REACTOME_GAMMA_CARBOXYLATION_HYPUSINE_FO | 1.161425 | 0.274366 | 0.774163 | REACTOME_GAMMA_CAI | 0.897177 | 0.594929 | 0.979173 |
| REACTOME_CHOLESTER | 1.470458 | 0.064706 | 0.504456 | REACTOME_CHOLESTEROL_BIOSYNTHESIS | 1.202061 | 0.236074 | 0.774362 | REACTOME_CHOLESTERO | 1.797441 | 0.003685 | 0.040753 |
| REACTOME_MET_PROM | 0.552062 | 0.972187 | 1 | REACTOME_MET_PROMOTES_CELL_MOTILITY | 1.164901 | 0.269517 | 0.775396 | REACTOME_MET_PROMC | -0.82128 | 0.713875 | 0.843575 |
| BIOCARTA_DEATH_PATH | 0.996131 | 0.452142 | 1 | BIOCARTA_DEATH_PATHWAY | 1.166813 | 0.268749 | 0.775542 | BIOCARTA_DEATH_PATHV | -0.76263 | 0.803507 | 0.898723 |
| REACTOME_RAC2_GTPA | -1.06084 | 0.358859 | 1 | REACTOME_RAC2_GTPASE_CYCLE | 1.142292 | 0.253408 | 0.775703 | REACTOME_RAC2_GTPAS | 0.720551 | 0.917091 | 1 |
| REACTOME_AMINO_AC | -0.67795 | 0.894796 | 1 | REACTOME_AMINO_ACID_TRANSPORT_ACROSS_THE_ | 1.156877 | 0.280625 | 0.776682 | REACTOME_AMINO_ACID | 0.979931 | 0.486387 | 0.87531 |
| REACTOME_TGF_BETA_ | -0.66294 | 0.8995 | 1 | REACTOME_TGF_BETA_RECEPTOR_SIGNALING_ACTIV | 1.199251 | 0.239082 | 0.776706 | REACTOME_TGF_BETA_RI | 0.713344 | 0.838455 | 0.999582 |
| WP_FOCAL_ADHESION | -0.8904 | 0.68697 | 1 | WP_FOCAL_ADHESION | 1.134401 | 0.240139 | 0.778022 | WP_FOCAL_ADHESION | -1.16873 | 0.145599 | 0.356819 |
| REACTOME_DRUG_ADM | 1.015296 | 0.419837 | 1 | REACTOME_DRUG_ADME | 1.196619 | 0.209171 | 0.778433 | REACTOME_DRUG_ADME | -1.0166 | 0.412371 | 0.562015 |
| REACTOME_DARPP_32_ | 1.146204 | 0.282562 | 0.860536 | REACTOME_DARPP_32_EVENTS | 1.139622 | 0.302182 | 0.778565 | REACTOME_DARPP_32_E | -1.05316 | 0.385825 | 0.510005 |
| REACTOME_CELL_DEATI | -0.92322 | 0.579021 | 1 | REACTOME_CELL_DEATH_SIGNALLING_VIA_NRAGE_NI | 1.167757 | 0.231624 | 0.779039 | REACTOME_CELL_DEATH_ | -0.91758 | 0.599566 | 0.709675 |
| REACTOME_DNA_DOUB | -1.67355 | 0.008642 | 0.403302 | REACTOME_DNA_DOUBLE_STRAND_BREAK_REPAIR | 1.179483 | 0.22152 | 0.779692 | REACTOME_DNA_DOUBLI | -0.9283 | 0.583897 | 0.695333 |
| REACTOME_SIGNALING_ | -1.08059 | 0.29218 | 1 | REACTOME_SIGNALING_BY_INTERLEUKINS | 1.142834 | 0.213154 | 0.779957 | REACTOME_SIGNALING_E | -1.11487 | 0.195933 | 0.423386 |
| REACTOME_PROCESSIN | 1.313819 | 0.133703 | 0.713082 | REACTOME_PROCESSING_OF_CAPPED_INTRONLESS_P | 1.135508 | 0.293419 | 0.78026 | REACTOME_PROCESSING_ | 1.792162 | 0.002378 | 0.040062 |
| REACTOME_ANCHORIN | -0.91458 | 0.607029 | 1 | REACTOME_ANCHORING_OF_THE_BASAL_BODY_TO_T | 1.202485 | 0.200665 | 0.780654 | REACTOME_ANCHORING_ | 1.311633 | 0.106065 | 0.385027 |
| REACTOME_SEROTONIN | 0.607298 | 0.941034 | 1 | REACTOME_SEROTONIN_NEUROTRANSMITTER_RELEA | -1.09073 | 0.331738 | 0.78112 | REACTOME_SEROTONIN_ | -1.60638 | 0.027714 | 0.075622 |
| REACTOME_BILE_ACID_ | 1.363989 | 0.113932 | 0.666119 | REACTOME_BILE_ACID_AND_BILE_SALT_METABOLISM | 1.157196 | 0.279342 | 0.782049 | REACTOME_BILE_ACID_A | 0.879976 | 0.62914 | 0.967698 |
| WP_PPAR_SIGNALING_I | 1.687301 | 0.011696 | 0.16634 | WP_PPAR_SIGNALING_PATHWAY | 1.143962 | 0.268174 | 0.78224 | WP_PPAR_SIGNALING_PA | 0.986777 | 0.47785 | 0.875082 |
| REACTOME_APOPTOTIC | -0.96546 | 0.508311 | 1 | REACTOME_APOPTOTIC_EXECUTION_PHASE | 1.168433 | 0.251041 | 0.783519 | REACTOME_APOPTOTIC_I | -0.84144 | 0.728466 | 0.817334 |
| REACTOME_TNF_SIGNA | 0.847164 | 0.714576 | 1 | REACTOME_TNF_SIGNALING | 1.172371 | 0.228559 | 0.783543 | REACTOME_TNF_SIGNALI | 0.970728 | 0.504649 | 0.888411 |
| REACTOME_CLASS_I_MI | -0.91139 | 0.691683 | 1 | REACTOME_CLASS_I_MHC_MEDIATED_ANTIGEN_PRO | 1.170206 | 0.155433 | 0.784134 | REACTOME_CLASS_I_MH | 0.868515 | 0.784258 | 0.96065 |
| REACTOME_SIALIC_ACIE | 1.180043 | 0.254581 | 0.831697 | REACTOME_SIALIC_ACID_METABOLISM | 1.135716 | 0.303678 | 0.78554 | REACTOME_SIALIC_ACID_ | 0.826449 | 0.692107 | 0.976499 |
| WP_SPHINGOLIPID_ME | 1.61796 | 0.026882 | 0.225063 | WP_SPHINGOLIPID_METABOLISM_OVERVIEW | -1.0911 | 0.340179 | 0.785743 | WP_SPHINGOLIPID_META | 1.684823 | 0.011356 | 0.088699 |
| REACTOME_MYD88_INE | -1.08996 | 0.307868 | 1 | REACTOME_MYD88_INDEPENDENT_TLR4_CASCADE | 1.126315 | 0.277731 | 0.786949 | REACTOME_MYD88_INDE | 0.656644 | 0.963319 | 0.973237 |
| REACTOME_BASE_EXCIS | -1.46728 | 0.066066 | 0.608694 | REACTOME_BASE_EXCISION_REPAIR | 1.173325 | 0.266775 | 0.787293 | REACTOME_BASE_EXCISI | 0.701088 | 0.855167 | 0.990372 |
| REACTOME_INTRA_GOL | 1.119019 | 0.210023 | 0.887324 | REACTOME_INTRA_GOLGI_AND_RETROGRADE_GOLGI | 1.174981 | 0.186549 | 0.788533 | REACTOME_INTRA_GOLG | 1.034175 | 0.399511 | 0.80991 |
| REACTOME_NEUROTRAI | 1.195844 | 0.14598 | 0.810632 | REACTOME_NEUROTRANSMITTER_RECEPTORS_AND_F | -1.09569 | 0.254258 | 0.788815 | REACTOME_NEUROTRANS | -1.42746 | 0.013519 | 0.143895 |
| REACTOME_A_TETRASA | 0.58303 | 0.95656 | 1 | REACTOME_A_TETRASACCHARIDE_LINKER_SEQUENCE | 1.12353 | 0.312861 | 0.790354 | REACTOME_A_TETRASAC | 0.846002 | 0.664325 | 0.961933 |
| REACTOME_METABOLIS | 1.257107 | 0.191737 | 0.736206 | REACTOME_METABOLISM_OF_PORPHYRINS | 1.121634 | 0.323057 | 0.790763 | REACTOME_METABOLISM | 0.923839 | 0.567536 | 0.959675 |
| WP_SPHINGOLIPID_ME | 1.617602 | 0.029221 | 0.218299 | WP_SPHINGOLIPID_METABOLISM_INTEGRATED_PATH | -1.09135 | 0.330977 | 0.790874 | WP_SPHINGOLIPID_META | 1.682079 | 0.014452 | 0.086526 |
| REACTOME_MAJOR_PA | 1.760818 | 5.05E-04 | 0.119708 | REACTOME_MAJOR_PATHWAY_OF_RRNA_PROCESSIN | 1.1176 | 0.259011 | 0.792479 | REACTOME_MAJOR_PATH | 2.142849 | 0 | 1.83E-04 |
| REACTOME_NUCLEOTID | 0.705401 | 0.880157 | 1 | REACTOME_NUCLEOTIDE_BINDING_DOMAIN_LEUCINE | 1.115702 | 0.312169 | 0.79284 | REACTOME_NUCLEOTIDE_ | -0.69758 | 0.897547 | 0.93881 |
| WP_NEURAL_CREST_DII | -1.57099 | 0.026425 | 0.582001 | WP_NEURAL_CREST_DIFFERENTIATION | -1.0924 | 0.318346 | 0.793453 | WP_NEURAL_CREST_DIFF | -0.81547 | 0.771701 | 0.841556 |
| BIOCARTA_FAS_PATHW | -1.09824 | 0.333457 | 1 | BIOCARTA_FAS_PATHWAY | 1.118698 | 0.315298 | 0.794564 | BIOCARTA_FAS_PATHWAY | -1.10435 | 0.315584 | 0.438704 |
| REACTOME_BRANCHED_ | -0.98142 | 0.479554 | 1 | REACTOME_BRANCHED_CHAIN_AMINO_ACID_CATABO | -1.07918 | 0.3482 | 0.801251 | REACTOME_BRANCHED_C | -1.22322 | 0.199948 | 0.299738 |
| WP_SPINAL_CORD_INJU | -1.3047 | 0.115529 | 0.849956 | WP_SPINAL_CORD_INJURY | -1.06419 | 0.340537 | 0.804743 | WP_SPINAL_CORD_INJUR | -1.21455 | 0.160524 | 0.307766 |
| REACTOME_SIGNALING_ | 0.86742 | 0.629118 | 1 | REACTOME_SIGNALING_BY_ERBB4 | -1.07922 | 0.338298 | 0.807004 | REACTOME_SIGNALING_E | -1.28736 | 0.153289 | 0.250432 |
| REACTOME_RAS_PROCE | 1.376319 | 0.117088 | 0.651313 | REACTOME_RAS_PROCESSING | -1.06496 | 0.362909 | 0.807803 | REACTOME_RAS_PROCES | -1.33243 | 0.129132 | 0.211652 |
| REACTOME_DOWNSTRE | 0.746909 | 0.817712 | 1 | REACTOME_DOWNSTREAM_SIGNAL_TRANSDUCTION | -1.08078 | 0.339504 | 0.807901 | REACTOME_DOWNSTREA | -1.71553 | 0.01021 | 0.04941 |
| REACTOME_VEGFR2_MI | 1.217909 | 0.201771 | 0.806597 | REACTOME_VEGFR2_MEDIATED_VASCULAR_PERMEAE | -1.07006 | 0.350371 | 0.80791 | REACTOME_VEGFR2_MEI | -1.48577 | 0.053499 | 0.117461 |
| REACTOME_MTORC1_N | -0.93909 | 0.540939 | 1 | REACTOME_MTORC1_MEDIATED_SIGNALLING | -1.0735 | 0.344294 | 0.808183 | REACTOME_MTORC1_ME | -1.61613 | 0.021687 | 0.074524 |
| REACTOME_PLATELET_I | -0.88274 | 0.663407 | 1 | REACTOME_PLATELET_HOMEOSTASIS | -1.06789 | 0.331885 | 0.809427 | REACTOME_PLATELET_HC | -1.34326 | 0.071451 | 0.204312 |
| REACTOME_EFFECTS_OI | -0.83772 | 0.687165 | 1 | REACTOME_EFFECTS_OF_PIP2_HYDROLYSIS | -1.07132 | 0.354822 | 0.809522 | REACTOME_EFFECTS_OF_ | -1.51774 | 0.045192 | 0.109874 |
| BIOCARTA_TOLL_PATHV | -1.24788 | 0.197775 | 0.872559 | BIOCARTA_TOLL_PATHWAY | -1.06549 | 0.374046 | 0.811709 | BIOCARTA_TOLL_PATHW | 0.784748 | 0.749041 | 0.998379 |

| | | | | | | | | | | | |
|---|---|---|---|---|---|---|---|---|---|---|---|
| REACTOME_HATS_ACET | -0.92424 | 0.558538 | 1 | REACTOME_HATS_ACETYLATE_HISTONES | -1.0736 | 0.352743 | 0.813722 | REACTOME_HATS_ACETYI | 0.935814 | 0.548245 | 0.940096 |
| REACTOME_PYRUVATE_ | 1.590145 | 0.015304 | 0.256602 | REACTOME_PYRUVATE_METABOLISM_AND_CITRIC_A( | 1.105609 | 0.312047 | 0.814205 | REACTOME_PYRUVATE_M | -1.58021 | 0.014303 | 0.08627 |
| REACTOME_REGULATIO | 0.665462 | 0.920248 | 1 | REACTOME_REGULATION_OF_TP53_ACTIVITY_THROU( | -1.05907 | 0.36227 | 0.815872 | REACTOME_REGULATION | 1.029514 | 0.423462 | 0.810503 |
| REACTOME_BUTYRATE_ | 1.277572 | 0.176534 | 0.771937 | REACTOME_BUTYRATE_RESPONSE_FACTOR_1_BRF1_E | 1.106795 | 0.337664 | 0.816065 | REACTOME_BUTYRATE_R | 1.6344 | 0.017138 | 0.093821 |
| REACTOME_PROCESSIN( | -1.74344 | 0.007442 | 0.411722 | REACTOME_PROCESSING_OF_DNA_DOUBLE_STRAND_ | -1.04874 | 0.379032 | 0.822112 | REACTOME_PROCESSING_ | -0.99558 | 0.447717 | 0.587929 |
| BIOCARTA_BIOPEPTIDES | 1.332733 | 0.124695 | 0.695955 | BIOCARTA_BIOPEPTIDES_PATHWAY | -1.04908 | 0.375556 | 0.826469 | BIOCARTA_BIOPEPTIDES_ | -1.42073 | 0.072472 | 0.148162 |
| REACTOME_TP53_REGU | -1.17106 | 0.270225 | 0.89327 | REACTOME_TP53_REGULATES_TRANSCRIPTION_OF_CI | 1.099492 | 0.350799 | 0.828519 | REACTOME_TP53_REGUL | -0.66122 | 0.903452 | 0.958092 |
| REACTOME_TNFR1_IND | -0.5402 | 0.981908 | 1 | REACTOME_TNFR1_INDUCED_NFKAPPAB_SIGNALING_ | 1.088974 | 0.341104 | 0.828993 | REACTOME_TNFR1_INDU | 0.655952 | 0.909621 | 0.969631 |
| REACTOME_G2_M_DNA | -1.75926 | 0.004176 | 0.468303 | REACTOME_G2_M_DNA_DAMAGE_CHECKPOINT | -1.0494 | 0.375561 | 0.83104 | REACTOME_G2_M_DNA_ | -1.01798 | 0.413032 | 0.560677 |
| WP_PARKINSONS_DISEA | -0.9193 | 0.576774 | 1 | WP_PARKINSONS_DISEASE | 1.089349 | 0.35612 | 0.833314 | WP_PARKINSONS_DISEAS | -1.08707 | 0.33175 | 0.463141 |
| REACTOME_RNA_POLYN | -1.67673 | 0.00763 | 0.522329 | REACTOME_RNA_POLYMERASE_II_TRANSCRIBES_SNRI | 1.096175 | 0.314178 | 0.83356 | REACTOME_RNA_POLYMI | 1.079656 | 0.345208 | 0.736845 |
| REACTOME_RAP1_SIGN | -0.83059 | 0.684298 | 1 | REACTOME_RAP1_SIGNALLING | 1.090593 | 0.353098 | 0.834824 | REACTOME_RAP1_SIGNAI | -1.44689 | 0.08144 | 0.133244 |
| REACTOME_REGULATIO | -0.90322 | 0.584888 | 1 | REACTOME_REGULATION_OF_APC_C_ACTIVATORS_BE | -1.0497 | 0.386931 | 0.835593 | REACTOME_REGULATION | -1.10993 | 0.316812 | 0.429714 |
| REACTOME_PCP_CE_PA | 1.053595 | 0.351472 | 0.963622 | REACTOME_PCP_CE_PATHWAY | -1.05018 | 0.346167 | 0.839564 | REACTOME_PCP_CE_PATI | -1.92088 | 0 | 0.019841 |
| REACTOME_NRAGE_SIG | -1.06785 | 0.354317 | 1 | REACTOME_NRAGE_SIGNALS_DEATH_THROUGH_JNK | 1.090621 | 0.328269 | 0.840406 | REACTOME_NRAGE_SIGN | -1.16434 | 0.224683 | 0.360168 |
| REACTOME_INTERLEUKI | -1.36461 | 0.077395 | 0.786247 | REACTOME_INTERLEUKIN_17_SIGNALING | 1.091466 | 0.335468 | 0.843367 | REACTOME_INTERLEUKIN | -0.70856 | 0.928317 | 0.932968 |
| REACTOME_RUNX1_INT | -1.42453 | 0.074851 | 0.672641 | REACTOME_RUNX1_INTERACTS_WITH_CO_FACTORS_\ | 1.074507 | 0.366419 | 0.859106 | REACTOME_RUNX1_INTEI | 1.336965 | 0.126192 | 0.347235 |
| REACTOME_ADAPTIVE_I | -0.66494 | 1 | 1 | REACTOME_ADAPTIVE_IMMUNE_SYSTEM | 1.077419 | 0.286228 | 0.860933 | REACTOME_ADAPTIVE_IN | -1.09089 | 0.144397 | 0.460055 |
| REACTOME_REGULATIO | 0.831373 | 0.686149 | 1 | REACTOME_REGULATION_OF_TP53_DEGRADATION | 1.06879 | 0.375459 | 0.861068 | REACTOME_REGULATION | -0.73332 | 0.838528 | 0.91374 |
| REACTOME_BINDING_A | 1.312886 | 0.158933 | 0.704277 | REACTOME_BINDING_AND_UPTAKE_OF_LIGANDS_BY_ | 1.075583 | 0.370059 | 0.861298 | REACTOME_BINDING_AN | 1.1973 | 0.243072 | 0.513739 |
| REACTOME_DDX58_IFIH | 0.845499 | 0.675136 | 1 | REACTOME_DDX58_IFIH1_MEDIATED_INDUCTION_OF_ | 1.069669 | 0.370818 | 0.863763 | REACTOME_DDX58_IFIH1 | 1.215804 | 0.21503 | 0.494834 |
| WP_NOTCH_SIGNALING | -1.11973 | 0.310069 | 0.984549 | WP_NOTCH_SIGNALING_PATHWAY | 1.070563 | 0.371752 | 0.866515 | WP_NOTCH_SIGNALING_I | 0.895506 | 0.603488 | 0.971101 |
| REACTOME_SPHINGOLIF | 1.688962 | 0.018579 | 0.171061 | REACTOME_SPHINGOLIPID_DE_NOVO_BIOSYNTHESIS | -1.03272 | 0.403265 | 0.869918 | REACTOME_SPHINGOLIPI | 1.636504 | 0.020756 | 0.095334 |
| REACTOME_GLUCONEO | 0.706645 | 0.878478 | 1 | REACTOME_GLUCONEOGENESIS | -1.02458 | 0.406993 | 0.874796 | REACTOME_GLUCONEOG | -1.27604 | 0.150968 | 0.25733 |
| REACTOME_RND3_GTPA | 1.20022 | 0.213466 | 0.803594 | REACTOME_RND3_GTPASE_CYCLE | -1.01785 | 0.421068 | 0.875318 | REACTOME_RND3_GTPAS | 0.725217 | 0.851228 | 1 |
| REACTOME_NEUROTRAI | 0.671864 | 0.95693 | 1 | REACTOME_NEUROTRANSMITTER_RELEASE_CYCLE | -1.02275 | 0.413625 | 0.875463 | REACTOME_NEUROTRANS | -1.71296 | 0.00494 | 0.049147 |
| REACTOME_MUSCLE_C( | -0.92762 | 0.594425 | 1 | REACTOME_MUSCLE_CONTRACTION | -1.01939 | 0.40625 | 0.875605 | REACTOME_MUSCLE_CON | -1.42857 | 0.017537 | 0.144498 |
| REACTOME_MTOR_SIGN | -0.65629 | 0.943192 | 1 | REACTOME_MTOR_SIGNALLING | -1.02011 | 0.416287 | 0.878713 | REACTOME_MTOR_SIGNA | -1.51839 | 0.029031 | 0.110158 |
| REACTOME_CYTOPROTE | 0.784935 | 0.807435 | 1 | REACTOME_CYTOPROTECTION_BY_HMOX1 | -1.02497 | 0.414543 | 0.879177 | REACTOME_CYTOPROTEC | 0.931629 | 0.556875 | 0.945098 |
| REACTOME_TP53_REGU | 0.54457 | 0.998377 | 1 | REACTOME_TP53_REGULATES_METABOLIC_GENES | -1.02749 | 0.392303 | 0.882099 | REACTOME_TP53_REGUL | -1.31617 | 0.076797 | 0.224322 |
| REACTOME_RHO_GTPA | -0.90839 | 0.67681 | 1 | REACTOME_RHO_GTPASE_EFFECTORS | -1.0255 | 0.388646 | 0.883071 | REACTOME_RHO_GTPASE | -1.53049 | 9.99E-04 | 0.105585 |
| REACTOME_TRISTETRAF | 1.335204 | 0.136665 | 0.713821 | REACTOME_TRISTETRAPROLIN_TTP_ZFP36_BINDS_ANI | 1.060219 | 0.386088 | 0.883498 | REACTOME_TRISTETRAPR | 1.584885 | 0.028291 | 0.117078 |
| REACTOME_VISUAL_PH( | -1.06868 | 0.360336 | 1 | REACTOME_VISUAL_PHOTOTRANSDUCTION | -1.00733 | 0.433286 | 0.893713 | REACTOME_VISUAL_PHO | -0.78164 | 0.810141 | 0.884754 |
| WP_CALCIUM_REGULAT | -0.67956 | 0.982865 | 1 | WP_CALCIUM_REGULATION_IN_CARDIAC_CELLS | -1.00527 | 0.432664 | 0.895265 | WP_CALCIUM_REGULATI( | -1.30754 | 0.046465 | 0.231539 |
| REACTOME_UNBLOCKIN | -0.81244 | 0.730819 | 1 | REACTOME_UNBLOCKING_OF_NMDA_RECEPTORS_GLI | -1.00821 | 0.441774 | 0.896357 | REACTOME_UNBLOCKING | -0.87635 | 0.632664 | 0.772506 |
| REACTOME_GENERIC_TI | -1.21001 | 0.065 | 0.860776 | REACTOME_GENERIC_TRANSCRIPTION_PATHWAY | 1.048483 | 0.354968 | 0.899364 | REACTOME_GENERIC_TR/ | 1.148147 | 0.156065 | 0.589838 |
| REACTOME_RUNX1_RE( | -1.21125 | 0.226942 | 0.866715 | REACTOME_RUNX1_REGULATES_GENES_INVOLVED_IN | 1.053598 | 0.399477 | 0.899633 | REACTOME_RUNX1_REGU | 1.358499 | 0.120238 | 0.324123 |
| REACTOME_PLATELET_/ | -0.76745 | 0.805621 | 1 | REACTOME_PLATELET_AGGREGATION_PLUG_FORMAT | 1.049999 | 0.399286 | 0.900111 | REACTOME_PLATELET_AG | -1.08661 | 0.323529 | 0.462596 |
| REACTOME_SIGNALING_ | 1.007949 | 0.425431 | 1 | REACTOME_SIGNALING_BY_WNT | -1.00843 | 0.426074 | 0.901162 | REACTOME_SIGNALING_E | -1.50366 | 0.002782 | 0.114677 |
| REACTOME_REGULATIO | -0.79581 | 0.798053 | 1 | REACTOME_REGULATION_OF_PLK1_ACTIVITY_AT_G2_ | 1.014491 | 0.440693 | 0.901509 | REACTOME_REGULATION | 0.56191 | 0.987466 | 0.993537 |
| WP_FACTORS_AND_PAT | -0.78798 | 0.75085 | 1 | WP_FACTORS_AND_PATHWAYS_AFFECTING_INSULINL | 1.05097 | 0.401066 | 0.902581 | WP_FACTORS_AND_PATH | 0.896636 | 0.599772 | 0.974334 |
| REACTOME_RHO_GTPA | -0.96311 | 0.508756 | 1 | REACTOME_RHO_GTPASES_ACTIVATE_PKNS | 1.020649 | 0.435317 | 0.902668 | REACTOME_RHO_GTPASE | 0.85197 | 0.667264 | 0.975987 |
| BIOCARTA_MCALPAIN_F | 0.481211 | 0.988428 | 1 | BIOCARTA_MCALPAIN_PATHWAY | -0.99972 | 0.450342 | 0.902915 | BIOCARTA_MCALPAIN_PA | -1.61087 | 0.028773 | 0.076031 |
| REACTOME_FCERI_MED | -1.7272 | 0.008663 | 0.389204 | REACTOME_FCERI_MEDIATED_MAPK_ACTIVATION | 1.018715 | 0.438404 | 0.903517 | REACTOME_FCERI_MEDI/ | -1.61412 | 0.023126 | 0.074901 |
| REACTOME_MRNA_DEC | 1.412188 | 0.099674 | 0.586882 | REACTOME_MRNA_DECAY_BY_3_TO_5_EXORIBONUCI | 1.017066 | 0.440158 | 0.903658 | REACTOME_MRNA_DECA | 1.519548 | 0.043664 | 0.170309 |
| REACTOME_ANTIGEN_P | 1.276155 | 0.173514 | 0.765201 | REACTOME_ANTIGEN_PRESENTATION_FOLDING_ASSE | 1.021506 | 0.437805 | 0.905133 | REACTOME_ANTIGEN_PR | 1.408219 | 0.086928 | 0.29562 |
| REACTOME_MITOTIC_M | -0.85269 | 0.800533 | 1 | REACTOME_MITOTIC_METAPHASE_AND_ANAPHASE | -0.99738 | 0.46942 | 0.905308 | REACTOME_MITOTIC_ME | -0.93105 | 0.672515 | 0.696572 |
| BIOCARTA_IGF1_PATHW | 0.897797 | 0.58646 | 1 | BIOCARTA_IGF1_PATHWAY | -0.99576 | 0.455626 | 0.905383 | BIOCARTA_IGF1_PATHWA | -1.54735 | 0.037911 | 0.098028 |
| REACTOME_APOPTOTIC | -1.00774 | 0.442906 | 1 | REACTOME_APOPTOTIC_CLEAVAGE_OF_CELLULAR_PR | 1.014734 | 0.441507 | 0.905775 | REACTOME_APOPTOTIC_( | -0.90252 | 0.62102 | 0.727205 |
| REACTOME_FATTY_ACID | 1.658859 | 0.002422 | 0.177438 | REACTOME_FATTY_ACID_METABOLISM | -1.00017 | 0.459196 | 0.906919 | REACTOME_FATTY_ACID_ | -1.14545 | 0.181571 | 0.38395 |
| REACTOME_RESPONSE_ | 1.271366 | 0.10583 | 0.748117 | REACTOME_RESPONSE_TO_ELEVATED_PLATELET_CYT( | 1.021983 | 0.42435 | 0.908777 | REACTOME_RESPONSE_T( | 0.881531 | 0.683984 | 0.969821 |
| REACTOME_FORMATIOI | -1.25777 | 0.181951 | 0.900825 | REACTOME_FORMATION_OF_THE_EARLY_ELONGATIO | 1.008039 | 0.442934 | 0.911308 | REACTOME_FORMATION_ | 1.421671 | 0.077178 | 0.281096 |
| WP_IL3_SIGNALING_PA | -1.10984 | 0.281491 | 0.996316 | WP_IL3_SIGNALING_PATHWAY | 1.022009 | 0.423328 | 0.913908 | WP_IL3_SIGNALING_PATH | -1.23041 | 0.122449 | 0.295134 |
| REACTOME_ION_CHANI | 1.335759 | 0.058168 | 0.72584 | REACTOME_ION_CHANNEL_TRANSPORT | 1.00859 | 0.4375 | 0.914626 | REACTOME_ION_CHANNE | 0.902184 | 0.654209 | 0.978806 |
| REACTOME_CARGO_CO | 1.74242 | 0.010033 | 0.133342 | REACTOME_CARGO_CONCENTRATION_IN_THE_ER | 1.022906 | 0.433757 | 0.916275 | REACTOME_CARGO_CON | 1.435002 | 0.065866 | 0.267974 |
| REACTOME_METABOLIS | -1.17761 | 0.251134 | 0.88675 | REACTOME_METABOLISM_OF_FAT_SOLUBLE_VITAMIN | 1.031755 | 0.419924 | 0.92032 | REACTOME_METABOLISM | 1.149043 | 0.279128 | 0.592977 |
| REACTOME_PROTEIN_U | -0.72359 | 0.895968 | 1 | REACTOME_PROTEIN_UBIQUITINATION | 1.023206 | 0.421029 | 0.920605 | REACTOME_PROTEIN_UB | 0.838928 | 0.721598 | 0.966453 |
| REACTOME_TCF_DEPEN | -0.65347 | 0.989428 | 1 | REACTOME_TCF_DEPENDENT_SIGNALING_IN_RESPON | 1.04018 | 0.382851 | 0.920912 | REACTOME_TCF_DEPEND | -1.00043 | 0.45444 | 0.582273 |
| REACTOME_CD209_DC_ | 1.064415 | 0.377697 | 0.952603 | REACTOME_CD209_DC_SIGN_SIGNALING | 1.026042 | 0.430822 | 0.922343 | REACTOME_CD209_DC_S | 0.970597 | 0.495063 | 0.882476 |
| REACTOME_RECRUITME | -0.56707 | 0.992451 | 1 | REACTOME_RECRUITMENT_OF_NUMA_TO_MITOTIC_( | 1.03226 | 0.40769 | 0.924237 | REACTOME_RECRUITMEN | -0.77715 | 0.870644 | 0.88504 |
| REACTOME_REGULATEC | 1.146809 | 0.260562 | 0.866717 | REACTOME_REGULATED_NECROSIS | 1.023342 | 0.422256 | 0.925539 | REACTOME_REGULATED_ | -1.127 | 0.272345 | 0.409629 |
| REACTOME_METABOLIS | -0.6854 | 0.994646 | 1 | REACTOME_METABOLISM_OF_AMINO_ACIDS_AND_D | -0.98811 | 0.494061 | 0.925586 | REACTOME_METABOLISM | -1.95034 | 0 | 0.018042 |
| REACTOME_TRANSLESI( | -1.225 | 0.220349 | 0.855757 | REACTOME_TRANSLESION_SYNTHESIS_BY_Y_FAMILY_I | 1.026294 | 0.427563 | 0.926987 | REACTOME_TRANSLESION | -0.70832 | 0.862191 | 0.931117 |
| WP_IL5_SIGNALING_PA | -1.15737 | 0.237152 | 0.928797 | WP_IL5_SIGNALING_PATHWAY | 1.033022 | 0.405398 | 0.927341 | WP_IL5_SIGNALING_PATH | -1.17124 | 0.205033 | 0.354028 |
| REACTOME_M_PHASE | -0.74398 | 0.98263 | 1 | REACTOME_M_PHASE | -0.98562 | 0.49677 | 0.928527 | REACTOME_M_PHASE | -0.74574 | 0.996947 | 0.911879 |
| REACTOME_RHOG_GTP/ | -0.96356 | 0.502708 | 1 | REACTOME_RHOG_GTPASE_CYCLE | 1.035698 | 0.398214 | 0.929882 | REACTOME_RHOG_GTPAS | -0.93021 | 0.58634 | 0.694757 |
| REACTOME_TRANSPORT | 1.267156 | 0.175503 | 0.752489 | REACTOME_TRANSPORT_OF_VITAMINS_NUCLEOSIDES | 1.033851 | 0.417953 | 0.930267 | REACTOME_TRANSPORT_ | 1.250896 | 0.189944 | 0.45331 |
| REACTOME_PRESYNAPT | -0.74228 | 0.812917 | 1 | REACTOME_PRESYNAPTIC_FUNCTION_OF_KAINATE_RI | -0.95906 | 0.500533 | 0.931813 | REACTOME_PRESYNAPTIC | -1.28471 | 0.165115 | 0.250594 |
| REACTOME_PEROXISON | 1.867626 | 4.34E-03 | 0.068713 | REACTOME_PEROXISOMAL_LIPID_METABOLISM | 1.026402 | 0.425626 | 0.932136 | REACTOME_PEROXISOMA | 1.274199 | 0.170951 | 0.41593 |
| REACTOME_REGULATIO | -1.00168 | 0.444767 | 1 | REACTOME_REGULATION_OF_TP53_ACTIVITY_THROU( | -0.96019 | 0.513872 | 0.933316 | REACTOME_REGULATION | 0.999447 | 0.460441 | 0.851857 |
| REACTOME_RHO_GTPA | 1.084569 | 0.335118 | 0.922235 | REACTOME_RHO_GTPASES_ACTIVATE_WASPS_AND_W | -0.9604 | 0.518326 | 0.93771 | REACTOME_RHO_GTPASE | -1.98298 | 2.80E-04 | 0.015214 |
| BIOCARTA_INSULIN_PAT | 1.055936 | 0.378918 | 0.969431 | BIOCARTA_INSULIN_PATHWAY | -0.96167 | 0.494026 | 0.938736 | BIOCARTA_INSULIN_PATH | -1.58667 | 0.036999 | 0.084695 |
| REACTOME_G_PROTEIN | -0.64108 | 0.925713 | 1 | REACTOME_G_PROTEIN_BETA_GAMMA_SIGNALLING | -0.98091 | 0.460296 | 0.938934 | REACTOME_G_PROTEIN_I | -1.57919 | 0.029674 | 0.086189 |
| REACTOME_ION_HOME | 0.722564 | 0.891771 | 1 | REACTOME_ION_HOMEOSTASIS | -0.95362 | 0.527067 | 0.939206 | REACTOME_ION_HOMEO | 0.869035 | 0.655257 | 0.965042 |
| REACTOME_CELL_CYCLE | -1.16309 | 0.175436 | 0.915553 | REACTOME_CELL_CYCLE_CHECKPOINTS | -0.93947 | 0.621495 | 0.939359 | REACTOME_CELL_CYCLE_ | -0.97815 | 0.511859 | 0.616456 |
| REACTOME_MITOTIC_SI | -0.85981 | 0.705265 | 1 | REACTOME_MITOTIC_SPINDLE_CHECKPOINT | -0.9402 | 0.567225 | 0.941942 | REACTOME_MITOTIC_SPI | 0.726166 | 0.900287 | 1 |
| REACTOME_SIGNALING_ | -0.63999 | 0.944736 | 1 | REACTOME_SIGNALING_BY_PTK6 | -0.954 | 0.516532 | 0.942973 | REACTOME_SIGNALING_E | -1.15974 | 0.24234 | 0.365191 |
| REACTOME_SLC_MEDIA | -0.85392 | 0.776241 | 1 | REACTOME_SLC_MEDIATED_TRANSMEMBRANE_TRAN | -0.96184 | 0.540708 | 0.943312 | REACTOME_SLC_MEDIATI | 0.795354 | 0.859872 | 0.994866 |
| REACTOME_SYNAPTIC_/ | 1.128139 | 0.295514 | 0.870088 | REACTOME_SYNAPTIC_ADHESION_LIKE_MOLECULES | -0.96321 | 0.505907 | 0.943953 | REACTOME_SYNAPTIC_AI | -0.93733 | 0.52796 | 0.690006 |
| WP_CHEMOKINE_SIGNA | 0.768071 | 0.922796 | 1 | WP_CHEMOKINE_SIGNALING_PATHWAY | -0.93579 | 0.624209 | 0.945742 | WP_CHEMOKINE_SIGNAL | -1.74366 | 0 | 0.041793 |
| WP_MAPK_CASCADE | -0.88285 | 0.624052 | 1 | WP_MAPK_CASCADE | -0.94039 | 0.553763 | 0.946183 | WP_MAPK_CASCADE | -1.36594 | 0.100081 | 0.187677 |
| REACTOME_DNA_DOUB | -1.37521 | 0.097828 | 0.780452 | REACTOME_DNA_DOUBLE_STRAND_BREAK_RESPONSI | -0.96728 | 0.485454 | 0.946435 | REACTOME_DNA_DOUBLI | -1.28648 | 0.154883 | 0.249385 |
| REACTOME_CELL_CYCLE | -1.0075 | 0.437985 | 1 | REACTOME_CELL_CYCLE | -0.94963 | 0.672117 | 0.946786 | REACTOME_CELL_CYCLE | 0.658924 | 0.99954 | 0.984117 |
| WP_PURINE_METABOLI | -0.74671 | 0.933434 | 1 | WP_PURINE_METABOLISM | -0.96392 | 0.547537 | 0.946852 | WP_PURINE_METABOLISI | -0.92931 | 0.640176 | 0.694916 |
| REACTOME_SIGNALING_ | -1.21555 | 0.227332 | 0.86033 | REACTOME_SIGNALING_BY_HIPPO | 0.986394 | 0.477489 | 0.946892 | REACTOME_SIGNALING_E | 0.722659 | 0.822939 | 1 |
| REACTOME_ACTIVATION | -0.64756 | 0.967924 | 1 | REACTOME_ACTIVATION_OF_NF_KAPPAB_IN_B_CELLS | -0.96543 | 0.516364 | 0.947215 | REACTOME_ACTIVATION_ | -2.06293 | 0 | 0.008192 |
| REACTOME_SULFUR_AN | -0.77449 | 0.764312 | 1 | REACTOME_SULFUR_AMINO_ACID_METABOLISM | -0.9683 | 0.500655 | 0.948472 | REACTOME_SULFUR_AMI | -1.52013 | 0.044346 | 0.109716 |
| REACTOME_FC_EPSILON | -1.0537 | 0.360887 | 1 | REACTOME_FC_EPSILON_RECEPTOR_FCERI_SIGNALIN( | -0.94068 | 0.589879 | 0.950163 | REACTOME_FC_EPSILON_ | -1.99662 | 0 | 0.013375 |
| REACTOME_ADP_SIGNA | -0.7044 | 0.852341 | 1 | REACTOME_ADP_SIGNALLING_THROUGH_P2Y_PURIN( | -0.94218 | 0.523607 | 0.950392 | REACTOME_ADP_SIGNALI | -1.50296 | 0.056276 | 0.11444 |
| REACTOME_NCAM_SIGI | 1.054813 | 0.38288 | 0.96638 | REACTOME_NCAM_SIGNALING_FOR_NEURITE_OUT_G | 0.986605 | 0.473217 | 0.95131 | REACTOME_NCAM_SIGN/ | -0.87663 | 0.640434 | 0.773872 |
| REACTOME_PHASE_I_FL | 1.20989 | 0.194292 | 0.787715 | REACTOME_PHASE_I_FUNCTIONALIZATION_OF_COMP | -0.97552 | 0.492453 | 0.951553 | REACTOME_PHASE_I_FUN | -1.03725 | 0.388636 | 0.5308 |
| REACTOME_CELLULAR_I | -0.8305 | 0.925234 | 1 | REACTOME_CELLULAR_RESPONSES_TO_STIMULI | -0.97055 | 0.594439 | 0.951798 | REACTOME_CELLULAR_RE | 0.801582 | 0.929563 | 0.993687 |
| REACTOME_G_ALPHA_Z | -0.83549 | 0.702965 | 1 | REACTOME_G_ALPHA_Z_SIGNALLING_EVENTS | -0.94631 | 0.524618 | 0.952302 | REACTOME_G_ALPHA_Z_ | -0.98089 | 0.472586 | 0.612179 |
| REACTOME_SENSORY_P | -1.08702 | 0.339121 | 1 | REACTOME_SENSORY_PERCEPTION | -0.94287 | 0.541716 | 0.953205 | REACTOME_SENSORY_PE | -1.03891 | 0.389879 | 0.528874 |
| REACTOME_POSITIVE_E | 0.871203 | 0.634146 | 1 | REACTOME_POSITIVE_EPIGENETIC_REGULATION_OF_F | -0.96847 | 0.486908 | 0.953252 | REACTOME_POSITIVE_EPI | 1.232636 | 0.208557 | 0.470023 |
| REACTOME_SUMOYLATI | -1.25645 | 0.115391 | 0.891819 | REACTOME_SUMOYLATION | 0.982598 | 0.488331 | 0.953368 | REACTOME_SUMOYLATIC | 1.437667 | 0.025944 | 0.268679 |
| REACTOME_KINESINS | -0.69638 | 0.897501 | 1 | REACTOME_KINESINS | 0.987335 | 0.476255 | 0.954179 | REACTOME_KINESINS | 0.725173 | 0.856833 | 1 |

| | | | | | | | | | | | |
|---|---|---|---|---|---|---|---|---|---|---|---|
| REACTOME_CDC42_GTF | -1.27535 | 0.112855 | 0.867591 | REACTOME_CDC42_GTPASE_CYCLE | 0.977083 | 0.498869 | 0.955027 | REACTOME_CDC42_GTPA | 0.702843 | 0.928744 | 0.99302 |
| WP_CHOLESTEROL_MET | 1.705891 | 0.006996 | 0.163596 | WP_CHOLESTEROL_METABOLISM_WITH_BLOCH_AND_ | 0.980172 | 0.482615 | 0.955665 | WP_CHOLESTEROL_META | 1.652451 | 0.010295 | 0.092276 |
| WP_OXIDATIVE_STRESS_ | 1.663716 | 0.018255 | 0.177254 | WP_OXIDATIVE_STRESS_RESPONSE | 0.988468 | 0.470245 | 0.955814 | WP_OXIDATIVE_STRESS_F | 1.128535 | 0.303437 | 0.630594 |
| REACTOME_ANTIGEN_P | -1.00187 | 0.443673 | 1 | REACTOME_ANTIGEN_PROCESSING_UBIQUITINATION_ | 0.978383 | 0.519775 | 0.956128 | REACTOME_ANTIGEN_PR | 0.827202 | 0.848577 | 0.980266 |
| REACTOME_GABA_RECE | 1.099622 | 0.303229 | 0.904438 | REACTOME_GABA_RECEPTOR_ACTIVATION | -0.97068 | 0.494453 | 0.956818 | REACTOME_GABA_RECEP | -1.24755 | 0.13912 | 0.277825 |
| REACTOME_NEGATIVE_ | 0.897278 | 0.584938 | 1 | REACTOME_NEGATIVE_REGULATION_OF_FGFR3_SIGN | -0.94313 | 0.535615 | 0.957391 | REACTOME_NEGATIVE_RI | -1.12956 | 0.296504 | 0.407953 |
| REACTOME_G_BETA_GA | -0.67183 | 0.896584 | 1 | REACTOME_G_BETA_GAMMA_SIGNALLING_THROUGH | -0.972 | 0.485699 | 0.957876 | REACTOME_G_BETA_GAN | -1.34877 | 0.119349 | 0.200336 |
| REACTOME_MITOTIC_G | 0.558873 | 0.999513 | 1 | REACTOME_MITOTIC_G2_G2_M_PHASES | 0.989412 | 0.478784 | 0.95804 | REACTOME_MITOTIC_G2_ | -1.20306 | 0.106764 | 0.32006 |
| WP_COMPREHENSIVE_I | 0.909749 | 0.614059 | 1 | WP_COMPREHENSIVE_IL17A_SIGNALING | 0.991009 | 0.471639 | 0.958348 | WP_COMPREHENSIVE_IL1 | -1.06568 | 0.328649 | 0.490541 |
| REACTOME_SIGNALING_ | -0.61205 | 0.941144 | 1 | REACTOME_SIGNALING_BY_RETINOIC_ACID | -0.92538 | 0.559725 | 0.962784 | REACTOME_SIGNALING_E | -1.66798 | 0.015893 | 0.059687 |
| WP_MICRORNAS_IN_CA | -0.73513 | 0.89744 | 1 | WP_MICRORNAS_IN_CARDIOMYOCYTE_HYPERTROPHY | -0.92347 | 0.599804 | 0.963644 | WP_MICRORNAS_IN_CAR | -1.27134 | 0.099874 | 0.259188 |
| BIOCARTA_HDAC_PATH\ | -1.2305 | 0.209221 | 0.844724 | BIOCARTA_HDAC_PATHWAY | -0.92131 | 0.558771 | 0.965243 | BIOCARTA_HDAC_PATHW | -1.56334 | 0.036331 | 0.091573 |
| BIOCARTA_BARRESTIN_ | 1.164254 | 0.273835 | 0.845477 | BIOCARTA_BARRESTIN_SRC_PATHWAY | -0.92557 | 0.547421 | 0.967043 | BIOCARTA_BARRESTIN_SF | -1.29685 | 0.163433 | 0.243708 |
| REACTOME_RHOA_GTP/ | -1.14315 | 0.214624 | 0.937284 | REACTOME_RHOA_GTPASE_CYCLE | 0.966222 | 0.520855 | 0.967614 | REACTOME_RHOA_GTPAS | 0.691707 | 0.967932 | 0.977849 |
| WP_APOPTOSIS | 0.836903 | 0.70583 | 1 | WP_APOPTOSIS | 0.970802 | 0.499388 | 0.968731 | WP_APOPTOSIS | -0.67646 | 0.94343 | 0.949605 |
| REACTOME_RHO_GTPAS | -0.71519 | 0.844098 | 1 | REACTOME_RHO_GTPASES_ACTIVATE_PAKS | -0.92577 | 0.560574 | 0.971267 | REACTOME_RHO_GTPASE | -1.63366 | 0.018499 | 0.068601 |
| REACTOME_TRANSFERR | 1.236244 | 0.205908 | 0.77996 | REACTOME_TRANSFERRIN_ENDOCYTOSIS_AND_RECYC | 0.968192 | 0.506102 | 0.97154 | REACTOME_TRANSFERRIN | -0.81674 | 0.727724 | 0.843381 |
| REACTOME_ORGANELLE_ | -1.02307 | 0.40336 | 1 | REACTOME_ORGANELLE_BIOGENESIS_AND_MAINTEN/ | 0.966238 | 0.524062 | 0.972496 | REACTOME_ORGANELLE_ | -0.89165 | 0.743261 | 0.748519 |
| REACTOME_DNA_REPLI( | -0.93881 | 0.56058 | 1 | REACTOME_DNA_REPLICATION | -0.91661 | 0.622922 | 0.974454 | REACTOME_DNA_REPLICA | -1.5104 | 0.013243 | 0.111287 |
| REACTOME_REGULATIO | -0.78587 | 0.865457 | 1 | REACTOME_REGULATION_OF_TP53_ACTIVITY | -0.89609 | 0.697216 | 0.977975 | REACTOME_REGULATION | 1.001823 | 0.452358 | 0.851536 |
| BIOCARTA_PYK2_PATHW | 1.333528 | 0.128983 | 0.70614 | BIOCARTA_PYK2_PATHWAY | -0.8976 | 0.592396 | 0.978339 | BIOCARTA_PYK2_PATHW/ | -1.22872 | 0.198253 | 0.296366 |
| REACTOME_EPH_EPHRII | 1.228957 | 0.163987 | 0.796274 | REACTOME_EPH_EPHRIN_SIGNALING | -0.89804 | 0.64906 | 0.981586 | REACTOME_EPH_EPHRIN_ | -1.87254 | 6.30E-04 | 0.025136 |
| WP_MAPK_SIGNALING_ | -1.08337 | 0.304378 | 1 | WP_MAPK_SIGNALING_PATHWAY | -0.89868 | 0.716989 | 0.984345 | WP_MAPK_SIGNALING_P/ | -1.67868 | 0 | 0.059603 |
| REACTOME_DETOXIFICA | 1.807249 | 0.004008 | 0.1 | REACTOME_DETOXIFICATION_OF_REACTIVE_OXYGEN_ | -0.90013 | 0.607379 | 0.984812 | REACTOME_DETOXIFICAT | 0.750724 | 0.813608 | 1 |
| REACTOME_AMPLIFICAT | -0.89051 | 0.634482 | 1 | REACTOME_AMPLIFICATION_OF_SIGNAL_FROM_THE_ | -0.90803 | 0.61435 | 0.985288 | REACTOME_AMPLIFICATI | 0.886324 | 0.648195 | 0.964002 |
| REACTOME_RHO_GTPAS | -0.76221 | 0.890767 | 1 | REACTOME_RHO_GTPASES_ACTIVATE_FORMINS | -0.89132 | 0.708271 | 0.986799 | REACTOME_RHO_GTPASE | -0.94907 | 0.570489 | 0.669427 |
| REACTOME_SIGNALING_ | 0.600426 | 0.995454 | 1 | REACTOME_SIGNALING_BY_VEGF | -0.90589 | 0.656446 | 0.986813 | REACTOME_SIGNALING_E | -1.74426 | 7.19E-04 | 0.04225 |
| REACTOME_DEATH_REC | -0.90993 | 0.629792 | 1 | REACTOME_DEATH_RECEPTOR_SIGNALING | 0.957822 | 0.536576 | 0.987589 | REACTOME_DEATH_RECE | -0.81864 | 0.877668 | 0.844154 |
| REACTOME_INTRA_GOL | 1.093633 | 0.323529 | 0.903888 | REACTOME_INTRA_GOLGI_TRAFFIC | 0.950957 | 0.528873 | 0.988271 | REACTOME_INTRA_GOLG | 1.473208 | 0.049266 | 0.223789 |
| WP_WNT_SIGNALING_P | -1.48999 | 0.021545 | 0.646087 | WP_WNT_SIGNALING_PATHWAY_WP539 | 0.955822 | 0.531448 | 0.988578 | WP_WNT_SIGNALING_PA | 0.790422 | 0.819128 | 0.998563 |
| REACTOME_CELL_JUNC1 | -0.63359 | 0.970661 | 1 | REACTOME_CELL_JUNCTION_ORGANIZATION | 0.949087 | 0.534638 | 0.988775 | REACTOME_CELL_JUNCTI | -1.41215 | 0.045909 | 0.152628 |
| REACTOME_CARGO_TR/ | -0.92455 | 0.566562 | 1 | REACTOME_CARGO_TRAFFICKING_TO_THE_PERICILIAF | -0.90845 | 0.591398 | 0.988849 | REACTOME_CARGO_TRAF | -1.11143 | 0.297267 | 0.428352 |
| WP_TOLLLIKE_RECEPTO | -0.73077 | 0.821141 | 1 | WP_TOLLLIKE_RECEPTOR_SIGNALING | -0.90031 | 0.594595 | 0.988982 | WP_TOLLLIKE_RECEPTOR_ | -0.95954 | 0.505125 | 0.647583 |
| BIOCARTA_FCER1_PATH | -0.78789 | 0.7664 | 1 | BIOCARTA_FCER1_PATHWAY | -0.90341 | 0.590972 | 0.989345 | BIOCARTA_FCER1_PATHW | -1.74681 | 7.77E-03 | 0.042003 |
| REACTOME_E3_UBIQUI1 | -0.71635 | 0.868528 | 1 | REACTOME_E3_UBIQUITIN_LIGASES_UBIQUITINATE_T/ | 0.953278 | 0.521308 | 0.991169 | REACTOME_E3_UBIQUITII | -0.85804 | 0.685463 | 0.795423 |
| REACTOME_SIGNALING_ | 0.711184 | 0.866518 | 1 | REACTOME_SIGNALING_BY_SCF_KIT | -0.90103 | 0.608861 | 0.991602 | REACTOME_SIGNALING_E | -1.58553 | 0.028097 | 0.084603 |
| BIOCARTA_PDGF_PATHV | 0.527142 | 0.982475 | 1 | BIOCARTA_PDGF_PATHWAY | -0.90907 | 0.58489 | 0.991855 | BIOCARTA_PDGF_PATHW | -1.66402 | 0.017562 | 0.060167 |
| REACTOME_GLYCOSPHII | 1.57335 | 0.026368 | 0.279259 | REACTOME_GLYCOSPHINGOLIPID_METABOLISM | 0.95134 | 0.537338 | 0.992068 | REACTOME_GLYCOSPHIN( | 1.204262 | 0.214899 | 0.517826 |
| REACTOME_CELL_SURF/ | -0.6987 | 0.928101 | 1 | REACTOME_CELL_SURFACE_INTERACTIONS_AT_THE_V | 0.944961 | 0.538841 | 0.995946 | REACTOME_CELL_SURFAC | -1.2022 | 0.171165 | 0.319016 |
| REACTOME_G2_M_CHE | -1.28437 | 0.107199 | 0.875023 | REACTOME_G2_M_CHECKPOINTS | -0.88359 | 0.697535 | 0.998859 | REACTOME_G2_M_CHECI | -1.50029 | 0.015225 | 0.114845 |
| REACTOME_INSULIN_RE | 1.094706 | 0.342585 | 0.907382 | REACTOME_INSULIN_RECEPTOR_RECYCLING | 0.94216 | 0.534483 | 0.999123 | REACTOME_INSULIN_REC | -0.7928 | 0.755717 | 0.870008 |
| BIOCARTA_BAD_PATHW | 0.56533 | 0.969876 | 1 | BIOCARTA_BAD_PATHWAY | -0.41079 | 0.999206 | 0.999355 | BIOCARTA_BAD_PATHWA | -1.06197 | 0.364997 | 0.496524 |
| REACTOME_ADHERENS_ | 0.814554 | 0.732067 | 1 | REACTOME_ADHERENS_JUNCTIONS_INTERACTIONS | 0.329669 | 1 | 0.999959 | REACTOME_ADHERENS_J | -1.54641 | 0.028757 | 0.097897 |
| REACTOME_FCGAMMA_ | 0.7562 | 0.894885 | 1 | REACTOME_FCGAMMA_RECEPTOR_FCGR_DEPENDENT | -0.59476 | 0.997522 | 1 | REACTOME_FCGAMMA_F | -2.14294 | 0 | 0.006166 |
| REACTOME_GLI3_IS_PR( | -0.74052 | 0.878545 | 1 | REACTOME_GLI3_IS_PROCESSED_TO_GLI3R_BY_THE_F | -0.81029 | 0.805538 | 1 | REACTOME_GLI3_IS_PRO | -1.96759 | 0.00E+00 | 0.017226 |
| REACTOME_THE_ROLE_ | 0.784828 | 0.844309 | 1 | REACTOME_THE_ROLE_OF_GTSE1_IN_G2_M_PROGRE | 0.586109 | 0.988455 | 1 | REACTOME_THE_ROLE_O | -1.94319 | 0 | 0.017643 |
| REACTOME_DEGRADATI | 0.580991 | 0.990544 | 1 | REACTOME_DEGRADATION_OF_DVL | 0.455639 | 0.998692 | 1 | REACTOME_DEGRADATIO | -1.94671 | 0 | 0.017847 |
| REACTOME_REGULATIO | 0.731503 | 0.911214 | 1 | REACTOME_REGULATION_OF_RAS_BY_GAPS | -0.71285 | 0.941869 | 1 | REACTOME_REGULATION | -1.90886 | 6.47E-04 | 0.020957 |
| REACTOME_TCR_SIGNA | -0.65618 | 0.977073 | 1 | REACTOME_TCR_SIGNALING | -0.81509 | 0.855395 | 1 | REACTOME_TCR_SIGNALI | -1.89827 | 0 | 0.022855 |
| REACTOME_DEGRADATI | 0.579513 | 0.996083 | 1 | REACTOME_DEGRADATION_OF_BETA_CATENIN_BY_TI | 0.521602 | 0.997038 | 1 | REACTOME_DEGRADATIO | -1.88797 | 6.50E-04 | 0.023622 |
| REACTOME_HEDGEHOG_ | -0.93776 | 0.565751 | 1 | REACTOME_HEDGEHOG_OFF_STATE | 0.629237 | 0.984198 | 1 | REACTOME_HEDGEHOG_( | -1.88111 | 0 | 0.024527 |
| REACTOME_UBIQUITIN_ | 0.754439 | 0.852265 | 1 | REACTOME_UBIQUITIN_MEDIATED_DEGRADATION_OI | -0.68869 | 0.942105 | 1 | REACTOME_UBIQUITIN_N | -1.86122 | 0.002112 | 0.027307 |
| REACTOME_CDK_MEDI/ | 0.662233 | 0.963943 | 1 | REACTOME_CDK_MEDIATED_PHOSPHORYLATION_ANI | -0.79806 | 0.84575 | 1 | REACTOME_CDK_MEDIAT | -1.80962 | 9.55E-04 | 0.028097 |
| REACTOME_METABOLIS | 0.834734 | 0.721353 | 1 | REACTOME_METABOLISM_OF_POLYAMINES | -0.53162 | 0.99682 | 1 | REACTOME_METABOLISM | -1.819 | 0.002454 | 0.028603 |
| REACTOME_DEGRADATI | 0.714618 | 0.913083 | 1 | REACTOME_DEGRADATION_OF_AXIN | -0.67419 | 0.951709 | 1 | REACTOME_DEGRADATIO | -1.81294 | 0.002114 | 0.02873 |
| REACTOME_CROSS_PRE | 0.625914 | 0.971929 | 1 | REACTOME_CROSS_PRESENTATION_OF_SOLUBLE_EXC | -0.58256 | 0.988633 | 1 | REACTOME_CROSS_PRESI | -1.81363 | 0.002109 | 0.029099 |
| REACTOME_REGULATIO | 0.914599 | 0.603752 | 1 | REACTOME_REGULATION_OF_PTEN_STABILITY_AND_/ | -0.74432 | 0.911225 | 1 | REACTOME_REGULATION | -1.82163 | 9.64E-04 | 0.029104 |
| REACTOME_DOWNSTRE | -0.64807 | 0.976995 | 1 | REACTOME_DOWNSTREAM_TCR_SIGNALING | -0.86999 | 0.728088 | 1 | REACTOME_DOWNSTREA | -1.82329 | 3.34E-04 | 0.029291 |
| REACTOME_COSTIMULA | 1.070018 | 0.345083 | 0.946582 | REACTOME_COSTIMULATION_BY_THE_CD28_FAMILY | 0.442955 | 0.998497 | 1 | REACTOME_COSTIMULAT | -1.80361 | 0.001718 | 0.029299 |
| REACTOME_RHO_GTPAS | 0.700915 | 0.881548 | 1 | REACTOME_RHO_GTPASES_ACTIVATE_IQGAPS | 0.887917 | 0.6164 | 1 | REACTOME_RHO_GTPASE | -1.80023 | 0.002912 | 0.029758 |
| REACTOME_DECTIN_1_I | 0.665027 | 0.962634 | 1 | REACTOME_DECTIN_1_MEDIATED_NONCANONICAL_N | -0.75648 | 0.882372 | 1 | REACTOME_DECTIN_1_M | -1.79663 | 9.28E-04 | 0.030173 |
| REACTOME_HEDGEHOG | -0.61378 | 0.981028 | 1 | REACTOME_HEDGEHOG_ON_STATE | -0.53421 | 0.99935 | 1 | REACTOME_HEDGEHOG_( | -1.84279 | 9.53E-04 | 0.030266 |
| REACTOME_SMOOTH_N | -1.14688 | 0.28045 | 0.951772 | REACTOME_SMOOTH_MUSCLE_CONTRACTION | -0.66731 | 0.919059 | 1 | REACTOME_SMOOTH_MU | -1.82904 | 0.003957 | 0.030372 |
| REACTOME_CD28_CO_S | 0.865378 | 0.651007 | 1 | REACTOME_CD28_CO_STIMULATION | -0.56971 | 0.971823 | 1 | REACTOME_CD28_CO_ST | -1.83126 | 0.004088 | 0.030426 |
| REACTOME_SCF_SKP2_I | -0.62785 | 0.973209 | 1 | REACTOME_SCF_SKP2_MEDIATED_DEGRADATION_OF_ | -0.78396 | 0.839577 | 1 | REACTOME_SCF_SKP2_M | -1.78866 | 0.002124 | 0.031457 |
| REACTOME_MAPK6_MA | -0.63715 | 0.973994 | 1 | REACTOME_MAPK6_MAPK4_SIGNALING | -0.78614 | 0.857093 | 1 | REACTOME_MAPK6_MAP | -1.78454 | 0.001984 | 0.032068 |
| REACTOME_AUF1_HNRI | -0.62043 | 0.974219 | 1 | REACTOME_AUF1_HNRNP_D0_BINDS_AND_DESTABILI | -0.7584 | 0.880705 | 1 | REACTOME_AUF1_HNRNI | -1.76317 | 0.002781 | 0.03757 |
| REACTOME_APC_C_CDH | 0.614594 | 0.987591 | 1 | REACTOME_APC_C_CDH1_MEDIATED_DEGRADATION_ | -0.73813 | 0.923327 | 1 | REACTOME_APC_C_CDH1 | -1.73723 | 0.002247 | 0.042874 |
| WP_REGULATION_OF_A | 0.519717 | 1 | 1 | WP_REGULATION_OF_ACTIN_CYTOSKELETON | -0.87796 | 0.756779 | 1 | WP_REGULATION_OF_AC | -1.71951 | 4.02E-04 | 0.048493 |
| REACTOME_EPHRIN_SIG | 0.743133 | 0.810764 | 1 | REACTOME_EPHRIN_SIGNALING | -0.79252 | 0.750395 | 1 | REACTOME_EPHRIN_SIGN | -1.71099 | 0.015597 | 0.04926 |
| REACTOME_DEVELOPM | 0.947953 | 0.612894 | 1 | REACTOME_DEVELOPMENTAL_BIOLOGY | -0.69953 | 1 | 1 | REACTOME_DEVELOPMEI | -1.71302 | 0 | 0.049792 |
| REACTOME_ACTIVATION | -0.56922 | 0.993485 | 1 | REACTOME_ACTIVATION_OF_APC_C_AND_APC_C_CD( | -0.66634 | 0.968499 | 1 | REACTOME_ACTIVATION_ | -1.6964 | 0.002985 | 0.054167 |
| REACTOME_APC_C_MEI | -0.68952 | 0.942008 | 1 | REACTOME_APC_C_MEDIATED_DEGRADATION_OF_CE | -0.81424 | 0.815586 | 1 | REACTOME_APC_C_MEDI | -1.69034 | 0.002615 | 0.056254 |
| REACTOME_ASYMMETR | 0.665397 | 0.955146 | 1 | REACTOME_ASYMMETRIC_LOCALIZATION_OF_PCP_PR | -0.65949 | 0.971456 | 1 | REACTOME_ASYMMETRIC | -1.68591 | 0.005789 | 0.056822 |
| REACTOME_FCERI_MED | -0.88582 | 0.657294 | 1 | REACTOME_FCERI_MEDIATED_NF_KB_ACTIVATION | -0.82462 | 0.80068 | 1 | REACTOME_FCERI_MEDIA | -1.6776 | 0.002659 | 0.059384 |
| REACTOME_NERVOUS_! | 1.071644 | 0.271598 | 0.947872 | REACTOME_NERVOUS_SYSTEM_DEVELOPMENT | -0.6899 | 1 | 1 | REACTOME_NERVOUS_SY | -1.66914 | 0 | 0.059842 |
| REACTOME_MAPK_FAM | 0.613591 | 0.999735 | 1 | REACTOME_MAPK_FAMILY_SIGNALING_CASCADES | -0.81823 | 0.955556 | 1 | REACTOME_MAPK_FAMIL | -1.67336 | 0 | 0.05902 |
| REACTOME_DOPAMINE_ | 0.593153 | 0.953368 | 1 | REACTOME_DOPAMINE_NEUROTRANSMITTER_RELEAS | -0.78959 | 0.77055 | 1 | REACTOME_DOPAMINE_I | -1.6615 | 0.016698 | 0.05997 |
| REACTOME_NEGATIVE_ | -0.92101 | 0.56547 | 1 | REACTOME_NEGATIVE_REGULATION_OF_MAPK_PATH | -0.84779 | 0.703107 | 1 | REACTOME_NEGATIVE_RI | -1.67432 | 0.009796 | 0.060194 |
| REACTOME_STABILIZATI | 0.628983 | 0.970627 | 1 | REACTOME_STABILIZATION_OF_P53 | -0.57418 | 0.994138 | 1 | REACTOME_STABILIZATIO | -1.65519 | 0.004103 | 0.061569 |
| REACTOME_C_TYPE_LE( | 0.64354 | 0.987843 | 1 | REACTOME_C_TYPE_LECTIN_RECEPTORS_CLRS | -0.80695 | 0.883137 | 1 | REACTOME_C_TYPE_LECT | -1.65277 | 0.003712 | 0.061997 |
| REACTOME_ORC1_REM( | 0.605 | 0.980073 | 1 | REACTOME_ORC1_REMOVAL_FROM_CHROMATIN | -0.73437 | 0.901219 | 1 | REACTOME_ORC1_REMO' | -1.61879 | 0.011163 | 0.074498 |
| WP_EGFR1_SIGNALING_ | -0.85067 | 0.78002 | 1 | WP_EGFR1_SIGNALING_PATHWAY | -0.67877 | 0.996244 | 1 | WP_EGFR1_SIGNALING_P | -1.6082 | 0.00176 | 0.075306 |
| REACTOME_GLYOXYLAT | -0.74104 | 0.82056 | 1 | REACTOME_GLYOXYLATE_METABOLISM_AND_GLYCIN | 0.73573 | 0.818138 | 1 | REACTOME_GLYOXYLATE_ | -1.61004 | 0.024632 | 0.075741 |
| REACTOME_SWITCHING | -0.5573 | 0.994685 | 1 | REACTOME_SWITCHING_OF_ORIGINS_TO_A_POST_RE | -0.82065 | 0.806136 | 1 | REACTOME_SWITCHING_( | -1.59467 | 0.010161 | 0.081366 |
| BIOCARTA_MET_PATHW | 0.766321 | 0.803481 | 1 | BIOCARTA_MET_PATHWAY | -0.68317 | 0.90582 | 1 | BIOCARTA_MET_PATHWA | -1.59009 | 0.021225 | 0.083366 |
| BIOCARTA_ECM_PATHW | 0.651364 | 0.905551 | 1 | BIOCARTA_ECM_PATHWAY | -0.68014 | 0.880115 | 1 | BIOCARTA_ECM_PATHWA | -1.57517 | 0.032504 | 0.086543 |
| REACTOME_REGULATIO | -0.48192 | 0.990304 | 1 | REACTOME_REGULATION_OF_SIGNALING_BY_CBL | 0.795444 | 0.727185 | 1 | REACTOME_REGULATION | -1.57004 | 0.039813 | 0.088926 |
| REACTOME_TNFR2_NOI | 0.685843 | 0.952392 | 1 | REACTOME_TNFR2_NON_CANONICAL_NF_KB_PATHW | -0.6083 | 0.986243 | 1 | REACTOME_TNFR2_NON_ | -1.56632 | 0.01039 | 0.090452 |
| REACTOME_NUCLEAR_E | 0.68217 | 0.940292 | 1 | REACTOME_NUCLEAR_EVENTS_MEDIATED_BY_NFE2L2 | -0.60991 | 0.985471 | 1 | REACTOME_NUCLEAR_EV | -1.55353 | 0.015755 | 0.095452 |
| BIOCARTA_GH_PATHW/ | 0.836785 | 0.677061 | 1 | BIOCARTA_GH_PATHWAY | -0.61229 | 0.944134 | 1 | BIOCARTA_GH_PATHWAY | -1.54833 | 0.038711 | 0.098174 |
| REACTOME_G1_S_DNA_ | 0.684266 | 0.939291 | 1 | REACTOME_G1_S_DNA_DAMAGE_CHECKPOINTS | 0.49518 | 0.996347 | 1 | REACTOME_G1_S_DNA_D | -1.54367 | 0.019051 | 0.098919 |
| REACTOME_DAP12_SIGI | 0.99906 | 0.448067 | 1 | REACTOME_DAP12_SIGNALING | 0.607444 | 0.934359 | 1 | REACTOME_DAP12_SIGN/ | -1.5384 | 0.047656 | 0.100878 |
| REACTOME_AGGREPHA( | -0.98551 | 0.474301 | 1 | REACTOME_AGGREPHAGY | 0.764758 | 0.801604 | 1 | REACTOME_AGGREPHAGY | -1.53011 | 0.032846 | 0.105044 |
| REACTOME_KSRP_KHSR | 1.339832 | 0.140373 | 0.724816 | REACTOME_KSRP_KHSRP_BINDS_AND_DESTABILIZES_I | 0.810061 | 0.699705 | 1 | REACTOME_KSRP_KHSRP_ | 1.605728 | 0.027117 | 0.106779 |
| REACTOME_MITOCHON | -0.73271 | 0.812535 | 1 | REACTOME_MITOCHONDRIAL_CALCIUM_ION_TRANSP | -0.75474 | 0.796831 | 1 | REACTOME_MITOCHOND | -1.52203 | 0.051263 | 0.109134 |
| REACTOME_HSP90_CHA | -0.62347 | 0.971268 | 1 | REACTOME_HSP90_CHAPERONE_CYCLE_FOR_STEROIE | 0.668881 | 0.927939 | 1 | REACTOME_HSP90_CHAP | -1.51408 | 0.023072 | 0.110912 |

| | | | | | | | | | | | |
|---|---|---|---|---|---|---|---|---|---|---|---|
| REACTOME_GABA_SYNT | -0.66681 | 0.894592 | 1 | REACTOME_GABA_SYNTHESIS_RELEASE_REUPTAKE_AI | -0.87133 | 0.632212 | 1 | REACTOME_GABA_SYNTH | -1.51134 | 0.050362 | 0.11131 |
| REACTOME_CELL_CELL_ | 0.684072 | 0.915698 | 1 | REACTOME_CELL_CELL_JUNCTION_ORGANIZATION | 0.606299 | 0.958683 | 1 | REACTOME_CELL_CELL_JU | -1.51199 | 0.034597 | 0.111608 |
| BIOCARTA_BCR_PATHW | 1.137683 | 0.288996 | 0.867215 | BIOCARTA_BCR_PATHWAY | -0.84167 | 0.679195 | 1 | BIOCARTA_BCR_PATHWA | -1.50002 | 0.052508 | 0.114267 |
| BIOCARTA_IGF1R_PATH | 0.49427 | 0.988233 | 1 | BIOCARTA_IGF1R_PATHWAY | -0.56228 | 0.970233 | 1 | BIOCARTA_IGF1R_PATHW | -1.50086 | 0.048046 | 0.115185 |
| REACTOME_CTLA4_INHI | 1.272685 | 0.175242 | 0.754684 | REACTOME_CTLA4_INHIBITORY_SIGNALING | 0.412238 | 0.997093 | 1 | REACTOME_CTLA4_INHIB | -1.48453 | 0.056136 | 0.116948 |
| REACTOME_DAP12_INT | 1.06458 | 0.367064 | 0.959323 | REACTOME_DAP12_INTERACTIONS | 0.545814 | 0.96601 | 1 | REACTOME_DAP12_INTEF | -1.49002 | 0.060861 | 0.117156 |
| REACTOME_DNA_REPLI | 0.673755 | 0.96342 | 1 | REACTOME_DNA_REPLICATION_PRE_INITIATION | -0.70858 | 0.945946 | 1 | REACTOME_DNA_REPLICA | -1.48216 | 0.020649 | 0.117226 |
| WP_WNT_SIGNALING_P | -1.29395 | 0.14398 | 0.848239 | WP_WNT_SIGNALING_PATHWAY_WP403 | 0.665778 | 0.906304 | 1 | WP_WNT_SIGNALING_PA | -1.49061 | 0.041506 | 0.117492 |
| REACTOME_UCH_PROTI | 0.798368 | 0.832137 | 1 | REACTOME_UCH_PROTEINASES | -0.86736 | 0.719429 | 1 | REACTOME_UCH_PROTEI | -1.48665 | 0.019351 | 0.117552 |
| REACTOME_CYCLIN_A_ | 0.573978 | 0.994059 | 1 | REACTOME_CYCLIN_A_CDK2_ASSOCIATED_EVENTS_AT | 0.650607 | 0.95471 | 1 | REACTOME_CYCLIN_A_CD | -1.48463 | 0.027328 | 0.117561 |
| REACTOME_CLATHRIN_ | -0.97711 | 0.501618 | 1 | REACTOME_CLATHRIN_MEDIATED_ENDOCYTOSIS | 0.854705 | 0.738632 | 1 | REACTOME_CLATHRIN_M | -1.49238 | 0.007323 | 0.117656 |
| REACTOME_REGULATIO | 0.914543 | 0.583446 | 1 | REACTOME_REGULATION_OF_RUNX3_EXPRESSION_AN | -0.57435 | 0.992671 | 1 | REACTOME_REGULATION | -1.48253 | 0.030812 | 0.117684 |
| REACTOME_GAP_JUNCT | 0.64946 | 0.937734 | 1 | REACTOME_GAP_JUNCTION_TRAFFICKING_AND_REGU | 0.750063 | 0.81832 | 1 | REACTOME_GAP_JUNCTI | -1.49396 | 0.038835 | 0.11807 |
| REACTOME_RECOGNITI | -1.45863 | 0.070707 | 0.620041 | REACTOME_RECOGNITION_OF_DNA_DAMAGE_BY_PC | 0.837074 | 0.678613 | 1 | REACTOME_RECOGNITIO | -1.49256 | 0.062045 | 0.118291 |
| REACTOME_TRANSCRIP | -0.78041 | 0.827955 | 1 | REACTOME_TRANSCRIPTIONAL_REGULATION_BY_RUN | -0.58654 | 0.989974 | 1 | REACTOME_TRANSCRIPTI | -1.4784 | 0.02474 | 0.118506 |
| REACTOME_SPHINGOLI | 1.789229 | 0.001842 | 0.110671 | REACTOME_SPHINGOLIPID_METABOLISM | 0.823081 | 0.75104 | 1 | REACTOME_SPHINGOLIPI | 1.579735 | 0.013668 | 0.119143 |
| REACTOME_EGFR_DOW | -1.01579 | 0.435305 | 1 | REACTOME_EGFR_DOWNREGULATION | -0.85408 | 0.667555 | 1 | REACTOME_EGFR_DOWN | -1.47405 | 0.058421 | 0.120298 |
| REACTOME_CA_DEPENE | 0.569715 | 0.971466 | 1 | REACTOME_CA_DEPENDENT_EVENTS | -0.88386 | 0.628664 | 1 | REACTOME_CA_DEPENDE | -1.47437 | 0.053905 | 0.12075 |
| REACTOME_GAP_JUNCT | -0.76857 | 0.778661 | 1 | REACTOME_GAP_JUNCTION_ASSEMBLY | 0.901167 | 0.595406 | 1 | REACTOME_GAP_JUNCTI | -1.4669 | 0.064541 | 0.124281 |
| REACTOME_RUNX1_REC | 0.614079 | 0.983838 | 1 | REACTOME_RUNX1_REGULATES_TRANSCRIPTION_OF_ | -0.61886 | 0.984355 | 1 | REACTOME_RUNX1_REGU | -1.46773 | 0.027239 | 0.124372 |
| REACTOME_CELL_CELL_ | -0.5927 | 0.993029 | 1 | REACTOME_CELL_CELL_COMMUNICATION | 0.769424 | 0.843636 | 1 | REACTOME_CELL_CELL_C | -1.4644 | 0.026316 | 0.125441 |
| REACTOME_ASSEMBLY_ | 0.678233 | 0.958219 | 1 | REACTOME_ASSEMBLY_OF_THE_PRE_REPLICATIVE_CC | -0.69541 | 0.956317 | 1 | REACTOME_ASSEMBLY_O | -1.46003 | 0.028031 | 0.128065 |
| REACTOME_CARGO_REC | -1.23226 | 0.141955 | 0.849235 | REACTOME_CARGO_RECOGNITION_FOR_CLATHRIN_M | 0.892544 | 0.648764 | 1 | REACTOME_CARGO_RECC | -1.45592 | 0.021862 | 0.129025 |
| REACTOME_TOLL_LIKE_ | -1.33729 | 0.062468 | 0.791833 | REACTOME_TOLL_LIKE_RECEPTOR_CASCADES | 0.750835 | 0.895671 | 1 | REACTOME_TOLL_LIKE_R | -1.45139 | 0.018304 | 0.131907 |
| WP_INSULIN_SIGNALIN | -0.97756 | 0.493233 | 1 | WP_INSULIN_SIGNALING | 0.785295 | 0.872812 | 1 | WP_INSULIN_SIGNALING | -1.44001 | 0.006748 | 0.137256 |
| WP_ALPHA_6_BETA_4_ | -0.59087 | 0.987595 | 1 | WP_ALPHA_6_BETA_4_INTEGRIN_SIGNALING_PATHW | 0.927647 | 0.570634 | 1 | WP_ALPHA_6_BETA_4_IN | -1.42814 | 0.03947 | 0.144096 |
| REACTOME_MACROAUT | -0.82333 | 0.816569 | 1 | REACTOME_MACROAUTOPHAGY | -0.81135 | 0.880677 | 1 | REACTOME_MACROAUTC | -1.42982 | 0.013857 | 0.14424 |
| REACTOME_INTRACELLU | -0.63652 | 0.999013 | 1 | REACTOME_INTRACELLULAR_SIGNALING_BY_SECOND_ | -0.83918 | 0.898951 | 1 | REACTOME_INTRACELLUL | -1.42328 | 0.003595 | 0.14675 |
| REACTOME_GPVI_MEDI | -1.06387 | 0.375231 | 1 | REACTOME_GPVI_MEDIATED_ACTIVATION_CASCADE | 0.645463 | 0.915936 | 1 | REACTOME_GPVI_MEDIAT | -1.41378 | 0.078775 | 0.15192 |
| BIOCARTA_AT1R_PATHW | 1.342265 | 0.127562 | 0.730257 | BIOCARTA_AT1R_PATHWAY | -0.82097 | 0.714133 | 1 | BIOCARTA_AT1R_PATHW | -1.41542 | 0.088974 | 0.152094 |
| REACTOME_MAP2K_AN | 0.873013 | 0.652781 | 1 | REACTOME_MAP2K_AND_MAPK_ACTIVATION | 0.667654 | 0.917633 | 1 | REACTOME_MAP2K_AND | -1.40707 | 0.06302 | 0.156464 |
| REACTOME_TRANSCRIP | 0.766211 | 0.859553 | 1 | REACTOME_TRANSCRIPTIONAL_REGULATION_BY_RUN | -0.64861 | 0.976509 | 1 | REACTOME_TRANSCRIPTI | -1.39386 | 0.049621 | 0.165416 |
| REACTOME_NEUTROPH | 1.062684 | 0.270559 | 0.951827 | REACTOME_NEUTROPHIL_DEGRANULATION | -0.82512 | 0.97343 | 1 | REACTOME_NEUTROPHIL | -1.38858 | 7.53E-04 | 0.168596 |
| REACTOME_INTERLEUKI | -0.5653 | 0.973116 | 1 | REACTOME_INTERLEUKIN_3_INTERLEUKIN_5_AND_GM | 0.859028 | 0.660398 | 1 | REACTOME_INTERLEUKIN | -1.3888 | 0.077304 | 0.169252 |
| REACTOME_PTEN_REGU | -0.63259 | 0.992353 | 1 | REACTOME_PTEN_REGULATION | -0.81438 | 0.857972 | 1 | REACTOME_PTEN_REGUL | -1.38115 | 0.027499 | 0.174951 |
| REACTOME_G_BETA_GA | -0.81052 | 0.71282 | 1 | REACTOME_G_BETA_GAMMA_SIGNALLING_THROUGH | -0.78814 | 0.749611 | 1 | REACTOME_G_BETA_GAN | -1.3743 | 0.115375 | 0.180898 |
| REACTOME_PLATELET_ | 0.968173 | 0.535961 | 1 | REACTOME_PLATELET_ACTIVATION_SIGNALING_AND_ | 0.736904 | 0.955107 | 1 | REACTOME_PLATELET_AC | -1.36209 | 0.013103 | 0.190874 |
| REACTOME_SIGNALING_ | 0.782168 | 0.980239 | 1 | REACTOME_SIGNALING_BY_RECEPTOR_TYROSINE_KIN | -0.80444 | 0.983199 | 1 | REACTOME_SIGNALING_E | -1.35642 | 0.001314 | 0.195765 |
| REACTOME_HEMOSTAS | 0.832615 | 0.951269 | 1 | REACTOME_HEMOSTASIS | 0.840995 | 0.855465 | 1 | REACTOME_HEMOSTASIS | -1.34943 | 0.009076 | 0.200553 |
| REACTOME_RECEPTOR_ | 0.90576 | 0.582005 | 1 | REACTOME_RECEPTOR_TYPE_TYROSINE_PROTEIN_PH | -0.59942 | 0.952919 | 1 | REACTOME_RECEPTOR_T | -1.3469 | 0.115855 | 0.201376 |
| REACTOME_INTERLEUKI | -0.74928 | 0.907761 | 1 | REACTOME_INTERLEUKIN_1_FAMILY_SIGNALING | 0.669574 | 0.962733 | 1 | REACTOME_INTERLEUKIN | -1.33972 | 0.052971 | 0.206322 |
| WP_SPLICING_FACTOR_ | -1.24716 | 0.158781 | 0.862187 | WP_SPLICING_FACTOR_NOVA_REGULATED_SYNAPTIC_ | 0.926437 | 0.577155 | 1 | WP_SPLICING_FACTOR_N | -1.33638 | 0.088367 | 0.209161 |
| REACTOME_MHC_CLAS | 1.135414 | 0.227186 | 0.867303 | REACTOME_MHC_CLASS_II_ANTIGEN_PRESENTATION | 0.670607 | 0.964542 | 1 | REACTOME_MHC_CLASS_ | -1.33266 | 0.046857 | 0.212383 |
| REACTOME_DISASSEMB | -0.72615 | 0.833821 | 1 | REACTOME_DISASSEMBLY_OF_THE_DESTRUCTION_CC | 0.781509 | 0.755427 | 1 | REACTOME_DISASSEMBLY | -1.3228 | 0.123692 | 0.221738 |
| REACTOME_SIGNALING_ | 0.73016 | 0.858029 | 1 | REACTOME_SIGNALING_BY_ERBB2 | -0.74582 | 0.853558 | 1 | REACTOME_SIGNALING_E | -1.31784 | 0.110371 | 0.225517 |
| REACTOME_CARBOXYTE | -0.67842 | 0.870865 | 1 | REACTOME_CARBOXYTERMINAL_POST_TRANSLATION | 0.859187 | 0.653115 | 1 | REACTOME_CARBOXYTER | -1.3168 | 0.143072 | 0.225665 |
| REACTOME_SELECTIVE_ | -0.74707 | 0.879316 | 1 | REACTOME_SELECTIVE_AUTOPHAGY | 0.795832 | 0.787191 | 1 | REACTOME_SELECTIVE_A | -1.3128 | 0.081486 | 0.227291 |
| REACTOME_SIGNALING_ | -0.63662 | 0.99154 | 1 | REACTOME_SIGNALING_BY_HEDGEHOG | 0.728318 | 0.929529 | 1 | REACTOME_SIGNALING_E | -1.31157 | 0.045098 | 0.227763 |
| REACTOME_RIPK1_MED | 1.260382 | 0.178185 | 0.744963 | REACTOME_RIPK1_MEDIATED_REGULATED_NECROSIS | 0.854362 | 0.654911 | 1 | REACTOME_RIPK1_MEDI | -1.29647 | 0.148659 | 0.243089 |
| REACTOME_NETRIN_1_ | -0.49617 | 0.985518 | 1 | REACTOME_NETRIN_1_SIGNALING | 0.543622 | 0.969653 | 1 | REACTOME_NETRIN_1_SI | -1.29209 | 0.159216 | 0.246548 |
| BIOCARTA_EIF4_PATHW | 0.86008 | 0.637058 | 1 | BIOCARTA_EIF4_PATHWAY | -0.82783 | 0.697137 | 1 | BIOCARTA_EIF4_PATHWA | -1.28804 | 0.155757 | 0.250624 |
| REACTOME_INTEGRIN_ | 0.470785 | 0.997521 | 0.999348 | REACTOME_INTEGRIN_SIGNALING | 0.777168 | 0.775343 | 1 | REACTOME_INTEGRIN_SI | -1.2822 | 0.138777 | 0.251657 |
| BIOCARTA_RAS_PATHW | 0.638994 | 0.917236 | 1 | BIOCARTA_RAS_PATHWAY | -0.80089 | 0.734815 | 1 | BIOCARTA_RAS_PATHWA | -1.28055 | 0.166323 | 0.252664 |
| BIOCARTA_IL2RB_PATH | 0.709064 | 0.858305 | 1 | BIOCARTA_IL2RB_PATHWAY | 0.440472 | 0.995531 | 1 | BIOCARTA_IL2RB_PATHW | -1.27485 | 0.160582 | 0.257757 |
| REACTOME_G1_S_TRAN | 0.752272 | 0.885306 | 1 | REACTOME_G1_S_TRANSITION | 0.750935 | 0.859915 | 1 | REACTOME_G1_S_TRANS | -1.27153 | 0.092665 | 0.260015 |
| BIOCARTA_NFAT_PATH | 0.958494 | 0.57823 | 1 | BIOCARTA_NFAT_PATHWAY | 0.737781 | 0.836625 | 1 | BIOCARTA_NFAT_PATHW | -1.2718 | 0.145297 | 0.260722 |
| BIOCARTA_RAC1_PATH | -0.77187 | 0.769849 | 1 | BIOCARTA_RAC1_PATHWAY | -0.6124 | 0.940724 | 1 | BIOCARTA_RAC1_PATHW | -1.26683 | 0.175032 | 0.262942 |
| WP_IL7_SIGNALING_PA | -0.87124 | 0.656966 | 1 | WP_IL7_SIGNALING_PATHWAY | -0.67712 | 0.92994 | 1 | WP_IL7_SIGNALING_PATH | -1.2669 | 0.143292 | 0.263936 |
| WP_PROTEASOME_DEG | 0.958935 | 0.50624 | 1 | WP_PROTEASOME_DEGRADATION | 0.555621 | 0.987977 | 1 | WP_PROTEASOME_DEGR | -1.26396 | 0.124103 | 0.265583 |
| WP_EPO_RECEPTOR_SI | 0.570185 | 0.966874 | 1 | WP_EPO_RECEPTOR_SIGNALING | -0.44836 | 0.996841 | 1 | WP_EPO_RECEPTOR_SIGN | -1.25584 | 0.177081 | 0.271018 |
| REACTOME_SIGNALLING | 0.712369 | 0.857955 | 1 | REACTOME_SIGNALLING_TO_ERKS | -0.77098 | 0.787177 | 1 | REACTOME_SIGNALLING_ | -1.25625 | 0.179144 | 0.271584 |
| REACTOME_CELLULAR_ | 0.919685 | 0.58584 | 1 | REACTOME_CELLULAR_RESPONSE_TO_HYPOXIA | 0.817333 | 0.749495 | 1 | REACTOME_CELLULAR_RE | -1.25276 | 0.126679 | 0.272965 |
| WP_INTEGRINMEDIATEI | -0.7063 | 0.945404 | 1 | WP_INTEGRINMEDIATED_CELL_ADHESION | 0.748784 | 0.879053 | 1 | WP_INTEGRINMEDIATED_ | -1.25325 | 0.098712 | 0.273402 |
| REACTOME_ACTIVATION | 0.850026 | 0.662136 | 1 | REACTOME_ACTIVATION_OF_BH3_ONLY_PROTEINS | -0.84603 | 0.662214 | 1 | REACTOME_ACTIVATION_ | -1.24388 | 0.199289 | 0.280694 |
| REACTOME_PKMTS_ME | -1.43903 | 0.07211 | 0.679156 | REACTOME_PKMTS_METHYLATE_HISTONE_LYSINES | 0.886685 | 0.609186 | 1 | REACTOME_PKMTS_METI | 1.406814 | 0.088662 | 0.282338 |
| REACTOME_SIGNAL_TR | 0.998654 | 0.452941 | 1 | REACTOME_SIGNAL_TRANSDUCTION_BY_L1 | -0.70857 | 0.861192 | 1 | REACTOME_SIGNAL_TRAI | -1.24056 | 0.1946 | 0.284204 |
| WP_FIBRIN_COMPLEME | -1.1435 | 0.287749 | 0.945639 | WP_FIBRIN_COMPLEMENT_RECEPTOR_3_SIGNALING_ | 0.775005 | 0.777971 | 1 | WP_FIBRIN_COMPLEMEN | -1.23414 | 0.18667 | 0.290969 |
| BIOCARTA_GLEEVEC_PA | 0.702573 | 0.850109 | 1 | BIOCARTA_GLEEVEC_PATHWAY | -0.50433 | 0.986975 | 1 | BIOCARTA_GLEEVEC_PATI | -1.22701 | 0.203145 | 0.297658 |
| BIOCARTA_TNFR1_PATH | -0.82296 | 0.70808 | 1 | BIOCARTA_TNFR1_PATHWAY | -0.83261 | 0.694518 | 1 | BIOCARTA_TNFR1_PATHW | -1.22516 | 0.19746 | 0.298061 |
| REACTOME_RAF_ACTIV | 0.745501 | 0.842094 | 1 | REACTOME_RAF_ACTIVATION | -0.81168 | 0.766589 | 1 | REACTOME_RAF_ACTIVAT | -1.22587 | 0.179466 | 0.298174 |
| REACTOME_S_PHASE | -1.03929 | 0.381941 | 1 | REACTOME_S_PHASE | 0.774958 | 0.852129 | 1 | REACTOME_S_PHASE | -1.21799 | 0.124367 | 0.306268 |
| REACTOME_SIGNALING_ | 0.612224 | 0.981153 | 1 | REACTOME_SIGNALING_BY_MET | 0.75943 | 0.830641 | 1 | REACTOME_SIGNALING_E | -1.2148 | 0.156467 | 0.308552 |
| REACTOME_SIGNALING_ | -0.8983 | 0.798958 | 1 | REACTOME_SIGNALING_BY_RHO_GTPASES_MIRO_GTF | -0.80663 | 1 | 1 | REACTOME_SIGNALING_E | -1.20944 | 0.015625 | 0.314059 |
| BIOCARTA_KERATINOCY | 0.945053 | 0.529711 | 1 | BIOCARTA_KERATINOCYTE_PATHWAY | 0.530353 | 0.984497 | 1 | BIOCARTA_KERATINOCYTI | -1.2075 | 0.196345 | 0.314684 |
| REACTOME_SUMOYLATI | -1.08181 | 0.350574 | 1 | REACTOME_SUMOYLATION_OF_TRANSCRIPTION_COF | 0.761484 | 0.782167 | 1 | REACTOME_SUMOYLATIC | 1.369061 | 0.102482 | 0.318387 |
| REACTOME_P75_NTR_R | -1.07036 | 0.339626 | 1 | REACTOME_P75_NTR_RECEPTOR_MEDIATED_SIGNALL | 0.805694 | 0.777474 | 1 | REACTOME_P75_NTR_RE | -1.19819 | 0.164155 | 0.321549 |
| REACTOME_UB_SPECIFI | 0.974481 | 0.512989 | 1 | REACTOME_UB_SPECIFIC_PROCESSING_PROTEASES | -0.78711 | 0.920974 | 1 | REACTOME_UB_SPECIFIC_ | -1.1924 | 0.129412 | 0.329163 |
| WP_MYOMETRIAL_REL | -0.98713 | 0.471801 | 1 | WP_MYOMETRIAL_RELAXATION_AND_CONTRACTION_ | -0.77214 | 0.928829 | 1 | WP_MYOMETRIAL_RELAX | -1.1914 | 0.129082 | 0.329553 |
| REACTOME_NEGATIVE_ | 0.701313 | 0.940124 | 1 | REACTOME_NEGATIVE_REGULATION_OF_THE_PI3K_AI | -0.76239 | 0.895882 | 1 | REACTOME_NEGATIVE_RI | -1.18993 | 0.160782 | 0.330576 |
| REACTOME_RET_SIGNAI | -0.71855 | 0.86075 | 1 | REACTOME_RET_SIGNALING | 0.620656 | 0.941726 | 1 | REACTOME_RET_SIGNALI | -1.18304 | 0.224292 | 0.340049 |
| REACTOME_RHOU_GTP | 1.167351 | 0.240749 | 0.851422 | REACTOME_RHOU_GTPASE_CYCLE | -0.73218 | 0.882845 | 1 | REACTOME_RHOU_GTPAS | -1.17819 | 0.209296 | 0.346426 |
| REACTOME_PURINE_CA | 0.870606 | 0.624702 | 1 | REACTOME_PURINE_CATABOLISM | 0.766066 | 0.770551 | 1 | REACTOME_PURINE_CAT | -1.17268 | 0.254495 | 0.352919 |
| REACTOME_NEDDYLATI | -0.75528 | 0.953701 | 1 | REACTOME_NEDDYLATION | 0.891965 | 0.688273 | 1 | REACTOME_NEDDYLATIO | -1.17302 | 0.118365 | 0.353608 |
| REACTOME_SEALING_O | -0.84464 | 0.683593 | 1 | REACTOME_SEALING_OF_THE_NUCLEAR_ENVELOPE_N | 0.77964 | 0.765056 | 1 | REACTOME_SEALING_OF_ | -1.16477 | 0.248087 | 0.360751 |
| REACTOME_RHOV_GTP | 0.885422 | 0.626056 | 1 | REACTOME_RHOV_GTPASE_CYCLE | -0.67703 | 0.935977 | 1 | REACTOME_RHOV_GTPAS | -1.16532 | 0.227066 | 0.361082 |
| REACTOME_IRON_UPTA | 1.106731 | 0.303491 | 0.893893 | REACTOME_IRON_UPTAKE_AND_TRANSPORT | 0.690109 | 0.89441 | 1 | REACTOME_IRON_UPTAK | -1.16126 | 0.227418 | 0.36387 |
| BIOCARTA_CREB_PATHW | 1.13398 | 0.290801 | 0.864597 | BIOCARTA_CREB_PATHWAY | -0.58371 | 0.963554 | 1 | BIOCARTA_CREB_PATHW | -1.15848 | 0.264652 | 0.366052 |
| REACTOME_SEPARATIO | -0.80775 | 0.872026 | 1 | REACTOME_SEPARATION_OF_SISTER_CHROMATIDS | -0.81216 | 0.904389 | 1 | REACTOME_SEPARATION_ | -1.15313 | 0.163039 | 0.373592 |
| REACTOME_MITOTIC_G | 0.607267 | 0.994166 | 1 | REACTOME_MITOTIC_G1_PHASE_AND_G1_S_TRANSIT | 0.810749 | 0.789122 | 1 | REACTOME_MITOTIC_G1_ | -1.14518 | 0.212204 | 0.383175 |
| REACTOME_KEAP1_NFE | 0.759897 | 0.884518 | 1 | REACTOME_KEAP1_NFE2L2_PATHWAY | 0.882597 | 0.652584 | 1 | REACTOME_KEAP1_NFE2I | -1.14592 | 0.203395 | 0.38444 |
| REACTOME_O_LINKED_ | 1.470378 | 0.07101 | 0.49068 | REACTOME_O_LINKED_GLYCOSYLATION | 0.742875 | 0.811345 | 1 | REACTOME_O_LINKED_GI | 1.295996 | 0.157672 | 0.39348 |
| REACTOME_CLASS_I_PE | 1.284837 | 0.158656 | 0.757483 | REACTOME_CLASS_I_PEROXISOMAL_MEMBRANE_PRC | 0.712582 | 0.835988 | 1 | REACTOME_CLASS_I_PER | 1.290896 | 0.167965 | 0.399583 |
| REACTOME_MITOPHAG | 0.665902 | 0.913854 | 1 | REACTOME_MITOPHAGY | -0.77519 | 0.792098 | 1 | REACTOME_MITOPHAGY | -1.13281 | 0.276153 | 0.403425 |
| BIOCARTA_RACCYCD_P | 0.958567 | 0.497833 | 1 | BIOCARTA_RACCYCD_PATHWAY | 0.585192 | 0.950056 | 1 | BIOCARTA_RACCYCD_PAT | -1.12771 | 0.297004 | 0.409784 |
| BIOCARTA_EDG1_PATH | 1.275542 | 0.175379 | 0.75578 | BIOCARTA_EDG1_PATHWAY | 0.486495 | 0.98673 | 1 | BIOCARTA_EDG1_PATHW | -1.12229 | 0.306901 | 0.415382 |
| REACTOME_SIGNALING_ | 1.397431 | 0.105771 | 0.607538 | REACTOME_SIGNALING_BY_NOTCH | 0.856142 | 0.649106 | 1 | REACTOME_SIGNALING_E | 1.27117 | 0.181367 | 0.417662 |
| REACTOME_OVARIAN_T | 0.812478 | 0.713546 | 1 | REACTOME_OVARIAN_TUMOR_DOMAIN_PROTEASES | 0.735224 | 0.812781 | 1 | REACTOME_OVARIAN_TU | -1.11623 | 0.313715 | 0.423655 |

| | | | | | | | | | | | |
|---|---|---|---|---|---|---|---|---|---|---|---|
| BIOCARTA_PAR1_PATHV | 0.61563 | 0.927598 | 1 | BIOCARTA_PAR1_PATHWAY | 0.519022 | 0.977162 | 1 | BIOCARTA_PAR1_PATHW. | -1.11529 | 0.311541 | 0.423987 |
| BIOCARTA_MTOR_PATH | -0.68381 | 0.873654 | 1 | BIOCARTA_MTOR_PATHWAY | 0.493782 | 0.985015 | 1 | BIOCARTA_MTOR_PATHW | -1.10187 | 0.323049 | 0.441936 |
| WP_IL2_SIGNALING_PA | -1.00531 | 0.443154 | 1 | WP_IL2_SIGNALING_PATHWAY | 0.697745 | 0.922421 | 1 | WP_IL2_SIGNALING_PATH | -1.10022 | 0.288703 | 0.443631 |
| REACTOME_GENE_EXPR | -1.24182 | 0.026551 | 0.846214 | REACTOME_GENE_EXPRESSION_TRANSCRIPTION | 0.903882 | 0.753236 | 1 | REACTOME_GENE_EXPRE | 1.245406 | 0.029905 | 0.4506 |
| REACTOME_ABC_FAMIL | -0.81265 | 0.82285 | 1 | REACTOME_ABC_FAMILY_PROTEINS_MEDIATED_TRAN | 0.801231 | 0.808822 | 1 | REACTOME_ABC_FAMILY_ | 1.247527 | 0.136062 | 0.450789 |
| REACTOME_NEGATIVE_ | -0.84599 | 0.673304 | 1 | REACTOME_NEGATIVE_REGULATION_OF_MET_ACTIVIT | -0.57806 | 0.96124 | 1 | REACTOME_NEGATIVE_RI | -1.08731 | 0.345117 | 0.464114 |
| BIOCARTA_FMLP_PATH\ | 1.075289 | 0.359103 | 0.941757 | BIOCARTA_FMLP_PATHWAY | -0.76525 | 0.79535 | 1 | BIOCARTA_FMLP_PATHW | -1.08745 | 0.339108 | 0.465317 |
| REACTOME_COPI_INDEF | 0.958992 | 0.510417 | 1 | REACTOME_COPI_INDEPENDENT_GOLGI_TO_ER_RETR | 0.69041 | 0.906682 | 1 | REACTOME_COPI_INDEPE | -1.08153 | 0.315293 | 0.469495 |
| REACTOME_PROGRAMN | 0.84881 | 0.773359 | 1 | REACTOME_PROGRAMMED_CELL_DEATH | 0.893691 | 0.659248 | 1 | REACTOME_PROGRAMMI | -1.07794 | 0.283849 | 0.473545 |
| REACTOME_MYD88_MA | -1.42943 | 0.043285 | 0.697046 | REACTOME_MYD88_MAL_TIRAP_CASCADE_INITIATED_ | 0.832045 | 0.740367 | 1 | REACTOME_MYD88_MAL | -1.0786 | 0.305107 | 0.473745 |
| REACTOME_REGULATIO | 0.799841 | 0.846374 | 1 | REACTOME_REGULATION_OF_MRNA_STABILITY_BY_P | 0.658976 | 0.965052 | 1 | REACTOME_REGULATION | -1.07517 | 0.298898 | 0.475985 |
| REACTOME_G_ALPHA_1 | -0.95733 | 0.528676 | 1 | REACTOME_G_ALPHA_12_13_SIGNALLING_EVENTS | 0.749896 | 0.862378 | 1 | REACTOME_G_ALPHA_12 | -1.07525 | 0.32424 | 0.477327 |
| REACTOME_TOLL_LIKE_ | -1.23646 | 0.139099 | 0.843758 | REACTOME_TOLL_LIKE_RECEPTOR_4_TLR4_CASCADE | 0.879135 | 0.67481 | 1 | REACTOME_TOLL_LIKE_RI | -1.07124 | 0.311033 | 0.482438 |
| BIOCARTA_TFF_PATHW | 0.733245 | 0.811679 | 1 | BIOCARTA_TFF_PATHWAY | -0.69996 | 0.842602 | 1 | BIOCARTA_TFF_PATHWAY | -1.06807 | 0.372052 | 0.487271 |
| REACTOME_POST_TRAN | 1.770477 | 0.007579 | 0.11617 | REACTOME_POST_TRANSLATIONAL_MODIFICATION_S | -0.57346 | 0.982117 | 1 | REACTOME_POST_TRANS | 1.217399 | 0.205628 | 0.49631 |
| REACTOME_INTRINSIC_I | 0.662432 | 0.92662 | 1 | REACTOME_INTRINSIC_PATHWAY_FOR_APOPTOSIS | -0.79631 | 0.783378 | 1 | REACTOME_INTRINSIC_P/ | -1.05571 | 0.363253 | 0.507848 |
| REACTOME_TOLL_LIKE_ | -1.40472 | 0.049913 | 0.714341 | REACTOME_TOLL_LIKE_RECEPTOR_9_TLR9_CASCADE | 0.797737 | 0.789871 | 1 | REACTOME_TOLL_LIKE_RI | -1.05335 | 0.347826 | 0.511161 |
| REACTOME_RHO_GTPA | -0.8255 | 0.940907 | 1 | REACTOME_RHO_GTPASE_CYCLE | 0.612749 | 1 | 1 | REACTOME_RHO_GTPASE | -1.0476 | 0.293617 | 0.518517 |
| REACTOME_PEPTIDE_H | 1.779025 | 0.007021 | 0.114038 | REACTOME_PEPTIDE_HORMONE_METABOLISM | -0.7954 | 0.791139 | 1 | REACTOME_PEPTIDE_HOI | 1.197426 | 0.220616 | 0.518664 |
| REACTOME_SIGNALING_ | 0.538257 | 0.991691 | 1 | REACTOME_SIGNALING_BY_FGFR1 | -0.75265 | 0.845826 | 1 | REACTOME_SIGNALING_E | -1.04679 | 0.379272 | 0.518669 |
| REACTOME_JNK_C_JUN_ | -1.63817 | 0.020946 | 0.487219 | REACTOME_JNK_C_JUN_KINASES_PHOSPHORYLATION_ | 0.898144 | 0.599178 | 1 | REACTOME_JNK_C_JUN_I | -1.04477 | 0.397533 | 0.519789 |
| REACTOME_DEADENYLA | -1.2777 | 0.169881 | 0.872871 | REACTOME_DEADENYLATION_OF_MRNA | 0.894432 | 0.607015 | 1 | REACTOME_DEADENYLAT | -1.04525 | 0.395474 | 0.520329 |
| REACTOME_FATTY_ACY | 1.211891 | 0.217904 | 0.789754 | REACTOME_FATTY_ACYL_COA_BIOSYNTHESIS | 0.763411 | 0.784221 | 1 | REACTOME_FATTY_ACYL_ | 1.190089 | 0.240605 | 0.521112 |
| WP_ERBB_SIGNALING_F | -0.97145 | 0.485866 | 1 | WP_ERBB_SIGNALING_PATHWAY | 0.651182 | 0.921645 | 1 | WP_ERBB_SIGNALING_PA | -1.03687 | 0.388812 | 0.530007 |
| REACTOME_SEMAPHOR | 0.780956 | 0.825227 | 1 | REACTOME_SEMAPHORIN_INTERACTIONS | 0.577385 | 0.9835 | 1 | REACTOME_SEMAPHORIN | -1.02168 | 0.410914 | 0.557577 |
| WP_FOCAL_ADHESION_ | 0.614571 | 0.999493 | 1 | WP_FOCAL_ADHESION_PI3KAKTMTOR_SIGNALING_PA | -0.83257 | 0.904 | 1 | WP_FOCAL_ADHESION_P | -1.02016 | 0.402338 | 0.559192 |
| BIOCARTA_RHO_PATHW | -0.57817 | 0.95544 | 1 | BIOCARTA_RHO_PATHWAY | -0.66625 | 0.895495 | 1 | BIOCARTA_RHO_PATHWA | -1.01566 | 0.429597 | 0.562368 |
| REACTOME_AMINO_AC | -1.25182 | 0.159001 | 0.883273 | REACTOME_AMINO_ACIDS_REGULATE_MTORC1 | 0.86314 | 0.675255 | 1 | REACTOME_AMINO_ACID | -1.01169 | 0.424501 | 0.564411 |
| REACTOME_ROS_AND_I | 1.332718 | 0.139836 | 0.683336 | REACTOME_ROS_AND_RNS_PRODUCTION_IN_PHAGO | 0.937184 | 0.549936 | 1 | REACTOME_ROS_AND_RN | -1.01077 | 0.444217 | 0.564828 |
| REACTOME_CLASS_B_2_ | -1.03292 | 0.412817 | 1 | REACTOME_CLASS_B_2_SECRETIN_FAMILY_RECEPTOR | 0.662441 | 0.900157 | 1 | REACTOME_CLASS_B_2_S | -1.01195 | 0.428649 | 0.567076 |
| BIOCARTA_IGF1MTOR_I | 0.488188 | 0.985504 | 1 | BIOCARTA_IGF1MTOR_PATHWAY | -0.45347 | 0.995313 | 1 | BIOCARTA_IGF1MTOR_PA | -1.00815 | 0.441681 | 0.568926 |
| REACTOME_AZATHIOPR | 0.760074 | 0.793006 | 1 | REACTOME_AZATHIOPRINE_ADME | 0.654167 | 0.899711 | 1 | REACTOME_AZATHIOPRIN | -0.9974 | 0.455051 | 0.587245 |
| REACTOME_SIGNALING_ | -1.11263 | 0.205194 | 1 | REACTOME_SIGNALING_BY_GPCR | 0.831913 | 0.855422 | 1 | REACTOME_SIGNALING_E | -0.9963 | 0.482736 | 0.588025 |
| WP_ESC_PLURIPOTENC | -0.62228 | 0.97705 | 1 | WP_ESC_PLURIPOTENCY_PATHWAYS | 0.792406 | 0.795305 | 1 | WP_ESC_PLURIPOTENCY_ | -0.98876 | 0.466646 | 0.598053 |
| REACTOME_SIGNALING_ | 0.734178 | 0.95628 | 1 | REACTOME_SIGNALING_BY_NUCLEAR_RECEPTORS | 0.835573 | 0.77132 | 1 | REACTOME_SIGNALING_E | -0.97641 | 0.509331 | 0.616921 |
| REACTOME_EPH_EPHRII | 0.795049 | 0.751126 | 1 | REACTOME_EPH_EPHRIN_MEDIATED_REPULSION_OF_ | 0.792514 | 0.739489 | 1 | REACTOME_EPH_EPHRIN_ | -0.97712 | 0.478637 | 0.617024 |
| REACTOME_FACTORS_II | -0.98712 | 0.474034 | 1 | REACTOME_FACTORS_INVOLVED_IN_MEGAKARYOCYT | 0.93572 | 0.572702 | 1 | REACTOME_FACTORS_IN\ | -0.97403 | 0.50766 | 0.620498 |
| REACTOME_MEMBRANI | 0.893472 | 0.86209 | 1 | REACTOME_MEMBRANE_TRAFFICKING | 0.654551 | 0.999567 | 1 | REACTOME_MEMBRANE_ | -0.97093 | 0.613187 | 0.625745 |
| REACTOME_CELLULAR_I | 1.103424 | 0.250687 | 0.898311 | REACTOME_CELLULAR_RESPONSE_TO_CHEMICAL_STR | 0.858674 | 0.742027 | 1 | REACTOME_CELLULAR_RE | -0.96183 | 0.557443 | 0.644237 |
| REACTOME_DEUBIQUITI | 0.779774 | 0.951933 | 1 | REACTOME_DEUBIQUITINATION | 0.816254 | 0.839292 | 1 | REACTOME_DEUBIQUITIN | -0.94873 | 0.612352 | 0.668376 |
| REACTOME_DOWNREGU | -0.8264 | 0.688312 | 1 | REACTOME_DOWNREGULATION_OF_ERBB2_SIGNALIN | 0.659736 | 0.884238 | 1 | REACTOME_DOWNREGUI | -0.94371 | 0.529382 | 0.677723 |
| REACTOME_G_ALPHA_S | -1.17327 | 0.225289 | 0.894268 | REACTOME_G_ALPHA_S_SIGNALLING_EVENTS | -0.62753 | 0.985388 | 1 | REACTOME_G_ALPHA_S_ | -0.93671 | 0.567228 | 0.689504 |
| REACTOME_IRS_MEDIA | -0.78002 | 0.76485 | 1 | REACTOME_IRS_MEDIATED_SIGNALLING | 0.850168 | 0.66235 | 1 | REACTOME_IRS_MEDIATE | -0.93507 | 0.54215 | 0.691401 |
| REACTOME_DEADENYLA | 0.663959 | 0.96418 | 1 | REACTOME_DEADENYLATION_DEPENDENT_MRNA_DE | 0.836115 | 0.730554 | 1 | REACTOME_DEADENYLAT | 1.100959 | 0.322884 | 0.694713 |
| REACTOME_ADRENALIN | -1.05837 | 0.382358 | 1 | REACTOME_ADRENALINE_NORADRENALINE_INHIBITS_ | -0.75381 | 0.809711 | 1 | REACTOME_ADRENALINE_ | -0.93085 | 0.555613 | 0.695179 |
| REACTOME_VESICLE_MI | 0.98409 | 0.512284 | 1 | REACTOME_VESICLE_MEDIATED_TRANSPORT | 0.725017 | 0.993954 | 1 | REACTOME_VESICLE_MEI | -0.9312 | 0.790888 | 0.698107 |
| REACTOME_SYNTHESIS_ | 0.566111 | 0.989633 | 1 | REACTOME_SYNTHESIS_OF_PIPS_AT_THE_PLASMA_M | 0.604748 | 0.965262 | 1 | REACTOME_SYNTHESIS_C | -0.92511 | 0.574978 | 0.698745 |
| REACTOME_TRANSCRIP | -1.31769 | 0.074579 | 0.825776 | REACTOME_TRANSCRIPTIONAL_REGULATION_BY_RUN | 0.822318 | 0.787643 | 1 | REACTOME_TRANSCRIPTI | -0.92157 | 0.639084 | 0.70469 |
| REACTOME_INSULIN_RE | -0.65425 | 0.908101 | 1 | REACTOME_INSULIN_RECEPTOR_SIGNALLING_CASCAD | 0.721648 | 0.840877 | 1 | REACTOME_INSULIN_REC | -0.91897 | 0.571967 | 0.708505 |
| REACTOME_SIGNALING_ | -0.76204 | 0.787351 | 1 | REACTOME_SIGNALING_BY_TYPE_1_INSULIN_LIKE_GR | 0.845631 | 0.66456 | 1 | REACTOME_SIGNALING_E | -0.91667 | 0.58292 | 0.709885 |
| REACTOME_APOPTOSIS | 0.691634 | 0.96478 | 1 | REACTOME_APOPTOSIS | 0.897236 | 0.633785 | 1 | REACTOME_APOPTOSIS | -0.91408 | 0.642932 | 0.713686 |
| REACTOME_FLT3_SIGNA | 1.217079 | 0.219726 | 0.799663 | REACTOME_FLT3_SIGNALING | 0.700442 | 0.855327 | 1 | REACTOME_FLT3_SIGNAL | -0.90843 | 0.583333 | 0.722156 |
| REACTOME_GPER1_SIGI | -0.70047 | 0.867439 | 1 | REACTOME_GPER1_SIGNALING | 0.456204 | 0.993729 | 1 | REACTOME_GPER1_SIGN/ | -0.90688 | 0.589895 | 0.723571 |
| REACTOME_HOMOLOG | -1.57334 | 0.023081 | 0.618706 | REACTOME_HOMOLOGY_DIRECTED_REPAIR | 0.794416 | 0.756502 | 1 | REACTOME_HOMOLOGY_ | -0.90314 | 0.603709 | 0.727712 |
| REACTOME_G_ALPHA_I_ | 0.959551 | 0.540319 | 1 | REACTOME_G_ALPHA_I_SIGNALLING_EVENTS | 0.731146 | 0.931733 | 1 | REACTOME_G_ALPHA_I_S | -0.90325 | 0.719678 | 0.729317 |
| REACTOME_HSF1_DEPE | -1.00568 | 0.449871 | 1 | REACTOME_HSF1_DEPENDENT_TRANSACTIVATION | -0.70338 | 0.883206 | 1 | REACTOME_HSF1_DEPEN | -0.87351 | 0.636506 | 0.774397 |
| REACTOME_SIGNALING_ | 0.869032 | 0.665614 | 1 | REACTOME_SIGNALING_BY_INSULIN_RECEPTOR | 0.815699 | 0.746641 | 1 | REACTOME_SIGNALING_E | -0.87434 | 0.682678 | 0.774613 |
| REACTOME_PLASMA_LII | 0.864766 | 0.671973 | 1 | REACTOME_PLASMA_LIPOPROTEIN_ASSEMBLY_REMO | 0.91241 | 0.588587 | 1 | REACTOME_PLASMA_LIP( | -0.86729 | 0.678293 | 0.784994 |
| REACTOME_ASPARAGIN | 1.825187 | 0 | 0.090367 | REACTOME_ASPARAGINE_N_LINKED_GLYCOSYLATION | 0.784616 | 0.910182 | 1 | REACTOME_ASPARAGINE_ | 1.054548 | 0.349803 | 0.790148 |
| REACTOME_OXIDATIVE_ | -1.34215 | 0.1223 | 0.807455 | REACTOME_OXIDATIVE_STRESS_INDUCED_SENESCENC | 0.749913 | 0.8022 | 1 | REACTOME_OXIDATIVE_S | -0.86283 | 0.656399 | 0.791924 |
| REACTOME_METABOLIS | 1.67045 | 0 | 0.174818 | REACTOME_METABOLISM_OF_LIPIDS | 0.885565 | 0.778218 | 1 | REACTOME_METABOLISM | 1.048384 | 0.349063 | 0.794816 |
| WP_KIT_RECEPTOR_SIG | 0.662188 | 0.955632 | 1 | WP_KIT_RECEPTOR_SIGNALING_PATHWAY | 0.731171 | 0.862672 | 1 | WP_KIT_RECEPTOR_SIGN. | -0.85925 | 0.71634 | 0.794985 |
| REACTOME_BIOSYNTHE | 1.448956 | 0.054385 | 0.488764 | REACTOME_BIOSYNTHESIS_OF_THE_N_GLYCAN_PRECI | -0.85651 | 0.719751 | 1 | REACTOME_BIOSYNTHESI | 1.041261 | 0.397223 | 0.795801 |
| REACTOME_ANTIGEN_A | -0.92419 | 0.559495 | 1 | REACTOME_ANTIGEN_ACTIVATES_B_CELL_RECEPTOR_ | 0.749067 | 0.784307 | 1 | REACTOME_ANTIGEN_AC | -0.85497 | 0.660247 | 0.797607 |
| REACTOME_MITOCHON | 0.835358 | 0.790754 | 1 | REACTOME_MITOCHONDRIAL_TRANSLATION | -0.64565 | 0.991092 | 1 | REACTOME_MITOCHOND | -0.84837 | 0.809595 | 0.808223 |
| REACTOME_ESR_MEDIA | -0.67007 | 0.974098 | 1 | REACTOME_ESR_MEDIATED_SIGNALING | 0.909546 | 0.613593 | 1 | REACTOME_ESR_MEDIAT | -0.8446 | 0.789397 | 0.81339 |
| REACTOME_PROTEIN_L | 1.947594 | 2.37E-04 | 0.037819 | REACTOME_PROTEIN_LOCALIZATION | -0.67912 | 0.977609 | 1 | REACTOME_PROTEIN_LO | 1.017344 | 0.42644 | 0.819944 |
| REACTOME_RND2_GTP/ | 0.962967 | 0.492257 | 1 | REACTOME_RND2_GTPASE_CYCLE | -0.8379 | 0.732382 | 1 | REACTOME_RND2_GTPAS | -0.83658 | 0.731771 | 0.822237 |
| BIOCARTA_CERAMIDE_F | 0.816464 | 0.700258 | 1 | BIOCARTA_CERAMIDE_PATHWAY | 0.724536 | 0.827392 | 1 | BIOCARTA_CERAMIDE_PA | -0.83338 | 0.701849 | 0.826033 |
| REACTOME_DOWNSTRE | -0.64599 | 0.910297 | 1 | REACTOME_DOWNSTREAM_SIGNALING_OF_ACTIVATE | 0.835529 | 0.678056 | 1 | REACTOME_DOWNSTREA | -0.82818 | 0.696764 | 0.833478 |
| REACTOME_PEROXISON | 1.918114 | 0.001362 | 0.047512 | REACTOME_PEROXISOMAL_PROTEIN_IMPORT | 0.622917 | 0.963703 | 1 | REACTOME_PEROXISOMA | 1.009653 | 0.443082 | 0.835504 |
| REACTOME_ACTIVATION | -0.58064 | 0.97785 | 1 | REACTOME_ACTIVATION_OF_NMDA_RECEPTORS_AND | -0.66304 | 0.943287 | 1 | REACTOME_ACTIVATION_ | -0.81739 | 0.760826 | 0.844306 |
| REACTOME_RAC1_GTPA | -1.07163 | 0.31495 | 1 | REACTOME_RAC1_GTPASE_CYCLE | 0.899183 | 0.662084 | 1 | REACTOME_RAC1_GTPAS | -0.81924 | 0.900223 | 0.845122 |
| WP_EXERCISEINDUCED_ | -0.99247 | 0.461354 | 1 | WP_EXERCISEINDUCED_CIRCADIAN_REGULATION | 0.710951 | 0.862481 | 1 | WP_EXERCISEINDUCED_C | -0.79486 | 0.785115 | 0.868816 |
| WP_ALZHEIMERS_DISEA | -0.86644 | 0.679635 | 1 | WP_ALZHEIMERS_DISEASE | -0.78718 | 0.853846 | 1 | WP_ALZHEIMERS_DISEAS | -0.79687 | 0.83946 | 0.869791 |
| WP_IL1_SIGNALING_PA | 0.590219 | 0.96772 | 1 | WP_IL1_SIGNALING_PATHWAY | 0.928748 | 0.56356 | 1 | WP_IL1_SIGNALING_PATH | 0.983262 | 0.486524 | 0.872189 |
| REACTOME_EPIGENETIC | -1.18395 | 0.239603 | 0.881324 | REACTOME_EPIGENETIC_REGULATION_OF_GENE_EXPI | 0.836055 | 0.687314 | 1 | REACTOME_EPIGENETIC_ | 0.984991 | 0.486141 | 0.873727 |
| BIOCARTA_INTEGRIN_P/ | 0.453646 | 0.999335 | 0.997659 | BIOCARTA_INTEGRIN_PATHWAY | -0.68948 | 0.923845 | 1 | BIOCARTA_INTEGRIN_PAT | -0.77945 | 0.807081 | 0.885846 |
| REACTOME_SENESCENC | -0.85404 | 0.662283 | 1 | REACTOME_SENESCENCE_ASSOCIATED_SECRETORY_PI | 0.755204 | 0.796285 | 1 | REACTOME_SENESCENCE_ | -0.77754 | 0.78916 | 0.886542 |
| WP_SIGNAL_TRANSDUC | 0.847515 | 0.654997 | 1 | WP_SIGNAL_TRANSDUCTION_OF_S1P_RECEPTOR | -0.74859 | 0.80602 | 1 | WP_SIGNAL_TRANSDUCT | -0.7631 | 0.783957 | 0.900188 |
| REACTOME_PI_METABC | 0.753828 | 0.892157 | 1 | REACTOME_PI_METABOLISM | -0.80315 | 0.83666 | 1 | REACTOME_PI_METABOL | -0.76411 | 0.900968 | 0.900933 |
| REACTOME_GLUCAGON | 0.694586 | 0.871253 | 1 | REACTOME_GLUCAGON_TYPE_LIGAND_RECEPTORS | 0.5288 | 0.972702 | 1 | REACTOME_GLUCAGON_ | -0.73987 | 0.817387 | 0.910515 |
| WP_GPCRS_OTHER | -1.88948 | 0.002629 | 0.292559 | WP_GPCRS_OTHER | -0.83896 | 0.683311 | 1 | WP_GPCRS_OTHER | -0.74151 | 0.815938 | 0.910657 |
| BIOCARTA_ERK_PATHW | 0.887486 | 0.611882 | 1 | BIOCARTA_ERK_PATHWAY | 0.85884 | 0.645648 | 1 | BIOCARTA_ERK_PATHWA | -0.74793 | 0.813103 | 0.91134 |
| BIOCARTA_HIVNEF_PAT | -0.71711 | 0.886796 | 1 | BIOCARTA_HIVNEF_PATHWAY | 0.933615 | 0.56039 | 1 | BIOCARTA_HIVNEF_PATH | -0.73668 | 0.884138 | 0.912025 |
| BIOCARTA_STRESS_PATI | 0.939492 | 0.529336 | 1 | BIOCARTA_STRESS_PATHWAY | 0.664906 | 0.880526 | 1 | BIOCARTA_STRESS_PATH\ | -0.74161 | 0.815296 | 0.912617 |
| REACTOME_COPI_DEPEI | 1.176635 | 0.192117 | 0.835334 | REACTOME_COPI_DEPENDENT_GOLGI_TO_ER_RETRO( | 0.699993 | 0.927912 | 1 | REACTOME_COPI_DEPENI | 0.947428 | 0.544796 | 0.914793 |
| BIOCARTA_AKT_PATHW | -1.22002 | 0.23324 | 0.864733 | BIOCARTA_AKT_PATHWAY | -0.51006 | 0.978822 | 1 | BIOCARTA_AKT_PATHWA | -0.72605 | 0.832776 | 0.91953 |
| REACTOME_TRANSCRIP | -0.87446 | 0.764563 | 1 | REACTOME_TRANSCRIPTIONAL_REGULATION_BY_TP5 | 0.897931 | 0.688588 | 1 | REACTOME_TRANSCRIPTI | 0.947453 | 0.582533 | 0.921043 |
| REACTOME_NUCLEOTID | 1.206528 | 0.216338 | 0.790304 | REACTOME_NUCLEOTIDE_CATABOLISM | 0.69031 | 0.876547 | 1 | REACTOME_NUCLEOTIDE_ | -0.71619 | 0.875169 | 0.927627 |
| REACTOME_PLATELET_S | 0.707338 | 0.842071 | 1 | REACTOME_PLATELET_SENSITIZATION_BY_LDL | 0.928353 | 0.556082 | 1 | REACTOME_PLATELET_SE | -0.69373 | 0.872212 | 0.940027 |
| REACTOME_PHOSPHOLI | 1.218761 | 0.111765 | 0.813512 | REACTOME_PHOSPHOLIPID_METABOLISM | 0.659726 | 0.986449 | 1 | REACTOME_PHOSPHOLIP | -0.6834 | 0.995502 | 0.946341 |
| REACTOME_TRANSPORT | 1.721794 | 2.50E-04 | 0.150822 | REACTOME_TRANSPORT_TO_THE_GOLGI_AND_SUBSE | 0.855966 | 0.754569 | 1 | REACTOME_TRANSPORT_ | 0.923158 | 0.621305 | 0.955112 |
| REACTOME_VASOPRESS | -0.67041 | 0.895612 | 1 | REACTOME_VASOPRESSIN_REGULATES_RENAL_WATEI | -0.59649 | 0.96229 | 1 | REACTOME_VASOPRESSIN | -0.64664 | 0.93526 | 0.964917 |
| REACTOME_ACTIVATED_ | -1.13355 | 0.306494 | 0.966875 | REACTOME_ACTIVATED_TAK1_MEDIATES_P38_MAPK_ | 0.599396 | 0.937058 | 1 | REACTOME_ACTIVATED_T | 0.916561 | 0.569241 | 0.966461 |
| REACTOME_HEDGEHOG | 1.22715 | 0.165667 | 0.792827 | REACTOME_HEDGEHOG_LIGAND_BIOGENESIS | 0.569414 | 0.986945 | 1 | REACTOME_HEDGEHOG_ | 0.887593 | 0.6487 | 0.966745 |
| REACTOME_TAK1_DEPE | -0.90471 | 0.59704 | 1 | REACTOME_TAK1_DEPENDENT_IKK_AND_NF_KAPPA_I | -0.57712 | 0.973132 | 1 | REACTOME_TAK1_DEPEN | 0.841204 | 0.68501 | 0.966897 |

| | | | | | | | | | | | |
|---|---|---|---|---|---|---|---|---|---|---|---|
| REACTOME_COPII_MED | 1.704452 | 0.005495 | 0.157944 | REACTOME_COPII_MEDIATED_VESICLE_TRANSPORT | 0.83454 | 0.734082 | 1 | REACTOME_COPII_MEDIA | 0.894392 | 0.638881 | 0.967864 |
| WP_ONECARBON_META | 1.075653 | 0.336114 | 0.947915 | WP_ONECARBON_METABOLISM_AND_RELATED_PATH | 0.914469 | 0.5825 | 1 | WP_ONECARBON_METAB | 0.869749 | 0.662529 | 0.969058 |
| REACTOME_ER_TO_GOL | 1.645782 | 0.001007 | 0.189551 | REACTOME_ER_TO_GOLGI_ANTEROGRADE_TRANSPOI | 0.691849 | 0.966799 | 1 | REACTOME_ER_TO_GOLG | 0.889005 | 0.693548 | 0.969243 |
| REACTOME_TRANSPORT | 1.049065 | 0.285003 | 0.973169 | REACTOME_TRANSPORT_OF_SMALL_MOLECULES | 0.831931 | 0.878675 | 1 | REACTOME_TRANSPORT_ | 0.847413 | 0.869318 | 0.969659 |
| REACTOME_AQUAPORII | -0.704 | 0.871324 | 1 | REACTOME_AQUAPORIN_MEDIATED_TRANSPORT | -0.54249 | 0.983514 | 1 | REACTOME_AQUAPORIN_ | -0.62276 | 0.948848 | 0.971118 |
| REACTOME_SYNTHESIS_ | 1.018962 | 0.422458 | 1 | REACTOME_SYNTHESIS_OF_PA | -0.65223 | 0.918659 | 1 | REACTOME_SYNTHESIS_C | -0.62695 | 0.9453 | 0.971269 |
| WP_GLUTATHIONE_ANI | -0.60892 | 0.955405 | 1 | WP_GLUTATHIONE_AND_ONECARBON_METABOLISM | 0.802779 | 0.742339 | 1 | WP_GLUTATHIONE_AND_ | -0.61613 | 0.95887 | 0.971939 |
| REACTOME_CELLULAR_! | -1.49942 | 0.027968 | 0.635944 | REACTOME_CELLULAR_SENESCENCE | 0.860634 | 0.688413 | 1 | REACTOME_CELLULAR_SE | 0.83357 | 0.735852 | 0.972397 |
| REACTOME_RESOLUTIO | -1.00339 | 0.447523 | 1 | REACTOME_RESOLUTION_OF_SISTER_CHROMATID_CC | 0.818873 | 0.777716 | 1 | REACTOME_RESOLUTION_ | 0.676691 | 0.951768 | 0.974272 |
| REACTOME_ION_TRANS | 0.906301 | 0.594697 | 1 | REACTOME_ION_TRANSPORT_BY_P_TYPE_ATPASES | -0.69066 | 0.924623 | 1 | REACTOME_ION_TRANSP | 0.869884 | 0.661445 | 0.974468 |
| REACTOME_PINK1_PRKI | -0.73636 | 0.813889 | 1 | REACTOME_PINK1_PRKN_MEDIATED_MITOPHAGY | 0.677344 | 0.880814 | 1 | REACTOME_PINK1_PRKN_ | 0.872016 | 0.632958 | 0.975158 |
| REACTOME_INTERFERO | -0.81505 | 0.753668 | 1 | REACTOME_INTERFERON_SIGNALING | 0.862555 | 0.668269 | 1 | REACTOME_INTERFERON_ | 0.900713 | 0.620411 | 0.976305 |
| WP_SELENIUM_METAB( | 0.750757 | 0.832454 | 1 | WP_SELENIUM_METABOLISM_SELENOPROTEINS | 0.893085 | 0.613935 | 1 | WP_SELENIUM_METABOL | 0.638211 | 0.943004 | 0.977245 |
| WP_IL9_SIGNALING_PA | 0.988441 | 0.465012 | 1 | WP_IL9_SIGNALING_PATHWAY | 0.787568 | 0.749183 | 1 | WP_IL9_SIGNALING_PATH | 0.687282 | 0.869451 | 0.977718 |
| REACTOME_STIMULI_SE | 1.409929 | 0.060496 | 0.580665 | REACTOME_STIMULI_SENSING_CHANNELS | 0.937469 | 0.551958 | 1 | REACTOME_STIMULI_SEN | 0.823269 | 0.742993 | 0.977719 |
| REACTOME_CYTOSOLIC_ | 0.907406 | 0.576857 | 1 | REACTOME_CYTOSOLIC_SENSORS_OF_PATHOGEN_AS! | 0.716382 | 0.836901 | 1 | REACTOME_CYTOSOLIC_S | 0.907058 | 0.58888 | 0.978799 |
| REACTOME_SIGNALING_ | 1.215813 | 0.227293 | 0.785162 | REACTOME_SIGNALING_BY_ROBO_RECEPTORS | 0.861533 | 0.645449 | 1 | REACTOME_SIGNALING_E | 0.694485 | 0.854187 | 0.979498 |
| REACTOME_ANTIVIRAL_ | -0.58489 | 0.96 | 1 | REACTOME_ANTIVIRAL_MECHANISM_BY_IFN_STIMUL | 0.812853 | 0.718562 | 1 | REACTOME_ANTIVIRAL_M | 0.698727 | 0.863307 | 0.983957 |
| REACTOME_COPI_MEDI | 1.469403 | 0.023969 | 0.480327 | REACTOME_COPI_MEDIATED_ANTEROGRADE_TRANSF | -0.81356 | 0.863077 | 1 | REACTOME_COPI_MEDIA | 0.811659 | 0.802108 | 0.984898 |
| REACTOME_SYNTHESIS_ | 0.582894 | 0.967884 | 1 | REACTOME_SYNTHESIS_OF_ACTIVE_UBIQUITIN_ROLES | 0.557625 | 0.970114 | 1 | REACTOME_SYNTHESIS_C | 0.808881 | 0.738462 | 0.985054 |
| REACTOME_GOLGI_TO_ | 1.1919 | 0.156766 | 0.806713 | REACTOME_GOLGI_TO_ER_RETROGRADE_TRANSPORT | 0.72053 | 0.932429 | 1 | REACTOME_GOLGI_TO_EI | 0.801376 | 0.839722 | 0.988925 |
| WP_TRYPTOPHAN_MET. | 1.458146 | 0.066595 | 0.474887 | WP_TRYPTOPHAN_METABOLISM | -0.46552 | 0.994694 | 1 | WP_TRYPTOPHAN_METAI | 0.597962 | 0.957227 | 0.989448 |
| REACTOME_DOWNREGU | 0.717505 | 0.822957 | 1 | REACTOME_DOWNREGULATION_OF_SMAD2_3_SMAD | 0.894068 | 0.602968 | 1 | REACTOME_DOWNREGUL | -0.5415 | 0.973286 | 0.990385 |
| BIOCARTA_PTEN_PATH\ | 0.622268 | 0.92369 | 1 | BIOCARTA_PTEN_PATHWAY | 0.587093 | 0.939997 | 1 | BIOCARTA_PTEN_PATHW. | -0.54946 | 0.968774 | 0.990907 |
| REACTOME_ENDOSOM/ | -1.30297 | 0.134104 | 0.824212 | REACTOME_ENDOSOMAL_SORTING_COMPLEX_REQUI | 0.647166 | 0.919247 | 1 | REACTOME_ENDOSOMAL | -0.50405 | 0.993648 | 0.993963 |
| REACTOME_REGULATIO | -1.0683 | 0.36259 | 1 | REACTOME_REGULATION_OF_PTEN_GENE_TRANSCRIF | -0.75114 | 0.847062 | 1 | REACTOME_REGULATION | 0.711686 | 0.879405 | 0.996873 |
| WP_GPCRS_NONODOR/ | -1.86001 | 3.49E-04 | 0.215451 | WP_GPCRS_NONODORANT | 0.843061 | 0.726362 | 1 | WP_GPCRS_NONODORAN | 0.706964 | 0.921464 | 0.997616 |
| REACTOME_SYNTHESIS_ | 1.326578 | 0.144666 | 0.680284 | REACTOME_SYNTHESIS_OF_BILE_ACIDS_AND_BILE_SA | 0.869389 | 0.637364 | 1 | REACTOME_SYNTHESIS_C | 0.741838 | 0.808047 | 1 |
| BIOCARTA_HER2_PATH\ | 1.292741 | 0.166415 | 0.740504 | BIOCARTA_HER2_PATHWAY | 0.715297 | 0.834839 | 1 | BIOCARTA_HER2_PATHW. | 0.759411 | 0.788547 | 1 |
| REACTOME_GLYCEROPH | 1.294218 | 0.0873 | 0.74712 | REACTOME_GLYCEROPHOSPHOLIPID_BIOSYNTHESIS | 0.837987 | 0.755058 | 1 | REACTOME_GLYCEROPHC | 0.726448 | 0.922798 | 1 |
| REACTOME_MITOTIC_PI | -0.78471 | 0.889772 | 1 | REACTOME_MITOTIC_PROMETAPHASE | 0.909981 | 0.636092 | 1 | REACTOME_MITOTIC_PR( | 0.716807 | 0.948887 | 1 |
| REACTOME_RHOBTB1_( | -0.9683 | 0.496678 | 1 | REACTOME_RHOBTB1_GTPASE_CYCLE | -0.80111 | 0.751704 | 1 | REACTOME_RHOBTB1_GT | 0.774298 | 0.772559 | 1 |
| WP_ONECARBON_META | -0.87882 | 0.632428 | 1 | WP_ONECARBON_METABOLISM | 0.893307 | 0.604425 | 1 | WP_ONECARBON_METAE | 0.775845 | 0.777939 | 1 |

**GSEA cluster**

** NbClust results. Method=ward.D2, distance=euclidean*

| | | NES | | |
|---|---|---|---|---|
| cluster | ID | SYP MNK1 | SYP MNK2 | SYP DKO |
| 1 | REACTOME_CELLULAR_RESPONSE_TO_HEAT_STRESS | -0.7214 | -1.57444 | 1.05994 |
| 1 | REACTOME_DETOXIFICATION_OF_REACTIVE_OXYGEN_SPECIES | 1.807249 | -0.90013 | 0.750724 |
| 1 | REACTOME_GENE_SILENCING_BY_RNA | -0.78141 | -1.70913 | 1.365919 |
| 1 | REACTOME_GLUCOSE_METABOLISM | 0.742212 | -2.03181 | 0.721595 |
| 1 | REACTOME_GLYCOLYSIS | 0.679522 | -1.95653 | 0.956136 |
| 1 | REACTOME_MITOTIC_PROPHASE | 0.72832 | -1.84208 | 1.204012 |
| 1 | REACTOME_MRNA_SPLICING | -1.55241 | -1.835 | 1.350378 |
| 1 | REACTOME_MRNA_SPLICING_MINOR_PATHWAY | -0.88051 | -2.03613 | 0.700826 |
| 1 | REACTOME_NUCLEAR_ENVELOPE_BREAKDOWN | -0.51275 | -2.18728 | 1.407661 |
| 1 | REACTOME_NUCLEAR_PORE_COMPLEX_NPC_DISASSEMBLY | -0.69559 | -2.16179 | 1.636719 |
| 1 | REACTOME_PEPTIDE_HORMONE_METABOLISM | 1.779025 | -0.7954 | 1.197426 |
| 1 | REACTOME_POST_TRANSLATIONAL_MODIFICATION_SYNTHESIS_( | 1.770477 | -0.57346 | 1.217399 |
| 1 | REACTOME_POSTMITOTIC_NUCLEAR_PORE_COMPLEX_NPC_REFC | -0.59096 | -1.99266 | 1.048955 |
| 1 | REACTOME_PROCESSING_OF_CAPPED_INTRON_CONTAINING_PRI | -1.28226 | -1.91038 | 1.692861 |
| 1 | REACTOME_PROTEIN_LOCALIZATION | 1.947594 | -0.67912 | 1.017344 |
| 1 | REACTOME_REGULATION_OF_HSF1_MEDIATED_HEAT_SHOCK_RE | -0.62035 | -1.82768 | 1.201697 |
| 1 | REACTOME_SNRNP_ASSEMBLY | 0.991794 | -1.92618 | 1.352834 |
| 1 | REACTOME_SPHINGOLIPID_DE_NOVO_BIOSYNTHESIS | 1.688962 | -1.03272 | 1.636504 |
| 1 | REACTOME_SUMOYLATION_OF_CHROMATIN_ORGANIZATION_PR | -0.94157 | -1.72061 | 1.847096 |
| 1 | REACTOME_SUMOYLATION_OF_DNA_REPLICATION_PROTEINS | -1.18072 | -2.01883 | 1.379512 |
| 1 | REACTOME_SUMOYLATION_OF_RNA_BINDING_PROTEINS | -0.99988 | -2.0389 | 1.71632 |
| 1 | REACTOME_SUMOYLATION_OF_SUMOYLATION_PROTEINS | -1.20708 | -2.03173 | 1.487596 |
| 1 | REACTOME_SUMOYLATION_OF_UBIQUITINYLATION_PROTEINS | -1.24275 | -1.92209 | 1.414128 |
| 1 | REACTOME_TRANSCRIPTIONAL_REGULATION_BY_SMALL_RNAS | -0.55007 | -2.15233 | 1.430609 |
| 1 | REACTOME_TRANSPORT_OF_MATURE_MRNAS_DERIVED_FROM_ | 0.777301 | -2.21124 | 1.833126 |
| 1 | REACTOME_TRANSPORT_OF_MATURE_TRANSCRIPT_TO_CYTOPLA | -0.88811 | -2.3427 | 1.651372 |
| 1 | REACTOME_TRANSPORT_OF_THE_SLBP_DEPENDANT_MATURE_M | -0.78723 | -2.2161 | 1.46113 |
| 1 | WP_GPCRS_NONODORANT | -1.86001 | 0.843061 | 0.706964 |
| 1 | WP_SPHINGOLIPID_METABOLISM_INTEGRATED_PATHWAY | 1.617602 | -1.09135 | 1.682079 |
| 1 | WP_SPHINGOLIPID_METABOLISM_OVERVIEW | 1.61796 | -1.0911 | 1.684823 |
| 2 | REACTOME_CHAPERONIN_MEDIATED_PROTEIN_FOLDING | 0.782386 | -1.5436 | -1.82633 |
| 2 | REACTOME_COMPLEX_I_BIOGENESIS | -1.26704 | -2.02219 | -2.03005 |
| 2 | REACTOME_FATTY_ACID_METABOLISM | 1.658859 | -1.00017 | -1.14545 |
| 2 | REACTOME_GLUTATHIONE_CONJUGATION | 1.255178 | -1.74353 | -1.55866 |
| 2 | REACTOME_NEUREXINS_AND_NEUROLIGINS | 0.519758 | -1.79752 | -2.25592 |
| 2 | REACTOME_POTASSIUM_CHANNELS | -1.03822 | -1.62591 | -1.19951 |
| 2 | REACTOME_RAB_GERANYLGERANYLATION | -0.91277 | -2.03501 | -1.48063 |
| 2 | REACTOME_RAB_REGULATION_OF_TRAFFICKING | -0.72139 | -1.65534 | -1.44862 |
| 2 | REACTOME_REGULATION_OF_INSULIN_SECRETION | -0.65621 | -1.66055 | -1.65025 |
| 2 | REACTOME_RESPIRATORY_ELECTRON_TRANSPORT | -0.93888 | -1.89568 | -1.84939 |
| 2 | REACTOME_RESPIRATORY_ELECTRON_TRANSPORT_ATP_SYNTHES | -0.84537 | -1.73265 | -1.79349 |
| 2 | REACTOME_RHOBTB2_GTPASE_CYCLE | 0.557364 | -1.70872 | -1.2933 |
| 2 | REACTOME_SIGNAL_AMPLIFICATION | 1.161491 | -1.53827 | -1.81031 |
| 2 | REACTOME_TBC_RABGAPS | 0.971685 | -1.64493 | -1.45702 |
| 2 | REACTOME_THE_CITRIC_ACID_TCA_CYCLE_AND_RESPIRATORY_EI | 0.921426 | -1.58813 | -1.94134 |
| 2 | REACTOME_TRAFFICKING_OF_AMPA_RECEPTORS | 1.165314 | -1.67256 | -1.48981 |
| 2 | REACTOME_TRAFFICKING_OF_GLUR2_CONTAINING_AMPA_RECEI | 1.685841 | -1.63804 | -1.62217 |

| | | | | |
|---|---|---|---|---|
| 2 | REACTOME_VOLTAGE_GATED_POTASSIUM_CHANNELS | -0.90878 | -1.70201 | -1.31662 |
| 2 | WP_ELECTRON_TRANSPORT_CHAIN | 0.834097 | -1.94647 | -2.06865 |
| 2 | WP_G_PROTEIN_SIGNALING_PATHWAYS | 0.998573 | -1.56746 | -2.13382 |
| 2 | WP_METAPATHWAY_BIOTRANSFORMATION | 1.873613 | -1.34878 | -1.0075 |
| 2 | WP_OXIDATIVE_PHOSPHORYLATION | -1.04895 | -1.79167 | -1.73451 |
| 2 | WP_TCA_CYCLE | 0.983286 | -1.99699 | -2.23446 |
| 3 | REACTOME_ACTIVATION_OF_THE_MRNA_UPON_BINDING_OF_TH | 2.220735 | 1.536282 | 2.117987 |
| 3 | REACTOME_ASPARAGINE_N_LINKED_GLYCOSYLATION | 1.825187 | 0.784616 | 1.054548 |
| 3 | REACTOME_CARGO_CONCENTRATION_IN_THE_ER | 1.74242 | 1.022906 | 1.435002 |
| 3 | REACTOME_CITRIC_ACID_CYCLE_TCA_CYCLE | 1.825698 | 1.359805 | -1.23579 |
| 3 | REACTOME_COMPLEMENT_CASCADE | 1.159981 | 1.949624 | 1.264958 |
| 3 | REACTOME_COPII_MEDIATED_VESICLE_TRANSPORT | 1.704452 | 0.83454 | 0.894392 |
| 3 | REACTOME_ER_TO_GOLGI_ANTEROGRADE_TRANSPORT | 1.645782 | 0.691849 | 0.889005 |
| 3 | REACTOME_EUKARYOTIC_TRANSLATION_INITIATION | 2.592438 | 1.661947 | 2.485513 |
| 3 | REACTOME_FORMATION_OF_A_POOL_OF_FREE_40S_SUBUNITS | 2.717268 | 1.624374 | 2.660923 |
| 3 | REACTOME_MAJOR_PATHWAY_OF_RRNA_PROCESSING_IN_THE_ | 1.760818 | 1.1176 | 2.142849 |
| 3 | REACTOME_METABOLISM_OF_LIPIDS | 1.67045 | 0.885565 | 1.048384 |
| 3 | REACTOME_NONSENSE_MEDIATED_DECAY_NMD | 2.432419 | 1.195268 | 2.327559 |
| 3 | REACTOME_NONSENSE_MEDIATED_DECAY_NMD_INDEPENDENT_ | 2.607835 | 1.464736 | 2.493042 |
| 3 | REACTOME_PEROXISOMAL_LIPID_METABOLISM | 1.867626 | 1.026402 | 1.274199 |
| 3 | REACTOME_PEROXISOMAL_PROTEIN_IMPORT | 1.918114 | 0.622917 | 1.009653 |
| 3 | REACTOME_SPHINGOLIPID_METABOLISM | 1.789229 | 0.823081 | 1.579735 |
| 3 | REACTOME_TRANSLATION | 2.198588 | 1.134119 | 1.811309 |
| 3 | REACTOME_TRANSPORT_TO_THE_GOLGI_AND_SUBSEQUENT_MC | 1.721794 | 0.855966 | 0.923158 |
| 3 | WP_CHOLESTEROL_METABOLISM_WITH_BLOCH_AND_KANDUTSC | 1.705891 | 0.980172 | 1.652451 |
| 3 | WP_CYTOPLASMIC_RIBOSOMAL_PROTEINS | 2.706107 | 1.413896 | 2.519593 |
| 3 | WP_OXIDATIVE_STRESS_RESPONSE | 1.663716 | 0.988468 | 1.128535 |
| 3 | WP_PPAR_SIGNALING_PATHWAY | 1.687301 | 1.143962 | 0.986777 |

**GSEA cellular component ribosomes**

* GSEA mouse MSigDB canonical pathways: m5.go.cc.v2022.1.Mm.symbols

| cellular component | | MNK1KO CX | | MNK2 CX | | MNKDKO CX | | MNK1KO SYP | | MNK2 SYP | | MNKDKO SYP | |
|---|---|---|---|---|---|---|---|---|---|---|---|---|---|
| NAME | SIZE | NES | FDR q-val | NES | FDR q-val | NES | FDR q-val | NES | FDR q-val | NES | FDR q-val | NES | FDR q-val |
| GOCC_RIBOSOME | 193 | 1.52142 | 0.304023 | 1.130573 | 0.72783 | 1.634043 | 0.425109 | 2.155775 | 4.12E-04 | 0.911822 | 1 | 1.823913 | 0.012035 |
| GOCC_RIBOSOMAL_SUBUNIT | 171 | 1.528584 | 0.304863 | 1.193163 | 0.633838 | 1.732 | 0.520912 | 2.283716 | 3.43E-05 | 0.881306 | 1 | 1.889939 | 0.005655 |
| GOCC_SMALL_RIBOSOMAL_SUBUNIT | 71 | 1.417562 | 0.351503 | 0.930775 | 0.944072 | 1.531242 | 0.418086 | 2.228482 | 5.83E-05 | 0.679602 | 1 | 1.4866 | 0.119646 |
| GOCC_LARGE_RIBOSOMAL_SUBUNIT | 104 | 1.409506 | 0.329721 | 1.231165 | 0.60338 | 1.584168 | 0.426088 | 1.976255 | 0.006237 | 1.081417 | 0.812605 | 1.862165 | 0.007765 |
| GOCC_CYTOSOLIC_RIBOSOME | 87 | 1.864964 | 0.047406 | 1.297749 | 0.584419 | 1.718933 | 0.382586 | 2.657334 | 0 | 1.218762 | 0.579024 | 2.446244 | 0 |
| GOCC_CYTOSOLIC_SMALL_RIBOSOMAL | 38 | 1.657538 | 0.188184 | 0.850063 | 0.996257 | 1.617309 | 0.39772 | 2.376504 | 0 | 0.89848 | 1 | 1.895957 | 0.005527 |
| GOCC_CYTOSOLIC_LARGE_RIBOSOMAL | 48 | 1.732056 | 0.135769 | 1.558418 | 0.320795 | 1.489458 | 0.412936 | 2.366335 | 0 | 1.397891 | 0.309473 | 2.422273 | 0 |
